# Transcriptome and histone epigenome of *Plasmodium vivax* salivary-gland sporozoites point to tight regulatory control and potential mechanisms for liver-stage differentiation

**DOI:** 10.1101/145250

**Authors:** Vivax Sporozoite Consortium, Ivo Muller, Aaron R. Jex, Stefan H. I. Kappe, Sebastian A. Mikolajczak, Jetsumon Sattabongkot, Rapatbhorn Patrapuvich, Scott Lindner, Erika L. Flannery, Cristian Koepfli, Brendan Ansell, Anita Lerch, Samantha J Emery-Corbin, Sarah Charnaud, Jeffrey Smith, Nicolas Merrienne, Kristian E. Swearingen, Robert L. Moritz, Michaela Petter, Michael Duffy, Vorada Chuenchob

## Abstract

*Plasmodium vivax* is the key obstacle to malaria elimination in Asia and Latin America, largely attributed to its ability to form resilient hypnozoites (sleeper-cells) in the host liver that escape treatment and cause relapsing infections. The decision to form hypnozoites is made early in the liver infection and may already be set in sporozoites prior to invasion. To better understand these early stages of infection, we undertook a comprehensive transcriptomic and histone epigenetic characterization of *P. vivax* sporozoites. The salivary-gland sporozoite transcriptome is heavily composed of transcripts associated with functions needed for early infection of the vertebrate host and development within hepatocytes. Through comparisons to recently published proteome data for the *P. vivax* sporozoite, our study finds that although highly transcribed, these transcripts are not detectable as proteins and may be regulated through translational repression; a finding we test for a small subset of transcripts and proteins through immunofluorescent microscopy of sporozoites and liver stages in humanized mice. We identify differential transcription between the sporozoite and published transcriptomes of asexual blood-stages and mixed versus hypnozoite-enriched liver stages. These comparisons point to multiple layers of transcriptional, post-transcriptional and post-translational control that appear active in sporozoites and to a lesser extent hypnozoites, but largely absent in replicating liver schizonts or mixed blood-stages. Common transcripts up-regulated in sporozoites and hypnozoites compared to mixed (i.e., schizont) liver-stages identify genes linked to dormancy/persistence in bacteria, amoebae and plants. We also characterise histone epigenetic modifications in the *P. vivax* sporozoite and explore their role in regulating transcription. Collectively, these data support the hypothesis that the sporozoite as a tightly programmed stage primed to infect the human host and identifies potential mechanisms for hypnozoite-formation that may be further explored in liver stage models.

## INTRODUCTION

Malaria is among the most significant infectious diseases impacting humans globally, with 3.3 billion people at risk of infection, 381 million suspected clinical cases and up to ~660,000 deaths attributed to malaria in 2014 [1]. Two major parasitic species contribute to the vast majority of human malaria, *Plasmodium falciparum* and *P. vivax*. Historically, *P. falciparum* has attracted the majority of global attention, due to its higher contribution to morbidity and mortality. However, *P. vivax* is broadly distributed, more pathogenic than previously thought, and is recognised as the key obstacle to malaria elimination in the Asia-Pacific and Americas [2]. Unlike *P. falciparum, P. vivax* can establish long-lasting ‘sleeper-cells’ (= hypnozoites) in the host liver that emerge weeks, months or years after the primary infection (= relapsing malaria) [3]. Primaquine is the only approved drug that prevents relapse. However, the short half-life, long dosage regimens and incompatibility of primaquine with glucose-6-phosphate-dehydrogenase deficiency (which requires pre-screening of recipients [4]) makes it unsuitable for widespread use. As a consequence, *P. vivax* is overtaking *P. falciparum* as the primary cause of malaria in a number of co-endemic regions [5]. Developing new tools to diagnose, treat and/or prevent hypnozoite infections is considered one of the highest priorities in the malaria elimination research agenda [6].

When *Plasmodium* sporozoites are deposited by an infected mosquito, they likely traverse the skin cells, enter the blood-stream and are trafficked to the host liver, as has been shown in rodents [7]. The sporozoites’ journey from skin deposition to hepatocytes takes less than a few minutes [8]. Upon reaching the liver, sporozoites traverse Kupffer and endothelial cells to reach the parenchyma, moving through several hepatocytes before invading a final hepatocyte suitable for development [7, 9]. Within hepatocytes, these parasites replicate, and undergo further development and differentiation to produce merozoites that emerge from the liver and infect red blood cells. However, *P. vivax* sporozoites are able to commit to two distinct developmental fates within the hepatocyte: they either immediately continue development as replicating schizonts and establish a blood infection, or delay replication and persist as hypnozoites. Regulation of this major developmental fate decision is not understood and this represents a key gap in current knowledge of *P. vivax* biology and control.

Sporozoites prepare for mammalian host infection while still residing in the mosquito salivary glands. It has been hypothesized that *P. vivax* sporozoites exist within an inoculum as replicating ‘tachysporozoites’ and relapsing ‘bradysporozoites’ [10] and that these subpopulations may have distinct developmental fates as schizonts or hypnozoites, thus contributing to their relapse phenotype [10-12]. This observation is supported by the stability of different hypnozoite phenotypes (ratios of hypnozoite to schizont formation) in *P. vivax* infections of liver-chimeric mouse models [13]. To determine fates in the sporozoite stage control of protein expression must take place. Studies using rodent malaria parasites have identified genes [14] that are transcribed in sporozoites but translationally repressed (i.e., present as transcript but un- or under-represented as protein), via RNA-binding proteins [15], and ready for immediate translation after the parasites’ infection of the mammalian host cell [13, 16]. It is therefore also possible that translational repression (i.e., the blocking of translation of present and retained transcripts) and other mechanisms of epigenetic control may contribute to the *P. vivax* sporozoite fate decision and hypnozoite formation, persistence and activation. Supporting this hypothesis, histone methyltransferase inhibitors stimulate increased activation of *P. cynomolgi* hypnozoites to become schizonts in macaque hepatocytes [17, 18]. Epigenetic control of stage development is further evidenced in *Plasmodium* through chromatin structure controlling expression of PfAP2-G, a specific transcription factor that, in turn, regulates gametocyte (dimorphic sexual stages) development in blood-stages [19]. It is well documented that *P. vivax* hypnozoite activation patterns stratify with climate and geography [11] and recent modelling suggests transmission potential selects for hypnozoite phenotype [20]. Clearly the ability for *P. vivax* to dynamically regulate hypnozoite formation and relapse phenotypes in response to high or low transmission periods in different climate conditions would confer a significant evolutionary advantage.

Unfortunately, despite recent advances [21] current approaches for *in vitro P. vivax* culture do not support routine maintenance in the laboratory and tools to directly perturb gene function are not established. This renders studies on *P. vivax*, particularly its sporozoites and liver stages, exceedingly difficult. Although *in-vitro* liver stage assays and humanised mouse models are being developed [13], ‘omics analysis of *P. vivax* liver stage dormancy has until recently [22] been impossible and even now is in its early stages. Recent characterization [23] of liver-stage (hypnozoites and schizonts) of *P. cynomolgi* (a related and relapsing parasite in macaques) provides valuable insight, but investigations in *P. vivax* directly are clearly needed. The systems analysis of *P. vivax* sporozoites that reside in the mosquito salivary glands and are poised for transmission and liver infection offer a key opportunity to gain insight into *P. vivax* infection. *Plasmodium vivax* sporozoites have been explored previously by microarray [24] and most recently, in a single RNA-seq replicate [25] and a study on sporozoite activation {Roth, 2018 #66}. Epigenetic regulation in sporozoites has only been explored in *P. falciparum* [26, 27]. Here, we present a detailed characterization of the *P. vivax* sporozoite transcriptome and histone epigenome and use these data to better understand this key infective stage and the role of sporozoite programming in invasion and infection of the human host, and development within the host liver.

## RESULTS AND DISCUSSION

Mosquito infections were generated by membrane feeding of blood samples taken from *P. vivax* infected patients in western Thailand (n = 9). Approximately 3-15 million *P. vivax* sporozoites were harvested per isolate from *Anopheles dirus* salivary glands. Using RNA-seq, we detected transcription for 5,714 *P. vivax* genes (based on the *P. vivax* P01 gene models: [28]) and obtained a high degree of coverage (4,930 with a mean counts per million (CPM) ≥ 1.0; Figure S1 and Table S1 and S2). Among the most highly transcribed genes in the infectious sporozoite stage are *csp* (<u>c</u>ircumsporozoite <u>p</u>rotein), five *etramps* (<u>e</u>arly <u>tr</u>anscribed <u>m</u>embrane <u>p</u>roteins), including *uis3* (<u>u</u>p-regulated in infective sporozoites), *uis4* and *lsap-1* (liver stage <u>a</u>ssociated <u>p</u>rotein 1), a variety of genes involved in cell transversal and initiation of invasion, including *celtos* (<u>cel</u>l traversal protein for <u>o</u>okinetes and sporozoites), *gest* (<u>g</u>amete <u>e</u>gress and <u>s</u>porozoite traversal protein), *spect1* (<u>s</u>porozoite <u>p</u>rotein <u>e</u>ssential for <u>c</u>ell traversal) and *siap-1* (<u>s</u>porozoite invasion <u>a</u>ssociated <u>p</u>rotein), and genes associated with translational repression (*alba1, alba4* and *Puf2*). Collectively, these genes account for >1/3^rd^ of all transcripts in the sporozoite. Although we found only moderate agreement (R^2^ = 0.35; Figure S2) between our RNA-seq data and previous microarray data for *P. vivax* sporozoites and blood-stages [24], improved transcript detection and quantitation is expected with the increased technical resolution of RNA-seq over microarray. Supporting this, we find higher correlation between RNA-seq data from *P. vivax* and *P. falciparum* (single replicate sequenced herein for comparative purposes) sporozoite datasets (R^2^ = 0.42), compared to either species relative to published microarray data (Figure S2 and Table S3).

Although microarray supports the high transcription in sporozoites of genes such as *uis4, csp, celtos* and several other *etramps*, 27% and 16% of the most abundant 1% of transcribed genes in our sporozoite RNA-seq data are absent from the top decile or quartile respectively in the existing *P. vivax* sporozoite microarray data [24]. Among these are genes involved in early invasion/hepatocyte development, such as *lsap-1, celtos, gest* and *siap-1*, or translational repression (e.g., *alba-1* and *alba-4*); orthologs of these genes are also in the top percentile of transcripts in RNA-seq (see [26, 29]) and previous microarray data [30, 31] for human-infecting *P. falciparum* and murine-infecting *P. yoelii* sporozoites, suggesting many are indeed more abundant than previously characterized. A subset of representative transcripts, including *Pv_AP2-X* (PVP01_0733100), *d13, gest, g10* (PVP01_1011100), 40S ribosomal protein S27 (PVP01_1409300), *puf-2, zipco* and 14-3-3, were tested by qPCR for their transcript abundance relative to *celtos* and *sera* (Figure 1A and Table S4). This representative set differed markedly in their relative abundance between our RNAseq and previous microarray data [24]. To control for batch effects introduced by collection of the sporozoites used here for RNAseq, this testing was conducted in an additional six sample replicates representing four additional clinical *P. vivax* isolates (PvSPZ-Thai13-16; with PvSPZ-Thai16 tested in technical triplicate). The qPCR results agreed with the RNAseq data for these transcripts (Table S4).

**Fig. 1.**
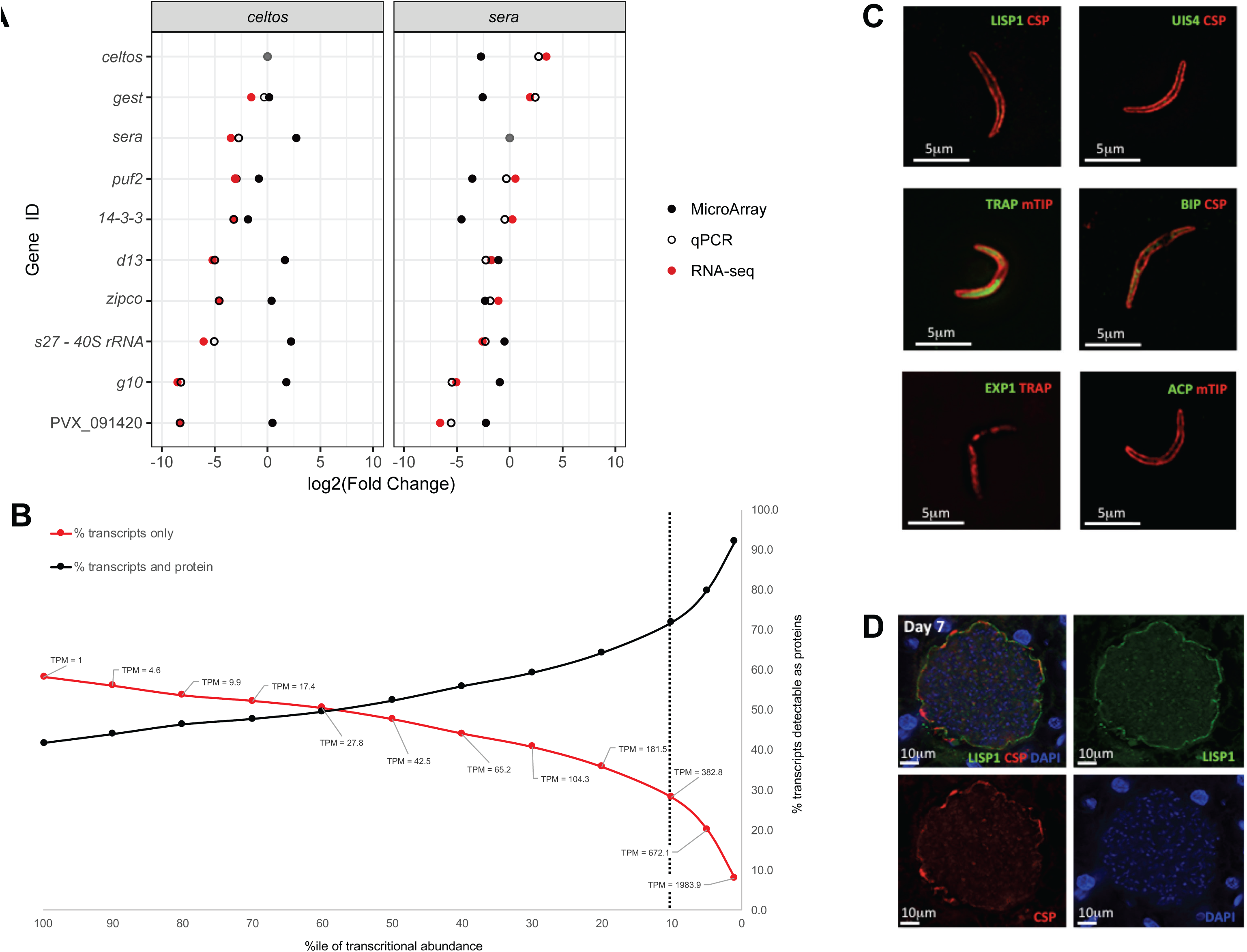
Transcriptional activity of the *P. vivax* sporozoite and evidence for translational repression. **a** Relative transcript abundance of key marker genes for sporozoites inferred by RNA-seq and qPCR (here) relative to previously published microarray data [24]; **b** Relative proportion of genes detectable as transcripts and proteins or transcripts only in RNA-seq and previously published proteomic data. Dashed line shows cut-off used in the current study for putatively repressed transcripts. Immunofluorescent staining of select proteins either known (UIS4) or predicted here (LISP1, EXP1 and ACP) to be translationally repressed in sporozoites in **c** sporozoite stages. CSP, mTIP as known positive controls and TRAP and BIP as exerpimental positive controls and **d** liver stages (schizonts) at 7 days post-infection in HuHep mice. Liver expression of EXP1 and ACP has been demonstrated by IFA in Mikolajczak et al [13], using the same antibodies as used here.

### *Transcription in* P. *vivax relative to other plasmodia* sporozoites

To gain insight into species-specific aspects of the *P.vivax* transcriptome, we qualitatively compared these data with available data for *P. falciparum* [27] and *P. yoelii* sporozoites (single replicate only) for 4,067 single-copy orthologs (SCO) (transcribed at ≥ 1 TPM in *P. vivax* infectious sporozoites) shared with *P. falciparum* and *P. yoelli* (Table S5). Genes highly transcribed in salivary-gland sporozoites of all three species include *celtos, gest, trap, siap1, spect1* and *puf2*. There are 696 *P. vivax* genes shared as orthologs between *P. vivax* P01 and *P. vivax* Sal1 lacking a defined SCO in *P. falciparum* or *P. yoelli* transcribed at a mean of ≥ 1 TPM in *P. vivax* salivary-gland sporozoites (Table S6). Prominent among these are *vir* (n=25) and *Pv-fam* (41 fam-e, 16 fam-b, 14 fam-a, 8 fam-d and 3 fam-h) genes, as well as hypothetical proteins or proteins of unknown function (n=212) and, interestingly, a number of ‘merozoite surface protein’ 3 and 7 homologs (n=5 of each). Both *msp3* and *msp7* have undergone significant expansion in *P. vivax* relative to *P. falciparum* and *P. yoelii* [32] and may have repurposed functions in sporozoites. In addition, there are 69 *P. vivax* P01 genes lacking a defined ortholog in *P. vivax* Sal1, *P. falciparum* or *P. yoelli* transcribed at ≥ 1 TPM in infectious *P. vivax* sporozoites; most of which are *Plasmodium* interspersed repeat (PIR) genes [32] found in telomeric regions of the P01 assembly and likely absent from the Sal1 assembly but present in the Sal1 genome, indicating the improved coverage of telomeric regions in P01 relative to Sal1.

### *P. vivax* sporozoites transcriptome compared with proteome

We compared relative protein abundance presented in a recently published *P. vivax* sporozoite proteome [33] to relative transcript abundance from the current study (Figure 1B and Table S7). The proteome study incorporated data from the same PvSPZ-Thai1 and PvSPZ-Thai5 isolates tested by RNAseq here. We identified 2,402 *P. vivax* genes transcribed in the sporozoite (CPM > 1) for which no protein expression was detected. Although many of these are lowly transcribed and likely below the detection sensitivity of LC-MS proteomics, others are among the most highly transcribed genes in the sporozoite, indicating these may be under translational repression.

Translational repression, the mechanism through which transcripts are held in stasis by RNA binding proteins, has been demonstrated to have important functional roles in the transition of *Plasmodium* spp. between the vertebrate to invertebrate host. More than 700 genes have been identified as translationally repressed in *Plasmodium berghei* (‘rodent malaria’) gametocytes based on DOZI (DEAD box RNA helicase “<u>d</u>evelopment <u>o</u>f <u>z</u>ygote inhibited”) pulldowns [34]. Translational repression mechanisms mediated through Puf-2 have been explored in sporozoites of several *Plasmodium* species and regulate some of the most abundant transcripts in the sporozoite, such as *uis-3* and *uis-4*. UIS3 and UIS4 are the best characterized proteins under translational repression by Puf-2 in sporozoites [35] and are essential for liver-stage development [14].

In considering genes that may be translationally repressed (i.e., transcribed but not translated) in the *P. vivax* sporozoite, we confine our observations to those transcripts representing the top decile of transcript abundance to ensure their lack of detection as proteins was not due to limitations in the detection sensitivity of the proteomic dataset. Approximately 1/3^rd^ of transcripts in the top decile of transcriptional abundance (n = 170 of 558) in *P. vivax* sporozoites were not detectable as peptides in multiple replicates (Figure 1B and Table S7). Of these 170 putatively repressed transcripts, 156 and 154 have orthologs in *P. falciparum* and *P. yoelii* respectively, with 89 and 118 of these also not detected as proteins in *P. falciparum* and *P. yoelii* salivary-gland sporozoites [36] despite being highly transcribed in these stages (see [26, 29]; Tables S3-S5), and 133 (78.2%) having no detectable sporozoite expression (>1 unique peptide count) in LC-MS data deposited for any species in PlasmoDB (Table S8). In contrast, 106 of these putatively repressed transcripts with orthologs in other *Plasmodium* species (Table S8) for which proteomic data is available in PlasmoDB, are detectable (>1 unique peptide count) by LC-MS methods in at least one other life-cycle stage, indicating against a technical issue (e.g., inability to be trypsin-digested) preventing their detection in the *P. vivax* sporozoite proteome [33]. In addition to *uis3* and *uis4*, genes involved in liver stage development and detectable as transcripts but not proteins in the *P. vivax* sporozoites include *lsap1* (liver stage <u>a</u>ssociated <u>p</u>rotein 1), *zipco* (<u>ZIP</u> domain-<u>co</u>ntaining protein), several other *etramps* (PVP01_1271000, PVP01_0422600, PVP01_0504800 and PVP01_0734800), *pv1* (<u>p</u>arasitophorous <u>v</u>acuole protein 1) and *lisp1* and *lisp2* (PVP01_1330800 and PVP01_0304700). Also notable among genes detectable as transcripts but not proteins in sporozoites is a putative ‘Yippee’ homolog (PVP01_0724100). Yippee is a DNA-binding protein that, in humans (YPEL3), suppresses cell growth [37] and is regulated through histone acetylation [38], making it noteworthy in the context of *P. vivax* hypnozoite developmental arrest.

Although verifying each putatively repressed transcript will require further empirical data, our system level approach is supported by immunofluorescent microscopy (Figure 1C) of UIS4, LISP1, EXP1 and ACP (PVP01_0416300). These represent one known and three putative (i.e., newly proposed here) translationally repressed genes in *P. vivax* sporozoites, and are compared to TRAP and BiP (which are both transcribed and expressed as protein in the *P. vivax* sporozoite; Table S8). The *lisp1* gene is an interesting find. In *P. berghei, lisp1* is essential for rupture of the PVM during liver stage development allowing release of the merozoite into the host blood stream. *Pv-lisp1* is ~350-fold and ~1,350-fold more highly transcribed in *P. vivax* sporozoites compared to sporozoites of either *P. falciparum* or *P. yoelii* (see Table S5). IFAs using LISP1 specific mAbs (Figure 1C) show that this protein is undetectable in sporozoites but clearly expressed at 7 days post-infection in liver schizonts.

### Up-regulated transcripts in *P. vivax* sporozoites relative to other life-cycle stages

Recently completed studies of the transcriptome of *P. vivax* for sporozoite activation [39], as well as, liver [22] and asexual blood-stages [40] support comparative transcriptomic study of sporozoites, their biology and transcriptional regulation over the *P. vivax* life-cycle. Recently published data for activate sporozoites from Roth et al [39] was significantly lower depth coverage, with ~0.03 to 0.6M reads mapped the *P. vivax* P01 coding domains; compared with 0.7 to 15.3 M, 2.4 to 10.6 M and 18.7 to 57.6M mapped reads for salivary sporozoite, liver-stages [22] and asexual blood stages [40] respectively. This lower coverage could not be compensated for through data normalization and therefore data from Roth et al [39] was not included in our quantitative analyses, although qualitatively, many of the highly transcribed genes in Roth et al [39] sporozoites were among the highly transcribed genes in salivary sporozoites from the present study. The remaining RNAseq data presents an analytical challenge in that each (sporozoites, liver-stages and blood stages) is produced in a separate study and may be influenced by technical batch effects that cannot be differentiated from biologically meaningful changes. To address this, we first examined *P. vivax* transcripts in a previous microarray study of multiple *P. vivax* life-cycle stages [24], including sporozoites and several blood-stages, to identify genes that may be transcriptionally stable across the life-cycle. We identified ~160 genes with low transcriptional variability between sporozoites and blood-stages that covered the breadth of transcript abundance levels in Westenberger et al [24]. These include genes typically associated with “house-keeping” functions, such as ribosomal proteins, histones, translation initiation complex proteins and various chaperones (see Figure S3 and Table S8). We assessed transcription of these 160 genes among the current and recently published RNA-seq data for *P. vivax* and all were of similarly low variability (Figure S4). This suggests that any batch effect between the studies is sufficiently lower than the biological differences between each life-cycle stage, allowing informative comparisons. We then combined all published RNAseq-based, transcriptomic data available for *P. vivax* [22, 39, 40] with the salivary sporozoite data generated here.

### P. vivax *sporozoite relative to blood-stage transcriptome*

To identify transcripts up-regulated in sporozoites, we first compared the *P. vivax* sporozoite transcriptome to RNA-seq data for *P. vivax* blood-stages [40] (Figure 2 and Figures S7-S10). We identified 1,672 up (Table S9); Interactive Glimma Plot - Supplementary Data 1) and 1,958 down-regulated (Table S9); Interactive Glimma Plot - Supplementary Data 1) transcripts (FDR ≤ 0.05; minimum 2-fold change in Counts per Million (CPM)) and explored patterns among these differentially transcribed genes (DTGs) by protein family (Figure 2C and Table S10) and Gene Ontology (GO) classifications (Table S11). RNA recognition motifs (RRM-1 and RRM-6) and helicase domains (Helicase-C and DEAD box helicases) are over-represented (p-value <0.05) among transcripts up-regulated in sporozoites, consistent with translational repression through ribonucleoprotein (RNP) granules [41]. Transcripts encoding nucleic acid binding domains, such as bromodomains (PF00439; which can also bind lysine-acetylated proteins), zinc fingers (PF13923) and EF hand domains (PF13499) are also enriched in sporozoites. Included among these proteins are a putative ApiAP2 transcription factor (PVP01_1211900) and a homologue of the *Drosophila* zinc-binding protein ‘Yippee’ (PVP01_0724100). Thrombospondin-1 like repeats (TSR: PF00090) and von Willebrand factor type A domains (PF00092) are enriched in sporozoites as well. In *P. falciparum* sporozoites, genes enriched in TSR domains are important in invasion of the mosquito salivary gland (e.g., *trap*) and secretory vesicles released by sporozoites upon entering the vertebrate host (e.g., *csp*) [42]. By comparison, genes up-regulated in blood-stages are enriched for *vir* gene domains (PF09687 and PF05796), Tryptophan-Threonine-rich *Plasmodium* antigens (PF12319; which are associated with merozoites [43]), markers of cell-division (PF02493; [44]) protein production/degradation (PF00112, PF10584, PF00152, PF09688 and PF00227) and ATP metabolism (PF08238 and PF12774). 47 of the 343 transcripts unique to *P. vivax* sporozoites relative to *P. falciparum* or *P. yoelii* are up-regulated in sporozoites compared to *P. vivax* blood stages. Nine of these are in the top decile of transcription, and include a Pv-fam-e (PVP01_0525200), a Pf-fam-b homolog (PVP01_0602000) and 7 proteins of unknown function. A further nine have an ortholog in *P. cynomolgi* (which also forms hypnozoites) but not the closely related *P. knowlesi* (which does not form hypnozoites) and include ‘*msp7*’-like (PVP01_1219600, PVP01_1220300 and PVP01_1219900), ‘*msp3*’-like (PVP01_1031300), Pv-fam-e genes (PVP01_0302100, PVP01_0524500 and PVP01_0523400), a serine-threonine protein kinase (PVP01_0207300) and a RecQ1 helicase homolog (PVP01_0717000). Notably, the *P. cynomolgi* ortholog of PVP01_0207300, PCYB_021650, is transcriptionally up-regulated in hypnozoites relative to replicating schizonts [23], indicating a target of significant interest when considering hypnozoite formation and/or biology and suggesting that the list here may contain other genes important in hypnozoite biology.

**Fig. 2.**
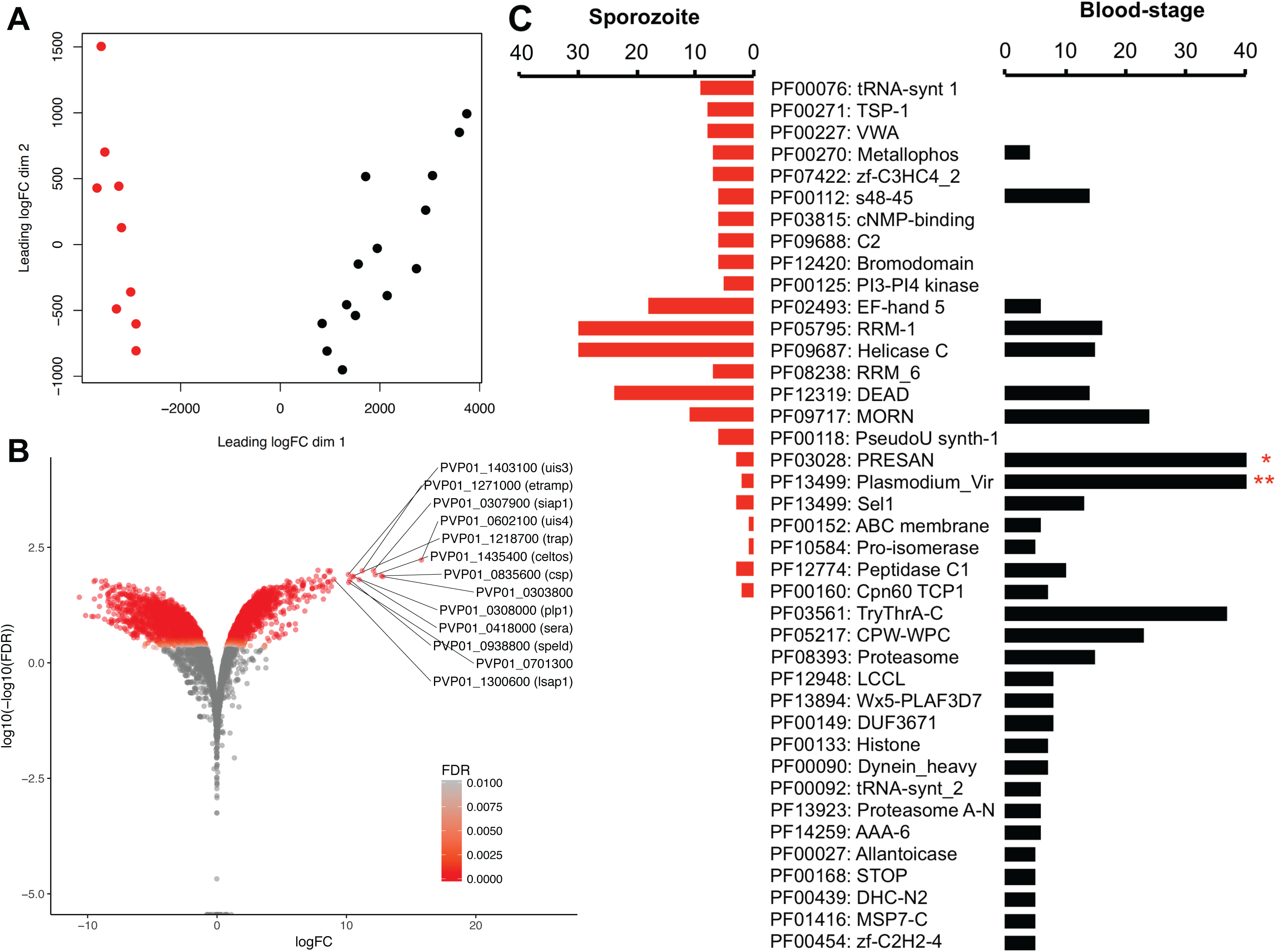
Differential transcription between *Plasmodium vivax* salivary-gland sporozoites and blood-stages. **A** BCV plot showing separation between blood-stage (black) and salivary-gland sporozoite (red) biological replicates. **B** Volcano plot of distribution of fold-changes (FC) in transcription between blood-stages and salivary-gland sporozoites relative to statistical significance threshold (False Discovery Rate (FDR) ≤ 0.05). Positive FC represents up-regulated transcription in the sporozoite stage. **C** Mirror plot showing pFam domains statistically significantly (FDR ≤ 0.05) over-represented in salivary-gland sporozoite up-regulated (red) or blood-stage up-regulated (black) transcripts. Scale bar truncated for presentation. * - 55 PRESAN domains are in this dataset. ** - 99 Vir domains are in this dataset.

### P. vivax *sporozoites are enriched in translational repressors*

In *Plasmodium*, translational repression regulates key life-cycle transitions coinciding with switching between the mosquito and the mammalian host (either as sporozoites or gametocytes) [41]. For example, although *uis4* is the most abundant transcript in the infectious sporozoite ([24, 31]; Table S2), UIS4 is translationally repressed in this stage [15] and only expressed after hepatocyte invasion [45]. In sporozoites, it is thought that PUF2 binds to mRNA transcripts and prevents their translation [29], and SAP1 stabilises the repressed transcripts and prevents their degradation [45]. Consistent with this, *Puf2* and *SAP1* (PVP01_0947600) are up-regulated in the sporozoite relative to blood-stages. Indeed, *Puf2* (PVP01_0526500) is among the top percentile of transcripts in salivary sporozoites and expressed at high levels in the proteome [33]. However, our data implicate other genes that may act in translational repression in *P. vivax* sporozoites, many of which are already known to be involved in translational repression in other *Plasmodium* stages and other protists [41]. Among these are *alba*-*2* and *alba-4*, both of which are among the top 2% of genes transcribed in sporozoites and ~14 to 20-fold more highly transcribed in sporozoites relative to blood-stages; ALBA-2 is in the top 100 most abundant proteins in the *P. vivax* sporozoite proteome [33]. In addition, *P. vivax* sporozoites are enriched for genes encoding RRM-6 RNA helicase domains. Intriguing among these are HoMu (homolog of Musashi; top decile of sporozoite proteins by abundance [33]) and ptbp (<u>p</u>olypyrimidine tract <u>b</u>inding <u>p</u>rotein). Musashi regulates eukaryotic stem cell differentiation through translational repression [46] and HoMu localizes with DOZI and CITH in *Plasmodium* gametocytes [47]. PTBP is linked to mRNA stability, splice regulation and translational initiation [48] and may perform a complementary role to SAP1.

### P. vivax *sporozoite relative to Plasmodium spp. liver stage transcriptomes*

New advances in *P. vivax* liver culture has allowed recent publication of mixed stage and hypnozoite-enriched transcriptomes [22]. This is an early, yet highly valuable, study and, due no doubt to the difficulty in generating the material, is limited to biological duplicates. Noting this, although we undertake differential transcriptomic studies of this dataset here, we recognize that additional biological replication is needed and have used a higher burden of significance (FDR ≤ 0.01 and ≥ 2-fold change) than used with blood-stages. Nevertheless, these comparisons identified 1,015 and 856 sporozoite up-regulated transcripts relative to mLS and HPZs respectively and 1,007 and 1,079 transcripts up-regulated in mLS and HPZs relative to sporozoites respectively (Figures 3 and S11-S13, Table S12 and S13 and Interactive Glimma Plot - Supplementary Data 1).

**Fig. 3.**
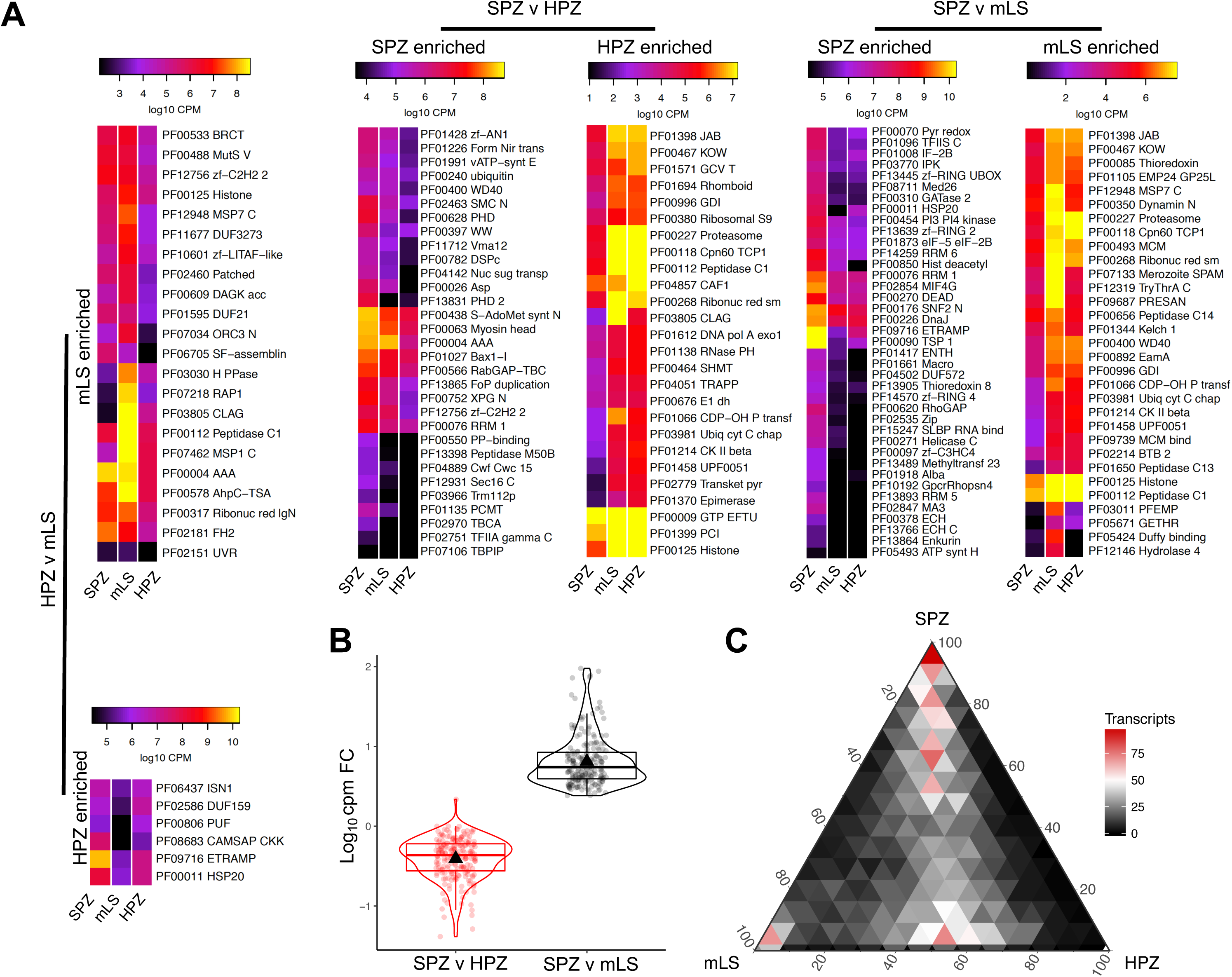
Differential transcription between *P. vivax* sporozoites (SPZ), mixed (mLS) and hypnozoite (HPZ) enriched liver stages (liver-stage data from Gural et al [22]). **A** Heatmap comparisons showing summed transcription of enriched Pfam domains in HPZ vs mLS (left), SPZ vs HPZ (top middle) and SPZ vs mLS (top right) comparisons. All Pfam domains statistically significantly enriched at p-value 0.05). All transcript data for stage up-regulated genes at FDR 0.01). **B** Violin box-plot showing relative fold-change differences between SPZ and HPZ compared with SPZ and mLS for genes down-regulated in mLS compared to SPZ, but not down-regulated in HPZ compared to SPZ. **C** Ternary heatmap summarizing relative transcript abundance in each of SPZ, mLS and HPZ stages.

Compared to mLS transcriptomes, sporozoites are enriched for many of the transcripts similarly up-regulated in comparison to blood-stages (e.g., *uis4, celtos, puf2, siap1* and *plp-1*). More broadly, SPZ up-regulated transcripts over-represent (p-value ≤ 0.05) Pfam domains (Figure 3A) associated with transcriptional regulation (PF00176, PF01096, PF01661 and PF08711), translational repression/regulation (PF00076, PF00279, PF01008, PF01873, PF01917, PF02847, PF02854, PF13893 and PF14259), DNA/RNA binding (PF0097, PF13445, PF13639, PF14570 and PF15247) and chromatin regulation (PF00271, PF00850 and PF13489). In contrast, the mLS transcriptome is enriched in genes involved in replication and merozoite formation [n = 14; including PVP01_0728900 (*msp1*), PVP01_0010670 (*msp3*) and PVP01_1446800 (*msp9*)], rhoptry function [n = 9; including PVP01_1469200 (*rnp3*), PVP01_1255000 (*rnp2*) and PVP01_1338500 (*rap1*)] and reticulocyte binding [n=10 including PVP01_0534300 (rbp2c), PVP01_1402400 (rbp2a), PVP01_0701100 (rbp1b) and PVP01_0800700 (rbp2b)]. These data are further enriched for Pfam domains associated with cell division (PF00493), merozoite formation (PF07133 and PF12984), proteasome function (PF00227, PF00400, PF00656, PF01344, PF01398 and PF03981), protein export / vesicle function (PF00350 and PF00996), membrane proteins (PF01105, PF03011, PF05424 and PF12139) and metabolism (PF00085, PF00118, PF00268, PF01066, PF01214 and PF01214). Collectively, in addition to markers consistent with sporozoite or merozoite formation, these data point towards the sporozoite stage as being highly regulated and controlled at transcriptional, translational and chromatin levels, with the mLS stages representing a release of this control allowing replication, protein turn-over, reconfiguration of the proteins on the plasma membrane and metabolic activity.

Comparison of sporozoites with HPZs does not indicate a similar release of control, or at least that any release is more specific than for mLS. The sporozoite is enriched, relative to HPZs, in genes such as PVP01_1258000 (*gest*), PVP01_0418000 (*sera*), PVP01_1435400 (*celtos*), PVP01_0835600 (*csp*) and PVP01_0602100 (*uis4*). At a broad level, sporozoite enriched Pfam domains include a smaller number associated with translational repression/regulation (PF00076) or DNA/RNA binding (PF01428 and PF12756). Interestingly, sporozoites are enriched in Pfam domains specifically associated with heterochromatin (H3K9me3) reading/interaction (PF02463, PF00628, PF13831 and PF13865). Ours (see below) and previous epigenetic studies of *Plasmodium* sporozoites [27] find dense heterochromatin in the telomeric to subtelomeric regions of the chromosome, which is more transcriptionally active in blood-stages [49]. Others have noted an up-regulation of methyl/acetyltransferases in *P. cynomolgi* HPZs [23] and/or shown methyltransferase inhibitors stimulate hypnozoite activation *in vitro* [17]. The potential that histone epigenetics of sporozoites has a role in or changes with liver-stage development and the formation of liver schizonts or HPZs is intriguing but requires detailed study of the chromatin of liver-stage parasites, which is not presently available for *P. vivax*. In contrast, HPZs were enriched, relative to sporozoites, for genes including histone proteins (PVP01_1138700, PVP01_1131700 and PVP01_0905900) and classic markers of metabolism (PVP01_MITO3300 and PVP01_MITO3400) and *lisp2*. Pfam data indicated– largely similar domain enrichment trends as were seen for the mLS stage relative to sporozoites, including a number of proteosomal (PF00227, PF00112, PF03981), vesicular transport (PF00996) and metabolic (PF00118, PF00268, PF01066, PF01214 a) associated functions. This supports HPZs being an arrested, rather than classically ‘dormant’, stage with active metabolism and protein turn-over. HPZs are also enriched for Pfams associated with mRNA/tRNA regulation and turnover (PF04857, PF01612, PF00009 and PF01138) and glycine metabolism (PF01571 and PF00464) and acetyl-CoA production (PF02779 and PF00676).

Finally, although not the focus of this study, we looked at differential transcription between mLS and HPZ stages using the Gural et al [22] data, but using the same approaches as employed here. In particular, we were interested in what these comparisons might provide in terms of sporozoite differentiation or development into liver schizonts or HPZs (Table S14). Among mLS up-regulated transcripts are genes associated with rhoptry function (n = 11; including PVP01_0107500, PVP01_1469200 and PVP01_1469200), cytoadherence to red-cells (PVP01_1401400 and PVP01_0734500), merozoite formation (PVP01_0728900 and PVP01_0612400) and exported proteins (n = 6; including PVP01_0504000, PVP01_0119200 and PVP01_0801600). Consistent with *P. cynomolgi* [23], HPZ up-regulated transcripts include several key sporozoite transcripts, specifically *uis4* (PVP01_0602100), *puf1* (PVP01_1015000) and *speld* (PVP01_0938800). At the Pfam domain level, mLS is enriched for metabolic (PF00317) and proteosomal (PF00112) domains also enriched in mLS or HPZs relative to sporozoites above, as well as domains associated with merozoite formation (PF12948, PF07462), rhoptry function (PF0712), DNA/RNA binding (PF12756, PF10601 and PF02151) and cell division, development and DNA replication (PF06705, PF00533, PF00488, PF02460, PF07034, PF02181). In contrast, HPZs are enriched in Pfam domains that overlap notably with key sporozoite markers, including *etramps* (PF09716) and *puf* proteins (PF00806), as well as domains associated with calcium (PF08683) and nucleotide metabolism (PF06437). These data largely indicate that the hypnozoite bears similarity both to the sporozoite and liver schizonts consistent with a stalled stage on the path to schizont development regulated by checkpoint signals that halt/restart normal schizont development, which has been proposed previously for this species [24].

With this is mind, we looked at transcripts that are differentially transcribed in mLS, but not HPZs, relative to SPZs. There are 107 transcripts down-regulated in mLS relative to SPZs that are transcribed at roughly similar levels in both SPZs and HPZs (Figure 3B). A common theme among many of these genes are their role in transcriptional, post-transcriptional, translational or post-translational regulation. Among transcriptional regulators are transcription factors including AP2-SP2 (PVP01_0303400) and three non-AP2-like transcription factors (PVP01_0306600, PVP01_0204300 and PVP01_1415800). Post-transcriptional controllers include several DNA/RNA-binding proteins (PVP01_1011000. PVP01_0932900, PVP01_0715300, PVP_1242600 and PVP01_0605200), RNA helicases (PVP01_1403600 and PVP01_1329800) and mRNA processing (PVP01_1443100 and PVP01_1458200) genes. Translational control includes several key regulators of translation initiation (PVP01_1467700), tRNA processing (PVP01_0318700 and PVP01_1017700) or ribosomal function/biogenesis (PVP01_1443700, PVP01_0421400, PVP01_1117200 and PVP01_0215100). Post-translational control includes two methyltransferases (PVP01_1428800 and PVP01_1465200), including CARM1, which methylates of H3R17 and, in mice, prevents differentiation in embryonic stem cells [50], and a putative histone methylation reading enzyme, EEML2 (PVP01_1014100). The remaining genes in this group have three noteworthy and largely overlapping themes: (1) an association with calcium binding, metabolism or signalling, (2) a role in organellar metabolism and (3) homologs in other organisms, including a variety of prokaryotes and eukaroytes, with key roles in germination, dormancy and persistent non-replicating stages. The latter most function is clearly intriguing in the context of HPZ formation and activation. These genes include a homolog of dihydrolipoamide acyltransferase (aka ‘sucB’), which is essential for growth in *Mycobacterium tuberculosis* [51] and a key regulator in persistent *Escherichia coli* stages [52]. Another example is gamete fusion factor HAP2, which, despite the name, has been shown to regulate dormancy in eukaryotes ranging from plants [53, 54] to amoebae [55].

In addition to data for *P. vivax*, two transcriptomic studies are now available for *P. cynomolgi* [27, 56] that compare mixed/schizont stage parasites with small-form “hypnozoites”. In comparing *P. cynomolgi* liver-stage RNA-seq and *P. vivax* liver-stage microarray data, Cubi et al [23] noted a moderate to good level of agreement (R^2^ = 0.50) as evidence of *P. cynomolgi* being predictive and representative of *P. vivax*. However, Voorberg van der Wel et al [56] explored congruence between their and the Cubi et al [23] studies and found generally good agreement among schizonts and overall relatively poor agreement among hypnozoites from each study. This highlights the complexity of these datasets and indicates caution in comparing the current data to *P. vivax*. ApiAP2 transcription factors feature prominently in each liver-stage transcriptomic study for *P. cynomolgi* [23, 56] and *P. vivax* [22]. Cubi et al [23] noted an ApiAP2 (dubbed “AP2-Q”; PCYB_102390) as transcriptionally up-regulated in *P. cynomolgi* hypnozoites and proposed this as a potential hypnozoite marker. We note that the *P. vivax* ortholog of *Pc*-AP2-Q (PVP01_1016100) is among the genes detectable as a transcript but not protein in *P. vivax* sporozoites. This may point to a translationally repressed signal in sporozoites to regulate hypnozoite formation. However, as *Pv*-AP2-Q is transcribed at an abundance (~50 TPM) at or below which ~50% of *P. vivax* genes are detectable as transcripts but not as proteins (Figure 1B), this could as likely result from LC-MS detection sensitivity. Further, although AP2-Q was reported as specific to hypnozoite forming *Plasmodium* species [23], it is indeed found in non-hypnozoite producing species, such as *P. knowlesi, P. gallinaceum* and *P. inui* [56]. Up-regulation of AP2-Q transcripts is not observed for hypnozoites in subsequent transcriptomic studies of *P. cynomolgi* [56] or *P. vivax* [22], nor do we see such an up-regulation here. Voorberg van der Wel et al [56] note transcription of a range of AP2s in *P. cynomolgi* liver stages, but do not find any to be up-regulated in hypnozoites. AP2s also feature among transcribed genes in *P. vivax* liver stages, with one, PVP01_0916300, significantly up-regulated in hypnozoites. We note that PVP01_0916300 is up-regulated in *P. vivax* sporozoites relative to blood-stages and found in the top quartile of transcripts by abundance (TPM = 104).

### Chromatin epigenetics in *P. vivax* sporozoites

As noted above, transcriptomic data for sporozoites, and their comparison with liver and blood-stages, implicate histone epigenetics as having an important role in sporozoite biology and liver stage differentiation. This concept has been alluded to in recent liver-stage studies of *P. cynomolgi* [17, 23] that propose methyltransferases as having a potential role in hypnozoite formation. No epigenetic data are currently available for any *P. vivax* life-cycle stage. Studies of *P. falciparum* blood-stages have identified the importance of histone modifications as a primary epigenetic regulator [57, 58] and characterized key markers of heterochromatin (H3K9me3) and euchromatin/transcriptional activation (H3K4me3 and H3K9ac). Recently, these marks have been explored with the maturation of *P. falciparum* sporozoites in the mosquito [26]. Here, we characterize major histone marks in *P. vivax* sporozoites and assess their relationship to transcript abundance.

### *Histone modifications in* P. vivax *sporozoites*

Using ChIP-seq, we identified 1,506, 1,999 and 5,262 ChIP-seq peaks stably represented in multiple *P. vivax* sporozoite replicates and associated with H3K9me3, H3K9ac and H3K4me3 histone marks respectively (Figure 4 and S14-S19). Peak width, spacing and stability differed with histone mark type (Figures S15 and S16). H3K4me3 covered the greatest breadth of the genome (36.0% of all bases) and was the most stable among replicates, with ~84% of bases associated with an H3K4me3 found in multiple biological replicates. By comparison H3K9me3 marks were least stable, with 46% of bases associated with this mark found in just one replicate. Consistent with observations in *P. falciparum*, H3K9me3 ‘heterochromatin’ marks primarily clustered in telomeric and subtelometric regions (Figure 4). In contrast, the ‘euchromatin’ / transcriptionally open histone marks, H3K4me3 and H3K9ac, were distributed in chromosome central regions and did not overlap with regions under H3K9me3 suppression. Both H3K9me3 and H3K4me3 marks were reasonably uniformly distributed (mean peak spacing ~500bp for each) within their respective regions of the genome. In contrast, H3K9ac peaks were spaced further apart (mean: ~2kb), but also with a greater variability in spacing (likely reflecting their association with promoter regions [59]; Figure S17 and S19). The instability of H3K9me3 may reflect its use in *Plasmodium* for regulating expression of contingency genes from multigene families, whose members have overlapping and redundant functions [49] and confer phenotypic plasticity [60].

**Fig. 4.**
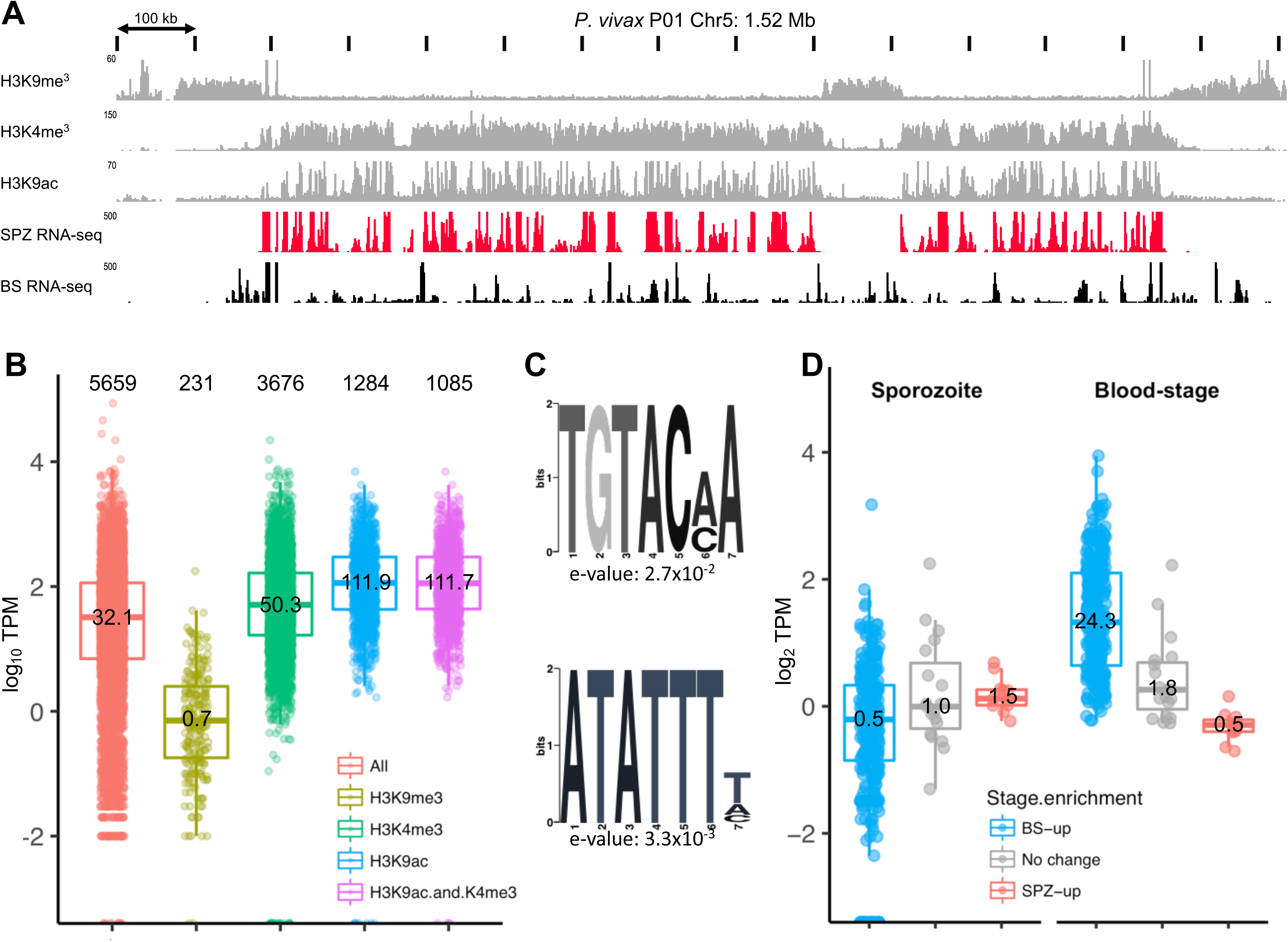
Histone epigenetics relative to transcriptional behaviour in salivary-gland sporozoites. **a** Representative H3K9me3, H3K4me3 and H3K9ac ChIP-seq data (grey) from a representative chromosome (*P. vivax* P01 Chr5) relative to mRNA transcription in salivary-gland sporozoites (black) and blood-stages (black). Small numbers to top left of each row show data range. **b** Salivary-gland sporozoite transcription relative to nearest stable histone epigenetic marks. Numbers at the top of the figure represent total genes included in each category. Numbers within in box plot represent mean transcription in transcripts per million (TPM). **c** Sequence motifs enriched within 1kb upstream of the Transcription Start Site of highly transcribed (top 10%) relative to lowly transcribed genes associated with H3K9ac marks in salivary-gland sporozoites. **d** Relative transcription of (sub)telomeric genes in *P. vivax* salivary-gland sporozoites and blood-stages categorized by gene sets up-regulated in blood-stages (blue), salivary sporozoites (red) or not stage enriched (grey). Numbers in each box show mean transcription in TPM.

### Genes under histone regulation

We explored an association between these histone marks and the transcriptional behaviour of protein coding genes (Figure 4 and S19 and Tables S15-S20). 485 coding genes stably intersected with an H3K9me3 mark; all are located near the ends of the chromosomal scaffolds (i.e., are (sub)telomeric). On average, these genes are transcribed at ~30 fold lower levels (mean 0.7 TPM) than genes not stably intersected by H3K9me3 marks. These data clearly support the function of this mark in transcriptional silencing. This is largely consistent with observations in *P. falciparum* sporozoites [26], however, in contrast to *P. falciparum* sporozoites where a single *var* gene was described to lack heterochromatin structure [27] we observe no genes within heterochromatin dense region that lacked a stable H3K9me3 signal or were transcribed at notable levels (i.e., above ~5 TPM). Whether this relates to differences in epigenetic control between the species is not clear. We note that (sub)telomeric genes are overall transcriptionally silent in *P. vivax* sporozoites relative to blood-stages (Figure 4 and Tables S21 and S22). Consistent with observations in *P. falciparum* [57], the bulk of these genes include complex protein families, such as *vir* and *Pv-fam* genes, which are so far described to function primarily in blood-stages. Also notable among the genes are several reticulocyte-binding proteins, including RBP2, 2a, 2b and 2c. This transcriptional silence in telomeric and subtelomeric regions was recently observed in *P. falciparum* sporozoites [27].

Outside of the telomeres and subtelomeres, H3K4me3 marks are stably associated with the intergenic regions of 3,676 genes. H3K9ac marks are also identified within 1kb of the transcriptional start site (TSS) of 1,284 coding genes, with 1085 of these stably marked also by H3K4me3 (Figure 4B). The average transcription of these genes is 50, 112 and 112 TPMs respectively (72, 160 and 160-fold higher than H3K9me3 marked genes). Gene-by-gene observations show that H3K9ac and H3K4me3 marks cluster densely in the 1000kb up and down-stream of the start and stop codon respectively of transcribed genes, but are much less dense within coding regions of these genes (Figure S15). This pattern directly correlates with transcription and contrasts H3K9me3 marks, which are distributed across the length of the gene at even density and are correlated with a lack of transcription. These data support the role of H3K9ac and H3K4me3 in transcriptional activation in *P. vivax*. The lower transcriptional abundance of H3K4me3 marked, compared with H3K9ac or H3K9ac and H3K4me3 marked genes suggest these marks work synergistically and that H3K9ac is possibly the better of the two, as a single mark indicator of transcriptional activity in *P. vivax*. This is consistent with recent observations in *P. falciparum* sporozoites [26].

Interestingly, H3K9ac-marked genes ranged in transcriptional activity from the most abundantly transcribed genes to many in the lower 50% and even lowest decile of transcription. This suggests more contributes to transcriptional activation in *P. vivax* sporozoites than, simply, gene accessibility through chromatin regulation. Specific activation by a transcription factor (e.g., ApiAP2s [61]) is the obvious candidate. To explore this, we compared upstream regions (within 1kb of the TSS or up to the 3’ end of the next gene upstream, whichever was less) of highly (top 10%) and lowly (bottom 10%) transcribed H3K9ac marked genes for over-represented sequence motifs in the highly expressed genes that might coincide with known ApiAP2 transcription factor binding sites [62]. We identified these based on the location of the nearest stable H3K9ac peak relative to the transcription start site for each gene (Figure S12). In most instances, these peaks were within 100bp of the TSS and, consistent with data from *P. falciparum* [59], *P. vivax* promoters appear to be no more than a few hundred to a maximum of 1000 bp upstream of the TSS. Exploring these regions, we identified two over-represented motifs: TGTACMA (e-value 2.7e^-2^) and ATATTTH (e-value 3.3e^-3^) (Fig. 2D). TGTAC is consistent with the known binding site for *Pf*-AP2-G, which regulates sexual differentiation in gametocytes [63], but its *P. vivax* ortholog (PVP01_1418100) is neither highly transcribed nor expressed in sporozoites. ATATTTH is similar to the binding motif for *Pf-*AP2-L (AATTTCC), a transcription factor that is important for liver stage development in *P. berghei* [64]. In contrast to AP2-G, *Pv*-AP2-L (PVX_081180) is in the top 10% of transcription and expression in *P. vivax* sporozoites and up-regulated relative to blood-stages. In *P. vivax* sporozoites, the ATATTTH motif is associated with a number of highly transcribed genes, including *lisp1* and *uis2-4*, known to be regulated by AP2-L in *P. berghei* [64] as well as many of the most highly transcribed, H3K9ac marked genes, including two *etramps* (PVP01_0734800 and PVP01_0504800), several RNA-binding proteins, including *Puf2, ddx5*, a putative ATP-dependent RNA helicase DBP1 (PVP01_1429700), and a putative *bax1* inhibitor (PVP01_1465600). Interestingly, a number of highly transcribed and translationally repressed genes associated with the ATATTTH motif, including *uis4, siap2* and *pv1*, are not stably marked by H3K9ac in all replicates (i.e., there is significant variation in the placement of the H3K9ac peak or their presence/absence among replicates for these genes). It may be that additional histone modifications, for example H3K27me, H3R17me3 or H2A or H4 modifications, are involved in regulating transcription of these genes. Certainly the incorporation of the H2A.Z histone variant, which is present in intergenic regions of *P. falciparum (*Petter et al 2011), and controls temperature responses in plants [65] is intriguing as a potential mark regulating sporozoite fate in *P. vivax* considering the association between hypnozoite activation rate and climate [11], as is H3R17me3 in consideration of the enrichment of markers/readers of this modification in HPZs noted above and the role of this mark in cell fate progression in other species [50].

## CONCLUSIONS

We provide the first comprehensive study of the transcriptome and epigenome of mature *Plasmodium vivax* sporozoites and undertake detailed comparisons with recently published proteomic data for *P. vivax* sporozoites [33] and transcriptomic data for *P. vivax* mixed and hypnozoite-enriched liver-stages [22] and mixed blood-stages [40]. These data support the proposal that the sporozoite is a highly-programmed stage that is primed for invasion of and development in the host hepatocyte. Cellular regulation, including at transcription, translational and epigenetic levels, appears to play a major role in shaping this stage (which continues on in some form in hypnozoites), and many of the genes proposed here as being under translational repression are involved in hepatocyte infection and early liver-stage development (Figure 5). We highlight a major role for RNA-binding proteins, including PUF2, ALBA2/4 and, intriguingly, ‘Homologue of Musashi’ (HoMu). We find that transcriptionally, the hypnozoite appears to be a transition point between the sporozoite and replicating schizonts, having many of the dominant sporozoite transcripts and retaining high transcription of a number of key regulatory pathways involved in transcription, translation and chromatin configuration (including histone arginine methylation). A consistent theme in the study is the prominence of a number of genes that have a role in numerous eukaryotic systems in cell fate determination and differentiation (e.g., HoMu, Yippee and CARM1) and overlap with dormancy and/or persistent cell states in bacteria, protists or higher eukaryotes (e.g., bacterial sucB and gamete fusion protein HAP2). These data do not point to one single programming switch for dormancy or liver developmental fate in *P. vivax*, but present a number of intriguing avenues for exploration in subsequent studies, particularly in model species such as *P. cynomolgi*. Our study contributes to understanding the early stages of hepatocyte infection and the developmental switch between liver trophozoite and hypnozoite formation. We also identify potential avenues for rationally prioritizing targets underpinning liver-stage differentiation for functional evaluation in humanized mouse and simian models for relapsing *Plasmodium* species and identifying novel avenues to understand and eradicate liver-stage infections.

**Fig. 5.**
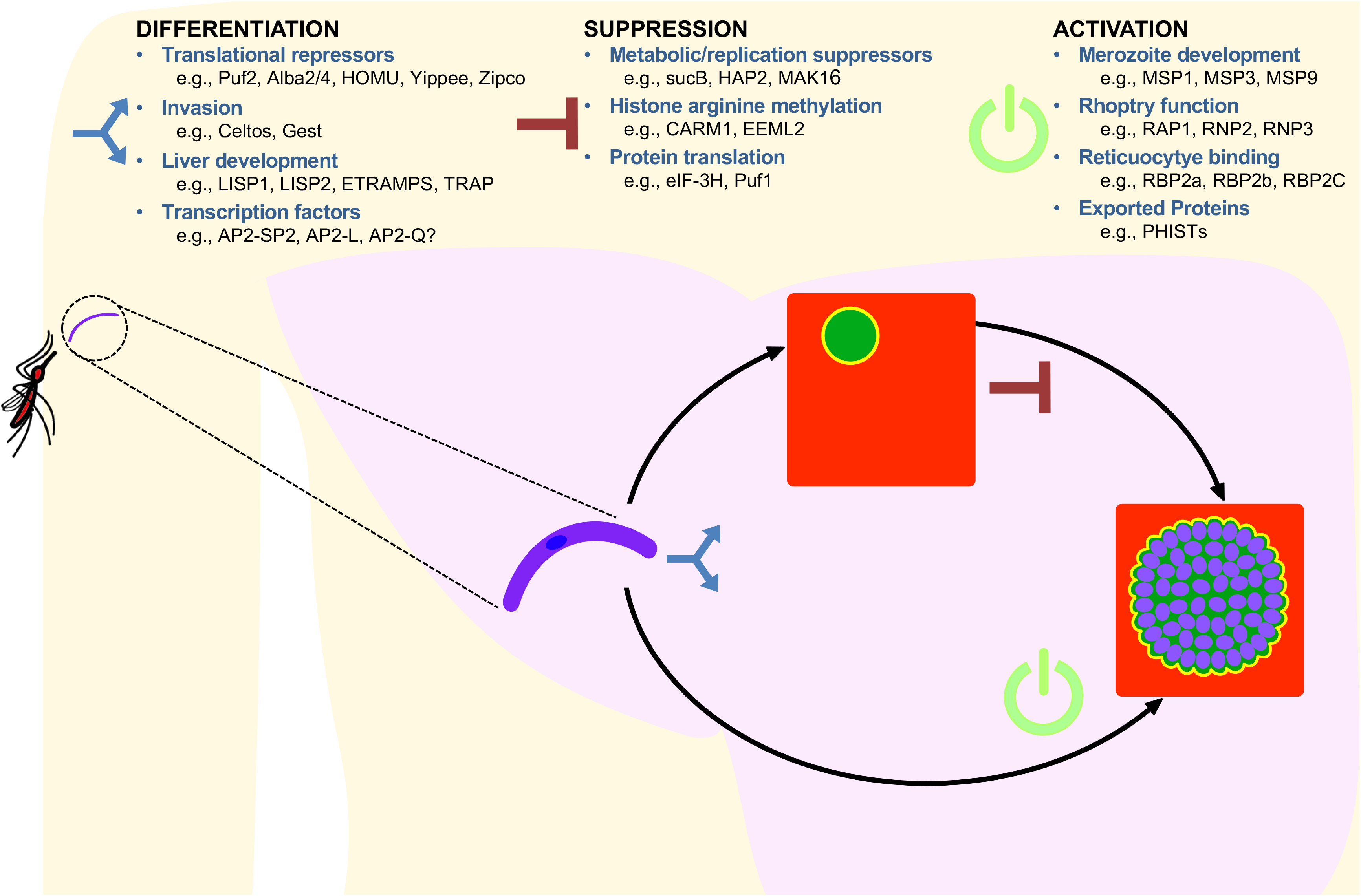
Schematic of potential mechanisms underpinning development in differentiation of *P. vivax* sporozoites during liver-stage infection as hypnozoites and schizonts. We suggest differentiation programming at different points in development; first, schizont or hypnozoite fate possibly encoded in the sporozoite as epigenetic signals or translationally repressed transcripts; secondly, suppression signals that halt progression of the hypnozoite to schizont stage and support persistence; and finally activation signals signified by a release in chromatin, (post)transcriptional and (post)translational control leading to up-regulation of replication, metabolic and protein export pathways.

## MATERIALS AND METHODS

### Ethics Statement

Collection of venous blood from human patients with naturally acquired vivax infection for the current study was approved by the Ethical Review Committee of the Faculty of Tropical Medicine, Mahidol University (Human Subjects Protocol number TMEC 11-033) with the informed written consent of each donor individual. All mouse tissue used in the current study was from preserved infected tissues generated previously [13]. All mouse infection work in [13] was carried out at the Centre for Infectious Diseases Research (CIDR) in Seattle, USA, under direct approval of the CIDR Institutional Animal Care and Use Committee (IACUC) and performed in strict accordance with the recommendations in the Guide for the Care and Use of Laboratory Animals of the National Institutes of Health, USA. The Centre for Infectious Disease Research Biomedical Research Institute has an Assurance from the Public Health Service (PHS Assurance number is A3640-01) through the Office of Laboratory Animal Welfare (OLAW) for work approved by its IACUC.

### Material collection, isolation and preparation

Nine field isolates (PvSpz-Thai 1 to 9), representing symptomatic blood-stage malaria infections were collected as venous blood (20 mL) from patients presenting at malaria clinics in Tak and Ubon Ratchatani provinces in Thailand. Each isolate was used to establish, infections in *Anopheles dirus* colonized at Mahidol University (Bangkok) by membrane feeding [13], after 14-16 days post blood feeding, ~3-15 million sporozoites were harvested per field isolate from the salivary glands of up to 1,000 of these mosquitoes as per [66] and shipped in preservative (trizol (RNA/DNA) or 1% paraformaldehyde (DNA for ChiP-seq) to the Walter and Eliza Hall Institute (WEHI).

### Transcriptomics sequencing and differential analysis

Upon arrival at WEHI, messenger RNAs were purified from an aliquot (~0.5-1 million sporozoites) of each *P. vivax* field isolate as per [40] and subjected to RNA-seq on Illumina NextSeq using TruSeq library construction chemistry as per the manufacturer’s instructions. Raw reads for each RNA-seq replicate are available through the Sequence Read Archive (XXX-XXX). Sequencing adaptors were removed and low quality reads trimmed and filtered using Trimmomatic v. 0.36 [67]. To remove host contaminants, processed reads were aligned, as single-end reads, to the *Anopholes dirus* wrari2 genome (VectorBase version W1) using Bowtie2 [68] (--very-sensitive preset). All non-host reads were then aligned to the manually curated transcripts of the *P. vivax* P01 genome (<u>http://www.genedb.org/Homepage/P*vivax*P01; [28]</u>) using RSEM [69] (pertinent settings: --bowtie2 --bowtie2-sensitivity-level very_sensitive --calc-ci --ci-memory 10240 --estimate-rspd --paired-end). Transcript abundance for each gene in each replicate was calculated by RSEM as raw count, posterior mean estimate expected counts (pme-EC) and transcripts per million (TPM).

Transcriptional abundance in *P. vivax* sporozoites was compared qualitatively (by ranked abundance) with previously published microarray data for *P. vivax* salivary-gland sporozoites [24]. As a further quality control, these RNA-seq data were compared also with previously published microarray data for *P. falciparum* salivary-gland sporozoites [30], as well as RNA-seq data from salivary-gland sporozoites generated here for *P. falciparum* (single replicate generated from *P. falciparum* 3D7 lab cultures isolated from *Anopholes stephensi* and processed as above) and previously published for *P. yoelii* [29]. RNA-seq data from these additional *Plasmodium* species were (re)analysed from raw reads and transcriptional abundance for each species was determined (raw counts and pme-EC and TPM data) as described above using gene models current as of 04-10-2016 (PlasmoDB release v29). Interspecific transcriptional behaviour was qualitatively compared by relative ranked abundance in each species using TPM data for single copy orthologs (SCOs; defined in PlasmoDB) only, shared between *P. vivax* and *P. faliciparum* or shared among *P. vivax, P. falciparum* and *P. yoelii*.

To define transcripts that were up-regulated in sporozoites, we remapped raw reads representing early (18-24 hours post-infection (HPI)), mid (30-40 HPI) and late (42-46 HPI) *P. vivax* blood-stage infections recently published by Zhu *et al* [40] to the *P. vivax* P01 transcripts using RSEM as above. All replicate data was assessed for mapping metrics, transcript saturation and other standard QC metrics using QualiMap v 2.1.3 [70]. Differential transcription between *P. vivax* salivary-gland sporozoites and mixed blood-stages [40] was assessed using pme-EC data in EdgeR [71] and limma [72] (differential transcription cut-off: ≥ 2-fold change in counts per million (CPM) and a False Discovery Rate (FDR) ≤ 0.05). Pearson Chi squared tests were used to detect over-represented Pfam domains and Gene Ontology (GO) terms among differentially transcribed genes in sporozoites (Bonferroni-corrected *p* < 0.05), based on gene annotations in PlasmoDB (release v29).

We also compared transcription of the sporozoite stages to recently published liver-stage data from Gural et al [22] as per the sporozoite to blood-stage comparisons above, with the following modifcations: (1) EC values were normalized using the ‘upper quartile’ method instead of TMM, (2) differential transcription was assessed using a quasi-likelihood generalize linear model (instead of a linear model) and (3) an FDR threshold for significance of ≤ 0.01 was used instead of ≤ 0.05. These differences related to specific attributes of the liver-stage dataset, particularly the small number of replicates (n = 2) per experiment condition. Data visualization and interactive R-shiny plots were produced in R using the ggplot[73], ggplot2 [74], gplots(heatmap.2) [75] and Glimma [76] packages.

### Assessment of Sporozoite RNA-seq transcriptome by selective RT-qPCR

Extracted RNA was DNase treated (Sigma D5307) as per manufacturer recommendations. RNA was quantified using the TapeStation High Sensitivity RNA kit (Agilent). Two intron-spanning primer pairs were designed per gene of interest using Primer3 and BLAST. Primer pairs were tested in two concentrations (0.75ng and 2.83ng per reaction) to determine efficiency and specificity. Product was run on a 1% agarose gel with ethidium bromide. Primer pairs indicating non-specific priming were removed. The resulting 11 primer pairs were used on four sporozoite samples; VUBR06, VUNL23, VUBR24, VTTY84. RNA was reverse transcribed (Sensifast, Bioline) and used at 0.75ng per reaction, run on a Roche LightCycler 480 II. Melt curves were assessed and products were run on a gel to ensure specificity again. Cp threshold was set automatically. ΔCp value was calculated as target gene – comparator gene (SERA and CelTOS were used). Data were log transformed and fold change calculated.

RT-qPCR Primers were as follows:

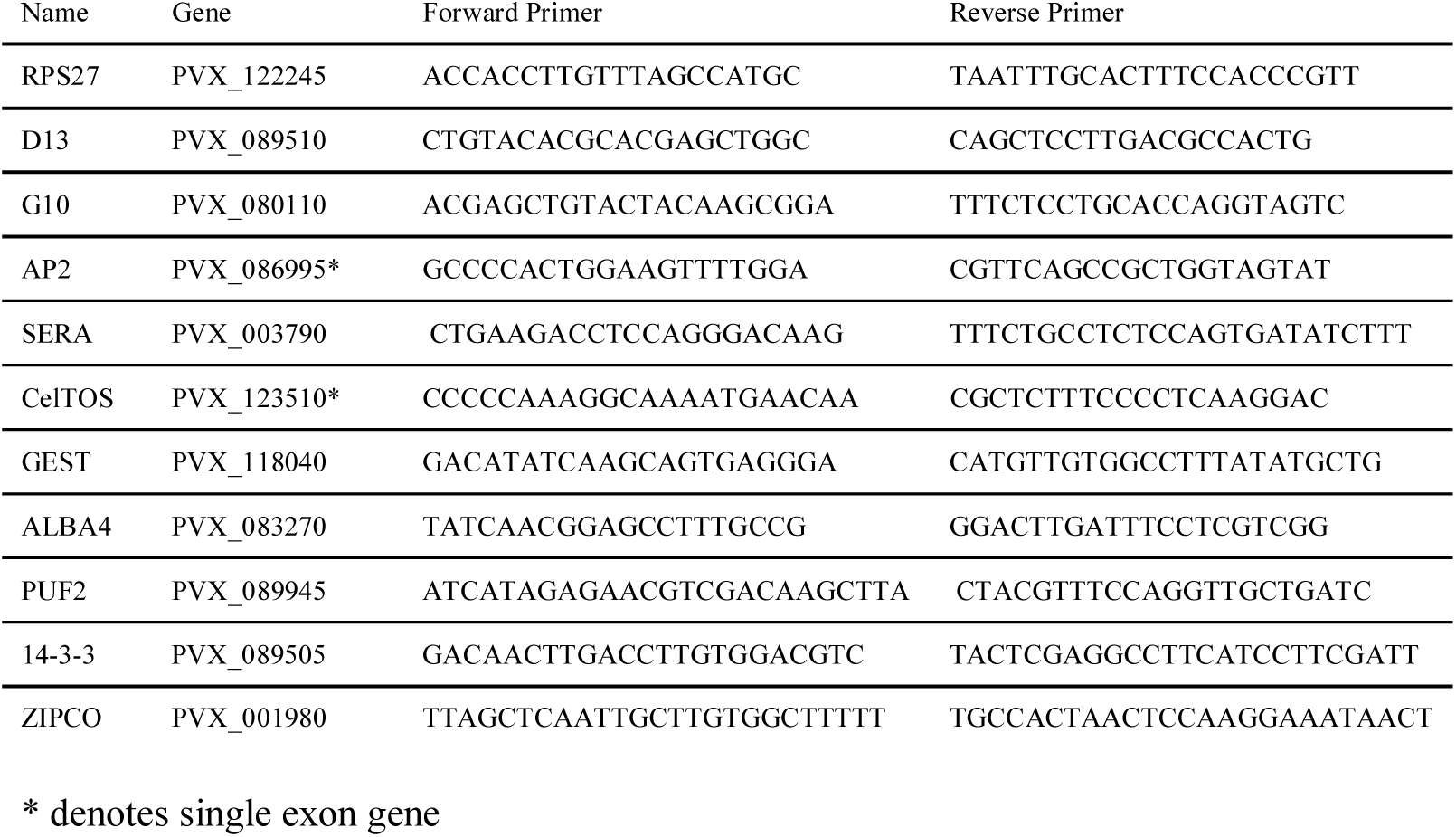

### Salivary-gland sporozoite and liver-stage immunofluorescence assays (IFAs)

IFAs were performed as per [13] using preserved, vivax infected mouse liver tissue generated previously for that study. In [13], female FRG [fumarylacetoacetate hydrolase (F), recombination activation gene 2 (R), interleukin-2 receptor subunit gamma (G)] triple KO mice engrafted with human hepatocytes (FRG KO huHep) were purchased from Yecuris Corporation (Oregon, USA). Mice were infected through intravenous injection into the tail with 3.5 × 10^5^ to 1 × 10^6^ sporozoites isolated from the salivary glands of infected mosquitoes in 100 µl of RPMI media. Liver stages for the current study were obtained from 10µm formalin fixed paraffin embedded day 7 liver stages generated previously [13] from FRG knockout huHep mice;[13] these were deparaffinized prior to staining. Fresh salivary-gland sporozoites were fixed in acetone per [13]. All cells were incubated twice for 3 minutes in Xylene, then 100% Ethanol, and finally once for 3 minutes each in 95%, 70%, and 50% Ethanol. The cells were rinsed in DI water and permeabilized immediately in 1XTBS, containing Triton X-100 and 30% hydrogen peroxide. The cells were blocked in 5% milk in 1XTBS. The hepatocytes were stained overnight with a rabbit polyclonal LISP1 antibody (A), a rabbit polyclonal UIS4 antibody (B), and a rabbit polyclonal BIP antibody (C) in blocking buffer. The cells were washed with 1XTBS and the primary antibodies were detected with goat anti-rabbit Alexa Fluor 488 antibody (Life Technologies). The cells were washed in 1XTBS. The hepatocytes were rinsed in KMNO4 and washed in 1XTBS. The cells were incubated in DAPI for 5 minutes.

### Histone ChIP sequencing and analysis

Aliquots of 2 – 6 million freshly isolated sporozoites were fixed with 1% paraformaldehyde for 10 min at 37°C and the reaction subsequently quenched by adding glycine to a final concentration of 125 mM. After three washes with PBS, sporozoite pellets were stored at −80°C and shipped to Australia. Nuclei were released from the sporozoites by dounce homogenization in lysis buffer (10 mM Hepes pH 7.9, 10 mM KCl, 0.1 mM EDTA, 0.1 mM EDTA, 1 mM DTT, 1x EDTA-free protease inhibitor cocktail (Roche), 0.25% NP40). Nuclei were pelleted by centrifugation at 21,000 g for 10 min at 4°C and resuspended in SDS lysis buffer (1% SDS, 10 mM EDTA, 50 mM Tris pH 8.1, 1x EDTA-free protease inhibitor cocktail). Chromatin was sheared into 200–1000 bp fragments by sonication for 16 cycles in 30 sec intervals (on/off, high setting) using a Bioruptor (Diagenode) and diluted 1:10 in ChIP dilution buffer (0.01% SDS, 1.1% Triton X-100, 1.2 mM EDTA, 16.7 mM Tris pH 8.1, 150 mM NaCl). Chromatin was precleared for 1 hour with protein A/G sepharose (4FastFlow, GE Healthcare) equilibrated in 0.1% BSA (Sigma-Aldrich, USA) in ChIP dilution buffer. Chromatin from 3 x 10^5^ nuclei was taken aside as input material. Chromatin from approximately 3 x 10^6^ sporozoite nuclei was used for each ChIP. ChIP was carried out over night at 4°C with 5 µg of antibody (H3K9me3 (Active Motif), H3K4me3 (Abcam), H3K9ac (Upstate), H4K16ac (Abcam)) and 10 µl each of equilibrated protein A and G sepharose beads (4FastFlow, GE Healthcare). After washes in low-salt, high-salt, LiCl, and TE buffers (EZ-ChIP Kit, Millipore), precipitated complexes were eluted in 1% SDS, 0.1 M NaHCO_3_. Cross-linking of the immune complexes and input material was reversed for 6 hours at 45°C after addition of 500 mM NaCl and 20 µg/ml of proteinase K (NEB). DNA was purified using the MinElute® PCR purification kit (Qiagen) and paired-end sequenced on Illumina NextSeq using TruSeq library construction chemistry as per the manufacturer’s instructions. Raw reads for each ChIP-seq replicate are available through the Sequence Read Archive (XXX-XXX).

Fastq files were checked for quality using fastqc (<u>http://www.bioinformatics.babraham.ac.uk/projects/fastqc/</u>) and adapter sequences were trimmed using cutadapt [77]. Paired end reads were mapped to the *P. vivax* P01 strain genome annotation using Bowtie2 [68]. The alignment files were converted to Bam format, sorted and indexed using Samtools [78]. ChIP peaks were called relative to input using MACS2[79] in paired end mode with a q value less than or equal to 0.01. Peaks and peak summits were converted to sorted BED files. Bedtools intersect[80] was used to identify genes that intersected H3K9me3 peaks and Bedtools closest was used to identify genes that were closest to and downstream of H3K9ac and H3K4me3 peak summits.

### Sequence motif analysis

Conserved sequence motifs were identified using the program DREME [81]. Only genes in the top decile of transcription showing no evidence of protein expression in multiple salivary-gland sporozoite replicates were considered as putatively translationally repressed (n = 170). We queried coding regions and regions upstream of the transcriptional start site (TSS) for each gene, defined by Zhu *et al* [40] and/or predicted here from all RNA-seq data using the Tuxedo suite [82], for enriched sequence motifs in comparison to 170 genes found to be in the top decile of both transcriptional and expressional abundance in the same sporozoite replicates. In searching for motifs associated with highly transcribed genes with stable H3K9ac marks within 1kb of the TSS (or up to the 3’ end of the next gene upstream), we compared H3K9ac marked genes in the top decile of transcription to the same number of H3K9ac marked genes in the bottom decile of transcription. In both instances, an e-value threshold of 0.05 was considered the minimum threshold for statistical significance.

## Supporting information

