## Supplementary material for "Transcriptome and histone epigenome of *Plasmodium vivax* salivary-gland sporozoites point to tight regulatory control and potential mechanisms for liver-stage differentiation"

**Supplementary Figure Legends:**

**Figure S1** Median and range in gene transcription (values in counts per million (CPM)) for each *P. vivax* salivary sporozoite isolate. **a** before and **b** following data normalization.

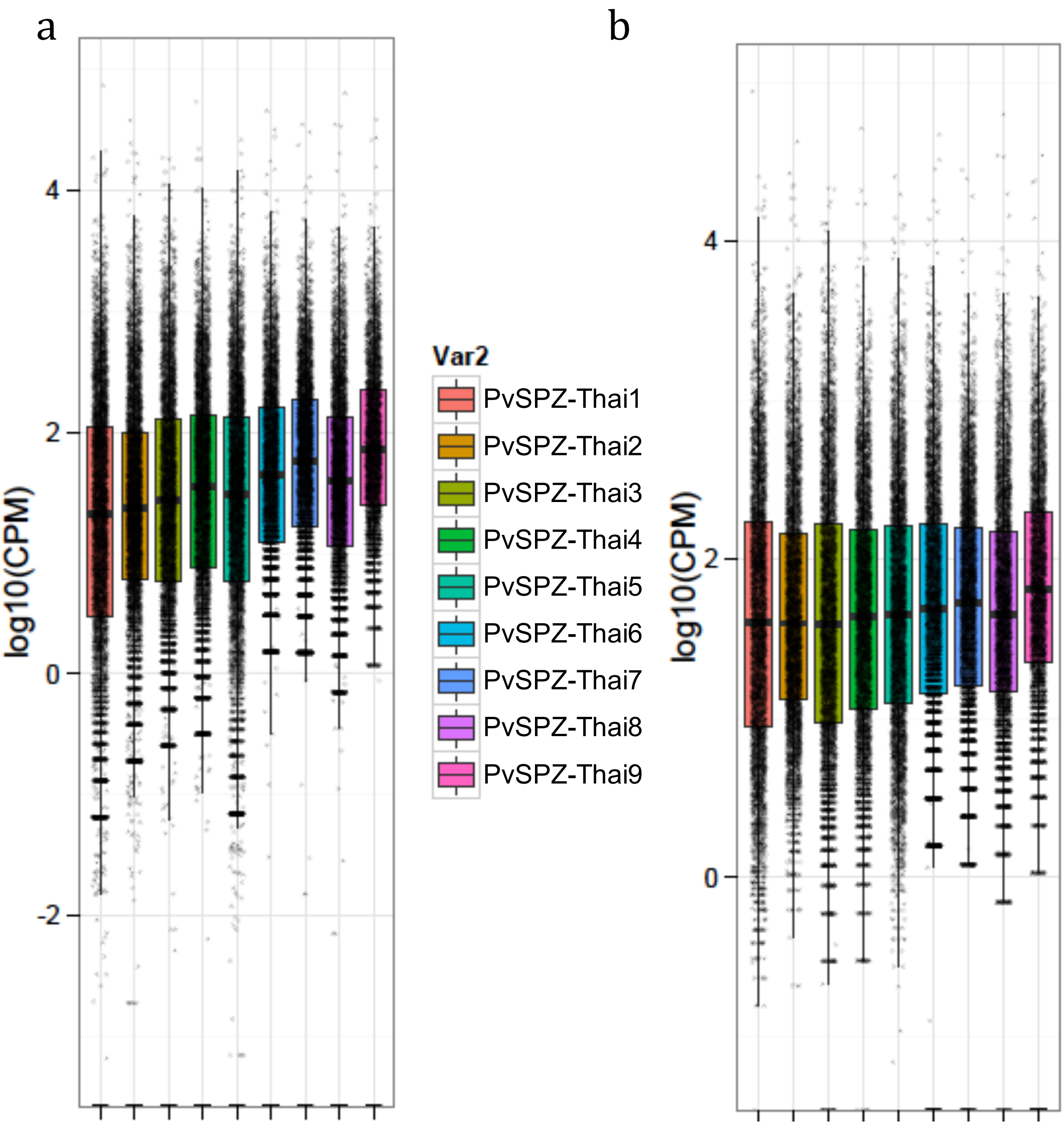

**Figure S2** Salivary sporozoite transcriptional correlation plots comparing **a** *P. vivax* microarray [1] and RNA-sequencing; **b** *P. vivax* and *P. falciparum* RNA sequencing; **c** *P. vivax* and *P. yoelli* RNA sequencing. All heatmaps based on single copy orthologous genes shared among all three species and represented on microarrays in previous publications. Data points represent transcription (TPKM or E for RNA-seq and microarray data respectively) as a proportion of total transcription (sum of TPKM or E for the data-set) for each experiment. Axes log10 transformed. $R^2$ values based on linear regression of log10 transformed data.

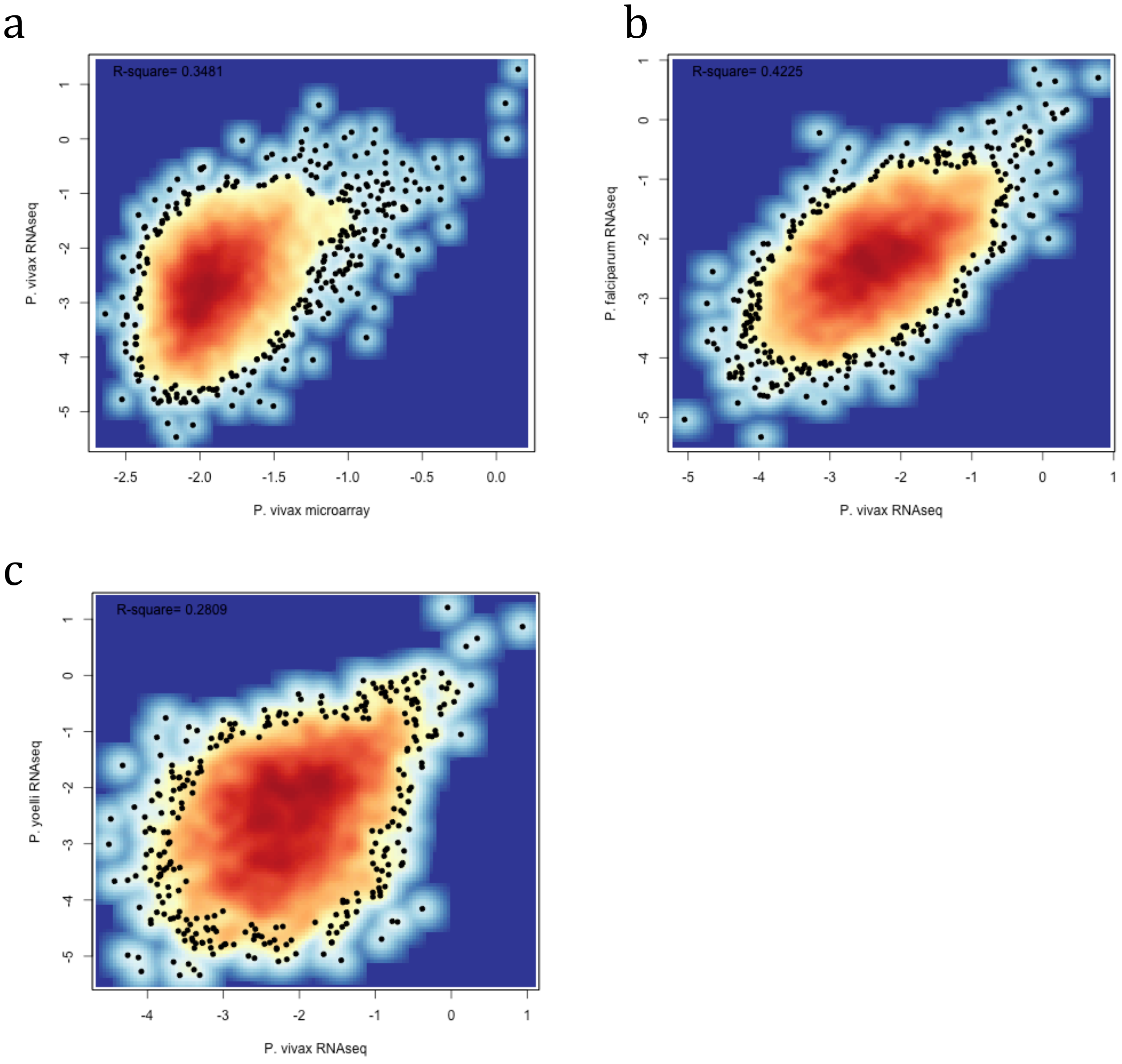

**Figure S3** Housekeeping genes for *P. vivax* inferred from Westenberger et al[1] microarray data. **a.** Scatterplot of median compared to median absolute deviation of expression of each *P. vivax* gene among all replicates and life-cycle stages in Westenberger et al[1] based on microarray data in log (left) and log(log) (right) transformation. **b.** Histogram showing frequency distribution of transcripts by abundance (per microarray data) for all data in Westenberger et al (left), compared to those transcripts with a MAD ≤ 0.07 (middle) and MAD ≤ 0.05 (right).

a

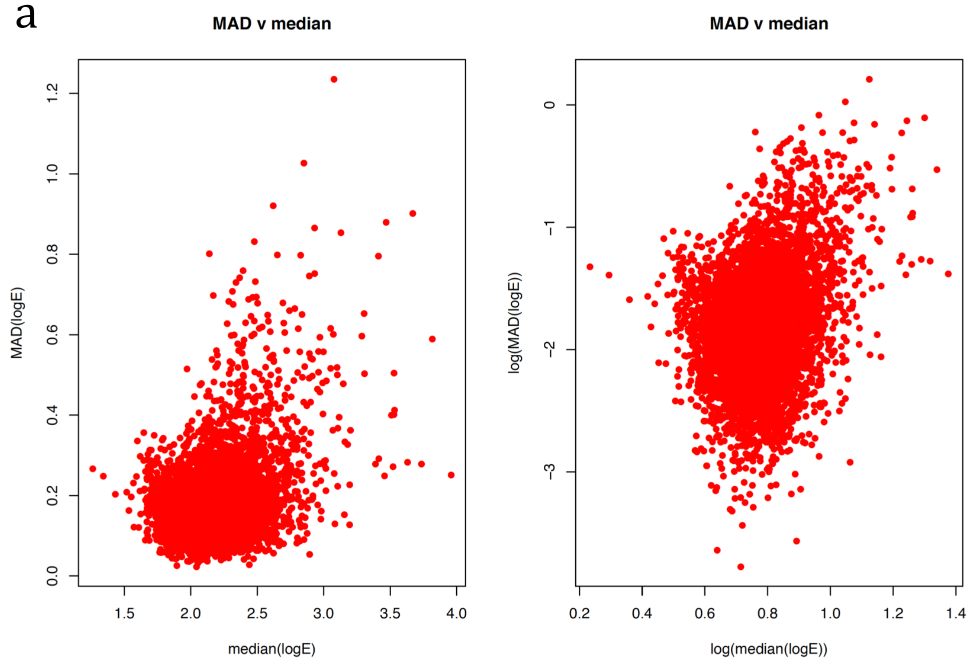


b

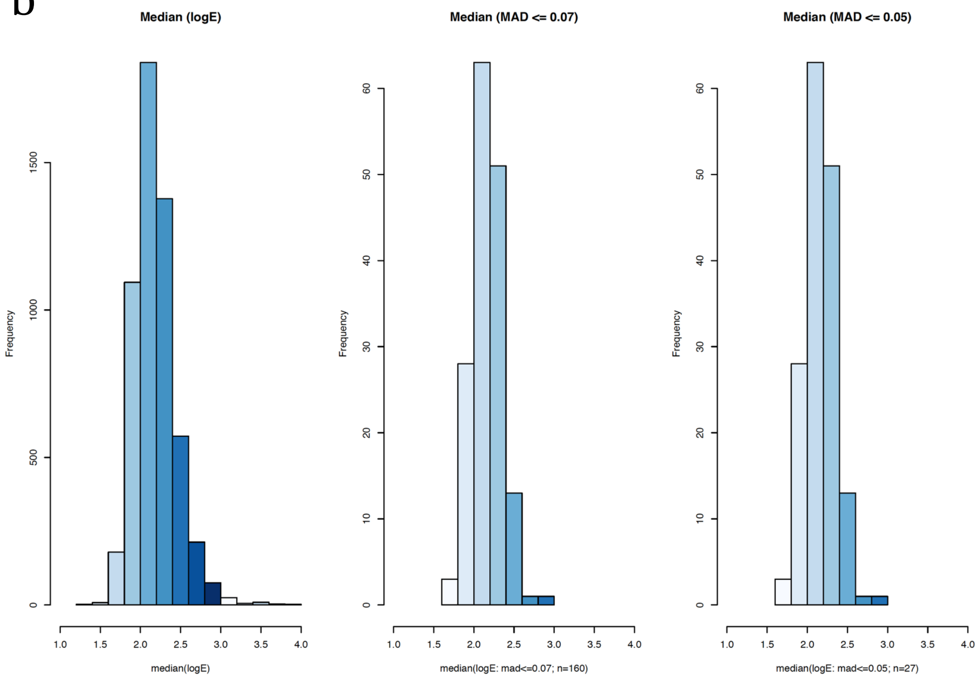

**Figure S4** MAD versus median transcription in sporozoite, blood-stage and liver-stage RNAseq data used in the current study (red dots), with low transcriptional-variation (i.e., 'housekeeping') genes inferred from Westenberger et al[1] microarray as having a MAD ≤ 0.07 in black.

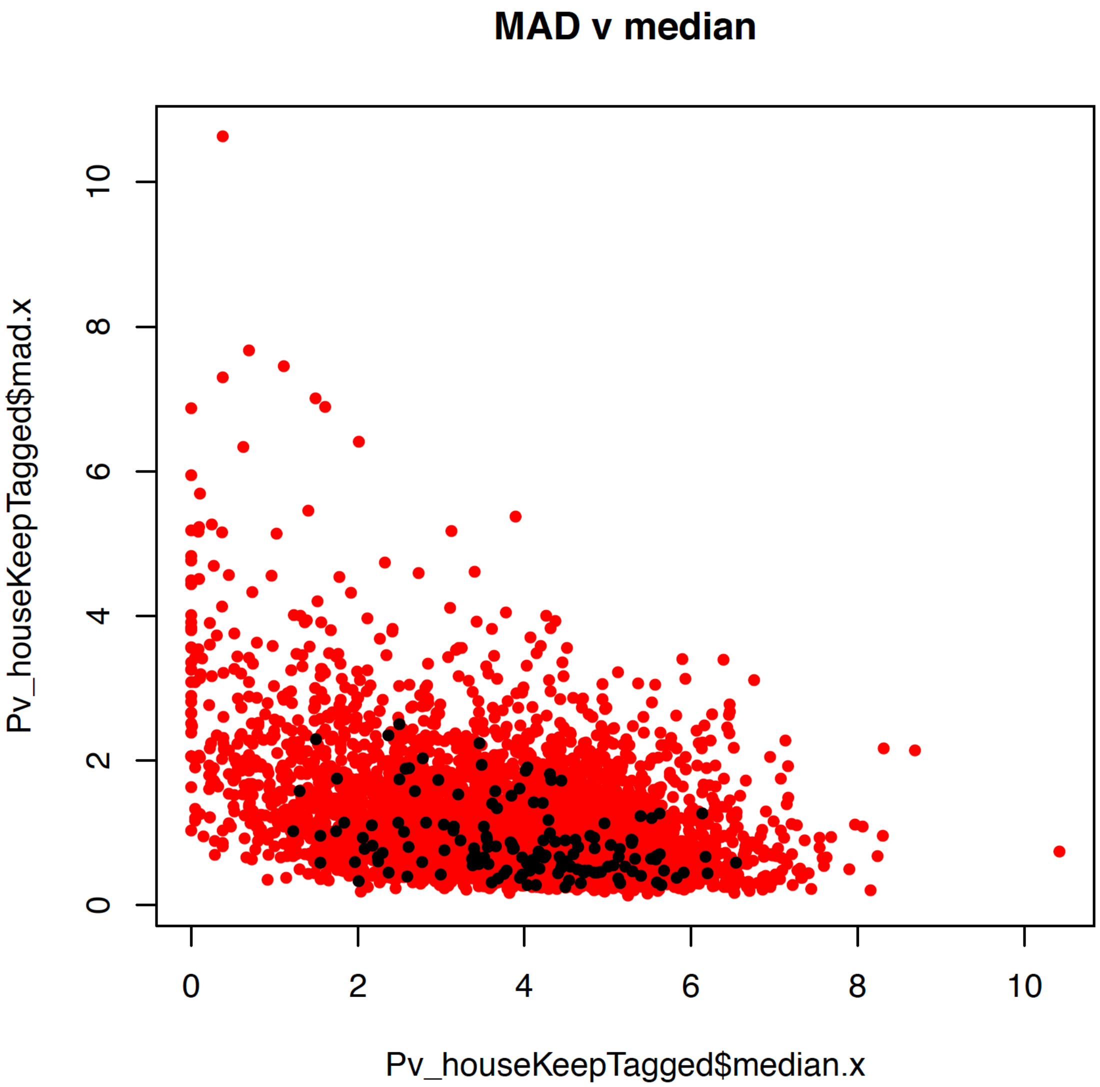

**Figure S5** Box and whisker plot of the mean of and range in abundance of each gene transcribed in *P. vivax* mixed salivary sporozoite, activated (RPMI + 3% BSA) sporozoite (source [2]), mixed and hypnozoite enriched liver-stage (source [4]), and asexual blood-stage (source [3]) isolates.

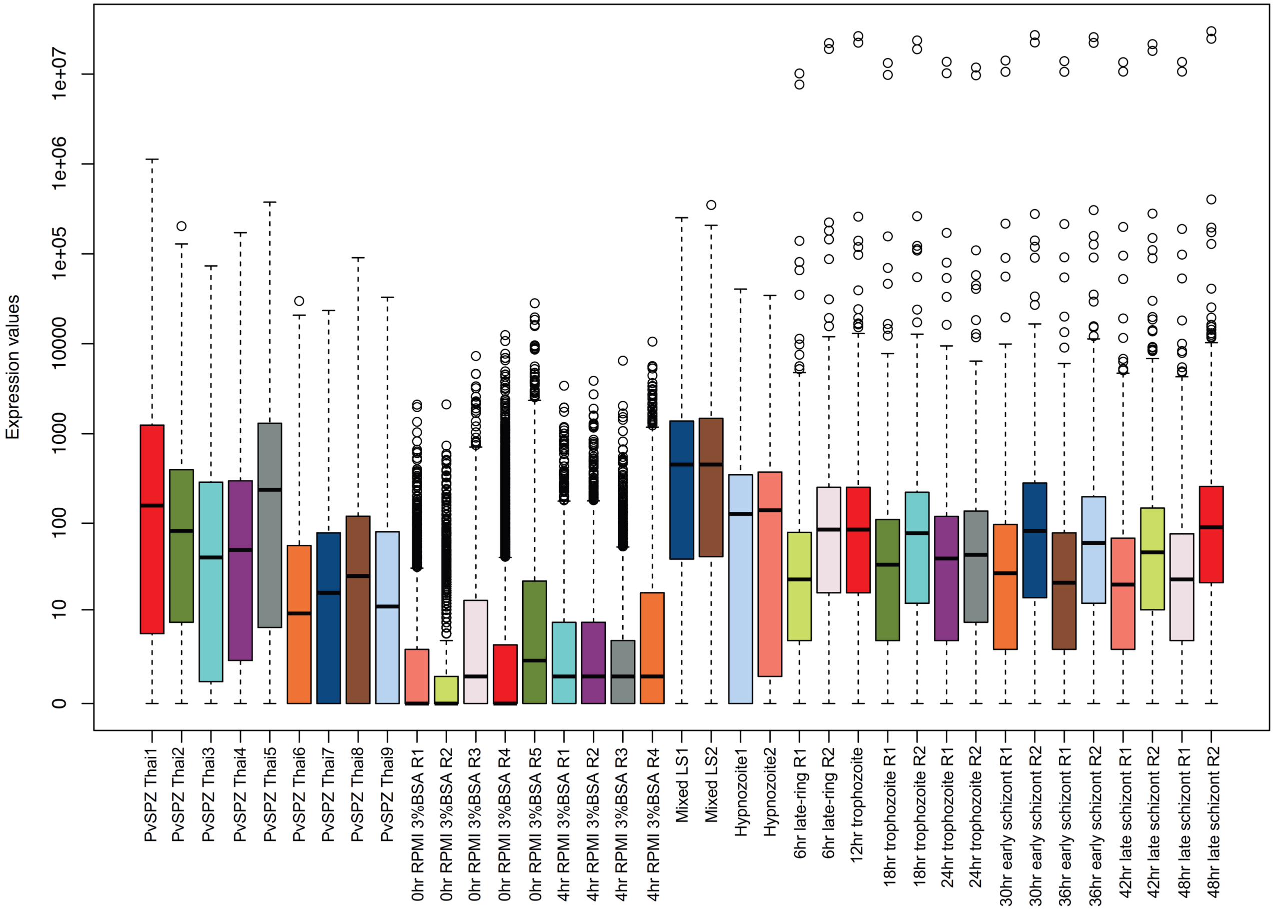

**Figure S6** Coding transcript coverage barplot by sample for (A) *P. vivax* mixed salivary sporozoite, (B) activated (RPMI + 3% BSA) sporozoite (source [2] (C) mixed and hypnozoite enriched liver-stage (source [4]), and (D) asexual blood-stage (source [3]) isolates.

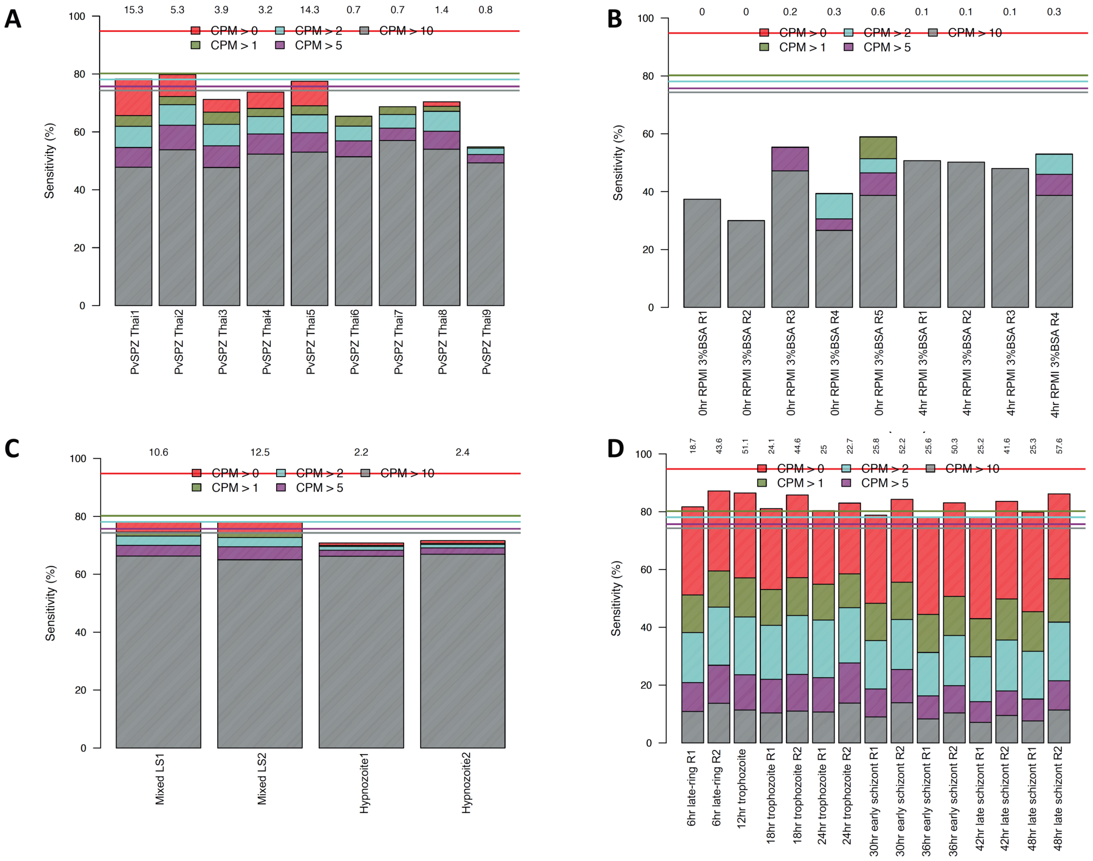

**Figure S7** Box and whisker plot of the mean of and range in abundance of each gene transcribed in *P. vivax* mixed blood-stage (source [3]) and salivary sporozoite isolates following TMM-normalization.

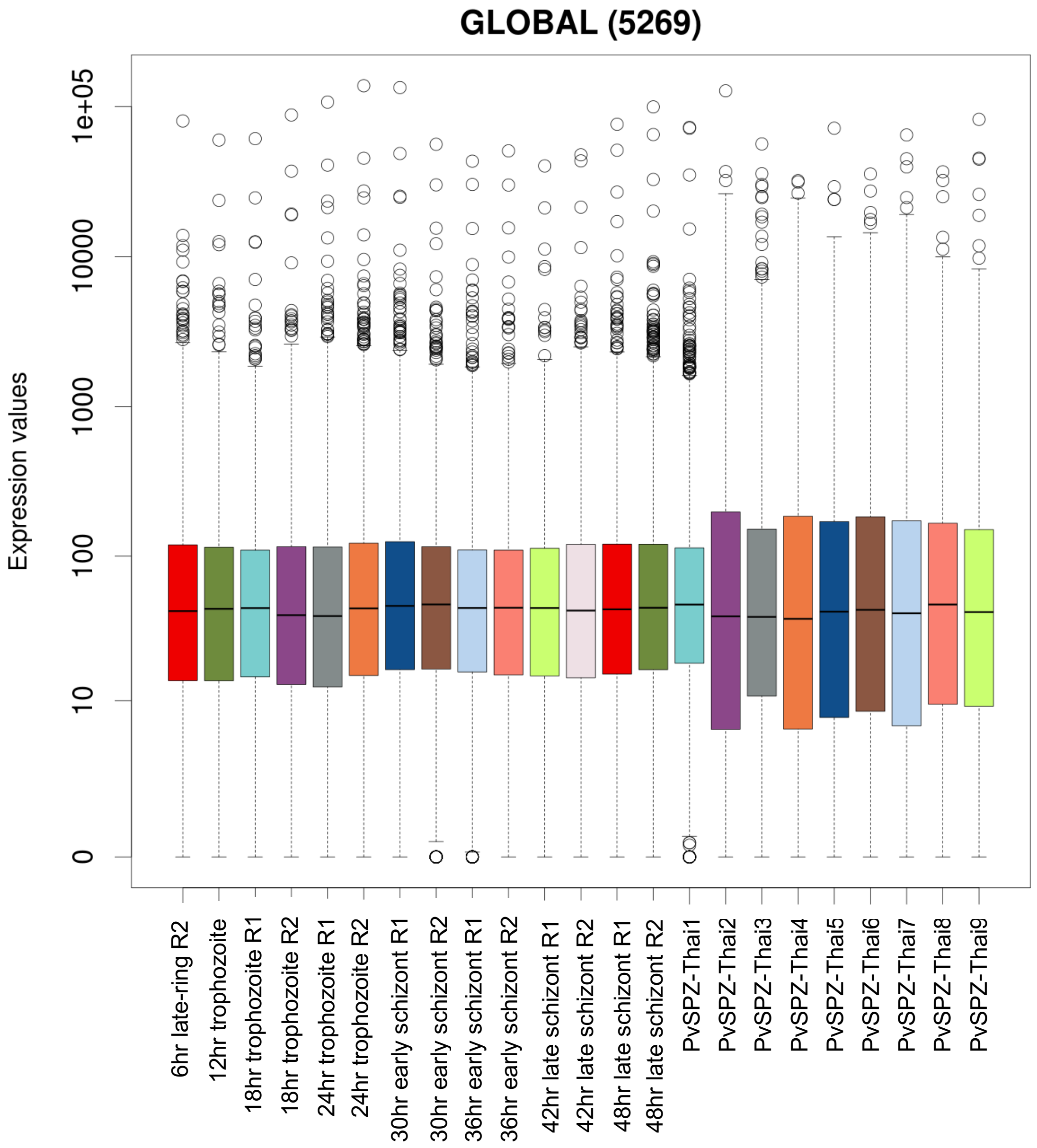

**Figure S8** Assessment of RNA-seq quantification bias by GC content in mixed *P. vivax* blood-stages[3] (red squares) and salivary sporozoites (blue circles).

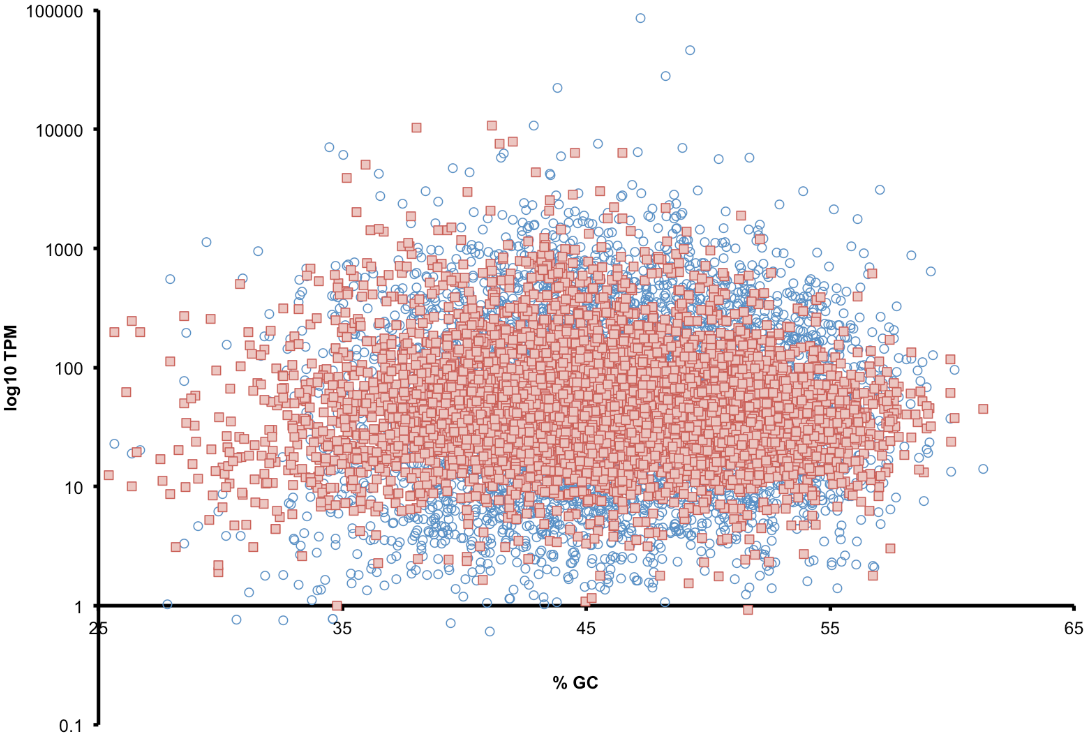

**Figure S9** Sporozoite enriched transcription in *P. vivax* relative to mixed blood stages. **a** pre-normalized BCV plot of pooled salivary sporozoite and blood-stage[3] RNA-sequencing data; **b** post-normalized BCV plot of pooled salivary sporozoite and blood-stage RNA-sequencing data

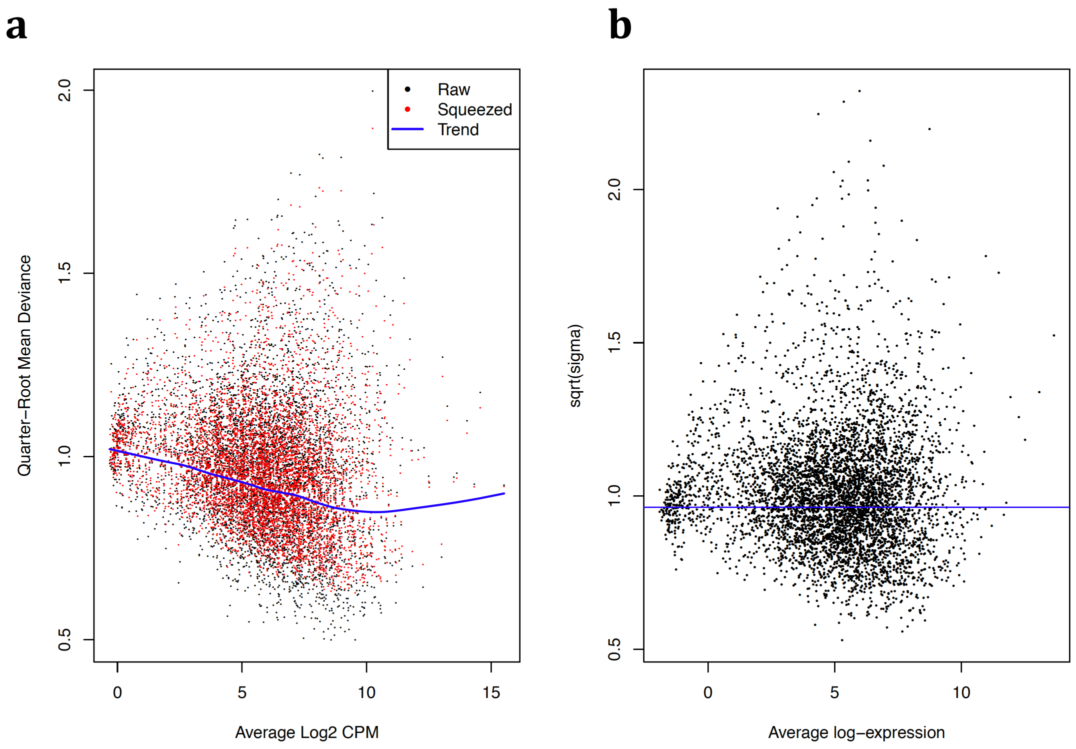

**Figure S10** MDS-plots for sporozoite vs blood-stage[3] comparisons by sample group (i.e., stage) and sequencing batch.

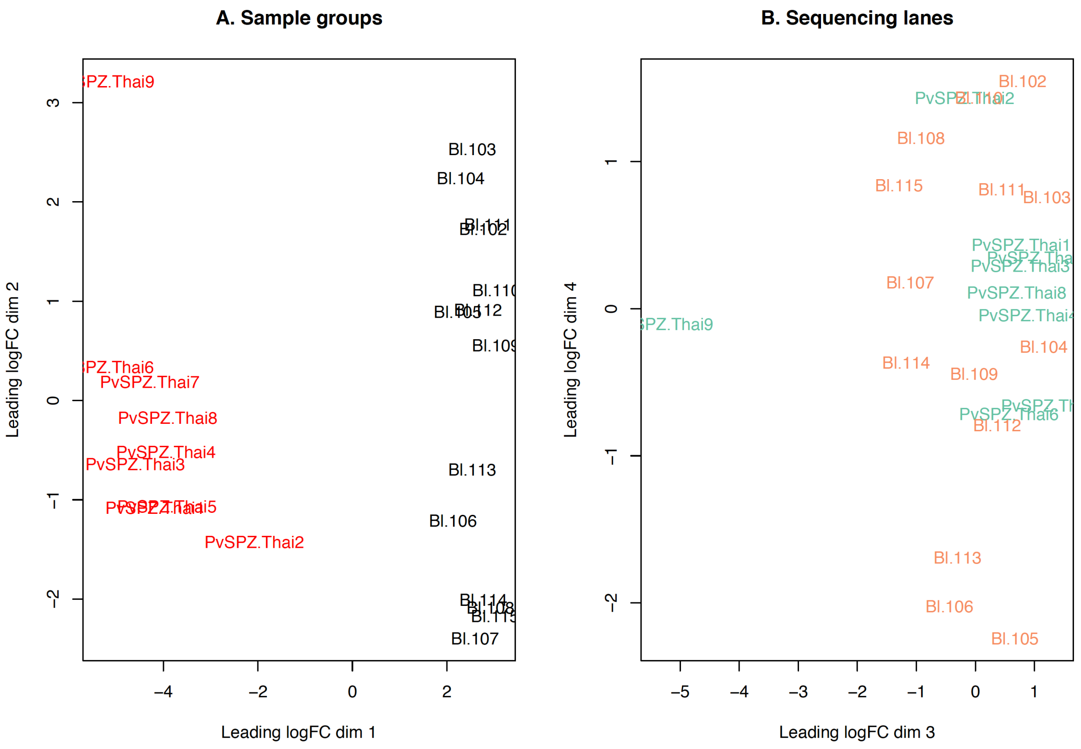

**Figure S11** Median and range in gene transcription (values in counts per million (CPM)) for each *P. vivax* salivary sporozoite isolate vs mixed liver-stage (MLS) and hypnozoite enriched liver stage data (HPZ) from Liver-stage data from Gural et al.[4] **a** before and **b** following data normalization

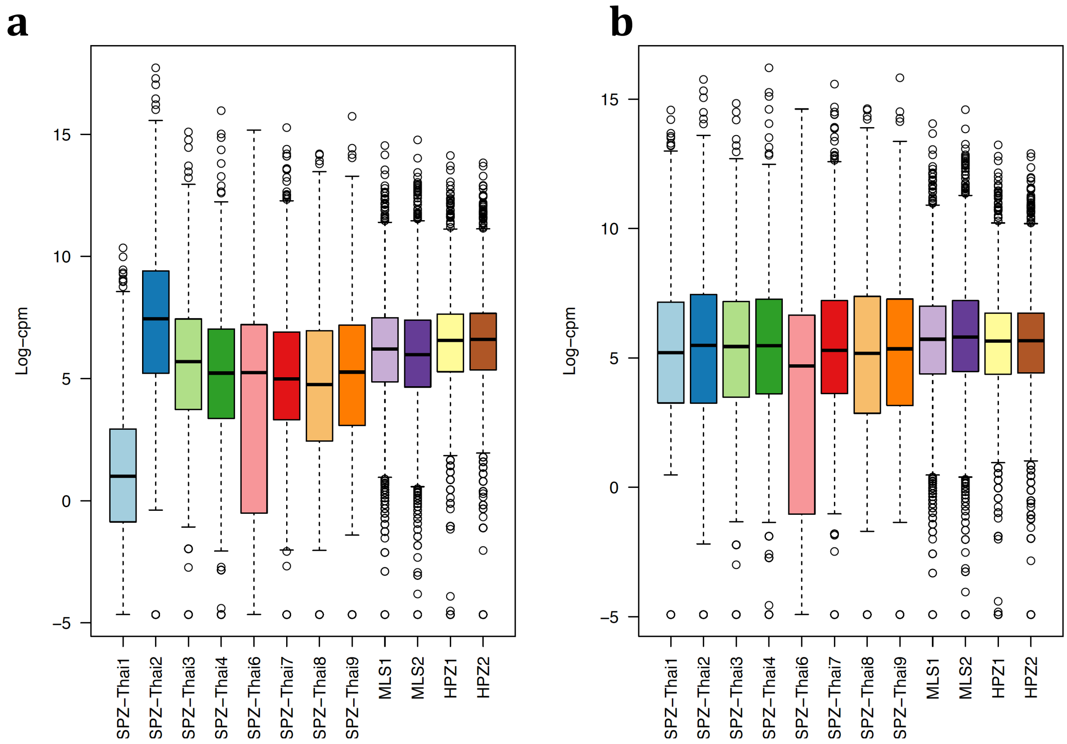

**Figure S12** Sporozoite enriched transcription in *P. vivax* relative to liver-stages (MLS and HPZ). **a** pre-normalized BCV plot of pooled salivary sporozoite and blood-stage RNA-sequencing data; **b** post-normalized BCV plot of pooled salivary sporozoite and liver-stage RNA-sequencing data. Liver-stage data from Gural et al.[4]

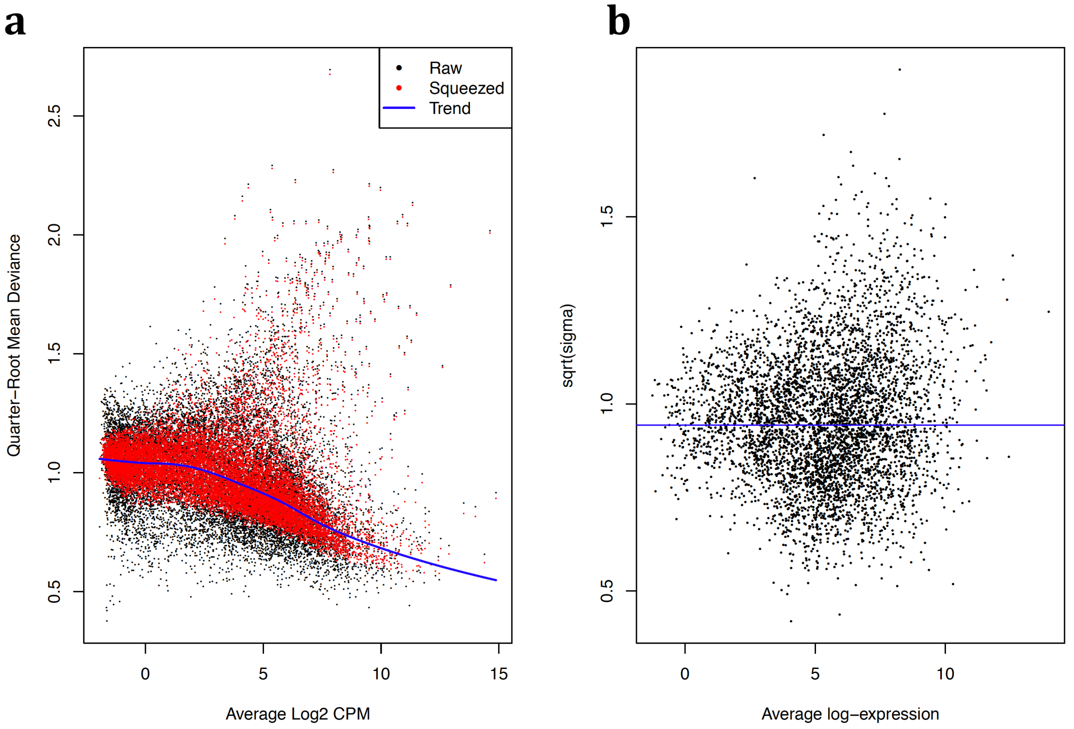

**Figure S13** MDS-plots for sporozoite vs liver-stage[4] comparisons by sample group (i.e., stage) and sequencing batch.

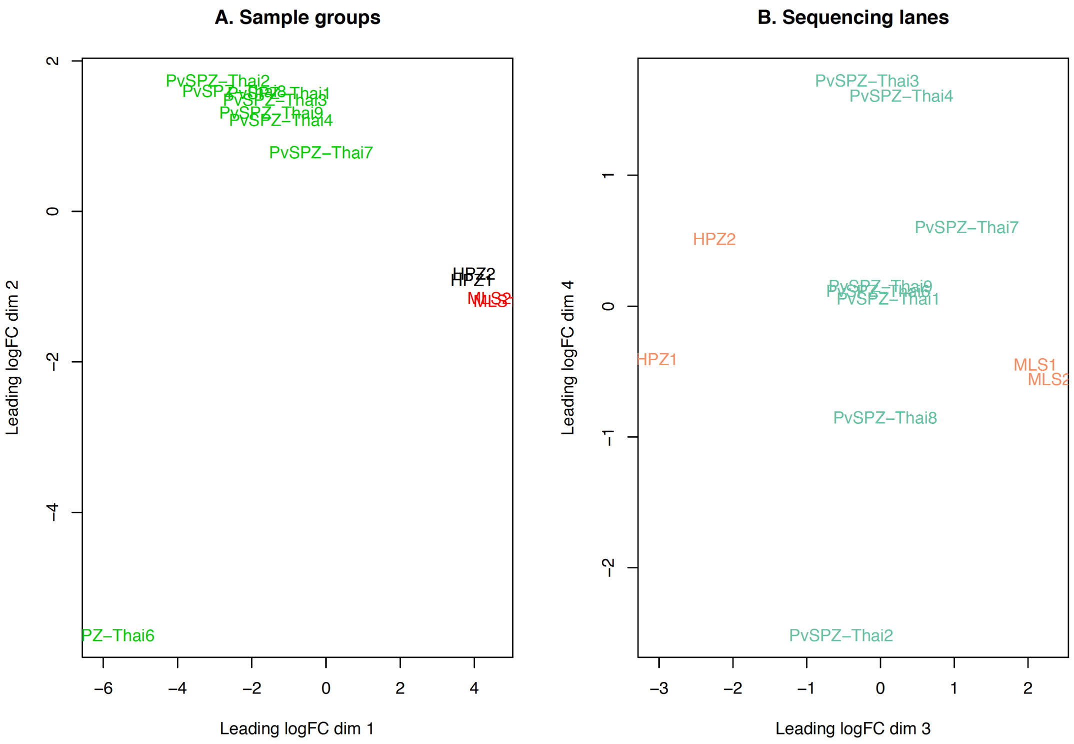

**Figure S14** Spearman's rank correlation matrix and self-organized heatmap of input normalized H3 histone modification ChIP-seq data by sporozoite isolate from the current study.

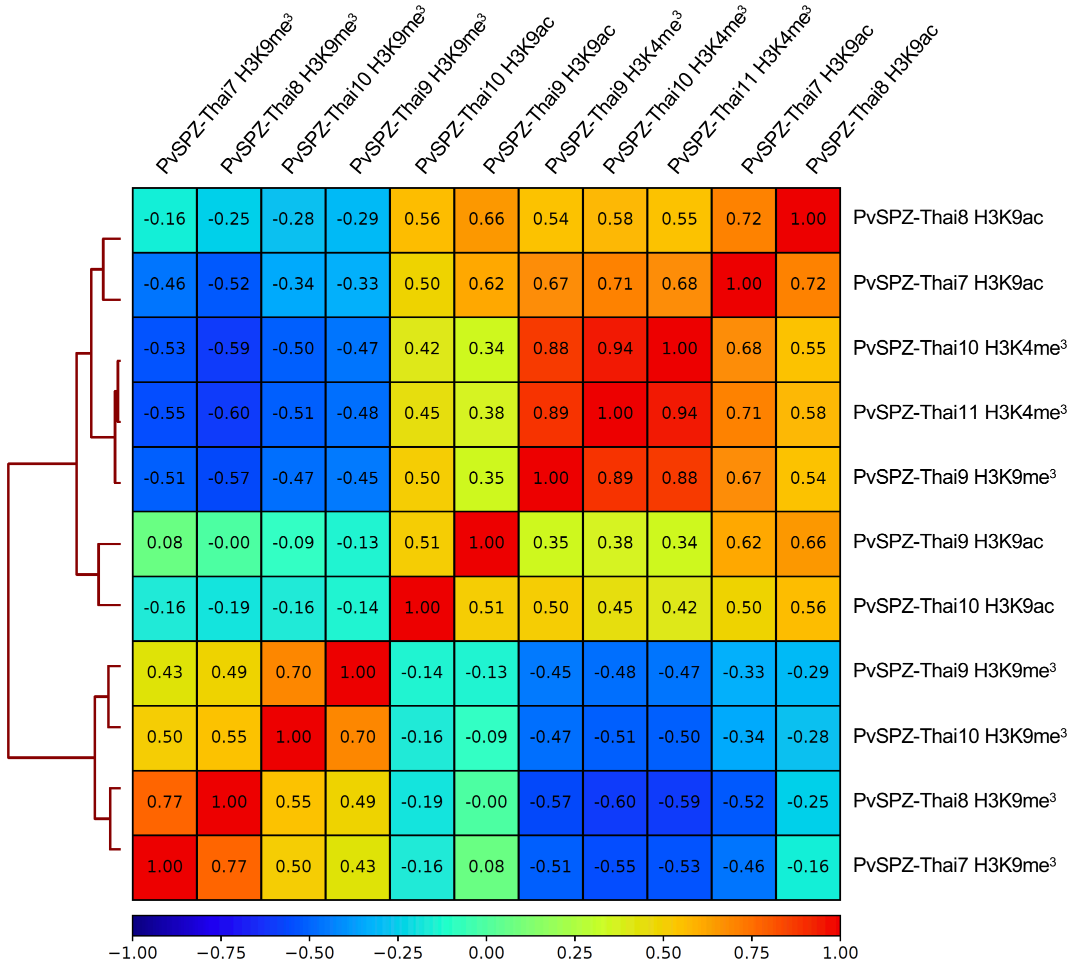

**Figure S15** Frequency distribution of peak widths associated with H3K9me3, H3K4me3 and H3K9ac histone modifications.

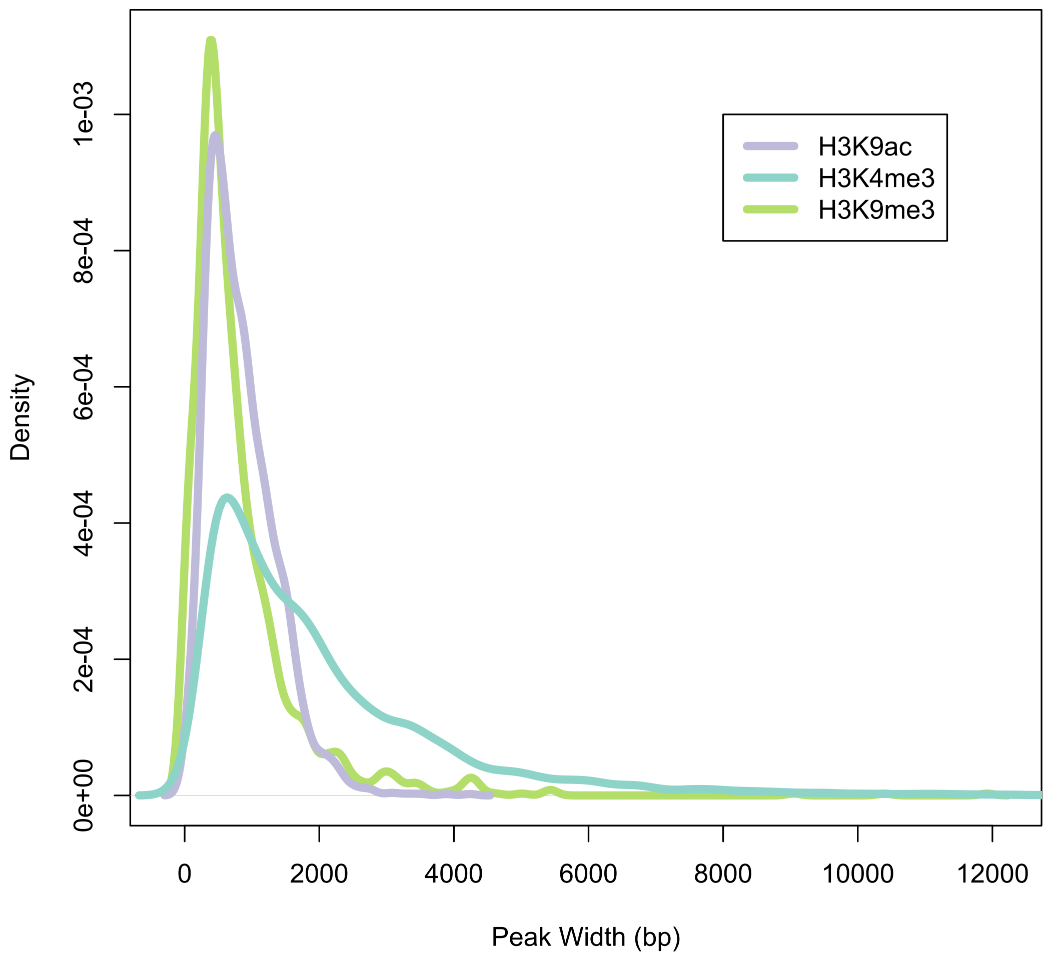

**Figure S16** Frequency distributions of spacing between peaks associated with **a** H3K9me3, **b** H3K4me3 and **c** H3K9ac histone modifications.

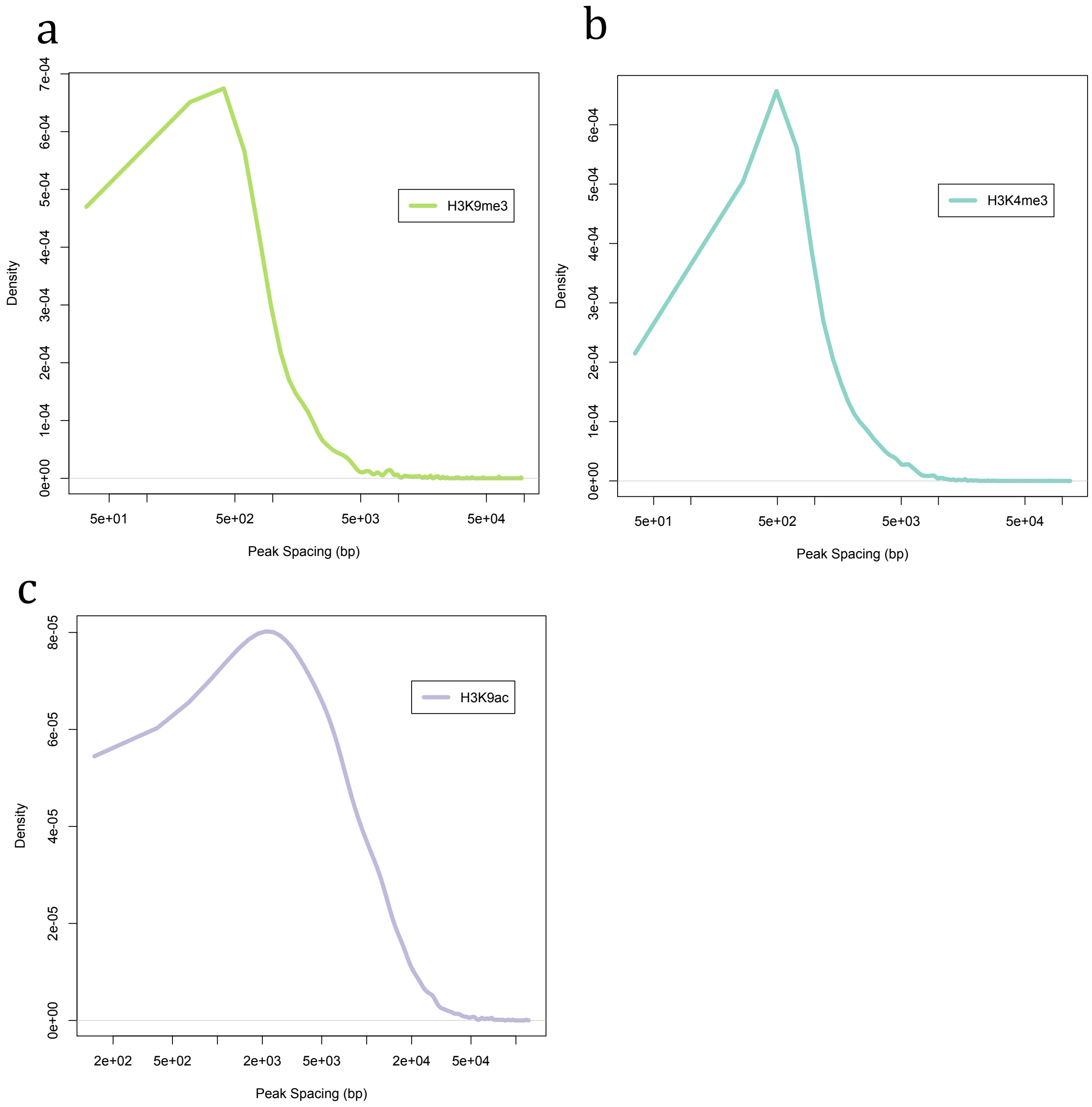

**Figure S17** Frequency distribution of distance of H3K4me3 and H3K9ac histone marks from the gene transcription start site.

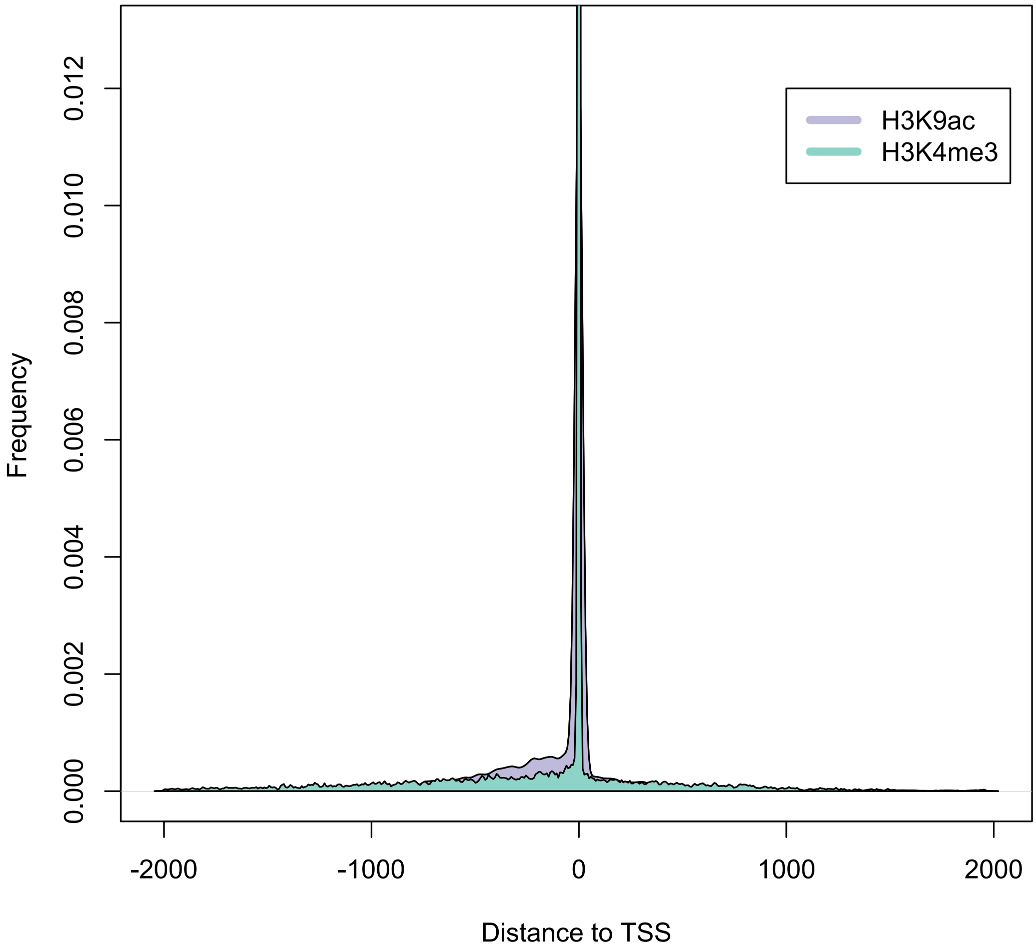

**Figure S18** Correlation matrix comparing major H3 histone marks by species and stage. *P. falciparum* data from [3,4]. Boxes marked with X show replicates with poor correlation with any other published data.

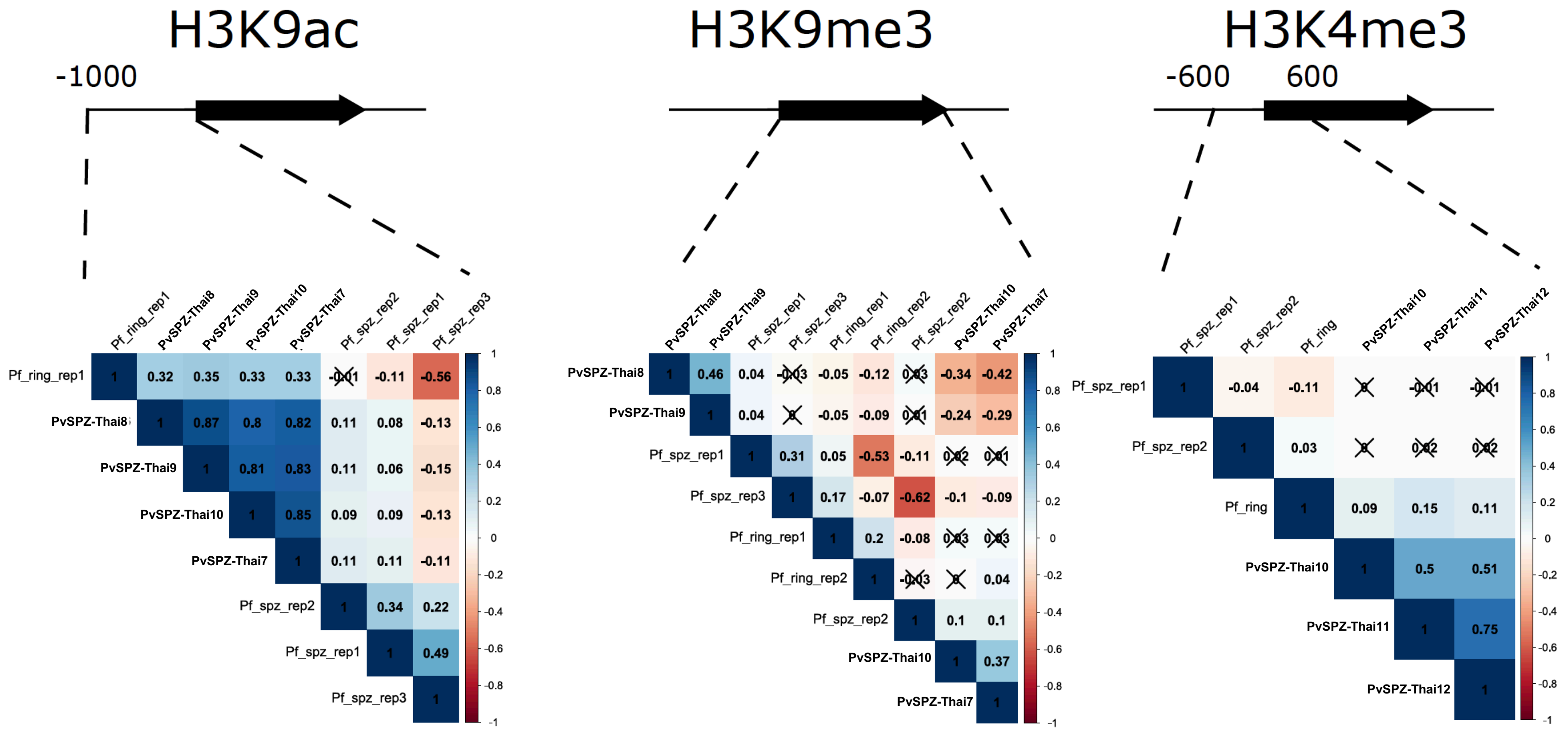

**Figure S19** Heatmap showing histone mark density in *P. vivax* sporozoites in regions up (-1kb), down (1kb) and within (ATG to stop) coding domains of each *P. vivax* coding gene. Each row represents a transcript. All transcripts are ranked (top to bottom) by TPM abundance. Data is a composite of mean density for each mark for each sample replicate subjected to ChiP-seq analyses in the current study.

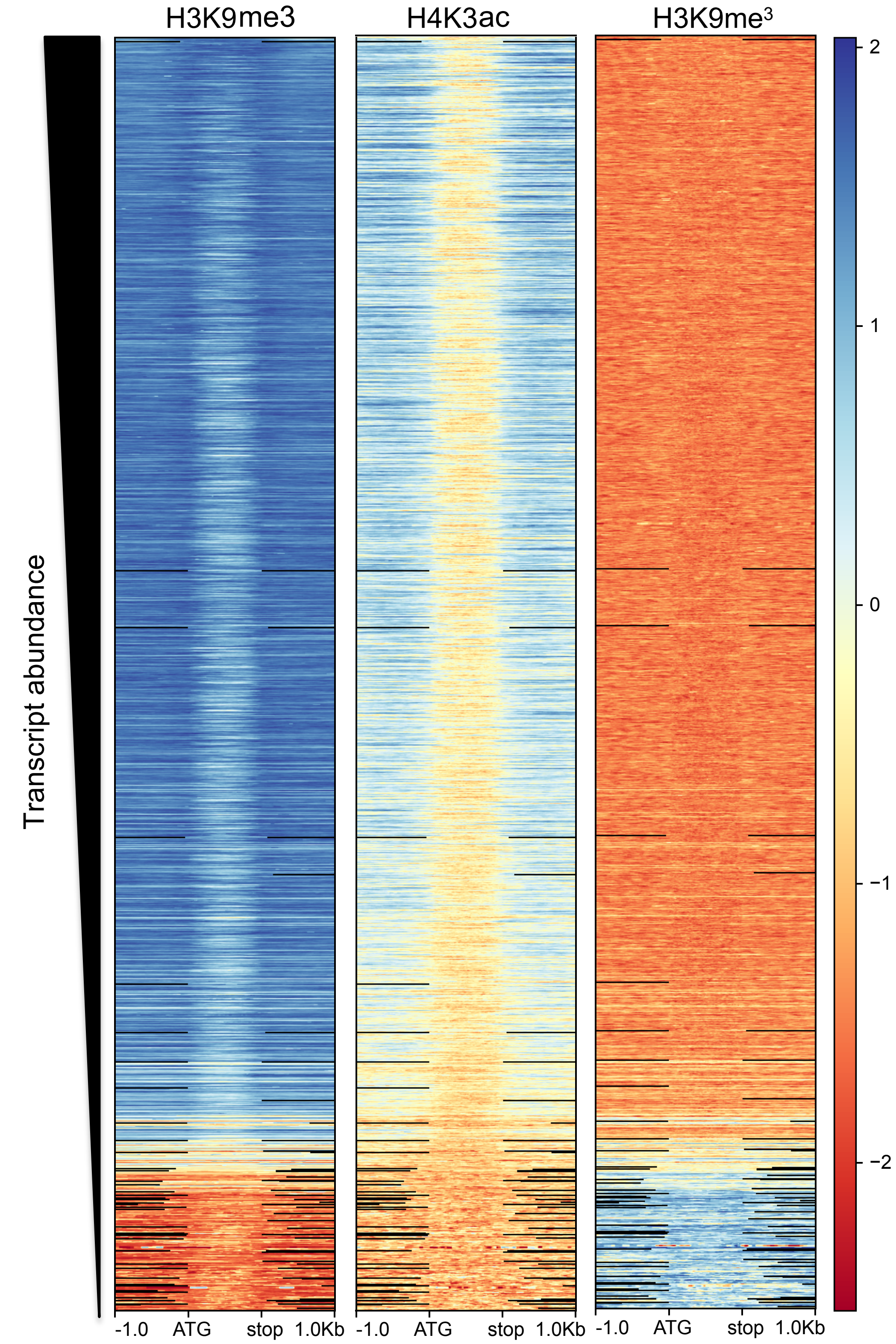

**Supplementary Table Legends:**

**Table S1** RNAseq and ChIPseq library and alignment metrics for *P. vivax* sporozoite data.

**Table S2** Transcript abundance (RNA-seq) per *P. vivax* salivary sporozoite replicate (presented in Counts per Million (CPM) and Transcripts per Million (TPM)).

**Table S3** Relative transcript abundance of genes transcribed in salivary sporozoites of *P. vivax* inferred by RNA-seq (here) compared to previously published microarray data[1]. TPM = transcripts per million; E = microarray expression. TPM proportion is the TPM for a given gene as a proportion of the sum of TPMs of all genes. E proportion is E for a given gene as a proportion of the sum of E of all genes.

**Table S4** Testing of additional *P. vivax* clinical isolates (PvSPZ-Thai13 to PvSPZ-Thai16) by qPCR for a representative panel of transcripts differing in our study compared with prior microarray[1] data, assessing abundance relative to *celtos* and *sera*.

**Table S5** Mean transcript abundance and relative transcript abundance rank comparison in salivary sporozoites (RNA-seq data only) of single copy orthologous genes shared among *P. vivax* P01, *P. falciparum* 3D7 and *P. yoelii* 17X (data from [5]). TPM = transcripts per million.

**Table S6** Mean transcript abundance (RNA-seq data only) in salivary sporozoites of *P. vivax* genes lacking orthologs in *P. falciparum* 3D7 or *P. yoelii* 17X. TPM = transcripts per million.

**Table S7** Comparison of relative mean transcription and protein expression of genes in *P. vivax* salivary sporozoites. TPM = transcripts per million; NSAF = normalize spectral abundance factor. Protein abundance rank based on mean ln[NSAF].

**Table S8** Independent LC-MS detection of orthologs of each *P. vivax* gene putatively defined as translationally repressed in sporozoites.

**Table S9** *P. vivax* transcripts statistically significantly (FDR ≤ 0.05) differentially transcribed in salivary sporozoites relative to mixed blood-stages (data from [6]). EC = expected count; CPM = counts per million; FDR = false discovery rate.

**Table S10** Protein family domain (Pfam) annotations for differentially enriched transcripts (FDR ≤ 0.05) between *P. vivax* salivary sporozoites and mixed blood-stages (data from [6]). EC = expected count; CPM = counts per million; FDR = false discovery rate.

**Table S11** Frequency counts of Gene Ontology terms associated with differentially enriched transcripts in *P. vivax* salivary sporozoites relative to mixed blood stages. BS = blood-stage; SPZ = salivary sporozoite.

**Table S12** *P. vivax* transcripts statistically significantly (FDR ≤ 0.01) differentially transcribed in salivary sporozoites relative to mixed liver-stages (data from [4]) . EC = expected count; CPM = counts per million; FDR = false discovery rate.

**Table S13** *P. vivax* transcripts statistically significantly (FDR ≤ 0.01) differentially transcribed in salivary sporozoites relative to hypnozoite-enriched liver-stages (data from [4]). EC = expected count; CPM = counts per million; FDR = false discovery rate.

**Table S14** *Plasmodium vivax* transcripts statistically significantly down-regulated (FDR ≤ 0.01) in mixed liver stages relative to sporozoites and hypnozoite-enriched liver-stages, but transcribed at similar levels in both sporozoites and hypnozoite-enriched liver-stages.

**Table S15** Summary of histone epigenetic marker data for H3K9me$^3$, H3K9ac and H3K4me$^3$ modifications, including information on the number and stability (i.e., representation among multiple replicates) of called peaks for each mark per genome per sample and intersecting with (H3K9me$^3$) or within 1kb of (H3K9ac and H3K4me$^3$) annotated protein coding genes in the *P. vivax* P01 genome annotation. Additionally, provides data on mean peak size and spacing, and the total proportion of the genome under each histone mark.

**Table S16** Transcriptional abundance of stably H3K9me$^3$ marked genes (in 3 or more replicates) in the *P. vivax* salivary sporozoite. TPM = transcripts per million.

**Table S17** Transcriptional abundance of stably H3K9ac marked genes (in 3 or more replicates) in the *P. vivax* salivary sporozoite. TPM = transcripts per million.

**Table S18** Transcriptional abundance of stably H3K4me$^3$ marked genes (in 2 or more replicates) in the *P. vivax* salivary sporozoite. TPM = transcripts per million.

**Table S19** Transcriptional abundance of *P. vivax* genes stably associated with both H3K9ac (3 or more replicates) and H3K4me$^3$ (2 or more replicates) histone marks in the salivary sporozoite. TPM = transcripts per million.

**Table S20** Relative transcriptional abundance of telomeric and subtelomeric (within genes (*P. vivax* P01 assembly) in *P. vivax* salivary sporozoites and mixed blood-stages (data from [6]). BS = blood-stage; SPZ = salivary sporozoite; EC = expected count; CPM = counts per million; TPM = transcripts per million; FDR = false discovery rate.

**Table S21** Relative transcriptional abundance of genes adjacent (within 50 kb) of the subtelomeres (*P. vivax* P01 assembly) in *P. vivax* salivary sporozoites and mixed blood-stages (data from [6]). BS = blood-stage; SPZ = salivary sporozoite; EC = expected count; CPM = counts per million; TPM = transcripts per million; FDR = false discovery rate.

**Table S22** Relative transcriptional abundance of non-telomeric (> 50kb away from the subtelomeres; *P. vivax* P01 assembly) in *P. vivax* salivary sporozoites and mixed blood-stages (data from [6]). BS = blood-stage; SPZ = salivary sporozoite; EC = expected count; CPM = counts per million; TPM = transcripts per million; FDR = false discovery rate.

1
