## Supplementary material for "Transcriptome and histone epigenome of *Plasmodium vivax* salivary-gland sporozoites point to tight regulatory control and potential mechanisms for liver-stage differentiation": glimma.min.js

```
(function t(e,n,i){function r(s,o){if(!n[s]){if(!e[s]){var l=typeof require=="function"&&require;if(!o&&l)return l(s,!0);if(a)return a(s,!0);var u=new Error("Cannot find module '"+s+"'");throw u.code="MODULE_NOT_FOUND",u}var f=n[s]={exports:{}};e[s][0].call(f.exports,function(t){var n=e[s][1][t];return r(n?n:t)},f,f.exports,t,e,n,i)}return n[s].exports}var a=typeof require=="function"&&require;for(var s=0;s<i.length;s++)r(i[s]);return r})({1:[function(t,e,n){t("../../js/transition.js");t("../../js/alert.js");t("../../js/button.js");t("../../js/carousel.js");t("../../js/collapse.js");t("../../js/dropdown.js");t("../../js/modal.js");t("../../js/tooltip.js");t("../../js/popover.js");t("../../js/scrollspy.js");t("../../js/tab.js");t("../../js/affix.js")},{"../../js/affix.js":2,"../../js/alert.js":3,"../../js/button.js":4,"../../js/carousel.js":5,"../../js/collapse.js":6,"../../js/dropdown.js":7,"../../js/modal.js":8,"../../js/popover.js":9,"../../js/scrollspy.js":10,"../../js/tab.js":11,"../../js/tooltip.js":12,"../../js/transition.js":13}],2:[function(t,e,n){+function(t){"use strict";var e=function(n,i){this.options=t.extend({},e.DEFAULTS,i);this.$target=t(this.options.target).on("scroll.bs.affix.data-api",t.proxy(this.checkPosition,this)).on("click.bs.affix.data-api",t.proxy(this.checkPositionWithEventLoop,this));this.$element=t(n);this.affixed=null;this.unpin=null;this.pinnedOffset=null;this.checkPosition()};e.VERSION="3.3.7";e.RESET="affix affix-top affix-bottom";e.DEFAULTS={offset:0,target:window};e.prototype.getState=function(t,e,n,i){var r=this.$target.scrollTop();var a=this.$element.offset();var s=this.$target.height();if(n!=null&&this.affixed=="top")return r<n?"top":false;if(this.affixed=="bottom"){if(n!=null)return r+this.unpin<=a.top?false:"bottom";return r+s<=t-i?false:"bottom"}var o=this.affixed==null;var l=o?r:a.top;var u=o?s:e;if(n!=null&&r<=n)return"top";if(i!=null&&l+u>=t-i)return"bottom";return false};e.prototype.getPinnedOffset=function(){if(this.pinnedOffset)return this.pinnedOffset;this.$element.removeClass(e.RESET).addClass("affix");var t=this.$target.scrollTop();var n=this.$element.offset();return this.pinnedOffset=n.top-t};e.prototype.checkPositionWithEventLoop=function(){setTimeout(t.proxy(this.checkPosition,this),1)};e.prototype.checkPosition=function(){if(!this.$element.is(":visible"))return;var n=this.$element.height();var i=this.options.offset;var r=i.top;var a=i.bottom;var s=Math.max(t(document).height(),t(document.body).height());if(typeof i!="object")a=r=i;if(typeof r=="function")r=i.top(this.$element);if(typeof a=="function")a=i.bottom(this.$element);var o=this.getState(s,n,r,a);if(this.affixed!=o){if(this.unpin!=null)this.$element.css("top","");var l="affix"+(o?"-"+o:"");var u=t.Event(l+".bs.affix");this.$element.trigger(u);if(u.isDefaultPrevented())return;this.affixed=o;this.unpin=o=="bottom"?this.getPinnedOffset():null;this.$element.removeClass(e.RESET).addClass(l).trigger(l.replace("affix","affixed")+".bs.affix")}if(o=="bottom"){this.$element.offset({top:s-n-a})}};function n(n){return this.each(function(){var i=t(this);var r=i.data("bs.affix");var a=typeof n=="object"&&n;if(!r)i.data("bs.affix",r=new e(this,a));if(typeof n=="string")r[n]()})}var i=t.fn.affix;t.fn.affix=n;t.fn.affix.Constructor=e;t.fn.affix.noConflict=function(){t.fn.affix=i;return this};t(window).on("load",function(){t('[data-spy="affix"]').each(function(){var e=t(this);var i=e.data();i.offset=i.offset||{};if(i.offsetBottom!=null)i.offset.bottom=i.offsetBottom;if(i.offsetTop!=null)i.offset.top=i.offsetTop;n.call(e,i)})})}(jQuery)},{}],3:[function(t,e,n){+function(t){"use strict";var e='[data-dismiss="alert"]';var n=function(n){t(n).on("click",e,this.close)};n.VERSION="3.3.7";n.TRANSITION_DURATION=150;n.prototype.close=function(e){var i=t(this);var r=i.attr("data-target");if(!r){r=i.attr("href");r=r&&r.replace(/.*(?=#[^\s]*$)/,"")}var a=t(r==="#"?[]:r);if(e)e.preventDefault();if(!a.length){a=i.closest(".alert")}a.trigger(e=t.Event("close.bs.alert"));if(e.isDefaultPrevented())return;a.removeClass("in");function s(){a.detach().trigger("closed.bs.alert").remove()}t.support.transition&&a.hasClass("fade")?a.one("bsTransitionEnd",s).emulateTransitionEnd(n.TRANSITION_DURATION):s()};function i(e){return this.each(function(){var i=t(this);var r=i.data("bs.alert");if(!r)i.data("bs.alert",r=new n(this));if(typeof e=="string")r[e].call(i)})}var r=t.fn.alert;t.fn.alert=i;t.fn.alert.Constructor=n;t.fn.alert.noConflict=function(){t.fn.alert=r;return this};t(document).on("click.bs.alert.data-api",e,n.prototype.close)}(jQuery)},{}],4:[function(t,e,n){+function(t){"use strict";var e=function(n,i){this.$element=t(n);this.options=t.extend({},e.DEFAULTS,i);this.isLoading=false};e.VERSION="3.3.7";e.DEFAULTS={loadingText:"loading..."};e.prototype.setState=function(e){var n="disabled";var i=this.$element;var r=i.is("input")?"val":"html";var a=i.data();e+="Text";if(a.resetText==null)i.data("resetText",i[r]());setTimeout(t.proxy(function(){i[r](a[e]==null?this.options[e]:a[e]);if(e=="loadingText"){this.isLoading=true;i.addClass(n).attr(n,n).prop(n,true)}else if(this.isLoading){this.isLoading=false;i.removeClass(n).removeAttr(n).prop(n,false)}},this),0)};e.prototype.toggle=function(){var t=true;var e=this.$element.closest('[data-toggle="buttons"]');if(e.length){var n=this.$element.find("input");if(n.prop("type")=="radio"){if(n.prop("checked"))t=false;e.find(".active").removeClass("active");this.$element.addClass("active")}else if(n.prop("type")=="checkbox"){if(n.prop("checked")!==this.$element.hasClass("active"))t=false;this.$element.toggleClass("active")}n.prop("checked",this.$element.hasClass("active"));if(t)n.trigger("change")}else{this.$element.attr("aria-pressed",!this.$element.hasClass("active"));this.$element.toggleClass("active")}};function n(n){return this.each(function(){var i=t(this);var r=i.data("bs.button");var a=typeof n=="object"&&n;if(!r)i.data("bs.button",r=new e(this,a));if(n=="toggle")r.toggle();else if(n)r.setState(n)})}var i=t.fn.button;t.fn.button=n;t.fn.button.Constructor=e;t.fn.button.noConflict=function(){t.fn.button=i;return this};t(document).on("click.bs.button.data-api",'[data-toggle^="button"]',function(e){var i=t(e.target).closest(".btn");n.call(i,"toggle");if(!t(e.target).is('input[type="radio"], input[type="checkbox"]')){e.preventDefault();if(i.is("input,button"))i.trigger("focus");else i.find("input:visible,button:visible").first().trigger("focus")}}).on("focus.bs.button.data-api blur.bs.button.data-api",'[data-toggle^="button"]',function(e){t(e.target).closest(".btn").toggleClass("focus",/^focus(in)?$/.test(e.type))})}(jQuery)},{}],5:[function(t,e,n){+function(t){"use strict";var e=function(e,n){this.$element=t(e);this.$indicators=this.$element.find(".carousel-indicators");this.options=n;this.paused=null;this.sliding=null;this.interval=null;this.$active=null;this.$items=null;this.options.keyboard&&this.$element.on("keydown.bs.carousel",t.proxy(this.keydown,this));this.options.pause=="hover"&&!("ontouchstart"in document.documentElement)&&this.$element.on("mouseenter.bs.carousel",t.proxy(this.pause,this)).on("mouseleave.bs.carousel",t.proxy(this.cycle,this))};e.VERSION="3.3.7";e.TRANSITION_DURATION=600;e.DEFAULTS={interval:5e3,pause:"hover",wrap:true,keyboard:true};e.prototype.keydown=function(t){if(/input|textarea/i.test(t.target.tagName))return;switch(t.which){case 37:this.prev();break;case 39:this.next();break;default:return}t.preventDefault()};e.prototype.cycle=function(e){e||(this.paused=false);this.interval&&clearInterval(this.interval);this.options.interval&&!this.paused&&(this.interval=setInterval(t.proxy(this.next,this),this.options.interval));return this};e.prototype.getItemIndex=function(t){this.$items=t.parent().children(".item");return this.$items.index(t||this.$active)};e.prototype.getItemForDirection=function(t,e){var n=this.getItemIndex(e);var i=t=="prev"&&n===0||t=="next"&&n==this.$items.length-1;if(i&&!this.options.wrap)return e;var r=t=="prev"?-1:1;var a=(n+r)%this.$items.length;return this.$items.eq(a)};e.prototype.to=function(t){var e=this;var n=this.getItemIndex(this.$active=this.$element.find(".item.active"));if(t>this.$items.length-1||t<0)return;if(this.sliding)return this.$element.one("slid.bs.carousel",function(){e.to(t)});if(n==t)return this.pause().cycle();return this.slide(t>n?"next":"prev",this.$items.eq(t))};e.prototype.pause=function(e){e||(this.paused=true);if(this.$element.find(".next, .prev").length&&t.support.transition){this.$element.trigger(t.support.transition.end);this.cycle(true)}this.interval=clearInterval(this.interval);return this};e.prototype.next=function(){if(this.sliding)return;return this.slide("next")};e.prototype.prev=function(){if(this.sliding)return;return this.slide("prev")};e.prototype.slide=function(n,i){var r=this.$element.find(".item.active");var a=i||this.getItemForDirection(n,r);var s=this.interval;var o=n=="next"?"left":"right";var l=this;if(a.hasClass("active"))return this.sliding=false;var u=a[0];var f=t.Event("slide.bs.carousel",{relatedTarget:u,direction:o});this.$element.trigger(f);if(f.isDefaultPrevented())return;this.sliding=true;s&&this.pause();if(this.$indicators.length){this.$indicators.find(".active").removeClass("active");var c=t(this.$indicators.children()[this.getItemIndex(a)]);c&&c.addClass("active")}var h=t.Event("slid.bs.carousel",{relatedTarget:u,direction:o});if(t.support.transition&&this.$element.hasClass("slide")){a.addClass(n);a[0].offsetWidth;r.addClass(o);a.addClass(o);r.one("bsTransitionEnd",function(){a.removeClass([n,o].join(" ")).addClass("active");r.removeClass(["active",o].join(" "));l.sliding=false;setTimeout(function(){l.$element.trigger(h)},0)}).emulateTransitionEnd(e.TRANSITION_DURATION)}else{r.removeClass("active");a.addClass("active");this.sliding=false;this.$element.trigger(h)}s&&this.cycle();return this};function n(n){return this.each(function(){var i=t(this);var r=i.data("bs.carousel");var a=t.extend({},e.DEFAULTS,i.data(),typeof n=="object"&&n);var s=typeof n=="string"?n:a.slide;if(!r)i.data("bs.carousel",r=new e(this,a));if(typeof n=="number")r.to(n);else if(s)r[s]();else if(a.interval)r.pause().cycle()})}var i=t.fn.carousel;t.fn.carousel=n;t.fn.carousel.Constructor=e;t.fn.carousel.noConflict=function(){t.fn.carousel=i;return this};var r=function(e){var i;var r=t(this);var a=t(r.attr("data-target")||(i=r.attr("href"))&&i.replace(/.*(?=#[^\s]+$)/,""));if(!a.hasClass("carousel"))return;var s=t.extend({},a.data(),r.data());var o=r.attr("data-slide-to");if(o)s.interval=false;n.call(a,s);if(o){a.data("bs.carousel").to(o)}e.preventDefault()};t(document).on("click.bs.carousel.data-api","[data-slide]",r).on("click.bs.carousel.data-api","[data-slide-to]",r);t(window).on("load",function(){t('[data-ride="carousel"]').each(function(){var e=t(this);n.call(e,e.data())})})}(jQuery)},{}],6:[function(t,e,n){+function(t){"use strict";var e=function(n,i){this.$element=t(n);this.options=t.extend({},e.DEFAULTS,i);this.$trigger=t('[data-toggle="collapse"][href="#'+n.id+'"],'+'[data-toggle="collapse"][data-target="#'+n.id+'"]');this.transitioning=null;if(this.options.parent){this.$parent=this.getParent()}else{this.addAriaAndCollapsedClass(this.$element,this.$trigger)}if(this.options.toggle)this.toggle()};e.VERSION="3.3.7";e.TRANSITION_DURATION=350;e.DEFAULTS={toggle:true};e.prototype.dimension=function(){var t=this.$element.hasClass("width");return t?"width":"height"};e.prototype.show=function(){if(this.transitioning||this.$element.hasClass("in"))return;var n;var r=this.$parent&&this.$parent.children(".panel").children(".in, .collapsing");if(r&&r.length){n=r.data("bs.collapse");if(n&&n.transitioning)return}var a=t.Event("show.bs.collapse");this.$element.trigger(a);if(a.isDefaultPrevented())return;if(r&&r.length){i.call(r,"hide");n||r.data("bs.collapse",null)}var s=this.dimension();this.$element.removeClass("collapse").addClass("collapsing")[s](0).attr("aria-expanded",true);this.$trigger.removeClass("collapsed").attr("aria-expanded",true);this.transitioning=1;var o=function(){this.$element.removeClass("collapsing").addClass("collapse in")[s]("");this.transitioning=0;this.$element.trigger("shown.bs.collapse")};if(!t.support.transition)return o.call(this);var l=t.camelCase(["scroll",s].join("-"));this.$element.one("bsTransitionEnd",t.proxy(o,this)).emulateTransitionEnd(e.TRANSITION_DURATION)[s](this.$element[0][l])};e.prototype.hide=function(){if(this.transitioning||!this.$element.hasClass("in"))return;var n=t.Event("hide.bs.collapse");this.$element.trigger(n);if(n.isDefaultPrevented())return;var i=this.dimension();this.$element[i](this.$element[i]())[0].offsetHeight;this.$element.addClass("collapsing").removeClass("collapse in").attr("aria-expanded",false);this.$trigger.addClass("collapsed").attr("aria-expanded",false);this.transitioning=1;var r=function(){this.transitioning=0;this.$element.removeClass("collapsing").addClass("collapse").trigger("hidden.bs.collapse")};if(!t.support.transition)return r.call(this);this.$element[i](0).one("bsTransitionEnd",t.proxy(r,this)).emulateTransitionEnd(e.TRANSITION_DURATION)};e.prototype.toggle=function(){this[this.$element.hasClass("in")?"hide":"show"]()};e.prototype.getParent=function(){return t(this.options.parent).find('[data-toggle="collapse"][data-parent="'+this.options.parent+'"]').each(t.proxy(function(e,i){var r=t(i);this.addAriaAndCollapsedClass(n(r),r)},this)).end()};e.prototype.addAriaAndCollapsedClass=function(t,e){var n=t.hasClass("in");t.attr("aria-expanded",n);e.toggleClass("collapsed",!n).attr("aria-expanded",n)};function n(e){var n;var i=e.attr("data-target")||(n=e.attr("href"))&&n.replace(/.*(?=#[^\s]+$)/,"");return t(i)}function i(n){return this.each(function(){var i=t(this);var r=i.data("bs.collapse");var a=t.extend({},e.DEFAULTS,i.data(),typeof n=="object"&&n);if(!r&&a.toggle&&/show|hide/.test(n))a.toggle=false;if(!r)i.data("bs.collapse",r=new e(this,a));if(typeof n=="string")r[n]()})}var r=t.fn.collapse;t.fn.collapse=i;t.fn.collapse.Constructor=e;t.fn.collapse.noConflict=function(){t.fn.collapse=r;return this};t(document).on("click.bs.collapse.data-api",'[data-toggle="collapse"]',function(e){var r=t(this);if(!r.attr("data-target"))e.preventDefault();var a=n(r);var s=a.data("bs.collapse");var o=s?"toggle":r.data();i.call(a,o)})}(jQuery)},{}],7:[function(t,e,n){+function(t){"use strict";var e=".dropdown-backdrop";var n='[data-toggle="dropdown"]';var i=function(e){t(e).on("click.bs.dropdown",this.toggle)};i.VERSION="3.3.7";function r(e){var n=e.attr("data-target");if(!n){n=e.attr("href");n=n&&/#[A-Za-z]/.test(n)&&n.replace(/.*(?=#[^\s]*$)/,"")}var i=n&&t(n);return i&&i.length?i:e.parent()}function a(i){if(i&&i.which===3)return;t(e).remove();t(n).each(function(){var e=t(this);var n=r(e);var a={relatedTarget:this};if(!n.hasClass("open"))return;if(i&&i.type=="click"&&/input|textarea/i.test(i.target.tagName)&&t.contains(n[0],i.target))return;n.trigger(i=t.Event("hide.bs.dropdown",a));if(i.isDefaultPrevented())return;e.attr("aria-expanded","false");n.removeClass("open").trigger(t.Event("hidden.bs.dropdown",a))})}i.prototype.toggle=function(e){var n=t(this);if(n.is(".disabled, :disabled"))return;var i=r(n);var s=i.hasClass("open");a();if(!s){if("ontouchstart"in document.documentElement&&!i.closest(".navbar-nav").length){t(document.createElement("div")).addClass("dropdown-backdrop").insertAfter(t(this)).on("click",a)}var o={relatedTarget:this};i.trigger(e=t.Event("show.bs.dropdown",o));if(e.isDefaultPrevented())return;n.trigger("focus").attr("aria-expanded","true");i.toggleClass("open").trigger(t.Event("shown.bs.dropdown",o))}return false};i.prototype.keydown=function(e){if(!/(38|40|27|32)/.test(e.which)||/input|textarea/i.test(e.target.tagName))return;var i=t(this);e.preventDefault();e.stopPropagation();if(i.is(".disabled, :disabled"))return;var a=r(i);var s=a.hasClass("open");if(!s&&e.which!=27||s&&e.which==27){if(e.which==27)a.find(n).trigger("focus");return i.trigger("click")}var o=" li:not(.disabled):visible a";var l=a.find(".dropdown-menu"+o);if(!l.length)return;var u=l.index(e.target);if(e.which==38&&u>0)u--;if(e.which==40&&u<l.length-1)u++;if(!~u)u=0;l.eq(u).trigger("focus")};function s(e){return this.each(function(){var n=t(this);var r=n.data("bs.dropdown");if(!r)n.data("bs.dropdown",r=new i(this));if(typeof e=="string")r[e].call(n)})}var o=t.fn.dropdown;t.fn.dropdown=s;t.fn.dropdown.Constructor=i;t.fn.dropdown.noConflict=function(){t.fn.dropdown=o;return this};t(document).on("click.bs.dropdown.data-api",a).on("click.bs.dropdown.data-api",".dropdown form",function(t){t.stopPropagation()}).on("click.bs.dropdown.data-api",n,i.prototype.toggle).on("keydown.bs.dropdown.data-api",n,i.prototype.keydown).on("keydown.bs.dropdown.data-api",".dropdown-menu",i.prototype.keydown)}(jQuery)},{}],8:[function(t,e,n){+function(t){"use strict";var e=function(e,n){this.options=n;this.$body=t(document.body);this.$element=t(e);this.$dialog=this.$element.find(".modal-dialog");this.$backdrop=null;this.isShown=null;this.originalBodyPad=null;this.scrollbarWidth=0;this.ignoreBackdropClick=false;if(this.options.remote){this.$element.find(".modal-content").load(this.options.remote,t.proxy(function(){this.$element.trigger("loaded.bs.modal")},this))}};e.VERSION="3.3.7";e.TRANSITION_DURATION=300;e.BACKDROP_TRANSITION_DURATION=150;e.DEFAULTS={backdrop:true,keyboard:true,show:true};e.prototype.toggle=function(t){return this.isShown?this.hide():this.show(t)};e.prototype.show=function(n){var i=this;var r=t.Event("show.bs.modal",{relatedTarget:n});this.$element.trigger(r);if(this.isShown||r.isDefaultPrevented())return;this.isShown=true;this.checkScrollbar();this.setScrollbar();this.$body.addClass("modal-open");this.escape();this.resize();this.$element.on("click.dismiss.bs.modal",'[data-dismiss="modal"]',t.proxy(this.hide,this));this.$dialog.on("mousedown.dismiss.bs.modal",function(){i.$element.one("mouseup.dismiss.bs.modal",function(e){if(t(e.target).is(i.$element))i.ignoreBackdropClick=true})});this.backdrop(function(){var r=t.support.transition&&i.$element.hasClass("fade");if(!i.$element.parent().length){i.$element.appendTo(i.$body)}i.$element.show().scrollTop(0);i.adjustDialog();if(r){i.$element[0].offsetWidth}i.$element.addClass("in");i.enforceFocus();var a=t.Event("shown.bs.modal",{relatedTarget:n});r?i.$dialog.one("bsTransitionEnd",function(){i.$element.trigger("focus").trigger(a)}).emulateTransitionEnd(e.TRANSITION_DURATION):i.$element.trigger("focus").trigger(a)})};e.prototype.hide=function(n){if(n)n.preventDefault();n=t.Event("hide.bs.modal");this.$element.trigger(n);if(!this.isShown||n.isDefaultPrevented())return;this.isShown=false;this.escape();this.resize();t(document).off("focusin.bs.modal");this.$element.removeClass("in").off("click.dismiss.bs.modal").off("mouseup.dismiss.bs.modal");this.$dialog.off("mousedown.dismiss.bs.modal");t.support.transition&&this.$element.hasClass("fade")?this.$element.one("bsTransitionEnd",t.proxy(this.hideModal,this)).emulateTransitionEnd(e.TRANSITION_DURATION):this.hideModal()};e.prototype.enforceFocus=function(){t(document).off("focusin.bs.modal").on("focusin.bs.modal",t.proxy(function(t){if(document!==t.target&&this.$element[0]!==t.target&&!this.$element.has(t.target).length){this.$element.trigger("focus")}},this))};e.prototype.escape=function(){if(this.isShown&&this.options.keyboard){this.$element.on("keydown.dismiss.bs.modal",t.proxy(function(t){t.which==27&&this.hide()},this))}else if(!this.isShown){this.$element.off("keydown.dismiss.bs.modal")}};e.prototype.resize=function(){if(this.isShown){t(window).on("resize.bs.modal",t.proxy(this.handleUpdate,this))}else{t(window).off("resize.bs.modal")}};e.prototype.hideModal=function(){var t=this;this.$element.hide();this.backdrop(function(){t.$body.removeClass("modal-open");t.resetAdjustments();t.resetScrollbar();t.$element.trigger("hidden.bs.modal")})};e.prototype.removeBackdrop=function(){this.$backdrop&&this.$backdrop.remove();this.$backdrop=null};e.prototype.backdrop=function(n){var i=this;var r=this.$element.hasClass("fade")?"fade":"";if(this.isShown&&this.options.backdrop){var a=t.support.transition&&r;this.$backdrop=t(document.createElement("div")).addClass("modal-backdrop "+r).appendTo(this.$body);this.$element.on("click.dismiss.bs.modal",t.proxy(function(t){if(this.ignoreBackdropClick){this.ignoreBackdropClick=false;return}if(t.target!==t.currentTarget)return;this.options.backdrop=="static"?this.$element[0].focus():this.hide()},this));if(a)this.$backdrop[0].offsetWidth;this.$backdrop.addClass("in");if(!n)return;a?this.$backdrop.one("bsTransitionEnd",n).emulateTransitionEnd(e.BACKDROP_TRANSITION_DURATION):n()}else if(!this.isShown&&this.$backdrop){this.$backdrop.removeClass("in");var s=function(){i.removeBackdrop();n&&n()};t.support.transition&&this.$element.hasClass("fade")?this.$backdrop.one("bsTransitionEnd",s).emulateTransitionEnd(e.BACKDROP_TRANSITION_DURATION):s()}else if(n){n()}};e.prototype.handleUpdate=function(){this.adjustDialog()};e.prototype.adjustDialog=function(){var t=this.$element[0].scrollHeight>document.documentElement.clientHeight;this.$element.css({paddingLeft:!this.bodyIsOverflowing&&t?this.scrollbarWidth:"",paddingRight:this.bodyIsOverflowing&&!t?this.scrollbarWidth:""})};e.prototype.resetAdjustments=function(){this.$element.css({paddingLeft:"",paddingRight:""})};e.prototype.checkScrollbar=function(){var t=window.innerWidth;if(!t){var e=document.documentElement.getBoundingClientRect();t=e.right-Math.abs(e.left)}this.bodyIsOverflowing=document.body.clientWidth<t;this.scrollbarWidth=this.measureScrollbar()};e.prototype.setScrollbar=function(){var t=parseInt(this.$body.css("padding-right")||0,10);this.originalBodyPad=document.body.style.paddingRight||"";if(this.bodyIsOverflowing)this.$body.css("padding-right",t+this.scrollbarWidth)};e.prototype.resetScrollbar=function(){this.$body.css("padding-right",this.originalBodyPad)};e.prototype.measureScrollbar=function(){var t=document.createElement("div");t.className="modal-scrollbar-measure";this.$body.append(t);var e=t.offsetWidth-t.clientWidth;this.$body[0].removeChild(t);return e};function n(n,i){return this.each(function(){var r=t(this);var a=r.data("bs.modal");var s=t.extend({},e.DEFAULTS,r.data(),typeof n=="object"&&n);if(!a)r.data("bs.modal",a=new e(this,s));if(typeof n=="string")a[n](i);else if(s.show)a.show(i)})}var i=t.fn.modal;t.fn.modal=n;t.fn.modal.Constructor=e;t.fn.modal.noConflict=function(){t.fn.modal=i;return this};t(document).on("click.bs.modal.data-api",'[data-toggle="modal"]',function(e){var i=t(this);var r=i.attr("href");var a=t(i.attr("data-target")||r&&r.replace(/.*(?=#[^\s]+$)/,""));var s=a.data("bs.modal")?"toggle":t.extend({remote:!/#/.test(r)&&r},a.data(),i.data());if(i.is("a"))e.preventDefault();a.one("show.bs.modal",function(t){if(t.isDefaultPrevented())return;a.one("hidden.bs.modal",function(){i.is(":visible")&&i.trigger("focus")})});n.call(a,s,this)})}(jQuery)},{}],9:[function(t,e,n){+function(t){"use strict";var e=function(t,e){this.init("popover",t,e)};if(!t.fn.tooltip)throw new Error("Popover requires tooltip.js");e.VERSION="3.3.7";e.DEFAULTS=t.extend({},t.fn.tooltip.Constructor.DEFAULTS,{placement:"right",trigger:"click",content:"",template:'<div class="popover" role="tooltip"><div class="arrow"></div><h3 class="popover-title"></h3><div class="popover-content"></div></div>'});e.prototype=t.extend({},t.fn.tooltip.Constructor.prototype);e.prototype.constructor=e;e.prototype.getDefaults=function(){return e.DEFAULTS};e.prototype.setContent=function(){var t=this.tip();var e=this.getTitle();var n=this.getContent();t.find(".popover-title")[this.options.html?"html":"text"](e);t.find(".popover-content").children().detach().end()[this.options.html?typeof n=="string"?"html":"append":"text"](n);t.removeClass("fade top bottom left right in");if(!t.find(".popover-title").html())t.find(".popover-title").hide()};e.prototype.hasContent=function(){return this.getTitle()||this.getContent()};e.prototype.getContent=function(){var t=this.$element;var e=this.options;return t.attr("data-content")||(typeof e.content=="function"?e.content.call(t[0]):e.content)};e.prototype.arrow=function(){return this.$arrow=this.$arrow||this.tip().find(".arrow")};function n(n){return this.each(function(){var i=t(this);var r=i.data("bs.popover");var a=typeof n=="object"&&n;if(!r&&/destroy|hide/.test(n))return;if(!r)i.data("bs.popover",r=new e(this,a));if(typeof n=="string")r[n]()})}var i=t.fn.popover;t.fn.popover=n;t.fn.popover.Constructor=e;t.fn.popover.noConflict=function(){t.fn.popover=i;return this}}(jQuery)},{}],10:[function(t,e,n){+function(t){"use strict";function e(n,i){this.$body=t(document.body);this.$scrollElement=t(n).is(document.body)?t(window):t(n);this.options=t.extend({},e.DEFAULTS,i);this.selector=(this.options.target||"")+" .nav li > a";this.offsets=[];this.targets=[];this.activeTarget=null;this.scrollHeight=0;this.$scrollElement.on("scroll.bs.scrollspy",t.proxy(this.process,this));this.refresh();this.process()}e.VERSION="3.3.7";e.DEFAULTS={offset:10};e.prototype.getScrollHeight=function(){return this.$scrollElement[0].scrollHeight||Math.max(this.$body[0].scrollHeight,document.documentElement.scrollHeight)};e.prototype.refresh=function(){var e=this;var n="offset";var i=0;this.offsets=[];this.targets=[];this.scrollHeight=this.getScrollHeight();if(!t.isWindow(this.$scrollElement[0])){n="position";i=this.$scrollElement.scrollTop()}this.$body.find(this.selector).map(function(){var e=t(this);var r=e.data("target")||e.attr("href");var a=/^#./.test(r)&&t(r);return a&&a.length&&a.is(":visible")&&[[a[n]().top+i,r]]||null}).sort(function(t,e){return t[0]-e[0]}).each(function(){e.offsets.push(this[0]);e.targets.push(this[1])})};e.prototype.process=function(){var t=this.$scrollElement.scrollTop()+this.options.offset;var e=this.getScrollHeight();var n=this.options.offset+e-this.$scrollElement.height();var i=this.offsets;var r=this.targets;var a=this.activeTarget;var s;if(this.scrollHeight!=e){this.refresh()}if(t>=n){return a!=(s=r[r.length-1])&&this.activate(s)}if(a&&t<i[0]){this.activeTarget=null;return this.clear()}for(s=i.length;s--;){a!=r[s]&&t>=i[s]&&(i[s+1]===undefined||t<i[s+1])&&this.activate(r[s])}};e.prototype.activate=function(e){this.activeTarget=e;this.clear();var n=this.selector+'[data-target="'+e+'"],'+this.selector+'[href="'+e+'"]';var i=t(n).parents("li").addClass("active");if(i.parent(".dropdown-menu").length){i=i.closest("li.dropdown").addClass("active")}i.trigger("activate.bs.scrollspy")};e.prototype.clear=function(){t(this.selector).parentsUntil(this.options.target,".active").removeClass("active")};function n(n){return this.each(function(){var i=t(this);var r=i.data("bs.scrollspy");var a=typeof n=="object"&&n;if(!r)i.data("bs.scrollspy",r=new e(this,a));if(typeof n=="string")r[n]()})}var i=t.fn.scrollspy;t.fn.scrollspy=n;t.fn.scrollspy.Constructor=e;t.fn.scrollspy.noConflict=function(){t.fn.scrollspy=i;return this};t(window).on("load.bs.scrollspy.data-api",function(){t('[data-spy="scroll"]').each(function(){var e=t(this);n.call(e,e.data())})})}(jQuery)},{}],11:[function(t,e,n){+function(t){"use strict";var e=function(e){this.element=t(e)};e.VERSION="3.3.7";e.TRANSITION_DURATION=150;e.prototype.show=function(){var e=this.element;var n=e.closest("ul:not(.dropdown-menu)");var i=e.data("target");if(!i){i=e.attr("href");i=i&&i.replace(/.*(?=#[^\s]*$)/,"")}if(e.parent("li").hasClass("active"))return;var r=n.find(".active:last a");var a=t.Event("hide.bs.tab",{relatedTarget:e[0]});var s=t.Event("show.bs.tab",{relatedTarget:r[0]});r.trigger(a);e.trigger(s);if(s.isDefaultPrevented()||a.isDefaultPrevented())return;var o=t(i);this.activate(e.closest("li"),n);this.activate(o,o.parent(),function(){r.trigger({type:"hidden.bs.tab",relatedTarget:e[0]});e.trigger({type:"shown.bs.tab",relatedTarget:r[0]})})};e.prototype.activate=function(n,i,r){var a=i.find("> .active");var s=r&&t.support.transition&&(a.length&&a.hasClass("fade")||!!i.find("> .fade").length);function o(){a.removeClass("active").find("> .dropdown-menu > .active").removeClass("active").end().find('[data-toggle="tab"]').attr("aria-expanded",false);n.addClass("active").find('[data-toggle="tab"]').attr("aria-expanded",true);if(s){n[0].offsetWidth;n.addClass("in")}else{n.removeClass("fade")}if(n.parent(".dropdown-menu").length){n.closest("li.dropdown").addClass("active").end().find('[data-toggle="tab"]').attr("aria-expanded",true)}r&&r()}a.length&&s?a.one("bsTransitionEnd",o).emulateTransitionEnd(e.TRANSITION_DURATION):o();a.removeClass("in")};function n(n){return this.each(function(){var i=t(this);var r=i.data("bs.tab");if(!r)i.data("bs.tab",r=new e(this));if(typeof n=="string")r[n]()})}var i=t.fn.tab;t.fn.tab=n;t.fn.tab.Constructor=e;t.fn.tab.noConflict=function(){t.fn.tab=i;return this};var r=function(e){e.preventDefault();n.call(t(this),"show")};t(document).on("click.bs.tab.data-api",'[data-toggle="tab"]',r).on("click.bs.tab.data-api",'[data-toggle="pill"]',r)}(jQuery)},{}],12:[function(t,e,n){+function(t){"use strict";var e=function(t,e){this.type=null;this.options=null;this.enabled=null;this.timeout=null;this.hoverState=null;this.$element=null;this.inState=null;this.init("tooltip",t,e)};e.VERSION="3.3.7";e.TRANSITION_DURATION=150;e.DEFAULTS={animation:true,placement:"top",selector:false,template:'<div class="tooltip" role="tooltip"><div class="tooltip-arrow"></div><div class="tooltip-inner"></div></div>',trigger:"hover focus",title:"",delay:0,html:false,container:false,viewport:{selector:"body",padding:0}};e.prototype.init=function(e,n,i){this.enabled=true;this.type=e;this.$element=t(n);this.options=this.getOptions(i);this.$viewport=this.options.viewport&&t(t.isFunction(this.options.viewport)?this.options.viewport.call(this,this.$element):this.options.viewport.selector||this.options.viewport);this.inState={click:false,hover:false,focus:false};if(this.$element[0]instanceof document.constructor&&!this.options.selector){throw new Error("`selector` option must be specified when initializing "+this.type+" on the window.document object!")}var r=this.options.trigger.split(" ");for(var a=r.length;a--;){var s=r[a];if(s=="click"){this.$element.on("click."+this.type,this.options.selector,t.proxy(this.toggle,this))}else if(s!="manual"){var o=s=="hover"?"mouseenter":"focusin";var l=s=="hover"?"mouseleave":"focusout";this.$element.on(o+"."+this.type,this.options.selector,t.proxy(this.enter,this));this.$element.on(l+"."+this.type,this.options.selector,t.proxy(this.leave,this))}}this.options.selector?this._options=t.extend({},this.options,{trigger:"manual",selector:""}):this.fixTitle()};e.prototype.getDefaults=function(){return e.DEFAULTS};e.prototype.getOptions=function(e){e=t.extend({},this.getDefaults(),this.$element.data(),e);if(e.delay&&typeof e.delay=="number"){e.delay={show:e.delay,hide:e.delay}}return e};e.prototype.getDelegateOptions=function(){var e={};var n=this.getDefaults();this._options&&t.each(this._options,function(t,i){if(n[t]!=i)e[t]=i});return e};e.prototype.enter=function(e){var n=e instanceof this.constructor?e:t(e.currentTarget).data("bs."+this.type);if(!n){n=new this.constructor(e.currentTarget,this.getDelegateOptions());t(e.currentTarget).data("bs."+this.type,n)}if(e instanceof t.Event){n.inState[e.type=="focusin"?"focus":"hover"]=true}if(n.tip().hasClass("in")||n.hoverState=="in"){n.hoverState="in";return}clearTimeout(n.timeout);n.hoverState="in";if(!n.options.delay||!n.options.delay.show)return n.show();n.timeout=setTimeout(function(){if(n.hoverState=="in")n.show()},n.options.delay.show)};e.prototype.isInStateTrue=function(){for(var t in this.inState){if(this.inState[t])return true}return false};e.prototype.leave=function(e){var n=e instanceof this.constructor?e:t(e.currentTarget).data("bs."+this.type);if(!n){n=new this.constructor(e.currentTarget,this.getDelegateOptions());t(e.currentTarget).data("bs."+this.type,n)}
if(e instanceof t.Event){n.inState[e.type=="focusout"?"focus":"hover"]=false}if(n.isInStateTrue())return;clearTimeout(n.timeout);n.hoverState="out";if(!n.options.delay||!n.options.delay.hide)return n.hide();n.timeout=setTimeout(function(){if(n.hoverState=="out")n.hide()},n.options.delay.hide)};e.prototype.show=function(){var n=t.Event("show.bs."+this.type);if(this.hasContent()&&this.enabled){this.$element.trigger(n);var i=t.contains(this.$element[0].ownerDocument.documentElement,this.$element[0]);if(n.isDefaultPrevented()||!i)return;var r=this;var a=this.tip();var s=this.getUID(this.type);this.setContent();a.attr("id",s);this.$element.attr("aria-describedby",s);if(this.options.animation)a.addClass("fade");var o=typeof this.options.placement=="function"?this.options.placement.call(this,a[0],this.$element[0]):this.options.placement;var l=/\s?auto?\s?/i;var u=l.test(o);if(u)o=o.replace(l,"")||"top";a.detach().css({top:0,left:0,display:"block"}).addClass(o).data("bs."+this.type,this);this.options.container?a.appendTo(this.options.container):a.insertAfter(this.$element);this.$element.trigger("inserted.bs."+this.type);var f=this.getPosition();var c=a[0].offsetWidth;var h=a[0].offsetHeight;if(u){var d=o;var p=this.getPosition(this.$viewport);o=o=="bottom"&&f.bottom+h>p.bottom?"top":o=="top"&&f.top-h<p.top?"bottom":o=="right"&&f.right+c>p.width?"left":o=="left"&&f.left-c<p.left?"right":o;a.removeClass(d).addClass(o)}var g=this.getCalculatedOffset(o,f,c,h);this.applyPlacement(g,o);var v=function(){var t=r.hoverState;r.$element.trigger("shown.bs."+r.type);r.hoverState=null;if(t=="out")r.leave(r)};t.support.transition&&this.$tip.hasClass("fade")?a.one("bsTransitionEnd",v).emulateTransitionEnd(e.TRANSITION_DURATION):v()}};e.prototype.applyPlacement=function(e,n){var i=this.tip();var r=i[0].offsetWidth;var a=i[0].offsetHeight;var s=parseInt(i.css("margin-top"),10);var o=parseInt(i.css("margin-left"),10);if(isNaN(s))s=0;if(isNaN(o))o=0;e.top+=s;e.left+=o;t.offset.setOffset(i[0],t.extend({using:function(t){i.css({top:Math.round(t.top),left:Math.round(t.left)})}},e),0);i.addClass("in");var l=i[0].offsetWidth;var u=i[0].offsetHeight;if(n=="top"&&u!=a){e.top=e.top+a-u}var f=this.getViewportAdjustedDelta(n,e,l,u);if(f.left)e.left+=f.left;else e.top+=f.top;var c=/top|bottom/.test(n);var h=c?f.left*2-r+l:f.top*2-a+u;var d=c?"offsetWidth":"offsetHeight";i.offset(e);this.replaceArrow(h,i[0][d],c)};e.prototype.replaceArrow=function(t,e,n){this.arrow().css(n?"left":"top",50*(1-t/e)+"%").css(n?"top":"left","")};e.prototype.setContent=function(){var t=this.tip();var e=this.getTitle();t.find(".tooltip-inner")[this.options.html?"html":"text"](e);t.removeClass("fade in top bottom left right")};e.prototype.hide=function(n){var i=this;var r=t(this.$tip);var a=t.Event("hide.bs."+this.type);function s(){if(i.hoverState!="in")r.detach();if(i.$element){i.$element.removeAttr("aria-describedby").trigger("hidden.bs."+i.type)}n&&n()}this.$element.trigger(a);if(a.isDefaultPrevented())return;r.removeClass("in");t.support.transition&&r.hasClass("fade")?r.one("bsTransitionEnd",s).emulateTransitionEnd(e.TRANSITION_DURATION):s();this.hoverState=null;return this};e.prototype.fixTitle=function(){var t=this.$element;if(t.attr("title")||typeof t.attr("data-original-title")!="string"){t.attr("data-original-title",t.attr("title")||"").attr("title","")}};e.prototype.hasContent=function(){return this.getTitle()};e.prototype.getPosition=function(e){e=e||this.$element;var n=e[0];var i=n.tagName=="BODY";var r=n.getBoundingClientRect();if(r.width==null){r=t.extend({},r,{width:r.right-r.left,height:r.bottom-r.top})}var a=window.SVGElement&&n instanceof window.SVGElement;var s=i?{top:0,left:0}:a?null:e.offset();var o={scroll:i?document.documentElement.scrollTop||document.body.scrollTop:e.scrollTop()};var l=i?{width:t(window).width(),height:t(window).height()}:null;return t.extend({},r,o,l,s)};e.prototype.getCalculatedOffset=function(t,e,n,i){return t=="bottom"?{top:e.top+e.height,left:e.left+e.width/2-n/2}:t=="top"?{top:e.top-i,left:e.left+e.width/2-n/2}:t=="left"?{top:e.top+e.height/2-i/2,left:e.left-n}:{top:e.top+e.height/2-i/2,left:e.left+e.width}};e.prototype.getViewportAdjustedDelta=function(t,e,n,i){var r={top:0,left:0};if(!this.$viewport)return r;var a=this.options.viewport&&this.options.viewport.padding||0;var s=this.getPosition(this.$viewport);if(/right|left/.test(t)){var o=e.top-a-s.scroll;var l=e.top+a-s.scroll+i;if(o<s.top){r.top=s.top-o}else if(l>s.top+s.height){r.top=s.top+s.height-l}}else{var u=e.left-a;var f=e.left+a+n;if(u<s.left){r.left=s.left-u}else if(f>s.right){r.left=s.left+s.width-f}}return r};e.prototype.getTitle=function(){var t;var e=this.$element;var n=this.options;t=e.attr("data-original-title")||(typeof n.title=="function"?n.title.call(e[0]):n.title);return t};e.prototype.getUID=function(t){do t+=~~(Math.random()*1e6);while(document.getElementById(t));return t};e.prototype.tip=function(){if(!this.$tip){this.$tip=t(this.options.template);if(this.$tip.length!=1){throw new Error(this.type+" `template` option must consist of exactly 1 top-level element!")}}return this.$tip};e.prototype.arrow=function(){return this.$arrow=this.$arrow||this.tip().find(".tooltip-arrow")};e.prototype.enable=function(){this.enabled=true};e.prototype.disable=function(){this.enabled=false};e.prototype.toggleEnabled=function(){this.enabled=!this.enabled};e.prototype.toggle=function(e){var n=this;if(e){n=t(e.currentTarget).data("bs."+this.type);if(!n){n=new this.constructor(e.currentTarget,this.getDelegateOptions());t(e.currentTarget).data("bs."+this.type,n)}}if(e){n.inState.click=!n.inState.click;if(n.isInStateTrue())n.enter(n);else n.leave(n)}else{n.tip().hasClass("in")?n.leave(n):n.enter(n)}};e.prototype.destroy=function(){var t=this;clearTimeout(this.timeout);this.hide(function(){t.$element.off("."+t.type).removeData("bs."+t.type);if(t.$tip){t.$tip.detach()}t.$tip=null;t.$arrow=null;t.$viewport=null;t.$element=null})};function n(n){return this.each(function(){var i=t(this);var r=i.data("bs.tooltip");var a=typeof n=="object"&&n;if(!r&&/destroy|hide/.test(n))return;if(!r)i.data("bs.tooltip",r=new e(this,a));if(typeof n=="string")r[n]()})}var i=t.fn.tooltip;t.fn.tooltip=n;t.fn.tooltip.Constructor=e;t.fn.tooltip.noConflict=function(){t.fn.tooltip=i;return this}}(jQuery)},{}],13:[function(t,e,n){+function(t){"use strict";function e(){var t=document.createElement("bootstrap");var e={WebkitTransition:"webkitTransitionEnd",MozTransition:"transitionend",OTransition:"oTransitionEnd otransitionend",transition:"transitionend"};for(var n in e){if(t.style[n]!==undefined){return{end:e[n]}}}return false}t.fn.emulateTransitionEnd=function(e){var n=false;var i=this;t(this).one("bsTransitionEnd",function(){n=true});var r=function(){if(!n)t(i).trigger(t.support.transition.end)};setTimeout(r,e);return this};t(function(){t.support.transition=e();if(!t.support.transition)return;t.event.special.bsTransitionEnd={bindType:t.support.transition.end,delegateType:t.support.transition.end,handle:function(e){if(t(e.target).is(this))return e.handleObj.handler.apply(this,arguments)}}})}(jQuery)},{}],14:[function(t,e,n){!function(){var t={version:"3.5.12"};var n=[].slice,i=function(t){return n.call(t)};var r=this.document;function a(t){return t&&(t.ownerDocument||t.document||t).documentElement}function s(t){return t&&(t.ownerDocument&&t.ownerDocument.defaultView||t.document&&t||t.defaultView)}if(r){try{i(r.documentElement.childNodes)[0].nodeType}catch(t){i=function(t){var e=t.length,n=new Array(e);while(e--)n[e]=t[e];return n}}}if(!Date.now)Date.now=function(){return+new Date};if(r){try{r.createElement("DIV").style.setProperty("opacity",0,"")}catch(t){var o=this.Element.prototype,l=o.setAttribute,u=o.setAttributeNS,f=this.CSSStyleDeclaration.prototype,c=f.setProperty;o.setAttribute=function(t,e){l.call(this,t,e+"")};o.setAttributeNS=function(t,e,n){u.call(this,t,e,n+"")};f.setProperty=function(t,e,n){c.call(this,t,e+"",n)}}}t.ascending=h;function h(t,e){return t<e?-1:t>e?1:t>=e?0:NaN}t.descending=function(t,e){return e<t?-1:e>t?1:e>=t?0:NaN};t.min=function(t,e){var n=-1,i=t.length,r,a;if(arguments.length===1){while(++n<i)if((a=t[n])!=null&&a>=a){r=a;break}while(++n<i)if((a=t[n])!=null&&r>a)r=a}else{while(++n<i)if((a=e.call(t,t[n],n))!=null&&a>=a){r=a;break}while(++n<i)if((a=e.call(t,t[n],n))!=null&&r>a)r=a}return r};t.max=function(t,e){var n=-1,i=t.length,r,a;if(arguments.length===1){while(++n<i)if((a=t[n])!=null&&a>=a){r=a;break}while(++n<i)if((a=t[n])!=null&&a>r)r=a}else{while(++n<i)if((a=e.call(t,t[n],n))!=null&&a>=a){r=a;break}while(++n<i)if((a=e.call(t,t[n],n))!=null&&a>r)r=a}return r};t.extent=function(t,e){var n=-1,i=t.length,r,a,s;if(arguments.length===1){while(++n<i)if((a=t[n])!=null&&a>=a){r=s=a;break}while(++n<i)if((a=t[n])!=null){if(r>a)r=a;if(s<a)s=a}}else{while(++n<i)if((a=e.call(t,t[n],n))!=null&&a>=a){r=s=a;break}while(++n<i)if((a=e.call(t,t[n],n))!=null){if(r>a)r=a;if(s<a)s=a}}return[r,s]};function d(t){return t===null?NaN:+t}function p(t){return!isNaN(t)}t.sum=function(t,e){var n=0,i=t.length,r,a=-1;if(arguments.length===1){while(++a<i)if(p(r=+t[a]))n+=r}else{while(++a<i)if(p(r=+e.call(t,t[a],a)))n+=r}return n};t.mean=function(t,e){var n=0,i=t.length,r,a=-1,s=i;if(arguments.length===1){while(++a<i)if(p(r=d(t[a])))n+=r;else--s}else{while(++a<i)if(p(r=d(e.call(t,t[a],a))))n+=r;else--s}if(s)return n/s};t.quantile=function(t,e){var n=(t.length-1)*e+1,i=Math.floor(n),r=+t[i-1],a=n-i;return a?r+a*(t[i]-r):r};t.median=function(e,n){var i=[],r=e.length,a,s=-1;if(arguments.length===1){while(++s<r)if(p(a=d(e[s])))i.push(a)}else{while(++s<r)if(p(a=d(n.call(e,e[s],s))))i.push(a)}if(i.length)return t.quantile(i.sort(h),.5)};t.variance=function(t,e){var n=t.length,i=0,r,a,s=0,o=-1,l=0;if(arguments.length===1){while(++o<n){if(p(r=d(t[o]))){a=r-i;i+=a/++l;s+=a*(r-i)}}}else{while(++o<n){if(p(r=d(e.call(t,t[o],o)))){a=r-i;i+=a/++l;s+=a*(r-i)}}}if(l>1)return s/(l-1)};t.deviation=function(){var e=t.variance.apply(this,arguments);return e?Math.sqrt(e):e};function g(t){return{left:function(e,n,i,r){if(arguments.length<3)i=0;if(arguments.length<4)r=e.length;while(i<r){var a=i+r>>>1;if(t(e[a],n)<0)i=a+1;else r=a}return i},right:function(e,n,i,r){if(arguments.length<3)i=0;if(arguments.length<4)r=e.length;while(i<r){var a=i+r>>>1;if(t(e[a],n)>0)r=a;else i=a+1}return i}}}var v=g(h);t.bisectLeft=v.left;t.bisect=t.bisectRight=v.right;t.bisector=function(t){return g(t.length===1?function(e,n){return h(t(e),n)}:t)};t.shuffle=function(t,e,n){if((i=arguments.length)<3){n=t.length;if(i<2)e=0}var i=n-e,r,a;while(i){a=Math.random()*i--|0;r=t[i+e],t[i+e]=t[a+e],t[a+e]=r}return t};t.permute=function(t,e){var n=e.length,i=new Array(n);while(n--)i[n]=t[e[n]];return i};t.pairs=function(t){var e=0,n=t.length-1,i,r=t[0],a=new Array(n<0?0:n);while(e<n)a[e]=[i=r,r=t[++e]];return a};t.zip=function(){if(!(a=arguments.length))return[];for(var e=-1,n=t.min(arguments,m),i=new Array(n);++e<n;){for(var r=-1,a,s=i[e]=new Array(a);++r<a;){s[r]=arguments[r][e]}}return i};function m(t){return t.length}t.transpose=function(e){return t.zip.apply(t,e)};t.keys=function(t){var e=[];for(var n in t)e.push(n);return e};t.values=function(t){var e=[];for(var n in t)e.push(t[n]);return e};t.entries=function(t){var e=[];for(var n in t)e.push({key:n,value:t[n]});return e};t.merge=function(t){var e=t.length,n,i=-1,r=0,a,s;while(++i<e)r+=t[i].length;a=new Array(r);while(--e>=0){s=t[e];n=s.length;while(--n>=0){a[--r]=s[n]}}return a};var y=Math.abs;t.range=function(t,e,n){if(arguments.length<3){n=1;if(arguments.length<2){e=t;t=0}}if((e-t)/n===Infinity)throw new Error("infinite range");var i=[],r=b(y(n)),a=-1,s;t*=r,e*=r,n*=r;if(n<0)while((s=t+n*++a)>e)i.push(s/r);else while((s=t+n*++a)<e)i.push(s/r);return i};function b(t){var e=1;while(t*e%1)e*=10;return e}function w(t,e){for(var n in e){Object.defineProperty(t.prototype,n,{value:e[n],enumerable:false})}}t.map=function(t,e){var n=new _;if(t instanceof _){t.forEach(function(t,e){n.set(t,e)})}else if(Array.isArray(t)){var i=-1,r=t.length,a;if(arguments.length===1)while(++i<r)n.set(i,t[i]);else while(++i<r)n.set(e.call(t,a=t[i],i),a)}else{for(var s in t)n.set(s,t[s])}return n};function _(){this._=Object.create(null)}var x="__proto__",S="\0";w(_,{has:T,get:function(t){return this._[D(t)]},set:function(t,e){return this._[D(t)]=e},remove:k,keys:M,values:function(){var t=[];for(var e in this._)t.push(this._[e]);return t},entries:function(){var t=[];for(var e in this._)t.push({key:C(e),value:this._[e]});return t},size:I,empty:A,forEach:function(t){for(var e in this._)t.call(this,C(e),this._[e])}});function D(t){return(t+="")===x||t[0]===S?S+t:t}function C(t){return(t+="")[0]===S?t.slice(1):t}function T(t){return D(t)in this._}function k(t){return(t=D(t))in this._&&delete this._[t]}function M(){var t=[];for(var e in this._)t.push(C(e));return t}function I(){var t=0;for(var e in this._)++t;return t}function A(){for(var t in this._)return false;return true}t.nest=function(){var e={},n=[],i=[],r,a;function s(t,i,o){if(o>=n.length)return a?a.call(e,i):r?i.sort(r):i;var l=-1,u=i.length,f=n[o++],c,h,d,p=new _,g;while(++l<u){if(g=p.get(c=f(h=i[l]))){g.push(h)}else{p.set(c,[h])}}if(t){h=t();d=function(e,n){h.set(e,s(t,n,o))}}else{h={};d=function(e,n){h[e]=s(t,n,o)}}p.forEach(d);return h}function o(t,e){if(e>=n.length)return t;var r=[],a=i[e++];t.forEach(function(t,n){r.push({key:t,values:o(n,e)})});return a?r.sort(function(t,e){return a(t.key,e.key)}):r}e.map=function(t,e){return s(e,t,0)};e.entries=function(e){return o(s(t.map,e,0),0)};e.key=function(t){n.push(t);return e};e.sortKeys=function(t){i[n.length-1]=t;return e};e.sortValues=function(t){r=t;return e};e.rollup=function(t){a=t;return e};return e};t.set=function(t){var e=new N;if(t)for(var n=0,i=t.length;n<i;++n)e.add(t[n]);return e};function N(){this._=Object.create(null)}w(N,{has:T,add:function(t){this._[D(t+="")]=true;return t},remove:k,values:M,size:I,empty:A,forEach:function(t){for(var e in this._)t.call(this,C(e))}});t.behavior={};function P(t){return t}t.rebind=function(t,e){var n=1,i=arguments.length,r;while(++n<i)t[r=arguments[n]]=E(t,e,e[r]);return t};function E(t,e,n){return function(){var i=n.apply(e,arguments);return i===e?t:i}}function F(t,e){if(e in t)return e;e=e.charAt(0).toUpperCase()+e.slice(1);for(var n=0,i=R.length;n<i;++n){var r=R[n]+e;if(r in t)return r}}var R=["webkit","ms","moz","Moz","o","O"];function L(){}t.dispatch=function(){var t=new j,e=-1,n=arguments.length;while(++e<n)t[arguments[e]]=O(t);return t};function j(){}j.prototype.on=function(t,e){var n=t.indexOf("."),i="";if(n>=0){i=t.slice(n+1);t=t.slice(0,n)}if(t)return arguments.length<2?this[t].on(i):this[t].on(i,e);if(arguments.length===2){if(e==null)for(t in this){if(this.hasOwnProperty(t))this[t].on(i,null)}return this}};function O(t){var e=[],n=new _;function i(){var n=e,i=-1,r=n.length,a;while(++i<r)if(a=n[i].on)a.apply(this,arguments);return t}i.on=function(i,r){var a=n.get(i),s;if(arguments.length<2)return a&&a.on;if(a){a.on=null;e=e.slice(0,s=e.indexOf(a)).concat(e.slice(s+1));n.remove(i)}if(r)e.push(n.set(i,{on:r}));return t};return i}t.event=null;function H(){t.event.preventDefault()}function W(){var e=t.event,n;while(n=e.sourceEvent)e=n;return e}function z(e){var n=new j,i=0,r=arguments.length;while(++i<r)n[arguments[i]]=O(n);n.of=function(i,r){return function(a){try{var s=a.sourceEvent=t.event;a.target=e;t.event=a;n[a.type].apply(i,r)}finally{t.event=s}}};return n}t.requote=function(t){return t.replace(q,"\\$&")};var q=/[\\\^\$\*\+\?\|\[\]\(\)\.\{\}]/g;var B={}.__proto__?function(t,e){t.__proto__=e}:function(t,e){for(var n in e)t[n]=e[n]};function $(t){B(t,X);return t}var U=function(t,e){return e.querySelector(t)},Y=function(t,e){return e.querySelectorAll(t)},V=function(t,e){var n=t.matches||t[F(t,"matchesSelector")];V=function(t,e){return n.call(t,e)};return V(t,e)};if(typeof Sizzle==="function"){U=function(t,e){return Sizzle(t,e)[0]||null};Y=Sizzle;V=Sizzle.matchesSelector}t.selection=function(){return t.select(r.documentElement)};var X=t.selection.prototype=[];X.select=function(t){var e=[],n,i,r,a;t=J(t);for(var s=-1,o=this.length;++s<o;){e.push(n=[]);n.parentNode=(r=this[s]).parentNode;for(var l=-1,u=r.length;++l<u;){if(a=r[l]){n.push(i=t.call(a,a.__data__,l,s));if(i&&"__data__"in a)i.__data__=a.__data__}else{n.push(null)}}}return $(e)};function J(t){return typeof t==="function"?t:function(){return U(t,this)}}X.selectAll=function(t){var e=[],n,r;t=G(t);for(var a=-1,s=this.length;++a<s;){for(var o=this[a],l=-1,u=o.length;++l<u;){if(r=o[l]){e.push(n=i(t.call(r,r.__data__,l,a)));n.parentNode=r}}}return $(e)};function G(t){return typeof t==="function"?t:function(){return Y(t,this)}}var K={svg:"http://www.w3.org/2000/svg",xhtml:"http://www.w3.org/1999/xhtml",xlink:"http://www.w3.org/1999/xlink",xml:"http://www.w3.org/XML/1998/namespace",xmlns:"http://www.w3.org/2000/xmlns/"};t.ns={prefix:K,qualify:function(t){var e=t.indexOf(":"),n=t;if(e>=0&&(n=t.slice(0,e))!=="xmlns")t=t.slice(e+1);return K.hasOwnProperty(n)?{space:K[n],local:t}:t}};X.attr=function(e,n){if(arguments.length<2){if(typeof e==="string"){var i=this.node();e=t.ns.qualify(e);return e.local?i.getAttributeNS(e.space,e.local):i.getAttribute(e)}for(n in e)this.each(Z(n,e[n]));return this}return this.each(Z(e,n))};function Z(e,n){e=t.ns.qualify(e);function i(){this.removeAttribute(e)}function r(){this.removeAttributeNS(e.space,e.local)}function a(){this.setAttribute(e,n)}function s(){this.setAttributeNS(e.space,e.local,n)}function o(){var t=n.apply(this,arguments);if(t==null)this.removeAttribute(e);else this.setAttribute(e,t)}function l(){var t=n.apply(this,arguments);if(t==null)this.removeAttributeNS(e.space,e.local);else this.setAttributeNS(e.space,e.local,t)}return n==null?e.local?r:i:typeof n==="function"?e.local?l:o:e.local?s:a}function Q(t){return t.trim().replace(/\s+/g," ")}X.classed=function(t,e){if(arguments.length<2){if(typeof t==="string"){var n=this.node(),i=(t=et(t)).length,r=-1;if(e=n.classList){while(++r<i)if(!e.contains(t[r]))return false}else{e=n.getAttribute("class");while(++r<i)if(!tt(t[r]).test(e))return false}return true}for(e in t)this.each(nt(e,t[e]));return this}return this.each(nt(t,e))};function tt(e){return new RegExp("(?:^|\\s+)"+t.requote(e)+"(?:\\s+|$)","g")}function et(t){return(t+"").trim().split(/^|\s+/)}function nt(t,e){t=et(t).map(it);var n=t.length;function i(){var i=-1;while(++i<n)t[i](this,e)}function r(){var i=-1,r=e.apply(this,arguments);while(++i<n)t[i](this,r)}return typeof e==="function"?r:i}function it(t){var e=tt(t);return function(n,i){if(r=n.classList)return i?r.add(t):r.remove(t);var r=n.getAttribute("class")||"";if(i){e.lastIndex=0;if(!e.test(r))n.setAttribute("class",Q(r+" "+t))}else{n.setAttribute("class",Q(r.replace(e," ")))}}}X.style=function(t,e,n){var i=arguments.length;if(i<3){if(typeof t!=="string"){if(i<2)e="";for(n in t)this.each(rt(n,t[n],e));return this}if(i<2){var r=this.node();return s(r).getComputedStyle(r,null).getPropertyValue(t)}n=""}return this.each(rt(t,e,n))};function rt(t,e,n){function i(){this.style.removeProperty(t)}function r(){this.style.setProperty(t,e,n)}function a(){var i=e.apply(this,arguments);if(i==null)this.style.removeProperty(t);else this.style.setProperty(t,i,n)}return e==null?i:typeof e==="function"?a:r}X.property=function(t,e){if(arguments.length<2){if(typeof t==="string")return this.node()[t];for(e in t)this.each(at(e,t[e]));return this}return this.each(at(t,e))};function at(t,e){function n(){delete this[t]}function i(){this[t]=e}function r(){var n=e.apply(this,arguments);if(n==null)delete this[t];else this[t]=n}return e==null?n:typeof e==="function"?r:i}X.text=function(t){return arguments.length?this.each(typeof t==="function"?function(){var e=t.apply(this,arguments);this.textContent=e==null?"":e}:t==null?function(){this.textContent=""}:function(){this.textContent=t}):this.node().textContent};X.html=function(t){return arguments.length?this.each(typeof t==="function"?function(){var e=t.apply(this,arguments);this.innerHTML=e==null?"":e}:t==null?function(){this.innerHTML=""}:function(){this.innerHTML=t}):this.node().innerHTML};X.append=function(t){t=st(t);return this.select(function(){return this.appendChild(t.apply(this,arguments))})};function st(e){function n(){var t=this.ownerDocument,n=this.namespaceURI;return n?t.createElementNS(n,e):t.createElement(e)}function i(){return this.ownerDocument.createElementNS(e.space,e.local)}return typeof e==="function"?e:(e=t.ns.qualify(e)).local?i:n}X.insert=function(t,e){t=st(t);e=J(e);return this.select(function(){return this.insertBefore(t.apply(this,arguments),e.apply(this,arguments)||null)})};X.remove=function(){return this.each(ot)};function ot(){var t=this.parentNode;if(t)t.removeChild(this)}X.data=function(t,e){var n=-1,i=this.length,r,a;if(!arguments.length){t=new Array(i=(r=this[0]).length);while(++n<i){if(a=r[n]){t[n]=a.__data__}}return t}function s(t,n){var i,r=t.length,a=n.length,s=Math.min(r,a),f=new Array(a),c=new Array(a),h=new Array(r),d,p;if(e){var g=new _,v=new Array(r),m;for(i=-1;++i<r;){if(d=t[i]){if(g.has(m=e.call(d,d.__data__,i))){h[i]=d}else{g.set(m,d)}v[i]=m}}for(i=-1;++i<a;){if(!(d=g.get(m=e.call(n,p=n[i],i)))){c[i]=lt(p)}else if(d!==true){f[i]=d;d.__data__=p}g.set(m,true)}for(i=-1;++i<r;){if(i in v&&g.get(v[i])!==true){h[i]=t[i]}}}else{for(i=-1;++i<s;){d=t[i];p=n[i];if(d){d.__data__=p;f[i]=d}else{c[i]=lt(p)}}for(;i<a;++i){c[i]=lt(n[i])}for(;i<r;++i){h[i]=t[i]}}c.update=f;c.parentNode=f.parentNode=h.parentNode=t.parentNode;o.push(c);l.push(f);u.push(h)}var o=ht([]),l=$([]),u=$([]);if(typeof t==="function"){while(++n<i){s(r=this[n],t.call(r,r.parentNode.__data__,n))}}else{while(++n<i){s(r=this[n],t)}}l.enter=function(){return o};l.exit=function(){return u};return l};function lt(t){return{__data__:t}}X.datum=function(t){return arguments.length?this.property("__data__",t):this.property("__data__")};X.filter=function(t){var e=[],n,i,r;if(typeof t!=="function")t=ut(t);for(var a=0,s=this.length;a<s;a++){e.push(n=[]);n.parentNode=(i=this[a]).parentNode;for(var o=0,l=i.length;o<l;o++){if((r=i[o])&&t.call(r,r.__data__,o,a)){n.push(r)}}}return $(e)};function ut(t){return function(){return V(this,t)}}X.order=function(){for(var t=-1,e=this.length;++t<e;){for(var n=this[t],i=n.length-1,r=n[i],a;--i>=0;){if(a=n[i]){if(r&&r!==a.nextSibling)r.parentNode.insertBefore(a,r);r=a}}}return this};X.sort=function(t){t=ft.apply(this,arguments);for(var e=-1,n=this.length;++e<n;)this[e].sort(t);return this.order()};function ft(t){if(!arguments.length)t=h;return function(e,n){return e&&n?t(e.__data__,n.__data__):!e-!n}}X.each=function(t){return ct(this,function(e,n,i){t.call(e,e.__data__,n,i)})};function ct(t,e){for(var n=0,i=t.length;n<i;n++){for(var r=t[n],a=0,s=r.length,o;a<s;a++){if(o=r[a])e(o,a,n)}}return t}X.call=function(t){var e=i(arguments);t.apply(e[0]=this,e);return this};X.empty=function(){return!this.node()};X.node=function(){for(var t=0,e=this.length;t<e;t++){for(var n=this[t],i=0,r=n.length;i<r;i++){var a=n[i];if(a)return a}}return null};X.size=function(){var t=0;ct(this,function(){++t});return t};function ht(t){B(t,dt);return t}var dt=[];t.selection.enter=ht;t.selection.enter.prototype=dt;dt.append=X.append;dt.empty=X.empty;dt.node=X.node;dt.call=X.call;dt.size=X.size;dt.select=function(t){var e=[],n,i,r,a,s;for(var o=-1,l=this.length;++o<l;){r=(a=this[o]).update;e.push(n=[]);n.parentNode=a.parentNode;for(var u=-1,f=a.length;++u<f;){if(s=a[u]){n.push(r[u]=i=t.call(a.parentNode,s.__data__,u,o));i.__data__=s.__data__}else{n.push(null)}}}return $(e)};dt.insert=function(t,e){if(arguments.length<2)e=pt(this);return X.insert.call(this,t,e)};function pt(t){var e,n;return function(i,r,a){var s=t[a].update,o=s.length,l;if(a!=n)n=a,e=0;if(r>=e)e=r+1;while(!(l=s[e])&&++e<o);return l}}t.select=function(t){var e;if(typeof t==="string"){e=[U(t,r)];e.parentNode=r.documentElement}else{e=[t];e.parentNode=a(t)}return $([e])};t.selectAll=function(t){var e;if(typeof t==="string"){e=i(Y(t,r));e.parentNode=r.documentElement}else{e=i(t);e.parentNode=null}return $([e])};X.on=function(t,e,n){var i=arguments.length;if(i<3){if(typeof t!=="string"){if(i<2)e=false;for(n in t)this.each(gt(n,t[n],e));return this}if(i<2)return(i=this.node()["__on"+t])&&i._;n=false}return this.each(gt(t,e,n))};function gt(e,n,r){var a="__on"+e,s=e.indexOf("."),o=mt;if(s>0)e=e.slice(0,s);var l=vt.get(e);if(l)e=l,o=yt;function u(){var t=this[a];if(t){this.removeEventListener(e,t,t.$);delete this[a]}}function f(){var t=o(n,i(arguments));u.call(this);this.addEventListener(e,this[a]=t,t.$=r);t._=n}function c(){var n=new RegExp("^__on([^.]+)"+t.requote(e)+"$"),i;for(var r in this){if(i=r.match(n)){var a=this[r];this.removeEventListener(i[1],a,a.$);delete this[r]}}}return s?n?f:u:n?L:c}var vt=t.map({mouseenter:"mouseover",mouseleave:"mouseout"});if(r){vt.forEach(function(t){if("on"+t in r)vt.remove(t)})}function mt(e,n){return function(i){var r=t.event;t.event=i;n[0]=this.__data__;try{e.apply(this,n)}finally{t.event=r}}}function yt(t,e){var n=mt(t,e);return function(t){var e=this,i=t.relatedTarget;if(!i||i!==e&&!(i.compareDocumentPosition(e)&8)){n.call(e,t)}}}var bt,wt=0;function _t(e){var n=".dragsuppress-"+ ++wt,i="click"+n,r=t.select(s(e)).on("touchmove"+n,H).on("dragstart"+n,H).on("selectstart"+n,H);if(bt==null){bt="onselectstart"in e?false:F(e.style,"userSelect")}if(bt){var o=a(e).style,l=o[bt];o[bt]="none"}return function(t){r.on(n,null);if(bt)o[bt]=l;if(t){var e=function(){r.on(i,null)};r.on(i,function(){H();e()},true);setTimeout(e,0)}}}t.mouse=function(t){return St(t,W())};var xt=this.navigator&&/WebKit/.test(this.navigator.userAgent)?-1:0;function St(e,n){if(n.changedTouches)n=n.changedTouches[0];var i=e.ownerSVGElement||e;if(i.createSVGPoint){var r=i.createSVGPoint();if(xt<0){var a=s(e);if(a.scrollX||a.scrollY){i=t.select("body").append("svg").style({position:"absolute",top:0,left:0,margin:0,padding:0,border:"none"},"important");var o=i[0][0].getScreenCTM();xt=!(o.f||o.e);i.remove()}}if(xt)r.x=n.pageX,r.y=n.pageY;else r.x=n.clientX,r.y=n.clientY;r=r.matrixTransform(e.getScreenCTM().inverse());return[r.x,r.y]}var l=e.getBoundingClientRect();return[n.clientX-l.left-e.clientLeft,n.clientY-l.top-e.clientTop]}t.touch=function(t,e,n){if(arguments.length<3)n=e,e=W().changedTouches;if(e)for(var i=0,r=e.length,a;i<r;++i){if((a=e[i]).identifier===n){return St(t,a)}}};t.behavior.drag=function(){var e=z(a,"drag","dragstart","dragend"),n=null,i=o(L,t.mouse,s,"mousemove","mouseup"),r=o(Dt,t.touch,P,"touchmove","touchend");function a(){this.on("mousedown.drag",i).on("touchstart.drag",r)}function o(i,r,a,s,o){return function(){var l=this,u=t.event.target,f=l.parentNode,c=e.of(l,arguments),h=0,d=i(),p=".drag"+(d==null?"":"-"+d),g,v=t.select(a(u)).on(s+p,b).on(o+p,w),m=_t(u),y=r(f,d);if(n){g=n.apply(l,arguments);g=[g.x-y[0],g.y-y[1]]}else{g=[0,0]}c({type:"dragstart"});function b(){var t=r(f,d),e,n;if(!t)return;e=t[0]-y[0];n=t[1]-y[1];h|=e|n;y=t;c({type:"drag",x:t[0]+g[0],y:t[1]+g[1],dx:e,dy:n})}function w(){if(!r(f,d))return;v.on(s+p,null).on(o+p,null);m(h);c({type:"dragend"})}}}a.origin=function(t){if(!arguments.length)return n;n=t;return a};return t.rebind(a,e,"on")};function Dt(){return t.event.changedTouches[0].identifier}t.touches=function(t,e){if(arguments.length<2)e=W().touches;return e?i(e).map(function(e){var n=St(t,e);n.identifier=e.identifier;return n}):[]};var Ct=1e-6,Tt=Ct*Ct,kt=Math.PI,Mt=2*kt,It=Mt-Ct,At=kt/2,Nt=kt/180,Pt=180/kt;function Et(t){return t>0?1:t<0?-1:0}function Ft(t,e,n){return(e[0]-t[0])*(n[1]-t[1])-(e[1]-t[1])*(n[0]-t[0])}function Rt(t){return t>1?0:t<-1?kt:Math.acos(t)}function Lt(t){return t>1?At:t<-1?-At:Math.asin(t)}function jt(t){return((t=Math.exp(t))-1/t)/2}function Ot(t){return((t=Math.exp(t))+1/t)/2}function Ht(t){return((t=Math.exp(2*t))-1)/(t+1)}function Wt(t){return(t=Math.sin(t/2))*t}var zt=Math.SQRT2,qt=2,Bt=4;t.interpolateZoom=function(t,e){var n=t[0],i=t[1],r=t[2],a=e[0],s=e[1],o=e[2],l=a-n,u=s-i,f=l*l+u*u,c,h;if(f<Tt){h=Math.log(o/r)/zt;c=function(t){return[n+t*l,i+t*u,r*Math.exp(zt*t*h)]}}else{var d=Math.sqrt(f),p=(o*o-r*r+Bt*f)/(2*r*qt*d),g=(o*o-r*r-Bt*f)/(2*o*qt*d),v=Math.log(Math.sqrt(p*p+1)-p),m=Math.log(Math.sqrt(g*g+1)-g);h=(m-v)/zt;c=function(t){var e=t*h,a=Ot(v),s=r/(qt*d)*(a*Ht(zt*e+v)-jt(v));return[n+s*l,i+s*u,r*a/Ot(zt*e+v)]}}c.duration=h*1e3;return c};t.behavior.zoom=function(){var e={x:0,y:0,k:1},n,i,a,o=[960,500],l=$t,u=250,f=0,c="mousedown.zoom",h="mousemove.zoom",d="mouseup.zoom",p,g="touchstart.zoom",v,m=z(x,"zoomstart","zoom","zoomend"),y,b,w,_;if(!Yt){Yt="onwheel"in r?(Ut=function(){return-t.event.deltaY*(t.event.deltaMode?120:1)},"wheel"):"onmousewheel"in r?(Ut=function(){return t.event.wheelDelta},"mousewheel"):(Ut=function(){return-t.event.detail},"MozMousePixelScroll")}function x(t){t.on(c,P).on(Yt+".zoom",F).on("dblclick.zoom",R).on(g,E)}x.event=function(n){n.each(function(){var n=m.of(this,arguments),r=e;if(Al){t.select(this).transition().each("start.zoom",function(){e=this.__chart__||{x:0,y:0,k:1};I(n)}).tween("zoom:zoom",function(){var a=o[0],s=o[1],l=i?i[0]:a/2,u=i?i[1]:s/2,f=t.interpolateZoom([(l-e.x)/e.k,(u-e.y)/e.k,a/e.k],[(l-r.x)/r.k,(u-r.y)/r.k,a/r.k]);return function(t){var i=f(t),r=a/i[2];this.__chart__=e={x:l-i[0]*r,y:u-i[1]*r,k:r};A(n)}}).each("interrupt.zoom",function(){N(n)}).each("end.zoom",function(){N(n)})}else{this.__chart__=e;I(n);A(n);N(n)}})};x.translate=function(t){if(!arguments.length)return[e.x,e.y];e={x:+t[0],y:+t[1],k:e.k};M();return x};x.scale=function(t){if(!arguments.length)return e.k;e={x:e.x,y:e.y,k:null};C(+t);M();return x};x.scaleExtent=function(t){if(!arguments.length)return l;l=t==null?$t:[+t[0],+t[1]];return x};x.center=function(t){if(!arguments.length)return a;a=t&&[+t[0],+t[1]];return x};x.size=function(t){if(!arguments.length)return o;o=t&&[+t[0],+t[1]];return x};x.duration=function(t){if(!arguments.length)return u;u=+t;return x};x.x=function(t){if(!arguments.length)return b;b=t;y=t.copy();e={x:0,y:0,k:1};return x};x.y=function(t){if(!arguments.length)return _;_=t;w=t.copy();e={x:0,y:0,k:1};return x};function S(t){return[(t[0]-e.x)/e.k,(t[1]-e.y)/e.k]}function D(t){return[t[0]*e.k+e.x,t[1]*e.k+e.y]}function C(t){e.k=Math.max(l[0],Math.min(l[1],t))}function T(t,n){n=D(n);e.x+=t[0]-n[0];e.y+=t[1]-n[1]}function k(n,r,a,s){n.__chart__={x:e.x,y:e.y,k:e.k};C(Math.pow(2,s));T(i=r,a);n=t.select(n);if(u>0)n=n.transition().duration(u);n.call(x.event)}function M(){if(b)b.domain(y.range().map(function(t){return(t-e.x)/e.k}).map(y.invert));if(_)_.domain(w.range().map(function(t){return(t-e.y)/e.k}).map(w.invert))}function I(t){if(!f++)t({type:"zoomstart"})}function A(t){M();t({type:"zoom",scale:e.k,translate:[e.x,e.y]})}function N(t){if(!--f)t({type:"zoomend"}),i=null}function P(){var e=this,n=m.of(e,arguments),i=0,r=t.select(s(e)).on(h,l).on(d,u),a=S(t.mouse(e)),o=_t(e);Cl.call(e);I(n);function l(){i=1;T(t.mouse(e),a);A(n)}function u(){r.on(h,null).on(d,null);o(i);N(n)}}function E(){var n=this,i=m.of(n,arguments),r={},a=0,s,o=".zoom-"+t.event.changedTouches[0].identifier,l="touchmove"+o,u="touchend"+o,f=[],h=t.select(n),d=_t(n);y();I(i);h.on(c,null).on(g,y);function p(){var i=t.touches(n);s=e.k;i.forEach(function(t){if(t.identifier in r)r[t.identifier]=S(t)});return i}function y(){var i=t.event.target;t.select(i).on(l,b).on(u,w);f.push(i);var s=t.event.changedTouches;for(var o=0,c=s.length;o<c;++o){r[s[o].identifier]=null}var h=p(),d=Date.now();if(h.length===1){if(d-v<500){var g=h[0];k(n,g,r[g.identifier],Math.floor(Math.log(e.k)/Math.LN2)+1);H()}v=d}else if(h.length>1){var g=h[0],m=h[1],y=g[0]-m[0],_=g[1]-m[1];a=y*y+_*_}}function b(){var e=t.touches(n),o,l,u,f;Cl.call(n);for(var c=0,h=e.length;c<h;++c,f=null){u=e[c];if(f=r[u.identifier]){if(l)break;o=u,l=f}}if(f){var d=(d=u[0]-o[0])*d+(d=u[1]-o[1])*d,p=a&&Math.sqrt(d/a);o=[(o[0]+u[0])/2,(o[1]+u[1])/2];l=[(l[0]+f[0])/2,(l[1]+f[1])/2];C(p*s)}v=null;T(o,l);A(i)}
function w(){if(t.event.touches.length){var e=t.event.changedTouches;for(var n=0,a=e.length;n<a;++n){delete r[e[n].identifier]}for(var s in r){return void p()}}t.selectAll(f).on(o,null);h.on(c,P).on(g,E);d();N(i)}}function F(){var r=m.of(this,arguments);if(p)clearTimeout(p);else Cl.call(this),n=S(i=a||t.mouse(this)),I(r);p=setTimeout(function(){p=null;N(r)},50);H();C(Math.pow(2,Ut()*.002)*e.k);T(i,n);A(r)}function R(){var n=t.mouse(this),i=Math.log(e.k)/Math.LN2;k(this,n,S(n),t.event.shiftKey?Math.ceil(i)-1:Math.floor(i)+1)}return t.rebind(x,m,"on")};var $t=[0,Infinity],Ut,Yt;t.color=Vt;function Vt(){}Vt.prototype.toString=function(){return this.rgb()+""};t.hsl=Xt;function Xt(t,e,n){return this instanceof Xt?void(this.h=+t,this.s=+e,this.l=+n):arguments.length<2?t instanceof Xt?new Xt(t.h,t.s,t.l):ve(""+t,me,Xt):new Xt(t,e,n)}var Jt=Xt.prototype=new Vt;Jt.brighter=function(t){t=Math.pow(.7,arguments.length?t:1);return new Xt(this.h,this.s,this.l/t)};Jt.darker=function(t){t=Math.pow(.7,arguments.length?t:1);return new Xt(this.h,this.s,t*this.l)};Jt.rgb=function(){return Gt(this.h,this.s,this.l)};function Gt(t,e,n){var i,r;t=isNaN(t)?0:(t%=360)<0?t+360:t;e=isNaN(e)?0:e<0?0:e>1?1:e;n=n<0?0:n>1?1:n;r=n<=.5?n*(1+e):n+e-n*e;i=2*n-r;function a(t){if(t>360)t-=360;else if(t<0)t+=360;if(t<60)return i+(r-i)*t/60;if(t<180)return r;if(t<240)return i+(r-i)*(240-t)/60;return i}function s(t){return Math.round(a(t)*255)}return new ce(s(t+120),s(t),s(t-120))}t.hcl=Kt;function Kt(e,n,i){return this instanceof Kt?void(this.h=+e,this.c=+n,this.l=+i):arguments.length<2?e instanceof Kt?new Kt(e.h,e.c,e.l):e instanceof te?oe(e.l,e.a,e.b):oe((e=ye((e=t.rgb(e)).r,e.g,e.b)).l,e.a,e.b):new Kt(e,n,i)}var Zt=Kt.prototype=new Vt;Zt.brighter=function(t){return new Kt(this.h,this.c,Math.min(100,this.l+ee*(arguments.length?t:1)))};Zt.darker=function(t){return new Kt(this.h,this.c,Math.max(0,this.l-ee*(arguments.length?t:1)))};Zt.rgb=function(){return Qt(this.h,this.c,this.l).rgb()};function Qt(t,e,n){if(isNaN(t))t=0;if(isNaN(e))e=0;return new te(n,Math.cos(t*=Nt)*e,Math.sin(t)*e)}t.lab=te;function te(t,e,n){return this instanceof te?void(this.l=+t,this.a=+e,this.b=+n):arguments.length<2?t instanceof te?new te(t.l,t.a,t.b):t instanceof Kt?Qt(t.h,t.c,t.l):ye((t=ce(t)).r,t.g,t.b):new te(t,e,n)}var ee=18;var ne=.95047,ie=1,re=1.08883;var ae=te.prototype=new Vt;ae.brighter=function(t){return new te(Math.min(100,this.l+ee*(arguments.length?t:1)),this.a,this.b)};ae.darker=function(t){return new te(Math.max(0,this.l-ee*(arguments.length?t:1)),this.a,this.b)};ae.rgb=function(){return se(this.l,this.a,this.b)};function se(t,e,n){var i=(t+16)/116,r=i+e/500,a=i-n/200;r=le(r)*ne;i=le(i)*ie;a=le(a)*re;return new ce(fe(3.2404542*r-1.5371385*i-.4985314*a),fe(-.969266*r+1.8760108*i+.041556*a),fe(.0556434*r-.2040259*i+1.0572252*a))}function oe(t,e,n){return t>0?new Kt(Math.atan2(n,e)*Pt,Math.sqrt(e*e+n*n),t):new Kt(NaN,NaN,t)}function le(t){return t>.206893034?t*t*t:(t-4/29)/7.787037}function ue(t){return t>.008856?Math.pow(t,1/3):7.787037*t+4/29}function fe(t){return Math.round(255*(t<=.00304?12.92*t:1.055*Math.pow(t,1/2.4)-.055))}t.rgb=ce;function ce(t,e,n){return this instanceof ce?void(this.r=~~t,this.g=~~e,this.b=~~n):arguments.length<2?t instanceof ce?new ce(t.r,t.g,t.b):ve(""+t,ce,Gt):new ce(t,e,n)}function he(t){return new ce(t>>16,t>>8&255,t&255)}function de(t){return he(t)+""}var pe=ce.prototype=new Vt;pe.brighter=function(t){t=Math.pow(.7,arguments.length?t:1);var e=this.r,n=this.g,i=this.b,r=30;if(!e&&!n&&!i)return new ce(r,r,r);if(e&&e<r)e=r;if(n&&n<r)n=r;if(i&&i<r)i=r;return new ce(Math.min(255,e/t),Math.min(255,n/t),Math.min(255,i/t))};pe.darker=function(t){t=Math.pow(.7,arguments.length?t:1);return new ce(t*this.r,t*this.g,t*this.b)};pe.hsl=function(){return me(this.r,this.g,this.b)};pe.toString=function(){return"#"+ge(this.r)+ge(this.g)+ge(this.b)};function ge(t){return t<16?"0"+Math.max(0,t).toString(16):Math.min(255,t).toString(16)}function ve(t,e,n){var i=0,r=0,a=0,s,o,l;s=/([a-z]+)\((.*)\)/.exec(t=t.toLowerCase());if(s){o=s[2].split(",");switch(s[1]){case"hsl":{return n(parseFloat(o[0]),parseFloat(o[1])/100,parseFloat(o[2])/100)}case"rgb":{return e(we(o[0]),we(o[1]),we(o[2]))}}}if(l=_e.get(t)){return e(l.r,l.g,l.b)}if(t!=null&&t.charAt(0)==="#"&&!isNaN(l=parseInt(t.slice(1),16))){if(t.length===4){i=(l&3840)>>4;i=i>>4|i;r=l&240;r=r>>4|r;a=l&15;a=a<<4|a}else if(t.length===7){i=(l&16711680)>>16;r=(l&65280)>>8;a=l&255}}return e(i,r,a)}function me(t,e,n){var i=Math.min(t/=255,e/=255,n/=255),r=Math.max(t,e,n),a=r-i,s,o,l=(r+i)/2;if(a){o=l<.5?a/(r+i):a/(2-r-i);if(t==r)s=(e-n)/a+(e<n?6:0);else if(e==r)s=(n-t)/a+2;else s=(t-e)/a+4;s*=60}else{s=NaN;o=l>0&&l<1?0:s}return new Xt(s,o,l)}function ye(t,e,n){t=be(t);e=be(e);n=be(n);var i=ue((.4124564*t+.3575761*e+.1804375*n)/ne),r=ue((.2126729*t+.7151522*e+.072175*n)/ie),a=ue((.0193339*t+.119192*e+.9503041*n)/re);return te(116*r-16,500*(i-r),200*(r-a))}function be(t){return(t/=255)<=.04045?t/12.92:Math.pow((t+.055)/1.055,2.4)}function we(t){var e=parseFloat(t);return t.charAt(t.length-1)==="%"?Math.round(e*2.55):e}var _e=t.map({aliceblue:15792383,antiquewhite:16444375,aqua:65535,aquamarine:8388564,azure:15794175,beige:16119260,bisque:16770244,black:0,blanchedalmond:16772045,blue:255,blueviolet:9055202,brown:10824234,burlywood:14596231,cadetblue:6266528,chartreuse:8388352,chocolate:13789470,coral:16744272,cornflowerblue:6591981,cornsilk:16775388,crimson:14423100,cyan:65535,darkblue:139,darkcyan:35723,darkgoldenrod:12092939,darkgray:11119017,darkgreen:25600,darkgrey:11119017,darkkhaki:12433259,darkmagenta:9109643,darkolivegreen:5597999,darkorange:16747520,darkorchid:10040012,darkred:9109504,darksalmon:15308410,darkseagreen:9419919,darkslateblue:4734347,darkslategray:3100495,darkslategrey:3100495,darkturquoise:52945,darkviolet:9699539,deeppink:16716947,deepskyblue:49151,dimgray:6908265,dimgrey:6908265,dodgerblue:2003199,firebrick:11674146,floralwhite:16775920,forestgreen:2263842,fuchsia:16711935,gainsboro:14474460,ghostwhite:16316671,gold:16766720,goldenrod:14329120,gray:8421504,green:32768,greenyellow:11403055,grey:8421504,honeydew:15794160,hotpink:16738740,indianred:13458524,indigo:4915330,ivory:16777200,khaki:15787660,lavender:15132410,lavenderblush:16773365,lawngreen:8190976,lemonchiffon:16775885,lightblue:11393254,lightcoral:15761536,lightcyan:14745599,lightgoldenrodyellow:16448210,lightgray:13882323,lightgreen:9498256,lightgrey:13882323,lightpink:16758465,lightsalmon:16752762,lightseagreen:2142890,lightskyblue:8900346,lightslategray:7833753,lightslategrey:7833753,lightsteelblue:11584734,lightyellow:16777184,lime:65280,limegreen:3329330,linen:16445670,magenta:16711935,maroon:8388608,mediumaquamarine:6737322,mediumblue:205,mediumorchid:12211667,mediumpurple:9662683,mediumseagreen:3978097,mediumslateblue:8087790,mediumspringgreen:64154,mediumturquoise:4772300,mediumvioletred:13047173,midnightblue:1644912,mintcream:16121850,mistyrose:16770273,moccasin:16770229,navajowhite:16768685,navy:128,oldlace:16643558,olive:8421376,olivedrab:7048739,orange:16753920,orangered:16729344,orchid:14315734,palegoldenrod:15657130,palegreen:10025880,paleturquoise:11529966,palevioletred:14381203,papayawhip:16773077,peachpuff:16767673,peru:13468991,pink:16761035,plum:14524637,powderblue:11591910,purple:8388736,rebeccapurple:6697881,red:16711680,rosybrown:12357519,royalblue:4286945,saddlebrown:9127187,salmon:16416882,sandybrown:16032864,seagreen:3050327,seashell:16774638,sienna:10506797,silver:12632256,skyblue:8900331,slateblue:6970061,slategray:7372944,slategrey:7372944,snow:16775930,springgreen:65407,steelblue:4620980,tan:13808780,teal:32896,thistle:14204888,tomato:16737095,turquoise:4251856,violet:15631086,wheat:16113331,white:16777215,whitesmoke:16119285,yellow:16776960,yellowgreen:10145074});_e.forEach(function(t,e){_e.set(t,he(e))});function xe(t){return typeof t==="function"?t:function(){return t}}t.functor=xe;t.xhr=Se(P);function Se(t){return function(e,n,i){if(arguments.length===2&&typeof n==="function")i=n,n=null;return De(e,n,t,i)}}function De(e,n,r,a){var s={},o=t.dispatch("beforesend","progress","load","error"),l={},u=new XMLHttpRequest,f=null;if(this.XDomainRequest&&!("withCredentials"in u)&&/^(http(s)?:)?\/\//.test(e))u=new XDomainRequest;"onload"in u?u.onload=u.onerror=c:u.onreadystatechange=function(){u.readyState>3&&c()};function c(){var t=u.status,e;if(!t&&Te(u)||t>=200&&t<300||t===304){try{e=r.call(s,u)}catch(t){o.error.call(s,t);return}o.load.call(s,e)}else{o.error.call(s,u)}}u.onprogress=function(e){var n=t.event;t.event=e;try{o.progress.call(s,u)}finally{t.event=n}};s.header=function(t,e){t=(t+"").toLowerCase();if(arguments.length<2)return l[t];if(e==null)delete l[t];else l[t]=e+"";return s};s.mimeType=function(t){if(!arguments.length)return n;n=t==null?null:t+"";return s};s.responseType=function(t){if(!arguments.length)return f;f=t;return s};s.response=function(t){r=t;return s};["get","post"].forEach(function(t){s[t]=function(){return s.send.apply(s,[t].concat(i(arguments)))}});s.send=function(t,i,r){if(arguments.length===2&&typeof i==="function")r=i,i=null;u.open(t,e,true);if(n!=null&&!("accept"in l))l["accept"]=n+",*/*";if(u.setRequestHeader)for(var a in l)u.setRequestHeader(a,l[a]);if(n!=null&&u.overrideMimeType)u.overrideMimeType(n);if(f!=null)u.responseType=f;if(r!=null)s.on("error",r).on("load",function(t){r(null,t)});o.beforesend.call(s,u);u.send(i==null?null:i);return s};s.abort=function(){u.abort();return s};t.rebind(s,o,"on");return a==null?s:s.get(Ce(a))}function Ce(t){return t.length===1?function(e,n){t(e==null?n:null)}:t}function Te(t){var e=t.responseType;return e&&e!=="text"?t.response:t.responseText}t.dsv=function(t,e){var n=new RegExp('["'+t+"\n]"),i=t.charCodeAt(0);function r(t,n,i){if(arguments.length<3)i=n,n=null;var r=De(t,e,n==null?a:s(n),i);r.row=function(t){return arguments.length?r.response((n=t)==null?a:s(t)):n};return r}function a(t){return r.parse(t.responseText)}function s(t){return function(e){return r.parse(e.responseText,t)}}r.parse=function(t,e){var n;return r.parseRows(t,function(t,i){if(n)return n(t,i-1);var r=new Function("d","return {"+t.map(function(t,e){return JSON.stringify(t)+": d["+e+"]"}).join(",")+"}");n=e?function(t,n){return e(r(t),n)}:r})};r.parseRows=function(t,e){var n={},r={},a=[],s=t.length,o=0,l=0,u,f;function c(){if(o>=s)return r;if(f)return f=false,n;var e=o;if(t.charCodeAt(e)===34){var a=e;while(a++<s){if(t.charCodeAt(a)===34){if(t.charCodeAt(a+1)!==34)break;++a}}o=a+2;var l=t.charCodeAt(a+1);if(l===13){f=true;if(t.charCodeAt(a+2)===10)++o}else if(l===10){f=true}return t.slice(e+1,a).replace(/""/g,'"')}while(o<s){var l=t.charCodeAt(o++),u=1;if(l===10)f=true;else if(l===13){f=true;if(t.charCodeAt(o)===10)++o,++u}else if(l!==i)continue;return t.slice(e,o-u)}return t.slice(e)}while((u=c())!==r){var h=[];while(u!==n&&u!==r){h.push(u);u=c()}if(e&&(h=e(h,l++))==null)continue;a.push(h)}return a};r.format=function(e){if(Array.isArray(e[0]))return r.formatRows(e);var n=new N,i=[];e.forEach(function(t){for(var e in t){if(!n.has(e)){i.push(n.add(e))}}});return[i.map(l).join(t)].concat(e.map(function(e){return i.map(function(t){return l(e[t])}).join(t)})).join("\n")};r.formatRows=function(t){return t.map(o).join("\n")};function o(e){return e.map(l).join(t)}function l(t){return n.test(t)?'"'+t.replace(/\"/g,'""')+'"':t}return r};t.csv=t.dsv(",","text/csv");t.tsv=t.dsv("\t","text/tab-separated-values");var ke,Me,Ie,Ae,Ne=this[F(this,"requestAnimationFrame")]||function(t){setTimeout(t,17)};t.timer=function(){Pe.apply(this,arguments)};function Pe(t,e,n){var i=arguments.length;if(i<2)e=0;if(i<3)n=Date.now();var r=n+e,a={c:t,t:r,n:null};if(Me)Me.n=a;else ke=a;Me=a;if(!Ie){Ae=clearTimeout(Ae);Ie=1;Ne(Ee)}return a}function Ee(){var t=Fe(),e=Re()-t;if(e>24){if(isFinite(e)){clearTimeout(Ae);Ae=setTimeout(Ee,e)}Ie=0}else{Ie=1;Ne(Ee)}}t.timer.flush=function(){Fe();Re()};function Fe(){var t=Date.now(),e=ke;while(e){if(t>=e.t&&e.c(t-e.t))e.c=null;e=e.n}return t}function Re(){var t,e=ke,n=Infinity;while(e){if(e.c){if(e.t<n)n=e.t;e=(t=e).n}else{e=t?t.n=e.n:ke=e.n}}Me=t;return n}function Le(t,e){return e-(t?Math.ceil(Math.log(t)/Math.LN10):1)}t.round=function(t,e){return e?Math.round(t*(e=Math.pow(10,e)))/e:Math.round(t)};var je=["y","z","a","f","p","n","µ","m","","k","M","G","T","P","E","Z","Y"].map(Oe);t.formatPrefix=function(e,n){var i=0;if(e=+e){if(e<0)e*=-1;if(n)e=t.round(e,Le(e,n));i=1+Math.floor(1e-12+Math.log(e)/Math.LN10);i=Math.max(-24,Math.min(24,Math.floor((i-1)/3)*3))}return je[8+i/3]};function Oe(t,e){var n=Math.pow(10,y(8-e)*3);return{scale:e>8?function(t){return t/n}:function(t){return t*n},symbol:t}}function He(e){var n=e.decimal,i=e.thousands,r=e.grouping,a=e.currency,s=r&&i?function(t,e){var n=t.length,a=[],s=0,o=r[0],l=0;while(n>0&&o>0){if(l+o+1>e)o=Math.max(1,e-l);a.push(t.substring(n-=o,n+o));if((l+=o+1)>e)break;o=r[s=(s+1)%r.length]}return a.reverse().join(i)}:P;return function(e){var i=We.exec(e),r=i[1]||" ",o=i[2]||">",l=i[3]||"-",u=i[4]||"",f=i[5],c=+i[6],h=i[7],d=i[8],p=i[9],g=1,v="",m="",y=false,b=true;if(d)d=+d.substring(1);if(f||r==="0"&&o==="="){f=r="0";o="="}switch(p){case"n":h=true;p="g";break;case"%":g=100;m="%";p="f";break;case"p":g=100;m="%";p="r";break;case"b":case"o":case"x":case"X":if(u==="#")v="0"+p.toLowerCase();case"c":b=false;case"d":y=true;d=0;break;case"s":g=-1;p="r";break}if(u==="$")v=a[0],m=a[1];if(p=="r"&&!d)p="g";if(d!=null){if(p=="g")d=Math.max(1,Math.min(21,d));else if(p=="e"||p=="f")d=Math.max(0,Math.min(20,d))}p=ze.get(p)||qe;var w=f&&h;return function(e){var i=m;if(y&&e%1)return"";var a=e<0||e===0&&1/e<0?(e=-e,"-"):l==="-"?"":l;if(g<0){var u=t.formatPrefix(e,d);e=u.scale(e);i=u.symbol+m}else{e*=g}e=p(e,d);var _=e.lastIndexOf("."),x,S;if(_<0){var D=b?e.lastIndexOf("e"):-1;if(D<0)x=e,S="";else x=e.substring(0,D),S=e.substring(D)}else{x=e.substring(0,_);S=n+e.substring(_+1)}if(!f&&h)x=s(x,Infinity);var C=v.length+x.length+S.length+(w?0:a.length),T=C<c?new Array(C=c-C+1).join(r):"";if(w)x=s(T+x,T.length?c-S.length:Infinity);a+=v;e=x+S;return(o==="<"?a+e+T:o===">"?T+a+e:o==="^"?T.substring(0,C>>=1)+a+e+T.substring(C):a+(w?e:T+e))+i}}}var We=/(?:([^{])?([<>=^]))?([+\- ])?([$#])?(0)?(\d+)?(,)?(\.-?\d+)?([a-z%])?/i;var ze=t.map({b:function(t){return t.toString(2)},c:function(t){return String.fromCharCode(t)},o:function(t){return t.toString(8)},x:function(t){return t.toString(16)},X:function(t){return t.toString(16).toUpperCase()},g:function(t,e){return t.toPrecision(e)},e:function(t,e){return t.toExponential(e)},f:function(t,e){return t.toFixed(e)},r:function(e,n){return(e=t.round(e,Le(e,n))).toFixed(Math.max(0,Math.min(20,Le(e*(1+1e-15),n))))}});function qe(t){return t+""}var Be=t.time={},$e=Date;function Ue(){this._=new Date(arguments.length>1?Date.UTC.apply(this,arguments):arguments[0])}Ue.prototype={getDate:function(){return this._.getUTCDate()},getDay:function(){return this._.getUTCDay()},getFullYear:function(){return this._.getUTCFullYear()},getHours:function(){return this._.getUTCHours()},getMilliseconds:function(){return this._.getUTCMilliseconds()},getMinutes:function(){return this._.getUTCMinutes()},getMonth:function(){return this._.getUTCMonth()},getSeconds:function(){return this._.getUTCSeconds()},getTime:function(){return this._.getTime()},getTimezoneOffset:function(){return 0},valueOf:function(){return this._.valueOf()},setDate:function(){Ye.setUTCDate.apply(this._,arguments)},setDay:function(){Ye.setUTCDay.apply(this._,arguments)},setFullYear:function(){Ye.setUTCFullYear.apply(this._,arguments)},setHours:function(){Ye.setUTCHours.apply(this._,arguments)},setMilliseconds:function(){Ye.setUTCMilliseconds.apply(this._,arguments)},setMinutes:function(){Ye.setUTCMinutes.apply(this._,arguments)},setMonth:function(){Ye.setUTCMonth.apply(this._,arguments)},setSeconds:function(){Ye.setUTCSeconds.apply(this._,arguments)},setTime:function(){Ye.setTime.apply(this._,arguments)}};var Ye=Date.prototype;function Ve(t,e,n){function i(e){var n=t(e),i=a(n,1);return e-n<i-e?n:i}function r(n){e(n=t(new $e(n-1)),1);return n}function a(t,n){e(t=new $e(+t),n);return t}function s(t,i,a){var s=r(t),o=[];if(a>1){while(s<i){if(!(n(s)%a))o.push(new Date(+s));e(s,1)}}else{while(s<i)o.push(new Date(+s)),e(s,1)}return o}function o(t,e,n){try{$e=Ue;var i=new Ue;i._=t;return s(i,e,n)}finally{$e=Date}}t.floor=t;t.round=i;t.ceil=r;t.offset=a;t.range=s;var l=t.utc=Xe(t);l.floor=l;l.round=Xe(i);l.ceil=Xe(r);l.offset=Xe(a);l.range=o;return t}function Xe(t){return function(e,n){try{$e=Ue;var i=new Ue;i._=e;return t(i,n)._}finally{$e=Date}}}Be.year=Ve(function(t){t=Be.day(t);t.setMonth(0,1);return t},function(t,e){t.setFullYear(t.getFullYear()+e)},function(t){return t.getFullYear()});Be.years=Be.year.range;Be.years.utc=Be.year.utc.range;Be.day=Ve(function(t){var e=new $e(2e3,0);e.setFullYear(t.getFullYear(),t.getMonth(),t.getDate());return e},function(t,e){t.setDate(t.getDate()+e)},function(t){return t.getDate()-1});Be.days=Be.day.range;Be.days.utc=Be.day.utc.range;Be.dayOfYear=function(t){var e=Be.year(t);return Math.floor((t-e-(t.getTimezoneOffset()-e.getTimezoneOffset())*6e4)/864e5)};["sunday","monday","tuesday","wednesday","thursday","friday","saturday"].forEach(function(t,e){e=7-e;var n=Be[t]=Ve(function(t){(t=Be.day(t)).setDate(t.getDate()-(t.getDay()+e)%7);return t},function(t,e){t.setDate(t.getDate()+Math.floor(e)*7)},function(t){var n=Be.year(t).getDay();return Math.floor((Be.dayOfYear(t)+(n+e)%7)/7)-(n!==e)});Be[t+"s"]=n.range;Be[t+"s"].utc=n.utc.range;Be[t+"OfYear"]=function(t){var n=Be.year(t).getDay();return Math.floor((Be.dayOfYear(t)+(n+e)%7)/7)}});Be.week=Be.sunday;Be.weeks=Be.sunday.range;Be.weeks.utc=Be.sunday.utc.range;Be.weekOfYear=Be.sundayOfYear;function Je(e){var n=e.dateTime,i=e.date,r=e.time,a=e.periods,s=e.days,o=e.shortDays,l=e.months,u=e.shortMonths;function f(t){var e=t.length;function n(n){var i=[],r=-1,a=0,s,o,l;while(++r<e){if(t.charCodeAt(r)===37){i.push(t.slice(a,r));if((o=Ge[s=t.charAt(++r)])!=null)s=t.charAt(++r);if(l=_[s])s=l(n,o==null?s==="e"?" ":"0":o);i.push(s);a=r+1}}i.push(t.slice(a,r));return i.join("")}n.parse=function(e){var n={y:1900,m:0,d:1,H:0,M:0,S:0,L:0,Z:null},i=c(n,t,e,0);if(i!=e.length)return null;if("p"in n)n.H=n.H%12+n.p*12;var r=n.Z!=null&&$e!==Ue,a=new(r?Ue:$e);if("j"in n)a.setFullYear(n.y,0,n.j);else if("W"in n||"U"in n){if(!("w"in n))n.w="W"in n?1:0;a.setFullYear(n.y,0,1);a.setFullYear(n.y,0,"W"in n?(n.w+6)%7+n.W*7-(a.getDay()+5)%7:n.w+n.U*7-(a.getDay()+6)%7)}else a.setFullYear(n.y,n.m,n.d);a.setHours(n.H+(n.Z/100|0),n.M+n.Z%100,n.S,n.L);return r?a._:a};n.toString=function(){return t};return n}function c(t,e,n,i){var r,a,s,o=0,l=e.length,u=n.length;while(o<l){if(i>=u)return-1;r=e.charCodeAt(o++);if(r===37){s=e.charAt(o++);a=x[s in Ge?e.charAt(o++):s];if(!a||(i=a(t,n,i))<0)return-1}else if(r!=n.charCodeAt(i++)){return-1}}return i}f.utc=function(t){var e=f(t);function n(t){try{$e=Ue;var n=new $e;n._=t;return e(n)}finally{$e=Date}}n.parse=function(t){try{$e=Ue;var n=e.parse(t);return n&&n._}finally{$e=Date}};n.toString=e.toString;return n};f.multi=f.utc.multi=bn;var h=t.map(),d=tn(s),p=en(s),g=tn(o),v=en(o),m=tn(l),y=en(l),b=tn(u),w=en(u);a.forEach(function(t,e){h.set(t.toLowerCase(),e)});var _={a:function(t){return o[t.getDay()]},A:function(t){return s[t.getDay()]},b:function(t){return u[t.getMonth()]},B:function(t){return l[t.getMonth()]},c:f(n),d:function(t,e){return Qe(t.getDate(),e,2)},e:function(t,e){return Qe(t.getDate(),e,2)},H:function(t,e){return Qe(t.getHours(),e,2)},I:function(t,e){return Qe(t.getHours()%12||12,e,2)},j:function(t,e){return Qe(1+Be.dayOfYear(t),e,3)},L:function(t,e){return Qe(t.getMilliseconds(),e,3)},m:function(t,e){return Qe(t.getMonth()+1,e,2)},M:function(t,e){return Qe(t.getMinutes(),e,2)},p:function(t){return a[+(t.getHours()>=12)]},S:function(t,e){return Qe(t.getSeconds(),e,2)},U:function(t,e){return Qe(Be.sundayOfYear(t),e,2)},w:function(t){return t.getDay()},W:function(t,e){return Qe(Be.mondayOfYear(t),e,2)},x:f(i),X:f(r),y:function(t,e){return Qe(t.getFullYear()%100,e,2)},Y:function(t,e){return Qe(t.getFullYear()%1e4,e,4)},Z:mn,"%":function(){return"%"}};var x={a:S,A:D,b:C,B:T,c:k,d:cn,e:cn,H:dn,I:dn,j:hn,L:vn,m:fn,M:pn,p:A,S:gn,U:rn,w:nn,W:an,x:M,X:I,y:on,Y:sn,Z:ln,"%":yn};function S(t,e,n){g.lastIndex=0;var i=g.exec(e.slice(n));return i?(t.w=v.get(i[0].toLowerCase()),n+i[0].length):-1}function D(t,e,n){d.lastIndex=0;var i=d.exec(e.slice(n));return i?(t.w=p.get(i[0].toLowerCase()),n+i[0].length):-1}function C(t,e,n){b.lastIndex=0;var i=b.exec(e.slice(n));return i?(t.m=w.get(i[0].toLowerCase()),n+i[0].length):-1}function T(t,e,n){m.lastIndex=0;var i=m.exec(e.slice(n));return i?(t.m=y.get(i[0].toLowerCase()),n+i[0].length):-1}function k(t,e,n){return c(t,_.c.toString(),e,n)}function M(t,e,n){return c(t,_.x.toString(),e,n)}function I(t,e,n){return c(t,_.X.toString(),e,n)}function A(t,e,n){var i=h.get(e.slice(n,n+=2).toLowerCase());return i==null?-1:(t.p=i,n)}return f}var Ge={"-":"",_:" ",0:"0"},Ke=/^\s*\d+/,Ze=/^%/;function Qe(t,e,n){var i=t<0?"-":"",r=(i?-t:t)+"",a=r.length;return i+(a<n?new Array(n-a+1).join(e)+r:r)}function tn(e){return new RegExp("^(?:"+e.map(t.requote).join("|")+")","i")}function en(t){var e=new _,n=-1,i=t.length;while(++n<i)e.set(t[n].toLowerCase(),n);return e}function nn(t,e,n){Ke.lastIndex=0;var i=Ke.exec(e.slice(n,n+1));return i?(t.w=+i[0],n+i[0].length):-1}function rn(t,e,n){Ke.lastIndex=0;var i=Ke.exec(e.slice(n));return i?(t.U=+i[0],n+i[0].length):-1}function an(t,e,n){Ke.lastIndex=0;var i=Ke.exec(e.slice(n));return i?(t.W=+i[0],n+i[0].length):-1}function sn(t,e,n){Ke.lastIndex=0;var i=Ke.exec(e.slice(n,n+4));return i?(t.y=+i[0],n+i[0].length):-1}function on(t,e,n){Ke.lastIndex=0;var i=Ke.exec(e.slice(n,n+2));return i?(t.y=un(+i[0]),n+i[0].length):-1}function ln(t,e,n){return/^[+-]\d{4}$/.test(e=e.slice(n,n+5))?(t.Z=-e,n+5):-1}function un(t){return t+(t>68?1900:2e3)}function fn(t,e,n){Ke.lastIndex=0;var i=Ke.exec(e.slice(n,n+2));return i?(t.m=i[0]-1,n+i[0].length):-1}function cn(t,e,n){Ke.lastIndex=0;var i=Ke.exec(e.slice(n,n+2));return i?(t.d=+i[0],n+i[0].length):-1}function hn(t,e,n){Ke.lastIndex=0;var i=Ke.exec(e.slice(n,n+3));return i?(t.j=+i[0],n+i[0].length):-1}function dn(t,e,n){Ke.lastIndex=0;var i=Ke.exec(e.slice(n,n+2));return i?(t.H=+i[0],n+i[0].length):-1}function pn(t,e,n){Ke.lastIndex=0;var i=Ke.exec(e.slice(n,n+2));return i?(t.M=+i[0],n+i[0].length):-1}function gn(t,e,n){Ke.lastIndex=0;var i=Ke.exec(e.slice(n,n+2));return i?(t.S=+i[0],n+i[0].length):-1}function vn(t,e,n){Ke.lastIndex=0;var i=Ke.exec(e.slice(n,n+3));return i?(t.L=+i[0],n+i[0].length):-1}function mn(t){var e=t.getTimezoneOffset(),n=e>0?"-":"+",i=y(e)/60|0,r=y(e)%60;return n+Qe(i,"0",2)+Qe(r,"0",2)}function yn(t,e,n){Ze.lastIndex=0;var i=Ze.exec(e.slice(n,n+1));return i?n+i[0].length:-1}function bn(t){var e=t.length,n=-1;while(++n<e)t[n][0]=this(t[n][0]);return function(e){var n=0,i=t[n];while(!i[1](e))i=t[++n];return i[0](e)}}t.locale=function(t){return{numberFormat:He(t),timeFormat:Je(t)}};var wn=t.locale({decimal:".",thousands:",",grouping:[3],currency:["$",""],dateTime:"%a %b %e %X %Y",date:"%m/%d/%Y",time:"%H:%M:%S",periods:["AM","PM"],days:["Sunday","Monday","Tuesday","Wednesday","Thursday","Friday","Saturday"],shortDays:["Sun","Mon","Tue","Wed","Thu","Fri","Sat"],months:["January","February","March","April","May","June","July","August","September","October","November","December"],shortMonths:["Jan","Feb","Mar","Apr","May","Jun","Jul","Aug","Sep","Oct","Nov","Dec"]});t.format=wn.numberFormat;t.geo={};function _n(){}_n.prototype={s:0,t:0,add:function(t){Sn(t,this.t,xn);Sn(xn.s,this.s,this);if(this.s)this.t+=xn.t;else this.s=xn.t},reset:function(){this.s=this.t=0},valueOf:function(){return this.s}};var xn=new _n;function Sn(t,e,n){var i=n.s=t+e,r=i-t,a=i-r;n.t=t-a+(e-r)}t.geo.stream=function(t,e){if(t&&Cn.hasOwnProperty(t.type)){Cn[t.type](t,e)}else{Dn(t,e)}};function Dn(t,e){if(t&&Tn.hasOwnProperty(t.type)){Tn[t.type](t,e)}}var Cn={Feature:function(t,e){Dn(t.geometry,e)},FeatureCollection:function(t,e){var n=t.features,i=-1,r=n.length;while(++i<r)Dn(n[i].geometry,e)}};var Tn={Sphere:function(t,e){e.sphere()},Point:function(t,e){t=t.coordinates;e.point(t[0],t[1],t[2])},MultiPoint:function(t,e){var n=t.coordinates,i=-1,r=n.length;while(++i<r)t=n[i],e.point(t[0],t[1],t[2])},LineString:function(t,e){kn(t.coordinates,e,0)},MultiLineString:function(t,e){var n=t.coordinates,i=-1,r=n.length;while(++i<r)kn(n[i],e,0)},Polygon:function(t,e){Mn(t.coordinates,e)},MultiPolygon:function(t,e){var n=t.coordinates,i=-1,r=n.length;while(++i<r)Mn(n[i],e)},GeometryCollection:function(t,e){var n=t.geometries,i=-1,r=n.length;while(++i<r)Dn(n[i],e)}};function kn(t,e,n){var i=-1,r=t.length-n,a;e.lineStart();while(++i<r)a=t[i],e.point(a[0],a[1],a[2]);e.lineEnd()}function Mn(t,e){var n=-1,i=t.length;e.polygonStart();while(++n<i)kn(t[n],e,1);e.polygonEnd()}t.geo.area=function(e){In=0;t.geo.stream(e,Nn);return In};var In,An=new _n;var Nn={sphere:function(){In+=4*kt},point:L,lineStart:L,lineEnd:L,polygonStart:function(){An.reset();Nn.lineStart=Pn},polygonEnd:function(){var t=2*An;In+=t<0?4*kt+t:t;Nn.lineStart=Nn.lineEnd=Nn.point=L}};function Pn(){var t,e,n,i,r;Nn.point=function(s,o){Nn.point=a;n=(t=s)*Nt,i=Math.cos(o=(e=o)*Nt/2+kt/4),r=Math.sin(o)};function a(t,e){t*=Nt;e=e*Nt/2+kt/4;var a=t-n,s=a>=0?1:-1,o=s*a,l=Math.cos(e),u=Math.sin(e),f=r*u,c=i*l+f*Math.cos(o),h=f*s*Math.sin(o);An.add(Math.atan2(h,c));n=t,i=l,r=u}Nn.lineEnd=function(){a(t,e)}}function En(t){var e=t[0],n=t[1],i=Math.cos(n);return[i*Math.cos(e),i*Math.sin(e),Math.sin(n)]}function Fn(t,e){return t[0]*e[0]+t[1]*e[1]+t[2]*e[2]}function Rn(t,e){return[t[1]*e[2]-t[2]*e[1],t[2]*e[0]-t[0]*e[2],t[0]*e[1]-t[1]*e[0]]}function Ln(t,e){t[0]+=e[0];t[1]+=e[1];t[2]+=e[2]}function jn(t,e){return[t[0]*e,t[1]*e,t[2]*e]}function On(t){var e=Math.sqrt(t[0]*t[0]+t[1]*t[1]+t[2]*t[2]);t[0]/=e;t[1]/=e;t[2]/=e}function Hn(t){return[Math.atan2(t[1],t[0]),Lt(t[2])]}function Wn(t,e){return y(t[0]-e[0])<Ct&&y(t[1]-e[1])<Ct}t.geo.bounds=function(){var e,n,i,r,a,s,o,l,u,f,c;var h={point:d,lineStart:g,lineEnd:v,polygonStart:function(){h.point=m;h.lineStart=b;h.lineEnd=w;u=0;Nn.polygonStart()},polygonEnd:function(){Nn.polygonEnd();h.point=d;h.lineStart=g;h.lineEnd=v;if(An<0)e=-(i=180),n=-(r=90);else if(u>Ct)r=90;else if(u<-Ct)n=-90;c[0]=e,c[1]=i}};function d(t,a){f.push(c=[e=t,i=t]);if(a<n)n=a;if(a>r)r=a}function p(t,s){var o=En([t*Nt,s*Nt]);if(l){var u=Rn(l,o),f=[u[1],-u[0],0],c=Rn(f,u);On(c);c=Hn(c);var h=t-a,p=h>0?1:-1,g=c[0]*Pt*p,v=y(h)>180;if(v^(p*a<g&&g<p*t)){var m=c[1]*Pt;if(m>r)r=m}else if(g=(g+360)%360-180,v^(p*a<g&&g<p*t)){var m=-c[1]*Pt;if(m<n)n=m}else{if(s<n)n=s;if(s>r)r=s}if(v){if(t<a){if(_(e,t)>_(e,i))i=t}else{if(_(t,i)>_(e,i))e=t}}else{if(i>=e){if(t<e)e=t;if(t>i)i=t}else{if(t>a){if(_(e,t)>_(e,i))i=t}else{if(_(t,i)>_(e,i))e=t}}}}else{d(t,s)}l=o,a=t}function g(){h.point=p}function v(){c[0]=e,c[1]=i;h.point=d;l=null}function m(t,e){if(l){var n=t-a;u+=y(n)>180?n+(n>0?360:-360):n}else s=t,o=e;Nn.point(t,e);p(t,e)}function b(){Nn.lineStart()}function w(){m(s,o);Nn.lineEnd();if(y(u)>Ct)e=-(i=180);c[0]=e,c[1]=i;l=null}function _(t,e){return(e-=t)<0?e+360:e}function x(t,e){return t[0]-e[0]}function S(t,e){return e[0]<=e[1]?e[0]<=t&&t<=e[1]:t<e[0]||e[1]<t}return function(a){r=i=-(e=n=Infinity);f=[];t.geo.stream(a,h);var s=f.length;if(s){f.sort(x);for(var o=1,l=f[0],u,d=[l];o<s;++o){u=f[o];if(S(u[0],l)||S(u[1],l)){if(_(l[0],u[1])>_(l[0],l[1]))l[1]=u[1];if(_(u[0],l[1])>_(l[0],l[1]))l[0]=u[0]}else{d.push(l=u)}}var p=-Infinity,g;for(var s=d.length-1,o=0,l=d[s],u;o<=s;l=u,++o){u=d[o];if((g=_(l[1],u[0]))>p)p=g,e=u[0],i=l[1]}}f=c=null;return e===Infinity||n===Infinity?[[NaN,NaN],[NaN,NaN]]:[[e,n],[i,r]]}}();t.geo.centroid=function(e){zn=qn=Bn=$n=Un=Yn=Vn=Xn=Jn=Gn=Kn=0;t.geo.stream(e,Zn);var n=Jn,i=Gn,r=Kn,a=n*n+i*i+r*r;if(a<Tt){n=Yn,i=Vn,r=Xn;if(qn<Ct)n=Bn,i=$n,r=Un;a=n*n+i*i+r*r;if(a<Tt)return[NaN,NaN]}return[Math.atan2(i,n)*Pt,Lt(r/Math.sqrt(a))*Pt]};var zn,qn,Bn,$n,Un,Yn,Vn,Xn,Jn,Gn,Kn;var Zn={sphere:L,point:Qn,lineStart:ei,lineEnd:ni,polygonStart:function(){Zn.lineStart=ii},polygonEnd:function(){Zn.lineStart=ei}};function Qn(t,e){t*=Nt;var n=Math.cos(e*=Nt);ti(n*Math.cos(t),n*Math.sin(t),Math.sin(e))}function ti(t,e,n){++zn;Bn+=(t-Bn)/zn;$n+=(e-$n)/zn;Un+=(n-Un)/zn}function ei(){var t,e,n;Zn.point=function(r,a){r*=Nt;var s=Math.cos(a*=Nt);t=s*Math.cos(r);e=s*Math.sin(r);n=Math.sin(a);Zn.point=i;ti(t,e,n)};function i(i,r){i*=Nt;var a=Math.cos(r*=Nt),s=a*Math.cos(i),o=a*Math.sin(i),l=Math.sin(r),u=Math.atan2(Math.sqrt((u=e*l-n*o)*u+(u=n*s-t*l)*u+(u=t*o-e*s)*u),t*s+e*o+n*l);qn+=u;Yn+=u*(t+(t=s));Vn+=u*(e+(e=o));Xn+=u*(n+(n=l));ti(t,e,n)}}function ni(){Zn.point=Qn}function ii(){var t,e,n,i,r;Zn.point=function(s,o){t=s,e=o;Zn.point=a;s*=Nt;var l=Math.cos(o*=Nt);n=l*Math.cos(s);i=l*Math.sin(s);r=Math.sin(o);ti(n,i,r)};Zn.lineEnd=function(){a(t,e);Zn.lineEnd=ni;Zn.point=Qn};function a(t,e){t*=Nt;var a=Math.cos(e*=Nt),s=a*Math.cos(t),o=a*Math.sin(t),l=Math.sin(e),u=i*l-r*o,f=r*s-n*l,c=n*o-i*s,h=Math.sqrt(u*u+f*f+c*c),d=n*s+i*o+r*l,p=h&&-Rt(d)/h,g=Math.atan2(h,d);Jn+=p*u;Gn+=p*f;Kn+=p*c;qn+=g;Yn+=g*(n+(n=s));Vn+=g*(i+(i=o));Xn+=g*(r+(r=l));ti(n,i,r)}}function ri(t,e){function n(n,i){return n=t(n,i),e(n[0],n[1])}if(t.invert&&e.invert)n.invert=function(n,i){return n=e.invert(n,i),n&&t.invert(n[0],n[1])};return n}function ai(){return true}function si(t,e,n,i,r){var a=[],s=[];t.forEach(function(t){if((e=t.length-1)<=0)return;var e,n=t[0],i=t[e];if(Wn(n,i)){r.lineStart();for(var o=0;o<e;++o)r.point((n=t[o])[0],n[1]);r.lineEnd();return}var l=new li(n,t,null,true),u=new li(n,null,l,false);l.o=u;a.push(l);s.push(u);l=new li(i,t,null,false);u=new li(i,null,l,true);l.o=u;a.push(l);s.push(u)});s.sort(e);oi(a);oi(s);if(!a.length)return;for(var o=0,l=n,u=s.length;o<u;++o){s[o].e=l=!l}var f=a[0],c,h;while(1){var d=f,p=true;while(d.v)if((d=d.n)===f)return;c=d.z;r.lineStart();do{d.v=d.o.v=true;if(d.e){if(p){for(var o=0,u=c.length;o<u;++o)r.point((h=c[o])[0],h[1])}else{i(d.x,d.n.x,1,r)}d=d.n}else{if(p){c=d.p.z;for(var o=c.length-1;o>=0;--o)r.point((h=c[o])[0],h[1])}else{i(d.x,d.p.x,-1,r)}d=d.p}d=d.o;c=d.z;p=!p}while(!d.v);r.lineEnd()}}function oi(t){if(!(e=t.length))return;var e,n=0,i=t[0],r;while(++n<e){i.n=r=t[n];r.p=i;i=r}i.n=r=t[0];r.p=i}function li(t,e,n,i){this.x=t;this.z=e;this.o=n;this.e=i;this.v=false;this.n=this.p=null}function ui(e,n,i,r){return function(a,s){var o=n(s),l=a.invert(r[0],r[1]);var u={point:f,lineStart:h,lineEnd:d,polygonStart:function(){u.point=w;u.lineStart=_;u.lineEnd=x;p=[];y=[]},polygonEnd:function(){u.point=f;u.lineStart=h;u.lineEnd=d;p=t.merge(p);var e=mi(l,y);if(p.length){if(!m)s.polygonStart(),m=true;si(p,hi,e,i,s)}else if(e){if(!m)s.polygonStart(),m=true;s.lineStart();i(null,null,1,s);s.lineEnd()}if(m)s.polygonEnd(),m=false;p=y=null},sphere:function(){s.polygonStart();s.lineStart();i(null,null,1,s);s.lineEnd();s.polygonEnd()}};function f(t,n){var i=a(t,n);if(e(t=i[0],n=i[1]))s.point(t,n)}function c(t,e){var n=a(t,e);o.point(n[0],n[1])}function h(){u.point=c;o.lineStart()}function d(){u.point=f;o.lineEnd()}var p;var g=ci(),v=n(g),m=false,y,b;function w(t,e){b.push([t,e]);var n=a(t,e);v.point(n[0],n[1])}function _(){v.lineStart();b=[]}function x(){w(b[0][0],b[0][1]);v.lineEnd();var t=v.clean(),e=g.buffer(),n,i=e.length;b.pop();y.push(b);b=null;if(!i)return;if(t&1){n=e[0];var i=n.length-1,r=-1,a;if(i>0){if(!m)s.polygonStart(),m=true;s.lineStart();while(++r<i)s.point((a=n[r])[0],a[1]);s.lineEnd()}return}if(i>1&&t&2)e.push(e.pop().concat(e.shift()));p.push(e.filter(fi))}return u}}function fi(t){return t.length>1}function ci(){var t=[],e;return{lineStart:function(){t.push(e=[])},point:function(t,n){e.push([t,n])},lineEnd:L,buffer:function(){var n=t;t=[];e=null;return n},rejoin:function(){if(t.length>1)t.push(t.pop().concat(t.shift()))}}}function hi(t,e){
return((t=t.x)[0]<0?t[1]-At-Ct:At-t[1])-((e=e.x)[0]<0?e[1]-At-Ct:At-e[1])}var di=ui(ai,pi,vi,[-kt,-kt/2]);function pi(t){var e=NaN,n=NaN,i=NaN,r;return{lineStart:function(){t.lineStart();r=1},point:function(a,s){var o=a>0?kt:-kt,l=y(a-e);if(y(l-kt)<Ct){t.point(e,n=(n+s)/2>0?At:-At);t.point(i,n);t.lineEnd();t.lineStart();t.point(o,n);t.point(a,n);r=0}else if(i!==o&&l>=kt){if(y(e-i)<Ct)e-=i*Ct;if(y(a-o)<Ct)a-=o*Ct;n=gi(e,n,a,s);t.point(i,n);t.lineEnd();t.lineStart();t.point(o,n);r=0}t.point(e=a,n=s);i=o},lineEnd:function(){t.lineEnd();e=n=NaN},clean:function(){return 2-r}}}function gi(t,e,n,i){var r,a,s=Math.sin(t-n);return y(s)>Ct?Math.atan((Math.sin(e)*(a=Math.cos(i))*Math.sin(n)-Math.sin(i)*(r=Math.cos(e))*Math.sin(t))/(r*a*s)):(e+i)/2}function vi(t,e,n,i){var r;if(t==null){r=n*At;i.point(-kt,r);i.point(0,r);i.point(kt,r);i.point(kt,0);i.point(kt,-r);i.point(0,-r);i.point(-kt,-r);i.point(-kt,0);i.point(-kt,r)}else if(y(t[0]-e[0])>Ct){var a=t[0]<e[0]?kt:-kt;r=n*a/2;i.point(-a,r);i.point(0,r);i.point(a,r)}else{i.point(e[0],e[1])}}function mi(t,e){var n=t[0],i=t[1],r=[Math.sin(n),-Math.cos(n),0],a=0,s=0;An.reset();for(var o=0,l=e.length;o<l;++o){var u=e[o],f=u.length;if(!f)continue;var c=u[0],h=c[0],d=c[1]/2+kt/4,p=Math.sin(d),g=Math.cos(d),v=1;while(true){if(v===f)v=0;t=u[v];var m=t[0],y=t[1]/2+kt/4,b=Math.sin(y),w=Math.cos(y),_=m-h,x=_>=0?1:-1,S=x*_,D=S>kt,C=p*b;An.add(Math.atan2(C*x*Math.sin(S),g*w+C*Math.cos(S)));a+=D?_+x*Mt:_;if(D^h>=n^m>=n){var T=Rn(En(c),En(t));On(T);var k=Rn(r,T);On(k);var M=(D^_>=0?-1:1)*Lt(k[2]);if(i>M||i===M&&(T[0]||T[1])){s+=D^_>=0?1:-1}}if(!v++)break;h=m,p=b,g=w,c=t}}return(a<-Ct||a<Ct&&An<0)^s&1}function yi(t){var e=Math.cos(t),n=e>0,i=y(e)>Ct,r=er(t,6*Nt);return ui(a,s,r,n?[0,-t]:[-kt,t-kt]);function a(t,n){return Math.cos(t)*Math.cos(n)>e}function s(t){var e,r,s,u,f;return{lineStart:function(){u=s=false;f=1},point:function(c,h){var d=[c,h],p,g=a(c,h),v=n?g?0:l(c,h):g?l(c+(c<0?kt:-kt),h):0;if(!e&&(u=s=g))t.lineStart();if(g!==s){p=o(e,d);if(Wn(e,p)||Wn(d,p)){d[0]+=Ct;d[1]+=Ct;g=a(d[0],d[1])}}if(g!==s){f=0;if(g){t.lineStart();p=o(d,e);t.point(p[0],p[1])}else{p=o(e,d);t.point(p[0],p[1]);t.lineEnd()}e=p}else if(i&&e&&n^g){var m;if(!(v&r)&&(m=o(d,e,true))){f=0;if(n){t.lineStart();t.point(m[0][0],m[0][1]);t.point(m[1][0],m[1][1]);t.lineEnd()}else{t.point(m[1][0],m[1][1]);t.lineEnd();t.lineStart();t.point(m[0][0],m[0][1])}}}if(g&&(!e||!Wn(e,d))){t.point(d[0],d[1])}e=d,s=g,r=v},lineEnd:function(){if(s)t.lineEnd();e=null},clean:function(){return f|(u&&s)<<1}}}function o(t,n,i){var r=En(t),a=En(n);var s=[1,0,0],o=Rn(r,a),l=Fn(o,o),u=o[0],f=l-u*u;if(!f)return!i&&t;var c=e*l/f,h=-e*u/f,d=Rn(s,o),p=jn(s,c),g=jn(o,h);Ln(p,g);var v=d,m=Fn(p,v),b=Fn(v,v),w=m*m-b*(Fn(p,p)-1);if(w<0)return;var _=Math.sqrt(w),x=jn(v,(-m-_)/b);Ln(x,p);x=Hn(x);if(!i)return x;var S=t[0],D=n[0],C=t[1],T=n[1],k;if(D<S)k=S,S=D,D=k;var M=D-S,I=y(M-kt)<Ct,A=I||M<Ct;if(!I&&T<C)k=C,C=T,T=k;if(A?I?C+T>0^x[1]<(y(x[0]-S)<Ct?C:T):C<=x[1]&&x[1]<=T:M>kt^(S<=x[0]&&x[0]<=D)){var N=jn(v,(-m+_)/b);Ln(N,p);return[x,Hn(N)]}}function l(e,i){var r=n?t:kt-t,a=0;if(e<-r)a|=1;else if(e>r)a|=2;if(i<-r)a|=4;else if(i>r)a|=8;return a}}function bi(t,e,n,i){return function(r){var a=r.a,s=r.b,o=a.x,l=a.y,u=s.x,f=s.y,c=0,h=1,d=u-o,p=f-l,g;g=t-o;if(!d&&g>0)return;g/=d;if(d<0){if(g<c)return;if(g<h)h=g}else if(d>0){if(g>h)return;if(g>c)c=g}g=n-o;if(!d&&g<0)return;g/=d;if(d<0){if(g>h)return;if(g>c)c=g}else if(d>0){if(g<c)return;if(g<h)h=g}g=e-l;if(!p&&g>0)return;g/=p;if(p<0){if(g<c)return;if(g<h)h=g}else if(p>0){if(g>h)return;if(g>c)c=g}g=i-l;if(!p&&g<0)return;g/=p;if(p<0){if(g>h)return;if(g>c)c=g}else if(p>0){if(g<c)return;if(g<h)h=g}if(c>0)r.a={x:o+c*d,y:l+c*p};if(h<1)r.b={x:o+h*d,y:l+h*p};return r}}var wi=1e9;t.geo.clipExtent=function(){var t,e,n,i,r,a,s={stream:function(t){if(r)r.valid=false;r=a(t);r.valid=true;return r},extent:function(o){if(!arguments.length)return[[t,e],[n,i]];a=_i(t=+o[0][0],e=+o[0][1],n=+o[1][0],i=+o[1][1]);if(r)r.valid=false,r=null;return s}};return s.extent([[0,0],[960,500]])};function _i(e,n,i,r){return function(l){var u=l,f=ci(),c=bi(e,n,i,r),h,d,p;var g={point:b,lineStart:M,lineEnd:I,polygonStart:function(){l=f;h=[];d=[];k=true},polygonEnd:function(){l=u;h=t.merge(h);var n=v([e,r]),i=k&&n,a=h.length;if(i||a){l.polygonStart();if(i){l.lineStart();m(null,null,1,l);l.lineEnd()}if(a){si(h,s,n,m,l)}l.polygonEnd()}h=d=p=null}};function v(t){var e=0,n=d.length,i=t[1];for(var r=0;r<n;++r){for(var a=1,s=d[r],o=s.length,l=s[0],u;a<o;++a){u=s[a];if(l[1]<=i){if(u[1]>i&&Ft(l,u,t)>0)++e}else{if(u[1]<=i&&Ft(l,u,t)<0)--e}l=u}}return e!==0}function m(t,s,l,u){var f=0,c=0;if(t==null||(f=a(t,l))!==(c=a(s,l))||o(t,s)<0^l>0){do{u.point(f===0||f===3?e:i,f>1?r:n)}while((f=(f+l+4)%4)!==c)}else{u.point(s[0],s[1])}}function y(t,a){return e<=t&&t<=i&&n<=a&&a<=r}function b(t,e){if(y(t,e))l.point(t,e)}var w,_,x,S,D,C,T,k;function M(){g.point=A;if(d)d.push(p=[]);T=true;C=false;S=D=NaN}function I(){if(h){A(w,_);if(x&&C)f.rejoin();h.push(f.buffer())}g.point=b;if(C)l.lineEnd()}function A(t,e){t=Math.max(-wi,Math.min(wi,t));e=Math.max(-wi,Math.min(wi,e));var n=y(t,e);if(d)p.push([t,e]);if(T){w=t,_=e,x=n;T=false;if(n){l.lineStart();l.point(t,e)}}else{if(n&&C)l.point(t,e);else{var i={a:{x:S,y:D},b:{x:t,y:e}};if(c(i)){if(!C){l.lineStart();l.point(i.a.x,i.a.y)}l.point(i.b.x,i.b.y);if(!n)l.lineEnd();k=false}else if(n){l.lineStart();l.point(t,e);k=false}}}S=t,D=e,C=n}return g};function a(t,r){return y(t[0]-e)<Ct?r>0?0:3:y(t[0]-i)<Ct?r>0?2:1:y(t[1]-n)<Ct?r>0?1:0:r>0?3:2}function s(t,e){return o(t.x,e.x)}function o(t,e){var n=a(t,1),i=a(e,1);return n!==i?n-i:n===0?e[1]-t[1]:n===1?t[0]-e[0]:n===2?t[1]-e[1]:e[0]-t[0]}}function xi(t){var e=0,n=kt/3,i=Vi(t),r=i(e,n);r.parallels=function(t){if(!arguments.length)return[e/kt*180,n/kt*180];return i(e=t[0]*kt/180,n=t[1]*kt/180)};return r}function Si(t,e){var n=Math.sin(t),i=(n+Math.sin(e))/2,r=1+n*(2*i-n),a=Math.sqrt(r)/i;function s(t,e){var n=Math.sqrt(r-2*i*Math.sin(e))/i;return[n*Math.sin(t*=i),a-n*Math.cos(t)]}s.invert=function(t,e){var n=a-e;return[Math.atan2(t,n)/i,Lt((r-(t*t+n*n)*i*i)/(2*i))]};return s}(t.geo.conicEqualArea=function(){return xi(Si)}).raw=Si;t.geo.albers=function(){return t.geo.conicEqualArea().rotate([96,0]).center([-.6,38.7]).parallels([29.5,45.5]).scale(1070)};t.geo.albersUsa=function(){var e=t.geo.albers();var n=t.geo.conicEqualArea().rotate([154,0]).center([-2,58.5]).parallels([55,65]);var i=t.geo.conicEqualArea().rotate([157,0]).center([-3,19.9]).parallels([8,18]);var r,a={point:function(t,e){r=[t,e]}},s,o,l;function u(t){var e=t[0],n=t[1];r=null;(s(e,n),r)||(o(e,n),r)||l(e,n);return r}u.invert=function(t){var r=e.scale(),a=e.translate(),s=(t[0]-a[0])/r,o=(t[1]-a[1])/r;return(o>=.12&&o<.234&&s>=-.425&&s<-.214?n:o>=.166&&o<.234&&s>=-.214&&s<-.115?i:e).invert(t)};u.stream=function(t){var r=e.stream(t),a=n.stream(t),s=i.stream(t);return{point:function(t,e){r.point(t,e);a.point(t,e);s.point(t,e)},sphere:function(){r.sphere();a.sphere();s.sphere()},lineStart:function(){r.lineStart();a.lineStart();s.lineStart()},lineEnd:function(){r.lineEnd();a.lineEnd();s.lineEnd()},polygonStart:function(){r.polygonStart();a.polygonStart();s.polygonStart()},polygonEnd:function(){r.polygonEnd();a.polygonEnd();s.polygonEnd()}}};u.precision=function(t){if(!arguments.length)return e.precision();e.precision(t);n.precision(t);i.precision(t);return u};u.scale=function(t){if(!arguments.length)return e.scale();e.scale(t);n.scale(t*.35);i.scale(t);return u.translate(e.translate())};u.translate=function(t){if(!arguments.length)return e.translate();var r=e.scale(),f=+t[0],c=+t[1];s=e.translate(t).clipExtent([[f-.455*r,c-.238*r],[f+.455*r,c+.238*r]]).stream(a).point;o=n.translate([f-.307*r,c+.201*r]).clipExtent([[f-.425*r+Ct,c+.12*r+Ct],[f-.214*r-Ct,c+.234*r-Ct]]).stream(a).point;l=i.translate([f-.205*r,c+.212*r]).clipExtent([[f-.214*r+Ct,c+.166*r+Ct],[f-.115*r-Ct,c+.234*r-Ct]]).stream(a).point;return u};return u.scale(1070)};var Di,Ci,Ti={point:L,lineStart:L,lineEnd:L,polygonStart:function(){Ci=0;Ti.lineStart=ki},polygonEnd:function(){Ti.lineStart=Ti.lineEnd=Ti.point=L;Di+=y(Ci/2)}};function ki(){var t,e,n,i;Ti.point=function(a,s){Ti.point=r;t=n=a,e=i=s};function r(t,e){Ci+=i*t-n*e;n=t,i=e}Ti.lineEnd=function(){r(t,e)}}var Mi,Ii,Ai,Ni;var Pi={point:Ei,lineStart:L,lineEnd:L,polygonStart:L,polygonEnd:L};function Ei(t,e){if(t<Mi)Mi=t;if(t>Ai)Ai=t;if(e<Ii)Ii=e;if(e>Ni)Ni=e}function Fi(){var t=Ri(4.5),e=[];var n={point:i,lineStart:function(){n.point=r},lineEnd:s,polygonStart:function(){n.lineEnd=o},polygonEnd:function(){n.lineEnd=s;n.point=i},pointRadius:function(e){t=Ri(e);return n},result:function(){if(e.length){var t=e.join("");e=[];return t}}};function i(n,i){e.push("M",n,",",i,t)}function r(t,i){e.push("M",t,",",i);n.point=a}function a(t,n){e.push("L",t,",",n)}function s(){n.point=i}function o(){e.push("Z")}return n}function Ri(t){return"m0,"+t+"a"+t+","+t+" 0 1,1 0,"+-2*t+"a"+t+","+t+" 0 1,1 0,"+2*t+"z"}var Li={point:ji,lineStart:Oi,lineEnd:Hi,polygonStart:function(){Li.lineStart=Wi},polygonEnd:function(){Li.point=ji;Li.lineStart=Oi;Li.lineEnd=Hi}};function ji(t,e){Bn+=t;$n+=e;++Un}function Oi(){var t,e;Li.point=function(i,r){Li.point=n;ji(t=i,e=r)};function n(n,i){var r=n-t,a=i-e,s=Math.sqrt(r*r+a*a);Yn+=s*(t+n)/2;Vn+=s*(e+i)/2;Xn+=s;ji(t=n,e=i)}}function Hi(){Li.point=ji}function Wi(){var t,e,n,i;Li.point=function(a,s){Li.point=r;ji(t=n=a,e=i=s)};function r(t,e){var r=t-n,a=e-i,s=Math.sqrt(r*r+a*a);Yn+=s*(n+t)/2;Vn+=s*(i+e)/2;Xn+=s;s=i*t-n*e;Jn+=s*(n+t);Gn+=s*(i+e);Kn+=s*3;ji(n=t,i=e)}Li.lineEnd=function(){r(t,e)}}function zi(t){var e=4.5;var n={point:i,lineStart:function(){n.point=r},lineEnd:s,polygonStart:function(){n.lineEnd=o},polygonEnd:function(){n.lineEnd=s;n.point=i},pointRadius:function(t){e=t;return n},result:L};function i(n,i){t.moveTo(n+e,i);t.arc(n,i,e,0,Mt)}function r(e,i){t.moveTo(e,i);n.point=a}function a(e,n){t.lineTo(e,n)}function s(){n.point=i}function o(){t.closePath()}return n}function qi(t){var e=.5,n=Math.cos(30*Nt),i=16;function r(t){return(i?s:a)(t)}function a(e){return Ui(e,function(n,i){n=t(n,i);e.point(n[0],n[1])})}function s(e){var n,r,a,s,l,u,f,c,h,d,p,g,v;var m={point:y,lineStart:b,lineEnd:_,polygonStart:function(){e.polygonStart();m.lineStart=x},polygonEnd:function(){e.polygonEnd();m.lineStart=b}};function y(n,i){n=t(n,i);e.point(n[0],n[1])}function b(){h=NaN;m.point=w;e.lineStart()}function w(n,r){var a=En([n,r]),s=t(n,r);o(h,d,c,p,g,v,h=s[0],d=s[1],c=n,p=a[0],g=a[1],v=a[2],i,e);e.point(h,d)}function _(){m.point=y;e.lineEnd()}function x(){b();m.point=S;m.lineEnd=D}function S(t,e){w(n=t,r=e),a=h,s=d,l=p,u=g,f=v;m.point=w}function D(){o(h,d,c,p,g,v,a,s,n,l,u,f,i,e);m.lineEnd=_;_()}return m}function o(i,r,a,s,l,u,f,c,h,d,p,g,v,m){var b=f-i,w=c-r,_=b*b+w*w;if(_>4*e&&v--){var x=s+d,S=l+p,D=u+g,C=Math.sqrt(x*x+S*S+D*D),T=Math.asin(D/=C),k=y(y(D)-1)<Ct||y(a-h)<Ct?(a+h)/2:Math.atan2(S,x),M=t(k,T),I=M[0],A=M[1],N=I-i,P=A-r,E=w*N-b*P;if(E*E/_>e||y((b*N+w*P)/_-.5)>.3||s*d+l*p+u*g<n){o(i,r,a,s,l,u,I,A,k,x/=C,S/=C,D,v,m);m.point(I,A);o(I,A,k,x,S,D,f,c,h,d,p,g,v,m)}}}r.precision=function(t){if(!arguments.length)return Math.sqrt(e);i=(e=t*t)>0&&16;return r};return r}t.geo.path=function(){var e=4.5,n,i,r,a,s;function o(n){if(n){if(typeof e==="function")a.pointRadius(+e.apply(this,arguments));if(!s||!s.valid)s=r(a);t.geo.stream(n,s)}return a.result()}o.area=function(e){Di=0;t.geo.stream(e,r(Ti));return Di};o.centroid=function(e){Bn=$n=Un=Yn=Vn=Xn=Jn=Gn=Kn=0;t.geo.stream(e,r(Li));return Kn?[Jn/Kn,Gn/Kn]:Xn?[Yn/Xn,Vn/Xn]:Un?[Bn/Un,$n/Un]:[NaN,NaN]};o.bounds=function(e){Ai=Ni=-(Mi=Ii=Infinity);t.geo.stream(e,r(Pi));return[[Mi,Ii],[Ai,Ni]]};o.projection=function(t){if(!arguments.length)return n;r=(n=t)?t.stream||Bi(t):P;return l()};o.context=function(t){if(!arguments.length)return i;a=(i=t)==null?new Fi:new zi(t);if(typeof e!=="function")a.pointRadius(e);return l()};o.pointRadius=function(t){if(!arguments.length)return e;e=typeof t==="function"?t:(a.pointRadius(+t),+t);return o};function l(){s=null;return o}return o.projection(t.geo.albersUsa()).context(null)};function Bi(t){var e=qi(function(e,n){return t([e*Pt,n*Pt])});return function(t){return Xi(e(t))}}t.geo.transform=function(t){return{stream:function(e){var n=new $i(e);for(var i in t)n[i]=t[i];return n}}};function $i(t){this.stream=t}$i.prototype={point:function(t,e){this.stream.point(t,e)},sphere:function(){this.stream.sphere()},lineStart:function(){this.stream.lineStart()},lineEnd:function(){this.stream.lineEnd()},polygonStart:function(){this.stream.polygonStart()},polygonEnd:function(){this.stream.polygonEnd()}};function Ui(t,e){return{point:e,sphere:function(){t.sphere()},lineStart:function(){t.lineStart()},lineEnd:function(){t.lineEnd()},polygonStart:function(){t.polygonStart()},polygonEnd:function(){t.polygonEnd()}}}t.geo.projection=Yi;t.geo.projectionMutator=Vi;function Yi(t){return Vi(function(){return t})()}function Vi(e){var n,i,r,a=qi(function(t,e){t=n(t,e);return[t[0]*s+p,g-t[1]*s]}),s=150,o=480,l=250,u=0,f=0,c=0,h=0,d=0,p,g,v=di,m=P,y=null,b=null,w;function _(t){t=r(t[0]*Nt,t[1]*Nt);return[t[0]*s+p,g-t[1]*s]}function x(t){t=r.invert((t[0]-p)/s,(g-t[1])/s);return t&&[t[0]*Pt,t[1]*Pt]}_.stream=function(t){if(w)w.valid=false;w=Xi(v(i,a(m(t))));w.valid=true;return w};_.clipAngle=function(t){if(!arguments.length)return y;v=t==null?(y=t,di):yi((y=+t)*Nt);return D()};_.clipExtent=function(t){if(!arguments.length)return b;b=t;m=t?_i(t[0][0],t[0][1],t[1][0],t[1][1]):P;return D()};_.scale=function(t){if(!arguments.length)return s;s=+t;return S()};_.translate=function(t){if(!arguments.length)return[o,l];o=+t[0];l=+t[1];return S()};_.center=function(t){if(!arguments.length)return[u*Pt,f*Pt];u=t[0]%360*Nt;f=t[1]%360*Nt;return S()};_.rotate=function(t){if(!arguments.length)return[c*Pt,h*Pt,d*Pt];c=t[0]%360*Nt;h=t[1]%360*Nt;d=t.length>2?t[2]%360*Nt:0;return S()};t.rebind(_,a,"precision");function S(){r=ri(i=Ki(c,h,d),n);var t=n(u,f);p=o-t[0]*s;g=l+t[1]*s;return D()}function D(){if(w)w.valid=false,w=null;return _}return function(){n=e.apply(this,arguments);_.invert=n.invert&&x;return S()}}function Xi(t){return Ui(t,function(e,n){t.point(e*Nt,n*Nt)})}function Ji(t,e){return[t,e]}(t.geo.equirectangular=function(){return Yi(Ji)}).raw=Ji.invert=Ji;t.geo.rotation=function(t){t=Ki(t[0]%360*Nt,t[1]*Nt,t.length>2?t[2]*Nt:0);function e(e){e=t(e[0]*Nt,e[1]*Nt);return e[0]*=Pt,e[1]*=Pt,e}e.invert=function(e){e=t.invert(e[0]*Nt,e[1]*Nt);return e[0]*=Pt,e[1]*=Pt,e};return e};function Gi(t,e){return[t>kt?t-Mt:t<-kt?t+Mt:t,e]}Gi.invert=Ji;function Ki(t,e,n){return t?e||n?ri(Qi(t),tr(e,n)):Qi(t):e||n?tr(e,n):Gi}function Zi(t){return function(e,n){return e+=t,[e>kt?e-Mt:e<-kt?e+Mt:e,n]}}function Qi(t){var e=Zi(t);e.invert=Zi(-t);return e}function tr(t,e){var n=Math.cos(t),i=Math.sin(t),r=Math.cos(e),a=Math.sin(e);function s(t,e){var s=Math.cos(e),o=Math.cos(t)*s,l=Math.sin(t)*s,u=Math.sin(e),f=u*n+o*i;return[Math.atan2(l*r-f*a,o*n-u*i),Lt(f*r+l*a)]}s.invert=function(t,e){var s=Math.cos(e),o=Math.cos(t)*s,l=Math.sin(t)*s,u=Math.sin(e),f=u*r-l*a;return[Math.atan2(l*r+u*a,o*n+f*i),Lt(f*n-o*i)]};return s}t.geo.circle=function(){var t=[0,0],e,n=6,i;function r(){var e=typeof t==="function"?t.apply(this,arguments):t,n=Ki(-e[0]*Nt,-e[1]*Nt,0).invert,r=[];i(null,null,1,{point:function(t,e){r.push(t=n(t,e));t[0]*=Pt,t[1]*=Pt}});return{type:"Polygon",coordinates:[r]}}r.origin=function(e){if(!arguments.length)return t;t=e;return r};r.angle=function(t){if(!arguments.length)return e;i=er((e=+t)*Nt,n*Nt);return r};r.precision=function(t){if(!arguments.length)return n;i=er(e*Nt,(n=+t)*Nt);return r};return r.angle(90)};function er(t,e){var n=Math.cos(t),i=Math.sin(t);return function(r,a,s,o){var l=s*e;if(r!=null){r=nr(n,r);a=nr(n,a);if(s>0?r<a:r>a)r+=s*Mt}else{r=t+s*Mt;a=t-.5*l}for(var u,f=r;s>0?f>a:f<a;f-=l){o.point((u=Hn([n,-i*Math.cos(f),-i*Math.sin(f)]))[0],u[1])}}}function nr(t,e){var n=En(e);n[0]-=t;On(n);var i=Rt(-n[1]);return((-n[2]<0?-i:i)+2*Math.PI-Ct)%(2*Math.PI)}t.geo.distance=function(t,e){var n=(e[0]-t[0])*Nt,i=t[1]*Nt,r=e[1]*Nt,a=Math.sin(n),s=Math.cos(n),o=Math.sin(i),l=Math.cos(i),u=Math.sin(r),f=Math.cos(r),c;return Math.atan2(Math.sqrt((c=f*a)*c+(c=l*u-o*f*s)*c),o*u+l*f*s)};t.geo.graticule=function(){var e,n,i,r,a,s,o,l,u=10,f=u,c=90,h=360,d,p,g,v,m=2.5;function b(){return{type:"MultiLineString",coordinates:w()}}function w(){return t.range(Math.ceil(r/c)*c,i,c).map(g).concat(t.range(Math.ceil(l/h)*h,o,h).map(v)).concat(t.range(Math.ceil(n/u)*u,e,u).filter(function(t){return y(t%c)>Ct}).map(d)).concat(t.range(Math.ceil(s/f)*f,a,f).filter(function(t){return y(t%h)>Ct}).map(p))}b.lines=function(){return w().map(function(t){return{type:"LineString",coordinates:t}})};b.outline=function(){return{type:"Polygon",coordinates:[g(r).concat(v(o).slice(1),g(i).reverse().slice(1),v(l).reverse().slice(1))]}};b.extent=function(t){if(!arguments.length)return b.minorExtent();return b.majorExtent(t).minorExtent(t)};b.majorExtent=function(t){if(!arguments.length)return[[r,l],[i,o]];r=+t[0][0],i=+t[1][0];l=+t[0][1],o=+t[1][1];if(r>i)t=r,r=i,i=t;if(l>o)t=l,l=o,o=t;return b.precision(m)};b.minorExtent=function(t){if(!arguments.length)return[[n,s],[e,a]];n=+t[0][0],e=+t[1][0];s=+t[0][1],a=+t[1][1];if(n>e)t=n,n=e,e=t;if(s>a)t=s,s=a,a=t;return b.precision(m)};b.step=function(t){if(!arguments.length)return b.minorStep();return b.majorStep(t).minorStep(t)};b.majorStep=function(t){if(!arguments.length)return[c,h];c=+t[0],h=+t[1];return b};b.minorStep=function(t){if(!arguments.length)return[u,f];u=+t[0],f=+t[1];return b};b.precision=function(t){if(!arguments.length)return m;m=+t;d=ir(s,a,90);p=rr(n,e,m);g=ir(l,o,90);v=rr(r,i,m);return b};return b.majorExtent([[-180,-90+Ct],[180,90-Ct]]).minorExtent([[-180,-80-Ct],[180,80+Ct]])};function ir(e,n,i){var r=t.range(e,n-Ct,i).concat(n);return function(t){return r.map(function(e){return[t,e]})}}function rr(e,n,i){var r=t.range(e,n-Ct,i).concat(n);return function(t){return r.map(function(e){return[e,t]})}}function ar(t){return t.source}function sr(t){return t.target}t.geo.greatArc=function(){var e=ar,n,i=sr,r;function a(){return{type:"LineString",coordinates:[n||e.apply(this,arguments),r||i.apply(this,arguments)]}}a.distance=function(){return t.geo.distance(n||e.apply(this,arguments),r||i.apply(this,arguments))};a.source=function(t){if(!arguments.length)return e;e=t,n=typeof t==="function"?null:t;return a};a.target=function(t){if(!arguments.length)return i;i=t,r=typeof t==="function"?null:t;return a};a.precision=function(){return arguments.length?a:0};return a};t.geo.interpolate=function(t,e){return or(t[0]*Nt,t[1]*Nt,e[0]*Nt,e[1]*Nt)};function or(t,e,n,i){var r=Math.cos(e),a=Math.sin(e),s=Math.cos(i),o=Math.sin(i),l=r*Math.cos(t),u=r*Math.sin(t),f=s*Math.cos(n),c=s*Math.sin(n),h=2*Math.asin(Math.sqrt(Wt(i-e)+r*s*Wt(n-t))),d=1/Math.sin(h);var p=h?function(t){var e=Math.sin(t*=h)*d,n=Math.sin(h-t)*d,i=n*l+e*f,r=n*u+e*c,s=n*a+e*o;return[Math.atan2(r,i)*Pt,Math.atan2(s,Math.sqrt(i*i+r*r))*Pt]}:function(){return[t*Pt,e*Pt]};p.distance=h;return p}t.geo.length=function(e){lr=0;t.geo.stream(e,ur);return lr};var lr;var ur={sphere:L,point:L,lineStart:fr,lineEnd:L,polygonStart:L,polygonEnd:L};function fr(){var t,e,n;ur.point=function(r,a){t=r*Nt,e=Math.sin(a*=Nt),n=Math.cos(a);ur.point=i};ur.lineEnd=function(){ur.point=ur.lineEnd=L};function i(i,r){var a=Math.sin(r*=Nt),s=Math.cos(r),o=y((i*=Nt)-t),l=Math.cos(o);lr+=Math.atan2(Math.sqrt((o=s*Math.sin(o))*o+(o=n*a-e*s*l)*o),e*a+n*s*l);t=i,e=a,n=s}}function cr(t,e){function n(e,n){var i=Math.cos(e),r=Math.cos(n),a=t(i*r);return[a*r*Math.sin(e),a*Math.sin(n)]}n.invert=function(t,n){var i=Math.sqrt(t*t+n*n),r=e(i),a=Math.sin(r),s=Math.cos(r);return[Math.atan2(t*a,i*s),Math.asin(i&&n*a/i)]};return n}var hr=cr(function(t){return Math.sqrt(2/(1+t))},function(t){return 2*Math.asin(t/2)});(t.geo.azimuthalEqualArea=function(){return Yi(hr)}).raw=hr;var dr=cr(function(t){var e=Math.acos(t);return e&&e/Math.sin(e)},P);(t.geo.azimuthalEquidistant=function(){return Yi(dr)}).raw=dr;function pr(t,e){var n=Math.cos(t),i=function(t){return Math.tan(kt/4+t/2)},r=t===e?Math.sin(t):Math.log(n/Math.cos(e))/Math.log(i(e)/i(t)),a=n*Math.pow(i(t),r)/r;if(!r)return mr;function s(t,e){if(a>0){if(e<-At+Ct)e=-At+Ct}else{if(e>At-Ct)e=At-Ct}var n=a/Math.pow(i(e),r);return[n*Math.sin(r*t),a-n*Math.cos(r*t)]}s.invert=function(t,e){var n=a-e,i=Et(r)*Math.sqrt(t*t+n*n);return[Math.atan2(t,n)/r,2*Math.atan(Math.pow(a/i,1/r))-At]};return s}(t.geo.conicConformal=function(){return xi(pr)}).raw=pr;function gr(t,e){var n=Math.cos(t),i=t===e?Math.sin(t):(n-Math.cos(e))/(e-t),r=n/i+t;if(y(i)<Ct)return Ji;function a(t,e){var n=r-e;return[n*Math.sin(i*t),r-n*Math.cos(i*t)]}a.invert=function(t,e){var n=r-e;return[Math.atan2(t,n)/i,r-Et(i)*Math.sqrt(t*t+n*n)]};return a}(t.geo.conicEquidistant=function(){return xi(gr)}).raw=gr;var vr=cr(function(t){return 1/t},Math.atan);(t.geo.gnomonic=function(){return Yi(vr)}).raw=vr;function mr(t,e){return[t,Math.log(Math.tan(kt/4+e/2))]}mr.invert=function(t,e){return[t,2*Math.atan(Math.exp(e))-At]};function yr(t){var e=Yi(t),n=e.scale,i=e.translate,r=e.clipExtent,a;e.scale=function(){var t=n.apply(e,arguments);return t===e?a?e.clipExtent(null):e:t};e.translate=function(){var t=i.apply(e,arguments);return t===e?a?e.clipExtent(null):e:t};e.clipExtent=function(t){var s=r.apply(e,arguments);if(s===e){if(a=t==null){var o=kt*n(),l=i();r([[l[0]-o,l[1]-o],[l[0]+o,l[1]+o]])}}else if(a){s=null}return s};return e.clipExtent(null)}(t.geo.mercator=function(){return yr(mr)}).raw=mr;var br=cr(function(){return 1},Math.asin);(t.geo.orthographic=function(){return Yi(br)}).raw=br;var wr=cr(function(t){return 1/(1+t)},function(t){return 2*Math.atan(t)});(t.geo.stereographic=function(){return Yi(wr)}).raw=wr;function _r(t,e){return[Math.log(Math.tan(kt/4+e/2)),-t]}_r.invert=function(t,e){return[-e,2*Math.atan(Math.exp(t))-At]};(t.geo.transverseMercator=function(){var t=yr(_r),e=t.center,n=t.rotate;t.center=function(t){return t?e([-t[1],t[0]]):(t=e(),[t[1],-t[0]])};t.rotate=function(t){return t?n([t[0],t[1],t.length>2?t[2]+90:90]):(t=n(),[t[0],t[1],t[2]-90])};return n([0,0,90])}).raw=_r;t.geom={};function xr(t){return t[0]}function Sr(t){return t[1]}t.geom.hull=function(t){var e=xr,n=Sr;if(arguments.length)return i(t);function i(t){if(t.length<3)return[];var i=xe(e),r=xe(n),a,s=t.length,o=[],l=[];for(a=0;a<s;a++){o.push([+i.call(this,t[a],a),+r.call(this,t[a],a),a])}o.sort(Cr);for(a=0;a<s;a++)l.push([o[a][0],-o[a][1]]);var u=Dr(o),f=Dr(l);var c=f[0]===u[0],h=f[f.length-1]===u[u.length-1],d=[];for(a=u.length-1;a>=0;--a)d.push(t[o[u[a]][2]]);for(a=+c;a<f.length-h;++a)d.push(t[o[f[a]][2]]);return d}i.x=function(t){return arguments.length?(e=t,i):e};i.y=function(t){return arguments.length?(n=t,i):n};return i};function Dr(t){var e=t.length,n=[0,1],i=2;for(var r=2;r<e;r++){while(i>1&&Ft(t[n[i-2]],t[n[i-1]],t[r])<=0)--i;n[i++]=r}return n.slice(0,i)}function Cr(t,e){return t[0]-e[0]||t[1]-e[1]}t.geom.polygon=function(t){B(t,Tr);return t};var Tr=t.geom.polygon.prototype=[];Tr.area=function(){var t=-1,e=this.length,n,i=this[e-1],r=0;while(++t<e){n=i;i=this[t];r+=n[1]*i[0]-n[0]*i[1]}return r*.5};Tr.centroid=function(t){var e=-1,n=this.length,i=0,r=0,a,s=this[n-1],o;if(!arguments.length)t=-1/(6*this.area());while(++e<n){a=s;s=this[e];o=a[0]*s[1]-s[0]*a[1];i+=(a[0]+s[0])*o;r+=(a[1]+s[1])*o}return[i*t,r*t]};Tr.clip=function(t){var e,n=Ir(t),i=-1,r=this.length-Ir(this),a,s,o=this[r-1],l,u,f;while(++i<r){e=t.slice();t.length=0;l=this[i];u=e[(s=e.length-n)-1];a=-1;while(++a<s){f=e[a];if(kr(f,o,l)){if(!kr(u,o,l)){t.push(Mr(u,f,o,l))}t.push(f)}else if(kr(u,o,l)){t.push(Mr(u,f,o,l))}u=f}if(n)t.push(t[0]);o=l}return t};function kr(t,e,n){return(n[0]-e[0])*(t[1]-e[1])<(n[1]-e[1])*(t[0]-e[0])}function Mr(t,e,n,i){var r=t[0],a=n[0],s=e[0]-r,o=i[0]-a,l=t[1],u=n[1],f=e[1]-l,c=i[1]-u,h=(o*(l-u)-c*(r-a))/(c*s-o*f);return[r+h*s,l+h*f]}function Ir(t){var e=t[0],n=t[t.length-1];return!(e[0]-n[0]||e[1]-n[1])}var Ar,Nr,Pr,Er=[],Fr,Rr,Lr=[];function jr(){ra(this);this.edge=this.site=this.circle=null}function Or(t){var e=Er.pop()||new jr;e.site=t;return e}function Hr(t){Jr(t);Pr.remove(t);Er.push(t);ra(t)}function Wr(t){var e=t.circle,n=e.x,i=e.cy,r={x:n,y:i},a=t.P,s=t.N,o=[t];Hr(t);var l=a;while(l.circle&&y(n-l.circle.x)<Ct&&y(i-l.circle.cy)<Ct){a=l.P;o.unshift(l);Hr(l);l=a}o.unshift(l);Jr(l);var u=s;while(u.circle&&y(n-u.circle.x)<Ct&&y(i-u.circle.cy)<Ct){s=u.N;o.push(u);Hr(u);u=s}o.push(u);Jr(u);var f=o.length,c;for(c=1;c<f;++c){u=o[c];l=o[c-1];ea(u.edge,l.site,u.site,r)}l=o[0];u=o[f-1];u.edge=Qr(l.site,u.site,null,r);Xr(l);Xr(u)}function zr(t){var e=t.x,n=t.y,i,r,a,s,o=Pr._;while(o){a=qr(o,n)-e;if(a>Ct)o=o.L;else{s=e-Br(o,n);if(s>Ct){if(!o.R){i=o;break}o=o.R}else{if(a>-Ct){i=o.P;r=o}else if(s>-Ct){i=o;r=o.N}else{i=r=o}break}}}var l=Or(t);Pr.insert(i,l);if(!i&&!r)return;if(i===r){Jr(i);r=Or(i.site);Pr.insert(l,r);l.edge=r.edge=Qr(i.site,l.site);Xr(i);Xr(r);return}if(!r){l.edge=Qr(i.site,l.site);return}Jr(i);Jr(r);var u=i.site,f=u.x,c=u.y,h=t.x-f,d=t.y-c,p=r.site,g=p.x-f,v=p.y-c,m=2*(h*v-d*g),y=h*h+d*d,b=g*g+v*v,w={x:(v*y-d*b)/m+f,y:(h*b-g*y)/m+c};ea(r.edge,u,p,w);l.edge=Qr(u,t,null,w);r.edge=Qr(t,p,null,w);Xr(i);Xr(r)}function qr(t,e){var n=t.site,i=n.x,r=n.y,a=r-e;if(!a)return i;var s=t.P;if(!s)return-Infinity;n=s.site;var o=n.x,l=n.y,u=l-e;if(!u)return o;var f=o-i,c=1/a-1/u,h=f/u;if(c)return(-h+Math.sqrt(h*h-2*c*(f*f/(-2*u)-l+u/2+r-a/2)))/c+i;return(i+o)/2}function Br(t,e){var n=t.N;if(n)return qr(n,e);var i=t.site;return i.y===e?i.x:Infinity}function $r(t){this.site=t;this.edges=[]}$r.prototype.prepare=function(){var t=this.edges,e=t.length,n;while(e--){n=t[e].edge;if(!n.b||!n.a)t.splice(e,1)}t.sort(Yr);return t.length};function Ur(t){var e=t[0][0],n=t[1][0],i=t[0][1],r=t[1][1],a,s,o,l,u=Nr,f=u.length,c,h,d,p,g,v;while(f--){c=u[f];if(!c||!c.prepare())continue;d=c.edges;p=d.length;h=0;while(h<p){v=d[h].end(),o=v.x,l=v.y;g=d[++h%p].start(),a=g.x,s=g.y;if(y(o-a)>Ct||y(l-s)>Ct){d.splice(h,0,new na(ta(c.site,v,y(o-e)<Ct&&r-l>Ct?{x:e,y:y(a-e)<Ct?s:r}:y(l-r)<Ct&&n-o>Ct?{x:y(s-r)<Ct?a:n,y:r}:y(o-n)<Ct&&l-i>Ct?{x:n,y:y(a-n)<Ct?s:i}:y(l-i)<Ct&&o-e>Ct?{x:y(s-i)<Ct?a:e,y:i}:null),c.site,null));++p}}}}function Yr(t,e){return e.angle-t.angle}function Vr(){ra(this);this.x=this.y=this.arc=this.site=this.cy=null}function Xr(t){var e=t.P,n=t.N;if(!e||!n)return;var i=e.site,r=t.site,a=n.site;if(i===a)return;var s=r.x,o=r.y,l=i.x-s,u=i.y-o,f=a.x-s,c=a.y-o;var h=2*(l*c-u*f);if(h>=-Tt)return;var d=l*l+u*u,p=f*f+c*c,g=(c*d-u*p)/h,v=(l*p-f*d)/h,c=v+o;var m=Lr.pop()||new Vr;m.arc=t;m.site=r;m.x=g+s;m.y=c+Math.sqrt(g*g+v*v);m.cy=c;t.circle=m;var y=null,b=Rr._;while(b){if(m.y<b.y||m.y===b.y&&m.x<=b.x){if(b.L)b=b.L;else{y=b.P;break}}else{if(b.R)b=b.R;else{y=b;break}}}Rr.insert(y,m);if(!y)Fr=m}function Jr(t){var e=t.circle;if(e){if(!e.P)Fr=e.N;Rr.remove(e);Lr.push(e);ra(e);t.circle=null}}function Gr(t){var e=Ar,n=bi(t[0][0],t[0][1],t[1][0],t[1][1]),i=e.length,r;while(i--){r=e[i];if(!Kr(r,t)||!n(r)||y(r.a.x-r.b.x)<Ct&&y(r.a.y-r.b.y)<Ct){r.a=r.b=null;e.splice(i,1)}}}function Kr(t,e){var n=t.b;if(n)return true;var i=t.a,r=e[0][0],a=e[1][0],s=e[0][1],o=e[1][1],l=t.l,u=t.r,f=l.x,c=l.y,h=u.x,d=u.y,p=(f+h)/2,g=(c+d)/2,v,m;if(d===c){if(p<r||p>=a)return;if(f>h){if(!i)i={x:p,y:s};else if(i.y>=o)return;n={x:p,y:o}}else{if(!i)i={x:p,y:o};else if(i.y<s)return;n={x:p,y:s}}}else{v=(f-h)/(d-c);m=g-v*p;if(v<-1||v>1){if(f>h){if(!i)i={x:(s-m)/v,y:s};else if(i.y>=o)return;n={x:(o-m)/v,y:o}}else{if(!i)i={x:(o-m)/v,y:o};else if(i.y<s)return;n={x:(s-m)/v,y:s}}}else{if(c<d){if(!i)i={x:r,y:v*r+m};else if(i.x>=a)return;n={x:a,y:v*a+m}}else{if(!i)i={x:a,y:v*a+m};else if(i.x<r)return;n={x:r,y:v*r+m}}}}t.a=i;t.b=n;return true}function Zr(t,e){this.l=t;this.r=e;this.a=this.b=null}function Qr(t,e,n,i){var r=new Zr(t,e);Ar.push(r);if(n)ea(r,t,e,n);if(i)ea(r,e,t,i);Nr[t.i].edges.push(new na(r,t,e));Nr[e.i].edges.push(new na(r,e,t));return r}function ta(t,e,n){var i=new Zr(t,null);i.a=e;i.b=n;Ar.push(i);return i}function ea(t,e,n,i){if(!t.a&&!t.b){t.a=i;t.l=e;t.r=n}else if(t.l===n){t.b=i}else{t.a=i}}function na(t,e,n){var i=t.a,r=t.b;this.edge=t;this.site=e;this.angle=n?Math.atan2(n.y-e.y,n.x-e.x):t.l===e?Math.atan2(r.x-i.x,i.y-r.y):Math.atan2(i.x-r.x,r.y-i.y)}na.prototype={start:function(){return this.edge.l===this.site?this.edge.a:this.edge.b},end:function(){return this.edge.l===this.site?this.edge.b:this.edge.a}};function ia(){this._=null}function ra(t){t.U=t.C=t.L=t.R=t.P=t.N=null}ia.prototype={insert:function(t,e){var n,i,r;if(t){e.P=t;e.N=t.N;if(t.N)t.N.P=e;t.N=e;if(t.R){t=t.R;while(t.L)t=t.L;t.L=e}else{t.R=e}n=t}else if(this._){t=oa(this._);e.P=null;e.N=t;t.P=t.L=e;n=t}else{e.P=e.N=null;this._=e;n=null}e.L=e.R=null;e.U=n;e.C=true;t=e;while(n&&n.C){i=n.U;if(n===i.L){r=i.R;if(r&&r.C){n.C=r.C=false;i.C=true;t=i}else{if(t===n.R){aa(this,n);t=n;n=t.U}n.C=false;i.C=true;sa(this,i)}}else{r=i.L;if(r&&r.C){n.C=r.C=false;i.C=true;t=i}else{if(t===n.L){sa(this,n);t=n;n=t.U}n.C=false;i.C=true;aa(this,i)}}n=t.U}this._.C=false},remove:function(t){if(t.N)t.N.P=t.P;if(t.P)t.P.N=t.N;t.N=t.P=null;var e=t.U,n,i=t.L,r=t.R,a,s;if(!i)a=r;else if(!r)a=i;else a=oa(r);if(e){if(e.L===t)e.L=a;else e.R=a}else{this._=a}if(i&&r){s=a.C;a.C=t.C;a.L=i;i.U=a;if(a!==r){e=a.U;a.U=t.U;t=a.R;e.L=t;a.R=r;r.U=a}else{a.U=e;e=a;t=a.R}}else{s=t.C;t=a}if(t)t.U=e;if(s)return;if(t&&t.C){t.C=false;return}do{if(t===this._)break;if(t===e.L){n=e.R;if(n.C){n.C=false;e.C=true;aa(this,e);n=e.R}if(n.L&&n.L.C||n.R&&n.R.C){if(!n.R||!n.R.C){n.L.C=false;n.C=true;sa(this,n);n=e.R}n.C=e.C;e.C=n.R.C=false;aa(this,e);t=this._;break}}else{n=e.L;if(n.C){n.C=false;e.C=true;sa(this,e);n=e.L}if(n.L&&n.L.C||n.R&&n.R.C){if(!n.L||!n.L.C){n.R.C=false;n.C=true;aa(this,n);n=e.L}n.C=e.C;e.C=n.L.C=false;sa(this,e);t=this._;break}}n.C=true;t=e;e=e.U}while(!t.C);if(t)t.C=false}};function aa(t,e){var n=e,i=e.R,r=n.U;if(r){if(r.L===n)r.L=i;else r.R=i}else{t._=i}i.U=r;n.U=i;n.R=i.L;if(n.R)n.R.U=n;i.L=n}function sa(t,e){var n=e,i=e.L,r=n.U;if(r){if(r.L===n)r.L=i;else r.R=i}else{t._=i}i.U=r;n.U=i;n.L=i.R;if(n.L)n.L.U=n;i.R=n}function oa(t){while(t.L)t=t.L;return t}function la(t,e){var n=t.sort(ua).pop(),i,r,a;Ar=[];Nr=new Array(t.length);Pr=new ia;Rr=new ia;while(true){a=Fr;if(n&&(!a||n.y<a.y||n.y===a.y&&n.x<a.x)){if(n.x!==i||n.y!==r){Nr[n.i]=new $r(n);zr(n);i=n.x,r=n.y}n=t.pop()}else if(a){Wr(a.arc)}else{break}}if(e)Gr(e),Ur(e);var s={cells:Nr,edges:Ar};Pr=Rr=Ar=Nr=null;return s}function ua(t,e){return e.y-t.y||e.x-t.x}t.geom.voronoi=function(t){var e=xr,n=Sr,i=e,r=n,a=fa;if(t)return s(t);function s(t){var e=new Array(t.length),n=a[0][0],i=a[0][1],r=a[1][0],s=a[1][1];la(o(t),a).cells.forEach(function(a,o){var l=a.edges,u=a.site,f=e[o]=l.length?l.map(function(t){var e=t.start();return[e.x,e.y]}):u.x>=n&&u.x<=r&&u.y>=i&&u.y<=s?[[n,s],[r,s],[r,i],[n,i]]:[];f.point=t[o]});return e}function o(t){return t.map(function(t,e){return{x:Math.round(i(t,e)/Ct)*Ct,y:Math.round(r(t,e)/Ct)*Ct,i:e}})}s.links=function(t){return la(o(t)).edges.filter(function(t){return t.l&&t.r}).map(function(e){return{source:t[e.l.i],target:t[e.r.i]}})};s.triangles=function(t){var e=[];la(o(t)).cells.forEach(function(n,i){var r=n.site,a=n.edges.sort(Yr),s=-1,o=a.length,l,u,f=a[o-1].edge,c=f.l===r?f.r:f.l;while(++s<o){l=f;u=c;f=a[s].edge;c=f.l===r?f.r:f.l;if(i<u.i&&i<c.i&&ca(r,u,c)<0){e.push([t[i],t[u.i],t[c.i]])}}});return e};s.x=function(t){return arguments.length?(i=xe(e=t),s):e};s.y=function(t){return arguments.length?(r=xe(n=t),s):n};s.clipExtent=function(t){if(!arguments.length)return a===fa?null:a;a=t==null?fa:t;return s};s.size=function(t){if(!arguments.length)return a===fa?null:a&&a[1];return s.clipExtent(t&&[[0,0],t])};return s};var fa=[[-1e6,-1e6],[1e6,1e6]];function ca(t,e,n){return(t.x-n.x)*(e.y-t.y)-(t.x-e.x)*(n.y-t.y)}t.geom.delaunay=function(e){return t.geom.voronoi().triangles(e)};t.geom.quadtree=function(t,e,n,i,r){var a=xr,s=Sr,o;if(o=arguments.length){a=ha;s=da;if(o===3){r=n;i=e;n=e=0}return l(t)}
function l(t){var l,u=xe(a),f=xe(s),c,h,d,p,g,v,m,b;if(e!=null){g=e,v=n,m=i,b=r}else{m=b=-(g=v=Infinity);c=[],h=[];p=t.length;if(o)for(d=0;d<p;++d){l=t[d];if(l.x<g)g=l.x;if(l.y<v)v=l.y;if(l.x>m)m=l.x;if(l.y>b)b=l.y;c.push(l.x);h.push(l.y)}else for(d=0;d<p;++d){var w=+u(l=t[d],d),_=+f(l,d);if(w<g)g=w;if(_<v)v=_;if(w>m)m=w;if(_>b)b=_;c.push(w);h.push(_)}}var x=m-g,S=b-v;if(x>S)b=v+x;else m=g+S;function D(t,e,n,i,r,a,s,o){if(isNaN(n)||isNaN(i))return;if(t.leaf){var l=t.x,u=t.y;if(l!=null){if(y(l-n)+y(u-i)<.01){C(t,e,n,i,r,a,s,o)}else{var f=t.point;t.x=t.y=t.point=null;C(t,f,l,u,r,a,s,o);C(t,e,n,i,r,a,s,o)}}else{t.x=n,t.y=i,t.point=e}}else{C(t,e,n,i,r,a,s,o)}}function C(t,e,n,i,r,a,s,o){var l=(r+s)*.5,u=(a+o)*.5,f=n>=l,c=i>=u,h=c<<1|f;t.leaf=false;t=t.nodes[h]||(t.nodes[h]=pa());if(f)r=l;else s=l;if(c)a=u;else o=u;D(t,e,n,i,r,a,s,o)}var T=pa();T.add=function(t){D(T,t,+u(t,++d),+f(t,d),g,v,m,b)};T.visit=function(t){ga(t,T,g,v,m,b)};T.find=function(t){return va(T,t[0],t[1],g,v,m,b)};d=-1;if(e==null){while(++d<p){D(T,t[d],c[d],h[d],g,v,m,b)}--d}else t.forEach(T.add);c=h=t=l=null;return T}l.x=function(t){return arguments.length?(a=t,l):a};l.y=function(t){return arguments.length?(s=t,l):s};l.extent=function(t){if(!arguments.length)return e==null?null:[[e,n],[i,r]];if(t==null)e=n=i=r=null;else e=+t[0][0],n=+t[0][1],i=+t[1][0],r=+t[1][1];return l};l.size=function(t){if(!arguments.length)return e==null?null:[i-e,r-n];if(t==null)e=n=i=r=null;else e=n=0,i=+t[0],r=+t[1];return l};return l};function ha(t){return t.x}function da(t){return t.y}function pa(){return{leaf:true,nodes:[],point:null,x:null,y:null}}function ga(t,e,n,i,r,a){if(!t(e,n,i,r,a)){var s=(n+r)*.5,o=(i+a)*.5,l=e.nodes;if(l[0])ga(t,l[0],n,i,s,o);if(l[1])ga(t,l[1],s,i,r,o);if(l[2])ga(t,l[2],n,o,s,a);if(l[3])ga(t,l[3],s,o,r,a)}}function va(t,e,n,i,r,a,s){var o=Infinity,l;(function t(u,f,c,h,d){if(f>a||c>s||h<i||d<r)return;if(p=u.point){var p,g=e-u.x,v=n-u.y,m=g*g+v*v;if(m<o){var y=Math.sqrt(o=m);i=e-y,r=n-y;a=e+y,s=n+y;l=p}}var b=u.nodes,w=(f+h)*.5,_=(c+d)*.5,x=e>=w,S=n>=_;for(var D=S<<1|x,C=D+4;D<C;++D){if(u=b[D&3])switch(D&3){case 0:t(u,f,c,w,_);break;case 1:t(u,w,c,h,_);break;case 2:t(u,f,_,w,d);break;case 3:t(u,w,_,h,d);break}}})(t,i,r,a,s);return l}t.interpolateRgb=ma;function ma(e,n){e=t.rgb(e);n=t.rgb(n);var i=e.r,r=e.g,a=e.b,s=n.r-i,o=n.g-r,l=n.b-a;return function(t){return"#"+ge(Math.round(i+s*t))+ge(Math.round(r+o*t))+ge(Math.round(a+l*t))}}t.interpolateObject=ya;function ya(t,e){var n={},i={},r;for(r in t){if(r in e){n[r]=Sa(t[r],e[r])}else{i[r]=t[r]}}for(r in e){if(!(r in t)){i[r]=e[r]}}return function(t){for(r in n)i[r]=n[r](t);return i}}t.interpolateNumber=ba;function ba(t,e){t=+t,e=+e;return function(n){return t*(1-n)+e*n}}t.interpolateString=wa;function wa(t,e){var n=_a.lastIndex=xa.lastIndex=0,i,r,a,s=-1,o=[],l=[];t=t+"",e=e+"";while((i=_a.exec(t))&&(r=xa.exec(e))){if((a=r.index)>n){a=e.slice(n,a);if(o[s])o[s]+=a;else o[++s]=a}if((i=i[0])===(r=r[0])){if(o[s])o[s]+=r;else o[++s]=r}else{o[++s]=null;l.push({i:s,x:ba(i,r)})}n=xa.lastIndex}if(n<e.length){a=e.slice(n);if(o[s])o[s]+=a;else o[++s]=a}return o.length<2?l[0]?(e=l[0].x,function(t){return e(t)+""}):function(){return e}:(e=l.length,function(t){for(var n=0,i;n<e;++n)o[(i=l[n]).i]=i.x(t);return o.join("")})}var _a=/[-+]?(?:\d+\.?\d*|\.?\d+)(?:[eE][-+]?\d+)?/g,xa=new RegExp(_a.source,"g");t.interpolate=Sa;function Sa(e,n){var i=t.interpolators.length,r;while(--i>=0&&!(r=t.interpolators[i](e,n)));return r}t.interpolators=[function(t,e){var n=typeof e;return(n==="string"?_e.has(e.toLowerCase())||/^(#|rgb\(|hsl\()/i.test(e)?ma:wa:e instanceof Vt?ma:Array.isArray(e)?Da:n==="object"&&isNaN(e)?ya:ba)(t,e)}];t.interpolateArray=Da;function Da(t,e){var n=[],i=[],r=t.length,a=e.length,s=Math.min(t.length,e.length),o;for(o=0;o<s;++o)n.push(Sa(t[o],e[o]));for(;o<r;++o)i[o]=t[o];for(;o<a;++o)i[o]=e[o];return function(t){for(o=0;o<s;++o)i[o]=n[o](t);return i}}var Ca=function(){return P};var Ta=t.map({linear:Ca,poly:Fa,quad:function(){return Na},cubic:function(){return Pa},sin:function(){return Ra},exp:function(){return La},circle:function(){return ja},elastic:Oa,back:Ha,bounce:function(){return Wa}});var ka=t.map({in:P,out:Ia,"in-out":Aa,"out-in":function(t){return Aa(Ia(t))}});t.ease=function(t){var e=t.indexOf("-"),i=e>=0?t.slice(0,e):t,r=e>=0?t.slice(e+1):"in";i=Ta.get(i)||Ca;r=ka.get(r)||P;return Ma(r(i.apply(null,n.call(arguments,1))))};function Ma(t){return function(e){return e<=0?0:e>=1?1:t(e)}}function Ia(t){return function(e){return 1-t(1-e)}}function Aa(t){return function(e){return.5*(e<.5?t(2*e):2-t(2-2*e))}}function Na(t){return t*t}function Pa(t){return t*t*t}function Ea(t){if(t<=0)return 0;if(t>=1)return 1;var e=t*t,n=e*t;return 4*(t<.5?n:3*(t-e)+n-.75)}function Fa(t){return function(e){return Math.pow(e,t)}}function Ra(t){return 1-Math.cos(t*At)}function La(t){return Math.pow(2,10*(t-1))}function ja(t){return 1-Math.sqrt(1-t*t)}function Oa(t,e){var n;if(arguments.length<2)e=.45;if(arguments.length)n=e/Mt*Math.asin(1/t);else t=1,n=e/4;return function(i){return 1+t*Math.pow(2,-10*i)*Math.sin((i-n)*Mt/e)}}function Ha(t){if(!t)t=1.70158;return function(e){return e*e*((t+1)*e-t)}}function Wa(t){return t<1/2.75?7.5625*t*t:t<2/2.75?7.5625*(t-=1.5/2.75)*t+.75:t<2.5/2.75?7.5625*(t-=2.25/2.75)*t+.9375:7.5625*(t-=2.625/2.75)*t+.984375}t.interpolateHcl=za;function za(e,n){e=t.hcl(e);n=t.hcl(n);var i=e.h,r=e.c,a=e.l,s=n.h-i,o=n.c-r,l=n.l-a;if(isNaN(o))o=0,r=isNaN(r)?n.c:r;if(isNaN(s))s=0,i=isNaN(i)?n.h:i;else if(s>180)s-=360;else if(s<-180)s+=360;return function(t){return Qt(i+s*t,r+o*t,a+l*t)+""}}t.interpolateHsl=qa;function qa(e,n){e=t.hsl(e);n=t.hsl(n);var i=e.h,r=e.s,a=e.l,s=n.h-i,o=n.s-r,l=n.l-a;if(isNaN(o))o=0,r=isNaN(r)?n.s:r;if(isNaN(s))s=0,i=isNaN(i)?n.h:i;else if(s>180)s-=360;else if(s<-180)s+=360;return function(t){return Gt(i+s*t,r+o*t,a+l*t)+""}}t.interpolateLab=Ba;function Ba(e,n){e=t.lab(e);n=t.lab(n);var i=e.l,r=e.a,a=e.b,s=n.l-i,o=n.a-r,l=n.b-a;return function(t){return se(i+s*t,r+o*t,a+l*t)+""}}t.interpolateRound=$a;function $a(t,e){e-=t;return function(n){return Math.round(t+e*n)}}t.transform=function(e){var n=r.createElementNS(t.ns.prefix.svg,"g");return(t.transform=function(t){if(t!=null){n.setAttribute("transform",t);var e=n.transform.baseVal.consolidate()}return new Ua(e?e.matrix:Ja)})(e)};function Ua(t){var e=[t.a,t.b],n=[t.c,t.d],i=Va(e),r=Ya(e,n),a=Va(Xa(n,e,-r))||0;if(e[0]*n[1]<n[0]*e[1]){e[0]*=-1;e[1]*=-1;i*=-1;r*=-1}this.rotate=(i?Math.atan2(e[1],e[0]):Math.atan2(-n[0],n[1]))*Pt;this.translate=[t.e,t.f];this.scale=[i,a];this.skew=a?Math.atan2(r,a)*Pt:0}Ua.prototype.toString=function(){return"translate("+this.translate+")rotate("+this.rotate+")skewX("+this.skew+")scale("+this.scale+")"};function Ya(t,e){return t[0]*e[0]+t[1]*e[1]}function Va(t){var e=Math.sqrt(Ya(t,t));if(e){t[0]/=e;t[1]/=e}return e}function Xa(t,e,n){t[0]+=n*e[0];t[1]+=n*e[1];return t}var Ja={a:1,b:0,c:0,d:1,e:0,f:0};t.interpolateTransform=es;function Ga(t){return t.length?t.pop()+",":""}function Ka(t,e,n,i){if(t[0]!==e[0]||t[1]!==e[1]){var r=n.push("translate(",null,",",null,")");i.push({i:r-4,x:ba(t[0],e[0])},{i:r-2,x:ba(t[1],e[1])})}else if(e[0]||e[1]){n.push("translate("+e+")")}}function Za(t,e,n,i){if(t!==e){if(t-e>180)e+=360;else if(e-t>180)t+=360;i.push({i:n.push(Ga(n)+"rotate(",null,")")-2,x:ba(t,e)})}else if(e){n.push(Ga(n)+"rotate("+e+")")}}function Qa(t,e,n,i){if(t!==e){i.push({i:n.push(Ga(n)+"skewX(",null,")")-2,x:ba(t,e)})}else if(e){n.push(Ga(n)+"skewX("+e+")")}}function ts(t,e,n,i){if(t[0]!==e[0]||t[1]!==e[1]){var r=n.push(Ga(n)+"scale(",null,",",null,")");i.push({i:r-4,x:ba(t[0],e[0])},{i:r-2,x:ba(t[1],e[1])})}else if(e[0]!==1||e[1]!==1){n.push(Ga(n)+"scale("+e+")")}}function es(e,n){var i=[],r=[];e=t.transform(e),n=t.transform(n);Ka(e.translate,n.translate,i,r);Za(e.rotate,n.rotate,i,r);Qa(e.skew,n.skew,i,r);ts(e.scale,n.scale,i,r);e=n=null;return function(t){var e=-1,n=r.length,a;while(++e<n)i[(a=r[e]).i]=a.x(t);return i.join("")}}function ns(t,e){e=(e-=t=+t)||1/e;return function(n){return(n-t)/e}}function is(t,e){e=(e-=t=+t)||1/e;return function(n){return Math.max(0,Math.min(1,(n-t)/e))}}t.layout={};t.layout.bundle=function(){return function(t){var e=[],n=-1,i=t.length;while(++n<i)e.push(rs(t[n]));return e}};function rs(t){var e=t.source,n=t.target,i=ss(e,n),r=[e];while(e!==i){e=e.parent;r.push(e)}var a=r.length;while(n!==i){r.splice(a,0,n);n=n.parent}return r}function as(t){var e=[],n=t.parent;while(n!=null){e.push(t);t=n;n=n.parent}e.push(t);return e}function ss(t,e){if(t===e)return t;var n=as(t),i=as(e),r=n.pop(),a=i.pop(),s=null;while(r===a){s=r;r=n.pop();a=i.pop()}return s}t.layout.chord=function(){var e={},n,i,r,a,s=0,o,l,u;function f(){var e={},f=[],h=t.range(a),d=[],p,g,v,m,y;n=[];i=[];p=0,m=-1;while(++m<a){g=0,y=-1;while(++y<a){g+=r[m][y]}f.push(g);d.push(t.range(a));p+=g}if(o){h.sort(function(t,e){return o(f[t],f[e])})}if(l){d.forEach(function(t,e){t.sort(function(t,n){return l(r[e][t],r[e][n])})})}p=(Mt-s*a)/p;g=0,m=-1;while(++m<a){v=g,y=-1;while(++y<a){var b=h[m],w=d[b][y],_=r[b][w],x=g,S=g+=_*p;e[b+"-"+w]={index:b,subindex:w,startAngle:x,endAngle:S,value:_}}i[b]={index:b,startAngle:v,endAngle:g,value:f[b]};g+=s}m=-1;while(++m<a){y=m-1;while(++y<a){var D=e[m+"-"+y],C=e[y+"-"+m];if(D.value||C.value){n.push(D.value<C.value?{source:C,target:D}:{source:D,target:C})}}}if(u)c()}function c(){n.sort(function(t,e){return u((t.source.value+t.target.value)/2,(e.source.value+e.target.value)/2)})}e.matrix=function(t){if(!arguments.length)return r;a=(r=t)&&r.length;n=i=null;return e};e.padding=function(t){if(!arguments.length)return s;s=t;n=i=null;return e};e.sortGroups=function(t){if(!arguments.length)return o;o=t;n=i=null;return e};e.sortSubgroups=function(t){if(!arguments.length)return l;l=t;n=null;return e};e.sortChords=function(t){if(!arguments.length)return u;u=t;if(n)c();return e};e.chords=function(){if(!n)f();return n};e.groups=function(){if(!i)f();return i};return e};t.layout.force=function(){var e={},n=t.dispatch("start","tick","end"),i,r=[1,1],a,s,o=.9,l=hs,u=ds,f=-30,c=ps,h=.1,d=.64,p=[],g=[],v,m,y;function b(t){return function(e,n,i,r){if(e.point!==t){var a=e.cx-t.x,s=e.cy-t.y,o=r-n,l=a*a+s*s;if(o*o/d<l){if(l<c){var u=e.charge/l;t.px-=a*u;t.py-=s*u}return true}if(e.point&&l&&l<c){var u=e.pointCharge/l;t.px-=a*u;t.py-=s*u}}return!e.charge}}e.tick=function(){if((s*=.99)<.005){i=null;n.end({type:"end",alpha:s=0});return true}var e=p.length,a=g.length,l,u,c,d,w,_,x,S,D;for(u=0;u<a;++u){c=g[u];d=c.source;w=c.target;S=w.x-d.x;D=w.y-d.y;if(_=S*S+D*D){_=s*m[u]*((_=Math.sqrt(_))-v[u])/_;S*=_;D*=_;w.x-=S*(x=d.weight+w.weight?d.weight/(d.weight+w.weight):.5);w.y-=D*x;d.x+=S*(x=1-x);d.y+=D*x}}if(x=s*h){S=r[0]/2;D=r[1]/2;u=-1;if(x)while(++u<e){c=p[u];c.x+=(S-c.x)*x;c.y+=(D-c.y)*x}}if(f){cs(l=t.geom.quadtree(p),s,y);u=-1;while(++u<e){if(!(c=p[u]).fixed){l.visit(b(c))}}}u=-1;while(++u<e){c=p[u];if(c.fixed){c.x=c.px;c.y=c.py}else{c.x-=(c.px-(c.px=c.x))*o;c.y-=(c.py-(c.py=c.y))*o}}n.tick({type:"tick",alpha:s})};e.nodes=function(t){if(!arguments.length)return p;p=t;return e};e.links=function(t){if(!arguments.length)return g;g=t;return e};e.size=function(t){if(!arguments.length)return r;r=t;return e};e.linkDistance=function(t){if(!arguments.length)return l;l=typeof t==="function"?t:+t;return e};e.distance=e.linkDistance;e.linkStrength=function(t){if(!arguments.length)return u;u=typeof t==="function"?t:+t;return e};e.friction=function(t){if(!arguments.length)return o;o=+t;return e};e.charge=function(t){if(!arguments.length)return f;f=typeof t==="function"?t:+t;return e};e.chargeDistance=function(t){if(!arguments.length)return Math.sqrt(c);c=t*t;return e};e.gravity=function(t){if(!arguments.length)return h;h=+t;return e};e.theta=function(t){if(!arguments.length)return Math.sqrt(d);d=t*t;return e};e.alpha=function(t){if(!arguments.length)return s;t=+t;if(s){if(t>0){s=t}else{i.c=null,i.t=NaN,i=null;n.end({type:"end",alpha:s=0})}}else if(t>0){n.start({type:"start",alpha:s=t});i=Pe(e.tick)}return e};e.start=function(){var t,n=p.length,i=g.length,a=r[0],s=r[1],o,c;for(t=0;t<n;++t){(c=p[t]).index=t;c.weight=0}for(t=0;t<i;++t){c=g[t];if(typeof c.source=="number")c.source=p[c.source];if(typeof c.target=="number")c.target=p[c.target];++c.source.weight;++c.target.weight}for(t=0;t<n;++t){c=p[t];if(isNaN(c.x))c.x=h("x",a);if(isNaN(c.y))c.y=h("y",s);if(isNaN(c.px))c.px=c.x;if(isNaN(c.py))c.py=c.y}v=[];if(typeof l==="function")for(t=0;t<i;++t)v[t]=+l.call(this,g[t],t);else for(t=0;t<i;++t)v[t]=l;m=[];if(typeof u==="function")for(t=0;t<i;++t)m[t]=+u.call(this,g[t],t);else for(t=0;t<i;++t)m[t]=u;y=[];if(typeof f==="function")for(t=0;t<n;++t)y[t]=+f.call(this,p[t],t);else for(t=0;t<n;++t)y[t]=f;function h(e,r){if(!o){o=new Array(n);for(l=0;l<n;++l){o[l]=[]}for(l=0;l<i;++l){var a=g[l];o[a.source.index].push(a.target);o[a.target.index].push(a.source)}}var s=o[t],l=-1,u=s.length,f;while(++l<u)if(!isNaN(f=s[l][e]))return f;return Math.random()*r}return e.resume()};e.resume=function(){return e.alpha(.1)};e.stop=function(){return e.alpha(0)};e.drag=function(){if(!a)a=t.behavior.drag().origin(P).on("dragstart.force",os).on("drag.force",w).on("dragend.force",ls);if(!arguments.length)return a;this.on("mouseover.force",us).on("mouseout.force",fs).call(a)};function w(n){n.px=t.event.x,n.py=t.event.y;e.resume()}return t.rebind(e,n,"on")};function os(t){t.fixed|=2}function ls(t){t.fixed&=~6}function us(t){t.fixed|=4;t.px=t.x,t.py=t.y}function fs(t){t.fixed&=~4}function cs(t,e,n){var i=0,r=0;t.charge=0;if(!t.leaf){var a=t.nodes,s=a.length,o=-1,l;while(++o<s){l=a[o];if(l==null)continue;cs(l,e,n);t.charge+=l.charge;i+=l.charge*l.cx;r+=l.charge*l.cy}}if(t.point){if(!t.leaf){t.point.x+=Math.random()-.5;t.point.y+=Math.random()-.5}var u=e*n[t.point.index];t.charge+=t.pointCharge=u;i+=u*t.point.x;r+=u*t.point.y}t.cx=i/t.charge;t.cy=r/t.charge}var hs=20,ds=1,ps=Infinity;t.layout.hierarchy=function(){var t=ws,e=ys,n=bs;function i(r){var a=[r],s=[],o;r.depth=0;while((o=a.pop())!=null){s.push(o);if((u=e.call(i,o,o.depth))&&(l=u.length)){var l,u,f;while(--l>=0){a.push(f=u[l]);f.parent=o;f.depth=o.depth+1}if(n)o.value=0;o.children=u}else{if(n)o.value=+n.call(i,o,o.depth)||0;delete o.children}}ms(r,function(e){var i,r;if(t&&(i=e.children))i.sort(t);if(n&&(r=e.parent))r.value+=e.value});return s}i.sort=function(e){if(!arguments.length)return t;t=e;return i};i.children=function(t){if(!arguments.length)return e;e=t;return i};i.value=function(t){if(!arguments.length)return n;n=t;return i};i.revalue=function(t){if(n){vs(t,function(t){if(t.children)t.value=0});ms(t,function(t){var e;if(!t.children)t.value=+n.call(i,t,t.depth)||0;if(e=t.parent)e.value+=t.value})}return t};return i};function gs(e,n){t.rebind(e,n,"sort","children","value");e.nodes=e;e.links=_s;return e}function vs(t,e){var n=[t];while((t=n.pop())!=null){e(t);if((r=t.children)&&(i=r.length)){var i,r;while(--i>=0)n.push(r[i])}}}function ms(t,e){var n=[t],i=[];while((t=n.pop())!=null){i.push(t);if((s=t.children)&&(a=s.length)){var r=-1,a,s;while(++r<a)n.push(s[r])}}while((t=i.pop())!=null){e(t)}}function ys(t){return t.children}function bs(t){return t.value}function ws(t,e){return e.value-t.value}function _s(e){return t.merge(e.map(function(t){return(t.children||[]).map(function(e){return{source:t,target:e}})}))}t.layout.partition=function(){var e=t.layout.hierarchy(),n=[1,1];function i(t,e,n,r){var a=t.children;t.x=e;t.y=t.depth*r;t.dx=n;t.dy=r;if(a&&(o=a.length)){var s=-1,o,l,u;n=t.value?n/t.value:0;while(++s<o){i(l=a[s],e,u=l.value*n,r);e+=u}}}function r(t){var e=t.children,n=0;if(e&&(a=e.length)){var i=-1,a;while(++i<a)n=Math.max(n,r(e[i]))}return 1+n}function a(t,a){var s=e.call(this,t,a);i(s[0],0,n[0],n[1]/r(s[0]));return s}a.size=function(t){if(!arguments.length)return n;n=t;return a};return gs(a,e)};t.layout.pie=function(){var e=Number,n=xs,i=0,r=Mt,a=0;function s(o){var l=o.length,u=o.map(function(t,n){return+e.call(s,t,n)}),f=+(typeof i==="function"?i.apply(this,arguments):i),c=(typeof r==="function"?r.apply(this,arguments):r)-f,h=Math.min(Math.abs(c)/l,+(typeof a==="function"?a.apply(this,arguments):a)),d=h*(c<0?-1:1),p=t.sum(u),g=p?(c-l*d)/p:0,v=t.range(l),m=[],y;if(n!=null)v.sort(n===xs?function(t,e){return u[e]-u[t]}:function(t,e){return n(o[t],o[e])});v.forEach(function(t){m[t]={data:o[t],value:y=u[t],startAngle:f,endAngle:f+=y*g+d,padAngle:h}});return m}s.value=function(t){if(!arguments.length)return e;e=t;return s};s.sort=function(t){if(!arguments.length)return n;n=t;return s};s.startAngle=function(t){if(!arguments.length)return i;i=t;return s};s.endAngle=function(t){if(!arguments.length)return r;r=t;return s};s.padAngle=function(t){if(!arguments.length)return a;a=t;return s};return s};var xs={};t.layout.stack=function(){var e=P,n=Ms,i=Is,r=Cs,a=Ss,s=Ds;function o(l,u){if(!(g=l.length))return l;var f=l.map(function(t,n){return e.call(o,t,n)});var c=f.map(function(t){return t.map(function(t,e){return[a.call(o,t,e),s.call(o,t,e)]})});var h=n.call(o,c,u);f=t.permute(f,h);c=t.permute(c,h);var d=i.call(o,c,u);var p=f[0].length,g,v,m,y;for(m=0;m<p;++m){r.call(o,f[0][m],y=d[m],c[0][m][1]);for(v=1;v<g;++v){r.call(o,f[v][m],y+=c[v-1][m][1],c[v][m][1])}}return l}o.values=function(t){if(!arguments.length)return e;e=t;return o};o.order=function(t){if(!arguments.length)return n;n=typeof t==="function"?t:Ts.get(t)||Ms;return o};o.offset=function(t){if(!arguments.length)return i;i=typeof t==="function"?t:ks.get(t)||Is;return o};o.x=function(t){if(!arguments.length)return a;a=t;return o};o.y=function(t){if(!arguments.length)return s;s=t;return o};o.out=function(t){if(!arguments.length)return r;r=t;return o};return o};function Ss(t){return t.x}function Ds(t){return t.y}function Cs(t,e,n){t.y0=e;t.y=n}var Ts=t.map({"inside-out":function(e){var n=e.length,i,r,a=e.map(As),s=e.map(Ns),o=t.range(n).sort(function(t,e){return a[t]-a[e]}),l=0,u=0,f=[],c=[];for(i=0;i<n;++i){r=o[i];if(l<u){l+=s[r];f.push(r)}else{u+=s[r];c.push(r)}}return c.reverse().concat(f)},reverse:function(e){return t.range(e.length).reverse()},default:Ms});var ks=t.map({silhouette:function(t){var e=t.length,n=t[0].length,i=[],r=0,a,s,o,l=[];for(s=0;s<n;++s){for(a=0,o=0;a<e;a++)o+=t[a][s][1];if(o>r)r=o;i.push(o)}for(s=0;s<n;++s){l[s]=(r-i[s])/2}return l},wiggle:function(t){var e=t.length,n=t[0],i=n.length,r,a,s,o,l,u,f,c,h,d=[];d[0]=c=h=0;for(a=1;a<i;++a){for(r=0,o=0;r<e;++r)o+=t[r][a][1];for(r=0,l=0,f=n[a][0]-n[a-1][0];r<e;++r){for(s=0,u=(t[r][a][1]-t[r][a-1][1])/(2*f);s<r;++s){u+=(t[s][a][1]-t[s][a-1][1])/f}l+=u*t[r][a][1]}d[a]=c-=o?l/o*f:0;if(c<h)h=c}for(a=0;a<i;++a)d[a]-=h;return d},expand:function(t){var e=t.length,n=t[0].length,i=1/e,r,a,s,o=[];for(a=0;a<n;++a){for(r=0,s=0;r<e;r++)s+=t[r][a][1];if(s)for(r=0;r<e;r++)t[r][a][1]/=s;else for(r=0;r<e;r++)t[r][a][1]=i}for(a=0;a<n;++a)o[a]=0;return o},zero:Is});function Ms(e){return t.range(e.length)}function Is(t){var e=-1,n=t[0].length,i=[];while(++e<n)i[e]=0;return i}function As(t){var e=1,n=0,i=t[0][1],r,a=t.length;for(;e<a;++e){if((r=t[e][1])>i){n=e;i=r}}return n}function Ns(t){return t.reduce(Ps,0)}function Ps(t,e){return t+e[1]}t.layout.histogram=function(){var e=true,n=Number,i=Rs,r=Es;function a(a,s){var o=[],l=a.map(n,this),u=i.call(this,l,s),f=r.call(this,u,l,s),c,s=-1,h=l.length,d=f.length-1,p=e?1:1/h,g;while(++s<d){c=o[s]=[];c.dx=f[s+1]-(c.x=f[s]);c.y=0}if(d>0){s=-1;while(++s<h){g=l[s];if(g>=u[0]&&g<=u[1]){c=o[t.bisect(f,g,1,d)-1];c.y+=p;c.push(a[s])}}}return o}a.value=function(t){if(!arguments.length)return n;n=t;return a};a.range=function(t){if(!arguments.length)return i;i=xe(t);return a};a.bins=function(t){if(!arguments.length)return r;r=typeof t==="number"?function(e){return Fs(e,t)}:xe(t);return a};a.frequency=function(t){if(!arguments.length)return e;e=!!t;return a};return a};function Es(t,e){return Fs(t,Math.ceil(Math.log(e.length)/Math.LN2+1))}function Fs(t,e){var n=-1,i=+t[0],r=(t[1]-i)/e,a=[];while(++n<=e)a[n]=r*n+i;return a}function Rs(e){return[t.min(e),t.max(e)]}t.layout.pack=function(){var e=t.layout.hierarchy().sort(Ls),n=0,i=[1,1],r;function a(t,a){var s=e.call(this,t,a),o=s[0],l=i[0],u=i[1],f=r==null?Math.sqrt:typeof r==="function"?r:function(){return r};o.x=o.y=0;ms(o,function(t){t.r=+f(t.value)});ms(o,Ws);if(n){var c=n*(r?1:Math.max(2*o.r/l,2*o.r/u))/2;ms(o,function(t){t.r+=c});ms(o,Ws);ms(o,function(t){t.r-=c})}Bs(o,l/2,u/2,r?1:1/Math.max(2*o.r/l,2*o.r/u));return s}a.size=function(t){if(!arguments.length)return i;i=t;return a};a.radius=function(t){if(!arguments.length)return r;r=t==null||typeof t==="function"?t:+t;return a};a.padding=function(t){if(!arguments.length)return n;n=+t;return a};return gs(a,e)};function Ls(t,e){return t.value-e.value}function js(t,e){var n=t._pack_next;t._pack_next=e;e._pack_prev=t;e._pack_next=n;n._pack_prev=e}function Os(t,e){t._pack_next=e;e._pack_prev=t}function Hs(t,e){var n=e.x-t.x,i=e.y-t.y,r=t.r+e.r;return.999*r*r>n*n+i*i}function Ws(t){if(!(e=t.children)||!(h=e.length))return;var e,n=Infinity,i=-Infinity,r=Infinity,a=-Infinity,s,o,l,u,f,c,h;function d(t){n=Math.min(t.x-t.r,n);i=Math.max(t.x+t.r,i);r=Math.min(t.y-t.r,r);a=Math.max(t.y+t.r,a)}e.forEach(zs);s=e[0];s.x=-s.r;s.y=0;d(s);if(h>1){o=e[1];o.x=o.r;o.y=0;d(o);if(h>2){l=e[2];$s(s,o,l);d(l);js(s,l);s._pack_prev=l;js(l,o);o=s._pack_next;for(u=3;u<h;u++){$s(s,o,l=e[u]);var p=0,g=1,v=1;for(f=o._pack_next;f!==o;f=f._pack_next,g++){if(Hs(f,l)){p=1;break}}if(p==1){for(c=s._pack_prev;c!==f._pack_prev;c=c._pack_prev,v++){if(Hs(c,l)){break}}}if(p){if(g<v||g==v&&o.r<s.r)Os(s,o=f);else Os(s=c,o);u--}else{js(s,l);o=l;d(l)}}}}var m=(n+i)/2,y=(r+a)/2,b=0;for(u=0;u<h;u++){l=e[u];l.x-=m;l.y-=y;b=Math.max(b,l.r+Math.sqrt(l.x*l.x+l.y*l.y))}t.r=b;e.forEach(qs)}function zs(t){t._pack_next=t._pack_prev=t}function qs(t){delete t._pack_next;delete t._pack_prev}function Bs(t,e,n,i){var r=t.children;t.x=e+=i*t.x;t.y=n+=i*t.y;t.r*=i;if(r){var a=-1,s=r.length;while(++a<s)Bs(r[a],e,n,i)}}function $s(t,e,n){var i=t.r+n.r,r=e.x-t.x,a=e.y-t.y;if(i&&(r||a)){var s=e.r+n.r,o=r*r+a*a;s*=s;i*=i;var l=.5+(i-s)/(2*o),u=Math.sqrt(Math.max(0,2*s*(i+o)-(i-=o)*i-s*s))/(2*o);n.x=t.x+l*r+u*a;n.y=t.y+l*a-u*r}else{n.x=t.x+i;n.y=t.y}}t.layout.tree=function(){var e=t.layout.hierarchy().sort(null).value(null),n=Us,i=[1,1],r=null;function a(t,a){var u=e.call(this,t,a),c=u[0],h=s(c);ms(h,o),h.parent.m=-h.z;vs(h,l);if(r)vs(c,f);else{var d=c,p=c,g=c;vs(c,function(t){if(t.x<d.x)d=t;if(t.x>p.x)p=t;if(t.depth>g.depth)g=t});var v=n(d,p)/2-d.x,m=i[0]/(p.x+n(p,d)/2+v),y=i[1]/(g.depth||1);vs(c,function(t){t.x=(t.x+v)*m;t.y=t.depth*y})}return u}function s(t){var e={A:null,children:[t]},n=[e],i;while((i=n.pop())!=null){for(var r=i.children,a,s=0,o=r.length;s<o;++s){n.push((r[s]=a={_:r[s],parent:i,children:(a=r[s].children)&&a.slice()||[],A:null,a:null,z:0,m:0,c:0,s:0,t:null,i:s}).a=a)}}return e.children[0]}function o(t){var e=t.children,i=t.parent.children,r=t.i?i[t.i-1]:null;if(e.length){Js(t);var a=(e[0].z+e[e.length-1].z)/2;if(r){t.z=r.z+n(t._,r._);t.m=t.z-a}else{t.z=a}}else if(r){t.z=r.z+n(t._,r._)}t.parent.A=u(t,r,t.parent.A||i[0])}function l(t){t._.x=t.z+t.parent.m;t.m+=t.parent.m}function u(t,e,i){if(e){var r=t,a=t,s=e,o=r.parent.children[0],l=r.m,u=a.m,f=s.m,c=o.m,h;while(s=Vs(s),r=Ys(r),s&&r){o=Ys(o);a=Vs(a);a.a=t;h=s.z+f-r.z-l+n(s._,r._);if(h>0){Xs(Gs(s,t,i),t,h);l+=h;u+=h}f+=s.m;l+=r.m;c+=o.m;u+=a.m}if(s&&!Vs(a)){a.t=s;a.m+=f-u}if(r&&!Ys(o)){o.t=r;o.m+=l-c;i=t}}return i}function f(t){t.x*=i[0];t.y=t.depth*i[1]}a.separation=function(t){if(!arguments.length)return n;n=t;return a};a.size=function(t){if(!arguments.length)return r?null:i;r=(i=t)==null?f:null;return a};a.nodeSize=function(t){if(!arguments.length)return r?i:null;r=(i=t)==null?null:f;return a};return gs(a,e)};function Us(t,e){return t.parent==e.parent?1:2}function Ys(t){var e=t.children;return e.length?e[0]:t.t}function Vs(t){var e=t.children,n;return(n=e.length)?e[n-1]:t.t}function Xs(t,e,n){var i=n/(e.i-t.i);e.c-=i;e.s+=n;t.c+=i;e.z+=n;e.m+=n}function Js(t){var e=0,n=0,i=t.children,r=i.length,a;while(--r>=0){a=i[r];a.z+=e;a.m+=e;e+=a.s+(n+=a.c)}}function Gs(t,e,n){return t.a.parent===e.parent?t.a:n}t.layout.cluster=function(){var e=t.layout.hierarchy().sort(null).value(null),n=Us,i=[1,1],r=false;function a(t,a){var s=e.call(this,t,a),o=s[0],l,u=0;ms(o,function(t){var e=t.children;if(e&&e.length){t.x=Zs(e);t.y=Ks(e)}else{t.x=l?u+=n(t,l):0;t.y=0;l=t}});var f=Qs(o),c=to(o),h=f.x-n(f,c)/2,d=c.x+n(c,f)/2;ms(o,r?function(t){t.x=(t.x-o.x)*i[0];t.y=(o.y-t.y)*i[1]}:function(t){t.x=(t.x-h)/(d-h)*i[0];t.y=(1-(o.y?t.y/o.y:1))*i[1]});return s}a.separation=function(t){if(!arguments.length)return n;n=t;return a};a.size=function(t){if(!arguments.length)return r?null:i;r=(i=t)==null;return a};a.nodeSize=function(t){if(!arguments.length)return r?i:null;r=(i=t)!=null;return a};return gs(a,e)};function Ks(e){return 1+t.max(e,function(t){return t.y})}function Zs(t){return t.reduce(function(t,e){return t+e.x},0)/t.length}function Qs(t){var e=t.children;return e&&e.length?Qs(e[0]):t}function to(t){var e=t.children,n;return e&&(n=e.length)?to(e[n-1]):t}t.layout.treemap=function(){var e=t.layout.hierarchy(),n=Math.round,i=[1,1],r=null,a=eo,s=false,o,l="squarify",u=.5*(1+Math.sqrt(5));function f(t,e){var n=-1,i=t.length,r,a;while(++n<i){a=(r=t[n]).value*(e<0?0:e);r.area=isNaN(a)||a<=0?0:a}}function c(t){var e=t.children;if(e&&e.length){var n=a(t),i=[],r=e.slice(),s,o=Infinity,u,h=l==="slice"?n.dx:l==="dice"?n.dy:l==="slice-dice"?t.depth&1?n.dy:n.dx:Math.min(n.dx,n.dy),g;f(r,n.dx*n.dy/t.value);i.area=0;while((g=r.length)>0){i.push(s=r[g-1]);i.area+=s.area;if(l!=="squarify"||(u=d(i,h))<=o){r.pop();o=u}else{i.area-=i.pop().area;p(i,h,n,false);h=Math.min(n.dx,n.dy);i.length=i.area=0;o=Infinity}}if(i.length){p(i,h,n,true);i.length=i.area=0}e.forEach(c)}}function h(t){var e=t.children;if(e&&e.length){var n=a(t),i=e.slice(),r,s=[];f(i,n.dx*n.dy/t.value);s.area=0;while(r=i.pop()){s.push(r);s.area+=r.area;if(r.z!=null){p(s,r.z?n.dx:n.dy,n,!i.length);s.length=s.area=0}}e.forEach(h)}}function d(t,e){var n=t.area,i,r=0,a=Infinity,s=-1,o=t.length;while(++s<o){if(!(i=t[s].area))continue;if(i<a)a=i;if(i>r)r=i}n*=n;e*=e;return n?Math.max(e*r*u/n,n/(e*a*u)):Infinity}function p(t,e,i,r){var a=-1,s=t.length,o=i.x,l=i.y,u=e?n(t.area/e):0,f;if(e==i.dx){if(r||u>i.dy)u=i.dy;while(++a<s){f=t[a];f.x=o;f.y=l;f.dy=u;o+=f.dx=Math.min(i.x+i.dx-o,u?n(f.area/u):0)}f.z=true;f.dx+=i.x+i.dx-o;i.y+=u;i.dy-=u}else{if(r||u>i.dx)u=i.dx;while(++a<s){f=t[a];f.x=o;f.y=l;f.dx=u;l+=f.dy=Math.min(i.y+i.dy-l,u?n(f.area/u):0)}f.z=false;f.dy+=i.y+i.dy-l;i.x+=u;i.dx-=u}}function g(t){var n=o||e(t),r=n[0];r.x=r.y=0;if(r.value)r.dx=i[0],r.dy=i[1];else r.dx=r.dy=0;if(o)e.revalue(r);f([r],r.dx*r.dy/r.value);(o?h:c)(r);if(s)o=n;return n}g.size=function(t){if(!arguments.length)return i;i=t;return g};g.padding=function(t){if(!arguments.length)return r;function e(e){var n=t.call(g,e,e.depth);return n==null?eo(e):no(e,typeof n==="number"?[n,n,n,n]:n)}function n(e){return no(e,t)}var i;a=(r=t)==null?eo:(i=typeof t)==="function"?e:i==="number"?(t=[t,t,t,t],n):n;return g};g.round=function(t){if(!arguments.length)return n!=Number;n=t?Math.round:Number;return g};g.sticky=function(t){if(!arguments.length)return s;s=t;o=null;return g};g.ratio=function(t){if(!arguments.length)return u;u=t;return g};g.mode=function(t){if(!arguments.length)return l;l=t+"";return g};return gs(g,e)};function eo(t){return{x:t.x,y:t.y,dx:t.dx,dy:t.dy}}function no(t,e){var n=t.x+e[3],i=t.y+e[0],r=t.dx-e[1]-e[3],a=t.dy-e[0]-e[2];if(r<0){n+=r/2;r=0}if(a<0){i+=a/2;a=0}return{x:n,y:i,dx:r,dy:a}}t.random={normal:function(t,e){var n=arguments.length;if(n<2)e=1;if(n<1)t=0;return function(){var n,i,r;do{n=Math.random()*2-1;i=Math.random()*2-1;r=n*n+i*i}while(!r||r>1);return t+e*n*Math.sqrt(-2*Math.log(r)/r)}},logNormal:function(){var e=t.random.normal.apply(t,arguments);return function(){return Math.exp(e())}},bates:function(e){var n=t.random.irwinHall(e);return function(){return n()/e}},irwinHall:function(t){return function(){for(var e=0,n=0;n<t;n++)e+=Math.random();return e}}};t.scale={};function io(t){var e=t[0],n=t[t.length-1];return e<n?[e,n]:[n,e]}function ro(t){return t.rangeExtent?t.rangeExtent():io(t.range())}function ao(t,e,n,i){var r=n(t[0],t[1]),a=i(e[0],e[1]);return function(t){return a(r(t))}}function so(t,e){var n=0,i=t.length-1,r=t[n],a=t[i],s;if(a<r){s=n,n=i,i=s;s=r,r=a,a=s}t[n]=e.floor(r);t[i]=e.ceil(a);return t}function oo(t){return t?{floor:function(e){return Math.floor(e/t)*t},ceil:function(e){return Math.ceil(e/t)*t}}:lo}var lo={floor:P,ceil:P};function uo(e,n,i,r){var a=[],s=[],o=0,l=Math.min(e.length,n.length)-1;if(e[l]<e[0]){e=e.slice().reverse();n=n.slice().reverse()}while(++o<=l){a.push(i(e[o-1],e[o]));s.push(r(n[o-1],n[o]))}return function(n){var i=t.bisect(e,n,1,l)-1;return s[i](a[i](n))}}t.scale.linear=function(){return fo([0,1],[0,1],Sa,false)};function fo(t,e,n,i){var r,a;function s(){var s=Math.min(t.length,e.length)>2?uo:ao,l=i?is:ns;r=s(t,e,l,n);a=s(e,t,l,Sa);return o}function o(t){return r(t)}o.invert=function(t){return a(t)};o.domain=function(e){if(!arguments.length)return t;t=e.map(Number);return s()};o.range=function(t){if(!arguments.length)return e;e=t;return s()};o.rangeRound=function(t){return o.range(t).interpolate($a)};o.clamp=function(t){if(!arguments.length)return i;i=t;return s()};o.interpolate=function(t){if(!arguments.length)return n;n=t;return s()};o.ticks=function(e){return go(t,e)};o.tickFormat=function(e,n){return vo(t,e,n)};o.nice=function(e){ho(t,e);return s()};o.copy=function(){return fo(t,e,n,i)};return s()}function co(e,n){return t.rebind(e,n,"range","rangeRound","interpolate","clamp")}function ho(t,e){so(t,oo(po(t,e)[2]));so(t,oo(po(t,e)[2]));return t}function po(t,e){if(e==null)e=10;var n=io(t),i=n[1]-n[0],r=Math.pow(10,Math.floor(Math.log(i/e)/Math.LN10)),a=e/i*r;if(a<=.15)r*=10;else if(a<=.35)r*=5;else if(a<=.75)r*=2;n[0]=Math.ceil(n[0]/r)*r;n[1]=Math.floor(n[1]/r)*r+r*.5;n[2]=r;return n}function go(e,n){return t.range.apply(t,po(e,n))}function vo(e,n,i){var r=po(e,n);if(i){var a=We.exec(i);a.shift();if(a[8]==="s"){var s=t.formatPrefix(Math.max(y(r[0]),y(r[1])));if(!a[7])a[7]="."+yo(s.scale(r[2]));a[8]="f";i=t.format(a.join(""));return function(t){return i(s.scale(t))+s.symbol}}if(!a[7])a[7]="."+bo(a[8],r);i=a.join("")}else{i=",."+yo(r[2])+"f"}return t.format(i)}var mo={s:1,g:1,p:1,r:1,e:1};function yo(t){return-Math.floor(Math.log(t)/Math.LN10+.01)}function bo(t,e){var n=yo(e[2]);return t in mo?Math.abs(n-yo(Math.max(y(e[0]),y(e[1]))))+ +(t!=="e"):n-(t==="%")*2}t.scale.log=function(){return wo(t.scale.linear().domain([0,1]),10,true,[1,10])};function wo(e,n,i,r){function a(t){return(i?Math.log(t<0?0:t):-Math.log(t>0?0:-t))/Math.log(n)}function s(t){return i?Math.pow(n,t):-Math.pow(n,-t)}function o(t){return e(a(t))}o.invert=function(t){return s(e.invert(t))};o.domain=function(t){if(!arguments.length)return r;i=t[0]>=0;e.domain((r=t.map(Number)).map(a));return o};o.base=function(t){if(!arguments.length)return n;n=+t;e.domain(r.map(a));return o};o.nice=function(){var t=so(r.map(a),i?Math:xo);e.domain(t);r=t.map(s);return o};o.ticks=function(){var t=io(r),e=[],o=t[0],l=t[1],u=Math.floor(a(o)),f=Math.ceil(a(l)),c=n%1?2:n;if(isFinite(f-u)){if(i){for(;u<f;u++)for(var h=1;h<c;h++)e.push(s(u)*h);e.push(s(u))}else{e.push(s(u));for(;u++<f;)for(var h=c-1;h>0;h--)e.push(s(u)*h)}for(u=0;e[u]<o;u++){}for(f=e.length;e[f-1]>l;f--){}e=e.slice(u,f)}return e};o.tickFormat=function(e,i){if(!arguments.length)return _o;if(arguments.length<2)i=_o;else if(typeof i!=="function")i=t.format(i);var r=Math.max(1,n*e/o.ticks().length);return function(t){var e=t/s(Math.round(a(t)));if(e*n<n-.5)e*=n;return e<=r?i(t):""}};o.copy=function(){return wo(e.copy(),n,i,r)};return co(o,e)}var _o=t.format(".0e"),xo={floor:function(t){return-Math.ceil(-t)},ceil:function(t){return-Math.floor(-t)}};t.scale.pow=function(){return So(t.scale.linear(),1,[0,1])};function So(t,e,n){var i=Do(e),r=Do(1/e);function a(e){
return t(i(e))}a.invert=function(e){return r(t.invert(e))};a.domain=function(e){if(!arguments.length)return n;t.domain((n=e.map(Number)).map(i));return a};a.ticks=function(t){return go(n,t)};a.tickFormat=function(t,e){return vo(n,t,e)};a.nice=function(t){return a.domain(ho(n,t))};a.exponent=function(s){if(!arguments.length)return e;i=Do(e=s);r=Do(1/e);t.domain(n.map(i));return a};a.copy=function(){return So(t.copy(),e,n)};return co(a,t)}function Do(t){return function(e){return e<0?-Math.pow(-e,t):Math.pow(e,t)}}t.scale.sqrt=function(){return t.scale.pow().exponent(.5)};t.scale.ordinal=function(){return Co([],{t:"range",a:[[]]})};function Co(e,n){var i,r,a;function s(t){return r[((i.get(t)||(n.t==="range"?i.set(t,e.push(t)):NaN))-1)%r.length]}function o(n,i){return t.range(e.length).map(function(t){return n+i*t})}s.domain=function(t){if(!arguments.length)return e;e=[];i=new _;var r=-1,a=t.length,o;while(++r<a)if(!i.has(o=t[r]))i.set(o,e.push(o));return s[n.t].apply(s,n.a)};s.range=function(t){if(!arguments.length)return r;r=t;a=0;n={t:"range",a:arguments};return s};s.rangePoints=function(t,i){if(arguments.length<2)i=0;var l=t[0],u=t[1],f=e.length<2?(l=(l+u)/2,0):(u-l)/(e.length-1+i);r=o(l+f*i/2,f);a=0;n={t:"rangePoints",a:arguments};return s};s.rangeRoundPoints=function(t,i){if(arguments.length<2)i=0;var l=t[0],u=t[1],f=e.length<2?(l=u=Math.round((l+u)/2),0):(u-l)/(e.length-1+i)|0;r=o(l+Math.round(f*i/2+(u-l-(e.length-1+i)*f)/2),f);a=0;n={t:"rangeRoundPoints",a:arguments};return s};s.rangeBands=function(t,i,l){if(arguments.length<2)i=0;if(arguments.length<3)l=i;var u=t[1]<t[0],f=t[u-0],c=t[1-u],h=(c-f)/(e.length-i+2*l);r=o(f+h*l,h);if(u)r.reverse();a=h*(1-i);n={t:"rangeBands",a:arguments};return s};s.rangeRoundBands=function(t,i,l){if(arguments.length<2)i=0;if(arguments.length<3)l=i;var u=t[1]<t[0],f=t[u-0],c=t[1-u],h=Math.floor((c-f)/(e.length-i+2*l));r=o(f+Math.round((c-f-(e.length-i)*h)/2),h);if(u)r.reverse();a=Math.round(h*(1-i));n={t:"rangeRoundBands",a:arguments};return s};s.rangeBand=function(){return a};s.rangeExtent=function(){return io(n.a[0])};s.copy=function(){return Co(e,n)};return s.domain(e)}t.scale.category10=function(){return t.scale.ordinal().range(To)};t.scale.category20=function(){return t.scale.ordinal().range(ko)};t.scale.category20b=function(){return t.scale.ordinal().range(Mo)};t.scale.category20c=function(){return t.scale.ordinal().range(Io)};var To=[2062260,16744206,2924588,14034728,9725885,9197131,14907330,8355711,12369186,1556175].map(de);var ko=[2062260,11454440,16744206,16759672,2924588,10018698,14034728,16750742,9725885,12955861,9197131,12885140,14907330,16234194,8355711,13092807,12369186,14408589,1556175,10410725].map(de);var Mo=[3750777,5395619,7040719,10264286,6519097,9216594,11915115,13556636,9202993,12426809,15186514,15190932,8666169,11356490,14049643,15177372,8077683,10834324,13528509,14589654].map(de);var Io=[3244733,7057110,10406625,13032431,15095053,16616764,16625259,16634018,3253076,7652470,10607003,13101504,7695281,10394312,12369372,14342891,6513507,9868950,12434877,14277081].map(de);t.scale.quantile=function(){return Ao([],[])};function Ao(e,n){var i;function r(){var r=0,s=n.length;i=[];while(++r<s)i[r-1]=t.quantile(e,r/s);return a}function a(e){if(!isNaN(e=+e))return n[t.bisect(i,e)]}a.domain=function(t){if(!arguments.length)return e;e=t.map(d).filter(p).sort(h);return r()};a.range=function(t){if(!arguments.length)return n;n=t;return r()};a.quantiles=function(){return i};a.invertExtent=function(t){t=n.indexOf(t);return t<0?[NaN,NaN]:[t>0?i[t-1]:e[0],t<i.length?i[t]:e[e.length-1]]};a.copy=function(){return Ao(e,n)};return r()}t.scale.quantize=function(){return No(0,1,[0,1])};function No(t,e,n){var i,r;function a(e){return n[Math.max(0,Math.min(r,Math.floor(i*(e-t))))]}function s(){i=n.length/(e-t);r=n.length-1;return a}a.domain=function(n){if(!arguments.length)return[t,e];t=+n[0];e=+n[n.length-1];return s()};a.range=function(t){if(!arguments.length)return n;n=t;return s()};a.invertExtent=function(e){e=n.indexOf(e);e=e<0?NaN:e/i+t;return[e,e+1/i]};a.copy=function(){return No(t,e,n)};return s()}t.scale.threshold=function(){return Po([.5],[0,1])};function Po(e,n){function i(i){if(i<=i)return n[t.bisect(e,i)]}i.domain=function(t){if(!arguments.length)return e;e=t;return i};i.range=function(t){if(!arguments.length)return n;n=t;return i};i.invertExtent=function(t){t=n.indexOf(t);return[e[t-1],e[t]]};i.copy=function(){return Po(e,n)};return i}t.scale.identity=function(){return Eo([0,1])};function Eo(t){function e(t){return+t}e.invert=e;e.domain=e.range=function(n){if(!arguments.length)return t;t=n.map(e);return e};e.ticks=function(e){return go(t,e)};e.tickFormat=function(e,n){return vo(t,e,n)};e.copy=function(){return Eo(t)};return e}t.svg={};function Fo(){return 0}t.svg.arc=function(){var t=Lo,e=jo,n=Fo,i=Ro,r=Oo,a=Ho,s=Wo;function o(){var o=Math.max(0,+t.apply(this,arguments)),u=Math.max(0,+e.apply(this,arguments)),f=r.apply(this,arguments)-At,c=a.apply(this,arguments)-At,h=Math.abs(c-f),d=f>c?0:1;if(u<o)p=u,u=o,o=p;if(h>=It)return l(u,d)+(o?l(o,1-d):"")+"Z";var p,g,v,m,y=0,b=0,w,_,x,S,D,C,T,k,M=[];if(m=(+s.apply(this,arguments)||0)/2){v=i===Ro?Math.sqrt(o*o+u*u):+i.apply(this,arguments);if(!d)b*=-1;if(u)b=Lt(v/u*Math.sin(m));if(o)y=Lt(v/o*Math.sin(m))}if(u){w=u*Math.cos(f+b);_=u*Math.sin(f+b);x=u*Math.cos(c-b);S=u*Math.sin(c-b);var I=Math.abs(c-f-2*b)<=kt?0:1;if(b&&zo(w,_,x,S)===d^I){var A=(f+c)/2;w=u*Math.cos(A);_=u*Math.sin(A);x=S=null}}else{w=_=0}if(o){D=o*Math.cos(c-y);C=o*Math.sin(c-y);T=o*Math.cos(f+y);k=o*Math.sin(f+y);var N=Math.abs(f-c+2*y)<=kt?0:1;if(y&&zo(D,C,T,k)===1-d^N){var P=(f+c)/2;D=o*Math.cos(P);C=o*Math.sin(P);T=k=null}}else{D=C=0}if(h>Ct&&(p=Math.min(Math.abs(u-o)/2,+n.apply(this,arguments)))>.001){g=o<u^d?0:1;var E=p,F=p;if(h<kt){var R=T==null?[D,C]:x==null?[w,_]:Mr([w,_],[T,k],[x,S],[D,C]),L=w-R[0],j=_-R[1],O=x-R[0],H=S-R[1],W=1/Math.sin(Math.acos((L*O+j*H)/(Math.sqrt(L*L+j*j)*Math.sqrt(O*O+H*H)))/2),z=Math.sqrt(R[0]*R[0]+R[1]*R[1]);F=Math.min(p,(o-z)/(W-1));E=Math.min(p,(u-z)/(W+1))}if(x!=null){var q=qo(T==null?[D,C]:[T,k],[w,_],u,E,d),B=qo([x,S],[D,C],u,E,d);if(p===E){M.push("M",q[0],"A",E,",",E," 0 0,",g," ",q[1],"A",u,",",u," 0 ",1-d^zo(q[1][0],q[1][1],B[1][0],B[1][1]),",",d," ",B[1],"A",E,",",E," 0 0,",g," ",B[0])}else{M.push("M",q[0],"A",E,",",E," 0 1,",g," ",B[0])}}else{M.push("M",w,",",_)}if(T!=null){var $=qo([w,_],[T,k],o,-F,d),U=qo([D,C],x==null?[w,_]:[x,S],o,-F,d);if(p===F){M.push("L",U[0],"A",F,",",F," 0 0,",g," ",U[1],"A",o,",",o," 0 ",d^zo(U[1][0],U[1][1],$[1][0],$[1][1]),",",1-d," ",$[1],"A",F,",",F," 0 0,",g," ",$[0])}else{M.push("L",U[0],"A",F,",",F," 0 0,",g," ",$[0])}}else{M.push("L",D,",",C)}}else{M.push("M",w,",",_);if(x!=null)M.push("A",u,",",u," 0 ",I,",",d," ",x,",",S);M.push("L",D,",",C);if(T!=null)M.push("A",o,",",o," 0 ",N,",",1-d," ",T,",",k)}M.push("Z");return M.join("")}function l(t,e){return"M0,"+t+"A"+t+","+t+" 0 1,"+e+" 0,"+-t+"A"+t+","+t+" 0 1,"+e+" 0,"+t}o.innerRadius=function(e){if(!arguments.length)return t;t=xe(e);return o};o.outerRadius=function(t){if(!arguments.length)return e;e=xe(t);return o};o.cornerRadius=function(t){if(!arguments.length)return n;n=xe(t);return o};o.padRadius=function(t){if(!arguments.length)return i;i=t==Ro?Ro:xe(t);return o};o.startAngle=function(t){if(!arguments.length)return r;r=xe(t);return o};o.endAngle=function(t){if(!arguments.length)return a;a=xe(t);return o};o.padAngle=function(t){if(!arguments.length)return s;s=xe(t);return o};o.centroid=function(){var n=(+t.apply(this,arguments)+ +e.apply(this,arguments))/2,i=(+r.apply(this,arguments)+ +a.apply(this,arguments))/2-At;return[Math.cos(i)*n,Math.sin(i)*n]};return o};var Ro="auto";function Lo(t){return t.innerRadius}function jo(t){return t.outerRadius}function Oo(t){return t.startAngle}function Ho(t){return t.endAngle}function Wo(t){return t&&t.padAngle}function zo(t,e,n,i){return(t-n)*e-(e-i)*t>0?0:1}function qo(t,e,n,i,r){var a=t[0]-e[0],s=t[1]-e[1],o=(r?i:-i)/Math.sqrt(a*a+s*s),l=o*s,u=-o*a,f=t[0]+l,c=t[1]+u,h=e[0]+l,d=e[1]+u,p=(f+h)/2,g=(c+d)/2,v=h-f,m=d-c,y=v*v+m*m,b=n-i,w=f*d-h*c,_=(m<0?-1:1)*Math.sqrt(Math.max(0,b*b*y-w*w)),x=(w*m-v*_)/y,S=(-w*v-m*_)/y,D=(w*m+v*_)/y,C=(-w*v+m*_)/y,T=x-p,k=S-g,M=D-p,I=C-g;if(T*T+k*k>M*M+I*I)x=D,S=C;return[[x-l,S-u],[x*n/b,S*n/b]]}function Bo(t){var e=xr,n=Sr,i=ai,r=Uo,a=r.key,s=.7;function o(a){var o=[],l=[],u=-1,f=a.length,c,h=xe(e),d=xe(n);function p(){o.push("M",r(t(l),s))}while(++u<f){if(i.call(this,c=a[u],u)){l.push([+h.call(this,c,u),+d.call(this,c,u)])}else if(l.length){p();l=[]}}if(l.length)p();return o.length?o.join(""):null}o.x=function(t){if(!arguments.length)return e;e=t;return o};o.y=function(t){if(!arguments.length)return n;n=t;return o};o.defined=function(t){if(!arguments.length)return i;i=t;return o};o.interpolate=function(t){if(!arguments.length)return a;if(typeof t==="function")a=r=t;else a=(r=$o.get(t)||Uo).key;return o};o.tension=function(t){if(!arguments.length)return s;s=t;return o};return o}t.svg.line=function(){return Bo(P)};var $o=t.map({linear:Uo,"linear-closed":Yo,step:Vo,"step-before":Xo,"step-after":Jo,basis:el,"basis-open":nl,"basis-closed":il,bundle:rl,cardinal:Zo,"cardinal-open":Go,"cardinal-closed":Ko,monotone:dl});$o.forEach(function(t,e){e.key=t;e.closed=/-closed$/.test(t)});function Uo(t){return t.length>1?t.join("L"):t+"Z"}function Yo(t){return t.join("L")+"Z"}function Vo(t){var e=0,n=t.length,i=t[0],r=[i[0],",",i[1]];while(++e<n)r.push("H",(i[0]+(i=t[e])[0])/2,"V",i[1]);if(n>1)r.push("H",i[0]);return r.join("")}function Xo(t){var e=0,n=t.length,i=t[0],r=[i[0],",",i[1]];while(++e<n)r.push("V",(i=t[e])[1],"H",i[0]);return r.join("")}function Jo(t){var e=0,n=t.length,i=t[0],r=[i[0],",",i[1]];while(++e<n)r.push("H",(i=t[e])[0],"V",i[1]);return r.join("")}function Go(t,e){return t.length<4?Uo(t):t[1]+Qo(t.slice(1,-1),tl(t,e))}function Ko(t,e){return t.length<3?Yo(t):t[0]+Qo((t.push(t[0]),t),tl([t[t.length-2]].concat(t,[t[1]]),e))}function Zo(t,e){return t.length<3?Uo(t):t[0]+Qo(t,tl(t,e))}function Qo(t,e){if(e.length<1||t.length!=e.length&&t.length!=e.length+2){return Uo(t)}var n=t.length!=e.length,i="",r=t[0],a=t[1],s=e[0],o=s,l=1;if(n){i+="Q"+(a[0]-s[0]*2/3)+","+(a[1]-s[1]*2/3)+","+a[0]+","+a[1];r=t[1];l=2}if(e.length>1){o=e[1];a=t[l];l++;i+="C"+(r[0]+s[0])+","+(r[1]+s[1])+","+(a[0]-o[0])+","+(a[1]-o[1])+","+a[0]+","+a[1];for(var u=2;u<e.length;u++,l++){a=t[l];o=e[u];i+="S"+(a[0]-o[0])+","+(a[1]-o[1])+","+a[0]+","+a[1]}}if(n){var f=t[l];i+="Q"+(a[0]+o[0]*2/3)+","+(a[1]+o[1]*2/3)+","+f[0]+","+f[1]}return i}function tl(t,e){var n=[],i=(1-e)/2,r,a=t[0],s=t[1],o=1,l=t.length;while(++o<l){r=a;a=s;s=t[o];n.push([i*(s[0]-r[0]),i*(s[1]-r[1])])}return n}function el(t){if(t.length<3)return Uo(t);var e=1,n=t.length,i=t[0],r=i[0],a=i[1],s=[r,r,r,(i=t[1])[0]],o=[a,a,a,i[1]],l=[r,",",a,"L",al(ll,s),",",al(ll,o)];t.push(t[n-1]);while(++e<=n){i=t[e];s.shift();s.push(i[0]);o.shift();o.push(i[1]);ul(l,s,o)}t.pop();l.push("L",i);return l.join("")}function nl(t){if(t.length<4)return Uo(t);var e=[],n=-1,i=t.length,r,a=[0],s=[0];while(++n<3){r=t[n];a.push(r[0]);s.push(r[1])}e.push(al(ll,a)+","+al(ll,s));--n;while(++n<i){r=t[n];a.shift();a.push(r[0]);s.shift();s.push(r[1]);ul(e,a,s)}return e.join("")}function il(t){var e,n=-1,i=t.length,r=i+4,a,s=[],o=[];while(++n<4){a=t[n%i];s.push(a[0]);o.push(a[1])}e=[al(ll,s),",",al(ll,o)];--n;while(++n<r){a=t[n%i];s.shift();s.push(a[0]);o.shift();o.push(a[1]);ul(e,s,o)}return e.join("")}function rl(t,e){var n=t.length-1;if(n){var i=t[0][0],r=t[0][1],a=t[n][0]-i,s=t[n][1]-r,o=-1,l,u;while(++o<=n){l=t[o];u=o/n;l[0]=e*l[0]+(1-e)*(i+u*a);l[1]=e*l[1]+(1-e)*(r+u*s)}}return el(t)}function al(t,e){return t[0]*e[0]+t[1]*e[1]+t[2]*e[2]+t[3]*e[3]}var sl=[0,2/3,1/3,0],ol=[0,1/3,2/3,0],ll=[0,1/6,2/3,1/6];function ul(t,e,n){t.push("C",al(sl,e),",",al(sl,n),",",al(ol,e),",",al(ol,n),",",al(ll,e),",",al(ll,n))}function fl(t,e){return(e[1]-t[1])/(e[0]-t[0])}function cl(t){var e=0,n=t.length-1,i=[],r=t[0],a=t[1],s=i[0]=fl(r,a);while(++e<n){i[e]=(s+(s=fl(r=a,a=t[e+1])))/2}i[e]=s;return i}function hl(t){var e=[],n,i,r,a,s=cl(t),o=-1,l=t.length-1;while(++o<l){n=fl(t[o],t[o+1]);if(y(n)<Ct){s[o]=s[o+1]=0}else{i=s[o]/n;r=s[o+1]/n;a=i*i+r*r;if(a>9){a=n*3/Math.sqrt(a);s[o]=a*i;s[o+1]=a*r}}}o=-1;while(++o<=l){a=(t[Math.min(l,o+1)][0]-t[Math.max(0,o-1)][0])/(6*(1+s[o]*s[o]));e.push([a||0,s[o]*a||0])}return e}function dl(t){return t.length<3?Uo(t):t[0]+Qo(t,hl(t))}t.svg.line.radial=function(){var t=Bo(pl);t.radius=t.x,delete t.x;t.angle=t.y,delete t.y;return t};function pl(t){var e,n=-1,i=t.length,r,a;while(++n<i){e=t[n];r=e[0];a=e[1]-At;e[0]=r*Math.cos(a);e[1]=r*Math.sin(a)}return t}function gl(t){var e=xr,n=xr,i=0,r=Sr,a=ai,s=Uo,o=s.key,l=s,u="L",f=.7;function c(o){var c=[],h=[],d=[],p=-1,g=o.length,v,m=xe(e),y=xe(i),b=e===n?function(){return _}:xe(n),w=i===r?function(){return x}:xe(r),_,x;function S(){c.push("M",s(t(d),f),u,l(t(h.reverse()),f),"Z")}while(++p<g){if(a.call(this,v=o[p],p)){h.push([_=+m.call(this,v,p),x=+y.call(this,v,p)]);d.push([+b.call(this,v,p),+w.call(this,v,p)])}else if(h.length){S();h=[];d=[]}}if(h.length)S();return c.length?c.join(""):null}c.x=function(t){if(!arguments.length)return n;e=n=t;return c};c.x0=function(t){if(!arguments.length)return e;e=t;return c};c.x1=function(t){if(!arguments.length)return n;n=t;return c};c.y=function(t){if(!arguments.length)return r;i=r=t;return c};c.y0=function(t){if(!arguments.length)return i;i=t;return c};c.y1=function(t){if(!arguments.length)return r;r=t;return c};c.defined=function(t){if(!arguments.length)return a;a=t;return c};c.interpolate=function(t){if(!arguments.length)return o;if(typeof t==="function")o=s=t;else o=(s=$o.get(t)||Uo).key;l=s.reverse||s;u=s.closed?"M":"L";return c};c.tension=function(t){if(!arguments.length)return f;f=t;return c};return c}Xo.reverse=Jo;Jo.reverse=Xo;t.svg.area=function(){return gl(P)};t.svg.area.radial=function(){var t=gl(pl);t.radius=t.x,delete t.x;t.innerRadius=t.x0,delete t.x0;t.outerRadius=t.x1,delete t.x1;t.angle=t.y,delete t.y;t.startAngle=t.y0,delete t.y0;t.endAngle=t.y1,delete t.y1;return t};t.svg.chord=function(){var t=ar,e=sr,n=vl,i=Oo,r=Ho;function a(n,i){var r=s(this,t,n,i),a=s(this,e,n,i);return"M"+r.p0+l(r.r,r.p1,r.a1-r.a0)+(o(r,a)?u(r.r,r.p1,r.r,r.p0):u(r.r,r.p1,a.r,a.p0)+l(a.r,a.p1,a.a1-a.a0)+u(a.r,a.p1,r.r,r.p0))+"Z"}function s(t,e,a,s){var o=e.call(t,a,s),l=n.call(t,o,s),u=i.call(t,o,s)-At,f=r.call(t,o,s)-At;return{r:l,a0:u,a1:f,p0:[l*Math.cos(u),l*Math.sin(u)],p1:[l*Math.cos(f),l*Math.sin(f)]}}function o(t,e){return t.a0==e.a0&&t.a1==e.a1}function l(t,e,n){return"A"+t+","+t+" 0 "+ +(n>kt)+",1 "+e}function u(t,e,n,i){return"Q 0,0 "+i}a.radius=function(t){if(!arguments.length)return n;n=xe(t);return a};a.source=function(e){if(!arguments.length)return t;t=xe(e);return a};a.target=function(t){if(!arguments.length)return e;e=xe(t);return a};a.startAngle=function(t){if(!arguments.length)return i;i=xe(t);return a};a.endAngle=function(t){if(!arguments.length)return r;r=xe(t);return a};return a};function vl(t){return t.radius}t.svg.diagonal=function(){var t=ar,e=sr,n=ml;function i(i,r){var a=t.call(this,i,r),s=e.call(this,i,r),o=(a.y+s.y)/2,l=[a,{x:a.x,y:o},{x:s.x,y:o},s];l=l.map(n);return"M"+l[0]+"C"+l[1]+" "+l[2]+" "+l[3]}i.source=function(e){if(!arguments.length)return t;t=xe(e);return i};i.target=function(t){if(!arguments.length)return e;e=xe(t);return i};i.projection=function(t){if(!arguments.length)return n;n=t;return i};return i};function ml(t){return[t.x,t.y]}t.svg.diagonal.radial=function(){var e=t.svg.diagonal(),n=ml,i=e.projection;e.projection=function(t){return arguments.length?i(yl(n=t)):n};return e};function yl(t){return function(){var e=t.apply(this,arguments),n=e[0],i=e[1]-At;return[n*Math.cos(i),n*Math.sin(i)]}}t.svg.symbol=function(){var t=wl,e=bl;function n(n,i){return(xl.get(t.call(this,n,i))||_l)(e.call(this,n,i))}n.type=function(e){if(!arguments.length)return t;t=xe(e);return n};n.size=function(t){if(!arguments.length)return e;e=xe(t);return n};return n};function bl(){return 64}function wl(){return"circle"}function _l(t){var e=Math.sqrt(t/kt);return"M0,"+e+"A"+e+","+e+" 0 1,1 0,"+-e+"A"+e+","+e+" 0 1,1 0,"+e+"Z"}var xl=t.map({circle:_l,cross:function(t){var e=Math.sqrt(t/5)/2;return"M"+-3*e+","+-e+"H"+-e+"V"+-3*e+"H"+e+"V"+-e+"H"+3*e+"V"+e+"H"+e+"V"+3*e+"H"+-e+"V"+e+"H"+-3*e+"Z"},diamond:function(t){var e=Math.sqrt(t/(2*Dl)),n=e*Dl;return"M0,"+-e+"L"+n+",0"+" 0,"+e+" "+-n+",0"+"Z"},square:function(t){var e=Math.sqrt(t)/2;return"M"+-e+","+-e+"L"+e+","+-e+" "+e+","+e+" "+-e+","+e+"Z"},"triangle-down":function(t){var e=Math.sqrt(t/Sl),n=e*Sl/2;return"M0,"+n+"L"+e+","+-n+" "+-e+","+-n+"Z"},"triangle-up":function(t){var e=Math.sqrt(t/Sl),n=e*Sl/2;return"M0,"+-n+"L"+e+","+n+" "+-e+","+n+"Z"}});t.svg.symbolTypes=xl.keys();var Sl=Math.sqrt(3),Dl=Math.tan(30*Nt);X.transition=function(t){var e=Al||++Il,n=Fl(t),i=[],r,a,s=Nl||{time:Date.now(),ease:Ea,delay:0,duration:250};for(var o=-1,l=this.length;++o<l;){i.push(r=[]);for(var u=this[o],f=-1,c=u.length;++f<c;){if(a=u[f])Rl(a,f,n,e,s);r.push(a)}}return kl(i,n,e)};X.interrupt=function(t){return this.each(t==null?Cl:Tl(Fl(t)))};var Cl=Tl(Fl());function Tl(t){return function(){var e,n,i;if((e=this[t])&&(i=e[n=e.active])){i.timer.c=null;i.timer.t=NaN;if(--e.count)delete e[n];else delete this[t];e.active+=.5;i.event&&i.event.interrupt.call(this,this.__data__,i.index)}}}function kl(t,e,n){B(t,Ml);t.namespace=e;t.id=n;return t}var Ml=[],Il=0,Al,Nl;Ml.call=X.call;Ml.empty=X.empty;Ml.node=X.node;Ml.size=X.size;t.transition=function(e,n){return e&&e.transition?Al?e.transition(n):e:t.selection().transition(e)};t.transition.prototype=Ml;Ml.select=function(t){var e=this.id,n=this.namespace,i=[],r,a,s;t=J(t);for(var o=-1,l=this.length;++o<l;){i.push(r=[]);for(var u=this[o],f=-1,c=u.length;++f<c;){if((s=u[f])&&(a=t.call(s,s.__data__,f,o))){if("__data__"in s)a.__data__=s.__data__;Rl(a,f,n,e,s[n][e]);r.push(a)}else{r.push(null)}}}return kl(i,n,e)};Ml.selectAll=function(t){var e=this.id,n=this.namespace,i=[],r,a,s,o,l;t=G(t);for(var u=-1,f=this.length;++u<f;){for(var c=this[u],h=-1,d=c.length;++h<d;){if(s=c[h]){l=s[n][e];a=t.call(s,s.__data__,h,u);i.push(r=[]);for(var p=-1,g=a.length;++p<g;){if(o=a[p])Rl(o,p,n,e,l);r.push(o)}}}}return kl(i,n,e)};Ml.filter=function(t){var e=[],n,i,r;if(typeof t!=="function")t=ut(t);for(var a=0,s=this.length;a<s;a++){e.push(n=[]);for(var i=this[a],o=0,l=i.length;o<l;o++){if((r=i[o])&&t.call(r,r.__data__,o,a)){n.push(r)}}}return kl(e,this.namespace,this.id)};Ml.tween=function(t,e){var n=this.id,i=this.namespace;if(arguments.length<2)return this.node()[i][n].tween.get(t);return ct(this,e==null?function(e){e[i][n].tween.remove(t)}:function(r){r[i][n].tween.set(t,e)})};function Pl(t,e,n,i){var r=t.id,a=t.namespace;return ct(t,typeof n==="function"?function(t,s,o){t[a][r].tween.set(e,i(n.call(t,t.__data__,s,o)))}:(n=i(n),function(t){t[a][r].tween.set(e,n)}))}Ml.attr=function(e,n){if(arguments.length<2){for(n in e)this.attr(n,e[n]);return this}var i=e=="transform"?es:Sa,r=t.ns.qualify(e);function a(){this.removeAttribute(r)}function s(){this.removeAttributeNS(r.space,r.local)}function o(t){return t==null?a:(t+="",function(){var e=this.getAttribute(r),n;return e!==t&&(n=i(e,t),function(t){this.setAttribute(r,n(t))})})}function l(t){return t==null?s:(t+="",function(){var e=this.getAttributeNS(r.space,r.local),n;return e!==t&&(n=i(e,t),function(t){this.setAttributeNS(r.space,r.local,n(t))})})}return Pl(this,"attr."+e,n,r.local?l:o)};Ml.attrTween=function(e,n){var i=t.ns.qualify(e);function r(t,e){var r=n.call(this,t,e,this.getAttribute(i));return r&&function(t){this.setAttribute(i,r(t))}}function a(t,e){var r=n.call(this,t,e,this.getAttributeNS(i.space,i.local));return r&&function(t){this.setAttributeNS(i.space,i.local,r(t))}}return this.tween("attr."+e,i.local?a:r)};Ml.style=function(t,e,n){var i=arguments.length;if(i<3){if(typeof t!=="string"){if(i<2)e="";for(n in t)this.style(n,t[n],e);return this}n=""}function r(){this.style.removeProperty(t)}function a(e){return e==null?r:(e+="",function(){var i=s(this).getComputedStyle(this,null).getPropertyValue(t),r;return i!==e&&(r=Sa(i,e),function(e){this.style.setProperty(t,r(e),n)})})}return Pl(this,"style."+t,e,a)};Ml.styleTween=function(t,e,n){if(arguments.length<3)n="";function i(i,r){var a=e.call(this,i,r,s(this).getComputedStyle(this,null).getPropertyValue(t));return a&&function(e){this.style.setProperty(t,a(e),n)}}return this.tween("style."+t,i)};Ml.text=function(t){return Pl(this,"text",t,El)};function El(t){if(t==null)t="";return function(){this.textContent=t}}Ml.remove=function(){var t=this.namespace;return this.each("end.transition",function(){var e;if(this[t].count<2&&(e=this.parentNode))e.removeChild(this)})};Ml.ease=function(e){var n=this.id,i=this.namespace;if(arguments.length<1)return this.node()[i][n].ease;if(typeof e!=="function")e=t.ease.apply(t,arguments);return ct(this,function(t){t[i][n].ease=e})};Ml.delay=function(t){var e=this.id,n=this.namespace;if(arguments.length<1)return this.node()[n][e].delay;return ct(this,typeof t==="function"?function(i,r,a){i[n][e].delay=+t.call(i,i.__data__,r,a)}:(t=+t,function(i){i[n][e].delay=t}))};Ml.duration=function(t){var e=this.id,n=this.namespace;if(arguments.length<1)return this.node()[n][e].duration;return ct(this,typeof t==="function"?function(i,r,a){i[n][e].duration=Math.max(1,t.call(i,i.__data__,r,a))}:(t=Math.max(1,t),function(i){i[n][e].duration=t}))};Ml.each=function(e,n){var i=this.id,r=this.namespace;if(arguments.length<2){var a=Nl,s=Al;try{Al=i;ct(this,function(t,n,a){Nl=t[r][i];e.call(t,t.__data__,n,a)})}finally{Nl=a;Al=s}}else{ct(this,function(a){var s=a[r][i];(s.event||(s.event=t.dispatch("start","end","interrupt"))).on(e,n)})}return this};Ml.transition=function(){var t=this.id,e=++Il,n=this.namespace,i=[],r,a,s,o;for(var l=0,u=this.length;l<u;l++){i.push(r=[]);for(var a=this[l],f=0,c=a.length;f<c;f++){if(s=a[f]){o=s[n][t];Rl(s,f,n,e,{time:o.time,ease:o.ease,delay:o.delay+o.duration,duration:o.duration})}r.push(s)}}return kl(i,n,e)};function Fl(t){return t==null?"__transition__":"__transition_"+t+"__"}function Rl(t,e,n,i,r){var a=t[n]||(t[n]={active:0,count:0}),s=a[i],o,l,u,f,c;function h(t){var e=s.delay;l.t=e+o;if(e<=t)return d(t-e);l.c=d}function d(n){var r=a.active,h=a[r];if(h){h.timer.c=null;h.timer.t=NaN;--a.count;delete a[r];h.event&&h.event.interrupt.call(t,t.__data__,h.index)}for(var d in a){if(+d<i){var g=a[d];g.timer.c=null;g.timer.t=NaN;--a.count;delete a[d]}}l.c=p;Pe(function(){if(l.c&&p(n||1)){l.c=null;l.t=NaN}return 1},0,o);a.active=i;s.event&&s.event.start.call(t,t.__data__,e);c=[];s.tween.forEach(function(n,i){if(i=i.call(t,t.__data__,e)){c.push(i)}});f=s.ease;u=s.duration}function p(r){var o=r/u,l=f(o),h=c.length;while(h>0){c[--h].call(t,l)}if(o>=1){s.event&&s.event.end.call(t,t.__data__,e);if(--a.count)delete a[i];else delete t[n];return 1}}if(!s){o=r.time;l=Pe(h,0,o);s=a[i]={tween:new _,time:o,timer:l,delay:r.delay,duration:r.duration,ease:r.ease,index:e};r=null;++a.count}}t.svg.axis=function(){var e=t.scale.linear(),n=Ll,r=6,a=6,s=3,o=[10],l=null,u;function f(i){i.each(function(){var i=t.select(this);var f=this.__chart__||e,c=this.__chart__=e.copy();var h=l==null?c.ticks?c.ticks.apply(c,o):c.domain():l,d=u==null?c.tickFormat?c.tickFormat.apply(c,o):P:u,p=i.selectAll(".tick").data(h,c),g=p.enter().insert("g",".domain").attr("class","tick").style("opacity",Ct),v=t.transition(p.exit()).style("opacity",Ct).remove(),m=t.transition(p.order()).style("opacity",1),y=Math.max(r,0)+s,b;var w=ro(c),_=i.selectAll(".domain").data([0]),x=(_.enter().append("path").attr("class","domain"),t.transition(_));g.append("line");g.append("text");var S=g.select("line"),D=m.select("line"),C=p.select("text").text(d),T=g.select("text"),k=m.select("text"),M=n==="top"||n==="left"?-1:1,I,A,N,E;if(n==="bottom"||n==="top"){b=Ol,I="x",N="y",A="x2",E="y2";C.attr("dy",M<0?"0em":".71em").style("text-anchor","middle");x.attr("d","M"+w[0]+","+M*a+"V0H"+w[1]+"V"+M*a)}else{b=Hl,I="y",N="x",A="y2",E="x2";C.attr("dy",".32em").style("text-anchor",M<0?"end":"start");x.attr("d","M"+M*a+","+w[0]+"H0V"+w[1]+"H"+M*a)}S.attr(E,M*r);T.attr(N,M*y);D.attr(A,0).attr(E,M*r);k.attr(I,0).attr(N,M*y);if(c.rangeBand){var F=c,R=F.rangeBand()/2;f=c=function(t){return F(t)+R}}else if(f.rangeBand){f=c}else{v.call(b,c,f)}g.call(b,f,c);m.call(b,c,c)})}f.scale=function(t){if(!arguments.length)return e;e=t;return f};f.orient=function(t){if(!arguments.length)return n;n=t in jl?t+"":Ll;return f};f.ticks=function(){if(!arguments.length)return o;o=i(arguments);return f};f.tickValues=function(t){if(!arguments.length)return l;l=t;return f};f.tickFormat=function(t){if(!arguments.length)return u;u=t;return f};f.tickSize=function(t){var e=arguments.length;if(!e)return r;r=+t;a=+arguments[e-1];return f};f.innerTickSize=function(t){if(!arguments.length)return r;r=+t;return f};f.outerTickSize=function(t){if(!arguments.length)return a;a=+t;return f};f.tickPadding=function(t){if(!arguments.length)return s;s=+t;return f};f.tickSubdivide=function(){return arguments.length&&f};return f};var Ll="bottom",jl={top:1,right:1,bottom:1,left:1};function Ol(t,e,n){t.attr("transform",function(t){var i=e(t);return"translate("+(isFinite(i)?i:n(t))+",0)"})}function Hl(t,e,n){t.attr("transform",function(t){var i=e(t);return"translate(0,"+(isFinite(i)?i:n(t))+")"})}t.svg.brush=function(){var e=z(h,"brushstart","brush","brushend"),n=null,i=null,r=[0,0],a=[0,0],o,l,u=true,f=true,c=zl[0];function h(e){e.each(function(){var e=t.select(this).style("pointer-events","all").style("-webkit-tap-highlight-color","rgba(0,0,0,0)").on("mousedown.brush",v).on("touchstart.brush",v);var r=e.selectAll(".background").data([0]);r.enter().append("rect").attr("class","background").style("visibility","hidden").style("cursor","crosshair");e.selectAll(".extent").data([0]).enter().append("rect").attr("class","extent").style("cursor","move");var a=e.selectAll(".resize").data(c,P);a.exit().remove();a.enter().append("g").attr("class",function(t){return"resize "+t}).style("cursor",function(t){return Wl[t]}).append("rect").attr("x",function(t){return/[ew]$/.test(t)?-3:null}).attr("y",function(t){return/^[ns]/.test(t)?-3:null}).attr("width",6).attr("height",6).style("visibility","hidden");a.style("display",h.empty()?"none":null);var s=t.transition(e),o=t.transition(r),l;if(n){l=ro(n);o.attr("x",l[0]).attr("width",l[1]-l[0]);p(s)}if(i){l=ro(i);o.attr("y",l[0]).attr("height",l[1]-l[0]);g(s)}d(s)})}h.event=function(n){n.each(function(){var n=e.of(this,arguments),i={x:r,y:a,i:o,j:l},s=this.__chart__||i;this.__chart__=i;if(Al){t.select(this).transition().each("start.brush",function(){o=s.i;l=s.j;r=s.x;a=s.y;n({type:"brushstart"})}).tween("brush:brush",function(){var t=Da(r,i.x),e=Da(a,i.y);o=l=null;return function(s){r=i.x=t(s);a=i.y=e(s);n({type:"brush",mode:"resize"})}}).each("end.brush",function(){o=i.i;l=i.j;n({type:"brush",mode:"resize"});n({type:"brushend"})})}else{n({type:"brushstart"});n({type:"brush",mode:"resize"});n({type:"brushend"})}})};function d(t){t.selectAll(".resize").attr("transform",function(t){return"translate("+r[+/e$/.test(t)]+","+a[+/^s/.test(t)]+")"})}function p(t){t.select(".extent").attr("x",r[0]);t.selectAll(".extent,.n>rect,.s>rect").attr("width",r[1]-r[0])}function g(t){t.select(".extent").attr("y",a[0]);t.selectAll(".extent,.e>rect,.w>rect").attr("height",a[1]-a[0])}function v(){var c=this,v=t.select(t.event.target),m=e.of(c,arguments),y=t.select(c),b=v.datum(),w=!/^(n|s)$/.test(b)&&n,_=!/^(e|w)$/.test(b)&&i,x=v.classed("extent"),S=_t(c),D,C=t.mouse(c),T;var k=t.select(s(c)).on("keydown.brush",A).on("keyup.brush",N);if(t.event.changedTouches){k.on("touchmove.brush",P).on("touchend.brush",F)}else{k.on("mousemove.brush",P).on("mouseup.brush",F)}y.interrupt().selectAll("*").interrupt();if(x){C[0]=r[0]-C[0];C[1]=a[0]-C[1]}else if(b){var M=+/w$/.test(b),I=+/^n/.test(b);T=[r[1-M]-C[0],a[1-I]-C[1]];C[0]=r[M];C[1]=a[I]}else if(t.event.altKey)D=C.slice();y.style("pointer-events","none").selectAll(".resize").style("display",null);t.select("body").style("cursor",v.style("cursor"));m({type:"brushstart"});P();function A(){if(t.event.keyCode==32){if(!x){D=null;C[0]-=r[1];C[1]-=a[1];x=2}H()}}function N(){if(t.event.keyCode==32&&x==2){C[0]+=r[1];C[1]+=a[1];x=0;H()}}function P(){var e=t.mouse(c),s=false;if(T){e[0]+=T[0];e[1]+=T[1]}if(!x){if(t.event.altKey){if(!D)D=[(r[0]+r[1])/2,(a[0]+a[1])/2];C[0]=r[+(e[0]<D[0])];C[1]=a[+(e[1]<D[1])]}else D=null}if(w&&E(e,n,0)){p(y);s=true}if(_&&E(e,i,1)){g(y);s=true}if(s){d(y);m({type:"brush",mode:x?"move":"resize"})}}function E(t,e,n){var i=ro(e),s=i[0],c=i[1],h=C[n],d=n?a:r,p=d[1]-d[0],g,v;if(x){s-=h;c-=p+h}g=(n?f:u)?Math.max(s,Math.min(c,t[n])):t[n];if(x){v=(g+=h)+p}else{if(D)h=Math.max(s,Math.min(c,2*D[n]-g));if(h<g){v=g;g=h}else{v=h}}if(d[0]!=g||d[1]!=v){if(n)l=null;else o=null;d[0]=g;d[1]=v;return true}}function F(){P();y.style("pointer-events","all").selectAll(".resize").style("display",h.empty()?"none":null);t.select("body").style("cursor",null);k.on("mousemove.brush",null).on("mouseup.brush",null).on("touchmove.brush",null).on("touchend.brush",null).on("keydown.brush",null).on("keyup.brush",null);S();m({type:"brushend"})}}h.x=function(t){if(!arguments.length)return n;n=t;c=zl[!n<<1|!i];return h};h.y=function(t){if(!arguments.length)return i;i=t;c=zl[!n<<1|!i];return h};h.clamp=function(t){if(!arguments.length)return n&&i?[u,f]:n?u:i?f:null;if(n&&i)u=!!t[0],f=!!t[1];else if(n)u=!!t;else if(i)f=!!t;return h};h.extent=function(t){var e,s,u,f,c;if(!arguments.length){if(n){if(o){e=o[0],s=o[1]}else{e=r[0],s=r[1];if(n.invert)e=n.invert(e),s=n.invert(s);if(s<e)c=e,e=s,s=c}}if(i){if(l){u=l[0],f=l[1]}else{u=a[0],f=a[1];if(i.invert)u=i.invert(u),f=i.invert(f);if(f<u)c=u,u=f,f=c}}return n&&i?[[e,u],[s,f]]:n?[e,s]:i&&[u,f]}if(n){e=t[0],s=t[1];if(i)e=e[0],s=s[0];o=[e,s];if(n.invert)e=n(e),s=n(s);if(s<e)c=e,e=s,s=c;if(e!=r[0]||s!=r[1])r=[e,s]}if(i){u=t[0],f=t[1];if(n)u=u[1],f=f[1];l=[u,f];if(i.invert)u=i(u),f=i(f);if(f<u)c=u,u=f,f=c;if(u!=a[0]||f!=a[1])a=[u,f]}return h};h.clear=function(){if(!h.empty()){r=[0,0],a=[0,0];o=l=null}return h};h.empty=function(){return!!n&&r[0]==r[1]||!!i&&a[0]==a[1]};return t.rebind(h,e,"on")};var Wl={n:"ns-resize",e:"ew-resize",s:"ns-resize",w:"ew-resize",nw:"nwse-resize",ne:"nesw-resize",se:"nwse-resize",sw:"nesw-resize"};var zl=[["n","e","s","w","nw","ne","se","sw"],["e","w"],["n","s"],[]];var ql=Be.format=wn.timeFormat;var Bl=ql.utc;var $l=Bl("%Y-%m-%dT%H:%M:%S.%LZ");ql.iso=Date.prototype.toISOString&&+new Date("2000-01-01T00:00:00.000Z")?Ul:$l;function Ul(t){return t.toISOString()}Ul.parse=function(t){var e=new Date(t);return isNaN(e)?null:e};Ul.toString=$l.toString;Be.second=Ve(function(t){return new $e(Math.floor(t/1e3)*1e3)},function(t,e){t.setTime(t.getTime()+Math.floor(e)*1e3)},function(t){return t.getSeconds()});Be.seconds=Be.second.range;Be.seconds.utc=Be.second.utc.range;Be.minute=Ve(function(t){return new $e(Math.floor(t/6e4)*6e4)},function(t,e){t.setTime(t.getTime()+Math.floor(e)*6e4)},function(t){return t.getMinutes()});Be.minutes=Be.minute.range;Be.minutes.utc=Be.minute.utc.range;Be.hour=Ve(function(t){var e=t.getTimezoneOffset()/60;return new $e((Math.floor(t/36e5-e)+e)*36e5)},function(t,e){t.setTime(t.getTime()+Math.floor(e)*36e5)},function(t){return t.getHours()});Be.hours=Be.hour.range;Be.hours.utc=Be.hour.utc.range;Be.month=Ve(function(t){t=Be.day(t);t.setDate(1);return t},function(t,e){t.setMonth(t.getMonth()+e)},function(t){return t.getMonth()});Be.months=Be.month.range;Be.months.utc=Be.month.utc.range;function Yl(e,n,i){function r(t){return e(t)}r.invert=function(t){return Vl(e.invert(t))};r.domain=function(t){if(!arguments.length)return e.domain().map(Vl);e.domain(t);return r};function a(e,i){var r=e[1]-e[0],a=r/i,s=t.bisect(Xl,a);return s==Xl.length?[n.year,po(e.map(function(t){return t/31536e6}),i)[2]]:!s?[Kl,po(e,i)[2]]:n[a/Xl[s-1]<Xl[s]/a?s-1:s]}
r.nice=function(t,e){var n=r.domain(),i=io(n),s=t==null?a(i,10):typeof t==="number"&&a(i,t);if(s)t=s[0],e=s[1];function o(n){return!isNaN(n)&&!t.range(n,Vl(+n+1),e).length}return r.domain(so(n,e>1?{floor:function(e){while(o(e=t.floor(e)))e=Vl(e-1);return e},ceil:function(e){while(o(e=t.ceil(e)))e=Vl(+e+1);return e}}:t))};r.ticks=function(t,e){var n=io(r.domain()),i=t==null?a(n,10):typeof t==="number"?a(n,t):!t.range&&[{range:t},e];if(i)t=i[0],e=i[1];return t.range(n[0],Vl(+n[1]+1),e<1?1:e)};r.tickFormat=function(){return i};r.copy=function(){return Yl(e.copy(),n,i)};return co(r,e)}function Vl(t){return new Date(t)}var Xl=[1e3,5e3,15e3,3e4,6e4,3e5,9e5,18e5,36e5,108e5,216e5,432e5,864e5,1728e5,6048e5,2592e6,7776e6,31536e6];var Jl=[[Be.second,1],[Be.second,5],[Be.second,15],[Be.second,30],[Be.minute,1],[Be.minute,5],[Be.minute,15],[Be.minute,30],[Be.hour,1],[Be.hour,3],[Be.hour,6],[Be.hour,12],[Be.day,1],[Be.day,2],[Be.week,1],[Be.month,1],[Be.month,3],[Be.year,1]];var Gl=ql.multi([[".%L",function(t){return t.getMilliseconds()}],[":%S",function(t){return t.getSeconds()}],["%I:%M",function(t){return t.getMinutes()}],["%I %p",function(t){return t.getHours()}],["%a %d",function(t){return t.getDay()&&t.getDate()!=1}],["%b %d",function(t){return t.getDate()!=1}],["%B",function(t){return t.getMonth()}],["%Y",ai]]);var Kl={range:function(e,n,i){return t.range(Math.ceil(e/i)*i,+n,i).map(Vl)},floor:P,ceil:P};Jl.year=Be.year;Be.scale=function(){return Yl(t.scale.linear(),Jl,Gl)};var Zl=Jl.map(function(t){return[t[0].utc,t[1]]});var Ql=Bl.multi([[".%L",function(t){return t.getUTCMilliseconds()}],[":%S",function(t){return t.getUTCSeconds()}],["%I:%M",function(t){return t.getUTCMinutes()}],["%I %p",function(t){return t.getUTCHours()}],["%a %d",function(t){return t.getUTCDay()&&t.getUTCDate()!=1}],["%b %d",function(t){return t.getUTCDate()!=1}],["%B",function(t){return t.getUTCMonth()}],["%Y",ai]]);Zl.year=Be.year.utc;Be.scale.utc=function(){return Yl(t.scale.linear(),Zl,Ql)};t.text=Se(function(t){return t.responseText});t.json=function(t,e){return De(t,"application/json",tu,e)};function tu(t){return JSON.parse(t.responseText)}t.html=function(t,e){return De(t,"text/html",eu,e)};function eu(t){var e=r.createRange();e.selectNode(r.body);return e.createContextualFragment(t.responseText)}t.xml=Se(function(t){return t.responseXML});if(typeof define==="function"&&define.amd)this.d3=t,define(t);else if(typeof e==="object"&&e.exports)e.exports=t;else this.d3=t}()},{}],15:[function(t,e,n){(function(i){if(typeof define==="function"&&define.amd){define(["jquery","datatables.net"],function(t){return i(t,window,document)})}else if(typeof n==="object"){e.exports=function(e,n){if(!e){e=window}if(!n||!n.fn.dataTable){n=t("datatables.net")(e,n).$}return i(n,e,e.document)}}else{i(jQuery,window,document)}})(function(t,e,n,i){"use strict";var r=t.fn.dataTable;var a=function(e,r){if(!(this instanceof a)){alert("Scroller warning: Scroller must be initialised with the 'new' keyword.");return}if(r===i){r={}}this.s={dt:t.fn.dataTable.Api(e).settings()[0],tableTop:0,tableBottom:0,redrawTop:0,redrawBottom:0,autoHeight:true,viewportRows:0,stateTO:null,drawTO:null,heights:{jump:null,page:null,virtual:null,scroll:null,row:null,viewport:null},topRowFloat:0,scrollDrawDiff:null,loaderVisible:false};this.s=t.extend(this.s,a.oDefaults,r);this.s.heights.row=this.s.rowHeight;this.dom={force:n.createElement("div"),scroller:null,table:null,loader:null};if(this.s.dt.oScroller){return}this.s.dt.oScroller=this;this._fnConstruct()};t.extend(a.prototype,{fnRowToPixels:function(t,e,n){var r;if(n){r=this._domain("virtualToPhysical",t*this.s.heights.row)}else{var a=t-this.s.baseRowTop;r=this.s.baseScrollTop+a*this.s.heights.row}return e||e===i?parseInt(r,10):r},fnPixelsToRow:function(t,e,n){var r=t-this.s.baseScrollTop;var a=n?this._domain("physicalToVirtual",t)/this.s.heights.row:r/this.s.heights.row+this.s.baseRowTop;return e||e===i?parseInt(a,10):a},fnScrollToRow:function(e,n){var i=this;var r=false;var a=this.fnRowToPixels(e);var s=(this.s.displayBuffer-1)/2*this.s.viewportRows;var o=e-s;if(o<0){o=0}if((a>this.s.redrawBottom||a<this.s.redrawTop)&&this.s.dt._iDisplayStart!==o){r=true;a=this.fnRowToPixels(e,false,true)}if(typeof n=="undefined"||n){this.s.ani=r;t(this.dom.scroller).animate({scrollTop:a},function(){setTimeout(function(){i.s.ani=false},25)})}else{t(this.dom.scroller).scrollTop(a)}},fnMeasure:function(e){if(this.s.autoHeight){this._fnCalcRowHeight()}var n=this.s.heights;if(n.row){n.viewport=t(this.dom.scroller).height();this.s.viewportRows=parseInt(n.viewport/n.row,10)+1;this.s.dt._iDisplayLength=this.s.viewportRows*this.s.displayBuffer}if(e===i||e){this.s.dt.oInstance.fnDraw(false)}},fnPageInfo:function(){var t=this.s.dt,e=this.dom.scroller.scrollTop,n=t.fnRecordsDisplay(),i=Math.ceil(this.fnPixelsToRow(e+this.s.heights.viewport,false,this.s.ani));return{start:Math.floor(this.fnPixelsToRow(e,false,this.s.ani)),end:n<i?n-1:i-1}},_fnConstruct:function(){var n=this;if(!this.s.dt.oFeatures.bPaginate){this.s.dt.oApi._fnLog(this.s.dt,0,"Pagination must be enabled for Scroller");return}this.dom.force.style.position="relative";this.dom.force.style.top="0px";this.dom.force.style.left="0px";this.dom.force.style.width="1px";this.dom.scroller=t("div."+this.s.dt.oClasses.sScrollBody,this.s.dt.nTableWrapper)[0];this.dom.scroller.appendChild(this.dom.force);this.dom.scroller.style.position="relative";this.dom.table=t(">table",this.dom.scroller)[0];this.dom.table.style.position="absolute";this.dom.table.style.top="0px";this.dom.table.style.left="0px";t(this.s.dt.nTableWrapper).addClass("DTS");if(this.s.loadingIndicator){this.dom.loader=t('<div class="dataTables_processing DTS_Loading">'+this.s.dt.oLanguage.sLoadingRecords+"</div>").css("display","none");t(this.dom.scroller.parentNode).css("position","relative").append(this.dom.loader)}if(this.s.heights.row&&this.s.heights.row!="auto"){this.s.autoHeight=false}this.fnMeasure(false);this.s.ingnoreScroll=true;this.s.stateSaveThrottle=this.s.dt.oApi._fnThrottle(function(){n.s.dt.oApi._fnSaveState(n.s.dt)},500);t(this.dom.scroller).on("scroll.DTS",function(t){n._fnScroll.call(n)});t(this.dom.scroller).on("touchstart.DTS",function(){n._fnScroll.call(n)});this.s.dt.aoDrawCallback.push({fn:function(){if(n.s.dt.bInitialised){n._fnDrawCallback.call(n)}},sName:"Scroller"});t(e).on("resize.DTS",function(){n.fnMeasure(false);n._fnInfo()});var i=true;this.s.dt.oApi._fnCallbackReg(this.s.dt,"aoStateSaveParams",function(t,e){if(i&&n.s.dt.oLoadedState){e.iScroller=n.s.dt.oLoadedState.iScroller;e.iScrollerTopRow=n.s.dt.oLoadedState.iScrollerTopRow;i=false}else{e.iScroller=n.dom.scroller.scrollTop;e.iScrollerTopRow=n.s.topRowFloat}},"Scroller_State");if(this.s.dt.oLoadedState){this.s.topRowFloat=this.s.dt.oLoadedState.iScrollerTopRow||0}t(this.s.dt.nTable).one("init.dt",function(){n.fnMeasure()});this.s.dt.aoDestroyCallback.push({sName:"Scroller",fn:function(){t(e).off("resize.DTS");t(n.dom.scroller).off("touchstart.DTS scroll.DTS");t(n.s.dt.nTableWrapper).removeClass("DTS");t("div.DTS_Loading",n.dom.scroller.parentNode).remove();t(n.s.dt.nTable).off("init.dt");n.dom.table.style.position="";n.dom.table.style.top="";n.dom.table.style.left=""}})},_fnScroll:function(){var e=this,n=this.s.heights,i=this.dom.scroller.scrollTop,r;if(this.s.skip){return}if(this.s.ingnoreScroll){return}if(this.s.dt.bFiltered||this.s.dt.bSorted){this.s.lastScrollTop=0;return}this._fnInfo();clearTimeout(this.s.stateTO);this.s.stateTO=setTimeout(function(){e.s.dt.oApi._fnSaveState(e.s.dt)},250);if(i<this.s.redrawTop||i>this.s.redrawBottom){var a=Math.ceil((this.s.displayBuffer-1)/2*this.s.viewportRows);if(Math.abs(i-this.s.lastScrollTop)>n.viewport||this.s.ani){r=parseInt(this._domain("physicalToVirtual",i)/n.row,10)-a;this.s.topRowFloat=this._domain("physicalToVirtual",i)/n.row}else{r=this.fnPixelsToRow(i)-a;this.s.topRowFloat=this.fnPixelsToRow(i,false)}if(r<=0){r=0}else if(r+this.s.dt._iDisplayLength>this.s.dt.fnRecordsDisplay()){r=this.s.dt.fnRecordsDisplay()-this.s.dt._iDisplayLength;if(r<0){r=0}}else if(r%2!==0){r++}if(r!=this.s.dt._iDisplayStart){this.s.tableTop=t(this.s.dt.nTable).offset().top;this.s.tableBottom=t(this.s.dt.nTable).height()+this.s.tableTop;var s=function(){if(e.s.scrollDrawReq===null){e.s.scrollDrawReq=i}e.s.dt._iDisplayStart=r;e.s.dt.oApi._fnDraw(e.s.dt)};if(this.s.dt.oFeatures.bServerSide){clearTimeout(this.s.drawTO);this.s.drawTO=setTimeout(s,this.s.serverWait)}else{s()}if(this.dom.loader&&!this.s.loaderVisible){this.dom.loader.css("display","block");this.s.loaderVisible=true}}}else{this.s.topRowFloat=this._domain("physicalToVirtual",i)/n.row}this.s.lastScrollTop=i;this.s.stateSaveThrottle()},_domain:function(t,e){var n=this.s.heights;var i;if(n.virtual===n.scroll){return e}var r=(n.scroll-n.viewport)/2;var a=(n.virtual-n.viewport)/2;i=a/(r*r);if(t==="virtualToPhysical"){if(e<a){return Math.pow(e/i,.5)}else{e=a*2-e;return e<0?n.scroll:r*2-Math.pow(e/i,.5)}}else if(t==="physicalToVirtual"){if(e<r){return e*e*i}else{e=r*2-e;return e<0?n.virtual:a*2-e*e*i}}},_fnDrawCallback:function(){var e=this,n=this.s.heights,i=this.dom.scroller.scrollTop,r=i,a=i+n.viewport,s=t(this.s.dt.nTable).height(),o=this.s.dt._iDisplayStart,l=this.s.dt._iDisplayLength,u=this.s.dt.fnRecordsDisplay();this.s.skip=true;this._fnScrollForce();if(o===0){i=this.s.topRowFloat*n.row}else if(o+l>=u){i=n.scroll-(u-this.s.topRowFloat)*n.row}else{i=this._domain("virtualToPhysical",this.s.topRowFloat*n.row)}this.dom.scroller.scrollTop=i;this.s.baseScrollTop=i;this.s.baseRowTop=this.s.topRowFloat;var f=i-(this.s.topRowFloat-o)*n.row;if(o===0){f=0}else if(o+l>=u){f=n.scroll-s}this.dom.table.style.top=f+"px";this.s.tableTop=f;this.s.tableBottom=s+this.s.tableTop;var c=(i-this.s.tableTop)*this.s.boundaryScale;this.s.redrawTop=i-c;this.s.redrawBottom=i+c;this.s.skip=false;if(this.s.dt.oFeatures.bStateSave&&this.s.dt.oLoadedState!==null&&typeof this.s.dt.oLoadedState.iScroller!="undefined"){var h=(this.s.dt.sAjaxSource||e.s.dt.ajax)&&!this.s.dt.oFeatures.bServerSide?true:false;if(h&&this.s.dt.iDraw==2||!h&&this.s.dt.iDraw==1){setTimeout(function(){t(e.dom.scroller).scrollTop(e.s.dt.oLoadedState.iScroller);e.s.redrawTop=e.s.dt.oLoadedState.iScroller-n.viewport/2;setTimeout(function(){e.s.ingnoreScroll=false},0)},0)}}else{e.s.ingnoreScroll=false}if(this.s.dt.oFeatures.bInfo){setTimeout(function(){e._fnInfo.call(e)},0)}if(this.dom.loader&&this.s.loaderVisible){this.dom.loader.css("display","none");this.s.loaderVisible=false}},_fnScrollForce:function(){var t=this.s.heights;var e=1e6;t.virtual=t.row*this.s.dt.fnRecordsDisplay();t.scroll=t.virtual;if(t.scroll>e){t.scroll=e}this.dom.force.style.height=t.scroll>this.s.heights.row?t.scroll+"px":this.s.heights.row+"px"},_fnCalcRowHeight:function(){var e=this.s.dt;var n=e.nTable;var i=n.cloneNode(false);var r=t("<tbody/>").appendTo(i);var a=t('<div class="'+e.oClasses.sWrapper+' DTS">'+'<div class="'+e.oClasses.sScrollWrapper+'">'+'<div class="'+e.oClasses.sScrollBody+'"></div>'+"</div>"+"</div>");t("tbody tr:lt(4)",n).clone().appendTo(r);while(t("tr",r).length<3){r.append("<tr><td>&nbsp;</td></tr>")}t("div."+e.oClasses.sScrollBody,a).append(i);var s=this.s.dt.nHolding||n.parentNode;if(!t(s).is(":visible")){s="body"}a.appendTo(s);this.s.heights.row=t("tr",r).eq(1).outerHeight();a.remove()},_fnInfo:function(){if(!this.s.dt.oFeatures.bInfo){return}var e=this.s.dt,n=e.oLanguage,i=this.dom.scroller.scrollTop,r=Math.floor(this.fnPixelsToRow(i,false,this.s.ani)+1),a=e.fnRecordsTotal(),s=e.fnRecordsDisplay(),o=Math.ceil(this.fnPixelsToRow(i+this.s.heights.viewport,false,this.s.ani)),l=s<o?s:o,u=e.fnFormatNumber(r),f=e.fnFormatNumber(l),c=e.fnFormatNumber(a),h=e.fnFormatNumber(s),d;if(e.fnRecordsDisplay()===0&&e.fnRecordsDisplay()==e.fnRecordsTotal()){d=n.sInfoEmpty+n.sInfoPostFix}else if(e.fnRecordsDisplay()===0){d=n.sInfoEmpty+" "+n.sInfoFiltered.replace("_MAX_",c)+n.sInfoPostFix}else if(e.fnRecordsDisplay()==e.fnRecordsTotal()){d=n.sInfo.replace("_START_",u).replace("_END_",f).replace("_MAX_",c).replace("_TOTAL_",h)+n.sInfoPostFix}else{d=n.sInfo.replace("_START_",u).replace("_END_",f).replace("_MAX_",c).replace("_TOTAL_",h)+" "+n.sInfoFiltered.replace("_MAX_",e.fnFormatNumber(e.fnRecordsTotal()))+n.sInfoPostFix}var p=n.fnInfoCallback;if(p){d=p.call(e.oInstance,e,r,l,a,s,d)}var g=e.aanFeatures.i;if(typeof g!="undefined"){for(var v=0,m=g.length;v<m;v++){t(g[v]).html(d)}}t(e.nTable).triggerHandler("info.dt")}});a.defaults={trace:false,rowHeight:"auto",serverWait:200,displayBuffer:9,boundaryScale:.5,loadingIndicator:false};a.oDefaults=a.defaults;a.version="1.4.2";if(typeof t.fn.dataTable=="function"&&typeof t.fn.dataTableExt.fnVersionCheck=="function"&&t.fn.dataTableExt.fnVersionCheck("1.10.0")){t.fn.dataTableExt.aoFeatures.push({fnInit:function(t){var e=t.oInit;var n=e.scroller||e.oScroller||{};new a(t,n)},cFeature:"S",sFeature:"Scroller"})}else{alert("Warning: Scroller requires DataTables 1.10.0 or greater - www.datatables.net/download")}t(n).on("preInit.dt.dtscroller",function(e,n){if(e.namespace!=="dt"){return}var i=n.oInit.scroller;var s=r.defaults.scroller;if(i||s){var o=t.extend({},i,s);if(i!==false){new a(n,o)}}});t.fn.dataTable.Scroller=a;t.fn.DataTable.Scroller=a;var s=t.fn.dataTable.Api;s.register("scroller()",function(){return this});s.register("scroller().rowToPixels()",function(t,e,n){var i=this.context;if(i.length&&i[0].oScroller){return i[0].oScroller.fnRowToPixels(t,e,n)}});s.register("scroller().pixelsToRow()",function(t,e,n){var i=this.context;if(i.length&&i[0].oScroller){return i[0].oScroller.fnPixelsToRow(t,e,n)}});s.register("scroller().scrollToRow()",function(t,e){this.iterator("table",function(n){if(n.oScroller){n.oScroller.fnScrollToRow(t,e)}});return this});s.register("row().scrollTo()",function(t){var e=this;this.iterator("row",function(n,i){if(n.oScroller){var r=e.rows({order:"applied",search:"applied"}).indexes().indexOf(i);n.oScroller.fnScrollToRow(r,t)}});return this});s.register("scroller.measure()",function(t){this.iterator("table",function(e){if(e.oScroller){e.oScroller.fnMeasure(t)}});return this});s.register("scroller.page()",function(){var t=this.context;if(t.length&&t[0].oScroller){return t[0].oScroller.fnPageInfo()}});return a})},{"datatables.net":19}],16:[function(t,e,n){(function(i){if(typeof define==="function"&&define.amd){define(["jquery","datatables.net"],function(t){return i(t,window,document)})}else if(typeof n==="object"){e.exports=function(e,n){if(!e){e=window}if(!n||!n.fn.dataTable){n=t("datatables.net")(e,n).$}return i(n,e,e.document)}}else{i(jQuery,window,document)}})(function(t,e,n,i){"use strict";var r=t.fn.dataTable;r.select={};r.select.version="1.1.2";r.select.init=function(e){var n=e.settings()[0];var a=n.oInit.select;var s=r.defaults.select;var o=a===i?s:a;var l="row";var u="api";var f=false;var c=true;var h="td, th";var d="selected";n._select={};if(o===true){u="os"}else if(typeof o==="string"){u=o}else if(t.isPlainObject(o)){if(o.blurable!==i){f=o.blurable}if(o.info!==i){c=o.info}if(o.items!==i){l=o.items}if(o.style!==i){u=o.style}if(o.selector!==i){h=o.selector}if(o.className!==i){d=o.className}}e.select.selector(h);e.select.items(l);e.select.style(u);e.select.blurable(f);e.select.info(c);n._select.className=d;t.fn.dataTable.ext.order["select-checkbox"]=function(e,n){return this.api().column(n,{order:"index"}).nodes().map(function(n){if(e._select.items==="row"){return t(n).parent().hasClass(e._select.className)}else if(e._select.items==="cell"){return t(n).hasClass(e._select.className)}return false})};if(t(e.table().node()).hasClass("selectable")){e.select.style("os")}};function a(t,e,n){var i;var r;var a;var s=function(e,n){if(e>n){var i=n;n=e;e=i}var r=false;return t.columns(":visible").indexes().filter(function(t){if(t===e){r=true}if(t===n){r=false;return true}return r})};var o=function(e,n){var i=t.rows({search:"applied"}).indexes();if(i.indexOf(e)>i.indexOf(n)){var r=n;n=e;e=r}var a=false;return i.filter(function(t){if(t===e){a=true}if(t===n){a=false;return true}return a})};if(!t.cells({selected:true}).any()&&!n){r=s(0,e.column);a=o(0,e.row)}else{r=s(n.column,e.column);a=o(n.row,e.row)}i=t.cells(a,r).flatten();if(!t.cells(e,{selected:true}).any()){t.cells(i).select()}else{t.cells(i).deselect()}}function s(e){var n=e.settings()[0];var i=n._select.selector;t(e.table().body()).off("mousedown.dtSelect",i).off("mouseup.dtSelect",i).off("click.dtSelect",i);t("body").off("click.dtSelect")}function o(n){var i=t(n.table().body());var r=n.settings()[0];var a=r._select.selector;i.on("mousedown.dtSelect",a,function(t){if(t.shiftKey){i.css("-moz-user-select","none").one("selectstart.dtSelect",a,function(){return false})}}).on("mouseup.dtSelect",a,function(t){i.css("-moz-user-select","")}).on("click.dtSelect",a,function(i){var r=n.select.items();var a;if(e.getSelection&&e.getSelection().toString()){return}var s=n.settings()[0];if(t(i.target).closest("div.dataTables_wrapper")[0]!=n.table().container()){return}var o=t(i.target).closest("td, th");var l=n.cell(o).index();if(!n.cell(o).any()){return}if(r==="row"){a=l.row;d(i,n,s,"row",a)}else if(r==="column"){a=n.cell(o).index().column;d(i,n,s,"column",a)}else if(r==="cell"){a=n.cell(o).index();d(i,n,s,"cell",a)}s._select_lastCell=l});t("body").on("click.dtSelect",function(e){if(r._select.blurable){if(t(e.target).parents().filter(n.table().container()).length){return}if(t(e.target).parents("div.DTE").length){return}h(r,true)}})}function l(e,n,i,r){if(r&&!e.flatten().length){return}i.unshift(e);t(e.table().node()).triggerHandler(n+".dt",i)}function u(e){var n=e.settings()[0];if(!n._select.info||!n.aanFeatures.i){return}var i=t('<span class="select-info"/>');var r=function(n,r){i.append(t('<span class="select-item"/>').append(e.i18n("select."+n+"s",{_:"%d "+n+"s selected",0:"",1:"1 "+n+" selected"},r)))};r("row",e.rows({selected:true}).flatten().length);r("column",e.columns({selected:true}).flatten().length);r("cell",e.cells({selected:true}).flatten().length);t.each(n.aanFeatures.i,function(e,n){n=t(n);var r=n.children("span.select-info");if(r.length){r.remove()}if(i.text()!==""){n.append(i)}})}function f(e){var n=new r.Api(e);e.aoRowCreatedCallback.push({fn:function(n,i,r){var a,s;var o=e.aoData[r];if(o._select_selected){t(n).addClass(e._select.className)}for(a=0,s=e.aoColumns.length;a<s;a++){if(e.aoColumns[a]._select_selected||o._selected_cells&&o._selected_cells[a]){t(o.anCells[a]).addClass(e._select.className)}}},sName:"select-deferRender"});n.on("preXhr.dt.dtSelect",function(){var t=n.rows({selected:true}).ids(true).filter(function(t){return t!==i});var e=n.cells({selected:true}).eq(0).map(function(t){var e=n.row(t.row).id(true);return e?{row:e,column:t.column}:i}).filter(function(t){return t!==i});n.one("draw.dt.dtSelect",function(){n.rows(t).select();if(e.any()){e.each(function(t){n.cells(t.row,t.column).select()})}})});n.on("draw.dtSelect.dt select.dtSelect.dt deselect.dtSelect.dt",function(){u(n)});n.on("destroy.dtSelect",function(){s(n);n.off(".dtSelect")})}function c(e,n,i,r){var a=e[n+"s"]({search:"applied"}).indexes();var s=t.inArray(r,a);var o=t.inArray(i,a);if(!e[n+"s"]({selected:true}).any()&&s===-1){a.splice(t.inArray(i,a)+1,a.length)}else{if(s>o){var l=o;o=s;s=l}a.splice(o+1,a.length);a.splice(0,s)}if(!e[n](i,{selected:true}).any()){e[n+"s"](a).select()}else{a.splice(t.inArray(i,a),1);e[n+"s"](a).deselect()}}function h(t,e){if(e||t._select.style==="single"){var n=new r.Api(t);n.rows({selected:true}).deselect();n.columns({selected:true}).deselect();n.cells({selected:true}).deselect()}}function d(t,e,n,i,r){var s=e.select.style();var o=e[i](r,{selected:true}).any();if(s==="os"){if(t.ctrlKey||t.metaKey){e[i](r).select(!o)}else if(t.shiftKey){if(i==="cell"){a(e,r,n._select_lastCell||null)}else{c(e,i,r,n._select_lastCell?n._select_lastCell[i]:null)}}else{var l=e[i+"s"]({selected:true});if(o&&l.flatten().length===1){e[i](r).deselect()}else{l.deselect();e[i](r).select()}}}else{e[i](r).select(!o)}}t.each([{type:"row",prop:"aoData"},{type:"column",prop:"aoColumns"}],function(t,e){r.ext.selector[e.type].push(function(t,n,r){var a=n.selected;var s;var o=[];if(a===i){return r}for(var l=0,u=r.length;l<u;l++){s=t[e.prop][r[l]];if(a===true&&s._select_selected===true||a===false&&!s._select_selected){o.push(r[l])}}return o})});r.ext.selector.cell.push(function(t,e,n){var r=e.selected;var a;var s=[];if(r===i){return n}for(var o=0,l=n.length;o<l;o++){a=t.aoData[n[o].row];if(r===true&&a._selected_cells&&a._selected_cells[n[o].column]===true||r===false&&(!a._selected_cells||!a._selected_cells[n[o].column])){s.push(n[o])}}return s});var p=r.Api.register;var g=r.Api.registerPlural;p("select()",function(){return this.iterator("table",function(t){r.select.init(new r.Api(t))})});p("select.blurable()",function(t){if(t===i){return this.context[0]._select.blurable}return this.iterator("table",function(e){e._select.blurable=t})});p("select.info()",function(t){if(u===i){return this.context[0]._select.info}return this.iterator("table",function(e){e._select.info=t})});p("select.items()",function(t){if(t===i){return this.context[0]._select.items}return this.iterator("table",function(e){e._select.items=t;l(new r.Api(e),"selectItems",[t])})});p("select.style()",function(t){if(t===i){return this.context[0]._select.style}return this.iterator("table",function(e){e._select.style=t;if(!e._select_init){f(e)}var n=new r.Api(e);s(n);if(t!=="api"){o(n)}l(new r.Api(e),"selectStyle",[t])})});p("select.selector()",function(t){if(t===i){return this.context[0]._select.selector}return this.iterator("table",function(e){s(new r.Api(e));e._select.selector=t;if(e._select.style!=="api"){o(new r.Api(e))}})});g("rows().select()","row().select()",function(e){var n=this;if(e===false){return this.deselect()}this.iterator("row",function(e,n){h(e);e.aoData[n]._select_selected=true;t(e.aoData[n].nTr).addClass(e._select.className)});this.iterator("table",function(t,e){l(n,"select",["row",n[e]],true)});return this});g("columns().select()","column().select()",function(e){var n=this;if(e===false){return this.deselect()}this.iterator("column",function(e,n){h(e);e.aoColumns[n]._select_selected=true;var i=new r.Api(e).column(n);t(i.header()).addClass(e._select.className);t(i.footer()).addClass(e._select.className);i.nodes().to$().addClass(e._select.className)});this.iterator("table",function(t,e){l(n,"select",["column",n[e]],true)});return this});g("cells().select()","cell().select()",function(e){var n=this;if(e===false){return this.deselect()}this.iterator("cell",function(e,n,r){h(e);var a=e.aoData[n];if(a._selected_cells===i){a._selected_cells=[]}a._selected_cells[r]=true;if(a.anCells){t(a.anCells[r]).addClass(e._select.className)}});this.iterator("table",function(t,e){l(n,"select",["cell",n[e]],true)});return this});g("rows().deselect()","row().deselect()",function(){var e=this;this.iterator("row",function(e,n){e.aoData[n]._select_selected=false;t(e.aoData[n].nTr).removeClass(e._select.className)});this.iterator("table",function(t,n){l(e,"deselect",["row",e[n]],true)});return this});g("columns().deselect()","column().deselect()",function(){var e=this;this.iterator("column",function(e,n){e.aoColumns[n]._select_selected=false;var i=new r.Api(e);var a=i.column(n);t(a.header()).removeClass(e._select.className);t(a.footer()).removeClass(e._select.className);i.cells(null,n).indexes().each(function(n){var i=e.aoData[n.row];var r=i._selected_cells;if(i.anCells&&(!r||!r[n.column])){t(i.anCells[n.column]).removeClass(e._select.className)}})});this.iterator("table",function(t,n){l(e,"deselect",["column",e[n]],true)});return this});g("cells().deselect()","cell().deselect()",function(){var e=this;this.iterator("cell",function(e,n,i){var r=e.aoData[n];r._selected_cells[i]=false;if(r.anCells&&!e.aoColumns[i]._select_selected){t(r.anCells[i]).removeClass(e._select.className)}});this.iterator("table",function(t,n){l(e,"deselect",["cell",e[n]],true)});return this});function v(t,e){return function(n){return n.i18n("buttons."+t,e)}}t.extend(r.ext.buttons,{selected:{text:v("selected","Selected"),className:"buttons-selected",init:function(t,e,n){var i=this;t.on("draw.dt.DT select.dt.DT deselect.dt.DT",function(){var t=i.rows({selected:true}).any()||i.columns({selected:true}).any()||i.cells({selected:true}).any();i.enable(t)});this.disable()}},selectedSingle:{text:v("selectedSingle","Selected single"),className:"buttons-selected-single",init:function(t,e,n){var i=this;t.on("draw.dt.DT select.dt.DT deselect.dt.DT",function(){var e=t.rows({selected:true}).flatten().length+t.columns({selected:true}).flatten().length+t.cells({selected:true}).flatten().length;i.enable(e===1)});this.disable()}},selectAll:{text:v("selectAll","Select all"),className:"buttons-select-all",action:function(){var t=this.select.items();this[t+"s"]().select()}},selectNone:{text:v("selectNone","Deselect all"),className:"buttons-select-none",action:function(){h(this.settings()[0],true)},init:function(t,e,n){var i=this;t.on("draw.dt.DT select.dt.DT deselect.dt.DT",function(){var e=t.rows({selected:true}).flatten().length+t.columns({selected:true}).flatten().length+t.cells({selected:true}).flatten().length;i.enable(e>0)});this.disable()}}});t.each(["Row","Column","Cell"],function(t,e){var n=e.toLowerCase();r.ext.buttons["select"+e+"s"]={text:v("select"+e+"s","Select "+n+"s"),className:"buttons-select-"+n+"s",action:function(){this.select.items(n)},init:function(t,e,i){var r=this;t.on("selectItems.dt.DT",function(t,e,i){r.active(i===n)})}}});t(n).on("preInit.dt.dtSelect",function(t,e){if(t.namespace!=="dt"){return}r.select.init(new r.Api(e))});return r.select})},{"datatables.net":17}],17:[function(t,e,n){(function(i){"use strict";if(typeof define==="function"&&define.amd){define(["jquery"],function(t){return i(t,window,document)})}else if(typeof n==="object"){e.exports=function(e,n){if(!e){e=window}if(!n){n=typeof window!=="undefined"?t("jquery"):t("jquery")(e)}return i(n,e,e.document)}}else{i(jQuery,window,document)}})(function(t,e,n,i){"use strict";var r;var a;var s;var o;var l;var u={};var f=/[\r\n]/g;var c=/<.*?>/g;var h=/^[\w\+\-]/;var d=/[\w\+\-]$/;var p=new RegExp("(\\"+["/",".","*","+","?","|","(",")","[","]","{","}","\\","$","^","-"].join("|\\")+")","g");var g=/[',$£€¥%\u2009\u202F\u20BD\u20a9\u20BArfk]/gi;var v=function(t){return!t||t===true||t==="-"?true:false};var m=function(t){var e=parseInt(t,10);return!isNaN(e)&&isFinite(t)?e:null};var y=function(t,e){if(!u[e]){u[e]=new RegExp(kt(e),"g")}return typeof t==="string"&&e!=="."?t.replace(/\./g,"").replace(u[e],"."):t};var b=function(t,e,n){var i=typeof t==="string";if(v(t)){return true}if(e&&i){t=y(t,e)}if(n&&i){t=t.replace(g,"")}return!isNaN(parseFloat(t))&&isFinite(t)};var w=function(t){return v(t)||typeof t==="string"};var _=function(t,e,n){if(v(t)){return true}var i=w(t);return!i?null:b(T(t),e,n)?true:null};var x=function(t,e,n){var r=[];var a=0,s=t.length;if(n!==i){for(;a<s;a++){if(t[a]&&t[a][e]){r.push(t[a][e][n])}}}else{for(;a<s;a++){if(t[a]){r.push(t[a][e])}}}return r};var S=function(t,e,n,r){var a=[];var s=0,o=e.length;if(r!==i){for(;s<o;s++){if(t[e[s]][n]){a.push(t[e[s]][n][r])}}}else{for(;s<o;s++){a.push(t[e[s]][n])}}return a};var D=function(t,e){var n=[];var r;if(e===i){e=0;r=t}else{r=e;e=t}for(var a=e;a<r;a++){n.push(a)}return n};var C=function(t){var e=[];for(var n=0,i=t.length;n<i;n++){if(t[n]){e.push(t[n])}}return e};var T=function(t){return t.replace(c,"")};var k=function(t){var e=[],n,i,r=t.length,a,s=0;t:for(i=0;i<r;i++){n=t[i];for(a=0;a<s;a++){if(e[a]===n){continue t}}e.push(n);s++}return e};function M(e){var n="a aa ai ao as b fn i m o s ",i,r,a={};t.each(e,function(t,s){i=t.match(/^([^A-Z]+?)([A-Z])/);if(i&&n.indexOf(i[1]+" ")!==-1){r=t.replace(i[0],i[2].toLowerCase());a[r]=t;if(i[1]==="o"){M(e[t])}}});e._hungarianMap=a}function I(e,n,r){if(!e._hungarianMap){M(e)}var a;t.each(n,function(s,o){a=e._hungarianMap[s];if(a!==i&&(r||n[a]===i)){if(a.charAt(0)==="o"){if(!n[a]){n[a]={}}t.extend(true,n[a],n[s]);I(e[a],n[a],r)}else{n[a]=n[s]}}})}function A(t){var e=r.defaults.oLanguage;var n=t.sZeroRecords;if(!t.sEmptyTable&&n&&e.sEmptyTable==="No data available in table"){ce(t,t,"sZeroRecords","sEmptyTable")}if(!t.sLoadingRecords&&n&&e.sLoadingRecords==="Loading..."){ce(t,t,"sZeroRecords","sLoadingRecords")}if(t.sInfoThousands){t.sThousands=t.sInfoThousands}var i=t.sDecimal;if(i){$e(i)}}var N=function(t,e,n){if(t[e]!==i){t[n]=t[e]}};function P(t){N(t,"ordering","bSort");N(t,"orderMulti","bSortMulti");N(t,"orderClasses","bSortClasses");N(t,"orderCellsTop","bSortCellsTop");N(t,"order","aaSorting");N(t,"orderFixed","aaSortingFixed");N(t,"paging","bPaginate");N(t,"pagingType","sPaginationType");N(t,"pageLength","iDisplayLength");N(t,"searching","bFilter");if(typeof t.sScrollX==="boolean"){t.sScrollX=t.sScrollX?"100%":""}if(typeof t.scrollX==="boolean"){t.scrollX=t.scrollX?"100%":""}var e=t.aoSearchCols;if(e){for(var n=0,i=e.length;n<i;n++){if(e[n]){I(r.models.oSearch,e[n])}}}}function E(e){N(e,"orderable","bSortable");N(e,"orderData","aDataSort");N(e,"orderSequence","asSorting");N(e,"orderDataType","sortDataType");var n=e.aDataSort;if(n&&!t.isArray(n)){e.aDataSort=[n]}}function F(e){if(!r.__browser){var n={};r.__browser=n;var i=t("<div/>").css({position:"fixed",top:0,left:0,height:1,width:1,overflow:"hidden"}).append(t("<div/>").css({position:"absolute",top:1,left:1,width:100,overflow:"scroll"}).append(t("<div/>").css({width:"100%",height:10}))).appendTo("body");var a=i.children();var s=a.children();n.barWidth=a[0].offsetWidth-a[0].clientWidth;n.bScrollOversize=s[0].offsetWidth===100&&a[0].clientWidth!==100;n.bScrollbarLeft=Math.round(s.offset().left)!==1;n.bBounding=i[0].getBoundingClientRect().width?true:false;i.remove()}t.extend(e.oBrowser,r.__browser);e.oScroll.iBarWidth=r.__browser.barWidth}function R(t,e,n,r,a,s){var o=r,l,u=false;if(n!==i){l=n;u=true}while(o!==a){if(!t.hasOwnProperty(o)){continue}l=u?e(l,t[o],o,t):t[o];u=true;o+=s}return l}function L(e,i){var a=r.defaults.column;var s=e.aoColumns.length;var o=t.extend({},r.models.oColumn,a,{nTh:i?i:n.createElement("th"),sTitle:a.sTitle?a.sTitle:i?i.innerHTML:"",aDataSort:a.aDataSort?a.aDataSort:[s],mData:a.mData?a.mData:s,idx:s});e.aoColumns.push(o);var l=e.aoPreSearchCols;l[s]=t.extend({},r.models.oSearch,l[s]);j(e,s,t(i).data())}function j(e,n,a){var s=e.aoColumns[n];var o=e.oClasses;var l=t(s.nTh);if(!s.sWidthOrig){s.sWidthOrig=l.attr("width")||null;var u=(l.attr("style")||"").match(/width:\s*(\d+[pxem%]+)/);if(u){s.sWidthOrig=u[1]}}if(a!==i&&a!==null){E(a);I(r.defaults.column,a);if(a.mDataProp!==i&&!a.mData){a.mData=a.mDataProp}if(a.sType){s._sManualType=a.sType}if(a.className&&!a.sClass){a.sClass=a.className}t.extend(s,a);ce(s,a,"sWidth","sWidthOrig");if(a.iDataSort!==i){s.aDataSort=[a.iDataSort]}ce(s,a,"aDataSort")}var f=s.mData;var c=tt(f);var h=s.mRender?tt(s.mRender):null;var d=function(t){return typeof t==="string"&&t.indexOf("@")!==-1};s._bAttrSrc=t.isPlainObject(f)&&(d(f.sort)||d(f.type)||d(f.filter));s._setter=null;s.fnGetData=function(t,e,n){var r=c(t,e,i,n);return h&&e?h(r,e,t,n):r};s.fnSetData=function(t,e,n){return et(f)(t,e,n)};if(typeof f!=="number"){e._rowReadObject=true}if(!e.oFeatures.bSort){s.bSortable=false;l.addClass(o.sSortableNone)}var p=t.inArray("asc",s.asSorting)!==-1;var g=t.inArray("desc",s.asSorting)!==-1;if(!s.bSortable||!p&&!g){s.sSortingClass=o.sSortableNone;s.sSortingClassJUI=""}else if(p&&!g){s.sSortingClass=o.sSortableAsc;s.sSortingClassJUI=o.sSortJUIAscAllowed}else if(!p&&g){s.sSortingClass=o.sSortableDesc;s.sSortingClassJUI=o.sSortJUIDescAllowed}else{s.sSortingClass=o.sSortable;s.sSortingClassJUI=o.sSortJUI}}function O(t){if(t.oFeatures.bAutoWidth!==false){var e=t.aoColumns;Xt(t);for(var n=0,i=e.length;n<i;n++){e[n].nTh.style.width=e[n].sWidth}}var r=t.oScroll;if(r.sY!==""||r.sX!==""){Ut(t)}ge(t,null,"column-sizing",[t])}function H(t,e){var n=q(t,"bVisible")
;return typeof n[e]==="number"?n[e]:null}function W(e,n){var i=q(e,"bVisible");var r=t.inArray(n,i);return r!==-1?r:null}function z(e){return t(x(e.aoColumns,"nTh")).filter(":visible").length}function q(e,n){var i=[];t.map(e.aoColumns,function(t,e){if(t[n]){i.push(e)}});return i}function B(t){var e=t.aoColumns;var n=t.aoData;var a=r.ext.type.detect;var s,o,l,u,f,c;var h,d,p,g;for(s=0,o=e.length;s<o;s++){h=e[s];g=[];if(!h.sType&&h._sManualType){h.sType=h._sManualType}else if(!h.sType){for(l=0,u=a.length;l<u;l++){for(f=0,c=n.length;f<c;f++){if(g[f]===i){g[f]=J(t,f,s,"type")}p=a[l](g[f],t);if(!p&&l!==a.length-1){break}if(p==="html"){break}}if(p){h.sType=p;break}}if(!h.sType){h.sType="string"}}}}function $(e,n,r,a){var s,o,l,u,f,c,h;var d=e.aoColumns;if(n){for(s=n.length-1;s>=0;s--){h=n[s];var p=h.targets!==i?h.targets:h.aTargets;if(!t.isArray(p)){p=[p]}for(l=0,u=p.length;l<u;l++){if(typeof p[l]==="number"&&p[l]>=0){while(d.length<=p[l]){L(e)}a(p[l],h)}else if(typeof p[l]==="number"&&p[l]<0){a(d.length+p[l],h)}else if(typeof p[l]==="string"){for(f=0,c=d.length;f<c;f++){if(p[l]=="_all"||t(d[f].nTh).hasClass(p[l])){a(f,h)}}}}}}if(r){for(s=0,o=r.length;s<o;s++){a(s,r[s])}}}function U(e,n,a,s){var o=e.aoData.length;var l=t.extend(true,{},r.models.oRow,{src:a?"dom":"data",idx:o});l._aData=n;e.aoData.push(l);var u,f;var c=e.aoColumns;for(var h=0,d=c.length;h<d;h++){c[h].sType=null}e.aiDisplayMaster.push(o);var p=e.rowIdFn(n);if(p!==i){e.aIds[p]=l}if(a||!e.oFeatures.bDeferRender){ot(e,o,a,s)}return o}function Y(e,n){var i;if(!(n instanceof t)){n=t(n)}return n.map(function(t,n){i=st(e,n);return U(e,i.data,n,i.cells)})}function V(t,e){return e._DT_RowIndex!==i?e._DT_RowIndex:null}function X(e,n,i){return t.inArray(i,e.aoData[n].anCells)}function J(t,e,n,r){var a=t.iDraw;var s=t.aoColumns[n];var o=t.aoData[e]._aData;var l=s.sDefaultContent;var u=s.fnGetData(o,r,{settings:t,row:e,col:n});if(u===i){if(t.iDrawError!=a&&l===null){fe(t,0,"Requested unknown parameter "+(typeof s.mData=="function"?"{function}":"'"+s.mData+"'")+" for row "+e+", column "+n,4);t.iDrawError=a}return l}if((u===o||u===null)&&l!==null&&r!==i){u=l}else if(typeof u==="function"){return u.call(o)}if(u===null&&r=="display"){return""}return u}function G(t,e,n,i){var r=t.aoColumns[n];var a=t.aoData[e]._aData;r.fnSetData(a,i,{settings:t,row:e,col:n})}var K=/\[.*?\]$/;var Z=/\(\)$/;function Q(e){return t.map(e.match(/(\\.|[^\.])+/g)||[""],function(t){return t.replace(/\\./g,".")})}function tt(e){if(t.isPlainObject(e)){var n={};t.each(e,function(t,e){if(e){n[t]=tt(e)}});return function(t,e,r,a){var s=n[e]||n._;return s!==i?s(t,e,r,a):t}}else if(e===null){return function(t){return t}}else if(typeof e==="function"){return function(t,n,i,r){return e(t,n,i,r)}}else if(typeof e==="string"&&(e.indexOf(".")!==-1||e.indexOf("[")!==-1||e.indexOf("(")!==-1)){var r=function(e,n,a){var s,o,l,u;if(a!==""){var f=Q(a);for(var c=0,h=f.length;c<h;c++){s=f[c].match(K);o=f[c].match(Z);if(s){f[c]=f[c].replace(K,"");if(f[c]!==""){e=e[f[c]]}l=[];f.splice(0,c+1);u=f.join(".");if(t.isArray(e)){for(var d=0,p=e.length;d<p;d++){l.push(r(e[d],n,u))}}var g=s[0].substring(1,s[0].length-1);e=g===""?l:l.join(g);break}else if(o){f[c]=f[c].replace(Z,"");e=e[f[c]]();continue}if(e===null||e[f[c]]===i){return i}e=e[f[c]]}}return e};return function(t,n){return r(t,n,e)}}else{return function(t,n){return t[e]}}}function et(e){if(t.isPlainObject(e)){return et(e._)}else if(e===null){return function(){}}else if(typeof e==="function"){return function(t,n,i){e(t,"set",n,i)}}else if(typeof e==="string"&&(e.indexOf(".")!==-1||e.indexOf("[")!==-1||e.indexOf("(")!==-1)){var n=function(e,r,a){var s=Q(a),o;var l=s[s.length-1];var u,f,c,h;for(var d=0,p=s.length-1;d<p;d++){u=s[d].match(K);f=s[d].match(Z);if(u){s[d]=s[d].replace(K,"");e[s[d]]=[];o=s.slice();o.splice(0,d+1);h=o.join(".");if(t.isArray(r)){for(var g=0,v=r.length;g<v;g++){c={};n(c,r[g],h);e[s[d]].push(c)}}else{e[s[d]]=r}return}else if(f){s[d]=s[d].replace(Z,"");e=e[s[d]](r)}if(e[s[d]]===null||e[s[d]]===i){e[s[d]]={}}e=e[s[d]]}if(l.match(Z)){e=e[l.replace(Z,"")](r)}else{e[l.replace(K,"")]=r}};return function(t,i){return n(t,i,e)}}else{return function(t,n){t[e]=n}}}function nt(t){return x(t.aoData,"_aData")}function it(t){t.aoData.length=0;t.aiDisplayMaster.length=0;t.aiDisplay.length=0;t.aIds={}}function rt(t,e,n){var r=-1;for(var a=0,s=t.length;a<s;a++){if(t[a]==e){r=a}else if(t[a]>e){t[a]--}}if(r!=-1&&n===i){t.splice(r,1)}}function at(t,e,n,r){var a=t.aoData[e];var s,o;var l=function(n,i){while(n.childNodes.length){n.removeChild(n.firstChild)}n.innerHTML=J(t,e,i,"display")};if(n==="dom"||(!n||n==="auto")&&a.src==="dom"){a._aData=st(t,a,r,r===i?i:a._aData).data}else{var u=a.anCells;if(u){if(r!==i){l(u[r],r)}else{for(s=0,o=u.length;s<o;s++){l(u[s],s)}}}}a._aSortData=null;a._aFilterData=null;var f=t.aoColumns;if(r!==i){f[r].sType=null}else{for(s=0,o=f.length;s<o;s++){f[s].sType=null}lt(t,a)}}function st(e,n,r,a){var s=[],o=n.firstChild,l,u,f,c=0,h,d=e.aoColumns,p=e._rowReadObject;a=a!==i?a:p?{}:[];var g=function(t,e){if(typeof t==="string"){var n=t.indexOf("@");if(n!==-1){var i=t.substring(n+1);var r=et(t);r(a,e.getAttribute(i))}}};var v=function(e){if(r===i||r===c){u=d[c];h=t.trim(e.innerHTML);if(u&&u._bAttrSrc){var n=et(u.mData._);n(a,h);g(u.mData.sort,e);g(u.mData.type,e);g(u.mData.filter,e)}else{if(p){if(!u._setter){u._setter=et(u.mData)}u._setter(a,h)}else{a[c]=h}}}c++};if(o){while(o){l=o.nodeName.toUpperCase();if(l=="TD"||l=="TH"){v(o);s.push(o)}o=o.nextSibling}}else{s=n.anCells;for(var m=0,y=s.length;m<y;m++){v(s[m])}}var b=n.firstChild?n:n.nTr;if(b){var w=b.getAttribute("id");if(w){et(e.rowId)(a,w)}}return{data:a,cells:s}}function ot(e,i,r,a){var s=e.aoData[i],o=s._aData,l=[],u,f,c,h,d;if(s.nTr===null){u=r||n.createElement("tr");s.nTr=u;s.anCells=l;u._DT_RowIndex=i;lt(e,s);for(h=0,d=e.aoColumns.length;h<d;h++){c=e.aoColumns[h];f=r?a[h]:n.createElement(c.sCellType);f._DT_CellIndex={row:i,column:h};l.push(f);if((!r||c.mRender||c.mData!==h)&&(!t.isPlainObject(c.mData)||c.mData._!==h+".display")){f.innerHTML=J(e,i,h,"display")}if(c.sClass){f.className+=" "+c.sClass}if(c.bVisible&&!r){u.appendChild(f)}else if(!c.bVisible&&r){f.parentNode.removeChild(f)}if(c.fnCreatedCell){c.fnCreatedCell.call(e.oInstance,f,J(e,i,h),o,i,h)}}ge(e,"aoRowCreatedCallback",null,[u,o,i])}s.nTr.setAttribute("role","row")}function lt(e,n){var i=n.nTr;var r=n._aData;if(i){var a=e.rowIdFn(r);if(a){i.id=a}if(r.DT_RowClass){var s=r.DT_RowClass.split(" ");n.__rowc=n.__rowc?k(n.__rowc.concat(s)):s;t(i).removeClass(n.__rowc.join(" ")).addClass(r.DT_RowClass)}if(r.DT_RowAttr){t(i).attr(r.DT_RowAttr)}if(r.DT_RowData){t(i).data(r.DT_RowData)}}}function ut(e){var n,i,r,a,s;var o=e.nTHead;var l=e.nTFoot;var u=t("th, td",o).length===0;var f=e.oClasses;var c=e.aoColumns;if(u){a=t("<tr/>").appendTo(o)}for(n=0,i=c.length;n<i;n++){s=c[n];r=t(s.nTh).addClass(s.sClass);if(u){r.appendTo(a)}if(e.oFeatures.bSort){r.addClass(s.sSortingClass);if(s.bSortable!==false){r.attr("tabindex",e.iTabIndex).attr("aria-controls",e.sTableId);re(e,s.nTh,n)}}if(s.sTitle!=r[0].innerHTML){r.html(s.sTitle)}me(e,"header")(e,r,s,f)}if(u){pt(e.aoHeader,o)}t(o).find(">tr").attr("role","row");t(o).find(">tr>th, >tr>td").addClass(f.sHeaderTH);t(l).find(">tr>th, >tr>td").addClass(f.sFooterTH);if(l!==null){var h=e.aoFooter[0];for(n=0,i=h.length;n<i;n++){s=c[n];s.nTf=h[n].cell;if(s.sClass){t(s.nTf).addClass(s.sClass)}}}}function ft(e,n,r){var a,s,o,l,u,f,c,h;var d=[];var p=[];var g=e.aoColumns.length;var v,m;if(!n){return}if(r===i){r=false}for(a=0,s=n.length;a<s;a++){d[a]=n[a].slice();d[a].nTr=n[a].nTr;for(o=g-1;o>=0;o--){if(!e.aoColumns[o].bVisible&&!r){d[a].splice(o,1)}}p.push([])}for(a=0,s=d.length;a<s;a++){h=d[a].nTr;if(h){while(c=h.firstChild){h.removeChild(c)}}for(o=0,l=d[a].length;o<l;o++){v=1;m=1;if(p[a][o]===i){h.appendChild(d[a][o].cell);p[a][o]=1;while(d[a+v]!==i&&d[a][o].cell==d[a+v][o].cell){p[a+v][o]=1;v++}while(d[a][o+m]!==i&&d[a][o].cell==d[a][o+m].cell){for(u=0;u<v;u++){p[a+u][o+m]=1}m++}t(d[a][o].cell).attr("rowspan",v).attr("colspan",m)}}}}function ct(e){var n=ge(e,"aoPreDrawCallback","preDraw",[e]);if(t.inArray(false,n)!==-1){Bt(e,false);return}var r,a,s;var o=[];var l=0;var u=e.asStripeClasses;var f=u.length;var c=e.aoOpenRows.length;var h=e.oLanguage;var d=e.iInitDisplayStart;var p=ye(e)=="ssp";var g=e.aiDisplay;e.bDrawing=true;if(d!==i&&d!==-1){e._iDisplayStart=p?d:d>=e.fnRecordsDisplay()?0:d;e.iInitDisplayStart=-1}var v=e._iDisplayStart;var m=e.fnDisplayEnd();if(e.bDeferLoading){e.bDeferLoading=false;e.iDraw++;Bt(e,false)}else if(!p){e.iDraw++}else if(!e.bDestroying&&!mt(e)){return}if(g.length!==0){var y=p?0:v;var b=p?e.aoData.length:m;for(var w=y;w<b;w++){var _=g[w];var x=e.aoData[_];if(x.nTr===null){ot(e,_)}var S=x.nTr;if(f!==0){var D=u[l%f];if(x._sRowStripe!=D){t(S).removeClass(x._sRowStripe).addClass(D);x._sRowStripe=D}}ge(e,"aoRowCallback",null,[S,x._aData,l,w]);o.push(S);l++}}else{var C=h.sZeroRecords;if(e.iDraw==1&&ye(e)=="ajax"){C=h.sLoadingRecords}else if(h.sEmptyTable&&e.fnRecordsTotal()===0){C=h.sEmptyTable}o[0]=t("<tr/>",{class:f?u[0]:""}).append(t("<td />",{valign:"top",colSpan:z(e),class:e.oClasses.sRowEmpty}).html(C))[0]}ge(e,"aoHeaderCallback","header",[t(e.nTHead).children("tr")[0],nt(e),v,m,g]);ge(e,"aoFooterCallback","footer",[t(e.nTFoot).children("tr")[0],nt(e),v,m,g]);var T=t(e.nTBody);T.children().detach();T.append(t(o));ge(e,"aoDrawCallback","draw",[e]);e.bSorted=false;e.bFiltered=false;e.bDrawing=false}function ht(t,e){var n=t.oFeatures,i=n.bSort,r=n.bFilter;if(i){ee(t)}if(r){xt(t,t.oPreviousSearch)}else{t.aiDisplay=t.aiDisplayMaster.slice()}if(e!==true){t._iDisplayStart=0}t._drawHold=e;ct(t);t._drawHold=false}function dt(e){var n=e.oClasses;var i=t(e.nTable);var a=t("<div/>").insertBefore(i);var s=e.oFeatures;var o=t("<div/>",{id:e.sTableId+"_wrapper",class:n.sWrapper+(e.nTFoot?"":" "+n.sNoFooter)});e.nHolding=a[0];e.nTableWrapper=o[0];e.nTableReinsertBefore=e.nTable.nextSibling;var l=e.sDom.split("");var u,f,c,h,d,p;for(var g=0;g<l.length;g++){u=null;f=l[g];if(f=="<"){c=t("<div/>")[0];h=l[g+1];if(h=="'"||h=='"'){d="";p=2;while(l[g+p]!=h){d+=l[g+p];p++}if(d=="H"){d=n.sJUIHeader}else if(d=="F"){d=n.sJUIFooter}if(d.indexOf(".")!=-1){var v=d.split(".");c.id=v[0].substr(1,v[0].length-1);c.className=v[1]}else if(d.charAt(0)=="#"){c.id=d.substr(1,d.length-1)}else{c.className=d}g+=p}o.append(c);o=t(c)}else if(f==">"){o=o.parent()}else if(f=="l"&&s.bPaginate&&s.bLengthChange){u=Ht(e)}else if(f=="f"&&s.bFilter){u=_t(e)}else if(f=="r"&&s.bProcessing){u=qt(e)}else if(f=="t"){u=$t(e)}else if(f=="i"&&s.bInfo){u=Et(e)}else if(f=="p"&&s.bPaginate){u=Wt(e)}else if(r.ext.feature.length!==0){var m=r.ext.feature;for(var y=0,b=m.length;y<b;y++){if(f==m[y].cFeature){u=m[y].fnInit(e);break}}}if(u){var w=e.aanFeatures;if(!w[f]){w[f]=[]}w[f].push(u);o.append(u)}}a.replaceWith(o);e.nHolding=null}function pt(e,n){var i=t(n).children("tr");var r,a;var s,o,l,u,f,c,h,d,p;var g;var v=function(t,e,n){var i=t[e];while(i[n]){n++}return n};e.splice(0,e.length);for(s=0,u=i.length;s<u;s++){e.push([])}for(s=0,u=i.length;s<u;s++){r=i[s];h=0;a=r.firstChild;while(a){if(a.nodeName.toUpperCase()=="TD"||a.nodeName.toUpperCase()=="TH"){d=a.getAttribute("colspan")*1;p=a.getAttribute("rowspan")*1;d=!d||d===0||d===1?1:d;p=!p||p===0||p===1?1:p;c=v(e,s,h);g=d===1?true:false;for(l=0;l<d;l++){for(o=0;o<p;o++){e[s+o][c+l]={cell:a,unique:g};e[s+o].nTr=r}}}a=a.nextSibling}}}function gt(t,e,n){var i=[];if(!n){n=t.aoHeader;if(e){n=[];pt(n,e)}}for(var r=0,a=n.length;r<a;r++){for(var s=0,o=n[r].length;s<o;s++){if(n[r][s].unique&&(!i[s]||!t.bSortCellsTop)){i[s]=n[r][s].cell}}}return i}function vt(e,n,i){ge(e,"aoServerParams","serverParams",[n]);if(n&&t.isArray(n)){var r={};var a=/(.*?)\[\]$/;t.each(n,function(t,e){var n=e.name.match(a);if(n){var i=n[0];if(!r[i]){r[i]=[]}r[i].push(e.value)}else{r[e.name]=e.value}});n=r}var s;var o=e.ajax;var l=e.oInstance;var u=function(t){ge(e,null,"xhr",[e,t,e.jqXHR]);i(t)};if(t.isPlainObject(o)&&o.data){s=o.data;var f=t.isFunction(s)?s(n,e):s;n=t.isFunction(s)&&f?f:t.extend(true,n,f);delete o.data}var c={data:n,success:function(t){var n=t.error||t.sError;if(n){fe(e,0,n)}e.json=t;u(t)},dataType:"json",cache:false,type:e.sServerMethod,error:function(n,i,r){var a=ge(e,null,"xhr",[e,null,e.jqXHR]);if(t.inArray(true,a)===-1){if(i=="parsererror"){fe(e,0,"Invalid JSON response",1)}else if(n.readyState===4){fe(e,0,"Ajax error",7)}}Bt(e,false)}};e.oAjaxData=n;ge(e,null,"preXhr",[e,n]);if(e.fnServerData){e.fnServerData.call(l,e.sAjaxSource,t.map(n,function(t,e){return{name:e,value:t}}),u,e)}else if(e.sAjaxSource||typeof o==="string"){e.jqXHR=t.ajax(t.extend(c,{url:o||e.sAjaxSource}))}else if(t.isFunction(o)){e.jqXHR=o.call(l,n,u,e)}else{e.jqXHR=t.ajax(t.extend(c,o));o.data=s}}function mt(t){if(t.bAjaxDataGet){t.iDraw++;Bt(t,true);vt(t,yt(t),function(e){bt(t,e)});return false}return true}function yt(e){var n=e.aoColumns,i=n.length,a=e.oFeatures,s=e.oPreviousSearch,o=e.aoPreSearchCols,l,u=[],f,c,h,d=te(e),p=e._iDisplayStart,g=a.bPaginate!==false?e._iDisplayLength:-1;var v=function(t,e){u.push({name:t,value:e})};v("sEcho",e.iDraw);v("iColumns",i);v("sColumns",x(n,"sName").join(","));v("iDisplayStart",p);v("iDisplayLength",g);var m={draw:e.iDraw,columns:[],order:[],start:p,length:g,search:{value:s.sSearch,regex:s.bRegex}};for(l=0;l<i;l++){c=n[l];h=o[l];f=typeof c.mData=="function"?"function":c.mData;m.columns.push({data:f,name:c.sName,searchable:c.bSearchable,orderable:c.bSortable,search:{value:h.sSearch,regex:h.bRegex}});v("mDataProp_"+l,f);if(a.bFilter){v("sSearch_"+l,h.sSearch);v("bRegex_"+l,h.bRegex);v("bSearchable_"+l,c.bSearchable)}if(a.bSort){v("bSortable_"+l,c.bSortable)}}if(a.bFilter){v("sSearch",s.sSearch);v("bRegex",s.bRegex)}if(a.bSort){t.each(d,function(t,e){m.order.push({column:e.col,dir:e.dir});v("iSortCol_"+t,e.col);v("sSortDir_"+t,e.dir)});v("iSortingCols",d.length)}var y=r.ext.legacy.ajax;if(y===null){return e.sAjaxSource?u:m}return y?u:m}function bt(t,e){var n=function(t,n){return e[t]!==i?e[t]:e[n]};var r=wt(t,e);var a=n("sEcho","draw");var s=n("iTotalRecords","recordsTotal");var o=n("iTotalDisplayRecords","recordsFiltered");if(a){if(a*1<t.iDraw){return}t.iDraw=a*1}it(t);t._iRecordsTotal=parseInt(s,10);t._iRecordsDisplay=parseInt(o,10);for(var l=0,u=r.length;l<u;l++){U(t,r[l])}t.aiDisplay=t.aiDisplayMaster.slice();t.bAjaxDataGet=false;ct(t);if(!t._bInitComplete){jt(t,e)}t.bAjaxDataGet=true;Bt(t,false)}function wt(e,n){var r=t.isPlainObject(e.ajax)&&e.ajax.dataSrc!==i?e.ajax.dataSrc:e.sAjaxDataProp;if(r==="data"){return n.aaData||n[r]}return r!==""?tt(r)(n):n}function _t(e){var i=e.oClasses;var r=e.sTableId;var a=e.oLanguage;var s=e.oPreviousSearch;var o=e.aanFeatures;var l='<input type="search" class="'+i.sFilterInput+'"/>';var u=a.sSearch;u=u.match(/_INPUT_/)?u.replace("_INPUT_",l):u+l;var f=t("<div/>",{id:!o.f?r+"_filter":null,class:i.sFilter}).append(t("<label/>").append(u));var c=function(){var t=o.f;var n=!this.value?"":this.value;if(n!=s.sSearch){xt(e,{sSearch:n,bRegex:s.bRegex,bSmart:s.bSmart,bCaseInsensitive:s.bCaseInsensitive});e._iDisplayStart=0;ct(e)}};var h=e.searchDelay!==null?e.searchDelay:ye(e)==="ssp"?400:0;var d=t("input",f).val(s.sSearch).attr("placeholder",a.sSearchPlaceholder).bind("keyup.DT search.DT input.DT paste.DT cut.DT",h?Jt(c,h):c).bind("keypress.DT",function(t){if(t.keyCode==13){return false}}).attr("aria-controls",r);t(e.nTable).on("search.dt.DT",function(t,i){if(e===i){try{if(d[0]!==n.activeElement){d.val(s.sSearch)}}catch(t){}}});return f[0]}function xt(t,e,n){var r=t.oPreviousSearch;var a=t.aoPreSearchCols;var s=function(t){r.sSearch=t.sSearch;r.bRegex=t.bRegex;r.bSmart=t.bSmart;r.bCaseInsensitive=t.bCaseInsensitive};var o=function(t){return t.bEscapeRegex!==i?!t.bEscapeRegex:t.bRegex};B(t);if(ye(t)!="ssp"){Ct(t,e.sSearch,n,o(e),e.bSmart,e.bCaseInsensitive);s(e);for(var l=0;l<a.length;l++){Dt(t,a[l].sSearch,l,o(a[l]),a[l].bSmart,a[l].bCaseInsensitive)}St(t)}else{s(e)}t.bFiltered=true;ge(t,null,"search",[t])}function St(e){var n=r.ext.search;var i=e.aiDisplay;var a,s;for(var o=0,l=n.length;o<l;o++){var u=[];for(var f=0,c=i.length;f<c;f++){s=i[f];a=e.aoData[s];if(n[o](e,a._aFilterData,s,a._aData,f)){u.push(s)}}i.length=0;t.merge(i,u)}}function Dt(t,e,n,i,r,a){if(e===""){return}var s;var o=t.aiDisplay;var l=Tt(e,i,r,a);for(var u=o.length-1;u>=0;u--){s=t.aoData[o[u]]._aFilterData[n];if(!l.test(s)){o.splice(u,1)}}}function Ct(t,e,n,i,a,s){var o=Tt(e,i,a,s);var l=t.oPreviousSearch.sSearch;var u=t.aiDisplayMaster;var f,c,h;if(r.ext.search.length!==0){n=true}c=At(t);if(e.length<=0){t.aiDisplay=u.slice()}else{if(c||n||l.length>e.length||e.indexOf(l)!==0||t.bSorted){t.aiDisplay=u.slice()}f=t.aiDisplay;for(h=f.length-1;h>=0;h--){if(!o.test(t.aoData[f[h]]._sFilterRow)){f.splice(h,1)}}}}function Tt(e,n,i,r){e=n?e:kt(e);if(i){var a=t.map(e.match(/"[^"]+"|[^ ]+/g)||[""],function(t){if(t.charAt(0)==='"'){var e=t.match(/^"(.*)"$/);t=e?e[1]:t}return t.replace('"',"")});e="^(?=.*?"+a.join(")(?=.*?")+").*$"}return new RegExp(e,r?"i":"")}function kt(t){return t.replace(p,"\\$1")}var Mt=t("<div>")[0];var It=Mt.textContent!==i;function At(t){var e=t.aoColumns;var n;var i,a,s,o,l,u,f;var c=r.ext.type.search;var h=false;for(i=0,s=t.aoData.length;i<s;i++){f=t.aoData[i];if(!f._aFilterData){l=[];for(a=0,o=e.length;a<o;a++){n=e[a];if(n.bSearchable){u=J(t,i,a,"filter");if(c[n.sType]){u=c[n.sType](u)}if(u===null){u=""}if(typeof u!=="string"&&u.toString){u=u.toString()}}else{u=""}if(u.indexOf&&u.indexOf("&")!==-1){Mt.innerHTML=u;u=It?Mt.textContent:Mt.innerText}if(u.replace){u=u.replace(/[\r\n]/g,"")}l.push(u)}f._aFilterData=l;f._sFilterRow=l.join("  ");h=true}}return h}function Nt(t){return{search:t.sSearch,smart:t.bSmart,regex:t.bRegex,caseInsensitive:t.bCaseInsensitive}}function Pt(t){return{sSearch:t.search,bSmart:t.smart,bRegex:t.regex,bCaseInsensitive:t.caseInsensitive}}function Et(e){var n=e.sTableId,i=e.aanFeatures.i,r=t("<div/>",{class:e.oClasses.sInfo,id:!i?n+"_info":null});if(!i){e.aoDrawCallback.push({fn:Ft,sName:"information"});r.attr("role","status").attr("aria-live","polite");t(e.nTable).attr("aria-describedby",n+"_info")}return r[0]}function Ft(e){var n=e.aanFeatures.i;if(n.length===0){return}var i=e.oLanguage,r=e._iDisplayStart+1,a=e.fnDisplayEnd(),s=e.fnRecordsTotal(),o=e.fnRecordsDisplay(),l=o?i.sInfo:i.sInfoEmpty;if(o!==s){l+=" "+i.sInfoFiltered}l+=i.sInfoPostFix;l=Rt(e,l);var u=i.fnInfoCallback;if(u!==null){l=u.call(e.oInstance,e,r,a,s,o,l)}t(n).html(l)}function Rt(t,e){var n=t.fnFormatNumber,i=t._iDisplayStart+1,r=t._iDisplayLength,a=t.fnRecordsDisplay(),s=r===-1;return e.replace(/_START_/g,n.call(t,i)).replace(/_END_/g,n.call(t,t.fnDisplayEnd())).replace(/_MAX_/g,n.call(t,t.fnRecordsTotal())).replace(/_TOTAL_/g,n.call(t,a)).replace(/_PAGE_/g,n.call(t,s?1:Math.ceil(i/r))).replace(/_PAGES_/g,n.call(t,s?1:Math.ceil(a/r)))}function Lt(t){var e,n,i=t.iInitDisplayStart;var r=t.aoColumns,a;var s=t.oFeatures;var o=t.bDeferLoading;if(!t.bInitialised){setTimeout(function(){Lt(t)},200);return}dt(t);ut(t);ft(t,t.aoHeader);ft(t,t.aoFooter);Bt(t,true);if(s.bAutoWidth){Xt(t)}for(e=0,n=r.length;e<n;e++){a=r[e];if(a.sWidth){a.nTh.style.width=Qt(a.sWidth)}}ge(t,null,"preInit",[t]);ht(t);var l=ye(t);if(l!="ssp"||o){if(l=="ajax"){vt(t,[],function(n){var r=wt(t,n);for(e=0;e<r.length;e++){U(t,r[e])}t.iInitDisplayStart=i;ht(t);Bt(t,false);jt(t,n)},t)}else{Bt(t,false);jt(t)}}}function jt(t,e){t._bInitComplete=true;if(e||t.oInit.aaData){O(t)}ge(t,null,"plugin-init",[t,e]);ge(t,"aoInitComplete","init",[t,e])}function Ot(t,e){var n=parseInt(e,10);t._iDisplayLength=n;ve(t);ge(t,null,"length",[t,n])}function Ht(e){var n=e.oClasses,i=e.sTableId,r=e.aLengthMenu,a=t.isArray(r[0]),s=a?r[0]:r,o=a?r[1]:r;var l=t("<select/>",{name:i+"_length","aria-controls":i,class:n.sLengthSelect});for(var u=0,f=s.length;u<f;u++){l[0][u]=new Option(o[u],s[u])}var c=t("<div><label/></div>").addClass(n.sLength);if(!e.aanFeatures.l){c[0].id=i+"_length"}c.children().append(e.oLanguage.sLengthMenu.replace("_MENU_",l[0].outerHTML));t("select",c).val(e._iDisplayLength).bind("change.DT",function(n){Ot(e,t(this).val());ct(e)});t(e.nTable).bind("length.dt.DT",function(n,i,r){if(e===i){t("select",c).val(r)}});return c[0]}function Wt(e){var n=e.sPaginationType,i=r.ext.pager[n],a=typeof i==="function",s=function(t){ct(t)},o=t("<div/>").addClass(e.oClasses.sPaging+n)[0],l=e.aanFeatures;if(!a){i.fnInit(e,o,s)}if(!l.p){o.id=e.sTableId+"_paginate";e.aoDrawCallback.push({fn:function(t){if(a){var e=t._iDisplayStart,n=t._iDisplayLength,r=t.fnRecordsDisplay(),o=n===-1,u=o?0:Math.ceil(e/n),f=o?1:Math.ceil(r/n),c=i(u,f),h,d;for(h=0,d=l.p.length;h<d;h++){me(t,"pageButton")(t,l.p[h],h,c,u,f)}}else{i.fnUpdate(t,s)}},sName:"pagination"})}return o}function zt(t,e,n){var i=t._iDisplayStart,r=t._iDisplayLength,a=t.fnRecordsDisplay();if(a===0||r===-1){i=0}else if(typeof e==="number"){i=e*r;if(i>a){i=0}}else if(e=="first"){i=0}else if(e=="previous"){i=r>=0?i-r:0;if(i<0){i=0}}else if(e=="next"){if(i+r<a){i+=r}}else if(e=="last"){i=Math.floor((a-1)/r)*r}else{fe(t,0,"Unknown paging action: "+e,5)}var s=t._iDisplayStart!==i;t._iDisplayStart=i;if(s){ge(t,null,"page",[t]);if(n){ct(t)}}return s}function qt(e){return t("<div/>",{id:!e.aanFeatures.r?e.sTableId+"_processing":null,class:e.oClasses.sProcessing}).html(e.oLanguage.sProcessing).insertBefore(e.nTable)[0]}function Bt(e,n){if(e.oFeatures.bProcessing){t(e.aanFeatures.r).css("display",n?"block":"none")}ge(e,null,"processing",[e,n])}function $t(e){var n=t(e.nTable);n.attr("role","grid");var i=e.oScroll;if(i.sX===""&&i.sY===""){return e.nTable}var r=i.sX;var a=i.sY;var s=e.oClasses;var o=n.children("caption");var l=o.length?o[0]._captionSide:null;var u=t(n[0].cloneNode(false));var f=t(n[0].cloneNode(false));var c=n.children("tfoot");var h="<div/>";var d=function(t){return!t?null:Qt(t)};if(!c.length){c=null}var p=t(h,{class:s.sScrollWrapper}).append(t(h,{class:s.sScrollHead}).css({overflow:"hidden",position:"relative",border:0,width:r?d(r):"100%"}).append(t(h,{class:s.sScrollHeadInner}).css({"box-sizing":"content-box",width:i.sXInner||"100%"}).append(u.removeAttr("id").css("margin-left",0).append(l==="top"?o:null).append(n.children("thead"))))).append(t(h,{class:s.sScrollBody}).css({position:"relative",overflow:"auto",width:d(r)}).append(n));if(c){p.append(t(h,{class:s.sScrollFoot}).css({overflow:"hidden",border:0,width:r?d(r):"100%"}).append(t(h,{class:s.sScrollFootInner}).append(f.removeAttr("id").css("margin-left",0).append(l==="bottom"?o:null).append(n.children("tfoot")))))}var g=p.children();var v=g[0];var m=g[1];var y=c?g[2]:null;if(r){t(m).on("scroll.DT",function(t){var e=this.scrollLeft;v.scrollLeft=e;if(c){y.scrollLeft=e}})}t(m).css(a&&i.bCollapse?"max-height":"height",a);e.nScrollHead=v;e.nScrollBody=m;e.nScrollFoot=y;e.aoDrawCallback.push({fn:Ut,sName:"scrolling"});return p[0]}function Ut(e){var n=e.oScroll,r=n.sX,a=n.sXInner,s=n.sY,o=n.iBarWidth,l=t(e.nScrollHead),u=l[0].style,f=l.children("div"),c=f[0].style,h=f.children("table"),d=e.nScrollBody,p=t(d),g=d.style,v=t(e.nScrollFoot),m=v.children("div"),y=m.children("table"),b=t(e.nTHead),w=t(e.nTable),_=w[0],S=_.style,D=e.nTFoot?t(e.nTFoot):null,C=e.oBrowser,T=C.bScrollOversize,k=x(e.aoColumns,"nTh"),M,I,A,N,P,E,F=[],R=[],L=[],j=[],W,z,q,B=function(t){var e=t.style;e.paddingTop="0";e.paddingBottom="0";e.borderTopWidth="0";e.borderBottomWidth="0";e.height=0};var $=d.scrollHeight>d.clientHeight;if(e.scrollBarVis!==$&&e.scrollBarVis!==i){e.scrollBarVis=$;O(e);return}else{e.scrollBarVis=$}w.children("thead, tfoot").remove();if(D){E=D.clone().prependTo(w);I=D.find("tr");N=E.find("tr")}P=b.clone().prependTo(w);M=b.find("tr");A=P.find("tr");P.find("th, td").removeAttr("tabindex");if(!r){g.width="100%";l[0].style.width="100%"}t.each(gt(e,P),function(t,n){W=H(e,t);n.style.width=e.aoColumns[W].sWidth});if(D){Yt(function(t){t.style.width=""},N)}q=w.outerWidth();if(r===""){S.width="100%";if(T&&(w.find("tbody").height()>d.offsetHeight||p.css("overflow-y")=="scroll")){S.width=Qt(w.outerWidth()-o)}q=w.outerWidth()}else if(a!==""){S.width=Qt(a);q=w.outerWidth()}Yt(B,A);Yt(function(e){L.push(e.innerHTML);F.push(Qt(t(e).css("width")))},A);Yt(function(e,n){if(t.inArray(e,k)!==-1){e.style.width=F[n]}},M);t(A).height(0);if(D){Yt(B,N);Yt(function(e){j.push(e.innerHTML);R.push(Qt(t(e).css("width")))},N);Yt(function(t,e){t.style.width=R[e]},I);t(N).height(0)}Yt(function(t,e){t.innerHTML='<div class="dataTables_sizing" style="height:0;overflow:hidden;">'+L[e]+"</div>";t.style.width=F[e]},A);if(D){Yt(function(t,e){t.innerHTML='<div class="dataTables_sizing" style="height:0;overflow:hidden;">'+j[e]+"</div>";t.style.width=R[e]},N)}if(w.outerWidth()<q){z=d.scrollHeight>d.offsetHeight||p.css("overflow-y")=="scroll"?q+o:q;if(T&&(d.scrollHeight>d.offsetHeight||p.css("overflow-y")=="scroll")){S.width=Qt(z-o)}if(r===""||a!==""){fe(e,1,"Possible column misalignment",6)}}else{z="100%"}g.width=Qt(z);u.width=Qt(z);if(D){e.nScrollFoot.style.width=Qt(z)}if(!s){if(T){g.height=Qt(_.offsetHeight+o)}}var U=w.outerWidth();h[0].style.width=Qt(U);c.width=Qt(U);var Y=w.height()>d.clientHeight||p.css("overflow-y")=="scroll";var V="padding"+(C.bScrollbarLeft?"Left":"Right");c[V]=Y?o+"px":"0px";if(D){y[0].style.width=Qt(U);m[0].style.width=Qt(U);m[0].style[V]=Y?o+"px":"0px"}w.children("colgroup").insertBefore(w.children("thead"));p.scroll();if((e.bSorted||e.bFiltered)&&!e._drawHold){d.scrollTop=0}}function Yt(t,e,n){var i=0,r=0,a=e.length;var s,o;while(r<a){s=e[r].firstChild;o=n?n[r].firstChild:null;while(s){if(s.nodeType===1){if(n){t(s,o,i)}else{t(s,i)}i++}s=s.nextSibling;o=n?o.nextSibling:null}r++}}var Vt=/<.*?>/g;function Xt(n){var i=n.nTable,r=n.aoColumns,a=n.oScroll,s=a.sY,o=a.sX,l=a.sXInner,u=r.length,f=q(n,"bVisible"),c=t("th",n.nTHead),h=i.getAttribute("width"),d=i.parentNode,p=false,g,v,m,y,b,w=n.oBrowser,_=w.bScrollOversize;var x=i.style.width;if(x&&x.indexOf("%")!==-1){h=x}for(g=0;g<f.length;g++){v=r[f[g]];if(v.sWidth!==null){v.sWidth=Gt(v.sWidthOrig,d);p=true}}if(_||!p&&!o&&!s&&u==z(n)&&u==c.length){for(g=0;g<u;g++){var S=H(n,g);if(S!==null){r[S].sWidth=Qt(c.eq(g).width())}}}else{var D=t(i).clone().css("visibility","hidden").removeAttr("id");D.find("tbody tr").remove();var C=t("<tr/>").appendTo(D.find("tbody"));D.find("thead, tfoot").remove();D.append(t(n.nTHead).clone()).append(t(n.nTFoot).clone());D.find("tfoot th, tfoot td").css("width","");c=gt(n,D.find("thead")[0]);for(g=0;g<f.length;g++){v=r[f[g]];c[g].style.width=v.sWidthOrig!==null&&v.sWidthOrig!==""?Qt(v.sWidthOrig):"";if(v.sWidthOrig&&o){t(c[g]).append(t("<div/>").css({width:v.sWidthOrig,margin:0,padding:0,border:0,height:1}))}}if(n.aoData.length){for(g=0;g<f.length;g++){m=f[g];v=r[m];t(Kt(n,m)).clone(false).append(v.sContentPadding).appendTo(C)}}t("[name]",D).removeAttr("name");var T=t("<div/>").css(o||s?{position:"absolute",top:0,left:0,height:1,right:0,overflow:"hidden"}:{}).append(D).appendTo(d);if(o&&l){D.width(l)}else if(o){D.css("width","auto");D.removeAttr("width");if(D.width()<d.clientWidth&&h){D.width(d.clientWidth)}}else if(s){D.width(d.clientWidth)}else if(h){D.width(h)}var k=0;for(g=0;g<f.length;g++){var M=t(c[g]);var I=M.outerWidth()-M.width();var A=w.bBounding?Math.ceil(c[g].getBoundingClientRect().width):M.outerWidth();k+=A;r[f[g]].sWidth=Qt(A-I)}i.style.width=Qt(k);T.remove()}if(h){i.style.width=Qt(h)}if((h||o)&&!n._reszEvt){var N=function(){t(e).bind("resize.DT-"+n.sInstance,Jt(function(){O(n)}))};if(_){setTimeout(N,1e3)}else{N()}n._reszEvt=true}}function Jt(t,e){var n=e!==i?e:200,r,a;return function(){var e=this,s=+new Date,o=arguments;if(r&&s<r+n){clearTimeout(a);a=setTimeout(function(){r=i;t.apply(e,o)},n)}else{r=s;t.apply(e,o)}}}function Gt(e,i){if(!e){return 0}var r=t("<div/>").css("width",Qt(e)).appendTo(i||n.body);var a=r[0].offsetWidth;r.remove();return a}function Kt(e,n){var i=Zt(e,n);if(i<0){return null}var r=e.aoData[i];return!r.nTr?t("<td/>").html(J(e,i,n,"display"))[0]:r.anCells[n]}function Zt(t,e){var n,i=-1,r=-1;for(var a=0,s=t.aoData.length;a<s;a++){n=J(t,a,e,"display")+"";n=n.replace(Vt,"");n=n.replace(/&nbsp;/g," ");if(n.length>i){i=n.length;r=a}}return r}function Qt(t){if(t===null){return"0px"}if(typeof t=="number"){return t<0?"0px":t+"px"}return t.match(/\d$/)?t+"px":t}function te(e){var n,a,s,o,l=[],u=[],f=e.aoColumns,c,h,d,p,g=e.aaSortingFixed,v=t.isPlainObject(g),m=[],y=function(e){if(e.length&&!t.isArray(e[0])){m.push(e)}else{t.merge(m,e)}};if(t.isArray(g)){y(g)}if(v&&g.pre){y(g.pre)}y(e.aaSorting);if(v&&g.post){y(g.post)}for(n=0;n<m.length;n++){p=m[n][0];c=f[p].aDataSort;for(s=0,o=c.length;s<o;s++){h=c[s];d=f[h].sType||"string";if(m[n]._idx===i){m[n]._idx=t.inArray(m[n][1],f[h].asSorting)}l.push({src:p,col:h,dir:m[n][1],index:m[n]._idx,type:d,formatter:r.ext.type.order[d+"-pre"]})}}return l}function ee(t){var e,n,i,a,s,o,l,u,f,c=[],h=r.ext.type.order,d=t.aoData,p=t.aoColumns,g,v,m,y,b,w=0,_,x=t.aiDisplayMaster,S;B(t);S=te(t);for(e=0,n=S.length;e<n;e++){_=S[e];if(_.formatter){w++}se(t,_.col)}if(ye(t)!="ssp"&&S.length!==0){for(e=0,i=x.length;e<i;e++){c[x[e]]=e}if(w===S.length){x.sort(function(t,e){var n,i,r,a,s,o=S.length,l=d[t]._aSortData,u=d[e]._aSortData;for(r=0;r<o;r++){s=S[r];n=l[s.col];i=u[s.col];a=n<i?-1:n>i?1:0;if(a!==0){return s.dir==="asc"?a:-a}}n=c[t];i=c[e];return n<i?-1:n>i?1:0})}else{x.sort(function(t,e){var n,i,r,a,s,o,l,u=S.length,f=d[t]._aSortData,p=d[e]._aSortData;for(r=0;r<u;r++){o=S[r];n=f[o.col];i=p[o.col];l=h[o.type+"-"+o.dir]||h["string-"+o.dir];s=l(n,i);if(s!==0){return s}}n=c[t];i=c[e];return n<i?-1:n>i?1:0})}}t.bSorted=true}function ne(t){var e;var n;var i=t.aoColumns;var r=te(t);var a=t.oLanguage.oAria;for(var s=0,o=i.length;s<o;s++){var l=i[s];var u=l.asSorting;var f=l.sTitle.replace(/<.*?>/g,"");var c=l.nTh;c.removeAttribute("aria-sort");if(l.bSortable){if(r.length>0&&r[0].col==s){c.setAttribute("aria-sort",r[0].dir=="asc"?"ascending":"descending");n=u[r[0].index+1]||u[0]}else{n=u[0]}e=f+(n==="asc"?a.sSortAscending:a.sSortDescending)}else{e=f}c.setAttribute("aria-label",e)}}function ie(e,n,r,a){var s=e.aoColumns[n];var o=e.aaSorting;var l=s.asSorting;var u;var f=function(e,n){var r=e._idx;if(r===i){r=t.inArray(e[1],l)}return r+1<l.length?r+1:n?null:0};if(typeof o[0]==="number"){o=e.aaSorting=[o]}if(r&&e.oFeatures.bSortMulti){var c=t.inArray(n,x(o,"0"));if(c!==-1){u=f(o[c],true);if(u===null&&o.length===1){u=0}if(u===null){o.splice(c,1)}else{o[c][1]=l[u];o[c]._idx=u}}else{o.push([n,l[0],0]);o[o.length-1]._idx=0}}else if(o.length&&o[0][0]==n){u=f(o[0]);o.length=1;o[0][1]=l[u];o[0]._idx=u}else{o.length=0;o.push([n,l[0]]);o[0]._idx=0}ht(e);if(typeof a=="function"){a(e)}}function re(t,e,n,i){var r=t.aoColumns[n];de(e,{},function(e){if(r.bSortable===false){return}if(t.oFeatures.bProcessing){Bt(t,true);setTimeout(function(){ie(t,n,e.shiftKey,i);if(ye(t)!=="ssp"){Bt(t,false)}},0)}else{ie(t,n,e.shiftKey,i)}})}function ae(e){var n=e.aLastSort;var i=e.oClasses.sSortColumn;var r=te(e);var a=e.oFeatures;var s,o,l;if(a.bSort&&a.bSortClasses){for(s=0,o=n.length;s<o;s++){l=n[s].src;t(x(e.aoData,"anCells",l)).removeClass(i+(s<2?s+1:3))}for(s=0,o=r.length;s<o;s++){l=r[s].src;t(x(e.aoData,"anCells",l)).addClass(i+(s<2?s+1:3))}}e.aLastSort=r}function se(t,e){var n=t.aoColumns[e];var i=r.ext.order[n.sSortDataType];var a;if(i){a=i.call(t.oInstance,t,e,W(t,e))}var s,o;var l=r.ext.type.order[n.sType+"-pre"];for(var u=0,f=t.aoData.length;u<f;u++){s=t.aoData[u];if(!s._aSortData){s._aSortData=[]}if(!s._aSortData[e]||i){o=i?a[u]:J(t,u,e,"sort");s._aSortData[e]=l?l(o):o}}}function oe(e){if(!e.oFeatures.bStateSave||e.bDestroying){return}var n={time:+new Date,start:e._iDisplayStart,length:e._iDisplayLength,order:t.extend(true,[],e.aaSorting),search:Nt(e.oPreviousSearch),columns:t.map(e.aoColumns,function(t,n){return{visible:t.bVisible,search:Nt(e.aoPreSearchCols[n])}})};ge(e,"aoStateSaveParams","stateSaveParams",[e,n]);e.oSavedState=n
;e.fnStateSaveCallback.call(e.oInstance,e,n)}function le(e,n){var r,a;var s=e.aoColumns;if(!e.oFeatures.bStateSave){return}var o=e.fnStateLoadCallback.call(e.oInstance,e);if(!o||!o.time){return}var l=ge(e,"aoStateLoadParams","stateLoadParams",[e,o]);if(t.inArray(false,l)!==-1){return}var u=e.iStateDuration;if(u>0&&o.time<+new Date-u*1e3){return}if(s.length!==o.columns.length){return}e.oLoadedState=t.extend(true,{},o);if(o.start!==i){e._iDisplayStart=o.start;e.iInitDisplayStart=o.start}if(o.length!==i){e._iDisplayLength=o.length}if(o.order!==i){e.aaSorting=[];t.each(o.order,function(t,n){e.aaSorting.push(n[0]>=s.length?[0,n[1]]:n)})}if(o.search!==i){t.extend(e.oPreviousSearch,Pt(o.search))}for(r=0,a=o.columns.length;r<a;r++){var f=o.columns[r];if(f.visible!==i){s[r].bVisible=f.visible}if(f.search!==i){t.extend(e.aoPreSearchCols[r],Pt(f.search))}}ge(e,"aoStateLoaded","stateLoaded",[e,o])}function ue(e){var n=r.settings;var i=t.inArray(e,x(n,"nTable"));return i!==-1?n[i]:null}function fe(t,n,i,a){i="DataTables warning: "+(t?"table id="+t.sTableId+" - ":"")+i;if(a){i+=". For more information about this error, please see "+"http://datatables.net/tn/"+a}if(!n){var s=r.ext;var o=s.sErrMode||s.errMode;if(t){ge(t,null,"error",[t,a,i])}if(o=="alert"){alert(i)}else if(o=="throw"){throw new Error(i)}else if(typeof o=="function"){o(t,a,i)}}else if(e.console&&console.log){console.log(i)}}function ce(e,n,r,a){if(t.isArray(r)){t.each(r,function(i,r){if(t.isArray(r)){ce(e,n,r[0],r[1])}else{ce(e,n,r)}});return}if(a===i){a=r}if(n[r]!==i){e[a]=n[r]}}function he(e,n,i){var r;for(var a in n){if(n.hasOwnProperty(a)){r=n[a];if(t.isPlainObject(r)){if(!t.isPlainObject(e[a])){e[a]={}}t.extend(true,e[a],r)}else if(i&&a!=="data"&&a!=="aaData"&&t.isArray(r)){e[a]=r.slice()}else{e[a]=r}}}return e}function de(e,n,i){t(e).bind("click.DT",n,function(t){e.blur();i(t)}).bind("keypress.DT",n,function(t){if(t.which===13){t.preventDefault();i(t)}}).bind("selectstart.DT",function(){return false})}function pe(t,e,n,i){if(n){t[e].push({fn:n,sName:i})}}function ge(e,n,i,r){var a=[];if(n){a=t.map(e[n].slice().reverse(),function(t,n){return t.fn.apply(e.oInstance,r)})}if(i!==null){var s=t.Event(i+".dt");t(e.nTable).trigger(s,r);a.push(s.result)}return a}function ve(t){var e=t._iDisplayStart,n=t.fnDisplayEnd(),i=t._iDisplayLength;if(e>=n){e=n-i}e-=e%i;if(i===-1||e<0){e=0}t._iDisplayStart=e}function me(e,n){var i=e.renderer;var a=r.ext.renderer[n];if(t.isPlainObject(i)&&i[n]){return a[i[n]]||a._}else if(typeof i==="string"){return a[i]||a._}return a._}function ye(t){if(t.oFeatures.bServerSide){return"ssp"}else if(t.ajax||t.sAjaxSource){return"ajax"}return"dom"}r=function(e){this.$=function(t,e){return this.api(true).$(t,e)};this._=function(t,e){return this.api(true).rows(t,e).data()};this.api=function(t){return t?new s(ue(this[a.iApiIndex])):new s(this)};this.fnAddData=function(e,n){var r=this.api(true);var a=t.isArray(e)&&(t.isArray(e[0])||t.isPlainObject(e[0]))?r.rows.add(e):r.row.add(e);if(n===i||n){r.draw()}return a.flatten().toArray()};this.fnAdjustColumnSizing=function(t){var e=this.api(true).columns.adjust();var n=e.settings()[0];var r=n.oScroll;if(t===i||t){e.draw(false)}else if(r.sX!==""||r.sY!==""){Ut(n)}};this.fnClearTable=function(t){var e=this.api(true).clear();if(t===i||t){e.draw()}};this.fnClose=function(t){this.api(true).row(t).child.hide()};this.fnDeleteRow=function(t,e,n){var r=this.api(true);var a=r.rows(t);var s=a.settings()[0];var o=s.aoData[a[0][0]];a.remove();if(e){e.call(this,s,o)}if(n===i||n){r.draw()}return o};this.fnDestroy=function(t){this.api(true).destroy(t)};this.fnDraw=function(t){this.api(true).draw(t)};this.fnFilter=function(t,e,n,r,a,s){var o=this.api(true);if(e===null||e===i){o.search(t,n,r,s)}else{o.column(e).search(t,n,r,s)}o.draw()};this.fnGetData=function(t,e){var n=this.api(true);if(t!==i){var r=t.nodeName?t.nodeName.toLowerCase():"";return e!==i||r=="td"||r=="th"?n.cell(t,e).data():n.row(t).data()||null}return n.data().toArray()};this.fnGetNodes=function(t){var e=this.api(true);return t!==i?e.row(t).node():e.rows().nodes().flatten().toArray()};this.fnGetPosition=function(t){var e=this.api(true);var n=t.nodeName.toUpperCase();if(n=="TR"){return e.row(t).index()}else if(n=="TD"||n=="TH"){var i=e.cell(t).index();return[i.row,i.columnVisible,i.column]}return null};this.fnIsOpen=function(t){return this.api(true).row(t).child.isShown()};this.fnOpen=function(t,e,n){return this.api(true).row(t).child(e,n).show().child()[0]};this.fnPageChange=function(t,e){var n=this.api(true).page(t);if(e===i||e){n.draw(false)}};this.fnSetColumnVis=function(t,e,n){var r=this.api(true).column(t).visible(e);if(n===i||n){r.columns.adjust().draw()}};this.fnSettings=function(){return ue(this[a.iApiIndex])};this.fnSort=function(t){this.api(true).order(t).draw()};this.fnSortListener=function(t,e,n){this.api(true).order.listener(t,e,n)};this.fnUpdate=function(t,e,n,r,a){var s=this.api(true);if(n===i||n===null){s.row(e).data(t)}else{s.cell(e,n).data(t)}if(a===i||a){s.columns.adjust()}if(r===i||r){s.draw()}return 0};this.fnVersionCheck=a.fnVersionCheck;var n=this;var o=e===i;var l=this.length;if(o){e={}}this.oApi=this.internal=a.internal;for(var u in r.ext.internal){if(u){this[u]=Ye(u)}}this.each(function(){var a={};var s=l>1?he(a,e,true):e;var u=0,f,c,h,d,p;var g=this.getAttribute("id");var v=false;var m=r.defaults;var y=t(this);if(this.nodeName.toLowerCase()!="table"){fe(null,0,"Non-table node initialisation ("+this.nodeName+")",2);return}P(m);E(m.column);I(m,m,true);I(m.column,m.column,true);I(m,t.extend(s,y.data()));var b=r.settings;for(u=0,f=b.length;u<f;u++){var w=b[u];if(w.nTable==this||w.nTHead.parentNode==this||w.nTFoot&&w.nTFoot.parentNode==this){var _=s.bRetrieve!==i?s.bRetrieve:m.bRetrieve;var x=s.bDestroy!==i?s.bDestroy:m.bDestroy;if(o||_){return w.oInstance}else if(x){w.oInstance.fnDestroy();break}else{fe(w,0,"Cannot reinitialise DataTable",3);return}}if(w.sTableId==this.id){b.splice(u,1);break}}if(g===null||g===""){g="DataTables_Table_"+r.ext._unique++;this.id=g}var S=t.extend(true,{},r.models.oSettings,{sDestroyWidth:y[0].style.width,sInstance:g,sTableId:g});S.nTable=this;S.oApi=n.internal;S.oInit=s;b.push(S);S.oInstance=n.length===1?n:y.dataTable();P(s);if(s.oLanguage){A(s.oLanguage)}if(s.aLengthMenu&&!s.iDisplayLength){s.iDisplayLength=t.isArray(s.aLengthMenu[0])?s.aLengthMenu[0][0]:s.aLengthMenu[0]}s=he(t.extend(true,{},m),s);ce(S.oFeatures,s,["bPaginate","bLengthChange","bFilter","bSort","bSortMulti","bInfo","bProcessing","bAutoWidth","bSortClasses","bServerSide","bDeferRender"]);ce(S,s,["asStripeClasses","ajax","fnServerData","fnFormatNumber","sServerMethod","aaSorting","aaSortingFixed","aLengthMenu","sPaginationType","sAjaxSource","sAjaxDataProp","iStateDuration","sDom","bSortCellsTop","iTabIndex","fnStateLoadCallback","fnStateSaveCallback","renderer","searchDelay","rowId",["iCookieDuration","iStateDuration"],["oSearch","oPreviousSearch"],["aoSearchCols","aoPreSearchCols"],["iDisplayLength","_iDisplayLength"],["bJQueryUI","bJUI"]]);ce(S.oScroll,s,[["sScrollX","sX"],["sScrollXInner","sXInner"],["sScrollY","sY"],["bScrollCollapse","bCollapse"]]);ce(S.oLanguage,s,"fnInfoCallback");pe(S,"aoDrawCallback",s.fnDrawCallback,"user");pe(S,"aoServerParams",s.fnServerParams,"user");pe(S,"aoStateSaveParams",s.fnStateSaveParams,"user");pe(S,"aoStateLoadParams",s.fnStateLoadParams,"user");pe(S,"aoStateLoaded",s.fnStateLoaded,"user");pe(S,"aoRowCallback",s.fnRowCallback,"user");pe(S,"aoRowCreatedCallback",s.fnCreatedRow,"user");pe(S,"aoHeaderCallback",s.fnHeaderCallback,"user");pe(S,"aoFooterCallback",s.fnFooterCallback,"user");pe(S,"aoInitComplete",s.fnInitComplete,"user");pe(S,"aoPreDrawCallback",s.fnPreDrawCallback,"user");S.rowIdFn=tt(s.rowId);F(S);var D=S.oClasses;if(s.bJQueryUI){t.extend(D,r.ext.oJUIClasses,s.oClasses);if(s.sDom===m.sDom&&m.sDom==="lfrtip"){S.sDom='<"H"lfr>t<"F"ip>'}if(!S.renderer){S.renderer="jqueryui"}else if(t.isPlainObject(S.renderer)&&!S.renderer.header){S.renderer.header="jqueryui"}}else{t.extend(D,r.ext.classes,s.oClasses)}y.addClass(D.sTable);if(S.iInitDisplayStart===i){S.iInitDisplayStart=s.iDisplayStart;S._iDisplayStart=s.iDisplayStart}if(s.iDeferLoading!==null){S.bDeferLoading=true;var C=t.isArray(s.iDeferLoading);S._iRecordsDisplay=C?s.iDeferLoading[0]:s.iDeferLoading;S._iRecordsTotal=C?s.iDeferLoading[1]:s.iDeferLoading}var T=S.oLanguage;t.extend(true,T,s.oLanguage);if(T.sUrl!==""){t.ajax({dataType:"json",url:T.sUrl,success:function(e){A(e);I(m.oLanguage,e);t.extend(true,T,e);Lt(S)},error:function(){Lt(S)}});v=true}if(s.asStripeClasses===null){S.asStripeClasses=[D.sStripeOdd,D.sStripeEven]}var k=S.asStripeClasses;var M=y.children("tbody").find("tr").eq(0);if(t.inArray(true,t.map(k,function(t,e){return M.hasClass(t)}))!==-1){t("tbody tr",this).removeClass(k.join(" "));S.asDestroyStripes=k.slice()}var N=[];var R;var O=this.getElementsByTagName("thead");if(O.length!==0){pt(S.aoHeader,O[0]);N=gt(S)}if(s.aoColumns===null){R=[];for(u=0,f=N.length;u<f;u++){R.push(null)}}else{R=s.aoColumns}for(u=0,f=R.length;u<f;u++){L(S,N?N[u]:null)}$(S,s.aoColumnDefs,R,function(t,e){j(S,t,e)});if(M.length){var H=function(t,e){return t.getAttribute("data-"+e)!==null?e:null};t(M[0]).children("th, td").each(function(t,e){var n=S.aoColumns[t];if(n.mData===t){var r=H(e,"sort")||H(e,"order");var a=H(e,"filter")||H(e,"search");if(r!==null||a!==null){n.mData={_:t+".display",sort:r!==null?t+".@data-"+r:i,type:r!==null?t+".@data-"+r:i,filter:a!==null?t+".@data-"+a:i};j(S,t)}}})}var W=S.oFeatures;if(s.bStateSave){W.bStateSave=true;le(S,s);pe(S,"aoDrawCallback",oe,"state_save")}if(s.aaSorting===i){var z=S.aaSorting;for(u=0,f=z.length;u<f;u++){z[u][1]=S.aoColumns[u].asSorting[0]}}ae(S);if(W.bSort){pe(S,"aoDrawCallback",function(){if(S.bSorted){var e=te(S);var n={};t.each(e,function(t,e){n[e.src]=e.dir});ge(S,null,"order",[S,e,n]);ne(S)}})}pe(S,"aoDrawCallback",function(){if(S.bSorted||ye(S)==="ssp"||W.bDeferRender){ae(S)}},"sc");var q=y.children("caption").each(function(){this._captionSide=y.css("caption-side")});var B=y.children("thead");if(B.length===0){B=t("<thead/>").appendTo(this)}S.nTHead=B[0];var V=y.children("tbody");if(V.length===0){V=t("<tbody/>").appendTo(this)}S.nTBody=V[0];var X=y.children("tfoot");if(X.length===0&&q.length>0&&(S.oScroll.sX!==""||S.oScroll.sY!=="")){X=t("<tfoot/>").appendTo(this)}if(X.length===0||X.children().length===0){y.addClass(D.sNoFooter)}else if(X.length>0){S.nTFoot=X[0];pt(S.aoFooter,S.nTFoot)}if(s.aaData){for(u=0;u<s.aaData.length;u++){U(S,s.aaData[u])}}else if(S.bDeferLoading||ye(S)=="dom"){Y(S,t(S.nTBody).children("tr"))}S.aiDisplay=S.aiDisplayMaster.slice();S.bInitialised=true;if(v===false){Lt(S)}});n=null;return this};var be=[];var we=Array.prototype;var _e=function(e){var n,i;var a=r.settings;var s=t.map(a,function(t,e){return t.nTable});if(!e){return[]}else if(e.nTable&&e.oApi){return[e]}else if(e.nodeName&&e.nodeName.toLowerCase()==="table"){n=t.inArray(e,s);return n!==-1?[a[n]]:null}else if(e&&typeof e.settings==="function"){return e.settings().toArray()}else if(typeof e==="string"){i=t(e)}else if(e instanceof t){i=e}if(i){return i.map(function(e){n=t.inArray(this,s);return n!==-1?a[n]:null}).toArray()}};s=function(e,n){if(!(this instanceof s)){return new s(e,n)}var i=[];var r=function(t){var e=_e(t);if(e){i=i.concat(e)}};if(t.isArray(e)){for(var a=0,o=e.length;a<o;a++){r(e[a])}}else{r(e)}this.context=k(i);if(n){t.merge(this,n)}this.selector={rows:null,cols:null,opts:null};s.extend(this,this,be)};r.Api=s;t.extend(s.prototype,{any:function(){return this.count()!==0},concat:we.concat,context:[],count:function(){return this.flatten().length},each:function(t){for(var e=0,n=this.length;e<n;e++){t.call(this,this[e],e,this)}return this},eq:function(t){var e=this.context;return e.length>t?new s(e[t],this[t]):null},filter:function(t){var e=[];if(we.filter){e=we.filter.call(this,t,this)}else{for(var n=0,i=this.length;n<i;n++){if(t.call(this,this[n],n,this)){e.push(this[n])}}}return new s(this.context,e)},flatten:function(){var t=[];return new s(this.context,t.concat.apply(t,this.toArray()))},join:we.join,indexOf:we.indexOf||function(t,e){for(var n=e||0,i=this.length;n<i;n++){if(this[n]===t){return n}}return-1},iterator:function(t,e,n,r){var a=[],o,l,u,f,c,h=this.context,d,p,g,v=this.selector;if(typeof t==="string"){r=n;n=e;e=t;t=false}for(l=0,u=h.length;l<u;l++){var m=new s(h[l]);if(e==="table"){o=n.call(m,h[l],l);if(o!==i){a.push(o)}}else if(e==="columns"||e==="rows"){o=n.call(m,h[l],this[l],l);if(o!==i){a.push(o)}}else if(e==="column"||e==="column-rows"||e==="row"||e==="cell"){p=this[l];if(e==="column-rows"){d=ke(h[l],v.opts)}for(f=0,c=p.length;f<c;f++){g=p[f];if(e==="cell"){o=n.call(m,h[l],g.row,g.column,l,f)}else{o=n.call(m,h[l],g,l,f,d)}if(o!==i){a.push(o)}}}}if(a.length||r){var y=new s(h,t?a.concat.apply([],a):a);var b=y.selector;b.rows=v.rows;b.cols=v.cols;b.opts=v.opts;return y}return this},lastIndexOf:we.lastIndexOf||function(t,e){return this.indexOf.apply(this.toArray.reverse(),arguments)},length:0,map:function(t){var e=[];if(we.map){e=we.map.call(this,t,this)}else{for(var n=0,i=this.length;n<i;n++){e.push(t.call(this,this[n],n))}}return new s(this.context,e)},pluck:function(t){return this.map(function(e){return e[t]})},pop:we.pop,push:we.push,reduce:we.reduce||function(t,e){return R(this,t,e,0,this.length,1)},reduceRight:we.reduceRight||function(t,e){return R(this,t,e,this.length-1,-1,-1)},reverse:we.reverse,selector:null,shift:we.shift,sort:we.sort,splice:we.splice,toArray:function(){return we.slice.call(this)},to$:function(){return t(this)},toJQuery:function(){return t(this)},unique:function(){return new s(this.context,k(this))},unshift:we.unshift});s.extend=function(e,n,i){if(!i.length||!n||!(n instanceof s)&&!n.__dt_wrapper){return}var r,a,o,l,u,f,c=function(t,e,n){return function(){var i=e.apply(t,arguments);s.extend(i,i,n.methodExt);return i}};for(r=0,a=i.length;r<a;r++){u=i[r];n[u.name]=typeof u.val==="function"?c(e,u.val,u):t.isPlainObject(u.val)?{}:u.val;n[u.name].__dt_wrapper=true;s.extend(e,n[u.name],u.propExt)}};s.register=o=function(e,n){if(t.isArray(e)){for(var i=0,r=e.length;i<r;i++){s.register(e[i],n)}return}var a,o,l=e.split("."),u=be,f,c;var h=function(t,e){for(var n=0,i=t.length;n<i;n++){if(t[n].name===e){return t[n]}}return null};for(a=0,o=l.length;a<o;a++){c=l[a].indexOf("()")!==-1;f=c?l[a].replace("()",""):l[a];var d=h(u,f);if(!d){d={name:f,val:{},methodExt:[],propExt:[]};u.push(d)}if(a===o-1){d.val=n}else{u=c?d.methodExt:d.propExt}}};s.registerPlural=l=function(e,n,r){s.register(e,r);s.register(n,function(){var e=r.apply(this,arguments);if(e===this){return this}else if(e instanceof s){return e.length?t.isArray(e[0])?new s(e.context,e[0]):e[0]:i}return e})};var xe=function(e,n){if(typeof e==="number"){return[n[e]]}var i=t.map(n,function(t,e){return t.nTable});return t(i).filter(e).map(function(e){var r=t.inArray(this,i);return n[r]}).toArray()};o("tables()",function(t){return t?new s(xe(t,this.context)):this});o("table()",function(t){var e=this.tables(t);var n=e.context;return n.length?new s(n[0]):e});l("tables().nodes()","table().node()",function(){return this.iterator("table",function(t){return t.nTable},1)});l("tables().body()","table().body()",function(){return this.iterator("table",function(t){return t.nTBody},1)});l("tables().header()","table().header()",function(){return this.iterator("table",function(t){return t.nTHead},1)});l("tables().footer()","table().footer()",function(){return this.iterator("table",function(t){return t.nTFoot},1)});l("tables().containers()","table().container()",function(){return this.iterator("table",function(t){return t.nTableWrapper},1)});o("draw()",function(t){return this.iterator("table",function(e){if(t==="page"){ct(e)}else{if(typeof t==="string"){t=t==="full-hold"?false:true}ht(e,t===false)}})});o("page()",function(t){if(t===i){return this.page.info().page}return this.iterator("table",function(e){zt(e,t)})});o("page.info()",function(t){if(this.context.length===0){return i}var e=this.context[0],n=e._iDisplayStart,r=e.oFeatures.bPaginate?e._iDisplayLength:-1,a=e.fnRecordsDisplay(),s=r===-1;return{page:s?0:Math.floor(n/r),pages:s?1:Math.ceil(a/r),start:n,end:e.fnDisplayEnd(),length:r,recordsTotal:e.fnRecordsTotal(),recordsDisplay:a,serverSide:ye(e)==="ssp"}});o("page.len()",function(t){if(t===i){return this.context.length!==0?this.context[0]._iDisplayLength:i}return this.iterator("table",function(e){Ot(e,t)})});var Se=function(t,e,n){if(n){var i=new s(t);i.one("draw",function(){n(i.ajax.json())})}if(ye(t)=="ssp"){ht(t,e)}else{Bt(t,true);var r=t.jqXHR;if(r&&r.readyState!==4){r.abort()}vt(t,[],function(n){it(t);var i=wt(t,n);for(var r=0,a=i.length;r<a;r++){U(t,i[r])}ht(t,e);Bt(t,false)})}};o("ajax.json()",function(){var t=this.context;if(t.length>0){return t[0].json}});o("ajax.params()",function(){var t=this.context;if(t.length>0){return t[0].oAjaxData}});o("ajax.reload()",function(t,e){return this.iterator("table",function(n){Se(n,e===false,t)})});o("ajax.url()",function(e){var n=this.context;if(e===i){if(n.length===0){return i}n=n[0];return n.ajax?t.isPlainObject(n.ajax)?n.ajax.url:n.ajax:n.sAjaxSource}return this.iterator("table",function(n){if(t.isPlainObject(n.ajax)){n.ajax.url=e}else{n.ajax=e}})});o("ajax.url().load()",function(t,e){return this.iterator("table",function(n){Se(n,e===false,t)})});var De=function(e,n,r,s,o){var l=[],u,f,c,h,d,p,g=typeof n;if(!n||g==="string"||g==="function"||n.length===i){n=[n]}for(c=0,h=n.length;c<h;c++){f=n[c]&&n[c].split?n[c].split(","):[n[c]];for(d=0,p=f.length;d<p;d++){u=r(typeof f[d]==="string"?t.trim(f[d]):f[d]);if(u&&u.length){l=l.concat(u)}}}var v=a.selector[e];if(v.length){for(c=0,h=v.length;c<h;c++){l=v[c](s,o,l)}}return k(l)};var Ce=function(e){if(!e){e={}}if(e.filter&&e.search===i){e.search=e.filter}return t.extend({search:"none",order:"current",page:"all"},e)};var Te=function(t){for(var e=0,n=t.length;e<n;e++){if(t[e].length>0){t[0]=t[e];t[0].length=1;t.length=1;t.context=[t.context[e]];return t}}t.length=0;return t};var ke=function(e,n){var i,r,a,s=[],o=e.aiDisplay,l=e.aiDisplayMaster;var u=n.search,f=n.order,c=n.page;if(ye(e)=="ssp"){return u==="removed"?[]:D(0,l.length)}else if(c=="current"){for(i=e._iDisplayStart,r=e.fnDisplayEnd();i<r;i++){s.push(o[i])}}else if(f=="current"||f=="applied"){s=u=="none"?l.slice():u=="applied"?o.slice():t.map(l,function(e,n){return t.inArray(e,o)===-1?e:null})}else if(f=="index"||f=="original"){for(i=0,r=e.aoData.length;i<r;i++){if(u=="none"){s.push(i)}else{a=t.inArray(i,o);if(a===-1&&u=="removed"||a>=0&&u=="applied"){s.push(i)}}}}return s};var Me=function(e,n,r){var a=function(n){var a=m(n);var s,o;if(a!==null&&!r){return[a]}var l=ke(e,r);if(a!==null&&t.inArray(a,l)!==-1){return[a]}else if(!n){return l}if(typeof n==="function"){return t.map(l,function(t){var i=e.aoData[t];return n(t,i._aData,i.nTr)?t:null})}var u=C(S(e.aoData,l,"nTr"));if(n.nodeName){if(n._DT_RowIndex!==i){return[n._DT_RowIndex]}else if(n._DT_CellIndex){return[n._DT_CellIndex.row]}else{var f=t(n).closest("*[data-dt-row]");return f.length?[f.data("dt-row")]:[]}}if(typeof n==="string"&&n.charAt(0)==="#"){var c=e.aIds[n.replace(/^#/,"")];if(c!==i){return[c.idx]}}return t(u).filter(n).map(function(){return this._DT_RowIndex}).toArray()};return De("row",n,a,e,r)};o("rows()",function(e,n){if(e===i){e=""}else if(t.isPlainObject(e)){n=e;e=""}n=Ce(n);var r=this.iterator("table",function(t){return Me(t,e,n)},1);r.selector.rows=e;r.selector.opts=n;return r});o("rows().nodes()",function(){return this.iterator("row",function(t,e){return t.aoData[e].nTr||i},1)});o("rows().data()",function(){return this.iterator(true,"rows",function(t,e){return S(t.aoData,e,"_aData")},1)});l("rows().cache()","row().cache()",function(t){return this.iterator("row",function(e,n){var i=e.aoData[n];return t==="search"?i._aFilterData:i._aSortData},1)});l("rows().invalidate()","row().invalidate()",function(t){return this.iterator("row",function(e,n){at(e,n,t)})});l("rows().indexes()","row().index()",function(){return this.iterator("row",function(t,e){return e},1)});l("rows().ids()","row().id()",function(t){var e=[];var n=this.context;for(var i=0,r=n.length;i<r;i++){for(var a=0,o=this[i].length;a<o;a++){var l=n[i].rowIdFn(n[i].aoData[this[i][a]]._aData);e.push((t===true?"#":"")+l)}}return new s(n,e)});l("rows().remove()","row().remove()",function(){var t=this;this.iterator("row",function(e,n,r){var a=e.aoData;var s=a[n];var o,l,u,f;var c,h;a.splice(n,1);for(o=0,l=a.length;o<l;o++){c=a[o];h=c.anCells;if(c.nTr!==null){c.nTr._DT_RowIndex=o}if(h!==null){for(u=0,f=h.length;u<f;u++){h[u]._DT_CellIndex.row=o}}}rt(e.aiDisplayMaster,n);rt(e.aiDisplay,n);rt(t[r],n,false);ve(e);var d=e.rowIdFn(s._aData);if(d!==i){delete e.aIds[d]}});this.iterator("table",function(t){for(var e=0,n=t.aoData.length;e<n;e++){t.aoData[e].idx=e}});return this});o("rows.add()",function(e){var n=this.iterator("table",function(t){var n,i,r;var a=[];for(i=0,r=e.length;i<r;i++){n=e[i];if(n.nodeName&&n.nodeName.toUpperCase()==="TR"){a.push(Y(t,n)[0])}else{a.push(U(t,n))}}return a},1);var i=this.rows(-1);i.pop();t.merge(i,n);return i});o("row()",function(t,e){return Te(this.rows(t,e))});o("row().data()",function(t){var e=this.context;if(t===i){return e.length&&this.length?e[0].aoData[this[0]]._aData:i}e[0].aoData[this[0]]._aData=t;at(e[0],this[0],"data");return this});o("row().node()",function(){var t=this.context;return t.length&&this.length?t[0].aoData[this[0]].nTr||null:null});o("row.add()",function(e){if(e instanceof t&&e.length){e=e[0]}var n=this.iterator("table",function(t){if(e.nodeName&&e.nodeName.toUpperCase()==="TR"){return Y(t,e)[0]}return U(t,e)});return this.row(n[0])});var Ie=function(e,n,i,r){var a=[];var s=function(n,i){if(t.isArray(n)||n instanceof t){for(var r=0,o=n.length;r<o;r++){s(n[r],i)}return}if(n.nodeName&&n.nodeName.toLowerCase()==="tr"){a.push(n)}else{var l=t("<tr><td/></tr>").addClass(i);t("td",l).addClass(i).html(n)[0].colSpan=z(e);a.push(l[0])}};s(i,r);if(n._details){n._details.remove()}n._details=t(a);if(n._detailsShow){n._details.insertAfter(n.nTr)}};var Ae=function(t,e){var n=t.context;if(n.length){var r=n[0].aoData[e!==i?e:t[0]];if(r&&r._details){r._details.remove();r._detailsShow=i;r._details=i}}};var Ne=function(t,e){var n=t.context;if(n.length&&t.length){var i=n[0].aoData[t[0]];if(i._details){i._detailsShow=e;if(e){i._details.insertAfter(i.nTr)}else{i._details.detach()}Pe(n[0])}}};var Pe=function(t){var e=new s(t);var n=".dt.DT_details";var i="draw"+n;var r="column-visibility"+n;var a="destroy"+n;var o=t.aoData;e.off(i+" "+r+" "+a);if(x(o,"_details").length>0){e.on(i,function(n,i){if(t!==i){return}e.rows({page:"current"}).eq(0).each(function(t){var e=o[t];if(e._detailsShow){e._details.insertAfter(e.nTr)}})});e.on(r,function(e,n,i,r){if(t!==n){return}var a,s=z(n);for(var l=0,u=o.length;l<u;l++){a=o[l];if(a._details){a._details.children("td[colspan]").attr("colspan",s)}}});e.on(a,function(n,i){if(t!==i){return}for(var r=0,a=o.length;r<a;r++){if(o[r]._details){Ae(e,r)}}})}};var Ee="";var Fe=Ee+"row().child";var Re=Fe+"()";o(Re,function(t,e){var n=this.context;if(t===i){return n.length&&this.length?n[0].aoData[this[0]]._details:i}else if(t===true){this.child.show()}else if(t===false){Ae(this)}else if(n.length&&this.length){Ie(n[0],n[0].aoData[this[0]],t,e)}return this});o([Fe+".show()",Re+".show()"],function(t){Ne(this,true);return this});o([Fe+".hide()",Re+".hide()"],function(){Ne(this,false);return this});o([Fe+".remove()",Re+".remove()"],function(){Ae(this);return this});o(Fe+".isShown()",function(){var t=this.context;if(t.length&&this.length){return t[0].aoData[this[0]]._detailsShow||false}return false});var Le=/^(.+):(name|visIdx|visible)$/;var je=function(t,e,n,i,r){var a=[];for(var s=0,o=r.length;s<o;s++){a.push(J(t,r[s],e))}return a};var Oe=function(e,n,i){var r=e.aoColumns,a=x(r,"sName"),s=x(r,"nTh");var o=function(n){var o=m(n);if(n===""){return D(r.length)}if(o!==null){return[o>=0?o:r.length+o]}if(typeof n==="function"){var l=ke(e,i);return t.map(r,function(t,i){return n(i,je(e,i,0,0,l),s[i])?i:null})}var u=typeof n==="string"?n.match(Le):"";if(u){switch(u[2]){case"visIdx":case"visible":var f=parseInt(u[1],10);if(f<0){var c=t.map(r,function(t,e){return t.bVisible?e:null});return[c[c.length+f]]}return[H(e,f)];case"name":return t.map(a,function(t,e){return t===u[1]?e:null});default:return[]}}if(n.nodeName&&n._DT_CellIndex){return[n._DT_CellIndex.column]}var h=t(s).filter(n).map(function(){return t.inArray(this,s)}).toArray();if(h.length||!n.nodeName){return h}var d=t(n).closest("*[data-dt-column]");return d.length?[d.data("dt-column")]:[]};return De("column",n,o,e,i)};var He=function(e,n,r,a){var s=e.aoColumns,o=s[n],l=e.aoData,u,f,c,h,d;if(r===i){return o.bVisible}if(o.bVisible===r){return}if(r){var p=t.inArray(true,x(s,"bVisible"),n+1);for(c=0,h=l.length;c<h;c++){d=l[c].nTr;f=l[c].anCells;if(d){d.insertBefore(f[n],f[p]||null)}}}else{t(x(e.aoData,"anCells",n)).detach()}o.bVisible=r;ft(e,e.aoHeader);ft(e,e.aoFooter);if(a===i||a){O(e)}ge(e,null,"column-visibility",[e,n,r,a]);oe(e)};o("columns()",function(e,n){if(e===i){e=""}else if(t.isPlainObject(e)){n=e;e=""}n=Ce(n);var r=this.iterator("table",function(t){return Oe(t,e,n)},1);r.selector.cols=e;r.selector.opts=n;return r});l("columns().header()","column().header()",function(t,e){return this.iterator("column",function(t,e){return t.aoColumns[e].nTh},1)});l("columns().footer()","column().footer()",function(t,e){return this.iterator("column",function(t,e){return t.aoColumns[e].nTf},1)});l("columns().data()","column().data()",function(){return this.iterator("column-rows",je,1)});l("columns().dataSrc()","column().dataSrc()",function(){return this.iterator("column",function(t,e){return t.aoColumns[e].mData},1)});l("columns().cache()","column().cache()",function(t){return this.iterator("column-rows",function(e,n,i,r,a){return S(e.aoData,a,t==="search"?"_aFilterData":"_aSortData",n)},1)});l("columns().nodes()","column().nodes()",function(){return this.iterator("column-rows",function(t,e,n,i,r){return S(t.aoData,r,"anCells",e)},1)});l("columns().visible()","column().visible()",function(t,e){return this.iterator("column",function(n,r){if(t===i){return n.aoColumns[r].bVisible}He(n,r,t,e)})});l("columns().indexes()","column().index()",function(t){return this.iterator("column",function(e,n){return t==="visible"?W(e,n):n},1)});o("columns.adjust()",function(){return this.iterator("table",function(t){O(t)},1)});o("column.index()",function(t,e){if(this.context.length!==0){var n=this.context[0];if(t==="fromVisible"||t==="toData"){return H(n,e)}else if(t==="fromData"||t==="toVisible"){return W(n,e)}}});o("column()",function(t,e){return Te(this.columns(t,e))});var We=function(e,n,r){var a=e.aoData;var s=ke(e,r);var o=C(S(a,s,"anCells"));var l=t([].concat.apply([],o));var u;var f=e.aoColumns.length;var c,h,d,p,g,v;var m=function(n){var r=typeof n==="function";if(n===null||n===i||r){c=[];for(h=0,d=s.length;h<d;h++){u=s[h];for(p=0;p<f;p++){g={row:u,column:p};if(r){v=a[u];if(n(g,J(e,u,p),v.anCells?v.anCells[p]:null)){c.push(g)}}else{c.push(g)}}}return c}if(t.isPlainObject(n)){return[n]}var o=l.filter(n).map(function(t,e){return{row:e._DT_CellIndex.row,column:e._DT_CellIndex.column}}).toArray();if(o.length||!n.nodeName){return o}v=t(n).closest("*[data-dt-row]");return v.length?[{row:v.data("dt-row"),column:v.data("dt-column")}]:[]};return De("cell",n,m,e,r)};o("cells()",function(e,n,r){if(t.isPlainObject(e)){if(e.row===i){r=e;e=null}else{r=n;n=null}}if(t.isPlainObject(n)){r=n;n=null}if(n===null||n===i){return this.iterator("table",function(t){return We(t,e,Ce(r))})}var a=this.columns(n,r);var s=this.rows(e,r);var o,l,u,f,c;var h=this.iterator("table",function(t,e){o=[];for(l=0,u=s[e].length;l<u;l++){for(f=0,c=a[e].length;f<c;f++){o.push({row:s[e][l],column:a[e][f]})}}return o},1);t.extend(h.selector,{cols:n,rows:e,opts:r});return h});l("cells().nodes()","cell().node()",function(){return this.iterator("cell",function(t,e,n){var r=t.aoData[e];return r&&r.anCells?r.anCells[n]:i},1)});o("cells().data()",function(){return this.iterator("cell",function(t,e,n){return J(t,e,n)},1)});l("cells().cache()","cell().cache()",function(t){t=t==="search"?"_aFilterData":"_aSortData";return this.iterator("cell",function(e,n,i){return e.aoData[n][t][i]},1)});l("cells().render()","cell().render()",function(t){return this.iterator("cell",function(e,n,i){return J(e,n,i,t)},1)});l("cells().indexes()","cell().index()",function(){return this.iterator("cell",function(t,e,n){return{row:e,column:n,columnVisible:W(t,n)}},1)});l("cells().invalidate()","cell().invalidate()",function(t){return this.iterator("cell",function(e,n,i){at(e,n,t,i)})});o("cell()",function(t,e,n){return Te(this.cells(t,e,n))});o("cell().data()",function(t){var e=this.context;var n=this[0];if(t===i){return e.length&&n.length?J(e[0],n[0].row,n[0].column):i}G(e[0],n[0].row,n[0].column,t);at(e[0],n[0].row,"data",n[0].column);return this});o("order()",function(e,n){var r=this.context;if(e===i){return r.length!==0?r[0].aaSorting:i}if(typeof e==="number"){e=[[e,n]]}else if(!t.isArray(e[0])){e=Array.prototype.slice.call(arguments)}return this.iterator("table",function(t){t.aaSorting=e.slice()})});o("order.listener()",function(t,e,n){return this.iterator("table",function(i){re(i,t,e,n)})});o("order.fixed()",function(e){if(!e){var n=this.context;var r=n.length?n[0].aaSortingFixed:i;return t.isArray(r)?{pre:r}:r}return this.iterator("table",function(n){n.aaSortingFixed=t.extend(true,{},e)})});o(["columns().order()","column().order()"],function(e){var n=this;return this.iterator("table",function(i,r){var a=[];t.each(n[r],function(t,n){a.push([n,e])});i.aaSorting=a})});o("search()",function(e,n,r,a){var s=this.context;if(e===i){return s.length!==0?s[0].oPreviousSearch.sSearch:i}return this.iterator("table",function(i){if(!i.oFeatures.bFilter){return}xt(i,t.extend({},i.oPreviousSearch,{sSearch:e+"",bRegex:n===null?false:n,bSmart:r===null?true:r,bCaseInsensitive:a===null?true:a}),1)})});l("columns().search()","column().search()",function(e,n,r,a){return this.iterator("column",function(s,o){var l=s.aoPreSearchCols;if(e===i){return l[o].sSearch}if(!s.oFeatures.bFilter){return}t.extend(l[o],{sSearch:e+"",bRegex:n===null?false:n,bSmart:r===null?true:r,bCaseInsensitive:a===null?true:a});xt(s,s.oPreviousSearch,1)})});o("state()",function(){return this.context.length?this.context[0].oSavedState:null});o("state.clear()",function(){return this.iterator("table",function(t){t.fnStateSaveCallback.call(t.oInstance,t,{})})});o("state.loaded()",function(){return this.context.length?this.context[0].oLoadedState:null});o("state.save()",function(){return this.iterator("table",function(t){oe(t)})});r.versionCheck=r.fnVersionCheck=function(t){var e=r.version.split(".");var n=t.split(".");var i,a;for(var s=0,o=n.length;s<o;s++){i=parseInt(e[s],10)||0;a=parseInt(n[s],10)||0;if(i===a){continue}return i>a}return true};r.isDataTable=r.fnIsDataTable=function(e){var n=t(e).get(0);var i=false;t.each(r.settings,function(e,r){var a=r.nScrollHead?t("table",r.nScrollHead)[0]:null;var s=r.nScrollFoot?t("table",r.nScrollFoot)[0]:null;if(r.nTable===n||a===n||s===n){i=true}});return i};r.tables=r.fnTables=function(e){var n=false;if(t.isPlainObject(e)){n=e.api;e=e.visible}var i=t.map(r.settings,function(n){if(!e||e&&t(n.nTable).is(":visible")){return n.nTable}});return n?new s(i):i};r.util={throttle:Jt,escapeRegex:kt};r.camelToHungarian=I;o("$()",function(e,n){var i=this.rows(n).nodes(),r=t(i);return t([].concat(r.filter(e).toArray(),r.find(e).toArray()))});t.each(["on","one","off"],function(e,n){o(n+"()",function(){var e=Array.prototype.slice.call(arguments);if(!e[0].match(/\.dt\b/)){e[0]+=".dt"}var i=t(this.tables().nodes());i[n].apply(i,e);return this})});o("clear()",function(){return this.iterator("table",function(t){it(t)})});o("settings()",function(){
return new s(this.context,this.context)});o("init()",function(){var t=this.context;return t.length?t[0].oInit:null});o("data()",function(){return this.iterator("table",function(t){return x(t.aoData,"_aData")}).flatten()});o("destroy()",function(n){n=n||false;return this.iterator("table",function(i){var a=i.nTableWrapper.parentNode;var o=i.oClasses;var l=i.nTable;var u=i.nTBody;var f=i.nTHead;var c=i.nTFoot;var h=t(l);var d=t(u);var p=t(i.nTableWrapper);var g=t.map(i.aoData,function(t){return t.nTr});var v,m;i.bDestroying=true;ge(i,"aoDestroyCallback","destroy",[i]);if(!n){new s(i).columns().visible(true)}p.unbind(".DT").find(":not(tbody *)").unbind(".DT");t(e).unbind(".DT-"+i.sInstance);if(l!=f.parentNode){h.children("thead").detach();h.append(f)}if(c&&l!=c.parentNode){h.children("tfoot").detach();h.append(c)}i.aaSorting=[];i.aaSortingFixed=[];ae(i);t(g).removeClass(i.asStripeClasses.join(" "));t("th, td",f).removeClass(o.sSortable+" "+o.sSortableAsc+" "+o.sSortableDesc+" "+o.sSortableNone);if(i.bJUI){t("th span."+o.sSortIcon+", td span."+o.sSortIcon,f).detach();t("th, td",f).each(function(){var e=t("div."+o.sSortJUIWrapper,this);t(this).append(e.contents());e.detach()})}d.children().detach();d.append(g);var y=n?"remove":"detach";h[y]();p[y]();if(!n&&a){a.insertBefore(l,i.nTableReinsertBefore);h.css("width",i.sDestroyWidth).removeClass(o.sTable);m=i.asDestroyStripes.length;if(m){d.children().each(function(e){t(this).addClass(i.asDestroyStripes[e%m])})}}var b=t.inArray(i,r.settings);if(b!==-1){r.settings.splice(b,1)}})});t.each(["column","row","cell"],function(t,e){o(e+"s().every()",function(t){var n=this.selector.opts;var r=this;return this.iterator(e,function(a,s,o,l,u){t.call(r[e](s,e==="cell"?o:n,e==="cell"?n:i),s,o,l,u)})})});o("i18n()",function(e,n,r){var a=this.context[0];var s=tt(e)(a.oLanguage);if(s===i){s=n}if(r!==i&&t.isPlainObject(s)){s=s[r]!==i?s[r]:s._}return s.replace("%d",r)});r.version="1.10.11";r.settings=[];r.models={};r.models.oSearch={bCaseInsensitive:true,sSearch:"",bRegex:false,bSmart:true};r.models.oRow={nTr:null,anCells:null,_aData:[],_aSortData:null,_aFilterData:null,_sFilterRow:null,_sRowStripe:"",src:null,idx:-1};r.models.oColumn={idx:null,aDataSort:null,asSorting:null,bSearchable:null,bSortable:null,bVisible:null,_sManualType:null,_bAttrSrc:false,fnCreatedCell:null,fnGetData:null,fnSetData:null,mData:null,mRender:null,nTh:null,nTf:null,sClass:null,sContentPadding:null,sDefaultContent:null,sName:null,sSortDataType:"std",sSortingClass:null,sSortingClassJUI:null,sTitle:null,sType:null,sWidth:null,sWidthOrig:null};r.defaults={aaData:null,aaSorting:[[0,"asc"]],aaSortingFixed:[],ajax:null,aLengthMenu:[10,25,50,100],aoColumns:null,aoColumnDefs:null,aoSearchCols:[],asStripeClasses:null,bAutoWidth:true,bDeferRender:false,bDestroy:false,bFilter:true,bInfo:true,bJQueryUI:false,bLengthChange:true,bPaginate:true,bProcessing:false,bRetrieve:false,bScrollCollapse:false,bServerSide:false,bSort:true,bSortMulti:true,bSortCellsTop:false,bSortClasses:true,bStateSave:false,fnCreatedRow:null,fnDrawCallback:null,fnFooterCallback:null,fnFormatNumber:function(t){return t.toString().replace(/\B(?=(\d{3})+(?!\d))/g,this.oLanguage.sThousands)},fnHeaderCallback:null,fnInfoCallback:null,fnInitComplete:null,fnPreDrawCallback:null,fnRowCallback:null,fnServerData:null,fnServerParams:null,fnStateLoadCallback:function(t){try{return JSON.parse((t.iStateDuration===-1?sessionStorage:localStorage).getItem("DataTables_"+t.sInstance+"_"+location.pathname))}catch(t){}},fnStateLoadParams:null,fnStateLoaded:null,fnStateSaveCallback:function(t,e){try{(t.iStateDuration===-1?sessionStorage:localStorage).setItem("DataTables_"+t.sInstance+"_"+location.pathname,JSON.stringify(e))}catch(t){}},fnStateSaveParams:null,iStateDuration:7200,iDeferLoading:null,iDisplayLength:10,iDisplayStart:0,iTabIndex:0,oClasses:{},oLanguage:{oAria:{sSortAscending:": activate to sort column ascending",sSortDescending:": activate to sort column descending"},oPaginate:{sFirst:"First",sLast:"Last",sNext:"Next",sPrevious:"Previous"},sEmptyTable:"No data available in table",sInfo:"Showing _START_ to _END_ of _TOTAL_ entries",sInfoEmpty:"Showing 0 to 0 of 0 entries",sInfoFiltered:"(filtered from _MAX_ total entries)",sInfoPostFix:"",sDecimal:"",sThousands:",",sLengthMenu:"Show _MENU_ entries",sLoadingRecords:"Loading...",sProcessing:"Processing...",sSearch:"Search:",sSearchPlaceholder:"",sUrl:"",sZeroRecords:"No matching records found"},oSearch:t.extend({},r.models.oSearch),sAjaxDataProp:"data",sAjaxSource:null,sDom:"lfrtip",searchDelay:null,sPaginationType:"simple_numbers",sScrollX:"",sScrollXInner:"",sScrollY:"",sServerMethod:"GET",renderer:null,rowId:"DT_RowId"};M(r.defaults);r.defaults.column={aDataSort:null,iDataSort:-1,asSorting:["asc","desc"],bSearchable:true,bSortable:true,bVisible:true,fnCreatedCell:null,mData:null,mRender:null,sCellType:"td",sClass:"",sContentPadding:"",sDefaultContent:null,sName:"",sSortDataType:"std",sTitle:null,sType:null,sWidth:null};M(r.defaults.column);r.models.oSettings={oFeatures:{bAutoWidth:null,bDeferRender:null,bFilter:null,bInfo:null,bLengthChange:null,bPaginate:null,bProcessing:null,bServerSide:null,bSort:null,bSortMulti:null,bSortClasses:null,bStateSave:null},oScroll:{bCollapse:null,iBarWidth:0,sX:null,sXInner:null,sY:null},oLanguage:{fnInfoCallback:null},oBrowser:{bScrollOversize:false,bScrollbarLeft:false,bBounding:false,barWidth:0},ajax:null,aanFeatures:[],aoData:[],aiDisplay:[],aiDisplayMaster:[],aIds:{},aoColumns:[],aoHeader:[],aoFooter:[],oPreviousSearch:{},aoPreSearchCols:[],aaSorting:null,aaSortingFixed:[],asStripeClasses:null,asDestroyStripes:[],sDestroyWidth:0,aoRowCallback:[],aoHeaderCallback:[],aoFooterCallback:[],aoDrawCallback:[],aoRowCreatedCallback:[],aoPreDrawCallback:[],aoInitComplete:[],aoStateSaveParams:[],aoStateLoadParams:[],aoStateLoaded:[],sTableId:"",nTable:null,nTHead:null,nTFoot:null,nTBody:null,nTableWrapper:null,bDeferLoading:false,bInitialised:false,aoOpenRows:[],sDom:null,searchDelay:null,sPaginationType:"two_button",iStateDuration:0,aoStateSave:[],aoStateLoad:[],oSavedState:null,oLoadedState:null,sAjaxSource:null,sAjaxDataProp:null,bAjaxDataGet:true,jqXHR:null,json:i,oAjaxData:i,fnServerData:null,aoServerParams:[],sServerMethod:null,fnFormatNumber:null,aLengthMenu:null,iDraw:0,bDrawing:false,iDrawError:-1,_iDisplayLength:10,_iDisplayStart:0,_iRecordsTotal:0,_iRecordsDisplay:0,bJUI:null,oClasses:{},bFiltered:false,bSorted:false,bSortCellsTop:null,oInit:null,aoDestroyCallback:[],fnRecordsTotal:function(){return ye(this)=="ssp"?this._iRecordsTotal*1:this.aiDisplayMaster.length},fnRecordsDisplay:function(){return ye(this)=="ssp"?this._iRecordsDisplay*1:this.aiDisplay.length},fnDisplayEnd:function(){var t=this._iDisplayLength,e=this._iDisplayStart,n=e+t,i=this.aiDisplay.length,r=this.oFeatures,a=r.bPaginate;if(r.bServerSide){return a===false||t===-1?e+i:Math.min(e+t,this._iRecordsDisplay)}else{return!a||n>i||t===-1?i:n}},oInstance:null,sInstance:null,iTabIndex:0,nScrollHead:null,nScrollFoot:null,aLastSort:[],oPlugins:{},rowIdFn:null,rowId:null};r.ext=a={buttons:{},classes:{},builder:"-source-",errMode:"alert",feature:[],search:[],selector:{cell:[],column:[],row:[]},internal:{},legacy:{ajax:null},pager:{},renderer:{pageButton:{},header:{}},order:{},type:{detect:[],search:{},order:{}},_unique:0,fnVersionCheck:r.fnVersionCheck,iApiIndex:0,oJUIClasses:{},sVersion:r.version};t.extend(a,{afnFiltering:a.search,aTypes:a.type.detect,ofnSearch:a.type.search,oSort:a.type.order,afnSortData:a.order,aoFeatures:a.feature,oApi:a.internal,oStdClasses:a.classes,oPagination:a.pager});t.extend(r.ext.classes,{sTable:"dataTable",sNoFooter:"no-footer",sPageButton:"paginate_button",sPageButtonActive:"current",sPageButtonDisabled:"disabled",sStripeOdd:"odd",sStripeEven:"even",sRowEmpty:"dataTables_empty",sWrapper:"dataTables_wrapper",sFilter:"dataTables_filter",sInfo:"dataTables_info",sPaging:"dataTables_paginate paging_",sLength:"dataTables_length",sProcessing:"dataTables_processing",sSortAsc:"sorting_asc",sSortDesc:"sorting_desc",sSortable:"sorting",sSortableAsc:"sorting_asc_disabled",sSortableDesc:"sorting_desc_disabled",sSortableNone:"sorting_disabled",sSortColumn:"sorting_",sFilterInput:"",sLengthSelect:"",sScrollWrapper:"dataTables_scroll",sScrollHead:"dataTables_scrollHead",sScrollHeadInner:"dataTables_scrollHeadInner",sScrollBody:"dataTables_scrollBody",sScrollFoot:"dataTables_scrollFoot",sScrollFootInner:"dataTables_scrollFootInner",sHeaderTH:"",sFooterTH:"",sSortJUIAsc:"",sSortJUIDesc:"",sSortJUI:"",sSortJUIAscAllowed:"",sSortJUIDescAllowed:"",sSortJUIWrapper:"",sSortIcon:"",sJUIHeader:"",sJUIFooter:""});(function(){var e="";e="";var n=e+"ui-state-default";var i=e+"css_right ui-icon ui-icon-";var a=e+"fg-toolbar ui-toolbar ui-widget-header ui-helper-clearfix";t.extend(r.ext.oJUIClasses,r.ext.classes,{sPageButton:"fg-button ui-button "+n,sPageButtonActive:"ui-state-disabled",sPageButtonDisabled:"ui-state-disabled",sPaging:"dataTables_paginate fg-buttonset ui-buttonset fg-buttonset-multi "+"ui-buttonset-multi paging_",sSortAsc:n+" sorting_asc",sSortDesc:n+" sorting_desc",sSortable:n+" sorting",sSortableAsc:n+" sorting_asc_disabled",sSortableDesc:n+" sorting_desc_disabled",sSortableNone:n+" sorting_disabled",sSortJUIAsc:i+"triangle-1-n",sSortJUIDesc:i+"triangle-1-s",sSortJUI:i+"carat-2-n-s",sSortJUIAscAllowed:i+"carat-1-n",sSortJUIDescAllowed:i+"carat-1-s",sSortJUIWrapper:"DataTables_sort_wrapper",sSortIcon:"DataTables_sort_icon",sScrollHead:"dataTables_scrollHead "+n,sScrollFoot:"dataTables_scrollFoot "+n,sHeaderTH:n,sFooterTH:n,sJUIHeader:a+" ui-corner-tl ui-corner-tr",sJUIFooter:a+" ui-corner-bl ui-corner-br"})})();var ze=r.ext.pager;function qe(t,e){var n=[],i=ze.numbers_length,r=Math.floor(i/2),a=1;if(e<=i){n=D(0,e)}else if(t<=r){n=D(0,i-2);n.push("ellipsis");n.push(e-1)}else if(t>=e-1-r){n=D(e-(i-2),e);n.splice(0,0,"ellipsis");n.splice(0,0,0)}else{n=D(t-r+2,t+r-1);n.push("ellipsis");n.push(e-1);n.splice(0,0,"ellipsis");n.splice(0,0,0)}n.DT_el="span";return n}t.extend(ze,{simple:function(t,e){return["previous","next"]},full:function(t,e){return["first","previous","next","last"]},numbers:function(t,e){return[qe(t,e)]},simple_numbers:function(t,e){return["previous",qe(t,e),"next"]},full_numbers:function(t,e){return["first","previous",qe(t,e),"next","last"]},_numbers:qe,numbers_length:7});t.extend(true,r.ext.renderer,{pageButton:{_:function(e,i,r,a,s,o){var l=e.oClasses;var u=e.oLanguage.oPaginate;var f=e.oLanguage.oAria.paginate||{};var c,h,d=0;var p=function(n,i){var a,g,v,m;var y=function(t){zt(e,t.data.action,true)};for(a=0,g=i.length;a<g;a++){m=i[a];if(t.isArray(m)){var b=t("<"+(m.DT_el||"div")+"/>").appendTo(n);p(b,m)}else{c=null;h="";switch(m){case"ellipsis":n.append('<span class="ellipsis">&#x2026;</span>');break;case"first":c=u.sFirst;h=m+(s>0?"":" "+l.sPageButtonDisabled);break;case"previous":c=u.sPrevious;h=m+(s>0?"":" "+l.sPageButtonDisabled);break;case"next":c=u.sNext;h=m+(s<o-1?"":" "+l.sPageButtonDisabled);break;case"last":c=u.sLast;h=m+(s<o-1?"":" "+l.sPageButtonDisabled);break;default:c=m+1;h=s===m?l.sPageButtonActive:"";break}if(c!==null){v=t("<a>",{class:l.sPageButton+" "+h,"aria-controls":e.sTableId,"aria-label":f[m],"data-dt-idx":d,tabindex:e.iTabIndex,id:r===0&&typeof m==="string"?e.sTableId+"_"+m:null}).html(c).appendTo(n);de(v,{action:m},y);d++}}}};var g;try{g=t(i).find(n.activeElement).data("dt-idx")}catch(t){}p(t(i).empty(),a);if(g){t(i).find("[data-dt-idx="+g+"]").focus()}}}});t.extend(r.ext.type.detect,[function(t,e){var n=e.oLanguage.sDecimal;return b(t,n)?"num"+n:null},function(t,e){if(t&&!(t instanceof Date)&&(!h.test(t)||!d.test(t))){return null}var n=Date.parse(t);return n!==null&&!isNaN(n)||v(t)?"date":null},function(t,e){var n=e.oLanguage.sDecimal;return b(t,n,true)?"num-fmt"+n:null},function(t,e){var n=e.oLanguage.sDecimal;return _(t,n)?"html-num"+n:null},function(t,e){var n=e.oLanguage.sDecimal;return _(t,n,true)?"html-num-fmt"+n:null},function(t,e){return v(t)||typeof t==="string"&&t.indexOf("<")!==-1?"html":null}]);t.extend(r.ext.type.search,{html:function(t){return v(t)?t:typeof t==="string"?t.replace(f," ").replace(c,""):""},string:function(t){return v(t)?t:typeof t==="string"?t.replace(f," "):t}});var Be=function(t,e,n,i){if(t!==0&&(!t||t==="-")){return-Infinity}if(e){t=y(t,e)}if(t.replace){if(n){t=t.replace(n,"")}if(i){t=t.replace(i,"")}}return t*1};function $e(e){t.each({num:function(t){return Be(t,e)},"num-fmt":function(t){return Be(t,e,g)},"html-num":function(t){return Be(t,e,c)},"html-num-fmt":function(t){return Be(t,e,c,g)}},function(t,n){a.type.order[t+e+"-pre"]=n;if(t.match(/^html\-/)){a.type.search[t+e]=a.type.search.html}})}t.extend(a.type.order,{"date-pre":function(t){return Date.parse(t)||0},"html-pre":function(t){return v(t)?"":t.replace?t.replace(/<.*?>/g,"").toLowerCase():t+""},"string-pre":function(t){return v(t)?"":typeof t==="string"?t.toLowerCase():!t.toString?"":t.toString()},"string-asc":function(t,e){return t<e?-1:t>e?1:0},"string-desc":function(t,e){return t<e?1:t>e?-1:0}});$e("");t.extend(true,r.ext.renderer,{header:{_:function(e,n,i,r){t(e.nTable).on("order.dt.DT",function(t,a,s,o){if(e!==a){return}var l=i.idx;n.removeClass(i.sSortingClass+" "+r.sSortAsc+" "+r.sSortDesc).addClass(o[l]=="asc"?r.sSortAsc:o[l]=="desc"?r.sSortDesc:i.sSortingClass)})},jqueryui:function(e,n,i,r){t("<div/>").addClass(r.sSortJUIWrapper).append(n.contents()).append(t("<span/>").addClass(r.sSortIcon+" "+i.sSortingClassJUI)).appendTo(n);t(e.nTable).on("order.dt.DT",function(t,a,s,o){if(e!==a){return}var l=i.idx;n.removeClass(r.sSortAsc+" "+r.sSortDesc).addClass(o[l]=="asc"?r.sSortAsc:o[l]=="desc"?r.sSortDesc:i.sSortingClass);n.find("span."+r.sSortIcon).removeClass(r.sSortJUIAsc+" "+r.sSortJUIDesc+" "+r.sSortJUI+" "+r.sSortJUIAscAllowed+" "+r.sSortJUIDescAllowed).addClass(o[l]=="asc"?r.sSortJUIAsc:o[l]=="desc"?r.sSortJUIDesc:i.sSortingClassJUI)})}}});var Ue=function(t){return typeof t==="string"?t.replace(/</g,"&lt;").replace(/>/g,"&gt;").replace(/"/g,"&quot;"):t};r.render={number:function(t,e,n,i,r){return{display:function(a){if(typeof a!=="number"&&typeof a!=="string"){return a}var s=a<0?"-":"";var o=parseFloat(a);if(isNaN(o)){return Ue(a)}a=Math.abs(o);var l=parseInt(a,10);var u=n?e+(a-l).toFixed(n).substring(2):"";return s+(i||"")+l.toString().replace(/\B(?=(\d{3})+(?!\d))/g,t)+u+(r||"")}}},text:function(){return{display:Ue}}};function Ye(t){return function(){var e=[ue(this[r.ext.iApiIndex])].concat(Array.prototype.slice.call(arguments));return r.ext.internal[t].apply(this,e)}}t.extend(r.ext.internal,{_fnExternApiFunc:Ye,_fnBuildAjax:vt,_fnAjaxUpdate:mt,_fnAjaxParameters:yt,_fnAjaxUpdateDraw:bt,_fnAjaxDataSrc:wt,_fnAddColumn:L,_fnColumnOptions:j,_fnAdjustColumnSizing:O,_fnVisibleToColumnIndex:H,_fnColumnIndexToVisible:W,_fnVisbleColumns:z,_fnGetColumns:q,_fnColumnTypes:B,_fnApplyColumnDefs:$,_fnHungarianMap:M,_fnCamelToHungarian:I,_fnLanguageCompat:A,_fnBrowserDetect:F,_fnAddData:U,_fnAddTr:Y,_fnNodeToDataIndex:V,_fnNodeToColumnIndex:X,_fnGetCellData:J,_fnSetCellData:G,_fnSplitObjNotation:Q,_fnGetObjectDataFn:tt,_fnSetObjectDataFn:et,_fnGetDataMaster:nt,_fnClearTable:it,_fnDeleteIndex:rt,_fnInvalidate:at,_fnGetRowElements:st,_fnCreateTr:ot,_fnBuildHead:ut,_fnDrawHead:ft,_fnDraw:ct,_fnReDraw:ht,_fnAddOptionsHtml:dt,_fnDetectHeader:pt,_fnGetUniqueThs:gt,_fnFeatureHtmlFilter:_t,_fnFilterComplete:xt,_fnFilterCustom:St,_fnFilterColumn:Dt,_fnFilter:Ct,_fnFilterCreateSearch:Tt,_fnEscapeRegex:kt,_fnFilterData:At,_fnFeatureHtmlInfo:Et,_fnUpdateInfo:Ft,_fnInfoMacros:Rt,_fnInitialise:Lt,_fnInitComplete:jt,_fnLengthChange:Ot,_fnFeatureHtmlLength:Ht,_fnFeatureHtmlPaginate:Wt,_fnPageChange:zt,_fnFeatureHtmlProcessing:qt,_fnProcessingDisplay:Bt,_fnFeatureHtmlTable:$t,_fnScrollDraw:Ut,_fnApplyToChildren:Yt,_fnCalculateColumnWidths:Xt,_fnThrottle:Jt,_fnConvertToWidth:Gt,_fnGetWidestNode:Kt,_fnGetMaxLenString:Zt,_fnStringToCss:Qt,_fnSortFlatten:te,_fnSort:ee,_fnSortAria:ne,_fnSortListener:ie,_fnSortAttachListener:re,_fnSortingClasses:ae,_fnSortData:se,_fnSaveState:oe,_fnLoadState:le,_fnSettingsFromNode:ue,_fnLog:fe,_fnMap:ce,_fnBindAction:de,_fnCallbackReg:pe,_fnCallbackFire:ge,_fnLengthOverflow:ve,_fnRenderer:me,_fnDataSource:ye,_fnRowAttributes:lt,_fnCalculateEnd:function(){}});t.fn.dataTable=r;r.$=t;t.fn.dataTableSettings=r.settings;t.fn.dataTableExt=r.ext;t.fn.DataTable=function(e){return t(this).dataTable(e).api()};t.each(r,function(e,n){t.fn.DataTable[e]=n});return t.fn.dataTable})},{jquery:18}],18:[function(t,e,n){(function(t,n){"use strict";if(typeof e==="object"&&typeof e.exports==="object"){e.exports=t.document?n(t,true):function(t){if(!t.document){throw new Error("jQuery requires a window with a document")}return n(t)}}else{n(t)}})(typeof window!=="undefined"?window:this,function(t,e){"use strict";var n=[];var i=t.document;var r=Object.getPrototypeOf;var a=n.slice;var s=n.concat;var o=n.push;var l=n.indexOf;var u={};var f=u.toString;var c=u.hasOwnProperty;var h=c.toString;var d=h.call(Object);var p={};function g(t,e){e=e||i;var n=e.createElement("script");n.text=t;e.head.appendChild(n).parentNode.removeChild(n)}var v="3.1.1",m=function(t,e){return new m.fn.init(t,e)},y=/^[\s\uFEFF\xA0]+|[\s\uFEFF\xA0]+$/g,b=/^-ms-/,w=/-([a-z])/g,_=function(t,e){return e.toUpperCase()};m.fn=m.prototype={jquery:v,constructor:m,length:0,toArray:function(){return a.call(this)},get:function(t){if(t==null){return a.call(this)}return t<0?this[t+this.length]:this[t]},pushStack:function(t){var e=m.merge(this.constructor(),t);e.prevObject=this;return e},each:function(t){return m.each(this,t)},map:function(t){return this.pushStack(m.map(this,function(e,n){return t.call(e,n,e)}))},slice:function(){return this.pushStack(a.apply(this,arguments))},first:function(){return this.eq(0)},last:function(){return this.eq(-1)},eq:function(t){var e=this.length,n=+t+(t<0?e:0);return this.pushStack(n>=0&&n<e?[this[n]]:[])},end:function(){return this.prevObject||this.constructor()},push:o,sort:n.sort,splice:n.splice};m.extend=m.fn.extend=function(){var t,e,n,i,r,a,s=arguments[0]||{},o=1,l=arguments.length,u=false;if(typeof s==="boolean"){u=s;s=arguments[o]||{};o++}if(typeof s!=="object"&&!m.isFunction(s)){s={}}if(o===l){s=this;o--}for(;o<l;o++){if((t=arguments[o])!=null){for(e in t){n=s[e];i=t[e];if(s===i){continue}if(u&&i&&(m.isPlainObject(i)||(r=m.isArray(i)))){if(r){r=false;a=n&&m.isArray(n)?n:[]}else{a=n&&m.isPlainObject(n)?n:{}}s[e]=m.extend(u,a,i)}else if(i!==undefined){s[e]=i}}}}return s};m.extend({expando:"jQuery"+(v+Math.random()).replace(/\D/g,""),isReady:true,error:function(t){throw new Error(t)},noop:function(){},isFunction:function(t){return m.type(t)==="function"},isArray:Array.isArray,isWindow:function(t){return t!=null&&t===t.window},isNumeric:function(t){var e=m.type(t);return(e==="number"||e==="string")&&!isNaN(t-parseFloat(t))},isPlainObject:function(t){var e,n;if(!t||f.call(t)!=="[object Object]"){return false}e=r(t);if(!e){return true}n=c.call(e,"constructor")&&e.constructor;return typeof n==="function"&&h.call(n)===d},isEmptyObject:function(t){var e;for(e in t){return false}return true},type:function(t){if(t==null){return t+""}return typeof t==="object"||typeof t==="function"?u[f.call(t)]||"object":typeof t},globalEval:function(t){g(t)},camelCase:function(t){return t.replace(b,"ms-").replace(w,_)},nodeName:function(t,e){return t.nodeName&&t.nodeName.toLowerCase()===e.toLowerCase()},each:function(t,e){var n,i=0;if(x(t)){n=t.length;for(;i<n;i++){if(e.call(t[i],i,t[i])===false){break}}}else{for(i in t){if(e.call(t[i],i,t[i])===false){break}}}return t},trim:function(t){return t==null?"":(t+"").replace(y,"")},makeArray:function(t,e){var n=e||[];if(t!=null){if(x(Object(t))){m.merge(n,typeof t==="string"?[t]:t)}else{o.call(n,t)}}return n},inArray:function(t,e,n){return e==null?-1:l.call(e,t,n)},merge:function(t,e){var n=+e.length,i=0,r=t.length;for(;i<n;i++){t[r++]=e[i]}t.length=r;return t},grep:function(t,e,n){var i,r=[],a=0,s=t.length,o=!n;for(;a<s;a++){i=!e(t[a],a);if(i!==o){r.push(t[a])}}return r},map:function(t,e,n){var i,r,a=0,o=[];if(x(t)){i=t.length;for(;a<i;a++){r=e(t[a],a,n);if(r!=null){o.push(r)}}}else{for(a in t){r=e(t[a],a,n);if(r!=null){o.push(r)}}}return s.apply([],o)},guid:1,proxy:function(t,e){var n,i,r;if(typeof e==="string"){n=t[e];e=t;t=n}if(!m.isFunction(t)){return undefined}i=a.call(arguments,2);r=function(){return t.apply(e||this,i.concat(a.call(arguments)))};r.guid=t.guid=t.guid||m.guid++;return r},now:Date.now,support:p});if(typeof Symbol==="function"){m.fn[Symbol.iterator]=n[Symbol.iterator]}m.each("Boolean Number String Function Array Date RegExp Object Error Symbol".split(" "),function(t,e){u["[object "+e+"]"]=e.toLowerCase()});function x(t){var e=!!t&&"length"in t&&t.length,n=m.type(t);if(n==="function"||m.isWindow(t)){return false}return n==="array"||e===0||typeof e==="number"&&e>0&&e-1 in t}var S=function(t){var e,n,i,r,a,s,o,l,u,f,c,h,d,p,g,v,m,y,b,w="sizzle"+1*new Date,_=t.document,x=0,S=0,D=st(),C=st(),T=st(),k=function(t,e){if(t===e){c=true}return 0},M={}.hasOwnProperty,I=[],A=I.pop,N=I.push,P=I.push,E=I.slice,F=function(t,e){var n=0,i=t.length;for(;n<i;n++){if(t[n]===e){return n}}return-1},R="checked|selected|async|autofocus|autoplay|controls|defer|disabled|hidden|ismap|loop|multiple|open|readonly|required|scoped",L="[\\x20\\t\\r\\n\\f]",j="(?:\\\\.|[\\w-]|[^\0-\\xa0])+",O="\\["+L+"*("+j+")(?:"+L+"*([*^$|!~]?=)"+L+"*(?:'((?:\\\\.|[^\\\\'])*)'|\"((?:\\\\.|[^\\\\\"])*)\"|("+j+"))|)"+L+"*\\]",H=":("+j+")(?:\\(("+"('((?:\\\\.|[^\\\\'])*)'|\"((?:\\\\.|[^\\\\\"])*)\")|"+"((?:\\\\.|[^\\\\()[\\]]|"+O+")*)|"+".*"+")\\)|)",W=new RegExp(L+"+","g"),z=new RegExp("^"+L+"+|((?:^|[^\\\\])(?:\\\\.)*)"+L+"+$","g"),q=new RegExp("^"+L+"*,"+L+"*"),B=new RegExp("^"+L+"*([>+~]|"+L+")"+L+"*"),$=new RegExp("="+L+"*([^\\]'\"]*?)"+L+"*\\]","g"),U=new RegExp(H),Y=new RegExp("^"+j+"$"),V={ID:new RegExp("^#("+j+")"),CLASS:new RegExp("^\\.("+j+")"),TAG:new RegExp("^("+j+"|[*])"),ATTR:new RegExp("^"+O),PSEUDO:new RegExp("^"+H),CHILD:new RegExp("^:(only|first|last|nth|nth-last)-(child|of-type)(?:\\("+L+"*(even|odd|(([+-]|)(\\d*)n|)"+L+"*(?:([+-]|)"+L+"*(\\d+)|))"+L+"*\\)|)","i"),bool:new RegExp("^(?:"+R+")$","i"),needsContext:new RegExp("^"+L+"*[>+~]|:(even|odd|eq|gt|lt|nth|first|last)(?:\\("+L+"*((?:-\\d)?\\d*)"+L+"*\\)|)(?=[^-]|$)","i")},X=/^(?:input|select|textarea|button)$/i,J=/^h\d$/i,G=/^[^{]+\{\s*\[native \w/,K=/^(?:#([\w-]+)|(\w+)|\.([\w-]+))$/,Z=/[+~]/,Q=new RegExp("\\\\([\\da-f]{1,6}"+L+"?|("+L+")|.)","ig"),tt=function(t,e,n){var i="0x"+e-65536;return i!==i||n?e:i<0?String.fromCharCode(i+65536):String.fromCharCode(i>>10|55296,i&1023|56320)},et=/([\0-\x1f\x7f]|^-?\d)|^-$|[^\0-\x1f\x7f-\uFFFF\w-]/g,nt=function(t,e){if(e){if(t==="\0"){return"�"}return t.slice(0,-1)+"\\"+t.charCodeAt(t.length-1).toString(16)+" "}return"\\"+t},it=function(){h()},rt=yt(function(t){return t.disabled===true&&("form"in t||"label"in t)},{dir:"parentNode",next:"legend"});try{P.apply(I=E.call(_.childNodes),_.childNodes);I[_.childNodes.length].nodeType}catch(t){P={apply:I.length?function(t,e){N.apply(t,E.call(e))}:function(t,e){var n=t.length,i=0;while(t[n++]=e[i++]){}t.length=n-1}}}function at(t,e,i,r){var a,o,u,f,c,p,m,y=e&&e.ownerDocument,x=e?e.nodeType:9;i=i||[];if(typeof t!=="string"||!t||x!==1&&x!==9&&x!==11){return i}if(!r){if((e?e.ownerDocument||e:_)!==d){h(e)}e=e||d;if(g){if(x!==11&&(c=K.exec(t))){if(a=c[1]){if(x===9){if(u=e.getElementById(a)){if(u.id===a){i.push(u);return i}}else{return i}}else{if(y&&(u=y.getElementById(a))&&b(e,u)&&u.id===a){i.push(u);return i}}}else if(c[2]){P.apply(i,e.getElementsByTagName(t));return i}else if((a=c[3])&&n.getElementsByClassName&&e.getElementsByClassName){P.apply(i,e.getElementsByClassName(a));return i}}if(n.qsa&&!T[t+" "]&&(!v||!v.test(t))){if(x!==1){y=e;m=t}else if(e.nodeName.toLowerCase()!=="object"){if(f=e.getAttribute("id")){f=f.replace(et,nt)}else{e.setAttribute("id",f=w)}p=s(t);o=p.length;while(o--){p[o]="#"+f+" "+mt(p[o])}m=p.join(",");y=Z.test(t)&&gt(e.parentNode)||e}if(m){try{P.apply(i,y.querySelectorAll(m));return i}catch(t){}finally{if(f===w){e.removeAttribute("id")}}}}}}return l(t.replace(z,"$1"),e,i,r)}function st(){var t=[];function e(n,r){if(t.push(n+" ")>i.cacheLength){delete e[t.shift()]}return e[n+" "]=r}return e}function ot(t){t[w]=true;return t}function lt(t){var e=d.createElement("fieldset");try{return!!t(e)}catch(t){return false}finally{if(e.parentNode){e.parentNode.removeChild(e)}e=null}}function ut(t,e){var n=t.split("|"),r=n.length;while(r--){i.attrHandle[n[r]]=e}}function ft(t,e){var n=e&&t,i=n&&t.nodeType===1&&e.nodeType===1&&t.sourceIndex-e.sourceIndex;if(i){return i}if(n){while(n=n.nextSibling){if(n===e){return-1}}}return t?1:-1}function ct(t){return function(e){var n=e.nodeName.toLowerCase();return n==="input"&&e.type===t}}function ht(t){return function(e){var n=e.nodeName.toLowerCase();return(n==="input"||n==="button")&&e.type===t}}function dt(t){return function(e){if("form"in e){if(e.parentNode&&e.disabled===false){if("label"in e){if("label"in e.parentNode){return e.parentNode.disabled===t}else{return e.disabled===t}}return e.isDisabled===t||e.isDisabled!==!t&&rt(e)===t}return e.disabled===t}else if("label"in e){return e.disabled===t}return false}}function pt(t){return ot(function(e){e=+e;return ot(function(n,i){var r,a=t([],n.length,e),s=a.length;while(s--){if(n[r=a[s]]){n[r]=!(i[r]=n[r])}}})})}function gt(t){return t&&typeof t.getElementsByTagName!=="undefined"&&t}n=at.support={};a=at.isXML=function(t){var e=t&&(t.ownerDocument||t).documentElement;return e?e.nodeName!=="HTML":false};h=at.setDocument=function(t){var e,r,s=t?t.ownerDocument||t:_;if(s===d||s.nodeType!==9||!s.documentElement){return d}d=s;p=d.documentElement;g=!a(d);if(_!==d&&(r=d.defaultView)&&r.top!==r){if(r.addEventListener){r.addEventListener("unload",it,false)}else if(r.attachEvent){r.attachEvent("onunload",it)}}n.attributes=lt(function(t){t.className="i";return!t.getAttribute("className")});n.getElementsByTagName=lt(function(t){t.appendChild(d.createComment(""));return!t.getElementsByTagName("*").length});n.getElementsByClassName=G.test(d.getElementsByClassName);n.getById=lt(function(t){p.appendChild(t).id=w;return!d.getElementsByName||!d.getElementsByName(w).length});if(n.getById){i.filter["ID"]=function(t){var e=t.replace(Q,tt);return function(t){return t.getAttribute("id")===e}};i.find["ID"]=function(t,e){if(typeof e.getElementById!=="undefined"&&g){var n=e.getElementById(t);return n?[n]:[]}}}else{i.filter["ID"]=function(t){var e=t.replace(Q,tt);return function(t){var n=typeof t.getAttributeNode!=="undefined"&&t.getAttributeNode("id");return n&&n.value===e}};i.find["ID"]=function(t,e){if(typeof e.getElementById!=="undefined"&&g){var n,i,r,a=e.getElementById(t);if(a){n=a.getAttributeNode("id");if(n&&n.value===t){return[a]}r=e.getElementsByName(t);i=0;while(a=r[i++]){n=a.getAttributeNode("id");if(n&&n.value===t){return[a]}}}return[]}}}i.find["TAG"]=n.getElementsByTagName?function(t,e){if(typeof e.getElementsByTagName!=="undefined"){return e.getElementsByTagName(t)}else if(n.qsa){return e.querySelectorAll(t)}}:function(t,e){var n,i=[],r=0,a=e.getElementsByTagName(t);if(t==="*"){while(n=a[r++]){if(n.nodeType===1){i.push(n)}}return i}return a};i.find["CLASS"]=n.getElementsByClassName&&function(t,e){if(typeof e.getElementsByClassName!=="undefined"&&g){return e.getElementsByClassName(t)}};m=[];v=[];if(n.qsa=G.test(d.querySelectorAll)){lt(function(t){p.appendChild(t).innerHTML="<a id='"+w+"'></a>"+"<select id='"+w+"-\r\\' msallowcapture=''>"+"<option selected=''></option></select>";if(t.querySelectorAll("[msallowcapture^='']").length){v.push("[*^$]="+L+"*(?:''|\"\")")}if(!t.querySelectorAll("[selected]").length){v.push("\\["+L+"*(?:value|"+R+")")}if(!t.querySelectorAll("[id~="+w+"-]").length){v.push("~=")}if(!t.querySelectorAll(":checked").length){v.push(":checked")}if(!t.querySelectorAll("a#"+w+"+*").length){v.push(".#.+[+~]")}});lt(function(t){t.innerHTML="<a href='' disabled='disabled'></a>"+"<select disabled='disabled'><option/></select>";var e=d.createElement("input");e.setAttribute("type","hidden");t.appendChild(e).setAttribute("name","D");if(t.querySelectorAll("[name=d]").length){v.push("name"+L+"*[*^$|!~]?=")}if(t.querySelectorAll(":enabled").length!==2){v.push(":enabled",":disabled")}p.appendChild(t).disabled=true;if(t.querySelectorAll(":disabled").length!==2){v.push(":enabled",":disabled")}t.querySelectorAll("*,:x");v.push(",.*:")})}if(n.matchesSelector=G.test(y=p.matches||p.webkitMatchesSelector||p.mozMatchesSelector||p.oMatchesSelector||p.msMatchesSelector)){lt(function(t){n.disconnectedMatch=y.call(t,"*");y.call(t,"[s!='']:x");m.push("!=",H)})}v=v.length&&new RegExp(v.join("|"));m=m.length&&new RegExp(m.join("|"));e=G.test(p.compareDocumentPosition);b=e||G.test(p.contains)?function(t,e){var n=t.nodeType===9?t.documentElement:t,i=e&&e.parentNode;return t===i||!!(i&&i.nodeType===1&&(n.contains?n.contains(i):t.compareDocumentPosition&&t.compareDocumentPosition(i)&16))}:function(t,e){if(e){while(e=e.parentNode){if(e===t){return true}}}return false};k=e?function(t,e){if(t===e){c=true;return 0}var i=!t.compareDocumentPosition-!e.compareDocumentPosition;if(i){return i}i=(t.ownerDocument||t)===(e.ownerDocument||e)?t.compareDocumentPosition(e):1;if(i&1||!n.sortDetached&&e.compareDocumentPosition(t)===i){if(t===d||t.ownerDocument===_&&b(_,t)){return-1}if(e===d||e.ownerDocument===_&&b(_,e)){return 1}return f?F(f,t)-F(f,e):0}return i&4?-1:1}:function(t,e){if(t===e){c=true;return 0}var n,i=0,r=t.parentNode,a=e.parentNode,s=[t],o=[e];if(!r||!a){return t===d?-1:e===d?1:r?-1:a?1:f?F(f,t)-F(f,e):0}else if(r===a){return ft(t,e)}n=t;while(n=n.parentNode){s.unshift(n)}n=e;while(n=n.parentNode){o.unshift(n)}while(s[i]===o[i]){i++}return i?ft(s[i],o[i]):s[i]===_?-1:o[i]===_?1:0};return d};at.matches=function(t,e){return at(t,null,null,e)};at.matchesSelector=function(t,e){if((t.ownerDocument||t)!==d){h(t)}e=e.replace($,"='$1']");if(n.matchesSelector&&g&&!T[e+" "]&&(!m||!m.test(e))&&(!v||!v.test(e))){try{var i=y.call(t,e);if(i||n.disconnectedMatch||t.document&&t.document.nodeType!==11){return i}}catch(t){}}return at(e,d,null,[t]).length>0};at.contains=function(t,e){if((t.ownerDocument||t)!==d){h(t)}return b(t,e)};at.attr=function(t,e){if((t.ownerDocument||t)!==d){h(t)}var r=i.attrHandle[e.toLowerCase()],a=r&&M.call(i.attrHandle,e.toLowerCase())?r(t,e,!g):undefined;return a!==undefined?a:n.attributes||!g?t.getAttribute(e):(a=t.getAttributeNode(e))&&a.specified?a.value:null};at.escape=function(t){return(t+"").replace(et,nt)};at.error=function(t){throw new Error("Syntax error, unrecognized expression: "+t)};at.uniqueSort=function(t){var e,i=[],r=0,a=0;c=!n.detectDuplicates;f=!n.sortStable&&t.slice(0);t.sort(k);if(c){while(e=t[a++]){if(e===t[a]){r=i.push(a)}}while(r--){t.splice(i[r],1)}}f=null;return t};r=at.getText=function(t){var e,n="",i=0,a=t.nodeType;if(!a){while(e=t[i++]){n+=r(e)}}else if(a===1||a===9||a===11){if(typeof t.textContent==="string"){return t.textContent}else{for(t=t.firstChild;t;t=t.nextSibling){n+=r(t)}}}else if(a===3||a===4){return t.nodeValue}return n};i=at.selectors={cacheLength:50,createPseudo:ot,match:V,attrHandle:{},find:{},relative:{">":{dir:"parentNode",first:true}," ":{dir:"parentNode"},"+":{dir:"previousSibling",first:true},"~":{dir:"previousSibling"}},preFilter:{ATTR:function(t){t[1]=t[1].replace(Q,tt);t[3]=(t[3]||t[4]||t[5]||"").replace(Q,tt);if(t[2]==="~="){t[3]=" "+t[3]+" "}return t.slice(0,4)},CHILD:function(t){t[1]=t[1].toLowerCase();if(t[1].slice(0,3)==="nth"){if(!t[3]){at.error(t[0])}t[4]=+(t[4]?t[5]+(t[6]||1):2*(t[3]==="even"||t[3]==="odd"));t[5]=+(t[7]+t[8]||t[3]==="odd")}else if(t[3]){at.error(t[0])}return t},PSEUDO:function(t){var e,n=!t[6]&&t[2];if(V["CHILD"].test(t[0])){return null}if(t[3]){t[2]=t[4]||t[5]||""}else if(n&&U.test(n)&&(e=s(n,true))&&(e=n.indexOf(")",n.length-e)-n.length)){t[0]=t[0].slice(0,e)
;t[2]=n.slice(0,e)}return t.slice(0,3)}},filter:{TAG:function(t){var e=t.replace(Q,tt).toLowerCase();return t==="*"?function(){return true}:function(t){return t.nodeName&&t.nodeName.toLowerCase()===e}},CLASS:function(t){var e=D[t+" "];return e||(e=new RegExp("(^|"+L+")"+t+"("+L+"|$)"))&&D(t,function(t){return e.test(typeof t.className==="string"&&t.className||typeof t.getAttribute!=="undefined"&&t.getAttribute("class")||"")})},ATTR:function(t,e,n){return function(i){var r=at.attr(i,t);if(r==null){return e==="!="}if(!e){return true}r+="";return e==="="?r===n:e==="!="?r!==n:e==="^="?n&&r.indexOf(n)===0:e==="*="?n&&r.indexOf(n)>-1:e==="$="?n&&r.slice(-n.length)===n:e==="~="?(" "+r.replace(W," ")+" ").indexOf(n)>-1:e==="|="?r===n||r.slice(0,n.length+1)===n+"-":false}},CHILD:function(t,e,n,i,r){var a=t.slice(0,3)!=="nth",s=t.slice(-4)!=="last",o=e==="of-type";return i===1&&r===0?function(t){return!!t.parentNode}:function(e,n,l){var u,f,c,h,d,p,g=a!==s?"nextSibling":"previousSibling",v=e.parentNode,m=o&&e.nodeName.toLowerCase(),y=!l&&!o,b=false;if(v){if(a){while(g){h=e;while(h=h[g]){if(o?h.nodeName.toLowerCase()===m:h.nodeType===1){return false}}p=g=t==="only"&&!p&&"nextSibling"}return true}p=[s?v.firstChild:v.lastChild];if(s&&y){h=v;c=h[w]||(h[w]={});f=c[h.uniqueID]||(c[h.uniqueID]={});u=f[t]||[];d=u[0]===x&&u[1];b=d&&u[2];h=d&&v.childNodes[d];while(h=++d&&h&&h[g]||(b=d=0)||p.pop()){if(h.nodeType===1&&++b&&h===e){f[t]=[x,d,b];break}}}else{if(y){h=e;c=h[w]||(h[w]={});f=c[h.uniqueID]||(c[h.uniqueID]={});u=f[t]||[];d=u[0]===x&&u[1];b=d}if(b===false){while(h=++d&&h&&h[g]||(b=d=0)||p.pop()){if((o?h.nodeName.toLowerCase()===m:h.nodeType===1)&&++b){if(y){c=h[w]||(h[w]={});f=c[h.uniqueID]||(c[h.uniqueID]={});f[t]=[x,b]}if(h===e){break}}}}}b-=r;return b===i||b%i===0&&b/i>=0}}},PSEUDO:function(t,e){var n,r=i.pseudos[t]||i.setFilters[t.toLowerCase()]||at.error("unsupported pseudo: "+t);if(r[w]){return r(e)}if(r.length>1){n=[t,t,"",e];return i.setFilters.hasOwnProperty(t.toLowerCase())?ot(function(t,n){var i,a=r(t,e),s=a.length;while(s--){i=F(t,a[s]);t[i]=!(n[i]=a[s])}}):function(t){return r(t,0,n)}}return r}},pseudos:{not:ot(function(t){var e=[],n=[],i=o(t.replace(z,"$1"));return i[w]?ot(function(t,e,n,r){var a,s=i(t,null,r,[]),o=t.length;while(o--){if(a=s[o]){t[o]=!(e[o]=a)}}}):function(t,r,a){e[0]=t;i(e,null,a,n);e[0]=null;return!n.pop()}}),has:ot(function(t){return function(e){return at(t,e).length>0}}),contains:ot(function(t){t=t.replace(Q,tt);return function(e){return(e.textContent||e.innerText||r(e)).indexOf(t)>-1}}),lang:ot(function(t){if(!Y.test(t||"")){at.error("unsupported lang: "+t)}t=t.replace(Q,tt).toLowerCase();return function(e){var n;do{if(n=g?e.lang:e.getAttribute("xml:lang")||e.getAttribute("lang")){n=n.toLowerCase();return n===t||n.indexOf(t+"-")===0}}while((e=e.parentNode)&&e.nodeType===1);return false}}),target:function(e){var n=t.location&&t.location.hash;return n&&n.slice(1)===e.id},root:function(t){return t===p},focus:function(t){return t===d.activeElement&&(!d.hasFocus||d.hasFocus())&&!!(t.type||t.href||~t.tabIndex)},enabled:dt(false),disabled:dt(true),checked:function(t){var e=t.nodeName.toLowerCase();return e==="input"&&!!t.checked||e==="option"&&!!t.selected},selected:function(t){if(t.parentNode){t.parentNode.selectedIndex}return t.selected===true},empty:function(t){for(t=t.firstChild;t;t=t.nextSibling){if(t.nodeType<6){return false}}return true},parent:function(t){return!i.pseudos["empty"](t)},header:function(t){return J.test(t.nodeName)},input:function(t){return X.test(t.nodeName)},button:function(t){var e=t.nodeName.toLowerCase();return e==="input"&&t.type==="button"||e==="button"},text:function(t){var e;return t.nodeName.toLowerCase()==="input"&&t.type==="text"&&((e=t.getAttribute("type"))==null||e.toLowerCase()==="text")},first:pt(function(){return[0]}),last:pt(function(t,e){return[e-1]}),eq:pt(function(t,e,n){return[n<0?n+e:n]}),even:pt(function(t,e){var n=0;for(;n<e;n+=2){t.push(n)}return t}),odd:pt(function(t,e){var n=1;for(;n<e;n+=2){t.push(n)}return t}),lt:pt(function(t,e,n){var i=n<0?n+e:n;for(;--i>=0;){t.push(i)}return t}),gt:pt(function(t,e,n){var i=n<0?n+e:n;for(;++i<e;){t.push(i)}return t})}};i.pseudos["nth"]=i.pseudos["eq"];for(e in{radio:true,checkbox:true,file:true,password:true,image:true}){i.pseudos[e]=ct(e)}for(e in{submit:true,reset:true}){i.pseudos[e]=ht(e)}function vt(){}vt.prototype=i.filters=i.pseudos;i.setFilters=new vt;s=at.tokenize=function(t,e){var n,r,a,s,o,l,u,f=C[t+" "];if(f){return e?0:f.slice(0)}o=t;l=[];u=i.preFilter;while(o){if(!n||(r=q.exec(o))){if(r){o=o.slice(r[0].length)||o}l.push(a=[])}n=false;if(r=B.exec(o)){n=r.shift();a.push({value:n,type:r[0].replace(z," ")});o=o.slice(n.length)}for(s in i.filter){if((r=V[s].exec(o))&&(!u[s]||(r=u[s](r)))){n=r.shift();a.push({value:n,type:s,matches:r});o=o.slice(n.length)}}if(!n){break}}return e?o.length:o?at.error(t):C(t,l).slice(0)};function mt(t){var e=0,n=t.length,i="";for(;e<n;e++){i+=t[e].value}return i}function yt(t,e,n){var i=e.dir,r=e.next,a=r||i,s=n&&a==="parentNode",o=S++;return e.first?function(e,n,r){while(e=e[i]){if(e.nodeType===1||s){return t(e,n,r)}}return false}:function(e,n,l){var u,f,c,h=[x,o];if(l){while(e=e[i]){if(e.nodeType===1||s){if(t(e,n,l)){return true}}}}else{while(e=e[i]){if(e.nodeType===1||s){c=e[w]||(e[w]={});f=c[e.uniqueID]||(c[e.uniqueID]={});if(r&&r===e.nodeName.toLowerCase()){e=e[i]||e}else if((u=f[a])&&u[0]===x&&u[1]===o){return h[2]=u[2]}else{f[a]=h;if(h[2]=t(e,n,l)){return true}}}}}return false}}function bt(t){return t.length>1?function(e,n,i){var r=t.length;while(r--){if(!t[r](e,n,i)){return false}}return true}:t[0]}function wt(t,e,n){var i=0,r=e.length;for(;i<r;i++){at(t,e[i],n)}return n}function _t(t,e,n,i,r){var a,s=[],o=0,l=t.length,u=e!=null;for(;o<l;o++){if(a=t[o]){if(!n||n(a,i,r)){s.push(a);if(u){e.push(o)}}}}return s}function xt(t,e,n,i,r,a){if(i&&!i[w]){i=xt(i)}if(r&&!r[w]){r=xt(r,a)}return ot(function(a,s,o,l){var u,f,c,h=[],d=[],p=s.length,g=a||wt(e||"*",o.nodeType?[o]:o,[]),v=t&&(a||!e)?_t(g,h,t,o,l):g,m=n?r||(a?t:p||i)?[]:s:v;if(n){n(v,m,o,l)}if(i){u=_t(m,d);i(u,[],o,l);f=u.length;while(f--){if(c=u[f]){m[d[f]]=!(v[d[f]]=c)}}}if(a){if(r||t){if(r){u=[];f=m.length;while(f--){if(c=m[f]){u.push(v[f]=c)}}r(null,m=[],u,l)}f=m.length;while(f--){if((c=m[f])&&(u=r?F(a,c):h[f])>-1){a[u]=!(s[u]=c)}}}}else{m=_t(m===s?m.splice(p,m.length):m);if(r){r(null,s,m,l)}else{P.apply(s,m)}}})}function St(t){var e,n,r,a=t.length,s=i.relative[t[0].type],o=s||i.relative[" "],l=s?1:0,f=yt(function(t){return t===e},o,true),c=yt(function(t){return F(e,t)>-1},o,true),h=[function(t,n,i){var r=!s&&(i||n!==u)||((e=n).nodeType?f(t,n,i):c(t,n,i));e=null;return r}];for(;l<a;l++){if(n=i.relative[t[l].type]){h=[yt(bt(h),n)]}else{n=i.filter[t[l].type].apply(null,t[l].matches);if(n[w]){r=++l;for(;r<a;r++){if(i.relative[t[r].type]){break}}return xt(l>1&&bt(h),l>1&&mt(t.slice(0,l-1).concat({value:t[l-2].type===" "?"*":""})).replace(z,"$1"),n,l<r&&St(t.slice(l,r)),r<a&&St(t=t.slice(r)),r<a&&mt(t))}h.push(n)}}return bt(h)}function Dt(t,e){var n=e.length>0,r=t.length>0,a=function(a,s,o,l,f){var c,p,v,m=0,y="0",b=a&&[],w=[],_=u,S=a||r&&i.find["TAG"]("*",f),D=x+=_==null?1:Math.random()||.1,C=S.length;if(f){u=s===d||s||f}for(;y!==C&&(c=S[y])!=null;y++){if(r&&c){p=0;if(!s&&c.ownerDocument!==d){h(c);o=!g}while(v=t[p++]){if(v(c,s||d,o)){l.push(c);break}}if(f){x=D}}if(n){if(c=!v&&c){m--}if(a){b.push(c)}}}m+=y;if(n&&y!==m){p=0;while(v=e[p++]){v(b,w,s,o)}if(a){if(m>0){while(y--){if(!(b[y]||w[y])){w[y]=A.call(l)}}}w=_t(w)}P.apply(l,w);if(f&&!a&&w.length>0&&m+e.length>1){at.uniqueSort(l)}}if(f){x=D;u=_}return b};return n?ot(a):a}o=at.compile=function(t,e){var n,i=[],r=[],a=T[t+" "];if(!a){if(!e){e=s(t)}n=e.length;while(n--){a=St(e[n]);if(a[w]){i.push(a)}else{r.push(a)}}a=T(t,Dt(r,i));a.selector=t}return a};l=at.select=function(t,e,n,r){var a,l,u,f,c,h=typeof t==="function"&&t,d=!r&&s(t=h.selector||t);n=n||[];if(d.length===1){l=d[0]=d[0].slice(0);if(l.length>2&&(u=l[0]).type==="ID"&&e.nodeType===9&&g&&i.relative[l[1].type]){e=(i.find["ID"](u.matches[0].replace(Q,tt),e)||[])[0];if(!e){return n}else if(h){e=e.parentNode}t=t.slice(l.shift().value.length)}a=V["needsContext"].test(t)?0:l.length;while(a--){u=l[a];if(i.relative[f=u.type]){break}if(c=i.find[f]){if(r=c(u.matches[0].replace(Q,tt),Z.test(l[0].type)&&gt(e.parentNode)||e)){l.splice(a,1);t=r.length&&mt(l);if(!t){P.apply(n,r);return n}break}}}}(h||o(t,d))(r,e,!g,n,!e||Z.test(t)&&gt(e.parentNode)||e);return n};n.sortStable=w.split("").sort(k).join("")===w;n.detectDuplicates=!!c;h();n.sortDetached=lt(function(t){return t.compareDocumentPosition(d.createElement("fieldset"))&1});if(!lt(function(t){t.innerHTML="<a href='#'></a>";return t.firstChild.getAttribute("href")==="#"})){ut("type|href|height|width",function(t,e,n){if(!n){return t.getAttribute(e,e.toLowerCase()==="type"?1:2)}})}if(!n.attributes||!lt(function(t){t.innerHTML="<input/>";t.firstChild.setAttribute("value","");return t.firstChild.getAttribute("value")===""})){ut("value",function(t,e,n){if(!n&&t.nodeName.toLowerCase()==="input"){return t.defaultValue}})}if(!lt(function(t){return t.getAttribute("disabled")==null})){ut(R,function(t,e,n){var i;if(!n){return t[e]===true?e.toLowerCase():(i=t.getAttributeNode(e))&&i.specified?i.value:null}})}return at}(t);m.find=S;m.expr=S.selectors;m.expr[":"]=m.expr.pseudos;m.uniqueSort=m.unique=S.uniqueSort;m.text=S.getText;m.isXMLDoc=S.isXML;m.contains=S.contains;m.escapeSelector=S.escape;var D=function(t,e,n){var i=[],r=n!==undefined;while((t=t[e])&&t.nodeType!==9){if(t.nodeType===1){if(r&&m(t).is(n)){break}i.push(t)}}return i};var C=function(t,e){var n=[];for(;t;t=t.nextSibling){if(t.nodeType===1&&t!==e){n.push(t)}}return n};var T=m.expr.match.needsContext;var k=/^<([a-z][^\/\0>:\x20\t\r\n\f]*)[\x20\t\r\n\f]*\/?>(?:<\/\1>|)$/i;var M=/^.[^:#\[\.,]*$/;function I(t,e,n){if(m.isFunction(e)){return m.grep(t,function(t,i){return!!e.call(t,i,t)!==n})}if(e.nodeType){return m.grep(t,function(t){return t===e!==n})}if(typeof e!=="string"){return m.grep(t,function(t){return l.call(e,t)>-1!==n})}if(M.test(e)){return m.filter(e,t,n)}e=m.filter(e,t);return m.grep(t,function(t){return l.call(e,t)>-1!==n&&t.nodeType===1})}m.filter=function(t,e,n){var i=e[0];if(n){t=":not("+t+")"}if(e.length===1&&i.nodeType===1){return m.find.matchesSelector(i,t)?[i]:[]}return m.find.matches(t,m.grep(e,function(t){return t.nodeType===1}))};m.fn.extend({find:function(t){var e,n,i=this.length,r=this;if(typeof t!=="string"){return this.pushStack(m(t).filter(function(){for(e=0;e<i;e++){if(m.contains(r[e],this)){return true}}}))}n=this.pushStack([]);for(e=0;e<i;e++){m.find(t,r[e],n)}return i>1?m.uniqueSort(n):n},filter:function(t){return this.pushStack(I(this,t||[],false))},not:function(t){return this.pushStack(I(this,t||[],true))},is:function(t){return!!I(this,typeof t==="string"&&T.test(t)?m(t):t||[],false).length}});var A,N=/^(?:\s*(<[\w\W]+>)[^>]*|#([\w-]+))$/,P=m.fn.init=function(t,e,n){var r,a;if(!t){return this}n=n||A;if(typeof t==="string"){if(t[0]==="<"&&t[t.length-1]===">"&&t.length>=3){r=[null,t,null]}else{r=N.exec(t)}if(r&&(r[1]||!e)){if(r[1]){e=e instanceof m?e[0]:e;m.merge(this,m.parseHTML(r[1],e&&e.nodeType?e.ownerDocument||e:i,true));if(k.test(r[1])&&m.isPlainObject(e)){for(r in e){if(m.isFunction(this[r])){this[r](e[r])}else{this.attr(r,e[r])}}}return this}else{a=i.getElementById(r[2]);if(a){this[0]=a;this.length=1}return this}}else if(!e||e.jquery){return(e||n).find(t)}else{return this.constructor(e).find(t)}}else if(t.nodeType){this[0]=t;this.length=1;return this}else if(m.isFunction(t)){return n.ready!==undefined?n.ready(t):t(m)}return m.makeArray(t,this)};P.prototype=m.fn;A=m(i);var E=/^(?:parents|prev(?:Until|All))/,F={children:true,contents:true,next:true,prev:true};m.fn.extend({has:function(t){var e=m(t,this),n=e.length;return this.filter(function(){var t=0;for(;t<n;t++){if(m.contains(this,e[t])){return true}}})},closest:function(t,e){var n,i=0,r=this.length,a=[],s=typeof t!=="string"&&m(t);if(!T.test(t)){for(;i<r;i++){for(n=this[i];n&&n!==e;n=n.parentNode){if(n.nodeType<11&&(s?s.index(n)>-1:n.nodeType===1&&m.find.matchesSelector(n,t))){a.push(n);break}}}}return this.pushStack(a.length>1?m.uniqueSort(a):a)},index:function(t){if(!t){return this[0]&&this[0].parentNode?this.first().prevAll().length:-1}if(typeof t==="string"){return l.call(m(t),this[0])}return l.call(this,t.jquery?t[0]:t)},add:function(t,e){return this.pushStack(m.uniqueSort(m.merge(this.get(),m(t,e))))},addBack:function(t){return this.add(t==null?this.prevObject:this.prevObject.filter(t))}});function R(t,e){while((t=t[e])&&t.nodeType!==1){}return t}m.each({parent:function(t){var e=t.parentNode;return e&&e.nodeType!==11?e:null},parents:function(t){return D(t,"parentNode")},parentsUntil:function(t,e,n){return D(t,"parentNode",n)},next:function(t){return R(t,"nextSibling")},prev:function(t){return R(t,"previousSibling")},nextAll:function(t){return D(t,"nextSibling")},prevAll:function(t){return D(t,"previousSibling")},nextUntil:function(t,e,n){return D(t,"nextSibling",n)},prevUntil:function(t,e,n){return D(t,"previousSibling",n)},siblings:function(t){return C((t.parentNode||{}).firstChild,t)},children:function(t){return C(t.firstChild)},contents:function(t){return t.contentDocument||m.merge([],t.childNodes)}},function(t,e){m.fn[t]=function(n,i){var r=m.map(this,e,n);if(t.slice(-5)!=="Until"){i=n}if(i&&typeof i==="string"){r=m.filter(i,r)}if(this.length>1){if(!F[t]){m.uniqueSort(r)}if(E.test(t)){r.reverse()}}return this.pushStack(r)}});var L=/[^\x20\t\r\n\f]+/g;function j(t){var e={};m.each(t.match(L)||[],function(t,n){e[n]=true});return e}m.Callbacks=function(t){t=typeof t==="string"?j(t):m.extend({},t);var e,n,i,r,a=[],s=[],o=-1,l=function(){r=t.once;i=e=true;for(;s.length;o=-1){n=s.shift();while(++o<a.length){if(a[o].apply(n[0],n[1])===false&&t.stopOnFalse){o=a.length;n=false}}}if(!t.memory){n=false}e=false;if(r){if(n){a=[]}else{a=""}}},u={add:function(){if(a){if(n&&!e){o=a.length-1;s.push(n)}(function e(n){m.each(n,function(n,i){if(m.isFunction(i)){if(!t.unique||!u.has(i)){a.push(i)}}else if(i&&i.length&&m.type(i)!=="string"){e(i)}})})(arguments);if(n&&!e){l()}}return this},remove:function(){m.each(arguments,function(t,e){var n;while((n=m.inArray(e,a,n))>-1){a.splice(n,1);if(n<=o){o--}}});return this},has:function(t){return t?m.inArray(t,a)>-1:a.length>0},empty:function(){if(a){a=[]}return this},disable:function(){r=s=[];a=n="";return this},disabled:function(){return!a},lock:function(){r=s=[];if(!n&&!e){a=n=""}return this},locked:function(){return!!r},fireWith:function(t,n){if(!r){n=n||[];n=[t,n.slice?n.slice():n];s.push(n);if(!e){l()}}return this},fire:function(){u.fireWith(this,arguments);return this},fired:function(){return!!i}};return u};function O(t){return t}function H(t){throw t}function W(t,e,n){var i;try{if(t&&m.isFunction(i=t.promise)){i.call(t).done(e).fail(n)}else if(t&&m.isFunction(i=t.then)){i.call(t,e,n)}else{e.call(undefined,t)}}catch(t){n.call(undefined,t)}}m.extend({Deferred:function(e){var n=[["notify","progress",m.Callbacks("memory"),m.Callbacks("memory"),2],["resolve","done",m.Callbacks("once memory"),m.Callbacks("once memory"),0,"resolved"],["reject","fail",m.Callbacks("once memory"),m.Callbacks("once memory"),1,"rejected"]],i="pending",r={state:function(){return i},always:function(){a.done(arguments).fail(arguments);return this},catch:function(t){return r.then(null,t)},pipe:function(){var t=arguments;return m.Deferred(function(e){m.each(n,function(n,i){var r=m.isFunction(t[i[4]])&&t[i[4]];a[i[1]](function(){var t=r&&r.apply(this,arguments);if(t&&m.isFunction(t.promise)){t.promise().progress(e.notify).done(e.resolve).fail(e.reject)}else{e[i[0]+"With"](this,r?[t]:arguments)}})});t=null}).promise()},then:function(e,i,r){var a=0;function s(e,n,i,r){return function(){var o=this,l=arguments,u=function(){var t,u;if(e<a){return}t=i.apply(o,l);if(t===n.promise()){throw new TypeError("Thenable self-resolution")}u=t&&(typeof t==="object"||typeof t==="function")&&t.then;if(m.isFunction(u)){if(r){u.call(t,s(a,n,O,r),s(a,n,H,r))}else{a++;u.call(t,s(a,n,O,r),s(a,n,H,r),s(a,n,O,n.notifyWith))}}else{if(i!==O){o=undefined;l=[t]}(r||n.resolveWith)(o,l)}},f=r?u:function(){try{u()}catch(t){if(m.Deferred.exceptionHook){m.Deferred.exceptionHook(t,f.stackTrace)}if(e+1>=a){if(i!==H){o=undefined;l=[t]}n.rejectWith(o,l)}}};if(e){f()}else{if(m.Deferred.getStackHook){f.stackTrace=m.Deferred.getStackHook()}t.setTimeout(f)}}}return m.Deferred(function(t){n[0][3].add(s(0,t,m.isFunction(r)?r:O,t.notifyWith));n[1][3].add(s(0,t,m.isFunction(e)?e:O));n[2][3].add(s(0,t,m.isFunction(i)?i:H))}).promise()},promise:function(t){return t!=null?m.extend(t,r):r}},a={};m.each(n,function(t,e){var s=e[2],o=e[5];r[e[1]]=s.add;if(o){s.add(function(){i=o},n[3-t][2].disable,n[0][2].lock)}s.add(e[3].fire);a[e[0]]=function(){a[e[0]+"With"](this===a?undefined:this,arguments);return this};a[e[0]+"With"]=s.fireWith});r.promise(a);if(e){e.call(a,a)}return a},when:function(t){var e=arguments.length,n=e,i=Array(n),r=a.call(arguments),s=m.Deferred(),o=function(t){return function(n){i[t]=this;r[t]=arguments.length>1?a.call(arguments):n;if(!--e){s.resolveWith(i,r)}}};if(e<=1){W(t,s.done(o(n)).resolve,s.reject);if(s.state()==="pending"||m.isFunction(r[n]&&r[n].then)){return s.then()}}while(n--){W(r[n],o(n),s.reject)}return s.promise()}});var z=/^(Eval|Internal|Range|Reference|Syntax|Type|URI)Error$/;m.Deferred.exceptionHook=function(e,n){if(t.console&&t.console.warn&&e&&z.test(e.name)){t.console.warn("jQuery.Deferred exception: "+e.message,e.stack,n)}};m.readyException=function(e){t.setTimeout(function(){throw e})};var q=m.Deferred();m.fn.ready=function(t){q.then(t).catch(function(t){m.readyException(t)});return this};m.extend({isReady:false,readyWait:1,holdReady:function(t){if(t){m.readyWait++}else{m.ready(true)}},ready:function(t){if(t===true?--m.readyWait:m.isReady){return}m.isReady=true;if(t!==true&&--m.readyWait>0){return}q.resolveWith(i,[m])}});m.ready.then=q.then;function B(){i.removeEventListener("DOMContentLoaded",B);t.removeEventListener("load",B);m.ready()}if(i.readyState==="complete"||i.readyState!=="loading"&&!i.documentElement.doScroll){t.setTimeout(m.ready)}else{i.addEventListener("DOMContentLoaded",B);t.addEventListener("load",B)}var $=function(t,e,n,i,r,a,s){var o=0,l=t.length,u=n==null;if(m.type(n)==="object"){r=true;for(o in n){$(t,e,o,n[o],true,a,s)}}else if(i!==undefined){r=true;if(!m.isFunction(i)){s=true}if(u){if(s){e.call(t,i);e=null}else{u=e;e=function(t,e,n){return u.call(m(t),n)}}}if(e){for(;o<l;o++){e(t[o],n,s?i:i.call(t[o],o,e(t[o],n)))}}}if(r){return t}if(u){return e.call(t)}return l?e(t[0],n):a};var U=function(t){return t.nodeType===1||t.nodeType===9||!+t.nodeType};function Y(){this.expando=m.expando+Y.uid++}Y.uid=1;Y.prototype={cache:function(t){var e=t[this.expando];if(!e){e={};if(U(t)){if(t.nodeType){t[this.expando]=e}else{Object.defineProperty(t,this.expando,{value:e,configurable:true})}}}return e},set:function(t,e,n){var i,r=this.cache(t);if(typeof e==="string"){r[m.camelCase(e)]=n}else{for(i in e){r[m.camelCase(i)]=e[i]}}return r},get:function(t,e){return e===undefined?this.cache(t):t[this.expando]&&t[this.expando][m.camelCase(e)]},access:function(t,e,n){if(e===undefined||e&&typeof e==="string"&&n===undefined){return this.get(t,e)}this.set(t,e,n);return n!==undefined?n:e},remove:function(t,e){var n,i=t[this.expando];if(i===undefined){return}if(e!==undefined){if(m.isArray(e)){e=e.map(m.camelCase)}else{e=m.camelCase(e);e=e in i?[e]:e.match(L)||[]}n=e.length;while(n--){delete i[e[n]]}}if(e===undefined||m.isEmptyObject(i)){if(t.nodeType){t[this.expando]=undefined}else{delete t[this.expando]}}},hasData:function(t){var e=t[this.expando];return e!==undefined&&!m.isEmptyObject(e)}};var V=new Y;var X=new Y;var J=/^(?:\{[\w\W]*\}|\[[\w\W]*\])$/,G=/[A-Z]/g;function K(t){if(t==="true"){return true}if(t==="false"){return false}if(t==="null"){return null}if(t===+t+""){return+t}if(J.test(t)){return JSON.parse(t)}return t}function Z(t,e,n){var i;if(n===undefined&&t.nodeType===1){i="data-"+e.replace(G,"-$&").toLowerCase();n=t.getAttribute(i);if(typeof n==="string"){try{n=K(n)}catch(t){}X.set(t,e,n)}else{n=undefined}}return n}m.extend({hasData:function(t){return X.hasData(t)||V.hasData(t)},data:function(t,e,n){return X.access(t,e,n)},removeData:function(t,e){X.remove(t,e)},_data:function(t,e,n){return V.access(t,e,n)},_removeData:function(t,e){V.remove(t,e)}});m.fn.extend({data:function(t,e){var n,i,r,a=this[0],s=a&&a.attributes;if(t===undefined){if(this.length){r=X.get(a);if(a.nodeType===1&&!V.get(a,"hasDataAttrs")){n=s.length;while(n--){if(s[n]){i=s[n].name;if(i.indexOf("data-")===0){i=m.camelCase(i.slice(5));Z(a,i,r[i])}}}V.set(a,"hasDataAttrs",true)}}return r}if(typeof t==="object"){return this.each(function(){X.set(this,t)})}return $(this,function(e){var n;if(a&&e===undefined){n=X.get(a,t);if(n!==undefined){return n}n=Z(a,t);if(n!==undefined){return n}return}this.each(function(){X.set(this,t,e)})},null,e,arguments.length>1,null,true)},removeData:function(t){return this.each(function(){X.remove(this,t)})}});m.extend({queue:function(t,e,n){var i;if(t){e=(e||"fx")+"queue";i=V.get(t,e);if(n){if(!i||m.isArray(n)){i=V.access(t,e,m.makeArray(n))}else{i.push(n)}}return i||[]}},dequeue:function(t,e){e=e||"fx";var n=m.queue(t,e),i=n.length,r=n.shift(),a=m._queueHooks(t,e),s=function(){m.dequeue(t,e)};if(r==="inprogress"){r=n.shift();i--}if(r){if(e==="fx"){n.unshift("inprogress")}delete a.stop;r.call(t,s,a)}if(!i&&a){a.empty.fire()}},_queueHooks:function(t,e){var n=e+"queueHooks";return V.get(t,n)||V.access(t,n,{empty:m.Callbacks("once memory").add(function(){V.remove(t,[e+"queue",n])})})}});m.fn.extend({queue:function(t,e){var n=2;if(typeof t!=="string"){e=t;t="fx";n--}if(arguments.length<n){return m.queue(this[0],t)}return e===undefined?this:this.each(function(){var n=m.queue(this,t,e);m._queueHooks(this,t);if(t==="fx"&&n[0]!=="inprogress"){m.dequeue(this,t)}})},dequeue:function(t){return this.each(function(){m.dequeue(this,t)})},clearQueue:function(t){return this.queue(t||"fx",[])},promise:function(t,e){var n,i=1,r=m.Deferred(),a=this,s=this.length,o=function(){if(!--i){r.resolveWith(a,[a])}};if(typeof t!=="string"){e=t;t=undefined}t=t||"fx";while(s--){n=V.get(a[s],t+"queueHooks");if(n&&n.empty){i++;n.empty.add(o)}}o();return r.promise(e)}});var Q=/[+-]?(?:\d*\.|)\d+(?:[eE][+-]?\d+|)/.source;var tt=new RegExp("^(?:([+-])=|)("+Q+")([a-z%]*)$","i");var et=["Top","Right","Bottom","Left"];var nt=function(t,e){t=e||t;return t.style.display==="none"||t.style.display===""&&m.contains(t.ownerDocument,t)&&m.css(t,"display")==="none"};var it=function(t,e,n,i){var r,a,s={};for(a in e){s[a]=t.style[a];t.style[a]=e[a]}r=n.apply(t,i||[]);for(a in e){t.style[a]=s[a]}return r};function rt(t,e,n,i){var r,a=1,s=20,o=i?function(){return i.cur()}:function(){return m.css(t,e,"")},l=o(),u=n&&n[3]||(m.cssNumber[e]?"":"px"),f=(m.cssNumber[e]||u!=="px"&&+l)&&tt.exec(m.css(t,e));if(f&&f[3]!==u){u=u||f[3];n=n||[];f=+l||1;do{a=a||".5";f=f/a;m.style(t,e,f+u)}while(a!==(a=o()/l)&&a!==1&&--s)}if(n){f=+f||+l||0;r=n[1]?f+(n[1]+1)*n[2]:+n[2];if(i){i.unit=u;i.start=f;i.end=r}}return r}var at={};function st(t){var e,n=t.ownerDocument,i=t.nodeName,r=at[i];if(r){return r}e=n.body.appendChild(n.createElement(i));r=m.css(e,"display");e.parentNode.removeChild(e);if(r==="none"){r="block"}at[i]=r;return r}function ot(t,e){var n,i,r=[],a=0,s=t.length;for(;a<s;a++){i=t[a];if(!i.style){continue}n=i.style.display;if(e){if(n==="none"){r[a]=V.get(i,"display")||null;if(!r[a]){i.style.display=""}}if(i.style.display===""&&nt(i)){r[a]=st(i)}}else{if(n!=="none"){r[a]="none";V.set(i,"display",n)}}}for(a=0;a<s;a++){if(r[a]!=null){t[a].style.display=r[a]}}return t}m.fn.extend({show:function(){return ot(this,true)},hide:function(){return ot(this)},toggle:function(t){if(typeof t==="boolean"){return t?this.show():this.hide()}return this.each(function(){if(nt(this)){m(this).show()}else{m(this).hide()}})}});var lt=/^(?:checkbox|radio)$/i;var ut=/<([a-z][^\/\0>\x20\t\r\n\f]+)/i;var ft=/^$|\/(?:java|ecma)script/i;var ct={option:[1,"<select multiple='multiple'>","</select>"],thead:[1,"<table>","</table>"],col:[2,"<table><colgroup>","</colgroup></table>"],tr:[2,"<table><tbody>","</tbody></table>"],td:[3,"<table><tbody><tr>","</tr></tbody></table>"],_default:[0,"",""]};ct.optgroup=ct.option;ct.tbody=ct.tfoot=ct.colgroup=ct.caption=ct.thead;ct.th=ct.td;function ht(t,e){var n;if(typeof t.getElementsByTagName!=="undefined"){n=t.getElementsByTagName(e||"*")}else if(typeof t.querySelectorAll!=="undefined"){n=t.querySelectorAll(e||"*")}else{n=[]}if(e===undefined||e&&m.nodeName(t,e)){return m.merge([t],n)}return n}function dt(t,e){var n=0,i=t.length;for(;n<i;n++){V.set(t[n],"globalEval",!e||V.get(e[n],"globalEval"))}}var pt=/<|&#?\w+;/;function gt(t,e,n,i,r){var a,s,o,l,u,f,c=e.createDocumentFragment(),h=[],d=0,p=t.length;for(;d<p;d++){a=t[d];if(a||a===0){if(m.type(a)==="object"){m.merge(h,a.nodeType?[a]:a)}else if(!pt.test(a)){h.push(e.createTextNode(a))}else{s=s||c.appendChild(e.createElement("div"));o=(ut.exec(a)||["",""])[1].toLowerCase();l=ct[o]||ct._default;s.innerHTML=l[1]+m.htmlPrefilter(a)+l[2];f=l[0];while(f--){s=s.lastChild}m.merge(h,s.childNodes);s=c.firstChild;s.textContent=""}}}c.textContent="";d=0;while(a=h[d++]){if(i&&m.inArray(a,i)>-1){if(r){r.push(a)}continue}u=m.contains(a.ownerDocument,a);s=ht(c.appendChild(a),"script");if(u){dt(s)}if(n){f=0;while(a=s[f++]){if(ft.test(a.type||"")){n.push(a)}}}}return c}(function(){var t=i.createDocumentFragment(),e=t.appendChild(i.createElement("div")),n=i.createElement("input");n.setAttribute("type","radio");n.setAttribute("checked","checked");n.setAttribute("name","t");e.appendChild(n);p.checkClone=e.cloneNode(true).cloneNode(true).lastChild.checked;e.innerHTML="<textarea>x</textarea>";p.noCloneChecked=!!e.cloneNode(true).lastChild.defaultValue})();var vt=i.documentElement;var mt=/^key/,yt=/^(?:mouse|pointer|contextmenu|drag|drop)|click/,bt=/^([^.]*)(?:\.(.+)|)/;function wt(){return true}function _t(){return false}function xt(){try{return i.activeElement}catch(t){}}function St(t,e,n,i,r,a){var s,o;if(typeof e==="object"){if(typeof n!=="string"){i=i||n;n=undefined}for(o in e){St(t,o,n,i,e[o],a)}return t}if(i==null&&r==null){r=n;i=n=undefined}else if(r==null){if(typeof n==="string"){r=i;i=undefined}else{r=i;i=n;n=undefined}}if(r===false){r=_t}else if(!r){return t}if(a===1){s=r;r=function(t){m().off(t);return s.apply(this,arguments)};r.guid=s.guid||(s.guid=m.guid++)}return t.each(function(){m.event.add(this,e,r,i,n)})}m.event={global:{},add:function(t,e,n,i,r){var a,s,o,l,u,f,c,h,d,p,g,v=V.get(t);if(!v){return}if(n.handler){a=n;n=a.handler;r=a.selector}if(r){m.find.matchesSelector(vt,r)}if(!n.guid){n.guid=m.guid++}if(!(l=v.events)){l=v.events={}}if(!(s=v.handle)){s=v.handle=function(e){return typeof m!=="undefined"&&m.event.triggered!==e.type?m.event.dispatch.apply(t,arguments):undefined}}e=(e||"").match(L)||[""];u=e.length;while(u--){o=bt.exec(e[u])||[];d=g=o[1];p=(o[2]||"").split(".").sort();if(!d){continue}c=m.event.special[d]||{};d=(r?c.delegateType:c.bindType)||d;c=m.event.special[d]||{};f=m.extend({type:d,origType:g,data:i,handler:n,guid:n.guid,selector:r,needsContext:r&&m.expr.match.needsContext.test(r),namespace:p.join(".")},a);if(!(h=l[d])){h=l[d]=[];h.delegateCount=0;if(!c.setup||c.setup.call(t,i,p,s)===false){if(t.addEventListener){t.addEventListener(d,s)}}}if(c.add){c.add.call(t,f);if(!f.handler.guid){f.handler.guid=n.guid}}if(r){h.splice(h.delegateCount++,0,f)}else{h.push(f)}m.event.global[d]=true}},remove:function(t,e,n,i,r){var a,s,o,l,u,f,c,h,d,p,g,v=V.hasData(t)&&V.get(t);if(!v||!(l=v.events)){return}e=(e||"").match(L)||[""];u=e.length;while(u--){o=bt.exec(e[u])||[];d=g=o[1];p=(o[2]||"").split(".").sort();if(!d){for(d in l){m.event.remove(t,d+e[u],n,i,true)}continue}c=m.event.special[d]||{};d=(i?c.delegateType:c.bindType)||d;h=l[d]||[];o=o[2]&&new RegExp("(^|\\.)"+p.join("\\.(?:.*\\.|)")+"(\\.|$)");s=a=h.length;while(a--){f=h[a];if((r||g===f.origType)&&(!n||n.guid===f.guid)&&(!o||o.test(f.namespace))&&(!i||i===f.selector||i==="**"&&f.selector)){h.splice(a,1);if(f.selector){h.delegateCount--}if(c.remove){c.remove.call(t,f)}}}if(s&&!h.length){if(!c.teardown||c.teardown.call(t,p,v.handle)===false){m.removeEvent(t,d,v.handle)}delete l[d]}}if(m.isEmptyObject(l)){V.remove(t,"handle events")}},dispatch:function(t){var e=m.event.fix(t);var n,i,r,a,s,o,l=new Array(arguments.length),u=(V.get(this,"events")||{})[e.type]||[],f=m.event.special[e.type]||{};l[0]=e;for(n=1;n<arguments.length;n++){l[n]=arguments[n]}e.delegateTarget=this;if(f.preDispatch&&f.preDispatch.call(this,e)===false){return}o=m.event.handlers.call(this,e,u);n=0;while((a=o[n++])&&!e.isPropagationStopped()){e.currentTarget=a.elem;i=0;while((s=a.handlers[i++])&&!e.isImmediatePropagationStopped()){if(!e.rnamespace||e.rnamespace.test(s.namespace)){e.handleObj=s;e.data=s.data;r=((m.event.special[s.origType]||{}).handle||s.handler).apply(a.elem,l);if(r!==undefined){if((e.result=r)===false){e.preventDefault();e.stopPropagation()}}}}}if(f.postDispatch){f.postDispatch.call(this,e)}return e.result},handlers:function(t,e){var n,i,r,a,s,o=[],l=e.delegateCount,u=t.target;if(l&&u.nodeType&&!(t.type==="click"&&t.button>=1)){for(;u!==this;u=u.parentNode||this){if(u.nodeType===1&&!(t.type==="click"&&u.disabled===true)){a=[];s={};for(n=0;n<l;n++){i=e[n];r=i.selector+" ";if(s[r]===undefined){s[r]=i.needsContext?m(r,this).index(u)>-1:m.find(r,this,null,[u]).length}if(s[r]){a.push(i)}}if(a.length){o.push({elem:u,handlers:a})}}}}u=this;if(l<e.length){o.push({elem:u,handlers:e.slice(l)})}return o},addProp:function(t,e){Object.defineProperty(m.Event.prototype,t,{enumerable:true,configurable:true,get:m.isFunction(e)?function(){if(this.originalEvent){return e(this.originalEvent)}}:function(){if(this.originalEvent){return this.originalEvent[t]}},set:function(e){Object.defineProperty(this,t,{enumerable:true,configurable:true,writable:true,value:e})}})},fix:function(t){return t[m.expando]?t:new m.Event(t)},special:{load:{noBubble:true},focus:{trigger:function(){if(this!==xt()&&this.focus){this.focus();return false}},delegateType:"focusin"},blur:{trigger:function(){if(this===xt()&&this.blur){this.blur();return false}},delegateType:"focusout"},click:{trigger:function(){if(this.type==="checkbox"&&this.click&&m.nodeName(this,"input")){this.click();return false}},_default:function(t){return m.nodeName(t.target,"a")}},beforeunload:{postDispatch:function(t){if(t.result!==undefined&&t.originalEvent){t.originalEvent.returnValue=t.result}}}}};m.removeEvent=function(t,e,n){if(t.removeEventListener){t.removeEventListener(e,n)}};m.Event=function(t,e){if(!(this instanceof m.Event)){return new m.Event(t,e)}if(t&&t.type){this.originalEvent=t;this.type=t.type;this.isDefaultPrevented=t.defaultPrevented||t.defaultPrevented===undefined&&t.returnValue===false?wt:_t;this.target=t.target&&t.target.nodeType===3?t.target.parentNode:t.target;this.currentTarget=t.currentTarget;this.relatedTarget=t.relatedTarget}else{this.type=t}if(e){m.extend(this,e)}this.timeStamp=t&&t.timeStamp||m.now();this[m.expando]=true};m.Event.prototype={constructor:m.Event,isDefaultPrevented:_t,isPropagationStopped:_t,isImmediatePropagationStopped:_t,isSimulated:false,preventDefault:function(){var t=this.originalEvent;this.isDefaultPrevented=wt;if(t&&!this.isSimulated){t.preventDefault()}},stopPropagation:function(){var t=this.originalEvent;this.isPropagationStopped=wt;if(t&&!this.isSimulated){t.stopPropagation()}},stopImmediatePropagation:function(){var t=this.originalEvent;this.isImmediatePropagationStopped=wt;if(t&&!this.isSimulated){t.stopImmediatePropagation()}this.stopPropagation()}};m.each({altKey:true,bubbles:true,cancelable:true,changedTouches:true,ctrlKey:true,detail:true,eventPhase:true,metaKey:true,pageX:true,pageY:true,shiftKey:true,
view:true,char:true,charCode:true,key:true,keyCode:true,button:true,buttons:true,clientX:true,clientY:true,offsetX:true,offsetY:true,pointerId:true,pointerType:true,screenX:true,screenY:true,targetTouches:true,toElement:true,touches:true,which:function(t){var e=t.button;if(t.which==null&&mt.test(t.type)){return t.charCode!=null?t.charCode:t.keyCode}if(!t.which&&e!==undefined&&yt.test(t.type)){if(e&1){return 1}if(e&2){return 3}if(e&4){return 2}return 0}return t.which}},m.event.addProp);m.each({mouseenter:"mouseover",mouseleave:"mouseout",pointerenter:"pointerover",pointerleave:"pointerout"},function(t,e){m.event.special[t]={delegateType:e,bindType:e,handle:function(t){var n,i=this,r=t.relatedTarget,a=t.handleObj;if(!r||r!==i&&!m.contains(i,r)){t.type=a.origType;n=a.handler.apply(this,arguments);t.type=e}return n}}});m.fn.extend({on:function(t,e,n,i){return St(this,t,e,n,i)},one:function(t,e,n,i){return St(this,t,e,n,i,1)},off:function(t,e,n){var i,r;if(t&&t.preventDefault&&t.handleObj){i=t.handleObj;m(t.delegateTarget).off(i.namespace?i.origType+"."+i.namespace:i.origType,i.selector,i.handler);return this}if(typeof t==="object"){for(r in t){this.off(r,e,t[r])}return this}if(e===false||typeof e==="function"){n=e;e=undefined}if(n===false){n=_t}return this.each(function(){m.event.remove(this,t,n,e)})}});var Dt=/<(?!area|br|col|embed|hr|img|input|link|meta|param)(([a-z][^\/\0>\x20\t\r\n\f]*)[^>]*)\/>/gi,Ct=/<script|<style|<link/i,Tt=/checked\s*(?:[^=]|=\s*.checked.)/i,kt=/^true\/(.*)/,Mt=/^\s*<!(?:\[CDATA\[|--)|(?:\]\]|--)>\s*$/g;function It(t,e){if(m.nodeName(t,"table")&&m.nodeName(e.nodeType!==11?e:e.firstChild,"tr")){return t.getElementsByTagName("tbody")[0]||t}return t}function At(t){t.type=(t.getAttribute("type")!==null)+"/"+t.type;return t}function Nt(t){var e=kt.exec(t.type);if(e){t.type=e[1]}else{t.removeAttribute("type")}return t}function Pt(t,e){var n,i,r,a,s,o,l,u;if(e.nodeType!==1){return}if(V.hasData(t)){a=V.access(t);s=V.set(e,a);u=a.events;if(u){delete s.handle;s.events={};for(r in u){for(n=0,i=u[r].length;n<i;n++){m.event.add(e,r,u[r][n])}}}}if(X.hasData(t)){o=X.access(t);l=m.extend({},o);X.set(e,l)}}function Et(t,e){var n=e.nodeName.toLowerCase();if(n==="input"&&lt.test(t.type)){e.checked=t.checked}else if(n==="input"||n==="textarea"){e.defaultValue=t.defaultValue}}function Ft(t,e,n,i){e=s.apply([],e);var r,a,o,l,u,f,c=0,h=t.length,d=h-1,v=e[0],y=m.isFunction(v);if(y||h>1&&typeof v==="string"&&!p.checkClone&&Tt.test(v)){return t.each(function(r){var a=t.eq(r);if(y){e[0]=v.call(this,r,a.html())}Ft(a,e,n,i)})}if(h){r=gt(e,t[0].ownerDocument,false,t,i);a=r.firstChild;if(r.childNodes.length===1){r=a}if(a||i){o=m.map(ht(r,"script"),At);l=o.length;for(;c<h;c++){u=r;if(c!==d){u=m.clone(u,true,true);if(l){m.merge(o,ht(u,"script"))}}n.call(t[c],u,c)}if(l){f=o[o.length-1].ownerDocument;m.map(o,Nt);for(c=0;c<l;c++){u=o[c];if(ft.test(u.type||"")&&!V.access(u,"globalEval")&&m.contains(f,u)){if(u.src){if(m._evalUrl){m._evalUrl(u.src)}}else{g(u.textContent.replace(Mt,""),f)}}}}}}return t}function Rt(t,e,n){var i,r=e?m.filter(e,t):t,a=0;for(;(i=r[a])!=null;a++){if(!n&&i.nodeType===1){m.cleanData(ht(i))}if(i.parentNode){if(n&&m.contains(i.ownerDocument,i)){dt(ht(i,"script"))}i.parentNode.removeChild(i)}}return t}m.extend({htmlPrefilter:function(t){return t.replace(Dt,"<$1></$2>")},clone:function(t,e,n){var i,r,a,s,o=t.cloneNode(true),l=m.contains(t.ownerDocument,t);if(!p.noCloneChecked&&(t.nodeType===1||t.nodeType===11)&&!m.isXMLDoc(t)){s=ht(o);a=ht(t);for(i=0,r=a.length;i<r;i++){Et(a[i],s[i])}}if(e){if(n){a=a||ht(t);s=s||ht(o);for(i=0,r=a.length;i<r;i++){Pt(a[i],s[i])}}else{Pt(t,o)}}s=ht(o,"script");if(s.length>0){dt(s,!l&&ht(t,"script"))}return o},cleanData:function(t){var e,n,i,r=m.event.special,a=0;for(;(n=t[a])!==undefined;a++){if(U(n)){if(e=n[V.expando]){if(e.events){for(i in e.events){if(r[i]){m.event.remove(n,i)}else{m.removeEvent(n,i,e.handle)}}}n[V.expando]=undefined}if(n[X.expando]){n[X.expando]=undefined}}}}});m.fn.extend({detach:function(t){return Rt(this,t,true)},remove:function(t){return Rt(this,t)},text:function(t){return $(this,function(t){return t===undefined?m.text(this):this.empty().each(function(){if(this.nodeType===1||this.nodeType===11||this.nodeType===9){this.textContent=t}})},null,t,arguments.length)},append:function(){return Ft(this,arguments,function(t){if(this.nodeType===1||this.nodeType===11||this.nodeType===9){var e=It(this,t);e.appendChild(t)}})},prepend:function(){return Ft(this,arguments,function(t){if(this.nodeType===1||this.nodeType===11||this.nodeType===9){var e=It(this,t);e.insertBefore(t,e.firstChild)}})},before:function(){return Ft(this,arguments,function(t){if(this.parentNode){this.parentNode.insertBefore(t,this)}})},after:function(){return Ft(this,arguments,function(t){if(this.parentNode){this.parentNode.insertBefore(t,this.nextSibling)}})},empty:function(){var t,e=0;for(;(t=this[e])!=null;e++){if(t.nodeType===1){m.cleanData(ht(t,false));t.textContent=""}}return this},clone:function(t,e){t=t==null?false:t;e=e==null?t:e;return this.map(function(){return m.clone(this,t,e)})},html:function(t){return $(this,function(t){var e=this[0]||{},n=0,i=this.length;if(t===undefined&&e.nodeType===1){return e.innerHTML}if(typeof t==="string"&&!Ct.test(t)&&!ct[(ut.exec(t)||["",""])[1].toLowerCase()]){t=m.htmlPrefilter(t);try{for(;n<i;n++){e=this[n]||{};if(e.nodeType===1){m.cleanData(ht(e,false));e.innerHTML=t}}e=0}catch(t){}}if(e){this.empty().append(t)}},null,t,arguments.length)},replaceWith:function(){var t=[];return Ft(this,arguments,function(e){var n=this.parentNode;if(m.inArray(this,t)<0){m.cleanData(ht(this));if(n){n.replaceChild(e,this)}}},t)}});m.each({appendTo:"append",prependTo:"prepend",insertBefore:"before",insertAfter:"after",replaceAll:"replaceWith"},function(t,e){m.fn[t]=function(t){var n,i=[],r=m(t),a=r.length-1,s=0;for(;s<=a;s++){n=s===a?this:this.clone(true);m(r[s])[e](n);o.apply(i,n.get())}return this.pushStack(i)}});var Lt=/^margin/;var jt=new RegExp("^("+Q+")(?!px)[a-z%]+$","i");var Ot=function(e){var n=e.ownerDocument.defaultView;if(!n||!n.opener){n=t}return n.getComputedStyle(e)};(function(){function e(){if(!l){return}l.style.cssText="box-sizing:border-box;"+"position:relative;display:block;"+"margin:auto;border:1px;padding:1px;"+"top:1%;width:50%";l.innerHTML="";vt.appendChild(o);var e=t.getComputedStyle(l);n=e.top!=="1%";s=e.marginLeft==="2px";r=e.width==="4px";l.style.marginRight="50%";a=e.marginRight==="4px";vt.removeChild(o);l=null}var n,r,a,s,o=i.createElement("div"),l=i.createElement("div");if(!l.style){return}l.style.backgroundClip="content-box";l.cloneNode(true).style.backgroundClip="";p.clearCloneStyle=l.style.backgroundClip==="content-box";o.style.cssText="border:0;width:8px;height:0;top:0;left:-9999px;"+"padding:0;margin-top:1px;position:absolute";o.appendChild(l);m.extend(p,{pixelPosition:function(){e();return n},boxSizingReliable:function(){e();return r},pixelMarginRight:function(){e();return a},reliableMarginLeft:function(){e();return s}})})();function Ht(t,e,n){var i,r,a,s,o=t.style;n=n||Ot(t);if(n){s=n.getPropertyValue(e)||n[e];if(s===""&&!m.contains(t.ownerDocument,t)){s=m.style(t,e)}if(!p.pixelMarginRight()&&jt.test(s)&&Lt.test(e)){i=o.width;r=o.minWidth;a=o.maxWidth;o.minWidth=o.maxWidth=o.width=s;s=n.width;o.width=i;o.minWidth=r;o.maxWidth=a}}return s!==undefined?s+"":s}function Wt(t,e){return{get:function(){if(t()){delete this.get;return}return(this.get=e).apply(this,arguments)}}}var zt=/^(none|table(?!-c[ea]).+)/,qt={position:"absolute",visibility:"hidden",display:"block"},Bt={letterSpacing:"0",fontWeight:"400"},$t=["Webkit","Moz","ms"],Ut=i.createElement("div").style;function Yt(t){if(t in Ut){return t}var e=t[0].toUpperCase()+t.slice(1),n=$t.length;while(n--){t=$t[n]+e;if(t in Ut){return t}}}function Vt(t,e,n){var i=tt.exec(e);return i?Math.max(0,i[2]-(n||0))+(i[3]||"px"):e}function Xt(t,e,n,i,r){var a,s=0;if(n===(i?"border":"content")){a=4}else{a=e==="width"?1:0}for(;a<4;a+=2){if(n==="margin"){s+=m.css(t,n+et[a],true,r)}if(i){if(n==="content"){s-=m.css(t,"padding"+et[a],true,r)}if(n!=="margin"){s-=m.css(t,"border"+et[a]+"Width",true,r)}}else{s+=m.css(t,"padding"+et[a],true,r);if(n!=="padding"){s+=m.css(t,"border"+et[a]+"Width",true,r)}}}return s}function Jt(t,e,n){var i,r=true,a=Ot(t),s=m.css(t,"boxSizing",false,a)==="border-box";if(t.getClientRects().length){i=t.getBoundingClientRect()[e]}if(i<=0||i==null){i=Ht(t,e,a);if(i<0||i==null){i=t.style[e]}if(jt.test(i)){return i}r=s&&(p.boxSizingReliable()||i===t.style[e]);i=parseFloat(i)||0}return i+Xt(t,e,n||(s?"border":"content"),r,a)+"px"}m.extend({cssHooks:{opacity:{get:function(t,e){if(e){var n=Ht(t,"opacity");return n===""?"1":n}}}},cssNumber:{animationIterationCount:true,columnCount:true,fillOpacity:true,flexGrow:true,flexShrink:true,fontWeight:true,lineHeight:true,opacity:true,order:true,orphans:true,widows:true,zIndex:true,zoom:true},cssProps:{float:"cssFloat"},style:function(t,e,n,i){if(!t||t.nodeType===3||t.nodeType===8||!t.style){return}var r,a,s,o=m.camelCase(e),l=t.style;e=m.cssProps[o]||(m.cssProps[o]=Yt(o)||o);s=m.cssHooks[e]||m.cssHooks[o];if(n!==undefined){a=typeof n;if(a==="string"&&(r=tt.exec(n))&&r[1]){n=rt(t,e,r);a="number"}if(n==null||n!==n){return}if(a==="number"){n+=r&&r[3]||(m.cssNumber[o]?"":"px")}if(!p.clearCloneStyle&&n===""&&e.indexOf("background")===0){l[e]="inherit"}if(!s||!("set"in s)||(n=s.set(t,n,i))!==undefined){l[e]=n}}else{if(s&&"get"in s&&(r=s.get(t,false,i))!==undefined){return r}return l[e]}},css:function(t,e,n,i){var r,a,s,o=m.camelCase(e);e=m.cssProps[o]||(m.cssProps[o]=Yt(o)||o);s=m.cssHooks[e]||m.cssHooks[o];if(s&&"get"in s){r=s.get(t,true,n)}if(r===undefined){r=Ht(t,e,i)}if(r==="normal"&&e in Bt){r=Bt[e]}if(n===""||n){a=parseFloat(r);return n===true||isFinite(a)?a||0:r}return r}});m.each(["height","width"],function(t,e){m.cssHooks[e]={get:function(t,n,i){if(n){return zt.test(m.css(t,"display"))&&(!t.getClientRects().length||!t.getBoundingClientRect().width)?it(t,qt,function(){return Jt(t,e,i)}):Jt(t,e,i)}},set:function(t,n,i){var r,a=i&&Ot(t),s=i&&Xt(t,e,i,m.css(t,"boxSizing",false,a)==="border-box",a);if(s&&(r=tt.exec(n))&&(r[3]||"px")!=="px"){t.style[e]=n;n=m.css(t,e)}return Vt(t,n,s)}}});m.cssHooks.marginLeft=Wt(p.reliableMarginLeft,function(t,e){if(e){return(parseFloat(Ht(t,"marginLeft"))||t.getBoundingClientRect().left-it(t,{marginLeft:0},function(){return t.getBoundingClientRect().left}))+"px"}});m.each({margin:"",padding:"",border:"Width"},function(t,e){m.cssHooks[t+e]={expand:function(n){var i=0,r={},a=typeof n==="string"?n.split(" "):[n];for(;i<4;i++){r[t+et[i]+e]=a[i]||a[i-2]||a[0]}return r}};if(!Lt.test(t)){m.cssHooks[t+e].set=Vt}});m.fn.extend({css:function(t,e){return $(this,function(t,e,n){var i,r,a={},s=0;if(m.isArray(e)){i=Ot(t);r=e.length;for(;s<r;s++){a[e[s]]=m.css(t,e[s],false,i)}return a}return n!==undefined?m.style(t,e,n):m.css(t,e)},t,e,arguments.length>1)}});function Gt(t,e,n,i,r){return new Gt.prototype.init(t,e,n,i,r)}m.Tween=Gt;Gt.prototype={constructor:Gt,init:function(t,e,n,i,r,a){this.elem=t;this.prop=n;this.easing=r||m.easing._default;this.options=e;this.start=this.now=this.cur();this.end=i;this.unit=a||(m.cssNumber[n]?"":"px")},cur:function(){var t=Gt.propHooks[this.prop];return t&&t.get?t.get(this):Gt.propHooks._default.get(this)},run:function(t){var e,n=Gt.propHooks[this.prop];if(this.options.duration){this.pos=e=m.easing[this.easing](t,this.options.duration*t,0,1,this.options.duration)}else{this.pos=e=t}this.now=(this.end-this.start)*e+this.start;if(this.options.step){this.options.step.call(this.elem,this.now,this)}if(n&&n.set){n.set(this)}else{Gt.propHooks._default.set(this)}return this}};Gt.prototype.init.prototype=Gt.prototype;Gt.propHooks={_default:{get:function(t){var e;if(t.elem.nodeType!==1||t.elem[t.prop]!=null&&t.elem.style[t.prop]==null){return t.elem[t.prop]}e=m.css(t.elem,t.prop,"");return!e||e==="auto"?0:e},set:function(t){if(m.fx.step[t.prop]){m.fx.step[t.prop](t)}else if(t.elem.nodeType===1&&(t.elem.style[m.cssProps[t.prop]]!=null||m.cssHooks[t.prop])){m.style(t.elem,t.prop,t.now+t.unit)}else{t.elem[t.prop]=t.now}}}};Gt.propHooks.scrollTop=Gt.propHooks.scrollLeft={set:function(t){if(t.elem.nodeType&&t.elem.parentNode){t.elem[t.prop]=t.now}}};m.easing={linear:function(t){return t},swing:function(t){return.5-Math.cos(t*Math.PI)/2},_default:"swing"};m.fx=Gt.prototype.init;m.fx.step={};var Kt,Zt,Qt=/^(?:toggle|show|hide)$/,te=/queueHooks$/;function ee(){if(Zt){t.requestAnimationFrame(ee);m.fx.tick()}}function ne(){t.setTimeout(function(){Kt=undefined});return Kt=m.now()}function ie(t,e){var n,i=0,r={height:t};e=e?1:0;for(;i<4;i+=2-e){n=et[i];r["margin"+n]=r["padding"+n]=t}if(e){r.opacity=r.width=t}return r}function re(t,e,n){var i,r=(oe.tweeners[e]||[]).concat(oe.tweeners["*"]),a=0,s=r.length;for(;a<s;a++){if(i=r[a].call(n,e,t)){return i}}}function ae(t,e,n){var i,r,a,s,o,l,u,f,c="width"in e||"height"in e,h=this,d={},p=t.style,g=t.nodeType&&nt(t),v=V.get(t,"fxshow");if(!n.queue){s=m._queueHooks(t,"fx");if(s.unqueued==null){s.unqueued=0;o=s.empty.fire;s.empty.fire=function(){if(!s.unqueued){o()}}}s.unqueued++;h.always(function(){h.always(function(){s.unqueued--;if(!m.queue(t,"fx").length){s.empty.fire()}})})}for(i in e){r=e[i];if(Qt.test(r)){delete e[i];a=a||r==="toggle";if(r===(g?"hide":"show")){if(r==="show"&&v&&v[i]!==undefined){g=true}else{continue}}d[i]=v&&v[i]||m.style(t,i)}}l=!m.isEmptyObject(e);if(!l&&m.isEmptyObject(d)){return}if(c&&t.nodeType===1){n.overflow=[p.overflow,p.overflowX,p.overflowY];u=v&&v.display;if(u==null){u=V.get(t,"display")}f=m.css(t,"display");if(f==="none"){if(u){f=u}else{ot([t],true);u=t.style.display||u;f=m.css(t,"display");ot([t])}}if(f==="inline"||f==="inline-block"&&u!=null){if(m.css(t,"float")==="none"){if(!l){h.done(function(){p.display=u});if(u==null){f=p.display;u=f==="none"?"":f}}p.display="inline-block"}}}if(n.overflow){p.overflow="hidden";h.always(function(){p.overflow=n.overflow[0];p.overflowX=n.overflow[1];p.overflowY=n.overflow[2]})}l=false;for(i in d){if(!l){if(v){if("hidden"in v){g=v.hidden}}else{v=V.access(t,"fxshow",{display:u})}if(a){v.hidden=!g}if(g){ot([t],true)}h.done(function(){if(!g){ot([t])}V.remove(t,"fxshow");for(i in d){m.style(t,i,d[i])}})}l=re(g?v[i]:0,i,h);if(!(i in v)){v[i]=l.start;if(g){l.end=l.start;l.start=0}}}}function se(t,e){var n,i,r,a,s;for(n in t){i=m.camelCase(n);r=e[i];a=t[n];if(m.isArray(a)){r=a[1];a=t[n]=a[0]}if(n!==i){t[i]=a;delete t[n]}s=m.cssHooks[i];if(s&&"expand"in s){a=s.expand(a);delete t[i];for(n in a){if(!(n in t)){t[n]=a[n];e[n]=r}}}else{e[i]=r}}}function oe(t,e,n){var i,r,a=0,s=oe.prefilters.length,o=m.Deferred().always(function(){delete l.elem}),l=function(){if(r){return false}var e=Kt||ne(),n=Math.max(0,u.startTime+u.duration-e),i=n/u.duration||0,a=1-i,s=0,l=u.tweens.length;for(;s<l;s++){u.tweens[s].run(a)}o.notifyWith(t,[u,a,n]);if(a<1&&l){return n}else{o.resolveWith(t,[u]);return false}},u=o.promise({elem:t,props:m.extend({},e),opts:m.extend(true,{specialEasing:{},easing:m.easing._default},n),originalProperties:e,originalOptions:n,startTime:Kt||ne(),duration:n.duration,tweens:[],createTween:function(e,n){var i=m.Tween(t,u.opts,e,n,u.opts.specialEasing[e]||u.opts.easing);u.tweens.push(i);return i},stop:function(e){var n=0,i=e?u.tweens.length:0;if(r){return this}r=true;for(;n<i;n++){u.tweens[n].run(1)}if(e){o.notifyWith(t,[u,1,0]);o.resolveWith(t,[u,e])}else{o.rejectWith(t,[u,e])}return this}}),f=u.props;se(f,u.opts.specialEasing);for(;a<s;a++){i=oe.prefilters[a].call(u,t,f,u.opts);if(i){if(m.isFunction(i.stop)){m._queueHooks(u.elem,u.opts.queue).stop=m.proxy(i.stop,i)}return i}}m.map(f,re,u);if(m.isFunction(u.opts.start)){u.opts.start.call(t,u)}m.fx.timer(m.extend(l,{elem:t,anim:u,queue:u.opts.queue}));return u.progress(u.opts.progress).done(u.opts.done,u.opts.complete).fail(u.opts.fail).always(u.opts.always)}m.Animation=m.extend(oe,{tweeners:{"*":[function(t,e){var n=this.createTween(t,e);rt(n.elem,t,tt.exec(e),n);return n}]},tweener:function(t,e){if(m.isFunction(t)){e=t;t=["*"]}else{t=t.match(L)}var n,i=0,r=t.length;for(;i<r;i++){n=t[i];oe.tweeners[n]=oe.tweeners[n]||[];oe.tweeners[n].unshift(e)}},prefilters:[ae],prefilter:function(t,e){if(e){oe.prefilters.unshift(t)}else{oe.prefilters.push(t)}}});m.speed=function(t,e,n){var r=t&&typeof t==="object"?m.extend({},t):{complete:n||!n&&e||m.isFunction(t)&&t,duration:t,easing:n&&e||e&&!m.isFunction(e)&&e};if(m.fx.off||i.hidden){r.duration=0}else{if(typeof r.duration!=="number"){if(r.duration in m.fx.speeds){r.duration=m.fx.speeds[r.duration]}else{r.duration=m.fx.speeds._default}}}if(r.queue==null||r.queue===true){r.queue="fx"}r.old=r.complete;r.complete=function(){if(m.isFunction(r.old)){r.old.call(this)}if(r.queue){m.dequeue(this,r.queue)}};return r};m.fn.extend({fadeTo:function(t,e,n,i){return this.filter(nt).css("opacity",0).show().end().animate({opacity:e},t,n,i)},animate:function(t,e,n,i){var r=m.isEmptyObject(t),a=m.speed(e,n,i),s=function(){var e=oe(this,m.extend({},t),a);if(r||V.get(this,"finish")){e.stop(true)}};s.finish=s;return r||a.queue===false?this.each(s):this.queue(a.queue,s)},stop:function(t,e,n){var i=function(t){var e=t.stop;delete t.stop;e(n)};if(typeof t!=="string"){n=e;e=t;t=undefined}if(e&&t!==false){this.queue(t||"fx",[])}return this.each(function(){var e=true,r=t!=null&&t+"queueHooks",a=m.timers,s=V.get(this);if(r){if(s[r]&&s[r].stop){i(s[r])}}else{for(r in s){if(s[r]&&s[r].stop&&te.test(r)){i(s[r])}}}for(r=a.length;r--;){if(a[r].elem===this&&(t==null||a[r].queue===t)){a[r].anim.stop(n);e=false;a.splice(r,1)}}if(e||!n){m.dequeue(this,t)}})},finish:function(t){if(t!==false){t=t||"fx"}return this.each(function(){var e,n=V.get(this),i=n[t+"queue"],r=n[t+"queueHooks"],a=m.timers,s=i?i.length:0;n.finish=true;m.queue(this,t,[]);if(r&&r.stop){r.stop.call(this,true)}for(e=a.length;e--;){if(a[e].elem===this&&a[e].queue===t){a[e].anim.stop(true);a.splice(e,1)}}for(e=0;e<s;e++){if(i[e]&&i[e].finish){i[e].finish.call(this)}}delete n.finish})}});m.each(["toggle","show","hide"],function(t,e){var n=m.fn[e];m.fn[e]=function(t,i,r){return t==null||typeof t==="boolean"?n.apply(this,arguments):this.animate(ie(e,true),t,i,r)}});m.each({slideDown:ie("show"),slideUp:ie("hide"),slideToggle:ie("toggle"),fadeIn:{opacity:"show"},fadeOut:{opacity:"hide"},fadeToggle:{opacity:"toggle"}},function(t,e){m.fn[t]=function(t,n,i){return this.animate(e,t,n,i)}});m.timers=[];m.fx.tick=function(){var t,e=0,n=m.timers;Kt=m.now();for(;e<n.length;e++){t=n[e];if(!t()&&n[e]===t){n.splice(e--,1)}}if(!n.length){m.fx.stop()}Kt=undefined};m.fx.timer=function(t){m.timers.push(t);if(t()){m.fx.start()}else{m.timers.pop()}};m.fx.interval=13;m.fx.start=function(){if(!Zt){Zt=t.requestAnimationFrame?t.requestAnimationFrame(ee):t.setInterval(m.fx.tick,m.fx.interval)}};m.fx.stop=function(){if(t.cancelAnimationFrame){t.cancelAnimationFrame(Zt)}else{t.clearInterval(Zt)}Zt=null};m.fx.speeds={slow:600,fast:200,_default:400};m.fn.delay=function(e,n){e=m.fx?m.fx.speeds[e]||e:e;n=n||"fx";return this.queue(n,function(n,i){var r=t.setTimeout(n,e);i.stop=function(){t.clearTimeout(r)}})};(function(){var t=i.createElement("input"),e=i.createElement("select"),n=e.appendChild(i.createElement("option"));t.type="checkbox";p.checkOn=t.value!=="";p.optSelected=n.selected;t=i.createElement("input");t.value="t";t.type="radio";p.radioValue=t.value==="t"})();var le,ue=m.expr.attrHandle;m.fn.extend({attr:function(t,e){return $(this,m.attr,t,e,arguments.length>1)},removeAttr:function(t){return this.each(function(){m.removeAttr(this,t)})}});m.extend({attr:function(t,e,n){var i,r,a=t.nodeType;if(a===3||a===8||a===2){return}if(typeof t.getAttribute==="undefined"){return m.prop(t,e,n)}if(a!==1||!m.isXMLDoc(t)){r=m.attrHooks[e.toLowerCase()]||(m.expr.match.bool.test(e)?le:undefined)}if(n!==undefined){if(n===null){m.removeAttr(t,e);return}if(r&&"set"in r&&(i=r.set(t,n,e))!==undefined){return i}t.setAttribute(e,n+"");return n}if(r&&"get"in r&&(i=r.get(t,e))!==null){return i}i=m.find.attr(t,e);return i==null?undefined:i},attrHooks:{type:{set:function(t,e){if(!p.radioValue&&e==="radio"&&m.nodeName(t,"input")){var n=t.value;t.setAttribute("type",e);if(n){t.value=n}return e}}}},removeAttr:function(t,e){var n,i=0,r=e&&e.match(L);if(r&&t.nodeType===1){while(n=r[i++]){t.removeAttribute(n)}}}});le={set:function(t,e,n){if(e===false){m.removeAttr(t,n)}else{t.setAttribute(n,n)}return n}};m.each(m.expr.match.bool.source.match(/\w+/g),function(t,e){var n=ue[e]||m.find.attr;ue[e]=function(t,e,i){var r,a,s=e.toLowerCase();if(!i){a=ue[s];ue[s]=r;r=n(t,e,i)!=null?s:null;ue[s]=a}return r}});var fe=/^(?:input|select|textarea|button)$/i,ce=/^(?:a|area)$/i;m.fn.extend({prop:function(t,e){return $(this,m.prop,t,e,arguments.length>1)},removeProp:function(t){return this.each(function(){delete this[m.propFix[t]||t]})}});m.extend({prop:function(t,e,n){var i,r,a=t.nodeType;if(a===3||a===8||a===2){return}if(a!==1||!m.isXMLDoc(t)){e=m.propFix[e]||e;r=m.propHooks[e]}if(n!==undefined){if(r&&"set"in r&&(i=r.set(t,n,e))!==undefined){return i}return t[e]=n}if(r&&"get"in r&&(i=r.get(t,e))!==null){return i}return t[e]},propHooks:{tabIndex:{get:function(t){var e=m.find.attr(t,"tabindex");if(e){return parseInt(e,10)}if(fe.test(t.nodeName)||ce.test(t.nodeName)&&t.href){return 0}return-1}}},propFix:{for:"htmlFor",class:"className"}});if(!p.optSelected){m.propHooks.selected={get:function(t){var e=t.parentNode;if(e&&e.parentNode){e.parentNode.selectedIndex}return null},set:function(t){var e=t.parentNode;if(e){e.selectedIndex;if(e.parentNode){e.parentNode.selectedIndex}}}}}m.each(["tabIndex","readOnly","maxLength","cellSpacing","cellPadding","rowSpan","colSpan","useMap","frameBorder","contentEditable"],function(){m.propFix[this.toLowerCase()]=this});function he(t){var e=t.match(L)||[];return e.join(" ")}function de(t){return t.getAttribute&&t.getAttribute("class")||""}m.fn.extend({addClass:function(t){var e,n,i,r,a,s,o,l=0;if(m.isFunction(t)){return this.each(function(e){m(this).addClass(t.call(this,e,de(this)))})}if(typeof t==="string"&&t){e=t.match(L)||[];while(n=this[l++]){r=de(n);i=n.nodeType===1&&" "+he(r)+" ";if(i){s=0;while(a=e[s++]){if(i.indexOf(" "+a+" ")<0){i+=a+" "}}o=he(i);if(r!==o){n.setAttribute("class",o)}}}}return this},removeClass:function(t){var e,n,i,r,a,s,o,l=0;if(m.isFunction(t)){return this.each(function(e){m(this).removeClass(t.call(this,e,de(this)))})}if(!arguments.length){return this.attr("class","")}if(typeof t==="string"&&t){e=t.match(L)||[];while(n=this[l++]){r=de(n);i=n.nodeType===1&&" "+he(r)+" ";if(i){s=0;while(a=e[s++]){while(i.indexOf(" "+a+" ")>-1){i=i.replace(" "+a+" "," ")}}o=he(i);if(r!==o){n.setAttribute("class",o)}}}}return this},toggleClass:function(t,e){var n=typeof t;if(typeof e==="boolean"&&n==="string"){return e?this.addClass(t):this.removeClass(t)}if(m.isFunction(t)){return this.each(function(n){m(this).toggleClass(t.call(this,n,de(this),e),e)})}return this.each(function(){var e,i,r,a;if(n==="string"){i=0;r=m(this);a=t.match(L)||[];while(e=a[i++]){if(r.hasClass(e)){r.removeClass(e)}else{r.addClass(e)}}}else if(t===undefined||n==="boolean"){e=de(this);if(e){V.set(this,"__className__",e)}if(this.setAttribute){this.setAttribute("class",e||t===false?"":V.get(this,"__className__")||"")}}})},hasClass:function(t){var e,n,i=0;e=" "+t+" ";while(n=this[i++]){if(n.nodeType===1&&(" "+he(de(n))+" ").indexOf(e)>-1){return true}}return false}});var pe=/\r/g;m.fn.extend({val:function(t){var e,n,i,r=this[0];if(!arguments.length){if(r){e=m.valHooks[r.type]||m.valHooks[r.nodeName.toLowerCase()];if(e&&"get"in e&&(n=e.get(r,"value"))!==undefined){return n}n=r.value;if(typeof n==="string"){return n.replace(pe,"")}return n==null?"":n}return}i=m.isFunction(t);return this.each(function(n){var r;if(this.nodeType!==1){return}if(i){r=t.call(this,n,m(this).val())}else{r=t}if(r==null){r=""}else if(typeof r==="number"){r+=""}else if(m.isArray(r)){r=m.map(r,function(t){return t==null?"":t+""})}e=m.valHooks[this.type]||m.valHooks[this.nodeName.toLowerCase()];if(!e||!("set"in e)||e.set(this,r,"value")===undefined){this.value=r}})}});m.extend({valHooks:{option:{get:function(t){var e=m.find.attr(t,"value");return e!=null?e:he(m.text(t))}},select:{get:function(t){var e,n,i,r=t.options,a=t.selectedIndex,s=t.type==="select-one",o=s?null:[],l=s?a+1:r.length;if(a<0){i=l}else{i=s?a:0}for(;i<l;i++){n=r[i];if((n.selected||i===a)&&!n.disabled&&(!n.parentNode.disabled||!m.nodeName(n.parentNode,"optgroup"))){e=m(n).val();if(s){return e}o.push(e)}}return o},set:function(t,e){var n,i,r=t.options,a=m.makeArray(e),s=r.length;while(s--){i=r[s];if(i.selected=m.inArray(m.valHooks.option.get(i),a)>-1){n=true}}if(!n){t.selectedIndex=-1}return a}}}});m.each(["radio","checkbox"],function(){m.valHooks[this]={set:function(t,e){if(m.isArray(e)){return t.checked=m.inArray(m(t).val(),e)>-1}}};if(!p.checkOn){m.valHooks[this].get=function(t){return t.getAttribute("value")===null?"on":t.value}}});var ge=/^(?:focusinfocus|focusoutblur)$/;m.extend(m.event,{trigger:function(e,n,r,a){var s,o,l,u,f,h,d,p=[r||i],g=c.call(e,"type")?e.type:e,v=c.call(e,"namespace")?e.namespace.split("."):[];o=l=r=r||i;if(r.nodeType===3||r.nodeType===8){return}if(ge.test(g+m.event.triggered)){return}if(g.indexOf(".")>-1){v=g.split(".");g=v.shift();v.sort()}f=g.indexOf(":")<0&&"on"+g;e=e[m.expando]?e:new m.Event(g,typeof e==="object"&&e);e.isTrigger=a?2:3;e.namespace=v.join(".");e.rnamespace=e.namespace?new RegExp("(^|\\.)"+v.join("\\.(?:.*\\.|)")+"(\\.|$)"):null;e.result=undefined;if(!e.target){e.target=r}n=n==null?[e]:m.makeArray(n,[e]);d=m.event.special[g]||{};if(!a&&d.trigger&&d.trigger.apply(r,n)===false){return}if(!a&&!d.noBubble&&!m.isWindow(r)){u=d.delegateType||g;if(!ge.test(u+g)){o=o.parentNode}for(;o;o=o.parentNode){p.push(o);l=o}if(l===(r.ownerDocument||i)){p.push(l.defaultView||l.parentWindow||t)}}s=0;while((o=p[s++])&&!e.isPropagationStopped()){e.type=s>1?u:d.bindType||g;h=(V.get(o,"events")||{})[e.type]&&V.get(o,"handle");if(h){h.apply(o,n)}h=f&&o[f];if(h&&h.apply&&U(o)){e.result=h.apply(o,n);if(e.result===false){e.preventDefault()}}}e.type=g;if(!a&&!e.isDefaultPrevented()){if((!d._default||d._default.apply(p.pop(),n)===false)&&U(r)){if(f&&m.isFunction(r[g])&&!m.isWindow(r)){l=r[f];if(l){r[f]=null}m.event.triggered=g;r[g]();m.event.triggered=undefined;if(l){r[f]=l}}}}return e.result},simulate:function(t,e,n){var i=m.extend(new m.Event,n,{type:t,isSimulated:true});m.event.trigger(i,null,e)}});m.fn.extend({trigger:function(t,e){return this.each(function(){m.event.trigger(t,e,this)})},triggerHandler:function(t,e){var n=this[0];if(n){return m.event.trigger(t,e,n,true)}}});m.each(("blur focus focusin focusout resize scroll click dblclick "+"mousedown mouseup mousemove mouseover mouseout mouseenter mouseleave "+"change select submit keydown keypress keyup contextmenu").split(" "),function(t,e){m.fn[e]=function(t,n){return arguments.length>0?this.on(e,null,t,n):this.trigger(e)}});m.fn.extend({hover:function(t,e){return this.mouseenter(t).mouseleave(e||t)}});p.focusin="onfocusin"in t;if(!p.focusin){m.each({focus:"focusin",blur:"focusout"},function(t,e){var n=function(t){m.event.simulate(e,t.target,m.event.fix(t))};m.event.special[e]={setup:function(){var i=this.ownerDocument||this,r=V.access(i,e);if(!r){i.addEventListener(t,n,true)}V.access(i,e,(r||0)+1)},teardown:function(){var i=this.ownerDocument||this,r=V.access(i,e)-1;if(!r){i.removeEventListener(t,n,true);V.remove(i,e)}else{V.access(i,e,r)}}}})}var ve=t.location;var me=m.now();var ye=/\?/;m.parseXML=function(e){var n;if(!e||typeof e!=="string"){return null}try{n=(new t.DOMParser).parseFromString(e,"text/xml")}catch(t){n=undefined}if(!n||n.getElementsByTagName("parsererror").length){m.error("Invalid XML: "+e)}return n};var be=/\[\]$/,we=/\r?\n/g,_e=/^(?:submit|button|image|reset|file)$/i,xe=/^(?:input|select|textarea|keygen)/i;function Se(t,e,n,i){var r;if(m.isArray(e)){m.each(e,function(e,r){if(n||be.test(t)){i(t,r)}else{Se(t+"["+(typeof r==="object"&&r!=null?e:"")+"]",r,n,i)}})}else if(!n&&m.type(e)==="object"){for(r in e){Se(t+"["+r+"]",e[r],n,i)}}else{i(t,e)}}m.param=function(t,e){var n,i=[],r=function(t,e){var n=m.isFunction(e)?e():e;i[i.length]=encodeURIComponent(t)+"="+encodeURIComponent(n==null?"":n)};if(m.isArray(t)||t.jquery&&!m.isPlainObject(t)){m.each(t,function(){r(this.name,this.value)})}else{for(n in t){Se(n,t[n],e,r)}}return i.join("&")};m.fn.extend({serialize:function(){return m.param(this.serializeArray())},serializeArray:function(){return this.map(function(){var t=m.prop(this,"elements");return t?m.makeArray(t):this}).filter(function(){var t=this.type;return this.name&&!m(this).is(":disabled")&&xe.test(this.nodeName)&&!_e.test(t)&&(this.checked||!lt.test(t))}).map(function(t,e){var n=m(this).val();if(n==null){return null}if(m.isArray(n)){return m.map(n,function(t){return{name:e.name,value:t.replace(we,"\r\n")}})}return{name:e.name,value:n.replace(we,"\r\n")}}).get()}});var De=/%20/g,Ce=/#.*$/,Te=/([?&])_=[^&]*/,ke=/^(.*?):[ \t]*([^\r\n]*)$/gm,Me=/^(?:about|app|app-storage|.+-extension|file|res|widget):$/,Ie=/^(?:GET|HEAD)$/,Ae=/^\/\//,Ne={},Pe={},Ee="*/".concat("*"),Fe=i.createElement("a");Fe.href=ve.href;function Re(t){return function(e,n){if(typeof e!=="string"){n=e;e="*"}var i,r=0,a=e.toLowerCase().match(L)||[];if(m.isFunction(n)){while(i=a[r++]){if(i[0]==="+"){i=i.slice(1)||"*";(t[i]=t[i]||[]).unshift(n)}else{(t[i]=t[i]||[]).push(n)}}}}}function Le(t,e,n,i){var r={},a=t===Pe;function s(o){var l;r[o]=true;m.each(t[o]||[],function(t,o){var u=o(e,n,i);if(typeof u==="string"&&!a&&!r[u]){e.dataTypes.unshift(u);s(u);return false}else if(a){return!(l=u)}});return l}return s(e.dataTypes[0])||!r["*"]&&s("*")}function je(t,e){var n,i,r=m.ajaxSettings.flatOptions||{};for(n in e){if(e[n]!==undefined){(r[n]?t:i||(i={}))[n]=e[n]}}if(i){m.extend(true,t,i)}return t}function Oe(t,e,n){var i,r,a,s,o=t.contents,l=t.dataTypes;while(l[0]==="*"){l.shift();if(i===undefined){i=t.mimeType||e.getResponseHeader("Content-Type")}}if(i){for(r in o){if(o[r]&&o[r].test(i)){l.unshift(r);break}}}if(l[0]in n){a=l[0]}else{for(r in n){if(!l[0]||t.converters[r+" "+l[0]]){a=r;break}if(!s){s=r}}a=a||s}if(a){if(a!==l[0]){l.unshift(a)}return n[a]}}function He(t,e,n,i){var r,a,s,o,l,u={},f=t.dataTypes.slice();if(f[1]){for(s in t.converters){u[s.toLowerCase()]=t.converters[s]}}a=f.shift();while(a){if(t.responseFields[a]){n[t.responseFields[a]]=e}if(!l&&i&&t.dataFilter){e=t.dataFilter(e,t.dataType)}l=a;a=f.shift();if(a){if(a==="*"){a=l}else if(l!=="*"&&l!==a){s=u[l+" "+a]||u["* "+a];if(!s){for(r in u){o=r.split(" ");if(o[1]===a){s=u[l+" "+o[0]]||u["* "+o[0]];if(s){if(s===true){s=u[r]}else if(u[r]!==true){a=o[0];f.unshift(o[1])}break}}}}if(s!==true){if(s&&t.throws){e=s(e)}else{try{e=s(e)}catch(t){return{state:"parsererror",error:s?t:"No conversion from "+l+" to "+a}}}}}}}return{state:"success",data:e}}m.extend({active:0,lastModified:{},etag:{},ajaxSettings:{url:ve.href,type:"GET",isLocal:Me.test(ve.protocol),global:true,processData:true,async:true,contentType:"application/x-www-form-urlencoded; charset=UTF-8",accepts:{"*":Ee,text:"text/plain",html:"text/html",xml:"application/xml, text/xml",json:"application/json, text/javascript"},contents:{xml:/\bxml\b/,html:/\bhtml/,json:/\bjson\b/},responseFields:{xml:"responseXML",text:"responseText",json:"responseJSON"},converters:{"* text":String,"text html":true,"text json":JSON.parse,"text xml":m.parseXML},flatOptions:{url:true,context:true}},ajaxSetup:function(t,e){return e?je(je(t,m.ajaxSettings),e):je(m.ajaxSettings,t)},ajaxPrefilter:Re(Ne),ajaxTransport:Re(Pe),ajax:function(e,n){if(typeof e==="object"){n=e;e=undefined}n=n||{};var r,a,s,o,l,u,f,c,h,d,p=m.ajaxSetup({},n),g=p.context||p,v=p.context&&(g.nodeType||g.jquery)?m(g):m.event,y=m.Deferred(),b=m.Callbacks("once memory"),w=p.statusCode||{},_={},x={},S="canceled",D={readyState:0,getResponseHeader:function(t){var e;if(f){if(!o){o={};while(e=ke.exec(s)){
```
