## Supplementary material for "Transcriptome and histone epigenome of *Plasmodium vivax* salivary-gland sporozoites point to tight regulatory control and potential mechanisms for liver-stage differentiation": Pv-HPZ-v-mLS-DTGs.js

```
glimma.layout.setupGrid(d3.select(".container"), "md", [1,2]);
glimma.storage.chartData.push(glimma.transform.toRowMajor({"GeneID":["PVP01_0000030","PVP01_0000040","PVP01_0000050","PVP01_0000080","PVP01_0000110","PVP01_0000120","PVP01_0000130","PVP01_0000140","PVP01_0000150","PVP01_0000160","PVP01_0000170","PVP01_0000180","PVP01_0000190","PVP01_0000200","PVP01_0000210","PVP01_0000220","PVP01_0000300","PVP01_0000480","PVP01_0000670","PVP01_0000740","PVP01_0000790","PVP01_0001030","PVP01_0001090","PVP01_0001310","PVP01_0001430","PVP01_0001440","PVP01_0001480","PVP01_0001550","PVP01_0001640","PVP01_0001660","PVP01_0001670","PVP01_0001900","PVP01_0001910","PVP01_0001940","PVP01_0001970","PVP01_0002020","PVP01_0002050","PVP01_0002120","PVP01_0002200","PVP01_0002210","PVP01_0002330","PVP01_0002370","PVP01_0002390","PVP01_0002470","PVP01_0002530","PVP01_0002540","PVP01_0002750","PVP01_0002810","PVP01_0002820","PVP01_0003010","PVP01_0003050","PVP01_0003150","PVP01_0003210","PVP01_0003350","PVP01_0003360","PVP01_0003370","PVP01_0003380","PVP01_0003400","PVP01_0003410","PVP01_0003420","PVP01_0003440","PVP01_0003530","PVP01_0003640","PVP01_0003690","PVP01_0003730","PVP01_0003800","PVP01_0003990","PVP01_0004040","PVP01_0004160","PVP01_0004170","PVP01_0004180","PVP01_0004190","PVP01_0004210","PVP01_0004220","PVP01_0004230","PVP01_0004240","PVP01_0004250","PVP01_0004260","PVP01_0004270","PVP01_0004280","PVP01_0004290","PVP01_0004300","PVP01_0004310","PVP01_0004320","PVP01_0004330","PVP01_0004350","PVP01_0004370","PVP01_0004380","PVP01_0004390","PVP01_0004400","PVP01_0004410","PVP01_0004640","PVP01_0004670","PVP01_0004910","PVP01_0004920","PVP01_0004950","PVP01_0004960","PVP01_0005010","PVP01_0005100","PVP01_0005160","PVP01_0005210","PVP01_0005300","PVP01_0005310","PVP01_0005420","PVP01_0005430","PVP01_0005540","PVP01_0005550","PVP01_0005640","PVP01_0005710","PVP01_0005740","PVP01_0005760","PVP01_0005800","PVP01_0005820","PVP01_0005870","PVP01_0005910","PVP01_0006000","PVP01_0006040","PVP01_0006080","PVP01_0006260","PVP01_0006650","PVP01_0006660","PVP01_0006670","PVP01_0006680","PVP01_0006700","PVP01_0006860","PVP01_0006890","PVP01_0007080","PVP01_0007160","PVP01_0007240","PVP01_0007290","PVP01_0007310","PVP01_0007320","PVP01_0007340","PVP01_0007610","PVP01_0007620","PVP01_0007690","PVP01_0007700","PVP01_0007720","PVP01_0007850","PVP01_0007900","PVP01_0007910","PVP01_0007920","PVP01_0008190","PVP01_0008430","PVP01_0008480","PVP01_0008490","PVP01_0008600","PVP01_0008700","PVP01_0008770","PVP01_0008830","PVP01_0008990","PVP01_0009010","PVP01_0009170","PVP01_0009180","PVP01_0009210","PVP01_0009370","PVP01_0009410","PVP01_0009470","PVP01_0009540","PVP01_0009620","PVP01_0009710","PVP01_0009720","PVP01_0009730","PVP01_0009740","PVP01_0009760","PVP01_0009770","PVP01_0009820","PVP01_0009850","PVP01_0009860","PVP01_0009870","PVP01_0009900","PVP01_0009940","PVP01_0009980","PVP01_0010020","PVP01_0010030","PVP01_0010070","PVP01_0010100","PVP01_0010120","PVP01_0010140","PVP01_0010170","PVP01_0010180","PVP01_0010200","PVP01_0010210","PVP01_0010220","PVP01_0010230","PVP01_0010240","PVP01_0010340","PVP01_0010380","PVP01_0010440","PVP01_0010470","PVP01_0010500","PVP01_0010540","PVP01_0010570","PVP01_0010600","PVP01_0010610","PVP01_0010640","PVP01_0010650","PVP01_0010670","PVP01_0010770","PVP01_0010840","PVP01_0010870","PVP01_0010900","PVP01_0010910","PVP01_0010960","PVP01_0010970","PVP01_0010990","PVP01_0011020","PVP01_0011030","PVP01_0011070","PVP01_0102100","PVP01_0102200","PVP01_0102300","PVP01_0102400","PVP01_0102500","PVP01_0102600","PVP01_0102700","PVP01_0102800","PVP01_0103100","PVP01_0103200","PVP01_0103300","PVP01_0103400","PVP01_0103500","PVP01_0103600","PVP01_0103700","PVP01_0103800","PVP01_0103900","PVP01_0104000","PVP01_0104100","PVP01_0104200","PVP01_0104400","PVP01_0104500","PVP01_0104600","PVP01_0104800","PVP01_0104900","PVP01_0105000","PVP01_0105100","PVP01_0105200","PVP01_0105300","PVP01_0105400","PVP01_0105500","PVP01_0105600","PVP01_0105700","PVP01_0105800","PVP01_0105900","PVP01_0106000","PVP01_0106100","PVP01_0106200","PVP01_0106300","PVP01_0106400","PVP01_0106500","PVP01_0106600","PVP01_0106700","PVP01_0106800","PVP01_0106900","PVP01_0107400","PVP01_0107600","PVP01_0107700","PVP01_0107800","PVP01_0107900","PVP01_0108000","PVP01_0108100","PVP01_0108200","PVP01_0108300","PVP01_0108400","PVP01_0108500","PVP01_0108600","PVP01_0108700","PVP01_0108800","PVP01_0108900","PVP01_0109000","PVP01_0109100","PVP01_0109200","PVP01_0109300","PVP01_0109400","PVP01_0109500","PVP01_0109600","PVP01_0109700","PVP01_0109800","PVP01_0109900","PVP01_0110000","PVP01_0110100","PVP01_0110200","PVP01_0110300","PVP01_0110400","PVP01_0110500","PVP01_0110600","PVP01_0110700","PVP01_0110800","PVP01_0110900","PVP01_0111000","PVP01_0111100","PVP01_0111200","PVP01_0111300","PVP01_0111400","PVP01_0111500","PVP01_0111600","PVP01_0111700","PVP01_0111800","PVP01_0111900","PVP01_0112000","PVP01_0112100","PVP01_0112200","PVP01_0112300","PVP01_0112400","PVP01_0112500","PVP01_0112600","PVP01_0112700","PVP01_0112800","PVP01_0112900","PVP01_0113000","PVP01_0113100","PVP01_0113200","PVP01_0113300","PVP01_0113400","PVP01_0113500","PVP01_0113600","PVP01_0113700","PVP01_0113800","PVP01_0113900","PVP01_0114000","PVP01_0114100","PVP01_0114200","PVP01_0114300","PVP01_0114400","PVP01_0114500","PVP01_0114600","PVP01_0114700","PVP01_0114800","PVP01_0114900","PVP01_0115000","PVP01_0115100","PVP01_0115200","PVP01_0115300","PVP01_0115400","PVP01_0115500","PVP01_0115600","PVP01_0115700","PVP01_0115800","PVP01_0115900","PVP01_0116000","PVP01_0116100","PVP01_0116200","PVP01_0116300","PVP01_0116400","PVP01_0116500","PVP01_0116600","PVP01_0116700","PVP01_0116800","PVP01_0116900","PVP01_0117100","PVP01_0117200","PVP01_0117300","PVP01_0117400","PVP01_0117500","PVP01_0117700","PVP01_0117800","PVP01_0117900","PVP01_0118000","PVP01_0118100","PVP01_0118200","PVP01_0118300","PVP01_0118400","PVP01_0118500","PVP01_0118600","PVP01_0118700","PVP01_0118800","PVP01_0118900","PVP01_0119300","PVP01_0119400","PVP01_0119500","PVP01_0119600","PVP01_0119700","PVP01_0119800","PVP01_0119900","PVP01_0120000","PVP01_0120100","PVP01_0120500","PVP01_0120800","PVP01_0121000","PVP01_0121700","PVP01_0200200","PVP01_0200700","PVP01_0201000","PVP01_0201100","PVP01_0201200","PVP01_0201300","PVP01_0201400","PVP01_0201500","PVP01_0201600","PVP01_0201700","PVP01_0201800","PVP01_0201900","PVP01_0202000","PVP01_0202100","PVP01_0202200","PVP01_0202400","PVP01_0202500","PVP01_0202600","PVP01_0203000","PVP01_0203100","PVP01_0203200","PVP01_0203300","PVP01_0203400","PVP01_0203500","PVP01_0203600","PVP01_0203700","PVP01_0203800","PVP01_0203900","PVP01_0204000","PVP01_0204100","PVP01_0204200","PVP01_0204300","PVP01_0204400","PVP01_0204500","PVP01_0204600","PVP01_0204700","PVP01_0204800","PVP01_0204900","PVP01_0205000","PVP01_0205100","PVP01_0205200","PVP01_0205300","PVP01_0205400","PVP01_0205500","PVP01_0205600","PVP01_0205700","PVP01_0205800","PVP01_0205900","PVP01_0206000","PVP01_0206100","PVP01_0206200","PVP01_0206300","PVP01_0206400","PVP01_0206500","PVP01_0206600","PVP01_0206700","PVP01_0206800","PVP01_0206900","PVP01_0207000","PVP01_0207100","PVP01_0207200","PVP01_0207300","PVP01_0207500","PVP01_0207600","PVP01_0207700","PVP01_0207800","PVP01_0207900","PVP01_0208000","PVP01_0208100","PVP01_0208200","PVP01_0208300","PVP01_0208400","PVP01_0208500","PVP01_0208600","PVP01_0208700","PVP01_0208800","PVP01_0208900","PVP01_0209000","PVP01_0209100","PVP01_0209200","PVP01_0209300","PVP01_0209400","PVP01_0209500","PVP01_0209600","PVP01_0209700","PVP01_0209900","PVP01_0210000","PVP01_0210100","PVP01_0210200","PVP01_0210300","PVP01_0210400","PVP01_0210600","PVP01_0210700","PVP01_0210800","PVP01_0210900","PVP01_0211000","PVP01_0211100","PVP01_0211200","PVP01_0211300","PVP01_0211400","PVP01_0211500","PVP01_0211600","PVP01_0211700","PVP01_0211800","PVP01_0211900","PVP01_0212000","PVP01_0212100","PVP01_0212200","PVP01_0212300","PVP01_0212400","PVP01_0212500","PVP01_0212600","PVP01_0212700","PVP01_0212800","PVP01_0212900","PVP01_0213000","PVP01_0213100","PVP01_0213200","PVP01_0213300","PVP01_0213400","PVP01_0213500","PVP01_0213600","PVP01_0213700","PVP01_0213800","PVP01_0213900","PVP01_0214000","PVP01_0214100","PVP01_0214200","PVP01_0214300","PVP01_0214500","PVP01_0214600","PVP01_0214700","PVP01_0214800","PVP01_0214900","PVP01_0215000","PVP01_0215100","PVP01_0215200","PVP01_0215300","PVP01_0215400","PVP01_0215500","PVP01_0215600","PVP01_0215700","PVP01_0215800","PVP01_0215900","PVP01_0216000","PVP01_0216100","PVP01_0216200","PVP01_0216500","PVP01_0216600","PVP01_0216700","PVP01_0216800","PVP01_0216900","PVP01_0217000","PVP01_0217100","PVP01_0217200","PVP01_0217300","PVP01_0217400","PVP01_0217500","PVP01_0218200","PVP01_0218300","PVP01_0218900","PVP01_0219400","PVP01_0220100","PVP01_0220200","PVP01_0221100","PVP01_0221300","PVP01_0221700","PVP01_0300100","PVP01_0300600","PVP01_0300700","PVP01_0300800","PVP01_0301100","PVP01_0301200","PVP01_0301500","PVP01_0301900","PVP01_0302000","PVP01_0302100","PVP01_0302300","PVP01_0302400","PVP01_0302500","PVP01_0302600","PVP01_0302700","PVP01_0302800","PVP01_0302900","PVP01_0303000","PVP01_0303100","PVP01_0303200","PVP01_0303300","PVP01_0303400","PVP01_0303500","PVP01_0303600","PVP01_0303700","PVP01_0303800","PVP01_0303900","PVP01_0304000","PVP01_0304100","PVP01_0304200","PVP01_0304400","PVP01_0304500","PVP01_0304600","PVP01_0304700","PVP01_0304800","PVP01_0304900","PVP01_0305000","PVP01_0305100","PVP01_0305200","PVP01_0305300","PVP01_0305400","PVP01_0305500","PVP01_0305600","PVP01_0305700","PVP01_0305800","PVP01_0305900","PVP01_0306000","PVP01_0306100","PVP01_0306200","PVP01_0306300","PVP01_0306400","PVP01_0306500","PVP01_0306600","PVP01_0306700","PVP01_0306800","PVP01_0306900","PVP01_0307000","PVP01_0307100","PVP01_0307200","PVP01_0307300","PVP01_0307400","PVP01_0307500","PVP01_0307600","PVP01_0307700","PVP01_0307800","PVP01_0307900","PVP01_0308000","PVP01_0308100","PVP01_0308200","PVP01_0308300","PVP01_0308400","PVP01_0308500","PVP01_0308600","PVP01_0308700","PVP01_0308800","PVP01_0308900","PVP01_0309000","PVP01_0309200","PVP01_0309300","PVP01_0309600","PVP01_0309700","PVP01_0309800","PVP01_0309900","PVP01_0310000","PVP01_0310100","PVP01_0310200","PVP01_0310300","PVP01_0310400","PVP01_0310500","PVP01_0310600","PVP01_0310700","PVP01_0311000","PVP01_0311100","PVP01_0311200","PVP01_0311300","PVP01_0311500","PVP01_0311600","PVP01_0311700","PVP01_0311800","PVP01_0311900","PVP01_0312000","PVP01_0312100","PVP01_0312200","PVP01_0312300","PVP01_0312400","PVP01_0312500","PVP01_0312600","PVP01_0312700","PVP01_0312800","PVP01_0312900","PVP01_0313000","PVP01_0313100","PVP01_0313200","PVP01_0313300","PVP01_0313400","PVP01_0313500","PVP01_0313600","PVP01_0313700","PVP01_0313800","PVP01_0313900","PVP01_0314000","PVP01_0314100","PVP01_0314200","PVP01_0314300","PVP01_0314400","PVP01_0314500","PVP01_0314600","PVP01_0314700","PVP01_0314800","PVP01_0314900","PVP01_0315000","PVP01_0315100","PVP01_0315200","PVP01_0315300","PVP01_0315400","PVP01_0315500","PVP01_0315600","PVP01_0315700","PVP01_0315800","PVP01_0315900","PVP01_0316000","PVP01_0316100","PVP01_0316200","PVP01_0316300","PVP01_0316400","PVP01_0316500","PVP01_0316600","PVP01_0316700","PVP01_0316800","PVP01_0316900","PVP01_0317000","PVP01_0317100","PVP01_0317200","PVP01_0317300","PVP01_0317400","PVP01_0317500","PVP01_0317600","PVP01_0317700","PVP01_0317800","PVP01_0318000","PVP01_0318100","PVP01_0318200","PVP01_0318300","PVP01_0318400","PVP01_0318500","PVP01_0318600","PVP01_0318700","PVP01_0318800","PVP01_0318900","PVP01_0319000","PVP01_0319100","PVP01_0319200","PVP01_0319300","PVP01_0319400","PVP01_0319600","PVP01_0319700","PVP01_0319800","PVP01_0319900","PVP01_0320000","PVP01_0320100","PVP01_0320200","PVP01_0320300","PVP01_0320400","PVP01_0320500","PVP01_0320600","PVP01_0320700","PVP01_0320800","PVP01_0320900","PVP01_0321000","PVP01_0321100","PVP01_0400900","PVP01_0403600","PVP01_0403700","PVP01_0403900","PVP01_0404000","PVP01_0404100","PVP01_0404200","PVP01_0404300","PVP01_0404400","PVP01_0404500","PVP01_0404600","PVP01_0404700","PVP01_0404800","PVP01_0404900","PVP01_0405000","PVP01_0405100","PVP01_0405200","PVP01_0405300","PVP01_0405400","PVP01_0405500","PVP01_0405600","PVP01_0405700","PVP01_0405800","PVP01_0405900","PVP01_0406300","PVP01_0406400","PVP01_0406500","PVP01_0406600","PVP01_0406700","PVP01_0406800","PVP01_0406900","PVP01_0407000","PVP01_0407100","PVP01_0407200","PVP01_0407300","PVP01_0407400","PVP01_0407500","PVP01_0407600","PVP01_0407700","PVP01_0407800","PVP01_0407900","PVP01_0408000","PVP01_0408100","PVP01_0408200","PVP01_0408300","PVP01_0408400","PVP01_0408500","PVP01_0408600","PVP01_0408700","PVP01_0408800","PVP01_0408900","PVP01_0409000","PVP01_0409100","PVP01_0409200","PVP01_0409300","PVP01_0409400","PVP01_0409500","PVP01_0409600","PVP01_0409700","PVP01_0409800","PVP01_0409900","PVP01_0410000","PVP01_0410100","PVP01_0410200","PVP01_0410300","PVP01_0410400","PVP01_0410500","PVP01_0410600","PVP01_0410700","PVP01_0410800","PVP01_0410900","PVP01_0411000","PVP01_0411100","PVP01_0411200","PVP01_0411400","PVP01_0411500","PVP01_0411600","PVP01_0411700","PVP01_0411800","PVP01_0411900","PVP01_0412000","PVP01_0412100","PVP01_0412200","PVP01_0412300","PVP01_0412400","PVP01_0412500","PVP01_0412600","PVP01_0412700","PVP01_0412800","PVP01_0412900","PVP01_0413000","PVP01_0413100","PVP01_0413200","PVP01_0413300","PVP01_0413400","PVP01_0413500","PVP01_0413600","PVP01_0413700","PVP01_0413800","PVP01_0413900","PVP01_0414000","PVP01_0414100","PVP01_0414200","PVP01_0414300","PVP01_0414400","PVP01_0414500","PVP01_0414600","PVP01_0414700","PVP01_0414800","PVP01_0414900","PVP01_0415000","PVP01_0415100","PVP01_0415200","PVP01_0415300","PVP01_0415400","PVP01_0415500","PVP01_0415600","PVP01_0415700","PVP01_0415800","PVP01_0415900","PVP01_0416000","PVP01_0416100","PVP01_0416200","PVP01_0416300","PVP01_0416400","PVP01_0416500","PVP01_0416600","PVP01_0416700","PVP01_0416800","PVP01_0416900","PVP01_0417000","PVP01_0417100","PVP01_0417200","PVP01_0417400","PVP01_0417500","PVP01_0417700","PVP01_0417800","PVP01_0417900","PVP01_0418000","PVP01_0418100","PVP01_0418200","PVP01_0418300","PVP01_0418400","PVP01_0418500","PVP01_0418600","PVP01_0418800","PVP01_0418900","PVP01_0419000","PVP01_0419100","PVP01_0419200","PVP01_0419300","PVP01_0419400","PVP01_0419500","PVP01_0419600","PVP01_0419700","PVP01_0419800","PVP01_0419900","PVP01_0420000","PVP01_0420100","PVP01_0420200","PVP01_0420300","PVP01_0420400","PVP01_0420500","PVP01_0420600","PVP01_0420700","PVP01_0420800","PVP01_0420900","PVP01_0421100","PVP01_0421200","PVP01_0421300","PVP01_0421400","PVP01_0421500","PVP01_0421700","PVP01_0421800","PVP01_0421900","PVP01_0422000","PVP01_0422100","PVP01_0422200","PVP01_0422300","PVP01_0422400","PVP01_0422500","PVP01_0422600","PVP01_0422700","PVP01_0422800","PVP01_0422900","PVP01_0423000","PVP01_0423100","PVP01_0423200","PVP01_0423300","PVP01_0423400","PVP01_0423500","PVP01_0423600","PVP01_0423800","PVP01_0423900","PVP01_0424300","PVP01_0424400","PVP01_0424500","PVP01_0500700","PVP01_0502300","PVP01_0502400","PVP01_0502900","PVP01_0503300","PVP01_0503400","PVP01_0503500","PVP01_0503600","PVP01_0503700","PVP01_0503800","PVP01_0503900","PVP01_0504100","PVP01_0504200","PVP01_0504300","PVP01_0504400","PVP01_0504800","PVP01_0504900","PVP01_0505000","PVP01_0505100","PVP01_0505200","PVP01_0505300","PVP01_0505400","PVP01_0505500","PVP01_0505600","PVP01_0505700","PVP01_0505800","PVP01_0505900","PVP01_0506000","PVP01_0506100","PVP01_0506200","PVP01_0506300","PVP01_0506400","PVP01_0506500","PVP01_0506600","PVP01_0506700","PVP01_0506800","PVP01_0506900","PVP01_0507000","PVP01_0507100","PVP01_0507200","PVP01_0507300","PVP01_0507400","PVP01_0507500","PVP01_0507600","PVP01_0507700","PVP01_0507800","PVP01_0507900","PVP01_0508000","PVP01_0508100","PVP01_0508200","PVP01_0508300","PVP01_0508400","PVP01_0508500","PVP01_0508600","PVP01_0508700","PVP01_0508800","PVP01_0508900","PVP01_0509000","PVP01_0509100","PVP01_0509200","PVP01_0509300","PVP01_0509400","PVP01_0509500","PVP01_0509600","PVP01_0509700","PVP01_0509800","PVP01_0509900","PVP01_0510000","PVP01_0510100","PVP01_0510200","PVP01_0510300","PVP01_0510400","PVP01_0510500","PVP01_0510600","PVP01_0510700","PVP01_0510800","PVP01_0510900","PVP01_0511000","PVP01_0511100","PVP01_0511200","PVP01_0511300","PVP01_0511500","PVP01_0511600","PVP01_0511700","PVP01_0511800","PVP01_0511900","PVP01_0512000","PVP01_0512100","PVP01_0512200","PVP01_0512300","PVP01_0512400","PVP01_0512500","PVP01_0512600","PVP01_0512700","PVP01_0512800","PVP01_0512900","PVP01_0513000","PVP01_0513100","PVP01_0513200","PVP01_0513300","PVP01_0513400","PVP01_0513500","PVP01_0513600","PVP01_0513700","PVP01_0513800","PVP01_0513900","PVP01_0514000","PVP01_0514100","PVP01_0514200","PVP01_0514300","PVP01_0514400","PVP01_0514500","PVP01_0514600","PVP01_0514700","PVP01_0514800","PVP01_0514900","PVP01_0515000","PVP01_0515100","PVP01_0515200","PVP01_0515300","PVP01_0515400","PVP01_0515500","PVP01_0515600","PVP01_0515700","PVP01_0515800","PVP01_0515900","PVP01_0516000","PVP01_0516100","PVP01_0516200","PVP01_0516300","PVP01_0516400","PVP01_0516500","PVP01_0516600","PVP01_0516700","PVP01_0516800","PVP01_0516900","PVP01_0517000","PVP01_0517100","PVP01_0517200","PVP01_0517300","PVP01_0517400","PVP01_0517500","PVP01_0517700","PVP01_0517800","PVP01_0517900","PVP01_0518000","PVP01_0518100","PVP01_0518200","PVP01_0518300","PVP01_0518400","PVP01_0518500","PVP01_0518600","PVP01_0518700","PVP01_0518800","PVP01_0518900","PVP01_0519000","PVP01_0519100","PVP01_0519200","PVP01_0519300","PVP01_0519400","PVP01_0519500","PVP01_0519600","PVP01_0519700","PVP01_0519800","PVP01_0519900","PVP01_0520000","PVP01_0520100","PVP01_0520200","PVP01_0520300","PVP01_0520400","PVP01_0520500","PVP01_0520600","PVP01_0520700","PVP01_0520800","PVP01_0520900","PVP01_0521000","PVP01_0521100","PVP01_0521200","PVP01_0521300","PVP01_0521400","PVP01_0521500","PVP01_0521600","PVP01_0521700","PVP01_0521800","PVP01_0521900","PVP01_0522000","PVP01_0522100","PVP01_0522200","PVP01_0522300","PVP01_0522400","PVP01_0522500","PVP01_0522600","PVP01_0522700","PVP01_0522800","PVP01_0522900","PVP01_0523000","PVP01_0523100","PVP01_0523200","PVP01_0523300","PVP01_0523400","PVP01_0523500","PVP01_0523600","PVP01_0523700","PVP01_0523800","PVP01_0523900","PVP01_0524000","PVP01_0524100","PVP01_0524200","PVP01_0524300","PVP01_0524500","PVP01_0524600","PVP01_0524700","PVP01_0524900","PVP01_0525000","PVP01_0525100","PVP01_0525200","PVP01_0525300","PVP01_0525400","PVP01_0525500","PVP01_0525600","PVP01_0525700","PVP01_0525800","PVP01_0525900","PVP01_0526000","PVP01_0526100","PVP01_0526200","PVP01_0526300","PVP01_0526400","PVP01_0526500","PVP01_0526600","PVP01_0526700","PVP01_0526800","PVP01_0526900","PVP01_0527000","PVP01_0527100","PVP01_0527200","PVP01_0527300","PVP01_0527400","PVP01_0527500","PVP01_0527600","PVP01_0527700","PVP01_0527800","PVP01_0527900","PVP01_0528000","PVP01_0528100","PVP01_0528200","PVP01_0528300","PVP01_0528400","PVP01_0528500","PVP01_0528600","PVP01_0528700","PVP01_0528800","PVP01_0528900","PVP01_0529000","PVP01_0529100","PVP01_0529200","PVP01_0529300","PVP01_0529400","PVP01_0529500","PVP01_0529600","PVP01_0529700","PVP01_0529800","PVP01_0529900","PVP01_0530000","PVP01_0530100","PVP01_0530200","PVP01_0530300","PVP01_0530400","PVP01_0530500","PVP01_0530600","PVP01_0530700","PVP01_0530800","PVP01_0530900","PVP01_0531000","PVP01_0531100","PVP01_0531200","PVP01_0531300","PVP01_0531400","PVP01_0531500","PVP01_0531600","PVP01_0531700","PVP01_0531800","PVP01_0531900","PVP01_0532100","PVP01_0532200","PVP01_0532300","PVP01_0532400","PVP01_0532500","PVP01_0532600","PVP01_0532700","PVP01_0532800","PVP01_0532900","PVP01_0533000","PVP01_0533100","PVP01_0533200","PVP01_0533300","PVP01_0533700","PVP01_0534000","PVP01_0534100","PVP01_0534200","PVP01_0534400","PVP01_0534600","PVP01_0534800","PVP01_0534900","PVP01_0535400","PVP01_0600100","PVP01_0600300","PVP01_0600400","PVP01_0600500","PVP01_0600600","PVP01_0600700","PVP01_0600800","PVP01_0600900","PVP01_0601000","PVP01_0601100","PVP01_0601200","PVP01_0601300","PVP01_0601400","PVP01_0601500","PVP01_0601600","PVP01_0601700","PVP01_0601800","PVP01_0601900","PVP01_0602000","PVP01_0602200","PVP01_0602300","PVP01_0602400","PVP01_0602500","PVP01_0602600","PVP01_0602700","PVP01_0602800","PVP01_0602900","PVP01_0603000","PVP01_0603100","PVP01_0603200","PVP01_0603300","PVP01_0603400","PVP01_0603500","PVP01_0603600","PVP01_0603700","PVP01_0603800","PVP01_0603900","PVP01_0604000","PVP01_0604100","PVP01_0604200","PVP01_0604300","PVP01_0604400","PVP01_0604500","PVP01_0604600","PVP01_0604700","PVP01_0604800","PVP01_0604900","PVP01_0605000","PVP01_0605100","PVP01_0605200","PVP01_0605300","PVP01_0605400","PVP01_0605500","PVP01_0605600","PVP01_0605700","PVP01_0605800","PVP01_0605900","PVP01_0606000","PVP01_0606100","PVP01_0606200","PVP01_0606300","PVP01_0606400","PVP01_0606500","PVP01_0606600","PVP01_0606700","PVP01_0606800","PVP01_0606900","PVP01_0607000","PVP01_0607100","PVP01_0607200","PVP01_0607300","PVP01_0607400","PVP01_0607500","PVP01_0607600","PVP01_0607700","PVP01_0607800","PVP01_0607900","PVP01_0608000","PVP01_0608100","PVP01_0608200","PVP01_0608300","PVP01_0608400","PVP01_0608500","PVP01_0608600","PVP01_0608700","PVP01_0608800","PVP01_0608900","PVP01_0609000","PVP01_0609200","PVP01_0609300","PVP01_0609400","PVP01_0609500","PVP01_0609600","PVP01_0609700","PVP01_0609800","PVP01_0609900","PVP01_0610000","PVP01_0610100","PVP01_0610200","PVP01_0610300","PVP01_0610400","PVP01_0610500","PVP01_0610600","PVP01_0610700","PVP01_0610800","PVP01_0610900","PVP01_0611000","PVP01_0611100","PVP01_0611200","PVP01_0611300","PVP01_0611400","PVP01_0611500","PVP01_0611600","PVP01_0611700","PVP01_0611800","PVP01_0611900","PVP01_0612000","PVP01_0612100","PVP01_0612200","PVP01_0612300","PVP01_0612500","PVP01_0612600","PVP01_0612700","PVP01_0612800","PVP01_0612900","PVP01_0613000","PVP01_0613100","PVP01_0613200","PVP01_0613300","PVP01_0613400","PVP01_0613500","PVP01_0613600","PVP01_0613700","PVP01_0613800","PVP01_0613900","PVP01_0614000","PVP01_0614100","PVP01_0614200","PVP01_0614300","PVP01_0614400","PVP01_0614500","PVP01_0614600","PVP01_0614700","PVP01_0614800","PVP01_0614900","PVP01_0615100","PVP01_0615200","PVP01_0615300","PVP01_0615400","PVP01_0615500","PVP01_0615600","PVP01_0615700","PVP01_0615800","PVP01_0615900","PVP01_0616000","PVP01_0616100","PVP01_0616200","PVP01_0616300","PVP01_0616400","PVP01_0616500","PVP01_0616600","PVP01_0616700","PVP01_0616800","PVP01_0616900","PVP01_0617000","PVP01_0617100","PVP01_0617200","PVP01_0617300","PVP01_0617400","PVP01_0617500","PVP01_0617600","PVP01_0617700","PVP01_0617800","PVP01_0617900","PVP01_0618000","PVP01_0618100","PVP01_0618200","PVP01_0618300","PVP01_0618400","PVP01_0618500","PVP01_0618600","PVP01_0618700","PVP01_0618800","PVP01_0618900","PVP01_0619000","PVP01_0619100","PVP01_0619200","PVP01_0619300","PVP01_0619400","PVP01_0619500","PVP01_0619600","PVP01_0619700","PVP01_0619800","PVP01_0619900","PVP01_0620000","PVP01_0620100","PVP01_0620200","PVP01_0620300","PVP01_0620400","PVP01_0620500","PVP01_0620600","PVP01_0620700","PVP01_0620900","PVP01_0621000","PVP01_0621100","PVP01_0621200","PVP01_0621300","PVP01_0621400","PVP01_0621500","PVP01_0621600","PVP01_0621700","PVP01_0621800","PVP01_0621900","PVP01_0622000","PVP01_0622100","PVP01_0622200","PVP01_0622300","PVP01_0622400","PVP01_0622500","PVP01_0622600","PVP01_0622700","PVP01_0623000","PVP01_0623100","PVP01_0623200","PVP01_0623300","PVP01_0623400","PVP01_0623500","PVP01_0623600","PVP01_0623700","PVP01_0623800","PVP01_0623900","PVP01_0624000","PVP01_0624100","PVP01_0624300","PVP01_0624400","PVP01_0624500","PVP01_0700100","PVP01_0700200","PVP01_0700300","PVP01_0700400","PVP01_0700500","PVP01_0700600","PVP01_0700700","PVP01_0700800","PVP01_0700900","PVP01_0701000","PVP01_0701100","PVP01_0701200","PVP01_0701300","PVP01_0701400","PVP01_0701500","PVP01_0701600","PVP01_0701700","PVP01_0701800","PVP01_0701900","PVP01_0702000","PVP01_0702100","PVP01_0702200","PVP01_0702300","PVP01_0702400","PVP01_0702500","PVP01_0702600","PVP01_0702700","PVP01_0702800","PVP01_0702900","PVP01_0703000","PVP01_0703100","PVP01_0703200","PVP01_0703300","PVP01_0703400","PVP01_0703500","PVP01_0703600","PVP01_0703700","PVP01_0703900","PVP01_0704000","PVP01_0704100","PVP01_0704200","PVP01_0704300","PVP01_0704400","PVP01_0704500","PVP01_0704600","PVP01_0704700","PVP01_0704800","PVP01_0704900","PVP01_0705000","PVP01_0705100","PVP01_0705200","PVP01_0705300","PVP01_0705400","PVP01_0705500","PVP01_0705600","PVP01_0705700","PVP01_0705800","PVP01_0706000","PVP01_0706100","PVP01_0706200","PVP01_0706300","PVP01_0706400","PVP01_0706500","PVP01_0706600","PVP01_0706700","PVP01_0706800","PVP01_0706900","PVP01_0707000","PVP01_0707100","PVP01_0707200","PVP01_0707300","PVP01_0707400","PVP01_0707500","PVP01_0707600","PVP01_0707700","PVP01_0707800","PVP01_0707900","PVP01_0708000","PVP01_0708100","PVP01_0708200","PVP01_0708300","PVP01_0708400","PVP01_0708500","PVP01_0708600","PVP01_0708700","PVP01_0708800","PVP01_0708900","PVP01_0709100","PVP01_0709200","PVP01_0709300","PVP01_0709400","PVP01_0709500","PVP01_0709600","PVP01_0709700","PVP01_0709800","PVP01_0709900","PVP01_0710000","PVP01_0710100","PVP01_0710200","PVP01_0710300","PVP01_0710400","PVP01_0710500","PVP01_0710600","PVP01_0710700","PVP01_0710800","PVP01_0710900","PVP01_0711000","PVP01_0711100","PVP01_0711200","PVP01_0711300","PVP01_0711400","PVP01_0711500","PVP01_0711600","PVP01_0711700","PVP01_0711800","PVP01_0711900","PVP01_0712000","PVP01_0712100","PVP01_0712200","PVP01_0712300","PVP01_0712400","PVP01_0712500","PVP01_0712600","PVP01_0712700","PVP01_0712800","PVP01_0712900","PVP01_0713000","PVP01_0713100","PVP01_0713200","PVP01_0713300","PVP01_0713400","PVP01_0713500","PVP01_0713600","PVP01_0713700","PVP01_0713800","PVP01_0713900","PVP01_0714000","PVP01_0714100","PVP01_0714200","PVP01_0714300","PVP01_0714400","PVP01_0714500","PVP01_0714600","PVP01_0714700","PVP01_0714800","PVP01_0714900","PVP01_0715000","PVP01_0715100","PVP01_0715200","PVP01_0715300","PVP01_0715400","PVP01_0715500","PVP01_0715600","PVP01_0715700","PVP01_0715800","PVP01_0715900","PVP01_0716000","PVP01_0716100","PVP01_0716200","PVP01_0716300","PVP01_0716400","PVP01_0716500","PVP01_0716600","PVP01_0716700","PVP01_0716800","PVP01_0717000","PVP01_0717100","PVP01_0717200","PVP01_0717300","PVP01_0717400","PVP01_0717500","PVP01_0717600","PVP01_0717700","PVP01_0717800","PVP01_0717900","PVP01_0718000","PVP01_0718100","PVP01_0718200","PVP01_0718300","PVP01_0718400","PVP01_0718500","PVP01_0718600","PVP01_0718700","PVP01_0718800","PVP01_0718900","PVP01_0719000","PVP01_0719100","PVP01_0719200","PVP01_0719300","PVP01_0719400","PVP01_0719500","PVP01_0719600","PVP01_0719700","PVP01_0719800","PVP01_0719900","PVP01_0720000","PVP01_0720100","PVP01_0720200","PVP01_0720300","PVP01_0720400","PVP01_0720500","PVP01_0720600","PVP01_0720700","PVP01_0720800","PVP01_0720900","PVP01_0721000","PVP01_0721100","PVP01_0721200","PVP01_0721300","PVP01_0721400","PVP01_0721500","PVP01_0721600","PVP01_0721700","PVP01_0721800","PVP01_0721900","PVP01_0722000","PVP01_0722100","PVP01_0722200","PVP01_0722300","PVP01_0722400","PVP01_0722500","PVP01_0722600","PVP01_0722700","PVP01_0722800","PVP01_0722900","PVP01_0723000","PVP01_0723200","PVP01_0723300","PVP01_0723400","PVP01_0723500","PVP01_0723700","PVP01_0723800","PVP01_0723900","PVP01_0724000","PVP01_0724100","PVP01_0724200","PVP01_0724300","PVP01_0724400","PVP01_0724500","PVP01_0724600","PVP01_0724700","PVP01_0724800","PVP01_0724900","PVP01_0725000","PVP01_0725100","PVP01_0725200","PVP01_0725300","PVP01_0725400","PVP01_0725500","PVP01_0725600","PVP01_0725700","PVP01_0725800","PVP01_0725900","PVP01_0726000","PVP01_0726100","PVP01_0726200","PVP01_0726300","PVP01_0726400","PVP01_0726500","PVP01_0726600","PVP01_0726700","PVP01_0726800","PVP01_0726900","PVP01_0727000","PVP01_0727100","PVP01_0727200","PVP01_0727300","PVP01_0727400","PVP01_0727500","PVP01_0727600","PVP01_0727700","PVP01_0727800","PVP01_0728000","PVP01_0728100","PVP01_0728200","PVP01_0728300","PVP01_0728400","PVP01_0728500","PVP01_0728600","PVP01_0728700","PVP01_0728800","PVP01_0729000","PVP01_0729100","PVP01_0729200","PVP01_0729300","PVP01_0729400","PVP01_0729500","PVP01_0729600","PVP01_0729700","PVP01_0729800","PVP01_0729900","PVP01_0730000","PVP01_0730100","PVP01_0730200","PVP01_0730300","PVP01_0730400","PVP01_0730500","PVP01_0730600","PVP01_0730700","PVP01_0730900","PVP01_0731000","PVP01_0731100","PVP01_0731200","PVP01_0731300","PVP01_0731400","PVP01_0731500","PVP01_0731600","PVP01_0731700","PVP01_0731800","PVP01_0731900","PVP01_0732000","PVP01_0732100","PVP01_0732200","PVP01_0732300","PVP01_0732400","PVP01_0732500","PVP01_0732600","PVP01_0732700","PVP01_0732800","PVP01_0732900","PVP01_0733000","PVP01_0733100","PVP01_0733200","PVP01_0733300","PVP01_0733400","PVP01_0733500","PVP01_0733600","PVP01_0733700","PVP01_0733800","PVP01_0733900","PVP01_0734000","PVP01_0734100","PVP01_0734200","PVP01_0734400","PVP01_0734600","PVP01_0734700","PVP01_0734800","PVP01_0734900","PVP01_0735100","PVP01_0735200","PVP01_0735300","PVP01_0735800","PVP01_0735900","PVP01_0800100","PVP01_0800200","PVP01_0800300","PVP01_0800400","PVP01_0800500","PVP01_0800600","PVP01_0800700","PVP01_0800900","PVP01_0801000","PVP01_0801100","PVP01_0801200","PVP01_0801300","PVP01_0801400","PVP01_0801500","PVP01_0801700","PVP01_0801800","PVP01_0802100","PVP01_0802200","PVP01_0802300","PVP01_0802400","PVP01_0802500","PVP01_0802600","PVP01_0802700","PVP01_0802800","PVP01_0802900","PVP01_0803000","PVP01_0803100","PVP01_0803200","PVP01_0803300","PVP01_0803400","PVP01_0803500","PVP01_0803600","PVP01_0803700","PVP01_0803800","PVP01_0803900","PVP01_0804000","PVP01_0804100","PVP01_0804200","PVP01_0804300","PVP01_0804400","PVP01_0804500","PVP01_0804600","PVP01_0804700","PVP01_0804800","PVP01_0804900","PVP01_0805000","PVP01_0805100","PVP01_0805200","PVP01_0805300","PVP01_0805400","PVP01_0805500","PVP01_0805600","PVP01_0805700","PVP01_0805800","PVP01_0805900","PVP01_0806000","PVP01_0806100","PVP01_0806200","PVP01_0806300","PVP01_0806400","PVP01_0806500","PVP01_0806600","PVP01_0806700","PVP01_0806800","PVP01_0806900","PVP01_0807000","PVP01_0807100","PVP01_0807200","PVP01_0807300","PVP01_0807400","PVP01_0807500","PVP01_0807600","PVP01_0807800","PVP01_0807900","PVP01_0808000","PVP01_0808100","PVP01_0808200","PVP01_0808300","PVP01_0808400","PVP01_0808500","PVP01_0808600","PVP01_0808700","PVP01_0808800","PVP01_0808900","PVP01_0809000","PVP01_0809100","PVP01_0809200","PVP01_0809300","PVP01_0809400","PVP01_0809500","PVP01_0809600","PVP01_0809700","PVP01_0809800","PVP01_0809900","PVP01_0810000","PVP01_0810100","PVP01_0810200","PVP01_0810300","PVP01_0810400","PVP01_0810500","PVP01_0810600","PVP01_0810700","PVP01_0810800","PVP01_0810900","PVP01_0811000","PVP01_0811100","PVP01_0811200","PVP01_0811300","PVP01_0811400","PVP01_0811500","PVP01_0811600","PVP01_0811700","PVP01_0811800","PVP01_0811900","PVP01_0812000","PVP01_0812100","PVP01_0812200","PVP01_0812400","PVP01_0812500","PVP01_0812600","PVP01_0812700","PVP01_0812800","PVP01_0812900","PVP01_0813000","PVP01_0813100","PVP01_0813200","PVP01_0813300","PVP01_0813400","PVP01_0813500","PVP01_0813600","PVP01_0813700","PVP01_0813800","PVP01_0814000","PVP01_0814100","PVP01_0814200","PVP01_0814300","PVP01_0814400","PVP01_0814500","PVP01_0814600","PVP01_0814700","PVP01_0814800","PVP01_0814900","PVP01_0815100","PVP01_0815200","PVP01_0815300","PVP01_0815400","PVP01_0815500","PVP01_0815600","PVP01_0815700","PVP01_0815800","PVP01_0815900","PVP01_0816000","PVP01_0816100","PVP01_0816200","PVP01_0816300","PVP01_0816400","PVP01_0816500","PVP01_0816600","PVP01_0816700","PVP01_0816900","PVP01_0817000","PVP01_0817100","PVP01_0817200","PVP01_0817300","PVP01_0817400","PVP01_0817500","PVP01_0817600","PVP01_0817700","PVP01_0817800","PVP01_0817900","PVP01_0818000","PVP01_0818100","PVP01_0818200","PVP01_0818300","PVP01_0818400","PVP01_0818500","PVP01_0818600","PVP01_0818700","PVP01_0818800","PVP01_0818900","PVP01_0819000","PVP01_0819100","PVP01_0819200","PVP01_0819400","PVP01_0819500","PVP01_0819600","PVP01_0819700","PVP01_0819800","PVP01_0819900","PVP01_0820000","PVP01_0820100","PVP01_0820200","PVP01_0820300","PVP01_0820400","PVP01_0820500","PVP01_0820600","PVP01_0820700","PVP01_0820800","PVP01_0820900","PVP01_0821000","PVP01_0821100","PVP01_0821200","PVP01_0821300","PVP01_0821400","PVP01_0821500","PVP01_0821600","PVP01_0821700","PVP01_0821800","PVP01_0821900","PVP01_0822000","PVP01_0822100","PVP01_0822200","PVP01_0822300","PVP01_0822400","PVP01_0822500","PVP01_0822600","PVP01_0822700","PVP01_0822800","PVP01_0822900","PVP01_0823000","PVP01_0823100","PVP01_0823200","PVP01_0823300","PVP01_0823400","PVP01_0823500","PVP01_0823600","PVP01_0823700","PVP01_0823800","PVP01_0823900","PVP01_0824000","PVP01_0824100","PVP01_0824200","PVP01_0824300","PVP01_0824400","PVP01_0824500","PVP01_0824600","PVP01_0824700","PVP01_0824800","PVP01_0824900","PVP01_0825000","PVP01_0825100","PVP01_0825200","PVP01_0825300","PVP01_0825400","PVP01_0825500","PVP01_0825600","PVP01_0825700","PVP01_0825800","PVP01_0825900","PVP01_0826000","PVP01_0826100","PVP01_0826200","PVP01_0826300","PVP01_0826400","PVP01_0826500","PVP01_0826600","PVP01_0826700","PVP01_0826800","PVP01_0826900","PVP01_0827000","PVP01_0827100","PVP01_0827200","PVP01_0827300","PVP01_0827400","PVP01_0827700","PVP01_0827800","PVP01_0827900","PVP01_0828000","PVP01_0828100","PVP01_0828200","PVP01_0828300","PVP01_0828400","PVP01_0828500","PVP01_0828600","PVP01_0828700","PVP01_0828800","PVP01_0828900","PVP01_0829000","PVP01_0829100","PVP01_0829200","PVP01_0829300","PVP01_0829400","PVP01_0829500","PVP01_0829600","PVP01_0829700","PVP01_0829800","PVP01_0829900","PVP01_0830000","PVP01_0830100","PVP01_0830200","PVP01_0830300","PVP01_0830400","PVP01_0830500","PVP01_0830600","PVP01_0830700","PVP01_0830900","PVP01_0831000","PVP01_0831100","PVP01_0831200","PVP01_0831300","PVP01_0831400","PVP01_0831500","PVP01_0831600","PVP01_0831700","PVP01_0831900","PVP01_0832000","PVP01_0832100","PVP01_0832200","PVP01_0832300","PVP01_0832400","PVP01_0832500","PVP01_0832600","PVP01_0832700","PVP01_0832800","PVP01_0832900","PVP01_0833000","PVP01_0833100","PVP01_0833200","PVP01_0833400","PVP01_0833500","PVP01_0833600","PVP01_0833700","PVP01_0833800","PVP01_0833900","PVP01_0834000","PVP01_0834100","PVP01_0834200","PVP01_0834300","PVP01_0834400","PVP01_0834500","PVP01_0834600","PVP01_0834700","PVP01_0834800","PVP01_0834900","PVP01_0835000","PVP01_0835100","PVP01_0835200","PVP01_0835300","PVP01_0835400","PVP01_0835500","PVP01_0835600","PVP01_0835700","PVP01_0835800","PVP01_0835900","PVP01_0836000","PVP01_0836100","PVP01_0836200","PVP01_0836300","PVP01_0836400","PVP01_0836500","PVP01_0836600","PVP01_0836700","PVP01_0836800","PVP01_0836900","PVP01_0837000","PVP01_0837100","PVP01_0837200","PVP01_0837300","PVP01_0837400","PVP01_0837500","PVP01_0837600","PVP01_0837700","PVP01_0837800","PVP01_0837900","PVP01_0838000","PVP01_0838100","PVP01_0838200","PVP01_0838300","PVP01_0838600","PVP01_0838700","PVP01_0838800","PVP01_0838900","PVP01_0839000","PVP01_0839100","PVP01_0839200","PVP01_0839400","PVP01_0839700","PVP01_0840700","PVP01_0841000","PVP01_0900700","PVP01_0901000","PVP01_0901100","PVP01_0901200","PVP01_0901300","PVP01_0901800","PVP01_0901900","PVP01_0902000","PVP01_0902100","PVP01_0902400","PVP01_0903400","PVP01_0903500","PVP01_0903600","PVP01_0903700","PVP01_0903800","PVP01_0903900","PVP01_0904200","PVP01_0904300","PVP01_0904400","PVP01_0904500","PVP01_0904600","PVP01_0904700","PVP01_0904800","PVP01_0904900","PVP01_0905000","PVP01_0905100","PVP01_0905200","PVP01_0905300","PVP01_0905400","PVP01_0905500","PVP01_0905600","PVP01_0905700","PVP01_0905800","PVP01_0905900","PVP01_0906000","PVP01_0906100","PVP01_0906200","PVP01_0906300","PVP01_0906400","PVP01_0906500","PVP01_0906600","PVP01_0906700","PVP01_0906800","PVP01_0906900","PVP01_0907000","PVP01_0907100","PVP01_0907200","PVP01_0907300","PVP01_0907400","PVP01_0907500","PVP01_0907600","PVP01_0907700","PVP01_0907800","PVP01_0907900","PVP01_0908000","PVP01_0908100","PVP01_0908200","PVP01_0908300","PVP01_0908400","PVP01_0908500","PVP01_0908600","PVP01_0908700","PVP01_0908800","PVP01_0908900","PVP01_0909000","PVP01_0909100","PVP01_0909200","PVP01_0909300","PVP01_0909400","PVP01_0909500","PVP01_0909600","PVP01_0909700","PVP01_0909900","PVP01_0910000","PVP01_0910200","PVP01_0910700","PVP01_0910800","PVP01_0910900","PVP01_0911000","PVP01_0911100","PVP01_0911200","PVP01_0911300","PVP01_0911400","PVP01_0911500","PVP01_0911600","PVP01_0911700","PVP01_0911800","PVP01_0911900","PVP01_0912000","PVP01_0912100","PVP01_0912200","PVP01_0912300","PVP01_0912400","PVP01_0912500","PVP01_0912600","PVP01_0912700","PVP01_0912800","PVP01_0912900","PVP01_0913000","PVP01_0913100","PVP01_0913200","PVP01_0913300","PVP01_0913400","PVP01_0913500","PVP01_0913600","PVP01_0913700","PVP01_0913900","PVP01_0914000","PVP01_0914100","PVP01_0914200","PVP01_0914300","PVP01_0914400","PVP01_0914500","PVP01_0914600","PVP01_0914700","PVP01_0914800","PVP01_0914900","PVP01_0915000","PVP01_0915100","PVP01_0915200","PVP01_0915300","PVP01_0915400","PVP01_0915500","PVP01_0915600","PVP01_0915700","PVP01_0915800","PVP01_0915900","PVP01_0916100","PVP01_0916200","PVP01_0916300","PVP01_0916400","PVP01_0916500","PVP01_0916700","PVP01_0916800","PVP01_0916900","PVP01_0917000","PVP01_0917100","PVP01_0917200","PVP01_0917300","PVP01_0917500","PVP01_0917600","PVP01_0917700","PVP01_0917800","PVP01_0917900","PVP01_0918000","PVP01_0918100","PVP01_0918200","PVP01_0918300","PVP01_0918400","PVP01_0918500","PVP01_0918600","PVP01_0918700","PVP01_0918800","PVP01_0918900","PVP01_0919000","PVP01_0919100","PVP01_0919200","PVP01_0919300","PVP01_0919400","PVP01_0919500","PVP01_0919600","PVP01_0919700","PVP01_0919800","PVP01_0919900","PVP01_0920000","PVP01_0920100","PVP01_0920200","PVP01_0920300","PVP01_0920400","PVP01_0920500","PVP01_0920600","PVP01_0920700","PVP01_0920800","PVP01_0920900","PVP01_0921000","PVP01_0921100","PVP01_0921200","PVP01_0921300","PVP01_0921400","PVP01_0921500","PVP01_0921600","PVP01_0921700","PVP01_0921800","PVP01_0921900","PVP01_0922000","PVP01_0922100","PVP01_0922200","PVP01_0922300","PVP01_0922400","PVP01_0922500","PVP01_0922600","PVP01_0922700","PVP01_0922800","PVP01_0922900","PVP01_0923000","PVP01_0923100","PVP01_0923200","PVP01_0923300","PVP01_0923400","PVP01_0923500","PVP01_0923600","PVP01_0923700","PVP01_0923800","PVP01_0923900","PVP01_0924000","PVP01_0924100","PVP01_0924200","PVP01_0924300","PVP01_0924400","PVP01_0924500","PVP01_0924600","PVP01_0924700","PVP01_0924800","PVP01_0924900","PVP01_0925000","PVP01_0925100","PVP01_0925200","PVP01_0925300","PVP01_0925400","PVP01_0925500","PVP01_0925600","PVP01_0925700","PVP01_0925800","PVP01_0925900","PVP01_0926000","PVP01_0926100","PVP01_0926200","PVP01_0926300","PVP01_0926400","PVP01_0926600","PVP01_0926700","PVP01_0926800","PVP01_0926900","PVP01_0927000","PVP01_0927100","PVP01_0927200","PVP01_0927300","PVP01_0927400","PVP01_0927500","PVP01_0927600","PVP01_0927700","PVP01_0927800","PVP01_0927900","PVP01_0928000","PVP01_0928100","PVP01_0928200","PVP01_0928300","PVP01_0928400","PVP01_0928500","PVP01_0928600","PVP01_0928700","PVP01_0928800","PVP01_0928900","PVP01_0929000","PVP01_0929100","PVP01_0929200","PVP01_0929300","PVP01_0929400","PVP01_0929500","PVP01_0929600","PVP01_0929700","PVP01_0929800","PVP01_0929900","PVP01_0930000","PVP01_0930100","PVP01_0930200","PVP01_0930300","PVP01_0930400","PVP01_0930500","PVP01_0930600","PVP01_0930700","PVP01_0930800","PVP01_0931200","PVP01_0931300","PVP01_0931400","PVP01_0931500","PVP01_0931600","PVP01_0931700","PVP01_0931800","PVP01_0932000","PVP01_0932100","PVP01_0932200","PVP01_0932300","PVP01_0932400","PVP01_0932500","PVP01_0932600","PVP01_0932700","PVP01_0932800","PVP01_0932900","PVP01_0933000","PVP01_0933100","PVP01_0933200","PVP01_0933300","PVP01_0933400","PVP01_0933500","PVP01_0933600","PVP01_0933700","PVP01_0933800","PVP01_0933900","PVP01_0934000","PVP01_0934100","PVP01_0934200","PVP01_0934300","PVP01_0934400","PVP01_0934500","PVP01_0934600","PVP01_0934700","PVP01_0934800","PVP01_0934900","PVP01_0935000","PVP01_0935100","PVP01_0935200","PVP01_0935300","PVP01_0935400","PVP01_0935500","PVP01_0935600","PVP01_0935700","PVP01_0935800","PVP01_0936000","PVP01_0936100","PVP01_0936200","PVP01_0936300","PVP01_0936400","PVP01_0936500","PVP01_0936600","PVP01_0936700","PVP01_0936800","PVP01_0936900","PVP01_0937000","PVP01_0937100","PVP01_0937200","PVP01_0937300","PVP01_0937400","PVP01_0937500","PVP01_0937600","PVP01_0937700","PVP01_0937800","PVP01_0938000","PVP01_0938100","PVP01_0938200","PVP01_0938300","PVP01_0938400","PVP01_0938500","PVP01_0938600","PVP01_0938700","PVP01_0938900","PVP01_0939000","PVP01_0939100","PVP01_0939200","PVP01_0939300","PVP01_0939400","PVP01_0939500","PVP01_0939600","PVP01_0939700","PVP01_0939800","PVP01_0939900","PVP01_0940000","PVP01_0940100","PVP01_0940200","PVP01_0940400","PVP01_0940500","PVP01_0940600","PVP01_0940700","PVP01_0940800","PVP01_0940900","PVP01_0941000","PVP01_0941100","PVP01_0941200","PVP01_0941300","PVP01_0941400","PVP01_0941500","PVP01_0941600","PVP01_0941700","PVP01_0941800","PVP01_0941900","PVP01_0942000","PVP01_0942100","PVP01_0942200","PVP01_0942300","PVP01_0942400","PVP01_0942500","PVP01_0942600","PVP01_0942700","PVP01_0942800","PVP01_0942900","PVP01_0943000","PVP01_0943100","PVP01_0943200","PVP01_0943300","PVP01_0943400","PVP01_0943500","PVP01_0943600","PVP01_0943700","PVP01_0943800","PVP01_0943900","PVP01_0944000","PVP01_0944100","PVP01_0944200","PVP01_0944300","PVP01_0944400","PVP01_0944500","PVP01_0944600","PVP01_0944700","PVP01_0944800","PVP01_0944900","PVP01_0945000","PVP01_0945100","PVP01_0945200","PVP01_0945300","PVP01_0945400","PVP01_0945500","PVP01_0945600","PVP01_0945700","PVP01_0945800","PVP01_0946000","PVP01_0946100","PVP01_0946200","PVP01_0946300","PVP01_0946400","PVP01_0946500","PVP01_0946600","PVP01_0946700","PVP01_0946800","PVP01_0946900","PVP01_0947000","PVP01_0947100","PVP01_0947200","PVP01_0947300","PVP01_0947400","PVP01_0947500","PVP01_0947600","PVP01_0947700","PVP01_0947800","PVP01_0947900","PVP01_0948000","PVP01_0948100","PVP01_0948200","PVP01_0948300","PVP01_0948400","PVP01_0948500","PVP01_0948600","PVP01_0948700","PVP01_0948800","PVP01_0948900","PVP01_0949000","PVP01_0949200","PVP01_0949300","PVP01_0949400","PVP01_0949600","PVP01_0950000","PVP01_0950200","PVP01_1000700","PVP01_1001900","PVP01_1002400","PVP01_1002500","PVP01_1002600","PVP01_1002700","PVP01_1002800","PVP01_1003000","PVP01_1003100","PVP01_1003200","PVP01_1003300","PVP01_1003400","PVP01_1003500","PVP01_1003600","PVP01_1003700","PVP01_1003800","PVP01_1003900","PVP01_1004000","PVP01_1004100","PVP01_1004200","PVP01_1004400","PVP01_1004500","PVP01_1004600","PVP01_1004700","PVP01_1004800","PVP01_1004900","PVP01_1005000","PVP01_1005100","PVP01_1005200","PVP01_1005300","PVP01_1005400","PVP01_1005500","PVP01_1005600","PVP01_1005700","PVP01_1005800","PVP01_1005900","PVP01_1006200","PVP01_1006300","PVP01_1006400","PVP01_1006500","PVP01_1006600","PVP01_1006700","PVP01_1006800","PVP01_1006900","PVP01_1007000","PVP01_1007100","PVP01_1007200","PVP01_1007300","PVP01_1007400","PVP01_1007500","PVP01_1007600","PVP01_1007700","PVP01_1007800","PVP01_1007900","PVP01_1008000","PVP01_1008100","PVP01_1008200","PVP01_1008300","PVP01_1008400","PVP01_1008500","PVP01_1008600","PVP01_1008700","PVP01_1008800","PVP01_1008900","PVP01_1009000","PVP01_1009100","PVP01_1009200","PVP01_1009300","PVP01_1009400","PVP01_1009500","PVP01_1009600","PVP01_1009700","PVP01_1009800","PVP01_1009900","PVP01_1010000","PVP01_1010100","PVP01_1010200","PVP01_1010300","PVP01_1010400","PVP01_1010500","PVP01_1010600","PVP01_1010700","PVP01_1010800","PVP01_1010900","PVP01_1011000","PVP01_1011100","PVP01_1011200","PVP01_1011300","PVP01_1011400","PVP01_1011500","PVP01_1011600","PVP01_1011700","PVP01_1011800","PVP01_1011900","PVP01_1012000","PVP01_1012100","PVP01_1012200","PVP01_1012300","PVP01_1012400","PVP01_1012500","PVP01_1012600","PVP01_1012700","PVP01_1012800","PVP01_1012900","PVP01_1013000","PVP01_1013100","PVP01_1013200","PVP01_1013300","PVP01_1013400","PVP01_1013500","PVP01_1013600","PVP01_1013700","PVP01_1013800","PVP01_1013900","PVP01_1014000","PVP01_1014100","PVP01_1014200","PVP01_1014300","PVP01_1014400","PVP01_1014500","PVP01_1014600","PVP01_1014700","PVP01_1014800","PVP01_1014900","PVP01_1015000","PVP01_1015100","PVP01_1015300","PVP01_1015400","PVP01_1015500","PVP01_1015600","PVP01_1015700","PVP01_1015800","PVP01_1015900","PVP01_1016000","PVP01_1016100","PVP01_1016200","PVP01_1016300","PVP01_1016400","PVP01_1016500","PVP01_1016600","PVP01_1016700","PVP01_1016800","PVP01_1016900","PVP01_1017000","PVP01_1017100","PVP01_1017200","PVP01_1017300","PVP01_1017400","PVP01_1017500","PVP01_1017600","PVP01_1017700","PVP01_1017800","PVP01_1017900","PVP01_1018000","PVP01_1018100","PVP01_1018200","PVP01_1018300","PVP01_1018400","PVP01_1018500","PVP01_1018600","PVP01_1018700","PVP01_1018800","PVP01_1018900","PVP01_1019000","PVP01_1019100","PVP01_1019200","PVP01_1019300","PVP01_1019400","PVP01_1019600","PVP01_1019700","PVP01_1019800","PVP01_1019900","PVP01_1020000","PVP01_1020100","PVP01_1020200","PVP01_1020300","PVP01_1020400","PVP01_1020500","PVP01_1020600","PVP01_1020700","PVP01_1020800","PVP01_1020900","PVP01_1021000","PVP01_1021100","PVP01_1021200","PVP01_1021300","PVP01_1021400","PVP01_1021500","PVP01_1021600","PVP01_1021700","PVP01_1021800","PVP01_1021900","PVP01_1022000","PVP01_1022100","PVP01_1022200","PVP01_1022300","PVP01_1022400","PVP01_1022500","PVP01_1022600","PVP01_1022700","PVP01_1022800","PVP01_1022900","PVP01_1023000","PVP01_1023200","PVP01_1023300","PVP01_1023500","PVP01_1023600","PVP01_1023700","PVP01_1023800","PVP01_1023900","PVP01_1024000","PVP01_1024100","PVP01_1024200","PVP01_1024300","PVP01_1024400","PVP01_1024500","PVP01_1024600","PVP01_1024700","PVP01_1024800","PVP01_1024900","PVP01_1025000","PVP01_1025100","PVP01_1025200","PVP01_1025300","PVP01_1025400","PVP01_1025600","PVP01_1025700","PVP01_1025800","PVP01_1025900","PVP01_1026000","PVP01_1026100","PVP01_1026200","PVP01_1026300","PVP01_1026400","PVP01_1026500","PVP01_1026600","PVP01_1026700","PVP01_1026800","PVP01_1026900","PVP01_1027000","PVP01_1027100","PVP01_1027200","PVP01_1027300","PVP01_1027400","PVP01_1027500","PVP01_1027600","PVP01_1027700","PVP01_1027800","PVP01_1027900","PVP01_1028000","PVP01_1028100","PVP01_1028200","PVP01_1028300","PVP01_1028400","PVP01_1028600","PVP01_1028700","PVP01_1028800","PVP01_1028900","PVP01_1029000","PVP01_1029100","PVP01_1029200","PVP01_1029300","PVP01_1029400","PVP01_1029500","PVP01_1029600","PVP01_1029700","PVP01_1029800","PVP01_1029900","PVP01_1030000","PVP01_1030100","PVP01_1030200","PVP01_1030300","PVP01_1030400","PVP01_1030500","PVP01_1030600","PVP01_1030700","PVP01_1030800","PVP01_1030900","PVP01_1031000","PVP01_1031100","PVP01_1031200","PVP01_1031300","PVP01_1031400","PVP01_1031500","PVP01_1031600","PVP01_1031700","PVP01_1031800","PVP01_1031900","PVP01_1032000","PVP01_1032100","PVP01_1032200","PVP01_1032300","PVP01_1032400","PVP01_1032500","PVP01_1032600","PVP01_1032700","PVP01_1032800","PVP01_1032900","PVP01_1033000","PVP01_1033100","PVP01_1033200","PVP01_1033300","PVP01_1033500","PVP01_1033600","PVP01_1033700","PVP01_1033800","PVP01_1033900","PVP01_1034000","PVP01_1034100","PVP01_1034200","PVP01_1034300","PVP01_1034400","PVP01_1034500","PVP01_1035200","PVP01_1100400","PVP01_1100500","PVP01_1101000","PVP01_1101200","PVP01_1101300","PVP01_1101400","PVP01_1101500","PVP01_1101600","PVP01_1101700","PVP01_1102300","PVP01_1102400","PVP01_1102500","PVP01_1102600","PVP01_1102700","PVP01_1102800","PVP01_1102900","PVP01_1103000","PVP01_1103100","PVP01_1103200","PVP01_1103300","PVP01_1103400","PVP01_1103500","PVP01_1103600","PVP01_1103700","PVP01_1103800","PVP01_1103900","PVP01_1104000","PVP01_1104100","PVP01_1104200","PVP01_1104300","PVP01_1104400","PVP01_1104500","PVP01_1104600","PVP01_1104700","PVP01_1104800","PVP01_1104900","PVP01_1105000","PVP01_1105100","PVP01_1105200","PVP01_1105300","PVP01_1105400","PVP01_1105500","PVP01_1105600","PVP01_1105700","PVP01_1105800","PVP01_1105900","PVP01_1106000","PVP01_1106100","PVP01_1106200","PVP01_1106300","PVP01_1106400","PVP01_1106500","PVP01_1106600","PVP01_1106700","PVP01_1106800","PVP01_1106900","PVP01_1107000","PVP01_1107100","PVP01_1107200","PVP01_1107300","PVP01_1107400","PVP01_1107500","PVP01_1107600","PVP01_1107700","PVP01_1107800","PVP01_1107900","PVP01_1108000","PVP01_1108100","PVP01_1108200","PVP01_1108300","PVP01_1108400","PVP01_1108500","PVP01_1108600","PVP01_1108700","PVP01_1108800","PVP01_1108900","PVP01_1109000","PVP01_1109100","PVP01_1109200","PVP01_1109300","PVP01_1109400","PVP01_1109500","PVP01_1109600","PVP01_1109700","PVP01_1109800","PVP01_1109900","PVP01_1110000","PVP01_1110100","PVP01_1110300","PVP01_1110400","PVP01_1110500","PVP01_1110600","PVP01_1110700","PVP01_1110800","PVP01_1110900","PVP01_1111000","PVP01_1111100","PVP01_1111200","PVP01_1111300","PVP01_1111400","PVP01_1111500","PVP01_1111600","PVP01_1111700","PVP01_1111800","PVP01_1111900","PVP01_1112000","PVP01_1112100","PVP01_1112200","PVP01_1112300","PVP01_1112400","PVP01_1112500","PVP01_1112600","PVP01_1112700","PVP01_1112800","PVP01_1112900","PVP01_1113000","PVP01_1113100","PVP01_1113200","PVP01_1113300","PVP01_1113400","PVP01_1113500","PVP01_1113600","PVP01_1113700","PVP01_1113800","PVP01_1113900","PVP01_1114000","PVP01_1114100","PVP01_1114200","PVP01_1114300","PVP01_1114400","PVP01_1114500","PVP01_1114600","PVP01_1114700","PVP01_1114800","PVP01_1114900","PVP01_1115000","PVP01_1115100","PVP01_1115200","PVP01_1115300","PVP01_1115400","PVP01_1115500","PVP01_1115600","PVP01_1115700","PVP01_1115800","PVP01_1115900","PVP01_1116000","PVP01_1116100","PVP01_1116200","PVP01_1116300","PVP01_1116400","PVP01_1116600","PVP01_1116700","PVP01_1116800","PVP01_1116900","PVP01_1117000","PVP01_1117100","PVP01_1117200","PVP01_1117300","PVP01_1117400","PVP01_1117500","PVP01_1117600","PVP01_1117700","PVP01_1117800","PVP01_1117900","PVP01_1118000","PVP01_1118100","PVP01_1118200","PVP01_1118300","PVP01_1118400","PVP01_1118500","PVP01_1118600","PVP01_1118700","PVP01_1118800","PVP01_1118900","PVP01_1119000","PVP01_1119200","PVP01_1119300","PVP01_1119400","PVP01_1119500","PVP01_1119600","PVP01_1119700","PVP01_1119800","PVP01_1119900","PVP01_1120000","PVP01_1120100","PVP01_1120200","PVP01_1120300","PVP01_1120400","PVP01_1120500","PVP01_1120600","PVP01_1120700","PVP01_1120800","PVP01_1120900","PVP01_1121000","PVP01_1121100","PVP01_1121200","PVP01_1121300","PVP01_1121400","PVP01_1121500","PVP01_1121600","PVP01_1121700","PVP01_1121800","PVP01_1121900","PVP01_1122000","PVP01_1122100","PVP01_1122200","PVP01_1122300","PVP01_1122400","PVP01_1122500","PVP01_1122600","PVP01_1122700","PVP01_1122800","PVP01_1122900","PVP01_1123000","PVP01_1123100","PVP01_1123200","PVP01_1123300","PVP01_1123400","PVP01_1123500","PVP01_1123600","PVP01_1123700","PVP01_1123800","PVP01_1123900","PVP01_1124000","PVP01_1124100","PVP01_1124200","PVP01_1124300","PVP01_1124400","PVP01_1124500","PVP01_1124600","PVP01_1124700","PVP01_1124800","PVP01_1124900","PVP01_1125000","PVP01_1125100","PVP01_1125200","PVP01_1125300","PVP01_1125400","PVP01_1125500","PVP01_1125600","PVP01_1125700","PVP01_1125800","PVP01_1125900","PVP01_1126000","PVP01_1126100","PVP01_1126200","PVP01_1126300","PVP01_1126400","PVP01_1126500","PVP01_1126600","PVP01_1126700","PVP01_1126800","PVP01_1126900","PVP01_1127000","PVP01_1127100","PVP01_1127200","PVP01_1127300","PVP01_1127400","PVP01_1127500","PVP01_1127600","PVP01_1127700","PVP01_1127800","PVP01_1128000","PVP01_1128100","PVP01_1128200","PVP01_1128300","PVP01_1128400","PVP01_1128500","PVP01_1128800","PVP01_1128900","PVP01_1129000","PVP01_1129100","PVP01_1129200","PVP01_1129300","PVP01_1129400","PVP01_1129500","PVP01_1129600","PVP01_1129700","PVP01_1129800","PVP01_1129900","PVP01_1130000","PVP01_1130100","PVP01_1130200","PVP01_1130300","PVP01_1130400","PVP01_1130500","PVP01_1130600","PVP01_1130700","PVP01_1130800","PVP01_1130900","PVP01_1131000","PVP01_1131100","PVP01_1131200","PVP01_1131300","PVP01_1131400","PVP01_1131500","PVP01_1131600","PVP01_1131700","PVP01_1131800","PVP01_1131900","PVP01_1132000","PVP01_1132100","PVP01_1132200","PVP01_1132300","PVP01_1132400","PVP01_1132500","PVP01_1132600","PVP01_1132700","PVP01_1132800","PVP01_1132900","PVP01_1133000","PVP01_1133100","PVP01_1133200","PVP01_1133300","PVP01_1133400","PVP01_1133500","PVP01_1133600","PVP01_1133700","PVP01_1133800","PVP01_1133900","PVP01_1134000","PVP01_1134100","PVP01_1134200","PVP01_1134300","PVP01_1134400","PVP01_1134500","PVP01_1134600","PVP01_1134700","PVP01_1134800","PVP01_1134900","PVP01_1135000","PVP01_1135100","PVP01_1135300","PVP01_1135400","PVP01_1135500","PVP01_1135600","PVP01_1135700","PVP01_1135800","PVP01_1135900","PVP01_1136000","PVP01_1136100","PVP01_1136200","PVP01_1136300","PVP01_1136400","PVP01_1136500","PVP01_1136600","PVP01_1136700","PVP01_1136800","PVP01_1136900","PVP01_1137000","PVP01_1137100","PVP01_1137200","PVP01_1137300","PVP01_1137400","PVP01_1137500","PVP01_1137600","PVP01_1137700","PVP01_1137800","PVP01_1137900","PVP01_1138000","PVP01_1138100","PVP01_1138200","PVP01_1138300","PVP01_1138400","PVP01_1138500","PVP01_1138600","PVP01_1138700","PVP01_1138800","PVP01_1138900","PVP01_1139000","PVP01_1139100","PVP01_1139300","PVP01_1139400","PVP01_1139500","PVP01_1139600","PVP01_1139700","PVP01_1139800","PVP01_1139900","PVP01_1140000","PVP01_1140100","PVP01_1140200","PVP01_1140300","PVP01_1140400","PVP01_1140500","PVP01_1140600","PVP01_1140700","PVP01_1140800","PVP01_1140900","PVP01_1141000","PVP01_1141100","PVP01_1141200","PVP01_1141300","PVP01_1141400","PVP01_1141500","PVP01_1141600","PVP01_1141700","PVP01_1141800","PVP01_1141900","PVP01_1142000","PVP01_1142100","PVP01_1142200","PVP01_1142300","PVP01_1142400","PVP01_1142500","PVP01_1142600","PVP01_1142700","PVP01_1142800","PVP01_1142900","PVP01_1143000","PVP01_1143100","PVP01_1143200","PVP01_1143300","PVP01_1143400","PVP01_1143500","PVP01_1143600","PVP01_1143700","PVP01_1143800","PVP01_1143900","PVP01_1144000","PVP01_1144100","PVP01_1144200","PVP01_1144300","PVP01_1144400","PVP01_1144500","PVP01_1144600","PVP01_1144700","PVP01_1144900","PVP01_1145000","PVP01_1145100","PVP01_1145200","PVP01_1145300","PVP01_1145400","PVP01_1145500","PVP01_1145600","PVP01_1145700","PVP01_1145800","PVP01_1145900","PVP01_1146000","PVP01_1146100","PVP01_1146200","PVP01_1146300","PVP01_1146400","PVP01_1146500","PVP01_1146600","PVP01_1146700","PVP01_1146800","PVP01_1146900","PVP01_1147000","PVP01_1147100","PVP01_1147200","PVP01_1147300","PVP01_1147400","PVP01_1147500","PVP01_1147600","PVP01_1147700","PVP01_1147800","PVP01_1147900","PVP01_1148200","PVP01_1148700","PVP01_1148800","PVP01_1149000","PVP01_1149100","PVP01_1149400","PVP01_1149500","PVP01_1200200","PVP01_1200800","PVP01_1201100","PVP01_1201300","PVP01_1201400","PVP01_1201500","PVP01_1201600","PVP01_1201700","PVP01_1201800","PVP01_1201900","PVP01_1202000","PVP01_1202100","PVP01_1202200","PVP01_1202300","PVP01_1202400","PVP01_1202500","PVP01_1202600","PVP01_1202700","PVP01_1202800","PVP01_1202900","PVP01_1203000","PVP01_1203100","PVP01_1203200","PVP01_1203300","PVP01_1203500","PVP01_1203600","PVP01_1203700","PVP01_1203800","PVP01_1203900","PVP01_1204000","PVP01_1204100","PVP01_1204200","PVP01_1204400","PVP01_1204500","PVP01_1204600","PVP01_1204700","PVP01_1204800","PVP01_1204900","PVP01_1205000","PVP01_1205100","PVP01_1205200","PVP01_1205300","PVP01_1205400","PVP01_1205500","PVP01_1205600","PVP01_1205700","PVP01_1205800","PVP01_1205900","PVP01_1206000","PVP01_1206100","PVP01_1206300","PVP01_1206400","PVP01_1206500","PVP01_1206600","PVP01_1206700","PVP01_1206800","PVP01_1206900","PVP01_1207000","PVP01_1207100","PVP01_1207200","PVP01_1207300","PVP01_1207400","PVP01_1207500","PVP01_1207600","PVP01_1207700","PVP01_1207800","PVP01_1207900","PVP01_1208000","PVP01_1208100","PVP01_1208200","PVP01_1208300","PVP01_1208400","PVP01_1208500","PVP01_1208600","PVP01_1208700","PVP01_1208800","PVP01_1208900","PVP01_1209000","PVP01_1209100","PVP01_1209300","PVP01_1209400","PVP01_1209500","PVP01_1209600","PVP01_1209700","PVP01_1209800","PVP01_1209900","PVP01_1210000","PVP01_1210100","PVP01_1210200","PVP01_1210300","PVP01_1210400","PVP01_1210500","PVP01_1210600","PVP01_1210700","PVP01_1210800","PVP01_1210900","PVP01_1211000","PVP01_1211100","PVP01_1211200","PVP01_1211300","PVP01_1211400","PVP01_1211500","PVP01_1211600","PVP01_1211700","PVP01_1211800","PVP01_1211900","PVP01_1212000","PVP01_1212100","PVP01_1212200","PVP01_1212300","PVP01_1212400","PVP01_1212500","PVP01_1212600","PVP01_1212700","PVP01_1212800","PVP01_1212900","PVP01_1213000","PVP01_1213100","PVP01_1213200","PVP01_1213300","PVP01_1213400","PVP01_1213600","PVP01_1213900","PVP01_1214000","PVP01_1214100","PVP01_1214200","PVP01_1214300","PVP01_1214400","PVP01_1214500","PVP01_1214600","PVP01_1214700","PVP01_1214800","PVP01_1214900","PVP01_1215000","PVP01_1215200","PVP01_1215300","PVP01_1215400","PVP01_1215500","PVP01_1215800","PVP01_1215900","PVP01_1216000","PVP01_1216100","PVP01_1216200","PVP01_1216300","PVP01_1216400","PVP01_1216500","PVP01_1216600","PVP01_1216700","PVP01_1216800","PVP01_1216900","PVP01_1217000","PVP01_1217200","PVP01_1217300","PVP01_1217400","PVP01_1217500","PVP01_1217600","PVP01_1217700","PVP01_1217800","PVP01_1217900","PVP01_1218000","PVP01_1218100","PVP01_1218200","PVP01_1218300","PVP01_1218400","PVP01_1218500","PVP01_1218600","PVP01_1218700","PVP01_1218800","PVP01_1218900","PVP01_1219000","PVP01_1219100","PVP01_1219200","PVP01_1219300","PVP01_1219400","PVP01_1219500","PVP01_1219700","PVP01_1219800","PVP01_1219900","PVP01_1220100","PVP01_1220200","PVP01_1220400","PVP01_1220500","PVP01_1220600","PVP01_1220700","PVP01_1220800","PVP01_1220900","PVP01_1221000","PVP01_1221100","PVP01_1221200","PVP01_1221300","PVP01_1221400","PVP01_1221500","PVP01_1221600","PVP01_1221700","PVP01_1221800","PVP01_1221900","PVP01_1222000","PVP01_1222100","PVP01_1222200","PVP01_1222300","PVP01_1222400","PVP01_1222500","PVP01_1222600","PVP01_1222700","PVP01_1222800","PVP01_1222900","PVP01_1223000","PVP01_1223100","PVP01_1223200","PVP01_1223300","PVP01_1223400","PVP01_1223500","PVP01_1223600","PVP01_1223700","PVP01_1223800","PVP01_1223900","PVP01_1224000","PVP01_1224100","PVP01_1224200","PVP01_1224300","PVP01_1224400","PVP01_1224500","PVP01_1224600","PVP01_1224700","PVP01_1224800","PVP01_1224900","PVP01_1225000","PVP01_1225100","PVP01_1225200","PVP01_1225300","PVP01_1225400","PVP01_1225500","PVP01_1225600","PVP01_1225700","PVP01_1225800","PVP01_1225900","PVP01_1226000","PVP01_1226100","PVP01_1226200","PVP01_1226300","PVP01_1226400","PVP01_1226600","PVP01_1226700","PVP01_1226800","PVP01_1226900","PVP01_1227000","PVP01_1227100","PVP01_1227200","PVP01_1227300","PVP01_1227400","PVP01_1227500","PVP01_1227600","PVP01_1227700","PVP01_1227800","PVP01_1227900","PVP01_1228000","PVP01_1228100","PVP01_1228200","PVP01_1228300","PVP01_1228400","PVP01_1228500","PVP01_1228600","PVP01_1228700","PVP01_1228800","PVP01_1228900","PVP01_1229000","PVP01_1229100","PVP01_1229200","PVP01_1229300","PVP01_1229400","PVP01_1229500","PVP01_1229600","PVP01_1229700","PVP01_1229800","PVP01_1229900","PVP01_1230000","PVP01_1230100","PVP01_1230200","PVP01_1230300","PVP01_1230400","PVP01_1230500","PVP01_1230600","PVP01_1230700","PVP01_1230800","PVP01_1231000","PVP01_1231100","PVP01_1231200","PVP01_1231300","PVP01_1231400","PVP01_1231500","PVP01_1231600","PVP01_1231700","PVP01_1231800","PVP01_1231900","PVP01_1232000","PVP01_1232100","PVP01_1232200","PVP01_1232300","PVP01_1232400","PVP01_1232500","PVP01_1232600","PVP01_1232700","PVP01_1232800","PVP01_1232900","PVP01_1233000","PVP01_1233100","PVP01_1233200","PVP01_1233300","PVP01_1233400","PVP01_1233600","PVP01_1233700","PVP01_1233800","PVP01_1233900","PVP01_1234000","PVP01_1234100","PVP01_1234200","PVP01_1234300","PVP01_1234400","PVP01_1234500","PVP01_1234600","PVP01_1234700","PVP01_1234800","PVP01_1234900","PVP01_1235000","PVP01_1235100","PVP01_1235200","PVP01_1235300","PVP01_1235500","PVP01_1235600","PVP01_1235700","PVP01_1235800","PVP01_1235900","PVP01_1236000","PVP01_1236100","PVP01_1236200","PVP01_1236300","PVP01_1236400","PVP01_1236500","PVP01_1236600","PVP01_1236700","PVP01_1236800","PVP01_1236900","PVP01_1237000","PVP01_1237100","PVP01_1237200","PVP01_1237300","PVP01_1237400","PVP01_1237500","PVP01_1237600","PVP01_1237700","PVP01_1237800","PVP01_1237900","PVP01_1238000","PVP01_1238100","PVP01_1238200","PVP01_1238300","PVP01_1238400","PVP01_1238500","PVP01_1238600","PVP01_1238700","PVP01_1238900","PVP01_1239000","PVP01_1239100","PVP01_1239200","PVP01_1239300","PVP01_1239400","PVP01_1239500","PVP01_1239600","PVP01_1239700","PVP01_1239800","PVP01_1239900","PVP01_1240000","PVP01_1240100","PVP01_1240200","PVP01_1240300","PVP01_1240400","PVP01_1240500","PVP01_1240600","PVP01_1240700","PVP01_1240800","PVP01_1240900","PVP01_1241000","PVP01_1241100","PVP01_1241200","PVP01_1241300","PVP01_1241400","PVP01_1241500","PVP01_1241600","PVP01_1241700","PVP01_1241800","PVP01_1241900","PVP01_1242000","PVP01_1242100","PVP01_1242200","PVP01_1242300","PVP01_1242400","PVP01_1242500","PVP01_1242600","PVP01_1242700","PVP01_1242800","PVP01_1243000","PVP01_1243100","PVP01_1243200","PVP01_1243300","PVP01_1243400","PVP01_1243500","PVP01_1243600","PVP01_1243700","PVP01_1243800","PVP01_1243900","PVP01_1244000","PVP01_1244100","PVP01_1244200","PVP01_1244300","PVP01_1244400","PVP01_1244500","PVP01_1244600","PVP01_1244700","PVP01_1244800","PVP01_1244900","PVP01_1245000","PVP01_1245100","PVP01_1245200","PVP01_1245300","PVP01_1245400","PVP01_1245500","PVP01_1245600","PVP01_1245700","PVP01_1245800","PVP01_1245900","PVP01_1246000","PVP01_1246100","PVP01_1246200","PVP01_1246300","PVP01_1246400","PVP01_1246500","PVP01_1246600","PVP01_1246700","PVP01_1246800","PVP01_1246900","PVP01_1247000","PVP01_1247100","PVP01_1247200","PVP01_1247300","PVP01_1247400","PVP01_1247500","PVP01_1247600","PVP01_1247700","PVP01_1247800","PVP01_1247900","PVP01_1248000","PVP01_1248100","PVP01_1248200","PVP01_1248300","PVP01_1248400","PVP01_1248500","PVP01_1248600","PVP01_1248700","PVP01_1248800","PVP01_1248900","PVP01_1249100","PVP01_1249200","PVP01_1249300","PVP01_1249400","PVP01_1249500","PVP01_1249600","PVP01_1249700","PVP01_1249800","PVP01_1249900","PVP01_1250000","PVP01_1250200","PVP01_1250300","PVP01_1250400","PVP01_1250500","PVP01_1250600","PVP01_1250700","PVP01_1250800","PVP01_1250900","PVP01_1251000","PVP01_1251100","PVP01_1251200","PVP01_1251300","PVP01_1251400","PVP01_1251500","PVP01_1251700","PVP01_1251800","PVP01_1251900","PVP01_1252000","PVP01_1252100","PVP01_1252200","PVP01_1252300","PVP01_1252400","PVP01_1252500","PVP01_1252600","PVP01_1252700","PVP01_1252800","PVP01_1252900","PVP01_1253000","PVP01_1253100","PVP01_1253200","PVP01_1253300","PVP01_1253400","PVP01_1253500","PVP01_1253600","PVP01_1253700","PVP01_1253800","PVP01_1253900","PVP01_1254000","PVP01_1254100","PVP01_1254200","PVP01_1254300","PVP01_1254400","PVP01_1254500","PVP01_1254600","PVP01_1254700","PVP01_1254800","PVP01_1254900","PVP01_1255100","PVP01_1255200","PVP01_1255300","PVP01_1255400","PVP01_1255500","PVP01_1255600","PVP01_1255700","PVP01_1255800","PVP01_1255900","PVP01_1256000","PVP01_1256100","PVP01_1256200","PVP01_1256300","PVP01_1256400","PVP01_1256500","PVP01_1256600","PVP01_1256700","PVP01_1256800","PVP01_1256900","PVP01_1257000","PVP01_1257100","PVP01_1257200","PVP01_1257300","PVP01_1257400","PVP01_1257500","PVP01_1257600","PVP01_1257700","PVP01_1257800","PVP01_1257900","PVP01_1258000","PVP01_1258100","PVP01_1258200","PVP01_1258300","PVP01_1258400","PVP01_1258500","PVP01_1258600","PVP01_1258700","PVP01_1258800","PVP01_1258900","PVP01_1259000","PVP01_1259100","PVP01_1259200","PVP01_1259300","PVP01_1259400","PVP01_1259500","PVP01_1259600","PVP01_1259700","PVP01_1259800","PVP01_1259900","PVP01_1260000","PVP01_1260100","PVP01_1260200","PVP01_1260300","PVP01_1260400","PVP01_1260500","PVP01_1260600","PVP01_1260700","PVP01_1260800","PVP01_1260900","PVP01_1261100","PVP01_1261200","PVP01_1261300","PVP01_1261400","PVP01_1261500","PVP01_1261600","PVP01_1261700","PVP01_1261800","PVP01_1261900","PVP01_1262000","PVP01_1262100","PVP01_1262200","PVP01_1262300","PVP01_1262400","PVP01_1262500","PVP01_1262600","PVP01_1262700","PVP01_1262800","PVP01_1262900","PVP01_1263000","PVP01_1263100","PVP01_1263200","PVP01_1263300","PVP01_1263400","PVP01_1263500","PVP01_1263600","PVP01_1263700","PVP01_1263800","PVP01_1263900","PVP01_1264000","PVP01_1264100","PVP01_1264200","PVP01_1264300","PVP01_1264400","PVP01_1264500","PVP01_1264600","PVP01_1264700","PVP01_1264800","PVP01_1264900","PVP01_1265000","PVP01_1265100","PVP01_1265200","PVP01_1265300","PVP01_1265400","PVP01_1265500","PVP01_1265600","PVP01_1265700","PVP01_1265800","PVP01_1265900","PVP01_1266000","PVP01_1266100","PVP01_1266200","PVP01_1266300","PVP01_1266400","PVP01_1266500","PVP01_1266600","PVP01_1266700","PVP01_1266800","PVP01_1266900","PVP01_1267100","PVP01_1267200","PVP01_1267300","PVP01_1267400","PVP01_1267500","PVP01_1267600","PVP01_1267700","PVP01_1267800","PVP01_1267900","PVP01_1268000","PVP01_1268100","PVP01_1268200","PVP01_1268300","PVP01_1268400","PVP01_1268500","PVP01_1268600","PVP01_1268900","PVP01_1269000","PVP01_1269100","PVP01_1269200","PVP01_1269300","PVP01_1269400","PVP01_1269500","PVP01_1269600","PVP01_1269700","PVP01_1269800","PVP01_1269900","PVP01_1270000","PVP01_1270100","PVP01_1270200","PVP01_1270300","PVP01_1270400","PVP01_1270500","PVP01_1270600","PVP01_1270700","PVP01_1270800","PVP01_1270900","PVP01_1271000","PVP01_1271100","PVP01_1271200","PVP01_1271300","PVP01_1271400","PVP01_1271500","PVP01_1271600","PVP01_1271900","PVP01_1272500","PVP01_1272600","PVP01_1273000","PVP01_1274300","PVP01_1300400","PVP01_1300500","PVP01_1300600","PVP01_1300700","PVP01_1300800","PVP01_1300900","PVP01_1301000","PVP01_1301100","PVP01_1301200","PVP01_1301300","PVP01_1301500","PVP01_1301600","PVP01_1301700","PVP01_1301800","PVP01_1301900","PVP01_1302000","PVP01_1302100","PVP01_1302200","PVP01_1302300","PVP01_1302400","PVP01_1302500","PVP01_1302600","PVP01_1302700","PVP01_1302800","PVP01_1302900","PVP01_1303000","PVP01_1303100","PVP01_1303200","PVP01_1303300","PVP01_1303400","PVP01_1303500","PVP01_1303600","PVP01_1303700","PVP01_1303800","PVP01_1303900","PVP01_1304000","PVP01_1304100","PVP01_1304200","PVP01_1304300","PVP01_1304400","PVP01_1304500","PVP01_1304600","PVP01_1304700","PVP01_1304800","PVP01_1304900","PVP01_1305000","PVP01_1305100","PVP01_1305300","PVP01_1305400","PVP01_1305500","PVP01_1305600","PVP01_1305700","PVP01_1305800","PVP01_1305900","PVP01_1306000","PVP01_1306100","PVP01_1306200","PVP01_1306300","PVP01_1306400","PVP01_1306500","PVP01_1306600","PVP01_1306700","PVP01_1306800","PVP01_1306900","PVP01_1307000","PVP01_1307100","PVP01_1307200","PVP01_1307300","PVP01_1307400","PVP01_1307500","PVP01_1307600","PVP01_1307700","PVP01_1307800","PVP01_1307900","PVP01_1308000","PVP01_1308100","PVP01_1308200","PVP01_1308300","PVP01_1308400","PVP01_1308500","PVP01_1308600","PVP01_1308700","PVP01_1308800","PVP01_1308900","PVP01_1309000","PVP01_1309100","PVP01_1309200","PVP01_1309300","PVP01_1309400","PVP01_1309500","PVP01_1309600","PVP01_1309700","PVP01_1309800","PVP01_1309900","PVP01_1310000","PVP01_1310100","PVP01_1310200","PVP01_1310300","PVP01_1310400","PVP01_1310500","PVP01_1310600","PVP01_1310700","PVP01_1310800","PVP01_1310900","PVP01_1311000","PVP01_1311100","PVP01_1311200","PVP01_1311300","PVP01_1311400","PVP01_1311500","PVP01_1311600","PVP01_1311700","PVP01_1311800","PVP01_1311900","PVP01_1312000","PVP01_1312100","PVP01_1312200","PVP01_1312400","PVP01_1312500","PVP01_1312600","PVP01_1312700","PVP01_1312800","PVP01_1312900","PVP01_1313000","PVP01_1313100","PVP01_1313200","PVP01_1313300","PVP01_1313400","PVP01_1313500","PVP01_1313600","PVP01_1313700","PVP01_1313800","PVP01_1313900","PVP01_1314000","PVP01_1314100","PVP01_1314200","PVP01_1314300","PVP01_1314400","PVP01_1314500","PVP01_1314600","PVP01_1314700","PVP01_1314800","PVP01_1314900","PVP01_1315000","PVP01_1315200","PVP01_1315300","PVP01_1315400","PVP01_1315500","PVP01_1315600","PVP01_1315800","PVP01_1315900","PVP01_1316000","PVP01_1316100","PVP01_1316200","PVP01_1316300","PVP01_1316400","PVP01_1316500","PVP01_1316600","PVP01_1316700","PVP01_1316800","PVP01_1316900","PVP01_1317000","PVP01_1317100","PVP01_1317200","PVP01_1317300","PVP01_1317400","PVP01_1317500","PVP01_1317600","PVP01_1317700","PVP01_1317800","PVP01_1317900","PVP01_1318000","PVP01_1318100","PVP01_1318200","PVP01_1318300","PVP01_1318400","PVP01_1318500","PVP01_1318600","PVP01_1318700","PVP01_1318800","PVP01_1318900","PVP01_1319000","PVP01_1319100","PVP01_1319200","PVP01_1319300","PVP01_1319400","PVP01_1319500","PVP01_1319600","PVP01_1319700","PVP01_1319800","PVP01_1319900","PVP01_1320000","PVP01_1320100","PVP01_1320200","PVP01_1320300","PVP01_1320400","PVP01_1320500","PVP01_1320600","PVP01_1320700","PVP01_1320800","PVP01_1320900","PVP01_1321000","PVP01_1321200","PVP01_1321300","PVP01_1321400","PVP01_1321500","PVP01_1321600","PVP01_1321700","PVP01_1321800","PVP01_1321900","PVP01_1322000","PVP01_1322100","PVP01_1322200","PVP01_1322300","PVP01_1322400","PVP01_1322500","PVP01_1322600","PVP01_1322700","PVP01_1322800","PVP01_1322900","PVP01_1323000","PVP01_1323100","PVP01_1323200","PVP01_1323300","PVP01_1323400","PVP01_1323500","PVP01_1323600","PVP01_1323700","PVP01_1323800","PVP01_1323900","PVP01_1324000","PVP01_1324100","PVP01_1324700","PVP01_1324800","PVP01_1325200","PVP01_1325300","PVP01_1325400","PVP01_1325500","PVP01_1325600","PVP01_1325700","PVP01_1325800","PVP01_1325900","PVP01_1326000","PVP01_1326100","PVP01_1326200","PVP01_1326300","PVP01_1326400","PVP01_1326500","PVP01_1326600","PVP01_1326700","PVP01_1326800","PVP01_1326900","PVP01_1327000","PVP01_1327100","PVP01_1327200","PVP01_1327300","PVP01_1327400","PVP01_1327500","PVP01_1327600","PVP01_1327700","PVP01_1327800","PVP01_1327900","PVP01_1328000","PVP01_1328100","PVP01_1328200","PVP01_1328300","PVP01_1328400","PVP01_1328500","PVP01_1328600","PVP01_1328700","PVP01_1328800","PVP01_1328900","PVP01_1329000","PVP01_1329100","PVP01_1329200","PVP01_1329300","PVP01_1329400","PVP01_1329500","PVP01_1329600","PVP01_1329700","PVP01_1329800","PVP01_1329900","PVP01_1330000","PVP01_1330600","PVP01_1330700","PVP01_1330800","PVP01_1330900","PVP01_1331000","PVP01_1331100","PVP01_1331200","PVP01_1331300","PVP01_1331400","PVP01_1331500","PVP01_1331600","PVP01_1331700","PVP01_1331800","PVP01_1331900","PVP01_1332000","PVP01_1332100","PVP01_1332200","PVP01_1332300","PVP01_1332400","PVP01_1332500","PVP01_1332600","PVP01_1332700","PVP01_1332800","PVP01_1332900","PVP01_1333000","PVP01_1333100","PVP01_1333200","PVP01_1333300","PVP01_1333400","PVP01_1333500","PVP01_1333600","PVP01_1333700","PVP01_1333800","PVP01_1333900","PVP01_1334000","PVP01_1334100","PVP01_1334200","PVP01_1334300","PVP01_1334400","PVP01_1334500","PVP01_1334600","PVP01_1334700","PVP01_1334800","PVP01_1334900","PVP01_1335000","PVP01_1335100","PVP01_1335200","PVP01_1335300","PVP01_1335400","PVP01_1335500","PVP01_1335600","PVP01_1335700","PVP01_1335800","PVP01_1335900","PVP01_1336000","PVP01_1336100","PVP01_1336200","PVP01_1336300","PVP01_1336400","PVP01_1336500","PVP01_1336600","PVP01_1336700","PVP01_1336800","PVP01_1336900","PVP01_1337000","PVP01_1337100","PVP01_1337200","PVP01_1337300","PVP01_1337400","PVP01_1337500","PVP01_1337600","PVP01_1337700","PVP01_1337800","PVP01_1337900","PVP01_1338000","PVP01_1338100","PVP01_1338200","PVP01_1338300","PVP01_1338400","PVP01_1338600","PVP01_1338700","PVP01_1338800","PVP01_1338900","PVP01_1339000","PVP01_1339100","PVP01_1339200","PVP01_1339300","PVP01_1339400","PVP01_1339500","PVP01_1339600","PVP01_1339700","PVP01_1339800","PVP01_1339900","PVP01_1340000","PVP01_1340100","PVP01_1340200","PVP01_1340300","PVP01_1340400","PVP01_1340500","PVP01_1340600","PVP01_1340700","PVP01_1340800","PVP01_1340900","PVP01_1341000","PVP01_1341100","PVP01_1341200","PVP01_1341300","PVP01_1341400","PVP01_1341500","PVP01_1341600","PVP01_1341700","PVP01_1341800","PVP01_1341900","PVP01_1342000","PVP01_1342100","PVP01_1342200","PVP01_1342300","PVP01_1342400","PVP01_1342500","PVP01_1342600","PVP01_1342700","PVP01_1342800","PVP01_1342900","PVP01_1343000","PVP01_1343100","PVP01_1343200","PVP01_1343300","PVP01_1343400","PVP01_1343500","PVP01_1343600","PVP01_1343700","PVP01_1343800","PVP01_1343900","PVP01_1344000","PVP01_1344100","PVP01_1344200","PVP01_1344300","PVP01_1344400","PVP01_1344500","PVP01_1344600","PVP01_1344700","PVP01_1344800","PVP01_1344900","PVP01_1345000","PVP01_1345100","PVP01_1345200","PVP01_1345300","PVP01_1345400","PVP01_1345500","PVP01_1345600","PVP01_1345700","PVP01_1345800","PVP01_1345900","PVP01_1346000","PVP01_1346100","PVP01_1346200","PVP01_1346400","PVP01_1346500","PVP01_1346600","PVP01_1346700","PVP01_1346800","PVP01_1346900","PVP01_1347000","PVP01_1347100","PVP01_1347200","PVP01_1347300","PVP01_1347500","PVP01_1347700","PVP01_1347800","PVP01_1400100","PVP01_1400400","PVP01_1400600","PVP01_1400700","PVP01_1400900","PVP01_1401000","PVP01_1401100","PVP01_1401200","PVP01_1401600","PVP01_1401700","PVP01_1401800","PVP01_1401900","PVP01_1402000","PVP01_1402100","PVP01_1402200","PVP01_1402300","PVP01_1402400","PVP01_1402500","PVP01_1402600","PVP01_1402700","PVP01_1402800","PVP01_1402900","PVP01_1403000","PVP01_1403100","PVP01_1403200","PVP01_1403300","PVP01_1403400","PVP01_1403500","PVP01_1403600","PVP01_1403700","PVP01_1403800","PVP01_1403900","PVP01_1404000","PVP01_1404100","PVP01_1404200","PVP01_1404300","PVP01_1404400","PVP01_1404500","PVP01_1404600","PVP01_1404700","PVP01_1404800","PVP01_1404900","PVP01_1405000","PVP01_1405100","PVP01_1405200","PVP01_1405300","PVP01_1405500","PVP01_1405600","PVP01_1405700","PVP01_1405800","PVP01_1405900","PVP01_1406000","PVP01_1406100","PVP01_1406200","PVP01_1406300","PVP01_1406400","PVP01_1406500","PVP01_1406600","PVP01_1406700","PVP01_1406800","PVP01_1406900","PVP01_1407000","PVP01_1407100","PVP01_1407200","PVP01_1407300","PVP01_1407400","PVP01_1407500","PVP01_1407600","PVP01_1407700","PVP01_1407800","PVP01_1407900","PVP01_1408000","PVP01_1408100","PVP01_1408200","PVP01_1408300","PVP01_1408400","PVP01_1408500","PVP01_1408600","PVP01_1408700","PVP01_1408800","PVP01_1408900","PVP01_1409100","PVP01_1409200","PVP01_1409300","PVP01_1409400","PVP01_1409500","PVP01_1409600","PVP01_1409700","PVP01_1409800","PVP01_1409900","PVP01_1410000","PVP01_1410100","PVP01_1410200","PVP01_1410300","PVP01_1410400","PVP01_1410500","PVP01_1410600","PVP01_1410700","PVP01_1410800","PVP01_1410900","PVP01_1411000","PVP01_1411100","PVP01_1411200","PVP01_1411300","PVP01_1411400","PVP01_1411500","PVP01_1411600","PVP01_1411700","PVP01_1411800","PVP01_1411900","PVP01_1412000","PVP01_1412100","PVP01_1412200","PVP01_1412300","PVP01_1412400","PVP01_1412500","PVP01_1412600","PVP01_1412700","PVP01_1412800","PVP01_1412900","PVP01_1413000","PVP01_1413100","PVP01_1413200","PVP01_1413300","PVP01_1413400","PVP01_1413450","PVP01_1413500","PVP01_1413600","PVP01_1413700","PVP01_1413800","PVP01_1413900","PVP01_1414000","PVP01_1414100","PVP01_1414200","PVP01_1414300","PVP01_1414400","PVP01_1414600","PVP01_1414700","PVP01_1414800","PVP01_1414900","PVP01_1415000","PVP01_1415100","PVP01_1415200","PVP01_1415300","PVP01_1415400","PVP01_1415500","PVP01_1415600","PVP01_1415700","PVP01_1415800","PVP01_1415900","PVP01_1416000","PVP01_1416100","PVP01_1416200","PVP01_1416300","PVP01_1416400","PVP01_1416500","PVP01_1416600","PVP01_1416700","PVP01_1416800","PVP01_1416900","PVP01_1417000","PVP01_1417100","PVP01_1417200","PVP01_1417300","PVP01_1417400","PVP01_1417500","PVP01_1417600","PVP01_1417700","PVP01_1417800","PVP01_1417900","PVP01_1418000","PVP01_1418100","PVP01_1418200","PVP01_1418400","PVP01_1418500","PVP01_1418600","PVP01_1418700","PVP01_1418800","PVP01_1418900","PVP01_1419000","PVP01_1419100","PVP01_1419200","PVP01_1419300","PVP01_1419400","PVP01_1419500","PVP01_1419600","PVP01_1419700","PVP01_1419800","PVP01_1419900","PVP01_1420000","PVP01_1420100","PVP01_1420200","PVP01_1420300","PVP01_1420400","PVP01_1420500","PVP01_1420600","PVP01_1420700","PVP01_1420800","PVP01_1420900","PVP01_1421000","PVP01_1421100","PVP01_1421200","PVP01_1421300","PVP01_1421400","PVP01_1421600","PVP01_1421700","PVP01_1421800","PVP01_1421900","PVP01_1422000","PVP01_1422100","PVP01_1422200","PVP01_1422300","PVP01_1422400","PVP01_1422500","PVP01_1422600","PVP01_1422700","PVP01_1422800","PVP01_1422900","PVP01_1423000","PVP01_1423100","PVP01_1423200","PVP01_1423300","PVP01_1423400","PVP01_1423500","PVP01_1423600","PVP01_1423700","PVP01_1424100","PVP01_1424200","PVP01_1424300","PVP01_1424400","PVP01_1424500","PVP01_1424600","PVP01_1424700","PVP01_1424800","PVP01_1424900","PVP01_1425000","PVP01_1425100","PVP01_1425200","PVP01_1425300","PVP01_1425400","PVP01_1425500","PVP01_1425600","PVP01_1425700","PVP01_1425800","PVP01_1425900","PVP01_1426000","PVP01_1426100","PVP01_1426200","PVP01_1426300","PVP01_1426400","PVP01_1426500","PVP01_1426600","PVP01_1426700","PVP01_1426800","PVP01_1426900","PVP01_1427000","PVP01_1427100","PVP01_1427200","PVP01_1427300","PVP01_1427400","PVP01_1427500","PVP01_1427600","PVP01_1427700","PVP01_1427800","PVP01_1427900","PVP01_1428000","PVP01_1428100","PVP01_1428200","PVP01_1428300","PVP01_1428400","PVP01_1428500","PVP01_1428600","PVP01_1428800","PVP01_1428900","PVP01_1429000","PVP01_1429100","PVP01_1429200","PVP01_1429300","PVP01_1429400","PVP01_1429500","PVP01_1429600","PVP01_1429700","PVP01_1429800","PVP01_1429900","PVP01_1430000","PVP01_1430100","PVP01_1430200","PVP01_1430300","PVP01_1430400","PVP01_1430500","PVP01_1430600","PVP01_1430700","PVP01_1430800","PVP01_1430900","PVP01_1431000","PVP01_1431100","PVP01_1431200","PVP01_1431300","PVP01_1431400","PVP01_1431500","PVP01_1431600","PVP01_1431700","PVP01_1431800","PVP01_1432000","PVP01_1432100","PVP01_1432200","PVP01_1432300","PVP01_1432400","PVP01_1432500","PVP01_1432600","PVP01_1432700","PVP01_1432800","PVP01_1432900","PVP01_1433000","PVP01_1433100","PVP01_1433200","PVP01_1433300","PVP01_1433400","PVP01_1433500","PVP01_1433600","PVP01_1433700","PVP01_1433800","PVP01_1433900","PVP01_1434000","PVP01_1434100","PVP01_1434200","PVP01_1434300","PVP01_1434400","PVP01_1434500","PVP01_1434600","PVP01_1434700","PVP01_1434800","PVP01_1434900","PVP01_1435000","PVP01_1435100","PVP01_1435200","PVP01_1435300","PVP01_1435400","PVP01_1435500","PVP01_1435700","PVP01_1435800","PVP01_1435900","PVP01_1436000","PVP01_1436100","PVP01_1436200","PVP01_1436300","PVP01_1436400","PVP01_1436500","PVP01_1436600","PVP01_1436700","PVP01_1436800","PVP01_1436900","PVP01_1437000","PVP01_1437100","PVP01_1437200","PVP01_1437300","PVP01_1437400","PVP01_1437500","PVP01_1437600","PVP01_1437700","PVP01_1437800","PVP01_1437900","PVP01_1438000","PVP01_1438100","PVP01_1438200","PVP01_1438300","PVP01_1438400","PVP01_1438500","PVP01_1438600","PVP01_1438700","PVP01_1438800","PVP01_1438900","PVP01_1439000","PVP01_1439200","PVP01_1439300","PVP01_1439400","PVP01_1439500","PVP01_1439600","PVP01_1439700","PVP01_1439800","PVP01_1439900","PVP01_1440000","PVP01_1440100","PVP01_1440300","PVP01_1440400","PVP01_1440500","PVP01_1440600","PVP01_1440700","PVP01_1440800","PVP01_1440900","PVP01_1441000","PVP01_1441100","PVP01_1441200","PVP01_1441300","PVP01_1441400","PVP01_1441500","PVP01_1441600","PVP01_1441700","PVP01_1441800","PVP01_1441900","PVP01_1442000","PVP01_1442100","PVP01_1442200","PVP01_1442300","PVP01_1442400","PVP01_1442500","PVP01_1442600","PVP01_1442700","PVP01_1442800","PVP01_1442900","PVP01_1443000","PVP01_1443100","PVP01_1443200","PVP01_1443300","PVP01_1443400","PVP01_1443500","PVP01_1443600","PVP01_1443700","PVP01_1443800","PVP01_1443900","PVP01_1444000","PVP01_1444100","PVP01_1444200","PVP01_1444300","PVP01_1444400","PVP01_1444500","PVP01_1444600","PVP01_1444700","PVP01_1444800","PVP01_1444900","PVP01_1445000","PVP01_1445100","PVP01_1445200","PVP01_1445300","PVP01_1445400","PVP01_1445500","PVP01_1445600","PVP01_1445700","PVP01_1445800","PVP01_1445900","PVP01_1446000","PVP01_1446100","PVP01_1446200","PVP01_1446300","PVP01_1446400","PVP01_1446500","PVP01_1446600","PVP01_1446700","PVP01_1446800","PVP01_1446900","PVP01_1447000","PVP01_1447100","PVP01_1447200","PVP01_1447300","PVP01_1447400","PVP01_1447500","PVP01_1447600","PVP01_1447700","PVP01_1447800","PVP01_1447900","PVP01_1448000","PVP01_1448100","PVP01_1448200","PVP01_1448300","PVP01_1448400","PVP01_1448500","PVP01_1448600","PVP01_1448700","PVP01_1448800","PVP01_1448900","PVP01_1449000","PVP01_1449100","PVP01_1449200","PVP01_1449300","PVP01_1449400","PVP01_1449600","PVP01_1449700","PVP01_1449800","PVP01_1449900","PVP01_1450000","PVP01_1450100","PVP01_1450200","PVP01_1450300","PVP01_1450400","PVP01_1450500","PVP01_1450600","PVP01_1450700","PVP01_1450800","PVP01_1450900","PVP01_1451000","PVP01_1451100","PVP01_1451200","PVP01_1451300","PVP01_1451400","PVP01_1451500","PVP01_1451600","PVP01_1451700","PVP01_1451800","PVP01_1451900","PVP01_1452000","PVP01_1452100","PVP01_1452200","PVP01_1452300","PVP01_1452400","PVP01_1452500","PVP01_1452600","PVP01_1452700","PVP01_1452800","PVP01_1452900","PVP01_1453000","PVP01_1453100","PVP01_1453200","PVP01_1453300","PVP01_1453400","PVP01_1453500","PVP01_1453600","PVP01_1453700","PVP01_1453800","PVP01_1453900","PVP01_1454000","PVP01_1454100","PVP01_1454200","PVP01_1454300","PVP01_1454400","PVP01_1454500","PVP01_1454600","PVP01_1454800","PVP01_1454900","PVP01_1455000","PVP01_1455100","PVP01_1455200","PVP01_1455300","PVP01_1455400","PVP01_1455500","PVP01_1455600","PVP01_1455700","PVP01_1455800","PVP01_1455900","PVP01_1456000","PVP01_1456100","PVP01_1456200","PVP01_1456400","PVP01_1456500","PVP01_1456600","PVP01_1456700","PVP01_1456800","PVP01_1456900","PVP01_1457000","PVP01_1457100","PVP01_1457200","PVP01_1457300","PVP01_1457400","PVP01_1457500","PVP01_1457600","PVP01_1457700","PVP01_1457800","PVP01_1457900","PVP01_1458000","PVP01_1458100","PVP01_1458200","PVP01_1458300","PVP01_1458400","PVP01_1458500","PVP01_1458600","PVP01_1458700","PVP01_1458800","PVP01_1458900","PVP01_1459000","PVP01_1459100","PVP01_1459200","PVP01_1459300","PVP01_1459400","PVP01_1459500","PVP01_1459600","PVP01_1459700","PVP01_1459800","PVP01_1459900","PVP01_1460000","PVP01_1460100","PVP01_1460200","PVP01_1460300","PVP01_1460400","PVP01_1460500","PVP01_1460600","PVP01_1460700","PVP01_1460800","PVP01_1460900","PVP01_1461000","PVP01_1461100","PVP01_1461200","PVP01_1461300","PVP01_1461500","PVP01_1461600","PVP01_1461700","PVP01_1461800","PVP01_1461900","PVP01_1462000","PVP01_1462100","PVP01_1462200","PVP01_1462300","PVP01_1462400","PVP01_1462500","PVP01_1462600","PVP01_1462700","PVP01_1462800","PVP01_1462900","PVP01_1463000","PVP01_1463100","PVP01_1463200","PVP01_1463300","PVP01_1463400","PVP01_1463500","PVP01_1463600","PVP01_1463700","PVP01_1463800","PVP01_1463900","PVP01_1464000","PVP01_1464100","PVP01_1464200","PVP01_1464300","PVP01_1464400","PVP01_1464500","PVP01_1464600","PVP01_1464700","PVP01_1464800","PVP01_1464900","PVP01_1465000","PVP01_1465100","PVP01_1465200","PVP01_1465300","PVP01_1465400","PVP01_1465500","PVP01_1465600","PVP01_1465700","PVP01_1465800","PVP01_1465900","PVP01_1466000","PVP01_1466100","PVP01_1466200","PVP01_1466300","PVP01_1466400","PVP01_1466500","PVP01_1466600","PVP01_1466700","PVP01_1466800","PVP01_1466900","PVP01_1467000","PVP01_1467100","PVP01_1467200","PVP01_1467300","PVP01_1467400","PVP01_1467500","PVP01_1467600","PVP01_1467700","PVP01_1467800","PVP01_1467900","PVP01_1468000","PVP01_1468100","PVP01_1468200","PVP01_1468300","PVP01_1468400","PVP01_1468500","PVP01_1468600","PVP01_1468700","PVP01_1468800","PVP01_1468900","PVP01_1469300","PVP01_1469400","PVP01_1469600","PVP01_1469700","PVP01_1469800","PVP01_1469900","PVP01_1470000","PVP01_1470100","PVP01_1470200","PVP01_1470300","PVP01_1470400","PVP01_1470500","PVP01_1470600","PVP01_1470700","PVP01_1470800","PVP01_1470900","PVP01_1471100","PVP01_1471300","PVP01_1471400","PVP01_1471500","PVP01_1471600","PVP01_1471700","PVP01_1471800","PVP01_1471900","PVP01_1472100","PVP01_1472200","PVP01_1472400","PVP01_API01200","PVP01_API01400","PVP01_API01700","PVP01_API02600","PVP01_API02800","PVP01_API03500","PVP01_API04100","PVP01_API04200","PVP01_API04300","PVP01_API04600","PVP01_MIT02700","PVP01_MIT03300","PVP01_MIT03400","PVP01_0000100","PVP01_0104300","PVP01_0104700","PVP01_0107500","PVP01_0117000","PVP01_0117600","PVP01_0119200","PVP01_0202300","PVP01_0207400","PVP01_0210500","PVP01_0214400","PVP01_0304300","PVP01_0309100","PVP01_0311400","PVP01_0317900","PVP01_0319500","PVP01_0411300","PVP01_0417300","PVP01_0417600","PVP01_0418700","PVP01_0421000","PVP01_0504000","PVP01_0511400","PVP01_0517600","PVP01_0532000","PVP01_0534300","PVP01_0602100","PVP01_0609100","PVP01_0612400","PVP01_0615000","PVP01_0620800","PVP01_0703800","PVP01_0705900","PVP01_0709000","PVP01_0723100","PVP01_0723600","PVP01_0727900","PVP01_0728900","PVP01_0730800","PVP01_0734300","PVP01_0734500","PVP01_0735000","PVP01_0801600","PVP01_0807700","PVP01_0812300","PVP01_0813900","PVP01_0815000","PVP01_0816800","PVP01_0819300","PVP01_0913800","PVP01_0916000","PVP01_0916600","PVP01_0917400","PVP01_0926500","PVP01_0931900","PVP01_0935900","PVP01_0938800","PVP01_0940300","PVP01_0945900","PVP01_1002900","PVP01_1015200","PVP01_1025500","PVP01_1028500","PVP01_1033400","PVP01_1110200","PVP01_1119100","PVP01_1135200","PVP01_1139200","PVP01_1144800","PVP01_1204300","PVP01_1206200","PVP01_1209200","PVP01_1215100","PVP01_1219600","PVP01_1220000","PVP01_1220300","PVP01_1230900","PVP01_1233500","PVP01_1235400","PVP01_1238800","PVP01_1242900","PVP01_1249000","PVP01_1250100","PVP01_1251600","PVP01_1255000","PVP01_1267000","PVP01_1301400","PVP01_1305200","PVP01_1312300","PVP01_1315100","PVP01_1315700","PVP01_1321100","PVP01_1338500","PVP01_1401300","PVP01_1401400","PVP01_1405400","PVP01_1409000","PVP01_1414500","PVP01_1418300","PVP01_1421500","PVP01_1428700","PVP01_1439100","PVP01_1440200","PVP01_1449500","PVP01_1454700","PVP01_1456300","PVP01_1469200","PVP01_1469500"],"Row.names":["PVP01_0000030","PVP01_0000040","PVP01_0000050","PVP01_0000080","PVP01_0000110","PVP01_0000120","PVP01_0000130","PVP01_0000140","PVP01_0000150","PVP01_0000160","PVP01_0000170","PVP01_0000180","PVP01_0000190","PVP01_0000200","PVP01_0000210","PVP01_0000220","PVP01_0000300","PVP01_0000480","PVP01_0000670","PVP01_0000740","PVP01_0000790","PVP01_0001030","PVP01_0001090","PVP01_0001310","PVP01_0001430","PVP01_0001440","PVP01_0001480","PVP01_0001550","PVP01_0001640","PVP01_0001660","PVP01_0001670","PVP01_0001900","PVP01_0001910","PVP01_0001940","PVP01_0001970","PVP01_0002020","PVP01_0002050","PVP01_0002120","PVP01_0002200","PVP01_0002210","PVP01_0002330","PVP01_0002370","PVP01_0002390","PVP01_0002470","PVP01_0002530","PVP01_0002540","PVP01_0002750","PVP01_0002810","PVP01_0002820","PVP01_0003010","PVP01_0003050","PVP01_0003150","PVP01_0003210","PVP01_0003350","PVP01_0003360","PVP01_0003370","PVP01_0003380","PVP01_0003400","PVP01_0003410","PVP01_0003420","PVP01_0003440","PVP01_0003530","PVP01_0003640","PVP01_0003690","PVP01_0003730","PVP01_0003800","PVP01_0003990","PVP01_0004040","PVP01_0004160","PVP01_0004170","PVP01_0004180","PVP01_0004190","PVP01_0004210","PVP01_0004220","PVP01_0004230","PVP01_0004240","PVP01_0004250","PVP01_0004260","PVP01_0004270","PVP01_0004280","PVP01_0004290","PVP01_0004300","PVP01_0004310","PVP01_0004320","PVP01_0004330","PVP01_0004350","PVP01_0004370","PVP01_0004380","PVP01_0004390","PVP01_0004400","PVP01_0004410","PVP01_0004640","PVP01_0004670","PVP01_0004910","PVP01_0004920","PVP01_0004950","PVP01_0004960","PVP01_0005010","PVP01_0005100","PVP01_0005160","PVP01_0005210","PVP01_0005300","PVP01_0005310","PVP01_0005420","PVP01_0005430","PVP01_0005540","PVP01_0005550","PVP01_0005640","PVP01_0005710","PVP01_0005740","PVP01_0005760","PVP01_0005800","PVP01_0005820","PVP01_0005870","PVP01_0005910","PVP01_0006000","PVP01_0006040","PVP01_0006080","PVP01_0006260","PVP01_0006650","PVP01_0006660","PVP01_0006670","PVP01_0006680","PVP01_0006700","PVP01_0006860","PVP01_0006890","PVP01_0007080","PVP01_0007160","PVP01_0007240","PVP01_0007290","PVP01_0007310","PVP01_0007320","PVP01_0007340","PVP01_0007610","PVP01_0007620","PVP01_0007690","PVP01_0007700","PVP01_0007720","PVP01_0007850","PVP01_0007900","PVP01_0007910","PVP01_0007920","PVP01_0008190","PVP01_0008430","PVP01_0008480","PVP01_0008490","PVP01_0008600","PVP01_0008700","PVP01_0008770","PVP01_0008830","PVP01_0008990","PVP01_0009010","PVP01_0009170","PVP01_0009180","PVP01_0009210","PVP01_0009370","PVP01_0009410","PVP01_0009470","PVP01_0009540","PVP01_0009620","PVP01_0009710","PVP01_0009720","PVP01_0009730","PVP01_0009740","PVP01_0009760","PVP01_0009770","PVP01_0009820","PVP01_0009850","PVP01_0009860","PVP01_0009870","PVP01_0009900","PVP01_0009940","PVP01_0009980","PVP01_0010020","PVP01_0010030","PVP01_0010070","PVP01_0010100","PVP01_0010120","PVP01_0010140","PVP01_0010170","PVP01_0010180","PVP01_0010200","PVP01_0010210","PVP01_0010220","PVP01_0010230","PVP01_0010240","PVP01_0010340","PVP01_0010380","PVP01_0010440","PVP01_0010470","PVP01_0010500","PVP01_0010540","PVP01_0010570","PVP01_0010600","PVP01_0010610","PVP01_0010640","PVP01_0010650","PVP01_0010670","PVP01_0010770","PVP01_0010840","PVP01_0010870","PVP01_0010900","PVP01_0010910","PVP01_0010960","PVP01_0010970","PVP01_0010990","PVP01_0011020","PVP01_0011030","PVP01_0011070","PVP01_0102100","PVP01_0102200","PVP01_0102300","PVP01_0102400","PVP01_0102500","PVP01_0102600","PVP01_0102700","PVP01_0102800","PVP01_0103100","PVP01_0103200","PVP01_0103300","PVP01_0103400","PVP01_0103500","PVP01_0103600","PVP01_0103700","PVP01_0103800","PVP01_0103900","PVP01_0104000","PVP01_0104100","PVP01_0104200","PVP01_0104400","PVP01_0104500","PVP01_0104600","PVP01_0104800","PVP01_0104900","PVP01_0105000","PVP01_0105100","PVP01_0105200","PVP01_0105300","PVP01_0105400","PVP01_0105500","PVP01_0105600","PVP01_0105700","PVP01_0105800","PVP01_0105900","PVP01_0106000","PVP01_0106100","PVP01_0106200","PVP01_0106300","PVP01_0106400","PVP01_0106500","PVP01_0106600","PVP01_0106700","PVP01_0106800","PVP01_0106900","PVP01_0107400","PVP01_0107600","PVP01_0107700","PVP01_0107800","PVP01_0107900","PVP01_0108000","PVP01_0108100","PVP01_0108200","PVP01_0108300","PVP01_0108400","PVP01_0108500","PVP01_0108600","PVP01_0108700","PVP01_0108800","PVP01_0108900","PVP01_0109000","PVP01_0109100","PVP01_0109200","PVP01_0109300","PVP01_0109400","PVP01_0109500","PVP01_0109600","PVP01_0109700","PVP01_0109800","PVP01_0109900","PVP01_0110000","PVP01_0110100","PVP01_0110200","PVP01_0110300","PVP01_0110400","PVP01_0110500","PVP01_0110600","PVP01_0110700","PVP01_0110800","PVP01_0110900","PVP01_0111000","PVP01_0111100","PVP01_0111200","PVP01_0111300","PVP01_0111400","PVP01_0111500","PVP01_0111600","PVP01_0111700","PVP01_0111800","PVP01_0111900","PVP01_0112000","PVP01_0112100","PVP01_0112200","PVP01_0112300","PVP01_0112400","PVP01_0112500","PVP01_0112600","PVP01_0112700","PVP01_0112800","PVP01_0112900","PVP01_0113000","PVP01_0113100","PVP01_0113200","PVP01_0113300","PVP01_0113400","PVP01_0113500","PVP01_0113600","PVP01_0113700","PVP01_0113800","PVP01_0113900","PVP01_0114000","PVP01_0114100","PVP01_0114200","PVP01_0114300","PVP01_0114400","PVP01_0114500","PVP01_0114600","PVP01_0114700","PVP01_0114800","PVP01_0114900","PVP01_0115000","PVP01_0115100","PVP01_0115200","PVP01_0115300","PVP01_0115400","PVP01_0115500","PVP01_0115600","PVP01_0115700","PVP01_0115800","PVP01_0115900","PVP01_0116000","PVP01_0116100","PVP01_0116200","PVP01_0116300","PVP01_0116400","PVP01_0116500","PVP01_0116600","PVP01_0116700","PVP01_0116800","PVP01_0116900","PVP01_0117100","PVP01_0117200","PVP01_0117300","PVP01_0117400","PVP01_0117500","PVP01_0117700","PVP01_0117800","PVP01_0117900","PVP01_0118000","PVP01_0118100","PVP01_0118200","PVP01_0118300","PVP01_0118400","PVP01_0118500","PVP01_0118600","PVP01_0118700","PVP01_0118800","PVP01_0118900","PVP01_0119300","PVP01_0119400","PVP01_0119500","PVP01_0119600","PVP01_0119700","PVP01_0119800","PVP01_0119900","PVP01_0120000","PVP01_0120100","PVP01_0120500","PVP01_0120800","PVP01_0121000","PVP01_0121700","PVP01_0200200","PVP01_0200700","PVP01_0201000","PVP01_0201100","PVP01_0201200","PVP01_0201300","PVP01_0201400","PVP01_0201500","PVP01_0201600","PVP01_0201700","PVP01_0201800","PVP01_0201900","PVP01_0202000","PVP01_0202100","PVP01_0202200","PVP01_0202400","PVP01_0202500","PVP01_0202600","PVP01_0203000","PVP01_0203100","PVP01_0203200","PVP01_0203300","PVP01_0203400","PVP01_0203500","PVP01_0203600","PVP01_0203700","PVP01_0203800","PVP01_0203900","PVP01_0204000","PVP01_0204100","PVP01_0204200","PVP01_0204300","PVP01_0204400","PVP01_0204500","PVP01_0204600","PVP01_0204700","PVP01_0204800","PVP01_0204900","PVP01_0205000","PVP01_0205100","PVP01_0205200","PVP01_0205300","PVP01_0205400","PVP01_0205500","PVP01_0205600","PVP01_0205700","PVP01_0205800","PVP01_0205900","PVP01_0206000","PVP01_0206100","PVP01_0206200","PVP01_0206300","PVP01_0206400","PVP01_0206500","PVP01_0206600","PVP01_0206700","PVP01_0206800","PVP01_0206900","PVP01_0207000","PVP01_0207100","PVP01_0207200","PVP01_0207300","PVP01_0207500","PVP01_0207600","PVP01_0207700","PVP01_0207800","PVP01_0207900","PVP01_0208000","PVP01_0208100","PVP01_0208200","PVP01_0208300","PVP01_0208400","PVP01_0208500","PVP01_0208600","PVP01_0208700","PVP01_0208800","PVP01_0208900","PVP01_0209000","PVP01_0209100","PVP01_0209200","PVP01_0209300","PVP01_0209400","PVP01_0209500","PVP01_0209600","PVP01_0209700","PVP01_0209900","PVP01_0210000","PVP01_0210100","PVP01_0210200","PVP01_0210300","PVP01_0210400","PVP01_0210600","PVP01_0210700","PVP01_0210800","PVP01_0210900","PVP01_0211000","PVP01_0211100","PVP01_0211200","PVP01_0211300","PVP01_0211400","PVP01_0211500","PVP01_0211600","PVP01_0211700","PVP01_0211800","PVP01_0211900","PVP01_0212000","PVP01_0212100","PVP01_0212200","PVP01_0212300","PVP01_0212400","PVP01_0212500","PVP01_0212600","PVP01_0212700","PVP01_0212800","PVP01_0212900","PVP01_0213000","PVP01_0213100","PVP01_0213200","PVP01_0213300","PVP01_0213400","PVP01_0213500","PVP01_0213600","PVP01_0213700","PVP01_0213800","PVP01_0213900","PVP01_0214000","PVP01_0214100","PVP01_0214200","PVP01_0214300","PVP01_0214500","PVP01_0214600","PVP01_0214700","PVP01_0214800","PVP01_0214900","PVP01_0215000","PVP01_0215100","PVP01_0215200","PVP01_0215300","PVP01_0215400","PVP01_0215500","PVP01_0215600","PVP01_0215700","PVP01_0215800","PVP01_0215900","PVP01_0216000","PVP01_0216100","PVP01_0216200","PVP01_0216500","PVP01_0216600","PVP01_0216700","PVP01_0216800","PVP01_0216900","PVP01_0217000","PVP01_0217100","PVP01_0217200","PVP01_0217300","PVP01_0217400","PVP01_0217500","PVP01_0218200","PVP01_0218300","PVP01_0218900","PVP01_0219400","PVP01_0220100","PVP01_0220200","PVP01_0221100","PVP01_0221300","PVP01_0221700","PVP01_0300100","PVP01_0300600","PVP01_0300700","PVP01_0300800","PVP01_0301100","PVP01_0301200","PVP01_0301500","PVP01_0301900","PVP01_0302000","PVP01_0302100","PVP01_0302300","PVP01_0302400","PVP01_0302500","PVP01_0302600","PVP01_0302700","PVP01_0302800","PVP01_0302900","PVP01_0303000","PVP01_0303100","PVP01_0303200","PVP01_0303300","PVP01_0303400","PVP01_0303500","PVP01_0303600","PVP01_0303700","PVP01_0303800","PVP01_0303900","PVP01_0304000","PVP01_0304100","PVP01_0304200","PVP01_0304400","PVP01_0304500","PVP01_0304600","PVP01_0304700","PVP01_0304800","PVP01_0304900","PVP01_0305000","PVP01_0305100","PVP01_0305200","PVP01_0305300","PVP01_0305400","PVP01_0305500","PVP01_0305600","PVP01_0305700","PVP01_0305800","PVP01_0305900","PVP01_0306000","PVP01_0306100","PVP01_0306200","PVP01_0306300","PVP01_0306400","PVP01_0306500","PVP01_0306600","PVP01_0306700","PVP01_0306800","PVP01_0306900","PVP01_0307000","PVP01_0307100","PVP01_0307200","PVP01_0307300","PVP01_0307400","PVP01_0307500","PVP01_0307600","PVP01_0307700","PVP01_0307800","PVP01_0307900","PVP01_0308000","PVP01_0308100","PVP01_0308200","PVP01_0308300","PVP01_0308400","PVP01_0308500","PVP01_0308600","PVP01_0308700","PVP01_0308800","PVP01_0308900","PVP01_0309000","PVP01_0309200","PVP01_0309300","PVP01_0309600","PVP01_0309700","PVP01_0309800","PVP01_0309900","PVP01_0310000","PVP01_0310100","PVP01_0310200","PVP01_0310300","PVP01_0310400","PVP01_0310500","PVP01_0310600","PVP01_0310700","PVP01_0311000","PVP01_0311100","PVP01_0311200","PVP01_0311300","PVP01_0311500","PVP01_0311600","PVP01_0311700","PVP01_0311800","PVP01_0311900","PVP01_0312000","PVP01_0312100","PVP01_0312200","PVP01_0312300","PVP01_0312400","PVP01_0312500","PVP01_0312600","PVP01_0312700","PVP01_0312800","PVP01_0312900","PVP01_0313000","PVP01_0313100","PVP01_0313200","PVP01_0313300","PVP01_0313400","PVP01_0313500","PVP01_0313600","PVP01_0313700","PVP01_0313800","PVP01_0313900","PVP01_0314000","PVP01_0314100","PVP01_0314200","PVP01_0314300","PVP01_0314400","PVP01_0314500","PVP01_0314600","PVP01_0314700","PVP01_0314800","PVP01_0314900","PVP01_0315000","PVP01_0315100","PVP01_0315200","PVP01_0315300","PVP01_0315400","PVP01_0315500","PVP01_0315600","PVP01_0315700","PVP01_0315800","PVP01_0315900","PVP01_0316000","PVP01_0316100","PVP01_0316200","PVP01_0316300","PVP01_0316400","PVP01_0316500","PVP01_0316600","PVP01_0316700","PVP01_0316800","PVP01_0316900","PVP01_0317000","PVP01_0317100","PVP01_0317200","PVP01_0317300","PVP01_0317400","PVP01_0317500","PVP01_0317600","PVP01_0317700","PVP01_0317800","PVP01_0318000","PVP01_0318100","PVP01_0318200","PVP01_0318300","PVP01_0318400","PVP01_0318500","PVP01_0318600","PVP01_0318700","PVP01_0318800","PVP01_0318900","PVP01_0319000","PVP01_0319100","PVP01_0319200","PVP01_0319300","PVP01_0319400","PVP01_0319600","PVP01_0319700","PVP01_0319800","PVP01_0319900","PVP01_0320000","PVP01_0320100","PVP01_0320200","PVP01_0320300","PVP01_0320400","PVP01_0320500","PVP01_0320600","PVP01_0320700","PVP01_0320800","PVP01_0320900","PVP01_0321000","PVP01_0321100","PVP01_0400900","PVP01_0403600","PVP01_0403700","PVP01_0403900","PVP01_0404000","PVP01_0404100","PVP01_0404200","PVP01_0404300","PVP01_0404400","PVP01_0404500","PVP01_0404600","PVP01_0404700","PVP01_0404800","PVP01_0404900","PVP01_0405000","PVP01_0405100","PVP01_0405200","PVP01_0405300","PVP01_0405400","PVP01_0405500","PVP01_0405600","PVP01_0405700","PVP01_0405800","PVP01_0405900","PVP01_0406300","PVP01_0406400","PVP01_0406500","PVP01_0406600","PVP01_0406700","PVP01_0406800","PVP01_0406900","PVP01_0407000","PVP01_0407100","PVP01_0407200","PVP01_0407300","PVP01_0407400","PVP01_0407500","PVP01_0407600","PVP01_0407700","PVP01_0407800","PVP01_0407900","PVP01_0408000","PVP01_0408100","PVP01_0408200","PVP01_0408300","PVP01_0408400","PVP01_0408500","PVP01_0408600","PVP01_0408700","PVP01_0408800","PVP01_0408900","PVP01_0409000","PVP01_0409100","PVP01_0409200","PVP01_0409300","PVP01_0409400","PVP01_0409500","PVP01_0409600","PVP01_0409700","PVP01_0409800","PVP01_0409900","PVP01_0410000","PVP01_0410100","PVP01_0410200","PVP01_0410300","PVP01_0410400","PVP01_0410500","PVP01_0410600","PVP01_0410700","PVP01_0410800","PVP01_0410900","PVP01_0411000","PVP01_0411100","PVP01_0411200","PVP01_0411400","PVP01_0411500","PVP01_0411600","PVP01_0411700","PVP01_0411800","PVP01_0411900","PVP01_0412000","PVP01_0412100","PVP01_0412200","PVP01_0412300","PVP01_0412400","PVP01_0412500","PVP01_0412600","PVP01_0412700","PVP01_0412800","PVP01_0412900","PVP01_0413000","PVP01_0413100","PVP01_0413200","PVP01_0413300","PVP01_0413400","PVP01_0413500","PVP01_0413600","PVP01_0413700","PVP01_0413800","PVP01_0413900","PVP01_0414000","PVP01_0414100","PVP01_0414200","PVP01_0414300","PVP01_0414400","PVP01_0414500","PVP01_0414600","PVP01_0414700","PVP01_0414800","PVP01_0414900","PVP01_0415000","PVP01_0415100","PVP01_0415200","PVP01_0415300","PVP01_0415400","PVP01_0415500","PVP01_0415600","PVP01_0415700","PVP01_0415800","PVP01_0415900","PVP01_0416000","PVP01_0416100","PVP01_0416200","PVP01_0416300","PVP01_0416400","PVP01_0416500","PVP01_0416600","PVP01_0416700","PVP01_0416800","PVP01_0416900","PVP01_0417000","PVP01_0417100","PVP01_0417200","PVP01_0417400","PVP01_0417500","PVP01_0417700","PVP01_0417800","PVP01_0417900","PVP01_0418000","PVP01_0418100","PVP01_0418200","PVP01_0418300","PVP01_0418400","PVP01_0418500","PVP01_0418600","PVP01_0418800","PVP01_0418900","PVP01_0419000","PVP01_0419100","PVP01_0419200","PVP01_0419300","PVP01_0419400","PVP01_0419500","PVP01_0419600","PVP01_0419700","PVP01_0419800","PVP01_0419900","PVP01_0420000","PVP01_0420100","PVP01_0420200","PVP01_0420300","PVP01_0420400","PVP01_0420500","PVP01_0420600","PVP01_0420700","PVP01_0420800","PVP01_0420900","PVP01_0421100","PVP01_0421200","PVP01_0421300","PVP01_0421400","PVP01_0421500","PVP01_0421700","PVP01_0421800","PVP01_0421900","PVP01_0422000","PVP01_0422100","PVP01_0422200","PVP01_0422300","PVP01_0422400","PVP01_0422500","PVP01_0422600","PVP01_0422700","PVP01_0422800","PVP01_0422900","PVP01_0423000","PVP01_0423100","PVP01_0423200","PVP01_0423300","PVP01_0423400","PVP01_0423500","PVP01_0423600","PVP01_0423800","PVP01_0423900","PVP01_0424300","PVP01_0424400","PVP01_0424500","PVP01_0500700","PVP01_0502300","PVP01_0502400","PVP01_0502900","PVP01_0503300","PVP01_0503400","PVP01_0503500","PVP01_0503600","PVP01_0503700","PVP01_0503800","PVP01_0503900","PVP01_0504100","PVP01_0504200","PVP01_0504300","PVP01_0504400","PVP01_0504800","PVP01_0504900","PVP01_0505000","PVP01_0505100","PVP01_0505200","PVP01_0505300","PVP01_0505400","PVP01_0505500","PVP01_0505600","PVP01_0505700","PVP01_0505800","PVP01_0505900","PVP01_0506000","PVP01_0506100","PVP01_0506200","PVP01_0506300","PVP01_0506400","PVP01_0506500","PVP01_0506600","PVP01_0506700","PVP01_0506800","PVP01_0506900","PVP01_0507000","PVP01_0507100","PVP01_0507200","PVP01_0507300","PVP01_0507400","PVP01_0507500","PVP01_0507600","PVP01_0507700","PVP01_0507800","PVP01_0507900","PVP01_0508000","PVP01_0508100","PVP01_0508200","PVP01_0508300","PVP01_0508400","PVP01_0508500","PVP01_0508600","PVP01_0508700","PVP01_0508800","PVP01_0508900","PVP01_0509000","PVP01_0509100","PVP01_0509200","PVP01_0509300","PVP01_0509400","PVP01_0509500","PVP01_0509600","PVP01_0509700","PVP01_0509800","PVP01_0509900","PVP01_0510000","PVP01_0510100","PVP01_0510200","PVP01_0510300","PVP01_0510400","PVP01_0510500","PVP01_0510600","PVP01_0510700","PVP01_0510800","PVP01_0510900","PVP01_0511000","PVP01_0511100","PVP01_0511200","PVP01_0511300","PVP01_0511500","PVP01_0511600","PVP01_0511700","PVP01_0511800","PVP01_0511900","PVP01_0512000","PVP01_0512100","PVP01_0512200","PVP01_0512300","PVP01_0512400","PVP01_0512500","PVP01_0512600","PVP01_0512700","PVP01_0512800","PVP01_0512900","PVP01_0513000","PVP01_0513100","PVP01_0513200","PVP01_0513300","PVP01_0513400","PVP01_0513500","PVP01_0513600","PVP01_0513700","PVP01_0513800","PVP01_0513900","PVP01_0514000","PVP01_0514100","PVP01_0514200","PVP01_0514300","PVP01_0514400","PVP01_0514500","PVP01_0514600","PVP01_0514700","PVP01_0514800","PVP01_0514900","PVP01_0515000","PVP01_0515100","PVP01_0515200","PVP01_0515300","PVP01_0515400","PVP01_0515500","PVP01_0515600","PVP01_0515700","PVP01_0515800","PVP01_0515900","PVP01_0516000","PVP01_0516100","PVP01_0516200","PVP01_0516300","PVP01_0516400","PVP01_0516500","PVP01_0516600","PVP01_0516700","PVP01_0516800","PVP01_0516900","PVP01_0517000","PVP01_0517100","PVP01_0517200","PVP01_0517300","PVP01_0517400","PVP01_0517500","PVP01_0517700","PVP01_0517800","PVP01_0517900","PVP01_0518000","PVP01_0518100","PVP01_0518200","PVP01_0518300","PVP01_0518400","PVP01_0518500","PVP01_0518600","PVP01_0518700","PVP01_0518800","PVP01_0518900","PVP01_0519000","PVP01_0519100","PVP01_0519200","PVP01_0519300","PVP01_0519400","PVP01_0519500","PVP01_0519600","PVP01_0519700","PVP01_0519800","PVP01_0519900","PVP01_0520000","PVP01_0520100","PVP01_0520200","PVP01_0520300","PVP01_0520400","PVP01_0520500","PVP01_0520600","PVP01_0520700","PVP01_0520800","PVP01_0520900","PVP01_0521000","PVP01_0521100","PVP01_0521200","PVP01_0521300","PVP01_0521400","PVP01_0521500","PVP01_0521600","PVP01_0521700","PVP01_0521800","PVP01_0521900","PVP01_0522000","PVP01_0522100","PVP01_0522200","PVP01_0522300","PVP01_0522400","PVP01_0522500","PVP01_0522600","PVP01_0522700","PVP01_0522800","PVP01_0522900","PVP01_0523000","PVP01_0523100","PVP01_0523200","PVP01_0523300","PVP01_0523400","PVP01_0523500","PVP01_0523600","PVP01_0523700","PVP01_0523800","PVP01_0523900","PVP01_0524000","PVP01_0524100","PVP01_0524200","PVP01_0524300","PVP01_0524500","PVP01_0524600","PVP01_0524700","PVP01_0524900","PVP01_0525000","PVP01_0525100","PVP01_0525200","PVP01_0525300","PVP01_0525400","PVP01_0525500","PVP01_0525600","PVP01_0525700","PVP01_0525800","PVP01_0525900","PVP01_0526000","PVP01_0526100","PVP01_0526200","PVP01_0526300","PVP01_0526400","PVP01_0526500","PVP01_0526600","PVP01_0526700","PVP01_0526800","PVP01_0526900","PVP01_0527000","PVP01_0527100","PVP01_0527200","PVP01_0527300","PVP01_0527400","PVP01_0527500","PVP01_0527600","PVP01_0527700","PVP01_0527800","PVP01_0527900","PVP01_0528000","PVP01_0528100","PVP01_0528200","PVP01_0528300","PVP01_0528400","PVP01_0528500","PVP01_0528600","PVP01_0528700","PVP01_0528800","PVP01_0528900","PVP01_0529000","PVP01_0529100","PVP01_0529200","PVP01_0529300","PVP01_0529400","PVP01_0529500","PVP01_0529600","PVP01_0529700","PVP01_0529800","PVP01_0529900","PVP01_0530000","PVP01_0530100","PVP01_0530200","PVP01_0530300","PVP01_0530400","PVP01_0530500","PVP01_0530600","PVP01_0530700","PVP01_0530800","PVP01_0530900","PVP01_0531000","PVP01_0531100","PVP01_0531200","PVP01_0531300","PVP01_0531400","PVP01_0531500","PVP01_0531600","PVP01_0531700","PVP01_0531800","PVP01_0531900","PVP01_0532100","PVP01_0532200","PVP01_0532300","PVP01_0532400","PVP01_0532500","PVP01_0532600","PVP01_0532700","PVP01_0532800","PVP01_0532900","PVP01_0533000","PVP01_0533100","PVP01_0533200","PVP01_0533300","PVP01_0533700","PVP01_0534000","PVP01_0534100","PVP01_0534200","PVP01_0534400","PVP01_0534600","PVP01_0534800","PVP01_0534900","PVP01_0535400","PVP01_0600100","PVP01_0600300","PVP01_0600400","PVP01_0600500","PVP01_0600600","PVP01_0600700","PVP01_0600800","PVP01_0600900","PVP01_0601000","PVP01_0601100","PVP01_0601200","PVP01_0601300","PVP01_0601400","PVP01_0601500","PVP01_0601600","PVP01_0601700","PVP01_0601800","PVP01_0601900","PVP01_0602000","PVP01_0602200","PVP01_0602300","PVP01_0602400","PVP01_0602500","PVP01_0602600","PVP01_0602700","PVP01_0602800","PVP01_0602900","PVP01_0603000","PVP01_0603100","PVP01_0603200","PVP01_0603300","PVP01_0603400","PVP01_0603500","PVP01_0603600","PVP01_0603700","PVP01_0603800","PVP01_0603900","PVP01_0604000","PVP01_0604100","PVP01_0604200","PVP01_0604300","PVP01_0604400","PVP01_0604500","PVP01_0604600","PVP01_0604700","PVP01_0604800","PVP01_0604900","PVP01_0605000","PVP01_0605100","PVP01_0605200","PVP01_0605300","PVP01_0605400","PVP01_0605500","PVP01_0605600","PVP01_0605700","PVP01_0605800","PVP01_0605900","PVP01_0606000","PVP01_0606100","PVP01_0606200","PVP01_0606300","PVP01_0606400","PVP01_0606500","PVP01_0606600","PVP01_0606700","PVP01_0606800","PVP01_0606900","PVP01_0607000","PVP01_0607100","PVP01_0607200","PVP01_0607300","PVP01_0607400","PVP01_0607500","PVP01_0607600","PVP01_0607700","PVP01_0607800","PVP01_0607900","PVP01_0608000","PVP01_0608100","PVP01_0608200","PVP01_0608300","PVP01_0608400","PVP01_0608500","PVP01_0608600","PVP01_0608700","PVP01_0608800","PVP01_0608900","PVP01_0609000","PVP01_0609200","PVP01_0609300","PVP01_0609400","PVP01_0609500","PVP01_0609600","PVP01_0609700","PVP01_0609800","PVP01_0609900","PVP01_0610000","PVP01_0610100","PVP01_0610200","PVP01_0610300","PVP01_0610400","PVP01_0610500","PVP01_0610600","PVP01_0610700","PVP01_0610800","PVP01_0610900","PVP01_0611000","PVP01_0611100","PVP01_0611200","PVP01_0611300","PVP01_0611400","PVP01_0611500","PVP01_0611600","PVP01_0611700","PVP01_0611800","PVP01_0611900","PVP01_0612000","PVP01_0612100","PVP01_0612200","PVP01_0612300","PVP01_0612500","PVP01_0612600","PVP01_0612700","PVP01_0612800","PVP01_0612900","PVP01_0613000","PVP01_0613100","PVP01_0613200","PVP01_0613300","PVP01_0613400","PVP01_0613500","PVP01_0613600","PVP01_0613700","PVP01_0613800","PVP01_0613900","PVP01_0614000","PVP01_0614100","PVP01_0614200","PVP01_0614300","PVP01_0614400","PVP01_0614500","PVP01_0614600","PVP01_0614700","PVP01_0614800","PVP01_0614900","PVP01_0615100","PVP01_0615200","PVP01_0615300","PVP01_0615400","PVP01_0615500","PVP01_0615600","PVP01_0615700","PVP01_0615800","PVP01_0615900","PVP01_0616000","PVP01_0616100","PVP01_0616200","PVP01_0616300","PVP01_0616400","PVP01_0616500","PVP01_0616600","PVP01_0616700","PVP01_0616800","PVP01_0616900","PVP01_0617000","PVP01_0617100","PVP01_0617200","PVP01_0617300","PVP01_0617400","PVP01_0617500","PVP01_0617600","PVP01_0617700","PVP01_0617800","PVP01_0617900","PVP01_0618000","PVP01_0618100","PVP01_0618200","PVP01_0618300","PVP01_0618400","PVP01_0618500","PVP01_0618600","PVP01_0618700","PVP01_0618800","PVP01_0618900","PVP01_0619000","PVP01_0619100","PVP01_0619200","PVP01_0619300","PVP01_0619400","PVP01_0619500","PVP01_0619600","PVP01_0619700","PVP01_0619800","PVP01_0619900","PVP01_0620000","PVP01_0620100","PVP01_0620200","PVP01_0620300","PVP01_0620400","PVP01_0620500","PVP01_0620600","PVP01_0620700","PVP01_0620900","PVP01_0621000","PVP01_0621100","PVP01_0621200","PVP01_0621300","PVP01_0621400","PVP01_0621500","PVP01_0621600","PVP01_0621700","PVP01_0621800","PVP01_0621900","PVP01_0622000","PVP01_0622100","PVP01_0622200","PVP01_0622300","PVP01_0622400","PVP01_0622500","PVP01_0622600","PVP01_0622700","PVP01_0623000","PVP01_0623100","PVP01_0623200","PVP01_0623300","PVP01_0623400","PVP01_0623500","PVP01_0623600","PVP01_0623700","PVP01_0623800","PVP01_0623900","PVP01_0624000","PVP01_0624100","PVP01_0624300","PVP01_0624400","PVP01_0624500","PVP01_0700100","PVP01_0700200","PVP01_0700300","PVP01_0700400","PVP01_0700500","PVP01_0700600","PVP01_0700700","PVP01_0700800","PVP01_0700900","PVP01_0701000","PVP01_0701100","PVP01_0701200","PVP01_0701300","PVP01_0701400","PVP01_0701500","PVP01_0701600","PVP01_0701700","PVP01_0701800","PVP01_0701900","PVP01_0702000","PVP01_0702100","PVP01_0702200","PVP01_0702300","PVP01_0702400","PVP01_0702500","PVP01_0702600","PVP01_0702700","PVP01_0702800","PVP01_0702900","PVP01_0703000","PVP01_0703100","PVP01_0703200","PVP01_0703300","PVP01_0703400","PVP01_0703500","PVP01_0703600","PVP01_0703700","PVP01_0703900","PVP01_0704000","PVP01_0704100","PVP01_0704200","PVP01_0704300","PVP01_0704400","PVP01_0704500","PVP01_0704600","PVP01_0704700","PVP01_0704800","PVP01_0704900","PVP01_0705000","PVP01_0705100","PVP01_0705200","PVP01_0705300","PVP01_0705400","PVP01_0705500","PVP01_0705600","PVP01_0705700","PVP01_0705800","PVP01_0706000","PVP01_0706100","PVP01_0706200","PVP01_0706300","PVP01_0706400","PVP01_0706500","PVP01_0706600","PVP01_0706700","PVP01_0706800","PVP01_0706900","PVP01_0707000","PVP01_0707100","PVP01_0707200","PVP01_0707300","PVP01_0707400","PVP01_0707500","PVP01_0707600","PVP01_0707700","PVP01_0707800","PVP01_0707900","PVP01_0708000","PVP01_0708100","PVP01_0708200","PVP01_0708300","PVP01_0708400","PVP01_0708500","PVP01_0708600","PVP01_0708700","PVP01_0708800","PVP01_0708900","PVP01_0709100","PVP01_0709200","PVP01_0709300","PVP01_0709400","PVP01_0709500","PVP01_0709600","PVP01_0709700","PVP01_0709800","PVP01_0709900","PVP01_0710000","PVP01_0710100","PVP01_0710200","PVP01_0710300","PVP01_0710400","PVP01_0710500","PVP01_0710600","PVP01_0710700","PVP01_0710800","PVP01_0710900","PVP01_0711000","PVP01_0711100","PVP01_0711200","PVP01_0711300","PVP01_0711400","PVP01_0711500","PVP01_0711600","PVP01_0711700","PVP01_0711800","PVP01_0711900","PVP01_0712000","PVP01_0712100","PVP01_0712200","PVP01_0712300","PVP01_0712400","PVP01_0712500","PVP01_0712600","PVP01_0712700","PVP01_0712800","PVP01_0712900","PVP01_0713000","PVP01_0713100","PVP01_0713200","PVP01_0713300","PVP01_0713400","PVP01_0713500","PVP01_0713600","PVP01_0713700","PVP01_0713800","PVP01_0713900","PVP01_0714000","PVP01_0714100","PVP01_0714200","PVP01_0714300","PVP01_0714400","PVP01_0714500","PVP01_0714600","PVP01_0714700","PVP01_0714800","PVP01_0714900","PVP01_0715000","PVP01_0715100","PVP01_0715200","PVP01_0715300","PVP01_0715400","PVP01_0715500","PVP01_0715600","PVP01_0715700","PVP01_0715800","PVP01_0715900","PVP01_0716000","PVP01_0716100","PVP01_0716200","PVP01_0716300","PVP01_0716400","PVP01_0716500","PVP01_0716600","PVP01_0716700","PVP01_0716800","PVP01_0717000","PVP01_0717100","PVP01_0717200","PVP01_0717300","PVP01_0717400","PVP01_0717500","PVP01_0717600","PVP01_0717700","PVP01_0717800","PVP01_0717900","PVP01_0718000","PVP01_0718100","PVP01_0718200","PVP01_0718300","PVP01_0718400","PVP01_0718500","PVP01_0718600","PVP01_0718700","PVP01_0718800","PVP01_0718900","PVP01_0719000","PVP01_0719100","PVP01_0719200","PVP01_0719300","PVP01_0719400","PVP01_0719500","PVP01_0719600","PVP01_0719700","PVP01_0719800","PVP01_0719900","PVP01_0720000","PVP01_0720100","PVP01_0720200","PVP01_0720300","PVP01_0720400","PVP01_0720500","PVP01_0720600","PVP01_0720700","PVP01_0720800","PVP01_0720900","PVP01_0721000","PVP01_0721100","PVP01_0721200","PVP01_0721300","PVP01_0721400","PVP01_0721500","PVP01_0721600","PVP01_0721700","PVP01_0721800","PVP01_0721900","PVP01_0722000","PVP01_0722100","PVP01_0722200","PVP01_0722300","PVP01_0722400","PVP01_0722500","PVP01_0722600","PVP01_0722700","PVP01_0722800","PVP01_0722900","PVP01_0723000","PVP01_0723200","PVP01_0723300","PVP01_0723400","PVP01_0723500","PVP01_0723700","PVP01_0723800","PVP01_0723900","PVP01_0724000","PVP01_0724100","PVP01_0724200","PVP01_0724300","PVP01_0724400","PVP01_0724500","PVP01_0724600","PVP01_0724700","PVP01_0724800","PVP01_0724900","PVP01_0725000","PVP01_0725100","PVP01_0725200","PVP01_0725300","PVP01_0725400","PVP01_0725500","PVP01_0725600","PVP01_0725700","PVP01_0725800","PVP01_0725900","PVP01_0726000","PVP01_0726100","PVP01_0726200","PVP01_0726300","PVP01_0726400","PVP01_0726500","PVP01_0726600","PVP01_0726700","PVP01_0726800","PVP01_0726900","PVP01_0727000","PVP01_0727100","PVP01_0727200","PVP01_0727300","PVP01_0727400","PVP01_0727500","PVP01_0727600","PVP01_0727700","PVP01_0727800","PVP01_0728000","PVP01_0728100","PVP01_0728200","PVP01_0728300","PVP01_0728400","PVP01_0728500","PVP01_0728600","PVP01_0728700","PVP01_0728800","PVP01_0729000","PVP01_0729100","PVP01_0729200","PVP01_0729300","PVP01_0729400","PVP01_0729500","PVP01_0729600","PVP01_0729700","PVP01_0729800","PVP01_0729900","PVP01_0730000","PVP01_0730100","PVP01_0730200","PVP01_0730300","PVP01_0730400","PVP01_0730500","PVP01_0730600","PVP01_0730700","PVP01_0730900","PVP01_0731000","PVP01_0731100","PVP01_0731200","PVP01_0731300","PVP01_0731400","PVP01_0731500","PVP01_0731600","PVP01_0731700","PVP01_0731800","PVP01_0731900","PVP01_0732000","PVP01_0732100","PVP01_0732200","PVP01_0732300","PVP01_0732400","PVP01_0732500","PVP01_0732600","PVP01_0732700","PVP01_0732800","PVP01_0732900","PVP01_0733000","PVP01_0733100","PVP01_0733200","PVP01_0733300","PVP01_0733400","PVP01_0733500","PVP01_0733600","PVP01_0733700","PVP01_0733800","PVP01_0733900","PVP01_0734000","PVP01_0734100","PVP01_0734200","PVP01_0734400","PVP01_0734600","PVP01_0734700","PVP01_0734800","PVP01_0734900","PVP01_0735100","PVP01_0735200","PVP01_0735300","PVP01_0735800","PVP01_0735900","PVP01_0800100","PVP01_0800200","PVP01_0800300","PVP01_0800400","PVP01_0800500","PVP01_0800600","PVP01_0800700","PVP01_0800900","PVP01_0801000","PVP01_0801100","PVP01_0801200","PVP01_0801300","PVP01_0801400","PVP01_0801500","PVP01_0801700","PVP01_0801800","PVP01_0802100","PVP01_0802200","PVP01_0802300","PVP01_0802400","PVP01_0802500","PVP01_0802600","PVP01_0802700","PVP01_0802800","PVP01_0802900","PVP01_0803000","PVP01_0803100","PVP01_0803200","PVP01_0803300","PVP01_0803400","PVP01_0803500","PVP01_0803600","PVP01_0803700","PVP01_0803800","PVP01_0803900","PVP01_0804000","PVP01_0804100","PVP01_0804200","PVP01_0804300","PVP01_0804400","PVP01_0804500","PVP01_0804600","PVP01_0804700","PVP01_0804800","PVP01_0804900","PVP01_0805000","PVP01_0805100","PVP01_0805200","PVP01_0805300","PVP01_0805400","PVP01_0805500","PVP01_0805600","PVP01_0805700","PVP01_0805800","PVP01_0805900","PVP01_0806000","PVP01_0806100","PVP01_0806200","PVP01_0806300","PVP01_0806400","PVP01_0806500","PVP01_0806600","PVP01_0806700","PVP01_0806800","PVP01_0806900","PVP01_0807000","PVP01_0807100","PVP01_0807200","PVP01_0807300","PVP01_0807400","PVP01_0807500","PVP01_0807600","PVP01_0807800","PVP01_0807900","PVP01_0808000","PVP01_0808100","PVP01_0808200","PVP01_0808300","PVP01_0808400","PVP01_0808500","PVP01_0808600","PVP01_0808700","PVP01_0808800","PVP01_0808900","PVP01_0809000","PVP01_0809100","PVP01_0809200","PVP01_0809300","PVP01_0809400","PVP01_0809500","PVP01_0809600","PVP01_0809700","PVP01_0809800","PVP01_0809900","PVP01_0810000","PVP01_0810100","PVP01_0810200","PVP01_0810300","PVP01_0810400","PVP01_0810500","PVP01_0810600","PVP01_0810700","PVP01_0810800","PVP01_0810900","PVP01_0811000","PVP01_0811100","PVP01_0811200","PVP01_0811300","PVP01_0811400","PVP01_0811500","PVP01_0811600","PVP01_0811700","PVP01_0811800","PVP01_0811900","PVP01_0812000","PVP01_0812100","PVP01_0812200","PVP01_0812400","PVP01_0812500","PVP01_0812600","PVP01_0812700","PVP01_0812800","PVP01_0812900","PVP01_0813000","PVP01_0813100","PVP01_0813200","PVP01_0813300","PVP01_0813400","PVP01_0813500","PVP01_0813600","PVP01_0813700","PVP01_0813800","PVP01_0814000","PVP01_0814100","PVP01_0814200","PVP01_0814300","PVP01_0814400","PVP01_0814500","PVP01_0814600","PVP01_0814700","PVP01_0814800","PVP01_0814900","PVP01_0815100","PVP01_0815200","PVP01_0815300","PVP01_0815400","PVP01_0815500","PVP01_0815600","PVP01_0815700","PVP01_0815800","PVP01_0815900","PVP01_0816000","PVP01_0816100","PVP01_0816200","PVP01_0816300","PVP01_0816400","PVP01_0816500","PVP01_0816600","PVP01_0816700","PVP01_0816900","PVP01_0817000","PVP01_0817100","PVP01_0817200","PVP01_0817300","PVP01_0817400","PVP01_0817500","PVP01_0817600","PVP01_0817700","PVP01_0817800","PVP01_0817900","PVP01_0818000","PVP01_0818100","PVP01_0818200","PVP01_0818300","PVP01_0818400","PVP01_0818500","PVP01_0818600","PVP01_0818700","PVP01_0818800","PVP01_0818900","PVP01_0819000","PVP01_0819100","PVP01_0819200","PVP01_0819400","PVP01_0819500","PVP01_0819600","PVP01_0819700","PVP01_0819800","PVP01_0819900","PVP01_0820000","PVP01_0820100","PVP01_0820200","PVP01_0820300","PVP01_0820400","PVP01_0820500","PVP01_0820600","PVP01_0820700","PVP01_0820800","PVP01_0820900","PVP01_0821000","PVP01_0821100","PVP01_0821200","PVP01_0821300","PVP01_0821400","PVP01_0821500","PVP01_0821600","PVP01_0821700","PVP01_0821800","PVP01_0821900","PVP01_0822000","PVP01_0822100","PVP01_0822200","PVP01_0822300","PVP01_0822400","PVP01_0822500","PVP01_0822600","PVP01_0822700","PVP01_0822800","PVP01_0822900","PVP01_0823000","PVP01_0823100","PVP01_0823200","PVP01_0823300","PVP01_0823400","PVP01_0823500","PVP01_0823600","PVP01_0823700","PVP01_0823800","PVP01_0823900","PVP01_0824000","PVP01_0824100","PVP01_0824200","PVP01_0824300","PVP01_0824400","PVP01_0824500","PVP01_0824600","PVP01_0824700","PVP01_0824800","PVP01_0824900","PVP01_0825000","PVP01_0825100","PVP01_0825200","PVP01_0825300","PVP01_0825400","PVP01_0825500","PVP01_0825600","PVP01_0825700","PVP01_0825800","PVP01_0825900","PVP01_0826000","PVP01_0826100","PVP01_0826200","PVP01_0826300","PVP01_0826400","PVP01_0826500","PVP01_0826600","PVP01_0826700","PVP01_0826800","PVP01_0826900","PVP01_0827000","PVP01_0827100","PVP01_0827200","PVP01_0827300","PVP01_0827400","PVP01_0827700","PVP01_0827800","PVP01_0827900","PVP01_0828000","PVP01_0828100","PVP01_0828200","PVP01_0828300","PVP01_0828400","PVP01_0828500","PVP01_0828600","PVP01_0828700","PVP01_0828800","PVP01_0828900","PVP01_0829000","PVP01_0829100","PVP01_0829200","PVP01_0829300","PVP01_0829400","PVP01_0829500","PVP01_0829600","PVP01_0829700","PVP01_0829800","PVP01_0829900","PVP01_0830000","PVP01_0830100","PVP01_0830200","PVP01_0830300","PVP01_0830400","PVP01_0830500","PVP01_0830600","PVP01_0830700","PVP01_0830900","PVP01_0831000","PVP01_0831100","PVP01_0831200","PVP01_0831300","PVP01_0831400","PVP01_0831500","PVP01_0831600","PVP01_0831700","PVP01_0831900","PVP01_0832000","PVP01_0832100","PVP01_0832200","PVP01_0832300","PVP01_0832400","PVP01_0832500","PVP01_0832600","PVP01_0832700","PVP01_0832800","PVP01_0832900","PVP01_0833000","PVP01_0833100","PVP01_0833200","PVP01_0833400","PVP01_0833500","PVP01_0833600","PVP01_0833700","PVP01_0833800","PVP01_0833900","PVP01_0834000","PVP01_0834100","PVP01_0834200","PVP01_0834300","PVP01_0834400","PVP01_0834500","PVP01_0834600","PVP01_0834700","PVP01_0834800","PVP01_0834900","PVP01_0835000","PVP01_0835100","PVP01_0835200","PVP01_0835300","PVP01_0835400","PVP01_0835500","PVP01_0835600","PVP01_0835700","PVP01_0835800","PVP01_0835900","PVP01_0836000","PVP01_0836100","PVP01_0836200","PVP01_0836300","PVP01_0836400","PVP01_0836500","PVP01_0836600","PVP01_0836700","PVP01_0836800","PVP01_0836900","PVP01_0837000","PVP01_0837100","PVP01_0837200","PVP01_0837300","PVP01_0837400","PVP01_0837500","PVP01_0837600","PVP01_0837700","PVP01_0837800","PVP01_0837900","PVP01_0838000","PVP01_0838100","PVP01_0838200","PVP01_0838300","PVP01_0838600","PVP01_0838700","PVP01_0838800","PVP01_0838900","PVP01_0839000","PVP01_0839100","PVP01_0839200","PVP01_0839400","PVP01_0839700","PVP01_0840700","PVP01_0841000","PVP01_0900700","PVP01_0901000","PVP01_0901100","PVP01_0901200","PVP01_0901300","PVP01_0901800","PVP01_0901900","PVP01_0902000","PVP01_0902100","PVP01_0902400","PVP01_0903400","PVP01_0903500","PVP01_0903600","PVP01_0903700","PVP01_0903800","PVP01_0903900","PVP01_0904200","PVP01_0904300","PVP01_0904400","PVP01_0904500","PVP01_0904600","PVP01_0904700","PVP01_0904800","PVP01_0904900","PVP01_0905000","PVP01_0905100","PVP01_0905200","PVP01_0905300","PVP01_0905400","PVP01_0905500","PVP01_0905600","PVP01_0905700","PVP01_0905800","PVP01_0905900","PVP01_0906000","PVP01_0906100","PVP01_0906200","PVP01_0906300","PVP01_0906400","PVP01_0906500","PVP01_0906600","PVP01_0906700","PVP01_0906800","PVP01_0906900","PVP01_0907000","PVP01_0907100","PVP01_0907200","PVP01_0907300","PVP01_0907400","PVP01_0907500","PVP01_0907600","PVP01_0907700","PVP01_0907800","PVP01_0907900","PVP01_0908000","PVP01_0908100","PVP01_0908200","PVP01_0908300","PVP01_0908400","PVP01_0908500","PVP01_0908600","PVP01_0908700","PVP01_0908800","PVP01_0908900","PVP01_0909000","PVP01_0909100","PVP01_0909200","PVP01_0909300","PVP01_0909400","PVP01_0909500","PVP01_0909600","PVP01_0909700","PVP01_0909900","PVP01_0910000","PVP01_0910200","PVP01_0910700","PVP01_0910800","PVP01_0910900","PVP01_0911000","PVP01_0911100","PVP01_0911200","PVP01_0911300","PVP01_0911400","PVP01_0911500","PVP01_0911600","PVP01_0911700","PVP01_0911800","PVP01_0911900","PVP01_0912000","PVP01_0912100","PVP01_0912200","PVP01_0912300","PVP01_0912400","PVP01_0912500","PVP01_0912600","PVP01_0912700","PVP01_0912800","PVP01_0912900","PVP01_0913000","PVP01_0913100","PVP01_0913200","PVP01_0913300","PVP01_0913400","PVP01_0913500","PVP01_0913600","PVP01_0913700","PVP01_0913900","PVP01_0914000","PVP01_0914100","PVP01_0914200","PVP01_0914300","PVP01_0914400","PVP01_0914500","PVP01_0914600","PVP01_0914700","PVP01_0914800","PVP01_0914900","PVP01_0915000","PVP01_0915100","PVP01_0915200","PVP01_0915300","PVP01_0915400","PVP01_0915500","PVP01_0915600","PVP01_0915700","PVP01_0915800","PVP01_0915900","PVP01_0916100","PVP01_0916200","PVP01_0916300","PVP01_0916400","PVP01_0916500","PVP01_0916700","PVP01_0916800","PVP01_0916900","PVP01_0917000","PVP01_0917100","PVP01_0917200","PVP01_0917300","PVP01_0917500","PVP01_0917600","PVP01_0917700","PVP01_0917800","PVP01_0917900","PVP01_0918000","PVP01_0918100","PVP01_0918200","PVP01_0918300","PVP01_0918400","PVP01_0918500","PVP01_0918600","PVP01_0918700","PVP01_0918800","PVP01_0918900","PVP01_0919000","PVP01_0919100","PVP01_0919200","PVP01_0919300","PVP01_0919400","PVP01_0919500","PVP01_0919600","PVP01_0919700","PVP01_0919800","PVP01_0919900","PVP01_0920000","PVP01_0920100","PVP01_0920200","PVP01_0920300","PVP01_0920400","PVP01_0920500","PVP01_0920600","PVP01_0920700","PVP01_0920800","PVP01_0920900","PVP01_0921000","PVP01_0921100","PVP01_0921200","PVP01_0921300","PVP01_0921400","PVP01_0921500","PVP01_0921600","PVP01_0921700","PVP01_0921800","PVP01_0921900","PVP01_0922000","PVP01_0922100","PVP01_0922200","PVP01_0922300","PVP01_0922400","PVP01_0922500","PVP01_0922600","PVP01_0922700","PVP01_0922800","PVP01_0922900","PVP01_0923000","PVP01_0923100","PVP01_0923200","PVP01_0923300","PVP01_0923400","PVP01_0923500","PVP01_0923600","PVP01_0923700","PVP01_0923800","PVP01_0923900","PVP01_0924000","PVP01_0924100","PVP01_0924200","PVP01_0924300","PVP01_0924400","PVP01_0924500","PVP01_0924600","PVP01_0924700","PVP01_0924800","PVP01_0924900","PVP01_0925000","PVP01_0925100","PVP01_0925200","PVP01_0925300","PVP01_0925400","PVP01_0925500","PVP01_0925600","PVP01_0925700","PVP01_0925800","PVP01_0925900","PVP01_0926000","PVP01_0926100","PVP01_0926200","PVP01_0926300","PVP01_0926400","PVP01_0926600","PVP01_0926700","PVP01_0926800","PVP01_0926900","PVP01_0927000","PVP01_0927100","PVP01_0927200","PVP01_0927300","PVP01_0927400","PVP01_0927500","PVP01_0927600","PVP01_0927700","PVP01_0927800","PVP01_0927900","PVP01_0928000","PVP01_0928100","PVP01_0928200","PVP01_0928300","PVP01_0928400","PVP01_0928500","PVP01_0928600","PVP01_0928700","PVP01_0928800","PVP01_0928900","PVP01_0929000","PVP01_0929100","PVP01_0929200","PVP01_0929300","PVP01_0929400","PVP01_0929500","PVP01_0929600","PVP01_0929700","PVP01_0929800","PVP01_0929900","PVP01_0930000","PVP01_0930100","PVP01_0930200","PVP01_0930300","PVP01_0930400","PVP01_0930500","PVP01_0930600","PVP01_0930700","PVP01_0930800","PVP01_0931200","PVP01_0931300","PVP01_0931400","PVP01_0931500","PVP01_0931600","PVP01_0931700","PVP01_0931800","PVP01_0932000","PVP01_0932100","PVP01_0932200","PVP01_0932300","PVP01_0932400","PVP01_0932500","PVP01_0932600","PVP01_0932700","PVP01_0932800","PVP01_0932900","PVP01_0933000","PVP01_0933100","PVP01_0933200","PVP01_0933300","PVP01_0933400","PVP01_0933500","PVP01_0933600","PVP01_0933700","PVP01_0933800","PVP01_0933900","PVP01_0934000","PVP01_0934100","PVP01_0934200","PVP01_0934300","PVP01_0934400","PVP01_0934500","PVP01_0934600","PVP01_0934700","PVP01_0934800","PVP01_0934900","PVP01_0935000","PVP01_0935100","PVP01_0935200","PVP01_0935300","PVP01_0935400","PVP01_0935500","PVP01_0935600","PVP01_0935700","PVP01_0935800","PVP01_0936000","PVP01_0936100","PVP01_0936200","PVP01_0936300","PVP01_0936400","PVP01_0936500","PVP01_0936600","PVP01_0936700","PVP01_0936800","PVP01_0936900","PVP01_0937000","PVP01_0937100","PVP01_0937200","PVP01_0937300","PVP01_0937400","PVP01_0937500","PVP01_0937600","PVP01_0937700","PVP01_0937800","PVP01_0938000","PVP01_0938100","PVP01_0938200","PVP01_0938300","PVP01_0938400","PVP01_0938500","PVP01_0938600","PVP01_0938700","PVP01_0938900","PVP01_0939000","PVP01_0939100","PVP01_0939200","PVP01_0939300","PVP01_0939400","PVP01_0939500","PVP01_0939600","PVP01_0939700","PVP01_0939800","PVP01_0939900","PVP01_0940000","PVP01_0940100","PVP01_0940200","PVP01_0940400","PVP01_0940500","PVP01_0940600","PVP01_0940700","PVP01_0940800","PVP01_0940900","PVP01_0941000","PVP01_0941100","PVP01_0941200","PVP01_0941300","PVP01_0941400","PVP01_0941500","PVP01_0941600","PVP01_0941700","PVP01_0941800","PVP01_0941900","PVP01_0942000","PVP01_0942100","PVP01_0942200","PVP01_0942300","PVP01_0942400","PVP01_0942500","PVP01_0942600","PVP01_0942700","PVP01_0942800","PVP01_0942900","PVP01_0943000","PVP01_0943100","PVP01_0943200","PVP01_0943300","PVP01_0943400","PVP01_0943500","PVP01_0943600","PVP01_0943700","PVP01_0943800","PVP01_0943900","PVP01_0944000","PVP01_0944100","PVP01_0944200","PVP01_0944300","PVP01_0944400","PVP01_0944500","PVP01_0944600","PVP01_0944700","PVP01_0944800","PVP01_0944900","PVP01_0945000","PVP01_0945100","PVP01_0945200","PVP01_0945300","PVP01_0945400","PVP01_0945500","PVP01_0945600","PVP01_0945700","PVP01_0945800","PVP01_0946000","PVP01_0946100","PVP01_0946200","PVP01_0946300","PVP01_0946400","PVP01_0946500","PVP01_0946600","PVP01_0946700","PVP01_0946800","PVP01_0946900","PVP01_0947000","PVP01_0947100","PVP01_0947200","PVP01_0947300","PVP01_0947400","PVP01_0947500","PVP01_0947600","PVP01_0947700","PVP01_0947800","PVP01_0947900","PVP01_0948000","PVP01_0948100","PVP01_0948200","PVP01_0948300","PVP01_0948400","PVP01_0948500","PVP01_0948600","PVP01_0948700","PVP01_0948800","PVP01_0948900","PVP01_0949000","PVP01_0949200","PVP01_0949300","PVP01_0949400","PVP01_0949600","PVP01_0950000","PVP01_0950200","PVP01_1000700","PVP01_1001900","PVP01_1002400","PVP01_1002500","PVP01_1002600","PVP01_1002700","PVP01_1002800","PVP01_1003000","PVP01_1003100","PVP01_1003200","PVP01_1003300","PVP01_1003400","PVP01_1003500","PVP01_1003600","PVP01_1003700","PVP01_1003800","PVP01_1003900","PVP01_1004000","PVP01_1004100","PVP01_1004200","PVP01_1004400","PVP01_1004500","PVP01_1004600","PVP01_1004700","PVP01_1004800","PVP01_1004900","PVP01_1005000","PVP01_1005100","PVP01_1005200","PVP01_1005300","PVP01_1005400","PVP01_1005500","PVP01_1005600","PVP01_1005700","PVP01_1005800","PVP01_1005900","PVP01_1006200","PVP01_1006300","PVP01_1006400","PVP01_1006500","PVP01_1006600","PVP01_1006700","PVP01_1006800","PVP01_1006900","PVP01_1007000","PVP01_1007100","PVP01_1007200","PVP01_1007300","PVP01_1007400","PVP01_1007500","PVP01_1007600","PVP01_1007700","PVP01_1007800","PVP01_1007900","PVP01_1008000","PVP01_1008100","PVP01_1008200","PVP01_1008300","PVP01_1008400","PVP01_1008500","PVP01_1008600","PVP01_1008700","PVP01_1008800","PVP01_1008900","PVP01_1009000","PVP01_1009100","PVP01_1009200","PVP01_1009300","PVP01_1009400","PVP01_1009500","PVP01_1009600","PVP01_1009700","PVP01_1009800","PVP01_1009900","PVP01_1010000","PVP01_1010100","PVP01_1010200","PVP01_1010300","PVP01_1010400","PVP01_1010500","PVP01_1010600","PVP01_1010700","PVP01_1010800","PVP01_1010900","PVP01_1011000","PVP01_1011100","PVP01_1011200","PVP01_1011300","PVP01_1011400","PVP01_1011500","PVP01_1011600","PVP01_1011700","PVP01_1011800","PVP01_1011900","PVP01_1012000","PVP01_1012100","PVP01_1012200","PVP01_1012300","PVP01_1012400","PVP01_1012500","PVP01_1012600","PVP01_1012700","PVP01_1012800","PVP01_1012900","PVP01_1013000","PVP01_1013100","PVP01_1013200","PVP01_1013300","PVP01_1013400","PVP01_1013500","PVP01_1013600","PVP01_1013700","PVP01_1013800","PVP01_1013900","PVP01_1014000","PVP01_1014100","PVP01_1014200","PVP01_1014300","PVP01_1014400","PVP01_1014500","PVP01_1014600","PVP01_1014700","PVP01_1014800","PVP01_1014900","PVP01_1015000","PVP01_1015100","PVP01_1015300","PVP01_1015400","PVP01_1015500","PVP01_1015600","PVP01_1015700","PVP01_1015800","PVP01_1015900","PVP01_1016000","PVP01_1016100","PVP01_1016200","PVP01_1016300","PVP01_1016400","PVP01_1016500","PVP01_1016600","PVP01_1016700","PVP01_1016800","PVP01_1016900","PVP01_1017000","PVP01_1017100","PVP01_1017200","PVP01_1017300","PVP01_1017400","PVP01_1017500","PVP01_1017600","PVP01_1017700","PVP01_1017800","PVP01_1017900","PVP01_1018000","PVP01_1018100","PVP01_1018200","PVP01_1018300","PVP01_1018400","PVP01_1018500","PVP01_1018600","PVP01_1018700","PVP01_1018800","PVP01_1018900","PVP01_1019000","PVP01_1019100","PVP01_1019200","PVP01_1019300","PVP01_1019400","PVP01_1019600","PVP01_1019700","PVP01_1019800","PVP01_1019900","PVP01_1020000","PVP01_1020100","PVP01_1020200","PVP01_1020300","PVP01_1020400","PVP01_1020500","PVP01_1020600","PVP01_1020700","PVP01_1020800","PVP01_1020900","PVP01_1021000","PVP01_1021100","PVP01_1021200","PVP01_1021300","PVP01_1021400","PVP01_1021500","PVP01_1021600","PVP01_1021700","PVP01_1021800","PVP01_1021900","PVP01_1022000","PVP01_1022100","PVP01_1022200","PVP01_1022300","PVP01_1022400","PVP01_1022500","PVP01_1022600","PVP01_1022700","PVP01_1022800","PVP01_1022900","PVP01_1023000","PVP01_1023200","PVP01_1023300","PVP01_1023500","PVP01_1023600","PVP01_1023700","PVP01_1023800","PVP01_1023900","PVP01_1024000","PVP01_1024100","PVP01_1024200","PVP01_1024300","PVP01_1024400","PVP01_1024500","PVP01_1024600","PVP01_1024700","PVP01_1024800","PVP01_1024900","PVP01_1025000","PVP01_1025100","PVP01_1025200","PVP01_1025300","PVP01_1025400","PVP01_1025600","PVP01_1025700","PVP01_1025800","PVP01_1025900","PVP01_1026000","PVP01_1026100","PVP01_1026200","PVP01_1026300","PVP01_1026400","PVP01_1026500","PVP01_1026600","PVP01_1026700","PVP01_1026800","PVP01_1026900","PVP01_1027000","PVP01_1027100","PVP01_1027200","PVP01_1027300","PVP01_1027400","PVP01_1027500","PVP01_1027600","PVP01_1027700","PVP01_1027800","PVP01_1027900","PVP01_1028000","PVP01_1028100","PVP01_1028200","PVP01_1028300","PVP01_1028400","PVP01_1028600","PVP01_1028700","PVP01_1028800","PVP01_1028900","PVP01_1029000","PVP01_1029100","PVP01_1029200","PVP01_1029300","PVP01_1029400","PVP01_1029500","PVP01_1029600","PVP01_1029700","PVP01_1029800","PVP01_1029900","PVP01_1030000","PVP01_1030100","PVP01_1030200","PVP01_1030300","PVP01_1030400","PVP01_1030500","PVP01_1030600","PVP01_1030700","PVP01_1030800","PVP01_1030900","PVP01_1031000","PVP01_1031100","PVP01_1031200","PVP01_1031300","PVP01_1031400","PVP01_1031500","PVP01_1031600","PVP01_1031700","PVP01_1031800","PVP01_1031900","PVP01_1032000","PVP01_1032100","PVP01_1032200","PVP01_1032300","PVP01_1032400","PVP01_1032500","PVP01_1032600","PVP01_1032700","PVP01_1032800","PVP01_1032900","PVP01_1033000","PVP01_1033100","PVP01_1033200","PVP01_1033300","PVP01_1033500","PVP01_1033600","PVP01_1033700","PVP01_1033800","PVP01_1033900","PVP01_1034000","PVP01_1034100","PVP01_1034200","PVP01_1034300","PVP01_1034400","PVP01_1034500","PVP01_1035200","PVP01_1100400","PVP01_1100500","PVP01_1101000","PVP01_1101200","PVP01_1101300","PVP01_1101400","PVP01_1101500","PVP01_1101600","PVP01_1101700","PVP01_1102300","PVP01_1102400","PVP01_1102500","PVP01_1102600","PVP01_1102700","PVP01_1102800","PVP01_1102900","PVP01_1103000","PVP01_1103100","PVP01_1103200","PVP01_1103300","PVP01_1103400","PVP01_1103500","PVP01_1103600","PVP01_1103700","PVP01_1103800","PVP01_1103900","PVP01_1104000","PVP01_1104100","PVP01_1104200","PVP01_1104300","PVP01_1104400","PVP01_1104500","PVP01_1104600","PVP01_1104700","PVP01_1104800","PVP01_1104900","PVP01_1105000","PVP01_1105100","PVP01_1105200","PVP01_1105300","PVP01_1105400","PVP01_1105500","PVP01_1105600","PVP01_1105700","PVP01_1105800","PVP01_1105900","PVP01_1106000","PVP01_1106100","PVP01_1106200","PVP01_1106300","PVP01_1106400","PVP01_1106500","PVP01_1106600","PVP01_1106700","PVP01_1106800","PVP01_1106900","PVP01_1107000","PVP01_1107100","PVP01_1107200","PVP01_1107300","PVP01_1107400","PVP01_1107500","PVP01_1107600","PVP01_1107700","PVP01_1107800","PVP01_1107900","PVP01_1108000","PVP01_1108100","PVP01_1108200","PVP01_1108300","PVP01_1108400","PVP01_1108500","PVP01_1108600","PVP01_1108700","PVP01_1108800","PVP01_1108900","PVP01_1109000","PVP01_1109100","PVP01_1109200","PVP01_1109300","PVP01_1109400","PVP01_1109500","PVP01_1109600","PVP01_1109700","PVP01_1109800","PVP01_1109900","PVP01_1110000","PVP01_1110100","PVP01_1110300","PVP01_1110400","PVP01_1110500","PVP01_1110600","PVP01_1110700","PVP01_1110800","PVP01_1110900","PVP01_1111000","PVP01_1111100","PVP01_1111200","PVP01_1111300","PVP01_1111400","PVP01_1111500","PVP01_1111600","PVP01_1111700","PVP01_1111800","PVP01_1111900","PVP01_1112000","PVP01_1112100","PVP01_1112200","PVP01_1112300","PVP01_1112400","PVP01_1112500","PVP01_1112600","PVP01_1112700","PVP01_1112800","PVP01_1112900","PVP01_1113000","PVP01_1113100","PVP01_1113200","PVP01_1113300","PVP01_1113400","PVP01_1113500","PVP01_1113600","PVP01_1113700","PVP01_1113800","PVP01_1113900","PVP01_1114000","PVP01_1114100","PVP01_1114200","PVP01_1114300","PVP01_1114400","PVP01_1114500","PVP01_1114600","PVP01_1114700","PVP01_1114800","PVP01_1114900","PVP01_1115000","PVP01_1115100","PVP01_1115200","PVP01_1115300","PVP01_1115400","PVP01_1115500","PVP01_1115600","PVP01_1115700","PVP01_1115800","PVP01_1115900","PVP01_1116000","PVP01_1116100","PVP01_1116200","PVP01_1116300","PVP01_1116400","PVP01_1116600","PVP01_1116700","PVP01_1116800","PVP01_1116900","PVP01_1117000","PVP01_1117100","PVP01_1117200","PVP01_1117300","PVP01_1117400","PVP01_1117500","PVP01_1117600","PVP01_1117700","PVP01_1117800","PVP01_1117900","PVP01_1118000","PVP01_1118100","PVP01_1118200","PVP01_1118300","PVP01_1118400","PVP01_1118500","PVP01_1118600","PVP01_1118700","PVP01_1118800","PVP01_1118900","PVP01_1119000","PVP01_1119200","PVP01_1119300","PVP01_1119400","PVP01_1119500","PVP01_1119600","PVP01_1119700","PVP01_1119800","PVP01_1119900","PVP01_1120000","PVP01_1120100","PVP01_1120200","PVP01_1120300","PVP01_1120400","PVP01_1120500","PVP01_1120600","PVP01_1120700","PVP01_1120800","PVP01_1120900","PVP01_1121000","PVP01_1121100","PVP01_1121200","PVP01_1121300","PVP01_1121400","PVP01_1121500","PVP01_1121600","PVP01_1121700","PVP01_1121800","PVP01_1121900","PVP01_1122000","PVP01_1122100","PVP01_1122200","PVP01_1122300","PVP01_1122400","PVP01_1122500","PVP01_1122600","PVP01_1122700","PVP01_1122800","PVP01_1122900","PVP01_1123000","PVP01_1123100","PVP01_1123200","PVP01_1123300","PVP01_1123400","PVP01_1123500","PVP01_1123600","PVP01_1123700","PVP01_1123800","PVP01_1123900","PVP01_1124000","PVP01_1124100","PVP01_1124200","PVP01_1124300","PVP01_1124400","PVP01_1124500","PVP01_1124600","PVP01_1124700","PVP01_1124800","PVP01_1124900","PVP01_1125000","PVP01_1125100","PVP01_1125200","PVP01_1125300","PVP01_1125400","PVP01_1125500","PVP01_1125600","PVP01_1125700","PVP01_1125800","PVP01_1125900","PVP01_1126000","PVP01_1126100","PVP01_1126200","PVP01_1126300","PVP01_1126400","PVP01_1126500","PVP01_1126600","PVP01_1126700","PVP01_1126800","PVP01_1126900","PVP01_1127000","PVP01_1127100","PVP01_1127200","PVP01_1127300","PVP01_1127400","PVP01_1127500","PVP01_1127600","PVP01_1127700","PVP01_1127800","PVP01_1128000","PVP01_1128100","PVP01_1128200","PVP01_1128300","PVP01_1128400","PVP01_1128500","PVP01_1128800","PVP01_1128900","PVP01_1129000","PVP01_1129100","PVP01_1129200","PVP01_1129300","PVP01_1129400","PVP01_1129500","PVP01_1129600","PVP01_1129700","PVP01_1129800","PVP01_1129900","PVP01_1130000","PVP01_1130100","PVP01_1130200","PVP01_1130300","PVP01_1130400","PVP01_1130500","PVP01_1130600","PVP01_1130700","PVP01_1130800","PVP01_1130900","PVP01_1131000","PVP01_1131100","PVP01_1131200","PVP01_1131300","PVP01_1131400","PVP01_1131500","PVP01_1131600","PVP01_1131700","PVP01_1131800","PVP01_1131900","PVP01_1132000","PVP01_1132100","PVP01_1132200","PVP01_1132300","PVP01_1132400","PVP01_1132500","PVP01_1132600","PVP01_1132700","PVP01_1132800","PVP01_1132900","PVP01_1133000","PVP01_1133100","PVP01_1133200","PVP01_1133300","PVP01_1133400","PVP01_1133500","PVP01_1133600","PVP01_1133700","PVP01_1133800","PVP01_1133900","PVP01_1134000","PVP01_1134100","PVP01_1134200","PVP01_1134300","PVP01_1134400","PVP01_1134500","PVP01_1134600","PVP01_1134700","PVP01_1134800","PVP01_1134900","PVP01_1135000","PVP01_1135100","PVP01_1135300","PVP01_1135400","PVP01_1135500","PVP01_1135600","PVP01_1135700","PVP01_1135800","PVP01_1135900","PVP01_1136000","PVP01_1136100","PVP01_1136200","PVP01_1136300","PVP01_1136400","PVP01_1136500","PVP01_1136600","PVP01_1136700","PVP01_1136800","PVP01_1136900","PVP01_1137000","PVP01_1137100","PVP01_1137200","PVP01_1137300","PVP01_1137400","PVP01_1137500","PVP01_1137600","PVP01_1137700","PVP01_1137800","PVP01_1137900","PVP01_1138000","PVP01_1138100","PVP01_1138200","PVP01_1138300","PVP01_1138400","PVP01_1138500","PVP01_1138600","PVP01_1138700","PVP01_1138800","PVP01_1138900","PVP01_1139000","PVP01_1139100","PVP01_1139300","PVP01_1139400","PVP01_1139500","PVP01_1139600","PVP01_1139700","PVP01_1139800","PVP01_1139900","PVP01_1140000","PVP01_1140100","PVP01_1140200","PVP01_1140300","PVP01_1140400","PVP01_1140500","PVP01_1140600","PVP01_1140700","PVP01_1140800","PVP01_1140900","PVP01_1141000","PVP01_1141100","PVP01_1141200","PVP01_1141300","PVP01_1141400","PVP01_1141500","PVP01_1141600","PVP01_1141700","PVP01_1141800","PVP01_1141900","PVP01_1142000","PVP01_1142100","PVP01_1142200","PVP01_1142300","PVP01_1142400","PVP01_1142500","PVP01_1142600","PVP01_1142700","PVP01_1142800","PVP01_1142900","PVP01_1143000","PVP01_1143100","PVP01_1143200","PVP01_1143300","PVP01_1143400","PVP01_1143500","PVP01_1143600","PVP01_1143700","PVP01_1143800","PVP01_1143900","PVP01_1144000","PVP01_1144100","PVP01_1144200","PVP01_1144300","PVP01_1144400","PVP01_1144500","PVP01_1144600","PVP01_1144700","PVP01_1144900","PVP01_1145000","PVP01_1145100","PVP01_1145200","PVP01_1145300","PVP01_1145400","PVP01_1145500","PVP01_1145600","PVP01_1145700","PVP01_1145800","PVP01_1145900","PVP01_1146000","PVP01_1146100","PVP01_1146200","PVP01_1146300","PVP01_1146400","PVP01_1146500","PVP01_1146600","PVP01_1146700","PVP01_1146800","PVP01_1146900","PVP01_1147000","PVP01_1147100","PVP01_1147200","PVP01_1147300","PVP01_1147400","PVP01_1147500","PVP01_1147600","PVP01_1147700","PVP01_1147800","PVP01_1147900","PVP01_1148200","PVP01_1148700","PVP01_1148800","PVP01_1149000","PVP01_1149100","PVP01_1149400","PVP01_1149500","PVP01_1200200","PVP01_1200800","PVP01_1201100","PVP01_1201300","PVP01_1201400","PVP01_1201500","PVP01_1201600","PVP01_1201700","PVP01_1201800","PVP01_1201900","PVP01_1202000","PVP01_1202100","PVP01_1202200","PVP01_1202300","PVP01_1202400","PVP01_1202500","PVP01_1202600","PVP01_1202700","PVP01_1202800","PVP01_1202900","PVP01_1203000","PVP01_1203100","PVP01_1203200","PVP01_1203300","PVP01_1203500","PVP01_1203600","PVP01_1203700","PVP01_1203800","PVP01_1203900","PVP01_1204000","PVP01_1204100","PVP01_1204200","PVP01_1204400","PVP01_1204500","PVP01_1204600","PVP01_1204700","PVP01_1204800","PVP01_1204900","PVP01_1205000","PVP01_1205100","PVP01_1205200","PVP01_1205300","PVP01_1205400","PVP01_1205500","PVP01_1205600","PVP01_1205700","PVP01_1205800","PVP01_1205900","PVP01_1206000","PVP01_1206100","PVP01_1206300","PVP01_1206400","PVP01_1206500","PVP01_1206600","PVP01_1206700","PVP01_1206800","PVP01_1206900","PVP01_1207000","PVP01_1207100","PVP01_1207200","PVP01_1207300","PVP01_1207400","PVP01_1207500","PVP01_1207600","PVP01_1207700","PVP01_1207800","PVP01_1207900","PVP01_1208000","PVP01_1208100","PVP01_1208200","PVP01_1208300","PVP01_1208400","PVP01_1208500","PVP01_1208600","PVP01_1208700","PVP01_1208800","PVP01_1208900","PVP01_1209000","PVP01_1209100","PVP01_1209300","PVP01_1209400","PVP01_1209500","PVP01_1209600","PVP01_1209700","PVP01_1209800","PVP01_1209900","PVP01_1210000","PVP01_1210100","PVP01_1210200","PVP01_1210300","PVP01_1210400","PVP01_1210500","PVP01_1210600","PVP01_1210700","PVP01_1210800","PVP01_1210900","PVP01_1211000","PVP01_1211100","PVP01_1211200","PVP01_1211300","PVP01_1211400","PVP01_1211500","PVP01_1211600","PVP01_1211700","PVP01_1211800","PVP01_1211900","PVP01_1212000","PVP01_1212100","PVP01_1212200","PVP01_1212300","PVP01_1212400","PVP01_1212500","PVP01_1212600","PVP01_1212700","PVP01_1212800","PVP01_1212900","PVP01_1213000","PVP01_1213100","PVP01_1213200","PVP01_1213300","PVP01_1213400","PVP01_1213600","PVP01_1213900","PVP01_1214000","PVP01_1214100","PVP01_1214200","PVP01_1214300","PVP01_1214400","PVP01_1214500","PVP01_1214600","PVP01_1214700","PVP01_1214800","PVP01_1214900","PVP01_1215000","PVP01_1215200","PVP01_1215300","PVP01_1215400","PVP01_1215500","PVP01_1215800","PVP01_1215900","PVP01_1216000","PVP01_1216100","PVP01_1216200","PVP01_1216300","PVP01_1216400","PVP01_1216500","PVP01_1216600","PVP01_1216700","PVP01_1216800","PVP01_1216900","PVP01_1217000","PVP01_1217200","PVP01_1217300","PVP01_1217400","PVP01_1217500","PVP01_1217600","PVP01_1217700","PVP01_1217800","PVP01_1217900","PVP01_1218000","PVP01_1218100","PVP01_1218200","PVP01_1218300","PVP01_1218400","PVP01_1218500","PVP01_1218600","PVP01_1218700","PVP01_1218800","PVP01_1218900","PVP01_1219000","PVP01_1219100","PVP01_1219200","PVP01_1219300","PVP01_1219400","PVP01_1219500","PVP01_1219700","PVP01_1219800","PVP01_1219900","PVP01_1220100","PVP01_1220200","PVP01_1220400","PVP01_1220500","PVP01_1220600","PVP01_1220700","PVP01_1220800","PVP01_1220900","PVP01_1221000","PVP01_1221100","PVP01_1221200","PVP01_1221300","PVP01_1221400","PVP01_1221500","PVP01_1221600","PVP01_1221700","PVP01_1221800","PVP01_1221900","PVP01_1222000","PVP01_1222100","PVP01_1222200","PVP01_1222300","PVP01_1222400","PVP01_1222500","PVP01_1222600","PVP01_1222700","PVP01_1222800","PVP01_1222900","PVP01_1223000","PVP01_1223100","PVP01_1223200","PVP01_1223300","PVP01_1223400","PVP01_1223500","PVP01_1223600","PVP01_1223700","PVP01_1223800","PVP01_1223900","PVP01_1224000","PVP01_1224100","PVP01_1224200","PVP01_1224300","PVP01_1224400","PVP01_1224500","PVP01_1224600","PVP01_1224700","PVP01_1224800","PVP01_1224900","PVP01_1225000","PVP01_1225100","PVP01_1225200","PVP01_1225300","PVP01_1225400","PVP01_1225500","PVP01_1225600","PVP01_1225700","PVP01_1225800","PVP01_1225900","PVP01_1226000","PVP01_1226100","PVP01_1226200","PVP01_1226300","PVP01_1226400","PVP01_1226600","PVP01_1226700","PVP01_1226800","PVP01_1226900","PVP01_1227000","PVP01_1227100","PVP01_1227200","PVP01_1227300","PVP01_1227400","PVP01_1227500","PVP01_1227600","PVP01_1227700","PVP01_1227800","PVP01_1227900","PVP01_1228000","PVP01_1228100","PVP01_1228200","PVP01_1228300","PVP01_1228400","PVP01_1228500","PVP01_1228600","PVP01_1228700","PVP01_1228800","PVP01_1228900","PVP01_1229000","PVP01_1229100","PVP01_1229200","PVP01_1229300","PVP01_1229400","PVP01_1229500","PVP01_1229600","PVP01_1229700","PVP01_1229800","PVP01_1229900","PVP01_1230000","PVP01_1230100","PVP01_1230200","PVP01_1230300","PVP01_1230400","PVP01_1230500","PVP01_1230600","PVP01_1230700","PVP01_1230800","PVP01_1231000","PVP01_1231100","PVP01_1231200","PVP01_1231300","PVP01_1231400","PVP01_1231500","PVP01_1231600","PVP01_1231700","PVP01_1231800","PVP01_1231900","PVP01_1232000","PVP01_1232100","PVP01_1232200","PVP01_1232300","PVP01_1232400","PVP01_1232500","PVP01_1232600","PVP01_1232700","PVP01_1232800","PVP01_1232900","PVP01_1233000","PVP01_1233100","PVP01_1233200","PVP01_1233300","PVP01_1233400","PVP01_1233600","PVP01_1233700","PVP01_1233800","PVP01_1233900","PVP01_1234000","PVP01_1234100","PVP01_1234200","PVP01_1234300","PVP01_1234400","PVP01_1234500","PVP01_1234600","PVP01_1234700","PVP01_1234800","PVP01_1234900","PVP01_1235000","PVP01_1235100","PVP01_1235200","PVP01_1235300","PVP01_1235500","PVP01_1235600","PVP01_1235700","PVP01_1235800","PVP01_1235900","PVP01_1236000","PVP01_1236100","PVP01_1236200","PVP01_1236300","PVP01_1236400","PVP01_1236500","PVP01_1236600","PVP01_1236700","PVP01_1236800","PVP01_1236900","PVP01_1237000","PVP01_1237100","PVP01_1237200","PVP01_1237300","PVP01_1237400","PVP01_1237500","PVP01_1237600","PVP01_1237700","PVP01_1237800","PVP01_1237900","PVP01_1238000","PVP01_1238100","PVP01_1238200","PVP01_1238300","PVP01_1238400","PVP01_1238500","PVP01_1238600","PVP01_1238700","PVP01_1238900","PVP01_1239000","PVP01_1239100","PVP01_1239200","PVP01_1239300","PVP01_1239400","PVP01_1239500","PVP01_1239600","PVP01_1239700","PVP01_1239800","PVP01_1239900","PVP01_1240000","PVP01_1240100","PVP01_1240200","PVP01_1240300","PVP01_1240400","PVP01_1240500","PVP01_1240600","PVP01_1240700","PVP01_1240800","PVP01_1240900","PVP01_1241000","PVP01_1241100","PVP01_1241200","PVP01_1241300","PVP01_1241400","PVP01_1241500","PVP01_1241600","PVP01_1241700","PVP01_1241800","PVP01_1241900","PVP01_1242000","PVP01_1242100","PVP01_1242200","PVP01_1242300","PVP01_1242400","PVP01_1242500","PVP01_1242600","PVP01_1242700","PVP01_1242800","PVP01_1243000","PVP01_1243100","PVP01_1243200","PVP01_1243300","PVP01_1243400","PVP01_1243500","PVP01_1243600","PVP01_1243700","PVP01_1243800","PVP01_1243900","PVP01_1244000","PVP01_1244100","PVP01_1244200","PVP01_1244300","PVP01_1244400","PVP01_1244500","PVP01_1244600","PVP01_1244700","PVP01_1244800","PVP01_1244900","PVP01_1245000","PVP01_1245100","PVP01_1245200","PVP01_1245300","PVP01_1245400","PVP01_1245500","PVP01_1245600","PVP01_1245700","PVP01_1245800","PVP01_1245900","PVP01_1246000","PVP01_1246100","PVP01_1246200","PVP01_1246300","PVP01_1246400","PVP01_1246500","PVP01_1246600","PVP01_1246700","PVP01_1246800","PVP01_1246900","PVP01_1247000","PVP01_1247100","PVP01_1247200","PVP01_1247300","PVP01_1247400","PVP01_1247500","PVP01_1247600","PVP01_1247700","PVP01_1247800","PVP01_1247900","PVP01_1248000","PVP01_1248100","PVP01_1248200","PVP01_1248300","PVP01_1248400","PVP01_1248500","PVP01_1248600","PVP01_1248700","PVP01_1248800","PVP01_1248900","PVP01_1249100","PVP01_1249200","PVP01_1249300","PVP01_1249400","PVP01_1249500","PVP01_1249600","PVP01_1249700","PVP01_1249800","PVP01_1249900","PVP01_1250000","PVP01_1250200","PVP01_1250300","PVP01_1250400","PVP01_1250500","PVP01_1250600","PVP01_1250700","PVP01_1250800","PVP01_1250900","PVP01_1251000","PVP01_1251100","PVP01_1251200","PVP01_1251300","PVP01_1251400","PVP01_1251500","PVP01_1251700","PVP01_1251800","PVP01_1251900","PVP01_1252000","PVP01_1252100","PVP01_1252200","PVP01_1252300","PVP01_1252400","PVP01_1252500","PVP01_1252600","PVP01_1252700","PVP01_1252800","PVP01_1252900","PVP01_1253000","PVP01_1253100","PVP01_1253200","PVP01_1253300","PVP01_1253400","PVP01_1253500","PVP01_1253600","PVP01_1253700","PVP01_1253800","PVP01_1253900","PVP01_1254000","PVP01_1254100","PVP01_1254200","PVP01_1254300","PVP01_1254400","PVP01_1254500","PVP01_1254600","PVP01_1254700","PVP01_1254800","PVP01_1254900","PVP01_1255100","PVP01_1255200","PVP01_1255300","PVP01_1255400","PVP01_1255500","PVP01_1255600","PVP01_1255700","PVP01_1255800","PVP01_1255900","PVP01_1256000","PVP01_1256100","PVP01_1256200","PVP01_1256300","PVP01_1256400","PVP01_1256500","PVP01_1256600","PVP01_1256700","PVP01_1256800","PVP01_1256900","PVP01_1257000","PVP01_1257100","PVP01_1257200","PVP01_1257300","PVP01_1257400","PVP01_1257500","PVP01_1257600","PVP01_1257700","PVP01_1257800","PVP01_1257900","PVP01_1258000","PVP01_1258100","PVP01_1258200","PVP01_1258300","PVP01_1258400","PVP01_1258500","PVP01_1258600","PVP01_1258700","PVP01_1258800","PVP01_1258900","PVP01_1259000","PVP01_1259100","PVP01_1259200","PVP01_1259300","PVP01_1259400","PVP01_1259500","PVP01_1259600","PVP01_1259700","PVP01_1259800","PVP01_1259900","PVP01_1260000","PVP01_1260100","PVP01_1260200","PVP01_1260300","PVP01_1260400","PVP01_1260500","PVP01_1260600","PVP01_1260700","PVP01_1260800","PVP01_1260900","PVP01_1261100","PVP01_1261200","PVP01_1261300","PVP01_1261400","PVP01_1261500","PVP01_1261600","PVP01_1261700","PVP01_1261800","PVP01_1261900","PVP01_1262000","PVP01_1262100","PVP01_1262200","PVP01_1262300","PVP01_1262400","PVP01_1262500","PVP01_1262600","PVP01_1262700","PVP01_1262800","PVP01_1262900","PVP01_1263000","PVP01_1263100","PVP01_1263200","PVP01_1263300","PVP01_1263400","PVP01_1263500","PVP01_1263600","PVP01_1263700","PVP01_1263800","PVP01_1263900","PVP01_1264000","PVP01_1264100","PVP01_1264200","PVP01_1264300","PVP01_1264400","PVP01_1264500","PVP01_1264600","PVP01_1264700","PVP01_1264800","PVP01_1264900","PVP01_1265000","PVP01_1265100","PVP01_1265200","PVP01_1265300","PVP01_1265400","PVP01_1265500","PVP01_1265600","PVP01_1265700","PVP01_1265800","PVP01_1265900","PVP01_1266000","PVP01_1266100","PVP01_1266200","PVP01_1266300","PVP01_1266400","PVP01_1266500","PVP01_1266600","PVP01_1266700","PVP01_1266800","PVP01_1266900","PVP01_1267100","PVP01_1267200","PVP01_1267300","PVP01_1267400","PVP01_1267500","PVP01_1267600","PVP01_1267700","PVP01_1267800","PVP01_1267900","PVP01_1268000","PVP01_1268100","PVP01_1268200","PVP01_1268300","PVP01_1268400","PVP01_1268500","PVP01_1268600","PVP01_1268900","PVP01_1269000","PVP01_1269100","PVP01_1269200","PVP01_1269300","PVP01_1269400","PVP01_1269500","PVP01_1269600","PVP01_1269700","PVP01_1269800","PVP01_1269900","PVP01_1270000","PVP01_1270100","PVP01_1270200","PVP01_1270300","PVP01_1270400","PVP01_1270500","PVP01_1270600","PVP01_1270700","PVP01_1270800","PVP01_1270900","PVP01_1271000","PVP01_1271100","PVP01_1271200","PVP01_1271300","PVP01_1271400","PVP01_1271500","PVP01_1271600","PVP01_1271900","PVP01_1272500","PVP01_1272600","PVP01_1273000","PVP01_1274300","PVP01_1300400","PVP01_1300500","PVP01_1300600","PVP01_1300700","PVP01_1300800","PVP01_1300900","PVP01_1301000","PVP01_1301100","PVP01_1301200","PVP01_1301300","PVP01_1301500","PVP01_1301600","PVP01_1301700","PVP01_1301800","PVP01_1301900","PVP01_1302000","PVP01_1302100","PVP01_1302200","PVP01_1302300","PVP01_1302400","PVP01_1302500","PVP01_1302600","PVP01_1302700","PVP01_1302800","PVP01_1302900","PVP01_1303000","PVP01_1303100","PVP01_1303200","PVP01_1303300","PVP01_1303400","PVP01_1303500","PVP01_1303600","PVP01_1303700","PVP01_1303800","PVP01_1303900","PVP01_1304000","PVP01_1304100","PVP01_1304200","PVP01_1304300","PVP01_1304400","PVP01_1304500","PVP01_1304600","PVP01_1304700","PVP01_1304800","PVP01_1304900","PVP01_1305000","PVP01_1305100","PVP01_1305300","PVP01_1305400","PVP01_1305500","PVP01_1305600","PVP01_1305700","PVP01_1305800","PVP01_1305900","PVP01_1306000","PVP01_1306100","PVP01_1306200","PVP01_1306300","PVP01_1306400","PVP01_1306500","PVP01_1306600","PVP01_1306700","PVP01_1306800","PVP01_1306900","PVP01_1307000","PVP01_1307100","PVP01_1307200","PVP01_1307300","PVP01_1307400","PVP01_1307500","PVP01_1307600","PVP01_1307700","PVP01_1307800","PVP01_1307900","PVP01_1308000","PVP01_1308100","PVP01_1308200","PVP01_1308300","PVP01_1308400","PVP01_1308500","PVP01_1308600","PVP01_1308700","PVP01_1308800","PVP01_1308900","PVP01_1309000","PVP01_1309100","PVP01_1309200","PVP01_1309300","PVP01_1309400","PVP01_1309500","PVP01_1309600","PVP01_1309700","PVP01_1309800","PVP01_1309900","PVP01_1310000","PVP01_1310100","PVP01_1310200","PVP01_1310300","PVP01_1310400","PVP01_1310500","PVP01_1310600","PVP01_1310700","PVP01_1310800","PVP01_1310900","PVP01_1311000","PVP01_1311100","PVP01_1311200","PVP01_1311300","PVP01_1311400","PVP01_1311500","PVP01_1311600","PVP01_1311700","PVP01_1311800","PVP01_1311900","PVP01_1312000","PVP01_1312100","PVP01_1312200","PVP01_1312400","PVP01_1312500","PVP01_1312600","PVP01_1312700","PVP01_1312800","PVP01_1312900","PVP01_1313000","PVP01_1313100","PVP01_1313200","PVP01_1313300","PVP01_1313400","PVP01_1313500","PVP01_1313600","PVP01_1313700","PVP01_1313800","PVP01_1313900","PVP01_1314000","PVP01_1314100","PVP01_1314200","PVP01_1314300","PVP01_1314400","PVP01_1314500","PVP01_1314600","PVP01_1314700","PVP01_1314800","PVP01_1314900","PVP01_1315000","PVP01_1315200","PVP01_1315300","PVP01_1315400","PVP01_1315500","PVP01_1315600","PVP01_1315800","PVP01_1315900","PVP01_1316000","PVP01_1316100","PVP01_1316200","PVP01_1316300","PVP01_1316400","PVP01_1316500","PVP01_1316600","PVP01_1316700","PVP01_1316800","PVP01_1316900","PVP01_1317000","PVP01_1317100","PVP01_1317200","PVP01_1317300","PVP01_1317400","PVP01_1317500","PVP01_1317600","PVP01_1317700","PVP01_1317800","PVP01_1317900","PVP01_1318000","PVP01_1318100","PVP01_1318200","PVP01_1318300","PVP01_1318400","PVP01_1318500","PVP01_1318600","PVP01_1318700","PVP01_1318800","PVP01_1318900","PVP01_1319000","PVP01_1319100","PVP01_1319200","PVP01_1319300","PVP01_1319400","PVP01_1319500","PVP01_1319600","PVP01_1319700","PVP01_1319800","PVP01_1319900","PVP01_1320000","PVP01_1320100","PVP01_1320200","PVP01_1320300","PVP01_1320400","PVP01_1320500","PVP01_1320600","PVP01_1320700","PVP01_1320800","PVP01_1320900","PVP01_1321000","PVP01_1321200","PVP01_1321300","PVP01_1321400","PVP01_1321500","PVP01_1321600","PVP01_1321700","PVP01_1321800","PVP01_1321900","PVP01_1322000","PVP01_1322100","PVP01_1322200","PVP01_1322300","PVP01_1322400","PVP01_1322500","PVP01_1322600","PVP01_1322700","PVP01_1322800","PVP01_1322900","PVP01_1323000","PVP01_1323100","PVP01_1323200","PVP01_1323300","PVP01_1323400","PVP01_1323500","PVP01_1323600","PVP01_1323700","PVP01_1323800","PVP01_1323900","PVP01_1324000","PVP01_1324100","PVP01_1324700","PVP01_1324800","PVP01_1325200","PVP01_1325300","PVP01_1325400","PVP01_1325500","PVP01_1325600","PVP01_1325700","PVP01_1325800","PVP01_1325900","PVP01_1326000","PVP01_1326100","PVP01_1326200","PVP01_1326300","PVP01_1326400","PVP01_1326500","PVP01_1326600","PVP01_1326700","PVP01_1326800","PVP01_1326900","PVP01_1327000","PVP01_1327100","PVP01_1327200","PVP01_1327300","PVP01_1327400","PVP01_1327500","PVP01_1327600","PVP01_1327700","PVP01_1327800","PVP01_1327900","PVP01_1328000","PVP01_1328100","PVP01_1328200","PVP01_1328300","PVP01_1328400","PVP01_1328500","PVP01_1328600","PVP01_1328700","PVP01_1328800","PVP01_1328900","PVP01_1329000","PVP01_1329100","PVP01_1329200","PVP01_1329300","PVP01_1329400","PVP01_1329500","PVP01_1329600","PVP01_1329700","PVP01_1329800","PVP01_1329900","PVP01_1330000","PVP01_1330600","PVP01_1330700","PVP01_1330800","PVP01_1330900","PVP01_1331000","PVP01_1331100","PVP01_1331200","PVP01_1331300","PVP01_1331400","PVP01_1331500","PVP01_1331600","PVP01_1331700","PVP01_1331800","PVP01_1331900","PVP01_1332000","PVP01_1332100","PVP01_1332200","PVP01_1332300","PVP01_1332400","PVP01_1332500","PVP01_1332600","PVP01_1332700","PVP01_1332800","PVP01_1332900","PVP01_1333000","PVP01_1333100","PVP01_1333200","PVP01_1333300","PVP01_1333400","PVP01_1333500","PVP01_1333600","PVP01_1333700","PVP01_1333800","PVP01_1333900","PVP01_1334000","PVP01_1334100","PVP01_1334200","PVP01_1334300","PVP01_1334400","PVP01_1334500","PVP01_1334600","PVP01_1334700","PVP01_1334800","PVP01_1334900","PVP01_1335000","PVP01_1335100","PVP01_1335200","PVP01_1335300","PVP01_1335400","PVP01_1335500","PVP01_1335600","PVP01_1335700","PVP01_1335800","PVP01_1335900","PVP01_1336000","PVP01_1336100","PVP01_1336200","PVP01_1336300","PVP01_1336400","PVP01_1336500","PVP01_1336600","PVP01_1336700","PVP01_1336800","PVP01_1336900","PVP01_1337000","PVP01_1337100","PVP01_1337200","PVP01_1337300","PVP01_1337400","PVP01_1337500","PVP01_1337600","PVP01_1337700","PVP01_1337800","PVP01_1337900","PVP01_1338000","PVP01_1338100","PVP01_1338200","PVP01_1338300","PVP01_1338400","PVP01_1338600","PVP01_1338700","PVP01_1338800","PVP01_1338900","PVP01_1339000","PVP01_1339100","PVP01_1339200","PVP01_1339300","PVP01_1339400","PVP01_1339500","PVP01_1339600","PVP01_1339700","PVP01_1339800","PVP01_1339900","PVP01_1340000","PVP01_1340100","PVP01_1340200","PVP01_1340300","PVP01_1340400","PVP01_1340500","PVP01_1340600","PVP01_1340700","PVP01_1340800","PVP01_1340900","PVP01_1341000","PVP01_1341100","PVP01_1341200","PVP01_1341300","PVP01_1341400","PVP01_1341500","PVP01_1341600","PVP01_1341700","PVP01_1341800","PVP01_1341900","PVP01_1342000","PVP01_1342100","PVP01_1342200","PVP01_1342300","PVP01_1342400","PVP01_1342500","PVP01_1342600","PVP01_1342700","PVP01_1342800","PVP01_1342900","PVP01_1343000","PVP01_1343100","PVP01_1343200","PVP01_1343300","PVP01_1343400","PVP01_1343500","PVP01_1343600","PVP01_1343700","PVP01_1343800","PVP01_1343900","PVP01_1344000","PVP01_1344100","PVP01_1344200","PVP01_1344300","PVP01_1344400","PVP01_1344500","PVP01_1344600","PVP01_1344700","PVP01_1344800","PVP01_1344900","PVP01_1345000","PVP01_1345100","PVP01_1345200","PVP01_1345300","PVP01_1345400","PVP01_1345500","PVP01_1345600","PVP01_1345700","PVP01_1345800","PVP01_1345900","PVP01_1346000","PVP01_1346100","PVP01_1346200","PVP01_1346400","PVP01_1346500","PVP01_1346600","PVP01_1346700","PVP01_1346800","PVP01_1346900","PVP01_1347000","PVP01_1347100","PVP01_1347200","PVP01_1347300","PVP01_1347500","PVP01_1347700","PVP01_1347800","PVP01_1400100","PVP01_1400400","PVP01_1400600","PVP01_1400700","PVP01_1400900","PVP01_1401000","PVP01_1401100","PVP01_1401200","PVP01_1401600","PVP01_1401700","PVP01_1401800","PVP01_1401900","PVP01_1402000","PVP01_1402100","PVP01_1402200","PVP01_1402300","PVP01_1402400","PVP01_1402500","PVP01_1402600","PVP01_1402700","PVP01_1402800","PVP01_1402900","PVP01_1403000","PVP01_1403100","PVP01_1403200","PVP01_1403300","PVP01_1403400","PVP01_1403500","PVP01_1403600","PVP01_1403700","PVP01_1403800","PVP01_1403900","PVP01_1404000","PVP01_1404100","PVP01_1404200","PVP01_1404300","PVP01_1404400","PVP01_1404500","PVP01_1404600","PVP01_1404700","PVP01_1404800","PVP01_1404900","PVP01_1405000","PVP01_1405100","PVP01_1405200","PVP01_1405300","PVP01_1405500","PVP01_1405600","PVP01_1405700","PVP01_1405800","PVP01_1405900","PVP01_1406000","PVP01_1406100","PVP01_1406200","PVP01_1406300","PVP01_1406400","PVP01_1406500","PVP01_1406600","PVP01_1406700","PVP01_1406800","PVP01_1406900","PVP01_1407000","PVP01_1407100","PVP01_1407200","PVP01_1407300","PVP01_1407400","PVP01_1407500","PVP01_1407600","PVP01_1407700","PVP01_1407800","PVP01_1407900","PVP01_1408000","PVP01_1408100","PVP01_1408200","PVP01_1408300","PVP01_1408400","PVP01_1408500","PVP01_1408600","PVP01_1408700","PVP01_1408800","PVP01_1408900","PVP01_1409100","PVP01_1409200","PVP01_1409300","PVP01_1409400","PVP01_1409500","PVP01_1409600","PVP01_1409700","PVP01_1409800","PVP01_1409900","PVP01_1410000","PVP01_1410100","PVP01_1410200","PVP01_1410300","PVP01_1410400","PVP01_1410500","PVP01_1410600","PVP01_1410700","PVP01_1410800","PVP01_1410900","PVP01_1411000","PVP01_1411100","PVP01_1411200","PVP01_1411300","PVP01_1411400","PVP01_1411500","PVP01_1411600","PVP01_1411700","PVP01_1411800","PVP01_1411900","PVP01_1412000","PVP01_1412100","PVP01_1412200","PVP01_1412300","PVP01_1412400","PVP01_1412500","PVP01_1412600","PVP01_1412700","PVP01_1412800","PVP01_1412900","PVP01_1413000","PVP01_1413100","PVP01_1413200","PVP01_1413300","PVP01_1413400","PVP01_1413450","PVP01_1413500","PVP01_1413600","PVP01_1413700","PVP01_1413800","PVP01_1413900","PVP01_1414000","PVP01_1414100","PVP01_1414200","PVP01_1414300","PVP01_1414400","PVP01_1414600","PVP01_1414700","PVP01_1414800","PVP01_1414900","PVP01_1415000","PVP01_1415100","PVP01_1415200","PVP01_1415300","PVP01_1415400","PVP01_1415500","PVP01_1415600","PVP01_1415700","PVP01_1415800","PVP01_1415900","PVP01_1416000","PVP01_1416100","PVP01_1416200","PVP01_1416300","PVP01_1416400","PVP01_1416500","PVP01_1416600","PVP01_1416700","PVP01_1416800","PVP01_1416900","PVP01_1417000","PVP01_1417100","PVP01_1417200","PVP01_1417300","PVP01_1417400","PVP01_1417500","PVP01_1417600","PVP01_1417700","PVP01_1417800","PVP01_1417900","PVP01_1418000","PVP01_1418100","PVP01_1418200","PVP01_1418400","PVP01_1418500","PVP01_1418600","PVP01_1418700","PVP01_1418800","PVP01_1418900","PVP01_1419000","PVP01_1419100","PVP01_1419200","PVP01_1419300","PVP01_1419400","PVP01_1419500","PVP01_1419600","PVP01_1419700","PVP01_1419800","PVP01_1419900","PVP01_1420000","PVP01_1420100","PVP01_1420200","PVP01_1420300","PVP01_1420400","PVP01_1420500","PVP01_1420600","PVP01_1420700","PVP01_1420800","PVP01_1420900","PVP01_1421000","PVP01_1421100","PVP01_1421200","PVP01_1421300","PVP01_1421400","PVP01_1421600","PVP01_1421700","PVP01_1421800","PVP01_1421900","PVP01_1422000","PVP01_1422100","PVP01_1422200","PVP01_1422300","PVP01_1422400","PVP01_1422500","PVP01_1422600","PVP01_1422700","PVP01_1422800","PVP01_1422900","PVP01_1423000","PVP01_1423100","PVP01_1423200","PVP01_1423300","PVP01_1423400","PVP01_1423500","PVP01_1423600","PVP01_1423700","PVP01_1424100","PVP01_1424200","PVP01_1424300","PVP01_1424400","PVP01_1424500","PVP01_1424600","PVP01_1424700","PVP01_1424800","PVP01_1424900","PVP01_1425000","PVP01_1425100","PVP01_1425200","PVP01_1425300","PVP01_1425400","PVP01_1425500","PVP01_1425600","PVP01_1425700","PVP01_1425800","PVP01_1425900","PVP01_1426000","PVP01_1426100","PVP01_1426200","PVP01_1426300","PVP01_1426400","PVP01_1426500","PVP01_1426600","PVP01_1426700","PVP01_1426800","PVP01_1426900","PVP01_1427000","PVP01_1427100","PVP01_1427200","PVP01_1427300","PVP01_1427400","PVP01_1427500","PVP01_1427600","PVP01_1427700","PVP01_1427800","PVP01_1427900","PVP01_1428000","PVP01_1428100","PVP01_1428200","PVP01_1428300","PVP01_1428400","PVP01_1428500","PVP01_1428600","PVP01_1428800","PVP01_1428900","PVP01_1429000","PVP01_1429100","PVP01_1429200","PVP01_1429300","PVP01_1429400","PVP01_1429500","PVP01_1429600","PVP01_1429700","PVP01_1429800","PVP01_1429900","PVP01_1430000","PVP01_1430100","PVP01_1430200","PVP01_1430300","PVP01_1430400","PVP01_1430500","PVP01_1430600","PVP01_1430700","PVP01_1430800","PVP01_1430900","PVP01_1431000","PVP01_1431100","PVP01_1431200","PVP01_1431300","PVP01_1431400","PVP01_1431500","PVP01_1431600","PVP01_1431700","PVP01_1431800","PVP01_1432000","PVP01_1432100","PVP01_1432200","PVP01_1432300","PVP01_1432400","PVP01_1432500","PVP01_1432600","PVP01_1432700","PVP01_1432800","PVP01_1432900","PVP01_1433000","PVP01_1433100","PVP01_1433200","PVP01_1433300","PVP01_1433400","PVP01_1433500","PVP01_1433600","PVP01_1433700","PVP01_1433800","PVP01_1433900","PVP01_1434000","PVP01_1434100","PVP01_1434200","PVP01_1434300","PVP01_1434400","PVP01_1434500","PVP01_1434600","PVP01_1434700","PVP01_1434800","PVP01_1434900","PVP01_1435000","PVP01_1435100","PVP01_1435200","PVP01_1435300","PVP01_1435400","PVP01_1435500","PVP01_1435700","PVP01_1435800","PVP01_1435900","PVP01_1436000","PVP01_1436100","PVP01_1436200","PVP01_1436300","PVP01_1436400","PVP01_1436500","PVP01_1436600","PVP01_1436700","PVP01_1436800","PVP01_1436900","PVP01_1437000","PVP01_1437100","PVP01_1437200","PVP01_1437300","PVP01_1437400","PVP01_1437500","PVP01_1437600","PVP01_1437700","PVP01_1437800","PVP01_1437900","PVP01_1438000","PVP01_1438100","PVP01_1438200","PVP01_1438300","PVP01_1438400","PVP01_1438500","PVP01_1438600","PVP01_1438700","PVP01_1438800","PVP01_1438900","PVP01_1439000","PVP01_1439200","PVP01_1439300","PVP01_1439400","PVP01_1439500","PVP01_1439600","PVP01_1439700","PVP01_1439800","PVP01_1439900","PVP01_1440000","PVP01_1440100","PVP01_1440300","PVP01_1440400","PVP01_1440500","PVP01_1440600","PVP01_1440700","PVP01_1440800","PVP01_1440900","PVP01_1441000","PVP01_1441100","PVP01_1441200","PVP01_1441300","PVP01_1441400","PVP01_1441500","PVP01_1441600","PVP01_1441700","PVP01_1441800","PVP01_1441900","PVP01_1442000","PVP01_1442100","PVP01_1442200","PVP01_1442300","PVP01_1442400","PVP01_1442500","PVP01_1442600","PVP01_1442700","PVP01_1442800","PVP01_1442900","PVP01_1443000","PVP01_1443100","PVP01_1443200","PVP01_1443300","PVP01_1443400","PVP01_1443500","PVP01_1443600","PVP01_1443700","PVP01_1443800","PVP01_1443900","PVP01_1444000","PVP01_1444100","PVP01_1444200","PVP01_1444300","PVP01_1444400","PVP01_1444500","PVP01_1444600","PVP01_1444700","PVP01_1444800","PVP01_1444900","PVP01_1445000","PVP01_1445100","PVP01_1445200","PVP01_1445300","PVP01_1445400","PVP01_1445500","PVP01_1445600","PVP01_1445700","PVP01_1445800","PVP01_1445900","PVP01_1446000","PVP01_1446100","PVP01_1446200","PVP01_1446300","PVP01_1446400","PVP01_1446500","PVP01_1446600","PVP01_1446700","PVP01_1446800","PVP01_1446900","PVP01_1447000","PVP01_1447100","PVP01_1447200","PVP01_1447300","PVP01_1447400","PVP01_1447500","PVP01_1447600","PVP01_1447700","PVP01_1447800","PVP01_1447900","PVP01_1448000","PVP01_1448100","PVP01_1448200","PVP01_1448300","PVP01_1448400","PVP01_1448500","PVP01_1448600","PVP01_1448700","PVP01_1448800","PVP01_1448900","PVP01_1449000","PVP01_1449100","PVP01_1449200","PVP01_1449300","PVP01_1449400","PVP01_1449600","PVP01_1449700","PVP01_1449800","PVP01_1449900","PVP01_1450000","PVP01_1450100","PVP01_1450200","PVP01_1450300","PVP01_1450400","PVP01_1450500","PVP01_1450600","PVP01_1450700","PVP01_1450800","PVP01_1450900","PVP01_1451000","PVP01_1451100","PVP01_1451200","PVP01_1451300","PVP01_1451400","PVP01_1451500","PVP01_1451600","PVP01_1451700","PVP01_1451800","PVP01_1451900","PVP01_1452000","PVP01_1452100","PVP01_1452200","PVP01_1452300","PVP01_1452400","PVP01_1452500","PVP01_1452600","PVP01_1452700","PVP01_1452800","PVP01_1452900","PVP01_1453000","PVP01_1453100","PVP01_1453200","PVP01_1453300","PVP01_1453400","PVP01_1453500","PVP01_1453600","PVP01_1453700","PVP01_1453800","PVP01_1453900","PVP01_1454000","PVP01_1454100","PVP01_1454200","PVP01_1454300","PVP01_1454400","PVP01_1454500","PVP01_1454600","PVP01_1454800","PVP01_1454900","PVP01_1455000","PVP01_1455100","PVP01_1455200","PVP01_1455300","PVP01_1455400","PVP01_1455500","PVP01_1455600","PVP01_1455700","PVP01_1455800","PVP01_1455900","PVP01_1456000","PVP01_1456100","PVP01_1456200","PVP01_1456400","PVP01_1456500","PVP01_1456600","PVP01_1456700","PVP01_1456800","PVP01_1456900","PVP01_1457000","PVP01_1457100","PVP01_1457200","PVP01_1457300","PVP01_1457400","PVP01_1457500","PVP01_1457600","PVP01_1457700","PVP01_1457800","PVP01_1457900","PVP01_1458000","PVP01_1458100","PVP01_1458200","PVP01_1458300","PVP01_1458400","PVP01_1458500","PVP01_1458600","PVP01_1458700","PVP01_1458800","PVP01_1458900","PVP01_1459000","PVP01_1459100","PVP01_1459200","PVP01_1459300","PVP01_1459400","PVP01_1459500","PVP01_1459600","PVP01_1459700","PVP01_1459800","PVP01_1459900","PVP01_1460000","PVP01_1460100","PVP01_1460200","PVP01_1460300","PVP01_1460400","PVP01_1460500","PVP01_1460600","PVP01_1460700","PVP01_1460800","PVP01_1460900","PVP01_1461000","PVP01_1461100","PVP01_1461200","PVP01_1461300","PVP01_1461500","PVP01_1461600","PVP01_1461700","PVP01_1461800","PVP01_1461900","PVP01_1462000","PVP01_1462100","PVP01_1462200","PVP01_1462300","PVP01_1462400","PVP01_1462500","PVP01_1462600","PVP01_1462700","PVP01_1462800","PVP01_1462900","PVP01_1463000","PVP01_1463100","PVP01_1463200","PVP01_1463300","PVP01_1463400","PVP01_1463500","PVP01_1463600","PVP01_1463700","PVP01_1463800","PVP01_1463900","PVP01_1464000","PVP01_1464100","PVP01_1464200","PVP01_1464300","PVP01_1464400","PVP01_1464500","PVP01_1464600","PVP01_1464700","PVP01_1464800","PVP01_1464900","PVP01_1465000","PVP01_1465100","PVP01_1465200","PVP01_1465300","PVP01_1465400","PVP01_1465500","PVP01_1465600","PVP01_1465700","PVP01_1465800","PVP01_1465900","PVP01_1466000","PVP01_1466100","PVP01_1466200","PVP01_1466300","PVP01_1466400","PVP01_1466500","PVP01_1466600","PVP01_1466700","PVP01_1466800","PVP01_1466900","PVP01_1467000","PVP01_1467100","PVP01_1467200","PVP01_1467300","PVP01_1467400","PVP01_1467500","PVP01_1467600","PVP01_1467700","PVP01_1467800","PVP01_1467900","PVP01_1468000","PVP01_1468100","PVP01_1468200","PVP01_1468300","PVP01_1468400","PVP01_1468500","PVP01_1468600","PVP01_1468700","PVP01_1468800","PVP01_1468900","PVP01_1469300","PVP01_1469400","PVP01_1469600","PVP01_1469700","PVP01_1469800","PVP01_1469900","PVP01_1470000","PVP01_1470100","PVP01_1470200","PVP01_1470300","PVP01_1470400","PVP01_1470500","PVP01_1470600","PVP01_1470700","PVP01_1470800","PVP01_1470900","PVP01_1471100","PVP01_1471300","PVP01_1471400","PVP01_1471500","PVP01_1471600","PVP01_1471700","PVP01_1471800","PVP01_1471900","PVP01_1472100","PVP01_1472200","PVP01_1472400","PVP01_API01200","PVP01_API01400","PVP01_API01700","PVP01_API02600","PVP01_API02800","PVP01_API03500","PVP01_API04100","PVP01_API04200","PVP01_API04300","PVP01_API04600","PVP01_MIT02700","PVP01_MIT03300","PVP01_MIT03400","PVP01_0000100","PVP01_0104300","PVP01_0104700","PVP01_0107500","PVP01_0117000","PVP01_0117600","PVP01_0119200","PVP01_0202300","PVP01_0207400","PVP01_0210500","PVP01_0214400","PVP01_0304300","PVP01_0309100","PVP01_0311400","PVP01_0317900","PVP01_0319500","PVP01_0411300","PVP01_0417300","PVP01_0417600","PVP01_0418700","PVP01_0421000","PVP01_0504000","PVP01_0511400","PVP01_0517600","PVP01_0532000","PVP01_0534300","PVP01_0602100","PVP01_0609100","PVP01_0612400","PVP01_0615000","PVP01_0620800","PVP01_0703800","PVP01_0705900","PVP01_0709000","PVP01_0723100","PVP01_0723600","PVP01_0727900","PVP01_0728900","PVP01_0730800","PVP01_0734300","PVP01_0734500","PVP01_0735000","PVP01_0801600","PVP01_0807700","PVP01_0812300","PVP01_0813900","PVP01_0815000","PVP01_0816800","PVP01_0819300","PVP01_0913800","PVP01_0916000","PVP01_0916600","PVP01_0917400","PVP01_0926500","PVP01_0931900","PVP01_0935900","PVP01_0938800","PVP01_0940300","PVP01_0945900","PVP01_1002900","PVP01_1015200","PVP01_1025500","PVP01_1028500","PVP01_1033400","PVP01_1110200","PVP01_1119100","PVP01_1135200","PVP01_1139200","PVP01_1144800","PVP01_1204300","PVP01_1206200","PVP01_1209200","PVP01_1215100","PVP01_1219600","PVP01_1220000","PVP01_1220300","PVP01_1230900","PVP01_1233500","PVP01_1235400","PVP01_1238800","PVP01_1242900","PVP01_1249000","PVP01_1250100","PVP01_1251600","PVP01_1255000","PVP01_1267000","PVP01_1301400","PVP01_1305200","PVP01_1312300","PVP01_1315100","PVP01_1315700","PVP01_1321100","PVP01_1338500","PVP01_1401300","PVP01_1401400","PVP01_1405400","PVP01_1409000","PVP01_1414500","PVP01_1418300","PVP01_1421500","PVP01_1428700","PVP01_1439100","PVP01_1440200","PVP01_1449500","PVP01_1454700","PVP01_1456300","PVP01_1469200","PVP01_1469500"],"logFC":[-7.461,-1.934,-3.312,-4.6,-3.867,3.735,-7.527,-9.994,-1.475,-6.649,-5.497,-6.912,-3.541,-12.82,-3.026,-8.008,-3.91,5.125e-15,5.125e-15,5.125e-15,5.125e-15,5.125e-15,-4.935,5.125e-15,-5.23,-8.626,5.125e-15,-6.448,5.125e-15,5.125e-15,-3.411,-1.698,5.125e-15,5.125e-15,5.125e-15,5.125e-15,1.03,5.125e-15,-1.459,5.125e-15,0.5561,5.125e-15,5.125e-15,5.125e-15,-0.2773,5.841,5.125e-15,5.125e-15,5.125e-15,5.125e-15,5.125e-15,-5.478,5.125e-15,-5.292,-5.045,-4.909,-10.01,-8.063,3.087,-1.891,-6.463,5.125e-15,0.4095,2.993,5.125e-15,5.125e-15,5.125e-15,-2.609,5.125e-15,5.125e-15,-1.44,5.125e-15,-3.71,-0.9055,-1.465,-3.009,-6.534,-2.435,-4.377,-7.929,-6.816,-6.785,-5.714,-7.626,-0.785,-6.269,-7.369,-6.277,-6.569,-5.259,-6.073,5.125e-15,5.125e-15,-4.066,-4.337,-1.155,-6.543,5.125e-15,5.125e-15,-0.8245,-0.4411,5.125e-15,5.125e-15,-3.28,-8.482,-0.4593,5.125e-15,5.125e-15,-4.496,-4.221,-7.759,5.125e-15,5.125e-15,0.6721,5.125e-15,5.125e-15,5.125e-15,5.125e-15,-7.778,-3.582,5.125e-15,5.125e-15,5.803,5.125e-15,5.125e-15,5.125e-15,-3.042,5.125e-15,-3.551,1.431,-8.367,-0.02539,-1.821,5.125e-15,-1.476,-0.4347,5.125e-15,4.712,5.125e-15,4.869,-0.6901,-0.04365,5.125e-15,6.61,5.125e-15,5.125e-15,5.125e-15,5.125e-15,5.125e-15,5.125e-15,5.125e-15,5.125e-15,5.125e-15,5.125e-15,1.125,0.5866,-5.38,5.125e-15,5.125e-15,5.125e-15,5.125e-15,-5.074,5.125e-15,2.002,5.125e-15,5.125e-15,5.125e-15,-0.584,-0.04233,-5.884,5.125e-15,5.125e-15,-3.537,-4.225,0.7421,5.364,5.125e-15,-0.8823,1.929,5.125e-15,-4.225,-5.065,-11.77,-3.439,-1.988,-3.756,-6.873,2.554,-0.3372,-5.319,-5.822,5.209,-6.038,4.96,3.672,-6.244,5.125e-15,-4.551,-7.441,2.248,-0.2724,-4.876,1.592,3.331,2.179,-7.773,-0.8035,5.125e-15,6.825,-9.242,-7.848,-9.5,-4.905,-0.6145,-6.898,-0.4365,-1.474,-0.04501,0.3792,-0.126,0.1778,1.566,1.501,0.4248,-1.913,-2.401,0.4761,0.6335,0.06302,-0.06753,-2.664,-1.284,-1.964,0.191,-1.887,-0.1427,0.02436,-2.147,0.9791,-1.112,-0.7284,-2.28,1.464,0.04085,1.15,-0.07386,0.274,-0.8497,-2.056,1.151,-0.358,-0.9728,2.431,2.132,-0.2549,0.8239,-1.657,0.7819,-0.919,0.2738,-0.3024,-0.37,1.339,0.9103,1.921,1.176,0.1373,0.9734,0.2664,-0.1439,0.4948,0.1154,-2.583,-0.163,1.501,0.2549,0.2134,-1.389,0.6408,0.4075,-1.501,0.8462,-0.7382,-1.694,-0.6844,-1.742,0.8156,0.6856,0.2217,0.1799,0.641,0.2316,0.9155,0.215,-1.509,0.3764,0.7674,1.123,-1.014,-1.051,-0.5159,-3.802,1.02,0.8095,0.5191,-0.9403,-2.973,-0.6938,0.5139,0.9801,-0.9927,1.648,0.0001807,0.4911,0.06203,-2.575,-1.306,0.269,1.357,-1.884,-0.2572,-0.003172,-0.7724,0.579,1.996,-1.125,1.133,-3.723,-1.907,-0.1073,0.2826,1.411,-1.798,1.909,-1.559,-1.691,0.4544,0.9436,-0.8654,-1.339,1.04,-0.1571,1.142,0.1422,-0.2176,0.404,0.08735,1.808,-0.4589,0.316,-0.6878,1.228,-1.048,-2.308,-1.765,-1.442,-0.03876,-0.2197,-2.525,0.4946,0.714,-1.218,0.3705,0.9862,0.6139,-1.589,-4.334,0.2285,2.019,1.2,3.795,1.782,-2.168,2.084,5.125e-15,5.125e-15,5.125e-15,5.125e-15,5.125e-15,5.125e-15,-1.459,-5.945,5.125e-15,0.3906,-1.099,-5.674,-7.501,-10.41,0.8138,-1.638,-10.04,-4.866,-1.598,1.166,-0.9312,-1.048,-3.138,1.489,1.525,-5.119,0.8187,0.2869,0.6575,-0.2244,0.4203,0.09356,-0.9942,-0.3762,-0.3299,-2.211,2.238,0.7038,-0.0612,-1.875,-0.2072,1.223,0.6655,0.8552,-0.03459,0.4618,-3.227,-0.06029,0.883,1.529,0.602,0.9576,1.218,0.7659,0.6453,-1.838,0.9992,-1.127,0.4807,-2.655,0.6202,-1.712,-0.02139,-0.1683,1.22,1.198,-1.682,1.746,-0.1969,1.029,-0.9776,-0.5971,1.394,-2.048,0.4132,1.046,-0.4749,0.8038,-1.432,-2.879,-0.3101,-1.618,-0.9606,-1.639,-0.105,0.7939,1.544,-2.279,-1.444,0.5569,0.9508,0.4847,3.336,1.474,0.6729,-0.1931,-2.188,-3.642,0.6784,1.428,0.2437,-0.3568,-0.5557,0.6616,-2.597,0.942,-1.799,0.3005,1.395,0.2057,0.7546,-0.6292,-1.426,-1.022,0.7909,0.4803,0.6496,1.412,-0.4341,0.3297,1.385,0.3295,0.2956,-0.2548,0.5967,0.2625,-0.7084,-1.713,0.1465,-2.544,-0.6288,0.7983,0.9413,1.202,0.3939,0.2347,0.3342,1.194,-1.484,1.585,0.5446,1.004,0.2437,0.7376,-0.2289,-7.825,1.721,0.166,0.9233,0.4259,-0.1495,-1.39,-0.2569,-8.14,-5.603,-4.062,-2.901,-8.57,-3.847,-9.553,-5.942,-2.191,-3.936,5.125e-15,5.125e-15,5.125e-15,-4.935,5.125e-15,-7.13,-2.596,5.125e-15,5.125e-15,5.125e-15,5.125e-15,-7.851,-10.15,-6.422,5.125e-15,5.125e-15,-3.546,-1.475,-0.3531,-4.637,0.1048,0.3308,1.032,0.7223,-0.1802,-0.04218,-2.112,0.6275,-1.985,-0.8234,0.05117,0.9263,1.599,0.048,0.2331,3.385,2.093,-1.98,-3.581,1.226,1.819,-1.04,-1.739,-1.654,-0.4027,0.5175,0.1709,-0.2869,1.251,-1.386,-0.7983,0.399,-2.729,0.2105,0.8376,-0.1397,-2.598,-0.3168,0.5648,-0.2919,1.198,0.5645,1.459,-0.961,0.07913,1.898,-1.955,-2.828,0.09214,-1.736,1.029,0.7757,-0.5999,1.212,-2.081,2.797,2.621,-3.43,-0.975,-0.5994,-1.68,-0.8311,1.142,0.6785,-0.4531,-1.272,0.3146,-0.4654,0.1318,-0.9692,0.2568,0.975,0.2688,0.7266,-1.642,0.4487,0.8596,-0.537,1.193,0.6347,0.7634,3.161,-0.9482,-0.9515,-1.351,-0.6439,0.6919,1.394,0.4756,0.06347,1.306,-0.4024,0.0794,1.142,1.335,2.086,-0.7125,-0.1347,1.105,-0.7984,-0.5395,-0.7592,-0.07246,-0.4892,-1.392,-0.6108,1.385,1.228,2.351,-0.2242,2.97,-0.03303,0.2981,1.238,0.09001,4.172,1.395,1.014,0.2797,0.09616,0.0174,0.2739,-1.138,-1.165,-0.207,0.5458,-1.238,0.1764,-0.3605,0.5207,0.4219,0.6752,0.5929,-0.2962,-2.042,0.02641,-0.005105,-0.6717,-0.3147,-0.02845,-0.8919,-0.3357,-1.051,0.315,1.476,0.1864,-0.1796,0.2257,-0.09423,-0.02913,-0.497,-1.074,-0.465,-1.7,1.067,-2.53,2.103,-1.742,2.232,-0.9282,-1.016,1.045,1.073,0.7427,-0.9921,-2.487,-1.037,1.383,-0.3012,-1.722,0.3276,-1.227,-1.177,1.777,2.055,0.6306,1.321,-1.819,1.069,-8.911,5.125e-15,-5.335,5.125e-15,-5.425,5.125e-15,-2.668,-9.273,-7.541,1.257,-5.981,-0.7914,1.392,1.371,-6.797,-6.696,0.8085,-3.156,-3.614,0.122,0.3949,-0.09223,-0.6721,0.4013,2.421,-1.009,0.9377,0.8251,0.0432,1.424,-1.919,1.922,0.192,0.4128,-0.01317,0.3534,0.05598,-2.146,0.2178,-1.341,-0.2486,0.1626,1.242,0.344,-0.09289,-1.263,-1.683,-0.8505,-0.2932,0.3059,2.025,0.3096,1.789,-2.77,0.04693,-1.031,-1.403,0.04712,-1.16,2.351,-1.736,-1.989,-3.833,-1.318,-0.61,-0.2227,0.04414,0.8212,1.537,1.652,-1.542,0.7787,0.9338,1.129,0.9467,0.844,2.669,1.642,2.469,-0.2458,1.692,-0.833,1.293,0.07102,-0.6747,-0.6237,-0.9087,-1.474,0.01242,-0.9421,1.355,-0.3894,-0.7893,0.3753,0.04703,0.4596,-1.531,-0.004779,0.09808,-0.5683,1.334,0.07022,-2.244,-1.318,-3.189,0.7142,0.8627,-0.448,-0.8214,-0.2883,-1.126,-0.4763,0.5695,-2.4,-0.7221,0.9396,0.2814,0.9279,-1.378,2.897,-1.52,0.3818,1.454,0.4344,-0.3198,-0.2068,1.421,0.3547,-1.676,-1.353,-0.642,-4.131,-1.432,-5.703,-4.829,-3.526,-5.198,0.01837,-2.313,-1.198,0.15,-1.327,-1.711,-3.559,0.5741,0.8417,1.352,-2.185,0.03268,-1.422,-0.8607,-0.6478,-0.2367,0.01895,-1.125,-0.2617,-1.187,-5.107,-6.597,-4.231,0.6857,0.8462,-0.7513,-0.189,-0.5847,-3.722,-0.8563,-0.5331,1.942,1.519,1.225,1.291,1.28,2.564,-0.05265,-0.1738,0.06121,-0.7747,0.7115,-0.6157,0.4415,-0.3126,-9.823,-4.68,-1.691,-8.479,-9.832,-2.622,-7.914,-8.486,-6.882,-4.953,-6.239,-4.453,5.125e-15,-5.376,5.185,1.979,5.125e-15,-5.246,-5.041,5.125e-15,-9.954,-8.97,-1.087,-8.685,-8.079,-11.22,-5.723,-3.299,-3.562,-3.165,-0.5837,2.197,0.3396,0.1967,0.7869,1.229,0.9005,-1.442,-5.556,-3.194,1.686,-2.999,1.857,0.6305,-1.61,-0.03222,2.132,2.145,-0.5014,-3.395,1.09,0.6919,0.1812,1.666,0.3161,2.849,1.229,-1.059,-0.1288,-0.1115,-0.7527,-0.7428,-0.6311,-1.881,1.134,-0.4383,1.516,-0.4825,0.5491,-6.3,-0.05903,-0.1521,0.8569,0.3887,0.2495,1.434,-1.445,0.2235,-1.058,-1.315,0.9607,1.773,1.205,0.02124,0.193,2.789,2.382,0.4362,-0.3768,0.9756,0.8721,3.433,1.048,-1.936,-0.8012,-0.3558,-1.301,0.4318,-0.6815,-1.649,-1.7,-0.1616,1.102,-0.1158,0.9822,0.9575,1.253,-1.517,-0.4796,1.058,0.9091,-4.14,0.602,0.1013,-1.445,-0.8731,0.2598,0.264,0.2812,-1.955,-1.799,0.5118,-2.385,0.2853,0.8928,-0.3932,-0.1665,2.322,-0.3475,-3.689,-0.2218,2.353,0.9624,1.252,0.9624,0.05957,0.1926,0.6583,-1.303,0.5803,-3.799,-3.854,-10.03,-8.894,-1.159,-0.2408,-5.88,-1.485,-1.188,0.1437,-1.586,0.835,3.161,-0.6653,-1.33,-0.3855,3.22,-1.86,-2.63,-0.5422,2.324,0.8835,0.8602,-0.728,1.345,1.997,0.5795,0.2338,0.4373,3.454,-0.2538,1.538,-0.2184,1.2,-0.2847,0.3478,-2.817,0.9362,0.3676,-1.255,-0.7956,-0.6565,-0.8139,-0.6936,-0.02264,1.432,-1.242,1.376,-1.266,0.1692,2.323,1.453,-1.56,0.2367,0.4478,-0.4986,-2.129,-0.8209,-0.5038,0.4826,-0.7164,0.03864,-2.353,-0.1902,0.3679,0.1108,0.115,0.9195,-1.911,1.051,-2.525,2.866,-1.829,-4.894,0.629,-7.444,-2.401,-5.701,0.1192,-10.9,-9.403,1.329,-3.429,-1.898,-10.96,-8.654,-1.881,-8.064,-4.129,-6.933,-11.05,-2.229,-2.27,-0.2231,0.5962,-0.3004,-1.119,-0.5942,-1.601,0.9545,0.8307,-0.94,0.7696,3.243,0.3094,0.4884,0.5241,-0.8242,0.7963,2.661,2.218,1,0.9346,0.299,-0.6672,0.1002,-1.269,1.027,0.8959,-0.9303,0.2361,2.496,-2.736,0.3316,-0.2257,-1.997,0.7248,-0.2061,0.6669,0.6638,-3.073,0.5096,-1.293,-0.6021,1.023,0.4986,0.4702,-1.374,-0.4087,-1.392,-1.643,0.06029,-1.117,1.599,-0.491,-1.369,-0.1422,0.0259,-0.06598,-0.5666,-0.4114,0.7246,0.2527,-0.2954,0.35,-0.1539,-0.3887,-3.55,-2.86,-0.896,-0.2218,-3.944,-1.968,-1.224,-3.693,-9.602,-0.5008,-9.961,1.338,-1.198,-1.192,-7.839,-12.88,-2.55,-7.175,-4.703,-8.449,5.125e-15,5.125e-15,-2.342,5.125e-15,-2.724,-3.936,-5.307,-6.762,-4.96,0.4264,-5.318,1.253,-6.167,-4.799,-1.793,-5.76,-1.886,-0.5182,-2.907,-4.82,-1.675,1.787,0.7507,0.5889,-3.995,0.4265,-2.309,0.338,-1.58,-0.6859,0.7742,-1.189,-1.243,1.08,1.314,-2.905,-1.263,-0.9998,0.4442,1.236,-2.544,-0.4548,-0.2315,-2.232,-2.536,-7.295,1.947,-0.162,-1.04,-2.458,-2.064,0.7788,-0.937,0.6947,2.28,-0.4956,0.5837,0.175,1.084,0.4347,0.8808,0.7277,-1.106,0.2626,-0.6018,0.1346,-2.065,0.2102,-0.01124,0.2533,-2.558,-1.594,-2.115,0.7672,1.085,0.3496,0.3329,-1.366,-0.1049,0.5125,-1.396,0.6515,0.7452,0.3412,0.3268,1.952,0.6112,-0.582,-0.04801,0.1411,2.01,-4.088,0.9538,1.706,0.02633,-0.2294,1.42,0.595,1.152,0.3469,-1.044,0.3179,-2.288,-1.845,-0.4573,-0.4178,-0.6113,2.042,-0.6604,0.2012,0.2533,-1.711,0.00692,-1.109,1.467,0.7363,1.45,4.743,-1.552,-0.4895,1.236,0.7965,0.4755,1.062,0.5571,0.3423,1.05,-1.212,-0.2724,0.6647,-1.225,-0.1601,-0.134,0.9397,1.459,-1.452,-2.11,-1.9,0.3181,-0.3422,1.635,1.543,-0.4089,0.8281,-0.4942,2.017,1,2.673,0.1317,0.145,-0.5778,-3.656,-0.8592,0.05295,-0.06938,-0.484,-1.288,-1.517,2.019,2.123,0.1521,-2.51,-1.217,-2.575,-1.627,-0.3698,-0.2161,-0.07033,1.684,1.759,-0.7281,2.32,0.6454,0.1963,-0.06897,1.022,0.3085,0.901,0.8436,1.425,-0.7104,-1.595,0.4476,2.159,-0.2245,-0.09015,-0.893,0.5488,-1.273,-0.6127,-1.856,-0.804,2.923,0.3802,-0.12,-2.325,-2.125,0.2459,-2.516,0.7873,-0.0629,-4.158,-5.679,-6.234,-2.196,-4.497,-5.557,1.202,0.8762,-0.3907,-0.8833,-0.6499,0.7334,1.394,0.1386,-1.013,-2.791,-1.488,0.6709,2.228,0.1227,-0.7466,1.079,2.076,-0.2148,-4.896,1.403,-8.086,-5.755,-7.242,-4.34,-8.981,-7.454,-6.439,-9.466,-8.219,-6.054,-5.383,-2.996,-7.923,2.088,-6.902,-6.445,-9.452,-6.513,-1.082,-1.561,-0.3122,-4.837,-6.96,-2.015,-3.397,2.425,0.7283,0.2167,-1.181,-0.9216,0.3063,0.2273,-2.032,-0.7348,1.258,-0.005001,0.1064,0.07977,-4.966,0.4778,-0.1865,0.2167,-3.098,-0.1186,0.5088,-0.2843,0.4848,-0.7979,1.037,-1.413,-2.686,1.83,0.3018,-0.2387,0.1709,0.8316,0.2749,-0.2906,1.901,0.7361,1.192,0.846,1.054,1.192,1.496,0.6489,-0.6807,-0.4228,-0.6973,1.4,-2.602,0.5033,0.336,0.08562,1.699,-1.256,0.004061,-1.114,-0.4522,-1.965,-1.775,0.7394,1.169,1.006,0.5402,0.2798,1.913,0.3619,0.9397,-1.017,-0.1099,-0.7563,-2.406,0.1551,1.321,1.876,0.315,-1.123,-0.1973,0.3553,-2.341,2.124,1.614,-0.4202,0.6055,-1.821,0.62,0.02304,1.041,0.08621,-1.839,0.1144,1.15,0.4737,-0.4622,1.351,2.817,0.292,-0.6509,0.9872,-0.8175,0.4614,-0.6108,0.2152,0.3833,-1.319,-0.09187,1.605,0.2108,0.5633,0.408,-0.3106,0.7162,1.402,0.2521,0.6255,-0.2639,-2.583,-0.8499,-0.06488,-0.3125,-1.219,-1.859,-0.06297,-0.1765,-0.2542,1.048,-0.8667,1.99,0.7763,-0.06158,0.8963,0.7709,-2.423,0.9242,1.603,-1.627,-1.056,0.6499,0.2551,-1.427,1.897,2.295,-1.764,-0.5472,-1.4,-3.942,-0.8497,-3.137,-0.8229,0.9053,0.1128,-1.143,-2.728,0.3969,0.4661,0.3702,2.072,-0.3303,-2.576,-0.275,0.09669,0.4063,-1.198,-0.02404,-0.05482,0.01764,-0.01578,-0.1065,1.63,-0.9482,-2.305,-0.6538,0.7445,0.1837,2.586,2.146,-0.3753,-0.535,-2.603,1.036,1.041,1.069,0.5292,0.8735,0.8818,1.127,-0.7429,0.32,-0.02018,-0.1504,0.8495,-0.8035,0.7369,-2.056,-1.225,2.931,1.389,0.2045,0.06606,-0.3331,-0.5121,-0.01979,0.2793,-0.5743,-1.436,0.599,-0.4742,-5.642,0.5224,1.178,0.5649,-0.8118,-2.54,0.5424,0.8917,0.08175,-1.044,-1.768,0.2326,2.04,-0.9545,1.557,-0.06928,-0.1377,0.5417,1.17,-0.8887,-0.9461,0.3013,0.152,-0.171,-1.257,-1.13,1.468,-0.9411,-1.455,2.242,-1.573,-0.6659,-0.4403,-1.266,-0.7269,1.853,1.366,-0.7867,-6.382,-2.583,-0.4842,-0.3827,-2.463,-0.6712,1.38,-0.2512,1.084,0.05272,1.223,0.6474,0.2595,-0.2979,-0.1467,0.07542,0.5383,-2.042,1.096,-2.855,0.7845,-0.09925,-0.4124,-0.5892,0.2066,-0.5558,2.491,0.4975,-0.7694,-2.398,0.03447,1.58,-0.2602,-3.025,-0.3814,0.5927,-0.7349,-0.06765,-0.4153,0.08091,-0.9573,-0.3427,0.5892,-0.1865,-0.3816,-2.035,-1.538,-2.261,0.05425,-1.007,0.7668,1.54,-1.457,0.3419,0.2804,0.5543,-0.3385,0.2106,-0.3685,-1.619,0.559,2.762,0.5134,0.6111,0.3679,0.1663,0.00547,0.8916,0.8734,-0.7724,-2.22,-1.898,-2.527,0.4445,0.2349,-1.193,0.01717,-0.07793,2.582,-0.7903,0.2752,-4.216,-6.483,-4.624,-8.211,0.9373,-3.084,5.125e-15,5.125e-15,5.125e-15,-2.892,-6.329,1.691,-6.636,-6.845,-5.554,-6.497,-8.765,-7.247,-1.896,-1.185,-0.6014,-6.317,-4.016,-3.022,-4.066,1.411,0.2696,0.3139,-0.1612,-0.1522,0.1068,0.4013,-0.9905,-1.345,-0.2803,0.1354,-3.379,0.363,-5.342,1.446,0.06221,1.95,0.08248,0.003618,-1.168,0.5808,-1.461,0.2029,-0.3016,0.4564,-0.9543,-0.04161,0.2739,-0.5464,2.651,3.385,-0.9002,0.196,1.49,-3.307,-0.3812,0.6022,2.088,0.08716,2.385,1.625,1.377,1.089,1.978,-0.1671,-1.809,0.2573,0.9013,-1.001,-3.427,0.8643,0.7485,0.1069,-1.375,2.497,0.3061,-2.016,-1.089,-0.8383,-0.8465,-0.4624,3.547,-0.1194,1.149,1.302,-12.45,-1.009,-0.3028,1.291,0.7014,1.35,-0.03746,0.4822,1.117,1.818,-6.476,1.06,0.7656,1.344,0.5564,0.1418,0.1201,-1.141,-0.1767,2.44,1.068,0.6231,0.6934,2.077,0.1909,1.231,0.0606,-0.4965,-0.09766,1.745,1.129,-0.9702,-0.8368,0.01342,-0.9478,-1.649,-0.05341,-1.059,1.335,0.7687,-0.1002,0.2003,-0.1156,0.1345,1.113,-1.949,0.1879,1.275,-1.672,-0.2611,-0.9365,0.4881,1.265,-3.741,1.756,0.5381,0.9441,-1.073,-2.1,0.3498,2.453,1.704,-0.6494,0.534,0.7896,-0.07566,1.122,1.186,-0.1602,-0.05702,0.461,-0.5895,-2.931,-0.7273,-0.7111,-1.025,-0.03808,-1.264,0.5756,-0.5083,-1.483,-1.024,-0.1946,0.1225,-0.8315,1.238,0.6136,0.2639,-0.9481,-3.492,-1.141,0.4833,-0.8575,-1.445,-0.6965,-0.9418,-0.5599,-4.545,-0.9897,1.054,-0.107,0.898,0.3438,-1.765,-0.6881,1.092,3.943,1.035,0.9877,0.5211,1.24,1.157,1.326,-0.6843,-0.2611,0.3517,-0.3821,0.7586,0.3987,1.54,-2.071,-2.389,1.408,-0.5389,-1.074,0.9638,0.1385,-0.7519,0.9593,1.898,1.737,0.4121,0.3383,0.1934,-1.278,-1.236,-2.001,-1.385,0.9171,0.1388,-0.5127,0.1077,-3.52,-1.507,-1.28,0.1081,-0.0718,-0.765,-1,-0.2372,1.246,-0.02294,0.8127,-0.2658,-0.2832,0.06747,0.7872,3.588,0.07105,0.04887,-0.793,-2.935,0.8457,-0.2303,0.3638,-0.6542,0.0296,-0.1716,-0.2531,-0.2684,0.2359,-0.3594,-0.3098,0.5824,-1.739,0.269,0.3494,-1.459,-0.8876,-0.0322,0.5097,1.355,-0.6047,0.3873,-2.254,-0.5487,-0.8889,-0.1789,1.443,0.7042,-0.003614,0.6873,1.45,-1.648,1.622,0.1508,-0.1337,0.6114,2.049,-1.418,0.08335,-0.1552,0.1203,0.1111,-3.009,-1.87,0.43,2.152,0.4159,2.975,0.2291,0.5418,-0.02667,-1.103,0.7032,-0.9538,-0.6918,-0.5056,-0.2332,0.4464,-0.3656,0.2539,-0.2368,-0.8314,-0.06551,-2.12,0.6936,1.977,0.1656,-1.526,1.031,-0.3863,0.8474,-2.436,0.1428,0.1618,-0.1041,-0.2966,0.6821,-0.03573,-0.1858,-0.2255,0.8852,2.261,-0.307,-0.4792,0.293,3.286,-0.1487,-0.3607,-2.309,0.6743,-0.5721,-1.595,-2.327,-0.15,-0.327,0.474,0.348,2.082,2.325,0.1046,1.239,0.2373,-2.356,2.386,-0.2607,0.3508,0.9677,0.5906,-1.358,-2.589,0.6808,-0.4368,0.9214,-0.3125,-0.124,-0.01318,-1.058,1.131,0.2161,1.312,0.2477,-4.483,-10.68,-5.826,-11.12,-7.583,-7.015,-7.092,1.313,-8.074,-4.888,5.125e-15,-6.973,5.125e-15,5.125e-15,-0.2572,-6.265,-7.227,-7.085,-7.093,-5.825,-5.041,-2.818,-3.742,0.2258,-3.026,-5.618,-5.184,-1.459,-1.713,-4.621,-0.1998,1.178,1.128,1.057,1.11,0.2572,-1.556,-0.4616,-0.02528,-0.3475,-0.0709,-1.076,-6.846,1.388,0.3119,0.1387,-2.467,-0.454,-0.9066,-0.309,-2.929,-0.1109,-0.2813,-1.363,0.109,0.07186,2.035,0.5665,-0.2184,0.3891,0.1182,-0.009895,0.01437,0.4377,0.1359,-0.06798,0.5818,0.4649,-1.433,0.7493,-0.9602,-0.919,0.7164,0.9949,0.5855,-2.112,2.254,0.4304,-0.1134,1.868,1.143,0.9382,-1.024,-0.6157,-0.2159,0.9717,1.02,0.4746,4.095,0.483,-0.255,1.753,-1.259,-0.5485,0.5986,-0.464,-0.4507,0.3215,-0.5787,-3.099,-0.1409,1.115,0.5807,1.346,-0.4552,0.9688,1.095,-0.4506,0.8042,0.647,0.3346,-0.3764,-0.9489,-0.657,-0.0602,1.036,-0.3554,-0.1342,0.65,0.9226,-1.377,-0.9665,1.117,-1.211,-0.006374,0.3531,0.8395,0.5843,-1.274,0.4647,1.269,-0.2804,-2.305,0.4305,1.388,1.323,-0.01107,-0.4381,-3.58,0.2833,-0.9796,-2.248,-1.773,1.184,2.133,-0.7169,0.2352,-2.836,-1.094,-2.739,0.14,-1.189,0.8125,-0.3512,-2.516,-0.8001,0.5383,0.8584,-1.083,-1.682,1.081,1.532,-0.08844,-0.3416,-1.335,-1.44,-1.235,0.809,1.382,-0.2067,0.614,-0.02872,-1.681,2.653,1.74,0.4873,-0.02635,-0.4557,0.5264,-1.072,0.4577,0.3506,-2.284,-2.763,-1.635,-1.589,0.6837,-0.7781,-0.2463,-1.693,-0.8964,1.299,-0.9853,-2.688,0.5521,-2.744,-1.226,1.018,-0.5733,-0.6694,-0.1346,-1.152,1.057,2.516,-0.8855,-0.5271,0.4292,1.338,-0.1615,-0.06645,-1.002,1.167,1.07,0.7264,1.608,-0.4732,-2.173,-0.5091,-0.5227,-0.6229,-1.283,0.6901,1.031,-0.6417,1.058,-1.807,-0.8206,-0.6174,-1.618,-2.792,0.478,-0.5424,1.009,-0.4529,-1.65,-0.1081,0.5381,-0.4276,0.7358,0.1317,0.009983,0.2357,0.2817,-0.6083,-0.2966,0.3735,0.6433,-0.03822,1.674,1.143,0.2432,-0.5908,-1.761,1.822,1.194,-1.118,0.1023,1.722,0.4356,0.7468,-0.8124,-1.028,-5.754,-0.1164,-2.375,-2.073,0.4094,0.243,0.2186,2.031,-0.2912,-0.2987,-0.001833,0.8221,-3.071,0.6377,1.166,-0.1612,-1.846,1.41,-0.145,0.1956,0.3282,0.04169,1.427,0.1314,0.1412,-0.8958,0.008728,-0.2574,-1.048,-3.247,0.3306,-0.2951,-0.6174,-2.031,0.5621,2.866,2.107,3.797,0.1003,-0.2483,-1.677,1.967,0.4342,0.1782,-0.2818,1.795,0.4801,0.7413,0.594,0.9815,0.4093,0.6286,-0.9825,-1.515,-4.182,0.7385,0.1034,0.7935,-0.8114,0.03812,1.447,0.106,0.9074,-1.314,-0.8709,-1.109,1.644,0.8711,-0.5145,2.226,1.166,-0.4052,0.206,-1.838,-2.32,1.843,1.336,0.8363,1.869,-0.1372,-3.43,2.096,-1.725,0.2303,0.2172,-1.005,1.578,-2.085,0.2123,0.09141,0.136,-0.7327,0.4778,0.4127,0.6627,0.2199,3.302,-0.3079,0.9848,-3.349,0.7847,1.366,2.118,-1.514,-0.4578,1.488,-1.07,-0.9946,3.026,0.4575,1.336,3.683,1.797,-0.1479,-0.8598,1.207,0.5961,0.9034,1.511,-5.578,0.499,-1.648,-0.8257,-0.3324,-0.6831,0.09639,1.563,0.6683,-1.789,-0.2029,0.2393,0.4814,1.019,-1.798,-0.5089,1.957,-1.048,1.212,-0.6124,0.06598,0.08936,0.1793,1.057,1.041,-0.7862,0.3689,-3.857,-0.7847,0.6948,0.7196,0.438,4.619,2.657,-0.5652,1.15,0.1457,1.238,2.532,-0.1891,1.391,1.167,-0.04877,0.0411,2.077,-0.7031,1.593,-1.25,0.9252,-1.947,-0.6247,-0.2707,-0.5607,0.4607,1.896,1.101,2.64,-1.304,0.3105,1.567,-2.107,2.88,1.99,0.3704,0.7343,3.902,-0.6416,-1.072,-0.1241,0.2123,0.4487,1.421,0.9352,-1.001,0.3259,-0.1115,-8.1,5.125e-15,-5.764,-7.835,-2.096,-6.897,-5.852,-0.5242,-2.402,5.125e-15,-4.858,0.416,-0.4206,-0.7055,1.175,-0.3692,1.087,2.344,1.327,0.4316,0.7106,0.1788,-2.4,-1.012,0.5564,2.977,-0.2988,-0.7434,2.102,-1.091,-2.42,2.604,0.371,-0.9958,1.264,0.5798,1.338,0.6814,-0.03666,-2.249,-2.182,-0.165,-1.145,0.1794,-0.6263,0.4931,0.6485,-0.04691,0.7836,0.3158,0.6858,-1.246,-0.8085,0.07918,-1.372,2.106,0.3379,-0.2405,-0.3073,0.8936,1.794,1.032,0.8702,-2.995,1.429,0.9016,0.1644,2.922,-0.4636,-2.023,-0.4382,0.2335,0.5616,0.1406,1.916,0.4767,-0.2175,-0.3038,-0.8458,0.8526,2.829,1.544,0.6599,0.9931,0.5372,0.9947,-1.023,-7.729,0.4111,0.338,0.834,0.3109,1.093,1.477,1.475,1.463,-1.785,-0.4184,-2.43,0.5185,-0.6462,-2.344,0.06023,1.512,0.3353,0.1852,0.316,0.8508,0.8049,0.825,0.8325,3.02,0.5645,0.6805,0.658,0.4761,1.173,3.199,0.5307,0.7996,0.9063,1.382,0.2408,0.7473,0.1738,-1.321,0.738,-2.636,0.6311,-0.3208,-1.095,0.949,-0.2486,3.423,2.902,-0.6864,-1.045,-2.53,-0.2556,-0.4196,0.1154,0.1529,-1.378,0.9566,-0.1181,0.6808,1.676,3.783,-0.5893,0.2931,0.2031,1.131,-0.3055,-0.1958,-2.763,-0.8952,0.5474,-0.1753,-1.352,1.05,0.3908,1.286,0.5925,0.7621,-5.112,1.902,0.3646,-1.823,-1,-7.693,1.773,1.007,-0.7196,-0.2527,0.9138,0.433,1.12,0.5861,-0.9223,0.06956,-0.9438,0.349,-0.6761,1.806,-1.285,-0.8439,1.37,1.03,-0.4757,-4.67,1.915,-0.3637,1.238,-0.8148,0.05675,0.4613,-2.106,0.6078,-0.1014,0.76,-0.4188,-0.04061,1.758,-1.838,-1.167,-3.819,-1.098,0.6084,0.3917,-0.5007,0.5466,-2.275,0.289,-3.756,-0.2798,0.0495,2.973,0.8742,-2.684,1.819,0.6096,-0.878,0.3565,0.2605,-0.009745,-0.6412,0.9549,1.557,-0.2202,-0.08428,-4.324,0.004074,0.1562,-0.1526,-0.3242,-1.172,-0.6331,-0.9338,-1.307,0.168,-0.9044,-0.2296,-0.1665,-3.451,-2.473,0.08948,-3.023,-0.2207,0.7111,-4.016,-1.356,-1.273,1.844,-0.1974,-0.4914,-0.2316,-0.5961,0.2518,-1.225,-0.6775,-0.6443,0.6387,-1.268,0.1433,-2.117,-1.34,-0.02638,0.9263,-3.789,-1.652,-1.016,0.4426,2.434,0.7355,-6.47,-2.165,0.3193,-1.337,0.1155,-0.8785,-0.02309,0.5078,-2.443,-0.1297,-0.5471,-0.01697,2.069,-3.614,-3.003,-5.963,-5.222,-5.085,-6.584,-6.625,-8.564,-0.6315,0.4203,-0.2714,1.113,-0.05478,0.3432,2.907,2.051,0.3646,0.284,1.519,0.7751,0.4398,0.5524,0.351,0.5613,-8.9,-7.599,-4.471,-6.758,-3.943,-10.48,-3.369,-9.583,-0.4413,-5.689,-5.6,-5.804,-6.518,-3.551,5.125e-15,-7.861,-9.503,-4.875,-6.897,-6.581,-4.617,0.8536,-0.4664,0.8743,0.4245,0.578,-0.8972,1.142,0.7995,-0.1541,1.132,2.154,1.419,-0.1468,-1.91,0.06637,-1.968,-0.467,-2.276,0.9376,1.351,0.2706,0.8317,1.214,0.7224,1.225,-0.5297,-2.033,-0.7068,-1.11,0.01729,1.05,-0.7567,-1.724,-1.776,-0.8266,0.5638,-0.1709,-0.04306,1.562,0.1225,-2.544,-0.4206,0.7215,0.3054,0.1198,1.325,1.204,0.1919,0.1712,1.047,0.6437,-0.3773,1.5,0.4727,0.1219,-1.089,-4.122,-0.355,-1.269,0.4955,0.7421,0.1939,-0.5923,-0.2628,1.43,0.3859,0.2342,1.334,1.66,-0.8307,-0.7929,0.1855,0.6953,0.222,1.243,1.553,1.13,0.1584,0.07026,0.5245,0.7383,-1.351,-2.362,1.231,-1.434,-1.059,1.839,0.4969,-0.5439,-0.04244,2.046,1.418,0.5474,-0.08001,-0.3209,0.7628,0.4046,-3.4,-1.92,-3.566,0.6081,-0.4942,1.172,-0.2692,0.5393,0.4571,-0.0566,0.3101,-0.7847,-0.3369,0.7355,-2.915,-0.2513,0.3058,0.7911,-0.7824,1.495,0.8575,0.3461,-0.2471,0.2235,-1.503,-0.5685,0.8181,0.429,-0.2131,-1.397,-3.943,0.2257,-2.347,1.065,-0.03419,0.1486,1.562,-1.056,1.369,0.8859,0.2946,-1.165,0.7366,-1.543,-0.6745,0.7242,-0.9785,0.498,2.296,1.979,1.663,1.375,-0.4265,0.01551,0.2329,0.4078,0.5616,0.05647,1.781,-0.2312,1.202,0.5163,-0.9914,-3.375,0.1162,0.1415,1.281,-0.8923,0.5857,-0.05745,0.04154,0.2056,0.9425,-1.117,0.7944,-1.436,1.78,0.5449,-1,1.062,-1.25,0.1769,0.1375,1.039,1.497,1.906,-0.5482,0.5774,1.439,-1.541,0.1757,-3.235,1.846,0.8055,0.07096,-0.3907,2.159,1.092,2.959,-0.9975,-1.118,-0.4375,0.727,1.117,-0.6742,3.894,1.986,4.204,-0.2365,0.5276,0.7657,-0.8957,-1.802,1.359,0.07262,1.352,0.6984,0.2125,-1.718,-0.1962,-0.1066,0.6278,-1.404,0.8325,-0.4299,0.8736,-1.436,-0.8335,0.7353,0.5614,0.3021,0.2969,-0.8104,-0.0932,-1.727,-0.07233,-0.5023,1.035,1.594,-0.2909,-0.003724,0.2738,1.971,-0.6265,-0.3079,0.3286,0.1425,-0.4367,0.4637,0.3757,-0.7655,0.5337,1.54,0.2706,-0.5782,0.645,1.048,2.922,0.4477,1.76,-3.653,-0.3006,-0.3883,-1.152,0.2639,-1.129,0.1035,1.148,0.9055,0.2628,-0.07817,0.3663,0.3824,0.05121,0.2388,0.5095,-0.769,-0.5302,0.6352,-1.822,-0.8362,0.2807,0.1206,1.116,0.7011,-0.6687,-0.2493,0.4649,1.489,-1.147,-0.8192,-1.109,-3.09,-1.897,-1.453,1.596,0.5284,-0.3266,-0.02308,-0.3984,0.9782,-0.4105,-1.395,-1.83,-2.235,0.5294,1.529,-0.166,0.3769,-1.106,-1.634,-0.4711,-0.1298,-0.2668,0.2197,-0.697,-0.6363,0.2332,-3.248,-1.441,-0.2005,0.2743,-1.974,0.8026,-1.332,-1.724,-3.978,0.8066,-1.401,0.234,-1.404,-3.94,0.1538,0.07469,1.791,0.2042,0.5569,-4.598,0.01967,0.5139,-0.4352,-1.318,-2.68,0.1994,-2.036,-0.1929,0.8585,-1.523,0.6642,-2.171,-1.799,0.3885,0.133,1.247,-0.6681,-1.255,-1,0.4803,-0.7386,1.402,-0.2592,0.8123,-0.6516,0.4778,0.5408,-3.139,1.252,-0.8911,-0.7992,-0.2763,-1.229,-1.839,0.2436,1.816,0.7019,1.253,-0.2266,-0.5213,0.09033,0.9804,0.5301,-0.0005684,0.5866,0.2328,-0.004959,-4.118,-2.13,0.0282,-0.03513,1.398,0.5323,1.173,-0.2473,1.296,-1.68,-0.3956,-0.07506,0.6494,-0.4396,-0.1757,0.8318,-1.747,-1.62,-1.861,2.28,0.2097,-0.934,-0.04042,-2.577,-0.1236,0.1272,0.9767,2.134,0.7299,0.03054,0.5368,-2.117,2.657,-0.5901,-1.267,-2.75,-0.1526,-0.9813,0.2569,-1.519,0.04012,0.8771,1.34,1.942,0.3124,0.004487,0.6153,0.619,1.803,-1.077,1.054,1.273,0.6988,0.7023,0.9663,0.8134,-3.403,-7.642,-3.794,-4.362,-9.426,-8.925,-6.067,-6.272,-8.574,-5.015,-5.384,5.125e-15,-4.095,5.125e-15,-4.477,-4.614,-0.9859,5.125e-15,5.124,6.929,-5.176,-7.154,-8.647,-8.742,-8.934,-9.406,-1.002,0.1968,0.5601,2.46,0.7625,-0.3245,1.176,1.404,0.8397,0.4421,0.1013,-2.793,-6.731,1.045,0.09596,-1.615,-0.1751,0.1793,-2.838,0.9321,1.661,1.342,-0.3304,1.586,0.2803,1.54,-0.9137,1.326,1.832,1.291,-0.7163,-0.5824,-1.465,0.5556,-2.349,-0.4332,-0.4522,3.654,0.09604,2.475,-0.02838,-1.291,0.6008,-1.519,-0.1317,-1.953,0.4034,0.8269,1.816,0.02172,3.168,-0.6132,0.2704,-1.362,0.1899,-0.4575,1.635,1.514,2.496,1.181,-0.292,0.1093,-1.065,0.8718,3.441,-0.8177,0.3885,-0.1483,0.6543,0.1766,-0.2584,-0.6268,-0.6598,1.649,-0.1937,0.5101,-1.706,1.621,0.3311,1.288,0.6848,1.093,-3.335,-0.559,-1.025,1.516,0.1997,-2.158,-2.011,0.3959,-0.3947,0.2522,-0.4918,0.3938,-2.433,0.2807,-0.06772,0.1568,0.63,-1.23,2.207,1.011,1.377,1.162,0.2365,-0.003025,-0.07982,1.968,-4.551,0.3858,2.972,-0.1993,0.2592,-0.4518,1.42,0.9751,0.3065,3.263,1.824,0.6131,-1.517,0.3952,1.303,0.7448,0.4708,0.9516,0.5087,-0.7716,0.1461,-0.2978,-0.5982,-2.487,0.3294,0.5888,2.576,0.02856,-0.2085,-0.9174,-0.3761,0.8838,-2.533,-2.884,-1.685,-0.3139,-0.1764,-0.008156,1.117,0.6304,-0.08566,0.05728,-0.6801,1.298,-0.06582,-0.8136,-0.4775,0.6207,-0.03257,0.4059,0.5169,-0.06672,-0.4035,0.2498,-2.95,-0.4744,-1.492,0.156,-4.43,-5.557,1.239,1.214,-8.243,-0.5401,1.056,-2.715,-0.326,1.166,1.506,-0.8524,0.5435,2.366,0.04444,-1.007,-0.2641,1.684,0.02961,-0.906,-0.3058,-0.09107,-0.5765,-0.1315,-0.3424,-0.6879,-4.572,2.375,0.02814,0.1981,0.07548,-0.6873,-1.341,2.052,-0.4599,-0.1805,0.8778,-2.269,-0.3452,-0.2705,0.2125,0.02364,-1.228,-0.1031,-0.2118,-0.7454,0.9314,0.1047,-0.1065,-1.7,-0.7815,-0.1566,0.1281,-1.355,0.8505,-1.277,-0.9609,-0.7831,-1.143,0.6857,1.797,0.7453,-1.885,-1.987,0.02856,-0.9287,0.7635,0.8893,0.4788,-1.584,-0.7912,-1.113,1.931,3.453,1.453,1.093,-0.5368,-1.297,-2.133,-1.313,-0.8961,1.344,-1.647,-3.583,0.4766,-0.1614,0.2049,-2.384,-1.637,-1.296,-0.1065,-0.2809,-2.631,0.1234,2.665,1.203,-0.9303,-0.08228,0.1662,-1.024,0.5322,-0.6647,0.5634,0.001675,-1.177,-1.484,0.2832,-0.9468,-0.6553,0.7868,-2.355,-0.1209,0.07946,0.1433,0.03459,1.299,2.509,2.297,0.5951,-0.1194,-0.9047,0.1414,-1.221,-0.5353,-0.556,-0.1243,-1.555,-0.9114,0.3941,1.641,-0.8626,2.694,0.7143,-1.039,-0.7801,-0.1012,-0.3894,0.2774,-0.3591,-0.3213,-1.588,-0.4534,0.2091,-1.326,-1.006,0.3515,0.2919,1.333,1.207,1.044,0.2392,0.967,0.559,-0.3539,-1.829,1.404,-0.307,0.2589,0.2765,0.3609,-0.2058,-0.3219,-0.4414,0.1028,1.372,0.4846,-0.4609,-0.5958,0.9044,-0.149,1.506,1.688,0.5609,-2.684,-0.2949,0.1152,0.7764,-0.009729,0.362,0.719,-0.05324,-2.465,-0.206,-0.5893,-1.734,-2.13,-0.5098,2.369,1.262,-0.3787,-0.21,0.5934,1.402,0.7592,0.498,-0.1174,0.8798,1.401,1.177,3.267,-0.8173,-1.521,0.4349,-0.5987,0.2966,-0.2334,1.948,-1.107,-1.002,-0.9081,0.6909,0.6735,1.022,0.7778,0.03803,0.2893,2.781,-1.64,-0.2253,0.271,0.5444,1.32,2.194,1.373,-1.131,0.3493,0.9265,0.6581,-1.642,-0.5072,0.7144,0.5955,0.3361,0.8624,0.7387,1.247,0.4914,-0.5174,1.202,-0.33,-0.3324,-0.7183,-0.08561,-1.58,-0.1882,1.077,0.6162,0.666,1.719,-2.097,-0.2588,-0.1978,0.9789,1.676,0.1798,0.07645,0.2661,-0.7662,0.3093,0.3807,-0.2681,0.8606,1.411,0.9929,-0.6679,-0.9837,-0.5869,1.203,-1.41,-0.152,-0.1882,0.2616,0.2058,0.6176,1.328,-0.8615,0.5662,0.7304,0.0007823,0.6953,0.8874,0.1987,0.7562,1.416,2.794,-2.65,0.08783,0.636,-0.3257,0.1662,-0.608,-0.404,0.2607,-2.135,0.9043,-1.774,0.6291,-0.5653,0.5244,0.9901,0.9269,2.042,-3.621,-0.3197,2.619,-0.6532,-2.491,-0.6264,0.7238,-0.1426,1.626,2.992,3.341,1.881,0.1225,0.6907,1.457,-0.09029,-0.7083,-0.06623,0.5719,-2.294,1.157,1.103,0.3336,0.2645,-1.498,-0.4763,0.3933,0.1681,-0.8208,1.544,0.8904,1.122,1.217,-0.6924,-0.2965,-0.5474,-0.9548,0.7264,1.218,1.295,1.022,0.08817,-0.7671,2.512,0.7483,-0.5565,0.8904,0.3416,-1.4,-0.8682,1.576,0.7253,-0.3597,1.074,-0.9989,-0.3656,0.3296,-0.9725,-0.2749,-0.5269,-0.3875,-0.9372,-0.8046,-3.088,-1.183,0.367,0.2026,0.9686,-1.547,-1.095,0.5579,1.595,0.8333,-0.1398,1.056,0.6742,-0.6974,0.1487,0.004731,0.9653,1.015,0.8187,-0.6434,3.269,1.741,0.06501,1.072,0.2808,0.2509,0.3097,-0.6072,0.6071,-1.588,-1.075,-0.9962,-0.01048,-1.072,0.5908,0.9877,0.2143,1.259,0.7252,0.238,-0.527,0.4021,1.477,0.6175,-0.5273,0.5057,0.6469,-0.6405,1.002,0.3133,-1.921,1.464,0.5193,1.73,1.437,-1.502,0.9375,1.494,-0.5144,0.1627,1.422,-1.004,-0.8836,1.396,-0.3859,0.7508,-0.2863,0.6202,-1.444,2.21,0.5909,0.5656,1.281,1.062,0.9348,0.3135,0.8377,1.218,1.612,-1.273,-0.0723,-0.3443,-2.291,-0.4418,-0.5085,2.545,0.1313,-0.5458,0.3989,2.55,1.728,-1.632,-0.4243,1.041,-1.753,1.608,0.3175,1.394,-0.0926,-0.9611,-0.6473,-1.559,2.073,0.1456,0.05595,1.358,-1.056,-0.2555,-0.594,1.581,-1.676,-1.449,-0.4163,1.298,-0.4411,0.9023,0.431,-0.05568,0.8601,0.6112,0.07604,1.226,0.6456,0.7614,-0.007295,2.227,1.678,-1.301,1.851,1.585,2.723,1.186,2.029,-3.565,-6.903,-6.303,-7.863,-10.9,-7.465,-2.167,-6.778,-0.8967,5.125e-15,5.125e-15,5.125e-15,5.125e-15,1.568,1.211,0.8616,2.377,2.706,1.015,0.289,1.878,0.9348,0.7177,1.046,2.793,-1.078,1.39,1.747,-0.2109,-0.4263,1.771,-0.5897,-0.7395,0.5259,1.069,1.197,-0.3627,-0.6269,-1.477,1.426,1.633,-2.117,0.9948,-0.0772,-0.7671,2.286,-0.3668,-1.418,2.375,-1.277,0.7498,-0.7654,-0.2503,0.9608,0.6016,-0.5015,-0.9723,-0.2678,0.5365,-1.511,-0.7815,-1.693,0.7121,-0.2726,-1.57,1.656,-1.744,-0.4354,0.7012,5.125e-15,1.476,-0.07795,0.2189,1.163,2.089,-0.6142,-0.8744,0.2805,2.273,1.072,-0.6526,0.2561,1.044,1.261,-0.2737,-1.157,0.7558,-0.2097,-0.2934,-0.4901,-1.437,2.484,1.872,0.6441,-0.8271,-0.6647,-1.441,-0.783,-0.8453,1.12,2.167,1.728,1.03,0.2728,-0.1885,1.37,1.603,2.114,-3.918,0.4507,-2.11,-0.3268,-0.5819,-0.841,-0.4257,-0.481,-2.16,1.837,0.7849,0.09339,-0.0395,1.235,0.2405,0.006982,0.4718,-1.074,-0.4616,-5.098,0.4203,-1.058,-0.04627,0.9257,0.9068,-2.136,-0.04526,0.153,-1.849,-3.251,-0.7632,-1.483,0.5939,0.8825,-1.725,-2.623,0.2935,0.2429,-0.3441,1.894,0.5106,0.4048,1.062,1.593,1.932,0.2587,-0.698,0.8439,-1.628,0.6857,0.9453,-2.836,1.146,-0.5199,-3.075,0.6276,-0.7597,0.9447,-0.5587,1.194,-4.677,-0.7131,-3.487,0.3689,-0.4651,3.123,-0.1012,-0.4893,0.4388,0.4906,0.1815,-0.3054,0.5225,-0.7685,-1.977,-1.002,2.611,1.725,1.243,0.148,0.1193,-0.129,0.151,1.31,0.03843,0.1818,-3.179,-0.778,-1.273,-1.307,-1.253,0.4237,-0.6957,0.7854,-0.1579,1.877,0.7912,-0.1063,1.919,-1.384,-1.196,-1.038,-0.454,-1.273,0.7759,-0.9953,0.4696,-0.3872,0.01432,1.275,0.4083,-1.394,-0.5037,-0.7286,0.2179,-0.3597,0.4173,0.2901,-0.9378,-0.1348,-0.3978,1.738,-3.312,1.655,1.114,-0.7758,-0.07872,0.7014,0.2625,-1.387,4.75,-1.142,-1.003,0.05636,-0.6907,-0.02437,-0.08308,0.9369,-1.219,1.471,0.523,0.714,-3.237,-0.1408,-0.2239,0.01368,0.6489,0.3626,0.1405,-1.02,-0.7124,1.018,0.2008,-1.369,0.9825,0.2384,0.3401,1.545,0.7158,0.2964,0.358,1.178,0.396,-1.889,-0.007082,0.1906,-0.06882,-0.9227,-0.4451,-1.532,-0.2979,-1.312,-1.132,0.4654,-1.187,-0.6292,0.746,-0.4747,-0.646,-1.544,-1.809,0.3922,0.3038,1.134,-1.141,1.556,0.7963,0.6826,-0.7697,1.196,0.8162,-0.9077,0.02571,-0.8754,1.674,-0.4467,-0.4563,-0.9621,0.2867,-0.02732,0.8507,0.7497,1.353,1.25,0.5095,0.9469,1.825,0.1571,-2.427,-0.4146,0.07603,-0.0271,-1.887,-1.273,0.206,-0.5494,-0.4185,0.04975,0.217,-0.4416,1.657,0.6633,1.022,-0.1523,-1.097,0.9335,0.2156,-0.3137,-0.9302,0.8038,-0.4136,0.1346,-0.4417,0.9695,0.7328,1.105,1.005,2.091,2.797,3.331,0.06845,0.238,0.8685,0.8542,2.163,-0.3846,-0.2711,0.3284,-0.6682,-4.377,-0.02159,-0.05464,0.2993,0.3694,0.8367,-0.3451,0.2291,-1.056,-1.03,0.5224,0.6799,-2.959,-0.3469,0.2425,1.55,-0.4539,-0.2499,0.3573,0.3624,0.1727,-0.03541,1.175,-2.01,-0.5334,-1.403,-1.906,-0.02964,1.084,-0.8094,0.02439,0.09406,-6.662,-1.023,0.3291,0.2441,-0.3468,3.308,1.07,0.3154,1.989,-0.02691,-0.8712,1.286,-0.2893,-1.04,1.174,3.152,0.6095,-3.47,0.06332,-0.5819,1.415,0.8654,-0.3591,-0.4332,0.3294,0.5729,1.494,1.156,0.4189,-1.207,-1.626,0.8967,0.6766,0.1454,-0.7802,-1.378,0.7795,-0.3469,1.107,-0.6492,1.42,0.05293,0.2669,0.5574,0.7057,0.9708,0.7771,0.3021,1.914,-1.307,-0.3752,-1.028,2.576,1.578,1.045,0.16,2.527,0.2202,-2.095,-0.6609,0.2301,-0.6918,-1.191,0.1878,0.5978,0.1011,-0.2827,-1.215,2.081,5.125e-15,5.125e-15,5.125e-15,5.125e-15,-3.518,-5.991,-5.237,-10.3,-9.22,-11.14,-8.308,-8.206,-6.557,-7.837,-10.27,-13.43,-9.446,-9.275,-8.104,-8.018,-7.018,-7.612,-8.1,-10.12,-12.62,-10.53,-2.19,-4.748,1.535,0.3119,2.009,0.3392,-0.1507,1.015,0.3566,2.082,0.7971,0.5964,0.3246,0.1082,0.7071,-1.59,-0.004716,-0.1,-1.342,-2.191,-0.5106,0.6888,-0.382,-0.2614,0.6524,1.099,0.6648,0.9847,0.3278,-0.173,-0.481,0.441,1.16,0.9364,0.01731,0.8156,0.4557,4.139,0.5644,-1.153,0.5059,-0.3263,2.001,-0.7826,1.556,-2.174,0.5997,0.2512,0.3868,1.292,1.094,0.3474,1.353,0.5672,-0.06228,-0.4893,-1.189,1.283,0.6695,-3.023,-0.247,-0.2662,-0.6466,-0.5489,0.2355,0.3321,-1.79,-0.2412,-0.6493,0.5657,0.08058,-1.656,0.645,0.538,0.7966,1.32,-0.666,1.369,1.123,0.5816,1.029,-4.288,0.673,1.117,1.064,0.7459,-2.996,1.083,0.6043,3.016,-0.3699,0.5808,1.037,-0.4966,-0.5297,-0.6007,1.672,-0.8878,-0.7409,-1.746,-2.198,-0.5218,0.3249,1.188,5.125e-15,-5.182,-0.08102,-1.135,-0.5054,0.9434,-1.533,1.333,0.7395,0.4047,-0.8772,0.5893,-0.3956,0.3516,1.123,1.983,-0.3025,0.949,0.1674,-1.224,0.5123,1.282,2.793,2.36,-1.26,-1.172,-0.08676,-2.97,0.4226,0.4608,-1.037,-0.1445,0.06407,0.2232,1.307,0.009985,-2.29,2.835,1.351,0.2892,-1.382,-0.09932,0.6881,-0.3767,-0.426,-1.201,-0.1422,-0.6903,0.9514,-0.3513,1.148,0.2137,-0.1991,-3.424,0.04397,0.2344,-0.7681,0.5324,1.738,-0.002064,-2.783,-1.441,1.878,1.304,-0.7587,-0.3473,1.196,0.331,0.7323,1.964,-0.1448,1.509,-0.3037,-0.9193,0.5797,1.086,-1.246,0.3305,-0.7171,0.8349,0.8945,-2.733,-2.846,-0.0569,-0.0822,-0.4052,-3.148,2.527,-1.782,1.285,0.1832,0.4795,-1.866,0.1417,-2.366,1.372,-1.08,-0.8222,2.391,-0.2811,0.03051,-0.1054,-4.471,-1.021,0.7904,0.8224,0.7374,1.011,1.075,0.1467,0.07796,0.3359,-0.9423,1.669,0.5601,1.724,0.136,1.736,-1.085,0.2263,0.7511,-0.4739,2.045,-0.06669,0.4185,1.966,1.143,0.767,0.725,-0.8549,1.189,-1.049,-1.075,0.5848,-0.2922,0.07231,-0.7019,-0.08287,-0.2195,2.403,-1.406,0.1798,-0.4909,1.166,-0.6355,1.029,-0.844,1.134,0.26,1.359,0.1238,1.215,-0.8414,-2.643,0.1215,1.071,0.419,0.5704,0.5183,-0.4642,0.07264,0.7172,-3.584,-0.01037,0.979,-0.2286,0.002682,0.8699,0.5993,-1.26,1.443,0.6991,0.04439,-0.5784,-0.1986,0.3894,0.554,-0.4979,0.8049,0.4065,3.07,1.846,-0.3741,1.984,0.7618,-0.1432,0.2381,-2.011,1.715,-0.1152,0.365,0.7438,0.3855,0.8944,2.17,-0.8782,1.355,1.874,1.575,1.918,-0.516,1.969,0.7059,0.9799,1.315,0.2094,0.1748,0.4391,1.571,0.9414,0.5766,2.102,2.518,1.592,0.3777,0.9764,0.9193,0.08749,0.7495,0.7261,0.01148,1.023,-0.3196,-0.4372,-0.3989,-1.198,-1.005,0.02152,-1.518,-0.5213,-1.517,1.045,1.491,-0.5454,1.633,0.03039,-0.3094,0.328,0.09027,-4.7,-0.01242,4.935,0.766,-2.136,-3.333,-0.5609,0.8929,-0.5207,-2.576,-2.447,-7.624,-1.696,-3.802,2.58,0.9584,0.7933,0.1302,-0.6578,1.151,0.1273,0.04773,-0.9391,2.332,-0.5409,-3.831,3.66,2.404,-1.359,-2.433,-1.372,0.07161,-1.882,-0.08613,-0.09904,1.304,0.5858,0.8817,1.356,-3.746,-1.118,-1.627,0.943,-1.162,1.622,-0.9727,1.069,0.5014,1.859,0.235,-0.6184,-0.5508,0.3893,1.175,0.2442,-1.17,-0.1334,-0.8028,1.752,0.6719,-0.6885,0.7289,-0.2317,0.9163,0.1054,0.7791,-1.215,4.176,-2.311,0.746,-2.213,-1.764,-0.9524,-0.3012,-2.771,-0.8362,-1.05,-0.1344,0.421,0.8056,0.8211,0.0645,-0.06416,1.111,-3.809,-0.838,-1.393,1.167,0.2276,-1.104,-2.624,-0.2233,0.08087,-0.08987,0.9001,-1.291,-1.881,2.353,-0.3097,1.058,-0.8208,1.347,0.03527,1.548,0.9729,-2.025,0.6156,1.302,0.3152,-0.6209,0.9924,-1.175,0.8847,-0.4157,1.737,0.06879,0.6173,0.4773,0.527,-0.02848,0.1292,-0.8458,-0.6461,-2.327,-1.58,-1.447,-0.002399,0.258,0.1657,-2.322,-1.61,0.1281,-0.3642,0.06107,-0.3489,-1.363,0.6082,-0.8224,-0.2366,-1.545,3.023,0.5497,-2.616,0.6694,0.4511,-0.5385,0.7418,-2.007,-3.281,-0.2313,-0.141,-2.247,-0.2521,1.228,2.622,-0.04521,0.1631,-1.531,1.096,-0.3787,-0.2992,3.623,-0.1028,0.3998,-1.818,0.09905,1.294,0.8896,0.2227,-0.6175,-1.703,1.172,-0.2119,1.088,0.4425,1.184,1.729,0.5019,-1.877,-0.9674,-0.9275,-0.02558,0.1923,-1.362,0.616,1.494,-2.361,0.2547,-6.696,-0.3605,1.112,-0.6169,-0.8527,0.2214,0.8488,0.6119,-1.675,0.3535,0.5801,-0.2831,-0.682,0.3911,0.7823,0.05997,1.712,-6.668,0.3102,1.657,1.041,0.3319,0.3147,0.03516,-0.7774,-1.133,-0.2311,0.1922,0.6362,2.457,-0.7376,-0.8911,-1.552,-1.928,-1.068,-2.013,0.4272,0.9165,0.5383,0.09176,1.077,0.0783,0.9017,-0.4092,0.5592,0.7659,-1.147,1.614,2.392,0.2124,0.6953,0.09049,-1.061,-0.2579,0.8471,0.253,-1.642,0.1163,-0.6196,-1.843,0.84,-0.1132,1.123,0.01969,0.2195,-0.7296,0.8529,0.8884,0.3411,0.1859,-2.779,-1.248,-2.07,2.965,0.8273,1.455,0.6184,0.1737,0.2321,2.487,-0.9285,0.9316,-0.6938,0.4415,-1.163,1.286,1.394,-0.2849,-0.007096,-1.939,0.5637,-0.8409,-4.853,-1.5,0.4524,0.4273,0.8662,-1.438,0.006079,2.249,0.8017,1.258,-1.027,-0.6602,0.1084,2.762,0.6085,4.176,1.353,0.9902,1.124,0.01428,-1.358,-0.9705,-9.116,-6.49,-9.752,-9.627,-3.202,-6.618,-9.672,-4.779,-7.482,-7.289,-7.1,-8.555,-5.963,-9.637,-0.4503,-7.101,-9.83,-4.884,-7.118,-8.357,-4.017,-7.501,-6.647,-9.92,-7.946,-1.937,5.125e-15,5.623,-5.55,1.473,0.3728,0.2461,0.04789,1.412,-6.062,-0.2329,0.1679,0.9137,0.875,1.291,-10.27,-4.16,-4.745,-7.159,-3.423,-3.814,-5.453,-6.219,-4.053,-3.494,-3.918,-5.691,-2.476,-3.321,-4.041,-2.858,3.404,-6.648,-5.841,-3.136,-4.529,-8.436,-3.894,-5.759,-4.21,-7.32,3.399,-2.959,-3.626,-5.369,-4.458,-7.539,-3.091,-2.571,-3.866,-2.829,-5.999,-5.468,-2.314,-6.178,-4.908,-7.6,-5.144,-3.049,-4.494,-4.207,-6.068,-3.082,-2.517,-4.469,-5.044,-4.881,-3.58,-3.663,-2.712,-2.83,3.554,3.078,-5.037,-3.058,4.432,-5.222,-2.779,-8.792,-4.168,-3.846,-4.345,-4.615,-4.952,-4.556,-3.273,-4.752,-2.353,-4.683,-5.299,-7.207,-5.399,2.372,-3.631,2.088,-4.16,-2.221,-4.465,-5.403,-4.522,-2.871,3.68,2.152,-3.048,-3.934,-3.281,-3.355,-6.712,-8.31,-7.888,2.865,-5.096,-3.973,-2.695,-3.771,-4.592,-5.497,-2.408,-9.814,4.295,4.107,-7.56,-6.699],"logCPM":[0.2335,0.8914,-0.909,-1.044,3.163,0.6133,5.003,2.106,1.12,6.624,0.944,-0.1883,1.833,4.629,-1.05,0.4613,-1.034,-1.221,-1.113,-1.154,-0.8025,-0.5506,-1.087,-0.8912,4.231,5.846,-0.4062,-0.5014,-1.269,-1.082,-1.037,-1.018,-0.8162,-0.4439,-0.3194,-1.205,5.047,-1.333,-0.6471,-1.063,-0.8348,-0.4968,-1.243,-1.325,4.202,-0.7709,-0.6452,-0.6829,-1.271,-0.9345,-1.305,-0.8035,-0.477,-0.8719,-0.6047,-1.128,2.171,0.5397,0.4155,-0.783,-0.4198,-0.7101,-0.06131,-0.5802,-0.5516,-1.319,-1.205,0.7128,-1.325,-1.031,-0.6486,-1.2,4.859,4.004,7.292,4.419,-0.2541,0.05658,-1.063,0.7089,3.866,-0.1543,-0.4205,0.4831,0.8699,-0.1943,0.1023,0.05213,-0.2669,3.389,-0.5134,-1.082,-0.6948,-0.9267,-0.1904,-0.6631,-0.1987,-1.139,-1.238,-0.8462,-0.6023,-0.7576,-1.405,-1.111,0.8912,1.358,-0.1736,-1.264,0.5787,0.7213,0.2753,-0.9965,-0.6714,0.7875,-0.7072,-1.029,-0.8471,-0.8726,0.3965,-0.1988,-0.7048,-0.551,-0.9555,-1.18,-0.9394,-0.8912,-0.6887,-0.176,-0.919,-1.01,0.7173,-1.227,-0.7852,-1.107,-1.042,-0.4406,-1.168,0.09461,-1.041,-1.188,-0.1335,1.254,-0.6179,-0.7133,-0.799,-1.319,-1.325,-1.405,-0.7855,-0.97,-0.3408,-0.4832,-1.294,0.2788,0.2009,0.5138,-0.02345,-1.019,-1.405,-1.04,-0.9892,0.9867,-0.9012,2.021,-1.214,-1.389,-1.061,-0.004036,2.9,0.3679,-0.1631,-0.484,-0.7328,-0.7121,0.2777,1.72,-1.294,-0.6928,0.5968,-0.5475,-0.2769,2.635,3.691,3.512,-1.055,-0.2817,-0.2713,-1.017,1.501,-0.9366,-0.7041,-0.4596,-0.6979,-1.31,-1.422,-0.6012,-0.406,6.243,0.5244,10.03,4.756,-1.066,-0.7452,3.143,5.357,2.96,2.894,-1.322,-0.407,1.489,5.436,5.915,1.124,5.629,1.73,1.548,2.064,1.297,5.732,5.346,8.008,5.571,4.384,9.505,5.624,3.914,5.214,8.249,5.372,9.581,7.531,7.692,9.62,6.055,8.267,5.312,5.342,6.881,5.048,6.251,8.939,8.391,5.735,5.951,1.803,4.829,8.226,7.331,0.9543,3.065,6.467,7.129,4.275,3.248,6.765,5.882,5.94,3.723,7.764,6.145,7.112,6.5,5.176,3.633,6.39,4.351,12.91,6.354,4.469,6.233,9.297,5.317,5.782,8.06,7.465,6.491,8.249,4.562,9.335,9.322,7.931,3.286,8.203,8.325,7.499,10.11,4.061,5.415,5.86,5.897,6.496,4.386,4.269,7.186,4.136,4.22,3.575,5.728,7.672,6.997,8.224,8.205,6.167,7.425,4.795,5.996,5.361,7.027,6.017,4.467,4.534,3.532,6.249,4.068,5.947,11.66,6.469,8.254,6.517,9.883,4.625,7.821,6.771,7.03,6.635,4.252,4.399,9.387,6.297,6.71,2.168,3.375,7.21,4.455,2.918,9.749,6.029,6.685,5.872,8.769,6.066,8.668,6.911,6.211,4.663,6.862,5.842,6.271,4.813,6.912,8.411,4.553,9.915,4.956,5.534,5.253,6.531,5.036,8.186,7.543,9.797,9.578,6.076,5.788,5.157,4.996,1.67,0.4254,3.378,4.533,2.296,5.005,0.4293,0.6483,-0.7163,-1.076,-1.052,-0.9892,-0.2229,-1.034,-1.134,0.1661,-1.045,3.891,5.787,-0.546,3.617,2.56,1.102,4.077,3.08,3.104,3.098,2.269,2.475,2.379,4.42,3.992,3.375,-1.061,4.812,0.7968,8.992,3.426,4.563,5.149,5.883,6.601,2.784,3.553,5.101,7.522,6.588,9.602,7.88,7.631,4.193,5.447,2.604,3.811,3.916,7.624,7.522,1.716,4.469,6.381,4.836,3.676,6.075,5.842,7.104,5.345,5.331,6.543,2.988,10.44,4.069,5.5,5.1,6.137,6.579,10.07,5.403,3.214,5.154,6.952,5.641,4.128,6.241,3.602,4.943,4.739,7.111,5.256,5.124,5.824,7.026,7.492,6.323,4.696,3.672,5.512,4.219,7.163,2.441,9.817,4.604,5.142,4.02,6.417,10.81,11.06,7.682,6.404,6.67,5.723,5.319,6.221,8.041,5.109,7.831,6,4.422,7.397,7.181,6.807,9.258,9.458,8.081,5.276,2.524,5.559,5.538,6.388,9.489,5.941,5.439,3.948,4.615,3.907,6.68,6.968,7.765,6.57,3.547,4.026,3.313,2.965,10.08,4.941,4.272,2.65,5.959,4.855,7.962,7.001,9.028,3.731,6.638,1.583,8.809,0.833,5.021,6.408,11.06,4.321,9.493,0.7227,3.016,4.527,2.873,1.395,3.758,1.785,1.264,-0.7617,-0.3822,-0.7949,-0.8444,-1.107,-1.062,-0.196,-0.1219,-0.867,0.008575,-1.038,-1.186,0.2391,0.5017,2.557,1.408,-0.9892,-1.205,-0.286,0.2555,3.083,5.513,6.26,3.005,3.266,9.017,9.31,5.007,6.48,5.247,7.157,8.954,5.613,5.666,3.747,7.495,9.743,10.2,10.1,8.126,6.064,9.258,8.13,8.149,6.371,10.9,8.6,4.288,6.034,11.96,4.691,7.215,10.36,7.748,4.228,4.503,7.576,4.071,6.27,5.286,6.982,5.151,5.072,5.491,6.746,1.489,3.964,3.189,5.795,6.125,7.409,4.536,5.706,4.705,5.701,4.476,9.026,13.09,12.8,4.66,6.882,7.345,8.554,8.506,5.287,6.163,5.328,8.339,6.083,4.886,6.902,9.52,5.967,5.045,7.667,6.892,8.212,6.499,5.154,3.587,5.922,5.367,8.608,2.365,6.474,6.331,7.947,2.933,3.747,4.17,8.777,7.482,3.094,7.713,4.441,7.065,4.936,4.03,6.908,4.641,4.955,9.024,6.912,8.061,7.657,11.15,7.657,5.979,6.143,7.801,6.287,6.416,6.281,3.011,4.387,7.616,5.221,5.581,9.19,6.207,5.183,5.229,4.514,5.287,7.699,7.711,6.365,5.807,6.323,4.025,7.43,3.998,4.018,9.362,4.571,6.031,8.023,1.862,4.816,5.804,8.075,5.908,8.476,6.15,5.793,7.302,3.389,4.683,6.079,7.3,8.863,4.755,6.192,6.631,7.232,4.153,5.96,6.614,5.69,4.625,3.364,9.108,8.794,3.867,4.377,5.811,5.823,5.253,7.853,7.51,6.889,5.218,4.108,9.402,6.824,5.569,5.271,7.018,6.801,2.606,7.423,3.752,-0.7945,-0.9865,-1.227,-0.8058,-0.9701,-1.2,1.406,2.522,0.5532,0.1818,1.461,1.537,1.363,0.9126,8.474,0.7756,3.739,3.951,10.24,5.608,6.425,4.426,6.228,5.044,7.569,6.595,4.864,5.487,3.596,7.413,4.583,5.472,5.537,6.567,5.958,10.68,9.634,3.875,2.778,6.136,7.291,7.606,8.11,8.008,7.904,8.785,7.704,6.918,5.854,5.547,4.711,4.274,5.138,5.967,8.191,8.078,7.937,8.551,4.166,6.607,6.346,6.806,6.463,3.319,8.091,5.667,6.171,4.463,3.535,7.671,8.8,9.098,7.529,6.079,5.335,4.613,1.631,5.075,6.294,3.008,4.664,4.149,4.267,9.602,8.59,9.385,6.518,5.225,5.254,5.064,4.646,5.763,7.457,5.664,6.479,6.457,4.595,5.286,4.401,4.341,8.701,5.916,8.701,5.67,5.011,3.121,4.42,7.714,5.644,5.231,9.458,8.648,6.845,5.978,4.834,4.844,6.122,8.985,5.293,7.065,6.26,6.499,5.616,6.706,8.548,8.123,6.565,9.013,7.707,6.856,7.693,8.343,8.386,6.805,6.286,7.72,3.176,6.61,10.97,5.059,9.815,4.865,5.114,4.805,2.291,7.29,8.859,4.079,6.476,8.303,2.473,5.517,4.643,4.695,7.531,6.546,3.731,0.86,2.144,3.67,4.465,7.61,4.562,7.723,-0.1508,4.767,6.423,7.814,7.353,9.769,6.463,1.665,4.891,4.993,7.278,8.608,6.378,4.391,3.281,5.585,7.305,6.487,4.064,8.125,0.9202,4.546,2.325,3.998,6.578,6.233,4.524,-0.0695,-0.6377,1.749,0.5699,-1.183,3.19,-1.135,-0.8169,-0.5826,-1.152,2.021,1.576,2.879,3.407,2.826,3.438,3.14,3.25,2.561,1.735,9.154,10.66,8.87,7.501,3.813,6.944,8.105,5.377,4.813,10.14,7.614,6.223,7.408,5.356,8.804,7.261,5.93,2.973,7.734,6.699,2.026,5.388,5.786,4.414,7.803,6.72,5.504,8.232,5.753,8.201,6.85,7.281,4.364,8.53,9.053,9.161,3.658,4.936,2.438,2.726,3.947,9.051,5.55,5.624,4.02,4.798,10.2,5.933,9.669,6.8,7.61,6.934,7.423,5.03,6.737,3.992,2.391,5.875,8.71,5.668,5.235,3.553,5.738,9.597,6.538,5.156,6.074,6.155,8.769,7.885,5.18,7.308,3.828,6.386,5.263,8.568,4.169,5.403,4.665,6.303,4.653,5.413,6.419,6.565,8.36,7.239,5.522,5.942,5.869,4.982,7.296,5.158,7.358,4.587,3.768,7.876,2.966,8.584,5.976,8.332,7.645,3.252,4.406,3.092,4.263,6.517,14.91,7.605,4.914,2.989,1.435,3.312,2.191,4.507,1.582,4.414,1.331,4.592,9.833,9.739,7.641,4.48,3.202,10.18,8.873,5.524,5.191,3.345,5.258,2.782,2.799,5.893,9.818,9.09,3.582,3.643,4.133,3.928,7.398,4.593,6.005,6.671,6.128,3.064,7.317,4.993,5.048,5.477,5.504,4.904,3.996,5.301,6.668,6,5.006,7.795,9.143,4.794,3.842,2.793,8.359,7.715,8.39,8.359,3.465,8.217,6.965,4.84,6.471,4.492,5.535,5.654,3.444,6.664,5.484,7.682,7.251,5.653,5.623,3.52,3.153,3.513,3.03,2.573,0.7383,4.123,3.384,2.852,0.01928,2.865,4.046,2.371,3.607,2.279,2.944,2.551,1.61,4.286,3.427,0.1291,3.161,7.497,4.06,2.98,4.871,3.439,6.792,7.647,5.192,5.263,2.81,6.694,6.872,3.344,11.02,9.798,9.213,8.762,6.465,7.014,5.42,7.677,5.635,6.798,4.01,6.844,8.603,4.274,5.112,6.68,5.924,4.847,8.148,4.001,5.769,6.594,4.78,7.706,5.909,7.033,6.632,7.048,10.17,10.49,9.014,4.426,4.751,10.69,3.445,3.024,6.345,8.034,7.158,1.953,4.79,6.537,7.071,5.867,6.348,8.799,4.397,5.218,7.334,6.091,5.169,4.242,3.236,7.544,5.373,8.615,9.224,6.606,3.955,2.228,2.78,1.823,4.014,2.403,4.376,1.555,2.602,0.7006,4.778,-1.178,0.1361,-1.099,4.447,-1.181,-0.8574,-1.325,-1.122,-1.118,-0.9769,-0.4607,0.1144,1.002,2.9,2.116,2.915,5.655,-0.04987,6.805,3.406,4.111,2.819,5.341,6.137,3.914,6.947,7.526,10.71,7.117,7.192,9.019,4.681,8.64,6.638,6.964,5.523,6.329,6.174,6.116,8.992,8.181,5.219,4.084,7.065,7.566,5.43,5.52,6.207,7.766,2.076,7.703,4.646,6.499,7.978,6.621,3.577,5.32,7.255,2.615,5.716,4.965,6.698,5.866,7.608,6.455,7.413,6.502,4.899,8.671,4.651,5.212,5.187,8.475,5.228,8.641,9.904,6.447,6.255,4.166,7.218,8.947,8.201,8.269,8.474,8.828,4.803,4.817,4.709,7.276,4.819,2.168,4.383,4.447,3.604,3.786,4.58,6.357,10.57,4.204,4.645,3.803,5.454,4.357,5.114,8.138,3.759,7.446,7.688,5.343,7.217,6.251,8.019,0.2897,4.377,3.46,6.729,6.116,2.095,7.023,6.181,5.799,5.012,10.36,6.912,4.232,5.48,7.908,7.745,4.981,2.397,4.787,5.4,8.41,5.819,5.99,6.595,5.664,5.185,5.019,9.245,5.617,6.627,4.447,4.463,10.5,7.538,6.241,3.426,7.096,3.043,7.09,10.45,5.905,4.281,8.508,9.408,9.572,4.18,6.304,6.249,4.684,6.933,4.831,8.318,3.792,6.578,5.289,8.747,7.282,7.62,6.333,6.6,4.903,4.167,5.809,5.38,6.195,4.053,7.398,6.926,8.167,5.281,7.909,6.025,11.31,5.487,7.097,7.998,6.701,7.744,9.669,5.628,7.692,7.288,2.385,4.105,3.483,5.486,4.714,4.436,4.98,6.929,8.447,6.027,7.739,5.865,6.514,5.535,4.085,4.076,4.461,3.536,5.875,5.474,10.28,5.519,5.42,5.09,6.819,6.971,6.704,9.147,5.326,5.171,4.88,6.457,5.898,2.601,3.923,3.668,3.729,0.8394,-0.1203,0.02446,3.715,1.323,5.951,6.29,1.658,1.282,-0.4155,0.1425,-0.4194,2.647,7.946,0.2375,2.271,1.613,6.886,0.08755,3.394,3.48,2.436,0.3118,6.315,8.454,7.596,6.17,6.344,7.788,10.43,9.27,10.53,6.062,9.125,4.64,6.053,6.89,8.011,3.468,4.392,4.627,4.573,7.129,5.769,5.886,6.345,4.661,10.67,7.275,5.093,4.584,4.27,5.899,9.159,7.817,6.708,6.525,3.638,3.891,5.597,5.102,3.498,10.23,4.891,3.191,4.328,5.854,6.153,6.352,6.2,5.932,5.559,6.192,6.01,4.903,7.858,5.588,6.886,4.778,5.019,6.215,5.886,4.847,5.335,6.448,6.774,7.51,7.036,5.179,8.633,1.379,7.863,6.658,3.938,6.724,3.684,8.249,9.8,7.056,7.072,6.224,4.632,4.462,7.383,4.066,5.765,3.987,8.592,3.809,3.904,5.1,6.075,5.414,11.96,8.426,6.541,4.637,5.364,8.794,4.359,2.536,4.41,5.536,3.774,4.907,6.665,6.941,3.398,5.248,6.279,7.318,4.355,4.883,5.106,9.572,5.459,5.529,5.14,8.373,5.417,9.376,5.769,9.867,8.702,4.572,4.819,6.719,8.482,4.824,5.063,5.031,2.359,3.769,8.564,8.94,6.84,8.008,7.813,4.172,4.215,8.191,1.237,6.44,8.449,4.471,3.741,4.736,3.085,7.2,5.604,4.208,5.731,9.898,9.007,8.334,7.139,6.61,5.346,6.516,8.882,6.219,9.976,8.869,7.78,3.848,6.345,5.606,8.784,4.774,5.206,8.919,6.67,7.78,8.017,6.635,2.804,7.145,3.222,6.044,7.203,6.184,4.387,6.38,4.704,3.767,6.856,5.217,4.122,5.05,6.669,6.907,5.821,5.118,8.605,9.424,10.16,3.575,4.014,7.417,8.129,9.175,7.983,4.593,7.17,4.818,5.396,4.582,7.442,4.852,5.531,2.715,4.463,6.182,6.066,5.913,6.2,6.245,8.358,6.358,5.905,0.3767,3.685,4.54,7.239,7.705,10.15,3.717,5.471,6.384,6.347,7.678,4.618,8.026,3.914,6.138,7.194,4.784,3.704,5.391,5.193,4.403,5.164,5.8,4.694,7.247,6.643,1.878,8.777,4.736,5.669,7.132,4.397,6.827,5.583,9.09,6.21,2.989,0.9769,4.818,4.434,5.226,7.095,3.434,6.539,2.982,3.337,5.587,5.531,3.384,3.627,6.162,7.689,7.5,11.11,9.236,7.138,6.219,4.699,9.421,1.466,7.391,5.908,9.713,6.188,6.383,4.955,7.237,3.212,6.038,5.929,3.941,2.558,8.801,6.302,5.33,4.409,4.377,6.183,9.118,4.677,5.811,5.969,2.059,3.481,3.944,5.592,7.257,4.791,6.318,4.62,5.963,5.949,2.968,4.077,3.95,8.238,7.538,8.979,7.086,7.641,3.371,1.405,4.562,2.318,4.589,3.33,4.316,5.815,4.024,5.519,3.513,1.198,3.732,-1.166,-1.218,-0.3903,3.516,-0.3931,0.8305,0.2322,-0.2245,-0.7174,5.277,1.613,5.764,7.544,5.374,2.552,-0.3163,4.843,3.707,5.37,4.544,9.541,6.761,5.391,5.713,9.215,4.378,5.764,2.103,4.161,5.956,10.25,5.939,6.101,7.005,5.978,2.15,7.368,6.469,8.627,8.293,7.758,4.583,3.181,2.908,4.796,5.607,6.653,1.403,3.647,6.331,5.827,5.986,4.679,4.984,6.285,0.529,2.711,4.536,0.3706,4.791,4.556,4.277,4.836,9.34,6.705,4.756,2.17,10.14,5.559,6.939,4.985,5.083,10.43,6.602,6.543,9.781,4.492,2.338,8.106,9.573,8.947,8.964,8.892,1.618,4.372,5.775,4.479,4.139,2.795,5.794,5.811,5.947,8.294,5.343,4.744,9.229,7.88,6.049,7.115,7.623,5.81,4.11,3.481,1.246,5.804,4.879,5.015,3.185,2.591,7.113,3.499,6.204,6.51,9.467,6.39,7.996,9.456,6.494,7.513,5.031,5.26,5.459,3.178,5.093,6.367,8.839,7.217,9.645,6.74,6.727,6.783,9.113,5.791,7.296,4.64,9.277,3.617,7.482,5.748,8.51,4.07,9.999,10.15,3.628,5.151,4.36,7.799,8.226,5.196,5.901,3.594,6.33,5.242,7.642,9.453,7.611,2.741,7.797,4.999,9.477,6.382,6.352,6.165,5.979,7.19,3.773,7.084,2.34,8.734,7.901,6.949,6.698,5.282,5.563,4.896,7.404,7.92,10.34,7.509,7.262,6.311,5.138,6.705,6.76,9.119,4.866,6.172,3.453,4.68,7.44,5.141,7.091,3.512,7.331,5.371,4.428,5.94,6.851,6.374,6.516,7.308,4.812,6.299,4.355,6.816,6.599,4.693,6.303,9.166,5.773,6.205,9.787,4.188,3.687,4.133,3.571,8.028,6.843,5.965,5.576,9.845,8.095,10.48,7.064,8.196,7.121,3.354,8.609,7.297,4.187,7.948,6.558,4.888,4.961,6.525,5.435,6.755,4.392,2.788,4.819,6.664,4.84,6.163,4.649,7.42,5.918,10.05,8.408,8.362,6.261,4.375,6.495,4.618,4.174,7.643,5.585,3.525,4.395,4.232,5.996,5.675,4.401,5.788,4.966,3.538,6.57,4.988,4.228,6.153,5.172,3.307,6.213,2.066,5.093,5.755,5.89,5.457,3.115,5.86,8.876,5.274,5.24,5.241,5.444,4.827,5.2,6.855,5.369,5.007,6.938,9.288,5.036,3.792,4.248,5.828,5.666,7.148,3.952,4.152,6.468,4.906,6.918,7.101,1.968,6.893,5.903,4.104,5.649,8.087,6.623,5.982,7.363,4.636,6.762,7.082,7.653,6.259,5.695,8.953,6.479,6.394,6.756,5.406,3.427,6.362,5.015,5.109,4.149,8.219,5.012,3.089,4.748,5.09,6.675,9.034,4.615,8.282,6.819,7.829,6.696,6.353,7.603,6.094,14.19,5.087,6.659,5.714,6.935,7.066,6.359,2.164,5.172,6.697,4.391,5.984,6.899,7.831,6.614,4.444,7.554,8.659,8.013,2.763,6.491,10.54,6.884,6.962,5.266,2.86,3.153,3.087,0.1806,-0.1605,-0.1313,1.274,0.5407,-1.07,-0.851,-0.1846,-0.936,-1.18,-0.94,-0.05853,-0.06275,1.87,-0.1456,-0.793,0.1674,-0.5127,-0.6933,-0.2115,-1.103,-0.8784,-0.8847,0.2962,3.304,4.469,8.033,5.999,4.429,5.341,5.268,7.891,2.683,6.229,6.354,10.42,4.702,6.544,5.98,5.998,2.868,4.305,8.746,10.6,10.46,1.835,6.742,7.693,5.174,8.942,5.276,8.361,2.482,5.932,5.634,7.038,5.48,2.939,5.155,6.356,4.847,4.389,1.179,3.019,10.28,6.745,12.61,5.898,3.826,6.987,5.799,8.701,6.138,9.002,5.916,2.856,3.054,5.566,4.592,7.999,7.612,2.686,5.97,6.493,3.381,6.023,4.896,2.062,6.403,4.904,6.614,11.51,7.498,4.379,5.645,2.44,6.42,5.945,6.158,6.418,6.925,5.166,4.852,4.856,7.601,10.12,4.707,5.878,8.009,5.871,3.959,3.88,3.548,7.354,6.592,7.411,6.474,7.308,5.323,6.857,6.777,3.551,3.849,5.727,4.2,4.825,3.415,9.443,8.112,4.176,3.772,2.04,8.476,4.285,5.087,3.792,1.83,8.494,6.694,8.203,5.832,6.537,9.497,7.816,1.238,1.726,5.321,6.209,6.136,9.057,9.18,6.131,4.724,8.604,7.363,5.122,5.85,3.74,7.765,5.775,7.257,4.737,4.543,8.739,5.93,4.981,7.598,7.528,8.005,4.397,3.505,3.421,3.913,6.076,7.514,8.641,5.412,5.028,7.295,5.395,6.752,8.958,7.45,5.077,7.836,8.377,8.632,7.021,5.742,1.759,3.07,8.5,8.274,2.073,6.839,4.786,5.757,8.355,6.967,3.131,4.254,3.551,4.367,5.076,7.575,7.034,6.69,7.015,6.55,10.93,8.378,11.18,8.251,5.281,7.298,7.39,8.708,7.115,3.413,7.304,4.959,5.508,3.949,4.327,8.882,2.933,5.145,6.876,4.364,4.414,11.64,5.592,5.564,5.775,7.248,7.482,8.495,4.62,3.422,6.621,3.58,11.14,7.362,7.894,8.275,5.879,6.769,7.322,7.486,7.266,3.765,5.543,6.177,5.843,4.936,3.517,5.9,6.648,3.005,9.673,7.852,10.23,6.232,6.741,5.538,4.985,6.792,6.302,4.195,4.792,5.86,6.691,10.13,8.985,5.885,4.21,3.586,3.327,4.068,8.408,5.568,6.611,9.188,5.774,6.735,4.503,6.743,2.632,3.902,3.735,6.102,5.583,5.144,2.282,6.787,5.493,7.607,7.469,9.178,5.193,6.163,6.24,9.855,6.915,6.069,5.869,8.79,3.977,3.152,8.246,6.128,5.606,10.76,4.777,5.404,5.941,6.725,6.635,11.67,5.749,9.097,8.197,4.01,8.365,7.999,6.378,6.141,4.369,3.384,6.435,6.106,4.635,1.827,5.362,7.076,2.444,2.899,5.978,7.6,4.465,7.103,7.419,6.756,8.173,5.381,4.649,6.558,8.232,5.719,7.122,4.139,5.685,5.036,3.746,3.567,3.306,4.812,7.555,1.919,2.117,3.692,7.765,10.04,4.394,7.142,10.98,5.76,6.04,6.83,3.319,7.514,5.307,5.257,0.6893,4.618,6.078,3.655,5.598,5.207,8.029,8.949,9.041,6.164,7.047,4.126,4.281,6.419,4.622,3.591,5.414,5.565,9.565,1.996,3.312,7.918,5.689,10.38,5.863,5.272,6.492,5.236,4.831,8.106,7.245,6.338,6.032,7.516,5.698,4.71,3.001,4.344,8.014,3.996,6.614,3.766,5.471,3.481,4.286,6.033,6.369,4.64,4.921,8.395,6.068,8.201,4.738,7.364,5.701,5.473,5.07,5.627,1.704,5.415,4.417,2.309,2.866,2.704,8.25,4.399,5.505,4.257,11.82,1.069,7.302,7.844,4.688,6.097,8.449,6.423,8.693,6.116,7.674,4.813,1.807,0.4207,0.3368,3.909,0.7663,-0.09173,0.7734,-1.093,0.3691,-0.9727,-1.141,9.154,6.349,5.079,10.97,9.772,2.98,2.296,7.873,6.963,4.813,6.1,6.726,6.557,5.237,7.92,8.845,5.691,5.101,3.8,9.612,5.097,3.72,8.641,1.63,4.974,2.947,2.603,4.437,7.246,7.684,8.494,8.435,4.925,5.569,3.903,8.925,6.39,6.363,7.304,5.982,6.664,7.386,5.263,7.128,7.832,9.038,6.253,6.679,6.957,4.566,5.81,3.408,8.611,4.239,4.393,5.875,6.265,5.857,7.936,7.505,7.815,5.628,4.794,7.355,3.897,4.396,4.738,6.594,6.246,6.608,5.213,9.336,6.685,7.818,6.775,4.633,0.9542,10.85,5.216,6.013,6.19,8.9,9.06,4.902,3.091,7.963,4.398,9.554,6.369,4.944,4.998,7.282,7.726,6.547,6.455,3.744,5.102,5.803,8.004,3.235,5.348,7.126,6.592,6.826,6.382,5.741,3.793,4.488,3.354,4.455,9.341,7.565,6.178,7.022,9.812,6.477,8.854,5.911,4.379,9.244,5.271,7.665,3.262,0.5856,8.625,5.325,2.902,6.905,5.524,7.589,5.676,7.304,7.004,5.201,8.226,3.075,2.505,8.188,4.886,5.543,4.469,5.953,6.82,6.359,9.682,6.847,5.704,8.204,7.049,5.66,4.788,3.776,2.69,7.074,7.23,4.959,4.538,8.525,4.571,4.775,6.431,5.336,6.324,7.336,5.24,2.905,5.101,7.698,7.195,5.271,3.56,4.512,4.414,5.971,6.314,6.456,7.538,6.701,5.579,5.338,4.255,6.73,4.303,5.174,8.497,3.793,2.837,6.855,5.025,4.576,6.523,4.538,10.43,2.459,3.808,11.25,11.31,9.86,9.929,2.881,7.244,2.323,-0.5344,9.604,5.676,3.575,6.6,7.99,8.626,5.35,8.769,4.48,5.702,4.588,8.715,6.628,2.957,6.44,8.024,8.365,5.436,6.18,5.756,7.594,5.619,6.297,8.528,9.408,6.453,6.357,6.772,5.159,4.707,3.481,3.883,4.741,8.354,1.669,7.256,8.975,5.179,5.193,11.12,3.745,5.087,6.288,5.165,5.806,4.902,5.196,5.879,7.372,6.135,7.972,7.999,7.173,5.657,2.469,6.527,6.018,5.245,1.417,5.424,1.161,8.368,5.553,7.84,5.685,6.459,8.179,6.666,6.844,8.163,4.399,4.753,5.104,8.082,4.916,5.136,7.533,5.749,5.526,6.516,4.326,7.614,6.733,6.29,3.367,3.404,3.868,2.935,2.514,8.446,7.855,6.188,5.391,6.248,5.36,5.845,3.181,1.367,5.827,-0.5714,7.532,6.163,2.552,-0.2882,1.755,0.7686,5.504,-0.6448,-0.2334,-0.4631,-0.9949,-0.9279,0.7312,1.932,3.499,1.882,-0.07521,2.742,3.798,6.855,7.312,8.595,5.048,5.391,3.211,5.748,8.945,6.571,4.511,3.495,9.093,8.292,5.521,8.916,6.236,2.007,4.022,3.234,8.397,5.048,8.162,6.462,3.45,2.736,5.062,5.565,8.826,4.676,4.808,5.313,4.518,4.761,8.786,6.065,6.437,3.097,7.38,6.767,5.808,4.222,5.193,3.858,6.647,3.78,4.067,4.712,5.272,6.623,3.704,5.882,5.486,7.586,7.668,9.691,4.733,7.958,6.718,6.445,4.958,5.03,6.951,6.749,3.522,4.91,6.962,4.167,3.87,6.796,9.005,5.474,5.802,9.568,4.978,5.713,4.412,5.427,6.32,5.701,4.604,4.859,1.027,2.075,10.76,10.09,5.328,6.711,11.44,8.201,9.094,5.938,4.823,4.417,6.84,5.06,7.676,3.348,8.473,8.263,2.906,9.894,7.134,6.858,6.842,8.066,5.231,6.363,6.001,5.643,6.012,4.156,5.096,7.121,8.328,7.207,4.081,7.103,7.476,2.603,8.884,7.195,1.788,5.372,6.955,11.48,4.114,11.33,2.306,5.225,5.146,5.948,5.472,3.886,5.169,6.161,4.632,3.041,8.162,5.392,6.119,6.052,6.999,9.928,5.238,5.457,8.333,7.104,5.633,8.292,8.349,5.917,4.829,5.251,5.526,4.559,7.432,4.612,7.478,9.1,7.411,6.251,3.897,5.333,9.143,5.592,6.691,1.569,4.779,4.42,6.4,9.039,10.62,6.364,7.758,7.7,13.62,7.028,8.798,3.842,4.863,4.052,4.244,2.309,4.402,3.968,6.747,10.12,11.57,3.667,7.163,7.306,8.966,5.524,6.714,2.637,9.376,3.783,9.002,2.709,4.88,9.662,7.957,3.133,5.742,7.92,6.919,4.897,3.694,8.934,2.281,4.958,2.978,6.736,5.769,6.584,6.363,5.962,5.087,8.9,2.227,6.77,3.823,10.54,3.891,6.269,2.043,6.349,3.503,9.625,4.311,6.337,6.478,7.058,6.83,4.481,4.517,6.72,10.37,6.173,5.652,9.934,4.949,4.876,6.202,5.155,5.832,6.343,4.792,-0.1024,6.43,6.458,7.043,6.698,2.632,5.444,5.124,7.519,5.953,5.669,4.213,6.316,9.53,7.717,8.471,4.931,5.398,6.017,7.512,8.18,7.371,9.009,10.63,7.391,7.7,6.681,10.2,4.2,4.479,4.075,4.688,7.653,4.496,6.199,4.504,3.586,6.122,8.983,10.59,7.878,7.092,7.729,5.855,6.924,5.955,5.848,4.954,10.46,6.855,5.937,7.913,5.737,4.173,4.138,4.886,6.256,7.437,6.742,9.415,6.039,5.485,5.388,5.402,6.831,4.79,6.034,6.384,6.517,4.035,5.944,7.959,6.36,6.027,7.211,8.843,6.486,5.582,4.14,6.108,1.607,6.275,4.191,6.833,6.431,6.146,4.053,2.39,4.623,3.737,6.845,6.983,0.3132,4.511,1.821,10.74,4.7,7.212,3.469,8.072,5.606,5.109,3.504,6.421,9.946,8.207,2.532,3.404,10.57,7.573,7.36,3.92,6.034,6.277,6.166,7.326,6.515,5.094,7.627,10.06,4.794,4.486,7.206,6.206,5.279,4.742,4.993,4.628,1.213,6.72,5.244,5.495,4.096,7.423,7.719,4.988,7.557,6.94,6.72,3.606,7.886,5.293,5.704,6.445,9.013,2.922,3.899,5.112,7.939,8.794,6.209,5.152,4.873,7.54,5.176,3.899,7.426,6.234,9.548,4.187,1.96,4.861,4.428,5.76,11.51,4.063,8.011,1.992,4.391,1.949,5.377,3.787,9.539,5.885,6.615,5.871,4.769,7.299,6.521,7.016,7.153,6.6,8.553,4.774,5.994,6.401,8.243,5.926,6.635,4.391,0.2436,1.858,3.446,1.96,4.767,6.845,3.968,0.8829,0.3518,-0.8686,-1.04,0.07887,-0.6,-0.8321,-0.8271,-0.5751,-0.3915,-0.1634,-0.447,-0.7435,7.004,4.184,2.284,1.643,2.736,6.863,7.912,8.082,4.287,7.833,10.57,6.561,8.246,7.915,8.171,8.538,8.685,6.1,5.432,8.274,5.554,6.617,2.264,6.98,9.093,5.378,4.95,5.342,5.536,5.789,3.988,7.204,0.1589,3.214,5.894,6.561,6.474,9.279,3.776,7.538,7.289,6.592,7.439,4.351,6.659,4.744,5.5,6.452,6.898,9.036,3.34,7.168,6.473,3.661,8.786,2.449,10.51,4.431,5.578,7.269,5.369,3.141,4.544,5.806,4.659,4.259,7.048,9.718,8.047,3.593,8.597,5.429,6.018,7.031,5.746,4.891,5.975,3.36,4.252,5.124,4.169,8.307,6.112,6.179,5.732,4.918,9.491,7.682,8.101,7.241,6.406,5.209,7.698,7.396,5.25,5.513,9.361,9.067,6.322,6.495,7.449,11.16,7.102,3.124,11.13,10.51,5.422,5.859,6.977,8.973,5.903,4.758,2.827,0.7966,4.63,3.727,8.512,6.896,9.861,4.655,4.299,5.686,6,4.968,7.35,1.086,5.518,4.543,4.746,2.313,8.212,4.997,6.549,5.792,3.572,1.828,6.292,5.95,5.115,2.402,7.398,7.952,6.663,5.59,4.165,5.945,5.704,7.281,5.574,6.197,7.101,3.477,5.164,9.018,9.06,7.955,3.963,5.866,6.668,9.056,5.732,8.875,12.46,6.299,5.706,5,4.387,4.26,4.624,6.196,0.6223,7.594,6.413,5.644,2.705,4.395,5.104,5.873,5.007,6.004,7.453,5.599,4.054,7.445,4.28,6.063,7.34,9.065,9.64,6.936,7.643,5.999,6.088,3.905,3.687,5.429,3.177,7.294,5.605,5.016,6.379,5.411,5.965,6.888,6.073,5.593,6.682,5.981,6.862,8.865,5.418,5.632,6.928,6.476,7.775,6.934,4.707,3.552,4.585,6.382,6.65,8.463,7.003,10.71,5.813,3.865,8.209,6.622,6.974,6.914,2.676,7.453,8.759,7.203,8.815,6.746,9.152,5.753,4.865,5.712,8.714,8.314,11.04,4.443,3.07,7.368,5.2,6.879,7.889,8.812,3.973,5.503,5.057,3.328,5.396,5.376,6.271,11.9,7.649,4.692,3.411,4.375,5.775,7.386,2.736,4.99,5.631,1.311,8.328,4.355,6.114,4.162,5.947,4.14,9.924,9.704,6.401,5.185,4.606,5.062,5.937,7.107,6.397,6.407,2.884,6.347,4.729,1.404,3.538,3.573,7.346,9.143,9.396,7.188,7.251,8.812,5.252,7.17,8.987,7.606,8.609,4.716,3.706,7.752,8.501,9.086,8.147,3.031,3.917,3.92,5.834,5.484,6.474,6.198,9.143,7.855,6.611,7.143,8.409,2.859,3.986,3.412,3.465,4.763,9.467,7.121,4.284,5.514,7.542,5.559,3.866,8.162,3.497,6.65,7.135,7.565,3.541,6.194,5.706,4.478,7.034,6.582,6.557,7.316,6.845,5.738,3.852,6.675,6.93,9.028,7.267,5.582,6.503,5.776,7.374,10.95,8.526,2.295,2.747,8.561,5.87,4.961,5.179,7.539,6.237,6.12,8.063,6.823,4.592,4.602,2.599,9.07,8.307,3.509,9.54,3.288,7.548,4.874,2.024,2.427,4.784,4.585,7.203,6.398,5.994,6.651,6.334,0.05197,9.954,12.47,8.653,4.115,7.641,7.1,4.028,3.837,3.288,5.524,3.435,5.807,4.022,6.213,6.625,6.788,5.61,5.813,10.77,6.294,4,1.684,7.137,10.67,3.011,6.736,4.044,7.337,8.153,3.278,8.69,4.651,8.157,6.881,6.028,7.784,5.567,5.283,5.883,7.107,5.711,7.616,3.816,5.004,3.17,6.37,8.985,4.789,5.899,9.718,4.241,7.753,7.003,7.79,4.221,5.585,5.26,8.106,4.661,5.871,3.792,8.62,9.269,5.021,9.622,4.832,2.75,9.325,5.694,8.118,8.699,7.993,7.18,6.043,8.106,4.813,7.908,2.184,8.695,5.712,7.642,3.727,6.65,5.932,4.14,6.418,3.336,5.848,10.38,6.622,6.484,7.706,5.604,5.936,2.526,2.427,3.133,2.978,7.304,5.822,8.573,9.018,4.554,7.742,2.55,6.495,9.025,8.619,9.665,6.616,5.471,7.073,0.5304,7.146,4.206,5.337,5.874,4.721,5.057,6.924,3.779,7.079,5.202,5.137,4.826,6.937,9.228,6.426,4.022,4.678,5.176,6.937,9.12,9.776,7.965,5.349,5.903,10.21,6.204,5.269,8.059,6.02,2.907,6.655,6.22,5.275,4.67,5.855,6.406,6.3,5.642,6.861,2.95,7.478,11.37,1.867,5.267,6.041,4.516,5.872,6.43,6.78,10.21,4.343,7.199,7.303,5.149,6.671,1.878,1.323,4.039,3.072,9.009,6.268,7.091,8.802,5.043,8.55,7.944,8.824,4.718,7.987,8.593,6.255,9.918,7.088,5.705,9.434,5.378,9.672,6.589,5.691,6.599,6.321,4.979,10.34,3.61,0.821,5.839,6.163,8.55,2.994,8.195,2.584,8.304,8.638,7.633,6.635,4.732,4.714,3.214,6.033,7.691,6.194,7.771,4.313,7.076,8.541,5.625,9.247,7.709,5.876,6.238,5.757,4.61,7.918,6.026,8.262,8.448,5.542,7.217,7.334,6.924,5.79,4.13,5.991,7.238,6.016,4.524,5.248,5.615,5.091,7.514,7.134,6.691,3.686,6.608,8.637,6.598,8.671,5.414,7.012,6.165,9.015,7.014,5.147,5.038,4.506,8.428,7.433,3.466,6.22,6.562,9.642,8.872,8.509,5.165,5.35,5.395,3.187,6.474,5.696,4.424,7.579,5.177,9.149,5.126,6.564,8.207,4.932,10.85,4.536,2.096,4.87,1.258,2.938,0.2456,-1.336,-0.3075,1.899,-1.167,-1.135,-1.096,0.4836,9.126,8.246,8.653,3.712,8.076,5.578,6.106,8.06,5.947,7.334,3.996,7.542,10.98,7.547,3.351,7.657,5.66,4.411,3.484,7.106,6.105,4.816,2.748,8.727,6.971,3.799,5.793,6.329,6.371,2.164,4.066,3.353,1.911,0.2956,8.717,4.977,6.423,4.994,5.689,9.51,5.42,3.809,9.057,8.824,5.17,9.515,10.5,8.374,4.571,6.879,10.36,7.256,4.929,8.259,8.623,5.936,1.116,11.05,5.27,5.552,3.488,7.35,3.734,2.438,8.36,5.246,3.199,5.131,6.268,2.121,5.449,10.28,8.137,4.455,9.339,4.176,7.62,6.318,11.05,5.132,4.515,5.028,6.799,4.849,7.867,7.975,5.338,5.744,6.552,6.037,4.598,7.502,8.745,5.119,7.122,4.121,5.461,5.711,8.473,3.647,5.558,7.119,10.29,8.498,6.316,6.628,6.108,6.463,10.14,3.822,5.896,5.484,7.993,4.57,4.486,3.833,8.732,3.805,3.599,4.976,6.989,4.215,3.659,8.854,6.12,6.896,6.719,5.071,4.772,6.994,7.696,5.089,4.453,10.61,5.21,4.481,4.845,5.825,6.207,3.215,6.975,9.019,9.437,7.888,5.283,4.183,8.78,8.445,3.91,7.126,9.913,5.18,6.435,7.222,1.371,4.758,6.826,7.704,5.609,5.302,6.995,7.012,8.725,6.601,6.413,5.712,6.894,7.214,5.152,6.098,6.584,7.707,3.5,4.814,5.493,3.65,5.991,4.681,5.032,4.53,7.427,6.704,6.293,8.672,4.298,6.628,4.007,4.008,4.416,4.378,2.347,3.016,5.445,2.646,4.205,2.19,4.015,5.974,7.683,6.226,6.969,5.437,7.018,4.042,1.505,8.36,7.548,7.847,9.275,4.377,8.914,6.304,4.497,8.918,7.528,6.524,5.256,-0.5858,4.887,6.031,5.583,7.503,5.673,5.938,7.836,3.081,0.657,4.497,5.704,5.921,8.762,8.784,7.572,11.22,7.693,6.15,6.982,9.775,5.946,6.153,6.463,7.404,5.329,9.033,5.559,9.944,6.181,4.895,7.551,6.268,2.463,3.922,3.021,4.847,2.608,5.101,3.793,6.416,9.953,5.029,6.169,4.431,6.87,5.922,7.007,5.577,6.336,8.518,4.269,4.111,7.693,9.031,6.983,5.884,3.264,9.763,6.788,8.168,6.405,7.426,5.926,4.787,4.744,11.88,7.774,3.745,8.26,6.739,10.02,6.984,10.11,5.748,11.44,4.19,6.019,4.206,5.267,0.9584,3.653,8.716,5.105,5.306,6.827,6.493,1.984,3.427,5.744,7.538,5.607,7.6,10.96,7.369,2.68,7.465,3.682,5.506,4.574,6.787,6.386,10.64,7.541,8.722,8.792,6.934,4.973,6.152,8.013,9.211,5.679,3.989,4.313,3.317,5.575,1.512,2.445,6.356,5.217,4.816,4.051,7.661,8.076,5.047,2.446,5.768,6.027,2.375,4.51,6.323,5.974,6.638,6.155,7.252,6.828,5.52,7.402,10.36,3.785,5.945,8.013,1.97,8.491,6.381,4.623,4.78,8.857,8.298,5.041,8.012,6.904,8.797,7.442,5.89,5.701,5.282,4.298,3.514,1.357,7.796,7.617,4.826,7.474,4.756,4.439,9.284,9.093,7.625,6.135,5.906,9.852,8.128,7.753,6.228,4.073,7.15,4.464,6.141,6.197,5.068,5.292,10.96,6.654,3.461,6.389,9.546,9.32,6.706,8.803,5.628,7.753,6.817,3.553,6.975,7.646,9.779,8.736,10.68,4.43,6.756,4.478,4.633,3.793,6.384,5.345,6.768,6.238,6.512,3.982,2.262,4.211,1.568,3.503,6.683,6.452,5.668,7.818,5.964,6.811,8.283,7.268,5.103,6.625,8.641,2.637,8.729,8.368,2.348,-0.3133,-1.231,-1.253,0.09074,-0.7202,0.8534,2.329,1.427,3.06,3.131,2.865,2.936,1.171,2.457,5.35,4.055,1.696,4.872,7.108,4.451,5.616,4.136,2.118,4.588,2.955,1.733,7.234,10.83,4.543,2.216,8.757,8.112,9.154,8.221,5.488,5.77,6.65,3.976,4.424,10.23,7.861,3.655,6.003,9.283,0.434,9.186,7.216,6.976,6.159,7.708,7.563,4.801,11.5,4.866,8.947,5.03,3.181,8.703,4.053,6.016,5.755,3.349,1.115,6.678,4.119,3.266,7.038,4.685,6.837,5.526,6.181,6.47,6.53,6.352,4.609,6.766,4.459,5.67,6.867,6.316,6.715,7.803,7.472,5.221,7.491,3.829,8.193,5.95,8.787,6.036,5.635,2.832,4.723,7.333,7.068,6.656,7.269,7.059,6.997,4.781,4.284,6.997,4.995,3.656,4.853,6.087,4.129,3.949,5.616,4.605,3.863,8.808,6.055,3.478,2.834,3.828,4.014,5.161,7.671,8.788,10.32,3.95,12.24,7.491,7.171,7.064,4.617,4.692,4.439,2.175,6.974,6.322,7.009,12.05,3.707,5.65,5.204,5.222,6.428,7.495,4.969,5.606,3.861,4.477,7.746,6.786,6.258,5.824,5.16,5.211,6.325,2.734,7.162,6.243,4.827,5.878,4.816,9.862,4.466,5.728,4.649,6.197,8.068,4.245,9.496,4.921,3.262,5.608,8.739,6.531,5.192,6.675,8.233,6.06,7.483,6.457,6.768,4.242,9.413,4.044,6.147,6.405,4.083,5.61,4.812,9.574,7.711,0.4922,6.48,7.936,8.102,3.347,8.641,8.096,5.754,7.512,8.864,1.871,7.022,6.498,4.844,9.756,5.574,5.28,5.896,3.786,5.245,6.866,6.414,3.018,1.591,7.874,1.909,9.095,7.363,7.523,6.843,8.487,4.182,8.082,9.519,4.12,8.68,6.651,4.048,9.217,6.908,6.056,9.261,6.591,4.98,-0.3449,9.532,6.455,1.4,2.347,4.734,3.653,7.739,5.269,6.413,7.189,6.027,6.62,9.287,5.969,8.873,7.873,9.925,8.369,10.08,4.294,7.225,6.523,4.664,5.898,5.574,6.785,7.685,9.22,6.001,8.872,9.909,3.766,5.542,4.841,7.206,10.01,9.425,2.581,4.473,7.826,4.98,7.847,5.412,7.514,7.862,9.073,9.516,7.716,5.87,7.621,6.95,7.687,2.674,11.51,4.058,7.449,6.994,6.693,3.601,5.261,10.01,4.708,2.171,7.658,5.495,8.815,6.761,6.224,6.728,6.152,6.661,5.152,10.19,7.599,10.31,5.194,2.435,10.59,5.887,4.135,4.906,8.508,7.301,4.757,8.342,3.596,5.481,3.862,3.661,6.614,7.132,2.656,6.187,5.476,3.625,3.449,5.16,7.596,1.078,4.684,7.025,2.33,9.558,5.012,4.797,8.421,5.653,7.824,4.744,12.37,10.03,8.643,3.992,3.374,6.722,5.407,9.795,7.187,4.009,8.347,4.446,7.386,8.37,5.699,10.76,7.727,8.192,6.602,10.16,4.299,3.546,5.818,9.478,9.048,11.46,3.804,-0.005438,4.87,2.956,5.82,6.198,6.774,5.988,6.143,4.955,7.131,8.73,0.8933,2.737,5.794,4.334,5.337,4.804,6.927,6.728,4.619,6.864,8.541,4.925,4.673,6.765,5.885,4.609,5.866,4.599,9.319,6.369,8.522,10.64,6.675,4.022,3.497,3.303,4.997,1.843,5.133,10.2,9.533,7.251,6.371,4.156,5.421,4.151,5.789,6.799,7.09,8.026,7.679,5.343,6.89,6.609,5.569,6.402,5.869,6.461,8.471,5.717,5.812,6.612,5.756,5.299,6.153,6.462,6.445,5.412,6.354,7.098,5.835,5.273,4.505,8.679,11.05,7.614,5.916,3.387,5.311,4.164,8.161,4.26,5.279,6.097,5.245,8.196,7.928,5.346,6.76,9.039,6.808,5.976,6.546,2.74,5.263,6.302,3.824,6.084,4.606,4.442,6.246,3.976,5.202,5.309,7.634,7.732,5.907,6.836,6.031,4.936,9.834,3.969,8.16,2.712,5.729,6.352,6.923,7.365,5.684,3.415,7.417,8.559,6.38,7.868,4.809,5.233,5.82,4.874,7.563,5.028,5.868,7.999,5.599,6.502,8.515,5.551,8.517,6.795,5.773,2.675,5.066,4.893,5.707,5.796,7.378,5.172,7.647,7.618,8.742,6.96,7.136,6.24,7.314,8.195,5.851,5.245,7.304,8.71,6.959,7.794,3.328,6.369,5.422,8.85,7.258,8.475,8.118,4.23,8.238,8.419,7.088,6.507,2.463,4.328,8.868,8.444,9.296,4.281,7.893,4.651,5.818,7.647,8.114,8.589,4.622,7.288,5.006,1.981,7.622,5.845,8.155,9.504,8.951,7.666,7.848,8.321,4.705,5.705,8.197,5.795,7.462,8.295,9.268,4.781,-0.2875,4.814,2.704,3.872,6.313,4.412,5.695,6.775,2.452,5.206,5.075,3.39,0.583,3.829,7.576,9.684,5.904,5.162,9.202,4.099,9.101,6.616,9.021,8.631,9.081,6.923,6.252,6.119,5.849,7.498,5.159,3.433,5.423,6.747,5.789,7.919,4.955,6.811,7.267,10.69,5.763,7.65,7.66,4.898,3.339,3.514,5.422,8.849,6.56,7.574,6.246,6.304,5.709,8.792,6.447,5.439,3.25,4.303,4.325,6.303,7.661,4.494,3.095,10.24,5.238,7.213,1.699,8.017,6.575,6.687,3.199,9.338,5.094,5.11,7.366,8.274,10.2,4.867,5.633,7.15,8.105,4.944,6.499,4.231,8.667,5.522,7.228,9.49,4.169,6.322,3.744,5.335,2.766,5.546,8.232,6.127,9.239,1.385,5.501,1.885,4.258,5.274,-0.1346,2.226,2.589,1.175,0.5323,-0.05413,1.183,-0.1004,4.283,-0.2792,3.4,1.926,-0.7782,7.292,0.7088,0.1006,0.6227,-0.312,3.915,0.4019,-0.7813,-1.322,-1.125,-0.8562,-0.06173,1.968,5.26,1.741,-0.4989,-0.6876,2.307,0.8779,5.838,10.06,9.26,6.033,8.884,7.836,8.528,7.478,6.833,7.221,4.622,7.977,8.196,9.642,7.282,9.402,6.849,7.641,7.991,7.506,7.905,8.282,8.2,10.14,6.4,7.153,7.458,7.658,7.431,14.25,11.19,10.77,7.163,9.342,9.932,9.712,8.585,6.096,9.976,9.638,11.67,7.674,7.449,9.439,4.913,6.853,11.44,5.546,8.593,8.041,7.535,8.69,8.988,9.91,7.648,9.692,7.397,8.374,9.213,10.17,8.605,8.739,10.53,6.241,6.258,8.974,8.773,13.59,7.619,9.386,6.864,8.095,9.263,8.816,6.63,8.984,8.048,6.535,6.34,7.098,8.336,5.703,10.4,7.1,9.31,8.785,6.601,9.569,9.017,7.514,8.002,8.546,9.724,6.586,8.521,9.416,5.985,9.571,10.66,6.045,7.015,8.022,6.283,9.399,9.257,8.79,3.605,7.247,7.773,10.28,9.496],"F":[5.08,0.8665,1.126,2.129,3.808,1.743,10.87,6.983,0.1791,12.32,2.02,5.44,2.505,20.6,0.9292,9.29,1.734,0,0,0,0,0,3.333,0,6.004,11.51,0,3.826,0,0,1.581,0.3875,0,0,0,0,0.5456,0,0.3904,0,0.178,0,0,0,0.04539,4.095,0,0,0,0,0,3.579,0,3.343,4.003,2.584,6.639,6.686,1.543,0.4074,4.998,0,0.03893,1.609,0,0,0,2.026,0,0,0.3512,0,5.002,0.315,1.659,2.409,4.7,0.9224,2.112,5.198,11.76,4.635,3.381,5.088,0.1142,3.542,5.53,3.462,4.042,4.298,4.004,0,0,2.101,2.057,0.2994,3.979,0,0,0.1686,0.04314,0,0,1.225,6.646,0.06869,0,0,2.257,2.975,6.3,0,0,0.08607,0,0,0,0,6.807,2.476,0,0,5.767,0,0,0,0.9133,0,1.302,0.4047,9.665,0.004498,0.387,0,0.3245,0.04667,0,3.883,0,4.232,0.1177,0.0002564,0,4.709,0,0,0,0,0,0,0,0,0,0,0.2797,0.08417,1.815,0,0,0,0,5.942,0,1.775,0,0,0,0.05858,0.0006673,4.456,0,0,1.286,1.97,0.128,3.815,0,0.1495,0.6413,0,2.222,7.778,18.59,3.127,0.5215,1.428,5.02,0.8659,0.02103,4.148,4.064,7.535,3.927,3.456,3.303,4.177,0,19.82,6.149,9.254,0.08435,3.538,0.722,2.408,8.341,9.974,0.2597,0,5.392,6.849,10.81,18.17,2.498,0.3501,9.939,0.03448,0.5138,0.0004967,0.4028,0.03703,0.09024,2.559,1.696,0.6237,2.241,1.388,0.1606,0.9224,0.003033,0.03197,11.9,4.838,11.03,0.02838,6.863,0.02519,0.0004736,4.77,1.116,2.239,2.601,16.68,2.311,0.00151,0.3045,0.003662,0.4047,2.332,1.042,0.4466,0.3095,3.877,2.255,1.572,0.2089,0.9693,7.204,0.3207,2.938,0.2046,0.1715,0.2929,4.386,0.4339,4.656,1.575,0.04518,2.275,0.1046,0.0311,0.4779,0.01336,11.29,0.1331,7.952,0.2098,0.1497,1.064,2.788,0.5238,6.604,0.3616,1.502,8.27,0.7537,4.737,0.5123,0.5567,0.06625,0.07009,0.8441,0.0673,1.022,0.1198,1.502,0.1383,0.382,3.131,4.018,3.508,1.357,9.188,2.056,2.657,0.3528,2.825,11.41,1.462,0.4221,0.5239,1.065,1.371,6.208e-08,0.1778,0.01238,18.84,5.024,0.3718,6.184,13.46,0.09499,2.207e-05,0.748,0.8543,4.562,1.659,1.397,15.61,9.359,0.01299,0.027,0.9568,5.871,3.303,0.9951,13.47,0.3996,3.032,1.359,6.036,2.151,0.1169,3.519,0.05756,0.0353,0.5267,0.02432,7.376,0.3226,0.2897,2.929,1.972,1.784,2.949,2.45,2.245,0.004855,0.02531,7.786,0.9906,1.066,11.18,0.2535,1.813,0.6566,2.439,1.827,0.01094,2.182,1.613,3.654,1.294,0.4829,0.8027,0,0,0,0,0,0,0.3376,2.638,0,0.08792,1.063,3.112,12.5,9.823,0.1446,1.401,14.45,6.129,0.8781,0.3264,0.2017,0.1728,3.568,2.123,1.109,2.858,0.4516,0.02266,2.49,0.01746,0.2442,0.01043,1.118,0.3576,0.03551,2.359,3.918,1.644,0.0117,17.59,0.2038,6.565,0.2179,1.439,0.0008205,0.1568,5.087,0.0136,2.971,0.6961,0.367,2.478,1.223,0.4533,0.8935,6.305,2.82,2.445,0.2291,19.31,0.1706,9.216,0.0003357,0.01657,2.074,3.187,2.642,5.852,0.04331,0.5051,0.9977,0.6738,5.167,2.682,0.2524,0.3909,0.1304,0.8593,7.136,13.17,0.1928,3.645,3.046,7.308,0.01999,0.7883,1.031,6.764,1.433,0.8209,0.4253,1.283,7.75,2.104,0.3213,0.05514,23.05,8.633,0.6666,1.835,0.1653,0.2386,0.4924,0.8822,22.9,1.813,10.26,0.2803,2.411,0.03592,1.229,1.343,9.169,7.775,1.771,0.1985,0.1248,4.452,0.3192,0.1775,7.514,0.2001,0.07815,0.04062,0.1608,0.03233,1.246,7.437,0.1139,5.551,0.2511,0.2605,0.4353,0.7371,0.837,0.08833,0.09338,0.7078,2.383,4.338,0.7397,2.52,0.2951,0.2809,0.1844,9.853,6.064,0.005773,1.086,0.1655,0.06242,0.9647,0.2266,6.195,6.82,7.212,2.502,4.74,2.522,9.365,3.146,0.5942,1.362,0,0,0,2.96,0,5.349,0.7905,0,0,0,0,5.368,5.667,6.675,0,0,1.294,0.3104,0.06332,12.94,0.0222,0.05003,0.5877,2.282,0.08012,0.001525,5.396,0.3636,7.365,2.46,0.00639,1.579,1.782,0.007506,0.18,20.75,10.93,14.25,19.98,4.244,13.31,2.227,6.4,5.844,0.5121,0.3037,0.04726,0.3784,1.161,4.284,2.058,0.3623,3.436,0.0346,2.735,0.009196,12.45,0.1891,0.3063,0.09324,2.224,0.6129,4.907,0.2767,0.00344,2.74,6.489,10.84,0.036,1.676,2.603,0.6622,0.3786,2.102,8.239,7.743,14.23,7.082,1.038,1.2,11.85,3.247,2.825,1.699,0.1653,5.782,0.2917,0.1171,0.06016,4.663,0.1796,1.718,0.2709,1.983,10.32,0.6824,0.4783,0.1821,1.791,0.6572,2.548,2.154,2.617,1.653,4.2,0.1704,0.2867,1.254,1.28,0.02158,0.6792,0.9161,0.009796,3.509,1.489,4.867,1.697,0.01946,1.445,2.234,0.9762,1.821,0.01573,0.374,6.415,0.7558,1.702,6.981,10.2,0.05137,9.952,0.0006183,0.08789,4.12,0.01877,17.55,3.531,2.606,0.07982,0.01409,0.0002913,0.1653,2.022,4.819,0.114,0.5752,4.291,0.04556,0.5866,0.3175,0.06086,0.8474,0.222,0.1062,22.11,0.0002246,2.712e-05,1.097,0.2498,0.002139,3.778,0.2507,1.947,0.09308,1.779,0.02612,0.02642,0.1279,0.02758,0.001094,0.9655,3.096,0.5956,1.625,1.001,7.136,9.217,2.195,1.353,2.216,7.344,1.287,1.582,1.129,2.152,6.869,2.576,10.06,0.2781,2.606,0.09556,2.971,4.125,6.523,3.58,1.284,5.117,0.9568,4.734,13.45,0,2.267,0,3.228,0,0.8364,13.27,10.07,0.2461,2.567,0.1489,0.5716,0.3685,4.501,19.34,0.121,3.665,5.827,0.06866,0.4614,0.02406,0.3301,0.1254,4.318,2.661,3.057,0.9202,0.003511,1.215,12.49,2.335,0.0588,0.3693,0.0005453,0.435,0.0206,14.39,0.02387,0.4956,0.1955,0.08804,3.717,0.2421,0.01384,9.266,11.38,2.938,0.2486,0.1503,8.54,0.1628,2.316,4.427,0.005356,5.416,11.43,0.008224,4.084,6.126,7.757,3.008,16.56,2.106,0.1299,0.05508,0.005559,1.731,2.291,1.401,10.45,3.119,4.859,3.109,2.611,1.255,7.897,0.6918,3.806,0.1316,1.12,0.9641,1.084,0.005132,0.829,1.492,4.232,5.057,0.0002886,0.7386,3.333,0.1082,0.4919,0.4942,0.003861,0.7249,6.105,2.049e-05,0.006398,0.2289,1.468,0.01366,10.75,8.081,9.3,0.7951,0.2645,0.2053,1.293,0.06672,1.602,1.239,1.883,19.13,0.4925,1.468,0.1134,2.148,8.859,17.45,5.888,0.6153,4.788,0.4158,0.249,0.1031,6.122,0.4583,6.55,5.732,0.9878,13.07,1.759,22.34,22.02,9.885,14.51,0.0001158,9.608,1.219,0.01815,3.903,2.142,8.847,0.1933,0.2488,5.215,10.19,0.000981,6.575,5.1,0.1778,0.1241,0.000578,0.7886,0.3013,2.362,6.691,3.701,2.829,0.3002,0.4771,0.7871,0.0437,1.653,1.726,1.124,1.073,8.339,5.372,9.405,2.2,0.5304,12.59,0.002046,0.1192,0.006885,1.649,0.3661,0.1736,0.4972,0.2262,8.438,6.536,5.584,7.491,10.98,1.832,12.79,6.741,10.54,5.495,3.766,1.955,0,4.812,5.316,1.7,0,2.908,2.333,0,9.704,5.105,0.3706,9.662,9.405,18.36,6.571,2.769,2.706,1.823,0.935,17.32,0.4018,0.1673,0.3951,2.685,4.334,3.747,10.71,14.11,9.979,14.01,17.1,0.3286,13.4,0.003132,9.225,1.462,0.73,15.67,0.3096,1.003,0.02991,4.79,0.3536,18.3,2.908,6.379,0.04086,0.02088,2.542,1.533,0.3434,11.41,3.282,0.5185,1.487,0.1699,0.07472,2.977,0.001189,0.1159,1.16,0.3995,0.02376,3.173,12.23,0.1234,6.255,3.403,4.648,7.793,5.859,0.000759,0.1538,3.481,1.771,0.4342,0.5162,0.9887,0.5114,4.357,0.8804,22,0.99,0.252,2.721,0.4117,0.9903,10.28,1.85,0.04202,0.9381,0.02887,0.7155,4.937,0.8269,1.585,0.2402,3.025,1.36,19.56,0.8194,0.0373,7.529,1.92,0.05497,0.1742,0.1582,1.941,11.54,0.4878,11.18,0.08268,0.5191,0.3956,0.0189,6.114,0.3215,18.26,0.05377,1.694,0.468,0.9516,0.6206,0.006756,0.101,1.171,1.299,0.09605,1.998,2.116,6.261,16.48,0.4191,0.02585,4.594,2.457,7.535,0.08194,8.417,0.4866,3.334,2.692,3.079,0.1118,6.695,1.057,9.036,0.1657,1.928,0.5313,3.541,0.969,0.7363,1.303,0.1798,0.02525,0.5415,9.35,0.1577,3.724,0.1066,1.136,0.3069,0.07976,6.167,2.117,0.1296,0.753,0.3337,0.314,0.567,0.3665,0.0006886,2.579,3.417,1.077,0.7436,0.01449,16.91,2.181,9.359,0.2719,0.0792,0.6214,12.75,0.5108,0.6543,0.3241,0.9373,0.002988,3.042,0.05554,0.1362,0.02396,0.04938,2.481,8.439,0.7045,1.563,2.133,0.9074,7.211,0.0997,22.62,2.409,6.509,0.002333,10.01,12.92,0.7525,3.33,0.8238,10.2,11.73,0.9331,6.994,2.635,4.073,12.84,8.475,2.54,0.01717,0.2681,0.05583,3.976,2.048,2.433,2.363,0.2816,2.447,1.456,2.369,0.1629,0.5138,1.245,1.158,0.7928,17.41,9.036,5.631,2.312,0.3032,0.4207,0.01925,1.913,0.5543,0.696,1.588,0.1468,5.345,14.13,0.1033,0.1192,9.858,0.8502,0.2013,1.388,1.364,10.67,1.062,5.461,1.303,5.673,0.2604,0.1603,3.018,0.07633,0.697,4.353,0.008338,0.6989,0.7254,0.2078,4.43,0.04417,0.0006446,0.005786,1.338,0.117,0.8596,0.1771,0.179,0.2264,0.01829,0.09146,20.8,6.326,4.131,0.176,10.76,1.642,0.3938,3.467,5.996,0.1397,7.835,0.5612,0.2951,0.4271,4.745,9.235,0.8161,5.465,2.827,18.76,0,0,0.721,0,0.879,1.596,3.17,4.99,2.732,0.05374,5.203,0.6117,18.65,2.037,1.357,6.612,1.519,0.07426,7.128,10.79,2.238,7.718,1.118,1.691,21.67,0.4424,19.91,0.07436,2.958,0.8207,0.8617,1.401,3.695,2.228,3.791,13.82,6.551,0.738,0.118,4.543,14.11,0.5136,0.1009,8.425,15.85,7.007,10.57,0.01184,4.047,19.82,13.32,0.4001,1.243,1.445,1.212,0.4172,0.4778,0.08719,1.663,0.692,1.918,0.8654,3.275,0.08801,0.6142,0.01091,4.552,0.04874,0.0003068,0.1501,18.12,14.98,3.838,1.691,0.4544,0.1274,0.3285,3.57,0.04488,1.034,4.213,0.6511,0.3451,0.1231,0.2534,2.453,0.1274,0.3905,0.001761,0.01093,1.846,1.912,2.342,7.521,0.0004992,0.04643,2.555,0.8374,1.608,0.155,4.278,0.06283,7.089,20.87,0.3798,0.6304,1.078,15.49,0.09591,0.03718,0.02853,8.208,0.000146,0.4898,8.59,1.05,4.746,5.299,4.803,0.8521,0.7199,0.8987,0.5698,3.039,0.2195,0.04085,1.659,1.426,0.237,0.4134,2.072,0.06253,0.01949,1.178,2.677,9.027,3.087,2.055,0.06527,0.1567,6.057,12.79,0.3569,0.5299,0.6403,1.401,4.199,13.34,0.03476,0.01019,2.029,15.08,2.541,0.001685,0.004157,0.3811,0.9329,3.335,2.577,4.614,0.01406,10.06,2.417,20.35,7.408,0.4882,0.07226,0.01413,1.69,3.066,1.071,9.307,1.557,0.02274,0.0156,2.133,0.4724,0.9928,1.636,3.773,3.398,1.909,0.4997,15.52,0.09377,0.01838,4.45,0.3871,2.181,1.784,1.187,0.4792,4.801,0.06361,0.01541,3.618,2.94,0.1518,23.09,2.069,0.01343,20.09,15.73,19.9,2.503,5.901,5.333,0.8819,1.864,0.3659,4.281,0.8967,0.4121,1.967,0.03619,1.854,5.481,8.272,0.5651,3.598,0.01548,0.6469,2.069,1.958,0.0206,5.27,0.7068,6.143,3.141,6.541,5.282,7.243,19.87,10.45,6.74,6.689,3.223,2.689,0.7162,12.33,4.019,5.217,8.017,8.351,6.661,0.328,0.8134,0.04969,4.922,4.38,2.252,11.54,7.499,0.896,0.04942,8.689,3.74,0.1234,0.2991,4.805,3.262,0.8879,3.394e-05,0.02488,0.0395,6.294,0.1422,0.01613,0.02806,5.974,0.01121,0.4301,0.2186,0.1535,3.038,3.317,3.052,1.862,2.128,0.2216,0.2762,0.04368,0.6562,0.1871,0.0428,2.101,0.4031,1.42,0.4225,6.848,0.7028,0.9855,0.3156,0.7779,0.3952,2.038,6.028,7.208,0.2179,0.3783,0.004351,3.408,4.314,4.745e-05,3.981,0.2708,2.827,6.498,1.517,2.061,1.499,1.295,0.2486,12.49,0.4324,1.627,5.632,0.002506,1.576,3.694,0.01481,3.95,2.29,0.2088,6.397,0.1352,0.4609,6.994,2.69,3.524,0.7557,0.2698,7.29,0.2789,0.00275,0.7404,0.004594,2.84,0.03239,2.356,0.6625,0.422,6.244,7.813,0.1518,2.494,1.22,0.2099,0.293,0.5736,0.02119,0.1863,2.052,0.03152,1.415,0.06688,0.843,0.7857,0.05201,0.4592,1.154,0.07399,0.987,0.02987,5.4,4.201,0.003563,0.1078,2.268,10.97,0.008813,0.02255,0.04255,2.946,3.719,3.051,0.7233,0.003066,0.2667,0.3327,11.5,4,6.024,6.523,4.985,0.1803,0.04638,2.143,0.6349,15.68,8.521,0.3798,1.154,8.489,0.4005,8.279,0.8978,0.4525,0.03165,1.672,8.148,0.4588,0.7387,0.2398,8.799,0.3486,18.48,0.1463,0.03167,0.2318,7.359,0.0002994,0.001989,0.000921,0.0003386,0.009648,6.204,4.431,7.81,2.05,1.917,0.0692,3.124,9.192,0.0563,0.5029,14.5,1.22,0.8283,2.741,0.5457,0.4418,2.058,1.392,0.5606,0.08269,0.001289,0.1016,1.906,0.9791,2.243,15.78,6.874,5.201,1.831,0.0577,0.005404,0.3747,1.106,0.0002004,0.333,0.2507,1.567,0.198,0.2163,14.2,0.3853,0.5099,0.2185,1.392,9.929,0.4663,2.123,0.01128,1.723,7.022,0.1492,0.6508,0.4259,2.006,0.008253,0.04651,1.125,1.014,0.9447,2.099,0.1906,0.07285,0.038,4.093,0.7013,2.994,2.297,2.297,2.434,5.29,0.4268,0.1627,1.459,0.5497,4.603,4.072,0.9865,8.765,13.65,0.1869,0.2627,13.03,0.2094,3.68,0.07559,4.176,0.007625,0.829,0.1048,0.07597,0.1117,0.03157,0.01682,0.1604,10.12,0.3672,4.638,0.4321,0.01302,0.06164,0.1745,0.1426,1.689,8.446,0.3432,2.497,11.38,0.0009041,2.475,0.2528,1.103,0.6516,0.7837,2.864,0.00522,0.6309,0.01115,1.748,0.06119,1.093,0.0538,0.09499,1.439,6.4,13.83,0.004187,0.9967,0.6419,5.751,7.672,0.09591,0.207,0.4144,0.04624,0.03268,0.1277,6.253,0.9849,9.407,0.902,0.3201,0.3595,0.115,9.921e-06,0.3515,0.4392,2.239,6.868,14.29,7.664,0.3488,0.01824,0.3806,0.0001605,0.002242,4.434,0.2231,0.04621,15.7,10.41,19.12,18.11,0.172,3.73,0,0,0,1.738,4.395,0.391,4.282,5.409,3.212,6.496,6.914,6.783,3.505,0.7733,0.121,3.131,6.826,2.469,10.93,2.168,0.3699,0.2097,0.01766,0.03549,0.03344,0.1448,1.958,0.5318,0.1131,0.03762,15.97,0.2442,12.76,4.229,0.006463,1.207,0.009864,5.168e-05,4.12,0.7412,7.691,0.04765,0.03015,0.075,1.452,0.002959,0.1045,0.05326,4.35,20.26,0.6421,0.08865,1.329,5.053,0.2648,0.05654,1.072,0.003933,0.93,1.972,2.155,0.682,3.234,0.1111,11.33,0.05016,0.2682,4.964,7.396,2.604,1.215,0.02508,9.292,10.35,0.1531,22.11,0.8491,0.2276,2.331,1.089,20.12,0.07744,2.292,0.5025,11.38,1.551,0.0593,0.6638,0.1951,5.291,0.003504,0.6441,5.569,2.437,10.95,5.669,1.574,5.92,0.9386,0.06935,0.03788,1.106,0.01456,1.158,0.8863,0.3496,0.4633,2.494,0.01572,2.616,0.001614,0.1862,0.009282,14.97,3.462,6.3,2.387,0.0005016,3.5,2.297,0.002589,0.9673,0.6736,0.3062,0.01197,0.1365,0.06274,0.06404,1.684,5.089,0.05747,5.243,5.593,0.1231,0.6396,0.8239,1.063,18.84,7.645,1.211,0.5668,5.386,10.85,0.09847,8.55,2.494,2.063,0.9742,0.5239,0.004725,0.5316,1.792,0.01632,0.007909,0.2711,0.9098,2.879,2.395,0.376,4.949,0.002518,1.587,1.061,0.2741,5,0.3558,0.0404,0.003891,4.077,4.97,1.056,0.1833,1.504,21.88,1.98,0.599,3.764,6.101,2.469,3.33,0.3748,9.487,2.218,3.958,0.02734,0.6392,0.2446,1.618,0.6174,2.686,18.89,0.9193,0.7162,0.477,1.977,1.307,5.185,1.953,0.224,0.3006,0.3903,0.4498,0.1284,1.657,11.39,12.76,1.631,0.5252,1.923,1.483,0.04731,2.316,0.4752,1.928,3.108,0.08142,0.5263,0.09073,1.953,1.92,16.8,8.522,1.575,0.03812,0.976,0.02629,4.008,11.73,2.939,0.01299,0.02258,2.046,0.8185,0.05714,3.714,0.0004143,1.485,0.03522,0.02773,0.002266,1.085,5.442,0.003087,0.002287,2.103,8.169,0.9457,0.1868,0.4252,1.308,0.001142,0.08949,0.0414,0.06744,0.01184,0.1765,0.06105,0.2424,3.923,0.1225,0.2867,1.656,1.748,0.0009927,0.1655,6.627,0.4466,0.1003,6.505,0.2264,0.4198,0.113,0.5348,0.834,2.883e-05,0.6608,3.186,1.228,6.084,0.05528,0.03408,0.7947,7.563,3.194,0.008164,0.01279,0.02129,0.01857,4.374,9.554,0.8569,1.869,0.0795,7.959,0.07272,0.2227,0.00123,0.6549,0.5969,1.364,0.3215,0.6303,0.1537,0.03793,0.3632,0.1786,0.0735,0.699,0.0244,7.36,0.8158,8.443,0.009661,3.12,1.679,0.4918,0.8531,8.239,0.04732,0.02957,0.02258,0.2094,1.085,0.000544,0.07887,0.03777,0.928,3.905,0.358,0.193,0.04821,4.545,0.01587,0.2857,22.5,0.4721,0.9755,6.571,17.81,0.05284,0.2571,0.4687,0.1543,9.085,7.295,0.008801,3.578,0.1168,9.139,15.76,0.03316,0.2555,2.32,0.4507,3.364,18.79,2.255,0.5563,0.6939,0.2729,0.07146,0.0007119,0.3125,2.559,0.1812,5.061,0.1971,10.53,13.2,9.513,11.96,8.285,6.812,8.211,0.2513,7.006,2.776,0,6.603,0,0,0.05335,3.056,5.281,8.299,4.91,3.475,1.969,1.105,1.386,0.009116,1.093,3.225,3.076,0.4989,1.319,5.986,0.1605,1.654,0.786,0.7784,1.135,0.1406,1.011,0.6207,0.0007607,0.5727,0.003764,1.184,19.87,1.586,0.07095,0.01836,6.177,0.07637,0.4418,0.02762,11.13,0.01417,0.09358,8.961,0.0251,0.0326,1.251,0.6651,0.1387,0.5346,0.02939,6.278e-05,0.0002723,0.7219,0.02036,0.005902,0.08199,0.1313,11.29,1.652,3.722,1.626,0.3467,1.659,0.4884,6.171,13.58,1.198,0.03543,1.267,0.4822,1.18,1.039,0.4336,0.1716,0.3848,2.853,0.4916,6.787,0.2168,0.03646,1.062,5.188,0.2003,1.104,1.152,0.903,0.04854,0.1721,2.821,0.07658,3.82,1.136,4.956,0.7647,1.87,2.397,0.3375,3.649,1.349,0.1117,0.2866,3.059,1.085,0.002254,0.3469,0.09598,0.06757,1.249,2.499,5.871,1.16,1.529,3.233,4.715e-05,0.04884,0.442,0.2851,0.9852,0.2644,0.8216,0.2609,8.667,0.09792,0.659,0.4516,0.0007521,0.1228,7.249,0.02592,0.2396,13.65,3.311,4.907,6.247,0.5147,0.1386,22.6,0.3235,1.493,0.01676,2.449,1.572,0.4296,15.29,2.03,0.2462,2.763,4.284,1.806,3.023,1.472,0.02025,0.1397,4.491,2.743,1.689,2.953,4.125,0.0602,0.6033,0.002374,7.674,5.582,1.361,0.09763,0.0005727,0.4819,0.9715,6.72,0.3523,0.2853,4.874,3.433,7.688,14.75,1.772,1.133,0.1114,8.132,2.175,5.366,1.778,1.508,0.12,11.1,6.314,0.2291,0.4481,0.3086,0.0168,4.766,1.672,2.745,0.4077,0.08476,0.1078,2.218,0.01979,0.007323,2.686,6.185,3.122,1.089,10.1,1.136,21.82,0.4818,0.4488,0.911,5.668,2.229,0.5684,1.417,1.86,4.812,0.4101,0.4018,5.159,3.868,0.4871,0.8411,0.6478,0.09134,17.42,0.02651,0.6325,0.3359,1.234,0.09189,0.0003415,0.05851,0.02699,0.7952,0.04349,0.5862,1.145,0.004265,9.29,1.564,0.1145,1.317,4.087,7.538,0.8043,0.9592,0.00752,2.819,0.246,0.1848,0.9608,1.804,8.795,0.1002,19.72,15.81,0.4523,0.1741,0.07327,3.144,0.06492,0.1569,2.294e-06,0.5422,18.75,0.3951,1.918,0.003688,3.695,2.433,0.01024,0.03033,0.06555,0.006584,2.799,0.009615,0.05197,1.582,0.00016,0.08439,2.587,1.475,0.07754,0.06766,0.7909,7.932,0.548,1.752,7.437,6.651,0.02887,0.29,12.28,8.952,0.2265,0.08835,0.3781,9.47,0.4882,1.402,0.9586,0.9783,0.1201,2.577,0.771,1.749,21.74,0.4195,0.00898,1.14,0.9783,0.003501,11.07,0.0189,1.897,6.201,0.3432,5.003,9.161,0.7086,0.3759,2.179,1.014,0.1294,0.03726,1.578,0.9141,4.822,5.668,0.2333,1.243,0.01813,18.34,2.484,4.808,0.1288,0.138,3.831,2.004,2.094,0.07879,0.01174,0.0538,1.45,0.1381,0.3813,0.7145,0.02072,5.753,0.03376,1.324,20.44,0.1916,0.4363,2.573,6.192,1.446,1.726,3.514,5.382,6.518,0.2698,3.784,6.565,16.48,0.01553,1.243,0.338,0.2583,1.733,0.707,15.99,0.2923,3.968,3.008,0.7408,1.136,0.03328,0.955,0.5394,4.92,0.03276,0.04163,0.5828,1.725,9.335,0.0645,2.293,2.843,4.038,1.412,0.0121,0.00575,0.02478,0.9716,0.6598,0.7509,0.1014,10.65,0.9229,2.268,0.8014,0.2663,7.259,4.219,0.8974,1.522,0.05129,0.7408,9.813,0.01366,1.066,3.214,0.007065,0.002409,5.469,2.149,7.219,7.03,0.9877,12.1,0.5478,0.1239,0.4602,0.1878,0.8369,1.799,3.873,0.6212,0.04039,0.8767,19.13,8.336,5.362,0.1337,1.963,3.761,1.079,5.338,0.01595,0.03902,0.9508,6.763,0.9803,1.153,0.3698,0.01159,8.193,0,3.253,9.367,1.255,4.817,4.172,0.09656,0.8784,0,2.559,0.8197,0.6478,0.3685,5.074,0.8212,0.4941,2.032,4.172,0.4308,0.5416,0.0265,7.691,1.248,0.1745,9.325,0.4309,0.4725,6.247,0.7837,15.55,3.953,0.06015,5.673,0.3027,0.1641,0.6435,0.1881,0.0008154,10.23,13.69,0.1717,8.014,0.01478,0.612,0.1392,2.261,0.004525,1.078,0.4284,0.08542,0.8768,1.134,0.008095,5.198,19.38,0.5586,0.05096,0.2758,2.915,2.452,2.399,0.4134,7.567,1.373,1.182,0.0879,20.63,0.3955,8.326,0.3059,0.04169,0.5867,0.01461,13.54,0.121,0.03606,0.08354,1.039,2.314,8.816,2.595,1.106,1.098,0.4734,1.828,0.7038,5.21,1.002,0.07164,1.117,0.115,3.665,6.698,2.515,0.9047,10.15,0.2529,15.64,0.8493,0.676,5.941,0.002966,8.501,0.2565,0.06311,0.0884,0.8525,0.8405,2.839,0.2552,21.43,1.081,0.5221,0.3771,0.5185,3.606,8.54,0.1302,0.2868,0.8761,11.77,0.09894,1.466,0.07816,6.328,0.6532,16.07,0.3329,0.0706,4.729,1.285,0.2943,7.192,1.179,2.226,1.456,1.989,0.2034,0.1436,0.04773,0.03229,6.14,2.445,0.01572,0.8205,0.8535,2.941,1.613,0.1046,0.06573,1.851,0.07584,0.02466,20.86,3.981,1.046,0.03372,4.924,2.496,0.2491,1.965,0.2289,0.2337,17.74,6.449,0.1282,1.765,5.166,10.17,4.384,2.585,0.9524,0.2036,3.809,0.313,0.3837,0.6747,3.536,0.01727,0.6474,0.07998,0.6341,3.172,2.431,2.528,1.096,1.379,0.2899,14.22,7.383,0.1027,5.879,0.418,0.006515,0.5328,1.616,0.1027,0.02351,0.738,0.1071,0.001667,1.576,14.06,0.756,6.043,3.276,1.56,0.4446,1.053,0.1541,9.238,0.03557,1.237,0.3129,0.0039,5.639,1.532,10.71,8.031,0.4508,4.802,0.09218,0.1457,0.0001101,1.513,1.501,0.8413,0.1592,0.02535,21.55,3.572e-05,0.03251,0.04045,0.1346,1.33,0.4814,1.153,4.03,0.04752,2.327,0.1658,0.05405,4.857,2.882,0.007638,8.171,0.1597,0.1525,18.23,11.26,2.249,3.911,0.1699,0.09696,0.0901,0.9863,0.08962,3.428,0.6913,0.2843,0.8707,5.683,0.05423,9.338,4.462,0.003341,1.415,3.206,7.624,0.9631,0.09092,1.228,0.4791,4.138,6.649,0.1052,10.67,0.01352,0.8112,0.0008811,0.8277,3.893,0.0548,0.1318,0.0001824,6.767,9.515,14.16,14.5,7.709,12.72,10.53,16.79,13.26,0.666,0.8039,0.1966,0.5814,0.001131,0.08671,3.251,1.569,0.2907,0.06373,4.298,0.783,0.4564,0.3092,0.162,0.1508,7.101,7.739,2.014,9.552,10.06,9.184,1.536,9.941,0.02528,14.19,3.103,2.725,4.539,1.402,0,4.899,9.573,4.415,5.308,4.606,4.629,0.4356,0.4725,1.31,0.6379,0.6611,1.677,0.7602,1.589,0.1383,3.388,4.358,1.205,0.09246,12.13,0.007698,20.48,0.334,1.182,0.5491,0.7044,0.496,0.6978,5.748,0.756,0.7129,0.1433,7.365,0.4763,5.39,0.0003664,1.307,1.321,0.8307,1.605,4.065,0.5441,0.05834,0.0007924,7.587,0.02656,3.902,0.1131,0.3941,0.05567,0.03465,0.8224,1.15,0.03833,0.03531,2.572,0.2568,0.2441,3.789,0.8085,0.05928,5.279,13.58,0.5212,2.58,0.6212,0.4234,0.05172,1.118,0.2272,1.094,0.1076,0.2383,1.031,1.215,1.051,1.551,0.08563,1.095,0.1025,2.156,1.93,1.113,0.01895,0.01722,0.2674,0.4408,2.193,1.276,0.3322,9.708,8.155,3.023,0.6529,1.014,0.01208,4.07,2.311,0.1986,0.006239,0.2807,0.3367,0.3824,3.547,8.592,12.74,0.1516,0.7464,6.536,0.1737,1.021,0.5203,0.006059,0.06809,0.4967,0.1343,0.784,6.787,0.07039,0.309,1.448,2.746,1.087,0.9217,0.1632,0.02073,0.2859,6.029,0.07871,0.4205,0.06189,0.07962,1.916,18.15,0.02646,5.38,1.477,0.002972,0.03,1.62,0.9483,2.773,1.04,0.05871,5.359,1.348,1.845,0.9112,2.159,3.273,0.4821,11.14,8.269,6.456,2.526,0.7906,0.0003282,0.08716,0.2113,0.3502,0.002518,2.443,0.2391,1.927,1.057,7.002,12.21,0.01655,0.02004,1.352,4.307,0.7592,0.003253,0.0003355,0.03572,0.6476,1.121,3.686,12.46,10.09,0.7781,1.305,6.006,1.388,0.07779,0.01216,0.9402,0.9499,2.82,0.08324,0.2048,1.208,3.171,0.07855,18.23,1.818,1.908,0.01096,0.5273,3.345,1.345,3.005,2.059,0.6054,0.5944,0.1669,0.9695,1.958,19.98,1.571,22.35,0.08834,0.4053,0.4413,0.4007,15.03,0.4293,0.003946,0.5315,1.375,0.09333,4.893,0.119,0.006805,0.3454,3.451,0.2035,0.3555,0.643,5.748,0.3244,0.5587,0.07197,0.196,0.04663,3.354,0.005828,3.895,0.02008,0.7423,1.564,1.701,0.05411,3.004e-05,0.4484,7.268,0.4965,0.3291,0.159,0.02943,0.4429,0.3239,0.3868,1.804,0.2265,0.3881,0.1766,0.3082,1.102,3.533,1.954,0.2025,2.109,15.97,0.1833,0.2654,0.8494,0.1617,6.078,0.03972,1.518,1.134,0.07873,0.0208,0.5681,0.514,0.008762,0.3113,1.144,1.89,1.39,0.8649,18.3,0.598,0.08126,0.01736,2.026,0.6662,0.7642,0.05681,0.2483,1.916,3.679,2.43,1.325,13.13,5.217,8.361,5.958,1.036,0.1653,0.000796,0.1057,0.5819,0.7189,2.14,7.026,5.348,0.3361,2.653,0.0387,0.1554,4.393,1.21,1.268,0.04809,0.1207,0.0401,0.8275,1.168,0.06505,14.26,3.719,0.09112,0.03694,7.604,1.795,2.702,2.858,11.98,3.589,3.925,0.04437,1.055,10.21,0.007048,0.01442,2.656,0.1307,0.776,15.02,0.0003315,0.0601,0.1912,1.066,9.248,0.0798,0.8381,0.02289,0.2392,5.635,0.3347,12.18,1.558,0.08216,0.01047,1.448,0.2116,1.335,6.01,1.376,0.1901,1.041,0.06465,4.011,1.647,0.08214,0.3999,8.717,1.394,1.708,1.533,0.1643,3.681,10.88,0.05131,2.787,1.038,2.995,0.02987,0.2951,0.005253,1.098,0.06605,4.399e-07,0.7082,0.07695,2.625e-05,17.78,11.02,0.000656,0.00459,2.256,0.4491,0.4941,0.2808,2.491,2.836,0.3009,0.02337,0.1276,0.1316,0.07172,3.771,8.434,2.438,2.149,2.119,0.1991,1.538,0.001723,16.79,0.02565,0.08386,0.5946,1.547,1.038,0.0008567,0.9007,20.81,2.298,1.064,0.6998,3.415,0.00564,0.5137,0.03872,7.425,0.003794,1.986,5.68,6.848,0.396,5.657e-05,1.64,1.927,8.599,4.849,0.688,4.194,0.6784,2.503,1.239,1.063,3.039,6.795,3.259,4.222,6.2,14.19,17.04,14.35,8.779,4.904,3.626,0,1.982,0,2.085,2.323,0.2111,0,3.242,6.625,2.556,6.169,14.89,5.258,8.989,12.31,0.8267,0.1598,0.759,3.073,2.423,0.5228,4.828,8.188,2.089,1.116,0.04989,15.57,18.48,0.6627,0.03619,4.19,0.08604,0.009564,15.86,2.546,4.639,2.345,0.08457,5.669,0.1337,1.468,1.519,0.313,1.81,2.706,1.031,0.7066,9.296,0.1704,11.62,0.2451,0.3349,13.06,0.01326,16.41,0.001252,3.835,0.4875,7.952,0.06278,1.485,0.3765,1.969,2.096,0.003042,1.885,2.075,0.05094,1.639,0.02814,0.1345,1.305,3.595,7.491,1.05,0.1261,0.05359,2.712,1.975,5.477,3.628,0.1687,0.05425,1.605,0.08331,0.04889,0.5984,0.2413,1.312,0.04163,0.1002,15.67,4.537,0.2283,1.618,0.8368,3.53,19.22,1.393,2.503,8.417,0.0836,10.02,11.75,0.2803,0.3324,0.2879,1.399,0.1288,6.406,0.2522,0.01729,0.07195,0.1729,4.207,8.876,0.8292,3.399,2.988,0.06581,2.71e-05,0.005542,1.796,2.832,0.1049,3.279,0.1751,0.1632,0.7725,0.9389,0.4016,0.2304,13.45,6.299,2.181,0.4355,0.2826,1.341,0.3761,0.07788,5.745,0.5055,2.012,0.0214,0.03942,0.1203,9.374,0.3429,0.8575,2.271,0.002494,0.07329,1.293,0.1453,0.5112,4.308,3.236,4.081,0.06504,0.05014,0.0002036,0.6376,0.9169,0.03659,0.02322,1.143,0.731,0.003702,1.785,1.501,0.6175,0.008044,0.3304,0.8131,0.01256,0.3327,0.03925,5.821,0.08025,4.734,0.005207,15.92,23.12,1.443,0.4587,16.14,0.2842,0.9044,4.252,0.08898,6.902,2.406,0.5096,1.629,3.188,0.005858,3.62,0.2337,7.549,0.003023,2.752,0.2196,0.02166,0.348,0.01044,0.1709,0.1322,11.49,12.59,0.000663,0.07648,0.01074,0.6529,4.823,8.699,0.3343,0.07825,2.143,9.501,0.4712,0.09998,0.07798,0.000858,1.74,0.03456,0.08814,0.7269,0.3404,0.008118,0.009231,1.285,3.101,0.07913,0.04649,1.574,0.3825,5.619,3.55,2.45,4.409,0.1699,11.67,2.526,12.88,15.83,0.001118,4.434,0.9615,1.119,0.3925,5.495,2.784,3.71,2.534,3.158,8.912,2.136,1.29,4.225,14.17,1.02,1.125,3.422,0.9619,19.93,0.2058,0.04257,0.07149,12.86,3.222,0.7444,0.008364,0.1654,18.28,0.005264,8.248,1.118,0.1946,0.006098,0.03374,4.025,0.2771,0.6833,0.2084,1.084e-05,4.837,5.896,0.1659,0.7941,0.8135,0.9589,5.883,0.03347,0.003814,0.007691,0.003139,3.019,1.145,2.999,0.3567,0.04431,2.431,0.06462,5.308,0.7631,1.324,0.02583,8.461,4.205,0.6451,5.978,0.7249,3.058,3.041,2.991,2.803,0.05674,0.07699,0.03048,0.07097,0.178,2.629,0.3108,0.1216,9.294,2.813,0.499,0.3517,11.29,0.7269,0.7334,0.0323,0.7401,0.4736,0.7331,3.395,1.163,0.1475,0.3556,0.1457,0.0651,0.2464,0.04465,0.4851,0.0287,9.236,0.1194,0.3142,0.7611,0.3947,0.04258,5.872,4.462,0.9195,5.28,0.04845,0.004299,0.5739,0.0002036,0.3293,1.121,0.002766,2.466,0.08216,0.9353,7.128,6.326,0.1003,2.911,6.466,0.327,0.03201,0.7757,8.908,1.684,0.6226,0.0827,3.056,1.307,1.12,2.831,1.237,3.735,0.07082,0.8873,0.04044,0.1268,3.492,0.3607,0.3119,0.9062,0.3828,1.456,2.95,1.549,0.00181,0.1215,1.438,13.78,0.2108,0.1461,0.2742,8.158,12.46,1.146,0.8843,0.04265,1.868,0.1315,2.788,0.1628,0.9318,0.7914,0.1818,1.043,1.434,2.794,0.2789,0.1469,0.4626,0.1904,0.4443,0.1721,0.02097,2.534,0.1213,2.87,0.1651,2.282,3.462,3.991,0.06743,0.05824,5.283,6.357,0.06663,0.0179,0.06417,0.5652,0.1409,0.08626,0.05079,0.4292,4.521,3.295,0.3838,1.963,1.57,0.7231,8.462,0.1179,0.1395,0.04491,0.1037,0.6135,4.456,0.3601,0.3547,0.5505,2.4e-06,3.806,1.361,0.08426,0.4028,0.7304,18.35,3.981,0.01378,2.118,0.3531,0.05964,0.4674,0.4232,0.05271,3.559,0.2175,10.8,0.2949,0.9716,0.1797,2.968,1.761,1.798,8.34,0.04221,12.25,1.449,15.89,1.406,1.67,0.05037,6.926,4.096,2.191,0.9932,0.005808,0.8666,1.912,0.02922,2.198,0.003004,0.9553,1.36,3.577,2.611,0.3752,0.2337,5.31,0.374,0.189,0.005426,2.63,1.526,0.6744,0.9609,1.279,0.2285,0.07651,0.1637,1.353,0.7062,0.9505,1.939,1.14,0.02101,0.9567,3.65,0.3098,0.221,1.618,0.3271,1.858,1.695,5.681,1.977,0.8327,2.963,1.638,0.5912,0.3573,0.3285,0.2769,0.4156,0.1263,0.3807,1.338,17.73,0.8499,0.2359,0.1137,0.3335,2.878,0.6031,0.07338,4.342,0.942,0.01174,1.328,0.2498,0.7954,0.04212,1.478e-05,1.61,1.835,0.6976,0.5828,1.547,0.7253,0.001066,0.384,0.3844,0.139,0.08692,1.006,0.2573,9.938,2.333,3.278,6.417e-05,4.497,1.48,1.868,0.1885,3.755,0.4834,0.186,0.3964,0.8891,3.226,0.6158,0.6674,0.851,0.4634,1.093,0.5599,0.02104,7.776,4.234,0.7344,0.7035,3.438,1.22,1.998,9.651,1.114,0.02666,1.033,0.5485,0.3158,2.104,0.6102,1.903,0.0695,0.2612,6.364,11.6,0.332,1.74,6.399,1.379,1.879,0.2888,0.8846,7.764,7.626,5.929,0.03006,0.1431,13.4,0.3039,0.2884,3.672,0.00486,0.2066,0.2169,3.472,3.852,1.187,0.153,1.078,7.375,6.371,0.3467,1.725,0.01544,1.536,1.127,9.722,6.078,0.03002,0.004147,7.179,2.925,0.02968,0.1881,1.939,8.799,4.781,0.09991,4.709,0.3367,5.005,0.9503,0.01134,0.579,0.398,0.004027,0.5602,1.132,1.478,7.116e-05,7.534,3.092,6.329,2.12,1.476,13.15,1.436,14.25,9.488,9.323,9.205,7.283,9.422,5.645,0.6491,7.456,0.2049,0,0,0,0,6.595,2.739,3.336,4.339,6.755,1.842,0.2622,13.85,1.879,1.573,1.038,22.4,6.194,3.928,0.9613,0.05474,0.2433,2.534,0.1811,1.274,0.731,1.877,0.5993,0.7375,0.6006,0.6975,4.093,8.63,8.286,0.1567,0.002462,0.1997,1.021,0.02719,5.757,7.598,1.279,0.5927,0.9495,0.3053,0.9365,0.2957,0.7235,3.298,0.07145,1.226,10.01,3.423,2.384,0.8459,0.2542,6.806,5.095,9.482,1.043,0.426,0,7.108,0.004544,0.03299,0.46,14.46,0.254,0.2679,0.339,7.029,0.4082,0.6751,0.1881,0.3247,2.116,0.4658,6.259,0.393,0.1055,0.1048,0.4908,3.588,13.25,5.488,0.2535,0.3902,0.6101,1.901,2.371,2.586,2.259,8.427,5.533,2.406,0.05667,0.1121,7.329,2.068,10.35,4.826,0.3314,4.382,0.5589,0.3617,1.004,0.7847,0.8826,5.81,6.479,2.262,0.01667,0.004503,9.485,0.02529,0.000143,0.3729,2.774,0.1354,7.031,0.1624,7.752,0.002301,0.343,1.288,10.97,0.002739,0.0119,6.325,7.041,1.328,2.803,0.2592,0.6776,2.856,10.79,0.07427,0.04777,0.5263,6.505,0.307,0.1301,0.5782,2.717,1.43,0.1844,1.939,5.815,5.099,0.7572,0.5607,18.73,9.836,0.1194,19.9,1.092,1.116,2.56,1.21,0.3565,9.981,0.6893,12.71,0.3812,0.2443,19.9,0.04029,0.6884,0.5166,0.874,0.06543,0.2237,0.1767,0.91,9.562,2.09,14.78,2.255,0.8374,0.04208,0.004035,0.03194,0.02527,2.11,0.001619,0.1111,17.72,1.585,5.563,1.011,4.35,0.07735,0.3585,0.8438,0.03366,0.8514,0.268,0.02033,1.161,1.49,0.4364,1.366,0.5847,6.149,0.9443,3.255,0.5062,0.1683,0.0001274,0.4295,0.157,6.781,1.221,3.204,0.02392,0.8484,0.6007,0.06261,4.334,0.04926,0.6037,2.463,1.346,4.905,2.315,0.8769,0.0106,0.9815,0.1815,8.801,8.534,0.2448,0.6479,0.006962,0.3414,0.003058,0.03071,3.605,7.24,8.458,0.6724,0.9633,19.36,0.03235,0.1062,0.0004499,1.242,0.3345,0.0855,2.119,2.803,3.825,0.03112,5.398,2.844,0.01831,0.05393,1.149,0.399,0.03529,0.1329,1.115,0.443,13.33,2.349e-05,0.05524,0.004174,1.404,0.5432,8.633,0.1553,6.465,3.467,0.2038,1.454,1.396,3.542,0.6476,0.342,1.595,16.99,0.4687,0.4749,2.931,2.444,2.756,0.6899,0.4344,2.048,4.55,0.6521,2.672,0.0004863,1.944,11.99,0.5251,0.7092,3.21,0.05699,0.0009518,0.5675,0.3075,0.3063,0.5786,0.3875,0.8199,2.908,0.05313,15.78,0.04845,0.003336,0.000657,2.564,4.099,0.1967,2.016,0.5548,0.0007476,0.1317,0.2245,6.489,0.3198,1.767,0.01574,3.401,2.299,0.09021,0.3064,3.431,0.7371,0.3549,0.05421,0.6137,1.344,0.2354,0.9519,0.5329,7.154,1.678,3.163,0.01557,0.05252,0.5924,0.5144,18.64,0.6196,0.07071,0.04552,0.5272,7.495,0.0001959,0.001773,0.216,0.2477,0.8299,0.2103,0.1812,2.589,0.7409,0.8572,2.786,2.63,0.3087,0.0758,0.8048,1.123,0.175,0.09286,0.08372,0.0976,0.005487,1.698,11.39,0.5644,11.64,4.9,0.00122,1.626,0.6244,0.0004114,0.006793,4.769,2.533,0.2316,0.06802,0.3862,5.973,1.696,0.2389,17.91,0.002166,1.426,2.329,0.4249,5.539,2.853,17.86,0.3173,19.47,0.003803,0.3492,5.747,1.534,0.3202,1.523,0.34,0.3027,2.189,10.38,1.258,2.238,7.921,2.163,2.351,0.09625,0.232,3.34,2.558,0.3771,3.148,1.47,1.89,0.009647,0.05023,0.3678,0.4308,2.478,0.6806,0.1587,10.28,3.089,0.05756,0.3205,4.621,0.7359,0.6447,0.08997,6.798,0.08399,5.699,0.4073,0.1062,2.992,4.138,0.03281,0.616,0.03347,0.04313,7.722,7.236,0,0,0,0,0.8151,3.537,2.025,9.412,8.659,12.15,5.477,15.21,5.791,4.846,9.278,7.145,19.76,7.687,18.54,3.286,7.517,22.52,4.882,18.19,11.87,8.633,0.8176,15.66,12.74,0.07348,1.175,0.5024,0.08029,4.068,0.3674,4.145,0.5635,0.8724,0.05959,0.008275,1.192,2.583,1.046e-05,0.01947,2.589,0.8708,1.351,1.904,0.6218,0.2452,1.522,2.708,0.3418,4.155,0.07855,0.07438,0.3781,0.1008,1.819,0.4069,0.000344,1.358,0.115,3.659,0.5415,1.031,0.1434,0.194,3.202,0.7645,4.411,4.277,0.4092,0.2092,0.3986,1.227,1.38,0.1836,3.356,0.6242,0.01007,0.4029,3.234,3.905,0.3718,15.47,0.02559,0.08267,0.8815,0.7611,0.05679,0.1101,1.15,0.05322,0.6224,1.031,0.01331,4.965,1.701,1.086,0.5339,1.588,1.943,3.274,0.7683,0.3317,2.348,7.709,0.3076,1.019,0.7193,0.4485,15.2,0.8176,0.212,3.167,0.05176,0.2905,1.105,0.8809,1.793,0.8018,1.311,2.251,1.1,5.496,11.25,0.1942,0.05316,1.198,0,8.84,0.01659,3.162,0.8704,0.6091,1.769,4.265,0.7834,0.6871,2.494,0.2618,0.1206,0.03961,0.7642,10.39,0.3628,1.064,0.0264,2.178,0.6057,2.983,2.855,16.08,1.131,1.116,0.02011,7.698,1.146,0.1504,0.9977,0.01665,0.005342,0.1965,1.402,0.0003645,2.992,2.253,3.875,0.4881,1.982,0.0042,0.9978,0.3155,0.4625,3.247,0.02018,1.11,0.777,0.4719,1.073,0.06222,0.05004,3.515,0.002939,0.07937,1.903,1.139,0.5953,7.184e-06,14.12,2.329,1.12,7.498,2.429,0.09923,4.289,0.2093,0.1627,10.09,0.02595,1.551,0.4861,1.465,0.5934,1.227,0.7739,0.06595,0.9104,2.611,0.3418,1.491,8.318,0.001226,0.02583,0.3395,19.62,16.99,10.2,0.7951,0.1746,0.8182,1.719,0.0255,13.31,1.341,2.699,1.691,7.482,0.3051,0.003173,0.008681,2.485,5.308,0.7282,0.1198,0.2724,1.266,0.5074,0.08696,0.006503,0.2242,2.602,7.883,0.6557,13.36,0.01112,3.886,4.398,0.2214,1.571,1.006,3.103,0.006837,0.2354,2.043,1.826,0.7026,0.6563,1.408,3.158,0.8289,6.093,1.395,0.03427,0.01013,0.5871,0.008677,0.1554,9.687,0.5209,0.02398,0.6338,2.759,2.161,1.604,2.146,2.911,0.4118,4.117,0.04338,4.984,2.856,7.301,0.05387,0.4132,0.3847,0.2482,0.865,0.1586,0.013,0.366,13.8,0.0003861,1.216,0.02121,2.829e-05,0.9877,2.155,4.605,2.713,0.6664,0.004613,1.154,0.06015,0.381,0.5491,0.5893,0.9656,0.06025,17.87,7.676,0.1967,2.138,1.026,0.0496,0.03117,12.55,1.986,0.01045,0.06936,0.1759,0.2782,3.838,1.927,1.869,1.248,1.447,0.9182,3.052,1.707,1.052,0.7659,1.921,0.331,0.2369,0.04696,0.1344,10.94,1.68,1.019,3.928,3.402,8.637,0.7933,0.365,0.6879,0.0117,0.953,0.8591,0.0001419,1.077,0.3722,0.1616,0.3306,2.745,1.337,0.001752,6.446,0.5746,3.948,6.791,1.754,0.1876,6.824,0.003976,0.3695,0.3313,0.0031,1.841,0.0001164,5.801,0.6905,3.95,9.419,0.7722,2.345,0.2036,6.215,12.5,3.927,1.007,12.63,5.209,0.5166,0.505,0.0761,1.702,1.019,0.05114,0.007309,0.5837,5.605,0.3054,7.908,6.726,13.61,1.243,11.06,2.054,0.01974,19.56,0.02037,0.004905,0.7433,0.1328,0.5493,0.3337,8.938,4.952,10.33,2.358,4.157,1.645,1.454,0.5129,0.6459,5.974,0.1687,2.129,0.722,0.194,3.762,0.1112,2.172,0.03373,1.971,4.677,0.7392,0.2706,0.853,0.1318,1.322,0.008741,1.22,3.271,16.58,10.11,0.974,10.48,5.644,2.195,0.04909,15.72,3.431,1.243,0.02044,0.06819,0.8182,0.6166,0.004417,0.004035,1.879,9.132,0.8829,7.556,4.038,0.1287,1.83,20.3,0.1066,0.02111,0.01659,0.2473,1.161,7.628,2.896,0.1546,0.7532,0.4595,1.232,0.0008737,3.768,2.187,17.61,1.949,3.776,0.1356,1.023,0.8163,3.172,0.4874,0.4405,1.224,0.005889,1.171,0.779,0.5146,0.002406,0.01014,3.307,0.8507,12.16,7.421,1.294,1.351e-05,0.06408,0.02015,4.355,2.219,0.0399,0.3329,0.006684,0.2337,7.382,0.5334,2.222,0.08804,5.309,1.783,0.2485,6.463,0.5515,0.2838,0.7297,0.6142,5.431,17.64,0.2299,0.1026,13.21,0.152,3.983,16.04,0.006811,0.03778,8.898,4.54,0.4366,0.415,9.965,0.01877,0.2836,7.045,0.05112,7.222,2.652,0.02924,2.391,10.81,4.31,0.1835,0.3832,0.2812,2.017,20.09,1.249,1.81,3.62,0.6365,0.0008353,0.1164,4.975,1.783,1.401,19.05,0.1126,7.474,0.4624,3.551,2.189,2.316,0.2336,2.6,1.323,9.571,0.1909,0.1763,0.2667,0.9563,0.3657,4.05,0.02205,3.049,5.223,0.129,0.9748,0.9345,0.3985,0.1114,0.001929,0.7463,0.4116,0.05488,0.02966,0.1844,1.719,0.2764,2.547,7.932,8.458,1.677,16.57,0.1562,4.888,0.6657,0.01586,4.773,0.01843,1.372,0.3611,0.4843,0.886,3.372,3.414,2.482,0.06231,0.1682,0.01532,5.301,0.08175,2.957,0.3708,8.406,0.04781,1.972,7.098,0.5497,0.006009,0.6883,0.0006699,0.11,0.3766,1.875,2.004,0.2523,0.0964,12.58,2.389,5.701,5.857,0.8554,1.681,0.8502,0.1477,0.02831,2.563,2.516,1.104,1.917,0.05105,5.875,3.319,7.531,0.03016,0.0002396,5.104,0.196,1.039,11.23,10.18,0.2841,0.3955,2.039,5.031,1.7e-05,14.37,0.8391,8.86,1.9,1.487,0.0614,6.761,0.9779,9.44,3.837,0.4315,1.617,0.0009173,4.054,6.73,6.376,18.82,7.918,13.61,3.661,3.949,6.976,5.116,5.649,3.733,5.355,5.574,2.699,12.53,0.03867,1.809,9.828,2.582,16.97,9.768,2.448,5.378,5.666,3.225,6.649,0.5725,0,3.959,4.544,0.358,0.05322,0.0473,0.0008554,0.4524,3.806,0.02172,0.005276,0.4022,0.6907,1.565,27.2,40.53,34.74,107.1,31.61,28.29,30.42,26.98,42.12,51.11,23.43,54.19,24.59,26.36,36.77,30.22,37.88,25.72,43.72,45.08,33.14,31.8,23.86,39.47,43.32,28.28,50.92,39.37,41.07,35.35,39.05,33.36,37.76,24.44,25.85,31.98,34.26,60.25,26.84,55.22,27.58,23.98,27.98,26.42,23.33,39.85,40.32,25.38,27.25,66.51,67.2,32.34,51.1,27.15,24.2,23.47,27.23,24.66,45.6,38.28,31.92,49.04,26.41,29.09,28.23,27.27,32.34,35.95,46.36,68.47,57.64,28.77,23.65,51.28,30.22,72.38,41.12,31.46,24.18,25.78,26.5,30.85,32.94,23.51,47.21,41.79,33.31,23.39,32.49,27.43,27.07,54.49,73.19,25.02,34.15,24.28,28.21,26.07,35.15,35.07,32.58,25.77,33,28.68,28.09,24.66,106.8,33.39],"PValue":[0.03996,0.366,0.3058,0.1656,0.06911,0.2057,0.004661,0.01873,0.6783,0.002979,0.1762,0.03437,0.1335,0.0004134,0.3507,0.008308,0.2081,1,1,1,1,1,0.08834,1,0.02643,0.003806,1,0.08774,1,1,0.2283,0.5432,1,1,1,1,0.4711,1,0.5417,1,0.6792,1,1,1,0.834,0.06161,1,1,1,1,1,0.07844,1,0.08791,0.06306,0.1482,0.02133,0.02096,0.2325,0.5331,0.04137,1,0.8461,0.2231,1,1,1,0.1743,1,1,0.5625,1,0.04023,0.5825,0.2164,0.1406,0.04704,0.3514,0.1672,0.03801,0.003529,0.04841,0.08627,0.03982,0.7435,0.07981,0.03312,0.08296,0.06314,0.05504,0.06426,1,1,0.1683,0.1711,0.5919,0.065,1,1,0.6873,0.8381,1,1,0.2862,0.02128,0.7967,1,1,0.1542,0.1056,0.03758,1,1,0.7731,1,1,1,1,0.02001,0.1356,1,1,0.03007,1,1,1,0.3547,1,0.2721,0.5339,0.007344,0.9474,0.5434,1,0.5775,0.8318,1,0.0667,1,0.05789,0.7364,0.9874,1,0.06327,1,1,1,1,1,1,1,1,1,1,0.6043,0.7755,0.1983,1,1,1,1,0.0271,1,0.2018,1,1,1,0.8119,0.9797,0.05236,1,1,0.275,0.1812,0.7253,0.06888,1,0.7042,0.4352,1,0.1572,0.01333,0.0006484,0.09644,0.4816,0.251,0.05678,0.3773,0.8866,0.06015,0.0625,0.01459,0.08439,0.1017,0.08834,0.07674,1,0.0004211,0.02581,0.007914,0.7753,0.07999,0.4083,0.1407,0.01088,0.006217,0.6174,1,0.03506,0.0197,0.004738,0.0006223,0.134,0.5625,0.00629,0.8551,0.484,0.9825,0.5348,0.8499,0.7678,0.1296,0.2116,0.4415,0.1543,0.2564,0.694,0.3514,0.9568,0.8604,0.003381,0.04322,0.004421,0.8684,0.01881,0.8759,0.9829,0.04452,0.3069,0.1544,0.1267,0.0008988,0.1484,0.9695,0.5888,0.9525,0.5339,0.1467,0.3229,0.5137,0.5859,0.0669,0.1531,0.2283,0.6539,0.3398,0.01652,0.5792,0.1062,0.6573,0.6844,0.596,0.05287,0.5197,0.04684,0.2278,0.8344,0.1514,0.7506,0.8623,0.4995,0.9094,0.004079,0.7201,0.01251,0.6532,0.704,0.318,0.1148,0.4799,0.0208,0.5562,0.2385,0.01116,0.3984,0.04518,0.4847,0.4666,0.8002,0.7947,0.3722,0.7987,0.3275,0.7339,0.2385,0.715,0.5454,0.09629,0.0626,0.07985,0.2616,0.008094,0.1713,0.123,0.561,0.1126,0.003932,0.2445,0.5253,0.4798,0.3177,0.2591,0.9998,0.679,0.9128,0.0005301,0.03985,0.5508,0.02458,0.002139,0.762,0.9963,0.4002,0.3693,0.04883,0.2164,0.2549,0.001188,0.007638,0.9107,0.8716,0.3428,0.0279,0.08832,0.3337,0.002134,0.5364,0.1012,0.2612,0.02609,0.1622,0.737,0.07942,0.8135,0.8534,0.4787,0.8781,0.01548,0.5781,0.598,0.1067,0.1797,0.2007,0.1056,0.1375,0.1539,0.9453,0.8756,0.0133,0.3347,0.3176,0.004218,0.6216,0.1973,0.4299,0.1384,0.197,0.918,0.1594,0.2226,0.07441,0.2724,0.498,0.3839,1,1,1,1,1,1,0.57,0.1256,1,0.7707,0.3183,0.0985,0.002826,0.006979,0.7089,0.2542,0.001803,0.02513,0.3629,0.5759,0.6595,0.6833,0.07754,0.1649,0.3082,0.131,0.5114,0.8823,0.1345,0.8966,0.628,0.9199,0.3063,0.5584,0.853,0.1445,0.06563,0.2184,0.9152,0.0007167,0.6578,0.02113,0.647,0.2481,0.9775,0.6974,0.03877,0.9086,0.1044,0.4167,0.5533,0.1354,0.2855,0.5106,0.3589,0.02342,0.1129,0.1379,0.6388,0.0004738,0.6852,0.008018,0.9856,0.8992,0.1695,0.09361,0.124,0.02812,0.8378,0.4877,0.333,0.424,0.03746,0.1214,0.6224,0.5408,0.7229,0.368,0.01695,0.002327,0.6666,0.07472,0.1005,0.01588,0.8894,0.388,0.3253,0.01955,0.2491,0.3786,0.5238,0.2743,0.01347,0.1666,0.5788,0.8174,0.0002072,0.009811,0.4265,0.1947,0.6898,0.632,0.4932,0.3618,0.0002139,0.1974,0.005665,0.6039,0.1404,0.8521,0.2843,0.2639,0.008147,0.01335,0.2024,0.662,0.7286,0.0513,0.5801,0.6792,0.0147,0.6608,0.7835,0.8429,0.6939,0.8596,0.2811,0.01513,0.7403,0.03185,0.6233,0.6169,0.519,0.4035,0.3741,0.7702,0.7639,0.4128,0.1426,0.05403,0.4027,0.1324,0.5946,0.6035,0.6735,0.00691,0.0258,0.9404,0.3132,0.6896,0.806,0.3409,0.6407,0.02535,0.01913,0.01646,0.1337,0.04624,0.1323,0.008104,0.09686,0.453,0.2617,1,1,1,0.1063,1,0.05082,0.3883,1,1,1,1,0.03541,0.03132,0.02024,1,1,0.2735,0.5858,0.8046,0.002487,0.8834,0.8259,0.4547,0.1508,0.7808,0.9693,0.03399,0.5551,0.01555,0.1367,0.9373,0.2273,0.201,0.9321,0.6772,0.0003413,0.00457,0.001711,0.0004056,0.0564,0.002231,0.1555,0.02255,0.02821,0.4848,0.5893,0.8307,0.5473,0.2975,0.05538,0.1711,0.5558,0.0827,0.8548,0.1181,0.9248,0.002869,0.6696,0.5878,0.7641,0.1557,0.4454,0.04193,0.6063,0.954,0.1178,0.02177,0.004695,0.852,0.2142,0.1266,0.428,0.5472,0.1668,0.01128,0.01351,0.001723,0.0173,0.3238,0.2898,0.003439,0.09081,0.1126,0.2113,0.6898,0.02895,0.5967,0.7368,0.8094,0.04668,0.6775,0.2089,0.61,0.1786,0.005549,0.4212,0.4993,0.6754,0.2,0.4297,0.1304,0.162,0.1257,0.2172,0.05754,0.6854,0.5998,0.2796,0.275,0.8851,0.4222,0.353,0.9224,0.07981,0.2404,0.04266,0.2115,0.8908,0.2472,0.1549,0.3381,0.1964,0.9018,0.5496,0.02241,0.3978,0.2109,0.01798,0.005765,0.8236,0.006264,0.9805,0.7708,0.05971,0.8928,0.0007227,0.07895,0.1264,0.7812,0.907,0.9866,0.6898,0.1747,0.04358,0.7401,0.4595,0.05521,0.8337,0.4551,0.5811,0.8083,0.3713,0.644,0.7489,0.0002529,0.9882,0.9959,0.3108,0.6241,0.9637,0.07012,0.6236,0.1823,0.7643,0.2013,0.8737,0.8729,0.7254,0.8702,0.974,0.3407,0.09801,0.4517,0.221,0.3324,0.01695,0.008015,0.1583,0.2621,0.1564,0.01567,0.2737,0.2269,0.3041,0.1622,0.01877,0.1285,0.006047,0.6053,0.1264,0.7613,0.1044,0.05959,0.02148,0.0771,0.2742,0.03829,0.3429,0.04525,0.002145,1,0.1722,1,0.09299,1,0.3752,0.002486,0.006022,0.6268,0.1304,0.7048,0.4608,0.5525,0.05135,0.0004712,0.7326,0.07402,0.02841,0.7967,0.5069,0.8787,0.5738,0.7279,0.05452,0.1228,0.09994,0.352,0.9535,0.2871,0.002832,0.1464,0.8115,0.5521,0.9817,0.5191,0.8877,0.001646,0.8792,0.4918,0.6644,0.7706,0.07219,0.6295,0.9078,0.007882,0.003962,0.1062,0.625,0.7035,0.01014,0.692,0.148,0.05189,0.9426,0.03371,0.003912,0.9289,0.06073,0.02516,0.01344,0.1025,0.0009269,0.1665,0.7234,0.8175,0.9415,0.2073,0.1501,0.2542,0.005321,0.09684,0.04283,0.09736,0.1261,0.2795,0.01277,0.418,0.06919,0.7217,0.306,0.3411,0.3136,0.9438,0.3764,0.24,0.05671,0.03928,0.9867,0.4031,0.08701,0.7465,0.4934,0.4924,0.9512,0.4074,0.02537,0.9964,0.9373,0.639,0.2435,0.9085,0.004831,0.01194,0.00779,0.3861,0.6142,0.6567,0.2726,0.7995,0.2241,0.2824,0.1893,0.0004941,0.4931,0.2436,0.7408,0.1625,0.009066,0.0007415,0.02771,0.4445,0.04418,0.5284,0.6247,0.7523,0.0252,0.5083,0.02125,0.02954,0.3354,0.002393,0.2038,0.000241,0.0002579,0.006403,0.001596,0.9915,0.00702,0.2863,0.8946,0.06608,0.1631,0.009104,0.6662,0.6249,0.03671,0.005787,0.9754,0.02105,0.03856,0.679,0.7293,0.9811,0.3879,0.5908,0.1443,0.02092,0.074,0.1137,0.5915,0.4998,0.3884,0.8371,0.2172,0.2091,0.3051,0.3161,0.01089,0.03433,0.007519,0.1578,0.4772,0.002748,0.9645,0.7345,0.9349,0.2178,0.5538,0.6826,0.4911,0.6409,0.01118,0.02137,0.03142,0.01552,0.004491,0.1951,0.002593,0.02064,0.005166,0.0326,0.07176,0.1828,1,0.04481,0.03618,0.2111,1,0.1092,0.148,1,0.007253,0.03953,0.5514,0.006896,0.007518,0.0006828,0.02108,0.116,0.1199,0.1961,0.3483,0.0007651,0.5353,0.6881,0.5387,0.1212,0.05413,0.07117,0.004895,0.001782,0.006207,0.001832,0.00081,0.5746,0.002175,0.9561,0.007992,0.2446,0.4058,0.001169,0.5858,0.3318,0.8649,0.04414,0.5606,0.0006021,0.1079,0.02273,0.8424,0.887,0.1308,0.234,0.5662,0.003931,0.08924,0.4821,0.2408,0.6858,0.7882,0.1055,0.9729,0.738,0.2978,0.5365,0.8795,0.09425,0.003064,0.73,0.02389,0.08408,0.04699,0.01326,0.02804,0.9784,0.7002,0.0809,0.2023,0.5195,0.483,0.3352,0.4851,0.05357,0.3623,0.0002593,0.3349,0.6226,0.1189,0.5304,0.3348,0.00562,0.193,0.8402,0.3475,0.8672,0.4104,0.04138,0.3769,0.2265,0.6309,0.1016,0.2609,0.0004476,0.3791,0.8493,0.01462,0.1853,0.8177,0.6821,0.6962,0.183,0.003776,0.4951,0.004219,0.7775,0.4819,0.5384,0.8924,0.02528,0.5787,0.0006076,0.8196,0.2119,0.5039,0.3441,0.4426,0.9355,0.7549,0.2955,0.2716,0.7607,0.1771,0.1655,0.0247,0.000946,0.5268,0.8743,0.04815,0.137,0.01459,0.7784,0.01059,0.4957,0.087,0.1208,0.09886,0.7426,0.02008,0.3195,0.008529,0.6895,0.1844,0.4768,0.07856,0.3399,0.4038,0.2708,0.6773,0.8758,0.4727,0.00766,0.6966,0.07193,0.7484,0.3026,0.5874,0.7813,0.02475,0.1654,0.7236,0.3986,0.5717,0.5832,0.4626,0.5536,0.9794,0.1282,0.08349,0.3151,0.4015,0.9057,0.0008484,0.1596,0.007637,0.6093,0.7821,0.4423,0.002624,0.4853,0.4307,0.5772,0.3477,0.9571,0.1007,0.8167,0.717,0.879,0.827,0.1352,0.01051,0.4139,0.2295,0.1639,0.3553,0.01647,0.7564,0.0002269,0.1406,0.0216,0.9621,0.006565,0.002497,0.3988,0.08715,0.3778,0.00618,0.003869,0.3487,0.01789,0.1245,0.06223,0.002805,0.01037,0.1309,0.8974,0.6119,0.8163,0.06385,0.172,0.1388,0.1442,0.6031,0.1377,0.2454,0.1437,0.6919,0.484,0.2813,0.2982,0.3867,0.0007493,0.008528,0.03081,0.1483,0.5897,0.526,0.8914,0.186,0.4676,0.4167,0.226,0.7067,0.03474,0.001773,0.7521,0.7345,0.006461,0.3705,0.6598,0.2563,0.2604,0.004953,0.3183,0.03308,0.2708,0.03028,0.6169,0.6943,0.102,0.7859,0.4163,0.05366,0.9284,0.4157,0.4072,0.6547,0.05183,0.8362,0.9801,0.9403,0.2647,0.7369,0.3679,0.6796,0.6779,0.6408,0.8941,0.7663,0.0003374,0.02322,0.05942,0.6805,0.00482,0.2186,0.5394,0.08147,0.02743,0.7136,0.01371,0.4649,0.5946,0.5229,0.04614,0.008461,0.3809,0.03401,0.1139,0.000539,1,1,0.4095,1,0.3637,0.2261,0.09573,0.04151,0.1196,0.8197,0.03794,0.4458,0.0005545,0.1745,0.2616,0.02074,0.236,0.7888,0.017,0.004769,0.1545,0.01363,0.3063,0.2123,0.0002783,0.5156,0.0004126,0.7887,0.1051,0.3787,0.3673,0.2542,0.07296,0.1554,0.06969,0.001932,0.02124,0.4032,0.7358,0.04926,0.001781,0.4841,0.7549,0.01056,0.001114,0.01856,0.005115,0.9147,0.06178,0.0004215,0.002225,0.5362,0.2817,0.2472,0.2875,0.5277,0.4995,0.7717,0.2159,0.418,0.1854,0.3664,0.08957,0.7706,0.4449,0.9182,0.04906,0.8281,0.9862,0.7036,0.0006286,0.001403,0.06815,0.2123,0.5101,0.7259,0.5747,0.07749,0.835,0.3247,0.0572,0.4318,0.5652,0.7303,0.6217,0.1373,0.7259,0.541,0.9671,0.9181,0.1935,0.1875,0.1459,0.01467,0.9825,0.8322,0.1299,0.374,0.2233,0.6991,0.05554,0.8053,0.01725,0.0003324,0.5466,0.4391,0.3149,0.001224,0.7609,0.8496,0.868,0.01141,0.9905,0.4943,0.009962,0.3211,0.04501,0.03541,0.04389,0.37,0.4089,0.3575,0.4615,0.1009,0.6458,0.8424,0.2164,0.2502,0.6331,0.5295,0.1697,0.8058,0.8907,0.2941,0.1217,0.008555,0.09841,0.1714,0.8017,0.6976,0.02587,0.002597,0.5587,0.4774,0.4356,0.2543,0.05757,0.00221,0.8545,0.9209,0.174,0.001367,0.1309,0.9678,0.9494,0.5459,0.3488,0.08693,0.1284,0.04771,0.9071,0.006049,0.14,0.0003732,0.01529,0.495,0.7916,0.9069,0.2123,0.0995,0.3163,0.007774,0.2304,0.8821,0.9022,0.1639,0.5019,0.3342,0.2195,0.07027,0.08427,0.1864,0.49,0.001215,0.7635,0.8939,0.05135,0.5428,0.1595,0.2007,0.2925,0.4989,0.04392,0.8042,0.9028,0.0757,0.1061,0.7021,0.0002057,0.17,0.9092,0.0003956,0.001149,0.0004136,0.1336,0.02756,0.03492,0.3619,0.1914,0.5539,0.05545,0.3581,0.5302,0.1803,0.8516,0.1925,0.0328,0.01115,0.4633,0.07645,0.9026,0.4333,0.17,0.1965,0.8877,0.03585,0.4132,0.02587,0.09713,0.02216,0.03567,0.017,0.0004161,0.005331,0.02053,0.02093,0.09325,0.1223,0.411,0.002976,0.06257,0.03772,0.01222,0.01143,0.02128,0.575,0.3808,0.8265,0.04166,0.05417,0.1533,0.003777,0.01478,0.3582,0.8269,0.009621,0.0714,0.7301,0.5922,0.04385,0.09014,0.3603,0.9954,0.8767,0.845,0.02352,0.7112,0.9006,0.8691,0.02675,0.917,0.5215,0.6465,0.7005,0.1009,0.08774,0.1002,0.1917,0.1644,0.6443,0.6066,0.8371,0.43,0.6712,0.8388,0.1669,0.5347,0.2511,0.5251,0.01892,0.4144,0.3359,0.5822,0.3911,0.5386,0.173,0.02617,0.01649,0.6471,0.5473,0.9482,0.08388,0.05462,0.9946,0.06371,0.6101,0.1125,0.0217,0.2363,0.1708,0.2389,0.2722,0.625,0.002833,0.5204,0.2208,0.03079,0.9607,0.2277,0.07297,0.9047,0.06465,0.1501,0.654,0.02257,0.7181,0.5071,0.01789,0.1209,0.07923,0.3978,0.6107,0.01599,0.6048,0.9588,0.4025,0.9468,0.1117,0.8595,0.1447,0.4279,0.5254,0.02399,0.01316,0.7021,0.1343,0.2859,0.6531,0.5959,0.4601,0.8861,0.6719,0.1717,0.8614,0.252,0.7993,0.3725,0.3888,0.8226,0.5079,0.299,0.7892,0.3356,0.865,0.03393,0.05753,0.9532,0.747,0.152,0.004506,0.9264,0.8826,0.8392,0.1058,0.07212,0.1003,0.4079,0.9565,0.6128,0.5723,0.003827,0.06315,0.02622,0.02148,0.04054,0.6769,0.8323,0.163,0.4375,0.001163,0.01021,0.5466,0.299,0.01032,0.536,0.01112,0.3577,0.5109,0.8611,0.2147,0.01166,0.5081,0.4031,0.6312,0.009257,0.5633,0.0005765,0.7072,0.861,0.6369,0.01558,0.9864,0.965,0.9762,0.9856,0.923,0.02438,0.0518,0.01318,0.1719,0.1856,0.7959,0.0966,0.008084,0.8155,0.4887,0.0016,0.2861,0.3765,0.1177,0.471,0.5159,0.1711,0.2557,0.4651,0.7775,0.9718,0.7542,0.1868,0.3375,0.1541,0.001134,0.01874,0.03692,0.1952,0.8133,0.9423,0.5492,0.3089,0.9889,0.5721,0.6236,0.2291,0.6624,0.6483,0.001735,0.5437,0.4857,0.6466,0.2557,0.006311,0.5047,0.1649,0.9168,0.2082,0.0177,0.7045,0.4319,0.5235,0.1763,0.9288,0.832,0.305,0.3292,0.3458,0.1671,0.6684,0.7908,0.8479,0.06047,0.4149,0.1032,0.1495,0.1495,0.1387,0.03555,0.523,0.6921,0.245,0.4694,0.04794,0.06105,0.3357,0.009913,0.002027,0.6714,0.6154,0.002419,0.6535,0.07348,0.787,0.05819,0.9315,0.3764,0.7505,0.7864,0.7426,0.8612,0.8985,0.6942,0.005929,0.5532,0.04722,0.5205,0.9106,0.8071,0.6818,0.7108,0.2125,0.01048,0.5663,0.1341,0.003968,0.9764,0.1356,0.6221,0.3106,0.4316,0.3894,0.1103,0.9433,0.4389,0.9172,0.2051,0.8078,0.3117,0.8196,0.762,0.2481,0.02255,0.001925,0.9492,0.3333,0.435,0.02931,0.01386,0.7609,0.6554,0.5291,0.8325,0.8589,0.7256,0.02391,0.3361,0.007514,0.3567,0.5796,0.5574,0.739,0.9975,0.5617,0.5172,0.1544,0.01878,0.001693,0.0139,0.5632,0.8943,0.5462,0.9901,0.9628,0.05174,0.6432,0.8326,0.00116,0.005401,0.0004953,0.0007246,0.684,0.07175,1,1,1,0.2064,0.05382,0.5408,0.05661,0.03481,0.09375,0.02171,0.01922,0.01956,0.07998,0.3925,0.7326,0.09757,0.01909,0.1361,0.004569,0.1607,0.5518,0.6533,0.896,0.853,0.8572,0.7087,0.1812,0.4766,0.7411,0.8487,0.001079,0.6281,0.002618,0.05678,0.9369,0.2885,0.9221,0.9944,0.05971,0.4023,0.01377,0.83,0.8644,0.7878,0.2461,0.9573,0.7508,0.8205,0.05373,0.0003809,0.4349,0.7698,0.2663,0.03936,0.614,0.8151,0.3162,0.9508,0.3495,0.1798,0.1619,0.4213,0.09141,0.7433,0.004025,0.8257,0.6117,0.0409,0.01536,0.1266,0.287,0.8762,0.007812,0.005497,0.7008,0.0002529,0.3708,0.6399,0.1468,0.3124,0.0003933,0.7844,0.15,0.4888,0.004297,0.2314,0.8108,0.4274,0.6647,0.03555,0.9535,0.4342,0.03162,0.1384,0.004534,0.03032,0.228,0.02735,0.3474,0.7957,0.8482,0.3089,0.9055,0.2981,0.3608,0.5628,0.506,0.1342,0.9018,0.1257,0.9685,0.672,0.9245,0.001408,0.08167,0.02347,0.1423,0.9824,0.08018,0.1495,0.9601,0.3403,0.4241,0.5878,0.9143,0.7167,0.8055,0.8035,0.2132,0.03874,0.8136,0.03627,0.0313,0.7303,0.4358,0.3778,0.3181,0.0005292,0.014,0.2878,0.4627,0.03413,0.00468,0.7578,0.0101,0.1342,0.1706,0.3386,0.4798,0.9461,0.4767,0.1997,0.9,0.9303,0.6099,0.3547,0.1095,0.1417,0.5486,0.04118,0.9606,0.2263,0.3187,0.6079,0.04026,0.5593,0.8433,0.951,0.06093,0.0408,0.3197,0.6744,0.2381,0.0002661,0.179,0.4505,0.07059,0.02541,0.1361,0.08716,0.5492,0.007314,0.1562,0.06441,0.8708,0.436,0.6278,0.2219,0.4437,0.1212,0.0005229,0.3522,0.4101,0.4999,0.1793,0.2701,0.03719,0.1818,0.6425,0.5912,0.5412,0.5122,0.7248,0.2166,0.00396,0.002619,0.2202,0.4793,0.1849,0.2413,0.8306,0.148,0.5007,0.1845,0.09738,0.7791,0.4789,0.7672,0.1817,0.1853,0.0008717,0.0102,0.2279,0.8477,0.3382,0.8733,0.06291,0.003558,0.1062,0.9107,0.8825,0.1722,0.3793,0.8142,0.0723,0.984,0.241,0.8535,0.8699,0.9626,0.3134,0.03335,0.9564,0.9625,0.1667,0.01157,0.3456,0.6715,0.5238,0.2699,0.9735,0.7688,0.8414,0.7985,0.9149,0.6801,0.808,0.6293,0.06546,0.731,0.5999,0.2168,0.2051,0.9753,0.6896,0.02062,0.5137,0.7556,0.02163,0.6408,0.5264,0.7412,0.4754,0.375,0.9958,0.4284,0.09364,0.2845,0.02559,0.8172,0.8559,0.3862,0.01444,0.09326,0.9292,0.9114,0.8859,0.8934,0.05315,0.007151,0.3686,0.1908,0.7817,0.01248,0.7909,0.6435,0.9725,0.4305,0.4513,0.2603,0.5787,0.4391,0.7003,0.8481,0.5554,0.6783,0.7898,0.4157,0.8779,0.01558,0.3801,0.01049,0.9229,0.09682,0.2138,0.4934,0.3697,0.01128,0.8306,0.8657,0.8825,0.6535,0.3134,0.9817,0.7825,0.8484,0.35,0.06602,0.5582,0.6664,0.829,0.04921,0.9013,0.6005,0.0002329,0.5021,0.3383,0.02108,0.0006791,0.8212,0.6192,0.5036,0.6997,0.008386,0.01596,0.9264,0.07718,0.7371,0.008231,0.001141,0.8578,0.6203,0.1476,0.5118,0.08571,0.0005362,0.153,0.4668,0.4174,0.6087,0.7927,0.9791,0.5841,0.1297,0.6761,0.03922,0.6631,0.005193,0.002532,0.007249,0.003612,0.01169,0.01998,0.012,0.6231,0.01857,0.1169,1,0.02163,1,1,0.8205,0.1013,0.05198,0.01886,0.05896,0.1009,0.1813,0.3091,0.2578,0.9261,0.3127,0.1119,0.1003,0.4911,0.2679,0.02663,0.6941,0.2171,0.3887,0.391,0.3029,0.7127,0.3299,0.4426,0.9783,0.4604,0.9519,0.2931,0.0004854,0.2264,0.7934,0.894,0.02465,0.7862,0.5161,0.8701,0.004293,0.9068,0.7637,0.008752,0.8762,0.859,0.2803,0.427,0.7146,0.4755,0.8661,0.9938,0.987,0.4083,0.8884,0.9397,0.7784,0.7219,0.004076,0.2174,0.07202,0.2209,0.5644,0.2165,0.4949,0.02471,0.002065,0.2902,0.8531,0.2772,0.4976,0.2937,0.3236,0.5198,0.6843,0.544,0.111,0.4935,0.01938,0.6479,0.851,0.3183,0.03713,0.6606,0.3093,0.2994,0.3564,0.8285,0.6839,0.1129,0.7856,0.06871,0.3027,0.04105,0.3951,0.1908,0.1415,0.5695,0.07458,0.2628,0.7427,0.5999,0.09984,0.3133,0.9627,0.5642,0.7608,0.7983,0.2807,0.1339,0.02791,0.2978,0.2345,0.09144,0.9946,0.828,0.5158,0.6009,0.336,0.6143,0.3784,0.6166,0.009693,0.7585,0.4291,0.5114,0.9785,0.7307,0.01623,0.8742,0.6312,0.002028,0.08797,0.04192,0.02397,0.4837,0.7146,0.0002279,0.5776,0.2399,0.8986,0.1375,0.2283,0.5217,0.001291,0.1738,0.6267,0.1164,0.05538,0.1982,0.1017,0.2431,0.8887,0.7136,0.05041,0.1176,0.2125,0.1054,0.05957,0.8094,0.4489,0.9618,0.01385,0.03144,0.2607,0.7588,0.9812,0.4977,0.3393,0.01988,0.5613,0.6007,0.04255,0.08283,0.01379,0.001492,0.2021,0.3033,0.7429,0.01172,0.1601,0.03443,0.2015,0.2375,0.7336,0.00433,0.02333,0.6388,0.513,0.5864,0.8985,0.04461,0.2148,0.1175,0.5323,0.7748,0.747,0.1563,0.8899,0.9329,0.1212,0.02457,0.0967,0.3126,0.005966,0.3027,0.0002696,0.4978,0.5126,0.3544,0.03033,0.1553,0.4621,0.2515,0.1919,0.04371,0.5312,0.5353,0.03759,0.06718,0.4955,0.373,0.4329,0.7664,0.0007469,0.8727,0.4383,0.5705,0.2834,0.7658,0.9855,0.812,0.8716,0.386,0.8375,0.4553,0.3007,0.9488,0.007816,0.2294,0.7395,0.2683,0.06065,0.01457,0.3834,0.3423,0.932,0.113,0.6268,0.6731,0.3419,0.1983,0.00981,0.7557,0.000431,0.001126,0.511,0.6822,0.7902,0.09566,0.8022,0.6974,0.9988,0.4724,0.0005403,0.5387,0.1854,0.9526,0.07296,0.1388,0.9207,0.864,0.8013,0.9364,0.1141,0.9231,0.8226,0.2269,0.9901,0.7752,0.1277,0.2426,0.7843,0.7982,0.3873,0.0126,0.4701,0.2047,0.01513,0.02043,0.8672,0.5978,0.003018,0.008779,0.6407,0.7702,0.5474,0.007354,0.495,0.254,0.3424,0.3376,0.7336,0.1284,0.3932,0.205,0.0002743,0.5265,0.9257,0.3019,0.3376,0.9536,0.004371,0.8924,0.1878,0.02442,0.5663,0.04022,0.00817,0.4126,0.5486,0.1597,0.3292,0.7238,0.8494,0.2274,0.3536,0.04353,0.03034,0.6357,0.2816,0.8946,0.0005969,0.135,0.04378,0.7244,0.7153,0.06837,0.1765,0.1676,0.7826,0.9151,0.8196,0.2464,0.7152,0.5458,0.4107,0.8874,0.02929,0.8566,0.2671,0.0003657,0.6675,0.5185,0.1287,0.0245,0.247,0.2079,0.07962,0.0342,0.02152,0.6107,0.0699,0.02112,0.0009462,0.9024,0.2817,0.5693,0.6183,0.207,0.4131,0.001074,0.5963,0.06409,0.1025,0.4024,0.3027,0.8576,0.3433,0.4735,0.04169,0.8587,0.841,0.4565,0.208,0.0077,0.8028,0.1499,0.1116,0.06205,0.2524,0.9138,0.9405,0.8769,0.3393,0.4288,0.3993,0.7544,0.00499,0.3513,0.152,0.3842,0.613,0.01618,0.05705,0.3578,0.2354,0.8238,0.4024,0.006558,0.9084,0.3175,0.09234,0.9341,0.9615,0.03297,0.1625,0.01643,0.01764,0.3354,0.003184,0.4702,0.7295,0.5074,0.6707,0.3741,0.199,0.06702,0.4424,0.8433,0.3633,0.000494,0.0109,0.03448,0.7195,0.1806,0.07067,0.3146,0.03484,0.9011,0.8459,0.3443,0.01956,0.3372,0.2993,0.5518,0.9156,0.01208,1,0.09187,0.007615,0.2796,0.04471,0.05951,0.7604,0.3629,1,0.15,0.379,0.4329,0.5525,0.039,0.3786,0.4924,0.1737,0.05829,0.5211,0.4727,0.8728,0.01377,0.2807,0.6818,0.007725,0.521,0.5019,0.02396,0.3894,0.001207,0.06456,0.8094,0.03028,0.5899,0.6909,0.4344,0.6704,0.9776,0.005719,0.002001,0.6842,0.01224,0.9048,0.4457,0.7141,0.1526,0.9472,0.3149,0.5223,0.7742,0.3633,0.303,0.9294,0.03697,0.0004664,0.4659,0.8243,0.6068,0.1075,0.1373,0.1414,0.5295,0.01442,0.2588,0.2934,0.7708,0.0003501,0.5385,0.01094,0.588,0.8408,0.4551,0.9053,0.002088,0.7325,0.8518,0.7763,0.3235,0.1481,0.009204,0.1271,0.309,0.3105,0.5015,0.1955,0.4141,0.03783,0.3319,0.7925,0.3067,0.739,0.074,0.02006,0.1327,0.356,0.005876,0.6221,0.001178,0.3707,0.4233,0.02711,0.9573,0.01028,0.6196,0.8049,0.7701,0.3698,0.3731,0.1118,0.6204,0.0002935,0.3142,0.4806,0.548,0.4821,0.07614,0.01014,0.723,0.5998,0.3635,0.003521,0.7573,0.2439,0.7835,0.0232,0.4311,0.001053,0.5721,0.7939,0.04534,0.2741,0.5951,0.01659,0.2939,0.1556,0.2455,0.1779,0.6582,0.7098,0.8299,0.8597,0.02501,0.1378,0.9018,0.3788,0.3696,0.1061,0.2226,0.7507,0.801,0.1929,0.7866,0.8772,0.0003326,0.06372,0.322,0.8567,0.04162,0.1341,0.6247,0.1805,0.639,0.6354,0.0006902,0.02212,0.7251,0.2031,0.03749,0.006253,0.05292,0.1279,0.3439,0.658,0.06907,0.5837,0.5446,0.4237,0.07876,0.8971,0.4331,0.781,0.4378,0.09433,0.1389,0.1318,0.311,0.2577,0.5978,0.001728,0.01544,0.7528,0.02781,0.5273,0.9367,0.4762,0.2222,0.7528,0.8801,0.4032,0.7478,0.968,0.2278,0.001804,0.3977,0.02602,0.08953,0.23,0.5146,0.3203,0.7,0.007956,0.8528,0.284,0.5838,0.951,0.0307,0.2341,0.004899,0.01216,0.5118,0.0439,0.7654,0.7078,0.9918,0.2368,0.2387,0.3729,0.6952,0.8755,0.000286,0.9953,0.8592,0.8432,0.7186,0.2661,0.498,0.2992,0.06227,0.8303,0.147,0.6894,0.8192,0.04285,0.1094,0.9315,0.01156,0.6948,0.7015,0.0006133,0.004114,0.1536,0.06583,0.6858,0.7596,0.768,0.3357,0.7686,0.08305,0.4182,0.6014,0.3649,0.03014,0.8189,0.007691,0.05107,0.9546,0.252,0.0927,0.01411,0.3413,0.767,0.2846,0.499,0.06041,0.02044,0.75,0.004964,0.9089,0.3814,0.9767,0.3767,0.0664,0.818,0.7214,0.9894,0.01953,0.007245,0.001754,0.001599,0.0138,0.00265,0.005188,0.0008742,0.002265,0.4267,0.3835,0.6635,0.4571,0.9736,0.7723,0.09064,0.2288,0.5974,0.804,0.05502,0.3896,0.5092,0.586,0.6928,0.703,0.01792,0.01365,0.1768,0.007677,0.006043,0.008606,0.2335,0.006718,0.8757,0.001743,0.09895,0.12,0.06721,0.2552,1,0.04316,0.007569,0.05217,0.03528,0.04901,0.04741,0.5188,0.5019,0.2696,0.4364,0.4283,0.2141,0.3964,0.2259,0.7149,0.08469,0.05355,0.289,0.7651,0.003158,0.9312,0.0003624,0.5715,0.2934,0.4696,0.4139,0.4916,0.4161,0.02935,0.3977,0.4112,0.7101,0.01554,0.5002,0.03408,0.985,0.2701,0.2677,0.3759,0.2237,0.06126,0.4716,0.8123,0.9779,0.01431,0.8726,0.06613,0.7411,0.5392,0.8165,0.8547,0.3782,0.2998,0.8473,0.8534,0.1287,0.6194,0.6281,0.06976,0.3822,0.8108,0.03572,0.002066,0.481,0.1281,0.4424,0.5247,0.823,0.3064,0.6402,0.3115,0.7472,0.6322,0.3253,0.287,0.3209,0.2312,0.7737,0.3112,0.7531,0.1618,0.1842,0.3074,0.8923,0.8973,0.6123,0.5164,0.1584,0.2756,0.5726,0.006791,0.01163,0.1017,0.4312,0.3292,0.9139,0.06111,0.1484,0.662,0.938,0.6037,0.57,0.5452,0.07835,0.009953,0.002632,0.7023,0.4007,0.02137,0.6825,0.3276,0.4813,0.9389,0.7975,0.4913,0.7189,0.3893,0.01937,0.7942,0.5862,0.2467,0.1174,0.3129,0.3516,0.6917,0.8874,0.6004,0.02616,0.7827,0.5261,0.8068,0.7815,0.1857,0.0006249,0.8729,0.03423,0.2422,0.9572,0.8647,0.2217,0.3449,0.1157,0.3233,0.8117,0.03454,0.263,0.1935,0.3543,0.1615,0.08967,0.4977,0.004269,0.01116,0.02206,0.132,0.3874,0.9858,0.7717,0.652,0.5625,0.9606,0.138,0.6317,0.1845,0.3195,0.01784,0.003084,0.8993,0.8892,0.2624,0.05481,0.3967,0.9552,0.9856,0.8525,0.433,0.3057,0.07326,0.002861,0.005975,0.3911,0.2704,0.02641,0.2563,0.784,0.9136,0.3469,0.3446,0.1129,0.7767,0.6571,0.2884,0.09436,0.7829,0.0006126,0.1967,0.1866,0.9179,0.4785,0.08652,0.2635,0.1027,0.171,0.4481,0.4522,0.6884,0.3398,0.1812,0.0004063,0.2284,0.0002406,0.7702,0.5335,0.5161,0.5359,0.001385,0.5218,0.9507,0.4767,0.2584,0.764,0.0422,0.7347,0.9353,0.5651,0.08212,0.6581,0.5596,0.4346,0.02935,0.5771,0.4658,0.792,0.664,0.8318,0.08614,0.9401,0.06634,0.8891,0.4019,0.2295,0.211,0.8191,0.9957,0.5128,0.01612,0.4914,0.5743,0.6955,0.866,0.5154,0.5773,0.5429,0.1984,0.6407,0.5423,0.68,0.5866,0.3097,0.07888,0.1816,0.6589,0.1662,0.001079,0.6744,0.6136,0.3707,0.693,0.02565,0.8446,0.2361,0.3031,0.7827,0.8872,0.4622,0.484,0.9266,0.5847,0.301,0.1885,0.2559,0.3665,0.000603,0.4508,0.7793,0.8969,0.1742,0.4266,0.3952,0.8147,0.6252,0.1857,0.0735,0.139,0.267,0.002354,0.03668,0.0108,0.02693,0.3243,0.6899,0.9778,0.7494,0.4569,0.4093,0.1633,0.01768,0.03469,0.5704,0.1233,0.8466,0.6988,0.05269,0.2879,0.2771,0.8293,0.7329,0.8439,0.3768,0.2962,0.802,0.001706,0.07212,0.7667,0.8501,0.01422,0.1995,0.1202,0.1107,0.003303,0.07678,0.06541,0.8359,0.32,0.005747,0.9342,0.9059,0.1231,0.7226,0.3917,0.001388,0.9857,0.8095,0.6679,0.3176,0.007931,0.7813,0.3738,0.8817,0.6316,0.03076,0.5711,0.003105,0.2303,0.7782,0.9198,0.2467,0.6518,0.2652,0.02637,0.2583,0.6688,0.3231,0.8026,0.06283,0.218,0.7782,0.5362,0.009525,0.2554,0.2102,0.234,0.6907,0.07343,0.00464,0.8237,0.1149,0.3238,0.1032,0.865,0.5946,0.9431,0.3107,0.8005,0.9995,0.4127,0.7851,0.996,0.000683,0.004436,0.9799,0.9468,0.153,0.5125,0.4924,0.6036,0.1345,0.112,0.5911,0.8804,0.7257,0.7216,0.7923,0.07034,0.01052,0.1384,0.1625,0.1652,0.6616,0.2332,0.9674,0.0008737,0.8748,0.7759,0.4521,0.2319,0.3238,0.977,0.357,0.0003367,0.1495,0.3179,0.4154,0.08358,0.9411,0.4841,0.8465,0.0152,0.9517,0.1783,0.03019,0.01892,0.5382,0.9941,0.2189,0.1845,0.009929,0.043,0.4193,0.05772,0.4225,0.1336,0.2825,0.3181,0.1009,0.02011,0.0903,0.05698,0.0253,0.001741,0.0008204,0.001666,0.009865,0.04198,0.07669,1,0.18,1,0.1698,0.1488,0.6527,1,0.09236,0.02145,0.1312,0.02574,0.001439,0.03707,0.009189,0.003263,0.377,0.6948,0.3968,0.09915,0.1395,0.4803,0.04341,0.01149,0.1681,0.3068,0.8261,0.001199,0.0005775,0.4278,0.8516,0.05782,0.7731,0.9233,0.001111,0.1305,0.04718,0.1456,0.775,0.03032,0.7195,0.2436,0.236,0.5837,0.1976,0.1199,0.3254,0.4132,0.0078,0.6853,0.003679,0.6274,0.571,0.0024,0.9098,0.0009626,0.9722,0.06823,0.4953,0.01252,0.8054,0.241,0.5483,0.1801,0.1674,0.9567,0.1891,0.1695,0.8243,0.2191,0.8689,0.7187,0.2705,0.07656,0.01483,0.3211,0.7273,0.8199,0.1195,0.1794,0.03285,0.07534,0.6869,0.8188,0.2237,0.7766,0.8279,0.4507,0.6301,0.2692,0.8409,0.7558,0.001169,0.04938,0.6394,0.2219,0.3742,0.079,0.0004837,0.2555,0.1336,0.01059,0.7763,0.006126,0.003544,0.6039,0.5724,0.5991,0.2545,0.7245,0.02249,0.6225,0.8971,0.792,0.6832,0.05736,0.009013,0.3763,0.08421,0.1035,0.8009,0.9959,0.9416,0.1993,0.1122,0.7503,0.08936,0.6813,0.6917,0.3927,0.3473,0.5354,0.6379,0.002145,0.02347,0.1596,0.5189,0.6025,0.2642,0.5485,0.7838,0.02939,0.4875,0.1756,0.8856,0.8452,0.7334,0.007599,0.5665,0.3685,0.1517,0.9608,0.7901,0.2726,0.7082,0.4851,0.05477,0.09133,0.06081,0.802,0.8257,0.9888,0.4365,0.3528,0.8508,0.8808,0.3013,0.4055,0.9523,0.2006,0.2385,0.4437,0.9297,0.5736,0.3808,0.9122,0.5723,0.8455,0.02848,0.7807,0.04524,0.9434,0.001094,0.0002044,0.2475,0.5081,0.001032,0.6014,0.356,0.05619,0.7694,0.01854,0.1408,0.4858,0.2204,0.09357,0.94,0.07561,0.6354,0.01451,0.9568,0.117,0.6458,0.8849,0.5637,0.9199,0.6849,0.721,0.003829,0.002755,0.9798,0.7857,0.9188,0.4312,0.0435,0.009585,0.5713,0.7833,0.163,0.007278,0.5025,0.756,0.7837,0.977,0.2061,0.8549,0.7705,0.4067,0.5679,0.9294,0.9247,0.274,0.09776,0.7822,0.8321,0.2281,0.5451,0.03095,0.07824,0.1375,0.05232,0.6858,0.003626,0.1319,0.002531,0.001121,0.9738,0.05174,0.3417,0.3061,0.54,0.03261,0.1151,0.07241,0.1314,0.09495,0.008901,0.1637,0.2732,0.05688,0.001752,0.328,0.3049,0.08329,0.3416,0.0004104,0.6563,0.8392,0.7927,0.00254,0.09196,0.4013,0.9283,0.6897,0.000605,0.9431,0.01125,0.3063,0.6651,0.9387,0.8566,0.06242,0.606,0.4209,0.6543,0.9974,0.04323,0.02762,0.6893,0.3863,0.3807,0.3423,0.02777,0.8572,0.9515,0.9312,0.956,0.1019,0.3009,0.1029,0.5589,0.836,0.1389,0.8026,0.03528,0.3955,0.2671,0.8744,0.01043,0.05742,0.4339,0.02671,0.4074,0.0999,0.1008,0.1034,0.1139,0.8148,0.785,0.8636,0.7934,0.6788,0.1249,0.585,0.732,0.007808,0.1133,0.4903,0.5616,0.004086,0.4067,0.4047,0.8597,0.4026,0.5014,0.4048,0.0844,0.2972,0.7061,0.5595,0.7078,0.8019,0.6265,0.8354,0.4963,0.8676,0.007962,0.7342,0.583,0.3962,0.5389,0.8392,0.02789,0.05107,0.3522,0.0357,0.8286,0.9486,0.4599,0.9888,0.5742,0.3059,0.9587,0.1363,0.7782,0.3482,0.017,0.02322,0.7556,0.1077,0.02197,0.5755,0.8603,0.3918,0.008915,0.2132,0.4418,0.7774,0.09999,0.2701,0.3061,0.1123,0.2828,0.07156,0.7936,0.3605,0.8432,0.7265,0.08049,0.5567,0.5844,0.3556,0.545,0.2454,0.1055,0.2316,0.9666,0.7321,0.2483,0.001951,0.6524,0.7075,0.6078,0.01161,0.002862,0.3007,0.3613,0.839,0.191,0.7217,0.1148,0.692,0.3491,0.3871,0.6757,0.3227,0.2488,0.1145,0.6048,0.7066,0.5064,0.6685,0.5148,0.6839,0.8867,0.1314,0.7322,0.11,0.69,0.1508,0.08165,0.06342,0.7985,0.8124,0.03567,0.02293,0.7997,0.8953,0.8033,0.4633,0.7124,0.7728,0.8246,0.5219,0.04974,0.08866,0.5445,0.1807,0.2286,0.4079,0.01042,0.7359,0.7138,0.8349,0.7517,0.4451,0.05122,0.557,0.56,0.4691,0.9988,0.06918,0.2608,0.7754,0.5348,0.4056,0.000595,0.06372,0.908,0.1653,0.5609,0.8102,0.5042,0.5248,0.8214,0.07789,0.6474,0.004756,0.5948,0.3392,0.6774,0.1046,0.2036,0.1991,0.01088,0.8399,0.003047,0.2466,0.001102,0.2534,0.215,0.8253,0.01837,0.06037,0.1586,0.3341,0.9402,0.366,0.1861,0.8665,0.1581,0.957,0.3432,0.261,0.07723,0.1261,0.549,0.6355,0.03526,0.5496,0.6696,0.9422,0.1248,0.235,0.4238,0.3419,0.275,0.6392,0.7857,0.6913,0.2622,0.4133,0.3444,0.1832,0.3019,0.8866,0.3429,0.07454,0.5857,0.6447,0.222,0.5755,0.1921,0.2117,0.03017,0.1792,0.3753,0.1049,0.2192,0.4534,0.5585,0.5747,0.6061,0.5285,0.727,0.5461,0.2648,0.0006913,0.3705,0.6339,0.7405,0.5718,0.1096,0.449,0.79,0.05392,0.3465,0.9151,0.2665,0.6242,0.386,0.84,0.997,0.2231,0.1948,0.4162,0.4566,0.2319,0.4072,0.9744,0.5444,0.5441,0.7143,0.772,0.331,0.619,0.006291,0.1466,0.08944,0.9937,0.05029,0.2417,0.191,0.6701,0.07089,0.4971,0.6722,0.538,0.36,0.0918,0.4443,0.4262,0.3703,0.506,0.3117,0.4654,0.8865,0.01334,0.05666,0.4044,0.4142,0.08265,0.286,0.1771,0.006922,0.3072,0.8724,0.3248,0.4699,0.5821,0.1666,0.4464,0.1871,0.7955,0.6164,0.02288,0.003703,0.5727,0.206,0.02255,0.2577,0.1898,0.5985,0.3612,0.0134,0.0141,0.02725,0.8646,0.7102,0.002173,0.5892,0.5988,0.07376,0.9453,0.6557,0.6478,0.08129,0.06769,0.2924,0.7009,0.315,0.01549,0.02281,0.5644,0.208,0.9027,0.2335,0.3045,0.00676,0.02566,0.8647,0.9495,0.01667,0.1069,0.8654,0.6704,0.1832,0.009257,0.04431,0.7561,0.04574,0.57,0.04018,0.3445,0.9165,0.458,0.5372,0.9502,0.4653,0.3034,0.2421,0.9934,0.01459,0.09818,0.02319,0.1651,0.2423,0.002341,0.2486,0.001714,0.007312,0.007732,0.008046,0.01675,0.007954,0.03159,0.4333,0.01572,0.657,1,1,1,1,0.02088,0.1178,0.08689,0.05401,0.01962,0.194,0.6158,0.001917,0.1897,0.2282,0.3238,0.0002381,0.02448,0.06531,0.3417,0.818,0.6287,0.1314,0.6762,0.2761,0.4054,0.19,0.4504,0.4034,0.4499,0.4162,0.06047,0.009823,0.01109,0.6976,0.9611,0.6611,0.3276,0.8711,0.02924,0.01425,0.2751,0.4528,0.3447,0.5884,0.3479,0.5943,0.4078,0.08855,0.7927,0.2849,0.006136,0.08324,0.1425,0.3716,0.6211,0.01923,0.03865,0.007326,0.3226,0.5234,1,0.01713,0.9471,0.8582,0.5075,0.001618,0.6213,0.612,0.5687,0.01765,0.5321,0.4236,0.6704,0.5769,0.1655,0.5049,0.02385,0.5398,0.7497,0.7504,0.4939,0.0768,0.002274,0.03271,0.6216,0.5412,0.4464,0.1874,0.1436,0.1278,0.1527,0.01055,0.03209,0.1408,0.8149,0.7422,0.01575,0.1701,0.005494,0.04455,0.573,0.05296,0.4658,0.5562,0.3315,0.3891,0.3618,0.02861,0.02186,0.1525,0.8989,0.9473,0.007318,0.8757,0.9906,0.5502,0.1157,0.7178,0.01764,0.6924,0.01346,0.9623,0.5665,0.2736,0.004513,0.9589,0.9145,0.02323,0.01757,0.2664,0.1139,0.6178,0.4228,0.1108,0.00478,0.7888,0.8298,0.4788,0.02163,0.5874,0.7231,0.4583,0.1192,0.2495,0.6734,0.1832,0.02855,0.03858,0.3974,0.4651,0.0005437,0.006508,0.7343,0.0004133,0.3119,0.3067,0.1295,0.288,0.559,0.006203,0.4189,0.002654,0.5458,0.628,0.0004131,0.8435,0.4192,0.4829,0.364,0.8014,0.6427,0.6799,0.3546,0.007131,0.1679,0.001481,0.153,0.374,0.8401,0.9502,0.8604,0.8757,0.1661,0.9684,0.7433,0.0006939,0.2265,0.03169,0.33,0.05373,0.7846,0.5579,0.3722,0.8568,0.3701,0.6119,0.8884,0.2976,0.2402,0.5185,0.26,0.4558,0.02493,0.3459,0.09046,0.4873,0.6872,0.9911,0.5217,0.6973,0.01942,0.2858,0.09281,0.8791,0.371,0.4499,0.8057,0.05412,0.8272,0.4488,0.1365,0.2644,0.04196,0.1481,0.3633,0.9193,0.3369,0.6758,0.009252,0.01016,0.6276,0.4329,0.9346,0.5674,0.9566,0.8631,0.07617,0.01629,0.01043,0.4245,0.3413,0.0004689,0.8596,0.7488,0.9833,0.2819,0.5712,0.7738,0.1652,0.114,0.06855,0.8622,0.03396,0.1115,0.8941,0.8194,0.2999,0.5367,0.8534,0.7203,0.307,0.5154,0.00222,0.9962,0.8172,0.9493,0.2537,0.472,0.009811,0.6988,0.02198,0.08148,0.6579,0.2458,0.2549,0.07853,0.433,0.567,0.2251,0.0008326,0.5036,0.5008,0.1066,0.1379,0.1168,0.4187,0.5194,0.1721,0.04911,0.4314,0.1221,0.9827,0.1827,0.003292,0.4793,0.4124,0.09251,0.8144,0.9758,0.4624,0.5871,0.5878,0.4581,0.5426,0.3789,0.1079,0.8207,0.001135,0.8286,0.9547,0.9799,0.1293,0.06029,0.6635,0.1752,0.4674,0.9785,0.7215,0.6422,0.02177,0.5797,0.2028,0.9018,0.08414,0.1494,0.7679,0.5877,0.08293,0.4035,0.5599,0.8189,0.4451,0.2636,0.6343,0.3441,0.4761,0.01683,0.2139,0.09474,0.9023,0.8217,0.4529,0.4838,0.0005547,0.4429,0.7938,0.8338,0.4785,0.01481,0.989,0.9669,0.6485,0.6256,0.3761,0.6528,0.6761,0.1276,0.4023,0.3686,0.115,0.1248,0.5863,0.7867,0.3832,0.3054,0.6814,0.7646,0.7761,0.7588,0.9419,0.2114,0.00396,0.4636,0.003659,0.04206,0.9726,0.2208,0.4412,0.9841,0.9354,0.04565,0.1314,0.637,0.7976,0.5432,0.02677,0.2116,0.6318,0.0006624,0.9635,0.2502,0.1469,0.5239,0.03201,0.111,0.0006712,0.5812,0.0004563,0.9516,0.563,0.02936,0.2337,0.5795,0.2354,0.5681,0.5899,0.1588,0.005441,0.2789,0.1545,0.01265,0.1611,0.1451,0.7605,0.6367,0.08671,0.1297,0.548,0.09545,0.2434,0.1886,0.923,0.8256,0.5529,0.5211,0.1354,0.4218,0.6957,0.005627,0.09835,0.8135,0.5793,0.04756,0.4039,0.434,0.7682,0.01929,0.7758,0.02995,0.5325,0.7489,0.1033,0.05922,0.8586,0.4442,0.8572,0.8381,0.01361,0.01632,1,1,1,1,0.3812,0.09839,0.1756,0.00798,0.01028,0.003421,0.03286,0.00132,0.02883,0.04415,0.008343,0.01863,0.000427,0.01446,0.000569,0.09432,0.01469,0.0002319,0.04238,0.0007102,0.003711,0.01037,0.3805,0.001172,0.002633,0.7899,0.2948,0.4888,0.7806,0.06116,0.5531,0.05904,0.464,0.3645,0.8103,0.9287,0.2915,0.128,0.9975,0.8908,0.1275,0.3649,0.2625,0.187,0.4421,0.6273,0.2355,0.1197,0.5671,0.05875,0.7829,0.7886,0.5474,0.755,0.1966,0.5328,0.9854,0.2613,0.739,0.07422,0.4727,0.3253,0.71,0.6656,0.09288,0.3951,0.05227,0.05556,0.5316,0.6537,0.5369,0.2846,0.2577,0.6742,0.08605,0.4413,0.9213,0.5348,0.09143,0.06602,0.5508,0.00123,0.8749,0.7775,0.3621,0.3962,0.8147,0.7445,0.2997,0.8205,0.4419,0.3253,0.9096,0.04089,0.211,0.3132,0.4758,0.226,0.1828,0.08962,0.394,0.5729,0.1454,0.01368,0.587,0.3281,0.4091,0.5128,0.001323,0.3796,0.6515,0.09455,0.823,0.5975,0.309,0.3622,0.1996,0.3841,0.2694,0.1534,0.3102,0.0326,0.00413,0.6655,0.8206,0.2903,1,0.009663,0.8991,0.09476,0.365,0.4468,0.2026,0.05585,0.3895,0.4196,0.1343,0.616,0.733,0.8448,0.3952,0.00543,0.5556,0.318,0.873,0.1598,0.448,0.1038,0.1109,0.001051,0.3037,0.3068,0.889,0.01373,0.3007,0.7034,0.333,0.899,0.9427,0.6636,0.254,0.985,0.1033,0.1532,0.06695,0.495,0.1787,0.9491,0.333,0.5823,0.5064,0.0908,0.8888,0.3082,0.3914,0.5022,0.316,0.8063,0.8259,0.07958,0.9575,0.7818,0.1871,0.3021,0.4519,0.9979,0.001775,0.1469,0.3061,0.01479,0.1391,0.7569,0.05525,0.6536,0.6921,0.005992,0.8741,0.2312,0.4959,0.2441,0.4526,0.2848,0.3923,0.8007,0.3545,0.1261,0.5671,0.2401,0.01097,0.9725,0.8744,0.5684,0.0004416,0.0008327,0.005773,0.386,0.6818,0.3794,0.2087,0.8752,0.002233,0.2642,0.1203,0.2122,0.01488,0.5885,0.9558,0.9269,0.1363,0.03529,0.4063,0.7339,0.609,0.2775,0.4867,0.7719,0.9367,0.6424,0.1267,0.01283,0.4302,0.002199,0.9174,0.06663,0.05258,0.6444,0.2285,0.331,0.09763,0.9351,0.6343,0.1726,0.1958,0.4145,0.43,0.2531,0.09497,0.3764,0.0255,0.2553,0.8555,0.9211,0.4549,0.927,0.6987,0.006838,0.4811,0.8789,0.4379,0.1166,0.1614,0.2238,0.1627,0.1077,0.5304,0.0598,0.8377,0.04055,0.1108,0.01592,0.8195,0.5296,0.544,0.6253,0.3664,0.6959,0.9107,0.5539,0.00194,0.9846,0.2868,0.8861,0.9958,0.3354,0.1619,0.0479,0.1195,0.4265,0.9467,0.299,0.8094,0.546,0.4697,0.4541,0.3407,0.8093,0.0006681,0.01385,0.6634,0.1634,0.3265,0.8266,0.8621,0.002783,0.1782,0.9199,0.7957,0.6806,0.6053,0.06816,0.1846,0.1909,0.2808,0.2468,0.3525,0.1002,0.2102,0.3206,0.3947,0.1852,0.5733,0.6332,0.8312,0.7188,0.004553,0.2136,0.3282,0.0653,0.0841,0.009796,0.3866,0.5544,0.4193,0.9152,0.3438,0.368,0.9906,0.3151,0.5505,0.6931,0.5734,0.1175,0.2648,0.9671,0.02214,0.4597,0.06471,0.01935,0.2044,0.6708,0.0191,0.9505,0.552,0.5731,0.9563,0.1954,0.9915,0.02872,0.4185,0.06464,0.007483,0.3928,0.1456,0.658,0.02428,0.002825,0.06659,0.331,0.002717,0.0368,0.4829,0.4878,0.7863,0.2108,0.3282,0.824,0.933,0.4562,0.03114,0.5883,0.01271,0.01984,0.002051,0.2818,0.004388,0.1714,0.8901,0.0004472,0.8883,0.9451,0.4016,0.7204,0.4696,0.5717,0.008821,0.04111,0.00553,0.1446,0.05869,0.2182,0.2458,0.4844,0.4336,0.02675,0.6868,0.1643,0.4083,0.6657,0.07065,0.7432,0.1604,0.8566,0.1798,0.0464,0.4029,0.6102,0.3697,0.7215,0.2674,0.9267,0.2861,0.08976,0.0009216,0.005949,0.3387,0.00528,0.03064,0.1583,0.8275,0.001153,0.0829,0.2817,0.8881,0.7974,0.3794,0.444,0.9479,0.9502,0.1898,0.008252,0.3617,0.01447,0.06204,0.7246,0.1953,0.0003772,0.7483,0.8863,0.8991,0.6259,0.2976,0.01409,0.1086,0.6994,0.3986,0.5078,0.2838,0.9768,0.07045,0.159,0.0007133,0.1822,0.07018,0.7176,0.3271,0.3799,0.09431,0.4953,0.5165,0.2854,0.9398,0.2956,0.3908,0.4837,0.9615,0.9211,0.08813,0.3703,0.003126,0.01522,0.2725,0.9971,0.8035,0.8889,0.05361,0.1562,0.8442,0.5722,0.9359,0.6355,0.01544,0.476,0.1559,0.7706,0.03526,0.2009,0.625,0.022,0.4687,0.6017,0.4059,0.4449,0.0335,0.0007073,0.6382,0.753,0.002295,0.7018,0.06365,0.00106,0.9353,0.8484,0.008943,0.04933,0.5184,0.5288,0.006235,0.8928,0.6018,0.01755,0.824,0.0164,0.1233,0.8664,0.142,0.004741,0.05473,0.6742,0.5448,0.6034,0.1752,0.0003961,0.2807,0.1976,0.07563,0.4369,0.9773,0.7375,0.04072,0.2009,0.2541,0.0005044,0.7417,0.01562,0.5064,0.0782,0.1589,0.1479,0.6355,0.1268,0.2673,0.007109,0.6681,0.6803,0.6128,0.343,0.554,0.0617,0.8839,0.1004,0.03762,0.7243,0.3385,0.3484,0.537,0.743,0.9655,0.4007,0.5304,0.8178,0.8655,0.6735,0.2087,0.6064,0.1305,0.01261,0.01043,0.214,0.000925,0.698,0.04227,0.4268,0.9014,0.04447,0.8937,0.2589,0.5565,0.4967,0.3609,0.08535,0.08361,0.1351,0.8061,0.6874,0.9031,0.03539,0.7787,0.1052,0.5513,0.01063,0.8297,0.1797,0.01719,0.4694,0.9392,0.4192,0.9797,0.7445,0.5482,0.1902,0.1765,0.6224,0.7603,0.002761,0.1422,0.02992,0.02807,0.3691,0.2135,0.3705,0.7059,0.8685,0.1294,0.1327,0.3093,0.1855,0.8242,0.02786,0.08763,0.01461,0.8644,0.9878,0.0385,0.664,0.3236,0.004153,0.005802,0.6015,0.5385,0.1729,0.03973,0.9968,0.001657,0.3735,0.009062,0.1875,0.2407,0.8075,0.01957,0.3377,0.00743,0.06819,0.5208,0.2221,0.9762,0.06157,0.01981,0.02362,0.0005325,0.01331,0.002051,0.07416,0.06591,0.01878,0.0383,0.03155,0.07288,0.0356,0.03253,0.1216,0.002804,0.8466,0.199,0.006967,0.1294,0.0008368,0.007102,0.1376,0.03526,0.03132,0.09316,0.03384,0.4613,1,0.08336,0.05038,0.5649,0.8205,0.8306,0.977,0.5187,0.08843,0.8862,0.9437,0.5351,0.4184,0.2292,9.083e-05,1.032e-05,2.466e-05,2.079e-08,4.131e-05,7.425e-05,5.067e-05,9.482e-05,8.256e-06,2.603e-06,0.0001915,1.819e-06,0.000151,0.0001067,1.797e-05,5.245e-05,1.519e-05,0.0001207,6.632e-06,5.54e-06,3.197e-05,3.999e-05,0.0001754,1.202e-05,7.007e-06,7.434e-05,2.663e-06,1.219e-05,9.553e-06,2.238e-05,1.278e-05,3.083e-05,1.546e-05,0.0001557,0.0001177,3.875e-05,2.665e-05,9.383e-07,9.731e-05,1.619e-06,8.467e-05,0.000171,7.854e-05,0.0001055,0.0001956,1.137e-05,1.063e-05,0.0001292,9.009e-05,5.002e-07,4.678e-07,3.652e-05,2.606e-06,9.167e-05,0.0001636,0.0001898,9.041e-05,0.000149,5.173e-06,1.431e-05,3.918e-05,3.345e-06,0.0001056,6.421e-05,7.499e-05,8.964e-05,3.646e-05,2.038e-05,4.69e-06,4.15e-07,1.24e-06,6.798e-05,0.0001831,2.55e-06,5.245e-05,2.894e-07,9.489e-06,4.237e-05,0.0001643,0.0001194,0.0001038,4.701e-05,3.305e-05,0.0001882,4.209e-06,8.642e-06,3.107e-05,0.0001931,3.559e-05,8.704e-05,9.313e-05,1.757e-06,2.691e-07,0.0001387,2.712e-05,0.0001609,7.535e-05,0.0001127,2.309e-05,2.339e-05,3.503e-05,0.0001195,3.268e-05,8.645e-05,7.694e-05,0.000149,2.131e-08,3.065e-05],"Contig":["Transfer.PvP01_00_1.final","Transfer.PvP01_00_1.final","Transfer.PvP01_00_1.final","Transfer.PvP01_00_1.final","Transfer.PvP01_00_1.final","Transfer.PvP01_00_1.final","Transfer.PvP01_00_1.final","Transfer.PvP01_00_1.final","Transfer.PvP01_00_1.final","Transfer.PvP01_00_1.final","Transfer.PvP01_00_1.final","Transfer.PvP01_00_1.final","Transfer.PvP01_00_1.final","Transfer.PvP01_00_1.final","Transfer.PvP01_00_1.final","Transfer.PvP01_00_1.final","Transfer.PvP01_00_1.final","Transfer.PvP01_00_1.final","Transfer.PvP01_00_1.final","Transfer.PvP01_00_1.final","Transfer.PvP01_00_1.final","Transfer.PvP01_00_2.final","Transfer.PvP01_00_2.final","Transfer.PvP01_00_2.final","Transfer.PvP01_00_2.final","Transfer.PvP01_00_2.final","Transfer.PvP01_00_2.final","Transfer.PvP01_00_2.final","Transfer.PvP01_00_3.final","Transfer.PvP01_00_3.final","Transfer.PvP01_00_3.final","Transfer.PvP01_00_3.final","Transfer.PvP01_00_3.final","Transfer.PvP01_00_3.final","Transfer.PvP01_00_3.final","Transfer.PvP01_00_3.final","Transfer.PvP01_00_4.final","Transfer.PvP01_00_4.final","Transfer.PvP01_00_4.final","Transfer.PvP01_00_4.final","Transfer.PvP01_00_4.final","Transfer.PvP01_00_4.final","Transfer.PvP01_00_4.final","Transfer.PvP01_00_5.final","Transfer.PvP01_00_5.final","Transfer.PvP01_00_5.final","Transfer.PvP01_00_6.final","Transfer.PvP01_00_6.final","Transfer.PvP01_00_6.final","Transfer.PvP01_00_6.final","Transfer.PvP01_00_7.final","Transfer.PvP01_00_7.final","Transfer.PvP01_00_7.final","Transfer.PvP01_00_8.final","Transfer.PvP01_00_8.final","Transfer.PvP01_00_8.final","Transfer.PvP01_00_8.final","Transfer.PvP01_00_8.final","Transfer.PvP01_00_8.final","Transfer.PvP01_00_8.final","Transfer.PvP01_00_8.final","Transfer.PvP01_00_8.final","Transfer.PvP01_00_8.final","Transfer.PvP01_00_9.final","Transfer.PvP01_00_9.final","Transfer.PvP01_00_9.final","Transfer.PvP01_00_10.final","Transfer.PvP01_00_10.final","Transfer.PvP01_00_10.final","Transfer.PvP01_00_10.final","Transfer.PvP01_00_10.final","Transfer.PvP01_00_10.final","Transfer.PvP01_00_11.final","Transfer.PvP01_00_11.final","Transfer.PvP01_00_11.final","Transfer.PvP01_00_11.final","Transfer.PvP01_00_11.final","Transfer.PvP01_00_11.final","Transfer.PvP01_00_11.final","Transfer.PvP01_00_11.final","Transfer.PvP01_00_11.final","Transfer.PvP01_00_11.final","Transfer.PvP01_00_11.final","Transfer.PvP01_00_11.final","Transfer.PvP01_00_11.final","Transfer.PvP01_00_11.final","Transfer.PvP01_00_11.final","Transfer.PvP01_00_11.final","Transfer.PvP01_00_11.final","Transfer.PvP01_00_11.final","Transfer.PvP01_00_11.final","Transfer.PvP01_00_12.final","Transfer.PvP01_00_12.final","Transfer.PvP01_00_13.final","Transfer.PvP01_00_13.final","Transfer.PvP01_00_14.final","Transfer.PvP01_00_14.final","Transfer.PvP01_00_14.final","Transfer.PvP01_00_14.final","Transfer.PvP01_00_14.final","Transfer.PvP01_00_15.final","Transfer.PvP01_00_15.final","Transfer.PvP01_00_15.final","Transfer.PvP01_00_15.final","Transfer.PvP01_00_15.final","Transfer.PvP01_00_16.final","Transfer.PvP01_00_16.final","Transfer.PvP01_00_16.final","Transfer.PvP01_00_17.final","Transfer.PvP01_00_17.final","Transfer.PvP01_00_17.final","Transfer.PvP01_00_17.final","Transfer.PvP01_00_17.final","Transfer.PvP01_00_17.final","Transfer.PvP01_00_17.final","Transfer.PvP01_00_18.final","Transfer.PvP01_00_18.final","Transfer.PvP01_00_18.final","Transfer.PvP01_00_19.final","Transfer.PvP01_00_22.final","Transfer.PvP01_00_22.final","Transfer.PvP01_00_22.final","Transfer.PvP01_00_22.final","Transfer.PvP01_00_22.final","Transfer.PvP01_00_23.final","Transfer.PvP01_00_23.final","Transfer.PvP01_00_24.final","Transfer.PvP01_00_25.final","Transfer.PvP01_00_26.final","Transfer.PvP01_00_26.final","Transfer.PvP01_00_26.final","Transfer.PvP01_00_26.final","Transfer.PvP01_00_26.final","Transfer.PvP01_00_29.final","Transfer.PvP01_00_29.final","Transfer.PvP01_00_29.final","Transfer.PvP01_00_30.final","Transfer.PvP01_00_30.final","Transfer.PvP01_00_31.final","Transfer.PvP01_00_31.final","Transfer.PvP01_00_31.final","Transfer.PvP01_00_32.final","Transfer.PvP01_00_34.final","Transfer.PvP01_00_37.final","Transfer.PvP01_00_38.final","Transfer.PvP01_00_38.final","Transfer.PvP01_00_39.final","Transfer.PvP01_00_41.final","Transfer.PvP01_00_42.final","Transfer.PvP01_00_42.final","Transfer.PvP01_00_45.final","Transfer.PvP01_00_45.final","Transfer.PvP01_00_48.final","Transfer.PvP01_00_48.final","Transfer.PvP01_00_48.final","Transfer.PvP01_00_51.final","Transfer.PvP01_00_52.final","Transfer.PvP01_00_53.final","Transfer.PvP01_00_55.final","Transfer.PvP01_00_57.final","Transfer.PvP01_00_61.final","Transfer.PvP01_00_61.final","Transfer.PvP01_00_61.final","Transfer.PvP01_00_63.final","Transfer.PvP01_00_64.final","Transfer.PvP01_00_64.final","Transfer.PvP01_00_66.final","Transfer.PvP01_00_67.final","Transfer.PvP01_00_67.final","Transfer.PvP01_00_67.final","Transfer.PvP01_00_69.final","Transfer.PvP01_00_71.final","Transfer.PvP01_00_74.final","Transfer.PvP01_00_77.final","Transfer.PvP01_00_77.final","Transfer.PvP01_00_79.final","Transfer.PvP01_00_81.final","Transfer.PvP01_00_82.final","Transfer.PvP01_00_83.final","Transfer.PvP01_00_86.final","Transfer.PvP01_00_88.final","Transfer.PvP01_00_89.final","Transfer.PvP01_00_89.final","Transfer.PvP01_00_89.final","Transfer.PvP01_00_90.final","Transfer.PvP01_00_90.final","Transfer.PvP01_00_104.final","Transfer.PvP01_00_108.final","Transfer.PvP01_00_119.final","Transfer.PvP01_00_124.final","Transfer.PvP01_00_129.final","Transfer.PvP01_00_138.final","Transfer.PvP01_00_143.final","Transfer.PvP01_00_150.final","Transfer.PvP01_00_152.final","Transfer.PvP01_00_157.final","Transfer.PvP01_00_158.final","Transfer.PvP01_00_161.final","Transfer.PvP01_00_180.final","Transfer.PvP01_00_199.final","Transfer.PvP01_00_203.final","Transfer.PvP01_00_208.final","Transfer.PvP01_00_209.final","Transfer.PvP01_00_217.final","Transfer.PvP01_00_221.final","Transfer.PvP01_00_223.final","Transfer.PvP01_00_227.final","Transfer.PvP01_00_233.final","Transfer.PvP01_00_300.final","PvP01_01_v1","PvP01_01_v1","PvP01_01_v1","PvP01_01_v1","PvP01_01_v1","PvP01_01_v1","PvP01_01_v1","PvP01_01_v1","PvP01_01_v1","PvP01_01_v1","PvP01_01_v1","PvP01_01_v1","PvP01_01_v1","PvP01_01_v1","PvP01_01_v1","PvP01_01_v1","PvP01_01_v1","PvP01_01_v1","PvP01_01_v1","PvP01_01_v1","PvP01_01_v1","PvP01_01_v1","PvP01_01_v1","PvP01_01_v1","PvP01_01_v1","PvP01_01_v1","PvP01_01_v1","PvP01_01_v1","PvP01_01_v1","PvP01_01_v1","PvP01_01_v1","PvP01_01_v1","PvP01_01_v1","PvP01_01_v1","PvP01_01_v1","PvP01_01_v1","PvP01_01_v1","PvP01_01_v1","PvP01_01_v1","PvP01_01_v1","PvP01_01_v1","PvP01_01_v1","PvP01_01_v1","PvP01_01_v1","PvP01_01_v1","PvP01_01_v1","PvP01_01_v1","PvP01_01_v1","PvP01_01_v1","PvP01_01_v1","PvP01_01_v1","PvP01_01_v1","PvP01_01_v1","PvP01_01_v1","PvP01_01_v1","PvP01_01_v1","PvP01_01_v1","PvP01_01_v1","PvP01_01_v1","PvP01_01_v1","PvP01_01_v1","PvP01_01_v1","PvP01_01_v1","PvP01_01_v1","PvP01_01_v1","PvP01_01_v1","PvP01_01_v1","PvP01_01_v1","PvP01_01_v1","PvP01_01_v1","PvP01_01_v1","PvP01_01_v1","PvP01_01_v1","PvP01_01_v1","PvP01_01_v1","PvP01_01_v1","PvP01_01_v1","PvP01_01_v1","PvP01_01_v1","PvP01_01_v1","PvP01_01_v1","PvP01_01_v1","PvP01_01_v1","PvP01_01_v1","PvP01_01_v1","PvP01_01_v1","PvP01_01_v1","PvP01_01_v1","PvP01_01_v1","PvP01_01_v1","PvP01_01_v1","PvP01_01_v1","PvP01_01_v1","PvP01_01_v1","PvP01_01_v1","PvP01_01_v1","PvP01_01_v1","PvP01_01_v1","PvP01_01_v1","PvP01_01_v1","PvP01_01_v1","PvP01_01_v1","PvP01_01_v1","PvP01_01_v1","PvP01_01_v1","PvP01_01_v1","PvP01_01_v1","PvP01_01_v1","PvP01_01_v1","PvP01_01_v1","PvP01_01_v1","PvP01_01_v1","PvP01_01_v1","PvP01_01_v1","PvP01_01_v1","PvP01_01_v1","PvP01_01_v1","PvP01_01_v1","PvP01_01_v1","PvP01_01_v1","PvP01_01_v1","PvP01_01_v1","PvP01_01_v1","PvP01_01_v1","PvP01_01_v1","PvP01_01_v1","PvP01_01_v1","PvP01_01_v1","PvP01_01_v1","PvP01_01_v1","PvP01_01_v1","PvP01_01_v1","PvP01_01_v1","PvP01_01_v1","PvP01_01_v1","PvP01_01_v1","PvP01_01_v1","PvP01_01_v1","PvP01_01_v1","PvP01_01_v1","PvP01_01_v1","PvP01_01_v1","PvP01_01_v1","PvP01_01_v1","PvP01_01_v1","PvP01_01_v1","PvP01_01_v1","PvP01_01_v1","PvP01_01_v1","PvP01_01_v1","PvP01_01_v1","PvP01_01_v1","PvP01_01_v1","PvP01_01_v1","PvP01_01_v1","PvP01_01_v1","PvP01_01_v1","PvP01_01_v1","PvP01_01_v1","PvP01_01_v1","PvP01_01_v1","PvP01_01_v1","PvP01_01_v1","PvP01_01_v1","PvP01_01_v1","PvP01_01_v1","PvP01_01_v1","PvP01_01_v1","PvP01_01_v1","PvP01_01_v1","PvP01_01_v1","PvP01_02_v1","PvP01_02_v1","PvP01_02_v1","PvP01_02_v1","PvP01_02_v1","PvP01_02_v1","PvP01_02_v1","PvP01_02_v1","PvP01_02_v1","PvP01_02_v1","PvP01_02_v1","PvP01_02_v1","PvP01_02_v1","PvP01_02_v1","PvP01_02_v1","PvP01_02_v1","PvP01_02_v1","PvP01_02_v1","PvP01_02_v1","PvP01_02_v1","PvP01_02_v1","PvP01_02_v1","PvP01_02_v1","PvP01_02_v1","PvP01_02_v1","PvP01_02_v1","PvP01_02_v1","PvP01_02_v1","PvP01_02_v1","PvP01_02_v1","PvP01_02_v1","PvP01_02_v1","PvP01_02_v1","PvP01_02_v1","PvP01_02_v1","PvP01_02_v1","PvP01_02_v1","PvP01_02_v1","PvP01_02_v1","PvP01_02_v1","PvP01_02_v1","PvP01_02_v1","PvP01_02_v1","PvP01_02_v1","PvP01_02_v1","PvP01_02_v1","PvP01_02_v1","PvP01_02_v1","PvP01_02_v1","PvP01_02_v1","PvP01_02_v1","PvP01_02_v1","PvP01_02_v1","PvP01_02_v1","PvP01_02_v1","PvP01_02_v1","PvP01_02_v1","PvP01_02_v1","PvP01_02_v1","PvP01_02_v1","PvP01_02_v1","PvP01_02_v1","PvP01_02_v1","PvP01_02_v1","PvP01_02_v1","PvP01_02_v1","PvP01_02_v1","PvP01_02_v1","PvP01_02_v1","PvP01_02_v1","PvP01_02_v1","PvP01_02_v1","PvP01_02_v1","PvP01_02_v1","PvP01_02_v1","PvP01_02_v1","PvP01_02_v1","PvP01_02_v1","PvP01_02_v1","PvP01_02_v1","PvP01_02_v1","PvP01_02_v1","PvP01_02_v1","PvP01_02_v1","PvP01_02_v1","PvP01_02_v1","PvP01_02_v1","PvP01_02_v1","PvP01_02_v1","PvP01_02_v1","PvP01_02_v1","PvP01_02_v1","PvP01_02_v1","PvP01_02_v1","PvP01_02_v1","PvP01_02_v1","PvP01_02_v1","PvP01_02_v1","PvP01_02_v1","PvP01_02_v1","PvP01_02_v1","PvP01_02_v1","PvP01_02_v1","PvP01_02_v1","PvP01_02_v1","PvP01_02_v1","PvP01_02_v1","PvP01_02_v1","PvP01_02_v1","PvP01_02_v1","PvP01_02_v1","PvP01_02_v1","PvP01_02_v1","PvP01_02_v1","PvP01_02_v1","PvP01_02_v1","PvP01_02_v1","PvP01_02_v1","PvP01_02_v1","PvP01_02_v1","PvP01_02_v1","PvP01_02_v1","PvP01_02_v1","PvP01_02_v1","PvP01_02_v1","PvP01_02_v1","PvP01_02_v1","PvP01_02_v1","PvP01_02_v1","PvP01_02_v1","PvP01_02_v1","PvP01_02_v1","PvP01_02_v1","PvP01_02_v1","PvP01_02_v1","PvP01_02_v1","PvP01_02_v1","PvP01_02_v1","PvP01_02_v1","PvP01_02_v1","PvP01_02_v1","PvP01_02_v1","PvP01_02_v1","PvP01_02_v1","PvP01_02_v1","PvP01_02_v1","PvP01_02_v1","PvP01_02_v1","PvP01_02_v1","PvP01_02_v1","PvP01_02_v1","PvP01_02_v1","PvP01_02_v1","PvP01_02_v1","PvP01_02_v1","PvP01_02_v1","PvP01_02_v1","PvP01_02_v1","PvP01_02_v1","PvP01_02_v1","PvP01_02_v1","PvP01_02_v1","PvP01_02_v1","PvP01_02_v1","PvP01_02_v1","PvP01_02_v1","PvP01_02_v1","PvP01_03_v1","PvP01_03_v1","PvP01_03_v1","PvP01_03_v1","PvP01_03_v1","PvP01_03_v1","PvP01_03_v1","PvP01_03_v1","PvP01_03_v1","PvP01_03_v1","PvP01_03_v1","PvP01_03_v1","PvP01_03_v1","PvP01_03_v1","PvP01_03_v1","PvP01_03_v1","PvP01_03_v1","PvP01_03_v1","PvP01_03_v1","PvP01_03_v1","PvP01_03_v1","PvP01_03_v1","PvP01_03_v1","PvP01_03_v1","PvP01_03_v1","PvP01_03_v1","PvP01_03_v1","PvP01_03_v1","PvP01_03_v1","PvP01_03_v1","PvP01_03_v1","PvP01_03_v1","PvP01_03_v1","PvP01_03_v1","PvP01_03_v1","PvP01_03_v1","PvP01_03_v1","PvP01_03_v1","PvP01_03_v1","PvP01_03_v1","PvP01_03_v1","PvP01_03_v1","PvP01_03_v1","PvP01_03_v1","PvP01_03_v1","PvP01_03_v1","PvP01_03_v1","PvP01_03_v1","PvP01_03_v1","PvP01_03_v1","PvP01_03_v1","PvP01_03_v1","PvP01_03_v1","PvP01_03_v1","PvP01_03_v1","PvP01_03_v1","PvP01_03_v1","PvP01_03_v1","PvP01_03_v1","PvP01_03_v1","PvP01_03_v1","PvP01_03_v1","PvP01_03_v1","PvP01_03_v1","PvP01_03_v1","PvP01_03_v1","PvP01_03_v1","PvP01_03_v1","PvP01_03_v1","PvP01_03_v1","PvP01_03_v1","PvP01_03_v1","PvP01_03_v1","PvP01_03_v1","PvP01_03_v1","PvP01_03_v1","PvP01_03_v1","PvP01_03_v1","PvP01_03_v1","PvP01_03_v1","PvP01_03_v1","PvP01_03_v1","PvP01_03_v1","PvP01_03_v1","PvP01_03_v1","PvP01_03_v1","PvP01_03_v1","PvP01_03_v1","PvP01_03_v1","PvP01_03_v1","PvP01_03_v1","PvP01_03_v1","PvP01_03_v1","PvP01_03_v1","PvP01_03_v1","PvP01_03_v1","PvP01_03_v1","PvP01_03_v1","PvP01_03_v1","PvP01_03_v1","PvP01_03_v1","PvP01_03_v1","PvP01_03_v1","PvP01_03_v1","PvP01_03_v1","PvP01_03_v1","PvP01_03_v1","PvP01_03_v1","PvP01_03_v1","PvP01_03_v1","PvP01_03_v1","PvP01_03_v1","PvP01_03_v1","PvP01_03_v1","PvP01_03_v1","PvP01_03_v1","PvP01_03_v1","PvP01_03_v1","PvP01_03_v1","PvP01_03_v1","PvP01_03_v1","PvP01_03_v1","PvP01_03_v1","PvP01_03_v1","PvP01_03_v1","PvP01_03_v1","PvP01_03_v1","PvP01_03_v1","PvP01_03_v1","PvP01_03_v1","PvP01_03_v1","PvP01_03_v1","PvP01_03_v1","PvP01_03_v1","PvP01_03_v1","PvP01_03_v1","PvP01_03_v1","PvP01_03_v1","PvP01_03_v1","PvP01_03_v1","PvP01_03_v1","PvP01_03_v1","PvP01_03_v1","PvP01_03_v1","PvP01_03_v1","PvP01_03_v1","PvP01_03_v1","PvP01_03_v1","PvP01_03_v1","PvP01_03_v1","PvP01_03_v1","PvP01_03_v1","PvP01_03_v1","PvP01_03_v1","PvP01_03_v1","PvP01_03_v1","PvP01_03_v1","PvP01_03_v1","PvP01_03_v1","PvP01_03_v1","PvP01_03_v1","PvP01_03_v1","PvP01_03_v1","PvP01_03_v1","PvP01_03_v1","PvP01_03_v1","PvP01_03_v1","PvP01_03_v1","PvP01_03_v1","PvP01_03_v1","PvP01_03_v1","PvP01_03_v1","PvP01_03_v1","PvP01_03_v1","PvP01_03_v1","PvP01_03_v1","PvP01_03_v1","PvP01_03_v1","PvP01_03_v1","PvP01_03_v1","PvP01_03_v1","PvP01_03_v1","PvP01_03_v1","PvP01_03_v1","PvP01_03_v1","PvP01_03_v1","PvP01_03_v1","PvP01_03_v1","PvP01_03_v1","PvP01_03_v1","PvP01_04_v1","PvP01_04_v1","PvP01_04_v1","PvP01_04_v1","PvP01_04_v1","PvP01_04_v1","PvP01_04_v1","PvP01_04_v1","PvP01_04_v1","PvP01_04_v1","PvP01_04_v1","PvP01_04_v1","PvP01_04_v1","PvP01_04_v1","PvP01_04_v1","PvP01_04_v1","PvP01_04_v1","PvP01_04_v1","PvP01_04_v1","PvP01_04_v1","PvP01_04_v1","PvP01_04_v1","PvP01_04_v1","PvP01_04_v1","PvP01_04_v1","PvP01_04_v1","PvP01_04_v1","PvP01_04_v1","PvP01_04_v1","PvP01_04_v1","PvP01_04_v1","PvP01_04_v1","PvP01_04_v1","PvP01_04_v1","PvP01_04_v1","PvP01_04_v1","PvP01_04_v1","PvP01_04_v1","PvP01_04_v1","PvP01_04_v1","PvP01_04_v1","PvP01_04_v1","PvP01_04_v1","PvP01_04_v1","PvP01_04_v1","PvP01_04_v1","PvP01_04_v1","PvP01_04_v1","PvP01_04_v1","PvP01_04_v1","PvP01_04_v1","PvP01_04_v1","PvP01_04_v1","PvP01_04_v1","PvP01_04_v1","PvP01_04_v1","PvP01_04_v1","PvP01_04_v1","PvP01_04_v1","PvP01_04_v1","PvP01_04_v1","PvP01_04_v1","PvP01_04_v1","PvP01_04_v1","PvP01_04_v1","PvP01_04_v1","PvP01_04_v1","PvP01_04_v1","PvP01_04_v1","PvP01_04_v1","PvP01_04_v1","PvP01_04_v1","PvP01_04_v1","PvP01_04_v1","PvP01_04_v1","PvP01_04_v1","PvP01_04_v1","PvP01_04_v1","PvP01_04_v1","PvP01_04_v1","PvP01_04_v1","PvP01_04_v1","PvP01_04_v1","PvP01_04_v1","PvP01_04_v1","PvP01_04_v1","PvP01_04_v1","PvP01_04_v1","PvP01_04_v1","PvP01_04_v1","PvP01_04_v1","PvP01_04_v1","PvP01_04_v1","PvP01_04_v1","PvP01_04_v1","PvP01_04_v1","PvP01_04_v1","PvP01_04_v1","PvP01_04_v1","PvP01_04_v1","PvP01_04_v1","PvP01_04_v1","PvP01_04_v1","PvP01_04_v1","PvP01_04_v1","PvP01_04_v1","PvP01_04_v1","PvP01_04_v1","PvP01_04_v1","PvP01_04_v1","PvP01_04_v1","PvP01_04_v1","PvP01_04_v1","PvP01_04_v1","PvP01_04_v1","PvP01_04_v1","PvP01_04_v1","PvP01_04_v1","PvP01_04_v1","PvP01_04_v1","PvP01_04_v1","PvP01_04_v1","PvP01_04_v1","PvP01_04_v1","PvP01_04_v1","PvP01_04_v1","PvP01_04_v1","PvP01_04_v1","PvP01_04_v1","PvP01_04_v1","PvP01_04_v1","PvP01_04_v1","PvP01_04_v1","PvP01_04_v1","PvP01_04_v1","PvP01_04_v1","PvP01_04_v1","PvP01_04_v1","PvP01_04_v1","PvP01_04_v1","PvP01_04_v1","PvP01_04_v1","PvP01_04_v1","PvP01_04_v1","PvP01_04_v1","PvP01_04_v1","PvP01_04_v1","PvP01_04_v1","PvP01_04_v1","PvP01_04_v1","PvP01_04_v1","PvP01_04_v1","PvP01_04_v1","PvP01_04_v1","PvP01_04_v1","PvP01_04_v1","PvP01_04_v1","PvP01_04_v1","PvP01_04_v1","PvP01_04_v1","PvP01_04_v1","PvP01_04_v1","PvP01_04_v1","PvP01_04_v1","PvP01_04_v1","PvP01_04_v1","PvP01_04_v1","PvP01_04_v1","PvP01_04_v1","PvP01_04_v1","PvP01_04_v1","PvP01_04_v1","PvP01_04_v1","PvP01_04_v1","PvP01_04_v1","PvP01_04_v1","PvP01_04_v1","PvP01_04_v1","PvP01_04_v1","PvP01_04_v1","PvP01_04_v1","PvP01_04_v1","PvP01_04_v1","PvP01_04_v1","PvP01_04_v1","PvP01_04_v1","PvP01_04_v1","PvP01_04_v1","PvP01_04_v1","PvP01_04_v1","PvP01_04_v1","PvP01_04_v1","PvP01_04_v1","PvP01_04_v1","PvP01_04_v1","PvP01_04_v1","PvP01_04_v1","PvP01_05_v1","PvP01_05_v1","PvP01_05_v1","PvP01_05_v1","PvP01_05_v1","PvP01_05_v1","PvP01_05_v1","PvP01_05_v1","PvP01_05_v1","PvP01_05_v1","PvP01_05_v1","PvP01_05_v1","PvP01_05_v1","PvP01_05_v1","PvP01_05_v1","PvP01_05_v1","PvP01_05_v1","PvP01_05_v1","PvP01_05_v1","PvP01_05_v1","PvP01_05_v1","PvP01_05_v1","PvP01_05_v1","PvP01_05_v1","PvP01_05_v1","PvP01_05_v1","PvP01_05_v1","PvP01_05_v1","PvP01_05_v1","PvP01_05_v1","PvP01_05_v1","PvP01_05_v1","PvP01_05_v1","PvP01_05_v1","PvP01_05_v1","PvP01_05_v1","PvP01_05_v1","PvP01_05_v1","PvP01_05_v1","PvP01_05_v1","PvP01_05_v1","PvP01_05_v1","PvP01_05_v1","PvP01_05_v1","PvP01_05_v1","PvP01_05_v1","PvP01_05_v1","PvP01_05_v1","PvP01_05_v1","PvP01_05_v1","PvP01_05_v1","PvP01_05_v1","PvP01_05_v1","PvP01_05_v1","PvP01_05_v1","PvP01_05_v1","PvP01_05_v1","PvP01_05_v1","PvP01_05_v1","PvP01_05_v1","PvP01_05_v1","PvP01_05_v1","PvP01_05_v1","PvP01_05_v1","PvP01_05_v1","PvP01_05_v1","PvP01_05_v1","PvP01_05_v1","PvP01_05_v1","PvP01_05_v1","PvP01_05_v1","PvP01_05_v1","PvP01_05_v1","PvP01_05_v1","PvP01_05_v1","PvP01_05_v1","PvP01_05_v1","PvP01_05_v1","PvP01_05_v1","PvP01_05_v1","PvP01_05_v1","PvP01_05_v1","PvP01_05_v1","PvP01_05_v1","PvP01_05_v1","PvP01_05_v1","PvP01_05_v1","PvP01_05_v1","PvP01_05_v1","PvP01_05_v1","PvP01_05_v1","PvP01_05_v1","PvP01_05_v1","PvP01_05_v1","PvP01_05_v1","PvP01_05_v1","PvP01_05_v1","PvP01_05_v1","PvP01_05_v1","PvP01_05_v1","PvP01_05_v1","PvP01_05_v1","PvP01_05_v1","PvP01_05_v1","PvP01_05_v1","PvP01_05_v1","PvP01_05_v1","PvP01_05_v1","PvP01_05_v1","PvP01_05_v1","PvP01_05_v1","PvP01_05_v1","PvP01_05_v1","PvP01_05_v1","PvP01_05_v1","PvP01_05_v1","PvP01_05_v1","PvP01_05_v1","PvP01_05_v1","PvP01_05_v1","PvP01_05_v1","PvP01_05_v1","PvP01_05_v1","PvP01_05_v1","PvP01_05_v1","PvP01_05_v1","PvP01_05_v1","PvP01_05_v1","PvP01_05_v1","PvP01_05_v1","PvP01_05_v1","PvP01_05_v1","PvP01_05_v1","PvP01_05_v1","PvP01_05_v1","PvP01_05_v1","PvP01_05_v1","PvP01_05_v1","PvP01_05_v1","PvP01_05_v1","PvP01_05_v1","PvP01_05_v1","PvP01_05_v1","PvP01_05_v1","PvP01_05_v1","PvP01_05_v1","PvP01_05_v1","PvP01_05_v1","PvP01_05_v1","PvP01_05_v1","PvP01_05_v1","PvP01_05_v1","PvP01_05_v1","PvP01_05_v1","PvP01_05_v1","PvP01_05_v1","PvP01_05_v1","PvP01_05_v1","PvP01_05_v1","PvP01_05_v1","PvP01_05_v1","PvP01_05_v1","PvP01_05_v1","PvP01_05_v1","PvP01_05_v1","PvP01_05_v1","PvP01_05_v1","PvP01_05_v1","PvP01_05_v1","PvP01_05_v1","PvP01_05_v1","PvP01_05_v1","PvP01_05_v1","PvP01_05_v1","PvP01_05_v1","PvP01_05_v1","PvP01_05_v1","PvP01_05_v1","PvP01_05_v1","PvP01_05_v1","PvP01_05_v1","PvP01_05_v1","PvP01_05_v1","PvP01_05_v1","PvP01_05_v1","PvP01_05_v1","PvP01_05_v1","PvP01_05_v1","PvP01_05_v1","PvP01_05_v1","PvP01_05_v1","PvP01_05_v1","PvP01_05_v1","PvP01_05_v1","PvP01_05_v1","PvP01_05_v1","PvP01_05_v1","PvP01_05_v1","PvP01_05_v1","PvP01_05_v1","PvP01_05_v1","PvP01_05_v1","PvP01_05_v1","PvP01_05_v1","PvP01_05_v1","PvP01_05_v1","PvP01_05_v1","PvP01_05_v1","PvP01_05_v1","PvP01_05_v1","PvP01_05_v1","PvP01_05_v1","PvP01_05_v1","PvP01_05_v1","PvP01_05_v1","PvP01_05_v1","PvP01_05_v1","PvP01_05_v1","PvP01_05_v1","PvP01_05_v1","PvP01_05_v1","PvP01_05_v1","PvP01_05_v1","PvP01_05_v1","PvP01_05_v1","PvP01_05_v1","PvP01_05_v1","PvP01_05_v1","PvP01_05_v1","PvP01_05_v1","PvP01_05_v1","PvP01_05_v1","PvP01_05_v1","PvP01_05_v1","PvP01_05_v1","PvP01_05_v1","PvP01_05_v1","PvP01_05_v1","PvP01_05_v1","PvP01_05_v1","PvP01_05_v1","PvP01_05_v1","PvP01_05_v1","PvP01_05_v1","PvP01_05_v1","PvP01_05_v1","PvP01_05_v1","PvP01_05_v1","PvP01_05_v1","PvP01_05_v1","PvP01_05_v1","PvP01_05_v1","PvP01_05_v1","PvP01_05_v1","PvP01_05_v1","PvP01_05_v1","PvP01_05_v1","PvP01_05_v1","PvP01_05_v1","PvP01_05_v1","PvP01_05_v1","PvP01_05_v1","PvP01_05_v1","PvP01_05_v1","PvP01_05_v1","PvP01_05_v1","PvP01_05_v1","PvP01_05_v1","PvP01_05_v1","PvP01_05_v1","PvP01_05_v1","PvP01_05_v1","PvP01_05_v1","PvP01_05_v1","PvP01_05_v1","PvP01_05_v1","PvP01_05_v1","PvP01_05_v1","PvP01_05_v1","PvP01_05_v1","PvP01_05_v1","PvP01_05_v1","PvP01_05_v1","PvP01_05_v1","PvP01_05_v1","PvP01_05_v1","PvP01_05_v1","PvP01_05_v1","PvP01_05_v1","PvP01_05_v1","PvP01_05_v1","PvP01_05_v1","PvP01_05_v1","PvP01_05_v1","PvP01_05_v1","PvP01_05_v1","PvP01_05_v1","PvP01_05_v1","PvP01_05_v1","PvP01_05_v1","PvP01_05_v1","PvP01_05_v1","PvP01_05_v1","PvP01_05_v1","PvP01_05_v1","PvP01_06_v1","PvP01_06_v1","PvP01_06_v1","PvP01_06_v1","PvP01_06_v1","PvP01_06_v1","PvP01_06_v1","PvP01_06_v1","PvP01_06_v1","PvP01_06_v1","PvP01_06_v1","PvP01_06_v1","PvP01_06_v1","PvP01_06_v1","PvP01_06_v1","PvP01_06_v1","PvP01_06_v1","PvP01_06_v1","PvP01_06_v1","PvP01_06_v1","PvP01_06_v1","PvP01_06_v1","PvP01_06_v1","PvP01_06_v1","PvP01_06_v1","PvP01_06_v1","PvP01_06_v1","PvP01_06_v1","PvP01_06_v1","PvP01_06_v1","PvP01_06_v1","PvP01_06_v1","PvP01_06_v1","PvP01_06_v1","PvP01_06_v1","PvP01_06_v1","PvP01_06_v1","PvP01_06_v1","PvP01_06_v1","PvP01_06_v1","PvP01_06_v1","PvP01_06_v1","PvP01_06_v1","PvP01_06_v1","PvP01_06_v1","PvP01_06_v1","PvP01_06_v1","PvP01_06_v1","PvP01_06_v1","PvP01_06_v1","PvP01_06_v1","PvP01_06_v1","PvP01_06_v1","PvP01_06_v1","PvP01_06_v1","PvP01_06_v1","PvP01_06_v1","PvP01_06_v1","PvP01_06_v1","PvP01_06_v1","PvP01_06_v1","PvP01_06_v1","PvP01_06_v1","PvP01_06_v1","PvP01_06_v1","PvP01_06_v1","PvP01_06_v1","PvP01_06_v1","PvP01_06_v1","PvP01_06_v1","PvP01_06_v1","PvP01_06_v1","PvP01_06_v1","PvP01_06_v1","PvP01_06_v1","PvP01_06_v1","PvP01_06_v1","PvP01_06_v1","PvP01_06_v1","PvP01_06_v1","PvP01_06_v1","PvP01_06_v1","PvP01_06_v1","PvP01_06_v1","PvP01_06_v1","PvP01_06_v1","PvP01_06_v1","PvP01_06_v1","PvP01_06_v1","PvP01_06_v1","PvP01_06_v1","PvP01_06_v1","PvP01_06_v1","PvP01_06_v1","PvP01_06_v1","PvP01_06_v1","PvP01_06_v1","PvP01_06_v1","PvP01_06_v1","PvP01_06_v1","PvP01_06_v1","PvP01_06_v1","PvP01_06_v1","PvP01_06_v1","PvP01_06_v1","PvP01_06_v1","PvP01_06_v1","PvP01_06_v1","PvP01_06_v1","PvP01_06_v1","PvP01_06_v1","PvP01_06_v1","PvP01_06_v1","PvP01_06_v1","PvP01_06_v1","PvP01_06_v1","PvP01_06_v1","PvP01_06_v1","PvP01_06_v1","PvP01_06_v1","PvP01_06_v1","PvP01_06_v1","PvP01_06_v1","PvP01_06_v1","PvP01_06_v1","PvP01_06_v1","PvP01_06_v1","PvP01_06_v1","PvP01_06_v1","PvP01_06_v1","PvP01_06_v1","PvP01_06_v1","PvP01_06_v1","PvP01_06_v1","PvP01_06_v1","PvP01_06_v1","PvP01_06_v1","PvP01_06_v1","PvP01_06_v1","PvP01_06_v1","PvP01_06_v1","PvP01_06_v1","PvP01_06_v1","PvP01_06_v1","PvP01_06_v1","PvP01_06_v1","PvP01_06_v1","PvP01_06_v1","PvP01_06_v1","PvP01_06_v1","PvP01_06_v1","PvP01_06_v1","PvP01_06_v1","PvP01_06_v1","PvP01_06_v1","PvP01_06_v1","PvP01_06_v1","PvP01_06_v1","PvP01_06_v1","PvP01_06_v1","PvP01_06_v1","PvP01_06_v1","PvP01_06_v1","PvP01_06_v1","PvP01_06_v1","PvP01_06_v1","PvP01_06_v1","PvP01_06_v1","PvP01_06_v1","PvP01_06_v1","PvP01_06_v1","PvP01_06_v1","PvP01_06_v1","PvP01_06_v1","PvP01_06_v1","PvP01_06_v1","PvP01_06_v1","PvP01_06_v1","PvP01_06_v1","PvP01_06_v1","PvP01_06_v1","PvP01_06_v1","PvP01_06_v1","PvP01_06_v1","PvP01_06_v1","PvP01_06_v1","PvP01_06_v1","PvP01_06_v1","PvP01_06_v1","PvP01_06_v1","PvP01_06_v1","PvP01_06_v1","PvP01_06_v1","PvP01_06_v1","PvP01_06_v1","PvP01_06_v1","PvP01_06_v1","PvP01_06_v1","PvP01_06_v1","PvP01_06_v1","PvP01_06_v1","PvP01_06_v1","PvP01_06_v1","PvP01_06_v1","PvP01_06_v1","PvP01_06_v1","PvP01_06_v1","PvP01_06_v1","PvP01_06_v1","PvP01_06_v1","PvP01_06_v1","PvP01_06_v1","PvP01_06_v1","PvP01_06_v1","PvP01_06_v1","PvP01_06_v1","PvP01_06_v1","PvP01_06_v1","PvP01_06_v1","PvP01_06_v1","PvP01_06_v1","PvP01_06_v1","PvP01_06_v1","PvP01_06_v1","PvP01_06_v1","PvP01_06_v1","PvP01_06_v1","PvP01_06_v1","PvP01_06_v1","PvP01_06_v1","PvP01_06_v1","PvP01_06_v1","PvP01_06_v1","PvP01_06_v1","PvP01_06_v1","PvP01_06_v1","PvP01_07_v1","PvP01_07_v1","PvP01_07_v1","PvP01_07_v1","PvP01_07_v1","PvP01_07_v1","PvP01_07_v1","PvP01_07_v1","PvP01_07_v1","PvP01_07_v1","PvP01_07_v1","PvP01_07_v1","PvP01_07_v1","PvP01_07_v1","PvP01_07_v1","PvP01_07_v1","PvP01_07_v1","PvP01_07_v1","PvP01_07_v1","PvP01_07_v1","PvP01_07_v1","PvP01_07_v1","PvP01_07_v1","PvP01_07_v1","PvP01_07_v1","PvP01_07_v1","PvP01_07_v1","PvP01_07_v1","PvP01_07_v1","PvP01_07_v1","PvP01_07_v1","PvP01_07_v1","PvP01_07_v1","PvP01_07_v1","PvP01_07_v1","PvP01_07_v1","PvP01_07_v1","PvP01_07_v1","PvP01_07_v1","PvP01_07_v1","PvP01_07_v1","PvP01_07_v1","PvP01_07_v1","PvP01_07_v1","PvP01_07_v1","PvP01_07_v1","PvP01_07_v1","PvP01_07_v1","PvP01_07_v1","PvP01_07_v1","PvP01_07_v1","PvP01_07_v1","PvP01_07_v1","PvP01_07_v1","PvP01_07_v1","PvP01_07_v1","PvP01_07_v1","PvP01_07_v1","PvP01_07_v1","PvP01_07_v1","PvP01_07_v1","PvP01_07_v1","PvP01_07_v1","PvP01_07_v1","PvP01_07_v1","PvP01_07_v1","PvP01_07_v1","PvP01_07_v1","PvP01_07_v1","PvP01_07_v1","PvP01_07_v1","PvP01_07_v1","PvP01_07_v1","PvP01_07_v1","PvP01_07_v1","PvP01_07_v1","PvP01_07_v1","PvP01_07_v1","PvP01_07_v1","PvP01_07_v1","PvP01_07_v1","PvP01_07_v1","PvP01_07_v1","PvP01_07_v1","PvP01_07_v1","PvP01_07_v1","PvP01_07_v1","PvP01_07_v1","PvP01_07_v1","PvP01_07_v1","PvP01_07_v1","PvP01_07_v1","PvP01_07_v1","PvP01_07_v1","PvP01_07_v1","PvP01_07_v1","PvP01_07_v1","PvP01_07_v1","PvP01_07_v1","PvP01_07_v1","PvP01_07_v1","PvP01_07_v1","PvP01_07_v1","PvP01_07_v1","PvP01_07_v1","PvP01_07_v1","PvP01_07_v1","PvP01_07_v1","PvP01_07_v1","PvP01_07_v1","PvP01_07_v1","PvP01_07_v1","PvP01_07_v1","PvP01_07_v1","PvP01_07_v1","PvP01_07_v1","PvP01_07_v1","PvP01_07_v1","PvP01_07_v1","PvP01_07_v1","PvP01_07_v1","PvP01_07_v1","PvP01_07_v1","PvP01_07_v1","PvP01_07_v1","PvP01_07_v1","PvP01_07_v1","PvP01_07_v1","PvP01_07_v1","PvP01_07_v1","PvP01_07_v1","PvP01_07_v1","PvP01_07_v1","PvP01_07_v1","PvP01_07_v1","PvP01_07_v1","PvP01_07_v1","PvP01_07_v1","PvP01_07_v1","PvP01_07_v1","PvP01_07_v1","PvP01_07_v1","PvP01_07_v1","PvP01_07_v1","PvP01_07_v1","PvP01_07_v1","PvP01_07_v1","PvP01_07_v1","PvP01_07_v1","PvP01_07_v1","PvP01_07_v1","PvP01_07_v1","PvP01_07_v1","PvP01_07_v1","PvP01_07_v1","PvP01_07_v1","PvP01_07_v1","PvP01_07_v1","PvP01_07_v1","PvP01_07_v1","PvP01_07_v1","PvP01_07_v1","PvP01_07_v1","PvP01_07_v1","PvP01_07_v1","PvP01_07_v1","PvP01_07_v1","PvP01_07_v1","PvP01_07_v1","PvP01_07_v1","PvP01_07_v1","PvP01_07_v1","PvP01_07_v1","PvP01_07_v1","PvP01_07_v1","PvP01_07_v1","PvP01_07_v1","PvP01_07_v1","PvP01_07_v1","PvP01_07_v1","PvP01_07_v1","PvP01_07_v1","PvP01_07_v1","PvP01_07_v1","PvP01_07_v1","PvP01_07_v1","PvP01_07_v1","PvP01_07_v1","PvP01_07_v1","PvP01_07_v1","PvP01_07_v1","PvP01_07_v1","PvP01_07_v1","PvP01_07_v1","PvP01_07_v1","PvP01_07_v1","PvP01_07_v1","PvP01_07_v1","PvP01_07_v1","PvP01_07_v1","PvP01_07_v1","PvP01_07_v1","PvP01_07_v1","PvP01_07_v1","PvP01_07_v1","PvP01_07_v1","PvP01_07_v1","PvP01_07_v1","PvP01_07_v1","PvP01_07_v1","PvP01_07_v1","PvP01_07_v1","PvP01_07_v1","PvP01_07_v1","PvP01_07_v1","PvP01_07_v1","PvP01_07_v1","PvP01_07_v1","PvP01_07_v1","PvP01_07_v1","PvP01_07_v1","PvP01_07_v1","PvP01_07_v1","PvP01_07_v1","PvP01_07_v1","PvP01_07_v1","PvP01_07_v1","PvP01_07_v1","PvP01_07_v1","PvP01_07_v1","PvP01_07_v1","PvP01_07_v1","PvP01_07_v1","PvP01_07_v1","PvP01_07_v1","PvP01_07_v1","PvP01_07_v1","PvP01_07_v1","PvP01_07_v1","PvP01_07_v1","PvP01_07_v1","PvP01_07_v1","PvP01_07_v1","PvP01_07_v1","PvP01_07_v1","PvP01_07_v1","PvP01_07_v1","PvP01_07_v1","PvP01_07_v1","PvP01_07_v1","PvP01_07_v1","PvP01_07_v1","PvP01_07_v1","PvP01_07_v1","PvP01_07_v1","PvP01_07_v1","PvP01_07_v1","PvP01_07_v1","PvP01_07_v1","PvP01_07_v1","PvP01_07_v1","PvP01_07_v1","PvP01_07_v1","PvP01_07_v1","PvP01_07_v1","PvP01_07_v1","PvP01_07_v1","PvP01_07_v1","PvP01_07_v1","PvP01_07_v1","PvP01_07_v1","PvP01_07_v1","PvP01_07_v1","PvP01_07_v1","PvP01_07_v1","PvP01_07_v1","PvP01_07_v1","PvP01_07_v1","PvP01_07_v1","PvP01_07_v1","PvP01_07_v1","PvP01_07_v1","PvP01_07_v1","PvP01_07_v1","PvP01_07_v1","PvP01_07_v1","PvP01_07_v1","PvP01_07_v1","PvP01_07_v1","PvP01_07_v1","PvP01_07_v1","PvP01_07_v1","PvP01_07_v1","PvP01_07_v1","PvP01_07_v1","PvP01_07_v1","PvP01_07_v1","PvP01_07_v1","PvP01_07_v1","PvP01_07_v1","PvP01_07_v1","PvP01_07_v1","PvP01_07_v1","PvP01_07_v1","PvP01_07_v1","PvP01_07_v1","PvP01_07_v1","PvP01_07_v1","PvP01_07_v1","PvP01_07_v1","PvP01_07_v1","PvP01_07_v1","PvP01_07_v1","PvP01_07_v1","PvP01_07_v1","PvP01_07_v1","PvP01_07_v1","PvP01_07_v1","PvP01_07_v1","PvP01_07_v1","PvP01_07_v1","PvP01_07_v1","PvP01_07_v1","PvP01_07_v1","PvP01_07_v1","PvP01_07_v1","PvP01_07_v1","PvP01_07_v1","PvP01_07_v1","PvP01_07_v1","PvP01_07_v1","PvP01_07_v1","PvP01_07_v1","PvP01_07_v1","PvP01_07_v1","PvP01_07_v1","PvP01_07_v1","PvP01_07_v1","PvP01_07_v1","PvP01_07_v1","PvP01_07_v1","PvP01_07_v1","PvP01_07_v1","PvP01_08_v1","PvP01_08_v1","PvP01_08_v1","PvP01_08_v1","PvP01_08_v1","PvP01_08_v1","PvP01_08_v1","PvP01_08_v1","PvP01_08_v1","PvP01_08_v1","PvP01_08_v1","PvP01_08_v1","PvP01_08_v1","PvP01_08_v1","PvP01_08_v1","PvP01_08_v1","PvP01_08_v1","PvP01_08_v1","PvP01_08_v1","PvP01_08_v1","PvP01_08_v1","PvP01_08_v1","PvP01_08_v1","PvP01_08_v1","PvP01_08_v1","PvP01_08_v1","PvP01_08_v1","PvP01_08_v1","PvP01_08_v1","PvP01_08_v1","PvP01_08_v1","PvP01_08_v1","PvP01_08_v1","PvP01_08_v1","PvP01_08_v1","PvP01_08_v1","PvP01_08_v1","PvP01_08_v1","PvP01_08_v1","PvP01_08_v1","PvP01_08_v1","PvP01_08_v1","PvP01_08_v1","PvP01_08_v1","PvP01_08_v1","PvP01_08_v1","PvP01_08_v1","PvP01_08_v1","PvP01_08_v1","PvP01_08_v1","PvP01_08_v1","PvP01_08_v1","PvP01_08_v1","PvP01_08_v1","PvP01_08_v1","PvP01_08_v1","PvP01_08_v1","PvP01_08_v1","PvP01_08_v1","PvP01_08_v1","PvP01_08_v1","PvP01_08_v1","PvP01_08_v1","PvP01_08_v1","PvP01_08_v1","PvP01_08_v1","PvP01_08_v1","PvP01_08_v1","PvP01_08_v1","PvP01_08_v1","PvP01_08_v1","PvP01_08_v1","PvP01_08_v1","PvP01_08_v1","PvP01_08_v1","PvP01_08_v1","PvP01_08_v1","PvP01_08_v1","PvP01_08_v1","PvP01_08_v1","PvP01_08_v1","PvP01_08_v1","PvP01_08_v1","PvP01_08_v1","PvP01_08_v1","PvP01_08_v1","PvP01_08_v1","PvP01_08_v1","PvP01_08_v1","PvP01_08_v1","PvP01_08_v1","PvP01_08_v1","PvP01_08_v1","PvP01_08_v1","PvP01_08_v1","PvP01_08_v1","PvP01_08_v1","PvP01_08_v1","PvP01_08_v1","PvP01_08_v1","PvP01_08_v1","PvP01_08_v1","PvP01_08_v1","PvP01_08_v1","PvP01_08_v1","PvP01_08_v1","PvP01_08_v1","PvP01_08_v1","PvP01_08_v1","PvP01_08_v1","PvP01_08_v1","PvP01_08_v1","PvP01_08_v1","PvP01_08_v1","PvP01_08_v1","PvP01_08_v1","PvP01_08_v1","PvP01_08_v1","PvP01_08_v1","PvP01_08_v1","PvP01_08_v1","PvP01_08_v1","PvP01_08_v1","PvP01_08_v1","PvP01_08_v1","PvP01_08_v1","PvP01_08_v1","PvP01_08_v1","PvP01_08_v1","PvP01_08_v1","PvP01_08_v1","PvP01_08_v1","PvP01_08_v1","PvP01_08_v1","PvP01_08_v1","PvP01_08_v1","PvP01_08_v1","PvP01_08_v1","PvP01_08_v1","PvP01_08_v1","PvP01_08_v1","PvP01_08_v1","PvP01_08_v1","PvP01_08_v1","PvP01_08_v1","PvP01_08_v1","PvP01_08_v1","PvP01_08_v1","PvP01_08_v1","PvP01_08_v1","PvP01_08_v1","PvP01_08_v1","PvP01_08_v1","PvP01_08_v1","PvP01_08_v1","PvP01_08_v1","PvP01_08_v1","PvP01_08_v1","PvP01_08_v1","PvP01_08_v1","PvP01_08_v1","PvP01_08_v1","PvP01_08_v1","PvP01_08_v1","PvP01_08_v1","PvP01_08_v1","PvP01_08_v1","PvP01_08_v1","PvP01_08_v1","PvP01_08_v1","PvP01_08_v1","PvP01_08_v1","PvP01_08_v1","PvP01_08_v1","PvP01_08_v1","PvP01_08_v1","PvP01_08_v1","PvP01_08_v1","PvP01_08_v1","PvP01_08_v1","PvP01_08_v1","PvP01_08_v1","PvP01_08_v1","PvP01_08_v1","PvP01_08_v1","PvP01_08_v1","PvP01_08_v1","PvP01_08_v1","PvP01_08_v1","PvP01_08_v1","PvP01_08_v1","PvP01_08_v1","PvP01_08_v1","PvP01_08_v1","PvP01_08_v1","PvP01_08_v1","PvP01_08_v1","PvP01_08_v1","PvP01_08_v1","PvP01_08_v1","PvP01_08_v1","PvP01_08_v1","PvP01_08_v1","PvP01_08_v1","PvP01_08_v1","PvP01_08_v1","PvP01_08_v1","PvP01_08_v1","PvP01_08_v1","PvP01_08_v1","PvP01_08_v1","PvP01_08_v1","PvP01_08_v1","PvP01_08_v1","PvP01_08_v1","PvP01_08_v1","PvP01_08_v1","PvP01_08_v1","PvP01_08_v1","PvP01_08_v1","PvP01_08_v1","PvP01_08_v1","PvP01_08_v1","PvP01_08_v1","PvP01_08_v1","PvP01_08_v1","PvP01_08_v1","PvP01_08_v1","PvP01_08_v1","PvP01_08_v1","PvP01_08_v1","PvP01_08_v1","PvP01_08_v1","PvP01_08_v1","PvP01_08_v1","PvP01_08_v1","PvP01_08_v1","PvP01_08_v1","PvP01_08_v1","PvP01_08_v1","PvP01_08_v1","PvP01_08_v1","PvP01_08_v1","PvP01_08_v1","PvP01_08_v1","PvP01_08_v1","PvP01_08_v1","PvP01_08_v1","PvP01_08_v1","PvP01_08_v1","PvP01_08_v1","PvP01_08_v1","PvP01_08_v1","PvP01_08_v1","PvP01_08_v1","PvP01_08_v1","PvP01_08_v1","PvP01_08_v1","PvP01_08_v1","PvP01_08_v1","PvP01_08_v1","PvP01_08_v1","PvP01_08_v1","PvP01_08_v1","PvP01_08_v1","PvP01_08_v1","PvP01_08_v1","PvP01_08_v1","PvP01_08_v1","PvP01_08_v1","PvP01_08_v1","PvP01_08_v1","PvP01_08_v1","PvP01_08_v1","PvP01_08_v1","PvP01_08_v1","PvP01_08_v1","PvP01_08_v1","PvP01_08_v1","PvP01_08_v1","PvP01_08_v1","PvP01_08_v1","PvP01_08_v1","PvP01_08_v1","PvP01_08_v1","PvP01_08_v1","PvP01_08_v1","PvP01_08_v1","PvP01_08_v1","PvP01_08_v1","PvP01_08_v1","PvP01_08_v1","PvP01_08_v1","PvP01_08_v1","PvP01_08_v1","PvP01_08_v1","PvP01_08_v1","PvP01_08_v1","PvP01_08_v1","PvP01_08_v1","PvP01_08_v1","PvP01_08_v1","PvP01_08_v1","PvP01_08_v1","PvP01_08_v1","PvP01_08_v1","PvP01_08_v1","PvP01_08_v1","PvP01_08_v1","PvP01_08_v1","PvP01_08_v1","PvP01_08_v1","PvP01_08_v1","PvP01_08_v1","PvP01_08_v1","PvP01_08_v1","PvP01_08_v1","PvP01_08_v1","PvP01_08_v1","PvP01_08_v1","PvP01_08_v1","PvP01_08_v1","PvP01_08_v1","PvP01_08_v1","PvP01_08_v1","PvP01_08_v1","PvP01_08_v1","PvP01_08_v1","PvP01_08_v1","PvP01_08_v1","PvP01_08_v1","PvP01_08_v1","PvP01_08_v1","PvP01_08_v1","PvP01_08_v1","PvP01_08_v1","PvP01_08_v1","PvP01_08_v1","PvP01_08_v1","PvP01_08_v1","PvP01_08_v1","PvP01_08_v1","PvP01_08_v1","PvP01_08_v1","PvP01_08_v1","PvP01_08_v1","PvP01_08_v1","PvP01_08_v1","PvP01_08_v1","PvP01_08_v1","PvP01_08_v1","PvP01_08_v1","PvP01_08_v1","PvP01_08_v1","PvP01_08_v1","PvP01_08_v1","PvP01_08_v1","PvP01_08_v1","PvP01_08_v1","PvP01_08_v1","PvP01_08_v1","PvP01_08_v1","PvP01_08_v1","PvP01_08_v1","PvP01_08_v1","PvP01_08_v1","PvP01_08_v1","PvP01_08_v1","PvP01_08_v1","PvP01_08_v1","PvP01_08_v1","PvP01_08_v1","PvP01_08_v1","PvP01_08_v1","PvP01_08_v1","PvP01_08_v1","PvP01_08_v1","PvP01_08_v1","PvP01_08_v1","PvP01_09_v1","PvP01_09_v1","PvP01_09_v1","PvP01_09_v1","PvP01_09_v1","PvP01_09_v1","PvP01_09_v1","PvP01_09_v1","PvP01_09_v1","PvP01_09_v1","PvP01_09_v1","PvP01_09_v1","PvP01_09_v1","PvP01_09_v1","PvP01_09_v1","PvP01_09_v1","PvP01_09_v1","PvP01_09_v1","PvP01_09_v1","PvP01_09_v1","PvP01_09_v1","PvP01_09_v1","PvP01_09_v1","PvP01_09_v1","PvP01_09_v1","PvP01_09_v1","PvP01_09_v1","PvP01_09_v1","PvP01_09_v1","PvP01_09_v1","PvP01_09_v1","PvP01_09_v1","PvP01_09_v1","PvP01_09_v1","PvP01_09_v1","PvP01_09_v1","PvP01_09_v1","PvP01_09_v1","PvP01_09_v1","PvP01_09_v1","PvP01_09_v1","PvP01_09_v1","PvP01_09_v1","PvP01_09_v1","PvP01_09_v1","PvP01_09_v1","PvP01_09_v1","PvP01_09_v1","PvP01_09_v1","PvP01_09_v1","PvP01_09_v1","PvP01_09_v1","PvP01_09_v1","PvP01_09_v1","PvP01_09_v1","PvP01_09_v1","PvP01_09_v1","PvP01_09_v1","PvP01_09_v1","PvP01_09_v1","PvP01_09_v1","PvP01_09_v1","PvP01_09_v1","PvP01_09_v1","PvP01_09_v1","PvP01_09_v1","PvP01_09_v1","PvP01_09_v1","PvP01_09_v1","PvP01_09_v1","PvP01_09_v1","PvP01_09_v1","PvP01_09_v1","PvP01_09_v1","PvP01_09_v1","PvP01_09_v1","PvP01_09_v1","PvP01_09_v1","PvP01_09_v1","PvP01_09_v1","PvP01_09_v1","PvP01_09_v1","PvP01_09_v1","PvP01_09_v1","PvP01_09_v1","PvP01_09_v1","PvP01_09_v1","PvP01_09_v1","PvP01_09_v1","PvP01_09_v1","PvP01_09_v1","PvP01_09_v1","PvP01_09_v1","PvP01_09_v1","PvP01_09_v1","PvP01_09_v1","PvP01_09_v1","PvP01_09_v1","PvP01_09_v1","PvP01_09_v1","PvP01_09_v1","PvP01_09_v1","PvP01_09_v1","PvP01_09_v1","PvP01_09_v1","PvP01_09_v1","PvP01_09_v1","PvP01_09_v1","PvP01_09_v1","PvP01_09_v1","PvP01_09_v1","PvP01_09_v1","PvP01_09_v1","PvP01_09_v1","PvP01_09_v1","PvP01_09_v1","PvP01_09_v1","PvP01_09_v1","PvP01_09_v1","PvP01_09_v1","PvP01_09_v1","PvP01_09_v1","PvP01_09_v1","PvP01_09_v1","PvP01_09_v1","PvP01_09_v1","PvP01_09_v1","PvP01_09_v1","PvP01_09_v1","PvP01_09_v1","PvP01_09_v1","PvP01_09_v1","PvP01_09_v1","PvP01_09_v1","PvP01_09_v1","PvP01_09_v1","PvP01_09_v1","PvP01_09_v1","PvP01_09_v1","PvP01_09_v1","PvP01_09_v1","PvP01_09_v1","PvP01_09_v1","PvP01_09_v1","PvP01_09_v1","PvP01_09_v1","PvP01_09_v1","PvP01_09_v1","PvP01_09_v1","PvP01_09_v1","PvP01_09_v1","PvP01_09_v1","PvP01_09_v1","PvP01_09_v1","PvP01_09_v1","PvP01_09_v1","PvP01_09_v1","PvP01_09_v1","PvP01_09_v1","PvP01_09_v1","PvP01_09_v1","PvP01_09_v1","PvP01_09_v1","PvP01_09_v1","PvP01_09_v1","PvP01_09_v1","PvP01_09_v1","PvP01_09_v1","PvP01_09_v1","PvP01_09_v1","PvP01_09_v1","PvP01_09_v1","PvP01_09_v1","PvP01_09_v1","PvP01_09_v1","PvP01_09_v1","PvP01_09_v1","PvP01_09_v1","PvP01_09_v1","PvP01_09_v1","PvP01_09_v1","PvP01_09_v1","PvP01_09_v1","PvP01_09_v1","PvP01_09_v1","PvP01_09_v1","PvP01_09_v1","PvP01_09_v1","PvP01_09_v1","PvP01_09_v1","PvP01_09_v1","PvP01_09_v1","PvP01_09_v1","PvP01_09_v1","PvP01_09_v1","PvP01_09_v1","PvP01_09_v1","PvP01_09_v1","PvP01_09_v1","PvP01_09_v1","PvP01_09_v1","PvP01_09_v1","PvP01_09_v1","PvP01_09_v1","PvP01_09_v1","PvP01_09_v1","PvP01_09_v1","PvP01_09_v1","PvP01_09_v1","PvP01_09_v1","PvP01_09_v1","PvP01_09_v1","PvP01_09_v1","PvP01_09_v1","PvP01_09_v1","PvP01_09_v1","PvP01_09_v1","PvP01_09_v1","PvP01_09_v1","PvP01_09_v1","PvP01_09_v1","PvP01_09_v1","PvP01_09_v1","PvP01_09_v1","PvP01_09_v1","PvP01_09_v1","PvP01_09_v1","PvP01_09_v1","PvP01_09_v1","PvP01_09_v1","PvP01_09_v1","PvP01_09_v1","PvP01_09_v1","PvP01_09_v1","PvP01_09_v1","PvP01_09_v1","PvP01_09_v1","PvP01_09_v1","PvP01_09_v1","PvP01_09_v1","PvP01_09_v1","PvP01_09_v1","PvP01_09_v1","PvP01_09_v1","PvP01_09_v1","PvP01_09_v1","PvP01_09_v1","PvP01_09_v1","PvP01_09_v1","PvP01_09_v1","PvP01_09_v1","PvP01_09_v1","PvP01_09_v1","PvP01_09_v1","PvP01_09_v1","PvP01_09_v1","PvP01_09_v1","PvP01_09_v1","PvP01_09_v1","PvP01_09_v1","PvP01_09_v1","PvP01_09_v1","PvP01_09_v1","PvP01_09_v1","PvP01_09_v1","PvP01_09_v1","PvP01_09_v1","PvP01_09_v1","PvP01_09_v1","PvP01_09_v1","PvP01_09_v1","PvP01_09_v1","PvP01_09_v1","PvP01_09_v1","PvP01_09_v1","PvP01_09_v1","PvP01_09_v1","PvP01_09_v1","PvP01_09_v1","PvP01_09_v1","PvP01_09_v1","PvP01_09_v1","PvP01_09_v1","PvP01_09_v1","PvP01_09_v1","PvP01_09_v1","PvP01_09_v1","PvP01_09_v1","PvP01_09_v1","PvP01_09_v1","PvP01_09_v1","PvP01_09_v1","PvP01_09_v1","PvP01_09_v1","PvP01_09_v1","PvP01_09_v1","PvP01_09_v1","PvP01_09_v1","PvP01_09_v1","PvP01_09_v1","PvP01_09_v1","PvP01_09_v1","PvP01_09_v1","PvP01_09_v1","PvP01_09_v1","PvP01_09_v1","PvP01_09_v1","PvP01_09_v1","PvP01_09_v1","PvP01_09_v1","PvP01_09_v1","PvP01_09_v1","PvP01_09_v1","PvP01_09_v1","PvP01_09_v1","PvP01_09_v1","PvP01_09_v1","PvP01_09_v1","PvP01_09_v1","PvP01_09_v1","PvP01_09_v1","PvP01_09_v1","PvP01_09_v1","PvP01_09_v1","PvP01_09_v1","PvP01_09_v1","PvP01_09_v1","PvP01_09_v1","PvP01_09_v1","PvP01_09_v1","PvP01_09_v1","PvP01_09_v1","PvP01_09_v1","PvP01_09_v1","PvP01_09_v1","PvP01_09_v1","PvP01_09_v1","PvP01_09_v1","PvP01_09_v1","PvP01_09_v1","PvP01_09_v1","PvP01_09_v1","PvP01_09_v1","PvP01_09_v1","PvP01_09_v1","PvP01_09_v1","PvP01_09_v1","PvP01_09_v1","PvP01_09_v1","PvP01_09_v1","PvP01_09_v1","PvP01_09_v1","PvP01_09_v1","PvP01_09_v1","PvP01_09_v1","PvP01_09_v1","PvP01_09_v1","PvP01_09_v1","PvP01_09_v1","PvP01_09_v1","PvP01_09_v1","PvP01_09_v1","PvP01_09_v1","PvP01_09_v1","PvP01_09_v1","PvP01_09_v1","PvP01_09_v1","PvP01_09_v1","PvP01_09_v1","PvP01_09_v1","PvP01_09_v1","PvP01_09_v1","PvP01_09_v1","PvP01_09_v1","PvP01_09_v1","PvP01_09_v1","PvP01_09_v1","PvP01_09_v1","PvP01_09_v1","PvP01_09_v1","PvP01_09_v1","PvP01_09_v1","PvP01_09_v1","PvP01_09_v1","PvP01_09_v1","PvP01_09_v1","PvP01_09_v1","PvP01_09_v1","PvP01_09_v1","PvP01_09_v1","PvP01_09_v1","PvP01_09_v1","PvP01_09_v1","PvP01_09_v1","PvP01_09_v1","PvP01_09_v1","PvP01_09_v1","PvP01_09_v1","PvP01_09_v1","PvP01_09_v1","PvP01_09_v1","PvP01_09_v1","PvP01_09_v1","PvP01_09_v1","PvP01_09_v1","PvP01_09_v1","PvP01_09_v1","PvP01_09_v1","PvP01_09_v1","PvP01_09_v1","PvP01_09_v1","PvP01_09_v1","PvP01_09_v1","PvP01_09_v1","PvP01_09_v1","PvP01_09_v1","PvP01_09_v1","PvP01_09_v1","PvP01_09_v1","PvP01_09_v1","PvP01_09_v1","PvP01_09_v1","PvP01_09_v1","PvP01_09_v1","PvP01_09_v1","PvP01_09_v1","PvP01_09_v1","PvP01_09_v1","PvP01_09_v1","PvP01_09_v1","PvP01_09_v1","PvP01_09_v1","PvP01_09_v1","PvP01_09_v1","PvP01_09_v1","PvP01_09_v1","PvP01_09_v1","PvP01_09_v1","PvP01_09_v1","PvP01_09_v1","PvP01_09_v1","PvP01_09_v1","PvP01_09_v1","PvP01_09_v1","PvP01_09_v1","PvP01_09_v1","PvP01_09_v1","PvP01_09_v1","PvP01_09_v1","PvP01_09_v1","PvP01_09_v1","PvP01_10_v1","PvP01_10_v1","PvP01_10_v1","PvP01_10_v1","PvP01_10_v1","PvP01_10_v1","PvP01_10_v1","PvP01_10_v1","PvP01_10_v1","PvP01_10_v1","PvP01_10_v1","PvP01_10_v1","PvP01_10_v1","PvP01_10_v1","PvP01_10_v1","PvP01_10_v1","PvP01_10_v1","PvP01_10_v1","PvP01_10_v1","PvP01_10_v1","PvP01_10_v1","PvP01_10_v1","PvP01_10_v1","PvP01_10_v1","PvP01_10_v1","PvP01_10_v1","PvP01_10_v1","PvP01_10_v1","PvP01_10_v1","PvP01_10_v1","PvP01_10_v1","PvP01_10_v1","PvP01_10_v1","PvP01_10_v1","PvP01_10_v1","PvP01_10_v1","PvP01_10_v1","PvP01_10_v1","PvP01_10_v1","PvP01_10_v1","PvP01_10_v1","PvP01_10_v1","PvP01_10_v1","PvP01_10_v1","PvP01_10_v1","PvP01_10_v1","PvP01_10_v1","PvP01_10_v1","PvP01_10_v1","PvP01_10_v1","PvP01_10_v1","PvP01_10_v1","PvP01_10_v1","PvP01_10_v1","PvP01_10_v1","PvP01_10_v1","PvP01_10_v1","PvP01_10_v1","PvP01_10_v1","PvP01_10_v1","PvP01_10_v1","PvP01_10_v1","PvP01_10_v1","PvP01_10_v1","PvP01_10_v1","PvP01_10_v1","PvP01_10_v1","PvP01_10_v1","PvP01_10_v1","PvP01_10_v1","PvP01_10_v1","PvP01_10_v1","PvP01_10_v1","PvP01_10_v1","PvP01_10_v1","PvP01_10_v1","PvP01_10_v1","PvP01_10_v1","PvP01_10_v1","PvP01_10_v1","PvP01_10_v1","PvP01_10_v1","PvP01_10_v1","PvP01_10_v1","PvP01_10_v1","PvP01_10_v1","PvP01_10_v1","PvP01_10_v1","PvP01_10_v1","PvP01_10_v1","PvP01_10_v1","PvP01_10_v1","PvP01_10_v1","PvP01_10_v1","PvP01_10_v1","PvP01_10_v1","PvP01_10_v1","PvP01_10_v1","PvP01_10_v1","PvP01_10_v1","PvP01_10_v1","PvP01_10_v1","PvP01_10_v1","PvP01_10_v1","PvP01_10_v1","PvP01_10_v1","PvP01_10_v1","PvP01_10_v1","PvP01_10_v1","PvP01_10_v1","PvP01_10_v1","PvP01_10_v1","PvP01_10_v1","PvP01_10_v1","PvP01_10_v1","PvP01_10_v1","PvP01_10_v1","PvP01_10_v1","PvP01_10_v1","PvP01_10_v1","PvP01_10_v1","PvP01_10_v1","PvP01_10_v1","PvP01_10_v1","PvP01_10_v1","PvP01_10_v1","PvP01_10_v1","PvP01_10_v1","PvP01_10_v1","PvP01_10_v1","PvP01_10_v1","PvP01_10_v1","PvP01_10_v1","PvP01_10_v1","PvP01_10_v1","PvP01_10_v1","PvP01_10_v1","PvP01_10_v1","PvP01_10_v1","PvP01_10_v1","PvP01_10_v1","PvP01_10_v1","PvP01_10_v1","PvP01_10_v1","PvP01_10_v1","PvP01_10_v1","PvP01_10_v1","PvP01_10_v1","PvP01_10_v1","PvP01_10_v1","PvP01_10_v1","PvP01_10_v1","PvP01_10_v1","PvP01_10_v1","PvP01_10_v1","PvP01_10_v1","PvP01_10_v1","PvP01_10_v1","PvP01_10_v1","PvP01_10_v1","PvP01_10_v1","PvP01_10_v1","PvP01_10_v1","PvP01_10_v1","PvP01_10_v1","PvP01_10_v1","PvP01_10_v1","PvP01_10_v1","PvP01_10_v1","PvP01_10_v1","PvP01_10_v1","PvP01_10_v1","PvP01_10_v1","PvP01_10_v1","PvP01_10_v1","PvP01_10_v1","PvP01_10_v1","PvP01_10_v1","PvP01_10_v1","PvP01_10_v1","PvP01_10_v1","PvP01_10_v1","PvP01_10_v1","PvP01_10_v1","PvP01_10_v1","PvP01_10_v1","PvP01_10_v1","PvP01_10_v1","PvP01_10_v1","PvP01_10_v1","PvP01_10_v1","PvP01_10_v1","PvP01_10_v1","PvP01_10_v1","PvP01_10_v1","PvP01_10_v1","PvP01_10_v1","PvP01_10_v1","PvP01_10_v1","PvP01_10_v1","PvP01_10_v1","PvP01_10_v1","PvP01_10_v1","PvP01_10_v1","PvP01_10_v1","PvP01_10_v1","PvP01_10_v1","PvP01_10_v1","PvP01_10_v1","PvP01_10_v1","PvP01_10_v1","PvP01_10_v1","PvP01_10_v1","PvP01_10_v1","PvP01_10_v1","PvP01_10_v1","PvP01_10_v1","PvP01_10_v1","PvP01_10_v1","PvP01_10_v1","PvP01_10_v1","PvP01_10_v1","PvP01_10_v1","PvP01_10_v1","PvP01_10_v1","PvP01_10_v1","PvP01_10_v1","PvP01_10_v1","PvP01_10_v1","PvP01_10_v1","PvP01_10_v1","PvP01_10_v1","PvP01_10_v1","PvP01_10_v1","PvP01_10_v1","PvP01_10_v1","PvP01_10_v1","PvP01_10_v1","PvP01_10_v1","PvP01_10_v1","PvP01_10_v1","PvP01_10_v1","PvP01_10_v1","PvP01_10_v1","PvP01_10_v1","PvP01_10_v1","PvP01_10_v1","PvP01_10_v1","PvP01_10_v1","PvP01_10_v1","PvP01_10_v1","PvP01_10_v1","PvP01_10_v1","PvP01_10_v1","PvP01_10_v1","PvP01_10_v1","PvP01_10_v1","PvP01_10_v1","PvP01_10_v1","PvP01_10_v1","PvP01_10_v1","PvP01_10_v1","PvP01_10_v1","PvP01_10_v1","PvP01_10_v1","PvP01_10_v1","PvP01_10_v1","PvP01_10_v1","PvP01_10_v1","PvP01_10_v1","PvP01_10_v1","PvP01_10_v1","PvP01_10_v1","PvP01_10_v1","PvP01_10_v1","PvP01_10_v1","PvP01_10_v1","PvP01_10_v1","PvP01_10_v1","PvP01_10_v1","PvP01_10_v1","PvP01_10_v1","PvP01_10_v1","PvP01_10_v1","PvP01_10_v1","PvP01_10_v1","PvP01_10_v1","PvP01_10_v1","PvP01_10_v1","PvP01_10_v1","PvP01_10_v1","PvP01_10_v1","PvP01_10_v1","PvP01_10_v1","PvP01_10_v1","PvP01_10_v1","PvP01_10_v1","PvP01_10_v1","PvP01_10_v1","PvP01_10_v1","PvP01_10_v1","PvP01_10_v1","PvP01_10_v1","PvP01_10_v1","PvP01_10_v1","PvP01_10_v1","PvP01_10_v1","PvP01_10_v1","PvP01_10_v1","PvP01_10_v1","PvP01_10_v1","PvP01_10_v1","PvP01_10_v1","PvP01_10_v1","PvP01_11_v1","PvP01_11_v1","PvP01_11_v1","PvP01_11_v1","PvP01_11_v1","PvP01_11_v1","PvP01_11_v1","PvP01_11_v1","PvP01_11_v1","PvP01_11_v1","PvP01_11_v1","PvP01_11_v1","PvP01_11_v1","PvP01_11_v1","PvP01_11_v1","PvP01_11_v1","PvP01_11_v1","PvP01_11_v1","PvP01_11_v1","PvP01_11_v1","PvP01_11_v1","PvP01_11_v1","PvP01_11_v1","PvP01_11_v1","PvP01_11_v1","PvP01_11_v1","PvP01_11_v1","PvP01_11_v1","PvP01_11_v1","PvP01_11_v1","PvP01_11_v1","PvP01_11_v1","PvP01_11_v1","PvP01_11_v1","PvP01_11_v1","PvP01_11_v1","PvP01_11_v1","PvP01_11_v1","PvP01_11_v1","PvP01_11_v1","PvP01_11_v1","PvP01_11_v1","PvP01_11_v1","PvP01_11_v1","PvP01_11_v1","PvP01_11_v1","PvP01_11_v1","PvP01_11_v1","PvP01_11_v1","PvP01_11_v1","PvP01_11_v1","PvP01_11_v1","PvP01_11_v1","PvP01_11_v1","PvP01_11_v1","PvP01_11_v1","PvP01_11_v1","PvP01_11_v1","PvP01_11_v1","PvP01_11_v1","PvP01_11_v1","PvP01_11_v1","PvP01_11_v1","PvP01_11_v1","PvP01_11_v1","PvP01_11_v1","PvP01_11_v1","PvP01_11_v1","PvP01_11_v1","PvP01_11_v1","PvP01_11_v1","PvP01_11_v1","PvP01_11_v1","PvP01_11_v1","PvP01_11_v1","PvP01_11_v1","PvP01_11_v1","PvP01_11_v1","PvP01_11_v1","PvP01_11_v1","PvP01_11_v1","PvP01_11_v1","PvP01_11_v1","PvP01_11_v1","PvP01_11_v1","PvP01_11_v1","PvP01_11_v1","PvP01_11_v1","PvP01_11_v1","PvP01_11_v1","PvP01_11_v1","PvP01_11_v1","PvP01_11_v1","PvP01_11_v1","PvP01_11_v1","PvP01_11_v1","PvP01_11_v1","PvP01_11_v1","PvP01_11_v1","PvP01_11_v1","PvP01_11_v1","PvP01_11_v1","PvP01_11_v1","PvP01_11_v1","PvP01_11_v1","PvP01_11_v1","PvP01_11_v1","PvP01_11_v1","PvP01_11_v1","PvP01_11_v1","PvP01_11_v1","PvP01_11_v1","PvP01_11_v1","PvP01_11_v1","PvP01_11_v1","PvP01_11_v1","PvP01_11_v1","PvP01_11_v1","PvP01_11_v1","PvP01_11_v1","PvP01_11_v1","PvP01_11_v1","PvP01_11_v1","PvP01_11_v1","PvP01_11_v1","PvP01_11_v1","PvP01_11_v1","PvP01_11_v1","PvP01_11_v1","PvP01_11_v1","PvP01_11_v1","PvP01_11_v1","PvP01_11_v1","PvP01_11_v1","PvP01_11_v1","PvP01_11_v1","PvP01_11_v1","PvP01_11_v1","PvP01_11_v1","PvP01_11_v1","PvP01_11_v1","PvP01_11_v1","PvP01_11_v1","PvP01_11_v1","PvP01_11_v1","PvP01_11_v1","PvP01_11_v1","PvP01_11_v1","PvP01_11_v1","PvP01_11_v1","PvP01_11_v1","PvP01_11_v1","PvP01_11_v1","PvP01_11_v1","PvP01_11_v1","PvP01_11_v1","PvP01_11_v1","PvP01_11_v1","PvP01_11_v1","PvP01_11_v1","PvP01_11_v1","PvP01_11_v1","PvP01_11_v1","PvP01_11_v1","PvP01_11_v1","PvP01_11_v1","PvP01_11_v1","PvP01_11_v1","PvP01_11_v1","PvP01_11_v1","PvP01_11_v1","PvP01_11_v1","PvP01_11_v1","PvP01_11_v1","PvP01_11_v1","PvP01_11_v1","PvP01_11_v1","PvP01_11_v1","PvP01_11_v1","PvP01_11_v1","PvP01_11_v1","PvP01_11_v1","PvP01_11_v1","PvP01_11_v1","PvP01_11_v1","PvP01_11_v1","PvP01_11_v1","PvP01_11_v1","PvP01_11_v1","PvP01_11_v1","PvP01_11_v1","PvP01_11_v1","PvP01_11_v1","PvP01_11_v1","PvP01_11_v1","PvP01_11_v1","PvP01_11_v1","PvP01_11_v1","PvP01_11_v1","PvP01_11_v1","PvP01_11_v1","PvP01_11_v1","PvP01_11_v1","PvP01_11_v1","PvP01_11_v1","PvP01_11_v1","PvP01_11_v1","PvP01_11_v1","PvP01_11_v1","PvP01_11_v1","PvP01_11_v1","PvP01_11_v1","PvP01_11_v1","PvP01_11_v1","PvP01_11_v1","PvP01_11_v1","PvP01_11_v1","PvP01_11_v1","PvP01_11_v1","PvP01_11_v1","PvP01_11_v1","PvP01_11_v1","PvP01_11_v1","PvP01_11_v1","PvP01_11_v1","PvP01_11_v1","PvP01_11_v1","PvP01_11_v1","PvP01_11_v1","PvP01_11_v1","PvP01_11_v1","PvP01_11_v1","PvP01_11_v1","PvP01_11_v1","PvP01_11_v1","PvP01_11_v1","PvP01_11_v1","PvP01_11_v1","PvP01_11_v1","PvP01_11_v1","PvP01_11_v1","PvP01_11_v1","PvP01_11_v1","PvP01_11_v1","PvP01_11_v1","PvP01_11_v1","PvP01_11_v1","PvP01_11_v1","PvP01_11_v1","PvP01_11_v1","PvP01_11_v1","PvP01_11_v1","PvP01_11_v1","PvP01_11_v1","PvP01_11_v1","PvP01_11_v1","PvP01_11_v1","PvP01_11_v1","PvP01_11_v1","PvP01_11_v1","PvP01_11_v1","PvP01_11_v1","PvP01_11_v1","PvP01_11_v1","PvP01_11_v1","PvP01_11_v1","PvP01_11_v1","PvP01_11_v1","PvP01_11_v1","PvP01_11_v1","PvP01_11_v1","PvP01_11_v1","PvP01_11_v1","PvP01_11_v1","PvP01_11_v1","PvP01_11_v1","PvP01_11_v1","PvP01_11_v1","PvP01_11_v1","PvP01_11_v1","PvP01_11_v1","PvP01_11_v1","PvP01_11_v1","PvP01_11_v1","PvP01_11_v1","PvP01_11_v1","PvP01_11_v1","PvP01_11_v1","PvP01_11_v1","PvP01_11_v1","PvP01_11_v1","PvP01_11_v1","PvP01_11_v1","PvP01_11_v1","PvP01_11_v1","PvP01_11_v1","PvP01_11_v1","PvP01_11_v1","PvP01_11_v1","PvP01_11_v1","PvP01_11_v1","PvP01_11_v1","PvP01_11_v1","PvP01_11_v1","PvP01_11_v1","PvP01_11_v1","PvP01_11_v1","PvP01_11_v1","PvP01_11_v1","PvP01_11_v1","PvP01_11_v1","PvP01_11_v1","PvP01_11_v1","PvP01_11_v1","PvP01_11_v1","PvP01_11_v1","PvP01_11_v1","PvP01_11_v1","PvP01_11_v1","PvP01_11_v1","PvP01_11_v1","PvP01_11_v1","PvP01_11_v1","PvP01_11_v1","PvP01_11_v1","PvP01_11_v1","PvP01_11_v1","PvP01_11_v1","PvP01_11_v1","PvP01_11_v1","PvP01_11_v1","PvP01_11_v1","PvP01_11_v1","PvP01_11_v1","PvP01_11_v1","PvP01_11_v1","PvP01_11_v1","PvP01_11_v1","PvP01_11_v1","PvP01_11_v1","PvP01_11_v1","PvP01_11_v1","PvP01_11_v1","PvP01_11_v1","PvP01_11_v1","PvP01_11_v1","PvP01_11_v1","PvP01_11_v1","PvP01_11_v1","PvP01_11_v1","PvP01_11_v1","PvP01_11_v1","PvP01_11_v1","PvP01_11_v1","PvP01_11_v1","PvP01_11_v1","PvP01_11_v1","PvP01_11_v1","PvP01_11_v1","PvP01_11_v1","PvP01_11_v1","PvP01_11_v1","PvP01_11_v1","PvP01_11_v1","PvP01_11_v1","PvP01_11_v1","PvP01_11_v1","PvP01_11_v1","PvP01_11_v1","PvP01_11_v1","PvP01_11_v1","PvP01_11_v1","PvP01_11_v1","PvP01_11_v1","PvP01_11_v1","PvP01_11_v1","PvP01_11_v1","PvP01_11_v1","PvP01_11_v1","PvP01_11_v1","PvP01_11_v1","PvP01_11_v1","PvP01_11_v1","PvP01_11_v1","PvP01_11_v1","PvP01_11_v1","PvP01_11_v1","PvP01_11_v1","PvP01_11_v1","PvP01_11_v1","PvP01_11_v1","PvP01_11_v1","PvP01_11_v1","PvP01_11_v1","PvP01_11_v1","PvP01_11_v1","PvP01_11_v1","PvP01_11_v1","PvP01_11_v1","PvP01_11_v1","PvP01_11_v1","PvP01_11_v1","PvP01_11_v1","PvP01_11_v1","PvP01_11_v1","PvP01_11_v1","PvP01_11_v1","PvP01_11_v1","PvP01_11_v1","PvP01_11_v1","PvP01_11_v1","PvP01_11_v1","PvP01_11_v1","PvP01_11_v1","PvP01_11_v1","PvP01_11_v1","PvP01_11_v1","PvP01_11_v1","PvP01_11_v1","PvP01_11_v1","PvP01_11_v1","PvP01_11_v1","PvP01_11_v1","PvP01_11_v1","PvP01_11_v1","PvP01_11_v1","PvP01_11_v1","PvP01_11_v1","PvP01_11_v1","PvP01_11_v1","PvP01_11_v1","PvP01_11_v1","PvP01_11_v1","PvP01_11_v1","PvP01_11_v1","PvP01_11_v1","PvP01_11_v1","PvP01_11_v1","PvP01_11_v1","PvP01_11_v1","PvP01_11_v1","PvP01_11_v1","PvP01_11_v1","PvP01_11_v1","PvP01_11_v1","PvP01_11_v1","PvP01_11_v1","PvP01_11_v1","PvP01_11_v1","PvP01_11_v1","PvP01_11_v1","PvP01_11_v1","PvP01_11_v1","PvP01_11_v1","PvP01_11_v1","PvP01_11_v1","PvP01_11_v1","PvP01_11_v1","PvP01_11_v1","PvP01_11_v1","PvP01_11_v1","PvP01_11_v1","PvP01_11_v1","PvP01_11_v1","PvP01_12_v1","PvP01_12_v1","PvP01_12_v1","PvP01_12_v1","PvP01_12_v1","PvP01_12_v1","PvP01_12_v1","PvP01_12_v1","PvP01_12_v1","PvP01_12_v1","PvP01_12_v1","PvP01_12_v1","PvP01_12_v1","PvP01_12_v1","PvP01_12_v1","PvP01_12_v1","PvP01_12_v1","PvP01_12_v1","PvP01_12_v1","PvP01_12_v1","PvP01_12_v1","PvP01_12_v1","PvP01_12_v1","PvP01_12_v1","PvP01_12_v1","PvP01_12_v1","PvP01_12_v1","PvP01_12_v1","PvP01_12_v1","PvP01_12_v1","PvP01_12_v1","PvP01_12_v1","PvP01_12_v1","PvP01_12_v1","PvP01_12_v1","PvP01_12_v1","PvP01_12_v1","PvP01_12_v1","PvP01_12_v1","PvP01_12_v1","PvP01_12_v1","PvP01_12_v1","PvP01_12_v1","PvP01_12_v1","PvP01_12_v1","PvP01_12_v1","PvP01_12_v1","PvP01_12_v1","PvP01_12_v1","PvP01_12_v1","PvP01_12_v1","PvP01_12_v1","PvP01_12_v1","PvP01_12_v1","PvP01_12_v1","PvP01_12_v1","PvP01_12_v1","PvP01_12_v1","PvP01_12_v1","PvP01_12_v1","PvP01_12_v1","PvP01_12_v1","PvP01_12_v1","PvP01_12_v1","PvP01_12_v1","PvP01_12_v1","PvP01_12_v1","PvP01_12_v1","PvP01_12_v1","PvP01_12_v1","PvP01_12_v1","PvP01_12_v1","PvP01_12_v1","PvP01_12_v1","PvP01_12_v1","PvP01_12_v1","PvP01_12_v1","PvP01_12_v1","PvP01_12_v1","PvP01_12_v1","PvP01_12_v1","PvP01_12_v1","PvP01_12_v1","PvP01_12_v1","PvP01_12_v1","PvP01_12_v1","PvP01_12_v1","PvP01_12_v1","PvP01_12_v1","PvP01_12_v1","PvP01_12_v1","PvP01_12_v1","PvP01_12_v1","PvP01_12_v1","PvP01_12_v1","PvP01_12_v1","PvP01_12_v1","PvP01_12_v1","PvP01_12_v1","PvP01_12_v1","PvP01_12_v1","PvP01_12_v1","PvP01_12_v1","PvP01_12_v1","PvP01_12_v1","PvP01_12_v1","PvP01_12_v1","PvP01_12_v1","PvP01_12_v1","PvP01_12_v1","PvP01_12_v1","PvP01_12_v1","PvP01_12_v1","PvP01_12_v1","PvP01_12_v1","PvP01_12_v1","PvP01_12_v1","PvP01_12_v1","PvP01_12_v1","PvP01_12_v1","PvP01_12_v1","PvP01_12_v1","PvP01_12_v1","PvP01_12_v1","PvP01_12_v1","PvP01_12_v1","PvP01_12_v1","PvP01_12_v1","PvP01_12_v1","PvP01_12_v1","PvP01_12_v1","PvP01_12_v1","PvP01_12_v1","PvP01_12_v1","PvP01_12_v1","PvP01_12_v1","PvP01_12_v1","PvP01_12_v1","PvP01_12_v1","PvP01_12_v1","PvP01_12_v1","PvP01_12_v1","PvP01_12_v1","PvP01_12_v1","PvP01_12_v1","PvP01_12_v1","PvP01_12_v1","PvP01_12_v1","PvP01_12_v1","PvP01_12_v1","PvP01_12_v1","PvP01_12_v1","PvP01_12_v1","PvP01_12_v1","PvP01_12_v1","PvP01_12_v1","PvP01_12_v1","PvP01_12_v1","PvP01_12_v1","PvP01_12_v1","PvP01_12_v1","PvP01_12_v1","PvP01_12_v1","PvP01_12_v1","PvP01_12_v1","PvP01_12_v1","PvP01_12_v1","PvP01_12_v1","PvP01_12_v1","PvP01_12_v1","PvP01_12_v1","PvP01_12_v1","PvP01_12_v1","PvP01_12_v1","PvP01_12_v1","PvP01_12_v1","PvP01_12_v1","PvP01_12_v1","PvP01_12_v1","PvP01_12_v1","PvP01_12_v1","PvP01_12_v1","PvP01_12_v1","PvP01_12_v1","PvP01_12_v1","PvP01_12_v1","PvP01_12_v1","PvP01_12_v1","PvP01_12_v1","PvP01_12_v1","PvP01_12_v1","PvP01_12_v1","PvP01_12_v1","PvP01_12_v1","PvP01_12_v1","PvP01_12_v1","PvP01_12_v1","PvP01_12_v1","PvP01_12_v1","PvP01_12_v1","PvP01_12_v1","PvP01_12_v1","PvP01_12_v1","PvP01_12_v1","PvP01_12_v1","PvP01_12_v1","PvP01_12_v1","PvP01_12_v1","PvP01_12_v1","PvP01_12_v1","PvP01_12_v1","PvP01_12_v1","PvP01_12_v1","PvP01_12_v1","PvP01_12_v1","PvP01_12_v1","PvP01_12_v1","PvP01_12_v1","PvP01_12_v1","PvP01_12_v1","PvP01_12_v1","PvP01_12_v1","PvP01_12_v1","PvP01_12_v1","PvP01_12_v1","PvP01_12_v1","PvP01_12_v1","PvP01_12_v1","PvP01_12_v1","PvP01_12_v1","PvP01_12_v1","PvP01_12_v1","PvP01_12_v1","PvP01_12_v1","PvP01_12_v1","PvP01_12_v1","PvP01_12_v1","PvP01_12_v1","PvP01_12_v1","PvP01_12_v1","PvP01_12_v1","PvP01_12_v1","PvP01_12_v1","PvP01_12_v1","PvP01_12_v1","PvP01_12_v1","PvP01_12_v1","PvP01_12_v1","PvP01_12_v1","PvP01_12_v1","PvP01_12_v1","PvP01_12_v1","PvP01_12_v1","PvP01_12_v1","PvP01_12_v1","PvP01_12_v1","PvP01_12_v1","PvP01_12_v1","PvP01_12_v1","PvP01_12_v1","PvP01_12_v1","PvP01_12_v1","PvP01_12_v1","PvP01_12_v1","PvP01_12_v1","PvP01_12_v1","PvP01_12_v1","PvP01_12_v1","PvP01_12_v1","PvP01_12_v1","PvP01_12_v1","PvP01_12_v1","PvP01_12_v1","PvP01_12_v1","PvP01_12_v1","PvP01_12_v1","PvP01_12_v1","PvP01_12_v1","PvP01_12_v1","PvP01_12_v1","PvP01_12_v1","PvP01_12_v1","PvP01_12_v1","PvP01_12_v1","PvP01_12_v1","PvP01_12_v1","PvP01_12_v1","PvP01_12_v1","PvP01_12_v1","PvP01_12_v1","PvP01_12_v1","PvP01_12_v1","PvP01_12_v1","PvP01_12_v1","PvP01_12_v1","PvP01_12_v1","PvP01_12_v1","PvP01_12_v1","PvP01_12_v1","PvP01_12_v1","PvP01_12_v1","PvP01_12_v1","PvP01_12_v1","PvP01_12_v1","PvP01_12_v1","PvP01_12_v1","PvP01_12_v1","PvP01_12_v1","PvP01_12_v1","PvP01_12_v1","PvP01_12_v1","PvP01_12_v1","PvP01_12_v1","PvP01_12_v1","PvP01_12_v1","PvP01_12_v1","PvP01_12_v1","PvP01_12_v1","PvP01_12_v1","PvP01_12_v1","PvP01_12_v1","PvP01_12_v1","PvP01_12_v1","PvP01_12_v1","PvP01_12_v1","PvP01_12_v1","PvP01_12_v1","PvP01_12_v1","PvP01_12_v1","PvP01_12_v1","PvP01_12_v1","PvP01_12_v1","PvP01_12_v1","PvP01_12_v1","PvP01_12_v1","PvP01_12_v1","PvP01_12_v1","PvP01_12_v1","PvP01_12_v1","PvP01_12_v1","PvP01_12_v1","PvP01_12_v1","PvP01_12_v1","PvP01_12_v1","PvP01_12_v1","PvP01_12_v1","PvP01_12_v1","PvP01_12_v1","PvP01_12_v1","PvP01_12_v1","PvP01_12_v1","PvP01_12_v1","PvP01_12_v1","PvP01_12_v1","PvP01_12_v1","PvP01_12_v1","PvP01_12_v1","PvP01_12_v1","PvP01_12_v1","PvP01_12_v1","PvP01_12_v1","PvP01_12_v1","PvP01_12_v1","PvP01_12_v1","PvP01_12_v1","PvP01_12_v1","PvP01_12_v1","PvP01_12_v1","PvP01_12_v1","PvP01_12_v1","PvP01_12_v1","PvP01_12_v1","PvP01_12_v1","PvP01_12_v1","PvP01_12_v1","PvP01_12_v1","PvP01_12_v1","PvP01_12_v1","PvP01_12_v1","PvP01_12_v1","PvP01_12_v1","PvP01_12_v1","PvP01_12_v1","PvP01_12_v1","PvP01_12_v1","PvP01_12_v1","PvP01_12_v1","PvP01_12_v1","PvP01_12_v1","PvP01_12_v1","PvP01_12_v1","PvP01_12_v1","PvP01_12_v1","PvP01_12_v1","PvP01_12_v1","PvP01_12_v1","PvP01_12_v1","PvP01_12_v1","PvP01_12_v1","PvP01_12_v1","PvP01_12_v1","PvP01_12_v1","PvP01_12_v1","PvP01_12_v1","PvP01_12_v1","PvP01_12_v1","PvP01_12_v1","PvP01_12_v1","PvP01_12_v1","PvP01_12_v1","PvP01_12_v1","PvP01_12_v1","PvP01_12_v1","PvP01_12_v1","PvP01_12_v1","PvP01_12_v1","PvP01_12_v1","PvP01_12_v1","PvP01_12_v1","PvP01_12_v1","PvP01_12_v1","PvP01_12_v1","PvP01_12_v1","PvP01_12_v1","PvP01_12_v1","PvP01_12_v1","PvP01_12_v1","PvP01_12_v1","PvP01_12_v1","PvP01_12_v1","PvP01_12_v1","PvP01_12_v1","PvP01_12_v1","PvP01_12_v1","PvP01_12_v1","PvP01_12_v1","PvP01_12_v1","PvP01_12_v1","PvP01_12_v1","PvP01_12_v1","PvP01_12_v1","PvP01_12_v1","PvP01_12_v1","PvP01_12_v1","PvP01_12_v1","PvP01_12_v1","PvP01_12_v1","PvP01_12_v1","PvP01_12_v1","PvP01_12_v1","PvP01_12_v1","PvP01_12_v1","PvP01_12_v1","PvP01_12_v1","PvP01_12_v1","PvP01_12_v1","PvP01_12_v1","PvP01_12_v1","PvP01_12_v1","PvP01_12_v1","PvP01_12_v1","PvP01_12_v1","PvP01_12_v1","PvP01_12_v1","PvP01_12_v1","PvP01_12_v1","PvP01_12_v1","PvP01_12_v1","PvP01_12_v1","PvP01_12_v1","PvP01_12_v1","PvP01_12_v1","PvP01_12_v1","PvP01_12_v1","PvP01_12_v1","PvP01_12_v1","PvP01_12_v1","PvP01_12_v1","PvP01_12_v1","PvP01_12_v1","PvP01_12_v1","PvP01_12_v1","PvP01_12_v1","PvP01_12_v1","PvP01_12_v1","PvP01_12_v1","PvP01_12_v1","PvP01_12_v1","PvP01_12_v1","PvP01_12_v1","PvP01_12_v1","PvP01_12_v1","PvP01_12_v1","PvP01_12_v1","PvP01_12_v1","PvP01_12_v1","PvP01_12_v1","PvP01_12_v1","PvP01_12_v1","PvP01_12_v1","PvP01_12_v1","PvP01_12_v1","PvP01_12_v1","PvP01_12_v1","PvP01_12_v1","PvP01_12_v1","PvP01_12_v1","PvP01_12_v1","PvP01_12_v1","PvP01_12_v1","PvP01_12_v1","PvP01_12_v1","PvP01_12_v1","PvP01_12_v1","PvP01_12_v1","PvP01_12_v1","PvP01_12_v1","PvP01_12_v1","PvP01_12_v1","PvP01_12_v1","PvP01_12_v1","PvP01_12_v1","PvP01_12_v1","PvP01_12_v1","PvP01_12_v1","PvP01_12_v1","PvP01_12_v1","PvP01_12_v1","PvP01_12_v1","PvP01_12_v1","PvP01_12_v1","PvP01_12_v1","PvP01_12_v1","PvP01_12_v1","PvP01_12_v1","PvP01_12_v1","PvP01_12_v1","PvP01_12_v1","PvP01_12_v1","PvP01_12_v1","PvP01_12_v1","PvP01_12_v1","PvP01_12_v1","PvP01_12_v1","PvP01_12_v1","PvP01_12_v1","PvP01_12_v1","PvP01_12_v1","PvP01_12_v1","PvP01_12_v1","PvP01_12_v1","PvP01_12_v1","PvP01_12_v1","PvP01_12_v1","PvP01_12_v1","PvP01_12_v1","PvP01_12_v1","PvP01_12_v1","PvP01_12_v1","PvP01_12_v1","PvP01_12_v1","PvP01_12_v1","PvP01_12_v1","PvP01_12_v1","PvP01_12_v1","PvP01_12_v1","PvP01_12_v1","PvP01_12_v1","PvP01_12_v1","PvP01_12_v1","PvP01_12_v1","PvP01_12_v1","PvP01_12_v1","PvP01_12_v1","PvP01_12_v1","PvP01_12_v1","PvP01_12_v1","PvP01_12_v1","PvP01_12_v1","PvP01_12_v1","PvP01_12_v1","PvP01_12_v1","PvP01_12_v1","PvP01_12_v1","PvP01_12_v1","PvP01_12_v1","PvP01_12_v1","PvP01_12_v1","PvP01_12_v1","PvP01_12_v1","PvP01_12_v1","PvP01_12_v1","PvP01_12_v1","PvP01_12_v1","PvP01_12_v1","PvP01_12_v1","PvP01_12_v1","PvP01_12_v1","PvP01_12_v1","PvP01_12_v1","PvP01_12_v1","PvP01_12_v1","PvP01_12_v1","PvP01_12_v1","PvP01_12_v1","PvP01_12_v1","PvP01_12_v1","PvP01_12_v1","PvP01_12_v1","PvP01_12_v1","PvP01_12_v1","PvP01_12_v1","PvP01_12_v1","PvP01_12_v1","PvP01_12_v1","PvP01_12_v1","PvP01_12_v1","PvP01_12_v1","PvP01_12_v1","PvP01_12_v1","PvP01_12_v1","PvP01_12_v1","PvP01_12_v1","PvP01_12_v1","PvP01_12_v1","PvP01_12_v1","PvP01_12_v1","PvP01_12_v1","PvP01_12_v1","PvP01_12_v1","PvP01_12_v1","PvP01_12_v1","PvP01_12_v1","PvP01_12_v1","PvP01_12_v1","PvP01_12_v1","PvP01_12_v1","PvP01_12_v1","PvP01_12_v1","PvP01_12_v1","PvP01_12_v1","PvP01_12_v1","PvP01_12_v1","PvP01_12_v1","PvP01_12_v1","PvP01_12_v1","PvP01_12_v1","PvP01_12_v1","PvP01_12_v1","PvP01_12_v1","PvP01_12_v1","PvP01_12_v1","PvP01_12_v1","PvP01_12_v1","PvP01_12_v1","PvP01_12_v1","PvP01_12_v1","PvP01_12_v1","PvP01_12_v1","PvP01_12_v1","PvP01_12_v1","PvP01_12_v1","PvP01_12_v1","PvP01_12_v1","PvP01_12_v1","PvP01_12_v1","PvP01_12_v1","PvP01_12_v1","PvP01_12_v1","PvP01_12_v1","PvP01_12_v1","PvP01_12_v1","PvP01_12_v1","PvP01_12_v1","PvP01_12_v1","PvP01_12_v1","PvP01_12_v1","PvP01_12_v1","PvP01_12_v1","PvP01_12_v1","PvP01_12_v1","PvP01_12_v1","PvP01_12_v1","PvP01_12_v1","PvP01_13_v1","PvP01_13_v1","PvP01_13_v1","PvP01_13_v1","PvP01_13_v1","PvP01_13_v1","PvP01_13_v1","PvP01_13_v1","PvP01_13_v1","PvP01_13_v1","PvP01_13_v1","PvP01_13_v1","PvP01_13_v1","PvP01_13_v1","PvP01_13_v1","PvP01_13_v1","PvP01_13_v1","PvP01_13_v1","PvP01_13_v1","PvP01_13_v1","PvP01_13_v1","PvP01_13_v1","PvP01_13_v1","PvP01_13_v1","PvP01_13_v1","PvP01_13_v1","PvP01_13_v1","PvP01_13_v1","PvP01_13_v1","PvP01_13_v1","PvP01_13_v1","PvP01_13_v1","PvP01_13_v1","PvP01_13_v1","PvP01_13_v1","PvP01_13_v1","PvP01_13_v1","PvP01_13_v1","PvP01_13_v1","PvP01_13_v1","PvP01_13_v1","PvP01_13_v1","PvP01_13_v1","PvP01_13_v1","PvP01_13_v1","PvP01_13_v1","PvP01_13_v1","PvP01_13_v1","PvP01_13_v1","PvP01_13_v1","PvP01_13_v1","PvP01_13_v1","PvP01_13_v1","PvP01_13_v1","PvP01_13_v1","PvP01_13_v1","PvP01_13_v1","PvP01_13_v1","PvP01_13_v1","PvP01_13_v1","PvP01_13_v1","PvP01_13_v1","PvP01_13_v1","PvP01_13_v1","PvP01_13_v1","PvP01_13_v1","PvP01_13_v1","PvP01_13_v1","PvP01_13_v1","PvP01_13_v1","PvP01_13_v1","PvP01_13_v1","PvP01_13_v1","PvP01_13_v1","PvP01_13_v1","PvP01_13_v1","PvP01_13_v1","PvP01_13_v1","PvP01_13_v1","PvP01_13_v1","PvP01_13_v1","PvP01_13_v1","PvP01_13_v1","PvP01_13_v1","PvP01_13_v1","PvP01_13_v1","PvP01_13_v1","PvP01_13_v1","PvP01_13_v1","PvP01_13_v1","PvP01_13_v1","PvP01_13_v1","PvP01_13_v1","PvP01_13_v1","PvP01_13_v1","PvP01_13_v1","PvP01_13_v1","PvP01_13_v1","PvP01_13_v1","PvP01_13_v1","PvP01_13_v1","PvP01_13_v1","PvP01_13_v1","PvP01_13_v1","PvP01_13_v1","PvP01_13_v1","PvP01_13_v1","PvP01_13_v1","PvP01_13_v1","PvP01_13_v1","PvP01_13_v1","PvP01_13_v1","PvP01_13_v1","PvP01_13_v1","PvP01_13_v1","PvP01_13_v1","PvP01_13_v1","PvP01_13_v1","PvP01_13_v1","PvP01_13_v1","PvP01_13_v1","PvP01_13_v1","PvP01_13_v1","PvP01_13_v1","PvP01_13_v1","PvP01_13_v1","PvP01_13_v1","PvP01_13_v1","PvP01_13_v1","PvP01_13_v1","PvP01_13_v1","PvP01_13_v1","PvP01_13_v1","PvP01_13_v1","PvP01_13_v1","PvP01_13_v1","PvP01_13_v1","PvP01_13_v1","PvP01_13_v1","PvP01_13_v1","PvP01_13_v1","PvP01_13_v1","PvP01_13_v1","PvP01_13_v1","PvP01_13_v1","PvP01_13_v1","PvP01_13_v1","PvP01_13_v1","PvP01_13_v1","PvP01_13_v1","PvP01_13_v1","PvP01_13_v1","PvP01_13_v1","PvP01_13_v1","PvP01_13_v1","PvP01_13_v1","PvP01_13_v1","PvP01_13_v1","PvP01_13_v1","PvP01_13_v1","PvP01_13_v1","PvP01_13_v1","PvP01_13_v1","PvP01_13_v1","PvP01_13_v1","PvP01_13_v1","PvP01_13_v1","PvP01_13_v1","PvP01_13_v1","PvP01_13_v1","PvP01_13_v1","PvP01_13_v1","PvP01_13_v1","PvP01_13_v1","PvP01_13_v1","PvP01_13_v1","PvP01_13_v1","PvP01_13_v1","PvP01_13_v1","PvP01_13_v1","PvP01_13_v1","PvP01_13_v1","PvP01_13_v1","PvP01_13_v1","PvP01_13_v1","PvP01_13_v1","PvP01_13_v1","PvP01_13_v1","PvP01_13_v1","PvP01_13_v1","PvP01_13_v1","PvP01_13_v1","PvP01_13_v1","PvP01_13_v1","PvP01_13_v1","PvP01_13_v1","PvP01_13_v1","PvP01_13_v1","PvP01_13_v1","PvP01_13_v1","PvP01_13_v1","PvP01_13_v1","PvP01_13_v1","PvP01_13_v1","PvP01_13_v1","PvP01_13_v1","PvP01_13_v1","PvP01_13_v1","PvP01_13_v1","PvP01_13_v1","PvP01_13_v1","PvP01_13_v1","PvP01_13_v1","PvP01_13_v1","PvP01_13_v1","PvP01_13_v1","PvP01_13_v1","PvP01_13_v1","PvP01_13_v1","PvP01_13_v1","PvP01_13_v1","PvP01_13_v1","PvP01_13_v1","PvP01_13_v1","PvP01_13_v1","PvP01_13_v1","PvP01_13_v1","PvP01_13_v1","PvP01_13_v1","PvP01_13_v1","PvP01_13_v1","PvP01_13_v1","PvP01_13_v1","PvP01_13_v1","PvP01_13_v1","PvP01_13_v1","PvP01_13_v1","PvP01_13_v1","PvP01_13_v1","PvP01_13_v1","PvP01_13_v1","PvP01_13_v1","PvP01_13_v1","PvP01_13_v1","PvP01_13_v1","PvP01_13_v1","PvP01_13_v1","PvP01_13_v1","PvP01_13_v1","PvP01_13_v1","PvP01_13_v1","PvP01_13_v1","PvP01_13_v1","PvP01_13_v1","PvP01_13_v1","PvP01_13_v1","PvP01_13_v1","PvP01_13_v1","PvP01_13_v1","PvP01_13_v1","PvP01_13_v1","PvP01_13_v1","PvP01_13_v1","PvP01_13_v1","PvP01_13_v1","PvP01_13_v1","PvP01_13_v1","PvP01_13_v1","PvP01_13_v1","PvP01_13_v1","PvP01_13_v1","PvP01_13_v1","PvP01_13_v1","PvP01_13_v1","PvP01_13_v1","PvP01_13_v1","PvP01_13_v1","PvP01_13_v1","PvP01_13_v1","PvP01_13_v1","PvP01_13_v1","PvP01_13_v1","PvP01_13_v1","PvP01_13_v1","PvP01_13_v1","PvP01_13_v1","PvP01_13_v1","PvP01_13_v1","PvP01_13_v1","PvP01_13_v1","PvP01_13_v1","PvP01_13_v1","PvP01_13_v1","PvP01_13_v1","PvP01_13_v1","PvP01_13_v1","PvP01_13_v1","PvP01_13_v1","PvP01_13_v1","PvP01_13_v1","PvP01_13_v1","PvP01_13_v1","PvP01_13_v1","PvP01_13_v1","PvP01_13_v1","PvP01_13_v1","PvP01_13_v1","PvP01_13_v1","PvP01_13_v1","PvP01_13_v1","PvP01_13_v1","PvP01_13_v1","PvP01_13_v1","PvP01_13_v1","PvP01_13_v1","PvP01_13_v1","PvP01_13_v1","PvP01_13_v1","PvP01_13_v1","PvP01_13_v1","PvP01_13_v1","PvP01_13_v1","PvP01_13_v1","PvP01_13_v1","PvP01_13_v1","PvP01_13_v1","PvP01_13_v1","PvP01_13_v1","PvP01_13_v1","PvP01_13_v1","PvP01_13_v1","PvP01_13_v1","PvP01_13_v1","PvP01_13_v1","PvP01_13_v1","PvP01_13_v1","PvP01_13_v1","PvP01_13_v1","PvP01_13_v1","PvP01_13_v1","PvP01_13_v1","PvP01_13_v1","PvP01_13_v1","PvP01_13_v1","PvP01_13_v1","PvP01_13_v1","PvP01_13_v1","PvP01_13_v1","PvP01_13_v1","PvP01_13_v1","PvP01_13_v1","PvP01_13_v1","PvP01_13_v1","PvP01_13_v1","PvP01_13_v1","PvP01_13_v1","PvP01_13_v1","PvP01_13_v1","PvP01_13_v1","PvP01_13_v1","PvP01_13_v1","PvP01_13_v1","PvP01_13_v1","PvP01_13_v1","PvP01_13_v1","PvP01_13_v1","PvP01_13_v1","PvP01_13_v1","PvP01_13_v1","PvP01_13_v1","PvP01_13_v1","PvP01_13_v1","PvP01_13_v1","PvP01_13_v1","PvP01_13_v1","PvP01_13_v1","PvP01_13_v1","PvP01_13_v1","PvP01_13_v1","PvP01_13_v1","PvP01_13_v1","PvP01_13_v1","PvP01_13_v1","PvP01_13_v1","PvP01_13_v1","PvP01_13_v1","PvP01_13_v1","PvP01_13_v1","PvP01_13_v1","PvP01_13_v1","PvP01_13_v1","PvP01_13_v1","PvP01_13_v1","PvP01_13_v1","PvP01_13_v1","PvP01_13_v1","PvP01_13_v1","PvP01_13_v1","PvP01_13_v1","PvP01_13_v1","PvP01_13_v1","PvP01_13_v1","PvP01_13_v1","PvP01_13_v1","PvP01_13_v1","PvP01_13_v1","PvP01_13_v1","PvP01_13_v1","PvP01_13_v1","PvP01_13_v1","PvP01_13_v1","PvP01_13_v1","PvP01_13_v1","PvP01_13_v1","PvP01_13_v1","PvP01_13_v1","PvP01_13_v1","PvP01_13_v1","PvP01_13_v1","PvP01_13_v1","PvP01_13_v1","PvP01_13_v1","PvP01_13_v1","PvP01_13_v1","PvP01_13_v1","PvP01_13_v1","PvP01_13_v1","PvP01_13_v1","PvP01_13_v1","PvP01_13_v1","PvP01_13_v1","PvP01_13_v1","PvP01_13_v1","PvP01_13_v1","PvP01_13_v1","PvP01_13_v1","PvP01_13_v1","PvP01_13_v1","PvP01_13_v1","PvP01_13_v1","PvP01_13_v1","PvP01_13_v1","PvP01_13_v1","PvP01_13_v1","PvP01_13_v1","PvP01_13_v1","PvP01_13_v1","PvP01_13_v1","PvP01_13_v1","PvP01_13_v1","PvP01_13_v1","PvP01_13_v1","PvP01_14_v1","PvP01_14_v1","PvP01_14_v1","PvP01_14_v1","PvP01_14_v1","PvP01_14_v1","PvP01_14_v1","PvP01_14_v1","PvP01_14_v1","PvP01_14_v1","PvP01_14_v1","PvP01_14_v1","PvP01_14_v1","PvP01_14_v1","PvP01_14_v1","PvP01_14_v1","PvP01_14_v1","PvP01_14_v1","PvP01_14_v1","PvP01_14_v1","PvP01_14_v1","PvP01_14_v1","PvP01_14_v1","PvP01_14_v1","PvP01_14_v1","PvP01_14_v1","PvP01_14_v1","PvP01_14_v1","PvP01_14_v1","PvP01_14_v1","PvP01_14_v1","PvP01_14_v1","PvP01_14_v1","PvP01_14_v1","PvP01_14_v1","PvP01_14_v1","PvP01_14_v1","PvP01_14_v1","PvP01_14_v1","PvP01_14_v1","PvP01_14_v1","PvP01_14_v1","PvP01_14_v1","PvP01_14_v1","PvP01_14_v1","PvP01_14_v1","PvP01_14_v1","PvP01_14_v1","PvP01_14_v1","PvP01_14_v1","PvP01_14_v1","PvP01_14_v1","PvP01_14_v1","PvP01_14_v1","PvP01_14_v1","PvP01_14_v1","PvP01_14_v1","PvP01_14_v1","PvP01_14_v1","PvP01_14_v1","PvP01_14_v1","PvP01_14_v1","PvP01_14_v1","PvP01_14_v1","PvP01_14_v1","PvP01_14_v1","PvP01_14_v1","PvP01_14_v1","PvP01_14_v1","PvP01_14_v1","PvP01_14_v1","PvP01_14_v1","PvP01_14_v1","PvP01_14_v1","PvP01_14_v1","PvP01_14_v1","PvP01_14_v1","PvP01_14_v1","PvP01_14_v1","PvP01_14_v1","PvP01_14_v1","PvP01_14_v1","PvP01_14_v1","PvP01_14_v1","PvP01_14_v1","PvP01_14_v1","PvP01_14_v1","PvP01_14_v1","PvP01_14_v1","PvP01_14_v1","PvP01_14_v1","PvP01_14_v1","PvP01_14_v1","PvP01_14_v1","PvP01_14_v1","PvP01_14_v1","PvP01_14_v1","PvP01_14_v1","PvP01_14_v1","PvP01_14_v1","PvP01_14_v1","PvP01_14_v1","PvP01_14_v1","PvP01_14_v1","PvP01_14_v1","PvP01_14_v1","PvP01_14_v1","PvP01_14_v1","PvP01_14_v1","PvP01_14_v1","PvP01_14_v1","PvP01_14_v1","PvP01_14_v1","PvP01_14_v1","PvP01_14_v1","PvP01_14_v1","PvP01_14_v1","PvP01_14_v1","PvP01_14_v1","PvP01_14_v1","PvP01_14_v1","PvP01_14_v1","PvP01_14_v1","PvP01_14_v1","PvP01_14_v1","PvP01_14_v1","PvP01_14_v1","PvP01_14_v1","PvP01_14_v1","PvP01_14_v1","PvP01_14_v1","PvP01_14_v1","PvP01_14_v1","PvP01_14_v1","PvP01_14_v1","PvP01_14_v1","PvP01_14_v1","PvP01_14_v1","PvP01_14_v1","PvP01_14_v1","PvP01_14_v1","PvP01_14_v1","PvP01_14_v1","PvP01_14_v1","PvP01_14_v1","PvP01_14_v1","PvP01_14_v1","PvP01_14_v1","PvP01_14_v1","PvP01_14_v1","PvP01_14_v1","PvP01_14_v1","PvP01_14_v1","PvP01_14_v1","PvP01_14_v1","PvP01_14_v1","PvP01_14_v1","PvP01_14_v1","PvP01_14_v1","PvP01_14_v1","PvP01_14_v1","PvP01_14_v1","PvP01_14_v1","PvP01_14_v1","PvP01_14_v1","PvP01_14_v1","PvP01_14_v1","PvP01_14_v1","PvP01_14_v1","PvP01_14_v1","PvP01_14_v1","PvP01_14_v1","PvP01_14_v1","PvP01_14_v1","PvP01_14_v1","PvP01_14_v1","PvP01_14_v1","PvP01_14_v1","PvP01_14_v1","PvP01_14_v1","PvP01_14_v1","PvP01_14_v1","PvP01_14_v1","PvP01_14_v1","PvP01_14_v1","PvP01_14_v1","PvP01_14_v1","PvP01_14_v1","PvP01_14_v1","PvP01_14_v1","PvP01_14_v1","PvP01_14_v1","PvP01_14_v1","PvP01_14_v1","PvP01_14_v1","PvP01_14_v1","PvP01_14_v1","PvP01_14_v1","PvP01_14_v1","PvP01_14_v1","PvP01_14_v1","PvP01_14_v1","PvP01_14_v1","PvP01_14_v1","PvP01_14_v1","PvP01_14_v1","PvP01_14_v1","PvP01_14_v1","PvP01_14_v1","PvP01_14_v1","PvP01_14_v1","PvP01_14_v1","PvP01_14_v1","PvP01_14_v1","PvP01_14_v1","PvP01_14_v1","PvP01_14_v1","PvP01_14_v1","PvP01_14_v1","PvP01_14_v1","PvP01_14_v1","PvP01_14_v1","PvP01_14_v1","PvP01_14_v1","PvP01_14_v1","PvP01_14_v1","PvP01_14_v1","PvP01_14_v1","PvP01_14_v1","PvP01_14_v1","PvP01_14_v1","PvP01_14_v1","PvP01_14_v1","PvP01_14_v1","PvP01_14_v1","PvP01_14_v1","PvP01_14_v1","PvP01_14_v1","PvP01_14_v1","PvP01_14_v1","PvP01_14_v1","PvP01_14_v1","PvP01_14_v1","PvP01_14_v1","PvP01_14_v1","PvP01_14_v1","PvP01_14_v1","PvP01_14_v1","PvP01_14_v1","PvP01_14_v1","PvP01_14_v1","PvP01_14_v1","PvP01_14_v1","PvP01_14_v1","PvP01_14_v1","PvP01_14_v1","PvP01_14_v1","PvP01_14_v1","PvP01_14_v1","PvP01_14_v1","PvP01_14_v1","PvP01_14_v1","PvP01_14_v1","PvP01_14_v1","PvP01_14_v1","PvP01_14_v1","PvP01_14_v1","PvP01_14_v1","PvP01_14_v1","PvP01_14_v1","PvP01_14_v1","PvP01_14_v1","PvP01_14_v1","PvP01_14_v1","PvP01_14_v1","PvP01_14_v1","PvP01_14_v1","PvP01_14_v1","PvP01_14_v1","PvP01_14_v1","PvP01_14_v1","PvP01_14_v1","PvP01_14_v1","PvP01_14_v1","PvP01_14_v1","PvP01_14_v1","PvP01_14_v1","PvP01_14_v1","PvP01_14_v1","PvP01_14_v1","PvP01_14_v1","PvP01_14_v1","PvP01_14_v1","PvP01_14_v1","PvP01_14_v1","PvP01_14_v1","PvP01_14_v1","PvP01_14_v1","PvP01_14_v1","PvP01_14_v1","PvP01_14_v1","PvP01_14_v1","PvP01_14_v1","PvP01_14_v1","PvP01_14_v1","PvP01_14_v1","PvP01_14_v1","PvP01_14_v1","PvP01_14_v1","PvP01_14_v1","PvP01_14_v1","PvP01_14_v1","PvP01_14_v1","PvP01_14_v1","PvP01_14_v1","PvP01_14_v1","PvP01_14_v1","PvP01_14_v1","PvP01_14_v1","PvP01_14_v1","PvP01_14_v1","PvP01_14_v1","PvP01_14_v1","PvP01_14_v1","PvP01_14_v1","PvP01_14_v1","PvP01_14_v1","PvP01_14_v1","PvP01_14_v1","PvP01_14_v1","PvP01_14_v1","PvP01_14_v1","PvP01_14_v1","PvP01_14_v1","PvP01_14_v1","PvP01_14_v1","PvP01_14_v1","PvP01_14_v1","PvP01_14_v1","PvP01_14_v1","PvP01_14_v1","PvP01_14_v1","PvP01_14_v1","PvP01_14_v1","PvP01_14_v1","PvP01_14_v1","PvP01_14_v1","PvP01_14_v1","PvP01_14_v1","PvP01_14_v1","PvP01_14_v1","PvP01_14_v1","PvP01_14_v1","PvP01_14_v1","PvP01_14_v1","PvP01_14_v1","PvP01_14_v1","PvP01_14_v1","PvP01_14_v1","PvP01_14_v1","PvP01_14_v1","PvP01_14_v1","PvP01_14_v1","PvP01_14_v1","PvP01_14_v1","PvP01_14_v1","PvP01_14_v1","PvP01_14_v1","PvP01_14_v1","PvP01_14_v1","PvP01_14_v1","PvP01_14_v1","PvP01_14_v1","PvP01_14_v1","PvP01_14_v1","PvP01_14_v1","PvP01_14_v1","PvP01_14_v1","PvP01_14_v1","PvP01_14_v1","PvP01_14_v1","PvP01_14_v1","PvP01_14_v1","PvP01_14_v1","PvP01_14_v1","PvP01_14_v1","PvP01_14_v1","PvP01_14_v1","PvP01_14_v1","PvP01_14_v1","PvP01_14_v1","PvP01_14_v1","PvP01_14_v1","PvP01_14_v1","PvP01_14_v1","PvP01_14_v1","PvP01_14_v1","PvP01_14_v1","PvP01_14_v1","PvP01_14_v1","PvP01_14_v1","PvP01_14_v1","PvP01_14_v1","PvP01_14_v1","PvP01_14_v1","PvP01_14_v1","PvP01_14_v1","PvP01_14_v1","PvP01_14_v1","PvP01_14_v1","PvP01_14_v1","PvP01_14_v1","PvP01_14_v1","PvP01_14_v1","PvP01_14_v1","PvP01_14_v1","PvP01_14_v1","PvP01_14_v1","PvP01_14_v1","PvP01_14_v1","PvP01_14_v1","PvP01_14_v1","PvP01_14_v1","PvP01_14_v1","PvP01_14_v1","PvP01_14_v1","PvP01_14_v1","PvP01_14_v1","PvP01_14_v1","PvP01_14_v1","PvP01_14_v1","PvP01_14_v1","PvP01_14_v1","PvP01_14_v1","PvP01_14_v1","PvP01_14_v1","PvP01_14_v1","PvP01_14_v1","PvP01_14_v1","PvP01_14_v1","PvP01_14_v1","PvP01_14_v1","PvP01_14_v1","PvP01_14_v1","PvP01_14_v1","PvP01_14_v1","PvP01_14_v1","PvP01_14_v1","PvP01_14_v1","PvP01_14_v1","PvP01_14_v1","PvP01_14_v1","PvP01_14_v1","PvP01_14_v1","PvP01_14_v1","PvP01_14_v1","PvP01_14_v1","PvP01_14_v1","PvP01_14_v1","PvP01_14_v1","PvP01_14_v1","PvP01_14_v1","PvP01_14_v1","PvP01_14_v1","PvP01_14_v1","PvP01_14_v1","PvP01_14_v1","PvP01_14_v1","PvP01_14_v1","PvP01_14_v1","PvP01_14_v1","PvP01_14_v1","PvP01_14_v1","PvP01_14_v1","PvP01_14_v1","PvP01_14_v1","PvP01_14_v1","PvP01_14_v1","PvP01_14_v1","PvP01_14_v1","PvP01_14_v1","PvP01_14_v1","PvP01_14_v1","PvP01_14_v1","PvP01_14_v1","PvP01_14_v1","PvP01_14_v1","PvP01_14_v1","PvP01_14_v1","PvP01_14_v1","PvP01_14_v1","PvP01_14_v1","PvP01_14_v1","PvP01_14_v1","PvP01_14_v1","PvP01_14_v1","PvP01_14_v1","PvP01_14_v1","PvP01_14_v1","PvP01_14_v1","PvP01_14_v1","PvP01_14_v1","PvP01_14_v1","PvP01_14_v1","PvP01_14_v1","PvP01_14_v1","PvP01_14_v1","PvP01_14_v1","PvP01_14_v1","PvP01_14_v1","PvP01_14_v1","PvP01_14_v1","PvP01_14_v1","PvP01_14_v1","PvP01_14_v1","PvP01_14_v1","PvP01_14_v1","PvP01_14_v1","PvP01_14_v1","PvP01_14_v1","PvP01_14_v1","PvP01_14_v1","PvP01_14_v1","PvP01_14_v1","PvP01_14_v1","PvP01_14_v1","PvP01_14_v1","PvP01_14_v1","PvP01_14_v1","PvP01_14_v1","PvP01_14_v1","PvP01_14_v1","PvP01_14_v1","PvP01_14_v1","PvP01_14_v1","PvP01_14_v1","PvP01_14_v1","PvP01_14_v1","PvP01_14_v1","PvP01_14_v1","PvP01_14_v1","PvP01_14_v1","PvP01_14_v1","PvP01_14_v1","PvP01_14_v1","PvP01_14_v1","PvP01_14_v1","PvP01_14_v1","PvP01_14_v1","PvP01_14_v1","PvP01_14_v1","PvP01_14_v1","PvP01_14_v1","PvP01_14_v1","PvP01_14_v1","PvP01_14_v1","PvP01_14_v1","PvP01_14_v1","PvP01_14_v1","PvP01_14_v1","PvP01_14_v1","PvP01_14_v1","PvP01_14_v1","PvP01_14_v1","PvP01_14_v1","PvP01_14_v1","PvP01_14_v1","PvP01_14_v1","PvP01_14_v1","PvP01_14_v1","PvP01_14_v1","PvP01_14_v1","PvP01_14_v1","PvP01_14_v1","PvP01_14_v1","PvP01_14_v1","PvP01_14_v1","PvP01_14_v1","PvP01_14_v1","PvP01_14_v1","PvP01_14_v1","PvP01_14_v1","PvP01_14_v1","PvP01_14_v1","PvP01_14_v1","PvP01_14_v1","PvP01_14_v1","PvP01_14_v1","PvP01_14_v1","PvP01_14_v1","PvP01_14_v1","PvP01_14_v1","PvP01_14_v1","PvP01_14_v1","PvP01_14_v1","PvP01_14_v1","PvP01_14_v1","PvP01_14_v1","PvP01_14_v1","PvP01_14_v1","PvP01_14_v1","PvP01_14_v1","PvP01_14_v1","PvP01_14_v1","PvP01_14_v1","PvP01_14_v1","PvP01_14_v1","PvP01_14_v1","PvP01_14_v1","PvP01_14_v1","PvP01_14_v1","PvP01_14_v1","PvP01_14_v1","PvP01_14_v1","PvP01_14_v1","PvP01_14_v1","PvP01_14_v1","PvP01_14_v1","PvP01_14_v1","PvP01_14_v1","PvP01_14_v1","PvP01_14_v1","PvP01_14_v1","PvP01_14_v1","PvP01_14_v1","PvP01_14_v1","PvP01_14_v1","PvP01_14_v1","PvP01_14_v1","PvP01_14_v1","PvP01_14_v1","PvP01_14_v1","PvP01_14_v1","PvP01_14_v1","PvP01_14_v1","PvP01_14_v1","PvP01_14_v1","PvP01_14_v1","PvP01_14_v1","PvP01_14_v1","PvP01_14_v1","PvP01_14_v1","PvP01_14_v1","PvP01_14_v1","PvP01_14_v1","PvP01_14_v1","PvP01_14_v1","PvP01_14_v1","PvP01_14_v1","PvP01_14_v1","PvP01_14_v1","PvP01_14_v1","PvP01_14_v1","PvP01_14_v1","PvP01_14_v1","PvP01_14_v1","PvP01_14_v1","PvP01_14_v1","PvP01_14_v1","PvP01_14_v1","PvP01_14_v1","PvP01_14_v1","PvP01_14_v1","PvP01_14_v1","PvP01_14_v1","PvP01_14_v1","PvP01_14_v1","PvP01_14_v1","PvP01_14_v1","PvP01_14_v1","PvP01_14_v1","PvP01_14_v1","PvP01_14_v1","PvP01_14_v1","PvP01_14_v1","PvP01_14_v1","PvP01_14_v1","PvP01_14_v1","PvP01_14_v1","PvP01_14_v1","PvP01_14_v1","PvP01_14_v1","PvP01_14_v1","PvP01_14_v1","PvP01_14_v1","PvP01_14_v1","PvP01_14_v1","PvP01_14_v1","PvP01_14_v1","PvP01_14_v1","PvP01_14_v1","PvP01_14_v1","PvP01_14_v1","PvP01_API_v1","PvP01_API_v1","PvP01_API_v1","PvP01_API_v1","PvP01_API_v1","PvP01_API_v1","PvP01_API_v1","PvP01_API_v1","PvP01_API_v1","PvP01_API_v1","PvP01_MIT_v1","PvP01_MIT_v1","PvP01_MIT_v1","Transfer.PvP01_00_1.final","PvP01_01_v1","PvP01_01_v1","PvP01_01_v1","PvP01_01_v1","PvP01_01_v1","PvP01_01_v1","PvP01_02_v1","PvP01_02_v1","PvP01_02_v1","PvP01_02_v1","PvP01_03_v1","PvP01_03_v1","PvP01_03_v1","PvP01_03_v1","PvP01_03_v1","PvP01_04_v1","PvP01_04_v1","PvP01_04_v1","PvP01_04_v1","PvP01_04_v1","PvP01_05_v1","PvP01_05_v1","PvP01_05_v1","PvP01_05_v1","PvP01_05_v1","PvP01_06_v1","PvP01_06_v1","PvP01_06_v1","PvP01_06_v1","PvP01_06_v1","PvP01_07_v1","PvP01_07_v1","PvP01_07_v1","PvP01_07_v1","PvP01_07_v1","PvP01_07_v1","PvP01_07_v1","PvP01_07_v1","PvP01_07_v1","PvP01_07_v1","PvP01_07_v1","PvP01_08_v1","PvP01_08_v1","PvP01_08_v1","PvP01_08_v1","PvP01_08_v1","PvP01_08_v1","PvP01_08_v1","PvP01_09_v1","PvP01_09_v1","PvP01_09_v1","PvP01_09_v1","PvP01_09_v1","PvP01_09_v1","PvP01_09_v1","PvP01_09_v1","PvP01_09_v1","PvP01_09_v1","PvP01_10_v1","PvP01_10_v1","PvP01_10_v1","PvP01_10_v1","PvP01_10_v1","PvP01_11_v1","PvP01_11_v1","PvP01_11_v1","PvP01_11_v1","PvP01_11_v1","PvP01_12_v1","PvP01_12_v1","PvP01_12_v1","PvP01_12_v1","PvP01_12_v1","PvP01_12_v1","PvP01_12_v1","PvP01_12_v1","PvP01_12_v1","PvP01_12_v1","PvP01_12_v1","PvP01_12_v1","PvP01_12_v1","PvP01_12_v1","PvP01_12_v1","PvP01_12_v1","PvP01_12_v1","PvP01_13_v1","PvP01_13_v1","PvP01_13_v1","PvP01_13_v1","PvP01_13_v1","PvP01_13_v1","PvP01_13_v1","PvP01_14_v1","PvP01_14_v1","PvP01_14_v1","PvP01_14_v1","PvP01_14_v1","PvP01_14_v1","PvP01_14_v1","PvP01_14_v1","PvP01_14_v1","PvP01_14_v1","PvP01_14_v1","PvP01_14_v1","PvP01_14_v1","PvP01_14_v1","PvP01_14_v1"],"Source":["EuPathDB","EuPathDB","EuPathDB","EuPathDB","EuPathDB","EuPathDB","EuPathDB","EuPathDB","EuPathDB","EuPathDB","EuPathDB","EuPathDB","EuPathDB","EuPathDB","EuPathDB","EuPathDB","EuPathDB","EuPathDB","EuPathDB","EuPathDB","EuPathDB","EuPathDB","EuPathDB","EuPathDB","EuPathDB","EuPathDB","EuPathDB","EuPathDB","EuPathDB","EuPathDB","EuPathDB","EuPathDB","EuPathDB","EuPathDB","EuPathDB","EuPathDB","EuPathDB","EuPathDB","EuPathDB","EuPathDB","EuPathDB","EuPathDB","EuPathDB","EuPathDB","EuPathDB","EuPathDB","EuPathDB","EuPathDB","EuPathDB","EuPathDB","EuPathDB","EuPathDB","EuPathDB","EuPathDB","EuPathDB","EuPathDB","EuPathDB","EuPathDB","EuPathDB","EuPathDB","EuPathDB","EuPathDB","EuPathDB","EuPathDB","EuPathDB","EuPathDB","EuPathDB","EuPathDB","EuPathDB","EuPathDB","EuPathDB","EuPathDB","EuPathDB","EuPathDB","EuPathDB","EuPathDB","EuPathDB","EuPathDB","EuPathDB","EuPathDB","EuPathDB","EuPathDB","EuPathDB","EuPathDB","EuPathDB","EuPathDB","EuPathDB","EuPathDB","EuPathDB","EuPathDB","EuPathDB","EuPathDB","EuPathDB","EuPathDB","EuPathDB","EuPathDB","EuPathDB","EuPathDB","EuPathDB","EuPathDB","EuPathDB","EuPathDB","EuPathDB","EuPathDB","EuPathDB","EuPathDB","EuPathDB","EuPathDB","EuPathDB","EuPathDB","EuPathDB","EuPathDB","EuPathDB","EuPathDB","EuPathDB","EuPathDB","EuPathDB","EuPathDB","EuPathDB","EuPathDB","EuPathDB","EuPathDB","EuPathDB","EuPathDB","EuPathDB","EuPathDB","EuPathDB","EuPathDB","EuPathDB","EuPathDB","EuPathDB","EuPathDB","EuPathDB","EuPathDB","EuPathDB","EuPathDB","EuPathDB","EuPathDB","EuPathDB","EuPathDB","EuPathDB","EuPathDB","EuPathDB","EuPathDB","EuPathDB","EuPathDB","EuPathDB","EuPathDB","EuPathDB","EuPathDB","EuPathDB","EuPathDB","EuPathDB","EuPathDB","EuPathDB","EuPathDB","EuPathDB","EuPathDB","EuPathDB","EuPathDB","EuPathDB","EuPathDB","EuPathDB","EuPathDB","EuPathDB","EuPathDB","EuPathDB","EuPathDB","EuPathDB","EuPathDB","EuPathDB","EuPathDB","EuPathDB","EuPathDB","EuPathDB","EuPathDB","EuPathDB","EuPathDB","EuPathDB","EuPathDB","EuPathDB","EuPathDB","EuPathDB","EuPathDB","EuPathDB","EuPathDB","EuPathDB","EuPathDB","EuPathDB","EuPathDB","EuPathDB","EuPathDB","EuPathDB","EuPathDB","EuPathDB","EuPathDB","EuPathDB","EuPathDB","EuPathDB","EuPathDB","EuPathDB","EuPathDB","EuPathDB","EuPathDB","EuPathDB","EuPathDB","EuPathDB","EuPathDB","EuPathDB","EuPathDB","EuPathDB","EuPathDB","EuPathDB","EuPathDB","EuPathDB","EuPathDB","EuPathDB","EuPathDB","EuPathDB","EuPathDB","EuPathDB","EuPathDB","EuPathDB","EuPathDB","EuPathDB","EuPathDB","EuPathDB","EuPathDB","EuPathDB","EuPathDB","EuPathDB","EuPathDB","EuPathDB","EuPathDB","EuPathDB","EuPathDB","EuPathDB","EuPathDB","EuPathDB","EuPathDB","EuPathDB","EuPathDB","EuPathDB","EuPathDB","EuPathDB","EuPathDB","EuPathDB","EuPathDB","EuPathDB","EuPathDB","EuPathDB","EuPathDB","EuPathDB","EuPathDB","EuPathDB","EuPathDB","EuPathDB","EuPathDB","EuPathDB","EuPathDB","EuPathDB","EuPathDB","EuPathDB","EuPathDB","EuPathDB","EuPathDB","EuPathDB","EuPathDB","EuPathDB","EuPathDB","EuPathDB","EuPathDB","EuPathDB","EuPathDB","EuPathDB","EuPathDB","EuPathDB","EuPathDB","EuPathDB","EuPathDB","EuPathDB","EuPathDB","EuPathDB","EuPathDB","EuPathDB","EuPathDB","EuPathDB","EuPathDB","EuPathDB","EuPathDB","EuPathDB","EuPathDB","EuPathDB","EuPathDB","EuPathDB","EuPathDB","EuPathDB","EuPathDB","EuPathDB","EuPathDB","EuPathDB","EuPathDB","EuPathDB","EuPathDB","EuPathDB","EuPathDB","EuPathDB","EuPathDB","EuPathDB","EuPathDB","EuPathDB","EuPathDB","EuPathDB","EuPathDB","EuPathDB","EuPathDB","EuPathDB","EuPathDB","EuPathDB","EuPathDB","EuPathDB","EuPathDB","EuPathDB","EuPathDB","EuPathDB","EuPathDB","EuPathDB","EuPathDB","EuPathDB","EuPathDB","EuPathDB","EuPathDB","EuPathDB","EuPathDB","EuPathDB","EuPathDB","EuPathDB","EuPathDB","EuPathDB","EuPathDB","EuPathDB","EuPathDB","EuPathDB","EuPathDB","EuPathDB","EuPathDB","EuPathDB","EuPathDB","EuPathDB","EuPathDB","EuPathDB","EuPathDB","EuPathDB","EuPathDB","EuPathDB","EuPathDB","EuPathDB","EuPathDB","EuPathDB","EuPathDB","EuPathDB","EuPathDB","EuPathDB","EuPathDB","EuPathDB","EuPathDB","EuPathDB","EuPathDB","EuPathDB","EuPathDB","EuPathDB","EuPathDB","EuPathDB","EuPathDB","EuPathDB","EuPathDB","EuPathDB","EuPathDB","EuPathDB","EuPathDB","EuPathDB","EuPathDB","EuPathDB","EuPathDB","EuPathDB","EuPathDB","EuPathDB","EuPathDB","EuPathDB","EuPathDB","EuPathDB","EuPathDB","EuPathDB","EuPathDB","EuPathDB","EuPathDB","EuPathDB","EuPathDB","EuPathDB","EuPathDB","EuPathDB","EuPathDB","EuPathDB","EuPathDB","EuPathDB","EuPathDB","EuPathDB","EuPathDB","EuPathDB","EuPathDB","EuPathDB","EuPathDB","EuPathDB","EuPathDB","EuPathDB","EuPathDB","EuPathDB","EuPathDB","EuPathDB","EuPathDB","EuPathDB","EuPathDB","EuPathDB","EuPathDB","EuPathDB","EuPathDB","EuPathDB","EuPathDB","EuPathDB","EuPathDB","EuPathDB","EuPathDB","EuPathDB","EuPathDB","EuPathDB","EuPathDB","EuPathDB","EuPathDB","EuPathDB","EuPathDB","EuPathDB","EuPathDB","EuPathDB","EuPathDB","EuPathDB","EuPathDB","EuPathDB","EuPathDB","EuPathDB","EuPathDB","EuPathDB","EuPathDB","EuPathDB","EuPathDB","EuPathDB","EuPathDB","EuPathDB","EuPathDB","EuPathDB","EuPathDB","EuPathDB","EuPathDB","EuPathDB","EuPathDB","EuPathDB","EuPathDB","EuPathDB","EuPathDB","Eu
```
