## Supplementary material for "Transcriptome and histone epigenome of *Plasmodium vivax* salivary-gland sporozoites point to tight regulatory control and potential mechanisms for liver-stage differentiation": Pv-SPZ-v-BS-DTGs.js

```
glimma.layout.setupGrid(d3.select(".container"), "md", [1,2]);
glimma.storage.chartData.push(glimma.transform.toRowMajor({"logFC":[-0.2437,0.07041,-0.7906,-1.868,-0.9006,-1.378,-1.057,0.6221,-0.5779,-0.8177,-1.175,-0.8003,-0.1452,-0.4046,-1.099,-0.2226,0.03712,-0.5423,-0.7507,-0.5852,-0.7404,-0.1816,-0.6285,-0.3188,0.04648,-0.08435,-0.938,-0.4779,0.3131,-0.3089,-1.478,-1.216,-1.009,-0.6125,0.02005,-0.7048,-0.8796,-0.2811,0.8094,0.03859,-0.8558,-1.265,-0.9504,-1.277,-0.3314,-0.5495,-0.8393,-1.007,-0.4137,-1.104,-0.2291,-0.03063,-0.1525,-0.9088,0.4208,-1.397,0.4183,-0.3273,0.06086,-0.5043,-0.2689,-1.126,-0.6369,-1.431,0.8809,-0.7275,-1.039,-0.7184,-0.7042,-0.02818,-0.4384,-0.4069,0.09384,-0.8099,-1.258,-0.4288,-0.09432,-1.118,-1.147,-0.5197,-1.352,-0.8173,-0.224,-0.3877,1.532,-0.8762,-0.9163,-0.335,-0.7211,-0.476,-0.1274,-0.1858,-0.3509,-0.6463,-0.03467,-0.9331,-1.032,0.5992,0.0009314,-0.9972,-0.5996,-0.8872,-1.239,-0.2207,-1.032,-0.5809,-0.3116,-0.3025,-0.7887,0.4583,-0.3605,0.09695,1.096,-0.2992,0.5505,-1.026,-0.6258,-0.8575,-1.462,-0.8012,-0.4159,-0.698,-1.079,1.046,-0.5639,-0.7941,0.142,1.232,-0.475,0.01917,0.3497,-1.667,-0.8181,-0.631,0.4188,-0.6212,0.2735,-0.1019,0.3613,-0.04348,-0.6955,-1.475,-0.4587,0.2541,0.04978,0.8923,-0.1158,0.4736,0.07029,0.4596,-0.3004,0.1293,0.6349,0.2029,0.2017,0.7192,0.465,0.296,-0.4941,-0.7244,-0.4794,0.07839,-0.9073,-0.2099,-0.4927,0.3525,0.4911,0.06868,-0.4305,0.1054,-0.452,0.8274,0.4463,0.4767,-0.3606,-0.3903,0.723,0.2691,-0.06335,-0.292,-0.5209,-0.7746,-1.148,-0.3585,-0.8879,0.7414,-0.4256,-0.8515,-0.6497,-0.8026,-0.4696,-0.01637,-0.1328,-0.03365,-0.6834,0.8897,-1.12,0.8795,0.712,-0.7038,-0.1712,0.6963,0.7261,0.4741,-0.7081,0.4642,0.3882,0.7469,0.01804,1.074,-0.4906,-0.5851,1.116,-0.1472,0.3203,0.4786,0.1977,0.9215,0.03777,1.062,-0.3628,-0.646,0.874,-0.09296,-0.7011,-0.6399,0.6813,0.5072,-1.009,-0.1151,-0.4824,0.8364,0.008176,-0.8405,-0.7403,0.3015,1.09,-0.8244,-0.5954,0.1816,-2.104,-1.612,-0.504,-1.172,-0.1333,-0.6525,-1.239,-1.499,0.6391,-0.5532,0.279,0.8664,-0.5353,0.5923,1.498,0.8277,0.2951,0.07695,0.8054,-0.08227,0.1307,-0.2149,0.9561,0.6045,-0.3351,0.3995,-0.5118,-0.4127,0.386,0.8233,0.5493,0.7951,0.544,0.1912,-0.339,-0.5831,0.1834,-0.4347,-0.4155,-0.8778,-0.7992,-0.4138,-0.08008,-0.3193,0.5427,-0.4675,0.3728,0.3201,-0.9978,-0.4216,0.9375,-0.4523,-1.423,0.3282,0.4179,-0.2982,-0.9706,-0.3012,0.3589,-0.5421,-0.255,0.02563,0.2149,-0.9669,0.8968,0.8831,-0.8652,0.5132,-1.504,-0.9225,-0.7381,-0.7306,-1.298,-0.9946,0.3844,-0.2863,0.4914,-0.2357,-0.0528,0.3696,0.782,0.3339,0.3166,-0.1309,0.7278,-0.3567,0.8025,-0.6664,0.7445,0.3935,-0.1563,0.2012,0.1585,0.1271,0.3806,0.6824,0.7883,1.216,-0.9494,-0.0983,0.4967,0.872,-0.1807,0.4123,0.8369,0.9102,-0.4912,1.315,-3.086,-2.526,0.2882,0.6155,-0.9946,-0.3561,0.2187,-0.4933,-0.8021,0.9329,0.5398,0.4016,0.7201,0.7719,0.6067,0.8007,0.3619,-0.9689,-0.4522,-0.5525,-1.383,-0.4835,-1.644,-2.629,0.4492,-0.2279,0.373,0.3703,0.8434,-0.4974,-0.8537,-0.7005,0.9516,-0.6614,-0.4682,-1.12,-0.09252,0.2646,0.03475,-0.5087,-0.2372,0.3136,0.3243,-0.7203,-0.2945,1.149,0.4107,0.9261,-0.2528,0.02076,-0.2573,0.5436,-0.4645,-0.4572,-0.3728,-1.606,-1.934,-2.059,0.08446,0.732,-0.2208,-0.07259,-0.683,-0.7737,0.9767,-0.01186,0.196,0.2465,-0.784,0.9216,-1.023,0.03302,-0.8837,0.08925,0.7163,-0.4199,0.7256,1.056,0.8783,-0.05966,-1.579,-2.774,0.323,0.005469,-0.8929,-0.5582,-0.7586,0.4807,0.4344,-0.5729,-0.4211,0.5948,-0.915,0.9846,0.02951,-0.8033,0.723,0.5174,0.8344,0.02145,0.4328,-1.034,-0.6069,-0.4796,0.5341,-0.6575,0.7112,0.8728,-0.4701,-0.627,-0.479,-0.9655,-0.7567,-0.5689,-1.237,-0.6672,-1.206,-0.3387,-1.115,-1.065,0.6287,-1.561,-1.744,0.3528,-1.934,0.9578,-0.1905,-0.9885,0.2255,0.8501,0.7436,-0.007634,0.5254,0.1581,-0.8313,-0.004362,-0.6185,0.9946,-0.1969,-0.6709,-0.7331,-0.8384,-0.3039,0.9332,-0.8186,-0.9024,-0.3553,0.8551,0.1144,0.7276,-0.5421,0.03876,0.7161,0.01362,0.3232,0.4889,-0.4236,-0.687,0.5625,0.05829,-1.333,0.6829,0.855,-0.8294,0.9075,0.1362,-0.7964,0.01809,-0.3301,-0.03978,-0.517,-0.2723,0.5358,-0.692,0.1943,-1.264,0.3703,1.289,0.5571,-0.07516,-0.6982,-0.687,-0.6357,-0.5026,0.07281,-0.952,0.806,0.6845,0.1127,0.4398,0.6893,0.2472,-1.846,0.8757,0.8246,0.2245,0.5229,0.3176,-1.389,-1.115,-0.8445,-0.8638,0.0531,0.5414,0.6012,0.8888,0.2675,-0.7396,0.3414,-1,-0.1403,-0.8158,-0.1792,0.5132,0.8915,-1.12,0.2196,0.3766,0.2172,-0.04333,0.2252,0.7258,0.7027,-0.7079,0.4982,0.6009,0.4933,0.4934,0.3968,0.913,0.5955,-0.5205,0.3805,0.3353,0.8464,0.1892,-0.8116,-0.2959,-0.2126,0.9994,-0.1546,-0.6292,-0.414,0.07569,-0.83,-1.48,0.2197,-1.347,-0.342,-0.182,-0.6048,-1.116,-0.05382,-0.2913,0.7991,-0.1095,-1.413,0.07026,-0.4369,-0.6571,-0.02642,-0.1202,0.7342,0.43,0.5814,0.07657,0.6998,-0.9589,0.6223,-0.9724,0.138,0.1306,0.3313,0.01824,-0.134,0.6961,-0.2699,0.2974,-1.114,1.023,-0.6254,-0.3233,-0.7066,0.8289,-0.08967,-0.01555,-0.9713,0.999,-0.09522,-0.6737,0.5124,-1.037,-0.8376,-0.08761,0.4045,-0.8903,0.1411,0.3252,0.8312,0.8998,-0.06009,-1.594,0.5169,0.2755,0.2601,0.1511,0.6324,0.8162,-0.003784,0.5749,-0.9004,-0.3361,-0.9336,-0.9828,1.01,0.1011,-1.147,-0.9531,-1.156,-0.4807,-1.131,-1.817,-1.412,-0.08879,0.03058,0.1133,-0.264,-0.558,0.2836,0.5743,0.06895,-0.8943,-1.004,0.6721,-0.8633,0.1982,-0.8957,0.9474,-0.486,1.201,0.3684,-0.01453,-0.2314,-1.448,-0.3825,0.7019,0.06245,-0.3373,-0.8358,-0.3685,-0.7678,0.8388,-0.1299,1.045,0.5181,-0.2097,-0.7482,-0.1266,0.3745,-1.902,0.07608,0.07711,-0.3767,-0.5508,0.8254,-0.1322,-0.1883,0.3004,0.7922,-0.8546,0.07766,0.09195,0.08044,0.3408,0.255,-0.6189,-0.194,-0.1045,0.6319,-0.7066,0.09924,0.05777,-0.4823,-0.1482,0.7578,-0.9636,0.9859,0.2387,0.4792,0.549,-0.4433,0.4236,0.6979,0.1556,0.1664,0.3912,-0.5713,-1.597,0.2256,-0.5997,0.8876,0.1377,0.4407,-0.07651,-0.4677,-0.07783,0.2449,0.6472,0.2676,-0.8894,0.05149,-0.5081,-0.2602,0.301,-0.2342,0.1059,0.2744,-0.8413,0.6268,0.5107,0.2915,0.3665,0.04379,-0.08221,0.4623,-0.5729,-0.8022,0.1749,-0.4276,0.3646,0.9069,-0.1233,0.214,-0.5779,-0.3301,0.9718,0.3132,-0.2494,0.2788,0.4664,-1.715,-0.3313,0.1606,-0.2489,-0.6272,-0.7097,-1.535,-1.111,1.076,0.8934,0.397,-0.4078,-0.7102,0.06193,0.2202,0.9694,0.7095,-0.8873,0.7536,0.7115,0.07708,0.9792,-0.125,-0.3118,0.7981,-0.03439,-0.1985,-0.3032,0.5023,-0.4718,-0.0751,0.2037,-0.1473,0.6765,0.8238,-0.7213,0.9207,0.7601,-0.3304,-0.149,0.4053,0.1299,0.02945,-0.8902,-0.3837,-0.04938,-0.3401,-0.2588,0.8654,0.3109,0.7951,-0.1699,0.6915,0.2927,-0.5309,-0.7952,-0.9783,0.6559,-0.199,-0.5694,0.3716,-0.9361,0.5513,0.5653,0.8451,-0.8863,0.3208,0.7041,-0.5647,-0.1403,1.026,0.6654,0.06436,0.8801,0.4582,-0.6854,0.4972,-0.29,0.7135,0.3654,-0.4434,-0.1852,0.9531,0.4281,-0.2266,0.4115,-0.3077,-0.1418,0.6409,0.7161,0.7151,1.181,-0.9851,-0.8489,0.8664,-0.1833,-0.8283,0.2714,0.3364,0.5212,-0.1698,-0.09658,0.003004,-1.431,-0.1618,-0.6455,-0.8985,0.1087,0.7503,0.523,0.5936,0.8584,0.9986,0.2691,0.456,0.3972,-1.9,-0.2885,0.2376,-0.5726,0.08328,0.8328,-0.7215,0.9673,0.01068,0.9345,-0.2685,0.1638,0.1988,-0.3961,-0.2923,-0.3414,-0.0634,-0.6178,0.2685,-1.471,-0.05539,0.2004,-0.5631,-0.03654,-0.2522,-0.74,0.3479,0.9293,-0.09166,-0.0001487,-0.05629,-0.8733,-0.6622,-0.03675,0.9429,-0.1698,0.9853,-0.737,-0.3492,0.5841,-0.549,-0.31,0.5313,-0.9922,0.824,0.6281,0.3588,-0.5959,-0.5624,0.6343,-0.2083,-0.9529,-0.6448,-0.3344,0.8991,-0.1489,0.3578,-0.9529,0.8348,0.1943,-0.5599,0.3624,0.6846,0.464,-0.2376,0.7611,0.5424,-0.2089,-0.8553,-0.6333,0.7353,0.5296,0.5018,-1.083,0.3447,-0.6809,0.9409,-0.4903,-0.6208,0.1098,0.5992,0.3307,-0.753,-0.4492,-0.7141,-0.535,-0.5144,-0.8733,0.7991,-0.1506,-0.6535,0.9274,0.1095,0.4609,-0.06374,-0.3948,0.4639,-0.8342,0.4936,0.7686,0.5611,-0.8432,-0.7366,-0.2151,-0.07544,-0.169,0.122,0.9556,-0.5887,-0.4768,1.219,-0.08293,-0.9942,0.6986,-0.3936,1.097,-0.1894,0.2448,-0.8097,-0.03999,0.5068,-0.5528,0.01958,-0.5545,-0.09862,0.9527,-0.4262,0.04982,0.06541,0.1726,0.6093,-0.1604,-0.4214,-0.9588,-0.4637,0.4899,-0.5886,-0.3947,-0.1503,-0.07423,0.65,-0.2007,-0.7102,0.1422,-0.2849,-1.235,-0.6381,0.1197,0.6458,-1.587,-0.2522,0.7823,-1.043,-1.621,0.5431,0.9488,0.4775,-0.7203,-1.095,-1.171,-1.281,-0.4008,0.02397,-0.1542,-1.075,-0.997,0.09644,-0.8752,0.2501,-0.6601,0.5594,-0.808,-0.398,0.1426,-0.005386,-0.2583,-0.08292,-0.04638,-0.8828,0.3311,0.8701,-0.4987,-0.4485,-0.5876,-0.01061,0.47,0.6338,-0.604,-1.029,0.8893,-0.1912,-0.3772,0.4615,-0.8664,0.1225,0.8473,1.116,-0.9992,-0.7242,0.002539,0.09639,0.3369,0.5029,0.4182,-0.5558,0.497,0.4577,0.615,0.4474,-0.1968,0.1671,-0.103,-0.5482,0.5892,-0.889,-0.01564,-1.064,0.1954,-0.7147,-0.05824,0.6819,-0.2113,0.4454,-0.0791,0.4842,-0.4909,-1.05,0.2456,1.226,0.4766,0.38,-0.05723,-0.678,-0.6391,0.1151,-1.023,0.04157,0.3185,0.4293,-0.8192,1.172,0.9696,0.9101,0.4759,0.1472,0.5217,0.02989,-0.3614,0.5089,-0.4861,-0.9715,0.4026,1.128,0.3839,-0.607,0.08554,1.012,0.5553,-0.6592,0.7671,-0.2299,0.8184,0.7501,-0.6407,0.01394,-0.01452,0.7995,-0.06593,0.5834,0.6996,-0.9849,-0.9331,0.6088,-0.07148,-0.1409,0.9751,0.5052,0.6908,-0.3413,-0.06585,0.914,0.2947,-0.9001,-0.1671,-0.1036,0.05053,-0.5734,-1.004,-0.7904,-0.06738,0.5331,-0.1703,0.1427,-0.4243,0.4478,0.6096,-0.3132,-0.2865,-0.9512,0.6125,-0.4657,0.02654,0.2307,0.2066,0.7013,-0.2036,-0.05773,-0.7129,0.5272,-1.042,0.2317,0.2298,-0.03864,-0.8925,0.7665,-0.9486,-0.6416,-1.333,-0.9306,-1.761,-0.2138,0.23,-0.1023,0.844,-0.2744,1.062,0.7509,0.2549,-0.84,0.1805,-0.3609,-0.5489,-0.7818,-0.4615,0.9336,0.2875,-0.5903,-0.5551,-0.9554,0.1653,-0.4837,-1.137,0.1707,0.5839,-0.2438,0.4002,0.07411,-1.152,0.73,0.1392,-0.8845,0.238,-0.231,0.8355,-0.3385,-0.6958,0.8376,-0.2949,-0.6391,-0.03427,0.1196,-0.6286,-0.7568,-0.4737,-0.998,-1.194,-0.2624,-0.1319,0.0807,-0.8263,-0.7338,-0.4253,-0.5051,0.166,-0.3493,-0.5711,-0.1715,0.1385,0.7598,1.563,-2.106,-0.7802,-1.755,1.639,0.8391,-0.5546,0.03076,-0.918,0.7359,-0.9806,0.6735,-0.4292,-0.4871,0.2077,-0.3634,-0.4535,0.8359,-0.7546,0.6281,-0.05913,-1.104,0.4141,-0.118,0.9003,-0.2626,-0.8904,-0.8697,-0.4373,0.1772,-0.5611,0.04806,-1.505,-0.8905,0.9976,-0.5059,0.7829,-0.8073,-0.4494,-0.802,-0.1427,-0.1359,0.4646,0.6658,-0.8178,-0.9477,0.2592,0.9228,-0.3153,-0.6739,0.4815,-0.07011,-1.016,-0.8486,-0.2353,-0.5904,-0.2936,-0.6902,1.685,0.4417,0.36,-0.6633,0.8004,1.111,-0.6807,-0.3539,-0.5057,-1.817,-0.6889,-0.2288,0.8365,-0.1136,0.9443,0.1739,0.4652,-0.1132,0.52,0.6379,-0.824,0.96,0.199,0.6589,0.409,-0.3948,-0.6667,0.526,0.3987,-0.4676,0.8297,0.1114,0.3743,0.3679,-0.5128,-0.3014,0.4633,-0.4153,-0.1351,0.7797,0.2493,0.8043,0.7304,0.9694,0.3299,0.2576,0.5097,-0.7883,0.05183,-1.04,0.5536,0.07732,-0.4372,0.9189,-0.8939,0.3313,-1.126,0.1696,0.1929,0.3837,0.2581,0.2043,0.001653,-0.06378,0.2101,-0.7827,-0.1021,-0.2274,-1.032,0.5171,0.9354,-0.08237,0.7834,-0.9248,0.6512,-0.9763,0.03132,-0.3342,0.5459,-0.987,0.4446,0.4414,0.6278,-0.3449,-0.7159,0.4716,0.3991,-1.312,-0.77,0.8315,-0.7702,0.1742,0.3652,0.6479,-0.5075,-0.3291,-0.8389,0.9178,0.281,-0.09477,0.1574,0.5802,0.6352,-0.4724,-0.7093,-0.4898,-0.05251,-0.5246,-1.137,-0.5354,-0.7898,0.7259,0.2539,-1.574,0.5288,0.1911,0.9883,-0.6794,-0.8512,0.56,-0.8442,-0.4396,0.9004,-0.8138,0.6381,-0.6463,0.5981,-0.6171,-0.5538,-2.411,-0.5881,0.7252,-0.4288,-0.4773,0.3356,0.8082,-0.7222,0.08979,-0.4827,-0.3632,-0.07609,-0.4607,-0.2406,0.3988,0.04626,0.4575,0.166,0.9906,-0.1824,0.9055,1.331,0.1841,0.2163,0.2146,0.1211,-0.1969,-0.9716,0.1862,-0.7085,-0.5111,0.7151,-0.7295,0.8775,-0.9653,-0.2766,-0.8529,0.878,0.2361,-0.905,-0.8161,-0.7808,-0.8094,-0.1461,0.2273,-0.4845,-0.2817,-0.3569,0.8556,0.3323,-0.3594,-0.04156,0.4798,0.2549,-0.1696,-0.7736,0.08114,0.6037,0.8369,-0.1287,0.1546,-0.09889,0.8166,0.0105,-0.4323,0.9628,-0.6878,0.3243,-0.7018,-0.2124,-0.2329,-0.1742,-0.254,0.1345,-0.8654,-0.3589,-0.867,-0.1352,1.146,0.4424,0.1234,0.9073,-0.4676,0.5193,0.2707,-0.9915,0.28,1.311,-0.9412,-0.428,-3.038,-0.09686,0.1169,-0.3291,0.06928,-0.3806,-0.2335,-0.5215,-0.838,0.06766,0.9078,0.5551,-0.6886,0.5363,0.5508,-0.6177,0.01121,-0.789,-0.6374,0.4102,-0.9681,-0.9355,-0.8061,-0.553,-0.9847,-1.042,-2.58,-0.8801,-1.102,-1.632,0.5376,0.1916,-0.8946,0.1453,-0.1068,-0.7163,-0.409,0.2047,0.1382,-0.1694,-0.8512,-0.9449,0.5446,-0.4765,-0.1432,-0.4739,-0.3027,1.044,-0.02319,-0.09812,0.3483,0.2331,0.9693,-0.5177,0.7404,-0.5489,0.7834,0.5522,-0.9757,-0.7213,0.6324,0.6771,0.1226,-0.7817,0.277,-0.1921,-0.8133,-0.9243,-1.13,0.3321,-0.4441,-0.5557,0.5477,0.2601,-1.415,0.4271,-0.141,0.1203,1.136,0.9759,0.7435,-1.319,-1.348,-0.1353,0.04593,-0.3038,-0.407,-0.2066,-0.7375,-0.4183,-0.1088,0.5865,-0.8244,-1.09,0.1101,-0.8335,-0.0566,-0.0772,-0.109,-0.162,-1.219,0.5734,0.2791,-0.1195,-0.6356,0.7276,-1.303,-0.8826,0.4938,0.04527,0.4694,-0.6369,-0.6781,-0.3569,0.364,0.536,-0.8301,0.08411,0.8819,0.7212,-0.9965,-0.3819,-0.292,-0.8171,0.6452,-0.2545,0.7288,0.6254,-0.6146,0.2192,0.7569,-0.3002,-0.3409,-0.9143,-0.4058,-0.3192,-0.8242,0.424,-0.08606,-0.6309,0.3596,-0.7959,-0.3243,-0.7527,1.276,0.4887,0.5986,-0.7308,-0.6275,0.3237,-1.296,-0.07217,1.32,-0.7942,-0.2584,-0.7226,0.8964,0.9929,0.1193,-0.6181,-0.8991,0.9431,0.9426,0.8751,0.212,-0.3859,-0.5607,0.3449,0.1124,-0.476,-0.9901,0.0347,-0.6375,0.2862,-0.2118,0.7031,0.9487,-0.2558,-0.6335,0.7128,0.4336,0.6631,0.8506,-0.2513,-0.6845,-0.6349,0.3256,-0.7748,-0.3912,0.713,-0.8865,0.9639,0.6168,0.7833,-0.9147,-0.3271,0.667,0.3498,-0.3568,-0.2467,-0.7504,0.5158,0.9723,-0.8246,0.6582,-0.8138,0.1363,0.363,0.3129,0.2402,0.4351,-0.5658,-0.6904,0.03292,0.6243,-0.492,-0.1656,0.2551,0.7383,0.2154,0.3373,0.9066,-0.5663,-0.2251,0.5142,-0.04885,-0.5306,0.5463,-0.6334,-1.313,-1.865,-0.4835,-0.4859,-0.4764,0.691,0.432,-0.8165,-0.6766,-1.54,-1.542,-4.327,-5.227,-6.511,-4.188,-4.87,-6.793,-7.072,-4.088,-7.139,-6.151,-7.535,-2.965,-5.154,-4.824,-2.652,-7.136,-9.02,-2.346,-2.783,-1.209,1.516,-3.874,-3.772,-2.27,1.986,-1.845,-2.054,-2.715,-2.39,-1.864,-1.978,-3.642,-1.681,-8.827,-4.553,-1.199,-3.057,-6.204,-2.532,-7.677,-6.804,-5.238,-2.387,-2.862,-2.682,-3.7,-3.07,-3.116,-4.346,-4.66,-5.064,-1.623,-2.385,-2.855,-2.429,-3.078,-4.745,-2.467,-1.705,-2.092,-1.862,-2.671,-3.134,-3.679,-3.063,-3.712,1.657,1.457,-5.291,-1.656,-1.561,1.84,2.115,-2.662,-1.448,-2.495,-2.019,-1.313,-2.947,-1.545,-4.278,-1.208,-1.3,-1.317,-1.854,-1.565,-4.103,-4.864,-1.47,-3.588,-1.375,-1.565,-2.663,-5.343,-3.332,-2.795,-4.887,-1.824,-4.843,-3.051,-2.872,-1.903,-2.993,-1.707,-4.441,1.513,-1.68,-1.867,1.789,-3.823,-2.198,-2.419,-3.21,-2.792,-2.217,8.864,-2.222,-1.664,1.061,2.638,-6.996,-2.798,-1.855,-3.077,2.672,-7.011,-6.39,-4.18,-2.373,1.143,1.838,3.375,-2.501,-4.89,-4.197,-3.606,-2.382,3.285,-3.44,2.528,-3.105,1.385,-1.509,1.701,-1.468,-3.017,-4.995,-3.585,1.027,-1.568,-3.945,-1.588,-3.404,-1.061,-3.409,2.256,1.52,1.681,1.329,-1.149,2.041,-1.794,-1.282,1.959,4.111,2.461,1.61,4.387,3.83,1.829,1.325,2.689,1.664,2.123,3.639,3.444,-2.695,1.925,-1.203,1.784,-3.292,-2.399,-1.206,2.286,4.232,1.88,-2.995,-2.889,1.577,-1.282,-1.706,-1.53,1.182,3.6,1.343,1.874,1.073,2.927,1.422,1.252,2.298,3.136,3.165,-1.353,1.722,1.293,-3.514,-2.83,3.963,-1.36,-3.236,3.371,-1.188,1.153,4.181,-3.26,-2.071,-1.373,2.533,2.367,1.629,4.538,-1.466,1.134,1.038,2.559,2.608,1.832,2.398,-1.767,-1.741,-6.675,-4.379,-2.898,-1.857,-2.22,-2.309,-1.651,-3.89,-2.57,-3.088,-7.156,-3.182,-7.815,-7.345,-4.02,-4.56,-5.383,-4.774,-6.873,-2.889,-7.741,-3.005,-4.24,-5.847,-4.484,-2.766,-5.533,-2.77,1.625,-1.462,-1.455,-3.183,-3.819,-2.379,1.462,-2.096,3.095,2.015,2.389,-1.361,-1.251,3.144,1.495,2.68,-1.707,-4.238,-2.534,1.521,-1.251,1.506,-2.637,-4.186,4.843,1.638,2.559,-3.125,2.007,-1.951,-1.219,-1.762,-2.301,4.138,-2.108,-7.173,4.598,-3.274,1.101,3.103,2.632,7.046,6.017,-2.58,4.498,1.28,1.242,-1.468,3.073,1.327,2.303,-1.35,-1.024,5.84,-4.107,-1.833,-2.523,-3.84,-4.226,-1.078,-4.171,-2.991,-3.586,1.377,2.852,5.602,-3.553,-4.444,-2.154,1.171,2.205,3.325,-2.677,1.291,1.011,1.822,-1.642,2.372,-2.754,-8.791,-6.577,-5.457,-5.467,-7.123,-7.208,1.444,-2.593,-2.35,-3.726,-1.893,-4.517,-12.47,-7.937,-2.576,-5.773,2.772,-3.467,2.559,2.072,5.881,-1.307,-3.041,1.877,4.675,1.561,8.041,12.6,9.025,3.147,4.085,-1.051,4.14,1.688,1.993,4.211,-5.77,-2.497,1.728,-1.942,3.046,1.02,1.023,1.712,-2.149,-1.604,2.88,-1.467,1.998,1.862,2.986,11.97,10.92,4.37,1.774,2.619,3.197,1.49,3.357,1.824,1.012,2.981,1.422,2.64,1.174,2.072,-2.345,-1.297,1.673,1.511,4.461,-3.19,-2.581,-1.772,-1.644,2.412,1.237,-2.247,1.365,2.145,-1.87,-3.114,3.405,3.493,2.408,2.499,4.843,-1.888,2.212,3.285,-1.367,-4.553,4.335,-1.771,4.508,1.602,3.986,6.664,1.81,-1.052,-2.6,-1.141,1.485,-1.874,-1.006,2.701,1.897,3.825,3.567,1.137,-1.361,1.213,2.427,1.666,-2.628,-1.984,-2.849,4.141,3.299,3.485,2.301,2.028,1.585,1.349,2.188,2.325,2.62,3.958,1.975,1.345,-2.079,3.035,2.785,3.368,3.411,5.651,-1.273,1.339,-2.426,2.351,-1.378,-2.095,-5.947,-3.062,-3.164,-3.614,-3.15,-3.82,-2.86,-3.253,-3.3,-7.349,-6.404,1.452,3.696,1.136,1.603,-3.161,-2.944,-4.198,1.277,1.104,-2.185,1.936,2.365,-1.359,-1.957,-2.721,3.047,4.441,3.864,-1.502,-2.991,4.107,3.773,3.259,2.691,2.268,1.209,1.62,2.734,-1.562,3.518,1.154,2.515,2.649,3.451,2.47,-4.172,-4.158,-2.577,3.326,2.44,1.405,-3.189,-1.081,1.835,3.692,3.758,2.202,2.844,-1.886,-2.842,-3.519,-2.271,-1.642,5.988,1.378,-2.681,-1.087,-1.533,1.412,2.422,1.918,1.621,-1.724,-3.12,-4.069,1.719,-1.922,2.97,-4.184,-1.559,-4.43,1.81,4.503,2.426,2.243,3.185,-3.081,2.549,4.438,3.505,3.221,1.014,3.684,3.066,-1.427,-4.565,-3.992,2.777,3.508,-4.236,-6.224,11.11,3.21,3.579,-4.324,-2.764,1.934,1.387,3.818,-1.371,1.326,-1.769,1.224,1.321,3.11,1.822,-3.975,-2.399,-1.414,-1.889,-1.566,-2.284,3.187,2.498,4.337,3.553,2.972,1.83,-3.782,2.199,3.752,-2.013,1.141,-8.863,-1.784,-3.214,-7.921,-2.077,-5.698,-6.973,-6.605,-6.513,-2.606,1.897,-2.839,-2.766,-2.636,-2.855,-2.999,-4.976,-2.249,-5.12,-6.325,-3.42,-7.057,-5.372,-6.435,-2.687,-4.917,-8.717,7.791,-3.306,-7.029,-1.448,9.026,4.454,1.393,1.421,4.645,-1.533,5.428,2.368,1.783,3.284,3.143,2.344,1.764,-2.663,-2.786,-2.778,-2.17,-2.054,-1.292,2.226,2.184,1.595,1.768,-3.051,2.599,4.861,-2.047,-2.266,-4.707,-6.494,-6,1.447,3.073,1.052,2.035,2.33,1.784,2.71,1.457,3.696,3.757,2.001,1.122,-2.074,-1.058,-4.309,1.318,1.645,1.536,6.371,2.212,2.962,1.107,1.68,4.041,-3.62,1.103,1.448,3.318,2.767,-1.698,-2.144,-1.809,1.346,1.618,-1.742,-1.685,1.57,-2.267,4.285,1.685,3.096,2.361,-3.621,-2.351,3.799,3.207,-4.294,-4.331,-4.482,-4.078,-1.845,-1.553,2.103,2.855,1.369,7.079,-1.368,-1.074,-2.494,-6.387,-3.146,2.172,5.672,3.544,-1.425,-3.508,-2.683,2.961,1.326,-2.595,1.232,-3.044,-3.87,-5.311,-4.445,-4.321,-4.582,-1.379,-1.63,1.864,-3.361,3.048,3.332,3.534,4.465,2.788,2.84,5.754,-1.387,-1.192,1.572,-2.31,1.652,-1.978,-3.073,-3.676,-3.178,-2.994,-5.596,-2.47,2.834,-2.125,-4.572,-4.079,-4.768,-4.983,-4.775,-4.863,-2.88,-1.748,-6.299,-2.23,2.234,-2.296,-2.377,-2.864,1.188,-1.765,-2.55,7.455,3.552,4.083,1.222,1.931,2.377,2.563,3.283,-3.948,-3.528,1.502,4.62,2.083,1.275,2.605,2.711,1.977,2.669,3.585,1.259,-1.175,-2.353,4.517,6.217,-1.166,-4.053,-3.194,-1.019,3.854,2.91,-2.487,3.454,1.879,6.058,-5.973,-4.258,-5.86,-7.967,-3.746,-7.158,-2.3,-7.002,-6.19,-5.043,-4.532,-4.967,-5.742,-4.136,-3.332,-5.786,-3.579,-4.54,-2.992,-5.757,-3.046,1.451,-4.897,-3.468,-4.871,-7.43,-1.369,4.149,2.39,15.75,3.936,2.895,2.587,4.841,4.191,1.238,2.999,2.336,2.296,1.591,1.643,1.716,-3.174,1.686,-1.224,3.414,-5.386,1.445,1.008,1.423,1.941,-1.813,1.301,1.637,1.484,6.194,1.092,-3.168,-1.632,-1.748,2.848,-1.399,2.9,-4.015,-1.149,-4.609,-2.893,8.76,3.422,2.3,5.171,-1.326,-1.545,-2.067,-2.664,-1.337,4.454,1.024,6.921,-1.138,-2.872,-4.356,-2.867,1.117,-2.895,-4.325,-2.075,2.347,1.747,2.519,7.892,4.092,-2.393,3.548,3.672,-3.253,-1.799,-3.006,-2.454,-1.461,2.552,3.032,-1.294,2.16,6.149,1.909,1.388,8.237,2.31,6.342,3.721,-3.277,-1.827,1.142,-5.299,-3.391,2.426,1.379,1.875,3.859,1.708,3.16,-1.428,-3.342,3.498,1.366,2.739,2.541,2.794,-1.662,1.559,2.254,2.053,-3.189,3.595,2.77,3.754,1.673,-2.009,-1.516,1.267,-3.365,-3.06,1.596,3.584,6.372,-3.871,2.719,-4.089,-4.515,-1.646,-1.354,-1.043,-4.025,-1.215,-5.271,1.848,3.255,-2.328,-3.842,-2.132,-2.16,-2.444,8.167,-4.893,-9.9,-5.834,-4.602,-6.868,-3.177,-3.231,-5.258,-3.58,-6.137,-5.599,-5.96,-4.396,-5.302,-1.967,-3.946,-1.959,-2.557,2.259,-3.237,-2.592,-2.804,-4.413,-5.228,9.381,1.72,-2.893,3.879,-1.785,-4.499,-3.241,-3.962,-3.104,-4.625,-4.426,-2.808,-4.219,-2.558,-4.079,-2.671,2.381,1.291,-3.525,-6.837,-3.107,1.265,2.527,-4.231,-2.973,-1.227,-1.822,-1.702,1.207,1.114,-2.533,-4.107,-2.668,2.238,1.564,2.09,1.845,2.24,1.39,-1.738,1.417,1.552,1.337,1.769,1.209,3.152,6.158,-5.424,2.435,-1.442,1.706,-2.759,1.793,1.05,1.635,-1.926,-1.58,3.318,-3.137,-1.288,2.664,4.414,3.741,2.653,-1.238,-2.687,2.652,-1.417,-4.235,-1.843,-5.13,1.368,1.023,1.372,1.636,3.95,4.993,-1.281,-1.487,-3.788,2.155,2.343,1.442,3.33,3.837,1.686,-1.407,-2.249,-1.451,2.685,5.857,1.54,5.392,1.482,-1.112,1.784,3.704,2.184,-1.338,-1.485,-7.253,-1.495,-4.956,3.432,2.151,-1.212,-3.22,6.861,2.951,1.368,7.27,3.975,-1.493,1.655,1.959,-2.633,3.094,3.808,1.843,-4.195,-3.024,-2.88,1.79,-3.962,-1.309,-1.432,1.465,3.799,1.818,-1.502,-5.824,1.351,3.962,1.445,1.972,-2.348,-3.031,-2.389,1.037,-1.359,-2.877,1.131,1.559,1.494,1.526,-1.787,-2.589,2.625,3.506,4.817,2.629,2.514,4.889,-1.061,1.487,-2.655,-1.468,-1.8,1.295,1.117,-3.153,-2.533,-2.35,-3.226,-1.418,2.497,2.873,2.62,-4.046,-2.265,-5.282,3.107,1.229,5.655,1.072,-1.73,-2.267,2.883,-1.752,-3.02,-5.79,-1.296,-1.604,1.128,4.795,7.724,-3.613,2.415,-1.233,-1.14,4.346,3.455,-1.618,-1.719,2.489,-3.541,2.224,2.32,1.746,1.766,-4.512,-1.039,-1.025,-1.497,1.247,1.013,2.046,-1.558,2.467,5.956,1.987,1.558,4.959,-5.176,-4.011,-3.682,-3.145,-2.243,-1.178,-4.103,-5.336,-2.951,2.53,-5.444,-8.747,-2.199,-2.159,-3.985,-3.176,-6.048,-3.497,-8.509,-4.809,-6.335,-4.801,-5.197,-3.812,-6.003,-6.834,-3.126,2.803,3.146,3.536,-1.215,-2.901,3.029,-2.244,-1.292,-2.503,1.447,2.079,4.886,-1.22,-4.154,-1.165,-2.141,-3.371,-1.591,1.426,2.276,-1.831,-3.434,-1.356,2.124,5.744,-3.126,-3.072,-4.693,-4.394,1.03,-1.865,2.246,-2.031,4.644,2.781,2.657,2.319,2.572,-2.607,3.342,-2.398,-2.423,3.901,8.061,-1.069,3.031,-7.521,-5.108,-4.607,-3.035,1.721,3.765,4.016,1.359,3.779,3.986,-1.675,-1.499,-1.604,-3.838,-3.174,-2.825,-1.274,-3.583,-2.4,-2.151,6.875,2.22,1.703,-1.54,1.702,1.456,-2.199,-2.466,-4.475,-4.478,-5.538,-1.923,3.046,1.799,4.887,-1.609,6.863,-2.017,3.824,3.763,2.516,-2.351,4.369,-2.004,2,2.896,-1.462,-3.659,-3.069,-2.164,-2.134,-4.284,4.049,1.036,-6.742,3.792,-2.531,2.218,-2.282,-3.949,2.671,3.098,3.162,3.088,-2.392,2.003,-1.955,1.146,2.686,-4.157,-4.117,2.846,3.092,-2.449,-4.274,-2.233,1.32,1.105,-4.079,-2.603,2.202,-2.108,2.046,1.789,1.288,-1.673,-1.07,-2.151,1.098,2.386,-1.27,1.067,-3.264,-2.477,-2.405,-2.502,-1.469,1.065,6.797,2.36,-1.052,-1.925,2.435,-1.326,3.882,1.546,2.132,-4.332,-5.284,-2.561,2.317,-2.808,1.628,7.295,4.01,2.293,1.164,1.187,-1.72,-4.134,-2.872,-2.695,1.852,1.319,1.552,-2.764,-3.96,5.679,-4.15,-2.18,-5.478,-1.817,-1.499,-1.084,2.752,4.672,-1.105,-2.607,-3.212,-1.498,-2.96,-3.594,-1.563,1.576,2.376,-2.153,1.563,-2.416,-4.055,-1.55,1.323,-2.733,2.844,-1.648,-1.874,2.548,-2.413,-1.552,2.342,2.907,2.798,-2.559,-1.38,-3.561,-4.058,2.241,-2.078,-2.818,-2.506,-1.66,2.03,-1.441,-1.842,1.378,-2.385,1.044,2.869,3.711,3.627,13.01,2.145,-1.311,-1.06,4.808,-3.144,1.089,1.107,3.159,-1.907,-2.246,2.407,-1.984,1.794,1.112,2.525,-6.884,-4.772,-8.384,-3.156,-4.166,-2.614,-1.602,-3.763,-2.816,-2.388,-6.417,-5.862,-5.063,-4.169,-1.504,-2.641,-4.194,1.396,-3.728,-2.605,-3.289,-4.47,-1.127,-1.435,2.483,-2.392,-5.646,2.826,2.939,-3.116,-3.6,3.884,-5.905,-5.974,-3.131,-3.646,-5.345,1.642,2.726,4.508,1.296,-1.624,1.264,-2.821,-2.304,-2.194,2.205,2.503,-1.463,-1.948,-1.692,2.982,-2.785,-5.233,-2.968,-5.153,-1.422,-5.499,-3.865,-3.696,3.2,2.775,-3.462,-1.869,2.288,2.474,5.733,-1.2,3.703,-2.799,-4.86,-3.108,1.445,1.276,4.194,2.972,2.087,2.868,-2.773,-4.476,-2.659,-2.812,-3.4,-2.223,1.678,5.586,1.75,-2.486,-3.681,-1.361,1.406,-1.824,-3.164,-4.748,-4.728,1.63,-4.978,-3.035,1.744,-4.892,6.035,-2.286,-4.553,-3.33,-2.225,1.822,5.607,-3.332,3.784,-2.057,1.431,-2.542,1.265,-1.825,2.174,3.147,3.687,-3.484,2.351,1.774,2.233,1.685,1.905,3.911,3.151,1.868,-2.265,-2.598,-2.312,4.396,-2.749,-1.748,-1.194,-1.119,-4.059,-5.533,-4.198,-3.472,1.566,2.495,6.101,1.033,3.819,1.906,-3.79,-3.869,-2.974,2.123,-2.973,2.191,-2.233,-2.314,3.578,-1.31,1.684,1.862,4.563,-1.77,2.078,1.946,4.354,2.746,3.193,2.605,1.693,7.074,2.157,3.114,3.192,2.399,1.325,-1.961,1.938,-2.374,-2.368,-1.454,-1.631,-3.83,-2.589,-1.042,1.594,-4.677,-1.408,-3.416,-1.572,-1.239,-3.721,-2.082,-1.171,1.999,1.247,-4.383,6.804,3.668,2.31,2.552,-4.199,2.739,3.832,2.083,2.945,-2.464,7.869,-2.243,-2.626,4.571,-2.41,1.648,1.701,2.171,2.845,-2.621,-3.243,3.126,-1.408,-3.116,-5.584,-2.557,-1.567,-1.205,-2.527,-4.118,1.14,-1.942,-5.315,3.193,-8.798,2.889,-2.88,-2.593,-4.101,10.52,-3.176,2.165,-2.003,1.353,-2.78,-1.009,3.39,-5.694,-3.648,-2.987,-1.835,7.365,4.282,-2.066,-1.825,-4.616,1.918,2.983,-4.108,-2.158,-2.134,1.032,4.772,-5.034,-1.956,-2.298,4.996,4.749,1.436,-5.285,-1.679,-1.921,-4.519,3.588,2.481,-4.464,-2.361,2.859,-1.423,-2.288,1.427,-2.161,3.89,-1.072,1.376,2.094,3.062,-1.883,3.027,-1.486,-1.278,-3.35,-1.404,-1.945,2.007,-1.419,-1.918,-1.491,7.167,-1.617,-1.203,1.865,-1.338,-1.543,4.141,2.146,5.603,1.039,-5.216,-5.854,-2.041,1.981,-3.469,-5.281,-5.876,-2.326,-4.02,-2,-2.853,-2.962,1.193,2.861,1.325,4.487,4.329,3.372,-2.875,-1.772,1.731,2.278,1.635,-1.306,-2.264,-2.802,-2.477,3.954,3.035,-2.697,-4.536,1.107,-1.648,1.775,-4.237,-2.487,-2.462,-3.279,1.828,4.274,1.125,1.067,-3.821,-2.17,1.442,3.435,-3.291,-4.552,-1.899,1.792,1.771,-1.371,3.014,1.994,-1.276,-1.905,-5.723,1.398,-1.937,1.426,1.288,6.215,-1.081,-1.82,-2.814,-2.231,3.998,-1.446,-2.042,-2.888,-3.596,1.269,2.558,3.156,7.573,1.993,-1.549,4.236,2.254,-1.097,2.241,-3.179,-1.679,1.84,-1.12,-4.337,-4.163,-1.847,3.501,-2.583,-1.292,-1.066,2.53,-4.656,2.771,1.438,2.805,-4.122,1.203,2.26,-2.71,1.549,1.837,1.996,-1.42,-2.722,-2.263,-3.627,2.65,-3.927,-2.564,-2.925,2.222,1.76,3.032,-2.968,-3.297,-1.389,-4.039,1.792,-3.08,2.273,2.362,1.909,1.402,1.609,2.983,2.048,-1.719,2.357,1.308,1.203,-4.497,-2.376,3.217,3.789,-1.572,1.171,1.024,3.847,2.596,-1.29,-1.336,9.109,4.231,-2.424,-4.897,-3.861,-3.816,2.849,4.249,-1.263,1.473,-1.761,2.586,2.99,-1.03,1.203,2.949,-1.694,1.364,3.966,3.512,4.259,-3.693,-1.433,-1.191,-2.13,3.307,4.214,3.447,1.803,-1.43,1.267,2.09,2.449,1.292,3.495,2.908,-1.925,-2.305,-6.253,-5.934,-3.925,3.039,3.045,1.034,-1.236,-1.938,4.376,1.815,-4.501,-4.241,2.85,-2.22,-2.713,-3.001,1.175,1.827,-2.562,-2.499,-1.275,-1.51,-1.808,4.533,2.021,-2.65,-2.813,-1.599,-5.042,-5.168,-7.752,-5.639,-6.089,-6.955,-3.44,-6.715,-7.987,-4.326,-8.02,-3.04,-2.402,-2.281,-1.414,-7.603,-7.351,-8.571,-4.356,-8.165,-2.896,2.302,-1.644,-2.069,2.611,3.845,2.312,2.857,-2.341,1.518,-1.468,2.732,-2.689,-3.23,-5.867,1.064,-2.543,1.13,-1.544,-2.276,1.612,4.686,-1.031,1.578,-1.536,-1.335,-2.955,-2.117,1.852,2.331,-1.555,-1.224,1.196,2.438,1.902,-1.927,2.89,-1.21,-1.459,-2.494,-2.396,-5.166,-1.535,1.892,-1.389,-3.228,2.219,-3.79,-1.574,4.27,-2.386,-2.149,-3.879,3.105,1.682,1.83,3.215,1.315,1.642,-1.357,2.036,-4.334,1.112,1.212,-7.049,-1.816,1.492,1.192,2.092,1.366,-1.57,-3.891,-5.024,-3.05,-1.365,2.211,-3.461,-1.373,2.688,-2.673,-2.517,-5.018,-1.618,-1.547,3.919,-2.619,-2.706,3.118,-1.581,-4.92,-2.372,-1.368,1.595,-1.72,-4.526,3.153,1.876,-1.177,2.556,6.96,2.576,-1.206,4.264,1,-2.058,-4.646,-5.859,1.345,-1.455,1.234,2.173,2.317,4.141,-4.808,-4.07,-1.935,1.866,3.756,-1.32,8.016,2.624,-2.961,-4.411,-3.095,-4.568,-1.876,-5.83,1.411,2.515,-1.095,-3.747,2.426,2.43,-2.233,2.447,3.213,-1.589,-1.471,-2.937,-3.76,-1.826,4.327,-1.875,5.063,-3.372,1.013,1.877,-1.371,-1.561,1.376,2.406,1.468,1.288,3.434,1.364,-2.307,1.35,1.65,1.293,2.116,-2.356,-1.968,-4.733,2.919,-5.446,-4.309,-4.983,-1.543,-1.025,-1.131,6.858,3.753,8.873,1.812,1.104,3.794,2.427,1.719,1.709,2.166,2.286,3.29,-2.6,-2.761,-1.723,-1.808,-3.379,-1.53,-2.251,-1.219,-1.552,-5.507,4.956,-2.342,1.278,1.105,1.668,-2.089,6.448,1.428,-2.202,1.729,2.075,1.203,1.456,-2.788,-3.191,-2.119,3.198,1.506,1.334,2.509,-4.898,3.528,-2.123,-1.47,4.107,-1.012,2.048,-2.685,-4.805,-3.092,-2.224,-1.672,2.048,-5.785,-4.664,-3.629,-3.615,-2.08,2.736,-2.48,-7.208,2.193,1.098,-3.126,-1.162,-2.052,-1.877,2.49,2.862,1.129,1.654,2.632,-1.538,-6.447,-3.639,-5.067,-2.322,-2.932,1.422,2.878,-3.509,-2.079,1.612,1.482,2.89,2.545,-1.531,-4.71,-1.478,2.035,-2.131,1.782,1.174,-3.936,-1.497,1.466,2.376,2.024,2.778,-2.683,-2.78,1.032,2.266,2.513,-1.636,-3.093,-2.361,-4.133,-1.735,1.765,-1.347,1.003,1.331,2.016,2.125,3.603,3.233,1.507,4.18,3.862,-2.021,-2.592,-5.036,-5.521,-5.094,-7.532,-5.312,-8.997,-6.019,-4.96,-3.106,-3.582,-4.537,-2.597,-4.214,-3.334,-2.22,-3.575,-6.231,-3.532,-9.832,-7.932,-2.648,2.912,1.555,3.296,3.428,3.929,3.782,2.253,3.817,1.166,-1.896,-2.923,-3.745,-1.821,4.388,2.261,2.387,-1.409,-1.304,-3.262,-2.193,3.612,1.826,1.372,-1.968,-2.778,-1.054,4.997,2.123,1.067,3.294,-1.941,1.21,4.098,2.213,1.327,-4.794,1.389,-2.699,-3.28,-3.804,-4.495,-1.006,1.267,-1.652,1.511,2.122,1.746,1.022,-1.987,-4.026,-2.025,-2.191,-2.378,-3.107,-3.793,5.038,-3.373,-2.543,-2.078,-2.017,2.24,1.632,-1.495,-1.511,4.198,-1.14,2.555,8.972,-3.25,1.402,1.798,2.977,-3.912,-3.963,-4.146,-3.004,-3.278,2.524,-2.137,1.625,-3.926,1.567,1.868,-2.203,-1.948,3.53,4.596,3.106,2.122,-3.118,-3.112,-5.428,-1.711,-1.454,-2.805,-2.24,-3.51,-1.999,-2.799,-2.866,-4.163,-1.107,2.928,3.361,3.831,-3.586,-4.619,3.675,2.578,3.342,3.809,10.83,-4.363,-4.781,1.376,3.02,2.464,-3.876,-6.076,-5.778,-1.802,1.849,2.076,3.195,-2.066,2.068,1.544,1.696,2.823,8.506,2.179,2.632,-2.83,-2.524,-1.444,2.866,2.262,-1.676,-1.341,-3.814,-2.716,1.932,1.6,3.94,1.929,-2.624,3.459,2.053,1.772,3.286,2.647,-1.925,6.384,-1.383,1.587,1.545,1.506,1.931,-1.642,4.71,1.857,-2.062,1.008,2.063,2.212,2.8,3.168,-2.173,-4.52,2.443,2.574,1.324,2.205,8.237,1.272,-1.24,-1.269,-3.668,1.091,-5.381,-2.591,-5.107,-1.965,2.802,3.517,-2.397,-1.752,-3.39,-5.988,-4.852,-2.92,3.059,-3.482,4.337,3.637,7.018,1.86,-1.316,1.139,3.468,4.807,-1.245,2.319,4.461,5.312,3.803,2.134,-1.976,-4.03,-1.213,-3.221,-4.313,3.718,2.204,2.464,1.054,1.931,3.33,2.707,-2.625,-4.223,-5.626,-1.766,-1.668,3.734,-4.225,-1.328,-3.546,-2.716,2.781,-1.726,2.98,4.331,1.767,1.574,1.885,2.445,-1.171,1.48,-2.156,-3.282,-1.126,-1.171,2.614,2.777,2.629,1.259,4.325,-1.428,3.133,4.128,-1.21,2.813,2.182,1.662,1.28,2.617,-3.686,1.314,-2.105,-4.642,3.147,-2.095,-1.017,-3.657,-3.213,-2.483,-1.924,1.9,2.279,-1.23,-2.445,-1.524,3.616,5.274,1.732,3.704,2.886,-1.36,-4.285,-1.268,1.718,1.916,-2.852,-3.569,-2.322,-2.021,-4.466,-3.14,-2.97,-4.551,-3.284,2.607,-2.642,2.588,-1.227,2.479,1.322,1.421,3.824,1.372,1.507,-2.125,3.954,-1.968,-2.104,3.481,4.18,3.011,2.988,3.261,-3.428,1.974,2.868,-1.006,-1.044,-3.286,-2.882,-2.799,2.568,3.467,1.056,3.295,2.414,3.466,3.03,-2.388,3.327,-4.464,-4.776,-5.19,-3.776,1.608,-2.387,-3.742,2.178,-1.349,-2.107,-4.012,4.643,1.377,-2.712,2.485,-2.863,1.505,2.501,-1.209,-3.6,-5.761,-2.859,-2.355,5.303,-3.116,1.013,3.84,2.066,1.419,-2.237,-2.101,-1.797,1.057,-3.737,-2.336,1.525,-2.815,-5.791,-3.592,-1.605,-1.064,-1.267,-4.354,1.895,-6.6,-2.054,-4.609,2.078,-3.751,2.124,3.692,1.706,3.884,-2.033,1.206,-3.074,-2.598,-1.438,2.065,-2.745,-1.209,1.281,6.965,1.354,-1.236,-1.491,-5.119,-1.815,4.675,-2.19,-2.825,-2.404,-3.007,3.591,2.589,-4.605,5.03,3.272,3.085,3.504,1.143,-3.455,1.618,2.677,-2.424,2.336,2.502,3.117,1.248,-1.676,-3.883,-1.785,1.081,1.136,-1.612,-3.249,2.406,3.737,1.358,1.327,3.948,-2.325,-6.886,-2.639,1.729,1.164,-3.97,-1.618,1.762,2.146,1.755,4.251,1.987,2.66,4.484,-2.738,-1.734,-1.918,-1.137,-3.045,-2.39,-2.937,-1.192,-3.713,-1.49,-1.622,-2.255,2.381,2.675,1.3,-1.367,-3.451,1.417,-3.338,-6.346,-4.083,-1.588,1.564,3.729,3.26,1.926,-3.412,-3.698,-3.391,-1.889,-1.199,-3.034,2.677,-4.675,2.667,4.95,8.074,4.19,11.78,-2.319,-4.922,-8.555,-6.538,-7.034,-3.192,-1.613,-2.64,-1.626,-5.635,-5.534,-7.271,-6.44,2.697,-1.844,1.627,9.422,7.643,3.415,-3.528,6.204,2.653,2.231,2.54,1.618,2.735,3.41,-1.693,-1.986,-1.188,-1.345,-1.241,1.527,-2.313,1.151,1.615,2.279,2.545,-3.654,-6.05,-4.428,-4.352,-4.115,3.533,1.778,-1.753,-1.135,-5.137,1.752,-1.818,3.386,4.154,3.519,3.28,2.018,8.138,1.43,-5.622,2.086,-1.722,-3.986,1.11,-1.821,3.422,-1.554,5.151,-2.309,6.659,1.519,-1.544,1.547,3.136,-3.337,2.117,5.212,-1.521,1.618,3.601,-1.293,3.221,-2.485,-2.892,1.659,-1.69,-2.695,2.218,2.571,-2.821,-2.176,1.416,2.825,-2.417,1.846,1.202,-3.343,-1.366,-1.344,-2.951,-1.695,-1.907,-3.486,-2.374,-2.488,-4.707,-2.816,1.953,-3.367,-3.38,-3.627,-2.428,6.392,2.017,2.226,1.292,3.227,2.775,-3.967,-3.261,-1.193,2.589,1.32,-1.222,3.718,1.693,3.923,1.353,-1.483,4.369,-1.455,-2.518,-3.213,-1.125,-1.133,2.022,1.854,5.261,-1.015,-2.593,-2.334,-1.063,-2.318,-4.896,-3.384,-1.223,-2.491,-1.966,-2.813,1.723,-2.75,1.917,2.446,-6.583,-2.045,2.737,2.568,1.864,-3.7,1.461,1.628,-3.07,-2.245,-3.129,-3.413,2.626,-2.965,-1.456,2.522,-7.428,-7.807,3.608,3.024,2.52,1.045,4.203,4.919,2.887,3.327,-1.632,-2.841,3.922,2.107,-2.686,-2.065,-2.18,-2.133,-3.557,-1.569,-1.549,2.549,-2.526,1.18,-1.946,1.092,1.501,-2.477,1.175,1.937,-1.578,1.687,2.917,2.242,1.716,1.674,-1.021,-9.449,-5.841,-4.421,-3.341,8.997,2.412,-1.242,2.601,1.119,4.434,-1.182,1.311,3.351,-2.557,-5.204,-3.82,-1.474,2.098,-1.717,-1.408,-3.105,-1.225,2.944,2.233,4.946,2.895,-2.576,2.692,2.009,-2.935,-1.771,2.454,5.179,3.178,-4.052,-4.431,-2.02,-2.669,-2.692,-3.379,1.695,1.233,-4.508,-3.358,-4.024,-5.393,-4.722,-1.532,1.315,1.329,1.298,-2.11,-4.503,3.99,-2.2,-2.031,-4.185,-2.826,-4.974,-2.87,-1.103,1.913,-2.098,1.212,-2.588,-3.536,-2.844,-2.007,-3.443,-1.422,7.264,-2.564,-3.96,8.771,2.748,2.512,1.086,3.257,3.425,2.082,-3.015,-1.539,-1.523,-3.229,-3.743,-1.242,2.782,1.495,3.494,3.296,1.641,2.548,-3.272,1.974,2.883,1.478,9.446,1.241,-1.556,1.383,1.948,1.095,-1.306,-2.159,-2.818,-1.371,-1.338,-3.026,3.458,1.164,-4.788,-4.481,-1.305,2.941,-1.674,1.002,1.188,3.157,8.814,3.253,-2.087,-1.607,-2.959,-3.417,-7.155,-5.423,-3.913,-3.81,-5.858,-6.681,-8.163,-7.671,-7.135,-5.598,-3.961,-8.627,-6.791,-4.574,-5.152,-5.021,-3.907,-7.603,-9.219,10.84,-1.986,-2.928,2.839,2.893,1.351,-2.352,-2.353,2.046,-2.75,-2.585,-3.506,3.963,1.206,4.172,3.265,5.863,2.748,6.322,1.534,2.798,-2.908,-1.739,-4.013,-4.92,-2.926,-1.827,-2.435,-1.598,-5.7,-1.262,-1.921,2.79,1.228,3.251,4.817,-1.749,-3.036,-2.483,-2.433,-1.44,-1.945,-4.871,2.488,-2.194,-2.585,1.462,-1.146,3.871,-3.051,-3.979,-2.5,-2.86,-2.727,-2.974,1.841,1.109,2.696,-1.415,-2.218,1.993,-1.599,-2.549,3.536,9.039,-1.58,3.24,1.081,-1.091,3.722,1.542,-1.356,-4.811,-4.284,-1.861,2.316,1.504,2.939,-1.837,-2.164,1.708,3.18,-2.912,-3.511,-2.238,-3.992,-1.232,-1.452,3.01,2.266,2.11,-4.367,1.381,2.733,-2.716,1.551,1.818,2.327,4.921,-3.705,-1.884,2.943,1.835,1.62,-1.202,3.934,1.877,-1.19,1.402,-1.659,-2.504,1.58,3.347,-1.141,1.209,-1.042,-3.281,-6.488,-2.191,-3.54,2.318,3.145,6.448,3.185,2.082,4.414,-4.297,-1.624,1.889,-3.727,3.868,3.278,-1.491,1.384,2.056,-4.109,-3.777,-2.019,-2.268,-3.727,-1.581,2.116,1.834,3.587,1.74,2.656,3.332,5.45,1.986,-3.209,-2.214,-1.635,-2.026,-3.485,2.526,5.753,3.818,-1.727,-1.172,-1.178,-1.782,2.713,8.174,-2.236,-1.09,-2.763,1.001,2.593,-3.392,3.361,1.476,-1.872,5.723,1.371,-3.549,2.158,-3.587,-1.13,-1.728,4.474,-2.726,2.922,-2.494,-1.082,2.264,2.961,1.306,-6.474,-1.05,6.171,2.142,-1.03,-1.014,-1.565,-3.635,-3.65,-5.83,4.412,4.529,-2.257,-4.604,-1.402,4.289,2.638,-2.399,3.402,1.89,3.581,1.854,2.782,-3.88,11.75,4.455,1.407,3.302,1.474,-1.908,4.472,-1.885,1.473,3.333,2.399,2.149,6.876,5.575,4.037,1.102,1.854,-1.793,3.052,-1.486,2.064,4.157,2.22,-1.147,-3.571,-1.52,-1.954,-3.468,1.167,2.959,1.586,8.647,-4.643,1.235,-5.505,-3.694,-3.217,-2.661,-2.624,1.054,3.51,-1.877,-4.647,-1.233,2.517,1.859,-1.33,2.241,1.937,-4.072,-5.712,-3.382,-5.234,-3.451,-2.105,2.329,2.467,-1.297,-1.501,-2.709,3.067,2.067,3.562,1.937,-2.635,1.229,-1.245,-1.826,-2.724,1.241,-2.342,-2.248,2.644,-1.09,1.419,-5.344,1.92,4.071,1.018,-1.444,2.026,-1.259,-1.084,-6.369,1.925,-2.572,1.221,1.445,-2.34,-2.414,-3.796,2.098,-2.465,2.651,1.062,3.646,1.903,-3.05,1.585,2.183,-2.703,-1.694,2.214,3.407,3.42,2.49,1.554,-2.922,-1.96,2.494,-1.179,1.103,2.503,2.579,2.46,-2.486,-5.356,-1.134,-2.268,2.695,2.723,3.318,2.06,1.018,1.744,1.79,-2.835,3.045,-1.243,-1.15,5.618,1.338,1.446,5.254,1.247,3.725,1.304,2.303,1.752,1.27,1.097,1.591,1.942,-1.881,-3.17,6.324,2.463,5.676,3.377,-3.255,-2.516,2.195,2.859,-3.888,-4.994,1.048,-2.806,1.615,1.468,3.658,2.939,2.507,-1.641,-5.438,2.914,1.597,2.908,3.074,-1.445,-3.128,-1.65,-3.625,-1.71,1.524,-3.72,-3.317,-3.818,-3.533,-2.548,-1.79,2.74,1.515,1.513,1.944,2.388,-2.274,-5.254,2.001,4.975,2.798,3.992,1.308,1.042,-2.138,1.545,2.144,1.65,2.19,2.273,1.843,1.691,-1.698,-1.874,-2.348,3.136,1.59,4.153,2.207,1.961,-1.429,2.705,2.798,1.565,-2.233,1.178,2.581,2.535,-2.069,-5.338,2.089,-2.274,-1.147,1.838,1.54,1.653,3.332,1.655,2.344,3.359,-1.364,1.9,1.652,-3.393,-4.817,-1.078,3.455,3.466,4.536,2.128,3.158,3.891,1.555,-3.347,-1.65,3.441,3.05,-4.374,-7.493,-7.597,-2.25,-5.101,-6.008,-8.308,-9.023,-3.702,-4.628,-2.123,-3.048,-4.043,-3.98,-8.173,-2.433,-3.774,-6.592,-3.658,-6.149,-5.34,-4.684,-3.905,-8.483,-2.859,-6.087,-2.359,-5.151,-7.166,2.582,-5.39,-2.26,-2.571,-4.028,-3.833],"logCPM":[-1.565,-1.285,-1.22,-0.2534,-1.199,-0.6265,-0.6491,-1.31,-1.503,-1.529,-1.135,-1.539,-1.375,-1.582,-0.5875,-1.529,-1.313,-1.306,-1.118,-1.13,-1.577,-1.747,4.022,-1.285,-1.158,-0.5198,-1.169,-0.9155,-0.2834,-1.497,-0.789,-1.285,-1.415,-1.648,-1.699,-1.6,-1.477,-1.431,-1.606,-1.489,-1.315,-0.7676,-1.119,-1.004,-1.631,-1.428,-1.317,-1.254,-1.135,-0.5,-0.9011,-1.566,-1.655,-1.458,-1.585,-0.5649,-1.65,-1.159,-1.805,-1.14,-1.594,-0.6839,0.8576,-0.6989,0.4001,-1.542,-1.13,-1.413,-1.424,-0.9674,-1.497,-1.768,-1.914,-1.53,-1.255,-1.766,-1.436,-0.2106,-0.7493,-1.375,-1.197,-1.533,-1.556,-1.784,-1.551,-1.092,-0.9057,-1.383,-1.577,-1.739,-1.505,-1.614,-1.62,-1.63,-1.838,-1.46,-1.21,-1.906,-1.618,-1.237,-1.306,-1.151,-1.275,-1.54,-0.7112,-1.666,1.309,-1.837,-0.7159,-1.729,-1.217,-1.188,-0.1612,-1.589,1.705,2.667,-0.6247,-1.01,-0.9612,-1.531,-1.401,-1.181,-1.004,-0.2829,-1.422,-1.358,1.231,5.068,8.287,8.876,7.566,5.285,9.752,5.439,4.689,6.488,5.154,8.3,2.156,5.874,7.135,6.316,5.833,7.934,5.863,6.849,6.563,5.729,9.095,4.978,6.844,3.943,5.317,4.181,6.088,7.629,6.975,6.367,7.654,4.223,3.972,5.826,6.196,6.25,7.538,9.044,5.684,7.218,7.098,5.847,4.997,6.915,5.021,6.853,5.655,4.815,8.171,6.48,5.139,1.053,-0.3543,-0.2601,-0.6395,-1.531,-0.7452,-1.294,3.328,4.557,-1.439,5.942,7.182,2.56,3.009,5.801,4.234,7.289,2.13,5.765,5.379,7.064,3.266,4.715,6.02,6.444,8.348,4.749,2.684,4.708,6.126,3.231,4.728,7.766,4.444,5.683,6.37,7.257,4.162,6.25,4.055,4.22,5.059,5.433,5.722,8.519,5.593,6.185,6.811,9.523,4.199,5.333,6.252,5.117,7.305,2.923,4.204,4.055,1.154,0.908,3.284,8.581,0.9386,0.1321,-1.719,-1.047,-0.3964,-1.367,-1.004,0.02985,5.316,3.306,7.209,5.484,3.153,6.304,4.592,5.566,8.026,5.395,3.757,5.23,7.546,6.178,4.262,4.492,5.462,5.774,5.383,5.552,4.148,6.184,5.35,6.219,4.686,7.109,5.405,7.059,6.728,6.769,8.386,6.608,8.134,4.686,4.096,4.904,7.753,2.732,5.315,7.675,5.203,4.72,3.978,8.677,2.573,5.048,7.163,5.86,5.346,6.485,6.142,4.11,4.266,5.775,8.122,6.004,7.713,3.948,7.398,6.829,2.255,-1.466,-1.306,-1.309,-1.21,-0.6547,-1.837,3.197,3.774,-1.225,7.081,5.136,6.19,7.39,6.744,4.904,8.076,5.894,6.408,4.828,4.781,3.435,3.104,6.088,8.843,6.813,5.185,4.69,6.092,4.161,4.8,7.981,4.583,6.282,4.668,5.511,6.987,5.167,5.118,6.292,5.305,6.306,6.518,3.367,2.323,3.948,6.793,8.56,0.4163,4.85,6.224,4.734,4.465,3.493,2.25,4.673,8.403,0.261,-1.234,-1.094,-0.07011,-1.402,-0.6249,1.088,2.699,3.969,4.659,4.791,7.041,5.682,8.075,7.611,3.942,3.172,6.721,2.087,7.957,3.119,2.53,5.18,5.89,6.373,5.745,8.184,2.573,3.184,4.376,5.816,5.931,6.495,5.983,7.068,1.791,3.14,6.523,0.08749,-0.2005,1.047,3.823,7.574,3.72,2.537,5.192,5.854,2.647,7.536,4.632,5.324,3.033,7.077,3.576,2.177,6.804,4.494,6.473,5.666,3.413,4.774,4.92,2.082,0.6807,0.6056,3.42,6.688,4.804,2.199,6.566,6.963,5.863,7.248,8.662,2.582,4.21,5.378,5.278,4.206,4.871,4.607,7.612,6.93,4.14,3.76,6.431,2.397,3.272,7.801,8.89,3.992,5.632,2.675,3.832,7.366,1.676,-1.667,-0.7405,-0.8659,-0.945,-1.202,-0.477,0.3746,2.702,1.261,2.485,4.876,3.215,4.437,5.938,6.746,5.316,3.38,7.566,5.726,5.435,6.628,3.689,2.952,5.529,6.054,6.957,5.14,9.137,7.982,4.966,4.191,4.823,3.708,3.929,11.19,3.941,4.923,4.342,5.193,7.074,7.633,5.182,7.164,7.876,4.761,6.678,6.668,1.969,3.854,9.948,4.62,5.802,6.092,6.561,4.764,4.722,5.943,6.123,3.637,2.217,7.035,5.602,7.097,3.832,2.687,4.005,3.021,6.182,5.669,3.287,6.751,7.611,6.095,6.636,3.492,8.782,6.01,5.162,2.264,2.012,5.431,10.09,5.208,4.119,4.513,0.07855,-1.347,-1.497,1.423,6.741,4.427,5.285,7.611,8.767,6.657,7.603,4.155,3.006,3.149,5.811,6.348,5.76,5.317,5.802,4.01,6.556,3.08,5.8,5.143,8.555,7.097,6.198,10.03,6.583,5.391,4.226,3.93,4.676,4.285,5.772,4.67,2.019,4.313,2.775,4.837,4.656,7.352,4.533,5.449,8.909,4.022,3.149,2.979,0.5562,2.928,2.186,3.616,5.099,7.833,5.075,6.748,5.768,7.383,3.445,5.925,6.942,3.141,6.12,6.76,3.689,6.724,3.647,6.835,5.112,5.746,9.523,4.266,8.032,3.798,7.093,6.166,6.435,5.941,6.158,6.268,7.654,2.522,3.28,5.488,4.367,1.972,6.872,0.3419,5.027,5.022,7.061,1.657,5.51,1.954,7.05,7.187,8.928,2.396,5.455,6.326,3.424,6.118,2.997,1.318,7.626,5.977,5.334,4.569,3.914,9.192,6.171,6.91,4.702,5.948,2.791,3.146,4.289,2.156,6.726,0.2916,-1.161,-1.554,-1.343,1.007,5.067,-0.7388,7.294,7.774,3.168,4.346,6.189,-0.09427,6.001,4.157,2.823,5.902,2.728,4.039,3.902,9.093,4.478,4.478,5.392,9.983,7.691,0.814,5.221,5.267,5.438,3.884,3.323,7.459,3.667,6.04,2.854,4.437,5.924,7.17,6.947,3.438,0.9805,6.612,8.655,10.05,7.914,3.67,7.775,6.634,2.86,6.085,6.585,4.619,7.007,1.552,4.845,7.627,6.848,5.543,4.459,8.32,5.627,7.574,6.777,7.32,6.04,4.491,5.678,4.036,7.833,6.255,8.058,6.769,6.76,7.072,6.575,4.965,3.522,6.355,4.799,3.218,4.128,4.218,5.383,5.454,5.811,6.309,6.421,1.926,4.574,6.011,3.158,6.988,5.557,5.7,4.158,5.376,6.849,4.579,7.127,4.205,7.852,6.655,5.881,7.481,5.046,8.702,6.223,2.75,4.915,9.23,7.151,8.021,6.358,5.209,7.41,5.217,6.449,6.042,7.192,8.439,5.389,3.918,-0.8234,-1.37,-1.678,-1.618,-0.5529,-1.389,-0.762,-1.358,1.412,1.897,5.68,8.288,5.451,5.573,5.411,6.575,2.632,5.053,6.556,6.693,8.922,6.082,5.717,2.461,7.248,2.806,5.344,7.384,2.872,5.716,6.309,7.408,1.204,6.301,5.557,6.366,5.935,4.028,3.674,4.904,7.742,5.457,7.645,6.222,8.229,7.864,6.604,6.168,9.486,6.318,6.632,5.024,5.554,3.277,7.633,4.435,4.933,4.253,4.188,4.139,8.862,5.501,5.305,4.547,6.808,7.999,7.309,8.276,4.442,5.293,3.519,6.046,6.65,6.495,8.528,5.48,6.899,7.585,8.505,3.115,7.163,4.146,6.488,5.055,5.791,2.967,3.225,7.552,5.502,6.644,7.489,6.991,3.036,5.246,3.903,2.565,5.472,6.322,6.604,5.495,6.014,5.232,6.458,6.643,5.87,4.83,8.181,6.272,5.494,5.041,6.888,7.386,6.13,5.599,5.283,3.822,3.124,4.567,5.016,5.997,3.649,3.905,9.283,5.807,5.303,6.853,7.975,4.795,6.771,5.635,5.253,4.412,3.417,4.079,4.676,7.416,-0.1674,4.848,6.334,4.248,5.377,9.554,5.3,6.881,2.795,3.198,5.263,5.801,3.737,3.585,3.649,0.1638,5.556,4.103,5.524,8.015,5.346,5.259,4.974,5.184,8.028,4.28,5.325,-0.7278,-1.667,4.528,4.1,2.238,3.391,5.309,5.425,7.49,5.833,6.406,5.56,6.087,3.507,5.368,3.202,3.375,6.243,4.348,7.787,5.614,4.626,2.385,5.609,6.986,5.701,0.5129,3.937,1.323,6.399,3.252,8.979,4.396,5.579,4.14,4.994,2.839,6.621,6.18,9.337,4.134,2.32,3.127,6.598,4.714,7.413,4.66,8.437,4.412,6.976,6.891,4.117,4.894,5.151,5.625,3.505,4.747,7.051,5.528,3.917,3.517,5.366,4.207,1.5,5.357,3.704,5.808,10.64,3.132,11.77,8.361,7.008,5.122,3.483,6.154,8.309,6.46,5.152,8.505,3.504,7.025,8.298,5.144,6.344,6.751,6.881,5.72,3.319,3.616,6.883,5.301,3.234,5.285,4.868,4.69,6.449,7.424,5.875,3.072,1.424,5.904,7.955,5.518,4.497,5.358,4.872,2.845,7.589,5.928,4.656,5.813,-0.6119,-0.7192,-0.7778,-1.609,-1.157,-1.302,-1.111,-0.7352,6.231,6.518,5.424,4.567,6.284,3.026,2.375,4.934,0.433,8.171,4.312,6.156,3.222,4.671,8.74,1.722,0.411,2.798,4.043,4.04,5.126,8.085,3.792,6.197,6.578,4.555,6.379,6.308,4.409,9.098,3.642,1.005,3.953,6.505,7.876,2.839,7.723,3.39,3.69,6.983,7.187,6.787,2.44,7.414,5.67,4.858,6.983,5.3,4.676,3.136,5.901,5.694,4.657,3.773,5.024,5,5.637,10.78,6.81,6.367,2.617,3.517,10.96,2.802,6.921,4.687,1.475,2.506,8.531,1.85,3.951,9.568,5.541,8.821,0.8304,6.498,4.942,6.713,6.094,5.209,2.86,10.5,2.676,5.66,2.481,7.331,3.666,3.964,5.884,6.423,4.697,6.666,7.072,5.798,3.815,7.216,7.209,4.492,6.625,6.386,5.46,6.907,5.897,7.192,3.982,4.259,5.779,7.714,6.198,5.649,5.263,2.653,5.112,5.637,6.006,5.541,0.789,3.652,3.794,2.573,6.809,0.0592,4.183,7.435,3.465,6.893,8.08,1.633,10.11,6.279,4.216,4.888,8.279,4.721,4.198,7.945,3.425,4.125,5.064,5.04,3.797,5.189,5.38,4.125,8.022,6.491,4.421,4.414,5.774,7.46,-1.257,-0.683,-1.456,-0.4965,-0.8389,-1.285,-1.519,5.735,7.921,3.292,5.003,8.095,5.261,6.037,5.839,4.022,3.949,3.136,4.848,7.815,4.977,5.614,2.9,6.176,3.861,6.398,3.936,9.795,2.213,8.341,7.022,5.393,8.387,5.128,6.339,3.609,4.176,6.641,5.806,6.736,4.663,5.511,5.407,1.027,7.853,1.83,5.894,6.026,6.069,0.1732,4.68,5.944,5.168,3.73,7.771,7.095,2.918,5.243,5.972,6.274,4.67,3.995,5.42,5.872,4.071,6.496,4.517,5.585,5.642,8.071,6.112,2.944,6.454,5.029,6.491,5.635,6.051,6.96,4.763,3.997,4.477,7.032,7.615,6.905,4.917,5.11,5.16,6.45,4.647,5.876,5.58,3.367,5.615,4.479,7.637,5.515,1.157,5.785,3.979,5.541,4.001,5.938,5.799,7.311,5.394,6.753,2.331,3.792,5.92,7.118,7.454,5.072,4.005,7.329,4.837,2.332,3.589,6.651,4.407,6.143,3.626,4.952,8.525,5.538,4.36,6.735,3.362,4.098,5.957,8.178,5.431,5.829,6.183,5.044,6.565,3.929,7.267,6.835,5.031,4.54,3.442,5.47,7.704,5.013,7.654,3.206,0.8078,7.317,8.784,7.554,7.308,8.132,5.021,4.68,4.578,8.55,0.9916,5.845,-1.08,3.497,3.448,5.983,8.419,5.741,5.097,7.983,2.379,6.635,6.269,5.218,0.6738,1.16,4.978,3.855,6.37,4.866,6.137,0.7171,4.51,9.018,1.387,8.151,5.779,2.103,8.093,6.577,6.083,6.086,6.931,4.26,8.747,7.312,5.396,5.122,7.683,4.987,6.767,6.116,3.588,5.695,6.376,4.591,6.128,7.568,2.383,5.737,3.259,3.741,4.189,-0.8555,5.79,5.858,7.1,3.802,6.021,4.932,6.929,-0.7908,4.377,9.808,3.224,9.443,8.842,4.756,6.651,6.434,5.367,8.213,5.118,1.96,5.165,3.2,7.998,4.889,5.174,5.074,4.466,3.579,5.745,4.138,5.772,4.205,5.514,7.117,7.004,10.07,8.45,6.305,5.775,6.648,1.868,9.149,2.879,6.321,6.412,7.807,3.247,5.895,4.273,6.599,9.416,4.689,6.712,10.18,0.04958,4.086,5.628,5.348,6.936,5.691,5.397,6.15,5.544,6.262,6.546,4.782,5.182,4.841,7.478,4.374,4.219,7.827,5.101,0.1054,6.997,6.036,3.925,7.375,6.793,5.363,5.748,5.304,5.883,2.212,5.478,-1.107,8.572,8.753,7.471,5.868,6.111,6.309,5.426,5.642,3.966,4.255,6.04,4.702,7.04,4.764,5.409,3.007,2.234,2.699,3.619,6.14,3.527,6.663,2.246,4.989,6.963,6.005,6.169,2.552,3.532,4.972,4.344,6.648,5.267,6.978,2.787,5.79,0.9661,5.191,5.77,5.98,6.477,3.008,7.478,0.8645,5.555,8.742,8.192,7.812,6.364,4.55,7.47,6.393,5.219,6.948,5.972,6.358,6.762,6.375,6.79,4.262,4.224,3.85,6.392,6.625,3.636,2.561,-0.1953,-1.355,-1.406,-1.382,1.377,-1.237,2.663,6.066,8.288,7.955,3.83,4.256,2.71,5.862,7.155,3.435,4.334,4.837,1.894,2.906,6.007,6.073,6.13,3.018,4.261,4.904,5.886,7.526,6.924,4.257,6.414,4.705,5.129,8.756,6.965,7.12,8.016,3.758,6.852,4.456,4.483,4.903,1.78,3.364,5.052,3.801,1.334,6.15,3.819,7.561,2.596,6.066,6.089,6.797,4.98,6.315,5.32,4.886,5.671,4.191,3.7,3.903,5.425,6.047,7.343,2.804,6.062,4.19,3.985,7.287,0.1952,4.043,5.495,7.291,2.921,8.556,5.197,4.499,5.441,5.952,3.984,2.812,3.947,6.401,0.1416,2.391,6.101,9.955,3.754,5.262,6.882,5.692,3.999,6.367,9.146,5.524,6.794,6.953,6.955,6.072,5.185,8.058,4.413,5.188,5.932,5.725,9.543,4.977,3.684,2.424,7.786,2.009,5.87,4.141,1.953,7.199,-0.774,4.521,6.604,3.14,5.595,8.728,3.385,4.485,6.399,4.034,6.643,4.103,6.209,5.255,3.227,6.99,9.885,5.727,6.107,6.733,4.958,6.808,5.954,5.542,6.241,5.577,6.367,6.194,5.615,5.571,4.772,4.288,5.843,8.467,5.959,6.743,2.889,5.247,6.089,6.489,5.593,4.536,5.108,7.792,5.495,5.549,3.344,3.84,2.7,0.1405,6.013,5.001,5.533,5.643,8.591,6.603,6.092,5,6.835,7.056,4.128,4.99,6.206,4.12,7.948,8.034,4.75,9.717,8.468,5.261,6.197,1.838,2.957,4.819,4.012,6.282,5.663,4.676,5.468,5.847,10.09,5.924,4.544,3.813,1.33,7.8,4.36,7.736,4.193,7.316,5.02,-0.004422,-0.3232,-1.28,-0.9567,-1.302,-1.409,-0.4736,-0.2966,-0.9038,-1.076,0.8056,1.311,1.406,2.7,5.774,2.541,2.82,6.447,1.307,3.823,5.026,4.596,0.4525,3.467,2.699,0.2051,3.818,5.941,-0.6112,-0.1608,-0.9725,-1.591,0.7678,1.96,0.04815,2.182,-0.3788,-0.756,-0.358,0.3731,-0.7151,-0.1979,0.1915,-0.8243,5.928,2.006,3.559,2.509,2.973,1.323,4.17,3.385,1.855,0.02113,0.774,0.05751,1.163,0.3366,0.1914,1.399,1.186,1.612,-0.562,0.7648,-0.1131,-0.3059,0.08764,0.8433,-0.5251,-0.9976,-0.5811,-0.8868,-0.0725,0.1482,0.747,0.002654,1.783,-1.182,-1.163,1.362,-0.718,-0.8932,-1.035,-1.264,0.1051,-0.1855,0.5997,0.208,-1.227,-0.2293,-1.09,1.481,-1.105,-1.245,-1.23,-0.4479,-0.677,1.377,1.092,-0.8819,2.174,-0.7608,-0.6204,0.67,1.477,-0.008342,-0.01047,2.91,-0.493,1.761,0.9435,1.268,-0.2243,0.5185,-0.9998,1.422,-1.562,-1.015,-0.6963,9.17,0.3066,0.06933,-0.5701,-0.06544,3.864,-0.2497,5.808,-0.318,-0.2154,3.857,3.676,5.477,1.412,-0.9003,0.3537,-1.15,2.743,2.568,1.088,5.418,4.952,7.042,4.149,4.161,8.81,3.15,4.854,5.654,8.073,6.812,7.066,5.274,5.874,4.76,8.351,9.002,3.488,2.804,4.021,7.059,0.1409,7.992,3.68,3.615,6.074,4.152,5.585,4.647,2.882,5.295,4.571,11.93,4.557,6.032,7.338,5.62,7.372,3.734,7.355,7.24,7.207,2.608,7.073,7.749,6.641,8.565,4.331,6.401,5.884,4.631,6.679,4.09,4.971,3.59,7.071,6.099,6.831,5.758,6.623,4.837,6.153,5.224,4.062,5.999,10.08,6.194,7.586,6.357,8.712,4.191,7.523,6.02,5.842,5.372,4.728,3.598,8.951,1.969,4.66,5.557,5.309,3.169,8.346,5.629,5.554,6.916,5.283,4.879,5.918,6.534,7.605,4.222,7.914,6.107,6.406,7.322,8.911,8.558,4.639,3.497,0.9994,7.57,8.835,5.915,5.769,2.214,0.7733,0.04703,-0.753,0.3294,-0.04019,5.188,6.344,1.35,4.803,4.691,6.106,4.477,4.694,3.196,4.594,0.6504,3.781,4.57,6.452,1.455,12.26,3.902,2.527,0.5951,8.501,2.77,4.086,5.472,5.115,2.782,7.105,7.238,8.415,7.238,6.809,3.723,3.249,6.357,4.161,5.605,4.941,3.47,8.065,6.725,6.498,10.23,4.017,4.062,7.762,6.462,4.586,3.981,3.937,5.477,7.429,5.669,3.258,3.623,1.687,7.39,7.494,3.741,5.605,9.471,6.853,7.36,4.747,5.884,4.786,7.232,3.61,6.842,5.808,8.921,7.218,2.494,5.54,6.603,4.616,3.216,4.141,3.959,6.25,7.203,6.228,2.819,3.274,1.59,8.889,6.858,3.684,1.343,6.922,7.267,5.837,7.664,3.271,5.976,8.616,3.939,5.7,10.07,0.6655,7.336,5.545,2.917,3.583,2.628,4.472,-1.577,-0.4442,-0.1799,0.4265,-0.5046,1.402,7.287,3.534,-0.2688,2.738,3.741,3.71,0.2624,7.895,6.331,6.309,4.824,8.322,3.37,6.98,5.441,3.283,4.381,6.559,4.403,8.828,10.03,3.334,5.924,8.488,4.486,4.651,6.836,3.05,5.778,4.286,5.153,6.028,1.035,3.365,1.867,6.663,6.701,3.145,7.912,6.338,6.905,2.143,6.754,7.588,7.187,8.089,4.726,8.619,4.545,8.193,5.307,6.54,6.525,7.661,6.78,3.699,5.435,4.739,6.616,2.886,4.038,3.085,4.087,7.822,7.08,2.673,4.009,6.29,6.762,4.891,7.58,5.438,7.167,6.708,8.754,6.813,5.331,6.271,6.732,6.138,3.832,4.881,5.626,4.729,3.911,5.841,5.44,4.176,5.128,6.171,7.203,5.784,7.136,6.615,3.488,2.301,7.824,5.662,6.704,5.929,7.697,5.754,6.913,4.686,3.834,4.46,6.026,7.531,6.819,5.832,6.505,3.437,6.002,4.732,3.799,7.581,8.369,3.488,6.911,6.448,5.891,3.698,7.937,1.916,6.23,6.093,3.123,6.298,-0.8035,-0.7602,1.885,0.4972,2.19,0.929,1.215,2.607,2.056,0.6257,2.163,7.319,3.982,3.172,8.631,5.168,5.627,3.542,5.257,8.363,7.293,4.269,3.143,6.979,3.454,5.621,5.781,5.595,4.603,8.561,7.939,3.087,3.219,5.66,6.496,6.453,6.643,7.902,7.255,5.24,4.142,5.651,3.482,6.024,7.188,6.916,7.211,2.942,7.658,6.604,2.906,6.674,4.386,5.805,2.414,8.462,8.047,7.449,6.084,5.372,6.343,0.2087,2.506,4.376,4.685,4.41,6.716,8.965,6.09,4.845,5.983,6.931,4.598,5.832,0.7695,3.49,3.828,3.573,7.756,5.246,7.347,1.359,4.333,5.24,8.793,6.792,5.184,3.963,7.7,7.42,5.31,4.579,7.052,6.924,6.231,7.462,6.413,7.407,5.342,5.229,4.312,4.602,4.116,6.328,4.896,3.708,8.209,3.826,1.225,7.637,6.91,7.082,2.423,4.882,5.305,4.527,6.935,5.342,3.192,2.777,3.726,3.794,3.139,6.513,5.357,6.334,9.113,5.856,5.695,8.56,5.25,4.04,6.424,6.985,7.454,6.661,7.613,4.267,1.163,4.924,3.7,3.141,3.2,4.242,2.746,0.8874,0.359,0.5433,0.5462,-0.411,-0.09566,-0.007783,1.841,-0.3877,3.281,3.168,1.422,5.673,2.907,4.969,1.52,3.535,4.019,4.759,13.65,5.335,9.402,5.662,6.697,6.881,5.529,5.766,5.995,7.471,6.707,5.365,5.885,7.744,6.495,4.886,4.09,8.336,7.598,2.654,6.334,5.355,6.905,7.52,5.254,5.98,3.908,7.108,6.829,10.4,3.467,5.463,5.031,4.956,2.161,7.713,4.448,3.603,9.409,5.418,8.56,6.148,5.862,5.215,6.263,4.46,2.536,5.41,5.595,5.18,9.014,6.074,5.444,4.753,6.334,5.268,4.53,6.794,5.029,3.57,5.86,5.238,7.218,5.378,6.29,4.681,6.853,4.197,5.318,3.298,7.241,2.78,6.583,5.283,7.021,6.339,2.802,4.127,13.07,6.125,4.988,3.074,2.55,3.321,1.613,3.128,8.994,8.686,9.687,5.239,5.587,1.061,6.556,3.725,2.854,4.834,7.209,7.597,3.239,3.297,6.178,5.627,5.598,6.121,4.884,5.618,3.759,2.946,4.667,6.38,4.159,4.182,6.96,8.518,3.442,2.259,6.839,6.086,6.377,7.245,1.175,5.252,5.105,4.973,5.893,7.33,6.454,6.829,3.849,3.153,1.747,0.6923,2.25,4.619,-1.055,0.1331,1.589,2.058,4.242,3.506,3.738,2.114,0.5013,1.615,3.96,1.181,6.308,1.6,4.342,2.878,7.258,4.248,4.192,6.924,8.166,7.327,8.497,5.875,6.033,4.33,6.288,5.298,5.978,6.038,5.902,5.508,4.777,5.525,5.699,9.365,9.301,7.325,10.38,3.321,2.68,5.979,3.888,6.337,7.173,6.768,5.539,5.623,5.103,3.153,5.887,7.96,6.166,8.101,2.443,5.183,3.561,1.791,4.168,4.06,4.077,3.202,2.025,1.205,2.792,1.962,1.372,0.7247,3.691,1.368,1.108,0.3996,1.949,1.289,1.561,5.095,2.99,8.896,4.88,3.065,5.235,5.887,5.364,8.936,5.686,6.131,6.841,6.762,6.505,6.041,5.352,5.264,8.37,7.686,6.648,7.284,7.455,2.748,6.399,4.119,7.467,6.489,4.52,6.523,7.142,6.52,5.342,7.373,4.162,8.055,4.374,5.238,5.553,7.175,6.546,8.711,6.817,7.142,2.743,5.884,4.028,6.704,7.566,6.443,4.035,8.164,3.326,1.686,3.782,2.14,5.811,7.131,3.684,4.289,9.018,3.994,5.817,0.8604,1.671,3.87,6.224,5.659,4.881,1.343,8.586,7.267,6.438,6.134,3.795,1.73,4.1,8.251,3.648,8.309,4.555,3.344,3.293,7.497,6.414,6.835,6.126,7.748,6.287,7.938,5.677,6.089,6.304,6.385,8.282,7.834,6.789,6.91,4.563,5.518,2.952,4.395,6.993,4.122,5.626,6.113,5.78,7.113,5.098,7.147,4.987,10.62,7.282,5.56,6.817,8.01,6.76,1.441,3.124,2.136,4.708,5.282,4.563,5.346,4.235,3.957,1.005,4.621,6.662,2.77,5.342,5.424,3.931,3.896,5.007,5.624,7.584,3.084,4.674,4.747,-0.7097,3.29,2.996,2.965,7.64,2.607,1.604,5.898,0.8752,2.17,3.183,0.8775,4.047,1.794,3.067,1.223,1.825,-0.3342,0.4056,-0.6566,5.577,-1.313,0.3211,3.797,1.648,2.034,4.963,2.133,9.275,8.547,8.736,5.596,6.1,5.761,5.877,3.539,2.61,2.546,7.269,3.218,5.916,6.508,3.204,9.635,6.807,6.999,5.356,2.559,5.566,8.142,6.459,6.711,6.816,3.543,5.555,4.998,9.378,3.593,1.573,3.498,4.543,8.853,5.108,4.818,6.993,4.52,4.216,5.094,4.23,4.805,5.773,6.569,6.16,3.961,4.344,6.693,3.431,2.454,7.438,6.211,8.064,6.884,3.897,3.436,7.157,4.389,4.953,4.194,9.867,6.88,5.45,5.209,5.586,7.852,4.399,5.447,4.291,6.087,6.547,4.593,5.961,6.694,2.477,7.179,5.159,4.768,5.358,7.745,4.249,5.113,8.571,6.89,6.168,4.591,4.863,4.225,7.434,6.078,6.071,5.385,7.433,3.703,6.968,4.795,7.639,4.834,2.993,4.348,5.432,8.995,6.769,6.22,5.977,8.622,5.448,8.716,8.356,2.484,6.773,4.775,4.275,8.228,6.696,4.86,6.374,6.519,2.699,5.385,3.893,5.644,2.815,4.493,4.442,6.262,6.861,9.429,4.639,7.283,7.062,7.395,1.563,6.022,4.13,4.804,4.265,5.142,2.996,5.172,5.287,5.494,5.243,7.987,7.408,2.037,1.775,2.972,7.668,5.88,6.362,7.928,2.925,4.991,6.364,5.35,9.073,5.705,6.317,5.674,6.352,4.462,5.012,5.437,6.288,5.446,8.029,3.941,4.849,2.692,5.75,5.304,6.503,5.596,2.72,5.611,5.877,3.867,3.373,6.927,4.709,1.824,7.462,5.309,6.977,8.737,8.31,5.651,4.15,5.573,8.268,4.645,5.762,6.127,5.231,8.083,5.144,4.315,5.585,3.071,4.329,4.343,6.044,4.631,6.303,5.103,4.097,7.156,4.637,8.393,6.331,5.377,3.546,0.8102,4.95,0.1847,7.99,4.802,5.551,4.486,3.77,2.355,7.732,6.481,-0.6774,-0.7074,1.655,0.5402,3.525,2.527,5.058,7.999,4.725,1.614,1.703,5.099,7.112,5.347,8.364,2.606,8.243,4.768,4.466,5.934,7.852,4.202,4.995,1.305,3.783,4.655,7.911,6.04,5.083,6.352,1.698,6.793,6.887,8.405,6.545,2.138,2.336,5.729,2.633,3.546,6.357,1.746,3.208,2.066,3.691,5.268,5.895,0.7353,7.962,5.943,3.871,4.37,9.634,6.79,10.02,4.226,2.663,7.748,4.217,9.52,6.837,4.261,4.294,3.327,3.75,4.857,6.544,7.235,5.216,5.307,5.849,6.319,4.675,1.519,4.539,4.386,3.395,2.27,2.92,4.405,6.136,5.923,4.878,7.323,9.945,5.551,7.037,3.921,5.159,5.297,3.723,4.621,6.44,7.501,6.391,7.44,5.667,5.703,6.877,7.062,7.572,4.778,2.859,7.963,3.594,4.307,6.489,5.196,4.778,2.304,5.749,3.262,8.852,5.888,6.809,5.649,7.859,5.95,5.383,1.883,7.231,7.647,6.784,5.775,5.604,4.993,4.679,6.643,7.489,9.26,5.416,3.039,4.977,7.801,6.482,3.122,4.901,6.796,4.699,6.116,4.95,5.622,4.293,5.067,5.589,3.601,5.79,2.779,7.04,5.994,8.2,6.8,9.579,3.041,4.832,3.194,6.468,5.524,9.687,7.019,7.261,6.877,4.387,7.565,4.358,4.8,5.042,5.992,4.74,2.124,4.473,3.028,5.325,6.562,5.578,6.541,7.493,6.076,6.45,5.023,4.051,4.822,5.761,5.064,4.294,5.46,5.102,3.733,3.934,5.171,3.903,4.398,4.122,5.635,4.662,7.827,2.892,6.033,3.777,4.334,6.036,6.758,4.307,7.973,8.773,2.862,2.815,5.278,6.239,4.625,4.9,6.611,2.009,5.94,5.737,4.908,4.101,3.266,7.434,6.106,6.519,5.013,5.938,6.244,1.799,6.891,4.427,3.068,5.852,5.486,4.103,7.285,4.087,5.06,6.248,3.598,7.988,5.681,6.094,3.332,7.09,4.087,6.118,6.445,3.79,3.826,6.342,3.814,6.497,6.563,4.093,6.839,2.5,9.734,6.16,5.683,4.558,5.228,4.328,0.4232,0.7868,-0.4304,-0.419,0.5578,0.8187,-0.5856,2.838,2.188,1.477,0.5108,-1.095,1.811,6.442,7.502,4.331,4.701,4.28,3.298,7.945,5.794,9.241,4.962,5.589,5.088,4.94,2.805,3.842,6.975,8.332,8.344,1.689,5.363,5.375,4.633,7.32,-0.04389,4.827,5.583,6.03,5.017,2.742,2.74,9.473,11.67,5.718,2.946,5.937,7.885,1.886,4.763,5.518,6.646,-1.155,4.696,1.129,5.94,5.374,10.43,1.915,4.499,5.772,6.622,7.319,3.452,4.653,3.763,2.122,3.571,6.49,5.962,5.195,5.079,6.167,3.666,6.785,3.279,3.504,4.983,4.653,4.906,2.735,6.617,7.232,3.704,2.474,2.525,7.94,2.747,5.583,2.624,3.845,8.104,6.162,7.273,4.76,4.573,6.535,5.899,2.1,2.666,5.839,8.465,6.503,4.859,6.965,5.978,6.746,4.424,8.001,7.002,6.554,5.966,6.295,3.266,5.086,6.697,6.443,8.582,4.455,6.014,7.091,7.975,7.818,1.661,2.932,6.508,2.899,6.329,7.822,6.674,3.978,2.997,2.073,2.984,6.472,5.476,7.819,8.25,9.39,6.525,6.022,5.094,5.188,7.692,2.957,3.62,3.832,4.228,10.13,6.227,6.555,6.818,6.426,3.6,5.416,6.031,9.153,6.326,6.891,6.407,5.245,2.371,2.117,8.42,6.52,9.361,5.171,5.812,5.409,4.527,3.16,9.883,4.295,3.707,4.307,4.001,7.876,4.426,-0.9861,6.487,4.864,7.277,3.111,3.965,3.459,7.631,5.636,4.589,2.166,5.777,8.464,3.941,5.676,8.683,5.04,6.773,6.841,5.982,6.584,6.868,6.838,9.418,5.408,7.975,7.62,7.342,5.968,6.642,5.755,1.212,5.202,2.321,3.6,1.304,2.071,5.581,5.644,6.69,7.818,5.259,3.595,6.821,10.5,5.924,4.208,5.5,1.786,4.568,2.135,2.347,7.143,9.565,4.402,7.015,9.341,6.488,4.104,7.748,3.785,4.883,5.457,5.605,3.603,3.472,7.108,7.848,5.999,6.321,4.005,4.388,7.122,4.846,3.293,8.55,3.087,8.095,5.167,4.601,6.698,4.779,4.312,6.405,5.725,6.529,5.602,6.292,4.398,3.085,7.372,5.248,4.055,4.813,5.835,7.421,4.507,8.419,3.22,7.472,5.167,1.779,5.552,2.286,1.598,2.641,6.151,2.65,7.852,-0.5018,8.142,6.874,2.186,5.687,6.274,5.225,5.384,6.87,7.45,4.349,1.269,-1.4,0.3315,2.704,2.088,-0.604,2.582,-0.7903,-0.2912,0.4713,-0.4401,7.7,5.654,8.446,7.662,8.828,4.037,0.3104,6.605,5.855,3.056,4.839,5.245,5.185,3.982,5.891,7.335,5.293,3.373,8.801,2.033,7.724,3.384,4.084,2.038,2.013,6.221,5.525,7.879,7.944,3.662,3.865,8.251,4.847,3.956,6.168,4.665,6.727,8.32,4.993,6.587,3.823,5.458,7.468,6.303,6.333,4.137,3.939,6.332,3.021,4.593,8.604,4.398,4.437,8.948,4.096,4.409,5.445,8.783,8.705,5.204,3.929,3.429,6.867,5.957,1.457,7.139,5.131,6.193,5.693,6.633,4.958,3.754,3.882,1.857,4.556,7.811,7.217,4.809,7.931,8.705,5.336,7.995,4.159,6.676,2.974,5.718,7.895,6.504,6.919,4.851,6.624,7.81,1.889,3.381,5.172,4.188,5.881,7.239,7.629,8.893,7.738,5.901,5.605,4.376,1.912,6.378,6.892,5.549,7.757,3.095,4.16,6.095,5.975,6.225,4.663,4.25,6.841,6.948,4.777,6.691,6.049,4.955,4.5,6.255,5.066,6.251,6.747,5.942,3.503,2.267,6.494,7.829,3.107,2.363,2.201,1.667,8.46,6.697,5.119,8.361,6.35,5.586,6.804,6.439,5.235,6.316,5.086,5.859,6.659,7.838,4.648,4.907,4.436,8.825,0.1317,7.641,3.178,9.619,4.336,4.829,5.176,5.216,6.103,8.609,6.438,6.657,7.499,6.745,3.21,4.059,3.923,6.918,4.192,7.385,4.887,6.236,5.916,5.961,4.423,4.509,4.054,4.65,4.393,3.567,5.775,2.883,2.196,2.847,3.1,2.398,8.266,5.7,4.892,4.289,5.855,3.009,3.637,2.01,5.616,1.778,4.643,6.822,1.404,2.714,3.93,1.586,7.614,-0.01154,-0.5786,-0.6333,-0.177,5.537,4.134,3.999,1.853,5.284,2.364,7.562,6.178,3.388,7.932,7.141,7.384,7.595,5.949,3.034,3.145,6.999,2.58,5.014,4.118,8.409,3.871,3.505,6.236,5.489,5.109,4.217,6.559,5.912,5.76,3.67,3.569,3.277,6.009,5.528,2.533,4.308,7.126,8.691,6.976,6.597,5.211,4.786,4.469,1.834,3.88,5.19,6.331,7.936,4.013,5.479,3.795,3.643,6.59,11.47,4.444,3.333,3.696,9.408,9.29,10.39,7.437,4.924,3.368,4.226,5.879,2.986,7.276,8.059,7.026,2.296,9.245,6.706,6.239,5.885,5.286,5.774,4.238,7.362,5.358,7.248,2.51,6.712,7.631,1.172,4.254,5.539,10.9,4.218,9.511,1.779,4.139,3.984,2.522,4.839,4.03,2.203,7.663,4.51,5.916,5.707,9.144,5.4,4.328,4.802,5.679,5.561,6.525,7.587,5.598,4.613,3.981,6.606,3.599,7.122,8.214,8.104,5.545,6.319,3.528,4.287,8.255,4.677,7.6,9.448,7.642,3.642,2.414,2.725,2.179,3.243,5.333,9.067,5.798,8.796,4.961,8.067,5.976,2.387,6.835,4.99,4.206,3.621,3.74,1.376,6.058,6.74,3.076,3.965,2.5,5.833,8.918,3.974,7.804,6.114,5.326,2.339,6.097,8.969,5.502,7.408,9.455,4.279,4.156,5.327,0.7063,6.552,5.288,5.404,5.063,4.589,4.838,6.046,5.332,4.077,6.093,5.908,3.892,4.247,5.49,6.522,6.488,8.432,9.77,6.857,5.066,8.782,3.857,4.664,4.051,4.282,6.279,5.083,4.111,5.708,8.547,8.513,5.789,7.41,7.255,5.568,6.573,5.373,6.74,5.035,5.766,8.976,4.507,6.43,5.413,7.128,4.186,6.467,6.804,5.811,5.597,8.238,8.349,7.432,5.784,2.94,4.584,6.661,5.492,4.825,3.668,3.027,6.825,2.899,10.02,2.244,2.973,3.032,0.5554,5.204,8.831,1.517,7.779,6.808,7.05,3.335,6.328,5.254,6.532,5.307,6.22,6.357,4.638,6.618,5.403,5.766,6.532,4.867,4.286,1.209,6.247,4.032,4.634,4.47,7.309,7.023,5.423,5.585,4.501,6.658,7.003,8.391,3.218,7.335,8.492,5.77,5.456,7.181,6.512,5.432,8.288,2.747,0.7559,3.557,10.7,7.065,2.446,5.402,2.986,5.258,3.563,8.934,5.435,5.6,6.792,5.829,6.255,5.425,5.092,8.125,2.581,6.563,5.998,3.156,1.351,3.216,4.288,4.504,5.102,7.251,4.38,1.287,-0.1384,0.1512,1.352,0.2163,1.34,-0.000806,-0.4816,0.769,4.29,2.608,6.214,5.29,3.392,6.534,7.124,9.044,5.064,6.151,5.891,7.348,7.07,8.429,5.673,0.5329,2.532,7.564,7.083,8.314,4.54,5.744,7.008,1.311,2.825,4.934,5.591,8.886,3.437,7.601,5.625,4.94,3.801,6.489,7.363,7.678,6.613,7.119,6.206,8.041,4.07,10.13,2.963,2.566,6.874,4.092,3.127,4.227,5.277,6.54,7.017,4.985,5.615,5.428,3.055,6.825,2.325,3.086,4.355,3.877,7.023,7.593,7.736,4.983,7.882,8.436,8.336,5.501,7.54,9.167,5.927,9.973,5.775,4.976,5.246,6.11,7.554,7.365,4.12,4.016,1.205,4.149,1.859,6.763,9.085,3.396,6.872,3.307,4.406,2.023,7.626,6.007,3.228,5.709,5.453,3.024,4.507,7.09,2.961,7.427,7.227,5.938,5.006,4.092,4.087,4.425,6.275,7.711,7.594,5.795,1.401,4.645,4.909,8.031,4.109,7.133,6.478,5.148,3.331,-0.9523,4.331,4.572,3.343,5.412,4.101,5.241,5.331,6.791,3.901,3.718,6.603,3.411,6.241,7.95,5.2,6.017,4.979,4.592,3.696,5.592,6.087,2.889,4.756,6.244,6.35,4.829,6.285,5.512,5.339,8.24,5.457,6.254,6.165,5.872,5.39,7.552,8.103,7.456,5.93,3.33,7.81,6.392,6.003,3.041,6.686,8.147,5.971,8.849,3.715,7.25,7.296,9.773,3.949,3.938,6.298,6.81,8.431,5.459,7.483,7.33,4.974,1.441,2.968,5.342,2.848,6.555,8.284,5.425,8.743,8.294,5.076,5.653,7.034,4.083,3.012,6.074,2.931,3.147,5.379,7.408,5.965,6.767,7.94,8.532,7.369,6.11,1.483,6.899,6.487,5.789,7.453,7.425,3.016,3.118,3.588,5.462,4.086,5.078,5.242,8.047,7.362,5.869,5.734,7.474,4.45,5.669,5.745,3.63,6.11,7.824,6.952,3.551,4.846,3.469,7.05,2.846,5.161,5.003,6.979,5.743,4.883,5.45,4.306,6.169,4.558,3.373,7.122,8.563,5.93,4.461,6.115,10.72,8.489,2.428,0.5427,6.719,4.727,6.303,5.544,5.325,7.623,3.539,2.669,8.757,8.262,4.517,8.093,3.643,7.596,5.766,3.989,2.611,3.692,6.299,5.361,5.097,5.003,0.4152,8.344,9.81,3.59,5.931,5.702,3.671,4.706,2.216,4.025,0.8599,3.933,3.471,5.999,5.358,8.976,5.015,2.405,2.201,8.329,9.531,1.634,7.166,4.059,7.734,4.435,7.328,6.096,4.569,5.089,6.542,5.072,3.726,2.947,4.102,6.809,3.282,4.01,8.102,2.944,6.322,6.26,3.543,5.604,2.751,5.159,4.438,7.469,7.783,3.819,7.965,4.754,7.027,6.63,6.463,6.56,5.377,6.169,2.275,6.927,4.432,7.515,4.206,5.585,2.262,1.252,8.334,8.189,5.796,6.259,6.482,4.86,5.02,4.748,2.267,2.5,3.463,4.232,6.388,6.03,2.314,7.182,7.086,7.713,8.988,5.953,6.986,6.775,6.354,3.173,4.453,4.8,3.936,5.783,2.897,6.76,3.45,3.549,6.605,5.704,6.567,3.781,5.046,8.323,9.103,6.999,3.437,4.973,8.359,5.501,5.596,4.111,3.338,6.074,5.295,6.14,7.156,6.883,7.572,4.046,3.625,6.522,4.159,7.176,8.043,7.702,3.082,2.155,1.747,7.331,5.466,6.375,6.3,7.111,6.691,6.419,5.83,2.022,9.104,4.876,5.13,8.244,4.041,8.234,5.491,5.178,6.214,3.419,10.07,1.835,0.223,6.121,4.654,6.803,1.625,7.206,6.641,3.168,3.962,1.302,4.166,7.107,6.862,3.422,7.353,5.172,3.62,5.984,4.64,6.917,6.266,5.871,6.772,7.897,3.785,1.753,4.863,5.792,4.267,3.657,5.681,4.144,6.836,8.02,7.428,8.155,5.025,6.567,8.054,4.966,3.039,3.863,6.742,5.561,7.916,7.206,7.477,3.894,4.964,5.515,1.235,5.888,4.671,5.733,7.78,7.769,1.769,3.477,2.212,4.196,2.053,4.141,5.425,2.961,3.307,0.1617,-0.8536,-0.1349,-0.8676,1.824,1.744,2.724,2.332,-1.03,-0.884,-1.52,3.592,3.819,7.014,5.943,2.674,3.883,6.541,5.873,5.791,9.911,5.817,3.271,6.726,6.227,3.098,3.819,6.73,2.753,8.149,3.045,5.256,5.429,6.008,3.53,2.977,2.667,1.149,7.269,5.247,3.814,6.561,5.914,8.196,4.192,6.488,8.662,6.574,8.699,8.007,6.834,4.656,4.842,6.461,2.108,2.078,4.453,1.483,8.788,8.089,6.847,7.625,7.73,3.513,3.872,7.129,6.444,7.333,4.66,4.024,4.266,6.987,6.908,3.74,2.797,5.909,5.764,7.834,3.857,5.887,5.071,8.989,2.504,5.589,4.759,6.723,4.349,7.999,3.391,2.452,4.976,3.986,3.569,8.815,5.892,6.49,3.424,4.165,6.545,4.882,10.01,4.345,4.011,4.444,2.059,6.575,8.492,7.034,2.952,7.089,7.257,2.926,7.808,4.694,5.375,6.765,6.892,6.082,6.495,7.016,5.904,7.316,5.655,4.174,3.838,3.994,5.003,4.377,5.911,5.672,6.224,5.352,7.525,4.058,4.96,5.446,3.574,3.648,5.304,1.838,5.327,3.057,5.517,6.475,7.502,6.123,5.866,3.486,6.851,6.299,8.23,2.28,7.937,8.293,5.374,0.9846,6.013,6.251,6.81,1.848,5.01,5.034,5.473,4.75,9.663,6.305,5.056,6.818,7.835,5.261,7.909,8.519,6.959,4.402,5.96,5.501,1.508,2.585,4.336,5.491,2.031,4.098,5.994,9.163,6.421,5.581,6.519,5.884,7.724,5.265,7.089,8.29,6.834,9.178,5.297,7.164,5.9,6.678,5.429,12.51,2.059,1.099,0.4288,6.553,6.732,8.002,5.174,9.55,7.976,5.586,10.95,4.408,3.378,4.443,1.778,4.144,7.931,5.481,4.033,6.531,3.724,6.244,10.1,4.989,9.349,7.605,7.429,6.063,4.807,5.669,6.718,6.738,4.131,1.478,3.734,3.133,4.22,4.288,3.602,6.98,7.395,4.975,2.93,4.997,2.913,1.364,4.577,5.503,5.129,6.363,6.213,3.812,8.501,5.304,6.903,3.407,3.748,3.621,7.932,6.64,8.072,5.446,4.809,4.906,3.223,3.179,2.152,8.684,7.94,1.219,5.728,8.806,4.498,6.554,5.303,5.425,8.478,6.584,6.622,8.396,3.46,6.524,3.769,5.078,5.498,9.822,2.598,7.941,7.823,4.886,6.683,2.79,7.01,8.185,7.952,5.683,6.276,2.999,2.886,5.438,5.974,5.182,3.133,2.606,3.87,2.407,4.541,4.838,5.018,6.499,5.107,7.27,6.999,7.145,5.575,7.943,7.324,2.937,-0.3626,-0.3096,-0.362,0.2076,1.399,2.856,2.517,1.137,0.7913,7.64,4.962,4.711,4.52,3.525,1.91,1.648,5.601,4.351,2.325,4.699,3.116,0.7385,5.117,5.827,4.98,3.129,2.243,6.541,7.539,5.819,3.455,4.142,9.017,6.529,9.251,1.033,7.359,5.912,4.375,6.125,7.543,5.993,8.324,3.739,7.634,3.302,8.187,3.337,4.722,1.365,6.158,3.274,2.976,4.952,5.892,5.566,5.287,5.457,5.499,3.982,3.051,7.072,4.944,4.843,3.391,3.246,6.302,5.875,7.2,3.694,2.562,2.682,6.982,4.892,1.467,0.6758,5.747,3.443,3.239,6.91,8.072,8.872,10.84,7.291,4.715,3.69,3.635,-0.02986,2.382,5.357,10.47,2.872,4.801,3.269,6.653,3.522,5.714,2.592,4.209,6.568,5.148,4.297,4.089,3.898,6.538,8.467,3.624,4.85,4.315,4.456,7.831,4.897,8.108,5.058,7.838,6.267,6.026,6.882,8.075,5.746,7.045,2.971,7.001,6.057,2.664,8.337,5.811,7.371,7.54,6.979,6.957,6.804,8.228,0.6961,5.132,4.765,4.223,3.28,6.416,6.334,1.719,4.853,5.51,1.15,7.898,5.365,3.419,6.89,7.127,7.504,8.447,6.771,8.201,6.424,4.451,7.799,5.168,9.1,4.164,3.372,0.7869,4.483,6.043,4.713,7.599,4.675,4.915,7.546,5.671,7.913,6.103,7.227,7.332,7.004,6.4,2.12,5.305,7.365,7.698,6.083,8.214,1.925,4.933,4.148,7.235,8.79,5.172,1.806,4.104,7.333,7.433,4.281,6.313,5.87,8.351,7.077,4.101,7.294,1.703,10.27,2.153,1.658,6.116,7.676,3.712,7.432,6.533,5.773,3.968,6.145,9.372,9.014,2.553,7.543,5.031,7.08,3.422,7.546,4.604,1.845,3.707,4.485,4.843,4.337,3.766,3.548,2.9,5.821,2.843,7.806,3.641,6.803,4.514,6.528,7.023,5.836,7.89,8.142,2.252,2.393,4.83,7.691,7.553,3.315,6.722,2.63,6.273,4.744,2.79,8.734,6.828,7.371,6.227,8.572,4.232,8.711,9.406,-1.21,3.956,3.704,5.191,5.466,5.309,5.69,4.721,3.421,4.538,5.389,8.067,2.576,1.532,5.936,4.363,4.061,5.994,7.237,8.576,6.298,7.929,4.586,8.528,5.859,7.65,9.853,4.735,3.037,2.675,2.69,2.758,3.53,9.278,6.328,5.931,3.114,3.786,4.578,7.438,5.955,6.135,7.361,5.559,7.954,6.463,6.091,5.326,5.692,4.91,3.938,7.257,5.384,5.741,8.058,9.116,7.112,3.12,7.277,5.286,4.72,5.978,7.367,7.929,8.785,5.958,6.835,4.291,5.322,5.78,3.715,6.602,6.507,8.177,7.458,7.57,5.806,5.925,6.09,3.579,6.751,7.05,5.147,6.872,3.809,5.346,4.693,6.536,8.867,5.515,5.838,7.592,4.32,1.949,4.059,6.555,6.407,6.485,4.236,6.513,6.906,8.526,6.418,6.676,6.408,6.857,6.253,5.007,6.022,5.997,7.117,4.866,5.991,7.207,6.857,7.458,7.407,3.109,7.691,8.05,6.174,0.9538,5.436,5.823,7.359,5.077,7.925,6.1,5.919,6.967,7.113,3.854,7.895,3.907,3.745,7.076,7.616,7.114,6.439,6.718,4.149,5.443,7.03,6.928,7.133,8.023,5.214,0.4846,5.233,2.509,4.315,3.628,6.066,6.379,1.732,4.748,3.906,-0.01682,6.389,9.068,5.244,8.319,8.081,6.218,8.89,7.708,5.928,7.744,5.23,5.159,7.326,1.765,4.681,7.059,3.989,5.902,4.314,7.091,6.899,3.985,2.669,4.71,7.451,5.666,5.828,5.431,5.646,6.09,7.838,5.137,4.485,4.714,2.95,5.29,6.654,2.871,5.179,9.246,5.655,8.268,4.829,6.149,2.603,7.905,4.257,3.628,5.756,2.349,9.448,5.136,9.294,3.683,7.632,6.94,3.926,7.338,3.262,5.03,1.534,4.778,2.216,5.558,6.567,7.875,6.402,3.003,9.144,0.3385,2.45,5.002,4.072,5.359,1.42,3.461,0.2737,0.6543,1.384,1.086,4.838,-0.05124,0.4745,2.314,1.162,1.868,2.483,2.313,1.061,4.614,-0.2797,1.655,-0.5784,1.528,4.542,-1.19,6.535,8.006,2.486,6.316,5.847],"cols":["#bebebe","#bebebe","#bebebe","#bebebe","#bebebe","#bebebe","#bebebe","#bebebe","#bebebe","#bebebe","#bebebe","#bebebe","#bebebe","#bebebe","#bebebe","#bebebe","#bebebe","#bebebe","#bebebe","#bebebe","#bebebe","#bebebe","#bebebe","#bebebe","#bebebe","#bebebe","#bebebe","#bebebe","#bebebe","#bebebe","#bebebe","#bebebe","#bebebe","#bebebe","#bebebe","#bebebe","#bebebe","#bebebe","#bebebe","#bebebe","#bebebe","#bebebe","#bebebe","#bebebe","#bebebe","#bebebe","#bebebe","#bebebe","#bebebe","#bebebe","#bebebe","#bebebe","#bebebe","#bebebe","#bebebe","#bebebe","#bebebe","#bebebe","#bebebe","#bebebe","#bebebe","#bebebe","#bebebe","#bebebe","#bebebe","#bebebe","#bebebe","#bebebe","#bebebe","#bebebe","#bebebe","#bebebe","#bebebe","#bebebe","#bebebe","#bebebe","#bebebe","#bebebe","#bebebe","#bebebe","#bebebe","#bebebe","#bebebe","#bebebe","#bebebe","#bebebe","#bebebe","#bebebe","#bebebe","#bebebe","#bebebe","#bebebe","#bebebe","#bebebe","#bebebe","#bebebe","#bebebe","#bebebe","#bebebe","#bebebe","#bebebe","#bebebe","#bebebe","#bebebe","#bebebe","#bebebe","#bebebe","#bebebe","#bebebe","#bebebe","#bebebe","#bebebe","#bebebe","#bebebe","#bebebe","#bebebe","#bebebe","#bebebe","#bebebe","#bebebe","#bebebe","#bebebe","#bebebe","#bebebe","#bebebe","#bebebe","#bebebe","#bebebe","#bebebe","#bebebe","#bebebe","#bebebe","#bebebe","#bebebe","#bebebe","#bebebe","#bebebe","#bebebe","#bebebe","#bebebe","#bebebe","#bebebe","#bebebe","#bebebe","#bebebe","#bebebe","#bebebe","#bebebe","#bebebe","#bebebe","#bebebe","#bebebe","#bebebe","#bebebe","#bebebe","#bebebe","#bebebe","#bebebe","#bebebe","#bebebe","#bebebe","#bebebe","#bebebe","#bebebe","#bebebe","#bebebe","#bebebe","#bebebe","#bebebe","#bebebe","#bebebe","#bebebe","#bebebe","#bebebe","#bebebe","#bebebe","#bebebe","#bebebe","#bebebe","#bebebe","#bebebe","#bebebe","#bebebe","#bebebe","#bebebe","#bebebe","#bebebe","#bebebe","#bebebe","#bebebe","#bebebe","#bebebe","#bebebe","#bebebe","#bebebe","#bebebe","#bebebe","#bebebe","#bebebe","#bebebe","#bebebe","#bebebe","#bebebe","#bebebe","#bebebe","#bebebe","#bebebe","#bebebe","#bebebe","#bebebe","#bebebe","#bebebe","#bebebe","#bebebe","#bebebe","#bebebe","#bebebe","#bebebe","#bebebe","#bebebe","#bebebe","#bebebe","#bebebe","#bebebe","#bebebe","#bebebe","#bebebe","#bebebe","#bebebe","#bebebe","#bebebe","#bebebe","#bebebe","#bebebe","#bebebe","#bebebe","#bebebe","#bebebe","#bebebe","#bebebe","#bebebe","#bebebe","#bebebe","#bebebe","#bebebe","#bebebe","#bebebe","#bebebe","#bebebe","#bebebe","#bebebe","#bebebe","#bebebe","#bebebe","#bebebe","#bebebe","#bebebe","#bebebe","#bebebe","#bebebe","#bebebe","#bebebe","#bebebe","#bebebe","#bebebe","#bebebe","#bebebe","#bebebe","#bebebe","#bebebe","#bebebe","#bebebe","#bebebe","#bebebe","#bebebe","#bebebe","#bebebe","#bebebe","#bebebe","#bebebe","#bebebe","#bebebe","#bebebe","#bebebe","#bebebe","#bebebe","#bebebe","#bebebe","#bebebe","#bebebe","#bebebe","#bebebe","#bebebe","#bebebe","#bebebe","#bebebe","#bebebe","#bebebe","#bebebe","#bebebe","#bebebe","#bebebe","#bebebe","#bebebe","#bebebe","#bebebe","#bebebe","#bebebe","#bebebe","#bebebe","#bebebe","#bebebe","#bebebe","#bebebe","#bebebe","#bebebe","#bebebe","#bebebe","#bebebe","#bebebe","#bebebe","#bebebe","#bebebe","#bebebe","#bebebe","#bebebe","#bebebe","#bebebe","#bebebe","#bebebe","#bebebe","#bebebe","#bebebe","#bebebe","#bebebe","#bebebe","#bebebe","#bebebe","#bebebe","#bebebe","#bebebe","#bebebe","#bebebe","#bebebe","#bebebe","#bebebe","#bebebe","#bebebe","#bebebe","#bebebe","#bebebe","#bebebe","#bebebe","#bebebe","#bebebe","#bebebe","#bebebe","#bebebe","#bebebe","#bebebe","#bebebe","#bebebe","#bebebe","#bebebe","#bebebe","#bebebe","#bebebe","#bebebe","#bebebe","#bebebe","#bebebe","#bebebe","#bebebe","#bebebe","#bebebe","#bebebe","#bebebe","#bebebe","#bebebe","#bebebe","#bebebe","#bebebe","#bebebe","#bebebe","#bebebe","#bebebe","#bebebe","#bebebe","#bebebe","#bebebe","#bebebe","#bebebe","#bebebe","#bebebe","#bebebe","#bebebe","#bebebe","#bebebe","#bebebe","#bebebe","#bebebe","#bebebe","#bebebe","#bebebe","#bebebe","#bebebe","#bebebe","#bebebe","#bebebe","#bebebe","#bebebe","#bebebe","#bebebe","#bebebe","#bebebe","#bebebe","#bebebe","#bebebe","#bebebe","#bebebe","#bebebe","#bebebe","#bebebe","#bebebe","#bebebe","#bebebe","#bebebe","#bebebe","#bebebe","#bebebe","#bebebe","#bebebe","#bebebe","#bebebe","#bebebe","#bebebe","#bebebe","#bebebe","#bebebe","#bebebe","#bebebe","#bebebe","#bebebe","#bebebe","#bebebe","#bebebe","#bebebe","#bebebe","#bebebe","#bebebe","#bebebe","#bebebe","#bebebe","#bebebe","#bebebe","#bebebe","#bebebe","#bebebe","#bebebe","#bebebe","#bebebe","#bebebe","#bebebe","#bebebe","#bebebe","#bebebe","#bebebe","#bebebe","#bebebe","#bebebe","#bebebe","#bebebe","#bebebe","#bebebe","#bebebe","#bebebe","#bebebe","#bebebe","#bebebe","#bebebe","#bebebe","#bebebe","#bebebe","#bebebe","#bebebe","#bebebe","#bebebe","#bebebe","#bebebe","#bebebe","#bebebe","#bebebe","#bebebe","#bebebe","#bebebe","#bebebe","#bebebe","#bebebe","#bebebe","#bebebe","#bebebe","#bebebe","#bebebe","#bebebe","#bebebe","#bebebe","#bebebe","#bebebe","#bebebe","#bebebe","#bebebe","#bebebe","#bebebe","#bebebe","#bebebe","#bebebe","#bebebe","#bebebe","#bebebe","#bebebe","#bebebe","#bebebe","#bebebe","#bebebe","#bebebe","#bebebe","#bebebe","#bebebe","#bebebe","#bebebe","#bebebe","#bebebe","#bebebe","#bebebe","#bebebe","#bebebe","#bebebe","#bebebe","#bebebe","#bebebe","#bebebe","#bebebe","#bebebe","#bebebe","#bebebe","#bebebe","#bebebe","#bebebe","#bebebe","#bebebe","#bebebe","#bebebe","#bebebe","#bebebe","#bebebe","#bebebe","#bebebe","#bebebe","#bebebe","#bebebe","#bebebe","#bebebe","#bebebe","#bebebe","#bebebe","#bebebe","#bebebe","#bebebe","#bebebe","#bebebe","#bebebe","#bebebe","#bebebe","#bebebe","#bebebe","#bebebe","#bebebe","#bebebe","#bebebe","#bebebe","#bebebe","#bebebe","#bebebe","#bebebe","#bebebe","#bebebe","#bebebe","#bebebe","#bebebe","#bebebe","#bebebe","#bebebe","#bebebe","#bebebe","#bebebe","#bebebe","#bebebe","#bebebe","#bebebe","#bebebe","#bebebe","#bebebe","#bebebe","#bebebe","#bebebe","#bebebe","#bebebe","#bebebe","#bebebe","#bebebe","#bebebe","#bebebe","#bebebe","#bebebe","#bebebe","#bebebe","#bebebe","#bebebe","#bebebe","#bebebe","#bebebe","#bebebe","#bebebe","#bebebe","#bebebe","#bebebe","#bebebe","#bebebe","#bebebe","#bebebe","#bebebe","#bebebe","#bebebe","#bebebe","#bebebe","#bebebe","#bebebe","#bebebe","#bebebe","#bebebe","#bebebe","#bebebe","#bebebe","#bebebe","#bebebe","#bebebe","#bebebe","#bebebe","#bebebe","#bebebe","#bebebe","#bebebe","#bebebe","#bebebe","#bebebe","#bebebe","#bebebe","#bebebe","#bebebe","#bebebe","#bebebe","#bebebe","#bebebe","#bebebe","#bebebe","#bebebe","#bebebe","#bebebe","#bebebe","#bebebe","#bebebe","#bebebe","#bebebe","#bebebe","#bebebe","#bebebe","#bebebe","#bebebe","#bebebe","#bebebe","#bebebe","#bebebe","#bebebe","#bebebe","#bebebe","#bebebe","#bebebe","#bebebe","#bebebe","#bebebe","#bebebe","#bebebe","#bebebe","#bebebe","#bebebe","#bebebe","#bebebe","#bebebe","#bebebe","#bebebe","#bebebe","#bebebe","#bebebe","#bebebe","#bebebe","#bebebe","#bebebe","#bebebe","#bebebe","#bebebe","#bebebe","#bebebe","#bebebe","#bebebe","#bebebe","#bebebe","#bebebe","#bebebe","#bebebe","#bebebe","#bebebe","#bebebe","#bebebe","#bebebe","#bebebe","#bebebe","#bebebe","#bebebe","#bebebe","#bebebe","#bebebe","#bebebe","#bebebe","#bebebe","#bebebe","#bebebe","#bebebe","#bebebe","#bebebe","#bebebe","#bebebe","#bebebe","#bebebe","#bebebe","#bebebe","#bebebe","#bebebe","#bebebe","#bebebe","#bebebe","#bebebe","#bebebe","#bebebe","#bebebe","#bebebe","#bebebe","#bebebe","#bebebe","#bebebe","#bebebe","#bebebe","#bebebe","#bebebe","#bebebe","#bebebe","#bebebe","#bebebe","#bebebe","#bebebe","#bebebe","#bebebe","#bebebe","#bebebe","#bebebe","#bebebe","#bebebe","#bebebe","#bebebe","#bebebe","#bebebe","#bebebe","#bebebe","#bebebe","#bebebe","#bebebe","#bebebe","#bebebe","#bebebe","#bebebe","#bebebe","#bebebe","#bebebe","#bebebe","#bebebe","#bebebe","#bebebe","#bebebe","#bebebe","#bebebe","#bebebe","#bebebe","#bebebe","#bebebe","#bebebe","#bebebe","#bebebe","#bebebe","#bebebe","#bebebe","#bebebe","#bebebe","#bebebe","#bebebe","#bebebe","#bebebe","#bebebe","#bebebe","#bebebe","#bebebe","#bebebe","#bebebe","#bebebe","#bebebe","#bebebe","#bebebe","#bebebe","#bebebe","#bebebe","#bebebe","#bebebe","#bebebe","#bebebe","#bebebe","#bebebe","#bebebe","#bebebe","#bebebe","#bebebe","#bebebe","#bebebe","#bebebe","#bebebe","#bebebe","#bebebe","#bebebe","#bebebe","#bebebe","#bebebe","#bebebe","#bebebe","#bebebe","#bebebe","#bebebe","#bebebe","#bebebe","#bebebe","#bebebe","#bebebe","#bebebe","#bebebe","#bebebe","#bebebe","#bebebe","#bebebe","#bebebe","#bebebe","#bebebe","#bebebe","#bebebe","#bebebe","#bebebe","#bebebe","#bebebe","#bebebe","#bebebe","#bebebe","#bebebe","#bebebe","#bebebe","#bebebe","#bebebe","#bebebe","#bebebe","#bebebe","#bebebe","#bebebe","#bebebe","#bebebe","#bebebe","#bebebe","#bebebe","#bebebe","#bebebe","#bebebe","#bebebe","#bebebe","#bebebe","#bebebe","#bebebe","#bebebe","#bebebe","#bebebe","#bebebe","#bebebe","#bebebe","#bebebe","#bebebe","#bebebe","#bebebe","#bebebe","#bebebe","#bebebe","#bebebe","#bebebe","#bebebe","#bebebe","#bebebe","#bebebe","#bebebe","#bebebe","#bebebe","#bebebe","#bebebe","#bebebe","#bebebe","#bebebe","#bebebe","#bebebe","#bebebe","#bebebe","#bebebe","#bebebe","#bebebe","#bebebe","#bebebe","#bebebe","#bebebe","#bebebe","#bebebe","#bebebe","#bebebe","#bebebe","#bebebe","#bebebe","#bebebe","#bebebe","#bebebe","#bebebe","#bebebe","#bebebe","#bebebe","#bebebe","#bebebe","#bebebe","#bebebe","#bebebe","#bebebe","#bebebe","#bebebe","#bebebe","#bebebe","#bebebe","#bebebe","#bebebe","#bebebe","#bebebe","#bebebe","#bebebe","#bebebe","#bebebe","#bebebe","#bebebe","#bebebe","#bebebe","#bebebe","#bebebe","#bebebe","#bebebe","#bebebe","#bebebe","#bebebe","#bebebe","#bebebe","#bebebe","#bebebe","#bebebe","#bebebe","#bebebe","#bebebe","#bebebe","#bebebe","#bebebe","#bebebe","#bebebe","#bebebe","#bebebe","#bebebe","#bebebe","#bebebe","#bebebe","#bebebe","#bebebe","#bebebe","#bebebe","#bebebe","#bebebe","#bebebe","#bebebe","#bebebe","#bebebe","#bebebe","#bebebe","#bebebe","#bebebe","#bebebe","#bebebe","#bebebe","#bebebe","#bebebe","#bebebe","#bebebe","#bebebe","#bebebe","#bebebe","#bebebe","#bebebe","#bebebe","#bebebe","#bebebe","#bebebe","#bebebe","#bebebe","#bebebe","#bebebe","#bebebe","#bebebe","#bebebe","#bebebe","#bebebe","#bebebe","#bebebe","#bebebe","#bebebe","#bebebe","#bebebe","#bebebe","#bebebe","#bebebe","#bebebe","#bebebe","#bebebe","#bebebe","#bebebe","#bebebe","#bebebe","#bebebe","#bebebe","#bebebe","#bebebe","#bebebe","#bebebe","#bebebe","#bebebe","#bebebe","#bebebe","#bebebe","#bebebe","#bebebe","#bebebe","#bebebe","#bebebe","#bebebe","#bebebe","#bebebe","#bebebe","#bebebe","#bebebe","#bebebe","#bebebe","#bebebe","#bebebe","#bebebe","#bebebe","#bebebe","#bebebe","#bebebe","#bebebe","#bebebe","#bebebe","#bebebe","#bebebe","#bebebe","#bebebe","#bebebe","#bebebe","#bebebe","#bebebe","#bebebe","#bebebe","#bebebe","#bebebe","#bebebe","#bebebe","#bebebe","#bebebe","#bebebe","#bebebe","#bebebe","#bebebe","#bebebe","#bebebe","#bebebe","#bebebe","#bebebe","#bebebe","#bebebe","#bebebe","#bebebe","#bebebe","#bebebe","#bebebe","#bebebe","#bebebe","#bebebe","#bebebe","#bebebe","#bebebe","#bebebe","#bebebe","#bebebe","#bebebe","#bebebe","#bebebe","#bebebe","#bebebe","#bebebe","#bebebe","#bebebe","#bebebe","#bebebe","#bebebe","#bebebe","#bebebe","#bebebe","#bebebe","#bebebe","#bebebe","#bebebe","#bebebe","#bebebe","#bebebe","#bebebe","#bebebe","#bebebe","#bebebe","#bebebe","#bebebe","#bebebe","#bebebe","#bebebe","#bebebe","#bebebe","#bebebe","#bebebe","#bebebe","#bebebe","#bebebe","#bebebe","#bebebe","#bebebe","#bebebe","#bebebe","#bebebe","#bebebe","#bebebe","#bebebe","#bebebe","#bebebe","#bebebe","#bebebe","#bebebe","#bebebe","#bebebe","#bebebe","#bebebe","#bebebe","#bebebe","#bebebe","#bebebe","#bebebe","#bebebe","#bebebe","#bebebe","#bebebe","#bebebe","#bebebe","#bebebe","#bebebe","#bebebe","#bebebe","#bebebe","#bebebe","#bebebe","#bebebe","#bebebe","#bebebe","#bebebe","#bebebe","#bebebe","#bebebe","#bebebe","#bebebe","#bebebe","#bebebe","#bebebe","#bebebe","#bebebe","#bebebe","#bebebe","#bebebe","#bebebe","#bebebe","#bebebe","#bebebe","#bebebe","#bebebe","#bebebe","#bebebe","#bebebe","#bebebe","#bebebe","#bebebe","#bebebe","#bebebe","#bebebe","#bebebe","#bebebe","#bebebe","#bebebe","#bebebe","#bebebe","#bebebe","#bebebe","#bebebe","#bebebe","#bebebe","#bebebe","#bebebe","#bebebe","#bebebe","#bebebe","#bebebe","#bebebe","#bebebe","#bebebe","#bebebe","#bebebe","#bebebe","#bebebe","#bebebe","#bebebe","#bebebe","#bebebe","#bebebe","#bebebe","#bebebe","#bebebe","#bebebe","#bebebe","#bebebe","#bebebe","#bebebe","#bebebe","#bebebe","#bebebe","#bebebe","#bebebe","#bebebe","#bebebe","#bebebe","#bebebe","#bebebe","#bebebe","#bebebe","#bebebe","#bebebe","#bebebe","#bebebe","#bebebe","#bebebe","#bebebe","#bebebe","#bebebe","#bebebe","#bebebe","#bebebe","#bebebe","#bebebe","#bebebe","#bebebe","#bebebe","#bebebe","#bebebe","#bebebe","#bebebe","#bebebe","#bebebe","#bebebe","#bebebe","#bebebe","#bebebe","#bebebe","#bebebe","#bebebe","#bebebe","#bebebe","#bebebe","#bebebe","#bebebe","#bebebe","#bebebe","#bebebe","#bebebe","#bebebe","#bebebe","#bebebe","#bebebe","#bebebe","#bebebe","#bebebe","#bebebe","#bebebe","#bebebe","#bebebe","#bebebe","#bebebe","#bebebe","#bebebe","#bebebe","#bebebe","#bebebe","#bebebe","#bebebe","#bebebe","#bebebe","#bebebe","#bebebe","#bebebe","#bebebe","#bebebe","#bebebe","#bebebe","#bebebe","#bebebe","#bebebe","#bebebe","#bebebe","#bebebe","#bebebe","#bebebe","#bebebe","#bebebe","#bebebe","#bebebe","#bebebe","#bebebe","#bebebe","#bebebe","#bebebe","#bebebe","#bebebe","#bebebe","#bebebe","#bebebe","#bebebe","#bebebe","#bebebe","#bebebe","#bebebe","#bebebe","#bebebe","#bebebe","#bebebe","#bebebe","#bebebe","#bebebe","#bebebe","#bebebe","#bebebe","#bebebe","#bebebe","#bebebe","#bebebe","#bebebe","#bebebe","#bebebe","#bebebe","#bebebe","#bebebe","#bebebe","#bebebe","#bebebe","#bebebe","#bebebe","#bebebe","#bebebe","#bebebe","#bebebe","#bebebe","#bebebe","#bebebe","#bebebe","#bebebe","#bebebe","#bebebe","#bebebe","#bebebe","#bebebe","#bebebe","#bebebe","#bebebe","#bebebe","#bebebe","#bebebe","#bebebe","#bebebe","#bebebe","#bebebe","#bebebe","#bebebe","#bebebe","#bebebe","#bebebe","#bebebe","#bebebe","#bebebe","#bebebe","#bebebe","#bebebe","#bebebe","#bebebe","#bebebe","#bebebe","#bebebe","#bebebe","#bebebe","#bebebe","#bebebe","#bebebe","#bebebe","#bebebe","#bebebe","#bebebe","#bebebe","#bebebe","#bebebe","#bebebe","#bebebe","#bebebe","#bebebe","#bebebe","#bebebe","#bebebe","#bebebe","#bebebe","#bebebe","#bebebe","#bebebe","#bebebe","#bebebe","#bebebe","#bebebe","#bebebe","#bebebe","#bebebe","#bebebe","#bebebe","#bebebe","#bebebe","#bebebe","#bebebe","#bebebe","#bebebe","#bebebe","#bebebe","#bebebe","#bebebe","#bebebe","#bebebe","#bebebe","#bebebe","#bebebe","#bebebe","#bebebe","#bebebe","#bebebe","#bebebe","#bebebe","#bebebe","#bebebe","#bebebe","#bebebe","#bebebe","#bebebe","#bebebe","#bebebe","#bebebe","#bebebe","#bebebe","#bebebe","#bebebe","#bebebe","#bebebe","#bebebe","#bebebe","#bebebe","#bebebe","#bebebe","#bebebe","#bebebe","#bebebe","#bebebe","#bebebe","#bebebe","#bebebe","#bebebe","#bebebe","#bebebe","#bebebe","#bebebe","#bebebe","#bebebe","#bebebe","#bebebe","#bebebe","#bebebe","#bebebe","#bebebe","#bebebe","#bebebe","#bebebe","#bebebe","#bebebe","#bebebe","#bebebe","#bebebe","#bebebe","#bebebe","#bebebe","#bebebe","#bebebe","#bebebe","#bebebe","#bebebe","#bebebe","#bebebe","#bebebe","#bebebe","#bebebe","#bebebe","#bebebe","#bebebe","#bebebe","#bebebe","#bebebe","#bebebe","#bebebe","#bebebe","#bebebe","#bebebe","#bebebe","#bebebe","#bebebe","#bebebe","#bebebe","#bebebe","#bebebe","#bebebe","#bebebe","#bebebe","#bebebe","#bebebe","#bebebe","#bebebe","#bebebe","#bebebe","#bebebe","#bebebe","#bebebe","#bebebe","#bebebe","#bebebe","#bebebe","#bebebe","#bebebe","#bebebe","#bebebe","#bebebe","#bebebe","#bebebe","#bebebe","#bebebe","#bebebe","#bebebe","#bebebe","#bebebe","#bebebe","#bebebe","#bebebe","#bebebe","#bebebe","#bebebe","#bebebe","#bebebe","#bebebe","#bebebe","#bebebe","#bebebe","#bebebe","#bebebe","#bebebe","#bebebe","#bebebe","#bebebe","#bebebe","#bebebe","#bebebe","#bebebe","#bebebe","#bebebe","#bebebe","#bebebe","#bebebe","#bebebe","#bebebe","#bebebe","#bebebe","#bebebe","#bebebe","#bebebe","#bebebe","#bebebe","#bebebe","#bebebe","#bebebe","#bebebe","#bebebe","#bebebe","#bebebe","#bebebe","#bebebe","#bebebe","#bebebe","#bebebe","#bebebe","#bebebe","#bebebe","#bebebe","#bebebe","#bebebe","#bebebe","#bebebe","#bebebe","#bebebe","#bebebe","#bebebe","#bebebe","#bebebe","#bebebe","#bebebe","#bebebe","#bebebe","#bebebe","#bebebe","#bebebe","#bebebe","#bebebe","#bebebe","#bebebe","#bebebe","#bebebe","#bebebe","#bebebe","#bebebe","#bebebe","#bebebe","#bebebe","#bebebe","#bebebe","#bebebe","#bebebe","#bebebe","#bebebe","#bebebe","#bebebe","#bebebe","#bebebe","#bebebe","#bebebe","#bebebe","#bebebe","#bebebe","#bebebe","#bebebe","#bebebe","#bebebe","#bebebe","#bebebe","#bebebe","#bebebe","#bebebe","#bebebe","#bebebe","#bebebe","#bebebe","#bebebe","#bebebe","#bebebe","#bebebe","#bebebe","#bebebe","#bebebe","#bebebe","#bebebe","#bebebe","#bebebe","#bebebe","#bebebe","#bebebe","#bebebe","#bebebe","#bebebe","#bebebe","#bebebe","#bebebe","#bebebe","#bebebe","#bebebe","#bebebe","#bebebe","#bebebe","#bebebe","#bebebe","#bebebe","#bebebe","#bebebe","#bebebe","#bebebe","#bebebe","#bebebe","#bebebe","#bebebe","#bebebe","#bebebe","#bebebe","#bebebe","#bebebe","#bebebe","#bebebe","#bebebe","#bebebe","#bebebe","#bebebe","#bebebe","#bebebe","#bebebe","#bebebe","#bebebe","#bebebe","#bebebe","#bebebe","#bebebe","#bebebe","#bebebe","#bebebe","#bebebe","#bebebe","#bebebe","#bebebe","#bebebe","#bebebe","#bebebe","#bebebe","#bebebe","#bebebe","#bebebe","#bebebe","#bebebe","#bebebe","#bebebe","#bebebe","#bebebe","#bebebe","#bebebe","#bebebe","#bebebe","#bebebe","#bebebe","#000000","#000000","#000000","#000000","#000000","#000000","#000000","#000000","#000000","#000000","#000000","#000000","#000000","#000000","#000000","#000000","#000000","#000000","#000000","#000000","#000000","#000000","#ff0000","#000000","#000000","#000000","#ff0000","#000000","#000000","#000000","#000000","#000000","#000000","#000000","#000000","#000000","#000000","#000000","#000000","#000000","#000000","#000000","#000000","#000000","#000000","#000000","#000000","#000000","#000000","#000000","#000000","#000000","#000000","#000000","#000000","#000000","#000000","#000000","#000000","#000000","#000000","#000000","#000000","#000000","#000000","#000000","#000000","#000000","#ff0000","#ff0000","#000000","#000000","#000000","#ff0000","#ff0000","#000000","#000000","#000000","#000000","#000000","#000000","#000000","#000000","#000000","#000000","#000000","#000000","#000000","#000000","#000000","#000000","#000000","#000000","#000000","#000000","#000000","#000000","#000000","#000000","#000000","#000000","#000000","#000000","#000000","#000000","#000000","#000000","#ff0000","#000000","#000000","#ff0000","#000000","#000000","#000000","#000000","#000000","#000000","#ff0000","#000000","#000000","#ff0000","#ff0000","#000000","#000000","#000000","#000000","#ff0000","#000000","#000000","#000000","#000000","#ff0000","#ff0000","#ff0000","#000000","#000000","#000000","#000000","#000000","#ff0000","#000000","#ff0000","#000000","#ff0000","#000000","#ff0000","#000000","#000000","#000000","#000000","#ff0000","#000000","#000000","#000000","#000000","#000000","#000000","#ff0000","#ff0000","#ff0000","#ff0000","#000000","#ff0000","#000000","#000000","#ff0000","#ff0000","#ff0000","#ff0000","#ff0000","#ff0000","#ff0000","#ff0000","#ff0000","#ff0000","#ff0000","#ff0000","#ff0000","#000000","#ff0000","#000000","#ff0000","#000000","#000000","#000000","#ff0000","#ff0000","#ff0000","#000000","#000000","#ff0000","#000000","#000000","#000000","#ff0000","#ff0000","#ff0000","#ff0000","#ff0000","#ff0000","#ff0000","#ff0000","#ff0000","#ff0000","#ff0000","#000000","#ff0000","#ff0000","#000000","#000000","#ff0000","#000000","#000000","#ff0000","#000000","#ff0000","#ff0000","#000000","#000000","#000000","#ff0000","#ff0000","#ff0000","#ff0000","#000000","#ff0000","#ff0000","#ff0000","#ff0000","#ff0000","#ff0000","#000000","#000000","#000000","#000000","#000000","#000000","#000000","#000000","#000000","#000000","#000000","#000000","#000000","#000000","#000000","#000000","#000000","#000000","#000000","#000000","#000000","#000000","#000000","#000000","#000000","#000000","#000000","#000000","#000000","#000000","#ff0000","#000000","#000000","#000000","#000000","#000000","#ff0000","#000000","#ff0000","#ff0000","#ff0000","#000000","#000000","#ff0000","#ff0000","#ff0000","#000000","#000000","#000000","#ff0000","#000000","#ff0000","#000000","#000000","#ff0000","#ff0000","#ff0000","#000000","#ff0000","#000000","#000000","#000000","#000000","#ff0000","#000000","#000000","#ff0000","#000000","#ff0000","#ff0000","#ff0000","#ff0000","#ff0000","#000000","#ff0000","#ff0000","#ff0000","#000000","#ff0000","#ff0000","#ff0000","#000000","#000000","#ff0000","#000000","#000000","#000000","#000000","#000000","#000000","#000000","#000000","#000000","#ff0000","#ff0000","#ff0000","#000000","#000000","#000000","#ff0000","#ff0000","#ff0000","#000000","#ff0000","#ff0000","#ff0000","#000000","#ff0000","#000000","#000000","#000000","#000000","#000000","#000000","#000000","#ff0000","#000000","#000000","#000000","#000000","#000000","#000000","#000000","#000000","#000000","#ff0000","#000000","#ff0000","#ff0000","#ff0000","#000000","#000000","#ff0000","#ff0000","#ff0000","#ff0000","#ff0000","#ff0000","#ff0000","#ff0000","#000000","#ff0000","#ff0000","#ff0000","#ff0000","#000000","#000000","#ff0000","#000000","#ff0000","#ff0000","#ff0000","#ff0000","#000000","#000000","#ff0000","#000000","#ff0000","#ff0000","#ff0000","#ff0000","#ff0000","#ff0000","#ff0000","#ff0000","#ff0000","#ff0000","#ff0000","#ff0000","#ff0000","#ff0000","#ff0000","#ff0000","#ff0000","#ff0000","#000000","#000000","#ff0000","#ff0000","#ff0000","#000000","#000000","#000000","#000000","#ff0000","#ff0000","#000000","#ff0000","#ff0000","#000000","#000000","#ff0000","#ff0000","#ff0000","#ff0000","#ff0000","#000000","#ff0000","#ff0000","#000000","#000000","#ff0000","#000000","#ff0000","#ff0000","#ff0000","#ff0000","#ff0000","#000000","#000000","#000000","#ff0000","#000000","#000000","#ff0000","#ff0000","#ff0000","#ff0000","#ff0000","#000000","#ff0000","#ff0000","#ff0000","#000000","#000000","#000000","#ff0000","#ff0000","#ff0000","#ff0000","#ff0000","#ff0000","#ff0000","#ff0000","#ff0000","#ff0000","#ff0000","#ff0000","#ff0000","#000000","#ff0000","#ff0000","#ff0000","#ff0000","#ff0000","#000000","#ff0000","#000000","#ff0000","#000000","#000000","#000000","#000000","#000000","#000000","#000000","#000000","#000000","#000000","#000000","#000000","#000000","#ff0000","#ff0000","#ff0000","#ff0000","#000000","#000000","#000000","#ff0000","#ff0000","#000000","#ff0000","#ff0000","#000000","#000000","#000000","#ff0000","#ff0000","#ff0000","#000000","#000000","#ff0000","#ff0000","#ff0000","#ff0000","#ff0000","#ff0000","#ff0000","#ff0000","#000000","#ff0000","#ff0000","#ff0000","#ff0000","#ff0000","#ff0000","#000000","#000000","#000000","#ff0000","#ff0000","#ff0000","#000000","#000000","#ff0000","#ff0000","#ff0000","#ff0000","#ff0000","#000000","#000000","#000000","#000000","#000000","#ff0000","#ff0000","#000000","#000000","#000000","#ff0000","#ff0000","#ff0000","#ff0000","#000000","#000000","#000000","#ff0000","#000000","#ff0000","#000000","#000000","#000000","#ff0000","#ff0000","#ff0000","#ff0000","#ff0000","#000000","#ff0000","#ff0000","#ff0000","#ff0000","#ff0000","#ff0000","#ff0000","#000000","#000000","#000000","#ff0000","#ff0000","#000000","#000000","#ff0000","#ff0000","#ff0000","#000000","#000000","#ff0000","#ff0000","#ff0000","#000000","#ff0000","#000000","#ff0000","#ff0000","#ff0000","#ff0000","#000000","#000000","#000000","#000000","#000000","#000000","#ff0000","#ff0000","#ff0000","#ff0000","#ff0000","#ff0000","#000000","#ff0000","#ff0000","#000000","#ff0000","#000000","#000000","#000000","#000000","#000000","#000000","#000000","#000000","#000000","#000000","#ff0000","#000000","#000000","#000000","#000000","#000000","#000000","#000000","#000000","#000000","#000000","#000000","#000000","#000000","#000000","#000000","#000000","#ff0000","#000000","#000000","#000000","#ff0000","#ff0000","#ff0000","#ff0000","#ff0000","#000000","#ff0000","#ff0000","#ff0000","#ff0000","#ff0000","#ff0000","#ff0000","#000000","#000000","#000000","#000000","#000000","#000000","#ff0000","#ff0000","#ff0000","#ff0000","#000000","#ff0000","#ff0000","#000000","#000000","#000000","#000000","#000000","#ff0000","#ff0000","#ff0000","#ff0000","#ff0000","#ff0000","#ff0000","#ff0000","#ff0000","#ff0000","#ff0000","#ff0000","#000000","#000000","#000000","#ff0000","#ff0000","#ff0000","#ff0000","#ff0000","#ff0000","#ff0000","#ff0000","#ff0000","#000000","#ff0000","#ff0000","#ff0000","#ff0000","#000000","#000000","#000000","#ff0000","#ff0000","#000000","#000000","#ff0000","#000000","#ff0000","#ff0000","#ff0000","#ff0000","#000000","#000000","#ff0000","#ff0000","#000000","#000000","#000000","#000000","#000000","#000000","#ff0000","#ff0000","#ff0000","#ff0000","#000000","#000000","#000000","#000000","#000000","#ff0000","#ff0000","#ff0000","#000000","#000000","#000000","#ff0000","#ff0000","#000000","#ff0000","#000000","#000000","#000000","#000000","#000000","#000000","#000000","#000000","#ff0000","#000000","#ff0000","#ff0000","#ff0000","#ff0000","#ff0000","#ff0000","#ff0000","#000000","#000000","#ff0000","#000000","#ff0000","#000000","#000000","#000000","#000000","#000000","#000000","#000000","#ff0000","#000000","#000000","#000000","#000000","#000000","#000000","#000000","#000000","#000000","#000000","#000000","#ff0000","#000000","#000000","#000000","#ff0000","#000000","#000000","#ff0000","#ff0000","#ff0000","#ff0000","#ff0000","#ff0000","#ff0000","#ff0000","#000000","#000000","#ff0000","#ff0000","#ff0000","#ff0000","#ff0000","#ff0000","#ff0000","#ff0000","#ff0000","#ff0000","#000000","#000000","#ff0000","#ff0000","#000000","#000000","#000000","#000000","#ff0000","#ff0000","#000000","#ff0000","#ff0000","#ff0000","#000000","#000000","#000000","#000000","#000000","#000000","#000000","#000000","#000000","#000000","#000000","#000000","#000000","#000000","#000000","#000000","#000000","#000000","#000000","#000000","#000000","#ff0000","#000000","#000000","#000000","#000000","#000000","#ff0000","#ff0000","#ff0000","#ff0000","#ff0000","#ff0000","#ff0000","#ff0000","#ff0000","#ff0000","#ff0000","#ff0000","#ff0000","#ff0000","#ff0000","#000000","#ff0000","#000000","#ff0000","#000000","#ff0000","#ff0000","#ff0000","#ff0000","#000000","#ff0000","#ff0000","#ff0000","#ff0000","#ff0000","#000000","#000000","#000000","#ff0000","#000000","#ff0000","#000000","#000000","#000000","#000000","#ff0000","#ff0000","#ff0000","#ff0000","#000000","#000000","#000000","#000000","#000000","#ff0000","#ff0000","#ff0000","#000000","#000000","#000000","#000000","#ff0000","#000000","#000000","#000000","#ff0000","#ff0000","#ff0000","#ff0000","#ff0000","#000000","#ff0000","#ff0000","#000000","#000000","#000000","#000000","#000000","#ff0000","#ff0000","#000000","#ff0000","#ff0000","#ff0000","#ff0000","#ff0000","#ff0000","#ff0000","#ff0000","#000000","#000000","#ff0000","#000000","#000000","#ff0000","#ff0000","#ff0000","#ff0000","#ff0000","#ff0000","#000000","#000000","#ff0000","#ff0000","#ff0000","#ff0000","#ff0000","#000000","#ff0000","#ff0000","#ff0000","#000000","#ff0000","#ff0000","#ff0000","#ff0000","#000000","#000000","#ff0000","#000000","#000000","#ff0000","#ff0000","#ff0000","#000000","#ff0000","#000000","#000000","#000000","#000000","#000000","#000000","#000000","#000000","#ff0000","#ff0000","#000000","#000000","#000000","#000000","#000000","#ff0000","#000000","#000000","#000000","#000000","#000000","#000000","#000000","#000000","#000000","#000000","#000000","#000000","#000000","#000000","#000000","#000000","#000000","#000000","#ff0000","#000000","#000000","#000000","#000000","#000000","#ff0000","#ff0000","#000000","#ff0000","#000000","#000000","#000000","#000000","#000000","#000000","#000000","#000000","#000000","#000000","#000000","#000000","#ff0000","#ff0000","#000000","#000000","#000000","#ff0000","#ff0000","#000000","#000000","#000000","#000000","#000000","#ff0000","#ff0000","#000000","#000000","#000000","#ff0000","#ff0000","#ff0000","#ff0000","#ff0000","#ff0000","#000000","#ff0000","#ff0000","#ff0000","#ff0000","#ff0000","#ff0000","#ff0000","#000000","#ff0000","#000000","#ff0000","#000000","#ff0000","#ff0000","#ff0000","#000000","#000000","#ff0000","#000000","#000000","#ff0000","#ff0000","#ff0000","#ff0000","#000000","#000000","#ff0000","#000000","#000000","#000000","#000000","#ff0000","#ff0000","#ff0000","#ff0000","#ff0000","#ff0000","#000000","#000000","#000000","#ff0000","#ff0000","#ff0000","#ff0000","#ff0000","#ff0000","#000000","#000000","#000000","#ff0000","#ff0000","#ff0000","#ff0000","#ff0000","#000000","#ff0000","#ff0000","#ff0000","#000000","#000000","#000000","#000000","#000000","#ff0000","#ff0000","#000000","#000000","#ff0000","#ff0000","#ff0000","#ff0000","#ff0000","#000000","#ff0000","#ff0000","#000000","#ff0000","#ff0000","#ff0000","#000000","#000000","#000000","#ff0000","#000000","#000000","#000000","#ff0000","#ff0000","#ff0000","#000000","#000000","#ff0000","#ff0000","#ff0000","#ff0000","#000000","#000000","#000000","#ff0000","#000000","#000000","#ff0000","#ff0000","#ff0000","#ff0000","#000000","#000000","#ff0000","#ff0000","#ff0000","#ff0000","#ff0000","#ff0000","#000000","#ff0000","#000000","#000000","#000000","#ff0000","#ff0000","#000000","#000000","#000000","#000000","#000000","#ff0000","#ff0000","#ff0000","#000000","#000000","#000000","#ff0000","#ff0000","#ff0000","#ff0000","#000000","#000000","#ff0000","#000000","#000000","#000000","#000000","#000000","#ff0000","#ff0000","#ff0000","#000000","#ff0000","#000000","#000000","#ff0000","#ff0000","#000000","#000000","#ff0000","#000000","#ff0000","#ff0000","#ff0000","#ff0000","#000000","#000000","#000000","#000000","#ff0000","#ff0000","#ff0000","#000000","#ff0000","#ff0000","#ff0000","#ff0000","#ff0000","#000000","#000000","#000000","#000000","#000000","#000000","#000000","#000000","#000000","#ff0000","#000000","#000000","#000000","#000000","#000000","#000000","#000000","#000000","#000000","#000000","#000000","#000000","#000000","#000000","#000000","#000000","#000000","#ff0000","#ff0000","#ff0000","#000000","#000000","#ff0000","#000000","#000000","#000000","#ff0000","#ff0000","#ff0000","#000000","#000000","#000000","#000000","#000000","#000000","#ff0000","#ff0000","#000000","#000000","#000000","#ff0000","#ff0000","#000000","#000000","#000000","#000000","#ff0000","#000000","#ff0000","#000000","#ff0000","#ff0000","#ff0000","#ff0000","#ff0000","#000000","#ff0000","#000000","#000000","#ff0000","#ff0000","#000000","#ff0000","#000000","#000000","#000000","#000000","#ff0000","#ff0000","#ff0000","#ff0000","#ff0000","#ff0000","#000000","#000000","#000000","#000000","#000000","#000000","#000000","#000000","#000000","#000000","#ff0000","#ff0000","#ff0000","#000000","#ff0000","#ff0000","#000000","#000000","#000000","#000000","#000000","#000000","#ff0000","#ff0000","#ff0000","#000000","#ff0000","#000000","#ff0000","#ff0000","#ff0000","#000000","#ff0000","#000000","#ff0000","#ff0000","#000000","#000000","#000000","#000000","#000000","#000000","#ff0000","#ff0000","#000000","#ff0000","#000000","#ff0000","#000000","#000000","#ff0000","#ff0000","#ff0000","#ff0000","#000000","#ff0000","#000000","#ff0000","#ff0000","#000000","#000000","#ff0000","#ff0000","#000000","#000000","#000000","#ff0000","#ff0000","#000000","#000000","#ff0000","#000000","#ff0000","#ff0000","#ff0000","#000000","#000000","#000000","#ff0000","#ff0000","#000000","#ff0000","#000000","#000000","#000000","#000000","#000000","#ff0000","#ff0000","#ff0000","#000000","#000000","#ff0000","#000000","#ff0000","#ff0000","#ff0000","#000000","#000000","#000000","#ff0000","#000000","#ff0000","#ff0000","#ff0000","#ff0000","#ff0000","#ff0000","#000000","#000000","#000000","#000000","#ff0000","#ff0000","#ff0000","#000000","#000000","#ff0000","#000000","#000000","#000000","#000000","#000000","#000000","#ff0000","#ff0000","#000000","#000000","#000000","#000000","#000000","#000000","#000000","#ff0000","#ff0000","#000000","#ff0000","#000000","#000000","#000000","#ff0000","#000000","#ff0000","#000000","#000000","#ff0000","#000000","#000000","#ff0000","#ff0000","#ff0000","#000000","#000000","#000000","#000000","#ff0000","#000000","#000000","#000000","#000000","#ff0000","#000000","#000000","#ff0000","#000000","#ff0000","#ff0000","#ff0000","#ff0000","#ff0000","#ff0000","#000000","#000000","#ff0000","#000000","#ff0000","#ff0000","#ff0000","#000000","#000000","#ff0000","#000000","#ff0000","#ff0000","#ff0000","#000000","#000000","#000000","#000000","#000000","#000000","#000000","#000000","#000000","#000000","#000000","#000000","#000000","#000000","#000000","#000000","#000000","#ff0000","#000000","#000000","#000000","#000000","#000000","#000000","#ff0000","#000000","#000000","#ff0000","#ff0000","#000000","#000000","#ff0000","#000000","#000000","#000000","#000000","#000000","#ff0000","#ff0000","#ff0000","#ff0000","#000000","#ff0000","#000000","#000000","#000000","#ff0000","#ff0000","#000000","#000000","#000000","#ff0000","#000000","#000000","#000000","#000000","#000000","#000000","#000000","#000000","#ff0000","#ff0000","#000000","#000000","#ff0000","#ff0000","#ff0000","#000000","#ff0000","#000000","#000000","#000000","#ff0000","#ff0000","#ff0000","#ff0000","#ff0000","#ff0000","#000000","#000000","#000000","#000000","#000000","#000000","#ff0000","#ff0000","#ff0000","#000000","#000000","#000000","#ff0000","#000000","#000000","#000000","#000000","#ff0000","#000000","#000000","#ff0000","#000000","#ff0000","#000000","#000000","#000000","#000000","#ff0000","#ff0000","#000000","#ff0000","#000000","#ff0000","#000000","#ff0000","#000000","#ff0000","#ff0000","#ff0000","#000000","#ff0000","#ff0000","#ff0000","#ff0000","#ff0000","#ff0000","#ff0000","#ff0000","#000000","#000000","#000000","#ff0000","#000000","#000000","#000000","#000000","#000000","#000000","#000000","#000000","#ff0000","#ff0000","#ff0000","#ff0000","#ff0000","#ff0000","#000000","#000000","#000000","#ff0000","#000000","#ff0000","#000000","#000000","#ff0000","#000000","#ff0000","#ff0000","#ff0000","#000000","#ff0000","#ff0000","#ff0000","#ff0000","#ff0000","#ff0000","#ff0000","#ff0000","#ff0000","#ff0000","#ff0000","#ff0000","#ff0000","#000000","#ff0000","#000000","#000000","#000000","#000000","#000000","#000000","#000000","#ff0000","#000000","#000000","#000000","#000000","#000000","#000000","#000000","#000000","#ff0000","#ff0000","#000000","#ff0000","#ff0000","#ff0000","#ff0000","#000000","#ff0000","#ff0000","#ff0000","#ff0000","#000000","#ff0000","#000000","#000000","#ff0000","#000000","#ff0000","#ff0000","#ff0000","#ff0000","#000000","#000000","#ff0000","#000000","#000000","#000000","#000000","#000000","#000000","#000000","#000000","#ff0000","#000000","#000000","#ff0000","#000000","#ff0000","#000000","#000000","#000000","#ff0000","#000000","#ff0000","#000000","#ff0000","#000000","#000000","#ff0000","#000000","#000000","#000000","#000000","#ff0000","#ff0000","#000000","#000000","#000000","#ff0000","#ff0000","#000000","#000000","#000000","#ff0000","#ff0000","#000000","#000000","#000000","#ff0000","#ff0000","#ff0000","#000000","#000000","#000000","#000000","#ff0000","#ff0000","#000000","#000000","#ff0000","#000000","#000000","#ff0000","#000000","#ff0000","#000000","#ff0000","#ff0000","#ff0000","#000000","#ff0000","#000000","#000000","#000000","#000000","#000000","#ff0000","#000000","#000000","#000000","#ff0000","#000000","#000000","#ff0000","#000000","#000000","#ff0000","#ff0000","#ff0000","#ff0000","#000000","#000000","#000000","#ff0000","#000000","#000000","#000000","#000000","#000000","#000000","#000000","#000000","#ff0000","#ff0000","#ff0000","#ff0000","#ff0000","#ff0000","#000000","#000000","#ff0000","#ff0000","#ff0000","#000000","#000000","#000000","#000000","#ff0000","#ff0000","#000000","#000000","#ff0000","#000000","#ff0000","#000000","#000000","#000000","#000000","#ff0000","#ff0000","#ff0000","#ff0000","#000000","#000000","#ff0000","#ff0000","#000000","#000000","#000000","#ff0000","#ff0000","#000000","#ff0000","#ff0000","#000000","#000000","#000000","#ff0000","#000000","#ff0000","#ff0000","#ff0000","#000000","#000000","#000000","#000000","#ff0000","#000000","#000000","#000000","#000000","#ff0000","#ff0000","#ff0000","#ff0000","#ff0000","#000000","#ff0000","#ff0000","#000000","#ff0000","#000000","#000000","#ff0000","#000000","#000000","#000000","#000000","#ff0000","#000000","#000000","#000000","#ff0000","#000000","#ff0000","#ff0000","#ff0000","#000000","#ff0000","#ff0000","#000000","#ff0000","#ff0000","#ff0000","#000000","#000000","#000000","#000000","#ff0000","#000000","#000000","#000000","#ff0000","#ff0000","#ff0000","#000000","#000000","#000000","#000000","#ff0000","#000000","#ff0000","#ff0000","#ff0000","#ff0000","#ff0000","#ff0000","#ff0000","#000000","#ff0000","#ff0000","#ff0000","#000000","#000000","#ff0000","#ff0000","#000000","#ff0000","#ff0000","#ff0000","#ff0000","#000000","#000000","#ff0000","#ff0000","#000000","#000000","#000000","#000000","#ff0000","#ff0000","#000000","#ff0000","#000000","#ff0000","#ff0000","#000000","#ff0000","#ff0000","#000000","#ff0000","#ff0000","#ff0000","#ff0000","#000000","#000000","#000000","#000000","#ff0000","#ff0000","#ff0000","#ff0000","#000000","#ff0000","#ff0000","#ff0000","#ff0000","#ff0000","#ff0000","#000000","#000000","#000000","#000000","#000000","#ff0000","#ff0000","#ff0000","#000000","#000000","#ff0000","#ff0000","#000000","#000000","#ff0000","#000000","#000000","#000000","#ff0000","#ff0000","#000000","#000000","#000000","#000000","#000000","#ff0000","#ff0000","#000000","#000000","#000000","#000000","#000000","#000000","#000000","#000000","#000000","#000000","#000000","#000000","#000000","#000000","#000000","#000000","#000000","#000000","#000000","#000000","#000000","#000000","#000000","#000000","#ff0000","#000000","#000000","#ff0000","#ff0000","#ff0000","#ff0000","#000000","#ff0000","#000000","#ff0000","#000000","#000000","#000000","#ff0000","#000000","#ff0000","#000000","#000000","#ff0000","#ff0000","#000000","#ff0000","#000000","#000000","#000000","#000000","#ff0000","#ff0000","#000000","#000000","#ff0000","#ff0000","#ff0000","#000000","#ff0000","#000000","#000000","#000000","#000000","#000000","#000000","#ff0000","#000000","#000000","#ff0000","#000000","#000000","#ff0000","#000000","#000000","#000000","#ff0000","#ff0000","#ff0000","#ff0000","#ff0000","#ff0000","#000000","#ff0000","#000000","#ff0000","#ff0000","#000000","#000000","#ff0000","#ff0000","#ff0000","#ff0000","#000000","#000000","#000000","#000000","#000000","#ff0000","#000000","#000000","#ff0000","#000000","#000000","#000000","#000000","#000000","#ff0000","#000000","#000000","#ff0000","#000000","#000000","#000000","#000000","#ff0000","#000000","#000000","#ff0000","#ff0000","#000000","#ff0000","#ff0000","#ff0000","#000000","#ff0000","#ff0000","#000000","#000000","#000000","#ff0000","#000000","#ff0000","#ff0000","#ff0000","#ff0000","#000000","#000000","#000000","#ff0000","#ff0000","#000000","#ff0000","#ff0000","#000000","#000000","#000000","#000000","#000000","#000000","#ff0000","#ff0000","#000000","#000000","#ff0000","#ff0000","#000000","#ff0000","#ff0000","#000000","#000000","#000000","#000000","#000000","#ff0000","#000000","#ff0000","#000000","#ff0000","#ff0000","#000000","#000000","#ff0000","#ff0000","#ff0000","#ff0000","#ff0000","#ff0000","#000000","#ff0000","#ff0000","#ff0000","#ff0000","#000000","#000000","#000000","#ff0000","#000000","#000000","#000000","#000000","#000000","#000000","#ff0000","#ff0000","#ff0000","#ff0000","#ff0000","#ff0000","#ff0000","#ff0000","#ff0000","#ff0000","#ff0000","#ff0000","#000000","#000000","#000000","#000000","#000000","#000000","#000000","#000000","#000000","#000000","#ff0000","#000000","#ff0000","#ff0000","#ff0000","#000000","#ff0000","#ff0000","#000000","#ff0000","#ff0000","#ff0000","#ff0000","#000000","#000000","#000000","#ff0000","#ff0000","#ff0000","#ff0000","#000000","#ff0000","#000000","#000000","#ff0000","#000000","#ff0000","#000000","#000000","#000000","#000000","#000000","#ff0000","#000000","#000000","#000000","#000000","#000000","#ff0000","#000000","#000000","#ff0000","#ff0000","#000000","#000000","#000000","#000000","#ff0000","#ff0000","#ff0000","#ff0000","#ff0000","#000000","#000000","#000000","#000000","#000000","#000000","#ff0000","#ff0000","#000000","#000000","#ff0000","#ff0000","#ff0000","#ff0000","#000000","#000000","#000000","#ff0000","#000000","#ff0000","#ff0000","#000000","#000000","#ff0000","#ff0000","#ff0000","#ff0000","#000000","#000000","#ff0000","#ff0000","#ff0000","#000000","#000000","#000000","#000000","#000000","#ff0000","#000000","#ff0000","#ff0000","#ff0000","#ff0000","#ff0000","#ff0000","#ff0000","#ff0000","#ff0000","#000000","#000000","#000000","#000000","#000000","#000000","#000000","#000000","#000000","#000000","#000000","#000000","#000000","#000000","#000000","#000000","#000000","#000000","#000000","#000000","#000000","#000000","#000000","#ff0000","#ff0000","#ff0000","#ff0000","#ff0000","#ff0000","#ff0000","#ff0000","#ff0000","#000000","#000000","#000000","#000000","#ff0000","#ff0000","#ff0000","#000000","#000000","#000000","#000000","#ff0000","#ff0000","#ff0000","#000000","#000000","#000000","#ff0000","#ff0000","#ff0000","#ff0000","#000000","#ff0000","#ff0000","#ff0000","#ff0000","#000000","#ff0000","#000000","#000000","#000000","#000000","#000000","#ff0000","#000000","#ff0000","#ff0000","#ff0000","#ff0000","#000000","#000000","#000000","#000000","#000000","#000000","#000000","#ff0000","#000000","#000000","#000000","#000000","#ff0000","#ff0000","#000000","#000000","#ff0000","#000000","#ff0000","#ff0000","#000000","#ff0000","#ff0000","#ff0000","#000000","#000000","#000000","#000000","#000000","#ff0000","#000000","#ff0000","#000000","#ff0000","#ff0000","#000000","#000000","#ff0000","#ff0000","#ff0000","#ff0000","#000000","#000000","#000000","#000000","#000000","#000000","#000000","#000000","#000000","#000000","#000000","#000000","#000000","#ff0000","#ff0000","#ff0000","#000000","#000000","#ff0000","#ff0000","#ff0000","#ff0000","#ff0000","#000000","#000000","#ff0000","#ff0000","#ff0000","#000000","#000000","#000000","#000000","#ff0000","#ff0000","#ff0000","#000000","#ff0000","#ff0000","#ff0000","#ff0000","#ff0000","#ff0000","#ff0000","#000000","#000000","#000000","#ff0000","#ff0000","#000000","#000000","#000000","#000000","#ff0000","#ff0000","#ff0000","#ff0000","#000000","#ff0000","#ff0000","#ff0000","#ff0000","#ff0000","#000000","#ff0000","#000000","#ff0000","#ff0000","#ff0000","#ff0000","#000000","#ff0000","#ff0000","#000000","#ff0000","#ff0000","#ff0000","#ff0000","#ff0000","#000000","#000000","#ff0000","#ff0000","#ff0000","#ff0000","#ff0000","#ff0000","#000000","#000000","#000000","#ff0000","#000000","#000000","#000000","#000000","#ff0000","#ff0000","#000000","#000000","#000000","#000000","#000000","#000000","#ff0000","#000000","#ff0000","#ff0000","#ff0000","#ff0000","#000000","#ff0000","#ff0000","#ff0000","#000000","#ff0000","#ff0000","#ff0000","#ff0000","#ff0000","#000000","#000000","#000000","#000000","#000000","#ff0000","#ff0000","#ff0000","#ff0000","#ff0000","#ff0000","#ff0000","#000000","#000000","#000000","#000000","#000000","#ff0000","#000000","#000000","#000000","#000000","#ff0000","#000000","#ff0000","#ff0000","#ff0000","#ff0000","#ff0000","#ff0000","#000000","#ff0000","#000000","#000000","#000000","#000000","#ff0000","#ff0000","#ff0000","#ff0000","#ff0000","#000000","#ff0000","#ff0000","#000000","#ff0000","#ff0000","#ff0000","#ff0000","#ff0000","#000000","#ff0000","#000000","#000000","#ff0000","#000000","#000000","#000000","#000000","#000000","#000000","#ff0000","#ff0000","#000000","#000000","#000000","#ff0000","#ff0000","#ff0000","#ff0000","#ff0000","#000000","#000000","#000000","#ff0000","#ff0000","#000000","#000000","#000000","#000000","#000000","#000000","#000000","#000000","#000000","#ff0000","#000000","#ff0000","#000000","#ff0000","#ff0000","#ff0000","#ff0000","#ff0000","#ff0000","#000000","#ff0000","#000000","#000000","#ff0000","#ff0000","#ff0000","#ff0000","#ff0000","#000000","#ff0000","#ff0000","#000000","#000000","#000000","#000000","#000000","#ff0000","#ff0000","#ff0000","#ff0000","#ff0000","#ff0000","#ff0000","#000000","#ff0000","#000000","#000000","#000000","#000000","#ff0000","#000000","#000000","#ff0000","#000000","#000000","#000000","#ff0000","#ff0000","#000000","#ff0000","#000000","#ff0000","#ff0000","#000000","#000000","#000000","#000000","#000000","#ff0000","#000000","#ff0000","#ff0000","#ff0000","#ff0000","#000000","#000000","#000000","#ff0000","#000000","#000000","#ff0000","#000000","#000000","#000000","#000000","#000000","#000000","#000000","#ff0000","#000000","#000000","#000000","#ff0000","#000000","#ff0000","#ff0000","#ff0000","#ff0000","#000000","#ff0000","#000000","#000000","#000000","#ff0000","#000000","#000000","#ff0000","#ff0000","#ff0000","#000000","#000000","#000000","#000000","#ff0000","#000000","#000000","#000000","#000000","#ff0000","#ff0000","#000000","#ff0000","#ff0000","#ff0000","#ff0000","#ff0000","#000000","#ff0000","#ff0000","#000000","#ff0000","#ff0000","#ff0000","#ff0000","#000000","#000000","#000000","#ff0000","#ff0000","#000000","#000000","#ff0000","#ff0000","#ff0000","#ff0000","#ff0000","#000000","#000000","#000000","#ff0000","#ff0000","#000000","#000000","#ff0000","#ff0000","#ff0000","#ff0000","#ff0000","#ff0000","#ff0000","#000000","#000000","#000000","#000000","#000000","#000000","#000000","#000000","#000000","#000000","#000000","#000000","#ff0000","#ff0000","#ff0000","#000000","#000000","#ff0000","#000000","#000000","#000000","#000000","#ff0000","#ff0000","#ff0000","#ff0000","#000000","#000000","#000000","#000000","#000000","#000000","#ff0000","#000000","#ff0000","#ff0000","#ff0000","#ff0000","#ff0000","#000000","#000000","#000000","#000000","#000000","#000000","#000000","#000000","#000000","#000000","#000000","#000000","#000000","#ff0000","#000000","#ff0000","#ff0000","#ff0000","#ff0000","#000000","#ff0000","#ff0000","#ff0000","#ff0000","#ff0000","#ff0000","#ff0000","#000000","#000000","#000000","#000000","#000000","#ff0000","#000000","#ff0000","#ff0000","#ff0000","#ff0000","#000000","#000000","#000000","#000000","#000000","#ff0000","#ff0000","#000000","#000000","#000000","#ff0000","#000000","#ff0000","#ff0000","#ff0000","#ff0000","#ff0000","#ff0000","#ff0000","#000000","#ff0000","#000000","#000000","#ff0000","#000000","#ff0000","#000000","#ff0000","#000000","#ff0000","#ff0000","#000000","#ff0000","#ff0000","#000000","#ff0000","#ff0000","#000000","#ff0000","#ff0000","#000000","#ff0000","#000000","#000000","#ff0000","#000000","#000000","#ff0000","#ff0000","#000000","#000000","#ff0000","#ff0000","#000000","#ff0000","#ff0000","#000000","#000000","#000000","#000000","#000000","#000000","#000000","#000000","#000000","#000000","#000000","#ff0000","#000000","#000000","#000000","#000000","#ff0000","#ff0000","#ff0000","#ff0000","#ff0000","#ff0000","#000000","#000000","#000000","#ff0000","#ff0000","#000000","#ff0000","#ff0000","#ff0000","#ff0000","#000000","#ff0000","#000000","#000000","#000000","#000000","#000000","#ff0000","#ff0000","#ff0000","#000000","#000000","#000000","#000000","#000000","#000000","#000000","#000000","#000000","#000000","#000000","#ff0000","#000000","#ff0000","#ff0000","#000000","#000000","#ff0000","#ff0000","#ff0000","#000000","#ff0000","#ff0000","#000000","#000000","#000000","#000000","#ff0000","#000000","#000000","#ff0000","#000000","#000000","#ff0000","#ff0000","#ff0000","#ff0000","#ff0000","#ff0000","#ff0000","#ff0000","#000000","#000000","#ff0000","#ff0000","#000000","#000000","#000000","#000000","#000000","#000000","#000000","#ff0000","#000000","#ff0000","#000000","#ff0000","#ff0000","#000000","#ff0000","#ff0000","#000000","#ff0000","#ff0000","#ff0000","#ff0000","#ff0000","#000000","#000000","#000000","#000000","#000000","#ff0000","#ff0000","#000000","#ff0000","#ff0000","#ff0000","#000000","#ff0000","#ff0000","#000000","#000000","#000000","#000000","#ff0000","#000000","#000000","#000000","#000000","#ff0000","#ff0000","#ff0000","#ff0000","#000000","#ff0000","#ff0000","#000000","#000000","#ff0000","#ff0000","#ff0000","#000000","#000000","#000000","#000000","#000000","#000000","#ff0000","#ff0000","#000000","#000000","#000000","#000000","#000000","#000000","#ff0000","#ff0000","#ff0000","#000000","#000000","#ff0000","#000000","#000000","#000000","#000000","#000000","#000000","#000000","#ff0000","#000000","#ff0000","#000000","#000000","#000000","#000000","#000000","#000000","#ff0000","#000000","#000000","#ff0000","#ff0000","#ff0000","#ff0000","#ff0000","#ff0000","#ff0000","#000000","#000000","#000000","#000000","#000000","#000000","#ff0000","#ff0000","#ff0000","#ff0000","#ff0000","#ff0000","#000000","#ff0000","#ff0000","#ff0000","#ff0000","#ff0000","#000000","#ff0000","#ff0000","#ff0000","#000000","#000000","#000000","#000000","#000000","#000000","#ff0000","#ff0000","#000000","#000000","#000000","#ff0000","#000000","#ff0000","#ff0000","#ff0000","#ff0000","#ff0000","#000000","#000000","#000000","#000000","#000000","#000000","#000000","#000000","#000000","#000000","#000000","#000000","#000000","#000000","#000000","#000000","#000000","#000000","#000000","#000000","#000000","#000000","#000000","#ff0000","#000000","#000000","#ff0000","#ff0000","#ff0000","#000000","#000000","#ff0000","#000000","#000000","#000000","#ff0000","#ff0000","#ff0000","#ff0000","#ff0000","#ff0000","#ff0000","#ff0000","#ff0000","#000000","#000000","#000000","#000000","#000000","#000000","#000000","#000000","#000000","#000000","#000000","#ff0000","#ff0000","#ff0000","#ff0000","#000000","#000000","#000000","#000000","#000000","#000000","#000000","#ff0000","#000000","#000000","#ff0000","#000000","#ff0000","#000000","#000000","#000000","#000000","#000000","#000000","#ff0000","#ff0000","#ff0000","#000000","#000000","#ff0000","#000000","#000000","#ff0000","#ff0000","#000000","#ff0000","#ff0000","#000000","#ff0000","#ff0000","#000000","#000000","#000000","#000000","#ff0000","#ff0000","#ff0000","#000000","#000000","#ff0000","#ff0000","#000000","#000000","#000000","#000000","#000000","#000000","#ff0000","#ff0000","#ff0000","#000000","#ff0000","#ff0000","#000000","#ff0000","#ff0000","#ff0000","#ff0000","#000000","#000000","#ff0000","#ff0000","#ff0000","#000000","#ff0000","#ff0000","#000000","#ff0000","#000000","#000000","#ff0000","#ff0000","#000000","#ff0000","#000000","#000000","#000000","#000000","#000000","#ff0000","#ff0000","#ff0000","#ff0000","#ff0000","#ff0000","#000000","#000000","#ff0000","#000000","#ff0000","#ff0000","#000000","#ff0000","#ff0000","#000000","#000000","#000000","#000000","#000000","#000000","#ff0000","#ff0000","#ff0000","#ff0000","#ff0000","#ff0000","#ff0000","#ff0000","#000000","#000000","#000000","#000000","#000000","#ff0000","#ff0000","#ff0000","#000000","#000000","#000000","#000000","#ff0000","#ff0000","#000000","#000000","#000000","#ff0000","#ff0000","#000000","#ff0000","#ff0000","#000000","#ff0000","#ff0000","#000000","#ff0000","#000000","#000000","#000000","#ff0000","#000000","#ff0000","#000000","#000000","#ff0000","#ff0000","#ff0000","#000000","#000000","#ff0000","#ff0000","#000000","#000000","#000000","#000000","#000000","#000000","#ff0000","#ff0000","#000000","#000000","#000000","#ff0000","#ff0000","#000000","#ff0000","#ff0000","#ff0000","#ff0000","#ff0000","#000000","#ff0000","#ff0000","#ff0000","#ff0000","#ff0000","#000000","#ff0000","#000000","#ff0000","#ff0000","#ff0000","#ff0000","#ff0000","#ff0000","#ff0000","#ff0000","#ff0000","#000000","#ff0000","#000000","#ff0000","#ff0000","#ff0000","#000000","#000000","#000000","#000000","#000000","#ff0000","#ff0000","#ff0000","#ff0000","#000000","#ff0000","#000000","#000000","#000000","#000000","#000000","#ff0000","#ff0000","#000000","#000000","#000000","#ff0000","#ff0000","#000000","#ff0000","#ff0000","#000000","#000000","#000000","#000000","#000000","#000000","#ff0000","#ff0000","#000000","#000000","#000000","#ff0000","#ff0000","#ff0000","#ff0000","#000000","#ff0000","#000000","#000000","#000000","#ff0000","#000000","#000000","#ff0000","#000000","#ff0000","#000000","#ff0000","#ff0000","#ff0000","#000000","#ff0000","#000000","#000000","#000000","#ff0000","#000000","#ff0000","#ff0000","#000000","#000000","#000000","#ff0000","#000000","#ff0000","#ff0000","#ff0000","#ff0000","#000000","#ff0000","#ff0000","#000000","#000000","#ff0000","#ff0000","#ff0000","#ff0000","#ff0000","#000000","#000000","#ff0000","#000000","#ff0000","#ff0000","#ff0000","#ff0000","#000000","#000000","#000000","#000000","#ff0000","#ff0000","#ff0000","#ff0000","#ff0000","#ff0000","#ff0000","#000000","#ff0000","#000000","#000000","#ff0000","#ff0000","#ff0000","#ff0000","#ff0000","#ff0000","#ff0000","#ff0000","#ff0000","#ff0000","#ff0000","#ff0000","#ff0000","#000000","#000000","#ff0000","#ff0000","#ff0000","#ff0000","#000000","#000000","#ff0000","#ff0000","#000000","#000000","#ff0000","#000000","#ff0000","#ff0000","#ff0000","#ff0000","#ff0000","#000000","#000000","#ff0000","#ff0000","#ff0000","#ff0000","#000000","#000000","#000000","#000000","#000000","#ff0000","#000000","#000000","#000000","#000000","#000000","#000000","#ff0000","#ff0000","#ff0000","#ff0000","#ff0000","#000000","#000000","#ff0000","#ff0000","#ff0000","#ff0000","#ff0000","#ff0000","#000000","#ff0000","#ff0000","#ff0000","#ff0000","#ff0000","#ff0000","#ff0000","#000000","#000000","#000000","#ff0000","#ff0000","#ff0000","#ff0000","#ff0000","#000000","#ff0000","#ff0000","#ff0000","#000000","#ff0000","#ff0000","#ff0000","#000000","#000000","#ff0000","#000000","#000000","#ff0000","#ff0000","#ff0000","#ff0000","#ff0000","#ff0000","#ff0000","#000000","#ff0000","#ff0000","#000000","#000000","#000000","#ff0000","#ff0000","#ff0000","#ff0000","#ff0000","#ff0000","#ff0000","#000000","#000000","#ff0000","#ff0000","#000000","#000000","#000000","#000000","#000000","#000000","#000000","#000000","#000000","#000000","#000000","#000000","#000000","#000000","#000000","#000000","#000000","#000000","#000000","#000000","#000000","#000000","#000000","#000000","#000000","#000000","#000000","#000000","#000000","#ff0000","#000000","#000000","#000000","#000000","#000000"],"PValue":[0.6421,0.8982,0.2852,0.0577,0.1325,0.06082,0.1202,0.2353,0.3133,0.1643,0.07726,0.1525,0.8148,0.5326,0.04816,0.6798,0.9361,0.3667,0.2453,0.3282,0.1899,0.7503,0.2875,0.5468,0.9313,0.9171,0.1718,0.4517,0.5545,0.6229,0.03939,0.04066,0.1108,0.2975,0.9737,0.2076,0.1844,0.663,0.2177,0.9533,0.1448,0.1011,0.1005,0.05288,0.5649,0.3859,0.2213,0.1313,0.5105,0.09645,0.6804,0.9531,0.7945,0.2193,0.4959,0.1064,0.4046,0.5533,0.9142,0.398,0.6199,0.2753,0.2803,0.0414,0.1448,0.2961,0.1495,0.2544,0.2013,0.9588,0.4656,0.4877,0.872,0.1808,0.06957,0.4612,0.8553,0.2135,0.09235,0.365,0.03788,0.1837,0.6716,0.5089,0.04369,0.1714,0.1751,0.5552,0.2679,0.4246,0.8168,0.782,0.5541,0.2471,0.9529,0.118,0.0692,0.3623,0.9988,0.08493,0.276,0.2569,0.05892,0.6785,0.1721,0.3521,0.5731,0.5871,0.2299,0.4537,0.568,0.8947,0.07615,0.6215,0.4489,0.3262,0.3606,0.149,0.05466,0.1752,0.4518,0.2651,0.08379,0.3637,0.366,0.2386,0.8498,0.07737,0.164,0.9712,0.1793,0.2552,0.001828,0.1909,0.2183,0.2882,0.5262,0.6879,0.5941,0.9065,0.009109,0.2664,0.3804,0.2128,0.8519,0.002326,0.7078,0.1423,0.8591,0.2623,0.2328,0.7841,0.1539,0.4988,0.5227,0.001426,0.03882,0.3732,0.08401,0.1473,0.2465,0.7251,0.001721,0.5263,0.1344,0.04095,0.08246,0.7757,0.11,0.7717,0.1546,0.001859,0.2985,0.2322,0.2283,0.5986,0.06696,0.3437,0.8676,0.6684,0.4511,0.2777,0.0643,0.5309,0.2247,0.1457,0.5374,0.3277,0.2414,0.06421,0.09085,0.971,0.8089,0.9087,0.3298,0.0008924,0.03607,0.1561,0.04577,0.239,0.7765,0.06445,0.01293,0.2,0.003595,0.248,0.4267,0.0157,0.9626,0.03991,0.405,0.01255,0.03675,0.7434,0.2788,0.2016,0.7606,0.02547,0.9373,0.04755,0.2676,0.1449,0.01623,0.6688,0.02612,0.05944,0.004417,0.009044,0.05714,0.7533,0.09088,0.01592,0.9785,0.07624,0.04162,0.4946,0.1048,0.2224,0.3782,0.6668,0.04402,0.04007,0.383,0.05599,0.8683,0.2806,0.04314,0.05086,0.06367,0.3243,0.421,0.05475,0.2629,0.1214,0.05421,0.3171,0.3212,0.8631,0.04783,0.7908,0.6075,0.6405,0.01834,0.08932,0.5355,0.11,0.309,0.1059,0.3593,0.02686,0.2179,0.01639,0.2568,0.3849,0.3246,0.01028,0.4241,0.2032,0.1669,0.05227,0.009652,0.2878,0.8476,0.4496,0.09419,0.3341,0.3204,0.3803,0.00604,0.1377,0.08267,0.008041,0.05203,0.2978,0.3085,0.4679,0.009352,0.3375,0.3652,0.3995,0.3471,0.9328,0.5782,0.001551,0.0004907,0.08618,0.01141,0.1652,0.214,0.1353,0.1569,0.2473,0.1176,0.06746,0.5139,0.6695,0.3476,0.7228,0.7764,0.2322,0.003503,0.1685,0.1654,0.7495,0.001908,0.2513,0.3379,0.0831,0.03038,0.1966,0.7613,0.5331,0.5284,0.6487,0.1434,0.0268,0.04444,0.1647,0.008338,0.7454,0.1092,0.06787,0.7586,0.2455,0.04864,0.0001322,0.1933,0.1213,0.0628,0.08152,0.7054,0.5403,0.4105,0.3913,0.3397,0.0861,0.1789,0.03198,0.02172,0.257,0.1571,0.2003,0.2612,0.002738,0.4393,0.3624,0.4997,0.4141,0.07727,0.3787,0.08974,0.04555,0.4311,0.7619,0.4545,0.3266,0.02011,0.092,0.03434,0.008819,0.01916,0.2331,0.1156,0.0502,0.7546,0.5708,0.9445,0.2005,0.4109,0.7253,0.3145,0.001493,0.5516,0.05431,0.4077,0.00248,0.4555,0.9666,0.4921,0.04818,0.3898,0.3603,0.237,0.09566,0.07993,0.04718,0.9315,0.001923,0.5739,0.8744,0.2188,0.06561,0.1357,0.9773,0.7142,0.4304,0.02366,0.0006658,0.03741,0.9441,0.001163,0.8471,0.01492,0.09115,0.2021,0.04589,0.009722,0.91,0.1078,0.03906,0.577,0.9918,0.00352,0.2584,0.00866,0.1384,0.08074,0.04513,0.3097,0.2924,0.08597,0.003474,0.9562,0.1186,0.03023,0.04568,0.004771,0.9502,0.3107,0.04107,0.07325,0.4225,0.3223,0.06226,0.00323,0.07451,0.2285,0.09513,0.6394,0.08346,0.326,0.3492,0.0479,0.3122,0.05128,0.5851,0.07638,0.05245,0.3184,0.1147,0.06271,0.5827,0.1419,0.1121,0.5909,0.006591,0.6823,0.09132,0.002331,0.9783,0.06865,0.6213,0.09907,0.9935,0.05619,0.01401,0.4721,0.05255,0.0139,0.006704,0.5129,0.0915,0.01272,0.04873,0.5346,5.354e-05,0.7373,0.00723,0.1903,0.9127,0.02966,0.968,0.2206,0.05683,0.2007,0.02147,0.2775,0.7858,0.03682,0.06674,0.1117,0.02231,0.06458,0.7401,0.01913,0.9588,0.3797,0.9592,0.06782,0.5562,0.2554,0.2676,0.5679,0.1429,0.4589,0.03874,0.2941,0.8573,0.2003,0.06049,0.2066,0.1357,0.7514,0.04605,0.001414,0.1353,0.5621,0.1371,0.4929,0.8125,0.1611,0.001613,4.056e-05,0.7526,0.3461,0.5926,0.1288,0.07645,0.1781,0.3096,0.9586,0.1632,0.07482,0.001227,0.3784,0.004898,0.09116,0.1189,0.733,0.152,0.5794,0.009548,0.002295,0.05787,0.58,0.2326,0.6147,0.9624,0.6686,0.01508,0.001212,0.1435,0.1594,0.003642,0.08135,0.1112,0.4669,0.009935,0.4352,0.3254,0.2573,0.3975,0.08523,0.6013,0.03725,0.5942,0.733,0.005678,0.7558,0.1148,0.1558,0.8706,0.1552,0.068,0.732,0.07766,0.5002,0.7452,0.05798,0.09128,0.8625,0.2687,0.0153,0.6283,0.03729,0.7793,0.119,0.2201,0.9285,0.8121,0.06007,0.307,0.06745,0.7986,0.0425,0.001372,0.001662,0.1019,0.6565,0.7474,0.352,0.941,0.5166,0.1134,0.3107,0.2889,0.05403,0.053,0.1755,0.2326,0.06016,0.1422,0.7654,0.9793,0.01168,0.02517,0.6794,0.3327,0.2622,0.09693,0.02916,0.7644,0.1216,0.1444,0.6998,0.3213,0.1347,0.008184,0.9186,0.06726,0.1137,0.5532,0.384,0.6403,0.1626,0.005825,0.9857,0.05539,0.06273,0.3925,0.06245,0.03846,0.1146,0.9029,0.2158,0.2527,0.09037,0.403,0.07393,0.07049,0.05499,0.886,0.9169,0.7464,0.5408,0.03271,0.4181,0.2452,0.7988,0.06221,0.1911,0.03665,0.1533,0.6734,0.07262,0.005649,0.2393,0.07485,0.3009,0.935,0.3736,0.2053,0.4595,0.1368,0.8859,0.4593,0.3603,0.2958,0.2385,0.02312,0.7978,0.2084,0.2006,0.4504,0.03794,0.7781,0.4028,0.1236,0.7118,0.766,0.6046,0.2406,0.0006023,0.7376,0.7012,0.3682,0.2011,0.1247,0.6997,0.882,0.8606,0.2239,0.2086,0.0461,0.7506,0.6999,0.0922,0.04507,0.7461,0.8211,0.06895,0.7645,0.1396,0.1381,0.0004031,0.5492,0.2527,0.1492,0.2295,0.04242,0.00748,0.6033,0.7533,0.3469,0.327,0.2193,0.5254,0.2528,0.2081,0.6832,0.2606,0.8678,0.08224,0.8845,0.395,0.1267,0.5989,0.01363,0.8905,0.1169,0.5639,0.5664,0.5127,0.8364,0.3392,0.05155,0.0305,0.2178,0.3408,0.254,0.9353,0.8185,0.1781,0.1549,0.1372,0.2841,0.1874,0.08719,0.0003192,0.7096,0.5862,0.08111,0.339,0.1023,0.1385,0.2794,0.4125,0.6934,0.05065,0.5727,0.7536,0.6764,0.395,0.3963,0.05256,0.0905,0.1985,0.06065,0.2431,0.2363,0.1393,0.8732,0.4356,0.00762,0.07441,0.05089,0.00181,0.04788,0.8626,0.02708,0.8406,0.4857,0.001944,0.9357,0.5648,0.4128,0.4075,0.2789,0.8714,0.2969,0.8335,0.001149,0.0009889,0.002221,0.0006708,0.02784,0.3227,0.7439,0.3193,0.5554,0.976,0.08856,0.3472,0.8072,0.3011,0.3029,0.06569,0.2291,0.004605,0.7342,0.05957,0.4458,0.0568,0.009925,0.009045,0.1964,0.7087,0.1808,0.09929,0.01073,0.1782,0.3055,0.001959,0.06795,0.4831,0.001493,0.3274,0.6773,0.0498,0.1389,0.8283,0.04556,0.004712,0.09001,0.08283,0.2852,0.0139,0.3997,0.1899,0.661,0.0002524,0.1973,0.4117,0.4008,0.4413,0.6959,0.08755,0.02707,0.01129,0.05299,0.03927,0.1064,0.09127,0.7568,0.09029,0.4252,0.1689,0.1842,0.7186,0.8831,0.9954,0.03902,0.7407,0.1308,0.01706,0.7206,0.005567,0.2537,0.08244,0.0008593,0.0006366,0.3798,0.1744,0.2665,0.04419,0.5606,0.6373,0.04921,0.8688,0.1257,0.008541,0.058,0.9715,0.0009303,0.5261,0.8334,0.5514,0.2357,0.2853,0.3621,0.9058,0.3073,0.5569,0.06519,0.9185,0.6439,0.01986,0.9596,0.6098,0.000134,0.4578,0.007556,0.8106,0.9998,0.8741,0.02542,0.1823,0.9051,0.08319,0.7866,0.0008815,0.05542,0.3875,0.005962,0.1791,0.3223,0.214,0.005637,0.01161,0.2245,0.2393,0.3177,0.3867,0.2334,0.6204,0.08082,0.3309,0.2498,0.005569,0.4376,0.3866,0.005662,0.0005789,0.4322,0.2461,0.2593,0.2686,0.3073,0.6049,0.0161,0.107,0.4909,0.06999,0.3236,0.08496,0.1686,0.186,0.1221,0.3929,0.2822,0.1007,0.2467,0.3765,0.8847,0.03914,0.4859,0.07316,0.4193,0.01438,0.182,0.05404,0.05228,0.1162,0.7614,0.04203,0.06784,0.7332,0.2415,0.73,0.1772,0.151,0.06366,0.3299,0.06239,0.05594,0.06053,0.07378,0.3994,0.8101,0.6808,0.7817,0.1066,0.04155,0.4231,0.04311,0.8444,0.03004,0.08638,0.05569,0.09093,0.34,0.2773,0.02008,0.9309,0.1673,0.1097,0.9701,0.05196,0.8079,0.005548,0.4409,0.8821,0.9141,0.6114,0.08101,0.4492,0.07793,0.0122,0.4186,0.2593,0.2426,0.2158,0.8128,0.7771,0.04551,0.6635,0.06084,0.6178,0.4944,0.03764,0.3618,0.8674,0.02484,0.1031,0.8124,0.4521,0.2663,0.1099,0.1345,0.004609,0.208,0.02701,0.0617,0.08057,0.1256,0.4976,0.97,0.7865,0.08388,0.0772,0.7784,0.02597,0.4728,0.08931,0.0817,0.1396,0.3747,0.6076,0.9943,0.4018,0.8474,0.8805,0.104,0.6655,0.001334,0.2223,0.4278,0.1847,0.9754,0.19,0.1244,0.05344,0.3053,0.003386,0.4353,0.4404,0.1041,0.03068,0.7881,0.003064,0.07413,0.1791,0.1803,0.994,0.5249,0.6581,0.2087,0.402,0.1433,0.04095,0.02511,0.176,0.2723,0.4655,0.5475,0.7876,0.251,0.0297,0.01926,0.9753,0.09801,0.5541,0.199,0.9023,0.1215,0.6679,0.2599,0.6554,0.1259,0.18,0.07583,0.6359,0.05653,0.1903,0.1871,0.9073,0.2507,0.3571,0.6014,0.08133,0.928,0.4293,0.2504,0.01338,0.1179,0.002278,0.002224,0.2575,0.7443,0.2636,0.9495,0.15,0.4372,0.3245,0.04189,0.07632,0.04839,0.2624,0.04103,0.6567,0.04942,0.125,0.3189,0.1159,0.5809,0.004061,0.0184,0.1445,0.9753,0.9597,0.03356,0.764,0.1605,0.06618,0.02976,0.008099,0.09882,0.7591,0.8262,4.23e-05,0.202,0.1803,0.3061,0.888,0.05302,0.5602,0.1502,0.6932,0.7746,0.9009,0.2758,0.08234,0.08383,0.8655,0.2571,0.6879,0.4375,0.3599,0.07332,0.1959,0.3797,0.4666,0.07928,0.1189,0.4151,0.9156,0.5913,0.4707,0.1822,0.5476,0.8869,0.03167,0.07421,0.1082,0.7404,0.5127,0.9089,0.1399,0.04545,0.003793,0.3047,0.1294,0.1491,0.04844,0.7305,0.6945,0.843,0.1813,0.3686,0.0629,0.4535,0.4124,0.05285,0.4841,0.3575,0.1999,0.1136,0.2425,0.02293,0.3438,0.216,0.07776,0.1555,0.6787,0.3911,0.05984,0.6623,0.01094,0.5558,0.06336,0.8266,0.09026,3.754e-05,0.6434,0.006166,0.6227,0.5621,0.01141,0.2275,0.1023,0.03777,0.3322,0.167,0.9422,0.6775,0.2007,0.007123,0.2799,0.02033,0.0389,0.4407,0.7498,0.7728,0.06481,0.1026,0.1293,0.3206,0.6871,0.1469,0.0282,0.4968,0.8181,0.4196,0.09636,0.08739,0.07452,0.2131,0.03667,0.006105,0.04055,0.9187,0.1013,0.08337,0.005866,0.01086,0.1442,0.1325,0.5684,0.2405,0.3782,0.04896,0.01997,0.001989,0.9021,0.04337,0.2169,0.7072,6.578e-05,0.5872,0.009437,0.05452,0.4771,0.6582,0.236,0.9202,0.03654,0.1739,0.0008966,0.1258,0.04343,0.08311,0.3556,0.05432,0.7652,0.7341,0.2574,0.1645,0.0109,0.005968,0.1748,0.002364,0.4554,0.1528,0.02506,0.8673,0.05657,0.02641,0.5225,0.244,0.5078,0.09343,0.06217,0.1287,0.101,0.09922,0.005128,0.06705,0.2467,0.3691,0.04734,0.07504,0.04281,0.3466,0.01212,0.5866,0.00187,0.5836,0.02122,0.708,0.1255,0.3736,0.07592,6.163e-05,0.5895,0.04988,0.4052,0.3659,0.008246,0.1195,0.2449,0.1352,0.002167,0.7843,0.712,0.5341,0.2049,0.5413,0.05699,0.4933,0.782,0.07486,0.5571,0.0006528,0.03848,0.02423,0.1797,0.579,0.01025,0.01624,0.9237,0.1532,0.3113,0.8485,0.3481,0.0396,0.02118,0.3058,0.1311,0.7315,0.3846,0.6126,0.5489,0.4742,0.9976,0.8753,0.5241,0.05898,0.7026,0.6105,0.1075,0.003893,0.002795,0.7985,0.001984,0.03418,0.3165,0.07082,0.906,0.3738,0.105,0.002074,0.3543,0.1282,0.0767,0.4738,0.05853,0.3018,0.546,0.2951,0.1916,0.01819,0.02424,0.6945,0.3965,0.05501,0.2685,0.2506,0.2714,0.05676,0.1568,0.7986,0.4636,0.09463,0.1746,0.3278,0.2115,0.1337,0.9051,0.281,0.07234,0.2689,0.07641,0.02239,0.4465,0.1094,0.1582,0.5669,0.005141,0.1991,0.05576,0.1721,0.08512,0.2356,0.0196,0.01879,0.002897,0.07867,0.09616,0.03112,0.06279,0.06249,0.07953,0.04433,0.4513,0.09638,0.5242,0.09873,0.07147,0.8904,0.06323,0.1684,0.8181,0.2695,0.344,0.459,0.9532,0.09139,0.5412,3.037e-05,0.6262,0.01854,0.05411,0.4452,0.5806,0.4089,0.7523,0.5248,0.001662,0.5661,0.1272,0.08173,0.06118,0.01875,0.08525,0.02156,0.2162,0.07386,0.02077,0.3537,0.04479,0.1026,0.06739,0.002765,0.7924,0.659,0.4225,0.3728,0.3206,0.001214,0.2905,0.1741,0.8896,0.1513,0.5714,0.7408,0.1203,0.7993,0.1938,5.301e-05,0.7472,0.7749,0.8342,0.01668,0.9797,0.1999,0.004295,0.1345,0.3198,0.26,0.5554,0.6105,0.602,0.4259,0.7808,0.03122,0.4235,0.09026,0.6471,0.05218,0.1149,0.8421,0.06785,0.5163,0.04875,0.4919,0.01372,0.4664,0.07875,0.009292,0.3079,0.04866,0.6237,0.6323,0.2022,0.8914,0.2753,0.2962,0.07051,0.05031,0.752,0.007129,0.06118,0.03336,0.01898,0.04441,0.3365,0.9848,0.02737,0.08115,0.2582,0.04493,0.1007,0.2628,0.3716,0.1934,0.1639,0.03993,0.1467,0.05814,0.0783,0.009178,0.5787,0.1289,0.7601,0.8177,0.02584,0.09661,0.6316,0.7329,0.5928,0.08081,0.03407,0.1369,0.1897,0.6683,0.4068,0.5702,0.04663,0.9459,0.6716,0.4111,0.6442,0.07638,0.5093,0.03686,0.07928,0.00937,0.005878,0.005211,0.1328,0.06302,0.02915,0.802,0.05058,0.7935,0.6754,0.1304,0.04421,0.04364,0.431,0.3634,0.1437,0.2404,0.4029,0.0405,0.1134,0.6984,0.7053,0.0436,0.0006383,0.002578,0.07096,0.1186,0.8263,0.9349,0.1897,0.3575,0.6196,0.03906,0.2064,0.7992,0.05534,0.2297,0.04476,0.8134,0.03118,0.916,0.7884,0.6765,0.7605,0.0709,0.463,0.6212,0.8426,0.205,0.0006366,0.045,0.03657,0.1172,0.8299,0.2272,0.04615,0.06395,0.445,0.3214,0.06984,0.006294,0.7464,0.04611,0.007065,0.01811,0.1094,0.3775,0.02779,0.04053,0.463,0.02741,0.02694,0.08692,0.4864,0.04941,0.6443,0.2058,0.03596,0.1138,0.63,0.08584,0.02204,0.8814,0.1066,0.185,0.1042,0.2663,0.003811,0.04154,0.522,0.004926,0.07615,0.0308,0.3556,0.05618,0.8708,0.05009,0.02968,0.1305,0.03481,0.02109,0.005535,0.6968,0.1451,0.01096,0.02256,0.0006761,0.01009,0.4025,0.3137,0.02796,0.3933,0.748,0.1623,0.007278,0.8673,0.01711,0.3233,0.6059,0.02082,0.03244,0.3815,0.2727,0.1163,0.1538,0.00208,0.02082,0.3495,0.2045,0.2274,0.6037,0.1756,0.2212,0.03366,0.02003,0.001506,0.008059,0.01873,0.001423,0.3846,0.04844,0.05451,0.362,0.5185,0.002248,0.382,0.007224,0.0009059,0.07196,0.04206,0.5659,0.2607,0.3108,0.6847,0.5694,0.09001,0.1749,0.9343,0.1257,0.216,0.771,0.5603,0.1106,0.5259,0.3548,0.007067,0.2538,0.254,0.1393,0.8786,0.09606,0.004658,0.04275,0.05802,0.04841,0.4005,0.4999,0.3804,0.2462,0.4094,0.1469,0.2312,0.02442,0.005347,5.304e-08,9.965e-12,7.753e-17,1.227e-10,9.816e-09,1.539e-08,5.595e-10,0.001282,1.207e-07,2.909e-05,3.669e-07,0.0002784,3.202e-09,4.251e-05,0.006187,4.574e-06,2.177e-08,0.004707,1.677e-05,0.02579,0.006686,5.945e-08,5.169e-07,0.0009679,0.001943,0.006235,0.03519,0.000765,2.021e-06,0.006724,0.004427,7.228e-07,0.007526,1.126e-15,1.555e-08,0.03043,0.001488,5.302e-14,0.001719,3.036e-09,2.264e-09,5.104e-11,0.0004118,0.0005853,0.0001198,1.926e-05,5.671e-05,0.001319,1.494e-07,1.76e-08,1.883e-06,0.01353,0.001829,0.0002002,0.0006062,2.769e-05,6.01e-09,0.0009297,0.009901,0.0008425,0.01113,0.0005605,0.0007565,1.678e-05,0.000739,1.448e-07,0.003359,0.009431,1.554e-09,0.008737,0.02991,0.0004932,2.306e-05,0.0005925,0.004994,0.001419,0.001333,0.03263,2.134e-05,0.02523,1.29e-06,0.03292,0.0343,0.03475,0.001894,0.01669,3.403e-07,5.627e-08,0.02377,4.182e-07,0.01737,0.007261,0.001188,8.721e-12,4.353e-05,0.0002132,7.911e-08,0.01789,7.573e-10,5.459e-06,2.25e-05,0.01097,4.684e-06,0.00898,6.305e-07,0.01641,0.02681,0.001757,2.736e-05,6.901e-05,0.0001419,0.00119,1.214e-05,8.649e-05,0.0003938,8.655e-16,0.0003473,0.0348,0.009411,4.422e-09,2.65e-16,0.002601,0.007795,0.002613,3.917e-06,3.029e-12,8.849e-11,3.655e-09,6.035e-08,2.809e-05,1.094e-08,1.835e-10,1.276e-06,5.158e-15,0.0004556,1.014e-06,0.000129,7.232e-19,1.823e-06,6.647e-06,6.961e-09,0.01163,0.002594,2.28e-05,4.107e-06,5.847e-07,6.208e-09,4.118e-07,0.001918,0.001528,2.352e-18,0.004039,5.958e-05,0.003249,2.419e-08,2.254e-07,1.608e-06,2.203e-05,0.009187,0.0003958,1.001e-06,5.717e-05,0.006156,1.872e-07,1.776e-13,5.558e-11,0.002415,1.65e-16,6.898e-13,3.919e-07,0.01052,6.167e-11,2.082e-05,1.258e-05,8.37e-09,3.215e-09,1.867e-06,1.577e-07,5.971e-05,9.048e-05,2.342e-06,1.373e-07,0.003514,7.246e-12,3.682e-07,2.002e-07,1.304e-08,2.215e-12,0.0008377,0.001086,5.079e-05,0.01094,0.0003216,7.783e-16,0.0005557,5.418e-09,0.0001302,1.742e-10,2.992e-05,0.0005613,2.956e-06,2.791e-11,1.371e-08,0.0001972,0.0002491,0.02648,1.771e-07,9.704e-05,9.455e-05,0.003685,1.627e-05,1.383e-13,5.65e-05,0.01163,3.811e-14,8.863e-09,0.0001111,6.952e-06,2.424e-05,5.099e-14,2.746e-05,2.284e-12,0.0001073,0.003838,2.092e-05,9.021e-10,7.262e-17,1.85e-06,0.009185,0.005159,0.001958,5.937e-08,1.571e-11,2.196e-08,0.009154,8.034e-05,0.01062,0.006299,3.318e-06,0.0003841,6.628e-07,6.895e-16,0.007738,2.981e-16,1.143e-14,8.553e-11,1.976e-10,1.555e-11,1.865e-07,3.617e-11,0.004043,4.685e-10,5.782e-07,0.0007673,1.625e-05,1.891e-13,0.0005646,2.149e-09,5.913e-07,9.106e-09,0.004139,0.000147,6.836e-07,2.214e-06,5.429e-05,2.213e-07,7.118e-09,1.348e-16,1.536e-11,8.075e-12,0.005328,0.001572,1.269e-14,0.007436,7.262e-11,0.0002939,6.728e-10,8.609e-06,4.102e-06,9.755e-06,2.76e-08,3.188e-06,1.593e-07,1.872e-13,0.0006205,5.515e-11,1.224e-06,7.589e-06,7.407e-07,0.0001085,0.0002239,0.000195,6.76e-09,0.008248,5.465e-14,6.072e-16,3.117e-05,0.005295,8.868e-17,0.002835,9.738e-13,4.855e-13,2.886e-05,1.431e-12,0.00181,0.003405,0.006163,1.701e-09,2.378e-07,5.814e-09,0.01654,0.0007065,4.363e-18,1.094e-07,2.2e-05,0.0001415,1.025e-09,8.216e-10,0.01804,2.683e-06,2.849e-05,1.952e-08,0.00645,1.374e-15,1.033e-09,2.83e-08,1.331e-09,6.714e-05,0.001623,8.03e-08,5.407e-19,1.369e-06,0.0001366,0.02605,0.001954,0.001491,1.159e-08,0.0001029,2.697e-18,5.033e-17,0.001006,7.165e-08,2.593e-14,2.596e-14,0.03053,9.987e-05,0.0004347,9.672e-05,0.004411,8.052e-05,1.123e-19,2.279e-16,1.912e-05,7.355e-10,0.002499,8.456e-08,0.0003076,6.646e-08,3.464e-16,0.005514,2.173e-07,2.074e-09,8.014e-15,6.292e-07,3.58e-18,2.644e-22,5.68e-17,3.818e-11,2.735e-05,0.00293,1.389e-15,0.003555,0.006879,1.048e-17,3.714e-06,2.098e-05,2.139e-10,8.748e-05,1.209e-09,0.003265,0.0009113,2.4e-06,7.619e-05,0.002221,9.132e-08,0.007604,1.039e-07,0.004622,3.699e-09,2.016e-18,4.531e-20,1.656e-09,1.192e-07,3.499e-14,6.27e-15,3.43e-10,6.112e-12,5.44e-07,0.03537,1.806e-14,0.0007218,1.323e-08,3.894e-06,4.944e-06,0.0001569,0.001802,8.022e-05,8.656e-05,9.393e-16,1.024e-05,1.193e-07,0.004348,0.003402,7.218e-09,0.0003894,0.0001745,0.0006788,7.864e-07,3.953e-06,4.099e-10,4.891e-15,5.587e-12,3.226e-11,4.491e-10,5.395e-12,2.795e-05,9.931e-08,2.853e-13,0.00854,6.353e-07,1.13e-10,3.645e-07,5.705e-15,4.353e-05,9.402e-10,3.983e-15,1.357e-05,0.007816,5.104e-06,0.001633,0.001131,7.125e-05,0.002038,1.551e-12,1.834e-06,1.05e-06,5.972e-11,0.001611,3.403e-05,1.82e-05,1.646e-12,8.081e-05,4.861e-07,0.002828,1.328e-07,1.333e-12,1.638e-11,5.959e-14,0.0005835,7.504e-10,2.639e-06,0.009005,1.018e-06,9.06e-08,2.178e-06,6.67e-11,2.281e-13,0.000346,1.851e-07,3.29e-10,1.427e-10,1.112e-09,5.019e-10,2.494e-13,0.007879,6.45e-05,7.701e-05,1.263e-07,0.02557,0.003783,3.683e-09,0.003208,0.0002399,3.271e-07,9.406e-05,4.59e-09,2.08e-06,0.0006978,1.311e-05,7.022e-08,3.575e-15,0.02845,2.167e-16,0.001817,8.042e-05,9.871e-08,7.771e-06,1.407e-15,1.548e-05,0.00908,3.674e-05,1.058e-08,3.216e-07,1.982e-05,3.945e-08,1.124e-06,3.441e-11,9.14e-20,3.181e-11,0.0003278,8.442e-06,1.531e-12,2.951e-11,1.145e-06,3.301e-07,1.494e-09,0.0009654,3.985e-05,6.885e-08,0.00278,9.969e-05,0.001854,2.792e-12,9.789e-15,7.975e-13,1.324e-07,5.515e-07,2.599e-10,0.001511,1.874e-06,3.698e-07,0.0002527,1.16e-06,0.0006507,7.92e-06,2.389e-14,3.577e-12,3.978e-10,9.779e-09,0.00588,4.419e-06,4.147e-08,0.0002437,0.00327,5.207e-13,8.513e-06,1.682e-06,0.0255,0.02272,2.482e-07,4.119e-08,3.07e-09,0.0106,0.0006723,4.335e-09,3.133e-07,1.007e-08,3.258e-05,2.528e-09,2.461e-05,0.000249,5.326e-11,3.073e-07,1.742e-16,1.404e-06,1.717e-09,5.693e-14,4.845e-05,3.493e-09,1.558e-11,9.984e-11,8.518e-11,0.0002676,2.31e-10,6.54e-06,3.415e-06,0.003414,0.02009,0.01076,3.062e-06,0.01863,1.018e-10,1.409e-16,8.287e-10,1.213e-11,2.292e-08,3.423e-07,5.875e-12,0.003606,2.939e-09,0.03167,7.74e-05,1.651e-05,0.0008689,1.278e-06,2.969e-09,0.002097,1.175e-07,1.888e-05,0.0006543,0.0001005,0.001843,2.37e-08,9.545e-13,7.49e-05,1.098e-13,6.876e-12,1.915e-12,9.12e-05,1.479e-08,2.556e-11,3.945e-11,2.723e-05,0.0001299,9.539e-12,0.008963,0.0001816,1.316e-09,0.006346,6.241e-11,0.0002067,0.0001002,0.00151,6.734e-06,0.003197,0.0001624,4.657e-05,5.019e-05,5.969e-06,3.57e-05,2.11e-08,0.01877,3.09e-11,7.165e-07,0.001426,4.212e-08,1.261e-09,6.124e-12,0.006385,2.039e-05,7.339e-17,9.568e-14,1.654e-08,3.964e-14,0.0001608,7.559e-22,4.085e-12,7.371e-05,0.002443,2.156e-13,9.372e-05,1.65e-10,6.58e-06,0.001442,4.139e-14,8.405e-14,4.651e-09,8.3e-05,8.936e-05,5.726e-11,3.218e-09,0.0001998,1.429e-05,0.003527,3.169e-11,2.587e-12,8.01e-05,2.482e-08,1.203e-07,1.589e-08,1.503e-15,2.873e-10,7.036e-06,1.406e-11,1.27e-09,1.449e-08,0.005924,2.109e-16,0.0004327,6.223e-05,8.312e-11,1.313e-05,8.984e-15,0.001691,1.032e-16,1.292e-12,1.413e-05,0.001907,0.001472,0.001117,1.477e-08,0.0258,6.756e-09,3.779e-05,4.266e-18,7.527e-05,7.947e-11,0.005428,0.002556,3.517e-16,9.093e-08,0.00515,3.18e-05,3.166e-15,2.125e-12,0.0004712,3.549e-07,0.004886,3.119e-07,1.451e-05,3.439e-07,0.003322,2.306e-05,1.624e-05,1.402e-08,0.0001134,1.565e-07,0.0001359,1.237e-06,1.362e-06,2.431e-12,5.513e-10,1.517e-09,2.64e-06,0.001167,2.81e-09,0.001304,0.006942,7.62e-10,1.059e-11,6.97e-10,2.049e-11,0.003208,0.02779,0.01817,6.665e-13,2.571e-07,0.001643,4.401e-20,3.373e-11,0.005771,4.449e-08,6.044e-06,4.225e-07,6.454e-05,4.268e-07,0.0005506,1.535e-07,1.369e-05,1.178e-11,8.136e-10,5.916e-11,8.885e-08,0.0001312,0.0006599,1.405e-05,1.884e-06,0.0004632,8.635e-13,2.328e-08,1.743e-13,2.873e-12,0.0001169,5.667e-08,0.001044,0.0005006,8.679e-06,4.249e-07,7.461e-08,0.002118,1.664e-08,1.595e-06,6.329e-06,2.991e-05,2.021e-07,7.558e-05,4.679e-07,0.02195,2.16e-05,0.0002152,1.039e-07,8.22e-07,2.371e-07,7.083e-05,1.874e-05,0.01702,3.977e-09,0.03116,7.72e-08,0.0003377,7.838e-05,6.283e-08,0.0006327,0.005652,0.003531,3.659e-16,2.028e-11,6.895e-17,0.0001822,4.1e-05,1.776e-06,2.079e-07,2.348e-16,2.836e-12,4.453e-09,0.004071,5.984e-08,2.828e-08,0.0001539,0.00235,6.455e-11,2.605e-10,1.526e-12,3.747e-15,0.004499,0.005447,0.0001614,7.094e-12,1.097e-10,0.003982,4.525e-08,1.323e-07,0.01449,2.624e-11,2.103e-10,3.772e-06,1.82e-05,2.045e-12,3.734e-16,2.261e-14,3.475e-07,4.093e-11,2.087e-11,2.97e-05,2.458e-09,0.003509,5.236e-14,1.288e-10,1.473e-07,0.0004343,0.004516,7.083e-09,2.683e-07,8.502e-08,8.433e-05,0.00536,1.014e-11,8.765e-05,7.778e-10,9.328e-07,0.03153,5.823e-10,0.006797,3.406e-12,8.11e-09,0.03497,3.896e-11,3.36e-05,1.291e-27,1.216e-15,0.0003995,3.171e-09,8.461e-16,6.406e-10,0.002824,3.965e-08,1.782e-10,2.958e-09,6.008e-05,2.487e-08,1.267e-05,7.826e-11,9.039e-06,0.03308,3.445e-09,5.224e-10,4.488e-06,0.0007482,0.00181,7.098e-07,0.0001028,2.285e-05,0.0001691,1.897e-06,1.696e-15,0.003057,1.415e-06,0.0002585,0.001342,6.397e-11,0.0001747,2.945e-11,1.086e-08,5.795e-06,1.55e-08,1.3e-07,1.196e-16,2.652e-13,2.548e-10,2.192e-10,0.002858,0.001765,0.0003119,3.526e-07,0.01844,1.381e-09,0.001855,6.611e-18,0.002349,5.303e-08,1.091e-17,1.264e-06,0.02121,9.499e-08,2.918e-09,0.0002179,2.218e-10,1.348e-08,1.268e-08,1.905e-13,1.817e-09,3.102e-06,9.808e-12,2.714e-10,6.852e-08,0.001621,0.0003372,8.479e-08,0.01555,2.486e-14,2.756e-05,0.0008624,0.0003154,7.647e-15,1.619e-06,6.002e-05,9.822e-17,3.357e-12,3.171e-12,3.482e-14,5.751e-08,1.043e-05,0.002697,2.851e-09,9.271e-07,1.055e-06,1.651e-07,8.541e-09,2.927e-12,0.0001022,1.918e-05,0.01055,3.786e-14,1.155e-07,0.0001633,8.121e-11,5.457e-07,2.13e-15,0.0001554,3.354e-05,1.057e-09,6.041e-11,0.003362,9.71e-12,1.354e-08,5.459e-19,3.314e-05,0.0009087,0.0005722,0.02019,4.429e-06,6.692e-09,9.705e-05,4.264e-11,6.776e-16,0.004368,0.0002845,0.0005394,2.655e-09,0.002514,0.001694,0.01775,6.881e-09,0.02222,1.239e-12,4.71e-05,1.726e-15,1.473e-05,1.549e-07,0.0004195,0.0006563,0.0007307,8.141e-15,2.362e-05,5.192e-10,3.749e-07,1.193e-06,9.403e-10,0.003016,0.02962,0.0005098,0.000745,1.282e-09,3.974e-09,2.42e-09,3.241e-07,1.043e-07,0.004279,5.606e-05,0.01221,1.399e-05,2.594e-05,1.657e-07,3.095e-05,0.0001485,4.579e-07,5.858e-05,9.6e-18,0.0007709,2e-09,1.028e-18,0.03092,3.447e-08,2.496e-08,8.026e-07,1.054e-05,2.15e-09,2.031e-08,8.287e-06,2.508e-07,2.069e-08,5.813e-10,2.074e-06,2.206e-12,1.057e-05,1.299e-05,0.005291,0.001337,0.001326,4.528e-14,8.042e-13,1.085e-07,9.733e-06,0.0003168,0.0006035,0.02067,1.393e-05,0.0003949,1.736e-10,5.053e-07,2.409e-07,2.494e-07,0.0007612,0.003725,1.032e-08,0.0003372,7.429e-05,0.0002906,0.0002182,0.0006127,6.176e-06,2.935e-05,1.376e-10,6.185e-15,1.258e-10,4.31e-07,0.009952,0.002536,1.83e-10,4.391e-09,0.0003579,4.792e-07,0.001155,0.01491,2.482e-11,1.015e-06,0.008578,2.29e-09,1.673e-17,2.153e-10,3.204e-10,0.007027,1.757e-07,2.702e-11,0.001461,1.115e-12,0.001209,8.474e-13,8.613e-06,0.003101,5.156e-06,2.337e-06,5.673e-06,5.917e-11,0.008212,0.008576,4.197e-07,5.576e-10,0.0002371,0.0002902,7.276e-15,3.672e-13,3.866e-07,0.0002288,1.027e-05,0.001206,1.987e-05,3.367e-19,0.0002998,2.832e-12,3.918e-06,0.0288,0.001701,8.749e-11,2.632e-08,8.015e-05,0.001169,1.922e-12,0.002595,4.211e-12,1.469e-10,5.075e-10,0.00555,6.745e-10,5.483e-11,6.73e-13,0.0003516,9.26e-13,3.997e-10,0.002087,4.217e-05,2.101e-06,0.0006579,7.419e-11,8.494e-15,6.571e-06,6.804e-09,2.052e-05,1.123e-05,0.0001297,2.417e-10,0.00193,0.008366,1.94e-05,6.879e-12,2.802e-09,0.003593,3.313e-09,0.0006605,7.479e-15,0.01523,2.818e-08,4.583e-06,3.757e-08,0.00496,0.00961,0.006864,5.07e-07,0.005115,6.278e-05,0.000192,4.531e-05,0.000157,6.081e-05,0.0008425,1.126e-05,2.416e-15,3.165e-10,1.545e-09,6.71e-17,0.02402,0.005021,5.421e-10,0.0009649,2.026e-08,0.003029,0.01208,1.102e-10,1.763e-07,1.122e-06,6.481e-09,0.0002522,4.409e-09,4.096e-07,1.835e-09,1.779e-10,9.898e-08,1.034e-11,2.029e-08,0.01234,3.541e-16,3.302e-05,0.0003252,0.0001818,7.855e-11,3.184e-05,4.467e-07,0.001079,0.007272,0.009248,4.017e-05,4.285e-12,4.785e-14,0.01068,8.044e-13,0.0006837,0.003214,4.821e-14,1.377e-18,0.0001988,1.556e-07,5.71e-09,2.339e-10,1.438e-08,3.188e-06,5.907e-05,2.205e-06,1.223e-09,0.00529,0.001195,3.918e-05,7.138e-05,0.0001694,4.033e-07,0.001628,5.749e-12,8.653e-12,1.009e-11,0.00811,3.934e-13,1.414e-10,3.023e-07,1.669e-07,0.0003939,0.002044,0.01234,0.02383,3.941e-05,0.004085,0.0005603,5.899e-15,8.902e-18,0.001947,0.005576,8.807e-06,0.0001065,0.0004409,1.557e-08,1.819e-08,9.288e-14,1.587e-13,2.665e-09,4.495e-08,5.89e-05,4.003e-10,8.453e-13,2.557e-05,1.841e-06,1.88e-12,5.294e-11,0.0209,6.392e-11,3.341e-12,2.865e-06,0.01297,9.21e-06,0.001264,2.221e-07,3.13e-08,0.003983,7.222e-05,0.004679,0.0005099,2.164e-08,3.663e-05,1.251e-05,3.459e-10,0.001439,3.09e-08,0.0005732,2.751e-05,5.702e-15,7.975e-07,7.502e-05,1.291e-09,5.67e-08,0.004089,2.262e-05,9.932e-08,0.002102,3.243e-14,1.43e-12,9.096e-08,1.155e-07,6.343e-09,4.791e-08,5.288e-12,0.0009912,8.566e-05,9.987e-16,2.601e-15,0.002192,1.367e-07,5.529e-11,1.543e-11,7.863e-12,0.0001934,3.562e-05,2.691e-15,1.056e-15,4.782e-05,5.271e-12,8.891e-15,0.0007443,0.008898,0.03184,5.244e-09,8.865e-08,5.456e-07,0.007221,7.703e-07,0.0006857,4.899e-05,5.709e-21,1.832e-08,1.366e-09,2.474e-07,1.071e-07,1.242e-07,5.911e-06,8.42e-08,2.002e-06,1.424e-09,8.787e-10,4.967e-05,7.559e-10,3.057e-09,6.986e-15,0.001841,8.524e-17,2.077e-10,8.486e-08,7.127e-13,1.343e-06,0.0002386,9.537e-18,0.0005609,0.0002907,7.682e-14,0.002379,7.15e-09,5.212e-08,0.0003203,0.0001788,5.934e-07,1.529e-11,0.0001305,3.293e-16,4.349e-19,4.415e-07,1.009e-09,2.527e-05,3.192e-08,3.758e-14,2.829e-08,1.395e-11,2.296e-08,0.0002277,3.516e-05,0.002501,4.313e-06,6.6e-12,6.032e-12,0.0004755,2.766e-12,4.285e-10,1.352e-09,3.251e-11,2.932e-08,0.0002505,0.01929,8.935e-09,9.559e-06,5.356e-07,6.929e-05,1.394e-06,9.821e-06,0.01628,0.0008927,0.01285,2.04e-05,0.00195,9.728e-06,0.0005139,3.743e-05,1.768e-05,7.632e-05,3.685e-05,4.487e-07,0.0007275,2.567e-06,9.608e-16,7.703e-11,0.00593,0.0001494,3.172e-13,0.003581,2.383e-10,3.433e-06,5.558e-06,1.009e-07,5.505e-09,9.126e-05,4.572e-05,9.978e-07,4.923e-05,6.222e-15,4.087e-16,9.555e-11,9.658e-06,0.0002494,3.408e-05,6.886e-09,6.001e-07,5.795e-08,1.995e-05,1.236e-05,6.604e-08,6.201e-05,1.819e-07,9.281e-18,8.64e-09,0.0001525,3.88e-12,0.000648,0.0001876,0.02969,1.411e-11,3.691e-14,0.003891,3.044e-08,8.034e-07,0.0001042,2.294e-05,0.0003074,3.101e-08,2.858e-06,0.0008302,0.0002906,1.558e-05,5.058e-07,6.666e-07,0.0002216,3.936e-06,1.176e-06,5.056e-08,6.235e-05,0.0003425,0.0002006,0.001424,0.0005607,6.723e-07,1.6e-09,1.883e-07,2.823e-06,0.001375,4.011e-08,1.863e-12,2.32e-08,6.388e-05,3.234e-07,3.084e-08,0.01467,3.865e-09,0.002891,0.001475,0.0002691,1.156e-05,0.000857,3.131e-09,4.881e-10,1.603e-08,3.141e-18,7.329e-08,0.02884,0.0002593,1.72e-11,4.588e-06,0.003417,0.00407,2.136e-14,2.368e-06,9.856e-06,7.137e-13,0.0004578,4.07e-09,8.595e-05,2.767e-11,1.361e-12,8.154e-10,1.458e-15,0.0003765,8.202e-05,0.006019,0.02526,4.348e-07,0.00284,0.01324,6.299e-09,8.63e-10,1.428e-08,2.011e-08,0.02953,7.998e-07,2.711e-09,0.0004222,4.149e-07,6.176e-05,1.829e-07,6.777e-10,0.001875,0.0004249,1.703e-14,6.097e-05,3.415e-10,1.556e-06,4.269e-06,6.437e-08,3.004e-09,3.783e-07,1.756e-09,4.539e-09,1.66e-05,0.0006327,1.754e-09,0.001186,3.294e-15,5.2e-10,0.0023,0.005527,0.007023,1.837e-06,0.00116,0.000421,5.865e-14,1.296e-13,8.944e-05,0.0006753,3.703e-05,2.729e-16,3.675e-05,2.154e-11,7.121e-09,1.263e-10,0.0322,1.512e-09,2.375e-08,2.138e-09,9.893e-11,4.176e-11,7.229e-09,0.01607,2.891e-12,5.124e-12,1.837e-15,0.01291,2.169e-12,5.851e-06,8.838e-11,8.26e-05,2.886e-05,0.0001522,1.582e-13,1.014e-07,7.276e-06,2.416e-09,2.673e-08,7.54e-09,6.007e-07,2.86e-05,7.942e-07,1.271e-05,0.005173,2.854e-17,2.304e-05,4.26e-05,1.738e-06,0.01062,1.28e-09,0.0006568,5.734e-05,2.296e-08,1.39e-08,0.0005105,6.78e-13,5.227e-11,3.804e-05,1.779e-11,1.041e-16,0.01267,4.107e-05,7.779e-06,5.259e-05,1.272e-05,1.227e-10,1.315e-06,6.312e-15,8.71e-07,0.0001869,1.978e-08,0.0002072,5.573e-05,4.848e-05,5.483e-11,1.685e-06,1.358e-07,2.972e-09,1.316e-06,2.419e-07,9.693e-11,1.378e-07,3.379e-11,1.994e-11,1.62e-07,2.656e-05,1.786e-05,5.793e-05,1.265e-11,3.396e-06,0.01358,0.0003784,0.005278,2.601e-05,6.505e-08,3.584e-09,8.347e-08,6.223e-05,8.464e-10,9.681e-17,0.03487,8.198e-19,4.406e-05,8.556e-11,1.627e-07,1.83e-07,5.168e-07,1.851e-08,3.443e-07,0.0004906,1.29e-05,1.273e-17,0.0001495,3.006e-06,1.19e-05,1.958e-13,0.0006388,0.001274,0.02639,2.096e-13,8.168e-09,3.064e-10,2.758e-08,0.003049,1.661e-12,0.001271,3.551e-16,3.278e-10,2.336e-14,0.001202,0.0001618,1.396e-05,1.753e-06,9.672e-06,0.0003532,0.002846,4.705e-08,1.609e-06,0.02675,1.407e-08,3.51e-09,0.009765,4.787e-09,0.0002902,0.0006182,4.66e-06,7.84e-06,0.01535,6.609e-06,0.001376,1.591e-07,2.681e-15,1.182e-15,3.803e-11,1.98e-10,2.542e-08,5.324e-12,2.902e-12,1.244e-06,6.163e-10,0.0003925,2.363e-08,2.489e-09,2.314e-08,2.421e-12,3.223e-05,1.849e-05,4.407e-05,1.669e-10,4.348e-08,2.468e-06,1.996e-11,5.671e-07,0.002622,0.008644,3.829e-10,3.49e-06,0.002612,0.004271,0.000158,8.731e-09,2.874e-06,5.327e-05,1.364e-07,3.718e-07,4.354e-17,5.683e-08,8.134e-07,4.958e-05,1.753e-07,2.806e-19,1.169e-07,0.0001405,0.002958,1.502e-07,3.288e-09,0.002968,1.075e-17,4.197e-13,1.436e-09,5.711e-11,0.0009662,2.792e-20,2.694e-13,0.0004114,0.0003821,3.435e-11,0.0003629,3.218e-15,4.685e-07,5.561e-05,3.883e-05,0.001086,2.208e-08,1.207e-08,0.0001162,1.875e-10,1.332e-12,1.159e-18,0.004908,1.389e-10,0.001296,0.001201,3.072e-12,4.895e-08,0.0002052,4.315e-07,2.742e-08,1.929e-12,0.002792,0.0002467,0.000189,3.803e-07,1.8e-08,0.02883,4.089e-06,1.305e-09,1.132e-09,0.005065,6.181e-09,2.662e-07,0.006208,6.714e-07,0.01973,0.0009412,0.001532,0.0296,0.001465,0.005259,1.588e-17,0.009671,0.001943,0.0009232,0.003512,0.0002268,5.919e-13,1.844e-08,3.771e-13,0.0002284,1.739e-07,2.066e-09,0.00295,0.0002844,0.0002674,5.044e-09,6.993e-09,0.001912,3.355e-06,0.005165,0.0004087,0.0001862,0.02363,2.982e-12,0.001001,3.602e-09,3.501e-20,1.077e-06,3.036e-05,0.0005389,1.519e-06,1.334e-10,0.0001115,0.01136,0.0001103,1.802e-07,6.545e-06,9.634e-09,1.483e-14,0.0002577,7.471e-14,0.007722,0.008237,1.976e-10,6.346e-07,5.854e-06,0.000199,6.522e-08,0.001435,1.873e-09,0.0001893,6.981e-06,1.977e-07,0.01831,0.0001698,1.928e-11,0.0001045,4.762e-07,6.14e-06,1.733e-07,1.825e-10,0.01585,0.001286,3.087e-06,0.004284,5.043e-07,1.336e-14,3.876e-06,3.974e-05,0.0003768,0.00463,4.141e-11,0.02244,7.415e-05,7.63e-08,0.0001189,7.885e-19,0.006583,0.0002252,1.275e-07,1.451e-10,7.722e-06,1.784e-08,1.147e-10,2.587e-16,2.242e-06,0.0005911,1.134e-11,2.142e-09,0.001465,1.197e-08,0.0001519,5.883e-05,5.423e-06,0.008655,6.856e-10,1.476e-08,0.001623,7.164e-14,2.398e-07,0.0003618,0.003315,1.269e-12,2.967e-08,2.271e-12,0.0005409,1.81e-12,2.346e-07,0.004638,5.248e-11,6.149e-11,1.462e-05,4.416e-06,3.244e-05,0.004289,0.0007273,0.002744,3.316e-09,1.1e-08,1.898e-08,3.246e-05,4.03e-06,3.748e-12,2.526e-07,2.52e-08,2.709e-06,4.007e-08,0.008956,1.339e-05,0.0003753,1.954e-05,9.589e-13,0.0001251,3.84e-07,4.225e-06,8.647e-07,4.613e-10,2.779e-06,0.001307,1.992e-10,1.298e-05,0.008148,6.243e-06,1.14e-06,2.293e-12,2.15e-08,0.02909,0.03166,0.001354,1.448e-13,1.464e-07,0.03196,0.002512,3.027e-22,5.862e-11,5.651e-05,3.726e-06,2.055e-05,4.364e-07,4.383e-14,1.345e-12,0.0004428,6.43e-08,8.674e-05,2.35e-10,1.201e-11,0.01177,0.0001163,1.589e-06,0.0008567,0.0004969,2.84e-08,8.103e-09,1.138e-12,3.359e-05,0.005673,0.0008976,0.0001027,2.72e-16,3.055e-11,3.839e-15,2.505e-06,0.0008314,5.386e-05,4.603e-06,1.72e-11,2.116e-05,4.294e-12,1.487e-07,2.327e-06,4.594e-06,7.661e-11,3.051e-10,1.138e-07,1.114e-08,2.817e-07,0.005416,0.0124,4.659e-05,1.086e-08,9.421e-09,7.237e-11,3.273e-09,0.0002115,0.01062,0.01836,0.00712,0.0003507,0.002004,0.001087,1.393e-05,0.02129,0.004701,0.0002949,2.529e-09,1.314e-06,4.766e-06,2.186e-06,0.008569,0.004163,9.693e-11,3.074e-08,1.237e-08,3.839e-06,5.491e-09,0.001222,2.998e-12,1.473e-17,6.956e-06,2.228e-16,1.454e-05,0.01374,0.008464,0.01765,2.699e-09,4.022e-11,3.749e-14,1.203e-07,4.445e-13,1.402e-06,1.125e-10,7.233e-08,4.027e-06,1.262e-12,9.019e-17,1.444e-08,2.548e-14,1.096e-07,0.003374,0.0124,9.487e-10,1.566e-07,5.818e-12,2.808e-15,9.926e-06,3.15e-06,0.01487,2.309e-07,0.0006686,4.795e-05,2.043e-09,0.0001573,3.079e-05,0.0028,0.012,0.0001279,0.0006696,5.551e-07,1.565e-07,0.002984,0.004717,1.396e-05,7.985e-13,9.144e-10,9.763e-06,4.219e-09,0.005009,0.0005777,0.0005866,0.0003478,6.061e-09,0.00728,0.0002142,0.0001668,3.503e-09,1.847e-05,6.335e-10,3.958e-06,3.652e-08,1.614e-06,3.08e-06,2.573e-06,9.916e-18,2.518e-10,1.946e-07,2.051e-06,0.000303,9.402e-05,0.0006168,5e-08,2.13e-09,0.0001368,0.009792,4.078e-13,0.001992,4.713e-08,4.46e-05,3.413e-07,6.382e-07,2.839e-06,1.847e-08,2.485e-10,3.188e-07,0.00405,1.709e-07,0.0001122,0.006676,2.963e-13,0.0005313,4.997e-05,3.603e-06,4.515e-06,0.002089,1.493e-06,0.0001657,2.12e-05,3.357e-09,0.001635,6.088e-11,2.17e-06,0.0009725,1.643e-08,0.0006263,2.654e-11,1.968e-14,1.659e-05,6.265e-05,4.382e-08,2.648e-18,2.222e-08,0.02582,1.228e-17,0.007391,3.372e-06,5.027e-09,7.957e-10,3.225e-06,0.0009081,8.73e-07,1.583e-13,5.358e-12,2.779e-08,5.88e-09,3.805e-07,0.0007994,1.193e-09,2.746e-09,0.008805,8.547e-25,5.506e-08,4.173e-05,2.53e-07,5.075e-06,1.243e-07,0.0001571,8.046e-12,5.316e-06,2.793e-10,2.195e-05,2.585e-06,4.223e-11,8.524e-07,0.001128,6.47e-07,1.299e-09,0.004397,0.003388,3.251e-07,1.451e-08,3.096e-07,8.456e-12,0.006816,2.531e-12,1.707e-05,0.002612,1.898e-11,0.007019,0.02573,2.136e-07,1.285e-05,0.005171,0.0002488,3.7e-18,0.002587,7.133e-08,1.32e-05,1.064e-05,0.00026,8.594e-09,0.0003787,5.446e-06,1.454e-10,4.503e-10,5.597e-09,1.005e-09,5.133e-10,3.581e-05,0.0006592,0.005154,2.389e-11,3.757e-15,2.72e-17,0.0001375,0.000101,8.084e-13,2.683e-10,2.404e-05,2.018e-07,4.505e-11,4.206e-08,6.477e-18,2.062e-05,5.916e-07,6.527e-05,8.386e-07,6.582e-07,0.001696,1.226e-06,0.01282,1.629e-06,4.503e-10,4.468e-13,3.704e-08,1.501e-05,0.009493,1.984e-07,0.0001491,9.736e-10,0.0001746,1.124e-08,3.864e-10,7.332e-08,2.171e-05,0.002017,6.501e-09,1.032e-08,4.253e-05,8.945e-10,0.001124,9.558e-05,0.005178,1.612e-07,2.246e-16,2.476e-05,0.008542,9.622e-15,0.00545,1.546e-07,0.03038,2.875e-10,2.831e-06,0.0001216,0.002307,6.201e-09,8.612e-11,1.029e-09,6.065e-10,4.209e-06,3.33e-05,4.898e-16,0.0002168,8.479e-10,8.393e-10,6.427e-05,6.257e-05,0.001191,0.004457,2.115e-05,9.349e-05,8.891e-09,0.0001388,7.886e-07,7.391e-09,8.957e-06,2.447e-11,2.025e-09,2.715e-10,2.705e-05,4.566e-05,0.00484,1.452e-05,1.671e-08,1.781e-05,0.00703,4.875e-06,2.595e-07,3.108e-07,0.001926,1.271e-10,0.0004132,1.399e-08,0.001524,6.505e-08,0.0003782,1.119e-10,0.01689,7.139e-05,7.456e-09,6.042e-07,2.057e-14,2.328e-05,1.386e-06,0.02745,4.785e-14,1.205e-09,0.0001951,1.908e-07,0.0005068,8.465e-06,0.00379,5.123e-11,0.0001777,0.01713,4.874e-06,2.565e-09,1.816e-09,8.12e-14,9.201e-11,2.367e-09,3.454e-11,1.753e-14,1.372e-05,0.01024,4.64e-11,1.687e-13,4.34e-12,3.146e-13,1.944e-05,1.126e-16,2.413e-12,8.961e-06,1.17e-05,1.127e-05,7.172e-09,2.999e-05,2.167e-06,4.307e-05,0.01577,3.662e-06,0.001527,0.01926,9.995e-16,3.151e-16,0.005592,2.555e-13,4.007e-09,1.401e-14,4.399e-13,4.78e-11,5.071e-11,1.946e-12,6.943e-17,0.0001646,0.001531,6.263e-05,1.617e-06,1.708e-07,1.88e-14,0.0001337,2.212e-08,0.0001111,2.8e-05,2.41e-05,1.847e-06,3.452e-12,2.909e-06,2.552e-07,0.0007517,8.889e-09,0.0151,2.041e-10,1.316e-06,0.01352,3.193e-08,1.357e-05,1.658e-05,1.41e-15,1.926e-05,7.854e-05,1.013e-07,3.587e-09,1.927e-06,0.000122,5.272e-07,1.357e-09,0.01716,5.993e-06,0.003349,2.642e-06,1.41e-08,0.0007665,0.0007992,5.276e-07,1.246e-09,3.002e-06,6.51e-06,1.962e-05,1.742e-07,2.316e-07,3.039e-16,7.468e-06,2.982e-12,3.772e-08,0.00338,2.775e-13,4.635e-07,0.00406,0.003697,3.987e-15,0.0001454,5.27e-07,6.867e-17,4.629e-06,0.001829,6.409e-06,9.84e-08,1.26e-08,3.202e-11,1.026e-08,7.736e-06,5.065e-08,0.0001554,4.366e-06,1.308e-06,1.01e-07,2.229e-05,2.97e-06,3.207e-05,0.0001514,3.56e-08,3.921e-16,8.228e-07,2.95e-09,1.542e-08,6.333e-06,3.857e-09,8.596e-05,0.02017,1.877e-08,5.455e-06,7.947e-10,0.001161,0.009153,0.002156,3.924e-07,0.002084,2.556e-13,4.592e-12,7e-11,3.532e-09,2.581e-12,7.913e-12,6.2e-15,1.581e-08,2.179e-18,1.349e-19,4.181e-10,0.001261,0.007429,0.001881,0.007109,5.175e-07,1.802e-12,5.153e-08,0.000841,0.001579,2.825e-07,5.448e-06,2.877e-06,1.118e-06,0.004552,5.012e-05,1.896e-11,2.712e-19,4.079e-08,2.742e-10,4.299e-08,1.972e-06,0.0002108,6.233e-05,4.792e-07,0.0001587,0.0005902,3.4e-09,7.421e-09,3.896e-07,3.965e-05,2.426e-08,4.941e-08,9.573e-07,2.521e-14,4.854e-08,0.001658,1.207e-05,3.47e-13,7.042e-09,8.533e-17,0.02382,0.003452,7.217e-07,7.891e-08,8.621e-10,0.01507,4.89e-15,4.917e-08,4.55e-05,0.007455,0.0001928,5.763e-07,7.901e-12,1.187e-08,4.383e-05,1.725e-07,3.643e-07,1.523e-09,2.747e-06,2.578e-08,5.217e-16,0.002039,0.01756,0.001719,2.698e-08,0.0005931,2.819e-13,4.852e-05,1.306e-09,0.0005782,7.853e-13,5.872e-16,2.396e-06,5.871e-05,7.987e-10,6.401e-14,1.623e-12,2.323e-07,3.599e-08,4.473e-06,8.64e-13,1.083e-12,3.824e-18,1.281e-05,0.0003365,2.041e-07,2.849e-17,8.602e-15,0.02552,3.747e-12,6.288e-12,3.89e-15,2.118e-15,1.263e-12,0.0001854,8.22e-07,0.02377,8.818e-09,3.229e-08,2.221e-07,4.359e-09,5.026e-12,7.956e-05,7.488e-09,4.812e-14,7.939e-13,2.538e-05,1.361e-07,7.982e-10,3.624e-05,7.09e-05,3.139e-18,3.081e-10,0.01406,4.161e-08,1.219e-05,2.402e-10,0.005103,4.251e-11,2.073e-13,1.976e-08,0.0001301,4.682e-07,1.838e-09,0.0176,0.000291,0.0281,3.835e-06,0.0006039,0.01155,1.131e-07,3.371e-07,3.783e-08,1.274e-06,4.912e-15,0.004308,1.948e-06,1.074e-13,0.008498,9.311e-08,2.64e-07,3.213e-06,8.84e-09,1.902e-05,6.305e-07,0.0002825,3.846e-07,1.131e-08,7.787e-11,0.001166,0.01582,1.694e-06,1.101e-05,0.001078,2.794e-05,3.522e-07,8.909e-07,0.004025,5.75e-07,0.02054,2.869e-16,3.434e-15,3.507e-06,1.536e-13,1.42e-12,0.006877,2.708e-11,0.01768,9.803e-05,0.001276,0.0001753,2.42e-09,0.001162,1.974e-05,2.228e-07,6.732e-08,6.927e-06,6.243e-06,1.212e-11,2.757e-12,1.076e-06,3.2e-08,0.01602,8.223e-14,0.004795,0.01783,2.063e-13,0.0001504,6.436e-06,7.588e-05,2.984e-09,5.056e-05,3.79e-05,1.499e-11,2.873e-13,1.009e-07,2.159e-05,1.087e-16,2.477e-06,4.3e-07,3.378e-11,0.01954,0.005386,8.832e-08,1.296e-08,8.634e-09,2.837e-13,3.949e-17,0.01569,2.745e-09,4.598e-05,7.64e-18,1.557e-11,3.162e-05,1.95e-11,1.08e-07,1.094e-11,1.128e-11,2.852e-07,0.002301,3.745e-06,1.037e-07,6.269e-07,0.03045,0.003617,9.338e-09,2.056e-14,0.01988,3.341e-11,1.29e-08,0.001611,0.0003939,3.92e-09,0.01118,2.61e-08,5.788e-10,3.929e-05,0.0001172,3.55e-17,0.0002519,0.004486,3.372e-12,6.694e-09,0.000171,2.987e-07,5.289e-05,5.276e-05,0.01719,2.959e-08,0.0001444,0.009733,5.71e-05,5.094e-12,5.892e-10,1.078e-05,0.00913,0.002928,0.001887,0.0001063,1.507e-10,1.033e-05,1.566e-11,1.242e-07,2.881e-10,9.924e-07,3.662e-10,4.768e-06,1.445e-20,0.000354,8.029e-05,3.675e-09,1.697e-06,2.96e-06,0.0003421,2.348e-06,0.00052,0.005512,5.068e-10,0.00118,0.01483,0.0008353,1.281e-08,3.54e-05,1.425e-12,1.539e-08,7.345e-05,0.000356,1.048e-06,4.77e-17,1.096e-09,4.472e-07,2.905e-17,3.343e-10,2.823e-08,4.7e-09,0.005575,1.471e-07,4.658e-10,2.211e-07,9.203e-05,2.333e-11,5.762e-08,2.877e-17,0.004153,6.031e-05,2.958e-10,0.0001338,0.004755,0.03346,0.009523,5.795e-09,1.517e-06,2.034e-13,0.01347,0.002621,5.545e-17,0.004726,1.527e-11,0.001398,0.0003821,0.0002757,4.915e-08,0.0003251,1.588e-05,1.615e-07,6.247e-07,1.076e-15,1.661e-07,1.261e-10,5.758e-13,1.053e-07,4.236e-05,8.074e-07,0.01519,0.000439,5.248e-06,3.452e-06,0.008668,0.0003,0.0004179,0.0002778,0.0002046,1.885e-07,3.151e-07,3.895e-06,0.008032,3.74e-09,0.0007646,6.215e-08,1.72e-14,9.823e-10,1.607e-05,5.5e-05,2.47e-15,1.126e-15,2.934e-08,3.29e-06,2.364e-08,2.34e-07,0.002263,0.003273,2.678e-07,1.329e-07,1.057e-08,1.924e-11,1.295e-09,2.239e-18,3.01e-08,3.506e-22,0.001216,0.0001043,1.908e-13,3.423e-07,8.571e-12,8.692e-05,0.03494,0.001583,0.01355,2.277e-11,9.033e-08,7.252e-13,5.013e-14,1.042e-06,0.009032,0.01944,3.062e-18,2.283e-17,8.874e-13,2.22e-07,1.283e-11,4.846e-08,2.328e-07,2.069e-09,0.0009754,1.558e-16,3.541e-10,0.001777,4.712e-07,0.01913,0.0004857,0.01229,3.528e-05,1.747e-05,9.172e-06,0.01158,2.541e-08,6.742e-09,1.54e-12,1.648e-09,4.102e-10,2.608e-08,5.214e-07,6.155e-15,0.007814,2.461e-05,0.0008418,2.857e-11,7.667e-09,0.0009169,3.627e-11,1.021e-11,4.176e-12,8.151e-14,7.601e-11,4.652e-17,0.001274,1.735e-09,2.093e-06,0.001622,1.614e-07,0.003274,0.00133,6.981e-16,1.169e-05,1.579e-11,9.528e-08,1.793e-16,0.0002521,0.00113,8.15e-06,1.283e-12,4.951e-08,7.107e-05,6.806e-15,0.00486,1.006e-05,6.754e-16,0.0006024,1.175e-08,1.452e-06,2.587e-05,6.616e-09,4.395e-05,1.837e-08,3.276e-08,4.244e-10,0.0001228,1.17e-06,0.00154,3.083e-07,9.862e-05,9.661e-10,0.007216,1.891e-05,0.004742,6.727e-05,5.183e-07,0.001739,0.02355,3.279e-12,2.317e-06,1.982e-06,4.25e-12,5.997e-08,1.528e-10,8.79e-11,2.392e-08,4.049e-06,0.001056,6.334e-11,1.104e-06,4.65e-06,0.02289,4.119e-12,2.708e-10,1.231e-08,1.441e-08,0.0143,5.661e-08,4.852e-06,0.0006169,3.33e-10,2.762e-07,6.888e-12,2.883e-05,0.0002372,8.068e-13,0.0226,2.692e-05,3.897e-05,0.01453,0.004567,0.007205,1.173e-08,7.476e-16,0.005495,6.194e-10,5.509e-07,0.009653,2.138e-07,9.706e-09,8.966e-07,0.002206,6.144e-07,2.249e-07,4.749e-08,4.366e-07,5.849e-07,2.105e-05,4.216e-10,1.062e-16,1.521e-05,9.731e-06,1.357e-10,2.063e-08,2.943e-07,0.004519,3.184e-08,3.667e-07,0.002032,1.625e-09,2.192e-08,7.464e-11,2.073e-05,0.0239,7.647e-11,3.026e-16,7.493e-15,2.772e-16,5.195e-10,2.718e-08,0.005837,1.828e-07,2.248e-11,3.004e-10,4.404e-16,0.000664,1.095e-07,1.952e-08,4.108e-06,5.985e-05,0.0002985,0.0002781,2.54e-07,4.995e-07,0.000696,4.314e-05,1.3e-14,1.247e-06,0.001593,5.437e-06,0.0001336,4.517e-06,3.309e-08,1.393e-05,2.638e-09,3.307e-05,1.163e-10,1.308e-10,7.672e-10,3.456e-06,0.004999,0.005064,2.335e-20,0.0005877,0.001361,0.006211,1.521e-20,1.202e-10,0.00338,0.0006101,0.0002781,4.853e-12,0.01265,4.269e-07,1.88e-07,2.261e-07,1.849e-12,2.084e-06,0.02716,3.645e-05,0.001407,0.001134,4.099e-10,0.002766,1.395e-13,6.889e-15,4.283e-12,1.054e-08,6.026e-08,1.827e-10,2.988e-10,2.43e-08,1.6e-07,1.905e-10,4.476e-12,2.21e-09,2.234e-07,1.392e-10,0.004378,4.02e-05,2.483e-05,3.041e-07,1.573e-06,0.0002649,9.193e-10,2.441e-06,1.99e-07,5.719e-06,1.391e-10,0.001559,0.0002624,0.0001073,7.168e-06,2.414e-07,6.082e-11,2.061e-20,6.246e-07,4.701e-09,8.177e-10,2.751e-06,5.771e-12,1.449e-13,0.006856,2.503e-08,2.566e-07,0.002146,7.569e-07,4.033e-06,9.753e-07,0.006224,1.717e-11,0.001663,3.08e-15,5.619e-11,1.431e-14,5.589e-18,4.004e-10,5.298e-10,0.0001361,3.755e-17,1.542e-13,2.821e-06,5.217e-07,0.0001417,0.013,9.807e-10,3.418e-09,0.0001911,4.094e-16,0.01295,1.6e-13,1.974e-15,0.00031,8.684e-09,1.384e-06,1.47e-09,1.38e-12,0.0002906,3.347e-13,2.752e-07,0.006093,0.0006011,7.953e-08,0.005422,0.02109,0.0005444,0.0001392,0.01677,0.02047,0.0005547,1.647e-06,0.0007037,9.625e-12,1.092e-11,0.003087,1.121e-15,4.133e-06,0.001039,0.0003013,7.814e-16,1.959e-16,0.0003343,0.01747,0.01306,0.0003063,1.629e-05,2.022e-12,0.0002236,0.00757,0.004016,3.413e-14,0.00132,5.331e-13,2.841e-16,0.0003404,1.769e-08,3.214e-05,1.095e-18,0.006535,0.003889,8.579e-05,0.002204,0.0002075,4.65e-09,2.13e-13,9.324e-22,0.003768,3.153e-05,1.423e-13,4.373e-10,8.875e-05,2.292e-06,1.016e-05,1.356e-09,0.0003715,4.182e-06,6.624e-05,6.543e-11,0.002473,5.723e-11,5.1e-10,6.773e-19,4.996e-09,3.072e-18,0.004934,7.859e-08,3.171e-05,0.0002707,9.036e-08,4.538e-08,3.027e-06,7.819e-06,0.0003959,0.004049,2.299e-14,0.005911,4.553e-05,5.453e-13,0.01627,7.348e-08,7.501e-10,0.00277,4.968e-05,3.866e-05,1.409e-06,0.002065,0.003742,4.313e-15,1.937e-08,1.51e-07,1.896e-06,0.002725,0.03495,2.179e-06,6.889e-05,8.018e-06,5.127e-06,0.001631,2.214e-06,2.112e-06,2.669e-08,2.009e-06,4.083e-09,2.746e-06,2.517e-09,5.518e-06,0.004623,0.0001424,2.386e-06,1.742e-17,0.00117,2.493e-15,0.0331,0.002931,9.631e-09,0.001692,0.01113,1.39e-10,4.777e-08,0.004926,3.491e-07,0.0005462,5.93e-06,1.763e-05,0.0009998,5.879e-07,2.736e-18,7.774e-07,8.97e-08,1.674e-05,2.03e-10,0.000204,0.004513,4.374e-14,2.403e-05,4.202e-10,1.326e-11,1.821e-05,6.661e-12,1.211e-06,1.215e-05,2.34e-07,1.627e-06,4.272e-16,1.47e-08,0.01587,1.127e-09,6.879e-07,0.00145,0.000437,2.36e-12,1.294e-10,0.001451,0.001025,0.0004501,1.907e-05,2.013e-06,8.326e-12,0.01235,0.002022,1.507e-05,4.616e-07,1.28e-10,0.0337,9.854e-10,2.06e-10,2.07e-09,1.162e-15,1.305e-12,1.756e-07,7.11e-16,1.127e-10,0.000445,6.34e-08,1.209e-12,4.272e-10,1.095e-14,0.004372,0.0002726,0.0002409,2.233e-07,7.675e-09,6.379e-05,1.225e-08,7.724e-10,3.537e-06,1.242e-07,0.0007407,3.949e-14,0.002434,9.577e-07,5.389e-07,1.06e-17,5.48e-09,1.312e-07,6.682e-05,0.0009093,2.446e-06,9.538e-13,2.314e-06,1.814e-09,1.984e-13,0.001336,0.0001034,0.0003009,1.592e-05,9.178e-12,1.228e-15,0.0001173,0.01901,2.698e-10,2.704e-05,6.219e-09,0.003757,1.853e-08,7.83e-08,4.037e-05,3.944e-12,0.0005398,1.241e-06,2.149e-08,1.052e-06,0.01956,0.0003002,7.684e-14,6.811e-06,7.886e-14,7.373e-10,0.00128,1.823e-06,1.413e-08,4.56e-05,1.077e-14,0.02219,9.483e-15,3.825e-05,0.0006873,0.02853,2.767e-05,8.604e-08,2.143e-07,4.237e-10,1.352e-15,1.072e-15,0.001997,3.371e-10,0.01111,2.403e-14,1.187e-07,0.003821,1.024e-15,0.002137,6.562e-11,2.839e-06,1.141e-08,4.366e-11,1.485e-15,1.999e-13,6.637e-07,4.33e-06,0.003587,0.0001001,1.549e-10,2.535e-06,0.005809,2.091e-14,3.3e-08,1.016e-07,4.146e-11,2.849e-13,7.562e-16,0.0002156,1.432e-07,1.104e-05,2.734e-14,0.01348,4.831e-07,5.993e-15,0.0005209,0.008859,2.087e-06,0.0003054,0.004728,0.001032,1.213e-05,3.418e-09,0.01726,1.807e-15,4.124e-11,0.0008985,2.404e-08,2.424e-10,2.463e-06,1.769e-05,2.185e-05,8.822e-05,3.061e-07,1.762e-08,7.863e-12,0.005913,7.29e-08,0.000512,0.0003633,1.488e-09,6.255e-11,6.072e-09,2.598e-13,7.587e-05,8.763e-10,3.624e-05,0.008026,3.594e-08,2.983e-08,0.005319,0.00283,4.646e-06,5.112e-12,1.299e-10,3.651e-14,3.176e-07,5.053e-08,1.531e-06,0.004702,0.001831,3.481e-09,0.0001626,3.514e-06,0.01607,5.282e-06,0.000368,0.0005731,5.114e-11,2.445e-05,2.645e-14,0.025,0.003317,0.0001793,0.02314,0.008773,0.007379,5.564e-08,1.784e-05,0.0002837,0.005264,2.933e-05,0.000541,2.725e-07,1.72e-05,0.0001109,1.27e-09,0.002856,1.392e-09,1.006e-07,1.04e-08,1.374e-05,2.253e-09,2.995e-05,0.0001549,1.412e-11,1.455e-11,6.641e-12,2.446e-08,0.005296,8.339e-07,7.09e-06,2.664e-05,0.001196,0.0006849,2.165e-09,5.441e-14,3.439e-09,0.0002629,4.886e-11,0.01636,9.377e-05,7.9e-10,3.748e-07,3.616e-08,0.0001313,4.833e-06,3.088e-06,4.001e-09,5.603e-07,3.463e-08,3.199e-05,0.000437,3.684e-18,1.815e-05,2.995e-06,2.878e-13,0.000218,2.243e-11,1.322e-06,3.505e-07,5.533e-05,0.004632,1.152e-05,1.893e-08,3.62e-06,0.001302,2.313e-06,2.994e-13,2.468e-09,6.321e-14,4.62e-16,4.606e-06,1.46e-09,0.0001164,1.224e-13,2.783e-07,8.867e-11,0.001156,5.461e-05,1.374e-08,0.0004898,8.322e-17,2.424e-13,6.081e-09,0.0009751,9.734e-10,2.548e-11,4.267e-06,1.258e-11,9.13e-16,0.0001939,1.605e-05,0.003199,1.727e-05,0.0003324,0.0001569,3.61e-08,1.189e-08,1.283e-08,2.597e-10,3.81e-07,0.002461,4.989e-07,5.599e-07,0.006076,4.927e-07,4.722e-08,1.343e-05,2.442e-15,5.158e-05,1.765e-13,3.717e-11,9.313e-11,0.0001763,0.009435,0.0005591,0.00401,1.765e-13,0.0008035,1.387e-10,5.623e-06,5.873e-10,0.002269,0.0003774,0.001909,8.584e-08,7.248e-12,0.00175,5.716e-13,4.73e-09,1.556e-10,0.001475,2.519e-09,2.107e-06,0.02171,0.0002559,0.003262,1.142e-13,2.375e-14,0.0004369,2.057e-09,5.236e-11,6.839e-06,9.89e-07,5.075e-06,4.433e-05,0.009437,1.112e-16,3.423e-05,0.002309,3.079e-07,0.01735,7.139e-12,3.319e-06,2.745e-05,4.929e-09,0.01125,6.935e-15,8.911e-09,1.084e-18,2.448e-05,2.957e-13,8.152e-10,2.014e-05,9.85e-10,0.0002401,6.456e-15,1.847e-16,0.01521,7.398e-14,1.228e-15,0.01284,0.0001239,3.813e-14,8.195e-13,5.069e-11,2.387e-05,0.000143,0.03326,0.002305,0.0001824,0.0009673,2.183e-09,0.001039,4.967e-06,4.997e-11,6.383e-06,6.631e-11,1.97e-07,1.143e-11,1.644e-06,2.049e-11,0.003506,1.337e-09,0.006084,4.299e-10,1.389e-12,6.556e-06,2.669e-09,3.393e-05,0.0001838,1.711e-10,1.088e-08],"Adj.PValue":[0.6827,0.9124,0.3306,0.07704,0.1651,0.08087,0.1509,0.2789,0.3601,0.2011,0.1006,0.1876,0.8382,0.5789,0.06537,0.7178,0.9444,0.4142,0.2893,0.3748,0.2296,0.7824,0.3331,0.5931,0.9409,0.9282,0.2097,0.4983,0.6001,0.6638,0.05429,0.05592,0.1399,0.3438,0.9771,0.2488,0.2234,0.7029,0.2598,0.9597,0.1789,0.1286,0.128,0.07117,0.6096,0.4332,0.2635,0.1637,0.5573,0.1231,0.7183,0.9597,0.8202,0.2614,0.5425,0.1349,0.4516,0.5991,0.9258,0.4454,0.6614,0.3202,0.3253,0.05686,0.1789,0.3424,0.1842,0.2985,0.242,0.9645,0.5118,0.5344,0.8894,0.2194,0.09145,0.5075,0.8754,0.2553,0.1181,0.4126,0.05236,0.2226,0.7105,0.5558,0.05975,0.2093,0.2133,0.6007,0.3122,0.4714,0.84,0.8091,0.5998,0.291,0.9597,0.1483,0.09098,0.4099,0.999,0.1096,0.3209,0.3012,0.07853,0.7168,0.21,0.3998,0.6168,0.63,0.273,0.5001,0.6121,0.909,0.09938,0.6626,0.4958,0.3729,0.4082,0.1837,0.07333,0.2134,0.4983,0.3095,0.1082,0.4111,0.4135,0.2823,0.87,0.1007,0.2008,0.975,0.2179,0.2994,0.00317,0.2306,0.2604,0.3338,0.5724,0.7252,0.6366,0.9193,0.01387,0.3107,0.4277,0.2546,0.872,0.003944,0.7439,0.1762,0.8789,0.3065,0.2761,0.8111,0.1892,0.5453,0.5695,0.002525,0.05359,0.421,0.1084,0.1817,0.2905,0.7606,0.002993,0.5725,0.1673,0.0563,0.1067,0.8041,0.139,0.8005,0.19,0.003215,0.3449,0.2755,0.2714,0.6412,0.08833,0.3913,0.8861,0.7078,0.4978,0.3227,0.08502,0.5772,0.2674,0.1799,0.5836,0.3744,0.2852,0.08491,0.1165,0.9749,0.8333,0.9212,0.3766,0.001636,0.05002,0.1916,0.06232,0.2827,0.8046,0.08519,0.01924,0.2408,0.005907,0.2919,0.4734,0.0231,0.9674,0.05497,0.452,0.01871,0.0509,0.7771,0.3239,0.2422,0.7907,0.03624,0.9455,0.06461,0.312,0.1789,0.02382,0.7079,0.03705,0.07918,0.007152,0.01378,0.0763,0.7844,0.1165,0.02341,0.9807,0.09948,0.05713,0.5412,0.1329,0.2648,0.4259,0.7065,0.06019,0.05517,0.4304,0.07493,0.8864,0.3256,0.05908,0.06867,0.08424,0.3712,0.4679,0.07344,0.307,0.1522,0.07281,0.3642,0.3682,0.8823,0.06497,0.8168,0.6495,0.6811,0.02668,0.1148,0.5816,0.139,0.3558,0.1343,0.4071,0.03803,0.26,0.02402,0.3011,0.4321,0.3714,0.0155,0.4709,0.2439,0.2041,0.07045,0.01461,0.3333,0.8681,0.4964,0.1204,0.3811,0.3676,0.4277,0.009482,0.171,0.1069,0.01237,0.07016,0.344,0.3553,0.5139,0.0142,0.3848,0.4127,0.4468,0.3946,0.9421,0.6219,0.002723,0.0009387,0.111,0.0171,0.2021,0.2558,0.1682,0.1926,0.2911,0.1479,0.08888,0.5606,0.7086,0.3951,0.7583,0.8046,0.2755,0.005776,0.2059,0.2022,0.7818,0.003292,0.2954,0.3852,0.1074,0.04263,0.2371,0.7912,0.5793,0.5746,0.689,0.1775,0.03798,0.06069,0.2015,0.0128,0.7786,0.1382,0.08934,0.7892,0.2895,0.06591,0.0002772,0.2334,0.1521,0.08321,0.1056,0.7417,0.5866,0.4575,0.4387,0.3869,0.1109,0.2175,0.04467,0.03118,0.3013,0.1927,0.2411,0.3054,0.004601,0.4861,0.4099,0.5461,0.4609,0.1006,0.4263,0.1153,0.06207,0.4778,0.7916,0.5009,0.3733,0.02903,0.1177,0.04776,0.01347,0.02776,0.2764,0.1455,0.06784,0.7856,0.6147,0.952,0.2412,0.4578,0.7607,0.3615,0.00263,0.5975,0.07293,0.4546,0.004191,0.5018,0.9713,0.5389,0.06537,0.4371,0.4081,0.2805,0.1222,0.1038,0.06414,0.941,0.003314,0.6176,0.8914,0.261,0.08664,0.1686,0.9798,0.7498,0.4772,0.03383,0.001241,0.05176,0.9518,0.002097,0.8679,0.02202,0.1168,0.2427,0.06247,0.01471,0.9222,0.1365,0.05387,0.6207,0.9931,0.005796,0.3025,0.01324,0.1717,0.1048,0.06154,0.3565,0.3385,0.1107,0.00573,0.9624,0.1491,0.04244,0.06222,0.007661,0.9572,0.3575,0.05643,0.09599,0.4694,0.3692,0.08264,0.005365,0.09743,0.2716,0.1215,0.6803,0.1078,0.3728,0.3967,0.06502,0.359,0.06921,0.6282,0.09959,0.07066,0.3656,0.1445,0.08315,0.6259,0.1758,0.1415,0.6337,0.01029,0.72,0.1169,0.003952,0.9806,0.0903,0.6626,0.1262,0.9945,0.07516,0.02074,0.5183,0.07077,0.02058,0.01046,0.5596,0.1171,0.01895,0.066,0.5807,0.00012,0.7716,0.0112,0.23,0.9247,0.04172,0.9725,0.2628,0.07593,0.2413,0.03086,0.3225,0.8126,0.05099,0.08807,0.141,0.03199,0.08534,0.7743,0.02773,0.9645,0.4273,0.9647,0.08933,0.6013,0.2996,0.312,0.6121,0.1768,0.5053,0.0535,0.3404,0.8773,0.2411,0.08048,0.2476,0.1686,0.7833,0.06268,0.002507,0.1682,0.6068,0.1703,0.5396,0.8362,0.1975,0.002826,9.287e-05,0.784,0.3937,0.6352,0.1608,0.09963,0.2166,0.3564,0.9645,0.1999,0.09781,0.002197,0.426,0.007854,0.1168,0.1493,0.7677,0.1871,0.6228,0.01446,0.0039,0.07723,0.6233,0.2759,0.6563,0.9674,0.7079,0.02225,0.002173,0.1776,0.1954,0.005978,0.1054,0.1404,0.5129,0.01501,0.4822,0.3722,0.3015,0.4449,0.1099,0.6438,0.05155,0.6366,0.7677,0.008953,0.7866,0.1446,0.1913,0.8885,0.1907,0.08948,0.7671,0.1011,0.5465,0.7786,0.07736,0.1169,0.882,0.313,0.02254,0.6692,0.0516,0.8069,0.1494,0.2623,0.9385,0.8361,0.07996,0.3539,0.08888,0.8238,0.05826,0.002437,0.0029,0.1296,0.697,0.7799,0.3997,0.949,0.5633,0.1431,0.3575,0.3345,0.07262,0.0713,0.2136,0.2759,0.08006,0.1761,0.7944,0.9812,0.01748,0.03585,0.7175,0.3795,0.3064,0.1236,0.04106,0.7939,0.1525,0.1786,0.7365,0.3683,0.1677,0.01258,0.9293,0.08868,0.1434,0.5991,0.4314,0.6811,0.1992,0.009175,0.9871,0.07423,0.08315,0.4399,0.08285,0.05314,0.1444,0.9164,0.2579,0.297,0.1159,0.4499,0.09679,0.09259,0.07375,0.9014,0.9281,0.7792,0.587,0.04564,0.465,0.2893,0.8239,0.08259,0.2308,0.05079,0.1885,0.7123,0.09521,0.008917,0.283,0.09781,0.3475,0.9439,0.4213,0.2462,0.5057,0.17,0.9014,0.5056,0.4081,0.3421,0.2823,0.03309,0.8235,0.2496,0.2412,0.4971,0.05243,0.8059,0.4499,0.1549,0.7475,0.7949,0.6467,0.2843,0.001134,0.7718,0.7376,0.4158,0.2417,0.1561,0.7365,0.8983,0.8802,0.2665,0.2498,0.06273,0.7826,0.7365,0.1179,0.06148,0.7792,0.8437,0.09067,0.7939,0.1731,0.1715,0.0007789,0.5952,0.297,0.1839,0.2726,0.05816,0.01156,0.6456,0.7844,0.3945,0.3737,0.2614,0.5719,0.297,0.2492,0.7208,0.3048,0.8862,0.1065,0.9004,0.4424,0.1583,0.6414,0.02021,0.9051,0.1471,0.6087,0.6108,0.5595,0.8579,0.3865,0.06955,0.04277,0.2599,0.388,0.2982,0.944,0.8411,0.2166,0.1903,0.1704,0.3294,0.2267,0.1122,0.0006275,0.7453,0.6293,0.1052,0.3863,0.13,0.1718,0.3245,0.4592,0.7307,0.06841,0.6164,0.7847,0.7151,0.4424,0.4437,0.07078,0.1161,0.2392,0.08066,0.287,0.2798,0.1728,0.8905,0.4824,0.01176,0.09734,0.06869,0.003141,0.06502,0.882,0.0383,0.8621,0.5325,0.003345,0.9442,0.6096,0.4595,0.4545,0.3239,0.889,0.3432,0.8552,0.002074,0.001799,0.003782,0.001249,0.03931,0.3696,0.7775,0.3664,0.6007,0.9787,0.1139,0.3946,0.8319,0.3477,0.3496,0.08672,0.2722,0.007434,0.7686,0.07933,0.4926,0.0759,0.01499,0.01378,0.2369,0.7445,0.2194,0.1264,0.01614,0.2166,0.3524,0.003365,0.08943,0.5299,0.00263,0.3741,0.7158,0.06736,0.1724,0.8504,0.06207,0.007581,0.1156,0.1071,0.3306,0.02058,0.447,0.2296,0.7011,0.0005048,0.2378,0.4586,0.448,0.488,0.7329,0.1126,0.0383,0.01694,0.0713,0.05413,0.1349,0.1169,0.7875,0.1159,0.4719,0.2063,0.2231,0.7542,0.8991,0.9959,0.05384,0.7746,0.1631,0.02495,0.7562,0.008802,0.298,0.1067,0.001577,0.001193,0.4273,0.2126,0.3108,0.06042,0.6055,0.6781,0.06661,0.8868,0.1573,0.01309,0.07737,0.9751,0.001698,0.5724,0.8552,0.5974,0.2792,0.3307,0.4097,0.9189,0.3541,0.602,0.0861,0.9293,0.6844,0.02871,0.9648,0.6517,0.0002806,0.5042,0.01167,0.8348,0.9998,0.8912,0.03617,0.2209,0.9184,0.1075,0.8131,0.001617,0.07424,0.4347,0.009367,0.2177,0.3692,0.2558,0.008901,0.01737,0.2672,0.283,0.3648,0.4339,0.2767,0.6618,0.1049,0.3777,0.294,0.008802,0.4843,0.4339,0.008932,0.001094,0.479,0.2902,0.3035,0.313,0.3541,0.647,0.02363,0.1355,0.5377,0.09196,0.3704,0.1096,0.2059,0.2252,0.153,0.4403,0.3273,0.1282,0.2906,0.4242,0.9004,0.05397,0.5326,0.0959,0.4662,0.02126,0.2207,0.07262,0.07045,0.1462,0.7912,0.05766,0.08933,0.7678,0.2852,0.7654,0.2155,0.186,0.08424,0.3766,0.08279,0.07488,0.08052,0.09663,0.4468,0.8343,0.7186,0.8091,0.1351,0.05704,0.47,0.05906,0.8653,0.04219,0.1112,0.07458,0.1166,0.3872,0.3223,0.029,0.9407,0.2044,0.1387,0.9742,0.07008,0.8324,0.008778,0.4875,0.8983,0.9258,0.653,0.1051,0.496,0.1014,0.01822,0.4655,0.3035,0.2864,0.2579,0.8363,0.8051,0.06203,0.7032,0.08087,0.6594,0.5411,0.05206,0.4095,0.886,0.03541,0.1309,0.8362,0.4985,0.3107,0.139,0.1675,0.007438,0.2492,0.03822,0.08195,0.1046,0.1572,0.5441,0.9742,0.8131,0.1083,0.1006,0.8061,0.03686,0.519,0.1148,0.1058,0.1731,0.4223,0.6495,0.995,0.4491,0.8681,0.8971,0.132,0.7053,0.002373,0.2647,0.4746,0.2237,0.9783,0.2297,0.1559,0.07185,0.3522,0.005598,0.4822,0.4872,0.1321,0.043,0.8143,0.005102,0.09703,0.2176,0.2188,0.9949,0.5715,0.6983,0.2498,0.4492,0.1774,0.0563,0.03577,0.2142,0.3169,0.5118,0.5937,0.814,0.2951,0.04175,0.02789,0.9783,0.1249,0.5998,0.2397,0.916,0.1523,0.7075,0.304,0.696,0.1574,0.2186,0.09903,0.6767,0.0756,0.2299,0.2265,0.9199,0.2948,0.4049,0.6438,0.1054,0.9382,0.4761,0.2946,0.01987,0.1482,0.003871,0.003784,0.3016,0.7777,0.3078,0.9567,0.1848,0.4841,0.3713,0.05748,0.09956,0.06565,0.3065,0.05639,0.697,0.06686,0.1565,0.3661,0.1459,0.6241,0.006614,0.02674,0.1787,0.9783,0.9648,0.04675,0.7936,0.1968,0.08735,0.04181,0.01246,0.1259,0.7896,0.8486,9.665e-05,0.2426,0.2189,0.3529,0.903,0.07131,0.6053,0.185,0.7307,0.8032,0.915,0.3208,0.1066,0.1082,0.8846,0.3013,0.7252,0.4843,0.4078,0.09605,0.2364,0.4273,0.5127,0.103,0.1493,0.4619,0.9271,0.634,0.5168,0.2209,0.5937,0.9021,0.04427,0.0971,0.137,0.7744,0.5595,0.9213,0.1734,0.06196,0.00621,0.3516,0.1615,0.1839,0.06567,0.7658,0.7316,0.864,0.2199,0.4161,0.08332,0.5,0.4592,0.07114,0.5309,0.4052,0.2407,0.1433,0.2864,0.03281,0.3913,0.258,0.1012,0.191,0.7169,0.4385,0.07967,0.7023,0.01644,0.6011,0.08387,0.8488,0.1159,8.656e-05,0.684,0.00966,0.6638,0.6068,0.0171,0.2705,0.13,0.05222,0.3791,0.2042,0.95,0.7159,0.2413,0.01106,0.3249,0.02932,0.05369,0.4874,0.7821,0.8016,0.08562,0.1303,0.1615,0.3677,0.7246,0.1813,0.03978,0.5434,0.8409,0.4665,0.123,0.1124,0.09743,0.2549,0.0508,0.009574,0.05578,0.9293,0.1288,0.1077,0.009235,0.01633,0.1783,0.1651,0.6125,0.2842,0.4259,0.06629,0.02885,0.003414,0.916,0.05938,0.2589,0.7434,0.0001447,0.63,0.01431,0.07316,0.5234,0.6983,0.2794,0.9306,0.05067,0.2121,0.001643,0.1573,0.05945,0.1074,0.4033,0.07293,0.7944,0.7686,0.3016,0.2013,0.01639,0.009373,0.213,0.004005,0.5018,0.188,0.03571,0.886,0.07564,0.03745,0.5694,0.2879,0.5547,0.1194,0.08255,0.1608,0.1285,0.1264,0.008189,0.08844,0.2906,0.4166,0.06433,0.09802,0.05866,0.3942,0.01812,0.6296,0.003232,0.6267,0.0305,0.7439,0.1571,0.4213,0.09912,0.0001365,0.6323,0.06745,0.4522,0.4135,0.01266,0.1501,0.2889,0.1682,0.003694,0.8112,0.7476,0.5803,0.2458,0.5873,0.07612,0.54,0.8091,0.09781,0.6021,0.001221,0.05316,0.03456,0.2183,0.6225,0.01547,0.02383,0.934,0.1884,0.358,0.8688,0.3955,0.05457,0.03047,0.3526,0.1635,0.7667,0.4319,0.6541,0.595,0.5203,0.998,0.8921,0.5709,0.07859,0.7389,0.6522,0.1361,0.006361,0.004687,0.8238,0.003408,0.04756,0.3636,0.09298,0.9189,0.4213,0.1332,0.003548,0.402,0.1602,0.09993,0.52,0.07802,0.3484,0.5922,0.3414,0.2313,0.02647,0.03457,0.7316,0.4438,0.07375,0.3129,0.2948,0.316,0.07587,0.1924,0.8238,0.5098,0.1209,0.2129,0.3744,0.253,0.1665,0.9184,0.326,0.09486,0.3131,0.0996,0.0321,0.4933,0.1384,0.194,0.6112,0.008208,0.2398,0.07466,0.21,0.1098,0.2791,0.02835,0.02726,0.004842,0.1023,0.1228,0.04359,0.08321,0.08288,0.1033,0.06058,0.498,0.123,0.5709,0.1258,0.09377,0.9051,0.08372,0.2058,0.8409,0.3137,0.3915,0.5054,0.9597,0.117,0.5873,7.13e-05,0.6671,0.02694,0.0727,0.492,0.6239,0.4559,0.7838,0.5715,0.0029,0.6106,0.159,0.1058,0.08128,0.02722,0.1099,0.03096,0.2582,0.09671,0.02993,0.4015,0.06114,0.1303,0.08883,0.004643,0.8183,0.6991,0.4694,0.4206,0.3677,0.002176,0.3363,0.2123,0.9045,0.1863,0.6152,0.7746,0.1509,0.824,0.2338,0.0001189,0.7799,0.8034,0.8558,0.02443,0.9815,0.2407,0.00697,0.1675,0.3669,0.3041,0.6007,0.6522,0.6444,0.4726,0.8084,0.04369,0.4704,0.1159,0.6874,0.07035,0.1448,0.8635,0.08933,0.5631,0.06602,0.5388,0.02033,0.5126,0.1024,0.01412,0.3547,0.06594,0.6646,0.6731,0.2428,0.9058,0.3202,0.3425,0.09259,0.06798,0.7837,0.01106,0.08128,0.04649,0.02754,0.06066,0.3836,0.9864,0.0387,0.1052,0.3023,0.06132,0.1282,0.307,0.4194,0.2334,0.2007,0.05498,0.1811,0.07752,0.1018,0.01396,0.6224,0.161,0.7905,0.8408,0.03669,0.1232,0.6725,0.7677,0.6354,0.1049,0.04742,0.17,0.2295,0.7078,0.4539,0.6141,0.0634,0.9533,0.7105,0.458,0.6845,0.09959,0.5562,0.05103,0.103,0.01423,0.009252,0.0083,0.1655,0.08346,0.04106,0.8267,0.06833,0.8193,0.7142,0.1627,0.06043,0.05971,0.4778,0.411,0.1777,0.2842,0.4499,0.05575,0.1431,0.7353,0.7417,0.05966,0.001195,0.004348,0.09312,0.149,0.8486,0.9439,0.2294,0.4052,0.6613,0.05387,0.2475,0.824,0.07416,0.2727,0.06112,0.8369,0.04365,0.9274,0.8144,0.7151,0.7907,0.09307,0.5093,0.6626,0.8638,0.2459,0.001193,0.06139,0.05069,0.1474,0.8519,0.2702,0.06276,0.08458,0.492,0.3683,0.09179,0.009843,0.7792,0.06273,0.01098,0.02637,0.1384,0.4253,0.03925,0.05577,0.5093,0.03874,0.03814,0.1119,0.5331,0.06686,0.6845,0.2468,0.04988,0.1435,0.6709,0.1106,0.03162,0.8978,0.1351,0.224,0.1322,0.3107,0.006238,0.05704,0.569,0.007892,0.09938,0.04315,0.4033,0.07516,0.8886,0.06772,0.04173,0.1628,0.04837,0.03035,0.00876,0.7337,0.1792,0.01647,0.03232,0.001258,0.01523,0.4497,0.3606,0.03946,0.4407,0.7805,0.1989,0.01127,0.886,0.02501,0.3702,0.6479,0.02997,0.0453,0.4288,0.3174,0.1464,0.1891,0.003557,0.02997,0.397,0.2454,0.2704,0.6459,0.2138,0.2635,0.04687,0.02893,0.002652,0.0124,0.02719,0.002521,0.4319,0.06567,0.07316,0.4097,0.5653,0.003825,0.4293,0.0112,0.001658,0.09438,0.05769,0.6105,0.3048,0.3576,0.7223,0.6134,0.1156,0.213,0.9435,0.1572,0.258,0.7999,0.6053,0.1398,0.5724,0.4026,0.01098,0.298,0.2982,0.1727,0.8953,0.1227,0.007504,0.05859,0.07738,0.06565,0.4478,0.5463,0.4277,0.2902,0.4563,0.1812,0.2745,0.03483,0.008491,2.092e-07,9.844e-11,4.604e-15,9.073e-10,4.584e-08,6.854e-08,3.51e-09,0.00229,4.434e-07,6.855e-05,1.211e-06,0.0005531,1.676e-08,9.704e-05,0.00969,1.235e-05,9.371e-08,0.007574,4.146e-05,0.03665,0.01043,2.325e-07,1.65e-06,0.001763,0.003344,0.009754,0.04884,0.001412,5.838e-06,0.01048,0.007166,2.238e-06,0.01162,3.597e-14,6.908e-08,0.0427,0.002624,9.622e-13,0.002991,1.596e-08,1.229e-08,4.219e-10,0.000795,0.001105,0.0002527,4.705e-05,0.0001266,0.002351,5.397e-07,7.759e-08,5.472e-06,0.02007,0.00317,0.0004067,0.00114,6.554e-05,2.963e-08,0.001698,0.01496,0.001548,0.0167,0.001063,0.001398,4.147e-05,0.001369,5.25e-07,0.00556,0.01431,8.726e-09,0.01335,0.04201,0.000943,5.538e-05,0.001117,0.007994,0.002516,0.002372,0.04555,5.161e-05,0.03592,3.854e-06,0.04593,0.04771,0.04832,0.003269,0.02444,1.133e-06,2.215e-07,0.03396,1.364e-06,0.02536,0.01125,0.002137,8.71e-11,9.913e-05,0.0004316,3.024e-07,0.02606,4.638e-09,1.451e-05,5.42e-05,0.01648,1.259e-05,0.0137,1.971e-06,0.02404,0.03798,0.003054,6.492e-05,0.0001515,0.0002958,0.002139,3.079e-05,0.0001867,0.0007625,2.974e-14,0.0006789,0.04837,0.01429,2.228e-08,1.187e-14,0.004382,0.01202,0.004398,1.073e-05,3.426e-11,6.723e-10,1.875e-08,2.353e-07,6.636e-05,5.054e-08,1.292e-09,3.815e-06,1.305e-13,0.0008739,3.081e-06,0.0002713,1.463e-16,5.321e-06,1.741e-05,3.368e-08,0.0174,0.004372,5.486e-05,1.119e-05,1.843e-06,3.044e-08,1.345e-06,0.003307,0.002686,3.381e-16,0.006589,0.0001324,0.005395,1.027e-07,7.787e-07,4.736e-06,5.311e-05,0.01396,0.0007655,3.042e-06,0.0001275,0.00965,6.585e-07,2.845e-12,4.511e-10,0.004088,8.194e-15,9.54e-12,1.283e-06,0.01586,4.911e-10,5.05e-05,3.183e-05,3.98e-08,1.681e-08,5.432e-06,5.66e-07,0.0001326,0.0001945,6.677e-06,4.988e-07,0.005787,7.387e-11,1.215e-06,6.999e-07,5.917e-08,2.63e-11,0.001541,0.001967,0.0001142,0.01645,0.0006317,2.719e-14,0.001055,2.693e-08,0.0002734,1.236e-09,7.036e-05,0.001063,8.277e-06,2.463e-10,6.199e-08,0.0004012,0.0004993,0.03754,6.261e-07,0.0002076,0.0002026,0.006048,4.036e-05,2.317e-12,0.0001262,0.0174,7.257e-13,4.181e-08,0.0002354,1.817e-05,5.802e-05,9.317e-13,6.51e-05,2.701e-11,0.0002278,0.006277,5.072e-05,5.38e-09,4.406e-15,5.384e-06,0.01396,0.008228,0.003365,2.323e-07,1.465e-10,9.441e-08,0.01393,0.0001742,0.01599,0.009848,9.205e-06,0.0007448,2.064e-06,2.495e-14,0.01193,1.272e-14,2.528e-13,6.547e-10,1.384e-09,1.457e-10,6.565e-07,3.102e-10,0.006593,2.99e-09,1.824e-06,0.001416,4.032e-05,3.012e-12,0.001069,1.173e-08,1.861e-06,4.278e-08,0.006732,0.0003059,2.123e-06,6.338e-06,0.0001216,7.686e-07,3.431e-08,6.88e-15,1.447e-10,8.139e-11,0.008463,0.00276,2.796e-13,0.01149,5.676e-10,0.0005813,4.167e-09,2.227e-05,1.118e-05,2.506e-05,1.153e-07,8.864e-06,5.711e-07,2.99e-12,0.001165,4.49e-10,3.676e-06,1.976e-05,2.293e-06,0.0002301,0.0004515,0.000397,3.283e-08,0.01266,9.854e-13,2.241e-14,7.309e-05,0.008417,5.047e-15,0.004747,1.276e-11,6.889e-12,6.804e-05,1.785e-11,0.003141,0.005623,0.009658,9.504e-09,8.164e-07,2.872e-08,0.02424,0.001311,4.674e-16,4.057e-07,5.306e-05,0.0002954,6.013e-09,4.954e-09,0.02627,7.573e-06,6.729e-05,8.492e-08,0.01008,4.218e-14,6.05e-09,1.177e-07,7.613e-09,0.0001475,0.002839,3.065e-07,1.193e-16,4.068e-06,0.0002856,0.03696,0.003359,0.002629,5.331e-08,0.000219,3.646e-16,3.313e-15,0.001828,2.762e-07,5.175e-13,5.175e-13,0.0428,0.0002133,0.0008364,0.000207,0.007144,0.0001745,3.408e-17,1.046e-14,4.677e-05,4.53e-09,0.004222,3.222e-07,0.0006058,2.573e-07,1.423e-14,0.008731,7.556e-07,1.138e-08,1.855e-13,1.969e-06,4.211e-16,3.831e-19,3.65e-15,3.254e-10,6.492e-05,0.004893,4.239e-14,0.005848,0.01071,9.314e-16,1.022e-05,5.086e-05,1.483e-09,0.0001885,7.017e-09,0.005418,0.001666,6.825e-06,0.000166,0.003782,3.443e-07,0.01173,3.87e-07,0.007457,1.892e-08,3.146e-16,1.547e-17,9.261e-09,4.39e-07,6.826e-13,1.522e-13,2.26e-09,6.372e-11,1.727e-06,0.04908,3.838e-13,0.001338,5.996e-08,1.068e-05,1.324e-05,0.0003247,0.003129,0.0001741,0.0001868,3.187e-14,2.621e-05,4.39e-07,0.007053,0.00562,3.468e-08,0.0007549,0.0003586,0.001263,2.429e-06,1.081e-05,2.665e-09,1.248e-13,5.892e-11,2.807e-10,2.883e-09,5.7e-11,6.611e-05,3.724e-07,4.247e-12,0.01309,1.983e-06,8.407e-10,1.205e-06,1.43e-13,9.913e-05,5.584e-09,1.032e-13,3.411e-05,0.01204,1.364e-05,0.002855,0.002043,0.0001561,0.003492,1.917e-11,5.349e-06,3.182e-06,4.791e-10,0.002823,7.908e-05,4.47e-05,2.026e-11,0.0001751,1.56e-06,0.004738,4.842e-07,1.694e-11,1.521e-10,1.064e-12,0.001102,4.606e-09,7.461e-06,0.01373,3.088e-06,3.423e-07,6.251e-06,5.235e-10,3.51e-12,0.0006766,6.521e-07,2.181e-09,1.028e-09,6.488e-09,3.195e-09,3.817e-12,0.01213,0.0001421,0.0001676,4.623e-07,0.03635,0.006196,1.886e-08,0.005331,0.0004823,1.092e-06,0.0002016,2.307e-08,6.001e-06,0.001295,3.303e-05,2.711e-07,9.526e-14,0.04014,1.021e-14,0.00315,0.0001743,3.709e-07,2.021e-05,4.256e-14,3.853e-05,0.01383,8.489e-05,4.906e-08,1.076e-06,4.829e-05,1.599e-07,3.392e-06,2.961e-10,2.937e-17,2.776e-10,0.0006432,2.187e-05,1.901e-11,2.592e-10,3.45e-06,1.101e-06,8.432e-09,0.00176,9.14e-05,2.66e-07,0.004665,0.000213,0.003208,3.224e-11,2.192e-13,1.085e-11,4.833e-07,1.747e-06,1.768e-09,0.002659,5.448e-06,1.219e-06,0.0005053,3.494e-06,0.001217,2.056e-05,4.869e-13,3.972e-11,2.603e-09,4.57e-08,0.009252,1.197e-05,1.673e-07,0.0004894,0.005425,7.369e-12,2.203e-05,4.93e-06,0.03627,0.03254,8.491e-07,1.663e-07,1.611e-08,0.01597,0.001252,2.193e-08,1.051e-06,4.697e-08,7.599e-05,1.357e-08,5.882e-05,0.0004993,4.355e-10,1.034e-06,8.571e-15,4.162e-06,9.582e-09,1.023e-12,0.0001095,1.804e-08,1.457e-10,7.503e-10,6.536e-10,0.0005331,1.593e-09,1.718e-05,9.451e-06,0.005637,0.029,0.01619,8.542e-06,0.02706,7.638e-10,7.126e-15,4.991e-09,1.169e-10,9.82e-08,1.138e-06,6.149e-11,0.005923,1.556e-08,0.04427,0.0001684,4.09e-05,0.001594,3.82e-06,1.567e-08,0.003581,4.334e-07,4.628e-05,0.001223,0.0002144,0.003189,1.009e-07,1.257e-11,0.0001635,1.862e-12,7.073e-11,2.319e-11,0.000196,6.597e-08,2.278e-10,3.352e-10,6.469e-05,0.0002731,9.492e-11,0.01367,0.000372,7.538e-09,0.00992,4.963e-10,0.0004191,0.0002138,0.002659,1.763e-05,0.005318,0.000335,0.0001054,0.0001129,1.578e-05,8.272e-05,9.112e-08,0.02724,2.706e-10,2.222e-06,0.002525,1.696e-07,7.299e-09,6.372e-11,0.009978,4.959e-05,4.406e-15,1.633e-12,7.312e-08,7.494e-13,0.0003319,6.882e-19,4.49e-11,0.0001611,0.004134,3.327e-12,0.0002011,1.177e-09,1.726e-05,0.00255,7.797e-13,1.444e-12,2.333e-08,0.0001796,0.0001923,4.62e-10,1.681e-08,0.0004061,3.572e-05,0.005806,2.77e-10,3.007e-11,0.000174,1.05e-07,4.423e-07,7.039e-08,4.463e-14,1.932e-09,1.837e-05,1.341e-10,7.334e-09,6.488e-08,0.009313,1.002e-14,0.0008332,0.0001376,6.387e-10,3.307e-05,2.037e-13,0.002949,5.632e-15,1.653e-11,3.533e-05,0.003292,0.002599,0.002021,6.592e-08,0.03665,3.283e-08,8.711e-05,4.661e-16,0.0001642,6.123e-10,0.008608,0.004312,1.426e-14,3.432e-07,0.008219,7.443e-05,8.604e-14,2.541e-11,0.000903,1.174e-06,0.007836,1.047e-06,3.625e-05,1.143e-06,0.005504,5.538e-05,4.03e-05,6.32e-08,0.0002399,5.623e-07,0.0002844,3.712e-06,4.049e-06,2.843e-11,3.466e-09,8.545e-09,7.461e-06,0.002103,1.492e-08,0.002326,0.0108,4.661e-09,1.036e-10,4.298e-09,1.86e-10,0.005331,0.03925,0.02645,9.288e-12,8.75e-07,0.00287,1.547e-17,2.916e-10,0.009096,1.786e-07,1.597e-05,1.376e-06,0.0001421,1.388e-06,0.001046,5.535e-07,3.438e-05,1.141e-10,4.928e-09,4.753e-10,3.366e-07,0.0002753,0.001232,3.515e-05,5.472e-06,0.0008879,1.148e-11,9.935e-08,2.818e-12,3.297e-11,0.0002469,2.227e-07,0.001894,0.0009565,2.243e-05,1.383e-06,2.864e-07,0.003615,7.35e-08,4.701e-06,1.668e-05,7.036e-05,7.054e-07,0.0001648,1.509e-06,0.0315,5.219e-05,0.0004353,3.87e-07,2.524e-06,8.145e-07,0.0001553,4.595e-05,0.02491,2.023e-08,0.04363,2.96e-07,0.0006612,0.0001705,2.446e-07,0.001186,0.008919,0.00581,1.459e-14,1.846e-10,4.262e-15,0.000373,9.383e-05,5.188e-06,7.249e-07,1.069e-14,3.262e-11,2.242e-08,0.006627,2.338e-07,1.177e-07,0.0003191,0.003982,5.096e-10,1.77e-09,1.899e-11,9.915e-14,0.007276,0.008636,0.0003331,7.271e-11,8.223e-10,0.006505,1.814e-07,4.831e-07,0.02143,2.335e-10,1.46e-09,1.037e-05,4.47e-05,2.45e-11,1.478e-14,4.68e-13,1.153e-06,3.466e-10,1.891e-10,6.99e-05,1.327e-08,0.005783,9.534e-13,9.442e-10,5.33e-07,0.0008361,0.007298,3.418e-08,9.103e-07,3.232e-07,0.0001824,0.00851,9.977e-11,0.0001888,4.742e-09,2.847e-06,0.04412,3.64e-09,0.01059,3.798e-11,3.863e-08,0.04855,3.316e-10,7.815e-05,7.051e-24,3.812e-14,0.0007723,1.661e-08,2.926e-14,3.973e-09,0.004732,1.606e-07,1.261e-09,1.563e-08,0.0001333,1.052e-07,3.205e-05,6.047e-10,2.332e-05,0.04613,1.782e-08,3.295e-09,1.214e-05,0.001384,0.003141,2.202e-06,0.0002189,5.497e-05,0.0003478,5.504e-06,5.009e-14,0.005092,4.19e-06,0.0005163,0.002385,5.057e-10,0.0003586,2.59e-10,5.03e-08,1.536e-05,6.891e-08,4.754e-07,6.165e-15,4.013e-12,1.738e-09,1.516e-09,0.004779,0.003067,0.0006138,1.167e-06,0.0268,7.852e-09,0.00321,6.688e-16,0.003981,2.092e-07,9.314e-16,3.783e-06,0.0305,3.576e-07,1.546e-08,0.0004402,1.532e-09,6.108e-08,5.779e-08,3.021e-12,1.008e-08,8.638e-06,9.707e-11,1.834e-09,2.649e-07,0.002838,0.0006606,3.228e-07,0.0229,5.029e-13,6.529e-05,0.001583,0.0006205,1.778e-13,4.757e-06,0.0001332,5.42e-15,3.758e-11,3.571e-11,6.819e-13,2.255e-07,2.665e-05,0.004534,1.512e-08,2.831e-06,3.192e-06,5.892e-07,4.058e-08,3.338e-11,0.0002179,4.689e-05,0.01591,7.257e-13,4.267e-07,0.0003365,6.249e-10,1.73e-06,6.094e-14,0.0003219,7.806e-05,6.18e-09,4.839e-10,0.005564,9.627e-11,6.126e-08,1.193e-16,7.721e-05,0.001662,0.001083,0.02913,1.199e-05,3.259e-08,0.0002076,3.573e-10,2.468e-14,0.007083,0.0005643,0.001027,1.421e-08,0.004244,0.002952,0.02587,3.335e-08,0.03187,1.604e-11,0.0001066,5.069e-14,3.673e-05,5.577e-07,0.0008089,0.001227,0.001354,1.877e-13,5.666e-05,3.284e-09,1.234e-06,3.586e-06,5.584e-09,0.005028,0.04167,0.0009734,0.001378,7.388e-09,2.023e-08,1.308e-08,1.083e-06,3.883e-07,0.006951,0.0001253,0.01824,3.5e-05,6.18e-05,5.91e-07,7.259e-05,0.0003088,1.479e-06,0.0001305,8.889e-16,0.001422,1.105e-08,1.869e-16,0.04331,1.41e-07,1.054e-07,2.472e-06,2.693e-05,1.173e-08,8.791e-08,2.148e-05,8.569e-07,8.942e-08,3.637e-09,5.985e-06,2.625e-11,2.699e-05,3.275e-05,0.008415,0.002378,0.002361,8.443e-13,1.088e-11,4.025e-07,2.501e-05,0.0006229,0.001136,0.02978,3.492e-05,0.0007641,1.233e-09,1.617e-06,8.263e-07,8.525e-07,0.001406,0.00611,4.803e-08,0.0006606,0.0001623,0.0005754,0.0004405,0.001151,1.63e-05,6.912e-05,1.001e-09,1.518e-13,9.29e-10,1.4e-06,0.01503,0.00428,1.29e-09,2.217e-08,0.0006979,1.54e-06,0.002085,0.02202,2.219e-10,3.082e-06,0.01313,1.243e-08,1.324e-15,1.491e-09,2.13e-09,0.01092,6.221e-07,2.397e-10,0.002581,1.454e-11,0.002169,1.132e-11,2.227e-05,0.00516,1.377e-05,6.667e-06,1.505e-05,4.753e-10,0.01262,0.01313,1.368e-06,3.501e-09,0.0004771,0.0005753,1.713e-13,5.321e-12,1.267e-06,0.0004605,2.627e-05,0.002163,4.839e-05,8.36e-17,0.0005924,3.262e-11,1.073e-05,0.0406,0.002962,6.675e-10,1.105e-07,0.000174,0.002105,2.323e-11,0.004372,4.601e-11,1.054e-09,3.224e-09,0.008779,4.173e-09,4.47e-10,9.355e-12,0.0006865,1.225e-11,2.611e-09,0.003568,9.64e-05,6.048e-06,0.001229,5.79e-10,1.95e-13,1.724e-05,3.301e-08,4.986e-05,2.862e-05,0.0002728,1.657e-09,0.003325,0.01283,4.737e-05,7.073e-11,1.489e-08,0.005905,1.725e-08,0.001232,1.749e-13,0.02245,1.175e-07,1.237e-05,1.526e-07,0.007942,0.01455,0.01069,1.621e-06,0.00817,0.0001385,0.0003915,0.0001029,0.0003247,0.0001348,0.001548,2.867e-05,6.875e-14,2.106e-09,8.686e-09,4.262e-15,0.03428,0.008031,3.412e-09,0.00176,8.783e-08,0.005049,0.01806,8.249e-10,6.237e-07,3.386e-06,3.167e-08,0.0005047,2.224e-08,1.338e-06,1.017e-08,1.26e-09,3.716e-07,1.014e-10,8.79e-08,0.01842,1.426e-14,7.698e-05,0.0006384,0.0003722,6.061e-10,7.45e-05,1.446e-06,0.001955,0.01126,0.01405,9.209e-05,4.654e-11,8.861e-13,0.01607,1.088e-11,0.001271,0.00534,8.867e-13,2.213e-16,0.0004043,5.601e-07,2.826e-08,1.611e-09,6.453e-08,8.864e-06,0.0001313,6.321e-06,7.094e-09,0.008415,0.002147,9.004e-05,0.0001563,0.0003485,1.318e-06,0.002847,6.052e-11,8.658e-11,9.947e-11,0.01247,5.67e-12,1.021e-09,1.019e-06,5.944e-07,0.0007625,0.003499,0.01842,0.03402,9.051e-05,0.006648,0.001063,1.471e-13,8.685e-16,0.003348,0.008809,2.275e-05,0.0002263,0.0008469,6.909e-08,7.989e-08,1.591e-12,2.588e-12,1.425e-08,1.803e-07,0.000131,2.611e-09,1.132e-11,6.098e-05,5.365e-06,2.283e-11,4.336e-10,0.03009,5.057e-10,3.748e-11,8.038e-06,0.01928,2.373e-05,0.002263,7.7e-07,1.289e-07,0.006505,0.000158,0.007535,0.0009734,9.324e-08,8.467e-05,3.166e-05,2.277e-09,0.002546,1.274e-07,0.001084,6.521e-05,1.43e-13,2.459e-06,0.0001637,7.43e-09,2.227e-07,0.006653,5.445e-05,3.724e-07,0.00359,6.395e-13,1.785e-11,3.432e-07,4.267e-07,3.102e-08,1.911e-07,5.62e-11,0.001803,0.0001852,3.33e-14,7.248e-14,0.003737,4.97e-07,4.495e-10,1.45e-10,7.984e-11,0.0003942,8.255e-05,7.425e-14,3.474e-14,0.0001081,5.613e-11,2.024e-13,0.001377,0.01358,0.04449,2.609e-08,3.361e-07,1.73e-06,0.01119,2.382e-06,0.001274,0.0001105,3.899e-18,8.038e-08,7.772e-09,8.467e-07,3.981e-07,4.554e-07,1.564e-05,3.21e-07,5.792e-06,8.08e-09,5.252e-09,0.0001119,4.635e-09,1.606e-08,1.652e-13,0.003189,4.907e-15,1.443e-09,3.228e-07,9.822e-12,3.995e-06,0.0004797,8.889e-16,0.001063,0.0005754,1.341e-12,0.004029,3.441e-08,2.059e-07,0.0006295,0.0003664,1.865e-06,1.443e-10,0.0002739,1.363e-14,1.033e-16,1.43e-06,5.927e-09,6.032e-05,1.311e-07,7.229e-13,1.177e-07,1.333e-10,9.824e-08,0.0004587,8.161e-05,0.004224,1.171e-05,6.841e-11,6.301e-11,0.0009109,3.201e-11,2.761e-09,7.72e-09,2.823e-10,1.217e-07,0.0005018,0.02793,4.201e-08,2.462e-05,1.702e-06,0.000152,4.137e-06,2.52e-05,0.02387,0.001636,0.01912,4.959e-05,0.003354,2.501e-05,0.0009799,8.636e-05,4.357e-05,0.0001662,8.508e-05,1.451e-06,0.001348,7.269e-06,3.24e-14,5.969e-10,0.009319,0.0003107,4.633e-12,0.00589,1.637e-09,9.496e-06,1.476e-05,3.775e-07,2.729e-08,0.000196,0.0001036,3.035e-06,0.000111,1.518e-13,1.586e-14,7.219e-10,2.486e-05,0.0004998,7.916e-05,3.335e-08,1.885e-06,2.269e-07,4.856e-05,3.131e-05,2.559e-07,0.0001373,6.422e-07,8.889e-16,4.094e-08,0.0003163,4.282e-11,0.001213,0.0003832,0.04174,1.342e-10,7.15e-13,0.00636,1.258e-07,2.473e-06,0.0002217,5.515e-05,0.0006056,1.278e-07,8.023e-06,0.001529,0.0005754,3.876e-05,1.618e-06,2.074e-06,0.0004473,1.077e-05,3.538e-06,2.001e-07,0.0001378,0.0006698,0.0004074,0.002522,0.001063,2.089e-06,8.973e-09,6.616e-07,7.941e-06,0.002442,1.622e-07,2.267e-11,9.91e-08,0.0001408,1.081e-06,1.273e-07,0.02168,1.972e-08,0.004832,0.002601,0.000536,2.94e-05,0.001574,1.641e-08,3.111e-09,7.094e-08,3.813e-16,2.819e-07,0.04064,0.0005177,1.59e-10,1.238e-05,0.005639,0.006627,4.437e-13,6.744e-06,2.528e-05,9.822e-12,0.0008778,2.065e-08,0.0001856,2.446e-10,1.721e-11,4.928e-09,4.376e-14,0.0007323,0.0001776,0.009452,0.03596,1.41e-06,0.004754,0.01967,3.084e-08,5.169e-09,6.414e-08,8.725e-08,0.04157,2.464e-06,1.445e-08,0.0008135,1.354e-06,0.0001368,6.449e-07,4.188e-09,0.00324,0.0008184,3.662e-13,0.0001351,2.253e-09,4.593e-06,1.159e-05,2.501e-07,1.581e-08,1.244e-06,9.77e-09,2.284e-08,4.106e-05,0.001186,9.766e-09,0.002134,8.864e-14,3.284e-09,0.003907,0.008749,0.01092,5.356e-06,0.002094,0.0008116,1.05e-12,2.178e-12,0.0001924,0.001257,8.547e-05,1.202e-14,8.489e-05,1.948e-10,3.431e-08,9.299e-10,0.04497,8.525e-09,1.011e-07,1.17e-08,7.445e-10,3.515e-10,3.471e-08,0.0236,3.311e-11,5.467e-11,5.339e-14,0.01921,2.588e-11,1.55e-05,6.723e-10,0.0001788,6.804e-05,0.0003157,2.588e-12,3.786e-07,1.896e-05,1.308e-08,1.12e-07,3.604e-08,1.886e-06,6.753e-05,2.45e-06,3.212e-05,0.008243,2.088e-15,5.537e-05,9.718e-05,5.083e-06,0.01599,7.385e-09,0.001227,0.0001278,9.824e-08,6.277e-08,0.0009741,9.401e-12,4.301e-10,8.761e-05,1.642e-10,5.632e-15,0.01888,9.396e-05,2.022e-05,0.0001181,3.213e-05,9.073e-10,3.92e-06,1.526e-13,2.666e-06,0.0003818,8.59e-08,0.0004201,0.0001246,0.0001095,4.47e-10,4.936e-06,4.947e-07,1.567e-08,3.92e-06,8.285e-07,7.304e-10,5e-07,2.916e-10,1.82e-10,5.789e-07,6.322e-05,4.393e-05,0.0001291,1.215e-10,9.403e-06,0.02014,0.0007349,0.008398,6.193e-05,2.524e-07,1.843e-08,3.184e-07,0.0001376,5.087e-09,5.396e-15,0.04845,1.544e-16,0.0001002,6.547e-10,5.81e-07,6.45e-07,1.65e-06,8.096e-08,1.143e-06,0.0009387,3.254e-05,1.054e-15,0.0003107,8.395e-06,3.023e-05,3.091e-12,0.001196,0.002278,0.03742,3.253e-12,3.887e-08,2.044e-09,1.153e-07,0.005079,2.039e-11,0.002273,1.426e-14,2.176e-09,4.798e-13,0.002157,0.0003338,3.494e-05,5.123e-06,2.489e-05,0.0006893,0.004762,1.883e-07,4.737e-06,0.03791,6.337e-08,1.809e-08,0.01476,2.392e-08,0.0005753,0.001161,1.253e-05,2.036e-05,0.02261,1.732e-05,0.002443,5.705e-07,7.425e-14,3.732e-14,3.246e-10,1.385e-09,1.071e-07,5.647e-11,3.317e-11,3.727e-06,3.835e-09,0.0007607,1.007e-07,1.341e-08,9.892e-08,2.838e-11,7.528e-05,4.536e-05,0.0001002,1.189e-09,1.748e-07,7.005e-06,1.82e-10,1.792e-06,0.00441,0.01323,2.511e-09,9.639e-06,0.004398,0.006941,0.0003264,4.13e-08,8.06e-06,0.0001195,4.963e-07,1.225e-06,2.974e-15,2.23e-07,2.501e-06,0.0001118,6.213e-07,7.299e-17,4.315e-07,0.0002933,0.004936,5.425e-07,1.714e-08,0.00495,9.314e-16,6.018e-12,8.136e-09,4.62e-10,0.001761,1.173e-17,4.066e-12,0.0007944,0.0007413,2.96e-10,0.000707,8.702e-14,1.509e-06,0.0001243,8.933e-05,0.001967,9.482e-08,5.522e-08,0.0002458,1.318e-09,1.694e-11,1.919e-16,0.007867,1.006e-09,0.002313,0.002157,3.468e-11,1.948e-07,0.0004162,1.401e-06,1.147e-07,2.326e-11,0.004684,0.0004951,0.0003859,1.25e-06,7.913e-08,0.04063,1.115e-05,7.485e-09,6.594e-09,0.008096,3.037e-08,9.043e-07,0.00972,2.088e-06,0.02853,0.001717,0.002691,0.04165,0.002587,0.008374,1.276e-15,0.01464,0.003345,0.001686,0.005786,0.000457,8.269e-12,8.077e-08,5.45e-12,0.0004599,6.171e-07,1.137e-08,0.004924,0.0005643,0.000533,2.512e-08,3.381e-08,0.003297,9.299e-06,0.008235,0.0007895,0.0003805,0.03378,3.387e-11,0.00182,1.849e-08,1.366e-17,3.256e-06,7.129e-05,0.001026,4.489e-06,9.728e-10,0.0002361,0.01704,0.000234,6.369e-07,1.719e-05,4.51e-08,3.202e-13,0.000515,1.312e-12,0.01191,0.01265,1.384e-09,1.982e-06,1.55e-05,0.0004047,2.529e-07,0.002539,1.036e-08,0.0003864,1.823e-05,6.924e-07,0.02664,0.0003491,1.767e-10,0.000222,1.532e-06,1.621e-05,6.155e-07,1.29e-09,0.02331,0.002296,8.602e-06,0.006958,1.615e-06,2.919e-13,1.063e-05,9.118e-05,0.0007326,0.007465,3.495e-10,0.03215,0.000162,2.927e-07,0.0002509,1.538e-16,0.01028,0.0004539,4.666e-07,1.044e-09,2.01e-05,7.847e-08,8.524e-10,1.168e-14,6.415e-06,0.001115,1.102e-10,1.171e-08,0.002587,5.481e-08,0.0003152,0.0001309,1.445e-05,0.01324,4.232e-09,6.592e-08,0.002839,1.267e-12,8.23e-07,0.0007052,0.005495,1.631e-11,1.229e-07,2.691e-11,0.001029,2.213e-11,8.065e-07,0.007474,4.305e-10,4.904e-10,3.647e-05,1.197e-05,7.573e-05,0.006963,0.001348,0.00461,1.725e-08,5.08e-08,8.273e-08,7.575e-05,1.101e-05,4.145e-11,8.624e-07,1.063e-07,7.644e-06,1.622e-07,0.01367,3.369e-05,0.0007301,4.769e-05,1.259e-11,0.0002633,1.26e-06,1.148e-05,2.648e-06,2.951e-09,7.825e-06,0.00233,1.392e-09,3.275e-05,0.01253,1.646e-05,3.436e-06,2.706e-11,9.272e-08,0.04098,0.04427,0.002406,2.399e-12,5.304e-07,0.04465,0.004243,3.831e-19,4.724e-10,0.0001262,1.025e-05,4.991e-05,1.415e-06,8.201e-13,1.705e-11,0.0008502,2.5e-07,0.0001871,1.617e-09,1.162e-10,0.0176,0.0002459,4.685e-06,0.001574,0.0009498,1.18e-07,3.863e-08,1.48e-11,7.815e-05,0.008946,0.001644,0.0002189,1.202e-14,2.679e-10,1.008e-13,7.101e-06,0.001531,0.0001207,1.24e-05,1.59e-10,5.122e-05,4.655e-11,5.375e-07,6.641e-06,1.239e-05,5.945e-10,2.038e-09,4.211e-07,5.139e-08,9.529e-07,0.008594,0.01849,0.0001054,5.03e-08,4.418e-08,5.664e-10,1.708e-08,0.0004283,0.016,0.0267,0.01105,0.000685,0.003437,0.001968,3.492e-05,0.03059,0.007568,0.0005831,1.357e-08,3.919e-06,1.28e-05,6.268e-06,0.01312,0.006767,7.304e-10,1.269e-07,5.644e-08,1.054e-05,2.725e-08,0.002189,3.398e-11,1.201e-15,1.817e-05,1.04e-14,3.63e-05,0.02035,0.01298,0.02574,1.44e-08,3.412e-10,7.229e-13,4.423e-07,6.34e-12,4.157e-06,8.394e-10,2.787e-07,1.101e-05,1.627e-11,5.079e-15,6.47e-08,5.118e-13,4.058e-07,0.005582,0.01849,5.628e-09,5.626e-07,6.101e-11,7.71e-14,2.545e-05,8.767e-06,0.02197,7.97e-07,0.001246,0.0001084,1.126e-08,0.0003252,7.225e-05,0.004694,0.01794,0.0002691,0.001248,1.757e-06,5.623e-07,0.004976,0.007586,3.494e-05,1.085e-11,5.447e-09,2.506e-05,2.136e-08,0.008012,0.001092,0.001107,0.0006796,2.986e-08,0.01127,0.0004334,0.0003433,1.807e-08,4.532e-05,3.933e-09,1.082e-05,1.486e-07,4.749e-06,8.588e-06,7.283e-06,9.029e-16,1.719e-09,6.823e-07,5.922e-06,0.0005977,0.0002016,0.001159,1.982e-07,1.167e-08,0.0002861,0.0148,5.863e-12,0.00342,1.885e-07,0.0001013,1.136e-06,1.991e-06,7.975e-06,8.085e-08,1.699e-09,1.067e-06,0.0066,6.08e-07,0.0002376,0.01042,4.362e-12,0.001012,0.0001125,9.93e-06,1.22e-05,0.00357,4.415e-06,0.0003412,5.13e-05,1.745e-08,0.002857,4.862e-10,6.232e-06,0.001771,7.266e-08,0.001175,2.357e-10,4.151e-13,4.106e-05,0.0001383,1.76e-07,3.646e-16,9.528e-08,0.03667,1.032e-15,0.01143,9.34e-06,2.506e-08,4.835e-09,8.956e-06,0.001661,2.67e-06,2.588e-12,5.673e-11,1.16e-07,2.902e-08,1.25e-06,0.001473,6.941e-09,1.461e-08,0.01345,2.335e-21,2.17e-07,9.543e-05,8.633e-07,1.357e-05,4.556e-07,0.0003249,8.125e-11,1.417e-05,1.881e-09,5.297e-05,7.314e-06,3.549e-10,2.612e-06,0.00204,2.017e-06,7.463e-09,0.007124,0.005598,1.086e-06,6.491e-08,1.04e-06,8.492e-11,0.01062,2.955e-11,4.215e-05,0.004398,1.746e-10,0.01092,0.03657,7.442e-07,3.244e-05,0.008243,0.0004991,4.211e-16,0.004363,2.752e-07,3.322e-05,2.717e-05,0.000519,4.079e-08,0.0007352,1.45e-05,1.045e-09,2.884e-09,2.772e-08,5.907e-09,3.253e-09,8.293e-05,0.001231,0.008223,2.143e-10,9.915e-14,2.064e-15,0.0002873,0.0002153,1.088e-11,1.821e-09,5.76e-05,7.05e-07,3.763e-10,1.695e-07,6.676e-16,5.008e-05,1.861e-06,0.0001437,2.571e-06,2.051e-06,0.002954,3.679e-06,0.01908,4.783e-06,2.884e-09,6.356e-12,1.506e-07,3.74e-05,0.01439,6.943e-07,0.00031,5.756e-09,0.0003586,5.181e-08,2.531e-09,2.819e-07,5.242e-05,0.003458,3.174e-08,4.803e-08,9.705e-05,5.341e-09,0.002033,0.0002047,0.008249,5.771e-07,1.04e-14,5.914e-05,0.01309,2.163e-13,0.008638,5.573e-07,0.04263,1.932e-09,7.959e-06,0.0002564,0.003915,3.044e-08,6.58e-10,6.032e-09,3.778e-09,1.145e-05,7.755e-05,1.845e-14,0.0004381,5.09e-09,5.05e-09,0.0001416,0.0001382,0.002141,0.007213,5.122e-05,0.0002007,4.187e-08,0.0002901,2.434e-06,3.545e-08,2.312e-05,2.191e-10,1.117e-08,1.834e-09,6.428e-05,0.0001035,0.007767,3.625e-05,7.375e-08,4.385e-05,0.01093,1.306e-05,8.827e-07,1.044e-06,0.003318,9.349e-10,0.0007974,6.312e-08,0.002682,2.524e-07,0.0007347,8.366e-10,0.02471,0.0001563,3.57e-08,1.896e-06,4.306e-13,5.588e-05,4.115e-06,0.03878,8.861e-13,7.003e-09,0.0003972,6.693e-07,0.000968,2.192e-05,0.006207,4.221e-10,0.0003645,0.02504,1.306e-05,1.375e-08,1.008e-08,1.408e-12,6.972e-10,1.283e-08,2.967e-10,3.74e-13,3.446e-05,0.01545,3.87e-10,2.736e-12,4.695e-11,4.608e-12,4.745e-05,5.858e-15,2.834e-11,2.312e-05,2.972e-05,2.868e-05,3.449e-08,7.047e-05,6.226e-06,9.82e-05,0.0232,1.008e-05,0.002686,0.02789,3.33e-14,1.314e-14,0.008832,3.89e-12,2.035e-08,3.05e-13,6.291e-12,3.981e-10,4.197e-10,2.341e-11,4.262e-15,0.0003391,0.002691,0.0001383,4.756e-06,6.078e-07,3.98e-13,0.0002801,9.492e-08,0.0002354,6.618e-05,5.771e-05,5.379e-06,3.841e-11,8.151e-06,8.697e-07,0.00139,4.187e-08,0.02228,1.422e-09,3.92e-06,0.02005,1.311e-07,3.411e-05,4.106e-05,4.256e-14,4.705e-05,0.0001707,3.785e-07,1.843e-08,5.588e-06,0.0002571,1.677e-06,7.731e-09,0.02508,1.584e-05,0.005545,7.463e-06,6.347e-08,0.001415,0.001473,1.678e-06,7.221e-09,8.389e-06,1.711e-05,4.784e-05,6.181e-07,7.987e-07,1.277e-14,1.945e-05,3.387e-11,1.531e-07,0.005589,4.176e-12,1.496e-06,0.006614,0.006065,1.032e-13,0.0003027,1.677e-06,4.262e-15,1.246e-05,0.00317,1.686e-05,3.7e-07,5.747e-08,2.79e-10,4.783e-08,2.012e-05,2.003e-07,0.0003219,1.184e-05,3.904e-06,3.775e-07,5.371e-05,8.309e-06,7.496e-05,0.0003143,1.454e-07,1.541e-14,2.525e-06,1.56e-08,6.86e-08,1.668e-05,1.969e-08,0.0001856,0.0291,8.197e-08,1.451e-05,4.835e-09,0.002094,0.01393,0.003677,1.284e-06,0.003563,3.89e-12,4.948e-11,5.487e-10,1.819e-08,3.007e-11,8.005e-11,1.518e-13,7.012e-08,3.306e-16,3.88e-17,2.713e-09,0.002258,0.01149,0.00325,0.01104,1.651e-06,2.207e-11,2.037e-07,0.001547,0.00277,9.552e-07,1.45e-05,8.063e-06,3.375e-06,0.007352,0.0001128,1.746e-10,7.299e-17,1.648e-07,1.849e-09,1.729e-07,5.713e-06,0.000427,0.0001378,1.54e-06,0.0003278,0.001113,1.766e-08,3.556e-08,1.276e-06,9.102e-05,1.029e-07,1.961e-07,2.919e-06,5.081e-13,1.933e-07,0.002895,3.063e-05,5.042e-12,3.401e-08,4.907e-15,0.03402,0.005695,2.236e-06,3.019e-07,5.169e-09,0.02224,1.248e-13,1.954e-07,0.0001033,0.01152,0.0003929,1.819e-06,8.005e-11,5.447e-08,9.978e-05,6.13e-07,1.205e-06,8.567e-09,7.744e-06,1.085e-07,1.952e-14,0.003492,0.02562,0.002991,1.13e-07,0.001118,4.231e-12,0.0001095,7.485e-09,0.001093,1.075e-11,2.182e-14,6.818e-06,0.0001307,4.843e-09,1.135e-12,2.001e-11,8.008e-07,1.467e-07,1.21e-05,1.148e-11,1.415e-11,4.263e-16,3.236e-05,0.0006596,7.119e-07,2.088e-15,1.966e-13,0.03629,4.145e-11,6.53e-11,1.017e-13,6.091e-14,1.627e-11,0.000379,2.524e-06,0.03396,4.167e-08,1.324e-07,7.7e-07,2.203e-08,5.395e-11,0.0001729,3.582e-08,8.867e-13,1.084e-11,6.055e-05,4.952e-07,4.843e-09,8.385e-05,0.0001554,3.813e-16,2.052e-09,0.0208,1.678e-07,3.088e-05,1.649e-09,0.008154,3.567e-10,3.227e-12,8.589e-08,0.0002733,1.509e-06,1.017e-08,0.02567,0.0005758,0.03966,1.053e-05,0.001136,0.0173,4.186e-07,1.124e-06,1.534e-07,3.812e-06,1.248e-13,0.00699,5.645e-06,1.828e-12,0.01303,3.508e-07,8.976e-07,8.929e-06,4.174e-08,4.657e-05,1.971e-06,0.000561,1.261e-06,5.207e-08,6.026e-10,0.002101,0.02327,4.961e-06,2.807e-05,0.001954,6.61e-05,1.167e-06,2.724e-06,0.006567,1.816e-06,0.02961,1.234e-14,9.195e-14,9.682e-06,2.535e-12,1.783e-11,0.01071,2.398e-10,0.02578,0.0002096,0.00228,0.0003598,1.308e-08,0.002095,4.812e-05,7.718e-07,2.605e-07,1.811e-05,1.646e-05,1.169e-10,3.197e-11,3.253e-06,1.313e-07,0.02354,1.417e-12,0.007698,0.02598,3.22e-12,0.0003123,1.693e-05,0.0001653,1.572e-08,0.0001137,8.734e-05,1.419e-10,4.262e-12,3.775e-07,5.218e-05,5.767e-15,7.025e-06,1.398e-06,2.916e-10,0.02828,0.008548,3.35e-07,5.886e-08,4.094e-08,4.247e-12,2.731e-15,0.02309,1.461e-08,0.0001041,7.588e-16,1.457e-10,7.407e-05,1.785e-10,4.011e-07,1.067e-10,1.099e-10,9.637e-07,0.003908,1.03e-05,3.867e-07,1.962e-06,0.04271,0.005939,4.383e-08,4.306e-13,0.02873,2.897e-10,5.861e-08,0.002823,0.0007625,1.998e-08,0.01678,1.097e-07,3.626e-09,9.025e-05,0.0002475,2.518e-15,0.0005044,0.007257,3.767e-11,3.259e-08,0.0003515,1.008e-06,0.0001187,0.0001185,0.02511,1.226e-07,0.0003007,0.01472,0.0001274,5.457e-11,3.674e-09,2.75e-05,0.0139,0.004892,0.003259,0.0002259,1.08e-09,2.643e-05,1.462e-10,4.554e-07,1.934e-09,3.02e-06,2.404e-09,1.28e-05,8.31e-18,0.0006906,0.0001742,1.883e-08,4.967e-06,8.285e-06,0.0006695,6.692e-06,0.0009912,0.00873,3.223e-09,0.002124,0.02192,0.001538,5.833e-08,8.208e-05,1.785e-11,6.854e-08,0.0001606,0.0006943,3.178e-06,3.178e-15,6.403e-09,1.446e-06,2.088e-15,2.211e-09,1.176e-07,2.354e-08,0.008809,5.327e-07,2.976e-09,7.685e-07,0.0001976,2.096e-10,2.258e-07,2.088e-15,0.006752,0.0001338,1.982e-09,0.0002804,0.007639,0.04662,0.01443,2.865e-08,4.485e-06,3.184e-12,0.01999,0.004409,3.606e-15,0.007598,1.443e-10,0.002481,0.0007413,0.0005486,1.954e-07,0.0006384,3.948e-05,5.773e-07,1.957e-06,3.498e-14,5.921e-07,9.296e-10,8.065e-12,3.917e-07,9.675e-05,2.483e-06,0.02239,0.0008435,1.401e-05,9.542e-06,0.01325,0.0005925,0.0008061,0.0005525,0.0004152,6.619e-07,1.056e-06,1.068e-05,0.01236,1.911e-08,0.001412,2.422e-07,3.684e-13,5.795e-09,3.991e-05,0.0001231,6.957e-14,3.597e-14,1.217e-07,9.134e-06,1.007e-07,8.051e-07,0.00385,0.005428,9.092e-07,4.843e-07,4.906e-08,1.767e-10,7.447e-09,3.306e-16,1.245e-07,3.831e-19,0.002179,0.0002217,3.021e-12,1.138e-06,8.591e-11,0.0001874,0.04853,0.002776,0.02009,2.049e-10,3.416e-07,9.954e-12,9.19e-13,3.161e-06,0.01377,0.02814,3.813e-16,1.757e-15,1.177e-11,7.7e-07,1.23e-10,1.931e-07,8.019e-07,1.137e-08,0.001775,7.808e-15,2.328e-09,0.003088,1.517e-06,0.02773,0.00093,0.01835,8.185e-05,4.31e-05,2.365e-05,0.01734,1.071e-07,3.28e-08,1.908e-11,9.223e-09,2.665e-09,1.097e-07,1.662e-06,1.518e-13,0.01204,5.882e-05,0.001548,2.517e-10,3.661e-08,0.001675,3.105e-10,1.003e-10,4.572e-11,1.409e-12,5.915e-10,3.138e-15,0.002278,9.67e-09,6.028e-06,0.002838,5.773e-07,0.005428,0.002369,2.509e-14,2.972e-05,1.47e-10,3.585e-07,8.743e-15,0.0005046,0.002043,2.113e-05,1.646e-11,1.964e-07,0.0001557,1.631e-13,0.007797,2.578e-05,2.468e-14,0.001134,5.395e-08,4.298e-06,6.165e-05,3.227e-08,0.0001,8.053e-08,1.342e-07,2.74e-09,0.0002586,3.522e-06,0.002706,1.037e-06,0.0002108,5.724e-09,0.01119,4.633e-05,0.00762,0.0001478,1.653e-06,0.003025,0.03367,3.686e-11,6.617e-06,5.739e-06,4.634e-11,2.342e-07,1.094e-09,6.697e-10,1.017e-07,1.105e-05,0.001915,5.022e-10,3.337e-06,1.251e-05,0.03277,4.518e-11,1.833e-09,5.621e-08,6.465e-08,0.02115,2.227e-07,1.301e-05,0.001159,2.205e-09,9.355e-07,7.073e-11,6.803e-05,0.0004771,1.088e-11,0.03237,6.403e-05,8.96e-05,0.02148,0.007375,0.01118,5.39e-08,2.652e-14,0.008706,3.85e-09,1.746e-06,0.01461,7.445e-07,4.54e-08,2.739e-06,0.003757,1.927e-06,7.776e-07,1.896e-07,1.415e-06,1.843e-06,5.099e-05,2.729e-09,5.685e-15,3.788e-05,2.501e-05,9.882e-10,8.924e-08,9.937e-07,0.007301,1.31e-07,1.211e-06,0.003483,9.104e-09,9.428e-08,5.817e-10,5.03e-05,0.03412,5.942e-10,1.277e-14,1.749e-13,1.211e-14,3.284e-09,1.138e-07,0.009192,6.449e-07,2.027e-10,2.008e-09,1.683e-14,0.001238,4.057e-07,8.492e-08,1.119e-05,0.0001329,0.00059,0.0005526,8.661e-07,1.6e-06,0.001292,9.833e-05,2.851e-13,3.736e-06,0.002793,1.448e-05,0.0002801,1.22e-05,1.354e-07,3.492e-05,1.413e-08,7.707e-05,8.635e-10,9.55e-10,4.688e-09,9.549e-06,0.007999,0.008096,1.063e-17,0.001109,0.002418,0.009723,8.31e-18,8.913e-10,0.005589,0.001147,0.0005526,5.219e-11,0.01885,1.388e-06,6.609e-07,7.808e-07,2.255e-11,6.007e-06,0.03842,8.432e-05,0.002495,0.002048,2.665e-09,0.004644,2.331e-12,1.644e-13,4.654e-11,4.897e-08,2.351e-07,1.29e-09,2e-09,1.03e-07,5.731e-07,1.338e-09,4.832e-11,1.203e-08,7.729e-07,1.006e-09,0.007095,9.212e-05,5.928e-05,1.025e-06,4.64e-06,0.0005281,5.471e-09,6.937e-06,6.96e-07,1.517e-05,1.006e-09,0.002736,0.0005236,0.0002278,1.869e-05,8.274e-07,4.862e-10,1.024e-17,1.957e-06,2.354e-08,4.936e-09,7.75e-06,6.062e-11,2.399e-12,0.01068,1.057e-07,8.739e-07,0.003662,2.341e-06,1.101e-05,2.972e-06,0.00974,1.59e-10,0.0029,8.412e-14,4.555e-10,3.101e-13,5.871e-16,2.611e-09,3.338e-09,0.0002848,2.63e-15,2.538e-12,7.939e-06,1.662e-06,0.0002956,0.01932,5.792e-09,1.772e-08,0.0003898,1.586e-14,0.01926,2.601e-12,5.705e-14,0.0006102,4.111e-08,4.113e-06,8.315e-09,1.741e-11,0.0005754,4.875e-12,9.325e-07,0.009556,0.001132,3.038e-07,0.008601,0.03034,0.001035,0.0002907,0.02455,0.02951,0.001053,4.831e-06,0.001306,9.56e-11,1.067e-10,0.005138,3.597e-14,1.125e-05,0.001885,0.0005943,2.719e-14,9.386e-15,0.0006554,0.02549,0.01941,0.0006036,4.037e-05,2.427e-11,0.0004511,0.01168,0.006554,6.706e-13,0.002353,7.526e-12,1.232e-14,0.0006662,7.785e-08,7.51e-05,1.869e-16,0.01021,0.006359,0.0001854,0.003756,0.0004204,2.333e-08,3.297e-12,7.276e-19,0.006173,7.389e-05,2.37e-12,2.81e-09,0.000191,6.555e-06,2.602e-05,7.731e-09,0.0007229,1.138e-05,0.0001457,5.158e-10,0.004182,4.62e-10,3.236e-09,1.423e-16,2.493e-08,3.813e-16,0.007902,3.009e-07,7.426e-05,0.0005389,3.416e-07,1.817e-07,8.45e-06,2.031e-05,0.0007655,0.0066,4.739e-13,0.009298,0.0001033,7.678e-12,0.02386,2.823e-07,4.606e-09,0.004648,0.0001119,8.897e-05,4.175e-06,0.003535,0.006136,1.112e-13,8.44e-08,5.447e-07,5.504e-06,0.00458,0.04853,6.251e-06,0.0001513,2.08e-05,1.37e-05,0.002851,6.338e-06,6.073e-06,1.12e-07,5.81e-06,2.069e-08,7.744e-06,1.354e-08,1.466e-05,0.007457,0.0002969,6.791e-06,1.359e-15,0.002105,6.984e-14,0.04616,0.004894,4.51e-08,0.002949,0.0167,1.006e-09,1.906e-07,0.007892,1.158e-06,0.001038,1.569e-05,4.346e-05,0.001818,1.851e-06,3.646e-16,2.402e-06,3.396e-07,4.14e-05,1.416e-09,0.0004141,0.007297,8.201e-13,5.76e-05,2.723e-09,1.269e-10,4.47e-05,6.878e-11,3.639e-06,3.08e-05,8.051e-07,4.78e-06,1.643e-14,6.57e-08,0.02333,6.573e-09,2.135e-06,0.002564,0.00084,2.778e-11,9.479e-10,0.002565,0.001862,0.0008636,4.668e-05,5.819e-06,8.377e-11,0.01842,0.003465,3.754e-05,1.49e-06,9.402e-10,0.04691,5.801e-09,1.434e-09,1.137e-08,3.692e-14,1.666e-11,6.22e-07,2.539e-14,8.401e-10,0.0008542,2.467e-07,1.569e-11,2.756e-09,2.432e-13,0.007087,0.0005425,0.0004837,7.729e-07,3.662e-08,0.0001407,5.599e-08,4.715e-09,9.755e-06,4.554e-07,0.001371,7.49e-13,0.00412,2.919e-06,1.712e-06,9.314e-16,2.722e-08,4.793e-07,0.0001469,0.001663,6.949e-06,1.257e-11,6.611e-06,1.008e-08,3.124e-12,0.002377,0.00022,0.0005939,3.958e-05,9.149e-11,3.812e-14,0.0002476,0.02756,1.828e-09,6.428e-05,3.047e-08,0.006158,8.096e-08,2.999e-07,9.248e-05,4.344e-11,0.001027,3.721e-06,9.272e-08,3.185e-06,0.0283,0.0005927,1.341e-12,1.783e-05,1.372e-12,4.536e-09,0.002286,5.321e-06,6.352e-08,0.0001034,2.401e-13,0.03184,2.141e-13,8.805e-05,0.001277,0.04023,6.552e-05,3.266e-07,7.456e-07,2.739e-09,4.174e-14,3.498e-14,0.003426,2.227e-09,0.01668,4.881e-13,4.375e-07,0.006251,3.391e-14,0.003647,5.165e-10,7.975e-06,5.254e-08,3.652e-10,4.432e-14,3.138e-12,2.066e-06,1.174e-05,0.005896,0.0002136,1.107e-09,7.183e-06,0.009153,4.359e-13,1.351e-07,3.793e-07,3.495e-10,4.247e-12,2.665e-14,0.0004358,5.193e-07,2.815e-05,5.412e-13,0.02,1.552e-06,1.488e-13,0.0009925,0.01353,6.014e-06,0.0006022,0.007599,0.001873,3.078e-05,1.772e-08,0.02521,5.278e-14,3.487e-10,0.001645,1.021e-07,1.66e-09,6.993e-06,4.358e-05,5.274e-05,0.00019,1.031e-06,7.761e-08,7.984e-11,0.009299,2.806e-07,0.0009766,0.0007075,8.406e-09,4.967e-10,2.989e-08,3.942e-12,0.0001653,5.243e-09,8.385e-05,0.01236,1.466e-07,1.234e-07,0.008452,0.004739,1.25e-05,5.466e-11,9.498e-10,7.097e-13,1.064e-06,2.001e-07,4.522e-06,0.007568,0.003171,1.799e-08,0.0003351,9.697e-06,0.0236,1.409e-05,0.0007164,0.001084,4.22e-10,5.849e-05,5.254e-13,0.03564,0.005496,0.0003673,0.03311,0.01341,0.01142,2.192e-07,4.39e-05,0.0005631,0.008379,6.909e-05,0.001029,9.24e-07,4.245e-05,0.0002351,7.334e-09,0.004777,7.904e-09,3.768e-07,4.837e-08,3.447e-05,1.225e-08,7.041e-05,0.000321,1.342e-10,1.38e-10,6.871e-11,1.036e-07,0.008417,2.558e-06,1.85e-05,6.339e-05,0.002149,0.001273,1.181e-08,9.843e-13,1.781e-08,0.0005243,4.063e-10,0.02399,0.0002011,4.811e-09,1.234e-06,1.472e-07,0.0002755,1.297e-05,8.602e-06,2.033e-08,1.771e-06,1.415e-07,7.482e-05,0.00084,4.211e-16,4.463e-05,8.374e-06,4.262e-12,0.0004403,2.026e-10,3.936e-06,1.162e-06,0.0001238,0.007466,2.931e-05,8.261e-08,9.972e-06,0.002322,6.611e-06,4.397e-12,1.331e-08,1.125e-12,1.753e-14,1.241e-05,8.264e-09,0.0002459,2.063e-12,9.421e-07,6.728e-10,0.002086,0.0001223,6.208e-08,0.0009375,4.889e-15,3.72e-12,2.99e-08,0.001775,5.756e-09,2.275e-10,1.159e-05,1.21e-10,3.117e-14,0.000395,3.987e-05,0.00532,4.262e-05,0.0006521,0.0003247,1.471e-07,5.447e-08,5.836e-08,1.768e-09,1.251e-06,0.004163,1.599e-06,1.771e-06,0.009535,1.581e-06,1.887e-07,3.377e-05,6.913e-14,0.0001159,2.836e-12,3.178e-10,7.047e-10,0.0003617,0.01431,0.001061,0.006546,2.836e-12,0.001481,1.006e-09,1.493e-05,3.667e-09,0.003858,0.0007335,0.003292,3.261e-07,7.387e-11,0.003043,8.028e-12,2.366e-08,1.111e-09,0.002601,1.354e-08,6.06e-06,0.03118,0.0005115,0.005415,1.932e-12,4.86e-13,0.00084,1.133e-08,4.302e-10,1.789e-05,3.012e-06,1.357e-05,0.0001007,0.01431,5.839e-15,7.947e-05,0.003918,1.036e-06,0.02534,7.303e-11,9.205e-06,6.51e-05,2.461e-08,0.01688,1.647e-13,4.193e-08,1.869e-16,5.856e-05,4.362e-12,4.928e-09,4.9e-05,5.801e-09,0.0004825,1.554e-13,8.932e-15,0.02242,1.304e-12,3.812e-14,0.01911,0.0002609,7.257e-13,1.1e-11,4.197e-10,5.725e-05,0.000298,0.04637,0.003913,0.0003732,0.001763,1.189e-08,0.001886,1.329e-05,4.149e-10,1.68e-05,5.213e-10,6.902e-07,1.11e-10,4.823e-06,1.86e-10,0.00578,7.643e-09,0.009545,2.766e-09,1.748e-11,1.721e-05,1.425e-08,7.887e-05,0.0003759,1.217e-09,5.031e-08],"GeneID":["PVP01_0000020","PVP01_0000030","PVP01_0000060","PVP01_0000210","PVP01_0000230","PVP01_0000240","PVP01_0000300","PVP01_0000670","PVP01_0000960","PVP01_0000980","PVP01_0001090","PVP01_0001110","PVP01_0001360","PVP01_0001450","PVP01_0001500","PVP01_0001510","PVP01_0001580","PVP01_0001640","PVP01_0001790","PVP01_0001850","PVP01_0001990","PVP01_0002000","PVP01_0002050","PVP01_0002160","PVP01_0002200","PVP01_0002210","PVP01_0002310","PVP01_0002330","PVP01_0002370","PVP01_0002390","PVP01_0002400","PVP01_0002430","PVP01_0002440","PVP01_0002490","PVP01_0002540","PVP01_0002740","PVP01_0002750","PVP01_0002760","PVP01_0002810","PVP01_0002820","PVP01_0002950","PVP01_0003060","PVP01_0003320","PVP01_0003330","PVP01_0003670","PVP01_0003700","PVP01_0003710","PVP01_0003820","PVP01_0003950","PVP01_0004100","PVP01_0004180","PVP01_0004450","PVP01_0004810","PVP01_0004880","PVP01_0004910","PVP01_0004960","PVP01_0005160","PVP01_0005170","PVP01_0005290","PVP01_0005420","PVP01_0005450","PVP01_0005550","PVP01_0005710","PVP01_0005720","PVP01_0005740","PVP01_0005750","PVP01_0005830","PVP01_0006010","PVP01_0006100","PVP01_0006260","PVP01_0006590","PVP01_0006680","PVP01_0006700","PVP01_0006770","PVP01_0007000","PVP01_0007050","PVP01_0007420","PVP01_0007660","PVP01_0007720","PVP01_0007800","PVP01_0007820","PVP01_0007860","PVP01_0007870","PVP01_0007880","PVP01_0007920","PVP01_0007950","PVP01_0008030","PVP01_0008040","PVP01_0008060","PVP01_0008080","PVP01_0008100","PVP01_0008190","PVP01_0008380","PVP01_0008650","PVP01_0008830","PVP01_0008920","PVP01_0008930","PVP01_0008990","PVP01_0009040","PVP01_0009220","PVP01_0009270","PVP01_0009420","PVP01_0009450","PVP01_0009460","PVP01_0009490","PVP01_0009510","PVP01_0009740","PVP01_0009920","PVP01_0009940","PVP01_0009960","PVP01_0009990","PVP01_0010170","PVP01_0010180","PVP01_0010190","PVP01_0010200","PVP01_0010220","PVP01_0010240","PVP01_0010290","PVP01_0010380","PVP01_0010500","PVP01_0010520","PVP01_0010690","PVP01_0010900","PVP01_0010960","PVP01_0010980","PVP01_0100200","PVP01_0102800","PVP01_0103800","PVP01_0104100","PVP01_0104300","PVP01_0104600","PVP01_0104700","PVP01_0104800","PVP01_0104900","PVP01_0105100","PVP01_0105500","PVP01_0105800","PVP01_0106200","PVP01_0106500","PVP01_0106600","PVP01_0107400","PVP01_0107500","PVP01_0107600","PVP01_0107900","PVP01_0108000","PVP01_0108100","PVP01_0108800","PVP01_0109000","PVP01_0109100","PVP01_0109200","PVP01_0109600","PVP01_0110700","PVP01_0110800","PVP01_0111300","PVP01_0111800","PVP01_0111900","PVP01_0112000","PVP01_0112300","PVP01_0112800","PVP01_0113000","PVP01_0113400","PVP01_0113500","PVP01_0114900","PVP01_0115000","PVP01_0115800","PVP01_0116000","PVP01_0116100","PVP01_0116300","PVP01_0116600","PVP01_0116700","PVP01_0116900","PVP01_0117100","PVP01_0117500","PVP01_0117600","PVP01_0117800","PVP01_0118000","PVP01_0118100","PVP01_0118500","PVP01_0118700","PVP01_0118900","PVP01_0119300","PVP01_0119800","PVP01_0120100","PVP01_0120400","PVP01_0120500","PVP01_0201000","PVP01_0202000","PVP01_0202300","PVP01_0203100","PVP01_0203800","PVP01_0203900","PVP01_0204000","PVP01_0204800","PVP01_0204900","PVP01_0205200","PVP01_0205300","PVP01_0205500","PVP01_0206100","PVP01_0206400","PVP01_0206500","PVP01_0206600","PVP01_0207000","PVP01_0207100","PVP01_0207200","PVP01_0207400","PVP01_0207500","PVP01_0207600","PVP01_0207700","PVP01_0208100","PVP01_0208200","PVP01_0208300","PVP01_0208500","PVP01_0208600","PVP01_0208800","PVP01_0208900","PVP01_0209000","PVP01_0209500","PVP01_0209600","PVP01_0210000","PVP01_0210100","PVP01_0211000","PVP01_0211100","PVP01_0211200","PVP01_0211300","PVP01_0211400","PVP01_0211600","PVP01_0212000","PVP01_0212200","PVP01_0212400","PVP01_0212600","PVP01_0212800","PVP01_0213000","PVP01_0213700","PVP01_0214100","PVP01_0214500","PVP01_0214900","PVP01_0215500","PVP01_0215700","PVP01_0216100","PVP01_0216200","PVP01_0216600","PVP01_0217300","PVP01_0218800","PVP01_0220300","PVP01_0221100","PVP01_0301400","PVP01_0301600","PVP01_0301900","PVP01_0302300","PVP01_0302800","PVP01_0303100","PVP01_0303400","PVP01_0303500","PVP01_0304000","PVP01_0304100","PVP01_0304300","PVP01_0304500","PVP01_0304700","PVP01_0304900","PVP01_0305000","PVP01_0305500","PVP01_0306000","PVP01_0306100","PVP01_0306300","PVP01_0307000","PVP01_0307400","PVP01_0307500","PVP01_0307600","PVP01_0307700","PVP01_0308200","PVP01_0308600","PVP01_0308700","PVP01_0308800","PVP01_0309300","PVP01_0309800","PVP01_0310200","PVP01_0311100","PVP01_0311200","PVP01_0311300","PVP01_0311400","PVP01_0312100","PVP01_0312400","PVP01_0312500","PVP01_0312800","PVP01_0313400","PVP01_0314100","PVP01_0314900","PVP01_0315300","PVP01_0315500","PVP01_0315700","PVP01_0316200","PVP01_0316400","PVP01_0316500","PVP01_0316600","PVP01_0316800","PVP01_0317200","PVP01_0318000","PVP01_0318100","PVP01_0318500","PVP01_0318900","PVP01_0319300","PVP01_0319400","PVP01_0319500","PVP01_0319700","PVP01_0319800","PVP01_0320200","PVP01_0320400","PVP01_0320700","PVP01_0321100","PVP01_0400300","PVP01_0403200","PVP01_0403500","PVP01_0403600","PVP01_0403800","PVP01_0404000","PVP01_0405200","PVP01_0405700","PVP01_0406100","PVP01_0406400","PVP01_0406600","PVP01_0407800","PVP01_0407900","PVP01_0408600","PVP01_0408900","PVP01_0409300","PVP01_0409800","PVP01_0410000","PVP01_0410600","PVP01_0411400","PVP01_0411500","PVP01_0411700","PVP01_0411800","PVP01_0412400","PVP01_0412600","PVP01_0412700","PVP01_0412900","PVP01_0413500","PVP01_0414300","PVP01_0414400","PVP01_0414700","PVP01_0414900","PVP01_0415200","PVP01_0415400","PVP01_0415600","PVP01_0415900","PVP01_0416000","PVP01_0416200","PVP01_0417100","PVP01_0417200","PVP01_0417600","PVP01_0417900","PVP01_0418300","PVP01_0418400","PVP01_0419000","PVP01_0419100","PVP01_0419200","PVP01_0420700","PVP01_0420800","PVP01_0420900","PVP01_0421700","PVP01_0421800","PVP01_0422200","PVP01_0422300","PVP01_0422400","PVP01_0422800","PVP01_0423600","PVP01_0423700","PVP01_0424000","PVP01_0424100","PVP01_0500900","PVP01_0502600","PVP01_0503400","PVP01_0505100","PVP01_0505500","PVP01_0506000","PVP01_0507000","PVP01_0507200","PVP01_0507300","PVP01_0507600","PVP01_0507800","PVP01_0508900","PVP01_0509100","PVP01_0510100","PVP01_0510300","PVP01_0510500","PVP01_0510600","PVP01_0510700","PVP01_0511200","PVP01_0511300","PVP01_0511400","PVP01_0511500","PVP01_0511700","PVP01_0512000","PVP01_0512400","PVP01_0512500","PVP01_0512700","PVP01_0512900","PVP01_0513300","PVP01_0513500","PVP01_0514000","PVP01_0514900","PVP01_0515100","PVP01_0515300","PVP01_0515800","PVP01_0516000","PVP01_0516400","PVP01_0516500","PVP01_0516800","PVP01_0516900","PVP01_0517000","PVP01_0517400","PVP01_0517700","PVP01_0518600","PVP01_0518700","PVP01_0518800","PVP01_0518900","PVP01_0519200","PVP01_0519300","PVP01_0519400","PVP01_0520800","PVP01_0521500","PVP01_0521600","PVP01_0521700","PVP01_0522000","PVP01_0522100","PVP01_0522300","PVP01_0522600","PVP01_0524500","PVP01_0524900","PVP01_0525000","PVP01_0525300","PVP01_0525700","PVP01_0526000","PVP01_0526100","PVP01_0526200","PVP01_0526300","PVP01_0527300","PVP01_0527400","PVP01_0527700","PVP01_0527800","PVP01_0527900","PVP01_0528100","PVP01_0528200","PVP01_0528400","PVP01_0528500","PVP01_0528700","PVP01_0528800","PVP01_0529000","PVP01_0529600","PVP01_0529700","PVP01_0530100","PVP01_0530400","PVP01_0530500","PVP01_0530700","PVP01_0531000","PVP01_0531100","PVP01_0531500","PVP01_0531700","PVP01_0531900","PVP01_0532000","PVP01_0532500","PVP01_0533500","PVP01_0533600","PVP01_0533900","PVP01_0534700","PVP01_0535400","PVP01_0600100","PVP01_0600600","PVP01_0600700","PVP01_0601300","PVP01_0601400","PVP01_0601600","PVP01_0601700","PVP01_0602600","PVP01_0603000","PVP01_0603100","PVP01_0603600","PVP01_0603700","PVP01_0603900","PVP01_0604000","PVP01_0604100","PVP01_0604700","PVP01_0605000","PVP01_0605300","PVP01_0605500","PVP01_0605900","PVP01_0606100","PVP01_0606200","PVP01_0606900","PVP01_0607600","PVP01_0608000","PVP01_0608100","PVP01_0608600","PVP01_0608700","PVP01_0608900","PVP01_0609100","PVP01_0609600","PVP01_0609700","PVP01_0609800","PVP01_0609900","PVP01_0610200","PVP01_0610300","PVP01_0610400","PVP01_0610500","PVP01_0610700","PVP01_0610900","PVP01_0611100","PVP01_0611200","PVP01_0612000","PVP01_0612100","PVP01_0612400","PVP01_0612700","PVP01_0613000","PVP01_0613100","PVP01_0613200","PVP01_0613400","PVP01_0613500","PVP01_0613800","PVP01_0614300","PVP01_0614400","PVP01_0614600","PVP01_0614700","PVP01_0614900","PVP01_0615000","PVP01_0615100","PVP01_0615500","PVP01_0615800","PVP01_0616200","PVP01_0616300","PVP01_0616400","PVP01_0617500","PVP01_0618600","PVP01_0618700","PVP01_0618900","PVP01_0619100","PVP01_0619700","PVP01_0620000","PVP01_0620100","PVP01_0620300","PVP01_0620700","PVP01_0620900","PVP01_0621100","PVP01_0621300","PVP01_0621900","PVP01_0622100","PVP01_0622200","PVP01_0623200","PVP01_0624600","PVP01_0624800","PVP01_0700700","PVP01_0701200","PVP01_0701400","PVP01_0701500","PVP01_0701600","PVP01_0702100","PVP01_0702400","PVP01_0702500","PVP01_0704000","PVP01_0704700","PVP01_0705000","PVP01_0705600","PVP01_0705700","PVP01_0705800","PVP01_0706100","PVP01_0706200","PVP01_0706400","PVP01_0706700","PVP01_0706900","PVP01_0707000","PVP01_0707800","PVP01_0707900","PVP01_0708200","PVP01_0708400","PVP01_0708700","PVP01_0708900","PVP01_0709100","PVP01_0709200","PVP01_0709300","PVP01_0709600","PVP01_0710000","PVP01_0710200","PVP01_0711000","PVP01_0711100","PVP01_0711500","PVP01_0711800","PVP01_0712300","PVP01_0712400","PVP01_0713100","PVP01_0713500","PVP01_0713600","PVP01_0713800","PVP01_0713900","PVP01_0714200","PVP01_0714900","PVP01_0715200","PVP01_0715700","PVP01_0715800","PVP01_0716100","PVP01_0716200","PVP01_0716400","PVP01_0716800","PVP01_0717000","PVP01_0717200","PVP01_0717500","PVP01_0717600","PVP01_0717800","PVP01_0718600","PVP01_0718700","PVP01_0719000","PVP01_0719100","PVP01_0719500","PVP01_0719700","PVP01_0719900","PVP01_0720100","PVP01_0720300","PVP01_0720400","PVP01_0720600","PVP01_0720900","PVP01_0721300","PVP01_0721800","PVP01_0721900","PVP01_0722400","PVP01_0722500","PVP01_0723000","PVP01_0723200","PVP01_0724300","PVP01_0724400","PVP01_0724700","PVP01_0725100","PVP01_0725300","PVP01_0725400","PVP01_0726000","PVP01_0726400","PVP01_0727100","PVP01_0727300","PVP01_0727400","PVP01_0727700","PVP01_0728000","PVP01_0728100","PVP01_0728400","PVP01_0728500","PVP01_0729100","PVP01_0729400","PVP01_0729500","PVP01_0729700","PVP01_0730000","PVP01_0730300","PVP01_0730400","PVP01_0730600","PVP01_0730700","PVP01_0730800","PVP01_0731000","PVP01_0731100","PVP01_0731200","PVP01_0731300","PVP01_0731500","PVP01_0731700","PVP01_0732300","PVP01_0732600","PVP01_0732800","PVP01_0732900","PVP01_0733000","PVP01_0734100","PVP01_0734200","PVP01_0734800","PVP01_0735100","PVP01_0735300","PVP01_0736500","PVP01_0737700","PVP01_0800100","PVP01_0801800","PVP01_0802000","PVP01_0803500","PVP01_0804100","PVP01_0804300","PVP01_0804600","PVP01_0804800","PVP01_0804900","PVP01_0805300","PVP01_0805400","PVP01_0805500","PVP01_0805600","PVP01_0805800","PVP01_0806100","PVP01_0806300","PVP01_0806500","PVP01_0806700","PVP01_0807000","PVP01_0807500","PVP01_0807800","PVP01_0808100","PVP01_0808700","PVP01_0808800","PVP01_0809200","PVP01_0809400","PVP01_0809600","PVP01_0809700","PVP01_0809900","PVP01_0810500","PVP01_0811200","PVP01_0811300","PVP01_0812300","PVP01_0813200","PVP01_0813300","PVP01_0813700","PVP01_0813800","PVP01_0814100","PVP01_0814200","PVP01_0814400","PVP01_0814700","PVP01_0815000","PVP01_0815100","PVP01_0815300","PVP01_0815900","PVP01_0816200","PVP01_0816700","PVP01_0816800","PVP01_0817000","PVP01_0817100","PVP01_0817400","PVP01_0818100","PVP01_0818500","PVP01_0818600","PVP01_0818900","PVP01_0819200","PVP01_0819300","PVP01_0820200","PVP01_0820500","PVP01_0820600","PVP01_0820800","PVP01_0821300","PVP01_0821400","PVP01_0821700","PVP01_0822300","PVP01_0822400","PVP01_0822600","PVP01_0822900","PVP01_0823100","PVP01_0823600","PVP01_0823800","PVP01_0823900","PVP01_0824000","PVP01_0824100","PVP01_0824800","PVP01_0825100","PVP01_0825300","PVP01_0825800","PVP01_0826000","PVP01_0826200","PVP01_0826300","PVP01_0826700","PVP01_0827800","PVP01_0828100","PVP01_0828200","PVP01_0828300","PVP01_0828500","PVP01_0828700","PVP01_0828800","PVP01_0829000","PVP01_0829100","PVP01_0830300","PVP01_0830500","PVP01_0831000","PVP01_0831100","PVP01_0831300","PVP01_0831700","PVP01_0832000","PVP01_0832100","PVP01_0832200","PVP01_0832300","PVP01_0832900","PVP01_0833000","PVP01_0833700","PVP01_0834300","PVP01_0834400","PVP01_0834700","PVP01_0835000","PVP01_0835100","PVP01_0835300","PVP01_0835900","PVP01_0836100","PVP01_0836400","PVP01_0836700","PVP01_0836800","PVP01_0837200","PVP01_0837300","PVP01_0837600","PVP01_0838000","PVP01_0839200","PVP01_0839400","PVP01_0839500","PVP01_0839800","PVP01_0841000","PVP01_0901200","PVP01_0901500","PVP01_0902100","PVP01_0903700","PVP01_0904700","PVP01_0904900","PVP01_0906400","PVP01_0906500","PVP01_0906800","PVP01_0906900","PVP01_0907000","PVP01_0907200","PVP01_0907500","PVP01_0908000","PVP01_0908400","PVP01_0908600","PVP01_0908700","PVP01_0908900","PVP01_0909100","PVP01_0909500","PVP01_0909600","PVP01_0909700","PVP01_0909900","PVP01_0910000","PVP01_0910200","PVP01_0910900","PVP01_0911300","PVP01_0911600","PVP01_0911700","PVP01_0911800","PVP01_0911900","PVP01_0912000","PVP01_0912200","PVP01_0912300","PVP01_0912400","PVP01_0912900","PVP01_0913000","PVP01_0913800","PVP01_0914100","PVP01_0916000","PVP01_0916700","PVP01_0917100","PVP01_0917200","PVP01_0917400","PVP01_0917600","PVP01_0917900","PVP01_0918000","PVP01_0918100","PVP01_0918200","PVP01_0918300","PVP01_0918700","PVP01_0919000","PVP01_0919400","PVP01_0919500","PVP01_0919700","PVP01_0920000","PVP01_0920100","PVP01_0920200","PVP01_0920400","PVP01_0920500","PVP01_0920700","PVP01_0921200","PVP01_0921700","PVP01_0922000","PVP01_0922100","PVP01_0922800","PVP01_0922900","PVP01_0923000","PVP01_0923100","PVP01_0923700","PVP01_0923800","PVP01_0923900","PVP01_0924000","PVP01_0924100","PVP01_0924300","PVP01_0924400","PVP01_0924500","PVP01_0925200","PVP01_0925700","PVP01_0925800","PVP01_0926300","PVP01_0926600","PVP01_0926900","PVP01_0927100","PVP01_0927200","PVP01_0927300","PVP01_0927400","PVP01_0927600","PVP01_0927700","PVP01_0928000","PVP01_0928100","PVP01_0928200","PVP01_0928300","PVP01_0928900","PVP01_0929000","PVP01_0929200","PVP01_0929600","PVP01_0929700","PVP01_0929900","PVP01_0930000","PVP01_0930600","PVP01_0930800","PVP01_0931300","PVP01_0932100","PVP01_0932200","PVP01_0932400","PVP01_0932600","PVP01_0933100","PVP01_0933400","PVP01_0933500","PVP01_0933700","PVP01_0934100","PVP01_0934300","PVP01_0934400","PVP01_0934500","PVP01_0935200","PVP01_0935700","PVP01_0935900","PVP01_0936000","PVP01_0936400","PVP01_0936500","PVP01_0936900","PVP01_0937100","PVP01_0937300","PVP01_0937700","PVP01_0938000","PVP01_0938400","PVP01_0938600","PVP01_0938700","PVP01_0938900","PVP01_0939000","PVP01_0939200","PVP01_0940600","PVP01_0940900","PVP01_0941100","PVP01_0941200","PVP01_0941400","PVP01_0941600","PVP01_0941700","PVP01_0941900","PVP01_0942200","PVP01_0942400","PVP01_0944100","PVP01_0944400","PVP01_0944600","PVP01_0944800","PVP01_0945000","PVP01_0945200","PVP01_0945400","PVP01_0945500","PVP01_0945800","PVP01_0946200","PVP01_0946300","PVP01_0946400","PVP01_0946700","PVP01_0947200","PVP01_0947300","PVP01_0948500","PVP01_0950100","PVP01_1001900","PVP01_1002600","PVP01_1004200","PVP01_1004600","PVP01_1005300","PVP01_1005900","PVP01_1006400","PVP01_1006600","PVP01_1006900","PVP01_1007100","PVP01_1007500","PVP01_1007600","PVP01_1007700","PVP01_1007800","PVP01_1007900","PVP01_1008100","PVP01_1008300","PVP01_1008400","PVP01_1008600","PVP01_1008900","PVP01_1009000","PVP01_1009200","PVP01_1009600","PVP01_1010100","PVP01_1010200","PVP01_1010400","PVP01_1011100","PVP01_1011200","PVP01_1011300","PVP01_1011400","PVP01_1011500","PVP01_1011800","PVP01_1012100","PVP01_1012400","PVP01_1012500","PVP01_1012700","PVP01_1013200","PVP01_1014300","PVP01_1014400","PVP01_1014600","PVP01_1015000","PVP01_1015500","PVP01_1015600","PVP01_1015700","PVP01_1016100","PVP01_1016200","PVP01_1016600","PVP01_1016900","PVP01_1017400","PVP01_1017500","PVP01_1017900","PVP01_1018400","PVP01_1019000","PVP01_1019600","PVP01_1020000","PVP01_1020100","PVP01_1020700","PVP01_1020900","PVP01_1021000","PVP01_1021200","PVP01_1021300","PVP01_1021500","PVP01_1021600","PVP01_1021800","PVP01_1021900","PVP01_1022000","PVP01_1022200","PVP01_1022400","PVP01_1022500","PVP01_1022800","PVP01_1023000","PVP01_1023600","PVP01_1023700","PVP01_1023800","PVP01_1023900","PVP01_1024400","PVP01_1024500","PVP01_1024600","PVP01_1024800","PVP01_1024900","PVP01_1025100","PVP01_1025200","PVP01_1025500","PVP01_1026200","PVP01_1026300","PVP01_1026400","PVP01_1026600","PVP01_1026700","PVP01_1027100","PVP01_1027300","PVP01_1027600","PVP01_1027800","PVP01_1028100","PVP01_1028200","PVP01_1028600","PVP01_1028900","PVP01_1029000","PVP01_1029100","PVP01_1029700","PVP01_1030500","PVP01_1030600","PVP01_1031000","PVP01_1031100","PVP01_1031300","PVP01_1031400","PVP01_1031500","PVP01_1031800","PVP01_1031900","PVP01_1032900","PVP01_1033000","PVP01_1034600","PVP01_1034800","PVP01_1034900","PVP01_1035000","PVP01_1035200","PVP01_1100800","PVP01_1101000","PVP01_1101100","PVP01_1102400","PVP01_1102500","PVP01_1102700","PVP01_1103000","PVP01_1103200","PVP01_1103300","PVP01_1103400","PVP01_1103700","PVP01_1104000","PVP01_1104300","PVP01_1104400","PVP01_1104600","PVP01_1104800","PVP01_1105500","PVP01_1105700","PVP01_1106000","PVP01_1106800","PVP01_1106900","PVP01_1107000","PVP01_1107100","PVP01_1107400","PVP01_1107600","PVP01_1107900","PVP01_1108500","PVP01_1108600","PVP01_1108800","PVP01_1108900","PVP01_1109400","PVP01_1109500","PVP01_1109600","PVP01_1110500","PVP01_1110600","PVP01_1111000","PVP01_1111100","PVP01_1111300","PVP01_1112100","PVP01_1112900","PVP01_1113500","PVP01_1113600","PVP01_1113700","PVP01_1113900","PVP01_1114200","PVP01_1114300","PVP01_1114500","PVP01_1115500","PVP01_1115600","PVP01_1115900","PVP01_1116600","PVP01_1117700","PVP01_1118000","PVP01_1118500","PVP01_1118600","PVP01_1118700","PVP01_1118800","PVP01_1119000","PVP01_1119500","PVP01_1119600","PVP01_1119800","PVP01_1120000","PVP01_1120300","PVP01_1120500","PVP01_1121100","PVP01_1121400","PVP01_1121500","PVP01_1121700","PVP01_1121900","PVP01_1122100","PVP01_1122300","PVP01_1122400","PVP01_1122500","PVP01_1122600","PVP01_1122700","PVP01_1123000","PVP01_1123500","PVP01_1123600","PVP01_1123900","PVP01_1124000","PVP01_1124100","PVP01_1124300","PVP01_1124400","PVP01_1124800","PVP01_1124900","PVP01_1125000","PVP01_1125100","PVP01_1125400","PVP01_1125900","PVP01_1126200","PVP01_1127100","PVP01_1127200","PVP01_1127300","PVP01_1127400","PVP01_1127800","PVP01_1128400","PVP01_1129000","PVP01_1129400","PVP01_1129700","PVP01_1130200","PVP01_1130900","PVP01_1131000","PVP01_1132300","PVP01_1132400","PVP01_1132700","PVP01_1132800","PVP01_1132900","PVP01_1133100","PVP01_1133200","PVP01_1133400","PVP01_1133500","PVP01_1133600","PVP01_1133800","PVP01_1133900","PVP01_1134000","PVP01_1134300","PVP01_1134500","PVP01_1135500","PVP01_1135800","PVP01_1135900","PVP01_1136100","PVP01_1136700","PVP01_1136800","PVP01_1136900","PVP01_1137100","PVP01_1137200","PVP01_1137600","PVP01_1137700","PVP01_1137800","PVP01_1138400","PVP01_1138500","PVP01_1139900","PVP01_1140300","PVP01_1140500","PVP01_1140700","PVP01_1141300","PVP01_1141500","PVP01_1141900","PVP01_1142000","PVP01_1142600","PVP01_1142700","PVP01_1143200","PVP01_1143400","PVP01_1143500","PVP01_1144100","PVP01_1144300","PVP01_1144500","PVP01_1144800","PVP01_1145500","PVP01_1145700","PVP01_1146400","PVP01_1146500","PVP01_1146900","PVP01_1148100","PVP01_1148200","PVP01_1148500","PVP01_1148700","PVP01_1149000","PVP01_1200200","PVP01_1200600","PVP01_1201900","PVP01_1202100","PVP01_1202200","PVP01_1203100","PVP01_1203300","PVP01_1203500","PVP01_1203600","PVP01_1204000","PVP01_1204100","PVP01_1204200","PVP01_1204600","PVP01_1205000","PVP01_1205500","PVP01_1206100","PVP01_1206300","PVP01_1206700","PVP01_1206900","PVP01_1207000","PVP01_1208000","PVP01_1208200","PVP01_1208400","PVP01_1208600","PVP01_1208700","PVP01_1209000","PVP01_1209200","PVP01_1209900","PVP01_1210000","PVP01_1210100","PVP01_1210200","PVP01_1210300","PVP01_1210700","PVP01_1210800","PVP01_1211100","PVP01_1211200","PVP01_1211300","PVP01_1211600","PVP01_1212100","PVP01_1213400","PVP01_1214000","PVP01_1214200","PVP01_1214300","PVP01_1214400","PVP01_1214600","PVP01_1214700","PVP01_1216100","PVP01_1216200","PVP01_1216800","PVP01_1217200","PVP01_1217500","PVP01_1217600","PVP01_1217700","PVP01_1218800","PVP01_1218900","PVP01_1219000","PVP01_1219100","PVP01_1219400","PVP01_1219600","PVP01_1219700","PVP01_1219900","PVP01_1220200","PVP01_1220300","PVP01_1221300","PVP01_1221800","PVP01_1221900","PVP01_1222400","PVP01_1223200","PVP01_1223300","PVP01_1223400","PVP01_1223800","PVP01_1223900","PVP01_1224000","PVP01_1224400","PVP01_1224500","PVP01_1224600","PVP01_1225500","PVP01_1225900","PVP01_1226100","PVP01_1226600","PVP01_1226800","PVP01_1227500","PVP01_1227600","PVP01_1227900","PVP01_1228000","PVP01_1228100","PVP01_1228200","PVP01_1228300","PVP01_1228400","PVP01_1229200","PVP01_1229500","PVP01_1229600","PVP01_1229900","PVP01_1230000","PVP01_1230100","PVP01_1230200","PVP01_1230500","PVP01_1230600","PVP01_1230900","PVP01_1231000","PVP01_1231100","PVP01_1231800","PVP01_1231900","PVP01_1232600","PVP01_1232700","PVP01_1232900","PVP01_1233100","PVP01_1235500","PVP01_1236000","PVP01_1236100","PVP01_1236800","PVP01_1237100","PVP01_1237200","PVP01_1237700","PVP01_1237900","PVP01_1238300","PVP01_1238400","PVP01_1238900","PVP01_1239300","PVP01_1239500","PVP01_1240000","PVP01_1240100","PVP01_1241300","PVP01_1241900","PVP01_1242300","PVP01_1242900","PVP01_1243500","PVP01_1243600","PVP01_1243900","PVP01_1244700","PVP01_1244800","PVP01_1244900","PVP01_1245500","PVP01_1245600","PVP01_1245800","PVP01_1246200","PVP01_1246500","PVP01_1247300","PVP01_1247800","PVP01_1247900","PVP01_1248700","PVP01_1248800","PVP01_1248900","PVP01_1249000","PVP01_1249200","PVP01_1249500","PVP01_1250200","PVP01_1250700","PVP01_1250900","PVP01_1252400","PVP01_1252600","PVP01_1252900","PVP01_1253000","PVP01_1253700","PVP01_1254200","PVP01_1254900","PVP01_1255100","PVP01_1255200","PVP01_1255700","PVP01_1256400","PVP01_1256600","PVP01_1256800","PVP01_1256900","PVP01_1257000","PVP01_1257600","PVP01_1257800","PVP01_1258100","PVP01_1258300","PVP01_1258900","PVP01_1259000","PVP01_1259200","PVP01_1259300","PVP01_1259500","PVP01_1260200","PVP01_1260500","PVP01_1260600","PVP01_1260800","PVP01_1262000","PVP01_1262800","PVP01_1262900","PVP01_1263300","PVP01_1263400","PVP01_1264000","PVP01_1264300","PVP01_1264500","PVP01_1264600","PVP01_1264700","PVP01_1264800","PVP01_1265000","PVP01_1265700","PVP01_1266400","PVP01_1267100","PVP01_1267200","PVP01_1267300","PVP01_1267400","PVP01_1267600","PVP01_1268000","PVP01_1268200","PVP01_1268600","PVP01_1269000","PVP01_1269100","PVP01_1270200","PVP01_1270600","PVP01_1271100","PVP01_1273000","PVP01_1301200","PVP01_1301500","PVP01_1301600","PVP01_1301700","PVP01_1302700","PVP01_1302800","PVP01_1303100","PVP01_1304000","PVP01_1304200","PVP01_1304600","PVP01_1304800","PVP01_1304900","PVP01_1305300","PVP01_1305600","PVP01_1305700","PVP01_1305900","PVP01_1306000","PVP01_1306100","PVP01_1306300","PVP01_1306400","PVP01_1306700","PVP01_1307000","PVP01_1307200","PVP01_1307300","PVP01_1307500","PVP01_1307800","PVP01_1308100","PVP01_1308300","PVP01_1308500","PVP01_1308900","PVP01_1309000","PVP01_1309100","PVP01_1309700","PVP01_1310200","PVP01_1310900","PVP01_1311000","PVP01_1311100","PVP01_1311300","PVP01_1311400","PVP01_1311500","PVP01_1312200","PVP01_1312300","PVP01_1312400","PVP01_1312900","PVP01_1313400","PVP01_1313900","PVP01_1314100","PVP01_1314300","PVP01_1314600","PVP01_1314900","PVP01_1315000","PVP01_1315400","PVP01_1315700","PVP01_1316100","PVP01_1316500","PVP01_1316600","PVP01_1316900","PVP01_1317000","PVP01_1317200","PVP01_1317400","PVP01_1317600","PVP01_1317800","PVP01_1317900","PVP01_1318000","PVP01_1318100","PVP01_1318500","PVP01_1318800","PVP01_1318900","PVP01_1319100","PVP01_1319700","PVP01_1320700","PVP01_1320900","PVP01_1321300","PVP01_1321600","PVP01_1321800","PVP01_1322300","PVP01_1322400","PVP01_1322600","PVP01_1322700","PVP01_1323000","PVP01_1323100","PVP01_1323400","PVP01_1323500","PVP01_1323700","PVP01_1324100","PVP01_1324200","PVP01_1324700","PVP01_1324800","PVP01_1325200","PVP01_1325800","PVP01_1325900","PVP01_1326000","PVP01_1326200","PVP01_1326400","PVP01_1327600","PVP01_1327900","PVP01_1328000","PVP01_1328100","PVP01_1328400","PVP01_1328800","PVP01_1329300","PVP01_1329400","PVP01_1330600","PVP01_1330700","PVP01_1331000","PVP01_1331400","PVP01_1331800","PVP01_1332700","PVP01_1332900","PVP01_1333100","PVP01_1333200","PVP01_1333300","PVP01_1333600","PVP01_1333700","PVP01_1333900","PVP01_1334000","PVP01_1334100","PVP01_1334200","PVP01_1334300","PVP01_1334500","PVP01_1335600","PVP01_1335700","PVP01_1335800","PVP01_1336000","PVP01_1337400","PVP01_1337500","PVP01_1337900","PVP01_1338100","PVP01_1338300","PVP01_1338400","PVP01_1338500","PVP01_1338600","PVP01_1339000","PVP01_1339100","PVP01_1339600","PVP01_1340500","PVP01_1340600","PVP01_1342200","PVP01_1342300","PVP01_1342600","PVP01_1342800","PVP01_1343100","PVP01_1343200","PVP01_1343500","PVP01_1343700","PVP01_1344200","PVP01_1344600","PVP01_1344800","PVP01_1344900","PVP01_1345700","PVP01_1346600","PVP01_1346900","PVP01_1347300","PVP01_1347500","PVP01_1400100","PVP01_1400800","PVP01_1401200","PVP01_1401500","PVP01_1402900","PVP01_1403000","PVP01_1403400","PVP01_1403700","PVP01_1403800","PVP01_1403900","PVP01_1404500","PVP01_1404600","PVP01_1405000","PVP01_1405600","PVP01_1406000","PVP01_1406500","PVP01_1406600","PVP01_1407000","PVP01_1407100","PVP01_1407600","PVP01_1407800","PVP01_1407900","PVP01_1408100","PVP01_1408200","PVP01_1408500","PVP01_1408600","PVP01_1408700","PVP01_1408800","PVP01_1408900","PVP01_1409000","PVP01_1409300","PVP01_1409400","PVP01_1410200","PVP01_1410300","PVP01_1410400","PVP01_1410500","PVP01_1410700","PVP01_1410800","PVP01_1411000","PVP01_1411100","PVP01_1411200","PVP01_1411300","PVP01_1411400","PVP01_1411500","PVP01_1412700","PVP01_1413100","PVP01_1413200","PVP01_1413700","PVP01_1413900","PVP01_1414300","PVP01_1414500","PVP01_1415100","PVP01_1415500","PVP01_1415600","PVP01_1415900","PVP01_1416000","PVP01_1416100","PVP01_1416200","PVP01_1417100","PVP01_1417200","PVP01_1417600","PVP01_1417900","PVP01_1418000","PVP01_1418600","PVP01_1418800","PVP01_1419000","PVP01_1419100","PVP01_1419300","PVP01_1419400","PVP01_1419800","PVP01_1420100","PVP01_1420500","PVP01_1420700","PVP01_1420800","PVP01_1421000","PVP01_1421300","PVP01_1421500","PVP01_1422300","PVP01_1422600","PVP01_1422900","PVP01_1423200","PVP01_1423700","PVP01_1424200","PVP01_1424800","PVP01_1425100","PVP01_1426100","PVP01_1426200","PVP01_1426700","PVP01_1426800","PVP01_1427100","PVP01_1428300","PVP01_1428600","PVP01_1429000","PVP01_1429200","PVP01_1429300","PVP01_1429900","PVP01_1430000","PVP01_1430100","PVP01_1430500","PVP01_1430700","PVP01_1430800","PVP01_1431100","PVP01_1431200","PVP01_1431300","PVP01_1431600","PVP01_1432000","PVP01_1432400","PVP01_1432500","PVP01_1432600","PVP01_1433000","PVP01_1433700","PVP01_1433800","PVP01_1434000","PVP01_1434200","PVP01_1434300","PVP01_1434800","PVP01_1436000","PVP01_1437500","PVP01_1437700","PVP01_1437900","PVP01_1438100","PVP01_1439600","PVP01_1440000","PVP01_1440300","PVP01_1440400","PVP01_1440700","PVP01_1440900","PVP01_1441000","PVP01_1441200","PVP01_1441700","PVP01_1442500","PVP01_1442900","PVP01_1443200","PVP01_1443300","PVP01_1443600","PVP01_1443800","PVP01_1443900","PVP01_1444200","PVP01_1444600","PVP01_1444900","PVP01_1445000","PVP01_1445300","PVP01_1445600","PVP01_1446100","PVP01_1446300","PVP01_1446400","PVP01_1446900","PVP01_1447000","PVP01_1447200","PVP01_1447700","PVP01_1447800","PVP01_1447900","PVP01_1448000","PVP01_1448200","PVP01_1448300","PVP01_1448700","PVP01_1448800","PVP01_1448900","PVP01_1449100","PVP01_1449300","PVP01_1449400","PVP01_1449500","PVP01_1449700","PVP01_1449900","PVP01_1450400","PVP01_1451100","PVP01_1451500","PVP01_1451600","PVP01_1452200","PVP01_1452300","PVP01_1452400","PVP01_1452700","PVP01_1453800","PVP01_1454200","PVP01_1454600","PVP01_1454900","PVP01_1455800","PVP01_1456500","PVP01_1456600","PVP01_1456900","PVP01_1457700","PVP01_1457900","PVP01_1458300","PVP01_1458700","PVP01_1459600","PVP01_1460300","PVP01_1460500","PVP01_1460900","PVP01_1461100","PVP01_1461900","PVP01_1462000","PVP01_1462300","PVP01_1462600","PVP01_1462700","PVP01_1463200","PVP01_1463300","PVP01_1463600","PVP01_1465000","PVP01_1465900","PVP01_1466000","PVP01_1467000","PVP01_1467300","PVP01_1467600","PVP01_1467900","PVP01_1468800","PVP01_1471100","PVP01_1472000","PVP01_1472300","PVP01_1472400","PVP01_MIT00800","PVP01_MIT01400","PVP01_MIT02200","PVP01_MIT02300","PVP01_MIT02800","PVP01_0000010","PVP01_0000040","PVP01_0000050","PVP01_0000070","PVP01_0000090","PVP01_0000100","PVP01_0000110","PVP01_0000120","PVP01_0000130","PVP01_0000140","PVP01_0000150","PVP01_0000160","PVP01_0000170","PVP01_0000180","PVP01_0000190","PVP01_0000200","PVP01_0000220","PVP01_0001430","PVP01_0001440","PVP01_0001470","PVP01_0001490","PVP01_0001600","PVP01_0001940","PVP01_0002090","PVP01_0002180","PVP01_0002240","PVP01_0002530","PVP01_0002710","PVP01_0002720","PVP01_0003140","PVP01_0003150","PVP01_0003290","PVP01_0003350","PVP01_0003980","PVP01_0004110","PVP01_0004210","PVP01_0004220","PVP01_0004230","PVP01_0004240","PVP01_0004250","PVP01_0004260","PVP01_0004280","PVP01_0004290","PVP01_0004300","PVP01_0004310","PVP01_0004320","PVP01_0004340","PVP01_0004350","PVP01_0004360","PVP01_0004370","PVP01_0004380","PVP01_0004390","PVP01_0004400","PVP01_0004410","PVP01_0004920","PVP01_0004950","PVP01_0005000","PVP01_0005010","PVP01_0005020","PVP01_0005030","PVP01_0005250","PVP01_0005430","PVP01_0005560","PVP01_0005570","PVP01_0005760","PVP01_0005800","PVP01_0005810","PVP01_0005820","PVP01_0005870","PVP01_0005910","PVP01_0005930","PVP01_0005940","PVP01_0005950","PVP01_0006660","PVP01_0006670","PVP01_0007080","PVP01_0007220","PVP01_0007240","PVP01_0007290","PVP01_0007650","PVP01_0007810","PVP01_0007830","PVP01_0008090","PVP01_0008110","PVP01_0008150","PVP01_0008500","PVP01_0008770","PVP01_0008820","PVP01_0009050","PVP01_0009070","PVP01_0009100","PVP01_0009180","PVP01_0009210","PVP01_0009370","PVP01_0009410","PVP01_0009430","PVP01_0009800","PVP01_0009810","PVP01_0009870","PVP01_0009950","PVP01_0009980","PVP01_0010020","PVP01_0010030","PVP01_0010070","PVP01_0010120","PVP01_0010130","PVP01_0010230","PVP01_0010440","PVP01_0010470","PVP01_0010510","PVP01_0010550","PVP01_0010570","PVP01_0010590","PVP01_0010600","PVP01_0010640","PVP01_0010670","PVP01_0010810","PVP01_0010840","PVP01_0010850","PVP01_0010860","PVP01_0010870","PVP01_0010970","PVP01_0010990","PVP01_0011020","PVP01_0011050","PVP01_0100800","PVP01_0102300","PVP01_0102600","PVP01_0102700","PVP01_0103100","PVP01_0103200","PVP01_0103300","PVP01_0103400","PVP01_0103500","PVP01_0103600","PVP01_0103700","PVP01_0103900","PVP01_0104000","PVP01_0104200","PVP01_0104400","PVP01_0104500","PVP01_0105000","PVP01_0105200","PVP01_0105300","PVP01_0105400","PVP01_0105600","PVP01_0105700","PVP01_0105900","PVP01_0106000","PVP01_0106100","PVP01_0106300","PVP01_0106400","PVP01_0106700","PVP01_0106800","PVP01_0106900","PVP01_0107700","PVP01_0107800","PVP01_0108200","PVP01_0108300","PVP01_0108400","PVP01_0108500","PVP01_0108600","PVP01_0108700","PVP01_0108900","PVP01_0109300","PVP01_0109400","PVP01_0109500","PVP01_0109700","PVP01_0109800","PVP01_0109900","PVP01_0110000","PVP01_0110100","PVP01_0110200","PVP01_0110300","PVP01_0110400","PVP01_0110500","PVP01_0110600","PVP01_0110900","PVP01_0111000","PVP01_0111100","PVP01_0111200","PVP01_0111400","PVP01_0111500","PVP01_0111600","PVP01_0111700","PVP01_0112100","PVP01_0112200","PVP01_0112400","PVP01_0112500","PVP01_0112600","PVP01_0112700","PVP01_0112900","PVP01_0113100","PVP01_0113200","PVP01_0113300","PVP01_0113600","PVP01_0113700","PVP01_0113800","PVP01_0113900","PVP01_0114000","PVP01_0114100","PVP01_0114200","PVP01_0114300","PVP01_0114400","PVP01_0114500","PVP01_0114600","PVP01_0114700","PVP01_0114800","PVP01_0115100","PVP01_0115200","PVP01_0115300","PVP01_0115400","PVP01_0115500","PVP01_0115600","PVP01_0115700","PVP01_0115900","PVP01_0116200","PVP01_0116400","PVP01_0116500","PVP01_0116800","PVP01_0117000","PVP01_0117200","PVP01_0117300","PVP01_0117400","PVP01_0117700","PVP01_0117900","PVP01_0118200","PVP01_0118300","PVP01_0118400","PVP01_0118600","PVP01_0118800","PVP01_0119000","PVP01_0119100","PVP01_0119200","PVP01_0119400","PVP01_0119500","PVP01_0119600","PVP01_0119900","PVP01_0121700","PVP01_0122100","PVP01_0122200","PVP01_0200200","PVP01_0201100","PVP01_0201200","PVP01_0201400","PVP01_0201500","PVP01_0201600","PVP01_0201700","PVP01_0201800","PVP01_0201900","PVP01_0202100","PVP01_0202200","PVP01_0202400","PVP01_0202500","PVP01_0202600","PVP01_0202700","PVP01_0202800","PVP01_0202900","PVP01_0203000","PVP01_0203200","PVP01_0203300","PVP01_0203400","PVP01_0203500","PVP01_0203600","PVP01_0203700","PVP01_0204100","PVP01_0204200","PVP01_0204300","PVP01_0204400","PVP01_0204500","PVP01_0204600","PVP01_0204700","PVP01_0205000","PVP01_0205100","PVP01_0205400","PVP01_0205600","PVP01_0205700","PVP01_0205800","PVP01_0205900","PVP01_0206000","PVP01_0206200","PVP01_0206300","PVP01_0206700","PVP01_0206800","PVP01_0206900","PVP01_0207300","PVP01_0207800","PVP01_0207900","PVP01_0208000","PVP01_0208400","PVP01_0208700","PVP01_0209100","PVP01_0209200","PVP01_0209300","PVP01_0209400","PVP01_0209700","PVP01_0209800","PVP01_0209900","PVP01_0210200","PVP01_0210300","PVP01_0210400","PVP01_0210500","PVP01_0210600","PVP01_0210700","PVP01_0210800","PVP01_0210900","PVP01_0211500","PVP01_0211700","PVP01_0211800","PVP01_0211900","PVP01_0212100","PVP01_0212300","PVP01_0212500","PVP01_0212700","PVP01_0212900","PVP01_0213100","PVP01_0213200","PVP01_0213300","PVP01_0213400","PVP01_0213500","PVP01_0213600","PVP01_0213800","PVP01_0213900","PVP01_0214000","PVP01_0214200","PVP01_0214300","PVP01_0214400","PVP01_0214600","PVP01_0214700","PVP01_0214800","PVP01_0215000","PVP01_0215100","PVP01_0215200","PVP01_0215300","PVP01_0215400","PVP01_0215600","PVP01_0215800","PVP01_0215900","PVP01_0216000","PVP01_0216500","PVP01_0216700","PVP01_0216800","PVP01_0216900","PVP01_0217000","PVP01_0217100","PVP01_0217200","PVP01_0219400","PVP01_0220100","PVP01_0220200","PVP01_0300200","PVP01_0300300","PVP01_0300600","PVP01_0300700","PVP01_0300800","PVP01_0300900","PVP01_0301500","PVP01_0302100","PVP01_0302400","PVP01_0302500","PVP01_0302600","PVP01_0302700","PVP01_0302900","PVP01_0303000","PVP01_0303200","PVP01_0303300","PVP01_0303600","PVP01_0303700","PVP01_0303800","PVP01_0304200","PVP01_0304400","PVP01_0304600","PVP01_0304800","PVP01_0305100","PVP01_0305200","PVP01_0305300","PVP01_0305400","PVP01_0305600","PVP01_0305700","PVP01_0305800","PVP01_0305900","PVP01_0306200","PVP01_0306400","PVP01_0306500","PVP01_0306600","PVP01_0306700","PVP01_0306800","PVP01_0306900","PVP01_0307100","PVP01_0307200","PVP01_0307300","PVP01_0307800","PVP01_0307900","PVP01_0308000","PVP01_0308100","PVP01_0308300","PVP01_0308400","PVP01_0308500","PVP01_0308900","PVP01_0309000","PVP01_0309100","PVP01_0309200","PVP01_0309600","PVP01_0309700","PVP01_0309900","PVP01_0310000","PVP01_0310100","PVP01_0310300","PVP01_0310400","PVP01_0310500","PVP01_0310600","PVP01_0310700","PVP01_0311000","PVP01_0311500","PVP01_0311600","PVP01_0311700","PVP01_0311800","PVP01_0311900","PVP01_0312000","PVP01_0312200","PVP01_0312300","PVP01_0312600","PVP01_0312700","PVP01_0312900","PVP01_0313000","PVP01_0313100","PVP01_0313200","PVP01_0313300","PVP01_0313500","PVP01_0313600","PVP01_0313700","PVP01_0313800","PVP01_0313900","PVP01_0314000","PVP01_0314200","PVP01_0314300","PVP01_0314400","PVP01_0314500","PVP01_0314600","PVP01_0314700","PVP01_0314800","PVP01_0315000","PVP01_0315100","PVP01_0315200","PVP01_0315400","PVP01_0315600","PVP01_0315800","PVP01_0315900","PVP01_0316000","PVP01_0316100","PVP01_0316300","PVP01_0316700","PVP01_0316900","PVP01_0317000","PVP01_0317100","PVP01_0317300","PVP01_0317400","PVP01_0317500","PVP01_0317600","PVP01_0317700","PVP01_0317800","PVP01_0317900","PVP01_0318200","PVP01_0318300","PVP01_0318400","PVP01_0318600","PVP01_0318700","PVP01_0318800","PVP01_0319000","PVP01_0319100","PVP01_0319200","PVP01_0319600","PVP01_0319900","PVP01_0320000","PVP01_0320100","PVP01_0320300","PVP01_0320500","PVP01_0320600","PVP01_0320800","PVP01_0320900","PVP01_0321000","PVP01_0403300","PVP01_0403400","PVP01_0403900","PVP01_0404200","PVP01_0404300","PVP01_0404400","PVP01_0404500","PVP01_0404600","PVP01_0404700","PVP01_0404800","PVP01_0404900","PVP01_0405000","PVP01_0405100","PVP01_0405300","PVP01_0405400","PVP01_0405500","PVP01_0405600","PVP01_0405800","PVP01_0405900","PVP01_0406200","PVP01_0406300","PVP01_0406500","PVP01_0406700","PVP01_0406800","PVP01_0406900","PVP01_0407000","PVP01_0407100","PVP01_0407200","PVP01_0407300","PVP01_0407400","PVP01_0407500","PVP01_0407600","PVP01_0407700","PVP01_0408000","PVP01_0408100","PVP01_0408200","PVP01_0408300","PVP01_0408400","PVP01_0408500","PVP01_0408700","PVP01_0408800","PVP01_0409000","PVP01_0409100","PVP01_0409200","PVP01_0409400","PVP01_0409500","PVP01_0409600","PVP01_0409700","PVP01_0409900","PVP01_0410100","PVP01_0410200","PVP01_0410300","PVP01_0410400","PVP01_0410500","PVP01_0410700","PVP01_0410800","PVP01_0410900","PVP01_0411000","PVP01_0411100","PVP01_0411200","PVP01_0411300","PVP01_0411600","PVP01_0411900","PVP01_0412000","PVP01_0412100","PVP01_0412200","PVP01_0412300","PVP01_0412500","PVP01_0412800","PVP01_0413000","PVP01_0413100","PVP01_0413200","PVP01_0413300","PVP01_0413400","PVP01_0413600","PVP01_0413700","PVP01_0413800","PVP01_0413900","PVP01_0414000","PVP01_0414100","PVP01_0414200","PVP01_0414500","PVP01_0414600","PVP01_0414800","PVP01_0415000","PVP01_0415100","PVP01_0415300","PVP01_0415500","PVP01_0415700","PVP01_0415800","PVP01_0416100","PVP01_0416300","PVP01_0416400","PVP01_0416500","PVP01_0416600","PVP01_0416700","PVP01_0416800","PVP01_0416900","PVP01_0417000","PVP01_0417300","PVP01_0417400","PVP01_0417500","PVP01_0417700","PVP01_0417800","PVP01_0418000","PVP01_0418100","PVP01_0418200","PVP01_0418500","PVP01_0418600","PVP01_0418700","PVP01_0418800","PVP01_0418900","PVP01_0419300","PVP01_0419400","PVP01_0419500","PVP01_0419600","PVP01_0419700","PVP01_0419800","PVP01_0419900","PVP01_0420000","PVP01_0420100","PVP01_0420200","PVP01_0420300","PVP01_0420400","PVP01_0420500","PVP01_0420600","PVP01_0421000","PVP01_0421100","PVP01_0421200","PVP01_0421300","PVP01_0421400","PVP01_0421500","PVP01_0421900","PVP01_0422000","PVP01_0422100","PVP01_0422500","PVP01_0422600","PVP01_0422700","PVP01_0422900","PVP01_0423000","PVP01_0423100","PVP01_0423200","PVP01_0423300","PVP01_0423400","PVP01_0423500","PVP01_0423800","PVP01_0423900","PVP01_0424200","PVP01_0424500","PVP01_0500400","PVP01_0502700","PVP01_0502800","PVP01_0502900","PVP01_0503000","PVP01_0503500","PVP01_0503600","PVP01_0503700","PVP01_0503800","PVP01_0503900","PVP01_0504000","PVP01_0504100","PVP01_0504200","PVP01_0504300","PVP01_0504400","PVP01_0504500","PVP01_0504600","PVP01_0504700","PVP01_0504800","PVP01_0504900","PVP01_0505000","PVP01_0505200","PVP01_0505300","PVP01_0505400","PVP01_0505600","PVP01_0505700","PVP01_0505800","PVP01_0505900","PVP01_0506100","PVP01_0506200","PVP01_0506300","PVP01_0506400","PVP01_0506500","PVP01_0506600","PVP01_0506700","PVP01_0506800","PVP01_0506900","PVP01_0507100","PVP01_0507400","PVP01_0507500","PVP01_0507700","PVP01_0507900","PVP01_0508000","PVP01_0508100","PVP01_0508200","PVP01_0508300","PVP01_0508400","PVP01_0508500","PVP01_0508600","PVP01_0508700","PVP01_0508800","PVP01_0509000","PVP01_0509200","PVP01_0509300","PVP01_0509400","PVP01_0509500","PVP01_0509600","PVP01_0509700","PVP01_0509800","PVP01_0509900","PVP01_0510000","PVP01_0510200","PVP01_0510400","PVP01_0510800","PVP01_0510900","PVP01_0511000","PVP01_0511100","PVP01_0511600","PVP01_0511800","PVP01_0511900","PVP01_0512100","PVP01_0512200","PVP01_0512300","PVP01_0512600","PVP01_0512800","PVP01_0513000","PVP01_0513100","PVP01_0513200","PVP01_0513400","PVP01_0513600","PVP01_0513700","PVP01_0513800","PVP01_0513900","PVP01_0514100","PVP01_0514200","PVP01_0514300","PVP01_0514400","PVP01_0514500","PVP01_0514600","PVP01_0514700","PVP01_0514800","PVP01_0515000","PVP01_0515200","PVP01_0515400","PVP01_0515500","PVP01_0515600","PVP01_0515700","PVP01_0515900","PVP01_0516100","PVP01_0516200","PVP01_0516300","PVP01_0516600","PVP01_0516700","PVP01_0517100","PVP01_0517200","PVP01_0517300","PVP01_0517500","PVP01_0517600","PVP01_0517800","PVP01_0517900","PVP01_0518000","PVP01_0518100","PVP01_0518200","PVP01_0518300","PVP01_0518400","PVP01_0518500","PVP01_0519000","PVP01_0519100","PVP01_0519500","PVP01_0519600","PVP01_0519700","PVP01_0519800","PVP01_0519900","PVP01_0520000","PVP01_0520100","PVP01_0520200","PVP01_0520300","PVP01_0520400","PVP01_0520500","PVP01_0520600","PVP01_0520700","PVP01_0520900","PVP01_0521000","PVP01_0521100","PVP01_0521200","PVP01_0521300","PVP01_0521400","PVP01_0521800","PVP01_0521900","PVP01_0522200","PVP01_0522400","PVP01_0522500","PVP01_0522700","PVP01_0522800","PVP01_0523000","PVP01_0523100","PVP01_0523200","PVP01_0523300","PVP01_0523400","PVP01_0523500","PVP01_0523700","PVP01_0523800","PVP01_0523900","PVP01_0524000","PVP01_0524100","PVP01_0524200","PVP01_0524300","PVP01_0524400","PVP01_0524600","PVP01_0524700","PVP01_0525100","PVP01_0525200","PVP01_0525400","PVP01_0525500","PVP01_0525600","PVP01_0525800","PVP01_0525900","PVP01_0526400","PVP01_0526500","PVP01_0526600","PVP01_0526700","PVP01_0526800","PVP01_0526900","PVP01_0527000","PVP01_0527100","PVP01_0527200","PVP01_0527500","PVP01_0527600","PVP01_0528000","PVP01_0528300","PVP01_0528600","PVP01_0528900","PVP01_0529100","PVP01_0529200","PVP01_0529300","PVP01_0529400","PVP01_0529500","PVP01_0529800","PVP01_0529900","PVP01_0530000","PVP01_0530200","PVP01_0530300","PVP01_0530600","PVP01_0530800","PVP01_0530900","PVP01_0531200","PVP01_0531300","PVP01_0531400","PVP01_0531600","PVP01_0531800","PVP01_0532100","PVP01_0532200","PVP01_0532300","PVP01_0532400","PVP01_0532600","PVP01_0532700","PVP01_0532800","PVP01_0532900","PVP01_0533000","PVP01_0533100","PVP01_0533200","PVP01_0533300","PVP01_0533400","PVP01_0533700","PVP01_0533800","PVP01_0534000","PVP01_0534200","PVP01_0534300","PVP01_0534400","PVP01_0534500","PVP01_0534600","PVP01_0534800","PVP01_0600400","PVP01_0600800","PVP01_0600900","PVP01_0601000","PVP01_0601200","PVP01_0601500","PVP01_0601800","PVP01_0601900","PVP01_0602000","PVP01_0602100","PVP01_0602200","PVP01_0602300","PVP01_0602400","PVP01_0602500","PVP01_0602700","PVP01_0602800","PVP01_0602900","PVP01_0603200","PVP01_0603300","PVP01_0603400","PVP01_0603500","PVP01_0603800","PVP01_0604200","PVP01_0604300","PVP01_0604400","PVP01_0604500","PVP01_0604600","PVP01_0604800","PVP01_0604900","PVP01_0605100","PVP01_0605200","PVP01_0605400","PVP01_0605600","PVP01_0605700","PVP01_0605800","PVP01_0606000","PVP01_0606300","PVP01_0606400","PVP01_0606500","PVP01_0606600","PVP01_0606700","PVP01_0606800","PVP01_0607000","PVP01_0607100","PVP01_0607200","PVP01_0607300","PVP01_0607400","PVP01_0607500","PVP01_0607700","PVP01_0607800","PVP01_0607900","PVP01_0608200","PVP01_0608300","PVP01_0608400","PVP01_0608500","PVP01_0608800","PVP01_0609000","PVP01_0609200","PVP01_0609300","PVP01_0609400","PVP01_0609500","PVP01_0610000","PVP01_0610100","PVP01_0610600","PVP01_0610800","PVP01_0611000","PVP01_0611300","PVP01_0611400","PVP01_0611500","PVP01_0611600","PVP01_0611700","PVP01_0611800","PVP01_0611900","PVP01_0612200","PVP01_0612300","PVP01_0612500","PVP01_0612600","PVP01_0612800","PVP01_0612900","PVP01_0613300","PVP01_0613600","PVP01_0613700","PVP01_0613900","PVP01_0614000","PVP01_0614100","PVP01_0614200","PVP01_0614500","PVP01_0614800","PVP01_0615200","PVP01_0615300","PVP01_0615400","PVP01_0615600","PVP01_0615700","PVP01_0615900","PVP01_0616000","PVP01_0616100","PVP01_0616500","PVP01_0616600","PVP01_0616700","PVP01_0616800","PVP01_0616900","PVP01_0617000","PVP01_0617100","PVP01_0617200","PVP01_0617300","PVP01_0617400","PVP01_0617600","PVP01_0617700","PVP01_0617800","PVP01_0617900","PVP01_0618000","PVP01_0618100","PVP01_0618200","PVP01_0618300","PVP01_0618400","PVP01_0618500","PVP01_0618800","PVP01_0619000","PVP01_0619200","PVP01_0619300","PVP01_0619400","PVP01_0619500","PVP01_0619600","PVP01_0619800","PVP01_0619900","PVP01_0620200","PVP01_0620400","PVP01_0620500","PVP01_0620600","PVP01_0620800","PVP01_0621000","PVP01_0621200","PVP01_0621400","PVP01_0621500","PVP01_0621600","PVP01_0621700","PVP01_0621800","PVP01_0622000","PVP01_0622300","PVP01_0622400","PVP01_0622500","PVP01_0622600","PVP01_0622700","PVP01_0622800","PVP01_0623000","PVP01_0623100","PVP01_0623300","PVP01_0623400","PVP01_0623500","PVP01_0623600","PVP01_0623700","PVP01_0623800","PVP01_0623900","PVP01_0624000","PVP01_0624100","PVP01_0624300","PVP01_0624400","PVP01_0624500","PVP01_0624700","PVP01_0624900","PVP01_0625000","PVP01_0700100","PVP01_0700300","PVP01_0700600","PVP01_0700800","PVP01_0700900","PVP01_0701000","PVP01_0701100","PVP01_0701300","PVP01_0701700","PVP01_0701800","PVP01_0701900","PVP01_0702000","PVP01_0702200","PVP01_0702300","PVP01_0702600","PVP01_0702700","PVP01_0702800","PVP01_0702900","PVP01_0703000","PVP01_0703100","PVP01_0703200","PVP01_0703300","PVP01_0703400","PVP01_0703500","PVP01_0703600","PVP01_0703700","PVP01_0703800","PVP01_0703900","PVP01_0704100","PVP01_0704200","PVP01_0704300","PVP01_0704400","PVP01_0704500","PVP01_0704600","PVP01_0704800","PVP01_0704900","PVP01_0705100","PVP01_0705200","PVP01_0705300","PVP01_0705400","PVP01_0705500","PVP01_0705900","PVP01_0706000","PVP01_0706300","PVP01_0706500","PVP01_0706600","PVP01_0706800","PVP01_0707100","PVP01_0707200","PVP01_0707300","PVP01_0707400","PVP01_0707500","PVP01_0707600","PVP01_0707700","PVP01_0708000","PVP01_0708100","PVP01_0708300","PVP01_0708500","PVP01_0708600","PVP01_0708800","PVP01_0709000","PVP01_0709400","PVP01_0709500","PVP01_0709700","PVP01_0709800","PVP01_0709900","PVP01_0710100","PVP01_0710300","PVP01_0710400","PVP01_0710500","PVP01_0710600","PVP01_0710700","PVP01_0710800","PVP01_0710900","PVP01_0711200","PVP01_0711300","PVP01_0711400","PVP01_0711600","PVP01_0711700","PVP01_0711900","PVP01_0712000","PVP01_0712100","PVP01_0712200","PVP01_0712500","PVP01_0712600","PVP01_0712700","PVP01_0712800","PVP01_0712900","PVP01_0713000","PVP01_0713200","PVP01_0713300","PVP01_0713400","PVP01_0713700","PVP01_0714000","PVP01_0714100","PVP01_0714300","PVP01_0714400","PVP01_0714500","PVP01_0714600","PVP01_0714700","PVP01_0714800","PVP01_0715000","PVP01_0715100","PVP01_0715300","PVP01_0715400","PVP01_0715500","PVP01_0715600","PVP01_0715900","PVP01_0716000","PVP01_0716300","PVP01_0716500","PVP01_0716600","PVP01_0716700","PVP01_0716900","PVP01_0717100","PVP01_0717300","PVP01_0717400","PVP01_0717700","PVP01_0717900","PVP01_0718000","PVP01_0718100","PVP01_0718200","PVP01_0718300","PVP01_0718400","PVP01_0718500","PVP01_0718800","PVP01_0718900","PVP01_0719200","PVP01_0719300","PVP01_0719400","PVP01_0719600","PVP01_0719800","PVP01_0720000","PVP01_0720200","PVP01_0720500","PVP01_0720700","PVP01_0720800","PVP01_0721000","PVP01_0721100","PVP01_0721200","PVP01_0721400","PVP01_0721500","PVP01_0721600","PVP01_0721700","PVP01_0722000","PVP01_0722100","PVP01_0722200","PVP01_0722300","PVP01_0722600","PVP01_0722700","PVP01_0722800","PVP01_0722900","PVP01_0723100","PVP01_0723300","PVP01_0723400","PVP01_0723500","PVP01_0723600","PVP01_0723700","PVP01_0723800","PVP01_0723900","PVP01_0724000","PVP01_0724100","PVP01_0724200","PVP01_0724500","PVP01_0724600","PVP01_0724800","PVP01_0724900","PVP01_0725000","PVP01_0725200","PVP01_0725500","PVP01_0725600","PVP01_0725700","PVP01_0725800","PVP01_0725900","PVP01_0726100","PVP01_0726200","PVP01_0726300","PVP01_0726500","PVP01_0726600","PVP01_0726700","PVP01_0726800","PVP01_0726900","PVP01_0727000","PVP01_0727200","PVP01_0727500","PVP01_0727600","PVP01_0727800","PVP01_0727900","PVP01_0728200","PVP01_0728300","PVP01_0728600","PVP01_0728700","PVP01_0728800","PVP01_0728900","PVP01_0729000","PVP01_0729200","PVP01_0729300","PVP01_0729600","PVP01_0729800","PVP01_0729900","PVP01_0730100","PVP01_0730200","PVP01_0730500","PVP01_0730900","PVP01_0731400","PVP01_0731600","PVP01_0731800","PVP01_0731900","PVP01_0732000","PVP01_0732100","PVP01_0732200","PVP01_0732400","PVP01_0732500","PVP01_0732700","PVP01_0733100","PVP01_0733200","PVP01_0733300","PVP01_0733400","PVP01_0733500","PVP01_0733600","PVP01_0733700","PVP01_0733800","PVP01_0733900","PVP01_0734000","PVP01_0734300","PVP01_0734400","PVP01_0734500","PVP01_0734600","PVP01_0734700","PVP01_0734900","PVP01_0735000","PVP01_0735200","PVP01_0735500","PVP01_0736200","PVP01_0800400","PVP01_0800600","PVP01_0800700","PVP01_0800900","PVP01_0801000","PVP01_0801100","PVP01_0801200","PVP01_0801300","PVP01_0801400","PVP01_0801500","PVP01_0801600","PVP01_0801700","PVP01_0801900","PVP01_0802100","PVP01_0802200","PVP01_0802300","PVP01_0802400","PVP01_0802500","PVP01_0802600","PVP01_0802700","PVP01_0802800","PVP01_0802900","PVP01_0803000","PVP01_0803100","PVP01_0803200","PVP01_0803300","PVP01_0803400","PVP01_0803600","PVP01_0803700","PVP01_0803800","PVP01_0803900","PVP01_0804000","PVP01_0804200","PVP01_0804400","PVP01_0804500","PVP01_0804700","PVP01_0805000","PVP01_0805100","PVP01_0805200","PVP01_0805700","PVP01_0805900","PVP01_0806000","PVP01_0806200","PVP01_0806400","PVP01_0806600","PVP01_0806800","PVP01_0806900","PVP01_0807100","PVP01_0807200","PVP01_0807300","PVP01_0807400","PVP01_0807600","PVP01_0807700","PVP01_0807900","PVP01_0808000","PVP01_0808200","PVP01_0808300","PVP01_0808400","PVP01_0808500","PVP01_0808600","PVP01_0808900","PVP01_0809000","PVP01_0809100","PVP01_0809300","PVP01_0809500","PVP01_0809800","PVP01_0810000","PVP01_0810100","PVP01_0810200","PVP01_0810300","PVP01_0810400","PVP01_0810600","PVP01_0810700","PVP01_0810800","PVP01_0810900","PVP01_0811000","PVP01_0811100","PVP01_0811400","PVP01_0811500","PVP01_0811600","PVP01_0811700","PVP01_0811800","PVP01_0811900","PVP01_0812000","PVP01_0812100","PVP01_0812200","PVP01_0812400","PVP01_0812500","PVP01_0812600","PVP01_0812700","PVP01_0812800","PVP01_0812900","PVP01_0813000","PVP01_0813100","PVP01_0813400","PVP01_0813500","PVP01_0813600","PVP01_0813900","PVP01_0814000","PVP01_0814300","PVP01_0814500","PVP01_0814600","PVP01_0814800","PVP01_0814900","PVP01_0815200","PVP01_0815400","PVP01_0815500","PVP01_0815600","PVP01_0815700","PVP01_0815800","PVP01_0816000","PVP01_0816100","PVP01_0816300","PVP01_0816400","PVP01_0816500","PVP01_0816600","PVP01_0816900","PVP01_0817200","PVP01_0817300","PVP01_0817500","PVP01_0817600","PVP01_0817700","PVP01_0817800","PVP01_0817900","PVP01_0818000","PVP01_0818200","PVP01_0818300","PVP01_0818400","PVP01_0818700","PVP01_0818800","PVP01_0819000","PVP01_0819100","PVP01_0819400","PVP01_0819500","PVP01_0819600","PVP01_0819700","PVP01_0819800","PVP01_0819900","PVP01_0820000","PVP01_0820100","PVP01_0820300","PVP01_0820400","PVP01_0820700","PVP01_0820900","PVP01_0821000","PVP01_0821100","PVP01_0821200","PVP01_0821500","PVP01_0821600","PVP01_0821800","PVP01_0821900","PVP01_0822000","PVP01_0822100","PVP01_0822200","PVP01_0822500","PVP01_0822700","PVP01_0822800","PVP01_0823000","PVP01_0823200","PVP01_0823300","PVP01_0823400","PVP01_0823500","PVP01_0823700","PVP01_0824200","PVP01_0824300","PVP01_0824400","PVP01_0824500","PVP01_0824600","PVP01_0824700","PVP01_0824900","PVP01_0825000","PVP01_0825200","PVP01_0825400","PVP01_0825500","PVP01_0825600","PVP01_0825700","PVP01_0825900","PVP01_0826100","PVP01_0826400","PVP01_0826500","PVP01_0826600","PVP01_0826800","PVP01_0826900","PVP01_0827000","PVP01_0827100","PVP01_0827200","PVP01_0827300","PVP01_0827400","PVP01_0827700","PVP01_0827900","PVP01_0828000","PVP01_0828400","PVP01_0828600","PVP01_0828900","PVP01_0829200","PVP01_0829300","PVP01_0829400","PVP01_0829500","PVP01_0829600","PVP01_0829700","PVP01_0829800","PVP01_0829900","PVP01_0830000","PVP01_0830100","PVP01_0830200","PVP01_0830400","PVP01_0830600","PVP01_0830700","PVP01_0830900","PVP01_0831200","PVP01_0831400","PVP01_0831500","PVP01_0831600","PVP01_0831800","PVP01_0831900","PVP01_0832400","PVP01_0832500","PVP01_0832600","PVP01_0832700","PVP01_0832800","PVP01_0833100","PVP01_0833200","PVP01_0833300","PVP01_0833400","PVP01_0833500","PVP01_0833600","PVP01_0833800","PVP01_0833900","PVP01_0834000","PVP01_0834100","PVP01_0834200","PVP01_0834500","PVP01_0834600","PVP01_0834800","PVP01_0834900","PVP01_0835200","PVP01_0835400","PVP01_0835500","PVP01_0835600","PVP01_0835700","PVP01_0835800","PVP01_0836000","PVP01_0836200","PVP01_0836300","PVP01_0836500","PVP01_0836600","PVP01_0836900","PVP01_0837000","PVP01_0837100","PVP01_0837400","PVP01_0837500","PVP01_0837700","PVP01_0837800","PVP01_0837900","PVP01_0838100","PVP01_0838200","PVP01_0838300","PVP01_0838500","PVP01_0838600","PVP01_0839300","PVP01_0839600","PVP01_0900600","PVP01_0900700","PVP01_0901400","PVP01_0901800","PVP01_0901900","PVP01_0902000","PVP01_0903000","PVP01_0903400","PVP01_0903600","PVP01_0903800","PVP01_0903900","PVP01_0904200","PVP01_0904300","PVP01_0904400","PVP01_0904500","PVP01_0904600","PVP01_0904800","PVP01_0905000","PVP01_0905100","PVP01_0905200","PVP01_0905300","PVP01_0905400","PVP01_0905500","PVP01_0905600","PVP01_0905700","PVP01_0905800","PVP01_0905900","PVP01_0906000","PVP01_0906100","PVP01_0906200","PVP01_0906300","PVP01_0906600","PVP01_0906700","PVP01_0907100","PVP01_0907300","PVP01_0907400","PVP01_0907600","PVP01_0907700","PVP01_0907800","PVP01_0907900","PVP01_0908100","PVP01_0908200","PVP01_0908300","PVP01_0908500","PVP01_0908800","PVP01_0909000","PVP01_0909200","PVP01_0909300","PVP01_0909400","PVP01_0910500","PVP01_0910700","PVP01_0910800","PVP01_0911000","PVP01_0911100","PVP01_0911200","PVP01_0911400","PVP01_0911500","PVP01_0912100","PVP01_0912500","PVP01_0912600","PVP01_0912700","PVP01_0912800","PVP01_0913100","PVP01_0913200","PVP01_0913300","PVP01_0913400","PVP01_0913500","PVP01_0913600","PVP01_0913700","PVP01_0913900","PVP01_0914000","PVP01_0914200","PVP01_0914300","PVP01_0914400","PVP01_0914500","PVP01_0914600","PVP01_0914700","PVP01_0914800","PVP01_0914900","PVP01_0915000","PVP01_0915100","PVP01_0915200","PVP01_0915300","PVP01_0915400","PVP01_0915500","PVP01_0915600","PVP01_0915700","PVP01_0915800","PVP01_0915900","PVP01_0916100","PVP01_0916200","PVP01_0916300","PVP01_0916400","PVP01_0916500","PVP01_0916600","PVP01_0916800","PVP01_0916900","PVP01_0917000","PVP01_0917300","PVP01_0917500","PVP01_0917700","PVP01_0917800","PVP01_0918400","PVP01_0918500","PVP01_0918600","PVP01_0918800","PVP01_0918900","PVP01_0919100","PVP01_0919200","PVP01_0919300","PVP01_0919600","PVP01_0919800","PVP01_0919900","PVP01_0920300","PVP01_0920600","PVP01_0920800","PVP01_0920900","PVP01_0921000","PVP01_0921100","PVP01_0921300","PVP01_0921400","PVP01_0921500","PVP01_0921600","PVP01_0921800","PVP01_0921900","PVP01_0922200","PVP01_0922300","PVP01_0922400","PVP01_0922500","PVP01_0922600","PVP01_0922700","PVP01_0923200","PVP01_0923300","PVP01_0923400","PVP01_0923500","PVP01_0923600","PVP01_0924200","PVP01_0924600","PVP01_0924700","PVP01_0924800","PVP01_0924900","PVP01_0925000","PVP01_0925100","PVP01_0925300","PVP01_0925400","PVP01_0925500","PVP01_0925600","PVP01_0925900","PVP01_0926000","PVP01_0926100","PVP01_0926200","PVP01_0926400","PVP01_0926500","PVP01_0926700","PVP01_0926800","PVP01_0927000","PVP01_0927500","PVP01_0927800","PVP01_0927900","PVP01_0928400","PVP01_0928500","PVP01_0928600","PVP01_0928700","PVP01_0928800","PVP01_0929100","PVP01_0929300","PVP01_0929400","PVP01_0929500","PVP01_0929800","PVP01_0930100","PVP01_0930200","PVP01_0930300","PVP01_0930400","PVP01_0930500","PVP01_0930700","PVP01_0931100","PVP01_0931200","PVP01_0931400","PVP01_0931500","PVP01_0931600","PVP01_0931700","PVP01_0931800","PVP01_0931900","PVP01_0932000","PVP01_0932300","PVP01_0932500","PVP01_0932700","PVP01_0932800","PVP01_0932900","PVP01_0933000","PVP01_0933200","PVP01_0933300","PVP01_0933600","PVP01_0933900","PVP01_0934000","PVP01_0934200","PVP01_0934600","PVP01_0934700","PVP01_0934800","PVP01_0934900","PVP01_0935000","PVP01_0935100","PVP01_0935300","PVP01_0935400","PVP01_0935500","PVP01_0935600","PVP01_0935800","PVP01_0936100","PVP01_0936200","PVP01_0936300","PVP01_0936600","PVP01_0936700","PVP01_0936800","PVP01_0937000","PVP01_0937200","PVP01_0937400","PVP01_0937500","PVP01_0937600","PVP01_0937800","PVP01_0937900","PVP01_0938100","PVP01_0938200","PVP01_0938300","PVP01_0938500","PVP01_0938800","PVP01_0939100","PVP01_0939300","PVP01_0939400","PVP01_0939500","PVP01_0939600","PVP01_0939700","PVP01_0939800","PVP01_0939900","PVP01_0940000","PVP01_0940100","PVP01_0940200","PVP01_0940300","PVP01_0940400","PVP01_0940500","PVP01_0940800","PVP01_0941000","PVP01_0941300","PVP01_0941500","PVP01_0941800","PVP01_0942000","PVP01_0942100","PVP01_0942300","PVP01_0942500","PVP01_0942600","PVP01_0942700","PVP01_0942800","PVP01_0942900","PVP01_0943000","PVP01_0943100","PVP01_0943200","PVP01_0943300","PVP01_0943400","PVP01_0943500","PVP01_0943600","PVP01_0943700","PVP01_0943800","PVP01_0943900","PVP01_0944000","PVP01_0944200","PVP01_0944300","PVP01_0944500","PVP01_0944700","PVP01_0944900","PVP01_0945100","PVP01_0945300","PVP01_0945600","PVP01_0945700","PVP01_0945900","PVP01_0946000","PVP01_0946100","PVP01_0946500","PVP01_0946600","PVP01_0946800","PVP01_0946900","PVP01_0947000","PVP01_0947100","PVP01_0947400","PVP01_0947500","PVP01_0947600","PVP01_0947700","PVP01_0947800","PVP01_0947900","PVP01_0948000","PVP01_0948100","PVP01_0948200","PVP01_0948300","PVP01_0948400","PVP01_0948600","PVP01_0948700","PVP01_0948800","PVP01_0948900","PVP01_0949000","PVP01_0949100","PVP01_0949200","PVP01_0949400","PVP01_0949500","PVP01_0949600","PVP01_0949700","PVP01_0949800","PVP01_0950000","PVP01_0950200","PVP01_1002400","PVP01_1002500","PVP01_1002700","PVP01_1002800","PVP01_1002900","PVP01_1003000","PVP01_1003100","PVP01_1003200","PVP01_1003300","PVP01_1003400","PVP01_1003500","PVP01_1003600","PVP01_1003700","PVP01_1003800","PVP01_1003900","PVP01_1004000","PVP01_1004100","PVP01_1004400","PVP01_1004500","PVP01_1004700","PVP01_1004800","PVP01_1004900","PVP01_1005000","PVP01_1005100","PVP01_1005200","PVP01_1005400","PVP01_1005500","PVP01_1005600","PVP01_1005700","PVP01_1005800","PVP01_1006200","PVP01_1006300","PVP01_1006500","PVP01_1006700","PVP01_1006800","PVP01_1007000","PVP01_1007200","PVP01_1007300","PVP01_1007400","PVP01_1008000","PVP01_1008200","PVP01_1008500","PVP01_1008700","PVP01_1008800","PVP01_1009100","PVP01_1009300","PVP01_1009400","PVP01_1009500","PVP01_1009700","PVP01_1009800","PVP01_1009900","PVP01_1010000","PVP01_1010300","PVP01_1010500","PVP01_1010600","PVP01_1010700","PVP01_1010800","PVP01_1010900","PVP01_1011000","PVP01_1011600","PVP01_1011700","PVP01_1011900","PVP01_1012000","PVP01_1012200","PVP01_1012300","PVP01_1012600","PVP01_1012800","PVP01_1012900","PVP01_1013000","PVP01_1013100","PVP01_1013300","PVP01_1013400","PVP01_1013500","PVP01_1013600","PVP01_1013700","PVP01_1013800","PVP01_1013900","PVP01_1014000","PVP01_1014100","PVP01_1014200","PVP01_1014500","PVP01_1014700","PVP01_1014800","PVP01_1014900","PVP01_1015100","PVP01_1015200","PVP01_1015300","PVP01_1015400","PVP01_1015800","PVP01_1015900","PVP01_1016000","PVP01_1016300","PVP01_1016400","PVP01_1016500","PVP01_1016700","PVP01_1016800","PVP01_1017000","PVP01_1017100","PVP01_1017200","PVP01_1017300","PVP01_1017600","PVP01_1017700","PVP01_1017800","PVP01_1018000","PVP01_1018100","PVP01_1018200","PVP01_1018300","PVP01_1018500","PVP01_1018600","PVP01_1018700","PVP01_1018800","PVP01_1018900","PVP01_1019100","PVP01_1019200","PVP01_1019300","PVP01_1019400","PVP01_1019700","PVP01_1019800","PVP01_1019900","PVP01_1020200","PVP01_1020300","PVP01_1020400","PVP01_1020500","PVP01_1020600","PVP01_1020800","PVP01_1021100","PVP01_1021400","PVP01_1021700","PVP01_1022100","PVP01_1022300","PVP01_1022600","PVP01_1022700","PVP01_1022900","PVP01_1023100","PVP01_1023200","PVP01_1023300","PVP01_1023500","PVP01_1024000","PVP01_1024100","PVP01_1024200","PVP01_1024300","PVP01_1024700","PVP01_1025000","PVP01_1025300","PVP01_1025400","PVP01_1025600","PVP01_1025700","PVP01_1025800","PVP01_1025900","PVP01_1026000","PVP01_1026100","PVP01_1026500","PVP01_1026800","PVP01_1026900","PVP01_1027000","PVP01_1027200","PVP01_1027400","PVP01_1027500","PVP01_1027700","PVP01_1027900","PVP01_1028000","PVP01_1028300","PVP01_1028400","PVP01_1028500","PVP01_1028700","PVP01_1028800","PVP01_1029200","PVP01_1029300","PVP01_1029400","PVP01_1029500","PVP01_1029600","PVP01_1029800","PVP01_1029900","PVP01_1030000","PVP01_1030100","PVP01_1030200","PVP01_1030300","PVP01_1030400","PVP01_1030700","PVP01_1030800","PVP01_1030900","PVP01_1031200","PVP01_1031600","PVP01_1031700","PVP01_1032000","PVP01_1032100","PVP01_1032200","PVP01_1032300","PVP01_1032400","PVP01_1032500","PVP01_1032600","PVP01_1032700","PVP01_1032800","PVP01_1033100","PVP01_1033200","PVP01_1033300","PVP01_1033400","PVP01_1033500","PVP01_1033600","PVP01_1033700","PVP01_1033800","PVP01_1033900","PVP01_1034000","PVP01_1034100","PVP01_1034200","PVP01_1034300","PVP01_1034400","PVP01_1034500","PVP01_1100300","PVP01_1100400","PVP01_1100500","PVP01_1101300","PVP01_1101400","PVP01_1101500","PVP01_1101600","PVP01_1101700","PVP01_1102300","PVP01_1102600","PVP01_1102800","PVP01_1102900","PVP01_1103100","PVP01_1103500","PVP01_1103600","PVP01_1103800","PVP01_1103900","PVP01_1104100","PVP01_1104200","PVP01_1104500","PVP01_1104700","PVP01_1104900","PVP01_1105000","PVP01_1105100","PVP01_1105200","PVP01_1105300","PVP01_1105400","PVP01_1105600","PVP01_1105800","PVP01_1105900","PVP01_1106100","PVP01_1106200","PVP01_1106300","PVP01_1106400","PVP01_1106500","PVP01_1106600","PVP01_1106700","PVP01_1107200","PVP01_1107300","PVP01_1107500","PVP01_1107700","PVP01_1107800","PVP01_1108000","PVP01_1108100","PVP01_1108200","PVP01_1108300","PVP01_1108400","PVP01_1108700","PVP01_1109000","PVP01_1109100","PVP01_1109200","PVP01_1109300","PVP01_1109700","PVP01_1109800","PVP01_1109900","PVP01_1110000","PVP01_1110100","PVP01_1110200","PVP01_1110300","PVP01_1110400","PVP01_1110700","PVP01_1110800","PVP01_1110900","PVP01_1111200","PVP01_1111400","PVP01_1111500","PVP01_1111600","PVP01_1111700","PVP01_1111800","PVP01_1111900","PVP01_1112000","PVP01_1112200","PVP01_1112300","PVP01_1112400","PVP01_1112500","PVP01_1112600","PVP01_1112700","PVP01_1112800","PVP01_1113000","PVP01_1113100","PVP01_1113200","PVP01_1113300","PVP01_1113400","PVP01_1113800","PVP01_1114000","PVP01_1114100","PVP01_1114400","PVP01_1114600","PVP01_1114700","PVP01_1114800","PVP01_1114900","PVP01_1115000","PVP01_1115100","PVP01_1115200","PVP01_1115300","PVP01_1115400","PVP01_1115700","PVP01_1115800","PVP01_1116000","PVP01_1116100","PVP01_1116200","PVP01_1116300","PVP01_1116400","PVP01_1116700","PVP01_1116800","PVP01_1116900","PVP01_1117000","PVP01_1117100","PVP01_1117200","PVP01_1117300","PVP01_1117400","PVP01_1117500","PVP01_1117600","PVP01_1117800","PVP01_1117900","PVP01_1118100","PVP01_1118200","PVP01_1118300","PVP01_1118400","PVP01_1118900","PVP01_1119100","PVP01_1119200","PVP01_1119300","PVP01_1119400","PVP01_1119700","PVP01_1119900","PVP01_1120100","PVP01_1120200","PVP01_1120400","PVP01_1120600","PVP01_1120700","PVP01_1120800","PVP01_1120900","PVP01_1121000","PVP01_1121200","PVP01_1121300","PVP01_1121600","PVP01_1121800","PVP01_1122000","PVP01_1122200","PVP01_1122800","PVP01_1122900","PVP01_1123100","PVP01_1123200","PVP01_1123300","PVP01_1123400","PVP01_1123700","PVP01_1123800","PVP01_1124200","PVP01_1124500","PVP01_1124600","PVP01_1124700","PVP01_1125200","PVP01_1125300","PVP01_1125500","PVP01_1125600","PVP01_1125700","PVP01_1125800","PVP01_1126000","PVP01_1126100","PVP01_1126300","PVP01_1126400","PVP01_1126500","PVP01_1126600","PVP01_1126700","PVP01_1126800","PVP01_1126900","PVP01_1127000","PVP01_1127500","PVP01_1127600","PVP01_1127700","PVP01_1128000","PVP01_1128100","PVP01_1128200","PVP01_1128300","PVP01_1128500","PVP01_1128800","PVP01_1128900","PVP01_1129100","PVP01_1129200","PVP01_1129300","PVP01_1129500","PVP01_1129600","PVP01_1129800","PVP01_1129900","PVP01_1130000","PVP01_1130100","PVP01_1130300","PVP01_1130400","PVP01_1130500","PVP01_1130600","PVP01_1130700","PVP01_1130800","PVP01_1131100","PVP01_1131200","PVP01_1131300","PVP01_1131400","PVP01_1131500","PVP01_1131600","PVP01_1131700","PVP01_1131800","PVP01_1131900","PVP01_1132000","PVP01_1132100","PVP01_1132200","PVP01_1132500","PVP01_1132600","PVP01_1133000","PVP01_1133300","PVP01_1133700","PVP01_1134100","PVP01_1134200","PVP01_1134400","PVP01_1134600","PVP01_1134700","PVP01_1134800","PVP01_1134900","PVP01_1135000","PVP01_1135100","PVP01_1135200","PVP01_1135300","PVP01_1135400","PVP01_1135600","PVP01_1135700","PVP01_1136000","PVP01_1136200","PVP01_1136300","PVP01_1136400","PVP01_1136500","PVP01_1136600","PVP01_1137000","PVP01_1137300","PVP01_1137400","PVP01_1137500","PVP01_1137900","PVP01_1138000","PVP01_1138100","PVP01_1138200","PVP01_1138300","PVP01_1138600","PVP01_1138700","PVP01_1138800","PVP01_1138900","PVP01_1139000","PVP01_1139100","PVP01_1139200","PVP01_1139300","PVP01_1139400","PVP01_1139500","PVP01_1139600","PVP01_1139700","PVP01_1139800","PVP01_1140000","PVP01_1140100","PVP01_1140200","PVP01_1140400","PVP01_1140600","PVP01_1140800","PVP01_1140900","PVP01_1141000","PVP01_1141100","PVP01_1141200","PVP01_1141400","PVP01_1141600","PVP01_1141700","PVP01_1141800","PVP01_1142100","PVP01_1142200","PVP01_1142300","PVP01_1142400","PVP01_1142500","PVP01_1142800","PVP01_1142900","PVP01_1143000","PVP01_1143100","PVP01_1143300","PVP01_1143600","PVP01_1143700","PVP01_1143800","PVP01_1143900","PVP01_1144000","PVP01_1144200","PVP01_1144400","PVP01_1144600","PVP01_1144700","PVP01_1144900","PVP01_1145000","PVP01_1145100","PVP01_1145200","PVP01_1145300","PVP01_1145400","PVP01_1145600","PVP01_1145800","PVP01_1145900","PVP01_1146000","PVP01_1146100","PVP01_1146200","PVP01_1146300","PVP01_1146600","PVP01_1146700","PVP01_1146800","PVP01_1147000","PVP01_1147100","PVP01_1147200","PVP01_1147300","PVP01_1147400","PVP01_1147500","PVP01_1147600","PVP01_1147700","PVP01_1147800","PVP01_1148400","PVP01_1148600","PVP01_1149100","PVP01_1149200","PVP01_1149500","PVP01_1200300","PVP01_1201000","PVP01_1201300","PVP01_1201400","PVP01_1201500","PVP01_1201600","PVP01_1201700","PVP01_1201800","PVP01_1202000","PVP01_1202300","PVP01_1202400","PVP01_1202500","PVP01_1202600","PVP01_1202700","PVP01_1202800","PVP01_1202900","PVP01_1203000","PVP01_1203200","PVP01_1203400","PVP01_1203700","PVP01_1203800","PVP01_1203900","PVP01_1204300","PVP01_1204400","PVP01_1204500","PVP01_1204700","PVP01_1204800","PVP01_1204900","PVP01_1205100","PVP01_1205200","PVP01_1205300","PVP01_1205400","PVP01_1205600","PVP01_1205700","PVP01_1205800","PVP01_1205900","PVP01_1206000","PVP01_1206200","PVP01_1206400","PVP01_1206500","PVP01_1206600","PVP01_1206800","PVP01_1207100","PVP01_1207200","PVP01_1207300","PVP01_1207400","PVP01_1207500","PVP01_1207600","PVP01_1207700","PVP01_1207800","PVP01_1207900","PVP01_1208100","PVP01_1208300","PVP01_1208500","PVP01_1208800","PVP01_1208900","PVP01_1209100","PVP01_1209300","PVP01_1209400","PVP01_1209500","PVP01_1209600","PVP01_1209700","PVP01_1209800","PVP01_1210400","PVP01_1210500","PVP01_1210600","PVP01_1210900","PVP01_1211000","PVP01_1211400","PVP01_1211500","PVP01_1211700","PVP01_1211800","PVP01_1211900","PVP01_1212000","PVP01_1212200","PVP01_1212300","PVP01_1212400","PVP01_1212500","PVP01_1212600","PVP01_1212700","PVP01_1212800","PVP01_1212900","PVP01_1213000","PVP01_1213100","PVP01_1213200","PVP01_1213300","PVP01_1213600","PVP01_1213900","PVP01_1214100","PVP01_1214500","PVP01_1214800","PVP01_1214900","PVP01_1215000","PVP01_1215100","PVP01_1215200","PVP01_1215300","PVP01_1215400","PVP01_1215500","PVP01_1215800","PVP01_1215900","PVP01_1216000","PVP01_1216300","PVP01_1216400","PVP01_1216500","PVP01_1216600","PVP01_1216700","PVP01_1216900","PVP01_1217000","PVP01_1217300","PVP01_1217400","PVP01_1217800","PVP01_1217900","PVP01_1218000","PVP01_1218100","PVP01_1218200","PVP01_1218300","PVP01_1218400","PVP01_1218500","PVP01_1218600","PVP01_1218700","PVP01_1219200","PVP01_1219300","PVP01_1219500","PVP01_1219800","PVP01_1220000","PVP01_1220100","PVP01_1220400","PVP01_1220500","PVP01_1220600","PVP01_1220700","PVP01_1220800","PVP01_1220900","PVP01_1221000","PVP01_1221100","PVP01_1221200","PVP01_1221400","PVP01_1221500","PVP01_1221600","PVP01_1221700","PVP01_1222000","PVP01_1222100","PVP01_1222200","PVP01_1222300","PVP01_1222500","PVP01_1222600","PVP01_1222700","PVP01_1222800","PVP01_1222900","PVP01_1223000","PVP01_1223100","PVP01_1223500","PVP01_1223600","PVP01_1223700","PVP01_1224100","PVP01_1224200","PVP01_1224300","PVP01_1224700","PVP01_1224800","PVP01_1224900","PVP01_1225000","PVP01_1225100","PVP01_1225200","PVP01_1225300","PVP01_1225400","PVP01_1225600","PVP01_1225700","PVP01_1225800","PVP01_1226000","PVP01_1226200","PVP01_1226300","PVP01_1226400","PVP01_1226700","PVP01_1226900","PVP01_1227000","PVP01_1227100","PVP01_1227200","PVP01_1227300","PVP01_1227400","PVP01_1227700","PVP01_1227800","PVP01_1228500","PVP01_1228600","PVP01_1228700","PVP01_1228800","PVP01_1228900","PVP01_1229000","PVP01_1229100","PVP01_1229300","PVP01_1229400","PVP01_1229700","PVP01_1229800","PVP01_1230300","PVP01_1230400","PVP01_1230700","PVP01_1230800","PVP01_1231200","PVP01_1231300","PVP01_1231400","PVP01_1231500","PVP01_1231600","PVP01_1231700","PVP01_1232000","PVP01_1232100","PVP01_1232200","PVP01_1232300","PVP01_1232400","PVP01_1232500","PVP01_1232800","PVP01_1233000","PVP01_1233200","PVP01_1233300","PVP01_1233400","PVP01_1233500","PVP01_1233600","PVP01_1233700","PVP01_1233800","PVP01_1233900","PVP01_1234000","PVP01_1234100","PVP01_1234200","PVP01_1234300","PVP01_1234400","PVP01_1234500","PVP01_1234600","PVP01_1234700","PVP01_1234800","PVP01_1234900","PVP01_1235000","PVP01_1235100","PVP01_1235200","PVP01_1235300","PVP01_1235400","PVP01_1235600","PVP01_1235700","PVP01_1235800","PVP01_1235900","PVP01_1236200","PVP01_1236300","PVP01_1236400","PVP01_1236500","PVP01_1236600","PVP01_1236700","PVP01_1236900","PVP01_1237000","PVP01_1237300","PVP01_1237400","PVP01_1237500","PVP01_1237600","PVP01_1237800","PVP01_1238000","PVP01_1238100","PVP01_1238200","PVP01_1238500","PVP01_1238600","PVP01_1238700","PVP01_1238800","PVP01_1239000","PVP01_1239100","PVP01_1239200","PVP01_1239400","PVP01_1239600","PVP01_1239700","PVP01_1239800","PVP01_1239900","PVP01_1240200","PVP01_1240300","PVP01_1240400","PVP01_1240500","PVP01_1240600","PVP01_1240700","PVP01_1240800","PVP01_1240900","PVP01_1241000","PVP01_1241100","PVP01_1241200","PVP01_1241400","PVP01_1241500","PVP01_1241600","PVP01_1241700","PVP01_1241800","PVP01_1242000","PVP01_1242100","PVP01_1242200","PVP01_1242400","PVP01_1242500","PVP01_1242600","PVP01_1242700","PVP01_1242800","PVP01_1243000","PVP01_1243100","PVP01_1243200","PVP01_1243300","PVP01_1243400","PVP01_1243700","PVP01_1243800","PVP01_1244000","PVP01_1244100","PVP01_1244200","PVP01_1244300","PVP01_1244400","PVP01_1244500","PVP01_1244600","PVP01_1245000","PVP01_1245100","PVP01_1245200","PVP01_1245300","PVP01_1245400","PVP01_1245700","PVP01_1245900","PVP01_1246000","PVP01_1246100","PVP01_1246300","PVP01_1246400","PVP01_1246600","PVP01_1246700","PVP01_1246800","PVP01_1246900","PVP01_1247000","PVP01_1247100","PVP01_1247200","PVP01_1247400","PVP01_1247500","PVP01_1247600","PVP01_1247700","PVP01_1248000","PVP01_1248100","PVP01_1248200","PVP01_1248300","PVP01_1248400","PVP01_1248500","PVP01_1248600","PVP01_1249100","PVP01_1249300","PVP01_1249400","PVP01_1249600","PVP01_1249700","PVP01_1249800","PVP01_1249900","PVP01_1250000","PVP01_1250100","PVP01_1250300","PVP01_1250400","PVP01_1250500","PVP01_1250600","PVP01_1250800","PVP01_1251000","PVP01_1251100","PVP01_1251200","PVP01_1251300","PVP01_1251400","PVP01_1251500","PVP01_1251600","PVP01_1251700","PVP01_1251800","PVP01_1251900","PVP01_1252000","PVP01_1252100","PVP01_1252200","PVP01_1252300","PVP01_1252500","PVP01_1252700","PVP01_1252800","PVP01_1253100","PVP01_1253200","PVP01_1253300","PVP01_1253400","PVP01_1253500","PVP01_1253600","PVP01_1253800","PVP01_1253900","PVP01_1254000","PVP01_1254100","PVP01_1254300","PVP01_1254400","PVP01_1254500","PVP01_1254600","PVP01_1254700","PVP01_1254800","PVP01_1255000","PVP01_1255300","PVP01_1255400","PVP01_1255500","PVP01_1255600","PVP01_1255800","PVP01_1255900","PVP01_1256000","PVP01_1256100","PVP01_1256200","PVP01_1256300","PVP01_1256500","PVP01_1256700","PVP01_1257100","PVP01_1257200","PVP01_1257300","PVP01_1257400","PVP01_1257500","PVP01_1257700","PVP01_1257900","PVP01_1258000","PVP01_1258200","PVP01_1258400","PVP01_1258500","PVP01_1258600","PVP01_1258700","PVP01_1258800","PVP01_1259100","PVP01_1259400","PVP01_1259600","PVP01_1259700","PVP01_1259800","PVP01_1259900","PVP01_1260000","PVP01_1260100","PVP01_1260300","PVP01_1260400","PVP01_1260700","PVP01_1260900","PVP01_1261000","PVP01_1261100","PVP01_1261200","PVP01_1261300","PVP01_1261400","PVP01_1261500","PVP01_1261600","PVP01_1261700","PVP01_1261800","PVP01_1261900","PVP01_1262100","PVP01_1262200","PVP01_1262300","PVP01_1262400","PVP01_1262500","PVP01_1262600","PVP01_1262700","PVP01_1263000","PVP01_1263100","PVP01_1263200","PVP01_1263500","PVP01_1263600","PVP01_1263700","PVP01_1263800","PVP01_1263900","PVP01_1264100","PVP01_1264200","PVP01_1264400","PVP01_1264900","PVP01_1265100","PVP01_1265200","PVP01_1265300","PVP01_1265400","PVP01_1265500","PVP01_1265600","PVP01_1265800","PVP01_1265900","PVP01_1266000","PVP01_1266100","PVP01_1266200","PVP01_1266300","PVP01_1266500","PVP01_1266600","PVP01_1266700","PVP01_1266800","PVP01_1266900","PVP01_1267000","PVP01_1267500","PVP01_1267700","PVP01_1267800","PVP01_1267900","PVP01_1268100","PVP01_1268300","PVP01_1268400","PVP01_1268500","PVP01_1268900","PVP01_1269200","PVP01_1269300","PVP01_1269400","PVP01_1269500","PVP01_1269600","PVP01_1269700","PVP01_1269800","PVP01_1269900","PVP01_1270000","PVP01_1270100","PVP01_1270300","PVP01_1270400","PVP01_1270500","PVP01_1270700","PVP01_1270800","PVP01_1270900","PVP01_1271000","PVP01_1271200","PVP01_1271300","PVP01_1271400","PVP01_1271500","PVP01_1271600","PVP01_1271700","PVP01_1271800","PVP01_1271900","PVP01_1272100","PVP01_1272200","PVP01_1272300","PVP01_1272400","PVP01_1272500","PVP01_1272600","PVP01_1273100","PVP01_1300500","PVP01_1300600","PVP01_1300700","PVP01_1300800","PVP01_1300900","PVP01_1301000","PVP01_1301100","PVP01_1301300","PVP01_1301400","PVP01_1301800","PVP01_1301900","PVP01_1302000","PVP01_1302100","PVP01_1302200","PVP01_1302300","PVP01_1302400","PVP01_1302500","PVP01_1302600","PVP01_1302900","PVP01_1303000","PVP01_1303200","PVP01_1303300","PVP01_1303400","PVP01_1303500","PVP01_1303600","PVP01_1303700","PVP01_1303800","PVP01_1303900","PVP01_1304100","PVP01_1304300","PVP01_1304400","PVP01_1304500","PVP01_1304700","PVP01_1305000","PVP01_1305100","PVP01_1305200","PVP01_1305400","PVP01_1305500","PVP01_1305800","PVP01_1306200","PVP01_1306500","PVP01_1306600","PVP01_1306800","PVP01_1306900","PVP01_1307100","PVP01_1307400","PVP01_1307600","PVP01_1307700","PVP01_1307900","PVP01_1308000","PVP01_1308200","PVP01_1308400","PVP01_1308600","PVP01_1308700","PVP01_1308800","PVP01_1309200","PVP01_1309300","PVP01_1309400","PVP01_1309500","PVP01_1309600","PVP01_1309800","PVP01_1309900","PVP01_1310000","PVP01_1310100","PVP01_1310300","PVP01_1310400","PVP01_1310500","PVP01_1310600","PVP01_1310700","PVP01_1310800","PVP01_1311200","PVP01_1311600","PVP01_1311700","PVP01_1311800","PVP01_1311900","PVP01_1312000","PVP01_1312100","PVP01_1312500","PVP01_1312600","PVP01_1312700","PVP01_1312800","PVP01_1313000","PVP01_1313100","PVP01_1313200","PVP01_1313300","PVP01_1313500","PVP01_1313600","PVP01_1313700","PVP01_1313800","PVP01_1314000","PVP01_1314200","PVP01_1314400","PVP01_1314500","PVP01_1314700","PVP01_1314800","PVP01_1315100","PVP01_1315200","PVP01_1315300","PVP01_1315500","PVP01_1315600","PVP01_1315800","PVP01_1315900","PVP01_1316000","PVP01_1316200","PVP01_1316300","PVP01_1316400","PVP01_1316700","PVP01_1316800","PVP01_1317100","PVP01_1317300","PVP01_1317500","PVP01_1317700","PVP01_1318200","PVP01_1318300","PVP01_1318400","PVP01_1318600","PVP01_1318700","PVP01_1319000","PVP01_1319200","PVP01_1319300","PVP01_1319400","PVP01_1319500","PVP01_1319600","PVP01_1319800","PVP01_1319900","PVP01_1320000","PVP01_1320100","PVP01_1320200","PVP01_1320300","PVP01_1320400","PVP01_1320500","PVP01_1320600","PVP01_1320800","PVP01_1321000","PVP01_1321100","PVP01_1321200","PVP01_1321400","PVP01_1321500","PVP01_1321700","PVP01_1321900","PVP01_1322000","PVP01_1322100","PVP01_1322200","PVP01_1322500","PVP01_1322800","PVP01_1322900","PVP01_1323200","PVP01_1323300","PVP01_1323600","PVP01_1323800","PVP01_1323900","PVP01_1324000","PVP01_1324600","PVP01_1324900","PVP01_1325300","PVP01_1325400","PVP01_1325500","PVP01_1325600","PVP01_1325700","PVP01_1326100","PVP01_1326300","PVP01_1326500","PVP01_1326600","PVP01_1326700","PVP01_1326800","PVP01_1326900","PVP01_1327000","PVP01_1327100","PVP01_1327200","PVP01_1327300","PVP01_1327400","PVP01_1327500","PVP01_1327700","PVP01_1327800","PVP01_1328200","PVP01_1328300","PVP01_1328500","PVP01_1328600","PVP01_1328700","PVP01_1328900","PVP01_1329000","PVP01_1329100","PVP01_1329200","PVP01_1329500","PVP01_1329600","PVP01_1329700","PVP01_1329800","PVP01_1329900","PVP01_1330000","PVP01_1330100","PVP01_1330200","PVP01_1330300","PVP01_1330400","PVP01_1330800","PVP01_1330900","PVP01_1331100","PVP01_1331200","PVP01_1331300","PVP01_1331500","PVP01_1331600","PVP01_1331700","PVP01_1331900","PVP01_1332000","PVP01_1332100","PVP01_1332200","PVP01_1332300","PVP01_1332400","PVP01_1332500","PVP01_1332600","PVP01_1332800","PVP01_1333000","PVP01_1333400","PVP01_1333500","PVP01_1333800","PVP01_1334400","PVP01_1334600","PVP01_1334700","PVP01_1334800","PVP01_1334900","PVP01_1335000","PVP01_1335100","PVP01_1335200","PVP01_1335300","PVP01_1335400","PVP01_1335500","PVP01_1335900","PVP01_1336100","PVP01_1336200","PVP01_1336300","PVP01_1336400","PVP01_1336500","PVP01_1336600","PVP01_1336700","PVP01_1336800","PVP01_1336900","PVP01_1337000","PVP01_1337100","PVP01_1337200","PVP01_1337300","PVP01_1337600","PVP01_1337700","PVP01_1337800","PVP01_1338000","PVP01_1338200","PVP01_1338700","PVP01_1338800","PVP01_1338900","PVP01_1339200","PVP01_1339300","PVP01_1339400","PVP01_1339500","PVP01_1339700","PVP01_1339800","PVP01_1339900","PVP01_1340000","PVP01_1340100","PVP01_1340200","PVP01_1340300","PVP01_1340400","PVP01_1340700","PVP01_1340800","PVP01_1340900","PVP01_1341000","PVP01_1341100","PVP01_1341200","PVP01_1341300","PVP01_1341400","PVP01_1341500","PVP01_1341600","PVP01_1341700","PVP01_1341800","PVP01_1341900","PVP01_1342000","PVP01_1342100","PVP01_1342400","PVP01_1342500","PVP01_1342700","PVP01_1342900","PVP01_1343000","PVP01_1343300","PVP01_1343400","PVP01_1343600","PVP01_1343800","PVP01_1343900","PVP01_1344000","PVP01_1344100","PVP01_1344300","PVP01_1344400","PVP01_1344500","PVP01_1344700","PVP01_1345000","PVP01_1345100","PVP01_1345200","PVP01_1345300","PVP01_1345400","PVP01_1345500","PVP01_1345600","PVP01_1345800","PVP01_1345900","PVP01_1346000","PVP01_1346100","PVP01_1346200","PVP01_1346400","PVP01_1346500","PVP01_1346700","PVP01_1346800","PVP01_1347000","PVP01_1347100","PVP01_1347200","PVP01_1347400","PVP01_1347700","PVP01_1400300","PVP01_1400400","PVP01_1400700","PVP01_1400900","PVP01_1401000","PVP01_1401100","PVP01_1401300","PVP01_1401400","PVP01_1401600","PVP01_1401700","PVP01_1401800","PVP01_1401900","PVP01_1402000","PVP01_1402100","PVP01_1402200","PVP01_1402300","PVP01_1402400","PVP01_1402500","PVP01_1402600","PVP01_1402700","PVP01_1402800","PVP01_1403100","PVP01_1403200","PVP01_1403300","PVP01_1403500","PVP01_1403600","PVP01_1404000","PVP01_1404100","PVP01_1404200","PVP01_1404300","PVP01_1404400","PVP01_1404700","PVP01_1404800","PVP01_1404900","PVP01_1405100","PVP01_1405200","PVP01_1405300","PVP01_1405400","PVP01_1405500","PVP01_1405700","PVP01_1405800","PVP01_1405900","PVP01_1406100","PVP01_1406200","PVP01_1406300","PVP01_1406400","PVP01_1406700","PVP01_1406800","PVP01_1406900","PVP01_1407200","PVP01_1407300","PVP01_1407400","PVP01_1407500","PVP01_1407700","PVP01_1408000","PVP01_1408300","PVP01_1408400","PVP01_1409100","PVP01_1409200","PVP01_1409500","PVP01_1409600","PVP01_1409700","PVP01_1409800","PVP01_1409900","PVP01_1410000","PVP01_1410100","PVP01_1410600","PVP01_1410900","PVP01_1411600","PVP01_1411700","PVP01_1411800","PVP01_1411900","PVP01_1412000","PVP01_1412100","PVP01_1412200","PVP01_1412300","PVP01_1412400","PVP01_1412500","PVP01_1412600","PVP01_1412800","PVP01_1412900","PVP01_1413000","PVP01_1413300","PVP01_1413400","PVP01_1413450","PVP01_1413500","PVP01_1413600","PVP01_1413800","PVP01_1414000","PVP01_1414100","PVP01_1414200","PVP01_1414400","PVP01_1414600","PVP01_1414700","PVP01_1414800","PVP01_1414900","PVP01_1415000","PVP01_1415200","PVP01_1415300","PVP01_1415400","PVP01_1415700","PVP01_1415800","PVP01_1416300","PVP01_1416400","PVP01_1416500","PVP01_1416600","PVP01_1416700","PVP01_1416800","PVP01_1416900","PVP01_1417000","PVP01_1417300","PVP01_1417400","PVP01_1417500","PVP01_1417700","PVP01_1417800","PVP01_1418100","PVP01_1418200","PVP01_1418300","PVP01_1418400","PVP01_1418500","PVP01_1418700","PVP01_1418900","PVP01_1419200","PVP01_1419500","PVP01_1419600","PVP01_1419700","PVP01_1419900","PVP01_1420000","PVP01_1420200","PVP01_1420300","PVP01_1420400","PVP01_1420600","PVP01_1420900","PVP01_1421100","PVP01_1421200","PVP01_1421400","PVP01_1421600","PVP01_1421700","PVP01_1421800","PVP01_1421900","PVP01_1422000","PVP01_1422100","PVP01_1422200","PVP01_1422400","PVP01_1422500","PVP01_1422700","PVP01_1422800","PVP01_1423000","PVP01_1423100","PVP01_1423300","PVP01_1423400","PVP01_1423500","PVP01_1423600","PVP01_1424100","PVP01_1424300","PVP01_1424400","PVP01_1424500","PVP01_1424600","PVP01_1424700","PVP01_1424900","PVP01_1425000","PVP01_1425200","PVP01_1425300","PVP01_1425400","PVP01_1425500","PVP01_1425600","PVP01_1425700","PVP01_1425800","PVP01_1425900","PVP01_1426000","PVP01_1426300","PVP01_1426400","PVP01_1426500","PVP01_1426600","PVP01_1426900","PVP01_1427000","PVP01_1427200","PVP01_1427300","PVP01_1427400","PVP01_1427500","PVP01_1427600","PVP01_1427700","PVP01_1427800","PVP01_1427900","PVP01_1428000","PVP01_1428100","PVP01_1428200","PVP01_1428400","PVP01_1428500","PVP01_1428700","PVP01_1428800","PVP01_1428900","PVP01_1429100","PVP01_1429400","PVP01_1429500","PVP01_1429600","PVP01_1429700","PVP01_1429800","PVP01_1430200","PVP01_1430300","PVP01_1430400","PVP01_1430600","PVP01_1430900","PVP01_1431000","PVP01_1431400","PVP01_1431500","PVP01_1431700","PVP01_1431800","PVP01_1431900","PVP01_1432100","PVP01_1432200","PVP01_1432300","PVP01_1432700","PVP01_1432800","PVP01_1432900","PVP01_1433100","PVP01_1433200","PVP01_1433300","PVP01_1433400","PVP01_1433500","PVP01_1433600","PVP01_1433900","PVP01_1434100","PVP01_1434400","PVP01_1434500","PVP01_1434600","PVP01_1434700","PVP01_1434900","PVP01_1435000","PVP01_1435100","PVP01_1435200","PVP01_1435300","PVP01_1435400","PVP01_1435500","PVP01_1435700","PVP01_1435800","PVP01_1435900","PVP01_1436100","PVP01_1436200","PVP01_1436300","PVP01_1436400","PVP01_1436500","PVP01_1436600","PVP01_1436700","PVP01_1436800","PVP01_1436900","PVP01_1437000","PVP01_1437100","PVP01_1437200","PVP01_1437300","PVP01_1437400","PVP01_1437600","PVP01_1437800","PVP01_1438000","PVP01_1438200","PVP01_1438300","PVP01_1438400","PVP01_1438500","PVP01_1438600","PVP01_1438700","PVP01_1438800","PVP01_1438900","PVP01_1439000","PVP01_1439100","PVP01_1439200","PVP01_1439300","PVP01_1439400","PVP01_1439500","PVP01_1439700","PVP01_1439800","PVP01_1439900","PVP01_1440100","PVP01_1440200","PVP01_1440500","PVP01_1440600","PVP01_1440800","PVP01_1441100","PVP01_1441300","PVP01_1441400","PVP01_1441500","PVP01_1441600","PVP01_1441800","PVP01_1441900","PVP01_1442000","PVP01_1442100","PVP01_1442200","PVP01_1442300","PVP01_1442400","PVP01_1442600","PVP01_1442700","PVP01_1442800","PVP01_1443000","PVP01_1443100","PVP01_1443400","PVP01_1443500","PVP01_1443700","PVP01_1444000","PVP01_1444100","PVP01_1444300","PVP01_1444400","PVP01_1444500","PVP01_1444700","PVP01_1444800","PVP01_1445100","PVP01_1445200","PVP01_1445400","PVP01_1445500","PVP01_1445700","PVP01_1445800","PVP01_1445900","PVP01_1446000","PVP01_1446200","PVP01_1446500","PVP01_1446600","PVP01_1446700","PVP01_1446800","PVP01_1447100","PVP01_1447300","PVP01_1447400","PVP01_1447500","PVP01_1447600","PVP01_1448100","PVP01_1448400","PVP01_1448500","PVP01_1448600","PVP01_1449000","PVP01_1449200","PVP01_1449600","PVP01_1449800","PVP01_1450000","PVP01_1450100","PVP01_1450200","PVP01_1450300","PVP01_1450500","PVP01_1450600","PVP01_1450700","PVP01_1450800","PVP01_1450900","PVP01_1451000","PVP01_1451200","PVP01_1451300","PVP01_1451400","PVP01_1451700","PVP01_1451800","PVP01_1451900","PVP01_1452000","PVP01_1452100","PVP01_1452500","PVP01_1452600","PVP01_1452800","PVP01_1452900","PVP01_1453000","PVP01_1453100","PVP01_1453200","PVP01_1453300","PVP01_1453400","PVP01_1453500","PVP01_1453600","PVP01_1453700","PVP01_1453900","PVP01_1454000","PVP01_1454100","PVP01_1454300","PVP01_1454400","PVP01_1454500","PVP01_1454700","PVP01_1454800","PVP01_1455000","PVP01_1455100","PVP01_1455200","PVP01_1455300","PVP01_1455400","PVP01_1455500","PVP01_1455600","PVP01_1455700","PVP01_1455900","PVP01_1456000","PVP01_1456100","PVP01_1456200","PVP01_1456300","PVP01_1456400","PVP01_1456700","PVP01_1456800","PVP01_1457000","PVP01_1457100","PVP01_1457200","PVP01_1457300","PVP01_1457400","PVP01_1457500","PVP01_1457600","PVP01_1457800","PVP01_1458000","PVP01_1458100","PVP01_1458200","PVP01_1458400","PVP01_1458500","PVP01_1458600","PVP01_1458800","PVP01_1458900","PVP01_1459000","PVP01_1459100","PVP01_1459200","PVP01_1459300","PVP01_1459400","PVP01_1459500","PVP01_1459700","PVP01_1459800","PVP01_1459900","PVP01_1460000","PVP01_1460100","PVP01_1460200","PVP01_1460400","PVP01_1460600","PVP01_1460700","PVP01_1460800","PVP01_1461000","PVP01_1461200","PVP01_1461300","PVP01_1461400","PVP01_1461500","PVP01_1461600","PVP01_1461700","PVP01_1461800","PVP01_1462100","PVP01_1462200","PVP01_1462400","PVP01_1462500","PVP01_1462800","PVP01_1462900","PVP01_1463000","PVP01_1463100","PVP01_1463400","PVP01_1463500","PVP01_1463700","PVP01_1463800","PVP01_1463900","PVP01_1464000","PVP01_1464100","PVP01_1464200","PVP01_1464300","PVP01_1464400","PVP01_1464500","PVP01_1464600","PVP01_1464700","PVP01_1464800","PVP01_1464900","PVP01_1465100","PVP01_1465200","PVP01_1465300","PVP01_1465400","PVP01_1465500","PVP01_1465600","PVP01_1465700","PVP01_1465800","PVP01_1466100","PVP01_1466200","PVP01_1466300","PVP01_1466400","PVP01_1466500","PVP01_1466600","PVP01_1466700","PVP01_1466800","PVP01_1466900","PVP01_1467100","PVP01_1467200","PVP01_1467400","PVP01_1467500","PVP01_1467700","PVP01_1467800","PVP01_1468000","PVP01_1468100","PVP01_1468200","PVP01_1468300","PVP01_1468400","PVP01_1468500","PVP01_1468600","PVP01_1468700","PVP01_1468900","PVP01_1469200","PVP01_1469300","PVP01_1469500","PVP01_1469600","PVP01_1469700","PVP01_1469800","PVP01_1469900","PVP01_1470000","PVP01_1470100","PVP01_1470200","PVP01_1470300","PVP01_1470400","PVP01_1470500","PVP01_1470600","PVP01_1470700","PVP01_1470800","PVP01_1471000","PVP01_1471200","PVP01_1471300","PVP01_1471500","PVP01_1471600","PVP01_1471700","PVP01_1471800","PVP01_1471900","PVP01_1472100","PVP01_1472200","PVP01_1472500","PVP01_API02800","PVP01_API04800","PVP01_API05500","PVP01_MIT00500","PVP01_MIT01200","PVP01_MIT02700","PVP01_MIT03300","PVP01_MIT03400"],"Row.names":["PVP01_0000020","PVP01_0000030","PVP01_0000060","PVP01_0000210","PVP01_0000230","PVP01_0000240","PVP01_0000300","PVP01_0000670","PVP01_0000960","PVP01_0000980","PVP01_0001090","PVP01_0001110","PVP01_0001360","PVP01_0001450","PVP01_0001500","PVP01_0001510","PVP01_0001580","PVP01_0001640","PVP01_0001790","PVP01_0001850","PVP01_0001990","PVP01_0002000","PVP01_0002050","PVP01_0002160","PVP01_0002200","PVP01_0002210","PVP01_0002310","PVP01_0002330","PVP01_0002370","PVP01_0002390","PVP01_0002400","PVP01_0002430","PVP01_0002440","PVP01_0002490","PVP01_0002540","PVP01_0002740","PVP01_0002750","PVP01_0002760","PVP01_0002810","PVP01_0002820","PVP01_0002950","PVP01_0003060","PVP01_0003320","PVP01_0003330","PVP01_0003670","PVP01_0003700","PVP01_0003710","PVP01_0003820","PVP01_0003950","PVP01_0004100","PVP01_0004180","PVP01_0004450","PVP01_0004810","PVP01_0004880","PVP01_0004910","PVP01_0004960","PVP01_0005160","PVP01_0005170","PVP01_0005290","PVP01_0005420","PVP01_0005450","PVP01_0005550","PVP01_0005710","PVP01_0005720","PVP01_0005740","PVP01_0005750","PVP01_0005830","PVP01_0006010","PVP01_0006100","PVP01_0006260","PVP01_0006590","PVP01_0006680","PVP01_0006700","PVP01_0006770","PVP01_0007000","PVP01_0007050","PVP01_0007420","PVP01_0007660","PVP01_0007720","PVP01_0007800","PVP01_0007820","PVP01_0007860","PVP01_0007870","PVP01_0007880","PVP01_0007920","PVP01_0007950","PVP01_0008030","PVP01_0008040","PVP01_0008060","PVP01_0008080","PVP01_0008100","PVP01_0008190","PVP01_0008380","PVP01_0008650","PVP01_0008830","PVP01_0008920","PVP01_0008930","PVP01_0008990","PVP01_0009040","PVP01_0009220","PVP01_0009270","PVP01_0009420","PVP01_0009450","PVP01_0009460","PVP01_0009490","PVP01_0009510","PVP01_0009740","PVP01_0009920","PVP01_0009940","PVP01_0009960","PVP01_0009990","PVP01_0010170","PVP01_0010180","PVP01_0010190","PVP01_0010200","PVP01_0010220","PVP01_0010240","PVP01_0010290","PVP01_0010380","PVP01_0010500","PVP01_0010520","PVP01_0010690","PVP01_0010900","PVP01_0010960","PVP01_0010980","PVP01_0100200","PVP01_0102800","PVP01_0103800","PVP01_0104100","PVP01_0104300","PVP01_0104600","PVP01_0104700","PVP01_0104800","PVP01_0104900","PVP01_0105100","PVP01_0105500","PVP01_0105800","PVP01_0106200","PVP01_0106500","PVP01_0106600","PVP01_0107400","PVP01_0107500","PVP01_0107600","PVP01_0107900","PVP01_0108000","PVP01_0108100","PVP01_0108800","PVP01_0109000","PVP01_0109100","PVP01_0109200","PVP01_0109600","PVP01_0110700","PVP01_0110800","PVP01_0111300","PVP01_0111800","PVP01_0111900","PVP01_0112000","PVP01_0112300","PVP01_0112800","PVP01_0113000","PVP01_0113400","PVP01_0113500","PVP01_0114900","PVP01_0115000","PVP01_0115800","PVP01_0116000","PVP01_0116100","PVP01_0116300","PVP01_0116600","PVP01_0116700","PVP01_0116900","PVP01_0117100","PVP01_0117500","PVP01_0117600","PVP01_0117800","PVP01_0118000","PVP01_0118100","PVP01_0118500","PVP01_0118700","PVP01_0118900","PVP01_0119300","PVP01_0119800","PVP01_0120100","PVP01_0120400","PVP01_0120500","PVP01_0201000","PVP01_0202000","PVP01_0202300","PVP01_0203100","PVP01_0203800","PVP01_0203900","PVP01_0204000","PVP01_0204800","PVP01_0204900","PVP01_0205200","PVP01_0205300","PVP01_0205500","PVP01_0206100","PVP01_0206400","PVP01_0206500","PVP01_0206600","PVP01_0207000","PVP01_0207100","PVP01_0207200","PVP01_0207400","PVP01_0207500","PVP01_0207600","PVP01_0207700","PVP01_0208100","PVP01_0208200","PVP01_0208300","PVP01_0208500","PVP01_0208600","PVP01_0208800","PVP01_0208900","PVP01_0209000","PVP01_0209500","PVP01_0209600","PVP01_0210000","PVP01_0210100","PVP01_0211000","PVP01_0211100","PVP01_0211200","PVP01_0211300","PVP01_0211400","PVP01_0211600","PVP01_0212000","PVP01_0212200","PVP01_0212400","PVP01_0212600","PVP01_0212800","PVP01_0213000","PVP01_0213700","PVP01_0214100","PVP01_0214500","PVP01_0214900","PVP01_0215500","PVP01_0215700","PVP01_0216100","PVP01_0216200","PVP01_0216600","PVP01_0217300","PVP01_0218800","PVP01_0220300","PVP01_0221100","PVP01_0301400","PVP01_0301600","PVP01_0301900","PVP01_0302300","PVP01_0302800","PVP01_0303100","PVP01_0303400","PVP01_0303500","PVP01_0304000","PVP01_0304100","PVP01_0304300","PVP01_0304500","PVP01_0304700","PVP01_0304900","PVP01_0305000","PVP01_0305500","PVP01_0306000","PVP01_0306100","PVP01_0306300","PVP01_0307000","PVP01_0307400","PVP01_0307500","PVP01_0307600","PVP01_0307700","PVP01_0308200","PVP01_0308600","PVP01_0308700","PVP01_0308800","PVP01_0309300","PVP01_0309800","PVP01_0310200","PVP01_0311100","PVP01_0311200","PVP01_0311300","PVP01_0311400","PVP01_0312100","PVP01_0312400","PVP01_0312500","PVP01_0312800","PVP01_0313400","PVP01_0314100","PVP01_0314900","PVP01_0315300","PVP01_0315500","PVP01_0315700","PVP01_0316200","PVP01_0316400","PVP01_0316500","PVP01_0316600","PVP01_0316800","PVP01_0317200","PVP01_0318000","PVP01_0318100","PVP01_0318500","PVP01_0318900","PVP01_0319300","PVP01_0319400","PVP01_0319500","PVP01_0319700","PVP01_0319800","PVP01_0320200","PVP01_0320400","PVP01_0320700","PVP01_0321100","PVP01_0400300","PVP01_0403200","PVP01_0403500","PVP01_0403600","PVP01_0403800","PVP01_0404000","PVP01_0405200","PVP01_0405700","PVP01_0406100","PVP01_0406400","PVP01_0406600","PVP01_0407800","PVP01_0407900","PVP01_0408600","PVP01_0408900","PVP01_0409300","PVP01_0409800","PVP01_0410000","PVP01_0410600","PVP01_0411400","PVP01_0411500","PVP01_0411700","PVP01_0411800","PVP01_0412400","PVP01_0412600","PVP01_0412700","PVP01_0412900","PVP01_0413500","PVP01_0414300","PVP01_0414400","PVP01_0414700","PVP01_0414900","PVP01_0415200","PVP01_0415400","PVP01_0415600","PVP01_0415900","PVP01_0416000","PVP01_0416200","PVP01_0417100","PVP01_0417200","PVP01_0417600","PVP01_0417900","PVP01_0418300","PVP01_0418400","PVP01_0419000","PVP01_0419100","PVP01_0419200","PVP01_0420700","PVP01_0420800","PVP01_0420900","PVP01_0421700","PVP01_0421800","PVP01_0422200","PVP01_0422300","PVP01_0422400","PVP01_0422800","PVP01_0423600","PVP01_0423700","PVP01_0424000","PVP01_0424100","PVP01_0500900","PVP01_0502600","PVP01_0503400","PVP01_0505100","PVP01_0505500","PVP01_0506000","PVP01_0507000","PVP01_0507200","PVP01_0507300","PVP01_0507600","PVP01_0507800","PVP01_0508900","PVP01_0509100","PVP01_0510100","PVP01_0510300","PVP01_0510500","PVP01_0510600","PVP01_0510700","PVP01_0511200","PVP01_0511300","PVP01_0511400","PVP01_0511500","PVP01_0511700","PVP01_0512000","PVP01_0512400","PVP01_0512500","PVP01_0512700","PVP01_0512900","PVP01_0513300","PVP01_0513500","PVP01_0514000","PVP01_0514900","PVP01_0515100","PVP01_0515300","PVP01_0515800","PVP01_0516000","PVP01_0516400","PVP01_0516500","PVP01_0516800","PVP01_0516900","PVP01_0517000","PVP01_0517400","PVP01_0517700","PVP01_0518600","PVP01_0518700","PVP01_0518800","PVP01_0518900","PVP01_0519200","PVP01_0519300","PVP01_0519400","PVP01_0520800","PVP01_0521500","PVP01_0521600","PVP01_0521700","PVP01_0522000","PVP01_0522100","PVP01_0522300","PVP01_0522600","PVP01_0524500","PVP01_0524900","PVP01_0525000","PVP01_0525300","PVP01_0525700","PVP01_0526000","PVP01_0526100","PVP01_0526200","PVP01_0526300","PVP01_0527300","PVP01_0527400","PVP01_0527700","PVP01_0527800","PVP01_0527900","PVP01_0528100","PVP01_0528200","PVP01_0528400","PVP01_0528500","PVP01_0528700","PVP01_0528800","PVP01_0529000","PVP01_0529600","PVP01_0529700","PVP01_0530100","PVP01_0530400","PVP01_0530500","PVP01_0530700","PVP01_0531000","PVP01_0531100","PVP01_0531500","PVP01_0531700","PVP01_0531900","PVP01_0532000","PVP01_0532500","PVP01_0533500","PVP01_0533600","PVP01_0533900","PVP01_0534700","PVP01_0535400","PVP01_0600100","PVP01_0600600","PVP01_0600700","PVP01_0601300","PVP01_0601400","PVP01_0601600","PVP01_0601700","PVP01_0602600","PVP01_0603000","PVP01_0603100","PVP01_0603600","PVP01_0603700","PVP01_0603900","PVP01_0604000","PVP01_0604100","PVP01_0604700","PVP01_0605000","PVP01_0605300","PVP01_0605500","PVP01_0605900","PVP01_0606100","PVP01_0606200","PVP01_0606900","PVP01_0607600","PVP01_0608000","PVP01_0608100","PVP01_0608600","PVP01_0608700","PVP01_0608900","PVP01_0609100","PVP01_0609600","PVP01_0609700","PVP01_0609800","PVP01_0609900","PVP01_0610200","PVP01_0610300","PVP01_0610400","PVP01_0610500","PVP01_0610700","PVP01_0610900","PVP01_0611100","PVP01_0611200","PVP01_0612000","PVP01_0612100","PVP01_0612400","PVP01_0612700","PVP01_0613000","PVP01_0613100","PVP01_0613200","PVP01_0613400","PVP01_0613500","PVP01_0613800","PVP01_0614300","PVP01_0614400","PVP01_0614600","PVP01_0614700","PVP01_0614900","PVP01_0615000","PVP01_0615100","PVP01_0615500","PVP01_0615800","PVP01_0616200","PVP01_0616300","PVP01_0616400","PVP01_0617500","PVP01_0618600","PVP01_0618700","PVP01_0618900","PVP01_0619100","PVP01_0619700","PVP01_0620000","PVP01_0620100","PVP01_0620300","PVP01_0620700","PVP01_0620900","PVP01_0621100","PVP01_0621300","PVP01_0621900","PVP01_0622100","PVP01_0622200","PVP01_0623200","PVP01_0624600","PVP01_0624800","PVP01_0700700","PVP01_0701200","PVP01_0701400","PVP01_0701500","PVP01_0701600","PVP01_0702100","PVP01_0702400","PVP01_0702500","PVP01_0704000","PVP01_0704700","PVP01_0705000","PVP01_0705600","PVP01_0705700","PVP01_0705800","PVP01_0706100","PVP01_0706200","PVP01_0706400","PVP01_0706700","PVP01_0706900","PVP01_0707000","PVP01_0707800","PVP01_0707900","PVP01_0708200","PVP01_0708400","PVP01_0708700","PVP01_0708900","PVP01_0709100","PVP01_0709200","PVP01_0709300","PVP01_0709600","PVP01_0710000","PVP01_0710200","PVP01_0711000","PVP01_0711100","PVP01_0711500","PVP01_0711800","PVP01_0712300","PVP01_0712400","PVP01_0713100","PVP01_0713500","PVP01_0713600","PVP01_0713800","PVP01_0713900","PVP01_0714200","PVP01_0714900","PVP01_0715200","PVP01_0715700","PVP01_0715800","PVP01_0716100","PVP01_0716200","PVP01_0716400","PVP01_0716800","PVP01_0717000","PVP01_0717200","PVP01_0717500","PVP01_0717600","PVP01_0717800","PVP01_0718600","PVP01_0718700","PVP01_0719000","PVP01_0719100","PVP01_0719500","PVP01_0719700","PVP01_0719900","PVP01_0720100","PVP01_0720300","PVP01_0720400","PVP01_0720600","PVP01_0720900","PVP01_0721300","PVP01_0721800","PVP01_0721900","PVP01_0722400","PVP01_0722500","PVP01_0723000","PVP01_0723200","PVP01_0724300","PVP01_0724400","PVP01_0724700","PVP01_0725100","PVP01_0725300","PVP01_0725400","PVP01_0726000","PVP01_0726400","PVP01_0727100","PVP01_0727300","PVP01_0727400","PVP01_0727700","PVP01_0728000","PVP01_0728100","PVP01_0728400","PVP01_0728500","PVP01_0729100","PVP01_0729400","PVP01_0729500","PVP01_0729700","PVP01_0730000","PVP01_0730300","PVP01_0730400","PVP01_0730600","PVP01_0730700","PVP01_0730800","PVP01_0731000","PVP01_0731100","PVP01_0731200","PVP01_0731300","PVP01_0731500","PVP01_0731700","PVP01_0732300","PVP01_0732600","PVP01_0732800","PVP01_0732900","PVP01_0733000","PVP01_0734100","PVP01_0734200","PVP01_0734800","PVP01_0735100","PVP01_0735300","PVP01_0736500","PVP01_0737700","PVP01_0800100","PVP01_0801800","PVP01_0802000","PVP01_0803500","PVP01_0804100","PVP01_0804300","PVP01_0804600","PVP01_0804800","PVP01_0804900","PVP01_0805300","PVP01_0805400","PVP01_0805500","PVP01_0805600","PVP01_0805800","PVP01_0806100","PVP01_0806300","PVP01_0806500","PVP01_0806700","PVP01_0807000","PVP01_0807500","PVP01_0807800","PVP01_0808100","PVP01_0808700","PVP01_0808800","PVP01_0809200","PVP01_0809400","PVP01_0809600","PVP01_0809700","PVP01_0809900","PVP01_0810500","PVP01_0811200","PVP01_0811300","PVP01_0812300","PVP01_0813200","PVP01_0813300","PVP01_0813700","PVP01_0813800","PVP01_0814100","PVP01_0814200","PVP01_0814400","PVP01_0814700","PVP01_0815000","PVP01_0815100","PVP01_0815300","PVP01_0815900","PVP01_0816200","PVP01_0816700","PVP01_0816800","PVP01_0817000","PVP01_0817100","PVP01_0817400","PVP01_0818100","PVP01_0818500","PVP01_0818600","PVP01_0818900","PVP01_0819200","PVP01_0819300","PVP01_0820200","PVP01_0820500","PVP01_0820600","PVP01_0820800","PVP01_0821300","PVP01_0821400","PVP01_0821700","PVP01_0822300","PVP01_0822400","PVP01_0822600","PVP01_0822900","PVP01_0823100","PVP01_0823600","PVP01_0823800","PVP01_0823900","PVP01_0824000","PVP01_0824100","PVP01_0824800","PVP01_0825100","PVP01_0825300","PVP01_0825800","PVP01_0826000","PVP01_0826200","PVP01_0826300","PVP01_0826700","PVP01_0827800","PVP01_0828100","PVP01_0828200","PVP01_0828300","PVP01_0828500","PVP01_0828700","PVP01_0828800","PVP01_0829000","PVP01_0829100","PVP01_0830300","PVP01_0830500","PVP01_0831000","PVP01_0831100","PVP01_0831300","PVP01_0831700","PVP01_0832000","PVP01_0832100","PVP01_0832200","PVP01_0832300","PVP01_0832900","PVP01_0833000","PVP01_0833700","PVP01_0834300","PVP01_0834400","PVP01_0834700","PVP01_0835000","PVP01_0835100","PVP01_0835300","PVP01_0835900","PVP01_0836100","PVP01_0836400","PVP01_0836700","PVP01_0836800","PVP01_0837200","PVP01_0837300","PVP01_0837600","PVP01_0838000","PVP01_0839200","PVP01_0839400","PVP01_0839500","PVP01_0839800","PVP01_0841000","PVP01_0901200","PVP01_0901500","PVP01_0902100","PVP01_0903700","PVP01_0904700","PVP01_0904900","PVP01_0906400","PVP01_0906500","PVP01_0906800","PVP01_0906900","PVP01_0907000","PVP01_0907200","PVP01_0907500","PVP01_0908000","PVP01_0908400","PVP01_0908600","PVP01_0908700","PVP01_0908900","PVP01_0909100","PVP01_0909500","PVP01_0909600","PVP01_0909700","PVP01_0909900","PVP01_0910000","PVP01_0910200","PVP01_0910900","PVP01_0911300","PVP01_0911600","PVP01_0911700","PVP01_0911800","PVP01_0911900","PVP01_0912000","PVP01_0912200","PVP01_0912300","PVP01_0912400","PVP01_0912900","PVP01_0913000","PVP01_0913800","PVP01_0914100","PVP01_0916000","PVP01_0916700","PVP01_0917100","PVP01_0917200","PVP01_0917400","PVP01_0917600","PVP01_0917900","PVP01_0918000","PVP01_0918100","PVP01_0918200","PVP01_0918300","PVP01_0918700","PVP01_0919000","PVP01_0919400","PVP01_0919500","PVP01_0919700","PVP01_0920000","PVP01_0920100","PVP01_0920200","PVP01_0920400","PVP01_0920500","PVP01_0920700","PVP01_0921200","PVP01_0921700","PVP01_0922000","PVP01_0922100","PVP01_0922800","PVP01_0922900","PVP01_0923000","PVP01_0923100","PVP01_0923700","PVP01_0923800","PVP01_0923900","PVP01_0924000","PVP01_0924100","PVP01_0924300","PVP01_0924400","PVP01_0924500","PVP01_0925200","PVP01_0925700","PVP01_0925800","PVP01_0926300","PVP01_0926600","PVP01_0926900","PVP01_0927100","PVP01_0927200","PVP01_0927300","PVP01_0927400","PVP01_0927600","PVP01_0927700","PVP01_0928000","PVP01_0928100","PVP01_0928200","PVP01_0928300","PVP01_0928900","PVP01_0929000","PVP01_0929200","PVP01_0929600","PVP01_0929700","PVP01_0929900","PVP01_0930000","PVP01_0930600","PVP01_0930800","PVP01_0931300","PVP01_0932100","PVP01_0932200","PVP01_0932400","PVP01_0932600","PVP01_0933100","PVP01_0933400","PVP01_0933500","PVP01_0933700","PVP01_0934100","PVP01_0934300","PVP01_0934400","PVP01_0934500","PVP01_0935200","PVP01_0935700","PVP01_0935900","PVP01_0936000","PVP01_0936400","PVP01_0936500","PVP01_0936900","PVP01_0937100","PVP01_0937300","PVP01_0937700","PVP01_0938000","PVP01_0938400","PVP01_0938600","PVP01_0938700","PVP01_0938900","PVP01_0939000","PVP01_0939200","PVP01_0940600","PVP01_0940900","PVP01_0941100","PVP01_0941200","PVP01_0941400","PVP01_0941600","PVP01_0941700","PVP01_0941900","PVP01_0942200","PVP01_0942400","PVP01_0944100","PVP01_0944400","PVP01_0944600","PVP01_0944800","PVP01_0945000","PVP01_0945200","PVP01_0945400","PVP01_0945500","PVP01_0945800","PVP01_0946200","PVP01_0946300","PVP01_0946400","PVP01_0946700","PVP01_0947200","PVP01_0947300","PVP01_0948500","PVP01_0950100","PVP01_1001900","PVP01_1002600","PVP01_1004200","PVP01_1004600","PVP01_1005300","PVP01_1005900","PVP01_1006400","PVP01_1006600","PVP01_1006900","PVP01_1007100","PVP01_1007500","PVP01_1007600","PVP01_1007700","PVP01_1007800","PVP01_1007900","PVP01_1008100","PVP01_1008300","PVP01_1008400","PVP01_1008600","PVP01_1008900","PVP01_1009000","PVP01_1009200","PVP01_1009600","PVP01_1010100","PVP01_1010200","PVP01_1010400","PVP01_1011100","PVP01_1011200","PVP01_1011300","PVP01_1011400","PVP01_1011500","PVP01_1011800","PVP01_1012100","PVP01_1012400","PVP01_1012500","PVP01_1012700","PVP01_1013200","PVP01_1014300","PVP01_1014400","PVP01_1014600","PVP01_1015000","PVP01_1015500","PVP01_1015600","PVP01_1015700","PVP01_1016100","PVP01_1016200","PVP01_1016600","PVP01_1016900","PVP01_1017400","PVP01_1017500","PVP01_1017900","PVP01_1018400","PVP01_1019000","PVP01_1019600","PVP01_1020000","PVP01_1020100","PVP01_1020700","PVP01_1020900","PVP01_1021000","PVP01_1021200","PVP01_1021300","PVP01_1021500","PVP01_1021600","PVP01_1021800","PVP01_1021900","PVP01_1022000","PVP01_1022200","PVP01_1022400","PVP01_1022500","PVP01_1022800","PVP01_1023000","PVP01_1023600","PVP01_1023700","PVP01_1023800","PVP01_1023900","PVP01_1024400","PVP01_1024500","PVP01_1024600","PVP01_1024800","PVP01_1024900","PVP01_1025100","PVP01_1025200","PVP01_1025500","PVP01_1026200","PVP01_1026300","PVP01_1026400","PVP01_1026600","PVP01_1026700","PVP01_1027100","PVP01_1027300","PVP01_1027600","PVP01_1027800","PVP01_1028100","PVP01_1028200","PVP01_1028600","PVP01_1028900","PVP01_1029000","PVP01_1029100","PVP01_1029700","PVP01_1030500","PVP01_1030600","PVP01_1031000","PVP01_1031100","PVP01_1031300","PVP01_1031400","PVP01_1031500","PVP01_1031800","PVP01_1031900","PVP01_1032900","PVP01_1033000","PVP01_1034600","PVP01_1034800","PVP01_1034900","PVP01_1035000","PVP01_1035200","PVP01_1100800","PVP01_1101000","PVP01_1101100","PVP01_1102400","PVP01_1102500","PVP01_1102700","PVP01_1103000","PVP01_1103200","PVP01_1103300","PVP01_1103400","PVP01_1103700","PVP01_1104000","PVP01_1104300","PVP01_1104400","PVP01_1104600","PVP01_1104800","PVP01_1105500","PVP01_1105700","PVP01_1106000","PVP01_1106800","PVP01_1106900","PVP01_1107000","PVP01_1107100","PVP01_1107400","PVP01_1107600","PVP01_1107900","PVP01_1108500","PVP01_1108600","PVP01_1108800","PVP01_1108900","PVP01_1109400","PVP01_1109500","PVP01_1109600","PVP01_1110500","PVP01_1110600","PVP01_1111000","PVP01_1111100","PVP01_1111300","PVP01_1112100","PVP01_1112900","PVP01_1113500","PVP01_1113600","PVP01_1113700","PVP01_1113900","PVP01_1114200","PVP01_1114300","PVP01_1114500","PVP01_1115500","PVP01_1115600","PVP01_1115900","PVP01_1116600","PVP01_1117700","PVP01_1118000","PVP01_1118500","PVP01_1118600","PVP01_1118700","PVP01_1118800","PVP01_1119000","PVP01_1119500","PVP01_1119600","PVP01_1119800","PVP01_1120000","PVP01_1120300","PVP01_1120500","PVP01_1121100","PVP01_1121400","PVP01_1121500","PVP01_1121700","PVP01_1121900","PVP01_1122100","PVP01_1122300","PVP01_1122400","PVP01_1122500","PVP01_1122600","PVP01_1122700","PVP01_1123000","PVP01_1123500","PVP01_1123600","PVP01_1123900","PVP01_1124000","PVP01_1124100","PVP01_1124300","PVP01_1124400","PVP01_1124800","PVP01_1124900","PVP01_1125000","PVP01_1125100","PVP01_1125400","PVP01_1125900","PVP01_1126200","PVP01_1127100","PVP01_1127200","PVP01_1127300","PVP01_1127400","PVP01_1127800","PVP01_1128400","PVP01_1129000","PVP01_1129400","PVP01_1129700","PVP01_1130200","PVP01_1130900","PVP01_1131000","PVP01_1132300","PVP01_1132400","PVP01_1132700","PVP01_1132800","PVP01_1132900","PVP01_1133100","PVP01_1133200","PVP01_1133400","PVP01_1133500","PVP01_1133600","PVP01_1133800","PVP01_1133900","PVP01_1134000","PVP01_1134300","PVP01_1134500","PVP01_1135500","PVP01_1135800","PVP01_1135900","PVP01_1136100","PVP01_1136700","PVP01_1136800","PVP01_1136900","PVP01_1137100","PVP01_1137200","PVP01_1137600","PVP01_1137700","PVP01_1137800","PVP01_1138400","PVP01_1138500","PVP01_1139900","PVP01_1140300","PVP01_1140500","PVP01_1140700","PVP01_1141300","PVP01_1141500","PVP01_1141900","PVP01_1142000","PVP01_1142600","PVP01_1142700","PVP01_1143200","PVP01_1143400","PVP01_1143500","PVP01_1144100","PVP01_1144300","PVP01_1144500","PVP01_1144800","PVP01_1145500","PVP01_1145700","PVP01_1146400","PVP01_1146500","PVP01_1146900","PVP01_1148100","PVP01_1148200","PVP01_1148500","PVP01_1148700","PVP01_1149000","PVP01_1200200","PVP01_1200600","PVP01_1201900","PVP01_1202100","PVP01_1202200","PVP01_1203100","PVP01_1203300","PVP01_1203500","PVP01_1203600","PVP01_1204000","PVP01_1204100","PVP01_1204200","PVP01_1204600","PVP01_1205000","PVP01_1205500","PVP01_1206100","PVP01_1206300","PVP01_1206700","PVP01_1206900","PVP01_1207000","PVP01_1208000","PVP01_1208200","PVP01_1208400","PVP01_1208600","PVP01_1208700","PVP01_1209000","PVP01_1209200","PVP01_1209900","PVP01_1210000","PVP01_1210100","PVP01_1210200","PVP01_1210300","PVP01_1210700","PVP01_1210800","PVP01_1211100","PVP01_1211200","PVP01_1211300","PVP01_1211600","PVP01_1212100","PVP01_1213400","PVP01_1214000","PVP01_1214200","PVP01_1214300","PVP01_1214400","PVP01_1214600","PVP01_1214700","PVP01_1216100","PVP01_1216200","PVP01_1216800","PVP01_1217200","PVP01_1217500","PVP01_1217600","PVP01_1217700","PVP01_1218800","PVP01_1218900","PVP01_1219000","PVP01_1219100","PVP01_1219400","PVP01_1219600","PVP01_1219700","PVP01_1219900","PVP01_1220200","PVP01_1220300","PVP01_1221300","PVP01_1221800","PVP01_1221900","PVP01_1222400","PVP01_1223200","PVP01_1223300","PVP01_1223400","PVP01_1223800","PVP01_1223900","PVP01_1224000","PVP01_1224400","PVP01_1224500","PVP01_1224600","PVP01_1225500","PVP01_1225900","PVP01_1226100","PVP01_1226600","PVP01_1226800","PVP01_1227500","PVP01_1227600","PVP01_1227900","PVP01_1228000","PVP01_1228100","PVP01_1228200","PVP01_1228300","PVP01_1228400","PVP01_1229200","PVP01_1229500","PVP01_1229600","PVP01_1229900","PVP01_1230000","PVP01_1230100","PVP01_1230200","PVP01_1230500","PVP01_1230600","PVP01_1230900","PVP01_1231000","PVP01_1231100","PVP01_1231800","PVP01_1231900","PVP01_1232600","PVP01_1232700","PVP01_1232900","PVP01_1233100","PVP01_1235500","PVP01_1236000","PVP01_1236100","PVP01_1236800","PVP01_1237100","PVP01_1237200","PVP01_1237700","PVP01_1237900","PVP01_1238300","PVP01_1238400","PVP01_1238900","PVP01_1239300","PVP01_1239500","PVP01_1240000","PVP01_1240100","PVP01_1241300","PVP01_1241900","PVP01_1242300","PVP01_1242900","PVP01_1243500","PVP01_1243600","PVP01_1243900","PVP01_1244700","PVP01_1244800","PVP01_1244900","PVP01_1245500","PVP01_1245600","PVP01_1245800","PVP01_1246200","PVP01_1246500","PVP01_1247300","PVP01_1247800","PVP01_1247900","PVP01_1248700","PVP01_1248800","PVP01_1248900","PVP01_1249000","PVP01_1249200","PVP01_1249500","PVP01_1250200","PVP01_1250700","PVP01_1250900","PVP01_1252400","PVP01_1252600","PVP01_1252900","PVP01_1253000","PVP01_1253700","PVP01_1254200","PVP01_1254900","PVP01_1255100","PVP01_1255200","PVP01_1255700","PVP01_1256400","PVP01_1256600","PVP01_1256800","PVP01_1256900","PVP01_1257000","PVP01_1257600","PVP01_1257800","PVP01_1258100","PVP01_1258300","PVP01_1258900","PVP01_1259000","PVP01_1259200","PVP01_1259300","PVP01_1259500","PVP01_1260200","PVP01_1260500","PVP01_1260600","PVP01_1260800","PVP01_1262000","PVP01_1262800","PVP01_1262900","PVP01_1263300","PVP01_1263400","PVP01_1264000","PVP01_1264300","PVP01_1264500","PVP01_1264600","PVP01_1264700","PVP01_1264800","PVP01_1265000","PVP01_1265700","PVP01_1266400","PVP01_1267100","PVP01_1267200","PVP01_1267300","PVP01_1267400","PVP01_1267600","PVP01_1268000","PVP01_1268200","PVP01_1268600","PVP01_1269000","PVP01_1269100","PVP01_1270200","PVP01_1270600","PVP01_1271100","PVP01_1273000","PVP01_1301200","PVP01_1301500","PVP01_1301600","PVP01_1301700","PVP01_1302700","PVP01_1302800","PVP01_1303100","PVP01_1304000","PVP01_1304200","PVP01_1304600","PVP01_1304800","PVP01_1304900","PVP01_1305300","PVP01_1305600","PVP01_1305700","PVP01_1305900","PVP01_1306000","PVP01_1306100","PVP01_1306300","PVP01_1306400","PVP01_1306700","PVP01_1307000","PVP01_1307200","PVP01_1307300","PVP01_1307500","PVP01_1307800","PVP01_1308100","PVP01_1308300","PVP01_1308500","PVP01_1308900","PVP01_1309000","PVP01_1309100","PVP01_1309700","PVP01_1310200","PVP01_1310900","PVP01_1311000","PVP01_1311100","PVP01_1311300","PVP01_1311400","PVP01_1311500","PVP01_1312200","PVP01_1312300","PVP01_1312400","PVP01_1312900","PVP01_1313400","PVP01_1313900","PVP01_1314100","PVP01_1314300","PVP01_1314600","PVP01_1314900","PVP01_1315000","PVP01_1315400","PVP01_1315700","PVP01_1316100","PVP01_1316500","PVP01_1316600","PVP01_1316900","PVP01_1317000","PVP01_1317200","PVP01_1317400","PVP01_1317600","PVP01_1317800","PVP01_1317900","PVP01_1318000","PVP01_1318100","PVP01_1318500","PVP01_1318800","PVP01_1318900","PVP01_1319100","PVP01_1319700","PVP01_1320700","PVP01_1320900","PVP01_1321300","PVP01_1321600","PVP01_1321800","PVP01_1322300","PVP01_1322400","PVP01_1322600","PVP01_1322700","PVP01_1323000","PVP01_1323100","PVP01_1323400","PVP01_1323500","PVP01_1323700","PVP01_1324100","PVP01_1324200","PVP01_1324700","PVP01_1324800","PVP01_1325200","PVP01_1325800","PVP01_1325900","PVP01_1326000","PVP01_1326200","PVP01_1326400","PVP01_1327600","PVP01_1327900","PVP01_1328000","PVP01_1328100","PVP01_1328400","PVP01_1328800","PVP01_1329300","PVP01_1329400","PVP01_1330600","PVP01_1330700","PVP01_1331000","PVP01_1331400","PVP01_1331800","PVP01_1332700","PVP01_1332900","PVP01_1333100","PVP01_1333200","PVP01_1333300","PVP01_1333600","PVP01_1333700","PVP01_1333900","PVP01_1334000","PVP01_1334100","PVP01_1334200","PVP01_1334300","PVP01_1334500","PVP01_1335600","PVP01_1335700","PVP01_1335800","PVP01_1336000","PVP01_1337400","PVP01_1337500","PVP01_1337900","PVP01_1338100","PVP01_1338300","PVP01_1338400","PVP01_1338500","PVP01_1338600","PVP01_1339000","PVP01_1339100","PVP01_1339600","PVP01_1340500","PVP01_1340600","PVP01_1342200","PVP01_1342300","PVP01_1342600","PVP01_1342800","PVP01_1343100","PVP01_1343200","PVP01_1343500","PVP01_1343700","PVP01_1344200","PVP01_1344600","PVP01_1344800","PVP01_1344900","PVP01_1345700","PVP01_1346600","PVP01_1346900","PVP01_1347300","PVP01_1347500","PVP01_1400100","PVP01_1400800","PVP01_1401200","PVP01_1401500","PVP01_1402900","PVP01_1403000","PVP01_1403400","PVP01_1403700","PVP01_1403800","PVP01_1403900","PVP01_1404500","PVP01_1404600","PVP01_1405000","PVP01_1405600","PVP01_1406000","PVP01_1406500","PVP01_1406600","PVP01_1407000","PVP01_1407100","PVP01_1407600","PVP01_1407800","PVP01_1407900","PVP01_1408100","PVP01_1408200","PVP01_1408500","PVP01_1408600","PVP01_1408700","PVP01_1408800","PVP01_1408900","PVP01_1409000","PVP01_1409300","PVP01_1409400","PVP01_1410200","PVP01_1410300","PVP01_1410400","PVP01_1410500","PVP01_1410700","PVP01_1410800","PVP01_1411000","PVP01_1411100","PVP01_1411200","PVP01_1411300","PVP01_1411400","PVP01_1411500","PVP01_1412700","PVP01_1413100","PVP01_1413200","PVP01_1413700","PVP01_1413900","PVP01_1414300","PVP01_1414500","PVP01_1415100","PVP01_1415500","PVP01_1415600","PVP01_1415900","PVP01_1416000","PVP01_1416100","PVP01_1416200","PVP01_1417100","PVP01_1417200","PVP01_1417600","PVP01_1417900","PVP01_1418000","PVP01_1418600","PVP01_1418800","PVP01_1419000","PVP01_1419100","PVP01_1419300","PVP01_1419400","PVP01_1419800","PVP01_1420100","PVP01_1420500","PVP01_1420700","PVP01_1420800","PVP01_1421000","PVP01_1421300","PVP01_1421500","PVP01_1422300","PVP01_1422600","PVP01_1422900","PVP01_1423200","PVP01_1423700","PVP01_1424200","PVP01_1424800","PVP01_1425100","PVP01_1426100","PVP01_1426200","PVP01_1426700","PVP01_1426800","PVP01_1427100","PVP01_1428300","PVP01_1428600","PVP01_1429000","PVP01_1429200","PVP01_1429300","PVP01_1429900","PVP01_1430000","PVP01_1430100","PVP01_1430500","PVP01_1430700","PVP01_1430800","PVP01_1431100","PVP01_1431200","PVP01_1431300","PVP01_1431600","PVP01_1432000","PVP01_1432400","PVP01_1432500","PVP01_1432600","PVP01_1433000","PVP01_1433700","PVP01_1433800","PVP01_1434000","PVP01_1434200","PVP01_1434300","PVP01_1434800","PVP01_1436000","PVP01_1437500","PVP01_1437700","PVP01_1437900","PVP01_1438100","PVP01_1439600","PVP01_1440000","PVP01_1440300","PVP01_1440400","PVP01_1440700","PVP01_1440900","PVP01_1441000","PVP01_1441200","PVP01_1441700","PVP01_1442500","PVP01_1442900","PVP01_1443200","PVP01_1443300","PVP01_1443600","PVP01_1443800","PVP01_1443900","PVP01_1444200","PVP01_1444600","PVP01_1444900","PVP01_1445000","PVP01_1445300","PVP01_1445600","PVP01_1446100","PVP01_1446300","PVP01_1446400","PVP01_1446900","PVP01_1447000","PVP01_1447200","PVP01_1447700","PVP01_1447800","PVP01_1447900","PVP01_1448000","PVP01_1448200","PVP01_1448300","PVP01_1448700","PVP01_1448800","PVP01_1448900","PVP01_1449100","PVP01_1449300","PVP01_1449400","PVP01_1449500","PVP01_1449700","PVP01_1449900","PVP01_1450400","PVP01_1451100","PVP01_1451500","PVP01_1451600","PVP01_1452200","PVP01_1452300","PVP01_1452400","PVP01_1452700","PVP01_1453800","PVP01_1454200","PVP01_1454600","PVP01_1454900","PVP01_1455800","PVP01_1456500","PVP01_1456600","PVP01_1456900","PVP01_1457700","PVP01_1457900","PVP01_1458300","PVP01_1458700","PVP01_1459600","PVP01_1460300","PVP01_1460500","PVP01_1460900","PVP01_1461100","PVP01_1461900","PVP01_1462000","PVP01_1462300","PVP01_1462600","PVP01_1462700","PVP01_1463200","PVP01_1463300","PVP01_1463600","PVP01_1465000","PVP01_1465900","PVP01_1466000","PVP01_1467000","PVP01_1467300","PVP01_1467600","PVP01_1467900","PVP01_1468800","PVP01_1471100","PVP01_1472000","PVP01_1472300","PVP01_1472400","PVP01_MIT00800","PVP01_MIT01400","PVP01_MIT02200","PVP01_MIT02300","PVP01_MIT02800","PVP01_0000010","PVP01_0000040","PVP01_0000050","PVP01_0000070","PVP01_0000090","PVP01_0000100","PVP01_0000110","PVP01_0000120","PVP01_0000130","PVP01_0000140","PVP01_0000150","PVP01_0000160","PVP01_0000170","PVP01_0000180","PVP01_0000190","PVP01_0000200","PVP01_0000220","PVP01_0001430","PVP01_0001440","PVP01_0001470","PVP01_0001490","PVP01_0001600","PVP01_0001940","PVP01_0002090","PVP01_0002180","PVP01_0002240","PVP01_0002530","PVP01_0002710","PVP01_0002720","PVP01_0003140","PVP01_0003150","PVP01_0003290","PVP01_0003350","PVP01_0003980","PVP01_0004110","PVP01_0004210","PVP01_0004220","PVP01_0004230","PVP01_0004240","PVP01_0004250","PVP01_0004260","PVP01_0004280","PVP01_0004290","PVP01_0004300","PVP01_0004310","PVP01_0004320","PVP01_0004340","PVP01_0004350","PVP01_0004360","PVP01_0004370","PVP01_0004380","PVP01_0004390","PVP01_0004400","PVP01_0004410","PVP01_0004920","PVP01_0004950","PVP01_0005000","PVP01_0005010","PVP01_0005020","PVP01_0005030","PVP01_0005250","PVP01_0005430","PVP01_0005560","PVP01_0005570","PVP01_0005760","PVP01_0005800","PVP01_0005810","PVP01_0005820","PVP01_0005870","PVP01_0005910","PVP01_0005930","PVP01_0005940","PVP01_0005950","PVP01_0006660","PVP01_0006670","PVP01_0007080","PVP01_0007220","PVP01_0007240","PVP01_0007290","PVP01_0007650","PVP01_0007810","PVP01_0007830","PVP01_0008090","PVP01_0008110","PVP01_0008150","PVP01_0008500","PVP01_0008770","PVP01_0008820","PVP01_0009050","PVP01_0009070","PVP01_0009100","PVP01_0009180","PVP01_0009210","PVP01_0009370","PVP01_0009410","PVP01_0009430","PVP01_0009800","PVP01_0009810","PVP01_0009870","PVP01_0009950","PVP01_0009980","PVP01_0010020","PVP01_0010030","PVP01_0010070","PVP01_0010120","PVP01_0010130","PVP01_0010230","PVP01_0010440","PVP01_0010470","PVP01_0010510","PVP01_0010550","PVP01_0010570","PVP01_0010590","PVP01_0010600","PVP01_0010640","PVP01_0010670","PVP01_0010810","PVP01_0010840","PVP01_0010850","PVP01_0010860","PVP01_0010870","PVP01_0010970","PVP01_0010990","PVP01_0011020","PVP01_0011050","PVP01_0100800","PVP01_0102300","PVP01_0102600","PVP01_0102700","PVP01_0103100","PVP01_0103200","PVP01_0103300","PVP01_0103400","PVP01_0103500","PVP01_0103600","PVP01_0103700","PVP01_0103900","PVP01_0104000","PVP01_0104200","PVP01_0104400","PVP01_0104500","PVP01_0105000","PVP01_0105200","PVP01_0105300","PVP01_0105400","PVP01_0105600","PVP01_0105700","PVP01_0105900","PVP01_0106000","PVP01_0106100","PVP01_0106300","PVP01_0106400","PVP01_0106700","PVP01_0106800","PVP01_0106900","PVP01_0107700","PVP01_0107800","PVP01_0108200","PVP01_0108300","PVP01_0108400","PVP01_0108500","PVP01_0108600","PVP01_0108700","PVP01_0108900","PVP01_0109300","PVP01_0109400","PVP01_0109500","PVP01_0109700","PVP01_0109800","PVP01_0109900","PVP01_0110000","PVP01_0110100","PVP01_0110200","PVP01_0110300","PVP01_0110400","PVP01_0110500","PVP01_0110600","PVP01_0110900","PVP01_0111000","PVP01_0111100","PVP01_0111200","PVP01_0111400","PVP01_0111500","PVP01_0111600","PVP01_0111700","PVP01_0112100","PVP01_0112200","PVP01_0112400","PVP01_0112500","PVP01_0112600","PVP01_0112700","PVP01_0112900","PVP01_0113100","PVP01_0113200","PVP01_0113300","PVP01_0113600","PVP01_0113700","PVP01_0113800","PVP01_0113900","PVP01_0114000","PVP01_0114100","PVP01_0114200","PVP01_0114300","PVP01_0114400","PVP01_0114500","PVP01_0114600","PVP01_0114700","PVP01_0114800","PVP01_0115100","PVP01_0115200","PVP01_0115300","PVP01_0115400","PVP01_0115500","PVP01_0115600","PVP01_0115700","PVP01_0115900","PVP01_0116200","PVP01_0116400","PVP01_0116500","PVP01_0116800","PVP01_0117000","PVP01_0117200","PVP01_0117300","PVP01_0117400","PVP01_0117700","PVP01_0117900","PVP01_0118200","PVP01_0118300","PVP01_0118400","PVP01_0118600","PVP01_0118800","PVP01_0119000","PVP01_0119100","PVP01_0119200","PVP01_0119400","PVP01_0119500","PVP01_0119600","PVP01_0119900","PVP01_0121700","PVP01_0122100","PVP01_0122200","PVP01_0200200","PVP01_0201100","PVP01_0201200","PVP01_0201400","PVP01_0201500","PVP01_0201600","PVP01_0201700","PVP01_0201800","PVP01_0201900","PVP01_0202100","PVP01_0202200","PVP01_0202400","PVP01_0202500","PVP01_0202600","PVP01_0202700","PVP01_0202800","PVP01_0202900","PVP01_0203000","PVP01_0203200","PVP01_0203300","PVP01_0203400","PVP01_0203500","PVP01_0203600","PVP01_0203700","PVP01_0204100","PVP01_0204200","PVP01_0204300","PVP01_0204400","PVP01_0204500","PVP01_0204600","PVP01_0204700","PVP01_0205000","PVP01_0205100","PVP01_0205400","PVP01_0205600","PVP01_0205700","PVP01_0205800","PVP01_0205900","PVP01_0206000","PVP01_0206200","PVP01_0206300","PVP01_0206700","PVP01_0206800","PVP01_0206900","PVP01_0207300","PVP01_0207800","PVP01_0207900","PVP01_0208000","PVP01_0208400","PVP01_0208700","PVP01_0209100","PVP01_0209200","PVP01_0209300","PVP01_0209400","PVP01_0209700","PVP01_0209800","PVP01_0209900","PVP01_0210200","PVP01_0210300","PVP01_0210400","PVP01_0210500","PVP01_0210600","PVP01_0210700","PVP01_0210800","PVP01_0210900","PVP01_0211500","PVP01_0211700","PVP01_0211800","PVP01_0211900","PVP01_0212100","PVP01_0212300","PVP01_0212500","PVP01_0212700","PVP01_0212900","PVP01_0213100","PVP01_0213200","PVP01_0213300","PVP01_0213400","PVP01_0213500","PVP01_0213600","PVP01_0213800","PVP01_0213900","PVP01_0214000","PVP01_0214200","PVP01_0214300","PVP01_0214400","PVP01_0214600","PVP01_0214700","PVP01_0214800","PVP01_0215000","PVP01_0215100","PVP01_0215200","PVP01_0215300","PVP01_0215400","PVP01_0215600","PVP01_0215800","PVP01_0215900","PVP01_0216000","PVP01_0216500","PVP01_0216700","PVP01_0216800","PVP01_0216900","PVP01_0217000","PVP01_0217100","PVP01_0217200","PVP01_0219400","PVP01_0220100","PVP01_0220200","PVP01_0300200","PVP01_0300300","PVP01_0300600","PVP01_0300700","PVP01_0300800","PVP01_0300900","PVP01_0301500","PVP01_0302100","PVP01_0302400","PVP01_0302500","PVP01_0302600","PVP01_0302700","PVP01_0302900","PVP01_0303000","PVP01_0303200","PVP01_0303300","PVP01_0303600","PVP01_0303700","PVP01_0303800","PVP01_0304200","PVP01_0304400","PVP01_0304600","PVP01_0304800","PVP01_0305100","PVP01_0305200","PVP01_0305300","PVP01_0305400","PVP01_0305600","PVP01_0305700","PVP01_0305800","PVP01_0305900","PVP01_0306200","PVP01_0306400","PVP01_0306500","PVP01_0306600","PVP01_0306700","PVP01_0306800","PVP01_0306900","PVP01_0307100","PVP01_0307200","PVP01_0307300","PVP01_0307800","PVP01_0307900","PVP01_0308000","PVP01_0308100","PVP01_0308300","PVP01_0308400","PVP01_0308500","PVP01_0308900","PVP01_0309000","PVP01_0309100","PVP01_0309200","PVP01_0309600","PVP01_0309700","PVP01_0309900","PVP01_0310000","PVP01_0310100","PVP01_0310300","PVP01_0310400","PVP01_0310500","PVP01_0310600","PVP01_0310700","PVP01_0311000","PVP01_0311500","PVP01_0311600","PVP01_0311700","PVP01_0311800","PVP01_0311900","PVP01_0312000","PVP01_0312200","PVP01_0312300","PVP01_0312600","PVP01_0312700","PVP01_0312900","PVP01_0313000","PVP01_0313100","PVP01_0313200","PVP01_0313300","PVP01_0313500","PVP01_0313600","PVP01_0313700","PVP01_0313800","PVP01_0313900","PVP01_0314000","PVP01_0314200","PVP01_0314300","PVP01_0314400","PVP01_0314500","PVP01_0314600","PVP01_0314700","PVP01_0314800","PVP01_0315000","PVP01_0315100","PVP01_0315200","PVP01_0315400","PVP01_0315600","PVP01_0315800","PVP01_0315900","PVP01_0316000","PVP01_0316100","PVP01_0316300","PVP01_0316700","PVP01_0316900","PVP01_0317000","PVP01_0317100","PVP01_0317300","PVP01_0317400","PVP01_0317500","PVP01_0317600","PVP01_0317700","PVP01_0317800","PVP01_0317900","PVP01_0318200","PVP01_0318300","PVP01_0318400","PVP01_0318600","PVP01_0318700","PVP01_0318800","PVP01_0319000","PVP01_0319100","PVP01_0319200","PVP01_0319600","PVP01_0319900","PVP01_0320000","PVP01_0320100","PVP01_0320300","PVP01_0320500","PVP01_0320600","PVP01_0320800","PVP01_0320900","PVP01_0321000","PVP01_0403300","PVP01_0403400","PVP01_0403900","PVP01_0404200","PVP01_0404300","PVP01_0404400","PVP01_0404500","PVP01_0404600","PVP01_0404700","PVP01_0404800","PVP01_0404900","PVP01_0405000","PVP01_0405100","PVP01_0405300","PVP01_0405400","PVP01_0405500","PVP01_0405600","PVP01_0405800","PVP01_0405900","PVP01_0406200","PVP01_0406300","PVP01_0406500","PVP01_0406700","PVP01_0406800","PVP01_0406900","PVP01_0407000","PVP01_0407100","PVP01_0407200","PVP01_0407300","PVP01_0407400","PVP01_0407500","PVP01_0407600","PVP01_0407700","PVP01_0408000","PVP01_0408100","PVP01_0408200","PVP01_0408300","PVP01_0408400","PVP01_0408500","PVP01_0408700","PVP01_0408800","PVP01_0409000","PVP01_0409100","PVP01_0409200","PVP01_0409400","PVP01_0409500","PVP01_0409600","PVP01_0409700","PVP01_0409900","PVP01_0410100","PVP01_0410200","PVP01_0410300","PVP01_0410400","PVP01_0410500","PVP01_0410700","PVP01_0410800","PVP01_0410900","PVP01_0411000","PVP01_0411100","PVP01_0411200","PVP01_0411300","PVP01_0411600","PVP01_0411900","PVP01_0412000","PVP01_0412100","PVP01_0412200","PVP01_0412300","PVP01_0412500","PVP01_0412800","PVP01_0413000","PVP01_0413100","PVP01_0413200","PVP01_0413300","PVP01_0413400","PVP01_0413600","PVP01_0413700","PVP01_0413800","PVP01_0413900","PVP01_0414000","PVP01_0414100","PVP01_0414200","PVP01_0414500","PVP01_0414600","PVP01_0414800","PVP01_0415000","PVP01_0415100","PVP01_0415300","PVP01_0415500","PVP01_0415700","PVP01_0415800","PVP01_0416100","PVP01_0416300","PVP01_0416400","PVP01_0416500","PVP01_0416600","PVP01_0416700","PVP01_0416800","PVP01_0416900","PVP01_0417000","PVP01_0417300","PVP01_0417400","PVP01_0417500","PVP01_0417700","PVP01_0417800","PVP01_0418000","PVP01_0418100","PVP01_0418200","PVP01_0418500","PVP01_0418600","PVP01_0418700","PVP01_0418800","PVP01_0418900","PVP01_0419300","PVP01_0419400","PVP01_0419500","PVP01_0419600","PVP01_0419700","PVP01_0419800","PVP01_0419900","PVP01_0420000","PVP01_0420100","PVP01_0420200","PVP01_0420300","PVP01_0420400","PVP01_0420500","PVP01_0420600","PVP01_0421000","PVP01_0421100","PVP01_0421200","PVP01_0421300","PVP01_0421400","PVP01_0421500","PVP01_0421900","PVP01_0422000","PVP01_0422100","PVP01_0422500","PVP01_0422600","PVP01_0422700","PVP01_0422900","PVP01_0423000","PVP01_0423100","PVP01_0423200","PVP01_0423300","PVP01_0423400","PVP01_0423500","PVP01_0423800","PVP01_0423900","PVP01_0424200","PVP01_0424500","PVP01_0500400","PVP01_0502700","PVP01_0502800","PVP01_0502900","PVP01_0503000","PVP01_0503500","PVP01_0503600","PVP01_0503700","PVP01_0503800","PVP01_0503900","PVP01_0504000","PVP01_0504100","PVP01_0504200","PVP01_0504300","PVP01_0504400","PVP01_0504500","PVP01_0504600","PVP01_0504700","PVP01_0504800","PVP01_0504900","PVP01_0505000","PVP01_0505200","PVP01_0505300","PVP01_0505400","PVP01_0505600","PVP01_0505700","PVP01_0505800","PVP01_0505900","PVP01_0506100","PVP01_0506200","PVP01_0506300","PVP01_0506400","PVP01_0506500","PVP01_0506600","PVP01_0506700","PVP01_0506800","PVP01_0506900","PVP01_0507100","PVP01_0507400","PVP01_0507500","PVP01_0507700","PVP01_0507900","PVP01_0508000","PVP01_0508100","PVP01_0508200","PVP01_0508300","PVP01_0508400","PVP01_0508500","PVP01_0508600","PVP01_0508700","PVP01_0508800","PVP01_0509000","PVP01_0509200","PVP01_0509300","PVP01_0509400","PVP01_0509500","PVP01_0509600","PVP01_0509700","PVP01_0509800","PVP01_0509900","PVP01_0510000","PVP01_0510200","PVP01_0510400","PVP01_0510800","PVP01_0510900","PVP01_0511000","PVP01_0511100","PVP01_0511600","PVP01_0511800","PVP01_0511900","PVP01_0512100","PVP01_0512200","PVP01_0512300","PVP01_0512600","PVP01_0512800","PVP01_0513000","PVP01_0513100","PVP01_0513200","PVP01_0513400","PVP01_0513600","PVP01_0513700","PVP01_0513800","PVP01_0513900","PVP01_0514100","PVP01_0514200","PVP01_0514300","PVP01_0514400","PVP01_0514500","PVP01_0514600","PVP01_0514700","PVP01_0514800","PVP01_0515000","PVP01_0515200","PVP01_0515400","PVP01_0515500","PVP01_0515600","PVP01_0515700","PVP01_0515900","PVP01_0516100","PVP01_0516200","PVP01_0516300","PVP01_0516600","PVP01_0516700","PVP01_0517100","PVP01_0517200","PVP01_0517300","PVP01_0517500","PVP01_0517600","PVP01_0517800","PVP01_0517900","PVP01_0518000","PVP01_0518100","PVP01_0518200","PVP01_0518300","PVP01_0518400","PVP01_0518500","PVP01_0519000","PVP01_0519100","PVP01_0519500","PVP01_0519600","PVP01_0519700","PVP01_0519800","PVP01_0519900","PVP01_0520000","PVP01_0520100","PVP01_0520200","PVP01_0520300","PVP01_0520400","PVP01_0520500","PVP01_0520600","PVP01_0520700","PVP01_0520900","PVP01_0521000","PVP01_0521100","PVP01_0521200","PVP01_0521300","PVP01_0521400","PVP01_0521800","PVP01_0521900","PVP01_0522200","PVP01_0522400","PVP01_0522500","PVP01_0522700","PVP01_0522800","PVP01_0523000","PVP01_0523100","PVP01_0523200","PVP01_0523300","PVP01_0523400","PVP01_0523500","PVP01_0523700","PVP01_0523800","PVP01_0523900","PVP01_0524000","PVP01_0524100","PVP01_0524200","PVP01_0524300","PVP01_0524400","PVP01_0524600","PVP01_0524700","PVP01_0525100","PVP01_0525200","PVP01_0525400","PVP01_0525500","PVP01_0525600","PVP01_0525800","PVP01_0525900","PVP01_0526400","PVP01_0526500","PVP01_0526600","PVP01_0526700","PVP01_0526800","PVP01_0526900","PVP01_0527000","PVP01_0527100","PVP01_0527200","PVP01_0527500","PVP01_0527600","PVP01_0528000","PVP01_0528300","PVP01_0528600","PVP01_0528900","PVP01_0529100","PVP01_0529200","PVP01_0529300","PVP01_0529400","PVP01_0529500","PVP01_0529800","PVP01_0529900","PVP01_0530000","PVP01_0530200","PVP01_0530300","PVP01_0530600","PVP01_0530800","PVP01_0530900","PVP01_0531200","PVP01_0531300","PVP01_0531400","PVP01_0531600","PVP01_0531800","PVP01_0532100","PVP01_0532200","PVP01_0532300","PVP01_0532400","PVP01_0532600","PVP01_0532700","PVP01_0532800","PVP01_0532900","PVP01_0533000","PVP01_0533100","PVP01_0533200","PVP01_0533300","PVP01_0533400","PVP01_0533700","PVP01_0533800","PVP01_0534000","PVP01_0534200","PVP01_0534300","PVP01_0534400","PVP01_0534500","PVP01_0534600","PVP01_0534800","PVP01_0600400","PVP01_0600800","PVP01_0600900","PVP01_0601000","PVP01_0601200","PVP01_0601500","PVP01_0601800","PVP01_0601900","PVP01_0602000","PVP01_0602100","PVP01_0602200","PVP01_0602300","PVP01_0602400","PVP01_0602500","PVP01_0602700","PVP01_0602800","PVP01_0602900","PVP01_0603200","PVP01_0603300","PVP01_0603400","PVP01_0603500","PVP01_0603800","PVP01_0604200","PVP01_0604300","PVP01_0604400","PVP01_0604500","PVP01_0604600","PVP01_0604800","PVP01_0604900","PVP01_0605100","PVP01_0605200","PVP01_0605400","PVP01_0605600","PVP01_0605700","PVP01_0605800","PVP01_0606000","PVP01_0606300","PVP01_0606400","PVP01_0606500","PVP01_0606600","PVP01_0606700","PVP01_0606800","PVP01_0607000","PVP01_0607100","PVP01_0607200","PVP01_0607300","PVP01_0607400","PVP01_0607500","PVP01_0607700","PVP01_0607800","PVP01_0607900","PVP01_0608200","PVP01_0608300","PVP01_0608400","PVP01_0608500","PVP01_0608800","PVP01_0609000","PVP01_0609200","PVP01_0609300","PVP01_0609400","PVP01_0609500","PVP01_0610000","PVP01_0610100","PVP01_0610600","PVP01_0610800","PVP01_0611000","PVP01_0611300","PVP01_0611400","PVP01_0611500","PVP01_0611600","PVP01_0611700","PVP01_0611800","PVP01_0611900","PVP01_0612200","PVP01_0612300","PVP01_0612500","PVP01_0612600","PVP01_0612800","PVP01_0612900","PVP01_0613300","PVP01_0613600","PVP01_0613700","PVP01_0613900","PVP01_0614000","PVP01_0614100","PVP01_0614200","PVP01_0614500","PVP01_0614800","PVP01_0615200","PVP01_0615300","PVP01_0615400","PVP01_0615600","PVP01_0615700","PVP01_0615900","PVP01_0616000","PVP01_0616100","PVP01_0616500","PVP01_0616600","PVP01_0616700","PVP01_0616800","PVP01_0616900","PVP01_0617000","PVP01_0617100","PVP01_0617200","PVP01_0617300","PVP01_0617400","PVP01_0617600","PVP01_0617700","PVP01_0617800","PVP01_0617900","PVP01_0618000","PVP01_0618100","PVP01_0618200","PVP01_0618300","PVP01_0618400","PVP01_0618500","PVP01_0618800","PVP01_0619000","PVP01_0619200","PVP01_0619300","PVP01_0619400","PVP01_0619500","PVP01_0619600","PVP01_0619800","PVP01_0619900","PVP01_0620200","PVP01_0620400","PVP01_0620500","PVP01_0620600","PVP01_0620800","PVP01_0621000","PVP01_0621200","PVP01_0621400","PVP01_0621500","PVP01_0621600","PVP01_0621700","PVP01_0621800","PVP01_0622000","PVP01_0622300","PVP01_0622400","PVP01_0622500","PVP01_0622600","PVP01_0622700","PVP01_0622800","PVP01_0623000","PVP01_0623100","PVP01_0623300","PVP01_0623400","PVP01_0623500","PVP01_0623600","PVP01_0623700","PVP01_0623800","PVP01_0623900","PVP01_0624000","PVP01_0624100","PVP01_0624300","PVP01_0624400","PVP01_0624500","PVP01_0624700","PVP01_0624900","PVP01_0625000","PVP01_0700100","PVP01_0700300","PVP01_0700600","PVP01_0700800","PVP01_0700900","PVP01_0701000","PVP01_0701100","PVP01_0701300","PVP01_0701700","PVP01_0701800","PVP01_0701900","PVP01_0702000","PVP01_0702200","PVP01_0702300","PVP01_0702600","PVP01_0702700","PVP01_0702800","PVP01_0702900","PVP01_0703000","PVP01_0703100","PVP01_0703200","PVP01_0703300","PVP01_0703400","PVP01_0703500","PVP01_0703600","PVP01_0703700","PVP01_0703800","PVP01_0703900","PVP01_0704100","PVP01_0704200","PVP01_0704300","PVP01_0704400","PVP01_0704500","PVP01_0704600","PVP01_0704800","PVP01_0704900","PVP01_0705100","PVP01_0705200","PVP01_0705300","PVP01_0705400","PVP01_0705500","PVP01_0705900","PVP01_0706000","PVP01_0706300","PVP01_0706500","PVP01_0706600","PVP01_0706800","PVP01_0707100","PVP01_0707200","PVP01_0707300","PVP01_0707400","PVP01_0707500","PVP01_0707600","PVP01_0707700","PVP01_0708000","PVP01_0708100","PVP01_0708300","PVP01_0708500","PVP01_0708600","PVP01_0708800","PVP01_0709000","PVP01_0709400","PVP01_0709500","PVP01_0709700","PVP01_0709800","PVP01_0709900","PVP01_0710100","PVP01_0710300","PVP01_0710400","PVP01_0710500","PVP01_0710600","PVP01_0710700","PVP01_0710800","PVP01_0710900","PVP01_0711200","PVP01_0711300","PVP01_0711400","PVP01_0711600","PVP01_0711700","PVP01_0711900","PVP01_0712000","PVP01_0712100","PVP01_0712200","PVP01_0712500","PVP01_0712600","PVP01_0712700","PVP01_0712800","PVP01_0712900","PVP01_0713000","PVP01_0713200","PVP01_0713300","PVP01_0713400","PVP01_0713700","PVP01_0714000","PVP01_0714100","PVP01_0714300","PVP01_0714400","PVP01_0714500","PVP01_0714600","PVP01_0714700","PVP01_0714800","PVP01_0715000","PVP01_0715100","PVP01_0715300","PVP01_0715400","PVP01_0715500","PVP01_0715600","PVP01_0715900","PVP01_0716000","PVP01_0716300","PVP01_0716500","PVP01_0716600","PVP01_0716700","PVP01_0716900","PVP01_0717100","PVP01_0717300","PVP01_0717400","PVP01_0717700","PVP01_0717900","PVP01_0718000","PVP01_0718100","PVP01_0718200","PVP01_0718300","PVP01_0718400","PVP01_0718500","PVP01_0718800","PVP01_0718900","PVP01_0719200","PVP01_0719300","PVP01_0719400","PVP01_0719600","PVP01_0719800","PVP01_0720000","PVP01_0720200","PVP01_0720500","PVP01_0720700","PVP01_0720800","PVP01_0721000","PVP01_0721100","PVP01_0721200","PVP01_0721400","PVP01_0721500","PVP01_0721600","PVP01_0721700","PVP01_0722000","PVP01_0722100","PVP01_0722200","PVP01_0722300","PVP01_0722600","PVP01_0722700","PVP01_0722800","PVP01_0722900","PVP01_0723100","PVP01_0723300","PVP01_0723400","PVP01_0723500","PVP01_0723600","PVP01_0723700","PVP01_0723800","PVP01_0723900","PVP01_0724000","PVP01_0724100","PVP01_0724200","PVP01_0724500","PVP01_0724600","PVP01_0724800","PVP01_0724900","PVP01_0725000","PVP01_0725200","PVP01_0725500","PVP01_0725600","PVP01_0725700","PVP01_0725800","PVP01_0725900","PVP01_0726100","PVP01_0726200","PVP01_0726300","PVP01_0726500","PVP01_0726600","PVP01_0726700","PVP01_0726800","PVP01_0726900","PVP01_0727000","PVP01_0727200","PVP01_0727500","PVP01_0727600","PVP01_0727800","PVP01_0727900","PVP01_0728200","PVP01_0728300","PVP01_0728600","PVP01_0728700","PVP01_0728800","PVP01_0728900","PVP01_0729000","PVP01_0729200","PVP01_0729300","PVP01_0729600","PVP01_0729800","PVP01_0729900","PVP01_0730100","PVP01_0730200","PVP01_0730500","PVP01_0730900","PVP01_0731400","PVP01_0731600","PVP01_0731800","PVP01_0731900","PVP01_0732000","PVP01_0732100","PVP01_0732200","PVP01_0732400","PVP01_0732500","PVP01_0732700","PVP01_0733100","PVP01_0733200","PVP01_0733300","PVP01_0733400","PVP01_0733500","PVP01_0733600","PVP01_0733700","PVP01_0733800","PVP01_0733900","PVP01_0734000","PVP01_0734300","PVP01_0734400","PVP01_0734500","PVP01_0734600","PVP01_0734700","PVP01_0734900","PVP01_0735000","PVP01_0735200","PVP01_0735500","PVP01_0736200","PVP01_0800400","PVP01_0800600","PVP01_0800700","PVP01_0800900","PVP01_0801000","PVP01_0801100","PVP01_0801200","PVP01_0801300","PVP01_0801400","PVP01_0801500","PVP01_0801600","PVP01_0801700","PVP01_0801900","PVP01_0802100","PVP01_0802200","PVP01_0802300","PVP01_0802400","PVP01_0802500","PVP01_0802600","PVP01_0802700","PVP01_0802800","PVP01_0802900","PVP01_0803000","PVP01_0803100","PVP01_0803200","PVP01_0803300","PVP01_0803400","PVP01_0803600","PVP01_0803700","PVP01_0803800","PVP01_0803900","PVP01_0804000","PVP01_0804200","PVP01_0804400","PVP01_0804500","PVP01_0804700","PVP01_0805000","PVP01_0805100","PVP01_0805200","PVP01_0805700","PVP01_0805900","PVP01_0806000","PVP01_0806200","PVP01_0806400","PVP01_0806600","PVP01_0806800","PVP01_0806900","PVP01_0807100","PVP01_0807200","PVP01_0807300","PVP01_0807400","PVP01_0807600","PVP01_0807700","PVP01_0807900","PVP01_0808000","PVP01_0808200","PVP01_0808300","PVP01_0808400","PVP01_0808500","PVP01_0808600","PVP01_0808900","PVP01_0809000","PVP01_0809100","PVP01_0809300","PVP01_0809500","PVP01_0809800","PVP01_0810000","PVP01_0810100","PVP01_0810200","PVP01_0810300","PVP01_0810400","PVP01_0810600","PVP01_0810700","PVP01_0810800","PVP01_0810900","PVP01_0811000","PVP01_0811100","PVP01_0811400","PVP01_0811500","PVP01_0811600","PVP01_0811700","PVP01_0811800","PVP01_0811900","PVP01_0812000","PVP01_0812100","PVP01_0812200","PVP01_0812400","PVP01_0812500","PVP01_0812600","PVP01_0812700","PVP01_0812800","PVP01_0812900","PVP01_0813000","PVP01_0813100","PVP01_0813400","PVP01_0813500","PVP01_0813600","PVP01_0813900","PVP01_0814000","PVP01_0814300","PVP01_0814500","PVP01_0814600","PVP01_0814800","PVP01_0814900","PVP01_0815200","PVP01_0815400","PVP01_0815500","PVP01_0815600","PVP01_0815700","PVP01_0815800","PVP01_0816000","PVP01_0816100","PVP01_0816300","PVP01_0816400","PVP01_0816500","PVP01_0816600","PVP01_0816900","PVP01_0817200","PVP01_0817300","PVP01_0817500","PVP01_0817600","PVP01_0817700","PVP01_0817800","PVP01_0817900","PVP01_0818000","PVP01_0818200","PVP01_0818300","PVP01_0818400","PVP01_0818700","PVP01_0818800","PVP01_0819000","PVP01_0819100","PVP01_0819400","PVP01_0819500","PVP01_0819600","PVP01_0819700","PVP01_0819800","PVP01_0819900","PVP01_0820000","PVP01_0820100","PVP01_0820300","PVP01_0820400","PVP01_0820700","PVP01_0820900","PVP01_0821000","PVP01_0821100","PVP01_0821200","PVP01_0821500","PVP01_0821600","PVP01_0821800","PVP01_0821900","PVP01_0822000","PVP01_0822100","PVP01_0822200","PVP01_0822500","PVP01_0822700","PVP01_0822800","PVP01_0823000","PVP01_0823200","PVP01_0823300","PVP01_0823400","PVP01_0823500","PVP01_0823700","PVP01_0824200","PVP01_0824300","PVP01_0824400","PVP01_0824500","PVP01_0824600","PVP01_0824700","PVP01_0824900","PVP01_0825000","PVP01_0825200","PVP01_0825400","PVP01_0825500","PVP01_0825600","PVP01_0825700","PVP01_0825900","PVP01_0826100","PVP01_0826400","PVP01_0826500","PVP01_0826600","PVP01_0826800","PVP01_0826900","PVP01_0827000","PVP01_0827100","PVP01_0827200","PVP01_0827300","PVP01_0827400","PVP01_0827700","PVP01_0827900","PVP01_0828000","PVP01_0828400","PVP01_0828600","PVP01_0828900","PVP01_0829200","PVP01_0829300","PVP01_0829400","PVP01_0829500","PVP01_0829600","PVP01_0829700","PVP01_0829800","PVP01_0829900","PVP01_0830000","PVP01_0830100","PVP01_0830200","PVP01_0830400","PVP01_0830600","PVP01_0830700","PVP01_0830900","PVP01_0831200","PVP01_0831400","PVP01_0831500","PVP01_0831600","PVP01_0831800","PVP01_0831900","PVP01_0832400","PVP01_0832500","PVP01_0832600","PVP01_0832700","PVP01_0832800","PVP01_0833100","PVP01_0833200","PVP01_0833300","PVP01_0833400","PVP01_0833500","PVP01_0833600","PVP01_0833800","PVP01_0833900","PVP01_0834000","PVP01_0834100","PVP01_0834200","PVP01_0834500","PVP01_0834600","PVP01_0834800","PVP01_0834900","PVP01_0835200","PVP01_0835400","PVP01_0835500","PVP01_0835600","PVP01_0835700","PVP01_0835800","PVP01_0836000","PVP01_0836200","PVP01_0836300","PVP01_0836500","PVP01_0836600","PVP01_0836900","PVP01_0837000","PVP01_0837100","PVP01_0837400","PVP01_0837500","PVP01_0837700","PVP01_0837800","PVP01_0837900","PVP01_0838100","PVP01_0838200","PVP01_0838300","PVP01_0838500","PVP01_0838600","PVP01_0839300","PVP01_0839600","PVP01_0900600","PVP01_0900700","PVP01_0901400","PVP01_0901800","PVP01_0901900","PVP01_0902000","PVP01_0903000","PVP01_0903400","PVP01_0903600","PVP01_0903800","PVP01_0903900","PVP01_0904200","PVP01_0904300","PVP01_0904400","PVP01_0904500","PVP01_0904600","PVP01_0904800","PVP01_0905000","PVP01_0905100","PVP01_0905200","PVP01_0905300","PVP01_0905400","PVP01_0905500","PVP01_0905600","PVP01_0905700","PVP01_0905800","PVP01_0905900","PVP01_0906000","PVP01_0906100","PVP01_0906200","PVP01_0906300","PVP01_0906600","PVP01_0906700","PVP01_0907100","PVP01_0907300","PVP01_0907400","PVP01_0907600","PVP01_0907700","PVP01_0907800","PVP01_0907900","PVP01_0908100","PVP01_0908200","PVP01_0908300","PVP01_0908500","PVP01_0908800","PVP01_0909000","PVP01_0909200","PVP01_0909300","PVP01_0909400","PVP01_0910500","PVP01_0910700","PVP01_0910800","PVP01_0911000","PVP01_0911100","PVP01_0911200","PVP01_0911400","PVP01_0911500","PVP01_0912100","PVP01_0912500","PVP01_0912600","PVP01_0912700","PVP01_0912800","PVP01_0913100","PVP01_0913200","PVP01_0913300","PVP01_0913400","PVP01_0913500","PVP01_0913600","PVP01_0913700","PVP01_0913900","PVP01_0914000","PVP01_0914200","PVP01_0914300","PVP01_0914400","PVP01_0914500","PVP01_0914600","PVP01_0914700","PVP01_0914800","PVP01_0914900","PVP01_0915000","PVP01_0915100","PVP01_0915200","PVP01_0915300","PVP01_0915400","PVP01_0915500","PVP01_0915600","PVP01_0915700","PVP01_0915800","PVP01_0915900","PVP01_0916100","PVP01_0916200","PVP01_0916300","PVP01_0916400","PVP01_0916500","PVP01_0916600","PVP01_0916800","PVP01_0916900","PVP01_0917000","PVP01_0917300","PVP01_0917500","PVP01_0917700","PVP01_0917800","PVP01_0918400","PVP01_0918500","PVP01_0918600","PVP01_0918800","PVP01_0918900","PVP01_0919100","PVP01_0919200","PVP01_0919300","PVP01_0919600","PVP01_0919800","PVP01_0919900","PVP01_0920300","PVP01_0920600","PVP01_0920800","PVP01_0920900","PVP01_0921000","PVP01_0921100","PVP01_0921300","PVP01_0921400","PVP01_0921500","PVP01_0921600","PVP01_0921800","PVP01_0921900","PVP01_0922200","PVP01_0922300","PVP01_0922400","PVP01_0922500","PVP01_0922600","PVP01_0922700","PVP01_0923200","PVP01_0923300","PVP01_0923400","PVP01_0923500","PVP01_0923600","PVP01_0924200","PVP01_0924600","PVP01_0924700","PVP01_0924800","PVP01_0924900","PVP01_0925000","PVP01_0925100","PVP01_0925300","PVP01_0925400","PVP01_0925500","PVP01_0925600","PVP01_0925900","PVP01_0926000","PVP01_0926100","PVP01_0926200","PVP01_0926400","PVP01_0926500","PVP01_0926700","PVP01_0926800","PVP01_0927000","PVP01_0927500","PVP01_0927800","PVP01_0927900","PVP01_0928400","PVP01_0928500","PVP01_0928600","PVP01_0928700","PVP01_0928800","PVP01_0929100","PVP01_0929300","PVP01_0929400","PVP01_0929500","PVP01_0929800","PVP01_0930100","PVP01_0930200","PVP01_0930300","PVP01_0930400","PVP01_0930500","PVP01_0930700","PVP01_0931100","PVP01_0931200","PVP01_0931400","PVP01_0931500","PVP01_0931600","PVP01_0931700","PVP01_0931800","PVP01_0931900","PVP01_0932000","PVP01_0932300","PVP01_0932500","PVP01_0932700","PVP01_0932800","PVP01_0932900","PVP01_0933000","PVP01_0933200","PVP01_0933300","PVP01_0933600","PVP01_0933900","PVP01_0934000","PVP01_0934200","PVP01_0934600","PVP01_0934700","PVP01_0934800","PVP01_0934900","PVP01_0935000","PVP01_0935100","PVP01_0935300","PVP01_0935400","PVP01_0935500","PVP01_0935600","PVP01_0935800","PVP01_0936100","PVP01_0936200","PVP01_0936300","PVP01_0936600","PVP01_0936700","PVP01_0936800","PVP01_0937000","PVP01_0937200","PVP01_0937400","PVP01_0937500","PVP01_0937600","PVP01_0937800","PVP01_0937900","PVP01_0938100","PVP01_0938200","PVP01_0938300","PVP01_0938500","PVP01_0938800","PVP01_0939100","PVP01_0939300","PVP01_0939400","PVP01_0939500","PVP01_0939600","PVP01_0939700","PVP01_0939800","PVP01_0939900","PVP01_0940000","PVP01_0940100","PVP01_0940200","PVP01_0940300","PVP01_0940400","PVP01_0940500","PVP01_0940800","PVP01_0941000","PVP01_0941300","PVP01_0941500","PVP01_0941800","PVP01_0942000","PVP01_0942100","PVP01_0942300","PVP01_0942500","PVP01_0942600","PVP01_0942700","PVP01_0942800","PVP01_0942900","PVP01_0943000","PVP01_0943100","PVP01_0943200","PVP01_0943300","PVP01_0943400","PVP01_0943500","PVP01_0943600","PVP01_0943700","PVP01_0943800","PVP01_0943900","PVP01_0944000","PVP01_0944200","PVP01_0944300","PVP01_0944500","PVP01_0944700","PVP01_0944900","PVP01_0945100","PVP01_0945300","PVP01_0945600","PVP01_0945700","PVP01_0945900","PVP01_0946000","PVP01_0946100","PVP01_0946500","PVP01_0946600","PVP01_0946800","PVP01_0946900","PVP01_0947000","PVP01_0947100","PVP01_0947400","PVP01_0947500","PVP01_0947600","PVP01_0947700","PVP01_0947800","PVP01_0947900","PVP01_0948000","PVP01_0948100","PVP01_0948200","PVP01_0948300","PVP01_0948400","PVP01_0948600","PVP01_0948700","PVP01_0948800","PVP01_0948900","PVP01_0949000","PVP01_0949100","PVP01_0949200","PVP01_0949400","PVP01_0949500","PVP01_0949600","PVP01_0949700","PVP01_0949800","PVP01_0950000","PVP01_0950200","PVP01_1002400","PVP01_1002500","PVP01_1002700","PVP01_1002800","PVP01_1002900","PVP01_1003000","PVP01_1003100","PVP01_1003200","PVP01_1003300","PVP01_1003400","PVP01_1003500","PVP01_1003600","PVP01_1003700","PVP01_1003800","PVP01_1003900","PVP01_1004000","PVP01_1004100","PVP01_1004400","PVP01_1004500","PVP01_1004700","PVP01_1004800","PVP01_1004900","PVP01_1005000","PVP01_1005100","PVP01_1005200","PVP01_1005400","PVP01_1005500","PVP01_1005600","PVP01_1005700","PVP01_1005800","PVP01_1006200","PVP01_1006300","PVP01_1006500","PVP01_1006700","PVP01_1006800","PVP01_1007000","PVP01_1007200","PVP01_1007300","PVP01_1007400","PVP01_1008000","PVP01_1008200","PVP01_1008500","PVP01_1008700","PVP01_1008800","PVP01_1009100","PVP01_1009300","PVP01_1009400","PVP01_1009500","PVP01_1009700","PVP01_1009800","PVP01_1009900","PVP01_1010000","PVP01_1010300","PVP01_1010500","PVP01_1010600","PVP01_1010700","PVP01_1010800","PVP01_1010900","PVP01_1011000","PVP01_1011600","PVP01_1011700","PVP01_1011900","PVP01_1012000","PVP01_1012200","PVP01_1012300","PVP01_1012600","PVP01_1012800","PVP01_1012900","PVP01_1013000","PVP01_1013100","PVP01_1013300","PVP01_1013400","PVP01_1013500","PVP01_1013600","PVP01_1013700","PVP01_1013800","PVP01_1013900","PVP01_1014000","PVP01_1014100","PVP01_1014200","PVP01_1014500","PVP01_1014700","PVP01_1014800","PVP01_1014900","PVP01_1015100","PVP01_1015200","PVP01_1015300","PVP01_1015400","PVP01_1015800","PVP01_1015900","PVP01_1016000","PVP01_1016300","PVP01_1016400","PVP01_1016500","PVP01_1016700","PVP01_1016800","PVP01_1017000","PVP01_1017100","PVP01_1017200","PVP01_1017300","PVP01_1017600","PVP01_1017700","PVP01_1017800","PVP01_1018000","PVP01_1018100","PVP01_1018200","PVP01_1018300","PVP01_1018500","PVP01_1018600","PVP01_1018700","PVP01_1018800","PVP01_1018900","PVP01_1019100","PVP01_1019200","PVP01_1019300","PVP01_1019400","PVP01_1019700","PVP01_1019800","PVP01_1019900","PVP01_1020200","PVP01_1020300","PVP01_1020400","PVP01_1020500","PVP01_1020600","PVP01_1020800","PVP01_1021100","PVP01_1021400","PVP01_1021700","PVP01_1022100","PVP01_1022300","PVP01_1022600","PVP01_1022700","PVP01_1022900","PVP01_1023100","PVP01_1023200","PVP01_1023300","PVP01_1023500","PVP01_1024000","PVP01_1024100","PVP01_1024200","PVP01_1024300","PVP01_1024700","PVP01_1025000","PVP01_1025300","PVP01_1025400","PVP01_1025600","PVP01_1025700","PVP01_1025800","PVP01_1025900","PVP01_1026000","PVP01_1026100","PVP01_1026500","PVP01_1026800","PVP01_1026900","PVP01_1027000","PVP01_1027200","PVP01_1027400","PVP01_1027500","PVP01_1027700","PVP01_1027900","PVP01_1028000","PVP01_1028300","PVP01_1028400","PVP01_1028500","PVP01_1028700","PVP01_1028800","PVP01_1029200","PVP01_1029300","PVP01_1029400","PVP01_1029500","PVP01_1029600","PVP01_1029800","PVP01_1029900","PVP01_1030000","PVP01_1030100","PVP01_1030200","PVP01_1030300","PVP01_1030400","PVP01_1030700","PVP01_1030800","PVP01_1030900","PVP01_1031200","PVP01_1031600","PVP01_1031700","PVP01_1032000","PVP01_1032100","PVP01_1032200","PVP01_1032300","PVP01_1032400","PVP01_1032500","PVP01_1032600","PVP01_1032700","PVP01_1032800","PVP01_1033100","PVP01_1033200","PVP01_1033300","PVP01_1033400","PVP01_1033500","PVP01_1033600","PVP01_1033700","PVP01_1033800","PVP01_1033900","PVP01_1034000","PVP01_1034100","PVP01_1034200","PVP01_1034300","PVP01_1034400","PVP01_1034500","PVP01_1100300","PVP01_1100400","PVP01_1100500","PVP01_1101300","PVP01_1101400","PVP01_1101500","PVP01_1101600","PVP01_1101700","PVP01_1102300","PVP01_1102600","PVP01_1102800","PVP01_1102900","PVP01_1103100","PVP01_1103500","PVP01_1103600","PVP01_1103800","PVP01_1103900","PVP01_1104100","PVP01_1104200","PVP01_1104500","PVP01_1104700","PVP01_1104900","PVP01_1105000","PVP01_1105100","PVP01_1105200","PVP01_1105300","PVP01_1105400","PVP01_1105600","PVP01_1105800","PVP01_1105900","PVP01_1106100","PVP01_1106200","PVP01_1106300","PVP01_1106400","PVP01_1106500","PVP01_1106600","PVP01_1106700","PVP01_1107200","PVP01_1107300","PVP01_1107500","PVP01_1107700","PVP01_1107800","PVP01_1108000","PVP01_1108100","PVP01_1108200","PVP01_1108300","PVP01_1108400","PVP01_1108700","PVP01_1109000","PVP01_1109100","PVP01_1109200","PVP01_1109300","PVP01_1109700","PVP01_1109800","PVP01_1109900","PVP01_1110000","PVP01_1110100","PVP01_1110200","PVP01_1110300","PVP01_1110400","PVP01_1110700","PVP01_1110800","PVP01_1110900","PVP01_1111200","PVP01_1111400","PVP01_1111500","PVP01_1111600","PVP01_1111700","PVP01_1111800","PVP01_1111900","PVP01_1112000","PVP01_1112200","PVP01_1112300","PVP01_1112400","PVP01_1112500","PVP01_1112600","PVP01_1112700","PVP01_1112800","PVP01_1113000","PVP01_1113100","PVP01_1113200","PVP01_1113300","PVP01_1113400","PVP01_1113800","PVP01_1114000","PVP01_1114100","PVP01_1114400","PVP01_1114600","PVP01_1114700","PVP01_1114800","PVP01_1114900","PVP01_1115000","PVP01_1115100","PVP01_1115200","PVP01_1115300","PVP01_1115400","PVP01_1115700","PVP01_1115800","PVP01_1116000","PVP01_1116100","PVP01_1116200","PVP01_1116300","PVP01_1116400","PVP01_1116700","PVP01_1116800","PVP01_1116900","PVP01_1117000","PVP01_1117100","PVP01_1117200","PVP01_1117300","PVP01_1117400","PVP01_1117500","PVP01_1117600","PVP01_1117800","PVP01_1117900","PVP01_1118100","PVP01_1118200","PVP01_1118300","PVP01_1118400","PVP01_1118900","PVP01_1119100","PVP01_1119200","PVP01_1119300","PVP01_1119400","PVP01_1119700","PVP01_1119900","PVP01_1120100","PVP01_1120200","PVP01_1120400","PVP01_1120600","PVP01_1120700","PVP01_1120800","PVP01_1120900","PVP01_1121000","PVP01_1121200","PVP01_1121300","PVP01_1121600","PVP01_1121800","PVP01_1122000","PVP01_1122200","PVP01_1122800","PVP01_1122900","PVP01_1123100","PVP01_1123200","PVP01_1123300","PVP01_1123400","PVP01_1123700","PVP01_1123800","PVP01_1124200","PVP01_1124500","PVP01_1124600","PVP01_1124700","PVP01_1125200","PVP01_1125300","PVP01_1125500","PVP01_1125600","PVP01_1125700","PVP01_1125800","PVP01_1126000","PVP01_1126100","PVP01_1126300","PVP01_1126400","PVP01_1126500","PVP01_1126600","PVP01_1126700","PVP01_1126800","PVP01_1126900","PVP01_1127000","PVP01_1127500","PVP01_1127600","PVP01_1127700","PVP01_1128000","PVP01_1128100","PVP01_1128200","PVP01_1128300","PVP01_1128500","PVP01_1128800","PVP01_1128900","PVP01_1129100","PVP01_1129200","PVP01_1129300","PVP01_1129500","PVP01_1129600","PVP01_1129800","PVP01_1129900","PVP01_1130000","PVP01_1130100","PVP01_1130300","PVP01_1130400","PVP01_1130500","PVP01_1130600","PVP01_1130700","PVP01_1130800","PVP01_1131100","PVP01_1131200","PVP01_1131300","PVP01_1131400","PVP01_1131500","PVP01_1131600","PVP01_1131700","PVP01_1131800","PVP01_1131900","PVP01_1132000","PVP01_1132100","PVP01_1132200","PVP01_1132500","PVP01_1132600","PVP01_1133000","PVP01_1133300","PVP01_1133700","PVP01_1134100","PVP01_1134200","PVP01_1134400","PVP01_1134600","PVP01_1134700","PVP01_1134800","PVP01_1134900","PVP01_1135000","PVP01_1135100","PVP01_1135200","PVP01_1135300","PVP01_1135400","PVP01_1135600","PVP01_1135700","PVP01_1136000","PVP01_1136200","PVP01_1136300","PVP01_1136400","PVP01_1136500","PVP01_1136600","PVP01_1137000","PVP01_1137300","PVP01_1137400","PVP01_1137500","PVP01_1137900","PVP01_1138000","PVP01_1138100","PVP01_1138200","PVP01_1138300","PVP01_1138600","PVP01_1138700","PVP01_1138800","PVP01_1138900","PVP01_1139000","PVP01_1139100","PVP01_1139200","PVP01_1139300","PVP01_1139400","PVP01_1139500","PVP01_1139600","PVP01_1139700","PVP01_1139800","PVP01_1140000","PVP01_1140100","PVP01_1140200","PVP01_1140400","PVP01_1140600","PVP01_1140800","PVP01_1140900","PVP01_1141000","PVP01_1141100","PVP01_1141200","PVP01_1141400","PVP01_1141600","PVP01_1141700","PVP01_1141800","PVP01_1142100","PVP01_1142200","PVP01_1142300","PVP01_1142400","PVP01_1142500","PVP01_1142800","PVP01_1142900","PVP01_1143000","PVP01_1143100","PVP01_1143300","PVP01_1143600","PVP01_1143700","PVP01_1143800","PVP01_1143900","PVP01_1144000","PVP01_1144200","PVP01_1144400","PVP01_1144600","PVP01_1144700","PVP01_1144900","PVP01_1145000","PVP01_1145100","PVP01_1145200","PVP01_1145300","PVP01_1145400","PVP01_1145600","PVP01_1145800","PVP01_1145900","PVP01_1146000","PVP01_1146100","PVP01_1146200","PVP01_1146300","PVP01_1146600","PVP01_1146700","PVP01_1146800","PVP01_1147000","PVP01_1147100","PVP01_1147200","PVP01_1147300","PVP01_1147400","PVP01_1147500","PVP01_1147600","PVP01_1147700","PVP01_1147800","PVP01_1148400","PVP01_1148600","PVP01_1149100","PVP01_1149200","PVP01_1149500","PVP01_1200300","PVP01_1201000","PVP01_1201300","PVP01_1201400","PVP01_1201500","PVP01_1201600","PVP01_1201700","PVP01_1201800","PVP01_1202000","PVP01_1202300","PVP01_1202400","PVP01_1202500","PVP01_1202600","PVP01_1202700","PVP01_1202800","PVP01_1202900","PVP01_1203000","PVP01_1203200","PVP01_1203400","PVP01_1203700","PVP01_1203800","PVP01_1203900","PVP01_1204300","PVP01_1204400","PVP01_1204500","PVP01_1204700","PVP01_1204800","PVP01_1204900","PVP01_1205100","PVP01_1205200","PVP01_1205300","PVP01_1205400","PVP01_1205600","PVP01_1205700","PVP01_1205800","PVP01_1205900","PVP01_1206000","PVP01_1206200","PVP01_1206400","PVP01_1206500","PVP01_1206600","PVP01_1206800","PVP01_1207100","PVP01_1207200","PVP01_1207300","PVP01_1207400","PVP01_1207500","PVP01_1207600","PVP01_1207700","PVP01_1207800","PVP01_1207900","PVP01_1208100","PVP01_1208300","PVP01_1208500","PVP01_1208800","PVP01_1208900","PVP01_1209100","PVP01_1209300","PVP01_1209400","PVP01_1209500","PVP01_1209600","PVP01_1209700","PVP01_1209800","PVP01_1210400","PVP01_1210500","PVP01_1210600","PVP01_1210900","PVP01_1211000","PVP01_1211400","PVP01_1211500","PVP01_1211700","PVP01_1211800","PVP01_1211900","PVP01_1212000","PVP01_1212200","PVP01_1212300","PVP01_1212400","PVP01_1212500","PVP01_1212600","PVP01_1212700","PVP01_1212800","PVP01_1212900","PVP01_1213000","PVP01_1213100","PVP01_1213200","PVP01_1213300","PVP01_1213600","PVP01_1213900","PVP01_1214100","PVP01_1214500","PVP01_1214800","PVP01_1214900","PVP01_1215000","PVP01_1215100","PVP01_1215200","PVP01_1215300","PVP01_1215400","PVP01_1215500","PVP01_1215800","PVP01_1215900","PVP01_1216000","PVP01_1216300","PVP01_1216400","PVP01_1216500","PVP01_1216600","PVP01_1216700","PVP01_1216900","PVP01_1217000","PVP01_1217300","PVP01_1217400","PVP01_1217800","PVP01_1217900","PVP01_1218000","PVP01_1218100","PVP01_1218200","PVP01_1218300","PVP01_1218400","PVP01_1218500","PVP01_1218600","PVP01_1218700","PVP01_1219200","PVP01_1219300","PVP01_1219500","PVP01_1219800","PVP01_1220000","PVP01_1220100","PVP01_1220400","PVP01_1220500","PVP01_1220600","PVP01_1220700","PVP01_1220800","PVP01_1220900","PVP01_1221000","PVP01_1221100","PVP01_1221200","PVP01_1221400","PVP01_1221500","PVP01_1221600","PVP01_1221700","PVP01_1222000","PVP01_1222100","PVP01_1222200","PVP01_1222300","PVP01_1222500","PVP01_1222600","PVP01_1222700","PVP01_1222800","PVP01_1222900","PVP01_1223000","PVP01_1223100","PVP01_1223500","PVP01_1223600","PVP01_1223700","PVP01_1224100","PVP01_1224200","PVP01_1224300","PVP01_1224700","PVP01_1224800","PVP01_1224900","PVP01_1225000","PVP01_1225100","PVP01_1225200","PVP01_1225300","PVP01_1225400","PVP01_1225600","PVP01_1225700","PVP01_1225800","PVP01_1226000","PVP01_1226200","PVP01_1226300","PVP01_1226400","PVP01_1226700","PVP01_1226900","PVP01_1227000","PVP01_1227100","PVP01_1227200","PVP01_1227300","PVP01_1227400","PVP01_1227700","PVP01_1227800","PVP01_1228500","PVP01_1228600","PVP01_1228700","PVP01_1228800","PVP01_1228900","PVP01_1229000","PVP01_1229100","PVP01_1229300","PVP01_1229400","PVP01_1229700","PVP01_1229800","PVP01_1230300","PVP01_1230400","PVP01_1230700","PVP01_1230800","PVP01_1231200","PVP01_1231300","PVP01_1231400","PVP01_1231500","PVP01_1231600","PVP01_1231700","PVP01_1232000","PVP01_1232100","PVP01_1232200","PVP01_1232300","PVP01_1232400","PVP01_1232500","PVP01_1232800","PVP01_1233000","PVP01_1233200","PVP01_1233300","PVP01_1233400","PVP01_1233500","PVP01_1233600","PVP01_1233700","PVP01_1233800","PVP01_1233900","PVP01_1234000","PVP01_1234100","PVP01_1234200","PVP01_1234300","PVP01_1234400","PVP01_1234500","PVP01_1234600","PVP01_1234700","PVP01_1234800","PVP01_1234900","PVP01_1235000","PVP01_1235100","PVP01_1235200","PVP01_1235300","PVP01_1235400","PVP01_1235600","PVP01_1235700","PVP01_1235800","PVP01_1235900","PVP01_1236200","PVP01_1236300","PVP01_1236400","PVP01_1236500","PVP01_1236600","PVP01_1236700","PVP01_1236900","PVP01_1237000","PVP01_1237300","PVP01_1237400","PVP01_1237500","PVP01_1237600","PVP01_1237800","PVP01_1238000","PVP01_1238100","PVP01_1238200","PVP01_1238500","PVP01_1238600","PVP01_1238700","PVP01_1238800","PVP01_1239000","PVP01_1239100","PVP01_1239200","PVP01_1239400","PVP01_1239600","PVP01_1239700","PVP01_1239800","PVP01_1239900","PVP01_1240200","PVP01_1240300","PVP01_1240400","PVP01_1240500","PVP01_1240600","PVP01_1240700","PVP01_1240800","PVP01_1240900","PVP01_1241000","PVP01_1241100","PVP01_1241200","PVP01_1241400","PVP01_1241500","PVP01_1241600","PVP01_1241700","PVP01_1241800","PVP01_1242000","PVP01_1242100","PVP01_1242200","PVP01_1242400","PVP01_1242500","PVP01_1242600","PVP01_1242700","PVP01_1242800","PVP01_1243000","PVP01_1243100","PVP01_1243200","PVP01_1243300","PVP01_1243400","PVP01_1243700","PVP01_1243800","PVP01_1244000","PVP01_1244100","PVP01_1244200","PVP01_1244300","PVP01_1244400","PVP01_1244500","PVP01_1244600","PVP01_1245000","PVP01_1245100","PVP01_1245200","PVP01_1245300","PVP01_1245400","PVP01_1245700","PVP01_1245900","PVP01_1246000","PVP01_1246100","PVP01_1246300","PVP01_1246400","PVP01_1246600","PVP01_1246700","PVP01_1246800","PVP01_1246900","PVP01_1247000","PVP01_1247100","PVP01_1247200","PVP01_1247400","PVP01_1247500","PVP01_1247600","PVP01_1247700","PVP01_1248000","PVP01_1248100","PVP01_1248200","PVP01_1248300","PVP01_1248400","PVP01_1248500","PVP01_1248600","PVP01_1249100","PVP01_1249300","PVP01_1249400","PVP01_1249600","PVP01_1249700","PVP01_1249800","PVP01_1249900","PVP01_1250000","PVP01_1250100","PVP01_1250300","PVP01_1250400","PVP01_1250500","PVP01_1250600","PVP01_1250800","PVP01_1251000","PVP01_1251100","PVP01_1251200","PVP01_1251300","PVP01_1251400","PVP01_1251500","PVP01_1251600","PVP01_1251700","PVP01_1251800","PVP01_1251900","PVP01_1252000","PVP01_1252100","PVP01_1252200","PVP01_1252300","PVP01_1252500","PVP01_1252700","PVP01_1252800","PVP01_1253100","PVP01_1253200","PVP01_1253300","PVP01_1253400","PVP01_1253500","PVP01_1253600","PVP01_1253800","PVP01_1253900","PVP01_1254000","PVP01_1254100","PVP01_1254300","PVP01_1254400","PVP01_1254500","PVP01_1254600","PVP01_1254700","PVP01_1254800","PVP01_1255000","PVP01_1255300","PVP01_1255400","PVP01_1255500","PVP01_1255600","PVP01_1255800","PVP01_1255900","PVP01_1256000","PVP01_1256100","PVP01_1256200","PVP01_1256300","PVP01_1256500","PVP01_1256700","PVP01_1257100","PVP01_1257200","PVP01_1257300","PVP01_1257400","PVP01_1257500","PVP01_1257700","PVP01_1257900","PVP01_1258000","PVP01_1258200","PVP01_1258400","PVP01_1258500","PVP01_1258600","PVP01_1258700","PVP01_1258800","PVP01_1259100","PVP01_1259400","PVP01_1259600","PVP01_1259700","PVP01_1259800","PVP01_1259900","PVP01_1260000","PVP01_1260100","PVP01_1260300","PVP01_1260400","PVP01_1260700","PVP01_1260900","PVP01_1261000","PVP01_1261100","PVP01_1261200","PVP01_1261300","PVP01_1261400","PVP01_1261500","PVP01_1261600","PVP01_1261700","PVP01_1261800","PVP01_1261900","PVP01_1262100","PVP01_1262200","PVP01_1262300","PVP01_1262400","PVP01_1262500","PVP01_1262600","PVP01_1262700","PVP01_1263000","PVP01_1263100","PVP01_1263200","PVP01_1263500","PVP01_1263600","PVP01_1263700","PVP01_1263800","PVP01_1263900","PVP01_1264100","PVP01_1264200","PVP01_1264400","PVP01_1264900","PVP01_1265100","PVP01_1265200","PVP01_1265300","PVP01_1265400","PVP01_1265500","PVP01_1265600","PVP01_1265800","PVP01_1265900","PVP01_1266000","PVP01_1266100","PVP01_1266200","PVP01_1266300","PVP01_1266500","PVP01_1266600","PVP01_1266700","PVP01_1266800","PVP01_1266900","PVP01_1267000","PVP01_1267500","PVP01_1267700","PVP01_1267800","PVP01_1267900","PVP01_1268100","PVP01_1268300","PVP01_1268400","PVP01_1268500","PVP01_1268900","PVP01_1269200","PVP01_1269300","PVP01_1269400","PVP01_1269500","PVP01_1269600","PVP01_1269700","PVP01_1269800","PVP01_1269900","PVP01_1270000","PVP01_1270100","PVP01_1270300","PVP01_1270400","PVP01_1270500","PVP01_1270700","PVP01_1270800","PVP01_1270900","PVP01_1271000","PVP01_1271200","PVP01_1271300","PVP01_1271400","PVP01_1271500","PVP01_1271600","PVP01_1271700","PVP01_1271800","PVP01_1271900","PVP01_1272100","PVP01_1272200","PVP01_1272300","PVP01_1272400","PVP01_1272500","PVP01_1272600","PVP01_1273100","PVP01_1300500","PVP01_1300600","PVP01_1300700","PVP01_1300800","PVP01_1300900","PVP01_1301000","PVP01_1301100","PVP01_1301300","PVP01_1301400","PVP01_1301800","PVP01_1301900","PVP01_1302000","PVP01_1302100","PVP01_1302200","PVP01_1302300","PVP01_1302400","PVP01_1302500","PVP01_1302600","PVP01_1302900","PVP01_1303000","PVP01_1303200","PVP01_1303300","PVP01_1303400","PVP01_1303500","PVP01_1303600","PVP01_1303700","PVP01_1303800","PVP01_1303900","PVP01_1304100","PVP01_1304300","PVP01_1304400","PVP01_1304500","PVP01_1304700","PVP01_1305000","PVP01_1305100","PVP01_1305200","PVP01_1305400","PVP01_1305500","PVP01_1305800","PVP01_1306200","PVP01_1306500","PVP01_1306600","PVP01_1306800","PVP01_1306900","PVP01_1307100","PVP01_1307400","PVP01_1307600","PVP01_1307700","PVP01_1307900","PVP01_1308000","PVP01_1308200","PVP01_1308400","PVP01_1308600","PVP01_1308700","PVP01_1308800","PVP01_1309200","PVP01_1309300","PVP01_1309400","PVP01_1309500","PVP01_1309600","PVP01_1309800","PVP01_1309900","PVP01_1310000","PVP01_1310100","PVP01_1310300","PVP01_1310400","PVP01_1310500","PVP01_1310600","PVP01_1310700","PVP01_1310800","PVP01_1311200","PVP01_1311600","PVP01_1311700","PVP01_1311800","PVP01_1311900","PVP01_1312000","PVP01_1312100","PVP01_1312500","PVP01_1312600","PVP01_1312700","PVP01_1312800","PVP01_1313000","PVP01_1313100","PVP01_1313200","PVP01_1313300","PVP01_1313500","PVP01_1313600","PVP01_1313700","PVP01_1313800","PVP01_1314000","PVP01_1314200","PVP01_1314400","PVP01_1314500","PVP01_1314700","PVP01_1314800","PVP01_1315100","PVP01_1315200","PVP01_1315300","PVP01_1315500","PVP01_1315600","PVP01_1315800","PVP01_1315900","PVP01_1316000","PVP01_1316200","PVP01_1316300","PVP01_1316400","PVP01_1316700","PVP01_1316800","PVP01_1317100","PVP01_1317300","PVP01_1317500","PVP01_1317700","PVP01_1318200","PVP01_1318300","PVP01_1318400","PVP01_1318600","PVP01_1318700","PVP01_1319000","PVP01_1319200","PVP01_1319300","PVP01_1319400","PVP01_1319500","PVP01_1319600","PVP01_1319800","PVP01_1319900","PVP01_1320000","PVP01_1320100","PVP01_1320200","PVP01_1320300","PVP01_1320400","PVP01_1320500","PVP01_1320600","PVP01_1320800","PVP01_1321000","PVP01_1321100","PVP01_1321200","PVP01_1321400","PVP01_1321500","PVP01_1321700","PVP01_1321900","PVP01_1322000","PVP01_1322100","PVP01_1322200","PVP01_1322500","PVP01_1322800","PVP01_1322900","PVP01_1323200","PVP01_1323300","PVP01_1323600","PVP01_1323800","PVP01_1323900","PVP01_1324000","PVP01_1324600","PVP01_1324900","PVP01_1325300","PVP01_1325400","PVP01_1325500","PVP01_1325600","PVP01_1325700","PVP01_1326100","PVP01_1326300","PVP01_1326500","PVP01_1326600","PVP01_1326700","PVP01_1326800","PVP01_1326900","PVP01_1327000","PVP01_1327100","PVP01_1327200","PVP01_1327300","PVP01_1327400","PVP01_1327500","PVP01_1327700","PVP01_1327800","PVP01_1328200","PVP01_1328300","PVP01_1328500","PVP01_1328600","PVP01_1328700","PVP01_1328900","PVP01_1329000","PVP01_1329100","PVP01_1329200","PVP01_1329500","PVP01_1329600","PVP01_1329700","PVP01_1329800","PVP01_1329900","PVP01_1330000","PVP01_1330100","PVP01_1330200","PVP01_1330300","PVP01_1330400","PVP01_1330800","PVP01_1330900","PVP01_1331100","PVP01_1331200","PVP01_1331300","PVP01_1331500","PVP01_1331600","PVP01_1331700","PVP01_1331900","PVP01_1332000","PVP01_1332100","PVP01_1332200","PVP01_1332300","PVP01_1332400","PVP01_1332500","PVP01_1332600","PVP01_1332800","PVP01_1333000","PVP01_1333400","PVP01_1333500","PVP01_1333800","PVP01_1334400","PVP01_1334600","PVP01_1334700","PVP01_1334800","PVP01_1334900","PVP01_1335000","PVP01_1335100","PVP01_1335200","PVP01_1335300","PVP01_1335400","PVP01_1335500","PVP01_1335900","PVP01_1336100","PVP01_1336200","PVP01_1336300","PVP01_1336400","PVP01_1336500","PVP01_1336600","PVP01_1336700","PVP01_1336800","PVP01_1336900","PVP01_1337000","PVP01_1337100","PVP01_1337200","PVP01_1337300","PVP01_1337600","PVP01_1337700","PVP01_1337800","PVP01_1338000","PVP01_1338200","PVP01_1338700","PVP01_1338800","PVP01_1338900","PVP01_1339200","PVP01_1339300","PVP01_1339400","PVP01_1339500","PVP01_1339700","PVP01_1339800","PVP01_1339900","PVP01_1340000","PVP01_1340100","PVP01_1340200","PVP01_1340300","PVP01_1340400","PVP01_1340700","PVP01_1340800","PVP01_1340900","PVP01_1341000","PVP01_1341100","PVP01_1341200","PVP01_1341300","PVP01_1341400","PVP01_1341500","PVP01_1341600","PVP01_1341700","PVP01_1341800","PVP01_1341900","PVP01_1342000","PVP01_1342100","PVP01_1342400","PVP01_1342500","PVP01_1342700","PVP01_1342900","PVP01_1343000","PVP01_1343300","PVP01_1343400","PVP01_1343600","PVP01_1343800","PVP01_1343900","PVP01_1344000","PVP01_1344100","PVP01_1344300","PVP01_1344400","PVP01_1344500","PVP01_1344700","PVP01_1345000","PVP01_1345100","PVP01_1345200","PVP01_1345300","PVP01_1345400","PVP01_1345500","PVP01_1345600","PVP01_1345800","PVP01_1345900","PVP01_1346000","PVP01_1346100","PVP01_1346200","PVP01_1346400","PVP01_1346500","PVP01_1346700","PVP01_1346800","PVP01_1347000","PVP01_1347100","PVP01_1347200","PVP01_1347400","PVP01_1347700","PVP01_1400300","PVP01_1400400","PVP01_1400700","PVP01_1400900","PVP01_1401000","PVP01_1401100","PVP01_1401300","PVP01_1401400","PVP01_1401600","PVP01_1401700","PVP01_1401800","PVP01_1401900","PVP01_1402000","PVP01_1402100","PVP01_1402200","PVP01_1402300","PVP01_1402400","PVP01_1402500","PVP01_1402600","PVP01_1402700","PVP01_1402800","PVP01_1403100","PVP01_1403200","PVP01_1403300","PVP01_1403500","PVP01_1403600","PVP01_1404000","PVP01_1404100","PVP01_1404200","PVP01_1404300","PVP01_1404400","PVP01_1404700","PVP01_1404800","PVP01_1404900","PVP01_1405100","PVP01_1405200","PVP01_1405300","PVP01_1405400","PVP01_1405500","PVP01_1405700","PVP01_1405800","PVP01_1405900","PVP01_1406100","PVP01_1406200","PVP01_1406300","PVP01_1406400","PVP01_1406700","PVP01_1406800","PVP01_1406900","PVP01_1407200","PVP01_1407300","PVP01_1407400","PVP01_1407500","PVP01_1407700","PVP01_1408000","PVP01_1408300","PVP01_1408400","PVP01_1409100","PVP01_1409200","PVP01_1409500","PVP01_1409600","PVP01_1409700","PVP01_1409800","PVP01_1409900","PVP01_1410000","PVP01_1410100","PVP01_1410600","PVP01_1410900","PVP01_1411600","PVP01_1411700","PVP01_1411800","PVP01_1411900","PVP01_1412000","PVP01_1412100","PVP01_1412200","PVP01_1412300","PVP01_1412400","PVP01_1412500","PVP01_1412600","PVP01_1412800","PVP01_1412900","PVP01_1413000","PVP01_1413300","PVP01_1413400","PVP01_1413450","PVP01_1413500","PVP01_1413600","PVP01_1413800","PVP01_1414000","PVP01_1414100","PVP01_1414200","PVP01_1414400","PVP01_1414600","PVP01_1414700","PVP01_1414800","PVP01_1414900","PVP01_1415000","PVP01_1415200","PVP01_1415300","PVP01_1415400","PVP01_1415700","PVP01_1415800","PVP01_1416300","PVP01_1416400","PVP01_1416500","PVP01_1416600","PVP01_1416700","PVP01_1416800","PVP01_1416900","PVP01_1417000","PVP01_1417300","PVP01_1417400","PVP01_1417500","PVP01_1417700","PVP01_1417800","PVP01_1418100","PVP01_1418200","PVP01_1418300","PVP01_1418400","PVP01_1418500","PVP01_1418700","PVP01_1418900","PVP01_1419200","PVP01_1419500","PVP01_1419600","PVP01_1419700","PVP01_1419900","PVP01_1420000","PVP01_1420200","PVP01_1420300","PVP01_1420400","PVP01_1420600","PVP01_1420900","PVP01_1421100","PVP01_1421200","PVP01_1421400","PVP01_1421600","PVP01_1421700","PVP01_1421800","PVP01_1421900","PVP01_1422000","PVP01_1422100","PVP01_1422200","PVP01_1422400","PVP01_1422500","PVP01_1422700","PVP01_1422800","PVP01_1423000","PVP01_1423100","PVP01_1423300","PVP01_1423400","PVP01_1423500","PVP01_1423600","PVP01_1424100","PVP01_1424300","PVP01_1424400","PVP01_1424500","PVP01_1424600","PVP01_1424700","PVP01_1424900","PVP01_1425000","PVP01_1425200","PVP01_1425300","PVP01_1425400","PVP01_1425500","PVP01_1425600","PVP01_1425700","PVP01_1425800","PVP01_1425900","PVP01_1426000","PVP01_1426300","PVP01_1426400","PVP01_1426500","PVP01_1426600","PVP01_1426900","PVP01_1427000","PVP01_1427200","PVP01_1427300","PVP01_1427400","PVP01_1427500","PVP01_1427600","PVP01_1427700","PVP01_1427800","PVP01_1427900","PVP01_1428000","PVP01_1428100","PVP01_1428200","PVP01_1428400","PVP01_1428500","PVP01_1428700","PVP01_1428800","PVP01_1428900","PVP01_1429100","PVP01_1429400","PVP01_1429500","PVP01_1429600","PVP01_1429700","PVP01_1429800","PVP01_1430200","PVP01_1430300","PVP01_1430400","PVP01_1430600","PVP01_1430900","PVP01_1431000","PVP01_1431400","PVP01_1431500","PVP01_1431700","PVP01_1431800","PVP01_1431900","PVP01_1432100","PVP01_1432200","PVP01_1432300","PVP01_1432700","PVP01_1432800","PVP01_1432900","PVP01_1433100","PVP01_1433200","PVP01_1433300","PVP01_1433400","PVP01_1433500","PVP01_1433600","PVP01_1433900","PVP01_1434100","PVP01_1434400","PVP01_1434500","PVP01_1434600","PVP01_1434700","PVP01_1434900","PVP01_1435000","PVP01_1435100","PVP01_1435200","PVP01_1435300","PVP01_1435400","PVP01_1435500","PVP01_1435700","PVP01_1435800","PVP01_1435900","PVP01_1436100","PVP01_1436200","PVP01_1436300","PVP01_1436400","PVP01_1436500","PVP01_1436600","PVP01_1436700","PVP01_1436800","PVP01_1436900","PVP01_1437000","PVP01_1437100","PVP01_1437200","PVP01_1437300","PVP01_1437400","PVP01_1437600","PVP01_1437800","PVP01_1438000","PVP01_1438200","PVP01_1438300","PVP01_1438400","PVP01_1438500","PVP01_1438600","PVP01_1438700","PVP01_1438800","PVP01_1438900","PVP01_1439000","PVP01_1439100","PVP01_1439200","PVP01_1439300","PVP01_1439400","PVP01_1439500","PVP01_1439700","PVP01_1439800","PVP01_1439900","PVP01_1440100","PVP01_1440200","PVP01_1440500","PVP01_1440600","PVP01_1440800","PVP01_1441100","PVP01_1441300","PVP01_1441400","PVP01_1441500","PVP01_1441600","PVP01_1441800","PVP01_1441900","PVP01_1442000","PVP01_1442100","PVP01_1442200","PVP01_1442300","PVP01_1442400","PVP01_1442600","PVP01_1442700","PVP01_1442800","PVP01_1443000","PVP01_1443100","PVP01_1443400","PVP01_1443500","PVP01_1443700","PVP01_1444000","PVP01_1444100","PVP01_1444300","PVP01_1444400","PVP01_1444500","PVP01_1444700","PVP01_1444800","PVP01_1445100","PVP01_1445200","PVP01_1445400","PVP01_1445500","PVP01_1445700","PVP01_1445800","PVP01_1445900","PVP01_1446000","PVP01_1446200","PVP01_1446500","PVP01_1446600","PVP01_1446700","PVP01_1446800","PVP01_1447100","PVP01_1447300","PVP01_1447400","PVP01_1447500","PVP01_1447600","PVP01_1448100","PVP01_1448400","PVP01_1448500","PVP01_1448600","PVP01_1449000","PVP01_1449200","PVP01_1449600","PVP01_1449800","PVP01_1450000","PVP01_1450100","PVP01_1450200","PVP01_1450300","PVP01_1450500","PVP01_1450600","PVP01_1450700","PVP01_1450800","PVP01_1450900","PVP01_1451000","PVP01_1451200","PVP01_1451300","PVP01_1451400","PVP01_1451700","PVP01_1451800","PVP01_1451900","PVP01_1452000","PVP01_1452100","PVP01_1452500","PVP01_1452600","PVP01_1452800","PVP01_1452900","PVP01_1453000","PVP01_1453100","PVP01_1453200","PVP01_1453300","PVP01_1453400","PVP01_1453500","PVP01_1453600","PVP01_1453700","PVP01_1453900","PVP01_1454000","PVP01_1454100","PVP01_1454300","PVP01_1454400","PVP01_1454500","PVP01_1454700","PVP01_1454800","PVP01_1455000","PVP01_1455100","PVP01_1455200","PVP01_1455300","PVP01_1455400","PVP01_1455500","PVP01_1455600","PVP01_1455700","PVP01_1455900","PVP01_1456000","PVP01_1456100","PVP01_1456200","PVP01_1456300","PVP01_1456400","PVP01_1456700","PVP01_1456800","PVP01_1457000","PVP01_1457100","PVP01_1457200","PVP01_1457300","PVP01_1457400","PVP01_1457500","PVP01_1457600","PVP01_1457800","PVP01_1458000","PVP01_1458100","PVP01_1458200","PVP01_1458400","PVP01_1458500","PVP01_1458600","PVP01_1458800","PVP01_1458900","PVP01_1459000","PVP01_1459100","PVP01_1459200","PVP01_1459300","PVP01_1459400","PVP01_1459500","PVP01_1459700","PVP01_1459800","PVP01_1459900","PVP01_1460000","PVP01_1460100","PVP01_1460200","PVP01_1460400","PVP01_1460600","PVP01_1460700","PVP01_1460800","PVP01_1461000","PVP01_1461200","PVP01_1461300","PVP01_1461400","PVP01_1461500","PVP01_1461600","PVP01_1461700","PVP01_1461800","PVP01_1462100","PVP01_1462200","PVP01_1462400","PVP01_1462500","PVP01_1462800","PVP01_1462900","PVP01_1463000","PVP01_1463100","PVP01_1463400","PVP01_1463500","PVP01_1463700","PVP01_1463800","PVP01_1463900","PVP01_1464000","PVP01_1464100","PVP01_1464200","PVP01_1464300","PVP01_1464400","PVP01_1464500","PVP01_1464600","PVP01_1464700","PVP01_1464800","PVP01_1464900","PVP01_1465100","PVP01_1465200","PVP01_1465300","PVP01_1465400","PVP01_1465500","PVP01_1465600","PVP01_1465700","PVP01_1465800","PVP01_1466100","PVP01_1466200","PVP01_1466300","PVP01_1466400","PVP01_1466500","PVP01_1466600","PVP01_1466700","PVP01_1466800","PVP01_1466900","PVP01_1467100","PVP01_1467200","PVP01_1467400","PVP01_1467500","PVP01_1467700","PVP01_1467800","PVP01_1468000","PVP01_1468100","PVP01_1468200","PVP01_1468300","PVP01_1468400","PVP01_1468500","PVP01_1468600","PVP01_1468700","PVP01_1468900","PVP01_1469200","PVP01_1469300","PVP01_1469500","PVP01_1469600","PVP01_1469700","PVP01_1469800","PVP01_1469900","PVP01_1470000","PVP01_1470100","PVP01_1470200","PVP01_1470300","PVP01_1470400","PVP01_1470500","PVP01_1470600","PVP01_1470700","PVP01_1470800","PVP01_1471000","PVP01_1471200","PVP01_1471300","PVP01_1471500","PVP01_1471600","PVP01_1471700","PVP01_1471800","PVP01_1471900","PVP01_1472100","PVP01_1472200","PVP01_1472500","PVP01_API02800","PVP01_API04800","PVP01_API05500","PVP01_MIT00500","PVP01_MIT01200","PVP01_MIT02700","PVP01_MIT03300","PVP01_MIT03400"],"coefficients":[-0.2437,0.07041,-0.7906,-1.868,-0.9006,-1.378,-1.057,0.6221,-0.5779,-0.8177,-1.175,-0.8003,-0.1452,-0.4046,-1.099,-0.2226,0.03712,-0.5423,-0.7507,-0.5852,-0.7404,-0.1816,-0.6285,-0.3188,0.04648,-0.08435,-0.938,-0.4779,0.3131,-0.3089,-1.478,-1.216,-1.009,-0.6125,0.02005,-0.7048,-0.8796,-0.2811,0.8094,0.03859,-0.8558,-1.265,-0.9504,-1.277,-0.3314,-0.5495,-0.8393,-1.007,-0.4137,-1.104,-0.2291,-0.03063,-0.1525,-0.9088,0.4208,-1.397,0.4183,-0.3273,0.06086,-0.5043,-0.2689,-1.126,-0.6369,-1.431,0.8809,-0.7275,-1.039,-0.7184,-0.7042,-0.02818,-0.4384,-0.4069,0.09384,-0.8099,-1.258,-0.4288,-0.09432,-1.118,-1.147,-0.5197,-1.352,-0.8173,-0.224,-0.3877,1.532,-0.8762,-0.9163,-0.335,-0.7211,-0.476,-0.1274,-0.1858,-0.3509,-0.6463,-0.03467,-0.9331,-1.032,0.5992,0.0009314,-0.9972,-0.5996,-0.8872,-1.239,-0.2207,-1.032,-0.5809,-0.3116,-0.3025,-0.7887,0.4583,-0.3605,0.09695,1.096,-0.2992,0.5505,-1.026,-0.6258,-0.8575,-1.462,-0.8012,-0.4159,-0.698,-1.079,1.046,-0.5639,-0.7941,0.142,1.232,-0.475,0.01917,0.3497,-1.667,-0.8181,-0.631,0.4188,-0.6212,0.2735,-0.1019,0.3613,-0.04348,-0.6955,-1.475,-0.4587,0.2541,0.04978,0.8923,-0.1158,0.4736,0.07029,0.4596,-0.3004,0.1293,0.6349,0.2029,0.2017,0.7192,0.465,0.296,-0.4941,-0.7244,-0.4794,0.07839,-0.9073,-0.2099,-0.4927,0.3525,0.4911,0.06868,-0.4305,0.1054,-0.452,0.8274,0.4463,0.4767,-0.3606,-0.3903,0.723,0.2691,-0.06335,-0.292,-0.5209,-0.7746,-1.148,-0.3585,-0.8879,0.7414,-0.4256,-0.8515,-0.6497,-0.8026,-0.4696,-0.01637,-0.1328,-0.03365,-0.6834,0.8897,-1.12,0.8795,0.712,-0.7038,-0.1712,0.6963,0.7261,0.4741,-0.7081,0.4642,0.3882,0.7469,0.01804,1.074,-0.4906,-0.5851,1.116,-0.1472,0.3203,0.4786,0.1977,0.9215,0.03777,1.062,-0.3628,-0.646,0.874,-0.09296,-0.7011,-0.6399,0.6813,0.5072,-1.009,-0.1151,-0.4824,0.8364,0.008176,-0.8405,-0.7403,0.3015,1.09,-0.8244,-0.5954,0.1816,-2.104,-1.612,-0.504,-1.172,-0.1333,-0.6525,-1.239,-1.499,0.6391,-0.5532,0.279,0.8664,-0.5353,0.5923,1.498,0.8277,0.2951,0.07695,0.8054,-0.08227,0.1307,-0.2149,0.9561,0.6045,-0.3351,0.3995,-0.5118,-0.4127,0.386,0.8233,0.5493,0.7951,0.544,0.1912,-0.339,-0.5831,0.1834,-0.4347,-0.4155,-0.8778,-0.7992,-0.4138,-0.08008,-0.3193,0.5427,-0.4675,0.3728,0.3201,-0.9978,-0.4216,0.9375,-0.4523,-1.423,0.3282,0.4179,-0.2982,-0.9706,-0.3012,0.3589,-0.5421,-0.255,0.02563,0.2149,-0.9669,0.8968,0.8831,-0.8652,0.5132,-1.504,-0.9225,-0.7381,-0.7306,-1.298,-0.9946,0.3844,-0.2863,0.4914,-0.2357,-0.0528,0.3696,0.782,0.3339,0.3166,-0.1309,0.7278,-0.3567,0.8025,-0.6664,0.7445,0.3935,-0.1563,0.2012,0.1585,0.1271,0.3806,0.6824,0.7883,1.216,-0.9494,-0.0983,0.4967,0.872,-0.1807,0.4123,0.8369,0.9102,-0.4912,1.315,-3.086,-2.526,0.2882,0.6155,-0.9946,-0.3561,0.2187,-0.4933,-0.8021,0.9329,0.5398,0.4016,0.7201,0.7719,0.6067,0.8007,0.3619,-0.9689,-0.4522,-0.5525,-1.383,-0.4835,-1.644,-2.629,0.4492,-0.2279,0.373,0.3703,0.8434,-0.4974,-0.8537,-0.7005,0.9516,-0.6614,-0.4682,-1.12,-0.09252,0.2646,0.03475,-0.5087,-0.2372,0.3136,0.3243,-0.7203,-0.2945,1.149,0.4107,0.9261,-0.2528,0.02076,-0.2573,0.5436,-0.4645,-0.4572,-0.3728,-1.606,-1.934,-2.059,0.08446,0.732,-0.2208,-0.07259,-0.683,-0.7737,0.9767,-0.01186,0.196,0.2465,-0.784,0.9216,-1.023,0.03302,-0.8837,0.08925,0.7163,-0.4199,0.7256,1.056,0.8783,-0.05966,-1.579,-2.774,0.323,0.005469,-0.8929,-0.5582,-0.7586,0.4807,0.4344,-0.5729,-0.4211,0.5948,-0.915,0.9846,0.02951,-0.8033,0.723,0.5174,0.8344,0.02145,0.4328,-1.034,-0.6069,-0.4796,0.5341,-0.6575,0.7112,0.8728,-0.4701,-0.627,-0.479,-0.9655,-0.7567,-0.5689,-1.237,-0.6672,-1.206,-0.3387,-1.115,-1.065,0.6287,-1.561,-1.744,0.3528,-1.934,0.9578,-0.1905,-0.9885,0.2255,0.8501,0.7436,-0.007634,0.5254,0.1581,-0.8313,-0.004362,-0.6185,0.9946,-0.1969,-0.6709,-0.7331,-0.8384,-0.3039,0.9332,-0.8186,-0.9024,-0.3553,0.8551,0.1144,0.7276,-0.5421,0.03876,0.7161,0.01362,0.3232,0.4889,-0.4236,-0.687,0.5625,0.05829,-1.333,0.6829,0.855,-0.8294,0.9075,0.1362,-0.7964,0.01809,-0
```
