## Supplementary material for "Transcriptome and histone epigenome of *Plasmodium vivax* salivary-gland sporozoites point to tight regulatory control and potential mechanisms for liver-stage differentiation": Pv-SPZ-v-HPZ-DTGs.js

```
glimma.layout.setupGrid(d3.select(".container"), "md", [1,2]);
glimma.storage.chartData.push(glimma.transform.toRowMajor({"GeneID":["PVP01_0000030","PVP01_0000040","PVP01_0000050","PVP01_0000080","PVP01_0000110","PVP01_0000120","PVP01_0000130","PVP01_0000140","PVP01_0000150","PVP01_0000160","PVP01_0000170","PVP01_0000180","PVP01_0000190","PVP01_0000200","PVP01_0000210","PVP01_0000220","PVP01_0000300","PVP01_0000480","PVP01_0000670","PVP01_0000740","PVP01_0000790","PVP01_0001030","PVP01_0001090","PVP01_0001310","PVP01_0001430","PVP01_0001440","PVP01_0001480","PVP01_0001550","PVP01_0001640","PVP01_0001660","PVP01_0001670","PVP01_0001900","PVP01_0001910","PVP01_0001940","PVP01_0001970","PVP01_0002020","PVP01_0002050","PVP01_0002120","PVP01_0002200","PVP01_0002210","PVP01_0002330","PVP01_0002370","PVP01_0002390","PVP01_0002470","PVP01_0002530","PVP01_0002540","PVP01_0002750","PVP01_0002810","PVP01_0002820","PVP01_0003010","PVP01_0003050","PVP01_0003150","PVP01_0003210","PVP01_0003350","PVP01_0003360","PVP01_0003370","PVP01_0003380","PVP01_0003400","PVP01_0003410","PVP01_0003420","PVP01_0003440","PVP01_0003530","PVP01_0003640","PVP01_0003690","PVP01_0003730","PVP01_0003800","PVP01_0003990","PVP01_0004040","PVP01_0004160","PVP01_0004170","PVP01_0004180","PVP01_0004190","PVP01_0004210","PVP01_0004240","PVP01_0004250","PVP01_0004260","PVP01_0004270","PVP01_0004280","PVP01_0004290","PVP01_0004300","PVP01_0004310","PVP01_0004320","PVP01_0004330","PVP01_0004350","PVP01_0004370","PVP01_0004380","PVP01_0004390","PVP01_0004400","PVP01_0004410","PVP01_0004640","PVP01_0004670","PVP01_0004910","PVP01_0004920","PVP01_0004950","PVP01_0004960","PVP01_0005010","PVP01_0005100","PVP01_0005160","PVP01_0005210","PVP01_0005300","PVP01_0005310","PVP01_0005420","PVP01_0005430","PVP01_0005550","PVP01_0005640","PVP01_0005710","PVP01_0005740","PVP01_0005760","PVP01_0005800","PVP01_0005820","PVP01_0005870","PVP01_0005910","PVP01_0006000","PVP01_0006040","PVP01_0006080","PVP01_0006260","PVP01_0006650","PVP01_0006660","PVP01_0006670","PVP01_0006680","PVP01_0006700","PVP01_0006860","PVP01_0006890","PVP01_0007080","PVP01_0007160","PVP01_0007240","PVP01_0007290","PVP01_0007310","PVP01_0007320","PVP01_0007340","PVP01_0007610","PVP01_0007620","PVP01_0007690","PVP01_0007700","PVP01_0007850","PVP01_0007900","PVP01_0007910","PVP01_0007920","PVP01_0008190","PVP01_0008430","PVP01_0008480","PVP01_0008490","PVP01_0008600","PVP01_0008700","PVP01_0008770","PVP01_0008830","PVP01_0008990","PVP01_0009010","PVP01_0009170","PVP01_0009180","PVP01_0009210","PVP01_0009410","PVP01_0009470","PVP01_0009540","PVP01_0009620","PVP01_0009710","PVP01_0009720","PVP01_0009730","PVP01_0009740","PVP01_0009760","PVP01_0009770","PVP01_0009820","PVP01_0009850","PVP01_0009860","PVP01_0009870","PVP01_0009900","PVP01_0009940","PVP01_0009980","PVP01_0010020","PVP01_0010030","PVP01_0010070","PVP01_0010100","PVP01_0010120","PVP01_0010140","PVP01_0010170","PVP01_0010180","PVP01_0010200","PVP01_0010210","PVP01_0010220","PVP01_0010230","PVP01_0010240","PVP01_0010340","PVP01_0010380","PVP01_0010440","PVP01_0010470","PVP01_0010500","PVP01_0010570","PVP01_0010600","PVP01_0010610","PVP01_0010640","PVP01_0010650","PVP01_0010670","PVP01_0010770","PVP01_0010870","PVP01_0010900","PVP01_0010910","PVP01_0010960","PVP01_0010970","PVP01_0010990","PVP01_0011030","PVP01_0011070","PVP01_0102100","PVP01_0102200","PVP01_0102300","PVP01_0102400","PVP01_0102500","PVP01_0102600","PVP01_0102700","PVP01_0102800","PVP01_0103100","PVP01_0103300","PVP01_0103400","PVP01_0103500","PVP01_0103600","PVP01_0103800","PVP01_0103900","PVP01_0104100","PVP01_0104200","PVP01_0104600","PVP01_0104700","PVP01_0104800","PVP01_0104900","PVP01_0105000","PVP01_0105100","PVP01_0105200","PVP01_0105300","PVP01_0105400","PVP01_0105500","PVP01_0105600","PVP01_0105700","PVP01_0105800","PVP01_0106000","PVP01_0106200","PVP01_0106300","PVP01_0106400","PVP01_0106500","PVP01_0106800","PVP01_0106900","PVP01_0107400","PVP01_0107500","PVP01_0107600","PVP01_0107700","PVP01_0107800","PVP01_0107900","PVP01_0108000","PVP01_0108100","PVP01_0108200","PVP01_0108300","PVP01_0108400","PVP01_0108500","PVP01_0108600","PVP01_0108800","PVP01_0109000","PVP01_0109100","PVP01_0109200","PVP01_0109300","PVP01_0109600","PVP01_0109800","PVP01_0110000","PVP01_0110200","PVP01_0110500","PVP01_0110700","PVP01_0110800","PVP01_0110900","PVP01_0111100","PVP01_0111200","PVP01_0111300","PVP01_0111500","PVP01_0111600","PVP01_0111700","PVP01_0112000","PVP01_0112100","PVP01_0112300","PVP01_0112500","PVP01_0112700","PVP01_0113000","PVP01_0113100","PVP01_0113200","PVP01_0113300","PVP01_0113400","PVP01_0113500","PVP01_0114100","PVP01_0114200","PVP01_0114500","PVP01_0114600","PVP01_0114700","PVP01_0114900","PVP01_0115000","PVP01_0115100","PVP01_0115200","PVP01_0115400","PVP01_0115500","PVP01_0116100","PVP01_0116300","PVP01_0116500","PVP01_0116600","PVP01_0116700","PVP01_0116900","PVP01_0117300","PVP01_0117500","PVP01_0117700","PVP01_0117800","PVP01_0118000","PVP01_0118500","PVP01_0118600","PVP01_0118700","PVP01_0118800","PVP01_0118900","PVP01_0119200","PVP01_0119300","PVP01_0119400","PVP01_0119500","PVP01_0119600","PVP01_0119700","PVP01_0119800","PVP01_0119900","PVP01_0120000","PVP01_0120100","PVP01_0120500","PVP01_0120800","PVP01_0121000","PVP01_0121700","PVP01_0200200","PVP01_0200700","PVP01_0201000","PVP01_0201100","PVP01_0201300","PVP01_0201400","PVP01_0201500","PVP01_0201600","PVP01_0201700","PVP01_0201800","PVP01_0201900","PVP01_0202000","PVP01_0202100","PVP01_0202200","PVP01_0202400","PVP01_0203000","PVP01_0203100","PVP01_0203300","PVP01_0203400","PVP01_0203500","PVP01_0203800","PVP01_0203900","PVP01_0204000","PVP01_0204100","PVP01_0204300","PVP01_0204400","PVP01_0204800","PVP01_0204900","PVP01_0205000","PVP01_0205100","PVP01_0205200","PVP01_0205300","PVP01_0205500","PVP01_0205600","PVP01_0205800","PVP01_0206100","PVP01_0206200","PVP01_0206300","PVP01_0206400","PVP01_0206500","PVP01_0206600","PVP01_0206800","PVP01_0207000","PVP01_0207100","PVP01_0207200","PVP01_0207500","PVP01_0207600","PVP01_0207700","PVP01_0207800","PVP01_0207900","PVP01_0208000","PVP01_0208100","PVP01_0208200","PVP01_0208300","PVP01_0208400","PVP01_0208600","PVP01_0208700","PVP01_0208800","PVP01_0208900","PVP01_0209000","PVP01_0209100","PVP01_0209200","PVP01_0209300","PVP01_0209500","PVP01_0209600","PVP01_0210000","PVP01_0210100","PVP01_0210300","PVP01_0211000","PVP01_0211100","PVP01_0211200","PVP01_0211400","PVP01_0211500","PVP01_0211600","PVP01_0211700","PVP01_0211800","PVP01_0211900","PVP01_0212400","PVP01_0212500","PVP01_0212600","PVP01_0212700","PVP01_0212800","PVP01_0213000","PVP01_0213200","PVP01_0213300","PVP01_0213400","PVP01_0213600","PVP01_0213700","PVP01_0213800","PVP01_0213900","PVP01_0214100","PVP01_0214200","PVP01_0214800","PVP01_0214900","PVP01_0215000","PVP01_0215100","PVP01_0215300","PVP01_0215400","PVP01_0215500","PVP01_0215600","PVP01_0215700","PVP01_0215800","PVP01_0215900","PVP01_0216100","PVP01_0216500","PVP01_0216600","PVP01_0216700","PVP01_0216800","PVP01_0216900","PVP01_0217000","PVP01_0217100","PVP01_0217200","PVP01_0217300","PVP01_0217400","PVP01_0217500","PVP01_0218200","PVP01_0218300","PVP01_0218900","PVP01_0219400","PVP01_0220100","PVP01_0220200","PVP01_0221100","PVP01_0221300","PVP01_0221700","PVP01_0300100","PVP01_0300600","PVP01_0300700","PVP01_0300800","PVP01_0301100","PVP01_0301200","PVP01_0301500","PVP01_0301900","PVP01_0302000","PVP01_0302100","PVP01_0302300","PVP01_0302400","PVP01_0302500","PVP01_0302600","PVP01_0302800","PVP01_0302900","PVP01_0303100","PVP01_0303400","PVP01_0303500","PVP01_0304100","PVP01_0304400","PVP01_0304500","PVP01_0304600","PVP01_0304800","PVP01_0304900","PVP01_0305000","PVP01_0305200","PVP01_0305300","PVP01_0305500","PVP01_0305600","PVP01_0305700","PVP01_0305800","PVP01_0305900","PVP01_0306000","PVP01_0306100","PVP01_0306300","PVP01_0306400","PVP01_0306500","PVP01_0306600","PVP01_0306700","PVP01_0306800","PVP01_0306900","PVP01_0307000","PVP01_0307100","PVP01_0307300","PVP01_0307400","PVP01_0307500","PVP01_0307600","PVP01_0307700","PVP01_0308200","PVP01_0308600","PVP01_0308800","PVP01_0309200","PVP01_0309300","PVP01_0309700","PVP01_0309800","PVP01_0309900","PVP01_0310000","PVP01_0310200","PVP01_0310300","PVP01_0310400","PVP01_0310500","PVP01_0310600","PVP01_0311000","PVP01_0311200","PVP01_0311300","PVP01_0311400","PVP01_0311500","PVP01_0311600","PVP01_0311700","PVP01_0312000","PVP01_0312100","PVP01_0312200","PVP01_0312300","PVP01_0312400","PVP01_0312500","PVP01_0312700","PVP01_0312800","PVP01_0313500","PVP01_0313800","PVP01_0314000","PVP01_0314100","PVP01_0314200","PVP01_0314700","PVP01_0314900","PVP01_0315000","PVP01_0315100","PVP01_0315200","PVP01_0315300","PVP01_0315700","PVP01_0315900","PVP01_0316000","PVP01_0316200","PVP01_0316300","PVP01_0316500","PVP01_0316700","PVP01_0316800","PVP01_0317100","PVP01_0317200","PVP01_0317400","PVP01_0318000","PVP01_0318100","PVP01_0318400","PVP01_0318700","PVP01_0318900","PVP01_0319200","PVP01_0319300","PVP01_0319400","PVP01_0319600","PVP01_0319700","PVP01_0320200","PVP01_0320400","PVP01_0320500","PVP01_0320600","PVP01_0320700","PVP01_0320800","PVP01_0320900","PVP01_0321000","PVP01_0321100","PVP01_0400900","PVP01_0403600","PVP01_0403700","PVP01_0403900","PVP01_0404000","PVP01_0404100","PVP01_0404200","PVP01_0404300","PVP01_0404400","PVP01_0404500","PVP01_0404600","PVP01_0404700","PVP01_0404800","PVP01_0404900","PVP01_0405000","PVP01_0405100","PVP01_0405200","PVP01_0405300","PVP01_0405500","PVP01_0405600","PVP01_0405700","PVP01_0405900","PVP01_0406500","PVP01_0406600","PVP01_0406700","PVP01_0406900","PVP01_0407000","PVP01_0407300","PVP01_0407600","PVP01_0407700","PVP01_0407900","PVP01_0408200","PVP01_0408600","PVP01_0408800","PVP01_0408900","PVP01_0409000","PVP01_0409100","PVP01_0409200","PVP01_0409700","PVP01_0409800","PVP01_0409900","PVP01_0410000","PVP01_0410200","PVP01_0410400","PVP01_0410500","PVP01_0410600","PVP01_0410900","PVP01_0411200","PVP01_0411400","PVP01_0411600","PVP01_0411800","PVP01_0411900","PVP01_0412100","PVP01_0412200","PVP01_0412700","PVP01_0412800","PVP01_0412900","PVP01_0413000","PVP01_0413100","PVP01_0413200","PVP01_0413300","PVP01_0413400","PVP01_0413500","PVP01_0414000","PVP01_0414100","PVP01_0414300","PVP01_0414400","PVP01_0414600","PVP01_0414900","PVP01_0415400","PVP01_0415500","PVP01_0415600","PVP01_0415800","PVP01_0416100","PVP01_0416600","PVP01_0416900","PVP01_0417100","PVP01_0417200","PVP01_0417300","PVP01_0417400","PVP01_0417600","PVP01_0417700","PVP01_0417800","PVP01_0417900","PVP01_0418100","PVP01_0418300","PVP01_0418400","PVP01_0419000","PVP01_0419200","PVP01_0419300","PVP01_0419400","PVP01_0419500","PVP01_0419600","PVP01_0419700","PVP01_0420000","PVP01_0420100","PVP01_0420200","PVP01_0420500","PVP01_0420700","PVP01_0420900","PVP01_0421400","PVP01_0421500","PVP01_0421700","PVP01_0421800","PVP01_0421900","PVP01_0422100","PVP01_0422200","PVP01_0422300","PVP01_0422400","PVP01_0422500","PVP01_0422600","PVP01_0422700","PVP01_0422900","PVP01_0423000","PVP01_0423100","PVP01_0423200","PVP01_0423300","PVP01_0423400","PVP01_0423500","PVP01_0423600","PVP01_0423800","PVP01_0423900","PVP01_0424300","PVP01_0424400","PVP01_0500700","PVP01_0502300","PVP01_0502400","PVP01_0502900","PVP01_0503300","PVP01_0503400","PVP01_0503500","PVP01_0503600","PVP01_0503700","PVP01_0503800","PVP01_0503900","PVP01_0504000","PVP01_0504100","PVP01_0504200","PVP01_0504300","PVP01_0505000","PVP01_0505100","PVP01_0505400","PVP01_0505500","PVP01_0505700","PVP01_0505800","PVP01_0506000","PVP01_0506300","PVP01_0506400","PVP01_0506600","PVP01_0506700","PVP01_0506800","PVP01_0506900","PVP01_0507000","PVP01_0507300","PVP01_0507500","PVP01_0507600","PVP01_0507700","PVP01_0507800","PVP01_0508000","PVP01_0508200","PVP01_0508500","PVP01_0508600","PVP01_0508700","PVP01_0509100","PVP01_0509200","PVP01_0509400","PVP01_0509600","PVP01_0509800","PVP01_0509900","PVP01_0510000","PVP01_0510100","PVP01_0510300","PVP01_0510500","PVP01_0510800","PVP01_0510900","PVP01_0511200","PVP01_0511300","PVP01_0511400","PVP01_0511500","PVP01_0511700","PVP01_0511800","PVP01_0512100","PVP01_0512200","PVP01_0512400","PVP01_0512500","PVP01_0512600","PVP01_0512700","PVP01_0512800","PVP01_0513000","PVP01_0513300","PVP01_0513400","PVP01_0513500","PVP01_0513700","PVP01_0513900","PVP01_0514100","PVP01_0514400","PVP01_0514600","PVP01_0514800","PVP01_0514900","PVP01_0515100","PVP01_0515200","PVP01_0515300","PVP01_0515600","PVP01_0515700","PVP01_0515800","PVP01_0515900","PVP01_0516000","PVP01_0516100","PVP01_0516200","PVP01_0516300","PVP01_0516400","PVP01_0516500","PVP01_0516900","PVP01_0517000","PVP01_0517300","PVP01_0517400","PVP01_0517600","PVP01_0517700","PVP01_0517900","PVP01_0518300","PVP01_0518500","PVP01_0518600","PVP01_0518700","PVP01_0518800","PVP01_0518900","PVP01_0519100","PVP01_0519200","PVP01_0519400","PVP01_0519500","PVP01_0519600","PVP01_0520700","PVP01_0520800","PVP01_0521300","PVP01_0521400","PVP01_0521500","PVP01_0521600","PVP01_0521800","PVP01_0521900","PVP01_0522000","PVP01_0522100","PVP01_0522200","PVP01_0522300","PVP01_0522500","PVP01_0522600","PVP01_0522700","PVP01_0522900","PVP01_0523000","PVP01_0523100","PVP01_0523200","PVP01_0523300","PVP01_0523500","PVP01_0523600","PVP01_0523700","PVP01_0523800","PVP01_0523900","PVP01_0524000","PVP01_0524100","PVP01_0524200","PVP01_0524300","PVP01_0524600","PVP01_0524700","PVP01_0524900","PVP01_0525000","PVP01_0525100","PVP01_0525300","PVP01_0525400","PVP01_0525500","PVP01_0525700","PVP01_0526100","PVP01_0526200","PVP01_0526400","PVP01_0526900","PVP01_0527000","PVP01_0527100","PVP01_0527300","PVP01_0527400","PVP01_0527500","PVP01_0527700","PVP01_0527800","PVP01_0528000","PVP01_0528100","PVP01_0528200","PVP01_0528400","PVP01_0528500","PVP01_0528600","PVP01_0528700","PVP01_0528900","PVP01_0529000","PVP01_0529100","PVP01_0529200","PVP01_0529600","PVP01_0529700","PVP01_0529900","PVP01_0530000","PVP01_0530100","PVP01_0530400","PVP01_0530500","PVP01_0530600","PVP01_0530700","PVP01_0530800","PVP01_0531100","PVP01_0531200","PVP01_0531500","PVP01_0531700","PVP01_0531900","PVP01_0532000","PVP01_0532300","PVP01_0532500","PVP01_0532600","PVP01_0532700","PVP01_0532900","PVP01_0533000","PVP01_0533100","PVP01_0533200","PVP01_0533300","PVP01_0533700","PVP01_0534000","PVP01_0534100","PVP01_0534200","PVP01_0534300","PVP01_0534400","PVP01_0534600","PVP01_0534800","PVP01_0534900","PVP01_0535400","PVP01_0600100","PVP01_0600300","PVP01_0600400","PVP01_0600500","PVP01_0600600","PVP01_0600700","PVP01_0600900","PVP01_0601000","PVP01_0601100","PVP01_0601200","PVP01_0601300","PVP01_0601400","PVP01_0601500","PVP01_0601600","PVP01_0601700","PVP01_0601800","PVP01_0602600","PVP01_0603000","PVP01_0603100","PVP01_0603300","PVP01_0603600","PVP01_0603700","PVP01_0604100","PVP01_0604200","PVP01_0604400","PVP01_0604700","PVP01_0605000","PVP01_0605100","PVP01_0605200","PVP01_0605300","PVP01_0605500","PVP01_0606000","PVP01_0606200","PVP01_0606400","PVP01_0606500","PVP01_0606600","PVP01_0606800","PVP01_0607100","PVP01_0607200","PVP01_0607600","PVP01_0608000","PVP01_0608100","PVP01_0608300","PVP01_0608500","PVP01_0608600","PVP01_0608700","PVP01_0608800","PVP01_0608900","PVP01_0609200","PVP01_0609400","PVP01_0609600","PVP01_0609700","PVP01_0609800","PVP01_0609900","PVP01_0610100","PVP01_0610300","PVP01_0610400","PVP01_0610500","PVP01_0610600","PVP01_0610700","PVP01_0610800","PVP01_0610900","PVP01_0611100","PVP01_0611300","PVP01_0611400","PVP01_0611700","PVP01_0612000","PVP01_0612100","PVP01_0612600","PVP01_0612700","PVP01_0613000","PVP01_0613100","PVP01_0613200","PVP01_0613400","PVP01_0613500","PVP01_0613800","PVP01_0613900","PVP01_0614000","PVP01_0614200","PVP01_0614300","PVP01_0614400","PVP01_0614600","PVP01_0614900","PVP01_0615000","PVP01_0615100","PVP01_0615500","PVP01_0615700","PVP01_0615800","PVP01_0615900","PVP01_0616000","PVP01_0616200","PVP01_0616300","PVP01_0616400","PVP01_0616800","PVP01_0616900","PVP01_0617000","PVP01_0617100","PVP01_0617200","PVP01_0617300","PVP01_0617400","PVP01_0617500","PVP01_0617600","PVP01_0617700","PVP01_0617900","PVP01_0618100","PVP01_0618300","PVP01_0618600","PVP01_0618700","PVP01_0618900","PVP01_0619000","PVP01_0619100","PVP01_0619200","PVP01_0619300","PVP01_0619400","PVP01_0619500","PVP01_0619600","PVP01_0619700","PVP01_0619800","PVP01_0619900","PVP01_0620100","PVP01_0620200","PVP01_0620300","PVP01_0620400","PVP01_0620500","PVP01_0620600","PVP01_0620700","PVP01_0620900","PVP01_0621100","PVP01_0621200","PVP01_0621400","PVP01_0621800","PVP01_0621900","PVP01_0622100","PVP01_0622200","PVP01_0622700","PVP01_0623000","PVP01_0623100","PVP01_0623200","PVP01_0623300","PVP01_0623400","PVP01_0623500","PVP01_0623600","PVP01_0623700","PVP01_0623800","PVP01_0623900","PVP01_0624000","PVP01_0624100","PVP01_0624300","PVP01_0624400","PVP01_0624500","PVP01_0700200","PVP01_0700300","PVP01_0700400","PVP01_0700500","PVP01_0700600","PVP01_0700700","PVP01_0700800","PVP01_0700900","PVP01_0701000","PVP01_0701200","PVP01_0701300","PVP01_0701500","PVP01_0701600","PVP01_0701700","PVP01_0702000","PVP01_0702100","PVP01_0702200","PVP01_0702400","PVP01_0702500","PVP01_0703000","PVP01_0703600","PVP01_0703700","PVP01_0703900","PVP01_0704000","PVP01_0704100","PVP01_0704500","PVP01_0704600","PVP01_0704700","PVP01_0704800","PVP01_0704900","PVP01_0705000","PVP01_0705500","PVP01_0705600","PVP01_0705800","PVP01_0706000","PVP01_0706100","PVP01_0706200","PVP01_0706300","PVP01_0706400","PVP01_0706600","PVP01_0706700","PVP01_0706900","PVP01_0707000","PVP01_0707100","PVP01_0707200","PVP01_0707300","PVP01_0707400","PVP01_0707500","PVP01_0707800","PVP01_0708000","PVP01_0708100","PVP01_0708200","PVP01_0708300","PVP01_0708400","PVP01_0708500","PVP01_0708800","PVP01_0708900","PVP01_0709100","PVP01_0709200","PVP01_0709300","PVP01_0709500","PVP01_0709600","PVP01_0709700","PVP01_0709900","PVP01_0710000","PVP01_0710100","PVP01_0710200","PVP01_0710300","PVP01_0710700","PVP01_0711000","PVP01_0711100","PVP01_0711200","PVP01_0711400","PVP01_0711500","PVP01_0711900","PVP01_0712000","PVP01_0712100","PVP01_0712200","PVP01_0712300","PVP01_0712400","PVP01_0712700","PVP01_0712800","PVP01_0713000","PVP01_0713100","PVP01_0713200","PVP01_0713500","PVP01_0713600","PVP01_0713700","PVP01_0713900","PVP01_0714200","PVP01_0714300","PVP01_0714600","PVP01_0714900","PVP01_0715000","PVP01_0715100","PVP01_0715200","PVP01_0715300","PVP01_0715500","PVP01_0715600","PVP01_0715700","PVP01_0715800","PVP01_0716000","PVP01_0716100","PVP01_0716200","PVP01_0716400","PVP01_0716600","PVP01_0716800","PVP01_0717000","PVP01_0717200","PVP01_0717400","PVP01_0717500","PVP01_0717600","PVP01_0717800","PVP01_0718000","PVP01_0718100","PVP01_0718300","PVP01_0718600","PVP01_0718700","PVP01_0718800","PVP01_0719000","PVP01_0719100","PVP01_0719400","PVP01_0719500","PVP01_0719700","PVP01_0719900","PVP01_0720000","PVP01_0720100","PVP01_0720200","PVP01_0720300","PVP01_0720400","PVP01_0720800","PVP01_0720900","PVP01_0721100","PVP01_0721400","PVP01_0721500","PVP01_0721600","PVP01_0721700","PVP01_0721800","PVP01_0721900","PVP01_0722000","PVP01_0722100","PVP01_0722200","PVP01_0722300","PVP01_0722400","PVP01_0722500","PVP01_0722600","PVP01_0722700","PVP01_0722800","PVP01_0723000","PVP01_0723200","PVP01_0723300","PVP01_0723400","PVP01_0723500","PVP01_0723700","PVP01_0723800","PVP01_0724100","PVP01_0724300","PVP01_0724500","PVP01_0724600","PVP01_0724700","PVP01_0724800","PVP01_0724900","PVP01_0725000","PVP01_0725100","PVP01_0725200","PVP01_0725300","PVP01_0725400","PVP01_0725500","PVP01_0725700","PVP01_0725800","PVP01_0726400","PVP01_0726700","PVP01_0726900","PVP01_0727000","PVP01_0727100","PVP01_0727200","PVP01_0727300","PVP01_0727400","PVP01_0727500","PVP01_0727600","PVP01_0727800","PVP01_0728000","PVP01_0728100","PVP01_0728200","PVP01_0728300","PVP01_0728400","PVP01_0729100","PVP01_0729200","PVP01_0729300","PVP01_0729400","PVP01_0729500","PVP01_0729700","PVP01_0730000","PVP01_0730100","PVP01_0730200","PVP01_0730300","PVP01_0730400","PVP01_0730600","PVP01_0730700","PVP01_0731000","PVP01_0731100","PVP01_0731200","PVP01_0731300","PVP01_0731600","PVP01_0731700","PVP01_0731800","PVP01_0731900","PVP01_0732000","PVP01_0732100","PVP01_0732200","PVP01_0732300","PVP01_0732400","PVP01_0732600","PVP01_0732800","PVP01_0732900","PVP01_0733000","PVP01_0733100","PVP01_0733500","PVP01_0733700","PVP01_0733800","PVP01_0733900","PVP01_0734100","PVP01_0734200","PVP01_0734400","PVP01_0734700","PVP01_0735100","PVP01_0735200","PVP01_0735300","PVP01_0735800","PVP01_0735900","PVP01_0800100","PVP01_0800200","PVP01_0800300","PVP01_0800400","PVP01_0800500","PVP01_0800600","PVP01_0800700","PVP01_0800900","PVP01_0801000","PVP01_0801100","PVP01_0801400","PVP01_0801500","PVP01_0801600","PVP01_0801700","PVP01_0801800","PVP01_0802100","PVP01_0802300","PVP01_0802400","PVP01_0802500","PVP01_0802900","PVP01_0803000","PVP01_0803100","PVP01_0803400","PVP01_0803500","PVP01_0803600","PVP01_0803700","PVP01_0803900","PVP01_0804100","PVP01_0804400","PVP01_0804500","PVP01_0804600","PVP01_0804700","PVP01_0804800","PVP01_0804900","PVP01_0805000","PVP01_0805200","PVP01_0805300","PVP01_0805400","PVP01_0805500","PVP01_0805600","PVP01_0805700","PVP01_0805800","PVP01_0806000","PVP01_0806100","PVP01_0806200","PVP01_0806300","PVP01_0806400","PVP01_0806500","PVP01_0806700","PVP01_0807000","PVP01_0807200","PVP01_0807300","PVP01_0807600","PVP01_0807900","PVP01_0808000","PVP01_0808100","PVP01_0808400","PVP01_0808500","PVP01_0808600","PVP01_0808700","PVP01_0808800","PVP01_0809100","PVP01_0809200","PVP01_0809300","PVP01_0809400","PVP01_0809700","PVP01_0809900","PVP01_0810000","PVP01_0810100","PVP01_0810300","PVP01_0810400","PVP01_0810500","PVP01_0810600","PVP01_0811000","PVP01_0811100","PVP01_0811200","PVP01_0811300","PVP01_0811500","PVP01_0811700","PVP01_0811900","PVP01_0812000","PVP01_0812300","PVP01_0812400","PVP01_0812600","PVP01_0812800","PVP01_0813000","PVP01_0813200","PVP01_0813300","PVP01_0813600","PVP01_0813700","PVP01_0813800","PVP01_0814200","PVP01_0814300","PVP01_0814400","PVP01_0814800","PVP01_0815100","PVP01_0815300","PVP01_0815400","PVP01_0816200","PVP01_0816300","PVP01_0816700","PVP01_0816800","PVP01_0817100","PVP01_0817400","PVP01_0818100","PVP01_0818300","PVP01_0818600","PVP01_0818800","PVP01_0819000","PVP01_0819200","PVP01_0819300","PVP01_0819600","PVP01_0819700","PVP01_0819800","PVP01_0820000","PVP01_0820100","PVP01_0820200","PVP01_0820300","PVP01_0820400","PVP01_0820500","PVP01_0820600","PVP01_0820700","PVP01_0820800","PVP01_0820900","PVP01_0821000","PVP01_0821200","PVP01_0821300","PVP01_0821400","PVP01_0821500","PVP01_0821700","PVP01_0821800","PVP01_0822100","PVP01_0822300","PVP01_0822600","PVP01_0822700","PVP01_0822900","PVP01_0823100","PVP01_0823200","PVP01_0823400","PVP01_0823600","PVP01_0823700","PVP01_0824000","PVP01_0824100","PVP01_0824200","PVP01_0824300","PVP01_0824500","PVP01_0824600","PVP01_0824700","PVP01_0824800","PVP01_0824900","PVP01_0825000","PVP01_0825100","PVP01_0825200","PVP01_0825400","PVP01_0825600","PVP01_0825700","PVP01_0825800","PVP01_0826000","PVP01_0826100","PVP01_0826200","PVP01_0826300","PVP01_0826400","PVP01_0826500","PVP01_0826700","PVP01_0826900","PVP01_0827100","PVP01_0827700","PVP01_0827800","PVP01_0828000","PVP01_0828100","PVP01_0828200","PVP01_0828300","PVP01_0828700","PVP01_0829000","PVP01_0829100","PVP01_0829200","PVP01_0829300","PVP01_0829500","PVP01_0829700","PVP01_0829800","PVP01_0829900","PVP01_0830100","PVP01_0830300","PVP01_0830400","PVP01_0830500","PVP01_0830700","PVP01_0830900","PVP01_0831000","PVP01_0831100","PVP01_0831400","PVP01_0831700","PVP01_0831900","PVP01_0832000","PVP01_0832100","PVP01_0832200","PVP01_0832400","PVP01_0832500","PVP01_0832600","PVP01_0832800","PVP01_0832900","PVP01_0833000","PVP01_0833500","PVP01_0833600","PVP01_0833700","PVP01_0833800","PVP01_0834000","PVP01_0834200","PVP01_0834300","PVP01_0834400","PVP01_0834500","PVP01_0834600","PVP01_0834900","PVP01_0835000","PVP01_0835300","PVP01_0835500","PVP01_0835700","PVP01_0835800","PVP01_0836300","PVP01_0836400","PVP01_0836500","PVP01_0836600","PVP01_0836700","PVP01_0836800","PVP01_0837200","PVP01_0837300","PVP01_0837500","PVP01_0837600","PVP01_0837700","PVP01_0837800","PVP01_0837900","PVP01_0838000","PVP01_0838100","PVP01_0838200","PVP01_0838300","PVP01_0838600","PVP01_0838700","PVP01_0838800","PVP01_0838900","PVP01_0839000","PVP01_0839100","PVP01_0839200","PVP01_0839400","PVP01_0839700","PVP01_0840700","PVP01_0841000","PVP01_0900700","PVP01_0901000","PVP01_0901100","PVP01_0901200","PVP01_0901300","PVP01_0901800","PVP01_0901900","PVP01_0902000","PVP01_0902100","PVP01_0902400","PVP01_0903400","PVP01_0903500","PVP01_0903600","PVP01_0903700","PVP01_0903800","PVP01_0903900","PVP01_0904300","PVP01_0904600","PVP01_0904700","PVP01_0904900","PVP01_0905100","PVP01_0906000","PVP01_0906300","PVP01_0906400","PVP01_0906500","PVP01_0906700","PVP01_0906800","PVP01_0906900","PVP01_0907000","PVP01_0907100","PVP01_0907200","PVP01_0907300","PVP01_0907400","PVP01_0907500","PVP01_0907600","PVP01_0907700","PVP01_0907800","PVP01_0908000","PVP01_0908200","PVP01_0908400","PVP01_0908500","PVP01_0908600","PVP01_0908900","PVP01_0909100","PVP01_0909300","PVP01_0909500","PVP01_0909600","PVP01_0910000","PVP01_0910200","PVP01_0910900","PVP01_0911000","PVP01_0911300","PVP01_0911500","PVP01_0911600","PVP01_0911700","PVP01_0911800","PVP01_0911900","PVP01_0912000","PVP01_0912100","PVP01_0912200","PVP01_0912400","PVP01_0912900","PVP01_0913000","PVP01_0913100","PVP01_0913400","PVP01_0913500","PVP01_0913600","PVP01_0913800","PVP01_0913900","PVP01_0914000","PVP01_0914200","PVP01_0914300","PVP01_0914400","PVP01_0914600","PVP01_0914800","PVP01_0915100","PVP01_0915300","PVP01_0915400","PVP01_0915600","PVP01_0915800","PVP01_0916000","PVP01_0916300","PVP01_0916600","PVP01_0916800","PVP01_0916900","PVP01_0917000","PVP01_0917100","PVP01_0917200","PVP01_0917900","PVP01_0918000","PVP01_0918100","PVP01_0918200","PVP01_0918300","PVP01_0918400","PVP01_0918600","PVP01_0918700","PVP01_0918800","PVP01_0918900","PVP01_0919000","PVP01_0919100","PVP01_0919400","PVP01_0919500","PVP01_0919600","PVP01_0919700","PVP01_0919900","PVP01_0920100","PVP01_0920200","PVP01_0920400","PVP01_0920700","PVP01_0920800","PVP01_0921400","PVP01_0921500","PVP01_0921800","PVP01_0921900","PVP01_0922000","PVP01_0922100","PVP01_0922300","PVP01_0922400","PVP01_0922800","PVP01_0922900","PVP01_0923000","PVP01_0923100","PVP01_0923200","PVP01_0923300","PVP01_0923800","PVP01_0923900","PVP01_0924000","PVP01_0924300","PVP01_0924400","PVP01_0924600","PVP01_0924700","PVP01_0924800","PVP01_0925000","PVP01_0925100","PVP01_0925200","PVP01_0925400","PVP01_0925600","PVP01_0925700","PVP01_0925800","PVP01_0925900","PVP01_0926000","PVP01_0926300","PVP01_0926400","PVP01_0926500","PVP01_0926600","PVP01_0926800","PVP01_0926900","PVP01_0927100","PVP01_0927200","PVP01_0927400","PVP01_0927600","PVP01_0927700","PVP01_0927800","PVP01_0927900","PVP01_0928000","PVP01_0928100","PVP01_0928200","PVP01_0928300","PVP01_0928800","PVP01_0928900","PVP01_0929000","PVP01_0929100","PVP01_0929200","PVP01_0929300","PVP01_0929400","PVP01_0929600","PVP01_0929700","PVP01_0929900","PVP01_0930000","PVP01_0930200","PVP01_0930300","PVP01_0930400","PVP01_0930500","PVP01_0931200","PVP01_0931300","PVP01_0931400","PVP01_0931500","PVP01_0931600","PVP01_0931700","PVP01_0932000","PVP01_0932200","PVP01_0932300","PVP01_0932500","PVP01_0932600","PVP01_0932900","PVP01_0933100","PVP01_0933300","PVP01_0933400","PVP01_0933500","PVP01_0933700","PVP01_0934000","PVP01_0934300","PVP01_0934400","PVP01_0934500","PVP01_0934600","PVP01_0935000","PVP01_0935200","PVP01_0935400","PVP01_0935500","PVP01_0935700","PVP01_0935800","PVP01_0935900","PVP01_0936000","PVP01_0936100","PVP01_0936300","PVP01_0936400","PVP01_0936700","PVP01_0936800","PVP01_0937000","PVP01_0937300","PVP01_0937500","PVP01_0937700","PVP01_0937800","PVP01_0938100","PVP01_0938200","PVP01_0938600","PVP01_0938700","PVP01_0938900","PVP01_0939000","PVP01_0939100","PVP01_0939200","PVP01_0939300","PVP01_0939600","PVP01_0939700","PVP01_0940200","PVP01_0940500","PVP01_0940600","PVP01_0940700","PVP01_0940900","PVP01_0941100","PVP01_0941200","PVP01_0941300","PVP01_0941400","PVP01_0941600","PVP01_0941800","PVP01_0942000","PVP01_0942200","PVP01_0942300","PVP01_0942400","PVP01_0942600","PVP01_0942700","PVP01_0942800","PVP01_0942900","PVP01_0943100","PVP01_0943300","PVP01_0943400","PVP01_0943700","PVP01_0943800","PVP01_0944100","PVP01_0944300","PVP01_0944400","PVP01_0944600","PVP01_0944800","PVP01_0945100","PVP01_0945200","PVP01_0945300","PVP01_0945500","PVP01_0945600","PVP01_0945700","PVP01_0945900","PVP01_0946000","PVP01_0946100","PVP01_0946200","PVP01_0946300","PVP01_0946400","PVP01_0946500","PVP01_0946600","PVP01_0946700","PVP01_0946800","PVP01_0946900","PVP01_0947000","PVP01_0947100","PVP01_0947300","PVP01_0947400","PVP01_0947600","PVP01_0947800","PVP01_0947900","PVP01_0948100","PVP01_0948200","PVP01_0948300","PVP01_0948500","PVP01_0948600","PVP01_0948700","PVP01_0948800","PVP01_0948900","PVP01_0949000","PVP01_0949200","PVP01_0949400","PVP01_0949600","PVP01_0950000","PVP01_0950200","PVP01_1000700","PVP01_1001900","PVP01_1002500","PVP01_1002600","PVP01_1002700","PVP01_1003000","PVP01_1003200","PVP01_1003400","PVP01_1003500","PVP01_1003900","PVP01_1004100","PVP01_1004500","PVP01_1004800","PVP01_1004900","PVP01_1005100","PVP01_1005200","PVP01_1005300","PVP01_1005400","PVP01_1005600","PVP01_1005900","PVP01_1006200","PVP01_1006300","PVP01_1006400","PVP01_1006500","PVP01_1006600","PVP01_1006700","PVP01_1006800","PVP01_1007200","PVP01_1007500","PVP01_1007700","PVP01_1007800","PVP01_1007900","PVP01_1008000","PVP01_1008100","PVP01_1008200","PVP01_1008300","PVP01_1008600","PVP01_1008700","PVP01_1008900","PVP01_1009000","PVP01_1009100","PVP01_1009300","PVP01_1009400","PVP01_1009500","PVP01_1009600","PVP01_1009700","PVP01_1010100","PVP01_1010200","PVP01_1010300","PVP01_1010400","PVP01_1011000","PVP01_1011100","PVP01_1011300","PVP01_1011400","PVP01_1011500","PVP01_1011600","PVP01_1011800","PVP01_1012100","PVP01_1012300","PVP01_1012400","PVP01_1012500","PVP01_1012700","PVP01_1013100","PVP01_1013200","PVP01_1013300","PVP01_1013400","PVP01_1013700","PVP01_1014100","PVP01_1014300","PVP01_1014600","PVP01_1014800","PVP01_1015000","PVP01_1015100","PVP01_1015200","PVP01_1015400","PVP01_1015500","PVP01_1015600","PVP01_1015700","PVP01_1015900","PVP01_1016100","PVP01_1016200","PVP01_1016400","PVP01_1016500","PVP01_1016600","PVP01_1016900","PVP01_1017100","PVP01_1017200","PVP01_1017400","PVP01_1017700","PVP01_1017900","PVP01_1018000","PVP01_1018100","PVP01_1018200","PVP01_1018400","PVP01_1018500","PVP01_1018800","PVP01_1019000","PVP01_1019400","PVP01_1019600","PVP01_1019900","PVP01_1020000","PVP01_1020100","PVP01_1020200","PVP01_1020300","PVP01_1020500","PVP01_1020700","PVP01_1020900","PVP01_1021000","PVP01_1021100","PVP01_1021200","PVP01_1021300","PVP01_1021500","PVP01_1021600","PVP01_1021700","PVP01_1021800","PVP01_1021900","PVP01_1022000","PVP01_1022100","PVP01_1022200","PVP01_1022300","PVP01_1022400","PVP01_1022900","PVP01_1023000","PVP01_1023200","PVP01_1023300","PVP01_1023600","PVP01_1023700","PVP01_1024300","PVP01_1024400","PVP01_1024500","PVP01_1024800","PVP01_1025200","PVP01_1025300","PVP01_1025400","PVP01_1025700","PVP01_1025800","PVP01_1026300","PVP01_1026400","PVP01_1026500","PVP01_1026600","PVP01_1026700","PVP01_1026800","PVP01_1026900","PVP01_1027100","PVP01_1027300","PVP01_1027400","PVP01_1027600","PVP01_1027700","PVP01_1027800","PVP01_1027900","PVP01_1028000","PVP01_1028100","PVP01_1028200","PVP01_1028400","PVP01_1028600","PVP01_1029000","PVP01_1029100","PVP01_1029200","PVP01_1029300","PVP01_1029500","PVP01_1029700","PVP01_1029800","PVP01_1029900","PVP01_1030000","PVP01_1030100","PVP01_1030200","PVP01_1030300","PVP01_1030400","PVP01_1030500","PVP01_1030600","PVP01_1030700","PVP01_1030900","PVP01_1031000","PVP01_1031100","PVP01_1031200","PVP01_1031300","PVP01_1031400","PVP01_1031500","PVP01_1031600","PVP01_1031700","PVP01_1031800","PVP01_1031900","PVP01_1032000","PVP01_1032100","PVP01_1032200","PVP01_1032400","PVP01_1032500","PVP01_1032800","PVP01_1032900","PVP01_1033000","PVP01_1033300","PVP01_1033400","PVP01_1033500","PVP01_1033600","PVP01_1033700","PVP01_1033800","PVP01_1033900","PVP01_1034000","PVP01_1034100","PVP01_1034200","PVP01_1034300","PVP01_1034400","PVP01_1034500","PVP01_1035200","PVP01_1100400","PVP01_1100500","PVP01_1101000","PVP01_1101200","PVP01_1101300","PVP01_1101400","PVP01_1101500","PVP01_1101600","PVP01_1101700","PVP01_1102400","PVP01_1102500","PVP01_1102600","PVP01_1102700","PVP01_1102800","PVP01_1102900","PVP01_1103200","PVP01_1103400","PVP01_1103700","PVP01_1104000","PVP01_1104100","PVP01_1104200","PVP01_1104300","PVP01_1104400","PVP01_1104600","PVP01_1104800","PVP01_1104900","PVP01_1105300","PVP01_1105400","PVP01_1105500","PVP01_1105600","PVP01_1105800","PVP01_1106000","PVP01_1106200","PVP01_1106300","PVP01_1106400","PVP01_1106600","PVP01_1106700","PVP01_1107000","PVP01_1107100","PVP01_1107200","PVP01_1107300","PVP01_1107400","PVP01_1107600","PVP01_1107700","PVP01_1107900","PVP01_1108000","PVP01_1108100","PVP01_1108300","PVP01_1108500","PVP01_1108600","PVP01_1108800","PVP01_1108900","PVP01_1109000","PVP01_1109100","PVP01_1109200","PVP01_1109300","PVP01_1109400","PVP01_1109600","PVP01_1109900","PVP01_1110500","PVP01_1110600","PVP01_1110700","PVP01_1110900","PVP01_1111100","PVP01_1111200","PVP01_1111300","PVP01_1111400","PVP01_1111500","PVP01_1111600","PVP01_1111700","PVP01_1111800","PVP01_1112000","PVP01_1112100","PVP01_1112400","PVP01_1112600","PVP01_1112700","PVP01_1112800","PVP01_1112900","PVP01_1113000","PVP01_1113300","PVP01_1113500","PVP01_1113700","PVP01_1113900","PVP01_1114100","PVP01_1114200","PVP01_1114300","PVP01_1114500","PVP01_1114600","PVP01_1114700","PVP01_1114800","PVP01_1115000","PVP01_1115200","PVP01_1115300","PVP01_1115500","PVP01_1115600","PVP01_1116400","PVP01_1116600","PVP01_1116800","PVP01_1116900","PVP01_1117000","PVP01_1117200","PVP01_1117300","PVP01_1117500","PVP01_1117700","PVP01_1118100","PVP01_1118500","PVP01_1118600","PVP01_1118700","PVP01_1119000","PVP01_1119300","PVP01_1119500","PVP01_1119600","PVP01_1119700","PVP01_1119900","PVP01_1120000","PVP01_1120100","PVP01_1120300","PVP01_1120400","PVP01_1120500","PVP01_1120900","PVP01_1121000","PVP01_1121100","PVP01_1121300","PVP01_1121500","PVP01_1121600","PVP01_1121700","PVP01_1121800","PVP01_1121900","PVP01_1122100","PVP01_1122300","PVP01_1122400","PVP01_1122500","PVP01_1122600","PVP01_1122700","PVP01_1122800","PVP01_1122900","PVP01_1123000","PVP01_1123100","PVP01_1123200","PVP01_1123300","PVP01_1123400","PVP01_1123500","PVP01_1123600","PVP01_1123900","PVP01_1124000","PVP01_1124100","PVP01_1124300","PVP01_1124400","PVP01_1124600","PVP01_1124700","PVP01_1124800","PVP01_1124900","PVP01_1125000","PVP01_1125100","PVP01_1125200","PVP01_1125300","PVP01_1125400","PVP01_1125600","PVP01_1125700","PVP01_1125800","PVP01_1126000","PVP01_1126100","PVP01_1126200","PVP01_1126300","PVP01_1126500","PVP01_1126600","PVP01_1126800","PVP01_1126900","PVP01_1127000","PVP01_1127200","PVP01_1127300","PVP01_1127400","PVP01_1127500","PVP01_1127800","PVP01_1128000","PVP01_1128100","PVP01_1128400","PVP01_1128500","PVP01_1128800","PVP01_1128900","PVP01_1129000","PVP01_1129500","PVP01_1129600","PVP01_1129700","PVP01_1130100","PVP01_1130400","PVP01_1130600","PVP01_1130700","PVP01_1131300","PVP01_1131400","PVP01_1131900","PVP01_1132100","PVP01_1132300","PVP01_1132400","PVP01_1132500","PVP01_1132700","PVP01_1132800","PVP01_1132900","PVP01_1133000","PVP01_1133100","PVP01_1133200","PVP01_1133300","PVP01_1133400","PVP01_1133600","PVP01_1133800","PVP01_1133900","PVP01_1134100","PVP01_1134200","PVP01_1134300","PVP01_1134400","PVP01_1134500","PVP01_1134700","PVP01_1134800","PVP01_1135000","PVP01_1135100","PVP01_1135300","PVP01_1135500","PVP01_1135600","PVP01_1135700","PVP01_1135800","PVP01_1135900","PVP01_1136000","PVP01_1136100","PVP01_1136300","PVP01_1136400","PVP01_1136600","PVP01_1136700","PVP01_1136800","PVP01_1137100","PVP01_1137200","PVP01_1137300","PVP01_1137600","PVP01_1137700","PVP01_1137800","PVP01_1138200","PVP01_1138400","PVP01_1138500","PVP01_1139000","PVP01_1139100","PVP01_1139200","PVP01_1139300","PVP01_1139400","PVP01_1139700","PVP01_1140000","PVP01_1140300","PVP01_1140500","PVP01_1140600","PVP01_1140700","PVP01_1140800","PVP01_1140900","PVP01_1141000","PVP01_1141200","PVP01_1141500","PVP01_1141700","PVP01_1141800","PVP01_1141900","PVP01_1142000","PVP01_1142200","PVP01_1142300","PVP01_1142500","PVP01_1142600","PVP01_1143100","PVP01_1143200","PVP01_1143400","PVP01_1143500","PVP01_1143700","PVP01_1144100","PVP01_1144200","PVP01_1144300","PVP01_1144500","PVP01_1144700","PVP01_1144900","PVP01_1145000","PVP01_1145100","PVP01_1145200","PVP01_1145500","PVP01_1145600","PVP01_1145700","PVP01_1145800","PVP01_1146200","PVP01_1146400","PVP01_1146500","PVP01_1146600","PVP01_1146800","PVP01_1146900","PVP01_1147000","PVP01_1147100","PVP01_1147200","PVP01_1147300","PVP01_1147400","PVP01_1147500","PVP01_1147600","PVP01_1147700","PVP01_1147800","PVP01_1147900","PVP01_1148200","PVP01_1148700","PVP01_1148800","PVP01_1149000","PVP01_1149100","PVP01_1149400","PVP01_1149500","PVP01_1200200","PVP01_1200800","PVP01_1201100","PVP01_1201300","PVP01_1201400","PVP01_1201500","PVP01_1201600","PVP01_1201700","PVP01_1201800","PVP01_1201900","PVP01_1202000","PVP01_1202100","PVP01_1202200","PVP01_1202300","PVP01_1202500","PVP01_1202700","PVP01_1203100","PVP01_1203200","PVP01_1203300","PVP01_1203500","PVP01_1203600","PVP01_1203700","PVP01_1203800","PVP01_1204000","PVP01_1204200","PVP01_1204400","PVP01_1204500","PVP01_1204600","PVP01_1204700","PVP01_1204800","PVP01_1205000","PVP01_1205200","PVP01_1205400","PVP01_1205600","PVP01_1205800","PVP01_1205900","PVP01_1206100","PVP01_1206300","PVP01_1206400","PVP01_1206700","PVP01_1206900","PVP01_1207000","PVP01_1207100","PVP01_1207200","PVP01_1207800","PVP01_1207900","PVP01_1208000","PVP01_1208100","PVP01_1208200","PVP01_1208300","PVP01_1208600","PVP01_1208700","PVP01_1208800","PVP01_1208900","PVP01_1209000","PVP01_1209200","PVP01_1209400","PVP01_1210000","PVP01_1210100","PVP01_1210300","PVP01_1210400","PVP01_1210500","PVP01_1210700","PVP01_1211000","PVP01_1211100","PVP01_1211200","PVP01_1211300","PVP01_1211500","PVP01_1211600","PVP01_1211700","PVP01_1212100","PVP01_1212500","PVP01_1212600","PVP01_1212700","PVP01_1213000","PVP01_1213100","PVP01_1213300","PVP01_1214200","PVP01_1214300","PVP01_1214400","PVP01_1214500","PVP01_1214600","PVP01_1214800","PVP01_1214900","PVP01_1215000","PVP01_1215300","PVP01_1215400","PVP01_1215800","PVP01_1215900","PVP01_1216000","PVP01_1216100","PVP01_1216200","PVP01_1216300","PVP01_1216700","PVP01_1216800","PVP01_1216900","PVP01_1217000","PVP01_1217200","PVP01_1217400","PVP01_1217500","PVP01_1217600","PVP01_1217700","PVP01_1218000","PVP01_1218500","PVP01_1218800","PVP01_1218900","PVP01_1219000","PVP01_1219100","PVP01_1219200","PVP01_1219400","PVP01_1219500","PVP01_1219600","PVP01_1219800","PVP01_1219900","PVP01_1220100","PVP01_1220200","PVP01_1220600","PVP01_1220700","PVP01_1220800","PVP01_1220900","PVP01_1221000","PVP01_1221200","PVP01_1221300","PVP01_1221400","PVP01_1221700","PVP01_1221800","PVP01_1221900","PVP01_1222100","PVP01_1222200","PVP01_1222300","PVP01_1222400","PVP01_1222700","PVP01_1223400","PVP01_1223500","PVP01_1223800","PVP01_1223900","PVP01_1224000","PVP01_1224300","PVP01_1224400","PVP01_1224500","PVP01_1224600","PVP01_1224700","PVP01_1224800","PVP01_1225000","PVP01_1225200","PVP01_1225300","PVP01_1225500","PVP01_1225800","PVP01_1225900","PVP01_1226100","PVP01_1226700","PVP01_1226800","PVP01_1227300","PVP01_1227400","PVP01_1227500","PVP01_1227600","PVP01_1227900","PVP01_1228000","PVP01_1228100","PVP01_1228200","PVP01_1228400","PVP01_1228500","PVP01_1228800","PVP01_1229100","PVP01_1229400","PVP01_1229500","PVP01_1229600","PVP01_1229800","PVP01_1229900","PVP01_1230000","PVP01_1230100","PVP01_1230200","PVP01_1230500","PVP01_1230600","PVP01_1230700","PVP01_1230800","PVP01_1231000","PVP01_1231100","PVP01_1231600","PVP01_1231700","PVP01_1231800","PVP01_1231900","PVP01_1232400","PVP01_1232600","PVP01_1232700","PVP01_1233100","PVP01_1233800","PVP01_1233900","PVP01_1234000","PVP01_1234400","PVP01_1235000","PVP01_1235100","PVP01_1235200","PVP01_1235300","PVP01_1235400","PVP01_1235800","PVP01_1236000","PVP01_1236100","PVP01_1236200","PVP01_1236400","PVP01_1236700","PVP01_1236800","PVP01_1236900","PVP01_1237000","PVP01_1237100","PVP01_1237200","PVP01_1237300","PVP01_1237500","PVP01_1237600","PVP01_1237700","PVP01_1237800","PVP01_1237900","PVP01_1238000","PVP01_1238400","PVP01_1238500","PVP01_1238600","PVP01_1238900","PVP01_1239000","PVP01_1239100","PVP01_1239300","PVP01_1239400","PVP01_1239800","PVP01_1239900","PVP01_1240000","PVP01_1240100","PVP01_1240200","PVP01_1240300","PVP01_1240500","PVP01_1240600","PVP01_1240800","PVP01_1240900","PVP01_1241000","PVP01_1241100","PVP01_1241200","PVP01_1241300","PVP01_1241600","PVP01_1241900","PVP01_1242000","PVP01_1242300","PVP01_1242400","PVP01_1242600","PVP01_1242700","PVP01_1242800","PVP01_1242900","PVP01_1243000","PVP01_1243100","PVP01_1243200","PVP01_1243500","PVP01_1243600","PVP01_1243900","PVP01_1244400","PVP01_1244700","PVP01_1244800","PVP01_1244900","PVP01_1245100","PVP01_1245300","PVP01_1245400","PVP01_1245500","PVP01_1245600","PVP01_1245900","PVP01_1246000","PVP01_1246100","PVP01_1246300","PVP01_1246400","PVP01_1246500","PVP01_1246600","PVP01_1246700","PVP01_1246900","PVP01_1247000","PVP01_1247500","PVP01_1247600","PVP01_1247700","PVP01_1247800","PVP01_1247900","PVP01_1248200","PVP01_1248500","PVP01_1248600","PVP01_1249000","PVP01_1249100","PVP01_1249200","PVP01_1249500","PVP01_1249600","PVP01_1249700","PVP01_1250200","PVP01_1250300","PVP01_1250600","PVP01_1250900","PVP01_1251100","PVP01_1251500","PVP01_1251600","PVP01_1251700","PVP01_1251800","PVP01_1251900","PVP01_1252000","PVP01_1252200","PVP01_1252400","PVP01_1252800","PVP01_1252900","PVP01_1253000","PVP01_1253300","PVP01_1253600","PVP01_1253700","PVP01_1253800","PVP01_1254100","PVP01_1254300","PVP01_1254600","PVP01_1254900","PVP01_1255100","PVP01_1255200","PVP01_1255300","PVP01_1255400","PVP01_1255500","PVP01_1256000","PVP01_1256100","PVP01_1256200","PVP01_1256400","PVP01_1256500","PVP01_1256700","PVP01_1256800","PVP01_1256900","PVP01_1257000","PVP01_1257200","PVP01_1257300","PVP01_1257500","PVP01_1257600","PVP01_1257700","PVP01_1257900","PVP01_1258100","PVP01_1258200","PVP01_1258300","PVP01_1258400","PVP01_1258500","PVP01_1258700","PVP01_1258900","PVP01_1259000","PVP01_1259100","PVP01_1259200","PVP01_1259300","PVP01_1259400","PVP01_1259500","PVP01_1259600","PVP01_1260200","PVP01_1260800","PVP01_1260900","PVP01_1261100","PVP01_1261400","PVP01_1261500","PVP01_1261700","PVP01_1262000","PVP01_1262200","PVP01_1262300","PVP01_1262400","PVP01_1262500","PVP01_1262600","PVP01_1262800","PVP01_1262900","PVP01_1263000","PVP01_1263100","PVP01_1263400","PVP01_1263700","PVP01_1263800","PVP01_1263900","PVP01_1264000","PVP01_1264300","PVP01_1264400","PVP01_1264600","PVP01_1264700","PVP01_1264800","PVP01_1264900","PVP01_1265000","PVP01_1265100","PVP01_1265300","PVP01_1265400","PVP01_1265600","PVP01_1265700","PVP01_1265900","PVP01_1266200","PVP01_1266400","PVP01_1266500","PVP01_1266600","PVP01_1266700","PVP01_1267300","PVP01_1267400","PVP01_1267500","PVP01_1267600","PVP01_1268000","PVP01_1268100","PVP01_1268200","PVP01_1268300","PVP01_1268600","PVP01_1268900","PVP01_1269000","PVP01_1269200","PVP01_1269500","PVP01_1269800","PVP01_1270200","PVP01_1270300","PVP01_1270400","PVP01_1270600","PVP01_1270700","PVP01_1270800","PVP01_1270900","PVP01_1271100","PVP01_1271200","PVP01_1271300","PVP01_1271400","PVP01_1271500","PVP01_1271600","PVP01_1271900","PVP01_1272500","PVP01_1272600","PVP01_1273000","PVP01_1274300","PVP01_1300400","PVP01_1300500","PVP01_1300700","PVP01_1300900","PVP01_1301100","PVP01_1301200","PVP01_1301300","PVP01_1301400","PVP01_1301500","PVP01_1301600","PVP01_1301700","PVP01_1302100","PVP01_1302300","PVP01_1302500","PVP01_1302600","PVP01_1302700","PVP01_1302800","PVP01_1302900","PVP01_1303000","PVP01_1303100","PVP01_1303200","PVP01_1303400","PVP01_1303600","PVP01_1303900","PVP01_1304000","PVP01_1304200","PVP01_1304300","PVP01_1304500","PVP01_1304800","PVP01_1304900","PVP01_1305200","PVP01_1305300","PVP01_1305500","PVP01_1305700","PVP01_1305900","PVP01_1306000","PVP01_1306100","PVP01_1306300","PVP01_1306400","PVP01_1306600","PVP01_1306700","PVP01_1306800","PVP01_1306900","PVP01_1307000","PVP01_1307100","PVP01_1307200","PVP01_1307300","PVP01_1307500","PVP01_1307700","PVP01_1307800","PVP01_1308100","PVP01_1308300","PVP01_1308400","PVP01_1308500","PVP01_1308800","PVP01_1308900","PVP01_1309000","PVP01_1309100","PVP01_1309200","PVP01_1309400","PVP01_1309500","PVP01_1309700","PVP01_1309800","PVP01_1310000","PVP01_1310400","PVP01_1310500","PVP01_1310700","PVP01_1310800","PVP01_1311000","PVP01_1311100","PVP01_1311200","PVP01_1311300","PVP01_1311400","PVP01_1311500","PVP01_1311600","PVP01_1311900","PVP01_1312000","PVP01_1312100","PVP01_1312200","PVP01_1312400","PVP01_1312600","PVP01_1312900","PVP01_1313000","PVP01_1313100","PVP01_1313200","PVP01_1313300","PVP01_1313400","PVP01_1314100","PVP01_1314300","PVP01_1314600","PVP01_1314800","PVP01_1315200","PVP01_1315400","PVP01_1315500","PVP01_1316000","PVP01_1316100","PVP01_1316200","PVP01_1316300","PVP01_1316500","PVP01_1316600","PVP01_1316700","PVP01_1316900","PVP01_1317000","PVP01_1317500","PVP01_1317600","PVP01_1317700","PVP01_1317800","PVP01_1317900","PVP01_1318000","PVP01_1318100","PVP01_1318300","PVP01_1318500","PVP01_1318600","PVP01_1318800","PVP01_1318900","PVP01_1319000","PVP01_1319300","PVP01_1319500","PVP01_1319600","PVP01_1319700","PVP01_1319800","PVP01_1319900","PVP01_1320000","PVP01_1320100","PVP01_1320200","PVP01_1320300","PVP01_1320500","PVP01_1320600","PVP01_1320700","PVP01_1321500","PVP01_1321600","PVP01_1321700","PVP01_1321900","PVP01_1322000","PVP01_1322200","PVP01_1322300","PVP01_1322400","PVP01_1322600","PVP01_1322900","PVP01_1323000","PVP01_1323100","PVP01_1323200","PVP01_1323700","PVP01_1323800","PVP01_1323900","PVP01_1324100","PVP01_1324700","PVP01_1324800","PVP01_1325200","PVP01_1325600","PVP01_1325800","PVP01_1325900","PVP01_1326000","PVP01_1326200","PVP01_1326400","PVP01_1326600","PVP01_1326900","PVP01_1327200","PVP01_1327300","PVP01_1327500","PVP01_1327600","PVP01_1327900","PVP01_1328000","PVP01_1328100","PVP01_1328500","PVP01_1328600","PVP01_1328700","PVP01_1328800","PVP01_1328900","PVP01_1329000","PVP01_1329300","PVP01_1329400","PVP01_1329600","PVP01_1329700","PVP01_1329800","PVP01_1329900","PVP01_1330000","PVP01_1330800","PVP01_1331000","PVP01_1331100","PVP01_1331200","PVP01_1331800","PVP01_1331900","PVP01_1332200","PVP01_1332300","PVP01_1332400","PVP01_1332500","PVP01_1332700","PVP01_1332800","PVP01_1332900","PVP01_1333000","PVP01_1333100","PVP01_1333200","PVP01_1333700","PVP01_1333900","PVP01_1334100","PVP01_1334200","PVP01_1334300","PVP01_1334500","PVP01_1335100","PVP01_1335300","PVP01_1335600","PVP01_1335800","PVP01_1335900","PVP01_1336400","PVP01_1336500","PVP01_1336700","PVP01_1336900","PVP01_1337100","PVP01_1337200","PVP01_1337300","PVP01_1337400","PVP01_1337500","PVP01_1337600","PVP01_1337900","PVP01_1338100","PVP01_1338300","PVP01_1338400","PVP01_1338500","PVP01_1338600","PVP01_1338700","PVP01_1339000","PVP01_1339400","PVP01_1339600","PVP01_1339800","PVP01_1339900","PVP01_1340100","PVP01_1340200","PVP01_1340300","PVP01_1340400","PVP01_1340500","PVP01_1340600","PVP01_1340800","PVP01_1341300","PVP01_1341600","PVP01_1341700","PVP01_1341800","PVP01_1341900","PVP01_1342200","PVP01_1342300","PVP01_1342400","PVP01_1342600","PVP01_1342700","PVP01_1342800","PVP01_1343100","PVP01_1343300","PVP01_1343500","PVP01_1343600","PVP01_1344000","PVP01_1344200","PVP01_1344300","PVP01_1344500","PVP01_1344600","PVP01_1344700","PVP01_1344900","PVP01_1345000","PVP01_1345200","PVP01_1345300","PVP01_1345400","PVP01_1345500","PVP01_1345600","PVP01_1345700","PVP01_1345900","PVP01_1346200","PVP01_1346700","PVP01_1346800","PVP01_1346900","PVP01_1347100","PVP01_1347200","PVP01_1347300","PVP01_1347500","PVP01_1347700","PVP01_1347800","PVP01_1400100","PVP01_1400400","PVP01_1400600","PVP01_1400700","PVP01_1400900","PVP01_1401000","PVP01_1401100","PVP01_1401200","PVP01_1401600","PVP01_1401700","PVP01_1401800","PVP01_1401900","PVP01_1402000","PVP01_1402100","PVP01_1402200","PVP01_1402300","PVP01_1402400","PVP01_1402500","PVP01_1402600","PVP01_1402700","PVP01_1402800","PVP01_1402900","PVP01_1403000","PVP01_1403300","PVP01_1403400","PVP01_1403500","PVP01_1403600","PVP01_1403700","PVP01_1403900","PVP01_1404000","PVP01_1404100","PVP01_1404200","PVP01_1404300","PVP01_1404500","PVP01_1404600","PVP01_1404700","PVP01_1404800","PVP01_1405000","PVP01_1405100","PVP01_1405300","PVP01_1405500","PVP01_1405600","PVP01_1405800","PVP01_1406000","PVP01_1406100","PVP01_1406600","PVP01_1406700","PVP01_1406900","PVP01_1407000","PVP01_1407100","PVP01_1407400","PVP01_1407500","PVP01_1407600","PVP01_1407800","PVP01_1408000","PVP01_1408100","PVP01_1408200","PVP01_1408400","PVP01_1408500","PVP01_1408600","PVP01_1408700","PVP01_1408800","PVP01_1408900","PVP01_1409000","PVP01_1409100","PVP01_1409300","PVP01_1409400","PVP01_1409700","PVP01_1410000","PVP01_1410100","PVP01_1410200","PVP01_1410300","PVP01_1410400","PVP01_1410500","PVP01_1410700","PVP01_1410800","PVP01_1410900","PVP01_1411000","PVP01_1411200","PVP01_1411300","PVP01_1411400","PVP01_1412100","PVP01_1412400","PVP01_1412500","PVP01_1412900","PVP01_1413000","PVP01_1413100","PVP01_1413200","PVP01_1413300","PVP01_1413450","PVP01_1413700","PVP01_1414000","PVP01_1414200","PVP01_1414300","PVP01_1414400","PVP01_1414500","PVP01_1414900","PVP01_1415000","PVP01_1415100","PVP01_1415200","PVP01_1415300","PVP01_1415500","PVP01_1415600","PVP01_1415700","PVP01_1415800","PVP01_1415900","PVP01_1416000","PVP01_1416100","PVP01_1416200","PVP01_1416900","PVP01_1417100","PVP01_1417200","PVP01_1417300","PVP01_1417600","PVP01_1417700","PVP01_1417900","PVP01_1418000","PVP01_1418100","PVP01_1418200","PVP01_1418400","PVP01_1418600","PVP01_1418800","PVP01_1418900","PVP01_1419100","PVP01_1419300","PVP01_1419400","PVP01_1419500","PVP01_1419700","PVP01_1419800","PVP01_1420000","PVP01_1420100","PVP01_1420300","PVP01_1420500","PVP01_1420900","PVP01_1421000","PVP01_1421100","PVP01_1421200","PVP01_1421300","PVP01_1421400","PVP01_1421500","PVP01_1421800","PVP01_1422100","PVP01_1422200","PVP01_1422300","PVP01_1422600","PVP01_1422700","PVP01_1422900","PVP01_1423100","PVP01_1423200","PVP01_1423500","PVP01_1423700","PVP01_1424100","PVP01_1424200","PVP01_1424500","PVP01_1425100","PVP01_1425200","PVP01_1425300","PVP01_1425400","PVP01_1425500","PVP01_1425600","PVP01_1425800","PVP01_1426000","PVP01_1426100","PVP01_1426200","PVP01_1426700","PVP01_1426800","PVP01_1427000","PVP01_1427100","PVP01_1427600","PVP01_1427700","PVP01_1427800","PVP01_1428000","PVP01_1428100","PVP01_1428300","PVP01_1428400","PVP01_1428500","PVP01_1428600","PVP01_1428800","PVP01_1428900","PVP01_1429200","PVP01_1429300","PVP01_1429500","PVP01_1429700","PVP01_1429900","PVP01_1430000","PVP01_1430100","PVP01_1430300","PVP01_1430400","PVP01_1430500","PVP01_1430600","PVP01_1430700","PVP01_1430800","PVP01_1431200","PVP01_1431300","PVP01_1431500","PVP01_1431700","PVP01_1431800","PVP01_1432000","PVP01_1432100","PVP01_1432300","PVP01_1432400","PVP01_1432600","PVP01_1432700","PVP01_1432800","PVP01_1432900","PVP01_1433600","PVP01_1433700","PVP01_1433800","PVP01_1434000","PVP01_1434200","PVP01_1434400","PVP01_1434500","PVP01_1434600","PVP01_1434800","PVP01_1434900","PVP01_1435000","PVP01_1435100","PVP01_1435200","PVP01_1435300","PVP01_1435800","PVP01_1435900","PVP01_1436000","PVP01_1436300","PVP01_1436400","PVP01_1436600","PVP01_1436700","PVP01_1437100","PVP01_1437200","PVP01_1437300","PVP01_1437400","PVP01_1437600","PVP01_1438100","PVP01_1438200","PVP01_1438300","PVP01_1438400","PVP01_1438600","PVP01_1438700","PVP01_1438800","PVP01_1438900","PVP01_1439000","PVP01_1439400","PVP01_1439600","PVP01_1439700","PVP01_1440000","PVP01_1440100","PVP01_1440300","PVP01_1440600","PVP01_1440700","PVP01_1440800","PVP01_1440900","PVP01_1441000","PVP01_1441100","PVP01_1441200","PVP01_1441400","PVP01_1441700","PVP01_1441800","PVP01_1442000","PVP01_1442200","PVP01_1442300","PVP01_1442600","PVP01_1442900","PVP01_1443000","PVP01_1443100","PVP01_1443200","PVP01_1443600","PVP01_1443700","PVP01_1443800","PVP01_1443900","PVP01_1444000","PVP01_1444200","PVP01_1444400","PVP01_1444800","PVP01_1444900","PVP01_1445000","PVP01_1445200","PVP01_1445300","PVP01_1445400","PVP01_1445600","PVP01_1445700","PVP01_1446000","PVP01_1446100","PVP01_1446200","PVP01_1446300","PVP01_1446400","PVP01_1446600","PVP01_1446800","PVP01_1446900","PVP01_1447200","PVP01_1447300","PVP01_1447500","PVP01_1447600","PVP01_1447800","PVP01_1447900","PVP01_1448100","PVP01_1448200","PVP01_1448300","PVP01_1448400","PVP01_1448600","PVP01_1448700","PVP01_1448800","PVP01_1449100","PVP01_1449200","PVP01_1449300","PVP01_1449700","PVP01_1450000","PVP01_1450100","PVP01_1450200","PVP01_1450400","PVP01_1450500","PVP01_1451000","PVP01_1451100","PVP01_1451300","PVP01_1451500","PVP01_1451600","PVP01_1451700","PVP01_1451800","PVP01_1452100","PVP01_1452300","PVP01_1452400","PVP01_1452500","PVP01_1452700","PVP01_1452800","PVP01_1452900","PVP01_1453100","PVP01_1453200","PVP01_1453800","PVP01_1454000","PVP01_1454100","PVP01_1454200","PVP01_1454400","PVP01_1454500","PVP01_1454600","PVP01_1454800","PVP01_1454900","PVP01_1455100","PVP01_1455200","PVP01_1455400","PVP01_1455500","PVP01_1455700","PVP01_1456000","PVP01_1456200","PVP01_1456500","PVP01_1456700","PVP01_1456900","PVP01_1457000","PVP01_1457100","PVP01_1457400","PVP01_1457500","PVP01_1457900","PVP01_1458200","PVP01_1458700","PVP01_1458800","PVP01_1459100","PVP01_1459200","PVP01_1459300","PVP01_1459400","PVP01_1459500","PVP01_1459600","PVP01_1459700","PVP01_1460300","PVP01_1460500","PVP01_1460800","PVP01_1460900","PVP01_1461100","PVP01_1461700","PVP01_1461900","PVP01_1462000","PVP01_1462100","PVP01_1462300","PVP01_1462500","PVP01_1462600","PVP01_1462900","PVP01_1463000","PVP01_1463200","PVP01_1463300","PVP01_1463500","PVP01_1463600","PVP01_1463700","PVP01_1463800","PVP01_1464100","PVP01_1464300","PVP01_1464400","PVP01_1464500","PVP01_1464700","PVP01_1464900","PVP01_1465000","PVP01_1465100","PVP01_1465200","PVP01_1465500","PVP01_1465700","PVP01_1465800","PVP01_1465900","PVP01_1466000","PVP01_1466200","PVP01_1466300","PVP01_1466500","PVP01_1466600","PVP01_1467000","PVP01_1467100","PVP01_1467200","PVP01_1467300","PVP01_1467500","PVP01_1467600","PVP01_1467700","PVP01_1468100","PVP01_1468300","PVP01_1468400","PVP01_1468600","PVP01_1468800","PVP01_1469200","PVP01_1469300","PVP01_1469400","PVP01_1469500","PVP01_1469600","PVP01_1469700","PVP01_1469800","PVP01_1469900","PVP01_1470000","PVP01_1470100","PVP01_1470200","PVP01_1470300","PVP01_1470400","PVP01_1470500","PVP01_1470600","PVP01_1470700","PVP01_1470800","PVP01_1470900","PVP01_1471100","PVP01_1471300","PVP01_1471400","PVP01_1471500","PVP01_1471600","PVP01_1471700","PVP01_1471800","PVP01_1471900","PVP01_1472100","PVP01_1472200","PVP01_1472400","PVP01_API01200","PVP01_API01400","PVP01_API01700","PVP01_API04100","PVP01_API04200","PVP01_0000100","PVP01_0004220","PVP01_0004230","PVP01_0005540","PVP01_0007720","PVP01_0009370","PVP01_0010540","PVP01_0010840","PVP01_0011020","PVP01_0103200","PVP01_0103700","PVP01_0104000","PVP01_0104300","PVP01_0104400","PVP01_0104500","PVP01_0105900","PVP01_0106100","PVP01_0106600","PVP01_0106700","PVP01_0108700","PVP01_0108900","PVP01_0109400","PVP01_0109500","PVP01_0109700","PVP01_0109900","PVP01_0110100","PVP01_0110300","PVP01_0110400","PVP01_0110600","PVP01_0111000","PVP01_0111400","PVP01_0111800","PVP01_0111900","PVP01_0112200","PVP01_0112400","PVP01_0112600","PVP01_0112800","PVP01_0112900","PVP01_0113600","PVP01_0113700","PVP01_0113800","PVP01_0113900","PVP01_0114000","PVP01_0114300","PVP01_0114400","PVP01_0114800","PVP01_0115300","PVP01_0115600","PVP01_0115700","PVP01_0115800","PVP01_0115900","PVP01_0116000","PVP01_0116200","PVP01_0116400","PVP01_0116800","PVP01_0117000","PVP01_0117100","PVP01_0117200","PVP01_0117400","PVP01_0117600","PVP01_0117900","PVP01_0118100","PVP01_0118200","PVP01_0118300","PVP01_0118400","PVP01_0201200","PVP01_0202300","PVP01_0202500","PVP01_0202600","PVP01_0203200","PVP01_0203600","PVP01_0203700","PVP01_0204200","PVP01_0204500","PVP01_0204600","PVP01_0204700","PVP01_0205400","PVP01_0205700","PVP01_0205900","PVP01_0206000","PVP01_0206700","PVP01_0206900","PVP01_0207300","PVP01_0207400","PVP01_0208500","PVP01_0209400","PVP01_0209700","PVP01_0209900","PVP01_0210200","PVP01_0210400","PVP01_0210500","PVP01_0210600","PVP01_0210700","PVP01_0210800","PVP01_0210900","PVP01_0211300","PVP01_0212000","PVP01_0212100","PVP01_0212200","PVP01_0212300","PVP01_0212900","PVP01_0213100","PVP01_0213500","PVP01_0214000","PVP01_0214300","PVP01_0214400","PVP01_0214500","PVP01_0214600","PVP01_0214700","PVP01_0215200","PVP01_0216000","PVP01_0216200","PVP01_0302700","PVP01_0303000","PVP01_0303200","PVP01_0303300","PVP01_0303600","PVP01_0303700","PVP01_0303800","PVP01_0303900","PVP01_0304000","PVP01_0304200","PVP01_0304300","PVP01_0304700","PVP01_0305100","PVP01_0305400","PVP01_0306200","PVP01_0307200","PVP01_0307800","PVP01_0307900","PVP01_0308000","PVP01_0308100","PVP01_0308300","PVP01_0308400","PVP01_0308500","PVP01_0308700","PVP01_0308900","PVP01_0309000","PVP01_0309100","PVP01_0309600","PVP01_0310100","PVP01_0310700","PVP01_0311100","PVP01_0311800","PVP01_0311900","PVP01_0312600","PVP01_0312900","PVP01_0313000","PVP01_0313100","PVP01_0313200","PVP01_0313300","PVP01_0313400","PVP01_0313600","PVP01_0313700","PVP01_0313900","PVP01_0314300","PVP01_0314400","PVP01_0314500","PVP01_0314600","PVP01_0314800","PVP01_0315400","PVP01_0315500","PVP01_0315600","PVP01_0315800","PVP01_0316100","PVP01_0316400","PVP01_0316600","PVP01_0316900","PVP01_0317000","PVP01_0317300","PVP01_0317500","PVP01_0317600","PVP01_0317700","PVP01_0317800","PVP01_0317900","PVP01_0318200","PVP01_0318300","PVP01_0318500","PVP01_0318600","PVP01_0318800","PVP01_0319000","PVP01_0319100","PVP01_0319500","PVP01_0319800","PVP01_0319900","PVP01_0320000","PVP01_0320100","PVP01_0320300","PVP01_0405400","PVP01_0405800","PVP01_0406300","PVP01_0406400","PVP01_0406800","PVP01_0407100","PVP01_0407200","PVP01_0407400","PVP01_0407500","PVP01_0407800","PVP01_0408000","PVP01_0408100","PVP01_0408300","PVP01_0408400","PVP01_0408500","PVP01_0408700","PVP01_0409300","PVP01_0409400","PVP01_0409500","PVP01_0409600","PVP01_0410100","PVP01_0410300","PVP01_0410700","PVP01_0410800","PVP01_0411000","PVP01_0411100","PVP01_0411300","PVP01_0411500","PVP01_0411700","PVP01_0412000","PVP01_0412300","PVP01_0412400","PVP01_0412500","PVP01_0412600","PVP01_0413600","PVP01_0413700","PVP01_0413800","PVP01_0413900","PVP01_0414200","PVP01_0414500","PVP01_0414700","PVP01_0414800","PVP01_0415000","PVP01_0415100","PVP01_0415200","PVP01_0415300","PVP01_0415700","PVP01_0415900","PVP01_0416000","PVP01_0416200","PVP01_0416300","PVP01_0416400","PVP01_0416500","PVP01_0416700","PVP01_0416800","PVP01_0417000","PVP01_0417500","PVP01_0418000","PVP01_0418200","PVP01_0418500","PVP01_0418600","PVP01_0418700","PVP01_0418800","PVP01_0418900","PVP01_0419100","PVP01_0419800","PVP01_0419900","PVP01_0420300","PVP01_0420400","PVP01_0420600","PVP01_0420800","PVP01_0421000","PVP01_0421100","PVP01_0421200","PVP01_0421300","PVP01_0422000","PVP01_0422800","PVP01_0424500","PVP01_0504400","PVP01_0504800","PVP01_0504900","PVP01_0505200","PVP01_0505300","PVP01_0505600","PVP01_0505900","PVP01_0506100","PVP01_0506200","PVP01_0506500","PVP01_0507100","PVP01_0507200","PVP01_0507400","PVP01_0507900","PVP01_0508100","PVP01_0508300","PVP01_0508400","PVP01_0508800","PVP01_0508900","PVP01_0509000","PVP01_0509300","PVP01_0509500","PVP01_0509700","PVP01_0510200","PVP01_0510400","PVP01_0510600","PVP01_0510700","PVP01_0511000","PVP01_0511100","PVP01_0511600","PVP01_0511900","PVP01_0512000","PVP01_0512300","PVP01_0512900","PVP01_0513100","PVP01_0513200","PVP01_0513600","PVP01_0513800","PVP01_0514000","PVP01_0514200","PVP01_0514300","PVP01_0514500","PVP01_0514700","PVP01_0515000","PVP01_0515400","PVP01_0515500","PVP01_0516600","PVP01_0516700","PVP01_0516800","PVP01_0517100","PVP01_0517200","PVP01_0517500","PVP01_0517800","PVP01_0518000","PVP01_0518100","PVP01_0518200","PVP01_0518400","PVP01_0519000","PVP01_0519300","PVP01_0519700","PVP01_0519800","PVP01_0519900","PVP01_0520000","PVP01_0520100","PVP01_0520200","PVP01_0520300","PVP01_0520400","PVP01_0520500","PVP01_0520600","PVP01_0520900","PVP01_0521000","PVP01_0521100","PVP01_0521200","PVP01_0521700","PVP01_0522400","PVP01_0522800","PVP01_0523400","PVP01_0524500","PVP01_0525200","PVP01_0525600","PVP01_0525800","PVP01_0525900","PVP01_0526000","PVP01_0526300","PVP01_0526500","PVP01_0526600","PVP01_0526700","PVP01_0526800","PVP01_0527200","PVP01_0527600","PVP01_0527900","PVP01_0528300","PVP01_0528800","PVP01_0529300","PVP01_0529400","PVP01_0529500","PVP01_0529800","PVP01_0530200","PVP01_0530300","PVP01_0530900","PVP01_0531000","PVP01_0531300","PVP01_0531400","PVP01_0531600","PVP01_0531800","PVP01_0532100","PVP01_0532200","PVP01_0532400","PVP01_0532800","PVP01_0600800","PVP01_0601900","PVP01_0602000","PVP01_0602100","PVP01_0602200","PVP01_0602300","PVP01_0602400","PVP01_0602500","PVP01_0602700","PVP01_0602800","PVP01_0602900","PVP01_0603200","PVP01_0603400","PVP01_0603500","PVP01_0603800","PVP01_0603900","PVP01_0604000","PVP01_0604300","PVP01_0604500","PVP01_0604600","PVP01_0604800","PVP01_0604900","PVP01_0605400","PVP01_0605600","PVP01_0605700","PVP01_0605800","PVP01_0605900","PVP01_0606100","PVP01_0606300","PVP01_0606700","PVP01_0606900","PVP01_0607000","PVP01_0607300","PVP01_0607400","PVP01_0607500","PVP01_0607700","PVP01_0607800","PVP01_0607900","PVP01_0608200","PVP01_0608400","PVP01_0609000","PVP01_0609100","PVP01_0609300","PVP01_0609500","PVP01_0610000","PVP01_0610200","PVP01_0611000","PVP01_0611200","PVP01_0611500","PVP01_0611600","PVP01_0611800","PVP01_0611900","PVP01_0612200","PVP01_0612300","PVP01_0612400","PVP01_0612500","PVP01_0612800","PVP01_0612900","PVP01_0613300","PVP01_0613600","PVP01_0613700","PVP01_0614100","PVP01_0614500","PVP01_0614700","PVP01_0614800","PVP01_0615200","PVP01_0615300","PVP01_0615400","PVP01_0615600","PVP01_0616100","PVP01_0616500","PVP01_0616600","PVP01_0616700","PVP01_0617800","PVP01_0618000","PVP01_0618200","PVP01_0618400","PVP01_0618500","PVP01_0618800","PVP01_0620000","PVP01_0620800","PVP01_0621000","PVP01_0621300","PVP01_0621500","PVP01_0621600","PVP01_0621700","PVP01_0622000","PVP01_0622300","PVP01_0622400","PVP01_0622500","PVP01_0622600","PVP01_0700100","PVP01_0701100","PVP01_0701400","PVP01_0701800","PVP01_0701900","PVP01_0702300","PVP01_0702600","PVP01_0702700","PVP01_0702800","PVP01_0702900","PVP01_0703100","PVP01_0703200","PVP01_0703300","PVP01_0703400","PVP01_0703500","PVP01_0703800","PVP01_0704200","PVP01_0704300","PVP01_0704400","PVP01_0705100","PVP01_0705200","PVP01_0705300","PVP01_0705400","PVP01_0705700","PVP01_0705900","PVP01_0706500","PVP01_0706800","PVP01_0707600","PVP01_0707700","PVP01_0707900","PVP01_0708600","PVP01_0708700","PVP01_0709000","PVP01_0709400","PVP01_0709800","PVP01_0710400","PVP01_0710500","PVP01_0710600","PVP01_0710800","PVP01_0710900","PVP01_0711300","PVP01_0711600","PVP01_0711700","PVP01_0711800","PVP01_0712500","PVP01_0712600","PVP01_0712900","PVP01_0713300","PVP01_0713400","PVP01_0713800","PVP01_0714000","PVP01_0714100","PVP01_0714400","PVP01_0714500","PVP01_0714700","PVP01_0714800","PVP01_0715400","PVP01_0715900","PVP01_0716300","PVP01_0716500","PVP01_0716700","PVP01_0717100","PVP01_0717300","PVP01_0717700","PVP01_0717900","PVP01_0718200","PVP01_0718400","PVP01_0718500","PVP01_0718900","PVP01_0719200","PVP01_0719300","PVP01_0719600","PVP01_0719800","PVP01_0720500","PVP01_0720600","PVP01_0720700","PVP01_0721000","PVP01_0721200","PVP01_0721300","PVP01_0722900","PVP01_0723100","PVP01_0723600","PVP01_0723900","PVP01_0724000","PVP01_0724200","PVP01_0724400","PVP01_0725600","PVP01_0725900","PVP01_0726000","PVP01_0726100","PVP01_0726200","PVP01_0726300","PVP01_0726500","PVP01_0726600","PVP01_0726800","PVP01_0727700","PVP01_0727900","PVP01_0728500","PVP01_0728600","PVP01_0728700","PVP01_0728800","PVP01_0728900","PVP01_0729000","PVP01_0729600","PVP01_0729800","PVP01_0729900","PVP01_0730500","PVP01_0730800","PVP01_0730900","PVP01_0731400","PVP01_0731500","PVP01_0732500","PVP01_0732700","PVP01_0733200","PVP01_0733300","PVP01_0733400","PVP01_0733600","PVP01_0734000","PVP01_0734300","PVP01_0734500","PVP01_0734600","PVP01_0734800","PVP01_0734900","PVP01_0735000","PVP01_0801200","PVP01_0801300","PVP01_0802200","PVP01_0802600","PVP01_0802700","PVP01_0802800","PVP01_0803200","PVP01_0803300","PVP01_0803800","PVP01_0804000","PVP01_0804200","PVP01_0804300","PVP01_0805100","PVP01_0805900","PVP01_0806600","PVP01_0806800","PVP01_0806900","PVP01_0807100","PVP01_0807400","PVP01_0807500","PVP01_0807700","PVP01_0807800","PVP01_0808200","PVP01_0808300","PVP01_0808900","PVP01_0809000","PVP01_0809500","PVP01_0809600","PVP01_0809800","PVP01_0810200","PVP01_0810700","PVP01_0810800","PVP01_0810900","PVP01_0811400","PVP01_0811600","PVP01_0811800","PVP01_0812100","PVP01_0812200","PVP01_0812500","PVP01_0812700","PVP01_0812900","PVP01_0813100","PVP01_0813400","PVP01_0813500","PVP01_0813900","PVP01_0814000","PVP01_0814100","PVP01_0814500","PVP01_0814600","PVP01_0814700","PVP01_0814900","PVP01_0815000","PVP01_0815200","PVP01_0815500","PVP01_0815600","PVP01_0815700","PVP01_0815800","PVP01_0815900","PVP01_0816000","PVP01_0816100","PVP01_0816400","PVP01_0816500","PVP01_0816600","PVP01_0816900","PVP01_0817000","PVP01_0817200","PVP01_0817300","PVP01_0817500","PVP01_0817600","PVP01_0817700","PVP01_0817800","PVP01_0817900","PVP01_0818000","PVP01_0818200","PVP01_0818400","PVP01_0818500","PVP01_0818700","PVP01_0818900","PVP01_0819100","PVP01_0819400","PVP01_0819500","PVP01_0819900","PVP01_0821100","PVP01_0821600","PVP01_0821900","PVP01_0822000","PVP01_0822200","PVP01_0822400","PVP01_0822500","PVP01_0822800","PVP01_0823000","PVP01_0823300","PVP01_0823500","PVP01_0823800","PVP01_0823900","PVP01_0824400","PVP01_0825300","PVP01_0825500","PVP01_0825900","PVP01_0826600","PVP01_0826800","PVP01_0827000","PVP01_0827200","PVP01_0827300","PVP01_0827400","PVP01_0827900","PVP01_0828400","PVP01_0828500","PVP01_0828600","PVP01_0828800","PVP01_0828900","PVP01_0829400","PVP01_0829600","PVP01_0830000","PVP01_0830200","PVP01_0830600","PVP01_0831200","PVP01_0831300","PVP01_0831500","PVP01_0831600","PVP01_0832300","PVP01_0832700","PVP01_0833100","PVP01_0833200","PVP01_0833400","PVP01_0833900","PVP01_0834100","PVP01_0834700","PVP01_0834800","PVP01_0835100","PVP01_0835200","PVP01_0835400","PVP01_0835600","PVP01_0835900","PVP01_0836000","PVP01_0836100","PVP01_0836200","PVP01_0836900","PVP01_0837000","PVP01_0837100","PVP01_0837400","PVP01_0904200","PVP01_0904400","PVP01_0904500","PVP01_0904800","PVP01_0905000","PVP01_0905200","PVP01_0905300","PVP01_0905400","PVP01_0905500","PVP01_0905600","PVP01_0905700","PVP01_0905800","PVP01_0905900","PVP01_0906100","PVP01_0906200","PVP01_0906600","PVP01_0907900","PVP01_0908100","PVP01_0908300","PVP01_0908700","PVP01_0908800","PVP01_0909000","PVP01_0909200","PVP01_0909400","PVP01_0909700","PVP01_0909900","PVP01_0910700","PVP01_0910800","PVP01_0911100","PVP01_0911200","PVP01_0911400","PVP01_0912300","PVP01_0912500","PVP01_0912600","PVP01_0912700","PVP01_0912800","PVP01_0913200","PVP01_0913300","PVP01_0913700","PVP01_0914100","PVP01_0914500","PVP01_0914700","PVP01_0914900","PVP01_0915000","PVP01_0915200","PVP01_0915500","PVP01_0915700","PVP01_0915900","PVP01_0916100","PVP01_0916200","PVP01_0916400","PVP01_0916500","PVP01_0916700","PVP01_0917300","PVP01_0917400","PVP01_0917500","PVP01_0917600","PVP01_0917700","PVP01_0917800","PVP01_0918500","PVP01_0919200","PVP01_0919300","PVP01_0919800","PVP01_0920000","PVP01_0920300","PVP01_0920500","PVP01_0920600","PVP01_0920900","PVP01_0921000","PVP01_0921100","PVP01_0921200","PVP01_0921300","PVP01_0921600","PVP01_0921700","PVP01_0922200","PVP01_0922500","PVP01_0922600","PVP01_0922700","PVP01_0923400","PVP01_0923500","PVP01_0923600","PVP01_0923700","PVP01_0924100","PVP01_0924200","PVP01_0924500","PVP01_0924900","PVP01_0925300","PVP01_0925500","PVP01_0926100","PVP01_0926200","PVP01_0926700","PVP01_0927000","PVP01_0927300","PVP01_0927500","PVP01_0928400","PVP01_0928500","PVP01_0928600","PVP01_0928700","PVP01_0929500","PVP01_0929800","PVP01_0930100","PVP01_0930600","PVP01_0930700","PVP01_0930800","PVP01_0931800","PVP01_0931900","PVP01_0932100","PVP01_0932400","PVP01_0932700","PVP01_0932800","PVP01_0933000","PVP01_0933200","PVP01_0933600","PVP01_0933800","PVP01_0933900","PVP01_0934100","PVP01_0934200","PVP01_0934700","PVP01_0934800","PVP01_0934900","PVP01_0935100","PVP01_0935300","PVP01_0935600","PVP01_0936200","PVP01_0936500","PVP01_0936600","PVP01_0936900","PVP01_0937100","PVP01_0937200","PVP01_0937400","PVP01_0937600","PVP01_0938000","PVP01_0938300","PVP01_0938400","PVP01_0938500","PVP01_0938800","PVP01_0939400","PVP01_0939500","PVP01_0939800","PVP01_0939900","PVP01_0940000","PVP01_0940100","PVP01_0940300","PVP01_0940400","PVP01_0940800","PVP01_0941000","PVP01_0941500","PVP01_0941700","PVP01_0941900","PVP01_0942100","PVP01_0942500","PVP01_0943000","PVP01_0943200","PVP01_0943500","PVP01_0943600","PVP01_0943900","PVP01_0944000","PVP01_0944200","PVP01_0944500","PVP01_0944700","PVP01_0944900","PVP01_0945000","PVP01_0945400","PVP01_0945800","PVP01_0947200","PVP01_0947500","PVP01_0947700","PVP01_0948000","PVP01_0948400","PVP01_0949300","PVP01_1002400","PVP01_1002800","PVP01_1002900","PVP01_1003100","PVP01_1003300","PVP01_1003600","PVP01_1003700","PVP01_1003800","PVP01_1004000","PVP01_1004200","PVP01_1004400","PVP01_1004600","PVP01_1004700","PVP01_1005000","PVP01_1005500","PVP01_1005700","PVP01_1005800","PVP01_1006900","PVP01_1007000","PVP01_1007100","PVP01_1007300","PVP01_1007400","PVP01_1007600","PVP01_1008400","PVP01_1008500","PVP01_1008800","PVP01_1009200","PVP01_1009800","PVP01_1009900","PVP01_1010000","PVP01_1010500","PVP01_1010600","PVP01_1010700","PVP01_1010800","PVP01_1010900","PVP01_1011200","PVP01_1011700","PVP01_1011900","PVP01_1012000","PVP01_1012200","PVP01_1012600","PVP01_1012800","PVP01_1012900","PVP01_1013000","PVP01_1013500","PVP01_1013600","PVP01_1013800","PVP01_1013900","PVP01_1014000","PVP01_1014200","PVP01_1014400","PVP01_1014500","PVP01_1014700","PVP01_1014900","PVP01_1015300","PVP01_1015800","PVP01_1016000","PVP01_1016300","PVP01_1016700","PVP01_1016800","PVP01_1017000","PVP01_1017300","PVP01_1017500","PVP01_1017600","PVP01_1017800","PVP01_1018300","PVP01_1018600","PVP01_1018700","PVP01_1018900","PVP01_1019100","PVP01_1019200","PVP01_1019300","PVP01_1019700","PVP01_1019800","PVP01_1020400","PVP01_1020600","PVP01_1020800","PVP01_1021400","PVP01_1022500","PVP01_1022600","PVP01_1022700","PVP01_1022800","PVP01_1023500","PVP01_1023800","PVP01_1023900","PVP01_1024000","PVP01_1024100","PVP01_1024200","PVP01_1024600","PVP01_1024700","PVP01_1024900","PVP01_1025000","PVP01_1025100","PVP01_1025500","PVP01_1025600","PVP01_1025900","PVP01_1026000","PVP01_1026100","PVP01_1026200","PVP01_1027000","PVP01_1027200","PVP01_1027500","PVP01_1028300","PVP01_1028500","PVP01_1028700","PVP01_1028800","PVP01_1028900","PVP01_1029400","PVP01_1029600","PVP01_1030800","PVP01_1032300","PVP01_1032600","PVP01_1032700","PVP01_1033100","PVP01_1033200","PVP01_1102300","PVP01_1103000","PVP01_1103100","PVP01_1103300","PVP01_1103500","PVP01_1103600","PVP01_1103800","PVP01_1103900","PVP01_1104500","PVP01_1104700","PVP01_1105000","PVP01_1105100","PVP01_1105200","PVP01_1105700","PVP01_1105900","PVP01_1106100","PVP01_1106500","PVP01_1106800","PVP01_1106900","PVP01_1107500","PVP01_1107800","PVP01_1108200","PVP01_1108400","PVP01_1108700","PVP01_1109500","PVP01_1109700","PVP01_1109800","PVP01_1110000","PVP01_1110100","PVP01_1110200","PVP01_1110300","PVP01_1110400","PVP01_1110800","PVP01_1111000","PVP01_1111900","PVP01_1112200","PVP01_1112300","PVP01_1112500","PVP01_1113100","PVP01_1113200","PVP01_1113400","PVP01_1113600","PVP01_1113800","PVP01_1114000","PVP01_1114400","PVP01_1114900","PVP01_1115100","PVP01_1115400","PVP01_1115700","PVP01_1115800","PVP01_1115900","PVP01_1116000","PVP01_1116100","PVP01_1116200","PVP01_1116300","PVP01_1116700","PVP01_1117100","PVP01_1117400","PVP01_1117600","PVP01_1117800","PVP01_1117900","PVP01_1118000","PVP01_1118200","PVP01_1118300","PVP01_1118400","PVP01_1118800","PVP01_1118900","PVP01_1119100","PVP01_1119200","PVP01_1119400","PVP01_1119800","PVP01_1120200","PVP01_1120600","PVP01_1120700","PVP01_1120800","PVP01_1121200","PVP01_1121400","PVP01_1122000","PVP01_1122200","PVP01_1123700","PVP01_1123800","PVP01_1124200","PVP01_1124500","PVP01_1125500","PVP01_1125900","PVP01_1126400","PVP01_1126700","PVP01_1127100","PVP01_1127600","PVP01_1127700","PVP01_1128200","PVP01_1128300","PVP01_1129100","PVP01_1129200","PVP01_1129300","PVP01_1129400","PVP01_1129800","PVP01_1129900","PVP01_1130000","PVP01_1130200","PVP01_1130300","PVP01_1130500","PVP01_1130800","PVP01_1130900","PVP01_1131000","PVP01_1131100","PVP01_1131200","PVP01_1131500","PVP01_1131600","PVP01_1131700","PVP01_1131800","PVP01_1132000","PVP01_1132200","PVP01_1132600","PVP01_1133500","PVP01_1133700","PVP01_1134000","PVP01_1134600","PVP01_1134900","PVP01_1135200","PVP01_1135400","PVP01_1136200","PVP01_1136500","PVP01_1136900","PVP01_1137000","PVP01_1137400","PVP01_1137500","PVP01_1137900","PVP01_1138000","PVP01_1138100","PVP01_1138300","PVP01_1138600","PVP01_1138700","PVP01_1138800","PVP01_1138900","PVP01_1139500","PVP01_1139600","PVP01_1139800","PVP01_1139900","PVP01_1140100","PVP01_1140200","PVP01_1140400","PVP01_1141100","PVP01_1141300","PVP01_1141400","PVP01_1141600","PVP01_1142100","PVP01_1142400","PVP01_1142700","PVP01_1142800","PVP01_1142900","PVP01_1143000","PVP01_1143300","PVP01_1143600","PVP01_1143800","PVP01_1143900","PVP01_1144000","PVP01_1144400","PVP01_1144600","PVP01_1144800","PVP01_1145300","PVP01_1145400","PVP01_1145900","PVP01_1146000","PVP01_1146100","PVP01_1146300","PVP01_1146700","PVP01_1202400","PVP01_1202600","PVP01_1202800","PVP01_1202900","PVP01_1203000","PVP01_1203900","PVP01_1204100","PVP01_1204300","PVP01_1204900","PVP01_1205100","PVP01_1205300","PVP01_1205500","PVP01_1205700","PVP01_1206000","PVP01_1206200","PVP01_1206500","PVP01_1206600","PVP01_1206800","PVP01_1207300","PVP01_1207400","PVP01_1207500","PVP01_1207600","PVP01_1207700","PVP01_1208400","PVP01_1208500","PVP01_1209100","PVP01_1209300","PVP01_1209500","PVP01_1209600","PVP01_1209700","PVP01_1209800","PVP01_1209900","PVP01_1210200","PVP01_1210600","PVP01_1210800","PVP01_1210900","PVP01_1211400","PVP01_1211800","PVP01_1211900","PVP01_1212000","PVP01_1212200","PVP01_1212300","PVP01_1212400","PVP01_1212800","PVP01_1212900","PVP01_1213200","PVP01_1213400","PVP01_1213600","PVP01_1213900","PVP01_1214000","PVP01_1214100","PVP01_1214700","PVP01_1215100","PVP01_1215200","PVP01_1215500","PVP01_1216400","PVP01_1216500","PVP01_1216600","PVP01_1217300","PVP01_1217800","PVP01_1217900","PVP01_1218100","PVP01_1218200","PVP01_1218300","PVP01_1218400","PVP01_1218600","PVP01_1218700","PVP01_1219300","PVP01_1219700","PVP01_1220000","PVP01_1220300","PVP01_1220400","PVP01_1220500","PVP01_1221100","PVP01_1221500","PVP01_1221600","PVP01_1222000","PVP01_1222500","PVP01_1222600","PVP01_1222800","PVP01_1222900","PVP01_1223000","PVP01_1223100","PVP01_1223200","PVP01_1223300","PVP01_1223600","PVP01_1223700","PVP01_1224100","PVP01_1224200","PVP01_1224900","PVP01_1225100","PVP01_1225400","PVP01_1225600","PVP01_1225700","PVP01_1226000","PVP01_1226200","PVP01_1226300","PVP01_1226400","PVP01_1226600","PVP01_1226900","PVP01_1227000","PVP01_1227100","PVP01_1227200","PVP01_1227700","PVP01_1227800","PVP01_1228300","PVP01_1228600","PVP01_1228700","PVP01_1228900","PVP01_1229000","PVP01_1229200","PVP01_1229300","PVP01_1229700","PVP01_1230300","PVP01_1230400","PVP01_1230900","PVP01_1231200","PVP01_1231300","PVP01_1231400","PVP01_1231500","PVP01_1232000","PVP01_1232100","PVP01_1232200","PVP01_1232300","PVP01_1232500","PVP01_1232800","PVP01_1232900","PVP01_1233000","PVP01_1233200","PVP01_1233300","PVP01_1233400","PVP01_1233500","PVP01_1233600","PVP01_1233700","PVP01_1234100","PVP01_1234200","PVP01_1234300","PVP01_1234500","PVP01_1234600","PVP01_1234700","PVP01_1234800","PVP01_1234900","PVP01_1235500","PVP01_1235600","PVP01_1235700","PVP01_1235900","PVP01_1236300","PVP01_1236500","PVP01_1236600","PVP01_1237400","PVP01_1238100","PVP01_1238200","PVP01_1238300","PVP01_1238700","PVP01_1238800","PVP01_1239200","PVP01_1239500","PVP01_1239600","PVP01_1239700","PVP01_1240400","PVP01_1240700","PVP01_1241400","PVP01_1241500","PVP01_1241700","PVP01_1241800","PVP01_1242100","PVP01_1242200","PVP01_1242500","PVP01_1243300","PVP01_1243400","PVP01_1243700","PVP01_1243800","PVP01_1244000","PVP01_1244100","PVP01_1244200","PVP01_1244300","PVP01_1244500","PVP01_1244600","PVP01_1245000","PVP01_1245200","PVP01_1245700","PVP01_1245800","PVP01_1246200","PVP01_1246800","PVP01_1247100","PVP01_1247200","PVP01_1247300","PVP01_1247400","PVP01_1248000","PVP01_1248100","PVP01_1248300","PVP01_1248400","PVP01_1248700","PVP01_1248800","PVP01_1248900","PVP01_1249300","PVP01_1249400","PVP01_1249800","PVP01_1249900","PVP01_1250000","PVP01_1250100","PVP01_1250400","PVP01_1250500","PVP01_1250700","PVP01_1250800","PVP01_1251000","PVP01_1251200","PVP01_1251300","PVP01_1251400","PVP01_1252100","PVP01_1252300","PVP01_1252500","PVP01_1252600","PVP01_1252700","PVP01_1253100","PVP01_1253200","PVP01_1253400","PVP01_1253500","PVP01_1253900","PVP01_1254000","PVP01_1254200","PVP01_1254400","PVP01_1254500","PVP01_1254700","PVP01_1254800","PVP01_1255000","PVP01_1255600","PVP01_1255700","PVP01_1255800","PVP01_1255900","PVP01_1256300","PVP01_1256600","PVP01_1257100","PVP01_1257400","PVP01_1257800","PVP01_1258000","PVP01_1258600","PVP01_1258800","PVP01_1259700","PVP01_1259800","PVP01_1259900","PVP01_1260000","PVP01_1260100","PVP01_1260300","PVP01_1260400","PVP01_1260500","PVP01_1260600","PVP01_1260700","PVP01_1261200","PVP01_1261300","PVP01_1261600","PVP01_1261800","PVP01_1261900","PVP01_1262100","PVP01_1262700","PVP01_1263200","PVP01_1263300","PVP01_1263500","PVP01_1263600","PVP01_1264100","PVP01_1264200","PVP01_1264500","PVP01_1265200","PVP01_1265500","PVP01_1265800","PVP01_1266000","PVP01_1266100","PVP01_1266300","PVP01_1266800","PVP01_1266900","PVP01_1267000","PVP01_1267100","PVP01_1267200","PVP01_1267700","PVP01_1267800","PVP01_1267900","PVP01_1268400","PVP01_1268500","PVP01_1269100","PVP01_1269300","PVP01_1269400","PVP01_1269600","PVP01_1269700","PVP01_1269900","PVP01_1270000","PVP01_1270100","PVP01_1270500","PVP01_1271000","PVP01_1300600","PVP01_1300800","PVP01_1301000","PVP01_1301800","PVP01_1301900","PVP01_1302000","PVP01_1302200","PVP01_1302400","PVP01_1303300","PVP01_1303500","PVP01_1303700","PVP01_1303800","PVP01_1304100","PVP01_1304400","PVP01_1304600","PVP01_1304700","PVP01_1305000","PVP01_1305100","PVP01_1305400","PVP01_1305600","PVP01_1305800","PVP01_1306200","PVP01_1306500","PVP01_1307400","PVP01_1307600","PVP01_1307900","PVP01_1308000","PVP01_1308200","PVP01_1308600","PVP01_1308700","PVP01_1309300","PVP01_1309600","PVP01_1309900","PVP01_1310100","PVP01_1310200","PVP01_1310300","PVP01_1310600","PVP01_1310900","PVP01_1311700","PVP01_1311800","PVP01_1312300","PVP01_1312500","PVP01_1312700","PVP01_1312800","PVP01_1313500","PVP01_1313600","PVP01_1313700","PVP01_1313800","PVP01_1313900","PVP01_1314000","PVP01_1314200","PVP01_1314400","PVP01_1314500","PVP01_1314700","PVP01_1314900","PVP01_1315000","PVP01_1315100","PVP01_1315300","PVP01_1315600","PVP01_1315700","PVP01_1315800","PVP01_1315900","PVP01_1316400","PVP01_1316800","PVP01_1317100","PVP01_1317200","PVP01_1317300","PVP01_1317400","PVP01_1318200","PVP01_1318400","PVP01_1318700","PVP01_1319100","PVP01_1319200","PVP01_1319400","PVP01_1320400","PVP01_1320800","PVP01_1320900","PVP01_1321000","PVP01_1321100","PVP01_1321200","PVP01_1321300","PVP01_1321400","PVP01_1321800","PVP01_1322100","PVP01_1322500","PVP01_1322700","PVP01_1322800","PVP01_1323300","PVP01_1323400","PVP01_1323500","PVP01_1323600","PVP01_1324000","PVP01_1325300","PVP01_1325400","PVP01_1325500","PVP01_1325700","PVP01_1326100","PVP01_1326300","PVP01_1326500","PVP01_1326700","PVP01_1326800","PVP01_1327000","PVP01_1327100","PVP01_1327400","PVP01_1327700","PVP01_1327800","PVP01_1328200","PVP01_1328300","PVP01_1328400","PVP01_1329100","PVP01_1329200","PVP01_1329500","PVP01_1330600","PVP01_1330700","PVP01_1330900","PVP01_1331300","PVP01_1331400","PVP01_1331500","PVP01_1331600","PVP01_1331700","PVP01_1332000","PVP01_1332100","PVP01_1332600","PVP01_1333300","PVP01_1333400","PVP01_1333500","PVP01_1333600","PVP01_1333800","PVP01_1334000","PVP01_1334400","PVP01_1334600","PVP01_1334700","PVP01_1334800","PVP01_1334900","PVP01_1335000","PVP01_1335200","PVP01_1335400","PVP01_1335500","PVP01_1335700","PVP01_1336000","PVP01_1336100","PVP01_1336200","PVP01_1336300","PVP01_1336600","PVP01_1336800","PVP01_1337000","PVP01_1337700","PVP01_1337800","PVP01_1338000","PVP01_1338200","PVP01_1338800","PVP01_1338900","PVP01_1339100","PVP01_1339200","PVP01_1339300","PVP01_1339500","PVP01_1339700","PVP01_1340000","PVP01_1340700","PVP01_1340900","PVP01_1341000","PVP01_1341100","PVP01_1341200","PVP01_1341400","PVP01_1341500","PVP01_1342000","PVP01_1342100","PVP01_1342500","PVP01_1342900","PVP01_1343000","PVP01_1343200","PVP01_1343400","PVP01_1343700","PVP01_1343800","PVP01_1343900","PVP01_1344100","PVP01_1344400","PVP01_1344800","PVP01_1345100","PVP01_1345800","PVP01_1346000","PVP01_1346100","PVP01_1346400","PVP01_1346500","PVP01_1346600","PVP01_1347000","PVP01_1401300","PVP01_1401400","PVP01_1403100","PVP01_1403200","PVP01_1403800","PVP01_1404400","PVP01_1404900","PVP01_1405200","PVP01_1405400","PVP01_1405700","PVP01_1405900","PVP01_1406200","PVP01_1406300","PVP01_1406400","PVP01_1406500","PVP01_1406800","PVP01_1407200","PVP01_1407300","PVP01_1407700","PVP01_1407900","PVP01_1408300","PVP01_1409200","PVP01_1409500","PVP01_1409600","PVP01_1409800","PVP01_1409900","PVP01_1410600","PVP01_1411100","PVP01_1411500","PVP01_1411600","PVP01_1411700","PVP01_1411800","PVP01_1411900","PVP01_1412000","PVP01_1412200","PVP01_1412300","PVP01_1412600","PVP01_1412700","PVP01_1412800","PVP01_1413400","PVP01_1413500","PVP01_1413600","PVP01_1413800","PVP01_1413900","PVP01_1414100","PVP01_1414600","PVP01_1414700","PVP01_1414800","PVP01_1415400","PVP01_1416300","PVP01_1416400","PVP01_1416500","PVP01_1416600","PVP01_1416700","PVP01_1416800","PVP01_1417000","PVP01_1417400","PVP01_1417500","PVP01_1417800","PVP01_1418300","PVP01_1418500","PVP01_1418700","PVP01_1419000","PVP01_1419200","PVP01_1419600","PVP01_1419900","PVP01_1420200","PVP01_1420400","PVP01_1420600","PVP01_1420700","PVP01_1420800","PVP01_1421600","PVP01_1421700","PVP01_1421900","PVP01_1422000","PVP01_1422400","PVP01_1422500","PVP01_1422800","PVP01_1423000","PVP01_1423300","PVP01_1423400","PVP01_1423600","PVP01_1424300","PVP01_1424400","PVP01_1424600","PVP01_1424700","PVP01_1424800","PVP01_1424900","PVP01_1425000","PVP01_1425700","PVP01_1425900","PVP01_1426300","PVP01_1426400","PVP01_1426500","PVP01_1426600","PVP01_1426900","PVP01_1427200","PVP01_1427300","PVP01_1427400","PVP01_1427500","PVP01_1427900","PVP01_1428200","PVP01_1428700","PVP01_1429000","PVP01_1429100","PVP01_1429400","PVP01_1429600","PVP01_1429800","PVP01_1430200","PVP01_1430900","PVP01_1431000","PVP01_1431100","PVP01_1431400","PVP01_1431600","PVP01_1432200","PVP01_1432500","PVP01_1433000","PVP01_1433100","PVP01_1433200","PVP01_1433300","PVP01_1433400","PVP01_1433500","PVP01_1433900","PVP01_1434100","PVP01_1434300","PVP01_1434700","PVP01_1435400","PVP01_1435500","PVP01_1435700","PVP01_1436100","PVP01_1436200","PVP01_1436500","PVP01_1436800","PVP01_1436900","PVP01_1437000","PVP01_1437500","PVP01_1437700","PVP01_1437800","PVP01_1437900","PVP01_1438000","PVP01_1438500","PVP01_1439100","PVP01_1439200","PVP01_1439300","PVP01_1439500","PVP01_1439800","PVP01_1439900","PVP01_1440200","PVP01_1440400","PVP01_1440500","PVP01_1441300","PVP01_1441500","PVP01_1441600","PVP01_1441900","PVP01_1442100","PVP01_1442400","PVP01_1442500","PVP01_1442700","PVP01_1442800","PVP01_1443300","PVP01_1443400","PVP01_1443500","PVP01_1444100","PVP01_1444300","PVP01_1444500","PVP01_1444600","PVP01_1444700","PVP01_1445100","PVP01_1445500","PVP01_1445800","PVP01_1445900","PVP01_1446500","PVP01_1446700","PVP01_1447000","PVP01_1447100","PVP01_1447400","PVP01_1447700","PVP01_1448000","PVP01_1448500","PVP01_1448900","PVP01_1449000","PVP01_1449400","PVP01_1449500","PVP01_1449600","PVP01_1449800","PVP01_1449900","PVP01_1450300","PVP01_1450600","PVP01_1450700","PVP01_1450800","PVP01_1450900","PVP01_1451200","PVP01_1451400","PVP01_1451900","PVP01_1452000","PVP01_1452200","PVP01_1452600","PVP01_1453000","PVP01_1453300","PVP01_1453400","PVP01_1453500","PVP01_1453600","PVP01_1453700","PVP01_1453900","PVP01_1454300","PVP01_1454700","PVP01_1455000","PVP01_1455300","PVP01_1455600","PVP01_1455800","PVP01_1455900","PVP01_1456100","PVP01_1456300","PVP01_1456400","PVP01_1456600","PVP01_1456800","PVP01_1457200","PVP01_1457300","PVP01_1457600","PVP01_1457700","PVP01_1457800","PVP01_1458000","PVP01_1458100","PVP01_1458300","PVP01_1458400","PVP01_1458500","PVP01_1458600","PVP01_1458900","PVP01_1459000","PVP01_1459800","PVP01_1459900","PVP01_1460000","PVP01_1460100","PVP01_1460200","PVP01_1460400","PVP01_1460600","PVP01_1460700","PVP01_1461000","PVP01_1461200","PVP01_1461300","PVP01_1461500","PVP01_1461600","PVP01_1461800","PVP01_1462200","PVP01_1462400","PVP01_1462700","PVP01_1462800","PVP01_1463100","PVP01_1463400","PVP01_1463900","PVP01_1464000","PVP01_1464200","PVP01_1464600","PVP01_1464800","PVP01_1465300","PVP01_1465400","PVP01_1465600","PVP01_1466100","PVP01_1466400","PVP01_1466700","PVP01_1466800","PVP01_1466900","PVP01_1467400","PVP01_1467800","PVP01_1467900","PVP01_1468000","PVP01_1468200","PVP01_1468500","PVP01_1468700","PVP01_1468900","PVP01_API02600","PVP01_API02800","PVP01_API03500","PVP01_API04300","PVP01_API04600","PVP01_MIT02700","PVP01_MIT03300","PVP01_MIT03400"],"Row.names":["PVP01_0000030","PVP01_0000040","PVP01_0000050","PVP01_0000080","PVP01_0000110","PVP01_0000120","PVP01_0000130","PVP01_0000140","PVP01_0000150","PVP01_0000160","PVP01_0000170","PVP01_0000180","PVP01_0000190","PVP01_0000200","PVP01_0000210","PVP01_0000220","PVP01_0000300","PVP01_0000480","PVP01_0000670","PVP01_0000740","PVP01_0000790","PVP01_0001030","PVP01_0001090","PVP01_0001310","PVP01_0001430","PVP01_0001440","PVP01_0001480","PVP01_0001550","PVP01_0001640","PVP01_0001660","PVP01_0001670","PVP01_0001900","PVP01_0001910","PVP01_0001940","PVP01_0001970","PVP01_0002020","PVP01_0002050","PVP01_0002120","PVP01_0002200","PVP01_0002210","PVP01_0002330","PVP01_0002370","PVP01_0002390","PVP01_0002470","PVP01_0002530","PVP01_0002540","PVP01_0002750","PVP01_0002810","PVP01_0002820","PVP01_0003010","PVP01_0003050","PVP01_0003150","PVP01_0003210","PVP01_0003350","PVP01_0003360","PVP01_0003370","PVP01_0003380","PVP01_0003400","PVP01_0003410","PVP01_0003420","PVP01_0003440","PVP01_0003530","PVP01_0003640","PVP01_0003690","PVP01_0003730","PVP01_0003800","PVP01_0003990","PVP01_0004040","PVP01_0004160","PVP01_0004170","PVP01_0004180","PVP01_0004190","PVP01_0004210","PVP01_0004240","PVP01_0004250","PVP01_0004260","PVP01_0004270","PVP01_0004280","PVP01_0004290","PVP01_0004300","PVP01_0004310","PVP01_0004320","PVP01_0004330","PVP01_0004350","PVP01_0004370","PVP01_0004380","PVP01_0004390","PVP01_0004400","PVP01_0004410","PVP01_0004640","PVP01_0004670","PVP01_0004910","PVP01_0004920","PVP01_0004950","PVP01_0004960","PVP01_0005010","PVP01_0005100","PVP01_0005160","PVP01_0005210","PVP01_0005300","PVP01_0005310","PVP01_0005420","PVP01_0005430","PVP01_0005550","PVP01_0005640","PVP01_0005710","PVP01_0005740","PVP01_0005760","PVP01_0005800","PVP01_0005820","PVP01_0005870","PVP01_0005910","PVP01_0006000","PVP01_0006040","PVP01_0006080","PVP01_0006260","PVP01_0006650","PVP01_0006660","PVP01_0006670","PVP01_0006680","PVP01_0006700","PVP01_0006860","PVP01_0006890","PVP01_0007080","PVP01_0007160","PVP01_0007240","PVP01_0007290","PVP01_0007310","PVP01_0007320","PVP01_0007340","PVP01_0007610","PVP01_0007620","PVP01_0007690","PVP01_0007700","PVP01_0007850","PVP01_0007900","PVP01_0007910","PVP01_0007920","PVP01_0008190","PVP01_0008430","PVP01_0008480","PVP01_0008490","PVP01_0008600","PVP01_0008700","PVP01_0008770","PVP01_0008830","PVP01_0008990","PVP01_0009010","PVP01_0009170","PVP01_0009180","PVP01_0009210","PVP01_0009410","PVP01_0009470","PVP01_0009540","PVP01_0009620","PVP01_0009710","PVP01_0009720","PVP01_0009730","PVP01_0009740","PVP01_0009760","PVP01_0009770","PVP01_0009820","PVP01_0009850","PVP01_0009860","PVP01_0009870","PVP01_0009900","PVP01_0009940","PVP01_0009980","PVP01_0010020","PVP01_0010030","PVP01_0010070","PVP01_0010100","PVP01_0010120","PVP01_0010140","PVP01_0010170","PVP01_0010180","PVP01_0010200","PVP01_0010210","PVP01_0010220","PVP01_0010230","PVP01_0010240","PVP01_0010340","PVP01_0010380","PVP01_0010440","PVP01_0010470","PVP01_0010500","PVP01_0010570","PVP01_0010600","PVP01_0010610","PVP01_0010640","PVP01_0010650","PVP01_0010670","PVP01_0010770","PVP01_0010870","PVP01_0010900","PVP01_0010910","PVP01_0010960","PVP01_0010970","PVP01_0010990","PVP01_0011030","PVP01_0011070","PVP01_0102100","PVP01_0102200","PVP01_0102300","PVP01_0102400","PVP01_0102500","PVP01_0102600","PVP01_0102700","PVP01_0102800","PVP01_0103100","PVP01_0103300","PVP01_0103400","PVP01_0103500","PVP01_0103600","PVP01_0103800","PVP01_0103900","PVP01_0104100","PVP01_0104200","PVP01_0104600","PVP01_0104700","PVP01_0104800","PVP01_0104900","PVP01_0105000","PVP01_0105100","PVP01_0105200","PVP01_0105300","PVP01_0105400","PVP01_0105500","PVP01_0105600","PVP01_0105700","PVP01_0105800","PVP01_0106000","PVP01_0106200","PVP01_0106300","PVP01_0106400","PVP01_0106500","PVP01_0106800","PVP01_0106900","PVP01_0107400","PVP01_0107500","PVP01_0107600","PVP01_0107700","PVP01_0107800","PVP01_0107900","PVP01_0108000","PVP01_0108100","PVP01_0108200","PVP01_0108300","PVP01_0108400","PVP01_0108500","PVP01_0108600","PVP01_0108800","PVP01_0109000","PVP01_0109100","PVP01_0109200","PVP01_0109300","PVP01_0109600","PVP01_0109800","PVP01_0110000","PVP01_0110200","PVP01_0110500","PVP01_0110700","PVP01_0110800","PVP01_0110900","PVP01_0111100","PVP01_0111200","PVP01_0111300","PVP01_0111500","PVP01_0111600","PVP01_0111700","PVP01_0112000","PVP01_0112100","PVP01_0112300","PVP01_0112500","PVP01_0112700","PVP01_0113000","PVP01_0113100","PVP01_0113200","PVP01_0113300","PVP01_0113400","PVP01_0113500","PVP01_0114100","PVP01_0114200","PVP01_0114500","PVP01_0114600","PVP01_0114700","PVP01_0114900","PVP01_0115000","PVP01_0115100","PVP01_0115200","PVP01_0115400","PVP01_0115500","PVP01_0116100","PVP01_0116300","PVP01_0116500","PVP01_0116600","PVP01_0116700","PVP01_0116900","PVP01_0117300","PVP01_0117500","PVP01_0117700","PVP01_0117800","PVP01_0118000","PVP01_0118500","PVP01_0118600","PVP01_0118700","PVP01_0118800","PVP01_0118900","PVP01_0119200","PVP01_0119300","PVP01_0119400","PVP01_0119500","PVP01_0119600","PVP01_0119700","PVP01_0119800","PVP01_0119900","PVP01_0120000","PVP01_0120100","PVP01_0120500","PVP01_0120800","PVP01_0121000","PVP01_0121700","PVP01_0200200","PVP01_0200700","PVP01_0201000","PVP01_0201100","PVP01_0201300","PVP01_0201400","PVP01_0201500","PVP01_0201600","PVP01_0201700","PVP01_0201800","PVP01_0201900","PVP01_0202000","PVP01_0202100","PVP01_0202200","PVP01_0202400","PVP01_0203000","PVP01_0203100","PVP01_0203300","PVP01_0203400","PVP01_0203500","PVP01_0203800","PVP01_0203900","PVP01_0204000","PVP01_0204100","PVP01_0204300","PVP01_0204400","PVP01_0204800","PVP01_0204900","PVP01_0205000","PVP01_0205100","PVP01_0205200","PVP01_0205300","PVP01_0205500","PVP01_0205600","PVP01_0205800","PVP01_0206100","PVP01_0206200","PVP01_0206300","PVP01_0206400","PVP01_0206500","PVP01_0206600","PVP01_0206800","PVP01_0207000","PVP01_0207100","PVP01_0207200","PVP01_0207500","PVP01_0207600","PVP01_0207700","PVP01_0207800","PVP01_0207900","PVP01_0208000","PVP01_0208100","PVP01_0208200","PVP01_0208300","PVP01_0208400","PVP01_0208600","PVP01_0208700","PVP01_0208800","PVP01_0208900","PVP01_0209000","PVP01_0209100","PVP01_0209200","PVP01_0209300","PVP01_0209500","PVP01_0209600","PVP01_0210000","PVP01_0210100","PVP01_0210300","PVP01_0211000","PVP01_0211100","PVP01_0211200","PVP01_0211400","PVP01_0211500","PVP01_0211600","PVP01_0211700","PVP01_0211800","PVP01_0211900","PVP01_0212400","PVP01_0212500","PVP01_0212600","PVP01_0212700","PVP01_0212800","PVP01_0213000","PVP01_0213200","PVP01_0213300","PVP01_0213400","PVP01_0213600","PVP01_0213700","PVP01_0213800","PVP01_0213900","PVP01_0214100","PVP01_0214200","PVP01_0214800","PVP01_0214900","PVP01_0215000","PVP01_0215100","PVP01_0215300","PVP01_0215400","PVP01_0215500","PVP01_0215600","PVP01_0215700","PVP01_0215800","PVP01_0215900","PVP01_0216100","PVP01_0216500","PVP01_0216600","PVP01_0216700","PVP01_0216800","PVP01_0216900","PVP01_0217000","PVP01_0217100","PVP01_0217200","PVP01_0217300","PVP01_0217400","PVP01_0217500","PVP01_0218200","PVP01_0218300","PVP01_0218900","PVP01_0219400","PVP01_0220100","PVP01_0220200","PVP01_0221100","PVP01_0221300","PVP01_0221700","PVP01_0300100","PVP01_0300600","PVP01_0300700","PVP01_0300800","PVP01_0301100","PVP01_0301200","PVP01_0301500","PVP01_0301900","PVP01_0302000","PVP01_0302100","PVP01_0302300","PVP01_0302400","PVP01_0302500","PVP01_0302600","PVP01_0302800","PVP01_0302900","PVP01_0303100","PVP01_0303400","PVP01_0303500","PVP01_0304100","PVP01_0304400","PVP01_0304500","PVP01_0304600","PVP01_0304800","PVP01_0304900","PVP01_0305000","PVP01_0305200","PVP01_0305300","PVP01_0305500","PVP01_0305600","PVP01_0305700","PVP01_0305800","PVP01_0305900","PVP01_0306000","PVP01_0306100","PVP01_0306300","PVP01_0306400","PVP01_0306500","PVP01_0306600","PVP01_0306700","PVP01_0306800","PVP01_0306900","PVP01_0307000","PVP01_0307100","PVP01_0307300","PVP01_0307400","PVP01_0307500","PVP01_0307600","PVP01_0307700","PVP01_0308200","PVP01_0308600","PVP01_0308800","PVP01_0309200","PVP01_0309300","PVP01_0309700","PVP01_0309800","PVP01_0309900","PVP01_0310000","PVP01_0310200","PVP01_0310300","PVP01_0310400","PVP01_0310500","PVP01_0310600","PVP01_0311000","PVP01_0311200","PVP01_0311300","PVP01_0311400","PVP01_0311500","PVP01_0311600","PVP01_0311700","PVP01_0312000","PVP01_0312100","PVP01_0312200","PVP01_0312300","PVP01_0312400","PVP01_0312500","PVP01_0312700","PVP01_0312800","PVP01_0313500","PVP01_0313800","PVP01_0314000","PVP01_0314100","PVP01_0314200","PVP01_0314700","PVP01_0314900","PVP01_0315000","PVP01_0315100","PVP01_0315200","PVP01_0315300","PVP01_0315700","PVP01_0315900","PVP01_0316000","PVP01_0316200","PVP01_0316300","PVP01_0316500","PVP01_0316700","PVP01_0316800","PVP01_0317100","PVP01_0317200","PVP01_0317400","PVP01_0318000","PVP01_0318100","PVP01_0318400","PVP01_0318700","PVP01_0318900","PVP01_0319200","PVP01_0319300","PVP01_0319400","PVP01_0319600","PVP01_0319700","PVP01_0320200","PVP01_0320400","PVP01_0320500","PVP01_0320600","PVP01_0320700","PVP01_0320800","PVP01_0320900","PVP01_0321000","PVP01_0321100","PVP01_0400900","PVP01_0403600","PVP01_0403700","PVP01_0403900","PVP01_0404000","PVP01_0404100","PVP01_0404200","PVP01_0404300","PVP01_0404400","PVP01_0404500","PVP01_0404600","PVP01_0404700","PVP01_0404800","PVP01_0404900","PVP01_0405000","PVP01_0405100","PVP01_0405200","PVP01_0405300","PVP01_0405500","PVP01_0405600","PVP01_0405700","PVP01_0405900","PVP01_0406500","PVP01_0406600","PVP01_0406700","PVP01_0406900","PVP01_0407000","PVP01_0407300","PVP01_0407600","PVP01_0407700","PVP01_0407900","PVP01_0408200","PVP01_0408600","PVP01_0408800","PVP01_0408900","PVP01_0409000","PVP01_0409100","PVP01_0409200","PVP01_0409700","PVP01_0409800","PVP01_0409900","PVP01_0410000","PVP01_0410200","PVP01_0410400","PVP01_0410500","PVP01_0410600","PVP01_0410900","PVP01_0411200","PVP01_0411400","PVP01_0411600","PVP01_0411800","PVP01_0411900","PVP01_0412100","PVP01_0412200","PVP01_0412700","PVP01_0412800","PVP01_0412900","PVP01_0413000","PVP01_0413100","PVP01_0413200","PVP01_0413300","PVP01_0413400","PVP01_0413500","PVP01_0414000","PVP01_0414100","PVP01_0414300","PVP01_0414400","PVP01_0414600","PVP01_0414900","PVP01_0415400","PVP01_0415500","PVP01_0415600","PVP01_0415800","PVP01_0416100","PVP01_0416600","PVP01_0416900","PVP01_0417100","PVP01_0417200","PVP01_0417300","PVP01_0417400","PVP01_0417600","PVP01_0417700","PVP01_0417800","PVP01_0417900","PVP01_0418100","PVP01_0418300","PVP01_0418400","PVP01_0419000","PVP01_0419200","PVP01_0419300","PVP01_0419400","PVP01_0419500","PVP01_0419600","PVP01_0419700","PVP01_0420000","PVP01_0420100","PVP01_0420200","PVP01_0420500","PVP01_0420700","PVP01_0420900","PVP01_0421400","PVP01_0421500","PVP01_0421700","PVP01_0421800","PVP01_0421900","PVP01_0422100","PVP01_0422200","PVP01_0422300","PVP01_0422400","PVP01_0422500","PVP01_0422600","PVP01_0422700","PVP01_0422900","PVP01_0423000","PVP01_0423100","PVP01_0423200","PVP01_0423300","PVP01_0423400","PVP01_0423500","PVP01_0423600","PVP01_0423800","PVP01_0423900","PVP01_0424300","PVP01_0424400","PVP01_0500700","PVP01_0502300","PVP01_0502400","PVP01_0502900","PVP01_0503300","PVP01_0503400","PVP01_0503500","PVP01_0503600","PVP01_0503700","PVP01_0503800","PVP01_0503900","PVP01_0504000","PVP01_0504100","PVP01_0504200","PVP01_0504300","PVP01_0505000","PVP01_0505100","PVP01_0505400","PVP01_0505500","PVP01_0505700","PVP01_0505800","PVP01_0506000","PVP01_0506300","PVP01_0506400","PVP01_0506600","PVP01_0506700","PVP01_0506800","PVP01_0506900","PVP01_0507000","PVP01_0507300","PVP01_0507500","PVP01_0507600","PVP01_0507700","PVP01_0507800","PVP01_0508000","PVP01_0508200","PVP01_0508500","PVP01_0508600","PVP01_0508700","PVP01_0509100","PVP01_0509200","PVP01_0509400","PVP01_0509600","PVP01_0509800","PVP01_0509900","PVP01_0510000","PVP01_0510100","PVP01_0510300","PVP01_0510500","PVP01_0510800","PVP01_0510900","PVP01_0511200","PVP01_0511300","PVP01_0511400","PVP01_0511500","PVP01_0511700","PVP01_0511800","PVP01_0512100","PVP01_0512200","PVP01_0512400","PVP01_0512500","PVP01_0512600","PVP01_0512700","PVP01_0512800","PVP01_0513000","PVP01_0513300","PVP01_0513400","PVP01_0513500","PVP01_0513700","PVP01_0513900","PVP01_0514100","PVP01_0514400","PVP01_0514600","PVP01_0514800","PVP01_0514900","PVP01_0515100","PVP01_0515200","PVP01_0515300","PVP01_0515600","PVP01_0515700","PVP01_0515800","PVP01_0515900","PVP01_0516000","PVP01_0516100","PVP01_0516200","PVP01_0516300","PVP01_0516400","PVP01_0516500","PVP01_0516900","PVP01_0517000","PVP01_0517300","PVP01_0517400","PVP01_0517600","PVP01_0517700","PVP01_0517900","PVP01_0518300","PVP01_0518500","PVP01_0518600","PVP01_0518700","PVP01_0518800","PVP01_0518900","PVP01_0519100","PVP01_0519200","PVP01_0519400","PVP01_0519500","PVP01_0519600","PVP01_0520700","PVP01_0520800","PVP01_0521300","PVP01_0521400","PVP01_0521500","PVP01_0521600","PVP01_0521800","PVP01_0521900","PVP01_0522000","PVP01_0522100","PVP01_0522200","PVP01_0522300","PVP01_0522500","PVP01_0522600","PVP01_0522700","PVP01_0522900","PVP01_0523000","PVP01_0523100","PVP01_0523200","PVP01_0523300","PVP01_0523500","PVP01_0523600","PVP01_0523700","PVP01_0523800","PVP01_0523900","PVP01_0524000","PVP01_0524100","PVP01_0524200","PVP01_0524300","PVP01_0524600","PVP01_0524700","PVP01_0524900","PVP01_0525000","PVP01_0525100","PVP01_0525300","PVP01_0525400","PVP01_0525500","PVP01_0525700","PVP01_0526100","PVP01_0526200","PVP01_0526400","PVP01_0526900","PVP01_0527000","PVP01_0527100","PVP01_0527300","PVP01_0527400","PVP01_0527500","PVP01_0527700","PVP01_0527800","PVP01_0528000","PVP01_0528100","PVP01_0528200","PVP01_0528400","PVP01_0528500","PVP01_0528600","PVP01_0528700","PVP01_0528900","PVP01_0529000","PVP01_0529100","PVP01_0529200","PVP01_0529600","PVP01_0529700","PVP01_0529900","PVP01_0530000","PVP01_0530100","PVP01_0530400","PVP01_0530500","PVP01_0530600","PVP01_0530700","PVP01_0530800","PVP01_0531100","PVP01_0531200","PVP01_0531500","PVP01_0531700","PVP01_0531900","PVP01_0532000","PVP01_0532300","PVP01_0532500","PVP01_0532600","PVP01_0532700","PVP01_0532900","PVP01_0533000","PVP01_0533100","PVP01_0533200","PVP01_0533300","PVP01_0533700","PVP01_0534000","PVP01_0534100","PVP01_0534200","PVP01_0534300","PVP01_0534400","PVP01_0534600","PVP01_0534800","PVP01_0534900","PVP01_0535400","PVP01_0600100","PVP01_0600300","PVP01_0600400","PVP01_0600500","PVP01_0600600","PVP01_0600700","PVP01_0600900","PVP01_0601000","PVP01_0601100","PVP01_0601200","PVP01_0601300","PVP01_0601400","PVP01_0601500","PVP01_0601600","PVP01_0601700","PVP01_0601800","PVP01_0602600","PVP01_0603000","PVP01_0603100","PVP01_0603300","PVP01_0603600","PVP01_0603700","PVP01_0604100","PVP01_0604200","PVP01_0604400","PVP01_0604700","PVP01_0605000","PVP01_0605100","PVP01_0605200","PVP01_0605300","PVP01_0605500","PVP01_0606000","PVP01_0606200","PVP01_0606400","PVP01_0606500","PVP01_0606600","PVP01_0606800","PVP01_0607100","PVP01_0607200","PVP01_0607600","PVP01_0608000","PVP01_0608100","PVP01_0608300","PVP01_0608500","PVP01_0608600","PVP01_0608700","PVP01_0608800","PVP01_0608900","PVP01_0609200","PVP01_0609400","PVP01_0609600","PVP01_0609700","PVP01_0609800","PVP01_0609900","PVP01_0610100","PVP01_0610300","PVP01_0610400","PVP01_0610500","PVP01_0610600","PVP01_0610700","PVP01_0610800","PVP01_0610900","PVP01_0611100","PVP01_0611300","PVP01_0611400","PVP01_0611700","PVP01_0612000","PVP01_0612100","PVP01_0612600","PVP01_0612700","PVP01_0613000","PVP01_0613100","PVP01_0613200","PVP01_0613400","PVP01_0613500","PVP01_0613800","PVP01_0613900","PVP01_0614000","PVP01_0614200","PVP01_0614300","PVP01_0614400","PVP01_0614600","PVP01_0614900","PVP01_0615000","PVP01_0615100","PVP01_0615500","PVP01_0615700","PVP01_0615800","PVP01_0615900","PVP01_0616000","PVP01_0616200","PVP01_0616300","PVP01_0616400","PVP01_0616800","PVP01_0616900","PVP01_0617000","PVP01_0617100","PVP01_0617200","PVP01_0617300","PVP01_0617400","PVP01_0617500","PVP01_0617600","PVP01_0617700","PVP01_0617900","PVP01_0618100","PVP01_0618300","PVP01_0618600","PVP01_0618700","PVP01_0618900","PVP01_0619000","PVP01_0619100","PVP01_0619200","PVP01_0619300","PVP01_0619400","PVP01_0619500","PVP01_0619600","PVP01_0619700","PVP01_0619800","PVP01_0619900","PVP01_0620100","PVP01_0620200","PVP01_0620300","PVP01_0620400","PVP01_0620500","PVP01_0620600","PVP01_0620700","PVP01_0620900","PVP01_0621100","PVP01_0621200","PVP01_0621400","PVP01_0621800","PVP01_0621900","PVP01_0622100","PVP01_0622200","PVP01_0622700","PVP01_0623000","PVP01_0623100","PVP01_0623200","PVP01_0623300","PVP01_0623400","PVP01_0623500","PVP01_0623600","PVP01_0623700","PVP01_0623800","PVP01_0623900","PVP01_0624000","PVP01_0624100","PVP01_0624300","PVP01_0624400","PVP01_0624500","PVP01_0700200","PVP01_0700300","PVP01_0700400","PVP01_0700500","PVP01_0700600","PVP01_0700700","PVP01_0700800","PVP01_0700900","PVP01_0701000","PVP01_0701200","PVP01_0701300","PVP01_0701500","PVP01_0701600","PVP01_0701700","PVP01_0702000","PVP01_0702100","PVP01_0702200","PVP01_0702400","PVP01_0702500","PVP01_0703000","PVP01_0703600","PVP01_0703700","PVP01_0703900","PVP01_0704000","PVP01_0704100","PVP01_0704500","PVP01_0704600","PVP01_0704700","PVP01_0704800","PVP01_0704900","PVP01_0705000","PVP01_0705500","PVP01_0705600","PVP01_0705800","PVP01_0706000","PVP01_0706100","PVP01_0706200","PVP01_0706300","PVP01_0706400","PVP01_0706600","PVP01_0706700","PVP01_0706900","PVP01_0707000","PVP01_0707100","PVP01_0707200","PVP01_0707300","PVP01_0707400","PVP01_0707500","PVP01_0707800","PVP01_0708000","PVP01_0708100","PVP01_0708200","PVP01_0708300","PVP01_0708400","PVP01_0708500","PVP01_0708800","PVP01_0708900","PVP01_0709100","PVP01_0709200","PVP01_0709300","PVP01_0709500","PVP01_0709600","PVP01_0709700","PVP01_0709900","PVP01_0710000","PVP01_0710100","PVP01_0710200","PVP01_0710300","PVP01_0710700","PVP01_0711000","PVP01_0711100","PVP01_0711200","PVP01_0711400","PVP01_0711500","PVP01_0711900","PVP01_0712000","PVP01_0712100","PVP01_0712200","PVP01_0712300","PVP01_0712400","PVP01_0712700","PVP01_0712800","PVP01_0713000","PVP01_0713100","PVP01_0713200","PVP01_0713500","PVP01_0713600","PVP01_0713700","PVP01_0713900","PVP01_0714200","PVP01_0714300","PVP01_0714600","PVP01_0714900","PVP01_0715000","PVP01_0715100","PVP01_0715200","PVP01_0715300","PVP01_0715500","PVP01_0715600","PVP01_0715700","PVP01_0715800","PVP01_0716000","PVP01_0716100","PVP01_0716200","PVP01_0716400","PVP01_0716600","PVP01_0716800","PVP01_0717000","PVP01_0717200","PVP01_0717400","PVP01_0717500","PVP01_0717600","PVP01_0717800","PVP01_0718000","PVP01_0718100","PVP01_0718300","PVP01_0718600","PVP01_0718700","PVP01_0718800","PVP01_0719000","PVP01_0719100","PVP01_0719400","PVP01_0719500","PVP01_0719700","PVP01_0719900","PVP01_0720000","PVP01_0720100","PVP01_0720200","PVP01_0720300","PVP01_0720400","PVP01_0720800","PVP01_0720900","PVP01_0721100","PVP01_0721400","PVP01_0721500","PVP01_0721600","PVP01_0721700","PVP01_0721800","PVP01_0721900","PVP01_0722000","PVP01_0722100","PVP01_0722200","PVP01_0722300","PVP01_0722400","PVP01_0722500","PVP01_0722600","PVP01_0722700","PVP01_0722800","PVP01_0723000","PVP01_0723200","PVP01_0723300","PVP01_0723400","PVP01_0723500","PVP01_0723700","PVP01_0723800","PVP01_0724100","PVP01_0724300","PVP01_0724500","PVP01_0724600","PVP01_0724700","PVP01_0724800","PVP01_0724900","PVP01_0725000","PVP01_0725100","PVP01_0725200","PVP01_0725300","PVP01_0725400","PVP01_0725500","PVP01_0725700","PVP01_0725800","PVP01_0726400","PVP01_0726700","PVP01_0726900","PVP01_0727000","PVP01_0727100","PVP01_0727200","PVP01_0727300","PVP01_0727400","PVP01_0727500","PVP01_0727600","PVP01_0727800","PVP01_0728000","PVP01_0728100","PVP01_0728200","PVP01_0728300","PVP01_0728400","PVP01_0729100","PVP01_0729200","PVP01_0729300","PVP01_0729400","PVP01_0729500","PVP01_0729700","PVP01_0730000","PVP01_0730100","PVP01_0730200","PVP01_0730300","PVP01_0730400","PVP01_0730600","PVP01_0730700","PVP01_0731000","PVP01_0731100","PVP01_0731200","PVP01_0731300","PVP01_0731600","PVP01_0731700","PVP01_0731800","PVP01_0731900","PVP01_0732000","PVP01_0732100","PVP01_0732200","PVP01_0732300","PVP01_0732400","PVP01_0732600","PVP01_0732800","PVP01_0732900","PVP01_0733000","PVP01_0733100","PVP01_0733500","PVP01_0733700","PVP01_0733800","PVP01_0733900","PVP01_0734100","PVP01_0734200","PVP01_0734400","PVP01_0734700","PVP01_0735100","PVP01_0735200","PVP01_0735300","PVP01_0735800","PVP01_0735900","PVP01_0800100","PVP01_0800200","PVP01_0800300","PVP01_0800400","PVP01_0800500","PVP01_0800600","PVP01_0800700","PVP01_0800900","PVP01_0801000","PVP01_0801100","PVP01_0801400","PVP01_0801500","PVP01_0801600","PVP01_0801700","PVP01_0801800","PVP01_0802100","PVP01_0802300","PVP01_0802400","PVP01_0802500","PVP01_0802900","PVP01_0803000","PVP01_0803100","PVP01_0803400","PVP01_0803500","PVP01_0803600","PVP01_0803700","PVP01_0803900","PVP01_0804100","PVP01_0804400","PVP01_0804500","PVP01_0804600","PVP01_0804700","PVP01_0804800","PVP01_0804900","PVP01_0805000","PVP01_0805200","PVP01_0805300","PVP01_0805400","PVP01_0805500","PVP01_0805600","PVP01_0805700","PVP01_0805800","PVP01_0806000","PVP01_0806100","PVP01_0806200","PVP01_0806300","PVP01_0806400","PVP01_0806500","PVP01_0806700","PVP01_0807000","PVP01_0807200","PVP01_0807300","PVP01_0807600","PVP01_0807900","PVP01_0808000","PVP01_0808100","PVP01_0808400","PVP01_0808500","PVP01_0808600","PVP01_0808700","PVP01_0808800","PVP01_0809100","PVP01_0809200","PVP01_0809300","PVP01_0809400","PVP01_0809700","PVP01_0809900","PVP01_0810000","PVP01_0810100","PVP01_0810300","PVP01_0810400","PVP01_0810500","PVP01_0810600","PVP01_0811000","PVP01_0811100","PVP01_0811200","PVP01_0811300","PVP01_0811500","PVP01_0811700","PVP01_0811900","PVP01_0812000","PVP01_0812300","PVP01_0812400","PVP01_0812600","PVP01_0812800","PVP01_0813000","PVP01_0813200","PVP01_0813300","PVP01_0813600","PVP01_0813700","PVP01_0813800","PVP01_0814200","PVP01_0814300","PVP01_0814400","PVP01_0814800","PVP01_0815100","PVP01_0815300","PVP01_0815400","PVP01_0816200","PVP01_0816300","PVP01_0816700","PVP01_0816800","PVP01_0817100","PVP01_0817400","PVP01_0818100","PVP01_0818300","PVP01_0818600","PVP01_0818800","PVP01_0819000","PVP01_0819200","PVP01_0819300","PVP01_0819600","PVP01_0819700","PVP01_0819800","PVP01_0820000","PVP01_0820100","PVP01_0820200","PVP01_0820300","PVP01_0820400","PVP01_0820500","PVP01_0820600","PVP01_0820700","PVP01_0820800","PVP01_0820900","PVP01_0821000","PVP01_0821200","PVP01_0821300","PVP01_0821400","PVP01_0821500","PVP01_0821700","PVP01_0821800","PVP01_0822100","PVP01_0822300","PVP01_0822600","PVP01_0822700","PVP01_0822900","PVP01_0823100","PVP01_0823200","PVP01_0823400","PVP01_0823600","PVP01_0823700","PVP01_0824000","PVP01_0824100","PVP01_0824200","PVP01_0824300","PVP01_0824500","PVP01_0824600","PVP01_0824700","PVP01_0824800","PVP01_0824900","PVP01_0825000","PVP01_0825100","PVP01_0825200","PVP01_0825400","PVP01_0825600","PVP01_0825700","PVP01_0825800","PVP01_0826000","PVP01_0826100","PVP01_0826200","PVP01_0826300","PVP01_0826400","PVP01_0826500","PVP01_0826700","PVP01_0826900","PVP01_0827100","PVP01_0827700","PVP01_0827800","PVP01_0828000","PVP01_0828100","PVP01_0828200","PVP01_0828300","PVP01_0828700","PVP01_0829000","PVP01_0829100","PVP01_0829200","PVP01_0829300","PVP01_0829500","PVP01_0829700","PVP01_0829800","PVP01_0829900","PVP01_0830100","PVP01_0830300","PVP01_0830400","PVP01_0830500","PVP01_0830700","PVP01_0830900","PVP01_0831000","PVP01_0831100","PVP01_0831400","PVP01_0831700","PVP01_0831900","PVP01_0832000","PVP01_0832100","PVP01_0832200","PVP01_0832400","PVP01_0832500","PVP01_0832600","PVP01_0832800","PVP01_0832900","PVP01_0833000","PVP01_0833500","PVP01_0833600","PVP01_0833700","PVP01_0833800","PVP01_0834000","PVP01_0834200","PVP01_0834300","PVP01_0834400","PVP01_0834500","PVP01_0834600","PVP01_0834900","PVP01_0835000","PVP01_0835300","PVP01_0835500","PVP01_0835700","PVP01_0835800","PVP01_0836300","PVP01_0836400","PVP01_0836500","PVP01_0836600","PVP01_0836700","PVP01_0836800","PVP01_0837200","PVP01_0837300","PVP01_0837500","PVP01_0837600","PVP01_0837700","PVP01_0837800","PVP01_0837900","PVP01_0838000","PVP01_0838100","PVP01_0838200","PVP01_0838300","PVP01_0838600","PVP01_0838700","PVP01_0838800","PVP01_0838900","PVP01_0839000","PVP01_0839100","PVP01_0839200","PVP01_0839400","PVP01_0839700","PVP01_0840700","PVP01_0841000","PVP01_0900700","PVP01_0901000","PVP01_0901100","PVP01_0901200","PVP01_0901300","PVP01_0901800","PVP01_0901900","PVP01_0902000","PVP01_0902100","PVP01_0902400","PVP01_0903400","PVP01_0903500","PVP01_0903600","PVP01_0903700","PVP01_0903800","PVP01_0903900","PVP01_0904300","PVP01_0904600","PVP01_0904700","PVP01_0904900","PVP01_0905100","PVP01_0906000","PVP01_0906300","PVP01_0906400","PVP01_0906500","PVP01_0906700","PVP01_0906800","PVP01_0906900","PVP01_0907000","PVP01_0907100","PVP01_0907200","PVP01_0907300","PVP01_0907400","PVP01_0907500","PVP01_0907600","PVP01_0907700","PVP01_0907800","PVP01_0908000","PVP01_0908200","PVP01_0908400","PVP01_0908500","PVP01_0908600","PVP01_0908900","PVP01_0909100","PVP01_0909300","PVP01_0909500","PVP01_0909600","PVP01_0910000","PVP01_0910200","PVP01_0910900","PVP01_0911000","PVP01_0911300","PVP01_0911500","PVP01_0911600","PVP01_0911700","PVP01_0911800","PVP01_0911900","PVP01_0912000","PVP01_0912100","PVP01_0912200","PVP01_0912400","PVP01_0912900","PVP01_0913000","PVP01_0913100","PVP01_0913400","PVP01_0913500","PVP01_0913600","PVP01_0913800","PVP01_0913900","PVP01_0914000","PVP01_0914200","PVP01_0914300","PVP01_0914400","PVP01_0914600","PVP01_0914800","PVP01_0915100","PVP01_0915300","PVP01_0915400","PVP01_0915600","PVP01_0915800","PVP01_0916000","PVP01_0916300","PVP01_0916600","PVP01_0916800","PVP01_0916900","PVP01_0917000","PVP01_0917100","PVP01_0917200","PVP01_0917900","PVP01_0918000","PVP01_0918100","PVP01_0918200","PVP01_0918300","PVP01_0918400","PVP01_0918600","PVP01_0918700","PVP01_0918800","PVP01_0918900","PVP01_0919000","PVP01_0919100","PVP01_0919400","PVP01_0919500","PVP01_0919600","PVP01_0919700","PVP01_0919900","PVP01_0920100","PVP01_0920200","PVP01_0920400","PVP01_0920700","PVP01_0920800","PVP01_0921400","PVP01_0921500","PVP01_0921800","PVP01_0921900","PVP01_0922000","PVP01_0922100","PVP01_0922300","PVP01_0922400","PVP01_0922800","PVP01_0922900","PVP01_0923000","PVP01_0923100","PVP01_0923200","PVP01_0923300","PVP01_0923800","PVP01_0923900","PVP01_0924000","PVP01_0924300","PVP01_0924400","PVP01_0924600","PVP01_0924700","PVP01_0924800","PVP01_0925000","PVP01_0925100","PVP01_0925200","PVP01_0925400","PVP01_0925600","PVP01_0925700","PVP01_0925800","PVP01_0925900","PVP01_0926000","PVP01_0926300","PVP01_0926400","PVP01_0926500","PVP01_0926600","PVP01_0926800","PVP01_0926900","PVP01_0927100","PVP01_0927200","PVP01_0927400","PVP01_0927600","PVP01_0927700","PVP01_0927800","PVP01_0927900","PVP01_0928000","PVP01_0928100","PVP01_0928200","PVP01_0928300","PVP01_0928800","PVP01_0928900","PVP01_0929000","PVP01_0929100","PVP01_0929200","PVP01_0929300","PVP01_0929400","PVP01_0929600","PVP01_0929700","PVP01_0929900","PVP01_0930000","PVP01_0930200","PVP01_0930300","PVP01_0930400","PVP01_0930500","PVP01_0931200","PVP01_0931300","PVP01_0931400","PVP01_0931500","PVP01_0931600","PVP01_0931700","PVP01_0932000","PVP01_0932200","PVP01_0932300","PVP01_0932500","PVP01_0932600","PVP01_0932900","PVP01_0933100","PVP01_0933300","PVP01_0933400","PVP01_0933500","PVP01_0933700","PVP01_0934000","PVP01_0934300","PVP01_0934400","PVP01_0934500","PVP01_0934600","PVP01_0935000","PVP01_0935200","PVP01_0935400","PVP01_0935500","PVP01_0935700","PVP01_0935800","PVP01_0935900","PVP01_0936000","PVP01_0936100","PVP01_0936300","PVP01_0936400","PVP01_0936700","PVP01_0936800","PVP01_0937000","PVP01_0937300","PVP01_0937500","PVP01_0937700","PVP01_0937800","PVP01_0938100","PVP01_0938200","PVP01_0938600","PVP01_0938700","PVP01_0938900","PVP01_0939000","PVP01_0939100","PVP01_0939200","PVP01_0939300","PVP01_0939600","PVP01_0939700","PVP01_0940200","PVP01_0940500","PVP01_0940600","PVP01_0940700","PVP01_0940900","PVP01_0941100","PVP01_0941200","PVP01_0941300","PVP01_0941400","PVP01_0941600","PVP01_0941800","PVP01_0942000","PVP01_0942200","PVP01_0942300","PVP01_0942400","PVP01_0942600","PVP01_0942700","PVP01_0942800","PVP01_0942900","PVP01_0943100","PVP01_0943300","PVP01_0943400","PVP01_0943700","PVP01_0943800","PVP01_0944100","PVP01_0944300","PVP01_0944400","PVP01_0944600","PVP01_0944800","PVP01_0945100","PVP01_0945200","PVP01_0945300","PVP01_0945500","PVP01_0945600","PVP01_0945700","PVP01_0945900","PVP01_0946000","PVP01_0946100","PVP01_0946200","PVP01_0946300","PVP01_0946400","PVP01_0946500","PVP01_0946600","PVP01_0946700","PVP01_0946800","PVP01_0946900","PVP01_0947000","PVP01_0947100","PVP01_0947300","PVP01_0947400","PVP01_0947600","PVP01_0947800","PVP01_0947900","PVP01_0948100","PVP01_0948200","PVP01_0948300","PVP01_0948500","PVP01_0948600","PVP01_0948700","PVP01_0948800","PVP01_0948900","PVP01_0949000","PVP01_0949200","PVP01_0949400","PVP01_0949600","PVP01_0950000","PVP01_0950200","PVP01_1000700","PVP01_1001900","PVP01_1002500","PVP01_1002600","PVP01_1002700","PVP01_1003000","PVP01_1003200","PVP01_1003400","PVP01_1003500","PVP01_1003900","PVP01_1004100","PVP01_1004500","PVP01_1004800","PVP01_1004900","PVP01_1005100","PVP01_1005200","PVP01_1005300","PVP01_1005400","PVP01_1005600","PVP01_1005900","PVP01_1006200","PVP01_1006300","PVP01_1006400","PVP01_1006500","PVP01_1006600","PVP01_1006700","PVP01_1006800","PVP01_1007200","PVP01_1007500","PVP01_1007700","PVP01_1007800","PVP01_1007900","PVP01_1008000","PVP01_1008100","PVP01_1008200","PVP01_1008300","PVP01_1008600","PVP01_1008700","PVP01_1008900","PVP01_1009000","PVP01_1009100","PVP01_1009300","PVP01_1009400","PVP01_1009500","PVP01_1009600","PVP01_1009700","PVP01_1010100","PVP01_1010200","PVP01_1010300","PVP01_1010400","PVP01_1011000","PVP01_1011100","PVP01_1011300","PVP01_1011400","PVP01_1011500","PVP01_1011600","PVP01_1011800","PVP01_1012100","PVP01_1012300","PVP01_1012400","PVP01_1012500","PVP01_1012700","PVP01_1013100","PVP01_1013200","PVP01_1013300","PVP01_1013400","PVP01_1013700","PVP01_1014100","PVP01_1014300","PVP01_1014600","PVP01_1014800","PVP01_1015000","PVP01_1015100","PVP01_1015200","PVP01_1015400","PVP01_1015500","PVP01_1015600","PVP01_1015700","PVP01_1015900","PVP01_1016100","PVP01_1016200","PVP01_1016400","PVP01_1016500","PVP01_1016600","PVP01_1016900","PVP01_1017100","PVP01_1017200","PVP01_1017400","PVP01_1017700","PVP01_1017900","PVP01_1018000","PVP01_1018100","PVP01_1018200","PVP01_1018400","PVP01_1018500","PVP01_1018800","PVP01_1019000","PVP01_1019400","PVP01_1019600","PVP01_1019900","PVP01_1020000","PVP01_1020100","PVP01_1020200","PVP01_1020300","PVP01_1020500","PVP01_1020700","PVP01_1020900","PVP01_1021000","PVP01_1021100","PVP01_1021200","PVP01_1021300","PVP01_1021500","PVP01_1021600","PVP01_1021700","PVP01_1021800","PVP01_1021900","PVP01_1022000","PVP01_1022100","PVP01_1022200","PVP01_1022300","PVP01_1022400","PVP01_1022900","PVP01_1023000","PVP01_1023200","PVP01_1023300","PVP01_1023600","PVP01_1023700","PVP01_1024300","PVP01_1024400","PVP01_1024500","PVP01_1024800","PVP01_1025200","PVP01_1025300","PVP01_1025400","PVP01_1025700","PVP01_1025800","PVP01_1026300","PVP01_1026400","PVP01_1026500","PVP01_1026600","PVP01_1026700","PVP01_1026800","PVP01_1026900","PVP01_1027100","PVP01_1027300","PVP01_1027400","PVP01_1027600","PVP01_1027700","PVP01_1027800","PVP01_1027900","PVP01_1028000","PVP01_1028100","PVP01_1028200","PVP01_1028400","PVP01_1028600","PVP01_1029000","PVP01_1029100","PVP01_1029200","PVP01_1029300","PVP01_1029500","PVP01_1029700","PVP01_1029800","PVP01_1029900","PVP01_1030000","PVP01_1030100","PVP01_1030200","PVP01_1030300","PVP01_1030400","PVP01_1030500","PVP01_1030600","PVP01_1030700","PVP01_1030900","PVP01_1031000","PVP01_1031100","PVP01_1031200","PVP01_1031300","PVP01_1031400","PVP01_1031500","PVP01_1031600","PVP01_1031700","PVP01_1031800","PVP01_1031900","PVP01_1032000","PVP01_1032100","PVP01_1032200","PVP01_1032400","PVP01_1032500","PVP01_1032800","PVP01_1032900","PVP01_1033000","PVP01_1033300","PVP01_1033400","PVP01_1033500","PVP01_1033600","PVP01_1033700","PVP01_1033800","PVP01_1033900","PVP01_1034000","PVP01_1034100","PVP01_1034200","PVP01_1034300","PVP01_1034400","PVP01_1034500","PVP01_1035200","PVP01_1100400","PVP01_1100500","PVP01_1101000","PVP01_1101200","PVP01_1101300","PVP01_1101400","PVP01_1101500","PVP01_1101600","PVP01_1101700","PVP01_1102400","PVP01_1102500","PVP01_1102600","PVP01_1102700","PVP01_1102800","PVP01_1102900","PVP01_1103200","PVP01_1103400","PVP01_1103700","PVP01_1104000","PVP01_1104100","PVP01_1104200","PVP01_1104300","PVP01_1104400","PVP01_1104600","PVP01_1104800","PVP01_1104900","PVP01_1105300","PVP01_1105400","PVP01_1105500","PVP01_1105600","PVP01_1105800","PVP01_1106000","PVP01_1106200","PVP01_1106300","PVP01_1106400","PVP01_1106600","PVP01_1106700","PVP01_1107000","PVP01_1107100","PVP01_1107200","PVP01_1107300","PVP01_1107400","PVP01_1107600","PVP01_1107700","PVP01_1107900","PVP01_1108000","PVP01_1108100","PVP01_1108300","PVP01_1108500","PVP01_1108600","PVP01_1108800","PVP01_1108900","PVP01_1109000","PVP01_1109100","PVP01_1109200","PVP01_1109300","PVP01_1109400","PVP01_1109600","PVP01_1109900","PVP01_1110500","PVP01_1110600","PVP01_1110700","PVP01_1110900","PVP01_1111100","PVP01_1111200","PVP01_1111300","PVP01_1111400","PVP01_1111500","PVP01_1111600","PVP01_1111700","PVP01_1111800","PVP01_1112000","PVP01_1112100","PVP01_1112400","PVP01_1112600","PVP01_1112700","PVP01_1112800","PVP01_1112900","PVP01_1113000","PVP01_1113300","PVP01_1113500","PVP01_1113700","PVP01_1113900","PVP01_1114100","PVP01_1114200","PVP01_1114300","PVP01_1114500","PVP01_1114600","PVP01_1114700","PVP01_1114800","PVP01_1115000","PVP01_1115200","PVP01_1115300","PVP01_1115500","PVP01_1115600","PVP01_1116400","PVP01_1116600","PVP01_1116800","PVP01_1116900","PVP01_1117000","PVP01_1117200","PVP01_1117300","PVP01_1117500","PVP01_1117700","PVP01_1118100","PVP01_1118500","PVP01_1118600","PVP01_1118700","PVP01_1119000","PVP01_1119300","PVP01_1119500","PVP01_1119600","PVP01_1119700","PVP01_1119900","PVP01_1120000","PVP01_1120100","PVP01_1120300","PVP01_1120400","PVP01_1120500","PVP01_1120900","PVP01_1121000","PVP01_1121100","PVP01_1121300","PVP01_1121500","PVP01_1121600","PVP01_1121700","PVP01_1121800","PVP01_1121900","PVP01_1122100","PVP01_1122300","PVP01_1122400","PVP01_1122500","PVP01_1122600","PVP01_1122700","PVP01_1122800","PVP01_1122900","PVP01_1123000","PVP01_1123100","PVP01_1123200","PVP01_1123300","PVP01_1123400","PVP01_1123500","PVP01_1123600","PVP01_1123900","PVP01_1124000","PVP01_1124100","PVP01_1124300","PVP01_1124400","PVP01_1124600","PVP01_1124700","PVP01_1124800","PVP01_1124900","PVP01_1125000","PVP01_1125100","PVP01_1125200","PVP01_1125300","PVP01_1125400","PVP01_1125600","PVP01_1125700","PVP01_1125800","PVP01_1126000","PVP01_1126100","PVP01_1126200","PVP01_1126300","PVP01_1126500","PVP01_1126600","PVP01_1126800","PVP01_1126900","PVP01_1127000","PVP01_1127200","PVP01_1127300","PVP01_1127400","PVP01_1127500","PVP01_1127800","PVP01_1128000","PVP01_1128100","PVP01_1128400","PVP01_1128500","PVP01_1128800","PVP01_1128900","PVP01_1129000","PVP01_1129500","PVP01_1129600","PVP01_1129700","PVP01_1130100","PVP01_1130400","PVP01_1130600","PVP01_1130700","PVP01_1131300","PVP01_1131400","PVP01_1131900","PVP01_1132100","PVP01_1132300","PVP01_1132400","PVP01_1132500","PVP01_1132700","PVP01_1132800","PVP01_1132900","PVP01_1133000","PVP01_1133100","PVP01_1133200","PVP01_1133300","PVP01_1133400","PVP01_1133600","PVP01_1133800","PVP01_1133900","PVP01_1134100","PVP01_1134200","PVP01_1134300","PVP01_1134400","PVP01_1134500","PVP01_1134700","PVP01_1134800","PVP01_1135000","PVP01_1135100","PVP01_1135300","PVP01_1135500","PVP01_1135600","PVP01_1135700","PVP01_1135800","PVP01_1135900","PVP01_1136000","PVP01_1136100","PVP01_1136300","PVP01_1136400","PVP01_1136600","PVP01_1136700","PVP01_1136800","PVP01_1137100","PVP01_1137200","PVP01_1137300","PVP01_1137600","PVP01_1137700","PVP01_1137800","PVP01_1138200","PVP01_1138400","PVP01_1138500","PVP01_1139000","PVP01_1139100","PVP01_1139200","PVP01_1139300","PVP01_1139400","PVP01_1139700","PVP01_1140000","PVP01_1140300","PVP01_1140500","PVP01_1140600","PVP01_1140700","PVP01_1140800","PVP01_1140900","PVP01_1141000","PVP01_1141200","PVP01_1141500","PVP01_1141700","PVP01_1141800","PVP01_1141900","PVP01_1142000","PVP01_1142200","PVP01_1142300","PVP01_1142500","PVP01_1142600","PVP01_1143100","PVP01_1143200","PVP01_1143400","PVP01_1143500","PVP01_1143700","PVP01_1144100","PVP01_1144200","PVP01_1144300","PVP01_1144500","PVP01_1144700","PVP01_1144900","PVP01_1145000","PVP01_1145100","PVP01_1145200","PVP01_1145500","PVP01_1145600","PVP01_1145700","PVP01_1145800","PVP01_1146200","PVP01_1146400","PVP01_1146500","PVP01_1146600","PVP01_1146800","PVP01_1146900","PVP01_1147000","PVP01_1147100","PVP01_1147200","PVP01_1147300","PVP01_1147400","PVP01_1147500","PVP01_1147600","PVP01_1147700","PVP01_1147800","PVP01_1147900","PVP01_1148200","PVP01_1148700","PVP01_1148800","PVP01_1149000","PVP01_1149100","PVP01_1149400","PVP01_1149500","PVP01_1200200","PVP01_1200800","PVP01_1201100","PVP01_1201300","PVP01_1201400","PVP01_1201500","PVP01_1201600","PVP01_1201700","PVP01_1201800","PVP01_1201900","PVP01_1202000","PVP01_1202100","PVP01_1202200","PVP01_1202300","PVP01_1202500","PVP01_1202700","PVP01_1203100","PVP01_1203200","PVP01_1203300","PVP01_1203500","PVP01_1203600","PVP01_1203700","PVP01_1203800","PVP01_1204000","PVP01_1204200","PVP01_1204400","PVP01_1204500","PVP01_1204600","PVP01_1204700","PVP01_1204800","PVP01_1205000","PVP01_1205200","PVP01_1205400","PVP01_1205600","PVP01_1205800","PVP01_1205900","PVP01_1206100","PVP01_1206300","PVP01_1206400","PVP01_1206700","PVP01_1206900","PVP01_1207000","PVP01_1207100","PVP01_1207200","PVP01_1207800","PVP01_1207900","PVP01_1208000","PVP01_1208100","PVP01_1208200","PVP01_1208300","PVP01_1208600","PVP01_1208700","PVP01_1208800","PVP01_1208900","PVP01_1209000","PVP01_1209200","PVP01_1209400","PVP01_1210000","PVP01_1210100","PVP01_1210300","PVP01_1210400","PVP01_1210500","PVP01_1210700","PVP01_1211000","PVP01_1211100","PVP01_1211200","PVP01_1211300","PVP01_1211500","PVP01_1211600","PVP01_1211700","PVP01_1212100","PVP01_1212500","PVP01_1212600","PVP01_1212700","PVP01_1213000","PVP01_1213100","PVP01_1213300","PVP01_1214200","PVP01_1214300","PVP01_1214400","PVP01_1214500","PVP01_1214600","PVP01_1214800","PVP01_1214900","PVP01_1215000","PVP01_1215300","PVP01_1215400","PVP01_1215800","PVP01_1215900","PVP01_1216000","PVP01_1216100","PVP01_1216200","PVP01_1216300","PVP01_1216700","PVP01_1216800","PVP01_1216900","PVP01_1217000","PVP01_1217200","PVP01_1217400","PVP01_1217500","PVP01_1217600","PVP01_1217700","PVP01_1218000","PVP01_1218500","PVP01_1218800","PVP01_1218900","PVP01_1219000","PVP01_1219100","PVP01_1219200","PVP01_1219400","PVP01_1219500","PVP01_1219600","PVP01_1219800","PVP01_1219900","PVP01_1220100","PVP01_1220200","PVP01_1220600","PVP01_1220700","PVP01_1220800","PVP01_1220900","PVP01_1221000","PVP01_1221200","PVP01_1221300","PVP01_1221400","PVP01_1221700","PVP01_1221800","PVP01_1221900","PVP01_1222100","PVP01_1222200","PVP01_1222300","PVP01_1222400","PVP01_1222700","PVP01_1223400","PVP01_1223500","PVP01_1223800","PVP01_1223900","PVP01_1224000","PVP01_1224300","PVP01_1224400","PVP01_1224500","PVP01_1224600","PVP01_1224700","PVP01_1224800","PVP01_1225000","PVP01_1225200","PVP01_1225300","PVP01_1225500","PVP01_1225800","PVP01_1225900","PVP01_1226100","PVP01_1226700","PVP01_1226800","PVP01_1227300","PVP01_1227400","PVP01_1227500","PVP01_1227600","PVP01_1227900","PVP01_1228000","PVP01_1228100","PVP01_1228200","PVP01_1228400","PVP01_1228500","PVP01_1228800","PVP01_1229100","PVP01_1229400","PVP01_1229500","PVP01_1229600","PVP01_1229800","PVP01_1229900","PVP01_1230000","PVP01_1230100","PVP01_1230200","PVP01_1230500","PVP01_1230600","PVP01_1230700","PVP01_1230800","PVP01_1231000","PVP01_1231100","PVP01_1231600","PVP01_1231700","PVP01_1231800","PVP01_1231900","PVP01_1232400","PVP01_1232600","PVP01_1232700","PVP01_1233100","PVP01_1233800","PVP01_1233900","PVP01_1234000","PVP01_1234400","PVP01_1235000","PVP01_1235100","PVP01_1235200","PVP01_1235300","PVP01_1235400","PVP01_1235800","PVP01_1236000","PVP01_1236100","PVP01_1236200","PVP01_1236400","PVP01_1236700","PVP01_1236800","PVP01_1236900","PVP01_1237000","PVP01_1237100","PVP01_1237200","PVP01_1237300","PVP01_1237500","PVP01_1237600","PVP01_1237700","PVP01_1237800","PVP01_1237900","PVP01_1238000","PVP01_1238400","PVP01_1238500","PVP01_1238600","PVP01_1238900","PVP01_1239000","PVP01_1239100","PVP01_1239300","PVP01_1239400","PVP01_1239800","PVP01_1239900","PVP01_1240000","PVP01_1240100","PVP01_1240200","PVP01_1240300","PVP01_1240500","PVP01_1240600","PVP01_1240800","PVP01_1240900","PVP01_1241000","PVP01_1241100","PVP01_1241200","PVP01_1241300","PVP01_1241600","PVP01_1241900","PVP01_1242000","PVP01_1242300","PVP01_1242400","PVP01_1242600","PVP01_1242700","PVP01_1242800","PVP01_1242900","PVP01_1243000","PVP01_1243100","PVP01_1243200","PVP01_1243500","PVP01_1243600","PVP01_1243900","PVP01_1244400","PVP01_1244700","PVP01_1244800","PVP01_1244900","PVP01_1245100","PVP01_1245300","PVP01_1245400","PVP01_1245500","PVP01_1245600","PVP01_1245900","PVP01_1246000","PVP01_1246100","PVP01_1246300","PVP01_1246400","PVP01_1246500","PVP01_1246600","PVP01_1246700","PVP01_1246900","PVP01_1247000","PVP01_1247500","PVP01_1247600","PVP01_1247700","PVP01_1247800","PVP01_1247900","PVP01_1248200","PVP01_1248500","PVP01_1248600","PVP01_1249000","PVP01_1249100","PVP01_1249200","PVP01_1249500","PVP01_1249600","PVP01_1249700","PVP01_1250200","PVP01_1250300","PVP01_1250600","PVP01_1250900","PVP01_1251100","PVP01_1251500","PVP01_1251600","PVP01_1251700","PVP01_1251800","PVP01_1251900","PVP01_1252000","PVP01_1252200","PVP01_1252400","PVP01_1252800","PVP01_1252900","PVP01_1253000","PVP01_1253300","PVP01_1253600","PVP01_1253700","PVP01_1253800","PVP01_1254100","PVP01_1254300","PVP01_1254600","PVP01_1254900","PVP01_1255100","PVP01_1255200","PVP01_1255300","PVP01_1255400","PVP01_1255500","PVP01_1256000","PVP01_1256100","PVP01_1256200","PVP01_1256400","PVP01_1256500","PVP01_1256700","PVP01_1256800","PVP01_1256900","PVP01_1257000","PVP01_1257200","PVP01_1257300","PVP01_1257500","PVP01_1257600","PVP01_1257700","PVP01_1257900","PVP01_1258100","PVP01_1258200","PVP01_1258300","PVP01_1258400","PVP01_1258500","PVP01_1258700","PVP01_1258900","PVP01_1259000","PVP01_1259100","PVP01_1259200","PVP01_1259300","PVP01_1259400","PVP01_1259500","PVP01_1259600","PVP01_1260200","PVP01_1260800","PVP01_1260900","PVP01_1261100","PVP01_1261400","PVP01_1261500","PVP01_1261700","PVP01_1262000","PVP01_1262200","PVP01_1262300","PVP01_1262400","PVP01_1262500","PVP01_1262600","PVP01_1262800","PVP01_1262900","PVP01_1263000","PVP01_1263100","PVP01_1263400","PVP01_1263700","PVP01_1263800","PVP01_1263900","PVP01_1264000","PVP01_1264300","PVP01_1264400","PVP01_1264600","PVP01_1264700","PVP01_1264800","PVP01_1264900","PVP01_1265000","PVP01_1265100","PVP01_1265300","PVP01_1265400","PVP01_1265600","PVP01_1265700","PVP01_1265900","PVP01_1266200","PVP01_1266400","PVP01_1266500","PVP01_1266600","PVP01_1266700","PVP01_1267300","PVP01_1267400","PVP01_1267500","PVP01_1267600","PVP01_1268000","PVP01_1268100","PVP01_1268200","PVP01_1268300","PVP01_1268600","PVP01_1268900","PVP01_1269000","PVP01_1269200","PVP01_1269500","PVP01_1269800","PVP01_1270200","PVP01_1270300","PVP01_1270400","PVP01_1270600","PVP01_1270700","PVP01_1270800","PVP01_1270900","PVP01_1271100","PVP01_1271200","PVP01_1271300","PVP01_1271400","PVP01_1271500","PVP01_1271600","PVP01_1271900","PVP01_1272500","PVP01_1272600","PVP01_1273000","PVP01_1274300","PVP01_1300400","PVP01_1300500","PVP01_1300700","PVP01_1300900","PVP01_1301100","PVP01_1301200","PVP01_1301300","PVP01_1301400","PVP01_1301500","PVP01_1301600","PVP01_1301700","PVP01_1302100","PVP01_1302300","PVP01_1302500","PVP01_1302600","PVP01_1302700","PVP01_1302800","PVP01_1302900","PVP01_1303000","PVP01_1303100","PVP01_1303200","PVP01_1303400","PVP01_1303600","PVP01_1303900","PVP01_1304000","PVP01_1304200","PVP01_1304300","PVP01_1304500","PVP01_1304800","PVP01_1304900","PVP01_1305200","PVP01_1305300","PVP01_1305500","PVP01_1305700","PVP01_1305900","PVP01_1306000","PVP01_1306100","PVP01_1306300","PVP01_1306400","PVP01_1306600","PVP01_1306700","PVP01_1306800","PVP01_1306900","PVP01_1307000","PVP01_1307100","PVP01_1307200","PVP01_1307300","PVP01_1307500","PVP01_1307700","PVP01_1307800","PVP01_1308100","PVP01_1308300","PVP01_1308400","PVP01_1308500","PVP01_1308800","PVP01_1308900","PVP01_1309000","PVP01_1309100","PVP01_1309200","PVP01_1309400","PVP01_1309500","PVP01_1309700","PVP01_1309800","PVP01_1310000","PVP01_1310400","PVP01_1310500","PVP01_1310700","PVP01_1310800","PVP01_1311000","PVP01_1311100","PVP01_1311200","PVP01_1311300","PVP01_1311400","PVP01_1311500","PVP01_1311600","PVP01_1311900","PVP01_1312000","PVP01_1312100","PVP01_1312200","PVP01_1312400","PVP01_1312600","PVP01_1312900","PVP01_1313000","PVP01_1313100","PVP01_1313200","PVP01_1313300","PVP01_1313400","PVP01_1314100","PVP01_1314300","PVP01_1314600","PVP01_1314800","PVP01_1315200","PVP01_1315400","PVP01_1315500","PVP01_1316000","PVP01_1316100","PVP01_1316200","PVP01_1316300","PVP01_1316500","PVP01_1316600","PVP01_1316700","PVP01_1316900","PVP01_1317000","PVP01_1317500","PVP01_1317600","PVP01_1317700","PVP01_1317800","PVP01_1317900","PVP01_1318000","PVP01_1318100","PVP01_1318300","PVP01_1318500","PVP01_1318600","PVP01_1318800","PVP01_1318900","PVP01_1319000","PVP01_1319300","PVP01_1319500","PVP01_1319600","PVP01_1319700","PVP01_1319800","PVP01_1319900","PVP01_1320000","PVP01_1320100","PVP01_1320200","PVP01_1320300","PVP01_1320500","PVP01_1320600","PVP01_1320700","PVP01_1321500","PVP01_1321600","PVP01_1321700","PVP01_1321900","PVP01_1322000","PVP01_1322200","PVP01_1322300","PVP01_1322400","PVP01_1322600","PVP01_1322900","PVP01_1323000","PVP01_1323100","PVP01_1323200","PVP01_1323700","PVP01_1323800","PVP01_1323900","PVP01_1324100","PVP01_1324700","PVP01_1324800","PVP01_1325200","PVP01_1325600","PVP01_1325800","PVP01_1325900","PVP01_1326000","PVP01_1326200","PVP01_1326400","PVP01_1326600","PVP01_1326900","PVP01_1327200","PVP01_1327300","PVP01_1327500","PVP01_1327600","PVP01_1327900","PVP01_1328000","PVP01_1328100","PVP01_1328500","PVP01_1328600","PVP01_1328700","PVP01_1328800","PVP01_1328900","PVP01_1329000","PVP01_1329300","PVP01_1329400","PVP01_1329600","PVP01_1329700","PVP01_1329800","PVP01_1329900","PVP01_1330000","PVP01_1330800","PVP01_1331000","PVP01_1331100","PVP01_1331200","PVP01_1331800","PVP01_1331900","PVP01_1332200","PVP01_1332300","PVP01_1332400","PVP01_1332500","PVP01_1332700","PVP01_1332800","PVP01_1332900","PVP01_1333000","PVP01_1333100","PVP01_1333200","PVP01_1333700","PVP01_1333900","PVP01_1334100","PVP01_1334200","PVP01_1334300","PVP01_1334500","PVP01_1335100","PVP01_1335300","PVP01_1335600","PVP01_1335800","PVP01_1335900","PVP01_1336400","PVP01_1336500","PVP01_1336700","PVP01_1336900","PVP01_1337100","PVP01_1337200","PVP01_1337300","PVP01_1337400","PVP01_1337500","PVP01_1337600","PVP01_1337900","PVP01_1338100","PVP01_1338300","PVP01_1338400","PVP01_1338500","PVP01_1338600","PVP01_1338700","PVP01_1339000","PVP01_1339400","PVP01_1339600","PVP01_1339800","PVP01_1339900","PVP01_1340100","PVP01_1340200","PVP01_1340300","PVP01_1340400","PVP01_1340500","PVP01_1340600","PVP01_1340800","PVP01_1341300","PVP01_1341600","PVP01_1341700","PVP01_1341800","PVP01_1341900","PVP01_1342200","PVP01_1342300","PVP01_1342400","PVP01_1342600","PVP01_1342700","PVP01_1342800","PVP01_1343100","PVP01_1343300","PVP01_1343500","PVP01_1343600","PVP01_1344000","PVP01_1344200","PVP01_1344300","PVP01_1344500","PVP01_1344600","PVP01_1344700","PVP01_1344900","PVP01_1345000","PVP01_1345200","PVP01_1345300","PVP01_1345400","PVP01_1345500","PVP01_1345600","PVP01_1345700","PVP01_1345900","PVP01_1346200","PVP01_1346700","PVP01_1346800","PVP01_1346900","PVP01_1347100","PVP01_1347200","PVP01_1347300","PVP01_1347500","PVP01_1347700","PVP01_1347800","PVP01_1400100","PVP01_1400400","PVP01_1400600","PVP01_1400700","PVP01_1400900","PVP01_1401000","PVP01_1401100","PVP01_1401200","PVP01_1401600","PVP01_1401700","PVP01_1401800","PVP01_1401900","PVP01_1402000","PVP01_1402100","PVP01_1402200","PVP01_1402300","PVP01_1402400","PVP01_1402500","PVP01_1402600","PVP01_1402700","PVP01_1402800","PVP01_1402900","PVP01_1403000","PVP01_1403300","PVP01_1403400","PVP01_1403500","PVP01_1403600","PVP01_1403700","PVP01_1403900","PVP01_1404000","PVP01_1404100","PVP01_1404200","PVP01_1404300","PVP01_1404500","PVP01_1404600","PVP01_1404700","PVP01_1404800","PVP01_1405000","PVP01_1405100","PVP01_1405300","PVP01_1405500","PVP01_1405600","PVP01_1405800","PVP01_1406000","PVP01_1406100","PVP01_1406600","PVP01_1406700","PVP01_1406900","PVP01_1407000","PVP01_1407100","PVP01_1407400","PVP01_1407500","PVP01_1407600","PVP01_1407800","PVP01_1408000","PVP01_1408100","PVP01_1408200","PVP01_1408400","PVP01_1408500","PVP01_1408600","PVP01_1408700","PVP01_1408800","PVP01_1408900","PVP01_1409000","PVP01_1409100","PVP01_1409300","PVP01_1409400","PVP01_1409700","PVP01_1410000","PVP01_1410100","PVP01_1410200","PVP01_1410300","PVP01_1410400","PVP01_1410500","PVP01_1410700","PVP01_1410800","PVP01_1410900","PVP01_1411000","PVP01_1411200","PVP01_1411300","PVP01_1411400","PVP01_1412100","PVP01_1412400","PVP01_1412500","PVP01_1412900","PVP01_1413000","PVP01_1413100","PVP01_1413200","PVP01_1413300","PVP01_1413450","PVP01_1413700","PVP01_1414000","PVP01_1414200","PVP01_1414300","PVP01_1414400","PVP01_1414500","PVP01_1414900","PVP01_1415000","PVP01_1415100","PVP01_1415200","PVP01_1415300","PVP01_1415500","PVP01_1415600","PVP01_1415700","PVP01_1415800","PVP01_1415900","PVP01_1416000","PVP01_1416100","PVP01_1416200","PVP01_1416900","PVP01_1417100","PVP01_1417200","PVP01_1417300","PVP01_1417600","PVP01_1417700","PVP01_1417900","PVP01_1418000","PVP01_1418100","PVP01_1418200","PVP01_1418400","PVP01_1418600","PVP01_1418800","PVP01_1418900","PVP01_1419100","PVP01_1419300","PVP01_1419400","PVP01_1419500","PVP01_1419700","PVP01_1419800","PVP01_1420000","PVP01_1420100","PVP01_1420300","PVP01_1420500","PVP01_1420900","PVP01_1421000","PVP01_1421100","PVP01_1421200","PVP01_1421300","PVP01_1421400","PVP01_1421500","PVP01_1421800","PVP01_1422100","PVP01_1422200","PVP01_1422300","PVP01_1422600","PVP01_1422700","PVP01_1422900","PVP01_1423100","PVP01_1423200","PVP01_1423500","PVP01_1423700","PVP01_1424100","PVP01_1424200","PVP01_1424500","PVP01_1425100","PVP01_1425200","PVP01_1425300","PVP01_1425400","PVP01_1425500","PVP01_1425600","PVP01_1425800","PVP01_1426000","PVP01_1426100","PVP01_1426200","PVP01_1426700","PVP01_1426800","PVP01_1427000","PVP01_1427100","PVP01_1427600","PVP01_1427700","PVP01_1427800","PVP01_1428000","PVP01_1428100","PVP01_1428300","PVP01_1428400","PVP01_1428500","PVP01_1428600","PVP01_1428800","PVP01_1428900","PVP01_1429200","PVP01_1429300","PVP01_1429500","PVP01_1429700","PVP01_1429900","PVP01_1430000","PVP01_1430100","PVP01_1430300","PVP01_1430400","PVP01_1430500","PVP01_1430600","PVP01_1430700","PVP01_1430800","PVP01_1431200","PVP01_1431300","PVP01_1431500","PVP01_1431700","PVP01_1431800","PVP01_1432000","PVP01_1432100","PVP01_1432300","PVP01_1432400","PVP01_1432600","PVP01_1432700","PVP01_1432800","PVP01_1432900","PVP01_1433600","PVP01_1433700","PVP01_1433800","PVP01_1434000","PVP01_1434200","PVP01_1434400","PVP01_1434500","PVP01_1434600","PVP01_1434800","PVP01_1434900","PVP01_1435000","PVP01_1435100","PVP01_1435200","PVP01_1435300","PVP01_1435800","PVP01_1435900","PVP01_1436000","PVP01_1436300","PVP01_1436400","PVP01_1436600","PVP01_1436700","PVP01_1437100","PVP01_1437200","PVP01_1437300","PVP01_1437400","PVP01_1437600","PVP01_1438100","PVP01_1438200","PVP01_1438300","PVP01_1438400","PVP01_1438600","PVP01_1438700","PVP01_1438800","PVP01_1438900","PVP01_1439000","PVP01_1439400","PVP01_1439600","PVP01_1439700","PVP01_1440000","PVP01_1440100","PVP01_1440300","PVP01_1440600","PVP01_1440700","PVP01_1440800","PVP01_1440900","PVP01_1441000","PVP01_1441100","PVP01_1441200","PVP01_1441400","PVP01_1441700","PVP01_1441800","PVP01_1442000","PVP01_1442200","PVP01_1442300","PVP01_1442600","PVP01_1442900","PVP01_1443000","PVP01_1443100","PVP01_1443200","PVP01_1443600","PVP01_1443700","PVP01_1443800","PVP01_1443900","PVP01_1444000","PVP01_1444200","PVP01_1444400","PVP01_1444800","PVP01_1444900","PVP01_1445000","PVP01_1445200","PVP01_1445300","PVP01_1445400","PVP01_1445600","PVP01_1445700","PVP01_1446000","PVP01_1446100","PVP01_1446200","PVP01_1446300","PVP01_1446400","PVP01_1446600","PVP01_1446800","PVP01_1446900","PVP01_1447200","PVP01_1447300","PVP01_1447500","PVP01_1447600","PVP01_1447800","PVP01_1447900","PVP01_1448100","PVP01_1448200","PVP01_1448300","PVP01_1448400","PVP01_1448600","PVP01_1448700","PVP01_1448800","PVP01_1449100","PVP01_1449200","PVP01_1449300","PVP01_1449700","PVP01_1450000","PVP01_1450100","PVP01_1450200","PVP01_1450400","PVP01_1450500","PVP01_1451000","PVP01_1451100","PVP01_1451300","PVP01_1451500","PVP01_1451600","PVP01_1451700","PVP01_1451800","PVP01_1452100","PVP01_1452300","PVP01_1452400","PVP01_1452500","PVP01_1452700","PVP01_1452800","PVP01_1452900","PVP01_1453100","PVP01_1453200","PVP01_1453800","PVP01_1454000","PVP01_1454100","PVP01_1454200","PVP01_1454400","PVP01_1454500","PVP01_1454600","PVP01_1454800","PVP01_1454900","PVP01_1455100","PVP01_1455200","PVP01_1455400","PVP01_1455500","PVP01_1455700","PVP01_1456000","PVP01_1456200","PVP01_1456500","PVP01_1456700","PVP01_1456900","PVP01_1457000","PVP01_1457100","PVP01_1457400","PVP01_1457500","PVP01_1457900","PVP01_1458200","PVP01_1458700","PVP01_1458800","PVP01_1459100","PVP01_1459200","PVP01_1459300","PVP01_1459400","PVP01_1459500","PVP01_1459600","PVP01_1459700","PVP01_1460300","PVP01_1460500","PVP01_1460800","PVP01_1460900","PVP01_1461100","PVP01_1461700","PVP01_1461900","PVP01_1462000","PVP01_1462100","PVP01_1462300","PVP01_1462500","PVP01_1462600","PVP01_1462900","PVP01_1463000","PVP01_1463200","PVP01_1463300","PVP01_1463500","PVP01_1463600","PVP01_1463700","PVP01_1463800","PVP01_1464100","PVP01_1464300","PVP01_1464400","PVP01_1464500","PVP01_1464700","PVP01_1464900","PVP01_1465000","PVP01_1465100","PVP01_1465200","PVP01_1465500","PVP01_1465700","PVP01_1465800","PVP01_1465900","PVP01_1466000","PVP01_1466200","PVP01_1466300","PVP01_1466500","PVP01_1466600","PVP01_1467000","PVP01_1467100","PVP01_1467200","PVP01_1467300","PVP01_1467500","PVP01_1467600","PVP01_1467700","PVP01_1468100","PVP01_1468300","PVP01_1468400","PVP01_1468600","PVP01_1468800","PVP01_1469200","PVP01_1469300","PVP01_1469400","PVP01_1469500","PVP01_1469600","PVP01_1469700","PVP01_1469800","PVP01_1469900","PVP01_1470000","PVP01_1470100","PVP01_1470200","PVP01_1470300","PVP01_1470400","PVP01_1470500","PVP01_1470600","PVP01_1470700","PVP01_1470800","PVP01_1470900","PVP01_1471100","PVP01_1471300","PVP01_1471400","PVP01_1471500","PVP01_1471600","PVP01_1471700","PVP01_1471800","PVP01_1471900","PVP01_1472100","PVP01_1472200","PVP01_1472400","PVP01_API01200","PVP01_API01400","PVP01_API01700","PVP01_API04100","PVP01_API04200","PVP01_0000100","PVP01_0004220","PVP01_0004230","PVP01_0005540","PVP01_0007720","PVP01_0009370","PVP01_0010540","PVP01_0010840","PVP01_0011020","PVP01_0103200","PVP01_0103700","PVP01_0104000","PVP01_0104300","PVP01_0104400","PVP01_0104500","PVP01_0105900","PVP01_0106100","PVP01_0106600","PVP01_0106700","PVP01_0108700","PVP01_0108900","PVP01_0109400","PVP01_0109500","PVP01_0109700","PVP01_0109900","PVP01_0110100","PVP01_0110300","PVP01_0110400","PVP01_0110600","PVP01_0111000","PVP01_0111400","PVP01_0111800","PVP01_0111900","PVP01_0112200","PVP01_0112400","PVP01_0112600","PVP01_0112800","PVP01_0112900","PVP01_0113600","PVP01_0113700","PVP01_0113800","PVP01_0113900","PVP01_0114000","PVP01_0114300","PVP01_0114400","PVP01_0114800","PVP01_0115300","PVP01_0115600","PVP01_0115700","PVP01_0115800","PVP01_0115900","PVP01_0116000","PVP01_0116200","PVP01_0116400","PVP01_0116800","PVP01_0117000","PVP01_0117100","PVP01_0117200","PVP01_0117400","PVP01_0117600","PVP01_0117900","PVP01_0118100","PVP01_0118200","PVP01_0118300","PVP01_0118400","PVP01_0201200","PVP01_0202300","PVP01_0202500","PVP01_0202600","PVP01_0203200","PVP01_0203600","PVP01_0203700","PVP01_0204200","PVP01_0204500","PVP01_0204600","PVP01_0204700","PVP01_0205400","PVP01_0205700","PVP01_0205900","PVP01_0206000","PVP01_0206700","PVP01_0206900","PVP01_0207300","PVP01_0207400","PVP01_0208500","PVP01_0209400","PVP01_0209700","PVP01_0209900","PVP01_0210200","PVP01_0210400","PVP01_0210500","PVP01_0210600","PVP01_0210700","PVP01_0210800","PVP01_0210900","PVP01_0211300","PVP01_0212000","PVP01_0212100","PVP01_0212200","PVP01_0212300","PVP01_0212900","PVP01_0213100","PVP01_0213500","PVP01_0214000","PVP01_0214300","PVP01_0214400","PVP01_0214500","PVP01_0214600","PVP01_0214700","PVP01_0215200","PVP01_0216000","PVP01_0216200","PVP01_0302700","PVP01_0303000","PVP01_0303200","PVP01_0303300","PVP01_0303600","PVP01_0303700","PVP01_0303800","PVP01_0303900","PVP01_0304000","PVP01_0304200","PVP01_0304300","PVP01_0304700","PVP01_0305100","PVP01_0305400","PVP01_0306200","PVP01_0307200","PVP01_0307800","PVP01_0307900","PVP01_0308000","PVP01_0308100","PVP01_0308300","PVP01_0308400","PVP01_0308500","PVP01_0308700","PVP01_0308900","PVP01_0309000","PVP01_0309100","PVP01_0309600","PVP01_0310100","PVP01_0310700","PVP01_0311100","PVP01_0311800","PVP01_0311900","PVP01_0312600","PVP01_0312900","PVP01_0313000","PVP01_0313100","PVP01_0313200","PVP01_0313300","PVP01_0313400","PVP01_0313600","PVP01_0313700","PVP01_0313900","PVP01_0314300","PVP01_0314400","PVP01_0314500","PVP01_0314600","PVP01_0314800","PVP01_0315400","PVP01_0315500","PVP01_0315600","PVP01_0315800","PVP01_0316100","PVP01_0316400","PVP01_0316600","PVP01_0316900","PVP01_0317000","PVP01_0317300","PVP01_0317500","PVP01_0317600","PVP01_0317700","PVP01_0317800","PVP01_0317900","PVP01_0318200","PVP01_0318300","PVP01_0318500","PVP01_0318600","PVP01_0318800","PVP01_0319000","PVP01_0319100","PVP01_0319500","PVP01_0319800","PVP01_0319900","PVP01_0320000","PVP01_0320100","PVP01_0320300","PVP01_0405400","PVP01_0405800","PVP01_0406300","PVP01_0406400","PVP01_0406800","PVP01_0407100","PVP01_0407200","PVP01_0407400","PVP01_0407500","PVP01_0407800","PVP01_0408000","PVP01_0408100","PVP01_0408300","PVP01_0408400","PVP01_0408500","PVP01_0408700","PVP01_0409300","PVP01_0409400","PVP01_0409500","PVP01_0409600","PVP01_0410100","PVP01_0410300","PVP01_0410700","PVP01_0410800","PVP01_0411000","PVP01_0411100","PVP01_0411300","PVP01_0411500","PVP01_0411700","PVP01_0412000","PVP01_0412300","PVP01_0412400","PVP01_0412500","PVP01_0412600","PVP01_0413600","PVP01_0413700","PVP01_0413800","PVP01_0413900","PVP01_0414200","PVP01_0414500","PVP01_0414700","PVP01_0414800","PVP01_0415000","PVP01_0415100","PVP01_0415200","PVP01_0415300","PVP01_0415700","PVP01_0415900","PVP01_0416000","PVP01_0416200","PVP01_0416300","PVP01_0416400","PVP01_0416500","PVP01_0416700","PVP01_0416800","PVP01_0417000","PVP01_0417500","PVP01_0418000","PVP01_0418200","PVP01_0418500","PVP01_0418600","PVP01_0418700","PVP01_0418800","PVP01_0418900","PVP01_0419100","PVP01_0419800","PVP01_0419900","PVP01_0420300","PVP01_0420400","PVP01_0420600","PVP01_0420800","PVP01_0421000","PVP01_0421100","PVP01_0421200","PVP01_0421300","PVP01_0422000","PVP01_0422800","PVP01_0424500","PVP01_0504400","PVP01_0504800","PVP01_0504900","PVP01_0505200","PVP01_0505300","PVP01_0505600","PVP01_0505900","PVP01_0506100","PVP01_0506200","PVP01_0506500","PVP01_0507100","PVP01_0507200","PVP01_0507400","PVP01_0507900","PVP01_0508100","PVP01_0508300","PVP01_0508400","PVP01_0508800","PVP01_0508900","PVP01_0509000","PVP01_0509300","PVP01_0509500","PVP01_0509700","PVP01_0510200","PVP01_0510400","PVP01_0510600","PVP01_0510700","PVP01_0511000","PVP01_0511100","PVP01_0511600","PVP01_0511900","PVP01_0512000","PVP01_0512300","PVP01_0512900","PVP01_0513100","PVP01_0513200","PVP01_0513600","PVP01_0513800","PVP01_0514000","PVP01_0514200","PVP01_0514300","PVP01_0514500","PVP01_0514700","PVP01_0515000","PVP01_0515400","PVP01_0515500","PVP01_0516600","PVP01_0516700","PVP01_0516800","PVP01_0517100","PVP01_0517200","PVP01_0517500","PVP01_0517800","PVP01_0518000","PVP01_0518100","PVP01_0518200","PVP01_0518400","PVP01_0519000","PVP01_0519300","PVP01_0519700","PVP01_0519800","PVP01_0519900","PVP01_0520000","PVP01_0520100","PVP01_0520200","PVP01_0520300","PVP01_0520400","PVP01_0520500","PVP01_0520600","PVP01_0520900","PVP01_0521000","PVP01_0521100","PVP01_0521200","PVP01_0521700","PVP01_0522400","PVP01_0522800","PVP01_0523400","PVP01_0524500","PVP01_0525200","PVP01_0525600","PVP01_0525800","PVP01_0525900","PVP01_0526000","PVP01_0526300","PVP01_0526500","PVP01_0526600","PVP01_0526700","PVP01_0526800","PVP01_0527200","PVP01_0527600","PVP01_0527900","PVP01_0528300","PVP01_0528800","PVP01_0529300","PVP01_0529400","PVP01_0529500","PVP01_0529800","PVP01_0530200","PVP01_0530300","PVP01_0530900","PVP01_0531000","PVP01_0531300","PVP01_0531400","PVP01_0531600","PVP01_0531800","PVP01_0532100","PVP01_0532200","PVP01_0532400","PVP01_0532800","PVP01_0600800","PVP01_0601900","PVP01_0602000","PVP01_0602100","PVP01_0602200","PVP01_0602300","PVP01_0602400","PVP01_0602500","PVP01_0602700","PVP01_0602800","PVP01_0602900","PVP01_0603200","PVP01_0603400","PVP01_0603500","PVP01_0603800","PVP01_0603900","PVP01_0604000","PVP01_0604300","PVP01_0604500","PVP01_0604600","PVP01_0604800","PVP01_0604900","PVP01_0605400","PVP01_0605600","PVP01_0605700","PVP01_0605800","PVP01_0605900","PVP01_0606100","PVP01_0606300","PVP01_0606700","PVP01_0606900","PVP01_0607000","PVP01_0607300","PVP01_0607400","PVP01_0607500","PVP01_0607700","PVP01_0607800","PVP01_0607900","PVP01_0608200","PVP01_0608400","PVP01_0609000","PVP01_0609100","PVP01_0609300","PVP01_0609500","PVP01_0610000","PVP01_0610200","PVP01_0611000","PVP01_0611200","PVP01_0611500","PVP01_0611600","PVP01_0611800","PVP01_0611900","PVP01_0612200","PVP01_0612300","PVP01_0612400","PVP01_0612500","PVP01_0612800","PVP01_0612900","PVP01_0613300","PVP01_0613600","PVP01_0613700","PVP01_0614100","PVP01_0614500","PVP01_0614700","PVP01_0614800","PVP01_0615200","PVP01_0615300","PVP01_0615400","PVP01_0615600","PVP01_0616100","PVP01_0616500","PVP01_0616600","PVP01_0616700","PVP01_0617800","PVP01_0618000","PVP01_0618200","PVP01_0618400","PVP01_0618500","PVP01_0618800","PVP01_0620000","PVP01_0620800","PVP01_0621000","PVP01_0621300","PVP01_0621500","PVP01_0621600","PVP01_0621700","PVP01_0622000","PVP01_0622300","PVP01_0622400","PVP01_0622500","PVP01_0622600","PVP01_0700100","PVP01_0701100","PVP01_0701400","PVP01_0701800","PVP01_0701900","PVP01_0702300","PVP01_0702600","PVP01_0702700","PVP01_0702800","PVP01_0702900","PVP01_0703100","PVP01_0703200","PVP01_0703300","PVP01_0703400","PVP01_0703500","PVP01_0703800","PVP01_0704200","PVP01_0704300","PVP01_0704400","PVP01_0705100","PVP01_0705200","PVP01_0705300","PVP01_0705400","PVP01_0705700","PVP01_0705900","PVP01_0706500","PVP01_0706800","PVP01_0707600","PVP01_0707700","PVP01_0707900","PVP01_0708600","PVP01_0708700","PVP01_0709000","PVP01_0709400","PVP01_0709800","PVP01_0710400","PVP01_0710500","PVP01_0710600","PVP01_0710800","PVP01_0710900","PVP01_0711300","PVP01_0711600","PVP01_0711700","PVP01_0711800","PVP01_0712500","PVP01_0712600","PVP01_0712900","PVP01_0713300","PVP01_0713400","PVP01_0713800","PVP01_0714000","PVP01_0714100","PVP01_0714400","PVP01_0714500","PVP01_0714700","PVP01_0714800","PVP01_0715400","PVP01_0715900","PVP01_0716300","PVP01_0716500","PVP01_0716700","PVP01_0717100","PVP01_0717300","PVP01_0717700","PVP01_0717900","PVP01_0718200","PVP01_0718400","PVP01_0718500","PVP01_0718900","PVP01_0719200","PVP01_0719300","PVP01_0719600","PVP01_0719800","PVP01_0720500","PVP01_0720600","PVP01_0720700","PVP01_0721000","PVP01_0721200","PVP01_0721300","PVP01_0722900","PVP01_0723100","PVP01_0723600","PVP01_0723900","PVP01_0724000","PVP01_0724200","PVP01_0724400","PVP01_0725600","PVP01_0725900","PVP01_0726000","PVP01_0726100","PVP01_0726200","PVP01_0726300","PVP01_0726500","PVP01_0726600","PVP01_0726800","PVP01_0727700","PVP01_0727900","PVP01_0728500","PVP01_0728600","PVP01_0728700","PVP01_0728800","PVP01_0728900","PVP01_0729000","PVP01_0729600","PVP01_0729800","PVP01_0729900","PVP01_0730500","PVP01_0730800","PVP01_0730900","PVP01_0731400","PVP01_0731500","PVP01_0732500","PVP01_0732700","PVP01_0733200","PVP01_0733300","PVP01_0733400","PVP01_0733600","PVP01_0734000","PVP01_0734300","PVP01_0734500","PVP01_0734600","PVP01_0734800","PVP01_0734900","PVP01_0735000","PVP01_0801200","PVP01_0801300","PVP01_0802200","PVP01_0802600","PVP01_0802700","PVP01_0802800","PVP01_0803200","PVP01_0803300","PVP01_0803800","PVP01_0804000","PVP01_0804200","PVP01_0804300","PVP01_0805100","PVP01_0805900","PVP01_0806600","PVP01_0806800","PVP01_0806900","PVP01_0807100","PVP01_0807400","PVP01_0807500","PVP01_0807700","PVP01_0807800","PVP01_0808200","PVP01_0808300","PVP01_0808900","PVP01_0809000","PVP01_0809500","PVP01_0809600","PVP01_0809800","PVP01_0810200","PVP01_0810700","PVP01_0810800","PVP01_0810900","PVP01_0811400","PVP01_0811600","PVP01_0811800","PVP01_0812100","PVP01_0812200","PVP01_0812500","PVP01_0812700","PVP01_0812900","PVP01_0813100","PVP01_0813400","PVP01_0813500","PVP01_0813900","PVP01_0814000","PVP01_0814100","PVP01_0814500","PVP01_0814600","PVP01_0814700","PVP01_0814900","PVP01_0815000","PVP01_0815200","PVP01_0815500","PVP01_0815600","PVP01_0815700","PVP01_0815800","PVP01_0815900","PVP01_0816000","PVP01_0816100","PVP01_0816400","PVP01_0816500","PVP01_0816600","PVP01_0816900","PVP01_0817000","PVP01_0817200","PVP01_0817300","PVP01_0817500","PVP01_0817600","PVP01_0817700","PVP01_0817800","PVP01_0817900","PVP01_0818000","PVP01_0818200","PVP01_0818400","PVP01_0818500","PVP01_0818700","PVP01_0818900","PVP01_0819100","PVP01_0819400","PVP01_0819500","PVP01_0819900","PVP01_0821100","PVP01_0821600","PVP01_0821900","PVP01_0822000","PVP01_0822200","PVP01_0822400","PVP01_0822500","PVP01_0822800","PVP01_0823000","PVP01_0823300","PVP01_0823500","PVP01_0823800","PVP01_0823900","PVP01_0824400","PVP01_0825300","PVP01_0825500","PVP01_0825900","PVP01_0826600","PVP01_0826800","PVP01_0827000","PVP01_0827200","PVP01_0827300","PVP01_0827400","PVP01_0827900","PVP01_0828400","PVP01_0828500","PVP01_0828600","PVP01_0828800","PVP01_0828900","PVP01_0829400","PVP01_0829600","PVP01_0830000","PVP01_0830200","PVP01_0830600","PVP01_0831200","PVP01_0831300","PVP01_0831500","PVP01_0831600","PVP01_0832300","PVP01_0832700","PVP01_0833100","PVP01_0833200","PVP01_0833400","PVP01_0833900","PVP01_0834100","PVP01_0834700","PVP01_0834800","PVP01_0835100","PVP01_0835200","PVP01_0835400","PVP01_0835600","PVP01_0835900","PVP01_0836000","PVP01_0836100","PVP01_0836200","PVP01_0836900","PVP01_0837000","PVP01_0837100","PVP01_0837400","PVP01_0904200","PVP01_0904400","PVP01_0904500","PVP01_0904800","PVP01_0905000","PVP01_0905200","PVP01_0905300","PVP01_0905400","PVP01_0905500","PVP01_0905600","PVP01_0905700","PVP01_0905800","PVP01_0905900","PVP01_0906100","PVP01_0906200","PVP01_0906600","PVP01_0907900","PVP01_0908100","PVP01_0908300","PVP01_0908700","PVP01_0908800","PVP01_0909000","PVP01_0909200","PVP01_0909400","PVP01_0909700","PVP01_0909900","PVP01_0910700","PVP01_0910800","PVP01_0911100","PVP01_0911200","PVP01_0911400","PVP01_0912300","PVP01_0912500","PVP01_0912600","PVP01_0912700","PVP01_0912800","PVP01_0913200","PVP01_0913300","PVP01_0913700","PVP01_0914100","PVP01_0914500","PVP01_0914700","PVP01_0914900","PVP01_0915000","PVP01_0915200","PVP01_0915500","PVP01_0915700","PVP01_0915900","PVP01_0916100","PVP01_0916200","PVP01_0916400","PVP01_0916500","PVP01_0916700","PVP01_0917300","PVP01_0917400","PVP01_0917500","PVP01_0917600","PVP01_0917700","PVP01_0917800","PVP01_0918500","PVP01_0919200","PVP01_0919300","PVP01_0919800","PVP01_0920000","PVP01_0920300","PVP01_0920500","PVP01_0920600","PVP01_0920900","PVP01_0921000","PVP01_0921100","PVP01_0921200","PVP01_0921300","PVP01_0921600","PVP01_0921700","PVP01_0922200","PVP01_0922500","PVP01_0922600","PVP01_0922700","PVP01_0923400","PVP01_0923500","PVP01_0923600","PVP01_0923700","PVP01_0924100","PVP01_0924200","PVP01_0924500","PVP01_0924900","PVP01_0925300","PVP01_0925500","PVP01_0926100","PVP01_0926200","PVP01_0926700","PVP01_0927000","PVP01_0927300","PVP01_0927500","PVP01_0928400","PVP01_0928500","PVP01_0928600","PVP01_0928700","PVP01_0929500","PVP01_0929800","PVP01_0930100","PVP01_0930600","PVP01_0930700","PVP01_0930800","PVP01_0931800","PVP01_0931900","PVP01_0932100","PVP01_0932400","PVP01_0932700","PVP01_0932800","PVP01_0933000","PVP01_0933200","PVP01_0933600","PVP01_0933800","PVP01_0933900","PVP01_0934100","PVP01_0934200","PVP01_0934700","PVP01_0934800","PVP01_0934900","PVP01_0935100","PVP01_0935300","PVP01_0935600","PVP01_0936200","PVP01_0936500","PVP01_0936600","PVP01_0936900","PVP01_0937100","PVP01_0937200","PVP01_0937400","PVP01_0937600","PVP01_0938000","PVP01_0938300","PVP01_0938400","PVP01_0938500","PVP01_0938800","PVP01_0939400","PVP01_0939500","PVP01_0939800","PVP01_0939900","PVP01_0940000","PVP01_0940100","PVP01_0940300","PVP01_0940400","PVP01_0940800","PVP01_0941000","PVP01_0941500","PVP01_0941700","PVP01_0941900","PVP01_0942100","PVP01_0942500","PVP01_0943000","PVP01_0943200","PVP01_0943500","PVP01_0943600","PVP01_0943900","PVP01_0944000","PVP01_0944200","PVP01_0944500","PVP01_0944700","PVP01_0944900","PVP01_0945000","PVP01_0945400","PVP01_0945800","PVP01_0947200","PVP01_0947500","PVP01_0947700","PVP01_0948000","PVP01_0948400","PVP01_0949300","PVP01_1002400","PVP01_1002800","PVP01_1002900","PVP01_1003100","PVP01_1003300","PVP01_1003600","PVP01_1003700","PVP01_1003800","PVP01_1004000","PVP01_1004200","PVP01_1004400","PVP01_1004600","PVP01_1004700","PVP01_1005000","PVP01_1005500","PVP01_1005700","PVP01_1005800","PVP01_1006900","PVP01_1007000","PVP01_1007100","PVP01_1007300","PVP01_1007400","PVP01_1007600","PVP01_1008400","PVP01_1008500","PVP01_1008800","PVP01_1009200","PVP01_1009800","PVP01_1009900","PVP01_1010000","PVP01_1010500","PVP01_1010600","PVP01_1010700","PVP01_1010800","PVP01_1010900","PVP01_1011200","PVP01_1011700","PVP01_1011900","PVP01_1012000","PVP01_1012200","PVP01_1012600","PVP01_1012800","PVP01_1012900","PVP01_1013000","PVP01_1013500","PVP01_1013600","PVP01_1013800","PVP01_1013900","PVP01_1014000","PVP01_1014200","PVP01_1014400","PVP01_1014500","PVP01_1014700","PVP01_1014900","PVP01_1015300","PVP01_1015800","PVP01_1016000","PVP01_1016300","PVP01_1016700","PVP01_1016800","PVP01_1017000","PVP01_1017300","PVP01_1017500","PVP01_1017600","PVP01_1017800","PVP01_1018300","PVP01_1018600","PVP01_1018700","PVP01_1018900","PVP01_1019100","PVP01_1019200","PVP01_1019300","PVP01_1019700","PVP01_1019800","PVP01_1020400","PVP01_1020600","PVP01_1020800","PVP01_1021400","PVP01_1022500","PVP01_1022600","PVP01_1022700","PVP01_1022800","PVP01_1023500","PVP01_1023800","PVP01_1023900","PVP01_1024000","PVP01_1024100","PVP01_1024200","PVP01_1024600","PVP01_1024700","PVP01_1024900","PVP01_1025000","PVP01_1025100","PVP01_1025500","PVP01_1025600","PVP01_1025900","PVP01_1026000","PVP01_1026100","PVP01_1026200","PVP01_1027000","PVP01_1027200","PVP01_1027500","PVP01_1028300","PVP01_1028500","PVP01_1028700","PVP01_1028800","PVP01_1028900","PVP01_1029400","PVP01_1029600","PVP01_1030800","PVP01_1032300","PVP01_1032600","PVP01_1032700","PVP01_1033100","PVP01_1033200","PVP01_1102300","PVP01_1103000","PVP01_1103100","PVP01_1103300","PVP01_1103500","PVP01_1103600","PVP01_1103800","PVP01_1103900","PVP01_1104500","PVP01_1104700","PVP01_1105000","PVP01_1105100","PVP01_1105200","PVP01_1105700","PVP01_1105900","PVP01_1106100","PVP01_1106500","PVP01_1106800","PVP01_1106900","PVP01_1107500","PVP01_1107800","PVP01_1108200","PVP01_1108400","PVP01_1108700","PVP01_1109500","PVP01_1109700","PVP01_1109800","PVP01_1110000","PVP01_1110100","PVP01_1110200","PVP01_1110300","PVP01_1110400","PVP01_1110800","PVP01_1111000","PVP01_1111900","PVP01_1112200","PVP01_1112300","PVP01_1112500","PVP01_1113100","PVP01_1113200","PVP01_1113400","PVP01_1113600","PVP01_1113800","PVP01_1114000","PVP01_1114400","PVP01_1114900","PVP01_1115100","PVP01_1115400","PVP01_1115700","PVP01_1115800","PVP01_1115900","PVP01_1116000","PVP01_1116100","PVP01_1116200","PVP01_1116300","PVP01_1116700","PVP01_1117100","PVP01_1117400","PVP01_1117600","PVP01_1117800","PVP01_1117900","PVP01_1118000","PVP01_1118200","PVP01_1118300","PVP01_1118400","PVP01_1118800","PVP01_1118900","PVP01_1119100","PVP01_1119200","PVP01_1119400","PVP01_1119800","PVP01_1120200","PVP01_1120600","PVP01_1120700","PVP01_1120800","PVP01_1121200","PVP01_1121400","PVP01_1122000","PVP01_1122200","PVP01_1123700","PVP01_1123800","PVP01_1124200","PVP01_1124500","PVP01_1125500","PVP01_1125900","PVP01_1126400","PVP01_1126700","PVP01_1127100","PVP01_1127600","PVP01_1127700","PVP01_1128200","PVP01_1128300","PVP01_1129100","PVP01_1129200","PVP01_1129300","PVP01_1129400","PVP01_1129800","PVP01_1129900","PVP01_1130000","PVP01_1130200","PVP01_1130300","PVP01_1130500","PVP01_1130800","PVP01_1130900","PVP01_1131000","PVP01_1131100","PVP01_1131200","PVP01_1131500","PVP01_1131600","PVP01_1131700","PVP01_1131800","PVP01_1132000","PVP01_1132200","PVP01_1132600","PVP01_1133500","PVP01_1133700","PVP01_1134000","PVP01_1134600","PVP01_1134900","PVP01_1135200","PVP01_1135400","PVP01_1136200","PVP01_1136500","PVP01_1136900","PVP01_1137000","PVP01_1137400","PVP01_1137500","PVP01_1137900","PVP01_1138000","PVP01_1138100","PVP01_1138300","PVP01_1138600","PVP01_1138700","PVP01_1138800","PVP01_1138900","PVP01_1139500","PVP01_1139600","PVP01_1139800","PVP01_1139900","PVP01_1140100","PVP01_1140200","PVP01_1140400","PVP01_1141100","PVP01_1141300","PVP01_1141400","PVP01_1141600","PVP01_1142100","PVP01_1142400","PVP01_1142700","PVP01_1142800","PVP01_1142900","PVP01_1143000","PVP01_1143300","PVP01_1143600","PVP01_1143800","PVP01_1143900","PVP01_1144000","PVP01_1144400","PVP01_1144600","PVP01_1144800","PVP01_1145300","PVP01_1145400","PVP01_1145900","PVP01_1146000","PVP01_1146100","PVP01_1146300","PVP01_1146700","PVP01_1202400","PVP01_1202600","PVP01_1202800","PVP01_1202900","PVP01_1203000","PVP01_1203900","PVP01_1204100","PVP01_1204300","PVP01_1204900","PVP01_1205100","PVP01_1205300","PVP01_1205500","PVP01_1205700","PVP01_1206000","PVP01_1206200","PVP01_1206500","PVP01_1206600","PVP01_1206800","PVP01_1207300","PVP01_1207400","PVP01_1207500","PVP01_1207600","PVP01_1207700","PVP01_1208400","PVP01_1208500","PVP01_1209100","PVP01_1209300","PVP01_1209500","PVP01_1209600","PVP01_1209700","PVP01_1209800","PVP01_1209900","PVP01_1210200","PVP01_1210600","PVP01_1210800","PVP01_1210900","PVP01_1211400","PVP01_1211800","PVP01_1211900","PVP01_1212000","PVP01_1212200","PVP01_1212300","PVP01_1212400","PVP01_1212800","PVP01_1212900","PVP01_1213200","PVP01_1213400","PVP01_1213600","PVP01_1213900","PVP01_1214000","PVP01_1214100","PVP01_1214700","PVP01_1215100","PVP01_1215200","PVP01_1215500","PVP01_1216400","PVP01_1216500","PVP01_1216600","PVP01_1217300","PVP01_1217800","PVP01_1217900","PVP01_1218100","PVP01_1218200","PVP01_1218300","PVP01_1218400","PVP01_1218600","PVP01_1218700","PVP01_1219300","PVP01_1219700","PVP01_1220000","PVP01_1220300","PVP01_1220400","PVP01_1220500","PVP01_1221100","PVP01_1221500","PVP01_1221600","PVP01_1222000","PVP01_1222500","PVP01_1222600","PVP01_1222800","PVP01_1222900","PVP01_1223000","PVP01_1223100","PVP01_1223200","PVP01_1223300","PVP01_1223600","PVP01_1223700","PVP01_1224100","PVP01_1224200","PVP01_1224900","PVP01_1225100","PVP01_1225400","PVP01_1225600","PVP01_1225700","PVP01_1226000","PVP01_1226200","PVP01_1226300","PVP01_1226400","PVP01_1226600","PVP01_1226900","PVP01_1227000","PVP01_1227100","PVP01_1227200","PVP01_1227700","PVP01_1227800","PVP01_1228300","PVP01_1228600","PVP01_1228700","PVP01_1228900","PVP01_1229000","PVP01_1229200","PVP01_1229300","PVP01_1229700","PVP01_1230300","PVP01_1230400","PVP01_1230900","PVP01_1231200","PVP01_1231300","PVP01_1231400","PVP01_1231500","PVP01_1232000","PVP01_1232100","PVP01_1232200","PVP01_1232300","PVP01_1232500","PVP01_1232800","PVP01_1232900","PVP01_1233000","PVP01_1233200","PVP01_1233300","PVP01_1233400","PVP01_1233500","PVP01_1233600","PVP01_1233700","PVP01_1234100","PVP01_1234200","PVP01_1234300","PVP01_1234500","PVP01_1234600","PVP01_1234700","PVP01_1234800","PVP01_1234900","PVP01_1235500","PVP01_1235600","PVP01_1235700","PVP01_1235900","PVP01_1236300","PVP01_1236500","PVP01_1236600","PVP01_1237400","PVP01_1238100","PVP01_1238200","PVP01_1238300","PVP01_1238700","PVP01_1238800","PVP01_1239200","PVP01_1239500","PVP01_1239600","PVP01_1239700","PVP01_1240400","PVP01_1240700","PVP01_1241400","PVP01_1241500","PVP01_1241700","PVP01_1241800","PVP01_1242100","PVP01_1242200","PVP01_1242500","PVP01_1243300","PVP01_1243400","PVP01_1243700","PVP01_1243800","PVP01_1244000","PVP01_1244100","PVP01_1244200","PVP01_1244300","PVP01_1244500","PVP01_1244600","PVP01_1245000","PVP01_1245200","PVP01_1245700","PVP01_1245800","PVP01_1246200","PVP01_1246800","PVP01_1247100","PVP01_1247200","PVP01_1247300","PVP01_1247400","PVP01_1248000","PVP01_1248100","PVP01_1248300","PVP01_1248400","PVP01_1248700","PVP01_1248800","PVP01_1248900","PVP01_1249300","PVP01_1249400","PVP01_1249800","PVP01_1249900","PVP01_1250000","PVP01_1250100","PVP01_1250400","PVP01_1250500","PVP01_1250700","PVP01_1250800","PVP01_1251000","PVP01_1251200","PVP01_1251300","PVP01_1251400","PVP01_1252100","PVP01_1252300","PVP01_1252500","PVP01_1252600","PVP01_1252700","PVP01_1253100","PVP01_1253200","PVP01_1253400","PVP01_1253500","PVP01_1253900","PVP01_1254000","PVP01_1254200","PVP01_1254400","PVP01_1254500","PVP01_1254700","PVP01_1254800","PVP01_1255000","PVP01_1255600","PVP01_1255700","PVP01_1255800","PVP01_1255900","PVP01_1256300","PVP01_1256600","PVP01_1257100","PVP01_1257400","PVP01_1257800","PVP01_1258000","PVP01_1258600","PVP01_1258800","PVP01_1259700","PVP01_1259800","PVP01_1259900","PVP01_1260000","PVP01_1260100","PVP01_1260300","PVP01_1260400","PVP01_1260500","PVP01_1260600","PVP01_1260700","PVP01_1261200","PVP01_1261300","PVP01_1261600","PVP01_1261800","PVP01_1261900","PVP01_1262100","PVP01_1262700","PVP01_1263200","PVP01_1263300","PVP01_1263500","PVP01_1263600","PVP01_1264100","PVP01_1264200","PVP01_1264500","PVP01_1265200","PVP01_1265500","PVP01_1265800","PVP01_1266000","PVP01_1266100","PVP01_1266300","PVP01_1266800","PVP01_1266900","PVP01_1267000","PVP01_1267100","PVP01_1267200","PVP01_1267700","PVP01_1267800","PVP01_1267900","PVP01_1268400","PVP01_1268500","PVP01_1269100","PVP01_1269300","PVP01_1269400","PVP01_1269600","PVP01_1269700","PVP01_1269900","PVP01_1270000","PVP01_1270100","PVP01_1270500","PVP01_1271000","PVP01_1300600","PVP01_1300800","PVP01_1301000","PVP01_1301800","PVP01_1301900","PVP01_1302000","PVP01_1302200","PVP01_1302400","PVP01_1303300","PVP01_1303500","PVP01_1303700","PVP01_1303800","PVP01_1304100","PVP01_1304400","PVP01_1304600","PVP01_1304700","PVP01_1305000","PVP01_1305100","PVP01_1305400","PVP01_1305600","PVP01_1305800","PVP01_1306200","PVP01_1306500","PVP01_1307400","PVP01_1307600","PVP01_1307900","PVP01_1308000","PVP01_1308200","PVP01_1308600","PVP01_1308700","PVP01_1309300","PVP01_1309600","PVP01_1309900","PVP01_1310100","PVP01_1310200","PVP01_1310300","PVP01_1310600","PVP01_1310900","PVP01_1311700","PVP01_1311800","PVP01_1312300","PVP01_1312500","PVP01_1312700","PVP01_1312800","PVP01_1313500","PVP01_1313600","PVP01_1313700","PVP01_1313800","PVP01_1313900","PVP01_1314000","PVP01_1314200","PVP01_1314400","PVP01_1314500","PVP01_1314700","PVP01_1314900","PVP01_1315000","PVP01_1315100","PVP01_1315300","PVP01_1315600","PVP01_1315700","PVP01_1315800","PVP01_1315900","PVP01_1316400","PVP01_1316800","PVP01_1317100","PVP01_1317200","PVP01_1317300","PVP01_1317400","PVP01_1318200","PVP01_1318400","PVP01_1318700","PVP01_1319100","PVP01_1319200","PVP01_1319400","PVP01_1320400","PVP01_1320800","PVP01_1320900","PVP01_1321000","PVP01_1321100","PVP01_1321200","PVP01_1321300","PVP01_1321400","PVP01_1321800","PVP01_1322100","PVP01_1322500","PVP01_1322700","PVP01_1322800","PVP01_1323300","PVP01_1323400","PVP01_1323500","PVP01_1323600","PVP01_1324000","PVP01_1325300","PVP01_1325400","PVP01_1325500","PVP01_1325700","PVP01_1326100","PVP01_1326300","PVP01_1326500","PVP01_1326700","PVP01_1326800","PVP01_1327000","PVP01_1327100","PVP01_1327400","PVP01_1327700","PVP01_1327800","PVP01_1328200","PVP01_1328300","PVP01_1328400","PVP01_1329100","PVP01_1329200","PVP01_1329500","PVP01_1330600","PVP01_1330700","PVP01_1330900","PVP01_1331300","PVP01_1331400","PVP01_1331500","PVP01_1331600","PVP01_1331700","PVP01_1332000","PVP01_1332100","PVP01_1332600","PVP01_1333300","PVP01_1333400","PVP01_1333500","PVP01_1333600","PVP01_1333800","PVP01_1334000","PVP01_1334400","PVP01_1334600","PVP01_1334700","PVP01_1334800","PVP01_1334900","PVP01_1335000","PVP01_1335200","PVP01_1335400","PVP01_1335500","PVP01_1335700","PVP01_1336000","PVP01_1336100","PVP01_1336200","PVP01_1336300","PVP01_1336600","PVP01_1336800","PVP01_1337000","PVP01_1337700","PVP01_1337800","PVP01_1338000","PVP01_1338200","PVP01_1338800","PVP01_1338900","PVP01_1339100","PVP01_1339200","PVP01_1339300","PVP01_1339500","PVP01_1339700","PVP01_1340000","PVP01_1340700","PVP01_1340900","PVP01_1341000","PVP01_1341100","PVP01_1341200","PVP01_1341400","PVP01_1341500","PVP01_1342000","PVP01_1342100","PVP01_1342500","PVP01_1342900","PVP01_1343000","PVP01_1343200","PVP01_1343400","PVP01_1343700","PVP01_1343800","PVP01_1343900","PVP01_1344100","PVP01_1344400","PVP01_1344800","PVP01_1345100","PVP01_1345800","PVP01_1346000","PVP01_1346100","PVP01_1346400","PVP01_1346500","PVP01_1346600","PVP01_1347000","PVP01_1401300","PVP01_1401400","PVP01_1403100","PVP01_1403200","PVP01_1403800","PVP01_1404400","PVP01_1404900","PVP01_1405200","PVP01_1405400","PVP01_1405700","PVP01_1405900","PVP01_1406200","PVP01_1406300","PVP01_1406400","PVP01_1406500","PVP01_1406800","PVP01_1407200","PVP01_1407300","PVP01_1407700","PVP01_1407900","PVP01_1408300","PVP01_1409200","PVP01_1409500","PVP01_1409600","PVP01_1409800","PVP01_1409900","PVP01_1410600","PVP01_1411100","PVP01_1411500","PVP01_1411600","PVP01_1411700","PVP01_1411800","PVP01_1411900","PVP01_1412000","PVP01_1412200","PVP01_1412300","PVP01_1412600","PVP01_1412700","PVP01_1412800","PVP01_1413400","PVP01_1413500","PVP01_1413600","PVP01_1413800","PVP01_1413900","PVP01_1414100","PVP01_1414600","PVP01_1414700","PVP01_1414800","PVP01_1415400","PVP01_1416300","PVP01_1416400","PVP01_1416500","PVP01_1416600","PVP01_1416700","PVP01_1416800","PVP01_1417000","PVP01_1417400","PVP01_1417500","PVP01_1417800","PVP01_1418300","PVP01_1418500","PVP01_1418700","PVP01_1419000","PVP01_1419200","PVP01_1419600","PVP01_1419900","PVP01_1420200","PVP01_1420400","PVP01_1420600","PVP01_1420700","PVP01_1420800","PVP01_1421600","PVP01_1421700","PVP01_1421900","PVP01_1422000","PVP01_1422400","PVP01_1422500","PVP01_1422800","PVP01_1423000","PVP01_1423300","PVP01_1423400","PVP01_1423600","PVP01_1424300","PVP01_1424400","PVP01_1424600","PVP01_1424700","PVP01_1424800","PVP01_1424900","PVP01_1425000","PVP01_1425700","PVP01_1425900","PVP01_1426300","PVP01_1426400","PVP01_1426500","PVP01_1426600","PVP01_1426900","PVP01_1427200","PVP01_1427300","PVP01_1427400","PVP01_1427500","PVP01_1427900","PVP01_1428200","PVP01_1428700","PVP01_1429000","PVP01_1429100","PVP01_1429400","PVP01_1429600","PVP01_1429800","PVP01_1430200","PVP01_1430900","PVP01_1431000","PVP01_1431100","PVP01_1431400","PVP01_1431600","PVP01_1432200","PVP01_1432500","PVP01_1433000","PVP01_1433100","PVP01_1433200","PVP01_1433300","PVP01_1433400","PVP01_1433500","PVP01_1433900","PVP01_1434100","PVP01_1434300","PVP01_1434700","PVP01_1435400","PVP01_1435500","PVP01_1435700","PVP01_1436100","PVP01_1436200","PVP01_1436500","PVP01_1436800","PVP01_1436900","PVP01_1437000","PVP01_1437500","PVP01_1437700","PVP01_1437800","PVP01_1437900","PVP01_1438000","PVP01_1438500","PVP01_1439100","PVP01_1439200","PVP01_1439300","PVP01_1439500","PVP01_1439800","PVP01_1439900","PVP01_1440200","PVP01_1440400","PVP01_1440500","PVP01_1441300","PVP01_1441500","PVP01_1441600","PVP01_1441900","PVP01_1442100","PVP01_1442400","PVP01_1442500","PVP01_1442700","PVP01_1442800","PVP01_1443300","PVP01_1443400","PVP01_1443500","PVP01_1444100","PVP01_1444300","PVP01_1444500","PVP01_1444600","PVP01_1444700","PVP01_1445100","PVP01_1445500","PVP01_1445800","PVP01_1445900","PVP01_1446500","PVP01_1446700","PVP01_1447000","PVP01_1447100","PVP01_1447400","PVP01_1447700","PVP01_1448000","PVP01_1448500","PVP01_1448900","PVP01_1449000","PVP01_1449400","PVP01_1449500","PVP01_1449600","PVP01_1449800","PVP01_1449900","PVP01_1450300","PVP01_1450600","PVP01_1450700","PVP01_1450800","PVP01_1450900","PVP01_1451200","PVP01_1451400","PVP01_1451900","PVP01_1452000","PVP01_1452200","PVP01_1452600","PVP01_1453000","PVP01_1453300","PVP01_1453400","PVP01_1453500","PVP01_1453600","PVP01_1453700","PVP01_1453900","PVP01_1454300","PVP01_1454700","PVP01_1455000","PVP01_1455300","PVP01_1455600","PVP01_1455800","PVP01_1455900","PVP01_1456100","PVP01_1456300","PVP01_1456400","PVP01_1456600","PVP01_1456800","PVP01_1457200","PVP01_1457300","PVP01_1457600","PVP01_1457700","PVP01_1457800","PVP01_1458000","PVP01_1458100","PVP01_1458300","PVP01_1458400","PVP01_1458500","PVP01_1458600","PVP01_1458900","PVP01_1459000","PVP01_1459800","PVP01_1459900","PVP01_1460000","PVP01_1460100","PVP01_1460200","PVP01_1460400","PVP01_1460600","PVP01_1460700","PVP01_1461000","PVP01_1461200","PVP01_1461300","PVP01_1461500","PVP01_1461600","PVP01_1461800","PVP01_1462200","PVP01_1462400","PVP01_1462700","PVP01_1462800","PVP01_1463100","PVP01_1463400","PVP01_1463900","PVP01_1464000","PVP01_1464200","PVP01_1464600","PVP01_1464800","PVP01_1465300","PVP01_1465400","PVP01_1465600","PVP01_1466100","PVP01_1466400","PVP01_1466700","PVP01_1466800","PVP01_1466900","PVP01_1467400","PVP01_1467800","PVP01_1467900","PVP01_1468000","PVP01_1468200","PVP01_1468500","PVP01_1468700","PVP01_1468900","PVP01_API02600","PVP01_API02800","PVP01_API03500","PVP01_API04300","PVP01_API04600","PVP01_MIT02700","PVP01_MIT03300","PVP01_MIT03400"],"logFC":[3.536,0.1259,4.02,2.586,-2.099,-3.165,4.358,4.769,7.214,-0.2274,6.87,2.082,0.7422,4.83,3.705,1.569,3.466,3.404,3.972,3.664,4.314,4.754,1.011,4.295,-2.038,2.115,5.18,0,3.348,3.845,3.694,4.016,4.492,5.102,5.189,3.482,-2.463,3.203,4.788,4.019,3.909,4.899,3.612,3.054,-0.9026,-2.541,4.596,5.005,3.52,4.306,3.359,2.456,5.066,2.367,1.466,0,5.37,2.57,-5.457,4.555,2.085,4.738,-5.159,-5.155,4.794,3.318,3.482,-4.232,3.054,4.031,4.768,3.738,-2.263,-1.973,3.419,2.765,2.776,4.919,-0.9614,3.313,4.156,4.571,-7.249,4.3,2.574,4.933,3.246,-1.373,3.029,3.807,4.739,3.789,5.287,-2.83,3.782,3.684,3.649,4.357,-3.66,4.631,2.866,3.365,3.694,5.629,3.285,6.562,6.655,0,4.05,4.916,-1.354,4.554,3.998,4.398,4.301,3.328,-0.5697,4.639,4.819,-3.987,3.561,4.188,4.295,4.528,5.497,3.975,-3.763,1.397,3.497,4.555,4.03,3.971,0.8023,3.787,3.947,-2.775,5.468,0.6643,4.805,-6.61,4.487,3.318,3.054,2.866,4.501,4.085,5.258,5.022,3.253,6.21,-4.891,5.412,4.309,2.866,3.801,4.073,-2.042,4.113,-3.355,3.605,2.955,4.124,4.119,-0.9808,5.909,5.57,5.038,4.351,4.121,-0.5319,-5.043,3.253,-1.666,-4.437,4.811,5.071,4.406,5.936,1.973,3.883,5.124,0,-5.614,-1.663,1.423,2.088,0,-4.96,-2.614,0,5.18,-1.893,5.076,0.8031,1.62,-0.08672,2.904,-0.6843,4.759,3.284,-3.874,4.221,0.8344,2.122,1.453,0.1361,-0.3151,-4.329,2.118,-4.735,1.913,1.172,1.45,-2.405,2.065,-1.125,0.3357,-1.646,0.7947,-1.327,1.173,-0.7951,2.064,0.08145,-2.379,1.998,-1.67,-0.3226,1.154,0.9637,-1.054,-2.714,-0.3226,1.468,-2.441,-1.414,-3.19,-3.201,0.1591,1.294,0.6659,0.01144,-3.232,1.588,-0.6244,1.529,1.762,-0.125,-1.428,1.623,-0.8478,0.7697,-1.169,0.2356,-0.2898,0.3474,-0.4048,3.004,1.335,-0.3098,2.38,0.4842,1.013,2.439,1.27,-1.313,-0.5994,-2.022,-0.8603,-1.482,0.9212,1.011,1.161,-1.75,2.65,-1.167,1.301,-0.0961,1.831,-0.5902,-0.443,1.367,1.712,2.268,-0.519,0.1974,-0.003305,0.07317,-3.758,0.08353,0.8961,-1.025,-0.1018,0.8707,-1.078,0.5554,-0.537,-0.366,1.823,2.993,0.2404,1.59,-0.07946,1.814,-0.9098,0.649,1.302,7.714,1.497,-2.283,-0.9274,-2.045,-1.986,-3.518,6.213,-3.548,4.697,3.916,3.895,4.073,5.74,4.102,3.705,5.37,4.073,-0.5425,3.658,0.969,6.029,-1.119,0.5026,8.459,2.341,2.049,-2.482,-2.517,-2.265,-1.533,0,-4.384,0.4508,-2.319,-0.1892,1.42,0.5237,0.09297,1.31,-1.34,-1.812,0.84,0.6313,-2.654,1.808,0.9744,-1.212,1.31,-2.099,1.184,1.429,1.088,1.27,1.42,0.6972,-2.039,0.08651,0.4327,1.752,-0.463,0.05432,0.2727,1.567,0.7025,-1.925,1.276,1.196,0.5752,0.4703,1.408,-1.222,0.4949,-0.6631,0.9175,1.136,-0.08188,-2.241,0.9106,-1.03,-1,1.302,-0.3026,-1.017,0.2535,0.5295,-0.8643,1.411,-0.2283,-0.161,-2.324,1.436,-2.186,-0.9558,-1.009,-0.8825,-0.3999,-0.5316,-2.871,-2.939,-3.314,1.237,-1.053,-1.659,-2.746,-1.887,-0.7317,0.9339,-1.758,0.6835,1.244,-3.089,-0.4623,7.145,1.948,-1.858,-0.04001,-2.392,-1.392,4.229,-0.3292,0.1846,0.2582,6.166,-0.801,5.012,7.184,4.705,4.97,4.525,4.394,4.03,1.361,5.677,0,4.211,5.789,4.051,3.757,6.267,3.881,6.95,0.2897,4.073,3.482,5.202,5.989,0.275,3.859,-0.7129,-2.168,-2.457,1.102,-2.475,-0.7949,1.849,1.962,-2.038,1.095,1.098,1.437,2.046,0.3157,0.5855,-1.461,-1.261,0.6454,0.02166,-3.327,-2.427,0.5592,-3.302,0.8712,-1.005,1.007,-0.6934,1.08,1.678,-2.921,-2.092,1.57,0.293,1.254,0.939,0.7541,1.755,-0.8988,-0.6104,2.689,1.489,0.4503,1.841,1.31,0.7914,-0.2782,1.101,0.5228,0.1078,1.186,-1.114,1.911,0.9225,-2.505,1.955,1.234,0.7043,-0.5638,-2.473,-1.802,-3.261,-0.1915,1.79,1.318,-1.404,-0.1061,-1.785,-1.292,0.1039,0.3235,2.842,-0.5887,-1.228,1.137,0.7775,-0.9469,0.4893,1.428,0.6028,0.8775,1.17,1.652,0.7519,1.496,1.701,1.385,-0.9069,1.531,-0.2425,0.7378,-0.05836,0.4125,2.237,1.33,2.607,0.5812,-1.456,0.3497,0.7881,1.975,2.474,1.661,-1.019,2.004,0.404,0.4942,-0.7444,0.8431,4.624,4.305,0,3.497,2.584,4.505,3.252,1.326,3.349,-2.853,5.55,-0.6562,-2.231,-4.74,6.415,-0.7461,-2.133,1.24,3.906,0.9888,-0.2249,-0.6223,-2.581,-0.2581,-0.03941,-2.791,0.2932,-1.308,1.317,-2.9,-0.8804,0.9569,1.248,0.6286,0.177,1.019,1.156,4.423,1.749,0.6605,-0.5959,-0.2069,2.502,-3.349,0.3786,1.452,0.3758,0.6359,0.9145,0.1242,-4.67,0.6663,-3.279,-1.444,-1.41,-0.06293,-0.2966,-0.3023,-0.1125,-0.3474,0.6419,1.423,1.226,1.299,1.48,-1.136,0.5171,-1.498,-1.677,0.5767,-0.6985,0.8985,-1.044,1.356,1.239,0.8613,0.1032,-1.011,-1.068,-0.7654,2.197,-0.1654,-2.63,-0.08511,1.094,3.054,-0.8698,-1.857,-0.9215,0.3607,-1.528,0.9726,-1.484,4.032,0.5486,6.483,8.291,-1.063,-1.303,5.337,0.4741,1.694,-0.5714,-1.438,0.2285,1.495,0.4405,-0.934,-0.5367,-0.819,1.967,5.83,3.176,4.025,4.565,2.987,1.606,1.636,-0.1575,-2.658,4.746,4.33,7.902,6.281,-3.898,3.523,2.954,4.286,3.765,3.67,6.033,-3.251,3.213,2.445,7.553,0.112,0.8927,-1.6,0.3672,-0.507,0.4633,-1.942,0.9071,2.456,1.028,2.346,-0.702,0.2808,2.021,1.9,-0.5665,0.3048,-2.259,1.409,-0.06144,1.164,-0.9295,0.5888,0.9875,1.277,0.4137,-0.8131,8.86,-0.6844,-2.045,-0.2192,0.437,1.593,1.536,-0.2752,1.315,-0.7373,-0.9617,-1.12,-0.9538,0.7419,0.8503,1.107,1.403,-0.998,0.8577,2.701,-0.6386,0.7888,-0.1911,-0.9787,-2.035,0.2939,-0.6364,1.061,-0.5012,-1.253,-0.147,-0.3426,-0.2672,-0.9808,-2.689,0.9254,1.503,-2.938,-0.2968,-2.253,0.01152,-2.62,-3.845,0.1118,-0.6432,5.351,3.826,-0.3104,-2.647,3.063,-0.3964,-1.119,-1.857,-0.4182,-1.127,1.224,2.048,-3.225,-2.249,1.362,-0.3688,0.8477,2.111,-1.441,0.2088,-1.17,-2.645,1.11,-0.2834,3.55,-0.8525,0.2848,1.338,-0.2638,0.6851,-0.06508,-1.652,-0.2011,3.254,0.3304,-0.1949,1.322,-1.336,0.5477,-0.9206,-3.035,-3.129,-3.299,-4.309,-2.485,-2.03,-2.204,4.731,3.127,-1.577,-1.393,0.06018,5.303,-0.3722,3.482,-0.6831,4.514,6.714,2.587,-3.389,-2.704,0.3053,-2.563,-1.171,-0.5196,2.048,1.194,0.8964,0.5233,0.3522,-1.437,0.06745,-3,2.097,0.1303,0.6269,-0.7334,-1.255,1.449,0.442,-1.36,0.1387,2.391,0.4276,-0.6007,-2.321,-0.8396,-2.008,0.07442,0.06585,-1.789,-0.6816,1.967,-2.07,0.8891,-0.8223,-0.1791,-1.356,-1.725,0.923,0.4199,-0.9103,1.695,5.005,6.834,-0.7309,-1.251,-3.18,5.094,7.337,3.469,3.941,1.819,-2.249,-0.3998,3.561,4.356,2.866,3.568,3.586,3.548,4.5,4.738,6.875,2.811,-0.9712,-0.5563,5.415,3.179,-0.3298,-2.585,-2.398,2.077,-0.9766,-1.549,0.9426,2.544,0.7905,1.97,1.456,0.3913,2.145,0.5533,8.079,1.081,0.4582,-0.06178,1.667,0.4131,0.7037,2.52,0.5187,-2.882,0.9219,0.2908,0.7317,-1.476,0.3403,-0.1719,-0.3292,0.2431,1.203,-3.993,-0.8815,-2.451,-1.042,-0.4987,-0.3228,-2.193,-0.2465,-0.3727,-0.9194,0.8151,-2.226,0.6063,-0.06279,1.371,1.243,-0.852,-3.49,-0.8247,1.577,1.468,1.376,4.313,-3.33,-1.74,-3.092,-1.803,1.526,1.352,-0.8624,-0.7408,-1.074,0.2482,-2.655,0.5423,-0.9163,-1.077,-0.2887,-2.434,-0.8691,2.165,0.8462,-0.1372,-1.4,2.473,2.447,-2.754,-1.955,-0.1239,0.4264,2.17,-0.6083,0.9114,-1.576,-1.407,1.927,-0.4714,-2.187,1.453,1.17,-1.896,0.5711,0.7132,1.275,0.5014,2.112,1.241,-0.05067,-1.727,-2.252,-1.368,-3.024,-1.596,0.7071,1.766,1.576,0.229,1.228,2.067,-1.825,1.266,-1.292,0.5078,0.9468,-0.2819,-1.341,-1.358,0.007229,-0.1479,-1.161,-0.9608,-2.641,-0.4076,-2.63,5.09,4.96,2.589,0.5221,4.578,-0.3807,-1.768,4.312,6.328,3.702,5.645,5.156,-0.6756,4.884,1.595,3.77,0.8194,-4.911,-0.7017,-0.5906,-0.8759,5.089,0.6584,1.199,-0.3674,1.134,0.4989,-0.838,-0.5333,-1.04,-1.394,-0.3622,0.662,0.08738,1.035,-3.077,-1.254,0.4095,-1.259,-1.316,-2.207,-1.369,0.7404,-1.709,0.3134,-0.9379,-0.1574,2.254,-1.744,-0.9453,1.455,-2.064,-0.7498,0.6546,-1.373,0.5212,-0.09371,0.3255,0.2648,0.9777,1.386,1.063,-0.1326,1.125,1.916,-2.097,-0.3393,0.5414,1.133,-0.5749,0.5741,-0.5343,-0.1773,-2.294,-0.5276,-2.662,-2.621,-0.3114,-0.8808,-0.07901,0.2823,-0.8598,2.559,0.7955,-1.467,-0.7929,0.1505,-0.5825,1.527,1.292,1.142,-0.4902,-0.9133,0.782,-0.9732,1.454,-1.402,0.8454,0.168,0.833,1.513,-1.496,1.678,-0.6076,-0.1366,-2.533,-1.457,1.049,-3.384,1.242,0.2136,-1.306,-0.1226,-1.401,-1.246,-0.0854,-1.567,-1.004,0.5381,-0.892,0.1843,0.4851,0.7022,-0.6518,-2.066,1.106,-1.448,-0.4443,-1.07,0.9547,-0.8384,1.426,0.0448,0.8683,0.2639,-0.9983,0.8144,0.8598,-2.302,1.02,0.5188,-0.6545,1.027,1.201,-0.6689,0.6404,-3.208,-0.2533,-2.503,-1.868,0.9049,3.468,1.557,-0.1533,-1.154,-1.746,0.7613,2.119,0.9416,0.4011,-0.9566,0.6024,0.4626,-0.2825,0.9202,1.235,1.904,0.9251,1.677,1.607,0.947,1.506,-0.8374,1.071,-0.136,-0.7102,0.2523,-0.1575,1.252,-0.1371,-1.753,-0.8437,1.527,0.6631,2.101,-0.1735,-3.106,-2.361,-1.14,-0.2254,1.136,1.666,-1.683,-1.883,-1.265,0.6375,-1.58,-2.849,-3.163,1.832,-1.599,-2.184,0.02162,7.51,-1.079,0.1097,-0.5512,1.635,2.357,0.7793,-1.872,-2.025,0.8305,0.3467,1.369,0.5112,2.063,1.01,2.144,-2.508,0.225,-0.7091,0.0901,-0.4245,2.01,-0.2408,-0.2171,-1.35,-3.195,-1.483,2.372,-3.116,-3.457,-2.229,1.383,-1.26,2.414,2.391,-2.963,-1.17,3.573,3.467,5.174,-1.745,3.097,-1.538,5.18,2.083,2.953,0.01411,6.489,0.5898,-1.761,3.671,0.6229,-0.5599,-0.2245,1.563,-0.154,1.337,-1.475,-1.726,-2.552,0.7534,-0.3376,0.04656,0.1188,0.09957,-2.824,-1.179,-0.7289,-3.334,-3.245,-0.5944,-1.051,0.7414,-2.183,1.095,-1.292,1.152,-1.408,-1.367,-0.07358,0.395,-0.462,-1.016,-1.234,-0.251,-1.675,-0.9523,1.068,-1.065,1.963,0.7068,1.454,-1.808,-1.706,-1.135,-0.5845,-0.6194,0.4883,-2.735,7.119,-0.7416,-0.6463,-0.6496,0.6696,-1.018,3.133,-1.113,-0.4749,0.7986,-1.09,-0.3738,0.1331,-1.361,-2.956,-3.493,-1.055,-0.1402,-1.646,-0.3815,-0.4728,0.2138,2.585,-2.494,-3.395,-0.5333,0.6558,-0.6386,2.368,0.5233,-1.16,-1.281,0.006723,1.464,0.2773,-1.898,-1.45,0.1019,-1.636,3.051,-0.5748,1.1,0.8136,1.152,-0.2825,2.205,1.18,0.09212,-2.172,-0.788,-0.3716,0.6805,-1.968,0.1992,1.563,0.1976,-0.7652,1.935,-2.163,0.896,0.5745,1.617,1.503,1.009,0.6719,-1.215,-0.7838,0.1546,-0.791,1.557,0.565,-0.1211,-0.6275,-0.3257,2.328,-0.4531,1.294,-0.3358,1.541,2.828,-0.6321,-0.9974,0.822,-2.185,1.549,2.448,-2.391,-4.043,-2.123,2.385,2.74,-2.817,0.2315,1.64,1.772,1.633,1.045,0.2798,-1.125,-0.3212,2.777,0.4324,-2.484,-1.245,-0.07945,3.869,-1.579,-2.428,1.357,1.433,0.1832,1.179,-1.058,2.867,1.586,0.5532,0.6117,0.9025,-3.202,-0.1404,-1.988,1.529,5.211,-0.4546,1.579,0.6032,-0.9052,-1.125,1.985,-0.09131,-2.842,-0.777,0.2718,0.2198,2.401,0.2345,-3.288,0.9008,1.031,1.77,0.4717,-0.5791,0.7126,-2.988,-0.2913,-1.435,-2.417,-1.719,-1.678,0.675,-1.678,1.679,1.592,0.9539,0.7992,-0.6656,-0.1861,-2.447,0.3242,-1.407,0.8002,-0.4195,1.299,1.167,-0.2206,-0.429,-1.395,-1.441,1.455,-0.6931,1.333,1.936,6.635,5.209,5.465,1.564,1.352,1.025,-4.028,2.251,1.474,4.193,1.352,4.232,3.855,4.281,4.79,0,-2.752,0,0,5.735,1.033,4.415,-6.319,3.467,0,2.584,6.106,0.514,4.308,0.7604,-2.167,-0.626,0.3172,0.3101,-1.472,-2.651,1,-1.01,-0.5008,0.8936,-1.014,-0.2927,-0.0006867,1.1,-0.3386,-0.02997,1.237,-0.4743,-1.14,0.291,-2.717,0.2603,-1.447,1.345,-1.349,2.188,-0.9217,-2.136,-0.6507,0.6318,-0.6249,-0.6476,0.1673,0.1124,-1.482,-0.01956,-2.582,0.05126,0.7132,-0.346,-0.8206,0.8797,0.6143,-1.58,0.8744,0.8152,-0.3598,-2.873,-0.2458,-0.3947,0.8287,1.122,1.738,0.63,-2.166,-3.484,-2.857,-0.6015,0.8557,-2.409,0.1806,1.124,1.123,-0.5758,0.7985,-1.294,-0.4144,-2.547,0.6204,-1.318,0.6196,-0.4935,-0.4503,0.8596,0.3602,-0.6251,-0.8999,-0.981,-1.775,-0.3056,-0.4582,-0.163,-1.11,1.774,1.784,3.246,-3.126,-0.3123,0.6374,-0.5612,1.859,0.8601,-0.2396,1.884,-1.323,-2.922,-1.865,-1.793,-0.7976,-1.424,-1.938,-2.106,-1.964,0.3664,0.5088,1.739,0.54,0.2986,-0.1047,1.438,0.726,-0.403,-0.311,-1.565,-0.7138,-1.121,-0.6579,-0.6355,0.9924,-1.572,-0.7631,-0.9158,-0.08422,1.231,1.163,-1.577,-0.5642,1.879,-0.3926,1.286,-1.053,0.1807,2.126,1.586,-2.666,-0.6573,0.3779,2.189,-1.237,-1.374,-1.481,0.3097,-0.7443,-0.175,0.5415,0.5879,-0.6584,2.265,-1.372,0.5453,-1.231,1.377,1.491,-2.377,-0.6327,-0.5813,0.7642,-1.284,0.04597,-0.5607,0.1582,2.142,-2.378,1.934,1.556,1.068,3.555,0.957,1.854,1.051,1.109,-0.2805,-0.3592,-0.3107,-1.488,-0.9664,0.005512,-0.9553,-1.417,0.4828,0.4177,-1.012,-1.486,1.758,-0.5909,1.207,0.09748,-2.374,-0.8327,-0.6592,-3.518,-1.622,-0.6613,0.1111,-1.313,-1.426,2.427,1.053,-2.511,-0.7588,3.76,0.09538,0.6732,-3.217,-3.616,-0.02741,-2.121,-0.8137,-1.614,0.1091,1.939,-1.178,0.1675,2.142,0.6107,1.947,1.037,-0.07261,1.886,-1.274,-0.6723,-0.5797,-0.2193,1.561,-1.921,-0.531,0.0698,0.6602,-1.332,-2.035,1.535,-1.059,-1.536,-2.352,-1.763,-1.492,1.954,0.6358,0.4766,0.6115,0.4884,1.274,0.7663,1.572,1.407,-0.2258,-1.015,-0.8073,0.3269,-1.766,-3.762,-1.739,2.506,-1.083,0.1984,-1.213,0.9304,-0.749,1.378,1.445,1.055,-2.072,1.208,-0.7653,-0.05192,-0.1697,1.022,7.397,6.427,5.919,1.579,3.361,6.478,3.887,1.282,4.151,0,0.04296,1.306,0.3247,-0.3349,-0.3183,-1.446,-1.674,0.2652,-2.453,1.504,0.9731,0.9736,-3.265,-3.55,-2.441,1.877,0.4108,0.2788,-1.626,0.2293,-0.1653,1.882,-0.2131,-1.369,-2.46,-0.4594,-0.9591,-2.39,-1.258,0.8116,1.747,-1.82,0.7636,0.8763,2.065,-1.46,0.4049,-0.9457,-1.218,-2.029,0.1442,2.544,-0.5851,-0.5854,-1.051,-1.379,-1.763,5.94,0.9078,-2.188,-1.017,-1.825,-0.1616,0.5197,0.5321,-1.378,0.6222,-1.773,-1.936,-1.054,-2.338,-0.5457,0.6058,-2.399,-2.131,1.858,-0.5893,-0.9272,-1.115,-0.745,-0.03451,-0.6904,0.05911,4.127,-1.345,-0.8515,1.11,1.043,-0.4606,-3.719,0.1929,1.362,-0.8488,-0.758,0.7725,0.9489,1.877,0.2567,-2.182,-3.231,-0.05675,1.151,-0.2805,1.116,0.666,0.09703,-0.08703,2.587,-0.9881,1.009,-1.637,-1.011,2.274,2.694,-1.027,0.461,-0.3463,1.837,-0.3563,2.247,-0.6504,1.882,1.909,-2.705,-0.3274,-2.823,1.396,-1.122,3.075,-2.174,0.5639,-3.382,4.782,-0.5584,0.004795,3.639,0.9402,1.528,0.7703,-0.712,0.8139,2.279,-1.619,1.648,-0.2896,-0.1446,-0.4295,1.045,0.02778,0.413,-0.1185,1.633,-0.08509,1.027,-0.9485,1.301,-1.454,-1.733,1.084,0.8763,-0.4384,1.418,-0.2117,1.524,2.555,0.8992,1.605,-2.565,6.902,2.611,1.858,1.141,-1.621,-1.439,1.589,1.253,0.3775,1.3,-3.162,-0.03523,-1.727,0.6293,1.608,3.508,1.887,2.13,1.448,3.645,1.275,-0.9451,-0.04289,0.3758,-3.044,-1.682,-1.628,0.1989,-0.2227,-1.406,-1.638,-1.436,5.267,2.651,4.416,-0.6397,-0.9972,5.462,1.192,4.447,0.1368,1.379,3.287,4.696,0,3.774,4.233,5.185,6.123,-0.01668,2.411,4.336,1.421,-0.05762,-1.774,1.474,1.675,-0.1154,-1.97,0.2454,-2.616,-1.438,-0.9532,0.1393,-1.429,-0.9726,-1.266,0.0243,0.5538,-1.623,-1.788,0.1703,4.156,0.5425,-0.4105,-2.408,0.1394,0.2791,-1.94,-1.811,1.196,-0.6195,-1.621,0.6764,-2.61,-0.2852,1.065,1.276,0.5404,1.062,-1.138,-2.043,0.1603,-0.3572,-0.5402,-0.511,-2.491,-2.526,-1.697,1.296,0.2365,1.057,2.315,-0.134,0.4477,-0.5972,1.057,-0.9218,0.3911,-0.872,3.249,-0.6136,-0.9601,-0.8223,1.441,1.4,1.608,-2.886,0.228,1.87,-0.7394,1.33,-1.553,-0.915,0.8927,0.9014,1.372,-1.728,1.025,0.1365,1.673,-2.021,-2.748,-1.84,0.4002,-2.766,-2.052,-0.1346,-1.197,-2.62,0.3498,1.583,-1.559,0.5257,1.232,-0.9737,2.23,-1.159,-0.06694,-0.1378,-0.09917,2.429,-0.9442,1.635,1.4,-0.4766,0.9901,1.051,-1.511,0.1481,0.5259,1.401,-2.879,-3.511,-2.977,-0.2739,0.3379,-2.376,0.004149,-0.3065,-1.222,-2.312,-2.948,-1.899,-0.2257,-2.152,-2.112,0.6495,-1.457,-3.505,-0.6276,2.407,-0.7954,-2.433,-1.429,0.08794,0.5208,0.8628,-0.8657,2.109,1.449,1.104,-0.158,1.824,-2.428,0.6064,-1.779,-1.907,-1.506,0.1981,-2.094,-1.726,2.187,1.426,-0.5718,-2.698,0.2766,0.01698,-0.4491,1.18,0.1759,-0.3916,1.201,-1.041,0.9761,1.071,-1.626,0.4369,0.7415,-0.06755,1.477,-1.414,-1.237,-0.2613,-0.3722,1.097,0.4864,0.3655,0.711,-0.3975,-2.262,-1.608,-1.239,-1.958,-0.5446,0.3005,0.9898,-0.004475,-1.147,-0.1936,2.12,0.4968,1.79,-1.631,-1.165,-0.06687,0.8822,1.104,0.1571,0.3444,1.873,1.438,-0.286,2.213,-0.6988,-2.865,0.3177,2.533,2.894,0.7989,1.2,-1.448,-1.595,1.633,-3.654,1.475,-1.54,0.9811,-0.4699,-1.569,-0.5614,-0.2406,-1.208,-0.7086,-0.5933,1.464,2.893,-0.03393,-1.853,-0.7023,-1.421,-2.29,0.1163,-0.629,1.493,3.495,1.228,-1.151,0.5402,-1.515,-2.876,0.4413,-4.094,1.212,0.5043,-1.193,1.435,0.4682,0.7938,2.059,0.6569,-1.098,1.2,-1.41,-0.8731,0.1056,1.884,0.3431,1.074,1.629,-0.8054,-0.4439,0.08352,-1.325,0.01084,0.8573,1.946,-2.154,-0.9126,0.1963,0.9424,-1.724,1.06,0.7396,-1.801,0.6852,-1.925,-1.548,-0.004745,0.2085,2.077,0.1378,0.9107,6.444,5.34,-0.2079,0.868,2.066,-1.451,2.09,4.048,5.703,4.974,3.781,3.679,4.917,5.217,0.668,-4.748,3.494,0.971,2.704,7.881,6.331,8.281,2.417,1.354,-0.299,0.4172,0.6582,1.173,1.768,4.525,-1.919,-0.4305,-0.4768,-0.9938,-3.652,1.56,0.4514,-1.998,1.624,-1.739,-1.392,-0.9423,-2.688,-1.016,0.7981,-2.316,-1.363,2.989,1.53,0.1002,0.1594,0.9969,-0.5666,-0.553,-0.751,0.5845,-1.027,-1.029,0.1193,0.4634,-1.995,-0.5557,0.8533,-2.228,0.4888,1.834,0.9096,-0.3729,0.3283,0.5043,-1.484,-0.5321,-1.68,1.384,-0.8424,1.049,0.0811,-0.6999,0.2907,0.01107,1.038,-2.138,-1.264,-3.464,-0.1937,-0.1984,2.932,-3.165,0.7471,-1.161,-0.7842,-0.2982,1.235,0.8373,-1.146,0.8048,-2.127,-1.845,-0.1254,1.017,-3.142,0.3466,1.733,-0.1616,-0.5149,0.2928,-1.953,-1.247,-2.317,-1.291,1.787,0.07611,-0.6332,-1.611,0.3278,1.495,0.5506,-1.38,1.288,-0.3728,0.3349,-0.4377,-0.438,-2.144,1.162,3.285,0.8097,-2.725,5.943,0.9879,2.465,0.8461,0.2877,-2.217,0.08298,0.1433,0.1268,0.9577,0.8328,0.882,-0.9312,-1.854,-0.9428,-0.1199,-2.199,1.613,0.6635,-0.03474,0.2734,0.9037,1.772,0.1381,0.3087,0.8961,2.24,3.94,-0.5152,-0.615,2.227,1.099,0.3996,1.033,-0.004817,0.4824,-0.7741,-2.656,0.6157,0.06707,0.3056,3.814,1.233,0.1538,1.942,-2.179,0.9392,0.3602,0.1195,-0.7031,-1.431,-1.57,0.02053,0.9895,-0.6378,2.514,-1.59,-0.5767,0.133,-0.8423,-0.3575,-1.339,1.429,-1.237,-0.7971,-2.142,-0.4078,0.5982,-0.08986,0.865,1.872,-1.298,-3.602,-1.242,1.706,1.312,-0.3155,-1.66,-2.097,1.46,-2.611,-0.4204,-1.237,-3.143,-2.747,1.297,-3.013,1.821,1.749,-3.031,-0.8472,0.8468,-0.4735,-2.474,-1.359,-2.355,-0.4666,-0.5246,1.372,2.208,1.759,2.435,-0.3874,-1.576,0.07285,-0.8605,1.719,1.074,0.9026,-0.5629,3.318,-3.589,-1.217,-1.531,-1.519,-0.7964,-2.093,3.169,-1.472,-1.014,1.127,-0.3632,-1.587,-0.9975,1.442,1.101,-2.094,-2.02,-0.8118,-0.9555,-0.6173,-0.9105,-1.151,-0.9913,-0.991,-0.6957,-0.5431,2.496,0.08152,0.9758,0.9492,1.878,-1.212,-1.395,-0.2781,0.239,-2.245,0.4186,-2.343,-0.9253,-2.788,-0.7506,2.745,0.4475,0.7695,-1.84,-1.215,-0.5446,0.08141,-2.118,-1.414,1.954,0.6822,3.732,0.7781,0.1949,-1.439,1.768,0.7326,-0.2945,0.6938,0.3394,-0.4415,0.5982,-0.2104,0.1587,1.155,-0.2568,-3.985,0.5573,-1.062,0.1769,-0.5303,-1.059,0.2348,-1.634,-3.584,0.7153,1.074,-2.249,-1.156,-1.626,-0.7578,-0.2833,1.246,-2.514,-3.11,1.243,0.7899,-0.3012,-0.861,-1.188,0.458,-0.07107,1.327,-0.4777,-2.636,-0.9696,-1.491,0.1711,-1.167,1.372,-1.013,-1.433,-1.714,-1.744,1.253,-0.07107,-0.9045,0.6386,-1.677,-1.321,-0.5914,0.05622,-2.526,-2.404,-0.9839,-1.101,2.034,0.9401,1.059,2.083,-0.5061,-1.267,1.741,-1.722,-3.606,-1.339,0.3187,-0.7608,0.6252,1.335,-0.2341,-1.475,-3.106,-0.6468,0.8488,-0.5438,0.06585,-0.09427,-0.8062,-0.1022,-0.6629,0.07331,-1.232,-1.042,-0.7672,1.186,-1.737,0.7034,1.176,-2.732,1.674,-1.611,-3.411,-0.8946,-1.502,-1.457,1.564,-1.147,-1.34,-1.202,-0.1828,-1.992,0.1673,-1.526,-2.203,1.599,1.19,-2.623,-1.133,0.2134,1.717,1.154,-2.704,2.053,1.565,1.218,3.623,1.269,6.526,5.901,3.628,2.866,1.022,-1.643,3.72,3.523,3.628,6.433,2.225,-0.2136,-0.7701,-0.6228,-0.9251,-1.227,-1.387,0.312,-0.4028,-1.845,-0.4259,0.2612,1.941,0.898,-0.4818,-3.172,0.9639,-0.06223,4.84,1.096,-1.501,-4.327,-3.587,-1.276,0.3011,0.6395,-0.3574,1.771,0.5379,-0.7114,1.179,0.5574,-1.373,-0.0787,1.33,-1.816,7.159,-0.03883,-1.764,-1.779,1.062,-0.9118,-1.622,0.6714,-0.9153,-1.759,-2.664,-0.467,-0.4677,-0.8268,-1.71,-1.132,-2.337,-1.466,-0.955,-0.849,1.486,-0.7081,-0.2275,-1.601,-1.633,0.3522,-1.326,-1.069,-1.537,-1.323,0.5623,1.373,-0.6491,-0.7688,-1.642,-0.247,0.9657,0.01818,1.71,-2.496,-1.057,-1.14,0.521,-0.5435,-1.728,-2.082,-0.6385,0.7089,-0.8727,-1.438,0.395,-2.252,-3.435,0.4243,-1.161,-0.4984,-0.009683,0.4911,-1.588,1.288,-4.055,2.437,-0.1601,0.8536,2.057,1.205,0.8825,-0.9158,-0.02127,1.949,1.822,1.469,-0.03342,-1.649,-2.85,2.101,-1.657,-0.9575,1.906,1.875,-0.4105,0.2065,-1.937,0.00185,0.002726,-2.878,-2.049,-1.003,0.8706,-1.09,-0.2032,-2.492,-2.825,1.588,0.5955,1.017,0.1629,-1.279,-2.196,1.262,4.657,-0.5427,-1.704,-1.72,-2.518,0.2507,-0.1423,0.05924,-1.058,-0.8353,0.8363,1.845,1.476,-0.7355,-0.6886,0.8553,1.148,-0.8965,0.882,-1.13,-1.254,-2.318,0.2207,-1.876,-0.5144,1.928,0.3255,0.405,0.7527,2.024,0.5936,1.181,-0.4427,0.1324,0.9952,1.25,0.971,1.629,-1.76,0.9893,-1.626,-1.299,1.216,-1.664,2.34,-1.547,-0.4237,2.506,-1.151,-0.25,-1.387,0.8099,-0.8036,-1.541,-0.1614,1.54,-1.311,-1.302,0.1988,-0.02559,-1.323,0.9697,0.6849,-1.013,-1.34,-0.5623,0.2972,1.252,-1.604,-3.182,-2.118,-0.09916,0.01499,-0.1517,-0.4607,0.3003,-1.181,4.683,-1.783,-2.775,-1.394,1.077,-1.316,-0.678,-0.9128,-1.021,-1.448,-2.112,-2.682,7.148,-1.079,-0.9337,-1.022,-0.0296,-1.435,-0.2311,0.8512,0.7033,-2.288,-0.3155,-0.5391,-0.9402,-0.9079,0.2703,-0.5798,0.4801,0.977,0.4401,-0.6279,-2.435,1.121,-0.6623,1.003,-1.649,-0.1065,-0.492,-1.223,1.008,-3.134,-2.038,-1.674,0.6293,-1.634,0.3424,0.6635,-0.7263,-0.6512,0.228,-0.6824,1.662,8.494,5.211,3.601,3.541,5.903,0,7.087,4.203,3.624,4.216,3.639,-0.7501,2.487,6.404,6.033,8.192,0.5127,5.708,1.668,3.887,0.9662,1.832,5.147,1.831,7.772,7.513,7.848,0.4827,-2.932,-0.289,1.557,1.443,-0.1393,-2.419,0.1037,-2.615,-1.462,0.5897,-0.5446,-0.4951,-1.434,-0.9452,-1.144,-0.5442,1.537,0.4925,-1.877,0.3626,-1.097,-1.835,-2.955,-4.265,-2.256,-1.842,-1.037,-0.7216,-1.474,-0.4893,-0.36,0.778,-0.3342,-0.3011,0.735,-1.153,-0.01482,-0.7424,-1.356,1.059,1.287,-0.8455,-0.9317,0.2508,1.022,1.048,-1.519,1.317,0.6901,1.439,-2.181,0.8882,-0.2259,-0.03204,-0.3668,0.3787,-1.453,-1.673,3.888,1.411,0.003426,-1.701,-1.252,0.4363,-1.153,-2.921,8.296,1.032,-1.609,0.5484,-0.8612,0.7789,0.3915,-2.697,0.335,1.14,-0.5578,0.5666,-1.34,0.337,-0.09372,0.09913,0.7464,2.039,0.4182,-0.8254,-1.198,-1.289,1.955,1.688,3.75,0.1754,-0.06814,-0.2604,0.9864,-0.0514,-0.9282,-2.108,-0.594,0.07341,-1.257,-0.412,-0.451,1.185,-0.2375,-0.4632,0.9296,0.8193,0.8488,-1.417,1.576,-0.3488,2.152,-0.1618,0.5867,0.9568,0.1421,1.596,0.7043,0.4284,0.5301,2.071,0.8281,-0.7206,-0.2047,-0.08415,2.236,1.263,-0.6,4.973,1.621,-0.04598,-0.0743,-0.7071,-1.147,0.6206,4.471,0.9038,1.141,0.1222,-2.382,-1.031,0.7357,0.4605,0.7736,-1.699,-1.688,1.51,-1.622,-1.571,-1.678,0.4701,1.124,-0.9988,1.74,0.3934,-0.773,-0.3406,1.486,1.188,-0.4249,-0.9278,-1.318,1.512,1.396,2.736,1.043,-0.1616,-1.739,-0.06865,-0.2271,2.163,0.8799,0.9863,0.3517,-2.128,0.8191,-0.5076,-1.697,-1.131,-2.739,-1.132,0.5404,-1.234,-2.428,-3.073,-0.3086,1.52,0.7562,0.001047,-1.461,0.5751,1.056,-0.3629,1.138,0.501,1.677,-0.1345,0.7818,-0.2368,0.226,0.2529,1.577,0.8181,1.367,-1.304,0.5072,0.5422,2.809,5.367,-2.455,-2.154,-1.432,-2.063,1.829,0.502,-0.2143,5.98,1.159,-1.241,-0.7457,0.2301,-2.296,-0.8456,-1.307,3.061,2.238,1.512,0.09838,-0.2934,-0.6584,0.2653,-1.435,-1.95,-0.3728,-0.8952,1.164,1.365,-2.798,1.545,-0.003023,-0.02043,1.846,1.12,1.542,-0.6698,-0.241,0.7492,-1.572,0.4753,1.758,1.322,0.3413,0.5114,0.7422,-2.274,2.486,0.8257,-1.62,-0.7937,-1.29,1.57,-1.944,1.594,0.5649,0.5768,-0.3525,-1.032,-0.7619,3.271,-1.902,1.944,0.4773,-1.691,-2.738,-0.7925,-0.4352,0.5482,1.684,-1.013,-2.421,0.1176,1.43,0.7594,0.1288,-1.778,2.714,1.444,-1.63,1.374,0.9287,1.83,1.491,0.3183,1.959,0.3389,-3.233,2.871,2.226,-0.8468,2.443,1.462,-1.203,-0.4056,-1.279,0.5707,-0.399,0.6837,-0.1447,0.6379,-1.44,0.9671,0.8896,1.005,0.9553,-0.09157,-1.709,0.3807,1.03,-0.5697,0.09869,1.348,1.109,-0.6337,7.991,0.4426,1.426,-0.6968,1.198,-1.275,2.603,-0.6432,-3.775,-0.5748,-0.7199,0.7337,-3.41,-0.6965,1.8,-0.1716,-0.69,1.296,0.4458,0.3425,-0.6906,-1.28,1.389,1.43,1.436,1.253,0.2469,0.1179,1.84,-0.5306,0.6833,-2.955,2.134,1.241,1.673,-1.082,2.239,-0.7574,0.2955,-1.287,2.058,-1.705,-1.312,0.8519,-1.214,-0.1645,0.5476,3.445,1.22,1.711,-0.2219,1.342,1.157,0.7946,1.6,-1.526,0.7671,0.3768,1.59,0.4573,-1.34,-1.345,-1.379,4.182,2.403,0.06666,4.364,2.687,-2.237,3.954,6.835,-1.27,6.619,5.383,2.576,5.384,4.739,3.625,-2.425,9.556,3.981,3.745,-0.8521,1.343,1.779,5.317,2.366,9.788,0,4.335,3.284,-5.623,1.359,-7.087,-6.37,0,6.38,-5.867,-4.571,-5.375,-6.759,-5.353,-5.796,5.263,-4.255,-2.948,-5.148,-3.916,2.574,2.159,-1.874,-5.998,-4.037,-1.842,-2.51,2.558,-2.068,2.724,2.17,3.689,3.894,2.904,3.582,3.432,5.763,-2.055,2.842,2.086,2.072,6.244,1.604,-1.714,2.368,-2.292,5.445,2.198,2.902,1.7,5.587,4.78,2.692,4.049,2.848,3.507,-2.236,2.084,4.103,3.942,4.742,-4.462,-2.867,4.986,2.822,2.982,5.726,4.382,2.392,3.806,1.867,4.366,3.584,-3.408,4.364,-4.679,-2.817,-8.154,-2.797,-2.615,-5.363,5.409,3.552,2.611,3.973,2.555,-4.057,3.629,4.306,-4.756,4.308,2.988,1.965,3.155,-3.602,1.789,-2.927,5.983,3.23,8.719,4.883,-3.621,3.187,3.981,2.407,4.221,2.369,2.381,3.378,-3.937,-3.387,-3.874,2.752,6.22,-2.654,-3.938,-7.911,3.529,1.97,-1.386,4.946,-3.419,2.44,2.873,3.384,4.938,7.971,4.56,-1.588,5.14,2.321,-5.753,3.993,6.214,4.523,2.029,4.248,10.47,9.151,5.947,3.064,3.645,3.831,1.757,3.584,2.953,2.75,3.8,3.546,3.314,2.155,4.321,2.203,-1.784,3.052,2.626,2.976,1.891,6.115,2.04,4.511,2.234,-3.521,3.813,2.105,2.939,2.825,-2.886,-2.829,-1.888,2.117,2.615,3.469,2.966,3.369,2.997,3.64,-3.436,-3.856,4.888,2.53,4.212,4.969,1.872,3.131,5.039,7.304,6.268,7.4,4.495,3.547,2.226,2.305,2.478,5.237,5.126,4.095,-5.768,3.074,1.798,5.137,-1.866,-4.081,3.923,4.929,1.898,2.652,3.768,2.582,3.326,1.918,2.865,2.244,3.917,2.299,5.129,-2.209,3.871,-5.29,1.591,4.133,2.763,3.335,-2.241,-3.804,-3.179,5.812,2.58,3.595,2.288,-4.04,-3.466,-3.014,-4.356,2.321,-5.829,3.257,-4.083,2.348,4.688,1.879,4.933,4.831,4.49,-1.477,-1.996,2.153,4.764,3.056,5.16,3.082,-2.746,4.735,10.99,3.885,-4.103,-4.935,5.208,1.842,5.027,2.745,6.347,9.947,-3.028,-2.947,1.835,3.445,5.931,4.804,2.316,3.218,4.418,7.026,-7.293,3.174,2.424,4.379,-1.869,2.413,6.57,1.367,5.594,2.532,-1.535,2.725,2.84,4.874,-3.112,4.134,-3.256,-2.861,2.601,-2.531,-1.854,3.925,4.495,1.711,-4.05,-2.011,-3.578,-3.24,3.189,3.153,7.167,3.613,-3.359,3.04,5.524,2.023,3.06,-1.86,2.627,2.75,-3.053,3.233,4.983,5.628,-5.713,4.037,2.606,2.624,4.275,2.54,1.322,4.751,-4.429,-4.359,4.477,6.494,5.554,-4.082,3.662,2.067,-3.098,-3.574,-4.974,-3.497,-2.758,-5.298,-2.919,-3.128,2.99,-3.998,2.57,3.835,4.6,3.006,2.364,-2.865,-3.046,4.628,8.403,3.727,-3.424,1.674,-2.732,-2.33,2.285,6.84,2.825,3.646,4.819,2.383,-4.022,-2.798,4.204,2.253,3.383,5.051,3.114,3.584,3.803,7.237,-2.327,3.428,2.36,3.864,-3.414,3.212,2.299,5.071,-4.565,-6.197,8.233,3.738,2.412,5.387,4.772,3.7,2.473,5.427,5.925,4.148,4.725,2.394,4.49,2.562,2.377,3.881,2.089,5.441,4.72,-4.613,3.706,2.43,2.726,1.978,2.932,3.055,3.198,1.968,4.063,3.522,2.789,4.225,-5.465,-3.792,3.649,2.382,2.046,4.215,-2.412,-4.266,10.82,4.398,6.145,-2.876,-2.391,2.821,-6.885,2.087,2.6,2.353,4.852,-1.732,4.234,2.555,1.639,-4.693,-3.358,-2.663,-4.15,4.136,5.828,3.419,3.331,-1.579,4.428,2.962,7.414,4.148,-2.98,-3.768,3.547,3.295,3.047,2.435,2.062,2.902,2.175,1.684,5.136,2.742,-3.496,-3.362,1.942,-5.299,-3.822,-6.576,3.987,-4.33,-5.123,-3.718,-10.45,-4.921,-4.03,-2.355,-3.033,3.902,-2.882,6.826,-3.252,-5.407,-5.826,-5.6,-2.288,-2.943,-4.393,2.037,-3.297,3.411,-5.056,-3.036,-1.241,-3.697,-7.405,-3.911,1.724,2.474,2.516,-2.533,2.598,5.007,2.561,-4.003,3.613,1.879,2.424,2.025,3.533,4.286,1.86,-2.359,4.309,-3.252,-3.972,1.698,-3.184,4.921,-2.697,3.77,5.193,3.388,1.45,-2.457,-2.653,3.225,3.523,4.231,2.05,4.228,-5.824,-3.928,4.085,-2.352,5.827,3.942,6.403,3.791,1.557,-1.965,2.251,-4.765,-4.196,-3.561,-4.989,-2.366,3.674,3.226,2.339,-4.287,3.737,1.746,3.438,3.322,5.522,4.769,-3.003,-2.356,-1.745,-3.272,3.671,7.928,5.349,-4.014,-2.99,-5.867,3.707,4.566,3.133,-2.417,2.082,3.689,5.455,7.86,-2.18,2.829,3.437,5.18,-3.055,-4.025,2.569,3.597,3.147,2.566,2.819,2.289,2.156,5.086,3.783,5.327,-6.84,3.41,-3.841,-2.759,4.487,9.605,6.196,-4.116,-6.115,1.838,2.61,-3.42,-2.115,6.013,-1.97,-3.503,2.867,1.784,-2.767,3.027,-5.385,3.129,-4.342,4.572,2.242,2.993,-2.371,6.018,2.203,3.36,3.582,-4.136,-5.165,1.863,-2.77,2.879,2.285,-4.528,-3.356,-5.035,-3.584,4.772,1.405,2.344,-2.6,-2.84,-3.966,2.227,5.055,-3.122,5.685,6.073,2.835,-3.995,-4.163,6.012,2.009,2.838,4.05,2.206,-4.061,-4.384,-3.178,-4.721,-2.312,-4.328,3.636,-3.937,5.376,-2.89,2.031,-2.493,-4.648,-2.527,4.392,3.52,2.733,3.334,-2.056,4.304,-3.52,4.16,1.977,-4.275,-1.947,3.776,-2.335,-3.285,-4.259,-3.585,3.102,3.024,-4.702,-3.679,2.397,-3.093,4.828,5.95,-1.927,2.723,-1.607,1.894,2.352,-2.583,4.588,2.134,2.867,2.319,3.182,-3.152,3.201,-2.603,-4.62,-1.746,4.294,-2.376,2.315,3.689,-4.197,-2.444,4.106,-3.699,-3.589,2.223,2.185,3.064,-2.321,2.201,2.786,-3.208,-1.791,-3.168,-2.373,2.942,3.038,-3.686,4.877,2.875,2.264,6.174,-1.768,-1.755,2.697,3.136,1.756,-1.602,-2.9,1.586,-5.163,-3.88,-6.653,-2.099,1.966,-4.76,12.22,4.879,-3.836,-3.787,5.501,-5.577,-5.592,-2.932,-6.661,2.666,4.07,5.42,-3.169,2.184,3.188,-5.132,-5.355,-5.108,-1.81,2.418,-4.015,-6.902,2.374,4.32,-6.918,-3.315,1.67,4.834,-3.264,2.614,-6.32,-3.032,2.618,-1.916,-3.067,-2.906,2.96,2.422,-4.48,-3.693,-4.887,3.465,-3.318,-4.493,-5.893,4.471,2.573,1.825,4.201,4.636,2.131,-2.911,3.927,2.394,2.941,3.467,2.745,2.546,3.664,3.25,4.244,2.638,3.501,2.135,2.819,3.279,5.626,2.306,-1.771,-4.663,-4.613,-4.787,4.482,3.261,5.889,3.641,1.671,1.434,-2.318,3.77,-3.079,6.415,3.928,-3.233,3.646,3.031,2.72,2.478,9.184,4.291,4.011,5.587,-4.638,-2.24,-2.349,-2.938,-5.944,-2.705,-3.182,3.291,2.567,2.571,2.19,4.154,-4.057,1.95,-1.95,-8.828,3.291,-3.947,6.838,-2.051,1.963,-2.559,2.073,1.927,-2.804,-3.791,1.693,-4.788,3.587,-2.519,-4.357,1.84,-3.976,-1.524,-2.65,-2.019,-5.857,7.208,-1.666,1.223,2.962,-4.408,-5.343,-2.259,5.267,1.711,-3.085,-4.786,3.528,2.215,-2.949,-2.771,4.051,5.105,-4.773,-4.716,4.129,-2.179,2.434,-2.102,2.569,-4.006,4.816,-5.707,-2.121,1.394,1.95,-3.962,-6.349,-4.718,6.29,-5.062,1.674,2.944,3.982,-5.729,2.271,-2.706,-3.531,-3.681,2.042,-2.467,-3.267,-5.861,-4.134,-3.542,3.579,1.529,-4.26,-1.924,-2.452,-1.489,1.443,-2.637,-1.884,-3.199,-2.135,-6.23,-3.462,-2.346,-3.085,-5.394,1.977,-3.399,-3.793,-3.444,-4.042,-4.129,2.986,5.009,1.812,-2.528,1.558,-2.07,2.456,-3.938,-4.618,-5.339,2.397,-2.902,-3.919,2.785,2.922,-4.508,1.632,2.455,3.458,1.778,1.914,-1.926,-2.746,4.144,-2.724,5.443,3.611,3.028,-3.329,3.508,4.007,10.78,2.176,2.762,2.165,2.776,2.651,2.361,3.371,-3.579,4.084,4.215,3.614,7.145,2.385,-2.108,3.553,-1.98,2.93,4.773,-2.197,2.879,3.466,3.686,1.775,1.994,4.284,5.682,5.528,5.694,3.155,4.069,2.352,-5.578,4.321,3.131,2.404,2.966,3.704,3.873,1.402,-7.346,-4.885,-3.199,-3.834,-2.001,6.462,-3.529,-2.517,-4.863,-3.018,1.438,-3.043,2.668,3.371,3.631,-2.126,1.531,-4.133,-5.51,1.678,-3.169,2.531,2.666,-3.434,-4.021,-5.681,-3.332,-3.35,4.236,2.399,-2.486,-4.373,-2.572,-3.417,-3.677,-5.644,-1.368,6.052,-3.986,-3.798,4.137,-3.638,-6.222,3.435,-3.852,2.023,-3.411,-6.238,-2.246,-2.443,2.771,-5.449,1.34,-2.328,6.853,3.007,-3.378,-3.406,2.882,-3.057,-3.5,3.199,-3.245,1.728,5.469,3.124,-2.157,-4.631,-5.924,-3.088,-3.608,1.514,2.762,-2.763,1.639,4.751,-4.227,-2.642,1.642,7.174,-4.209,-5.638,-5.215,-5.296,3.208,-5.565,3.6,-3.713,-6.108,-2.329,3.95,2.298,1.944,4.078,2.052,-2.373,-2.039,-4.288,-4.576,-4.444,3.925,3.327,7.605,-2.227,2.47,2.45,2.018,3.492,2.761,4.299,-2.286,-1.992,-2.333,-1.972,-3.044,-1.739,-2.154,-5.931,6.651,3.702,1.751,5.9,1.694,3.842,-3.612,-1.752,3.205,4.654,3.666,-1.67,-4.372,2.751,-2.085,4.593,-8.02,-6.442,-5.464,-7.445,3.879,-5.152,-8.449,2.531,2.303,4.34,2.245,3.274,4.087,-4.4,-4.177,-4.261,-4.148,4.277,3.639,2.025,-2.524,3.296,-2.117,1.644,3.322,-2.633,3.624,3.549,2.707,-4.232,-5.892,5.491,2.032,4.088,4.802,-1.981,2.3,1.919,2.023,3.025,2.616,3.302,1.777,3.132,4.108,3.827,4.6,-2.186,4.683,-3.956,2.345,3.192,2.921,-2.877,2.479,3.781,3.939,3.081,2.499,2.257,-4.087,-6.883,-3.26,-4.446,4.03,2.761,-2.888,-5.525,-3.384,-4.206,-3.772,-4.456,3.565,-3.15,-2.699,-2.13,-2.724,2.352,-1.843,5.672,-2.908,2.18,6.614,-3.787,-1.694,-4.262,-3.392,-1.489,-2.725,1.996,-5.386,-4.813,-1.974,4.675,2.136,-3.296,-2.588,-2.809,-3.559,-3.56,1.758,2.101,-7.794,-5.245,3.769,2.241,2.96,9.447,-4.171,-2.551,2.69,6.399,-4.023,-6.933,2.071,3.087,5.489,3.394,6.011,-2.923,-1.843,-2.452,-3.817,2.224,2.388,-1.97,4.299,3.334,-3.143,2.926,3.408,3.746,3.264,1.768,2.057,2.841,3.614,-3.019,1.702,-2.439,2.467,2.507,4.329,-3.218,3.335,4.233,3.867,6.68,4.499,-3.643,-4.679,3.113,-5.327,-3.435,4.454,5.685,4.225,-2.736,-6.667,-4.054,-3.31,2.154,3.389,5.523,4.006,2.402,4.02,2.427,4.242,-3.925,2.644,4.538,3.984,3.126,1.827,-3.138,-4.077,4.057,2.199,2.613,1.619,2.507,2.7,-2.131,4.907,-2.48,-4.238,2.201,2.001,4.667,-4.575,-2.002,2.258,-3.508,4.145,4.685,4.583,-2.164,1.664,2.974,2.362,2.607,-3.251,2.11,-3.175,-4.062,4.922,4.261,1.886,-3.609,-3.06,-2.85,-2.478,-6.038,-4.611,-4.58,-4.232,3.728,-4.765,3.214,1.956,2.421,-1.743,-2.898,2.43,1.491,-5.848,2.839,1.817,-5.131,-3.314,2.087,2.914,-2.877,-4.427,-4.3,2.158,2.017,-3.165,-4.249,-4.699,-1.982,-2.006,-3.532,-1.808,-3.989,-4.565,3.721,2.231,-3.177,-4.981,-3.63,4.097,-1.274,-5.176,3.241,1.734,-2.15,3.388,-4.563,-3.65,-2.909,-5.147,-4.139,-3.749,-3.31,-3.289,-3.03,-2.044,2.883,-2.407,3.923,1.572,-3.918,3.325,-3.988,11.39,-6.11,4.854,-4.385,2.255,3.18,-4.527,4.54,2.879,2.775,2.01,-4.292,2.168,-1.85,-3.186,2.214,-2.059,-3.623,-4.14,2.459,2.417,-1.633,-4.167,-4.827,-3.568,-3.172,-2.902,2.678,3.154,-2.029,-3.848,-4.597,-4.297,-2.803,-2.458,4.836,-2.19,-1.479,2.221,-2.506,-3.108,-7.352,-4.338,-2.172,3.382,2.237,-4.021,-4.117,-4.54,-2.131,-3.952,2.382,3.255,1.954,1.664,-2.439,-2.259,4.047,2.472,-2.725,-4.658,2.489,-2.355,-4.994,-4.124,5.191,-3.668,2.167,-3.989,2.297,-2.853,4.926,4.831,2.191,2.343,5.824,-4.983,-2.096,3.383,-1.203,3.158,4.966,-2.235,1.824,3.206,2.905,-3.431,4.297,10.36,1.992,-1.497,-3.846,-3.059,2.491,1.593,-4.472,-2.008,-2.252,-4.298,-3.332,-2.875,2.455,-3.057,3.991,-3.11,-4.014,-5.126,-1.562,2.476,6.781,2.535,3.004,1.916,1.547,-5.105,2.12,5.256,2.489,2.228,3.9,-1.866,4.097,-3.539,-2.801,1.849,2.687,5.288,-4.296,2.469,3.612,-2.816,2.849,2.726,-1.675,-6.346,-1.397,-5.846,2.375,1.735,-3.23,-4.304,-2.149,-1.577,2.316,3.44,6.352,3.922,2.77,4.694,4.628,4.265,2.711,-3.338,3.236,-4.481,-3.706,-4.284,-2.514,5.032,-2.337,2.068,3.031,1.663,-1.708,3.591,-3.351,-3.28,1.902,2.962,-1.595,3.854,-1.533,2.749,-4.496,-4.823,-3.764,2.3,1.596,2.431,-1.919,2.692,-1.988,3.804,-3.707,2.376,2.204,-3.318,-1.91,4.268,-6.91,-5.208,4.054,-1.385,-3.268,-3.118,-3.815,-4.348,-3.986,-6.055,-2.208,-4.253,2.795,-2.897,-4.746,-3.434,-1.209,-5.868,-1.815,2.473,-2.45,-3.947,4.377,-3.995,4.524,1.818,2.853,3.122,3.734,-3.851,-3.969,3.754,1.861,2.568,-3.045,2.096,2.662,1.512,2.23,5.497,-3.535,-2.911,-2.57,3.108,-4.01,-4.178,2.53,-1.571,-2.663,4.175,5.737,-3.321,6.855,-3.734,-3.191,-2.871,3.267,2.997,1.925,4.457,2.007,-2.229,-3.684,-5.235,-1.951,-2.251,-4.074,-5.81,2.838,-3.171,2.426,-3.436,-3.727,-3.219,-3.558,-4.784,-3.688,-3.263,-3.17,-3.268,3.087,-4.082,-7.035,-6.613,-2.861,-4.871,3.742,-4.788,-2.957,-3.913,13.22,-3.347,3.07,-2.895,-2.678,-3.019,-3.175,-4.605,-2.704,1.729,-3.779,-3.297,-3.073,-5.816,-1.96,1.924,2.063,-2.993,3.043,1.685,3.867,-3.186,-2.28,3.214,3.105,2.099,-2.588,-4.883,-3.714,-2.951,-1.783,-1.738,-6.08,-2.771,-5.932,3.576,3.349,3.444,-3.312,2.42,-3.448,3.309,2.442,-2.239,-8.416,-2.802,-3.556,-4.494,-3.686,5.605,3.031,-2.86,-2.289,-5.511,-2.845,-2.913,3.099,2.689,-3.853,-2.508,5.682,-2.88,-3.023,-1.806,-2.559,5.642,-5.364,-5.499,-4.61,1.734,-2.263,-2.129,-2.429,-1.65,5.709,-4.317,-4.153,-3.595,-5.874,-4.587,2.426,1.619,-4.085,-4.744,-4.781,1.507,10.86,4.771,1.541,-3.079,2.638,2.301,4.55,4.693,3.779,-3.142,-1.562,2.119,-1.302,2.446,-2.247,8.772,-3.725,2.094,-4.817,-3.839,-3.558,4.043,-1.787,-1.905,2.734,1.628,3.571,-5.208,-7.871,3.056,1.923,-1.948,-3.385,2.773,3.328,4.16,1.927,-1.936,-2.617,2.88,2.614,-4.959,2.521,4.719,4.58,4.033,-2.871,3.595,3.249,3.07,2.77,2.632,4.973,2.863,3.519,-3.619,9.404,3.572,2.648,-6.297,-4.217,2.67,3.412,5.033,2.879,-2.445,4.294,3.677,5.462,3.512,-5.158,2.087,3.841,2.598,1.719,4.575,-3.143,3.126,3.766,3.189,3.971,2.28,3.462,-1.502,-4.281,5.738,3.962,4.299,2.895,-2.297,-4.215,-2.141,2.04,-2.054,2.031,3.044,2.49,3.219,-2.585,-4.92,2.397,2.602,3.397,-2.79,-2.776,-4.344,-3.506,-3.798,-5.76,1.716,1.893,2.762,1.699,-2.803,3.609,5.079,3.815,2.465,-4.043,2.342,3.006,-3.038,2.871,-3.122,2.559,3.125,4.936,2.879,2.827,-3.512,2.321,-1.78,4.18,3.034,-4.101,3.516,-4.168,3.052,2.786,3.984,2.423,-3.799,2.643,1.984,-6.346,-8.645,-6.732,-9.279,-7.819,-5.396,-7.241,-6.939],"logCPM":[0.2335,0.8914,-0.909,-1.044,3.163,0.6133,5.003,2.106,1.12,6.624,0.944,-0.1883,1.833,4.629,-1.05,0.4613,-1.034,-1.221,-1.113,-1.154,-0.8025,-0.5506,-1.087,-0.8912,4.231,5.846,-0.4062,-0.5014,-1.269,-1.082,-1.037,-1.018,-0.8162,-0.4439,-0.3194,-1.205,5.047,-1.333,-0.6471,-1.063,-0.8348,-0.4968,-1.243,-1.325,4.202,-0.7709,-0.6452,-0.6829,-1.271,-0.9345,-1.305,-0.8035,-0.477,-0.8719,-0.6047,-1.128,2.171,0.5397,0.4155,-0.783,-0.4198,-0.7101,-0.06131,-0.5802,-0.5516,-1.319,-1.205,0.7128,-1.325,-1.031,-0.6486,-1.2,4.859,4.419,-0.2541,0.05658,-1.063,0.7089,3.866,-0.1543,-0.4205,0.4831,0.8699,-0.1943,0.1023,0.05213,-0.2669,3.389,-0.5134,-1.082,-0.6948,-0.9267,-0.1904,-0.6631,-0.1987,-1.139,-1.238,-0.8462,-0.6023,-0.7576,-1.405,-1.111,0.8912,-0.1736,-1.264,0.5787,0.7213,0.2753,-0.9965,-0.6714,0.7875,-0.7072,-1.029,-0.8471,-0.8726,0.3965,-0.1988,-0.7048,-0.551,-0.9555,-1.18,-0.9394,-0.8912,-0.6887,-0.176,-0.919,-1.01,0.7173,-1.227,-0.7852,-1.107,-1.042,-0.4406,-1.168,-1.041,-1.188,-0.1335,1.254,-0.6179,-0.7133,-0.799,-1.319,-1.325,-1.405,-0.7855,-0.97,-0.3408,-0.4832,-1.294,0.2788,0.2009,-0.02345,-1.019,-1.405,-1.04,-0.9892,0.9867,-0.9012,2.021,-1.214,-1.389,-1.061,-0.004036,2.9,0.3679,-0.1631,-0.484,-0.7328,-0.7121,0.2777,1.72,-1.294,-0.6928,0.5968,-0.5475,-0.2769,2.635,3.691,3.512,-1.055,-0.2817,-0.2713,-1.017,1.501,-0.9366,-0.7041,-0.6979,-1.31,-1.422,-0.6012,-0.406,6.243,0.5244,4.756,-1.066,-0.7452,3.143,5.357,2.96,-1.322,-0.407,1.489,5.436,5.915,1.124,5.629,1.73,1.548,2.064,1.297,5.346,8.008,5.571,4.384,5.624,3.914,8.249,5.372,7.692,7.836,9.62,6.055,8.267,5.312,5.342,6.881,5.048,6.251,8.939,8.391,5.735,1.803,8.226,7.331,0.9543,3.065,4.275,3.248,6.765,8.528,5.882,5.94,3.723,7.764,6.145,7.112,6.5,5.176,3.633,6.39,4.351,6.354,6.233,9.297,5.317,5.782,6.491,4.562,9.322,3.286,7.499,4.061,5.415,5.86,6.496,4.386,4.269,4.136,4.22,3.575,6.997,8.224,6.167,4.795,5.361,4.467,4.534,3.532,6.249,4.068,5.947,4.625,7.821,6.635,4.252,4.399,6.297,6.71,2.168,3.375,4.455,2.918,6.066,6.911,4.663,6.862,5.842,4.813,4.553,4.956,5.534,5.253,5.036,6.076,5.788,5.157,4.996,1.67,7.221,0.4254,3.378,4.533,2.296,5.005,0.4293,0.6483,-0.7163,-1.076,-1.052,-0.9892,-0.2229,-1.034,-1.134,0.1661,-1.045,3.891,-0.546,3.617,2.56,1.102,4.077,3.08,3.104,3.098,2.269,2.475,2.379,3.375,-1.061,0.7968,8.992,3.426,5.883,6.601,2.784,3.553,7.522,6.588,4.193,5.447,2.604,3.811,3.916,7.624,1.716,4.469,4.836,5.842,7.104,5.345,5.331,6.543,2.988,4.069,5.1,6.137,6.579,5.403,3.214,5.154,6.952,5.641,4.128,6.241,3.602,4.943,4.739,5.256,5.124,5.824,7.026,7.492,6.323,4.696,3.672,4.219,7.163,4.604,5.142,6.417,5.723,5.319,6.221,5.109,7.831,6,4.422,7.397,7.181,5.276,2.524,5.559,5.538,6.388,5.941,3.948,4.615,3.907,6.968,7.765,6.57,3.547,3.313,2.965,5.959,4.855,7.962,7.001,3.731,6.638,1.583,8.809,0.833,5.021,6.408,4.321,0.7227,3.016,4.527,2.873,1.395,3.758,1.785,1.264,-0.7617,-0.3822,-0.7949,-0.8444,-1.107,-1.062,-0.196,-0.1219,-0.867,0.008575,-1.038,-1.186,0.2391,0.5017,2.557,1.408,-0.9892,-1.205,-0.286,0.2555,3.083,5.513,6.26,3.005,3.266,9.017,5.007,6.48,7.157,5.666,3.747,6.064,8.13,8.149,6.371,8.6,4.288,6.034,4.691,7.215,7.748,4.228,4.503,7.576,4.071,6.27,5.286,5.151,5.072,5.491,6.746,1.489,3.964,3.189,5.795,6.125,4.536,5.706,4.705,5.701,4.476,6.882,5.287,5.328,4.886,6.902,5.967,5.045,7.667,6.892,6.499,5.154,3.587,5.922,5.367,2.365,6.331,7.947,6.849,2.933,3.747,4.17,3.094,7.713,4.441,7.065,4.936,4.03,4.641,4.955,5.979,6.287,6.281,3.011,4.387,6.207,5.229,4.514,5.287,7.699,7.711,4.025,3.998,4.018,4.571,6.031,1.862,5.804,8.075,6.15,5.793,3.389,4.755,6.192,4.153,5.69,3.364,3.867,4.377,5.811,5.823,5.253,4.108,6.824,5.569,5.271,7.018,6.801,2.606,7.423,3.752,-0.7945,-0.9865,-1.227,-0.8058,-0.9701,-1.2,1.406,2.522,0.5532,0.1818,1.461,1.537,1.363,0.9126,8.474,0.7756,3.739,3.951,5.608,6.425,4.426,5.044,4.864,5.487,3.596,4.583,5.472,5.958,3.875,2.778,7.291,8.008,6.918,5.547,4.711,4.274,5.138,5.967,4.166,6.607,6.346,6.806,3.319,5.667,6.171,4.463,8.8,6.079,5.335,1.631,6.294,3.008,4.149,4.267,5.225,5.254,5.064,4.646,5.763,7.457,5.664,6.479,6.457,8.701,5.916,5.67,5.011,4.42,5.231,4.834,4.844,6.122,5.293,6.499,6.565,6.856,8.343,8.386,7.905,6.805,8.282,7.72,3.176,6.61,5.059,4.865,5.114,4.079,8.303,2.473,5.517,4.643,4.695,7.531,0.86,2.144,3.67,4.562,-0.1508,6.423,6.463,1.665,4.891,4.993,7.278,6.378,4.391,3.281,5.585,7.305,6.487,4.064,0.9202,4.546,2.325,3.998,6.578,6.233,4.524,-0.0695,-0.6377,1.749,0.5699,-1.183,-1.135,-0.8169,-0.5826,-1.152,2.021,1.576,2.879,3.407,2.826,3.438,3.14,6.4,3.25,2.561,1.735,7.501,3.813,5.377,4.813,7.614,6.223,5.356,5.93,2.973,6.699,2.026,5.388,5.786,4.414,5.504,5.753,8.201,6.85,7.281,8.53,9.161,2.438,2.726,3.947,4.02,4.798,5.933,6.8,6.934,7.423,5.03,6.737,2.391,8.71,3.553,5.738,5.156,6.074,7.153,6.155,7.885,5.18,6.386,5.263,4.169,5.403,4.665,6.303,4.653,6.419,7.239,5.522,5.942,4.982,5.158,4.587,2.966,5.976,7.645,3.252,3.092,4.263,6.517,4.914,2.989,1.435,3.312,2.191,4.507,1.582,4.414,1.331,4.592,4.48,3.202,5.524,5.191,7.458,5.258,2.799,3.582,4.133,3.928,7.398,4.593,6.005,6.128,3.064,4.993,5.048,5.477,3.842,2.793,3.465,8.217,6.965,4.84,4.492,5.535,5.654,3.444,6.664,5.484,7.251,5.653,5.623,3.153,3.513,3.03,2.573,0.7383,3.384,2.852,0.01928,2.865,4.046,2.371,3.607,2.279,2.944,1.61,4.286,3.427,0.1291,3.161,4.06,2.98,4.871,6.792,2.81,6.694,3.344,6.465,7.014,5.42,5.635,6.798,4.01,8.603,4.274,6.68,5.924,4.847,4.001,5.769,6.594,4.78,5.909,7.033,6.632,7.048,4.426,4.751,3.445,3.024,6.345,1.953,4.79,6.537,7.071,5.867,4.397,5.218,5.169,3.236,5.373,7.658,6.606,2.228,2.78,1.823,2.403,4.376,1.555,2.602,0.7006,4.778,-1.178,0.1361,-1.099,7.431,4.447,-1.181,-0.8574,-1.325,-1.122,-1.118,-0.9769,-0.4607,0.1144,1.002,2.9,2.915,5.655,-0.04987,6.805,3.406,4.111,2.819,5.341,6.137,3.914,4.681,5.523,6.329,6.116,5.219,4.084,5.52,6.207,2.076,6.499,3.577,5.32,7.255,2.615,4.965,7.413,4.899,4.651,5.212,5.187,5.228,6.447,6.255,8.201,4.803,4.817,7.276,2.168,4.383,4.447,3.604,3.786,6.357,4.204,3.803,5.454,4.357,5.114,3.759,7.688,5.343,7.217,6.251,8.019,0.2897,4.377,6.729,2.095,7.023,5.012,4.232,5.48,2.397,4.787,5.819,5.99,6.595,5.185,5.019,6.627,4.447,4.463,7.538,6.241,3.426,3.043,5.905,7.163,4.281,4.18,6.249,4.684,6.933,4.831,3.792,6.578,5.289,6.333,6.6,4.903,4.167,5.809,5.38,6.195,4.053,7.398,6.926,5.281,6.025,5.487,6.701,7.744,5.628,7.692,7.288,2.385,4.105,3.483,5.486,4.714,4.436,4.98,6.929,6.027,7.739,5.865,6.514,5.535,4.085,4.076,4.461,5.875,5.474,5.519,6.971,6.704,5.326,5.171,3.923,3.668,3.729,0.8394,-0.1203,0.02446,3.715,1.323,5.951,6.29,1.658,1.282,-0.4155,0.1425,-0.4194,2.647,0.2375,2.271,1.613,6.886,0.08755,3.394,3.48,2.436,0.3118,8.454,7.596,6.344,7.788,10.43,6.062,9.125,4.64,6.89,8.011,7.129,7.275,5.093,4.584,4.27,5.899,6.525,3.638,3.891,5.597,5.102,3.498,5.854,6.153,6.2,5.932,5.559,6.192,6.01,4.903,5.588,6.886,5.019,6.215,5.886,4.847,5.335,6.448,6.774,5.179,1.379,7.863,6.658,3.938,6.724,3.684,7.056,7.072,6.224,4.632,4.462,4.066,5.765,3.987,3.809,3.904,5.1,6.075,5.414,4.637,4.359,2.536,4.41,3.774,4.907,5.248,6.279,7.318,4.355,4.883,5.106,5.529,5.14,5.417,9.376,5.769,4.572,4.819,6.719,4.824,2.359,3.769,6.84,4.172,4.215,8.191,1.237,6.44,4.471,3.741,4.736,3.085,5.604,4.208,5.731,9.007,7.139,5.346,6.516,6.219,8.869,7.78,3.848,5.606,4.774,5.206,6.67,6.635,2.804,7.145,6.044,7.203,6.38,4.704,6.856,4.122,5.05,6.669,6.907,5.821,5.118,3.575,4.014,8.129,4.593,7.17,4.818,5.396,4.582,7.442,4.852,5.531,2.715,4.463,6.182,6.066,5.913,6.2,6.245,6.358,5.905,0.3767,3.685,4.54,7.239,7.705,5.471,6.347,4.618,8.026,3.914,6.138,7.194,4.784,3.704,5.391,5.193,4.403,5.164,4.694,7.247,7.132,5.583,6.21,2.989,0.9769,4.818,4.434,5.226,7.095,3.434,2.982,3.337,5.587,5.531,3.384,3.627,7.138,6.219,4.699,9.421,1.466,5.908,6.383,4.955,7.237,3.212,6.038,3.941,2.558,6.302,5.33,4.409,4.377,4.677,5.811,5.969,2.059,3.481,3.944,5.592,7.257,4.791,4.62,5.949,2.968,4.077,3.95,7.086,3.371,1.405,4.562,4.589,3.33,4.316,4.024,1.198,3.732,-1.166,-1.218,-0.3903,3.516,-0.3931,0.8305,0.2322,-0.2245,-0.7174,5.277,1.613,5.764,7.544,-0.3163,4.843,6.853,3.707,5.37,4.544,6.761,5.391,5.713,2.103,4.161,5.956,6.101,7.005,5.978,2.15,6.469,8.293,3.181,2.908,4.796,5.607,6.653,1.403,3.647,5.827,5.986,4.679,4.984,6.285,0.529,2.711,0.3706,4.791,4.556,4.277,4.836,9.34,4.756,5.559,4.985,5.083,6.543,4.492,2.338,8.106,8.964,8.892,1.618,4.372,5.775,2.795,5.794,5.811,5.947,4.744,7.88,6.049,7.115,5.81,4.11,3.481,1.246,3.185,2.591,7.113,3.499,6.51,6.39,9.456,6.494,5.546,5.26,3.178,6.367,7.217,6.74,6.727,5.791,7.296,4.64,7.482,5.748,8.51,3.628,4.36,8.226,5.196,2.741,7.797,6.352,7.535,7.19,2.34,4.896,7.92,7.262,5.138,6.76,4.866,8.69,4.68,7.44,5.141,3.512,7.331,5.371,4.428,5.94,6.851,6.374,6.516,7.308,4.812,6.299,6.816,6.599,4.693,6.303,5.773,6.205,3.687,3.571,5.965,5.576,8.095,7.064,8.196,3.354,7.297,4.187,4.888,4.961,6.525,5.435,4.392,2.788,4.819,6.664,4.84,6.163,4.649,7.42,10.05,8.362,6.261,4.375,4.618,4.174,7.643,5.585,3.525,4.395,5.996,4.401,4.966,4.228,6.153,3.307,6.213,2.066,5.093,3.115,5.274,5.24,5.241,5.444,5.2,5.369,5.007,6.938,5.036,4.248,5.828,5.666,3.952,4.152,6.468,4.906,1.968,4.104,5.649,8.087,6.623,5.982,4.636,6.762,7.082,6.259,5.695,8.953,5.406,3.427,6.362,5.015,4.149,5.012,3.089,4.748,5.09,6.675,8.282,6.819,6.353,6.094,5.087,6.659,2.164,5.172,6.697,4.391,5.984,6.899,7.554,8.659,2.763,6.491,10.54,6.884,6.962,5.266,2.86,3.153,3.087,0.1806,-0.1605,-0.1313,1.274,0.5407,-1.07,-0.851,-0.1846,-0.936,-1.18,-0.94,-0.05853,-0.06275,1.87,-0.1456,-0.793,0.1674,-0.5127,-0.6933,-0.2115,-1.103,-0.8784,-0.8847,0.2962,3.304,4.469,8.033,4.429,7.891,2.683,6.354,4.702,1.835,5.174,8.942,5.276,2.482,5.932,5.634,7.038,5.48,2.939,5.155,6.356,4.847,4.389,1.179,3.019,6.745,5.898,6.987,5.799,8.701,5.916,3.054,4.592,7.612,2.686,3.381,6.023,6.403,4.904,7.498,5.645,2.44,6.42,5.945,6.158,6.418,6.925,5.166,4.856,8.009,5.871,3.959,7.354,6.592,7.411,8.988,7.308,5.323,6.777,3.551,3.849,4.2,3.415,4.176,2.04,8.476,5.087,1.83,9.91,5.832,7.648,1.238,1.726,5.321,6.209,6.136,7.363,5.122,5.85,3.74,7.765,5.775,4.737,4.543,8.739,5.93,4.981,7.598,4.397,3.505,3.421,3.913,7.514,5.412,5.028,5.395,7.45,5.077,1.759,3.07,2.073,6.839,4.786,5.757,6.967,3.131,5.076,7.575,7.034,6.69,7.015,6.55,5.281,7.298,7.39,3.413,7.304,5.508,3.949,4.327,2.933,5.145,6.876,4.414,5.592,5.564,5.775,7.248,7.482,3.422,6.621,7.397,3.58,7.362,7.894,5.879,6.769,7.486,3.765,5.543,6.177,5.843,4.936,3.517,5.9,6.648,6.232,6.741,5.538,4.985,6.792,6.302,4.195,5.86,6.691,8.985,5.885,3.586,3.327,4.068,8.408,5.774,6.735,4.503,6.743,2.632,3.902,6.102,5.144,2.282,5.493,7.607,5.193,6.24,6.915,6.069,5.869,3.977,6.128,4.777,5.404,5.941,6.725,9.097,4.01,7.999,6.378,4.369,3.384,9.213,6.435,6.106,1.827,5.362,2.899,5.978,4.465,6.756,5.381,6.558,8.232,7.122,4.139,3.567,3.306,4.812,7.555,1.919,2.117,3.692,4.394,7.142,3.319,5.307,5.257,0.6893,6.078,5.598,5.207,8.029,8.949,6.164,4.126,6.419,3.591,5.414,5.565,1.996,3.312,7.918,5.689,5.863,6.492,5.236,7.245,6.338,5.698,3.001,4.344,3.996,3.766,4.286,6.033,6.369,4.921,8.395,6.068,8.739,4.738,7.364,5.701,5.473,5.07,5.627,1.704,5.415,4.417,2.309,2.866,2.704,4.399,5.505,11.82,7.302,7.844,6.097,8.449,6.423,6.116,7.674,4.813,1.807,0.4207,0.3368,3.909,-0.09173,0.7734,-1.093,0.3691,-0.9727,-1.141,6.349,5.079,10.97,2.98,7.873,4.813,6.1,7.92,5.691,9.612,8.641,1.63,2.947,2.603,4.437,7.246,8.494,5.569,3.903,8.925,6.39,6.363,7.304,5.982,6.664,7.832,6.679,4.566,5.81,3.408,8.611,4.239,4.393,5.875,7.936,7.505,5.628,4.794,7.355,4.396,4.738,6.594,6.246,6.608,7.818,6.775,4.633,0.9542,9.06,4.902,7.963,4.398,9.554,6.369,4.998,6.547,3.744,5.102,5.803,3.235,6.826,6.382,5.741,3.793,4.455,7.022,6.477,4.379,5.271,3.262,0.5856,6.241,5.325,2.902,6.905,5.524,5.676,7.004,5.201,3.075,2.505,8.188,4.469,6.82,6.359,6.847,7.049,4.788,3.776,2.69,7.074,4.959,4.538,4.775,5.336,2.905,5.101,5.271,3.56,4.512,4.414,5.971,6.456,6.701,5.338,4.255,6.73,4.303,5.174,3.793,2.837,6.855,5.025,4.576,6.523,4.538,10.43,2.459,3.808,2.881,7.244,2.323,-0.5344,5.676,3.575,4.48,5.702,4.588,2.957,5.436,6.18,5.756,5.619,6.297,6.772,5.159,4.707,3.481,3.883,4.741,8.354,7.256,5.179,5.193,3.745,5.087,6.288,5.165,5.806,4.902,5.196,7.372,6.135,5.657,2.469,6.527,6.018,1.417,1.161,8.368,5.553,7.84,5.685,6.459,8.179,6.666,6.844,8.163,4.399,5.104,8.082,4.916,5.136,7.533,5.749,5.526,6.516,4.326,7.614,6.733,6.29,3.367,3.404,2.935,2.514,6.188,5.391,6.248,3.181,8.773,1.367,5.827,-0.5714,7.532,6.163,2.552,-0.2882,1.755,0.7686,5.504,-0.6448,-0.2334,-0.4631,-0.9949,-0.9279,0.7312,1.932,3.499,1.882,-0.07521,2.742,6.855,7.312,8.595,5.048,5.391,3.211,6.571,3.495,5.521,2.007,4.022,3.234,8.397,5.048,6.462,2.736,5.062,4.808,5.313,4.518,4.761,6.065,3.097,6.767,5.808,4.222,3.858,6.647,4.712,5.272,6.623,3.704,5.882,7.586,7.668,4.733,7.958,6.718,4.958,6.951,6.749,4.91,6.962,4.167,3.87,6.796,9.005,5.474,9.568,4.412,4.859,1.027,2.075,10.09,6.711,11.44,8.201,9.094,5.938,4.823,4.417,6.84,7.676,3.348,2.906,7.134,6.858,6.842,8.066,5.231,5.643,4.156,7.121,7.207,7.103,7.476,2.603,7.195,1.788,5.372,6.955,4.114,2.306,5.225,5.948,5.472,6.119,6.052,9.928,5.238,5.457,7.104,5.633,8.349,4.829,7.432,7.411,6.251,3.897,5.592,1.569,4.42,6.4,9.039,6.364,7.758,7.7,7.028,8.798,3.842,2.309,4.402,3.968,10.12,3.667,7.163,7.306,8.966,5.524,2.637,3.783,9.002,2.709,4.88,9.662,7.957,3.133,5.742,7.92,6.919,4.897,3.694,8.934,2.281,6.736,5.769,6.584,5.962,5.087,2.227,6.77,3.823,10.54,3.891,6.269,2.043,6.349,3.503,4.311,6.337,6.478,6.83,4.481,4.517,6.72,6.173,5.652,4.949,4.876,6.202,5.832,6.343,4.792,-0.1024,7.043,6.698,2.632,7.519,5.953,5.669,4.213,6.316,5.398,6.017,7.512,10.63,6.681,4.2,4.479,4.504,3.586,7.092,5.855,5.955,5.848,4.954,6.855,5.937,7.913,5.737,4.173,4.138,4.886,6.256,6.742,6.039,5.485,5.402,6.831,4.79,6.034,6.384,4.035,5.944,6.36,6.027,7.211,6.486,5.582,4.14,6.108,1.607,6.275,4.191,6.431,6.146,2.39,4.623,3.737,0.3132,4.511,1.821,7.212,3.469,8.072,6.421,8.207,2.532,3.92,6.034,6.864,6.277,6.166,5.094,4.794,6.206,4.742,4.993,4.628,1.213,6.72,5.244,4.096,4.988,6.94,6.72,3.606,7.886,5.704,6.445,2.922,3.899,5.152,4.873,5.176,3.899,6.234,4.861,4.428,5.76,4.063,1.992,4.391,1.949,5.377,3.787,6.615,5.871,4.769,7.299,6.6,4.774,5.994,6.401,5.926,6.635,4.391,0.2436,1.858,3.446,1.96,4.767,6.845,3.968,0.8829,0.3518,-0.8686,-1.04,0.07887,-0.6,-0.8321,-0.8271,-0.5751,-0.3915,-0.1634,-0.447,-0.7435,7.004,4.184,2.284,1.643,2.736,6.863,7.912,8.082,4.287,7.833,6.561,7.915,6.1,5.432,8.274,5.554,6.617,2.264,6.98,5.378,5.342,5.536,5.789,3.988,7.204,0.1589,5.894,6.474,3.776,7.289,7.439,4.351,4.744,5.5,6.452,3.34,6.473,3.661,8.786,2.449,3.141,4.544,5.806,4.659,4.259,7.048,3.593,8.597,5.429,6.018,7.031,6.63,5.975,6.112,6.179,4.918,9.491,7.682,7.241,7.698,7.396,5.25,5.513,9.067,6.322,6.495,3.124,5.859,6.977,8.973,2.827,0.7966,3.727,5.686,6,4.968,7.35,1.086,4.543,4.746,2.313,4.997,6.549,3.572,1.828,6.292,5.95,5.115,2.402,5.59,4.165,5.945,5.704,7.281,6.197,7.101,3.477,5.164,7.955,5.732,6.299,5.706,5,4.387,4.26,6.196,0.6223,8.048,6.413,5.644,2.705,4.395,5.007,6.004,7.453,5.599,4.054,4.28,6.063,7.34,6.936,7.643,5.999,3.905,3.687,5.429,3.177,5.016,6.682,5.981,5.418,5.632,6.928,6.934,4.707,3.552,4.585,6.382,6.65,7.003,5.813,3.865,6.622,2.676,7.453,7.203,4.865,5.712,3.07,7.368,5.2,6.879,3.973,5.503,5.057,3.328,5.376,6.271,4.692,5.775,4.99,5.631,1.311,4.355,6.114,4.162,5.947,4.14,6.401,5.185,4.606,5.062,5.937,7.107,4.729,1.404,3.538,3.573,7.251,5.252,7.17,4.716,3.031,3.917,3.92,6.198,2.859,3.986,3.412,3.465,5.703,4.284,7.542,5.559,3.866,3.497,7.565,3.541,6.194,5.706,4.478,7.034,6.582,7.316,6.845,5.738,3.852,6.675,6.93,6.503,5.776,7.374,8.526,2.295,2.747,5.87,4.961,6.12,8.063,6.823,4.592,4.602,2.599,8.307,3.509,3.288,7.548,4.874,2.024,2.427,4.784,6.398,6.334,0.05197,8.653,4.115,7.1,4.028,3.837,7.1,3.288,5.524,3.435,6.213,6.625,5.813,7.137,6.736,4.044,7.337,3.278,4.651,8.157,6.881,6.028,5.283,5.883,7.107,7.616,3.816,5.004,3.17,6.37,4.789,5.899,7.79,4.221,5.585,5.26,8.106,3.792,5.021,9.622,9.31,5.694,8.118,7.18,6.043,8.106,8.695,5.712,6.65,6.418,5.848,7.706,6.601,5.604,5.936,2.526,2.427,2.978,5.822,7.742,2.55,6.495,9.665,7.073,0.5304,7.146,5.874,5.057,7.079,4.826,6.937,9.228,6.426,4.022,4.678,7.965,5.349,5.903,6.204,5.269,6.02,2.907,6.655,6.22,4.67,5.855,6.3,5.642,6.861,7.478,1.867,5.267,6.041,4.516,5.872,6.78,4.343,7.199,7.303,5.149,6.671,1.878,1.323,4.039,5.043,8.593,6.255,9.918,9.434,5.378,6.589,6.321,10.34,3.61,0.821,5.839,6.163,2.994,8.195,2.584,8.304,6.635,3.214,6.033,7.691,6.194,7.076,8.541,9.247,7.709,5.876,6.238,5.757,4.61,6.026,8.262,5.542,7.217,6.924,5.991,6.016,4.524,5.248,5.615,3.686,6.608,8.637,6.598,6.165,9.015,7.014,5.147,8.428,7.433,3.466,6.562,8.509,5.395,4.424,7.579,5.177,5.126,6.564,8.207,4.932,4.536,2.096,4.87,1.258,2.938,0.2456,-1.336,-0.3075,1.899,-1.167,-1.135,-1.096,0.4836,8.246,3.712,5.578,6.106,8.06,7.514,5.947,7.334,3.996,3.351,5.66,3.484,7.106,6.105,4.816,2.748,8.727,6.971,3.799,6.329,2.164,1.911,0.2956,4.977,6.423,5.689,3.809,9.057,8.002,9.515,8.374,6.879,7.256,4.929,8.259,5.936,1.116,5.27,5.552,3.488,7.35,3.734,2.438,8.36,5.246,5.131,2.121,5.449,4.455,4.176,7.62,6.318,4.515,5.028,6.799,4.849,7.867,5.338,5.744,6.037,4.598,8.745,5.461,5.711,3.647,5.558,10.29,8.498,6.316,6.628,6.108,6.463,10.14,5.484,7.993,4.57,4.486,3.833,3.805,6.989,4.215,3.659,8.854,6.12,6.896,4.453,5.21,5.825,3.215,9.437,5.283,4.183,7.126,9.913,5.18,6.435,1.371,4.758,6.826,5.609,5.302,6.413,5.712,6.894,7.214,5.152,6.098,6.584,3.5,5.493,3.65,4.681,5.032,4.53,6.293,4.298,6.628,4.007,4.008,4.416,4.378,2.347,3.016,5.445,4.205,2.19,4.015,4.042,1.505,8.36,7.847,9.275,8.914,6.304,4.497,7.528,-0.5858,4.887,6.031,5.583,3.081,0.657,4.497,5.921,8.762,8.784,7.572,6.982,5.946,6.153,6.463,5.329,5.559,6.181,6.268,3.021,4.847,5.101,3.793,5.029,6.169,4.431,5.577,6.336,8.518,4.269,4.111,7.693,5.884,3.264,6.788,8.168,6.405,7.426,5.926,11.88,3.745,8.26,6.739,4.19,6.019,0.9584,3.653,8.716,5.105,6.827,6.493,1.984,3.427,5.744,7.538,2.68,3.682,4.574,6.787,6.386,7.541,8.013,5.679,3.317,1.512,2.445,7.661,8.076,2.446,6.027,4.51,6.323,5.974,6.638,6.155,7.252,7.402,3.785,8.013,1.97,9.416,8.491,6.381,8.857,6.904,7.442,5.701,5.282,3.514,1.357,7.796,7.617,4.826,7.474,4.439,5.906,7.753,6.228,4.073,7.15,6.197,5.068,5.292,6.654,3.461,6.389,6.706,5.628,6.817,3.553,8.736,4.43,6.756,4.633,3.793,6.384,6.768,6.238,3.982,2.262,4.211,1.568,3.503,6.683,5.668,6.811,6.625,8.641,2.637,8.368,2.348,-0.3133,-1.231,-1.253,0.09074,-0.7202,0.8534,2.329,1.427,3.06,3.131,2.865,2.936,1.171,2.457,5.35,4.055,1.696,4.872,7.108,4.451,5.616,4.136,2.118,4.588,2.955,1.733,7.234,2.216,8.757,8.112,9.154,8.221,5.77,6.65,3.976,4.424,10.23,3.655,6.003,9.283,0.434,7.216,6.976,7.708,7.563,4.801,4.866,5.03,3.181,3.349,1.115,4.119,3.266,7.038,5.526,6.181,6.47,6.352,6.766,4.459,5.67,6.316,6.715,7.803,7.472,5.221,7.491,6.045,3.829,5.95,8.787,2.832,7.068,6.656,7.269,7.059,6.997,4.781,6.997,4.995,3.656,4.853,4.129,3.949,5.616,3.828,7.671,8.788,7.491,7.171,7.064,4.617,4.692,2.175,7.009,5.65,5.222,6.428,7.495,7.015,4.477,7.746,6.786,6.258,5.824,5.211,6.325,2.734,7.162,6.243,4.827,5.878,4.816,4.245,4.921,3.262,5.608,5.192,6.675,6.06,7.483,6.457,6.768,4.242,4.044,6.405,4.083,4.812,7.711,0.4922,6.48,8.102,3.347,8.096,5.754,8.864,7.022,5.574,5.28,5.896,3.786,5.245,6.866,6.283,1.591,9.095,7.363,7.523,4.182,8.082,4.12,6.651,4.048,6.056,6.591,4.98,-0.3449,1.4,6.413,7.189,6.027,6.62,9.287,5.969,7.873,8.369,10.08,4.294,5.574,6.785,9.22,6.001,4.841,7.206,10.01,2.581,4.473,4.98,7.847,5.412,7.514,7.862,9.073,5.87,7.621,7.687,11.51,7.449,6.994,6.693,5.261,10.01,4.708,2.171,7.658,5.495,6.728,6.152,5.152,7.599,10.31,5.194,2.435,5.887,4.135,8.508,7.301,4.757,8.342,2.656,6.187,5.476,3.449,7.596,4.684,7.025,2.33,5.012,4.797,8.421,5.653,7.824,4.744,3.992,3.374,6.722,7.187,4.009,4.446,7.386,7.727,8.192,6.602,10.16,3.546,3.804,-0.005438,4.87,2.956,6.198,6.774,5.988,6.143,4.955,0.8933,5.794,4.334,6.927,6.728,4.619,4.925,4.673,6.765,5.885,4.609,5.866,4.599,6.369,6.675,4.022,3.303,1.843,5.133,7.251,5.421,4.151,5.789,6.799,5.343,6.89,6.609,5.569,6.402,6.461,5.717,5.299,6.153,6.462,5.412,6.354,7.098,5.273,4.505,7.614,5.916,3.387,5.311,4.164,4.26,6.097,5.245,5.346,6.76,6.808,5.976,2.74,5.263,3.824,6.084,4.606,4.442,3.976,5.202,5.309,5.907,6.836,6.031,3.969,5.729,6.352,6.923,5.684,3.415,4.809,5.233,4.874,5.028,5.868,7.999,5.599,5.551,6.795,5.773,2.675,4.893,5.707,5.796,5.172,7.647,7.314,5.851,5.245,7.304,6.959,7.794,3.328,6.369,5.422,7.258,8.475,4.23,8.238,7.088,4.328,8.444,4.281,4.651,7.647,8.114,8.589,5.006,1.981,9.504,7.848,5.795,7.462,4.781,-0.2875,4.814,2.704,3.872,6.313,4.412,3.39,3.829,5.904,5.162,4.099,9.081,6.252,6.119,5.849,5.159,5.423,6.747,4.955,6.811,10.69,5.763,7.66,4.898,3.339,3.514,6.56,6.246,6.304,5.709,6.447,3.25,4.303,4.325,6.303,3.095,5.238,7.213,1.699,8.017,6.687,3.199,5.094,5.11,4.867,5.633,7.15,8.105,6.499,4.231,8.667,4.169,3.744,5.335,5.546,6.127,10.28,1.385,5.501,9.496,1.885,4.258,5.274,-0.1346,2.226,2.589,1.175,0.5323,-0.05413,1.183,-0.1004,4.283,-0.2792,3.4,1.926,-0.7782,7.292,0.7088,0.1006,0.6227,-0.312,3.915,0.4019,-0.7813,-1.322,-1.125,-0.8562,-0.06173,-0.4989,-0.6876,6.033,4.004,7.292,1.358,0.09461,0.5138,-0.4596,10.03,2.894,5.732,9.505,5.214,8.884,9.581,7.531,5.951,4.829,6.467,7.129,12.91,4.469,8.06,7.465,8.249,9.335,7.931,8.203,8.325,10.11,5.897,7.186,5.728,7.672,8.205,7.425,5.996,7.027,6.017,11.66,6.469,8.254,6.517,9.883,6.771,7.03,9.387,7.21,9.749,6.029,6.685,5.872,8.769,8.668,6.211,6.271,7.478,6.912,8.411,9.915,6.833,6.531,8.186,7.543,9.797,9.578,5.787,4.622,4.42,3.992,4.812,4.563,5.149,5.101,9.602,7.88,7.631,7.522,6.381,3.676,6.075,10.44,5.5,10.07,7.977,7.111,5.512,2.441,9.817,4.02,10.81,8.196,11.06,7.682,6.404,6.67,8.041,6.807,9.258,9.458,8.081,9.489,5.439,6.68,4.026,10.08,9.642,4.941,4.272,2.65,9.028,11.06,9.493,9.31,5.247,8.954,5.613,7.495,9.743,10.2,10.1,8.126,9.258,7.282,10.9,11.96,10.36,6.982,7.409,9.026,13.09,12.8,4.66,7.345,8.554,8.506,6.163,8.339,6.083,9.402,9.52,8.212,8.608,6.474,8.777,7.482,6.908,9.024,6.912,8.061,7.657,11.15,7.657,6.143,7.801,6.416,7.616,5.221,5.581,9.19,5.183,6.365,5.807,6.323,7.43,9.362,8.023,4.816,5.908,8.476,7.302,4.683,6.079,7.3,8.863,7.641,6.631,7.232,5.96,6.614,4.625,9.108,8.794,7.991,7.853,7.51,6.889,5.218,9.402,10.24,6.228,7.569,6.595,7.413,5.537,6.567,10.68,9.634,6.136,7.606,8.11,7.904,8.785,7.704,5.854,8.191,8.078,7.937,8.551,6.463,8.091,3.535,7.671,9.098,7.529,7.506,4.613,5.075,4.664,9.602,8.59,9.385,6.518,4.595,5.286,4.401,4.341,8.701,3.121,7.714,5.644,9.458,8.648,6.845,5.978,8.985,7.065,6.26,5.616,6.706,8.548,8.123,9.013,7.707,7.693,6.286,10.97,9.815,4.805,2.291,8.2,7.29,8.859,6.476,6.546,3.731,4.465,7.61,7.723,4.767,10.14,7.814,7.353,9.769,8.608,8.125,3.19,9.154,10.66,8.87,6.944,8.105,10.14,7.408,8.804,7.261,7.734,7.803,6.72,8.232,4.364,9.053,3.658,4.936,9.051,5.55,5.624,10.2,9.669,7.61,3.992,5.875,5.668,5.235,9.597,6.538,8.769,7.308,3.828,8.568,5.413,6.565,8.36,5.869,7.296,7.358,3.768,7.876,8.584,8.332,4.406,14.91,7.605,9.833,9.739,7.641,10.18,8.873,3.345,2.782,5.893,9.818,9.09,3.643,6.671,7.317,5.504,4.904,3.996,5.301,6.668,6,5.006,7.795,9.143,4.794,8.359,7.715,8.39,8.359,6.471,7.682,3.52,4.123,2.551,7.497,3.439,7.647,5.192,5.263,6.872,11.02,9.798,9.213,8.762,7.677,6.844,5.112,8.148,7.706,10.17,10.49,9.014,10.69,8.034,7.158,6.348,8.799,7.334,6.091,4.242,7.544,8.615,9.224,3.955,4.014,2.116,6.947,7.526,14.25,10.71,7.117,7.192,9.019,8.64,6.638,6.964,6.174,8.992,8.181,7.065,7.566,5.43,7.766,7.703,4.646,7.978,6.621,5.716,6.698,5.866,7.608,6.455,6.502,8.671,8.475,8.641,9.904,4.166,7.218,8.947,8.269,8.474,8.828,4.709,4.819,4.58,11.19,10.57,4.645,8.138,7.446,3.46,6.116,6.181,5.799,10.36,6.912,7.908,7.745,10.77,4.981,5.4,8.41,5.664,9.245,5.617,10.5,7.096,7.09,10.45,8.508,9.408,9.572,6.304,8.318,8.747,7.282,7.62,8.167,7.909,11.31,7.097,7.998,9.669,8.447,9.342,3.536,10.28,5.42,5.09,6.819,9.147,4.88,6.457,5.898,2.601,7.946,6.315,6.17,9.27,10.53,6.053,3.468,4.392,4.627,4.573,5.769,5.886,6.345,4.661,10.67,9.932,9.159,7.817,6.708,10.23,4.891,3.191,4.328,6.352,9.712,7.858,4.778,7.51,7.036,8.633,8.249,9.8,8.585,7.383,8.592,11.96,8.426,6.541,5.364,8.794,5.536,6.665,6.941,3.398,9.572,5.459,8.373,9.867,8.702,8.482,5.063,5.031,8.564,8.94,8.008,7.813,8.449,7.2,9.898,8.334,6.61,8.882,9.976,6.345,8.784,8.919,7.78,8.017,3.222,6.184,4.387,3.767,5.217,8.605,9.424,10.16,7.417,9.175,7.983,8.358,6.096,9.976,10.15,3.717,6.384,7.678,5.8,6.643,1.878,8.777,4.736,5.669,4.397,6.827,9.09,6.539,9.638,6.162,7.689,7.5,11.11,11.67,9.236,7.391,9.713,6.188,5.929,7.674,8.801,6.183,9.118,6.318,5.963,8.238,7.538,8.979,7.641,2.318,7.449,9.439,5.815,5.519,3.513,4.913,5.374,2.552,9.541,9.215,4.378,5.764,10.25,5.939,7.368,8.627,7.758,4.583,6.331,4.536,6.705,2.17,10.14,6.939,10.43,6.602,11.44,9.781,9.573,8.947,4.479,4.139,8.294,5.343,9.229,7.623,5.804,4.879,5.015,6.204,9.467,7.996,7.513,5.031,5.459,5.093,8.839,9.645,6.783,9.113,8.593,9.277,3.617,4.07,9.999,10.15,5.151,8.041,7.799,5.901,3.594,6.33,5.242,7.642,9.453,7.611,4.999,9.477,6.382,6.165,5.979,3.773,7.084,8.734,7.901,6.949,6.698,5.282,5.563,7.404,10.34,7.509,6.311,6.705,9.119,6.172,3.453,7.091,4.355,9.166,9.787,4.188,4.133,8.028,6.843,9.845,10.48,7.121,8.609,7.948,6.558,6.755,5.918,8.408,6.495,4.232,5.675,5.788,3.538,6.57,4.988,5.172,5.755,5.89,5.457,5.86,8.876,4.827,6.855,9.288,3.792,7.148,6.918,7.101,6.893,5.903,7.363,7.653,6.479,6.394,6.756,5.109,8.219,9.034,4.615,7.829,6.696,7.603,14.19,5.714,6.935,7.066,6.359,7.831,6.614,4.444,8.013,5.999,5.341,5.268,6.229,10.42,6.544,5.98,5.998,2.868,4.305,8.746,10.6,10.46,6.742,7.693,8.361,10.28,12.61,3.826,6.138,9.002,2.856,5.566,7.999,5.97,6.493,4.896,2.062,6.614,11.51,4.379,4.852,7.601,10.12,4.707,5.878,3.88,3.548,6.474,6.857,5.727,4.825,9.443,8.112,3.772,4.285,3.792,8.494,6.694,8.203,6.537,9.497,7.816,9.057,9.692,9.18,6.131,4.724,8.604,7.257,7.528,8.005,6.076,8.641,7.295,6.752,8.958,7.836,8.377,8.632,7.021,5.742,8.5,8.274,8.355,4.254,3.551,4.367,10.93,8.378,11.18,8.251,8.708,7.115,4.959,8.882,4.364,11.64,8.495,4.62,11.14,8.275,7.322,7.266,3.005,9.673,7.852,10.23,4.792,10.13,4.21,5.568,6.611,9.188,3.735,8.374,5.583,6.787,7.469,9.178,6.163,9.855,8.79,3.152,8.246,5.606,10.76,6.635,11.67,5.749,8.197,8.365,6.141,4.635,7.076,2.444,7.6,7.103,7.419,8.173,4.649,5.719,5.685,5.036,3.746,10.17,7.765,10.04,10.98,5.76,6.04,6.83,8.605,7.514,4.618,3.655,9.041,7.047,4.281,4.622,9.565,10.38,5.272,4.831,8.106,6.032,7.516,4.71,8.014,6.614,5.471,3.481,4.64,8.201,8.25,4.257,1.069,4.688,8.693,0.7663,9.154,9.772,10.53,2.296,6.963,6.726,6.557,5.237,8.845,5.101,3.8,5.097,3.72,4.974,7.684,8.435,4.925,7.386,5.263,7.128,9.038,6.253,6.957,6.265,5.857,7.815,3.897,5.213,9.336,6.685,10.85,5.216,6.013,6.19,8.9,3.091,4.944,7.282,7.726,6.455,8.004,5.348,7.126,6.592,4.488,3.354,9.341,7.565,6.178,9.812,8.854,5.911,9.244,7.665,8.625,7.589,7.304,8.226,4.886,5.543,5.953,9.682,5.704,8.204,5.66,7.23,8.525,4.571,6.431,6.324,7.336,5.24,7.698,7.195,6.314,7.538,5.579,8.497,11.25,11.31,9.86,9.929,9.604,6.6,7.99,8.626,5.35,8.769,8.715,6.628,6.44,8.024,8.365,6.258,7.594,8.528,9.408,6.453,6.357,1.669,8.975,11.12,5.879,8.974,7.972,7.999,7.173,5.245,5.424,4.753,3.868,8.446,7.855,5.36,5.845,3.798,5.748,8.945,4.511,9.093,8.292,8.916,6.236,8.162,3.45,5.565,8.826,4.676,8.786,6.437,7.38,5.193,3.78,4.067,5.486,9.691,6.445,5.03,3.522,5.802,4.978,5.713,5.427,6.32,13.59,5.701,4.604,10.76,5.328,5.06,8.473,8.263,9.894,6.363,6.001,6.012,5.096,8.328,4.081,8.884,11.48,11.33,5.146,3.886,5.169,6.161,4.632,3.041,8.162,5.392,6.999,8.333,8.292,5.917,5.251,5.526,4.559,4.612,7.478,9.1,5.333,9.143,7.619,6.691,4.779,10.62,13.62,4.863,4.052,4.244,6.747,11.57,6.714,9.376,4.958,2.978,6.363,8.9,9.625,7.058,10.37,9.934,5.155,6.43,6.458,5.444,5.124,9.53,7.717,8.471,4.931,8.18,7.371,9.009,7.391,7.7,10.2,4.075,4.688,7.653,4.496,6.199,6.122,8.983,10.59,7.878,7.729,6.924,10.46,7.437,9.415,5.388,6.517,7.959,9.386,8.843,6.833,4.053,6.845,6.983,10.74,4.7,5.606,5.109,3.504,9.946,3.404,10.57,7.573,7.36,7.326,6.515,7.627,10.06,4.486,7.206,5.279,5.495,7.423,7.719,7.557,5.293,9.013,5.112,7.939,8.794,6.209,7.54,7.426,9.548,4.187,1.96,11.51,8.011,8.095,9.539,5.885,6.521,7.016,7.153,8.553,8.243,10.57,8.246,8.171,8.538,8.685,9.093,4.95,9.263,3.214,6.561,9.279,7.538,6.592,6.659,8.816,6.898,9.036,7.168,10.51,4.431,5.578,7.269,5.369,9.718,8.047,5.746,4.891,3.36,4.252,5.124,4.169,8.307,5.732,8.101,6.406,5.209,9.361,7.449,11.16,7.102,11.13,10.51,5.422,5.903,4.758,4.63,8.512,6.896,9.861,4.655,4.299,5.518,8.984,8.212,5.792,7.398,7.952,6.663,5.574,9.018,9.06,3.963,5.866,6.668,9.056,8.875,12.46,4.624,7.594,6.535,6.34,5.104,5.873,7.445,9.065,9.64,6.088,7.294,5.605,6.379,5.411,5.965,6.888,6.073,5.593,6.862,8.865,6.476,7.775,8.463,10.71,8.209,6.974,6.914,8.759,8.815,6.746,9.152,5.753,8.714,8.314,11.04,4.443,7.889,8.812,5.396,11.9,7.649,3.411,4.375,7.386,2.736,8.328,9.924,9.704,7.098,6.397,6.407,2.884,6.347,7.346,9.143,9.396,7.188,8.812,8.987,7.606,8.609,3.706,7.752,8.501,8.336,9.086,8.147,5.834,5.484,6.474,9.143,7.855,6.611,7.143,8.409,4.763,9.467,7.121,5.514,8.162,6.65,7.135,6.557,9.028,7.267,5.582,10.95,10.4,8.561,5.179,7.539,6.237,9.07,9.54,4.585,7.203,5.994,6.651,9.954,12.47,7.641,5.807,4.022,6.788,5.61,10.77,6.294,4,1.684,10.67,3.011,8.153,8.69,7.784,5.567,5.711,8.985,9.718,4.241,7.753,7.003,4.661,5.871,8.62,9.269,4.832,2.75,9.325,8.699,7.993,4.813,7.908,2.184,8.785,7.642,3.727,5.932,4.14,3.336,10.38,6.622,6.484,3.133,7.304,8.573,9.018,4.554,9.025,8.619,6.616,5.471,4.206,5.337,4.721,6.924,3.779,5.202,5.137,9.569,5.176,6.937,9.12,9.776,10.21,8.059,5.275,6.406,2.95,11.37,6.43,10.21,3.072,9.009,6.268,7.091,8.802,8.55,7.944,8.824,4.718,7.987,7.088,5.705,9.672,5.691,6.599,4.979,8.55,8.638,7.633,4.732,4.714,7.771,4.313,5.625,7.918,8.448,7.334,5.79,4.13,7.238,5.091,7.514,9.017,7.134,6.691,8.671,5.414,7.012,5.038,4.506,6.22,9.642,8.872,5.165,5.35,3.187,6.474,5.696,9.149,10.85,9.126,8.653,8.076,7.542,10.98,7.547,7.657,4.411,5.793,6.371,4.066,3.353,8.717,4.994,9.51,5.42,8.824,5.17,10.5,4.571,10.36,8.623,11.05,3.199,6.268,10.28,8.137,9.339,11.05,5.132,7.975,6.552,7.502,5.119,7.122,4.121,8.473,7.119,3.822,5.896,8.546,8.732,3.599,4.976,6.719,5.071,4.772,6.994,7.696,5.089,10.61,4.481,4.845,6.207,6.975,9.019,9.724,7.888,8.78,6.586,8.445,3.91,7.222,7.704,6.995,7.012,8.725,6.601,7.707,4.814,5.991,7.427,6.704,8.672,2.646,5.974,7.683,6.226,8.521,6.969,5.437,7.018,7.548,4.377,8.918,6.524,5.256,7.503,5.673,5.938,7.836,5.704,11.22,7.693,6.15,9.775,7.404,9.033,9.944,4.895,7.551,2.463,3.922,2.608,6.416,9.953,6.87,5.922,7.007,9.031,6.983,9.763,4.787,4.744,7.774,10.02,6.984,10.11,5.748,11.44,4.206,5.267,5.306,5.607,7.6,10.96,7.369,7.465,5.506,10.64,8.722,8.792,6.934,4.973,6.152,9.211,3.989,4.313,5.575,6.356,5.217,4.816,4.051,5.047,5.768,2.375,6.828,5.52,10.36,5.945,4.623,4.78,8.298,5.041,8.012,8.797,5.89,4.298,4.756,9.284,9.093,7.625,6.135,9.852,8.128,4.464,6.141,10.96,9.546,9.32,8.803,7.753,6.975,7.646,9.779,10.68,4.478,5.345,6.512,6.452,7.818,5.964,8.283,7.268,5.103,8.729,5.985,9.571,10.83,4.543,5.488,7.861,9.186,6.159,10.66,11.5,8.947,8.703,4.053,6.016,5.755,6.678,4.685,6.837,6.53,4.609,6.867,8.193,6.036,5.635,4.723,7.333,4.284,6.087,4.605,3.863,8.808,6.055,3.478,2.834,4.014,5.161,10.32,3.95,12.24,4.439,6.974,6.322,12.05,3.707,5.204,4.969,5.606,3.861,5.16,9.862,4.466,5.728,4.649,6.197,8.068,9.496,8.739,6.531,8.233,8.022,9.413,6.147,5.61,9.574,7.936,8.641,7.512,1.871,6.498,4.844,9.756,6.414,3.018,7.874,1.909,6.843,8.487,9.519,8.68,9.217,6.908,9.261,9.532,6.455,2.347,4.734,3.653,7.739,5.269,8.873,9.925,7.225,6.523,4.664,5.898,7.685,8.872,9.909,3.766,5.542,9.425,7.826,9.399,9.516,7.716,6.95,2.674,4.058,3.601,8.815,6.761,6.224,6.661,10.19,10.59,4.906,3.596,5.481,3.862,3.661,6.614,7.132,3.625,5.16,1.078,9.558,12.37,10.03,8.643,5.407,9.795,8.347,8.37,5.699,10.76,4.299,5.818,9.478,9.048,11.46,5.82,9.257,7.131,8.73,2.737,5.337,4.804,8.79,6.864,8.541,9.319,8.522,10.64,3.497,4.997,10.2,9.533,6.371,4.156,7.09,8.026,7.679,5.869,8.471,5.812,6.612,5.756,6.445,5.835,8.679,11.05,8.161,5.279,8.196,7.928,9.039,6.546,6.302,6.246,7.634,7.732,4.936,3.605,9.834,8.16,2.712,7.365,7.417,8.559,6.38,7.868,5.82,7.563,6.502,8.515,8.517,5.066,7.378,7.618,8.742,6.96,7.136,6.24,8.195,8.71,7.247,8.85,8.118,8.419,6.507,2.463,8.868,7.773,9.296,7.893,5.818,4.622,7.288,7.622,5.845,8.155,8.951,7.666,8.321,4.705,5.705,8.197,8.295,9.268,5.695,6.775,2.452,5.206,5.075,0.583,7.576,9.684,9.202,9.101,6.616,9.021,8.631,6.923,7.498,3.433,5.789,7.919,7.267,7.65,5.422,8.849,7.574,8.792,5.439,7.661,4.494,10.24,6.575,9.338,7.366,8.274,10.2,4.944,5.522,7.228,9.49,6.322,2.766,8.232,9.239,1.968,5.26,1.741,2.307,0.8779,5.838,10.06,9.26],"F":[1.564,0.005865,1.722,0.9152,2.43,3.115,5.248,2.004,2.778,0.03432,3.245,0.8101,0.1821,2.796,1.429,0.7557,1.562,1.385,1.835,1.783,1.935,2.147,0.3761,1.703,2.037,1.264,1.991,0,1.317,1.597,1.984,1.608,2.011,2.595,2.383,1.404,6.478,1.325,2.706,1.921,1.701,3.195,1.357,1.234,0.8556,1.696,1.955,2.226,1.314,2.133,1.331,1.053,1.956,0.9967,0.5049,0,2.387,1.052,10.14,1.803,0.843,2.056,10.13,9.19,2.615,1.234,1.404,9.367,1.234,1.992,2.461,1.543,4.313,2.193,1.775,1.405,1.115,2.576,0.4614,1.526,2.204,2.39,14.05,2.09,1.031,2.569,1.366,0.6204,1.374,1.736,2.067,2.063,3.212,2.968,1.755,1.536,1.365,2.046,4.877,2.188,1.059,1.414,1.764,2.096,1.326,4.727,7.155,0,1.815,2.395,0.6373,2.277,1.783,2.399,1.716,1.832,0.1013,2.614,2.932,5.763,1.434,1.666,1.703,1.961,2.094,1.741,4.797,0.5462,1.475,1.806,1.693,1.601,0.2146,1.62,1.579,2.464,2.676,0.08459,2.389,10.22,1.769,1.234,1.234,1.059,2.181,1.64,2.324,1.953,1.28,3.28,9.743,2.028,1.618,1.059,1.744,1.625,1.746,1.862,11.09,1.712,1.094,1.76,2.781,0.6381,4.943,3.086,2.484,2.025,2.12,0.1088,9.702,1.28,0.8683,7.098,2.175,3.37,7.026,5.477,1.469,1.635,2.688,0,7.523,0.9567,0.5615,0.8287,0,6.275,2.563,0,1.999,8.333,3.596,1.043,0.6915,0.003441,2.43,1.564,5.03,1.227,4.631,1.863,0.2416,1.547,0.3494,0.02702,0.03149,6.454,1.301,10.34,10.37,5.3,2.991,9.279,3.306,0.6024,0.398,3.982,2.727,6.782,5.698,0.8688,10.37,0.01292,9.176,5.359,6.373,0.3206,8.924,4.548,2.272,3.515,0.9374,9.121,2.897,1.298,9.072,8.023,0.1272,7.668,0.9323,0.0005721,11.59,11.32,1.841,5.63,8.257,0.06411,2.042,4.519,1.491,2.075,3.774,0.1685,0.1393,0.3454,0.8911,5.413,7.439,0.08118,10.42,0.2724,1.696,9.082,4.451,4.024,0.7645,5.456,1.283,2.718,3.849,7.239,3.681,7.824,11.41,1.39,2.462,0.007895,7.888,0.4414,1.067,3.524,8.025,7.358,0.6117,0.06738,4.775e-05,0.009539,9.814,0.005464,1.095,0.7982,0.03403,2.961,1.573,1.474,1.569,0.3432,5.536,5.679,0.07383,3.597,0.005324,7.511,2.811,1.075,2.264,5.727,4.232,2.043,0.8859,9.555,2.463,11.29,3.077,5.058,2.095,1.866,1.866,1.625,2.561,2.01,1.497,2.465,1.94,0.2898,1.668,0.373,4.028,0.4921,0.2043,11.54,2.137,1.798,3.057,2.943,1.598,2.228,0,9.977,1.771,3.728,0.06736,6.654,0.1319,0.007055,7.614,10.53,3.185,1.996,0.3957,10.68,2.297,4.936,0.8319,2.32,7.475,3.736,7.652,3.18,2.14,8.089,0.3131,6.031,0.01692,0.6418,3.741,0.4046,0.002266,0.121,5.951,1.972,4.916,3.215,0.7041,0.2818,0.4446,4.682,5.521,0.5308,2.568,3.368,3.177,0.01365,4.566,0.8263,5.132,1.512,2.272,0.225,3.507,0.1583,0.8445,2.756,8.722,0.2661,0.05422,9.382,5.854,8.316,0.4847,4.27,2.356,0.4396,0.8908,10.56,8.172,10.78,5.464,10.74,5.059,9.828,3.486,0.4892,1.37,10.87,1.7,5.272,10.41,1.276,9.189,9.909,1.364,0.00332,10.72,1.869,2.257,0.04695,0.02856,0.03399,2.968,0.2261,3.31,4.783,2.236,2.25,1.974,1.738,1.693,0.4287,3.09,0,1.922,2.707,1.585,1.592,2.989,1.789,3.17,0.02191,1.625,1.404,2.745,3.226,0.05851,11.09,1.797,4.286,6.896,7.321,10.72,1.456,8.381,8.721,6.027,3.086,7.047,5.629,11.13,0.4825,0.5734,6.555,2.26,1.402,0.001705,11.53,9.371,1.825,10.65,2.209,3.438,1.534,1.333,3.089,8.496,5.115,4.747,2.569,0.2376,3.238,0.7187,2.047,4.276,1.526,0.9438,8.735,6.369,0.245,2.257,7.866,2.436,0.2354,6.207,1.539,0.06217,1.248,1.428,5.757,1.959,3.234,8.557,4.848,2.033,0.2232,7.533,4.245,9.078,0.3411,6.163,6.347,3.22,0.02199,6.612,3.764,0.03451,0.326,11.51,0.335,2.758,4.543,1.315,1.536,0.7857,4.245,1.939,1.586,2.177,1.183,0.5154,3.494,1.175,6.148,3.729,6.694,0.1738,0.6653,0.007031,1.005,3.434,5.298,2.283,0.5968,5.617,0.3845,1.97,5.765,6.189,10.59,4.121,4.45,0.8011,1.104,0.2905,4.269,5.115,1.943,0,1.376,1.044,2.046,1.26,0.5141,2.928,2.57,2.499,0.1754,3.004,8.837,4.477,0.6504,1.627,0.8737,7.697,4.023,0.235,0.4907,11.23,0.1503,0.004676,10.02,0.08909,5.081,8.01,8.782,0.3859,4.251,3.337,1.671,0.1096,2.439,1.393,11.3,9.25,0.7766,1.661,0.05712,9.778,8.105,0.6179,7.121,0.2168,3.075,3.478,0.04309,11.68,1.405,9.235,2.633,3.79,0.01188,0.123,0.2849,0.01461,0.1606,2.114,4.607,6.939,6.073,7.828,5.467,0.4234,6.697,5.552,0.6302,1.436,1.625,5.03,5.845,4.847,3.825,0.04047,1.655,2.055,0.9017,7.203,0.08427,10.16,0.003976,3.213,7.91,1.032,5.508,1.387,1.38,1.862,2.912,6.696,9.75,1.959,3.935,11.02,1.316,3.852,3.531,1.272,4.91,0.1843,8.414,0.05943,11.36,0.8081,1.14,0.2236,3.035,10.79,4.21,4.059,2.306,3.356,2.79,0.9291,0.5046,0.01273,3.582,2.639,2.079,10.76,6.892,5.612,1.489,1.24,1.971,1.607,1.664,2.838,6.789,1.989,1.349,9.793,0.005104,0.7287,1.37,0.05065,0.08682,1.389,4.791,2.154,3.331,5.409,11.28,0.7164,0.2672,1.705,7.127,0.1447,0.3002,9.409,4.729,0.01201,4.507,2.61,2.308,3.794,7.46,0.699,0.2879,6.121,0.2757,3.152,0.1265,0.7083,6.544,7.965,0.5213,3.85,3.942,0.5692,8.417,0.7304,0.6485,2.037,2.763,5.056,3.993,4.134,5.353,1.523,0.6717,0.03241,1.235,8.569,0.3652,1.17,3.517,1.094,2.378,0.08783,0.1058,0.2203,1.752,9.95,3.194,3.178,6.052,0.09061,6.949,0.000402,11.04,8.741,0.003064,0.1187,2.221,4.663,0.04979,6.419,1.762,0.3035,1.615,2.655,0.2218,1.79,3.51,7.29,8.525,4.259,1.316,0.1049,2.891,5.115,9.622,0.1511,2.023,9.554,1.477,0.3264,5.984,0.6475,0.04888,5.926,0.3504,0.5203,0.009486,9.653,0.1323,6.493,0.2556,0.06265,8.614,9.967,1.089,0.4012,5.623,5.604,6.17,8.946,5.464,1.589,1.427,2.235,2.113,2.074,1.163,0.001368,2.861,0.06295,2.045,0.1489,2.198,5.66,3.977,8.08,11.49,0.463,5.493,7.109,0.1232,6.333,5.352,2.274,1.093,0.6403,3.669,0.008825,10.09,9.686,0.07014,0.5511,0.8825,6.833,7.221,0.4786,10.93,0.0941,8.514,1.13,0.6523,7.938,0.5672,2.899,0.01478,0.002028,5.383,1.978,10.17,8.214,0.769,1.978,0.09682,2.071,4.989,3.755,0.2296,0.3915,1.093,2.082,4.422,0.3014,0.5938,6.251,2.547,3.349,1.452,2.225,0.7044,6.912,0.07198,1.709,1.93,1.049,1.58,1.504,1.501,2.659,3.044,5.302,2.576,0.6744,0.3495,2.772,4.593,0.04462,6.024,3.183,4.929,1.03,3.758,0.809,7.32,2.193,10.64,2.064,0.1386,10.21,0.831,9.083,6.105,0.2086,0.0088,10.56,0.06535,1.009,11.69,0.5088,10.39,1.364,0.1426,1.802,3.826,0.5119,0.09574,0.2788,0.05721,4.626,11.15,1.598,9.335,1.076,0.2079,0.4549,6.872,0.1313,0.5536,1.874,1.217,6.176,3.513,0.01152,8.922,5.99,5.195,5.084,1.101,9.352,1.129,10.33,6.389,11.1,8.423,6.88,9.764,2.825,3.407,3.233,1.305,2.752,0.05759,9.241,0.5807,8.479,4.53,0.1075,4.449,2.695,9.591,0.4911,0.01832,6.037,3.985,10.16,10.75,4.561,0.0441,0.4565,8.532,1.825,0.7337,4.985,7.596,8.647,1.421,5.622,8.97,3.86,8.801,0.9288,0.5754,4.02,0.8461,6.828,2.854,0.01974,2.025,7.603,2.264,8.468,5.202,0.5264,2.714,7.947,0.2737,6.848,7.5,4.165,1.589,1.722,0.1413,0.2924,0.3242,8.059,7.372,0.0001532,0.02843,3.128,1.319,6.294,0.07618,5.283,3.11,2.715,1.299,0.1418,2.473,0.1151,1.933,1.832,4.704,1.565,3.221,1.96,0.1475,3.244,0.8248,1.771,0.228,11.24,0.2952,0.3043,0.329,2.871,0.7576,2.745,0.2381,11.17,1.661,1.551,2.979,1.13,8.079,1.366,0.4628,0.03799,2.332,5.473,1.971,0.6156,7.312,1.63,6.087,2.651,0.8093,3.365,0.2536,3.498,0.129,6.988,5.119,5.391,1.629,10.58,2.832,2.045,2.743,0.8768,0.03974,0.2499,0.1629,5.928,8.002,2.906,0.005773,4.792,3.131,5.357,0.4502,0.3003,6.055,2.082,0.6789,0.3157,0.07284,7.864,1.103,10.49,9.834,0.09915,1.225,0.02488,0.2235,1.481,9.393,0.285,5.608,0.5055,0.04483,0.8775,8.007,10.49,0.9535,0.3691,0.91,0.3738,1.518,2.32,4.123,1.584,0.03172,0.6478,8.067,3.527,1.196,0.3595,0.07469,5.658,2.849,1.654,4.459,6.77,0.08968,1.914,0.01613,2.044,3.851,0.006585,11.68,2.277,1.464,3.15,0.169,0.6792,1.005,3.892,4.381,4.931,3.367,0.8191,3.35,2.605,0.6515,5.753,0.00561,2.551,0.2628,3.515,2.536,1.066,8.678,4.546,1.793,1.991,2.227,1.368,0.7637,0.7411,11.04,0.4529,9.658,5.384,0.6382,11.39,1.946,0.05423,0.9164,4.078,1.773,9.064,1.974,0.658,2.781,1.262,0.8824,0.02135,0.5611,1.776,7.565,2.937,4.278,6.888,1.623,7.775,0.6936,2.282,0.07932,0.9859,0.05171,0.08977,2.027,0.02552,5.538,1.803,6.765,1.522,6.265,0.1347,11.37,2.739,2.687,0.1047,2.569,10.32,3.021,2.158,1.854,0.4214,6.303,6.026,10.45,8.99,3.743,9.956,0.002787,6.25,4.766,0.06938,0.8835,6.539,3.334,2.751,4.46,2.903,2.887,0.2595,2.456,0.4282,4.181,3.718,7.353,5.054,0.05745,0.8228,0.03186,0.9654,6.904,0.0823,0.3213,1.121,9.618,2.435,8.541,6.666,6.351,5.402,1.884,1.05,4.062,2.363,3.497,1.107,1.45,1.565,2.347,1.351,1.471,0.6278,3.13,0.8203,1.241,6.556e-05,4.532,0.09432,6.289,1.395,0.296,0.7721,0.02535,2.594,0.0436,5.029,2.805,8.845,3.891,1.169,0.3896,0.002109,0.04667,0.02602,5.554,10.11,2.065,7.673,7.9,0.9793,3.426,1.102,1.626,1.151,2.497,4.153,2.332,1.832,0.01591,0.03593,0.09591,0.324,2.213,0.1221,3.183,1.445,6.226,1.562,3.491,1.581,6.02,10.5,4.087,0.7614,1.943,3.614,0.6331,4.627,4.254,1.495,0.2872,2.202,1.623,5.221,3.975,6.182,1.299,2.774,5.69,0.2019,0.01287,0.7382,11.34,10.81,3.612,0.01395,5.099,0.6799,1.316,0.1972,11.14,11.47,9.424,0.5793,2.937,0.9854,9.127,0.8513,4.489,2.251,0.0001195,7.311,0.4972,5.653,3.606,0.05619,4.386,3.712,2.602,1.686,2.933,4.247,0.0337,8.817,9.744,0.05103,5.13,3.991,0.186,3.224,10.01,0.142,4.896,0.04504,1.824,6.035,9.387,3.446,2.043,10.89,7.084,3.775,0.5171,2.21,3.097,0.09195,0.9426,5.627,0.7654,0.05859,0.3885,0.0846,7.635,0.4475,10.22,0.3728,11.34,3.31,1.281,1.987,0.7981,9.731,7.574,5.339,5.773,11.49,4.478,11.47,4.39,10.14,0.07909,11.47,5.205,10.83,4.621,0.1561,1.497,0.1599,1.765,0.3838,7.952,2.053,0.01724,11.55,4.536,8.032,3.377,1.436,0.1842,0.5001,3.446,4.184,6.105,0.9665,1.067,1.878,11.43,0.0481,4.252,8.711,11.32,0.3655,4.408,0.4069,0.7932,2.811,7.212,0.009082,3.101,1.383,0.116,0.4258,11.54,0.1454,7.803,1.591,2.367,4.685,0.5075,1.343,1.724,7.822,0.3216,2.882,9.936,4.789,3.036,0.3339,3.898,7.779,9.645,3.427,2.191,0.9842,0.08424,9.893,0.07732,7.739,2.304,0.3844,4.211,5.773,0.2241,1.447,1.012,8.079,10.65,2.552,7.541,3.093,6.131,8.857,3.444,0.7456,0.5179,0.4853,4.86,0.8547,0.4377,1.734,0.5079,2.23,1.444,2.312,2.157,0,2.509,0,0,2.71,0.2039,2.038,11.07,1.512,0,1.077,5.417,0.1837,6.095,3.333,5.971,1.446,0.06613,0.1747,3.069,4.031,1.637,9.265,0.8994,0.3654,3.883,0.4122,2.666e-06,3.472,0.1209,0.001891,7.739,0.4194,3.024,0.03109,9.213,0.3105,7.738,4.073,4.889,8.351,4.183,3.431,0.7362,2.148,0.278,0.3789,0.04074,0.0668,2.781,0.002734,7.081,0.001357,2.828,0.6287,4.044,3.09,2.045,9.68,1.795,3.266,0.5475,10.59,0.3739,0.7806,2.911,7.567,4.743,0.7278,10.97,9.925,10.75,0.3867,0.5444,6.237,0.01332,10.53,1.168,0.1416,3.368,4.666,0.5324,3.457,0.1231,2.774,0.9966,0.9988,1.283,0.6984,0.5205,0.4399,3.759,2.071,8.327,0.1749,1.631,0.09708,3.168,6.281,3.569,5.295,8.413,0.1328,2.09,0.9099,9.86,0.5305,0.3622,8.179,0.7137,6.936,1.51,8.639,0.7765,3.55,11.4,4.286,9.787,0.1545,0.638,10.23,2.015,0.3812,0.03299,4.433,1.796,0.1478,0.5583,7.238,0.5449,2.426,0.4085,1.495,3.92,2.093,2.318,3.296,0.02103,4.677,9.689,1.606,1.182,10.68,0.1274,6.128,5.91,0.06236,10.33,4.452,8.507,0.5876,0.1553,5.668,3.684,1.184,6.129,0.2548,2.63,0.1477,0.6646,0.4179,0.5781,10.44,2.384,1.011,2.758,0.3528,3.294,5.555,0.5441,0.3529,3.17,6.107,0.00703,0.6833,0.09381,0.8769,8.133,9.454,5.408,0.3839,7.623,3.666,10.54,4.22,5.245,0.277,0.5566,0.1652,3.422,1.305,4.063e-05,3.009,5.713,0.8151,0.1199,6.637,3.979,1.863,0.4582,6.563,0.01182,10.11,0.2173,1.142,9.86,4.881,0.4651,0.05681,2.713,6.732,9.316,4.155,7.808,0.6373,5.003,0.01988,1.43,6.683,6.547,0.0007465,7.248,3.676,2.995,0.01326,7.749,0.5894,0.09431,3.948,0.644,7.015,6.613,0.02081,1.788,4.9,0.5683,1.457,0.1339,0.7755,4.648,1.275,0.02174,1.76,2.567,7.311,2.253,1.787,7.043,5.178,4.096,4.976,2.312,0.3374,0.8211,1.625,0.4876,9.475,2.513,7.899,3.039,0.2866,2.628,1.946,0.239,5.4,7.214,8.941,4.507,0.7815,0.02535,1.005,1.409,1.436,9.146,7.159,7.256,7.446,9.296,3.592,0.005062,0.1641,1.341,7.623,3.713,3.745,0.6722,1.59,5.439,1.612,0.3505,1.79,0,0.01074,1.686,0.6079,0.07953,0.4142,4.281,4.488,0.1326,10.58,8.559,7.673,0.2555,8.08,10.74,7.355,9.473,1.606,0.187,2.896,0.4415,0.09163,7.688,0.3211,0.6461,7.209,1.701,4.845,9.417,6.774,0.5186,3.704,4.519,1.233,3.521,11.06,6.483,0.4624,1.183,10.89,6.208,0.02934,10.58,1.904,0.7478,3.325,6.755,4.123,3.682,3.704,11.57,6.332,9.359,0.1254,1.273,0.4948,8.187,0.5002,7.322,9.747,0.748,9.812,1.173,1.447,11.53,9.769,10.93,0.7256,1.055,3.302,0.7179,0.0002929,1.804,0.007476,5.462,10.59,1.052,2.32,4.076,0.404,9.256,0.01472,11.39,1.895,0.6503,2.641,4.411,9.945,0.1242,6.25,8.753,0.00483,1.734,0.07288,2.514,1.194,0.004616,0.02412,5.477,1.152,1.965,5.337,2.793,3.622,10.16,4.122,0.2459,0.7929,2.639,0.4281,2.295,0.2035,9.862,5.745,9.229,0.1805,8.958,11.29,1.28,4.977,4.874,0.9157,9.958,2.064,0.8496,2.702e-05,9.575,2.653,3.476,0.3079,1.902,1.255,10.43,4.986,4.14,0.438,0.06616,0.1487,0.7759,0.001164,0.2625,0.0748,4.816,0.01663,1.786,0.6464,3.767,11.17,8.248,3.768,1.64,0.2235,9.535,0.194,5.005,1.965,3.33,3.105,3.039,5.057,11.51,4.375,10.88,5.111,4.173,5.298,6.762,0.1557,7.288,9.211,0.0034,5.147,1.041,1.93,4.774,2.925,1.946,1.593,3.715,3.635,7.34,0.007882,0.1025,7.184,2.417,2.04,0.1197,0.1074,8.839,2.468,2.187,3.168,1.658,2.205,0.221,1.427,3.064,0.2713,2.8,0.003717,1.555,1.424,2.118,0,1.712,1.873,2.691,4.891,0.0001015,1.004,2.528,0.7284,0.01164,10.72,9.967,7.04,0.0455,4.559,0.2517,8.731,6.825,0.3913,0.01926,1.536,11.61,3.035,0.001374,0.2306,9.576,7.517,0.105,4.7,0.233,0.4888,4.986,0.05387,0.08006,4.741,3.791,4.644,0.6867,6.055,1.594,8.696,0.2311,5.656,8.644,0.4305,6.408,3.922,6.453,0.1285,0.7027,0.3612,1.939,7.553,6.062,8.621,5.56,0.2147,3.192,5.557,0.03604,0.07281,0.1369,11.4,4.053,0.7985,9.112,11.15,0.7774,1.104,1.157,7.358,6.002,1.186,7.078,0.3934,10.24,3.391,5.837,8.706,1.773,1.014,3.778,11.2,7.42,1.978,0.009816,9.773,1.931,9.885,2.253,0.2438,8.073,8.835,0.07468,3.59,11.41,0.3749,11.09,9.068,0.9538,5.007,2.367,7.853,3.134,0.03233,0.03943,0.0195,6.728,3.564,0.6517,1.904,0.3546,8.041,5.143,11.53,0.04561,0.3749,6.37,10.98,6.802,11.42,0.07513,0.443,6.513,8.184e-05,0.3404,9.632,8.498,7.176,3.48,0.2617,2.943,7.109,2.673,6.734,11.05,1.126,10.34,1.639,9.211,1.941,0.06002,0.09648,2.994,2.756,9.159,1.631,1.469,0.01196,7.89,10.24,1.563,2.877,7.513,0.9746,0.131,4.617,3.866,7.677,10.16,0.8385,10.59,0.07492,0.000994,0.687,2.389,0.07092,0.3721,4.514,5.404,4.114,1.258,0.808,0.7658,2.599,0.001931,4.133,7.663,4.993,0.07343,0.5398,1.875,1.199,0.8607,3.236,0.5746,8.871,5.08,3.276,6.809,0.7855,0.3395,2.107,4.766e-05,1.605,0.2625,6.032,0.8196,4.607,6.148,2.954,0.01006,1.202,0.7904,0.1098,0.3057,7.353,7.798,0.1619,9.063,1.581,8.319,0.3209,11.21,8.996,0.8673,4.008,3.216,2.671,2.768,7.843,7.264,3.979,3.365,0.3452,1.061,0.545,0.05951,0.5227,0.5375,0.1968,7.778,4.542,0.001008,5.846,5.176,1.313,3.772,0.02915,1.514,2.968,11.17,4.347,2.097,0.8496,4.755,11.05,0.3393,7.715,2.686,0.7828,2.782,2.21,0.3932,1.433,1.878,2.887,2.308,3.731,1.132,0.9205,0.01151,1.956,0.3196,1.662,5.629,2.25,0.3048,0.03454,1.208,8.521e-05,0.5268,1.113,5.033,0.8651,0.1569,4.064,11.23,6.274,2.213,4.016,1.818,10.31,6.197,5.855e-05,0.02058,0.836,0.007918,0.3208,3.51,6.993,0.04741,0.5284,0.8364,0.746,0.8495,1.584,3.861,2.34,1.752,1.776,2.547,2.235,0.08571,7.895,1.472,0.2485,2.355,4.757,5.464,10.64,5.533,9.95,0.3622,0.1484,2.659,6.669,11.63,11.37,4.511,1.233,0.6469,5.008,8.197,6.985,0.5411,6.113,7.918,10.02,2.347,2.953,2.533,3.132,1.902,6.019,4.585,11.45,4.319,0.02446,0.09366,1.873,0.2255,1.532,0.6655,3.241,0.4209,0.9793,0.03654,0.4323,6.097,0.7785,4.621,5.553,1.958,4.639,2.861,0.8821,0.2788,0.5789,7.614,1.01,9.791,7.58,2.594,3.674,0.02425,2.672,0.2319,0.0004168,8.626,7.66,3.533,11.01,0.1143,0.1466,10.62,10.35,0.1279,1.064,2.661,0.2226,4.125,5.81,0.4526,0.7536,6.21,2.319,0.04993,4.879,9.064,0.06082,6.282,0.1346,1.123,0.05026,7.725,1.908,8.03,1.361,5.966,0.01472,2.128,2.604,0.3803,7.17,0.7229,10.97,6.213,0.475,0.1073,0.2443,0.716,1.881,5.751,11.69,0.9051,4.86,10.67,0.877,5.738,5.287,0.1407,6.896,0.006829,0.09511,0.09186,4.403,3.349,2.574,1.62,3.992,2.324,0.006487,8.134,7.918,1.806,0.002639,0.198,1.76,7.641,0.03931,0.05796,0.8353,4.732,6.743,1.463,0.5774,3.121,6.439,0.08707,5.651,0.000143,0.4983,1.814,4.654,2.442,0.013,0.6421,8.581,2.888,0.07236,1.694,8.695,2.014,0.2464,0.04699,1.135,3.076,1.034,0.0008156,5.299,0.6896,11.01,3.194,1.591,0.05728,1.127,0.4061,5.257,3.101,5.236,0.2234,5.728,0.2816,1.405,0.02174,3.861,4.226,1.581,10.83,1.569,10.13,1.172,0.1134,2.973,7.002,6.144,8.598,1.587,5.406,10.28,6.673,11.36,9.656,6.054,8.198,9.422,2.45,2.749,1.122,10.2,1.935,3.634,0.3559,1.012,1.109,11.02,10.39,10.07,0.09559,2.707,0.01914,0.9534,9.31,9.473,4.588,0.3748,9.433,8.469,4.64,1.655,2.008,2.614,8.64,3.169,1.272,2.062,4.985,0.3208,0.9673,3.598,2.517,4.678,5.521,5.654,2.259,0.564,1.455,0.4473,4.493,3.993,4.711,1.482,1.446,7.398,0.03597,0.5781,1.559,4.214,2.748,5.444,0.2634,0.2696,9.266,0.3894,6.588,1.083,9.396,2.336,7.116,0.6187,3.343,4.35,6.683,0.8218,0.02737,9.299,6.603,9.809,4.614,8.434,1.547,0.1275,4.984,10.06,2.811,0.108,2.151,0.1354,0.6643,1.692,0.08098,0.09721,4.984,0.05732,7.684,0.1767,1.922,0.1433,0.1322,5.607,0.2853,6.322,4.606,2.918,1.239,4.814,3.711,6.068,1.127,0.3603,3.405,8.359,11.35,4.707,2.137,0.5716,3.467,4.333,1.034,0.002837,8.522,0.5828,6.107,3.531,2.625,0.08002,6.968,3.1,0.4342,6.948,7.853,3.539,2.552,0.01342,0.961,1.047,9.924,3.431,0.8545,0.0008553,3.197,2.883,1.227,9.508,9.66,5.071,5.062,7.381,0.674,9.99,10.1,3.26,5.449,7.477,0.3191,0.2487,0.9908,1.313,0.209,4.171,8.112,0.8128,4.189,1.728,0.02173,0.03738,6.319,0.06815,0.9554,0.01856,8.274,2.506,3.198,7.15,7.086,2,2.091,10.8,2.121,6.796,11.22,1.216,3.919,7.257,5.276,6.554,4.456,10.82,0.1464,3.544,0.1441,10.55,5.572,5.634,6.973,9.849,3.129,0.1175,4.29,1.192,9.412,10.13,3.292,1.78,3.643,0.7183,5.814,3.33,1.862,1.057,0.4792,1.304,1.647,1.489,1.529,2.704,11.03,0.06352,1.903,2.107,6.46,8.533,7.922,0.4591,0.2635,2.22,0.4098,0.05468,10.7,3.011,0.6602,8.928,7.268,0.009592,6.952,5.594,0.6797,8.198,4.79,4.522,0.1121,0.9753,0.1744,11.2,2.358,3.805,10.63,0.7726,10.26,0.01952,7.685,5.658,7.602,0.001804,4.182,2.14,5.562,0.9996,1.755,2.825,2.238,9.534,4.383,0.5073,0.2574,1.463,11.68,4.219,6.821,2.342,2.283,1.205,11.14,1.614,0.1665,11.41,3.903,0.7644,5.382,2.189,4.778,4.687,1.789,3.346,1.497,3.87,9.94,0.2901,8.319,0.0008827,8.963,8.08,0.6819,2.177,0.4303,1.316,7.698,4.366,1.395,0.5687,3.132,3.156,0.4568,5.385,10.22,2.245,4.014,0.2693,0.0003667,1.005,9.385,6.439,8.614,4.036,0.05721,2.891,5.709,7.125,2.177,3.614,0.0004713,7.144,10.68,5.923,0.001427,10.07,4.675,5.809,6.713,1.794,11.67,2.613,0.8083,0.02847,5.476,7.496e-06,1.584e-05,4.386,3.568,3.261,0.8575,0.6652,0.08664,7.852,4.397,3.028,2.518,8.713,0.2726,10.56,7.201,5.748,2.649,0.9491,10.77,8.418,6.685,0.01822,0.02147,0.004001,10.54,5.529,4.146,7.966,4.593,2.006,1.988,2.636,3.697,5.406,3.291,1.17,2.292,11.25,0.06191,3.233,0.6866,3.978,0.2823,0.9569,2.254,4.647,0.5444,6.681,0.2733,0.01889,4.195,10.76,3.068,6.44,7.213,4.747,5.003,10.41,1.459,3.692,7.996,0.7689,0.115,10.49,2.263,0.2221,10.67,0.2613,0.6536,4.054,0.03264,0.9171,3.657,2.316,0.1061,0.0007106,8.831,3.913,0.9904,0.9903,0.8097,0.1836,0.5881,8.787,2.059,9.835,5.312,0.03837,0.0006505,0.04505,0.6359,0.477,8.15,6.603,8.047,5.496,9.291,8.693,9.085,2.63,2.976,2.725,5.633,8.568,11.24,5.878,5.255,3.362,2.158,0.004519,5.866,0.1278,2.182,1.522,8.997,0.3219,1.485,3.29,3.651,0.3528,0.5317,0.3524,2.092,0.7939,3.108,4.586,4.499,0.7415,4.792,6.199,0.01597,1.104,4.831,4.226,8.341,2.469,4.22,0.1771,3.082,0.6272,1.109,1.856,1.28,0.2638,0.4324,6.287,10.89,2.792,1.616,1.464,2.168,0,3.699,1.935,1.814,1.987,1.548,0.1959,1.339,3.772,3.92,2.941,0.0754,3.656,1.436,1.252,0.2783,2.478,2.661,0.8958,5.14,5.169,8.84,0.3246,5.504,0.6077,10.94,10.89,0.09153,10.74,0.04175,7.951,2.847,1.231,0.2372,0.8113,5.816,0.2874,9.767,2.168,10.91,0.831,5.401,0.1458,3.593,3.403,10.05,10.25,8.042,3.716,3.569,1.738,4.035,0.4669,0.5795,1.021,0.2827,0.2857,2.016,4.132,0.000826,2.375,2.884,3.028,3.272,0.5258,3.301,0.2466,0.542,4.901,9.035,4.362,2.844,10.16,8.097,4.9,0.1517,0.001022,0.2217,0.114,2.726,4.531,5.578,9.317,0.0001205,11.39,5.552,0.7234,1.742,8.964,8.399,3.688,3.892,0.6428,5.559,2.851,0.4803,9.341,0.4865,6.99,0.642,0.4453,7.795,0.3243,0.00591,0.04943,0.5866,4.124,0.7019,1.193,2.244,1.912,1.51,7.889,5.828,0.102,0.01907,0.2593,1.346,0.009968,1.343,7.349,0.7676,0.002948,4.235,1.157,0.06899,3.19,0.1083,0.1229,5.155,0.7856,1.952,10.69,5.566,0.3607,5.811,0.02171,0.3058,2.279,0.09536,0.7159,2.742,0.5723,0.9667,2.556,5.067,0.4713,0.1796,0.008402,8.486,7.243,0.4842,3.307,0.5949,0.006758,0.02651,2.609,5.098,2.683,10.86,4.376,4.974,0.1056,9.254,2.33,0.9826,0.73,0.6575,6.648,7.001,9.451,1.341,3.496,11.33,1.781,2.661,5.515,8.734,1.428,3.653,0.7926,10.39,4.183,0.9877,1.143,8.02,3.804,9.137,10.63,0.6041,0.1677,7.784,0.01045,0.1984,8.384,1.964,3.224,0.2831,3.251,2.291,0.6139,10.05,5.686,8.524,7.861,0.1918,6.932,8.498,7.614,1.021,4.617,1.673,3.99e-07,6.211,0.3401,7.112,0.4248,5.419,0.36,1.384,0.0242,1.335,0.09941,0.08281,0.08308,6.596,2.775,5.212,5.66,2.431,0.2741,3.274,2.567,9.257,3.095,3.618,8.457,10.13,1.12,0.05665,2.87,1.953,2.51,4.387,0.3244,8.418,0.8481,3.626,10.19,3.868,1.853,0.03982,0.0987,0.8497,0.2971,1.932,2.872,0.04307,1.088,4.949,3.82,7.429,7.892,2.693e-05,0.0008546,11.69,3.186,4.942,1.478,0.1546,0.4632,3.724,0.6854,8.724,4.771,0.3135,0.9049,1.942,5.623,7.936,1.094,1.915,1.415,2.855,3.085,5.61,4.109,1.168,0.7516,0.4446,6.227,0.3137,7.792,4.11,7.374,0.2338,3.783,10.83,1.822,0.7519,1.022,4.867,4.957,7.521,0.02698,8.246,2.839,0.07694,3.704,5.146,6.357,3.749,2.213,2.934,10.33,5.141,0.2243,7.282,0.4048,4.866,9.256,7.177,1.781,7.654,4.008,7.3,0.9207,4.319,1.901,0.8176,3.146,0.03177,1.053,6.978,6.78,4.965,0.8213,8.032,0.04368,8.125,1.562,0.7958,0.4167,0.04834,6.635,7.929,1.205,11.12,0.9504,9.419,1.752,4.633,3.326,1.252,0.9595,11.16,0.4972,3.288,0.8711,11.15,0.4309,9.689,0.07183,0.7094,6.69,0.6442,0.2781,1.27,4.194,3.491,0.9342,3.293,8.762,0.3088,0.07708,9.205,0.3796,0.316,10.14,3.825,5.311,7.642,5.952,9.158,0.7773,0.169,2.54,11.3,2.591,4.151,4.125,0.7005,0.197,1.784,4.507,2.886,2.277,0.1124,5.161,4.999,2.277,10.16,5.842,4.869,0.2176,2.309,0.68,4.409,7.674,11.27,1.782,4.431,0.008404,2.043,1.548,4.071,1.842,4.164,0.7486,5.019,2.484,1.07,2.797,2.047,2.59,1.95,3.432,2.077,1.829,0.6279,0.5238,0.6759,3.35,1.129,3.372,0,2.27,1.227,7.774,0.5343,13.02,14.95,0,14.1,24.78,35.14,17.22,17.44,12.57,21.36,43.75,14.27,50.92,194.1,23.06,22,38.17,13.05,64.36,22.99,16.11,54.06,17.26,12.53,40.16,20.31,42.64,95.78,27.14,34.39,35.04,40.41,18.22,22.35,13.18,20.24,21.87,13.4,18.46,19.52,17.12,71.56,17.1,45.68,12.62,94.94,24.8,20.26,20.58,16.58,59.04,19.59,17.86,28.67,49.57,89.27,119.6,41.25,65.39,24.79,57.85,40.4,40.39,20.99,18.41,17.32,34.96,95.4,21.87,17.54,16.42,16.37,72.4,22.37,16.74,50.55,119.8,58.09,33.9,56.72,20.03,26.91,27.96,53.55,26.72,33,30.29,16.77,14.5,12.94,21.66,12.77,134.4,53.16,33.85,29.45,26.24,28.96,55.09,22.39,73.77,48.27,18.43,45.72,29.51,60.89,13.06,43.94,52.11,23.27,27.28,48.58,59.94,13.01,12.51,49.2,23.02,24.55,21.56,37.03,65.7,94.13,47.74,18.95,61.62,15.13,153.8,66.41,88.91,17.48,20.82,32.65,64.84,107.3,19.88,32.63,55.92,65.11,14.28,45.48,27.27,36.24,67.95,48.84,48.46,16.1,93.36,30.24,20.99,34.04,25.51,29.51,13.03,41.49,16.99,17.09,27.6,26.86,38.07,12.17,12.52,15.99,17.73,44.42,13.6,15.21,33.96,21.99,52.42,12.05,24.99,60.96,23.72,23.6,17.05,17.89,48.99,57.21,11.71,27.79,20.24,48.53,25.15,96.26,125.8,51.47,14.01,33.32,21.13,21.51,42.4,69.25,51.46,25.94,14.13,77.33,14.87,111.2,92.53,66.75,13.89,19.12,27.48,43.94,46.51,18.34,14.15,30.23,85.23,22.44,65.65,12.27,15.5,29.75,14.76,86.93,20.6,44.57,12.68,21.4,29.36,45.95,28.29,64.4,14.56,30.8,16.92,13.43,34.04,29.57,22.96,20.71,27.73,34.36,108.7,15.76,20.01,92.94,46.65,17.54,17.48,13.26,45.86,30.5,51.86,31.66,14.85,18.14,62.54,31.81,20.94,17.01,115.1,12.3,47.24,27.44,54.32,28.54,12.95,25.75,20.08,18.46,54.56,46.95,14.98,67.25,31.22,71.25,42.27,28.18,25.88,58.43,12.62,35.7,47.61,12.98,129.2,21.5,13.17,28.47,22.6,114.1,17.46,40.22,15.04,12.41,37.19,21.1,17.85,85.82,98.41,18.75,17.38,18.42,28.64,15.13,64.48,15.9,76.8,20.28,18.75,52.46,34.66,17.79,36.13,13.61,28.54,17.64,12.83,27.07,25.81,39.41,33.06,39.87,20.38,41.23,63.68,25.03,14.16,33.8,12.51,21.32,12.71,143.8,43.62,12.35,21.51,19.2,19.43,12.69,26.05,18.47,13.62,43.73,23.78,27.21,21.18,20.06,25.24,14.94,72.2,45.6,16.49,33.64,12.52,12.03,12.01,24.42,15,20.51,15.14,29.06,14.98,52.84,18.35,57.73,33.2,37.02,65.9,14.34,33.56,27.8,37.92,73.55,54.85,20.33,31.16,22.81,14.58,48.28,17.61,29.17,18.85,20.87,31.61,74,18.18,39.23,12.74,34.58,13.2,117.4,92.04,22.23,16.74,92.92,32.28,27.63,28.25,12.77,32.44,30.65,19.7,33.96,12.89,63.35,57.81,19.83,47.24,22.4,13.8,13.4,13.29,35.55,26.36,12.75,25.51,29.23,25.76,97.16,23.78,32.22,37.71,26.24,19.68,35.29,12.45,26.66,17.97,86.58,73.65,14.9,46.96,12.57,36.57,15.75,14.67,14.96,40.03,20.87,40.67,19.94,13,32.61,23.21,47.27,39.47,67.97,20.45,27.27,29.22,20.54,35.75,56.02,48.23,53.51,16.12,34,41.68,31.89,34.38,33.18,11.77,60.06,13.85,12.89,118.5,32.89,66.46,14.88,25.31,43.3,32.5,156.6,56.3,40.37,63.38,53.34,61.79,51.32,19.95,19.32,25.73,80.91,23.49,11.85,13.89,26.85,39.33,50.09,17.65,49.06,26.44,23.83,20.73,55.39,80.35,18.73,18.14,14.52,42.59,24.21,15.66,31.59,19.81,17.27,27.27,70.21,40.15,72.42,64.69,17.8,28.45,25.31,35.58,32.16,15.03,20.15,96.73,34.2,40.73,13.47,12.27,23.01,38.25,78.7,71.61,25.16,13.68,14.96,11.71,22.68,57.32,41.1,22.77,45.86,61.82,43.65,44.22,19.81,78.24,48.12,22.54,18.29,15.5,37.14,20.48,18.17,43.49,20.92,29.86,12.72,53.7,42.13,29.35,53.89,44.47,16.03,18.69,23.69,105.3,72.43,14.29,26.84,18.79,23.6,20.88,14.01,50.91,26.8,33.48,30.12,26.62,64.84,26.26,16.84,17.19,71.33,37.87,52.73,28.33,36.23,52.28,113.4,22.18,52.61,40.04,35.05,26.1,27.27,29.41,16.12,20.64,31.8,56.63,40.98,27.55,25.43,48.26,16.5,20.67,26.01,19.19,19.92,21.34,21.07,21.89,22.22,18.08,43.34,14.3,37.89,26.91,14.72,18.21,20.87,29.82,36.82,12.96,89.06,20.62,47.27,21.32,85.31,32.98,56.99,24.36,23.21,22.54,19.38,12.55,45.09,20.73,33.94,21.58,50.67,20.46,98.85,17.55,24.8,12.21,14.7,17.26,19.54,72.59,33.6,82.32,76.81,29.65,22.97,23.59,124.6,12.84,13.94,24.82,27.86,28.78,17.28,28.03,29.21,26.54,51.68,33.42,24.37,107.4,30.34,15.91,13.56,14.81,14.05,100.2,40.46,22.9,29.45,14.25,35.48,68.27,46.63,24.11,46.31,17.31,31.89,23.02,11.86,33.82,19.57,16.77,39.1,24.17,30.97,29.96,49.08,86.05,48.95,16.68,43.07,21.85,15.41,14.4,14.64,63.42,16.28,12,14.57,23.39,13.24,39.37,17.02,33.28,13.04,24.13,18.11,15.06,31.39,43.37,18.02,70.48,13.05,47.58,14.2,15.75,26.48,30.25,13.13,27.5,24.54,13.03,50.55,13.78,34.51,44.12,30.09,70.82,20.65,12.39,60.21,14.65,12.5,14.29,31.47,18.79,14.44,14.43,13.09,67.63,22.79,78.7,25.79,22.06,49.12,78.99,18.31,22.1,28.7,26.11,24.12,35.49,26.02,98.92,48.84,85.03,91.57,14.3,16.47,66.84,20.51,79.9,63.03,17.97,14.52,19.08,29.18,19.92,87,39.37,47.36,20.02,62.29,22.37,15.3,24.41,14.34,24.12,16.44,16.62,21.62,30.35,11.82,14.72,18.34,15.61,34.53,25.56,155.4,69.32,43.07,23.44,14.25,75.91,48.59,17.14,15.06,54.2,16.89,26.15,33.58,19.03,42.85,13.09,32.14,97.27,13.99,35.59,14.68,28.04,20.31,40.06,26.29,22.64,23.3,13.13,27.63,35.68,43.96,127.8,63.46,12.45,12.64,18.51,27.5,12.89,189,48.76,22.99,53.08,33.37,30.4,16.68,23.6,119.4,56.5,92.99,36.58,14.84,14.2,25.87,39.45,39.91,16.34,40.23,15.07,13.37,26.31,70.68,42.23,21.82,20.73,77.03,71.34,25.69,49.09,20.26,25.46,22.64,18.86,18.67,23.27,14.49,11.81,15.71,23.14,22.19,98.5,16.03,16.62,12.87,32.55,13.26,27.65,94.78,15.16,13.87,50.62,34.28,75.56,22.75,69.5,19.85,14.62,15.15,80.84,20.43,21.97,12.45,45.3,78.32,34.04,28.9,18.87,14.44,31.67,12.27,20.58,82.38,37.66,23.56,12.75,11.77,21.36,31.95,22.1,47.27,33.5,12.51,16.7,54.66,68.76,25.87,13.81,22.23,32.81,16.31,23.97,19.1,14.78,45.08,15.6,13.05,38.55,18.75,17.57,12.9,15.83,12.05,13.24,12.71,26.02,60.65,16.85,59.16,13.28,12.76,51.83,68.29,27.9,15.36,49.76,29.93,110.5,15.66,15.36,17.22,15.7,24.24,13.11,23.47,23.17,37.73,20.66,25.7,41.49,30.23,86.85,31,23.3,36.18,13.55,32.25,55.66,13.99,14.86,13.16,18.97,25.72,12.51,114.6,14.07,26.46,38.14,23.16,76.94,21.42,14.45,26.32,25.46,14.06,32.38,22.66,40.83,36.4,13.14,39.89,34.53,139.4,18.69,37.89,48.62,15.87,15.11,49.68,11.87,55.3,43.46,22.05,12.81,16.99,24.21,61.7,31.2,33.22,24.61,25.88,18.22,17.2,103.5,43.45,14.16,35.71,29.28,35.79,12.32,43.21,44.4,13.58,22.74,17.54,25.25,26.91,17.15,29.48,48.03,15.63,19.72,33.2,39.86,71.21,13.94,12.05,17.44,57.82,15.82,25.7,42.76,15.54,82.14,25.97,31.08,19.05,41.52,71.95,16.62,17.39,22.25,31.12,35.38,24.13,47.76,12.27,54.31,33.07,24.9,74.64,26.83,43.9,28.76,37.81,14.96,17.58,59.92,14.95,13.48,19.32,30.44,13.56,19.52,45.15,12.8,16.7,21.02,13.66,26.24,17.22,41.93,55.59,15.48,54.52,43.49,15.01,49.4,54.17,16.4,37.62,11.74,59.36,13.49,18.59,43.16,37.43,12.16,21.34,176.3,33.33,27.85,46.21,79.03,21.4,73.37,26.13,21.75,20.86,34.06,25.94,31.26,13.48,89.38,17.36,17.25,20.05,35.76,44.52,30.08,68.9,41.03,51.58,14.06,24.08,22.55,26.55,38.53,40.76,94.43,12.52,13.09,15.24,13.15,17.85,16.8,34.46,79.53,49.64,53.53,14.21,15.75,13.17,78.5,23.01,13.56,29.42,40.61,71.54,16.95,33.48,11.82,17.51,44.92,79.78,46.25,58.18,40.41,84.8,29.61,122.1,43.88,23.77,36.38,14.16,26.94,51.2,36.51,79.11,22.17,50.98,21.57,33.29,18.18,20.03,44.94,20.76,18.82,32.02,13.86,57.36,34.46,41.03,23.82,24.22,114.4,15.12,39.26,63.76,18.32,17.09,19.49,24.78,40.48,38.47,53.86,16.85,57.52,74.13,30.97,53.67,12.92,80.46,18.67,12.73,46.39,20.77,28.45,20.74,85.34,51.67,35.29,16.2,32.58,24.46,74.77,17.53,26.58,35.88,21.66,46.34,44.01,13.12,18.93,33.33,16.05,69.42,21.1,68.18,34.93,32.17,28.47,21.47,89.91,51.95,15.83,60.01,25.22,16.46,32.98,17.09,18.69,37.58,18.09,28.55,20.72,14,85.97,34.6,23.02,42.42,27.91,41.94,17.69,19.09,36.8,43.68,48.01,36.83,38.34,69.33,96.22,12.75,13.89,11.96,66.12,33.26,73.57,28.22,32.79,65.32,30.16,19.27,44.15,13.02,23.1,42.95,15.89,14.59,12.13,33.02,43.71,24.64,29.28,58.66,37.21,37.94,15.63,17.37,39.43,53.7,26.33,18.9,20.3,15.62,31.47,49.88,15.89,28.66,53.41,26.26,50.58,43.47,12.29,33.06,29.44,18.58,22.72,65.81,86.48,31.99,35.59,50.6,12.54,60.8,16.88,34.1,75.69,53.89,28.04,75.41,27.67,37.52,15.66,46.07,49.85,90.86,46.42,22.69,35.63,37.4,23.1,30.7,21.21,13.44,29.01,51.96,13.79,115.3,13.6,58.66,32.77,11.99,51.18,72.99,20.22,12.91,25.37,38.26,117.4,75.53,20.81,16.38,27.61,11.88,18.55,17.99,17.02,55.11,43.68,105.6,65.9,20.99,29.25,12.35,27.44,17.86,138.3,52.07,23.64,11.83,52.51,15.14,26.52,23.92,37.19,14.11,16.96,22.63,13.12,34.31,37.8,20.73,24.73,25.87,20.17,70.89,12.3,15.28,107.1,28.75,16.19,16.33,32.37,11.86,16.94,24.56,17.04,13.34,15.73,17.31,44.52,16.39,76.74,15.06,51.69,54.33,13.43,36.87,23.22,12.63,22.82,19.1,29.18,24.4,15.69,47.5,19.31,40.07,15.31,69.7,13.73,17.33,24.64,11.74,90.83,13.94,27.18,23.87,12.01,38.75,41.05,36.68,13.56,28.63,22.97,39.39,48.41,36.09,17.03,16.23,22.49,21.93,16.49,19.92,31.32,13.77,67.23,40.39,18.54,29.72,21.83,19.13,25.97,23.13,14.45,16.84,42.11,58.23,12.85,20.16,11.79,53.76,16.52,31.44,111.5,25.15,14.36,23.9,19.74,28.95,47.56,32.21,27.47,70.37,29.7,26.69,38.5,16.32,25.01,85.32,24.75,39.57,12.93,15.87,12.84,34.13,80.25,14.48,19.05,38.92,14.66,22.31,20.51,11.72,63.62,30.33,26.72,34.96,21.46,16.87,89.67,17.93,19.11,34.49,84.4,18.27,25.3,70.03,12.86,24.28,47.61,16.86,14.98,20.72,28.15,20.93,42.34,42.3,24.91,18.58,13.53,57.47,28.16,22.79,16.72,12.72,13.59,28.83,20.67,17.45,11.9,16.91,64.3,23.9,27.05,60.91,12.92,27.48,68.13,14.37,24.93,13.25,23.51,22.5,20.61,27.46,16.82,22.65,40.56,18.48,37.96,14.81,31.27,14.09,16.15,77.92,12.81,19.52,48.96,26.3,49.78,26.9,12.47,87.03,13.33,33.58,32,14.32,19.01,67.53,18.65,12.56,29.06,25.96,139.2,58.58,20.55,40.05,55.3,69.57,44.19,17.84,31.48,12.75,13.41,14.83,36.9,80.61,18.56,13.98,36.57,22.37,16.37,67.91,26.07,21.32,14.37,23.52,22.13,36.23,15.3,28.6,33.49,26.77,28.05,15.84,15,44.52,23.17,22.06,19.52,38.88,57.4,19.95,22.76,26.65,14.94,50.62,37.17,44.16,28.21,12,20.88,16.28,21.9,37.18,39.64,27.23,23.35,26.99,50.64,44.9,33.89,15.92,11.86,86.17,19.59,41.9,17.06,22.48,12.49,82.74,85.84,12.15,16.54,50.5,68.78,29.5,34.51,107.2,33.7,52.27,61.61,26.91,14.08,12.52,18.05,80.37,18.59,20.03,25.3,12.06,47.96,34.61,44.78,14.21,13.62,83.6,15.78,18.61,177.4,22.22,22.27,18.48,55.58,33.87,15.24,74.07,11.97,14.03,13.55,63.64,15.59,17.59,30.31,85.23,28.63,16.04,12.99,29.48,30.2,21.85,24.32,71.62,26.94,50.89,13.67,18.22,19.17,25.35,50.07,32.25,14.6,51.76,29.35,23.5,53.77,28.06,94.28,60.01,33.06,12.09,36.84,14.48,16.31,13.98,22.22,23.93,21.46,21.58,15.7,87.77,30.2,16.36,29.49,20.14,21.34,19.47,52.95,29.19,15.91,33.91,20.48,23.71,42.84,14.37,36.4,12.98,33.03,22.37,29.63,18.7,23.59,37.65,37.8,41.42,29.76,15.57,63.97,42.12,34.75,11.98,49.04,20.51,11.84,173.7,30.34,33.4,41.03,26.37,14.36,34.94,24.22,34.85,52.96,32.13,12.53,24.81,46.03,46.03,39.46,14.85,38.37,28.87,20.84,45.94,31.61,22.58,30.95,12.35,41.87,13.35,54.29,23.01,25.77,21.53,34.71,13.86,12.49,16.18,15.48,42.02,12.6,15.78,40,67.4,16.92,34.19,12.49,22.14,34.75,26.14,50.32,17.07,12.58,22.7,12.31,20.67,12.14,54.17,30.15,11.98,15.98,17.89,21.79,92.99,19.73,23.11,16.66,12.88,71.95,24.7,74.21,38.96,18.32,23.61,15.87,25.32,61.83,37.35,13.91,12.31,22.96,21.66,12.15,63.88,13.57,43.17,87.7,15.77,26.81,50.1,32.5,32.12,17.01,17.43,15.37,40.08,62.64,23.37,28.53,28.82,19.56,30.74,70.92,36.38,23.51,51.6,27.27,11.74,14.42,25.08,95.27,39.82,45.01,13.02,26.66,31.86,19.02,51.23,49.69,26.48,51.66,21.31,32.57,20.36,45.95,17.56,12.39,37.36,28.29,80.85,34.79,13.56,24.16,34.51,17.66,24.74,28.45,45.61,25.38,37.5,20.88,26.47,21.64,49.84,69.56,25.16,20.04,11.83,26.59,24.45,19.05,11.95,14.99,35.94,21.34,37.89,23.49,88.62,23.75,17.89,16.51,11.76,45.12,112.9,45.17,35.12,16.53,26.24,36.69,12.62,41.77,13.67,18.22,12.88,73.96,14.21,21.49,56.32,16.39,17.83,28.63,75.55,17.7,13.47,34.32,34.32,27.03,88.7,26.83,41.36,16.17,29.08,91.88,90.51],"PValue":[0.2307,0.9399,0.2096,0.3542,0.139,0.09707,0.03619,0.1777,0.1167,0.8554,0.09221,0.3826,0.6754,0.1157,0.2508,0.3987,0.231,0.2594,0.1975,0.2036,0.1865,0.1655,0.5491,0.2134,0.1731,0.2778,0.1806,1,0.2708,0.2274,0.1798,0.2245,0.1786,0.1301,0.1455,0.2563,0.02186,0.2695,0.1212,0.1879,0.2122,0.09606,0.264,0.2859,0.369,0.2129,0.1843,0.1585,0.2714,0.1667,0.2683,0.3215,0.1842,0.3343,0.4878,1,0.1437,0.3216,0.005888,0.1997,0.3734,0.1741,0.005907,0.00809,0.1287,0.2858,0.2563,0.007615,0.2859,0.1804,0.1381,0.235,0.05465,0.1584,0.203,0.2535,0.308,0.1298,0.5069,0.2362,0.1588,0.1434,0.004876,0.1693,0.3264,0.1303,0.2612,0.4426,0.2598,0.2093,0.173,0.1719,0.09245,0.1046,0.2055,0.2361,0.2627,0.1736,0.04249,0.1618,0.3214,0.2532,0.2044,0.1703,0.2692,0.0465,0.01756,1,0.1998,0.1446,0.4366,0.154,0.2036,0.1442,0.2118,0.1964,0.7545,0.1287,0.1094,0.03011,0.2514,0.2182,0.2134,0.1822,0.1704,0.2073,0.04401,0.5116,0.2437,0.1994,0.2147,0.2255,0.6496,0.2243,0.2299,0.1378,0.1231,0.775,0.145,0.0134,0.2052,0.2858,0.2859,0.3214,0.1624,0.2216,0.1502,0.1845,0.2774,0.09214,0.006712,0.1754,0.2245,0.3214,0.2083,0.2237,0.2054,0.1944,0.004349,0.2123,0.3136,0.2064,0.1152,0.4363,0.04237,0.1013,0.1379,0.1756,0.1665,0.7459,0.006804,0.2774,0.3656,0.0172,0.1629,0.08673,0.01767,0.03384,0.2435,0.2209,0.1224,1,0.02353,0.3429,0.5096,0.3774,1,0.03786,0.1294,1,0.1797,0.01091,0.07781,0.3227,0.4334,0.954,0.139,0.2295,0.03974,0.287,0.04848,0.1929,0.6299,0.2319,0.5629,0.8715,0.8614,0.02207,0.2712,0.005522,0.005467,0.03541,0.1034,0.007848,0.08822,0.4492,0.5372,0.06367,0.1185,0.01941,0.02996,0.3654,0.005455,0.9109,0.008126,0.03453,0.0228,0.5793,0.008865,0.04914,0.1517,0.07958,0.3476,0.008283,0.1085,0.2717,0.008424,0.0122,0.7261,0.01388,0.3489,0.9812,0.00372,0.004046,0.1941,0.03082,0.01121,0.8034,0.1726,0.04978,0.2401,0.1695,0.07024,0.687,0.714,0.5651,0.3595,0.03375,0.01512,0.7794,0.005374,0.609,0.2116,0.008394,0.05133,0.06245,0.3951,0.03314,0.2743,0.1191,0.06779,0.0163,0.07344,0.01311,0.003934,0.256,0.1366,0.9303,0.01281,0.5161,0.3174,0.07921,0.01219,0.01558,0.4458,0.7986,0.9946,0.9234,0.006555,0.942,0.3112,0.3851,0.856,0.105,0.2281,0.2427,0.2287,0.5663,0.03205,0.0302,0.7894,0.07648,0.9428,0.01472,0.1135,0.3155,0.1523,0.03056,0.0567,0.1725,0.3609,0.007147,0.1365,0.004076,0.1003,0.03927,0.1704,0.194,0.1939,0.2237,0.1324,0.1786,0.2404,0.1377,0.1859,0.5979,0.2165,0.5502,0.06355,0.4933,0.6575,0.00409,0.1635,0.1991,0.09996,0.1059,0.2247,0.1554,1,0.006212,0.2023,0.07182,0.7986,0.0204,0.7213,0.9341,0.01417,0.005186,0.09371,0.1773,0.5384,0.004943,0.1495,0.0414,0.3755,0.1477,0.01491,0.07153,0.01397,0.09394,0.1633,0.01191,0.5837,0.02615,0.8982,0.435,0.07138,0.5339,0.9626,0.7325,0.027,0.1798,0.04177,0.09231,0.414,0.603,0.5146,0.04629,0.03226,0.477,0.129,0.08553,0.09406,0.9085,0.04875,0.3771,0.03804,0.237,0.1517,0.6418,0.07991,0.6961,0.372,0.1167,0.00951,0.6131,0.8189,0.007578,0.0281,0.01098,0.4965,0.05572,0.1448,0.5169,0.3596,0.00513,0.01156,0.004795,0.03304,0.004851,0.03925,0.006526,0.08069,0.4945,0.2593,0.004656,0.2112,0.03583,0.005386,0.2756,0.008592,0.006351,0.2603,0.9548,0.004883,0.1908,0.1542,0.8313,0.868,0.8561,0.1059,0.641,0.08932,0.04538,0.156,0.1549,0.1824,0.2091,0.2147,0.5238,0.1011,1,0.1863,0.1227,0.2291,0.2282,0.1063,0.2014,0.0957,0.8842,0.2237,0.2563,0.1188,0.09309,0.812,0.004342,0.1992,0.05533,0.01857,0.0158,0.004874,0.2454,0.01072,0.009513,0.02618,0.09847,0.01753,0.03083,0.004294,0.4975,0.4601,0.02121,0.1526,0.2541,0.9676,0.003792,0.007604,0.1959,0.004989,0.1571,0.08264,0.2338,0.2656,0.09832,0.0103,0.03832,0.04499,0.1289,0.6327,0.09125,0.4093,0.1722,0.05558,0.2349,0.3461,0.009467,0.02283,0.6275,0.1529,0.01291,0.1385,0.6343,0.02435,0.2331,0.8063,0.2808,0.2499,0.02924,0.1811,0.0914,0.01008,0.04303,0.1735,0.6431,0.0146,0.05638,0.008405,0.5675,0.02479,0.02303,0.09208,0.884,0.02074,0.07057,0.855,0.5761,0.003811,0.571,0.1166,0.04925,0.2686,0.2334,0.3888,0.05638,0.1832,0.2263,0.1599,0.2933,0.4834,0.08038,0.2948,0.02494,0.07176,0.02009,0.6824,0.4269,0.9342,0.3313,0.0828,0.03544,0.1507,0.4513,0.03099,0.5441,0.1799,0.02914,0.02452,0.005081,0.05969,0.05136,0.3843,0.3094,0.5975,0.05576,0.03832,0.1856,1,0.2608,0.3234,0.1751,0.2797,0.5238,0.1068,0.1289,0.1352,0.6811,0.1027,0.009137,0.05189,0.4321,0.2207,0.3641,0.01374,0.06248,0.6346,0.4939,0.004156,0.7035,0.9463,0.00612,0.7693,0.03888,0.01225,0.009314,0.5434,0.05623,0.08685,0.2149,0.745,0.1384,0.2555,0.004071,0.007925,0.3915,0.2161,0.8142,0.006634,0.01184,0.4436,0.01704,0.6479,0.09904,0.08101,0.8382,0.003617,0.2536,0.007965,0.1246,0.06971,0.9146,0.7305,0.601,0.9053,0.694,0.1657,0.04786,0.01828,0.02571,0.01309,0.03299,0.5247,0.02006,0.03184,0.4391,0.2486,0.2209,0.03975,0.0282,0.04305,0.06856,0.8431,0.2169,0.1713,0.3567,0.01652,0.7754,0.005846,0.9505,0.09236,0.01271,0.3252,0.03243,0.2565,0.2577,0.1917,0.1076,0.02008,0.006697,0.1811,0.06632,0.004792,0.2686,0.0677,0.08024,0.2763,0.04188,0.6735,0.0106,0.8106,0.003994,0.3823,0.3018,0.6429,0.1011,0.004768,0.05865,0.06143,0.1501,0.08603,0.1147,0.3497,0.4887,0.9116,0.07705,0.1255,0.1704,0.005634,0.01938,0.03202,0.243,0.2833,0.1811,0.2262,0.217,0.1132,0.01936,0.178,0.2628,0.007047,0.9439,0.4062,0.2593,0.8249,0.7721,0.2561,0.04412,0.162,0.08711,0.03381,0.004088,0.4101,0.6124,0.2104,0.01701,0.7088,0.5915,0.007509,0.04534,0.9141,0.05006,0.1262,0.1486,0.06957,0.015,0.4157,0.5991,0.0261,0.6069,0.09525,0.7268,0.4127,0.0213,0.01245,0.4809,0.06776,0.06488,0.4618,0.01059,0.4056,0.4327,0.1732,0.1163,0.03931,0.06336,0.05932,0.03462,0.2354,0.4247,0.8594,0.2832,0.01004,0.5543,0.2958,0.07949,0.3114,0.143,0.7708,0.7492,0.6452,0.2046,0.006266,0.09325,0.094,0.02592,0.7674,0.0182,0.9843,0.004416,0.009447,0.9566,0.735,0.1573,0.04668,0.8263,0.02238,0.2034,0.5895,0.2224,0.1232,0.6442,0.2001,0.07976,0.01599,0.01019,0.05601,0.2686,0.7503,0.1088,0.03832,0.006989,0.7028,0.1745,0.007151,0.2422,0.5759,0.02665,0.433,0.8279,0.02729,0.5624,0.4814,0.9236,0.006916,0.7209,0.02173,0.6202,0.8056,0.009876,0.006232,0.3125,0.5356,0.03091,0.03115,0.02471,0.008796,0.03303,0.2259,0.25,0.1561,0.1658,0.1695,0.2972,0.971,0.1119,0.8052,0.1723,0.7048,0.1594,0.03141,0.06383,0.01195,0.003834,0.5062,0.03263,0.01712,0.7302,0.02315,0.03463,0.1514,0.3117,0.4356,0.07387,0.9263,0.005982,0.006841,0.7946,0.4688,0.3618,0.01903,0.01641,0.4992,0.00457,0.7631,0.01023,0.3038,0.4314,0.01258,0.4625,0.1084,0.9048,0.9647,0.03418,0.1791,0.005822,0.01139,0.3938,0.1791,0.7598,0.1698,0.04046,0.07089,0.6385,0.5405,0.3117,0.1701,0.05317,0.5908,0.4524,0.02393,0.1318,0.08765,0.2474,0.157,0.4148,0.01846,0.792,0.2127,0.1869,0.3222,0.2298,0.2393,0.2399,0.1243,0.1019,0.03639,0.1285,0.4239,0.5628,0.1171,0.04815,0.8354,0.02622,0.09377,0.04152,0.3255,0.07079,0.382,0.01581,0.1585,0.005011,0.1704,0.7147,0.005758,0.3758,0.008901,0.02537,0.6541,0.9265,0.005137,0.8016,0.3304,0.003609,0.4861,0.005436,0.2603,0.7108,0.1986,0.06854,0.4849,0.7611,0.6049,0.8141,0.04747,0.004259,0.2247,0.007698,0.3154,0.6547,0.5099,0.01875,0.722,0.4679,0.1904,0.2865,0.02466,0.07964,0.9159,0.008869,0.02658,0.03702,0.03883,0.3099,0.007654,0.3041,0.005531,0.02265,0.004331,0.01057,0.01869,0.006666,0.1126,0.08391,0.09146,0.2705,0.117,0.8135,0.007949,0.4573,0.01036,0.04955,0.7474,0.05139,0.1206,0.007061,0.4937,0.8941,0.02608,0.06358,0.005847,0.004837,0.04885,0.8364,0.5091,0.01017,0.1959,0.4046,0.04053,0.01426,0.009762,0.2509,0.03092,0.008723,0.06745,0.009251,0.3498,0.4594,0.06255,0.3716,0.01907,0.111,0.89,0.1743,0.01422,0.1523,0.0104,0.03691,0.4788,0.1193,0.01254,0.6082,0.01892,0.01478,0.05848,0.2258,0.2084,0.712,0.5963,0.5771,0.01204,0.01551,0.9903,0.8683,0.09643,0.2681,0.02352,0.7861,0.03567,0.0986,0.1207,0.2727,0.7116,0.1371,0.7389,0.1839,0.1964,0.04697,0.2305,0.09332,0.1823,0.7061,0.09226,0.3775,0.2035,0.6401,0.004149,0.5945,0.589,0.5744,0.1113,0.3972,0.1175,0.6324,0.004232,0.2162,0.2313,0.104,0.3039,0.01195,0.26,0.5063,0.848,0.1466,0.03291,0.1799,0.4444,0.01585,0.2203,0.02556,0.1234,0.3819,0.08564,0.6216,0.08024,0.7243,0.01793,0.03825,0.03406,0.2204,0.005107,0.1122,0.1723,0.1176,0.3633,0.8446,0.6241,0.692,0.02726,0.01229,0.108,0.9404,0.0441,0.09629,0.03456,0.512,0.5914,0.02589,0.1687,0.4223,0.5821,0.7908,0.01292,0.3096,0.005252,0.006512,0.757,0.285,0.8767,0.6429,0.2416,0.00755,0.6009,0.0311,0.4875,0.835,0.3631,0.01227,0.005259,0.3437,0.5522,0.3546,0.5497,0.2361,0.1477,0.05962,0.2266,0.8609,0.4329,0.012,0.07913,0.2906,0.5574,0.7882,0.03046,0.1112,0.2171,0.05116,0.0195,0.7685,0.1859,0.9006,0.1724,0.06774,0.9364,0.003621,0.1512,0.2443,0.09534,0.6865,0.4222,0.3314,0.06643,0.05298,0.0415,0.08555,0.3792,0.08628,0.1265,0.4316,0.02929,0.9412,0.1302,0.6154,0.07957,0.1313,0.3176,0.009657,0.04919,0.1997,0.1778,0.1555,0.2596,0.3954,0.4023,0.004406,0.5108,0.006905,0.03417,0.4363,0.003952,0.1825,0.8189,0.353,0.06089,0.2021,0.008446,0.1795,0.4294,0.1153,0.2781,0.3618,0.8857,0.4649,0.2017,0.01443,0.1063,0.05552,0.01864,0.2212,0.01335,0.4174,0.1508,0.7819,0.3358,0.823,0.7684,0.1741,0.8751,0.03202,0.1984,0.01954,0.2355,0.0238,0.7186,0.003982,0.1178,0.1211,0.7505,0.129,0.005556,0.1018,0.1617,0.1926,0.5256,0.02343,0.02619,0.005317,0.008664,0.07129,0.006255,0.9586,0.0248,0.04461,0.7957,0.3615,0.02134,0.08697,0.1171,0.05112,0.1082,0.1091,0.6176,0.1371,0.5224,0.05807,0.07215,0.01562,0.03934,0.8137,0.3781,0.8606,0.3408,0.01852,0.778,0.5788,0.3058,0.006998,0.1386,0.01013,0.02031,0.02299,0.03391,0.1892,0.3211,0.06133,0.1442,0.08029,0.3088,0.2489,0.232,0.1483,0.2625,0.2443,0.44,0.09764,0.3797,0.2831,0.9936,0.05064,0.7629,0.02356,0.2563,0.594,0.3929,0.8755,0.1273,0.8373,0.03977,0.1138,0.009109,0.06645,0.2959,0.5415,0.9639,0.8317,0.8739,0.03182,0.005944,0.1703,0.01386,0.01276,0.3374,0.08312,0.3098,0.2208,0.2997,0.1341,0.05882,0.1467,0.1951,0.9012,0.8521,0.7609,0.5773,0.1567,0.7314,0.09376,0.2471,0.02417,0.2297,0.08052,0.227,0.02626,0.005242,0.06065,0.3961,0.1828,0.07586,0.4381,0.04745,0.05733,0.2396,0.5995,0.1577,0.2212,0.03661,0.06389,0.0246,0.2714,0.1156,0.03007,0.6593,0.9111,0.4032,0.004012,0.004752,0.07592,0.9075,0.03859,0.422,0.2686,0.663,0.004272,0.00386,0.007471,0.4579,0.1063,0.336,0.008265,0.3702,0.05047,0.1534,0.9914,0.01586,0.4911,0.03053,0.07614,0.8157,0.05286,0.07237,0.1267,0.2129,0.1065,0.05633,0.8567,0.009199,0.00671,0.8242,0.03807,0.0634,0.6721,0.09185,0.006149,0.7113,0.04214,0.8347,0.196,0.0261,0.007565,0.08232,0.1726,0.004623,0.01729,0.07022,0.4827,0.157,0.09794,0.7657,0.3464,0.03085,0.3949,0.8119,0.5421,0.775,0.01406,0.5133,0.005727,0.5502,0.004023,0.08802,0.2748,0.1782,0.3852,0.00674,0.01438,0.03482,0.02905,0.003833,0.05072,0.00386,0.05276,0.005895,0.7822,0.003853,0.03687,0.004707,0.04757,0.6981,0.2392,0.6947,0.2052,0.5445,0.01251,0.1715,0.8972,0.003766,0.0494,0.01215,0.08513,0.2486,0.6736,0.4899,0.0823,0.05798,0.02538,0.3405,0.3173,0.1899,0.003903,0.8292,0.05619,0.009548,0.004046,0.5541,0.05235,0.5328,0.3866,0.1134,0.01647,0.9253,0.09772,0.2571,0.7379,0.5235,0.003779,0.7081,0.01321,0.2256,0.1439,0.04622,0.4867,0.2639,0.2081,0.01312,0.5787,0.1094,0.006296,0.04416,0.101,0.5716,0.06623,0.01333,0.006936,0.08308,0.1586,0.3362,0.7754,0.006386,0.7846,0.01353,0.149,0.5442,0.05726,0.02905,0.6425,0.2468,0.3297,0.01195,0.00499,0.1301,0.01455,0.09814,0.02599,0.009074,0.08369,0.4018,0.5238,0.5238,0.0428,0.3702,0.5238,0.2096,0.5238,0.158,0.2499,0.1496,0.1631,1,0.1489,1,1,0.1209,0.6577,0.1744,0.009304,0.2382,1,0.3161,0.0347,0.674,0.02547,0.08702,0.02678,0.247,0.8004,0.6816,0.09934,0.06223,0.2193,0.007884,0.3574,0.5542,0.06669,0.5301,0.9987,0.08125,0.7327,0.9659,0.01353,0.5266,0.1017,0.8623,0.008027,0.5853,0.01353,0.06103,0.04226,0.01084,0.05801,0.08292,0.4038,0.1626,0.6054,0.5471,0.8426,0.7994,0.1152,0.959,0.01731,0.9711,0.1124,0.4397,0.06187,0.09831,0.1723,0.006854,0.1994,0.08995,0.4703,0.005092,0.5497,0.3903,0.1077,0.01442,0.04506,0.4064,0.004507,0.006318,0.004842,0.543,0.4715,0.02406,0.9096,0.005185,0.2962,0.7117,0.08552,0.04662,0.4764,0.08185,0.7304,0.1156,0.3333,0.3328,0.2743,0.4159,0.4813,0.5168,0.07075,0.1698,0.01093,0.6815,0.2202,0.7595,0.0945,0.02364,0.07752,0.03548,0.0106,0.7204,0.168,0.3546,0.006457,0.4771,0.5559,0.01153,0.4109,0.01829,0.2372,0.00979,0.3915,0.07822,0.003945,0.05532,0.006613,0.6996,0.4364,0.005711,0.1754,0.5458,0.8582,0.05176,0.1993,0.7058,0.466,0.0163,0.4713,0.1393,0.532,0.2396,0.06555,0.1677,0.1478,0.08864,0.8865,0.0464,0.006835,0.2236,0.2933,0.004943,0.726,0.02514,0.02746,0.806,0.005542,0.0513,0.01026,0.4547,0.6989,0.03034,0.07334,0.2929,0.02513,0.6207,0.1248,0.7059,0.4272,0.5274,0.4583,0.005342,0.1425,0.33,0.1166,0.5633,0.0887,0.0318,0.4717,0.561,0.09439,0.02535,0.9342,0.4209,0.7634,0.3633,0.01172,0.007396,0.03382,0.5445,0.01412,0.07398,0.005179,0.05702,0.03625,0.606,0.4667,0.6899,0.08328,0.2705,0.995,0.1024,0.02977,0.3803,0.7338,0.02054,0.06376,0.1916,0.5083,0.02114,0.9148,0.005946,0.6475,0.3015,0.006456,0.04241,0.5052,0.8147,0.1195,0.0198,0.007749,0.05876,0.01319,0.4366,0.04021,0.8897,0.2495,0.02017,0.02128,0.9785,0.01624,0.07363,0.1032,0.9098,0.01348,0.4541,0.7628,0.0647,0.4343,0.01775,0.02073,0.8871,0.2003,0.04206,0.4621,0.2453,0.7194,0.3918,0.04699,0.2757,0.8847,0.2037,0.1291,0.01586,0.1532,0.2004,0.01756,0.03729,0.06038,0.04068,0.1483,0.5696,0.3786,0.2209,0.4952,0.007343,0.1329,0.01276,0.1009,0.5999,0.1249,0.1825,0.6317,0.03394,0.01645,0.008813,0.05005,0.3901,0.8755,0.3312,0.2529,0.2486,0.008212,0.0168,0.0162,0.01508,0.007802,0.07666,0.9442,0.6909,0.2643,0.0148,0.07504,0.0725,0.4246,0.227,0.03438,0.224,0.5623,0.2028,1,0.9188,0.2129,0.4472,0.7816,0.5292,0.05545,0.05047,0.7206,0.005099,0.01007,0.01386,0.6203,0.01195,0.004853,0.0156,0.007349,0.2235,0.6714,0.1086,0.5161,0.7661,0.01378,0.579,0.4345,0.01649,0.211,0.04309,0.007487,0.01947,0.482,0.07262,0.04978,0.2836,0.07937,0.004386,0.02182,0.5064,0.2932,0.004624,0.02435,0.8662,0.005107,0.187,0.4002,0.08738,0.01962,0.05964,0.07466,0.07265,0.003743,0.02316,0.007637,0.7279,0.2762,0.4921,0.0115,0.4898,0.0158,0.006702,0.4002,0.006561,0.2953,0.2469,0.003784,0.006653,0.004561,0.4072,0.32,0.08839,0.4096,0.9866,0.1984,0.9322,0.03306,0.005096,0.3207,0.1476,0.06094,0.5342,0.007909,0.905,0.003959,0.188,0.4321,0.1241,0.05227,0.006277,0.7292,0.02394,0.009408,0.9455,0.2068,0.7907,0.1328,0.2911,0.9467,0.8786,0.03285,0.2994,0.1805,0.03486,0.1146,0.07554,0.00585,0.05965,0.6269,0.3867,0.1242,0.5224,0.1497,0.6581,0.006452,0.02938,0.007982,0.6767,0.00876,0.004083,0.275,0.04067,0.04253,0.3531,0.00625,0.1718,0.3706,0.9959,0.007099,0.1233,0.0811,0.5868,0.1872,0.2795,0.005356,0.04051,0.05916,0.5177,0.8004,0.705,0.3917,0.9732,0.6156,0.788,0.04363,0.899,0.2005,0.4334,0.07048,0.004231,0.01125,0.07043,0.2189,0.6429,0.007197,0.6656,0.04019,0.1805,0.08717,0.09756,0.1009,0.04034,0.003807,0.05314,0.004633,0.03838,0.05828,0.03544,0.01957,0.6984,0.016,0.008032,0.9542,0.03779,0.3231,0.1841,0.04467,0.107,0.1825,0.2254,0.07226,0.07508,0.01569,0.9304,0.7531,0.01664,0.14,0.1728,0.734,0.7475,0.009129,0.1361,0.159,0.0958,0.2167,0.1587,0.6452,0.2499,0.1009,0.6098,0.1155,0.9522,0.2307,0.2517,0.1666,1,0.2108,0.1932,0.1222,0.04331,0.9921,0.3316,0.1331,0.4063,0.9154,0.004884,0.006232,0.01758,0.8338,0.04891,0.6228,0.00948,0.01909,0.5406,0.8914,0.2334,0.003695,0.1011,0.9709,0.6377,0.007096,0.01469,0.7502,0.04593,0.636,0.4947,0.04052,0.8195,0.7809,0.04509,0.06969,0.04708,0.4197,0.02589,0.2253,0.009597,0.6374,0.03049,0.009773,0.5213,0.02248,0.06548,0.02208,0.7248,0.4145,0.5564,0.1833,0.01449,0.02581,0.009852,0.03173,0.6495,0.09337,0.03178,0.8519,0.7908,0.7163,0.003939,0.06161,0.385,0.008308,0.004256,0.3913,0.3094,0.2983,0.01558,0.02645,0.2927,0.01733,0.5396,0.005703,0.08456,0.0283,0.009561,0.202,0.3292,0.07012,0.004201,0.01523,0.1791,0.9223,0.006645,0.1841,0.006403,0.1535,0.6283,0.01198,0.009141,0.7882,0.07673,0.003936,0.5491,0.004345,0.008435,0.3436,0.04015,0.1439,0.01297,0.09612,0.8596,0.8452,0.8907,0.01983,0.07771,0.4316,0.187,0.56,0.01212,0.03786,0.003793,0.8336,0.5491,0.02282,0.004498,0.01926,0.003919,0.7876,0.5153,0.02156,0.9929,0.5679,0.006965,0.01029,0.01669,0.08094,0.6161,0.1059,0.01712,0.122,0.01978,0.004394,0.3047,0.005509,0.2191,0.008031,0.183,0.8096,0.7602,0.1032,0.1168,0.008176,0.2201,0.2435,0.9143,0.0128,0.0057,0.2296,0.1096,0.01471,0.3385,0.7222,0.04765,0.06725,0.01384,0.005857,0.3737,0.005082,0.7879,0.9752,0.4196,0.1421,0.7935,0.5506,0.04989,0.03388,0.05989,0.2789,0.3823,0.3947,0.1269,0.9655,0.05935,0.01391,0.04039,0.7899,0.4734,0.1902,0.2901,0.3676,0.09132,0.4597,0.009029,0.0389,0.08952,0.01921,0.3889,0.5684,0.1664,0.9946,0.2237,0.6156,0.02614,0.379,0.04787,0.02494,0.1053,0.9214,0.2894,0.3874,0.7448,0.5881,0.01562,0.01324,0.6929,0.00845,0.227,0.01096,0.5791,0.00418,0.008645,0.3658,0.0629,0.09225,0.1221,0.1161,0.01302,0.01614,0.06377,0.08566,0.5652,0.3186,0.4713,0.8104,0.4804,0.4743,0.6634,0.01334,0.04928,0.9751,0.02819,0.03732,0.269,0.07033,0.8666,0.2367,0.1046,0.004234,0.0538,0.1673,0.3707,0.04483,0.004396,0.5685,0.01365,0.1212,0.3896,0.1152,0.157,0.5396,0.2491,0.1899,0.1091,0.1486,0.07171,0.3035,0.3519,0.9159,0.1815,0.5798,0.216,0.03083,0.1535,0.5887,0.855,0.2884,0.9928,0.4787,0.3074,0.03969,0.3664,0.6973,0.06129,0.004159,0.02371,0.1567,0.06267,0.1967,0.005578,0.02445,0.994,0.8878,0.3753,0.9302,0.5791,0.08105,0.0179,0.8304,0.478,0.3752,0.4008,0.3716,0.2292,0.06865,0.1489,0.2059,0.203,0.1318,0.1576,0.7738,0.01342,0.2442,0.6256,0.1448,0.0459,0.03403,0.00538,0.0321,0.006266,0.5559,0.7053,0.1229,0.02029,0.003671,0.00398,0.04997,0.2836,0.4332,0.04012,0.01145,0.01795,0.4728,0.02529,0.01267,0.006127,0.1455,0.1054,0.1315,0.09623,0.1873,0.02627,0.04834,0.00388,0.0545,0.8777,0.7636,0.1904,0.6414,0.2341,0.4268,0.09111,0.5259,0.3374,0.8509,0.5204,0.02545,0.3909,0.04756,0.03182,0.1812,0.04719,0.1105,0.3619,0.6049,0.458,0.01417,0.3301,0.006606,0.01434,0.1272,0.07369,0.8782,0.1221,0.6368,0.984,0.009836,0.01393,0.0789,0.004453,0.7398,0.7069,0.005047,0.005496,0.7253,0.3179,0.1228,0.6436,0.05957,0.02862,0.5109,0.3984,0.02433,0.1477,0.8261,0.04244,0.008447,0.8084,0.02363,0.7186,0.3053,0.8255,0.0136,0.1865,0.01216,0.2608,0.02685,0.905,0.1644,0.1266,0.5463,0.01673,0.408,0.004506,0.02429,0.5008,0.7476,0.628,0.4102,0.1895,0.02931,0.003605,0.3559,0.04279,0.004968,0.3632,0.02947,0.0356,0.7127,0.01858,0.9352,0.7618,0.7658,0.05247,0.08636,0.1286,0.2216,0.06339,0.1474,0.9368,0.01172,0.01267,0.1981,0.9597,0.6625,0.2036,0.01403,0.8454,0.8129,0.3746,0.04527,0.01971,0.2444,0.4586,0.09676,0.0222,0.7718,0.03055,0.9906,0.4906,0.1972,0.04687,0.1381,0.9107,0.4349,0.009991,0.109,0.7914,0.2119,0.009601,0.1754,0.6265,0.8312,0.3029,0.099,0.3247,0.9776,0.03542,0.4188,0.004451,0.09327,0.2256,0.814,0.3044,0.5332,0.03606,0.09776,0.03638,0.643,0.02959,0.6031,0.2535,0.8847,0.0674,0.05686,0.227,0.004721,0.2288,0.00591,0.2954,0.7408,0.1043,0.01783,0.02497,0.009934,0.2263,0.03385,0.005632,0.02025,0.003993,0.00691,0.0259,0.01145,0.007476,0.1375,0.1172,0.3056,0.005774,0.1837,0.07511,0.5593,0.3297,0.3082,0.004444,0.005424,0.006029,0.7613,0.1198,0.8917,0.3437,0.007766,0.007348,0.04828,0.5492,0.007447,0.01039,0.04716,0.2169,0.176,0.1259,0.009785,0.09443,0.2765,0.1706,0.04053,0.5791,0.3403,0.07643,0.1326,0.04639,0.03226,0.03052,0.1527,0.4638,0.2457,0.5133,0.05037,0.06334,0.0457,0.2415,0.247,0.01535,0.852,0.4584,0.2301,0.05718,0.1173,0.03331,0.615,0.6109,0.007881,0.5416,0.02094,0.3138,0.007541,0.1463,0.01708,0.4433,0.08661,0.05374,0.02018,0.3784,0.8707,0.007793,0.02081,0.006567,0.04772,0.01052,0.2319,0.7258,0.04054,0.006052,0.1134,0.7468,0.1623,0.7179,0.4273,0.2122,0.7797,0.7593,0.04055,0.8139,0.01381,0.6799,0.1851,0.7101,0.7211,0.03111,0.6007,0.02326,0.04789,0.1074,0.2824,0.04367,0.07239,0.02575,0.3045,0.5569,0.08398,0.01081,0.004008,0.04578,0.1635,0.4609,0.08148,0.05416,0.3247,0.9582,0.0102,0.4565,0.02535,0.07895,0.1251,0.781,0.01807,0.0978,0.5195,0.01821,0.01297,0.07866,0.1301,0.9092,0.3418,0.3217,0.006321,0.08293,0.3693,0.977,0.09314,0.1093,0.2848,0.007262,0.006901,0.03905,0.03921,0.01545,0.424,0.006184,0.00597,0.09022,0.03324,0.01491,0.5801,0.6249,0.3347,0.269,0.6538,0.05834,0.01181,0.381,0.05783,0.2075,0.8847,0.8492,0.02329,0.7975,0.3432,0.8934,0.01114,0.1334,0.09306,0.01686,0.01727,0.1768,0.1679,0.004757,0.165,0.01931,0.004171,0.2868,0.06558,0.01619,0.03577,0.02122,0.05121,0.004726,0.7071,0.07846,0.7093,0.005154,0.03157,0.03077,0.01804,0.00648,0.09637,0.7363,0.05524,0.2914,0.0075,0.005916,0.08883,0.2012,0.07478,0.4095,0.02951,0.08847,0.1929,0.3206,0.5238,0.2707,0.2208,0.243,0.2371,0.1229,0.004429,0.8043,0.1871,0.1664,0.02202,0.01016,0.01265,0.5079,0.6149,0.1561,0.5313,0.8181,0.00491,0.1023,0.4287,0.00885,0.01612,0.9232,0.01818,0.03128,0.4221,0.01145,0.04414,0.04971,0.7422,0.3384,0.6819,0.004191,0.1446,0.06921,0.005023,0.3927,0.005667,0.8907,0.0138,0.03047,0.01571,0.9667,0.05802,0.1632,0.0317,0.3326,0.2042,0.1126,0.1546,0.007199,0.05294,0.4868,0.619,0.2444,0.003616,0.05704,0.01912,0.1459,0.1507,0.2889,0.004274,0.2225,0.6888,0.003933,0.06608,0.3952,0.0342,0.1589,0.04436,0.0462,0.2001,0.08647,0.2392,0.06712,0.006288,0.5977,0.01096,0.9767,0.008746,0.01195,0.4213,0.1599,0.5214,0.2686,0.01373,0.05335,0.2552,0.462,0.09624,0.09508,0.509,0.03415,0.005727,0.1539,0.06274,0.6111,0.985,0.3313,0.007569,0.0222,0.009876,0.06209,0.8141,0.1088,0.02983,0.01702,0.1599,0.07586,0.983,0.0169,0.004944,0.02732,0.9703,0.006025,0.04643,0.02862,0.01994,0.1996,0.003624,0.1259,0.3822,0.8682,0.03287,0.9979,0.9969,0.05286,0.07755,0.09019,0.3685,0.4269,0.7724,0.01298,0.05261,0.1014,0.1325,0.009538,0.6089,0.005139,0.01654,0.02935,0.1249,0.3448,0.004808,0.01058,0.02016,0.8944,0.8854,0.9504,0.005178,0.03214,0.059,0.01245,0.04817,0.1763,0.1781,0.1244,0.07288,0.03386,0.08885,0.2959,0.15,0.004133,0.8067,0.09147,0.4198,0.06379,0.6026,0.3428,0.1532,0.04701,0.4715,0.02019,0.6084,0.8924,0.05769,0.004823,0.09938,0.02219,0.01646,0.04498,0.04021,0.00539,0.245,0.07304,0.01232,0.3938,0.739,0.005257,0.1524,0.6439,0.004959,0.6163,0.4309,0.06157,0.8589,0.3528,0.0743,0.1479,0.7489,0.9791,0.009153,0.06579,0.3348,0.3348,0.3818,0.6741,0.4546,0.009295,0.171,0.006509,0.03523,0.8472,0.98,0.8346,0.4371,0.4999,0.01165,0.02082,0.01209,0.03259,0.007815,0.009607,0.008385,0.1248,0.1042,0.1187,0.03078,0.01004,0.004148,0.02876,0.0361,0.08579,0.1616,0.9473,0.02796,0.7255,0.1594,0.2356,0.008643,0.5785,0.241,0.08888,0.0745,0.561,0.4767,0.5612,0.1678,0.3864,0.09738,0.04832,0.05024,0.4022,0.0441,0.02443,0.901,0.3093,0.04334,0.05685,0.01088,0.1361,0.05702,0.6796,0.0987,0.4402,0.3083,0.1923,0.275,0.6147,0.5204,0.02358,0.005406,0.1174,0.2249,0.2468,0.1621,1,0.07407,0.185,0.1985,0.1795,0.2318,0.6641,0.2645,0.07156,0.06678,0.1092,0.7872,0.07558,0.2486,0.2845,0.6052,0.1355,0.1227,0.3673,0.03896,0.03848,0.009662,0.5769,0.03248,0.4473,0.004546,0.004627,0.7662,0.004845,0.8407,0.01252,0.1113,0.2839,0.633,0.3814,0.02854,0.5994,0.006657,0.1607,0.004602,0.3758,0.03393,0.7077,0.07663,0.08405,0.006068,0.005689,0.01211,0.07221,0.0775,0.2064,0.06211,0.5044,0.4578,0.3276,0.6024,0.6005,0.1752,0.05939,0.9774,0.1433,0.1092,0.1014,0.0897,0.4791,0.08841,0.6264,0.4725,0.04204,0.00853,0.05345,0.1115,0.005849,0.01187,0.04205,0.7022,0.9749,0.6442,0.7401,0.1186,0.04952,0.03149,0.007747,0.9914,0.00396,0.03183,0.4078,0.2059,0.008742,0.01193,0.0732,0.06641,0.4347,0.03174,0.1111,0.4984,0.007683,0.4957,0.01792,0.435,0.5143,0.01325,0.5771,0.9397,0.8269,0.4551,0.0596,0.4148,0.2913,0.154,0.1861,0.2373,0.0128,0.0284,0.7536,0.8919,0.6177,0.2633,0.9217,0.2639,0.01564,0.3942,0.9574,0.05664,0.2983,0.7962,0.09344,0.7464,0.7306,0.03766,0.3888,0.1818,0.004931,0.03166,0.5567,0.02859,0.8847,0.588,0.1511,0.7615,0.4102,0.1176,0.4606,0.3404,0.1298,0.03912,0.5024,0.6775,0.9281,0.01033,0.01627,0.4967,0.08945,0.452,0.9355,0.8727,0.1262,0.03861,0.1213,0.004665,0.05311,0.04072,0.7495,0.007914,0.1468,0.3366,0.4058,0.4296,0.02045,0.01784,0.007403,0.2642,0.08032,0.004026,0.2011,0.1228,0.03234,0.00947,0.2499,0.07445,0.3868,0.005434,0.05801,0.3354,0.3013,0.01221,0.06926,0.008237,0.005021,0.4486,0.6877,0.01331,0.9199,0.6621,0.01071,0.1805,0.09187,0.6021,0.09063,0.15,0.445,0.006072,0.03012,0.0102,0.01294,0.6674,0.01832,0.01029,0.01417,0.3275,0.04765,0.2146,0.9995,0.02431,0.5681,0.0171,0.524,0.03367,0.5571,0.257,0.8784,0.2653,0.7567,0.7773,0.7769,0.02087,0.1156,0.03675,0.03043,0.1389,0.6079,0.0896,0.1304,0.007905,0.09805,0.07572,0.01044,0.005899,0.3059,0.815,0.1114,0.1817,0.1331,0.05283,0.5771,0.01059,0.3711,0.07543,0.005792,0.06717,0.1927,0.8444,0.7575,0.3706,0.5934,0.1839,0.1099,0.8383,0.3127,0.04117,0.06872,0.01517,0.01279,0.9959,0.977,0.003607,0.09364,0.0413,0.242,0.6995,0.5061,0.07193,0.4202,0.009504,0.04451,0.5834,0.3559,0.1829,0.03091,0.01259,0.3115,0.1858,0.2519,0.1109,0.09856,0.03107,0.06001,0.2962,0.3991,0.5146,0.02415,0.5833,0.01327,0.05999,0.01549,0.6354,0.06994,0.004713,0.1962,0.399,0.3273,0.04268,0.04103,0.01466,0.8716,0.01125,0.1118,0.7851,0.07264,0.03781,0.02293,0.0711,0.1567,0.1065,0.005539,0.0379,0.6423,0.01604,0.5338,0.04269,0.007909,0.01669,0.2011,0.01396,0.06292,0.01593,0.3519,0.05449,0.1874,0.3796,0.09554,0.8608,0.3204,0.018,0.01942,0.04089,0.3785,0.01216,0.8371,0.01175,0.2297,0.3859,0.5279,0.8288,0.02055,0.01262,0.289,0.004651,0.3444,0.007482,0.2046,0.0473,0.08733,0.2811,0.3422,0.004251,0.4911,0.08897,0.3648,0.00426,0.5211,0.006834,0.7922,0.4123,0.02012,0.4342,0.6054,0.2767,0.0577,0.08053,0.3486,0.08877,0.009378,0.5862,0.7849,0.008046,0.5467,0.582,0.005894,0.06857,0.03524,0.01402,0.027,0.008177,0.3913,0.6866,0.131,0.004066,0.1274,0.05885,0.05958,0.4152,0.6633,0.2007,0.05006,0.1091,0.1512,0.7419,0.03756,0.04029,0.1512,0.005847,0.02823,0.04263,0.6473,0.1486,0.422,0.05231,0.01386,0.004108,0.2023,0.05181,0.9281,0.1738,0.2317,0.06108,0.1953,0.05972,0.4,0.041,0.1364,0.3177,0.1157,0.1735,0.1275,0.1821,0.08415,0.1706,0.1967,0.44,0.5238,0.4233,0.0876,0.3051,0.08666,1,0.1531,0.287,0.02462,0.5238,0.006037,0.004089,1,0.001788,0.0001453,2.314e-05,0.0007858,0.0007433,0.002771,0.0002976,6.613e-06,0.001704,2.663e-06,3.053e-10,0.0002067,0.0002593,1.456e-05,0.002405,6.175e-07,0.0002101,0.001041,1.845e-06,0.0007778,0.002804,1.087e-05,0.0003764,7.686e-06,4.476e-08,9.189e-05,2.609e-05,2.35e-05,1.05e-05,0.0006143,0.0002406,0.002317,0.0003827,0.0002666,0.002175,0.0005792,0.0004511,0.0008048,3.115e-07,0.0008098,5.124e-06,0.002732,4.753e-08,0.0001448,0.0003809,0.0003547,0.0009219,1.066e-06,0.0004437,0.0006708,6.93e-05,3.134e-06,7.208e-08,9.718e-09,9.314e-06,5.578e-07,0.000145,1.212e-06,1.051e-05,1.052e-05,0.0003234,0.0005865,0.0007665,2.382e-05,4.602e-08,0.0002663,0.0007249,0.0009608,0.0009743,2.889e-07,0.0002392,0.0008855,2.784e-06,9.584e-09,1.18e-06,2.824e-05,1.37e-06,0.0004017,9.604e-05,7.885e-05,1.956e-06,9.959e-05,3.27e-05,5.184e-05,0.000879,0.0016,0.002482,0.0002792,0.002608,4.274e-09,2.048e-06,2.846e-05,6.012e-05,0.0001092,6.564e-05,1.642e-06,0.0002386,2.555e-07,3.682e-06,0.0005835,5.091e-06,5.948e-05,8.779e-07,0.002399,6.446e-06,2.313e-06,0.000198,8.953e-05,3.545e-06,9.69e-07,0.002436,0.002819,3.283e-06,0.0002085,0.0001523,0.0002849,1.727e-05,5.413e-07,5.04e-08,3.937e-06,0.0005154,8.139e-07,0.001347,1.639e-09,5.05e-07,7.404e-08,0.0007364,0.0003361,3.468e-05,5.887e-07,2.053e-08,0.000415,3.478e-05,1.497e-06,5.732e-07,0.001698,5.253e-06,8.971e-05,1.949e-05,4.359e-07,3.428e-06,3.596e-06,0.001045,5.327e-08,5.23e-05,0.0003234,2.759e-05,0.0001258,5.952e-05,0.002418,9.006e-06,0.0008313,0.0008116,8.433e-05,9.684e-05,1.477e-05,0.003123,0.002808,0.001074,0.0006919,6.045e-06,0.002056,0.00132,2.793e-05,0.0002597,2.231e-06,0.003231,0.0001394,8.714e-07,0.0001803,0.000185,0.0008196,0.0006653,3.369e-06,1.299e-06,0.003589,8.138e-05,0.0003825,3.565e-06,0.000135,4.328e-08,6.819e-09,2.496e-06,0.001829,3.103e-05,0.0003134,0.0002886,7.938e-06,3.856e-07,2.497e-06,0.0001156,0.00177,1.875e-07,0.001446,1.613e-08,5.661e-08,4.887e-07,0.001896,0.0004958,8.617e-05,6.448e-06,4.597e-06,0.0005964,0.00176,5.243e-05,9.831e-08,0.0002358,5.436e-07,0.003024,0.00122,5.703e-05,0.001491,8.614e-08,0.0003524,5.922e-06,0.002682,0.0002953,6.108e-05,4.948e-06,7.423e-05,6.15e-07,0.001571,4.742e-05,0.0008456,0.002156,2.757e-05,5.889e-05,0.0002113,0.0003441,8.235e-05,2.622e-05,1.884e-08,0.001141,0.0004028,5.493e-08,4.519e-06,0.000725,0.0007356,0.002264,5.002e-06,4.996e-05,2.382e-06,4.09e-05,0.001452,0.000626,7.41e-07,3.992e-05,0.000327,0.0008266,1.268e-08,0.003003,4.189e-06,8.694e-05,1.792e-06,8.869e-05,0.002476,0.0001201,0.0003968,0.0005792,1.744e-06,4.351e-06,0.001405,4.656e-07,4.416e-05,3.207e-07,8.087e-06,7.577e-05,0.0001171,1.138e-06,0.002731,2.12e-05,4.001e-06,0.002457,5.642e-09,0.0002892,0.002324,7.176e-05,0.000228,1.345e-08,0.0007391,1.078e-05,0.001382,0.002904,1.684e-05,0.0003153,0.0006713,9.391e-08,3.725e-08,0.0005409,0.0007553,0.0005855,6.964e-05,0.001346,6.102e-07,0.001098,1.962e-07,0.0003789,0.0005408,2.221e-06,2.497e-05,0.0006812,1.982e-05,0.00205,7.091e-05,0.0007077,0.002563,9.308e-05,0.0001186,1.212e-05,3.238e-05,1.135e-05,0.000371,9.348e-06,6.603e-07,0.0001384,0.001755,2.867e-05,0.002822,0.0003003,0.002658,2.644e-09,6.727e-06,0.002957,0.0002882,0.0004859,0.0004613,0.002676,0.0001132,0.0005777,0.002042,6.626e-06,0.000178,9.08e-05,0.0003099,0.0003987,0.0001328,0.001417,2.941e-07,5.172e-06,0.0009442,2.945e-05,0.002806,0.003254,0.00356,0.0001564,0.001397,0.0003598,0.001344,6.446e-05,0.001402,2.124e-06,0.0005953,1.227e-06,3.166e-05,1.73e-05,5.306e-07,0.001671,2.984e-05,8.131e-05,1.51e-05,2.605e-07,1.688e-06,0.0003751,4.455e-05,0.0002179,0.001565,3.676e-06,0.0007134,6.327e-05,0.0005282,0.0003322,4.128e-05,2.503e-07,0.0006204,1.244e-05,0.00288,2.531e-05,0.002303,1.106e-08,5.867e-08,0.0002469,0.0008858,5.502e-08,3.689e-05,8.379e-05,7.471e-05,0.002614,3.59e-05,4.868e-05,0.0004329,2.797e-05,0.002521,6.827e-07,1.216e-06,0.0004202,4.191e-06,0.0002379,0.001943,0.002177,0.002247,2.171e-05,0.0001066,0.002627,0.0001258,6.255e-05,0.0001199,4.063e-08,0.0001782,3.726e-05,1.559e-05,0.0001092,0.0004346,2.26e-05,0.002866,0.0001007,0.0007478,8.85e-08,2.582e-07,0.001433,4.342e-06,0.002768,1.853e-05,0.001143,0.001528,0.001412,1.109e-05,0.0003322,1.012e-05,0.0004095,0.00244,3.49e-05,0.0002004,4.176e-06,1.201e-05,4.35e-07,0.000365,8.962e-05,6.266e-05,0.0003573,2.102e-05,1.48e-06,3.703e-06,1.966e-06,0.001039,2.775e-05,8.781e-06,3.933e-05,2.614e-05,3.174e-05,0.003521,9.567e-07,0.001916,0.002519,1.035e-08,3.328e-05,5.027e-07,0.00144,0.0001309,7.021e-06,3.555e-05,1.443e-09,1.434e-06,1.056e-05,6.805e-07,2.003e-06,3.155e-05,2.538e-06,0.0004089,0.0004731,0.0001205,1.39e-07,0.0001893,0.003439,0.001891,9.713e-05,1.226e-05,2.943e-06,0.0007061,3.338e-06,0.000105,0.0001763,0.000343,1.587e-06,1.457e-07,0.0005433,0.0006268,0.001588,7.738e-06,0.0001632,0.00117,4.139e-05,0.0004221,0.0007756,8.977e-05,3.525e-07,1.089e-05,2.883e-07,5.974e-07,0.0006806,7.21e-05,0.0001309,2.159e-05,3.759e-05,0.001386,0.0003906,4.187e-08,2.689e-05,1.003e-05,0.002131,0.003029,0.0002089,1.437e-05,1.671e-07,3.101e-07,0.0001347,0.002009,0.00141,0.003586,0.0002241,1.283e-06,9.519e-06,0.0002201,5.001e-06,7.974e-07,6.701e-06,6.205e-06,0.0004222,1.736e-07,3.75e-06,0.0002308,0.0006032,0.001221,1.698e-05,0.0003628,0.0006211,6.844e-06,0.0003286,5.597e-05,0.002648,1.924e-06,8.243e-06,6.128e-05,1.881e-06,6.007e-06,0.001062,0.0005487,0.0001816,2.347e-08,2.88e-07,0.001693,9.724e-05,0.0005352,0.0001848,0.0003313,0.00203,2.667e-06,9.805e-05,3.02e-05,5.346e-05,0.0001015,5.885e-07,0.0001088,0.0008636,0.0007912,3.182e-07,1.521e-05,2.15e-06,7.361e-05,1.952e-05,2.268e-06,1.401e-08,0.0002491,2.181e-06,1.106e-05,2.348e-05,0.0001122,8.965e-05,6.06e-05,0.001039,0.0003499,3.997e-05,1.384e-06,9.678e-06,8.518e-05,0.0001279,3.686e-06,0.000942,0.0003475,0.00014,0.0004879,0.0004115,0.000299,0.0003178,0.0002653,0.0002474,0.0006351,6.987e-06,0.001689,1.516e-05,9.597e-05,0.001504,0.0006159,0.0003318,5.635e-05,1.784e-05,0.002469,7.323e-08,0.0003515,4.178e-06,0.0003008,9.771e-08,3.279e-05,1.33e-06,0.0001583,0.0002005,0.0002309,0.0004662,0.002786,5.53e-06,0.000343,2.803e-05,0.0002839,2.744e-06,0.0003644,3.612e-08,0.0007242,0.0001448,0.003083,0.001514,0.0007768,0.0004493,2.839e-07,2.965e-05,1.24e-07,1.961e-07,5.804e-05,0.0002108,0.0001853,7.264e-09,0.002558,0.001867,0.0001443,8.036e-05,6.785e-05,0.0007742,7.784e-05,6.281e-05,0.000103,2.434e-06,3.052e-05,0.0001579,2.042e-08,5.143e-05,0.001098,0.002077,0.00147,0.001813,3.296e-08,1.043e-05,0.000214,6.015e-05,0.001714,2.195e-05,4.226e-07,4.528e-06,0.0001666,4.718e-06,0.0007686,3.937e-05,0.0002085,0.003428,2.861e-05,0.000446,0.0008794,1.269e-05,0.0001645,4.606e-05,5.492e-05,3.332e-06,9.226e-08,3.386e-06,0.000899,7.246e-06,0.0002678,0.001252,0.001643,0.00154,6.778e-07,0.0009954,0.003278,0.00157,0.0001933,0.002275,1.22e-05,0.0008254,3.123e-05,0.002414,0.0001659,0.0006314,0.001375,4.286e-05,6.954e-06,0.0006452,3.44e-07,0.002406,4.015e-06,0.001739,0.001145,0.0001043,5.217e-05,0.002351,8.594e-05,0.0001527,0.002421,2.784e-06,0.001951,2.557e-05,6.29e-06,5.367e-05,3.333e-07,0.0003491,0.002919,9.422e-07,0.001534,0.002828,0.001696,4.225e-05,0.0005357,0.001624,0.001631,0.002381,4.492e-07,0.0002189,1.67e-07,0.0001192,0.000256,3.313e-06,2.743e-07,0.000601,0.0002535,6.882e-05,0.000112,0.0002063,2.31e-05,0.000114,3.595e-08,3.431e-06,9.987e-08,6.071e-08,0.001688,0.0009499,4.847e-07,0.0003602,1.511e-07,7.05e-07,0.0006522,0.00159,0.0005009,6.315e-05,0.0004114,8.567e-08,1.219e-05,4.126e-06,0.0004027,7.599e-07,0.0002393,0.001289,0.0001566,0.001673,0.0001661,0.0009574,0.0009128,0.0002815,5.134e-05,0.003466,0.001507,0.0005962,0.001187,2.549e-05,0.0001246,1.526e-09,3.83e-07,7.249e-06,0.000191,0.001711,2.118e-07,3.54e-06,0.0008012,0.001375,1.816e-06,0.0008538,0.000111,2.974e-05,0.0005063,7.471e-06,0.002378,3.772e-05,4.031e-08,0.001839,2.157e-05,0.001521,7.773e-05,0.0003767,1.104e-05,0.0001081,0.0002259,0.0001967,0.002348,8.378e-05,2.126e-05,6.425e-06,6.086e-09,6.752e-07,0.002868,0.002715,0.000573,8.587e-05,0.002523,3.704e-10,3.465e-06,0.00021,2.065e-06,3.078e-05,5.085e-05,0.0009002,0.0002221,9.828e-09,1.404e-06,5.473e-08,1.848e-05,0.001457,0.001738,0.0001172,1.206e-05,1.127e-05,0.0009799,1.077e-05,0.001371,0.002193,0.0001076,3.378e-07,8.133e-06,0.0002697,0.0003426,1.924e-07,3.181e-07,0.0001214,3.325e-06,0.0003808,0.0001271,0.000226,0.0005271,0.0005515,0.0001979,0.001603,0.003478,0.001155,0.0002033,0.0002488,3.702e-08,0.001064,0.0009139,0.002536,3.524e-05,0.002267,8.361e-05,4.81e-08,0.001336,0.001902,2.763e-06,2.655e-05,2.184e-07,0.000221,3.766e-07,0.0004185,0.001549,0.00134,1.399e-07,0.0003662,0.0002609,0.002872,5.384e-06,1.724e-07,2.758e-05,6.642e-05,0.0005256,0.001625,4.084e-05,0.00303,0.0003547,1.233e-07,1.569e-05,0.0001865,0.002625,0.003516,0.0002978,3.899e-05,0.0002539,4.173e-06,3.01e-05,0.002817,0.0008944,1.723e-06,4.038e-07,0.0001173,0.001936,0.0002469,3.374e-05,0.0009876,0.0001713,0.0004982,0.001482,5.542e-06,0.001191,0.002407,1.375e-05,0.0005409,0.0007194,0.00251,0.00112,0.003228,0.002277,0.00266,0.000114,9.001e-07,0.0008606,1.052e-06,0.002252,0.002619,2.392e-06,4.219e-07,7.982e-05,0.001266,3.065e-06,5.523e-05,1.676e-08,0.00117,0.001268,0.0007859,0.001159,0.0001622,0.002367,0.0001899,0.0002021,1.553e-05,0.0003477,0.0001212,9.011e-06,5.237e-05,8.672e-08,4.58e-05,0.0001968,1.966e-05,0.002082,3.707e-05,1.541e-06,0.001839,0.001449,0.002332,0.0005139,0.0001207,0.002819,1.304e-08,0.001802,0.0001047,1.46e-05,0.0002026,1.938e-07,0.0003468,0.001621,0.0001075,0.0001269,0.001808,3.623e-05,0.0002252,9.889e-06,1.902e-05,0.002347,1.131e-05,2.552e-05,3.305e-09,0.0005489,1.516e-05,3.525e-06,0.001108,0.001357,3.095e-06,0.003412,1.605e-06,6.871e-06,0.0002564,0.002582,0.0008317,0.0001633,8.077e-07,4.428e-05,3.153e-05,0.0001505,0.000117,0.000614,0.0007899,2.634e-08,6.884e-06,0.001759,2.116e-05,6.204e-05,2.092e-05,0.002978,7.115e-06,6.057e-06,0.00207,0.0002211,0.0007248,0.0001323,9.601e-05,0.0007987,5.98e-05,3.792e-06,0.001179,0.0004311,3.161e-05,1.136e-05,3.218e-07,0.001869,0.003235,0.0007433,1.215e-06,0.001124,0.0001213,7.559e-06,0.001208,1.258e-07,0.0001151,4.52e-05,0.0005038,8.977e-06,3.007e-07,0.0009135,0.000752,0.0002458,4.488e-05,2.228e-05,0.0001658,3.923e-06,0.003023,1.794e-06,3.235e-05,0.000142,2.367e-07,9.754e-05,6.476e-06,6.817e-05,1.536e-05,0.00141,0.0007187,9.716e-07,0.001415,0.002126,0.0004729,5.051e-05,0.00208,0.0004512,5.491e-06,0.00259,0.0008957,0.0003213,0.002019,0.0001092,0.0007848,8.471e-06,1.553e-06,0.001227,1.752e-06,6.847e-06,0.001391,3.202e-06,1.822e-06,0.0009671,1.579e-05,0.003552,1.031e-06,0.002118,0.0005622,7.156e-06,1.625e-05,0.003124,0.0002991,6.133e-10,3.096e-05,8.048e-05,4.779e-06,1.625e-07,0.0002954,2.649e-07,0.0001114,0.0002735,0.0003326,2.751e-05,0.0001156,4.383e-05,0.002128,7.147e-08,0.0007586,0.0007795,0.0003996,2.101e-05,5.965e-06,5.377e-05,3.983e-07,9.617e-06,2.462e-06,0.001806,0.0001676,0.0002304,0.0001028,1.379e-05,9.993e-06,4.93e-08,0.002813,0.002381,0.001307,0.002335,0.0006727,0.0008725,2.577e-05,1.558e-07,3.108e-06,1.961e-06,0.001732,0.001144,0.002322,1.698e-07,0.0002089,0.002077,6.044e-05,1.021e-05,3.123e-07,0.0008403,3.022e-05,0.003466,0.0007304,5.657e-06,1.527e-07,4.757e-06,1.169e-06,1.049e-05,1.017e-07,5.844e-05,8.414e-09,6.497e-06,0.0001785,1.907e-05,0.001757,9.545e-05,2.578e-06,1.869e-05,1.614e-07,0.0002498,2.644e-06,0.0002848,3.118e-05,0.0006194,0.0004012,5.639e-06,0.0003407,0.0005318,3.853e-05,0.001911,1.278e-06,2.578e-05,9.613e-06,0.0001765,0.0001628,1.32e-08,0.001351,1.239e-05,6.55e-07,0.000599,0.0008108,0.0004545,0.0001453,1.039e-05,1.391e-05,1.889e-06,0.000861,1.256e-06,2.475e-07,4.609e-05,1.929e-06,0.002501,1.442e-07,0.0005516,0.002643,4.675e-06,0.0003399,7.212e-05,0.0003421,9.752e-08,2.435e-06,2.259e-05,0.001017,3.505e-05,0.0001551,2.339e-07,0.0007276,0.0001023,2.062e-05,0.0002792,4.704e-06,6.38e-06,0.002358,0.0005178,3.098e-05,0.001058,3.795e-07,0.0003157,4.265e-07,2.394e-05,3.755e-05,7.185e-05,0.000291,6.871e-08,2.358e-06,0.001121,9.623e-07,0.0001332,0.0009513,3.281e-05,0.0008101,0.0005492,1.59e-05,0.0006337,7.075e-05,0.0003436,0.001834,9.278e-08,2.522e-05,0.0002087,7.926e-06,7.96e-05,8.466e-06,0.0006982,0.0004994,1.787e-05,6.671e-06,3.803e-06,1.779e-05,1.419e-05,3.826e-07,4.34e-08,0.002627,0.001894,0.003327,5.196e-07,3.131e-05,2.602e-07,7.517e-05,3.387e-05,5.612e-07,5.307e-05,0.0004787,6.262e-06,0.002425,0.0002051,7.369e-06,0.001102,0.00156,0.003157,3.262e-05,6.647e-06,0.0001495,6.206e-05,1.11e-06,1.681e-05,1.506e-05,0.001181,0.0007571,1.208e-05,1.924e-06,0.0001072,0.0005216,0.0003775,0.001184,4.231e-05,3.021e-06,0.001102,6.936e-05,1.989e-06,0.0001086,2.775e-06,6.864e-06,0.003011,3.239e-05,6.021e-05,0.0005636,0.0002222,5.355e-07,8.922e-08,3.873e-05,2.158e-05,2.769e-06,0.002792,8.86e-07,0.0008557,2.733e-05,2.16e-07,1.881e-06,7.77e-05,2.212e-07,8.325e-05,1.603e-05,0.00117,4.871e-06,3.031e-06,6.4e-08,4.651e-06,0.0002234,2.144e-05,1.633e-05,0.0002051,4.83e-05,0.0003077,0.002153,6.512e-05,2.354e-06,0.001948,1.253e-08,0.002057,1.11e-06,3.399e-05,0.003294,2.581e-06,2.738e-07,0.0003848,0.002506,0.0001293,1.436e-05,1.103e-08,2.19e-07,0.0003369,0.0009699,8.415e-05,0.003406,0.0005674,0.0006488,0.0008255,1.638e-06,6.673e-06,2.298e-08,5.304e-07,0.0003235,6.232e-05,0.002954,8.681e-05,0.0006698,3.498e-09,2.325e-06,0.0001835,0.003451,2.208e-06,0.001345,0.0001034,0.0001732,1.686e-05,0.00178,0.0008382,0.0002264,0.002355,2.643e-05,1.537e-05,0.0003427,0.0001469,0.0001173,0.0003889,3.314e-07,0.003,0.001296,2.081e-08,6.829e-05,0.001021,0.0009826,3.629e-05,0.003426,0.0008421,0.000152,0.0008217,0.002211,0.001151,0.0007669,5.968e-06,0.0009676,1.973e-07,0.001374,2.432e-06,1.79e-06,0.002157,1.769e-05,0.0002,0.002721,0.0002175,0.0004976,6.318e-05,0.000157,0.00116,4.053e-06,0.0004735,1.102e-05,0.001284,3.696e-07,0.001983,0.0007642,0.0001497,0.003549,6.416e-08,0.001865,9.123e-05,0.0001748,0.003269,1.334e-05,9.593e-06,1.822e-05,0.002077,6.97e-05,0.0002108,1.215e-05,3.616e-06,1.996e-05,0.0008242,0.001008,0.0002335,0.0002631,0.0009452,0.0004119,4.337e-05,0.001959,4.666e-07,1.052e-05,0.0005683,5.728e-05,0.000269,0.0004949,0.0001151,0.0002038,0.001623,0.0008627,8.271e-06,1.163e-06,0.00255,0.0003897,0.0035,1.911e-06,0.000937,4.251e-05,1.575e-08,0.0001352,0.001663,0.0001739,0.0004294,6.58e-05,4.023e-06,3.731e-05,8.638e-05,3.475e-07,5.752e-05,0.0001002,1.386e-05,0.0009855,0.0001389,9.766e-08,0.0001463,1.185e-05,0.002492,0.001109,0.002554,2.718e-05,1.468e-07,0.00161,0.0005043,1.301e-05,0.00153,0.0002425,0.0003597,0.003574,6.644e-07,5.148e-05,9.951e-05,2.381e-05,0.0002917,0.0008574,6.995e-08,0.0006593,0.0004973,2.565e-05,1.05e-07,0.0006064,0.0001312,3.587e-07,0.002542,0.0001607,4.001e-06,0.00086,0.001405,0.0003431,7.622e-05,0.0003277,8.011e-06,1.111e-05,0.0001418,0.0005629,0.002099,1.262e-06,7.604e-05,0.000219,0.0008908,0.002646,0.00206,6.726e-05,0.0003473,0.000741,0.003385,0.000849,6.208e-07,0.0001739,9.355e-05,8.76e-07,0.002497,8.616e-05,4.284e-07,0.001656,0.0001412,0.002272,0.0001882,0.0002328,0.0003518,8.648e-05,0.0008686,0.0002257,1.027e-05,0.0005769,1.501e-05,0.001469,4.375e-05,0.001791,0.001032,1.783e-07,0.002577,0.0004514,3.381e-06,0.0001079,3.055e-06,9.625e-05,0.002853,8.547e-08,0.00222,2.973e-05,3.862e-05,0.00168,0.0005087,4.534e-07,0.0005541,0.002775,6.451e-05,0.0001151,3.342e-09,1.12e-06,0.0003566,1.105e-05,1.605e-06,3.741e-07,6.23e-06,0.0006742,4.223e-05,0.002627,0.002167,0.001461,1.762e-05,1.425e-07,0.0005664,0.001846,1.852e-05,0.0002396,0.0009747,4.373e-07,0.0001128,0.0003008,0.001658,0.0001881,0.000252,1.951e-05,0.001287,7.015e-05,3.016e-05,9.857e-05,7.754e-05,0.001117,0.001396,5.961e-06,0.0002021,0.000256,0.0004514,1.31e-05,1.272e-06,0.0004092,0.0002203,0.0001009,0.001418,2.763e-06,1.69e-05,6.26e-06,7.528e-05,0.003285,0.0003312,0.0009955,0.0002647,1.688e-05,1.172e-05,9.032e-05,0.0001948,9.454e-05,2.754e-06,5.669e-06,2.825e-05,0.001095,0.003421,9.141e-08,0.0004439,8.51e-06,0.0008174,0.000234,0.002834,1.198e-07,9.377e-08,0.003142,0.0009319,2.803e-06,4.028e-07,5.96e-05,2.559e-05,2.068e-08,2.917e-05,2.269e-06,8.146e-07,9.597e-05,0.001796,0.00281,0.0006395,1.453e-07,0.0005615,0.0004013,0.0001313,0.00322,3.829e-06,2.516e-05,5.761e-06,0.001734,0.002043,1.118e-07,0.001135,0.0005587,5.878e-10,0.0002473,0.0002444,0.0005763,1.554e-06,2.835e-05,0.001308,2.488e-07,0.003316,0.001822,0.002086,6.634e-07,0.001192,0.0007164,5.167e-05,9.838e-08,6.969e-05,0.00106,0.002446,5.983e-05,5.269e-05,0.0002676,0.0001597,3.098e-07,9.55e-05,2.674e-06,0.002014,0.0006135,0.0004894,0.0001299,2.949e-06,3.705e-05,0.001554,2.41e-06,6.123e-05,0.0001889,1.907e-06,7.749e-05,8.943e-08,9.624e-07,3.237e-05,0.003191,1.777e-05,0.001607,0.0009887,0.001849,0.0002474,0.0001728,0.0002913,0.0002842,0.001157,8.078e-08,5.263e-05,0.0009771,5.976e-05,0.0003914,0.0002995,0.0004563,2.097e-06,6.297e-05,0.001097,2.818e-05,0.0003626,0.0001809,7.481e-06,0.001656,1.902e-05,0.002454,3.255e-05,0.0002395,5.827e-05,0.0005474,0.0001853,1.572e-05,1.536e-05,9.106e-06,5.692e-05,0.001199,6.419e-07,8.254e-06,2.462e-05,0.003303,3.345e-06,0.00036,0.00344,6.837e-10,5.139e-05,3.06e-05,9.612e-06,0.0001065,0.00166,2.389e-05,0.0001629,2.425e-05,2.094e-06,3.778e-05,0.002805,0.0001445,4.891e-06,4.893e-06,1.204e-05,0.001452,1.412e-05,6.672e-05,0.0003346,4.953e-06,4.127e-05,0.0002287,4.619e-05,0.002958,8.541e-06,0.002204,1.798e-06,0.0002092,0.0001195,0.0002869,2.48e-05,0.001912,0.002833,0.001022,0.001227,8.372e-06,0.002743,0.001133,1.113e-05,4.591e-07,0.0008454,2.695e-05,0.002831,0.0002517,2.461e-05,0.0001113,2.862e-06,0.0008156,0.002758,0.0002234,0.002993,0.0003475,0.003148,1.822e-06,5.313e-05,0.003304,0.001076,0.0006651,0.0002711,5.474e-08,0.0004298,0.0002049,0.0009052,0.002529,3.007e-07,0.0001479,2.457e-07,1.294e-05,0.0005993,0.0001845,0.001108,0.0001306,7.971e-07,1.646e-05,0.001881,0.002986,0.0002112,0.0002793,0.00314,6.474e-07,0.002073,7.151e-06,8.119e-08,0.001137,9.793e-05,2.941e-06,3.554e-05,3.788e-05,0.0008282,0.0007454,0.001263,1.101e-05,7.333e-07,0.0001937,8.878e-05,6.736e-05,0.0004468,4.797e-05,3.303e-07,1.908e-05,0.0001885,2.458e-06,8.968e-05,0.003547,0.001636,0.000137,4.643e-08,1.143e-05,5.59e-06,0.002426,0.0001008,3.953e-05,0.0005069,2.566e-06,3.091e-06,0.0001043,2.439e-06,0.0003015,3.512e-05,0.0003721,4.944e-06,0.0007209,0.00292,1.643e-05,7.42e-05,1.398e-07,2.445e-05,0.002077,0.0001647,2.56e-05,0.0007039,0.0001466,7.205e-05,5.171e-06,0.0001292,1.607e-05,0.0003314,0.0001043,0.0002799,3.036e-06,3.747e-07,0.0001349,0.0004005,0.003455,0.000102,0.0001554,0.0005044,0.003331,0.001399,2.041e-05,0.0002996,1.517e-05,0.0001889,7.571e-08,0.0001792,0.0006657,0.0009383,0.003528,5.507e-06,1.444e-08,5.474e-06,2.321e-05,0.0009356,0.0001092,1.819e-05,0.002728,8.661e-06,0.002013,0.0006135,0.00253,2.513e-07,0.001733,0.0002894,1.432e-06,0.0009682,0.0006752,6.969e-05,2.185e-07,0.0006965,0.002135,2.639e-05,2.639e-05,9.388e-05,7.525e-08,9.756e-05,0.0001425,0.003255,6.425e-05,5.933e-08,6.57e-08],"Contig":["Transfer.PvP01_00_1.final","Transfer.PvP01_00_1.final","Transfer.PvP01_00_1.final","Transfer.PvP01_00_1.final","Transfer.PvP01_00_1.final","Transfer.PvP01_00_1.final","Transfer.PvP01_00_1.final","Transfer.PvP01_00_1.final","Transfer.PvP01_00_1.final","Transfer.PvP01_00_1.final","Transfer.PvP01_00_1.final","Transfer.PvP01_00_1.final","Transfer.PvP01_00_1.final","Transfer.PvP01_00_1.final","Transfer.PvP01_00_1.final","Transfer.PvP01_00_1.final","Transfer.PvP01_00_1.final","Transfer.PvP01_00_1.final","Transfer.PvP01_00_1.final","Transfer.PvP01_00_1.final","Transfer.PvP01_00_1.final","Transfer.PvP01_00_2.final","Transfer.PvP01_00_2.final","Transfer.PvP01_00_2.final","Transfer.PvP01_00_2.final","Transfer.PvP01_00_2.final","Transfer.PvP01_00_2.final","Transfer.PvP01_00_2.final","Transfer.PvP01_00_3.final","Transfer.PvP01_00_3.final","Transfer.PvP01_00_3.final","Transfer.PvP01_00_3.final","Transfer.PvP01_00_3.final","Transfer.PvP01_00_3.final","Transfer.PvP01_00_3.final","Transfer.PvP01_00_3.final","Transfer.PvP01_00_4.final","Transfer.PvP01_00_4.final","Transfer.PvP01_00_4.final","Transfer.PvP01_00_4.final","Transfer.PvP01_00_4.final","Transfer.PvP01_00_4.final","Transfer.PvP01_00_4.final","Transfer.PvP01_00_5.final","Transfer.PvP01_00_5.final","Transfer.PvP01_00_5.final","Transfer.PvP01_00_6.final","Transfer.PvP01_00_6.final","Transfer.PvP01_00_6.final","Transfer.PvP01_00_6.final","Transfer.PvP01_00_7.final","Transfer.PvP01_00_7.final","Transfer.PvP01_00_7.final","Transfer.PvP01_00_8.final","Transfer.PvP01_00_8.final","Transfer.PvP01_00_8.final","Transfer.PvP01_00_8.final","Transfer.PvP01_00_8.final","Transfer.PvP01_00_8.final","Transfer.PvP01_00_8.final","Transfer.PvP01_00_8.final","Transfer.PvP01_00_8.final","Transfer.PvP01_00_8.final","Transfer.PvP01_00_9.final","Transfer.PvP01_00_9.final","Transfer.PvP01_00_9.final","Transfer.PvP01_00_10.final","Transfer.PvP01_00_10.final","Transfer.PvP01_00_10.final","Transfer.PvP01_00_10.final","Transfer.PvP01_00_10.final","Transfer.PvP01_00_10.final","Transfer.PvP01_00_11.final","Transfer.PvP01_00_11.final","Transfer.PvP01_00_11.final","Transfer.PvP01_00_11.final","Transfer.PvP01_00_11.final","Transfer.PvP01_00_11.final","Transfer.PvP01_00_11.final","Transfer.PvP01_00_11.final","Transfer.PvP01_00_11.final","Transfer.PvP01_00_11.final","Transfer.PvP01_00_11.final","Transfer.PvP01_00_11.final","Transfer.PvP01_00_11.final","Transfer.PvP01_00_11.final","Transfer.PvP01_00_11.final","Transfer.PvP01_00_11.final","Transfer.PvP01_00_11.final","Transfer.PvP01_00_12.final","Transfer.PvP01_00_12.final","Transfer.PvP01_00_13.final","Transfer.PvP01_00_13.final","Transfer.PvP01_00_14.final","Transfer.PvP01_00_14.final","Transfer.PvP01_00_14.final","Transfer.PvP01_00_14.final","Transfer.PvP01_00_14.final","Transfer.PvP01_00_15.final","Transfer.PvP01_00_15.final","Transfer.PvP01_00_15.final","Transfer.PvP01_00_15.final","Transfer.PvP01_00_15.final","Transfer.PvP01_00_16.final","Transfer.PvP01_00_16.final","Transfer.PvP01_00_17.final","Transfer.PvP01_00_17.final","Transfer.PvP01_00_17.final","Transfer.PvP01_00_17.final","Transfer.PvP01_00_17.final","Transfer.PvP01_00_17.final","Transfer.PvP01_00_17.final","Transfer.PvP01_00_18.final","Transfer.PvP01_00_18.final","Transfer.PvP01_00_18.final","Transfer.PvP01_00_19.final","Transfer.PvP01_00_22.final","Transfer.PvP01_00_22.final","Transfer.PvP01_00_22.final","Transfer.PvP01_00_22.final","Transfer.PvP01_00_22.final","Transfer.PvP01_00_23.final","Transfer.PvP01_00_23.final","Transfer.PvP01_00_24.final","Transfer.PvP01_00_25.final","Transfer.PvP01_00_26.final","Transfer.PvP01_00_26.final","Transfer.PvP01_00_26.final","Transfer.PvP01_00_26.final","Transfer.PvP01_00_26.final","Transfer.PvP01_00_29.final","Transfer.PvP01_00_29.final","Transfer.PvP01_00_29.final","Transfer.PvP01_00_30.final","Transfer.PvP01_00_31.final","Transfer.PvP01_00_31.final","Transfer.PvP01_00_31.final","Transfer.PvP01_00_32.final","Transfer.PvP01_00_34.final","Transfer.PvP01_00_37.final","Transfer.PvP01_00_38.final","Transfer.PvP01_00_38.final","Transfer.PvP01_00_39.final","Transfer.PvP01_00_41.final","Transfer.PvP01_00_42.final","Transfer.PvP01_00_42.final","Transfer.PvP01_00_45.final","Transfer.PvP01_00_45.final","Transfer.PvP01_00_48.final","Transfer.PvP01_00_48.final","Transfer.PvP01_00_48.final","Transfer.PvP01_00_52.final","Transfer.PvP01_00_53.final","Transfer.PvP01_00_55.final","Transfer.PvP01_00_57.final","Transfer.PvP01_00_61.final","Transfer.PvP01_00_61.final","Transfer.PvP01_00_61.final","Transfer.PvP01_00_63.final","Transfer.PvP01_00_64.final","Transfer.PvP01_00_64.final","Transfer.PvP01_00_66.final","Transfer.PvP01_00_67.final","Transfer.PvP01_00_67.final","Transfer.PvP01_00_67.final","Transfer.PvP01_00_69.final","Transfer.PvP01_00_71.final","Transfer.PvP01_00_74.final","Transfer.PvP01_00_77.final","Transfer.PvP01_00_77.final","Transfer.PvP01_00_79.final","Transfer.PvP01_00_81.final","Transfer.PvP01_00_82.final","Transfer.PvP01_00_83.final","Transfer.PvP01_00_86.final","Transfer.PvP01_00_88.final","Transfer.PvP01_00_89.final","Transfer.PvP01_00_89.final","Transfer.PvP01_00_89.final","Transfer.PvP01_00_90.final","Transfer.PvP01_00_90.final","Transfer.PvP01_00_104.final","Transfer.PvP01_00_108.final","Transfer.PvP01_00_119.final","Transfer.PvP01_00_124.final","Transfer.PvP01_00_129.final","Transfer.PvP01_00_143.final","Transfer.PvP01_00_150.final","Transfer.PvP01_00_152.final","Transfer.PvP01_00_157.final","Transfer.PvP01_00_158.final","Transfer.PvP01_00_161.final","Transfer.PvP01_00_180.final","Transfer.PvP01_00_203.final","Transfer.PvP01_00_208.final","Transfer.PvP01_00_209.final","Transfer.PvP01_00_217.final","Transfer.PvP01_00_221.final","Transfer.PvP01_00_223.final","Transfer.PvP01_00_233.final","Transfer.PvP01_00_300.final","PvP01_01_v1","PvP01_01_v1","PvP01_01_v1","PvP01_01_v1","PvP01_01_v1","PvP01_01_v1","PvP01_01_v1","PvP01_01_v1","PvP01_01_v1","PvP01_01_v1","PvP01_01_v1","PvP01_01_v1","PvP01_01_v1","PvP01_01_v1","PvP01_01_v1","PvP01_01_v1","PvP01_01_v1","PvP01_01_v1","PvP01_01_v1","PvP01_01_v1","PvP01_01_v1","PvP01_01_v1","PvP01_01_v1","PvP01_01_v1","PvP01_01_v1","PvP01_01_v1","PvP01_01_v1","PvP01_01_v1","PvP01_01_v1","PvP01_01_v1","PvP01_01_v1","PvP01_01_v1","PvP01_01_v1","PvP01_01_v1","PvP01_01_v1","PvP01_01_v1","PvP01_01_v1","PvP01_01_v1","PvP01_01_v1","PvP01_01_v1","PvP01_01_v1","PvP01_01_v1","PvP01_01_v1","PvP01_01_v1","PvP01_01_v1","PvP01_01_v1","PvP01_01_v1","PvP01_01_v1","PvP01_01_v1","PvP01_01_v1","PvP01_01_v1","PvP01_01_v1","PvP01_01_v1","PvP01_01_v1","PvP01_01_v1","PvP01_01_v1","PvP01_01_v1","PvP01_01_v1","PvP01_01_v1","PvP01_01_v1","PvP01_01_v1","PvP01_01_v1","PvP01_01_v1","PvP01_01_v1","PvP01_01_v1","PvP01_01_v1","PvP01_01_v1","PvP01_01_v1","PvP01_01_v1","PvP01_01_v1","PvP01_01_v1","PvP01_01_v1","PvP01_01_v1","PvP01_01_v1","PvP01_01_v1","PvP01_01_v1","PvP01_01_v1","PvP01_01_v1","PvP01_01_v1","PvP01_01_v1","PvP01_01_v1","PvP01_01_v1","PvP01_01_v1","PvP01_01_v1","PvP01_01_v1","PvP01_01_v1","PvP01_01_v1","PvP01_01_v1","PvP01_01_v1","PvP01_01_v1","PvP01_01_v1","PvP01_01_v1","PvP01_01_v1","PvP01_01_v1","PvP01_01_v1","PvP01_01_v1","PvP01_01_v1","PvP01_01_v1","PvP01_01_v1","PvP01_01_v1","PvP01_01_v1","PvP01_01_v1","PvP01_01_v1","PvP01_01_v1","PvP01_01_v1","PvP01_01_v1","PvP01_01_v1","PvP01_01_v1","PvP01_01_v1","PvP01_01_v1","PvP01_01_v1","PvP01_01_v1","PvP01_01_v1","PvP01_01_v1","PvP01_01_v1","PvP01_01_v1","PvP01_01_v1","PvP01_01_v1","PvP01_01_v1","PvP01_01_v1","PvP01_01_v1","PvP01_02_v1","PvP01_02_v1","PvP01_02_v1","PvP01_02_v1","PvP01_02_v1","PvP01_02_v1","PvP01_02_v1","PvP01_02_v1","PvP01_02_v1","PvP01_02_v1","PvP01_02_v1","PvP01_02_v1","PvP01_02_v1","PvP01_02_v1","PvP01_02_v1","PvP01_02_v1","PvP01_02_v1","PvP01_02_v1","PvP01_02_v1","PvP01_02_v1","PvP01_02_v1","PvP01_02_v1","PvP01_02_v1","PvP01_02_v1","PvP01_02_v1","PvP01_02_v1","PvP01_02_v1","PvP01_02_v1","PvP01_02_v1","PvP01_02_v1","PvP01_02_v1","PvP01_02_v1","PvP01_02_v1","PvP01_02_v1","PvP01_02_v1","PvP01_02_v1","PvP01_02_v1","PvP01_02_v1","PvP01_02_v1","PvP01_02_v1","PvP01_02_v1","PvP01_02_v1","PvP01_02_v1","PvP01_02_v1","PvP01_02_v1","PvP01_02_v1","PvP01_02_v1","PvP01_02_v1","PvP01_02_v1","PvP01_02_v1","PvP01_02_v1","PvP01_02_v1","PvP01_02_v1","PvP01_02_v1","PvP01_02_v1","PvP01_02_v1","PvP01_02_v1","PvP01_02_v1","PvP01_02_v1","PvP01_02_v1","PvP01_02_v1","PvP01_02_v1","PvP01_02_v1","PvP01_02_v1","PvP01_02_v1","PvP01_02_v1","PvP01_02_v1","PvP01_02_v1","PvP01_02_v1","PvP01_02_v1","PvP01_02_v1","PvP01_02_v1","PvP01_02_v1","PvP01_02_v1","PvP01_02_v1","PvP01_02_v1","PvP01_02_v1","PvP01_02_v1","PvP01_02_v1","PvP01_02_v1","PvP01_02_v1","PvP01_02_v1","PvP01_02_v1","PvP01_02_v1","PvP01_02_v1","PvP01_02_v1","PvP01_02_v1","PvP01_02_v1","PvP01_02_v1","PvP01_02_v1","PvP01_02_v1","PvP01_02_v1","PvP01_02_v1","PvP01_02_v1","PvP01_02_v1","PvP01_02_v1","PvP01_02_v1","PvP01_02_v1","PvP01_02_v1","PvP01_02_v1","PvP01_02_v1","PvP01_02_v1","PvP01_02_v1","PvP01_02_v1","PvP01_02_v1","PvP01_02_v1","PvP01_02_v1","PvP01_02_v1","PvP01_02_v1","PvP01_02_v1","PvP01_02_v1","PvP01_02_v1","PvP01_02_v1","PvP01_02_v1","PvP01_02_v1","PvP01_02_v1","PvP01_02_v1","PvP01_02_v1","PvP01_02_v1","PvP01_02_v1","PvP01_02_v1","PvP01_02_v1","PvP01_02_v1","PvP01_02_v1","PvP01_03_v1","PvP01_03_v1","PvP01_03_v1","PvP01_03_v1","PvP01_03_v1","PvP01_03_v1","PvP01_03_v1","PvP01_03_v1","PvP01_03_v1","PvP01_03_v1","PvP01_03_v1","PvP01_03_v1","PvP01_03_v1","PvP01_03_v1","PvP01_03_v1","PvP01_03_v1","PvP01_03_v1","PvP01_03_v1","PvP01_03_v1","PvP01_03_v1","PvP01_03_v1","PvP01_03_v1","PvP01_03_v1","PvP01_03_v1","PvP01_03_v1","PvP01_03_v1","PvP01_03_v1","PvP01_03_v1","PvP01_03_v1","PvP01_03_v1","PvP01_03_v1","PvP01_03_v1","PvP01_03_v1","PvP01_03_v1","PvP01_03_v1","PvP01_03_v1","PvP01_03_v1","PvP01_03_v1","PvP01_03_v1","PvP01_03_v1","PvP01_03_v1","PvP01_03_v1","PvP01_03_v1","PvP01_03_v1","PvP01_03_v1","PvP01_03_v1","PvP01_03_v1","PvP01_03_v1","PvP01_03_v1","PvP01_03_v1","PvP01_03_v1","PvP01_03_v1","PvP01_03_v1","PvP01_03_v1","PvP01_03_v1","PvP01_03_v1","PvP01_03_v1","PvP01_03_v1","PvP01_03_v1","PvP01_03_v1","PvP01_03_v1","PvP01_03_v1","PvP01_03_v1","PvP01_03_v1","PvP01_03_v1","PvP01_03_v1","PvP01_03_v1","PvP01_03_v1","PvP01_03_v1","PvP01_03_v1","PvP01_03_v1","PvP01_03_v1","PvP01_03_v1","PvP01_03_v1","PvP01_03_v1","PvP01_03_v1","PvP01_03_v1","PvP01_03_v1","PvP01_03_v1","PvP01_03_v1","PvP01_03_v1","PvP01_03_v1","PvP01_03_v1","PvP01_03_v1","PvP01_03_v1","PvP01_03_v1","PvP01_03_v1","PvP01_03_v1","PvP01_03_v1","PvP01_03_v1","PvP01_03_v1","PvP01_03_v1","PvP01_03_v1","PvP01_03_v1","PvP01_03_v1","PvP01_03_v1","PvP01_03_v1","PvP01_03_v1","PvP01_03_v1","PvP01_03_v1","PvP01_03_v1","PvP01_03_v1","PvP01_03_v1","PvP01_03_v1","PvP01_03_v1","PvP01_03_v1","PvP01_03_v1","PvP01_03_v1","PvP01_03_v1","PvP01_03_v1","PvP01_03_v1","PvP01_03_v1","PvP01_03_v1","PvP01_03_v1","PvP01_03_v1","PvP01_03_v1","PvP01_03_v1","PvP01_03_v1","PvP01_03_v1","PvP01_04_v1","PvP01_04_v1","PvP01_04_v1","PvP01_04_v1","PvP01_04_v1","PvP01_04_v1","PvP01_04_v1","PvP01_04_v1","PvP01_04_v1","PvP01_04_v1","PvP01_04_v1","PvP01_04_v1","PvP01_04_v1","PvP01_04_v1","PvP01_04_v1","PvP01_04_v1","PvP01_04_v1","PvP01_04_v1","PvP01_04_v1","PvP01_04_v1","PvP01_04_v1","PvP01_04_v1","PvP01_04_v1","PvP01_04_v1","PvP01_04_v1","PvP01_04_v1","PvP01_04_v1","PvP01_04_v1","PvP01_04_v1","PvP01_04_v1","PvP01_04_v1","PvP01_04_v1","PvP01_04_v1","PvP01_04_v1","PvP01_04_v1","PvP01_04_v1","PvP01_04_v1","PvP01_04_v1","PvP01_04_v1","PvP01_04_v1","PvP01_04_v1","PvP01_04_v1","PvP01_04_v1","PvP01_04_v1","PvP01_04_v1","PvP01_04_v1","PvP01_04_v1","PvP01_04_v1","PvP01_04_v1","PvP01_04_v1","PvP01_04_v1","PvP01_04_v1","PvP01_04_v1","PvP01_04_v1","PvP01_04_v1","PvP01_04_v1","PvP01_04_v1","PvP01_04_v1","PvP01_04_v1","PvP01_04_v1","PvP01_04_v1","PvP01_04_v1","PvP01_04_v1","PvP01_04_v1","PvP01_04_v1","PvP01_04_v1","PvP01_04_v1","PvP01_04_v1","PvP01_04_v1","PvP01_04_v1","PvP01_04_v1","PvP01_04_v1","PvP01_04_v1","PvP01_04_v1","PvP01_04_v1","PvP01_04_v1","PvP01_04_v1","PvP01_04_v1","PvP01_04_v1","PvP01_04_v1","PvP01_04_v1","PvP01_04_v1","PvP01_04_v1","PvP01_04_v1","PvP01_04_v1","PvP01_04_v1","PvP01_04_v1","PvP01_04_v1","PvP01_04_v1","PvP01_04_v1","PvP01_04_v1","PvP01_04_v1","PvP01_04_v1","PvP01_04_v1","PvP01_04_v1","PvP01_04_v1","PvP01_04_v1","PvP01_04_v1","PvP01_04_v1","PvP01_04_v1","PvP01_04_v1","PvP01_04_v1","PvP01_04_v1","PvP01_04_v1","PvP01_04_v1","PvP01_04_v1","PvP01_04_v1","PvP01_04_v1","PvP01_04_v1","PvP01_04_v1","PvP01_04_v1","PvP01_04_v1","PvP01_04_v1","PvP01_04_v1","PvP01_04_v1","PvP01_04_v1","PvP01_04_v1","PvP01_04_v1","PvP01_04_v1","PvP01_04_v1","PvP01_04_v1","PvP01_04_v1","PvP01_04_v1","PvP01_04_v1","PvP01_05_v1","PvP01_05_v1","PvP01_05_v1","PvP01_05_v1","PvP01_05_v1","PvP01_05_v1","PvP01_05_v1","PvP01_05_v1","PvP01_05_v1","PvP01_05_v1","PvP01_05_v1","PvP01_05_v1","PvP01_05_v1","PvP01_05_v1","PvP01_05_v1","PvP01_05_v1","PvP01_05_v1","PvP01_05_v1","PvP01_05_v1","PvP01_05_v1","PvP01_05_v1","PvP01_05_v1","PvP01_05_v1","PvP01_05_v1","PvP01_05_v1","PvP01_05_v1","PvP01_05_v1","PvP01_05_v1","PvP01_05_v1","PvP01_05_v1","PvP01_05_v1","PvP01_05_v1","PvP01_05_v1","PvP01_05_v1","PvP01_05_v1","PvP01_05_v1","PvP01_05_v1","PvP01_05_v1","PvP01_05_v1","PvP01_05_v1","PvP01_05_v1","PvP01_05_v1","PvP01_05_v1","PvP01_05_v1","PvP01_05_v1","PvP01_05_v1","PvP01_05_v1","PvP01_05_v1","PvP01_05_v1","PvP01_05_v1","PvP01_05_v1","PvP01_05_v1","PvP01_05_v1","PvP01_05_v1","PvP01_05_v1","PvP01_05_v1","PvP01_05_v1","PvP01_05_v1","PvP01_05_v1","PvP01_05_v1","PvP01_05_v1","PvP01_05_v1","PvP01_05_v1","PvP01_05_v1","PvP01_05_v1","PvP01_05_v1","PvP01_05_v1","PvP01_05_v1","PvP01_05_v1","PvP01_05_v1","PvP01_05_v1","PvP01_05_v1","PvP01_05_v1","PvP01_05_v1","PvP01_05_v1","PvP01_05_v1","PvP01_05_v1","PvP01_05_v1","PvP01_05_v1","PvP01_05_v1","PvP01_05_v1","PvP01_05_v1","PvP01_05_v1","PvP01_05_v1","PvP01_05_v1","PvP01_05_v1","PvP01_05_v1","PvP01_05_v1","PvP01_05_v1","PvP01_05_v1","PvP01_05_v1","PvP01_05_v1","PvP01_05_v1","PvP01_05_v1","PvP01_05_v1","PvP01_05_v1","PvP01_05_v1","PvP01_05_v1","PvP01_05_v1","PvP01_05_v1","PvP01_05_v1","PvP01_05_v1","PvP01_05_v1","PvP01_05_v1","PvP01_05_v1","PvP01_05_v1","PvP01_05_v1","PvP01_05_v1","PvP01_05_v1","PvP01_05_v1","PvP01_05_v1","PvP01_05_v1","PvP01_05_v1","PvP01_05_v1","PvP01_05_v1","PvP01_05_v1","PvP01_05_v1","PvP01_05_v1","PvP01_05_v1","PvP01_05_v1","PvP01_05_v1","PvP01_05_v1","PvP01_05_v1","PvP01_05_v1","PvP01_05_v1","PvP01_05_v1","PvP01_05_v1","PvP01_05_v1","PvP01_05_v1","PvP01_05_v1","PvP01_05_v1","PvP01_05_v1","PvP01_05_v1","PvP01_05_v1","PvP01_05_v1","PvP01_05_v1","PvP01_05_v1","PvP01_05_v1","PvP01_05_v1","PvP01_05_v1","PvP01_05_v1","PvP01_05_v1","PvP01_05_v1","PvP01_05_v1","PvP01_05_v1","PvP01_05_v1","PvP01_05_v1","PvP01_05_v1","PvP01_05_v1","PvP01_05_v1","PvP01_05_v1","PvP01_05_v1","PvP01_05_v1","PvP01_05_v1","PvP01_05_v1","PvP01_05_v1","PvP01_05_v1","PvP01_05_v1","PvP01_05_v1","PvP01_05_v1","PvP01_05_v1","PvP01_05_v1","PvP01_05_v1","PvP01_05_v1","PvP01_05_v1","PvP01_05_v1","PvP01_05_v1","PvP01_05_v1","PvP01_05_v1","PvP01_05_v1","PvP01_05_v1","PvP01_05_v1","PvP01_05_v1","PvP01_05_v1","PvP01_05_v1","PvP01_05_v1","PvP01_05_v1","PvP01_05_v1","PvP01_05_v1","PvP01_05_v1","PvP01_05_v1","PvP01_05_v1","PvP01_05_v1","PvP01_05_v1","PvP01_05_v1","PvP01_05_v1","PvP01_05_v1","PvP01_05_v1","PvP01_05_v1","PvP01_05_v1","PvP01_05_v1","PvP01_05_v1","PvP01_05_v1","PvP01_05_v1","PvP01_05_v1","PvP01_05_v1","PvP01_05_v1","PvP01_05_v1","PvP01_05_v1","PvP01_05_v1","PvP01_05_v1","PvP01_06_v1","PvP01_06_v1","PvP01_06_v1","PvP01_06_v1","PvP01_06_v1","PvP01_06_v1","PvP01_06_v1","PvP01_06_v1","PvP01_06_v1","PvP01_06_v1","PvP01_06_v1","PvP01_06_v1","PvP01_06_v1","PvP01_06_v1","PvP01_06_v1","PvP01_06_v1","PvP01_06_v1","PvP01_06_v1","PvP01_06_v1","PvP01_06_v1","PvP01_06_v1","PvP01_06_v1","PvP01_06_v1","PvP01_06_v1","PvP01_06_v1","PvP01_06_v1","PvP01_06_v1","PvP01_06_v1","PvP01_06_v1","PvP01_06_v1","PvP01_06_v1","PvP01_06_v1","PvP01_06_v1","PvP01_06_v1","PvP01_06_v1","PvP01_06_v1","PvP01_06_v1","PvP01_06_v1","PvP01_06_v1","PvP01_06_v1","PvP01_06_v1","PvP01_06_v1","PvP01_06_v1","PvP01_06_v1","PvP01_06_v1","PvP01_06_v1","PvP01_06_v1","PvP01_06_v1","PvP01_06_v1","PvP01_06_v1","PvP01_06_v1","PvP01_06_v1","PvP01_06_v1","PvP01_06_v1","PvP01_06_v1","PvP01_06_v1","PvP01_06_v1","PvP01_06_v1","PvP01_06_v1","PvP01_06_v1","PvP01_06_v1","PvP01_06_v1","PvP01_06_v1","PvP01_06_v1","PvP01_06_v1","PvP01_06_v1","PvP01_06_v1","PvP01_06_v1","PvP01_06_v1","PvP01_06_v1","PvP01_06_v1","PvP01_06_v1","PvP01_06_v1","PvP01_06_v1","PvP01_06_v1","PvP01_06_v1","PvP01_06_v1","PvP01_06_v1","PvP01_06_v1","PvP01_06_v1","PvP01_06_v1","PvP01_06_v1","PvP01_06_v1","PvP01_06_v1","PvP01_06_v1","PvP01_06_v1","PvP01_06_v1","PvP01_06_v1","PvP01_06_v1","PvP01_06_v1","PvP01_06_v1","PvP01_06_v1","PvP01_06_v1","PvP01_06_v1","PvP01_06_v1","PvP01_06_v1","PvP01_06_v1","PvP01_06_v1","PvP01_06_v1","PvP01_06_v1","PvP01_06_v1","PvP01_06_v1","PvP01_06_v1","PvP01_06_v1","PvP01_06_v1","PvP01_06_v1","PvP01_06_v1","PvP01_06_v1","PvP01_06_v1","PvP01_06_v1","PvP01_06_v1","PvP01_06_v1","PvP01_06_v1","PvP01_06_v1","PvP01_06_v1","PvP01_06_v1","PvP01_06_v1","PvP01_06_v1","PvP01_06_v1","PvP01_06_v1","PvP01_06_v1","PvP01_06_v1","PvP01_06_v1","PvP01_06_v1","PvP01_06_v1","PvP01_06_v1","PvP01_06_v1","PvP01_06_v1","PvP01_06_v1","PvP01_06_v1","PvP01_06_v1","PvP01_06_v1","PvP01_06_v1","PvP01_06_v1","PvP01_06_v1","PvP01_06_v1","PvP01_06_v1","PvP01_06_v1","PvP01_06_v1","PvP01_06_v1","PvP01_06_v1","PvP01_06_v1","PvP01_06_v1","PvP01_06_v1","PvP01_06_v1","PvP01_06_v1","PvP01_06_v1","PvP01_06_v1","PvP01_06_v1","PvP01_06_v1","PvP01_07_v1","PvP01_07_v1","PvP01_07_v1","PvP01_07_v1","PvP01_07_v1","PvP01_07_v1","PvP01_07_v1","PvP01_07_v1","PvP01_07_v1","PvP01_07_v1","PvP01_07_v1","PvP01_07_v1","PvP01_07_v1","PvP01_07_v1","PvP01_07_v1","PvP01_07_v1","PvP01_07_v1","PvP01_07_v1","PvP01_07_v1","PvP01_07_v1","PvP01_07_v1","PvP01_07_v1","PvP01_07_v1","PvP01_07_v1","PvP01_07_v1","PvP01_07_v1","PvP01_07_v1","PvP01_07_v1","PvP01_07_v1","PvP01_07_v1","PvP01_07_v1","PvP01_07_v1","PvP01_07_v1","PvP01_07_v1","PvP01_07_v1","PvP01_07_v1","PvP01_07_v1","PvP01_07_v1","PvP01_07_v1","PvP01_07_v1","PvP01_07_v1","PvP01_07_v1","PvP01_07_v1","PvP01_07_v1","PvP01_07_v1","PvP01_07_v1","PvP01_07_v1","PvP01_07_v1","PvP01_07_v1","PvP01_07_v1","PvP01_07_v1","PvP01_07_v1","PvP01_07_v1","PvP01_07_v1","PvP01_07_v1","PvP01_07_v1","PvP01_07_v1","PvP01_07_v1","PvP01_07_v1","PvP01_07_v1","PvP01_07_v1","PvP01_07_v1","PvP01_07_v1","PvP01_07_v1","PvP01_07_v1","PvP01_07_v1","PvP01_07_v1","PvP01_07_v1","PvP01_07_v1","PvP01_07_v1","PvP01_07_v1","PvP01_07_v1","PvP01_07_v1","PvP01_07_v1","PvP01_07_v1","PvP01_07_v1","PvP01_07_v1","PvP01_07_v1","PvP01_07_v1","PvP01_07_v1","PvP01_07_v1","PvP01_07_v1","PvP01_07_v1","PvP01_07_v1","PvP01_07_v1","PvP01_07_v1","PvP01_07_v1","PvP01_07_v1","PvP01_07_v1","PvP01_07_v1","PvP01_07_v1","PvP01_07_v1","PvP01_07_v1","PvP01_07_v1","PvP01_07_v1","PvP01_07_v1","PvP01_07_v1","PvP01_07_v1","PvP01_07_v1","PvP01_07_v1","PvP01_07_v1","PvP01_07_v1","PvP01_07_v1","PvP01_07_v1","PvP01_07_v1","PvP01_07_v1","PvP01_07_v1","PvP01_07_v1","PvP01_07_v1","PvP01_07_v1","PvP01_07_v1","PvP01_07_v1","PvP01_07_v1","PvP01_07_v1","PvP01_07_v1","PvP01_07_v1","PvP01_07_v1","PvP01_07_v1","PvP01_07_v1","PvP01_07_v1","PvP01_07_v1","PvP01_07_v1","PvP01_07_v1","PvP01_07_v1","PvP01_07_v1","PvP01_07_v1","PvP01_07_v1","PvP01_07_v1","PvP01_07_v1","PvP01_07_v1","PvP01_07_v1","PvP01_07_v1","PvP01_07_v1","PvP01_07_v1","PvP01_07_v1","PvP01_07_v1","PvP01_07_v1","PvP01_07_v1","PvP01_07_v1","PvP01_07_v1","PvP01_07_v1","PvP01_07_v1","PvP01_07_v1","PvP01_07_v1","PvP01_07_v1","PvP01_07_v1","PvP01_07_v1","PvP01_07_v1","PvP01_07_v1","PvP01_07_v1","PvP01_07_v1","PvP01_07_v1","PvP01_07_v1","PvP01_07_v1","PvP01_07_v1","PvP01_07_v1","PvP01_07_v1","PvP01_07_v1","PvP01_07_v1","PvP01_07_v1","PvP01_07_v1","PvP01_07_v1","PvP01_07_v1","PvP01_07_v1","PvP01_07_v1","PvP01_07_v1","PvP01_07_v1","PvP01_07_v1","PvP01_07_v1","PvP01_07_v1","PvP01_07_v1","PvP01_07_v1","PvP01_07_v1","PvP01_07_v1","PvP01_07_v1","PvP01_07_v1","PvP01_07_v1","PvP01_07_v1","PvP01_07_v1","PvP01_07_v1","PvP01_07_v1","PvP01_07_v1","PvP01_07_v1","PvP01_07_v1","PvP01_07_v1","PvP01_07_v1","PvP01_07_v1","PvP01_07_v1","PvP01_07_v1","PvP01_07_v1","PvP01_07_v1","PvP01_07_v1","PvP01_07_v1","PvP01_07_v1","PvP01_07_v1","PvP01_07_v1","PvP01_07_v1","PvP01_07_v1","PvP01_07_v1","PvP01_07_v1","PvP01_07_v1","PvP01_07_v1","PvP01_07_v1","PvP01_07_v1","PvP01_07_v1","PvP01_07_v1","PvP01_07_v1","PvP01_07_v1","PvP01_07_v1","PvP01_07_v1","PvP01_07_v1","PvP01_07_v1","PvP01_07_v1","PvP01_07_v1","PvP01_07_v1","PvP01_07_v1","PvP01_07_v1","PvP01_07_v1","PvP01_07_v1","PvP01_07_v1","PvP01_07_v1","PvP01_07_v1","PvP01_07_v1","PvP01_07_v1","PvP01_07_v1","PvP01_07_v1","PvP01_07_v1","PvP01_07_v1","PvP01_07_v1","PvP01_07_v1","PvP01_08_v1","PvP01_08_v1","PvP01_08_v1","PvP01_08_v1","PvP01_08_v1","PvP01_08_v1","PvP01_08_v1","PvP01_08_v1","PvP01_08_v1","PvP01_08_v1","PvP01_08_v1","PvP01_08_v1","PvP01_08_v1","PvP01_08_v1","PvP01_08_v1","PvP01_08_v1","PvP01_08_v1","PvP01_08_v1","PvP01_08_v1","PvP01_08_v1","PvP01_08_v1","PvP01_08_v1","PvP01_08_v1","PvP01_08_v1","PvP01_08_v1","PvP01_08_v1","PvP01_08_v1","PvP01_08_v1","PvP01_08_v1","PvP01_08_v1","PvP01_08_v1","PvP01_08_v1","PvP01_08_v1","PvP01_08_v1","PvP01_08_v1","PvP01_08_v1","PvP01_08_v1","PvP01_08_v1","PvP01_08_v1","PvP01_08_v1","PvP01_08_v1","PvP01_08_v1","PvP01_08_v1","PvP01_08_v1","PvP01_08_v1","PvP01_08_v1","PvP01_08_v1","PvP01_08_v1","PvP01_08_v1","PvP01_08_v1","PvP01_08_v1","PvP01_08_v1","PvP01_08_v1","PvP01_08_v1","PvP01_08_v1","PvP01_08_v1","PvP01_08_v1","PvP01_08_v1","PvP01_08_v1","PvP01_08_v1","PvP01_08_v1","PvP01_08_v1","PvP01_08_v1","PvP01_08_v1","PvP01_08_v1","PvP01_08_v1","PvP01_08_v1","PvP01_08_v1","PvP01_08_v1","PvP01_08_v1","PvP01_08_v1","PvP01_08_v1","PvP01_08_v1","PvP01_08_v1","PvP01_08_v1","PvP01_08_v1","PvP01_08_v1","PvP01_08_v1","PvP01_08_v1","PvP01_08_v1","PvP01_08_v1","PvP01_08_v1","PvP01_08_v1","PvP01_08_v1","PvP01_08_v1","PvP01_08_v1","PvP01_08_v1","PvP01_08_v1","PvP01_08_v1","PvP01_08_v1","PvP01_08_v1","PvP01_08_v1","PvP01_08_v1","PvP01_08_v1","PvP01_08_v1","PvP01_08_v1","PvP01_08_v1","PvP01_08_v1","PvP01_08_v1","PvP01_08_v1","PvP01_08_v1","PvP01_08_v1","PvP01_08_v1","PvP01_08_v1","PvP01_08_v1","PvP01_08_v1","PvP01_08_v1","PvP01_08_v1","PvP01_08_v1","PvP01_08_v1","PvP01_08_v1","PvP01_08_v1","PvP01_08_v1","PvP01_08_v1","PvP01_08_v1","PvP01_08_v1","PvP01_08_v1","PvP01_08_v1","PvP01_08_v1","PvP01_08_v1","PvP01_08_v1","PvP01_08_v1","PvP01_08_v1","PvP01_08_v1","PvP01_08_v1","PvP01_08_v1","PvP01_08_v1","PvP01_08_v1","PvP01_08_v1","PvP01_08_v1","PvP01_08_v1","PvP01_08_v1","PvP01_08_v1","PvP01_08_v1","PvP01_08_v1","PvP01_08_v1","PvP01_08_v1","PvP01_08_v1","PvP01_08_v1","PvP01_08_v1","PvP01_08_v1","PvP01_08_v1","PvP01_08_v1","PvP01_08_v1","PvP01_08_v1","PvP01_08_v1","PvP01_08_v1","PvP01_08_v1","PvP01_08_v1","PvP01_08_v1","PvP01_08_v1","PvP01_08_v1","PvP01_08_v1","PvP01_08_v1","PvP01_08_v1","PvP01_08_v1","PvP01_08_v1","PvP01_08_v1","PvP01_08_v1","PvP01_08_v1","PvP01_08_v1","PvP01_08_v1","PvP01_08_v1","PvP01_08_v1","PvP01_08_v1","PvP01_08_v1","PvP01_08_v1","PvP01_08_v1","PvP01_08_v1","PvP01_08_v1","PvP01_08_v1","PvP01_08_v1","PvP01_08_v1","PvP01_08_v1","PvP01_08_v1","PvP01_08_v1","PvP01_08_v1","PvP01_08_v1","PvP01_08_v1","PvP01_08_v1","PvP01_08_v1","PvP01_08_v1","PvP01_08_v1","PvP01_08_v1","PvP01_08_v1","PvP01_08_v1","PvP01_08_v1","PvP01_08_v1","PvP01_08_v1","PvP01_08_v1","PvP01_08_v1","PvP01_08_v1","PvP01_08_v1","PvP01_08_v1","PvP01_08_v1","PvP01_08_v1","PvP01_08_v1","PvP01_08_v1","PvP01_08_v1","PvP01_08_v1","PvP01_08_v1","PvP01_08_v1","PvP01_08_v1","PvP01_08_v1","PvP01_08_v1","PvP01_08_v1","PvP01_08_v1","PvP01_08_v1","PvP01_08_v1","PvP01_08_v1","PvP01_08_v1","PvP01_08_v1","PvP01_08_v1","PvP01_08_v1","PvP01_08_v1","PvP01_08_v1","PvP01_08_v1","PvP01_08_v1","PvP01_08_v1","PvP01_08_v1","PvP01_08_v1","PvP01_08_v1","PvP01_08_v1","PvP01_08_v1","PvP01_08_v1","PvP01_08_v1","PvP01_08_v1","PvP01_08_v1","PvP01_08_v1","PvP01_08_v1","PvP01_08_v1","PvP01_08_v1","PvP01_08_v1","PvP01_08_v1","PvP01_08_v1","PvP01_08_v1","PvP01_08_v1","PvP01_08_v1","PvP01_08_v1","PvP01_08_v1","PvP01_08_v1","PvP01_08_v1","PvP01_08_v1","PvP01_08_v1","PvP01_08_v1","PvP01_09_v1","PvP01_09_v1","PvP01_09_v1","PvP01_09_v1","PvP01_09_v1","PvP01_09_v1","PvP01_09_v1","PvP01_09_v1","PvP01_09_v1","PvP01_09_v1","PvP01_09_v1","PvP01_09_v1","PvP01_09_v1","PvP01_09_v1","PvP01_09_v1","PvP01_09_v1","PvP01_09_v1","PvP01_09_v1","PvP01_09_v1","PvP01_09_v1","PvP01_09_v1","PvP01_09_v1","PvP01_09_v1","PvP01_09_v1","PvP01_09_v1","PvP01_09_v1","PvP01_09_v1","PvP01_09_v1","PvP01_09_v1","PvP01_09_v1","PvP01_09_v1","PvP01_09_v1","PvP01_09_v1","PvP01_09_v1","PvP01_09_v1","PvP01_09_v1","PvP01_09_v1","PvP01_09_v1","PvP01_09_v1","PvP01_09_v1","PvP01_09_v1","PvP01_09_v1","PvP01_09_v1","PvP01_09_v1","PvP01_09_v1","PvP01_09_v1","PvP01_09_v1","PvP01_09_v1","PvP01_09_v1","PvP01_09_v1","PvP01_09_v1","PvP01_09_v1","PvP01_09_v1","PvP01_09_v1","PvP01_09_v1","PvP01_09_v1","PvP01_09_v1","PvP01_09_v1","PvP01_09_v1","PvP01_09_v1","PvP01_09_v1","PvP01_09_v1","PvP01_09_v1","PvP01_09_v1","PvP01_09_v1","PvP01_09_v1","PvP01_09_v1","PvP01_09_v1","PvP01_09_v1","PvP01_09_v1","PvP01_09_v1","PvP01_09_v1","PvP01_09_v1","PvP01_09_v1","PvP01_09_v1","PvP01_09_v1","PvP01_09_v1","PvP01_09_v1","PvP01_09_v1","PvP01_09_v1","PvP01_09_v1","PvP01_09_v1","PvP01_09_v1","PvP01_09_v1","PvP01_09_v1","PvP01_09_v1","PvP01_09_v1","PvP01_09_v1","PvP01_09_v1","PvP01_09_v1","PvP01_09_v1","PvP01_09_v1","PvP01_09_v1","PvP01_09_v1","PvP01_09_v1","PvP01_09_v1","PvP01_09_v1","PvP01_09_v1","PvP01_09_v1","PvP01_09_v1","PvP01_09_v1","PvP01_09_v1","PvP01_09_v1","PvP01_09_v1","PvP01_09_v1","PvP01_09_v1","PvP01_09_v1","PvP01_09_v1","PvP01_09_v1","PvP01_09_v1","PvP01_09_v1","PvP01_09_v1","PvP01_09_v1","PvP01_09_v1","PvP01_09_v1","PvP01_09_v1","PvP01_09_v1","PvP01_09_v1","PvP01_09_v1","PvP01_09_v1","PvP01_09_v1","PvP01_09_v1","PvP01_09_v1","PvP01_09_v1","PvP01_09_v1","PvP01_09_v1","PvP01_09_v1","PvP01_09_v1","PvP01_09_v1","PvP01_09_v1","PvP01_09_v1","PvP01_09_v1","PvP01_09_v1","PvP01_09_v1","PvP01_09_v1","PvP01_09_v1","PvP01_09_v1","PvP01_09_v1","PvP01_09_v1","PvP01_09_v1","PvP01_09_v1","PvP01_09_v1","PvP01_09_v1","PvP01_09_v1","PvP01_09_v1","PvP01_09_v1","PvP01_09_v1","PvP01_09_v1","PvP01_09_v1","PvP01_09_v1","PvP01_09_v1","PvP01_09_v1","PvP01_09_v1","PvP01_09_v1","PvP01_09_v1","PvP01_09_v1","PvP01_09_v1","PvP01_09_v1","PvP01_09_v1","PvP01_09_v1","PvP01_09_v1","PvP01_09_v1","PvP01_09_v1","PvP01_09_v1","PvP01_09_v1","PvP01_09_v1","PvP01_09_v1","PvP01_09_v1","PvP01_09_v1","PvP01_09_v1","PvP01_09_v1","PvP01_09_v1","PvP01_09_v1","PvP01_09_v1","PvP01_09_v1","PvP01_09_v1","PvP01_09_v1","PvP01_09_v1","PvP01_09_v1","PvP01_09_v1","PvP01_09_v1","PvP01_09_v1","PvP01_09_v1","PvP01_09_v1","PvP01_09_v1","PvP01_09_v1","PvP01_09_v1","PvP01_09_v1","PvP01_09_v1","PvP01_09_v1","PvP01_09_v1","PvP01_09_v1","PvP01_09_v1","PvP01_09_v1","PvP01_09_v1","PvP01_09_v1","PvP01_09_v1","PvP01_09_v1","PvP01_09_v1","PvP01_09_v1","PvP01_09_v1","PvP01_09_v1","PvP01_09_v1","PvP01_09_v1","PvP01_09_v1","PvP01_09_v1","PvP01_09_v1","PvP01_09_v1","PvP01_09_v1","PvP01_09_v1","PvP01_09_v1","PvP01_09_v1","PvP01_09_v1","PvP01_09_v1","PvP01_09_v1","PvP01_09_v1","PvP01_09_v1","PvP01_09_v1","PvP01_09_v1","PvP01_09_v1","PvP01_09_v1","PvP01_09_v1","PvP01_09_v1","PvP01_09_v1","PvP01_09_v1","PvP01_09_v1","PvP01_09_v1","PvP01_09_v1","PvP01_09_v1","PvP01_09_v1","PvP01_09_v1","PvP01_09_v1","PvP01_09_v1","PvP01_09_v1","PvP01_09_v1","PvP01_09_v1","PvP01_09_v1","PvP01_09_v1","PvP01_09_v1","PvP01_09_v1","PvP01_09_v1","PvP01_09_v1","PvP01_09_v1","PvP01_09_v1","PvP01_09_v1","PvP01_09_v1","PvP01_09_v1","PvP01_09_v1","PvP01_09_v1","PvP01_09_v1","PvP01_09_v1","PvP01_09_v1","PvP01_09_v1","PvP01_09_v1","PvP01_09_v1","PvP01_09_v1","PvP01_09_v1","PvP01_09_v1","PvP01_09_v1","PvP01_09_v1","PvP01_09_v1","PvP01_09_v1","PvP01_09_v1","PvP01_09_v1","PvP01_09_v1","PvP01_09_v1","PvP01_09_v1","PvP01_09_v1","PvP01_09_v1","PvP01_09_v1","PvP01_09_v1","PvP01_09_v1","PvP01_09_v1","PvP01_09_v1","PvP01_09_v1","PvP01_09_v1","PvP01_09_v1","PvP01_09_v1","PvP01_09_v1","PvP01_09_v1","PvP01_09_v1","PvP01_09_v1","PvP01_09_v1","PvP01_09_v1","PvP01_09_v1","PvP01_09_v1","PvP01_09_v1","PvP01_09_v1","PvP01_09_v1","PvP01_09_v1","PvP01_09_v1","PvP01_10_v1","PvP01_10_v1","PvP01_10_v1","PvP01_10_v1","PvP01_10_v1","PvP01_10_v1","PvP01_10_v1","PvP01_10_v1","PvP01_10_v1","PvP01_10_v1","PvP01_10_v1","PvP01_10_v1","PvP01_10_v1","PvP01_10_v1","PvP01_10_v1","PvP01_10_v1","PvP01_10_v1","PvP01_10_v1","PvP01_10_v1","PvP01_10_v1","PvP01_10_v1","PvP01_10_v1","PvP01_10_v1","PvP01_10_v1","PvP01_10_v1","PvP01_10_v1","PvP01_10_v1","PvP01_10_v1","PvP01_10_v1","PvP01_10_v1","PvP01_10_v1","PvP01_10_v1","PvP01_10_v1","PvP01_10_v1","PvP01_10_v1","PvP01_10_v1","PvP01_10_v1","PvP01_10_v1","PvP01_10_v1","PvP01_10_v1","PvP01_10_v1","PvP01_10_v1","PvP01_10_v1","PvP01_10_v1","PvP01_10_v1","PvP01_10_v1","PvP01_10_v1","PvP01_10_v1","PvP01_10_v1","PvP01_10_v1","PvP01_10_v1","PvP01_10_v1","PvP01_10_v1","PvP01_10_v1","PvP01_10_v1","PvP01_10_v1","PvP01_10_v1","PvP01_10_v1","PvP01_10_v1","PvP01_10_v1","PvP01_10_v1","PvP01_10_v1","PvP01_10_v1","PvP01_10_v1","PvP01_10_v1","PvP01_10_v1","PvP01_10_v1","PvP01_10_v1","PvP01_10_v1","PvP01_10_v1","PvP01_10_v1","PvP01_10_v1","PvP01_10_v1","PvP01_10_v1","PvP01_10_v1","PvP01_10_v1","PvP01_10_v1","PvP01_10_v1","PvP01_10_v1","PvP01_10_v1","PvP01_10_v1","PvP01_10_v1","PvP01_10_v1","PvP01_10_v1","PvP01_10_v1","PvP01_10_v1","PvP01_10_v1","PvP01_10_v1","PvP01_10_v1","PvP01_10_v1","PvP01_10_v1","PvP01_10_v1","PvP01_10_v1","PvP01_10_v1","PvP01_10_v1","PvP01_10_v1","PvP01_10_v1","PvP01_10_v1","PvP01_10_v1","PvP01_10_v1","PvP01_10_v1","PvP01_10_v1","PvP01_10_v1","PvP01_10_v1","PvP01_10_v1","PvP01_10_v1","PvP01_10_v1","PvP01_10_v1","PvP01_10_v1","PvP01_10_v1","PvP01_10_v1","PvP01_10_v1","PvP01_10_v1","PvP01_10_v1","PvP01_10_v1","PvP01_10_v1","PvP01_10_v1","PvP01_10_v1","PvP01_10_v1","PvP01_10_v1","PvP01_10_v1","PvP01_10_v1","PvP01_10_v1","PvP01_10_v1","PvP01_10_v1","PvP01_10_v1","PvP01_10_v1","PvP01_10_v1","PvP01_10_v1","PvP01_10_v1","PvP01_10_v1","PvP01_10_v1","PvP01_10_v1","PvP01_10_v1","PvP01_10_v1","PvP01_10_v1","PvP01_10_v1","PvP01_10_v1","PvP01_10_v1","PvP01_10_v1","PvP01_10_v1","PvP01_10_v1","PvP01_10_v1","PvP01_10_v1","PvP01_10_v1","PvP01_10_v1","PvP01_10_v1","PvP01_10_v1","PvP01_10_v1","PvP01_10_v1","PvP01_10_v1","PvP01_10_v1","PvP01_10_v1","PvP01_10_v1","PvP01_10_v1","PvP01_10_v1","PvP01_10_v1","PvP01_10_v1","PvP01_10_v1","PvP01_10_v1","PvP01_10_v1","PvP01_10_v1","PvP01_10_v1","PvP01_10_v1","PvP01_10_v1","PvP01_10_v1","PvP01_10_v1","PvP01_10_v1","PvP01_10_v1","PvP01_10_v1","PvP01_10_v1","PvP01_10_v1","PvP01_10_v1","PvP01_10_v1","PvP01_10_v1","PvP01_10_v1","PvP01_10_v1","PvP01_10_v1","PvP01_10_v1","PvP01_10_v1","PvP01_10_v1","PvP01_10_v1","PvP01_10_v1","PvP01_10_v1","PvP01_10_v1","PvP01_10_v1","PvP01_10_v1","PvP01_10_v1","PvP01_10_v1","PvP01_10_v1","PvP01_10_v1","PvP01_10_v1","PvP01_10_v1","PvP01_10_v1","PvP01_10_v1","PvP01_10_v1","PvP01_10_v1","PvP01_10_v1","PvP01_10_v1","PvP01_10_v1","PvP01_10_v1","PvP01_10_v1","PvP01_10_v1","PvP01_10_v1","PvP01_11_v1","PvP01_11_v1","PvP01_11_v1","PvP01_11_v1","PvP01_11_v1","PvP01_11_v1","PvP01_11_v1","PvP01_11_v1","PvP01_11_v1","PvP01_11_v1","PvP01_11_v1","PvP01_11_v1","PvP01_11_v1","PvP01_11_v1","PvP01_11_v1","PvP01_11_v1","PvP01_11_v1","PvP01_11_v1","PvP01_11_v1","PvP01_11_v1","PvP01_11_v1","PvP01_11_v1","PvP01_11_v1","PvP01_11_v1","PvP01_11_v1","PvP01_11_v1","PvP01_11_v1","PvP01_11_v1","PvP01_11_v1","PvP01_11_v1","PvP01_11_v1","PvP01_11_v1","PvP01_11_v1","PvP01_11_v1","PvP01_11_v1","PvP01_11_v1","PvP01_11_v1","PvP01_11_v1","PvP01_11_v1","PvP01_11_v1","PvP01_11_v1","PvP01_11_v1","PvP01_11_v1","PvP01_11_v1","PvP01_11_v1","PvP01_11_v1","PvP01_11_v1","PvP01_11_v1","PvP01_11_v1","PvP01_11_v1","PvP01_11_v1","PvP01_11_v1","PvP01_11_v1","PvP01_11_v1","PvP01_11_v1","PvP01_11_v1","PvP01_11_v1","PvP01_11_v1","PvP01_11_v1","PvP01_11_v1","PvP01_11_v1","PvP01_11_v1","PvP01_11_v1","PvP01_11_v1","PvP01_11_v1","PvP01_11_v1","PvP01_11_v1","PvP01_11_v1","PvP01_11_v1","PvP01_11_v1","PvP01_11_v1","PvP01_11_v1","PvP01_11_v1","PvP01_11_v1","PvP01_11_v1","PvP01_11_v1","PvP01_11_v1","PvP01_11_v1","PvP01_11_v1","PvP01_11_v1","PvP01_11_v1","PvP01_11_v1","PvP01_11_v1","PvP01_11_v1","PvP01_11_v1","PvP01_11_v1","PvP01_11_v1","PvP01_11_v1","PvP01_11_v1","PvP01_11_v1","PvP01_11_v1","PvP01_11_v1","PvP01_11_v1","PvP01_11_v1","PvP01_11_v1","PvP01_11_v1","PvP01_11_v1","PvP01_11_v1","PvP01_11_v1","PvP01_11_v1","PvP01_11_v1","PvP01_11_v1","PvP01_11_v1","PvP01_11_v1","PvP01_11_v1","PvP01_11_v1","PvP01_11_v1","PvP01_11_v1","PvP01_11_v1","PvP01_11_v1","PvP01_11_v1","PvP01_11_v1","PvP01_11_v1","PvP01_11_v1","PvP01_11_v1","PvP01_11_v1","PvP01_11_v1","PvP01_11_v1","PvP01_11_v1","PvP01_11_v1","PvP01_11_v1","PvP01_11_v1","PvP01_11_v1","PvP01_11_v1","PvP01_11_v1","PvP01_11_v1","PvP01_11_v1","PvP01_11_v1","PvP01_11_v1","PvP01_11_v1","PvP01_11_v1","PvP01_11_v1","PvP01_11_v1","PvP01_11_v1","PvP01_11_v1","PvP01_11_v1","PvP01_11_v1","PvP01_11_v1","PvP01_11_v1","PvP01_11_v1","PvP01_11_v1","PvP01_11_v1","PvP01_11_v1","PvP01_11_v1","PvP01_11_v1","PvP01_11_v1","PvP01_11_v1","PvP01_11_v1","PvP01_11_v1","PvP01_11_v1","PvP01_11_v1","PvP01_11_v1","PvP01_11_v1","PvP01_11_v1","PvP01_11_v1","PvP01_11_v1","PvP01_11_v1","PvP01_11_v1","PvP01_11_v1","PvP01_11_v1","PvP01_11_v1","PvP01_11_v1","PvP01_11_v1","PvP01_11_v1","PvP01_11_v1","PvP01_11_v1","PvP01_11_v1","PvP01_11_v1","PvP01_11_v1","PvP01_11_v1","PvP01_11_v1","PvP01_11_v1","PvP01_11_v1","PvP01_11_v1","PvP01_11_v1","PvP01_11_v1","PvP01_11_v1","PvP01_11_v1","PvP01_11_v1","PvP01_11_v1","PvP01_11_v1","PvP01_11_v1","PvP01_11_v1","PvP01_11_v1","PvP01_11_v1","PvP01_11_v1","PvP01_11_v1","PvP01_11_v1","PvP01_11_v1","PvP01_11_v1","PvP01_11_v1","PvP01_11_v1","PvP01_11_v1","PvP01_11_v1","PvP01_11_v1","PvP01_11_v1","PvP01_11_v1","PvP01_11_v1","PvP01_11_v1","PvP01_11_v1","PvP01_11_v1","PvP01_11_v1","PvP01_11_v1","PvP01_11_v1","PvP01_11_v1","PvP01_11_v1","PvP01_11_v1","PvP01_11_v1","PvP01_11_v1","PvP01_11_v1","PvP01_11_v1","PvP01_11_v1","PvP01_11_v1","PvP01_11_v1","PvP01_11_v1","PvP01_11_v1","PvP01_11_v1","PvP01_11_v1","PvP01_11_v1","PvP01_11_v1","PvP01_11_v1","PvP01_11_v1","PvP01_11_v1","PvP01_11_v1","PvP01_11_v1","PvP01_11_v1","PvP01_11_v1","PvP01_11_v1","PvP01_11_v1","PvP01_11_v1","PvP01_11_v1","PvP01_11_v1","PvP01_11_v1","PvP01_11_v1","PvP01_11_v1","PvP01_11_v1","PvP01_11_v1","PvP01_11_v1","PvP01_11_v1","PvP01_11_v1","PvP01_11_v1","PvP01_11_v1","PvP01_11_v1","PvP01_11_v1","PvP01_11_v1","PvP01_11_v1","PvP01_11_v1","PvP01_11_v1","PvP01_11_v1","PvP01_11_v1","PvP01_11_v1","PvP01_11_v1","PvP01_11_v1","PvP01_11_v1","PvP01_11_v1","PvP01_11_v1","PvP01_11_v1","PvP01_11_v1","PvP01_11_v1","PvP01_11_v1","PvP01_11_v1","PvP01_11_v1","PvP01_11_v1","PvP01_11_v1","PvP01_11_v1","PvP01_11_v1","PvP01_11_v1","PvP01_11_v1","PvP01_11_v1","PvP01_11_v1","PvP01_11_v1","PvP01_11_v1","PvP01_11_v1","PvP01_11_v1","PvP01_11_v1","PvP01_11_v1","PvP01_11_v1","PvP01_11_v1","PvP01_11_v1","PvP01_11_v1","PvP01_11_v1","PvP01_11_v1","PvP01_11_v1","PvP01_11_v1","PvP01_11_v1","PvP01_11_v1","PvP01_11_v1","PvP01_11_v1","PvP01_11_v1","PvP01_11_v1","PvP01_11_v1","PvP01_11_v1","PvP01_11_v1","PvP01_11_v1","PvP01_11_v1","PvP01_11_v1","PvP01_11_v1","PvP01_11_v1","PvP01_11_v1","PvP01_11_v1","PvP01_11_v1","PvP01_11_v1","PvP01_12_v1","PvP01_12_v1","PvP01_12_v1","PvP01_12_v1","PvP01_12_v1","PvP01_12_v1","PvP01_12_v1","PvP01_12_v1","PvP01_12_v1","PvP01_12_v1","PvP01_12_v1","PvP01_12_v1","PvP01_12_v1","PvP01_12_v1","PvP01_12_v1","PvP01_12_v1","PvP01_12_v1","PvP01_12_v1","PvP01_12_v1","PvP01_12_v1","PvP01_12_v1","PvP01_12_v1","PvP01_12_v1","PvP01_12_v1","PvP01_12_v1","PvP01_12_v1","PvP01_12_v1","PvP01_12_v1","PvP01_12_v1","PvP01_12_v1","PvP01_12_v1","PvP01_12_v1","PvP01_12_v1","PvP01_12_v1","PvP01_12_v1","PvP01_12_v1","PvP01_12_v1","PvP01_12_v1","PvP01_12_v1","PvP01_12_v1","PvP01_12_v1","PvP01_12_v1","PvP01_12_v1","PvP01_12_v1","PvP01_12_v1","PvP01_12_v1","PvP01_12_v1","PvP01_12_v1","PvP01_12_v1","PvP01_12_v1","PvP01_12_v1","PvP01_12_v1","PvP01_12_v1","PvP01_12_v1","PvP01_12_v1","PvP01_12_v1","PvP01_12_v1","PvP01_12_v1","PvP01_12_v1","PvP01_12_v1","PvP01_12_v1","PvP01_12_v1","PvP01_12_v1","PvP01_12_v1","PvP01_12_v1","PvP01_12_v1","PvP01_12_v1","PvP01_12_v1","PvP01_12_v1","PvP01_12_v1","PvP01_12_v1","PvP01_12_v1","PvP01_12_v1","PvP01_12_v1","PvP01_12_v1","PvP01_12_v1","PvP01_12_v1","PvP01_12_v1","PvP01_12_v1","PvP01_12_v1","PvP01_12_v1","PvP01_12_v1","PvP01_12_v1","PvP01_12_v1","PvP01_12_v1","PvP01_12_v1","PvP01_12_v1","PvP01_12_v1","PvP01_12_v1","PvP01_12_v1","PvP01_12_v1","PvP01_12_v1","PvP01_12_v1","PvP01_12_v1","PvP01_12_v1","PvP01_12_v1","PvP01_12_v1","PvP01_12_v1","PvP01_12_v1","PvP01_12_v1","PvP01_12_v1","PvP01_12_v1","PvP01_12_v1","PvP01_12_v1","PvP01_12_v1","PvP01_12_v1","PvP01_12_v1","PvP01_12_v1","PvP01_12_v1","PvP01_12_v1","PvP01_12_v1","PvP01_12_v1","PvP01_12_v1","PvP01_12_v1","PvP01_12_v1","PvP01_12_v1","PvP01_12_v1","PvP01_12_v1","PvP01_12_v1","PvP01_12_v1","PvP01_12_v1","PvP01_12_v1","PvP01_12_v1","PvP01_12_v1","PvP01_12_v1","PvP01_12_v1","PvP01_12_v1","PvP01_12_v1","PvP01_12_v1","PvP01_12_v1","PvP01_12_v1","PvP01_12_v1","PvP01_12_v1","PvP01_12_v1","PvP01_12_v1","PvP01_12_v1","PvP01_12_v1","PvP01_12_v1","PvP01_12_v1","PvP01_12_v1","PvP01_12_v1","PvP01_12_v1","PvP01_12_v1","PvP01_12_v1","PvP01_12_v1","PvP01_12_v1","PvP01_12_v1","PvP01_12_v1","PvP01_12_v1","PvP01_12_v1","PvP01_12_v1","PvP01_12_v1","PvP01_12_v1","PvP01_12_v1","PvP01_12_v1","PvP01_12_v1","PvP01_12_v1","PvP01_12_v1","PvP01_12_v1","PvP01_12_v1","PvP01_12_v1","PvP01_12_v1","PvP01_12_v1","PvP01_12_v1","PvP01_12_v1","PvP01_12_v1","PvP01_12_v1","PvP01_12_v1","PvP01_12_v1","PvP01_12_v1","PvP01_12_v1","PvP01_12_v1","PvP01_12_v1","PvP01_12_v1","PvP01_12_v1","PvP01_12_v1","PvP01_12_v1","PvP01_12_v1","PvP01_12_v1","PvP01_12_v1","PvP01_12_v1","PvP01_12_v1","PvP01_12_v1","PvP01_12_v1","PvP01_12_v1","PvP01_12_v1","PvP01_12_v1","PvP01_12_v1","PvP01_12_v1","PvP01_12_v1","PvP01_12_v1","PvP01_12_v1","PvP01_12_v1","PvP01_12_v1","PvP01_12_v1","PvP01_12_v1","PvP01_12_v1","PvP01_12_v1","PvP01_12_v1","PvP01_12_v1","PvP01_12_v1","PvP01_12_v1","PvP01_12_v1","PvP01_12_v1","PvP01_12_v1","PvP01_12_v1","PvP01_12_v1","PvP01_12_v1","PvP01_12_v1","PvP01_12_v1","PvP01_12_v1","PvP01_12_v1","PvP01_12_v1","PvP01_12_v1","PvP01_12_v1","PvP01_12_v1","PvP01_12_v1","PvP01_12_v1","PvP01_12_v1","PvP01_12_v1","PvP01_12_v1","PvP01_12_v1","PvP01_12_v1","PvP01_12_v1","PvP01_12_v1","PvP01_12_v1","PvP01_12_v1","PvP01_12_v1","PvP01_12_v1","PvP01_12_v1","PvP01_12_v1","PvP01_12_v1","PvP01_12_v1","PvP01_12_v1","PvP01_12_v1","PvP01_12_v1","PvP01_12_v1","PvP01_12_v1","PvP01_12_v1","PvP01_12_v1","PvP01_12_v1","PvP01_12_v1","PvP01_12_v1","PvP01_12_v1","PvP01_12_v1","PvP01_12_v1","PvP01_12_v1","PvP01_12_v1","PvP01_12_v1","PvP01_12_v1","PvP01_12_v1","PvP01_12_v1","PvP01_12_v1","PvP01_12_v1","PvP01_12_v1","PvP01_12_v1","PvP01_12_v1","PvP01_12_v1","PvP01_12_v1","PvP01_12_v1","PvP01_12_v1","PvP01_12_v1","PvP01_12_v1","PvP01_12_v1","PvP01_12_v1","PvP01_12_v1","PvP01_12_v1","PvP01_12_v1","PvP01_12_v1","PvP01_12_v1","PvP01_12_v1","PvP01_12_v1","PvP01_12_v1","PvP01_12_v1","PvP01_12_v1","PvP01_12_v1","PvP01_12_v1","PvP01_12_v1","PvP01_12_v1","PvP01_12_v1","PvP01_12_v1","PvP01_12_v1","PvP01_12_v1","PvP01_12_v1","PvP01_12_v1","PvP01_12_v1","PvP01_12_v1","PvP01_12_v1","PvP01_12_v1","PvP01_12_v1","PvP01_12_v1","PvP01_12_v1","PvP01_12_v1","PvP01_12_v1","PvP01_12_v1","PvP01_12_v1","PvP01_12_v1","PvP01_12_v1","PvP01_12_v1","PvP01_12_v1","PvP01_12_v1","PvP01_12_v1","PvP01_12_v1","PvP01_12_v1","PvP01_12_v1","PvP01_12_v1","PvP01_12_v1","PvP01_12_v1","PvP01_12_v1","PvP01_12_v1","PvP01_12_v1","PvP01_12_v1","PvP01_12_v1","PvP01_12_v1","PvP01_12_v1","PvP01_12_v1","PvP01_12_v1","PvP01_12_v1","PvP01_12_v1","PvP01_12_v1","PvP01_12_v1","PvP01_12_v1","PvP01_12_v1","PvP01_12_v1","PvP01_12_v1","PvP01_12_v1","PvP01_12_v1","PvP01_12_v1","PvP01_12_v1","PvP01_12_v1","PvP01_12_v1","PvP01_12_v1","PvP01_12_v1","PvP01_12_v1","PvP01_12_v1","PvP01_12_v1","PvP01_12_v1","PvP01_12_v1","PvP01_12_v1","PvP01_12_v1","PvP01_12_v1","PvP01_12_v1","PvP01_12_v1","PvP01_12_v1","PvP01_12_v1","PvP01_12_v1","PvP01_12_v1","PvP01_12_v1","PvP01_12_v1","PvP01_12_v1","PvP01_12_v1","PvP01_12_v1","PvP01_12_v1","PvP01_12_v1","PvP01_12_v1","PvP01_12_v1","PvP01_12_v1","PvP01_12_v1","PvP01_12_v1","PvP01_12_v1","PvP01_12_v1","PvP01_12_v1","PvP01_12_v1","PvP01_12_v1","PvP01_12_v1","PvP01_12_v1","PvP01_12_v1","PvP01_12_v1","PvP01_12_v1","PvP01_12_v1","PvP01_12_v1","PvP01_12_v1","PvP01_12_v1","PvP01_12_v1","PvP01_12_v1","PvP01_12_v1","PvP01_12_v1","PvP01_12_v1","PvP01_12_v1","PvP01_12_v1","PvP01_12_v1","PvP01_12_v1","PvP01_12_v1","PvP01_12_v1","PvP01_12_v1","PvP01_12_v1","PvP01_12_v1","PvP01_12_v1","PvP01_12_v1","PvP01_12_v1","PvP01_12_v1","PvP01_12_v1","PvP01_12_v1","PvP01_12_v1","PvP01_12_v1","PvP01_12_v1","PvP01_12_v1","PvP01_12_v1","PvP01_12_v1","PvP01_12_v1","PvP01_12_v1","PvP01_12_v1","PvP01_12_v1","PvP01_12_v1","PvP01_12_v1","PvP01_12_v1","PvP01_12_v1","PvP01_12_v1","PvP01_12_v1","PvP01_12_v1","PvP01_12_v1","PvP01_12_v1","PvP01_12_v1","PvP01_13_v1","PvP01_13_v1","PvP01_13_v1","PvP01_13_v1","PvP01_13_v1","PvP01_13_v1","PvP01_13_v1","PvP01_13_v1","PvP01_13_v1","PvP01_13_v1","PvP01_13_v1","PvP01_13_v1","PvP01_13_v1","PvP01_13_v1","PvP01_13_v1","PvP01_13_v1","PvP01_13_v1","PvP01_13_v1","PvP01_13_v1","PvP01_13_v1","PvP01_13_v1","PvP01_13_v1","PvP01_13_v1","PvP01_13_v1","PvP01_13_v1","PvP01_13_v1","PvP01_13_v1","PvP01_13_v1","PvP01_13_v1","PvP01_13_v1","PvP01_13_v1","PvP01_13_v1","PvP01_13_v1","PvP01_13_v1","PvP01_13_v1","PvP01_13_v1","PvP01_13_v1","PvP01_13_v1","PvP01_13_v1","PvP01_13_v1","PvP01_13_v1","PvP01_13_v1","PvP01_13_v1","PvP01_13_v1","PvP01_13_v1","PvP01_13_v1","PvP01_13_v1","PvP01_13_v1","PvP01_13_v1","PvP01_13_v1","PvP01_13_v1","PvP01_13_v1","PvP01_13_v1","PvP01_13_v1","PvP01_13_v1","PvP01_13_v1","PvP01_13_v1","PvP01_13_v1","PvP01_13_v1","PvP01_13_v1","PvP01_13_v1","PvP01_13_v1","PvP01_13_v1","PvP01_13_v1","PvP01_13_v1","PvP01_13_v1","PvP01_13_v1","PvP01_13_v1","PvP01_13_v1","PvP01_13_v1","PvP01_13_v1","PvP01_13_v1","PvP01_13_v1","PvP01_13_v1","PvP01_13_v1","PvP01_13_v1","PvP01_13_v1","PvP01_13_v1","PvP01_13_v1","PvP01_13_v1","PvP01_13_v1","PvP01_13_v1","PvP01_13_v1","PvP01_13_v1","PvP01_13_v1","PvP01_13_v1","PvP01_13_v1","PvP01_13_v1","PvP01_13_v1","PvP01_13_v1","PvP01_13_v1","PvP01_13_v1","PvP01_13_v1","PvP01_13_v1","PvP01_13_v1","PvP01_13_v1","PvP01_13_v1","PvP01_13_v1","PvP01_13_v1","PvP01_13_v1","PvP01_13_v1","PvP01_13_v1","PvP01_13_v1","PvP01_13_v1","PvP01_13_v1","PvP01_13_v1","PvP01_13_v1","PvP01_13_v1","PvP01_13_v1","PvP01_13_v1","PvP01_13_v1","PvP01_13_v1","PvP01_13_v1","PvP01_13_v1","PvP01_13_v1","PvP01_13_v1","PvP01_13_v1","PvP01_13_v1","PvP01_13_v1","PvP01_13_v1","PvP01_13_v1","PvP01_13_v1","PvP01_13_v1","PvP01_13_v1","PvP01_13_v1","PvP01_13_v1","PvP01_13_v1","PvP01_13_v1","PvP01_13_v1","PvP01_13_v1","PvP01_13_v1","PvP01_13_v1","PvP01_13_v1","PvP01_13_v1","PvP01_13_v1","PvP01_13_v1","PvP01_13_v1","PvP01_13_v1","PvP01_13_v1","PvP01_13_v1","PvP01_13_v1","PvP01_13_v1","PvP01_13_v1","PvP01_13_v1","PvP01_13_v1","PvP01_13_v1","PvP01_13_v1","PvP01_13_v1","PvP01_13_v1","PvP01_13_v1","PvP01_13_v1","PvP01_13_v1","PvP01_13_v1","PvP01_13_v1","PvP01_13_v1","PvP01_13_v1","PvP01_13_v1","PvP01_13_v1","PvP01_13_v1","PvP01_13_v1","PvP01_13_v1","PvP01_13_v1","PvP01_13_v1","PvP01_13_v1","PvP01_13_v1","PvP01_13_v1","PvP01_13_v1","PvP01_13_v1","PvP01_13_v1","PvP01_13_v1","PvP01_13_v1","PvP01_13_v1","PvP01_13_v1","PvP01_13_v1","PvP01_13_v1","PvP01_13_v1","PvP01_13_v1","PvP01_13_v1","PvP01_13_v1","PvP01_13_v1","PvP01_13_v1","PvP01_13_v1","PvP01_13_v1","PvP01_13_v1","PvP01_13_v1","PvP01_13_v1","PvP01_13_v1","PvP01_13_v1","PvP01_13_v1","PvP01_13_v1","PvP01_13_v1","PvP01_13_v1","PvP01_13_v1","PvP01_13_v1","PvP01_13_v1","PvP01_13_v1","PvP01_13_v1","PvP01_13_v1","PvP01_13_v1","PvP01_13_v1","PvP01_13_v1","PvP01_13_v1","PvP01_13_v1","PvP01_13_v1","PvP01_13_v1","PvP01_13_v1","PvP01_13_v1","PvP01_13_v1","PvP01_13_v1","PvP01_13_v1","PvP01_13_v1","PvP01_13_v1","PvP01_13_v1","PvP01_13_v1","PvP01_13_v1","PvP01_13_v1","PvP01_13_v1","PvP01_13_v1","PvP01_13_v1","PvP01_13_v1","PvP01_13_v1","PvP01_13_v1","PvP01_13_v1","PvP01_13_v1","PvP01_13_v1","PvP01_13_v1","PvP01_13_v1","PvP01_13_v1","PvP01_13_v1","PvP01_13_v1","PvP01_13_v1","PvP01_13_v1","PvP01_13_v1","PvP01_13_v1","PvP01_13_v1","PvP01_13_v1","PvP01_13_v1","PvP01_13_v1","PvP01_13_v1","PvP01_13_v1","PvP01_13_v1","PvP01_13_v1","PvP01_13_v1","PvP01_13_v1","PvP01_13_v1","PvP01_13_v1","PvP01_13_v1","PvP01_13_v1","PvP01_13_v1","PvP01_13_v1","PvP01_13_v1","PvP01_13_v1","PvP01_13_v1","PvP01_13_v1","PvP01_13_v1","PvP01_13_v1","PvP01_13_v1","PvP01_13_v1","PvP01_13_v1","PvP01_13_v1","PvP01_13_v1","PvP01_13_v1","PvP01_13_v1","PvP01_13_v1","PvP01_13_v1","PvP01_13_v1","PvP01_13_v1","PvP01_13_v1","PvP01_13_v1","PvP01_13_v1","PvP01_13_v1","PvP01_13_v1","PvP01_13_v1","PvP01_14_v1","PvP01_14_v1","PvP01_14_v1","PvP01_14_v1","PvP01_14_v1","PvP01_14_v1","PvP01_14_v1","PvP01_14_v1","PvP01_14_v1","PvP01_14_v1","PvP01_14_v1","PvP01_14_v1","PvP01_14_v1","PvP01_14_v1","PvP01_14_v1","PvP01_14_v1","PvP01_14_v1","PvP01_14_v1","PvP01_14_v1","PvP01_14_v1","PvP01_14_v1","PvP01_14_v1","PvP01_14_v1","PvP01_14_v1","PvP01_14_v1","PvP01_14_v1","PvP01_14_v1","PvP01_14_v1","PvP01_14_v1","PvP01_14_v1","PvP01_14_v1","PvP01_14_v1","PvP01_14_v1","PvP01_14_v1","PvP01_14_v1","PvP01_14_v1","PvP01_14_v1","PvP01_14_v1","PvP01_14_v1","PvP01_14_v1","PvP01_14_v1","PvP01_14_v1","PvP01_14_v1","PvP01_14_v1","PvP01_14_v1","PvP01_14_v1","PvP01_14_v1","PvP01_14_v1","PvP01_14_v1","PvP01_14_v1","PvP01_14_v1","PvP01_14_v1","PvP01_14_v1","PvP01_14_v1","PvP01_14_v1","PvP01_14_v1","PvP01_14_v1","PvP01_14_v1","PvP01_14_v1","PvP01_14_v1","PvP01_14_v1","PvP01_14_v1","PvP01_14_v1","PvP01_14_v1","PvP01_14_v1","PvP01_14_v1","PvP01_14_v1","PvP01_14_v1","PvP01_14_v1","PvP01_14_v1","PvP01_14_v1","PvP01_14_v1","PvP01_14_v1","PvP01_14_v1","PvP01_14_v1","PvP01_14_v1","PvP01_14_v1","PvP01_14_v1","PvP01_14_v1","PvP01_14_v1","PvP01_14_v1","PvP01_14_v1","PvP01_14_v1","PvP01_14_v1","PvP01_14_v1","PvP01_14_v1","PvP01_14_v1","PvP01_14_v1","PvP01_14_v1","PvP01_14_v1","PvP01_14_v1","PvP01_14_v1","PvP01_14_v1","PvP01_14_v1","PvP01_14_v1","PvP01_14_v1","PvP01_14_v1","PvP01_14_v1","PvP01_14_v1","PvP01_14_v1","PvP01_14_v1","PvP01_14_v1","PvP01_14_v1","PvP01_14_v1","PvP01_14_v1","PvP01_14_v1","PvP01_14_v1","PvP01_14_v1","PvP01_14_v1","PvP01_14_v1","PvP01_14_v1","PvP01_14_v1","PvP01_14_v1","PvP01_14_v1","PvP01_14_v1","PvP01_14_v1","PvP01_14_v1","PvP01_14_v1","PvP01_14_v1","PvP01_14_v1","PvP01_14_v1","PvP01_14_v1","PvP01_14_v1","PvP01_14_v1","PvP01_14_v1","PvP01_14_v1","PvP01_14_v1","PvP01_14_v1","PvP01_14_v1","PvP01_14_v1","PvP01_14_v1","PvP01_14_v1","PvP01_14_v1","PvP01_14_v1","PvP01_14_v1","PvP01_14_v1","PvP01_14_v1","PvP01_14_v1","PvP01_14_v1","PvP01_14_v1","PvP01_14_v1","PvP01_14_v1","PvP01_14_v1","PvP01_14_v1","PvP01_14_v1","PvP01_14_v1","PvP01_14_v1","PvP01_14_v1","PvP01_14_v1","PvP01_14_v1","PvP01_14_v1","PvP01_14_v1","PvP01_14_v1","PvP01_14_v1","PvP01_14_v1","PvP01_14_v1","PvP01_14_v1","PvP01_14_v1","PvP01_14_v1","PvP01_14_v1","PvP01_14_v1","PvP01_14_v1","PvP01_14_v1","PvP01_14_v1","PvP01_14_v1","PvP01_14_v1","PvP01_14_v1","PvP01_14_v1","PvP01_14_v1","PvP01_14_v1","PvP01_14_v1","PvP01_14_v1","PvP01_14_v1","PvP01_14_v1","PvP01_14_v1","PvP01_14_v1","PvP01_14_v1","PvP01_14_v1","PvP01_14_v1","PvP01_14_v1","PvP01_14_v1","PvP01_14_v1","PvP01_14_v1","PvP01_14_v1","PvP01_14_v1","PvP01_14_v1","PvP01_14_v1","PvP01_14_v1","PvP01_14_v1","PvP01_14_v1","PvP01_14_v1","PvP01_14_v1","PvP01_14_v1","PvP01_14_v1","PvP01_14_v1","PvP01_14_v1","PvP01_14_v1","PvP01_14_v1","PvP01_14_v1","PvP01_14_v1","PvP01_14_v1","PvP01_14_v1","PvP01_14_v1","PvP01_14_v1","PvP01_14_v1","PvP01_14_v1","PvP01_14_v1","PvP01_14_v1","PvP01_14_v1","PvP01_14_v1","PvP01_14_v1","PvP01_14_v1","PvP01_14_v1","PvP01_14_v1","PvP01_14_v1","PvP01_14_v1","PvP01_14_v1","PvP01_14_v1","PvP01_14_v1","PvP01_14_v1","PvP01_14_v1","PvP01_14_v1","PvP01_14_v1","PvP01_14_v1","PvP01_14_v1","PvP01_14_v1","PvP01_14_v1","PvP01_14_v1","PvP01_14_v1","PvP01_14_v1","PvP01_14_v1","PvP01_14_v1","PvP01_14_v1","PvP01_14_v1","PvP01_14_v1","PvP01_14_v1","PvP01_14_v1","PvP01_14_v1","PvP01_14_v1","PvP01_14_v1","PvP01_14_v1","PvP01_14_v1","PvP01_14_v1","PvP01_14_v1","PvP01_14_v1","PvP01_14_v1","PvP01_14_v1","PvP01_14_v1","PvP01_14_v1","PvP01_14_v1","PvP01_14_v1","PvP01_14_v1","PvP01_14_v1","PvP01_14_v1","PvP01_14_v1","PvP01_14_v1","PvP01_14_v1","PvP01_14_v1","PvP01_14_v1","PvP01_14_v1","PvP01_14_v1","PvP01_14_v1","PvP01_14_v1","PvP01_14_v1","PvP01_14_v1","PvP01_14_v1","PvP01_14_v1","PvP01_14_v1","PvP01_14_v1","PvP01_14_v1","PvP01_14_v1","PvP01_14_v1","PvP01_14_v1","PvP01_14_v1","PvP01_14_v1","PvP01_14_v1","PvP01_14_v1","PvP01_14_v1","PvP01_14_v1","PvP01_14_v1","PvP01_14_v1","PvP01_14_v1","PvP01_14_v1","PvP01_14_v1","PvP01_14_v1","PvP01_14_v1","PvP01_14_v1","PvP01_14_v1","PvP01_14_v1","PvP01_14_v1","PvP01_14_v1","PvP01_14_v1","PvP01_14_v1","PvP01_14_v1","PvP01_14_v1","PvP01_14_v1","PvP01_14_v1","PvP01_14_v1","PvP01_14_v1","PvP01_14_v1","PvP01_14_v1","PvP01_14_v1","PvP01_14_v1","PvP01_14_v1","PvP01_14_v1","PvP01_14_v1","PvP01_14_v1","PvP01_14_v1","PvP01_14_v1","PvP01_14_v1","PvP01_14_v1","PvP01_14_v1","PvP01_14_v1","PvP01_14_v1","PvP01_14_v1","PvP01_14_v1","PvP01_14_v1","PvP01_14_v1","PvP01_14_v1","PvP01_14_v1","PvP01_14_v1","PvP01_14_v1","PvP01_14_v1","PvP01_14_v1","PvP01_14_v1","PvP01_14_v1","PvP01_14_v1","PvP01_14_v1","PvP01_14_v1","PvP01_14_v1","PvP01_14_v1","PvP01_14_v1","PvP01_14_v1","PvP01_14_v1","PvP01_14_v1","PvP01_14_v1","PvP01_14_v1","PvP01_14_v1","PvP01_14_v1","PvP01_14_v1","PvP01_14_v1","PvP01_14_v1","PvP01_14_v1","PvP01_14_v1","PvP01_14_v1","PvP01_14_v1","PvP01_14_v1","PvP01_14_v1","PvP01_14_v1","PvP01_14_v1","PvP01_14_v1","PvP01_14_v1","PvP01_14_v1","PvP01_14_v1","PvP01_14_v1","PvP01_14_v1","PvP01_14_v1","PvP01_14_v1","PvP01_14_v1","PvP01_14_v1","PvP01_14_v1","PvP01_14_v1","PvP01_14_v1","PvP01_14_v1","PvP01_14_v1","PvP01_14_v1","PvP01_14_v1","PvP01_14_v1","PvP01_14_v1","PvP01_14_v1","PvP01_14_v1","PvP01_14_v1","PvP01_14_v1","PvP01_14_v1","PvP01_14_v1","PvP01_14_v1","PvP01_14_v1","PvP01_14_v1","PvP01_14_v1","PvP01_14_v1","PvP01_14_v1","PvP01_14_v1","PvP01_14_v1","PvP01_14_v1","PvP01_14_v1","PvP01_14_v1","PvP01_14_v1","PvP01_14_v1","PvP01_14_v1","PvP01_14_v1","PvP01_14_v1","PvP01_14_v1","PvP01_14_v1","PvP01_14_v1","PvP01_14_v1","PvP01_14_v1","PvP01_14_v1","PvP01_14_v1","PvP01_14_v1","PvP01_14_v1","PvP01_14_v1","PvP01_14_v1","PvP01_14_v1","PvP01_14_v1","PvP01_14_v1","PvP01_14_v1","PvP01_14_v1","PvP01_14_v1","PvP01_14_v1","PvP01_14_v1","PvP01_14_v1","PvP01_14_v1","PvP01_14_v1","PvP01_14_v1","PvP01_14_v1","PvP01_14_v1","PvP01_14_v1","PvP01_14_v1","PvP01_14_v1","PvP01_14_v1","PvP01_14_v1","PvP01_14_v1","PvP01_14_v1","PvP01_14_v1","PvP01_14_v1","PvP01_14_v1","PvP01_14_v1","PvP01_14_v1","PvP01_14_v1","PvP01_14_v1","PvP01_14_v1","PvP01_14_v1","PvP01_14_v1","PvP01_14_v1","PvP01_14_v1","PvP01_14_v1","PvP01_14_v1","PvP01_14_v1","PvP01_API_v1","PvP01_API_v1","PvP01_API_v1","PvP01_API_v1","PvP01_API_v1","Transfer.PvP01_00_1.final","Transfer.PvP01_00_11.final","Transfer.PvP01_00_11.final","Transfer.PvP01_00_16.final","Transfer.PvP01_00_30.final","Transfer.PvP01_00_51.final","Transfer.PvP01_00_138.final","Transfer.PvP01_00_199.final","Transfer.PvP01_00_227.final","PvP01_01_v1","PvP01_01_v1","PvP01_01_v1","PvP01_01_v1","PvP01_01_v1","PvP01_01_v1","PvP01_01_v1","PvP01_01_v1","PvP01_01_v1","PvP01_01_v1","PvP01_01_v1","PvP01_01_v1","PvP01_01_v1","PvP01_01_v1","PvP01_01_v1","PvP01_01_v1","PvP01_01_v1","PvP01_01_v1","PvP01_01_v1","PvP01_01_v1","PvP01_01_v1","PvP01_01_v1","PvP01_01_v1","PvP01_01_v1","PvP01_01_v1","PvP01_01_v1","PvP01_01_v1","PvP01_01_v1","PvP01_01_v1","PvP01_01_v1","PvP01_01_v1","PvP01_01_v1","PvP01_01_v1","PvP01_01_v1","PvP01_01_v1","PvP01_01_v1","PvP01_01_v1","PvP01_01_v1","PvP01_01_v1","PvP01_01_v1","PvP01_01_v1","PvP01_01_v1","PvP01_01_v1","PvP01_01_v1","PvP01_01_v1","PvP01_01_v1","PvP01_01_v1","PvP01_01_v1","PvP01_01_v1","PvP01_01_v1","PvP01_01_v1","PvP01_01_v1","PvP01_01_v1","PvP01_01_v1","PvP01_01_v1","PvP01_01_v1","PvP01_02_v1","PvP01_02_v1","PvP01_02_v1","PvP01_02_v1","PvP01_02_v1","PvP01_02_v1","PvP01_02_v1","PvP01_02_v1","PvP01_02_v1","PvP01_02_v1","PvP01_02_v1","PvP01_02_v1","PvP01_02_v1","PvP01_02_v1","PvP01_02_v1","PvP01_02_v1","PvP01_02_v1","PvP01_02_v1","PvP01_02_v1","PvP01_02_v1","PvP01_02_v1","PvP01_02_v1","PvP01_02_v1","PvP01_02_v1","PvP01_02_v1","PvP01_02_v1","PvP01_02_v1","PvP01_02_v1","PvP01_02_v1","PvP01_02_v1","PvP01_02_v1","PvP01_02_v1","PvP01_02_v1","PvP01_02_v1","PvP01_02_v1","PvP01_02_v1","PvP01_02_v1","PvP01_02_v1","PvP01_02_v1","PvP01_02_v1","PvP01_02_v1","PvP01_02_v1","PvP01_02_v1","PvP01_02_v1","PvP01_02_v1","PvP01_02_v1","PvP01_02_v1","PvP01_03_v1","PvP01_03_v1","PvP01_03_v1","PvP01_03_v1","PvP01_03_v1","PvP01_03_v1","PvP01_03_v1","PvP01_03_v1","PvP01_03_v1","PvP01_03_v1","PvP01_03_v1","PvP01_03_v1","PvP01_03_v1","PvP01_03_v1","PvP01_03_v1","PvP01_03_v1","PvP01_03_v1","PvP01_03_v1","PvP01_03_v1","PvP01_03_v1","PvP01_03_v1","PvP01_03_v1","PvP01_03_v1","PvP01_03_v1","PvP01_03_v1","PvP01_03_v1","PvP01_03_v1","PvP01_03_v1","PvP01_03_v1","PvP01_03_v1","PvP01_03_v1","PvP01_03_v1","PvP01_03_v1","PvP01_03_v1","PvP01_03_v1","PvP01_03_v1","PvP01_03_v1","PvP01_03_v1","PvP01_03_v1","PvP01_03_v1","PvP01_03_v1","PvP01_03_v1","PvP01_03_v1","PvP01_03_v1","PvP01_03_v1","PvP01_03_v1","PvP01_03_v1","PvP01_03_v1","PvP01_03_v1","PvP01_03_v1","PvP01_03_v1","PvP01_03_v1","PvP01_03_v1","PvP01_03_v1","PvP01_03_v1","PvP01_03_v1","PvP01_03_v1","PvP01_03_v1","PvP01_03_v1","PvP01_03_v1","PvP01_03_v1","PvP01_03_v1","PvP01_03_v1","PvP01_03_v1","PvP01_03_v1","PvP01_03_v1","PvP01_03_v1","PvP01_03_v1","PvP01_03_v1","PvP01_03_v1","PvP01_03_v1","PvP01_03_v1","PvP01_03_v1","PvP01_03_v1","PvP01_03_v1","PvP01_03_v1","PvP01_04_v1","PvP01_04_v1","PvP01_04_v1","PvP01_04_v1","PvP01_04_v1","PvP01_04_v1","PvP01_04_v1","PvP01_04_v1","PvP01_04_v1","PvP01_04_v1","PvP01_04_v1","PvP01_04_v1","PvP01_04_v1","PvP01_04_v1","PvP01_04_v1","PvP01_04_v1","PvP01_04_v1","PvP01_04_v1","PvP01_04_v1","PvP01_04_v1","PvP01_04_v1","PvP01_04_v1","PvP01_04_v1","PvP01_04_v1","PvP01_04_v1","PvP01_04_v1","PvP01_04_v1","PvP01_04_v1","PvP01_04_v1","PvP01_04_v1","PvP01_04_v1","PvP01_04_v1","PvP01_04_v1","PvP01_04_v1","PvP01_04_v1","PvP01_04_v1","PvP01_04_v1","PvP01_04_v1","PvP01_04_v1","PvP01_04_v1","PvP01_04_v1","PvP01_04_v1","PvP01_04_v1","PvP01_04_v1","PvP01_04_v1","PvP01_04_v1","PvP01_04_v1","PvP01_04_v1","PvP01_04_v1","PvP01_04_v1","PvP01_04_v1","PvP01_04_v1","PvP01_04_v1","PvP01_04_v1","PvP01_04_v1","PvP01_04_v1","PvP01_04_v1","PvP01_04_v1","PvP01_04_v1","PvP01_04_v1","PvP01_04_v1","PvP01_04_v1","PvP01_04_v1","PvP01_04_v1","PvP01_04_v1","PvP01_04_v1","PvP01_04_v1","PvP01_04_v1","PvP01_04_v1","PvP01_04_v1","PvP01_04_v1","PvP01_04_v1","PvP01_04_v1","PvP01_04_v1","PvP01_04_v1","PvP01_04_v1","PvP01_04_v1","PvP01_04_v1","PvP01_05_v1","PvP01_05_v1","PvP01_05_v1","PvP01_05_v1","PvP01_05_v1","PvP01_05_v1","PvP01_05_v1","PvP01_05_v1","PvP01_05_v1","PvP01_05_v1","PvP01_05_v1","PvP01_05_v1","PvP01_05_v1","PvP01_05_v1","PvP01_05_v1","PvP01_05_v1","PvP01_05_v1","PvP01_05_v1","PvP01_05_v1","PvP01_05_v1","PvP01_05_v1","PvP01_05_v1","PvP01_05_v1","PvP01_05_v1","PvP01_05_v1","PvP01_05_v1","PvP01_05_v1","PvP01_05_v1","PvP01_05_v1","PvP01_05_v1","PvP01_05_v1","PvP01_05_v1","PvP01_05_v1","PvP01_05_v1","PvP01_05_v1","PvP01_05_v1","PvP01_05_v1","PvP01_05_v1","PvP01_05_v1","PvP01_05_v1","PvP01_05_v1","PvP01_05_v1","PvP01_05_v1","PvP01_05_v1","PvP01_05_v1","PvP01_05_v1","PvP01_05_v1","PvP01_05_v1","PvP01_05_v1","PvP01_05_v1","PvP01_05_v1","PvP01_05_v1","PvP01_05_v1","PvP01_05_v1","PvP01_05_v1","PvP01_05_v1","PvP01_05_v1","PvP01_05_v1","PvP01_05_v1","PvP01_05_v1","PvP01_05_v1","PvP01_05_v1","PvP01_05_v1","PvP01_05_v1","PvP01_05_v1","PvP01_05_v1","PvP01_05_v1","PvP01_05_v1","PvP01_05_v1","PvP01_05_v1","PvP01_05_v1","PvP01_05_v1","PvP01_05_v1","PvP01_05_v1","PvP01_05_v1","PvP01_05_v1","PvP01_05_v1","PvP01_05_v1","PvP01_05_v1","PvP01_05_v1","PvP01_05_v1","PvP01_05_v1","PvP01_05_v1","PvP01_05_v1","PvP01_05_v1","PvP01_05_v1","PvP01_05_v1","PvP01_05_v1","PvP01_05_v1","PvP01_05_v1","PvP01_05_v1","PvP01_05_v1","PvP01_05_v1","PvP01_05_v1","PvP01_05_v1","PvP01_05_v1","PvP01_05_v1","PvP01_05_v1","PvP01_05_v1","PvP01_05_v1","PvP01_05_v1","PvP01_05_v1","PvP01_05_v1","PvP01_05_v1","PvP01_05_v1","PvP01_05_v1","PvP01_05_v1","PvP01_05_v1","PvP01_05_v1","PvP01_06_v1","PvP01_06_v1","PvP01_06_v1","PvP01_06_v1","PvP01_06_v1","PvP01_06_v1","PvP01_06_v1","PvP01_06_v1","PvP01_06_v1","PvP01_06_v1","PvP01_06_v1","PvP01_06_v1","PvP01_06_v1","PvP01_06_v1","PvP01_06_v1","PvP01_06_v1","PvP01_06_v1","PvP01_06_v1","PvP01_06_v1","PvP01_06_v1","PvP01_06_v1","PvP01_06_v1","PvP01_06_v1","PvP01_06_v1","PvP01_06_v1","PvP01_06_v1","PvP01_06_v1","PvP01_06_v1","PvP01_06_v1","PvP01_06_v1","PvP01_06_v1","PvP01_06_v1","PvP01_06_v1","PvP01_06_v1","PvP01_06_v1","PvP01_06_v1","PvP01_06_v1","PvP01_06_v1","PvP01_06_v1","PvP01_06_v1","PvP01_06_v1","PvP01_06_v1","PvP01_06_v1","PvP01_06_v1","PvP01_06_v1","PvP01_06_v1","PvP01_06_v1","PvP01_06_v1","PvP01_06_v1","PvP01_06_v1","PvP01_06_v1","PvP01_06_v1","PvP01_06_v1","PvP01_06_v1","PvP01_06_v1","PvP01_06_v1","PvP01_06_v1","PvP01_06_v1","PvP01_06_v1","PvP01_06_v1","PvP01_06_v1","PvP01_06_v1","PvP01_06_v1","PvP01_06_v1","PvP01_06_v1","PvP01_06_v1","PvP01_06_v1","PvP01_06_v1","PvP01_06_v1","PvP01_06_v1","PvP01_06_v1","PvP01_06_v1","PvP01_06_v1","PvP01_06_v1","PvP01_06_v1","PvP01_06_v1","PvP01_06_v1","PvP01_06_v1","PvP01_06_v1","PvP01_06_v1","PvP01_06_v1","PvP01_06_v1","PvP01_06_v1","PvP01_06_v1","PvP01_06_v1","PvP01_06_v1","PvP01_06_v1","PvP01_06_v1","PvP01_06_v1","PvP01_06_v1","PvP01_06_v1","PvP01_07_v1","PvP01_07_v1","PvP01_07_v1","PvP01_07_v1","PvP01_07_v1","PvP01_07_v1","PvP01_07_v1","PvP01_07_v1","PvP01_07_v1","PvP01_07_v1","PvP01_07_v1","PvP01_07_v1","PvP01_07_v1","PvP01_07_v1","PvP01_07_v1","PvP01_07_v1","PvP01_07_v1","PvP01_07_v1","PvP01_07_v1","PvP01_07_v1","PvP01_07_v1","PvP01_07_v1","PvP01_07_v1","PvP01_07_v1","PvP01_07_v1","PvP01_07_v1","PvP01_07_v1","PvP01_07_v1","PvP01_07_v1","PvP01_07_v1","PvP01_07_v1","PvP01_07_v1","PvP01_07_v1","PvP01_07_v1","PvP01_07_v1","PvP01_07_v1","PvP01_07_v1","PvP01_07_v1","PvP01_07_v1","PvP01_07_v1","PvP01_07_v1","PvP01_07_v1","PvP01_07_v1","PvP01_07_v1","PvP01_07_v1","PvP01_07_v1","PvP01_07_v1","PvP01_07_v1","PvP01_07_v1","PvP01_07_v1","PvP01_07_v1","PvP01_07_v1","PvP01_07_v1","PvP01_07_v1","PvP01_07_v1","PvP01_07_v1","PvP01_07_v1","PvP01_07_v1","PvP01_07_v1","PvP01_07_v1","PvP01_07_v1","PvP01_07_v1","PvP01_07_v1","PvP01_07_v1","PvP01_07_v1","PvP01_07_v1","PvP01_07_v1","PvP01_07_v1","PvP01_07_v1","PvP01_07_v1","PvP01_07_v1","PvP01_07_v1","PvP01_07_v1","PvP01_07_v1","PvP01_07_v1","PvP01_07_v1","PvP01_07_v1","PvP01_07_v1","PvP01_07_v1","PvP01_07_v1","PvP01_07_v1","PvP01_07_v1","PvP01_07_v1","PvP01_07_v1","PvP01_07_v1","PvP01_07_v1","PvP01_07_v1","PvP01_07_v1","PvP01_07_v1","PvP01_07_v1","PvP01_07_v1","PvP01_07_v1","PvP01_07_v1","PvP01_07_v1","PvP01_07_v1","PvP01_07_v1","PvP01_07_v1","PvP01_07_v1","PvP01_07_v1","PvP01_07_v1","PvP01_07_v1","PvP01_07_v1","PvP01_07_v1","PvP01_07_v1","PvP01_07_v1","PvP01_07_v1","PvP01_07_v1","PvP01_07_v1","PvP01_07_v1","PvP01_07_v1","PvP01_07_v1","PvP01_07_v1","PvP01_07_v1","PvP01_07_v1","PvP01_07_v1","PvP01_07_v1","PvP01_07_v1","PvP01_07_v1","PvP01_07_v1","PvP01_07_v1","PvP01_07_v1","PvP01_07_v1","PvP01_07_v1","PvP01_07_v1","PvP01_08_v1","PvP01_08_v1","PvP01_08_v1","PvP01_08_v1","PvP01_08_v1","PvP01_08_v1","PvP01_08_v1","PvP01_08_v1","PvP01_08_v1","PvP01_08_v1","PvP01_08_v1","PvP01_08_v1","PvP01_08_v1","PvP01_08_v1","PvP01_08_v1","PvP01_08_v1","PvP01_08_v1","PvP01_08_v1","PvP01_08_v1","PvP01_08_v1","PvP01_08_v1","PvP01_08_v1","PvP01_08_v1","PvP01_08_v1","PvP01_08_v1","PvP01_08_v1","PvP01_08_v1","PvP01_08_v1","PvP01_08_v1","PvP01_08_v1","PvP01_08_v1","PvP01_08_v1","PvP01_08_v1","PvP01_08_v1","PvP01_08_v1","PvP01_08_v1","PvP01_08_v1","PvP01_08_v1","PvP01_08_v1","PvP01_08_v1","PvP01_08_v1","PvP01_08_v1","PvP01_08_v1","PvP01_08_v1","PvP01_08_v1","PvP01_08_v1","PvP01_08_v1","PvP01_08_v1","PvP01_08_v1","PvP01_08_v1","PvP01_08_v1","PvP01_08_v1","PvP01_08_v1","PvP01_08_v1","PvP01_08_v1","PvP01_08_v1","PvP01_08_v1","PvP01_08_v1","PvP01_08_v1","PvP01_08_v1","PvP01_08_v1","PvP01_08_v1","PvP01_08_v1","PvP01_08_v1","PvP01_08_v1","PvP01_08_v1","PvP01_08_v1","PvP01_08_v1","PvP01_08_v1","PvP01_08_v1","PvP01_08_v1","PvP01_08_v1","PvP01_08_v1","PvP01_08_v1","PvP01_08_v1","PvP01_08_v1","PvP01_08_v1","PvP01_08_v1","PvP01_08_v1","PvP01_08_v1","PvP01_08_v1","PvP01_08_v1","PvP01_08_v1","PvP01_08_v1","PvP01_08_v1","PvP01_08_v1","PvP01_08_v1","PvP01_08_v1","PvP01_08_v1","PvP01_08_v1","PvP01_08_v1","PvP01_08_v1","PvP01_08_v1","PvP01_08_v1","PvP01_08_v1","PvP01_08_v1","PvP01_08_v1","PvP01_08_v1","PvP01_08_v1","PvP01_08_v1","PvP01_08_v1","PvP01_08_v1","PvP01_08_v1","PvP01_08_v1","PvP01_08_v1","PvP01_08_v1","PvP01_08_v1","PvP01_08_v1","PvP01_08_v1","PvP01_08_v1","PvP01_08_v1","PvP01_08_v1","PvP01_08_v1","PvP01_08_v1","PvP01_08_v1","PvP01_08_v1","PvP01_08_v1","PvP01_08_v1","PvP01_08_v1","PvP01_08_v1","PvP01_08_v1","PvP01_08_v1","PvP01_08_v1","PvP01_08_v1","PvP01_08_v1","PvP01_08_v1","PvP01_08_v1","PvP01_08_v1","PvP01_08_v1","PvP01_08_v1","PvP01_08_v1","PvP01_08_v1","PvP01_08_v1","PvP01_08_v1","PvP01_08_v1","PvP01_08_v1","PvP01_08_v1","PvP01_08_v1","PvP01_08_v1","PvP01_08_v1","PvP01_08_v1","PvP01_09_v1","PvP01_09_v1","PvP01_09_v1","PvP01_09_v1","PvP01_09_v1","PvP01_09_v1","PvP01_09_v1","PvP01_09_v1","PvP01_09_v1","PvP01_09_v1","PvP01_09_v1","PvP01_09_v1","PvP01_09_v1","PvP01_09_v1","PvP01_09_v1","PvP01_09_v1","PvP01_09_v1","PvP01_09_v1","PvP01_09_v1","PvP01_09_v1","PvP01_09_v1","PvP01_09_v1","PvP01_09_v1","PvP01_09_v1","PvP01_09_v1","PvP01_09_v1","PvP01_09_v1","PvP01_09_v1","PvP01_09_v1","PvP01_09_v1","PvP01_09_v1","PvP01_09_v1","PvP01_09_v1","PvP01_09_v1","PvP01_09_v1","PvP01_09_v1","PvP01_09_v1","PvP01_09_v1","PvP01_09_v1","PvP01_09_v1","PvP01_09_v1","PvP01_09_v1","PvP01_09_v1","PvP01_09_v1","PvP01_09_v1","PvP01_09_v1","PvP01_09_v1","PvP01_09_v1","PvP01_09_v1","PvP01_09_v1","PvP01_09_v1","PvP01_09_v1","PvP01_09_v1","PvP01_09_v1","PvP01_09_v1","PvP01_09_v1","PvP01_09_v1","PvP01_09_v1","PvP01_09_v1","PvP01_09_v1","PvP01_09_v1","PvP01_09_v1","PvP01_09_v1","PvP01_09_v1","PvP01_09_v1","PvP01_09_v1","PvP01_09_v1","PvP01_09_v1","PvP01_09_v1","PvP01_09_v1","PvP01_09_v1","PvP01_09_v1","PvP01_09_v1","PvP01_09_v1","PvP01_09_v1","PvP01_09_v1","PvP01_09_v1","PvP01_09_v1","PvP01_09_v1","PvP01_09_v1","PvP01_09_v1","PvP01_09_v1","PvP01_09_v1","PvP01_09_v1","PvP01_09_v1","PvP01_09_v1","PvP01_09_v1","PvP01_09_v1","PvP01_09_v1","PvP01_09_v1","PvP01_09_v1","PvP01_09_v1","PvP01_09_v1","PvP01_09_v1","PvP01_09_v1","PvP01_09_v1","PvP01_09_v1","PvP01_09_v1","PvP01_09_v1","PvP01_09_v1","PvP01_09_v1","PvP01_09_v1","PvP01_09_v1","PvP01_09_v1","PvP01_09_v1","PvP01_09_v1","PvP01_09_v1","PvP01_09_v1","PvP01_09_v1","PvP01_09_v1","PvP01_09_v1","PvP01_09_v1","PvP01_09_v1","PvP01_09_v1","PvP01_09_v1","PvP01_09_v1","PvP01_09_v1","PvP01_09_v1","PvP01_09_v1","PvP01_09_v1","PvP01_09_v1","PvP01_09_v1","PvP01_09_v1","PvP01_09_v1","PvP01_09_v1","PvP01_09_v1","PvP01_09_v1","PvP01_09_v1","PvP01_09_v1","PvP01_09_v1","PvP01_09_v1","PvP01_09_v1","PvP01_09_v1","PvP01_09_v1","PvP01_09_v1","PvP01_09_v1","PvP01_09_v1","PvP01_09_v1","PvP01_09_v1","PvP01_09_v1","PvP01_09_v1","PvP01_09_v1","PvP01_09_v1","PvP01_09_v1","PvP01_09_v1","PvP01_09_v1","PvP01_09_v1","PvP01_09_v1","PvP01_09_v1","PvP01_09_v1","PvP01_09_v1","PvP01_09_v1","PvP01_09_v1","PvP01_09_v1","PvP01_09_v1","PvP01_09_v1","PvP01_09_v1","PvP01_09_v1","PvP01_09_v1","PvP01_09_v1","PvP01_09_v1","PvP01_09_v1","PvP01_09_v1","PvP01_09_v1","PvP01_09_v1","PvP01_09_v1","PvP01_09_v1","PvP01_09_v1","PvP01_09_v1","PvP01_09_v1","PvP01_10_v1","PvP01_10_v1","PvP01_10_v1","PvP01_10_v1","PvP01_10_v1","PvP01_10_v1","PvP01_10_v1","PvP01_10_v1","PvP01_10_v1","PvP01_10_v1","PvP01_10_v1","PvP01_10_v1","PvP01_10_v1","PvP01_10_v1","PvP01_10_v1","PvP01_10_v1","PvP01_10_v1","PvP01_10_v1","PvP01_10_v1","PvP01_10_v1","PvP01_10_v1","PvP01_10_v1","PvP01_10_v1","PvP01_10_v1","PvP01_10_v1","PvP01_10_v1","PvP01_10_v1","PvP01_10_v1","PvP01_10_v1","PvP01_10_v1","PvP01_10_v1","PvP01_10_v1","PvP01_10_v1","PvP01_10_v1","PvP01_10_v1","PvP01_10_v1","PvP01_10_v1","PvP01_10_v1","PvP01_10_v1","PvP01_10_v1","PvP01_10_v1","PvP01_10_v1","PvP01_10_v1","PvP01_10_v1","PvP01_10_v1","PvP01_10_v1","PvP01_10_v1","PvP01_10_v1","PvP01_10_v1","PvP01_10_v1","PvP01_10_v1","PvP01_10_v1","PvP01_10_v1","PvP01_10_v1","PvP01_10_v1","PvP01_10_v1","PvP01_10_v1","PvP01_10_v1","PvP01_10_v1","PvP01_10_v1","PvP01_10_v1","PvP01_10_v1","PvP01_10_v1","PvP01_10_v1","PvP01_10_v1","PvP01_10_v1","PvP01_10_v1","PvP01_10_v1","PvP01_10_v1","PvP01_10_v1","PvP01_10_v1","PvP01_10_v1","PvP01_10_v1","PvP01_10_v1","PvP01_10_v1","PvP01_10_v1","PvP01_10_v1","PvP01_10_v1","PvP01_10_v1","PvP01_10_v1","PvP01_10_v1","PvP01_10_v1","PvP01_10_v1","PvP01_10_v1","PvP01_10_v1","PvP01_10_v1","PvP01_10_v1","PvP01_10_v1","PvP01_10_v1","PvP01_10_v1","PvP01_10_v1","PvP01_10_v1","PvP01_10_v1","PvP01_10_v1","PvP01_10_v1","PvP01_10_v1","PvP01_10_v1","PvP01_10_v1","PvP01_10_v1","PvP01_10_v1","PvP01_10_v1","PvP01_10_v1","PvP01_10_v1","PvP01_10_v1","PvP01_10_v1","PvP01_10_v1","PvP01_10_v1","PvP01_10_v1","PvP01_10_v1","PvP01_10_v1","PvP01_10_v1","PvP01_10_v1","PvP01_10_v1","PvP01_10_v1","PvP01_10_v1","PvP01_11_v1","PvP01_11_v1","PvP01_11_v1","PvP01_11_v1","PvP01_11_v1","PvP01_11_v1","PvP01_11_v1","PvP01_11_v1","PvP01_11_v1","PvP01_11_v1","PvP01_11_v1","PvP01_11_v1","PvP01_11_v1","PvP01_11_v1","PvP01_11_v1","PvP01_11_v1","PvP01_11_v1","PvP01_11_v1","PvP01_11_v1","PvP01_11_v1","PvP01_11_v1","PvP01_11_v1","PvP01_11_v1","PvP01_11_v1","PvP01_11_v1","PvP01_11_v1","PvP01_11_v1","PvP01_11_v1","PvP01_11_v1","PvP01_11_v1","PvP01_11_v1","PvP01_11_v1","PvP01_11_v1","PvP01_11_v1","PvP01_11_v1","PvP01_11_v1","PvP01_11_v1","PvP01_11_v1","PvP01_11_v1","PvP01_11_v1","PvP01_11_v1","PvP01_11_v1","PvP01_11_v1","PvP01_11_v1","PvP01_11_v1","PvP01_11_v1","PvP01_11_v1","PvP01_11_v1","PvP01_11_v1","PvP01_11_v1","PvP01_11_v1","PvP01_11_v1","PvP01_11_v1","PvP01_11_v1","PvP01_11_v1","PvP01_11_v1","PvP01_11_v1","PvP01_11_v1","PvP01_11_v1","PvP01_11_v1","PvP01_11_v1","PvP01_11_v1","PvP01_11_v1","PvP01_11_v1","PvP01_11_v1","PvP01_11_v1","PvP01_11_v1","PvP01_11_v1","PvP01_11_v1","PvP01_11_v1","PvP01_11_v1","PvP01_11_v1","PvP01_11_v1","PvP01_11_v1","PvP01_11_v1","PvP01_11_v1","PvP01_11_v1","PvP01_11_v1","PvP01_11_v1","PvP01_11_v1","PvP01_11_v1","PvP01_11_v1","PvP01_11_v1","PvP01_11_v1","PvP01_11_v1","PvP01_11_v1","PvP01_11_v1","PvP01_11_v1","PvP01_11_v1","PvP01_11_v1","PvP01_11_v1","PvP01_11_v1","PvP01_11_v1","PvP01_11_v1","PvP01_11_v1","PvP01_11_v1","PvP01_11_v1","PvP01_11_v1","PvP01_11_v1","PvP01_11_v1","PvP01_11_v1","PvP01_11_v1","PvP01_11_v1","PvP01_11_v1","PvP01_11_v1","PvP01_11_v1","PvP01_11_v1","PvP01_11_v1","PvP01_11_v1","PvP01_11_v1","PvP01_11_v1","PvP01_11_v1","PvP01_11_v1","PvP01_11_v1","PvP01_11_v1","PvP01_11_v1","PvP01_11_v1","PvP01_11_v1","PvP01_11_v1","PvP01_11_v1","PvP01_11_v1","PvP01_11_v1","PvP01_11_v1","PvP01_11_v1","PvP01_11_v1","PvP01_11_v1","PvP01_11_v1","PvP01_11_v1","PvP01_11_v1","PvP01_11_v1","PvP01_11_v1","PvP01_11_v1","PvP01_11_v1","PvP01_11_v1","PvP01_11_v1","PvP01_11_v1","PvP01_11_v1","PvP01_11_v1","PvP01_11_v1","PvP01_11_v1","PvP01_11_v1","PvP01_11_v1","PvP01_11_v1","PvP01_11_v1","PvP01_11_v1","PvP01_11_v1","PvP01_11_v1","PvP01_11_v1","PvP01_11_v1","PvP01_11_v1","PvP01_11_v1","PvP01_11_v1","PvP01_11_v1","PvP01_11_v1","PvP01_11_v1","PvP01_11_v1","PvP01_11_v1","PvP01_11_v1","PvP01_11_v1","PvP01_11_v1","PvP01_11_v1","PvP01_11_v1","PvP01_11_v1","PvP01_11_v1","PvP01_11_v1","PvP01_11_v1","PvP01_11_v1","PvP01_12_v1","PvP01_12_v1","PvP01_12_v1","PvP01_12_v1","PvP01_12_v1","PvP01_12_v1","PvP01_12_v1","PvP01_12_v1","PvP01_12_v1","PvP01_12_v1","PvP01_12_v1","PvP01_12_v1","PvP01_12_v1","PvP01_12_v1","PvP01_12_v1","PvP01_12_v1","PvP01_12_v1","PvP01_12_v1","PvP01_12_v1","PvP01_12_v1","PvP01_12_v1","PvP01_12_v1","PvP01_12_v1","PvP01_12_v1","PvP01_12_v1","PvP01_12_v1","PvP01_12_v1","PvP01_12_v1","PvP01_12_v1","PvP01_12_v1","PvP01_12_v1","PvP01_12_v1","PvP01_12_v1","PvP01_12_v1","PvP01_12_v1","PvP01_12_v1","PvP01_12_v1","PvP01_12_v1","PvP01_12_v1","PvP01_12_v1","PvP01_12_v1","PvP01_12_v1","PvP01_12_v1","PvP01_12_v1","PvP01_12_v1","PvP01_12_v1","PvP01_12_v1","PvP01_12_v1","PvP01_12_v1","PvP01_12_v1","PvP01_12_v1","PvP01_12_v1","PvP01_12_v1","PvP01_12_v1","PvP01_12_v1","PvP01_12_v1","PvP01_12_v1","PvP01_12_v1","PvP01_12_v1","PvP01_12_v1","PvP01_12_v1","PvP01_12_v1","PvP01_12_v1","PvP01_12_v1","PvP01_12_v1","PvP01_12_v1","PvP01_12_v1","PvP01_12_v1","PvP01_12_v1","PvP01_12_v1","PvP01_12_v1","PvP01_12_v1","PvP01_12_v1","PvP01_12_v1","PvP01_12_v1","PvP01_12_v1","PvP01_12_v1","PvP01_12_v1","PvP01_12_v1","PvP01_12_v1","PvP01_12_v1","PvP01_12_v1","PvP01_12_v1","PvP01_12_v1","PvP01_12_v1","PvP01_12_v1","PvP01_12_v1","PvP01_12_v1","PvP01_12_v1","PvP01_12_v1","PvP01_12_v1","PvP01_12_v1","PvP01_12_v1","PvP01_12_v1","PvP01_12_v1","PvP01_12_v1","PvP01_12_v1","PvP01_12_v1","PvP01_12_v1","PvP01_12_v1","PvP01_12_v1","PvP01_12_v1","PvP01_12_v1","PvP01_12_v1","PvP01_12_v1","PvP01_12_v1","PvP01_12_v1","PvP01_12_v1","PvP01_12_v1","PvP01_12_v1","PvP01_12_v1","PvP01_12_v1","PvP01_12_v1","PvP01_12_v1","PvP01_12_v1","PvP01_12_v1","PvP01_12_v1","PvP01_12_v1","PvP01_12_v1","PvP01_12_v1","PvP01_12_v1","PvP01_12_v1","PvP01_12_v1","PvP01_12_v1","PvP01_12_v1","PvP01_12_v1","PvP01_12_v1","PvP01_12_v1","PvP01_12_v1","PvP01_12_v1","PvP01_12_v1","PvP01_12_v1","PvP01_12_v1","PvP01_12_v1","PvP01_12_v1","PvP01_12_v1","PvP01_12_v1","PvP01_12_v1","PvP01_12_v1","PvP01_12_v1","PvP01_12_v1","PvP01_12_v1","PvP01_12_v1","PvP01_12_v1","PvP01_12_v1","PvP01_12_v1","PvP01_12_v1","PvP01_12_v1","PvP01_12_v1","PvP01_12_v1","PvP01_12_v1","PvP01_12_v1","PvP01_12_v1","PvP01_12_v1","PvP01_12_v1","PvP01_12_v1","PvP01_12_v1","PvP01_12_v1","PvP01_12_v1","PvP01_12_v1","PvP01_12_v1","PvP01_12_v1","PvP01_12_v1","PvP01_12_v1","PvP01_12_v1","PvP01_12_v1","PvP01_12_v1","PvP01_12_v1","PvP01_12_v1","PvP01_12_v1","PvP01_12_v1","PvP01_12_v1","PvP01_12_v1","PvP01_12_v1","PvP01_12_v1","PvP01_12_v1","PvP01_12_v1","PvP01_12_v1","PvP01_12_v1","PvP01_12_v1","PvP01_12_v1","PvP01_12_v1","PvP01_12_v1","PvP01_12_v1","PvP01_12_v1","PvP01_12_v1","PvP01_12_v1","PvP01_12_v1","PvP01_12_v1","PvP01_12_v1","PvP01_12_v1","PvP01_12_v1","PvP01_12_v1","PvP01_12_v1","PvP01_12_v1","PvP01_12_v1","PvP01_12_v1","PvP01_12_v1","PvP01_12_v1","PvP01_12_v1","PvP01_12_v1","PvP01_12_v1","PvP01_12_v1","PvP01_12_v1","PvP01_12_v1","PvP01_12_v1","PvP01_12_v1","PvP01_12_v1","PvP01_12_v1","PvP01_12_v1","PvP01_12_v1","PvP01_12_v1","PvP01_12_v1","PvP01_12_v1","PvP01_12_v1","PvP01_12_v1","PvP01_12_v1","PvP01_12_v1","PvP01_12_v1","PvP01_12_v1","PvP01_12_v1","PvP01_12_v1","PvP01_12_v1","PvP01_12_v1","PvP01_12_v1","PvP01_12_v1","PvP01_12_v1","PvP01_12_v1","PvP01_12_v1","PvP01_12_v1","PvP01_12_v1","PvP01_12_v1","PvP01_12_v1","PvP01_12_v1","PvP01_12_v1","PvP01_12_v1","PvP01_12_v1","PvP01_12_v1","PvP01_12_v1","PvP01_12_v1","PvP01_12_v1","PvP01_12_v1","PvP01_12_v1","PvP01_12_v1","PvP01_12_v1","PvP01_12_v1","PvP01_12_v1","PvP01_12_v1","PvP01_12_v1","PvP01_12_v1","PvP01_12_v1","PvP01_12_v1","PvP01_12_v1","PvP01_12_v1","PvP01_12_v1","PvP01_12_v1","PvP01_12_v1","PvP01_12_v1","PvP01_12_v1","PvP01_12_v1","PvP01_12_v1","PvP01_12_v1","PvP01_12_v1","PvP01_12_v1","PvP01_12_v1","PvP01_12_v1","PvP01_12_v1","PvP01_12_v1","PvP01_12_v1","PvP01_12_v1","PvP01_12_v1","PvP01_12_v1","PvP01_12_v1","PvP01_12_v1","PvP01_12_v1","PvP01_12_v1","PvP01_12_v1","PvP01_12_v1","PvP01_12_v1","PvP01_12_v1","PvP01_12_v1","PvP01_12_v1","PvP01_12_v1","PvP01_12_v1","PvP01_12_v1","PvP01_12_v1","PvP01_12_v1","PvP01_12_v1","PvP01_13_v1","PvP01_13_v1","PvP01_13_v1","PvP01_13_v1","PvP01_13_v1","PvP01_13_v1","PvP01_13_v1","PvP01_13_v1","PvP01_13_v1","PvP01_13_v1","PvP01_13_v1","PvP01_13_v1","PvP01_13_v1","PvP01_13_v1","PvP01_13_v1","PvP01_13_v1","PvP01_13_v1","PvP01_13_v1","PvP01_13_v1","PvP01_13_v1","PvP01_13_v1","PvP01_13_v1","PvP01_13_v1","PvP01_13_v1","PvP01_13_v1","PvP01_13_v1","PvP01_13_v1","PvP01_13_v1","PvP01_13_v1","PvP01_13_v1","PvP01_13_v1","PvP01_13_v1","PvP01_13_v1","PvP01_13_v1","PvP01_13_v1","PvP01_13_v1","PvP01_13_v1","PvP01_13_v1","PvP01_13_v1","PvP01_13_v1","PvP01_13_v1","PvP01_13_v1","PvP01_13_v1","PvP01_13_v1","PvP01_13_v1","PvP01_13_v1","PvP01_13_v1","PvP01_13_v1","PvP01_13_v1","PvP01_13_v1","PvP01_13_v1","PvP01_13_v1","PvP01_13_v1","PvP01_13_v1","PvP01_13_v1","PvP01_13_v1","PvP01_13_v1","PvP01_13_v1","PvP01_13_v1","PvP01_13_v1","PvP01_13_v1","PvP01_13_v1","PvP01_13_v1","PvP01_13_v1","PvP01_13_v1","PvP01_13_v1","PvP01_13_v1","PvP01_13_v1","PvP01_13_v1","PvP01_13_v1","PvP01_13_v1","PvP01_13_v1","PvP01_13_v1","PvP01_13_v1","PvP01_13_v1","PvP01_13_v1","PvP01_13_v1","PvP01_13_v1","PvP01_13_v1","PvP01_13_v1","PvP01_13_v1","PvP01_13_v1","PvP01_13_v1","PvP01_13_v1","PvP01_13_v1","PvP01_13_v1","PvP01_13_v1","PvP01_13_v1","PvP01_13_v1","PvP01_13_v1","PvP01_13_v1","PvP01_13_v1","PvP01_13_v1","PvP01_13_v1","PvP01_13_v1","PvP01_13_v1","PvP01_13_v1","PvP01_13_v1","PvP01_13_v1","PvP01_13_v1","PvP01_13_v1","PvP01_13_v1","PvP01_13_v1","PvP01_13_v1","PvP01_13_v1","PvP01_13_v1","PvP01_13_v1","PvP01_13_v1","PvP01_13_v1","PvP01_13_v1","PvP01_13_v1","PvP01_13_v1","PvP01_13_v1","PvP01_13_v1","PvP01_13_v1","PvP01_13_v1","PvP01_13_v1","PvP01_13_v1","PvP01_13_v1","PvP01_13_v1","PvP01_13_v1","PvP01_13_v1","PvP01_13_v1","PvP01_13_v1","PvP01_13_v1","PvP01_13_v1","PvP01_13_v1","PvP01_13_v1","PvP01_13_v1","PvP01_13_v1","PvP01_13_v1","PvP01_13_v1","PvP01_13_v1","PvP01_13_v1","PvP01_13_v1","PvP01_13_v1","PvP01_13_v1","PvP01_13_v1","PvP01_13_v1","PvP01_13_v1","PvP01_13_v1","PvP01_13_v1","PvP01_13_v1","PvP01_13_v1","PvP01_13_v1","PvP01_13_v1","PvP01_13_v1","PvP01_13_v1","PvP01_13_v1","PvP01_13_v1","PvP01_13_v1","PvP01_13_v1","PvP01_13_v1","PvP01_13_v1","PvP01_13_v1","PvP01_13_v1","PvP01_13_v1","PvP01_13_v1","PvP01_13_v1","PvP01_13_v1","PvP01_13_v1","PvP01_13_v1","PvP01_13_v1","PvP01_13_v1","PvP01_13_v1","PvP01_13_v1","PvP01_13_v1","PvP01_13_v1","PvP01_13_v1","PvP01_13_v1","PvP01_13_v1","PvP01_13_v1","PvP01_13_v1","PvP01_13_v1","PvP01_13_v1","PvP01_13_v1","PvP01_13_v1","PvP01_13_v1","PvP01_13_v1","PvP01_13_v1","PvP01_13_v1","PvP01_13_v1","PvP01_13_v1","PvP01_13_v1","PvP01_13_v1","PvP01_13_v1","PvP01_14_v1","PvP01_14_v1","PvP01_14_v1","PvP01_14_v1","PvP01_14_v1","PvP01_14_v1","PvP01_14_v1","PvP01_14_v1","PvP01_14_v1","PvP01_14_v1","PvP01_14_v1","PvP01_14_v1","PvP01_14_v1","PvP01_14_v1","PvP01_14_v1","PvP01_14_v1","PvP01_14_v1","PvP01_14_v1","PvP01_14_v1","PvP01_14_v1","PvP01_14_v1","PvP01_14_v1","PvP01_14_v1","PvP01_14_v1","PvP01_14_v1","PvP01_14_v1","PvP01_14_v1","PvP01_14_v1","PvP01_14_v1","PvP01_14_v1","PvP01_14_v1","PvP01_14_v1","PvP01_14_v1","PvP01_14_v1","PvP01_14_v1","PvP01_14_v1","PvP01_14_v1","PvP01_14_v1","PvP01_14_v1","PvP01_14_v1","PvP01_14_v1","PvP01_14_v1","PvP01_14_v1","PvP01_14_v1","PvP01_14_v1","PvP01_14_v1","PvP01_14_v1","PvP01_14_v1","PvP01_14_v1","PvP01_14_v1","PvP01_14_v1","PvP01_14_v1","PvP01_14_v1","PvP01_14_v1","PvP01_14_v1","PvP01_14_v1","PvP01_14_v1","PvP01_14_v1","PvP01_14_v1","PvP01_14_v1","PvP01_14_v1","PvP01_14_v1","PvP01_14_v1","PvP01_14_v1","PvP01_14_v1","PvP01_14_v1","PvP01_14_v1","PvP01_14_v1","PvP01_14_v1","PvP01_14_v1","PvP01_14_v1","PvP01_14_v1","PvP01_14_v1","PvP01_14_v1","PvP01_14_v1","PvP01_14_v1","PvP01_14_v1","PvP01_14_v1","PvP01_14_v1","PvP01_14_v1","PvP01_14_v1","PvP01_14_v1","PvP01_14_v1","PvP01_14_v1","PvP01_14_v1","PvP01_14_v1","PvP01_14_v1","PvP01_14_v1","PvP01_14_v1","PvP01_14_v1","PvP01_14_v1","PvP01_14_v1","PvP01_14_v1","PvP01_14_v1","PvP01_14_v1","PvP01_14_v1","PvP01_14_v1","PvP01_14_v1","PvP01_14_v1","PvP01_14_v1","PvP01_14_v1","PvP01_14_v1","PvP01_14_v1","PvP01_14_v1","PvP01_14_v1","PvP01_14_v1","PvP01_14_v1","PvP01_14_v1","PvP01_14_v1","PvP01_14_v1","PvP01_14_v1","PvP01_14_v1","PvP01_14_v1","PvP01_14_v1","PvP01_14_v1","PvP01_14_v1","PvP01_14_v1","PvP01_14_v1","PvP01_14_v1","PvP01_14_v1","PvP01_14_v1","PvP01_14_v1","PvP01_14_v1","PvP01_14_v1","PvP01_14_v1","PvP01_14_v1","PvP01_14_v1","PvP01_14_v1","PvP01_14_v1","PvP01_14_v1","PvP01_14_v1","PvP01_14_v1","PvP01_14_v1","PvP01_14_v1","PvP01_14_v1","PvP01_14_v1","PvP01_14_v1","PvP01_14_v1","PvP01_14_v1","PvP01_14_v1","PvP01_14_v1","PvP01_14_v1","PvP01_14_v1","PvP01_14_v1","PvP01_14_v1","PvP01_14_v1","PvP01_14_v1","PvP01_14_v1","PvP01_14_v1","PvP01_14_v1","PvP01_14_v1","PvP01_14_v1","PvP01_14_v1","PvP01_14_v1","PvP01_14_v1","PvP01_14_v1","PvP01_14_v1","PvP01_14_v1","PvP01_14_v1","PvP01_14_v1","PvP01_14_v1","PvP01_14_v1","PvP01_14_v1","PvP01_14_v1","PvP01_14_v1","PvP01_14_v1","PvP01_14_v1","PvP01_14_v1","PvP01_14_v1","PvP01_14_v1","PvP01_14_v1","PvP01_14_v1","PvP01_14_v1","PvP01_14_v1","PvP01_14_v1","PvP01_14_v1","PvP01_14_v1","PvP01_14_v1","PvP01_14_v1","PvP01_14_v1","PvP01_14_v1","PvP01_14_v1","PvP01_14_v1","PvP01_14_v1","PvP01_14_v1","PvP01_14_v1","PvP01_14_v1","PvP01_14_v1","PvP01_14_v1","PvP01_14_v1","PvP01_14_v1","PvP01_14_v1","PvP01_14_v1","PvP01_14_v1","PvP01_14_v1","PvP01_14_v1","PvP01_14_v1","PvP01_14_v1","PvP01_14_v1","PvP01_14_v1","PvP01_14_v1","PvP01_14_v1","PvP01_14_v1","PvP01_14_v1","PvP01_14_v1","PvP01_14_v1","PvP01_14_v1","PvP01_14_v1","PvP01_14_v1","PvP01_14_v1","PvP01_14_v1","PvP01_14_v1","PvP01_14_v1","PvP01_14_v1","PvP01_14_v1","PvP01_14_v1","PvP01_14_v1","PvP01_14_v1","PvP01_14_v1","PvP01_14_v1","PvP01_14_v1","PvP01_14_v1","PvP01_14_v1","PvP01_14_v1","PvP01_14_v1","PvP01_14_v1","PvP01_14_v1","PvP01_14_v1","PvP01_14_v1","PvP01_14_v1","PvP01_14_v1","PvP01_14_v1","PvP01_14_v1","PvP01_14_v1","PvP01_14_v1","PvP01_14_v1","PvP01_14_v1","PvP01_14_v1","PvP01_14_v1","PvP01_14_v1","PvP01_14_v1","PvP01_14_v1","PvP01_14_v1","PvP01_14_v1","PvP01_14_v1","PvP01_14_v1","PvP01_14_v1","PvP01_14_v1","PvP01_14_v1","PvP01_14_v1","PvP01_14_v1","PvP01_14_v1","PvP01_14_v1","PvP01_14_v1","PvP01_14_v1","PvP01_14_v1","PvP01_14_v1","PvP01_14_v1","PvP01_14_v1","PvP01_14_v1","PvP01_14_v1","PvP01_14_v1","PvP01_14_v1","PvP01_14_v1","PvP01_14_v1","PvP01_14_v1","PvP01_14_v1","PvP01_14_v1","PvP01_14_v1","PvP01_14_v1","PvP01_API_v1","PvP01_API_v1","PvP01_API_v1","PvP01_API_v1","PvP01_API_v1","PvP01_MIT_v1","PvP01_MIT_v1","PvP01_MIT_v1"],"Source":["EuPathDB","EuPathDB","EuPathDB","EuPathDB","EuPathDB","EuPathDB","EuPathDB","EuPathDB","EuPathDB","EuPathDB","EuPathDB","EuPathDB","EuPathDB","EuPathDB","EuPathDB","EuPathDB","EuPathDB","EuPathDB","EuPathDB","EuPathDB","EuPathDB","EuPathDB","EuPathDB","EuPathDB","EuPathDB","EuPathDB","EuPathDB","EuPathDB","EuPathDB","EuPathDB","EuPathDB","EuPathDB","EuPathDB","EuPathDB","EuPathDB","EuPathDB","EuPathDB","EuPathDB","EuPathDB","EuPathDB","EuPathDB","EuPathDB","EuPathDB","EuPathDB","EuPathDB","EuPathDB","EuPathDB","EuPathDB","EuPathDB","EuPathDB","EuPathDB","EuPathDB","EuPathDB","EuPathDB","EuPathDB","EuPathDB","EuPathDB","EuPathDB","EuPathDB","EuPathDB","EuPathDB","EuPathDB","EuPathDB","EuPathDB","EuPathDB","EuPathDB","EuPathDB","EuPathDB","EuPathDB","EuPathDB","EuPathDB","EuPathDB","EuPathDB","EuPathDB","EuPathDB","EuPathDB","EuPathDB","EuPathDB","EuPathDB","EuPathDB","EuPathDB","EuPathDB","EuPathDB","EuPathDB","EuPathDB","EuPathDB","EuPathDB","EuPathDB","EuPathDB","EuPathDB","EuPathDB","EuPathDB","EuPathDB","EuPathDB","EuPathDB","EuPathDB","EuPathDB","EuPathDB","EuPathDB","EuPathDB","EuPathDB","EuPathDB","EuPathDB","EuPathDB","EuPathDB","EuPathDB","EuPathDB","EuPathDB","EuPathDB","EuPathDB","EuPathDB","EuPathDB","EuPathDB","EuPathDB","EuPathDB","EuPathDB","EuPathDB","EuPathDB","EuPathDB","EuPathDB","EuPathDB","EuPathDB","EuPathDB","EuPathDB","EuPathDB","EuPathDB","EuPathDB","EuPathDB","EuPathDB","EuPathDB","EuPathDB","EuPathDB","EuPathDB","EuPathDB","EuPathDB","EuPathDB","EuPathDB","EuPathDB","EuPathDB","EuPathDB","EuPathDB","EuPathDB","EuPathDB","EuPathDB","EuPathDB","EuPathDB","EuPathDB","EuPathDB","EuPathDB","EuPathDB","EuPathDB","EuPathDB","EuPathDB","EuPathDB","EuPathDB","EuPathDB","EuPathDB","EuPathDB","EuPathDB","EuPathDB","EuPathDB","EuPathDB","EuPathDB","EuPathDB","EuPathDB","EuPathDB","EuPathDB","EuPathDB","EuPathDB","EuPathDB","EuPathDB","EuPathDB","EuPathDB","EuPathDB","EuPathDB","EuPathDB","EuPathDB","EuPathDB","EuPathDB","EuPathDB","EuPathDB","EuPathDB","EuPathDB","EuPathDB","EuPathDB","EuPathDB","EuPathDB","EuPathDB","EuPathDB","EuPathDB","EuPathDB","EuPathDB","EuPathDB","EuPathDB","EuPathDB","EuPathDB","EuPathDB","EuPathDB","EuPathDB","EuPathDB","EuPathDB","EuPathDB","EuPathDB","EuPathDB","EuPathDB","EuPathDB","EuPathDB","EuPathDB","EuPathDB","EuPathDB","EuPathDB","EuPathDB","EuPathDB","EuPathDB","EuPathDB","EuPathDB","EuPathDB","EuPathDB","EuPathDB","EuPathDB","EuPathDB","EuPathDB","EuPathDB","EuPathDB","EuPathDB","EuPathDB","EuPathDB","EuPathDB","EuPathDB","EuPathDB","EuPathDB","EuPathDB","EuPathDB","EuPathDB","EuPathDB","EuPathDB","EuPathDB","EuPathDB","EuPathDB","EuPathDB","EuPathDB","EuPathDB","EuPathDB","EuPathDB","EuPathDB","EuPathDB","EuPathDB","EuPathDB","EuPathDB","EuPathDB","EuPathDB","EuPathDB","EuPathDB","EuPathDB","EuPathDB","EuPathDB","EuPathDB","EuPathDB","EuPathDB","EuPathDB","EuPathDB","EuPathDB","EuPathDB","EuPathDB","EuPathDB","EuPathDB","EuPathDB","EuPathDB","EuPathDB","EuPathDB","EuPathDB","EuPathDB","EuPathDB","EuPathDB","EuPathDB","EuPathDB","EuPathDB","EuPathDB","EuPathDB","EuPathDB","EuPathDB","EuPathDB","EuPathDB","EuPathDB","EuPathDB","EuPathDB","EuPathDB","EuPathDB","EuPathDB","EuPathDB","EuPathDB","EuPathDB","EuPathDB","EuPathDB","EuPathDB","EuPathDB","EuPathDB","EuPathDB","EuPathDB","EuPathDB","EuPathDB","EuPathDB","EuPathDB","EuPathDB","EuPathDB","EuPathDB","EuPathDB","EuPathDB","EuPathDB","EuPathDB","EuPathDB","EuPathDB","EuPathDB","EuPathDB","EuPathDB","EuPathDB","EuPathDB","EuPathDB","EuPathDB","EuPathDB","EuPathDB","EuPathDB","EuPathDB","EuPathDB","EuPathDB","EuPathDB","EuPathDB","EuPathDB","EuPathDB","EuPathDB","EuPathDB","EuPathDB","EuPathDB","EuPathDB","EuPathDB","EuPathDB","EuPathDB","EuPathDB","EuPathDB","EuPathDB","EuPathDB","EuPathDB","EuPathDB","EuPathDB","EuPathDB","EuPathDB","EuPathDB","EuPathDB","EuPathDB","EuPathDB","EuPathDB","EuPathDB","EuPathDB","EuPathDB","EuPathDB","EuPathDB","EuPathDB","EuPathDB","EuPathDB","EuPathDB","EuPathDB","EuPathDB","EuPathDB","EuPathDB","EuPathDB","EuPathDB","EuPathDB","EuPathDB","EuPathDB","EuPathDB","EuPathDB","EuPath
```
