## Supplementary material for "Transcriptome and histone epigenome of *Plasmodium vivax* salivary-gland sporozoites point to tight regulatory control and potential mechanisms for liver-stage differentiation": Pv-SPZ-v-liver-DTGs.js

```
glimma.layout.setupGrid(d3.select(".container"), "md", [1,2]);
glimma.storage.chartData.push(glimma.transform.toRowMajor({"GeneID":["PVP01_0000030","PVP01_0000040","PVP01_0000050","PVP01_0000080","PVP01_0000100","PVP01_0000120","PVP01_0000130","PVP01_0000140","PVP01_0000150","PVP01_0000170","PVP01_0000180","PVP01_0000190","PVP01_0000200","PVP01_0000210","PVP01_0000220","PVP01_0000300","PVP01_0000480","PVP01_0000670","PVP01_0000740","PVP01_0000790","PVP01_0001030","PVP01_0001090","PVP01_0001310","PVP01_0001440","PVP01_0001480","PVP01_0001550","PVP01_0001640","PVP01_0001660","PVP01_0001670","PVP01_0001900","PVP01_0001910","PVP01_0001940","PVP01_0001970","PVP01_0002020","PVP01_0002050","PVP01_0002120","PVP01_0002200","PVP01_0002210","PVP01_0002330","PVP01_0002370","PVP01_0002390","PVP01_0002470","PVP01_0002530","PVP01_0002540","PVP01_0002750","PVP01_0002810","PVP01_0002820","PVP01_0003010","PVP01_0003050","PVP01_0003150","PVP01_0003210","PVP01_0003350","PVP01_0003360","PVP01_0003370","PVP01_0003380","PVP01_0003400","PVP01_0003410","PVP01_0003420","PVP01_0003440","PVP01_0003530","PVP01_0003690","PVP01_0003730","PVP01_0003800","PVP01_0003990","PVP01_0004160","PVP01_0004170","PVP01_0004180","PVP01_0004190","PVP01_0004240","PVP01_0004250","PVP01_0004260","PVP01_0004270","PVP01_0004280","PVP01_0004300","PVP01_0004310","PVP01_0004320","PVP01_0004350","PVP01_0004370","PVP01_0004380","PVP01_0004390","PVP01_0004400","PVP01_0004410","PVP01_0004640","PVP01_0004670","PVP01_0004910","PVP01_0004920","PVP01_0004950","PVP01_0004960","PVP01_0005010","PVP01_0005100","PVP01_0005160","PVP01_0005210","PVP01_0005300","PVP01_0005310","PVP01_0005420","PVP01_0005430","PVP01_0005550","PVP01_0005640","PVP01_0005710","PVP01_0005740","PVP01_0005760","PVP01_0005800","PVP01_0005820","PVP01_0005870","PVP01_0005910","PVP01_0006000","PVP01_0006040","PVP01_0006080","PVP01_0006260","PVP01_0006650","PVP01_0006660","PVP01_0006670","PVP01_0006680","PVP01_0006700","PVP01_0006860","PVP01_0006890","PVP01_0007080","PVP01_0007160","PVP01_0007240","PVP01_0007290","PVP01_0007310","PVP01_0007320","PVP01_0007340","PVP01_0007610","PVP01_0007620","PVP01_0007690","PVP01_0007700","PVP01_0007720","PVP01_0007850","PVP01_0007900","PVP01_0007910","PVP01_0007920","PVP01_0008190","PVP01_0008430","PVP01_0008480","PVP01_0008490","PVP01_0008600","PVP01_0008700","PVP01_0008770","PVP01_0008830","PVP01_0008990","PVP01_0009010","PVP01_0009170","PVP01_0009180","PVP01_0009210","PVP01_0009410","PVP01_0009470","PVP01_0009540","PVP01_0009620","PVP01_0009710","PVP01_0009730","PVP01_0009740","PVP01_0009760","PVP01_0009770","PVP01_0009820","PVP01_0009850","PVP01_0009860","PVP01_0009870","PVP01_0009900","PVP01_0009940","PVP01_0009980","PVP01_0010020","PVP01_0010030","PVP01_0010070","PVP01_0010100","PVP01_0010120","PVP01_0010140","PVP01_0010170","PVP01_0010180","PVP01_0010200","PVP01_0010210","PVP01_0010220","PVP01_0010230","PVP01_0010240","PVP01_0010340","PVP01_0010380","PVP01_0010440","PVP01_0010470","PVP01_0010500","PVP01_0010540","PVP01_0010570","PVP01_0010600","PVP01_0010610","PVP01_0010640","PVP01_0010650","PVP01_0010770","PVP01_0010870","PVP01_0010900","PVP01_0010910","PVP01_0010960","PVP01_0010970","PVP01_0010990","PVP01_0011030","PVP01_0011070","PVP01_0102100","PVP01_0102200","PVP01_0102300","PVP01_0102400","PVP01_0102500","PVP01_0102600","PVP01_0102700","PVP01_0102800","PVP01_0103400","PVP01_0103600","PVP01_0103800","PVP01_0103900","PVP01_0104100","PVP01_0104200","PVP01_0104300","PVP01_0104600","PVP01_0104800","PVP01_0104900","PVP01_0105000","PVP01_0105100","PVP01_0105300","PVP01_0105400","PVP01_0105500","PVP01_0105600","PVP01_0105700","PVP01_0105800","PVP01_0106000","PVP01_0106200","PVP01_0106300","PVP01_0106400","PVP01_0106500","PVP01_0106800","PVP01_0106900","PVP01_0107400","PVP01_0107600","PVP01_0107700","PVP01_0107900","PVP01_0108000","PVP01_0108100","PVP01_0108300","PVP01_0108400","PVP01_0108600","PVP01_0108800","PVP01_0109000","PVP01_0109100","PVP01_0109200","PVP01_0109300","PVP01_0109600","PVP01_0109800","PVP01_0110200","PVP01_0110700","PVP01_0110800","PVP01_0111200","PVP01_0111300","PVP01_0111600","PVP01_0111700","PVP01_0112000","PVP01_0112100","PVP01_0112500","PVP01_0112700","PVP01_0113000","PVP01_0113100","PVP01_0113200","PVP01_0113400","PVP01_0113500","PVP01_0114100","PVP01_0114600","PVP01_0114700","PVP01_0114900","PVP01_0115000","PVP01_0115200","PVP01_0115400","PVP01_0115500","PVP01_0116100","PVP01_0116500","PVP01_0116600","PVP01_0116700","PVP01_0116900","PVP01_0117500","PVP01_0117700","PVP01_0117800","PVP01_0118000","PVP01_0118600","PVP01_0118700","PVP01_0118800","PVP01_0118900","PVP01_0119200","PVP01_0119300","PVP01_0119400","PVP01_0119500","PVP01_0119600","PVP01_0119700","PVP01_0119800","PVP01_0119900","PVP01_0120000","PVP01_0120100","PVP01_0120500","PVP01_0120800","PVP01_0121000","PVP01_0121700","PVP01_0200200","PVP01_0200700","PVP01_0201000","PVP01_0201100","PVP01_0201300","PVP01_0201400","PVP01_0201500","PVP01_0201600","PVP01_0201700","PVP01_0201800","PVP01_0201900","PVP01_0202000","PVP01_0202100","PVP01_0202200","PVP01_0202300","PVP01_0202400","PVP01_0203000","PVP01_0203100","PVP01_0203400","PVP01_0203500","PVP01_0203800","PVP01_0203900","PVP01_0204000","PVP01_0204100","PVP01_0204800","PVP01_0204900","PVP01_0205000","PVP01_0205200","PVP01_0205300","PVP01_0205500","PVP01_0205600","PVP01_0205800","PVP01_0206100","PVP01_0206300","PVP01_0206400","PVP01_0206500","PVP01_0206600","PVP01_0206800","PVP01_0207000","PVP01_0207100","PVP01_0207200","PVP01_0207400","PVP01_0207500","PVP01_0207600","PVP01_0207700","PVP01_0207800","PVP01_0208100","PVP01_0208200","PVP01_0208300","PVP01_0208400","PVP01_0208600","PVP01_0208700","PVP01_0208800","PVP01_0209000","PVP01_0209100","PVP01_0209200","PVP01_0209300","PVP01_0209500","PVP01_0209600","PVP01_0210000","PVP01_0210100","PVP01_0210300","PVP01_0211000","PVP01_0211100","PVP01_0211200","PVP01_0211400","PVP01_0211500","PVP01_0211600","PVP01_0211700","PVP01_0212400","PVP01_0212500","PVP01_0212600","PVP01_0212700","PVP01_0212800","PVP01_0213000","PVP01_0213300","PVP01_0213600","PVP01_0213700","PVP01_0214100","PVP01_0214200","PVP01_0214800","PVP01_0214900","PVP01_0215000","PVP01_0215300","PVP01_0215400","PVP01_0215500","PVP01_0215700","PVP01_0215800","PVP01_0216100","PVP01_0216500","PVP01_0216600","PVP01_0216700","PVP01_0216800","PVP01_0216900","PVP01_0217000","PVP01_0217100","PVP01_0217200","PVP01_0217300","PVP01_0217400","PVP01_0217500","PVP01_0218200","PVP01_0218300","PVP01_0218900","PVP01_0219400","PVP01_0220100","PVP01_0220200","PVP01_0221100","PVP01_0221300","PVP01_0221700","PVP01_0300100","PVP01_0300600","PVP01_0300700","PVP01_0300800","PVP01_0301100","PVP01_0301200","PVP01_0301500","PVP01_0301900","PVP01_0302000","PVP01_0302100","PVP01_0302300","PVP01_0302400","PVP01_0302500","PVP01_0303100","PVP01_0303500","PVP01_0304100","PVP01_0304300","PVP01_0304500","PVP01_0304600","PVP01_0304800","PVP01_0304900","PVP01_0305000","PVP01_0305200","PVP01_0305300","PVP01_0305500","PVP01_0305800","PVP01_0306000","PVP01_0306100","PVP01_0306300","PVP01_0306400","PVP01_0306500","PVP01_0306700","PVP01_0306800","PVP01_0306900","PVP01_0307000","PVP01_0307100","PVP01_0307300","PVP01_0307400","PVP01_0307600","PVP01_0307700","PVP01_0308800","PVP01_0309200","PVP01_0309700","PVP01_0309800","PVP01_0310000","PVP01_0310200","PVP01_0310300","PVP01_0310400","PVP01_0310600","PVP01_0311000","PVP01_0311200","PVP01_0311300","PVP01_0311400","PVP01_0311500","PVP01_0311600","PVP01_0311700","PVP01_0312000","PVP01_0312100","PVP01_0312400","PVP01_0312500","PVP01_0312700","PVP01_0312800","PVP01_0313500","PVP01_0313800","PVP01_0314100","PVP01_0314200","PVP01_0314900","PVP01_0315000","PVP01_0315100","PVP01_0315200","PVP01_0315300","PVP01_0315700","PVP01_0315900","PVP01_0316000","PVP01_0316200","PVP01_0316300","PVP01_0316500","PVP01_0316700","PVP01_0316800","PVP01_0317100","PVP01_0317200","PVP01_0317400","PVP01_0318000","PVP01_0318100","PVP01_0318400","PVP01_0318900","PVP01_0319200","PVP01_0319300","PVP01_0319400","PVP01_0319600","PVP01_0319700","PVP01_0320400","PVP01_0320500","PVP01_0320700","PVP01_0320800","PVP01_0320900","PVP01_0321100","PVP01_0400900","PVP01_0403600","PVP01_0403700","PVP01_0403900","PVP01_0404000","PVP01_0404100","PVP01_0404200","PVP01_0404300","PVP01_0404400","PVP01_0404500","PVP01_0404600","PVP01_0404700","PVP01_0404800","PVP01_0404900","PVP01_0405100","PVP01_0405200","PVP01_0405300","PVP01_0405500","PVP01_0405600","PVP01_0405700","PVP01_0405900","PVP01_0406500","PVP01_0406600","PVP01_0406700","PVP01_0406900","PVP01_0407000","PVP01_0407700","PVP01_0407900","PVP01_0408200","PVP01_0408600","PVP01_0408800","PVP01_0408900","PVP01_0409000","PVP01_0409700","PVP01_0409900","PVP01_0410000","PVP01_0410400","PVP01_0410600","PVP01_0410800","PVP01_0411400","PVP01_0411500","PVP01_0411600","PVP01_0411800","PVP01_0411900","PVP01_0412100","PVP01_0412200","PVP01_0412700","PVP01_0412800","PVP01_0412900","PVP01_0413000","PVP01_0413100","PVP01_0413200","PVP01_0413300","PVP01_0413500","PVP01_0414300","PVP01_0414400","PVP01_0414600","PVP01_0414900","PVP01_0415200","PVP01_0415400","PVP01_0415500","PVP01_0415600","PVP01_0416600","PVP01_0416900","PVP01_0417100","PVP01_0417400","PVP01_0417800","PVP01_0417900","PVP01_0418300","PVP01_0419000","PVP01_0419200","PVP01_0419300","PVP01_0419400","PVP01_0419500","PVP01_0419700","PVP01_0420000","PVP01_0420200","PVP01_0420500","PVP01_0420700","PVP01_0420900","PVP01_0421500","PVP01_0421700","PVP01_0421800","PVP01_0422100","PVP01_0422200","PVP01_0422300","PVP01_0422400","PVP01_0422600","PVP01_0422700","PVP01_0422900","PVP01_0423000","PVP01_0423100","PVP01_0423200","PVP01_0423300","PVP01_0423600","PVP01_0423800","PVP01_0424300","PVP01_0424400","PVP01_0500700","PVP01_0502300","PVP01_0502400","PVP01_0502900","PVP01_0503300","PVP01_0503400","PVP01_0503600","PVP01_0503700","PVP01_0503800","PVP01_0503900","PVP01_0504100","PVP01_0504200","PVP01_0504300","PVP01_0505000","PVP01_0505100","PVP01_0505200","PVP01_0505400","PVP01_0505500","PVP01_0505800","PVP01_0506000","PVP01_0506300","PVP01_0506400","PVP01_0506600","PVP01_0506700","PVP01_0506800","PVP01_0507300","PVP01_0507500","PVP01_0507600","PVP01_0507700","PVP01_0507800","PVP01_0508000","PVP01_0508200","PVP01_0508500","PVP01_0508600","PVP01_0508700","PVP01_0509100","PVP01_0509200","PVP01_0509400","PVP01_0509600","PVP01_0509900","PVP01_0510000","PVP01_0510100","PVP01_0510200","PVP01_0510300","PVP01_0510800","PVP01_0510900","PVP01_0511200","PVP01_0511300","PVP01_0511400","PVP01_0511500","PVP01_0511700","PVP01_0511800","PVP01_0512100","PVP01_0512200","PVP01_0512400","PVP01_0512500","PVP01_0512700","PVP01_0512800","PVP01_0513000","PVP01_0513300","PVP01_0513400","PVP01_0513500","PVP01_0513700","PVP01_0513900","PVP01_0514100","PVP01_0514600","PVP01_0514800","PVP01_0514900","PVP01_0515100","PVP01_0515200","PVP01_0515300","PVP01_0515700","PVP01_0515800","PVP01_0515900","PVP01_0516000","PVP01_0516100","PVP01_0516200","PVP01_0516300","PVP01_0516400","PVP01_0516500","PVP01_0516900","PVP01_0517000","PVP01_0517100","PVP01_0517300","PVP01_0517400","PVP01_0517700","PVP01_0517900","PVP01_0518300","PVP01_0518400","PVP01_0518500","PVP01_0518600","PVP01_0518700","PVP01_0519100","PVP01_0519200","PVP01_0519500","PVP01_0519600","PVP01_0520700","PVP01_0520800","PVP01_0521300","PVP01_0521400","PVP01_0521600","PVP01_0521800","PVP01_0522000","PVP01_0522100","PVP01_0522200","PVP01_0522300","PVP01_0522600","PVP01_0522700","PVP01_0522900","PVP01_0523000","PVP01_0523300","PVP01_0523400","PVP01_0523700","PVP01_0523800","PVP01_0523900","PVP01_0524000","PVP01_0524100","PVP01_0524200","PVP01_0524300","PVP01_0524500","PVP01_0524600","PVP01_0524700","PVP01_0524900","PVP01_0525000","PVP01_0525100","PVP01_0525300","PVP01_0525700","PVP01_0526100","PVP01_0526400","PVP01_0527300","PVP01_0527400","PVP01_0527500","PVP01_0527700","PVP01_0527800","PVP01_0528100","PVP01_0528200","PVP01_0528400","PVP01_0528600","PVP01_0528700","PVP01_0528900","PVP01_0529000","PVP01_0529100","PVP01_0529200","PVP01_0529600","PVP01_0529700","PVP01_0529900","PVP01_0530000","PVP01_0530100","PVP01_0530400","PVP01_0530500","PVP01_0531100","PVP01_0531200","PVP01_0531500","PVP01_0531700","PVP01_0531800","PVP01_0532300","PVP01_0532500","PVP01_0532600","PVP01_0532700","PVP01_0532900","PVP01_0533000","PVP01_0533100","PVP01_0533300","PVP01_0533700","PVP01_0534000","PVP01_0534100","PVP01_0534200","PVP01_0534600","PVP01_0534800","PVP01_0534900","PVP01_0535400","PVP01_0600100","PVP01_0600300","PVP01_0600400","PVP01_0600500","PVP01_0600600","PVP01_0600700","PVP01_0600900","PVP01_0601100","PVP01_0601200","PVP01_0601300","PVP01_0601500","PVP01_0601600","PVP01_0602600","PVP01_0603000","PVP01_0603100","PVP01_0603600","PVP01_0603700","PVP01_0604200","PVP01_0604400","PVP01_0604700","PVP01_0605000","PVP01_0605100","PVP01_0605300","PVP01_0605500","PVP01_0606200","PVP01_0606500","PVP01_0606600","PVP01_0606800","PVP01_0607200","PVP01_0607600","PVP01_0608000","PVP01_0608100","PVP01_0608300","PVP01_0608600","PVP01_0608800","PVP01_0608900","PVP01_0609200","PVP01_0609400","PVP01_0609600","PVP01_0609700","PVP01_0609800","PVP01_0609900","PVP01_0610100","PVP01_0610200","PVP01_0610300","PVP01_0610400","PVP01_0610600","PVP01_0610700","PVP01_0610800","PVP01_0610900","PVP01_0611100","PVP01_0611300","PVP01_0612000","PVP01_0612100","PVP01_0612400","PVP01_0612600","PVP01_0612700","PVP01_0613000","PVP01_0613100","PVP01_0613200","PVP01_0613400","PVP01_0613500","PVP01_0613800","PVP01_0614000","PVP01_0614200","PVP01_0614300","PVP01_0614400","PVP01_0614600","PVP01_0614900","PVP01_0615000","PVP01_0615100","PVP01_0615500","PVP01_0615800","PVP01_0615900","PVP01_0616000","PVP01_0616200","PVP01_0616300","PVP01_0616400","PVP01_0616900","PVP01_0617000","PVP01_0617100","PVP01_0617400","PVP01_0617500","PVP01_0617900","PVP01_0618100","PVP01_0618300","PVP01_0618600","PVP01_0618700","PVP01_0619000","PVP01_0619100","PVP01_0619200","PVP01_0619400","PVP01_0619500","PVP01_0619600","PVP01_0619700","PVP01_0619800","PVP01_0620100","PVP01_0620300","PVP01_0620500","PVP01_0620600","PVP01_0620700","PVP01_0620900","PVP01_0621100","PVP01_0621800","PVP01_0621900","PVP01_0622100","PVP01_0622200","PVP01_0622700","PVP01_0623000","PVP01_0623100","PVP01_0623200","PVP01_0623300","PVP01_0623400","PVP01_0623500","PVP01_0623600","PVP01_0623900","PVP01_0624000","PVP01_0624100","PVP01_0624300","PVP01_0624400","PVP01_0624500","PVP01_0700200","PVP01_0700300","PVP01_0700400","PVP01_0700500","PVP01_0700700","PVP01_0700800","PVP01_0700900","PVP01_0701000","PVP01_0701200","PVP01_0701500","PVP01_0701600","PVP01_0701700","PVP01_0702100","PVP01_0702200","PVP01_0702500","PVP01_0703000","PVP01_0703600","PVP01_0703700","PVP01_0704000","PVP01_0704100","PVP01_0704500","PVP01_0704600","PVP01_0704700","PVP01_0704800","PVP01_0704900","PVP01_0705000","PVP01_0705100","PVP01_0705500","PVP01_0705600","PVP01_0705800","PVP01_0705900","PVP01_0706000","PVP01_0706100","PVP01_0706200","PVP01_0706300","PVP01_0706400","PVP01_0706600","PVP01_0706700","PVP01_0707000","PVP01_0707100","PVP01_0707200","PVP01_0707300","PVP01_0707800","PVP01_0708000","PVP01_0708100","PVP01_0708200","PVP01_0708300","PVP01_0708400","PVP01_0708500","PVP01_0708800","PVP01_0708900","PVP01_0709000","PVP01_0709100","PVP01_0709200","PVP01_0709300","PVP01_0709500","PVP01_0709900","PVP01_0710000","PVP01_0710100","PVP01_0710200","PVP01_0710300","PVP01_0710700","PVP01_0711100","PVP01_0711200","PVP01_0711400","PVP01_0711500","PVP01_0711800","PVP01_0711900","PVP01_0712200","PVP01_0712300","PVP01_0712400","PVP01_0712700","PVP01_0713000","PVP01_0713100","PVP01_0713200","PVP01_0713500","PVP01_0713600","PVP01_0713900","PVP01_0714200","PVP01_0714300","PVP01_0714600","PVP01_0714900","PVP01_0715000","PVP01_0715100","PVP01_0715200","PVP01_0715500","PVP01_0715600","PVP01_0715700","PVP01_0715800","PVP01_0716000","PVP01_0716100","PVP01_0716600","PVP01_0716800","PVP01_0717000","PVP01_0717200","PVP01_0717400","PVP01_0717600","PVP01_0717800","PVP01_0718000","PVP01_0718100","PVP01_0718600","PVP01_0718700","PVP01_0719000","PVP01_0719100","PVP01_0719400","PVP01_0719500","PVP01_0719700","PVP01_0719800","PVP01_0719900","PVP01_0720000","PVP01_0720100","PVP01_0720200","PVP01_0720300","PVP01_0720400","PVP01_0720800","PVP01_0720900","PVP01_0721100","PVP01_0721500","PVP01_0721800","PVP01_0722000","PVP01_0722100","PVP01_0722200","PVP01_0722300","PVP01_0722400","PVP01_0722500","PVP01_0722600","PVP01_0722700","PVP01_0722800","PVP01_0723000","PVP01_0723100","PVP01_0723200","PVP01_0723300","PVP01_0723400","PVP01_0723500","PVP01_0723800","PVP01_0724100","PVP01_0724500","PVP01_0724600","PVP01_0724700","PVP01_0724800","PVP01_0724900","PVP01_0725000","PVP01_0725100","PVP01_0725200","PVP01_0725300","PVP01_0725400","PVP01_0725700","PVP01_0726400","PVP01_0726700","PVP01_0726900","PVP01_0727000","PVP01_0727100","PVP01_0727200","PVP01_0727300","PVP01_0727400","PVP01_0727600","PVP01_0727800","PVP01_0728100","PVP01_0728200","PVP01_0728300","PVP01_0729100","PVP01_0729200","PVP01_0729300","PVP01_0729400","PVP01_0729500","PVP01_0729700","PVP01_0730000","PVP01_0730100","PVP01_0730200","PVP01_0730300","PVP01_0730400","PVP01_0730600","PVP01_0730700","PVP01_0731000","PVP01_0731100","PVP01_0731200","PVP01_0731300","PVP01_0731600","PVP01_0731700","PVP01_0731900","PVP01_0732000","PVP01_0732100","PVP01_0732200","PVP01_0732300","PVP01_0732600","PVP01_0732800","PVP01_0732900","PVP01_0733000","PVP01_0733100","PVP01_0733500","PVP01_0733700","PVP01_0733900","PVP01_0734100","PVP01_0734200","PVP01_0734300","PVP01_0734400","PVP01_0734700","PVP01_0735000","PVP01_0735100","PVP01_0735200","PVP01_0735300","PVP01_0735800","PVP01_0735900","PVP01_0800100","PVP01_0800200","PVP01_0800300","PVP01_0800400","PVP01_0800500","PVP01_0800600","PVP01_0800700","PVP01_0800900","PVP01_0801000","PVP01_0801400","PVP01_0801500","PVP01_0801700","PVP01_0801800","PVP01_0802100","PVP01_0802400","PVP01_0802900","PVP01_0803000","PVP01_0803100","PVP01_0803500","PVP01_0803600","PVP01_0803700","PVP01_0804100","PVP01_0804200","PVP01_0804500","PVP01_0804600","PVP01_0804700","PVP01_0804800","PVP01_0804900","PVP01_0805000","PVP01_0805200","PVP01_0805300","PVP01_0805400","PVP01_0805600","PVP01_0805700","PVP01_0805800","PVP01_0806000","PVP01_0806100","PVP01_0806200","PVP01_0806300","PVP01_0806400","PVP01_0806500","PVP01_0806700","PVP01_0807000","PVP01_0807200","PVP01_0807500","PVP01_0807900","PVP01_0808000","PVP01_0808100","PVP01_0808400","PVP01_0808500","PVP01_0808600","PVP01_0808700","PVP01_0808800","PVP01_0809100","PVP01_0809200","PVP01_0809300","PVP01_0809400","PVP01_0809600","PVP01_0809700","PVP01_0809900","PVP01_0810000","PVP01_0810100","PVP01_0810300","PVP01_0810400","PVP01_0810500","PVP01_0810600","PVP01_0811000","PVP01_0811200","PVP01_0811300","PVP01_0811500","PVP01_0811700","PVP01_0811800","PVP01_0811900","PVP01_0812000","PVP01_0812300","PVP01_0812600","PVP01_0812800","PVP01_0813000","PVP01_0813200","PVP01_0813300","PVP01_0813600","PVP01_0813700","PVP01_0813800","PVP01_0814400","PVP01_0814700","PVP01_0814800","PVP01_0815000","PVP01_0815100","PVP01_0815300","PVP01_0815400","PVP01_0816200","PVP01_0816300","PVP01_0816700","PVP01_0816800","PVP01_0817100","PVP01_0817400","PVP01_0818100","PVP01_0818300","PVP01_0818600","PVP01_0819000","PVP01_0819200","PVP01_0819300","PVP01_0819700","PVP01_0820000","PVP01_0820100","PVP01_0820300","PVP01_0820500","PVP01_0820800","PVP01_0820900","PVP01_0821000","PVP01_0821300","PVP01_0821400","PVP01_0821500","PVP01_0821700","PVP01_0821800","PVP01_0822100","PVP01_0822300","PVP01_0822600","PVP01_0822700","PVP01_0822900","PVP01_0823100","PVP01_0823400","PVP01_0823600","PVP01_0823700","PVP01_0824000","PVP01_0824300","PVP01_0824500","PVP01_0824700","PVP01_0825100","PVP01_0825700","PVP01_0825800","PVP01_0826000","PVP01_0826100","PVP01_0826200","PVP01_0826300","PVP01_0826500","PVP01_0826700","PVP01_0827100","PVP01_0827700","PVP01_0827800","PVP01_0828000","PVP01_0828100","PVP01_0828200","PVP01_0828300","PVP01_0828700","PVP01_0829000","PVP01_0829100","PVP01_0829300","PVP01_0829700","PVP01_0829900","PVP01_0830300","PVP01_0830400","PVP01_0830500","PVP01_0830700","PVP01_0830900","PVP01_0831000","PVP01_0831100","PVP01_0831400","PVP01_0831700","PVP01_0831900","PVP01_0832000","PVP01_0832100","PVP01_0832200","PVP01_0832400","PVP01_0832500","PVP01_0832600","PVP01_0832900","PVP01_0833000","PVP01_0833500","PVP01_0833600","PVP01_0833700","PVP01_0833800","PVP01_0834000","PVP01_0834200","PVP01_0834300","PVP01_0834400","PVP01_0834500","PVP01_0835000","PVP01_0835300","PVP01_0835500","PVP01_0835700","PVP01_0835900","PVP01_0836100","PVP01_0836300","PVP01_0836400","PVP01_0836500","PVP01_0836600","PVP01_0836700","PVP01_0836800","PVP01_0837200","PVP01_0837300","PVP01_0837500","PVP01_0837600","PVP01_0837800","PVP01_0838000","PVP01_0838100","PVP01_0838200","PVP01_0838300","PVP01_0838600","PVP01_0838700","PVP01_0838800","PVP01_0838900","PVP01_0839000","PVP01_0839100","PVP01_0839200","PVP01_0839400","PVP01_0839700","PVP01_0840700","PVP01_0841000","PVP01_0900700","PVP01_0901000","PVP01_0901100","PVP01_0901200","PVP01_0901300","PVP01_0901800","PVP01_0901900","PVP01_0902000","PVP01_0902100","PVP01_0902400","PVP01_0903400","PVP01_0903500","PVP01_0903600","PVP01_0903700","PVP01_0903800","PVP01_0903900","PVP01_0904300","PVP01_0904600","PVP01_0904700","PVP01_0904900","PVP01_0905100","PVP01_0906000","PVP01_0906300","PVP01_0906500","PVP01_0906700","PVP01_0906800","PVP01_0906900","PVP01_0907000","PVP01_0907100","PVP01_0907200","PVP01_0907300","PVP01_0907500","PVP01_0907600","PVP01_0907700","PVP01_0907800","PVP01_0908000","PVP01_0908500","PVP01_0908600","PVP01_0908900","PVP01_0909100","PVP01_0909300","PVP01_0909500","PVP01_0909600","PVP01_0910000","PVP01_0910200","PVP01_0910900","PVP01_0911000","PVP01_0911300","PVP01_0911600","PVP01_0911700","PVP01_0911800","PVP01_0911900","PVP01_0912100","PVP01_0912200","PVP01_0912400","PVP01_0912900","PVP01_0913000","PVP01_0913400","PVP01_0913500","PVP01_0913900","PVP01_0914000","PVP01_0914400","PVP01_0914600","PVP01_0914800","PVP01_0915000","PVP01_0915100","PVP01_0915300","PVP01_0915600","PVP01_0915800","PVP01_0916300","PVP01_0916700","PVP01_0916800","PVP01_0916900","PVP01_0917000","PVP01_0917100","PVP01_0917200","PVP01_0917900","PVP01_0918000","PVP01_0918100","PVP01_0918200","PVP01_0918300","PVP01_0918400","PVP01_0918700","PVP01_0918800","PVP01_0918900","PVP01_0919000","PVP01_0919600","PVP01_0919700","PVP01_0919900","PVP01_0920100","PVP01_0920400","PVP01_0920700","PVP01_0920800","PVP01_0921400","PVP01_0921500","PVP01_0921800","PVP01_0922000","PVP01_0922100","PVP01_0922300","PVP01_0922400","PVP01_0922800","PVP01_0922900","PVP01_0923000","PVP01_0923100","PVP01_0923300","PVP01_0923800","PVP01_0923900","PVP01_0924000","PVP01_0924100","PVP01_0924300","PVP01_0924400","PVP01_0924700","PVP01_0924800","PVP01_0925000","PVP01_0925100","PVP01_0925200","PVP01_0925400","PVP01_0925600","PVP01_0925700","PVP01_0925800","PVP01_0926300","PVP01_0926400","PVP01_0926500","PVP01_0926600","PVP01_0926900","PVP01_0927100","PVP01_0927400","PVP01_0927600","PVP01_0927700","PVP01_0927800","PVP01_0928000","PVP01_0928100","PVP01_0928300","PVP01_0928800","PVP01_0928900","PVP01_0929000","PVP01_0929100","PVP01_0929200","PVP01_0929400","PVP01_0929600","PVP01_0929700","PVP01_0929900","PVP01_0930000","PVP01_0930100","PVP01_0930200","PVP01_0930300","PVP01_0930400","PVP01_0930500","PVP01_0931300","PVP01_0931400","PVP01_0931500","PVP01_0931600","PVP01_0931700","PVP01_0932300","PVP01_0932600","PVP01_0933100","PVP01_0933400","PVP01_0933500","PVP01_0933700","PVP01_0934000","PVP01_0934300","PVP01_0934400","PVP01_0934500","PVP01_0935000","PVP01_0935200","PVP01_0935400","PVP01_0935500","PVP01_0935700","PVP01_0935800","PVP01_0935900","PVP01_0936000","PVP01_0936300","PVP01_0936400","PVP01_0936700","PVP01_0936800","PVP01_0937000","PVP01_0937300","PVP01_0937500","PVP01_0937700","PVP01_0938100","PVP01_0938200","PVP01_0938600","PVP01_0938700","PVP01_0938900","PVP01_0939000","PVP01_0939100","PVP01_0939200","PVP01_0939300","PVP01_0939500","PVP01_0939600","PVP01_0940200","PVP01_0940500","PVP01_0940600","PVP01_0940700","PVP01_0940900","PVP01_0941100","PVP01_0941200","PVP01_0941300","PVP01_0941400","PVP01_0941600","PVP01_0941800","PVP01_0942200","PVP01_0942300","PVP01_0942400","PVP01_0942600","PVP01_0942700","PVP01_0942800","PVP01_0942900","PVP01_0943100","PVP01_0943300","PVP01_0943400","PVP01_0943700","PVP01_0944100","PVP01_0944300","PVP01_0944400","PVP01_0944600","PVP01_0944800","PVP01_0945100","PVP01_0945200","PVP01_0945300","PVP01_0945500","PVP01_0945600","PVP01_0945800","PVP01_0945900","PVP01_0946000","PVP01_0946200","PVP01_0946300","PVP01_0946400","PVP01_0946500","PVP01_0946600","PVP01_0946700","PVP01_0946900","PVP01_0947000","PVP01_0947100","PVP01_0947200","PVP01_0947400","PVP01_0947800","PVP01_0947900","PVP01_0948100","PVP01_0948300","PVP01_0948500","PVP01_0948600","PVP01_0948700","PVP01_0948800","PVP01_0948900","PVP01_0949000","PVP01_0949200","PVP01_0949400","PVP01_0949600","PVP01_0950000","PVP01_0950200","PVP01_1000700","PVP01_1001900","PVP01_1002500","PVP01_1002600","PVP01_1002700","PVP01_1003000","PVP01_1003200","PVP01_1003400","PVP01_1003500","PVP01_1003900","PVP01_1004200","PVP01_1004500","PVP01_1004800","PVP01_1004900","PVP01_1005100","PVP01_1005300","PVP01_1005400","PVP01_1005600","PVP01_1005700","PVP01_1005900","PVP01_1006200","PVP01_1006300","PVP01_1006400","PVP01_1006600","PVP01_1006700","PVP01_1007200","PVP01_1007500","PVP01_1007700","PVP01_1007800","PVP01_1007900","PVP01_1008000","PVP01_1008100","PVP01_1008200","PVP01_1008300","PVP01_1008600","PVP01_1008900","PVP01_1009000","PVP01_1009100","PVP01_1009300","PVP01_1009400","PVP01_1009600","PVP01_1009700","PVP01_1009800","PVP01_1010100","PVP01_1010200","PVP01_1010300","PVP01_1010400","PVP01_1011100","PVP01_1011600","PVP01_1011800","PVP01_1012100","PVP01_1012300","PVP01_1012400","PVP01_1012500","PVP01_1012700","PVP01_1012800","PVP01_1013100","PVP01_1013200","PVP01_1013300","PVP01_1013400","PVP01_1013700","PVP01_1014300","PVP01_1014600","PVP01_1014800","PVP01_1015000","PVP01_1015100","PVP01_1015400","PVP01_1015500","PVP01_1015700","PVP01_1015900","PVP01_1016200","PVP01_1016400","PVP01_1016500","PVP01_1016900","PVP01_1017100","PVP01_1017200","PVP01_1017900","PVP01_1018000","PVP01_1018100","PVP01_1018400","PVP01_1018500","PVP01_1019000","PVP01_1019400","PVP01_1019600","PVP01_1019900","PVP01_1020000","PVP01_1020100","PVP01_1020200","PVP01_1020500","PVP01_1020800","PVP01_1020900","PVP01_1021000","PVP01_1021100","PVP01_1021200","PVP01_1021300","PVP01_1021500","PVP01_1021600","PVP01_1022000","PVP01_1022100","PVP01_1022200","PVP01_1022300","PVP01_1022400","PVP01_1022900","PVP01_1023000","PVP01_1023300","PVP01_1023600","PVP01_1023700","PVP01_1023900","PVP01_1024400","PVP01_1024500","PVP01_1024800","PVP01_1025200","PVP01_1025300","PVP01_1025800","PVP01_1026300","PVP01_1026400","PVP01_1026500","PVP01_1026600","PVP01_1026700","PVP01_1026800","PVP01_1026900","PVP01_1027100","PVP01_1027300","PVP01_1027400","PVP01_1027600","PVP01_1027700","PVP01_1027900","PVP01_1028000","PVP01_1028100","PVP01_1028200","PVP01_1028400","PVP01_1028600","PVP01_1029100","PVP01_1029200","PVP01_1029300","PVP01_1029500","PVP01_1029700","PVP01_1029800","PVP01_1029900","PVP01_1030000","PVP01_1030100","PVP01_1030200","PVP01_1030300","PVP01_1030500","PVP01_1030900","PVP01_1031100","PVP01_1031200","PVP01_1031300","PVP01_1031400","PVP01_1031500","PVP01_1031600","PVP01_1031700","PVP01_1031800","PVP01_1031900","PVP01_1032000","PVP01_1032100","PVP01_1032200","PVP01_1032400","PVP01_1032500","PVP01_1032800","PVP01_1032900","PVP01_1033000","PVP01_1033300","PVP01_1033500","PVP01_1033600","PVP01_1033700","PVP01_1034000","PVP01_1034100","PVP01_1034200","PVP01_1034300","PVP01_1034400","PVP01_1034500","PVP01_1035200","PVP01_1100400","PVP01_1100500","PVP01_1101000","PVP01_1101200","PVP01_1101300","PVP01_1101400","PVP01_1101500","PVP01_1101600","PVP01_1101700","PVP01_1102400","PVP01_1102500","PVP01_1102800","PVP01_1102900","PVP01_1103200","PVP01_1103400","PVP01_1103700","PVP01_1104000","PVP01_1104100","PVP01_1104200","PVP01_1104300","PVP01_1104400","PVP01_1104600","PVP01_1104800","PVP01_1105300","PVP01_1105400","PVP01_1105500","PVP01_1105600","PVP01_1105800","PVP01_1106000","PVP01_1106200","PVP01_1106300","PVP01_1106400","PVP01_1106600","PVP01_1106700","PVP01_1107000","PVP01_1107100","PVP01_1107200","PVP01_1107300","PVP01_1107400","PVP01_1107900","PVP01_1108000","PVP01_1108300","PVP01_1108500","PVP01_1108600","PVP01_1108800","PVP01_1108900","PVP01_1109000","PVP01_1109100","PVP01_1109300","PVP01_1109400","PVP01_1109600","PVP01_1110500","PVP01_1110600","PVP01_1110700","PVP01_1110900","PVP01_1111100","PVP01_1111200","PVP01_1111300","PVP01_1111500","PVP01_1111600","PVP01_1111700","PVP01_1111800","PVP01_1112100","PVP01_1112400","PVP01_1112600","PVP01_1112800","PVP01_1113300","PVP01_1113500","PVP01_1113700","PVP01_1113900","PVP01_1114100","PVP01_1114200","PVP01_1114300","PVP01_1114500","PVP01_1114600","PVP01_1114700","PVP01_1114800","PVP01_1115000","PVP01_1115200","PVP01_1115500","PVP01_1115600","PVP01_1116600","PVP01_1116800","PVP01_1116900","PVP01_1117300","PVP01_1117700","PVP01_1118100","PVP01_1118600","PVP01_1119000","PVP01_1119300","PVP01_1119500","PVP01_1119600","PVP01_1119800","PVP01_1120100","PVP01_1120300","PVP01_1120900","PVP01_1121100","PVP01_1121300","PVP01_1121500","PVP01_1121600","PVP01_1121700","PVP01_1121900","PVP01_1122100","PVP01_1122300","PVP01_1122400","PVP01_1122500","PVP01_1122600","PVP01_1122700","PVP01_1122800","PVP01_1122900","PVP01_1123000","PVP01_1123200","PVP01_1123300","PVP01_1123400","PVP01_1123500","PVP01_1123600","PVP01_1123900","PVP01_1124000","PVP01_1124100","PVP01_1124300","PVP01_1124400","PVP01_1124600","PVP01_1124800","PVP01_1124900","PVP01_1125000","PVP01_1125100","PVP01_1125200","PVP01_1125300","PVP01_1125400","PVP01_1125600","PVP01_1125700","PVP01_1126000","PVP01_1126100","PVP01_1126200","PVP01_1126300","PVP01_1126500","PVP01_1126600","PVP01_1126800","PVP01_1126900","PVP01_1127000","PVP01_1127200","PVP01_1127300","PVP01_1127400","PVP01_1127500","PVP01_1127800","PVP01_1128100","PVP01_1128400","PVP01_1128800","PVP01_1128900","PVP01_1129000","PVP01_1129500","PVP01_1129600","PVP01_1129700","PVP01_1130100","PVP01_1130400","PVP01_1130700","PVP01_1130900","PVP01_1131300","PVP01_1131400","PVP01_1131900","PVP01_1132100","PVP01_1132300","PVP01_1132400","PVP01_1132500","PVP01_1132700","PVP01_1132800","PVP01_1132900","PVP01_1133000","PVP01_1133100","PVP01_1133200","PVP01_1133300","PVP01_1133400","PVP01_1133600","PVP01_1133800","PVP01_1133900","PVP01_1134100","PVP01_1134200","PVP01_1134300","PVP01_1134400","PVP01_1134500","PVP01_1134700","PVP01_1134800","PVP01_1135100","PVP01_1135300","PVP01_1135500","PVP01_1135600","PVP01_1135700","PVP01_1135800","PVP01_1135900","PVP01_1136100","PVP01_1136300","PVP01_1136600","PVP01_1136700","PVP01_1136800","PVP01_1136900","PVP01_1137100","PVP01_1137200","PVP01_1137300","PVP01_1137600","PVP01_1137700","PVP01_1137800","PVP01_1138400","PVP01_1138500","PVP01_1139000","PVP01_1139100","PVP01_1139300","PVP01_1139700","PVP01_1140000","PVP01_1140300","PVP01_1140500","PVP01_1140700","PVP01_1140800","PVP01_1140900","PVP01_1141000","PVP01_1141200","PVP01_1141500","PVP01_1141700","PVP01_1141800","PVP01_1141900","PVP01_1142000","PVP01_1142300","PVP01_1142500","PVP01_1142600","PVP01_1143100","PVP01_1143200","PVP01_1143400","PVP01_1143500","PVP01_1143700","PVP01_1144100","PVP01_1144200","PVP01_1144300","PVP01_1144500","PVP01_1144700","PVP01_1144900","PVP01_1145000","PVP01_1145100","PVP01_1145200","PVP01_1145500","PVP01_1145700","PVP01_1146400","PVP01_1146600","PVP01_1146800","PVP01_1146900","PVP01_1147000","PVP01_1147100","PVP01_1147200","PVP01_1147300","PVP01_1147400","PVP01_1147500","PVP01_1147700","PVP01_1147800","PVP01_1147900","PVP01_1148200","PVP01_1148700","PVP01_1148800","PVP01_1149000","PVP01_1149100","PVP01_1149400","PVP01_1149500","PVP01_1200200","PVP01_1200800","PVP01_1201100","PVP01_1201300","PVP01_1201400","PVP01_1201500","PVP01_1201600","PVP01_1201700","PVP01_1201800","PVP01_1201900","PVP01_1202100","PVP01_1202200","PVP01_1203100","PVP01_1203200","PVP01_1203300","PVP01_1203500","PVP01_1203600","PVP01_1203700","PVP01_1203800","PVP01_1204000","PVP01_1204100","PVP01_1204200","PVP01_1204600","PVP01_1204700","PVP01_1204800","PVP01_1205000","PVP01_1205200","PVP01_1205400","PVP01_1205600","PVP01_1205900","PVP01_1206100","PVP01_1206300","PVP01_1206400","PVP01_1206700","PVP01_1206900","PVP01_1207000","PVP01_1207100","PVP01_1207200","PVP01_1207800","PVP01_1207900","PVP01_1208000","PVP01_1208100","PVP01_1208200","PVP01_1208300","PVP01_1208600","PVP01_1208700","PVP01_1208800","PVP01_1208900","PVP01_1209000","PVP01_1209400","PVP01_1210000","PVP01_1210100","PVP01_1210300","PVP01_1210700","PVP01_1211000","PVP01_1211200","PVP01_1211300","PVP01_1211500","PVP01_1211600","PVP01_1212500","PVP01_1212600","PVP01_1213000","PVP01_1213100","PVP01_1213300","PVP01_1214200","PVP01_1214300","PVP01_1214600","PVP01_1214800","PVP01_1214900","PVP01_1215000","PVP01_1215300","PVP01_1215400","PVP01_1215900","PVP01_1216000","PVP01_1216100","PVP01_1216200","PVP01_1216300","PVP01_1216800","PVP01_1217000","PVP01_1217200","PVP01_1217400","PVP01_1217500","PVP01_1217600","PVP01_1217700","PVP01_1218000","PVP01_1218500","PVP01_1218800","PVP01_1218900","PVP01_1219000","PVP01_1219100","PVP01_1219200","PVP01_1219400","PVP01_1219500","PVP01_1219800","PVP01_1219900","PVP01_1220000","PVP01_1220100","PVP01_1220200","PVP01_1220600","PVP01_1220700","PVP01_1220900","PVP01_1221200","PVP01_1221300","PVP01_1221400","PVP01_1221700","PVP01_1221800","PVP01_1221900","PVP01_1222100","PVP01_1222200","PVP01_1222300","PVP01_1222400","PVP01_1223500","PVP01_1223800","PVP01_1223900","PVP01_1224000","PVP01_1224400","PVP01_1224500","PVP01_1224600","PVP01_1224700","PVP01_1224800","PVP01_1225000","PVP01_1225200","PVP01_1225300","PVP01_1225500","PVP01_1225800","PVP01_1226100","PVP01_1226700","PVP01_1226800","PVP01_1227200","PVP01_1227300","PVP01_1227500","PVP01_1227600","PVP01_1228000","PVP01_1228100","PVP01_1228200","PVP01_1228400","PVP01_1228500","PVP01_1228800","PVP01_1229100","PVP01_1229400","PVP01_1229500","PVP01_1229600","PVP01_1229800","PVP01_1229900","PVP01_1230000","PVP01_1230200","PVP01_1230600","PVP01_1230700","PVP01_1230800","PVP01_1231000","PVP01_1231100","PVP01_1231600","PVP01_1231700","PVP01_1231800","PVP01_1231900","PVP01_1232400","PVP01_1232600","PVP01_1232700","PVP01_1233100","PVP01_1233200","PVP01_1233800","PVP01_1234000","PVP01_1235000","PVP01_1235100","PVP01_1235200","PVP01_1235300","PVP01_1235400","PVP01_1235800","PVP01_1236000","PVP01_1236100","PVP01_1236400","PVP01_1236900","PVP01_1237000","PVP01_1237100","PVP01_1237200","PVP01_1237500","PVP01_1237700","PVP01_1237800","PVP01_1237900","PVP01_1238000","PVP01_1238400","PVP01_1238900","PVP01_1239000","PVP01_1239100","PVP01_1239300","PVP01_1239400","PVP01_1240100","PVP01_1240300","PVP01_1240600","PVP01_1240800","PVP01_1240900","PVP01_1241000","PVP01_1241100","PVP01_1241200","PVP01_1241300","PVP01_1241900","PVP01_1242000","PVP01_1242300","PVP01_1242400","PVP01_1242700","PVP01_1243000","PVP01_1243100","PVP01_1243200","PVP01_1243500","PVP01_1243600","PVP01_1243900","PVP01_1244400","PVP01_1244700","PVP01_1244800","PVP01_1244900","PVP01_1245100","PVP01_1245300","PVP01_1245400","PVP01_1245500","PVP01_1245600","PVP01_1245800","PVP01_1245900","PVP01_1246000","PVP01_1246300","PVP01_1246400","PVP01_1246500","PVP01_1246600","PVP01_1246700","PVP01_1246900","PVP01_1247000","PVP01_1247500","PVP01_1247600","PVP01_1247700","PVP01_1247800","PVP01_1247900","PVP01_1248200","PVP01_1248500","PVP01_1249000","PVP01_1249100","PVP01_1249200","PVP01_1249500","PVP01_1250200","PVP01_1250300","PVP01_1250600","PVP01_1250700","PVP01_1250900","PVP01_1251100","PVP01_1251300","PVP01_1251500","PVP01_1251700","PVP01_1251900","PVP01_1252000","PVP01_1252200","PVP01_1252400","PVP01_1252800","PVP01_1252900","PVP01_1253000","PVP01_1253300","PVP01_1253600","PVP01_1253700","PVP01_1253800","PVP01_1254100","PVP01_1254300","PVP01_1254900","PVP01_1255100","PVP01_1255200","PVP01_1255300","PVP01_1255400","PVP01_1256000","PVP01_1256200","PVP01_1256400","PVP01_1256700","PVP01_1256800","PVP01_1257000","PVP01_1257500","PVP01_1257600","PVP01_1257700","PVP01_1257900","PVP01_1258100","PVP01_1258200","PVP01_1258300","PVP01_1258400","PVP01_1258500","PVP01_1258700","PVP01_1258900","PVP01_1259000","PVP01_1259100","PVP01_1259200","PVP01_1259300","PVP01_1259400","PVP01_1259500","PVP01_1259600","PVP01_1260200","PVP01_1260800","PVP01_1261100","PVP01_1261200","PVP01_1261400","PVP01_1261500","PVP01_1261700","PVP01_1262000","PVP01_1262300","PVP01_1262400","PVP01_1262600","PVP01_1262800","PVP01_1262900","PVP01_1263000","PVP01_1263100","PVP01_1263400","PVP01_1263800","PVP01_1263900","PVP01_1264000","PVP01_1264300","PVP01_1264400","PVP01_1264600","PVP01_1264700","PVP01_1264800","PVP01_1264900","PVP01_1265000","PVP01_1265100","PVP01_1265300","PVP01_1265400","PVP01_1265700","PVP01_1265900","PVP01_1266400","PVP01_1266500","PVP01_1266700","PVP01_1267300","PVP01_1267500","PVP01_1267800","PVP01_1268000","PVP01_1268100","PVP01_1268200","PVP01_1268300","PVP01_1268600","PVP01_1269000","PVP01_1269200","PVP01_1270200","PVP01_1270300","PVP01_1270600","PVP01_1270700","PVP01_1270900","PVP01_1271100","PVP01_1271200","PVP01_1271300","PVP01_1271400","PVP01_1271500","PVP01_1271600","PVP01_1271900","PVP01_1272500","PVP01_1272600","PVP01_1273000","PVP01_1274300","PVP01_1300400","PVP01_1300500","PVP01_1300900","PVP01_1301000","PVP01_1301100","PVP01_1301200","PVP01_1301300","PVP01_1301400","PVP01_1301500","PVP01_1301600","PVP01_1301700","PVP01_1301800","PVP01_1302100","PVP01_1302300","PVP01_1302500","PVP01_1302700","PVP01_1302800","PVP01_1302900","PVP01_1303000","PVP01_1303100","PVP01_1303200","PVP01_1303600","PVP01_1303900","PVP01_1304000","PVP01_1304200","PVP01_1304300","PVP01_1304500","PVP01_1304800","PVP01_1305300","PVP01_1305500","PVP01_1305700","PVP01_1306000","PVP01_1306100","PVP01_1306300","PVP01_1306600","PVP01_1306700","PVP01_1306800","PVP01_1307000","PVP01_1307100","PVP01_1307200","PVP01_1307300","PVP01_1307700","PVP01_1307800","PVP01_1308100","PVP01_1308300","PVP01_1308700","PVP01_1308800","PVP01_1308900","PVP01_1309000","PVP01_1309100","PVP01_1309400","PVP01_1309500","PVP01_1309700","PVP01_1309800","PVP01_1310000","PVP01_1310400","PVP01_1310700","PVP01_1311000","PVP01_1311100","PVP01_1311200","PVP01_1311300","PVP01_1311500","PVP01_1311900","PVP01_1312000","PVP01_1312300","PVP01_1312400","PVP01_1312600","PVP01_1313100","PVP01_1313300","PVP01_1313400","PVP01_1313900","PVP01_1314100","PVP01_1314300","PVP01_1314600","PVP01_1314800","PVP01_1315200","PVP01_1315400","PVP01_1315500","PVP01_1315700","PVP01_1316100","PVP01_1316500","PVP01_1316600","PVP01_1316700","PVP01_1316900","PVP01_1317000","PVP01_1317600","PVP01_1317700","PVP01_1317800","PVP01_1317900","PVP01_1318000","PVP01_1318100","PVP01_1318300","PVP01_1318600","PVP01_1318900","PVP01_1319000","PVP01_1319200","PVP01_1319500","PVP01_1319600","PVP01_1319700","PVP01_1319900","PVP01_1320000","PVP01_1320100","PVP01_1320200","PVP01_1320300","PVP01_1320400","PVP01_1320500","PVP01_1320600","PVP01_1320700","PVP01_1321500","PVP01_1321600","PVP01_1321700","PVP01_1321900","PVP01_1322000","PVP01_1322200","PVP01_1322400","PVP01_1322600","PVP01_1322900","PVP01_1323000","PVP01_1323100","PVP01_1323700","PVP01_1323800","PVP01_1323900","PVP01_1324100","PVP01_1324800","PVP01_1325800","PVP01_1325900","PVP01_1326000","PVP01_1326200","PVP01_1326400","PVP01_1326600","PVP01_1327200","PVP01_1327300","PVP01_1327600","PVP01_1327900","PVP01_1328000","PVP01_1328100","PVP01_1328500","PVP01_1328600","PVP01_1328700","PVP01_1328800","PVP01_1328900","PVP01_1329000","PVP01_1329300","PVP01_1329400","PVP01_1329600","PVP01_1329900","PVP01_1330000","PVP01_1330800","PVP01_1331000","PVP01_1331200","PVP01_1331400","PVP01_1331800","PVP01_1332200","PVP01_1332300","PVP01_1332500","PVP01_1332700","PVP01_1332900","PVP01_1333000","PVP01_1333100","PVP01_1333200","PVP01_1333700","PVP01_1333900","PVP01_1334000","PVP01_1334100","PVP01_1334200","PVP01_1334300","PVP01_1334500","PVP01_1335100","PVP01_1335300","PVP01_1335600","PVP01_1335800","PVP01_1335900","PVP01_1336700","PVP01_1337100","PVP01_1337200","PVP01_1337300","PVP01_1337400","PVP01_1337500","PVP01_1337600","PVP01_1337900","PVP01_1338100","PVP01_1338300","PVP01_1338400","PVP01_1338600","PVP01_1339000","PVP01_1339400","PVP01_1339800","PVP01_1340200","PVP01_1340400","PVP01_1340500","PVP01_1340600","PVP01_1340800","PVP01_1341300","PVP01_1341600","PVP01_1341800","PVP01_1342200","PVP01_1342300","PVP01_1342400","PVP01_1342600","PVP01_1342700","PVP01_1342800","PVP01_1343100","PVP01_1343300","PVP01_1343500","PVP01_1343600","PVP01_1344200","PVP01_1344300","PVP01_1344500","PVP01_1344600","PVP01_1344700","PVP01_1344900","PVP01_1345300","PVP01_1345400","PVP01_1345500","PVP01_1345600","PVP01_1345700","PVP01_1345900","PVP01_1346200","PVP01_1346700","PVP01_1346800","PVP01_1346900","PVP01_1347300","PVP01_1347500","PVP01_1347700","PVP01_1347800","PVP01_1400100","PVP01_1400400","PVP01_1400600","PVP01_1400700","PVP01_1400900","PVP01_1401000","PVP01_1401100","PVP01_1401200","PVP01_1401300","PVP01_1401600","PVP01_1401700","PVP01_1401800","PVP01_1401900","PVP01_1402000","PVP01_1402100","PVP01_1402200","PVP01_1402300","PVP01_1402400","PVP01_1402500","PVP01_1402600","PVP01_1402700","PVP01_1402800","PVP01_1403000","PVP01_1403300","PVP01_1403400","PVP01_1403700","PVP01_1403900","PVP01_1404000","PVP01_1404100","PVP01_1404200","PVP01_1404300","PVP01_1404500","PVP01_1404600","PVP01_1404800","PVP01_1405000","PVP01_1405100","PVP01_1405500","PVP01_1405600","PVP01_1405800","PVP01_1406000","PVP01_1406100","PVP01_1406200","PVP01_1406700","PVP01_1407000","PVP01_1407100","PVP01_1407400","PVP01_1407600","PVP01_1407800","PVP01_1408000","PVP01_1408100","PVP01_1408200","PVP01_1408400","PVP01_1408500","PVP01_1408600","PVP01_1408700","PVP01_1408800","PVP01_1408900","PVP01_1409000","PVP01_1409100","PVP01_1409300","PVP01_1409400","PVP01_1409700","PVP01_1410200","PVP01_1410300","PVP01_1410500","PVP01_1410700","PVP01_1410800","PVP01_1410900","PVP01_1411000","PVP01_1411200","PVP01_1411300","PVP01_1411400","PVP01_1412100","PVP01_1412500","PVP01_1413100","PVP01_1413200","PVP01_1413300","PVP01_1413700","PVP01_1414200","PVP01_1414300","PVP01_1414400","PVP01_1414900","PVP01_1415100","PVP01_1415200","PVP01_1415300","PVP01_1415500","PVP01_1415600","PVP01_1415700","PVP01_1415900","PVP01_1416000","PVP01_1416100","PVP01_1416900","PVP01_1417100","PVP01_1417200","PVP01_1417700","PVP01_1417900","PVP01_1418000","PVP01_1418100","PVP01_1418200","PVP01_1418300","PVP01_1418400","PVP01_1418600","PVP01_1418800","PVP01_1418900","PVP01_1419100","PVP01_1419300","PVP01_1419400","PVP01_1419500","PVP01_1419700","PVP01_1419800","PVP01_1420000","PVP01_1420100","PVP01_1420300","PVP01_1420500","PVP01_1420700","PVP01_1420900","PVP01_1421000","PVP01_1421200","PVP01_1421300","PVP01_1421400","PVP01_1421800","PVP01_1422100","PVP01_1422200","PVP01_1422300","PVP01_1422600","PVP01_1422700","PVP01_1422900","PVP01_1423200","PVP01_1424100","PVP01_1424200","PVP01_1424500","PVP01_1425100","PVP01_1425200","PVP01_1425300","PVP01_1425400","PVP01_1425800","PVP01_1426100","PVP01_1426200","PVP01_1426700","PVP01_1426800","PVP01_1427000","PVP01_1427100","PVP01_1427700","PVP01_1428000","PVP01_1428100","PVP01_1428300","PVP01_1428400","PVP01_1428500","PVP01_1428900","PVP01_1429000","PVP01_1429200","PVP01_1429300","PVP01_1429700","PVP01_1429900","PVP01_1430000","PVP01_1430300","PVP01_1430600","PVP01_1430700","PVP01_1430800","PVP01_1431100","PVP01_1431200","PVP01_1431300","PVP01_1431700","PVP01_1431800","PVP01_1432000","PVP01_1432100","PVP01_1432400","PVP01_1432600","PVP01_1432700","PVP01_1432800","PVP01_1433600","PVP01_1433800","PVP01_1434000","PVP01_1434200","PVP01_1434300","PVP01_1434500","PVP01_1434600","PVP01_1434800","PVP01_1434900","PVP01_1435100","PVP01_1435300","PVP01_1435800","PVP01_1435900","PVP01_1436000","PVP01_1436300","PVP01_1436400","PVP01_1436600","PVP01_1436700","PVP01_1437100","PVP01_1437200","PVP01_1437600","PVP01_1437700","PVP01_1438100","PVP01_1438200","PVP01_1438400","PVP01_1438900","PVP01_1439000","PVP01_1439300","PVP01_1439400","PVP01_1439600","PVP01_1439700","PVP01_1440000","PVP01_1440100","PVP01_1440300","PVP01_1440600","PVP01_1440700","PVP01_1440900","PVP01_1441100","PVP01_1441200","PVP01_1441400","PVP01_1441700","PVP01_1441800","PVP01_1442000","PVP01_1442200","PVP01_1442900","PVP01_1443000","PVP01_1443200","PVP01_1443600","PVP01_1443800","PVP01_1443900","PVP01_1444000","PVP01_1444200","PVP01_1444400","PVP01_1444800","PVP01_1444900","PVP01_1445000","PVP01_1445200","PVP01_1445300","PVP01_1445400","PVP01_1445600","PVP01_1445700","PVP01_1446000","PVP01_1446100","PVP01_1446200","PVP01_1446300","PVP01_1446400","PVP01_1446600","PVP01_1446900","PVP01_1447200","PVP01_1447300","PVP01_1447500","PVP01_1447600","PVP01_1447800","PVP01_1447900","PVP01_1448100","PVP01_1448300","PVP01_1448400","PVP01_1448700","PVP01_1448800","PVP01_1449100","PVP01_1449200","PVP01_1449700","PVP01_1450000","PVP01_1450200","PVP01_1450400","PVP01_1450500","PVP01_1451000","PVP01_1451300","PVP01_1451500","PVP01_1451600","PVP01_1451800","PVP01_1452100","PVP01_1452400","PVP01_1452500","PVP01_1452700","PVP01_1452900","PVP01_1453200","PVP01_1453800","PVP01_1454000","PVP01_1454100","PVP01_1454200","PVP01_1454400","PVP01_1454500","PVP01_1454600","PVP01_1454800","PVP01_1454900","PVP01_1455400","PVP01_1455500","PVP01_1455700","PVP01_1456000","PVP01_1456500","PVP01_1456700","PVP01_1456900","PVP01_1457000","PVP01_1457400","PVP01_1457500","PVP01_1457900","PVP01_1458700","PVP01_1458800","PVP01_1459100","PVP01_1459200","PVP01_1459300","PVP01_1459400","PVP01_1459500","PVP01_1459600","PVP01_1459700","PVP01_1460500","PVP01_1460700","PVP01_1460800","PVP01_1460900","PVP01_1461100","PVP01_1461900","PVP01_1462000","PVP01_1462100","PVP01_1462300","PVP01_1462400","PVP01_1462500","PVP01_1462600","PVP01_1462900","PVP01_1463200","PVP01_1463300","PVP01_1463500","PVP01_1463600","PVP01_1463700","PVP01_1463800","PVP01_1464100","PVP01_1464500","PVP01_1464700","PVP01_1464900","PVP01_1465000","PVP01_1465100","PVP01_1465500","PVP01_1465700","PVP01_1465800","PVP01_1465900","PVP01_1466000","PVP01_1466100","PVP01_1466300","PVP01_1466500","PVP01_1466600","PVP01_1467000","PVP01_1467300","PVP01_1467600","PVP01_1468100","PVP01_1468400","PVP01_1468600","PVP01_1469300","PVP01_1469400","PVP01_1469600","PVP01_1469700","PVP01_1469900","PVP01_1470000","PVP01_1470100","PVP01_1470200","PVP01_1470300","PVP01_1470400","PVP01_1470500","PVP01_1470600","PVP01_1470700","PVP01_1470800","PVP01_1470900","PVP01_1471100","PVP01_1471300","PVP01_1471500","PVP01_1471600","PVP01_1471700","PVP01_1471800","PVP01_1471900","PVP01_1472100","PVP01_1472200","PVP01_1472400","PVP01_API01200","PVP01_API01400","PVP01_API01700","PVP01_API04100","PVP01_API04200","PVP01_0000110","PVP01_0000160","PVP01_0001430","PVP01_0003640","PVP01_0004040","PVP01_0004210","PVP01_0004220","PVP01_0004230","PVP01_0004290","PVP01_0004330","PVP01_0005540","PVP01_0009370","PVP01_0009720","PVP01_0010670","PVP01_0010840","PVP01_0011020","PVP01_0103100","PVP01_0103200","PVP01_0103300","PVP01_0103500","PVP01_0103700","PVP01_0104000","PVP01_0104400","PVP01_0104500","PVP01_0104700","PVP01_0105200","PVP01_0105900","PVP01_0106100","PVP01_0106600","PVP01_0106700","PVP01_0107500","PVP01_0107800","PVP01_0108200","PVP01_0108500","PVP01_0108700","PVP01_0108900","PVP01_0109400","PVP01_0109500","PVP01_0109700","PVP01_0109900","PVP01_0110000","PVP01_0110100","PVP01_0110300","PVP01_0110400","PVP01_0110500","PVP01_0110600","PVP01_0110900","PVP01_0111000","PVP01_0111100","PVP01_0111400","PVP01_0111500","PVP01_0111800","PVP01_0111900","PVP01_0112200","PVP01_0112300","PVP01_0112400","PVP01_0112600","PVP01_0112800","PVP01_0112900","PVP01_0113300","PVP01_0113600","PVP01_0113700","PVP01_0113800","PVP01_0113900","PVP01_0114000","PVP01_0114200","PVP01_0114300","PVP01_0114400","PVP01_0114500","PVP01_0114800","PVP01_0115100","PVP01_0115300","PVP01_0115600","PVP01_0115700","PVP01_0115800","PVP01_0115900","PVP01_0116000","PVP01_0116200","PVP01_0116300","PVP01_0116400","PVP01_0116800","PVP01_0117000","PVP01_0117100","PVP01_0117200","PVP01_0117300","PVP01_0117400","PVP01_0117600","PVP01_0117900","PVP01_0118100","PVP01_0118200","PVP01_0118300","PVP01_0118400","PVP01_0118500","PVP01_0201200","PVP01_0202500","PVP01_0202600","PVP01_0203200","PVP01_0203300","PVP01_0203600","PVP01_0203700","PVP01_0204200","PVP01_0204300","PVP01_0204400","PVP01_0204500","PVP01_0204600","PVP01_0204700","PVP01_0205100","PVP01_0205400","PVP01_0205700","PVP01_0205900","PVP01_0206000","PVP01_0206200","PVP01_0206700","PVP01_0206900","PVP01_0207300","PVP01_0207900","PVP01_0208000","PVP01_0208500","PVP01_0208900","PVP01_0209400","PVP01_0209700","PVP01_0209900","PVP01_0210200","PVP01_0210400","PVP01_0210500","PVP01_0210600","PVP01_0210700","PVP01_0210800","PVP01_0210900","PVP01_0211300","PVP01_0211800","PVP01_0211900","PVP01_0212000","PVP01_0212100","PVP01_0212200","PVP01_0212300","PVP01_0212900","PVP01_0213100","PVP01_0213200","PVP01_0213400","PVP01_0213500","PVP01_0213800","PVP01_0213900","PVP01_0214000","PVP01_0214300","PVP01_0214400","PVP01_0214500","PVP01_0214600","PVP01_0214700","PVP01_0215100","PVP01_0215200","PVP01_0215600","PVP01_0215900","PVP01_0216000","PVP01_0216200","PVP01_0302600","PVP01_0302700","PVP01_0302800","PVP01_0302900","PVP01_0303000","PVP01_0303200","PVP01_0303300","PVP01_0303400","PVP01_0303600","PVP01_0303700","PVP01_0303800","PVP01_0303900","PVP01_0304000","PVP01_0304200","PVP01_0304400","PVP01_0304700","PVP01_0305100","PVP01_0305400","PVP01_0305600","PVP01_0305700","PVP01_0305900","PVP01_0306200","PVP01_0306600","PVP01_0307200","PVP01_0307500","PVP01_0307800","PVP01_0307900","PVP01_0308000","PVP01_0308100","PVP01_0308200","PVP01_0308300","PVP01_0308400","PVP01_0308500","PVP01_0308600","PVP01_0308700","PVP01_0308900","PVP01_0309000","PVP01_0309100","PVP01_0309300","PVP01_0309600","PVP01_0309900","PVP01_0310100","PVP01_0310500","PVP01_0310700","PVP01_0311100","PVP01_0311800","PVP01_0311900","PVP01_0312200","PVP01_0312300","PVP01_0312600","PVP01_0312900","PVP01_0313000","PVP01_0313100","PVP01_0313200","PVP01_0313300","PVP01_0313400","PVP01_0313600","PVP01_0313700","PVP01_0313900","PVP01_0314000","PVP01_0314300","PVP01_0314400","PVP01_0314500","PVP01_0314600","PVP01_0314700","PVP01_0314800","PVP01_0315400","PVP01_0315500","PVP01_0315600","PVP01_0315800","PVP01_0316100","PVP01_0316400","PVP01_0316600","PVP01_0316900","PVP01_0317000","PVP01_0317300","PVP01_0317500","PVP01_0317600","PVP01_0317700","PVP01_0317800","PVP01_0317900","PVP01_0318200","PVP01_0318300","PVP01_0318500","PVP01_0318600","PVP01_0318700","PVP01_0318800","PVP01_0319000","PVP01_0319100","PVP01_0319500","PVP01_0319800","PVP01_0319900","PVP01_0320000","PVP01_0320100","PVP01_0320200","PVP01_0320300","PVP01_0320600","PVP01_0321000","PVP01_0405000","PVP01_0405400","PVP01_0405800","PVP01_0406300","PVP01_0406400","PVP01_0406800","PVP01_0407100","PVP01_0407200","PVP01_0407300","PVP01_0407400","PVP01_0407500","PVP01_0407600","PVP01_0407800","PVP01_0408000","PVP01_0408100","PVP01_0408300","PVP01_0408400","PVP01_0408500","PVP01_0408700","PVP01_0409100","PVP01_0409200","PVP01_0409300","PVP01_0409400","PVP01_0409500","PVP01_0409600","PVP01_0409800","PVP01_0410100","PVP01_0410200","PVP01_0410300","PVP01_0410500","PVP01_0410700","PVP01_0410900","PVP01_0411000","PVP01_0411100","PVP01_0411200","PVP01_0411300","PVP01_0411700","PVP01_0412000","PVP01_0412300","PVP01_0412400","PVP01_0412500","PVP01_0412600","PVP01_0413400","PVP01_0413600","PVP01_0413700","PVP01_0413800","PVP01_0413900","PVP01_0414000","PVP01_0414100","PVP01_0414200","PVP01_0414500","PVP01_0414700","PVP01_0414800","PVP01_0415000","PVP01_0415100","PVP01_0415300","PVP01_0415700","PVP01_0415800","PVP01_0415900","PVP01_0416000","PVP01_0416100","PVP01_0416200","PVP01_0416300","PVP01_0416400","PVP01_0416500","PVP01_0416700","PVP01_0416800","PVP01_0417000","PVP01_0417200","PVP01_0417300","PVP01_0417500","PVP01_0417600","PVP01_0417700","PVP01_0418000","PVP01_0418100","PVP01_0418200","PVP01_0418400","PVP01_0418500","PVP01_0418600","PVP01_0418700","PVP01_0418800","PVP01_0418900","PVP01_0419100","PVP01_0419600","PVP01_0419800","PVP01_0419900","PVP01_0420100","PVP01_0420300","PVP01_0420400","PVP01_0420600","PVP01_0420800","PVP01_0421000","PVP01_0421100","PVP01_0421200","PVP01_0421300","PVP01_0421400","PVP01_0421900","PVP01_0422000","PVP01_0422500","PVP01_0422800","PVP01_0423400","PVP01_0423500","PVP01_0423900","PVP01_0424500","PVP01_0503500","PVP01_0504000","PVP01_0504400","PVP01_0504800","PVP01_0504900","PVP01_0505300","PVP01_0505600","PVP01_0505700","PVP01_0505900","PVP01_0506100","PVP01_0506200","PVP01_0506500","PVP01_0506900","PVP01_0507000","PVP01_0507100","PVP01_0507200","PVP01_0507400","PVP01_0507900","PVP01_0508100","PVP01_0508300","PVP01_0508400","PVP01_0508800","PVP01_0508900","PVP01_0509000","PVP01_0509300","PVP01_0509500","PVP01_0509700","PVP01_0509800","PVP01_0510400","PVP01_0510500","PVP01_0510600","PVP01_0510700","PVP01_0511000","PVP01_0511100","PVP01_0511600","PVP01_0511900","PVP01_0512000","PVP01_0512300","PVP01_0512600","PVP01_0512900","PVP01_0513100","PVP01_0513200","PVP01_0513600","PVP01_0513800","PVP01_0514000","PVP01_0514200","PVP01_0514300","PVP01_0514400","PVP01_0514500","PVP01_0514700","PVP01_0515000","PVP01_0515400","PVP01_0515500","PVP01_0515600","PVP01_0516600","PVP01_0516700","PVP01_0516800","PVP01_0517200","PVP01_0517500","PVP01_0517600","PVP01_0517800","PVP01_0518000","PVP01_0518100","PVP01_0518200","PVP01_0518800","PVP01_0518900","PVP01_0519000","PVP01_0519300","PVP01_0519400","PVP01_0519700","PVP01_0519800","PVP01_0519900","PVP01_0520000","PVP01_0520100","PVP01_0520200","PVP01_0520300","PVP01_0520400","PVP01_0520500","PVP01_0520600","PVP01_0520900","PVP01_0521000","PVP01_0521100","PVP01_0521200","PVP01_0521500","PVP01_0521700","PVP01_0521900","PVP01_0522400","PVP01_0522500","PVP01_0522800","PVP01_0523100","PVP01_0523200","PVP01_0523500","PVP01_0523600","PVP01_0525200","PVP01_0525400","PVP01_0525500","PVP01_0525600","PVP01_0525800","PVP01_0525900","PVP01_0526000","PVP01_0526200","PVP01_0526300","PVP01_0526500","PVP01_0526600","PVP01_0526700","PVP01_0526800","PVP01_0526900","PVP01_0527000","PVP01_0527100","PVP01_0527200","PVP01_0527600","PVP01_0527900","PVP01_0528000","PVP01_0528300","PVP01_0528500","PVP01_0528800","PVP01_0529300","PVP01_0529400","PVP01_0529500","PVP01_0529800","PVP01_0530200","PVP01_0530300","PVP01_0530600","PVP01_0530700","PVP01_0530800","PVP01_0530900","PVP01_0531000","PVP01_0531300","PVP01_0531400","PVP01_0531600","PVP01_0531900","PVP01_0532000","PVP01_0532100","PVP01_0532200","PVP01_0532400","PVP01_0532800","PVP01_0533200","PVP01_0534300","PVP01_0534400","PVP01_0600800","PVP01_0601000","PVP01_0601400","PVP01_0601700","PVP01_0601800","PVP01_0601900","PVP01_0602000","PVP01_0602100","PVP01_0602200","PVP01_0602300","PVP01_0602400","PVP01_0602500","PVP01_0602700","PVP01_0602800","PVP01_0602900","PVP01_0603200","PVP01_0603300","PVP01_0603400","PVP01_0603500","PVP01_0603800","PVP01_0603900","PVP01_0604000","PVP01_0604100","PVP01_0604300","PVP01_0604500","PVP01_0604600","PVP01_0604800","PVP01_0604900","PVP01_0605200","PVP01_0605400","PVP01_0605600","PVP01_0605700","PVP01_0605800","PVP01_0605900","PVP01_0606000","PVP01_0606100","PVP01_0606300","PVP01_0606400","PVP01_0606700","PVP01_0606900","PVP01_0607000","PVP01_0607100","PVP01_0607300","PVP01_0607400","PVP01_0607500","PVP01_0607700","PVP01_0607800","PVP01_0607900","PVP01_0608200","PVP01_0608400","PVP01_0608500","PVP01_0608700","PVP01_0609000","PVP01_0609100","PVP01_0609300","PVP01_0609500","PVP01_0610000","PVP01_0610500","PVP01_0611000","PVP01_0611200","PVP01_0611400","PVP01_0611500","PVP01_0611600","PVP01_0611700","PVP01_0611800","PVP01_0611900","PVP01_0612200","PVP01_0612300","PVP01_0612500","PVP01_0612800","PVP01_0612900","PVP01_0613300","PVP01_0613600","PVP01_0613700","PVP01_0613900","PVP01_0614100","PVP01_0614500","PVP01_0614700","PVP01_0614800","PVP01_0615200","PVP01_0615300","PVP01_0615400","PVP01_0615600","PVP01_0615700","PVP01_0616100","PVP01_0616500","PVP01_0616600","PVP01_0616700","PVP01_0616800","PVP01_0617200","PVP01_0617300","PVP01_0617600","PVP01_0617700","PVP01_0617800","PVP01_0618000","PVP01_0618200","PVP01_0618400","PVP01_0618500","PVP01_0618800","PVP01_0618900","PVP01_0619300","PVP01_0619900","PVP01_0620000","PVP01_0620200","PVP01_0620400","PVP01_0620800","PVP01_0621000","PVP01_0621200","PVP01_0621300","PVP01_0621400","PVP01_0621500","PVP01_0621600","PVP01_0621700","PVP01_0622000","PVP01_0622300","PVP01_0622400","PVP01_0622500","PVP01_0622600","PVP01_0623700","PVP01_0623800","PVP01_0700100","PVP01_0700600","PVP01_0701100","PVP01_0701300","PVP01_0701400","PVP01_0701800","PVP01_0701900","PVP01_0702000","PVP01_0702300","PVP01_0702400","PVP01_0702600","PVP01_0702700","PVP01_0702800","PVP01_0702900","PVP01_0703100","PVP01_0703200","PVP01_0703300","PVP01_0703400","PVP01_0703500","PVP01_0703800","PVP01_0703900","PVP01_0704200","PVP01_0704300","PVP01_0704400","PVP01_0705200","PVP01_0705300","PVP01_0705400","PVP01_0705700","PVP01_0706500","PVP01_0706800","PVP01_0706900","PVP01_0707400","PVP01_0707500","PVP01_0707600","PVP01_0707700","PVP01_0707900","PVP01_0708600","PVP01_0708700","PVP01_0709400","PVP01_0709600","PVP01_0709700","PVP01_0709800","PVP01_0710400","PVP01_0710500","PVP01_0710600","PVP01_0710800","PVP01_0710900","PVP01_0711000","PVP01_0711300","PVP01_0711600","PVP01_0711700","PVP01_0712000","PVP01_0712100","PVP01_0712500","PVP01_0712600","PVP01_0712800","PVP01_0712900","PVP01_0713300","PVP01_0713400","PVP01_0713700","PVP01_0713800","PVP01_0714000","PVP01_0714100","PVP01_0714400","PVP01_0714500","PVP01_0714700","PVP01_0714800","PVP01_0715300","PVP01_0715400","PVP01_0715900","PVP01_0716200","PVP01_0716300","PVP01_0716400","PVP01_0716500","PVP01_0716700","PVP01_0717100","PVP01_0717300","PVP01_0717500","PVP01_0717700","PVP01_0717900","PVP01_0718200","PVP01_0718300","PVP01_0718400","PVP01_0718500","PVP01_0718800","PVP01_0718900","PVP01_0719200","PVP01_0719300","PVP01_0719600","PVP01_0720500","PVP01_0720600","PVP01_0720700","PVP01_0721000","PVP01_0721200","PVP01_0721300","PVP01_0721400","PVP01_0721600","PVP01_0721700","PVP01_0721900","PVP01_0722900","PVP01_0723600","PVP01_0723700","PVP01_0723900","PVP01_0724000","PVP01_0724200","PVP01_0724300","PVP01_0724400","PVP01_0725500","PVP01_0725600","PVP01_0725800","PVP01_0725900","PVP01_0726000","PVP01_0726100","PVP01_0726200","PVP01_0726300","PVP01_0726500","PVP01_0726600","PVP01_0726800","PVP01_0727500","PVP01_0727700","PVP01_0727900","PVP01_0728000","PVP01_0728400","PVP01_0728500","PVP01_0728600","PVP01_0728700","PVP01_0728800","PVP01_0728900","PVP01_0729000","PVP01_0729600","PVP01_0729800","PVP01_0729900","PVP01_0730500","PVP01_0730800","PVP01_0730900","PVP01_0731400","PVP01_0731500","PVP01_0731800","PVP01_0732400","PVP01_0732500","PVP01_0732700","PVP01_0733200","PVP01_0733300","PVP01_0733400","PVP01_0733600","PVP01_0733800","PVP01_0734000","PVP01_0734500","PVP01_0734600","PVP01_0734800","PVP01_0734900","PVP01_0801100","PVP01_0801200","PVP01_0801300","PVP01_0801600","PVP01_0802200","PVP01_0802300","PVP01_0802500","PVP01_0802600","PVP01_0802700","PVP01_0802800","PVP01_0803200","PVP01_0803300","PVP01_0803400","PVP01_0803800","PVP01_0803900","PVP01_0804000","PVP01_0804300","PVP01_0804400","PVP01_0805100","PVP01_0805500","PVP01_0805900","PVP01_0806600","PVP01_0806800","PVP01_0806900","PVP01_0807100","PVP01_0807300","PVP01_0807400","PVP01_0807600","PVP01_0807700","PVP01_0807800","PVP01_0808200","PVP01_0808300","PVP01_0808900","PVP01_0809000","PVP01_0809500","PVP01_0809800","PVP01_0810200","PVP01_0810700","PVP01_0810800","PVP01_0810900","PVP01_0811100","PVP01_0811400","PVP01_0811600","PVP01_0812100","PVP01_0812200","PVP01_0812400","PVP01_0812500","PVP01_0812700","PVP01_0812900","PVP01_0813100","PVP01_0813400","PVP01_0813500","PVP01_0813900","PVP01_0814000","PVP01_0814100","PVP01_0814200","PVP01_0814300","PVP01_0814500","PVP01_0814600","PVP01_0814900","PVP01_0815200","PVP01_0815500","PVP01_0815600","PVP01_0815700","PVP01_0815800","PVP01_0815900","PVP01_0816000","PVP01_0816100","PVP01_0816400","PVP01_0816500","PVP01_0816600","PVP01_0816900","PVP01_0817000","PVP01_0817200","PVP01_0817300","PVP01_0817500","PVP01_0817600","PVP01_0817700","PVP01_0817800","PVP01_0817900","PVP01_0818000","PVP01_0818200","PVP01_0818400","PVP01_0818500","PVP01_0818700","PVP01_0818800","PVP01_0818900","PVP01_0819100","PVP01_0819400","PVP01_0819500","PVP01_0819600","PVP01_0819800","PVP01_0819900","PVP01_0820200","PVP01_0820400","PVP01_0820600","PVP01_0820700","PVP01_0821100","PVP01_0821200","PVP01_0821600","PVP01_0821900","PVP01_0822000","PVP01_0822200","PVP01_0822400","PVP01_0822500","PVP01_0822800","PVP01_0823000","PVP01_0823200","PVP01_0823300","PVP01_0823500","PVP01_0823800","PVP01_0823900","PVP01_0824100","PVP01_0824200","PVP01_0824400","PVP01_0824600","PVP01_0824800","PVP01_0824900","PVP01_0825000","PVP01_0825200","PVP01_0825300","PVP01_0825400","PVP01_0825500","PVP01_0825600","PVP01_0825900","PVP01_0826400","PVP01_0826600","PVP01_0826800","PVP01_0826900","PVP01_0827000","PVP01_0827200","PVP01_0827300","PVP01_0827400","PVP01_0827900","PVP01_0828400","PVP01_0828500","PVP01_0828600","PVP01_0828800","PVP01_0828900","PVP01_0829200","PVP01_0829400","PVP01_0829500","PVP01_0829600","PVP01_0829800","PVP01_0830000","PVP01_0830100","PVP01_0830200","PVP01_0830600","PVP01_0831200","PVP01_0831300","PVP01_0831500","PVP01_0831600","PVP01_0832300","PVP01_0832700","PVP01_0832800","PVP01_0833100","PVP01_0833200","PVP01_0833400","PVP01_0833900","PVP01_0834100","PVP01_0834600","PVP01_0834700","PVP01_0834800","PVP01_0834900","PVP01_0835100","PVP01_0835200","PVP01_0835400","PVP01_0835600","PVP01_0835800","PVP01_0836000","PVP01_0836200","PVP01_0836900","PVP01_0837000","PVP01_0837100","PVP01_0837400","PVP01_0837700","PVP01_0837900","PVP01_0904200","PVP01_0904400","PVP01_0904500","PVP01_0904800","PVP01_0905000","PVP01_0905200","PVP01_0905300","PVP01_0905400","PVP01_0905500","PVP01_0905600","PVP01_0905700","PVP01_0905800","PVP01_0905900","PVP01_0906100","PVP01_0906200","PVP01_0906400","PVP01_0906600","PVP01_0907400","PVP01_0907900","PVP01_0908100","PVP01_0908200","PVP01_0908300","PVP01_0908400","PVP01_0908700","PVP01_0908800","PVP01_0909000","PVP01_0909200","PVP01_0909400","PVP01_0909700","PVP01_0909900","PVP01_0910700","PVP01_0910800","PVP01_0911100","PVP01_0911200","PVP01_0911400","PVP01_0911500","PVP01_0912000","PVP01_0912300","PVP01_0912500","PVP01_0912600","PVP01_0912700","PVP01_0912800","PVP01_0913100","PVP01_0913200","PVP01_0913300","PVP01_0913600","PVP01_0913700","PVP01_0913800","PVP01_0914100","PVP01_0914200","PVP01_0914300","PVP01_0914500","PVP01_0914700","PVP01_0914900","PVP01_0915200","PVP01_0915400","PVP01_0915500","PVP01_0915700","PVP01_0915900","PVP01_0916000","PVP01_0916100","PVP01_0916200","PVP01_0916400","PVP01_0916500","PVP01_0916600","PVP01_0917300","PVP01_0917400","PVP01_0917500","PVP01_0917600","PVP01_0917700","PVP01_0917800","PVP01_0918500","PVP01_0918600","PVP01_0919100","PVP01_0919200","PVP01_0919300","PVP01_0919400","PVP01_0919500","PVP01_0919800","PVP01_0920000","PVP01_0920200","PVP01_0920300","PVP01_0920500","PVP01_0920600","PVP01_0920900","PVP01_0921000","PVP01_0921100","PVP01_0921200","PVP01_0921300","PVP01_0921600","PVP01_0921700","PVP01_0921900","PVP01_0922200","PVP01_0922500","PVP01_0922600","PVP01_0922700","PVP01_0923200","PVP01_0923400","PVP01_0923500","PVP01_0923600","PVP01_0923700","PVP01_0924200","PVP01_0924500","PVP01_0924600","PVP01_0924900","PVP01_0925300","PVP01_0925500","PVP01_0925900","PVP01_0926000","PVP01_0926100","PVP01_0926200","PVP01_0926700","PVP01_0926800","PVP01_0927000","PVP01_0927200","PVP01_0927300","PVP01_0927500","PVP01_0927900","PVP01_0928200","PVP01_0928400","PVP01_0928500","PVP01_0928600","PVP01_0928700","PVP01_0929300","PVP01_0929500","PVP01_0929800","PVP01_0930600","PVP01_0930700","PVP01_0930800","PVP01_0931200","PVP01_0931800","PVP01_0931900","PVP01_0932000","PVP01_0932100","PVP01_0932200","PVP01_0932400","PVP01_0932500","PVP01_0932700","PVP01_0932800","PVP01_0932900","PVP01_0933000","PVP01_0933200","PVP01_0933300","PVP01_0933600","PVP01_0933800","PVP01_0933900","PVP01_0934100","PVP01_0934200","PVP01_0934600","PVP01_0934700","PVP01_0934800","PVP01_0934900","PVP01_0935100","PVP01_0935300","PVP01_0935600","PVP01_0936100","PVP01_0936200","PVP01_0936500","PVP01_0936600","PVP01_0936900","PVP01_0937100","PVP01_0937200","PVP01_0937400","PVP01_0937600","PVP01_0937800","PVP01_0938000","PVP01_0938300","PVP01_0938400","PVP01_0938500","PVP01_0938800","PVP01_0939400","PVP01_0939700","PVP01_0939800","PVP01_0939900","PVP01_0940000","PVP01_0940100","PVP01_0940300","PVP01_0940400","PVP01_0940800","PVP01_0941000","PVP01_0941500","PVP01_0941700","PVP01_0941900","PVP01_0942000","PVP01_0942100","PVP01_0942500","PVP01_0943000","PVP01_0943200","PVP01_0943500","PVP01_0943600","PVP01_0943800","PVP01_0943900","PVP01_0944000","PVP01_0944200","PVP01_0944500","PVP01_0944700","PVP01_0944900","PVP01_0945000","PVP01_0945400","PVP01_0945700","PVP01_0946100","PVP01_0946800","PVP01_0947300","PVP01_0947500","PVP01_0947600","PVP01_0947700","PVP01_0948000","PVP01_0948200","PVP01_0948400","PVP01_0949300","PVP01_1002400","PVP01_1002800","PVP01_1002900","PVP01_1003100","PVP01_1003300","PVP01_1003600","PVP01_1003700","PVP01_1003800","PVP01_1004000","PVP01_1004100","PVP01_1004400","PVP01_1004600","PVP01_1004700","PVP01_1005000","PVP01_1005200","PVP01_1005500","PVP01_1005800","PVP01_1006500","PVP01_1006800","PVP01_1006900","PVP01_1007000","PVP01_1007100","PVP01_1007300","PVP01_1007400","PVP01_1007600","PVP01_1008400","PVP01_1008500","PVP01_1008700","PVP01_1008800","PVP01_1009200","PVP01_1009500","PVP01_1009900","PVP01_1010000","PVP01_1010500","PVP01_1010600","PVP01_1010700","PVP01_1010800","PVP01_1010900","PVP01_1011000","PVP01_1011200","PVP01_1011300","PVP01_1011400","PVP01_1011500","PVP01_1011700","PVP01_1011900","PVP01_1012000","PVP01_1012200","PVP01_1012600","PVP01_1012900","PVP01_1013000","PVP01_1013500","PVP01_1013600","PVP01_1013800","PVP01_1013900","PVP01_1014000","PVP01_1014100","PVP01_1014200","PVP01_1014400","PVP01_1014500","PVP01_1014700","PVP01_1014900","PVP01_1015200","PVP01_1015300","PVP01_1015600","PVP01_1015800","PVP01_1016000","PVP01_1016100","PVP01_1016300","PVP01_1016600","PVP01_1016700","PVP01_1016800","PVP01_1017000","PVP01_1017300","PVP01_1017400","PVP01_1017500","PVP01_1017600","PVP01_1017700","PVP01_1017800","PVP01_1018200","PVP01_1018300","PVP01_1018600","PVP01_1018700","PVP01_1018800","PVP01_1018900","PVP01_1019100","PVP01_1019200","PVP01_1019300","PVP01_1019700","PVP01_1019800","PVP01_1020300","PVP01_1020400","PVP01_1020600","PVP01_1020700","PVP01_1021400","PVP01_1021700","PVP01_1021800","PVP01_1021900","PVP01_1022500","PVP01_1022600","PVP01_1022700","PVP01_1022800","PVP01_1023200","PVP01_1023500","PVP01_1023800","PVP01_1024000","PVP01_1024100","PVP01_1024200","PVP01_1024300","PVP01_1024600","PVP01_1024700","PVP01_1024900","PVP01_1025000","PVP01_1025100","PVP01_1025400","PVP01_1025500","PVP01_1025600","PVP01_1025700","PVP01_1025900","PVP01_1026000","PVP01_1026100","PVP01_1026200","PVP01_1027000","PVP01_1027200","PVP01_1027500","PVP01_1027800","PVP01_1028300","PVP01_1028500","PVP01_1028700","PVP01_1028800","PVP01_1028900","PVP01_1029000","PVP01_1029400","PVP01_1029600","PVP01_1030400","PVP01_1030600","PVP01_1030700","PVP01_1030800","PVP01_1031000","PVP01_1032300","PVP01_1032600","PVP01_1032700","PVP01_1033100","PVP01_1033200","PVP01_1033400","PVP01_1033800","PVP01_1033900","PVP01_1102300","PVP01_1102600","PVP01_1102700","PVP01_1103000","PVP01_1103100","PVP01_1103300","PVP01_1103500","PVP01_1103600","PVP01_1103800","PVP01_1103900","PVP01_1104500","PVP01_1104700","PVP01_1104900","PVP01_1105000","PVP01_1105100","PVP01_1105200","PVP01_1105700","PVP01_1105900","PVP01_1106100","PVP01_1106500","PVP01_1106800","PVP01_1106900","PVP01_1107500","PVP01_1107600","PVP01_1107700","PVP01_1107800","PVP01_1108100","PVP01_1108200","PVP01_1108400","PVP01_1108700","PVP01_1109200","PVP01_1109500","PVP01_1109700","PVP01_1109800","PVP01_1109900","PVP01_1110000","PVP01_1110100","PVP01_1110200","PVP01_1110300","PVP01_1110400","PVP01_1110800","PVP01_1111000","PVP01_1111400","PVP01_1111900","PVP01_1112000","PVP01_1112200","PVP01_1112300","PVP01_1112500","PVP01_1112700","PVP01_1112900","PVP01_1113000","PVP01_1113100","PVP01_1113200","PVP01_1113400","PVP01_1113600","PVP01_1113800","PVP01_1114000","PVP01_1114400","PVP01_1114900","PVP01_1115100","PVP01_1115300","PVP01_1115400","PVP01_1115700","PVP01_1115800","PVP01_1115900","PVP01_1116000","PVP01_1116100","PVP01_1116200","PVP01_1116300","PVP01_1116400","PVP01_1116700","PVP01_1117000","PVP01_1117100","PVP01_1117200","PVP01_1117400","PVP01_1117500","PVP01_1117600","PVP01_1117800","PVP01_1117900","PVP01_1118000","PVP01_1118200","PVP01_1118300","PVP01_1118400","PVP01_1118500","PVP01_1118700","PVP01_1118800","PVP01_1118900","PVP01_1119100","PVP01_1119200","PVP01_1119400","PVP01_1119700","PVP01_1119900","PVP01_1120000","PVP01_1120200","PVP01_1120400","PVP01_1120500","PVP01_1120600","PVP01_1120700","PVP01_1120800","PVP01_1121000","PVP01_1121200","PVP01_1121400","PVP01_1121800","PVP01_1122000","PVP01_1122200","PVP01_1123100","PVP01_1123700","PVP01_1123800","PVP01_1124200","PVP01_1124500","PVP01_1124700","PVP01_1125500","PVP01_1125800","PVP01_1125900","PVP01_1126400","PVP01_1126700","PVP01_1127100","PVP01_1127600","PVP01_1127700","PVP01_1128000","PVP01_1128200","PVP01_1128300","PVP01_1128500","PVP01_1129100","PVP01_1129200","PVP01_1129300","PVP01_1129400","PVP01_1129800","PVP01_1129900","PVP01_1130000","PVP01_1130200","PVP01_1130300","PVP01_1130500","PVP01_1130600","PVP01_1130800","PVP01_1131000","PVP01_1131100","PVP01_1131200","PVP01_1131500","PVP01_1131600","PVP01_1131700","PVP01_1131800","PVP01_1132000","PVP01_1132200","PVP01_1132600","PVP01_1133500","PVP01_1133700","PVP01_1134000","PVP01_1134600","PVP01_1134900","PVP01_1135000","PVP01_1135200","PVP01_1135400","PVP01_1136000","PVP01_1136200","PVP01_1136400","PVP01_1136500","PVP01_1137000","PVP01_1137400","PVP01_1137500","PVP01_1137900","PVP01_1138000","PVP01_1138100","PVP01_1138200","PVP01_1138300","PVP01_1138600","PVP01_1138700","PVP01_1138800","PVP01_1138900","PVP01_1139200","PVP01_1139400","PVP01_1139500","PVP01_1139600","PVP01_1139800","PVP01_1139900","PVP01_1140100","PVP01_1140200","PVP01_1140400","PVP01_1140600","PVP01_1141100","PVP01_1141300","PVP01_1141400","PVP01_1141600","PVP01_1142100","PVP01_1142200","PVP01_1142400","PVP01_1142700","PVP01_1142800","PVP01_1142900","PVP01_1143000","PVP01_1143300","PVP01_1143600","PVP01_1143800","PVP01_1143900","PVP01_1144000","PVP01_1144400","PVP01_1144600","PVP01_1144800","PVP01_1145300","PVP01_1145400","PVP01_1145600","PVP01_1145800","PVP01_1145900","PVP01_1146000","PVP01_1146100","PVP01_1146200","PVP01_1146300","PVP01_1146500","PVP01_1146700","PVP01_1147600","PVP01_1202000","PVP01_1202300","PVP01_1202400","PVP01_1202500","PVP01_1202600","PVP01_1202700","PVP01_1202800","PVP01_1202900","PVP01_1203000","PVP01_1203900","PVP01_1204300","PVP01_1204400","PVP01_1204500","PVP01_1204900","PVP01_1205100","PVP01_1205300","PVP01_1205500","PVP01_1205700","PVP01_1205800","PVP01_1206000","PVP01_1206200","PVP01_1206500","PVP01_1206600","PVP01_1206800","PVP01_1207300","PVP01_1207400","PVP01_1207500","PVP01_1207600","PVP01_1207700","PVP01_1208400","PVP01_1208500","PVP01_1209100","PVP01_1209200","PVP01_1209300","PVP01_1209500","PVP01_1209600","PVP01_1209700","PVP01_1209800","PVP01_1209900","PVP01_1210200","PVP01_1210400","PVP01_1210500","PVP01_1210600","PVP01_1210800","PVP01_1210900","PVP01_1211100","PVP01_1211400","PVP01_1211700","PVP01_1211800","PVP01_1211900","PVP01_1212000","PVP01_1212100","PVP01_1212200","PVP01_1212300","PVP01_1212400","PVP01_1212700","PVP01_1212800","PVP01_1212900","PVP01_1213200","PVP01_1213400","PVP01_1213600","PVP01_1213900","PVP01_1214000","PVP01_1214100","PVP01_1214400","PVP01_1214500","PVP01_1214700","PVP01_1215100","PVP01_1215200","PVP01_1215500","PVP01_1215800","PVP01_1216400","PVP01_1216500","PVP01_1216600","PVP01_1216700","PVP01_1216900","PVP01_1217300","PVP01_1217800","PVP01_1217900","PVP01_1218100","PVP01_1218200","PVP01_1218300","PVP01_1218400","PVP01_1218600","PVP01_1218700","PVP01_1219300","PVP01_1219600","PVP01_1219700","PVP01_1220300","PVP01_1220400","PVP01_1220500","PVP01_1220800","PVP01_1221000","PVP01_1221100","PVP01_1221500","PVP01_1221600","PVP01_1222000","PVP01_1222500","PVP01_1222600","PVP01_1222700","PVP01_1222800","PVP01_1222900","PVP01_1223000","PVP01_1223100","PVP01_1223200","PVP01_1223300","PVP01_1223400","PVP01_1223600","PVP01_1223700","PVP01_1224100","PVP01_1224200","PVP01_1224300","PVP01_1224900","PVP01_1225100","PVP01_1225400","PVP01_1225600","PVP01_1225700","PVP01_1225900","PVP01_1226000","PVP01_1226200","PVP01_1226300","PVP01_1226400","PVP01_1226600","PVP01_1226900","PVP01_1227000","PVP01_1227100","PVP01_1227400","PVP01_1227700","PVP01_1227800","PVP01_1227900","PVP01_1228300","PVP01_1228600","PVP01_1228700","PVP01_1228900","PVP01_1229000","PVP01_1229200","PVP01_1229300","PVP01_1229700","PVP01_1230100","PVP01_1230300","PVP01_1230400","PVP01_1230500","PVP01_1230900","PVP01_1231200","PVP01_1231300","PVP01_1231400","PVP01_1231500","PVP01_1232000","PVP01_1232100","PVP01_1232200","PVP01_1232300","PVP01_1232500","PVP01_1232800","PVP01_1232900","PVP01_1233000","PVP01_1233300","PVP01_1233400","PVP01_1233500","PVP01_1233600","PVP01_1233700","PVP01_1233900","PVP01_1234100","PVP01_1234200","PVP01_1234300","PVP01_1234400","PVP01_1234500","PVP01_1234600","PVP01_1234700","PVP01_1234800","PVP01_1234900","PVP01_1235500","PVP01_1235600","PVP01_1235700","PVP01_1235900","PVP01_1236200","PVP01_1236300","PVP01_1236500","PVP01_1236600","PVP01_1236700","PVP01_1236800","PVP01_1237300","PVP01_1237400","PVP01_1237600","PVP01_1238100","PVP01_1238200","PVP01_1238300","PVP01_1238500","PVP01_1238600","PVP01_1238700","PVP01_1238800","PVP01_1239200","PVP01_1239500","PVP01_1239600","PVP01_1239700","PVP01_1239800","PVP01_1239900","PVP01_1240000","PVP01_1240200","PVP01_1240400","PVP01_1240500","PVP01_1240700","PVP01_1241400","PVP01_1241500","PVP01_1241600","PVP01_1241700","PVP01_1241800","PVP01_1242100","PVP01_1242200","PVP01_1242500","PVP01_1242600","PVP01_1242800","PVP01_1242900","PVP01_1243300","PVP01_1243400","PVP01_1243700","PVP01_1243800","PVP01_1244000","PVP01_1244100","PVP01_1244200","PVP01_1244300","PVP01_1244500","PVP01_1244600","PVP01_1245000","PVP01_1245200","PVP01_1245700","PVP01_1246100","PVP01_1246200","PVP01_1246800","PVP01_1247100","PVP01_1247200","PVP01_1247300","PVP01_1247400","PVP01_1248000","PVP01_1248100","PVP01_1248300","PVP01_1248400","PVP01_1248600","PVP01_1248700","PVP01_1248800","PVP01_1248900","PVP01_1249300","PVP01_1249400","PVP01_1249600","PVP01_1249700","PVP01_1249800","PVP01_1249900","PVP01_1250000","PVP01_1250100","PVP01_1250400","PVP01_1250500","PVP01_1250800","PVP01_1251000","PVP01_1251200","PVP01_1251400","PVP01_1251600","PVP01_1251800","PVP01_1252100","PVP01_1252300","PVP01_1252500","PVP01_1252600","PVP01_1252700","PVP01_1253100","PVP01_1253200","PVP01_1253400","PVP01_1253500","PVP01_1253900","PVP01_1254000","PVP01_1254200","PVP01_1254400","PVP01_1254500","PVP01_1254600","PVP01_1254700","PVP01_1254800","PVP01_1255000","PVP01_1255500","PVP01_1255600","PVP01_1255700","PVP01_1255800","PVP01_1255900","PVP01_1256100","PVP01_1256300","PVP01_1256500","PVP01_1256600","PVP01_1256900","PVP01_1257100","PVP01_1257200","PVP01_1257300","PVP01_1257400","PVP01_1257800","PVP01_1258000","PVP01_1258600","PVP01_1258800","PVP01_1259700","PVP01_1259800","PVP01_1259900","PVP01_1260000","PVP01_1260100","PVP01_1260300","PVP01_1260400","PVP01_1260500","PVP01_1260600","PVP01_1260700","PVP01_1260900","PVP01_1261300","PVP01_1261600","PVP01_1261800","PVP01_1261900","PVP01_1262100","PVP01_1262200","PVP01_1262500","PVP01_1262700","PVP01_1263200","PVP01_1263300","PVP01_1263500","PVP01_1263600","PVP01_1263700","PVP01_1264100","PVP01_1264200","PVP01_1264500","PVP01_1265200","PVP01_1265500","PVP01_1265600","PVP01_1265800","PVP01_1266000","PVP01_1266100","PVP01_1266200","PVP01_1266300","PVP01_1266600","PVP01_1266800","PVP01_1266900","PVP01_1267000","PVP01_1267100","PVP01_1267200","PVP01_1267400","PVP01_1267600","PVP01_1267700","PVP01_1267900","PVP01_1268400","PVP01_1268500","PVP01_1268900","PVP01_1269100","PVP01_1269300","PVP01_1269400","PVP01_1269500","PVP01_1269600","PVP01_1269700","PVP01_1269800","PVP01_1269900","PVP01_1270000","PVP01_1270100","PVP01_1270400","PVP01_1270500","PVP01_1270800","PVP01_1271000","PVP01_1300600","PVP01_1300700","PVP01_1300800","PVP01_1301900","PVP01_1302000","PVP01_1302200","PVP01_1302400","PVP01_1302600","PVP01_1303300","PVP01_1303400","PVP01_1303500","PVP01_1303700","PVP01_1303800","PVP01_1304100","PVP01_1304400","PVP01_1304600","PVP01_1304700","PVP01_1304900","PVP01_1305000","PVP01_1305100","PVP01_1305200","PVP01_1305400","PVP01_1305600","PVP01_1305800","PVP01_1305900","PVP01_1306200","PVP01_1306400","PVP01_1306500","PVP01_1306900","PVP01_1307400","PVP01_1307500","PVP01_1307600","PVP01_1307900","PVP01_1308000","PVP01_1308200","PVP01_1308400","PVP01_1308500","PVP01_1308600","PVP01_1309200","PVP01_1309300","PVP01_1309600","PVP01_1309900","PVP01_1310100","PVP01_1310200","PVP01_1310300","PVP01_1310500","PVP01_1310600","PVP01_1310800","PVP01_1310900","PVP01_1311400","PVP01_1311600","PVP01_1311700","PVP01_1311800","PVP01_1312100","PVP01_1312200","PVP01_1312500","PVP01_1312700","PVP01_1312800","PVP01_1312900","PVP01_1313000","PVP01_1313200","PVP01_1313500","PVP01_1313600","PVP01_1313700","PVP01_1313800","PVP01_1314000","PVP01_1314200","PVP01_1314400","PVP01_1314500","PVP01_1314700","PVP01_1314900","PVP01_1315000","PVP01_1315100","PVP01_1315300","PVP01_1315600","PVP01_1315800","PVP01_1315900","PVP01_1316000","PVP01_1316200","PVP01_1316300","PVP01_1316400","PVP01_1316800","PVP01_1317100","PVP01_1317200","PVP01_1317300","PVP01_1317400","PVP01_1317500","PVP01_1318200","PVP01_1318400","PVP01_1318500","PVP01_1318700","PVP01_1318800","PVP01_1319100","PVP01_1319300","PVP01_1319400","PVP01_1319800","PVP01_1320800","PVP01_1320900","PVP01_1321000","PVP01_1321100","PVP01_1321200","PVP01_1321300","PVP01_1321400","PVP01_1321800","PVP01_1322100","PVP01_1322300","PVP01_1322500","PVP01_1322700","PVP01_1322800","PVP01_1323200","PVP01_1323300","PVP01_1323400","PVP01_1323500","PVP01_1323600","PVP01_1324000","PVP01_1324700","PVP01_1325200","PVP01_1325300","PVP01_1325400","PVP01_1325500","PVP01_1325600","PVP01_1325700","PVP01_1326100","PVP01_1326300","PVP01_1326500","PVP01_1326700","PVP01_1326800","PVP01_1326900","PVP01_1327000","PVP01_1327100","PVP01_1327400","PVP01_1327500","PVP01_1327700","PVP01_1327800","PVP01_1328200","PVP01_1328300","PVP01_1328400","PVP01_1329100","PVP01_1329200","PVP01_1329500","PVP01_1329700","PVP01_1329800","PVP01_1330600","PVP01_1330700","PVP01_1330900","PVP01_1331100","PVP01_1331300","PVP01_1331500","PVP01_1331600","PVP01_1331700","PVP01_1331900","PVP01_1332000","PVP01_1332100","PVP01_1332400","PVP01_1332600","PVP01_1332800","PVP01_1333300","PVP01_1333400","PVP01_1333500","PVP01_1333600","PVP01_1333800","PVP01_1334400","PVP01_1334600","PVP01_1334700","PVP01_1334800","PVP01_1334900","PVP01_1335000","PVP01_1335200","PVP01_1335400","PVP01_1335500","PVP01_1335700","PVP01_1336000","PVP01_1336100","PVP01_1336200","PVP01_1336300","PVP01_1336400","PVP01_1336500","PVP01_1336600","PVP01_1336800","PVP01_1336900","PVP01_1337000","PVP01_1337700","PVP01_1337800","PVP01_1338000","PVP01_1338200","PVP01_1338500","PVP01_1338700","PVP01_1338800","PVP01_1338900","PVP01_1339100","PVP01_1339200","PVP01_1339300","PVP01_1339500","PVP01_1339600","PVP01_1339700","PVP01_1339900","PVP01_1340000","PVP01_1340100","PVP01_1340300","PVP01_1340700","PVP01_1340900","PVP01_1341000","PVP01_1341100","PVP01_1341200","PVP01_1341400","PVP01_1341500","PVP01_1341700","PVP01_1341900","PVP01_1342000","PVP01_1342100","PVP01_1342500","PVP01_1342900","PVP01_1343000","PVP01_1343200","PVP01_1343400","PVP01_1343700","PVP01_1343800","PVP01_1343900","PVP01_1344000","PVP01_1344100","PVP01_1344400","PVP01_1344800","PVP01_1345000","PVP01_1345100","PVP01_1345200","PVP01_1345800","PVP01_1346000","PVP01_1346100","PVP01_1346400","PVP01_1346500","PVP01_1346600","PVP01_1347000","PVP01_1347100","PVP01_1347200","PVP01_1401400","PVP01_1402900","PVP01_1403100","PVP01_1403200","PVP01_1403500","PVP01_1403600","PVP01_1403800","PVP01_1404400","PVP01_1404700","PVP01_1404900","PVP01_1405200","PVP01_1405300","PVP01_1405400","PVP01_1405700","PVP01_1405900","PVP01_1406300","PVP01_1406400","PVP01_1406500","PVP01_1406600","PVP01_1406800","PVP01_1406900","PVP01_1407200","PVP01_1407300","PVP01_1407500","PVP01_1407700","PVP01_1407900","PVP01_1408300","PVP01_1409200","PVP01_1409500","PVP01_1409600","PVP01_1409800","PVP01_1409900","PVP01_1410000","PVP01_1410100","PVP01_1410400","PVP01_1410600","PVP01_1411100","PVP01_1411500","PVP01_1411600","PVP01_1411700","PVP01_1411800","PVP01_1411900","PVP01_1412000","PVP01_1412200","PVP01_1412300","PVP01_1412400","PVP01_1412600","PVP01_1412700","PVP01_1412800","PVP01_1412900","PVP01_1413000","PVP01_1413400","PVP01_1413450","PVP01_1413500","PVP01_1413600","PVP01_1413800","PVP01_1413900","PVP01_1414000","PVP01_1414100","PVP01_1414500","PVP01_1414600","PVP01_1414700","PVP01_1414800","PVP01_1415000","PVP01_1415400","PVP01_1415800","PVP01_1416200","PVP01_1416300","PVP01_1416400","PVP01_1416500","PVP01_1416600","PVP01_1416700","PVP01_1416800","PVP01_1417000","PVP01_1417300","PVP01_1417400","PVP01_1417500","PVP01_1417600","PVP01_1417800","PVP01_1418500","PVP01_1418700","PVP01_1419000","PVP01_1419200","PVP01_1419600","PVP01_1419900","PVP01_1420200","PVP01_1420400","PVP01_1420600","PVP01_1420800","PVP01_1421100","PVP01_1421500","PVP01_1421600","PVP01_1421700","PVP01_1421900","PVP01_1422000","PVP01_1422400","PVP01_1422500","PVP01_1422800","PVP01_1423000","PVP01_1423100","PVP01_1423300","PVP01_1423400","PVP01_1423500","PVP01_1423600","PVP01_1423700","PVP01_1424300","PVP01_1424400","PVP01_1424600","PVP01_1424700","PVP01_1424800","PVP01_1424900","PVP01_1425000","PVP01_1425500","PVP01_1425600","PVP01_1425700","PVP01_1425900","PVP01_1426000","PVP01_1426300","PVP01_1426400","PVP01_1426500","PVP01_1426600","PVP01_1426900","PVP01_1427200","PVP01_1427300","PVP01_1427400","PVP01_1427500","PVP01_1427600","PVP01_1427800","PVP01_1427900","PVP01_1428200","PVP01_1428600","PVP01_1428700","PVP01_1428800","PVP01_1429100","PVP01_1429400","PVP01_1429500","PVP01_1429600","PVP01_1429800","PVP01_1430100","PVP01_1430200","PVP01_1430400","PVP01_1430500","PVP01_1430900","PVP01_1431000","PVP01_1431400","PVP01_1431500","PVP01_1431600","PVP01_1432200","PVP01_1432300","PVP01_1432500","PVP01_1432900","PVP01_1433000","PVP01_1433100","PVP01_1433200","PVP01_1433300","PVP01_1433400","PVP01_1433500","PVP01_1433700","PVP01_1433900","PVP01_1434100","PVP01_1434400","PVP01_1434700","PVP01_1435000","PVP01_1435200","PVP01_1435400","PVP01_1435500","PVP01_1435700","PVP01_1436100","PVP01_1436200","PVP01_1436500","PVP01_1436800","PVP01_1436900","PVP01_1437000","PVP01_1437300","PVP01_1437400","PVP01_1437500","PVP01_1437800","PVP01_1437900","PVP01_1438000","PVP01_1438300","PVP01_1438500","PVP01_1438600","PVP01_1438700","PVP01_1438800","PVP01_1439100","PVP01_1439200","PVP01_1439500","PVP01_1439800","PVP01_1439900","PVP01_1440200","PVP01_1440400","PVP01_1440500","PVP01_1440800","PVP01_1441000","PVP01_1441300","PVP01_1441500","PVP01_1441600","PVP01_1441900","PVP01_1442100","PVP01_1442300","PVP01_1442400","PVP01_1442500","PVP01_1442600","PVP01_1442700","PVP01_1442800","PVP01_1443100","PVP01_1443300","PVP01_1443400","PVP01_1443500","PVP01_1443700","PVP01_1444100","PVP01_1444300","PVP01_1444500","PVP01_1444600","PVP01_1444700","PVP01_1445100","PVP01_1445500","PVP01_1445800","PVP01_1445900","PVP01_1446500","PVP01_1446700","PVP01_1446800","PVP01_1447000","PVP01_1447100","PVP01_1447400","PVP01_1447700","PVP01_1448000","PVP01_1448200","PVP01_1448500","PVP01_1448600","PVP01_1448900","PVP01_1449000","PVP01_1449300","PVP01_1449400","PVP01_1449500","PVP01_1449600","PVP01_1449800","PVP01_1449900","PVP01_1450100","PVP01_1450300","PVP01_1450600","PVP01_1450700","PVP01_1450800","PVP01_1450900","PVP01_1451100","PVP01_1451200","PVP01_1451400","PVP01_1451700","PVP01_1451900","PVP01_1452000","PVP01_1452200","PVP01_1452300","PVP01_1452600","PVP01_1452800","PVP01_1453000","PVP01_1453100","PVP01_1453300","PVP01_1453400","PVP01_1453500","PVP01_1453600","PVP01_1453700","PVP01_1453900","PVP01_1454300","PVP01_1454700","PVP01_1455000","PVP01_1455100","PVP01_1455200","PVP01_1455300","PVP01_1455600","PVP01_1455800","PVP01_1455900","PVP01_1456100","PVP01_1456200","PVP01_1456300","PVP01_1456400","PVP01_1456600","PVP01_1456800","PVP01_1457100","PVP01_1457200","PVP01_1457300","PVP01_1457600","PVP01_1457700","PVP01_1457800","PVP01_1458000","PVP01_1458100","PVP01_1458200","PVP01_1458300","PVP01_1458400","PVP01_1458500","PVP01_1458600","PVP01_1458900","PVP01_1459000","PVP01_1459800","PVP01_1459900","PVP01_1460000","PVP01_1460100","PVP01_1460200","PVP01_1460300","PVP01_1460400","PVP01_1460600","PVP01_1461000","PVP01_1461200","PVP01_1461300","PVP01_1461500","PVP01_1461600","PVP01_1461700","PVP01_1461800","PVP01_1462200","PVP01_1462700","PVP01_1462800","PVP01_1463000","PVP01_1463100","PVP01_1463400","PVP01_1463900","PVP01_1464000","PVP01_1464200","PVP01_1464300","PVP01_1464400","PVP01_1464600","PVP01_1464800","PVP01_1465200","PVP01_1465300","PVP01_1465400","PVP01_1465600","PVP01_1466200","PVP01_1466400","PVP01_1466700","PVP01_1466800","PVP01_1466900","PVP01_1467100","PVP01_1467200","PVP01_1467400","PVP01_1467500","PVP01_1467700","PVP01_1467800","PVP01_1467900","PVP01_1468000","PVP01_1468200","PVP01_1468300","PVP01_1468500","PVP01_1468700","PVP01_1468800","PVP01_1468900","PVP01_1469200","PVP01_1469500","PVP01_1469800","PVP01_1471400","PVP01_API02600","PVP01_API02800","PVP01_API03500","PVP01_API04300","PVP01_API04600","PVP01_MIT02700","PVP01_MIT03300","PVP01_MIT03400"],"Row.names":["PVP01_0000030","PVP01_0000040","PVP01_0000050","PVP01_0000080","PVP01_0000100","PVP01_0000120","PVP01_0000130","PVP01_0000140","PVP01_0000150","PVP01_0000170","PVP01_0000180","PVP01_0000190","PVP01_0000200","PVP01_0000210","PVP01_0000220","PVP01_0000300","PVP01_0000480","PVP01_0000670","PVP01_0000740","PVP01_0000790","PVP01_0001030","PVP01_0001090","PVP01_0001310","PVP01_0001440","PVP01_0001480","PVP01_0001550","PVP01_0001640","PVP01_0001660","PVP01_0001670","PVP01_0001900","PVP01_0001910","PVP01_0001940","PVP01_0001970","PVP01_0002020","PVP01_0002050","PVP01_0002120","PVP01_0002200","PVP01_0002210","PVP01_0002330","PVP01_0002370","PVP01_0002390","PVP01_0002470","PVP01_0002530","PVP01_0002540","PVP01_0002750","PVP01_0002810","PVP01_0002820","PVP01_0003010","PVP01_0003050","PVP01_0003150","PVP01_0003210","PVP01_0003350","PVP01_0003360","PVP01_0003370","PVP01_0003380","PVP01_0003400","PVP01_0003410","PVP01_0003420","PVP01_0003440","PVP01_0003530","PVP01_0003690","PVP01_0003730","PVP01_0003800","PVP01_0003990","PVP01_0004160","PVP01_0004170","PVP01_0004180","PVP01_0004190","PVP01_0004240","PVP01_0004250","PVP01_0004260","PVP01_0004270","PVP01_0004280","PVP01_0004300","PVP01_0004310","PVP01_0004320","PVP01_0004350","PVP01_0004370","PVP01_0004380","PVP01_0004390","PVP01_0004400","PVP01_0004410","PVP01_0004640","PVP01_0004670","PVP01_0004910","PVP01_0004920","PVP01_0004950","PVP01_0004960","PVP01_0005010","PVP01_0005100","PVP01_0005160","PVP01_0005210","PVP01_0005300","PVP01_0005310","PVP01_0005420","PVP01_0005430","PVP01_0005550","PVP01_0005640","PVP01_0005710","PVP01_0005740","PVP01_0005760","PVP01_0005800","PVP01_0005820","PVP01_0005870","PVP01_0005910","PVP01_0006000","PVP01_0006040","PVP01_0006080","PVP01_0006260","PVP01_0006650","PVP01_0006660","PVP01_0006670","PVP01_0006680","PVP01_0006700","PVP01_0006860","PVP01_0006890","PVP01_0007080","PVP01_0007160","PVP01_0007240","PVP01_0007290","PVP01_0007310","PVP01_0007320","PVP01_0007340","PVP01_0007610","PVP01_0007620","PVP01_0007690","PVP01_0007700","PVP01_0007720","PVP01_0007850","PVP01_0007900","PVP01_0007910","PVP01_0007920","PVP01_0008190","PVP01_0008430","PVP01_0008480","PVP01_0008490","PVP01_0008600","PVP01_0008700","PVP01_0008770","PVP01_0008830","PVP01_0008990","PVP01_0009010","PVP01_0009170","PVP01_0009180","PVP01_0009210","PVP01_0009410","PVP01_0009470","PVP01_0009540","PVP01_0009620","PVP01_0009710","PVP01_0009730","PVP01_0009740","PVP01_0009760","PVP01_0009770","PVP01_0009820","PVP01_0009850","PVP01_0009860","PVP01_0009870","PVP01_0009900","PVP01_0009940","PVP01_0009980","PVP01_0010020","PVP01_0010030","PVP01_0010070","PVP01_0010100","PVP01_0010120","PVP01_0010140","PVP01_0010170","PVP01_0010180","PVP01_0010200","PVP01_0010210","PVP01_0010220","PVP01_0010230","PVP01_0010240","PVP01_0010340","PVP01_0010380","PVP01_0010440","PVP01_0010470","PVP01_0010500","PVP01_0010540","PVP01_0010570","PVP01_0010600","PVP01_0010610","PVP01_0010640","PVP01_0010650","PVP01_0010770","PVP01_0010870","PVP01_0010900","PVP01_0010910","PVP01_0010960","PVP01_0010970","PVP01_0010990","PVP01_0011030","PVP01_0011070","PVP01_0102100","PVP01_0102200","PVP01_0102300","PVP01_0102400","PVP01_0102500","PVP01_0102600","PVP01_0102700","PVP01_0102800","PVP01_0103400","PVP01_0103600","PVP01_0103800","PVP01_0103900","PVP01_0104100","PVP01_0104200","PVP01_0104300","PVP01_0104600","PVP01_0104800","PVP01_0104900","PVP01_0105000","PVP01_0105100","PVP01_0105300","PVP01_0105400","PVP01_0105500","PVP01_0105600","PVP01_0105700","PVP01_0105800","PVP01_0106000","PVP01_0106200","PVP01_0106300","PVP01_0106400","PVP01_0106500","PVP01_0106800","PVP01_0106900","PVP01_0107400","PVP01_0107600","PVP01_0107700","PVP01_0107900","PVP01_0108000","PVP01_0108100","PVP01_0108300","PVP01_0108400","PVP01_0108600","PVP01_0108800","PVP01_0109000","PVP01_0109100","PVP01_0109200","PVP01_0109300","PVP01_0109600","PVP01_0109800","PVP01_0110200","PVP01_0110700","PVP01_0110800","PVP01_0111200","PVP01_0111300","PVP01_0111600","PVP01_0111700","PVP01_0112000","PVP01_0112100","PVP01_0112500","PVP01_0112700","PVP01_0113000","PVP01_0113100","PVP01_0113200","PVP01_0113400","PVP01_0113500","PVP01_0114100","PVP01_0114600","PVP01_0114700","PVP01_0114900","PVP01_0115000","PVP01_0115200","PVP01_0115400","PVP01_0115500","PVP01_0116100","PVP01_0116500","PVP01_0116600","PVP01_0116700","PVP01_0116900","PVP01_0117500","PVP01_0117700","PVP01_0117800","PVP01_0118000","PVP01_0118600","PVP01_0118700","PVP01_0118800","PVP01_0118900","PVP01_0119200","PVP01_0119300","PVP01_0119400","PVP01_0119500","PVP01_0119600","PVP01_0119700","PVP01_0119800","PVP01_0119900","PVP01_0120000","PVP01_0120100","PVP01_0120500","PVP01_0120800","PVP01_0121000","PVP01_0121700","PVP01_0200200","PVP01_0200700","PVP01_0201000","PVP01_0201100","PVP01_0201300","PVP01_0201400","PVP01_0201500","PVP01_0201600","PVP01_0201700","PVP01_0201800","PVP01_0201900","PVP01_0202000","PVP01_0202100","PVP01_0202200","PVP01_0202300","PVP01_0202400","PVP01_0203000","PVP01_0203100","PVP01_0203400","PVP01_0203500","PVP01_0203800","PVP01_0203900","PVP01_0204000","PVP01_0204100","PVP01_0204800","PVP01_0204900","PVP01_0205000","PVP01_0205200","PVP01_0205300","PVP01_0205500","PVP01_0205600","PVP01_0205800","PVP01_0206100","PVP01_0206300","PVP01_0206400","PVP01_0206500","PVP01_0206600","PVP01_0206800","PVP01_0207000","PVP01_0207100","PVP01_0207200","PVP01_0207400","PVP01_0207500","PVP01_0207600","PVP01_0207700","PVP01_0207800","PVP01_0208100","PVP01_0208200","PVP01_0208300","PVP01_0208400","PVP01_0208600","PVP01_0208700","PVP01_0208800","PVP01_0209000","PVP01_0209100","PVP01_0209200","PVP01_0209300","PVP01_0209500","PVP01_0209600","PVP01_0210000","PVP01_0210100","PVP01_0210300","PVP01_0211000","PVP01_0211100","PVP01_0211200","PVP01_0211400","PVP01_0211500","PVP01_0211600","PVP01_0211700","PVP01_0212400","PVP01_0212500","PVP01_0212600","PVP01_0212700","PVP01_0212800","PVP01_0213000","PVP01_0213300","PVP01_0213600","PVP01_0213700","PVP01_0214100","PVP01_0214200","PVP01_0214800","PVP01_0214900","PVP01_0215000","PVP01_0215300","PVP01_0215400","PVP01_0215500","PVP01_0215700","PVP01_0215800","PVP01_0216100","PVP01_0216500","PVP01_0216600","PVP01_0216700","PVP01_0216800","PVP01_0216900","PVP01_0217000","PVP01_0217100","PVP01_0217200","PVP01_0217300","PVP01_0217400","PVP01_0217500","PVP01_0218200","PVP01_0218300","PVP01_0218900","PVP01_0219400","PVP01_0220100","PVP01_0220200","PVP01_0221100","PVP01_0221300","PVP01_0221700","PVP01_0300100","PVP01_0300600","PVP01_0300700","PVP01_0300800","PVP01_0301100","PVP01_0301200","PVP01_0301500","PVP01_0301900","PVP01_0302000","PVP01_0302100","PVP01_0302300","PVP01_0302400","PVP01_0302500","PVP01_0303100","PVP01_0303500","PVP01_0304100","PVP01_0304300","PVP01_0304500","PVP01_0304600","PVP01_0304800","PVP01_0304900","PVP01_0305000","PVP01_0305200","PVP01_0305300","PVP01_0305500","PVP01_0305800","PVP01_0306000","PVP01_0306100","PVP01_0306300","PVP01_0306400","PVP01_0306500","PVP01_0306700","PVP01_0306800","PVP01_0306900","PVP01_0307000","PVP01_0307100","PVP01_0307300","PVP01_0307400","PVP01_0307600","PVP01_0307700","PVP01_0308800","PVP01_0309200","PVP01_0309700","PVP01_0309800","PVP01_0310000","PVP01_0310200","PVP01_0310300","PVP01_0310400","PVP01_0310600","PVP01_0311000","PVP01_0311200","PVP01_0311300","PVP01_0311400","PVP01_0311500","PVP01_0311600","PVP01_0311700","PVP01_0312000","PVP01_0312100","PVP01_0312400","PVP01_0312500","PVP01_0312700","PVP01_0312800","PVP01_0313500","PVP01_0313800","PVP01_0314100","PVP01_0314200","PVP01_0314900","PVP01_0315000","PVP01_0315100","PVP01_0315200","PVP01_0315300","PVP01_0315700","PVP01_0315900","PVP01_0316000","PVP01_0316200","PVP01_0316300","PVP01_0316500","PVP01_0316700","PVP01_0316800","PVP01_0317100","PVP01_0317200","PVP01_0317400","PVP01_0318000","PVP01_0318100","PVP01_0318400","PVP01_0318900","PVP01_0319200","PVP01_0319300","PVP01_0319400","PVP01_0319600","PVP01_0319700","PVP01_0320400","PVP01_0320500","PVP01_0320700","PVP01_0320800","PVP01_0320900","PVP01_0321100","PVP01_0400900","PVP01_0403600","PVP01_0403700","PVP01_0403900","PVP01_0404000","PVP01_0404100","PVP01_0404200","PVP01_0404300","PVP01_0404400","PVP01_0404500","PVP01_0404600","PVP01_0404700","PVP01_0404800","PVP01_0404900","PVP01_0405100","PVP01_0405200","PVP01_0405300","PVP01_0405500","PVP01_0405600","PVP01_0405700","PVP01_0405900","PVP01_0406500","PVP01_0406600","PVP01_0406700","PVP01_0406900","PVP01_0407000","PVP01_0407700","PVP01_0407900","PVP01_0408200","PVP01_0408600","PVP01_0408800","PVP01_0408900","PVP01_0409000","PVP01_0409700","PVP01_0409900","PVP01_0410000","PVP01_0410400","PVP01_0410600","PVP01_0410800","PVP01_0411400","PVP01_0411500","PVP01_0411600","PVP01_0411800","PVP01_0411900","PVP01_0412100","PVP01_0412200","PVP01_0412700","PVP01_0412800","PVP01_0412900","PVP01_0413000","PVP01_0413100","PVP01_0413200","PVP01_0413300","PVP01_0413500","PVP01_0414300","PVP01_0414400","PVP01_0414600","PVP01_0414900","PVP01_0415200","PVP01_0415400","PVP01_0415500","PVP01_0415600","PVP01_0416600","PVP01_0416900","PVP01_0417100","PVP01_0417400","PVP01_0417800","PVP01_0417900","PVP01_0418300","PVP01_0419000","PVP01_0419200","PVP01_0419300","PVP01_0419400","PVP01_0419500","PVP01_0419700","PVP01_0420000","PVP01_0420200","PVP01_0420500","PVP01_0420700","PVP01_0420900","PVP01_0421500","PVP01_0421700","PVP01_0421800","PVP01_0422100","PVP01_0422200","PVP01_0422300","PVP01_0422400","PVP01_0422600","PVP01_0422700","PVP01_0422900","PVP01_0423000","PVP01_0423100","PVP01_0423200","PVP01_0423300","PVP01_0423600","PVP01_0423800","PVP01_0424300","PVP01_0424400","PVP01_0500700","PVP01_0502300","PVP01_0502400","PVP01_0502900","PVP01_0503300","PVP01_0503400","PVP01_0503600","PVP01_0503700","PVP01_0503800","PVP01_0503900","PVP01_0504100","PVP01_0504200","PVP01_0504300","PVP01_0505000","PVP01_0505100","PVP01_0505200","PVP01_0505400","PVP01_0505500","PVP01_0505800","PVP01_0506000","PVP01_0506300","PVP01_0506400","PVP01_0506600","PVP01_0506700","PVP01_0506800","PVP01_0507300","PVP01_0507500","PVP01_0507600","PVP01_0507700","PVP01_0507800","PVP01_0508000","PVP01_0508200","PVP01_0508500","PVP01_0508600","PVP01_0508700","PVP01_0509100","PVP01_0509200","PVP01_0509400","PVP01_0509600","PVP01_0509900","PVP01_0510000","PVP01_0510100","PVP01_0510200","PVP01_0510300","PVP01_0510800","PVP01_0510900","PVP01_0511200","PVP01_0511300","PVP01_0511400","PVP01_0511500","PVP01_0511700","PVP01_0511800","PVP01_0512100","PVP01_0512200","PVP01_0512400","PVP01_0512500","PVP01_0512700","PVP01_0512800","PVP01_0513000","PVP01_0513300","PVP01_0513400","PVP01_0513500","PVP01_0513700","PVP01_0513900","PVP01_0514100","PVP01_0514600","PVP01_0514800","PVP01_0514900","PVP01_0515100","PVP01_0515200","PVP01_0515300","PVP01_0515700","PVP01_0515800","PVP01_0515900","PVP01_0516000","PVP01_0516100","PVP01_0516200","PVP01_0516300","PVP01_0516400","PVP01_0516500","PVP01_0516900","PVP01_0517000","PVP01_0517100","PVP01_0517300","PVP01_0517400","PVP01_0517700","PVP01_0517900","PVP01_0518300","PVP01_0518400","PVP01_0518500","PVP01_0518600","PVP01_0518700","PVP01_0519100","PVP01_0519200","PVP01_0519500","PVP01_0519600","PVP01_0520700","PVP01_0520800","PVP01_0521300","PVP01_0521400","PVP01_0521600","PVP01_0521800","PVP01_0522000","PVP01_0522100","PVP01_0522200","PVP01_0522300","PVP01_0522600","PVP01_0522700","PVP01_0522900","PVP01_0523000","PVP01_0523300","PVP01_0523400","PVP01_0523700","PVP01_0523800","PVP01_0523900","PVP01_0524000","PVP01_0524100","PVP01_0524200","PVP01_0524300","PVP01_0524500","PVP01_0524600","PVP01_0524700","PVP01_0524900","PVP01_0525000","PVP01_0525100","PVP01_0525300","PVP01_0525700","PVP01_0526100","PVP01_0526400","PVP01_0527300","PVP01_0527400","PVP01_0527500","PVP01_0527700","PVP01_0527800","PVP01_0528100","PVP01_0528200","PVP01_0528400","PVP01_0528600","PVP01_0528700","PVP01_0528900","PVP01_0529000","PVP01_0529100","PVP01_0529200","PVP01_0529600","PVP01_0529700","PVP01_0529900","PVP01_0530000","PVP01_0530100","PVP01_0530400","PVP01_0530500","PVP01_0531100","PVP01_0531200","PVP01_0531500","PVP01_0531700","PVP01_0531800","PVP01_0532300","PVP01_0532500","PVP01_0532600","PVP01_0532700","PVP01_0532900","PVP01_0533000","PVP01_0533100","PVP01_0533300","PVP01_0533700","PVP01_0534000","PVP01_0534100","PVP01_0534200","PVP01_0534600","PVP01_0534800","PVP01_0534900","PVP01_0535400","PVP01_0600100","PVP01_0600300","PVP01_0600400","PVP01_0600500","PVP01_0600600","PVP01_0600700","PVP01_0600900","PVP01_0601100","PVP01_0601200","PVP01_0601300","PVP01_0601500","PVP01_0601600","PVP01_0602600","PVP01_0603000","PVP01_0603100","PVP01_0603600","PVP01_0603700","PVP01_0604200","PVP01_0604400","PVP01_0604700","PVP01_0605000","PVP01_0605100","PVP01_0605300","PVP01_0605500","PVP01_0606200","PVP01_0606500","PVP01_0606600","PVP01_0606800","PVP01_0607200","PVP01_0607600","PVP01_0608000","PVP01_0608100","PVP01_0608300","PVP01_0608600","PVP01_0608800","PVP01_0608900","PVP01_0609200","PVP01_0609400","PVP01_0609600","PVP01_0609700","PVP01_0609800","PVP01_0609900","PVP01_0610100","PVP01_0610200","PVP01_0610300","PVP01_0610400","PVP01_0610600","PVP01_0610700","PVP01_0610800","PVP01_0610900","PVP01_0611100","PVP01_0611300","PVP01_0612000","PVP01_0612100","PVP01_0612400","PVP01_0612600","PVP01_0612700","PVP01_0613000","PVP01_0613100","PVP01_0613200","PVP01_0613400","PVP01_0613500","PVP01_0613800","PVP01_0614000","PVP01_0614200","PVP01_0614300","PVP01_0614400","PVP01_0614600","PVP01_0614900","PVP01_0615000","PVP01_0615100","PVP01_0615500","PVP01_0615800","PVP01_0615900","PVP01_0616000","PVP01_0616200","PVP01_0616300","PVP01_0616400","PVP01_0616900","PVP01_0617000","PVP01_0617100","PVP01_0617400","PVP01_0617500","PVP01_0617900","PVP01_0618100","PVP01_0618300","PVP01_0618600","PVP01_0618700","PVP01_0619000","PVP01_0619100","PVP01_0619200","PVP01_0619400","PVP01_0619500","PVP01_0619600","PVP01_0619700","PVP01_0619800","PVP01_0620100","PVP01_0620300","PVP01_0620500","PVP01_0620600","PVP01_0620700","PVP01_0620900","PVP01_0621100","PVP01_0621800","PVP01_0621900","PVP01_0622100","PVP01_0622200","PVP01_0622700","PVP01_0623000","PVP01_0623100","PVP01_0623200","PVP01_0623300","PVP01_0623400","PVP01_0623500","PVP01_0623600","PVP01_0623900","PVP01_0624000","PVP01_0624100","PVP01_0624300","PVP01_0624400","PVP01_0624500","PVP01_0700200","PVP01_0700300","PVP01_0700400","PVP01_0700500","PVP01_0700700","PVP01_0700800","PVP01_0700900","PVP01_0701000","PVP01_0701200","PVP01_0701500","PVP01_0701600","PVP01_0701700","PVP01_0702100","PVP01_0702200","PVP01_0702500","PVP01_0703000","PVP01_0703600","PVP01_0703700","PVP01_0704000","PVP01_0704100","PVP01_0704500","PVP01_0704600","PVP01_0704700","PVP01_0704800","PVP01_0704900","PVP01_0705000","PVP01_0705100","PVP01_0705500","PVP01_0705600","PVP01_0705800","PVP01_0705900","PVP01_0706000","PVP01_0706100","PVP01_0706200","PVP01_0706300","PVP01_0706400","PVP01_0706600","PVP01_0706700","PVP01_0707000","PVP01_0707100","PVP01_0707200","PVP01_0707300","PVP01_0707800","PVP01_0708000","PVP01_0708100","PVP01_0708200","PVP01_0708300","PVP01_0708400","PVP01_0708500","PVP01_0708800","PVP01_0708900","PVP01_0709000","PVP01_0709100","PVP01_0709200","PVP01_0709300","PVP01_0709500","PVP01_0709900","PVP01_0710000","PVP01_0710100","PVP01_0710200","PVP01_0710300","PVP01_0710700","PVP01_0711100","PVP01_0711200","PVP01_0711400","PVP01_0711500","PVP01_0711800","PVP01_0711900","PVP01_0712200","PVP01_0712300","PVP01_0712400","PVP01_0712700","PVP01_0713000","PVP01_0713100","PVP01_0713200","PVP01_0713500","PVP01_0713600","PVP01_0713900","PVP01_0714200","PVP01_0714300","PVP01_0714600","PVP01_0714900","PVP01_0715000","PVP01_0715100","PVP01_0715200","PVP01_0715500","PVP01_0715600","PVP01_0715700","PVP01_0715800","PVP01_0716000","PVP01_0716100","PVP01_0716600","PVP01_0716800","PVP01_0717000","PVP01_0717200","PVP01_0717400","PVP01_0717600","PVP01_0717800","PVP01_0718000","PVP01_0718100","PVP01_0718600","PVP01_0718700","PVP01_0719000","PVP01_0719100","PVP01_0719400","PVP01_0719500","PVP01_0719700","PVP01_0719800","PVP01_0719900","PVP01_0720000","PVP01_0720100","PVP01_0720200","PVP01_0720300","PVP01_0720400","PVP01_0720800","PVP01_0720900","PVP01_0721100","PVP01_0721500","PVP01_0721800","PVP01_0722000","PVP01_0722100","PVP01_0722200","PVP01_0722300","PVP01_0722400","PVP01_0722500","PVP01_0722600","PVP01_0722700","PVP01_0722800","PVP01_0723000","PVP01_0723100","PVP01_0723200","PVP01_0723300","PVP01_0723400","PVP01_0723500","PVP01_0723800","PVP01_0724100","PVP01_0724500","PVP01_0724600","PVP01_0724700","PVP01_0724800","PVP01_0724900","PVP01_0725000","PVP01_0725100","PVP01_0725200","PVP01_0725300","PVP01_0725400","PVP01_0725700","PVP01_0726400","PVP01_0726700","PVP01_0726900","PVP01_0727000","PVP01_0727100","PVP01_0727200","PVP01_0727300","PVP01_0727400","PVP01_0727600","PVP01_0727800","PVP01_0728100","PVP01_0728200","PVP01_0728300","PVP01_0729100","PVP01_0729200","PVP01_0729300","PVP01_0729400","PVP01_0729500","PVP01_0729700","PVP01_0730000","PVP01_0730100","PVP01_0730200","PVP01_0730300","PVP01_0730400","PVP01_0730600","PVP01_0730700","PVP01_0731000","PVP01_0731100","PVP01_0731200","PVP01_0731300","PVP01_0731600","PVP01_0731700","PVP01_0731900","PVP01_0732000","PVP01_0732100","PVP01_0732200","PVP01_0732300","PVP01_0732600","PVP01_0732800","PVP01_0732900","PVP01_0733000","PVP01_0733100","PVP01_0733500","PVP01_0733700","PVP01_0733900","PVP01_0734100","PVP01_0734200","PVP01_0734300","PVP01_0734400","PVP01_0734700","PVP01_0735000","PVP01_0735100","PVP01_0735200","PVP01_0735300","PVP01_0735800","PVP01_0735900","PVP01_0800100","PVP01_0800200","PVP01_0800300","PVP01_0800400","PVP01_0800500","PVP01_0800600","PVP01_0800700","PVP01_0800900","PVP01_0801000","PVP01_0801400","PVP01_0801500","PVP01_0801700","PVP01_0801800","PVP01_0802100","PVP01_0802400","PVP01_0802900","PVP01_0803000","PVP01_0803100","PVP01_0803500","PVP01_0803600","PVP01_0803700","PVP01_0804100","PVP01_0804200","PVP01_0804500","PVP01_0804600","PVP01_0804700","PVP01_0804800","PVP01_0804900","PVP01_0805000","PVP01_0805200","PVP01_0805300","PVP01_0805400","PVP01_0805600","PVP01_0805700","PVP01_0805800","PVP01_0806000","PVP01_0806100","PVP01_0806200","PVP01_0806300","PVP01_0806400","PVP01_0806500","PVP01_0806700","PVP01_0807000","PVP01_0807200","PVP01_0807500","PVP01_0807900","PVP01_0808000","PVP01_0808100","PVP01_0808400","PVP01_0808500","PVP01_0808600","PVP01_0808700","PVP01_0808800","PVP01_0809100","PVP01_0809200","PVP01_0809300","PVP01_0809400","PVP01_0809600","PVP01_0809700","PVP01_0809900","PVP01_0810000","PVP01_0810100","PVP01_0810300","PVP01_0810400","PVP01_0810500","PVP01_0810600","PVP01_0811000","PVP01_0811200","PVP01_0811300","PVP01_0811500","PVP01_0811700","PVP01_0811800","PVP01_0811900","PVP01_0812000","PVP01_0812300","PVP01_0812600","PVP01_0812800","PVP01_0813000","PVP01_0813200","PVP01_0813300","PVP01_0813600","PVP01_0813700","PVP01_0813800","PVP01_0814400","PVP01_0814700","PVP01_0814800","PVP01_0815000","PVP01_0815100","PVP01_0815300","PVP01_0815400","PVP01_0816200","PVP01_0816300","PVP01_0816700","PVP01_0816800","PVP01_0817100","PVP01_0817400","PVP01_0818100","PVP01_0818300","PVP01_0818600","PVP01_0819000","PVP01_0819200","PVP01_0819300","PVP01_0819700","PVP01_0820000","PVP01_0820100","PVP01_0820300","PVP01_0820500","PVP01_0820800","PVP01_0820900","PVP01_0821000","PVP01_0821300","PVP01_0821400","PVP01_0821500","PVP01_0821700","PVP01_0821800","PVP01_0822100","PVP01_0822300","PVP01_0822600","PVP01_0822700","PVP01_0822900","PVP01_0823100","PVP01_0823400","PVP01_0823600","PVP01_0823700","PVP01_0824000","PVP01_0824300","PVP01_0824500","PVP01_0824700","PVP01_0825100","PVP01_0825700","PVP01_0825800","PVP01_0826000","PVP01_0826100","PVP01_0826200","PVP01_0826300","PVP01_0826500","PVP01_0826700","PVP01_0827100","PVP01_0827700","PVP01_0827800","PVP01_0828000","PVP01_0828100","PVP01_0828200","PVP01_0828300","PVP01_0828700","PVP01_0829000","PVP01_0829100","PVP01_0829300","PVP01_0829700","PVP01_0829900","PVP01_0830300","PVP01_0830400","PVP01_0830500","PVP01_0830700","PVP01_0830900","PVP01_0831000","PVP01_0831100","PVP01_0831400","PVP01_0831700","PVP01_0831900","PVP01_0832000","PVP01_0832100","PVP01_0832200","PVP01_0832400","PVP01_0832500","PVP01_0832600","PVP01_0832900","PVP01_0833000","PVP01_0833500","PVP01_0833600","PVP01_0833700","PVP01_0833800","PVP01_0834000","PVP01_0834200","PVP01_0834300","PVP01_0834400","PVP01_0834500","PVP01_0835000","PVP01_0835300","PVP01_0835500","PVP01_0835700","PVP01_0835900","PVP01_0836100","PVP01_0836300","PVP01_0836400","PVP01_0836500","PVP01_0836600","PVP01_0836700","PVP01_0836800","PVP01_0837200","PVP01_0837300","PVP01_0837500","PVP01_0837600","PVP01_0837800","PVP01_0838000","PVP01_0838100","PVP01_0838200","PVP01_0838300","PVP01_0838600","PVP01_0838700","PVP01_0838800","PVP01_0838900","PVP01_0839000","PVP01_0839100","PVP01_0839200","PVP01_0839400","PVP01_0839700","PVP01_0840700","PVP01_0841000","PVP01_0900700","PVP01_0901000","PVP01_0901100","PVP01_0901200","PVP01_0901300","PVP01_0901800","PVP01_0901900","PVP01_0902000","PVP01_0902100","PVP01_0902400","PVP01_0903400","PVP01_0903500","PVP01_0903600","PVP01_0903700","PVP01_0903800","PVP01_0903900","PVP01_0904300","PVP01_0904600","PVP01_0904700","PVP01_0904900","PVP01_0905100","PVP01_0906000","PVP01_0906300","PVP01_0906500","PVP01_0906700","PVP01_0906800","PVP01_0906900","PVP01_0907000","PVP01_0907100","PVP01_0907200","PVP01_0907300","PVP01_0907500","PVP01_0907600","PVP01_0907700","PVP01_0907800","PVP01_0908000","PVP01_0908500","PVP01_0908600","PVP01_0908900","PVP01_0909100","PVP01_0909300","PVP01_0909500","PVP01_0909600","PVP01_0910000","PVP01_0910200","PVP01_0910900","PVP01_0911000","PVP01_0911300","PVP01_0911600","PVP01_0911700","PVP01_0911800","PVP01_0911900","PVP01_0912100","PVP01_0912200","PVP01_0912400","PVP01_0912900","PVP01_0913000","PVP01_0913400","PVP01_0913500","PVP01_0913900","PVP01_0914000","PVP01_0914400","PVP01_0914600","PVP01_0914800","PVP01_0915000","PVP01_0915100","PVP01_0915300","PVP01_0915600","PVP01_0915800","PVP01_0916300","PVP01_0916700","PVP01_0916800","PVP01_0916900","PVP01_0917000","PVP01_0917100","PVP01_0917200","PVP01_0917900","PVP01_0918000","PVP01_0918100","PVP01_0918200","PVP01_0918300","PVP01_0918400","PVP01_0918700","PVP01_0918800","PVP01_0918900","PVP01_0919000","PVP01_0919600","PVP01_0919700","PVP01_0919900","PVP01_0920100","PVP01_0920400","PVP01_0920700","PVP01_0920800","PVP01_0921400","PVP01_0921500","PVP01_0921800","PVP01_0922000","PVP01_0922100","PVP01_0922300","PVP01_0922400","PVP01_0922800","PVP01_0922900","PVP01_0923000","PVP01_0923100","PVP01_0923300","PVP01_0923800","PVP01_0923900","PVP01_0924000","PVP01_0924100","PVP01_0924300","PVP01_0924400","PVP01_0924700","PVP01_0924800","PVP01_0925000","PVP01_0925100","PVP01_0925200","PVP01_0925400","PVP01_0925600","PVP01_0925700","PVP01_0925800","PVP01_0926300","PVP01_0926400","PVP01_0926500","PVP01_0926600","PVP01_0926900","PVP01_0927100","PVP01_0927400","PVP01_0927600","PVP01_0927700","PVP01_0927800","PVP01_0928000","PVP01_0928100","PVP01_0928300","PVP01_0928800","PVP01_0928900","PVP01_0929000","PVP01_0929100","PVP01_0929200","PVP01_0929400","PVP01_0929600","PVP01_0929700","PVP01_0929900","PVP01_0930000","PVP01_0930100","PVP01_0930200","PVP01_0930300","PVP01_0930400","PVP01_0930500","PVP01_0931300","PVP01_0931400","PVP01_0931500","PVP01_0931600","PVP01_0931700","PVP01_0932300","PVP01_0932600","PVP01_0933100","PVP01_0933400","PVP01_0933500","PVP01_0933700","PVP01_0934000","PVP01_0934300","PVP01_0934400","PVP01_0934500","PVP01_0935000","PVP01_0935200","PVP01_0935400","PVP01_0935500","PVP01_0935700","PVP01_0935800","PVP01_0935900","PVP01_0936000","PVP01_0936300","PVP01_0936400","PVP01_0936700","PVP01_0936800","PVP01_0937000","PVP01_0937300","PVP01_0937500","PVP01_0937700","PVP01_0938100","PVP01_0938200","PVP01_0938600","PVP01_0938700","PVP01_0938900","PVP01_0939000","PVP01_0939100","PVP01_0939200","PVP01_0939300","PVP01_0939500","PVP01_0939600","PVP01_0940200","PVP01_0940500","PVP01_0940600","PVP01_0940700","PVP01_0940900","PVP01_0941100","PVP01_0941200","PVP01_0941300","PVP01_0941400","PVP01_0941600","PVP01_0941800","PVP01_0942200","PVP01_0942300","PVP01_0942400","PVP01_0942600","PVP01_0942700","PVP01_0942800","PVP01_0942900","PVP01_0943100","PVP01_0943300","PVP01_0943400","PVP01_0943700","PVP01_0944100","PVP01_0944300","PVP01_0944400","PVP01_0944600","PVP01_0944800","PVP01_0945100","PVP01_0945200","PVP01_0945300","PVP01_0945500","PVP01_0945600","PVP01_0945800","PVP01_0945900","PVP01_0946000","PVP01_0946200","PVP01_0946300","PVP01_0946400","PVP01_0946500","PVP01_0946600","PVP01_0946700","PVP01_0946900","PVP01_0947000","PVP01_0947100","PVP01_0947200","PVP01_0947400","PVP01_0947800","PVP01_0947900","PVP01_0948100","PVP01_0948300","PVP01_0948500","PVP01_0948600","PVP01_0948700","PVP01_0948800","PVP01_0948900","PVP01_0949000","PVP01_0949200","PVP01_0949400","PVP01_0949600","PVP01_0950000","PVP01_0950200","PVP01_1000700","PVP01_1001900","PVP01_1002500","PVP01_1002600","PVP01_1002700","PVP01_1003000","PVP01_1003200","PVP01_1003400","PVP01_1003500","PVP01_1003900","PVP01_1004200","PVP01_1004500","PVP01_1004800","PVP01_1004900","PVP01_1005100","PVP01_1005300","PVP01_1005400","PVP01_1005600","PVP01_1005700","PVP01_1005900","PVP01_1006200","PVP01_1006300","PVP01_1006400","PVP01_1006600","PVP01_1006700","PVP01_1007200","PVP01_1007500","PVP01_1007700","PVP01_1007800","PVP01_1007900","PVP01_1008000","PVP01_1008100","PVP01_1008200","PVP01_1008300","PVP01_1008600","PVP01_1008900","PVP01_1009000","PVP01_1009100","PVP01_1009300","PVP01_1009400","PVP01_1009600","PVP01_1009700","PVP01_1009800","PVP01_1010100","PVP01_1010200","PVP01_1010300","PVP01_1010400","PVP01_1011100","PVP01_1011600","PVP01_1011800","PVP01_1012100","PVP01_1012300","PVP01_1012400","PVP01_1012500","PVP01_1012700","PVP01_1012800","PVP01_1013100","PVP01_1013200","PVP01_1013300","PVP01_1013400","PVP01_1013700","PVP01_1014300","PVP01_1014600","PVP01_1014800","PVP01_1015000","PVP01_1015100","PVP01_1015400","PVP01_1015500","PVP01_1015700","PVP01_1015900","PVP01_1016200","PVP01_1016400","PVP01_1016500","PVP01_1016900","PVP01_1017100","PVP01_1017200","PVP01_1017900","PVP01_1018000","PVP01_1018100","PVP01_1018400","PVP01_1018500","PVP01_1019000","PVP01_1019400","PVP01_1019600","PVP01_1019900","PVP01_1020000","PVP01_1020100","PVP01_1020200","PVP01_1020500","PVP01_1020800","PVP01_1020900","PVP01_1021000","PVP01_1021100","PVP01_1021200","PVP01_1021300","PVP01_1021500","PVP01_1021600","PVP01_1022000","PVP01_1022100","PVP01_1022200","PVP01_1022300","PVP01_1022400","PVP01_1022900","PVP01_1023000","PVP01_1023300","PVP01_1023600","PVP01_1023700","PVP01_1023900","PVP01_1024400","PVP01_1024500","PVP01_1024800","PVP01_1025200","PVP01_1025300","PVP01_1025800","PVP01_1026300","PVP01_1026400","PVP01_1026500","PVP01_1026600","PVP01_1026700","PVP01_1026800","PVP01_1026900","PVP01_1027100","PVP01_1027300","PVP01_1027400","PVP01_1027600","PVP01_1027700","PVP01_1027900","PVP01_1028000","PVP01_1028100","PVP01_1028200","PVP01_1028400","PVP01_1028600","PVP01_1029100","PVP01_1029200","PVP01_1029300","PVP01_1029500","PVP01_1029700","PVP01_1029800","PVP01_1029900","PVP01_1030000","PVP01_1030100","PVP01_1030200","PVP01_1030300","PVP01_1030500","PVP01_1030900","PVP01_1031100","PVP01_1031200","PVP01_1031300","PVP01_1031400","PVP01_1031500","PVP01_1031600","PVP01_1031700","PVP01_1031800","PVP01_1031900","PVP01_1032000","PVP01_1032100","PVP01_1032200","PVP01_1032400","PVP01_1032500","PVP01_1032800","PVP01_1032900","PVP01_1033000","PVP01_1033300","PVP01_1033500","PVP01_1033600","PVP01_1033700","PVP01_1034000","PVP01_1034100","PVP01_1034200","PVP01_1034300","PVP01_1034400","PVP01_1034500","PVP01_1035200","PVP01_1100400","PVP01_1100500","PVP01_1101000","PVP01_1101200","PVP01_1101300","PVP01_1101400","PVP01_1101500","PVP01_1101600","PVP01_1101700","PVP01_1102400","PVP01_1102500","PVP01_1102800","PVP01_1102900","PVP01_1103200","PVP01_1103400","PVP01_1103700","PVP01_1104000","PVP01_1104100","PVP01_1104200","PVP01_1104300","PVP01_1104400","PVP01_1104600","PVP01_1104800","PVP01_1105300","PVP01_1105400","PVP01_1105500","PVP01_1105600","PVP01_1105800","PVP01_1106000","PVP01_1106200","PVP01_1106300","PVP01_1106400","PVP01_1106600","PVP01_1106700","PVP01_1107000","PVP01_1107100","PVP01_1107200","PVP01_1107300","PVP01_1107400","PVP01_1107900","PVP01_1108000","PVP01_1108300","PVP01_1108500","PVP01_1108600","PVP01_1108800","PVP01_1108900","PVP01_1109000","PVP01_1109100","PVP01_1109300","PVP01_1109400","PVP01_1109600","PVP01_1110500","PVP01_1110600","PVP01_1110700","PVP01_1110900","PVP01_1111100","PVP01_1111200","PVP01_1111300","PVP01_1111500","PVP01_1111600","PVP01_1111700","PVP01_1111800","PVP01_1112100","PVP01_1112400","PVP01_1112600","PVP01_1112800","PVP01_1113300","PVP01_1113500","PVP01_1113700","PVP01_1113900","PVP01_1114100","PVP01_1114200","PVP01_1114300","PVP01_1114500","PVP01_1114600","PVP01_1114700","PVP01_1114800","PVP01_1115000","PVP01_1115200","PVP01_1115500","PVP01_1115600","PVP01_1116600","PVP01_1116800","PVP01_1116900","PVP01_1117300","PVP01_1117700","PVP01_1118100","PVP01_1118600","PVP01_1119000","PVP01_1119300","PVP01_1119500","PVP01_1119600","PVP01_1119800","PVP01_1120100","PVP01_1120300","PVP01_1120900","PVP01_1121100","PVP01_1121300","PVP01_1121500","PVP01_1121600","PVP01_1121700","PVP01_1121900","PVP01_1122100","PVP01_1122300","PVP01_1122400","PVP01_1122500","PVP01_1122600","PVP01_1122700","PVP01_1122800","PVP01_1122900","PVP01_1123000","PVP01_1123200","PVP01_1123300","PVP01_1123400","PVP01_1123500","PVP01_1123600","PVP01_1123900","PVP01_1124000","PVP01_1124100","PVP01_1124300","PVP01_1124400","PVP01_1124600","PVP01_1124800","PVP01_1124900","PVP01_1125000","PVP01_1125100","PVP01_1125200","PVP01_1125300","PVP01_1125400","PVP01_1125600","PVP01_1125700","PVP01_1126000","PVP01_1126100","PVP01_1126200","PVP01_1126300","PVP01_1126500","PVP01_1126600","PVP01_1126800","PVP01_1126900","PVP01_1127000","PVP01_1127200","PVP01_1127300","PVP01_1127400","PVP01_1127500","PVP01_1127800","PVP01_1128100","PVP01_1128400","PVP01_1128800","PVP01_1128900","PVP01_1129000","PVP01_1129500","PVP01_1129600","PVP01_1129700","PVP01_1130100","PVP01_1130400","PVP01_1130700","PVP01_1130900","PVP01_1131300","PVP01_1131400","PVP01_1131900","PVP01_1132100","PVP01_1132300","PVP01_1132400","PVP01_1132500","PVP01_1132700","PVP01_1132800","PVP01_1132900","PVP01_1133000","PVP01_1133100","PVP01_1133200","PVP01_1133300","PVP01_1133400","PVP01_1133600","PVP01_1133800","PVP01_1133900","PVP01_1134100","PVP01_1134200","PVP01_1134300","PVP01_1134400","PVP01_1134500","PVP01_1134700","PVP01_1134800","PVP01_1135100","PVP01_1135300","PVP01_1135500","PVP01_1135600","PVP01_1135700","PVP01_1135800","PVP01_1135900","PVP01_1136100","PVP01_1136300","PVP01_1136600","PVP01_1136700","PVP01_1136800","PVP01_1136900","PVP01_1137100","PVP01_1137200","PVP01_1137300","PVP01_1137600","PVP01_1137700","PVP01_1137800","PVP01_1138400","PVP01_1138500","PVP01_1139000","PVP01_1139100","PVP01_1139300","PVP01_1139700","PVP01_1140000","PVP01_1140300","PVP01_1140500","PVP01_1140700","PVP01_1140800","PVP01_1140900","PVP01_1141000","PVP01_1141200","PVP01_1141500","PVP01_1141700","PVP01_1141800","PVP01_1141900","PVP01_1142000","PVP01_1142300","PVP01_1142500","PVP01_1142600","PVP01_1143100","PVP01_1143200","PVP01_1143400","PVP01_1143500","PVP01_1143700","PVP01_1144100","PVP01_1144200","PVP01_1144300","PVP01_1144500","PVP01_1144700","PVP01_1144900","PVP01_1145000","PVP01_1145100","PVP01_1145200","PVP01_1145500","PVP01_1145700","PVP01_1146400","PVP01_1146600","PVP01_1146800","PVP01_1146900","PVP01_1147000","PVP01_1147100","PVP01_1147200","PVP01_1147300","PVP01_1147400","PVP01_1147500","PVP01_1147700","PVP01_1147800","PVP01_1147900","PVP01_1148200","PVP01_1148700","PVP01_1148800","PVP01_1149000","PVP01_1149100","PVP01_1149400","PVP01_1149500","PVP01_1200200","PVP01_1200800","PVP01_1201100","PVP01_1201300","PVP01_1201400","PVP01_1201500","PVP01_1201600","PVP01_1201700","PVP01_1201800","PVP01_1201900","PVP01_1202100","PVP01_1202200","PVP01_1203100","PVP01_1203200","PVP01_1203300","PVP01_1203500","PVP01_1203600","PVP01_1203700","PVP01_1203800","PVP01_1204000","PVP01_1204100","PVP01_1204200","PVP01_1204600","PVP01_1204700","PVP01_1204800","PVP01_1205000","PVP01_1205200","PVP01_1205400","PVP01_1205600","PVP01_1205900","PVP01_1206100","PVP01_1206300","PVP01_1206400","PVP01_1206700","PVP01_1206900","PVP01_1207000","PVP01_1207100","PVP01_1207200","PVP01_1207800","PVP01_1207900","PVP01_1208000","PVP01_1208100","PVP01_1208200","PVP01_1208300","PVP01_1208600","PVP01_1208700","PVP01_1208800","PVP01_1208900","PVP01_1209000","PVP01_1209400","PVP01_1210000","PVP01_1210100","PVP01_1210300","PVP01_1210700","PVP01_1211000","PVP01_1211200","PVP01_1211300","PVP01_1211500","PVP01_1211600","PVP01_1212500","PVP01_1212600","PVP01_1213000","PVP01_1213100","PVP01_1213300","PVP01_1214200","PVP01_1214300","PVP01_1214600","PVP01_1214800","PVP01_1214900","PVP01_1215000","PVP01_1215300","PVP01_1215400","PVP01_1215900","PVP01_1216000","PVP01_1216100","PVP01_1216200","PVP01_1216300","PVP01_1216800","PVP01_1217000","PVP01_1217200","PVP01_1217400","PVP01_1217500","PVP01_1217600","PVP01_1217700","PVP01_1218000","PVP01_1218500","PVP01_1218800","PVP01_1218900","PVP01_1219000","PVP01_1219100","PVP01_1219200","PVP01_1219400","PVP01_1219500","PVP01_1219800","PVP01_1219900","PVP01_1220000","PVP01_1220100","PVP01_1220200","PVP01_1220600","PVP01_1220700","PVP01_1220900","PVP01_1221200","PVP01_1221300","PVP01_1221400","PVP01_1221700","PVP01_1221800","PVP01_1221900","PVP01_1222100","PVP01_1222200","PVP01_1222300","PVP01_1222400","PVP01_1223500","PVP01_1223800","PVP01_1223900","PVP01_1224000","PVP01_1224400","PVP01_1224500","PVP01_1224600","PVP01_1224700","PVP01_1224800","PVP01_1225000","PVP01_1225200","PVP01_1225300","PVP01_1225500","PVP01_1225800","PVP01_1226100","PVP01_1226700","PVP01_1226800","PVP01_1227200","PVP01_1227300","PVP01_1227500","PVP01_1227600","PVP01_1228000","PVP01_1228100","PVP01_1228200","PVP01_1228400","PVP01_1228500","PVP01_1228800","PVP01_1229100","PVP01_1229400","PVP01_1229500","PVP01_1229600","PVP01_1229800","PVP01_1229900","PVP01_1230000","PVP01_1230200","PVP01_1230600","PVP01_1230700","PVP01_1230800","PVP01_1231000","PVP01_1231100","PVP01_1231600","PVP01_1231700","PVP01_1231800","PVP01_1231900","PVP01_1232400","PVP01_1232600","PVP01_1232700","PVP01_1233100","PVP01_1233200","PVP01_1233800","PVP01_1234000","PVP01_1235000","PVP01_1235100","PVP01_1235200","PVP01_1235300","PVP01_1235400","PVP01_1235800","PVP01_1236000","PVP01_1236100","PVP01_1236400","PVP01_1236900","PVP01_1237000","PVP01_1237100","PVP01_1237200","PVP01_1237500","PVP01_1237700","PVP01_1237800","PVP01_1237900","PVP01_1238000","PVP01_1238400","PVP01_1238900","PVP01_1239000","PVP01_1239100","PVP01_1239300","PVP01_1239400","PVP01_1240100","PVP01_1240300","PVP01_1240600","PVP01_1240800","PVP01_1240900","PVP01_1241000","PVP01_1241100","PVP01_1241200","PVP01_1241300","PVP01_1241900","PVP01_1242000","PVP01_1242300","PVP01_1242400","PVP01_1242700","PVP01_1243000","PVP01_1243100","PVP01_1243200","PVP01_1243500","PVP01_1243600","PVP01_1243900","PVP01_1244400","PVP01_1244700","PVP01_1244800","PVP01_1244900","PVP01_1245100","PVP01_1245300","PVP01_1245400","PVP01_1245500","PVP01_1245600","PVP01_1245800","PVP01_1245900","PVP01_1246000","PVP01_1246300","PVP01_1246400","PVP01_1246500","PVP01_1246600","PVP01_1246700","PVP01_1246900","PVP01_1247000","PVP01_1247500","PVP01_1247600","PVP01_1247700","PVP01_1247800","PVP01_1247900","PVP01_1248200","PVP01_1248500","PVP01_1249000","PVP01_1249100","PVP01_1249200","PVP01_1249500","PVP01_1250200","PVP01_1250300","PVP01_1250600","PVP01_1250700","PVP01_1250900","PVP01_1251100","PVP01_1251300","PVP01_1251500","PVP01_1251700","PVP01_1251900","PVP01_1252000","PVP01_1252200","PVP01_1252400","PVP01_1252800","PVP01_1252900","PVP01_1253000","PVP01_1253300","PVP01_1253600","PVP01_1253700","PVP01_1253800","PVP01_1254100","PVP01_1254300","PVP01_1254900","PVP01_1255100","PVP01_1255200","PVP01_1255300","PVP01_1255400","PVP01_1256000","PVP01_1256200","PVP01_1256400","PVP01_1256700","PVP01_1256800","PVP01_1257000","PVP01_1257500","PVP01_1257600","PVP01_1257700","PVP01_1257900","PVP01_1258100","PVP01_1258200","PVP01_1258300","PVP01_1258400","PVP01_1258500","PVP01_1258700","PVP01_1258900","PVP01_1259000","PVP01_1259100","PVP01_1259200","PVP01_1259300","PVP01_1259400","PVP01_1259500","PVP01_1259600","PVP01_1260200","PVP01_1260800","PVP01_1261100","PVP01_1261200","PVP01_1261400","PVP01_1261500","PVP01_1261700","PVP01_1262000","PVP01_1262300","PVP01_1262400","PVP01_1262600","PVP01_1262800","PVP01_1262900","PVP01_1263000","PVP01_1263100","PVP01_1263400","PVP01_1263800","PVP01_1263900","PVP01_1264000","PVP01_1264300","PVP01_1264400","PVP01_1264600","PVP01_1264700","PVP01_1264800","PVP01_1264900","PVP01_1265000","PVP01_1265100","PVP01_1265300","PVP01_1265400","PVP01_1265700","PVP01_1265900","PVP01_1266400","PVP01_1266500","PVP01_1266700","PVP01_1267300","PVP01_1267500","PVP01_1267800","PVP01_1268000","PVP01_1268100","PVP01_1268200","PVP01_1268300","PVP01_1268600","PVP01_1269000","PVP01_1269200","PVP01_1270200","PVP01_1270300","PVP01_1270600","PVP01_1270700","PVP01_1270900","PVP01_1271100","PVP01_1271200","PVP01_1271300","PVP01_1271400","PVP01_1271500","PVP01_1271600","PVP01_1271900","PVP01_1272500","PVP01_1272600","PVP01_1273000","PVP01_1274300","PVP01_1300400","PVP01_1300500","PVP01_1300900","PVP01_1301000","PVP01_1301100","PVP01_1301200","PVP01_1301300","PVP01_1301400","PVP01_1301500","PVP01_1301600","PVP01_1301700","PVP01_1301800","PVP01_1302100","PVP01_1302300","PVP01_1302500","PVP01_1302700","PVP01_1302800","PVP01_1302900","PVP01_1303000","PVP01_1303100","PVP01_1303200","PVP01_1303600","PVP01_1303900","PVP01_1304000","PVP01_1304200","PVP01_1304300","PVP01_1304500","PVP01_1304800","PVP01_1305300","PVP01_1305500","PVP01_1305700","PVP01_1306000","PVP01_1306100","PVP01_1306300","PVP01_1306600","PVP01_1306700","PVP01_1306800","PVP01_1307000","PVP01_1307100","PVP01_1307200","PVP01_1307300","PVP01_1307700","PVP01_1307800","PVP01_1308100","PVP01_1308300","PVP01_1308700","PVP01_1308800","PVP01_1308900","PVP01_1309000","PVP01_1309100","PVP01_1309400","PVP01_1309500","PVP01_1309700","PVP01_1309800","PVP01_1310000","PVP01_1310400","PVP01_1310700","PVP01_1311000","PVP01_1311100","PVP01_1311200","PVP01_1311300","PVP01_1311500","PVP01_1311900","PVP01_1312000","PVP01_1312300","PVP01_1312400","PVP01_1312600","PVP01_1313100","PVP01_1313300","PVP01_1313400","PVP01_1313900","PVP01_1314100","PVP01_1314300","PVP01_1314600","PVP01_1314800","PVP01_1315200","PVP01_1315400","PVP01_1315500","PVP01_1315700","PVP01_1316100","PVP01_1316500","PVP01_1316600","PVP01_1316700","PVP01_1316900","PVP01_1317000","PVP01_1317600","PVP01_1317700","PVP01_1317800","PVP01_1317900","PVP01_1318000","PVP01_1318100","PVP01_1318300","PVP01_1318600","PVP01_1318900","PVP01_1319000","PVP01_1319200","PVP01_1319500","PVP01_1319600","PVP01_1319700","PVP01_1319900","PVP01_1320000","PVP01_1320100","PVP01_1320200","PVP01_1320300","PVP01_1320400","PVP01_1320500","PVP01_1320600","PVP01_1320700","PVP01_1321500","PVP01_1321600","PVP01_1321700","PVP01_1321900","PVP01_1322000","PVP01_1322200","PVP01_1322400","PVP01_1322600","PVP01_1322900","PVP01_1323000","PVP01_1323100","PVP01_1323700","PVP01_1323800","PVP01_1323900","PVP01_1324100","PVP01_1324800","PVP01_1325800","PVP01_1325900","PVP01_1326000","PVP01_1326200","PVP01_1326400","PVP01_1326600","PVP01_1327200","PVP01_1327300","PVP01_1327600","PVP01_1327900","PVP01_1328000","PVP01_1328100","PVP01_1328500","PVP01_1328600","PVP01_1328700","PVP01_1328800","PVP01_1328900","PVP01_1329000","PVP01_1329300","PVP01_1329400","PVP01_1329600","PVP01_1329900","PVP01_1330000","PVP01_1330800","PVP01_1331000","PVP01_1331200","PVP01_1331400","PVP01_1331800","PVP01_1332200","PVP01_1332300","PVP01_1332500","PVP01_1332700","PVP01_1332900","PVP01_1333000","PVP01_1333100","PVP01_1333200","PVP01_1333700","PVP01_1333900","PVP01_1334000","PVP01_1334100","PVP01_1334200","PVP01_1334300","PVP01_1334500","PVP01_1335100","PVP01_1335300","PVP01_1335600","PVP01_1335800","PVP01_1335900","PVP01_1336700","PVP01_1337100","PVP01_1337200","PVP01_1337300","PVP01_1337400","PVP01_1337500","PVP01_1337600","PVP01_1337900","PVP01_1338100","PVP01_1338300","PVP01_1338400","PVP01_1338600","PVP01_1339000","PVP01_1339400","PVP01_1339800","PVP01_1340200","PVP01_1340400","PVP01_1340500","PVP01_1340600","PVP01_1340800","PVP01_1341300","PVP01_1341600","PVP01_1341800","PVP01_1342200","PVP01_1342300","PVP01_1342400","PVP01_1342600","PVP01_1342700","PVP01_1342800","PVP01_1343100","PVP01_1343300","PVP01_1343500","PVP01_1343600","PVP01_1344200","PVP01_1344300","PVP01_1344500","PVP01_1344600","PVP01_1344700","PVP01_1344900","PVP01_1345300","PVP01_1345400","PVP01_1345500","PVP01_1345600","PVP01_1345700","PVP01_1345900","PVP01_1346200","PVP01_1346700","PVP01_1346800","PVP01_1346900","PVP01_1347300","PVP01_1347500","PVP01_1347700","PVP01_1347800","PVP01_1400100","PVP01_1400400","PVP01_1400600","PVP01_1400700","PVP01_1400900","PVP01_1401000","PVP01_1401100","PVP01_1401200","PVP01_1401300","PVP01_1401600","PVP01_1401700","PVP01_1401800","PVP01_1401900","PVP01_1402000","PVP01_1402100","PVP01_1402200","PVP01_1402300","PVP01_1402400","PVP01_1402500","PVP01_1402600","PVP01_1402700","PVP01_1402800","PVP01_1403000","PVP01_1403300","PVP01_1403400","PVP01_1403700","PVP01_1403900","PVP01_1404000","PVP01_1404100","PVP01_1404200","PVP01_1404300","PVP01_1404500","PVP01_1404600","PVP01_1404800","PVP01_1405000","PVP01_1405100","PVP01_1405500","PVP01_1405600","PVP01_1405800","PVP01_1406000","PVP01_1406100","PVP01_1406200","PVP01_1406700","PVP01_1407000","PVP01_1407100","PVP01_1407400","PVP01_1407600","PVP01_1407800","PVP01_1408000","PVP01_1408100","PVP01_1408200","PVP01_1408400","PVP01_1408500","PVP01_1408600","PVP01_1408700","PVP01_1408800","PVP01_1408900","PVP01_1409000","PVP01_1409100","PVP01_1409300","PVP01_1409400","PVP01_1409700","PVP01_1410200","PVP01_1410300","PVP01_1410500","PVP01_1410700","PVP01_1410800","PVP01_1410900","PVP01_1411000","PVP01_1411200","PVP01_1411300","PVP01_1411400","PVP01_1412100","PVP01_1412500","PVP01_1413100","PVP01_1413200","PVP01_1413300","PVP01_1413700","PVP01_1414200","PVP01_1414300","PVP01_1414400","PVP01_1414900","PVP01_1415100","PVP01_1415200","PVP01_1415300","PVP01_1415500","PVP01_1415600","PVP01_1415700","PVP01_1415900","PVP01_1416000","PVP01_1416100","PVP01_1416900","PVP01_1417100","PVP01_1417200","PVP01_1417700","PVP01_1417900","PVP01_1418000","PVP01_1418100","PVP01_1418200","PVP01_1418300","PVP01_1418400","PVP01_1418600","PVP01_1418800","PVP01_1418900","PVP01_1419100","PVP01_1419300","PVP01_1419400","PVP01_1419500","PVP01_1419700","PVP01_1419800","PVP01_1420000","PVP01_1420100","PVP01_1420300","PVP01_1420500","PVP01_1420700","PVP01_1420900","PVP01_1421000","PVP01_1421200","PVP01_1421300","PVP01_1421400","PVP01_1421800","PVP01_1422100","PVP01_1422200","PVP01_1422300","PVP01_1422600","PVP01_1422700","PVP01_1422900","PVP01_1423200","PVP01_1424100","PVP01_1424200","PVP01_1424500","PVP01_1425100","PVP01_1425200","PVP01_1425300","PVP01_1425400","PVP01_1425800","PVP01_1426100","PVP01_1426200","PVP01_1426700","PVP01_1426800","PVP01_1427000","PVP01_1427100","PVP01_1427700","PVP01_1428000","PVP01_1428100","PVP01_1428300","PVP01_1428400","PVP01_1428500","PVP01_1428900","PVP01_1429000","PVP01_1429200","PVP01_1429300","PVP01_1429700","PVP01_1429900","PVP01_1430000","PVP01_1430300","PVP01_1430600","PVP01_1430700","PVP01_1430800","PVP01_1431100","PVP01_1431200","PVP01_1431300","PVP01_1431700","PVP01_1431800","PVP01_1432000","PVP01_1432100","PVP01_1432400","PVP01_1432600","PVP01_1432700","PVP01_1432800","PVP01_1433600","PVP01_1433800","PVP01_1434000","PVP01_1434200","PVP01_1434300","PVP01_1434500","PVP01_1434600","PVP01_1434800","PVP01_1434900","PVP01_1435100","PVP01_1435300","PVP01_1435800","PVP01_1435900","PVP01_1436000","PVP01_1436300","PVP01_1436400","PVP01_1436600","PVP01_1436700","PVP01_1437100","PVP01_1437200","PVP01_1437600","PVP01_1437700","PVP01_1438100","PVP01_1438200","PVP01_1438400","PVP01_1438900","PVP01_1439000","PVP01_1439300","PVP01_1439400","PVP01_1439600","PVP01_1439700","PVP01_1440000","PVP01_1440100","PVP01_1440300","PVP01_1440600","PVP01_1440700","PVP01_1440900","PVP01_1441100","PVP01_1441200","PVP01_1441400","PVP01_1441700","PVP01_1441800","PVP01_1442000","PVP01_1442200","PVP01_1442900","PVP01_1443000","PVP01_1443200","PVP01_1443600","PVP01_1443800","PVP01_1443900","PVP01_1444000","PVP01_1444200","PVP01_1444400","PVP01_1444800","PVP01_1444900","PVP01_1445000","PVP01_1445200","PVP01_1445300","PVP01_1445400","PVP01_1445600","PVP01_1445700","PVP01_1446000","PVP01_1446100","PVP01_1446200","PVP01_1446300","PVP01_1446400","PVP01_1446600","PVP01_1446900","PVP01_1447200","PVP01_1447300","PVP01_1447500","PVP01_1447600","PVP01_1447800","PVP01_1447900","PVP01_1448100","PVP01_1448300","PVP01_1448400","PVP01_1448700","PVP01_1448800","PVP01_1449100","PVP01_1449200","PVP01_1449700","PVP01_1450000","PVP01_1450200","PVP01_1450400","PVP01_1450500","PVP01_1451000","PVP01_1451300","PVP01_1451500","PVP01_1451600","PVP01_1451800","PVP01_1452100","PVP01_1452400","PVP01_1452500","PVP01_1452700","PVP01_1452900","PVP01_1453200","PVP01_1453800","PVP01_1454000","PVP01_1454100","PVP01_1454200","PVP01_1454400","PVP01_1454500","PVP01_1454600","PVP01_1454800","PVP01_1454900","PVP01_1455400","PVP01_1455500","PVP01_1455700","PVP01_1456000","PVP01_1456500","PVP01_1456700","PVP01_1456900","PVP01_1457000","PVP01_1457400","PVP01_1457500","PVP01_1457900","PVP01_1458700","PVP01_1458800","PVP01_1459100","PVP01_1459200","PVP01_1459300","PVP01_1459400","PVP01_1459500","PVP01_1459600","PVP01_1459700","PVP01_1460500","PVP01_1460700","PVP01_1460800","PVP01_1460900","PVP01_1461100","PVP01_1461900","PVP01_1462000","PVP01_1462100","PVP01_1462300","PVP01_1462400","PVP01_1462500","PVP01_1462600","PVP01_1462900","PVP01_1463200","PVP01_1463300","PVP01_1463500","PVP01_1463600","PVP01_1463700","PVP01_1463800","PVP01_1464100","PVP01_1464500","PVP01_1464700","PVP01_1464900","PVP01_1465000","PVP01_1465100","PVP01_1465500","PVP01_1465700","PVP01_1465800","PVP01_1465900","PVP01_1466000","PVP01_1466100","PVP01_1466300","PVP01_1466500","PVP01_1466600","PVP01_1467000","PVP01_1467300","PVP01_1467600","PVP01_1468100","PVP01_1468400","PVP01_1468600","PVP01_1469300","PVP01_1469400","PVP01_1469600","PVP01_1469700","PVP01_1469900","PVP01_1470000","PVP01_1470100","PVP01_1470200","PVP01_1470300","PVP01_1470400","PVP01_1470500","PVP01_1470600","PVP01_1470700","PVP01_1470800","PVP01_1470900","PVP01_1471100","PVP01_1471300","PVP01_1471500","PVP01_1471600","PVP01_1471700","PVP01_1471800","PVP01_1471900","PVP01_1472100","PVP01_1472200","PVP01_1472400","PVP01_API01200","PVP01_API01400","PVP01_API01700","PVP01_API04100","PVP01_API04200","PVP01_0000110","PVP01_0000160","PVP01_0001430","PVP01_0003640","PVP01_0004040","PVP01_0004210","PVP01_0004220","PVP01_0004230","PVP01_0004290","PVP01_0004330","PVP01_0005540","PVP01_0009370","PVP01_0009720","PVP01_0010670","PVP01_0010840","PVP01_0011020","PVP01_0103100","PVP01_0103200","PVP01_0103300","PVP01_0103500","PVP01_0103700","PVP01_0104000","PVP01_0104400","PVP01_0104500","PVP01_0104700","PVP01_0105200","PVP01_0105900","PVP01_0106100","PVP01_0106600","PVP01_0106700","PVP01_0107500","PVP01_0107800","PVP01_0108200","PVP01_0108500","PVP01_0108700","PVP01_0108900","PVP01_0109400","PVP01_0109500","PVP01_0109700","PVP01_0109900","PVP01_0110000","PVP01_0110100","PVP01_0110300","PVP01_0110400","PVP01_0110500","PVP01_0110600","PVP01_0110900","PVP01_0111000","PVP01_0111100","PVP01_0111400","PVP01_0111500","PVP01_0111800","PVP01_0111900","PVP01_0112200","PVP01_0112300","PVP01_0112400","PVP01_0112600","PVP01_0112800","PVP01_0112900","PVP01_0113300","PVP01_0113600","PVP01_0113700","PVP01_0113800","PVP01_0113900","PVP01_0114000","PVP01_0114200","PVP01_0114300","PVP01_0114400","PVP01_0114500","PVP01_0114800","PVP01_0115100","PVP01_0115300","PVP01_0115600","PVP01_0115700","PVP01_0115800","PVP01_0115900","PVP01_0116000","PVP01_0116200","PVP01_0116300","PVP01_0116400","PVP01_0116800","PVP01_0117000","PVP01_0117100","PVP01_0117200","PVP01_0117300","PVP01_0117400","PVP01_0117600","PVP01_0117900","PVP01_0118100","PVP01_0118200","PVP01_0118300","PVP01_0118400","PVP01_0118500","PVP01_0201200","PVP01_0202500","PVP01_0202600","PVP01_0203200","PVP01_0203300","PVP01_0203600","PVP01_0203700","PVP01_0204200","PVP01_0204300","PVP01_0204400","PVP01_0204500","PVP01_0204600","PVP01_0204700","PVP01_0205100","PVP01_0205400","PVP01_0205700","PVP01_0205900","PVP01_0206000","PVP01_0206200","PVP01_0206700","PVP01_0206900","PVP01_0207300","PVP01_0207900","PVP01_0208000","PVP01_0208500","PVP01_0208900","PVP01_0209400","PVP01_0209700","PVP01_0209900","PVP01_0210200","PVP01_0210400","PVP01_0210500","PVP01_0210600","PVP01_0210700","PVP01_0210800","PVP01_0210900","PVP01_0211300","PVP01_0211800","PVP01_0211900","PVP01_0212000","PVP01_0212100","PVP01_0212200","PVP01_0212300","PVP01_0212900","PVP01_0213100","PVP01_0213200","PVP01_0213400","PVP01_0213500","PVP01_0213800","PVP01_0213900","PVP01_0214000","PVP01_0214300","PVP01_0214400","PVP01_0214500","PVP01_0214600","PVP01_0214700","PVP01_0215100","PVP01_0215200","PVP01_0215600","PVP01_0215900","PVP01_0216000","PVP01_0216200","PVP01_0302600","PVP01_0302700","PVP01_0302800","PVP01_0302900","PVP01_0303000","PVP01_0303200","PVP01_0303300","PVP01_0303400","PVP01_0303600","PVP01_0303700","PVP01_0303800","PVP01_0303900","PVP01_0304000","PVP01_0304200","PVP01_0304400","PVP01_0304700","PVP01_0305100","PVP01_0305400","PVP01_0305600","PVP01_0305700","PVP01_0305900","PVP01_0306200","PVP01_0306600","PVP01_0307200","PVP01_0307500","PVP01_0307800","PVP01_0307900","PVP01_0308000","PVP01_0308100","PVP01_0308200","PVP01_0308300","PVP01_0308400","PVP01_0308500","PVP01_0308600","PVP01_0308700","PVP01_0308900","PVP01_0309000","PVP01_0309100","PVP01_0309300","PVP01_0309600","PVP01_0309900","PVP01_0310100","PVP01_0310500","PVP01_0310700","PVP01_0311100","PVP01_0311800","PVP01_0311900","PVP01_0312200","PVP01_0312300","PVP01_0312600","PVP01_0312900","PVP01_0313000","PVP01_0313100","PVP01_0313200","PVP01_0313300","PVP01_0313400","PVP01_0313600","PVP01_0313700","PVP01_0313900","PVP01_0314000","PVP01_0314300","PVP01_0314400","PVP01_0314500","PVP01_0314600","PVP01_0314700","PVP01_0314800","PVP01_0315400","PVP01_0315500","PVP01_0315600","PVP01_0315800","PVP01_0316100","PVP01_0316400","PVP01_0316600","PVP01_0316900","PVP01_0317000","PVP01_0317300","PVP01_0317500","PVP01_0317600","PVP01_0317700","PVP01_0317800","PVP01_0317900","PVP01_0318200","PVP01_0318300","PVP01_0318500","PVP01_0318600","PVP01_0318700","PVP01_0318800","PVP01_0319000","PVP01_0319100","PVP01_0319500","PVP01_0319800","PVP01_0319900","PVP01_0320000","PVP01_0320100","PVP01_0320200","PVP01_0320300","PVP01_0320600","PVP01_0321000","PVP01_0405000","PVP01_0405400","PVP01_0405800","PVP01_0406300","PVP01_0406400","PVP01_0406800","PVP01_0407100","PVP01_0407200","PVP01_0407300","PVP01_0407400","PVP01_0407500","PVP01_0407600","PVP01_0407800","PVP01_0408000","PVP01_0408100","PVP01_0408300","PVP01_0408400","PVP01_0408500","PVP01_0408700","PVP01_0409100","PVP01_0409200","PVP01_0409300","PVP01_0409400","PVP01_0409500","PVP01_0409600","PVP01_0409800","PVP01_0410100","PVP01_0410200","PVP01_0410300","PVP01_0410500","PVP01_0410700","PVP01_0410900","PVP01_0411000","PVP01_0411100","PVP01_0411200","PVP01_0411300","PVP01_0411700","PVP01_0412000","PVP01_0412300","PVP01_0412400","PVP01_0412500","PVP01_0412600","PVP01_0413400","PVP01_0413600","PVP01_0413700","PVP01_0413800","PVP01_0413900","PVP01_0414000","PVP01_0414100","PVP01_0414200","PVP01_0414500","PVP01_0414700","PVP01_0414800","PVP01_0415000","PVP01_0415100","PVP01_0415300","PVP01_0415700","PVP01_0415800","PVP01_0415900","PVP01_0416000","PVP01_0416100","PVP01_0416200","PVP01_0416300","PVP01_0416400","PVP01_0416500","PVP01_0416700","PVP01_0416800","PVP01_0417000","PVP01_0417200","PVP01_0417300","PVP01_0417500","PVP01_0417600","PVP01_0417700","PVP01_0418000","PVP01_0418100","PVP01_0418200","PVP01_0418400","PVP01_0418500","PVP01_0418600","PVP01_0418700","PVP01_0418800","PVP01_0418900","PVP01_0419100","PVP01_0419600","PVP01_0419800","PVP01_0419900","PVP01_0420100","PVP01_0420300","PVP01_0420400","PVP01_0420600","PVP01_0420800","PVP01_0421000","PVP01_0421100","PVP01_0421200","PVP01_0421300","PVP01_0421400","PVP01_0421900","PVP01_0422000","PVP01_0422500","PVP01_0422800","PVP01_0423400","PVP01_0423500","PVP01_0423900","PVP01_0424500","PVP01_0503500","PVP01_0504000","PVP01_0504400","PVP01_0504800","PVP01_0504900","PVP01_0505300","PVP01_0505600","PVP01_0505700","PVP01_0505900","PVP01_0506100","PVP01_0506200","PVP01_0506500","PVP01_0506900","PVP01_0507000","PVP01_0507100","PVP01_0507200","PVP01_0507400","PVP01_0507900","PVP01_0508100","PVP01_0508300","PVP01_0508400","PVP01_0508800","PVP01_0508900","PVP01_0509000","PVP01_0509300","PVP01_0509500","PVP01_0509700","PVP01_0509800","PVP01_0510400","PVP01_0510500","PVP01_0510600","PVP01_0510700","PVP01_0511000","PVP01_0511100","PVP01_0511600","PVP01_0511900","PVP01_0512000","PVP01_0512300","PVP01_0512600","PVP01_0512900","PVP01_0513100","PVP01_0513200","PVP01_0513600","PVP01_0513800","PVP01_0514000","PVP01_0514200","PVP01_0514300","PVP01_0514400","PVP01_0514500","PVP01_0514700","PVP01_0515000","PVP01_0515400","PVP01_0515500","PVP01_0515600","PVP01_0516600","PVP01_0516700","PVP01_0516800","PVP01_0517200","PVP01_0517500","PVP01_0517600","PVP01_0517800","PVP01_0518000","PVP01_0518100","PVP01_0518200","PVP01_0518800","PVP01_0518900","PVP01_0519000","PVP01_0519300","PVP01_0519400","PVP01_0519700","PVP01_0519800","PVP01_0519900","PVP01_0520000","PVP01_0520100","PVP01_0520200","PVP01_0520300","PVP01_0520400","PVP01_0520500","PVP01_0520600","PVP01_0520900","PVP01_0521000","PVP01_0521100","PVP01_0521200","PVP01_0521500","PVP01_0521700","PVP01_0521900","PVP01_0522400","PVP01_0522500","PVP01_0522800","PVP01_0523100","PVP01_0523200","PVP01_0523500","PVP01_0523600","PVP01_0525200","PVP01_0525400","PVP01_0525500","PVP01_0525600","PVP01_0525800","PVP01_0525900","PVP01_0526000","PVP01_0526200","PVP01_0526300","PVP01_0526500","PVP01_0526600","PVP01_0526700","PVP01_0526800","PVP01_0526900","PVP01_0527000","PVP01_0527100","PVP01_0527200","PVP01_0527600","PVP01_0527900","PVP01_0528000","PVP01_0528300","PVP01_0528500","PVP01_0528800","PVP01_0529300","PVP01_0529400","PVP01_0529500","PVP01_0529800","PVP01_0530200","PVP01_0530300","PVP01_0530600","PVP01_0530700","PVP01_0530800","PVP01_0530900","PVP01_0531000","PVP01_0531300","PVP01_0531400","PVP01_0531600","PVP01_0531900","PVP01_0532000","PVP01_0532100","PVP01_0532200","PVP01_0532400","PVP01_0532800","PVP01_0533200","PVP01_0534300","PVP01_0534400","PVP01_0600800","PVP01_0601000","PVP01_0601400","PVP01_0601700","PVP01_0601800","PVP01_0601900","PVP01_0602000","PVP01_0602100","PVP01_0602200","PVP01_0602300","PVP01_0602400","PVP01_0602500","PVP01_0602700","PVP01_0602800","PVP01_0602900","PVP01_0603200","PVP01_0603300","PVP01_0603400","PVP01_0603500","PVP01_0603800","PVP01_0603900","PVP01_0604000","PVP01_0604100","PVP01_0604300","PVP01_0604500","PVP01_0604600","PVP01_0604800","PVP01_0604900","PVP01_0605200","PVP01_0605400","PVP01_0605600","PVP01_0605700","PVP01_0605800","PVP01_0605900","PVP01_0606000","PVP01_0606100","PVP01_0606300","PVP01_0606400","PVP01_0606700","PVP01_0606900","PVP01_0607000","PVP01_0607100","PVP01_0607300","PVP01_0607400","PVP01_0607500","PVP01_0607700","PVP01_0607800","PVP01_0607900","PVP01_0608200","PVP01_0608400","PVP01_0608500","PVP01_0608700","PVP01_0609000","PVP01_0609100","PVP01_0609300","PVP01_0609500","PVP01_0610000","PVP01_0610500","PVP01_0611000","PVP01_0611200","PVP01_0611400","PVP01_0611500","PVP01_0611600","PVP01_0611700","PVP01_0611800","PVP01_0611900","PVP01_0612200","PVP01_0612300","PVP01_0612500","PVP01_0612800","PVP01_0612900","PVP01_0613300","PVP01_0613600","PVP01_0613700","PVP01_0613900","PVP01_0614100","PVP01_0614500","PVP01_0614700","PVP01_0614800","PVP01_0615200","PVP01_0615300","PVP01_0615400","PVP01_0615600","PVP01_0615700","PVP01_0616100","PVP01_0616500","PVP01_0616600","PVP01_0616700","PVP01_0616800","PVP01_0617200","PVP01_0617300","PVP01_0617600","PVP01_0617700","PVP01_0617800","PVP01_0618000","PVP01_0618200","PVP01_0618400","PVP01_0618500","PVP01_0618800","PVP01_0618900","PVP01_0619300","PVP01_0619900","PVP01_0620000","PVP01_0620200","PVP01_0620400","PVP01_0620800","PVP01_0621000","PVP01_0621200","PVP01_0621300","PVP01_0621400","PVP01_0621500","PVP01_0621600","PVP01_0621700","PVP01_0622000","PVP01_0622300","PVP01_0622400","PVP01_0622500","PVP01_0622600","PVP01_0623700","PVP01_0623800","PVP01_0700100","PVP01_0700600","PVP01_0701100","PVP01_0701300","PVP01_0701400","PVP01_0701800","PVP01_0701900","PVP01_0702000","PVP01_0702300","PVP01_0702400","PVP01_0702600","PVP01_0702700","PVP01_0702800","PVP01_0702900","PVP01_0703100","PVP01_0703200","PVP01_0703300","PVP01_0703400","PVP01_0703500","PVP01_0703800","PVP01_0703900","PVP01_0704200","PVP01_0704300","PVP01_0704400","PVP01_0705200","PVP01_0705300","PVP01_0705400","PVP01_0705700","PVP01_0706500","PVP01_0706800","PVP01_0706900","PVP01_0707400","PVP01_0707500","PVP01_0707600","PVP01_0707700","PVP01_0707900","PVP01_0708600","PVP01_0708700","PVP01_0709400","PVP01_0709600","PVP01_0709700","PVP01_0709800","PVP01_0710400","PVP01_0710500","PVP01_0710600","PVP01_0710800","PVP01_0710900","PVP01_0711000","PVP01_0711300","PVP01_0711600","PVP01_0711700","PVP01_0712000","PVP01_0712100","PVP01_0712500","PVP01_0712600","PVP01_0712800","PVP01_0712900","PVP01_0713300","PVP01_0713400","PVP01_0713700","PVP01_0713800","PVP01_0714000","PVP01_0714100","PVP01_0714400","PVP01_0714500","PVP01_0714700","PVP01_0714800","PVP01_0715300","PVP01_0715400","PVP01_0715900","PVP01_0716200","PVP01_0716300","PVP01_0716400","PVP01_0716500","PVP01_0716700","PVP01_0717100","PVP01_0717300","PVP01_0717500","PVP01_0717700","PVP01_0717900","PVP01_0718200","PVP01_0718300","PVP01_0718400","PVP01_0718500","PVP01_0718800","PVP01_0718900","PVP01_0719200","PVP01_0719300","PVP01_0719600","PVP01_0720500","PVP01_0720600","PVP01_0720700","PVP01_0721000","PVP01_0721200","PVP01_0721300","PVP01_0721400","PVP01_0721600","PVP01_0721700","PVP01_0721900","PVP01_0722900","PVP01_0723600","PVP01_0723700","PVP01_0723900","PVP01_0724000","PVP01_0724200","PVP01_0724300","PVP01_0724400","PVP01_0725500","PVP01_0725600","PVP01_0725800","PVP01_0725900","PVP01_0726000","PVP01_0726100","PVP01_0726200","PVP01_0726300","PVP01_0726500","PVP01_0726600","PVP01_0726800","PVP01_0727500","PVP01_0727700","PVP01_0727900","PVP01_0728000","PVP01_0728400","PVP01_0728500","PVP01_0728600","PVP01_0728700","PVP01_0728800","PVP01_0728900","PVP01_0729000","PVP01_0729600","PVP01_0729800","PVP01_0729900","PVP01_0730500","PVP01_0730800","PVP01_0730900","PVP01_0731400","PVP01_0731500","PVP01_0731800","PVP01_0732400","PVP01_0732500","PVP01_0732700","PVP01_0733200","PVP01_0733300","PVP01_0733400","PVP01_0733600","PVP01_0733800","PVP01_0734000","PVP01_0734500","PVP01_0734600","PVP01_0734800","PVP01_0734900","PVP01_0801100","PVP01_0801200","PVP01_0801300","PVP01_0801600","PVP01_0802200","PVP01_0802300","PVP01_0802500","PVP01_0802600","PVP01_0802700","PVP01_0802800","PVP01_0803200","PVP01_0803300","PVP01_0803400","PVP01_0803800","PVP01_0803900","PVP01_0804000","PVP01_0804300","PVP01_0804400","PVP01_0805100","PVP01_0805500","PVP01_0805900","PVP01_0806600","PVP01_0806800","PVP01_0806900","PVP01_0807100","PVP01_0807300","PVP01_0807400","PVP01_0807600","PVP01_0807700","PVP01_0807800","PVP01_0808200","PVP01_0808300","PVP01_0808900","PVP01_0809000","PVP01_0809500","PVP01_0809800","PVP01_0810200","PVP01_0810700","PVP01_0810800","PVP01_0810900","PVP01_0811100","PVP01_0811400","PVP01_0811600","PVP01_0812100","PVP01_0812200","PVP01_0812400","PVP01_0812500","PVP01_0812700","PVP01_0812900","PVP01_0813100","PVP01_0813400","PVP01_0813500","PVP01_0813900","PVP01_0814000","PVP01_0814100","PVP01_0814200","PVP01_0814300","PVP01_0814500","PVP01_0814600","PVP01_0814900","PVP01_0815200","PVP01_0815500","PVP01_0815600","PVP01_0815700","PVP01_0815800","PVP01_0815900","PVP01_0816000","PVP01_0816100","PVP01_0816400","PVP01_0816500","PVP01_0816600","PVP01_0816900","PVP01_0817000","PVP01_0817200","PVP01_0817300","PVP01_0817500","PVP01_0817600","PVP01_0817700","PVP01_0817800","PVP01_0817900","PVP01_0818000","PVP01_0818200","PVP01_0818400","PVP01_0818500","PVP01_0818700","PVP01_0818800","PVP01_0818900","PVP01_0819100","PVP01_0819400","PVP01_0819500","PVP01_0819600","PVP01_0819800","PVP01_0819900","PVP01_0820200","PVP01_0820400","PVP01_0820600","PVP01_0820700","PVP01_0821100","PVP01_0821200","PVP01_0821600","PVP01_0821900","PVP01_0822000","PVP01_0822200","PVP01_0822400","PVP01_0822500","PVP01_0822800","PVP01_0823000","PVP01_0823200","PVP01_0823300","PVP01_0823500","PVP01_0823800","PVP01_0823900","PVP01_0824100","PVP01_0824200","PVP01_0824400","PVP01_0824600","PVP01_0824800","PVP01_0824900","PVP01_0825000","PVP01_0825200","PVP01_0825300","PVP01_0825400","PVP01_0825500","PVP01_0825600","PVP01_0825900","PVP01_0826400","PVP01_0826600","PVP01_0826800","PVP01_0826900","PVP01_0827000","PVP01_0827200","PVP01_0827300","PVP01_0827400","PVP01_0827900","PVP01_0828400","PVP01_0828500","PVP01_0828600","PVP01_0828800","PVP01_0828900","PVP01_0829200","PVP01_0829400","PVP01_0829500","PVP01_0829600","PVP01_0829800","PVP01_0830000","PVP01_0830100","PVP01_0830200","PVP01_0830600","PVP01_0831200","PVP01_0831300","PVP01_0831500","PVP01_0831600","PVP01_0832300","PVP01_0832700","PVP01_0832800","PVP01_0833100","PVP01_0833200","PVP01_0833400","PVP01_0833900","PVP01_0834100","PVP01_0834600","PVP01_0834700","PVP01_0834800","PVP01_0834900","PVP01_0835100","PVP01_0835200","PVP01_0835400","PVP01_0835600","PVP01_0835800","PVP01_0836000","PVP01_0836200","PVP01_0836900","PVP01_0837000","PVP01_0837100","PVP01_0837400","PVP01_0837700","PVP01_0837900","PVP01_0904200","PVP01_0904400","PVP01_0904500","PVP01_0904800","PVP01_0905000","PVP01_0905200","PVP01_0905300","PVP01_0905400","PVP01_0905500","PVP01_0905600","PVP01_0905700","PVP01_0905800","PVP01_0905900","PVP01_0906100","PVP01_0906200","PVP01_0906400","PVP01_0906600","PVP01_0907400","PVP01_0907900","PVP01_0908100","PVP01_0908200","PVP01_0908300","PVP01_0908400","PVP01_0908700","PVP01_0908800","PVP01_0909000","PVP01_0909200","PVP01_0909400","PVP01_0909700","PVP01_0909900","PVP01_0910700","PVP01_0910800","PVP01_0911100","PVP01_0911200","PVP01_0911400","PVP01_0911500","PVP01_0912000","PVP01_0912300","PVP01_0912500","PVP01_0912600","PVP01_0912700","PVP01_0912800","PVP01_0913100","PVP01_0913200","PVP01_0913300","PVP01_0913600","PVP01_0913700","PVP01_0913800","PVP01_0914100","PVP01_0914200","PVP01_0914300","PVP01_0914500","PVP01_0914700","PVP01_0914900","PVP01_0915200","PVP01_0915400","PVP01_0915500","PVP01_0915700","PVP01_0915900","PVP01_0916000","PVP01_0916100","PVP01_0916200","PVP01_0916400","PVP01_0916500","PVP01_0916600","PVP01_0917300","PVP01_0917400","PVP01_0917500","PVP01_0917600","PVP01_0917700","PVP01_0917800","PVP01_0918500","PVP01_0918600","PVP01_0919100","PVP01_0919200","PVP01_0919300","PVP01_0919400","PVP01_0919500","PVP01_0919800","PVP01_0920000","PVP01_0920200","PVP01_0920300","PVP01_0920500","PVP01_0920600","PVP01_0920900","PVP01_0921000","PVP01_0921100","PVP01_0921200","PVP01_0921300","PVP01_0921600","PVP01_0921700","PVP01_0921900","PVP01_0922200","PVP01_0922500","PVP01_0922600","PVP01_0922700","PVP01_0923200","PVP01_0923400","PVP01_0923500","PVP01_0923600","PVP01_0923700","PVP01_0924200","PVP01_0924500","PVP01_0924600","PVP01_0924900","PVP01_0925300","PVP01_0925500","PVP01_0925900","PVP01_0926000","PVP01_0926100","PVP01_0926200","PVP01_0926700","PVP01_0926800","PVP01_0927000","PVP01_0927200","PVP01_0927300","PVP01_0927500","PVP01_0927900","PVP01_0928200","PVP01_0928400","PVP01_0928500","PVP01_0928600","PVP01_0928700","PVP01_0929300","PVP01_0929500","PVP01_0929800","PVP01_0930600","PVP01_0930700","PVP01_0930800","PVP01_0931200","PVP01_0931800","PVP01_0931900","PVP01_0932000","PVP01_0932100","PVP01_0932200","PVP01_0932400","PVP01_0932500","PVP01_0932700","PVP01_0932800","PVP01_0932900","PVP01_0933000","PVP01_0933200","PVP01_0933300","PVP01_0933600","PVP01_0933800","PVP01_0933900","PVP01_0934100","PVP01_0934200","PVP01_0934600","PVP01_0934700","PVP01_0934800","PVP01_0934900","PVP01_0935100","PVP01_0935300","PVP01_0935600","PVP01_0936100","PVP01_0936200","PVP01_0936500","PVP01_0936600","PVP01_0936900","PVP01_0937100","PVP01_0937200","PVP01_0937400","PVP01_0937600","PVP01_0937800","PVP01_0938000","PVP01_0938300","PVP01_0938400","PVP01_0938500","PVP01_0938800","PVP01_0939400","PVP01_0939700","PVP01_0939800","PVP01_0939900","PVP01_0940000","PVP01_0940100","PVP01_0940300","PVP01_0940400","PVP01_0940800","PVP01_0941000","PVP01_0941500","PVP01_0941700","PVP01_0941900","PVP01_0942000","PVP01_0942100","PVP01_0942500","PVP01_0943000","PVP01_0943200","PVP01_0943500","PVP01_0943600","PVP01_0943800","PVP01_0943900","PVP01_0944000","PVP01_0944200","PVP01_0944500","PVP01_0944700","PVP01_0944900","PVP01_0945000","PVP01_0945400","PVP01_0945700","PVP01_0946100","PVP01_0946800","PVP01_0947300","PVP01_0947500","PVP01_0947600","PVP01_0947700","PVP01_0948000","PVP01_0948200","PVP01_0948400","PVP01_0949300","PVP01_1002400","PVP01_1002800","PVP01_1002900","PVP01_1003100","PVP01_1003300","PVP01_1003600","PVP01_1003700","PVP01_1003800","PVP01_1004000","PVP01_1004100","PVP01_1004400","PVP01_1004600","PVP01_1004700","PVP01_1005000","PVP01_1005200","PVP01_1005500","PVP01_1005800","PVP01_1006500","PVP01_1006800","PVP01_1006900","PVP01_1007000","PVP01_1007100","PVP01_1007300","PVP01_1007400","PVP01_1007600","PVP01_1008400","PVP01_1008500","PVP01_1008700","PVP01_1008800","PVP01_1009200","PVP01_1009500","PVP01_1009900","PVP01_1010000","PVP01_1010500","PVP01_1010600","PVP01_1010700","PVP01_1010800","PVP01_1010900","PVP01_1011000","PVP01_1011200","PVP01_1011300","PVP01_1011400","PVP01_1011500","PVP01_1011700","PVP01_1011900","PVP01_1012000","PVP01_1012200","PVP01_1012600","PVP01_1012900","PVP01_1013000","PVP01_1013500","PVP01_1013600","PVP01_1013800","PVP01_1013900","PVP01_1014000","PVP01_1014100","PVP01_1014200","PVP01_1014400","PVP01_1014500","PVP01_1014700","PVP01_1014900","PVP01_1015200","PVP01_1015300","PVP01_1015600","PVP01_1015800","PVP01_1016000","PVP01_1016100","PVP01_1016300","PVP01_1016600","PVP01_1016700","PVP01_1016800","PVP01_1017000","PVP01_1017300","PVP01_1017400","PVP01_1017500","PVP01_1017600","PVP01_1017700","PVP01_1017800","PVP01_1018200","PVP01_1018300","PVP01_1018600","PVP01_1018700","PVP01_1018800","PVP01_1018900","PVP01_1019100","PVP01_1019200","PVP01_1019300","PVP01_1019700","PVP01_1019800","PVP01_1020300","PVP01_1020400","PVP01_1020600","PVP01_1020700","PVP01_1021400","PVP01_1021700","PVP01_1021800","PVP01_1021900","PVP01_1022500","PVP01_1022600","PVP01_1022700","PVP01_1022800","PVP01_1023200","PVP01_1023500","PVP01_1023800","PVP01_1024000","PVP01_1024100","PVP01_1024200","PVP01_1024300","PVP01_1024600","PVP01_1024700","PVP01_1024900","PVP01_1025000","PVP01_1025100","PVP01_1025400","PVP01_1025500","PVP01_1025600","PVP01_1025700","PVP01_1025900","PVP01_1026000","PVP01_1026100","PVP01_1026200","PVP01_1027000","PVP01_1027200","PVP01_1027500","PVP01_1027800","PVP01_1028300","PVP01_1028500","PVP01_1028700","PVP01_1028800","PVP01_1028900","PVP01_1029000","PVP01_1029400","PVP01_1029600","PVP01_1030400","PVP01_1030600","PVP01_1030700","PVP01_1030800","PVP01_1031000","PVP01_1032300","PVP01_1032600","PVP01_1032700","PVP01_1033100","PVP01_1033200","PVP01_1033400","PVP01_1033800","PVP01_1033900","PVP01_1102300","PVP01_1102600","PVP01_1102700","PVP01_1103000","PVP01_1103100","PVP01_1103300","PVP01_1103500","PVP01_1103600","PVP01_1103800","PVP01_1103900","PVP01_1104500","PVP01_1104700","PVP01_1104900","PVP01_1105000","PVP01_1105100","PVP01_1105200","PVP01_1105700","PVP01_1105900","PVP01_1106100","PVP01_1106500","PVP01_1106800","PVP01_1106900","PVP01_1107500","PVP01_1107600","PVP01_1107700","PVP01_1107800","PVP01_1108100","PVP01_1108200","PVP01_1108400","PVP01_1108700","PVP01_1109200","PVP01_1109500","PVP01_1109700","PVP01_1109800","PVP01_1109900","PVP01_1110000","PVP01_1110100","PVP01_1110200","PVP01_1110300","PVP01_1110400","PVP01_1110800","PVP01_1111000","PVP01_1111400","PVP01_1111900","PVP01_1112000","PVP01_1112200","PVP01_1112300","PVP01_1112500","PVP01_1112700","PVP01_1112900","PVP01_1113000","PVP01_1113100","PVP01_1113200","PVP01_1113400","PVP01_1113600","PVP01_1113800","PVP01_1114000","PVP01_1114400","PVP01_1114900","PVP01_1115100","PVP01_1115300","PVP01_1115400","PVP01_1115700","PVP01_1115800","PVP01_1115900","PVP01_1116000","PVP01_1116100","PVP01_1116200","PVP01_1116300","PVP01_1116400","PVP01_1116700","PVP01_1117000","PVP01_1117100","PVP01_1117200","PVP01_1117400","PVP01_1117500","PVP01_1117600","PVP01_1117800","PVP01_1117900","PVP01_1118000","PVP01_1118200","PVP01_1118300","PVP01_1118400","PVP01_1118500","PVP01_1118700","PVP01_1118800","PVP01_1118900","PVP01_1119100","PVP01_1119200","PVP01_1119400","PVP01_1119700","PVP01_1119900","PVP01_1120000","PVP01_1120200","PVP01_1120400","PVP01_1120500","PVP01_1120600","PVP01_1120700","PVP01_1120800","PVP01_1121000","PVP01_1121200","PVP01_1121400","PVP01_1121800","PVP01_1122000","PVP01_1122200","PVP01_1123100","PVP01_1123700","PVP01_1123800","PVP01_1124200","PVP01_1124500","PVP01_1124700","PVP01_1125500","PVP01_1125800","PVP01_1125900","PVP01_1126400","PVP01_1126700","PVP01_1127100","PVP01_1127600","PVP01_1127700","PVP01_1128000","PVP01_1128200","PVP01_1128300","PVP01_1128500","PVP01_1129100","PVP01_1129200","PVP01_1129300","PVP01_1129400","PVP01_1129800","PVP01_1129900","PVP01_1130000","PVP01_1130200","PVP01_1130300","PVP01_1130500","PVP01_1130600","PVP01_1130800","PVP01_1131000","PVP01_1131100","PVP01_1131200","PVP01_1131500","PVP01_1131600","PVP01_1131700","PVP01_1131800","PVP01_1132000","PVP01_1132200","PVP01_1132600","PVP01_1133500","PVP01_1133700","PVP01_1134000","PVP01_1134600","PVP01_1134900","PVP01_1135000","PVP01_1135200","PVP01_1135400","PVP01_1136000","PVP01_1136200","PVP01_1136400","PVP01_1136500","PVP01_1137000","PVP01_1137400","PVP01_1137500","PVP01_1137900","PVP01_1138000","PVP01_1138100","PVP01_1138200","PVP01_1138300","PVP01_1138600","PVP01_1138700","PVP01_1138800","PVP01_1138900","PVP01_1139200","PVP01_1139400","PVP01_1139500","PVP01_1139600","PVP01_1139800","PVP01_1139900","PVP01_1140100","PVP01_1140200","PVP01_1140400","PVP01_1140600","PVP01_1141100","PVP01_1141300","PVP01_1141400","PVP01_1141600","PVP01_1142100","PVP01_1142200","PVP01_1142400","PVP01_1142700","PVP01_1142800","PVP01_1142900","PVP01_1143000","PVP01_1143300","PVP01_1143600","PVP01_1143800","PVP01_1143900","PVP01_1144000","PVP01_1144400","PVP01_1144600","PVP01_1144800","PVP01_1145300","PVP01_1145400","PVP01_1145600","PVP01_1145800","PVP01_1145900","PVP01_1146000","PVP01_1146100","PVP01_1146200","PVP01_1146300","PVP01_1146500","PVP01_1146700","PVP01_1147600","PVP01_1202000","PVP01_1202300","PVP01_1202400","PVP01_1202500","PVP01_1202600","PVP01_1202700","PVP01_1202800","PVP01_1202900","PVP01_1203000","PVP01_1203900","PVP01_1204300","PVP01_1204400","PVP01_1204500","PVP01_1204900","PVP01_1205100","PVP01_1205300","PVP01_1205500","PVP01_1205700","PVP01_1205800","PVP01_1206000","PVP01_1206200","PVP01_1206500","PVP01_1206600","PVP01_1206800","PVP01_1207300","PVP01_1207400","PVP01_1207500","PVP01_1207600","PVP01_1207700","PVP01_1208400","PVP01_1208500","PVP01_1209100","PVP01_1209200","PVP01_1209300","PVP01_1209500","PVP01_1209600","PVP01_1209700","PVP01_1209800","PVP01_1209900","PVP01_1210200","PVP01_1210400","PVP01_1210500","PVP01_1210600","PVP01_1210800","PVP01_1210900","PVP01_1211100","PVP01_1211400","PVP01_1211700","PVP01_1211800","PVP01_1211900","PVP01_1212000","PVP01_1212100","PVP01_1212200","PVP01_1212300","PVP01_1212400","PVP01_1212700","PVP01_1212800","PVP01_1212900","PVP01_1213200","PVP01_1213400","PVP01_1213600","PVP01_1213900","PVP01_1214000","PVP01_1214100","PVP01_1214400","PVP01_1214500","PVP01_1214700","PVP01_1215100","PVP01_1215200","PVP01_1215500","PVP01_1215800","PVP01_1216400","PVP01_1216500","PVP01_1216600","PVP01_1216700","PVP01_1216900","PVP01_1217300","PVP01_1217800","PVP01_1217900","PVP01_1218100","PVP01_1218200","PVP01_1218300","PVP01_1218400","PVP01_1218600","PVP01_1218700","PVP01_1219300","PVP01_1219600","PVP01_1219700","PVP01_1220300","PVP01_1220400","PVP01_1220500","PVP01_1220800","PVP01_1221000","PVP01_1221100","PVP01_1221500","PVP01_1221600","PVP01_1222000","PVP01_1222500","PVP01_1222600","PVP01_1222700","PVP01_1222800","PVP01_1222900","PVP01_1223000","PVP01_1223100","PVP01_1223200","PVP01_1223300","PVP01_1223400","PVP01_1223600","PVP01_1223700","PVP01_1224100","PVP01_1224200","PVP01_1224300","PVP01_1224900","PVP01_1225100","PVP01_1225400","PVP01_1225600","PVP01_1225700","PVP01_1225900","PVP01_1226000","PVP01_1226200","PVP01_1226300","PVP01_1226400","PVP01_1226600","PVP01_1226900","PVP01_1227000","PVP01_1227100","PVP01_1227400","PVP01_1227700","PVP01_1227800","PVP01_1227900","PVP01_1228300","PVP01_1228600","PVP01_1228700","PVP01_1228900","PVP01_1229000","PVP01_1229200","PVP01_1229300","PVP01_1229700","PVP01_1230100","PVP01_1230300","PVP01_1230400","PVP01_1230500","PVP01_1230900","PVP01_1231200","PVP01_1231300","PVP01_1231400","PVP01_1231500","PVP01_1232000","PVP01_1232100","PVP01_1232200","PVP01_1232300","PVP01_1232500","PVP01_1232800","PVP01_1232900","PVP01_1233000","PVP01_1233300","PVP01_1233400","PVP01_1233500","PVP01_1233600","PVP01_1233700","PVP01_1233900","PVP01_1234100","PVP01_1234200","PVP01_1234300","PVP01_1234400","PVP01_1234500","PVP01_1234600","PVP01_1234700","PVP01_1234800","PVP01_1234900","PVP01_1235500","PVP01_1235600","PVP01_1235700","PVP01_1235900","PVP01_1236200","PVP01_1236300","PVP01_1236500","PVP01_1236600","PVP01_1236700","PVP01_1236800","PVP01_1237300","PVP01_1237400","PVP01_1237600","PVP01_1238100","PVP01_1238200","PVP01_1238300","PVP01_1238500","PVP01_1238600","PVP01_1238700","PVP01_1238800","PVP01_1239200","PVP01_1239500","PVP01_1239600","PVP01_1239700","PVP01_1239800","PVP01_1239900","PVP01_1240000","PVP01_1240200","PVP01_1240400","PVP01_1240500","PVP01_1240700","PVP01_1241400","PVP01_1241500","PVP01_1241600","PVP01_1241700","PVP01_1241800","PVP01_1242100","PVP01_1242200","PVP01_1242500","PVP01_1242600","PVP01_1242800","PVP01_1242900","PVP01_1243300","PVP01_1243400","PVP01_1243700","PVP01_1243800","PVP01_1244000","PVP01_1244100","PVP01_1244200","PVP01_1244300","PVP01_1244500","PVP01_1244600","PVP01_1245000","PVP01_1245200","PVP01_1245700","PVP01_1246100","PVP01_1246200","PVP01_1246800","PVP01_1247100","PVP01_1247200","PVP01_1247300","PVP01_1247400","PVP01_1248000","PVP01_1248100","PVP01_1248300","PVP01_1248400","PVP01_1248600","PVP01_1248700","PVP01_1248800","PVP01_1248900","PVP01_1249300","PVP01_1249400","PVP01_1249600","PVP01_1249700","PVP01_1249800","PVP01_1249900","PVP01_1250000","PVP01_1250100","PVP01_1250400","PVP01_1250500","PVP01_1250800","PVP01_1251000","PVP01_1251200","PVP01_1251400","PVP01_1251600","PVP01_1251800","PVP01_1252100","PVP01_1252300","PVP01_1252500","PVP01_1252600","PVP01_1252700","PVP01_1253100","PVP01_1253200","PVP01_1253400","PVP01_1253500","PVP01_1253900","PVP01_1254000","PVP01_1254200","PVP01_1254400","PVP01_1254500","PVP01_1254600","PVP01_1254700","PVP01_1254800","PVP01_1255000","PVP01_1255500","PVP01_1255600","PVP01_1255700","PVP01_1255800","PVP01_1255900","PVP01_1256100","PVP01_1256300","PVP01_1256500","PVP01_1256600","PVP01_1256900","PVP01_1257100","PVP01_1257200","PVP01_1257300","PVP01_1257400","PVP01_1257800","PVP01_1258000","PVP01_1258600","PVP01_1258800","PVP01_1259700","PVP01_1259800","PVP01_1259900","PVP01_1260000","PVP01_1260100","PVP01_1260300","PVP01_1260400","PVP01_1260500","PVP01_1260600","PVP01_1260700","PVP01_1260900","PVP01_1261300","PVP01_1261600","PVP01_1261800","PVP01_1261900","PVP01_1262100","PVP01_1262200","PVP01_1262500","PVP01_1262700","PVP01_1263200","PVP01_1263300","PVP01_1263500","PVP01_1263600","PVP01_1263700","PVP01_1264100","PVP01_1264200","PVP01_1264500","PVP01_1265200","PVP01_1265500","PVP01_1265600","PVP01_1265800","PVP01_1266000","PVP01_1266100","PVP01_1266200","PVP01_1266300","PVP01_1266600","PVP01_1266800","PVP01_1266900","PVP01_1267000","PVP01_1267100","PVP01_1267200","PVP01_1267400","PVP01_1267600","PVP01_1267700","PVP01_1267900","PVP01_1268400","PVP01_1268500","PVP01_1268900","PVP01_1269100","PVP01_1269300","PVP01_1269400","PVP01_1269500","PVP01_1269600","PVP01_1269700","PVP01_1269800","PVP01_1269900","PVP01_1270000","PVP01_1270100","PVP01_1270400","PVP01_1270500","PVP01_1270800","PVP01_1271000","PVP01_1300600","PVP01_1300700","PVP01_1300800","PVP01_1301900","PVP01_1302000","PVP01_1302200","PVP01_1302400","PVP01_1302600","PVP01_1303300","PVP01_1303400","PVP01_1303500","PVP01_1303700","PVP01_1303800","PVP01_1304100","PVP01_1304400","PVP01_1304600","PVP01_1304700","PVP01_1304900","PVP01_1305000","PVP01_1305100","PVP01_1305200","PVP01_1305400","PVP01_1305600","PVP01_1305800","PVP01_1305900","PVP01_1306200","PVP01_1306400","PVP01_1306500","PVP01_1306900","PVP01_1307400","PVP01_1307500","PVP01_1307600","PVP01_1307900","PVP01_1308000","PVP01_1308200","PVP01_1308400","PVP01_1308500","PVP01_1308600","PVP01_1309200","PVP01_1309300","PVP01_1309600","PVP01_1309900","PVP01_1310100","PVP01_1310200","PVP01_1310300","PVP01_1310500","PVP01_1310600","PVP01_1310800","PVP01_1310900","PVP01_1311400","PVP01_1311600","PVP01_1311700","PVP01_1311800","PVP01_1312100","PVP01_1312200","PVP01_1312500","PVP01_1312700","PVP01_1312800","PVP01_1312900","PVP01_1313000","PVP01_1313200","PVP01_1313500","PVP01_1313600","PVP01_1313700","PVP01_1313800","PVP01_1314000","PVP01_1314200","PVP01_1314400","PVP01_1314500","PVP01_1314700","PVP01_1314900","PVP01_1315000","PVP01_1315100","PVP01_1315300","PVP01_1315600","PVP01_1315800","PVP01_1315900","PVP01_1316000","PVP01_1316200","PVP01_1316300","PVP01_1316400","PVP01_1316800","PVP01_1317100","PVP01_1317200","PVP01_1317300","PVP01_1317400","PVP01_1317500","PVP01_1318200","PVP01_1318400","PVP01_1318500","PVP01_1318700","PVP01_1318800","PVP01_1319100","PVP01_1319300","PVP01_1319400","PVP01_1319800","PVP01_1320800","PVP01_1320900","PVP01_1321000","PVP01_1321100","PVP01_1321200","PVP01_1321300","PVP01_1321400","PVP01_1321800","PVP01_1322100","PVP01_1322300","PVP01_1322500","PVP01_1322700","PVP01_1322800","PVP01_1323200","PVP01_1323300","PVP01_1323400","PVP01_1323500","PVP01_1323600","PVP01_1324000","PVP01_1324700","PVP01_1325200","PVP01_1325300","PVP01_1325400","PVP01_1325500","PVP01_1325600","PVP01_1325700","PVP01_1326100","PVP01_1326300","PVP01_1326500","PVP01_1326700","PVP01_1326800","PVP01_1326900","PVP01_1327000","PVP01_1327100","PVP01_1327400","PVP01_1327500","PVP01_1327700","PVP01_1327800","PVP01_1328200","PVP01_1328300","PVP01_1328400","PVP01_1329100","PVP01_1329200","PVP01_1329500","PVP01_1329700","PVP01_1329800","PVP01_1330600","PVP01_1330700","PVP01_1330900","PVP01_1331100","PVP01_1331300","PVP01_1331500","PVP01_1331600","PVP01_1331700","PVP01_1331900","PVP01_1332000","PVP01_1332100","PVP01_1332400","PVP01_1332600","PVP01_1332800","PVP01_1333300","PVP01_1333400","PVP01_1333500","PVP01_1333600","PVP01_1333800","PVP01_1334400","PVP01_1334600","PVP01_1334700","PVP01_1334800","PVP01_1334900","PVP01_1335000","PVP01_1335200","PVP01_1335400","PVP01_1335500","PVP01_1335700","PVP01_1336000","PVP01_1336100","PVP01_1336200","PVP01_1336300","PVP01_1336400","PVP01_1336500","PVP01_1336600","PVP01_1336800","PVP01_1336900","PVP01_1337000","PVP01_1337700","PVP01_1337800","PVP01_1338000","PVP01_1338200","PVP01_1338500","PVP01_1338700","PVP01_1338800","PVP01_1338900","PVP01_1339100","PVP01_1339200","PVP01_1339300","PVP01_1339500","PVP01_1339600","PVP01_1339700","PVP01_1339900","PVP01_1340000","PVP01_1340100","PVP01_1340300","PVP01_1340700","PVP01_1340900","PVP01_1341000","PVP01_1341100","PVP01_1341200","PVP01_1341400","PVP01_1341500","PVP01_1341700","PVP01_1341900","PVP01_1342000","PVP01_1342100","PVP01_1342500","PVP01_1342900","PVP01_1343000","PVP01_1343200","PVP01_1343400","PVP01_1343700","PVP01_1343800","PVP01_1343900","PVP01_1344000","PVP01_1344100","PVP01_1344400","PVP01_1344800","PVP01_1345000","PVP01_1345100","PVP01_1345200","PVP01_1345800","PVP01_1346000","PVP01_1346100","PVP01_1346400","PVP01_1346500","PVP01_1346600","PVP01_1347000","PVP01_1347100","PVP01_1347200","PVP01_1401400","PVP01_1402900","PVP01_1403100","PVP01_1403200","PVP01_1403500","PVP01_1403600","PVP01_1403800","PVP01_1404400","PVP01_1404700","PVP01_1404900","PVP01_1405200","PVP01_1405300","PVP01_1405400","PVP01_1405700","PVP01_1405900","PVP01_1406300","PVP01_1406400","PVP01_1406500","PVP01_1406600","PVP01_1406800","PVP01_1406900","PVP01_1407200","PVP01_1407300","PVP01_1407500","PVP01_1407700","PVP01_1407900","PVP01_1408300","PVP01_1409200","PVP01_1409500","PVP01_1409600","PVP01_1409800","PVP01_1409900","PVP01_1410000","PVP01_1410100","PVP01_1410400","PVP01_1410600","PVP01_1411100","PVP01_1411500","PVP01_1411600","PVP01_1411700","PVP01_1411800","PVP01_1411900","PVP01_1412000","PVP01_1412200","PVP01_1412300","PVP01_1412400","PVP01_1412600","PVP01_1412700","PVP01_1412800","PVP01_1412900","PVP01_1413000","PVP01_1413400","PVP01_1413450","PVP01_1413500","PVP01_1413600","PVP01_1413800","PVP01_1413900","PVP01_1414000","PVP01_1414100","PVP01_1414500","PVP01_1414600","PVP01_1414700","PVP01_1414800","PVP01_1415000","PVP01_1415400","PVP01_1415800","PVP01_1416200","PVP01_1416300","PVP01_1416400","PVP01_1416500","PVP01_1416600","PVP01_1416700","PVP01_1416800","PVP01_1417000","PVP01_1417300","PVP01_1417400","PVP01_1417500","PVP01_1417600","PVP01_1417800","PVP01_1418500","PVP01_1418700","PVP01_1419000","PVP01_1419200","PVP01_1419600","PVP01_1419900","PVP01_1420200","PVP01_1420400","PVP01_1420600","PVP01_1420800","PVP01_1421100","PVP01_1421500","PVP01_1421600","PVP01_1421700","PVP01_1421900","PVP01_1422000","PVP01_1422400","PVP01_1422500","PVP01_1422800","PVP01_1423000","PVP01_1423100","PVP01_1423300","PVP01_1423400","PVP01_1423500","PVP01_1423600","PVP01_1423700","PVP01_1424300","PVP01_1424400","PVP01_1424600","PVP01_1424700","PVP01_1424800","PVP01_1424900","PVP01_1425000","PVP01_1425500","PVP01_1425600","PVP01_1425700","PVP01_1425900","PVP01_1426000","PVP01_1426300","PVP01_1426400","PVP01_1426500","PVP01_1426600","PVP01_1426900","PVP01_1427200","PVP01_1427300","PVP01_1427400","PVP01_1427500","PVP01_1427600","PVP01_1427800","PVP01_1427900","PVP01_1428200","PVP01_1428600","PVP01_1428700","PVP01_1428800","PVP01_1429100","PVP01_1429400","PVP01_1429500","PVP01_1429600","PVP01_1429800","PVP01_1430100","PVP01_1430200","PVP01_1430400","PVP01_1430500","PVP01_1430900","PVP01_1431000","PVP01_1431400","PVP01_1431500","PVP01_1431600","PVP01_1432200","PVP01_1432300","PVP01_1432500","PVP01_1432900","PVP01_1433000","PVP01_1433100","PVP01_1433200","PVP01_1433300","PVP01_1433400","PVP01_1433500","PVP01_1433700","PVP01_1433900","PVP01_1434100","PVP01_1434400","PVP01_1434700","PVP01_1435000","PVP01_1435200","PVP01_1435400","PVP01_1435500","PVP01_1435700","PVP01_1436100","PVP01_1436200","PVP01_1436500","PVP01_1436800","PVP01_1436900","PVP01_1437000","PVP01_1437300","PVP01_1437400","PVP01_1437500","PVP01_1437800","PVP01_1437900","PVP01_1438000","PVP01_1438300","PVP01_1438500","PVP01_1438600","PVP01_1438700","PVP01_1438800","PVP01_1439100","PVP01_1439200","PVP01_1439500","PVP01_1439800","PVP01_1439900","PVP01_1440200","PVP01_1440400","PVP01_1440500","PVP01_1440800","PVP01_1441000","PVP01_1441300","PVP01_1441500","PVP01_1441600","PVP01_1441900","PVP01_1442100","PVP01_1442300","PVP01_1442400","PVP01_1442500","PVP01_1442600","PVP01_1442700","PVP01_1442800","PVP01_1443100","PVP01_1443300","PVP01_1443400","PVP01_1443500","PVP01_1443700","PVP01_1444100","PVP01_1444300","PVP01_1444500","PVP01_1444600","PVP01_1444700","PVP01_1445100","PVP01_1445500","PVP01_1445800","PVP01_1445900","PVP01_1446500","PVP01_1446700","PVP01_1446800","PVP01_1447000","PVP01_1447100","PVP01_1447400","PVP01_1447700","PVP01_1448000","PVP01_1448200","PVP01_1448500","PVP01_1448600","PVP01_1448900","PVP01_1449000","PVP01_1449300","PVP01_1449400","PVP01_1449500","PVP01_1449600","PVP01_1449800","PVP01_1449900","PVP01_1450100","PVP01_1450300","PVP01_1450600","PVP01_1450700","PVP01_1450800","PVP01_1450900","PVP01_1451100","PVP01_1451200","PVP01_1451400","PVP01_1451700","PVP01_1451900","PVP01_1452000","PVP01_1452200","PVP01_1452300","PVP01_1452600","PVP01_1452800","PVP01_1453000","PVP01_1453100","PVP01_1453300","PVP01_1453400","PVP01_1453500","PVP01_1453600","PVP01_1453700","PVP01_1453900","PVP01_1454300","PVP01_1454700","PVP01_1455000","PVP01_1455100","PVP01_1455200","PVP01_1455300","PVP01_1455600","PVP01_1455800","PVP01_1455900","PVP01_1456100","PVP01_1456200","PVP01_1456300","PVP01_1456400","PVP01_1456600","PVP01_1456800","PVP01_1457100","PVP01_1457200","PVP01_1457300","PVP01_1457600","PVP01_1457700","PVP01_1457800","PVP01_1458000","PVP01_1458100","PVP01_1458200","PVP01_1458300","PVP01_1458400","PVP01_1458500","PVP01_1458600","PVP01_1458900","PVP01_1459000","PVP01_1459800","PVP01_1459900","PVP01_1460000","PVP01_1460100","PVP01_1460200","PVP01_1460300","PVP01_1460400","PVP01_1460600","PVP01_1461000","PVP01_1461200","PVP01_1461300","PVP01_1461500","PVP01_1461600","PVP01_1461700","PVP01_1461800","PVP01_1462200","PVP01_1462700","PVP01_1462800","PVP01_1463000","PVP01_1463100","PVP01_1463400","PVP01_1463900","PVP01_1464000","PVP01_1464200","PVP01_1464300","PVP01_1464400","PVP01_1464600","PVP01_1464800","PVP01_1465200","PVP01_1465300","PVP01_1465400","PVP01_1465600","PVP01_1466200","PVP01_1466400","PVP01_1466700","PVP01_1466800","PVP01_1466900","PVP01_1467100","PVP01_1467200","PVP01_1467400","PVP01_1467500","PVP01_1467700","PVP01_1467800","PVP01_1467900","PVP01_1468000","PVP01_1468200","PVP01_1468300","PVP01_1468500","PVP01_1468700","PVP01_1468800","PVP01_1468900","PVP01_1469200","PVP01_1469500","PVP01_1469800","PVP01_1471400","PVP01_API02600","PVP01_API02800","PVP01_API03500","PVP01_API04300","PVP01_API04600","PVP01_MIT02700","PVP01_MIT03300","PVP01_MIT03400"],"logFC":[-0.1944,-0.8413,2.363,0.286,1.244,-1.297,0.595,-0.2286,6.477,4.122,-1.374,-1.028,-1.577,2.192,-2.435,1.511,3.404,3.972,3.664,4.314,4.754,-1.457,4.295,-2.198,5.18,-3.224,3.348,3.845,1.988,3.166,4.492,5.102,5.189,3.482,-1.948,3.203,4.059,4.019,4.187,4.899,3.612,3.054,-1.041,0.3795,4.596,5.005,3.52,4.306,3.359,-0.2824,5.066,-0.2788,-1.056,-2.454,0.3658,-1.461,-3.914,3.61,-1.147,4.738,-3.659,4.794,3.318,3.482,3.054,4.031,4.048,3.738,-3.478,0.1526,1.547,0.5879,0.9548,-0.07939,1.299,0.758,1.166,-1.11,1.794,-0.03865,-4.003,-0.007531,3.807,4.739,1.756,3.119,-3.407,0.51,3.684,3.649,3.945,-3.881,4.631,2.866,1.724,-0.5467,5.629,3.285,4.314,4.545,-3.879,4.05,4.916,-1.018,4.554,3.998,4.398,4.301,-0.5611,-2.361,4.639,4.819,-1.086,3.561,4.188,4.295,3.007,5.497,2.199,-3.047,-2.786,3.484,3.645,4.03,3.233,0.5849,3.787,-4.403,3.947,-0.3408,5.123,0.6425,4.805,-3.305,4.487,3.318,3.054,2.866,4.501,4.085,5.258,5.022,3.253,6.21,-4.328,2.722,4.309,2.866,3.801,4.073,4.113,-2.354,3.605,2.955,4.124,3.827,-1.002,2.967,5.57,5.038,2.583,2.008,-0.1608,-2.361,3.253,-2.107,-3.472,4.811,2.959,1.874,0.05033,0.2539,2.889,3.246,-3.436,-4.337,-1.831,-1.236,-0.8228,-3.191,-3.019,-2.48,-0.7779,-3.122,5.18,1.356,0.6669,-0.8186,0.7095,4.57,0.4052,0.8726,3.284,-0.4615,-0.4001,-3.09,-2.628,-0.9991,-0.1711,-3.764,-4.547,1.381,1.261,-1.655,1.109,-2.325,0.6524,-1.614,0.4934,0.1529,0.1907,-0.6996,1.121,0.01012,0.9246,-1.18,-0.8784,0.7895,-0.1762,-0.3221,-2.139,-0.1856,1.043,-3.469,-0.8388,-1.974,-2.135,0.03168,1.078,-0.8173,1.128,-0.4875,1.378,0.5444,-0.9731,-0.2599,1.256,-1.241,0.483,-0.2321,-0.9442,-0.2773,2.309,0.1133,0.892,1.356,-1.197,-0.1417,-0.6721,-1.098,0.3955,0.7532,-1.491,1.164,-0.6767,0.8042,0.7281,-0.3446,-0.412,1.239,-1.082,0.7641,-0.9566,0.0195,0.789,1.851,-1.805,0.4184,-1.187,0.7573,-0.4933,-0.5954,1.839,-0.6422,0.8688,-0.1893,-0.4167,0.956,0.5075,5.547,-1.23,-2.169,0.08202,-1.445,-0.08851,-2.627,5.129,-2.506,4.697,3.916,3.895,4.073,5.74,4.102,2.976,2.398,4.073,-0.3472,0.8214,-2.782,0.8245,-0.712,-0.3164,3.439,-0.092,1.25,-1.898,-2.983,1.255,-2.789,-0.7702,-2.56,0.7795,-2.431,-0.6863,1.232,0.3587,-1.012,-1.479,1.268,0.614,0.1944,0.9442,-0.4474,1.611,-1.49,0.2649,0.5243,1.51,0.09281,1.007,-2.05,0.6964,1.032,0.911,0.9614,-0.5615,0.5687,-0.2161,1.268,1.483,1.719,0.3377,0.8722,-0.0315,-1.377,-0.3141,0.09776,1.084,0.3151,-1.469,0.1887,-0.751,0.6678,2.039,-0.3992,-1.196,-0.02428,0.8604,-0.3933,0.5115,-0.07801,0.5362,-1.946,-0.631,-0.3026,-1.1,-0.235,-0.3668,-2.64,0.3803,-0.9802,-1.416,-0.1309,0.1918,-0.9652,0.9558,-2.72,-0.5768,3.233,-1.775,0.4217,-2.087,0.1587,-3.131,-1.846,-1.192,1.881,-2.724,0.2354,4.213,3.61,3.002,4.525,4.394,4.03,-1.107,5.677,-3.565,2.913,5.789,4.051,3.757,6.267,-0.04485,1.874,-2.921,4.073,3.482,3.429,5.252,0.09844,1.54,-0.6606,-2.003,-1.941,0.8564,-1.239,-0.6956,-0.5249,0.9166,1.177,0.1143,0.8442,-1.375,-0.6359,-0.04764,0.2211,0.978,-0.4276,-1.164,0.8608,-0.09434,1.362,-3.401,-2.053,2.519,-0.6847,-0.1596,0.07087,1.269,-1.199,-0.004544,0.2237,1.608,0.9198,0.2093,0.8861,0.3322,1.616,-1.383,1.24,-0.9248,1.479,0.5584,-0.9564,-0.8858,-2.127,-1.105,-2.608,-0.3927,-0.7372,0.9372,-1.852,-0.7388,-0.2015,1.499,-0.6052,-1.079,0.8256,-0.9382,0.6263,0.8596,0.02007,0.9657,1.43,1.862,1.048,1.348,1.715,1.049,-1.064,1.363,-0.7681,1.476,-0.07293,0.164,1.387,3.723,1.104,-0.919,0.7211,0.292,0.7314,1.073,-0.1308,0.7193,1.155,-1.654,0.1691,4.305,-2.667,3.497,-0.1291,4.505,1.918,-3.31,-0.422,-2.224,2.56,-1.052,-1.535,-4.055,3.016,-1.728,-0.3381,2.099,1.186,-0.271,-0.9584,-1.371,0.1545,-0.01781,-2.079,1.254,-1.212,-1.551,1.038,1.202,0.482,1.19,1.174,2.05,1.836,-1.202,0.5854,0.4007,1.144,0.8203,0.5462,-0.9068,-3.849,0.5434,-2.433,-0.7976,-1.375,-0.05672,-0.7677,0.3753,-0.3072,-0.7421,0.8295,1.447,0.5337,-1.078,-1.141,-1.901,0.01352,0.679,-0.2287,1.039,-0.5802,1.039,-0.2178,-1.728,-0.218,-0.07592,-0.06291,-1.725,-0.9051,-0.06965,-1.852,0.8542,-1.475,0.4177,3.184,-0.7198,-1.397,3.476,0.2076,0.06868,-0.1557,0.2022,0.05321,-0.5782,-0.8446,-0.5982,0.9189,0.8366,-0.2142,-0.3508,1.676,-2.351,-2.607,1.627,2.104,3.593,-1.306,3.523,0.3313,1.765,3.765,-1.307,1.548,-1.129,-1.595,1.943,-2.75,-3.25,-1.414,-2.09,0.5616,-1.548,-1.254,0.1862,-0.3221,0.8466,-0.3868,1.347,3.093,0.2028,-0.02166,0.6507,0.5533,1.099,-0.9852,0.2125,0.6161,0.3369,0.1945,-0.5385,5.709,-0.7139,-1.92,0.4976,0.5488,0.9358,0.3272,1.326,-0.6408,-2.656,0.2293,0.7627,1.266,0.6724,0.4563,-0.5441,-0.7821,0.03336,1.851,-0.6965,1.28,0.4352,-1.737,0.8231,-0.1819,1.362,-0.9378,-1.124,-0.01497,-1.32,-0.01134,-0.8381,0.7517,1.392,-1.761,0.3293,-1.772,0.04131,-3.554,-1.788,-2.57,0.3332,-0.6214,-0.8898,-2.767,0.1224,-1.139,-0.7017,-0.2766,0.9897,-0.611,0.4825,0.7327,-2.063,-1.576,-3.084,1.652,-0.2518,1.066,0.09965,-0.5705,-0.2986,0.1847,2.917,-0.7679,0.5087,1.089,0.2747,0.1762,-0.1818,2.078,0.2353,-0.01093,-0.8761,-0.4076,-2.183,-1.602,-3.995,0.9063,-2.145,-0.7197,-1.575,-0.913,-3.107,-0.8888,-0.1778,4.076,-1.313,-0.5504,-2.748,1.047,1.186,1.452,-0.2542,-2.147,1.102,0.9906,0.5018,-1.77,-0.5669,-2.486,0.2484,1.875,-0.5676,0.4509,0.8044,-1.026,0.4706,0.8547,0.6824,-0.3513,-2.086,-1.044,-2.704,-0.7473,0.8651,-2.034,0.6834,-0.46,-0.004073,-1.551,1.437,-1.552,-1.522,-0.151,0.2043,1.854,-0.06172,-1.85,1.174,0.8983,2.194,0.3533,-0.5326,3.561,4.356,1.695,3.568,2.224,1.58,1.847,1.357,4.395,3.024,-0.3448,3.015,2.283,-3.21,-2.657,0.6236,1.112,1.949,0.1692,0.9564,0.6134,-0.5629,4.432,0.5615,0.8476,-0.5303,1.553,0.9955,0.65,-0.1104,0.3959,0.8583,0.7239,-0.8546,-0.003502,0.6156,1.367,-1.172,-0.9716,0.5063,0.1541,-2.18,0.4637,-0.07518,-0.3432,0.9886,-2.067,1.677,-0.3161,-0.2914,0.937,0.169,-3.82,-0.7241,0.7218,0.9134,-2.712,-1.342,-0.1741,-2.921,-1.278,1.858,0.7391,-0.9424,-0.2709,-0.3444,-0.7017,0.3712,-0.1449,-1.282,0.1254,-1.425,-0.8033,-0.519,0.9187,-0.1108,1.829,1.688,-1.744,-1.879,-1.379,-0.1822,-0.6434,1.753,-0.6963,-0.1487,-2.089,-1.445,1.284,-0.08423,1.162,0.4564,0.6042,-0.357,-2.655,0.09371,-2.834,-1.656,-0.4553,0.7038,0.6226,-0.01192,-1.852,-2.39,-1.741,-1.832,0.1562,-0.4994,-1.543,-0.8251,0.1533,-2.749,-2.855,-1.928,1.047,2.082,-1.032,-1.648,0.08755,-0.4203,2.219,0.6748,2.954,3.658,-4.637,1.433,-1.627,-0.9555,-2.437,-1.482,-0.7467,-3.294,1.609,-1.04,-0.259,0.544,0.03814,-0.9007,-0.4114,-0.3224,-0.8869,0.6056,0.3288,-0.3385,0.5604,-1.121,-1.461,-1.256,-1.001,1.337,-1.286,-0.7146,-0.02696,-1.149,0.5427,0.9278,0.9529,-1.493,-0.7773,1.498,-1.215,-0.7478,0.09747,-0.3664,0.276,0.91,0.7677,1.532,-0.1875,0.747,0.7123,-2.019,0.3211,1.479,1.034,-0.3973,0.5933,-0.5964,0.5279,0.6296,-1.991,-2.101,-0.2683,-1.8,-0.02179,0.8573,0.5486,0.3867,-1.237,-0.6853,0.3421,-2.382,-0.4771,0.9865,-0.1321,-0.2124,0.65,1.421,-1.558,0.236,0.07974,0.706,-0.5006,2.126,-0.2222,0.665,-2.208,-1.329,0.3358,-2.436,-0.06002,-2.006,-2.093,-1.826,-1.658,0.3672,0.7712,0.144,0.01918,0.3476,0.9054,-2.078,1.115,-1.501,0.3707,1.047,0.4547,-0.2227,-0.4333,0.7986,-0.7337,1.255,-1.802,0.4884,-2.142,1.01,0.4436,-0.2298,0.6253,2.667,0.02549,0.6735,-0.1136,1.204,-1.264,0.1079,-0.5652,-1.464,0.3554,0.8496,1.213,0.847,-0.9158,-0.2817,1.389,0.5789,0.7374,0.4429,2.014,0.8562,1.232,0.8615,0.8771,-1.402,1.805,-0.6065,-1.438,1.373,-0.9442,0.9195,-0.3573,0.08262,-0.5684,1.975,-0.1471,-2.495,-2.037,-1.01,-0.3743,1.063,-1.414,-1.335,1.03,-1.63,-3.055,0.6332,-1.582,-1.394,-0.1085,5.998,-0.7826,-0.09789,-0.5108,1.156,2.186,1.074,-2.063,-3.043,-0.3001,0.3738,0.8651,0.8946,2.234,1.15,-2.677,0.3303,-0.8934,-0.7194,-0.145,0.06475,-0.1339,-1.347,-2.749,-1.047,1.109,-2.998,-2.22,2.674,-1.655,0.3213,2.552,-0.8503,2.395,-2.495,-2.712,3.573,3.467,5.174,-3.191,-0.06783,-0.6921,1.862,-1.34,0.1763,-3.234,2.107,-3.034,0.5126,-1.385,-1.736,-0.4696,0.5516,-1.556,-3.224,0.6132,-0.2699,0.842,0.1307,-1.849,-0.4385,1.053,-3.016,-1.072,-1.072,0.8784,-2.456,2.42,-1.742,1.25,-0.6627,-0.2642,0.6961,0.5818,0.176,-0.4213,0.4374,-1.13,0.03653,0.9844,-0.9363,0.2499,1.081,-1.122,-2.251,-1.555,-1.008,-0.6791,1.063,-2.083,0.8966,-1.246,-0.2955,0.02535,0.6509,-0.777,-1.861,-0.105,-0.73,0.1972,1.077,-1.03,-0.9443,0.04473,-0.1409,-1.918,-0.4396,-0.1099,-1.695,0.1832,0.9201,-0.8912,0.2205,0.3379,-2.727,-0.5834,0.598,-0.08201,1.393,-0.3128,-1.291,-1.749,0.5463,0.959,-1.723,1.016,-0.598,0.3689,-1.241,1.586,-0.9385,0.4678,-0.7276,0.4099,-0.2213,1.635,0.7513,-0.3788,-0.2608,0.07747,-0.5781,0.7452,0.6915,-0.5047,-1.584,0.2324,0.8184,1.051,-1.015,-1.04,-0.08719,1.288,1.047,-0.05183,0.3215,-0.1197,1.689,-1.071,0.6017,-0.2664,1.068,-1.272,-0.9433,0.322,2.437,-2.524,-2.089,0.256,0.7177,0.2946,-1.252,-0.4554,2.895,0.2527,-0.954,0.05504,-1.595,-2.234,0.2298,0.9889,0.09377,1.9,-0.7057,2.044,1.519,0.8589,0.1932,-0.08488,0.5941,1.033,1.694,0.8741,-1.457,-0.7733,1.508,-0.4372,-2.619,-0.8954,-0.1439,0.1871,1.341,0.5813,-3.206,0.1376,1.547,-0.7461,-0.5077,1.054,-3.006,-0.3842,-1.548,-1.287,-1.958,-1.532,2.318,-1.752,0.1563,0.6357,-0.4916,0.9765,-1.148,1.519,0.1938,-1.232,1.284,-0.1242,0.62,-0.1275,-0.3768,-0.491,-1.924,-0.8757,-0.037,-0.3055,1.295,2.296,-0.09341,-2.228,-2.156,-2.521,-3.371,-1.786,-0.9695,4.193,-2.135,4.232,3.855,4.152,1.657,-3.614,-6.294,-3.546,-2.912,3.215,-0.3766,2.544,-6.206,1.954,-2.809,-0.007534,5.377,-0.3425,1.998,0.6605,-1.603,-0.4974,-0.4606,0.2974,-1.507,-2.806,0.8595,-0.4463,1.911,-0.7309,-0.4019,0.1939,1.159,-0.3436,-0.02278,-0.4063,-1.174,0.582,-2.484,0.635,-1.056,1.133,-0.9784,-1.565,-1.163,0.5239,-0.1391,1.4,0.4088,-0.517,-1.756,-0.2449,-1.498,0.6428,0.2116,-0.5303,0.3867,-1.096,0.6492,0.3408,-0.6883,-0.3129,-0.06968,1.255,1.188,-2.437,-1.239,1.49,1.269,-2.194,0.8423,-0.667,-1.066,-0.2272,1.155,-3.094,-0.7493,-1.248,0.025,-0.08723,-0.9917,0.01852,0.9009,0.1408,-0.9441,-1.152,-0.9234,-0.05372,0.5281,-1.213,-2.882,-0.3254,0.9006,-0.3323,-0.5216,0.1023,1.495,-2.667,-2.646,-1.356,-1.132,-1.491,-1.409,-0.8483,-1.295,0.2857,0.4756,1.238,0.8334,-0.3592,1.177,0.4146,1.03,0.1125,-0.6319,-1.124,-1.43,-2.054,-0.3965,0.7212,-1.799,-0.8172,-0.6467,-0.298,-1.436,-0.8684,0.04822,-0.5409,-1.073,0.7523,0.705,-2.069,-1.216,0.429,-1.019,-1.001,-0.2046,-0.5396,-0.05353,0.6508,1.604,-0.804,-1.373,-0.9901,-0.9124,1.297,0.5678,-1.644,-2.45,-0.5349,-0.4172,0.785,0.05033,-0.6894,-0.3658,0.5187,-2.213,2.501,1.007,1.14,-0.04043,0.01143,0.18,-1.98,-0.5971,0.05721,-0.5586,0.9365,-0.01775,-0.1898,-1.05,2.871,-0.007974,-0.2085,-0.1051,-1.993,0.2623,-2.583,-1.691,0.3865,0.2197,-0.5239,-1.32,0.6868,-2.272,0.8924,3.607,0.5878,-1.001,-2.825,-2.933,1.031,0.9936,-1.376,0.2278,0.03517,1.509,-0.575,0.6192,-0.6475,0.8602,1.123,0.6237,-0.4141,2.667,-0.5526,-0.339,0.2901,1.306,-0.9421,-1.055,0.676,0.6932,-1.242,-1.506,1.719,-1.176,-0.04254,-0.4346,-0.9172,2.573,1.331,1.06,0.5872,1.527,0.9228,0.7694,-0.9459,1.87,-1.327,-0.9426,0.0465,-1.536,-2.814,-1.189,-1.735,0.3537,-0.4295,0.8965,0.2459,1.125,0.5193,-1.966,-0.05472,-0.5523,-0.006696,0.9661,3.347,6.427,3.037,-2.339,-0.08726,3.552,3.625,0.08078,4.151,-2.429,-0.1674,0.9534,0.9123,0.2084,0.3454,-1.091,-1.585,1.754,-1.416,0.2941,0.4752,1.606,-2.596,-2.46,0.7521,0.3283,0.9561,-0.03429,-1.379,0.5536,-0.1888,-0.05518,-1.026,0.5933,-1.113,-1.493,-0.7423,1.247,0.2491,-1.105,1.214,0.9585,1.053,0.6857,-0.8753,-0.2601,-2.138,-0.007743,-0.1588,0.8292,-1.574,-0.7824,-0.8818,-2.275,2.075,-1.45,0.779,-0.6397,-1.211,0.7802,-1.348,-1.534,-0.6375,-0.5601,-2.009,-0.3076,1.192,-0.7996,-1.677,-0.2203,-1.088,-0.6402,0.9663,1.416,-0.4634,2.862,-1.061,1.187,-0.5197,-2.881,2.084,-0.2836,-0.8559,-0.6091,0.8998,-1.886,-2.85,1.333,-1.192,0.3062,0.6571,0.206,2.115,-0.8136,0.6714,-0.7344,2.959,1.749,-0.06988,0.2792,0.2725,1.429,-0.3279,1.194,-0.3465,-0.3477,-1.944,0.4775,-1.705,1.166,-1.901,-0.5734,2.904,-0.5337,1.491,1.587,1.071,1.523,1.549,-0.7099,0.892,1.332,-0.4044,-0.2279,-2.155,-0.1921,0.07252,-1.098,-0.2288,-0.3745,-0.7214,1.949,-1.194,1.185,-1.607,0.4718,0.5376,-0.7605,0.7839,-0.1401,0.6601,0.07312,1.097,-1.348,3.667,1.528,2.017,0.4722,-1.563,-1.878,1.577,-0.8441,0.9995,-0.8723,-1.373,0.8972,-0.6553,-1.162,-1.865,-0.637,0.959,-0.735,-0.1786,0.9325,-3.072,-0.2285,-0.6028,0.9582,0.1649,-1.186,-1.357,0.8166,-1.148,2.18,0.223,-0.4921,-0.344,-0.08384,-1.465,0.487,1.794,-3.259,1.998,4.233,1.255,1.372,-2.454,-1.038,1.046,-0.887,-0.2908,-1.336,-0.564,-1.399,0.8114,-1.907,-1.405,-2.091,0.6082,-0.7538,-0.8373,-0.85,0.3855,0.289,-1.263,-0.208,3.294,-0.3454,-0.1287,-2.429,0.2006,-0.993,-2.15,-1.658,1.256,-0.5235,-1.535,1.2,-2.288,-0.4738,-1.52,0.8841,-1.672,-0.1359,-0.4885,-0.3473,-0.3939,-1.825,-1.696,0.8994,0.3292,1.168,-0.8095,-0.733,0.01819,0.5278,-0.6734,0.1191,-0.8932,0.09546,-0.6864,-0.8623,1.28,-0.09236,-2.582,0.8139,-0.4698,-1.083,-0.5649,1.054,0.981,-1.3,1.198,0.01291,0.922,-2.305,-2.338,-1.626,-0.2981,-2.653,-0.1517,-1.123,0.01254,1.094,-1.31,-0.2861,-0.9547,-0.1825,-0.04109,-0.6514,1.656,1.871,-1.035,0.9241,-0.352,-0.09914,-3.785,0.4456,0.4257,-1.453,0.4069,-0.2711,-1.232,-1.468,-2.458,-0.4444,-1.788,-1.554,0.3124,0.49,-2.512,1.475,-0.5316,-2.051,-1.877,-0.8129,1.2,1.212,-0.7594,1.25,1.396,1.418,0.2583,-1.991,-0.1118,-2.196,-1.54,-1.225,0.3491,-1.946,-1.773,1.323,-0.05407,-1.901,0.1311,0.01512,0.5362,0.8671,0.3402,-0.3203,0.9823,-0.8531,0.5934,1.338,-0.8565,0.7594,1.394,-0.3494,-1.432,-0.8373,-0.2403,1.228,0.4473,0.5487,0.9657,-0.07989,-1.467,-1.434,-1.007,-1.213,-1.493,1.099,0.8265,-0.01602,-1.346,-0.3989,1.423,-0.4182,0.6721,-1.366,-0.401,-0.1499,1.071,0.2871,0.0922,0.211,1.524,1.12,-0.1694,0.589,-1.419,-2.728,-0.6693,2.032,-1.19,0.4994,-1.331,-2.297,-0.3375,-3.577,-0.6446,1.26,-1.312,-0.779,-0.8998,1.411,-2.226,-0.8051,-0.1641,0.3781,1.993,0.1603,-0.4621,-1.79,-2.051,0.3866,-0.07656,1.09,-1.029,1.167,-1.776,0.9315,-3.829,1.212,0.7976,-1.196,1.449,1.167,1.06,2.645,0.5333,1.002,-1.085,-1.093,-0.8248,3.024,-0.1239,1.054,1.567,-0.4404,-0.4286,0.3519,0.003502,-0.6229,-0.5176,1.87,-2.645,-0.7842,0.6349,-0.7529,-1.274,-1.576,-1.064,0.4019,-1.493,-1.743,-1.759,-1.27,1.731,0.8776,-2.268,-2.221,-3.958,-0.6021,4.048,3.655,4.974,1.543,1.372,4.424,5.217,3.23,-1.284,0.9059,-2.606,-1.619,3.509,1.865,3.578,1.916,-0.01898,1.647,1.159,-1.397,-0.3825,-1.284,-1.081,-3.562,0.1415,1.282,-1.514,-2.163,-0.6217,-1.399,-2.025,-0.3701,0.5069,-2.039,-1.58,1.578,0.08597,-0.4861,1.297,-1.543,-0.1396,0.157,0.5954,0.5564,-0.2112,0.8765,1.712,-1.404,-0.7017,0.908,-0.5077,0.0799,2.028,0.8354,-0.0457,0.1909,-0.6736,-0.3666,-1.337,0.5363,-0.998,0.4887,-0.1863,0.7924,-1.941,0.4947,0.3826,-2.181,-1.529,0.3254,-0.631,1.333,-1.905,1.456,-1.755,-1.61,0.129,0.6315,0.04754,0.4896,0.003122,-0.2205,1.581,-0.8054,-2.733,0.9449,-0.01211,-0.6373,-1.053,0.643,1.155,0.861,-1.121,1.255,-0.5745,0.4598,-1.913,-1.184,-2.066,0.5064,1.429,0.04034,-2.118,1.822,-0.3695,2.302,1.041,1.266,0.1655,-0.3765,0.9725,0.3797,0.7292,-1.219,-1.92,-1.114,-0.4638,1.102,-0.17,0.3796,0.9155,-0.2345,0.7744,0.9485,2.187,3.09,-0.5935,-1.292,2.653,0.6183,0.7424,-0.9474,0.927,-0.5347,-2.253,-0.9292,0.6134,0.03716,0.7853,0.8259,1.119,-1.941,0.8585,-0.4584,-0.02091,0.6295,-0.829,-2.035,0.1036,0.4773,-0.3717,-1.308,0.2746,-1.316,-0.6852,-0.9451,0.252,-0.5879,0.4572,-0.9936,-0.1102,0.3305,-0.152,0.08737,1.441,-2.579,-1.493,-1.421,1.916,0.2063,-1.54,-1.613,-0.3551,-1.909,-0.291,-1.099,-2.908,1.591,1.451,-2.579,-0.9217,-0.1931,-1.507,-2.297,-0.07839,-0.5295,0.1389,1.37,-0.6423,-0.3912,-0.1165,-0.9655,0.1377,-1.955,-1.313,-1.371,-0.9131,-1.119,2.616,-1.973,-1.468,-0.2185,-0.1965,-0.862,1.714,-1.408,-0.7808,-0.1541,-0.5814,-0.7938,-0.6935,-0.6216,-0.8607,-0.5859,1.706,-0.01256,1.284,1.809,0.8294,-1.342,-1.494,-0.9055,-0.1882,0.2772,0.5732,-2.153,-1.059,-2.358,-0.04517,2.411,-0.04435,0.6755,-1.709,-1.112,-0.2358,0.7456,-1.753,-0.9702,-0.4282,2.407,0.822,0.278,-0.1542,0.0201,1.189,-1.344,-1.471,0.8681,0.9854,0.9601,0.08743,1.239,-2.315,0.6185,-0.3332,0.4628,-1.677,-0.4804,0.367,-1.437,-3.5,0.3049,1.635,-2.595,-0.9781,-0.247,-0.2392,0.8622,-1.258,0.8087,0.06145,-0.3239,0.6228,-0.5573,-0.7412,-2.082,0.3546,-1.066,0.5984,-0.7337,-0.6352,-1.298,-1.814,1.781,-0.4198,-0.9021,1.121,-1.17,-0.9119,-0.9131,1.691,-1.656,-2.371,-0.6804,-0.8057,1.047,-1.22,1.178,1.819,0.2323,-1.014,-1.221,-3.449,1.051,0.1044,1.344,0.5835,0.2347,-1.393,0.0513,0.6559,-0.1684,-0.6563,1.011,-0.5234,0.5384,-0.132,0.5407,-1.075,-0.6232,0.03856,0.5499,-0.442,0.9218,2.95,-0.7474,-1.107,-0.8046,1.084,-1.47,-1.312,-0.5227,-0.7107,-2.12,-0.6705,-2.412,1.378,-1.136,1.327,2.079,-1.912,2.158,-0.5652,0.1713,-1.883,2.594,0.4521,-0.1049,1.783,-2.367,-2.091,3.72,3.523,3.628,6.433,0.9747,-1.086,-0.2626,-0.4783,0.01372,0.6132,-0.9194,0.6708,0.1204,-0.8623,-0.9712,-0.639,-0.03364,1.161,0.05279,-2.574,0.7825,-0.3757,4.102,-1.003,-3.184,-3.771,-0.08892,-0.3372,0.2568,-0.05656,-0.4431,0.7884,0.9135,0.7491,0.4576,-1.466,-0.07781,-1.655,-1.198,-1.219,-2.059,0.8117,0.2214,-2.142,0.1637,-0.08982,-0.9736,-1.299,-2.015,-1.879,-1.287,-1.569,-0.1479,0.8561,-1.086,-1.497,1.037,-1.1,-1.828,0.3218,0.2935,0.2696,-0.3763,-0.2667,0.2541,1.173,0.9667,-0.93,0.4978,-2.005,-0.9168,-1.254,1.143,-1.316,1.342,-1.721,-2.469,0.8462,-0.8176,-0.02573,0.2761,0.8049,-3.458,0.09813,-0.5167,1.038,1.824,0.9733,-1.068,0.24,1.565,0.8337,0.9686,0.8293,-2.79,-1.002,-0.9383,1.098,1.221,-1.037,0.4183,0.3946,-0.07624,-1.939,-1.653,-1.056,-3.336,0.1787,-1.688,-0.7224,-2.485,-2.187,1.792,0.3437,0.6529,-0.01695,-2.051,1.194,3,0.2848,-1.147,-0.143,-0.3203,-0.6437,-0.2861,-0.8769,1.405,-0.8474,-0.6818,1.037,0.638,-0.3873,-0.3577,-0.8957,0.8096,-1.88,-0.4191,1.894,0.1765,-0.2511,0.1866,2.257,-3.688e-05,0.8665,-0.7657,-0.6397,1.191,1.058,-0.9822,0.6044,-1.218,1.229,-0.758,-1.521,-0.8704,0.2013,-0.6777,-0.1714,0.6026,-0.7656,-1.554,-1.105,1.565,-1.532,-1.159,-0.9708,0.7096,-0.1018,-0.8565,1.037,1.17,-0.51,0.05873,1.103,-1.44,-2.145,0.05051,0.1997,0.2667,-0.6332,0.4148,-0.9201,3.203,-1.662,-2,0.8505,-0.5917,-1.179,-0.906,3.817,-0.7692,-0.9004,-0.203,-0.8997,0.4118,1.438,-1.984,0.1683,-0.5075,-1.087,0.435,-0.2933,1.227,0.3736,0.8884,-0.5552,-2.825,0.04766,1.029,-1.37,0.2463,-0.006601,-1.072,-2.552,-0.3862,1.418,-1.112,0.4224,0.7736,-0.6113,-0.3524,0.2786,-0.8237,5.211,3.601,3.541,4.144,-2.996,4.468,-0.9485,-0.9863,-1.355,-0.5146,-4.853,-0.7908,1.582,2.485,0.8971,1.475,-4.21,1.07,-2.384,-0.1217,-2.543,-1.974,1.097,-3.232,1.464,2.25,-1.891,-1.927,-0.1194,0.03901,-2.02,0.4019,-2.452,-1.408,0.9433,-0.5469,-0.5451,-2.04,-0.7997,-0.7352,1.042,-1.545,0.5265,-1.338,-1.615,-1.649,-2.195,-1.59,-1.2,0.05638,-0.1894,-0.1666,1.325,-0.1605,0.3754,0.7038,-1.397,-0.6092,-0.1009,-1.021,-0.452,-1.26,-0.969,-1.255,-0.02367,0.1272,0.4891,1.013,-1.783,0.5552,0.4588,0.5296,-0.07604,-1.765,-1.116,-1.114,3.703,-0.2614,-0.6629,-1.414,-2.758,0.4644,0.9181,-0.6589,0.3403,-2.136,0.9882,-0.08332,0.6503,-1.084,0.9777,1.303,0.1166,1.453,0.3748,-0.545,-2.434,3.373,0.5195,-0.2812,-0.8608,0.9152,-0.3965,0.3381,-0.4526,-1.534,-0.6936,-1.638,-1.14,-0.1458,0.4182,1.184,-0.9582,0.4757,0.5502,0.6457,1.014,-0.4347,-2.197,1.117,-0.05896,-0.7848,0.7519,0.5982,0.23,0.6632,0.2258,-1.044,2.714,0.9197,-1.654,0.6017,-0.6526,2.737,2.032,0.122,-0.5455,0.1273,-0.8672,0.3611,-0.1148,-1.36,-0.6479,1.098,1.055,0.2492,-1.73,-2.325,-1.481,-1.095,0.1524,1.638,0.5233,-1.126,-0.1653,-0.7613,1.398,-0.1657,-1.16,-0.2806,0.9288,-0.1603,-1.304,-1.407,0.2809,-0.2049,1.157,0.7373,0.7541,-1.925,-0.6947,-1.316,-1.202,-2.62,1.626,-1.751,-2.286,-0.5666,-3.796,1.246,0.6587,-1.374,0.7947,0.1078,1.552,2.165,0.3251,0.8255,-0.2311,0.7373,0.03426,1.378,0.05887,1.107,0.2695,-0.7458,2.854,3.017,0.3136,0.9485,-0.4746,0.8701,2.168,-0.7422,0.04871,-0.6806,-0.09876,-1.721,-1.315,-0.1416,0.3227,1.3,-0.9728,-1.344,0.2222,-1.484,-1.657,0.3055,0.8791,-2.263,0.9266,0.1742,1.242,0.9568,-0.7365,0.6351,0.405,-1.52,0.8648,1.151,0.1661,0.7143,-0.5952,0.266,-2.425,1.961,0.7585,-1.41,-0.3909,-0.8792,1.538,1.175,0.6787,0.02483,-0.4641,-0.9914,-0.3118,2.625,-0.7251,1.006,-2.101,-0.01847,0.05127,1.199,1.842,-1.978,0.152,0.998,0.1145,-1.713,1.991,-1.547,0.569,0.9927,1.522,0.6224,-0.4335,-1.721,1.563,-0.6212,0.4588,-0.5893,-0.4282,-1.198,-0.1948,-0.5884,0.5341,1.667,0.5865,-1.24,1.116,0.6466,0.4946,-1.488,0.09154,-1.033,0.1948,0.6669,-0.5064,4.643,0.01619,-1.038,1.394,-0.4191,-0.7312,-0.4881,-2.946,-0.05443,-0.554,0.891,-1.065,1.117,0.836,-0.7055,-0.4764,0.2413,0.622,-0.3076,-0.4737,-2.847,1.495,1.777,1.307,-0.5742,0.1761,0.9181,-0.1107,0.6267,-2.394,1.77,-0.989,1.615,0.7251,0.7091,-0.5596,-0.4611,-0.8464,0.505,-0.9928,-0.7458,-1.138,3.302,0.2504,1.993,0.004261,0.07543,-1.125,1.758,1.134,-0.778,-0.3763,-0.8418,-0.5122,-2.126,0.6447,1.999,-3.66,2.877,1.738,-0.9741,1.106,1.757,-1.194,-2.65,6.005,-0.9338,1.303,-2.835,-0.2295,1.567,-0.957,4.828,-3.973,3.367,3.284,-2.811,-1.416,-6.351,-5.664,-3.031,-4.033,-3.552,-4.653,-4.954,-5.536,-4.118,-6.319,-5.304,-4.37,-7.642,-5.605,-5.06,-4.579,-4.169,6.387,-4.656,-4.758,-2.759,1.85,2.233,-4.936,-3.678,2.125,-3.206,-3.699,-2.367,-5.977,-4.073,-2.021,-2.997,-2.285,-2.841,1.577,2.584,2.626,-1.934,2.643,2.921,3.796,4.214,1.539,2.154,3.213,2.585,2.037,4.892,2.55,-1.965,1.591,2.949,-2.777,2.648,1.565,4.343,1.672,2.008,-2.184,2.021,-2.035,1.831,4.157,1.545,3.036,2.379,4.645,1.711,4.394,2.982,3.266,2.187,-3.616,1.949,2.662,-2.009,2.556,3.67,3.272,4.664,1.442,-4.391,-1.963,3.274,2.98,2.638,2.437,5.202,2.475,2.372,2.544,2.114,4.723,2.975,1.999,-3.958,-6.248,-2.073,-7.745,-4.241,-2.587,-2.568,-4.244,1.662,-1.371,4.472,3.448,3.223,-2.423,4.415,3.034,-3.674,3.952,1.928,3.449,-4.84,5.181,1.399,-2.949,1.249,-1.143,2.015,-3.127,2.032,-2.59,4.889,1.483,6.897,5.222,-2.907,3.309,2.682,-2.221,1.814,2.093,3.509,1.858,2.777,4.071,-3.789,-2.999,-3.183,-3.741,-2.931,-3.061,-3.475,2.949,4.261,-2.537,-3.771,-7.314,1.746,3.651,2.809,-2.179,1.896,-1.515,1.463,4.856,-2.496,-1.851,-3.106,2.028,2.898,2.425,3.408,5.055,9.663,5.607,-2.578,5.753,2.008,-6.58,3.849,5.814,-4.692,-2.322,-3.372,4.806,2.407,2.075,2.143,3.207,11.87,10.46,4.232,2.202,2.765,2.805,3.415,2.06,2.096,2.948,3.11,1.512,1.376,3.315,1.235,2.725,2.508,3.695,1.681,4.559,2.234,1.83,1.889,-2.14,2.653,2.356,2.597,1.855,5.87,1.344,5.203,2.848,-3.633,4.327,4.433,2.15,5.025,3.523,1.644,-2.746,-2.932,-1.615,1.498,2.435,3.807,1.945,3.366,2.983,3.194,-3.278,-3.762,4.798,2.643,4.165,2.948,1.335,2.898,5.573,6.039,2.381,5.397,6.936,3.987,2.119,1.707,2.996,2.328,4.376,2.638,4.513,3.032,1.378,-4.094,4.156,-5.567,2.57,2.267,4.177,-1.659,-4.088,1.493,3.951,3.856,-2.791,1.774,3.274,3.94,1.951,2.485,1.493,3.018,3.038,1.773,1.729,3.216,2.322,4.549,-1.464,-2.868,-3.654,3.76,1.863,-4.464,1.025,4.599,3.327,1.388,5.036,-2.569,-3.595,5.474,2.268,3.141,1.551,1.455,-4.043,-3.417,-3.298,-3.689,1.515,-2.258,1.662,-5.398,2.846,-4.227,2.11,4.973,4.572,4.142,2.804,3.73,-1.286,1.966,-1.779,1.994,4.66,3.766,4.322,2.406,-4.812,-3.92,-4.09,2.972,-3.086,-5.229,10.39,3.129,3.221,-3.636,-3.816,-4.514,3.64,2.518,3.935,2.034,3.47,5.754,7.393,6.175,-2.605,-3.323,1.543,3.017,3.667,5.775,3.075,3.831,2.34,1.408,4.448,1.811,6.181,-3.599,-5.135,7.902,-6.303,-3.795,-3.325,2.882,3.522,4.548,2.863,4.973,1.871,2.295,4.789,2.516,-1.786,-2.169,2.242,2.883,4.265,4.344,-3.428,4.701,-2.498,-3.102,2.525,-2.103,-1.66,3.203,3.966,2.191,2.423,-1.793,-1.308,-3.09,-2.804,2.221,2.753,6.826,3.533,-2.808,3.519,-2.275,3.454,2.074,2.337,-1.719,1.727,1.557,-2.607,3.037,-2.773,6.144,3.784,-5.232,4.134,2.935,-3.271,2.03,4.347,1.747,4.086,-5.359,-1.656,-4.63,4.919,6.925,5.19,3.838,-1.568,4.431,1.924,-2.471,-2.914,-4.202,-5.372,-3.825,-3.165,-5.645,-2.93,-2.412,2.369,-3.31,3.731,4.561,3.819,3.124,-1.328,2.112,-2.011,-2.81,1.379,-2.521,-4.043,-5.746,-3.685,-4.88,2.613,-3.501,-2.406,-3.574,1.377,-3.532,-1.853,-1.641,2.67,6.994,3.07,3.908,4.407,2.446,2.525,2.006,2.883,-3.972,-2.35,1.632,2.836,-1.367,2.15,2.737,4.75,3.625,2.897,3.833,6.678,-1.366,1.896,-2.057,-2.36,3.145,2.486,3.717,-3.491,-3.155,-1.182,1.851,4.96,-5.549,-6.448,-3.776,-5.909,-4.624,5.574,-3.64,-3.528,-3.386,-2.386,4.632,2.788,7.086,5.067,1.702,2.686,4.272,5.135,3.806,5.112,2.934,2.627,3.038,1.931,2.995,2.609,1.861,2.029,4.173,5.694,-4.694,2.476,1.398,2.014,2.478,2.065,3.474,3.273,3.638,2.884,1.415,3.762,-2.814,3.516,1.51,3.428,-2.533,-4.923,-3.617,3.815,2.33,2.302,3.517,-2.241,-3.29,-3.688,-2.475,8.774,2.919,6.998,-2.991,-2.913,1.162,-6.758,2.091,2.109,2.968,3.078,6.685,4.076,-1.977,4.472,3.086,-4.414,-3.964,-2.799,-4.217,3.41,4.773,-2.496,4.236,3.084,-1.079,5.764,2.673,5.586,3.718,-3.015,-1.642,-2.707,2.259,2.481,2.862,2.061,-1.771,3.087,1.418,1.681,2.589,2.484,2.547,2.399,2.763,4.69,2.386,-2.654,1.698,1.484,1.196,-4.665,-5.725,-2.761,-1.536,1.5,-1.683,-4.932,-3.124,-6.507,3.243,-4.269,-5.497,-3.178,-9.41,-4.107,-4.987,-3.877,-5.452,-5.038,2.411,-1.991,-2.88,4.016,-1.854,-2.884,-1.341,4.343,-3.014,-5.5,-5.717,-5.659,-2.034,-3.085,-4.151,1.638,-7.066,-4.42,3.292,-4.97,-2.62,-3.101,-6.657,-3.586,1.375,1.888,-2.76,-2.355,1.248,1.526,3.554,5.188,2.052,-3.845,3.052,2.214,-1.438,-2.352,2.037,3.77,4.055,2.535,-2.213,3.984,3.052,-3.558,-4.631,1.652,1.809,1.496,5.047,-2.385,-2.265,3.345,4.264,3.357,2.038,1.016,-2.069,-2.683,2.013,3.985,3.417,1.522,2.39,3.346,-7.393,-1.511,-4.5,-2.368,4.283,-2.167,4.538,3.991,-1.251,6.375,3.783,1.083,-2.223,-2.292,2.623,2.499,-4.953,-3.678,-3.04,-4.552,4.043,2.198,1.727,-4.184,3.571,1.49,-3.218,-2.79,-2.586,3.231,2.916,4.108,1.869,5.04,-2.417,-2.829,1.757,-1.669,-2.386,-3.636,2.21,3.278,4.737,4.058,-4.256,-3.181,-6.202,4.397,5.108,1.704,2.113,-5.417,-2.692,-3.458,2.185,3.411,6.7,8.109,-4.914,2.444,3.246,4.812,-3.089,-4.119,1.412,2.828,3.917,1.837,2.422,3.391,3.076,2.473,1.77,3.976,2.834,5.549,-4.053,-6.879,-6.295,-4.867,2.175,5.499,-2.709,-4.709,-6.415,-3.132,1.972,1.494,-1.803,2.663,-3.22,-2.611,4.323,-1.788,-2.625,-3.461,-1.177,2.283,-2.665,-3.484,4.72,-3.02,-5.342,2.224,-3.892,4.071,2.674,1.507,2.306,-1.655,4.494,1.195,3.129,5.356,-4.288,-4.52,2.421,3.409,2.356,-3.994,-3.045,-4.688,-3.398,-3.832,5.644,1.87,-3.424,-2.52,-3.369,-3.582,2.327,5.122,-3.028,6.323,3.969,3.079,-3.363,-1.864,2.342,-3.691,5.476,4.064,1.881,-4.099,-3.823,-2.585,-4.801,-2.34,-4.097,3.341,-4.292,4.863,-2.909,2.319,-2.747,-5.16,-2.624,3.976,4.138,3.04,3.466,-2.53,2.558,-3.279,3.437,1.629,-4.555,-4.444,-2.442,3.723,-2.163,-4.167,-2.312,3.535,-3.741,2.555,1.559,1.487,1.678,-2.815,-1.819,2.565,2.648,-4.223,-2.811,2.566,-2.996,3.827,6.409,1.285,-1.873,1.969,-1.643,1.511,-2.303,2.172,2.758,-4.185,2.778,4.534,-2.781,1.244,-4.051,2.195,4.473,1.815,2.049,-2.639,1.997,2.493,3.14,2.738,-2.897,3.878,-2.905,-4.894,-1.748,4.637,-1.651,3.127,3.764,1.636,-4.155,-3.279,-2.383,-3.492,4.321,6.286,-3.491,-3.602,1.97,2.068,2.881,-2.194,3.19,2.593,2.194,-3.127,-1.843,-3.316,-1.931,2.788,1.499,1.883,-3.349,1.306,3.713,2.8,2.501,7.215,-2.394,-1.636,4.329,2.097,-1.82,-2.439,1.579,1.563,1.457,-4.574,-3.352,-6.098,-2.33,1.792,-5.298,8.798,5.573,-3.68,-3.718,4.268,-5.804,-6.046,-4.397,-6.717,-1.692,2.702,1.456,3.354,4.94,-1.906,-2.811,1.843,3.311,3.404,-4.198,-4.886,-5.416,-1.3,2.656,-4.143,-6.025,2.673,4.088,-6.757,-2.871,1.553,-2.768,2.072,5.158,-3.096,2.426,-2.903,-5.801,-3.209,1.29,1.93,-1.112,-2.522,-2.169,-3.307,-2.775,-2.673,2.82,-3.786,1.118,-3.912,-4.745,2.341,-1.724,-4.205,-3.901,-6.252,4.588,-2.855,1.649,2.411,3.378,1.731,-2.641,4.356,1.727,-2.495,2.081,2.926,2.627,3.11,4.116,2.518,2.01,2.034,2.522,2.433,3.449,2.515,2.655,1.686,3.468,2.786,4.254,1.693,-2.079,-2.346,-5.106,-4.876,-4.573,1.123,4.845,4.065,5.652,2.555,1.779,-1.789,-2.468,2.96,-2.575,5.59,1.599,1.229,3.933,-3.115,3.833,1.607,3.868,2.248,2.424,3.389,3.05,-1.887,6.307,4.233,2.824,4.551,2.116,-4.227,-1.658,-2.225,-5.878,-2.634,-1.732,-3.33,1.934,1.626,1.551,1.837,3.624,5.454,2.066,3.315,2.837,-3.84,1.809,2.006,-1.653,-8.624,3.605,-4.704,4.747,-1.823,-2.032,2.686,-2.506,1.416,1.372,-3.062,-2.271,-4.711,2.36,-4.37,1.872,-3.381,-4.242,1.338,-5.018,2.473,-1.456,-2.443,-1.687,-5.747,8.985,-2.423,-1.349,2.465,-2.895,-5.114,-1.591,6.806,2.609,-2.787,-4.031,3.362,2.264,-2.615,-2.169,-2.873,3.152,4.799,-4.728,-4.196,3.735,-2.988,-2.571,2.781,-1.883,2.286,-3.933,6.082,-5.802,-2.101,1.563,-1.2,3.826,2.37,-3.776,1.745,-4.398,-4.78,1.433,6.758,-6.11,1.882,2.759,2.453,-4.557,2.487,-3.906,-4.037,-3.403,1.893,-2.825,-3.812,-4.559,-3.948,-3.252,-3.209,2.488,-4.17,2.274,-3.083,-2.329,-2.412,-2.175,1.612,-2.757,-1.437,-1.738,-2.367,-1.679,-6.113,-3.224,2.121,-2.755,-4.897,2.183,-3.23,-3.376,-3.288,-3.495,1.646,-3.397,-1.909,-2.034,-1.377,2.663,5.04,2.568,-2.435,1.971,2.738,-3.598,-4.352,-4.939,3.088,-2.782,-3.545,1.945,2.124,1.603,-4.192,1.085,2.331,1.526,3.115,-1.472,1.835,1.225,1.521,-1.585,1.068,-2.599,4.246,-2.876,4.996,1.223,3.524,2.352,2.402,-3.133,-2.613,4.458,3.507,6.931,2.002,2.679,2.636,2.622,2.993,2.19,2.396,-1.653,2.949,-3.064,2.456,4.446,1.831,2.289,-2.914,3.065,7.45,2.581,-2.358,-3.237,3.413,-1.543,5.682,-1.892,2.44,3.818,3.145,4.164,1.665,1.952,2.123,2.203,3.071,5.366,-2.205,5.227,2.502,4.153,1.9,-5.222,3.643,3.032,-1.752,2.723,1.577,2.645,3.204,1.389,1.987,-7.125,-4.517,1.507,1.235,-3.435,-3.207,-3.534,-3.662,-1.818,6.604,-3.253,-2.341,-5.832,-4.018,-2.969,-4.436,1.686,1.964,-2.618,1.361,-1.966,2.595,2.417,2.647,-2.36,2.138,-3.52,-2.64,-5.864,1.124,-3.16,2.117,2.581,-2.653,-3.66,-5.019,-2.73,-2.6,1.302,1.337,3.691,-1.773,2.647,-2.389,-3.658,-2.113,-2.224,-2.795,-2.9,2.88,-5.565,-1.333,3.968,-3.724,-3.429,3.419,-2.719,4.272,-5.84,1.602,2.474,-5.635,1.776,1.735,1.559,-1.582,-3.256,-6.631,-1.878,-2.569,3.167,-4.701,1.452,-2.435,4.882,-3.226,3.539,-2.597,-3.933,3.566,-2.614,-3.353,2.617,-2.877,-3.392,2.09,1.674,6.459,2.063,2.91,2.238,-2.041,-4.351,-5.896,-2.198,-3.007,1.772,2.267,-1.826,2.499,-2.122,1.193,2.828,-4.256,-2.539,1.387,1.941,-1.239,7.705,1.489,-2.81,-3.689,-4.889,-4.262,-2.688,-6.067,1.591,-1.418,-5.019,3.102,2.288,-3.677,-5.432,-2.427,3.248,1.609,1.893,1.39,1.693,4.215,1.898,-2.141,-1.904,-4.577,1.265,-4.352,-3.563,-1.564,3.361,3.378,8.179,-1.775,2.661,2.476,2.138,3.108,2.496,3.389,-2.68,-2.225,-1.982,-2.307,-3.169,-2.313,-2.564,-6.486,5.106,2.975,2.015,6.389,1.141,3.606,-3.502,-1.852,3.607,1.867,2.481,4.07,1.512,-1.568,-2.769,-4.362,-1.985,3.831,-7.688,-6.375,-4.84,-7.779,-2.48,3.379,-4.451,-8.578,2.937,1.977,-2.937,4.121,3.894,1.845,2.659,3.167,-3.492,-3.826,-4.375,-2.831,-4.031,2.218,2.574,2.007,-1.876,-1.938,3.259,-2.205,2.06,2.449,-3.443,3.729,2.261,2.77,-3.744,-4.826,4.433,1.737,1.612,4.043,-1.961,1.613,1.217,2.302,2.226,2.333,1.641,2.487,1.322,2.967,-3.241,1.452,1.039,3.14,1.761,2.479,2.188,3.353,4.159,2.431,5.066,2.405,2.417,-1.599,-3.039,1.987,2.46,1.746,-3.103,4.815,3.716,2.145,3.18,3.015,2.7,1.951,-3.952,-7.564,-3.165,-4.675,3.497,3.197,-2.8,-2.048,-5.655,-3.714,-3.382,-3.869,-4.201,2.711,-2.506,1.93,-2.51,-2.979,-1.372,-2.624,-1.705,2.478,-2.48,-1.703,5.639,-2.829,-3.149,1.565,7.718,-3.282,3.05,-1.696,-4.302,-3.199,-1.588,-2.595,1.77,-4.676,-4.326,2.147,1.144,-1.776,3.498,2.612,-3.223,-3.291,-2.574,-2.914,-4.018,-2.141,-3.584,-3.717,1.715,2.129,-7.145,-5.278,3.362,2.002,2.943,9.65,-4.408,-1.18,-4.766,2.795,-4.293,-6.405,1.429,-2.643,2.343,2.954,6.331,3.348,3.725,-1.736,-2.185,-1.744,-2.414,-4.161,1.554,3.414,-2.2,1.523,3.165,3.162,-3.757,2.874,1.666,3.017,3.81,2.626,1.376,1.486,1.931,3.214,2.62,-3.004,1.238,-2.057,1.676,2.112,3.773,1.342,2.687,3.166,3.158,2.076,6.783,3.307,-4.291,-4.732,1.797,-5.265,-3.477,2.182,4.455,5.097,-1.319,1.525,-2.796,-6.628,-3.982,-3.293,2.094,2.937,5.594,3.395,2.124,3.564,2.624,5.063,3.001,4.019,5.17,2.736,1.777,-3.463,-3.298,-4.87,3.831,1.81,1.536,2.11,1.795,2.653,3.366,-1.852,4.73,-3.394,-4.392,-2.962,2.098,1.781,4.718,1.982,-2.771,1.6,-3.731,-3.816,-1.821,2.618,-3.535,2.105,1.465,3.279,5.729,5.214,-1.867,2.365,3.354,1.968,1.015,1.343,3.907,1.954,-1.977,2.308,-2.906,2.447,1.638,-2.786,-4.043,4.102,4.149,2.547,2.198,-2.585,-2.892,-4.43,-3.313,-2.682,-2.046,-5.415,-4.365,-4.839,-3.63,3.562,-5.124,3.753,2.289,2.911,-2.112,-3.281,2.926,1.197,-5.247,2.134,1.741,-5.562,-3.031,2.088,3.261,2.053,-2.498,-3.719,-2.903,2.476,1.854,-1.743,1.566,-3.035,-5.316,-4.247,-4.215,-2.289,-3.27,-2.968,-4.725,3.395,-3.49,-2.912,1.968,-4.041,-3.285,4.052,-1.628,-5.21,3.793,1.901,-2.899,3.15,-3.791,-3.205,-2.301,-5.295,-4.413,-1.633,-3.386,-2.701,-5.55,-2.736,-3.308,-1.599,3.054,-3.107,1.578,3.743,-1.688,1.389,1.189,-4.112,-3.104,-1.372,1.781,-3.503,10.85,-5.773,4.929,-3.849,2.396,3.305,-4.373,4.236,2.085,2.237,1.512,-4.297,1.632,2.527,-2.824,2.415,-1.751,-3.887,-3.816,1.42,-2.3,2.718,3.164,-1.89,-3.457,-5.329,-3.548,-3.712,-2.862,-2.606,3.288,3.118,-1.91,-2.25,-2.576,-4.531,-3.005,-4.097,-4.227,-2.282,-3.334,3.401,-1.386,-1.321,-1.503,-1.471,1.442,-3.035,-7.649,-3.548,-2.25,-1.523,3.833,2.453,1.162,-3.591,-3.811,-2.585,-3.926,-1.808,-3.572,2.556,1.731,3.414,4.269,2.738,2.831,2.095,3.508,3.167,-2.83,-3.773,1.572,3.202,1.912,-3.414,-5.033,-4.508,4.482,-3.293,2.042,-3.509,1.52,1.811,-2.987,1.614,4.171,3.985,2.054,-2.158,2.125,7.159,6.562,2.106,-4.447,-2.085,-1.968,3.246,-1.781,3.053,-1.955,-1.851,6.208,1.095,1.402,4.071,2.811,-2.63,5.354,8.399,-2.124,1.829,-1.743,-1.71,-1.595,1.583,-3.726,-3.056,-2.727,-3.606,1.064,-4.009,-1.554,-1.612,-1.751,-1.563,-2.993,-4.001,-2.89,-3.737,-2.91,3.819,-2.855,-3.812,-4.33,-1.432,2.127,4.814,1.72,1.586,2.12,-5.365,-1.547,-1.968,1.76,1.841,3.513,4.051,2.177,3.656,-1.647,1.451,5.403,-2.917,-1.575,-2.865,2.176,1.94,1.517,4.651,-2.285,2.242,2.975,-2.428,1.171,2.228,-1.44,-6.54,-2.094,-5.737,-1.071,1.906,1.536,-2.361,-2.107,-4.343,-1.799,-1.446,1.622,3.468,-1.071,1.305,5.742,4.658,3.031,2.202,3.075,4.952,4.335,2.354,-3.237,2.552,1.373,-4.362,-3.536,-4.136,-2.139,-2.316,4.087,-2.798,1.846,2.265,2.036,-1.946,2.686,1.402,1.538,-2.953,-2.938,2.5,-1.753,2.525,3.63,-1.762,2.268,2.326,-4.071,-4.448,2.761,-2.852,-2.6,1.663,1.699,2.156,-2.128,2.8,3.256,-3.599,2.219,1.739,-2.916,-2.117,4.047,-6.544,-4.656,5.1,-1.351,-3.149,-2.684,-3.388,1.379,1.06,-4.483,-4.32,-5.371,-6.065,-2.736,-4.768,3.135,-3.07,-4.75,-1.441,-4.567,-3.253,-1.226,-5.281,-2.82,1.771,-1.974,-2.465,-2.517,-3.934,-2.635,-1.59,6.031,-3.838,5.518,1.804,2.417,2.977,3.214,2.279,-2.05,-3.819,-4.26,3.537,2.439,2.777,-3.858,2.435,1.973,1.901,2.057,1.674,5.172,-3.402,-2.523,1.965,-3.224,-3.321,4.372,-5.058,-4.508,2.184,-2.167,-2.569,3.568,2.702,8.494,-7.265,6.753,7.623,-3.578,1.482,1.95,-2.15,-3.666,-2.105,3.011,2.867,1.863,3.358,4.95,1.921,-3.216,-5.226,-1.543,-2.727,-1.969,-2.832,-3.074,-6.202,-2.561,2.964,-2.525,2.709,-3.569,-3.609,-3.053,-3.679,-5.109,1.331,-1.479,1.708,-3.028,-2.748,-2.637,-2.895,1.589,-3.541,-6.733,-5.105,-2.571,-4.352,1.163,3.442,-3.952,-3.401,-2.071,-2.125,-3.319,8.296,10.63,-3.387,2.817,-2.423,-2.376,-2.012,-1.595,-2.724,-3.373,-4.43,1.326,-3.316,1.279,-2.31,1.94,-3.548,-3.816,-3.145,-5.784,-1.848,1.929,2.364,2.208,-3.684,3.7,2.855,3.691,-3.079,-2.258,2.83,1.713,2.751,-1.99,-4.517,-3.787,-1.935,2.694,-1.744,-1.321,-5.633,-4.194,-5.96,4.84,2.458,3.683,-3.241,-1.388,1.88,-3.859,3.432,3.168,1.278,1.932,-1.843,-8.047,-2.297,-3.019,-4.421,-3.647,1.482,4.539,6.473,3.144,1.517,-2.893,-2.08,-4.528,-2.274,-3.341,2.561,2.982,-4,-2.471,-2.05,1.4,6.884,-3.126,-1.421,-5.318,2.307,-2.497,4.321,1.547,-4.828,-5.214,-1.282,-4.252,1.391,3.225,2.033,-2.893,-2.718,2.064,-1.456,7.244,1.742,-3.325,-2.137,-3.295,-3.652,-5.692,-4.216,2.618,2.067,-1.673,-3.148,-3.785,1.873,1.612,1.841,1.426,12.12,5.567,1.729,-2.704,3.001,2.141,3.951,4.191,3.79,-2.063,1.03,-2.397,2.134,-1.457,2.61,-2.461,-1.864,-2.5,-3.729,1.548,6.023,-5.013,-5.665,-3.359,-3.161,2.839,-1.723,-1.881,2.79,3.342,1.518,1.664,2.63,-4.556,-7.431,-2.768,2.497,1.109,1.636,-2.529,-2.574,1.795,2.89,3.018,3.885,2.433,1.525,-1.6,-2.253,2.765,3.072,-2.871,1.639,3.333,4.162,4.066,-2.316,-3.849,2.899,3.832,1.758,2.725,1.692,1.789,5.646,-2.721,1.85,3.826,-1.324,-3.123,4.497,2.984,2.44,-5.429,1.739,-3.954,2.247,3.089,3.87,2.09,1.443,-2.316,3.133,1.647,3.502,4.78,3.101,1.841,-4.883,2.561,1.817,2.814,2.2,2.482,1.649,3.452,-3.269,4.438,4.314,5.336,3.062,1.017,1.536,2.725,2.61,-1.608,-3.737,6.33,1.245,6.015,4.55,2.411,-2.31,1.417,-3.468,-3.321,1.86,-1.497,1.723,3.154,2.914,1.732,2.382,-2.408,-4.63,2.256,2.993,3.427,-2.772,-3.165,-4.911,-3.622,-3.702,-3.092,-4.531,1.271,1.755,2.157,-2.534,3.655,5.618,1.335,4.266,1.892,2.387,2.476,1.676,-2.911,2.561,-3.112,2.669,3.551,1.685,1.843,3.547,1.844,2.367,2.914,-3.396,1.857,1.245,4.177,2.613,-6.527,2.766,1.555,1.59,-4.165,2.725,1.396,2.538,2.456,4.038,2.727,3.678,-3.304,2.65,-2.024,1.499,-5.159,-3.283,-3.838,-4.411,-6.16,-8.522,-6.708,-9.396,-7.736,-4.939,-6.803,-6.293],"logCPM":[0.2335,0.8914,-0.909,-1.044,6.033,0.6133,5.003,2.106,1.12,0.944,-0.1883,1.833,4.629,-1.05,0.4613,-1.034,-1.221,-1.113,-1.154,-0.8025,-0.5506,-1.087,-0.8912,5.846,-0.4062,-0.5014,-1.269,-1.082,-1.037,-1.018,-0.8162,-0.4439,-0.3194,-1.205,5.047,-1.333,-0.6471,-1.063,-0.8348,-0.4968,-1.243,-1.325,4.202,-0.7709,-0.6452,-0.6829,-1.271,-0.9345,-1.305,-0.8035,-0.477,-0.8719,-0.6047,-1.128,2.171,0.5397,0.4155,-0.783,-0.4198,-0.7101,-0.5802,-0.5516,-1.319,-1.205,-1.325,-1.031,-0.6486,-1.2,4.419,-0.2541,0.05658,-1.063,0.7089,-0.1543,-0.4205,0.4831,-0.1943,0.1023,0.05213,-0.2669,3.389,-0.5134,-1.082,-0.6948,-0.9267,-0.1904,-0.6631,-0.1987,-1.139,-1.238,-0.8462,-0.6023,-0.7576,-1.405,-1.111,0.8912,-0.1736,-1.264,0.5787,0.7213,0.2753,-0.9965,-0.6714,0.7875,-0.7072,-1.029,-0.8471,-0.8726,0.3965,-0.1988,-0.7048,-0.551,-0.9555,-1.18,-0.9394,-0.8912,-0.6887,-0.176,-0.919,-1.01,0.7173,-1.227,-0.7852,-1.107,-1.042,-0.4406,-1.168,0.09461,-1.041,-1.188,-0.1335,1.254,-0.6179,-0.7133,-0.799,-1.319,-1.325,-1.405,-0.7855,-0.97,-0.3408,-0.4832,-1.294,0.2788,0.2009,-0.02345,-1.019,-1.405,-1.04,-0.9892,-0.9012,2.021,-1.214,-1.389,-1.061,-0.004036,2.9,0.3679,-0.1631,-0.484,-0.7328,-0.7121,0.2777,1.72,-1.294,-0.6928,0.5968,-0.5475,-0.2769,2.635,3.691,3.512,-1.055,-0.2817,-0.2713,-1.017,1.501,-0.9366,-0.7041,-0.4596,-0.6979,-1.31,-1.422,-0.6012,-0.406,0.5244,4.756,-1.066,-0.7452,3.143,5.357,2.96,-1.322,-0.407,1.489,5.436,5.915,1.124,5.629,1.73,1.548,2.064,8.008,4.384,5.624,3.914,8.249,5.372,8.884,7.692,9.62,6.055,8.267,5.312,6.881,5.048,6.251,8.939,8.391,5.735,1.803,8.226,7.331,0.9543,3.065,4.275,3.248,6.765,5.882,5.94,7.764,6.145,7.112,5.176,3.633,4.351,6.354,6.233,9.297,5.317,5.782,6.491,4.562,3.286,4.061,5.415,4.386,4.269,4.22,3.575,6.997,8.224,4.795,5.361,4.467,4.534,3.532,4.068,5.947,4.625,4.252,4.399,6.297,6.71,3.375,4.455,2.918,6.066,4.663,6.862,5.842,4.813,4.956,5.534,5.253,5.036,5.788,5.157,4.996,1.67,7.221,0.4254,3.378,4.533,2.296,5.005,0.4293,0.6483,-0.7163,-1.076,-1.052,-0.9892,-0.2229,-1.034,-1.134,0.1661,-1.045,3.891,-0.546,3.617,2.56,1.102,4.077,3.08,3.104,3.098,2.269,2.475,4.622,2.379,3.375,-1.061,8.992,3.426,5.883,6.601,2.784,3.553,4.193,5.447,2.604,3.916,7.624,1.716,4.469,4.836,5.842,5.345,5.331,6.543,2.988,4.069,5.1,6.137,6.579,7.977,5.403,3.214,5.154,6.952,6.241,3.602,4.943,4.739,5.256,5.124,5.824,7.492,6.323,4.696,3.672,4.219,7.163,4.604,5.142,6.417,5.723,5.319,6.221,5.109,7.831,6,4.422,5.276,2.524,5.559,5.538,6.388,5.941,4.615,6.968,7.765,3.313,2.965,5.959,4.855,7.962,3.731,6.638,1.583,0.833,5.021,4.321,0.7227,3.016,4.527,2.873,1.395,3.758,1.785,1.264,-0.7617,-0.3822,-0.7949,-0.8444,-1.107,-1.062,-0.196,-0.1219,-0.867,0.008575,-1.038,-1.186,0.2391,0.5017,2.557,1.408,-0.9892,-1.205,-0.286,0.2555,3.083,5.513,6.26,3.005,3.266,7.157,3.747,6.064,7.282,8.149,6.371,8.6,4.288,6.034,4.691,7.215,7.748,7.576,6.27,5.286,5.151,5.072,5.491,1.489,3.964,3.189,5.795,6.125,4.536,5.706,5.701,4.476,5.328,4.886,5.967,5.045,6.892,6.499,5.154,3.587,5.367,2.365,6.331,7.947,6.849,2.933,3.747,4.17,3.094,7.713,4.936,4.03,4.641,4.955,5.979,6.287,3.011,4.387,5.229,4.514,5.287,7.699,7.711,4.025,3.998,4.018,4.571,6.031,1.862,5.804,8.075,6.15,5.793,3.389,4.755,6.192,4.153,3.364,3.867,4.377,5.811,5.823,5.253,6.824,5.569,7.018,6.801,2.606,3.752,-0.7945,-0.9865,-1.227,-0.8058,-0.9701,-1.2,1.406,2.522,0.5532,0.1818,1.461,1.537,1.363,0.9126,0.7756,3.739,3.951,5.608,6.425,4.426,5.044,4.864,5.487,3.596,4.583,5.472,2.778,7.291,8.008,6.918,5.547,4.711,4.274,4.166,6.346,6.806,5.667,4.463,7.671,5.335,4.613,1.631,6.294,3.008,4.149,4.267,5.225,5.254,5.064,4.646,5.763,7.457,5.664,6.457,5.67,5.011,4.42,5.231,6.845,4.834,4.844,6.122,6.565,6.856,8.343,6.805,3.176,6.61,4.865,4.079,8.303,2.473,5.517,4.643,7.531,0.86,3.67,4.562,-0.1508,6.423,1.665,4.891,4.993,6.378,4.391,3.281,5.585,6.487,4.064,0.9202,4.546,2.325,3.998,6.578,-0.0695,-0.6377,0.5699,-1.183,-1.135,-0.8169,-0.5826,-1.152,2.021,1.576,3.407,2.826,3.438,3.14,3.25,2.561,1.735,7.501,3.813,6.944,5.377,4.813,6.223,5.356,5.93,2.973,6.699,2.026,5.388,5.504,5.753,8.201,6.85,7.281,8.53,9.161,2.438,2.726,3.947,4.02,4.798,5.933,6.8,7.423,5.03,6.737,3.992,2.391,3.553,5.738,5.156,6.074,7.153,6.155,7.885,5.18,6.386,5.263,4.169,5.403,6.303,4.653,6.419,7.239,5.522,5.942,4.982,5.158,4.587,5.976,7.645,3.252,3.092,4.263,6.517,2.989,1.435,3.312,2.191,4.507,1.582,4.414,1.331,4.592,4.48,3.202,10.18,5.524,5.191,5.258,2.799,3.582,3.643,4.133,3.928,7.398,6.128,3.064,5.048,5.477,3.842,2.793,3.465,8.217,4.84,4.492,5.654,3.444,6.664,5.484,5.653,5.623,3.153,3.513,0.7383,4.123,0.01928,2.865,4.046,2.371,3.607,2.279,2.944,2.551,1.61,4.286,3.427,0.1291,3.161,4.06,6.792,2.81,3.344,5.635,6.798,4.01,8.603,4.274,5.924,4.847,4.001,6.594,4.78,5.909,7.033,6.632,7.048,4.426,4.751,3.445,3.024,6.345,1.953,4.79,4.397,5.218,5.169,3.236,7.544,6.606,2.228,2.78,1.823,2.403,4.376,1.555,0.7006,4.778,-1.178,0.1361,-1.099,-1.181,-0.8574,-1.325,-1.122,-1.118,-0.9769,-0.4607,0.1144,1.002,2.9,2.915,-0.04987,6.805,3.406,2.819,5.341,4.681,5.523,6.329,5.219,4.084,6.207,2.076,6.499,3.577,5.32,2.615,4.965,4.899,5.212,5.187,5.228,6.255,8.201,4.803,4.817,7.276,4.383,3.604,3.786,6.357,4.204,3.803,5.454,4.357,5.114,3.759,7.446,7.688,5.343,6.251,8.019,0.2897,4.377,6.729,2.095,4.232,5.48,10.77,2.397,4.787,5.819,5.99,6.595,5.185,5.019,6.627,4.463,7.538,6.241,3.426,3.043,5.905,7.163,4.281,4.18,4.684,6.933,4.831,3.792,6.578,5.289,6.6,4.903,4.167,6.195,4.053,5.281,6.025,5.487,6.701,7.744,7.692,7.288,2.385,3.483,5.486,4.714,4.436,4.98,6.027,5.865,5.535,4.085,4.076,4.461,5.875,6.971,6.704,5.326,5.171,3.923,3.668,3.729,0.8394,-0.1203,0.02446,3.715,1.323,1.658,1.282,-0.4155,0.1425,-0.4194,2.647,0.2375,2.271,1.613,6.886,3.394,3.48,2.436,0.3118,8.454,6.344,7.788,10.43,9.125,4.64,8.011,7.129,7.275,5.093,4.27,5.899,6.525,3.638,3.891,5.597,5.102,3.498,10.23,5.854,6.153,6.2,9.712,5.932,5.559,6.192,6.01,4.903,5.588,6.886,6.215,5.886,4.847,5.335,5.179,1.379,7.863,6.658,3.938,6.724,3.684,7.056,7.072,8.585,6.224,4.632,4.462,4.066,3.809,3.904,5.1,6.075,5.414,4.637,2.536,4.41,3.774,4.907,3.398,5.248,4.355,4.883,5.106,5.529,5.417,9.376,5.769,4.572,4.819,4.824,2.359,3.769,6.84,4.172,4.215,8.191,1.237,4.471,3.741,4.736,3.085,5.604,4.208,7.139,5.346,6.516,6.219,8.869,3.848,5.606,4.774,5.206,6.635,2.804,6.044,7.203,6.38,4.704,6.856,5.217,4.122,5.05,6.669,6.907,5.821,5.118,3.575,4.014,8.129,7.17,4.582,4.852,5.531,2.715,4.463,6.182,6.066,5.913,6.2,6.245,6.358,6.096,5.905,0.3767,3.685,4.54,7.705,5.471,4.618,8.026,3.914,6.138,7.194,4.784,3.704,5.391,5.193,4.403,4.694,7.132,5.583,6.21,2.989,0.9769,4.818,4.434,5.226,3.434,2.982,5.587,5.531,3.384,7.138,6.219,4.699,9.421,1.466,5.908,6.383,4.955,7.237,3.212,6.038,3.941,2.558,6.302,5.33,4.409,4.377,4.677,5.811,2.059,3.481,3.944,5.592,7.257,4.62,5.949,2.968,4.077,3.95,7.086,3.371,4.562,4.589,3.33,7.449,4.316,4.024,4.913,1.198,3.732,-1.166,-1.218,-0.3903,3.516,-0.3931,0.8305,0.2322,-0.2245,-0.7174,5.277,1.613,5.764,-0.3163,4.843,3.707,5.37,4.544,5.391,2.103,4.161,5.956,7.005,5.978,2.15,8.293,7.758,2.908,4.796,5.607,6.653,1.403,3.647,5.827,5.986,4.679,6.285,0.529,2.711,0.3706,4.791,4.556,4.277,4.836,9.34,4.756,5.559,4.985,6.602,4.492,2.338,8.106,8.964,8.892,1.618,4.372,5.775,2.795,5.794,5.811,5.947,5.343,4.744,7.88,6.049,7.115,5.81,4.11,3.481,1.246,3.185,7.113,3.499,6.51,6.39,7.996,9.456,6.494,5.546,3.178,6.367,7.217,6.74,6.727,5.791,7.296,4.64,8.51,10.15,3.628,8.041,4.36,8.226,5.196,2.741,7.797,6.352,7.535,7.19,2.34,4.896,7.92,7.262,6.76,4.866,8.69,7.44,3.512,7.331,4.428,6.851,7.308,4.812,6.299,6.599,4.693,6.303,5.773,6.205,3.687,3.571,5.965,5.576,8.095,7.064,3.354,7.297,4.187,4.888,5.435,4.392,4.819,4.649,6.261,4.375,4.618,4.174,7.643,5.585,4.395,5.996,4.966,4.228,6.153,3.307,6.213,2.066,5.093,3.115,5.274,5.24,5.444,5.369,6.938,4.248,5.828,5.666,3.952,4.152,6.468,4.906,1.968,4.104,5.649,8.087,6.623,5.982,4.636,6.762,7.082,5.695,8.953,5.406,3.427,6.362,5.015,4.149,5.012,3.089,4.748,5.09,6.819,6.353,6.094,5.087,5.714,7.066,2.164,5.172,6.697,4.391,5.984,6.899,7.554,8.659,2.763,6.491,6.884,5.266,2.86,3.153,3.087,0.1806,-0.1605,-0.1313,1.274,0.5407,-1.07,-0.851,-0.1846,-0.936,-1.18,-0.94,-0.05853,-0.06275,1.87,-0.1456,-0.793,0.1674,-0.5127,-0.6933,-0.2115,-1.103,-0.8784,-0.8847,0.2962,3.304,4.469,8.033,4.429,7.891,2.683,6.354,4.702,1.835,5.174,5.276,2.482,5.932,5.634,7.038,5.48,2.939,5.155,4.847,4.389,1.179,3.019,6.745,5.799,8.701,5.916,3.054,4.592,7.612,2.686,3.381,6.023,6.403,4.904,7.498,2.44,6.42,5.945,6.158,6.925,5.166,4.856,8.009,5.871,7.354,6.592,7.308,5.323,3.849,4.2,3.415,8.112,4.176,2.04,5.087,1.83,5.832,7.816,1.238,1.726,5.321,6.209,6.136,7.363,5.122,5.85,3.74,7.765,5.775,4.543,8.739,5.93,4.981,3.421,3.913,7.514,5.412,5.395,7.45,5.077,1.759,3.07,2.073,4.786,5.757,6.967,3.131,5.076,7.575,7.034,6.69,6.55,5.281,7.298,7.39,8.708,3.413,7.304,3.949,4.327,2.933,5.145,6.876,4.414,5.592,5.564,5.775,3.422,6.621,7.397,3.58,7.894,5.879,7.486,3.765,5.543,6.177,4.936,3.517,6.648,6.232,6.741,5.538,4.985,6.792,4.195,5.86,6.691,8.985,5.885,4.21,3.586,3.327,4.068,8.408,6.735,4.503,6.743,2.632,3.902,2.282,7.607,6.24,6.069,5.869,3.977,6.128,4.777,5.404,5.941,9.097,4.01,7.999,6.378,4.369,3.384,9.213,6.435,1.827,5.362,2.899,5.978,4.465,6.756,5.381,6.558,7.122,4.139,3.567,3.306,4.812,7.555,1.919,2.117,3.692,10.04,4.394,3.319,5.307,5.257,0.6893,6.078,5.598,5.207,8.029,8.949,6.164,4.126,3.591,5.414,5.565,1.996,3.312,7.918,5.689,5.863,6.492,5.236,7.245,5.698,3.001,4.344,3.996,3.766,4.286,6.033,6.369,4.921,8.395,8.201,8.739,4.738,5.701,5.473,5.07,5.627,1.704,5.415,2.309,2.866,2.704,8.25,5.505,7.302,7.844,6.097,6.423,6.116,7.674,4.813,1.807,0.4207,0.3368,3.909,-0.09173,0.7734,-1.093,0.3691,-0.9727,-1.141,6.349,5.079,10.97,2.98,7.873,4.813,6.1,7.92,5.101,9.612,8.641,1.63,2.947,4.437,7.246,8.494,8.435,5.569,3.903,8.925,6.39,7.304,5.982,7.832,6.679,4.566,5.81,3.408,8.611,4.239,4.393,5.875,7.936,5.628,4.794,7.355,4.396,4.738,6.246,6.608,5.213,7.818,6.775,4.633,0.9542,4.902,6.369,4.998,6.547,3.744,5.102,5.803,3.235,5.348,6.826,6.382,5.741,3.793,4.455,6.477,4.379,5.271,3.262,0.5856,5.325,2.902,5.524,5.676,5.201,3.075,2.505,4.469,6.82,6.359,4.788,3.776,2.69,4.959,4.538,5.336,2.905,5.101,5.271,3.56,4.512,4.414,6.456,5.579,5.338,4.255,6.73,4.303,5.174,3.793,2.837,6.523,4.538,10.43,2.459,3.808,2.881,7.244,-0.5344,5.676,3.575,7.99,5.702,4.588,2.957,5.436,6.18,6.297,6.772,5.159,4.707,3.481,3.883,4.741,8.354,7.256,5.179,5.193,3.745,5.087,5.165,5.806,4.902,5.196,7.372,6.135,2.469,6.527,6.018,1.417,1.161,8.368,5.553,7.84,5.685,6.459,8.179,6.844,5.104,4.916,5.136,7.533,5.749,5.526,6.516,4.326,7.614,6.733,6.29,3.367,3.404,2.935,2.514,6.188,5.391,6.248,3.181,1.367,5.827,-0.5714,2.552,-0.2882,1.755,0.7686,5.504,-0.6448,-0.2334,-0.4631,-0.9949,-0.9279,0.7312,1.932,3.499,1.882,-0.07521,2.742,6.855,7.312,5.391,3.211,6.571,3.495,5.521,2.007,4.022,3.234,8.397,5.048,6.462,2.736,4.808,5.313,4.518,4.761,6.065,3.097,6.767,5.808,4.222,3.858,6.647,4.712,5.272,6.623,3.704,5.882,4.733,7.958,4.958,6.951,6.749,4.91,6.962,4.167,3.87,9.005,5.474,9.568,4.859,1.027,2.075,10.09,6.711,11.44,8.201,5.938,4.823,4.417,6.84,3.348,2.906,7.134,6.842,5.643,4.156,7.121,7.207,7.103,7.476,2.603,7.195,1.788,5.372,6.955,4.114,2.306,5.948,5.472,6.052,9.928,5.238,5.633,4.829,7.432,6.251,5.592,1.569,4.42,6.4,10.62,7.7,7.028,2.309,3.968,10.12,3.667,7.163,7.306,5.524,2.637,3.783,9.002,2.709,4.88,9.662,7.957,3.133,5.742,6.919,4.897,3.694,8.934,2.281,6.736,5.769,6.584,5.962,5.087,2.227,3.823,10.54,3.891,6.269,2.043,6.349,3.503,4.311,6.337,6.83,4.481,4.517,6.72,6.173,5.652,4.949,4.876,6.202,5.832,6.343,4.792,-0.1024,7.043,2.632,7.519,5.669,4.213,6.316,5.398,6.017,7.512,10.63,6.681,4.479,4.688,4.504,3.586,7.092,5.855,5.955,5.848,4.954,6.855,5.937,7.913,5.737,4.173,4.138,4.886,6.256,6.742,6.039,5.485,5.402,6.831,4.79,6.034,6.384,4.035,5.944,6.027,7.211,6.486,5.582,4.14,6.108,1.607,4.191,6.431,2.39,4.623,3.737,6.845,0.3132,4.511,1.821,7.212,3.469,8.072,8.207,2.532,3.92,6.034,6.277,5.094,4.794,6.206,4.742,4.628,1.213,6.72,5.244,4.096,4.988,6.94,6.72,3.606,7.886,6.445,2.922,3.899,5.152,4.873,5.176,3.899,6.234,4.861,4.428,5.76,4.063,1.992,4.391,1.949,5.377,3.787,6.615,4.769,4.774,6.401,5.926,6.635,4.391,0.2436,1.858,3.446,1.96,4.767,3.968,0.8829,0.3518,-0.8686,-1.04,0.07887,-0.6,-0.8321,-0.8271,-0.5751,-0.3915,-0.1634,-0.447,-0.7435,7.004,4.184,2.284,1.643,2.736,6.863,8.082,4.287,6.1,5.432,8.274,5.554,6.617,2.264,6.98,5.378,4.95,5.342,3.988,7.204,0.1589,5.894,6.474,3.776,7.289,4.351,4.744,5.5,6.452,3.34,6.473,3.661,8.786,2.449,3.141,4.544,5.806,4.659,4.259,7.048,3.593,8.597,5.429,6.018,7.031,5.975,6.112,6.179,4.918,7.241,7.698,5.25,5.513,9.067,6.322,5.859,6.977,2.827,0.7966,3.727,5.686,6,1.086,4.543,4.746,2.313,4.997,6.549,1.828,6.292,5.95,5.115,2.402,4.165,5.704,7.281,6.197,7.101,3.477,5.164,7.955,5.732,6.299,5.706,5,4.387,4.26,6.196,0.6223,6.413,5.644,6.535,2.705,4.395,5.007,6.004,5.599,4.28,6.063,7.34,6.936,7.643,5.999,3.905,3.687,5.429,3.177,5.981,5.418,5.632,6.928,4.707,3.552,4.585,6.382,6.65,7.003,5.813,3.865,6.622,2.676,7.203,4.865,5.712,4.443,3.07,5.2,6.879,5.503,5.057,3.328,5.376,6.271,4.692,5.775,4.99,5.631,1.311,4.355,6.114,4.162,4.14,5.185,4.606,5.062,5.937,7.107,4.729,1.404,3.538,3.573,7.251,5.252,7.17,4.716,3.706,3.031,3.92,2.859,3.986,3.412,3.465,5.703,4.284,7.542,5.559,3.497,6.194,5.706,4.478,7.034,7.316,5.738,3.852,6.675,6.93,6.503,8.526,2.295,2.747,5.87,4.961,4.592,2.599,3.509,3.288,7.548,4.874,2.024,2.427,4.784,6.334,0.05197,8.653,4.115,4.028,3.288,5.524,3.435,6.213,6.625,5.813,7.137,6.736,4.044,7.337,3.278,4.651,8.157,6.881,6.028,5.567,5.283,5.883,7.616,3.816,5.004,3.17,6.37,4.789,5.899,7.79,4.221,5.585,5.26,8.106,3.792,5.021,9.31,5.694,8.118,7.18,8.695,5.712,6.65,5.932,6.418,5.848,6.622,7.706,5.604,2.526,2.427,2.978,5.822,7.742,2.55,6.495,9.665,7.073,0.5304,7.146,5.874,5.057,4.826,6.937,9.228,6.426,4.022,7.965,5.903,6.204,6.02,2.907,6.22,6.3,5.642,6.861,7.478,1.867,5.267,6.041,4.516,5.872,6.78,4.343,7.199,7.303,5.149,6.671,1.878,1.323,4.039,5.043,8.593,9.918,7.088,9.434,5.378,6.589,6.321,3.61,0.821,6.163,2.994,8.195,2.584,8.304,6.635,6.033,7.691,6.194,7.076,8.541,9.247,7.709,5.876,6.238,5.757,4.61,6.026,8.262,7.217,6.924,6.016,4.524,5.615,3.686,8.637,5.414,6.165,9.015,7.014,5.147,8.428,3.466,6.562,4.424,7.579,5.126,6.564,4.932,4.536,2.096,4.87,1.258,2.938,0.2456,-1.336,-0.3075,1.899,-1.167,-1.135,-1.096,0.4836,3.712,8.076,5.578,6.106,8.06,7.514,5.947,7.334,3.996,7.542,3.351,5.66,3.484,6.105,4.816,2.748,8.727,6.971,3.799,2.164,1.911,0.2956,4.977,6.423,5.689,3.809,9.515,8.374,6.879,4.929,8.259,5.936,5.27,5.552,3.488,3.734,2.438,8.36,5.246,2.121,5.449,4.455,4.176,5.132,4.515,5.028,6.799,4.849,5.338,5.744,6.037,4.598,8.745,5.461,3.647,10.29,8.498,6.316,6.628,6.463,5.484,7.993,8.546,3.833,3.805,3.659,6.12,6.896,7.696,4.453,5.21,5.825,3.215,9.437,5.283,4.183,6.586,9.913,1.371,4.758,6.826,5.609,5.302,5.712,6.894,7.214,5.152,6.098,6.584,3.5,3.65,5.032,4.53,6.704,4.298,6.628,4.007,4.416,4.378,2.347,3.016,5.445,2.646,4.205,2.19,4.015,4.042,1.505,8.36,7.847,9.275,8.914,4.497,7.528,-0.5858,4.887,6.031,3.081,0.657,4.497,5.921,8.784,5.946,6.153,6.463,5.329,5.559,6.181,3.021,4.847,3.793,5.029,6.169,4.431,5.577,6.336,8.518,4.269,4.111,7.693,5.884,3.264,6.788,7.426,5.926,11.88,3.745,6.739,6.984,4.19,0.9584,3.653,5.105,6.827,1.984,3.427,5.744,7.538,2.68,3.682,5.506,4.574,6.787,6.386,7.541,8.013,5.679,3.317,1.512,2.445,2.446,4.51,6.323,5.974,6.638,6.155,7.252,7.402,3.785,8.013,1.97,8.491,8.857,6.904,5.701,1.357,7.617,4.826,7.474,4.439,5.906,7.753,4.073,6.197,5.068,5.292,6.654,3.461,6.389,6.706,5.628,6.817,3.553,4.43,6.756,4.633,3.793,6.384,6.768,2.262,4.211,1.568,3.503,6.683,5.668,6.811,6.625,8.641,2.637,-0.3133,-1.231,-1.253,0.09074,-0.7202,0.8534,2.329,1.427,3.06,3.131,2.865,2.936,5.985,1.171,2.457,5.35,4.055,1.696,4.872,7.108,4.451,5.616,4.136,2.118,4.588,2.955,7.234,2.216,8.757,8.221,5.77,6.65,3.976,4.424,10.23,3.655,6.003,0.434,7.216,6.976,7.563,4.801,4.866,5.03,3.181,8.703,1.115,3.266,7.038,5.526,6.47,6.352,6.766,4.459,5.67,6.316,6.715,7.803,7.472,5.221,7.491,6.045,3.829,5.95,8.787,2.832,7.269,7.059,4.781,6.997,4.995,3.656,4.853,4.129,3.949,5.616,3.828,8.788,7.064,4.617,4.692,7.009,5.222,6.428,7.495,4.477,6.786,6.258,5.824,5.211,6.325,2.734,6.243,4.827,5.878,4.245,4.921,3.262,6.675,6.06,7.483,6.457,6.768,8.022,4.242,4.044,6.405,4.083,4.812,7.711,0.4922,6.48,8.102,3.347,8.096,5.754,8.864,7.022,4.844,5.574,5.28,3.786,5.245,6.866,1.591,9.095,7.363,7.523,4.182,8.082,4.12,4.048,4.98,-0.3449,1.4,6.413,7.189,6.027,6.62,7.873,10.08,4.294,5.574,6.785,9.22,6.001,7.206,2.581,4.473,4.98,7.847,5.412,9.073,9.516,5.87,7.621,11.51,7.449,6.994,5.261,2.171,7.658,5.495,6.224,6.728,6.152,7.599,10.31,5.194,2.435,4.135,8.508,7.301,4.757,2.656,5.476,3.449,7.596,1.078,7.025,2.33,5.012,4.797,5.653,4.744,3.992,3.374,6.722,7.187,4.009,4.446,7.386,7.727,8.192,3.546,5.818,3.804,-0.005438,2.956,6.143,4.955,8.73,0.8933,5.794,4.334,6.927,6.728,4.619,4.925,4.673,5.885,5.866,4.599,6.369,6.675,4.022,3.303,1.843,5.421,4.151,6.799,5.343,6.609,5.569,6.402,6.461,5.717,5.299,6.153,6.462,5.412,6.354,7.098,5.273,4.505,7.614,5.916,3.387,5.311,4.164,4.26,5.245,5.346,6.76,6.808,5.976,2.74,5.263,3.824,4.606,4.442,5.202,5.309,5.907,6.836,3.969,5.729,6.923,5.684,3.415,4.809,4.874,5.028,5.868,5.599,5.551,5.773,2.675,4.893,5.796,7.647,7.314,5.851,5.245,7.304,6.959,7.794,3.328,6.369,5.422,4.23,8.238,7.088,4.328,4.281,4.651,7.647,8.114,5.006,1.981,9.504,5.795,7.462,4.781,-0.2875,4.814,2.704,3.872,6.313,4.412,3.829,9.684,5.904,5.162,4.099,6.252,6.119,5.849,5.159,3.433,5.423,6.747,4.955,10.69,5.763,7.66,4.898,3.339,3.514,6.56,5.709,6.447,3.25,4.303,4.325,3.095,5.238,7.213,1.699,8.017,6.575,3.199,5.094,5.11,4.867,8.105,4.231,4.169,5.335,5.546,1.385,5.501,1.885,4.258,-0.1346,2.226,2.589,1.175,0.5323,-0.05413,1.183,-0.1004,4.283,-0.2792,3.4,1.926,-0.7782,0.7088,0.1006,0.6227,-0.312,3.915,0.4019,-0.7813,-1.322,-1.125,-0.8562,-0.06173,-0.4989,-0.6876,3.163,6.624,4.231,-0.06131,0.7128,4.859,4.004,7.292,3.866,0.8699,1.358,0.5138,0.9867,6.243,10.03,2.894,1.297,5.732,5.346,5.571,9.505,5.214,9.581,7.531,7.836,5.342,5.951,4.829,6.467,7.129,8.528,3.723,6.5,6.39,12.91,4.469,8.06,7.465,8.249,9.335,9.322,7.931,8.203,8.325,7.499,10.11,5.86,5.897,6.496,7.186,4.136,5.728,7.672,8.205,6.167,7.425,5.996,7.027,6.017,6.249,11.66,6.469,8.254,6.517,9.883,7.821,6.771,7.03,6.635,9.387,2.168,7.21,9.749,6.029,6.685,5.872,8.769,8.668,6.911,6.211,6.271,7.478,6.912,8.411,4.553,9.915,6.833,6.531,8.186,7.543,9.797,9.578,6.076,5.787,4.42,3.992,4.812,0.7968,4.563,5.149,5.101,7.522,6.588,9.602,7.88,7.631,3.811,7.522,6.381,3.676,6.075,7.104,10.44,5.5,10.07,5.641,4.128,7.111,7.026,5.512,2.441,9.817,4.02,10.81,8.196,11.06,7.682,6.404,6.67,8.041,7.397,7.181,6.807,9.258,9.458,8.081,9.489,5.439,3.948,3.907,6.68,6.57,3.547,4.026,10.08,9.642,4.941,4.272,2.65,7.001,9.028,8.809,6.408,11.06,9.493,9.017,9.31,5.007,6.48,5.247,8.954,5.613,5.666,7.495,9.743,10.2,10.1,8.126,9.258,8.13,10.9,11.96,10.36,4.228,4.503,4.071,6.982,6.746,7.409,4.705,9.026,13.09,12.8,4.66,6.882,7.345,8.554,8.506,5.287,6.163,8.339,6.083,9.402,6.902,9.52,7.667,8.212,5.922,8.608,6.474,8.777,7.482,4.441,7.065,6.908,9.024,6.912,8.061,7.657,11.15,7.657,6.143,7.801,6.416,6.281,7.616,5.221,5.581,9.19,6.207,5.183,6.365,5.807,6.323,7.43,9.362,8.023,4.816,5.908,8.476,7.302,4.683,6.079,7.3,8.863,7.641,6.631,7.232,5.96,6.614,5.69,4.625,9.108,8.794,7.991,7.853,7.51,6.889,5.218,4.108,9.402,5.271,7.423,8.474,10.24,6.228,7.569,6.595,7.413,5.537,6.567,5.958,10.68,9.634,3.875,6.136,7.606,8.11,7.904,8.785,7.704,5.854,5.138,5.967,8.191,8.078,7.937,8.551,6.607,6.463,3.319,8.091,6.171,3.535,8.8,9.098,7.529,6.079,7.506,5.075,4.664,9.602,8.59,9.385,6.518,6.479,4.595,5.286,4.401,4.341,8.701,5.916,8.701,3.121,7.714,5.644,9.458,8.648,5.978,8.985,5.293,7.065,6.26,6.499,5.616,6.706,8.548,8.123,9.013,7.707,7.693,8.386,7.905,6.286,8.282,7.72,10.97,5.059,9.815,5.114,4.805,2.291,8.2,7.29,8.859,6.476,4.695,6.546,3.731,2.144,4.465,7.61,7.723,4.767,10.14,7.814,7.353,9.769,6.463,7.278,8.608,7.305,8.125,6.233,4.524,1.749,3.19,2.879,6.4,9.154,10.66,8.87,8.105,10.14,7.614,7.408,8.804,7.261,7.734,5.786,4.414,7.803,6.72,8.232,4.364,9.053,3.658,4.936,9.051,5.55,5.624,10.2,9.669,7.61,6.934,5.875,8.71,5.668,5.235,9.597,6.538,8.769,7.308,3.828,8.568,4.665,5.413,6.565,8.36,5.869,7.296,7.358,3.768,7.876,2.966,8.584,8.332,4.406,14.91,7.605,4.914,9.833,9.739,7.641,8.873,3.345,7.458,2.782,5.893,9.818,9.09,4.593,6.005,6.671,7.317,4.993,5.504,4.904,3.996,5.301,6.668,6,5.006,7.795,9.143,4.794,8.359,7.715,8.39,8.359,6.965,6.471,5.535,7.682,7.251,3.52,3.03,2.573,3.384,2.852,7.497,2.98,4.871,3.439,7.647,5.192,5.263,6.694,6.872,11.02,9.798,9.213,8.762,6.465,7.014,5.42,7.677,6.844,5.112,6.68,8.148,5.769,7.706,10.17,10.49,9.014,10.69,8.034,7.158,6.537,7.071,5.867,6.348,8.799,7.334,6.091,4.242,5.373,7.658,8.615,9.224,3.955,4.014,2.602,7.431,4.447,2.116,5.655,4.111,6.137,3.914,6.947,7.526,14.25,10.71,7.117,7.192,9.019,8.64,6.638,6.964,6.174,6.116,8.992,8.181,7.065,7.566,5.43,5.52,7.766,7.703,4.646,7.978,6.621,7.255,5.716,6.698,5.866,7.608,6.455,7.413,6.502,8.671,4.651,8.475,8.641,9.904,6.447,4.166,7.218,8.947,8.269,8.474,8.828,4.709,4.819,2.168,4.447,4.58,11.19,10.57,4.645,8.138,7.217,3.46,6.116,7.023,6.181,5.799,5.012,10.36,6.912,7.908,7.745,4.981,5.4,8.41,5.664,9.245,5.617,4.447,10.5,7.096,7.09,10.45,8.508,9.408,9.572,6.304,6.249,8.318,8.747,7.282,7.62,6.333,5.809,5.38,7.398,6.926,8.167,7.909,11.31,7.097,7.998,9.669,5.628,4.105,6.929,8.447,7.739,6.514,9.342,3.536,5.474,10.28,5.519,5.42,5.09,6.819,9.147,4.88,6.457,5.898,2.601,5.951,6.29,7.946,0.08755,6.315,7.596,6.17,9.27,10.53,6.062,6.053,6.89,3.468,4.392,4.627,4.573,5.769,5.886,6.345,4.661,10.67,9.932,4.584,9.159,7.817,6.708,4.891,3.191,4.328,6.352,7.858,4.778,5.019,6.448,6.774,7.51,7.036,8.633,8.249,9.8,7.383,5.765,3.987,8.592,11.96,8.426,6.541,5.364,8.794,4.359,5.536,6.665,6.941,6.279,7.318,9.572,5.459,5.14,8.373,9.867,8.702,6.719,8.482,5.063,5.031,8.564,8.94,8.008,7.813,6.44,8.449,7.2,5.731,9.898,9.007,8.334,6.61,8.882,9.976,7.78,6.345,8.784,8.919,6.67,7.78,8.017,7.145,3.222,6.184,4.387,3.767,8.605,9.424,10.16,7.417,9.175,7.983,4.593,4.818,5.396,7.442,8.358,9.976,7.239,10.15,3.717,6.384,6.347,7.678,5.164,5.8,7.247,6.643,1.878,8.777,4.736,5.669,4.397,6.827,9.09,7.095,6.539,9.638,3.337,3.627,6.162,7.689,7.5,11.11,11.67,9.236,7.391,9.713,6.188,5.929,7.674,8.801,6.183,9.118,5.969,4.791,6.318,5.963,8.238,7.538,8.979,7.641,1.405,2.318,9.439,5.815,5.519,3.513,7.544,5.374,2.552,6.853,9.541,6.761,5.713,9.215,4.378,5.764,10.25,5.939,6.101,7.368,6.469,8.627,4.583,3.181,6.331,4.984,4.536,6.705,2.17,10.14,6.939,5.083,10.43,6.543,11.44,9.781,9.573,8.947,4.479,4.139,8.294,9.229,7.623,5.804,4.879,5.015,2.591,6.204,9.467,7.513,5.031,5.26,5.459,5.093,8.839,9.645,6.783,9.113,8.593,9.277,3.617,7.482,5.748,4.07,9.999,5.151,7.799,5.901,3.594,6.33,5.242,7.642,9.453,7.611,4.999,9.477,6.382,6.165,5.979,3.773,7.084,8.734,7.901,6.949,6.698,5.282,5.563,7.404,10.34,7.509,6.311,5.138,6.705,9.119,6.172,3.453,4.68,5.141,7.091,5.371,5.94,6.374,6.516,4.355,6.816,9.166,9.787,4.188,4.133,8.028,6.843,9.845,10.48,8.196,7.121,8.609,7.948,6.558,4.961,6.525,6.755,2.788,6.664,4.84,6.163,7.42,5.918,10.05,8.408,8.362,6.495,3.525,4.232,5.675,4.401,5.788,3.538,6.57,4.988,5.172,5.755,5.89,5.457,5.86,8.876,5.241,4.827,5.2,6.855,5.007,9.288,5.036,3.792,7.148,6.918,7.101,6.893,5.903,7.363,7.653,6.259,6.479,6.394,6.756,5.109,8.219,6.675,9.034,4.615,8.282,7.829,6.696,7.603,14.19,6.659,6.935,6.359,7.831,6.614,4.444,8.013,10.54,6.962,5.999,5.341,5.268,6.229,10.42,6.544,5.98,5.998,2.868,4.305,8.746,10.6,10.46,6.742,7.693,8.942,8.361,6.356,10.28,12.61,5.898,3.826,6.987,6.138,9.002,2.856,5.566,7.999,5.97,6.493,4.896,2.062,6.614,11.51,4.379,5.645,6.418,4.852,7.601,10.12,4.707,5.878,3.959,3.88,3.548,7.411,6.474,8.988,6.857,6.777,3.551,5.727,4.825,9.443,3.772,8.476,4.285,3.792,8.494,9.91,6.694,8.203,6.537,9.497,7.648,9.057,9.692,9.18,6.131,4.724,8.604,7.257,4.737,7.598,7.528,8.005,4.397,3.505,6.076,8.641,5.028,7.295,6.752,8.958,7.836,8.377,8.632,7.021,5.742,8.5,8.274,6.839,8.355,4.254,3.551,4.367,7.015,10.93,8.378,11.18,8.251,7.115,4.959,5.508,8.882,4.364,11.64,7.248,7.482,8.495,4.62,11.14,7.362,8.275,6.769,7.322,7.266,5.843,5.9,3.005,9.673,7.852,10.23,6.302,4.792,10.13,5.568,6.611,9.188,5.774,3.735,8.374,6.102,5.583,5.144,6.787,5.493,7.469,9.178,5.193,6.163,9.855,6.915,8.79,3.152,8.246,5.606,10.76,6.725,6.635,11.67,5.749,8.197,8.365,6.141,6.106,4.635,7.076,2.444,7.6,7.103,7.419,8.173,4.649,8.232,5.719,5.685,5.036,3.746,10.17,7.765,7.142,10.98,5.76,6.04,6.83,8.605,7.514,4.618,3.655,9.041,7.047,4.281,6.419,4.622,9.565,10.38,5.272,4.831,8.106,6.338,6.032,7.516,4.71,8.014,6.614,5.471,3.481,4.64,6.068,7.364,4.417,4.399,4.257,11.82,1.069,4.688,8.449,8.693,0.7663,9.154,9.772,10.53,2.296,6.963,6.726,6.557,5.237,8.845,5.691,3.8,5.097,3.72,4.974,2.603,7.684,4.925,6.363,6.664,7.386,5.263,7.128,9.038,6.253,6.957,6.265,5.857,7.505,7.815,3.897,6.594,9.336,6.685,10.85,5.216,6.013,6.19,8.9,9.06,3.091,7.963,4.398,9.554,4.944,7.282,7.726,6.455,8.004,7.126,6.592,4.488,3.354,9.341,7.565,6.178,7.022,9.812,8.854,5.911,9.244,7.665,6.241,8.625,6.905,7.589,7.304,7.004,8.226,8.188,4.886,5.543,5.953,9.682,6.847,5.704,8.204,7.049,5.66,7.074,7.23,8.525,4.571,4.775,6.431,6.324,7.336,5.24,7.698,7.195,5.971,6.314,7.538,6.701,8.497,6.855,5.025,4.576,11.25,11.31,9.86,9.929,2.323,9.604,6.6,8.626,5.35,8.769,4.48,8.715,6.628,6.44,8.024,8.365,5.756,6.258,7.594,5.619,8.528,9.408,6.453,6.357,1.669,8.975,11.12,6.288,5.879,8.974,7.972,7.999,7.173,5.657,5.245,5.424,6.666,8.163,4.399,4.753,8.082,3.868,8.446,7.855,5.36,5.845,8.773,7.532,6.163,3.798,8.595,5.048,5.748,8.945,4.511,9.093,8.292,8.916,6.236,8.162,3.45,5.062,5.565,8.826,4.676,8.786,6.437,7.38,5.193,3.78,4.067,5.486,7.586,7.668,9.691,6.718,6.445,5.03,3.522,6.796,5.802,4.978,5.713,4.412,5.427,6.32,13.59,5.701,4.604,10.76,5.328,9.094,5.06,7.676,8.473,8.263,9.894,6.858,8.066,5.231,6.363,6.001,6.012,5.096,8.328,4.081,8.884,11.48,11.33,5.225,5.146,3.886,5.169,6.161,4.632,3.041,8.162,5.392,6.119,6.999,5.457,8.333,7.104,8.292,8.349,5.917,5.251,5.526,4.559,4.612,7.478,9.1,7.411,3.897,5.333,9.143,7.619,6.691,4.779,9.039,6.364,7.758,13.62,8.798,3.842,4.863,4.052,4.244,4.402,6.747,11.57,8.966,6.714,9.376,7.92,4.958,2.978,6.363,8.9,6.77,9.625,6.478,7.058,10.37,9.934,5.155,6.43,6.458,6.698,5.444,5.124,5.953,9.53,7.717,8.471,4.931,8.18,7.371,9.009,7.391,7.7,10.2,4.2,4.075,7.653,4.496,6.199,6.122,8.983,10.59,7.878,7.729,6.924,10.46,7.437,9.415,5.388,6.517,7.959,6.36,9.386,8.843,6.275,6.833,6.146,4.053,6.983,10.74,4.7,5.606,5.109,3.504,6.421,9.946,3.404,10.57,7.573,7.36,6.864,6.166,7.326,6.515,7.627,10.06,4.486,7.206,5.279,4.993,5.495,7.423,7.719,7.557,5.293,5.704,9.013,5.112,7.939,8.794,6.209,7.54,7.426,9.548,4.187,1.96,11.51,8.011,8.095,9.539,5.885,5.871,7.299,6.521,7.016,7.153,6.6,8.553,5.994,8.243,6.845,7.912,7.833,10.57,6.561,8.246,7.915,8.171,8.538,8.685,9.093,9.263,5.536,5.789,3.214,6.561,9.279,7.538,6.592,7.439,6.659,8.816,6.898,9.036,7.168,10.51,4.431,5.578,7.269,5.369,9.718,8.047,5.746,6.63,4.891,3.36,4.252,5.124,4.169,8.307,5.732,9.491,7.682,8.101,6.406,5.209,7.396,9.361,6.495,7.449,11.16,7.102,3.124,11.13,10.51,5.422,8.973,5.903,4.758,4.63,8.512,6.896,9.861,4.655,4.299,4.968,7.35,5.518,8.984,8.212,5.792,3.572,7.398,7.952,6.663,5.59,5.945,5.574,9.018,9.06,3.963,5.866,6.668,9.056,8.875,12.46,4.624,8.048,7.594,6.34,5.104,5.873,7.453,4.054,7.445,9.065,9.64,6.088,7.294,5.605,5.016,6.379,5.411,5.965,6.888,6.073,5.593,6.682,6.862,8.865,6.476,7.775,6.934,8.463,10.71,8.209,6.974,6.914,7.453,8.759,8.815,6.746,9.152,5.753,8.714,8.314,11.04,7.368,7.889,8.812,3.973,5.396,11.9,7.649,3.411,4.375,7.386,2.736,8.328,5.947,9.924,9.704,6.401,7.098,6.397,6.407,2.884,6.347,7.346,9.143,9.396,7.188,8.812,8.987,7.606,8.609,7.752,8.501,8.336,9.086,8.147,3.917,5.834,5.484,6.474,6.198,9.143,7.855,6.611,7.143,8.409,4.763,9.467,7.121,5.514,3.866,8.162,6.65,7.135,7.565,3.541,6.582,6.557,6.845,9.028,7.267,5.582,5.776,7.374,10.95,10.4,8.561,5.179,7.539,6.237,6.12,8.063,6.823,4.602,9.07,8.307,9.54,4.585,7.203,6.398,5.994,6.651,9.954,12.47,7.641,7.1,3.837,7.1,5.807,4.022,6.788,5.61,10.77,6.294,4,1.684,10.67,3.011,8.153,8.69,7.784,7.107,5.711,8.985,9.718,4.241,7.753,7.003,4.661,5.871,8.62,9.269,9.622,4.832,2.75,9.325,8.699,7.993,6.043,8.106,4.813,7.908,2.184,8.785,7.642,3.727,4.14,3.336,10.38,6.484,6.601,5.936,3.133,7.304,8.573,9.018,4.554,9.025,8.619,6.616,5.471,4.206,5.337,4.721,6.924,3.779,7.079,5.202,5.137,9.569,4.678,5.176,6.937,9.12,9.776,5.349,10.21,5.269,8.059,6.655,5.275,4.67,5.855,6.406,2.95,11.37,6.43,10.21,3.072,9.009,6.268,7.091,8.802,8.55,7.944,8.824,4.718,7.987,6.255,5.705,9.672,5.691,6.599,4.979,10.34,5.839,8.55,8.638,7.633,4.732,4.714,3.214,7.771,4.313,5.625,7.918,8.448,5.542,7.334,5.79,4.13,5.991,7.238,5.248,5.091,7.514,9.017,7.134,6.691,6.608,6.598,8.671,7.012,5.038,4.506,7.433,6.22,9.642,8.872,8.509,5.165,5.35,5.395,3.187,6.474,5.696,5.177,9.149,8.207,10.85,9.126,8.246,8.653,10.98,7.547,7.657,4.411,7.106,5.793,6.329,6.371,4.066,3.353,8.717,4.994,9.51,5.42,9.057,8.824,5.17,8.002,10.5,4.571,10.36,7.256,8.623,1.116,11.05,7.35,3.199,5.131,6.268,10.28,8.137,9.339,7.62,6.318,11.05,7.867,7.975,6.552,7.502,5.119,7.122,4.121,5.711,8.473,5.558,7.119,6.108,10.14,3.822,5.896,4.57,4.486,8.732,3.599,4.976,6.989,4.215,8.854,6.719,5.071,4.772,6.994,5.089,10.61,4.481,4.845,6.207,6.975,9.019,9.724,7.888,8.78,8.445,3.91,7.126,5.18,6.435,7.222,7.704,6.995,7.012,8.725,6.601,6.413,7.707,4.814,5.493,5.991,4.681,7.427,6.293,8.672,4.008,5.974,7.683,6.226,8.521,6.969,5.437,7.018,7.548,4.377,6.304,8.918,6.524,5.256,5.583,7.503,5.673,5.938,7.836,5.704,8.762,7.572,11.22,7.693,6.15,6.982,9.775,7.404,9.033,9.944,4.895,7.551,6.268,2.463,3.922,2.608,5.101,6.416,9.953,6.87,5.922,7.007,9.031,6.983,9.763,8.168,6.405,4.787,4.744,7.774,8.26,10.02,10.11,5.748,11.44,6.019,4.206,5.267,8.716,5.306,6.493,5.607,7.6,10.96,7.369,7.465,10.64,8.722,8.792,6.934,4.973,6.152,9.211,3.989,4.313,5.575,6.356,5.217,4.816,4.051,7.661,8.076,5.047,5.768,6.027,2.375,6.828,5.52,10.36,5.945,9.416,6.381,4.623,4.78,8.298,5.041,8.012,8.797,7.442,5.89,5.282,4.298,3.514,7.796,4.756,9.284,9.093,7.625,6.135,9.852,8.128,6.228,7.15,4.464,6.141,10.96,9.546,9.32,8.803,7.753,6.975,7.646,9.779,8.736,10.68,4.478,5.345,6.238,6.512,3.982,6.452,7.818,5.964,8.283,7.268,5.103,8.729,8.368,2.348,9.571,1.733,10.83,4.543,8.112,9.154,5.488,7.861,9.283,9.186,6.159,7.708,10.66,11.5,8.947,4.053,6.016,5.755,3.349,6.678,4.119,4.685,6.837,6.181,6.53,4.609,6.867,8.193,6.036,5.635,4.723,7.333,7.068,6.656,6.997,4.284,6.087,4.605,3.863,8.808,6.055,3.478,2.834,4.014,5.161,7.671,10.32,3.95,12.24,7.491,7.171,4.439,2.175,6.974,6.322,12.05,3.707,5.65,5.204,7.015,4.969,5.606,3.861,7.746,5.16,7.162,4.816,9.862,4.466,5.728,4.649,6.197,8.068,9.496,5.608,8.739,6.531,5.192,8.233,9.413,6.147,5.61,9.574,7.936,8.641,7.512,1.871,6.498,9.756,5.896,6.283,6.414,3.018,7.874,1.909,6.843,8.487,9.519,8.68,6.651,9.217,6.908,6.056,9.261,6.591,9.532,6.455,2.347,4.734,3.653,7.739,5.269,9.287,5.969,8.873,9.925,8.369,7.225,6.523,4.664,5.898,7.685,8.872,9.909,3.766,5.542,4.841,10.01,9.425,7.826,7.514,9.399,7.862,7.716,6.95,7.687,2.674,4.058,6.693,3.601,10.01,4.708,8.815,6.761,6.661,5.152,10.19,10.59,5.887,4.906,8.342,3.596,5.481,3.862,3.661,6.614,7.132,6.187,3.625,5.16,4.684,9.558,8.421,7.824,12.37,10.03,8.643,5.407,9.795,8.347,8.37,5.699,10.76,6.602,10.16,4.299,9.478,9.048,11.46,4.87,5.82,6.198,6.774,5.988,9.257,7.131,2.737,5.337,4.804,8.79,6.864,8.541,6.765,4.609,9.319,8.522,10.64,3.497,4.997,5.133,10.2,9.533,7.251,6.371,4.156,5.789,7.09,8.026,7.679,6.89,5.869,8.471,5.812,6.612,5.756,6.445,5.835,8.679,11.05,8.161,5.279,6.097,8.196,7.928,9.039,6.546,6.302,6.084,6.246,3.976,7.634,7.732,6.031,4.936,3.605,9.834,8.16,2.712,6.352,7.365,7.417,8.559,6.38,7.868,5.233,5.82,7.563,7.999,6.502,8.515,8.517,6.795,5.066,5.707,7.378,5.172,7.618,8.742,6.96,7.136,6.24,8.195,8.71,7.247,8.85,7.258,8.475,8.118,8.419,6.507,2.463,8.868,8.444,7.773,9.296,7.893,5.818,8.589,4.622,7.288,7.622,5.845,8.155,8.951,7.666,7.848,8.321,4.705,5.705,8.197,8.295,9.268,5.695,6.775,2.452,5.206,5.075,3.39,0.583,7.576,9.202,9.101,6.616,9.021,8.631,9.081,6.923,7.498,5.789,7.919,6.811,7.267,7.65,5.422,8.849,7.574,6.246,6.304,8.792,5.439,6.303,7.661,4.494,10.24,6.687,9.338,7.366,8.274,10.2,5.633,7.15,4.944,6.499,8.667,5.522,7.228,9.49,6.322,3.744,2.766,8.232,6.127,9.239,10.28,9.496,5.274,7.292,1.968,5.26,1.741,2.307,0.8779,5.838,10.06,9.26],"F":[0.1516,0.4696,1.551,0.2283,1.573,0.6649,0.2841,0.1038,5.275,3.194,0.02906,0.6805,0.01766,1.289,0.007765,0.9089,3.616,4.64,4.596,4.885,5.331,0.01666,4.287,2.983,4.855,1.74e-07,3.428,4.091,1.541,2.36,5.018,6.341,5.82,3.649,5.645,3.438,4.59,4.865,4.839,7.881,3.465,3.264,1.801,0.5492,4.883,5.427,3.36,5.351,3.428,0.1403,4.788,0.1393,0.1253,1.806e-07,0.2055,0.02813,6.512,2.698,0.04119,5.08,5.748,6.477,3.178,3.649,3.264,5.055,4.187,3.927,9.668,0.2786,0.9929,0.3619,0.5473,0.1871,0.8427,0.4718,0.6743,0.04453,1.147,0.1811,8.251,0.2048,4.458,5.11,1.297,2.961,5.94,0.3549,3.959,3.476,3.944,7.137,5.428,2.775,1.034,0.1022,5.03,3.476,5.357,8.666,1.746e-07,4.612,5.868,0.5464,5.685,4.536,6.013,4.322,0.1241,2.838,6.497,7.251,0.1095,3.716,4.21,4.287,2.18,5.056,1.422,4.184,0.003239,3.731,2.749,4.265,2.489,0.2131,4.125,8.803,4.029,0.2809,5.493,0.1386,5.902,2.114e-07,4.42,3.178,3.264,2.775,5.446,4.165,5.653,4.789,3.342,7.75,9.509,1.455,4.035,2.775,4.497,4.125,4.74,7.3,4.397,2.86,4.426,5.246,1.044,3.56,7.428,6.086,1.869,1.454,0.01567,2.636,3.342,2.119,5.573,5.384,3.068,3.297,0.2216,0.05448,2.158,2.802,2.997e-07,5.704,1.739,0.02875,0.06251,6.672,1.367e-07,2.013e-07,0.002786,1.666e-07,4.874,1.065,1.274,0.06157,0.3673,9.268,0.8087,0.5181,3.169,0.2507,0.1008,6.618,4.435,0.3157,0.0737,6.804,8.817,1.122,10.77,6.133,1.948,4.068,2.485,5.68,1.883,0.1866,0.2934,1.059,6.22,0.000337,2.361,4.654,4.012,7.681,0.295,0.3165,3.178,0.5021,8.654,8.199,0.677,4.677,5.12,0.008579,4.066,5.141,10.83,1.782,8.364,1.908,1.411,0.2136,9.158,6.556,1.171,0.1459,4.739,0.6743,6.559,0.01735,1.531,5.172,5.076,0.06654,1.224,2.219,1.309,7.257,8.314,4.866,0.7052,1.775,0.7037,0.2384,1.494,5.261,4.414,1.624,7.077,0.001142,0.771,7.39,3.954,0.9133,2.97,4.631,2.129,1.504,4.629,0.95,2.099,0.05071,0.9005,3.925,0.6669,7.779,6.328,2.621,0.01042,6.803,0.005357,8.356,5.059,3.351,5.187,4.754,4.775,4.125,6.111,5.074,2.258,1.475,4.902,0.1894,0.4904,5.792,0.4536,0.3049,0.1485,5.703,0.006121,1.354,2.497,5.719,3.951,3.427,0.8201,2.264e-07,8.788,5.772,1.51,9.166,0.109,1.507,3.095,7.606,0.6535,0.05807,8.173,0.1676,6.072,5.359,0.3636,1.384,5.265,0.07064,1.108,8.777,1.746,5.896,2.011,7.404,0.97,0.3999,0.1341,7.263,7.563,2.452,0.1717,2.53,0.004675,10.79,0.392,0.07232,5.152,0.3255,2.76,0.06694,4.19,0.9349,9.158,0.6452,7.586,0.002518,3.717,0.8754,2.249,0.0506,0.9392,9.368,0.3266,0.5757,5.806,0.2442,0.6768,8.928,1,14.37,2.844,0.02392,0.1089,4.712,5.606,10.86,3.239,5.606,1.834,0.5919,6.934,0.2579,7.499,5.341,1.345,0.9535,4.302,0.2886,4.569,3.15,2.161,4.92,4.357,4.265,0.02648,7.398,3.751e-07,2.27,6.474,4.015,4.054,7.043,0.1831,0.7652,3.465,4.125,3.649,3.054,5.86,0.01296,4.415,2.447,5.224,6.013,3.645,3.16,2.549,1.867,4.338,7.438,0.1093,2.013,8.722,0.8549,0.01401,0.2922,9.228,1.038,7.226,2.009,0.03779,8.441,9.334,6.545,10.74,2.36,0.1059,0.007859,9.572,4.316,7.659e-05,0.1059,3.228,5.672,0.2108,7.339,0.9734,3.93,3.437,6.017,0.5914,9.454,1.889,7.328,0.906,7.782,2.301,7.768,2.396,1.307,2.599,10.45,1.837,0.2234,10.57,0.5717,3.249,2.577,2.358,2.184,2.925,0.00391,3.346,5.635,2.669,1.697,5.189,2.195,6.554,8.103,9.739,2.959,4.305,0.01831,0.2776,2.694,7.626,3.537,3.31,2.693,0.4947,1.658,8.521,0.1001,4.213,9.688,2.371,0.041,4.91,2.033e-07,3.56,0.1601,5.05,1.122,0.001461,0.05973,2.206,1.566,0.7421,2.01,8.123,2.915,1.53,0.1317,5.278,9.996,0.5633,1.9,4.341,0.0858,0.001589,8.002,2.525,6.618,1.939,8.716,5.528,1.722,7.578,5.624,7.047,9.181,3.346,1.226,1.18,3.189,7.672,1.353,2.855,10.05,1.639,6.813,1.182,5.439,0.01607,1.388,0.6829,0.1835,1.225,5.998,8.528,1.975,3.523,5.766,10.59,0.0006334,4.262,0.2383,3.764,2.359,9.559,0.3098,7.651,0.1645,0.005269,0.02082,6.569,2.098,0.09075,4.137,4.006,9.927,1.979,2.754,0.924,6.77,3.871,0.4285,0.004273,0.1388,0.07908,0.02088,0.6732,0.907,2.529,0.3178,0.6971,0.1786,0.01941,1.95,3.925,2.815,1.09,1.409,6.198,0.06233,3.892,0.2824,1.052,4.108,0.03212,0.7101,0.3909,0.9313,2.013,5.598,8.447,1.424,2.545,3.468,4.398,8.174,0.1706,0.1344,3.209,0.3404,9.39,6.479,0.1764,0.0003136,2.256,1.538,7.191,4.597,0.5365,2.696,1.014,0.269,0.1977,7.137,0.479,3.995,1.014,1.895,4.371,1.152,6.991,4.704,9.782,0.048,0.6477,3.115,2.271,0.886,1.614,3.785,0.01179,5.198,2.908,2.935,0.2644,6.208,4.633,0.1486,9.913,6.326,2.909,0.001492,2.693,0.0006385,1.984,3.766,4.96,2.95,0.1747,6.067,0.008615,9.613,1.434,3.191,0.1848,0.2374,0.6969,9.619,0.01845,4.24,0.9622,0.07925,14.65,0.7786,0.4559,1.976,4.689,2.947,9.187,3.365,0.07945,7.819,0.0587,0.7199,0.214,0.2197,8.358,0.829,0.2601,7.192,0.151,0.1136,0.1781,5.794,0.2225,0.0003203,6.401,1.121,3.666,2.097,9.717,1.335,1.932,0.05705,0.7965,1.016,8.853,0.5318,0.1184,8.351,1.35,0.1101,4.031,0.5895,0.7207,2.672,0.5698,5.361,0.7831,6.443,2.185,8.453,1.09,9.385,0.4248,7.466,0.8342,1.389,2.629,9.331,1.771,2.389,4.81,0.3532,9.036,1.399,7.583,2.639,0.5511,10.31,0.8148,0.9591,8.141e-05,4.14,9.674,6.009,1.772,0.01741,0.1743,1.096,0.003255,2.034,0.632,0.2083,1.429,0.3588,0.08739,4.428,4.848,0.9102,4.12,1.451,0.91,1.402,1.004,5.818,5.514,0.1283,2.35,4.964,7.439,5.472,0.955,1.948,8.588,0.1856,1.69,0.5718,1.614,7.864,3.065,1.183,1.114,1.417,3.411,1.359,0.03754,0.4454,4.266,3.809,4.064,4.958e-05,0.6025,10.41,4.507,1.453,0.322,0.1635,9.641,0.7233,0.03597,0.3966,3.077,7.582,9.489,1.777,0.4195,6.263,0.301,7.981,1.335,3.865,0.8437,9.78,7.307,0.3147,8.242,7.112,7.27,1.943,6.097,0.2696,0.4224,0.8329,0.477,0.3174,10,0.03241,2.114,3.612,1.3,1.008,0.01978,4.38,9.792,5.905,6.107,9.636,0.1499,3.28,4.354,1.406,0.2239,7.295,7.389,7.531,0.01492,6.03,1.208,1.284,1.666,7.118,0.01538,9.984,8.345,0.4164,0.8791,3.308,0.0004858,7.36,9.111,3.232,2.235,0.1582,1.265,5.465,2.399,0.04906,9.344,6.482,3.922,0.6801,1.48,0.06283,2.75,0.2186,0.09244,1.873,0.3876,2.475,2.454,7.516,1.094,1.677,0.05491,4.119,2.169,0.791,7.732,1.049,3.682,0.1916,4.825,0.01731,13.94,0.2653,1.758,1.61,2.926,0.4451,0.2098,1.946,8.821,3.08,2.765,2.122,4.319,2.79,8.981,0.003281,8.29,2.39,9.786,2.666,5.492,5.658,3.087,5.18,4.47,0.08253,0.7991,0.5556,3.15,2.217,10.17,0.01943,3.884,0.8992,7.114,0.6257,3.517,9.029,1.58,3.703,1.381,0.4586,1.419,8.367,9.473,0.1205,8.275,0.003132,3.319,0.8596,0.1234,5.953,0.5962,0.3845,9.243,0.9406,1.287,0.04235,0.07413,0.456,4.003,7.708,0.2287,0.01216,0.8253,0.5684,3.344,0.07539,2.737,5.978,3.561,0.32,3.091,0.01225,6.979,8.355,5.307,10.52,0.1955,5.06,0.121,0.003131,0.6067,2.838,6.372,8.868,5.464,0.8707,5.451,0.2794,0.2368,1.238,3.884,2.925,10.05,10.34,0.6247,10.6,7.863,2.269,0.3819,1.518,10.14,0.001725,1.413,0.1481,1.926,2.843,0.04357,0.3302,4.199,0.6994,3.079,5.598,4.808,3.977,0.5126,9.806,2.346,0.2301,0.2401,7.712,4.444,4.401,2.386,5.069,3.143,10.56,2.69,6.627,2.33,5.582,2.017,0.2913,0.02585,2.13,10.33,0.1596,9.883,2.882,3.241,0.4813,4.021,3.138,1.565,1.832,9.996,9.351,2.219,5.448,5.714,0.1182,9.979,3.833,0.09428,1.22,6.225,5.458,8.842,8.027,9.424,0.7245,0.5137,1.858,2.179,8.842,8.38,8.023,0.2085,2.104,3.601,0.1793,0.009594,0.2002,1.702,9.528,1.799,3.979,8.242,7.594,10.86,2.802,0.6005,8.305,0.694,8.309,3.374,9.335,3.761,4.064,5.726,6.652,0.2002,0.1844,1.358,0.03083,0.2372,6.442,1.578,4.952,0.2948,2.865,2.675,0.5052,0.8748,4.701,8.564,1.371,0.4049,3.689,0.07511,3.254,1.167,10.1,9.076,5.227,5.537,2.654,2.94,8.583,6.984,8.586,0.758,0.3457,0.1895,0.2293,0.01643,0.3798,0.5774,2.117,0.003174,9.412,1.865,0.124,6.198,6.584,10.54,2.249,9.461,6.973,4.863,3.701,0.3004,6.847,0.09437,0.005149,2.665,4.681,7.762,0.0095,4.038,0.3457,8.551,7.845,2.17,0.00248,0.01035,6.481,0.9424,0.01422,8.015,0.2451,14.17,7.724,0.3546,0.4534,8.092,1.121,4.252,0.02505,6.537,0.5601,8.583,6.484,3.195,5.999,6.764,4.218,0.8917,1.209,3.712,2.261,11.34,0.5747,4.609,1.026,0.03385,9.554,7.319,1.497,0.6702,0.0128,4.446,3.195,0.8936,1.239,7.141,0.5942,4.455,2.116,2.347,7.542,0.01731,7.149,4.324,0.0177,0.1534,0.01846,8.003,4.186,4.225,0.3814,1.09,8.521,2.769,0.2248,9.844,8.996,6.198,0.1637,3.979,0.2941,2.877,0.5309,3.528,0.2282,1.822,0.01361,6.896,9.515,0.1938,1.294,0.08205,2.155,2.342,4.433,10.2,3.903,0.1616,0.02898,2.608,2.7,9.036,1.439,3.282,2.022,8.043,0.3543,3.591,2.93,0.05702,0.5225,7.519,1.467,9.745,0.06986,8.875,2.375,1.649,6.333,10.62,0.9224,5.069,3.887,9.175,3.744,5.692,6.299,0.1741,2.458,0.8492,3.456,8.949,9.685,0.04798,8.91,9.84,0.05371,1.836,0.1374,1.087,3.086,2.943,4.402,0.01109,0.1721,0.9193,4.635,0.1473,0.01026,0.007465,0.004825,4.4,0.01598,0.03073,4.401,0.00754,5.617,3.653,5.204,0.8985,3.935e-07,11.44,3.363e-07,4.116e-07,2.334,0.04222,1.795,12.52,1.269,3.401e-07,0.1912,9.899,0.1503,3.305,4.417,4.636,1.45,0.2523,0.2737,4.848,6.222,2.165,1.151,2.643,3.1,1.283,0.3491,6.788,0.204,0.001822,0.4968,4.953,0.2088,10.59,3.027,4.502,4.628,7.407,2.619,3.85,2.579,0.02137,2.377,0.402,2.483,5.902,0.7258,2.113,4.018,0.3683,2.625,1.43,6.839,1.775,1.041,3.334,0.9998,0.03869,4.729,4.24,10.66,2.713,2.715,6.851,7.245,0.4642,0.8443,0.8051,0.2075,10.32,7.09,0.3419,3.778,0.002964,0.04905,10.37,0.0006231,5.277,0.0341,6.502,4.479,2.705,0.03528,1.59,5.878,9.575,0.2367,7.032,0.5061,0.3822,0.1058,9.829,4.513,7.716,1.157,2.516,5.886,8.659,0.9639,6.086,0.1612,0.9609,9.942,4.8,0.6563,5.474,1.048,9.161,0.0181,3.842,2.197,6.159,6.675,0.9189,3.744,4.097,4.227,2.545,0.4441,1.983,4.574,0.01551,0.3988,9.504,1.734,1.742,7.43,3.3,0.3403,3.801,0.9423,0.1968,2.17,0.02261,1.638,4.831,1.388,3.613,6.377,2.29,0.5638,0.946,9.74,8.281,0.6203,0.2898,5.808,0.0141,1.675,0.8786,0.1146,9.888,3.19,7.094,8.69,0.009275,0.0008984,0.08768,9.144,0.7649,0.007286,1.577,5.032,0.0003813,0.3468,2.942,7.915,0.0001209,0.3952,0.02344,2.037,0.2709,7.069,7.885,0.2359,0.3699,0.6341,8.666,3.236,8.894,1.234,9.059,1.216,6.162,6.896,5.677,1.602,16.67,4.315,0.08301,0.002333,9.019,0.2111,2.089,0.8484,2.159,4.665,4.395,1.152,5.967,0.6104,0.7891,0.371,1.021,1.584,8.321,3.243,3.353,3.383,5.724,4.983,6.116,0.001351,0.3467,2.77,6.846,2.39,6.587,2.596,7.364,9.176,6.846,6.34,9.149,7.077,4.223,0.008449,5.976,5.307,6.071,3.179,0.1348,0.1861,9.242,0.2296,8.047,3.267,9.606,0.02779,0.9845,0.0004168,2.129,4.653,8.72,2.793,2.91,0.1904,4.578,3.306,0.003659,4.524,1.743e-07,0.2772,1.669,7.714,0.04874,0.7568,3.631,5.934,8.43,8.698,0.6635,3.374,1.16,6.815,10.41,3.128,1.765,14.02,0.004922,3.077,4.225,0.1977,0.03504,0.5435,4.225,10.25,5.119,3.528,2.054,0.159,2.398,5.199,7.319,5.898,2.211,1.583,0.7244,9.927,0.0001443,0.2192,2.15,8.036,2.824,4.102,10.12,1.489,7.167,4.8,1.345,9.51,1.344,6.161,8.818,0.4177,2.352,10.21,0.5913,9.137,1.729,8.609,0.159,2.294,1.647,1.682,0.7599,0.7988,5.916,2.599,4.658,0.837,7.307,2.401,0.3227,1.32,3.187,2.44,6.645,9.088,4.056,2.268,0.4528,0.3406,0.2197,7.223,1.212,1.588,1.545,10.42,6.054,0.02792,0.1573,0.765,3.06,0.5911,1.353,0.09098,0.3332,5.807,2.59,4.66,1.681,5.318,1.778,1.986,1.247,3.801,9.715,5.966,6.24,2.013,3.012,2.613,5.065,1.41,0.2736,6.432,0.05134,0.01326,3.5,0.4656,0.5498,2.071,10.26,1.623,5.659,10.43,1.341,1.12,1.113,5.58,0.1391,0.308,0.04168,2.771,1.147,4.069,8.314,9.243,3.528,7.05,10.77,9.42,1.445,4.152,3.859,3.153,0.7962,0.8015,1.351,5.858,0.2279,3.788,6.872,0.2299,1.026,9.772,0.06062,0.3993,4.373,0.09424,9.436,2.506,0.5013,0.7658,1.538,0.2068,0.07261,0.1469,0.002399,3.846,0.3407,1.012,2.163e-07,1.301,4.723,0.6967,0.9929,4.101,0.3123,0.7175,0.5996,0.5015,8.848,1.856,3.3,4.425,6.414,9.812,3.063,0.5947,0.6292,13.34,2.055,0.5611,0.1117,5.434,0.2721,6.314,0.1748,0.07654,7.113,0.1869,1.87,8.463,4.643,9.04,0.7815,8.048,8.242,9.245,1.059,6.645,7.986,6.195,0.1593,2.162,0.238,1.845,5.628,3.789,4.973,0.6996,6.79,2.272,0.3613,0.0002137,5.276,3.347,0.1293,15.04,0.02878,0.8714,2.014,10.61,0.007836,7.712,8.031,2.14,3.929,0.7981,8.915,10.67,6.121,4.676,0.0001518,5.819,3.649,9.801,2.559,0.2443,10.18,0.1572,4.838,0.0008464,10.05,9.503,0.307,3.252,0.4028,0.005532,2.617,1.224,5.797,2.784,13.18,0.4503,0.02414,10.19,0.309,1.187,3.393,1.27,0.4343,3.355,2.392,8.494,1.692,2.906,5.473,1.102,1.02,7.555,8.247,1.141,9.495,5.108,9.111,0.832,9.967,3.322,6.507,2.735,4.151,0.0516,9.639,0.09739,6.452,7.074,0.9644,0.679,5.72,6.023,5.707,0.01163,7.238,0.029,0.001317,1.47,2.366,0.442,0.404,5.497,5.594,2.778,3.379,0.3257,3.841,1.2,0.4901,10.28,1.284,0.3633,4.107,1.748,3.251,10.09,0.03693,6.27,9.712,3.285,3.736,9.879,7.141,2.632,0.00102,3.432,1.862,5.41,1.071,1.332,6.374,0.5183,0.08464,3.045,0.1023,0.06409,0.198,9.247,8.777,0.09247,1.411,10.68,10.23,2.599,9.217,3.85,1.323,4.079,8.301,0.2554,9.718,1.008,9.478,1.105,1.711,1.419,6.776,2.821,1.113,0.02355,1.039,4.525,0.03691,3.518,3.184,4.265,0.53,0.01618,6.192,2.573,6.41,9.839,2.473,8.593,4.772,3.277,4.293,4.046,3.908,4.337,5.339,3.332,4.722,1.005,2.295,1.254,8.026,0.07384,2.843,9.486,1.043,0.4614,1.007,1.901e-05,0.4774,0.3757,1.919,10.76,0.9953,2.664,3.058,2.916,9.97,4.318,0.6778,1.951,0.01802,2.348,1.253,0.8952,0.5655,7.3,0.009215,7.062,0.0856,4.013,4.144,5.731,0.9405,0.8591,4.833,5.441,3.381,0.08946,0.4967,3.736,1.625,2.817,1.639,5.944,6.827,0.002339,3.492,2.091,3.439,1.574,7.891,9.268,10.1,0.1193,6.887,8.761,10.33,0.686,10.34,1.992,0.6252,1.376,6.62,9.365,8.259,0.03035,1.54,5.414,2.809,0.1528,0.04374,5.79,0.1755,0.06121,3.1,8.876,4.324,2.017,9.098,0.385,0.09295,10.11,4.269,0.02109,0.1467,2.283,0.7652,9.108,1.783,6.573,1.095,0.2002,9.073,9,1.162,0.8583,6.815,0.9836,0.1184,2.706,6.271,1.985,4.031,6.027,2.58,0.08593,3.444,0.002036,1.036,8.216e-05,0.3264,2.207,1.196,9.368,3.318,0.0006293,3.448,1.63,2.422,7.974,2.958,10.88,10.57,1.863,0.3416,7.997,8.803,2.53,0.7144,4.614,0.006991,4.316,2.948,0.2423,9.472,2.91,2.387,0.2134,1.411,7.965,1.269,3.129,4.405,6.241,5.114,0.1654,8.274,0.1065,0.6423,3.164,0.1965,0.5967,1.636,8.369,8.772,3.146,4.211,7.795,3.785,0.5001,9.765,3.032,1.354,10.48,0.7594,1.739,0.0165,2.184,3.312,1.134,9.809,2.982,0.724,0.002447,1.311,1.524,2.637,0.0346,2.282,0.3693,3.204,0.4063,4.424,2.483,3.921,0.1941,1.757,0.1103,1.717,0.0328,0.7589,0.1039,0.07402,4.805,8.927,3.215,3.17,4.201,0.0764,3.79,5.909,0.7664,6.355,1.223,6.497,10.18,8.629,10.6,9.208,4.583,0.2961,3.619,4.866,0.01583,1.662,0.02299,6.662,0.4352,0.2352,0.08329,1.901,0.03444,3.319,1.812,2.448,5.461,3.473,4.432,3.48,6.834,0.1876,0.01622,4.158,6.214,3.545,0.583,0.1409,0.2812,3.242,3.004,2.84,3.641,2.717,7.06,0.001443,1.712,9.073,1.677,5.179,9.451,5.465,0.1968,0.6125,1.231,7.823,2.24,9.207,0.0128,10.72,0.01085,4.524,5.482,8.555,0.2434,3.625,9.092,4.624,3.442,7.835,2.993,0.4349,0.2335,0.0008071,10.42,10.71,4.801,3.736,6.77,7.258,0.05005,1.929,3.39,0.3755,0.2822,1.614,2.247,1.739,1.166,7.221,5.735,0.9561,4.775,9.016,3.201,0.1841,0.4193,3.024,2.871,3.707,0.03798,0.7492,3.23,0.3006,2.305,8.312,0.5713,8.904,1.141,0.3533,1.994,6.574,5.647,8.664,0.7972,1.503,5.318,7.042,2.447,3.322,1.117,1.899,3.913,0.9028,7.764,10.91,10.33,10.9,10.67,0.2175,9.679,2.393,6.509,5.499,0.007073,7.343,0.4893,0.3329,5.581,0.007813,4.455,0.2603,3.793,6.445,4.141,2.97,0.05846,1.627,9.599,1.364,0.0122,2.904,1.503,2.34,10.23,2.11,2.951,1.646,4.802,9.374,6.481,3.041,3.773,5.784,4.132,9.484,7.747,4.865,6.88,6.194,6.469,10.56,0.7916,0.04557,3.415,2.846,0.2448,0.1975,0.9873,0.005919,3.179,4.215,3.892,3.977,6.354,1.97,3.491,0.3403,1.974,0.002107,2.89,5.18,3.491,0.03704,6.838,0.8792,1.518,0.001575,8.59,0.01239,7.895,8.567,0.5941,10.69,0.45,5.744,6.888,0.03118,0.2509,0.2822,0.00698,2.312,8.771,3.453,2.737,1.773,5.359,0.01212,5.4,1.406,2.844,4.25,7.052,0.191,4.119,0.09665,0.01499,3.223,7.969,7.139,5.83,6.559,6.644,0.1082,3.533,7.693,4.857,10.56,5.596,10.17,1.031,0.3039,0.3868,1.437,0.5569,0.2828,8.098,9.326,2.214,0.6794,5.816,1.86,10.32,5.547,3.978,8.198,4.439,7.054,14.66,3.023,0.001117,0.5954,4.493,8.093,0.0163,1.008,7.349,8.475,4.6,7.757,0.09795,8.806,4.53,4.877,1.361,6.069,3.567,2.69,6.022,2.179,8.657,0.1947,0.5542,0.02093,2.716,3.329,5.645,9.971,0.06775,2.531,1.866,10.86,3.701,6.788,1.481,6.551,0.005038,8.978,9.233,2.811,0.397,7.098,0.02421,0.05232,0.7519,0.1605,9.616,7.548,4.252,3.106,6.66,2.132,1.54,0.1736,1.765,1.338,4.728,0.7313,7.062,0.1432,0.6622,0.2537,10.34,3.869e-08,6.59,1.352,0.7841,10.36,5.222,3.231,3.234,4.128,2.646,7.222,4.559,0.3577,0.04046,1.184,0.1707,0.2601,0.9394,6.17,2.66,1.717,7.65,9.429,1.939,2.179,0.01884,5.524,7.807,4.775,0.3818,0.002747,0.9994,2.472,7.797,0.01633,0.1901,0.2226,1.966,1.531,7.515,7.169,10.22,3.92,9.786,3.176,7.87,3.261,4.788,3.541,2.593,0.3585,3.413,0.6345,10.23,9.53,0.2217,1.468,8.281,1.528,0.2151,3.667,0.5701,5.314,3.873,8.579,0.005905,8.865,6.303,0.1383,0.0003087,5.65,5.59,0.3139,1.458,2.09,1.614,2.588,2.076,0.5882,0.6627,1.018,6.813,4.137,3.754,2.707,1.192e-07,3.941,0.04742,0.05795,0.02609,0.04235,8.978,0.2902,3.561,1.777,0.4821,0.2542,5.915,0.6409,6.116,0.003188,3.939,6.527,0.3566,0.002566,0.5505,1.436,9.67,3.23,0.1678,0.01171,10.52,1.028,9.664,3.969,5.244,0.3868,1.595,2.218,7.244,6.435,6.047,5.305,0.5179,8.335,3.848,10.83,3.474,4.036,7.467,0.01608,0.1109,0.1998,4.874,0.1054,0.6903,3.227,9.416,2.419,0.06683,2.454,1.105,6.732,1.094,9.494,0.003799,0.01643,1.167,10.29,7.675,3.464,0.976,0.4458,0.01523,4.752,2.401,2.932,9.991,1.193,3.096,4.073,10.81,1.392,2.999,5.07,0.9873,8.443,9.463,0.02232,1.004,7.683,4.374,1.677,0.02641,4.111,0.9687,0.6913,10.55,6.964,1.464,0.5481,4.813,2.056,1.011,1.627,0.4879,5.529,1.688,2.68,5.305,0.2312,0.09527,5.657,3.016,0.1964,3.287,0.8676,4.788,1.454,9.785,5.304,0.01649,0.8806,0.8552,1.622,0.03271,4.234,0.2768,7.172,7.484,10.81,4.095,0.6745,0.9235,2.731,1.643,0.07813,2.449,0.1275,4.436,1.288,0.1595,4.186,1.404,3.67,6.187,0.1249,10.85,4.138,4.631,7.029,0.3296,9.474,4.261,8.323,0.2549,6.62,10.03,0.2396,2.822,0.2765,0.8643,0.2723,6.382,7.634,0.2829,0.2655,5.775,3.246,2.139,3.809,1.878,8.841,9.932,10.66,2.692,6.165,5.692,5.721,10.03,7.516,0.2163,8.248,1.099,0.05913,5.333,3.967,0.2268,2.583,0.1555,1.429,0.002635,9.274,0.02686,6.274,0.1201,4.155,6.341,2.218,0.08928,6.583,0.4666,4.335,1.226,1.659,0.005658,5.779,0.1037,6.654,3.295,0.06056,0.1821,10.41,1.851,5.821,0.3549,3.127,2.99,0.04552,2.986,6.654,3.881,0.1021,6.848,3.655,2.863,1.64,0.2435,5.184,3.752,7.256,0.1511,2.26,2.288,0.4541,9.022,9.733,1.624,2.143,0.5306,1.991,5.377,4.229,2.878,0.002524,1.267,8.921,0.08141,10.28,0.8377,1.696,9.466,0.001472,0.0164,7.899,10.34,7.03,0.07566,8.334,0.1026,5.053,5.649,5.011,0.7382,5.851,9.605,1.419,1.196,1.833,6.004,1.493,0.784,2.611,1.672,5.764,0.4066,2.923,3.375,5.873,1.549,7.756,1.767,6.661,2.014,9.056,0.01222,2.259,0.3138,3.111,1.223,10.64,0.002255,6.296,10.9,0.5265,0.092,0.8768,8.928,0.006899,3.069,2.206,1.639,10.36,4.219,2.073,0.5336,0.3293,1.524,0.3928,0.8429,10.74,7.17,2.489,4.971,2.996,0.2877,4.61,0.02589,0.4661,9.033,5.04,7.674,9.442,1.017,1.596,0.7087,0.2654,2.581,2.621,0.7176,6.927,7.595,8.095,0.2384,5.433,6.738e-05,0.03815,4.702,7.017,6.708,2.207,0.1021,1.315,0.08772,1.329,0.403,1.177,9.896,2.858,0.9183,0.05551,0.5798,0.9464,0.4161,3.291,3.892,0.05912,0.8049,0.002922,0.0122,1.102,0.06547,2.464,2.107e-07,3.297,3.169,3.483e-07,0.02154,12.32,14.22,1.346e-07,12.7,14.76,15.33,11.44,19.79,19.53,32.82,55.12,15.84,17.65,22.18,13.63,12.65,56.52,110.2,20.97,12.78,60.11,17.85,11.65,214.7,25.97,68.09,54.41,78.15,12.72,74,29.32,28.48,105.4,52.03,11.94,12.26,18.47,33.42,15.73,71.1,61.77,84.59,205.4,17.23,30.54,54.78,41.57,14.7,62.1,18.19,23.89,12.01,44.42,14.74,36.47,22.41,25.44,12.79,36.19,44.29,27.38,19.42,14.37,93.17,16.77,92,41.3,141.1,14.5,42.4,44.73,24.93,14.28,11.72,16.15,71.09,22.38,46.57,46.4,70.43,167.6,13.37,142.5,26.37,66.47,50.72,88.36,16.79,68.41,30.97,38.31,18.18,39.26,75.08,133.8,16.25,37.5,34.02,12.47,74.14,11.71,26.09,22.23,39.94,21.01,16.73,175.9,105,89.14,12.28,126.9,49.55,28.18,60.78,23.6,72.63,33.38,82.53,12.64,16.64,13.41,12.45,12.79,12.81,49.21,13.57,196.1,25.88,49.34,62.63,22.28,58,55.66,12.04,16.03,32.49,105.9,58.15,44.14,115.3,34.71,15.53,13.1,95.23,23.19,16.56,13.54,91.7,59,29.22,31.82,47.77,17.48,119.5,33.89,12.46,22.14,22.62,21.92,92.27,15.17,12.88,24.98,32.92,41.08,23.08,70.77,130.7,231.8,122.9,74.18,139.3,37.59,219.6,119.4,157.8,28.6,11.99,14.55,36.8,29.05,39.62,11.09,40.22,147.6,247.6,24.11,11.56,51.71,69.41,103.4,20.55,35.5,62.91,55.77,23.58,15.37,104,13.62,60.26,16.86,108.5,19.02,193.5,57.01,11.69,21.81,44.29,50.58,39.47,44.04,22.93,76.15,14.61,40.51,76.65,36.74,41.66,90.98,23.23,52.89,42.64,15.92,21.73,64.4,14.41,15.02,55.93,48.02,47.65,22.91,46.66,93.85,28.19,28.6,32.27,35.74,92.02,49.19,11.52,46.07,45.39,75.47,26.88,41.1,174.3,201.3,41.21,16.11,95.53,35.19,32.51,12.87,67.96,16.5,18.82,30.87,134.9,56.56,36.17,38.64,110.6,17.1,139.2,18.06,177.7,89,11.01,22.45,50.26,55.95,50.17,54.12,21.26,28.92,12.12,17.21,35.04,118.9,42.1,106.6,16.65,29.02,12.51,28.21,20.08,26.26,13.29,194.6,52.07,13.37,156.5,12.91,48.97,82.04,42.1,98.21,13.45,16.97,38.7,21.38,21.34,31.13,14.66,33.87,30.18,23.55,31.32,36.91,52.6,227.6,34.87,142.3,37.71,67.73,19.87,20.01,19.99,21.28,85.49,79.8,77.27,39.29,57.15,41.88,40.9,16.97,51.25,50.27,117.4,15.53,44.98,30.2,22.96,17.56,130.2,38.58,63.29,30.61,14.87,94.03,43.03,16.06,12.94,43.3,26.67,28.41,52.18,117.2,44.46,166.4,15.72,18.28,60.45,17.01,118.5,11.93,18,24.65,37.51,12.01,20.16,45.2,88.41,118.4,88.02,63,28.74,58.78,200.9,39.46,26.71,12.25,19.55,58.27,80.13,185.8,27.87,92.91,11.87,19.59,65.64,20.24,20.81,118.4,156.1,52.65,32.14,21.06,17.82,27.99,15.12,65.95,23.97,141.9,37.15,17.42,124.4,15.4,33.5,34.03,43.12,16.93,25.5,12.13,12.61,46.05,14.5,66.55,42.59,33.36,78.59,46.18,23.3,49.14,124.7,24.1,51.93,21.69,13.81,29.49,28.2,305.2,76.85,25.13,17.2,54.66,31.01,11.49,22.97,22.04,35.82,28.24,24.1,57.51,32.25,21.79,26.82,17.81,85.78,36.94,104.8,91,15.34,25.09,21.24,43.67,16.63,11.58,12.13,25.11,16.38,13.79,26.05,11.23,12.5,21.2,25.93,33.75,25.83,21.71,36.03,106.1,39.21,121.8,56.03,15.79,36.42,17.66,94.27,80.72,13.83,11.36,34.34,12.44,47.17,50.54,129.6,131.2,27.27,59.99,40.98,13.02,17.41,11.63,21.04,79.37,35.56,52.54,25.61,24.33,12.53,39.78,138.3,31.57,48.47,11.5,66.09,14.28,15.89,25.53,15.16,19.58,13.41,90.83,31.16,328.5,192.6,11.31,35.68,127,51.43,46.45,61.13,33.27,32.01,33.77,34.79,53.67,34.02,19.32,17.06,83.51,145.1,24.93,46.38,15.46,26.79,21.98,26.43,32.83,74.6,61.05,27.18,12.83,43.22,13.4,55.16,16.35,134.1,16.98,23.49,37.51,76.58,46.63,44.22,50.98,15.03,20.54,12.17,13.27,31.48,86.87,170,21.68,96.24,11.72,40.4,29.01,39.94,34.05,43.42,21.86,59.37,40.06,84.46,50.64,35.58,41.22,71.2,50.6,96.04,30.43,11.21,71.66,48.75,14.08,99.77,88.42,64.7,86.28,22.06,12.62,23.2,37.31,37.62,58.21,14.35,18.3,35.27,15.39,13.35,68.11,29.73,90.12,30.15,55.24,200.3,15.52,14.9,16.42,20.79,11.59,38.3,203,13.36,16.13,28.94,17.29,45.41,28.34,175.1,77.49,48.63,84.82,50.83,56.22,23.65,22.48,40.61,16.87,37.1,17.14,19.38,31.03,160.6,12.31,31.71,11.28,12.53,15.82,32.66,44.45,59.7,19.75,72.03,29.45,29.48,106.2,14.4,98.74,93.05,18.8,13.35,40.32,26.1,18.8,22.27,28.24,12.36,16.51,17.09,84.45,143.2,50.79,84.42,93.41,45.02,13.72,11.26,46.72,74.37,56.6,46.95,24.89,164.3,23.51,53.2,67.12,23.21,19.59,24.51,45.94,41.07,13.16,123.8,103.5,46.78,24.77,12.62,15.66,16.43,19.21,131.3,56.49,24.39,39.08,60.96,106.6,16.16,69.64,20.05,90.67,23.6,106.4,93.23,23.47,44.52,34.64,14.2,22.81,72.85,48.95,27.94,23.64,42.59,20.04,30.59,118,41.59,31.65,63.77,78.55,21.78,14.67,16.55,15,19.25,27.05,132.2,13.35,150.7,12.56,53.78,14.63,25.23,15.1,37.71,23.56,33.2,14.03,65.64,37.43,50.44,38.59,65.64,146.4,19.43,25.39,103.8,13.06,16.41,34.34,119.7,96.73,107.5,181.8,52.83,90.28,196.8,30.15,68.75,25.68,44.99,69.47,28.25,16.7,30.46,63.54,34,26.69,42.73,67.6,84.29,11.27,31.97,142.1,65.86,12.24,25.05,21.7,31.58,27.12,40.26,43.39,11.02,14.21,42.04,25.82,39.15,52.41,17.01,12.27,47.87,15.49,34.18,23.03,10.95,77.5,13.56,34.79,39.02,13.19,143.4,51.3,11.58,56.96,12.84,109.1,19.97,95.75,83.71,30.91,21.35,55.75,110.8,40.26,33.19,23.45,53.65,13.34,29.75,241.1,30.81,28.57,16.25,27.56,18.03,38.76,142.6,42,184.5,80.69,63.35,21.04,16.63,30.36,23.65,212.7,46.18,38.69,36.63,16.65,24.88,36.48,38.21,57.77,55.48,35.92,180,41.38,36.72,22.75,21.74,20.91,165.1,98.36,50.93,59.23,30.89,29.63,77.15,66.13,31.11,64.03,29.64,39.28,59.2,27.77,24.38,19.98,35.45,33.15,17.88,17.04,16.87,15.59,15.91,27.2,22.94,58.73,24.3,23.81,62.18,61.32,116.7,105,14.58,22.83,45.86,33.93,18.69,15.38,24.96,34.87,15.42,27.47,17.97,13.4,12.45,47.46,13.77,117.4,23.65,27.84,12.47,12.16,30.52,15.94,34.52,14.95,100.4,29.02,44.82,19.2,52.15,12.21,46.01,61.42,11.86,53.01,15.79,23.87,18.57,145.1,28.56,15,61.54,21.14,26.3,44.98,38.02,45,45.49,12.44,30.79,20.2,72.76,12.8,59.11,11.45,36.44,32.24,12.07,90.66,37.01,27.07,143.7,13.2,15.93,97.76,46.67,27.8,13.87,23.4,21.75,15.87,65.41,22.01,77.93,45.36,34.09,71.19,118.6,42.92,26.06,35.33,34.99,29.98,46.81,74.2,113.5,41.12,92.88,18.63,119.7,154.8,20.37,15.05,12.91,60.76,139.5,17.22,80.87,82.51,13.43,31.53,25.35,26.41,45.09,152.8,43.06,12,15.69,43.94,53.17,131.4,26.62,25.12,14.61,24.39,21.38,11.74,26.46,15.52,41.13,15.64,11.69,18.26,24.79,52.38,13.34,18.45,26.11,17.9,34.07,31.71,51.61,147.9,88.38,85.72,42.95,21.57,59.04,57.38,21.78,16.8,120.7,17.56,24.15,15.13,48.58,40.32,16.85,14.61,30.74,52.76,21.12,13.5,37.47,134.9,24,42.83,17.7,73.35,29.45,51.86,28.04,17.05,58.25,33.24,17.71,30.88,14.16,76.54,116.8,234,67.45,33.54,15.51,26.94,35.47,12.14,304.3,13.47,19.04,93.02,28.22,108.6,16.47,92.11,20.89,46.65,51.77,18.27,15.03,30.83,223.4,61.53,134,17.11,37.84,11.45,20.18,44.81,51.69,17.1,23.59,30.77,12.64,11.53,13.18,43.45,26.97,43.67,94.9,40.03,47.85,34.78,27.45,21.47,82.19,155.4,44.3,57.6,14.35,28.54,79.75,30.01,17.36,18.37,37.33,12.97,27.35,38.43,16.24,14.64,53.98,115.6,16.29,31.81,17.78,17.55,38.08,13.3,31.27,242.6,47.41,16.27,69.54,19.4,82.95,15.87,189.6,75.16,16.01,13.52,141.6,39.03,23.37,21.91,18.14,58.13,138,40.52,28.55,30.92,22.32,28.46,73.4,14.08,31.28,99.96,100.7,28.43,17.87,16.73,13.98,16.65,13.66,36.9,25.14,13.15,58.71,22.64,71.95,21.67,37.23,91.78,59.58,20.4,29.75,60.61,54.03,18.05,38.31,19.41,26,34.3,17.94,14.36,11.51,40.23,20.98,19.54,15.51,27.34,21.35,39.17,29.04,19.02,21.86,24.05,29.5,12.94,65.8,15.07,14.37,55.38,68.45,60.51,18.19,50.98,35.62,106.2,19.69,13.68,35.4,17.04,15.94,23.95,33.65,52.38,31.16,35.99,51.57,40.3,22.71,26.69,117,37.46,91.19,21.59,36.35,15.32,38.96,11.46,54.61,11.53,88.91,19.31,26.89,11.99,14.52,12.86,12.89,23.23,51.41,18.96,196.2,12.5,25.82,32.46,27.88,44.5,18.42,63.28,119.2,25.93,13,37.96,45.49,64.96,29.84,43.42,42.92,11.84,62.26,34.99,16.21,82.59,17.12,14.45,14.59,50.15,287.2,39.44,67.52,12.01,86.26,13.84,122.4,12.5,78.21,19.75,70.29,50.57,20.73,29.88,14.68,18.17,47.14,58.27,13.8,57.27,31.42,51.17,23.18,18.15,152.5,77.34,24.53,32.41,22.23,32.18,50.47,21.69,14.54,46.29,46.68,16.93,11.81,14.27,18.02,30.74,26.57,20.84,51.09,29.85,20.61,53.59,14.22,20.05,30.32,22.84,16.99,48.74,25.29,11.13,58.89,43.33,80.64,24.46,39.39,16.25,37.65,75.4,13.61,33.83,58.26,27.43,65.21,27.45,29.54,17.04,33.33,14.53,16.79,111.3,14.87,36.35,22.35,19.88,19.75,32.23,31.4,19.77,14.9,54.62,17.63,57.1,36.73,26.21,105.8,19.82,32.11,45.59,13.76,31.63,92.45,21.67,16.31,13.94,13.49,20.95,76.08,14.78,20.72,44.84,27.93,28.04,29.89,55,30.84,31.35,13.16,35.73,36.11,25.84,20.63,56.61,58.1,25.7,39.36,14.75,132.5,22.72,72.76,14.53,19.19,53.88,62.37,11.23,34.32,28.08,78.98,12.26,13.17,11.39,18.66,36.9,46.94,15.46,26.6,28.04,11.51,377,12.75,14.16,32.65,25.65,38.73,12.57,116.9,11.96,19.93,72.39,38.96,17.53,27.3,19.84,51.88,36.3,11.45,40.95,17.37,19.2,179,27.64,19.69,25.09,49.56,12.48,49.75,24.81,14.2,102.7,79.36,111.7,12.63,50.57,42.5,53.65,60.52,64.12,124.8,17.54,16.76,15.47,26.01,25.67,43.73,69.05,105.3,67.72,71.65,33.11,34.02,11.49,135.1,27.93,22.29,66.81,12.24,28.62,159.3,13.67,22.05,20.43,41.18,22.71,65.45,83.06,52.06,55.84,49.38,15.22,129.7,27.42,136.4,104.4,33.57,54.57,27.36,59.25,18.48,36.08,65.35,29.55,83.95,28.79,14.47,60.37,14.16,35.89,32.75,15.37,11.31,83.1,32.17,50.85,36.39,31.77,113.8,30.9,79.33,23.71,20.04,162.1,20.91,15.57,94.84,25.92,19.45,14.34,31.61,46.17,58.11,17.22,54.69,10.99,88.13,20.25,20.23,11.05,93.43,24.99,54.75,30.75,120.6,144.2,28.05,118.5,53.28,28.91,12.44,14.38,17.62,56.5,15.95,44.31,44.66,74.33,63.37,70.38,63.9,34.24,46.37,28.81,99.06,21.74,35.63,54.81,51.36,58.8,20.57,53.84,20.43,15.79,44.45,17.77,84.24,17.96,24.77,37.2,112.4,20.49,40.82,26.09,57.4,20.97,26.67,172.5,66.26,11.97,16.06,142.8,24.59,21.18,24.35,41.59,19.96,31.94,46.56,26.65,26.52,20.78,20.03,18.35,16.36,106.1,89.64,28.93,13.15,57.73,40.51,67.34,13.43,26.22,24.7,33.08,69.01,42.43,57.54,58.64,57.87,128.6,194.4,17.55,12.48,61.66,36.28,46.92,73.39,24.78,13.84,64.06,57.25,153.6,55.98,18.63,21.13,11.41,16.99,31.19,63.66,15.42,48.74,21.84,12.87,39.27,75.62,45.46,53.21,12.41,90.98,72.4,49.92,17.96,17.71,31.52,90.04,60.23,34.82,19.03,21.28,14.62,43.33,77.24,18.6,37.94,64.8,12.25,17.87,100.4,51.91,21.39,41.02,21.56,21.7,30.11,16.04,126,143.6,13.8,10.92,50.56,56.96,15.27,78.79,29.51,50.99,149.2,79.52,41.85,119.2,58.44,92.94,105.7,79.72,253.9,69.96,39.2,12.96,51.86,64.2,40.54,20.44,29.7,27.29,29.23,59.21,138.7,14.84,210.3,34.47,77.53,12.17,55.38,17.81,100.3,43.79,11.01,15.82,61.85,32.35,24.05,30.61,33.27,18.68,13.5,50.55,299.8,174.7,22.04,55.11,63,21.5,15.03,16.98,23.16,15.73,18.99,27.95,19.12,40.33,17.63,56.7,54.16,155,120.7,64.4,29.11,13.33,46.26,54.57,19.18,32.95,17.43,133.7,57.59,31.83,11.14,92.63,20.92,63.77,57.43,93.71,11.59,28.95,57.14,15.68,33.32,43.39,34.9,34.04,28.6,36.94,159.1,19.51,12.63,13.74,64.22,66.85,25.48,11.08,14.64,19.96,58.75,12.08,103.4,45.95,19.07,11.4,22.57,72.98,120.8,27.31,23.44,12.49,55.02,101.9,33.69,44.42,55.99,26.95,58.41,32.27,25.6,24.54,13.36,59.22,27.1,11.54,42.28,13.6,229.3,11.78,21.66,15.16,50.58,26.79,13.69,160.5,13.6,20.01,12.41,37.44,11.5,11.2,15.53,11.94,73.36,43.15,72.06,13.24,58.44,46.13,45.25,83.78,38.86,22.01,17.76,27.9,24.28,25.79,20.88,66.83,14.25,98.02,43.5,12.61,33.81,40.64,86.02,43.45,16.89,37.24,13.92,31.96,15.71,18.42,109.7,107.4,12.62,22.68,11.96,14.05,17.69,61,21.68,14.97,78.51,125.5,14.18,17.1,11.63,16.75,20.24,36.69,57.28,27.62,34.44,19.1,161.5,63.97,11.83,27.21,41.81,13.16,15.43,25.65,89.28,15.63,26.03,43.48,113.4,42.15,30.52,43.2,122.3,40.46,27.91,32.51,13.35,41.36,27.5,64.02,24.99,17.25,99.33,31.79,44.16,34.76,15.39,26,24.92,32.62,134.8,26.31,31.33,38.73,53.71,17.63,192.1,34.34,18.01,20.39,31.89,123.5,43.74,43.16,22.53,17.64,125,11.32,16.85,56.75,49.88,16.34,111.5,73.34,14.87,39.05,12.49,36.45,13.9,36.71,16.21,76.36,13.35,12.63,19.41,16.89,10.94,19.38,11.65,13.82,33.46,31.38,20.64,38.51,20.51,114.3,26.94,30.93,54.21,16.16,39.11,82.6,13.51,15.37,74.34,29.38,16.84,21.55,20.35,29.45,29.02,68.24,40.06,69.8,20.9,17.86,108.3,13.28,13.6,44.31,11.38,27.91,14.11,125.3,11.06,30.71,68,26.68,19.06,35.57,13.57,104.5,50.08,37.91,11.2,40.19,20.82,13.65,18.69,84.64,18.58,15.68,28.51,49.98,16.71,16.78,235.6,144.4,44.61,19.8,39.91,117.5,136.3,64.7,22.01,40.03,13.27,14.98,15.74,17.31,13.48,43.47,114.1,37.01,20.98,42.02,57.97,31.34,80.12,23.66,12.74,26.8,22.72,42.13,29.1,33.66,62.7,30.31,38.36,14.65,34.42,28.08,22.86,21.2,57.01,16.5,30.29,66.83,41.3,43.89,57.68,69.3,32.72,27.52,27.28,26.64,89.26,38.65,43.34,75.75,17.24,25.58,16.2,22.35,19.29,11.43,48.5,57.57,34.48,31.9,50.41,40.89,113.9,67.6,151.3,16.62,38.43,18.7,18.71,84.11,68.45,43.19,16.15,24.04,17.34,28.21,14.89,17.91,37.48,96.43,218.8,21.77,23.44,87.66,103,23.27,24.47,36.86,49.25,185.2,98.25,110.6,127.1,63.91,15.63,34.05,28.79,16.7,142.5,22.37,20.39,25.44,53.93,12.37,38.53,89.9,49.24,64.46,40.4,17.41,125.3,26.55,24.82,102.9,15.9,396.9,26.22,17.99,33.63,13.71,39.42,21.96,91.66,59.17,27.78,71.12,165.1,20.21,13.47,75.32,14.05,11.57,19.02,17.82,22.51,109.9,18.36,57.57,13.67,29.01,40.99,36.3,26.04,33.38,96.49,13.53,12.8,24.9,23.8,48.12,12.69,19.03,11.37,24.46,52.89,23.77,16.03,51.27,11.64,48.78,20.24,93.58,25.02,25.14,26.13,19.18,167.9,80.13,53.75,11.54,12.67,28.63,15.19,15.88,24.16,15.94,11.76,44.92,12.33,15.28,53.21,24.34,37.09,21.87,101.1,38.77,29.92,25.91,60.73,39.75,10.93,36.02,93.62,36.12,22,51.74,13.8,68.98,35.04,15.17,48.09,56.97,15.72,26.51,18.84,30.24,54.93,27.9,112.5,42.74,86.9,37.34,14.4,18.49,102,31.85,73.97,13.2,42.68,11.48,50.79,18.83,11.22,206.2,38.06,24.16,21.47,78.73,81.55,14.93,36.36,16.72,29.4,21.1,60.66,72.42,70.88,17.02,33.41,14.42,14.77,115.4,72.43,17.61,144.6,26.03,50.43,38.3,20.07,20.63,49.48,11.51,33.45,16.22,26.12,54.32,70.6,72.69,14.23,15.31,144.4,16.39,17.65,44.23,21,28.53,38.37,14.58,26.34,45.18,19.71,11.97,34.09,12.15,32.03,34.98,14.61,89.42,161.3,37.54,35.43,28.74,35.99,53.96,42.89,95.84,23.18,16.34,13.38,42.1,23.51,42.62,12.96,11.8,16.71,38.19,13.66,65.67,65.14,25.99,17.73,22.53,100.9,26.86,32.67,16.45,13.05,11.01,24.17,82.59,23.34,75.53,16.81,52.12,12.16,16.44,57.38,12.24,18.69,50.64,99.59,64.36,34.5,16.69,12.13,23.5,37.89,29.6,28.33,11.82,48.69,145.6,30.46,23.77,30.78,67.08,79.41,23.07,30.93,14.97,11.62,35.67,14.6,35.28,135.2,13.31,22.78,19.54,40.82,31.67,27.56,21.12,78.35,50.45,37.81,67.72,29.5,11.41,14.69,16.87,15.54,44.24,152.7,61.67,11.91,48.84,16.84,18.68,18.05,20.23,54.75,31.95,64.03,70.83,85.62,120.7,86.51,41.1,13.1,24.18,50.78,54.68,30.22,12.02,82.89,26.15,97.61,168.2,47.84,19.29,22.13,21,115.3,27.43,18.76,38.78,90.58,61.21,24.04,42.7,25.04,28.65,36.01,116.8,133,33.73,34.96,18.41,36.54,29.66,12.09,14.61,12.46,30.44,58.96,42.19,45.37,197.2,12.59,53.66,20.87,22.5,61.13,26.64,138.7,69.76,46.3,32.89,60.52,16.6,16.48,42.89,10.99,26.44,82.04,16.62,22.68,14.79,141,20.82,60.24,72.04,12.13,15.96,20.4,46.53,26.27,24.74,43.22,146.8,40.07,17.27,13.24,64.07,26.73,37.77,216,36.41,16.41,26.93,29.75,95.78,30.75,47.02,17.92,29.56,93.55,87.74],"PValue":[0.7026,0.5032,0.2325,0.6398,0.2282,0.4271,0.6015,0.7519,0.0368,0.0946,0.8881,0.4218,0.8961,0.2744,0.931,0.3559,0.0785,0.04959,0.05053,0.04469,0.03714,0.899,0.05785,0.1038,0.04525,0.9997,0.08581,0.06314,0.2339,0.1458,0.04225,0.02493,0.03052,0.0773,0.03063,0.0854,0.04936,0.04506,0.0443,0.01424,0.08433,0.09288,0.1987,0.4703,0.04472,0.03572,0.08868,0.03684,0.08581,0.7302,0.04655,0.7168,0.8095,0.9997,0.6569,0.8691,0.02157,0.1217,0.8697,0.04118,0.02935,0.02368,0.09683,0.0773,0.09288,0.04161,0.05908,0.06801,0.006883,0.6055,0.3342,0.5566,0.4711,0.6716,0.3735,0.5029,0.4247,0.8358,0.3016,0.6766,0.01123,0.6575,0.05366,0.04066,0.273,0.105,0.02713,0.5604,0.067,0.08387,0.06607,0.01694,0.03569,0.1185,0.3257,0.7538,0.04204,0.08389,0.03558,0.01025,0.9997,0.05019,0.02995,0.4707,0.03219,0.05187,0.0283,0.05696,0.7297,0.1119,0.02351,0.0178,0.786,0.07492,0.05985,0.05785,0.161,0.04159,0.2521,0.05799,0.9642,0.07296,0.1186,0.0584,0.1359,0.6507,0.06218,0.009245,0.06492,0.604,0.03363,0.7146,0.02954,0.9996,0.05455,0.09683,0.09288,0.1185,0.03544,0.06106,0.03261,0.04653,0.08944,0.01491,0.00726,0.2469,0.06475,0.1185,0.05275,0.06218,0.04751,0.01593,0.05512,0.1135,0.05443,0.03623,0.3224,0.07912,0.0167,0.0275,0.1921,0.2469,0.902,0.1244,0.08944,0.1653,0.03156,0.03634,0.1007,0.08859,0.6447,0.8185,0.1629,0.1153,0.9996,0.04167,0.2062,0.8958,0.8389,0.02026,0.9997,0.9997,0.9586,0.9997,0.04488,0.3189,0.2759,0.8291,0.5532,0.007876,0.3821,0.4822,0.09728,0.624,0.7554,0.0207,0.05171,0.5822,0.7896,0.01925,0.0092,0.3056,0.004804,0.02509,0.1823,0.06118,0.1349,0.03019,0.1893,0.6717,0.5957,0.319,0.02423,0.9856,0.1443,0.04688,0.06278,0.01382,0.5947,0.5817,0.09403,0.489,0.009738,0.01144,0.423,0.04641,0.03824,0.9274,0.06124,0.03789,0.004717,0.2009,0.01079,0.1865,0.2525,0.6503,0.008177,0.02121,0.2955,0.7076,0.04513,0.4239,0.02118,0.8969,0.2342,0.03739,0.03896,0.7998,0.2852,0.1562,0.2697,0.01619,0.01098,0.0427,0.4137,0.2018,0.4142,0.6321,0.2397,0.036,0.05221,0.2212,0.01733,0.9735,0.3932,0.0154,0.06452,0.3537,0.1045,0.04735,0.1643,0.2382,0.04739,0.3445,0.1671,0.8247,0.3571,0.06539,0.4264,0.01399,0.02321,0.1254,0.92,0.01926,0.9426,0.01082,0.04031,0.08624,0.03939,0.04723,0.04681,0.06218,0.02724,0.04127,0.1541,0.2437,0.04437,0.6693,0.4947,0.02882,0.5111,0.5886,0.7051,0.03086,0.9386,0.262,0.134,0.02971,0.06461,0.08307,0.3789,0.9996,0.009294,0.02906,0.2373,0.008156,0.7457,0.2377,0.09806,0.01421,0.431,0.8127,0.01155,0.6878,0.02571,0.03453,0.5551,0.2571,0.03594,0.7939,0.3085,0.009327,0.2054,0.02762,0.1757,0.01532,0.3396,0.5363,0.7191,0.01615,0.01443,0.1373,0.6842,0.1317,0.9464,0.004772,0.5403,0.7915,0.03771,0.5764,0.1165,0.7992,0.05782,0.3483,0.008178,0.4339,0.01431,0.9606,0.07218,0.3637,0.1536,0.8249,0.3472,0.007612,0.5758,0.4593,0.02866,0.628,0.423,0.008852,0.3325,0.001656,0.1115,0.8791,0.7458,0.04567,0.03113,0.004668,0.09121,0.0321,0.1948,0.4531,0.01831,0.6191,0.01478,0.03479,0.2635,0.3447,0.05491,0.5992,0.04982,0.09668,0.1626,0.04403,0.0561,0.0584,0.8894,0.01688,0.9995,0.1531,0.02371,0.06533,0.0642,0.01919,0.675,0.3958,0.08154,0.06218,0.0773,0.1014,0.02897,0.9108,0.05219,0.1377,0.03657,0.02634,0.07473,0.09486,0.1303,0.1911,0.05404,0.01513,0.7453,0.1756,0.009509,0.3692,0.9073,0.5964,0.007984,0.3239,0.01638,0.1759,0.8484,0.0105,0.007701,0.02129,0.004853,0.1444,0.7491,0.9305,0.007108,0.05457,0.9931,0.7492,0.0917,0.03029,0.6525,0.01569,0.3388,0.06524,0.08266,0.02629,0.4533,0.007396,0.1887,0.01576,0.3556,0.01332,0.1492,0.01338,0.1416,0.2701,0.1269,0.00533,0.1945,0.643,0.005121,0.4608,0.09074,0.1284,0.1446,0.1593,0.1069,0.9509,0.08646,0.03076,0.1222,0.2115,0.03712,0.1583,0.02122,0.01185,0.006721,0.1051,0.05485,0.8941,0.6056,0.1206,0.0141,0.07874,0.08803,0.1207,0.4922,0.2166,0.0102,0.7558,0.0572,0.006836,0.1436,0.8421,0.04421,0.9997,0.08061,0.6948,0.04169,0.3066,0.9754,0.8101,0.1573,0.2304,0.402,0.1759,0.01176,0.1088,0.2344,0.7216,0.03574,0.006173,0.464,0.1875,0.05394,0.7734,0.9687,0.01229,0.132,0.02069,0.1832,0.009529,0.03215,0.2083,0.01436,0.03089,0.01753,0.008115,0.08645,0.2848,0.2938,0.09351,0.01387,0.2622,0.1109,0.006067,0.2191,0.01918,0.2934,0.03339,0.9007,0.2563,0.421,0.6743,0.2851,0.02649,0.01018,0.1794,0.07928,0.02914,0.005084,0.9802,0.05593,0.6322,0.07057,0.1445,0.007137,0.5857,0.01397,0.6906,0.9431,0.8871,0.0211,0.1672,0.7672,0.05925,0.06297,0.006314,0.1791,0.1183,0.351,0.0195,0.06834,0.5222,0.9487,0.7145,0.7822,0.8869,0.4242,0.3554,0.1318,0.5815,0.4163,0.6787,0.891,0.1821,0.06542,0.1148,0.3134,0.2541,0.02532,0.8355,0.0691,0.6031,0.3216,0.06264,0.8602,0.4129,0.5408,0.3492,0.1768,0.03124,0.01048,0.2505,0.1306,0.08144,0.05258,0.01155,0.6852,0.7189,0.09258,0.5679,0.007558,0.02185,0.6802,0.9861,0.153,0.2331,0.0166,0.04807,0.4747,0.1205,0.3293,0.6113,0.6627,0.01768,0.499,0.0633,0.3293,0.188,0.05323,0.2993,0.01791,0.04584,0.006626,0.8294,0.433,0.09704,0.1517,0.3609,0.2225,0.06988,0.9149,0.03698,0.1079,0.1064,0.6143,0.02434,0.04731,0.7051,0.006344,0.02322,0.1078,0.9697,0.1207,0.9802,0.1785,0.0705,0.04097,0.1056,0.6817,0.02576,0.9272,0.00701,0.2489,0.0934,0.6735,0.6329,0.4164,0.006995,0.8937,0.05651,0.3415,0.782,0.001536,0.3909,0.5094,0.1793,0.04616,0.1057,0.008096,0.08567,0.7817,0.01314,0.8117,0.409,0.65,0.6457,0.01081,0.3764,0.6172,0.01659,0.7028,0.7406,0.6788,0.0288,0.6436,0.9859,0.02254,0.3057,0.07396,0.1673,0.00677,0.2653,0.184,0.8145,0.3856,0.3288,0.009085,0.4766,0.7357,0.01143,0.2627,0.7445,0.06225,0.4548,0.4095,0.1221,0.4615,0.0345,0.3896,0.02217,0.1592,0.01045,0.3122,0.00757,0.524,0.01497,0.3749,0.2562,0.1249,0.007708,0.2023,0.1422,0.04376,0.5608,0.008528,0.2545,0.01433,0.1242,0.4688,0.00557,0.3804,0.3423,0.9929,0.05916,0.006867,0.02638,0.2022,0.8967,0.6823,0.3121,0.9552,0.1734,0.4393,0.6548,0.2508,0.5583,0.7717,0.05437,0.04538,0.3555,0.0623,0.2474,0.3556,0.2552,0.3326,0.02946,0.03235,0.725,0.1465,0.0409,0.01512,0.03292,0.3433,0.1823,0.009967,0.6725,0.2124,0.4608,0.2225,0.01357,0.09953,0.2932,0.3072,0.2516,0.08372,0.2611,0.8489,0.5143,0.05583,0.0691,0.06128,0.9945,0.4492,0.00539,0.05007,0.246,0.5784,0.6914,0.006943,0.4079,0.852,0.538,0.09896,0.01433,0.007308,0.2016,0.5266,0.02381,0.591,0.01239,0.2652,0.06728,0.3723,0.00663,0.01589,0.5827,0.01127,0.01711,0.01611,0.1828,0.02545,0.6109,0.5252,0.3753,0.4999,0.5811,0.006161,0.8594,0.1657,0.0759,0.2714,0.3305,0.89,0.05301,0.006604,0.02751,0.02535,0.006957,0.7039,0.08932,0.05364,0.2534,0.6426,0.01596,0.01541,0.01461,0.9043,0.02616,0.2883,0.2742,0.2156,0.01707,0.9029,0.006197,0.01086,0.5281,0.3627,0.08811,0.9827,0.01557,0.008311,0.09152,0.1547,0.6962,0.2777,0.03302,0.1414,0.8276,0.007676,0.02183,0.06549,0.4228,0.2431,0.8056,0.1171,0.6469,0.7654,0.1917,0.5431,0.137,0.1385,0.01469,0.3125,0.2141,0.818,0.06113,0.1606,0.3872,0.01356,0.3224,0.07339,0.6676,0.04347,0.897,0.001868,0.6137,0.2039,0.2231,0.1069,0.5144,0.6532,0.1825,0.009186,0.09877,0.1162,0.1649,0.0545,0.1147,0.008692,0.955,0.01108,0.1421,0.006618,0.1225,0.03265,0.03046,0.09843,0.03726,0.05089,0.7777,0.3849,0.4671,0.09538,0.1563,0.005839,0.8909,0.06668,0.3574,0.01709,0.4408,0.07949,0.008549,0.2272,0.07267,0.2575,0.5081,0.2513,0.01078,0.007347,0.7331,0.01114,0.9561,0.08765,0.3679,0.7301,0.02698,0.4515,0.5441,0.007944,0.3469,0.2737,0.8396,0.789,0.5093,0.06304,0.01368,0.6391,0.9136,0.3774,0.4621,0.08654,0.7872,0.1179,0.02671,0.07783,0.5796,0.09823,0.9133,0.01799,0.01083,0.0353,0.005212,0.6644,0.03923,0.7326,0.9561,0.4477,0.1118,0.0228,0.009038,0.03304,0.3649,0.03321,0.6045,0.6333,0.2827,0.06668,0.1069,0.006073,0.005524,0.4411,0.005081,0.01293,0.1519,0.5455,0.2361,0.005894,0.9674,0.2523,0.7055,0.1846,0.1116,0.8373,0.5737,0.05758,0.4156,0.09885,0.03124,0.04379,0.06383,0.4846,0.006572,0.1456,0.6381,0.6309,0.01366,0.05151,0.0525,0.1424,0.03908,0.09568,0.00514,0.1209,0.02062,0.1468,0.03144,0.1752,0.597,0.8743,0.1642,0.005527,0.6949,0.006407,0.1093,0.09108,0.498,0.06252,0.09592,0.2293,0.1951,0.006172,0.007657,0.1562,0.03326,0.02976,0.7355,0.006635,0.06831,0.7628,0.2861,0.02418,0.03311,0.009119,0.01218,0.007469,0.4075,0.4841,0.1922,0.1597,0.009121,0.01073,0.01219,0.6542,0.1667,0.07633,0.6777,0.9232,0.6607,0.2108,0.007213,0.1989,0.06377,0.01127,0.01427,0.004676,0.114,0.4499,0.01102,0.4173,0.011,0.08526,0.0077,0.07337,0.06389,0.03168,0.02042,0.6611,0.6735,0.2625,0.8855,0.6334,0.02218,0.2287,0.04132,0.5953,0.1103,0.1219,0.4877,0.3638,0.04591,0.01005,0.2591,0.5338,0.07316,0.7876,0.09053,0.2963,0.005969,0.008411,0.03652,0.03204,0.1232,0.1061,0.009984,0.01796,0.009974,0.3971,0.5649,0.6693,0.6386,0.8996,0.5466,0.4586,0.1654,0.9558,0.0075,0.1913,0.7294,0.02444,0.02097,0.005179,0.1536,0.007379,0.01804,0.04274,0.07274,0.5919,0.01893,0.7627,0.9437,0.1225,0.04632,0.01342,0.9236,0.06203,0.5649,0.0101,0.01301,0.1605,0.9609,0.9202,0.02184,0.3464,0.9066,0.01223,0.6274,0.00175,0.0136,0.56,0.5105,0.01189,0.3058,0.05618,0.8763,0.02137,0.4653,0.009985,0.02181,0.09323,0.02648,0.01955,0.05708,0.3594,0.2882,0.07234,0.1525,0.004017,0.4596,0.04783,0.3266,0.8564,0.007149,0.01582,0.2392,0.4253,0.9114,0.05145,0.09324,0.3589,0.2824,0.01692,0.4523,0.05125,0.1655,0.1455,0.01455,0.897,0.01686,0.05437,0.8959,0.7006,0.8936,0.01228,0.05793,0.05689,0.5457,0.3123,0.01021,0.116,0.642,0.00649,0.008647,0.02444,0.6912,0.06377,0.5953,0.1096,0.477,0.08127,0.6394,0.1962,0.9086,0.01858,0.007244,0.6658,0.2724,0.7783,0.1619,0.1458,0.05176,0.005774,0.06608,0.6931,0.867,0.1263,0.1202,0.008527,0.2481,0.08925,0.1746,0.01211,0.5602,0.07669,0.1067,0.8144,0.4804,0.01468,0.2437,0.006708,0.795,0.009017,0.1432,0.2177,0.02315,0.00504,0.3514,0.03909,0.0666,0.00813,0.07125,0.03004,0.02347,0.6821,0.1369,0.3707,0.0819,0.008789,0.006842,0.8294,0.008908,0.006499,0.8197,0.1947,0.7158,0.313,0.09847,0.1059,0.05247,0.9175,0.6839,0.3532,0.04728,0.7067,0.9207,0.9458,0.9473,0.05252,0.9011,0.8861,0.05501,0.945,0.03307,0.07717,0.03792,0.3585,0.9995,0.008543,0.9996,0.9995,0.1479,0.8398,0.2007,0.006711,0.2781,0.9995,0.7047,0.006809,0.7034,0.08823,0.05212,0.04725,0.2464,0.6224,0.6082,0.04302,0.0242,0.161,0.2996,0.124,0.09781,0.2743,0.563,0.01937,0.6577,0.9665,0.4912,0.0411,0.654,0.005087,0.1015,0.05016,0.04743,0.0153,0.1256,0.06776,0.1283,0.8856,0.1431,0.5352,0.135,0.02755,0.4071,0.1657,0.06262,0.5526,0.1252,0.2495,0.01899,0.2018,0.3231,0.08698,0.3325,0.8466,0.04534,0.0565,0.004971,0.1194,0.1193,0.0189,0.01626,0.5056,0.3721,0.3831,0.655,0.005561,0.01725,0.567,0.07012,0.9573,0.8276,0.005464,0.9804,0.03576,0.8559,0.02166,0.05069,0.1199,0.8534,0.2257,0.02783,0.007099,0.6333,0.01763,0.4873,0.5453,0.7493,0.006522,0.04991,0.01364,0.2984,0.1327,0.02774,0.009721,0.3411,0.02557,0.6935,0.3419,0.006283,0.04395,0.43,0.0329,0.3216,0.008169,0.8947,0.06802,0.1581,0.02482,0.02024,0.3523,0.07125,0.06036,0.05684,0.1306,0.5148,0.1786,0.04858,0.9025,0.5368,0.007271,0.2069,0.2059,0.01517,0.08846,0.568,0.06935,0.3464,0.6634,0.1605,0.8824,0.2192,0.04335,0.2564,0.07588,0.02276,0.1501,0.4669,0.3455,0.006718,0.01112,0.4427,0.5979,0.02863,0.907,0.2144,0.3628,0.7394,0.006396,0.09346,0.01722,0.009617,0.9245,0.9765,0.771,0.008217,0.395,0.9331,0.2276,0.03972,0.9847,0.5643,0.106,0.01268,0.9914,0.5386,0.8803,0.1731,0.61,0.01738,0.01282,0.6339,0.5518,0.4378,0.009699,0.09135,0.008958,0.2834,0.008461,0.2869,0.0248,0.01857,0.03022,0.2242,0.0009027,0.05461,0.777,0.9621,0.008579,0.6522,0.1681,0.371,0.1615,0.04664,0.05266,0.2994,0.02684,0.4463,0.3878,0.5512,0.3277,0.2267,0.01096,0.09101,0.08615,0.08492,0.02964,0.04057,0.02526,0.9711,0.5644,0.1159,0.01894,0.1421,0.02095,0.1271,0.01555,0.008128,0.01894,0.02309,0.008203,0.01733,0.05695,0.9279,0.02674,0.0353,0.02573,0.09396,0.7184,0.6721,0.007946,0.6384,0.01209,0.08993,0.007027,0.8697,0.3362,0.984,0.1643,0.04801,0.01071,0.1159,0.1077,0.6689,0.04962,0.08949,0.9525,0.05213,0.9997,0.6059,0.2152,0.01365,0.8281,0.3974,0.07522,0.02719,0.01054,0.009589,0.4275,0.08529,0.2978,0.01917,0.005386,0.09643,0.203,0.001826,0.945,0.09892,0.05689,0.6626,0.8539,0.4727,0.05689,0.005686,0.03825,0.0791,0.1715,0.6954,0.1414,0.03696,0.01581,0.02759,0.1569,0.2267,0.4075,0.006315,0.9906,0.6461,0.1624,0.01214,0.1127,0.06022,0.005931,0.2416,0.01675,0.04395,0.2636,0.007256,0.2637,0.0248,0.009196,0.5274,0.1451,0.005747,0.4534,0.008238,0.2075,0.009895,0.6955,0.1498,0.218,0.2135,0.3965,0.385,0.0274,0.1269,0.04678,0.3741,0.01589,0.1413,0.5781,0.2678,0.09357,0.1383,0.02048,0.008377,0.0615,0.152,0.5108,0.5678,0.6457,0.0164,0.2875,0.226,0.2322,0.005375,0.0259,0.8694,0.697,0.395,0.09982,0.4534,0.2622,0.7669,0.572,0.02865,0.1275,0.04675,0.2136,0.03514,0.2014,0.1796,0.2809,0.06937,0.006774,0.02685,0.02403,0.1756,0.1023,0.126,0.03915,0.2528,0.6082,0.02227,0.8237,0.9098,0.08015,0.505,0.4694,0.1698,0.005669,0.2212,0.03045,0.00536,0.2642,0.306,0.3074,0.03147,0.7142,0.5867,0.8409,0.1159,0.3004,0.06237,0.01098,0.007945,0.07909,0.01751,0.004801,0.007481,0.2472,0.05883,0.06746,0.09523,0.3859,0.3842,0.2625,0.02805,0.6397,0.06979,0.01875,0.6382,0.3265,0.006647,0.8087,0.5366,0.05319,0.7629,0.007439,0.1334,0.49,0.3949,0.2344,0.656,0.7911,0.707,0.9616,0.06789,0.5683,0.3306,0.9996,0.2722,0.04786,0.4173,0.3352,0.06025,0.5842,0.4106,0.4503,0.4892,0.0091,0.1923,0.08846,0.05195,0.02242,0.006559,0.09965,0.4521,0.4395,0.002214,0.1714,0.4649,0.7427,0.03345,0.6092,0.02333,0.6816,0.7857,0.0171,0.6714,0.1907,0.01042,0.04709,0.008515,0.3901,0.01208,0.01127,0.007938,0.3192,0.02048,0.01236,0.02447,0.6952,0.1613,0.6324,0.1936,0.03085,0.06976,0.04075,0.4155,0.01935,0.1516,0.5564,0.9885,0.03576,0.08642,0.7239,0.001382,0.8675,0.3647,0.1755,0.005062,0.9306,0.01366,0.01216,0.1632,0.06527,0.3852,0.008891,0.004965,0.02521,0.04642,0.9903,0.0285,0.07457,0.006584,0.1299,0.628,0.00581,0.6971,0.04321,0.9772,0.006059,0.007273,0.5873,0.09061,0.5348,0.9417,0.1257,0.2854,0.02876,0.115,0.002319,0.512,0.8785,0.005789,0.5861,0.2924,0.08447,0.2767,0.5195,0.08607,0.1419,0.0103,0.2121,0.108,0.03291,0.3097,0.3279,0.01448,0.01125,0.3016,0.007294,0.03842,0.008312,0.3755,0.006231,0.08749,0.02162,0.118,0.05888,0.8232,0.006949,0.7591,0.02209,0.01735,0.341,0.4223,0.0297,0.02623,0.02985,0.9155,0.01631,0.8669,0.9715,0.2433,0.1439,0.5158,0.5342,0.03258,0.03128,0.1154,0.08508,0.5763,0.06805,0.2899,0.4941,0.005626,0.2742,0.5553,0.06008,0.2051,0.09062,0.005979,0.8501,0.02375,0.006781,0.0891,0.07154,0.006415,0.01691,0.1247,0.9749,0.08287,0.1916,0.0338,0.3165,0.2657,0.02278,0.4822,0.7749,0.1006,0.7533,0.8034,0.6624,0.007932,0.00933,0.7651,0.2526,0.004943,0.005724,0.1269,0.008014,0.06776,0.2673,0.06085,0.01103,0.6203,0.006769,0.3307,0.007335,0.3092,0.2097,0.2514,0.01946,0.1128,0.3075,0.88,0.3235,0.04966,0.8501,0.07948,0.09373,0.05587,0.4773,0.9004,0.0245,0.1287,0.02246,0.006501,0.1358,0.009951,0.04449,0.08946,0.05514,0.06179,0.06592,0.05404,0.03483,0.08708,0.04548,0.3313,0.1497,0.2796,0.01218,0.7894,0.1116,0.007316,0.3227,0.5069,0.331,0.9966,0.4997,0.5487,0.1854,0.00482,0.3336,0.1226,0.0999,0.1075,0.006226,0.05452,0.4227,0.182,0.9131,0.1454,0.2798,0.3594,0.4632,0.01593,0.9372,0.01743,0.8119,0.0654,0.06025,0.03161,0.3479,0.369,0.04442,0.03551,0.08628,0.7691,0.492,0.073,0.221,0.1144,0.2203,0.02801,0.01908,0.962,0.08048,0.1679,0.08259,0.2281,0.01279,0.007877,0.005972,0.7344,0.01864,0.009382,0.005535,0.42,0.005516,0.1776,0.4409,0.2583,0.02067,0.007621,0.0112,0.8639,0.2329,0.03374,0.1136,0.7011,0.837,0.02885,0.681,0.8078,0.0978,0.009013,0.05438,0.1751,0.008348,0.5439,0.7645,0.005936,0.05576,0.8864,0.7069,0.1507,0.3949,0.008321,0.2008,0.02106,0.3113,0.6607,0.008419,0.008635,0.2973,0.3683,0.01916,0.3364,0.7353,0.1199,0.02373,0.1784,0.06223,0.02619,0.1282,0.7733,0.08238,0.9646,0.3243,0.9929,0.5759,0.1573,0.2906,0.007613,0.08766,0.9803,0.08223,0.2202,0.1396,0.01241,0.1051,0.004643,0.005128,0.1916,0.5672,0.01231,0.009246,0.1317,0.4107,0.04771,0.9344,0.05457,0.1057,0.6293,0.00735,0.1078,0.1423,0.6505,0.2525,0.01246,0.2769,0.09635,0.05241,0.02403,0.03833,0.6898,0.01114,0.7485,0.4349,0.0947,0.6636,0.4513,0.2195,0.01077,0.009345,0.09554,0.05725,0.01325,0.06988,0.4898,0.006663,0.1012,0.2619,0.005281,0.3967,0.2062,0.8994,0.1593,0.08794,0.3032,0.006566,0.1039,0.4077,0.9612,0.2694,0.2353,0.1243,0.8548,0.1508,0.5521,0.09279,0.5331,0.05195,0.1351,0.06551,0.6655,0.2039,0.7442,0.209,0.8586,0.3968,0.7515,0.7891,0.04384,0.008855,0.09228,0.09441,0.05751,0.7859,0.06973,0.02747,0.3945,0.02296,0.2854,0.0217,0.005806,0.009824,0.005071,0.008041,0.04839,0.594,0.07566,0.04269,0.9015,0.216,0.8814,0.02034,0.519,0.6344,0.7767,0.1873,0.8551,0.08762,0.1974,0.1376,0.03308,0.08121,0.05178,0.08096,0.01903,0.6709,0.9003,0.05867,0.02429,0.07842,0.4565,0.7124,0.6034,0.09106,0.1027,0.1118,0.07485,0.1192,0.01744,0.9702,0.2096,0.00842,0.2141,0.03728,0.007402,0.03302,0.6634,0.4455,0.284,0.01312,0.1544,0.008042,0.9113,0.004882,0.9184,0.04968,0.03278,0.01008,0.6286,0.07543,0.008366,0.04751,0.08248,0.01306,0.1033,0.5192,0.6356,0.9777,0.005379,0.004904,0.04391,0.07152,0.0195,0.01618,0.8259,0.1843,0.08459,0.5488,0.6027,0.2224,0.1537,0.2062,0.2966,0.01641,0.02951,0.343,0.04442,0.008587,0.09295,0.6737,0.5267,0.1017,0.11,0.07253,0.848,0.3998,0.09159,0.5912,0.1489,0.01099,0.461,0.008927,0.3015,0.5607,0.1775,0.02105,0.03061,0.009704,0.3855,0.2383,0.03514,0.01757,0.1377,0.08749,0.3066,0.1875,0.06576,0.3565,0.0134,0.004592,0.00554,0.004605,0.004954,0.6474,0.006856,0.1418,0.0216,0.03255,0.934,0.01567,0.4945,0.5722,0.03146,0.9307,0.05125,0.617,0.06961,0.02215,0.05914,0.1045,0.8121,0.2208,0.007044,0.2602,0.9134,0.1081,0.2383,0.146,0.005717,0.1661,0.1055,0.2182,0.04391,0.007598,0.02184,0.1008,0.07029,0.02892,0.05939,0.007321,0.01349,0.0427,0.01869,0.02448,0.02194,0.00514,0.3871,0.8337,0.08358,0.1127,0.6281,0.6633,0.3365,0.9415,0.09399,0.05973,0.0691,0.06647,0.02482,0.1799,0.08052,0.5679,0.1796,0.964,0.1089,0.03727,0.0805,0.8499,0.019,0.3626,0.2361,0.9688,0.00996,0.9128,0.01277,0.01004,0.4523,0.004937,0.5121,0.0294,0.01863,0.8621,0.6234,0.6027,0.9345,0.1483,0.009347,0.08204,0.1179,0.2021,0.03453,0.9137,0.03394,0.2534,0.1115,0.05625,0.0175,0.6681,0.05973,0.76,0.9041,0.0919,0.01244,0.01693,0.02837,0.02117,0.02049,0.7465,0.07889,0.01376,0.04287,0.005135,0.03126,0.005835,0.3254,0.5892,0.5429,0.2485,0.4666,0.6023,0.01187,0.007722,0.1566,0.4222,0.02854,0.1919,0.005553,0.03191,0.0638,0.01145,0.0516,0.01749,0.001531,0.1017,0.9738,0.4518,0.05037,0.01189,0.9,0.3307,0.01564,0.01037,0.04801,0.01344,0.7584,0.009236,0.04955,0.04248,0.2608,0.02574,0.07758,0.1209,0.02624,0.1597,0.009727,0.6651,0.4676,0.8868,0.1193,0.0872,0.03063,0.006224,0.798,0.1316,0.1912,0.004671,0.07273,0.01937,0.2417,0.02124,0.9443,0.008699,0.00797,0.1148,0.5377,0.01719,0.8783,0.822,0.3989,0.6941,0.007002,0.01452,0.05619,0.09751,0.02036,0.164,0.2329,0.6826,0.203,0.2647,0.04536,0.4053,0.01743,0.7102,0.428,0.6215,0.005525,0.9998,0.02092,0.2624,0.3893,0.005474,0.0366,0.09154,0.0914,0.05951,0.1237,0.0164,0.04891,0.5583,0.8432,0.293,0.6851,0.6172,0.3471,0.02472,0.1229,0.209,0.01398,0.007459,0.1832,0.1597,0.8926,0.03222,0.01319,0.04444,0.5455,0.9589,0.3327,0.1359,0.01324,0.8999,0.6688,0.6435,0.1804,0.2342,0.0147,0.01673,0.005737,0.06555,0.006618,0.09409,0.0129,0.09019,0.04529,0.07857,0.1273,0.5579,0.08367,0.4376,0.00572,0.007208,0.6442,0.2437,0.01112,0.2347,0.6492,0.07393,0.4614,0.0352,0.06703,0.01,0.9397,0.009046,0.02344,0.715,0.9862,0.03056,0.03133,0.5832,0.2452,0.168,0.2225,0.1277,0.1693,0.4545,0.4278,0.3282,0.02089,0.06184,0.07362,0.1212,0.9997,0.06615,0.8306,0.8131,0.8739,0.8396,0.008701,0.5977,0.07783,0.2029,0.4983,0.6222,0.02741,0.4362,0.02526,0.9559,0.06498,0.02145,0.5589,0.9633,0.4698,0.2498,0.006878,0.09162,0.6876,0.9152,0.005198,0.3261,0.006891,0.06407,0.03626,0.5429,0.2251,0.1562,0.01627,0.02224,0.02597,0.03533,0.4823,0.0109,0.06782,0.004713,0.0812,0.0621,0.01496,0.9007,0.7435,0.661,0.04253,0.7497,0.4185,0.09175,0.00749,0.1398,0.7994,0.1372,0.3091,0.0198,0.3115,0.007295,0.9516,0.8996,0.2964,0.005614,0.01385,0.08158,0.3382,0.514,0.9034,0.04488,0.1412,0.1066,0.006183,0.2913,0.09802,0.06104,0.004751,0.2556,0.103,0.03907,0.3355,0.01049,0.007373,0.8831,0.3315,0.01381,0.05315,0.2141,0.873,0.05997,0.34,0.4182,0.005161,0.0181,0.2442,0.47,0.04369,0.1712,0.3299,0.2208,0.4951,0.03215,0.2126,0.1215,0.03533,0.6373,0.7616,0.03048,0.1021,0.6637,0.08903,0.3657,0.04417,0.2458,0.006619,0.03534,0.8995,0.3623,0.3691,0.2214,0.8588,0.05667,0.6061,0.01672,0.01487,0.004748,0.06042,0.4238,0.3512,0.1196,0.2185,0.7835,0.1376,0.7258,0.05167,0.2734,0.695,0.05793,0.2537,0.07384,0.02455,0.7285,0.004678,0.05922,0.04735,0.01765,0.574,0.007345,0.05597,0.01095,0.6207,0.02068,0.006112,0.6312,0.1128,0.6064,0.3667,0.6091,0.02271,0.01406,0.6023,0.6136,0.02903,0.09087,0.1634,0.0691,0.1898,0.009123,0.006305,0.004976,0.1208,0.02477,0.03004,0.02968,0.006103,0.01469,0.6483,0.01125,0.3103,0.811,0.03492,0.06412,0.6405,0.128,0.6987,0.2498,0.9597,0.007862,0.8719,0.02371,0.7335,0.05875,0.02308,0.1576,0.769,0.02098,0.5045,0.05409,0.2861,0.2165,0.941,0.02898,0.7517,0.0204,0.0887,0.8088,0.6754,0.005401,0.1929,0.02848,0.5599,0.09648,0.1034,0.8338,0.1036,0.0204,0.06678,0.7536,0.01893,0.07437,0.1105,0.2189,0.6285,0.0372,0.07099,0.0162,0.7027,0.1526,0.1503,0.5102,0.00857,0.006735,0.2211,0.163,0.4771,0.1778,0.03427,0.0568,0.1096,0.9606,0.2773,0.008872,0.7791,0.005633,0.3739,0.2116,0.007364,0.9699,0.8997,0.01276,0.005507,0.01764,0.7869,0.01091,0.753,0.03935,0.03057,0.04008,0.4032,0.02813,0.00703,0.2512,0.2907,0.195,0.02643,0.2398,0.3893,0.1261,0.2148,0.02916,0.5329,0.107,0.08522,0.02788,0.2316,0.01344,0.2028,0.02035,0.1755,0.008468,0.9134,0.1527,0.5833,0.09723,0.2854,0.005389,0.9627,0.0235,0.004617,0.4788,0.7659,0.3633,0.008851,0.9349,0.09935,0.1573,0.219,0.005487,0.05704,0.1696,0.4759,0.5742,0.2352,0.5399,0.3725,0.004857,0.01673,0.1349,0.04078,0.1031,0.5992,0.0478,0.8742,0.5048,0.008537,0.03957,0.01386,0.007425,0.3286,0.2249,0.4125,0.6136,0.1281,0.1254,0.4097,0.01836,0.01427,0.01188,0.6321,0.03347,0.9936,0.8477,0.04588,0.01773,0.01998,0.1572,0.7539,0.2688,0.7712,0.2663,0.5353,0.2955,0.006379,0.112,0.3534,0.817,0.4585,0.3464,0.5282,0.08885,0.06766,0.8113,0.3841,0.9661,0.9134,0.3108,0.8181,0.1378,0.9996,0.0899,0.09728,0.9995,0.9102,0.007011,0.004717,0.9997,0.002668,0.001491,0.001277,0.003891,0.0004243,0.0004506,3.366e-05,1.638e-06,0.001116,0.002503,0.0002495,0.002037,0.002707,1.4e-06,1.717e-08,0.0003249,0.002603,9.526e-07,0.0006717,0.003652,1.464e-10,0.0001151,4.301e-07,1.774e-06,1.749e-07,0.002648,2.505e-07,6.153e-05,7.172e-05,2.321e-08,2.334e-06,0.00334,0.003032,0.0005788,3.05e-05,0.001149,3.25e-07,8.016e-07,1.035e-07,2.022e-10,0.0007828,4.967e-05,1.701e-06,8.917e-06,0.001515,7.754e-07,0.0006192,0.000174,0.003274,6.046e-06,0.001499,1.881e-05,0.0002373,0.0001277,0.002592,1.966e-05,6.152e-06,8.781e-05,0.000462,0.001659,5.402e-08,0.0008801,5.882e-08,9.253e-06,3.035e-09,0.001598,7.939e-06,5.798e-06,0.000141,0.0017,0.003578,0.001029,3.251e-07,0.0002389,4.566e-06,4.663e-06,3.455e-07,8.846e-10,0.002192,2.83e-09,0.0001064,5.019e-07,2.727e-06,7.722e-08,0.0008743,4.171e-07,4.605e-05,1.424e-05,0.0006207,1.239e-05,2.278e-07,4.419e-09,0.001003,1.608e-05,2.767e-05,0.002853,2.473e-07,0.003586,0.0001124,0.0002465,1.124e-05,0.0003216,0.0008878,6.25e-10,2.388e-08,7.28e-08,0.003021,6.419e-09,3.142e-06,7.577e-05,8.88e-07,0.0001847,2.829e-07,3.071e-05,1.219e-07,0.002708,0.0009093,0.00217,0.002867,0.002595,0.002583,3.277e-06,0.00207,2.837e-10,0.000117,3.227e-06,7.339e-07,0.0002442,1.191e-06,1.541e-06,0.00324,0.001062,3.557e-05,2.256e-08,1.172e-06,6.273e-06,1.255e-08,2.476e-05,0.001213,0.002374,4.656e-08,0.0002012,0.0009279,0.002089,6.012e-08,1.071e-06,6.265e-05,3.985e-05,3.922e-06,0.0007365,9.726e-09,2.829e-05,0.002863,0.0002516,0.000227,0.0002635,5.766e-08,0.001333,0.002531,0.0001397,3.312e-05,9.543e-06,0.0002059,3.349e-07,5.191e-09,8.322e-11,7.999e-09,2.464e-07,3.324e-09,1.587e-05,1.241e-10,9.834e-09,1.363e-09,7.007e-05,0.00329,0.001579,1.787e-05,6.459e-05,1.176e-05,0.00434,1.078e-05,2.194e-09,5.123e-11,0.0001664,0.003751,2.424e-06,3.797e-07,2.659e-08,0.0003565,2.188e-05,7.141e-07,1.522e-06,0.0001856,0.001264,2.549e-08,0.002044,9.377e-07,0.000859,1.907e-08,0.0005075,3.131e-10,1.328e-06,0.003609,0.0002699,6.148e-06,2.776e-06,1.201e-05,6.359e-06,0.0002125,2.076e-07,0.001553,1.035e-05,1.987e-07,1.804e-05,8.805e-06,6.344e-08,0.0001997,2.112e-06,7.687e-06,0.001093,0.0002749,6.15e-07,0.001637,0.001388,1.495e-06,3.797e-06,3.979e-06,0.0002137,4.511e-06,5.14e-08,7.559e-05,7.006e-05,3.695e-05,2.107e-05,5.874e-08,3.284e-06,0.0038,4.866e-06,5.315e-06,2.2e-07,9.664e-05,9.519e-06,6.673e-10,2.34e-10,9.374e-06,0.001042,4.558e-08,2.298e-05,3.546e-05,0.002535,4.355e-07,0.0009429,0.0005318,4.689e-05,4.161e-09,1.395e-06,1.969e-05,1.356e-05,1.673e-08,0.0008094,3.33e-09,0.0006389,5.79e-10,7.358e-08,0.004453,0.0002351,2.883e-06,1.492e-06,2.916e-06,1.833e-06,0.0003047,6.621e-05,0.003168,0.0007875,2.353e-05,1.007e-08,8.283e-06,2.158e-08,0.0009066,6.493e-05,0.002815,7.527e-05,0.0003967,0.0001087,0.002246,3e-10,2.323e-06,0.002192,1.45e-09,0.002505,3.377e-06,1.269e-07,8.277e-06,3.775e-08,0.002147,0.0008351,1.345e-05,0.0002967,0.0002991,4.48e-05,0.001531,2.836e-05,5.287e-05,0.0001866,4.339e-05,1.758e-05,2.185e-06,9.516e-11,2.417e-05,2.847e-09,1.558e-05,4.449e-07,0.0004166,0.0004034,0.0004048,0.0003032,9.634e-08,1.524e-07,1.885e-07,1.234e-05,1.308e-06,8.539e-06,9.797e-06,0.0008362,2.561e-06,2.88e-06,1.104e-08,0.001213,5.614e-06,5.264e-05,0.0002114,0.0007216,5.354e-09,1.369e-05,6.867e-07,4.903e-05,0.001445,5.074e-08,1.009e-05,0.001191,0.002487,7.023e-06,0.0001005,7.266e-05,2.293e-06,1.116e-08,6.013e-06,9.306e-10,0.001151,0.0006056,9.194e-07,0.000828,1.033e-08,0.003351,0.000648,0.0002228,1.605e-05,0.003269,0.0003895,5.449e-06,7.695e-08,1.042e-08,7.925e-08,7.072e-07,6.837e-05,1.097e-06,2.379e-10,1.204e-05,9.976e-05,0.003041,0.0004484,1.158e-06,1.483e-07,4.203e-10,8.019e-05,5.505e-08,0.003417,0.000444,5.44e-07,0.0003824,0.0003366,1.042e-08,1.476e-09,2.173e-06,3.772e-05,0.0003185,0.0006774,7.84e-05,0.001353,5.278e-07,0.0001712,2.905e-09,1.695e-05,0.0007472,7.378e-09,0.001255,3.014e-05,2.764e-05,7.199e-06,0.0008453,0.0001261,0.003157,0.002733,4.878e-06,0.001601,4.984e-07,7.733e-06,3.082e-05,1.686e-07,4.802e-06,0.0001966,3.306e-06,7.23e-09,0.0001668,2.363e-06,0.000277,0.001937,5.975e-05,7.551e-05,1.08e-11,1.954e-07,0.0001357,0.000789,1.723e-06,4.577e-05,0.003839,0.0002109,0.0002568,2.079e-05,7.494e-05,0.000167,1.257e-06,3.705e-05,0.000271,9.774e-05,0.0006781,9.421e-08,1.75e-05,2.428e-08,6.334e-08,0.001275,0.0001368,0.0003061,6.68e-06,0.0009099,0.003732,0.003158,0.0001361,0.0009713,0.001948,0.0001132,0.004155,0.002827,0.0003086,0.0001159,2.891e-05,0.0001182,0.0002757,2.013e-05,2.225e-08,1.248e-05,8.55e-09,1.479e-06,0.001131,1.895e-05,0.0007039,4.99e-08,1.413e-07,0.001923,0.003996,2.628e-05,0.002873,4.228e-06,2.788e-06,5.536e-09,5.062e-09,8.978e-05,9.641e-07,9.683e-06,0.002426,0.0007493,0.003673,0.0003197,1.579e-07,2.167e-05,2.199e-06,0.0001235,0.0001592,0.002803,1.15e-05,3.498e-09,4.157e-05,3.592e-06,0.00382,5.208e-07,0.0017,0.001241,0.0001253,0.001338,0.0004451,0.002167,6.413e-08,4.456e-05,6.227e-12,3.228e-10,0.004054,2.126e-05,6.375e-09,2.506e-06,4.635e-06,8.567e-07,3.127e-05,3.861e-05,2.882e-05,2.448e-05,1.929e-06,2.768e-05,0.0004726,0.0008176,1.127e-07,2.479e-09,0.000141,4.675e-06,0.001233,9.829e-05,0.0002603,0.0001051,3.363e-05,2.376e-07,8.637e-07,9.132e-05,0.002568,7.099e-06,0.002176,1.629e-06,0.0009795,4.332e-09,0.0008333,0.0001893,1.607e-05,1.999e-07,4.531e-06,6.205e-06,2.644e-06,0.001384,0.0003573,0.00312,0.002259,5.396e-05,8.659e-08,7.973e-10,0.000278,4.334e-08,0.003569,1.051e-05,6.507e-05,1.123e-05,2.753e-05,6.912e-06,0.0002672,1.03e-06,1.104e-05,1.045e-07,2.754e-06,2.159e-05,9.354e-06,3.221e-07,2.767e-06,4.395e-08,5.055e-05,0.004177,3.089e-07,3.469e-06,0.001795,3.392e-08,7.689e-08,5.971e-07,9.061e-08,0.0002559,0.002732,0.000201,1.654e-05,1.579e-05,1.166e-06,0.001665,0.0006023,2.266e-05,0.001257,0.002205,4.293e-07,5.724e-05,6.763e-08,5.311e-05,1.614e-06,2.426e-10,0.001215,0.001436,0.0009619,0.0003383,0.003713,1.426e-05,2.203e-10,0.0022,0.001035,6.595e-05,0.0007722,5.304e-06,7.351e-05,6.442e-10,1.849e-07,3.519e-06,1.016e-07,2.693e-06,4.532e-05,0.0001829,0.0002337,1.02e-05,0.0008565,1.71e-05,0.000801,0.0004659,4.557e-05,1.201e-09,0.002988,4.057e-05,0.004096,0.002805,0.001122,3.459e-05,6.019e-06,9.94e-07,0.0004284,2.986e-07,6.016e-05,5.985e-05,2.203e-08,0.001644,3.641e-08,5.45e-08,0.0005349,0.002209,1.063e-05,0.0001122,0.0005345,0.0002447,7.493e-05,0.002949,0.0009381,0.0008105,1.046e-07,2.727e-09,2.705e-06,1.048e-07,5.309e-08,5.584e-06,0.001985,0.004117,4.48e-06,2.423e-07,1.388e-06,4.351e-06,0.0001422,1.02e-09,0.0001882,2.038e-06,4.715e-07,0.0002003,0.0004442,0.0001536,4.954e-06,9.565e-06,0.002332,7.609e-09,2.638e-08,4.443e-06,0.0001458,0.002724,0.00117,0.0009578,0.0004855,5.053e-09,1.406e-06,0.0001572,1.273e-05,8.717e-07,2.155e-08,0.001027,3.717e-07,0.0003995,6.489e-08,0.0001849,2.186e-08,5.378e-08,0.0001898,5.966e-06,2.507e-05,0.001735,0.000218,2.773e-07,3.385e-06,7.91e-05,0.0001833,7.733e-06,0.0004001,4.924e-05,1.066e-08,8.89e-06,4.098e-05,6.546e-07,1.692e-07,0.0002721,0.001527,0.0009288,0.001396,0.0004803,9.356e-05,4.808e-09,0.00221,1.899e-09,0.002779,1.905e-06,0.001544,0.0001329,0.00136,1.557e-05,0.0001866,3.165e-05,0.002016,5.443e-07,1.625e-05,2.823e-06,1.368e-05,5.441e-07,2.336e-09,0.0004611,0.0001289,2.587e-08,0.002397,0.0009626,2.629e-05,9.613e-09,4.187e-08,2.029e-08,4.914e-10,2.127e-06,6.682e-08,2.759e-10,5.313e-05,4.039e-07,0.0001218,5.603e-06,3.777e-07,7.472e-05,0.000896,5.035e-05,6.698e-07,2.775e-05,0.0001001,7.594e-06,4.504e-07,1.059e-07,0.004105,3.885e-05,2.88e-09,5.327e-07,0.00305,0.0001675,0.0002765,4.149e-05,9.222e-05,1.072e-05,6.937e-06,0.004435,0.00173,8.349e-06,0.0001184,1.259e-05,2.232e-06,0.000827,0.003023,3.873e-06,0.001226,2.699e-05,0.000208,0.004537,1.848e-07,0.00208,2.448e-05,1.283e-05,0.002314,2.695e-09,2.545e-06,0.003736,1.334e-06,0.002557,1.839e-08,0.0004067,4.487e-08,1.109e-07,4.655e-05,0.0002986,1.525e-06,1.652e-08,1.073e-05,3.167e-05,0.0001906,1.934e-06,0.002211,5.703e-05,6.227e-11,4.737e-05,7.055e-05,0.001005,8.496e-05,0.0006439,1.334e-05,2.816e-09,8.391e-06,4.411e-10,1.416e-07,6.825e-07,0.0003197,0.000912,5.117e-05,0.0001832,1.568e-10,4.798e-06,1.347e-05,1.837e-05,0.0009057,0.0001425,1.88e-05,1.446e-05,1.222e-06,1.572e-06,2.047e-05,5.293e-10,9.153e-06,1.811e-05,0.0002209,0.0002743,0.0003293,9.85e-10,3.738e-08,2.662e-06,1.045e-06,4.674e-05,5.824e-05,1.904e-07,5.191e-07,4.5e-05,6.378e-07,5.814e-05,1.235e-05,1.048e-06,8.165e-05,0.0001575,0.0004064,2.203e-05,3.189e-05,0.0006672,0.0008217,0.0008563,0.001194,0.001098,9.087e-05,0.000212,1.102e-06,0.0001602,0.000177,7.688e-07,8.395e-07,1.15e-08,2.392e-08,0.001565,0.0002172,5.001e-06,2.808e-05,0.0005491,0.001262,0.0001402,2.415e-05,0.001247,8.64e-05,0.0006525,0.002177,0.00287,4.074e-06,0.001959,1.106e-08,0.0001829,8.061e-05,0.00285,0.003132,4.981e-05,0.001087,2.554e-05,0.001414,3.238e-08,6.494e-05,5.729e-06,0.0004864,2.302e-06,0.00308,4.904e-06,8.305e-07,0.003425,2.083e-06,0.001132,0.000175,0.0005651,2.478e-09,7.069e-05,0.001395,8.209e-07,0.0003125,0.0001079,5.61e-06,1.487e-05,5.598e-06,5.246e-06,0.002873,4.749e-05,0.0003857,2.796e-07,0.002591,1.058e-06,0.003884,1.89e-05,3.712e-05,0.003213,6.496e-08,1.732e-05,9.307e-05,2.666e-09,0.002307,0.001091,3.896e-08,4.51e-06,8.128e-05,0.001906,0.0001927,0.0002734,0.001109,5.563e-07,0.0002589,1.782e-07,5.337e-06,2.737e-05,3.223e-07,2.015e-08,7.394e-06,0.0001129,2.246e-05,2.371e-05,7.166e-05,4.739e-06,2.459e-07,1.393e-08,9.493e-06,5.516e-08,0.0005565,9.629e-09,1.566e-09,0.0003714,0.001377,0.002508,8.899e-07,3.287e-09,0.0007857,1.395e-07,1.221e-07,0.002154,4.187e-05,0.0001299,0.0001056,5.533e-06,1.714e-09,7.259e-06,0.003278,0.001162,6.443e-06,2.044e-06,5.012e-09,0.0001014,0.0001359,0.001551,0.0001572,0.0002965,0.003554,0.0001046,0.001214,9.473e-06,0.001176,0.00361,0.0006086,0.0001452,2.241e-06,0.002213,0.0005804,0.0001119,0.0006642,2.744e-05,4.06e-05,2.454e-06,2.164e-09,7.712e-08,9.468e-08,7.366e-06,0.0002848,1.067e-06,1.274e-06,0.0002721,0.0008718,9.09e-09,0.0007224,0.0001651,0.001348,3.541e-06,1.064e-05,0.0008613,0.001551,4.79e-05,2.145e-06,0.000314,0.002112,1.615e-05,4.163e-09,0.0001702,7.485e-06,0.0006974,2.653e-07,6.012e-05,2.383e-06,7.765e-05,0.0008201,1.161e-06,3.143e-05,0.0006948,4.677e-05,0.001757,2.007e-07,1.143e-08,7.763e-11,4.571e-07,2.995e-05,0.00122,9.555e-05,2.196e-05,0.003146,1.106e-11,0.002133,0.000505,5.459e-08,7.517e-05,1.9e-08,0.0009493,5.833e-08,0.0003304,4.519e-06,2.409e-06,0.0006068,0.001384,6.008e-05,1.091e-10,8.214e-07,4.374e-09,0.0008074,1.529e-05,0.003877,0.0003884,5.736e-06,2.432e-06,0.0008096,0.0001853,4.765e-05,0.002713,0.003783,0.002316,6.882e-06,9.489e-05,6.679e-06,4.769e-08,1.109e-05,3.879e-06,2.452e-05,8.669e-05,0.0002909,1.253e-07,1.525e-09,6.143e-06,1.244e-06,0.001665,7.095e-05,1.53e-07,5.45e-05,0.0007576,0.0005922,1.65e-05,0.002466,8.84e-05,1.399e-05,0.001005,0.001541,1.864e-06,1.228e-08,0.0009938,3.991e-05,0.0006834,0.0007233,1.474e-05,0.002241,4.374e-05,5.954e-11,4.103e-06,0.0009998,3.753e-07,0.0004643,1.178e-07,0.001107,3.624e-10,2.262e-07,0.001069,0.0021,2.957e-09,1.28e-05,0.0001937,0.0002642,0.0006268,1.175e-06,3.547e-09,1.034e-05,7.074e-05,4.645e-05,0.0002418,7.192e-05,2.64e-07,0.001797,4.365e-05,3.346e-08,3.193e-08,7.232e-05,0.0006695,0.0008871,0.001846,0.000906,0.002019,1.761e-05,0.0001354,0.002336,1.104e-06,0.0002261,3.008e-07,0.0002786,1.675e-05,5.978e-08,1.007e-06,0.0003693,5.698e-05,9.037e-07,1.852e-06,0.0006395,1.425e-05,0.0004632,0.0001144,2.647e-05,0.0006568,0.001661,0.003812,1.077e-05,0.0003243,0.000449,0.001217,8.849e-05,0.0002989,1.255e-05,6.471e-05,0.0005076,0.0002671,0.0001685,5.961e-05,0.002487,5.359e-07,0.001369,0.001657,1.59e-06,4.155e-07,9.136e-07,0.0006192,2.645e-06,2.147e-05,2.211e-08,0.0004336,0.002008,2.223e-05,0.0008204,0.001088,0.0001721,2.938e-05,2.242e-06,4.459e-05,2.027e-05,2.464e-06,1.066e-05,0.0002229,0.0001002,1.132e-08,1.617e-05,6.247e-08,0.0002835,1.916e-05,0.001282,1.295e-05,0.00387,1.734e-06,0.003791,7.405e-08,0.0004742,9.644e-05,0.003294,0.001589,0.00254,0.002518,0.0001998,2.512e-06,0.0005147,2.827e-10,0.002828,0.0001184,3.578e-05,7.999e-05,5.983e-06,0.000586,6.873e-07,9.923e-09,0.000142,0.00244,1.502e-05,5.247e-06,5.815e-07,5.611e-05,6.911e-06,7.396e-06,0.003445,7.621e-07,2.371e-05,0.001015,1.213e-07,0.0008059,0.001623,0.00156,2.922e-06,1.7e-11,1.207e-05,4.538e-07,0.00327,9.071e-08,0.001922,8.268e-09,0.002825,1.741e-07,0.0004283,3.502e-07,2.778e-06,0.000343,5.572e-05,0.001524,0.0006223,4.247e-06,1.158e-06,0.001944,1.29e-06,4.265e-05,2.586e-06,0.0002016,0.0006254,1.745e-09,1.874e-07,0.0001529,3.609e-05,0.0002469,3.748e-05,2.81e-06,0.0002775,0.001581,4.73e-06,4.501e-06,0.0008436,0.00348,0.001702,0.0006449,4.793e-05,0.0001025,0.0003345,2.61e-06,5.603e-05,0.0003523,1.947e-06,0.001938,0.0003997,5.16e-05,0.0002164,0.0008319,3.473e-06,0.0001314,0.004295,1.084e-06,6.997e-06,1.421e-07,0.000155,1.216e-05,0.001004,1.572e-05,2.215e-07,0.002048,2.855e-05,1.159e-06,8.707e-05,5.675e-07,8.674e-05,5.922e-05,0.0008218,3.096e-05,0.001584,0.0008756,1.603e-08,0.001446,1.915e-05,0.0002407,0.0004152,0.0004282,3.719e-05,4.278e-05,0.0004265,0.001432,1.732e-06,0.0007095,1.315e-06,1.807e-05,0.0001097,2.265e-08,0.0004214,3.793e-05,5.18e-06,0.001966,4.115e-05,5.691e-08,0.0002784,0.0009885,0.001866,0.002118,0.0003264,2.088e-07,0.001482,0.0003434,5.718e-06,7.936e-05,7.767e-05,5.561e-05,1.66e-06,4.709e-05,4.315e-05,0.002329,2.108e-05,1.99e-05,0.000118,0.0003506,1.387e-06,1.179e-06,0.0001212,1.221e-05,0.001492,4.733e-09,0.0002221,2.797e-07,0.001587,0.0004879,1.883e-06,7.536e-07,0.004153,2.636e-05,7.712e-05,1.631e-07,0.003034,0.002322,0.003952,0.0005532,1.763e-05,4.353e-06,0.001233,0.0001019,7.775e-05,0.003812,2.211e-12,0.002625,0.001757,3.467e-05,0.0001224,1.339e-05,0.002765,1.139e-08,0.003317,0.0004109,2.89e-07,1.294e-05,0.0007265,8.919e-05,0.000419,2.377e-06,1.931e-05,0.003886,9.727e-06,0.0007561,0.0004867,5.512e-10,8.376e-05,0.000434,0.0001366,3.14e-06,0.002847,3.069e-06,0.0001445,0.001735,2.779e-08,1.581e-07,1.56e-08,0.002718,2.78e-06,7.835e-06,1.934e-06,9.126e-07,6.323e-07,7.208e-09,0.0007243,0.000882,0.001231,0.0001141,0.000122,6.63e-06,3.927e-07,2.345e-08,4.453e-07,3.09e-07,3.212e-05,2.768e-05,0.003838,4.116e-09,7.925e-05,0.0002438,4.86e-07,0.003055,6.99e-05,1.277e-09,0.002015,0.0002565,0.0003662,9.416e-06,0.0002227,5.546e-07,1.168e-07,2.326e-06,1.51e-06,3.211e-06,0.001315,5.481e-09,8.713e-05,3.85e-09,2.488e-08,2.976e-05,1.742e-06,8.819e-05,1.043e-06,0.000577,1.998e-05,5.601e-07,5.905e-05,1.088e-07,6.778e-05,0.001611,9.265e-07,0.001756,2.056e-05,3.405e-05,0.001266,0.004057,1.165e-07,3.758e-05,2.686e-06,1.903e-05,4.014e-05,1.37e-08,4.661e-05,1.585e-07,0.0001808,0.0004007,1.123e-09,0.0003293,0.001198,4.79e-08,0.0001162,0.0004589,0.001672,4.128e-05,4.807e-06,1.177e-06,0.0007861,1.718e-06,0.004478,7.859e-08,0.0003817,0.0003836,0.004401,5.3e-08,0.0001395,1.706e-06,4.784e-05,9.147e-09,2.6e-09,7.756e-05,1.035e-08,2.017e-06,6.632e-05,0.00288,0.001654,0.0007115,1.403e-06,0.001085,6.131e-06,5.855e-06,2.431e-07,6.813e-07,3.47e-07,6.464e-07,2.671e-05,4.683e-06,6.745e-05,3.559e-08,0.0002742,2.141e-05,1.694e-06,2.529e-06,1.093e-06,0.0003553,1.892e-06,0.0003669,0.001131,6.019e-06,0.0006859,1.064e-07,0.0006548,0.0001457,1.682e-05,1.492e-08,0.0003619,9.904e-06,0.0001124,1.272e-06,0.000325,0.0001004,7.193e-10,5.126e-07,0.003315,0.001055,2.788e-09,0.000151,0.0003101,0.0001586,8.89e-06,0.0004075,3.906e-05,4.569e-06,0.0001009,0.0001035,0.0003388,0.0004014,0.0005951,0.0009753,2.222e-08,7.01e-08,6.609e-05,0.00234,1.227e-06,1.035e-05,4.619e-07,0.002159,0.0001096,0.0001478,3.229e-05,3.942e-07,7.915e-06,1.253e-06,1.113e-06,1.209e-06,5.849e-09,3.022e-10,0.0007229,0.002839,8.104e-07,1.936e-05,4.365e-06,2.644e-07,0.0001455,0.001919,6.362e-07,1.293e-06,1.652e-09,1.487e-06,0.0005564,0.0003135,0.003932,0.0008312,4.432e-05,6.618e-07,0.001247,3.471e-06,0.0002682,0.002534,1.237e-05,2.172e-07,5.27e-06,2.035e-06,0.002902,6.342e-08,2.889e-07,3.006e-06,0.0006542,0.0006949,4.192e-05,6.805e-08,9.406e-07,2.434e-05,0.0005064,0.0003031,0.001549,6.991e-06,1.891e-07,0.0005602,1.506e-05,5.908e-07,0.003044,0.0006688,3.238e-08,2.369e-06,0.0002957,9.626e-06,0.0002853,0.0002768,5.356e-05,0.001061,6.735e-09,2.669e-09,0.00194,0.004581,2.781e-06,1.335e-06,0.001298,1.658e-07,5.956e-05,2.641e-06,2.038e-09,1.559e-07,8.572e-06,9.913e-09,1.137e-06,5.49e-08,2.278e-08,1.534e-07,4.243e-11,3.61e-07,1.249e-05,0.002466,2.383e-06,6.27e-07,1.031e-05,0.0003658,5.758e-05,8.938e-05,6.253e-05,1.047e-06,3.417e-09,0.001459,1.7e-10,2.576e-05,1.843e-07,0.003115,1.59e-06,0.0006784,3.26e-08,6.579e-06,0.004456,0.001123,7.95e-07,3.644e-05,0.0001687,4.903e-05,3.126e-05,0.0005495,0.002112,2.786e-06,1.234e-11,6.55e-10,0.0002572,1.638e-06,7.075e-07,0.0002887,0.001383,0.000833,0.0002027,0.001151,0.000511,7.902e-05,0.0004961,1.061e-05,0.0007086,1.374e-06,1.825e-06,1.549e-09,9.075e-09,6.148e-07,6.389e-05,0.002221,4.75e-06,1.742e-06,0.0004882,3.297e-05,0.0007459,4.425e-09,1.246e-06,3.978e-05,0.004277,5.616e-08,0.0003282,6.549e-07,1.268e-06,5.195e-08,0.003722,6.583e-05,1.308e-06,0.001165,3.101e-05,6.939e-06,2.404e-05,2.76e-05,7.015e-05,1.751e-05,1.284e-09,0.0004521,0.002723,0.001978,6.261e-07,4.841e-07,0.0001266,0.004362,0.00154,0.000408,1.1e-06,0.003208,2.659e-08,4.944e-06,0.0005019,0.00394,0.0002295,2.741e-07,9.037e-09,8.905e-05,0.0001912,0.002835,1.656e-06,2.926e-08,2.922e-05,6.045e-06,1.485e-06,9.528e-05,1.14e-06,3.694e-05,0.0001236,0.0001526,0.0022,1.045e-06,9.26e-05,0.003777,8.076e-06,0.002053,9.003e-11,0.003506,0.0002788,0.001337,2.776e-06,9.824e-05,0.002002,1.206e-09,0.002054,0.000403,0.002901,1.622e-05,0.00382,0.004199,0.001211,0.003344,2.65e-07,7.168e-06,2.978e-07,0.002279,1.136e-06,4.831e-06,5.416e-06,1.103e-07,1.314e-05,0.0002586,0.0006869,7.969e-05,0.0001609,0.0001192,0.0003314,4.85e-07,0.001711,3.827e-08,6.835e-06,0.002732,2.866e-05,1.015e-05,9.242e-08,6.885e-06,0.0008519,1.673e-05,0.00188,3.89e-05,0.001157,0.0005847,1.769e-08,2.046e-08,0.002725,0.0002241,0.003326,0.001812,0.0006987,8.679e-07,0.0002775,0.001406,1.698e-07,6.94e-09,0.001746,0.0008088,0.003677,0.0008844,0.000383,1.818e-05,1.289e-06,8.402e-05,2.588e-05,0.0004978,1.157e-09,6.416e-07,0.003456,9.064e-05,8.615e-06,0.002333,0.001246,0.0001224,7.203e-08,0.00118,0.0001136,6.858e-06,1.401e-08,8.22e-06,4.98e-05,7.117e-06,8.282e-09,1.042e-05,7.964e-05,3.545e-05,0.002205,9.176e-06,8.59e-05,6.385e-07,0.0001395,0.0007787,3.495e-08,4.003e-05,6.255e-06,2.46e-05,0.001257,0.0001144,0.0001415,3.483e-05,4.19e-09,0.0001076,4.329e-05,1.34e-05,1.922e-06,0.0009381,3.289e-10,2.63e-05,0.0006461,0.0003702,3.936e-05,7.741e-09,6.619e-06,7.154e-06,0.0002316,0.0007068,7.121e-09,0.004038,0.0008626,1.366e-06,3.021e-06,0.0009805,1.578e-08,4.353e-07,0.001445,1.277e-05,0.002838,1.887e-05,0.001888,1.814e-05,0.001016,2.038e-07,0.00221,0.002717,0.0004636,0.0008528,0.004558,0.0004664,0.003645,0.001932,3.03e-05,4.29e-05,0.0003494,1.383e-05,0.0003598,1.33e-08,9.544e-05,4.639e-05,1.815e-06,0.001028,1.266e-05,1.212e-07,0.002107,0.001263,2.43e-07,6.092e-05,0.0008628,0.0002859,0.0003735,6.012e-05,6.495e-05,4.237e-07,1.103e-05,3.663e-07,0.0003302,0.0006703,1.932e-08,0.002253,0.002056,6.136e-06,0.003965,7.955e-05,0.001783,6.995e-09,0.004386,4.822e-05,4.337e-07,0.0001004,0.0005024,2.162e-05,0.002074,2.472e-08,2.948e-06,1.511e-05,0.0042,1.083e-05,0.0003355,0.002029,0.0005485,1.03e-07,0.0005636,0.001164,7.134e-05,2.984e-06,0.0008932,0.0008762,7.383e-11,2.569e-09,5.891e-06,0.0004228,1.127e-05,1.095e-08,3.874e-09,5.968e-07,0.0002586,1.109e-05,0.002257,0.001403,0.001147,0.000768,0.002129,6.865e-06,1.343e-08,1.732e-05,0.0003243,8.372e-06,1.195e-06,4.322e-05,1.484e-07,0.0001824,0.002637,9.809e-05,0.0002224,8.239e-06,6.411e-05,2.936e-05,7.292e-07,5.166e-05,1.415e-05,0.001536,2.597e-05,7.715e-05,0.0002156,0.000309,1.327e-06,0.0009419,5.184e-05,4.851e-07,9.251e-06,6.49e-06,1.234e-06,3.837e-07,3.428e-05,8.558e-05,8.947e-05,0.000101,7.213e-08,1.355e-05,6.986e-06,2.149e-07,0.0007813,0.0001241,0.001017,0.0002406,0.0004762,0.003908,3.577e-06,1.248e-06,2.571e-05,3.929e-05,2.833e-06,9.81e-06,1.36e-08,4.506e-07,1.841e-09,0.0009137,1.399e-05,0.0005476,0.0005455,1.074e-07,4.156e-07,7.13e-06,0.001031,0.000169,0.0007611,7.535e-05,0.00144,0.0006618,1.613e-05,4.275e-08,1.271e-10,0.0002722,0.000191,8.143e-08,2.728e-08,0.0001979,0.0001547,1.772e-05,3.262e-06,4.303e-10,3.765e-08,1.675e-08,6.33e-09,6.458e-07,0.00118,2.754e-05,6.77e-05,0.0008949,2.823e-09,0.0002394,0.0003699,0.0001275,1.873e-06,0.00294,1.379e-05,6.873e-08,3.266e-06,6.111e-07,1.051e-05,0.0007482,7.005e-09,0.0001028,0.0002161,2.738e-08,0.001238,1.501e-12,0.0001095,0.0006495,2.949e-05,0.001994,1.21e-05,0.0002612,6.032e-08,1.051e-06,8.155e-05,3.242e-07,9.84e-10,0.0003851,0.002133,2.23e-07,0.00181,0.003737,0.0005078,0.0006772,0.0002325,1.746e-08,0.0005943,1.249e-06,0.002017,6.508e-05,9.67e-06,1.93e-05,0.0001134,3.073e-05,4.258e-08,0.002098,0.002588,0.0001419,0.0001775,3.751e-06,0.00267,0.0005067,0.003986,0.0001551,2.113e-06,0.0001784,0.001064,2.555e-06,0.003665,3.456e-06,0.0003825,5.244e-08,0.0001386,0.0001353,0.0001115,0.0006642,1.974e-09,1.483e-07,1.911e-06,0.00378,0.002692,6.973e-05,0.001325,0.001105,0.0001649,0.001089,0.003534,5.659e-06,0.002973,0.001294,2.036e-06,0.0001588,1.712e-05,0.0002666,3.095e-08,1.332e-05,5.531e-05,0.0001164,8.928e-07,1.153e-05,0.00457,2.017e-05,5.229e-08,1.987e-05,0.0002591,2.416e-06,0.001942,3.954e-07,2.351e-05,0.001332,3.765e-06,1.332e-06,0.001152,0.0001037,0.00053,5.232e-05,1.673e-06,7.981e-05,1.483e-08,7.583e-06,8.637e-08,1.648e-05,0.001642,0.0005751,2.925e-08,3.962e-05,2.511e-07,0.002306,7.646e-06,0.003843,2.707e-06,0.0005302,0.004171,1.963e-10,1.479e-05,0.0001647,0.0002909,1.666e-07,1.319e-07,0.001423,1.912e-05,0.0008903,6.067e-05,0.0003154,8.99e-07,2.884e-07,3.315e-07,0.0008252,3.058e-05,0.001633,0.001487,1.247e-08,2.879e-07,0.0007133,2.54e-09,0.0001136,2.826e-06,1.428e-05,0.0003979,0.0003508,3.172e-06,0.003806,3.035e-05,0.001012,0.0001118,1.793e-06,3.401e-07,2.815e-07,0.00172,0.001283,2.575e-09,0.0009686,0.0007053,6.197e-06,0.0003223,7.107e-05,1.412e-05,0.001566,0.000107,5.47e-06,0.0004321,0.003315,2.736e-05,0.003138,3.845e-05,2.375e-05,0.001553,7.126e-08,1.167e-09,1.599e-05,2.211e-05,6.839e-05,2.026e-05,1.867e-06,7.429e-06,4.458e-08,0.0002019,0.000981,0.002187,8.284e-06,0.0001883,7.706e-06,0.00247,0.003482,0.0008928,1.451e-05,0.002022,5.427e-07,5.713e-07,0.0001145,0.0006916,0.0002314,3.145e-08,9.697e-05,3.455e-05,0.0009545,0.002407,0.004453,0.0001646,1.213e-07,0.0001951,2.189e-07,0.0008698,2.311e-06,0.003132,0.0009559,1.274e-06,0.003055,0.0005492,2.753e-06,3.434e-08,6.172e-07,2.564e-05,0.000898,0.00316,0.0001889,1.516e-05,5.859e-05,7.366e-05,0.003464,3.493e-06,2.424e-09,5.037e-05,0.0001787,4.764e-05,4.735e-07,1.574e-07,0.0002065,4.641e-05,0.001406,0.003679,2.13e-05,0.001554,2.265e-05,4.094e-09,0.002234,0.0002194,0.000522,9.903e-06,4.089e-05,8.496e-05,0.000314,1.72e-07,2.818e-06,1.534e-05,4.454e-07,5.962e-05,0.003926,0.001519,0.0008563,0.00121,6.187e-06,1.726e-09,8.095e-07,0.003375,3.432e-06,0.0008646,0.00055,0.0006403,0.0003832,1.707e-06,3.893e-05,6.376e-07,3.331e-07,9.54e-08,9.115e-09,8.903e-08,9.524e-06,0.00237,0.000164,2.708e-06,1.72e-06,5.248e-05,0.003263,1.184e-07,0.0001111,3.938e-08,8.638e-10,3.887e-06,0.0004758,0.0002518,0.0003229,1.249e-08,8.71e-05,0.0005393,1.329e-05,6.532e-08,8.491e-07,0.0001687,7.618e-06,0.0001382,6.946e-05,2.021e-05,1.143e-08,4.592e-09,2.902e-05,2.382e-05,0.0005864,1.86e-05,5.793e-05,0.003197,0.00155,0.002856,5.049e-05,1.075e-06,8.179e-06,5.331e-06,2.723e-10,0.002751,1.932e-06,0.0003318,0.000233,8.562e-07,0.000101,3.422e-09,3.676e-07,4.727e-06,3.331e-05,9.123e-07,0.0009179,0.0009471,7.428e-06,0.004487,0.0001051,1.268e-07,0.0009122,0.000224,0.001477,3.048e-09,0.0003361,9.393e-07,2.984e-07,0.00316,0.001083,0.0003691,4.587e-06,0.0001086,0.0001467,7.098e-06,2.286e-09,1.102e-05,0.0007748,0.00228,6.351e-07,9.939e-05,1.543e-05,1.4e-10,1.898e-05,0.0009637,9.567e-05,5.703e-05,4.478e-08,4.787e-05,8.878e-05,0.002387,5.899e-05,5.254e-08,8.095e-08],"Contig":["Transfer.PvP01_00_1.final","Transfer.PvP01_00_1.final","Transfer.PvP01_00_1.final","Transfer.PvP01_00_1.final","Transfer.PvP01_00_1.final","Transfer.PvP01_00_1.final","Transfer.PvP01_00_1.final","Transfer.PvP01_00_1.final","Transfer.PvP01_00_1.final","Transfer.PvP01_00_1.final","Transfer.PvP01_00_1.final","Transfer.PvP01_00_1.final","Transfer.PvP01_00_1.final","Transfer.PvP01_00_1.final","Transfer.PvP01_00_1.final","Transfer.PvP01_00_1.final","Transfer.PvP01_00_1.final","Transfer.PvP01_00_1.final","Transfer.PvP01_00_1.final","Transfer.PvP01_00_1.final","Transfer.PvP01_00_2.final","Transfer.PvP01_00_2.final","Transfer.PvP01_00_2.final","Transfer.PvP01_00_2.final","Transfer.PvP01_00_2.final","Transfer.PvP01_00_2.final","Transfer.PvP01_00_3.final","Transfer.PvP01_00_3.final","Transfer.PvP01_00_3.final","Transfer.PvP01_00_3.final","Transfer.PvP01_00_3.final","Transfer.PvP01_00_3.final","Transfer.PvP01_00_3.final","Transfer.PvP01_00_3.final","Transfer.PvP01_00_4.final","Transfer.PvP01_00_4.final","Transfer.PvP01_00_4.final","Transfer.PvP01_00_4.final","Transfer.PvP01_00_4.final","Transfer.PvP01_00_4.final","Transfer.PvP01_00_4.final","Transfer.PvP01_00_5.final","Transfer.PvP01_00_5.final","Transfer.PvP01_00_5.final","Transfer.PvP01_00_6.final","Transfer.PvP01_00_6.final","Transfer.PvP01_00_6.final","Transfer.PvP01_00_6.final","Transfer.PvP01_00_7.final","Transfer.PvP01_00_7.final","Transfer.PvP01_00_7.final","Transfer.PvP01_00_8.final","Transfer.PvP01_00_8.final","Transfer.PvP01_00_8.final","Transfer.PvP01_00_8.final","Transfer.PvP01_00_8.final","Transfer.PvP01_00_8.final","Transfer.PvP01_00_8.final","Transfer.PvP01_00_8.final","Transfer.PvP01_00_8.final","Transfer.PvP01_00_9.final","Transfer.PvP01_00_9.final","Transfer.PvP01_00_9.final","Transfer.PvP01_00_10.final","Transfer.PvP01_00_10.final","Transfer.PvP01_00_10.final","Transfer.PvP01_00_10.final","Transfer.PvP01_00_10.final","Transfer.PvP01_00_11.final","Transfer.PvP01_00_11.final","Transfer.PvP01_00_11.final","Transfer.PvP01_00_11.final","Transfer.PvP01_00_11.final","Transfer.PvP01_00_11.final","Transfer.PvP01_00_11.final","Transfer.PvP01_00_11.final","Transfer.PvP01_00_11.final","Transfer.PvP01_00_11.final","Transfer.PvP01_00_11.final","Transfer.PvP01_00_11.final","Transfer.PvP01_00_11.final","Transfer.PvP01_00_11.final","Transfer.PvP01_00_12.final","Transfer.PvP01_00_12.final","Transfer.PvP01_00_13.final","Transfer.PvP01_00_13.final","Transfer.PvP01_00_14.final","Transfer.PvP01_00_14.final","Transfer.PvP01_00_14.final","Transfer.PvP01_00_14.final","Transfer.PvP01_00_14.final","Transfer.PvP01_00_15.final","Transfer.PvP01_00_15.final","Transfer.PvP01_00_15.final","Transfer.PvP01_00_15.final","Transfer.PvP01_00_15.final","Transfer.PvP01_00_16.final","Transfer.PvP01_00_16.final","Transfer.PvP01_00_17.final","Transfer.PvP01_00_17.final","Transfer.PvP01_00_17.final","Transfer.PvP01_00_17.final","Transfer.PvP01_00_17.final","Transfer.PvP01_00_17.final","Transfer.PvP01_00_17.final","Transfer.PvP01_00_18.final","Transfer.PvP01_00_18.final","Transfer.PvP01_00_18.final","Transfer.PvP01_00_19.final","Transfer.PvP01_00_22.final","Transfer.PvP01_00_22.final","Transfer.PvP01_00_22.final","Transfer.PvP01_00_22.final","Transfer.PvP01_00_22.final","Transfer.PvP01_00_23.final","Transfer.PvP01_00_23.final","Transfer.PvP01_00_24.final","Transfer.PvP01_00_25.final","Transfer.PvP01_00_26.final","Transfer.PvP01_00_26.final","Transfer.PvP01_00_26.final","Transfer.PvP01_00_26.final","Transfer.PvP01_00_26.final","Transfer.PvP01_00_29.final","Transfer.PvP01_00_29.final","Transfer.PvP01_00_29.final","Transfer.PvP01_00_30.final","Transfer.PvP01_00_30.final","Transfer.PvP01_00_31.final","Transfer.PvP01_00_31.final","Transfer.PvP01_00_31.final","Transfer.PvP01_00_32.final","Transfer.PvP01_00_34.final","Transfer.PvP01_00_37.final","Transfer.PvP01_00_38.final","Transfer.PvP01_00_38.final","Transfer.PvP01_00_39.final","Transfer.PvP01_00_41.final","Transfer.PvP01_00_42.final","Transfer.PvP01_00_42.final","Transfer.PvP01_00_45.final","Transfer.PvP01_00_45.final","Transfer.PvP01_00_48.final","Transfer.PvP01_00_48.final","Transfer.PvP01_00_48.final","Transfer.PvP01_00_52.final","Transfer.PvP01_00_53.final","Transfer.PvP01_00_55.final","Transfer.PvP01_00_57.final","Transfer.PvP01_00_61.final","Transfer.PvP01_00_61.final","Transfer.PvP01_00_63.final","Transfer.PvP01_00_64.final","Transfer.PvP01_00_64.final","Transfer.PvP01_00_66.final","Transfer.PvP01_00_67.final","Transfer.PvP01_00_67.final","Transfer.PvP01_00_67.final","Transfer.PvP01_00_69.final","Transfer.PvP01_00_71.final","Transfer.PvP01_00_74.final","Transfer.PvP01_00_77.final","Transfer.PvP01_00_77.final","Transfer.PvP01_00_79.final","Transfer.PvP01_00_81.final","Transfer.PvP01_00_82.final","Transfer.PvP01_00_83.final","Transfer.PvP01_00_86.final","Transfer.PvP01_00_88.final","Transfer.PvP01_00_89.final","Transfer.PvP01_00_89.final","Transfer.PvP01_00_89.final","Transfer.PvP01_00_90.final","Transfer.PvP01_00_90.final","Transfer.PvP01_00_104.final","Transfer.PvP01_00_108.final","Transfer.PvP01_00_119.final","Transfer.PvP01_00_124.final","Transfer.PvP01_00_129.final","Transfer.PvP01_00_138.final","Transfer.PvP01_00_143.final","Transfer.PvP01_00_150.final","Transfer.PvP01_00_152.final","Transfer.PvP01_00_157.final","Transfer.PvP01_00_158.final","Transfer.PvP01_00_180.final","Transfer.PvP01_00_203.final","Transfer.PvP01_00_208.final","Transfer.PvP01_00_209.final","Transfer.PvP01_00_217.final","Transfer.PvP01_00_221.final","Transfer.PvP01_00_223.final","Transfer.PvP01_00_233.final","Transfer.PvP01_00_300.final","PvP01_01_v1","PvP01_01_v1","PvP01_01_v1","PvP01_01_v1","PvP01_01_v1","PvP01_01_v1","PvP01_01_v1","PvP01_01_v1","PvP01_01_v1","PvP01_01_v1","PvP01_01_v1","PvP01_01_v1","PvP01_01_v1","PvP01_01_v1","PvP01_01_v1","PvP01_01_v1","PvP01_01_v1","PvP01_01_v1","PvP01_01_v1","PvP01_01_v1","PvP01_01_v1","PvP01_01_v1","PvP01_01_v1","PvP01_01_v1","PvP01_01_v1","PvP01_01_v1","PvP01_01_v1","PvP01_01_v1","PvP01_01_v1","PvP01_01_v1","PvP01_01_v1","PvP01_01_v1","PvP01_01_v1","PvP01_01_v1","PvP01_01_v1","PvP01_01_v1","PvP01_01_v1","PvP01_01_v1","PvP01_01_v1","PvP01_01_v1","PvP01_01_v1","PvP01_01_v1","PvP01_01_v1","PvP01_01_v1","PvP01_01_v1","PvP01_01_v1","PvP01_01_v1","PvP01_01_v1","PvP01_01_v1","PvP01_01_v1","PvP01_01_v1","PvP01_01_v1","PvP01_01_v1","PvP01_01_v1","PvP01_01_v1","PvP01_01_v1","PvP01_01_v1","PvP01_01_v1","PvP01_01_v1","PvP01_01_v1","PvP01_01_v1","PvP01_01_v1","PvP01_01_v1","PvP01_01_v1","PvP01_01_v1","PvP01_01_v1","PvP01_01_v1","PvP01_01_v1","PvP01_01_v1","PvP01_01_v1","PvP01_01_v1","PvP01_01_v1","PvP01_01_v1","PvP01_01_v1","PvP01_01_v1","PvP01_01_v1","PvP01_01_v1","PvP01_01_v1","PvP01_01_v1","PvP01_01_v1","PvP01_01_v1","PvP01_01_v1","PvP01_01_v1","PvP01_01_v1","PvP01_01_v1","PvP01_01_v1","PvP01_01_v1","PvP01_01_v1","PvP01_01_v1","PvP01_01_v1","PvP01_01_v1","PvP01_01_v1","PvP01_01_v1","PvP01_01_v1","PvP01_01_v1","PvP01_01_v1","PvP01_01_v1","PvP01_01_v1","PvP01_01_v1","PvP01_01_v1","PvP01_02_v1","PvP01_02_v1","PvP01_02_v1","PvP01_02_v1","PvP01_02_v1","PvP01_02_v1","PvP01_02_v1","PvP01_02_v1","PvP01_02_v1","PvP01_02_v1","PvP01_02_v1","PvP01_02_v1","PvP01_02_v1","PvP01_02_v1","PvP01_02_v1","PvP01_02_v1","PvP01_02_v1","PvP01_02_v1","PvP01_02_v1","PvP01_02_v1","PvP01_02_v1","PvP01_02_v1","PvP01_02_v1","PvP01_02_v1","PvP01_02_v1","PvP01_02_v1","PvP01_02_v1","PvP01_02_v1","PvP01_02_v1","PvP01_02_v1","PvP01_02_v1","PvP01_02_v1","PvP01_02_v1","PvP01_02_v1","PvP01_02_v1","PvP01_02_v1","PvP01_02_v1","PvP01_02_v1","PvP01_02_v1","PvP01_02_v1","PvP01_02_v1","PvP01_02_v1","PvP01_02_v1","PvP01_02_v1","PvP01_02_v1","PvP01_02_v1","PvP01_02_v1","PvP01_02_v1","PvP01_02_v1","PvP01_02_v1","PvP01_02_v1","PvP01_02_v1","PvP01_02_v1","PvP01_02_v1","PvP01_02_v1","PvP01_02_v1","PvP01_02_v1","PvP01_02_v1","PvP01_02_v1","PvP01_02_v1","PvP01_02_v1","PvP01_02_v1","PvP01_02_v1","PvP01_02_v1","PvP01_02_v1","PvP01_02_v1","PvP01_02_v1","PvP01_02_v1","PvP01_02_v1","PvP01_02_v1","PvP01_02_v1","PvP01_02_v1","PvP01_02_v1","PvP01_02_v1","PvP01_02_v1","PvP01_02_v1","PvP01_02_v1","PvP01_02_v1","PvP01_02_v1","PvP01_02_v1","PvP01_02_v1","PvP01_02_v1","PvP01_02_v1","PvP01_02_v1","PvP01_02_v1","PvP01_02_v1","PvP01_02_v1","PvP01_02_v1","PvP01_02_v1","PvP01_02_v1","PvP01_02_v1","PvP01_02_v1","PvP01_02_v1","PvP01_02_v1","PvP01_02_v1","PvP01_02_v1","PvP01_02_v1","PvP01_02_v1","PvP01_02_v1","PvP01_02_v1","PvP01_02_v1","PvP01_02_v1","PvP01_02_v1","PvP01_02_v1","PvP01_02_v1","PvP01_02_v1","PvP01_02_v1","PvP01_02_v1","PvP01_02_v1","PvP01_03_v1","PvP01_03_v1","PvP01_03_v1","PvP01_03_v1","PvP01_03_v1","PvP01_03_v1","PvP01_03_v1","PvP01_03_v1","PvP01_03_v1","PvP01_03_v1","PvP01_03_v1","PvP01_03_v1","PvP01_03_v1","PvP01_03_v1","PvP01_03_v1","PvP01_03_v1","PvP01_03_v1","PvP01_03_v1","PvP01_03_v1","PvP01_03_v1","PvP01_03_v1","PvP01_03_v1","PvP01_03_v1","PvP01_03_v1","PvP01_03_v1","PvP01_03_v1","PvP01_03_v1","PvP01_03_v1","PvP01_03_v1","PvP01_03_v1","PvP01_03_v1","PvP01_03_v1","PvP01_03_v1","PvP01_03_v1","PvP01_03_v1","PvP01_03_v1","PvP01_03_v1","PvP01_03_v1","PvP01_03_v1","PvP01_03_v1","PvP01_03_v1","PvP01_03_v1","PvP01_03_v1","PvP01_03_v1","PvP01_03_v1","PvP01_03_v1","PvP01_03_v1","PvP01_03_v1","PvP01_03_v1","PvP01_03_v1","PvP01_03_v1","PvP01_03_v1","PvP01_03_v1","PvP01_03_v1","PvP01_03_v1","PvP01_03_v1","PvP01_03_v1","PvP01_03_v1","PvP01_03_v1","PvP01_03_v1","PvP01_03_v1","PvP01_03_v1","PvP01_03_v1","PvP01_03_v1","PvP01_03_v1","PvP01_03_v1","PvP01_03_v1","PvP01_03_v1","PvP01_03_v1","PvP01_03_v1","PvP01_03_v1","PvP01_03_v1","PvP01_03_v1","PvP01_03_v1","PvP01_03_v1","PvP01_03_v1","PvP01_03_v1","PvP01_03_v1","PvP01_03_v1","PvP01_03_v1","PvP01_03_v1","PvP01_03_v1","PvP01_03_v1","PvP01_03_v1","PvP01_03_v1","PvP01_03_v1","PvP01_03_v1","PvP01_03_v1","PvP01_03_v1","PvP01_03_v1","PvP01_03_v1","PvP01_03_v1","PvP01_03_v1","PvP01_03_v1","PvP01_03_v1","PvP01_03_v1","PvP01_03_v1","PvP01_04_v1","PvP01_04_v1","PvP01_04_v1","PvP01_04_v1","PvP01_04_v1","PvP01_04_v1","PvP01_04_v1","PvP01_04_v1","PvP01_04_v1","PvP01_04_v1","PvP01_04_v1","PvP01_04_v1","PvP01_04_v1","PvP01_04_v1","PvP01_04_v1","PvP01_04_v1","PvP01_04_v1","PvP01_04_v1","PvP01_04_v1","PvP01_04_v1","PvP01_04_v1","PvP01_04_v1","PvP01_04_v1","PvP01_04_v1","PvP01_04_v1","PvP01_04_v1","PvP01_04_v1","PvP01_04_v1","PvP01_04_v1","PvP01_04_v1","PvP01_04_v1","PvP01_04_v1","PvP01_04_v1","PvP01_04_v1","PvP01_04_v1","PvP01_04_v1","PvP01_04_v1","PvP01_04_v1","PvP01_04_v1","PvP01_04_v1","PvP01_04_v1","PvP01_04_v1","PvP01_04_v1","PvP01_04_v1","PvP01_04_v1","PvP01_04_v1","PvP01_04_v1","PvP01_04_v1","PvP01_04_v1","PvP01_04_v1","PvP01_04_v1","PvP01_04_v1","PvP01_04_v1","PvP01_04_v1","PvP01_04_v1","PvP01_04_v1","PvP01_04_v1","PvP01_04_v1","PvP01_04_v1","PvP01_04_v1","PvP01_04_v1","PvP01_04_v1","PvP01_04_v1","PvP01_04_v1","PvP01_04_v1","PvP01_04_v1","PvP01_04_v1","PvP01_04_v1","PvP01_04_v1","PvP01_04_v1","PvP01_04_v1","PvP01_04_v1","PvP01_04_v1","PvP01_04_v1","PvP01_04_v1","PvP01_04_v1","PvP01_04_v1","PvP01_04_v1","PvP01_04_v1","PvP01_04_v1","PvP01_04_v1","PvP01_04_v1","PvP01_04_v1","PvP01_04_v1","PvP01_04_v1","PvP01_04_v1","PvP01_04_v1","PvP01_04_v1","PvP01_04_v1","PvP01_04_v1","PvP01_04_v1","PvP01_04_v1","PvP01_04_v1","PvP01_04_v1","PvP01_04_v1","PvP01_04_v1","PvP01_04_v1","PvP01_04_v1","PvP01_05_v1","PvP01_05_v1","PvP01_05_v1","PvP01_05_v1","PvP01_05_v1","PvP01_05_v1","PvP01_05_v1","PvP01_05_v1","PvP01_05_v1","PvP01_05_v1","PvP01_05_v1","PvP01_05_v1","PvP01_05_v1","PvP01_05_v1","PvP01_05_v1","PvP01_05_v1","PvP01_05_v1","PvP01_05_v1","PvP01_05_v1","PvP01_05_v1","PvP01_05_v1","PvP01_05_v1","PvP01_05_v1","PvP01_05_v1","PvP01_05_v1","PvP01_05_v1","PvP01_05_v1","PvP01_05_v1","PvP01_05_v1","PvP01_05_v1","PvP01_05_v1","PvP01_05_v1","PvP01_05_v1","PvP01_05_v1","PvP01_05_v1","PvP01_05_v1","PvP01_05_v1","PvP01_05_v1","PvP01_05_v1","PvP01_05_v1","PvP01_05_v1","PvP01_05_v1","PvP01_05_v1","PvP01_05_v1","PvP01_05_v1","PvP01_05_v1","PvP01_05_v1","PvP01_05_v1","PvP01_05_v1","PvP01_05_v1","PvP01_05_v1","PvP01_05_v1","PvP01_05_v1","PvP01_05_v1","PvP01_05_v1","PvP01_05_v1","PvP01_05_v1","PvP01_05_v1","PvP01_05_v1","PvP01_05_v1","PvP01_05_v1","PvP01_05_v1","PvP01_05_v1","PvP01_05_v1","PvP01_05_v1","PvP01_05_v1","PvP01_05_v1","PvP01_05_v1","PvP01_05_v1","PvP01_05_v1","PvP01_05_v1","PvP01_05_v1","PvP01_05_v1","PvP01_05_v1","PvP01_05_v1","PvP01_05_v1","PvP01_05_v1","PvP01_05_v1","PvP01_05_v1","PvP01_05_v1","PvP01_05_v1","PvP01_05_v1","PvP01_05_v1","PvP01_05_v1","PvP01_05_v1","PvP01_05_v1","PvP01_05_v1","PvP01_05_v1","PvP01_05_v1","PvP01_05_v1","PvP01_05_v1","PvP01_05_v1","PvP01_05_v1","PvP01_05_v1","PvP01_05_v1","PvP01_05_v1","PvP01_05_v1","PvP01_05_v1","PvP01_05_v1","PvP01_05_v1","PvP01_05_v1","PvP01_05_v1","PvP01_05_v1","PvP01_05_v1","PvP01_05_v1","PvP01_05_v1","PvP01_05_v1","PvP01_05_v1","PvP01_05_v1","PvP01_05_v1","PvP01_05_v1","PvP01_05_v1","PvP01_05_v1","PvP01_05_v1","PvP01_05_v1","PvP01_05_v1","PvP01_05_v1","PvP01_05_v1","PvP01_05_v1","PvP01_05_v1","PvP01_05_v1","PvP01_05_v1","PvP01_05_v1","PvP01_05_v1","PvP01_05_v1","PvP01_05_v1","PvP01_05_v1","PvP01_05_v1","PvP01_05_v1","PvP01_05_v1","PvP01_05_v1","PvP01_05_v1","PvP01_05_v1","PvP01_05_v1","PvP01_05_v1","PvP01_05_v1","PvP01_05_v1","PvP01_05_v1","PvP01_05_v1","PvP01_05_v1","PvP01_05_v1","PvP01_05_v1","PvP01_05_v1","PvP01_05_v1","PvP01_05_v1","PvP01_05_v1","PvP01_05_v1","PvP01_05_v1","PvP01_05_v1","PvP01_05_v1","PvP01_05_v1","PvP01_05_v1","PvP01_05_v1","PvP01_05_v1","PvP01_05_v1","PvP01_05_v1","PvP01_05_v1","PvP01_05_v1","PvP01_05_v1","PvP01_05_v1","PvP01_05_v1","PvP01_05_v1","PvP01_05_v1","PvP01_05_v1","PvP01_05_v1","PvP01_05_v1","PvP01_05_v1","PvP01_05_v1","PvP01_05_v1","PvP01_05_v1","PvP01_05_v1","PvP01_06_v1","PvP01_06_v1","PvP01_06_v1","PvP01_06_v1","PvP01_06_v1","PvP01_06_v1","PvP01_06_v1","PvP01_06_v1","PvP01_06_v1","PvP01_06_v1","PvP01_06_v1","PvP01_06_v1","PvP01_06_v1","PvP01_06_v1","PvP01_06_v1","PvP01_06_v1","PvP01_06_v1","PvP01_06_v1","PvP01_06_v1","PvP01_06_v1","PvP01_06_v1","PvP01_06_v1","PvP01_06_v1","PvP01_06_v1","PvP01_06_v1","PvP01_06_v1","PvP01_06_v1","PvP01_06_v1","PvP01_06_v1","PvP01_06_v1","PvP01_06_v1","PvP01_06_v1","PvP01_06_v1","PvP01_06_v1","PvP01_06_v1","PvP01_06_v1","PvP01_06_v1","PvP01_06_v1","PvP01_06_v1","PvP01_06_v1","PvP01_06_v1","PvP01_06_v1","PvP01_06_v1","PvP01_06_v1","PvP01_06_v1","PvP01_06_v1","PvP01_06_v1","PvP01_06_v1","PvP01_06_v1","PvP01_06_v1","PvP01_06_v1","PvP01_06_v1","PvP01_06_v1","PvP01_06_v1","PvP01_06_v1","PvP01_06_v1","PvP01_06_v1","PvP01_06_v1","PvP01_06_v1","PvP01_06_v1","PvP01_06_v1","PvP01_06_v1","PvP01_06_v1","PvP01_06_v1","PvP01_06_v1","PvP01_06_v1","PvP01_06_v1","PvP01_06_v1","PvP01_06_v1","PvP01_06_v1","PvP01_06_v1","PvP01_06_v1","PvP01_06_v1","PvP01_06_v1","PvP01_06_v1","PvP01_06_v1","PvP01_06_v1","PvP01_06_v1","PvP01_06_v1","PvP01_06_v1","PvP01_06_v1","PvP01_06_v1","PvP01_06_v1","PvP01_06_v1","PvP01_06_v1","PvP01_06_v1","PvP01_06_v1","PvP01_06_v1","PvP01_06_v1","PvP01_06_v1","PvP01_06_v1","PvP01_06_v1","PvP01_06_v1","PvP01_06_v1","PvP01_06_v1","PvP01_06_v1","PvP01_06_v1","PvP01_06_v1","PvP01_06_v1","PvP01_06_v1","PvP01_06_v1","PvP01_06_v1","PvP01_06_v1","PvP01_06_v1","PvP01_06_v1","PvP01_06_v1","PvP01_06_v1","PvP01_06_v1","PvP01_06_v1","PvP01_06_v1","PvP01_06_v1","PvP01_06_v1","PvP01_06_v1","PvP01_06_v1","PvP01_06_v1","PvP01_06_v1","PvP01_06_v1","PvP01_06_v1","PvP01_06_v1","PvP01_06_v1","PvP01_06_v1","PvP01_07_v1","PvP01_07_v1","PvP01_07_v1","PvP01_07_v1","PvP01_07_v1","PvP01_07_v1","PvP01_07_v1","PvP01_07_v1","PvP01_07_v1","PvP01_07_v1","PvP01_07_v1","PvP01_07_v1","PvP01_07_v1","PvP01_07_v1","PvP01_07_v1","PvP01_07_v1","PvP01_07_v1","PvP01_07_v1","PvP01_07_v1","PvP01_07_v1","PvP01_07_v1","PvP01_07_v1","PvP01_07_v1","PvP01_07_v1","PvP01_07_v1","PvP01_07_v1","PvP01_07_v1","PvP01_07_v1","PvP01_07_v1","PvP01_07_v1","PvP01_07_v1","PvP01_07_v1","PvP01_07_v1","PvP01_07_v1","PvP01_07_v1","PvP01_07_v1","PvP01_07_v1","PvP01_07_v1","PvP01_07_v1","PvP01_07_v1","PvP01_07_v1","PvP01_07_v1","PvP01_07_v1","PvP01_07_v1","PvP01_07_v1","PvP01_07_v1","PvP01_07_v1","PvP01_07_v1","PvP01_07_v1","PvP01_07_v1","PvP01_07_v1","PvP01_07_v1","PvP01_07_v1","PvP01_07_v1","PvP01_07_v1","PvP01_07_v1","PvP01_07_v1","PvP01_07_v1","PvP01_07_v1","PvP01_07_v1","PvP01_07_v1","PvP01_07_v1","PvP01_07_v1","PvP01_07_v1","PvP01_07_v1","PvP01_07_v1","PvP01_07_v1","PvP01_07_v1","PvP01_07_v1","PvP01_07_v1","PvP01_07_v1","PvP01_07_v1","PvP01_07_v1","PvP01_07_v1","PvP01_07_v1","PvP01_07_v1","PvP01_07_v1","PvP01_07_v1","PvP01_07_v1","PvP01_07_v1","PvP01_07_v1","PvP01_07_v1","PvP01_07_v1","PvP01_07_v1","PvP01_07_v1","PvP01_07_v1","PvP01_07_v1","PvP01_07_v1","PvP01_07_v1","PvP01_07_v1","PvP01_07_v1","PvP01_07_v1","PvP01_07_v1","PvP01_07_v1","PvP01_07_v1","PvP01_07_v1","PvP01_07_v1","PvP01_07_v1","PvP01_07_v1","PvP01_07_v1","PvP01_07_v1","PvP01_07_v1","PvP01_07_v1","PvP01_07_v1","PvP01_07_v1","PvP01_07_v1","PvP01_07_v1","PvP01_07_v1","PvP01_07_v1","PvP01_07_v1","PvP01_07_v1","PvP01_07_v1","PvP01_07_v1","PvP01_07_v1","PvP01_07_v1","PvP01_07_v1","PvP01_07_v1","PvP01_07_v1","PvP01_07_v1","PvP01_07_v1","PvP01_07_v1","PvP01_07_v1","PvP01_07_v1","PvP01_07_v1","PvP01_07_v1","PvP01_07_v1","PvP01_07_v1","PvP01_07_v1","PvP01_07_v1","PvP01_07_v1","PvP01_07_v1","PvP01_07_v1","PvP01_07_v1","PvP01_07_v1","PvP01_07_v1","PvP01_07_v1","PvP01_07_v1","PvP01_07_v1","PvP01_07_v1","PvP01_07_v1","PvP01_07_v1","PvP01_07_v1","PvP01_07_v1","PvP01_07_v1","PvP01_07_v1","PvP01_07_v1","PvP01_07_v1","PvP01_07_v1","PvP01_07_v1","PvP01_07_v1","PvP01_07_v1","PvP01_07_v1","PvP01_07_v1","PvP01_07_v1","PvP01_07_v1","PvP01_07_v1","PvP01_07_v1","PvP01_07_v1","PvP01_07_v1","PvP01_07_v1","PvP01_07_v1","PvP01_07_v1","PvP01_07_v1","PvP01_07_v1","PvP01_07_v1","PvP01_07_v1","PvP01_07_v1","PvP01_07_v1","PvP01_07_v1","PvP01_07_v1","PvP01_07_v1","PvP01_07_v1","PvP01_07_v1","PvP01_07_v1","PvP01_07_v1","PvP01_07_v1","PvP01_07_v1","PvP01_07_v1","PvP01_07_v1","PvP01_07_v1","PvP01_07_v1","PvP01_07_v1","PvP01_07_v1","PvP01_07_v1","PvP01_07_v1","PvP01_07_v1","PvP01_07_v1","PvP01_07_v1","PvP01_07_v1","PvP01_07_v1","PvP01_07_v1","PvP01_07_v1","PvP01_07_v1","PvP01_07_v1","PvP01_07_v1","PvP01_07_v1","PvP01_07_v1","PvP01_07_v1","PvP01_07_v1","PvP01_07_v1","PvP01_07_v1","PvP01_07_v1","PvP01_07_v1","PvP01_08_v1","PvP01_08_v1","PvP01_08_v1","PvP01_08_v1","PvP01_08_v1","PvP01_08_v1","PvP01_08_v1","PvP01_08_v1","PvP01_08_v1","PvP01_08_v1","PvP01_08_v1","PvP01_08_v1","PvP01_08_v1","PvP01_08_v1","PvP01_08_v1","PvP01_08_v1","PvP01_08_v1","PvP01_08_v1","PvP01_08_v1","PvP01_08_v1","PvP01_08_v1","PvP01_08_v1","PvP01_08_v1","PvP01_08_v1","PvP01_08_v1","PvP01_08_v1","PvP01_08_v1","PvP01_08_v1","PvP01_08_v1","PvP01_08_v1","PvP01_08_v1","PvP01_08_v1","PvP01_08_v1","PvP01_08_v1","PvP01_08_v1","PvP01_08_v1","PvP01_08_v1","PvP01_08_v1","PvP01_08_v1","PvP01_08_v1","PvP01_08_v1","PvP01_08_v1","PvP01_08_v1","PvP01_08_v1","PvP01_08_v1","PvP01_08_v1","PvP01_08_v1","PvP01_08_v1","PvP01_08_v1","PvP01_08_v1","PvP01_08_v1","PvP01_08_v1","PvP01_08_v1","PvP01_08_v1","PvP01_08_v1","PvP01_08_v1","PvP01_08_v1","PvP01_08_v1","PvP01_08_v1","PvP01_08_v1","PvP01_08_v1","PvP01_08_v1","PvP01_08_v1","PvP01_08_v1","PvP01_08_v1","PvP01_08_v1","PvP01_08_v1","PvP01_08_v1","PvP01_08_v1","PvP01_08_v1","PvP01_08_v1","PvP01_08_v1","PvP01_08_v1","PvP01_08_v1","PvP01_08_v1","PvP01_08_v1","PvP01_08_v1","PvP01_08_v1","PvP01_08_v1","PvP01_08_v1","PvP01_08_v1","PvP01_08_v1","PvP01_08_v1","PvP01_08_v1","PvP01_08_v1","PvP01_08_v1","PvP01_08_v1","PvP01_08_v1","PvP01_08_v1","PvP01_08_v1","PvP01_08_v1","PvP01_08_v1","PvP01_08_v1","PvP01_08_v1","PvP01_08_v1","PvP01_08_v1","PvP01_08_v1","PvP01_08_v1","PvP01_08_v1","PvP01_08_v1","PvP01_08_v1","PvP01_08_v1","PvP01_08_v1","PvP01_08_v1","PvP01_08_v1","PvP01_08_v1","PvP01_08_v1","PvP01_08_v1","PvP01_08_v1","PvP01_08_v1","PvP01_08_v1","PvP01_08_v1","PvP01_08_v1","PvP01_08_v1","PvP01_08_v1","PvP01_08_v1","PvP01_08_v1","PvP01_08_v1","PvP01_08_v1","PvP01_08_v1","PvP01_08_v1","PvP01_08_v1","PvP01_08_v1","PvP01_08_v1","PvP01_08_v1","PvP01_08_v1","PvP01_08_v1","PvP01_08_v1","PvP01_08_v1","PvP01_08_v1","PvP01_08_v1","PvP01_08_v1","PvP01_08_v1","PvP01_08_v1","PvP01_08_v1","PvP01_08_v1","PvP01_08_v1","PvP01_08_v1","PvP01_08_v1","PvP01_08_v1","PvP01_08_v1","PvP01_08_v1","PvP01_08_v1","PvP01_08_v1","PvP01_08_v1","PvP01_08_v1","PvP01_08_v1","PvP01_08_v1","PvP01_08_v1","PvP01_08_v1","PvP01_08_v1","PvP01_08_v1","PvP01_08_v1","PvP01_08_v1","PvP01_08_v1","PvP01_08_v1","PvP01_08_v1","PvP01_08_v1","PvP01_08_v1","PvP01_08_v1","PvP01_08_v1","PvP01_08_v1","PvP01_08_v1","PvP01_08_v1","PvP01_08_v1","PvP01_08_v1","PvP01_08_v1","PvP01_08_v1","PvP01_08_v1","PvP01_08_v1","PvP01_08_v1","PvP01_08_v1","PvP01_08_v1","PvP01_08_v1","PvP01_08_v1","PvP01_08_v1","PvP01_08_v1","PvP01_08_v1","PvP01_08_v1","PvP01_08_v1","PvP01_08_v1","PvP01_08_v1","PvP01_08_v1","PvP01_08_v1","PvP01_08_v1","PvP01_08_v1","PvP01_08_v1","PvP01_08_v1","PvP01_08_v1","PvP01_08_v1","PvP01_08_v1","PvP01_08_v1","PvP01_08_v1","PvP01_08_v1","PvP01_08_v1","PvP01_08_v1","PvP01_08_v1","PvP01_08_v1","PvP01_08_v1","PvP01_08_v1","PvP01_08_v1","PvP01_08_v1","PvP01_08_v1","PvP01_08_v1","PvP01_08_v1","PvP01_08_v1","PvP01_08_v1","PvP01_08_v1","PvP01_08_v1","PvP01_09_v1","PvP01_09_v1","PvP01_09_v1","PvP01_09_v1","PvP01_09_v1","PvP01_09_v1","PvP01_09_v1","PvP01_09_v1","PvP01_09_v1","PvP01_09_v1","PvP01_09_v1","PvP01_09_v1","PvP01_09_v1","PvP01_09_v1","PvP01_09_v1","PvP01_09_v1","PvP01_09_v1","PvP01_09_v1","PvP01_09_v1","PvP01_09_v1","PvP01_09_v1","PvP01_09_v1","PvP01_09_v1","PvP01_09_v1","PvP01_09_v1","PvP01_09_v1","PvP01_09_v1","PvP01_09_v1","PvP01_09_v1","PvP01_09_v1","PvP01_09_v1","PvP01_09_v1","PvP01_09_v1","PvP01_09_v1","PvP01_09_v1","PvP01_09_v1","PvP01_09_v1","PvP01_09_v1","PvP01_09_v1","PvP01_09_v1","PvP01_09_v1","PvP01_09_v1","PvP01_09_v1","PvP01_09_v1","PvP01_09_v1","PvP01_09_v1","PvP01_09_v1","PvP01_09_v1","PvP01_09_v1","PvP01_09_v1","PvP01_09_v1","PvP01_09_v1","PvP01_09_v1","PvP01_09_v1","PvP01_09_v1","PvP01_09_v1","PvP01_09_v1","PvP01_09_v1","PvP01_09_v1","PvP01_09_v1","PvP01_09_v1","PvP01_09_v1","PvP01_09_v1","PvP01_09_v1","PvP01_09_v1","PvP01_09_v1","PvP01_09_v1","PvP01_09_v1","PvP01_09_v1","PvP01_09_v1","PvP01_09_v1","PvP01_09_v1","PvP01_09_v1","PvP01_09_v1","PvP01_09_v1","PvP01_09_v1","PvP01_09_v1","PvP01_09_v1","PvP01_09_v1","PvP01_09_v1","PvP01_09_v1","PvP01_09_v1","PvP01_09_v1","PvP01_09_v1","PvP01_09_v1","PvP01_09_v1","PvP01_09_v1","PvP01_09_v1","PvP01_09_v1","PvP01_09_v1","PvP01_09_v1","PvP01_09_v1","PvP01_09_v1","PvP01_09_v1","PvP01_09_v1","PvP01_09_v1","PvP01_09_v1","PvP01_09_v1","PvP01_09_v1","PvP01_09_v1","PvP01_09_v1","PvP01_09_v1","PvP01_09_v1","PvP01_09_v1","PvP01_09_v1","PvP01_09_v1","PvP01_09_v1","PvP01_09_v1","PvP01_09_v1","PvP01_09_v1","PvP01_09_v1","PvP01_09_v1","PvP01_09_v1","PvP01_09_v1","PvP01_09_v1","PvP01_09_v1","PvP01_09_v1","PvP01_09_v1","PvP01_09_v1","PvP01_09_v1","PvP01_09_v1","PvP01_09_v1","PvP01_09_v1","PvP01_09_v1","PvP01_09_v1","PvP01_09_v1","PvP01_09_v1","PvP01_09_v1","PvP01_09_v1","PvP01_09_v1","PvP01_09_v1","PvP01_09_v1","PvP01_09_v1","PvP01_09_v1","PvP01_09_v1","PvP01_09_v1","PvP01_09_v1","PvP01_09_v1","PvP01_09_v1","PvP01_09_v1","PvP01_09_v1","PvP01_09_v1","PvP01_09_v1","PvP01_09_v1","PvP01_09_v1","PvP01_09_v1","PvP01_09_v1","PvP01_09_v1","PvP01_09_v1","PvP01_09_v1","PvP01_09_v1","PvP01_09_v1","PvP01_09_v1","PvP01_09_v1","PvP01_09_v1","PvP01_09_v1","PvP01_09_v1","PvP01_09_v1","PvP01_09_v1","PvP01_09_v1","PvP01_09_v1","PvP01_09_v1","PvP01_09_v1","PvP01_09_v1","PvP01_09_v1","PvP01_09_v1","PvP01_09_v1","PvP01_09_v1","PvP01_09_v1","PvP01_09_v1","PvP01_09_v1","PvP01_09_v1","PvP01_09_v1","PvP01_09_v1","PvP01_09_v1","PvP01_09_v1","PvP01_09_v1","PvP01_09_v1","PvP01_09_v1","PvP01_09_v1","PvP01_09_v1","PvP01_09_v1","PvP01_09_v1","PvP01_09_v1","PvP01_09_v1","PvP01_09_v1","PvP01_09_v1","PvP01_09_v1","PvP01_09_v1","PvP01_09_v1","PvP01_09_v1","PvP01_09_v1","PvP01_09_v1","PvP01_09_v1","PvP01_09_v1","PvP01_09_v1","PvP01_09_v1","PvP01_09_v1","PvP01_09_v1","PvP01_09_v1","PvP01_09_v1","PvP01_09_v1","PvP01_09_v1","PvP01_09_v1","PvP01_09_v1","PvP01_09_v1","PvP01_09_v1","PvP01_09_v1","PvP01_09_v1","PvP01_09_v1","PvP01_09_v1","PvP01_09_v1","PvP01_09_v1","PvP01_09_v1","PvP01_09_v1","PvP01_09_v1","PvP01_09_v1","PvP01_09_v1","PvP01_09_v1","PvP01_09_v1","PvP01_09_v1","PvP01_09_v1","PvP01_09_v1","PvP01_09_v1","PvP01_09_v1","PvP01_09_v1","PvP01_09_v1","PvP01_09_v1","PvP01_09_v1","PvP01_09_v1","PvP01_09_v1","PvP01_09_v1","PvP01_09_v1","PvP01_09_v1","PvP01_09_v1","PvP01_09_v1","PvP01_09_v1","PvP01_09_v1","PvP01_09_v1","PvP01_09_v1","PvP01_09_v1","PvP01_09_v1","PvP01_09_v1","PvP01_09_v1","PvP01_09_v1","PvP01_09_v1","PvP01_09_v1","PvP01_09_v1","PvP01_09_v1","PvP01_09_v1","PvP01_09_v1","PvP01_10_v1","PvP01_10_v1","PvP01_10_v1","PvP01_10_v1","PvP01_10_v1","PvP01_10_v1","PvP01_10_v1","PvP01_10_v1","PvP01_10_v1","PvP01_10_v1","PvP01_10_v1","PvP01_10_v1","PvP01_10_v1","PvP01_10_v1","PvP01_10_v1","PvP01_10_v1","PvP01_10_v1","PvP01_10_v1","PvP01_10_v1","PvP01_10_v1","PvP01_10_v1","PvP01_10_v1","PvP01_10_v1","PvP01_10_v1","PvP01_10_v1","PvP01_10_v1","PvP01_10_v1","PvP01_10_v1","PvP01_10_v1","PvP01_10_v1","PvP01_10_v1","PvP01_10_v1","PvP01_10_v1","PvP01_10_v1","PvP01_10_v1","PvP01_10_v1","PvP01_10_v1","PvP01_10_v1","PvP01_10_v1","PvP01_10_v1","PvP01_10_v1","PvP01_10_v1","PvP01_10_v1","PvP01_10_v1","PvP01_10_v1","PvP01_10_v1","PvP01_10_v1","PvP01_10_v1","PvP01_10_v1","PvP01_10_v1","PvP01_10_v1","PvP01_10_v1","PvP01_10_v1","PvP01_10_v1","PvP01_10_v1","PvP01_10_v1","PvP01_10_v1","PvP01_10_v1","PvP01_10_v1","PvP01_10_v1","PvP01_10_v1","PvP01_10_v1","PvP01_10_v1","PvP01_10_v1","PvP01_10_v1","PvP01_10_v1","PvP01_10_v1","PvP01_10_v1","PvP01_10_v1","PvP01_10_v1","PvP01_10_v1","PvP01_10_v1","PvP01_10_v1","PvP01_10_v1","PvP01_10_v1","PvP01_10_v1","PvP01_10_v1","PvP01_10_v1","PvP01_10_v1","PvP01_10_v1","PvP01_10_v1","PvP01_10_v1","PvP01_10_v1","PvP01_10_v1","PvP01_10_v1","PvP01_10_v1","PvP01_10_v1","PvP01_10_v1","PvP01_10_v1","PvP01_10_v1","PvP01_10_v1","PvP01_10_v1","PvP01_10_v1","PvP01_10_v1","PvP01_10_v1","PvP01_10_v1","PvP01_10_v1","PvP01_10_v1","PvP01_10_v1","PvP01_10_v1","PvP01_10_v1","PvP01_10_v1","PvP01_10_v1","PvP01_10_v1","PvP01_10_v1","PvP01_10_v1","PvP01_10_v1","PvP01_10_v1","PvP01_10_v1","PvP01_10_v1","PvP01_10_v1","PvP01_10_v1","PvP01_10_v1","PvP01_10_v1","PvP01_10_v1","PvP01_10_v1","PvP01_10_v1","PvP01_10_v1","PvP01_10_v1","PvP01_10_v1","PvP01_10_v1","PvP01_10_v1","PvP01_10_v1","PvP01_10_v1","PvP01_10_v1","PvP01_10_v1","PvP01_10_v1","PvP01_10_v1","PvP01_10_v1","PvP01_10_v1","PvP01_10_v1","PvP01_10_v1","PvP01_10_v1","PvP01_10_v1","PvP01_10_v1","PvP01_10_v1","PvP01_10_v1","PvP01_10_v1","PvP01_10_v1","PvP01_10_v1","PvP01_10_v1","PvP01_10_v1","PvP01_10_v1","PvP01_10_v1","PvP01_10_v1","PvP01_10_v1","PvP01_10_v1","PvP01_10_v1","PvP01_10_v1","PvP01_10_v1","PvP01_10_v1","PvP01_10_v1","PvP01_10_v1","PvP01_10_v1","PvP01_10_v1","PvP01_10_v1","PvP01_10_v1","PvP01_10_v1","PvP01_10_v1","PvP01_10_v1","PvP01_10_v1","PvP01_10_v1","PvP01_10_v1","PvP01_10_v1","PvP01_10_v1","PvP01_10_v1","PvP01_10_v1","PvP01_10_v1","PvP01_10_v1","PvP01_10_v1","PvP01_10_v1","PvP01_10_v1","PvP01_10_v1","PvP01_11_v1","PvP01_11_v1","PvP01_11_v1","PvP01_11_v1","PvP01_11_v1","PvP01_11_v1","PvP01_11_v1","PvP01_11_v1","PvP01_11_v1","PvP01_11_v1","PvP01_11_v1","PvP01_11_v1","PvP01_11_v1","PvP01_11_v1","PvP01_11_v1","PvP01_11_v1","PvP01_11_v1","PvP01_11_v1","PvP01_11_v1","PvP01_11_v1","PvP01_11_v1","PvP01_11_v1","PvP01_11_v1","PvP01_11_v1","PvP01_11_v1","PvP01_11_v1","PvP01_11_v1","PvP01_11_v1","PvP01_11_v1","PvP01_11_v1","PvP01_11_v1","PvP01_11_v1","PvP01_11_v1","PvP01_11_v1","PvP01_11_v1","PvP01_11_v1","PvP01_11_v1","PvP01_11_v1","PvP01_11_v1","PvP01_11_v1","PvP01_11_v1","PvP01_11_v1","PvP01_11_v1","PvP01_11_v1","PvP01_11_v1","PvP01_11_v1","PvP01_11_v1","PvP01_11_v1","PvP01_11_v1","PvP01_11_v1","PvP01_11_v1","PvP01_11_v1","PvP01_11_v1","PvP01_11_v1","PvP01_11_v1","PvP01_11_v1","PvP01_11_v1","PvP01_11_v1","PvP01_11_v1","PvP01_11_v1","PvP01_11_v1","PvP01_11_v1","PvP01_11_v1","PvP01_11_v1","PvP01_11_v1","PvP01_11_v1","PvP01_11_v1","PvP01_11_v1","PvP01_11_v1","PvP01_11_v1","PvP01_11_v1","PvP01_11_v1","PvP01_11_v1","PvP01_11_v1","PvP01_11_v1","PvP01_11_v1","PvP01_11_v1","PvP01_11_v1","PvP01_11_v1","PvP01_11_v1","PvP01_11_v1","PvP01_11_v1","PvP01_11_v1","PvP01_11_v1","PvP01_11_v1","PvP01_11_v1","PvP01_11_v1","PvP01_11_v1","PvP01_11_v1","PvP01_11_v1","PvP01_11_v1","PvP01_11_v1","PvP01_11_v1","PvP01_11_v1","PvP01_11_v1","PvP01_11_v1","PvP01_11_v1","PvP01_11_v1","PvP01_11_v1","PvP01_11_v1","PvP01_11_v1","PvP01_11_v1","PvP01_11_v1","PvP01_11_v1","PvP01_11_v1","PvP01_11_v1","PvP01_11_v1","PvP01_11_v1","PvP01_11_v1","PvP01_11_v1","PvP01_11_v1","PvP01_11_v1","PvP01_11_v1","PvP01_11_v1","PvP01_11_v1","PvP01_11_v1","PvP01_11_v1","PvP01_11_v1","PvP01_11_v1","PvP01_11_v1","PvP01_11_v1","PvP01_11_v1","PvP01_11_v1","PvP01_11_v1","PvP01_11_v1","PvP01_11_v1","PvP01_11_v1","PvP01_11_v1","PvP01_11_v1","PvP01_11_v1","PvP01_11_v1","PvP01_11_v1","PvP01_11_v1","PvP01_11_v1","PvP01_11_v1","PvP01_11_v1","PvP01_11_v1","PvP01_11_v1","PvP01_11_v1","PvP01_11_v1","PvP01_11_v1","PvP01_11_v1","PvP01_11_v1","PvP01_11_v1","PvP01_11_v1","PvP01_11_v1","PvP01_11_v1","PvP01_11_v1","PvP01_11_v1","PvP01_11_v1","PvP01_11_v1","PvP01_11_v1","PvP01_11_v1","PvP01_11_v1","PvP01_11_v1","PvP01_11_v1","PvP01_11_v1","PvP01_11_v1","PvP01_11_v1","PvP01_11_v1","PvP01_11_v1","PvP01_11_v1","PvP01_11_v1","PvP01_11_v1","PvP01_11_v1","PvP01_11_v1","PvP01_11_v1","PvP01_11_v1","PvP01_11_v1","PvP01_11_v1","PvP01_11_v1","PvP01_11_v1","PvP01_11_v1","PvP01_11_v1","PvP01_11_v1","PvP01_11_v1","PvP01_11_v1","PvP01_11_v1","PvP01_11_v1","PvP01_11_v1","PvP01_11_v1","PvP01_11_v1","PvP01_11_v1","PvP01_11_v1","PvP01_11_v1","PvP01_11_v1","PvP01_11_v1","PvP01_11_v1","PvP01_11_v1","PvP01_11_v1","PvP01_11_v1","PvP01_11_v1","PvP01_11_v1","PvP01_11_v1","PvP01_11_v1","PvP01_11_v1","PvP01_11_v1","PvP01_11_v1","PvP01_11_v1","PvP01_11_v1","PvP01_11_v1","PvP01_11_v1","PvP01_11_v1","PvP01_11_v1","PvP01_11_v1","PvP01_11_v1","PvP01_11_v1","PvP01_11_v1","PvP01_11_v1","PvP01_11_v1","PvP01_11_v1","PvP01_11_v1","PvP01_11_v1","PvP01_11_v1","PvP01_11_v1","PvP01_11_v1","PvP01_11_v1","PvP01_11_v1","PvP01_11_v1","PvP01_11_v1","PvP01_11_v1","PvP01_11_v1","PvP01_11_v1","PvP01_11_v1","PvP01_11_v1","PvP01_11_v1","PvP01_11_v1","PvP01_11_v1","PvP01_11_v1","PvP01_11_v1","PvP01_11_v1","PvP01_11_v1","PvP01_11_v1","PvP01_11_v1","PvP01_11_v1","PvP01_11_v1","PvP01_11_v1","PvP01_11_v1","PvP01_11_v1","PvP01_11_v1","PvP01_11_v1","PvP01_11_v1","PvP01_11_v1","PvP01_11_v1","PvP01_11_v1","PvP01_11_v1","PvP01_11_v1","PvP01_11_v1","PvP01_11_v1","PvP01_11_v1","PvP01_11_v1","PvP01_11_v1","PvP01_11_v1","PvP01_11_v1","PvP01_11_v1","PvP01_11_v1","PvP01_11_v1","PvP01_11_v1","PvP01_11_v1","PvP01_12_v1","PvP01_12_v1","PvP01_12_v1","PvP01_12_v1","PvP01_12_v1","PvP01_12_v1","PvP01_12_v1","PvP01_12_v1","PvP01_12_v1","PvP01_12_v1","PvP01_12_v1","PvP01_12_v1","PvP01_12_v1","PvP01_12_v1","PvP01_12_v1","PvP01_12_v1","PvP01_12_v1","PvP01_12_v1","PvP01_12_v1","PvP01_12_v1","PvP01_12_v1","PvP01_12_v1","PvP01_12_v1","PvP01_12_v1","PvP01_12_v1","PvP01_12_v1","PvP01_12_v1","PvP01_12_v1","PvP01_12_v1","PvP01_12_v1","PvP01_12_v1","PvP01_12_v1","PvP01_12_v1","PvP01_12_v1","PvP01_12_v1","PvP01_12_v1","PvP01_12_v1","PvP01_12_v1","PvP01_12_v1","PvP01_12_v1","PvP01_12_v1","PvP01_12_v1","PvP01_12_v1","PvP01_12_v1","PvP01_12_v1","PvP01_12_v1","PvP01_12_v1","PvP01_12_v1","PvP01_12_v1","PvP01_12_v1","PvP01_12_v1","PvP01_12_v1","PvP01_12_v1","PvP01_12_v1","PvP01_12_v1","PvP01_12_v1","PvP01_12_v1","PvP01_12_v1","PvP01_12_v1","PvP01_12_v1","PvP01_12_v1","PvP01_12_v1","PvP01_12_v1","PvP01_12_v1","PvP01_12_v1","PvP01_12_v1","PvP01_12_v1","PvP01_12_v1","PvP01_12_v1","PvP01_12_v1","PvP01_12_v1","PvP01_12_v1","PvP01_12_v1","PvP01_12_v1","PvP01_12_v1","PvP01_12_v1","PvP01_12_v1","PvP01_12_v1","PvP01_12_v1","PvP01_12_v1","PvP01_12_v1","PvP01_12_v1","PvP01_12_v1","PvP01_12_v1","PvP01_12_v1","PvP01_12_v1","PvP01_12_v1","PvP01_12_v1","PvP01_12_v1","PvP01_12_v1","PvP01_12_v1","PvP01_12_v1","PvP01_12_v1","PvP01_12_v1","PvP01_12_v1","PvP01_12_v1","PvP01_12_v1","PvP01_12_v1","PvP01_12_v1","PvP01_12_v1","PvP01_12_v1","PvP01_12_v1","PvP01_12_v1","PvP01_12_v1","PvP01_12_v1","PvP01_12_v1","PvP01_12_v1","PvP01_12_v1","PvP01_12_v1","PvP01_12_v1","PvP01_12_v1","PvP01_12_v1","PvP01_12_v1","PvP01_12_v1","PvP01_12_v1","PvP01_12_v1","PvP01_12_v1","PvP01_12_v1","PvP01_12_v1","PvP01_12_v1","PvP01_12_v1","PvP01_12_v1","PvP01_12_v1","PvP01_12_v1","PvP01_12_v1","PvP01_12_v1","PvP01_12_v1","PvP01_12_v1","PvP01_12_v1","PvP01_12_v1","PvP01_12_v1","PvP01_12_v1","PvP01_12_v1","PvP01_12_v1","PvP01_12_v1","PvP01_12_v1","PvP01_12_v1","PvP01_12_v1","PvP01_12_v1","PvP01_12_v1","PvP01_12_v1","PvP01_12_v1","PvP01_12_v1","PvP01_12_v1","PvP01_12_v1","PvP01_12_v1","PvP01_12_v1","PvP01_12_v1","PvP01_12_v1","PvP01_12_v1","PvP01_12_v1","PvP01_12_v1","PvP01_12_v1","PvP01_12_v1","PvP01_12_v1","PvP01_12_v1","PvP01_12_v1","PvP01_12_v1","PvP01_12_v1","PvP01_12_v1","PvP01_12_v1","PvP01_12_v1","PvP01_12_v1","PvP01_12_v1","PvP01_12_v1","PvP01_12_v1","PvP01_12_v1","PvP01_12_v1","PvP01_12_v1","PvP01_12_v1","PvP01_12_v1","PvP01_12_v1","PvP01_12_v1","PvP01_12_v1","PvP01_12_v1","PvP01_12_v1","PvP01_12_v1","PvP01_12_v1","PvP01_12_v1","PvP01_12_v1","PvP01_12_v1","PvP01_12_v1","PvP01_12_v1","PvP01_12_v1","PvP01_12_v1","PvP01_12_v1","PvP01_12_v1","PvP01_12_v1","PvP01_12_v1","PvP01_12_v1","PvP01_12_v1","PvP01_12_v1","PvP01_12_v1","PvP01_12_v1","PvP01_12_v1","PvP01_12_v1","PvP01_12_v1","PvP01_12_v1","PvP01_12_v1","PvP01_12_v1","PvP01_12_v1","PvP01_12_v1","PvP01_12_v1","PvP01_12_v1","PvP01_12_v1","PvP01_12_v1","PvP01_12_v1","PvP01_12_v1","PvP01_12_v1","PvP01_12_v1","PvP01_12_v1","PvP01_12_v1","PvP01_12_v1","PvP01_12_v1","PvP01_12_v1","PvP01_12_v1","PvP01_12_v1","PvP01_12_v1","PvP01_12_v1","PvP01_12_v1","PvP01_12_v1","PvP01_12_v1","PvP01_12_v1","PvP01_12_v1","PvP01_12_v1","PvP01_12_v1","PvP01_12_v1","PvP01_12_v1","PvP01_12_v1","PvP01_12_v1","PvP01_12_v1","PvP01_12_v1","PvP01_12_v1","PvP01_12_v1","PvP01_12_v1","PvP01_12_v1","PvP01_12_v1","PvP01_12_v1","PvP01_12_v1","PvP01_12_v1","PvP01_12_v1","PvP01_12_v1","PvP01_12_v1","PvP01_12_v1","PvP01_12_v1","PvP01_12_v1","PvP01_12_v1","PvP01_12_v1","PvP01_12_v1","PvP01_12_v1","PvP01_12_v1","PvP01_12_v1","PvP01_12_v1","PvP01_12_v1","PvP01_12_v1","PvP01_12_v1","PvP01_12_v1","PvP01_12_v1","PvP01_12_v1","PvP01_12_v1","PvP01_12_v1","PvP01_12_v1","PvP01_12_v1","PvP01_12_v1","PvP01_12_v1","PvP01_12_v1","PvP01_12_v1","PvP01_12_v1","PvP01_12_v1","PvP01_12_v1","PvP01_12_v1","PvP01_12_v1","PvP01_12_v1","PvP01_12_v1","PvP01_12_v1","PvP01_12_v1","PvP01_12_v1","PvP01_12_v1","PvP01_12_v1","PvP01_12_v1","PvP01_12_v1","PvP01_12_v1","PvP01_12_v1","PvP01_12_v1","PvP01_12_v1","PvP01_12_v1","PvP01_12_v1","PvP01_12_v1","PvP01_12_v1","PvP01_12_v1","PvP01_12_v1","PvP01_12_v1","PvP01_12_v1","PvP01_12_v1","PvP01_12_v1","PvP01_12_v1","PvP01_12_v1","PvP01_12_v1","PvP01_12_v1","PvP01_12_v1","PvP01_12_v1","PvP01_12_v1","PvP01_12_v1","PvP01_12_v1","PvP01_12_v1","PvP01_12_v1","PvP01_12_v1","PvP01_12_v1","PvP01_12_v1","PvP01_12_v1","PvP01_12_v1","PvP01_12_v1","PvP01_12_v1","PvP01_12_v1","PvP01_12_v1","PvP01_12_v1","PvP01_12_v1","PvP01_12_v1","PvP01_12_v1","PvP01_12_v1","PvP01_12_v1","PvP01_12_v1","PvP01_12_v1","PvP01_12_v1","PvP01_12_v1","PvP01_12_v1","PvP01_12_v1","PvP01_12_v1","PvP01_12_v1","PvP01_12_v1","PvP01_12_v1","PvP01_12_v1","PvP01_12_v1","PvP01_12_v1","PvP01_12_v1","PvP01_12_v1","PvP01_12_v1","PvP01_12_v1","PvP01_12_v1","PvP01_12_v1","PvP01_12_v1","PvP01_12_v1","PvP01_12_v1","PvP01_12_v1","PvP01_12_v1","PvP01_12_v1","PvP01_12_v1","PvP01_13_v1","PvP01_13_v1","PvP01_13_v1","PvP01_13_v1","PvP01_13_v1","PvP01_13_v1","PvP01_13_v1","PvP01_13_v1","PvP01_13_v1","PvP01_13_v1","PvP01_13_v1","PvP01_13_v1","PvP01_13_v1","PvP01_13_v1","PvP01_13_v1","PvP01_13_v1","PvP01_13_v1","PvP01_13_v1","PvP01_13_v1","PvP01_13_v1","PvP01_13_v1","PvP01_13_v1","PvP01_13_v1","PvP01_13_v1","PvP01_13_v1","PvP01_13_v1","PvP01_13_v1","PvP01_13_v1","PvP01_13_v1","PvP01_13_v1","PvP01_13_v1","PvP01_13_v1","PvP01_13_v1","PvP01_13_v1","PvP01_13_v1","PvP01_13_v1","PvP01_13_v1","PvP01_13_v1","PvP01_13_v1","PvP01_13_v1","PvP01_13_v1","PvP01_13_v1","PvP01_13_v1","PvP01_13_v1","PvP01_13_v1","PvP01_13_v1","PvP01_13_v1","PvP01_13_v1","PvP01_13_v1","PvP01_13_v1","PvP01_13_v1","PvP01_13_v1","PvP01_13_v1","PvP01_13_v1","PvP01_13_v1","PvP01_13_v1","PvP01_13_v1","PvP01_13_v1","PvP01_13_v1","PvP01_13_v1","PvP01_13_v1","PvP01_13_v1","PvP01_13_v1","PvP01_13_v1","PvP01_13_v1","PvP01_13_v1","PvP01_13_v1","PvP01_13_v1","PvP01_13_v1","PvP01_13_v1","PvP01_13_v1","PvP01_13_v1","PvP01_13_v1","PvP01_13_v1","PvP01_13_v1","PvP01_13_v1","PvP01_13_v1","PvP01_13_v1","PvP01_13_v1","PvP01_13_v1","PvP01_13_v1","PvP01_13_v1","PvP01_13_v1","PvP01_13_v1","PvP01_13_v1","PvP01_13_v1","PvP01_13_v1","PvP01_13_v1","PvP01_13_v1","PvP01_13_v1","PvP01_13_v1","PvP01_13_v1","PvP01_13_v1","PvP01_13_v1","PvP01_13_v1","PvP01_13_v1","PvP01_13_v1","PvP01_13_v1","PvP01_13_v1","PvP01_13_v1","PvP01_13_v1","PvP01_13_v1","PvP01_13_v1","PvP01_13_v1","PvP01_13_v1","PvP01_13_v1","PvP01_13_v1","PvP01_13_v1","PvP01_13_v1","PvP01_13_v1","PvP01_13_v1","PvP01_13_v1","PvP01_13_v1","PvP01_13_v1","PvP01_13_v1","PvP01_13_v1","PvP01_13_v1","PvP01_13_v1","PvP01_13_v1","PvP01_13_v1","PvP01_13_v1","PvP01_13_v1","PvP01_13_v1","PvP01_13_v1","PvP01_13_v1","PvP01_13_v1","PvP01_13_v1","PvP01_13_v1","PvP01_13_v1","PvP01_13_v1","PvP01_13_v1","PvP01_13_v1","PvP01_13_v1","PvP01_13_v1","PvP01_13_v1","PvP01_13_v1","PvP01_13_v1","PvP01_13_v1","PvP01_13_v1","PvP01_13_v1","PvP01_13_v1","PvP01_13_v1","PvP01_13_v1","PvP01_13_v1","PvP01_13_v1","PvP01_13_v1","PvP01_13_v1","PvP01_13_v1","PvP01_13_v1","PvP01_13_v1","PvP01_13_v1","PvP01_13_v1","PvP01_13_v1","PvP01_13_v1","PvP01_13_v1","PvP01_13_v1","PvP01_13_v1","PvP01_13_v1","PvP01_13_v1","PvP01_13_v1","PvP01_13_v1","PvP01_13_v1","PvP01_13_v1","PvP01_13_v1","PvP01_13_v1","PvP01_13_v1","PvP01_13_v1","PvP01_13_v1","PvP01_13_v1","PvP01_13_v1","PvP01_13_v1","PvP01_13_v1","PvP01_13_v1","PvP01_13_v1","PvP01_13_v1","PvP01_13_v1","PvP01_13_v1","PvP01_13_v1","PvP01_13_v1","PvP01_13_v1","PvP01_13_v1","PvP01_13_v1","PvP01_13_v1","PvP01_13_v1","PvP01_13_v1","PvP01_13_v1","PvP01_13_v1","PvP01_13_v1","PvP01_13_v1","PvP01_13_v1","PvP01_13_v1","PvP01_13_v1","PvP01_13_v1","PvP01_13_v1","PvP01_13_v1","PvP01_13_v1","PvP01_13_v1","PvP01_13_v1","PvP01_13_v1","PvP01_13_v1","PvP01_13_v1","PvP01_13_v1","PvP01_13_v1","PvP01_13_v1","PvP01_13_v1","PvP01_13_v1","PvP01_13_v1","PvP01_13_v1","PvP01_13_v1","PvP01_13_v1","PvP01_13_v1","PvP01_13_v1","PvP01_13_v1","PvP01_13_v1","PvP01_13_v1","PvP01_13_v1","PvP01_13_v1","PvP01_13_v1","PvP01_13_v1","PvP01_13_v1","PvP01_13_v1","PvP01_13_v1","PvP01_13_v1","PvP01_13_v1","PvP01_13_v1","PvP01_14_v1","PvP01_14_v1","PvP01_14_v1","PvP01_14_v1","PvP01_14_v1","PvP01_14_v1","PvP01_14_v1","PvP01_14_v1","PvP01_14_v1","PvP01_14_v1","PvP01_14_v1","PvP01_14_v1","PvP01_14_v1","PvP01_14_v1","PvP01_14_v1","PvP01_14_v1","PvP01_14_v1","PvP01_14_v1","PvP01_14_v1","PvP01_14_v1","PvP01_14_v1","PvP01_14_v1","PvP01_14_v1","PvP01_14_v1","PvP01_14_v1","PvP01_14_v1","PvP01_14_v1","PvP01_14_v1","PvP01_14_v1","PvP01_14_v1","PvP01_14_v1","PvP01_14_v1","PvP01_14_v1","PvP01_14_v1","PvP01_14_v1","PvP01_14_v1","PvP01_14_v1","PvP01_14_v1","PvP01_14_v1","PvP01_14_v1","PvP01_14_v1","PvP01_14_v1","PvP01_14_v1","PvP01_14_v1","PvP01_14_v1","PvP01_14_v1","PvP01_14_v1","PvP01_14_v1","PvP01_14_v1","PvP01_14_v1","PvP01_14_v1","PvP01_14_v1","PvP01_14_v1","PvP01_14_v1","PvP01_14_v1","PvP01_14_v1","PvP01_14_v1","PvP01_14_v1","PvP01_14_v1","PvP01_14_v1","PvP01_14_v1","PvP01_14_v1","PvP01_14_v1","PvP01_14_v1","PvP01_14_v1","PvP01_14_v1","PvP01_14_v1","PvP01_14_v1","PvP01_14_v1","PvP01_14_v1","PvP01_14_v1","PvP01_14_v1","PvP01_14_v1","PvP01_14_v1","PvP01_14_v1","PvP01_14_v1","PvP01_14_v1","PvP01_14_v1","PvP01_14_v1","PvP01_14_v1","PvP01_14_v1","PvP01_14_v1","PvP01_14_v1","PvP01_14_v1","PvP01_14_v1","PvP01_14_v1","PvP01_14_v1","PvP01_14_v1","PvP01_14_v1","PvP01_14_v1","PvP01_14_v1","PvP01_14_v1","PvP01_14_v1","PvP01_14_v1","PvP01_14_v1","PvP01_14_v1","PvP01_14_v1","PvP01_14_v1","PvP01_14_v1","PvP01_14_v1","PvP01_14_v1","PvP01_14_v1","PvP01_14_v1","PvP01_14_v1","PvP01_14_v1","PvP01_14_v1","PvP01_14_v1","PvP01_14_v1","PvP01_14_v1","PvP01_14_v1","PvP01_14_v1","PvP01_14_v1","PvP01_14_v1","PvP01_14_v1","PvP01_14_v1","PvP01_14_v1","PvP01_14_v1","PvP01_14_v1","PvP01_14_v1","PvP01_14_v1","PvP01_14_v1","PvP01_14_v1","PvP01_14_v1","PvP01_14_v1","PvP01_14_v1","PvP01_14_v1","PvP01_14_v1","PvP01_14_v1","PvP01_14_v1","PvP01_14_v1","PvP01_14_v1","PvP01_14_v1","PvP01_14_v1","PvP01_14_v1","PvP01_14_v1","PvP01_14_v1","PvP01_14_v1","PvP01_14_v1","PvP01_14_v1","PvP01_14_v1","PvP01_14_v1","PvP01_14_v1","PvP01_14_v1","PvP01_14_v1","PvP01_14_v1","PvP01_14_v1","PvP01_14_v1","PvP01_14_v1","PvP01_14_v1","PvP01_14_v1","PvP01_14_v1","PvP01_14_v1","PvP01_14_v1","PvP01_14_v1","PvP01_14_v1","PvP01_14_v1","PvP01_14_v1","PvP01_14_v1","PvP01_14_v1","PvP01_14_v1","PvP01_14_v1","PvP01_14_v1","PvP01_14_v1","PvP01_14_v1","PvP01_14_v1","PvP01_14_v1","PvP01_14_v1","PvP01_14_v1","PvP01_14_v1","PvP01_14_v1","PvP01_14_v1","PvP01_14_v1","PvP01_14_v1","PvP01_14_v1","PvP01_14_v1","PvP01_14_v1","PvP01_14_v1","PvP01_14_v1","PvP01_14_v1","PvP01_14_v1","PvP01_14_v1","PvP01_14_v1","PvP01_14_v1","PvP01_14_v1","PvP01_14_v1","PvP01_14_v1","PvP01_14_v1","PvP01_14_v1","PvP01_14_v1","PvP01_14_v1","PvP01_14_v1","PvP01_14_v1","PvP01_14_v1","PvP01_14_v1","PvP01_14_v1","PvP01_14_v1","PvP01_14_v1","PvP01_14_v1","PvP01_14_v1","PvP01_14_v1","PvP01_14_v1","PvP01_14_v1","PvP01_14_v1","PvP01_14_v1","PvP01_14_v1","PvP01_14_v1","PvP01_14_v1","PvP01_14_v1","PvP01_14_v1","PvP01_14_v1","PvP01_14_v1","PvP01_14_v1","PvP01_14_v1","PvP01_14_v1","PvP01_14_v1","PvP01_14_v1","PvP01_14_v1","PvP01_14_v1","PvP01_14_v1","PvP01_14_v1","PvP01_14_v1","PvP01_14_v1","PvP01_14_v1","PvP01_14_v1","PvP01_14_v1","PvP01_14_v1","PvP01_14_v1","PvP01_14_v1","PvP01_14_v1","PvP01_14_v1","PvP01_14_v1","PvP01_14_v1","PvP01_14_v1","PvP01_14_v1","PvP01_14_v1","PvP01_14_v1","PvP01_14_v1","PvP01_14_v1","PvP01_14_v1","PvP01_14_v1","PvP01_14_v1","PvP01_14_v1","PvP01_14_v1","PvP01_14_v1","PvP01_14_v1","PvP01_14_v1","PvP01_14_v1","PvP01_14_v1","PvP01_14_v1","PvP01_14_v1","PvP01_14_v1","PvP01_14_v1","PvP01_14_v1","PvP01_14_v1","PvP01_14_v1","PvP01_14_v1","PvP01_14_v1","PvP01_14_v1","PvP01_14_v1","PvP01_14_v1","PvP01_14_v1","PvP01_14_v1","PvP01_14_v1","PvP01_14_v1","PvP01_14_v1","PvP01_14_v1","PvP01_14_v1","PvP01_14_v1","PvP01_14_v1","PvP01_14_v1","PvP01_14_v1","PvP01_14_v1","PvP01_14_v1","PvP01_14_v1","PvP01_14_v1","PvP01_14_v1","PvP01_14_v1","PvP01_14_v1","PvP01_14_v1","PvP01_14_v1","PvP01_14_v1","PvP01_14_v1","PvP01_14_v1","PvP01_14_v1","PvP01_14_v1","PvP01_14_v1","PvP01_14_v1","PvP01_14_v1","PvP01_14_v1","PvP01_14_v1","PvP01_14_v1","PvP01_14_v1","PvP01_14_v1","PvP01_14_v1","PvP01_14_v1","PvP01_14_v1","PvP01_14_v1","PvP01_14_v1","PvP01_14_v1","PvP01_14_v1","PvP01_14_v1","PvP01_14_v1","PvP01_14_v1","PvP01_14_v1","PvP01_14_v1","PvP01_14_v1","PvP01_14_v1","PvP01_14_v1","PvP01_14_v1","PvP01_14_v1","PvP01_14_v1","PvP01_14_v1","PvP01_14_v1","PvP01_14_v1","PvP01_14_v1","PvP01_14_v1","PvP01_14_v1","PvP01_14_v1","PvP01_14_v1","PvP01_14_v1","PvP01_14_v1","PvP01_14_v1","PvP01_14_v1","PvP01_14_v1","PvP01_14_v1","PvP01_14_v1","PvP01_14_v1","PvP01_14_v1","PvP01_14_v1","PvP01_14_v1","PvP01_14_v1","PvP01_14_v1","PvP01_14_v1","PvP01_14_v1","PvP01_14_v1","PvP01_14_v1","PvP01_14_v1","PvP01_14_v1","PvP01_14_v1","PvP01_14_v1","PvP01_14_v1","PvP01_14_v1","PvP01_14_v1","PvP01_14_v1","PvP01_14_v1","PvP01_14_v1","PvP01_14_v1","PvP01_14_v1","PvP01_14_v1","PvP01_14_v1","PvP01_14_v1","PvP01_14_v1","PvP01_14_v1","PvP01_14_v1","PvP01_14_v1","PvP01_14_v1","PvP01_14_v1","PvP01_14_v1","PvP01_14_v1","PvP01_14_v1","PvP01_14_v1","PvP01_API_v1","PvP01_API_v1","PvP01_API_v1","PvP01_API_v1","PvP01_API_v1","Transfer.PvP01_00_1.final","Transfer.PvP01_00_1.final","Transfer.PvP01_00_2.final","Transfer.PvP01_00_8.final","Transfer.PvP01_00_10.final","Transfer.PvP01_00_11.final","Transfer.PvP01_00_11.final","Transfer.PvP01_00_11.final","Transfer.PvP01_00_11.final","Transfer.PvP01_00_11.final","Transfer.PvP01_00_16.final","Transfer.PvP01_00_51.final","Transfer.PvP01_00_61.final","Transfer.PvP01_00_161.final","Transfer.PvP01_00_199.final","Transfer.PvP01_00_227.final","PvP01_01_v1","PvP01_01_v1","PvP01_01_v1","PvP01_01_v1","PvP01_01_v1","PvP01_01_v1","PvP01_01_v1","PvP01_01_v1","PvP01_01_v1","PvP01_01_v1","PvP01_01_v1","PvP01_01_v1","PvP01_01_v1","PvP01_01_v1","PvP01_01_v1","PvP01_01_v1","PvP01_01_v1","PvP01_01_v1","PvP01_01_v1","PvP01_01_v1","PvP01_01_v1","PvP01_01_v1","PvP01_01_v1","PvP01_01_v1","PvP01_01_v1","PvP01_01_v1","PvP01_01_v1","PvP01_01_v1","PvP01_01_v1","PvP01_01_v1","PvP01_01_v1","PvP01_01_v1","PvP01_01_v1","PvP01_01_v1","PvP01_01_v1","PvP01_01_v1","PvP01_01_v1","PvP01_01_v1","PvP01_01_v1","PvP01_01_v1","PvP01_01_v1","PvP01_01_v1","PvP01_01_v1","PvP01_01_v1","PvP01_01_v1","PvP01_01_v1","PvP01_01_v1","PvP01_01_v1","PvP01_01_v1","PvP01_01_v1","PvP01_01_v1","PvP01_01_v1","PvP01_01_v1","PvP01_01_v1","PvP01_01_v1","PvP01_01_v1","PvP01_01_v1","PvP01_01_v1","PvP01_01_v1","PvP01_01_v1","PvP01_01_v1","PvP01_01_v1","PvP01_01_v1","PvP01_01_v1","PvP01_01_v1","PvP01_01_v1","PvP01_01_v1","PvP01_01_v1","PvP01_01_v1","PvP01_01_v1","PvP01_01_v1","PvP01_01_v1","PvP01_01_v1","PvP01_01_v1","PvP01_01_v1","PvP01_01_v1","PvP01_01_v1","PvP01_02_v1","PvP01_02_v1","PvP01_02_v1","PvP01_02_v1","PvP01_02_v1","PvP01_02_v1","PvP01_02_v1","PvP01_02_v1","PvP01_02_v1","PvP01_02_v1","PvP01_02_v1","PvP01_02_v1","PvP01_02_v1","PvP01_02_v1","PvP01_02_v1","PvP01_02_v1","PvP01_02_v1","PvP01_02_v1","PvP01_02_v1","PvP01_02_v1","PvP01_02_v1","PvP01_02_v1","PvP01_02_v1","PvP01_02_v1","PvP01_02_v1","PvP01_02_v1","PvP01_02_v1","PvP01_02_v1","PvP01_02_v1","PvP01_02_v1","PvP01_02_v1","PvP01_02_v1","PvP01_02_v1","PvP01_02_v1","PvP01_02_v1","PvP01_02_v1","PvP01_02_v1","PvP01_02_v1","PvP01_02_v1","PvP01_02_v1","PvP01_02_v1","PvP01_02_v1","PvP01_02_v1","PvP01_02_v1","PvP01_02_v1","PvP01_02_v1","PvP01_02_v1","PvP01_02_v1","PvP01_02_v1","PvP01_02_v1","PvP01_02_v1","PvP01_02_v1","PvP01_02_v1","PvP01_02_v1","PvP01_02_v1","PvP01_02_v1","PvP01_02_v1","PvP01_02_v1","PvP01_02_v1","PvP01_02_v1","PvP01_02_v1","PvP01_02_v1","PvP01_03_v1","PvP01_03_v1","PvP01_03_v1","PvP01_03_v1","PvP01_03_v1","PvP01_03_v1","PvP01_03_v1","PvP01_03_v1","PvP01_03_v1","PvP01_03_v1","PvP01_03_v1","PvP01_03_v1","PvP01_03_v1","PvP01_03_v1","PvP01_03_v1","PvP01_03_v1","PvP01_03_v1","PvP01_03_v1","PvP01_03_v1","PvP01_03_v1","PvP01_03_v1","PvP01_03_v1","PvP01_03_v1","PvP01_03_v1","PvP01_03_v1","PvP01_03_v1","PvP01_03_v1","PvP01_03_v1","PvP01_03_v1","PvP01_03_v1","PvP01_03_v1","PvP01_03_v1","PvP01_03_v1","PvP01_03_v1","PvP01_03_v1","PvP01_03_v1","PvP01_03_v1","PvP01_03_v1","PvP01_03_v1","PvP01_03_v1","PvP01_03_v1","PvP01_03_v1","PvP01_03_v1","PvP01_03_v1","PvP01_03_v1","PvP01_03_v1","PvP01_03_v1","PvP01_03_v1","PvP01_03_v1","PvP01_03_v1","PvP01_03_v1","PvP01_03_v1","PvP01_03_v1","PvP01_03_v1","PvP01_03_v1","PvP01_03_v1","PvP01_03_v1","PvP01_03_v1","PvP01_03_v1","PvP01_03_v1","PvP01_03_v1","PvP01_03_v1","PvP01_03_v1","PvP01_03_v1","PvP01_03_v1","PvP01_03_v1","PvP01_03_v1","PvP01_03_v1","PvP01_03_v1","PvP01_03_v1","PvP01_03_v1","PvP01_03_v1","PvP01_03_v1","PvP01_03_v1","PvP01_03_v1","PvP01_03_v1","PvP01_03_v1","PvP01_03_v1","PvP01_03_v1","PvP01_03_v1","PvP01_03_v1","PvP01_03_v1","PvP01_03_v1","PvP01_03_v1","PvP01_03_v1","PvP01_03_v1","PvP01_03_v1","PvP01_03_v1","PvP01_03_v1","PvP01_03_v1","PvP01_03_v1","PvP01_03_v1","PvP01_03_v1","PvP01_03_v1","PvP01_03_v1","PvP01_03_v1","PvP01_03_v1","PvP01_03_v1","PvP01_04_v1","PvP01_04_v1","PvP01_04_v1","PvP01_04_v1","PvP01_04_v1","PvP01_04_v1","PvP01_04_v1","PvP01_04_v1","PvP01_04_v1","PvP01_04_v1","PvP01_04_v1","PvP01_04_v1","PvP01_04_v1","PvP01_04_v1","PvP01_04_v1","PvP01_04_v1","PvP01_04_v1","PvP01_04_v1","PvP01_04_v1","PvP01_04_v1","PvP01_04_v1","PvP01_04_v1","PvP01_04_v1","PvP01_04_v1","PvP01_04_v1","PvP01_04_v1","PvP01_04_v1","PvP01_04_v1","PvP01_04_v1","PvP01_04_v1","PvP01_04_v1","PvP01_04_v1","PvP01_04_v1","PvP01_04_v1","PvP01_04_v1","PvP01_04_v1","PvP01_04_v1","PvP01_04_v1","PvP01_04_v1","PvP01_04_v1","PvP01_04_v1","PvP01_04_v1","PvP01_04_v1","PvP01_04_v1","PvP01_04_v1","PvP01_04_v1","PvP01_04_v1","PvP01_04_v1","PvP01_04_v1","PvP01_04_v1","PvP01_04_v1","PvP01_04_v1","PvP01_04_v1","PvP01_04_v1","PvP01_04_v1","PvP01_04_v1","PvP01_04_v1","PvP01_04_v1","PvP01_04_v1","PvP01_04_v1","PvP01_04_v1","PvP01_04_v1","PvP01_04_v1","PvP01_04_v1","PvP01_04_v1","PvP01_04_v1","PvP01_04_v1","PvP01_04_v1","PvP01_04_v1","PvP01_04_v1","PvP01_04_v1","PvP01_04_v1","PvP01_04_v1","PvP01_04_v1","PvP01_04_v1","PvP01_04_v1","PvP01_04_v1","PvP01_04_v1","PvP01_04_v1","PvP01_04_v1","PvP01_04_v1","PvP01_04_v1","PvP01_04_v1","PvP01_04_v1","PvP01_04_v1","PvP01_04_v1","PvP01_04_v1","PvP01_04_v1","PvP01_04_v1","PvP01_04_v1","PvP01_04_v1","PvP01_04_v1","PvP01_04_v1","PvP01_04_v1","PvP01_04_v1","PvP01_04_v1","PvP01_04_v1","PvP01_04_v1","PvP01_04_v1","PvP01_04_v1","PvP01_04_v1","PvP01_04_v1","PvP01_04_v1","PvP01_04_v1","PvP01_05_v1","PvP01_05_v1","PvP01_05_v1","PvP01_05_v1","PvP01_05_v1","PvP01_05_v1","PvP01_05_v1","PvP01_05_v1","PvP01_05_v1","PvP01_05_v1","PvP01_05_v1","PvP01_05_v1","PvP01_05_v1","PvP01_05_v1","PvP01_05_v1","PvP01_05_v1","PvP01_05_v1","PvP01_05_v1","PvP01_05_v1","PvP01_05_v1","PvP01_05_v1","PvP01_05_v1","PvP01_05_v1","PvP01_05_v1","PvP01_05_v1","PvP01_05_v1","PvP01_05_v1","PvP01_05_v1","PvP01_05_v1","PvP01_05_v1","PvP01_05_v1","PvP01_05_v1","PvP01_05_v1","PvP01_05_v1","PvP01_05_v1","PvP01_05_v1","PvP01_05_v1","PvP01_05_v1","PvP01_05_v1","PvP01_05_v1","PvP01_05_v1","PvP01_05_v1","PvP01_05_v1","PvP01_05_v1","PvP01_05_v1","PvP01_05_v1","PvP01_05_v1","PvP01_05_v1","PvP01_05_v1","PvP01_05_v1","PvP01_05_v1","PvP01_05_v1","PvP01_05_v1","PvP01_05_v1","PvP01_05_v1","PvP01_05_v1","PvP01_05_v1","PvP01_05_v1","PvP01_05_v1","PvP01_05_v1","PvP01_05_v1","PvP01_05_v1","PvP01_05_v1","PvP01_05_v1","PvP01_05_v1","PvP01_05_v1","PvP01_05_v1","PvP01_05_v1","PvP01_05_v1","PvP01_05_v1","PvP01_05_v1","PvP01_05_v1","PvP01_05_v1","PvP01_05_v1","PvP01_05_v1","PvP01_05_v1","PvP01_05_v1","PvP01_05_v1","PvP01_05_v1","PvP01_05_v1","PvP01_05_v1","PvP01_05_v1","PvP01_05_v1","PvP01_05_v1","PvP01_05_v1","PvP01_05_v1","PvP01_05_v1","PvP01_05_v1","PvP01_05_v1","PvP01_05_v1","PvP01_05_v1","PvP01_05_v1","PvP01_05_v1","PvP01_05_v1","PvP01_05_v1","PvP01_05_v1","PvP01_05_v1","PvP01_05_v1","PvP01_05_v1","PvP01_05_v1","PvP01_05_v1","PvP01_05_v1","PvP01_05_v1","PvP01_05_v1","PvP01_05_v1","PvP01_05_v1","PvP01_05_v1","PvP01_05_v1","PvP01_05_v1","PvP01_05_v1","PvP01_05_v1","PvP01_05_v1","PvP01_05_v1","PvP01_05_v1","PvP01_05_v1","PvP01_05_v1","PvP01_05_v1","PvP01_05_v1","PvP01_05_v1","PvP01_05_v1","PvP01_05_v1","PvP01_05_v1","PvP01_05_v1","PvP01_05_v1","PvP01_05_v1","PvP01_05_v1","PvP01_05_v1","PvP01_05_v1","PvP01_05_v1","PvP01_05_v1","PvP01_05_v1","PvP01_05_v1","PvP01_05_v1","PvP01_05_v1","PvP01_05_v1","PvP01_05_v1","PvP01_05_v1","PvP01_05_v1","PvP01_05_v1","PvP01_06_v1","PvP01_06_v1","PvP01_06_v1","PvP01_06_v1","PvP01_06_v1","PvP01_06_v1","PvP01_06_v1","PvP01_06_v1","PvP01_06_v1","PvP01_06_v1","PvP01_06_v1","PvP01_06_v1","PvP01_06_v1","PvP01_06_v1","PvP01_06_v1","PvP01_06_v1","PvP01_06_v1","PvP01_06_v1","PvP01_06_v1","PvP01_06_v1","PvP01_06_v1","PvP01_06_v1","PvP01_06_v1","PvP01_06_v1","PvP01_06_v1","PvP01_06_v1","PvP01_06_v1","PvP01_06_v1","PvP01_06_v1","PvP01_06_v1","PvP01_06_v1","PvP01_06_v1","PvP01_06_v1","PvP01_06_v1","PvP01_06_v1","PvP01_06_v1","PvP01_06_v1","PvP01_06_v1","PvP01_06_v1","PvP01_06_v1","PvP01_06_v1","PvP01_06_v1","PvP01_06_v1","PvP01_06_v1","PvP01_06_v1","PvP01_06_v1","PvP01_06_v1","PvP01_06_v1","PvP01_06_v1","PvP01_06_v1","PvP01_06_v1","PvP01_06_v1","PvP01_06_v1","PvP01_06_v1","PvP01_06_v1","PvP01_06_v1","PvP01_06_v1","PvP01_06_v1","PvP01_06_v1","PvP01_06_v1","PvP01_06_v1","PvP01_06_v1","PvP01_06_v1","PvP01_06_v1","PvP01_06_v1","PvP01_06_v1","PvP01_06_v1","PvP01_06_v1","PvP01_06_v1","PvP01_06_v1","PvP01_06_v1","PvP01_06_v1","PvP01_06_v1","PvP01_06_v1","PvP01_06_v1","PvP01_06_v1","PvP01_06_v1","PvP01_06_v1","PvP01_06_v1","PvP01_06_v1","PvP01_06_v1","PvP01_06_v1","PvP01_06_v1","PvP01_06_v1","PvP01_06_v1","PvP01_06_v1","PvP01_06_v1","PvP01_06_v1","PvP01_06_v1","PvP01_06_v1","PvP01_06_v1","PvP01_06_v1","PvP01_06_v1","PvP01_06_v1","PvP01_06_v1","PvP01_06_v1","PvP01_06_v1","PvP01_06_v1","PvP01_06_v1","PvP01_06_v1","PvP01_06_v1","PvP01_06_v1","PvP01_06_v1","PvP01_06_v1","PvP01_06_v1","PvP01_06_v1","PvP01_06_v1","PvP01_06_v1","PvP01_06_v1","PvP01_06_v1","PvP01_06_v1","PvP01_06_v1","PvP01_06_v1","PvP01_06_v1","PvP01_06_v1","PvP01_06_v1","PvP01_06_v1","PvP01_06_v1","PvP01_06_v1","PvP01_06_v1","PvP01_07_v1","PvP01_07_v1","PvP01_07_v1","PvP01_07_v1","PvP01_07_v1","PvP01_07_v1","PvP01_07_v1","PvP01_07_v1","PvP01_07_v1","PvP01_07_v1","PvP01_07_v1","PvP01_07_v1","PvP01_07_v1","PvP01_07_v1","PvP01_07_v1","PvP01_07_v1","PvP01_07_v1","PvP01_07_v1","PvP01_07_v1","PvP01_07_v1","PvP01_07_v1","PvP01_07_v1","PvP01_07_v1","PvP01_07_v1","PvP01_07_v1","PvP01_07_v1","PvP01_07_v1","PvP01_07_v1","PvP01_07_v1","PvP01_07_v1","PvP01_07_v1","PvP01_07_v1","PvP01_07_v1","PvP01_07_v1","PvP01_07_v1","PvP01_07_v1","PvP01_07_v1","PvP01_07_v1","PvP01_07_v1","PvP01_07_v1","PvP01_07_v1","PvP01_07_v1","PvP01_07_v1","PvP01_07_v1","PvP01_07_v1","PvP01_07_v1","PvP01_07_v1","PvP01_07_v1","PvP01_07_v1","PvP01_07_v1","PvP01_07_v1","PvP01_07_v1","PvP01_07_v1","PvP01_07_v1","PvP01_07_v1","PvP01_07_v1","PvP01_07_v1","PvP01_07_v1","PvP01_07_v1","PvP01_07_v1","PvP01_07_v1","PvP01_07_v1","PvP01_07_v1","PvP01_07_v1","PvP01_07_v1","PvP01_07_v1","PvP01_07_v1","PvP01_07_v1","PvP01_07_v1","PvP01_07_v1","PvP01_07_v1","PvP01_07_v1","PvP01_07_v1","PvP01_07_v1","PvP01_07_v1","PvP01_07_v1","PvP01_07_v1","PvP01_07_v1","PvP01_07_v1","PvP01_07_v1","PvP01_07_v1","PvP01_07_v1","PvP01_07_v1","PvP01_07_v1","PvP01_07_v1","PvP01_07_v1","PvP01_07_v1","PvP01_07_v1","PvP01_07_v1","PvP01_07_v1","PvP01_07_v1","PvP01_07_v1","PvP01_07_v1","PvP01_07_v1","PvP01_07_v1","PvP01_07_v1","PvP01_07_v1","PvP01_07_v1","PvP01_07_v1","PvP01_07_v1","PvP01_07_v1","PvP01_07_v1","PvP01_07_v1","PvP01_07_v1","PvP01_07_v1","PvP01_07_v1","PvP01_07_v1","PvP01_07_v1","PvP01_07_v1","PvP01_07_v1","PvP01_07_v1","PvP01_07_v1","PvP01_07_v1","PvP01_07_v1","PvP01_07_v1","PvP01_07_v1","PvP01_07_v1","PvP01_07_v1","PvP01_07_v1","PvP01_07_v1","PvP01_07_v1","PvP01_07_v1","PvP01_07_v1","PvP01_07_v1","PvP01_07_v1","PvP01_07_v1","PvP01_07_v1","PvP01_07_v1","PvP01_07_v1","PvP01_07_v1","PvP01_07_v1","PvP01_07_v1","PvP01_07_v1","PvP01_07_v1","PvP01_07_v1","PvP01_07_v1","PvP01_07_v1","PvP01_07_v1","PvP01_07_v1","PvP01_07_v1","PvP01_07_v1","PvP01_07_v1","PvP01_07_v1","PvP01_07_v1","PvP01_07_v1","PvP01_07_v1","PvP01_07_v1","PvP01_07_v1","PvP01_07_v1","PvP01_07_v1","PvP01_07_v1","PvP01_08_v1","PvP01_08_v1","PvP01_08_v1","PvP01_08_v1","PvP01_08_v1","PvP01_08_v1","PvP01_08_v1","PvP01_08_v1","PvP01_08_v1","PvP01_08_v1","PvP01_08_v1","PvP01_08_v1","PvP01_08_v1","PvP01_08_v1","PvP01_08_v1","PvP01_08_v1","PvP01_08_v1","PvP01_08_v1","PvP01_08_v1","PvP01_08_v1","PvP01_08_v1","PvP01_08_v1","PvP01_08_v1","PvP01_08_v1","PvP01_08_v1","PvP01_08_v1","PvP01_08_v1","PvP01_08_v1","PvP01_08_v1","PvP01_08_v1","PvP01_08_v1","PvP01_08_v1","PvP01_08_v1","PvP01_08_v1","PvP01_08_v1","PvP01_08_v1","PvP01_08_v1","PvP01_08_v1","PvP01_08_v1","PvP01_08_v1","PvP01_08_v1","PvP01_08_v1","PvP01_08_v1","PvP01_08_v1","PvP01_08_v1","PvP01_08_v1","PvP01_08_v1","PvP01_08_v1","PvP01_08_v1","PvP01_08_v1","PvP01_08_v1","PvP01_08_v1","PvP01_08_v1","PvP01_08_v1","PvP01_08_v1","PvP01_08_v1","PvP01_08_v1","PvP01_08_v1","PvP01_08_v1","PvP01_08_v1","PvP01_08_v1","PvP01_08_v1","PvP01_08_v1","PvP01_08_v1","PvP01_08_v1","PvP01_08_v1","PvP01_08_v1","PvP01_08_v1","PvP01_08_v1","PvP01_08_v1","PvP01_08_v1","PvP01_08_v1","PvP01_08_v1","PvP01_08_v1","PvP01_08_v1","PvP01_08_v1","PvP01_08_v1","PvP01_08_v1","PvP01_08_v1","PvP01_08_v1","PvP01_08_v1","PvP01_08_v1","PvP01_08_v1","PvP01_08_v1","PvP01_08_v1","PvP01_08_v1","PvP01_08_v1","PvP01_08_v1","PvP01_08_v1","PvP01_08_v1","PvP01_08_v1","PvP01_08_v1","PvP01_08_v1","PvP01_08_v1","PvP01_08_v1","PvP01_08_v1","PvP01_08_v1","PvP01_08_v1","PvP01_08_v1","PvP01_08_v1","PvP01_08_v1","PvP01_08_v1","PvP01_08_v1","PvP01_08_v1","PvP01_08_v1","PvP01_08_v1","PvP01_08_v1","PvP01_08_v1","PvP01_08_v1","PvP01_08_v1","PvP01_08_v1","PvP01_08_v1","PvP01_08_v1","PvP01_08_v1","PvP01_08_v1","PvP01_08_v1","PvP01_08_v1","PvP01_08_v1","PvP01_08_v1","PvP01_08_v1","PvP01_08_v1","PvP01_08_v1","PvP01_08_v1","PvP01_08_v1","PvP01_08_v1","PvP01_08_v1","PvP01_08_v1","PvP01_08_v1","PvP01_08_v1","PvP01_08_v1","PvP01_08_v1","PvP01_08_v1","PvP01_08_v1","PvP01_08_v1","PvP01_08_v1","PvP01_08_v1","PvP01_08_v1","PvP01_08_v1","PvP01_08_v1","PvP01_08_v1","PvP01_08_v1","PvP01_08_v1","PvP01_08_v1","PvP01_08_v1","PvP01_08_v1","PvP01_08_v1","PvP01_08_v1","PvP01_08_v1","PvP01_08_v1","PvP01_08_v1","PvP01_08_v1","PvP01_08_v1","PvP01_08_v1","PvP01_08_v1","PvP01_08_v1","PvP01_08_v1","PvP01_08_v1","PvP01_08_v1","PvP01_08_v1","PvP01_08_v1","PvP01_08_v1","PvP01_08_v1","PvP01_08_v1","PvP01_08_v1","PvP01_08_v1","PvP01_08_v1","PvP01_08_v1","PvP01_08_v1","PvP01_08_v1","PvP01_08_v1","PvP01_08_v1","PvP01_08_v1","PvP01_08_v1","PvP01_08_v1","PvP01_08_v1","PvP01_08_v1","PvP01_08_v1","PvP01_09_v1","PvP01_09_v1","PvP01_09_v1","PvP01_09_v1","PvP01_09_v1","PvP01_09_v1","PvP01_09_v1","PvP01_09_v1","PvP01_09_v1","PvP01_09_v1","PvP01_09_v1","PvP01_09_v1","PvP01_09_v1","PvP01_09_v1","PvP01_09_v1","PvP01_09_v1","PvP01_09_v1","PvP01_09_v1","PvP01_09_v1","PvP01_09_v1","PvP01_09_v1","PvP01_09_v1","PvP01_09_v1","PvP01_09_v1","PvP01_09_v1","PvP01_09_v1","PvP01_09_v1","PvP01_09_v1","PvP01_09_v1","PvP01_09_v1","PvP01_09_v1","PvP01_09_v1","PvP01_09_v1","PvP01_09_v1","PvP01_09_v1","PvP01_09_v1","PvP01_09_v1","PvP01_09_v1","PvP01_09_v1","PvP01_09_v1","PvP01_09_v1","PvP01_09_v1","PvP01_09_v1","PvP01_09_v1","PvP01_09_v1","PvP01_09_v1","PvP01_09_v1","PvP01_09_v1","PvP01_09_v1","PvP01_09_v1","PvP01_09_v1","PvP01_09_v1","PvP01_09_v1","PvP01_09_v1","PvP01_09_v1","PvP01_09_v1","PvP01_09_v1","PvP01_09_v1","PvP01_09_v1","PvP01_09_v1","PvP01_09_v1","PvP01_09_v1","PvP01_09_v1","PvP01_09_v1","PvP01_09_v1","PvP01_09_v1","PvP01_09_v1","PvP01_09_v1","PvP01_09_v1","PvP01_09_v1","PvP01_09_v1","PvP01_09_v1","PvP01_09_v1","PvP01_09_v1","PvP01_09_v1","PvP01_09_v1","PvP01_09_v1","PvP01_09_v1","PvP01_09_v1","PvP01_09_v1","PvP01_09_v1","PvP01_09_v1","PvP01_09_v1","PvP01_09_v1","PvP01_09_v1","PvP01_09_v1","PvP01_09_v1","PvP01_09_v1","PvP01_09_v1","PvP01_09_v1","PvP01_09_v1","PvP01_09_v1","PvP01_09_v1","PvP01_09_v1","PvP01_09_v1","PvP01_09_v1","PvP01_09_v1","PvP01_09_v1","PvP01_09_v1","PvP01_09_v1","PvP01_09_v1","PvP01_09_v1","PvP01_09_v1","PvP01_09_v1","PvP01_09_v1","PvP01_09_v1","PvP01_09_v1","PvP01_09_v1","PvP01_09_v1","PvP01_09_v1","PvP01_09_v1","PvP01_09_v1","PvP01_09_v1","PvP01_09_v1","PvP01_09_v1","PvP01_09_v1","PvP01_09_v1","PvP01_09_v1","PvP01_09_v1","PvP01_09_v1","PvP01_09_v1","PvP01_09_v1","PvP01_09_v1","PvP01_09_v1","PvP01_09_v1","PvP01_09_v1","PvP01_09_v1","PvP01_09_v1","PvP01_09_v1","PvP01_09_v1","PvP01_09_v1","PvP01_09_v1","PvP01_09_v1","PvP01_09_v1","PvP01_09_v1","PvP01_09_v1","PvP01_09_v1","PvP01_09_v1","PvP01_09_v1","PvP01_09_v1","PvP01_09_v1","PvP01_09_v1","PvP01_09_v1","PvP01_09_v1","PvP01_09_v1","PvP01_09_v1","PvP01_09_v1","PvP01_09_v1","PvP01_09_v1","PvP01_09_v1","PvP01_09_v1","PvP01_09_v1","PvP01_09_v1","PvP01_09_v1","PvP01_09_v1","PvP01_09_v1","PvP01_09_v1","PvP01_09_v1","PvP01_09_v1","PvP01_09_v1","PvP01_09_v1","PvP01_09_v1","PvP01_09_v1","PvP01_09_v1","PvP01_09_v1","PvP01_09_v1","PvP01_09_v1","PvP01_09_v1","PvP01_09_v1","PvP01_09_v1","PvP01_09_v1","PvP01_09_v1","PvP01_09_v1","PvP01_09_v1","PvP01_09_v1","PvP01_09_v1","PvP01_09_v1","PvP01_09_v1","PvP01_09_v1","PvP01_09_v1","PvP01_09_v1","PvP01_09_v1","PvP01_09_v1","PvP01_09_v1","PvP01_09_v1","PvP01_09_v1","PvP01_09_v1","PvP01_09_v1","PvP01_09_v1","PvP01_09_v1","PvP01_09_v1","PvP01_09_v1","PvP01_09_v1","PvP01_09_v1","PvP01_09_v1","PvP01_09_v1","PvP01_09_v1","PvP01_09_v1","PvP01_09_v1","PvP01_09_v1","PvP01_09_v1","PvP01_09_v1","PvP01_09_v1","PvP01_09_v1","PvP01_09_v1","PvP01_09_v1","PvP01_09_v1","PvP01_09_v1","PvP01_09_v1","PvP01_09_v1","PvP01_10_v1","PvP01_10_v1","PvP01_10_v1","PvP01_10_v1","PvP01_10_v1","PvP01_10_v1","PvP01_10_v1","PvP01_10_v1","PvP01_10_v1","PvP01_10_v1","PvP01_10_v1","PvP01_10_v1","PvP01_10_v1","PvP01_10_v1","PvP01_10_v1","PvP01_10_v1","PvP01_10_v1","PvP01_10_v1","PvP01_10_v1","PvP01_10_v1","PvP01_10_v1","PvP01_10_v1","PvP01_10_v1","PvP01_10_v1","PvP01_10_v1","PvP01_10_v1","PvP01_10_v1","PvP01_10_v1","PvP01_10_v1","PvP01_10_v1","PvP01_10_v1","PvP01_10_v1","PvP01_10_v1","PvP01_10_v1","PvP01_10_v1","PvP01_10_v1","PvP01_10_v1","PvP01_10_v1","PvP01_10_v1","PvP01_10_v1","PvP01_10_v1","PvP01_10_v1","PvP01_10_v1","PvP01_10_v1","PvP01_10_v1","PvP01_10_v1","PvP01_10_v1","PvP01_10_v1","PvP01_10_v1","PvP01_10_v1","PvP01_10_v1","PvP01_10_v1","PvP01_10_v1","PvP01_10_v1","PvP01_10_v1","PvP01_10_v1","PvP01_10_v1","PvP01_10_v1","PvP01_10_v1","PvP01_10_v1","PvP01_10_v1","PvP01_10_v1","PvP01_10_v1","PvP01_10_v1","PvP01_10_v1","PvP01_10_v1","PvP01_10_v1","PvP01_10_v1","PvP01_10_v1","PvP01_10_v1","PvP01_10_v1","PvP01_10_v1","PvP01_10_v1","PvP01_10_v1","PvP01_10_v1","PvP01_10_v1","PvP01_10_v1","PvP01_10_v1","PvP01_10_v1","PvP01_10_v1","PvP01_10_v1","PvP01_10_v1","PvP01_10_v1","PvP01_10_v1","PvP01_10_v1","PvP01_10_v1","PvP01_10_v1","PvP01_10_v1","PvP01_10_v1","PvP01_10_v1","PvP01_10_v1","PvP01_10_v1","PvP01_10_v1","PvP01_10_v1","PvP01_10_v1","PvP01_10_v1","PvP01_10_v1","PvP01_10_v1","PvP01_10_v1","PvP01_10_v1","PvP01_10_v1","PvP01_10_v1","PvP01_10_v1","PvP01_10_v1","PvP01_10_v1","PvP01_10_v1","PvP01_10_v1","PvP01_10_v1","PvP01_10_v1","PvP01_10_v1","PvP01_10_v1","PvP01_10_v1","PvP01_10_v1","PvP01_10_v1","PvP01_10_v1","PvP01_10_v1","PvP01_10_v1","PvP01_10_v1","PvP01_10_v1","PvP01_10_v1","PvP01_10_v1","PvP01_10_v1","PvP01_10_v1","PvP01_10_v1","PvP01_10_v1","PvP01_10_v1","PvP01_10_v1","PvP01_10_v1","PvP01_10_v1","PvP01_10_v1","PvP01_10_v1","PvP01_10_v1","PvP01_10_v1","PvP01_10_v1","PvP01_10_v1","PvP01_10_v1","PvP01_10_v1","PvP01_10_v1","PvP01_10_v1","PvP01_10_v1","PvP01_10_v1","PvP01_10_v1","PvP01_10_v1","PvP01_10_v1","PvP01_10_v1","PvP01_10_v1","PvP01_11_v1","PvP01_11_v1","PvP01_11_v1","PvP01_11_v1","PvP01_11_v1","PvP01_11_v1","PvP01_11_v1","PvP01_11_v1","PvP01_11_v1","PvP01_11_v1","PvP01_11_v1","PvP01_11_v1","PvP01_11_v1","PvP01_11_v1","PvP01_11_v1","PvP01_11_v1","PvP01_11_v1","PvP01_11_v1","PvP01_11_v1","PvP01_11_v1","PvP01_11_v1","PvP01_11_v1","PvP01_11_v1","PvP01_11_v1","PvP01_11_v1","PvP01_11_v1","PvP01_11_v1","PvP01_11_v1","PvP01_11_v1","PvP01_11_v1","PvP01_11_v1","PvP01_11_v1","PvP01_11_v1","PvP01_11_v1","PvP01_11_v1","PvP01_11_v1","PvP01_11_v1","PvP01_11_v1","PvP01_11_v1","PvP01_11_v1","PvP01_11_v1","PvP01_11_v1","PvP01_11_v1","PvP01_11_v1","PvP01_11_v1","PvP01_11_v1","PvP01_11_v1","PvP01_11_v1","PvP01_11_v1","PvP01_11_v1","PvP01_11_v1","PvP01_11_v1","PvP01_11_v1","PvP01_11_v1","PvP01_11_v1","PvP01_11_v1","PvP01_11_v1","PvP01_11_v1","PvP01_11_v1","PvP01_11_v1","PvP01_11_v1","PvP01_11_v1","PvP01_11_v1","PvP01_11_v1","PvP01_11_v1","PvP01_11_v1","PvP01_11_v1","PvP01_11_v1","PvP01_11_v1","PvP01_11_v1","PvP01_11_v1","PvP01_11_v1","PvP01_11_v1","PvP01_11_v1","PvP01_11_v1","PvP01_11_v1","PvP01_11_v1","PvP01_11_v1","PvP01_11_v1","PvP01_11_v1","PvP01_11_v1","PvP01_11_v1","PvP01_11_v1","PvP01_11_v1","PvP01_11_v1","PvP01_11_v1","PvP01_11_v1","PvP01_11_v1","PvP01_11_v1","PvP01_11_v1","PvP01_11_v1","PvP01_11_v1","PvP01_11_v1","PvP01_11_v1","PvP01_11_v1","PvP01_11_v1","PvP01_11_v1","PvP01_11_v1","PvP01_11_v1","PvP01_11_v1","PvP01_11_v1","PvP01_11_v1","PvP01_11_v1","PvP01_11_v1","PvP01_11_v1","PvP01_11_v1","PvP01_11_v1","PvP01_11_v1","PvP01_11_v1","PvP01_11_v1","PvP01_11_v1","PvP01_11_v1","PvP01_11_v1","PvP01_11_v1","PvP01_11_v1","PvP01_11_v1","PvP01_11_v1","PvP01_11_v1","PvP01_11_v1","PvP01_11_v1","PvP01_11_v1","PvP01_11_v1","PvP01_11_v1","PvP01_11_v1","PvP01_11_v1","PvP01_11_v1","PvP01_11_v1","PvP01_11_v1","PvP01_11_v1","PvP01_11_v1","PvP01_11_v1","PvP01_11_v1","PvP01_11_v1","PvP01_11_v1","PvP01_11_v1","PvP01_11_v1","PvP01_11_v1","PvP01_11_v1","PvP01_11_v1","PvP01_11_v1","PvP01_11_v1","PvP01_11_v1","PvP01_11_v1","PvP01_11_v1","PvP01_11_v1","PvP01_11_v1","PvP01_11_v1","PvP01_11_v1","PvP01_11_v1","PvP01_11_v1","PvP01_11_v1","PvP01_11_v1","PvP01_11_v1","PvP01_11_v1","PvP01_11_v1","PvP01_11_v1","PvP01_11_v1","PvP01_11_v1","PvP01_11_v1","PvP01_11_v1","PvP01_11_v1","PvP01_11_v1","PvP01_11_v1","PvP01_11_v1","PvP01_11_v1","PvP01_11_v1","PvP01_11_v1","PvP01_11_v1","PvP01_11_v1","PvP01_11_v1","PvP01_11_v1","PvP01_11_v1","PvP01_11_v1","PvP01_11_v1","PvP01_11_v1","PvP01_11_v1","PvP01_11_v1","PvP01_11_v1","PvP01_11_v1","PvP01_11_v1","PvP01_11_v1","PvP01_11_v1","PvP01_11_v1","PvP01_11_v1","PvP01_11_v1","PvP01_11_v1","PvP01_11_v1","PvP01_11_v1","PvP01_11_v1","PvP01_11_v1","PvP01_11_v1","PvP01_11_v1","PvP01_11_v1","PvP01_11_v1","PvP01_11_v1","PvP01_11_v1","PvP01_11_v1","PvP01_11_v1","PvP01_11_v1","PvP01_11_v1","PvP01_11_v1","PvP01_11_v1","PvP01_11_v1","PvP01_11_v1","PvP01_11_v1","PvP01_11_v1","PvP01_11_v1","PvP01_11_v1","PvP01_11_v1","PvP01_11_v1","PvP01_12_v1","PvP01_12_v1","PvP01_12_v1","PvP01_12_v1","PvP01_12_v1","PvP01_12_v1","PvP01_12_v1","PvP01_12_v1","PvP01_12_v1","PvP01_12_v1","PvP01_12_v1","PvP01_12_v1","PvP01_12_v1","PvP01_12_v1","PvP01_12_v1","PvP01_12_v1","PvP01_12_v1","PvP01_12_v1","PvP01_12_v1","PvP01_12_v1","PvP01_12_v1","PvP01_12_v1","PvP01_12_v1","PvP01_12_v1","PvP01_12_v1","PvP01_12_v1","PvP01_12_v1","PvP01_12_v1","PvP01_12_v1","PvP01_12_v1","PvP01_12_v1","PvP01_12_v1","PvP01_12_v1","PvP01_12_v1","PvP01_12_v1","PvP01_12_v1","PvP01_12_v1","PvP01_12_v1","PvP01_12_v1","PvP01_12_v1","PvP01_12_v1","PvP01_12_v1","PvP01_12_v1","PvP01_12_v1","PvP01_12_v1","PvP01_12_v1","PvP01_12_v1","PvP01_12_v1","PvP01_12_v1","PvP01_12_v1","PvP01_12_v1","PvP01_12_v1","PvP01_12_v1","PvP01_12_v1","PvP01_12_v1","PvP01_12_v1","PvP01_12_v1","PvP01_12_v1","PvP01_12_v1","PvP01_12_v1","PvP01_12_v1","PvP01_12_v1","PvP01_12_v1","PvP01_12_v1","PvP01_12_v1","PvP01_12_v1","PvP01_12_v1","PvP01_12_v1","PvP01_12_v1","PvP01_12_v1","PvP01_12_v1","PvP01_12_v1","PvP01_12_v1","PvP01_12_v1","PvP01_12_v1","PvP01_12_v1","PvP01_12_v1","PvP01_12_v1","PvP01_12_v1","PvP01_12_v1","PvP01_12_v1","PvP01_12_v1","PvP01_12_v1","PvP01_12_v1","PvP01_12_v1","PvP01_12_v1","PvP01_12_v1","PvP01_12_v1","PvP01_12_v1","PvP01_12_v1","PvP01_12_v1","PvP01_12_v1","PvP01_12_v1","PvP01_12_v1","PvP01_12_v1","PvP01_12_v1","PvP01_12_v1","PvP01_12_v1","PvP01_12_v1","PvP01_12_v1","PvP01_12_v1","PvP01_12_v1","PvP01_12_v1","PvP01_12_v1","PvP01_12_v1","PvP01_12_v1","PvP01_12_v1","PvP01_12_v1","PvP01_12_v1","PvP01_12_v1","PvP01_12_v1","PvP01_12_v1","PvP01_12_v1","PvP01_12_v1","PvP01_12_v1","PvP01_12_v1","PvP01_12_v1","PvP01_12_v1","PvP01_12_v1","PvP01_12_v1","PvP01_12_v1","PvP01_12_v1","PvP01_12_v1","PvP01_12_v1","PvP01_12_v1","PvP01_12_v1","PvP01_12_v1","PvP01_12_v1","PvP01_12_v1","PvP01_12_v1","PvP01_12_v1","PvP01_12_v1","PvP01_12_v1","PvP01_12_v1","PvP01_12_v1","PvP01_12_v1","PvP01_12_v1","PvP01_12_v1","PvP01_12_v1","PvP01_12_v1","PvP01_12_v1","PvP01_12_v1","PvP01_12_v1","PvP01_12_v1","PvP01_12_v1","PvP01_12_v1","PvP01_12_v1","PvP01_12_v1","PvP01_12_v1","PvP01_12_v1","PvP01_12_v1","PvP01_12_v1","PvP01_12_v1","PvP01_12_v1","PvP01_12_v1","PvP01_12_v1","PvP01_12_v1","PvP01_12_v1","PvP01_12_v1","PvP01_12_v1","PvP01_12_v1","PvP01_12_v1","PvP01_12_v1","PvP01_12_v1","PvP01_12_v1","PvP01_12_v1","PvP01_12_v1","PvP01_12_v1","PvP01_12_v1","PvP01_12_v1","PvP01_12_v1","PvP01_12_v1","PvP01_12_v1","PvP01_12_v1","PvP01_12_v1","PvP01_12_v1","PvP01_12_v1","PvP01_12_v1","PvP01_12_v1","PvP01_12_v1","PvP01_12_v1","PvP01_12_v1","PvP01_12_v1","PvP01_12_v1","PvP01_12_v1","PvP01_12_v1","PvP01_12_v1","PvP01_12_v1","PvP01_12_v1","PvP01_12_v1","PvP01_12_v1","PvP01_12_v1","PvP01_12_v1","PvP01_12_v1","PvP01_12_v1","PvP01_12_v1","PvP01_12_v1","PvP01_12_v1","PvP01_12_v1","PvP01_12_v1","PvP01_12_v1","PvP01_12_v1","PvP01_12_v1","PvP01_12_v1","PvP01_12_v1","PvP01_12_v1","PvP01_12_v1","PvP01_12_v1","PvP01_12_v1","PvP01_12_v1","PvP01_12_v1","PvP01_12_v1","PvP01_12_v1","PvP01_12_v1","PvP01_12_v1","PvP01_12_v1","PvP01_12_v1","PvP01_12_v1","PvP01_12_v1","PvP01_12_v1","PvP01_12_v1","PvP01_12_v1","PvP01_12_v1","PvP01_12_v1","PvP01_12_v1","PvP01_12_v1","PvP01_12_v1","PvP01_12_v1","PvP01_12_v1","PvP01_12_v1","PvP01_12_v1","PvP01_12_v1","PvP01_12_v1","PvP01_12_v1","PvP01_12_v1","PvP01_12_v1","PvP01_12_v1","PvP01_12_v1","PvP01_12_v1","PvP01_12_v1","PvP01_12_v1","PvP01_12_v1","PvP01_12_v1","PvP01_12_v1","PvP01_12_v1","PvP01_12_v1","PvP01_12_v1","PvP01_12_v1","PvP01_12_v1","PvP01_12_v1","PvP01_12_v1","PvP01_12_v1","PvP01_12_v1","PvP01_12_v1","PvP01_12_v1","PvP01_12_v1","PvP01_12_v1","PvP01_12_v1","PvP01_12_v1","PvP01_12_v1","PvP01_12_v1","PvP01_12_v1","PvP01_12_v1","PvP01_12_v1","PvP01_12_v1","PvP01_12_v1","PvP01_12_v1","PvP01_12_v1","PvP01_12_v1","PvP01_12_v1","PvP01_12_v1","PvP01_12_v1","PvP01_12_v1","PvP01_12_v1","PvP01_12_v1","PvP01_12_v1","PvP01_12_v1","PvP01_12_v1","PvP01_12_v1","PvP01_12_v1","PvP01_12_v1","PvP01_12_v1","PvP01_12_v1","PvP01_12_v1","PvP01_12_v1","PvP01_12_v1","PvP01_12_v1","PvP01_12_v1","PvP01_12_v1","PvP01_12_v1","PvP01_12_v1","PvP01_12_v1","PvP01_12_v1","PvP01_12_v1","PvP01_12_v1","PvP01_12_v1","PvP01_12_v1","PvP01_12_v1","PvP01_12_v1","PvP01_12_v1","PvP01_12_v1","PvP01_12_v1","PvP01_12_v1","PvP01_12_v1","PvP01_12_v1","PvP01_12_v1","PvP01_12_v1","PvP01_12_v1","PvP01_12_v1","PvP01_12_v1","PvP01_12_v1","PvP01_12_v1","PvP01_12_v1","PvP01_12_v1","PvP01_12_v1","PvP01_12_v1","PvP01_12_v1","PvP01_12_v1","PvP01_12_v1","PvP01_12_v1","PvP01_12_v1","PvP01_12_v1","PvP01_12_v1","PvP01_12_v1","PvP01_12_v1","PvP01_12_v1","PvP01_12_v1","PvP01_12_v1","PvP01_12_v1","PvP01_12_v1","PvP01_12_v1","PvP01_12_v1","PvP01_12_v1","PvP01_12_v1","PvP01_12_v1","PvP01_12_v1","PvP01_12_v1","PvP01_12_v1","PvP01_12_v1","PvP01_12_v1","PvP01_12_v1","PvP01_12_v1","PvP01_12_v1","PvP01_12_v1","PvP01_12_v1","PvP01_12_v1","PvP01_12_v1","PvP01_12_v1","PvP01_12_v1","PvP01_12_v1","PvP01_12_v1","PvP01_12_v1","PvP01_12_v1","PvP01_12_v1","PvP01_13_v1","PvP01_13_v1","PvP01_13_v1","PvP01_13_v1","PvP01_13_v1","PvP01_13_v1","PvP01_13_v1","PvP01_13_v1","PvP01_13_v1","PvP01_13_v1","PvP01_13_v1","PvP01_13_v1","PvP01_13_v1","PvP01_13_v1","PvP01_13_v1","PvP01_13_v1","PvP01_13_v1","PvP01_13_v1","PvP01_13_v1","PvP01_13_v1","PvP01_13_v1","PvP01_13_v1","PvP01_13_v1","PvP01_13_v1","PvP01_13_v1","PvP01_13_v1","PvP01_13_v1","PvP01_13_v1","PvP01_13_v1","PvP01_13_v1","PvP01_13_v1","PvP01_13_v1","PvP01_13_v1","PvP01_13_v1","PvP01_13_v1","PvP01_13_v1","PvP01_13_v1","PvP01_13_v1","PvP01_13_v1","PvP01_13_v1","PvP01_13_v1","PvP01_13_v1","PvP01_13_v1","PvP01_13_v1","PvP01_13_v1","PvP01_13_v1","PvP01_13_v1","PvP01_13_v1","PvP01_13_v1","PvP01_13_v1","PvP01_13_v1","PvP01_13_v1","PvP01_13_v1","PvP01_13_v1","PvP01_13_v1","PvP01_13_v1","PvP01_13_v1","PvP01_13_v1","PvP01_13_v1","PvP01_13_v1","PvP01_13_v1","PvP01_13_v1","PvP01_13_v1","PvP01_13_v1","PvP01_13_v1","PvP01_13_v1","PvP01_13_v1","PvP01_13_v1","PvP01_13_v1","PvP01_13_v1","PvP01_13_v1","PvP01_13_v1","PvP01_13_v1","PvP01_13_v1","PvP01_13_v1","PvP01_13_v1","PvP01_13_v1","PvP01_13_v1","PvP01_13_v1","PvP01_13_v1","PvP01_13_v1","PvP01_13_v1","PvP01_13_v1","PvP01_13_v1","PvP01_13_v1","PvP01_13_v1","PvP01_13_v1","PvP01_13_v1","PvP01_13_v1","PvP01_13_v1","PvP01_13_v1","PvP01_13_v1","PvP01_13_v1","PvP01_13_v1","PvP01_13_v1","PvP01_13_v1","PvP01_13_v1","PvP01_13_v1","PvP01_13_v1","PvP01_13_v1","PvP01_13_v1","PvP01_13_v1","PvP01_13_v1","PvP01_13_v1","PvP01_13_v1","PvP01_13_v1","PvP01_13_v1","PvP01_13_v1","PvP01_13_v1","PvP01_13_v1","PvP01_13_v1","PvP01_13_v1","PvP01_13_v1","PvP01_13_v1","PvP01_13_v1","PvP01_13_v1","PvP01_13_v1","PvP01_13_v1","PvP01_13_v1","PvP01_13_v1","PvP01_13_v1","PvP01_13_v1","PvP01_13_v1","PvP01_13_v1","PvP01_13_v1","PvP01_13_v1","PvP01_13_v1","PvP01_13_v1","PvP01_13_v1","PvP01_13_v1","PvP01_13_v1","PvP01_13_v1","PvP01_13_v1","PvP01_13_v1","PvP01_13_v1","PvP01_13_v1","PvP01_13_v1","PvP01_13_v1","PvP01_13_v1","PvP01_13_v1","PvP01_13_v1","PvP01_13_v1","PvP01_13_v1","PvP01_13_v1","PvP01_13_v1","PvP01_13_v1","PvP01_13_v1","PvP01_13_v1","PvP01_13_v1","PvP01_13_v1","PvP01_13_v1","PvP01_13_v1","PvP01_13_v1","PvP01_13_v1","PvP01_13_v1","PvP01_13_v1","PvP01_13_v1","PvP01_13_v1","PvP01_13_v1","PvP01_13_v1","PvP01_13_v1","PvP01_13_v1","PvP01_13_v1","PvP01_13_v1","PvP01_13_v1","PvP01_13_v1","PvP01_13_v1","PvP01_13_v1","PvP01_13_v1","PvP01_13_v1","PvP01_13_v1","PvP01_13_v1","PvP01_13_v1","PvP01_13_v1","PvP01_13_v1","PvP01_13_v1","PvP01_13_v1","PvP01_13_v1","PvP01_13_v1","PvP01_13_v1","PvP01_13_v1","PvP01_13_v1","PvP01_13_v1","PvP01_13_v1","PvP01_13_v1","PvP01_13_v1","PvP01_13_v1","PvP01_13_v1","PvP01_13_v1","PvP01_13_v1","PvP01_13_v1","PvP01_13_v1","PvP01_13_v1","PvP01_13_v1","PvP01_13_v1","PvP01_13_v1","PvP01_13_v1","PvP01_13_v1","PvP01_13_v1","PvP01_13_v1","PvP01_13_v1","PvP01_13_v1","PvP01_13_v1","PvP01_13_v1","PvP01_13_v1","PvP01_13_v1","PvP01_13_v1","PvP01_13_v1","PvP01_13_v1","PvP01_13_v1","PvP01_13_v1","PvP01_13_v1","PvP01_13_v1","PvP01_13_v1","PvP01_13_v1","PvP01_13_v1","PvP01_13_v1","PvP01_13_v1","PvP01_13_v1","PvP01_13_v1","PvP01_13_v1","PvP01_13_v1","PvP01_13_v1","PvP01_13_v1","PvP01_13_v1","PvP01_13_v1","PvP01_13_v1","PvP01_13_v1","PvP01_13_v1","PvP01_13_v1","PvP01_13_v1","PvP01_13_v1","PvP01_13_v1","PvP01_13_v1","PvP01_14_v1","PvP01_14_v1","PvP01_14_v1","PvP01_14_v1","PvP01_14_v1","PvP01_14_v1","PvP01_14_v1","PvP01_14_v1","PvP01_14_v1","PvP01_14_v1","PvP01_14_v1","PvP01_14_v1","PvP01_14_v1","PvP01_14_v1","PvP01_14_v1","PvP01_14_v1","PvP01_14_v1","PvP01_14_v1","PvP01_14_v1","PvP01_14_v1","PvP01_14_v1","PvP01_14_v1","PvP01_14_v1","PvP01_14_v1","PvP01_14_v1","PvP01_14_v1","PvP01_14_v1","PvP01_14_v1","PvP01_14_v1","PvP01_14_v1","PvP01_14_v1","PvP01_14_v1","PvP01_14_v1","PvP01_14_v1","PvP01_14_v1","PvP01_14_v1","PvP01_14_v1","PvP01_14_v1","PvP01_14_v1","PvP01_14_v1","PvP01_14_v1","PvP01_14_v1","PvP01_14_v1","PvP01_14_v1","PvP01_14_v1","PvP01_14_v1","PvP01_14_v1","PvP01_14_v1","PvP01_14_v1","PvP01_14_v1","PvP01_14_v1","PvP01_14_v1","PvP01_14_v1","PvP01_14_v1","PvP01_14_v1","PvP01_14_v1","PvP01_14_v1","PvP01_14_v1","PvP01_14_v1","PvP01_14_v1","PvP01_14_v1","PvP01_14_v1","PvP01_14_v1","PvP01_14_v1","PvP01_14_v1","PvP01_14_v1","PvP01_14_v1","PvP01_14_v1","PvP01_14_v1","PvP01_14_v1","PvP01_14_v1","PvP01_14_v1","PvP01_14_v1","PvP01_14_v1","PvP01_14_v1","PvP01_14_v1","PvP01_14_v1","PvP01_14_v1","PvP01_14_v1","PvP01_14_v1","PvP01_14_v1","PvP01_14_v1","PvP01_14_v1","PvP01_14_v1","PvP01_14_v1","PvP01_14_v1","PvP01_14_v1","PvP01_14_v1","PvP01_14_v1","PvP01_14_v1","PvP01_14_v1","PvP01_14_v1","PvP01_14_v1","PvP01_14_v1","PvP01_14_v1","PvP01_14_v1","PvP01_14_v1","PvP01_14_v1","PvP01_14_v1","PvP01_14_v1","PvP01_14_v1","PvP01_14_v1","PvP01_14_v1","PvP01_14_v1","PvP01_14_v1","PvP01_14_v1","PvP01_14_v1","PvP01_14_v1","PvP01_14_v1","PvP01_14_v1","PvP01_14_v1","PvP01_14_v1","PvP01_14_v1","PvP01_14_v1","PvP01_14_v1","PvP01_14_v1","PvP01_14_v1","PvP01_14_v1","PvP01_14_v1","PvP01_14_v1","PvP01_14_v1","PvP01_14_v1","PvP01_14_v1","PvP01_14_v1","PvP01_14_v1","PvP01_14_v1","PvP01_14_v1","PvP01_14_v1","PvP01_14_v1","PvP01_14_v1","PvP01_14_v1","PvP01_14_v1","PvP01_14_v1","PvP01_14_v1","PvP01_14_v1","PvP01_14_v1","PvP01_14_v1","PvP01_14_v1","PvP01_14_v1","PvP01_14_v1","PvP01_14_v1","PvP01_14_v1","PvP01_14_v1","PvP01_14_v1","PvP01_14_v1","PvP01_14_v1","PvP01_14_v1","PvP01_14_v1","PvP01_14_v1","PvP01_14_v1","PvP01_14_v1","PvP01_14_v1","PvP01_14_v1","PvP01_14_v1","PvP01_14_v1","PvP01_14_v1","PvP01_14_v1","PvP01_14_v1","PvP01_14_v1","PvP01_14_v1","PvP01_14_v1","PvP01_14_v1","PvP01_14_v1","PvP01_14_v1","PvP01_14_v1","PvP01_14_v1","PvP01_14_v1","PvP01_14_v1","PvP01_14_v1","PvP01_14_v1","PvP01_14_v1","PvP01_14_v1","PvP01_14_v1","PvP01_14_v1","PvP01_14_v1","PvP01_14_v1","PvP01_14_v1","PvP01_14_v1","PvP01_14_v1","PvP01_14_v1","PvP01_14_v1","PvP01_14_v1","PvP01_14_v1","PvP01_14_v1","PvP01_14_v1","PvP01_14_v1","PvP01_14_v1","PvP01_14_v1","PvP01_14_v1","PvP01_14_v1","PvP01_14_v1","PvP01_14_v1","PvP01_14_v1","PvP01_14_v1","PvP01_14_v1","PvP01_14_v1","PvP01_14_v1","PvP01_14_v1","PvP01_14_v1","PvP01_14_v1","PvP01_14_v1","PvP01_14_v1","PvP01_14_v1","PvP01_14_v1","PvP01_14_v1","PvP01_14_v1","PvP01_14_v1","PvP01_14_v1","PvP01_14_v1","PvP01_14_v1","PvP01_14_v1","PvP01_14_v1","PvP01_14_v1","PvP01_14_v1","PvP01_14_v1","PvP01_14_v1","PvP01_14_v1","PvP01_14_v1","PvP01_14_v1","PvP01_14_v1","PvP01_14_v1","PvP01_14_v1","PvP01_14_v1","PvP01_14_v1","PvP01_14_v1","PvP01_14_v1","PvP01_14_v1","PvP01_14_v1","PvP01_14_v1","PvP01_14_v1","PvP01_14_v1","PvP01_14_v1","PvP01_14_v1","PvP01_14_v1","PvP01_14_v1","PvP01_14_v1","PvP01_14_v1","PvP01_14_v1","PvP01_14_v1","PvP01_14_v1","PvP01_14_v1","PvP01_14_v1","PvP01_14_v1","PvP01_14_v1","PvP01_14_v1","PvP01_14_v1","PvP01_14_v1","PvP01_14_v1","PvP01_14_v1","PvP01_14_v1","PvP01_14_v1","PvP01_14_v1","PvP01_14_v1","PvP01_14_v1","PvP01_14_v1","PvP01_14_v1","PvP01_14_v1","PvP01_14_v1","PvP01_14_v1","PvP01_14_v1","PvP01_14_v1","PvP01_14_v1","PvP01_14_v1","PvP01_14_v1","PvP01_14_v1","PvP01_14_v1","PvP01_14_v1","PvP01_14_v1","PvP01_14_v1","PvP01_14_v1","PvP01_14_v1","PvP01_14_v1","PvP01_14_v1","PvP01_14_v1","PvP01_14_v1","PvP01_14_v1","PvP01_14_v1","PvP01_14_v1","PvP01_14_v1","PvP01_14_v1","PvP01_14_v1","PvP01_14_v1","PvP01_14_v1","PvP01_14_v1","PvP01_14_v1","PvP01_14_v1","PvP01_14_v1","PvP01_14_v1","PvP01_14_v1","PvP01_14_v1","PvP01_14_v1","PvP01_14_v1","PvP01_14_v1","PvP01_14_v1","PvP01_14_v1","PvP01_14_v1","PvP01_14_v1","PvP01_14_v1","PvP01_14_v1","PvP01_14_v1","PvP01_14_v1","PvP01_14_v1","PvP01_14_v1","PvP01_14_v1","PvP01_14_v1","PvP01_14_v1","PvP01_14_v1","PvP01_14_v1","PvP01_14_v1","PvP01_14_v1","PvP01_14_v1","PvP01_14_v1","PvP01_14_v1","PvP01_14_v1","PvP01_14_v1","PvP01_14_v1","PvP01_14_v1","PvP01_14_v1","PvP01_14_v1","PvP01_14_v1","PvP01_14_v1","PvP01_14_v1","PvP01_14_v1","PvP01_14_v1","PvP01_14_v1","PvP01_14_v1","PvP01_14_v1","PvP01_14_v1","PvP01_14_v1","PvP01_14_v1","PvP01_14_v1","PvP01_14_v1","PvP01_14_v1","PvP01_14_v1","PvP01_14_v1","PvP01_14_v1","PvP01_14_v1","PvP01_14_v1","PvP01_14_v1","PvP01_14_v1","PvP01_14_v1","PvP01_14_v1","PvP01_14_v1","PvP01_14_v1","PvP01_14_v1","PvP01_14_v1","PvP01_14_v1","PvP01_API_v1","PvP01_API_v1","PvP01_API_v1","PvP01_API_v1","PvP01_API_v1","PvP01_MIT_v1","PvP01_MIT_v1","PvP01_MIT_v1"],"Source":["EuPathDB","EuPathDB","EuPathDB","EuPathDB","EuPathDB","EuPathDB","EuPathDB","EuPathDB","EuPathDB","EuPathDB","EuPathDB","EuPathDB","EuPathDB","EuPathDB","EuPathDB","EuPathDB","EuPathDB","EuPathDB","EuPathDB","EuPathDB","EuPathDB","EuPathDB","EuPathDB","EuPathDB","EuPathDB","EuPathDB","EuPathDB","EuPathDB","EuPathDB","EuPathDB","EuPathDB","EuPathDB","EuPathDB","EuPathDB","EuPathDB","EuPathDB","EuPathDB","EuPathDB","EuPathDB","EuPathDB","EuPathDB","EuPathDB","EuPathDB","EuPathDB","EuPathDB","EuPathDB","EuPathDB","EuPathDB","EuPathDB","EuPathDB","EuPathDB","EuPathDB","EuPathDB","EuPathDB","EuPathDB","EuPathDB","EuPathDB","EuPathDB","EuPathDB","EuPathDB","EuPathDB","EuPathDB","EuPathDB","EuPathDB","EuPathDB","EuPathDB","EuPathDB","EuPathDB","EuPathDB","EuPathDB","EuPathDB","EuPathDB","EuPathDB","EuPathDB","EuPathDB","EuPathDB","EuPathDB","EuPathDB","EuPathDB","EuPathDB","EuPathDB","EuPathDB","EuPathDB","EuPathDB","EuPathDB","EuPathDB","EuPathDB","EuPathDB","EuPathDB","EuPathDB","EuPathDB","EuPathDB","EuPathDB","EuPathDB","EuPathDB","EuPathDB","EuPathDB","EuPathDB","EuPathDB","EuPathDB","EuPathDB","EuPathDB","EuPathDB","EuPathDB","EuPathDB","EuPathDB","EuPathDB","EuPathDB","EuPathDB","EuPathDB","EuPathDB","EuPathDB","EuPathDB","EuPathDB","EuPathDB","EuPathDB","EuPathDB","EuPathDB","EuPathDB","EuPathDB","EuPathDB","EuPathDB","EuPathDB","EuPathDB","EuPathDB","EuPathDB","EuPathDB","EuPathDB","EuPathDB","EuPathDB","EuPathDB","EuPathDB","EuPathDB","EuPathDB","EuPathDB","EuPathDB","EuPathDB","EuPathDB","EuPathDB","EuPathDB","EuPathDB","EuPathDB","EuPathDB","EuPathDB","EuPathDB","EuPathDB","EuPathDB","EuPathDB","EuPathDB","EuPathDB","EuPathDB","EuPathDB","EuPathDB","EuPathDB","EuPathDB","EuPathDB","EuPathDB","EuPathDB","EuPathDB","EuPathDB","EuPathDB","EuPathDB","EuPathDB","EuPathDB","EuPathDB","EuPathDB","EuPathDB","EuPathDB","EuPathDB","EuPathDB","EuPathDB","EuPathDB","EuPathDB","EuPathDB","EuPathDB","EuPathDB","EuPathDB","EuPathDB","EuPathDB","EuPathDB","EuPathDB","EuPathDB","EuPathDB","EuPathDB","EuPathDB","EuPathDB","EuPathDB","EuPathDB","EuPathDB","EuPathDB","EuPathDB","EuPathDB","EuPathDB","EuPathDB","EuPathDB","EuPathDB","EuPathDB","EuPathDB","EuPathDB","EuPathDB","EuPathDB","EuPathDB","EuPathDB","EuPathDB","EuPathDB","EuPathDB","EuPathDB","EuPathDB","EuPathDB","EuPathDB","EuPathDB","EuPathDB","EuPathDB","EuPathDB","EuPathDB","EuPathDB","EuPathDB","EuPathDB","EuPathDB","EuPathDB","EuPathDB","EuPathDB","EuPathDB","EuPathDB","EuPathDB","EuPathDB",
```
