## Supplementary material for "Transcriptome and histone epigenome of *Plasmodium vivax* salivary-gland sporozoites point to tight regulatory control and potential mechanisms for liver-stage differentiation": Pv-SPZ-v-mLS-DTGs.js

```
glimma.layout.setupGrid(d3.select(".container"), "md", [1,2]);
glimma.storage.chartData.push(glimma.transform.toRowMajor({"GeneID":["PVP01_0000030","PVP01_0000040","PVP01_0000050","PVP01_0000080","PVP01_0000120","PVP01_0000130","PVP01_0000140","PVP01_0000150","PVP01_0000170","PVP01_0000180","PVP01_0000190","PVP01_0000210","PVP01_0000300","PVP01_0000480","PVP01_0000670","PVP01_0000740","PVP01_0000790","PVP01_0001030","PVP01_0001090","PVP01_0001310","PVP01_0001480","PVP01_0001550","PVP01_0001640","PVP01_0001660","PVP01_0001670","PVP01_0001900","PVP01_0001910","PVP01_0001940","PVP01_0001970","PVP01_0002020","PVP01_0002050","PVP01_0002120","PVP01_0002200","PVP01_0002210","PVP01_0002330","PVP01_0002370","PVP01_0002390","PVP01_0002470","PVP01_0002530","PVP01_0002540","PVP01_0002750","PVP01_0002810","PVP01_0002820","PVP01_0003010","PVP01_0003050","PVP01_0003150","PVP01_0003210","PVP01_0003350","PVP01_0003360","PVP01_0003370","PVP01_0003380","PVP01_0003410","PVP01_0003420","PVP01_0003440","PVP01_0003530","PVP01_0003640","PVP01_0003690","PVP01_0003730","PVP01_0003800","PVP01_0003990","PVP01_0004160","PVP01_0004170","PVP01_0004180","PVP01_0004190","PVP01_0004250","PVP01_0004260","PVP01_0004270","PVP01_0004280","PVP01_0004300","PVP01_0004310","PVP01_0004320","PVP01_0004350","PVP01_0004370","PVP01_0004380","PVP01_0004390","PVP01_0004410","PVP01_0004640","PVP01_0004670","PVP01_0004910","PVP01_0004920","PVP01_0004950","PVP01_0004960","PVP01_0005010","PVP01_0005100","PVP01_0005160","PVP01_0005210","PVP01_0005300","PVP01_0005310","PVP01_0005420","PVP01_0005430","PVP01_0005550","PVP01_0005640","PVP01_0005710","PVP01_0005740","PVP01_0005800","PVP01_0005820","PVP01_0005870","PVP01_0005910","PVP01_0006000","PVP01_0006040","PVP01_0006080","PVP01_0006260","PVP01_0006650","PVP01_0006660","PVP01_0006670","PVP01_0006680","PVP01_0006700","PVP01_0006860","PVP01_0006890","PVP01_0007080","PVP01_0007160","PVP01_0007240","PVP01_0007290","PVP01_0007320","PVP01_0007340","PVP01_0007610","PVP01_0007620","PVP01_0007690","PVP01_0007700","PVP01_0007720","PVP01_0007850","PVP01_0007900","PVP01_0007910","PVP01_0007920","PVP01_0008190","PVP01_0008430","PVP01_0008480","PVP01_0008490","PVP01_0008600","PVP01_0008700","PVP01_0008770","PVP01_0008830","PVP01_0008990","PVP01_0009010","PVP01_0009170","PVP01_0009180","PVP01_0009210","PVP01_0009370","PVP01_0009410","PVP01_0009470","PVP01_0009540","PVP01_0009620","PVP01_0009710","PVP01_0009730","PVP01_0009740","PVP01_0009760","PVP01_0009770","PVP01_0009820","PVP01_0009850","PVP01_0009860","PVP01_0009870","PVP01_0009900","PVP01_0009940","PVP01_0009980","PVP01_0010020","PVP01_0010030","PVP01_0010070","PVP01_0010100","PVP01_0010120","PVP01_0010140","PVP01_0010170","PVP01_0010180","PVP01_0010200","PVP01_0010220","PVP01_0010230","PVP01_0010240","PVP01_0010340","PVP01_0010380","PVP01_0010440","PVP01_0010470","PVP01_0010500","PVP01_0010540","PVP01_0010570","PVP01_0010600","PVP01_0010610","PVP01_0010640","PVP01_0010650","PVP01_0010770","PVP01_0010870","PVP01_0010900","PVP01_0010910","PVP01_0010960","PVP01_0010970","PVP01_0010990","PVP01_0011030","PVP01_0011070","PVP01_0102100","PVP01_0102400","PVP01_0102500","PVP01_0102700","PVP01_0102800","PVP01_0103100","PVP01_0103300","PVP01_0103400","PVP01_0103500","PVP01_0103600","PVP01_0103800","PVP01_0103900","PVP01_0104100","PVP01_0104200","PVP01_0104600","PVP01_0104800","PVP01_0104900","PVP01_0105000","PVP01_0105100","PVP01_0105200","PVP01_0105300","PVP01_0105400","PVP01_0105500","PVP01_0105600","PVP01_0105700","PVP01_0105800","PVP01_0106000","PVP01_0106200","PVP01_0106300","PVP01_0106400","PVP01_0106500","PVP01_0106800","PVP01_0106900","PVP01_0107400","PVP01_0107600","PVP01_0107800","PVP01_0107900","PVP01_0108000","PVP01_0108100","PVP01_0108200","PVP01_0108300","PVP01_0108400","PVP01_0108600","PVP01_0108800","PVP01_0108900","PVP01_0109000","PVP01_0109100","PVP01_0109200","PVP01_0109600","PVP01_0109800","PVP01_0110100","PVP01_0110200","PVP01_0110400","PVP01_0110500","PVP01_0110700","PVP01_0110800","PVP01_0110900","PVP01_0111100","PVP01_0111200","PVP01_0111300","PVP01_0111600","PVP01_0111700","PVP01_0111900","PVP01_0112000","PVP01_0112100","PVP01_0112200","PVP01_0112300","PVP01_0112500","PVP01_0112700","PVP01_0112800","PVP01_0112900","PVP01_0113000","PVP01_0113100","PVP01_0113200","PVP01_0113300","PVP01_0113400","PVP01_0113500","PVP01_0113700","PVP01_0114100","PVP01_0114200","PVP01_0114600","PVP01_0114700","PVP01_0114800","PVP01_0115000","PVP01_0115100","PVP01_0115200","PVP01_0115300","PVP01_0115400","PVP01_0115500","PVP01_0116100","PVP01_0116500","PVP01_0116600","PVP01_0116700","PVP01_0116800","PVP01_0116900","PVP01_0117000","PVP01_0117500","PVP01_0117600","PVP01_0117700","PVP01_0117800","PVP01_0118000","PVP01_0118100","PVP01_0118500","PVP01_0118600","PVP01_0118700","PVP01_0118800","PVP01_0118900","PVP01_0119300","PVP01_0119400","PVP01_0119500","PVP01_0119600","PVP01_0119700","PVP01_0119800","PVP01_0119900","PVP01_0120000","PVP01_0120100","PVP01_0120500","PVP01_0120800","PVP01_0121000","PVP01_0121700","PVP01_0200200","PVP01_0200700","PVP01_0201000","PVP01_0201100","PVP01_0201300","PVP01_0201500","PVP01_0201600","PVP01_0201700","PVP01_0201800","PVP01_0201900","PVP01_0202000","PVP01_0202100","PVP01_0202200","PVP01_0202300","PVP01_0202400","PVP01_0202600","PVP01_0203000","PVP01_0203100","PVP01_0203300","PVP01_0203400","PVP01_0203500","PVP01_0203800","PVP01_0203900","PVP01_0204000","PVP01_0204100","PVP01_0204800","PVP01_0204900","PVP01_0205000","PVP01_0205100","PVP01_0205200","PVP01_0205300","PVP01_0205500","PVP01_0205600","PVP01_0205800","PVP01_0206100","PVP01_0206300","PVP01_0206400","PVP01_0206500","PVP01_0206600","PVP01_0206800","PVP01_0207000","PVP01_0207100","PVP01_0207200","PVP01_0207400","PVP01_0207500","PVP01_0207600","PVP01_0207700","PVP01_0207800","PVP01_0208100","PVP01_0208200","PVP01_0208300","PVP01_0208400","PVP01_0208500","PVP01_0208600","PVP01_0208700","PVP01_0208800","PVP01_0209000","PVP01_0209100","PVP01_0209200","PVP01_0209300","PVP01_0209400","PVP01_0209500","PVP01_0209600","PVP01_0209700","PVP01_0210000","PVP01_0210100","PVP01_0210200","PVP01_0210300","PVP01_0210500","PVP01_0210800","PVP01_0211000","PVP01_0211100","PVP01_0211200","PVP01_0211300","PVP01_0211400","PVP01_0211500","PVP01_0211600","PVP01_0211700","PVP01_0211800","PVP01_0212400","PVP01_0212500","PVP01_0212600","PVP01_0212700","PVP01_0212800","PVP01_0213000","PVP01_0213300","PVP01_0213400","PVP01_0213600","PVP01_0213700","PVP01_0214000","PVP01_0214100","PVP01_0214200","PVP01_0214800","PVP01_0214900","PVP01_0215000","PVP01_0215300","PVP01_0215400","PVP01_0215500","PVP01_0215700","PVP01_0215800","PVP01_0215900","PVP01_0216000","PVP01_0216100","PVP01_0216500","PVP01_0216800","PVP01_0216900","PVP01_0217000","PVP01_0217100","PVP01_0217200","PVP01_0217300","PVP01_0217400","PVP01_0217500","PVP01_0218200","PVP01_0218300","PVP01_0218900","PVP01_0219400","PVP01_0220200","PVP01_0221100","PVP01_0221300","PVP01_0221700","PVP01_0300100","PVP01_0300600","PVP01_0300700","PVP01_0301100","PVP01_0301200","PVP01_0301500","PVP01_0301900","PVP01_0302000","PVP01_0302100","PVP01_0302300","PVP01_0302400","PVP01_0302500","PVP01_0302800","PVP01_0303100","PVP01_0303500","PVP01_0304500","PVP01_0304600","PVP01_0304800","PVP01_0304900","PVP01_0305000","PVP01_0305200","PVP01_0305300","PVP01_0305500","PVP01_0305700","PVP01_0305800","PVP01_0306100","PVP01_0306300","PVP01_0306400","PVP01_0306500","PVP01_0306700","PVP01_0306800","PVP01_0306900","PVP01_0307000","PVP01_0307100","PVP01_0307300","PVP01_0307400","PVP01_0307500","PVP01_0307600","PVP01_0307700","PVP01_0307800","PVP01_0308100","PVP01_0308200","PVP01_0308800","PVP01_0309100","PVP01_0309200","PVP01_0309300","PVP01_0309700","PVP01_0309800","PVP01_0309900","PVP01_0310000","PVP01_0310200","PVP01_0310300","PVP01_0310400","PVP01_0310600","PVP01_0311000","PVP01_0311100","PVP01_0311200","PVP01_0311300","PVP01_0311500","PVP01_0311600","PVP01_0311700","PVP01_0312000","PVP01_0312100","PVP01_0312200","PVP01_0312400","PVP01_0312500","PVP01_0312700","PVP01_0312800","PVP01_0313400","PVP01_0313500","PVP01_0314100","PVP01_0314200","PVP01_0314900","PVP01_0315000","PVP01_0315100","PVP01_0315200","PVP01_0315300","PVP01_0315500","PVP01_0315600","PVP01_0315700","PVP01_0315900","PVP01_0316000","PVP01_0316200","PVP01_0316300","PVP01_0316400","PVP01_0316500","PVP01_0316700","PVP01_0316800","PVP01_0317100","PVP01_0317200","PVP01_0317400","PVP01_0317900","PVP01_0318000","PVP01_0318100","PVP01_0318200","PVP01_0318400","PVP01_0318900","PVP01_0319200","PVP01_0319300","PVP01_0319400","PVP01_0319500","PVP01_0319600","PVP01_0319700","PVP01_0319800","PVP01_0320200","PVP01_0320400","PVP01_0320500","PVP01_0320700","PVP01_0320900","PVP01_0400900","PVP01_0403600","PVP01_0403700","PVP01_0403900","PVP01_0404000","PVP01_0404100","PVP01_0404400","PVP01_0404500","PVP01_0404600","PVP01_0404700","PVP01_0404800","PVP01_0404900","PVP01_0405100","PVP01_0405200","PVP01_0405300","PVP01_0405500","PVP01_0405600","PVP01_0405700","PVP01_0405900","PVP01_0406500","PVP01_0406600","PVP01_0406700","PVP01_0406900","PVP01_0407000","PVP01_0407100","PVP01_0407600","PVP01_0407700","PVP01_0407800","PVP01_0407900","PVP01_0408200","PVP01_0408500","PVP01_0408600","PVP01_0408900","PVP01_0409000","PVP01_0409100","PVP01_0409200","PVP01_0409300","PVP01_0409700","PVP01_0409900","PVP01_0410000","PVP01_0410400","PVP01_0410600","PVP01_0410800","PVP01_0411400","PVP01_0411500","PVP01_0411600","PVP01_0411700","PVP01_0411800","PVP01_0411900","PVP01_0412100","PVP01_0412200","PVP01_0412600","PVP01_0412700","PVP01_0412800","PVP01_0412900","PVP01_0413000","PVP01_0413100","PVP01_0413200","PVP01_0413300","PVP01_0413500","PVP01_0414000","PVP01_0414200","PVP01_0414400","PVP01_0414600","PVP01_0414900","PVP01_0415200","PVP01_0415400","PVP01_0415500","PVP01_0415600","PVP01_0416000","PVP01_0416200","PVP01_0416300","PVP01_0416600","PVP01_0416900","PVP01_0417100","PVP01_0417500","PVP01_0417800","PVP01_0417900","PVP01_0418100","PVP01_0418300","PVP01_0419000","PVP01_0419100","PVP01_0419200","PVP01_0419300","PVP01_0419400","PVP01_0419500","PVP01_0419600","PVP01_0419700","PVP01_0420000","PVP01_0420100","PVP01_0420200","PVP01_0420300","PVP01_0420500","PVP01_0420600","PVP01_0420700","PVP01_0420900","PVP01_0421000","PVP01_0421500","PVP01_0421700","PVP01_0421800","PVP01_0421900","PVP01_0422100","PVP01_0422200","PVP01_0422300","PVP01_0422400","PVP01_0422500","PVP01_0422600","PVP01_0422700","PVP01_0422900","PVP01_0423100","PVP01_0423600","PVP01_0423800","PVP01_0424300","PVP01_0424400","PVP01_0500700","PVP01_0502300","PVP01_0502400","PVP01_0502900","PVP01_0503400","PVP01_0504200","PVP01_0504300","PVP01_0505000","PVP01_0505100","PVP01_0505200","PVP01_0505400","PVP01_0505500","PVP01_0505800","PVP01_0506000","PVP01_0506400","PVP01_0506600","PVP01_0506700","PVP01_0506800","PVP01_0506900","PVP01_0507300","PVP01_0507500","PVP01_0507600","PVP01_0507700","PVP01_0507800","PVP01_0508000","PVP01_0508200","PVP01_0508300","PVP01_0508500","PVP01_0508600","PVP01_0508700","PVP01_0508900","PVP01_0509000","PVP01_0509100","PVP01_0509200","PVP01_0509400","PVP01_0509600","PVP01_0509900","PVP01_0510000","PVP01_0510100","PVP01_0510200","PVP01_0510300","PVP01_0510400","PVP01_0510700","PVP01_0510800","PVP01_0510900","PVP01_0511100","PVP01_0511200","PVP01_0511300","PVP01_0511500","PVP01_0511700","PVP01_0511800","PVP01_0512000","PVP01_0512100","PVP01_0512200","PVP01_0512400","PVP01_0512500","PVP01_0512700","PVP01_0512800","PVP01_0512900","PVP01_0513000","PVP01_0513300","PVP01_0513400","PVP01_0513500","PVP01_0513600","PVP01_0513700","PVP01_0513800","PVP01_0513900","PVP01_0514000","PVP01_0514100","PVP01_0514200","PVP01_0514400","PVP01_0514600","PVP01_0514700","PVP01_0514800","PVP01_0514900","PVP01_0515100","PVP01_0515200","PVP01_0515300","PVP01_0515700","PVP01_0515800","PVP01_0515900","PVP01_0516000","PVP01_0516200","PVP01_0516300","PVP01_0516400","PVP01_0516500","PVP01_0516800","PVP01_0516900","PVP01_0517000","PVP01_0517100","PVP01_0517300","PVP01_0517400","PVP01_0517700","PVP01_0517900","PVP01_0518300","PVP01_0518400","PVP01_0518500","PVP01_0518600","PVP01_0518700","PVP01_0519100","PVP01_0519200","PVP01_0519400","PVP01_0519500","PVP01_0519600","PVP01_0520400","PVP01_0520500","PVP01_0520600","PVP01_0520700","PVP01_0520800","PVP01_0521300","PVP01_0521400","PVP01_0521600","PVP01_0521700","PVP01_0521800","PVP01_0522000","PVP01_0522100","PVP01_0522200","PVP01_0522300","PVP01_0522500","PVP01_0522600","PVP01_0522700","PVP01_0522800","PVP01_0522900","PVP01_0523000","PVP01_0523300","PVP01_0523700","PVP01_0524000","PVP01_0524200","PVP01_0524500","PVP01_0524600","PVP01_0524700","PVP01_0524900","PVP01_0525000","PVP01_0525200","PVP01_0525300","PVP01_0525400","PVP01_0525500","PVP01_0525700","PVP01_0525800","PVP01_0526000","PVP01_0526100","PVP01_0526400","PVP01_0526900","PVP01_0527300","PVP01_0527400","PVP01_0527500","PVP01_0527700","PVP01_0527800","PVP01_0527900","PVP01_0528000","PVP01_0528100","PVP01_0528200","PVP01_0528300","PVP01_0528400","PVP01_0528500","PVP01_0528600","PVP01_0528700","PVP01_0528900","PVP01_0529000","PVP01_0529100","PVP01_0529200","PVP01_0529600","PVP01_0529700","PVP01_0529800","PVP01_0529900","PVP01_0530000","PVP01_0530100","PVP01_0530400","PVP01_0530500","PVP01_0530700","PVP01_0530800","PVP01_0531100","PVP01_0531200","PVP01_0531500","PVP01_0531700","PVP01_0531800","PVP01_0532500","PVP01_0532600","PVP01_0532700","PVP01_0532900","PVP01_0533000","PVP01_0533100","PVP01_0533300","PVP01_0534000","PVP01_0534100","PVP01_0534200","PVP01_0534600","PVP01_0534800","PVP01_0534900","PVP01_0535400","PVP01_0600100","PVP01_0600300","PVP01_0600400","PVP01_0600500","PVP01_0600600","PVP01_0600700","PVP01_0600800","PVP01_0600900","PVP01_0601100","PVP01_0601200","PVP01_0601500","PVP01_0601600","PVP01_0602300","PVP01_0602600","PVP01_0603000","PVP01_0603100","PVP01_0603400","PVP01_0603500","PVP01_0603600","PVP01_0603700","PVP01_0603900","PVP01_0604000","PVP01_0604100","PVP01_0604200","PVP01_0604400","PVP01_0604700","PVP01_0604800","PVP01_0604900","PVP01_0605000","PVP01_0605100","PVP01_0605300","PVP01_0605400","PVP01_0605500","PVP01_0606100","PVP01_0606200","PVP01_0606400","PVP01_0606500","PVP01_0606600","PVP01_0606800","PVP01_0606900","PVP01_0607200","PVP01_0607600","PVP01_0608000","PVP01_0608100","PVP01_0608200","PVP01_0608300","PVP01_0608400","PVP01_0608500","PVP01_0608600","PVP01_0608700","PVP01_0608800","PVP01_0608900","PVP01_0609200","PVP01_0609400","PVP01_0609600","PVP01_0609700","PVP01_0609800","PVP01_0609900","PVP01_0610100","PVP01_0610200","PVP01_0610400","PVP01_0610500","PVP01_0610600","PVP01_0610700","PVP01_0610800","PVP01_0610900","PVP01_0611100","PVP01_0611300","PVP01_0612000","PVP01_0612100","PVP01_0612600","PVP01_0612700","PVP01_0613000","PVP01_0613100","PVP01_0613200","PVP01_0613400","PVP01_0613500","PVP01_0613700","PVP01_0613800","PVP01_0613900","PVP01_0614000","PVP01_0614200","PVP01_0614300","PVP01_0614400","PVP01_0614600","PVP01_0614700","PVP01_0614900","PVP01_0615100","PVP01_0615500","PVP01_0615700","PVP01_0615800","PVP01_0615900","PVP01_0616000","PVP01_0616100","PVP01_0616200","PVP01_0616400","PVP01_0616500","PVP01_0616600","PVP01_0616800","PVP01_0616900","PVP01_0617000","PVP01_0617100","PVP01_0617400","PVP01_0617500","PVP01_0617600","PVP01_0617900","PVP01_0618100","PVP01_0618300","PVP01_0618600","PVP01_0618700","PVP01_0618900","PVP01_0619000","PVP01_0619100","PVP01_0619200","PVP01_0619400","PVP01_0619500","PVP01_0619600","PVP01_0619700","PVP01_0619800","PVP01_0619900","PVP01_0620000","PVP01_0620100","PVP01_0620200","PVP01_0620900","PVP01_0621000","PVP01_0621100","PVP01_0621300","PVP01_0621800","PVP01_0622100","PVP01_0622200","PVP01_0622700","PVP01_0623100","PVP01_0623200","PVP01_0623300","PVP01_0623400","PVP01_0623600","PVP01_0623900","PVP01_0624000","PVP01_0624100","PVP01_0624300","PVP01_0624400","PVP01_0700200","PVP01_0700700","PVP01_0700800","PVP01_0701000","PVP01_0701400","PVP01_0701500","PVP01_0701600","PVP01_0701700","PVP01_0702200","PVP01_0702400","PVP01_0702500","PVP01_0702600","PVP01_0702700","PVP01_0703000","PVP01_0703200","PVP01_0703500","PVP01_0703600","PVP01_0703700","PVP01_0704000","PVP01_0704100","PVP01_0704400","PVP01_0704500","PVP01_0704600","PVP01_0704700","PVP01_0704800","PVP01_0704900","PVP01_0705000","PVP01_0705100","PVP01_0705200","PVP01_0705500","PVP01_0705600","PVP01_0705700","PVP01_0705800","PVP01_0705900","PVP01_0706000","PVP01_0706100","PVP01_0706200","PVP01_0706300","PVP01_0706400","PVP01_0706500","PVP01_0706600","PVP01_0706700","PVP01_0707000","PVP01_0707100","PVP01_0707200","PVP01_0707300","PVP01_0707500","PVP01_0707800","PVP01_0708000","PVP01_0708100","PVP01_0708200","PVP01_0708300","PVP01_0708400","PVP01_0708500","PVP01_0708800","PVP01_0708900","PVP01_0709000","PVP01_0709100","PVP01_0709200","PVP01_0709300","PVP01_0709500","PVP01_0709700","PVP01_0709900","PVP01_0710000","PVP01_0710200","PVP01_0710300","PVP01_0710700","PVP01_0711100","PVP01_0711200","PVP01_0711400","PVP01_0711500","PVP01_0711800","PVP01_0711900","PVP01_0712200","PVP01_0712300","PVP01_0712400","PVP01_0712700","PVP01_0713000","PVP01_0713100","PVP01_0713200","PVP01_0713500","PVP01_0713600","PVP01_0713800","PVP01_0713900","PVP01_0714000","PVP01_0714200","PVP01_0714300","PVP01_0714400","PVP01_0714600","PVP01_0714800","PVP01_0714900","PVP01_0715000","PVP01_0715100","PVP01_0715200","PVP01_0715500","PVP01_0715600","PVP01_0715800","PVP01_0716000","PVP01_0716100","PVP01_0716200","PVP01_0716600","PVP01_0716800","PVP01_0717000","PVP01_0717200","PVP01_0717400","PVP01_0717600","PVP01_0717800","PVP01_0718000","PVP01_0718100","PVP01_0718200","PVP01_0718600","PVP01_0718700","PVP01_0719000","PVP01_0719300","PVP01_0719400","PVP01_0719500","PVP01_0719700","PVP01_0719800","PVP01_0719900","PVP01_0720000","PVP01_0720100","PVP01_0720200","PVP01_0720300","PVP01_0720400","PVP01_0720600","PVP01_0720700","PVP01_0720800","PVP01_0720900","PVP01_0721100","PVP01_0721300","PVP01_0721400","PVP01_0721500","PVP01_0721800","PVP01_0721900","PVP01_0722100","PVP01_0722200","PVP01_0722300","PVP01_0722400","PVP01_0722500","PVP01_0722600","PVP01_0722700","PVP01_0722800","PVP01_0722900","PVP01_0723000","PVP01_0723100","PVP01_0723200","PVP01_0723300","PVP01_0723400","PVP01_0723500","PVP01_0723700","PVP01_0723800","PVP01_0724000","PVP01_0724100","PVP01_0724300","PVP01_0724500","PVP01_0724600","PVP01_0724700","PVP01_0724800","PVP01_0724900","PVP01_0725000","PVP01_0725100","PVP01_0725300","PVP01_0725400","PVP01_0725700","PVP01_0726000","PVP01_0726700","PVP01_0726900","PVP01_0727000","PVP01_0727100","PVP01_0727200","PVP01_0727300","PVP01_0727400","PVP01_0727500","PVP01_0727600","PVP01_0727700","PVP01_0727800","PVP01_0728100","PVP01_0728200","PVP01_0728300","PVP01_0729100","PVP01_0729200","PVP01_0729300","PVP01_0729400","PVP01_0729500","PVP01_0729700","PVP01_0730000","PVP01_0730100","PVP01_0730200","PVP01_0730300","PVP01_0730400","PVP01_0730600","PVP01_0730800","PVP01_0731100","PVP01_0731200","PVP01_0731300","PVP01_0731500","PVP01_0731600","PVP01_0731700","PVP01_0731800","PVP01_0731900","PVP01_0732000","PVP01_0732100","PVP01_0732200","PVP01_0732300","PVP01_0732600","PVP01_0732800","PVP01_0732900","PVP01_0733000","PVP01_0733100","PVP01_0733200","PVP01_0733500","PVP01_0733700","PVP01_0733900","PVP01_0734100","PVP01_0734200","PVP01_0734400","PVP01_0734800","PVP01_0734900","PVP01_0735000","PVP01_0735100","PVP01_0735300","PVP01_0735800","PVP01_0735900","PVP01_0800100","PVP01_0800200","PVP01_0800300","PVP01_0800400","PVP01_0800500","PVP01_0800600","PVP01_0800900","PVP01_0801400","PVP01_0801700","PVP01_0801800","PVP01_0802100","PVP01_0802300","PVP01_0802400","PVP01_0802500","PVP01_0802900","PVP01_0803000","PVP01_0803100","PVP01_0803300","PVP01_0803500","PVP01_0803600","PVP01_0803700","PVP01_0803900","PVP01_0804000","PVP01_0804100","PVP01_0804200","PVP01_0804400","PVP01_0804500","PVP01_0804600","PVP01_0804700","PVP01_0804800","PVP01_0804900","PVP01_0805000","PVP01_0805200","PVP01_0805300","PVP01_0805400","PVP01_0805600","PVP01_0805700","PVP01_0805800","PVP01_0806000","PVP01_0806100","PVP01_0806200","PVP01_0806300","PVP01_0806400","PVP01_0806500","PVP01_0806600","PVP01_0806700","PVP01_0806800","PVP01_0807000","PVP01_0807200","PVP01_0807300","PVP01_0807500","PVP01_0807600","PVP01_0807800","PVP01_0808000","PVP01_0808400","PVP01_0808500","PVP01_0808600","PVP01_0808800","PVP01_0809100","PVP01_0809200","PVP01_0809300","PVP01_0809400","PVP01_0809600","PVP01_0809700","PVP01_0809900","PVP01_0810000","PVP01_0810100","PVP01_0810300","PVP01_0810400","PVP01_0810500","PVP01_0810600","PVP01_0811000","PVP01_0811100","PVP01_0811200","PVP01_0811300","PVP01_0811500","PVP01_0811700","PVP01_0811800","PVP01_0811900","PVP01_0812000","PVP01_0812100","PVP01_0812300","PVP01_0812600","PVP01_0812700","PVP01_0812800","PVP01_0813000","PVP01_0813200","PVP01_0813300","PVP01_0813600","PVP01_0813700","PVP01_0813800","PVP01_0814100","PVP01_0814400","PVP01_0814700","PVP01_0814800","PVP01_0815100","PVP01_0815300","PVP01_0815400","PVP01_0815600","PVP01_0815700","PVP01_0816200","PVP01_0816700","PVP01_0817100","PVP01_0817400","PVP01_0818000","PVP01_0818100","PVP01_0818300","PVP01_0818500","PVP01_0818600","PVP01_0819000","PVP01_0819200","PVP01_0819700","PVP01_0820000","PVP01_0820100","PVP01_0820300","PVP01_0820500","PVP01_0820600","PVP01_0820700","PVP01_0820800","PVP01_0820900","PVP01_0821000","PVP01_0821100","PVP01_0821400","PVP01_0821500","PVP01_0821600","PVP01_0821700","PVP01_0821800","PVP01_0822100","PVP01_0822200","PVP01_0822300","PVP01_0822600","PVP01_0822700","PVP01_0822900","PVP01_0823100","PVP01_0823200","PVP01_0823400","PVP01_0823500","PVP01_0823700","PVP01_0823900","PVP01_0824000","PVP01_0824300","PVP01_0824500","PVP01_0824700","PVP01_0825000","PVP01_0825100","PVP01_0825200","PVP01_0825400","PVP01_0825700","PVP01_0825800","PVP01_0826000","PVP01_0826100","PVP01_0826200","PVP01_0826300","PVP01_0826400","PVP01_0826500","PVP01_0826600","PVP01_0826700","PVP01_0826900","PVP01_0827100","PVP01_0827200","PVP01_0827700","PVP01_0827800","PVP01_0828000","PVP01_0828100","PVP01_0828200","PVP01_0828300","PVP01_0828600","PVP01_0828700","PVP01_0829000","PVP01_0829100","PVP01_0829300","PVP01_0829700","PVP01_0829900","PVP01_0830200","PVP01_0830300","PVP01_0830400","PVP01_0830500","PVP01_0830700","PVP01_0830900","PVP01_0831000","PVP01_0831100","PVP01_0831200","PVP01_0831400","PVP01_0831700","PVP01_0831900","PVP01_0832000","PVP01_0832100","PVP01_0832200","PVP01_0832400","PVP01_0832500","PVP01_0832600","PVP01_0832800","PVP01_0833000","PVP01_0833500","PVP01_0833600","PVP01_0833700","PVP01_0833800","PVP01_0833900","PVP01_0834000","PVP01_0834200","PVP01_0834300","PVP01_0834400","PVP01_0834500","PVP01_0834600","PVP01_0834700","PVP01_0834900","PVP01_0835000","PVP01_0835300","PVP01_0835500","PVP01_0835700","PVP01_0835800","PVP01_0835900","PVP01_0836000","PVP01_0836100","PVP01_0836300","PVP01_0836400","PVP01_0836500","PVP01_0836600","PVP01_0836700","PVP01_0837100","PVP01_0837200","PVP01_0837300","PVP01_0837500","PVP01_0837600","PVP01_0837800","PVP01_0837900","PVP01_0838000","PVP01_0838200","PVP01_0838900","PVP01_0839100","PVP01_0839200","PVP01_0839700","PVP01_0840700","PVP01_0841000","PVP01_0900700","PVP01_0901200","PVP01_0901300","PVP01_0901800","PVP01_0901900","PVP01_0902000","PVP01_0902100","PVP01_0902400","PVP01_0903400","PVP01_0903500","PVP01_0903600","PVP01_0903700","PVP01_0903800","PVP01_0903900","PVP01_0904300","PVP01_0904600","PVP01_0904700","PVP01_0904900","PVP01_0905100","PVP01_0905700","PVP01_0906000","PVP01_0906300","PVP01_0906500","PVP01_0906700","PVP01_0906800","PVP01_0906900","PVP01_0907000","PVP01_0907100","PVP01_0907200","PVP01_0907300","PVP01_0907500","PVP01_0907600","PVP01_0907700","PVP01_0907800","PVP01_0908000","PVP01_0908300","PVP01_0908400","PVP01_0908500","PVP01_0908600","PVP01_0908900","PVP01_0909000","PVP01_0909100","PVP01_0909300","PVP01_0909500","PVP01_0909600","PVP01_0909700","PVP01_0910000","PVP01_0910200","PVP01_0910900","PVP01_0911000","PVP01_0911300","PVP01_0911500","PVP01_0911600","PVP01_0911700","PVP01_0911800","PVP01_0911900","PVP01_0912100","PVP01_0912200","PVP01_0912400","PVP01_0912900","PVP01_0913000","PVP01_0913100","PVP01_0913400","PVP01_0913500","PVP01_0913600","PVP01_0913700","PVP01_0913900","PVP01_0914000","PVP01_0914200","PVP01_0914300","PVP01_0914400","PVP01_0914500","PVP01_0914600","PVP01_0914800","PVP01_0915000","PVP01_0915100","PVP01_0915200","PVP01_0915300","PVP01_0915400","PVP01_0915600","PVP01_0915800","PVP01_0915900","PVP01_0916300","PVP01_0916700","PVP01_0916800","PVP01_0916900","PVP01_0917000","PVP01_0917100","PVP01_0917200","PVP01_0917300","PVP01_0917400","PVP01_0917600","PVP01_0917700","PVP01_0918000","PVP01_0918100","PVP01_0918200","PVP01_0918300","PVP01_0918400","PVP01_0918500","PVP01_0918700","PVP01_0918800","PVP01_0918900","PVP01_0919000","PVP01_0919100","PVP01_0919300","PVP01_0919500","PVP01_0919600","PVP01_0919700","PVP01_0919900","PVP01_0920100","PVP01_0920300","PVP01_0920400","PVP01_0920500","PVP01_0920700","PVP01_0920800","PVP01_0921100","PVP01_0921400","PVP01_0921500","PVP01_0921700","PVP01_0921800","PVP01_0922000","PVP01_0922100","PVP01_0922300","PVP01_0922400","PVP01_0922800","PVP01_0922900","PVP01_0923000","PVP01_0923100","PVP01_0923300","PVP01_0923800","PVP01_0923900","PVP01_0924000","PVP01_0924100","PVP01_0924300","PVP01_0924400","PVP01_0924500","PVP01_0924700","PVP01_0924800","PVP01_0924900","PVP01_0925100","PVP01_0925200","PVP01_0925300","PVP01_0925400","PVP01_0925600","PVP01_0925700","PVP01_0925800","PVP01_0925900","PVP01_0926300","PVP01_0926400","PVP01_0926600","PVP01_0926900","PVP01_0927100","PVP01_0927400","PVP01_0927600","PVP01_0927700","PVP01_0927800","PVP01_0928000","PVP01_0928100","PVP01_0928300","PVP01_0928400","PVP01_0928800","PVP01_0928900","PVP01_0929000","PVP01_0929100","PVP01_0929200","PVP01_0929300","PVP01_0929400","PVP01_0929700","PVP01_0929800","PVP01_0929900","PVP01_0930000","PVP01_0930100","PVP01_0930200","PVP01_0930300","PVP01_0930400","PVP01_0930500","PVP01_0930600","PVP01_0931300","PVP01_0931400","PVP01_0931500","PVP01_0931600","PVP01_0931700","PVP01_0931900","PVP01_0932000","PVP01_0932100","PVP01_0932200","PVP01_0932300","PVP01_0932600","PVP01_0933100","PVP01_0933400","PVP01_0933500","PVP01_0933600","PVP01_0933700","PVP01_0934000","PVP01_0934300","PVP01_0934400","PVP01_0934500","PVP01_0935000","PVP01_0935100","PVP01_0935200","PVP01_0935300","PVP01_0935400","PVP01_0935500","PVP01_0935700","PVP01_0935800","PVP01_0936000","PVP01_0936100","PVP01_0936300","PVP01_0936400","PVP01_0936600","PVP01_0936700","PVP01_0936800","PVP01_0936900","PVP01_0937000","PVP01_0937300","PVP01_0937400","PVP01_0937500","PVP01_0937700","PVP01_0937800","PVP01_0938000","PVP01_0938100","PVP01_0938200","PVP01_0938400","PVP01_0938600","PVP01_0938700","PVP01_0938900","PVP01_0939100","PVP01_0939200","PVP01_0939300","PVP01_0939500","PVP01_0939600","PVP01_0939900","PVP01_0940100","PVP01_0940200","PVP01_0940500","PVP01_0940600","PVP01_0940700","PVP01_0940800","PVP01_0940900","PVP01_0941000","PVP01_0941200","PVP01_0941300","PVP01_0941400","PVP01_0941600","PVP01_0941800","PVP01_0942200","PVP01_0942300","PVP01_0942400","PVP01_0942600","PVP01_0942700","PVP01_0942900","PVP01_0943100","PVP01_0943300","PVP01_0943400","PVP01_0943700","PVP01_0944100","PVP01_0944200","PVP01_0944300","PVP01_0944400","PVP01_0944600","PVP01_0944800","PVP01_0945100","PVP01_0945200","PVP01_0945300","PVP01_0945500","PVP01_0945600","PVP01_0945800","PVP01_0946000","PVP01_0946200","PVP01_0946300","PVP01_0946400","PVP01_0946500","PVP01_0946600","PVP01_0946700","PVP01_0946900","PVP01_0947000","PVP01_0947100","PVP01_0947200","PVP01_0947400","PVP01_0947700","PVP01_0947800","PVP01_0947900","PVP01_0948100","PVP01_0948300","PVP01_0948500","PVP01_0948600","PVP01_0948700","PVP01_0948800","PVP01_0948900","PVP01_0949000","PVP01_0949400","PVP01_0949600","PVP01_0950000","PVP01_0950200","PVP01_1000700","PVP01_1001900","PVP01_1002500","PVP01_1002600","PVP01_1002700","PVP01_1002900","PVP01_1003000","PVP01_1003100","PVP01_1003200","PVP01_1003400","PVP01_1003500","PVP01_1004200","PVP01_1004500","PVP01_1004800","PVP01_1004900","PVP01_1005000","PVP01_1005100","PVP01_1005200","PVP01_1005300","PVP01_1005400","PVP01_1005500","PVP01_1005600","PVP01_1005700","PVP01_1005900","PVP01_1006200","PVP01_1006300","PVP01_1006400","PVP01_1006600","PVP01_1006700","PVP01_1007500","PVP01_1007600","PVP01_1007700","PVP01_1007800","PVP01_1007900","PVP01_1008000","PVP01_1008100","PVP01_1008200","PVP01_1008300","PVP01_1008400","PVP01_1008600","PVP01_1008700","PVP01_1008900","PVP01_1009000","PVP01_1009100","PVP01_1009200","PVP01_1009300","PVP01_1009400","PVP01_1009500","PVP01_1009600","PVP01_1009700","PVP01_1009800","PVP01_1010100","PVP01_1010200","PVP01_1010300","PVP01_1010400","PVP01_1011100","PVP01_1011200","PVP01_1011600","PVP01_1011700","PVP01_1011800","PVP01_1012100","PVP01_1012300","PVP01_1012400","PVP01_1012500","PVP01_1012700","PVP01_1012800","PVP01_1013100","PVP01_1013200","PVP01_1013300","PVP01_1013400","PVP01_1013700","PVP01_1014200","PVP01_1014300","PVP01_1014400","PVP01_1014600","PVP01_1014700","PVP01_1014800","PVP01_1015000","PVP01_1015100","PVP01_1015400","PVP01_1015500","PVP01_1015700","PVP01_1015900","PVP01_1016000","PVP01_1016200","PVP01_1016300","PVP01_1016400","PVP01_1016500","PVP01_1016600","PVP01_1016900","PVP01_1017100","PVP01_1017400","PVP01_1017600","PVP01_1017900","PVP01_1018000","PVP01_1018100","PVP01_1018400","PVP01_1018500","PVP01_1018700","PVP01_1019000","PVP01_1019400","PVP01_1019600","PVP01_1019900","PVP01_1020000","PVP01_1020100","PVP01_1020200","PVP01_1020500","PVP01_1020700","PVP01_1020800","PVP01_1020900","PVP01_1021000","PVP01_1021100","PVP01_1021200","PVP01_1021300","PVP01_1021500","PVP01_1021600","PVP01_1021700","PVP01_1021800","PVP01_1022000","PVP01_1022100","PVP01_1022200","PVP01_1022300","PVP01_1022400","PVP01_1022900","PVP01_1023000","PVP01_1023200","PVP01_1023300","PVP01_1023600","PVP01_1023700","PVP01_1023800","PVP01_1023900","PVP01_1024100","PVP01_1024300","PVP01_1024400","PVP01_1024500","PVP01_1024800","PVP01_1024900","PVP01_1025100","PVP01_1025200","PVP01_1025300","PVP01_1025400","PVP01_1025500","PVP01_1025800","PVP01_1026000","PVP01_1026200","PVP01_1026300","PVP01_1026400","PVP01_1026600","PVP01_1026700","PVP01_1026900","PVP01_1027300","PVP01_1027400","PVP01_1027600","PVP01_1027700","PVP01_1027900","PVP01_1028000","PVP01_1028100","PVP01_1028200","PVP01_1028400","PVP01_1028500","PVP01_1028600","PVP01_1028700","PVP01_1029000","PVP01_1029100","PVP01_1029200","PVP01_1029300","PVP01_1029500","PVP01_1029700","PVP01_1029800","PVP01_1029900","PVP01_1030000","PVP01_1030100","PVP01_1030300","PVP01_1030500","PVP01_1030600","PVP01_1030900","PVP01_1031300","PVP01_1031800","PVP01_1031900","PVP01_1032000","PVP01_1032100","PVP01_1032200","PVP01_1032400","PVP01_1032500","PVP01_1032600","PVP01_1032800","PVP01_1032900","PVP01_1033000","PVP01_1033300","PVP01_1033500","PVP01_1033700","PVP01_1034100","PVP01_1034300","PVP01_1034500","PVP01_1035200","PVP01_1100400","PVP01_1100500","PVP01_1101000","PVP01_1101200","PVP01_1101300","PVP01_1101500","PVP01_1101600","PVP01_1101700","PVP01_1102400","PVP01_1102500","PVP01_1102800","PVP01_1102900","PVP01_1103200","PVP01_1103300","PVP01_1103400","PVP01_1103600","PVP01_1103700","PVP01_1104000","PVP01_1104100","PVP01_1104200","PVP01_1104300","PVP01_1104400","PVP01_1104600","PVP01_1104700","PVP01_1104800","PVP01_1105100","PVP01_1105300","PVP01_1105400","PVP01_1105500","PVP01_1105600","PVP01_1105800","PVP01_1106000","PVP01_1106200","PVP01_1106300","PVP01_1106400","PVP01_1106600","PVP01_1106700","PVP01_1106900","PVP01_1107000","PVP01_1107100","PVP01_1107200","PVP01_1107300","PVP01_1107400","PVP01_1107600","PVP01_1107700","PVP01_1108000","PVP01_1108300","PVP01_1108500","PVP01_1108600","PVP01_1108700","PVP01_1108800","PVP01_1108900","PVP01_1109000","PVP01_1109100","PVP01_1109300","PVP01_1109400","PVP01_1109600","PVP01_1109800","PVP01_1109900","PVP01_1110100","PVP01_1110200","PVP01_1110500","PVP01_1110600","PVP01_1110700","PVP01_1110900","PVP01_1111000","PVP01_1111100","PVP01_1111200","PVP01_1111300","PVP01_1111500","PVP01_1111600","PVP01_1111700","PVP01_1111800","PVP01_1112000","PVP01_1112100","PVP01_1112200","PVP01_1112400","PVP01_1112500","PVP01_1112700","PVP01_1112800","PVP01_1112900","PVP01_1113000","PVP01_1113300","PVP01_1113400","PVP01_1113500","PVP01_1113700","PVP01_1113900","PVP01_1114100","PVP01_1114200","PVP01_1114300","PVP01_1114500","PVP01_1114600","PVP01_1114700","PVP01_1114800","PVP01_1115000","PVP01_1115200","PVP01_1115500","PVP01_1115600","PVP01_1115700","PVP01_1116600","PVP01_1116800","PVP01_1116900","PVP01_1117300","PVP01_1117500","PVP01_1117700","PVP01_1118000","PVP01_1118100","PVP01_1118600","PVP01_1118700","PVP01_1118800","PVP01_1118900","PVP01_1119000","PVP01_1119100","PVP01_1119300","PVP01_1119400","PVP01_1119500","PVP01_1119600","PVP01_1119800","PVP01_1120000","PVP01_1120100","PVP01_1120300","PVP01_1120400","PVP01_1120500","PVP01_1120900","PVP01_1121000","PVP01_1121100","PVP01_1121300","PVP01_1121400","PVP01_1121500","PVP01_1121600","PVP01_1121700","PVP01_1121900","PVP01_1122100","PVP01_1122300","PVP01_1122400","PVP01_1122500","PVP01_1122600","PVP01_1122700","PVP01_1122800","PVP01_1122900","PVP01_1123100","PVP01_1123200","PVP01_1123300","PVP01_1123400","PVP01_1123600","PVP01_1123900","PVP01_1124000","PVP01_1124100","PVP01_1124300","PVP01_1124400","PVP01_1124600","PVP01_1124700","PVP01_1124800","PVP01_1124900","PVP01_1125000","PVP01_1125100","PVP01_1125200","PVP01_1125300","PVP01_1125400","PVP01_1125600","PVP01_1125700","PVP01_1125800","PVP01_1125900","PVP01_1126000","PVP01_1126100","PVP01_1126200","PVP01_1126300","PVP01_1126500","PVP01_1126600","PVP01_1126800","PVP01_1126900","PVP01_1127000","PVP01_1127100","PVP01_1127200","PVP01_1127300","PVP01_1127400","PVP01_1127500","PVP01_1127800","PVP01_1128100","PVP01_1128300","PVP01_1128800","PVP01_1128900","PVP01_1129000","PVP01_1129400","PVP01_1129500","PVP01_1129600","PVP01_1129700","PVP01_1130100","PVP01_1130400","PVP01_1130700","PVP01_1130800","PVP01_1130900","PVP01_1131000","PVP01_1131300","PVP01_1131400","PVP01_1132100","PVP01_1132300","PVP01_1132400","PVP01_1132500","PVP01_1132700","PVP01_1132800","PVP01_1132900","PVP01_1133000","PVP01_1133100","PVP01_1133200","PVP01_1133300","PVP01_1133400","PVP01_1133500","PVP01_1133600","PVP01_1133800","PVP01_1133900","PVP01_1134100","PVP01_1134200","PVP01_1134300","PVP01_1134400","PVP01_1134700","PVP01_1134800","PVP01_1135000","PVP01_1135100","PVP01_1135200","PVP01_1135500","PVP01_1135600","PVP01_1135700","PVP01_1135800","PVP01_1135900","PVP01_1136000","PVP01_1136100","PVP01_1136300","PVP01_1136600","PVP01_1136700","PVP01_1136800","PVP01_1136900","PVP01_1137100","PVP01_1137200","PVP01_1137300","PVP01_1137600","PVP01_1137700","PVP01_1137800","PVP01_1138400","PVP01_1138500","PVP01_1139000","PVP01_1139100","PVP01_1139300","PVP01_1139600","PVP01_1139700","PVP01_1140000","PVP01_1140300","PVP01_1140500","PVP01_1140600","PVP01_1140700","PVP01_1140800","PVP01_1140900","PVP01_1141000","PVP01_1141200","PVP01_1141300","PVP01_1141400","PVP01_1141500","PVP01_1141700","PVP01_1141800","PVP01_1141900","PVP01_1142000","PVP01_1142100","PVP01_1142300","PVP01_1142500","PVP01_1142600","PVP01_1142900","PVP01_1143100","PVP01_1143200","PVP01_1143400","PVP01_1143500","PVP01_1143600","PVP01_1143700","PVP01_1144000","PVP01_1144100","PVP01_1144200","PVP01_1144300","PVP01_1144500","PVP01_1144600","PVP01_1144700","PVP01_1144800","PVP01_1144900","PVP01_1145000","PVP01_1145100","PVP01_1145200","PVP01_1145500","PVP01_1145700","PVP01_1145800","PVP01_1146400","PVP01_1146600","PVP01_1146800","PVP01_1146900","PVP01_1147000","PVP01_1147200","PVP01_1147300","PVP01_1147400","PVP01_1147500","PVP01_1148200","PVP01_1148700","PVP01_1148800","PVP01_1149000","PVP01_1149100","PVP01_1149400","PVP01_1149500","PVP01_1200200","PVP01_1200800","PVP01_1201100","PVP01_1201300","PVP01_1201600","PVP01_1201700","PVP01_1201800","PVP01_1201900","PVP01_1202100","PVP01_1202200","PVP01_1202300","PVP01_1203000","PVP01_1203100","PVP01_1203200","PVP01_1203300","PVP01_1203600","PVP01_1203700","PVP01_1203800","PVP01_1204000","PVP01_1204100","PVP01_1204200","PVP01_1204300","PVP01_1204500","PVP01_1204600","PVP01_1204800","PVP01_1204900","PVP01_1205000","PVP01_1205100","PVP01_1205200","PVP01_1205400","PVP01_1205500","PVP01_1205600","PVP01_1205900","PVP01_1206100","PVP01_1206200","PVP01_1206300","PVP01_1206400","PVP01_1206700","PVP01_1206900","PVP01_1207000","PVP01_1207100","PVP01_1207200","PVP01_1207800","PVP01_1207900","PVP01_1208100","PVP01_1208200","PVP01_1208300","PVP01_1208600","PVP01_1208700","PVP01_1208800","PVP01_1208900","PVP01_1209000","PVP01_1209400","PVP01_1209600","PVP01_1210000","PVP01_1210100","PVP01_1210200","PVP01_1210300","PVP01_1210700","PVP01_1210800","PVP01_1211200","PVP01_1211300","PVP01_1211500","PVP01_1211600","PVP01_1212100","PVP01_1212200","PVP01_1212500","PVP01_1212600","PVP01_1213000","PVP01_1213100","PVP01_1213300","PVP01_1214200","PVP01_1214600","PVP01_1214700","PVP01_1214800","PVP01_1214900","PVP01_1215000","PVP01_1215300","PVP01_1215400","PVP01_1215800","PVP01_1215900","PVP01_1216000","PVP01_1216100","PVP01_1216200","PVP01_1216300","PVP01_1216700","PVP01_1216800","PVP01_1217200","PVP01_1217400","PVP01_1217500","PVP01_1217600","PVP01_1217700","PVP01_1218000","PVP01_1218500","PVP01_1218800","PVP01_1218900","PVP01_1219000","PVP01_1219100","PVP01_1219500","PVP01_1219900","PVP01_1220100","PVP01_1220200","PVP01_1220300","PVP01_1220600","PVP01_1220700","PVP01_1220900","PVP01_1221200","PVP01_1221300","PVP01_1221400","PVP01_1221700","PVP01_1221800","PVP01_1221900","PVP01_1222100","PVP01_1222200","PVP01_1222300","PVP01_1222400","PVP01_1222500","PVP01_1222600","PVP01_1222700","PVP01_1222800","PVP01_1223100","PVP01_1223400","PVP01_1223500","PVP01_1223600","PVP01_1223800","PVP01_1223900","PVP01_1224000","PVP01_1224300","PVP01_1224400","PVP01_1224500","PVP01_1224600","PVP01_1224700","PVP01_1224800","PVP01_1225000","PVP01_1225200","PVP01_1225300","PVP01_1225500","PVP01_1225600","PVP01_1225700","PVP01_1225800","PVP01_1226400","PVP01_1226600","PVP01_1226700","PVP01_1226800","PVP01_1226900","PVP01_1227200","PVP01_1227300","PVP01_1227500","PVP01_1227600","PVP01_1227900","PVP01_1228000","PVP01_1228100","PVP01_1228200","PVP01_1228300","PVP01_1228400","PVP01_1228500","PVP01_1228800","PVP01_1229100","PVP01_1229200","PVP01_1229400","PVP01_1229500","PVP01_1229600","PVP01_1229800","PVP01_1229900","PVP01_1230000","PVP01_1230100","PVP01_1230200","PVP01_1230600","PVP01_1230700","PVP01_1230800","PVP01_1230900","PVP01_1231000","PVP01_1231100","PVP01_1231600","PVP01_1231700","PVP01_1231800","PVP01_1231900","PVP01_1232400","PVP01_1232600","PVP01_1232700","PVP01_1233100","PVP01_1233200","PVP01_1233800","PVP01_1233900","PVP01_1234000","PVP01_1234500","PVP01_1234600","PVP01_1235000","PVP01_1235100","PVP01_1235200","PVP01_1235300","PVP01_1235500","PVP01_1235800","PVP01_1236000","PVP01_1236100","PVP01_1236200","PVP01_1236400","PVP01_1236500","PVP01_1236800","PVP01_1236900","PVP01_1237000","PVP01_1237100","PVP01_1237200","PVP01_1237500","PVP01_1237700","PVP01_1237800","PVP01_1237900","PVP01_1238000","PVP01_1238400","PVP01_1238500","PVP01_1238600","PVP01_1238900","PVP01_1239000","PVP01_1239100","PVP01_1239300","PVP01_1239400","PVP01_1239500","PVP01_1239900","PVP01_1240100","PVP01_1240300","PVP01_1240400","PVP01_1240600","PVP01_1240800","PVP01_1240900","PVP01_1241000","PVP01_1241100","PVP01_1241200","PVP01_1241300","PVP01_1241400","PVP01_1241900","PVP01_1242000","PVP01_1242300","PVP01_1242400","PVP01_1242700","PVP01_1243000","PVP01_1243100","PVP01_1243200","PVP01_1243500","PVP01_1243600","PVP01_1243800","PVP01_1243900","PVP01_1244300","PVP01_1244400","PVP01_1244700","PVP01_1244800","PVP01_1244900","PVP01_1245100","PVP01_1245300","PVP01_1245400","PVP01_1245500","PVP01_1245600","PVP01_1245800","PVP01_1245900","PVP01_1246000","PVP01_1246100","PVP01_1246300","PVP01_1246400","PVP01_1246500","PVP01_1246600","PVP01_1246700","PVP01_1246900","PVP01_1247000","PVP01_1247100","PVP01_1247500","PVP01_1247600","PVP01_1247700","PVP01_1247800","PVP01_1247900","PVP01_1248200","PVP01_1248500","PVP01_1248700","PVP01_1248800","PVP01_1249100","PVP01_1249200","PVP01_1249500","PVP01_1249600","PVP01_1249700","PVP01_1250000","PVP01_1250200","PVP01_1250300","PVP01_1250600","PVP01_1250700","PVP01_1250800","PVP01_1251100","PVP01_1251300","PVP01_1251500","PVP01_1251700","PVP01_1251900","PVP01_1252000","PVP01_1252100","PVP01_1252200","PVP01_1252400","PVP01_1252800","PVP01_1252900","PVP01_1253000","PVP01_1253300","PVP01_1253600","PVP01_1253700","PVP01_1253800","PVP01_1254100","PVP01_1254200","PVP01_1254300","PVP01_1254800","PVP01_1254900","PVP01_1255100","PVP01_1255200","PVP01_1255300","PVP01_1255400","PVP01_1255500","PVP01_1255700","PVP01_1256000","PVP01_1256200","PVP01_1256400","PVP01_1256600","PVP01_1256700","PVP01_1256800","PVP01_1256900","PVP01_1257000","PVP01_1257400","PVP01_1257500","PVP01_1257600","PVP01_1257700","PVP01_1257800","PVP01_1257900","PVP01_1258100","PVP01_1258200","PVP01_1258300","PVP01_1258400","PVP01_1258500","PVP01_1258700","PVP01_1258900","PVP01_1259000","PVP01_1259100","PVP01_1259200","PVP01_1259300","PVP01_1259400","PVP01_1259500","PVP01_1259600","PVP01_1259700","PVP01_1260200","PVP01_1260300","PVP01_1260400","PVP01_1260500","PVP01_1260700","PVP01_1260800","PVP01_1261100","PVP01_1261200","PVP01_1261300","PVP01_1261400","PVP01_1261500","PVP01_1261700","PVP01_1261800","PVP01_1262000","PVP01_1262200","PVP01_1262300","PVP01_1262400","PVP01_1262600","PVP01_1262800","PVP01_1262900","PVP01_1263000","PVP01_1263100","PVP01_1263400","PVP01_1263500","PVP01_1263800","PVP01_1263900","PVP01_1264000","PVP01_1264200","PVP01_1264300","PVP01_1264500","PVP01_1264600","PVP01_1264700","PVP01_1264800","PVP01_1264900","PVP01_1265000","PVP01_1265100","PVP01_1265300","PVP01_1265400","PVP01_1265600","PVP01_1265900","PVP01_1266000","PVP01_1266100","PVP01_1266400","PVP01_1266500","PVP01_1266700","PVP01_1266800","PVP01_1267100","PVP01_1267200","PVP01_1267300","PVP01_1267400","PVP01_1267500","PVP01_1267700","PVP01_1267800","PVP01_1268000","PVP01_1268100","PVP01_1268200","PVP01_1268300","PVP01_1269000","PVP01_1269100","PVP01_1269200","PVP01_1269500","PVP01_1269800","PVP01_1269900","PVP01_1270000","PVP01_1270200","PVP01_1270300","PVP01_1270500","PVP01_1270600","PVP01_1270700","PVP01_1270900","PVP01_1271100","PVP01_1271200","PVP01_1271400","PVP01_1271600","PVP01_1271900","PVP01_1272600","PVP01_1273000","PVP01_1274300","PVP01_1300400","PVP01_1300500","PVP01_1300900","PVP01_1301000","PVP01_1301100","PVP01_1301200","PVP01_1301300","PVP01_1301500","PVP01_1301600","PVP01_1301700","PVP01_1301800","PVP01_1302100","PVP01_1302300","PVP01_1302500","PVP01_1302600","PVP01_1302700","PVP01_1302800","PVP01_1302900","PVP01_1303000","PVP01_1303100","PVP01_1303200","PVP01_1303600","PVP01_1303900","PVP01_1304000","PVP01_1304200","PVP01_1304300","PVP01_1304500","PVP01_1304800","PVP01_1304900","PVP01_1305000","PVP01_1305300","PVP01_1305500","PVP01_1305600","PVP01_1305700","PVP01_1306000","PVP01_1306100","PVP01_1306300","PVP01_1306400","PVP01_1306600","PVP01_1306700","PVP01_1306800","PVP01_1307000","PVP01_1307100","PVP01_1307200","PVP01_1307300","PVP01_1307400","PVP01_1307700","PVP01_1307800","PVP01_1308100","PVP01_1308300","PVP01_1308700","PVP01_1308800","PVP01_1308900","PVP01_1309000","PVP01_1309100","PVP01_1309200","PVP01_1309300","PVP01_1309400","PVP01_1309500","PVP01_1309700","PVP01_1309800","PVP01_1310100","PVP01_1310400","PVP01_1310700","PVP01_1311000","PVP01_1311100","PVP01_1311200","PVP01_1311300","PVP01_1311400","PVP01_1311500","PVP01_1311900","PVP01_1312000","PVP01_1312300","PVP01_1312400","PVP01_1312500","PVP01_1312600","PVP01_1312700","PVP01_1312800","PVP01_1313000","PVP01_1313100","PVP01_1313300","PVP01_1313700","PVP01_1313900","PVP01_1314100","PVP01_1314600","PVP01_1314800","PVP01_1314900","PVP01_1315300","PVP01_1315400","PVP01_1315500","PVP01_1315600","PVP01_1315700","PVP01_1316100","PVP01_1316500","PVP01_1316600","PVP01_1316700","PVP01_1316800","PVP01_1316900","PVP01_1317000","PVP01_1317400","PVP01_1317600","PVP01_1317700","PVP01_1317800","PVP01_1317900","PVP01_1318000","PVP01_1318100","PVP01_1318300","PVP01_1318400","PVP01_1318500","PVP01_1318600","PVP01_1318800","PVP01_1318900","PVP01_1319000","PVP01_1319200","PVP01_1319300","PVP01_1319500","PVP01_1319700","PVP01_1319800","PVP01_1319900","PVP01_1320000","PVP01_1320100","PVP01_1320200","PVP01_1320300","PVP01_1320400","PVP01_1320500","PVP01_1320600","PVP01_1320700","PVP01_1321100","PVP01_1321300","PVP01_1321500","PVP01_1321600","PVP01_1321700","PVP01_1321900","PVP01_1322000","PVP01_1322200","PVP01_1322300","PVP01_1322400","PVP01_1322600","PVP01_1322700","PVP01_1322800","PVP01_1322900","PVP01_1323000","PVP01_1323100","PVP01_1323400","PVP01_1323500","PVP01_1323600","PVP01_1323700","PVP01_1323800","PVP01_1323900","PVP01_1324100","PVP01_1324700","PVP01_1324800","PVP01_1325700","PVP01_1325800","PVP01_1325900","PVP01_1326000","PVP01_1326200","PVP01_1326400","PVP01_1326600","PVP01_1327000","PVP01_1327100","PVP01_1327200","PVP01_1327300","PVP01_1327500","PVP01_1327600","PVP01_1327900","PVP01_1328000","PVP01_1328100","PVP01_1328300","PVP01_1328400","PVP01_1328500","PVP01_1328600","PVP01_1328700","PVP01_1328800","PVP01_1328900","PVP01_1329000","PVP01_1329300","PVP01_1329400","PVP01_1329600","PVP01_1329900","PVP01_1330000","PVP01_1330800","PVP01_1331000","PVP01_1331200","PVP01_1331400","PVP01_1331800","PVP01_1331900","PVP01_1332200","PVP01_1332300","PVP01_1332500","PVP01_1332600","PVP01_1332700","PVP01_1332900","PVP01_1333000","PVP01_1333100","PVP01_1333200","PVP01_1333300","PVP01_1333700","PVP01_1333900","PVP01_1334000","PVP01_1334100","PVP01_1334200","PVP01_1334300","PVP01_1334500","PVP01_1334800","PVP01_1335100","PVP01_1335300","PVP01_1335600","PVP01_1335800","PVP01_1335900","PVP01_1336000","PVP01_1336200","PVP01_1336500","PVP01_1336700","PVP01_1337100","PVP01_1337200","PVP01_1337300","PVP01_1337400","PVP01_1337500","PVP01_1337600","PVP01_1337900","PVP01_1338100","PVP01_1338300","PVP01_1338400","PVP01_1338600","PVP01_1339000","PVP01_1339400","PVP01_1339500","PVP01_1339800","PVP01_1340100","PVP01_1340200","PVP01_1340400","PVP01_1340500","PVP01_1340600","PVP01_1340800","PVP01_1341200","PVP01_1341300","PVP01_1341600","PVP01_1341800","PVP01_1342200","PVP01_1342300","PVP01_1342400","PVP01_1342600","PVP01_1342700","PVP01_1342800","PVP01_1343100","PVP01_1343300","PVP01_1343500","PVP01_1343600","PVP01_1343700","PVP01_1344200","PVP01_1344300","PVP01_1344500","PVP01_1344600","PVP01_1344700","PVP01_1344800","PVP01_1344900","PVP01_1345200","PVP01_1345300","PVP01_1345400","PVP01_1345500","PVP01_1345600","PVP01_1345700","PVP01_1345900","PVP01_1346200","PVP01_1346700","PVP01_1346800","PVP01_1346900","PVP01_1347300","PVP01_1347500","PVP01_1347700","PVP01_1347800","PVP01_1400100","PVP01_1400400","PVP01_1401000","PVP01_1401200","PVP01_1401300","PVP01_1401600","PVP01_1401700","PVP01_1401800","PVP01_1402000","PVP01_1402200","PVP01_1402500","PVP01_1402800","PVP01_1402900","PVP01_1403300","PVP01_1403400","PVP01_1403500","PVP01_1403700","PVP01_1403800","PVP01_1403900","PVP01_1404000","PVP01_1404100","PVP01_1404200","PVP01_1404300","PVP01_1404500","PVP01_1404600","PVP01_1404800","PVP01_1405000","PVP01_1405100","PVP01_1405500","PVP01_1405600","PVP01_1405800","PVP01_1405900","PVP01_1406000","PVP01_1406100","PVP01_1406200","PVP01_1406300","PVP01_1406500","PVP01_1406600","PVP01_1406700","PVP01_1406800","PVP01_1407000","PVP01_1407100","PVP01_1407200","PVP01_1407400","PVP01_1407600","PVP01_1407800","PVP01_1407900","PVP01_1408000","PVP01_1408100","PVP01_1408200","PVP01_1408400","PVP01_1408500","PVP01_1408600","PVP01_1408700","PVP01_1408800","PVP01_1409100","PVP01_1409300","PVP01_1409400","PVP01_1409700","PVP01_1410000","PVP01_1410100","PVP01_1410200","PVP01_1410300","PVP01_1410500","PVP01_1410600","PVP01_1410700","PVP01_1410800","PVP01_1410900","PVP01_1411000","PVP01_1411300","PVP01_1411400","PVP01_1411500","PVP01_1411600","PVP01_1411700","PVP01_1412000","PVP01_1412100","PVP01_1412200","PVP01_1412400","PVP01_1412500","PVP01_1412700","PVP01_1413200","PVP01_1413300","PVP01_1413450","PVP01_1413700","PVP01_1413900","PVP01_1414100","PVP01_1414200","PVP01_1414300","PVP01_1414400","PVP01_1414600","PVP01_1414900","PVP01_1415100","PVP01_1415200","PVP01_1415300","PVP01_1415500","PVP01_1415600","PVP01_1415700","PVP01_1415900","PVP01_1416000","PVP01_1416100","PVP01_1416900","PVP01_1417200","PVP01_1417600","PVP01_1417700","PVP01_1417900","PVP01_1418000","PVP01_1418100","PVP01_1418200","PVP01_1418300","PVP01_1418400","PVP01_1418600","PVP01_1418800","PVP01_1418900","PVP01_1419100","PVP01_1419300","PVP01_1419400","PVP01_1419500","PVP01_1419600","PVP01_1419700","PVP01_1419800","PVP01_1420000","PVP01_1420100","PVP01_1420200","PVP01_1420300","PVP01_1420400","PVP01_1420500","PVP01_1420700","PVP01_1420900","PVP01_1421000","PVP01_1421100","PVP01_1421200","PVP01_1421300","PVP01_1421400","PVP01_1421600","PVP01_1421800","PVP01_1422100","PVP01_1422200","PVP01_1422500","PVP01_1422600","PVP01_1422700","PVP01_1422900","PVP01_1423200","PVP01_1423300","PVP01_1423700","PVP01_1424100","PVP01_1424200","PVP01_1424400","PVP01_1424500","PVP01_1424700","PVP01_1424800","PVP01_1425100","PVP01_1425200","PVP01_1425300","PVP01_1425400","PVP01_1425800","PVP01_1426100","PVP01_1426200","PVP01_1426400","PVP01_1426600","PVP01_1426700","PVP01_1426800","PVP01_1427000","PVP01_1427100","PVP01_1427700","PVP01_1427800","PVP01_1428000","PVP01_1428100","PVP01_1428300","PVP01_1428400","PVP01_1428500","PVP01_1428900","PVP01_1429000","PVP01_1429200","PVP01_1429300","PVP01_1429700","PVP01_1429900","PVP01_1430000","PVP01_1430100","PVP01_1430300","PVP01_1430400","PVP01_1430600","PVP01_1430700","PVP01_1430800","PVP01_1431100","PVP01_1431200","PVP01_1431300","PVP01_1431500","PVP01_1431600","PVP01_1431700","PVP01_1431800","PVP01_1432000","PVP01_1432100","PVP01_1432400","PVP01_1432500","PVP01_1432600","PVP01_1432700","PVP01_1432800","PVP01_1433000","PVP01_1433300","PVP01_1433600","PVP01_1433800","PVP01_1433900","PVP01_1434000","PVP01_1434200","PVP01_1434300","PVP01_1434400","PVP01_1434500","PVP01_1434600","PVP01_1434800","PVP01_1434900","PVP01_1435100","PVP01_1435200","PVP01_1435300","PVP01_1435800","PVP01_1435900","PVP01_1436000","PVP01_1436300","PVP01_1436400","PVP01_1436600","PVP01_1436700","PVP01_1437100","PVP01_1437200","PVP01_1437500","PVP01_1437600","PVP01_1437700","PVP01_1438100","PVP01_1438200","PVP01_1438300","PVP01_1438400","PVP01_1438500","PVP01_1438800","PVP01_1438900","PVP01_1439000","PVP01_1439300","PVP01_1439400","PVP01_1439700","PVP01_1439800","PVP01_1440000","PVP01_1440100","PVP01_1440300","PVP01_1440600","PVP01_1440700","PVP01_1440800","PVP01_1440900","PVP01_1441200","PVP01_1441300","PVP01_1441400","PVP01_1441700","PVP01_1441800","PVP01_1442000","PVP01_1442200","PVP01_1442500","PVP01_1442800","PVP01_1442900","PVP01_1443000","PVP01_1443200","PVP01_1443600","PVP01_1443800","PVP01_1443900","PVP01_1444000","PVP01_1444100","PVP01_1444200","PVP01_1444300","PVP01_1444400","PVP01_1444500","PVP01_1444800","PVP01_1444900","PVP01_1445000","PVP01_1445100","PVP01_1445200","PVP01_1445300","PVP01_1445500","PVP01_1445600","PVP01_1445700","PVP01_1445800","PVP01_1446000","PVP01_1446100","PVP01_1446200","PVP01_1446300","PVP01_1446400","PVP01_1446600","PVP01_1446700","PVP01_1446900","PVP01_1447200","PVP01_1447300","PVP01_1447400","PVP01_1447500","PVP01_1447600","PVP01_1447800","PVP01_1447900","PVP01_1448000","PVP01_1448100","PVP01_1448200","PVP01_1448300","PVP01_1448400","PVP01_1448600","PVP01_1448700","PVP01_1448800","PVP01_1448900","PVP01_1449100","PVP01_1449200","PVP01_1449500","PVP01_1449700","PVP01_1450000","PVP01_1450200","PVP01_1450400","PVP01_1450500","PVP01_1450900","PVP01_1451000","PVP01_1451100","PVP01_1451200","PVP01_1451300","PVP01_1451400","PVP01_1451500","PVP01_1451600","PVP01_1451700","PVP01_1451800","PVP01_1452100","PVP01_1452300","PVP01_1452400","PVP01_1452500","PVP01_1452700","PVP01_1452800","PVP01_1452900","PVP01_1453000","PVP01_1453200","PVP01_1453300","PVP01_1453800","PVP01_1454000","PVP01_1454100","PVP01_1454200","PVP01_1454400","PVP01_1454500","PVP01_1454600","PVP01_1454800","PVP01_1454900","PVP01_1455100","PVP01_1455400","PVP01_1455500","PVP01_1455700","PVP01_1455900","PVP01_1456000","PVP01_1456500","PVP01_1456700","PVP01_1456900","PVP01_1457000","PVP01_1457200","PVP01_1457400","PVP01_1457500","PVP01_1457700","PVP01_1457900","PVP01_1458300","PVP01_1458700","PVP01_1458800","PVP01_1459100","PVP01_1459200","PVP01_1459300","PVP01_1459400","PVP01_1459500","PVP01_1459600","PVP01_1459700","PVP01_1460300","PVP01_1460400","PVP01_1460500","PVP01_1460600","PVP01_1460700","PVP01_1460800","PVP01_1460900","PVP01_1461000","PVP01_1461100","PVP01_1461700","PVP01_1461900","PVP01_1462000","PVP01_1462100","PVP01_1462200","PVP01_1462300","PVP01_1462400","PVP01_1462500","PVP01_1462600","PVP01_1462900","PVP01_1463300","PVP01_1463500","PVP01_1463600","PVP01_1463700","PVP01_1463800","PVP01_1464100","PVP01_1464400","PVP01_1464500","PVP01_1464600","PVP01_1464700","PVP01_1464800","PVP01_1464900","PVP01_1465000","PVP01_1465100","PVP01_1465500","PVP01_1465600","PVP01_1465700","PVP01_1465800","PVP01_1465900","PVP01_1466100","PVP01_1466300","PVP01_1466500","PVP01_1466600","PVP01_1466700","PVP01_1467000","PVP01_1467100","PVP01_1467300","PVP01_1467600","PVP01_1467800","PVP01_1468100","PVP01_1468400","PVP01_1468500","PVP01_1468600","PVP01_1468900","PVP01_1469300","PVP01_1469900","PVP01_1470000","PVP01_1470200","PVP01_1470300","PVP01_1470400","PVP01_1470500","PVP01_1470600","PVP01_1470800","PVP01_1470900","PVP01_1471300","PVP01_1471600","PVP01_1471700","PVP01_1471800","PVP01_1471900","PVP01_1472200","PVP01_1472400","PVP01_API01200","PVP01_API01400","PVP01_API01700","PVP01_API04100","PVP01_API04200","PVP01_0000100","PVP01_0000110","PVP01_0000160","PVP01_0000200","PVP01_0000220","PVP01_0001430","PVP01_0001440","PVP01_0003400","PVP01_0004040","PVP01_0004210","PVP01_0004220","PVP01_0004230","PVP01_0004240","PVP01_0004290","PVP01_0004330","PVP01_0004400","PVP01_0005540","PVP01_0005760","PVP01_0007310","PVP01_0009720","PVP01_0010210","PVP01_0010670","PVP01_0010840","PVP01_0011020","PVP01_0102200","PVP01_0102300","PVP01_0102600","PVP01_0103200","PVP01_0103700","PVP01_0104000","PVP01_0104300","PVP01_0104400","PVP01_0104500","PVP01_0104700","PVP01_0105900","PVP01_0106100","PVP01_0106600","PVP01_0106700","PVP01_0107500","PVP01_0107700","PVP01_0108500","PVP01_0108700","PVP01_0109300","PVP01_0109400","PVP01_0109500","PVP01_0109700","PVP01_0109900","PVP01_0110000","PVP01_0110300","PVP01_0110600","PVP01_0111000","PVP01_0111400","PVP01_0111500","PVP01_0111800","PVP01_0112400","PVP01_0112600","PVP01_0113600","PVP01_0113800","PVP01_0113900","PVP01_0114000","PVP01_0114300","PVP01_0114400","PVP01_0114500","PVP01_0114900","PVP01_0115600","PVP01_0115700","PVP01_0115800","PVP01_0115900","PVP01_0116000","PVP01_0116200","PVP01_0116300","PVP01_0116400","PVP01_0117100","PVP01_0117200","PVP01_0117300","PVP01_0117400","PVP01_0117900","PVP01_0118200","PVP01_0118300","PVP01_0118400","PVP01_0119200","PVP01_0201200","PVP01_0201400","PVP01_0202500","PVP01_0203200","PVP01_0203600","PVP01_0203700","PVP01_0204200","PVP01_0204300","PVP01_0204400","PVP01_0204500","PVP01_0204600","PVP01_0204700","PVP01_0205400","PVP01_0205700","PVP01_0205900","PVP01_0206000","PVP01_0206200","PVP01_0206700","PVP01_0206900","PVP01_0207300","PVP01_0207900","PVP01_0208000","PVP01_0208900","PVP01_0209900","PVP01_0210400","PVP01_0210600","PVP01_0210700","PVP01_0210900","PVP01_0211900","PVP01_0212000","PVP01_0212100","PVP01_0212200","PVP01_0212300","PVP01_0212900","PVP01_0213100","PVP01_0213200","PVP01_0213500","PVP01_0213800","PVP01_0213900","PVP01_0214300","PVP01_0214400","PVP01_0214500","PVP01_0214600","PVP01_0214700","PVP01_0215100","PVP01_0215200","PVP01_0215600","PVP01_0216200","PVP01_0216600","PVP01_0216700","PVP01_0220100","PVP01_0300800","PVP01_0302600","PVP01_0302700","PVP01_0302900","PVP01_0303000","PVP01_0303200","PVP01_0303300","PVP01_0303400","PVP01_0303600","PVP01_0303700","PVP01_0303800","PVP01_0303900","PVP01_0304000","PVP01_0304100","PVP01_0304200","PVP01_0304300","PVP01_0304400","PVP01_0304700","PVP01_0305100","PVP01_0305400","PVP01_0305600","PVP01_0305900","PVP01_0306000","PVP01_0306200","PVP01_0306600","PVP01_0307200","PVP01_0307900","PVP01_0308000","PVP01_0308300","PVP01_0308400","PVP01_0308500","PVP01_0308600","PVP01_0308700","PVP01_0308900","PVP01_0309000","PVP01_0309600","PVP01_0310100","PVP01_0310500","PVP01_0310700","PVP01_0311400","PVP01_0311800","PVP01_0311900","PVP01_0312300","PVP01_0312600","PVP01_0312900","PVP01_0313000","PVP01_0313100","PVP01_0313200","PVP01_0313300","PVP01_0313600","PVP01_0313700","PVP01_0313800","PVP01_0313900","PVP01_0314000","PVP01_0314300","PVP01_0314400","PVP01_0314500","PVP01_0314600","PVP01_0314700","PVP01_0314800","PVP01_0315400","PVP01_0315800","PVP01_0316100","PVP01_0316600","PVP01_0316900","PVP01_0317000","PVP01_0317300","PVP01_0317500","PVP01_0317600","PVP01_0317700","PVP01_0317800","PVP01_0318300","PVP01_0318500","PVP01_0318600","PVP01_0318700","PVP01_0318800","PVP01_0319000","PVP01_0319100","PVP01_0319900","PVP01_0320000","PVP01_0320100","PVP01_0320300","PVP01_0320600","PVP01_0320800","PVP01_0321000","PVP01_0321100","PVP01_0404200","PVP01_0404300","PVP01_0405000","PVP01_0405400","PVP01_0405800","PVP01_0406300","PVP01_0406400","PVP01_0406800","PVP01_0407200","PVP01_0407300","PVP01_0407400","PVP01_0407500","PVP01_0408000","PVP01_0408100","PVP01_0408300","PVP01_0408400","PVP01_0408700","PVP01_0408800","PVP01_0409400","PVP01_0409500","PVP01_0409600","PVP01_0409800","PVP01_0410100","PVP01_0410200","PVP01_0410300","PVP01_0410500","PVP01_0410700","PVP01_0410900","PVP01_0411000","PVP01_0411100","PVP01_0411200","PVP01_0411300","PVP01_0412000","PVP01_0412300","PVP01_0412400","PVP01_0412500","PVP01_0413400","PVP01_0413600","PVP01_0413700","PVP01_0413800","PVP01_0413900","PVP01_0414100","PVP01_0414300","PVP01_0414500","PVP01_0414700","PVP01_0414800","PVP01_0415000","PVP01_0415100","PVP01_0415300","PVP01_0415700","PVP01_0415800","PVP01_0415900","PVP01_0416100","PVP01_0416400","PVP01_0416500","PVP01_0416700","PVP01_0416800","PVP01_0417000","PVP01_0417200","PVP01_0417300","PVP01_0417400","PVP01_0417600","PVP01_0417700","PVP01_0418000","PVP01_0418200","PVP01_0418400","PVP01_0418500","PVP01_0418600","PVP01_0418700","PVP01_0418800","PVP01_0418900","PVP01_0419800","PVP01_0419900","PVP01_0420400","PVP01_0420800","PVP01_0421100","PVP01_0421200","PVP01_0421300","PVP01_0421400","PVP01_0422000","PVP01_0422800","PVP01_0423000","PVP01_0423200","PVP01_0423300","PVP01_0423400","PVP01_0423500","PVP01_0423900","PVP01_0424500","PVP01_0503300","PVP01_0503500","PVP01_0503600","PVP01_0503700","PVP01_0503800","PVP01_0503900","PVP01_0504000","PVP01_0504100","PVP01_0504400","PVP01_0504800","PVP01_0504900","PVP01_0505300","PVP01_0505600","PVP01_0505700","PVP01_0505900","PVP01_0506100","PVP01_0506200","PVP01_0506300","PVP01_0506500","PVP01_0507000","PVP01_0507100","PVP01_0507200","PVP01_0507400","PVP01_0507900","PVP01_0508100","PVP01_0508400","PVP01_0508800","PVP01_0509300","PVP01_0509500","PVP01_0509700","PVP01_0509800","PVP01_0510500","PVP01_0510600","PVP01_0511000","PVP01_0511400","PVP01_0511600","PVP01_0511900","PVP01_0512300","PVP01_0512600","PVP01_0513100","PVP01_0513200","PVP01_0514300","PVP01_0514500","PVP01_0515000","PVP01_0515400","PVP01_0515500","PVP01_0515600","PVP01_0516100","PVP01_0516600","PVP01_0516700","PVP01_0517200","PVP01_0517500","PVP01_0517600","PVP01_0517800","PVP01_0518000","PVP01_0518100","PVP01_0518200","PVP01_0518800","PVP01_0518900","PVP01_0519000","PVP01_0519300","PVP01_0519700","PVP01_0519800","PVP01_0519900","PVP01_0520000","PVP01_0520100","PVP01_0520200","PVP01_0520300","PVP01_0520900","PVP01_0521000","PVP01_0521100","PVP01_0521200","PVP01_0521500","PVP01_0521900","PVP01_0522400","PVP01_0523100","PVP01_0523200","PVP01_0523400","PVP01_0523500","PVP01_0523600","PVP01_0523800","PVP01_0523900","PVP01_0524100","PVP01_0524300","PVP01_0525100","PVP01_0525600","PVP01_0525900","PVP01_0526200","PVP01_0526300","PVP01_0526500","PVP01_0526600","PVP01_0526700","PVP01_0526800","PVP01_0527000","PVP01_0527100","PVP01_0527200","PVP01_0527600","PVP01_0528800","PVP01_0529300","PVP01_0529400","PVP01_0529500","PVP01_0530200","PVP01_0530300","PVP01_0530600","PVP01_0530900","PVP01_0531000","PVP01_0531300","PVP01_0531400","PVP01_0531600","PVP01_0531900","PVP01_0532000","PVP01_0532100","PVP01_0532200","PVP01_0532300","PVP01_0532400","PVP01_0532800","PVP01_0533200","PVP01_0533700","PVP01_0534300","PVP01_0534400","PVP01_0601000","PVP01_0601300","PVP01_0601400","PVP01_0601700","PVP01_0601800","PVP01_0601900","PVP01_0602000","PVP01_0602100","PVP01_0602200","PVP01_0602400","PVP01_0602500","PVP01_0602700","PVP01_0602800","PVP01_0602900","PVP01_0603200","PVP01_0603300","PVP01_0603800","PVP01_0604300","PVP01_0604500","PVP01_0604600","PVP01_0605200","PVP01_0605600","PVP01_0605700","PVP01_0605800","PVP01_0605900","PVP01_0606000","PVP01_0606300","PVP01_0606700","PVP01_0607000","PVP01_0607100","PVP01_0607300","PVP01_0607400","PVP01_0607500","PVP01_0607700","PVP01_0607800","PVP01_0607900","PVP01_0609000","PVP01_0609100","PVP01_0609300","PVP01_0609500","PVP01_0610000","PVP01_0610300","PVP01_0611000","PVP01_0611200","PVP01_0611400","PVP01_0611500","PVP01_0611600","PVP01_0611700","PVP01_0611800","PVP01_0611900","PVP01_0612200","PVP01_0612300","PVP01_0612400","PVP01_0612500","PVP01_0612800","PVP01_0612900","PVP01_0613300","PVP01_0613600","PVP01_0614100","PVP01_0614500","PVP01_0614800","PVP01_0615000","PVP01_0615200","PVP01_0615300","PVP01_0615400","PVP01_0615600","PVP01_0616300","PVP01_0616700","PVP01_0617200","PVP01_0617300","PVP01_0617700","PVP01_0617800","PVP01_0618000","PVP01_0618200","PVP01_0618400","PVP01_0618500","PVP01_0618800","PVP01_0619300","PVP01_0620300","PVP01_0620400","PVP01_0620500","PVP01_0620600","PVP01_0620700","PVP01_0620800","PVP01_0621200","PVP01_0621400","PVP01_0621500","PVP01_0621600","PVP01_0621700","PVP01_0621900","PVP01_0622000","PVP01_0622300","PVP01_0622400","PVP01_0622500","PVP01_0622600","PVP01_0623000","PVP01_0623500","PVP01_0623700","PVP01_0623800","PVP01_0624500","PVP01_0700100","PVP01_0700300","PVP01_0700400","PVP01_0700500","PVP01_0700600","PVP01_0700900","PVP01_0701100","PVP01_0701200","PVP01_0701300","PVP01_0701800","PVP01_0701900","PVP01_0702000","PVP01_0702100","PVP01_0702300","PVP01_0702800","PVP01_0702900","PVP01_0703100","PVP01_0703300","PVP01_0703400","PVP01_0703800","PVP01_0703900","PVP01_0704200","PVP01_0704300","PVP01_0705300","PVP01_0705400","PVP01_0706800","PVP01_0706900","PVP01_0707400","PVP01_0707600","PVP01_0707700","PVP01_0707900","PVP01_0708600","PVP01_0708700","PVP01_0709400","PVP01_0709600","PVP01_0709800","PVP01_0710100","PVP01_0710400","PVP01_0710500","PVP01_0710600","PVP01_0710800","PVP01_0710900","PVP01_0711000","PVP01_0711300","PVP01_0711600","PVP01_0711700","PVP01_0712000","PVP01_0712100","PVP01_0712500","PVP01_0712600","PVP01_0712800","PVP01_0712900","PVP01_0713300","PVP01_0713400","PVP01_0713700","PVP01_0714100","PVP01_0714500","PVP01_0714700","PVP01_0715300","PVP01_0715400","PVP01_0715700","PVP01_0715900","PVP01_0716300","PVP01_0716400","PVP01_0716500","PVP01_0716700","PVP01_0717100","PVP01_0717300","PVP01_0717500","PVP01_0717700","PVP01_0717900","PVP01_0718300","PVP01_0718400","PVP01_0718500","PVP01_0718800","PVP01_0718900","PVP01_0719100","PVP01_0719200","PVP01_0719600","PVP01_0720500","PVP01_0721000","PVP01_0721200","PVP01_0721600","PVP01_0721700","PVP01_0722000","PVP01_0723600","PVP01_0723900","PVP01_0724200","PVP01_0724400","PVP01_0725200","PVP01_0725500","PVP01_0725600","PVP01_0725800","PVP01_0725900","PVP01_0726100","PVP01_0726200","PVP01_0726300","PVP01_0726400","PVP01_0726500","PVP01_0726600","PVP01_0726800","PVP01_0727900","PVP01_0728000","PVP01_0728400","PVP01_0728500","PVP01_0728600","PVP01_0728700","PVP01_0728800","PVP01_0728900","PVP01_0729000","PVP01_0729600","PVP01_0729800","PVP01_0729900","PVP01_0730500","PVP01_0730700","PVP01_0730900","PVP01_0731000","PVP01_0731400","PVP01_0732400","PVP01_0732500","PVP01_0732700","PVP01_0733300","PVP01_0733400","PVP01_0733600","PVP01_0733800","PVP01_0734000","PVP01_0734300","PVP01_0734500","PVP01_0734600","PVP01_0734700","PVP01_0735200","PVP01_0800700","PVP01_0801000","PVP01_0801100","PVP01_0801200","PVP01_0801300","PVP01_0801500","PVP01_0801600","PVP01_0802200","PVP01_0802600","PVP01_0802700","PVP01_0802800","PVP01_0803200","PVP01_0803400","PVP01_0803800","PVP01_0804300","PVP01_0805100","PVP01_0805500","PVP01_0805900","PVP01_0806900","PVP01_0807100","PVP01_0807400","PVP01_0807700","PVP01_0807900","PVP01_0808100","PVP01_0808200","PVP01_0808300","PVP01_0808700","PVP01_0808900","PVP01_0809000","PVP01_0809500","PVP01_0809800","PVP01_0810200","PVP01_0810700","PVP01_0810800","PVP01_0810900","PVP01_0811400","PVP01_0811600","PVP01_0812200","PVP01_0812400","PVP01_0812500","PVP01_0812900","PVP01_0813100","PVP01_0813400","PVP01_0813500","PVP01_0813900","PVP01_0814000","PVP01_0814200","PVP01_0814300","PVP01_0814500","PVP01_0814600","PVP01_0814900","PVP01_0815000","PVP01_0815200","PVP01_0815500","PVP01_0815800","PVP01_0815900","PVP01_0816000","PVP01_0816100","PVP01_0816300","PVP01_0816400","PVP01_0816500","PVP01_0816600","PVP01_0816800","PVP01_0816900","PVP01_0817000","PVP01_0817200","PVP01_0817300","PVP01_0817500","PVP01_0817600","PVP01_0817700","PVP01_0817800","PVP01_0817900","PVP01_0818200","PVP01_0818400","PVP01_0818700","PVP01_0818800","PVP01_0818900","PVP01_0819100","PVP01_0819300","PVP01_0819400","PVP01_0819500","PVP01_0819600","PVP01_0819800","PVP01_0819900","PVP01_0820200","PVP01_0820400","PVP01_0821200","PVP01_0821300","PVP01_0821900","PVP01_0822000","PVP01_0822400","PVP01_0822500","PVP01_0822800","PVP01_0823000","PVP01_0823300","PVP01_0823600","PVP01_0823800","PVP01_0824100","PVP01_0824200","PVP01_0824400","PVP01_0824600","PVP01_0824800","PVP01_0824900","PVP01_0825300","PVP01_0825500","PVP01_0825600","PVP01_0825900","PVP01_0826800","PVP01_0827000","PVP01_0827300","PVP01_0827400","PVP01_0827900","PVP01_0828400","PVP01_0828500","PVP01_0828800","PVP01_0828900","PVP01_0829200","PVP01_0829400","PVP01_0829500","PVP01_0829600","PVP01_0829800","PVP01_0830000","PVP01_0830100","PVP01_0830600","PVP01_0831300","PVP01_0831500","PVP01_0831600","PVP01_0832300","PVP01_0832700","PVP01_0832900","PVP01_0833100","PVP01_0833200","PVP01_0833400","PVP01_0834100","PVP01_0834800","PVP01_0835100","PVP01_0835200","PVP01_0835400","PVP01_0835600","PVP01_0836200","PVP01_0836800","PVP01_0836900","PVP01_0837000","PVP01_0837400","PVP01_0837700","PVP01_0838100","PVP01_0838300","PVP01_0838600","PVP01_0838700","PVP01_0838800","PVP01_0839000","PVP01_0839400","PVP01_0901000","PVP01_0901100","PVP01_0904200","PVP01_0904400","PVP01_0904500","PVP01_0904800","PVP01_0905000","PVP01_0905200","PVP01_0905300","PVP01_0905400","PVP01_0905500","PVP01_0905600","PVP01_0905800","PVP01_0905900","PVP01_0906100","PVP01_0906200","PVP01_0906400","PVP01_0906600","PVP01_0907400","PVP01_0907900","PVP01_0908100","PVP01_0908200","PVP01_0908700","PVP01_0908800","PVP01_0909200","PVP01_0909400","PVP01_0909900","PVP01_0910700","PVP01_0910800","PVP01_0911100","PVP01_0911200","PVP01_0911400","PVP01_0912000","PVP01_0912300","PVP01_0912500","PVP01_0912600","PVP01_0912700","PVP01_0912800","PVP01_0913200","PVP01_0913300","PVP01_0913800","PVP01_0914100","PVP01_0914700","PVP01_0914900","PVP01_0915500","PVP01_0915700","PVP01_0916000","PVP01_0916100","PVP01_0916200","PVP01_0916400","PVP01_0916500","PVP01_0916600","PVP01_0917500","PVP01_0917800","PVP01_0917900","PVP01_0918600","PVP01_0919200","PVP01_0919400","PVP01_0919800","PVP01_0920000","PVP01_0920200","PVP01_0920600","PVP01_0920900","PVP01_0921000","PVP01_0921200","PVP01_0921300","PVP01_0921600","PVP01_0921900","PVP01_0922200","PVP01_0922500","PVP01_0922600","PVP01_0922700","PVP01_0923200","PVP01_0923400","PVP01_0923500","PVP01_0923600","PVP01_0923700","PVP01_0924200","PVP01_0924600","PVP01_0925000","PVP01_0925500","PVP01_0926000","PVP01_0926100","PVP01_0926200","PVP01_0926500","PVP01_0926700","PVP01_0926800","PVP01_0927000","PVP01_0927200","PVP01_0927300","PVP01_0927500","PVP01_0927900","PVP01_0928200","PVP01_0928500","PVP01_0928600","PVP01_0928700","PVP01_0929500","PVP01_0929600","PVP01_0930700","PVP01_0930800","PVP01_0931200","PVP01_0931800","PVP01_0932400","PVP01_0932500","PVP01_0932700","PVP01_0932800","PVP01_0932900","PVP01_0933000","PVP01_0933200","PVP01_0933300","PVP01_0933800","PVP01_0933900","PVP01_0934100","PVP01_0934200","PVP01_0934600","PVP01_0934700","PVP01_0934800","PVP01_0934900","PVP01_0935600","PVP01_0935900","PVP01_0936200","PVP01_0936500","PVP01_0937100","PVP01_0937200","PVP01_0937600","PVP01_0938300","PVP01_0938500","PVP01_0938800","PVP01_0939000","PVP01_0939400","PVP01_0939700","PVP01_0939800","PVP01_0940000","PVP01_0940300","PVP01_0940400","PVP01_0941100","PVP01_0941500","PVP01_0941700","PVP01_0941900","PVP01_0942000","PVP01_0942100","PVP01_0942500","PVP01_0942800","PVP01_0943000","PVP01_0943200","PVP01_0943500","PVP01_0943600","PVP01_0943800","PVP01_0943900","PVP01_0944000","PVP01_0944500","PVP01_0944700","PVP01_0944900","PVP01_0945000","PVP01_0945400","PVP01_0945700","PVP01_0945900","PVP01_0946100","PVP01_0946800","PVP01_0947300","PVP01_0947500","PVP01_0947600","PVP01_0948000","PVP01_0948200","PVP01_0948400","PVP01_0949200","PVP01_0949300","PVP01_1002400","PVP01_1002800","PVP01_1003300","PVP01_1003600","PVP01_1003700","PVP01_1003800","PVP01_1003900","PVP01_1004000","PVP01_1004100","PVP01_1004400","PVP01_1004600","PVP01_1004700","PVP01_1005800","PVP01_1006500","PVP01_1006800","PVP01_1006900","PVP01_1007000","PVP01_1007100","PVP01_1007200","PVP01_1007300","PVP01_1007400","PVP01_1008500","PVP01_1008800","PVP01_1009900","PVP01_1010000","PVP01_1010500","PVP01_1010600","PVP01_1010700","PVP01_1010800","PVP01_1010900","PVP01_1011000","PVP01_1011300","PVP01_1011400","PVP01_1011500","PVP01_1011900","PVP01_1012000","PVP01_1012200","PVP01_1012600","PVP01_1012900","PVP01_1013000","PVP01_1013500","PVP01_1013600","PVP01_1013800","PVP01_1013900","PVP01_1014000","PVP01_1014100","PVP01_1014500","PVP01_1014900","PVP01_1015200","PVP01_1015300","PVP01_1015600","PVP01_1015800","PVP01_1016100","PVP01_1016700","PVP01_1016800","PVP01_1017000","PVP01_1017200","PVP01_1017300","PVP01_1017500","PVP01_1017700","PVP01_1017800","PVP01_1018200","PVP01_1018300","PVP01_1018600","PVP01_1018800","PVP01_1018900","PVP01_1019100","PVP01_1019200","PVP01_1019300","PVP01_1019700","PVP01_1019800","PVP01_1020300","PVP01_1020400","PVP01_1020600","PVP01_1021400","PVP01_1021900","PVP01_1022500","PVP01_1022600","PVP01_1022700","PVP01_1022800","PVP01_1023500","PVP01_1024000","PVP01_1024200","PVP01_1024600","PVP01_1024700","PVP01_1025000","PVP01_1025600","PVP01_1025700","PVP01_1025900","PVP01_1026100","PVP01_1026500","PVP01_1026800","PVP01_1027000","PVP01_1027100","PVP01_1027200","PVP01_1027500","PVP01_1027800","PVP01_1028300","PVP01_1028800","PVP01_1028900","PVP01_1029400","PVP01_1029600","PVP01_1030200","PVP01_1030400","PVP01_1030700","PVP01_1030800","PVP01_1031000","PVP01_1031100","PVP01_1031200","PVP01_1031400","PVP01_1031500","PVP01_1031600","PVP01_1031700","PVP01_1032300","PVP01_1032700","PVP01_1033100","PVP01_1033200","PVP01_1033400","PVP01_1033600","PVP01_1033800","PVP01_1033900","PVP01_1034000","PVP01_1034200","PVP01_1034400","PVP01_1101400","PVP01_1102300","PVP01_1102600","PVP01_1102700","PVP01_1103000","PVP01_1103100","PVP01_1103500","PVP01_1103800","PVP01_1103900","PVP01_1104500","PVP01_1104900","PVP01_1105000","PVP01_1105200","PVP01_1105700","PVP01_1105900","PVP01_1106100","PVP01_1106500","PVP01_1106800","PVP01_1107500","PVP01_1107800","PVP01_1107900","PVP01_1108100","PVP01_1108200","PVP01_1108400","PVP01_1109200","PVP01_1109500","PVP01_1109700","PVP01_1110000","PVP01_1110300","PVP01_1110400","PVP01_1110800","PVP01_1111400","PVP01_1111900","PVP01_1112300","PVP01_1112600","PVP01_1113100","PVP01_1113200","PVP01_1113600","PVP01_1113800","PVP01_1114000","PVP01_1114400","PVP01_1114900","PVP01_1115100","PVP01_1115300","PVP01_1115400","PVP01_1115800","PVP01_1115900","PVP01_1116000","PVP01_1116100","PVP01_1116200","PVP01_1116300","PVP01_1116400","PVP01_1116700","PVP01_1117000","PVP01_1117100","PVP01_1117200","PVP01_1117400","PVP01_1117600","PVP01_1117800","PVP01_1117900","PVP01_1118200","PVP01_1118300","PVP01_1118400","PVP01_1118500","PVP01_1119200","PVP01_1119700","PVP01_1119900","PVP01_1120200","PVP01_1120600","PVP01_1120700","PVP01_1120800","PVP01_1121200","PVP01_1121800","PVP01_1122000","PVP01_1122200","PVP01_1123000","PVP01_1123500","PVP01_1123700","PVP01_1123800","PVP01_1124200","PVP01_1124500","PVP01_1125500","PVP01_1126400","PVP01_1126700","PVP01_1127600","PVP01_1127700","PVP01_1128000","PVP01_1128200","PVP01_1128400","PVP01_1128500","PVP01_1129100","PVP01_1129200","PVP01_1129300","PVP01_1129800","PVP01_1129900","PVP01_1130000","PVP01_1130200","PVP01_1130300","PVP01_1130500","PVP01_1130600","PVP01_1131100","PVP01_1131200","PVP01_1131500","PVP01_1131600","PVP01_1131700","PVP01_1131800","PVP01_1131900","PVP01_1132000","PVP01_1132200","PVP01_1132600","PVP01_1133700","PVP01_1134000","PVP01_1134500","PVP01_1134600","PVP01_1134900","PVP01_1135300","PVP01_1135400","PVP01_1136200","PVP01_1136400","PVP01_1136500","PVP01_1137000","PVP01_1137400","PVP01_1137500","PVP01_1137900","PVP01_1138000","PVP01_1138100","PVP01_1138200","PVP01_1138300","PVP01_1138600","PVP01_1138700","PVP01_1138800","PVP01_1138900","PVP01_1139200","PVP01_1139400","PVP01_1139500","PVP01_1139800","PVP01_1139900","PVP01_1140100","PVP01_1140200","PVP01_1140400","PVP01_1141100","PVP01_1141600","PVP01_1142200","PVP01_1142400","PVP01_1142700","PVP01_1142800","PVP01_1143000","PVP01_1143300","PVP01_1143800","PVP01_1143900","PVP01_1144400","PVP01_1145300","PVP01_1145400","PVP01_1145600","PVP01_1145900","PVP01_1146000","PVP01_1146100","PVP01_1146200","PVP01_1146300","PVP01_1146500","PVP01_1146700","PVP01_1147100","PVP01_1147600","PVP01_1147700","PVP01_1147800","PVP01_1147900","PVP01_1201400","PVP01_1201500","PVP01_1202000","PVP01_1202400","PVP01_1202500","PVP01_1202600","PVP01_1202700","PVP01_1202800","PVP01_1202900","PVP01_1203500","PVP01_1203900","PVP01_1204400","PVP01_1204700","PVP01_1205300","PVP01_1205700","PVP01_1205800","PVP01_1206000","PVP01_1206500","PVP01_1206600","PVP01_1206800","PVP01_1207300","PVP01_1207400","PVP01_1207500","PVP01_1207600","PVP01_1207700","PVP01_1208000","PVP01_1208400","PVP01_1208500","PVP01_1209100","PVP01_1209200","PVP01_1209300","PVP01_1209500","PVP01_1209700","PVP01_1209800","PVP01_1209900","PVP01_1210400","PVP01_1210500","PVP01_1210600","PVP01_1210900","PVP01_1211000","PVP01_1211100","PVP01_1211400","PVP01_1211700","PVP01_1211800","PVP01_1211900","PVP01_1212000","PVP01_1212300","PVP01_1212400","PVP01_1212700","PVP01_1212800","PVP01_1212900","PVP01_1213200","PVP01_1213400","PVP01_1213600","PVP01_1213900","PVP01_1214000","PVP01_1214100","PVP01_1214300","PVP01_1214400","PVP01_1214500","PVP01_1215100","PVP01_1215200","PVP01_1215500","PVP01_1216400","PVP01_1216500","PVP01_1216600","PVP01_1216900","PVP01_1217000","PVP01_1217300","PVP01_1217800","PVP01_1217900","PVP01_1218100","PVP01_1218200","PVP01_1218300","PVP01_1218400","PVP01_1218600","PVP01_1218700","PVP01_1219200","PVP01_1219300","PVP01_1219400","PVP01_1219600","PVP01_1219700","PVP01_1219800","PVP01_1220000","PVP01_1220400","PVP01_1220500","PVP01_1220800","PVP01_1221000","PVP01_1221100","PVP01_1221500","PVP01_1221600","PVP01_1222000","PVP01_1222900","PVP01_1223000","PVP01_1223200","PVP01_1223300","PVP01_1223700","PVP01_1224100","PVP01_1224200","PVP01_1224900","PVP01_1225100","PVP01_1225400","PVP01_1225900","PVP01_1226000","PVP01_1226100","PVP01_1226200","PVP01_1226300","PVP01_1227000","PVP01_1227100","PVP01_1227400","PVP01_1227700","PVP01_1227800","PVP01_1228600","PVP01_1228700","PVP01_1228900","PVP01_1229000","PVP01_1229300","PVP01_1229700","PVP01_1230300","PVP01_1230400","PVP01_1230500","PVP01_1231200","PVP01_1231300","PVP01_1231400","PVP01_1231500","PVP01_1232000","PVP01_1232100","PVP01_1232200","PVP01_1232300","PVP01_1232500","PVP01_1232800","PVP01_1232900","PVP01_1233000","PVP01_1233300","PVP01_1233400","PVP01_1233500","PVP01_1233600","PVP01_1233700","PVP01_1234100","PVP01_1234200","PVP01_1234300","PVP01_1234400","PVP01_1234700","PVP01_1234800","PVP01_1234900","PVP01_1235400","PVP01_1235600","PVP01_1235700","PVP01_1235900","PVP01_1236300","PVP01_1236600","PVP01_1236700","PVP01_1237300","PVP01_1237400","PVP01_1237600","PVP01_1238100","PVP01_1238200","PVP01_1238300","PVP01_1238700","PVP01_1238800","PVP01_1239200","PVP01_1239600","PVP01_1239700","PVP01_1239800","PVP01_1240000","PVP01_1240200","PVP01_1240500","PVP01_1240700","PVP01_1241500","PVP01_1241600","PVP01_1241700","PVP01_1241800","PVP01_1242100","PVP01_1242200","PVP01_1242500","PVP01_1242600","PVP01_1242800","PVP01_1242900","PVP01_1243300","PVP01_1243400","PVP01_1243700","PVP01_1244000","PVP01_1244100","PVP01_1244200","PVP01_1244500","PVP01_1244600","PVP01_1245000","PVP01_1245200","PVP01_1245700","PVP01_1246200","PVP01_1246800","PVP01_1247200","PVP01_1247300","PVP01_1247400","PVP01_1248000","PVP01_1248100","PVP01_1248300","PVP01_1248400","PVP01_1248600","PVP01_1248900","PVP01_1249000","PVP01_1249300","PVP01_1249400","PVP01_1249800","PVP01_1249900","PVP01_1250100","PVP01_1250400","PVP01_1250500","PVP01_1250900","PVP01_1251000","PVP01_1251200","PVP01_1251400","PVP01_1251600","PVP01_1251800","PVP01_1252300","PVP01_1252500","PVP01_1252600","PVP01_1252700","PVP01_1253100","PVP01_1253200","PVP01_1253400","PVP01_1253500","PVP01_1253900","PVP01_1254000","PVP01_1254400","PVP01_1254500","PVP01_1254600","PVP01_1254700","PVP01_1255000","PVP01_1255600","PVP01_1255800","PVP01_1255900","PVP01_1256100","PVP01_1256300","PVP01_1256500","PVP01_1257100","PVP01_1257200","PVP01_1257300","PVP01_1258000","PVP01_1258600","PVP01_1258800","PVP01_1259800","PVP01_1259900","PVP01_1260000","PVP01_1260100","PVP01_1260600","PVP01_1260900","PVP01_1261600","PVP01_1261900","PVP01_1262100","PVP01_1262500","PVP01_1262700","PVP01_1263200","PVP01_1263300","PVP01_1263600","PVP01_1263700","PVP01_1264100","PVP01_1264400","PVP01_1265200","PVP01_1265500","PVP01_1265700","PVP01_1265800","PVP01_1266200","PVP01_1266300","PVP01_1266600","PVP01_1266900","PVP01_1267000","PVP01_1267600","PVP01_1267900","PVP01_1268400","PVP01_1268500","PVP01_1268600","PVP01_1268900","PVP01_1269300","PVP01_1269400","PVP01_1269600","PVP01_1269700","PVP01_1270100","PVP01_1270400","PVP01_1270800","PVP01_1271000","PVP01_1271300","PVP01_1271500","PVP01_1272500","PVP01_1300600","PVP01_1300700","PVP01_1300800","PVP01_1301400","PVP01_1301900","PVP01_1302000","PVP01_1302200","PVP01_1302400","PVP01_1303300","PVP01_1303400","PVP01_1303500","PVP01_1303700","PVP01_1303800","PVP01_1304100","PVP01_1304400","PVP01_1304600","PVP01_1304700","PVP01_1305100","PVP01_1305200","PVP01_1305400","PVP01_1305800","PVP01_1305900","PVP01_1306200","PVP01_1306500","PVP01_1306900","PVP01_1307500","PVP01_1307600","PVP01_1307900","PVP01_1308000","PVP01_1308200","PVP01_1308400","PVP01_1308500","PVP01_1308600","PVP01_1309600","PVP01_1309900","PVP01_1310000","PVP01_1310200","PVP01_1310300","PVP01_1310500","PVP01_1310600","PVP01_1310800","PVP01_1310900","PVP01_1311600","PVP01_1311700","PVP01_1311800","PVP01_1312100","PVP01_1312200","PVP01_1312900","PVP01_1313200","PVP01_1313400","PVP01_1313500","PVP01_1313600","PVP01_1313800","PVP01_1314000","PVP01_1314200","PVP01_1314300","PVP01_1314400","PVP01_1314500","PVP01_1314700","PVP01_1315000","PVP01_1315100","PVP01_1315200","PVP01_1315800","PVP01_1315900","PVP01_1316000","PVP01_1316200","PVP01_1316300","PVP01_1316400","PVP01_1317100","PVP01_1317200","PVP01_1317300","PVP01_1317500","PVP01_1318200","PVP01_1318700","PVP01_1319100","PVP01_1319400","PVP01_1319600","PVP01_1320800","PVP01_1320900","PVP01_1321000","PVP01_1321200","PVP01_1321400","PVP01_1321800","PVP01_1322100","PVP01_1322500","PVP01_1323200","PVP01_1323300","PVP01_1324000","PVP01_1325200","PVP01_1325300","PVP01_1325400","PVP01_1325500","PVP01_1325600","PVP01_1326100","PVP01_1326300","PVP01_1326500","PVP01_1326700","PVP01_1326800","PVP01_1326900","PVP01_1327400","PVP01_1327700","PVP01_1327800","PVP01_1328200","PVP01_1329100","PVP01_1329200","PVP01_1329500","PVP01_1329700","PVP01_1329800","PVP01_1330600","PVP01_1330700","PVP01_1330900","PVP01_1331100","PVP01_1331300","PVP01_1331500","PVP01_1331600","PVP01_1331700","PVP01_1332000","PVP01_1332100","PVP01_1332400","PVP01_1332800","PVP01_1333400","PVP01_1333500","PVP01_1333600","PVP01_1333800","PVP01_1334400","PVP01_1334600","PVP01_1334700","PVP01_1334900","PVP01_1335000","PVP01_1335200","PVP01_1335400","PVP01_1335500","PVP01_1335700","PVP01_1336100","PVP01_1336300","PVP01_1336400","PVP01_1336600","PVP01_1336800","PVP01_1336900","PVP01_1337000","PVP01_1337700","PVP01_1337800","PVP01_1338000","PVP01_1338200","PVP01_1338500","PVP01_1338700","PVP01_1338800","PVP01_1338900","PVP01_1339100","PVP01_1339200","PVP01_1339300","PVP01_1339600","PVP01_1339700","PVP01_1339900","PVP01_1340000","PVP01_1340300","PVP01_1340700","PVP01_1340900","PVP01_1341000","PVP01_1341100","PVP01_1341400","PVP01_1341500","PVP01_1341700","PVP01_1341900","PVP01_1342000","PVP01_1342100","PVP01_1342500","PVP01_1342900","PVP01_1343000","PVP01_1343200","PVP01_1343400","PVP01_1343800","PVP01_1343900","PVP01_1344000","PVP01_1344100","PVP01_1344400","PVP01_1345000","PVP01_1345100","PVP01_1345800","PVP01_1346000","PVP01_1346100","PVP01_1346400","PVP01_1346500","PVP01_1346600","PVP01_1347000","PVP01_1347100","PVP01_1347200","PVP01_1400600","PVP01_1400700","PVP01_1400900","PVP01_1401100","PVP01_1401400","PVP01_1401900","PVP01_1402100","PVP01_1402300","PVP01_1402400","PVP01_1402600","PVP01_1402700","PVP01_1403000","PVP01_1403100","PVP01_1403200","PVP01_1403600","PVP01_1404400","PVP01_1404700","PVP01_1404900","PVP01_1405200","PVP01_1405300","PVP01_1405400","PVP01_1405700","PVP01_1406400","PVP01_1406900","PVP01_1407300","PVP01_1407500","PVP01_1407700","PVP01_1408300","PVP01_1408900","PVP01_1409000","PVP01_1409200","PVP01_1409500","PVP01_1409600","PVP01_1409800","PVP01_1409900","PVP01_1410400","PVP01_1411100","PVP01_1411200","PVP01_1411800","PVP01_1411900","PVP01_1412300","PVP01_1412600","PVP01_1412800","PVP01_1412900","PVP01_1413000","PVP01_1413100","PVP01_1413400","PVP01_1413500","PVP01_1413600","PVP01_1413800","PVP01_1414000","PVP01_1414500","PVP01_1414700","PVP01_1414800","PVP01_1415000","PVP01_1415400","PVP01_1415800","PVP01_1416200","PVP01_1416300","PVP01_1416400","PVP01_1416500","PVP01_1416600","PVP01_1416700","PVP01_1416800","PVP01_1417000","PVP01_1417100","PVP01_1417300","PVP01_1417400","PVP01_1417500","PVP01_1417800","PVP01_1418500","PVP01_1418700","PVP01_1419000","PVP01_1419200","PVP01_1419900","PVP01_1420600","PVP01_1420800","PVP01_1421500","PVP01_1421700","PVP01_1421900","PVP01_1422000","PVP01_1422300","PVP01_1422400","PVP01_1422800","PVP01_1423000","PVP01_1423100","PVP01_1423400","PVP01_1423500","PVP01_1423600","PVP01_1424300","PVP01_1424600","PVP01_1424900","PVP01_1425000","PVP01_1425500","PVP01_1425600","PVP01_1425700","PVP01_1425900","PVP01_1426000","PVP01_1426300","PVP01_1426500","PVP01_1426900","PVP01_1427200","PVP01_1427300","PVP01_1427400","PVP01_1427500","PVP01_1427600","PVP01_1427900","PVP01_1428200","PVP01_1428600","PVP01_1428700","PVP01_1428800","PVP01_1429100","PVP01_1429400","PVP01_1429500","PVP01_1429600","PVP01_1429800","PVP01_1430200","PVP01_1430500","PVP01_1430900","PVP01_1431000","PVP01_1431400","PVP01_1432200","PVP01_1432300","PVP01_1432900","PVP01_1433100","PVP01_1433200","PVP01_1433400","PVP01_1433500","PVP01_1433700","PVP01_1434100","PVP01_1434700","PVP01_1435000","PVP01_1435400","PVP01_1435500","PVP01_1435700","PVP01_1436100","PVP01_1436200","PVP01_1436500","PVP01_1436800","PVP01_1436900","PVP01_1437000","PVP01_1437300","PVP01_1437400","PVP01_1437800","PVP01_1437900","PVP01_1438000","PVP01_1438600","PVP01_1438700","PVP01_1439100","PVP01_1439200","PVP01_1439500","PVP01_1439600","PVP01_1439900","PVP01_1440200","PVP01_1440400","PVP01_1440500","PVP01_1441000","PVP01_1441100","PVP01_1441500","PVP01_1441600","PVP01_1441900","PVP01_1442100","PVP01_1442300","PVP01_1442400","PVP01_1442600","PVP01_1442700","PVP01_1443100","PVP01_1443300","PVP01_1443400","PVP01_1443500","PVP01_1443700","PVP01_1444600","PVP01_1444700","PVP01_1445400","PVP01_1445900","PVP01_1446500","PVP01_1446800","PVP01_1447000","PVP01_1447100","PVP01_1447700","PVP01_1448500","PVP01_1449000","PVP01_1449300","PVP01_1449400","PVP01_1449600","PVP01_1449800","PVP01_1449900","PVP01_1450100","PVP01_1450300","PVP01_1450600","PVP01_1450700","PVP01_1450800","PVP01_1451900","PVP01_1452000","PVP01_1452200","PVP01_1452600","PVP01_1453100","PVP01_1453400","PVP01_1453500","PVP01_1453600","PVP01_1453700","PVP01_1453900","PVP01_1454300","PVP01_1454700","PVP01_1455000","PVP01_1455200","PVP01_1455300","PVP01_1455600","PVP01_1455800","PVP01_1456100","PVP01_1456200","PVP01_1456300","PVP01_1456400","PVP01_1456600","PVP01_1456800","PVP01_1457100","PVP01_1457300","PVP01_1457600","PVP01_1457800","PVP01_1458000","PVP01_1458100","PVP01_1458200","PVP01_1458400","PVP01_1458500","PVP01_1458600","PVP01_1458900","PVP01_1459000","PVP01_1459800","PVP01_1459900","PVP01_1460000","PVP01_1460100","PVP01_1460200","PVP01_1461200","PVP01_1461300","PVP01_1461500","PVP01_1461600","PVP01_1461800","PVP01_1462700","PVP01_1462800","PVP01_1463000","PVP01_1463100","PVP01_1463200","PVP01_1463400","PVP01_1463900","PVP01_1464000","PVP01_1464200","PVP01_1464300","PVP01_1465200","PVP01_1465300","PVP01_1465400","PVP01_1466000","PVP01_1466200","PVP01_1466400","PVP01_1466800","PVP01_1466900","PVP01_1467200","PVP01_1467400","PVP01_1467500","PVP01_1467700","PVP01_1467900","PVP01_1468000","PVP01_1468200","PVP01_1468300","PVP01_1468700","PVP01_1468800","PVP01_1469200","PVP01_1469400","PVP01_1469500","PVP01_1469600","PVP01_1469700","PVP01_1469800","PVP01_1470100","PVP01_1470700","PVP01_1471100","PVP01_1471400","PVP01_1471500","PVP01_1472100","PVP01_API02600","PVP01_API02800","PVP01_API03500","PVP01_API04300","PVP01_API04600","PVP01_MIT02700","PVP01_MIT03300","PVP01_MIT03400"],"Row.names":["PVP01_0000030","PVP01_0000040","PVP01_0000050","PVP01_0000080","PVP01_0000120","PVP01_0000130","PVP01_0000140","PVP01_0000150","PVP01_0000170","PVP01_0000180","PVP01_0000190","PVP01_0000210","PVP01_0000300","PVP01_0000480","PVP01_0000670","PVP01_0000740","PVP01_0000790","PVP01_0001030","PVP01_0001090","PVP01_0001310","PVP01_0001480","PVP01_0001550","PVP01_0001640","PVP01_0001660","PVP01_0001670","PVP01_0001900","PVP01_0001910","PVP01_0001940","PVP01_0001970","PVP01_0002020","PVP01_0002050","PVP01_0002120","PVP01_0002200","PVP01_0002210","PVP01_0002330","PVP01_0002370","PVP01_0002390","PVP01_0002470","PVP01_0002530","PVP01_0002540","PVP01_0002750","PVP01_0002810","PVP01_0002820","PVP01_0003010","PVP01_0003050","PVP01_0003150","PVP01_0003210","PVP01_0003350","PVP01_0003360","PVP01_0003370","PVP01_0003380","PVP01_0003410","PVP01_0003420","PVP01_0003440","PVP01_0003530","PVP01_0003640","PVP01_0003690","PVP01_0003730","PVP01_0003800","PVP01_0003990","PVP01_0004160","PVP01_0004170","PVP01_0004180","PVP01_0004190","PVP01_0004250","PVP01_0004260","PVP01_0004270","PVP01_0004280","PVP01_0004300","PVP01_0004310","PVP01_0004320","PVP01_0004350","PVP01_0004370","PVP01_0004380","PVP01_0004390","PVP01_0004410","PVP01_0004640","PVP01_0004670","PVP01_0004910","PVP01_0004920","PVP01_0004950","PVP01_0004960","PVP01_0005010","PVP01_0005100","PVP01_0005160","PVP01_0005210","PVP01_0005300","PVP01_0005310","PVP01_0005420","PVP01_0005430","PVP01_0005550","PVP01_0005640","PVP01_0005710","PVP01_0005740","PVP01_0005800","PVP01_0005820","PVP01_0005870","PVP01_0005910","PVP01_0006000","PVP01_0006040","PVP01_0006080","PVP01_0006260","PVP01_0006650","PVP01_0006660","PVP01_0006670","PVP01_0006680","PVP01_0006700","PVP01_0006860","PVP01_0006890","PVP01_0007080","PVP01_0007160","PVP01_0007240","PVP01_0007290","PVP01_0007320","PVP01_0007340","PVP01_0007610","PVP01_0007620","PVP01_0007690","PVP01_0007700","PVP01_0007720","PVP01_0007850","PVP01_0007900","PVP01_0007910","PVP01_0007920","PVP01_0008190","PVP01_0008430","PVP01_0008480","PVP01_0008490","PVP01_0008600","PVP01_0008700","PVP01_0008770","PVP01_0008830","PVP01_0008990","PVP01_0009010","PVP01_0009170","PVP01_0009180","PVP01_0009210","PVP01_0009370","PVP01_0009410","PVP01_0009470","PVP01_0009540","PVP01_0009620","PVP01_0009710","PVP01_0009730","PVP01_0009740","PVP01_0009760","PVP01_0009770","PVP01_0009820","PVP01_0009850","PVP01_0009860","PVP01_0009870","PVP01_0009900","PVP01_0009940","PVP01_0009980","PVP01_0010020","PVP01_0010030","PVP01_0010070","PVP01_0010100","PVP01_0010120","PVP01_0010140","PVP01_0010170","PVP01_0010180","PVP01_0010200","PVP01_0010220","PVP01_0010230","PVP01_0010240","PVP01_0010340","PVP01_0010380","PVP01_0010440","PVP01_0010470","PVP01_0010500","PVP01_0010540","PVP01_0010570","PVP01_0010600","PVP01_0010610","PVP01_0010640","PVP01_0010650","PVP01_0010770","PVP01_0010870","PVP01_0010900","PVP01_0010910","PVP01_0010960","PVP01_0010970","PVP01_0010990","PVP01_0011030","PVP01_0011070","PVP01_0102100","PVP01_0102400","PVP01_0102500","PVP01_0102700","PVP01_0102800","PVP01_0103100","PVP01_0103300","PVP01_0103400","PVP01_0103500","PVP01_0103600","PVP01_0103800","PVP01_0103900","PVP01_0104100","PVP01_0104200","PVP01_0104600","PVP01_0104800","PVP01_0104900","PVP01_0105000","PVP01_0105100","PVP01_0105200","PVP01_0105300","PVP01_0105400","PVP01_0105500","PVP01_0105600","PVP01_0105700","PVP01_0105800","PVP01_0106000","PVP01_0106200","PVP01_0106300","PVP01_0106400","PVP01_0106500","PVP01_0106800","PVP01_0106900","PVP01_0107400","PVP01_0107600","PVP01_0107800","PVP01_0107900","PVP01_0108000","PVP01_0108100","PVP01_0108200","PVP01_0108300","PVP01_0108400","PVP01_0108600","PVP01_0108800","PVP01_0108900","PVP01_0109000","PVP01_0109100","PVP01_0109200","PVP01_0109600","PVP01_0109800","PVP01_0110100","PVP01_0110200","PVP01_0110400","PVP01_0110500","PVP01_0110700","PVP01_0110800","PVP01_0110900","PVP01_0111100","PVP01_0111200","PVP01_0111300","PVP01_0111600","PVP01_0111700","PVP01_0111900","PVP01_0112000","PVP01_0112100","PVP01_0112200","PVP01_0112300","PVP01_0112500","PVP01_0112700","PVP01_0112800","PVP01_0112900","PVP01_0113000","PVP01_0113100","PVP01_0113200","PVP01_0113300","PVP01_0113400","PVP01_0113500","PVP01_0113700","PVP01_0114100","PVP01_0114200","PVP01_0114600","PVP01_0114700","PVP01_0114800","PVP01_0115000","PVP01_0115100","PVP01_0115200","PVP01_0115300","PVP01_0115400","PVP01_0115500","PVP01_0116100","PVP01_0116500","PVP01_0116600","PVP01_0116700","PVP01_0116800","PVP01_0116900","PVP01_0117000","PVP01_0117500","PVP01_0117600","PVP01_0117700","PVP01_0117800","PVP01_0118000","PVP01_0118100","PVP01_0118500","PVP01_0118600","PVP01_0118700","PVP01_0118800","PVP01_0118900","PVP01_0119300","PVP01_0119400","PVP01_0119500","PVP01_0119600","PVP01_0119700","PVP01_0119800","PVP01_0119900","PVP01_0120000","PVP01_0120100","PVP01_0120500","PVP01_0120800","PVP01_0121000","PVP01_0121700","PVP01_0200200","PVP01_0200700","PVP01_0201000","PVP01_0201100","PVP01_0201300","PVP01_0201500","PVP01_0201600","PVP01_0201700","PVP01_0201800","PVP01_0201900","PVP01_0202000","PVP01_0202100","PVP01_0202200","PVP01_0202300","PVP01_0202400","PVP01_0202600","PVP01_0203000","PVP01_0203100","PVP01_0203300","PVP01_0203400","PVP01_0203500","PVP01_0203800","PVP01_0203900","PVP01_0204000","PVP01_0204100","PVP01_0204800","PVP01_0204900","PVP01_0205000","PVP01_0205100","PVP01_0205200","PVP01_0205300","PVP01_0205500","PVP01_0205600","PVP01_0205800","PVP01_0206100","PVP01_0206300","PVP01_0206400","PVP01_0206500","PVP01_0206600","PVP01_0206800","PVP01_0207000","PVP01_0207100","PVP01_0207200","PVP01_0207400","PVP01_0207500","PVP01_0207600","PVP01_0207700","PVP01_0207800","PVP01_0208100","PVP01_0208200","PVP01_0208300","PVP01_0208400","PVP01_0208500","PVP01_0208600","PVP01_0208700","PVP01_0208800","PVP01_0209000","PVP01_0209100","PVP01_0209200","PVP01_0209300","PVP01_0209400","PVP01_0209500","PVP01_0209600","PVP01_0209700","PVP01_0210000","PVP01_0210100","PVP01_0210200","PVP01_0210300","PVP01_0210500","PVP01_0210800","PVP01_0211000","PVP01_0211100","PVP01_0211200","PVP01_0211300","PVP01_0211400","PVP01_0211500","PVP01_0211600","PVP01_0211700","PVP01_0211800","PVP01_0212400","PVP01_0212500","PVP01_0212600","PVP01_0212700","PVP01_0212800","PVP01_0213000","PVP01_0213300","PVP01_0213400","PVP01_0213600","PVP01_0213700","PVP01_0214000","PVP01_0214100","PVP01_0214200","PVP01_0214800","PVP01_0214900","PVP01_0215000","PVP01_0215300","PVP01_0215400","PVP01_0215500","PVP01_0215700","PVP01_0215800","PVP01_0215900","PVP01_0216000","PVP01_0216100","PVP01_0216500","PVP01_0216800","PVP01_0216900","PVP01_0217000","PVP01_0217100","PVP01_0217200","PVP01_0217300","PVP01_0217400","PVP01_0217500","PVP01_0218200","PVP01_0218300","PVP01_0218900","PVP01_0219400","PVP01_0220200","PVP01_0221100","PVP01_0221300","PVP01_0221700","PVP01_0300100","PVP01_0300600","PVP01_0300700","PVP01_0301100","PVP01_0301200","PVP01_0301500","PVP01_0301900","PVP01_0302000","PVP01_0302100","PVP01_0302300","PVP01_0302400","PVP01_0302500","PVP01_0302800","PVP01_0303100","PVP01_0303500","PVP01_0304500","PVP01_0304600","PVP01_0304800","PVP01_0304900","PVP01_0305000","PVP01_0305200","PVP01_0305300","PVP01_0305500","PVP01_0305700","PVP01_0305800","PVP01_0306100","PVP01_0306300","PVP01_0306400","PVP01_0306500","PVP01_0306700","PVP01_0306800","PVP01_0306900","PVP01_0307000","PVP01_0307100","PVP01_0307300","PVP01_0307400","PVP01_0307500","PVP01_0307600","PVP01_0307700","PVP01_0307800","PVP01_0308100","PVP01_0308200","PVP01_0308800","PVP01_0309100","PVP01_0309200","PVP01_0309300","PVP01_0309700","PVP01_0309800","PVP01_0309900","PVP01_0310000","PVP01_0310200","PVP01_0310300","PVP01_0310400","PVP01_0310600","PVP01_0311000","PVP01_0311100","PVP01_0311200","PVP01_0311300","PVP01_0311500","PVP01_0311600","PVP01_0311700","PVP01_0312000","PVP01_0312100","PVP01_0312200","PVP01_0312400","PVP01_0312500","PVP01_0312700","PVP01_0312800","PVP01_0313400","PVP01_0313500","PVP01_0314100","PVP01_0314200","PVP01_0314900","PVP01_0315000","PVP01_0315100","PVP01_0315200","PVP01_0315300","PVP01_0315500","PVP01_0315600","PVP01_0315700","PVP01_0315900","PVP01_0316000","PVP01_0316200","PVP01_0316300","PVP01_0316400","PVP01_0316500","PVP01_0316700","PVP01_0316800","PVP01_0317100","PVP01_0317200","PVP01_0317400","PVP01_0317900","PVP01_0318000","PVP01_0318100","PVP01_0318200","PVP01_0318400","PVP01_0318900","PVP01_0319200","PVP01_0319300","PVP01_0319400","PVP01_0319500","PVP01_0319600","PVP01_0319700","PVP01_0319800","PVP01_0320200","PVP01_0320400","PVP01_0320500","PVP01_0320700","PVP01_0320900","PVP01_0400900","PVP01_0403600","PVP01_0403700","PVP01_0403900","PVP01_0404000","PVP01_0404100","PVP01_0404400","PVP01_0404500","PVP01_0404600","PVP01_0404700","PVP01_0404800","PVP01_0404900","PVP01_0405100","PVP01_0405200","PVP01_0405300","PVP01_0405500","PVP01_0405600","PVP01_0405700","PVP01_0405900","PVP01_0406500","PVP01_0406600","PVP01_0406700","PVP01_0406900","PVP01_0407000","PVP01_0407100","PVP01_0407600","PVP01_0407700","PVP01_0407800","PVP01_0407900","PVP01_0408200","PVP01_0408500","PVP01_0408600","PVP01_0408900","PVP01_0409000","PVP01_0409100","PVP01_0409200","PVP01_0409300","PVP01_0409700","PVP01_0409900","PVP01_0410000","PVP01_0410400","PVP01_0410600","PVP01_0410800","PVP01_0411400","PVP01_0411500","PVP01_0411600","PVP01_0411700","PVP01_0411800","PVP01_0411900","PVP01_0412100","PVP01_0412200","PVP01_0412600","PVP01_0412700","PVP01_0412800","PVP01_0412900","PVP01_0413000","PVP01_0413100","PVP01_0413200","PVP01_0413300","PVP01_0413500","PVP01_0414000","PVP01_0414200","PVP01_0414400","PVP01_0414600","PVP01_0414900","PVP01_0415200","PVP01_0415400","PVP01_0415500","PVP01_0415600","PVP01_0416000","PVP01_0416200","PVP01_0416300","PVP01_0416600","PVP01_0416900","PVP01_0417100","PVP01_0417500","PVP01_0417800","PVP01_0417900","PVP01_0418100","PVP01_0418300","PVP01_0419000","PVP01_0419100","PVP01_0419200","PVP01_0419300","PVP01_0419400","PVP01_0419500","PVP01_0419600","PVP01_0419700","PVP01_0420000","PVP01_0420100","PVP01_0420200","PVP01_0420300","PVP01_0420500","PVP01_0420600","PVP01_0420700","PVP01_0420900","PVP01_0421000","PVP01_0421500","PVP01_0421700","PVP01_0421800","PVP01_0421900","PVP01_0422100","PVP01_0422200","PVP01_0422300","PVP01_0422400","PVP01_0422500","PVP01_0422600","PVP01_0422700","PVP01_0422900","PVP01_0423100","PVP01_0423600","PVP01_0423800","PVP01_0424300","PVP01_0424400","PVP01_0500700","PVP01_0502300","PVP01_0502400","PVP01_0502900","PVP01_0503400","PVP01_0504200","PVP01_0504300","PVP01_0505000","PVP01_0505100","PVP01_0505200","PVP01_0505400","PVP01_0505500","PVP01_0505800","PVP01_0506000","PVP01_0506400","PVP01_0506600","PVP01_0506700","PVP01_0506800","PVP01_0506900","PVP01_0507300","PVP01_0507500","PVP01_0507600","PVP01_0507700","PVP01_0507800","PVP01_0508000","PVP01_0508200","PVP01_0508300","PVP01_0508500","PVP01_0508600","PVP01_0508700","PVP01_0508900","PVP01_0509000","PVP01_0509100","PVP01_0509200","PVP01_0509400","PVP01_0509600","PVP01_0509900","PVP01_0510000","PVP01_0510100","PVP01_0510200","PVP01_0510300","PVP01_0510400","PVP01_0510700","PVP01_0510800","PVP01_0510900","PVP01_0511100","PVP01_0511200","PVP01_0511300","PVP01_0511500","PVP01_0511700","PVP01_0511800","PVP01_0512000","PVP01_0512100","PVP01_0512200","PVP01_0512400","PVP01_0512500","PVP01_0512700","PVP01_0512800","PVP01_0512900","PVP01_0513000","PVP01_0513300","PVP01_0513400","PVP01_0513500","PVP01_0513600","PVP01_0513700","PVP01_0513800","PVP01_0513900","PVP01_0514000","PVP01_0514100","PVP01_0514200","PVP01_0514400","PVP01_0514600","PVP01_0514700","PVP01_0514800","PVP01_0514900","PVP01_0515100","PVP01_0515200","PVP01_0515300","PVP01_0515700","PVP01_0515800","PVP01_0515900","PVP01_0516000","PVP01_0516200","PVP01_0516300","PVP01_0516400","PVP01_0516500","PVP01_0516800","PVP01_0516900","PVP01_0517000","PVP01_0517100","PVP01_0517300","PVP01_0517400","PVP01_0517700","PVP01_0517900","PVP01_0518300","PVP01_0518400","PVP01_0518500","PVP01_0518600","PVP01_0518700","PVP01_0519100","PVP01_0519200","PVP01_0519400","PVP01_0519500","PVP01_0519600","PVP01_0520400","PVP01_0520500","PVP01_0520600","PVP01_0520700","PVP01_0520800","PVP01_0521300","PVP01_0521400","PVP01_0521600","PVP01_0521700","PVP01_0521800","PVP01_0522000","PVP01_0522100","PVP01_0522200","PVP01_0522300","PVP01_0522500","PVP01_0522600","PVP01_0522700","PVP01_0522800","PVP01_0522900","PVP01_0523000","PVP01_0523300","PVP01_0523700","PVP01_0524000","PVP01_0524200","PVP01_0524500","PVP01_0524600","PVP01_0524700","PVP01_0524900","PVP01_0525000","PVP01_0525200","PVP01_0525300","PVP01_0525400","PVP01_0525500","PVP01_0525700","PVP01_0525800","PVP01_0526000","PVP01_0526100","PVP01_0526400","PVP01_0526900","PVP01_0527300","PVP01_0527400","PVP01_0527500","PVP01_0527700","PVP01_0527800","PVP01_0527900","PVP01_0528000","PVP01_0528100","PVP01_0528200","PVP01_0528300","PVP01_0528400","PVP01_0528500","PVP01_0528600","PVP01_0528700","PVP01_0528900","PVP01_0529000","PVP01_0529100","PVP01_0529200","PVP01_0529600","PVP01_0529700","PVP01_0529800","PVP01_0529900","PVP01_0530000","PVP01_0530100","PVP01_0530400","PVP01_0530500","PVP01_0530700","PVP01_0530800","PVP01_0531100","PVP01_0531200","PVP01_0531500","PVP01_0531700","PVP01_0531800","PVP01_0532500","PVP01_0532600","PVP01_0532700","PVP01_0532900","PVP01_0533000","PVP01_0533100","PVP01_0533300","PVP01_0534000","PVP01_0534100","PVP01_0534200","PVP01_0534600","PVP01_0534800","PVP01_0534900","PVP01_0535400","PVP01_0600100","PVP01_0600300","PVP01_0600400","PVP01_0600500","PVP01_0600600","PVP01_0600700","PVP01_0600800","PVP01_0600900","PVP01_0601100","PVP01_0601200","PVP01_0601500","PVP01_0601600","PVP01_0602300","PVP01_0602600","PVP01_0603000","PVP01_0603100","PVP01_0603400","PVP01_0603500","PVP01_0603600","PVP01_0603700","PVP01_0603900","PVP01_0604000","PVP01_0604100","PVP01_0604200","PVP01_0604400","PVP01_0604700","PVP01_0604800","PVP01_0604900","PVP01_0605000","PVP01_0605100","PVP01_0605300","PVP01_0605400","PVP01_0605500","PVP01_0606100","PVP01_0606200","PVP01_0606400","PVP01_0606500","PVP01_0606600","PVP01_0606800","PVP01_0606900","PVP01_0607200","PVP01_0607600","PVP01_0608000","PVP01_0608100","PVP01_0608200","PVP01_0608300","PVP01_0608400","PVP01_0608500","PVP01_0608600","PVP01_0608700","PVP01_0608800","PVP01_0608900","PVP01_0609200","PVP01_0609400","PVP01_0609600","PVP01_0609700","PVP01_0609800","PVP01_0609900","PVP01_0610100","PVP01_0610200","PVP01_0610400","PVP01_0610500","PVP01_0610600","PVP01_0610700","PVP01_0610800","PVP01_0610900","PVP01_0611100","PVP01_0611300","PVP01_0612000","PVP01_0612100","PVP01_0612600","PVP01_0612700","PVP01_0613000","PVP01_0613100","PVP01_0613200","PVP01_0613400","PVP01_0613500","PVP01_0613700","PVP01_0613800","PVP01_0613900","PVP01_0614000","PVP01_0614200","PVP01_0614300","PVP01_0614400","PVP01_0614600","PVP01_0614700","PVP01_0614900","PVP01_0615100","PVP01_0615500","PVP01_0615700","PVP01_0615800","PVP01_0615900","PVP01_0616000","PVP01_0616100","PVP01_0616200","PVP01_0616400","PVP01_0616500","PVP01_0616600","PVP01_0616800","PVP01_0616900","PVP01_0617000","PVP01_0617100","PVP01_0617400","PVP01_0617500","PVP01_0617600","PVP01_0617900","PVP01_0618100","PVP01_0618300","PVP01_0618600","PVP01_0618700","PVP01_0618900","PVP01_0619000","PVP01_0619100","PVP01_0619200","PVP01_0619400","PVP01_0619500","PVP01_0619600","PVP01_0619700","PVP01_0619800","PVP01_0619900","PVP01_0620000","PVP01_0620100","PVP01_0620200","PVP01_0620900","PVP01_0621000","PVP01_0621100","PVP01_0621300","PVP01_0621800","PVP01_0622100","PVP01_0622200","PVP01_0622700","PVP01_0623100","PVP01_0623200","PVP01_0623300","PVP01_0623400","PVP01_0623600","PVP01_0623900","PVP01_0624000","PVP01_0624100","PVP01_0624300","PVP01_0624400","PVP01_0700200","PVP01_0700700","PVP01_0700800","PVP01_0701000","PVP01_0701400","PVP01_0701500","PVP01_0701600","PVP01_0701700","PVP01_0702200","PVP01_0702400","PVP01_0702500","PVP01_0702600","PVP01_0702700","PVP01_0703000","PVP01_0703200","PVP01_0703500","PVP01_0703600","PVP01_0703700","PVP01_0704000","PVP01_0704100","PVP01_0704400","PVP01_0704500","PVP01_0704600","PVP01_0704700","PVP01_0704800","PVP01_0704900","PVP01_0705000","PVP01_0705100","PVP01_0705200","PVP01_0705500","PVP01_0705600","PVP01_0705700","PVP01_0705800","PVP01_0705900","PVP01_0706000","PVP01_0706100","PVP01_0706200","PVP01_0706300","PVP01_0706400","PVP01_0706500","PVP01_0706600","PVP01_0706700","PVP01_0707000","PVP01_0707100","PVP01_0707200","PVP01_0707300","PVP01_0707500","PVP01_0707800","PVP01_0708000","PVP01_0708100","PVP01_0708200","PVP01_0708300","PVP01_0708400","PVP01_0708500","PVP01_0708800","PVP01_0708900","PVP01_0709000","PVP01_0709100","PVP01_0709200","PVP01_0709300","PVP01_0709500","PVP01_0709700","PVP01_0709900","PVP01_0710000","PVP01_0710200","PVP01_0710300","PVP01_0710700","PVP01_0711100","PVP01_0711200","PVP01_0711400","PVP01_0711500","PVP01_0711800","PVP01_0711900","PVP01_0712200","PVP01_0712300","PVP01_0712400","PVP01_0712700","PVP01_0713000","PVP01_0713100","PVP01_0713200","PVP01_0713500","PVP01_0713600","PVP01_0713800","PVP01_0713900","PVP01_0714000","PVP01_0714200","PVP01_0714300","PVP01_0714400","PVP01_0714600","PVP01_0714800","PVP01_0714900","PVP01_0715000","PVP01_0715100","PVP01_0715200","PVP01_0715500","PVP01_0715600","PVP01_0715800","PVP01_0716000","PVP01_0716100","PVP01_0716200","PVP01_0716600","PVP01_0716800","PVP01_0717000","PVP01_0717200","PVP01_0717400","PVP01_0717600","PVP01_0717800","PVP01_0718000","PVP01_0718100","PVP01_0718200","PVP01_0718600","PVP01_0718700","PVP01_0719000","PVP01_0719300","PVP01_0719400","PVP01_0719500","PVP01_0719700","PVP01_0719800","PVP01_0719900","PVP01_0720000","PVP01_0720100","PVP01_0720200","PVP01_0720300","PVP01_0720400","PVP01_0720600","PVP01_0720700","PVP01_0720800","PVP01_0720900","PVP01_0721100","PVP01_0721300","PVP01_0721400","PVP01_0721500","PVP01_0721800","PVP01_0721900","PVP01_0722100","PVP01_0722200","PVP01_0722300","PVP01_0722400","PVP01_0722500","PVP01_0722600","PVP01_0722700","PVP01_0722800","PVP01_0722900","PVP01_0723000","PVP01_0723100","PVP01_0723200","PVP01_0723300","PVP01_0723400","PVP01_0723500","PVP01_0723700","PVP01_0723800","PVP01_0724000","PVP01_0724100","PVP01_0724300","PVP01_0724500","PVP01_0724600","PVP01_0724700","PVP01_0724800","PVP01_0724900","PVP01_0725000","PVP01_0725100","PVP01_0725300","PVP01_0725400","PVP01_0725700","PVP01_0726000","PVP01_0726700","PVP01_0726900","PVP01_0727000","PVP01_0727100","PVP01_0727200","PVP01_0727300","PVP01_0727400","PVP01_0727500","PVP01_0727600","PVP01_0727700","PVP01_0727800","PVP01_0728100","PVP01_0728200","PVP01_0728300","PVP01_0729100","PVP01_0729200","PVP01_0729300","PVP01_0729400","PVP01_0729500","PVP01_0729700","PVP01_0730000","PVP01_0730100","PVP01_0730200","PVP01_0730300","PVP01_0730400","PVP01_0730600","PVP01_0730800","PVP01_0731100","PVP01_0731200","PVP01_0731300","PVP01_0731500","PVP01_0731600","PVP01_0731700","PVP01_0731800","PVP01_0731900","PVP01_0732000","PVP01_0732100","PVP01_0732200","PVP01_0732300","PVP01_0732600","PVP01_0732800","PVP01_0732900","PVP01_0733000","PVP01_0733100","PVP01_0733200","PVP01_0733500","PVP01_0733700","PVP01_0733900","PVP01_0734100","PVP01_0734200","PVP01_0734400","PVP01_0734800","PVP01_0734900","PVP01_0735000","PVP01_0735100","PVP01_0735300","PVP01_0735800","PVP01_0735900","PVP01_0800100","PVP01_0800200","PVP01_0800300","PVP01_0800400","PVP01_0800500","PVP01_0800600","PVP01_0800900","PVP01_0801400","PVP01_0801700","PVP01_0801800","PVP01_0802100","PVP01_0802300","PVP01_0802400","PVP01_0802500","PVP01_0802900","PVP01_0803000","PVP01_0803100","PVP01_0803300","PVP01_0803500","PVP01_0803600","PVP01_0803700","PVP01_0803900","PVP01_0804000","PVP01_0804100","PVP01_0804200","PVP01_0804400","PVP01_0804500","PVP01_0804600","PVP01_0804700","PVP01_0804800","PVP01_0804900","PVP01_0805000","PVP01_0805200","PVP01_0805300","PVP01_0805400","PVP01_0805600","PVP01_0805700","PVP01_0805800","PVP01_0806000","PVP01_0806100","PVP01_0806200","PVP01_0806300","PVP01_0806400","PVP01_0806500","PVP01_0806600","PVP01_0806700","PVP01_0806800","PVP01_0807000","PVP01_0807200","PVP01_0807300","PVP01_0807500","PVP01_0807600","PVP01_0807800","PVP01_0808000","PVP01_0808400","PVP01_0808500","PVP01_0808600","PVP01_0808800","PVP01_0809100","PVP01_0809200","PVP01_0809300","PVP01_0809400","PVP01_0809600","PVP01_0809700","PVP01_0809900","PVP01_0810000","PVP01_0810100","PVP01_0810300","PVP01_0810400","PVP01_0810500","PVP01_0810600","PVP01_0811000","PVP01_0811100","PVP01_0811200","PVP01_0811300","PVP01_0811500","PVP01_0811700","PVP01_0811800","PVP01_0811900","PVP01_0812000","PVP01_0812100","PVP01_0812300","PVP01_0812600","PVP01_0812700","PVP01_0812800","PVP01_0813000","PVP01_0813200","PVP01_0813300","PVP01_0813600","PVP01_0813700","PVP01_0813800","PVP01_0814100","PVP01_0814400","PVP01_0814700","PVP01_0814800","PVP01_0815100","PVP01_0815300","PVP01_0815400","PVP01_0815600","PVP01_0815700","PVP01_0816200","PVP01_0816700","PVP01_0817100","PVP01_0817400","PVP01_0818000","PVP01_0818100","PVP01_0818300","PVP01_0818500","PVP01_0818600","PVP01_0819000","PVP01_0819200","PVP01_0819700","PVP01_0820000","PVP01_0820100","PVP01_0820300","PVP01_0820500","PVP01_0820600","PVP01_0820700","PVP01_0820800","PVP01_0820900","PVP01_0821000","PVP01_0821100","PVP01_0821400","PVP01_0821500","PVP01_0821600","PVP01_0821700","PVP01_0821800","PVP01_0822100","PVP01_0822200","PVP01_0822300","PVP01_0822600","PVP01_0822700","PVP01_0822900","PVP01_0823100","PVP01_0823200","PVP01_0823400","PVP01_0823500","PVP01_0823700","PVP01_0823900","PVP01_0824000","PVP01_0824300","PVP01_0824500","PVP01_0824700","PVP01_0825000","PVP01_0825100","PVP01_0825200","PVP01_0825400","PVP01_0825700","PVP01_0825800","PVP01_0826000","PVP01_0826100","PVP01_0826200","PVP01_0826300","PVP01_0826400","PVP01_0826500","PVP01_0826600","PVP01_0826700","PVP01_0826900","PVP01_0827100","PVP01_0827200","PVP01_0827700","PVP01_0827800","PVP01_0828000","PVP01_0828100","PVP01_0828200","PVP01_0828300","PVP01_0828600","PVP01_0828700","PVP01_0829000","PVP01_0829100","PVP01_0829300","PVP01_0829700","PVP01_0829900","PVP01_0830200","PVP01_0830300","PVP01_0830400","PVP01_0830500","PVP01_0830700","PVP01_0830900","PVP01_0831000","PVP01_0831100","PVP01_0831200","PVP01_0831400","PVP01_0831700","PVP01_0831900","PVP01_0832000","PVP01_0832100","PVP01_0832200","PVP01_0832400","PVP01_0832500","PVP01_0832600","PVP01_0832800","PVP01_0833000","PVP01_0833500","PVP01_0833600","PVP01_0833700","PVP01_0833800","PVP01_0833900","PVP01_0834000","PVP01_0834200","PVP01_0834300","PVP01_0834400","PVP01_0834500","PVP01_0834600","PVP01_0834700","PVP01_0834900","PVP01_0835000","PVP01_0835300","PVP01_0835500","PVP01_0835700","PVP01_0835800","PVP01_0835900","PVP01_0836000","PVP01_0836100","PVP01_0836300","PVP01_0836400","PVP01_0836500","PVP01_0836600","PVP01_0836700","PVP01_0837100","PVP01_0837200","PVP01_0837300","PVP01_0837500","PVP01_0837600","PVP01_0837800","PVP01_0837900","PVP01_0838000","PVP01_0838200","PVP01_0838900","PVP01_0839100","PVP01_0839200","PVP01_0839700","PVP01_0840700","PVP01_0841000","PVP01_0900700","PVP01_0901200","PVP01_0901300","PVP01_0901800","PVP01_0901900","PVP01_0902000","PVP01_0902100","PVP01_0902400","PVP01_0903400","PVP01_0903500","PVP01_0903600","PVP01_0903700","PVP01_0903800","PVP01_0903900","PVP01_0904300","PVP01_0904600","PVP01_0904700","PVP01_0904900","PVP01_0905100","PVP01_0905700","PVP01_0906000","PVP01_0906300","PVP01_0906500","PVP01_0906700","PVP01_0906800","PVP01_0906900","PVP01_0907000","PVP01_0907100","PVP01_0907200","PVP01_0907300","PVP01_0907500","PVP01_0907600","PVP01_0907700","PVP01_0907800","PVP01_0908000","PVP01_0908300","PVP01_0908400","PVP01_0908500","PVP01_0908600","PVP01_0908900","PVP01_0909000","PVP01_0909100","PVP01_0909300","PVP01_0909500","PVP01_0909600","PVP01_0909700","PVP01_0910000","PVP01_0910200","PVP01_0910900","PVP01_0911000","PVP01_0911300","PVP01_0911500","PVP01_0911600","PVP01_0911700","PVP01_0911800","PVP01_0911900","PVP01_0912100","PVP01_0912200","PVP01_0912400","PVP01_0912900","PVP01_0913000","PVP01_0913100","PVP01_0913400","PVP01_0913500","PVP01_0913600","PVP01_0913700","PVP01_0913900","PVP01_0914000","PVP01_0914200","PVP01_0914300","PVP01_0914400","PVP01_0914500","PVP01_0914600","PVP01_0914800","PVP01_0915000","PVP01_0915100","PVP01_0915200","PVP01_0915300","PVP01_0915400","PVP01_0915600","PVP01_0915800","PVP01_0915900","PVP01_0916300","PVP01_0916700","PVP01_0916800","PVP01_0916900","PVP01_0917000","PVP01_0917100","PVP01_0917200","PVP01_0917300","PVP01_0917400","PVP01_0917600","PVP01_0917700","PVP01_0918000","PVP01_0918100","PVP01_0918200","PVP01_0918300","PVP01_0918400","PVP01_0918500","PVP01_0918700","PVP01_0918800","PVP01_0918900","PVP01_0919000","PVP01_0919100","PVP01_0919300","PVP01_0919500","PVP01_0919600","PVP01_0919700","PVP01_0919900","PVP01_0920100","PVP01_0920300","PVP01_0920400","PVP01_0920500","PVP01_0920700","PVP01_0920800","PVP01_0921100","PVP01_0921400","PVP01_0921500","PVP01_0921700","PVP01_0921800","PVP01_0922000","PVP01_0922100","PVP01_0922300","PVP01_0922400","PVP01_0922800","PVP01_0922900","PVP01_0923000","PVP01_0923100","PVP01_0923300","PVP01_0923800","PVP01_0923900","PVP01_0924000","PVP01_0924100","PVP01_0924300","PVP01_0924400","PVP01_0924500","PVP01_0924700","PVP01_0924800","PVP01_0924900","PVP01_0925100","PVP01_0925200","PVP01_0925300","PVP01_0925400","PVP01_0925600","PVP01_0925700","PVP01_0925800","PVP01_0925900","PVP01_0926300","PVP01_0926400","PVP01_0926600","PVP01_0926900","PVP01_0927100","PVP01_0927400","PVP01_0927600","PVP01_0927700","PVP01_0927800","PVP01_0928000","PVP01_0928100","PVP01_0928300","PVP01_0928400","PVP01_0928800","PVP01_0928900","PVP01_0929000","PVP01_0929100","PVP01_0929200","PVP01_0929300","PVP01_0929400","PVP01_0929700","PVP01_0929800","PVP01_0929900","PVP01_0930000","PVP01_0930100","PVP01_0930200","PVP01_0930300","PVP01_0930400","PVP01_0930500","PVP01_0930600","PVP01_0931300","PVP01_0931400","PVP01_0931500","PVP01_0931600","PVP01_0931700","PVP01_0931900","PVP01_0932000","PVP01_0932100","PVP01_0932200","PVP01_0932300","PVP01_0932600","PVP01_0933100","PVP01_0933400","PVP01_0933500","PVP01_0933600","PVP01_0933700","PVP01_0934000","PVP01_0934300","PVP01_0934400","PVP01_0934500","PVP01_0935000","PVP01_0935100","PVP01_0935200","PVP01_0935300","PVP01_0935400","PVP01_0935500","PVP01_0935700","PVP01_0935800","PVP01_0936000","PVP01_0936100","PVP01_0936300","PVP01_0936400","PVP01_0936600","PVP01_0936700","PVP01_0936800","PVP01_0936900","PVP01_0937000","PVP01_0937300","PVP01_0937400","PVP01_0937500","PVP01_0937700","PVP01_0937800","PVP01_0938000","PVP01_0938100","PVP01_0938200","PVP01_0938400","PVP01_0938600","PVP01_0938700","PVP01_0938900","PVP01_0939100","PVP01_0939200","PVP01_0939300","PVP01_0939500","PVP01_0939600","PVP01_0939900","PVP01_0940100","PVP01_0940200","PVP01_0940500","PVP01_0940600","PVP01_0940700","PVP01_0940800","PVP01_0940900","PVP01_0941000","PVP01_0941200","PVP01_0941300","PVP01_0941400","PVP01_0941600","PVP01_0941800","PVP01_0942200","PVP01_0942300","PVP01_0942400","PVP01_0942600","PVP01_0942700","PVP01_0942900","PVP01_0943100","PVP01_0943300","PVP01_0943400","PVP01_0943700","PVP01_0944100","PVP01_0944200","PVP01_0944300","PVP01_0944400","PVP01_0944600","PVP01_0944800","PVP01_0945100","PVP01_0945200","PVP01_0945300","PVP01_0945500","PVP01_0945600","PVP01_0945800","PVP01_0946000","PVP01_0946200","PVP01_0946300","PVP01_0946400","PVP01_0946500","PVP01_0946600","PVP01_0946700","PVP01_0946900","PVP01_0947000","PVP01_0947100","PVP01_0947200","PVP01_0947400","PVP01_0947700","PVP01_0947800","PVP01_0947900","PVP01_0948100","PVP01_0948300","PVP01_0948500","PVP01_0948600","PVP01_0948700","PVP01_0948800","PVP01_0948900","PVP01_0949000","PVP01_0949400","PVP01_0949600","PVP01_0950000","PVP01_0950200","PVP01_1000700","PVP01_1001900","PVP01_1002500","PVP01_1002600","PVP01_1002700","PVP01_1002900","PVP01_1003000","PVP01_1003100","PVP01_1003200","PVP01_1003400","PVP01_1003500","PVP01_1004200","PVP01_1004500","PVP01_1004800","PVP01_1004900","PVP01_1005000","PVP01_1005100","PVP01_1005200","PVP01_1005300","PVP01_1005400","PVP01_1005500","PVP01_1005600","PVP01_1005700","PVP01_1005900","PVP01_1006200","PVP01_1006300","PVP01_1006400","PVP01_1006600","PVP01_1006700","PVP01_1007500","PVP01_1007600","PVP01_1007700","PVP01_1007800","PVP01_1007900","PVP01_1008000","PVP01_1008100","PVP01_1008200","PVP01_1008300","PVP01_1008400","PVP01_1008600","PVP01_1008700","PVP01_1008900","PVP01_1009000","PVP01_1009100","PVP01_1009200","PVP01_1009300","PVP01_1009400","PVP01_1009500","PVP01_1009600","PVP01_1009700","PVP01_1009800","PVP01_1010100","PVP01_1010200","PVP01_1010300","PVP01_1010400","PVP01_1011100","PVP01_1011200","PVP01_1011600","PVP01_1011700","PVP01_1011800","PVP01_1012100","PVP01_1012300","PVP01_1012400","PVP01_1012500","PVP01_1012700","PVP01_1012800","PVP01_1013100","PVP01_1013200","PVP01_1013300","PVP01_1013400","PVP01_1013700","PVP01_1014200","PVP01_1014300","PVP01_1014400","PVP01_1014600","PVP01_1014700","PVP01_1014800","PVP01_1015000","PVP01_1015100","PVP01_1015400","PVP01_1015500","PVP01_1015700","PVP01_1015900","PVP01_1016000","PVP01_1016200","PVP01_1016300","PVP01_1016400","PVP01_1016500","PVP01_1016600","PVP01_1016900","PVP01_1017100","PVP01_1017400","PVP01_1017600","PVP01_1017900","PVP01_1018000","PVP01_1018100","PVP01_1018400","PVP01_1018500","PVP01_1018700","PVP01_1019000","PVP01_1019400","PVP01_1019600","PVP01_1019900","PVP01_1020000","PVP01_1020100","PVP01_1020200","PVP01_1020500","PVP01_1020700","PVP01_1020800","PVP01_1020900","PVP01_1021000","PVP01_1021100","PVP01_1021200","PVP01_1021300","PVP01_1021500","PVP01_1021600","PVP01_1021700","PVP01_1021800","PVP01_1022000","PVP01_1022100","PVP01_1022200","PVP01_1022300","PVP01_1022400","PVP01_1022900","PVP01_1023000","PVP01_1023200","PVP01_1023300","PVP01_1023600","PVP01_1023700","PVP01_1023800","PVP01_1023900","PVP01_1024100","PVP01_1024300","PVP01_1024400","PVP01_1024500","PVP01_1024800","PVP01_1024900","PVP01_1025100","PVP01_1025200","PVP01_1025300","PVP01_1025400","PVP01_1025500","PVP01_1025800","PVP01_1026000","PVP01_1026200","PVP01_1026300","PVP01_1026400","PVP01_1026600","PVP01_1026700","PVP01_1026900","PVP01_1027300","PVP01_1027400","PVP01_1027600","PVP01_1027700","PVP01_1027900","PVP01_1028000","PVP01_1028100","PVP01_1028200","PVP01_1028400","PVP01_1028500","PVP01_1028600","PVP01_1028700","PVP01_1029000","PVP01_1029100","PVP01_1029200","PVP01_1029300","PVP01_1029500","PVP01_1029700","PVP01_1029800","PVP01_1029900","PVP01_1030000","PVP01_1030100","PVP01_1030300","PVP01_1030500","PVP01_1030600","PVP01_1030900","PVP01_1031300","PVP01_1031800","PVP01_1031900","PVP01_1032000","PVP01_1032100","PVP01_1032200","PVP01_1032400","PVP01_1032500","PVP01_1032600","PVP01_1032800","PVP01_1032900","PVP01_1033000","PVP01_1033300","PVP01_1033500","PVP01_1033700","PVP01_1034100","PVP01_1034300","PVP01_1034500","PVP01_1035200","PVP01_1100400","PVP01_1100500","PVP01_1101000","PVP01_1101200","PVP01_1101300","PVP01_1101500","PVP01_1101600","PVP01_1101700","PVP01_1102400","PVP01_1102500","PVP01_1102800","PVP01_1102900","PVP01_1103200","PVP01_1103300","PVP01_1103400","PVP01_1103600","PVP01_1103700","PVP01_1104000","PVP01_1104100","PVP01_1104200","PVP01_1104300","PVP01_1104400","PVP01_1104600","PVP01_1104700","PVP01_1104800","PVP01_1105100","PVP01_1105300","PVP01_1105400","PVP01_1105500","PVP01_1105600","PVP01_1105800","PVP01_1106000","PVP01_1106200","PVP01_1106300","PVP01_1106400","PVP01_1106600","PVP01_1106700","PVP01_1106900","PVP01_1107000","PVP01_1107100","PVP01_1107200","PVP01_1107300","PVP01_1107400","PVP01_1107600","PVP01_1107700","PVP01_1108000","PVP01_1108300","PVP01_1108500","PVP01_1108600","PVP01_1108700","PVP01_1108800","PVP01_1108900","PVP01_1109000","PVP01_1109100","PVP01_1109300","PVP01_1109400","PVP01_1109600","PVP01_1109800","PVP01_1109900","PVP01_1110100","PVP01_1110200","PVP01_1110500","PVP01_1110600","PVP01_1110700","PVP01_1110900","PVP01_1111000","PVP01_1111100","PVP01_1111200","PVP01_1111300","PVP01_1111500","PVP01_1111600","PVP01_1111700","PVP01_1111800","PVP01_1112000","PVP01_1112100","PVP01_1112200","PVP01_1112400","PVP01_1112500","PVP01_1112700","PVP01_1112800","PVP01_1112900","PVP01_1113000","PVP01_1113300","PVP01_1113400","PVP01_1113500","PVP01_1113700","PVP01_1113900","PVP01_1114100","PVP01_1114200","PVP01_1114300","PVP01_1114500","PVP01_1114600","PVP01_1114700","PVP01_1114800","PVP01_1115000","PVP01_1115200","PVP01_1115500","PVP01_1115600","PVP01_1115700","PVP01_1116600","PVP01_1116800","PVP01_1116900","PVP01_1117300","PVP01_1117500","PVP01_1117700","PVP01_1118000","PVP01_1118100","PVP01_1118600","PVP01_1118700","PVP01_1118800","PVP01_1118900","PVP01_1119000","PVP01_1119100","PVP01_1119300","PVP01_1119400","PVP01_1119500","PVP01_1119600","PVP01_1119800","PVP01_1120000","PVP01_1120100","PVP01_1120300","PVP01_1120400","PVP01_1120500","PVP01_1120900","PVP01_1121000","PVP01_1121100","PVP01_1121300","PVP01_1121400","PVP01_1121500","PVP01_1121600","PVP01_1121700","PVP01_1121900","PVP01_1122100","PVP01_1122300","PVP01_1122400","PVP01_1122500","PVP01_1122600","PVP01_1122700","PVP01_1122800","PVP01_1122900","PVP01_1123100","PVP01_1123200","PVP01_1123300","PVP01_1123400","PVP01_1123600","PVP01_1123900","PVP01_1124000","PVP01_1124100","PVP01_1124300","PVP01_1124400","PVP01_1124600","PVP01_1124700","PVP01_1124800","PVP01_1124900","PVP01_1125000","PVP01_1125100","PVP01_1125200","PVP01_1125300","PVP01_1125400","PVP01_1125600","PVP01_1125700","PVP01_1125800","PVP01_1125900","PVP01_1126000","PVP01_1126100","PVP01_1126200","PVP01_1126300","PVP01_1126500","PVP01_1126600","PVP01_1126800","PVP01_1126900","PVP01_1127000","PVP01_1127100","PVP01_1127200","PVP01_1127300","PVP01_1127400","PVP01_1127500","PVP01_1127800","PVP01_1128100","PVP01_1128300","PVP01_1128800","PVP01_1128900","PVP01_1129000","PVP01_1129400","PVP01_1129500","PVP01_1129600","PVP01_1129700","PVP01_1130100","PVP01_1130400","PVP01_1130700","PVP01_1130800","PVP01_1130900","PVP01_1131000","PVP01_1131300","PVP01_1131400","PVP01_1132100","PVP01_1132300","PVP01_1132400","PVP01_1132500","PVP01_1132700","PVP01_1132800","PVP01_1132900","PVP01_1133000","PVP01_1133100","PVP01_1133200","PVP01_1133300","PVP01_1133400","PVP01_1133500","PVP01_1133600","PVP01_1133800","PVP01_1133900","PVP01_1134100","PVP01_1134200","PVP01_1134300","PVP01_1134400","PVP01_1134700","PVP01_1134800","PVP01_1135000","PVP01_1135100","PVP01_1135200","PVP01_1135500","PVP01_1135600","PVP01_1135700","PVP01_1135800","PVP01_1135900","PVP01_1136000","PVP01_1136100","PVP01_1136300","PVP01_1136600","PVP01_1136700","PVP01_1136800","PVP01_1136900","PVP01_1137100","PVP01_1137200","PVP01_1137300","PVP01_1137600","PVP01_1137700","PVP01_1137800","PVP01_1138400","PVP01_1138500","PVP01_1139000","PVP01_1139100","PVP01_1139300","PVP01_1139600","PVP01_1139700","PVP01_1140000","PVP01_1140300","PVP01_1140500","PVP01_1140600","PVP01_1140700","PVP01_1140800","PVP01_1140900","PVP01_1141000","PVP01_1141200","PVP01_1141300","PVP01_1141400","PVP01_1141500","PVP01_1141700","PVP01_1141800","PVP01_1141900","PVP01_1142000","PVP01_1142100","PVP01_1142300","PVP01_1142500","PVP01_1142600","PVP01_1142900","PVP01_1143100","PVP01_1143200","PVP01_1143400","PVP01_1143500","PVP01_1143600","PVP01_1143700","PVP01_1144000","PVP01_1144100","PVP01_1144200","PVP01_1144300","PVP01_1144500","PVP01_1144600","PVP01_1144700","PVP01_1144800","PVP01_1144900","PVP01_1145000","PVP01_1145100","PVP01_1145200","PVP01_1145500","PVP01_1145700","PVP01_1145800","PVP01_1146400","PVP01_1146600","PVP01_1146800","PVP01_1146900","PVP01_1147000","PVP01_1147200","PVP01_1147300","PVP01_1147400","PVP01_1147500","PVP01_1148200","PVP01_1148700","PVP01_1148800","PVP01_1149000","PVP01_1149100","PVP01_1149400","PVP01_1149500","PVP01_1200200","PVP01_1200800","PVP01_1201100","PVP01_1201300","PVP01_1201600","PVP01_1201700","PVP01_1201800","PVP01_1201900","PVP01_1202100","PVP01_1202200","PVP01_1202300","PVP01_1203000","PVP01_1203100","PVP01_1203200","PVP01_1203300","PVP01_1203600","PVP01_1203700","PVP01_1203800","PVP01_1204000","PVP01_1204100","PVP01_1204200","PVP01_1204300","PVP01_1204500","PVP01_1204600","PVP01_1204800","PVP01_1204900","PVP01_1205000","PVP01_1205100","PVP01_1205200","PVP01_1205400","PVP01_1205500","PVP01_1205600","PVP01_1205900","PVP01_1206100","PVP01_1206200","PVP01_1206300","PVP01_1206400","PVP01_1206700","PVP01_1206900","PVP01_1207000","PVP01_1207100","PVP01_1207200","PVP01_1207800","PVP01_1207900","PVP01_1208100","PVP01_1208200","PVP01_1208300","PVP01_1208600","PVP01_1208700","PVP01_1208800","PVP01_1208900","PVP01_1209000","PVP01_1209400","PVP01_1209600","PVP01_1210000","PVP01_1210100","PVP01_1210200","PVP01_1210300","PVP01_1210700","PVP01_1210800","PVP01_1211200","PVP01_1211300","PVP01_1211500","PVP01_1211600","PVP01_1212100","PVP01_1212200","PVP01_1212500","PVP01_1212600","PVP01_1213000","PVP01_1213100","PVP01_1213300","PVP01_1214200","PVP01_1214600","PVP01_1214700","PVP01_1214800","PVP01_1214900","PVP01_1215000","PVP01_1215300","PVP01_1215400","PVP01_1215800","PVP01_1215900","PVP01_1216000","PVP01_1216100","PVP01_1216200","PVP01_1216300","PVP01_1216700","PVP01_1216800","PVP01_1217200","PVP01_1217400","PVP01_1217500","PVP01_1217600","PVP01_1217700","PVP01_1218000","PVP01_1218500","PVP01_1218800","PVP01_1218900","PVP01_1219000","PVP01_1219100","PVP01_1219500","PVP01_1219900","PVP01_1220100","PVP01_1220200","PVP01_1220300","PVP01_1220600","PVP01_1220700","PVP01_1220900","PVP01_1221200","PVP01_1221300","PVP01_1221400","PVP01_1221700","PVP01_1221800","PVP01_1221900","PVP01_1222100","PVP01_1222200","PVP01_1222300","PVP01_1222400","PVP01_1222500","PVP01_1222600","PVP01_1222700","PVP01_1222800","PVP01_1223100","PVP01_1223400","PVP01_1223500","PVP01_1223600","PVP01_1223800","PVP01_1223900","PVP01_1224000","PVP01_1224300","PVP01_1224400","PVP01_1224500","PVP01_1224600","PVP01_1224700","PVP01_1224800","PVP01_1225000","PVP01_1225200","PVP01_1225300","PVP01_1225500","PVP01_1225600","PVP01_1225700","PVP01_1225800","PVP01_1226400","PVP01_1226600","PVP01_1226700","PVP01_1226800","PVP01_1226900","PVP01_1227200","PVP01_1227300","PVP01_1227500","PVP01_1227600","PVP01_1227900","PVP01_1228000","PVP01_1228100","PVP01_1228200","PVP01_1228300","PVP01_1228400","PVP01_1228500","PVP01_1228800","PVP01_1229100","PVP01_1229200","PVP01_1229400","PVP01_1229500","PVP01_1229600","PVP01_1229800","PVP01_1229900","PVP01_1230000","PVP01_1230100","PVP01_1230200","PVP01_1230600","PVP01_1230700","PVP01_1230800","PVP01_1230900","PVP01_1231000","PVP01_1231100","PVP01_1231600","PVP01_1231700","PVP01_1231800","PVP01_1231900","PVP01_1232400","PVP01_1232600","PVP01_1232700","PVP01_1233100","PVP01_1233200","PVP01_1233800","PVP01_1233900","PVP01_1234000","PVP01_1234500","PVP01_1234600","PVP01_1235000","PVP01_1235100","PVP01_1235200","PVP01_1235300","PVP01_1235500","PVP01_1235800","PVP01_1236000","PVP01_1236100","PVP01_1236200","PVP01_1236400","PVP01_1236500","PVP01_1236800","PVP01_1236900","PVP01_1237000","PVP01_1237100","PVP01_1237200","PVP01_1237500","PVP01_1237700","PVP01_1237800","PVP01_1237900","PVP01_1238000","PVP01_1238400","PVP01_1238500","PVP01_1238600","PVP01_1238900","PVP01_1239000","PVP01_1239100","PVP01_1239300","PVP01_1239400","PVP01_1239500","PVP01_1239900","PVP01_1240100","PVP01_1240300","PVP01_1240400","PVP01_1240600","PVP01_1240800","PVP01_1240900","PVP01_1241000","PVP01_1241100","PVP01_1241200","PVP01_1241300","PVP01_1241400","PVP01_1241900","PVP01_1242000","PVP01_1242300","PVP01_1242400","PVP01_1242700","PVP01_1243000","PVP01_1243100","PVP01_1243200","PVP01_1243500","PVP01_1243600","PVP01_1243800","PVP01_1243900","PVP01_1244300","PVP01_1244400","PVP01_1244700","PVP01_1244800","PVP01_1244900","PVP01_1245100","PVP01_1245300","PVP01_1245400","PVP01_1245500","PVP01_1245600","PVP01_1245800","PVP01_1245900","PVP01_1246000","PVP01_1246100","PVP01_1246300","PVP01_1246400","PVP01_1246500","PVP01_1246600","PVP01_1246700","PVP01_1246900","PVP01_1247000","PVP01_1247100","PVP01_1247500","PVP01_1247600","PVP01_1247700","PVP01_1247800","PVP01_1247900","PVP01_1248200","PVP01_1248500","PVP01_1248700","PVP01_1248800","PVP01_1249100","PVP01_1249200","PVP01_1249500","PVP01_1249600","PVP01_1249700","PVP01_1250000","PVP01_1250200","PVP01_1250300","PVP01_1250600","PVP01_1250700","PVP01_1250800","PVP01_1251100","PVP01_1251300","PVP01_1251500","PVP01_1251700","PVP01_1251900","PVP01_1252000","PVP01_1252100","PVP01_1252200","PVP01_1252400","PVP01_1252800","PVP01_1252900","PVP01_1253000","PVP01_1253300","PVP01_1253600","PVP01_1253700","PVP01_1253800","PVP01_1254100","PVP01_1254200","PVP01_1254300","PVP01_1254800","PVP01_1254900","PVP01_1255100","PVP01_1255200","PVP01_1255300","PVP01_1255400","PVP01_1255500","PVP01_1255700","PVP01_1256000","PVP01_1256200","PVP01_1256400","PVP01_1256600","PVP01_1256700","PVP01_1256800","PVP01_1256900","PVP01_1257000","PVP01_1257400","PVP01_1257500","PVP01_1257600","PVP01_1257700","PVP01_1257800","PVP01_1257900","PVP01_1258100","PVP01_1258200","PVP01_1258300","PVP01_1258400","PVP01_1258500","PVP01_1258700","PVP01_1258900","PVP01_1259000","PVP01_1259100","PVP01_1259200","PVP01_1259300","PVP01_1259400","PVP01_1259500","PVP01_1259600","PVP01_1259700","PVP01_1260200","PVP01_1260300","PVP01_1260400","PVP01_1260500","PVP01_1260700","PVP01_1260800","PVP01_1261100","PVP01_1261200","PVP01_1261300","PVP01_1261400","PVP01_1261500","PVP01_1261700","PVP01_1261800","PVP01_1262000","PVP01_1262200","PVP01_1262300","PVP01_1262400","PVP01_1262600","PVP01_1262800","PVP01_1262900","PVP01_1263000","PVP01_1263100","PVP01_1263400","PVP01_1263500","PVP01_1263800","PVP01_1263900","PVP01_1264000","PVP01_1264200","PVP01_1264300","PVP01_1264500","PVP01_1264600","PVP01_1264700","PVP01_1264800","PVP01_1264900","PVP01_1265000","PVP01_1265100","PVP01_1265300","PVP01_1265400","PVP01_1265600","PVP01_1265900","PVP01_1266000","PVP01_1266100","PVP01_1266400","PVP01_1266500","PVP01_1266700","PVP01_1266800","PVP01_1267100","PVP01_1267200","PVP01_1267300","PVP01_1267400","PVP01_1267500","PVP01_1267700","PVP01_1267800","PVP01_1268000","PVP01_1268100","PVP01_1268200","PVP01_1268300","PVP01_1269000","PVP01_1269100","PVP01_1269200","PVP01_1269500","PVP01_1269800","PVP01_1269900","PVP01_1270000","PVP01_1270200","PVP01_1270300","PVP01_1270500","PVP01_1270600","PVP01_1270700","PVP01_1270900","PVP01_1271100","PVP01_1271200","PVP01_1271400","PVP01_1271600","PVP01_1271900","PVP01_1272600","PVP01_1273000","PVP01_1274300","PVP01_1300400","PVP01_1300500","PVP01_1300900","PVP01_1301000","PVP01_1301100","PVP01_1301200","PVP01_1301300","PVP01_1301500","PVP01_1301600","PVP01_1301700","PVP01_1301800","PVP01_1302100","PVP01_1302300","PVP01_1302500","PVP01_1302600","PVP01_1302700","PVP01_1302800","PVP01_1302900","PVP01_1303000","PVP01_1303100","PVP01_1303200","PVP01_1303600","PVP01_1303900","PVP01_1304000","PVP01_1304200","PVP01_1304300","PVP01_1304500","PVP01_1304800","PVP01_1304900","PVP01_1305000","PVP01_1305300","PVP01_1305500","PVP01_1305600","PVP01_1305700","PVP01_1306000","PVP01_1306100","PVP01_1306300","PVP01_1306400","PVP01_1306600","PVP01_1306700","PVP01_1306800","PVP01_1307000","PVP01_1307100","PVP01_1307200","PVP01_1307300","PVP01_1307400","PVP01_1307700","PVP01_1307800","PVP01_1308100","PVP01_1308300","PVP01_1308700","PVP01_1308800","PVP01_1308900","PVP01_1309000","PVP01_1309100","PVP01_1309200","PVP01_1309300","PVP01_1309400","PVP01_1309500","PVP01_1309700","PVP01_1309800","PVP01_1310100","PVP01_1310400","PVP01_1310700","PVP01_1311000","PVP01_1311100","PVP01_1311200","PVP01_1311300","PVP01_1311400","PVP01_1311500","PVP01_1311900","PVP01_1312000","PVP01_1312300","PVP01_1312400","PVP01_1312500","PVP01_1312600","PVP01_1312700","PVP01_1312800","PVP01_1313000","PVP01_1313100","PVP01_1313300","PVP01_1313700","PVP01_1313900","PVP01_1314100","PVP01_1314600","PVP01_1314800","PVP01_1314900","PVP01_1315300","PVP01_1315400","PVP01_1315500","PVP01_1315600","PVP01_1315700","PVP01_1316100","PVP01_1316500","PVP01_1316600","PVP01_1316700","PVP01_1316800","PVP01_1316900","PVP01_1317000","PVP01_1317400","PVP01_1317600","PVP01_1317700","PVP01_1317800","PVP01_1317900","PVP01_1318000","PVP01_1318100","PVP01_1318300","PVP01_1318400","PVP01_1318500","PVP01_1318600","PVP01_1318800","PVP01_1318900","PVP01_1319000","PVP01_1319200","PVP01_1319300","PVP01_1319500","PVP01_1319700","PVP01_1319800","PVP01_1319900","PVP01_1320000","PVP01_1320100","PVP01_1320200","PVP01_1320300","PVP01_1320400","PVP01_1320500","PVP01_1320600","PVP01_1320700","PVP01_1321100","PVP01_1321300","PVP01_1321500","PVP01_1321600","PVP01_1321700","PVP01_1321900","PVP01_1322000","PVP01_1322200","PVP01_1322300","PVP01_1322400","PVP01_1322600","PVP01_1322700","PVP01_1322800","PVP01_1322900","PVP01_1323000","PVP01_1323100","PVP01_1323400","PVP01_1323500","PVP01_1323600","PVP01_1323700","PVP01_1323800","PVP01_1323900","PVP01_1324100","PVP01_1324700","PVP01_1324800","PVP01_1325700","PVP01_1325800","PVP01_1325900","PVP01_1326000","PVP01_1326200","PVP01_1326400","PVP01_1326600","PVP01_1327000","PVP01_1327100","PVP01_1327200","PVP01_1327300","PVP01_1327500","PVP01_1327600","PVP01_1327900","PVP01_1328000","PVP01_1328100","PVP01_1328300","PVP01_1328400","PVP01_1328500","PVP01_1328600","PVP01_1328700","PVP01_1328800","PVP01_1328900","PVP01_1329000","PVP01_1329300","PVP01_1329400","PVP01_1329600","PVP01_1329900","PVP01_1330000","PVP01_1330800","PVP01_1331000","PVP01_1331200","PVP01_1331400","PVP01_1331800","PVP01_1331900","PVP01_1332200","PVP01_1332300","PVP01_1332500","PVP01_1332600","PVP01_1332700","PVP01_1332900","PVP01_1333000","PVP01_1333100","PVP01_1333200","PVP01_1333300","PVP01_1333700","PVP01_1333900","PVP01_1334000","PVP01_1334100","PVP01_1334200","PVP01_1334300","PVP01_1334500","PVP01_1334800","PVP01_1335100","PVP01_1335300","PVP01_1335600","PVP01_1335800","PVP01_1335900","PVP01_1336000","PVP01_1336200","PVP01_1336500","PVP01_1336700","PVP01_1337100","PVP01_1337200","PVP01_1337300","PVP01_1337400","PVP01_1337500","PVP01_1337600","PVP01_1337900","PVP01_1338100","PVP01_1338300","PVP01_1338400","PVP01_1338600","PVP01_1339000","PVP01_1339400","PVP01_1339500","PVP01_1339800","PVP01_1340100","PVP01_1340200","PVP01_1340400","PVP01_1340500","PVP01_1340600","PVP01_1340800","PVP01_1341200","PVP01_1341300","PVP01_1341600","PVP01_1341800","PVP01_1342200","PVP01_1342300","PVP01_1342400","PVP01_1342600","PVP01_1342700","PVP01_1342800","PVP01_1343100","PVP01_1343300","PVP01_1343500","PVP01_1343600","PVP01_1343700","PVP01_1344200","PVP01_1344300","PVP01_1344500","PVP01_1344600","PVP01_1344700","PVP01_1344800","PVP01_1344900","PVP01_1345200","PVP01_1345300","PVP01_1345400","PVP01_1345500","PVP01_1345600","PVP01_1345700","PVP01_1345900","PVP01_1346200","PVP01_1346700","PVP01_1346800","PVP01_1346900","PVP01_1347300","PVP01_1347500","PVP01_1347700","PVP01_1347800","PVP01_1400100","PVP01_1400400","PVP01_1401000","PVP01_1401200","PVP01_1401300","PVP01_1401600","PVP01_1401700","PVP01_1401800","PVP01_1402000","PVP01_1402200","PVP01_1402500","PVP01_1402800","PVP01_1402900","PVP01_1403300","PVP01_1403400","PVP01_1403500","PVP01_1403700","PVP01_1403800","PVP01_1403900","PVP01_1404000","PVP01_1404100","PVP01_1404200","PVP01_1404300","PVP01_1404500","PVP01_1404600","PVP01_1404800","PVP01_1405000","PVP01_1405100","PVP01_1405500","PVP01_1405600","PVP01_1405800","PVP01_1405900","PVP01_1406000","PVP01_1406100","PVP01_1406200","PVP01_1406300","PVP01_1406500","PVP01_1406600","PVP01_1406700","PVP01_1406800","PVP01_1407000","PVP01_1407100","PVP01_1407200","PVP01_1407400","PVP01_1407600","PVP01_1407800","PVP01_1407900","PVP01_1408000","PVP01_1408100","PVP01_1408200","PVP01_1408400","PVP01_1408500","PVP01_1408600","PVP01_1408700","PVP01_1408800","PVP01_1409100","PVP01_1409300","PVP01_1409400","PVP01_1409700","PVP01_1410000","PVP01_1410100","PVP01_1410200","PVP01_1410300","PVP01_1410500","PVP01_1410600","PVP01_1410700","PVP01_1410800","PVP01_1410900","PVP01_1411000","PVP01_1411300","PVP01_1411400","PVP01_1411500","PVP01_1411600","PVP01_1411700","PVP01_1412000","PVP01_1412100","PVP01_1412200","PVP01_1412400","PVP01_1412500","PVP01_1412700","PVP01_1413200","PVP01_1413300","PVP01_1413450","PVP01_1413700","PVP01_1413900","PVP01_1414100","PVP01_1414200","PVP01_1414300","PVP01_1414400","PVP01_1414600","PVP01_1414900","PVP01_1415100","PVP01_1415200","PVP01_1415300","PVP01_1415500","PVP01_1415600","PVP01_1415700","PVP01_1415900","PVP01_1416000","PVP01_1416100","PVP01_1416900","PVP01_1417200","PVP01_1417600","PVP01_1417700","PVP01_1417900","PVP01_1418000","PVP01_1418100","PVP01_1418200","PVP01_1418300","PVP01_1418400","PVP01_1418600","PVP01_1418800","PVP01_1418900","PVP01_1419100","PVP01_1419300","PVP01_1419400","PVP01_1419500","PVP01_1419600","PVP01_1419700","PVP01_1419800","PVP01_1420000","PVP01_1420100","PVP01_1420200","PVP01_1420300","PVP01_1420400","PVP01_1420500","PVP01_1420700","PVP01_1420900","PVP01_1421000","PVP01_1421100","PVP01_1421200","PVP01_1421300","PVP01_1421400","PVP01_1421600","PVP01_1421800","PVP01_1422100","PVP01_1422200","PVP01_1422500","PVP01_1422600","PVP01_1422700","PVP01_1422900","PVP01_1423200","PVP01_1423300","PVP01_1423700","PVP01_1424100","PVP01_1424200","PVP01_1424400","PVP01_1424500","PVP01_1424700","PVP01_1424800","PVP01_1425100","PVP01_1425200","PVP01_1425300","PVP01_1425400","PVP01_1425800","PVP01_1426100","PVP01_1426200","PVP01_1426400","PVP01_1426600","PVP01_1426700","PVP01_1426800","PVP01_1427000","PVP01_1427100","PVP01_1427700","PVP01_1427800","PVP01_1428000","PVP01_1428100","PVP01_1428300","PVP01_1428400","PVP01_1428500","PVP01_1428900","PVP01_1429000","PVP01_1429200","PVP01_1429300","PVP01_1429700","PVP01_1429900","PVP01_1430000","PVP01_1430100","PVP01_1430300","PVP01_1430400","PVP01_1430600","PVP01_1430700","PVP01_1430800","PVP01_1431100","PVP01_1431200","PVP01_1431300","PVP01_1431500","PVP01_1431600","PVP01_1431700","PVP01_1431800","PVP01_1432000","PVP01_1432100","PVP01_1432400","PVP01_1432500","PVP01_1432600","PVP01_1432700","PVP01_1432800","PVP01_1433000","PVP01_1433300","PVP01_1433600","PVP01_1433800","PVP01_1433900","PVP01_1434000","PVP01_1434200","PVP01_1434300","PVP01_1434400","PVP01_1434500","PVP01_1434600","PVP01_1434800","PVP01_1434900","PVP01_1435100","PVP01_1435200","PVP01_1435300","PVP01_1435800","PVP01_1435900","PVP01_1436000","PVP01_1436300","PVP01_1436400","PVP01_1436600","PVP01_1436700","PVP01_1437100","PVP01_1437200","PVP01_1437500","PVP01_1437600","PVP01_1437700","PVP01_1438100","PVP01_1438200","PVP01_1438300","PVP01_1438400","PVP01_1438500","PVP01_1438800","PVP01_1438900","PVP01_1439000","PVP01_1439300","PVP01_1439400","PVP01_1439700","PVP01_1439800","PVP01_1440000","PVP01_1440100","PVP01_1440300","PVP01_1440600","PVP01_1440700","PVP01_1440800","PVP01_1440900","PVP01_1441200","PVP01_1441300","PVP01_1441400","PVP01_1441700","PVP01_1441800","PVP01_1442000","PVP01_1442200","PVP01_1442500","PVP01_1442800","PVP01_1442900","PVP01_1443000","PVP01_1443200","PVP01_1443600","PVP01_1443800","PVP01_1443900","PVP01_1444000","PVP01_1444100","PVP01_1444200","PVP01_1444300","PVP01_1444400","PVP01_1444500","PVP01_1444800","PVP01_1444900","PVP01_1445000","PVP01_1445100","PVP01_1445200","PVP01_1445300","PVP01_1445500","PVP01_1445600","PVP01_1445700","PVP01_1445800","PVP01_1446000","PVP01_1446100","PVP01_1446200","PVP01_1446300","PVP01_1446400","PVP01_1446600","PVP01_1446700","PVP01_1446900","PVP01_1447200","PVP01_1447300","PVP01_1447400","PVP01_1447500","PVP01_1447600","PVP01_1447800","PVP01_1447900","PVP01_1448000","PVP01_1448100","PVP01_1448200","PVP01_1448300","PVP01_1448400","PVP01_1448600","PVP01_1448700","PVP01_1448800","PVP01_1448900","PVP01_1449100","PVP01_1449200","PVP01_1449500","PVP01_1449700","PVP01_1450000","PVP01_1450200","PVP01_1450400","PVP01_1450500","PVP01_1450900","PVP01_1451000","PVP01_1451100","PVP01_1451200","PVP01_1451300","PVP01_1451400","PVP01_1451500","PVP01_1451600","PVP01_1451700","PVP01_1451800","PVP01_1452100","PVP01_1452300","PVP01_1452400","PVP01_1452500","PVP01_1452700","PVP01_1452800","PVP01_1452900","PVP01_1453000","PVP01_1453200","PVP01_1453300","PVP01_1453800","PVP01_1454000","PVP01_1454100","PVP01_1454200","PVP01_1454400","PVP01_1454500","PVP01_1454600","PVP01_1454800","PVP01_1454900","PVP01_1455100","PVP01_1455400","PVP01_1455500","PVP01_1455700","PVP01_1455900","PVP01_1456000","PVP01_1456500","PVP01_1456700","PVP01_1456900","PVP01_1457000","PVP01_1457200","PVP01_1457400","PVP01_1457500","PVP01_1457700","PVP01_1457900","PVP01_1458300","PVP01_1458700","PVP01_1458800","PVP01_1459100","PVP01_1459200","PVP01_1459300","PVP01_1459400","PVP01_1459500","PVP01_1459600","PVP01_1459700","PVP01_1460300","PVP01_1460400","PVP01_1460500","PVP01_1460600","PVP01_1460700","PVP01_1460800","PVP01_1460900","PVP01_1461000","PVP01_1461100","PVP01_1461700","PVP01_1461900","PVP01_1462000","PVP01_1462100","PVP01_1462200","PVP01_1462300","PVP01_1462400","PVP01_1462500","PVP01_1462600","PVP01_1462900","PVP01_1463300","PVP01_1463500","PVP01_1463600","PVP01_1463700","PVP01_1463800","PVP01_1464100","PVP01_1464400","PVP01_1464500","PVP01_1464600","PVP01_1464700","PVP01_1464800","PVP01_1464900","PVP01_1465000","PVP01_1465100","PVP01_1465500","PVP01_1465600","PVP01_1465700","PVP01_1465800","PVP01_1465900","PVP01_1466100","PVP01_1466300","PVP01_1466500","PVP01_1466600","PVP01_1466700","PVP01_1467000","PVP01_1467100","PVP01_1467300","PVP01_1467600","PVP01_1467800","PVP01_1468100","PVP01_1468400","PVP01_1468500","PVP01_1468600","PVP01_1468900","PVP01_1469300","PVP01_1469900","PVP01_1470000","PVP01_1470200","PVP01_1470300","PVP01_1470400","PVP01_1470500","PVP01_1470600","PVP01_1470800","PVP01_1470900","PVP01_1471300","PVP01_1471600","PVP01_1471700","PVP01_1471800","PVP01_1471900","PVP01_1472200","PVP01_1472400","PVP01_API01200","PVP01_API01400","PVP01_API01700","PVP01_API04100","PVP01_API04200","PVP01_0000100","PVP01_0000110","PVP01_0000160","PVP01_0000200","PVP01_0000220","PVP01_0001430","PVP01_0001440","PVP01_0003400","PVP01_0004040","PVP01_0004210","PVP01_0004220","PVP01_0004230","PVP01_0004240","PVP01_0004290","PVP01_0004330","PVP01_0004400","PVP01_0005540","PVP01_0005760","PVP01_0007310","PVP01_0009720","PVP01_0010210","PVP01_0010670","PVP01_0010840","PVP01_0011020","PVP01_0102200","PVP01_0102300","PVP01_0102600","PVP01_0103200","PVP01_0103700","PVP01_0104000","PVP01_0104300","PVP01_0104400","PVP01_0104500","PVP01_0104700","PVP01_0105900","PVP01_0106100","PVP01_0106600","PVP01_0106700","PVP01_0107500","PVP01_0107700","PVP01_0108500","PVP01_0108700","PVP01_0109300","PVP01_0109400","PVP01_0109500","PVP01_0109700","PVP01_0109900","PVP01_0110000","PVP01_0110300","PVP01_0110600","PVP01_0111000","PVP01_0111400","PVP01_0111500","PVP01_0111800","PVP01_0112400","PVP01_0112600","PVP01_0113600","PVP01_0113800","PVP01_0113900","PVP01_0114000","PVP01_0114300","PVP01_0114400","PVP01_0114500","PVP01_0114900","PVP01_0115600","PVP01_0115700","PVP01_0115800","PVP01_0115900","PVP01_0116000","PVP01_0116200","PVP01_0116300","PVP01_0116400","PVP01_0117100","PVP01_0117200","PVP01_0117300","PVP01_0117400","PVP01_0117900","PVP01_0118200","PVP01_0118300","PVP01_0118400","PVP01_0119200","PVP01_0201200","PVP01_0201400","PVP01_0202500","PVP01_0203200","PVP01_0203600","PVP01_0203700","PVP01_0204200","PVP01_0204300","PVP01_0204400","PVP01_0204500","PVP01_0204600","PVP01_0204700","PVP01_0205400","PVP01_0205700","PVP01_0205900","PVP01_0206000","PVP01_0206200","PVP01_0206700","PVP01_0206900","PVP01_0207300","PVP01_0207900","PVP01_0208000","PVP01_0208900","PVP01_0209900","PVP01_0210400","PVP01_0210600","PVP01_0210700","PVP01_0210900","PVP01_0211900","PVP01_0212000","PVP01_0212100","PVP01_0212200","PVP01_0212300","PVP01_0212900","PVP01_0213100","PVP01_0213200","PVP01_0213500","PVP01_0213800","PVP01_0213900","PVP01_0214300","PVP01_0214400","PVP01_0214500","PVP01_0214600","PVP01_0214700","PVP01_0215100","PVP01_0215200","PVP01_0215600","PVP01_0216200","PVP01_0216600","PVP01_0216700","PVP01_0220100","PVP01_0300800","PVP01_0302600","PVP01_0302700","PVP01_0302900","PVP01_0303000","PVP01_0303200","PVP01_0303300","PVP01_0303400","PVP01_0303600","PVP01_0303700","PVP01_0303800","PVP01_0303900","PVP01_0304000","PVP01_0304100","PVP01_0304200","PVP01_0304300","PVP01_0304400","PVP01_0304700","PVP01_0305100","PVP01_0305400","PVP01_0305600","PVP01_0305900","PVP01_0306000","PVP01_0306200","PVP01_0306600","PVP01_0307200","PVP01_0307900","PVP01_0308000","PVP01_0308300","PVP01_0308400","PVP01_0308500","PVP01_0308600","PVP01_0308700","PVP01_0308900","PVP01_0309000","PVP01_0309600","PVP01_0310100","PVP01_0310500","PVP01_0310700","PVP01_0311400","PVP01_0311800","PVP01_0311900","PVP01_0312300","PVP01_0312600","PVP01_0312900","PVP01_0313000","PVP01_0313100","PVP01_0313200","PVP01_0313300","PVP01_0313600","PVP01_0313700","PVP01_0313800","PVP01_0313900","PVP01_0314000","PVP01_0314300","PVP01_0314400","PVP01_0314500","PVP01_0314600","PVP01_0314700","PVP01_0314800","PVP01_0315400","PVP01_0315800","PVP01_0316100","PVP01_0316600","PVP01_0316900","PVP01_0317000","PVP01_0317300","PVP01_0317500","PVP01_0317600","PVP01_0317700","PVP01_0317800","PVP01_0318300","PVP01_0318500","PVP01_0318600","PVP01_0318700","PVP01_0318800","PVP01_0319000","PVP01_0319100","PVP01_0319900","PVP01_0320000","PVP01_0320100","PVP01_0320300","PVP01_0320600","PVP01_0320800","PVP01_0321000","PVP01_0321100","PVP01_0404200","PVP01_0404300","PVP01_0405000","PVP01_0405400","PVP01_0405800","PVP01_0406300","PVP01_0406400","PVP01_0406800","PVP01_0407200","PVP01_0407300","PVP01_0407400","PVP01_0407500","PVP01_0408000","PVP01_0408100","PVP01_0408300","PVP01_0408400","PVP01_0408700","PVP01_0408800","PVP01_0409400","PVP01_0409500","PVP01_0409600","PVP01_0409800","PVP01_0410100","PVP01_0410200","PVP01_0410300","PVP01_0410500","PVP01_0410700","PVP01_0410900","PVP01_0411000","PVP01_0411100","PVP01_0411200","PVP01_0411300","PVP01_0412000","PVP01_0412300","PVP01_0412400","PVP01_0412500","PVP01_0413400","PVP01_0413600","PVP01_0413700","PVP01_0413800","PVP01_0413900","PVP01_0414100","PVP01_0414300","PVP01_0414500","PVP01_0414700","PVP01_0414800","PVP01_0415000","PVP01_0415100","PVP01_0415300","PVP01_0415700","PVP01_0415800","PVP01_0415900","PVP01_0416100","PVP01_0416400","PVP01_0416500","PVP01_0416700","PVP01_0416800","PVP01_0417000","PVP01_0417200","PVP01_0417300","PVP01_0417400","PVP01_0417600","PVP01_0417700","PVP01_0418000","PVP01_0418200","PVP01_0418400","PVP01_0418500","PVP01_0418600","PVP01_0418700","PVP01_0418800","PVP01_0418900","PVP01_0419800","PVP01_0419900","PVP01_0420400","PVP01_0420800","PVP01_0421100","PVP01_0421200","PVP01_0421300","PVP01_0421400","PVP01_0422000","PVP01_0422800","PVP01_0423000","PVP01_0423200","PVP01_0423300","PVP01_0423400","PVP01_0423500","PVP01_0423900","PVP01_0424500","PVP01_0503300","PVP01_0503500","PVP01_0503600","PVP01_0503700","PVP01_0503800","PVP01_0503900","PVP01_0504000","PVP01_0504100","PVP01_0504400","PVP01_0504800","PVP01_0504900","PVP01_0505300","PVP01_0505600","PVP01_0505700","PVP01_0505900","PVP01_0506100","PVP01_0506200","PVP01_0506300","PVP01_0506500","PVP01_0507000","PVP01_0507100","PVP01_0507200","PVP01_0507400","PVP01_0507900","PVP01_0508100","PVP01_0508400","PVP01_0508800","PVP01_0509300","PVP01_0509500","PVP01_0509700","PVP01_0509800","PVP01_0510500","PVP01_0510600","PVP01_0511000","PVP01_0511400","PVP01_0511600","PVP01_0511900","PVP01_0512300","PVP01_0512600","PVP01_0513100","PVP01_0513200","PVP01_0514300","PVP01_0514500","PVP01_0515000","PVP01_0515400","PVP01_0515500","PVP01_0515600","PVP01_0516100","PVP01_0516600","PVP01_0516700","PVP01_0517200","PVP01_0517500","PVP01_0517600","PVP01_0517800","PVP01_0518000","PVP01_0518100","PVP01_0518200","PVP01_0518800","PVP01_0518900","PVP01_0519000","PVP01_0519300","PVP01_0519700","PVP01_0519800","PVP01_0519900","PVP01_0520000","PVP01_0520100","PVP01_0520200","PVP01_0520300","PVP01_0520900","PVP01_0521000","PVP01_0521100","PVP01_0521200","PVP01_0521500","PVP01_0521900","PVP01_0522400","PVP01_0523100","PVP01_0523200","PVP01_0523400","PVP01_0523500","PVP01_0523600","PVP01_0523800","PVP01_0523900","PVP01_0524100","PVP01_0524300","PVP01_0525100","PVP01_0525600","PVP01_0525900","PVP01_0526200","PVP01_0526300","PVP01_0526500","PVP01_0526600","PVP01_0526700","PVP01_0526800","PVP01_0527000","PVP01_0527100","PVP01_0527200","PVP01_0527600","PVP01_0528800","PVP01_0529300","PVP01_0529400","PVP01_0529500","PVP01_0530200","PVP01_0530300","PVP01_0530600","PVP01_0530900","PVP01_0531000","PVP01_0531300","PVP01_0531400","PVP01_0531600","PVP01_0531900","PVP01_0532000","PVP01_0532100","PVP01_0532200","PVP01_0532300","PVP01_0532400","PVP01_0532800","PVP01_0533200","PVP01_0533700","PVP01_0534300","PVP01_0534400","PVP01_0601000","PVP01_0601300","PVP01_0601400","PVP01_0601700","PVP01_0601800","PVP01_0601900","PVP01_0602000","PVP01_0602100","PVP01_0602200","PVP01_0602400","PVP01_0602500","PVP01_0602700","PVP01_0602800","PVP01_0602900","PVP01_0603200","PVP01_0603300","PVP01_0603800","PVP01_0604300","PVP01_0604500","PVP01_0604600","PVP01_0605200","PVP01_0605600","PVP01_0605700","PVP01_0605800","PVP01_0605900","PVP01_0606000","PVP01_0606300","PVP01_0606700","PVP01_0607000","PVP01_0607100","PVP01_0607300","PVP01_0607400","PVP01_0607500","PVP01_0607700","PVP01_0607800","PVP01_0607900","PVP01_0609000","PVP01_0609100","PVP01_0609300","PVP01_0609500","PVP01_0610000","PVP01_0610300","PVP01_0611000","PVP01_0611200","PVP01_0611400","PVP01_0611500","PVP01_0611600","PVP01_0611700","PVP01_0611800","PVP01_0611900","PVP01_0612200","PVP01_0612300","PVP01_0612400","PVP01_0612500","PVP01_0612800","PVP01_0612900","PVP01_0613300","PVP01_0613600","PVP01_0614100","PVP01_0614500","PVP01_0614800","PVP01_0615000","PVP01_0615200","PVP01_0615300","PVP01_0615400","PVP01_0615600","PVP01_0616300","PVP01_0616700","PVP01_0617200","PVP01_0617300","PVP01_0617700","PVP01_0617800","PVP01_0618000","PVP01_0618200","PVP01_0618400","PVP01_0618500","PVP01_0618800","PVP01_0619300","PVP01_0620300","PVP01_0620400","PVP01_0620500","PVP01_0620600","PVP01_0620700","PVP01_0620800","PVP01_0621200","PVP01_0621400","PVP01_0621500","PVP01_0621600","PVP01_0621700","PVP01_0621900","PVP01_0622000","PVP01_0622300","PVP01_0622400","PVP01_0622500","PVP01_0622600","PVP01_0623000","PVP01_0623500","PVP01_0623700","PVP01_0623800","PVP01_0624500","PVP01_0700100","PVP01_0700300","PVP01_0700400","PVP01_0700500","PVP01_0700600","PVP01_0700900","PVP01_0701100","PVP01_0701200","PVP01_0701300","PVP01_0701800","PVP01_0701900","PVP01_0702000","PVP01_0702100","PVP01_0702300","PVP01_0702800","PVP01_0702900","PVP01_0703100","PVP01_0703300","PVP01_0703400","PVP01_0703800","PVP01_0703900","PVP01_0704200","PVP01_0704300","PVP01_0705300","PVP01_0705400","PVP01_0706800","PVP01_0706900","PVP01_0707400","PVP01_0707600","PVP01_0707700","PVP01_0707900","PVP01_0708600","PVP01_0708700","PVP01_0709400","PVP01_0709600","PVP01_0709800","PVP01_0710100","PVP01_0710400","PVP01_0710500","PVP01_0710600","PVP01_0710800","PVP01_0710900","PVP01_0711000","PVP01_0711300","PVP01_0711600","PVP01_0711700","PVP01_0712000","PVP01_0712100","PVP01_0712500","PVP01_0712600","PVP01_0712800","PVP01_0712900","PVP01_0713300","PVP01_0713400","PVP01_0713700","PVP01_0714100","PVP01_0714500","PVP01_0714700","PVP01_0715300","PVP01_0715400","PVP01_0715700","PVP01_0715900","PVP01_0716300","PVP01_0716400","PVP01_0716500","PVP01_0716700","PVP01_0717100","PVP01_0717300","PVP01_0717500","PVP01_0717700","PVP01_0717900","PVP01_0718300","PVP01_0718400","PVP01_0718500","PVP01_0718800","PVP01_0718900","PVP01_0719100","PVP01_0719200","PVP01_0719600","PVP01_0720500","PVP01_0721000","PVP01_0721200","PVP01_0721600","PVP01_0721700","PVP01_0722000","PVP01_0723600","PVP01_0723900","PVP01_0724200","PVP01_0724400","PVP01_0725200","PVP01_0725500","PVP01_0725600","PVP01_0725800","PVP01_0725900","PVP01_0726100","PVP01_0726200","PVP01_0726300","PVP01_0726400","PVP01_0726500","PVP01_0726600","PVP01_0726800","PVP01_0727900","PVP01_0728000","PVP01_0728400","PVP01_0728500","PVP01_0728600","PVP01_0728700","PVP01_0728800","PVP01_0728900","PVP01_0729000","PVP01_0729600","PVP01_0729800","PVP01_0729900","PVP01_0730500","PVP01_0730700","PVP01_0730900","PVP01_0731000","PVP01_0731400","PVP01_0732400","PVP01_0732500","PVP01_0732700","PVP01_0733300","PVP01_0733400","PVP01_0733600","PVP01_0733800","PVP01_0734000","PVP01_0734300","PVP01_0734500","PVP01_0734600","PVP01_0734700","PVP01_0735200","PVP01_0800700","PVP01_0801000","PVP01_0801100","PVP01_0801200","PVP01_0801300","PVP01_0801500","PVP01_0801600","PVP01_0802200","PVP01_0802600","PVP01_0802700","PVP01_0802800","PVP01_0803200","PVP01_0803400","PVP01_0803800","PVP01_0804300","PVP01_0805100","PVP01_0805500","PVP01_0805900","PVP01_0806900","PVP01_0807100","PVP01_0807400","PVP01_0807700","PVP01_0807900","PVP01_0808100","PVP01_0808200","PVP01_0808300","PVP01_0808700","PVP01_0808900","PVP01_0809000","PVP01_0809500","PVP01_0809800","PVP01_0810200","PVP01_0810700","PVP01_0810800","PVP01_0810900","PVP01_0811400","PVP01_0811600","PVP01_0812200","PVP01_0812400","PVP01_0812500","PVP01_0812900","PVP01_0813100","PVP01_0813400","PVP01_0813500","PVP01_0813900","PVP01_0814000","PVP01_0814200","PVP01_0814300","PVP01_0814500","PVP01_0814600","PVP01_0814900","PVP01_0815000","PVP01_0815200","PVP01_0815500","PVP01_0815800","PVP01_0815900","PVP01_0816000","PVP01_0816100","PVP01_0816300","PVP01_0816400","PVP01_0816500","PVP01_0816600","PVP01_0816800","PVP01_0816900","PVP01_0817000","PVP01_0817200","PVP01_0817300","PVP01_0817500","PVP01_0817600","PVP01_0817700","PVP01_0817800","PVP01_0817900","PVP01_0818200","PVP01_0818400","PVP01_0818700","PVP01_0818800","PVP01_0818900","PVP01_0819100","PVP01_0819300","PVP01_0819400","PVP01_0819500","PVP01_0819600","PVP01_0819800","PVP01_0819900","PVP01_0820200","PVP01_0820400","PVP01_0821200","PVP01_0821300","PVP01_0821900","PVP01_0822000","PVP01_0822400","PVP01_0822500","PVP01_0822800","PVP01_0823000","PVP01_0823300","PVP01_0823600","PVP01_0823800","PVP01_0824100","PVP01_0824200","PVP01_0824400","PVP01_0824600","PVP01_0824800","PVP01_0824900","PVP01_0825300","PVP01_0825500","PVP01_0825600","PVP01_0825900","PVP01_0826800","PVP01_0827000","PVP01_0827300","PVP01_0827400","PVP01_0827900","PVP01_0828400","PVP01_0828500","PVP01_0828800","PVP01_0828900","PVP01_0829200","PVP01_0829400","PVP01_0829500","PVP01_0829600","PVP01_0829800","PVP01_0830000","PVP01_0830100","PVP01_0830600","PVP01_0831300","PVP01_0831500","PVP01_0831600","PVP01_0832300","PVP01_0832700","PVP01_0832900","PVP01_0833100","PVP01_0833200","PVP01_0833400","PVP01_0834100","PVP01_0834800","PVP01_0835100","PVP01_0835200","PVP01_0835400","PVP01_0835600","PVP01_0836200","PVP01_0836800","PVP01_0836900","PVP01_0837000","PVP01_0837400","PVP01_0837700","PVP01_0838100","PVP01_0838300","PVP01_0838600","PVP01_0838700","PVP01_0838800","PVP01_0839000","PVP01_0839400","PVP01_0901000","PVP01_0901100","PVP01_0904200","PVP01_0904400","PVP01_0904500","PVP01_0904800","PVP01_0905000","PVP01_0905200","PVP01_0905300","PVP01_0905400","PVP01_0905500","PVP01_0905600","PVP01_0905800","PVP01_0905900","PVP01_0906100","PVP01_0906200","PVP01_0906400","PVP01_0906600","PVP01_0907400","PVP01_0907900","PVP01_0908100","PVP01_0908200","PVP01_0908700","PVP01_0908800","PVP01_0909200","PVP01_0909400","PVP01_0909900","PVP01_0910700","PVP01_0910800","PVP01_0911100","PVP01_0911200","PVP01_0911400","PVP01_0912000","PVP01_0912300","PVP01_0912500","PVP01_0912600","PVP01_0912700","PVP01_0912800","PVP01_0913200","PVP01_0913300","PVP01_0913800","PVP01_0914100","PVP01_0914700","PVP01_0914900","PVP01_0915500","PVP01_0915700","PVP01_0916000","PVP01_0916100","PVP01_0916200","PVP01_0916400","PVP01_0916500","PVP01_0916600","PVP01_0917500","PVP01_0917800","PVP01_0917900","PVP01_0918600","PVP01_0919200","PVP01_0919400","PVP01_0919800","PVP01_0920000","PVP01_0920200","PVP01_0920600","PVP01_0920900","PVP01_0921000","PVP01_0921200","PVP01_0921300","PVP01_0921600","PVP01_0921900","PVP01_0922200","PVP01_0922500","PVP01_0922600","PVP01_0922700","PVP01_0923200","PVP01_0923400","PVP01_0923500","PVP01_0923600","PVP01_0923700","PVP01_0924200","PVP01_0924600","PVP01_0925000","PVP01_0925500","PVP01_0926000","PVP01_0926100","PVP01_0926200","PVP01_0926500","PVP01_0926700","PVP01_0926800","PVP01_0927000","PVP01_0927200","PVP01_0927300","PVP01_0927500","PVP01_0927900","PVP01_0928200","PVP01_0928500","PVP01_0928600","PVP01_0928700","PVP01_0929500","PVP01_0929600","PVP01_0930700","PVP01_0930800","PVP01_0931200","PVP01_0931800","PVP01_0932400","PVP01_0932500","PVP01_0932700","PVP01_0932800","PVP01_0932900","PVP01_0933000","PVP01_0933200","PVP01_0933300","PVP01_0933800","PVP01_0933900","PVP01_0934100","PVP01_0934200","PVP01_0934600","PVP01_0934700","PVP01_0934800","PVP01_0934900","PVP01_0935600","PVP01_0935900","PVP01_0936200","PVP01_0936500","PVP01_0937100","PVP01_0937200","PVP01_0937600","PVP01_0938300","PVP01_0938500","PVP01_0938800","PVP01_0939000","PVP01_0939400","PVP01_0939700","PVP01_0939800","PVP01_0940000","PVP01_0940300","PVP01_0940400","PVP01_0941100","PVP01_0941500","PVP01_0941700","PVP01_0941900","PVP01_0942000","PVP01_0942100","PVP01_0942500","PVP01_0942800","PVP01_0943000","PVP01_0943200","PVP01_0943500","PVP01_0943600","PVP01_0943800","PVP01_0943900","PVP01_0944000","PVP01_0944500","PVP01_0944700","PVP01_0944900","PVP01_0945000","PVP01_0945400","PVP01_0945700","PVP01_0945900","PVP01_0946100","PVP01_0946800","PVP01_0947300","PVP01_0947500","PVP01_0947600","PVP01_0948000","PVP01_0948200","PVP01_0948400","PVP01_0949200","PVP01_0949300","PVP01_1002400","PVP01_1002800","PVP01_1003300","PVP01_1003600","PVP01_1003700","PVP01_1003800","PVP01_1003900","PVP01_1004000","PVP01_1004100","PVP01_1004400","PVP01_1004600","PVP01_1004700","PVP01_1005800","PVP01_1006500","PVP01_1006800","PVP01_1006900","PVP01_1007000","PVP01_1007100","PVP01_1007200","PVP01_1007300","PVP01_1007400","PVP01_1008500","PVP01_1008800","PVP01_1009900","PVP01_1010000","PVP01_1010500","PVP01_1010600","PVP01_1010700","PVP01_1010800","PVP01_1010900","PVP01_1011000","PVP01_1011300","PVP01_1011400","PVP01_1011500","PVP01_1011900","PVP01_1012000","PVP01_1012200","PVP01_1012600","PVP01_1012900","PVP01_1013000","PVP01_1013500","PVP01_1013600","PVP01_1013800","PVP01_1013900","PVP01_1014000","PVP01_1014100","PVP01_1014500","PVP01_1014900","PVP01_1015200","PVP01_1015300","PVP01_1015600","PVP01_1015800","PVP01_1016100","PVP01_1016700","PVP01_1016800","PVP01_1017000","PVP01_1017200","PVP01_1017300","PVP01_1017500","PVP01_1017700","PVP01_1017800","PVP01_1018200","PVP01_1018300","PVP01_1018600","PVP01_1018800","PVP01_1018900","PVP01_1019100","PVP01_1019200","PVP01_1019300","PVP01_1019700","PVP01_1019800","PVP01_1020300","PVP01_1020400","PVP01_1020600","PVP01_1021400","PVP01_1021900","PVP01_1022500","PVP01_1022600","PVP01_1022700","PVP01_1022800","PVP01_1023500","PVP01_1024000","PVP01_1024200","PVP01_1024600","PVP01_1024700","PVP01_1025000","PVP01_1025600","PVP01_1025700","PVP01_1025900","PVP01_1026100","PVP01_1026500","PVP01_1026800","PVP01_1027000","PVP01_1027100","PVP01_1027200","PVP01_1027500","PVP01_1027800","PVP01_1028300","PVP01_1028800","PVP01_1028900","PVP01_1029400","PVP01_1029600","PVP01_1030200","PVP01_1030400","PVP01_1030700","PVP01_1030800","PVP01_1031000","PVP01_1031100","PVP01_1031200","PVP01_1031400","PVP01_1031500","PVP01_1031600","PVP01_1031700","PVP01_1032300","PVP01_1032700","PVP01_1033100","PVP01_1033200","PVP01_1033400","PVP01_1033600","PVP01_1033800","PVP01_1033900","PVP01_1034000","PVP01_1034200","PVP01_1034400","PVP01_1101400","PVP01_1102300","PVP01_1102600","PVP01_1102700","PVP01_1103000","PVP01_1103100","PVP01_1103500","PVP01_1103800","PVP01_1103900","PVP01_1104500","PVP01_1104900","PVP01_1105000","PVP01_1105200","PVP01_1105700","PVP01_1105900","PVP01_1106100","PVP01_1106500","PVP01_1106800","PVP01_1107500","PVP01_1107800","PVP01_1107900","PVP01_1108100","PVP01_1108200","PVP01_1108400","PVP01_1109200","PVP01_1109500","PVP01_1109700","PVP01_1110000","PVP01_1110300","PVP01_1110400","PVP01_1110800","PVP01_1111400","PVP01_1111900","PVP01_1112300","PVP01_1112600","PVP01_1113100","PVP01_1113200","PVP01_1113600","PVP01_1113800","PVP01_1114000","PVP01_1114400","PVP01_1114900","PVP01_1115100","PVP01_1115300","PVP01_1115400","PVP01_1115800","PVP01_1115900","PVP01_1116000","PVP01_1116100","PVP01_1116200","PVP01_1116300","PVP01_1116400","PVP01_1116700","PVP01_1117000","PVP01_1117100","PVP01_1117200","PVP01_1117400","PVP01_1117600","PVP01_1117800","PVP01_1117900","PVP01_1118200","PVP01_1118300","PVP01_1118400","PVP01_1118500","PVP01_1119200","PVP01_1119700","PVP01_1119900","PVP01_1120200","PVP01_1120600","PVP01_1120700","PVP01_1120800","PVP01_1121200","PVP01_1121800","PVP01_1122000","PVP01_1122200","PVP01_1123000","PVP01_1123500","PVP01_1123700","PVP01_1123800","PVP01_1124200","PVP01_1124500","PVP01_1125500","PVP01_1126400","PVP01_1126700","PVP01_1127600","PVP01_1127700","PVP01_1128000","PVP01_1128200","PVP01_1128400","PVP01_1128500","PVP01_1129100","PVP01_1129200","PVP01_1129300","PVP01_1129800","PVP01_1129900","PVP01_1130000","PVP01_1130200","PVP01_1130300","PVP01_1130500","PVP01_1130600","PVP01_1131100","PVP01_1131200","PVP01_1131500","PVP01_1131600","PVP01_1131700","PVP01_1131800","PVP01_1131900","PVP01_1132000","PVP01_1132200","PVP01_1132600","PVP01_1133700","PVP01_1134000","PVP01_1134500","PVP01_1134600","PVP01_1134900","PVP01_1135300","PVP01_1135400","PVP01_1136200","PVP01_1136400","PVP01_1136500","PVP01_1137000","PVP01_1137400","PVP01_1137500","PVP01_1137900","PVP01_1138000","PVP01_1138100","PVP01_1138200","PVP01_1138300","PVP01_1138600","PVP01_1138700","PVP01_1138800","PVP01_1138900","PVP01_1139200","PVP01_1139400","PVP01_1139500","PVP01_1139800","PVP01_1139900","PVP01_1140100","PVP01_1140200","PVP01_1140400","PVP01_1141100","PVP01_1141600","PVP01_1142200","PVP01_1142400","PVP01_1142700","PVP01_1142800","PVP01_1143000","PVP01_1143300","PVP01_1143800","PVP01_1143900","PVP01_1144400","PVP01_1145300","PVP01_1145400","PVP01_1145600","PVP01_1145900","PVP01_1146000","PVP01_1146100","PVP01_1146200","PVP01_1146300","PVP01_1146500","PVP01_1146700","PVP01_1147100","PVP01_1147600","PVP01_1147700","PVP01_1147800","PVP01_1147900","PVP01_1201400","PVP01_1201500","PVP01_1202000","PVP01_1202400","PVP01_1202500","PVP01_1202600","PVP01_1202700","PVP01_1202800","PVP01_1202900","PVP01_1203500","PVP01_1203900","PVP01_1204400","PVP01_1204700","PVP01_1205300","PVP01_1205700","PVP01_1205800","PVP01_1206000","PVP01_1206500","PVP01_1206600","PVP01_1206800","PVP01_1207300","PVP01_1207400","PVP01_1207500","PVP01_1207600","PVP01_1207700","PVP01_1208000","PVP01_1208400","PVP01_1208500","PVP01_1209100","PVP01_1209200","PVP01_1209300","PVP01_1209500","PVP01_1209700","PVP01_1209800","PVP01_1209900","PVP01_1210400","PVP01_1210500","PVP01_1210600","PVP01_1210900","PVP01_1211000","PVP01_1211100","PVP01_1211400","PVP01_1211700","PVP01_1211800","PVP01_1211900","PVP01_1212000","PVP01_1212300","PVP01_1212400","PVP01_1212700","PVP01_1212800","PVP01_1212900","PVP01_1213200","PVP01_1213400","PVP01_1213600","PVP01_1213900","PVP01_1214000","PVP01_1214100","PVP01_1214300","PVP01_1214400","PVP01_1214500","PVP01_1215100","PVP01_1215200","PVP01_1215500","PVP01_1216400","PVP01_1216500","PVP01_1216600","PVP01_1216900","PVP01_1217000","PVP01_1217300","PVP01_1217800","PVP01_1217900","PVP01_1218100","PVP01_1218200","PVP01_1218300","PVP01_1218400","PVP01_1218600","PVP01_1218700","PVP01_1219200","PVP01_1219300","PVP01_1219400","PVP01_1219600","PVP01_1219700","PVP01_1219800","PVP01_1220000","PVP01_1220400","PVP01_1220500","PVP01_1220800","PVP01_1221000","PVP01_1221100","PVP01_1221500","PVP01_1221600","PVP01_1222000","PVP01_1222900","PVP01_1223000","PVP01_1223200","PVP01_1223300","PVP01_1223700","PVP01_1224100","PVP01_1224200","PVP01_1224900","PVP01_1225100","PVP01_1225400","PVP01_1225900","PVP01_1226000","PVP01_1226100","PVP01_1226200","PVP01_1226300","PVP01_1227000","PVP01_1227100","PVP01_1227400","PVP01_1227700","PVP01_1227800","PVP01_1228600","PVP01_1228700","PVP01_1228900","PVP01_1229000","PVP01_1229300","PVP01_1229700","PVP01_1230300","PVP01_1230400","PVP01_1230500","PVP01_1231200","PVP01_1231300","PVP01_1231400","PVP01_1231500","PVP01_1232000","PVP01_1232100","PVP01_1232200","PVP01_1232300","PVP01_1232500","PVP01_1232800","PVP01_1232900","PVP01_1233000","PVP01_1233300","PVP01_1233400","PVP01_1233500","PVP01_1233600","PVP01_1233700","PVP01_1234100","PVP01_1234200","PVP01_1234300","PVP01_1234400","PVP01_1234700","PVP01_1234800","PVP01_1234900","PVP01_1235400","PVP01_1235600","PVP01_1235700","PVP01_1235900","PVP01_1236300","PVP01_1236600","PVP01_1236700","PVP01_1237300","PVP01_1237400","PVP01_1237600","PVP01_1238100","PVP01_1238200","PVP01_1238300","PVP01_1238700","PVP01_1238800","PVP01_1239200","PVP01_1239600","PVP01_1239700","PVP01_1239800","PVP01_1240000","PVP01_1240200","PVP01_1240500","PVP01_1240700","PVP01_1241500","PVP01_1241600","PVP01_1241700","PVP01_1241800","PVP01_1242100","PVP01_1242200","PVP01_1242500","PVP01_1242600","PVP01_1242800","PVP01_1242900","PVP01_1243300","PVP01_1243400","PVP01_1243700","PVP01_1244000","PVP01_1244100","PVP01_1244200","PVP01_1244500","PVP01_1244600","PVP01_1245000","PVP01_1245200","PVP01_1245700","PVP01_1246200","PVP01_1246800","PVP01_1247200","PVP01_1247300","PVP01_1247400","PVP01_1248000","PVP01_1248100","PVP01_1248300","PVP01_1248400","PVP01_1248600","PVP01_1248900","PVP01_1249000","PVP01_1249300","PVP01_1249400","PVP01_1249800","PVP01_1249900","PVP01_1250100","PVP01_1250400","PVP01_1250500","PVP01_1250900","PVP01_1251000","PVP01_1251200","PVP01_1251400","PVP01_1251600","PVP01_1251800","PVP01_1252300","PVP01_1252500","PVP01_1252600","PVP01_1252700","PVP01_1253100","PVP01_1253200","PVP01_1253400","PVP01_1253500","PVP01_1253900","PVP01_1254000","PVP01_1254400","PVP01_1254500","PVP01_1254600","PVP01_1254700","PVP01_1255000","PVP01_1255600","PVP01_1255800","PVP01_1255900","PVP01_1256100","PVP01_1256300","PVP01_1256500","PVP01_1257100","PVP01_1257200","PVP01_1257300","PVP01_1258000","PVP01_1258600","PVP01_1258800","PVP01_1259800","PVP01_1259900","PVP01_1260000","PVP01_1260100","PVP01_1260600","PVP01_1260900","PVP01_1261600","PVP01_1261900","PVP01_1262100","PVP01_1262500","PVP01_1262700","PVP01_1263200","PVP01_1263300","PVP01_1263600","PVP01_1263700","PVP01_1264100","PVP01_1264400","PVP01_1265200","PVP01_1265500","PVP01_1265700","PVP01_1265800","PVP01_1266200","PVP01_1266300","PVP01_1266600","PVP01_1266900","PVP01_1267000","PVP01_1267600","PVP01_1267900","PVP01_1268400","PVP01_1268500","PVP01_1268600","PVP01_1268900","PVP01_1269300","PVP01_1269400","PVP01_1269600","PVP01_1269700","PVP01_1270100","PVP01_1270400","PVP01_1270800","PVP01_1271000","PVP01_1271300","PVP01_1271500","PVP01_1272500","PVP01_1300600","PVP01_1300700","PVP01_1300800","PVP01_1301400","PVP01_1301900","PVP01_1302000","PVP01_1302200","PVP01_1302400","PVP01_1303300","PVP01_1303400","PVP01_1303500","PVP01_1303700","PVP01_1303800","PVP01_1304100","PVP01_1304400","PVP01_1304600","PVP01_1304700","PVP01_1305100","PVP01_1305200","PVP01_1305400","PVP01_1305800","PVP01_1305900","PVP01_1306200","PVP01_1306500","PVP01_1306900","PVP01_1307500","PVP01_1307600","PVP01_1307900","PVP01_1308000","PVP01_1308200","PVP01_1308400","PVP01_1308500","PVP01_1308600","PVP01_1309600","PVP01_1309900","PVP01_1310000","PVP01_1310200","PVP01_1310300","PVP01_1310500","PVP01_1310600","PVP01_1310800","PVP01_1310900","PVP01_1311600","PVP01_1311700","PVP01_1311800","PVP01_1312100","PVP01_1312200","PVP01_1312900","PVP01_1313200","PVP01_1313400","PVP01_1313500","PVP01_1313600","PVP01_1313800","PVP01_1314000","PVP01_1314200","PVP01_1314300","PVP01_1314400","PVP01_1314500","PVP01_1314700","PVP01_1315000","PVP01_1315100","PVP01_1315200","PVP01_1315800","PVP01_1315900","PVP01_1316000","PVP01_1316200","PVP01_1316300","PVP01_1316400","PVP01_1317100","PVP01_1317200","PVP01_1317300","PVP01_1317500","PVP01_1318200","PVP01_1318700","PVP01_1319100","PVP01_1319400","PVP01_1319600","PVP01_1320800","PVP01_1320900","PVP01_1321000","PVP01_1321200","PVP01_1321400","PVP01_1321800","PVP01_1322100","PVP01_1322500","PVP01_1323200","PVP01_1323300","PVP01_1324000","PVP01_1325200","PVP01_1325300","PVP01_1325400","PVP01_1325500","PVP01_1325600","PVP01_1326100","PVP01_1326300","PVP01_1326500","PVP01_1326700","PVP01_1326800","PVP01_1326900","PVP01_1327400","PVP01_1327700","PVP01_1327800","PVP01_1328200","PVP01_1329100","PVP01_1329200","PVP01_1329500","PVP01_1329700","PVP01_1329800","PVP01_1330600","PVP01_1330700","PVP01_1330900","PVP01_1331100","PVP01_1331300","PVP01_1331500","PVP01_1331600","PVP01_1331700","PVP01_1332000","PVP01_1332100","PVP01_1332400","PVP01_1332800","PVP01_1333400","PVP01_1333500","PVP01_1333600","PVP01_1333800","PVP01_1334400","PVP01_1334600","PVP01_1334700","PVP01_1334900","PVP01_1335000","PVP01_1335200","PVP01_1335400","PVP01_1335500","PVP01_1335700","PVP01_1336100","PVP01_1336300","PVP01_1336400","PVP01_1336600","PVP01_1336800","PVP01_1336900","PVP01_1337000","PVP01_1337700","PVP01_1337800","PVP01_1338000","PVP01_1338200","PVP01_1338500","PVP01_1338700","PVP01_1338800","PVP01_1338900","PVP01_1339100","PVP01_1339200","PVP01_1339300","PVP01_1339600","PVP01_1339700","PVP01_1339900","PVP01_1340000","PVP01_1340300","PVP01_1340700","PVP01_1340900","PVP01_1341000","PVP01_1341100","PVP01_1341400","PVP01_1341500","PVP01_1341700","PVP01_1341900","PVP01_1342000","PVP01_1342100","PVP01_1342500","PVP01_1342900","PVP01_1343000","PVP01_1343200","PVP01_1343400","PVP01_1343800","PVP01_1343900","PVP01_1344000","PVP01_1344100","PVP01_1344400","PVP01_1345000","PVP01_1345100","PVP01_1345800","PVP01_1346000","PVP01_1346100","PVP01_1346400","PVP01_1346500","PVP01_1346600","PVP01_1347000","PVP01_1347100","PVP01_1347200","PVP01_1400600","PVP01_1400700","PVP01_1400900","PVP01_1401100","PVP01_1401400","PVP01_1401900","PVP01_1402100","PVP01_1402300","PVP01_1402400","PVP01_1402600","PVP01_1402700","PVP01_1403000","PVP01_1403100","PVP01_1403200","PVP01_1403600","PVP01_1404400","PVP01_1404700","PVP01_1404900","PVP01_1405200","PVP01_1405300","PVP01_1405400","PVP01_1405700","PVP01_1406400","PVP01_1406900","PVP01_1407300","PVP01_1407500","PVP01_1407700","PVP01_1408300","PVP01_1408900","PVP01_1409000","PVP01_1409200","PVP01_1409500","PVP01_1409600","PVP01_1409800","PVP01_1409900","PVP01_1410400","PVP01_1411100","PVP01_1411200","PVP01_1411800","PVP01_1411900","PVP01_1412300","PVP01_1412600","PVP01_1412800","PVP01_1412900","PVP01_1413000","PVP01_1413100","PVP01_1413400","PVP01_1413500","PVP01_1413600","PVP01_1413800","PVP01_1414000","PVP01_1414500","PVP01_1414700","PVP01_1414800","PVP01_1415000","PVP01_1415400","PVP01_1415800","PVP01_1416200","PVP01_1416300","PVP01_1416400","PVP01_1416500","PVP01_1416600","PVP01_1416700","PVP01_1416800","PVP01_1417000","PVP01_1417100","PVP01_1417300","PVP01_1417400","PVP01_1417500","PVP01_1417800","PVP01_1418500","PVP01_1418700","PVP01_1419000","PVP01_1419200","PVP01_1419900","PVP01_1420600","PVP01_1420800","PVP01_1421500","PVP01_1421700","PVP01_1421900","PVP01_1422000","PVP01_1422300","PVP01_1422400","PVP01_1422800","PVP01_1423000","PVP01_1423100","PVP01_1423400","PVP01_1423500","PVP01_1423600","PVP01_1424300","PVP01_1424600","PVP01_1424900","PVP01_1425000","PVP01_1425500","PVP01_1425600","PVP01_1425700","PVP01_1425900","PVP01_1426000","PVP01_1426300","PVP01_1426500","PVP01_1426900","PVP01_1427200","PVP01_1427300","PVP01_1427400","PVP01_1427500","PVP01_1427600","PVP01_1427900","PVP01_1428200","PVP01_1428600","PVP01_1428700","PVP01_1428800","PVP01_1429100","PVP01_1429400","PVP01_1429500","PVP01_1429600","PVP01_1429800","PVP01_1430200","PVP01_1430500","PVP01_1430900","PVP01_1431000","PVP01_1431400","PVP01_1432200","PVP01_1432300","PVP01_1432900","PVP01_1433100","PVP01_1433200","PVP01_1433400","PVP01_1433500","PVP01_1433700","PVP01_1434100","PVP01_1434700","PVP01_1435000","PVP01_1435400","PVP01_1435500","PVP01_1435700","PVP01_1436100","PVP01_1436200","PVP01_1436500","PVP01_1436800","PVP01_1436900","PVP01_1437000","PVP01_1437300","PVP01_1437400","PVP01_1437800","PVP01_1437900","PVP01_1438000","PVP01_1438600","PVP01_1438700","PVP01_1439100","PVP01_1439200","PVP01_1439500","PVP01_1439600","PVP01_1439900","PVP01_1440200","PVP01_1440400","PVP01_1440500","PVP01_1441000","PVP01_1441100","PVP01_1441500","PVP01_1441600","PVP01_1441900","PVP01_1442100","PVP01_1442300","PVP01_1442400","PVP01_1442600","PVP01_1442700","PVP01_1443100","PVP01_1443300","PVP01_1443400","PVP01_1443500","PVP01_1443700","PVP01_1444600","PVP01_1444700","PVP01_1445400","PVP01_1445900","PVP01_1446500","PVP01_1446800","PVP01_1447000","PVP01_1447100","PVP01_1447700","PVP01_1448500","PVP01_1449000","PVP01_1449300","PVP01_1449400","PVP01_1449600","PVP01_1449800","PVP01_1449900","PVP01_1450100","PVP01_1450300","PVP01_1450600","PVP01_1450700","PVP01_1450800","PVP01_1451900","PVP01_1452000","PVP01_1452200","PVP01_1452600","PVP01_1453100","PVP01_1453400","PVP01_1453500","PVP01_1453600","PVP01_1453700","PVP01_1453900","PVP01_1454300","PVP01_1454700","PVP01_1455000","PVP01_1455200","PVP01_1455300","PVP01_1455600","PVP01_1455800","PVP01_1456100","PVP01_1456200","PVP01_1456300","PVP01_1456400","PVP01_1456600","PVP01_1456800","PVP01_1457100","PVP01_1457300","PVP01_1457600","PVP01_1457800","PVP01_1458000","PVP01_1458100","PVP01_1458200","PVP01_1458400","PVP01_1458500","PVP01_1458600","PVP01_1458900","PVP01_1459000","PVP01_1459800","PVP01_1459900","PVP01_1460000","PVP01_1460100","PVP01_1460200","PVP01_1461200","PVP01_1461300","PVP01_1461500","PVP01_1461600","PVP01_1461800","PVP01_1462700","PVP01_1462800","PVP01_1463000","PVP01_1463100","PVP01_1463200","PVP01_1463400","PVP01_1463900","PVP01_1464000","PVP01_1464200","PVP01_1464300","PVP01_1465200","PVP01_1465300","PVP01_1465400","PVP01_1466000","PVP01_1466200","PVP01_1466400","PVP01_1466800","PVP01_1466900","PVP01_1467200","PVP01_1467400","PVP01_1467500","PVP01_1467700","PVP01_1467900","PVP01_1468000","PVP01_1468200","PVP01_1468300","PVP01_1468700","PVP01_1468800","PVP01_1469200","PVP01_1469400","PVP01_1469500","PVP01_1469600","PVP01_1469700","PVP01_1469800","PVP01_1470100","PVP01_1470700","PVP01_1471100","PVP01_1471400","PVP01_1471500","PVP01_1472100","PVP01_API02600","PVP01_API02800","PVP01_API03500","PVP01_API04300","PVP01_API04600","PVP01_MIT02700","PVP01_MIT03300","PVP01_MIT03400"],"logFC":[-3.925,-1.809,0.7073,-2.014,0.5704,-3.169,-5.226,5.74,1.373,-4.83,-2.798,0.6786,-0.4437,3.404,3.972,3.664,4.314,4.754,-3.925,4.295,5.18,-6.448,3.348,3.845,0.283,2.317,4.492,5.102,5.189,3.482,-1.433,3.203,3.329,4.019,4.465,4.899,3.612,3.054,-1.18,3.3,4.596,5.005,3.52,4.306,3.359,-3.021,5.066,-2.925,-3.578,-4.909,-4.638,-2.37,2.665,-4.379,4.738,-4.749,-2.162,4.794,3.318,3.482,3.054,4.031,3.328,3.738,-3.114,0.3296,-1.601,-3.01,-3.472,-1.559,-3.055,-1.969,-4.795,-1.344,-3.323,-3.044,3.807,4.739,-0.2773,0.95,-3.984,-2.762,3.684,3.649,3.533,-4.101,4.631,2.866,0.08414,-4.788,5.629,3.285,2.066,2.434,4.05,4.916,-0.6816,4.554,3.998,4.398,4.301,-4.45,-4.152,4.639,4.819,1.816,3.561,4.188,4.295,1.486,5.497,0.4232,-2.331,3.471,2.734,4.03,2.495,0.3675,3.787,-2.047,3.947,2.094,4.778,0.6207,4.805,5.125e-15,4.487,3.318,3.054,2.866,4.501,4.085,5.258,5.022,3.253,6.21,-3.765,-4.767,0.0323,4.309,2.866,3.801,4.073,4.113,-1.353,3.605,2.955,4.124,3.535,-1.023,0.02525,5.57,5.038,0.8142,-0.1041,0.2102,0.3217,3.253,-2.548,-2.508,4.811,0.8463,-0.6589,-1.465,1.895,1.368,-6.873,-3.06,-2,-3.895,-3.734,-0.5872,-6.038,5.125e-15,1.058,-6.244,5.18,-2.365,0.5307,-3.257,1.506,6.235,1.495,-3.014,3.284,2.951,-5.021,-3.451,-0.4784,-4.765,0.644,-4.78,1.787,1.35,3.016,-0.9045,0.1523,-3.525,0.9691,-1.583,-0.4889,-0.7913,-0.6041,0.177,-0.06121,-2.355,-0.1491,-0.6906,-1.434,0.4253,-1.316,0.4101,-1.564,-0.04858,0.6185,-4.497,-0.2636,-0.7589,-1.069,-0.09579,1.49,-2.45,0.6687,-0.3506,1.227,1.392,1.214,-0.518,0.328,1.743,-1.801,-1.313,0.7304,-0.1744,-0.1498,1.615,1.403,0.5364,1.738,1.695,1.3,1.699,2.66,1.911,-1.081,0.3161,-0.4839,-0.7142,1.058,-0.1301,0.4952,2.442,2.182,-1.231,-0.323,1.674,-1.778,-0.1867,0.3078,1.552,1.831,-0.09905,-0.381,0.8924,1.11,1.709,-1.644,1.331,0.3258,-0.03417,-3.475,1.495,1.05,2.805,-2.584,0.9386,-1.296,0.9593,-0.4496,-1.059,-0.8249,1.563,0.6845,0.5679,-1.525,0.148,-0.2992,1.281,2.185,0.0764,1.263,-0.2872,3.38,-2.055,1.091,-0.845,1.809,-1.736,4.045,-1.464,4.697,3.916,3.895,4.073,5.74,4.102,2.246,-0.5747,4.073,-0.1519,-2.016,-4.38,-0.3051,-1.135,-1.581,-2.525,0.4512,-1.315,-3.448,-1.855,-3.312,-1.328,-0.007433,-5.119,-4.097,1.108,-2.544,-1.183,1.044,0.1938,-2.118,-1.146,1.695,0.5967,-2.192,-1.419,0.9141,0.3169,1.912,-0.8815,-0.6539,-0.03942,1.75,-1.235,1.317,-2.061,1.306,1.631,0.07003,-1.065,-0.66,1.083,-0.7049,0.9698,1.689,2.242,0.1002,1.274,0.5326,-1.471,-1.532,-1.123,-0.7219,1.031,0.712,-0.6967,0.8758,-0.5332,-0.4726,-2.651,2.336,2.776,-2.254,-0.4957,-0.2643,-2.193,-1.374,-0.3021,1.191,1.384,0.07769,-0.388,0.07224,1.234,-2.119,-1.706,-0.3062,0.4036,-1.317,-0.07019,-0.202,-2.342,-3.052,-0.4762,-0.9069,-3.076,-0.9455,0.4699,-0.5503,-0.1728,1.228,-2.351,-0.6913,-0.6799,-1.692,0.8833,-1.966,1.821,-2.782,-3.911,-2.643,-2.404,-4.648,-4.541,1.242,2.514,1.035,4.525,4.394,4.03,-3.574,5.677,1.616,5.789,4.051,3.757,6.267,-3.97,-3.203,4.073,3.482,1.656,4.514,-0.07813,-0.7782,-0.6082,-1.838,-1.425,-2.517,-0.136,-0.4391,0.3964,0.3074,-0.08706,1.103,-1.29,-0.01049,-0.7407,0.4206,-2.217,1.397,-1.322,0.7149,0.5048,1.644,-3.882,-2.013,3.467,-1.662,-1.574,-0.7973,1.783,2.531,-1.499,0.6013,2.166,2.517,1.714,-0.002848,0.2744,1.376,1.442,1.048,0.6968,1.369,1.249,0.5565,2.046,-1.651,1.557,0.6559,1.206,1.004,-0.1173,-1.208,-1.781,-0.4079,-1.954,-0.5939,1.869,-0.06994,1.98,-1.92,-0.1861,0.6485,-0.5069,-0.6218,-0.9302,0.8736,-0.9295,0.7632,0.2908,-0.5626,-1.343,0.8795,1.054,1.691,2.073,1.345,1.2,0.9238,1.728,0.713,-1.222,1.195,-1.294,2.214,0.9277,-0.08749,-0.08454,0.7975,0.5373,4.839,1.626,-0.3824,1.092,0.6898,-0.204,-0.5121,1.189,2.801,0.4844,0.7578,1.035,-2.564,4.305,-5.335,3.497,-2.842,4.505,0.5839,-1.596,-0.4301,-1.448,-0.8387,-3.369,-0.3822,-1.324,-1.916,0.2922,1.384,-0.3171,-1.294,-0.16,0.567,0.003789,-1.367,2.216,-1.116,-1.453,-2.682,-2.222,1.65,1.12,1.155,1.068,0.3353,1.329,2.944,1.653,1.796,1.213,3.011,-2.196,-1.331,0.4227,1.912,0.0491,0.9682,0.4275,-3.028,-1.335,0.4205,-1.587,-0.151,-1.339,0.8139,-0.05051,-1.239,1.053,-0.5019,-1.137,1.017,1.47,-0.2319,1.55,1.004,-0.7843,-2.125,-0.5497,-0.5213,0.2411,1.18,-0.1163,-1.095,-1.562,1.834,1.216,-0.5388,-2.444,1.208,-0.06674,-1.22,3.204,-2.58,-0.8888,1.323,-0.5,-2.176,0.7359,-1.465,2.907,0.2868,-0.1142,4.06,-0.3769,-2.182,-1.492,1.25,1.615,-0.05898,1.402,0.7087,1.126,0.1759,1.321,-0.3341,-0.2225,-1.152,-0.3775,1.655,-3.993,-1.503,-4.454,0.365,-1.493,-0.1225,0.905,1.286,3.523,-2.292,-0.7551,3.765,-2.937,-3.195,-3.672,0.66,-1.155,-0.6397,-0.5346,-3.1,-0.6531,-0.07156,4.165,-1.495,0.5231,0.9967,-2.078,1.168,1.035,-1.041,-0.1638,0.2447,-0.6034,-0.02467,-1.741,-0.264,2.559,-0.7434,-1.674,-1.465,-1.795,1.214,0.6606,0.2784,0.9295,1.337,-0.5443,-1.261,1.42,-1.575,-2.368,2.479,1.79,2.352,0.4945,-0.1943,-0.5662,-0.7909,1.001,-2.257,-0.7543,1.771,1.061,-2.496,1.352,0.2727,1.384,1.663,-1.374,-0.9936,0.117,-1.578,-2.297,0.8274,0.2446,0.3647,-0.6955,-2.16,-2.856,0.578,1.939,1.281,-0.585,0.9553,-1.291,0.07109,-3.264,-3.687,-4.497,-4.684,-1.469,-2.888,-2.818,-1.882,0.9536,-0.2843,1.304,0.6571,-0.8038,2.092,-0.5824,-0.9005,-0.9039,-2.085,1.942,-0.1349,1.285,-0.009536,0.0293,-2.297,-1.707,0.6528,-1.696,1.748,-2.621,2.284,-0.6833,0.7326,0.8396,-0.1358,1.861,0.4176,-0.1624,0.9009,0.1402,0.173,1.437,-0.4163,-1.363,-1.995,-3.446,-0.169,-3.68,-2.085,-0.2486,-1.838,-0.251,-2.253,-4.583,-4.812,-2.419,1.498,0.317,-3.612,-2.108,-0.8137,1.079,-1.376,-1.732,2.724,2.844,1.458,0.6513,-2.104,-1.201,-1.973,-1.902,1.167,0.3664,3.123,1.468,-0.4018,-1.48,-0.5475,1.167,-0.6927,0.8025,-0.6817,0.9372,-0.102,-1.85,2.21,-1.248,-3.4,-1.569,1.664,-2.28,1.824,-2.044,0.4777,-0.09777,0.1709,-1.745,-0.3385,-2.134,-1.997,-4.597,-3.126,0.6075,-2.449,-2.745,0.919,-3.234,-2.884,3.561,4.356,0.5244,3.568,0.8624,-0.388,-0.8068,-2.024,1.915,3.237,2.915,0.2816,0.616,1.386,-2.916,-0.8301,-0.2954,1.281,1.355,-0.452,1.585,1.299,0.4565,0.8355,1.337,1.634,1.914,-1.679,0.784,0.04169,1.247,0.3658,1.237,-0.9987,2.693,2.23,1.287,0.8616,0.7813,-2.747,-1.143,0.501,0.985,0.2309,1.108,-1.537,0.3222,0.9882,-2.071,1.53,-2.315,-3.382,-1.463,-2.499,-0.9011,1.511,0.631,-2.167,1.174,0.2223,0.233,1.162,-1.908,0.5325,-0.5201,0.9528,0.6314,1.19,-4.15,-0.6235,-0.1337,0.3591,-2.094,-0.9436,-2.749,-0.753,2.191,0.1266,-1.022,0.199,0.3848,3.718,-1.652,-2.337,0.2001,0.6265,-1.486,0.5395,-0.4165,-0.5789,-0.7374,0.9912,-0.0843,-1.884,1.185,0.9296,-0.7343,-1.646,-1.803,-0.7908,0.9718,1.668,1.953,-0.6786,2.596,0.1831,0.174,-1.991,1.384,-0.9948,1.996,-0.8817,1.05,0.4113,2.66,-0.03234,-0.6634,-3.583,1.555,-2.644,-1.716,-1.618,-0.3585,1.821,0.2261,1.016,1.165,-4.61,-2.16,0.5943,1.059,-1.006,-0.4897,1.267,-2.856,-1.227,-2.996,-0.7952,-4.654,-4.403,-5.153,-1.89,-2.352,0.2627,2.161,-2.018,-2.263,-0.9029,-1.871,-1.627,-0.1507,-0.04636,-0.4227,0.2176,-1.288,-0.2825,1.86,-2.775,-2.436,-1.779,1.24,1.124,-0.3777,0.5766,0.7113,-2.204,-0.9837,-1.606,-0.3059,-0.6333,1.933,-0.8635,-0.1877,-2.505,-0.3673,-1.361,1.027,1.243,-0.6179,-0.3482,-1.241,-0.6093,1.541,-0.365,1.26,-0.7458,-0.4597,-1.254,0.6457,1.495,1.27,1.666,2.002,-0.2425,0.3688,-0.4909,-1.942,0.9816,2.417,0.9357,-0.2197,-0.6922,-1.767,1.59,1.436,-1.688,-2.042,-1.58,-0.2252,0.03543,1.432,1.957,-0.02202,-1.006,-0.5777,0.5338,-1.58,-0.3717,0.8312,0.226,0.4885,0.5181,1.389,-1.714,-0.3734,-0.008522,0.5789,0.5831,0.4945,-1.681,2.574,0.1633,0.8012,1.467,0.9936,-1.883,-1.201,-0.3778,-1.487,-0.3336,-2.707,-2.251,-2.069,0.8199,-1.455,1.004,1.18,-0.146,0.2101,1.109,-2.09,1.123,-1.554,1.186,0.6088,1.138,1.748,-0.4902,-2.52,1.333,-0.4691,1.696,-1.239,0.1169,-1.982,1,0.3684,0.1949,0.2236,1.17,1.114,4.132,0.7199,0.7065,1.234,-3.228,0.02607,1.504,2.993,0.3691,0.02399,-1.181,-0.05055,-0.4203,1.484,1.293,-0.8749,2.394,-1.166,-0.5438,0.6952,1.757,-0.03435,2.792,1.834,0.7873,-1.832,0.788,1.908,0.776,0.2487,-1.967,2.539,-1.077,-2.165,2.494,0.5865,-0.5774,1.009,1.546,1.849,-0.1208,-1.883,-1.714,-0.8805,-0.5233,0.9894,1.742,-1.145,1.092,-0.7872,1.422,-1.679,-3.261,-0.566,-1.564,-0.6038,-0.2386,4.485,-0.4863,-0.3055,-0.4703,0.6772,2.015,1.369,-2.254,0.255,0.4009,0.3615,1.278,1.109,2.405,1.29,2.699,-2.846,0.4356,-1.078,-1.529,0.1345,0.3703,-0.05075,-1.344,-2.303,-0.6099,1.384,-0.1548,-2.881,-2.212,3.965,-2.051,2.689,-0.1373,1.394,-1.405,-2.026,3.573,3.467,5.174,-4.637,-3.232,0.1534,-1.456,-4.762,-2.601,-2.276,-2.646,-3.247,-2.502,1.257,1.651,-1.636,-1.879,-3.897,0.4731,-0.2022,-1.607,1.565,0.1618,-0.8741,-1.175,1.699,-0.1481,0.323,-3.635,-2.788,-1.549,-1.093,1.015,-2.729,3.745,-2.192,1.348,0.08233,-0.4548,0.9972,1.626,1.368,0.3912,1.126,-0.5857,1.025,0.9009,1.32,-0.8077,-3.441,-1.463,1.455,1.56,0.1265,-1.502,0.1868,-1.974,-0.7388,1.638,-1.432,-1.75,0.05516,0.7003,0.6322,-0.5359,-0.9524,-3.343,-0.3472,0.8694,1.355,-0.9699,-1.515,-0.04363,1.079,-0.8789,-3.302,0.176,-0.07955,-1.744,0.7479,0.435,-1.31,0.2272,1.396,-1.909,-2.06,-3.198,-0.6335,0.5402,0.4746,0.4189,-1.149,-1.421,-2.217,-2.73,0.8153,-0.09088,-1.548,0.2542,0.6359,-0.8463,-3.262,-1.992,0.1204,-0.1641,-0.3319,-0.16,0.8122,1.064,0.3225,1.281,-0.8496,0.2665,0.5265,1.291,1.185,-0.2441,-1.006,-0.1098,1.356,1.854,0.6274,1.43,-0.8159,-2.045,0.6166,1.018,2.028,1.529,0.01742,1.271,-1.942,0.08642,1.05,-1.689,-0.09083,-0.197,1.029,-0.6923,1.216,-0.8892,1.129,-0.178,2.425,-2.657,-2.056,-2.746,0.2804,0.8473,2.618,0.3906,0.3094,-1.378,-0.5897,3.012,0.07304,-2.794,-0.6628,1.128,0.1895,2.411,-1.611,-2.642,-2.041,-0.8973,0.5445,0.004317,2.622,-0.3536,-0.9253,1.22,1.452,1.165,-0.516,-0.02933,-0.3407,-3.283,2.521,1.808,1.145,-2.008,-0.4217,1.032,-0.7831,1.717,-2.395,-1.014,-0.5596,0.1543,0.2804,0.9282,-3.123,-0.6256,2.062,2.617,-0.4362,1.395,-3.024,-0.4771,-1.66,-1.488,-0.1564,-2.198,-1.385,3.961,-1.826,1.318,0.7287,1.02,-0.6413,0.4722,-0.3176,2.139,-2.342,-0.5288,-1.518,0.3409,0.06348,-1.056,1.768,0.1711,-0.05919,-1.978,-0.533,-0.553,-2.453,-0.3102,0.6191,1.581,-2.547,-0.617,-2.715,-3.413,4.193,4.232,3.855,4.024,-1.476,-7.093,-5.825,0.6942,-1.786,0.6731,-6.093,0.4406,-5.618,-2.599,4.647,-1.199,-0.3129,0.5606,-1.039,-0.3688,-1.238,0.2848,-1.543,3.035,-2.96,0.7189,-0.3918,2.928,-0.4476,-0.5111,0.3884,1.219,-0.3485,-0.01559,-0.3383,-1.208,0.8729,-2.252,1.01,-2.453,2.34,-0.7636,0.07686,-1.035,-3.264,-0.9934,-1.675,0.4159,0.3468,-0.7905,3.448,0.6504,-1.146,-2.03,-0.4702,-3.16,-3.047,0.5723,0.7692,-0.24,0.1591,-0.6112,0.4239,-0.1336,-1.017,-2.933,-0.38,0.2553,1.751,1.241,0.7713,1.747,-2.173,-3.131,-2.017,-2.483,-1.876,2.125,0.1168,-1.979,-3.092,1.504,1.113,-2.457,-1.555,1.217,0.8392,-0.2626,-3.641,-2.119,-1.178,-0.5696,0.319,1.474,0.6207,1.33,-2.372,-0.8226,1.442,0.9068,-0.9884,-1.323,1.059,-1.541,0.3508,1.219,-1.317,2.388,1.787,4.986,-2.638,-0.3386,1.164,-0.1035,1.38,-1.903,1.615,0.4441,1.105,1.238,-4.011,-2.369,1.08,-0.8469,-1.467,-1.558,-0.8803,0.4097,-0.6266,0.2049,0.4423,0.7366,1.368,-0.6137,0.9154,0.1032,0.3887,0.628,-0.9527,-1.26,-1.534,-1.739,2.151,-0.1575,0.45,-2.07,-2.025,-0.8712,-0.3777,-0.5118,1.967,-1.295,-1.173,-0.6892,-1.092,1.324,-0.1755,-1.472,-1.776,0.4802,-0.801,-0.6276,-0.7188,3.43,-0.3349,0.06798,0.76,2.619,-0.9497,1.966,-1.374,-0.5935,-1.075,1.216,-0.3552,-0.9394,-2.522,-0.4371,-0.2531,0.8058,-1.511,0.05469,-0.8181,-0.8898,-1.105,-2.048,0.5783,1.317,0.5359,2.118,3.934,1.057,1.229,0.1996,0.3821,-1.356,0.6708,-2.471,-0.2279,0.1089,-0.1618,1.39,0.7591,-0.4532,0.8175,0.6321,-0.6148,3.983,0.575,-0.3077,-2.168,-3.152,1.184,-3.951,-1.648,-1.759,0.1567,1.434,0.3283,0.8354,0.2651,-1.214,2.519,-1.388,0.3204,-2.033,-1.356,2.544,3.453,1.08,-2.433,-2.25,2.09,0.7647,-0.6323,-1.382,-0.9231,2.069,-0.03877,1.08,0.02847,-2.489,1.071,-3.275,1.11,0.2987,0.2109,-0.7557,3.449,-0.433,-0.09826,0.7996,1.052,0.03649,1.282,0.7262,-1.153,-0.9777,1.904,-0.8161,-1.664,2.267,0.8939,-0.3424,3.193,2.026,1.644,0.5628,2.566,0.5712,0.1446,2.332,-1.639,-1.078,-0.2339,-1.305,-1.866,-0.6385,-2.387,0.5089,0.3541,-0.1572,1.241,-2.447,0.8039,-0.01686,-1.86,0.6558,-1.053,0.1563,0.9104,-0.7026,6.427,0.1547,-3.536,0.6258,3.363,-1.12,4.151,-4.858,-0.3777,0.6006,1.5,0.9245,0.7518,-3.385,1.009,-0.7357,-1.495,-0.3652,-0.9158,-0.02274,2.238,-2.962,-1.927,-2.868,-2.478,-0.3725,1.397,0.2458,0.3835,-0.3474,-1.133,0.8778,-0.2122,0.1027,-0.683,-1.266,-0.9907,-0.5962,-0.2265,1.682,-1.248,-0.3904,1.665,1.041,-0.2769,0.04191,-1.898,0.9665,-0.805,0.6977,-2.985,-2.246,-0.1596,1.698,0.2675,2.244,-0.802,-0.5138,-0.3844,-2.786,-1.79,-0.7129,-2.666,1.038,2.34,-1.812,-1.043,0.9382,-0.9225,-1.132,-0.2213,0.9498,-1.68,-0.06959,1.779,0.7999,-1.224,1.463,0.1487,0.2852,-1.248,0.5369,-0.1657,2.678,2.867,-0.986,1.597,-1.271,1.263,0.536,-0.5787,-1.245,-2.042,3.976,0.7728,0.2817,-0.9537,1.496,1.677,1.543,-1.589,-2.469,1.515,-2.103,3.084,-0.05358,1.217,0.4991,1.643,-0.6391,0.3333,0.1686,3.644,2.218,-0.5863,0.8874,0.09734,0.8913,1.022,-0.2995,0.1412,-0.04255,1.781,2.669,-0.368,-1.065,-0.4414,-2.289,-0.7436,-1.627,-1.711,-3.093,1.026,-0.5089,2.978,-1.105,0.2453,-1.587,3.996,1.201,1.519,2.328,1.554,-0.03917,-0.7079,0.9701,2.126,0.4597,1.015,1.848,1.448,-0.5192,-0.3111,-1.429,0.1173,-0.3392,-1.358,2.871,-1.44,1.069,-1.481,-0.1407,0.1988,-1.083,0.15,0.187,-0.0684,1.587,2.45,-1.234,-0.753,0.5891,-0.1312,0.4323,0.446,2.177,-0.1964,-1.506,1.566,-2.066,1.17,2.034,-1.714,0.6432,-0.5248,-0.3143,1.489,-3.099,1.225,0.4225,-1.636,1.718,0.5524,-0.9662,-1.076,-3.634,-0.05517,-2.176,-0.3045,-2.313,-1.108,-6.518,0.2226,4.233,-2.676,-3.38,-4.486,-2.245,-3.195,-0.524,-0.8993,-1.013,-0.8277,1.377,-0.8894,-1.198,1.462,-1.371,-3.229,1.077,-0.0781,-0.702,-0.4341,0.7467,-2.908,0.02413,0.5688,-0.7376,-0.5864,2.432,-1.233,0.1532,-2.451,0.2619,-2.265,-2.36,-1.506,1.316,-2.127,-0.4276,-1.45,1.723,-1.966,-0.6624,1.538,1.398,0.7066,-1.301,-0.432,-0.6199,-2.943,-0.1543,-0.2768,-1.158,-0.8658,0.503,0.422,1.279,-2.123,3.445,-1.298,1.884,-1.485,-1.914,0.6335,-0.00184,-1.799,-0.4249,-0.1528,-0.9144,0.8045,-0.4127,-0.9023,1.12,1.805,-1.792,1.514,-2.278,1.529,1.6,-0.2001,1.787,-1.61,-1.252,-1.511,-2.023,1.207,0.5898,-0.8709,1.371,-0.1106,0.1706,-2.589,-1.929,-1.411,-0.9963,-2.54,-0.1688,-1.049,-1.816,-0.3247,0.6049,-1.061,0.4016,2.246,-0.7508,-1.307,-0.2981,0.01698,2.57,-1.482,0.7466,-0.3585,0.9048,1.677,-2.437,2.342,-1.594,0.2063,-0.9661,-0.8521,-0.7242,1.577,-2.741,-4.059,-2.4,1.165,0.5136,-0.0264,-0.5301,0.8096,-0.2356,-0.1524,0.01147,-3.017,-0.6631,-1.425,-0.9951,-0.02475,2.437,-1.519,2.17,-0.2678,-1.668,-2.325,1.879,1.561,-0.6532,0.3906,1.342,1.732,0.6745,1.395,-1.554,-0.83,-2.613,-1.172,-0.9446,0.5002,-1.798,-1.819,0.4596,1.354,1.442,0.4637,-1.105,-0.01437,0.01326,1.522,0.5538,0.5045,-0.2491,0.764,-1.909,-0.6653,0.2107,1.605,-0.08669,1.082,2.855,-2.683,-1.626,-1.413,-0.1083,-1.322,1.36,0.4082,0.7318,1.22,0.2377,-1.327,-2.165,-0.8762,-1.632,-0.7745,-0.4691,1.897,0.6633,-0.02756,-1.545,-0.6041,0.7253,-1.333,-0.4454,-1.101,0.3632,-0.2329,1.259,0.5874,-0.5299,0.02728,0.07757,1.176,0.8014,-0.05286,-1.035,-2.591,-1.656,1.201,1.17,0.308,-0.2012,-1.214,-2.999,-2.308,-3.5,1.549,0.2508,1.538,-1.055,-0.9965,-1.559,0.07052,-3.244,-0.9016,0.2652,-0.7076,1.094,0.3546,-0.2219,-2.159,-1.812,0.657,-1.646,1.446,0.9519,-0.9072,1.793,-2.037,-2.785,1.422,-3.564,1.212,1.091,-1.198,0.1587,1.509,1.463,1.866,1.326,3.232,0.4096,-1.228,0.8043,-0.7602,-1.313,1.575,-1.755,4.164,-0.5909,1.034,0.9725,1.505,-3.759,-0.07551,-0.4133,0.6203,1.332,1.442,-1.257,-0.8643,-1.893,1.793,-3.136,-0.6557,1.073,0.218,1.373,-0.7472,-1.227,-0.5813,0.8086,-3.195,-3.656,-3.451,-2.982,-3.585,-3.294,4.048,1.608,4.974,-0.6959,-0.9352,3.931,5.217,5.792,2.181,-1.682,-0.8618,-2.602,-1.124,1.415,0.2611,2.878,1.421,1.035,-2.206,-0.8741,-0.3345,-1.169,-3.473,-1.277,2.112,-0.8432,-2.328,0.1265,-1.458,0.1483,-1.362,-2.123,0.2755,1.629,0.2157,-1.761,0.5712,-1.796,1.626,0.07178,0.5088,-1.132,1.598,-2.519,0.2738,1.065,0.6063,2.14,0.6062,1.634,-0.8132,-0.8477,0.9626,1.213,-0.329,2.222,0.7613,0.2815,-0.1225,-2.557,0.1368,-0.201,-1.862,-0.995,0.02401,-0.614,0.6866,-0.3837,0.5465,-1.744,-2.834,0.95,1.183,0.9637,-1.197,-3.804,1.811,-0.4778,-2.664,-1.579,2.108,-1.382,-1.374,0.3833,0.2457,-3.44,-0.2515,-0.754,0.1678,0.07389,2.869,-2.329,-0.3635,0.1025,-0.1003,-0.6414,-0.4945,0.9582,0.815,1.171,-0.863,1.221,-0.7763,0.5846,-1.988,2.048,-1.511,-2.3,-0.8082,-1.727,2.139,1.793,2.449,0.1877,-0.8798,0.9873,-0.07326,0.5763,-1.508,-1.985,-1.285,-0.8077,1.439,-0.5483,-2.171,-1.645,0.883,1.433,1.541,2.03,-0.3053,0.4859,0.9274,1.56,-0.6072,1.24,1.001,2.133,2.24,-0.6718,-1.97,3.078,0.1378,0.9845,0.9142,1.085,0.7734,-1.675,1.372,-0.2952,0.8839,-1.287,0.797,1.16,-0.2313,2.502,0.3373,1.498,0.2954,0.2843,-1.703,0.7778,-1.277,-0.1613,0.482,1.962,-0.2273,-2.5,0.1867,-0.03498,-0.1056,1.85,-1.026,0.4162,-1.789,-1.013,-1.174,-0.5517,-0.9255,0.06159,1.711,0.1548,0.1874,0.0628,-0.2142,-0.6903,1.01,-1.232,-1.687,-3.325,-1.601,0.8734,1.607,2.519,0.728,-1.421,-1.13,-1.572,-1.207,-0.1616,-0.9609,-2.782,-3.069,1.56,-2.529,1.361,1.153,-2.127,-0.9962,0.08735,-1.654,-2.239,0.3098,-0.5343,-1.094,2.002,1.17,0.3051,-0.8973,0.7931,-0.3058,-1.07,-1.57,0.9562,0.8383,-0.3216,1.545,-1.096,-1.222,-1.03,-0.1451,2.062,-2.474,-1.922,-2.56,-0.0739,1.194,-0.7266,1.987,-0.7212,-0.6062,0.3092,-0.2523,-0.4366,-0.3958,-1.615,-0.2523,-3.029,-1.026,-0.6287,0.9153,-0.1066,1.592,2.668,-0.2191,-1.471,-1.593,-0.06764,-0.09835,0.3154,-1.979,0.7278,-1.962,-1.193,-1.928,0.6603,2.077,-0.5362,0.9036,0.5814,-1.579,-1.009,0.07303,1.41,-1.387,-0.5264,-2.12,-3.011,1.082,0.8659,0.3611,-2.047,1.364,-3.795,-1.041,0.3347,1.684,-0.8807,-1.947,2.178,-0.2602,1.322,0.01614,2.735,-0.644,-3.1,0.6798,0.3954,0.7488,-2.824,0.09796,0.4993,-1.241,-3.416,-0.1055,2.196,-1.692,-2.942,-2.092,-0.3305,0.2638,-0.1951,0.4787,-0.002711,-2.362,-1.154,0.3746,0.4241,0.2131,1.206,0.7876,-1.044,1.052,-1.005,0.237,-2.674,0.5382,-0.9646,-3.019,-0.1753,-0.4547,0.1624,-0.8809,-1.883,2.309,-0.7684,-0.8998,1.604,-0.6619,-0.5025,-1.235,3.325,-0.7848,-2.339,-3.313,-0.3768,1.29,1.7,1.014,1.096,-0.5103,1.154,-0.5905,-2.461,1.297,1.556,0.9707,-1.442,-0.7614,1.1,-0.72,-3.292,1.782,0.9695,2.062,-0.1676,0.7034,-1.312,-2.746,0.7495,0.463,0.207,-2.552,-1.378,-2.311,-0.2406,1.179,0.399,1.008,-0.9182,-0.2043,0.8443,-0.08668,-2.082,0.6676,-1.304,-4.466,4.225,0.1163,-1.319,-1.762,-0.5822,-1.162,-0.1076,-1.55,0.6031,0.6621,-0.4334,-1.284,0.1564,-1.239,-2.247,-2.62,-0.8734,1.158,1.134,-2.547,-3.313,-1.486,-1.14,2.441,1.08,3.005,-1.119,2.751,-2.348,-3.28,-1.337,-3.837,0.6991,-2.54,3.72,3.523,3.628,6.433,2.163,0.2675,0.2449,-0.3338,0.9525,-0.4519,1.03,0.6435,0.5343,-0.0974,-0.8522,-0.3285,1.202,1.424,0.5874,-1.975,0.6012,-0.6891,3.363,-0.5059,-2.041,-3.954,1.098,-0.9754,-0.1259,0.2442,1.269,1.325,-0.1749,0.3976,3.138,1.27,1.577,-0.4145,-1.115,7.159,-0.1168,-1.545,-0.616,-1.526,-2.496,0.952,1.358,-3.911,-1.621,0.7945,0.2881,-1.12,-0.3624,-1.693,-2.293,-1.62,-2.29,0.7034,0.9791,0.4123,1.94,-0.5707,-1.361,-1.829,-0.8751,-2.119,0.08133,-0.7864,1.188,0.0161,-1.548,-0.2865,0.4899,0.6358,-0.5574,-0.7199,0.5352,0.4747,-3.546,-1.101,-1.774,-1.929,-2.543,-2.449,-0.1681,-1.195,-1.19,-1.503,-1.303,0.9063,-0.4748,0.4469,0.1679,-1.364,1.119,-2.861,-2.24,-0.8732,1.769,1.222,1.592,-1.427,1.064,-1.221,0.5013,1.18,-0.1546,0.4679,1.692,-2.296,-1.501,-2.731,2.252,-0.3475,-0.9191,-0.4915,1.128,0.5678,0.6302,-2.633,0.7873,-0.1552,-1.001,-1.258,-1.109,-2.377,-0.5132,-2.286,-1.242,-0.5063,-1.205,-2.478,-1.55,1.997,0.09182,0.2886,-0.1968,-0.8622,-1.906,1.127,1.337,-1.492,1.344,1.112,-0.5894,-1.448,-1.315,0.9287,2.232,-0.8912,-1.145,-0.6314,-1.083,-0.9184,1.456,1.335,-0.9594,-0.6749,1.218,0.1278,0.122,-4.243,-3.366,0.4149,-0.5378,-1.96,1.398,-1.883,-0.3238,1.859,1.623,1.499,0.02761,-0.9072,-0.3794,2.489,-0.5937,0.5519,-1.089,-1.412,1.387,0.4875,-0.2042,0.2196,-0.8102,1.242,0.07928,-1.377,2.313,-0.1938,0.8263,-0.2042,-1.939,-0.0929,0.3953,-0.7275,-1.568,-2.048,1.027,1.59,-1.752,-0.3306,-0.6391,1.22,-0.1779,-0.3897,1.274,1.104,1.654,-0.007221,1.457,2.769,-1.316,-2.25,0.8679,-1.276,-2.173,0.2002,0.3844,0.685,-0.8058,0.5294,-0.6589,1.723,-1.54,-1.225,0.6235,-0.5053,-1.446,1.07,-0.364,-2.588,0.4858,-0.6047,-0.7783,-0.3764,-0.3648,1.982,1.055,2.025,-1.679,0.8757,-0.07481,-1.267,0.5997,-0.006877,1.974,-0.2298,1.337,-0.4825,-3.215,1.284,0.7577,1.056,-1.091,0.5992,0.4788,-2.134,-0.9211,-3.509,-3.066,0.9019,2.208,-0.5891,0.5023,0.8837,-0.4962,-0.05346,0.3291,-0.965,5.211,3.601,3.541,2.385,-5.991,1.849,-4.669,-4.069,-2.573,-1.433,-4.239,-5.242,-3.567,-4.131,-2.953,-3.012,5.658,-0.9229,0.0502,1.406,0.2173,-1.109,-1.622,0.7001,-2.29,-1.354,1.297,-0.5493,-0.5951,-3.136,-0.4553,-0.9262,1.591,-1.212,0.6904,1.834,-1.578,-1.394,-1.069,-2.748,-1.136,-2.499,-0.1253,-1.687,-1.337,-1.363,-2.073,0.8344,0.1104,0.0268,-1.878,1.872,0.01324,1.052,0.6727,-1.642,-1.204,0.5406,-0.6863,-1.093,-1.578,-0.2981,-0.7676,1.614,-1.439,-0.339,1.335,-1.384,-2.368,0.2222,1.144,1.091,0.2148,-0.78,-0.5557,-2.105,-2.522,0.0905,-3.597,3.518,-2.281,0.9148,-0.5262,-3.116,-1.675,-2.596,8.296,-0.103,-1.951,-1.345,1.288,-0.4566,-0.09829,-2.429,-1.575,0.837,0.3912,0.734,-0.8282,1.619,2.699,-0.5131,0.8664,0.3314,0.1085,4.79,3.651,0.8635,-0.4942,-1.461,0.8441,-0.7417,-1.009,0.02314,-0.9602,-0.7931,-3.35,-1.023,0.1204,1.287,1.183,0.3215,-1.679,1.415,0.1709,0.472,-1.392,1.18,-4.15,0.5473,-1.442,0.6569,0.2309,3.237,-1.408,0.9172,0.2397,-0.9034,-1.136,0.6221,0.02322,1.567,3.356,1.011,-2.587,1.288,1.34,1.294,-0.7053,0.5015,-1.448,2.444,-1.791,-2.481,0.2899,-1.017,0.9618,-0.5872,-0.1816,-0.3517,-0.337,-1.871,-1.702,-0.2644,1.461,1.649,-0.2753,-1.771,1.29,-3.028,-1.392,-0.512,-0.1654,2.152,0.6533,-0.447,0.4424,-1.182,1.607,0.0934,-1.392,-1.246,-2.073,1.386,0.8145,-0.1589,-0.8695,-0.6854,0.6305,-0.1827,1.964,-1.261,1.434,0.4884,1.157,-1.721,-0.8818,-2.333,-0.9354,-1.274,-2.501,-2.438,-3.844,2.711,-1.074,-2.211,-1.499,-0.8246,-2.812,2.226,1.736,1.316,-1.287,1.014,0.5785,1.714,2.603,2.653,0.7848,0.8693,-0.2253,1.249,-0.1843,1.179,-0.7003,0.846,-1.652,-0.003184,0.07076,2.899,0.6671,-2.467,2.781,-1.482,1.268,1.395,-0.735,-0.3536,-1.645,1.339,-2.88,-0.6155,-0.4276,-1.145,-1.785,1.024,2.52,-1.593,-1.652,0.3009,-2.03,0.1792,-1.534,-1.364,0.9837,0.2957,-1.763,0.3928,-1.729,1.856,0.3689,1.364,0.3717,-0.8032,1.124,1.511,-1.264,0.0607,-1.888,-1.467,1.254,0.5434,-0.7829,-0.9895,1.087,0.7572,-0.2102,-2.575,1.947,1.436,0.6913,-1.199,0.0119,-0.4687,1.506,-1.761,0.7565,0.7926,-0.5271,0.4462,-0.5758,-0.9509,0.1382,1.979,0.7512,0.4516,1.634,1.535,-2.512,-2.703,0.7555,0.5377,0.8378,1.85,1.999,-0.4094,-1.536,0.1864,1.237,0.1003,-1.648,1.3,1.267,1.442,-2.187,-1.464,1.972,-0.2359,1.057,1.465,1.552,0.9265,1.723,-1.206,-0.2096,0.2555,2.895,-0.3956,1.548,-0.5447,0.5598,0.02458,-0.4508,-1.116,-0.9603,-0.7777,0.3845,3.478,0.5351,-1.04,1.066,1.227,0.3378,1.081,-3.192,-1.267,-0.8471,-1.497,0.291,-0.01406,-2.721,-0.379,1.295,-0.9412,-0.4102,1.544,-1.379,1.589,0.4369,-4.065,-0.3329,-2.118,0.4659,-0.388,1.048,-2.774,-3.303,-1.434,0.825,0.3415,-0.128,-1.239,0.7482,-0.2628,1.374,0.03668,0.9016,0.07533,1.319,0.333,-1.651,1.601,2.125,1.178,0.2342,-0.003417,0.3093,0.5701,-1.832,1.405,2.014,-0.8961,2.157,0.991,0.8097,2.208,1.123,0.1677,0.7824,1.393,-0.3806,0.1582,-0.7721,-0.4947,3.16,-0.719,2.275,2.193,0.2305,1.769,-0.6437,-0.7245,2.025,3.139,1.81,-2.809,-0.2158,1.014,-4.934,-2.664,-2.836,-0.8636,-1.906,-4.524,-3.172,-1.224,-2.875,2.455,-1.139,-2.238,-2.183,-4.28,-0.1318,2.398,3.284,5.125e-15,-4.191,-5.614,-4.958,-6.062,-3.891,-5.966,-6.876,-7.985,-6.439,-7.268,-6.511,-5.493,-6.841,-5.973,-6.772,-6.036,-4.982,-7.778,-8.034,-6.632,-5.834,-7.759,-6.97,-7.116,-5.835,-6.445,7.511,-5.058,-7.014,-7.378,-7.213,-2.569,-4.724,-3.44,-1.587,2.092,-4.538,-6.071,-5.957,-4.11,-2.2,-3.483,-5.865,-1.646,3.544,2.695,-2.236,2.561,3.671,3.903,4.535,1.743,2.844,4.021,-1.876,3.057,-3.531,3.209,2.413,-2.654,2.87,3.171,3.057,3.703,4.007,3.271,4.264,-1.91,1.816,-1.782,3.028,3.238,2.603,4.585,2.013,-4.319,3.138,2.294,3.051,4.678,2.353,2.362,5.08,2.366,-3.956,-4.508,-6.532,-7.818,-7.335,-2.377,-2.521,-3.125,2.014,-1.402,3.534,3.345,3.835,4.856,3.513,-3.291,4.274,2.428,2.593,-4.924,6.054,2.096,-3.973,-1.624,2.274,3.795,5.076,5.561,3.431,2.191,1.778,2.796,1.347,3.172,4.763,-3.641,-3.126,-4.095,-4.203,-3.375,3.146,2.302,-2.42,-3.603,-6.717,2.248,3.772,3.669,-1.643,-5.932,-3.877,-7.13,-6.132,1.824,4.766,-2.907,-2.792,1.616,2.924,2.889,3.432,5.171,11.36,6.654,-3.568,-2.486,6.367,-3.37,2.917,-7.408,3.706,5.415,-6.056,-3.442,-1.726,5.088,3.137,2.121,13.27,11.77,2.465,1.965,3,2.631,2.435,2.312,3.267,2.831,1.904,3.104,4.077,-2.617,4.797,2.266,2.46,-2.497,2.254,2.086,2.217,1.819,5.625,5.896,3.462,2.674,-3.745,5.812,5.052,2.195,7.111,4.221,2.151,-2.607,-3.036,2.254,4.144,3.364,2.969,2.748,-3.121,-3.669,4.708,2.756,4.118,2.666,6.106,4.775,3.433,4.526,6.471,3.479,3.687,2.177,3.515,3.899,4.059,1.815,1.912,-4.286,-7.947,-4.193,-7.442,4.217,-5.367,2.065,2.736,3.217,-4.095,1.67,3.979,2.783,3.895,4.112,1.319,1.644,3.171,2.202,2.514,2.346,3.969,-2.332,-3.527,-3.959,3.649,2.274,-3.638,1.415,5.066,3.892,1.861,6.738,-4.012,5.137,1.956,2.687,1.685,-4.045,-3.368,-3.582,-3.022,-3.379,-2.672,-4.967,2.435,-4.371,1.871,5.258,4.211,3.453,4.253,2.97,2.693,4.557,4.477,3.484,1.729,-6.877,-6.771,-7.414,-2.633,-6.006,-7.828,9.79,2.558,-5.416,-3.528,-4.094,2.072,3.194,2.842,5.161,4.839,-3.699,2.589,6.746,3.835,4.444,2.985,4.479,5.336,-5.267,-6.309,-6.85,-7.04,-7.611,7.902,-5.314,-6.284,-4.338,-5.472,-5.634,-3.666,-5.611,-7.544,-4.899,2.59,4.621,4.718,3.313,3.376,2.714,3.224,3.984,2.5,2.413,-2.036,3.075,3.041,5.689,3.815,-3.743,5.268,-3.344,2.449,2.48,3.437,2.671,3.309,-1.497,-2.602,1.252,-2.491,6.485,3.452,3.998,-2.514,2.125,1.615,2.84,7.305,-4.75,4.23,3.264,-3.923,-5.068,1.436,4.419,3.421,-6.289,-4.535,-4.901,5.361,7.355,4.826,5.565,-1.695,5.2,1.782,-2.731,-4.829,-5.77,-4.153,-3.572,-5.992,-2.942,4.892,5.288,3.039,3.242,-2.393,-2.369,-2.754,-4.958,-8.193,-2.816,-4.886,-7.731,-6.17,-6.276,-4.822,-5.658,-4.341,-3.724,-4.333,-2.111,3.055,7.149,3.314,4.17,3.995,3.855,3.115,3.383,-3.921,2.047,2.09,4.449,4.137,3.863,6.119,-2.05,-2.393,2.861,2.613,3.569,-3.568,-4.585,-3.287,1.403,4.849,-3.524,-6.533,-6.698,-4.372,-5.54,-9.569,-8.848,-6.723,-6.09,-4.471,-5.796,-3.224,5.525,3.163,8.785,5.361,2.899,3.118,4.345,3.463,5.499,3.474,3.284,3.614,2.904,6.667,-4.775,2.362,2.153,4.016,3.49,4.079,3.248,3.461,3.511,2.631,-3.591,-4.38,-3.443,3.982,2.277,2.559,2.819,6.73,1.439,7.85,-3.106,-3.435,-1.238,-6.632,2.094,2.842,3.336,3.803,9.056,3.3,-2.221,4.71,3.617,-1.987,-4.135,-4.57,-2.935,-4.284,2.684,5.053,2.837,7.1,-3.203,2.384,3.759,3.288,-3.05,-2.634,2.677,-2.135,4.247,2.192,2.744,2.906,2.192,2.622,3.843,4.243,-3.056,-2.091,-7.504,-4.969,-3.488,-3.989,-7.954,-1.731,-2.007,-4.565,-2.427,-6.438,-2.939,2.499,-4.207,-5.87,-2.638,-8.372,-5.303,-3.818,-7.834,-8.207,-8.599,-2.833,-4.85,-5.681,-5.693,-5.994,-5.713,-6.045,-2.739,3.624,-2.727,4.129,-2.87,-1.268,-2.887,-5.594,-5.609,-5.719,-3.227,-3.908,-10.84,-5.763,3.173,-4.885,-5.909,-3.262,-2.986,-3.338,1.518,4.511,5.369,1.543,-3.688,2.49,2.004,-2.348,2.048,-2.719,4.007,3.824,3.211,-2.067,3.659,3.546,-3.863,-5.291,1.607,2.091,1.7,5.173,-2.072,-3.557,2.92,3.335,3.325,2.562,-2.714,4.447,2.604,3.537,2.464,-4.064,-8.962,-5.071,-3.732,4.482,-1.982,3.25,4.039,-1.85,6.348,3.775,-3.375,-2.619,2.996,3.572,-5.14,-1.735,-3.16,-4.116,4.411,-4.082,3.404,-3.077,-3.304,-4.084,2.693,5.311,-3.302,-1.593,-1.731,-3.019,-3.999,2.893,2.884,2.766,-4.498,-3.373,-1.8,-6.538,5.087,5.65,-8.416,-4.12,-3.752,2.288,3.133,7.945,8.358,-7.648,2.059,3.056,4.445,-3.122,-4.212,-4.06,2.059,-1.431,4.687,4.772,3.332,2.657,2.866,1.885,5.772,-4.65,-6.918,-2.768,-8.749,-6.975,-4.092,-4.254,-6.483,-6.658,-3.657,-5.301,-6.716,-3.393,-5.704,2.107,2.716,-3.019,-3.106,2.634,-5.296,-3.42,-2.564,6.412,-4.674,-5.298,3.571,3.106,1.618,2.969,-2.795,-1.431,2.897,7.13,-5.326,-4.439,-3.874,2.979,3.939,2.427,-3.46,-2.733,-4.342,-4.08,6.517,-4.248,-2.547,-3.898,2.427,5.19,-2.934,6.96,1.866,3.323,-3.734,3.219,-3.219,4.939,5.291,-2.018,1.556,-4.136,-4.882,-2.369,-3.867,3.046,-1.302,-4.648,4.351,-2.928,-2.269,2.607,-3.001,-5.672,-2.721,3.561,4.757,3.347,3.598,-3.005,-3.037,2.714,-4.835,-6.717,-2.936,3.669,-1.837,-1.991,-5.049,-2.656,5.507,-3.224,3.174,2.222,-2.855,-2.234,2.272,-3.743,2.736,-2.899,2.827,6.867,-1.819,-1.912,-1.679,-2.422,2.795,3.164,-4.326,3.172,6.327,-5.518,4.358,1.997,1.963,2.668,2.294,4.556,-3.207,-5.169,-1.75,4.981,3.938,3.839,2.661,-4.113,-3.357,-2.323,-4.996,4.536,7.362,-3.615,1.951,2.698,-2.067,4.179,2.4,-1.964,-3.046,-1.895,-3.464,2.635,-3.012,2.55,2.725,2.738,8.257,5.522,-1.422,2.437,-2.039,1.573,1.671,-4.045,-5.652,-6.02,-5.664,-6.067,-5.823,-5.621,-7.227,-9.837,-3.985,-2.823,-5.543,-2.561,1.618,-5.836,5.376,6.268,-3.524,-3.649,-6.031,-6.499,-5.861,-6.772,-2.374,2.738,1.674,2.638,4.46,-2.366,4.438,3.619,-4.417,-5.723,2.893,-4.27,-5.149,2.972,3.856,-6.596,2.226,-2.22,2.474,5.481,-2.929,2.238,-5.283,-3.387,-3.347,-3.127,-2.441,2.68,-4.131,-4.603,-4.246,-5.091,-3.309,-6.61,4.706,-5.295,2.12,4.785,-1.533,-3.215,2.912,4.437,2.29,1.474,2.21,2.655,2.392,1.809,4.117,2.294,2.882,-2.366,-2.922,-5.548,-5.14,-4.358,1.707,5.208,4.869,5.416,1.468,2.124,-3.372,-3.45,4.765,1.295,3.938,-2.997,-1.783,4.019,1.929,4.705,2.37,2.129,4.301,3.911,-2.293,4.174,1.636,3.515,-3.816,-2.525,-5.812,-2.564,-2.179,-3.478,4.678,7.352,1.942,2.477,3.821,-3.623,1.668,2.904,-8.419,3.92,-5.462,2.656,-2.229,-2.013,3.41,-2.453,-3.319,-1.624,-5.63,3.028,-4.244,-4.127,-6.061,-2.237,-5.637,10.76,-2.675,-3.18,-1.884,1.968,-4.885,8.345,3.507,-3.437,3.196,2.312,-2.281,-3.063,-2.974,2.253,-1.58,4.493,-4.683,-3.675,3.342,-4.917,-2.964,3.129,2.004,-3.86,7.348,-5.897,-2.08,2.359,-3.464,-2.173,5.146,3.81,-3.591,2.112,-4.843,1.657,7.225,-6.257,-7.158,2.09,2.575,2.702,-5.106,-4.543,-3.124,3.242,1.744,-3.196,-4.357,-3.257,-3.763,-4.08,2.666,-3.706,-2.733,-2.373,-2.861,1.646,1.781,-2.877,-2.599,-5.996,-2.425,-4.401,2.388,-3.061,-2.959,-3.133,-2.949,2.385,-2.802,-2.243,-2.591,5.07,3.324,-2.343,2.383,3.021,-3.258,-4.087,-4.539,3.779,-2.662,-3.171,2.032,-3.876,2.207,3.742,2.771,-1.6,1.893,2,-2.453,4.347,-3.029,-1.991,4.548,3.436,2.927,-2.938,-5.168,5.409,3.007,2.889,3.183,2.51,3.079,3.209,1.729,2.431,-2.295,2.527,-2.549,4.676,-3.124,2.516,7.754,2.777,-2.609,3.274,6.592,2.001,2.825,4.641,1.909,5.204,-2.79,4.761,4.237,-3.88,-2.61,-4.867,-2.382,2.965,2.934,-2.05,3.043,2.534,1.376,-6.903,-4.15,-2.317,1.761,-3.709,-3.216,-5.341,-2.374,-4.354,-3.198,-4.454,-5.178,-4.919,-3.491,6.746,-2.977,-2.166,-10.23,-4.948,-7.397,-4.94,-5.016,-5.135,-4.31,-4.891,-4.009,1.898,2.253,-2.218,1.284,2.522,1.663,-2.593,2.745,-3.656,-6.217,-3.151,1.704,2.495,-1.872,-3.299,-4.356,-1.85,3.147,-3.581,-2.407,2.894,-2.293,-2.528,-1.876,-2.173,-5.486,-3.462,-3.06,2.702,5.295,-5.459,-7.418,1.4,-3.101,-7.023,-2.695,3.562,-3.954,1.564,-2.541,2.911,-4.4,4.071,-4.461,4.25,-2.171,-3.206,2.035,-2.509,-4.163,2.453,2.822,7.448,2.895,2.697,-1.924,-4.07,-5.868,-2.406,2.031,1.771,-3.513,-4.285,1.785,2.831,8.236,-3.169,-4.14,-3.309,-6.837,-1.613,-4.473,2.603,3.577,-1.714,-3.641,-4.756,-2.525,2.546,1.488,4.352,1.744,-1.768,-4.866,1.789,-4.128,-2.176,-1.715,2.796,3.43,8.753,2.853,2.502,2.257,2.723,2.231,2.478,-3.098,-2.641,-3.294,-2.886,-2.974,-7.041,3.561,-2.442,2.249,2.279,6.878,3.371,-3.392,-2.14,-1.952,4.008,-3.179,4.473,-1.466,-5.068,-4.352,-1.886,3.069,-7.356,-6.309,-4.217,-8.113,-3.108,2.879,-3.75,-8.708,3.343,1.651,-5.244,4.747,3.449,2.045,2.248,-2.584,-3.475,-4.488,-3.915,1.99,-2.778,3.221,-2.293,2.475,-4.253,3.834,2.834,-3.256,3.374,3.283,-1.941,2.283,2.305,2.534,2.642,2.543,1.949,1.958,3.318,-5.564,-6.275,-5.404,-6.508,-6.466,-6.183,-5.943,1.551,2.978,2.348,3.181,2.608,3.574,4.21,-2.092,5.532,3.209,-1.856,1.727,-3.329,6.642,4.954,2.421,2.949,2.902,1.644,-3.817,-8.245,-3.07,-4.904,2.96,2.965,3.633,-2.712,-4.424,-5.784,-4.044,-3.966,-3.946,1.858,2.477,-4.178,-3.258,-2.524,-2.077,-2.711,2.604,-3.697,-1.563,5.605,-2.751,8.821,-2.776,3.168,-1.697,-4.342,-3.006,-1.688,-2.465,1.544,-3.966,-3.838,2.964,3.059,1.45,2.322,3.088,-3.15,-2.56,-3.018,-4.476,-4.851,-4.175,-3.874,1.672,2.158,-6.496,-5.311,2.955,1.763,2.927,9.853,-3.387,-4.645,-1.93,-3.521,-6.981,-2.272,-2.609,-4.563,-5.877,2.012,-3.07,2.614,2.822,7.173,3.303,-2.376,-4.505,4.439,-2.43,2.989,-4.371,2.823,2.626,3.874,1.987,2.829,3.586,-1.89,1.627,-2.99,1.716,3.216,2.069,2.038,2.099,6.885,2.114,-4.94,-4.785,-5.204,-3.518,4.456,4.508,-2.061,-2.857,-6.588,-3.911,-3.275,2.034,2.485,5.665,2.784,1.846,3.109,2.821,5.884,3.358,3.499,6.356,2.346,1.726,-3.459,-5.664,3.604,1.915,1.971,2.798,4.033,-2.171,4.554,-4.309,-4.545,1.995,4.769,2.668,2.352,-2.887,-5.158,-1.64,2.977,-3.561,2.412,6.773,5.845,3.066,3.733,2.217,1.782,4.495,-2.738,2.008,2.783,2.148,-2.397,-4.024,3.282,4.036,3.207,3.295,-3.151,-4.971,-5.25,-3.567,-2.514,-4.792,-4.12,-5.098,3.395,-5.483,4.291,2.622,3.4,-3.664,3.423,-4.645,1.429,1.665,-5.993,-2.748,2.088,3.609,2.153,-1.506,-1.539,2.794,1.691,-2.905,-6.383,-6.447,-2.571,-3.008,-3.282,-4.885,3.068,-3.804,-5.613,2.781,-2.939,4.007,-1.982,-5.243,4.344,2.067,-3.648,2.912,-3.019,-2.76,-5.443,-4.686,-2.111,-3.023,-7.811,-3.586,3.225,-3.808,2.366,3.563,-2.187,-4.306,-3.573,-1.774,10.3,-5.436,5.003,2.536,3.431,-4.218,3.932,-4.303,3.021,2.616,-4.15,-3.493,-3.26,2.978,3.911,-2.148,-5.831,-3.99,-3.855,2.115,3.897,3.082,-1.587,-2.471,-3.278,-3.898,-5.043,-4.211,1.965,-1.794,-2.962,-7.946,-2.757,-1.508,-2.975,4.284,2.668,-3.161,-3.506,-3.191,3.395,4.776,5.283,-5.034,-4.997,-5.756,3.522,3.436,2.526,2.453,2.969,3.862,-2.936,-2.888,3.915,2.729,-4.472,-5.071,-4.891,3.773,-2.918,1.917,-3.028,-3.121,2.689,3.415,1.918,-2.943,1.908,7.3,3.151,-2.411,-1.84,3.109,-2.359,2.948,-2.2,-2.57,7.45,4.935,2.717,1.722,6.411,6.44,-3.179,1.665,-2.164,-1.923,2.2,-3.606,-3.052,-2.958,-6.155,-2.68,-2.488,-1.636,-3.735,-3.704,-4.6,-2.764,3.647,2.289,-2.6,-3.61,-3.533,1.778,2.846,1.268,2.693,-5.625,-3.085,-2.348,2.233,1.561,5.612,2.127,3.411,1.696,6.708,-2.93,2.031,4.015,-1.664,2.015,2.338,-2.04,1.731,-6.734,-2.791,-5.628,1.437,-2.495,-4.383,3.496,1.773,5.132,5.393,3.293,2.559,5.277,4.406,1.998,-3.137,1.867,1.865,-3.988,-2.118,3.142,-3.26,2.409,-2.183,1.781,1.554,2.105,-2.555,-2.597,3.098,-2.206,2.087,3.407,-1.99,1.787,-3.646,-4.073,3.016,-3.814,1.802,1.881,-2.337,2.909,2.708,-3.491,2.062,-2.514,-2.323,3.826,-6.177,-4.104,6.145,-3.03,-2.961,2.46,-4.619,-4.654,-7.56,-6.076,-3.264,-5.283,3.475,-3.244,-8.106,-1.566,-4.389,-3.071,-1.244,-4.693,-3.825,-2.927,-2.479,-2.922,-3.922,-2.102,7.685,-3.68,6.513,1.791,2.833,2.694,3.855,-3.785,-3.788,-4.551,3.321,3.018,2.986,-4.671,2.773,2.291,1.883,2.228,4.848,-3.268,2.922,-3.877,5.636,-6.105,-4.839,1.839,-2.762,-2.475,2.961,3.743,8.494,-6.1,-5.596,-6.926,-8.956,-11.21,-8.934,-6.436,-6.052,-5.78,-8.294,-4.843,-4.265,8.39,-3.422,2.457,-4.461,-2.776,2.756,2.736,2.189,4.791,5.442,-5.218,-3.409,-6.593,-3.648,3.09,2.993,-1.963,-3.808,-3.702,-3.492,-2.887,-3.799,-5.433,1.977,-2.233,-3.909,-2.999,-6.431,-3.834,3.141,-3.845,-2.442,-2.998,-1.762,-2.725,8.042,-3.428,2.564,-3.142,-3.582,-3.571,-4.254,2.318,-3.927,2.459,-3.795,2.152,-3.318,-4.334,-3.218,-5.752,-1.737,1.934,-3.579,3.039,2.352,-4.375,2.666,3.516,-2.972,-2.236,2.445,3.403,-3.859,-2.087,-3.629,-5.186,-5.617,-5.989,-2.618,6.103,3.923,-3.17,-2.571,-4.27,4.627,3.028,1.421,-7.679,-4.348,-3.608,2.344,4.607,7.342,3.257,1.892,-2.927,-3.545,-3.768,2.024,3.274,-4.146,-2.435,-2.401,8.086,-3.371,-1.843,-7.614,2.874,-2.435,2.999,1.607,-4.293,-4.929,-3.893,3.715,2.333,-3.523,-3.008,8.779,2.665,-3.143,-3.71,-5.509,2.811,2.514,-2.112,-2.826,1.716,2.627,13.38,6.363,1.918,-2.33,3.364,1.981,3.352,3.688,3.801,-2.821,1.553,2.149,-1.611,2.774,-3.568,-5.396,3.274,-6.301,-6.513,-2.643,-2.764,1.635,-1.659,-1.857,5.172,2.502,1.699,1.689,-3.905,-6.99,-4.641,1.938,2.107,-3.11,2.046,3.008,2.709,3.609,3.021,2.649,3.531,-1.702,3.744,4.098,-5.753,2.202,4.415,2.68,6.32,4.134,-1.634,-2.627,2.397,2.232,-4.56,2.048,-3.69,1.824,2.766,2.707,3.328,4.099,2.69,-4.608,3.185,2.366,1.578,2.328,-3.395,5.749,4.862,7.484,2.153,2.183,3.17,1.759,-1.714,6.922,2.109,8.069,4.801,1.928,-2.323,1.725,-4.501,1.68,1.414,3.265,3.338,2.038,-2.231,-4.34,2.114,3.384,3.457,-2.755,-3.553,-5.478,-3.737,-3.606,2.615,-2.264,3.701,6.156,4.717,2.432,1.945,2.1,-2.785,-1.395,2.251,-3.102,2.778,3.978,2.129,2.676,3.001,-3.28,-1.327,1.942,4.173,-8.954,2.016,2.023,-4.162,3.849,2.025,2.126,4.092,3.031,5.766,2.657,-2.703,-8.939,-4.087,-6.632,-5.388,-6.94,-5.439,-6.049,-6.012,-5.849,-7.97,-7.014,-7.946,-5.973,-8.399,-6.684,-9.512,-7.652,-4.482,-6.366,-5.648],"logCPM":[0.2335,0.8914,-0.909,-1.044,0.6133,5.003,2.106,1.12,0.944,-0.1883,1.833,-1.05,-1.034,-1.221,-1.113,-1.154,-0.8025,-0.5506,-1.087,-0.8912,-0.4062,-0.5014,-1.269,-1.082,-1.037,-1.018,-0.8162,-0.4439,-0.3194,-1.205,5.047,-1.333,-0.6471,-1.063,-0.8348,-0.4968,-1.243,-1.325,4.202,-0.7709,-0.6452,-0.6829,-1.271,-0.9345,-1.305,-0.8035,-0.477,-0.8719,-0.6047,-1.128,2.171,0.4155,-0.783,-0.4198,-0.7101,-0.06131,-0.5802,-0.5516,-1.319,-1.205,-1.325,-1.031,-0.6486,-1.2,-0.2541,0.05658,-1.063,0.7089,-0.1543,-0.4205,0.4831,-0.1943,0.1023,0.05213,-0.2669,-0.5134,-1.082,-0.6948,-0.9267,-0.1904,-0.6631,-0.1987,-1.139,-1.238,-0.8462,-0.6023,-0.7576,-1.405,-1.111,0.8912,-0.1736,-1.264,0.5787,0.7213,-0.9965,-0.6714,0.7875,-0.7072,-1.029,-0.8471,-0.8726,0.3965,-0.1988,-0.7048,-0.551,-0.9555,-1.18,-0.9394,-0.8912,-0.6887,-0.176,-0.919,-1.01,-1.227,-0.7852,-1.107,-1.042,-0.4406,-1.168,0.09461,-1.041,-1.188,-0.1335,1.254,-0.6179,-0.7133,-0.799,-1.319,-1.325,-1.405,-0.7855,-0.97,-0.3408,-0.4832,-1.294,0.2788,0.2009,0.5138,-0.02345,-1.019,-1.405,-1.04,-0.9892,-0.9012,2.021,-1.214,-1.389,-1.061,-0.004036,2.9,0.3679,-0.1631,-0.484,-0.7328,-0.7121,0.2777,1.72,-1.294,-0.6928,0.5968,-0.5475,-0.2769,2.635,3.512,-1.055,-0.2817,-0.2713,-1.017,1.501,-0.9366,-0.7041,-0.4596,-0.6979,-1.31,-1.422,-0.6012,-0.406,0.5244,4.756,-1.066,-0.7452,3.143,5.357,2.96,-1.322,-0.407,1.489,1.124,5.629,1.548,2.064,1.297,5.346,8.008,5.571,4.384,5.624,3.914,8.249,5.372,7.692,9.62,6.055,8.267,5.312,5.342,6.881,5.048,6.251,8.939,8.391,5.735,1.803,8.226,7.331,0.9543,3.065,4.275,3.248,6.765,5.882,3.723,7.764,6.145,7.112,6.5,5.176,3.633,4.351,6.354,4.469,6.233,9.297,5.317,6.491,4.562,7.931,3.286,8.325,7.499,4.061,5.415,5.86,6.496,4.386,4.269,4.22,3.575,7.672,6.997,8.224,8.205,6.167,4.795,5.361,7.027,6.017,4.467,4.534,3.532,6.249,4.068,5.947,6.469,4.625,7.821,4.252,4.399,9.387,6.71,2.168,3.375,7.21,4.455,2.918,6.066,4.663,6.862,5.842,6.271,4.813,7.478,4.956,6.833,5.534,5.253,5.036,8.186,6.076,5.788,5.157,4.996,1.67,0.4254,3.378,4.533,2.296,5.005,0.4293,0.6483,-0.7163,-1.076,-1.052,-0.9892,-0.2229,-1.034,-1.134,0.1661,-1.045,3.891,-0.546,2.56,1.102,4.077,3.08,3.104,3.098,2.269,2.475,4.622,2.379,3.992,3.375,-1.061,0.7968,8.992,3.426,5.883,6.601,2.784,3.553,4.193,5.447,2.604,3.811,3.916,7.624,1.716,4.469,4.836,5.842,5.345,5.331,6.543,2.988,4.069,5.1,6.137,6.579,7.977,5.403,3.214,5.154,6.952,6.241,3.602,4.943,4.739,7.111,5.256,5.124,5.824,7.492,6.323,4.696,3.672,5.512,4.219,7.163,2.441,4.604,5.142,4.02,6.417,8.196,6.404,5.723,5.319,6.221,8.041,5.109,7.831,6,4.422,7.397,5.276,2.524,5.559,5.538,6.388,5.941,4.615,3.907,6.968,7.765,4.026,3.313,2.965,5.959,4.855,7.962,3.731,6.638,1.583,0.833,5.021,6.408,11.06,4.321,0.7227,2.873,1.395,3.758,1.785,1.264,-0.7617,-0.3822,-0.7949,-0.8444,-1.107,-1.062,-0.196,-0.867,0.008575,-1.038,-1.186,0.2391,0.5017,2.557,-0.9892,-1.205,-0.286,0.2555,3.083,5.513,6.26,3.005,3.266,5.007,7.157,3.747,8.149,6.371,8.6,4.288,6.034,4.691,7.215,7.748,4.503,7.576,5.286,5.151,5.072,5.491,1.489,3.964,3.189,5.795,6.125,4.536,5.706,4.705,5.701,4.476,9.026,4.66,6.882,5.328,9.402,4.886,6.902,5.967,5.045,7.667,6.892,6.499,5.154,3.587,5.367,2.365,6.474,6.331,7.947,2.933,3.747,4.17,3.094,7.713,4.441,4.936,4.03,4.641,4.955,7.657,5.979,3.011,4.387,5.229,4.514,5.287,7.699,7.711,5.807,6.323,4.025,3.998,4.018,4.571,6.031,8.023,1.862,5.804,8.075,6.15,5.793,3.389,7.641,4.755,6.192,6.631,4.153,3.364,3.867,4.377,5.811,7.991,5.823,5.253,7.853,4.108,6.824,5.569,7.018,2.606,-0.7945,-0.9865,-1.227,-0.8058,-0.9701,-1.2,0.5532,0.1818,1.461,1.537,1.363,0.9126,0.7756,3.739,3.951,5.608,6.425,4.426,5.044,4.864,5.487,3.596,4.583,5.472,5.537,3.875,2.778,6.136,7.291,8.008,7.704,6.918,4.711,4.274,5.138,5.967,8.191,4.166,6.346,6.806,5.667,4.463,7.671,5.335,4.613,1.631,5.075,6.294,3.008,4.149,4.267,6.518,5.225,5.254,5.064,4.646,5.763,7.457,5.664,6.457,8.701,8.701,5.011,4.42,5.231,6.845,4.834,4.844,6.122,6.26,5.616,6.706,6.565,6.856,8.343,6.286,3.176,6.61,5.059,4.865,4.079,6.476,8.303,2.473,5.517,4.643,4.695,7.531,0.86,2.144,3.67,4.465,4.562,7.723,-0.1508,6.423,10.14,1.665,4.891,4.993,7.278,6.378,4.391,3.281,5.585,7.305,6.487,4.064,0.9202,2.325,-0.0695,-0.6377,0.5699,-1.183,-1.135,-0.8169,-0.5826,-1.152,1.576,2.561,1.735,7.501,3.813,6.944,5.377,4.813,6.223,5.356,2.973,6.699,2.026,5.388,5.786,5.504,5.753,8.201,6.85,7.281,8.53,9.161,3.658,2.438,2.726,3.947,5.55,5.624,4.02,4.798,5.933,6.8,7.423,5.03,6.737,3.992,2.391,5.875,5.235,3.553,5.738,6.538,5.156,6.074,6.155,7.885,5.18,3.828,6.386,5.263,4.169,5.403,6.303,4.653,5.413,6.419,7.239,5.522,5.942,5.869,4.982,7.296,5.158,7.358,4.587,3.768,2.966,5.976,8.332,7.645,3.252,3.092,4.263,6.517,2.989,1.435,3.312,2.191,1.582,4.414,1.331,4.592,7.641,4.48,3.202,10.18,5.524,5.191,5.258,2.799,3.582,3.643,4.133,3.928,7.398,6.128,3.064,4.993,5.048,5.477,7.795,9.143,4.794,3.842,2.793,3.465,8.217,4.84,6.471,4.492,5.654,3.444,6.664,5.484,7.251,5.653,5.623,3.52,3.153,3.513,0.7383,0.01928,2.371,2.279,2.551,1.61,4.286,3.427,0.1291,7.497,4.06,2.98,4.871,6.792,7.647,5.263,2.81,3.344,6.465,5.635,6.798,4.01,8.603,4.274,5.112,6.68,5.924,4.847,8.148,4.001,5.769,6.594,4.78,5.909,7.033,6.632,7.048,4.426,4.751,10.69,3.445,3.024,6.345,1.953,4.79,7.071,5.867,4.397,5.218,5.169,3.236,7.544,2.228,2.78,1.823,2.403,4.376,1.555,0.7006,-1.178,0.1361,-1.099,-1.181,-0.8574,-1.325,-1.122,-1.118,-0.9769,-0.4607,0.1144,1.002,2.9,2.116,2.915,-0.04987,6.805,2.819,5.341,7.117,4.681,5.523,6.329,8.992,8.181,5.219,4.084,7.566,5.43,5.52,6.207,2.076,6.499,7.978,6.621,3.577,5.32,2.615,5.716,4.965,6.502,4.899,4.651,5.212,5.187,5.228,8.641,6.255,8.201,4.803,4.817,4.709,7.276,4.819,2.168,4.383,4.447,3.604,3.786,6.357,4.204,3.803,5.454,4.357,5.114,3.759,7.446,5.343,7.217,6.251,8.019,0.2897,4.377,6.729,2.095,4.232,5.48,2.397,4.787,5.819,5.99,6.595,5.185,5.019,5.617,6.627,4.447,4.463,7.538,6.241,3.426,3.043,7.09,5.905,4.281,4.18,6.249,4.684,6.933,4.831,8.318,3.792,5.289,8.747,7.282,6.333,6.6,4.903,4.167,6.195,4.053,7.398,5.281,6.025,5.487,6.701,7.744,5.628,7.692,7.288,2.385,3.483,5.486,4.714,4.436,4.98,6.929,8.447,6.027,7.739,4.461,3.536,5.875,10.28,6.971,5.326,5.171,3.923,3.729,0.8394,-0.1203,0.02446,1.323,1.658,1.282,-0.4155,0.1425,-0.4194,0.2375,3.394,3.48,0.3118,6.17,6.344,7.788,10.43,4.64,6.89,8.011,3.468,4.392,7.129,5.886,10.67,7.275,5.093,4.27,5.899,6.708,6.525,3.638,3.891,5.597,5.102,3.498,10.23,4.891,5.854,6.153,6.352,6.2,9.712,5.932,5.559,6.192,6.01,4.903,7.858,5.588,6.886,6.215,5.886,4.847,5.335,6.774,5.179,1.379,7.863,6.658,3.938,6.724,3.684,7.056,7.072,8.585,6.224,4.632,4.462,4.066,3.987,3.809,3.904,6.075,5.414,4.637,2.536,4.41,3.774,4.907,3.398,5.248,4.355,4.883,5.106,5.529,5.417,9.376,5.769,4.572,4.819,8.482,4.824,5.063,2.359,3.769,8.564,6.84,7.813,4.172,4.215,8.191,1.237,4.471,3.741,3.085,5.604,4.208,5.731,7.139,5.346,6.516,6.219,8.869,3.848,5.606,4.774,5.206,8.919,6.635,2.804,6.044,4.387,6.38,4.704,6.856,5.217,4.122,5.05,6.669,6.907,5.821,5.118,9.424,10.16,3.575,4.014,8.129,7.983,4.593,7.17,4.582,7.442,5.531,2.715,4.463,6.182,6.066,5.913,6.2,6.245,8.358,6.358,6.096,5.905,0.3767,3.685,4.54,7.239,7.705,3.717,5.471,6.347,4.618,8.026,3.914,6.138,7.194,4.784,3.704,5.193,4.403,4.694,1.878,5.583,6.21,2.989,0.9769,4.818,4.434,5.226,7.095,3.434,6.539,2.982,5.587,5.531,3.384,7.138,6.219,4.699,9.421,1.466,5.908,6.383,4.955,7.237,3.212,6.038,3.941,7.674,5.33,4.409,4.377,9.118,4.677,5.811,5.969,2.059,3.481,3.944,5.592,7.257,4.62,5.949,2.968,4.077,3.95,8.238,7.086,3.371,4.562,4.589,3.33,4.316,5.519,3.513,4.913,1.198,-1.166,-1.218,-0.3903,3.516,-0.3931,0.8305,0.2322,-0.2245,-0.7174,1.613,-0.3163,3.707,5.37,4.544,6.761,5.391,5.713,2.103,4.161,5.956,5.939,7.005,5.978,2.15,6.469,8.627,8.293,7.758,3.181,2.908,4.796,5.607,6.653,1.403,3.647,5.827,5.986,4.679,6.285,0.529,2.711,0.3706,4.791,4.556,4.277,4.836,9.34,6.705,4.756,2.17,5.559,4.985,5.083,6.602,6.543,9.781,2.338,8.964,8.892,1.618,5.775,2.795,5.794,5.811,5.947,5.343,4.744,7.88,6.049,7.115,5.81,4.11,3.481,1.246,3.185,2.591,7.113,3.499,6.51,6.39,7.996,9.456,6.494,7.513,5.546,3.178,5.093,6.367,7.217,6.74,6.727,5.791,7.296,4.64,3.617,8.51,10.15,3.628,4.36,8.226,5.196,3.594,6.33,2.741,6.352,7.19,2.34,5.563,4.896,7.92,7.509,7.262,6.76,4.866,7.44,3.512,7.331,4.428,6.851,6.374,6.516,7.308,4.812,6.299,4.355,4.693,6.303,9.166,5.773,6.205,3.687,4.133,3.571,5.965,5.576,8.095,7.064,8.196,3.354,8.609,4.187,6.558,4.888,5.435,4.392,4.819,6.163,4.649,7.42,10.05,6.261,4.375,4.618,4.174,7.643,5.585,3.525,4.395,4.232,5.996,4.401,4.966,3.538,4.228,6.153,3.307,6.213,2.066,5.093,5.457,3.115,5.274,5.24,5.444,5.369,6.938,3.792,4.248,5.828,5.666,3.952,4.152,6.468,4.906,6.918,1.968,4.104,5.649,8.087,6.623,5.982,4.636,6.762,7.082,6.259,8.953,5.406,3.427,6.362,5.015,5.109,4.149,5.012,3.089,4.748,5.09,6.675,9.034,8.282,6.819,6.353,6.094,5.087,6.659,5.714,6.935,7.066,2.164,5.172,6.697,4.391,5.984,4.444,7.554,8.659,2.763,6.491,6.884,6.962,5.266,3.153,1.274,-1.07,-0.851,-0.936,-1.18,-0.94,-0.05853,-0.1456,-0.793,0.1674,-0.5127,-0.6933,-0.2115,-1.103,-0.8784,-0.8847,0.2962,3.304,4.469,8.033,4.429,7.891,2.683,6.354,4.702,8.746,1.835,5.174,5.276,2.482,5.932,5.634,7.038,5.48,2.939,5.155,4.847,4.389,1.179,3.019,6.745,3.826,6.987,5.799,8.701,5.916,2.856,3.054,4.592,7.612,2.686,5.97,3.381,6.023,6.403,4.904,7.498,5.645,2.44,6.42,5.945,6.158,6.925,5.166,4.856,8.009,5.871,3.959,7.354,6.592,7.411,6.474,7.308,5.323,6.777,3.551,3.849,5.727,4.2,3.415,8.112,4.176,3.772,2.04,8.476,5.087,1.83,8.494,5.832,7.816,1.238,1.726,5.321,6.209,6.136,9.057,9.692,6.131,4.724,5.122,5.85,3.74,7.765,5.775,7.257,4.543,8.739,5.93,4.981,7.598,8.005,3.505,3.421,3.913,7.514,5.412,7.295,5.395,6.752,7.45,5.077,8.632,1.759,3.07,8.274,2.073,4.786,5.757,6.967,3.131,5.076,7.575,7.034,6.69,6.55,5.281,7.298,7.39,8.708,3.413,7.304,4.959,3.949,4.327,8.882,5.145,6.876,4.364,4.414,5.592,5.564,5.775,7.248,3.422,6.621,3.58,7.894,5.879,7.486,3.765,5.543,6.177,4.936,3.517,6.648,3.005,6.232,6.741,5.538,4.985,6.792,6.302,4.195,6.691,10.13,8.985,5.885,4.21,3.586,3.327,4.068,8.408,5.568,6.735,4.503,6.743,2.632,3.902,8.374,6.102,5.583,5.144,2.282,7.607,6.24,6.069,5.869,8.79,3.977,6.128,4.777,5.404,5.941,9.097,8.197,4.01,8.365,7.999,6.378,4.369,3.384,6.435,6.106,1.827,5.362,2.444,2.899,5.978,7.6,4.465,6.756,8.173,5.381,6.558,8.232,5.719,7.122,4.139,5.036,3.567,3.306,4.812,1.919,2.117,3.692,10.04,4.394,5.76,6.83,3.319,5.307,5.257,0.6893,4.618,6.078,3.655,5.207,8.029,8.949,6.164,4.126,3.591,5.414,5.565,1.996,3.312,5.689,5.863,6.492,5.236,7.245,5.698,4.71,3.001,4.344,3.996,3.766,4.286,6.033,6.369,4.921,8.395,8.201,4.738,5.701,5.473,5.07,5.627,1.704,5.415,2.309,2.866,2.704,8.25,5.505,1.069,7.302,7.844,6.097,6.423,6.116,7.674,4.813,1.807,0.4207,0.3368,-0.09173,0.7734,-1.093,0.3691,-0.9727,-1.141,6.349,5.079,10.97,10.53,2.98,2.296,7.873,4.813,6.1,5.101,9.612,8.641,1.63,4.974,2.947,2.603,4.437,7.246,7.684,8.494,8.435,5.569,3.903,8.925,6.39,7.304,5.982,6.679,6.957,4.566,5.81,3.408,8.611,4.239,4.393,5.875,6.265,7.936,7.505,5.628,4.794,7.355,3.897,4.396,4.738,6.594,6.246,6.608,5.213,7.818,6.775,4.633,0.9542,4.902,3.091,6.369,4.944,4.998,6.547,3.744,5.102,5.803,3.235,5.348,6.826,6.382,5.741,3.793,4.455,9.812,6.477,8.854,4.379,9.244,5.271,3.262,0.5856,5.325,2.902,5.524,5.676,7.304,5.201,8.226,3.075,2.505,8.188,4.469,6.82,6.847,8.204,4.788,3.776,2.69,4.959,4.538,4.571,5.336,2.905,5.101,5.271,3.56,4.512,4.414,6.456,6.701,5.579,5.338,4.255,6.73,4.303,5.174,3.793,2.837,6.855,5.025,6.523,4.538,10.43,2.459,3.808,2.881,7.244,2.323,-0.5344,5.676,3.575,6.6,7.99,5.35,4.48,5.702,4.588,2.957,6.44,8.365,5.436,6.18,5.756,6.258,6.297,9.408,6.357,6.772,5.159,3.481,3.883,8.354,5.179,5.193,3.745,5.087,5.165,5.806,4.902,5.196,7.372,8.974,6.135,7.972,5.657,2.469,6.527,6.018,1.417,1.161,8.368,5.553,7.84,5.685,8.179,6.844,8.163,5.104,7.533,7.614,6.733,6.29,3.367,3.404,2.935,2.514,8.446,6.188,5.391,6.248,3.181,1.367,-0.5714,-0.2882,0.7686,-0.6448,-0.2334,-0.4631,-0.9949,-0.9279,0.7312,1.932,1.882,-0.07521,2.742,6.855,7.312,5.391,3.211,6.571,4.511,3.495,8.292,5.521,2.007,4.022,3.234,8.397,5.048,6.462,3.45,2.736,8.826,4.808,5.313,4.518,4.761,6.065,3.097,6.767,5.808,4.222,3.858,6.647,4.067,4.712,5.272,6.623,3.704,5.882,7.586,7.668,7.958,4.958,6.951,6.749,3.522,4.91,6.962,4.167,3.87,9.005,5.474,9.568,5.713,4.412,6.32,13.59,4.859,1.027,2.075,10.09,5.328,6.711,11.44,8.201,5.938,4.823,4.417,6.84,7.676,3.348,8.473,2.906,9.894,6.858,6.842,8.066,5.231,5.643,6.012,4.156,7.121,7.207,7.103,7.476,2.603,7.195,1.788,5.372,6.955,4.114,2.306,5.948,5.472,3.886,6.052,9.928,5.238,5.633,8.349,4.829,4.559,7.432,6.251,3.897,5.333,9.143,5.592,7.619,1.569,4.779,4.42,6.4,10.62,7.758,7.7,7.028,8.798,3.842,2.309,4.402,3.968,10.12,11.57,3.667,7.163,7.306,5.524,2.637,3.783,9.002,2.709,4.88,9.662,7.957,3.133,7.92,6.919,4.897,3.694,2.281,6.736,5.769,6.584,5.962,5.087,2.227,6.77,3.823,10.54,3.891,6.269,2.043,6.349,3.503,4.311,6.337,6.478,7.058,6.83,4.481,4.517,6.72,6.173,5.652,4.949,4.876,6.202,5.155,5.832,6.343,4.792,-0.1024,7.043,2.632,5.124,5.669,4.213,6.316,4.931,5.398,6.017,7.512,10.63,6.681,4.479,4.075,4.688,7.653,4.504,3.586,5.855,5.955,5.848,4.954,6.855,5.937,7.913,5.737,4.173,4.138,4.886,6.256,7.437,6.742,6.039,5.485,5.402,6.831,4.79,6.034,4.035,5.944,6.36,6.027,9.386,6.486,5.582,4.14,6.108,1.607,6.275,4.191,6.431,2.39,4.623,3.737,6.845,0.3132,4.511,1.821,7.212,3.469,8.072,8.207,2.532,3.92,6.034,6.277,6.515,5.094,4.794,6.206,4.742,4.993,4.628,1.213,6.72,5.244,4.096,7.423,7.719,4.988,6.94,6.72,3.606,7.886,5.293,6.445,2.922,3.899,8.794,5.152,4.873,5.176,3.899,7.426,6.234,1.96,4.861,4.428,5.76,4.063,8.011,1.992,8.095,4.391,1.949,5.377,3.787,6.615,4.769,7.299,4.774,6.401,5.926,6.635,4.391,1.858,3.446,1.96,4.767,-0.8686,-1.04,0.07887,-0.6,-0.8321,-0.8271,-0.5751,-0.3915,-0.1634,-0.447,-0.7435,2.284,1.643,2.736,6.863,8.082,4.287,7.833,8.685,6.1,5.432,8.274,6.617,2.264,6.98,5.378,4.95,5.342,9.263,5.789,3.988,0.1589,3.214,5.894,6.561,6.474,3.776,7.538,7.289,4.351,4.744,8.816,5.5,6.452,3.34,6.473,3.661,8.786,2.449,3.141,4.544,4.659,4.259,7.048,3.593,8.597,5.429,6.018,7.031,5.975,4.252,6.112,6.179,5.732,4.918,7.241,6.406,5.25,5.513,9.067,6.322,3.124,11.13,5.859,6.977,2.827,0.7966,3.727,5.686,1.086,5.518,4.543,4.746,2.313,4.997,6.549,3.572,1.828,6.292,5.95,5.115,2.402,5.59,4.165,7.281,6.197,7.101,3.477,5.164,7.955,5.732,6.299,5.706,5,4.387,0.6223,5.644,2.705,4.395,6.34,5.007,6.004,5.599,4.28,6.063,7.34,6.936,7.643,5.999,3.905,3.687,5.429,3.177,7.294,5.605,5.016,6.379,6.888,6.682,5.981,6.862,5.418,5.632,6.928,6.934,4.707,3.552,4.585,6.382,6.65,7.003,5.813,3.865,6.622,6.974,6.914,2.676,9.152,5.753,4.865,5.712,8.714,4.443,3.07,5.2,6.879,3.973,5.503,5.057,3.328,5.396,5.376,6.271,4.692,5.775,7.386,4.99,5.631,1.311,4.355,6.114,4.162,5.947,4.14,5.185,4.606,5.062,7.098,5.937,7.107,4.729,1.404,3.538,3.573,7.251,5.252,7.17,4.716,3.706,3.031,3.917,3.92,9.143,7.855,2.859,3.986,3.412,3.465,4.763,4.284,7.542,5.559,3.866,3.497,6.65,3.541,6.194,5.706,4.478,7.034,7.316,5.738,3.852,6.675,6.93,6.503,5.776,7.374,8.526,2.295,2.747,5.87,4.961,5.179,8.063,4.592,2.599,9.07,3.509,3.288,7.548,4.874,2.024,2.427,4.784,4.585,6.334,0.05197,8.653,4.115,4.028,3.288,5.524,3.435,6.213,6.625,5.61,5.813,1.684,7.137,6.736,4.044,7.337,3.278,4.651,8.157,6.881,6.028,5.567,5.283,5.883,7.107,7.616,3.816,5.004,3.17,6.37,4.789,5.899,9.718,7.79,4.221,5.585,5.26,8.106,3.792,5.021,4.832,2.75,5.694,8.118,7.18,6.043,8.106,2.184,8.695,5.712,6.65,5.932,4.14,5.848,6.622,7.706,5.604,2.526,2.427,3.133,2.978,5.822,7.742,2.55,6.495,9.665,7.073,0.5304,7.146,5.874,4.721,5.057,5.137,4.826,6.937,9.228,6.426,4.022,4.678,6.937,7.965,5.903,6.204,8.059,6.02,2.907,6.655,6.22,6.406,6.3,5.642,6.861,2.95,7.478,1.867,5.267,6.041,4.516,5.872,6.78,4.343,7.199,7.303,5.149,6.671,1.878,1.323,4.039,3.072,5.043,8.55,7.944,8.824,7.987,8.593,9.918,7.088,5.705,9.434,5.378,6.589,5.691,6.321,10.34,3.61,0.821,6.163,2.994,8.195,2.584,8.304,6.635,4.732,6.033,7.691,6.194,4.313,7.076,5.625,9.247,7.709,5.876,6.238,5.757,4.61,6.026,8.262,5.542,6.924,5.79,4.13,6.016,4.524,5.615,5.091,7.134,6.691,3.686,6.608,8.637,8.671,5.414,6.165,9.015,7.014,5.147,3.466,6.22,6.562,8.509,5.395,3.187,6.474,4.424,7.579,9.149,5.126,6.564,4.932,4.536,2.096,1.258,0.2456,-1.336,1.899,-1.167,-1.135,-1.096,0.4836,3.712,8.076,5.578,6.106,8.06,5.947,7.334,3.996,7.542,3.351,5.66,3.484,7.106,6.105,4.816,2.748,8.727,6.971,3.799,2.164,1.911,0.2956,4.977,6.423,5.689,3.809,9.057,8.824,9.515,8.374,4.571,6.879,4.929,8.259,5.936,1.116,5.27,5.552,3.488,3.734,2.438,8.36,5.246,3.199,2.121,5.449,4.455,4.176,5.132,4.515,5.028,6.799,4.849,7.867,7.975,5.338,5.744,6.037,4.598,5.119,5.461,3.647,10.29,8.498,6.316,6.628,6.108,6.463,5.484,7.993,8.546,3.833,8.732,3.805,3.599,4.976,4.215,3.659,6.12,4.772,7.696,4.453,5.825,3.215,6.975,7.888,5.283,4.183,8.78,6.586,9.913,1.371,4.758,6.826,7.704,5.609,5.302,6.601,5.712,6.894,7.214,5.152,6.098,6.584,3.5,4.814,5.493,3.65,4.681,5.032,4.53,6.704,6.293,4.298,4.007,4.008,4.416,4.378,2.347,3.016,5.445,2.646,4.205,2.19,4.015,8.521,5.437,4.042,1.505,8.36,7.847,9.275,8.914,6.304,4.497,7.528,6.524,5.256,-0.5858,4.887,6.031,5.673,5.938,7.836,3.081,0.657,4.497,5.921,8.762,8.784,9.775,5.946,6.153,6.463,5.329,5.559,6.181,2.463,3.922,3.021,4.847,5.101,3.793,5.029,6.169,4.431,5.922,7.007,5.577,6.336,8.518,4.269,4.111,7.693,5.884,3.264,6.788,7.426,5.926,11.88,3.745,6.739,6.984,4.19,6.019,0.9584,3.653,5.105,5.306,6.827,1.984,3.427,5.744,7.538,5.607,2.68,3.682,5.506,4.574,6.787,6.386,7.541,6.934,8.013,5.679,3.317,1.512,2.445,6.356,4.816,8.076,2.446,4.51,6.323,5.974,6.638,6.155,7.252,7.402,3.785,8.013,1.97,8.491,8.857,6.904,8.797,5.701,3.514,1.357,7.617,4.826,7.474,4.439,6.135,5.906,7.753,4.073,6.197,5.068,5.292,6.654,3.461,6.389,6.706,5.628,6.817,3.553,6.975,4.43,6.756,4.633,3.793,6.384,5.345,6.768,3.982,2.262,4.211,1.568,3.503,6.683,5.668,6.811,6.625,8.641,2.637,-0.3133,-1.231,-1.253,0.09074,-0.7202,0.8534,3.131,2.936,5.985,1.171,2.457,5.35,1.696,7.108,4.136,2.955,1.733,2.216,8.757,8.112,8.221,5.488,5.77,6.65,3.976,4.424,10.23,3.655,6.003,0.434,7.216,6.976,7.563,4.801,4.866,8.947,5.03,3.181,8.703,4.053,5.755,3.349,1.115,6.678,3.266,7.038,4.685,5.526,6.47,6.352,4.609,6.766,4.459,5.67,6.316,6.715,7.803,7.472,5.221,3.829,5.95,8.787,2.832,7.068,6.656,7.269,7.059,4.781,4.284,6.997,4.995,3.656,4.853,3.949,5.616,4.605,3.863,8.808,2.834,3.828,4.014,7.671,8.788,3.95,4.617,4.692,2.175,7.009,3.707,5.204,5.222,6.428,7.495,4.969,4.477,6.786,6.258,5.824,5.211,6.325,2.734,6.243,4.827,5.878,4.245,3.262,5.192,6.675,6.06,7.483,6.457,6.768,8.022,4.242,4.044,6.405,4.083,4.812,7.711,0.4922,6.48,7.936,8.102,3.347,8.096,5.754,7.512,8.864,1.871,7.022,4.844,5.574,5.28,5.896,3.786,5.245,6.866,6.414,1.591,9.095,7.363,8.487,4.182,8.082,4.12,4.048,9.217,6.591,4.98,-0.3449,6.455,1.4,4.734,3.653,6.413,7.189,6.027,6.62,7.873,10.08,4.294,6.523,5.898,5.574,6.785,9.22,6.001,7.206,10.01,2.581,4.473,4.98,7.847,5.412,9.073,9.516,5.87,7.621,11.51,7.449,6.994,6.693,5.261,10.01,2.171,7.658,5.495,6.224,6.728,6.152,5.152,10.19,7.599,10.31,5.194,2.435,4.135,4.906,8.508,7.301,4.757,3.596,3.661,2.656,5.476,3.625,3.449,7.596,1.078,4.684,7.025,2.33,5.012,4.797,5.653,7.824,4.744,3.992,3.374,6.722,7.187,4.009,4.446,7.386,7.727,8.192,4.299,3.546,5.818,3.804,-0.005438,4.87,2.956,5.82,5.988,6.143,4.955,8.73,0.8933,4.334,5.337,6.927,6.728,4.619,4.925,4.673,6.765,5.885,4.599,9.319,6.369,6.675,4.022,3.303,1.843,9.533,4.156,5.421,4.151,6.799,5.343,6.609,5.569,6.402,5.869,6.461,8.471,5.717,5.812,5.299,6.153,6.462,6.445,5.412,6.354,5.835,5.273,4.505,8.679,7.614,5.916,3.387,5.311,4.164,4.26,5.279,5.245,5.346,6.76,9.039,6.808,5.976,2.74,5.263,6.302,3.824,6.084,4.606,4.442,3.976,5.202,5.309,7.634,5.907,6.836,3.605,3.969,5.729,6.923,5.684,3.415,7.868,4.809,5.233,5.82,4.874,7.563,5.028,5.868,7.999,5.599,5.551,6.795,5.773,2.675,4.893,5.707,5.796,7.378,7.647,7.618,7.314,5.851,5.245,7.304,6.959,7.794,3.328,6.369,5.422,7.258,4.23,8.238,7.088,2.463,4.328,4.281,4.651,7.647,8.114,4.622,5.006,1.981,5.845,9.504,8.321,5.795,7.462,4.781,-0.2875,4.814,2.704,3.872,6.313,4.412,3.39,0.583,3.829,7.576,9.684,5.904,5.162,9.202,4.099,9.081,6.252,6.119,5.849,7.498,5.159,3.433,5.423,6.747,4.955,5.763,7.66,4.898,3.339,3.514,6.56,6.304,5.709,8.792,6.447,5.439,3.25,4.303,4.325,3.095,10.24,5.238,7.213,1.699,6.575,3.199,5.094,5.11,7.366,4.867,5.633,8.105,4.231,5.522,4.169,5.335,2.766,5.546,9.239,1.385,-0.1346,2.226,1.175,0.5323,-0.05413,1.183,-0.1004,-0.2792,3.4,-0.7782,0.1006,0.6227,-0.312,3.915,-0.7813,-1.322,-1.125,-0.8562,-0.06173,-0.4989,-0.6876,6.033,3.163,6.624,4.629,0.4613,4.231,5.846,0.5397,0.7128,4.859,4.004,7.292,4.419,3.866,0.8699,3.389,1.358,0.2753,0.7173,0.9867,3.691,6.243,10.03,2.894,5.436,5.915,1.73,5.732,9.505,5.214,8.884,9.581,7.531,7.836,5.951,4.829,6.467,7.129,8.528,5.94,6.39,12.91,5.782,8.06,7.465,8.249,9.335,9.322,8.203,10.11,5.897,7.186,4.136,5.728,7.425,5.996,11.66,8.254,6.517,9.883,6.771,7.03,6.635,6.297,9.749,6.029,6.685,5.872,8.769,8.668,6.911,6.211,6.912,8.411,4.553,9.915,6.531,7.543,9.797,9.578,7.221,5.787,3.617,4.42,4.812,4.563,5.149,5.101,7.522,6.588,9.602,7.88,7.631,7.522,6.381,3.676,6.075,7.104,10.44,5.5,10.07,5.641,4.128,7.026,9.817,10.81,11.06,7.682,6.67,7.181,6.807,9.258,9.458,8.081,9.489,5.439,3.948,6.68,6.57,3.547,10.08,9.642,4.941,4.272,2.65,7.001,9.028,8.809,9.493,3.016,4.527,-0.1219,1.408,9.017,9.31,6.48,5.247,8.954,5.613,5.666,7.495,9.743,10.2,10.1,8.126,6.064,9.258,7.282,8.13,10.9,11.96,10.36,4.228,4.071,6.27,6.982,6.746,7.409,13.09,12.8,7.345,8.554,8.506,5.287,6.163,8.339,6.083,9.52,8.212,5.922,8.608,6.849,8.777,7.482,7.065,6.908,9.024,6.912,8.061,7.657,11.15,6.143,7.801,6.287,6.416,6.281,7.616,5.221,5.581,9.19,6.207,5.183,6.365,7.43,9.362,4.816,5.908,8.476,7.302,4.683,6.079,7.3,8.863,7.232,5.96,6.614,5.69,4.625,9.108,8.794,7.51,6.889,5.218,9.402,5.271,6.801,7.423,3.752,1.406,2.522,8.474,10.24,6.228,7.569,6.595,7.413,6.567,5.958,10.68,9.634,7.606,8.11,7.904,8.785,5.854,5.547,8.078,7.937,8.551,6.607,6.463,3.319,8.091,6.171,3.535,8.8,9.098,7.529,6.079,7.506,4.664,9.602,8.59,9.385,6.479,4.595,5.286,4.401,4.341,5.916,5.67,3.121,7.714,5.644,9.458,8.648,5.978,8.985,5.293,7.065,6.499,8.548,8.123,9.013,7.707,7.693,8.386,7.905,6.805,8.282,7.72,10.97,9.815,5.114,4.805,2.291,8.2,7.29,8.859,6.546,3.731,7.61,4.767,7.814,7.353,9.769,6.463,8.608,8.125,4.546,3.998,6.578,6.233,4.524,1.749,3.19,2.021,2.879,3.407,2.826,3.438,3.14,6.4,3.25,9.154,10.66,8.87,8.105,10.14,7.614,7.408,8.804,7.261,5.93,7.734,4.414,7.803,6.72,8.232,4.364,9.053,4.936,9.051,10.2,9.669,7.61,6.934,8.71,5.668,9.597,7.153,8.769,7.308,8.568,4.665,6.565,8.36,7.876,8.584,4.406,14.91,7.605,4.914,4.507,9.833,9.739,8.873,3.345,7.458,2.782,5.893,9.818,9.09,4.593,6.005,6.671,7.317,5.504,4.904,3.996,5.301,6.668,6,5.006,8.359,7.715,8.39,8.359,6.965,5.535,7.682,3.03,2.573,4.123,3.384,2.852,2.865,4.046,3.607,2.944,3.161,3.439,5.192,6.694,6.872,11.02,9.798,9.213,8.762,7.014,5.42,7.677,6.844,7.706,10.17,10.49,9.014,8.034,7.158,6.537,6.348,8.799,7.334,6.091,4.242,5.373,7.658,8.615,9.224,6.606,3.955,4.014,2.602,4.778,7.431,4.447,5.655,3.406,4.111,6.137,3.914,6.947,7.526,14.25,10.71,7.192,9.019,8.64,6.638,6.964,6.174,6.116,7.065,7.766,7.703,4.646,7.255,6.698,5.866,7.608,6.455,7.413,8.671,8.475,9.904,6.447,4.166,7.218,8.947,8.269,8.474,8.828,4.58,11.19,10.57,4.645,8.138,7.688,3.46,6.116,7.023,6.181,5.799,5.012,10.36,6.912,7.908,7.745,10.77,4.981,5.4,8.41,5.664,9.245,10.5,7.096,10.45,7.163,8.508,9.408,9.572,6.304,6.578,7.62,5.809,5.38,6.926,8.167,7.909,11.31,7.097,7.998,9.669,4.105,5.865,6.514,5.535,4.085,4.076,9.342,5.474,5.519,5.42,5.09,6.819,6.704,9.147,4.88,6.457,5.898,2.601,3.668,3.715,5.951,6.29,2.647,7.946,2.271,1.613,6.886,0.08755,2.436,6.315,8.454,7.596,9.27,10.53,6.062,9.125,6.053,4.627,4.573,5.769,6.345,4.661,9.932,4.584,9.159,7.817,3.191,4.328,4.778,5.019,6.448,7.51,7.036,8.633,8.249,9.8,7.383,5.765,8.592,5.1,11.96,8.426,6.541,5.364,8.794,4.359,5.536,6.665,6.941,6.279,7.318,9.572,5.459,5.14,8.373,9.867,8.702,6.719,5.031,8.94,8.008,6.44,8.449,4.736,7.2,9.898,9.007,8.334,6.61,8.882,9.976,7.78,6.345,8.784,6.67,7.78,8.017,7.145,3.222,7.203,6.184,3.767,8.605,7.417,9.175,4.818,5.396,4.852,9.976,10.15,6.384,7.678,5.391,5.164,5.8,7.247,6.643,8.777,4.736,5.669,7.132,4.397,6.827,9.09,9.638,3.337,3.627,6.162,7.689,7.5,11.11,11.67,9.236,7.391,9.713,6.188,5.929,2.558,8.801,6.302,6.183,4.791,6.318,5.963,7.538,8.979,7.641,1.405,2.318,7.449,9.439,5.815,4.024,3.732,5.277,5.764,7.544,5.374,2.552,4.843,6.853,9.541,9.215,4.378,5.764,10.25,6.101,7.368,4.583,6.331,4.984,4.536,10.14,6.939,10.43,11.44,4.492,8.106,9.573,8.947,4.372,4.479,4.139,8.294,9.229,7.623,5.804,4.879,5.015,6.204,9.467,5.031,5.26,5.459,8.839,9.645,6.783,9.113,8.593,9.277,7.482,5.748,4.07,9.999,5.151,8.041,7.799,5.901,5.242,7.642,9.453,7.611,7.797,4.999,9.477,6.382,7.535,6.165,5.979,3.773,7.084,8.734,7.901,6.949,6.698,5.282,7.404,10.34,6.311,5.138,6.705,9.119,8.69,6.172,3.453,4.68,5.141,7.091,5.371,5.94,6.816,6.599,9.787,4.188,8.028,6.843,9.845,10.48,7.121,7.297,7.948,4.961,6.525,6.755,2.788,6.664,4.84,5.918,8.408,8.362,6.495,5.675,5.788,6.57,4.988,5.172,5.755,5.89,5.86,8.876,5.241,4.827,5.2,6.855,5.007,9.288,5.036,7.148,7.101,6.893,5.903,7.363,7.653,5.695,6.479,6.394,6.756,8.219,4.615,7.829,6.696,7.603,14.19,6.359,6.899,7.831,6.614,8.013,10.54,2.86,3.087,0.1806,-0.1605,-0.1313,0.5407,-0.1846,-0.06275,1.87,5.999,5.341,5.268,6.229,10.42,6.544,5.98,5.998,2.868,4.305,10.6,10.46,6.742,7.693,8.942,8.361,6.356,10.28,12.61,5.898,6.138,9.002,5.566,7.999,6.493,4.896,2.062,6.614,11.51,4.379,6.418,4.852,7.601,10.12,4.707,5.878,3.88,3.548,8.988,6.857,4.825,9.443,4.285,3.792,9.91,6.694,8.203,6.537,9.497,7.648,9.18,8.604,7.363,4.737,7.528,4.397,6.076,8.641,5.028,8.958,7.836,8.377,7.021,5.742,8.5,6.839,8.355,4.254,3.551,4.367,7.015,10.93,8.378,11.18,8.251,7.115,5.508,2.933,11.64,7.482,8.495,4.62,7.397,11.14,7.362,8.275,6.769,7.322,7.266,5.843,5.9,9.673,7.852,10.23,4.792,5.86,6.611,9.188,5.774,3.735,6.787,5.493,7.469,9.178,5.193,6.163,9.855,6.915,3.152,8.246,5.606,10.76,6.725,6.635,11.67,5.749,6.141,9.213,4.635,7.076,7.103,7.419,4.649,5.685,3.746,10.17,7.555,7.765,7.142,10.98,6.04,8.605,7.514,5.598,9.041,7.047,4.281,6.419,4.622,9.565,7.918,10.38,5.272,4.831,8.106,6.338,6.032,7.516,8.014,6.614,5.471,3.481,4.64,6.068,8.739,7.364,4.417,4.399,4.257,11.82,4.688,8.449,8.693,3.909,0.7663,9.154,9.772,6.963,6.726,6.557,5.237,7.92,8.845,5.691,3.8,5.097,3.72,4.925,6.363,6.664,7.386,5.263,7.128,7.832,9.038,6.253,5.857,7.815,9.336,6.685,10.85,5.216,6.013,6.19,8.9,9.06,7.963,4.398,9.554,7.282,7.726,6.455,8.004,7.126,6.592,4.488,3.354,9.341,7.565,6.178,7.022,5.911,7.665,6.241,8.625,6.905,7.589,7.004,4.886,5.543,5.953,6.359,9.682,5.704,7.049,5.66,7.074,7.23,8.525,4.775,6.431,6.324,7.336,5.24,7.698,7.195,5.971,6.314,7.538,8.497,4.576,11.25,11.31,9.86,9.929,9.604,8.626,8.769,8.715,6.628,8.024,7.594,5.619,8.528,6.453,4.707,4.741,1.669,7.256,8.975,11.12,6.288,5.879,7.999,7.173,5.245,5.424,6.459,6.666,4.399,4.753,8.082,4.916,5.136,5.749,5.526,6.516,4.326,3.868,7.855,5.36,5.845,8.773,5.827,7.532,6.163,2.552,1.755,5.504,3.499,3.798,8.595,5.048,5.748,8.945,9.093,8.916,6.236,8.162,5.062,5.565,4.676,8.786,6.437,7.38,5.193,3.78,5.486,9.691,4.733,6.718,6.445,5.03,6.796,5.802,4.978,5.427,5.701,4.604,10.76,9.094,5.06,8.263,7.134,6.363,6.001,5.096,8.328,4.081,8.884,11.48,11.33,5.225,5.146,5.169,6.161,4.632,3.041,8.162,5.392,6.119,6.999,5.457,8.333,7.104,8.292,5.917,5.251,5.526,4.612,7.478,9.1,7.411,6.691,9.039,6.364,13.62,4.863,4.052,4.244,6.747,8.966,6.714,9.376,5.742,8.934,4.958,2.978,6.363,8.9,9.625,10.37,9.934,6.43,6.458,6.698,5.444,7.519,5.953,9.53,7.717,8.471,8.18,7.371,9.009,7.391,7.7,10.2,4.2,4.496,6.199,6.122,8.983,10.59,7.878,7.092,7.729,6.924,10.46,9.415,5.388,6.384,6.517,7.959,7.211,8.843,6.833,6.146,4.053,6.983,10.74,4.7,5.606,5.109,3.504,6.421,9.946,3.404,10.57,7.573,7.36,6.864,6.166,7.326,7.627,10.06,4.486,7.206,5.279,5.495,7.557,5.704,9.013,5.112,7.939,6.209,7.54,9.548,4.187,11.51,9.539,5.885,5.871,6.521,7.016,7.153,6.6,8.553,5.994,8.243,0.2436,6.845,3.968,0.8829,0.3518,7.004,4.184,7.912,10.57,6.561,8.246,7.915,8.171,8.538,5.554,9.093,5.536,7.204,9.279,6.592,7.439,6.659,6.898,9.036,7.168,10.51,4.431,5.578,7.269,5.369,5.806,9.718,8.047,5.746,6.63,4.891,3.36,5.124,4.169,8.307,9.491,7.682,8.101,5.209,7.698,7.396,9.361,6.495,7.449,11.16,7.102,10.51,5.422,8.973,5.903,4.758,4.63,8.512,6.896,9.861,4.655,4.299,6,4.968,7.35,8.984,8.212,5.792,7.398,7.952,6.663,5.945,5.704,5.574,9.018,9.06,3.963,5.866,6.668,9.056,8.875,12.46,4.26,4.624,6.196,8.048,7.594,6.413,6.535,5.104,5.873,7.453,4.054,7.445,9.065,9.64,6.088,5.411,5.965,6.073,5.593,8.865,6.476,7.775,8.463,10.71,8.209,7.453,8.759,7.203,8.815,6.746,8.314,11.04,7.368,7.889,8.812,11.9,7.649,3.411,4.375,2.736,8.328,9.924,9.704,6.401,6.397,6.407,2.884,6.347,7.346,9.143,9.396,7.188,8.812,8.987,7.606,8.609,7.752,8.501,8.336,9.086,8.147,5.834,5.484,6.474,6.198,6.611,7.143,8.409,5.703,9.467,7.121,5.514,8.162,7.135,7.565,6.582,6.557,6.845,9.028,7.267,5.582,10.95,10.4,8.561,7.539,6.237,6.12,6.823,4.602,8.307,9.54,7.203,6.398,5.994,6.651,9.954,12.47,7.641,7.1,3.837,7.1,5.807,4.022,6.788,10.77,6.294,4,10.67,3.011,8.153,8.69,7.784,5.711,8.985,4.241,7.753,7.003,4.661,5.871,8.62,9.269,9.622,9.325,9.31,8.699,7.993,4.813,7.908,8.785,7.642,3.727,6.418,3.336,10.38,6.484,6.601,5.936,7.304,8.573,9.018,4.554,9.025,8.619,6.616,5.471,4.206,5.337,6.924,3.779,7.079,5.202,9.569,5.176,9.12,9.776,5.349,10.21,5.269,5.275,4.67,5.855,11.37,6.43,10.21,9.009,6.268,7.091,8.802,4.718,6.255,9.672,6.599,4.979,5.839,8.55,8.638,7.633,4.714,3.214,7.771,8.541,7.918,8.448,7.217,7.334,5.991,7.238,5.248,7.514,9.017,6.598,7.012,5.038,4.506,8.428,7.433,9.642,8.872,5.165,5.35,5.696,5.177,8.207,10.85,4.87,2.938,-0.3075,9.126,8.246,8.653,7.514,10.98,7.547,7.657,4.411,5.793,6.329,6.371,4.066,3.353,8.717,4.994,9.51,5.42,5.17,8.002,10.5,10.36,7.256,8.623,11.05,7.35,5.131,6.268,10.28,8.137,9.339,7.62,6.318,11.05,6.552,7.502,8.745,7.122,4.121,5.711,8.473,5.558,7.119,10.14,3.822,5.896,4.57,4.486,6.989,8.854,6.896,6.719,5.071,6.994,5.089,10.61,5.21,4.481,4.845,6.207,9.019,9.724,9.437,8.445,3.91,7.126,5.18,6.435,7.222,6.995,7.012,8.725,6.413,7.707,5.991,7.427,8.672,6.628,5.974,7.683,6.226,6.969,7.018,7.548,4.377,8.918,5.583,7.503,5.704,7.572,11.22,7.693,6.15,6.982,7.404,9.033,9.944,4.895,7.551,6.268,2.608,6.416,9.953,6.87,9.031,6.983,9.763,8.168,6.405,4.787,4.744,7.774,8.26,10.02,10.11,5.748,11.44,4.206,5.267,8.716,6.493,7.6,10.96,7.369,7.465,10.64,8.722,8.792,4.973,6.152,9.211,3.989,4.313,5.575,5.217,4.051,7.661,5.047,5.768,6.027,2.375,6.828,5.52,10.36,5.945,9.416,6.381,4.623,4.78,8.298,5.041,8.012,7.442,5.89,5.282,4.298,7.796,4.756,9.284,9.093,7.625,9.852,8.128,6.228,7.15,4.464,6.141,10.96,9.546,9.32,8.803,7.753,7.646,9.779,8.736,10.68,4.478,6.238,6.512,6.452,7.818,5.964,8.283,7.268,5.103,8.729,8.368,2.348,2.329,1.427,3.06,2.865,9.571,4.055,4.872,4.451,5.616,2.118,4.588,7.234,10.83,4.543,9.154,7.861,9.283,9.186,6.159,7.708,10.66,11.5,6.016,4.119,6.837,6.181,6.53,6.867,7.491,6.045,8.193,6.036,5.635,4.723,7.333,6.997,6.087,4.129,6.055,3.478,5.161,10.32,12.24,7.491,7.171,7.064,4.439,6.974,6.322,12.05,5.65,7.015,5.606,3.861,7.746,5.16,7.162,4.816,9.862,4.466,5.728,4.649,6.197,8.068,9.496,4.921,5.608,8.739,6.531,8.233,9.413,6.147,5.61,9.574,8.641,6.498,9.756,6.283,3.018,7.874,1.909,7.523,6.843,9.519,8.68,6.651,6.908,6.056,9.261,9.532,2.347,7.739,5.269,9.287,5.969,8.873,9.925,8.369,7.225,4.664,7.685,8.872,9.909,3.766,5.542,4.841,9.425,7.826,7.514,9.399,7.862,7.716,6.95,7.687,2.674,4.058,3.601,4.708,8.815,6.761,6.661,10.59,5.887,8.342,5.481,3.862,6.614,7.132,6.187,5.16,9.558,8.421,12.37,10.03,8.643,5.407,9.795,8.347,8.37,5.699,10.76,6.602,10.16,9.478,9.048,11.46,6.198,6.774,9.257,7.131,2.737,5.794,4.804,8.79,6.864,8.541,4.609,5.866,8.522,10.64,3.497,4.997,5.133,10.2,7.251,6.371,5.789,7.09,8.026,7.679,6.89,6.612,5.756,7.098,11.05,8.161,6.097,8.196,7.928,6.546,6.246,7.732,6.031,4.936,9.834,8.16,2.712,6.352,7.365,7.417,8.559,6.38,6.502,8.515,8.517,5.066,5.172,8.742,6.96,7.136,6.24,8.195,8.71,7.247,8.85,8.475,8.118,8.419,6.507,8.868,8.444,7.773,9.296,7.893,5.818,8.589,7.288,7.622,8.155,8.951,7.666,7.848,4.705,5.705,8.197,8.295,9.268,5.695,6.775,2.452,5.206,5.075,9.101,6.616,9.021,8.631,6.923,5.789,7.919,6.811,7.267,10.69,7.65,5.422,8.849,7.574,6.246,6.303,7.661,4.494,8.017,6.687,9.338,8.274,10.2,7.15,4.944,6.499,8.667,7.228,9.49,6.322,3.744,8.232,6.127,10.28,5.501,9.496,1.885,4.258,5.274,2.589,4.283,1.926,7.292,0.7088,0.4019,1.968,5.26,1.741,2.307,0.8779,5.838,10.06,9.26],"F":[5.757,1.61,0.1133,1.247,0.07326,7.744,10.44,2.684,0.3402,9.767,4.059,0.09796,0.06092,2.558,3.184,3.198,3.245,3.461,6.487,2.86,3.097,11.99,2.451,2.808,0.0234,0.9397,3.314,4.061,3.707,2.57,2.027,2.509,2.167,3.316,3.474,5.086,2.45,2.377,1.524,2.198,3.195,3.503,2.393,3.577,2.474,3.654,3.07,3.396,6.579,7.847,8.11,1.86,1.08,8.197,3.313,8.894,1.684,4.2,2.305,2.57,2.377,3.435,1.985,2.746,4.22,0.03306,0.8386,3.385,4.745,0.8764,3.637,1.316,9.207,0.5897,3.983,3.63,3.064,3.332,0.02699,0.2448,6.779,2.765,2.748,2.453,2.165,6.619,3.559,2.095,0.001764,9.815,3.152,2.485,1.056,2.077,3.123,3.8,0.1554,3.735,3.084,3.99,2.88,9.932,8.454,4.253,4.699,1.469,2.602,2.827,2.86,0.4302,3.182,0.04258,1.997,2.608,1.131,2.918,1.088,0.05478,2.867,1.631,2.746,1.658,3.075,0.07721,3.83,0,2.917,2.305,2.377,2.095,3.591,2.812,3.593,3.076,2.408,4.757,5.872,10.19,0.0001966,2.712,2.095,3.081,2.789,3.185,1.601,3.092,2.141,3.005,2.785,0.7085,0.0002631,4.664,3.907,0.1555,0.002935,0.01645,0.02846,2.408,2.443,2.201,3.488,0.2216,0.3575,1.346,0.7083,0.4212,15.9,2.863,1.466,7.031,5.699,0.2405,12.19,0,0.6342,13.03,3.109,2.706,0.4796,4.9,0.8961,7.474,5.506,6.471,2.302,2.083,9.873,3.77,0.3657,7.962,0.1573,10.7,9.331,6.866,10.38,1.149,0.02417,7.095,3.031,3.684,1.24,3.415,0.4923,0.1011,0.007504,9.024,0.04114,0.9855,7.276,1.352,11.51,0.2849,1.089,0.02052,1.845,11.03,0.03975,0.4254,0.7708,0.04795,4.158,6.466,2.31,0.5631,3.81,5.486,5.019,0.2442,0.1923,9.259,9.368,4.858,1.509,0.05003,0.1185,1.963,7.94,0.2193,11.53,5.883,1.761,4.345,10.6,9.215,2.683,0.1894,0.3912,0.5853,6.161,0.0897,1.878,5.572,11.24,3.702,0.2666,10.89,9.939,0.03175,0.162,1.641,7.958,0.01178,0.787,3.452,2.447,8.03,7.05,2.618,0.2282,0.00212,8.453,1.443,2.96,8.187,5.942,2.503,2.347,4.162,1.095,4.883,1.865,10.49,0.4277,1.192,3.733,0.03907,0.07847,3.018,10.41,0.01753,3.753,0.1404,2.412,1.691,0.9392,1.457,1.268,2.506,2.204,0.7823,3.388,3.255,3.264,2.789,3.818,3.44,0.9574,0.08322,3.329,0.02181,1.39,10.55,0.03421,1.313,2.052,4.898,0.1142,0.7896,5.862,7.876,3.681,3.269,4.323e-05,8.709,8.886,9.734,4.598,3.003,3.828,0.01939,4.798,1.202,7.215,0.364,7.134,2.261,4.399,0.04784,4.575,1.176,1.493,0.004971,3.825,8.858,1.04,6.223,3.253,7.682,0.007712,6.872,0.8492,0.7871,0.9315,2.506,5.333,2.154,0.009265,2.927,1.534,7.636,9.025,3.413,2.629,2.676,0.9297,0.3818,1.593,0.3527,1.013,6.846,5.174,8.396,7.343,0.6223,0.5944,9.075,6.693,0.2449,3.898,9.72,0.01982,0.8576,0.02573,2.625,7.699,4.894,0.04659,0.5701,5.544,0.01302,0.1239,5.053,9.121,1.04,7.843,8.128,0.8038,0.1738,0.5912,0.08813,5.076,5.853,2.949,0.3135,1.153,1.428,7.044,11.36,8.401,6.495,5.016,1.849,9.766,11.5,0.4018,1.142,0.221,3.243,2.89,2.918,4.76,4.631,0.5498,4.032,2.727,2.826,4.318,5.963,3.405,2.789,2.57,0.5894,2.887,0.005055,0.9017,1.296,3.034,2.152,11.19,0.06097,0.2378,0.501,0.3279,0.0389,1.907,5.045,0.0001344,2.242,0.6095,7.76,10.09,6.203,0.8162,0.6039,6.645,9.547,4.407,9.714,9.805,7.475,0.6691,9.895,8.036,4.56,0.7818,11.39,6.034,4.128,1.053e-05,0.5201,1.369,9.393,4.144,1.296,9.264,7.927,1.561,3.298,3.357,5.126,0.1576,5.861,2.621,0.05345,1.126,3.74,0.1875,3.058,3.467,6.749,0.006824,5.704,7.795,0.06872,2.12,0.8989,0.3812,1.542,1.653,1.49,1.852,0.2089,1.992,6.56,3.185,2.26,4.269,1.783,1.527,2.371,6.9,1.26,1.819,7.027,4.325,5.662,4.886,3.566,0.01601,0.04557,2.512,0.2606,5.945,4.052,0.3445,3.386,2.969,0.1534,0.5646,4.682,7.734,1.079,1.802,4.809,4.198,3.262,6.933,2.515,2.976,3.365,0.07896,0.7592,0.04395,0.9651,0.3785,4.447,0.05173,0.6021,3.194,0.07608,7.491,0.4754,2.321,0.03397,0.6509,4.331e-05,2.171,3.863,3.644,8.723,7.491,2.861,10.98,5.701,2.908,6.48,0.4984,4.005,7.032,2.415,9.775,10.33,11.55,8.028,4.77,0.7713,4.517,0.01769,2.336,0.3423,4.836,2.296,0.5832,1.961,0.02482,3.423,2.315,0.007706,2.426,2.868,0.3093,1.907,5.041,4.931,0.2431,8.518,6.738,1.706,9.337,0.6758,1.726,0.152,2.716,0.05575,9.299,10.35,10.15,7.273,1.207,11.01,1.997,0.002476,5.499,8.683,10.71,1.298,7.942,2.99,4.069,1.743,6.563,6.113,0.5579,0.003817,5.675,0.1538,6.452,5.192,10.21,0.7011,0.02136,5.613,0.2437,3.699,0.0358,9.145,0.521,0.05958,1.126,0.6134,8.041,7.08,1.828,9.704,0.06508,0.7924,0.004044,0.3758,1.037,2.713,1.839,0.1608,2.849,2.858,5.721,6.208,2.749,1.584,1.3,0.9262,10.68,1.364,0.006895,5.485,7.061,0.1087,2.923,7.892,3.657,3.666,3.322,0.1998,0.2608,2.179,0.002648,3.948,0.02882,1.46,0.3311,8.694,10.81,2.4,3.183,1.577,0.244,5.022,4.009,2.105,1.457,0.9064,10.92,7.791,3.057,3.253,10.05,0.7333,0.1035,1.225,4.428,0.9631,8.083,2.166,2.951,0.8433,9.277,6.652,0.1915,4.05,7.948,9.144,1.46,0.05403,9.602,5.895,3.719,0.1732,0.4441,0.8586,6.156,11.46,1.321,7.773,2.396,0.197,0.797,2.11,0.01526,6.276,5.02,7.669,7.763,1.323,7.788,3.64,8.155,4.486,0.0947,0.8813,3.861,0.8661,3.947,0.8763,0.5614,0.6109,3.003,2.488,0.01376,6.249,0.0003267,0.001103,7.095,5.115,1.528,7.327,8.697,8.329,3.084,0.4143,0.3079,2.519,0.02324,11.1,0.3682,0.08649,0.7332,0.04753,0.04727,10.05,0.8725,8.72,5.088,6.93,0.01362,6.602,1.334,0.04476,1.641,0.04436,2.926,9.606,9.708,2.032,5.483,0.08574,9.327,6.761,3.838,9.361,9.387,2.388,2.14,10.96,7.441,2.105,8.438,3.292,4.126,6.277,3.47,0.5394,9.559,6.074,0.2566,9.793,1.381,3.026,2.641,2.871,1.088,5.059,0.01778,4.9,9.394,1.331,9.111,8.075,1.047,9.158,8.994,8.032,0.2389,0.02563,0.0847,3.61,0.391,2.497,2.558,7.421,4.591,0.1749,2.605,2.611,0.1926,4.741,3.288,3.101,3.223,0.05858,2.865,0.1703,0.04093,0.2316,1.775,1.005,3.344,4.003,0.04876,0.09037,1.146,4.913,1.212,0.2556,1.436,2.516,0.8596,6.099,9.534,0.2381,0.5991,5.861,8.552,8.518,10.2,0.2893,0.01063,7.681,0.6953,1.376,2.616,2.026,10.08,3.128,2.905,1.122,9.436,2.804,0.4143,3.175,0.2581,4.872,9.027,0.246,0.8558,9.018,7.155,7.43,7.999,4.744,9.804,0.8001,1.446,1.526,6.749,2.454,0.1828,0.1043,2.374,4.467,0.6318,0.8464,4.61,1.706,7.645,7.594,0.6197,0.08683,0.08281,4.141,2.292,5.43,1.532,5.322,0.0361,4.649,0.08358,0.2938,11.47,3.235,7.07,0.08405,3.226,9.048,0.3388,0.1076,2.473,1.92,0.6682,0.00694,11.5,1.13,1.851,0.6421,5.835,3.866,1.868,4.564,10.43,7.203,2.301,4.681,0.0547,0.1783,4.627,8.26,2.225,9.239,1.104,2.838,0.5784,10.12,0.002341,3.682,9.95,1.993,6.389,6.127,3.765,0.1548,10.3,0.325,4.835,6.244,11.5,5.828,1.282,8.582,3.396,0.5169,1.694,7.504,1.032,3.942,0.201,10.73,8.808,10.22,1.654,1.785,0.01891,0.6736,1.819,3.644,0.7495,1.282,8.729,0.03907,0.02222,1.355,0.04232,6.839,0.8237,1.957,9.981,8.976,10.3,9.935,5.422,0.3842,0.3307,1.79,9.465,4.348,2.532,0.09632,0.5237,4.659,0.7931,0.3642,6.409,0.3856,7.756,6.214,6.596,3.079,0.247,2.478,2.16,1.819,0.2772,6.011,2.818,1.184,6.442,1.708,4.473,3.258,11.15,9.059,0.02034,0.5773,0.2934,4.577,3.133,4.599,4.273,0.2911,3.498,8.659,2.078,3.816,4.089,5.971,3.329,0.05186,0.004952,4.89,5.185,0.0002502,2.529,0.2642,0.5388,2.74,0.3502,0.5366,0.0717,0.2171,0.1721,2.161,6.36,0.3706,8.45e-05,0.3284,2.522,0.3014,6.632,2.551,0.02367,2.004,6.847,6.285,3.006,1.904,0.2634,0.7774,0.2386,9.32,5.768,11.53,0.5386,9.998,4.778,4.159,0.1115,0.1334,2.362,4.534,5.122,3.952,4.656,2.733,3.624,1.991,0.7265,10.12,5.759,0.733,9.692,3.18,0.02226,6.318,4.405,0.9277,0.1584,0.12,7.496,7.992,10.78,0.7373,0.8954,8.673,11.26,0.004635,1.632,9.145,0.2941,0.0003484,1.777,0.008847,0.5217,4.557,6.028,2.314,10.56,6.013,1.131,1.94,0.6536,0.0009133,7.295,7.136,2.179,4.979,1.092,9.358,1.132,0.2562,4.354,10.38,5.62,10.73,3.682,0.4959,0.4848,2.011,1.569,5.103,0.06513,3.919,1.409,1.567,0.5935,2.011,11.21,1.337,4.353,0.3404,1.881,7.229,8.109,1.218,3.587,0.652,0.3519,4.11,0.9053,0.5723,0.6418,1.301,2.641,7.826,6.721,0.5578,0.3473,0.202,2.416,6.329,5.49,5.869,10.85,6.756,0.2121,2.008,11.28,0.09012,0.1803,0.01727,1.128,4.811,0.3759,9.565,0.05303,5.694,5.35,10.82,3.036,4.939,0.03892,2.04,3.105,1.58,2.628,2.87,3.652,10.94,4.314,0.005278,0.7925,9.59,2.42,2.366,2.084,7.11,11.12,2.391,7.356,3.528,10.67,9.746,0.4869,0.138,9.257,6.575,0.06851,0.4574,10.09,11.24,0.07915,0.6017,9.283,5.784,7.475,3.743,1.994,2.698,9.306,7.841,5.565,0.006679,0.6435,0.2215,0.8926,0.447,0.1815,2.034,0.3477,1.287,4.548,8.626,0.8784,8.135,3.15,6.056,6.911,0.04642,7.065,0.3197,2.556,5.225,6.019,1.17,9.343,0.00194,2.135,1.466,1.371,1.278,10.32,0.549,3.622,7.383,4.468,3.836,0.001439,0.3456,0.8525,9.721,0.08592,0.004521,5.794,2.239,1.947,11.16,0.2235,10.18,11.37,3.266,11.1,0.832,2.035,0.4689,0.3845,5.163,6.946,7.41,10.42,3.983,0.03574,3.68,0.09019,2.031,1.085,9.48,10.34,0.009361,0.0454,0.4384,0.01086,2.151,2.473,0.8283,11.27,4.93,0.3978,0.3321,5.103,1.423,0.174,1.829,0.08258,8.024,10.31,1.549,2.116,0.9564,5.946,0.4762,2.629,8.391,4.892,0.001196,1.23,7.982,0.005706,1.899,7.208,0.06178,0.1264,5.466,0.3451,10.75,1.574,6.182,0.04367,5.336,7.297,4.203,9.632,0.1164,3.461,10.05,0.7162,0.1922,2.328,0.5639,2.021,0.01162,10.33,0.5481,2.464,0.09503,5.691,4.771,9.171,5.562,2.043,0.2419,0.0001055,2.057,0.3554,2.413,1.004,5.282,3.948,0.7576,0.002082,0.5702,10.25,7.355,5.637,1.364,4.409,0.3659,2.263,0.7381,9.19,2.182,2.448,0.555,0.2123,0.2173,2.071,7.036,0.9557,8.152,9.117,0.7492,6.017,8.097,0.8884,3.972,5.065,0.03313,8.2,2.035,7.206,4.713,5.095,3.573,4.323,1.947,0.8074,0.2154,7.978,9.032,1.149,9.168,0.3256,0.003163,4.225,9.785,0.05956,0.01075,6.381,1.367,2.448,3.48,0.3284,1.706,10.27,9.518,0.3146,2.157,4.095,2.942,3.769,2.537,3.266,0.6229,15.64,10.74,0.102,1.053,0.1065,10.67,0.05045,9.919,2.533,4.906,1.279,0.06552,1.87,1.242,0.4868,1.264,0.1488,3.422,11.04,5.208,0.8909,0.5467,2.971,0.7092,1.302,0.8102,4.224,0.1305,0.000515,0.2114,3.451,0.2668,6.219,4.203,8.338,10.69,1.473,0.01409,5.376,8.266,0.6723,5.515,0.9633,0.07625,3.092,6.013,0.5774,8.211,5.527,1.684,10.95,6.757,1.87,2.669,0.3225,0.1471,1.307,0.4558,0.1008,4.776,11.15,0.9127,0.3,11.38,6.711,1.084,4.781,11.07,7.994,5.122,10.63,4.367,2.824,0.03878,4.109,6.903,0.7807,10.37,8.713,1.188,5.928,1.488,0.3711,7.631,2.087,2.197,1,0.3753,9.812,2.772,7.604,9.77,0.8162,7.155,0.7561,4.592,3.932,4.054,5.195,0.8563,4.471,4.594,10.44,11.43,10.21,5.914,0.1585,6.468,0.02931,2.609,3.714,10.17,1.134,3.208,5.659,8.005,4.473,6.94,0.2856,2.856,4.335,2.118,0.1222,0.8702,0.04959,0.4889,2.146,6.868,1.223,1.95,0.03981,0.8157,0.3147,5.696,5.013,2.794,6.271,11.3,0.08688,0.8773,5.532,3.643,3.08,0.5272,0.828,10.74,1.065,5.514,0.4135,6.388,2.849,0.07084,2.384,4.878,0.2485,1.48,0.229,1.585,8.614,0.5078,0.02164,1.279,6.193,1.251,8.3,2.413,0.5964,3.069,0.2823,0.2467,2.006,6.375,0.2568,0.06473,3.512,6.218,0.009984,1.516,3.424,0.3848,5.936,1.856,4.851,0.9012,9.323,3.527,4.423,5.661,0.132,0.5714,9.498,0.6776,10.26,0.06658,0.01575,0.07842,5.919,3.084,0.1614,3.975,2.083,0.6195,7.053,0.3741,0.1251,8.346,3.896,2.856,10.81,1.962,5.843,0.07336,1.638,0.4828,3.848,0.09106,4.786,9.937,10.59,0.4302,4.989,5.661,4.501,4.599,2.229,3.733,2.414,3.216,5.815,0.5579,2.949,3.357,3.02,0.001723,2.767,0.0003148,9.296,3.399,7.042,1.992,0.2116,0.3092,2.479,4.849,0.2313,0.03955,1.549,0.3955,0.001355,6.187,2.124,1.892,1.529,3.297,1.847,7.458,2.694,0.757,0.2311,5.254,2.819,8.269,1.394,9.984,2.118,0.1525,7.369,7.374,3.606,0.1336,2.835,1.64,1.075,4.378,0.1627,0.07109,0.1895,2.996,3.19,2.447,0.00217,5.921,2.181,2.377,0.1334,1.095,0.2689,5.316,0.007196,5.069,0.1427,1.973,0.4042,3.048,7.767,0.8836,0.4008,10.92,5.749,0.3493,9.099,3.451,1.029,3.536,0.3409,4.455,0.004848,1.158,9.01,2.634,6.926,7.648,0.5202,8.113,0.59,1.401,0.3191,1.348,5.895,0.1527,0.07166,0.1494,8.776,6.591,0.4946,0.1944,1.99,2.866,0.1787,5.187,4.883,0.3531,0.006157,11.44,2.445,0.8498,2.782,9.787,7.796,0.0379,5.389,0.356,7.332,1.293,0.7458,0.468,11.15,1.24,1.066,6.32,4.732,10.57,7.75,4.536,1.101,1.824,3.092,0.03008,3.431,4.823,0.01798,10.56,0.8928,2.989,10.4,0.04201,0.328,2.001,1.728,0.06515,5.748,1.416,2.403,1.248,2.475,2.961,1.437,0.6526,5.161,2.605,3.692,4.012,0.181,1.058,10.16,9.921,3.746,3.182,4.984,2.88,5.057,5.788,0.008648,0.6278,0.7361,2.582,0.4675,0.2397,0.04575,7.722,7.451,0.5453,2.387,0.01169,4.424,0.9396,0.3026,0.01268,0.0008157,9.019,10.17,0.2303,1.111,1.468,6.057,0.5228,2.642,11.23,8.356,0.2102,0.7057,6.84,4.56,0.1579,5.918,11.3,4.196,3.484,2.275,10.23,0.003784,1.892,1.754,9.402,0.8712,1.74,10.22,7.971,1.456,0.3157,2.066,0.02075,0.6328,5.001,10.73,1.592,2.665,5.919,0.07645,0.09428,1.487,0.1292,0.2138,0.01996,7.331,11.39,0.8285,2.955,0.4909,0.006316,0.06131,0.4641,5.797,0.3925,4.383,5.178,6.243,6.038,8.426,2.355,1.02,2.157,0.4416,1.385,7.531,0.8884,0.1081,11.38,7.172,0.5965,3.995,1.018,6.102,0.0007524,1.632,0.02042,1.859,0.3399,14.25,0.0122,3.164,2.546,6.778,8.552,2.08,6.52,1.029,2.522,3.957,0.7239,6.748,1.426,1.64,9.931,6.204,5.558,1.015,0.003926,5.865,0.3253,1.175,8.453,0.0004845,2.146,1.151,1.391,2.123,1.539,0.06336,5.238,0.1876,7.102,7.31,2.571,5.554,7.349,0.3219,4.793,8.894,4.762,1.318,11.03,10.22,3.001,2.457,1.016,2.199,9.879,0.02824,0.5534,1.465,0.6169,0.9433,0.6708,4.527,7.545,10.65,11.01,9.082,5.262,1.877,0.1329,4.027e-05,5.994,0.8121,0.1316,10.09,1.103,0.1919,1.42,4.69,9.425,2.384,7.412,4.305,9.205,7.858,0.2324,9.875,9.464,3.471,6.38,7.594,6.502,2.334,1.725,3.374,0.006808,0.1275,3.375,4.64,1.276,1.844,6.826,0.1187,2.725,4.448,0.3571,1.872,4.009,0.3373,7.971,1.264,2.6,0.6636,0.0005658,7.548,3.543,4.411,0.4801,2.731,0.7109,10.28,4.696,4.527,0.4165,4.453,1.729,0.8469,7.887,9.968,9.316,7.242,1.116,0.9988,0.002319,0.2721,2.784,0.1997,0.0295,8.598e-05,9.53,2.395,1.226,1.425,0.004279,11.05,1.858,8.747,0.1741,4.099,5.615,1.051,8.878,1.538,0.4087,1.435,3.324,0.1979,4.947,3.944,3.56,6.663,2.652,0.3676,0.8041,3.352,4.368,0.4424,9.314,8.033,0.4798,1.563,0.0002129,0.0006088,5.969,0.5825,0.5615,0.149,1.965,10.85,2.121,0.2157,2.636,0.002078,4.287,2.289,10.62,9.018,2.498,0.04435,4.596,2.796,0.8598,3.281,8.848,0.1891,3.392,11.23,2.277,7.065,1.222,0.3359,10.75,1.001,0.001825,3.062,2.709,0.8787,7.498,0.399,2.676,0.2374,0.126,2.33,1.876,0.2309,0.003391,0.01624,3.256,2.679,0.0054,3.18,6.756,11.21,3.101,1.924,0.3306,0.1389,2.218,10.52,9.153,7.289,7.989,0.08518,7.655,0.462,1.829,2.954,0.01162,4.738,0.8992,0.03592,2.485,0.8753,0.1045,0.4867,3.3,2.295,0.8667,5.525,6.668,2.752,1.277,7.816,9.07,10.38,3.077,5.892,2.699,3.388,2.833,0.05429,7.841,2.316,5.097,3.713,3.993,1.167,4.273,1.79,0.3101,2.212,9.301,4.011,7.003,1.088,1.573,3.797,4.936,10.03,0.01812,0.2657,1.776,0.859,8.343,1.339,3.57,3.685,1.007,11.32,0.4376,4.145,0.143,10.08,0.6218,3.925,0.7871,1.519,6.821,8.211,7.407,3.427,10.93,4.459,2.726,0.6874,3.696,0.1453,0.284,2.567,3.466,4.124,1.664,0.8822,0.2437,3.959,0.918,2.216,0.256,4.956,11.09,3.4,6.935,0.8452,0.7365,7.095,7.48,6.95,9.331,1.696,8.556,0.1125,6.897,0.02219,0.6166,5.031,0.1955,6.886,0.1521,3.358,1.13,8.321,4.882,0.01273,2.492,5.625,4.426,5.494,0.3369,1.05,3.481,1.188,0.2769,5.533,0.9043,1.898,5.811,1.093,0.9962,6.504,2.064,0.4608,0.03759,6.723,0.05317,0.1387,6.855,3.219,0.002245,2.494,1.231,0.5319,2.573,4.954,7.271,3.607,3.514,2.94,1.297,5.773,1.733,0.9559,2.897,8.694,4.305,2.469,1.25,0.4381,0.3205,11.06,0.03591,1.71,0.1395,0.02153,3.369,11.36,0.147,0.02528,0.02636,2.192,0.2181,2.988,2.365,3.007,4.068,5.696,2.187,0.3189,1.662,4.853,1.38,5.487,3.016,3.853,4.584,4.405,4.239,0.1632,5.061,4.68,0.02962,1.158,4.572,4.675,4.518,0.3279,2.103,1.265,7.96,10.23,3.078,6.456,8.615,10.27,0.2131,0.6111,1.856,6.149,0.8485,0.8274,1.038,4.401,2.812,2.546,6.886,5.375,0.1173,5.48,4.092,0.5927,4.488,9.069,3.566,0.2496,2.541,2.235,0.288,3.34,0.3981,4.622,0.249,5.681,0.05149,0.2458,5.13,1.424,3.87,0.08965,1.047,6.074,0.06756,2.92,0.06666,0.00774,0.01779,6.651,1.265,0.5429,5.675,3.566,5.36,0.8187,1.817,0.01109,0.7448,0.02329,0.0555,0.01679,0.1266,3.066,1.42,1.329,2.818,9.224,2.733,5.906,9.351,3.695,0.5263,2.151,1.873,7.2,1.645,0.227,3.179,7.987,8.533,7.834,6.694,3.648,3.937,4.432,3.459,0.03548,2.967,3.29,0.1416,1.055,0.9994,9.448,5.042,0.2171,0.5585,0.5131,0.3576,1.524,10.48,7.668,0.6889,0.05304,5.746,0.8192,1.277,4.505,0.03372,1.673,3.977,8.187,10.89,0.01283,0.4102,1.851,4.471,0.5734,0.2206,0.3238,0.03205,0.5953,0.5935,7.132,0.2797,5.983,3.361,1.967,1.289,0.06339,1.429,9.652,0.07812,4.171,7.274,0.01755,0.03239,0.4677,7.075,1.129,4.531,1.871,4.284,1.496,4.587,1.025,5.269,1.967,3.151,4.533,0.01369,6.772,3.77,0.8313,6.422,6.949,1.064,1.896,0.4289,10.71,6.378,7.773,7.22,0.1286,10.99,2.889,3.453,11.11,0.3238,7.446,0.001033,4.283,0.1626,5.764,0.2632,0.2212,2.371,4.775,0.04151,1.243,3.513,4.295,0.07163,4.415,4.915,8.617,5.77,0.2172,0.1195,0.1691,0.567,7.533e-06,6.335,5.087,0.4876,1.03,0.1851,8.7,2.933,0.7058,5.614,2.765,0.1933,9.293,0.7561,4.666,6.788,0.06358,0.08353,0.07364,1.907,4.214,7.469,1.727,0.9595,5.745,1.386,0.4536,4.036,1.899,0.2693,2.74,7.69,0.168,9.186,7.515,4.801,6.597,1.904,7.414,1.47,11.53,7.338,4.512,2.021,6.417,3.418,4.44,0.5164,4.614,8.088,0.3228,8.73,0.02659,1.656,3.258,8.044,0.9037,1.323,0.2273,9.144,11.43,10.38,0.5239,7.55,0.3011,3.081,4.462,0.08744,3.116,0.04602,10.53,0.73,2.009,11.18,9.482,0.02895,2.791,6.111,1.512,8.595,0.01719,8.314,0.9065,2.683,0.4839,4.086,0.1544,7.688,4.628,8.178,3.93,3.178,6.403,9.294,8.641,11.53,3.2,11.1,6.186,6.219,1.418,8.645,10.43,8.281,0.9293,6.436,0.1069,3.419,2.936,2.713,2.753,3.873,4.652,0.1153,0.1691,0.5859,5.305,0.7574,4.517,0.5868,1.387,0.005101,1.743,0.09538,4.606,7.05,0.8526,3.265,2.988,1.283,4.332,0.07017,1.695,6.088,2.45,1.4,0.04249,0.07586,6.214,8.262,0.2147,1.36,9.971,3.623,6.203,0.9615,1.997,10.57,0.01663,3.161,0.2283,3.031,4.545,5.461,3.611,11.36,1.522,1.237,0.08885,2.806,0.3666,3.423,6.194,7.075,10.16,2.813,4.91,0.4731,8.871,1.297,2.655,5.471,2.242,9.665,0.04012,1.489,3.887,0.001537,8.802,0.3943,0.604,1.459,2.197,0.8345,3.016,0.3649,10.49,3.525,8.211,3.738,10.68,10.79,0.08236,2.141,1.372,1.752,8.788,2.35,0.6224,0.1919,0.1184,10.64,4.759,4.224,6.478,1.873,5.086,5.662,3.715,10.41,3.106,6.675,0.2439,2.979,0.1037,0.7011,2.879,5.932,8.272,4.304,6.63,0.2557,1.66,0.8656,4.63,0.2963,0.2527,10.72,1.23,0.05326,0.4605,1.264,4.064,3.727,0.3675,3.348,3.724,2.565,6.192,7.819,1.23,4.516,0.06453,0.7813,0.4187,4.593,5.34,4.696,9.092,3.639,0.4646,3.179,1.145,7.998,8.542,5.769,2.942,0.2805,1.594,0.502,11.07,6.759,6.071,3.872,3.524,1.915,5.112,0.05366,0.08794,11.54,11.06,0.1305,0.3901,7.876,2.118,3.281,0.2667,3.801,9.299,11.29,0.002135,5.756,0.6731,6.681,0.6514,1.607,1.8,2.654,7.727,0.6865,0.08151,0.2617,1.148,1.524,0.04477,2.481,7.928,0.01058,0.371,0.06374,6.871,0.03014,0.06819,0.5376,4.232,6.498,3.869,0.9968,6.904,0.4529,0.5203,3.456,0.03526,0.6869,8.791,4.988,5.04,4.467e-05,0.6735,2.801,10.81,8.141,4.48,1.281,5.655,0.1507,0.4082,0.8189,2.043,1.439,2.39,1.441,5.883,0.9609,3.12,1.431,7.95,9.676,0.3149,10.51,0.08592,1.357,1.224,0.7682,0.337,9.175,2.226,10.41,4.648,3.227,0.01877,7.458,1.663,7.062e-05,4.798,0.1383,6.465,1.808,8.393,4.049,0.8035,5.296,2.583,0.4627,0.9182,10.33,2.657,10.65,6.064,0.8888,1.782,0.36,1.325,1.919,0.8446,0.008,0.5425,0.9124,4.336,2.92,2.665,0.7192,10.97,0.6153,7.699,8.174,10.2,0.7119,9.479,8.768,6.202,3.588,2.832,4.453,7.595,0.4693,0.01753,9.161,0.2124,2.318,4.555,1.755,6.026,2.434,5.367,0.2446,1.194,4.123,1.427,6.605,7.394,2.125,0.5089,10.27,7.878,1.906,2.899,7.355,4.903,7.084,0.006397,9.465,1.885,6.413,7.303,1.884,0.02203,0.00306,5.245,5.049,0.0004277,2.891,1.715,8.848,6.341,1.06,0.6884,0.9134,10.21,0.3765,0.4039,10.72,8.066,0.3683,9.714,3.06,10.59,0.3394,3.216,1.021,0.07074,0.7351,0.4438,5.722,10.57,0.02562,9.754,4.993,9.038,4.23,3.056,9.847,3.91,7.009,11.29,0.04354,5.217,8.403,3.206,1.491,0.05166,9.149,2.946,3.966,0.2785,0.7345,2.826,6.223,3.309,0.3331,0.9009,0.4495,0.01571,6.242,5.732,2.246,1.069,9.45,1.013,2.282,10.22,0.0007444,1.376,1.409,8.597,2.751,0.0921,0.4658,3.2,0.3322,6.412,0.8857,0.195,0.2767,11.28,3.598,10.51,1.244,2.796,1.12,0.147,11.38,1.942,0.7183,0.1596,5.555,0.5595,2.169,0.001787,10.76,5.668,7.374,7.336,1.63,5.608,7.608,0.6836,0.08215,4.7,1.254,7.867,5.559,0.2575,5.618,3.843,1.253,0.2071,0.9348,0.1512,9.321,8.041,0.1399,3.494,7.863,0.09739,7.784,7.138,5.255,2.715,0.9313,0.2418,8.452,3.797,0.784,1.021,10.61,7.192,0.04461,2.717,7.13,11.48,9.024,0.393,0.1625,1.784,1.11,0.8014,0.1283,7.225,7.502,4.82,0.8511,2.743,2.089,1.961,6.367,2.821,7.35,7.055,8.479,9.765,3.572,1.479,4.264,1.638,7.808,4.252,9.028,7.654,0.4572,4.76,1.002,0.9521,11.33,7.186,3.054,0.7324,1.637,0.09016,2.201,0.04745,3.925,2.519,2.162,4.311,1.03e-05,0.02127,3.541,0.09986,9.417,2.718,4.965,5.349,7.624,0.7198,0.4886,0.7848,2.071,9.859,2.951,1.232,1.897,4.208,1.641,7.537,3.357,3.709,0.2891,9.367,0.1378,2.252,1.346,0.254,0.5517,3.945,0.369,2.651,7.758,0.2656,4.584,0.3454,2.171,5.395,4.735,4.932,0.003387,11.41,3.228,4.281,1.006,1.313,3.731,2.868,1.612,0.1802,7.407,10.85,3.108,0.788,1.017,0.0002879,0.3449,2.91,9.397,1.061,2.242,0.7354,0.9995,1.226,5.267,0.009319,3.51,1.898,0.1759,5.512,2.087,8.968,10.62,1.348,1.009,4.654,9.662,6.583,0.2608,2.841,0.06761,7.047,0.0472,3.183,7.047,1.418,6.411,9.277,2.998,4.255,0.08336,3.757,7.02,5.575,1.752,5.871,6.312,0.016,0.1103,11.15,0.3694,7.806,0.7439,0.9188,0.002618,1.151,3.251,6.635,3.271,1.042,10.97,0.7565,3.508,8.153,1.202,1.102,5.154,6.825,4.304,0.7097,3.232,0.4101,0.0008838,9.744,0.4202,0.9056,4.678,0.9198,11.06,7.467,7.719,0.315,7.678,0.249,3.296,0.2863,0.9198,1.718,7.264,6.321,2.012,3.167,0.614,0.06571,4.312,3.557,0.09834,7.449,0.004651,1.787,0.01373,6.099,0.2323,2.474,4.541,1.872,2.331,0.3011,4.182e-05,0.1159,0.2277,3.654,1.862,10.61,4.018,10.31,2.18,1.38,4.367,2.185,0.03619,0.403,7.532,0.3145,0.1579,0.2759,0.8654,4.12,1.335,3.747,8.444,0.1148,8.499,1.83,1.214,9.274,10.17,8.784,7.208,0.1003,10.37,9.271,2.521,3.44,0.3075,1.065,8.406,3.871,0.4074,3.002,0.7131,0.4305,2.078,1.981,8.77,0.002954,1.248,2.302,0,8.287,8.683,9.839,11.82,21.16,22.21,42.05,48.48,22.99,30.05,26.39,12.67,25.81,32.2,32.38,59.01,15.6,44.82,17.3,18.53,20.46,20.39,24.79,28.81,29.77,104.3,69.93,20.35,30.77,45.64,33.66,38.11,165.2,17.74,14.81,36.23,83.77,158.6,63.71,23.96,23.73,108.4,255.1,14.55,16.44,18.79,19.41,36.52,47.24,46.65,120,11.94,24.17,24.48,15.02,25.26,18.15,26.81,27.05,47.65,27.79,52.75,33.69,53.08,19.66,27.77,19.84,20.01,20.21,12.05,33.11,20.55,26.15,85.76,13.41,112.5,29.56,37.96,13.15,30.73,20.61,25.88,43.43,49.49,50.92,38.73,33.64,44.12,60.27,15.87,15.57,17.19,16.27,11.63,65.26,53.37,61.98,76.33,33.37,17.59,36.17,19.15,24.59,28.68,52.98,14.37,23.24,17.19,32.61,70.94,16.99,35.66,32.78,12.24,13.5,39.46,18.18,29.33,76.03,25.26,12.76,90.01,36.5,15.37,54.39,11.93,19.18,22.87,35.95,14.91,66.52,27.59,18.04,22.75,18.22,17.05,19.37,18.05,46.97,23.44,15.07,12.2,22.52,16.68,38.13,70.31,142.1,79.9,107.1,24.58,82.57,61.63,38.19,235.1,59.46,74.32,38.2,11.7,12.17,20.93,24.17,22.56,84.96,144.4,23.18,20.74,45,16.98,25.04,22.78,32.36,43.22,17.91,12.89,66.47,40.95,108.9,31.88,18.76,43.46,20.88,17.56,18.36,12.24,37.22,25.11,55.89,15.83,30.59,32.98,57.45,13.23,43.65,29.59,14.08,14.33,51.78,26.77,28.76,12.31,24.98,39.56,19.44,21.42,16.62,20.67,47.65,21.7,26.87,29.64,26.2,17.6,82.9,86.7,70.52,17.17,12.73,28.82,13.81,12.29,18.79,15.05,36.98,12.99,74.32,72.43,45.14,13.64,28.71,39.69,112.1,12.32,94.61,28.43,34.88,31.47,13.89,14.54,16.82,12.66,43.01,23.31,45.98,30.37,33.34,11.57,14.29,15.56,14.13,13.58,116.7,35.18,12.6,120.3,47.87,39.12,17.87,40.94,12.32,31.03,16.01,19.43,16.11,57.01,16.22,16.98,13.11,31.9,23.45,128.4,16.48,57.2,38.38,25.46,18.59,43.42,54.35,29.55,12.22,90.34,92.6,99,19.01,143.8,85.84,56.89,16.61,51.99,15.46,11.82,28.16,30.56,20.33,43.09,15.91,41.43,12.09,74.58,33.35,109.1,12.71,32.05,51.18,16.19,31.06,16.92,35.53,31.73,14.59,23.47,18.88,12.48,17.12,19.14,13.26,19.72,91.42,15.1,20.44,69.86,65.35,59.48,18.69,29.48,55.32,80.9,21.18,14.48,24.28,17.88,34.09,62.98,80.94,25.85,56.92,17.35,33.79,41.95,66.38,39.88,28.72,15.66,14.67,13.21,25.64,69.18,19.09,79.9,13.63,19.47,12.47,22.22,43.01,23.38,42.68,29.29,25.95,24.74,14.72,67,21.01,24.71,79.59,27.17,16.72,169.9,36.26,22.53,13.71,35.99,14.97,14.92,23.44,34.71,26.41,23.54,54.89,24.31,67.18,23.81,39.16,51.56,36.04,21.13,30.98,14.58,43.82,12.99,22.76,30.25,18.18,26.25,16.66,16.33,12.94,18.02,39.75,25.33,24.13,56.15,23.61,70.68,25.54,38.22,19.97,65.04,62.76,23.76,17.43,61.65,84.56,32.12,19.39,20.71,15.54,36.56,20.92,26.39,20.79,39.69,73.68,13.48,69.95,23.52,36.31,45.35,12.25,12.8,133.5,64.82,67.75,22.19,19.11,43.89,18.4,60.8,20.5,220.6,108.4,21.79,41.69,21.3,21.46,35.37,23.28,24.68,38.48,25.6,92.77,21.3,19.23,15.57,21.77,43.95,38.48,17.6,20.18,29.23,46.97,25.77,15.6,26.46,43.12,24.44,28.65,19.2,14.34,14.06,101.1,17.65,101.4,18.74,34.32,16,35.74,21.99,32.21,16.28,22.74,35.82,47.75,34.6,30.44,25.5,43.65,58.23,42.2,35.06,48.32,22.94,67.82,39.37,39.59,19.34,37.87,16.98,25.84,28.15,18.77,30.61,11.69,40.44,20.76,37.97,18.98,49.55,90.83,14.76,13.42,75.75,42.79,14.69,13.39,302.9,14.05,17.22,32.57,12.53,151.1,14.72,27.3,38.29,81.52,26.01,40.32,17.76,11.63,70.1,47.36,42.88,16.77,16.09,14.82,17.16,17.45,19.13,43.48,19.02,17.72,20.54,88.09,21.77,18.39,23.66,28.71,36.8,52.18,59.8,21.01,184.2,19.53,49.68,75.39,28.29,16.72,24.69,18.66,13.26,63.66,78.69,17,61.4,36.05,20.8,26.43,25.87,13.86,42.98,27.39,37.08,15.23,76.41,15.98,49.07,71.19,12.29,13.9,17.16,24.78,21.78,24.56,53.4,37.68,24.55,19.92,12.37,81.16,19.64,39.6,20,24.33,130.2,71.65,37.46,50.9,13.73,34.12,49.99,35.71,23.39,18.27,39.42,69.08,32.7,26.61,21.21,14.24,24.36,20.55,70.58,48.97,38.74,15.11,18.41,24.23,36.38,84.35,55.31,15.48,13.13,17.97,35.83,19.98,14.57,19.28,33.82,43.34,15.29,36.86,42.4,87.5,185.3,23.64,15.07,20.34,56.01,61.85,57.24,325.6,21.52,43.94,91.9,23.32,57.81,13.05,14.36,11.6,47.68,26.86,38.65,20.81,13.77,18.49,46.4,11.98,28.36,37.04,211.7,105.9,14.91,17.73,20.67,19.63,29.95,32.72,25.71,13.3,101,26.66,23.41,17.19,41.42,12.89,38.24,36.13,15.57,61.39,25.16,29.03,62.09,35.18,16.88,30.58,11.95,12.87,45.32,63.36,15.19,26.86,12.86,42.3,73.18,23.01,19.74,14.04,38.12,26.74,151.2,34.27,12.07,28.65,22.6,75.62,29.54,108,12.82,38.13,50.4,27.62,13.97,95.95,35.46,14.07,15.34,29.96,31.27,28.02,41.32,25.66,14.58,34.08,80.31,31.29,33,24.29,20.25,21.86,16.51,73.94,63.27,31.73,33.44,32.38,50.23,24.24,59.03,53.04,42,30.66,32.12,16.39,28.73,19.25,35.61,19.17,13.73,17.52,49.3,25.23,24.63,15.39,37.31,42.91,38.64,59.24,14.77,13.46,24.05,12.23,20.67,23.38,13.32,18.27,14.85,69.05,59.39,15.39,14.24,18.56,14.06,67.19,26.71,41.45,13.17,30.44,35.03,33.47,15.03,41.85,12.69,16.21,29.55,81.7,18.23,48.4,13.08,21.83,23.66,36.1,21.7,11.89,22.07,14.78,61.1,29.06,19.83,26.84,19.14,17,87.29,72.24,12.17,32.86,24.39,12.94,13.61,11.99,18.96,19.65,15.43,20.83,14.12,14.88,16.87,37.51,40.91,11.77,56.81,39.73,15.75,71.94,42.99,26.26,18.73,26.74,27.69,46.09,104.1,101.8,57.94,51.17,13.38,43.56,69.74,22.47,50.13,81.41,55.38,77.66,19.58,21.68,16.69,28.89,73.71,36.26,16.29,20.23,39.19,74.31,17.9,12,17.53,18.27,106.9,47.15,14.97,25.94,23.19,13.97,144.9,60.38,83.81,84.98,46.51,111.1,14.42,72.46,16.91,30.08,25.92,15.39,14.32,16.88,13.4,47.56,11.98,12.17,50.43,11.65,15.2,15.77,66.58,32.65,16.32,23.09,16.99,44.13,79.69,114.4,14.19,25.15,37.84,14.41,126.5,11.82,49.22,19.7,15.73,61.45,12.59,64.61,12.46,20.44,39.29,14.33,15.82,115.1,13.29,47.96,24.88,30.88,37.96,35.61,19.2,19.85,33.47,20.68,21.57,31.8,34.17,33.64,16.66,27.73,70.89,93.39,48.58,12.51,15.21,19.52,62.52,20.77,33.35,17.22,31.7,31.28,65.97,88.59,37.81,22.71,25.85,152.8,32.81,60.84,22.09,25.8,63.98,125.9,64.71,19.64,69.68,22.08,12.74,32.43,14.56,18.05,12.74,65.61,32.94,17.62,13.86,46.18,28.06,47.64,13.66,76.61,66.67,25.17,12.31,18.93,65.45,32.84,13.51,15.73,26.26,19.29,49.93,16.39,40.53,22.65,25.51,24.52,44.15,18.49,81.26,54.62,11.64,12.86,18.3,18.71,26.94,14.17,12.97,16.19,13.86,17.17,27.52,14.83,49.32,16.17,19.22,15.36,25.86,55.39,31.05,46.45,38.35,12.39,29.74,24.63,57.67,20.58,55.83,14.74,41.21,17.81,42.67,20.65,27.23,32.53,25.59,16.3,18.88,85.16,25.18,56.24,12.82,26.87,27.15,28.21,39.54,15.43,15.67,13.05,14.97,28.27,15.75,24.8,89.72,13.32,20.88,29.43,54.45,43.51,49.96,13.11,26.96,22.79,45.38,17.97,15.85,23.92,16.33,26.24,17.82,46.59,12.62,19.52,155.1,23.97,59.96,42.99,77.22,30.44,31.71,31.2,15.84,29.28,16.22,26.31,27.9,15.97,14.8,13.3,16.94,58.86,39.24,23.54,20.92,18.59,11.97,38.81,32.35,11.71,12.45,12.94,13.8,53.68,21.27,25.92,15.32,16.9,35.71,19.62,18.87,27.36,18.91,12.46,121.8,12.86,35.76,42.45,12.47,15.04,26.57,13.04,20.22,15.56,11.84,24.94,12.77,30.36,19.87,21.46,32.52,54.9,72.22,25.55,21.96,14.09,22.52,17.34,18.75,11.69,46.12,28.54,19.7,22.72,14.69,20.43,15.84,13.54,45.44,24.85,15.95,38.81,22.81,34.71,126.9,12.52,14.44,73.91,16.73,28.7,16.35,17.89,23.57,13.25,44.42,20.64,36.59,24.92,12.67,14.46,20.16,32.28,30.1,28.18,19.72,82.13,21.74,34.61,11.78,38.34,53.38,16.02,19.64,28.27,34.17,38.53,23.25,28.87,209.8,18.68,15.37,18.65,124.8,17.47,48.65,15.75,20.07,28.29,20.99,12.98,40.68,13.16,14.79,98.43,13.14,14.8,45.87,13.02,36.44,14.26,11.66,40.95,43.3,62.72,30.52,23.47,32.13,26.33,28.89,40.35,17.56,24.89,21.13,50.22,68.79,106.2,21.39,19.1,24.46,22.4,19.32,64.61,20.29,17.25,17.2,41.48,20.83,96.19,12.84,52.2,33.37,14.13,24.61,68.78,44.67,35.42,47.54,17.82,53.7,15.85,127.8,68.61,13.53,131.9,17.71,26.2,12.63,20.04,12.12,54.51,24.66,45.53,17.74,17.07,43.51,24.81,37.9,37.76,62.92,44.25,13.97,56.61,36.55,17.61,19.68,17.3,31.24,38.83,20.13,19.7,12.37,56.28,13.79,57.09,35.26,20.19,20.5,16.16,29.94,12.72,45.87,22.59,44.21,22.45,70.81,77.29,14.9,69.67,24.8,12.62,16.83,38.82,35.94,59.68,24.45,33.05,20.75,18.93,21.4,102.5,15.48,32.27,12.28,22.52,33.35,40.68,73.33,48.06,19.13,37.08,12.65,24.21,20.7,78.02,101.7,27.47,20.38,48.19,33.72,35.24,15.07,88.66,46.27,86.27,13.21,12.03,16.61,34.38,13.33,24.52,30.41,11.59,15.78,13.27,13.75,19.57,15.97,29.4,63.26,21.01,41.5,32.54,66.71,37.42,16.72,21.13,17.53,38.59,31.4,49.27,25.69,25.53,68.35,101.8,19.02,15.88,16.46,85.08,101.5,11.9,22.34,42.93,54.99,25.23,13.98,41.76,28.54,93.18,29.38,21.71,60.01,38.35,19.18,37,48.52,27.81,39.08,39.17,16.98,32.68,56.83,27.47,14.53,25.88,16.62,31.98,22.29,12.98,17.78,52.7,13.52,22.9,34.7,17.89,23.89,66.03,63.05,23.86,39.15,49.63,11.76,59.66,15.41,20.89,78.83,31.18,18.02,51.27,35.51,59.42,67.55,34.25,174.4,29.31,20.63,43.87,70.94,19.69,12.48,19.01,34.9,96.85,21.73,104.2,43.09,67.62,27.86,53.59,39.44,17.05,28.46,46.18,13.16,20.34,26.28,16.47,193.5,106.3,45.56,39.61,14.51,15.77,15.11,26.56,12.03,27.01,15.66,30.38,42.96,58.35,60.9,50.33,30.56,14.85,103.8,61.78,17.17,21.04,90.87,41.8,29.25,45.63,20.01,41.1,39.09,64.13,27.86,39.12,22.23,11.84,17.91,33.33,17.47,20.24,99,11.58,11.67,31.2,44.07,11.99,13.69,68.97,183.2,42.02,12.24,18.07,19.92,33.15,111.8,76.88,22.9,33.31,52.89,34.79,37.92,36.11,17.13,70.44,15.39,12.57,13.61,52.8,24.9,13.58,25.67,349.2,19.65,29.46,30.81,15.3,78.78,16.22,32.99,11.71,12.93,35.81,33.3,38.37,34.84,26.14,34.33,39.55,22.7,18.63,41.32,88.79,28.67,50.89,25.34,63.4,39.49,37.26,13.81,27.12,13.65,75.63,56.4,13.71,19.69,15.95,44.31,24.73,96.93,27.47,17.18,26.22,54.69,12.63,14.44,44.43,100.4,39.79,16.27,27.84,55.01,13.29,37.84,80.39,20.06,12.3,18.38,33.65,22.22,32.28,20.45,50.07,29.24,22.36,14.65,30.05,27.41,85.39,21.23,16.6,40.62,18.87,21.73,19.82,20.51,43.03,52.55,15.26,52.99,24.45,113.5,36.21,18.97,19.17,61.53,54.89,21.85,20.16,24.69,81.58,39.44,25.45,14.95,73.82,33.06,22.86,18.32,13.62,31.95,36,11.92,57.43,11.7,28.83,39.21,25.29,11.99,39.82,21.43,45.83,13.71,56.29,11.68,16.41,21.87,29.14,15.74,19.74,17.73,60.38,27.21,49.98,21.97,16.92,12.22,56.9,21.08,33.31,13.19,74.95,34.55,16.62,53.23,15.38,14.06,24.77,13.2,12.64,96.57,59.29,31.43,13.49,18.92,70.08,27.1,16.27,104.7,92.73,27.29,13.88,67.2,73.25,26.67,15.74,12.86,12.78,12.95,25.59,40.39,37.82,42.13,27.87,21.69,15.93,12.25,14.73,13.05,32.2,32.63,13.26,30.14,26.99,13.91,22.04,19.26,14.16,93.05,18.62,28.92,35.55,25.13,22.95,50.8,15.78,14.85,22.92,43.3,30.42,27.83,50.99,17.89,12.97,29.37,42.09,54.12,53.75,27.65,53.92,41.18,71.28,57.26,319.2,13.25,29.2,12.66,12.64,56.77,95.74,26.15,17.58,17.31,22.32,22.03,26.47,70.09,141.5,11.89,43.38,41.66,29.12,60.69,28.67,45.42,89.08,75.02,66.76,146.9,42.87,25.73,13.57,15.11,67.75,15.87,26.96,60.18,28.82,106.1,46.28,26.27,48.01,11.71,49.87,23.76,14.57,17.48,15.26,25.63,54.04,182.7,62.99,45.2,19.5,49.1,50.85,13.23,37.4,229.3,18.65,27.25,45.74,24.12,42.64,29.67,20.17,63.81,98.03,63.4,19.6,106.5,28.18,33.07,18.37,15.25,48.27,34.41,26.48,17.43,27.98,91.01,17.79,22.74,18.2,13.44,42.74,32.31,22.49,92.69,24.9,36.37,15.52,13.21,77.21,63.27,24.8,16.52,58.08,20.94,12.01,17.43,48.91,21.57,31.4,34.86,16.48,37.88,17.41,86.2,23.31,16.53,17.25,21.2,36.84,44.06,17.36,46.03,25.3,15.33,21.95,51.85,39.45,46.4,87.25,22.03,77.16,24.26,33.44,80.61,50.52,27.18,36.16,98.96,26.34,36.78,13.71,41.42,163,29.16,29.71,11.6,47.76,45.99,12.26,27.75,14.69,59.68,26.46,43.93,14.63,23.42,14.13,72.67,64.21,20.5,226.7,20.28,34.56,11.81,11.98,13.57,37.47,22.67,17.41,37.53,79.52,66.3,94.04,18.59,70.4,23.07,30.95,16,34.35,22.14,14.59,19.87,35.01,50.71,99.91,24.87,18.91,18.37,17.26,22.28,19,50.82,29.91,19.53,23.29,19.46,25.37,25.7,61.78,14.67,82.69,28.79,16.06,12.87,21.49,16.88,21.9,13.15,18.18,13.92,21.09,14.05,60.49,36.34,18.44,14.15,64.71,12.97,29.03,44.98,30.56,26.56,19.12,19.71,13.25,65.4,16.26,53.2,23.04,51.65,16.28,21.63,80.21,13.84,11.94,15.34,14.83,16.53,15.52,54.28,19.34,16.99,20.95,21.86,64.34,26.34,36.37,11.86,27.43,16.47,18.34,58.61,64.27,75.13,69.91,12.41,24.74,34.78,15.22,23.43,48.13,37.22,73.32,95.02,17.85,13.96,17.55,163.7,12.69,15.17,51.04,40.67,17.63,15.31,20.8,17.62,75.79,71.65,24.59,33.78,18.97,30.48,22.08,44.3,24.04,24.49,115.8,32.65,12.67,22.15,21.81,94.1,12.13,30.56,34.82,18.93,38.09,13.8,17.62,46.19,11.92,14.81,18.32,74.05,88.8,22.96,13.5,16.42,42.87,28.21,18.48,78.84,25.63,17.36,34.76,34.69,458.9,26.5,107.2,12.66,31.62,25.75,22.18,23.39,17.69,65.93,25.28,21.62,24.33,84.57,26.72,43.62,15.71,20.46,74.25,63.53],"PValue":[0.03019,0.2231,0.7412,0.2821,0.7902,0.0135,0.005726,0.1226,0.5686,0.007105,0.06144,0.7587,0.8085,0.1326,0.09658,0.09589,0.09373,0.0845,0.02263,0.1135,0.1008,0.009109,0.1403,0.1165,0.8805,0.3481,0.09067,0.06399,0.07525,0.1318,0.1741,0.1361,0.1621,0.09058,0.08247,0.04107,0.1404,0.1459,0.2352,0.1594,0.09604,0.0828,0.1447,0.07996,0.1386,0.07566,0.1021,0.08565,0.02101,0.02414,0.01244,0.1919,0.3154,0.01206,0.09068,0.008957,0.2132,0.06013,0.1517,0.1318,0.1459,0.08554,0.1798,0.1202,0.05821,0.8581,0.3746,0.08612,0.04613,0.3643,0.07629,0.2697,0.008542,0.4547,0.06488,0.07655,0.1024,0.08988,0.8717,0.6276,0.01943,0.1175,0.1201,0.1401,0.1623,0.02068,0.08064,0.1703,0.9671,0.006997,0.09807,0.1378,0.3208,0.1706,0.09951,0.07209,0.6988,0.07427,0.1014,0.06609,0.1123,0.006737,0.01045,0.05872,0.04836,0.2447,0.1296,0.1154,0.1135,0.522,0.09665,0.8394,0.1771,0.1276,0.3048,0.1102,0.3138,0.818,0.1131,0.2202,0.1202,0.2178,0.1004,0.7847,0.07107,1,0.1102,0.1517,0.1459,0.1703,0.07944,0.1163,0.07938,0.1019,0.1436,0.04717,0.02789,0.005785,0.989,0.1224,0.1703,0.1016,0.1176,0.0965,0.2243,0.101,0.166,0.1055,0.115,0.4126,0.9873,0.04908,0.06863,0.699,0.9575,0.8996,0.8682,0.1436,0.138,0.1578,0.08339,0.6448,0.5585,0.2633,0.4135,0.5264,0.004381,0.1262,0.2439,0.0184,0.03091,0.6306,0.008744,1,0.4377,0.007391,0.1002,0.1212,0.4987,0.04314,0.3582,0.01492,0.03246,0.02192,0.152,0.1699,0.006865,0.0704,0.554,0.01247,0.697,0.004908,0.007708,0.0188,0.005438,0.3001,0.8784,0.01722,0.1013,0.07335,0.2823,0.08356,0.4932,0.7548,0.9321,0.008563,0.8419,0.3359,0.01608,0.2623,0.00381,0.601,0.3124,0.8879,0.1937,0.004421,0.8445,0.5237,0.3932,0.8295,0.05867,0.02197,0.1485,0.4641,0.06906,0.03273,0.03993,0.6281,0.667,0.007901,0.007614,0.04285,0.2374,0.8259,0.7352,0.1807,0.01257,0.646,0.003793,0.02777,0.2035,0.05385,0.005078,0.008021,0.1214,0.6693,0.5407,0.4556,0.0248,0.7685,0.1898,0.03158,0.004145,0.0727,0.6128,0.004632,0.00629,0.8609,0.6927,0.2188,0.01249,0.915,0.3884,0.08207,0.1377,0.01216,0.01751,0.1256,0.6394,0.9639,0.01045,0.2475,0.105,0.01149,0.02711,0.1336,0.1455,0.05856,0.3113,0.04237,0.1914,0.005258,0.5226,0.2914,0.07164,0.8459,0.783,0.1019,0.0054,0.8964,0.07096,0.7129,0.1417,0.2123,0.3472,0.2453,0.2771,0.1334,0.1588,0.3898,0.08746,0.09328,0.09287,0.1176,0.07147,0.08533,0.3437,0.777,0.08998,0.8845,0.2571,0.00553,0.8556,0.2691,0.173,0.04209,0.74,0.3877,0.02801,0.01287,0.07343,0.08985,0.9948,0.01928,0.008982,0.006732,0.04805,0.1027,0.06846,0.891,0.04398,0.2896,0.01645,0.5549,0.01696,0.1526,0.05256,0.8297,0.04855,0.2946,0.2399,0.9447,0.06856,0.009069,0.3233,0.02419,0.09057,0.01381,0.9311,0.01875,0.3708,0.3884,0.3491,0.1334,0.03491,0.1619,0.9245,0.1068,0.2338,0.01405,0.008559,0.08364,0.1248,0.1218,0.3496,0.5455,0.2254,0.5611,0.3295,0.01894,0.03736,0.01067,0.01567,0.442,0.4522,0.008413,0.0201,0.6276,0.06623,0.006763,0.8898,0.3684,0.8746,0.1251,0.01373,0.04217,0.8319,0.4614,0.03195,0.9106,0.7295,0.03935,0.008283,0.3234,0.01302,0.01174,0.3835,0.6824,0.4534,0.7705,0.03897,0.02811,0.1056,0.584,0.2993,0.2499,0.01755,0.003989,0.01065,0.02255,0.04,0.1945,0.00666,0.004143,0.5359,0.3026,0.6452,0.09384,0.1118,0.1102,0.04583,0.04979,0.4701,0.06484,0.1215,0.1154,0.05707,0.02779,0.08528,0.1176,0.1318,0.4548,0.1104,0.9442,0.3567,0.272,0.1011,0.1621,0.00421,0.8082,0.6325,0.4894,0.575,0.8462,0.1866,0.0395,0.9909,0.1542,0.4466,0.01342,0.005982,0.02439,0.38,0.4487,0.02048,0.007168,0.05237,0.006776,0.006574,0.01492,0.4256,0.006382,0.01214,0.04887,0.39,0.003956,0.02611,0.05948,0.9975,0.4814,0.2594,0.00755,0.05905,0.272,0.007888,0.01263,0.2299,0.08855,0.08598,0.03813,0.6967,0.02802,0.1254,0.8202,0.3047,0.07139,0.6709,0.09991,0.08146,0.01967,0.9352,0.02989,0.01325,0.7966,0.1651,0.3575,0.5458,0.2326,0.2172,0.2403,0.1928,0.6539,0.1777,0.02117,0.09369,0.1526,0.05575,0.2009,0.2348,0.1436,0.01854,0.2785,0.1966,0.01766,0.05436,0.03041,0.04232,0.07762,0.9009,0.8337,0.1329,0.6168,0.02707,0.06162,0.5656,0.08477,0.1045,0.7006,0.4636,0.04629,0.01355,0.3147,0.1986,0.04377,0.05761,0.09297,0.03115,0.1356,0.1055,0.08845,0.7826,0.3967,0.8368,0.3408,0.5472,0.05142,0.8232,0.4493,0.09325,0.7863,0.01483,0.5006,0.1476,0.8561,0.4319,0.9948,0.1604,0.06734,0.07475,0.009507,0.01483,0.1106,0.004491,0.02992,0.1079,0.02185,0.4905,0.063,0.01763,0.1402,0.00664,0.005532,0.003769,0.01217,0.04452,0.3931,0.04984,0.8959,0.1464,0.5668,0.04324,0.1496,0.4564,0.1808,0.8768,0.08326,0.1481,0.9312,0.1393,0.1101,0.586,0.1866,0.03956,0.0415,0.6288,0.01022,0.01975,0.2103,0.007695,0.4234,0.2078,0.7019,0.1192,0.8164,0.007794,0.005499,0.005859,0.01609,0.2885,0.004456,0.1772,0.9609,0.03256,0.009639,0.004892,0.2717,0.01256,0.1034,0.06115,0.2057,0.02115,0.02529,0.4662,0.9516,0.03122,0.7002,0.02209,0.03706,0.005746,0.4158,0.8857,0.03104,0.6284,0.07283,0.8524,0.008214,0.4811,0.8103,0.3047,0.4452,0.01211,0.01818,0.1956,0.007252,0.802,0.3877,0.9502,0.5493,0.325,0.1223,0.1956,0.6942,0.1141,0.112,0.02968,0.02434,0.1172,0.2267,0.2714,0.3505,0.004953,0.2604,0.9349,0.03275,0.01744,0.746,0.1071,0.01279,0.07429,0.07397,0.0875,0.661,0.6167,0.1597,0.9596,0.06469,0.8674,0.2461,0.5732,0.009604,0.004742,0.1413,0.0938,0.2276,0.6282,0.03988,0.06287,0.1666,0.2453,0.3555,0.004583,0.01327,0.09994,0.09055,0.00607,0.4047,0.7519,0.285,0.05187,0.3413,0.01193,0.1609,0.1055,0.3724,0.007852,0.02042,0.6676,0.06168,0.01253,0.008216,0.2448,0.8192,0.007035,0.02763,0.0721,0.6829,0.5149,0.3682,0.02485,0.003873,0.2676,0.01336,0.1416,0.6632,0.3855,0.1661,0.9033,0.02369,0.03991,0.01388,0.01407,0.2673,0.01329,0.07491,0.01163,0.05052,0.7623,0.3621,0.06739,0.3662,0.06472,0.3634,0.4648,0.4461,0.1028,0.1347,0.9081,0.02395,0.9858,0.9739,0.01721,0.03832,0.2346,0.01577,0.009592,0.01092,0.0986,0.5291,0.5868,0.1324,0.8808,0.004331,0.5527,0.7725,0.4047,0.8302,0.8307,0.006064,0.3644,0.009515,0.03876,0.01833,0.9086,0.02082,0.2655,0.8352,0.2188,0.8361,0.1069,0.007028,0.006791,0.175,0.03277,0.7735,0.007721,0.01957,0.06814,0.007631,0.007564,0.1422,0.1633,0.004523,0.01511,0.1665,0.01051,0.08879,0.05956,0.02368,0.08135,0.4735,0.007139,0.02569,0.6195,0.006601,0.2575,0.1015,0.1241,0.1099,0.3128,0.03925,0.8956,0.04206,0.007548,0.2658,0.00831,0.01197,0.3217,0.008179,0.008651,0.01216,0.6318,0.8749,0.7748,0.076,0.5408,0.1341,0.1297,0.01592,0.04934,0.6814,0.1265,0.1274,0.6672,0.04622,0.0903,0.1006,0.09476,0.8121,0.1132,0.6858,0.8425,0.6374,0.2031,0.3323,0.08656,0.06429,0.8281,0.7679,0.3005,0.04182,0.2876,0.6202,0.2486,0.1327,0.3679,0.02543,0.007199,0.6323,0.4504,0.02802,0.0101,0.01022,0.005782,0.5987,0.9192,0.01382,0.4169,0.2583,0.1258,0.1742,0.006005,0.09641,0.108,0.3056,0.00744,0.1139,0.5291,0.09414,0.6185,0.04257,0.008555,0.6268,0.3689,0.008579,0.01683,0.01517,0.0123,0.04503,0.006577,0.3846,0.247,0.2349,0.01966,0.1372,0.6748,0.751,0.1433,0.05096,0.4386,0.3715,0.04779,0.2104,0.014,0.01427,0.4429,0.7721,0.7773,0.05915,0.1499,0.03351,0.234,0.03507,0.8518,0.04697,0.7763,0.5954,0.003862,0.09138,0.01738,0.7757,0.09178,0.008493,0.5688,0.7472,0.1358,0.1853,0.4259,0.9347,0.003828,0.3038,0.1929,0.4349,0.02832,0.06724,0.191,0.04878,0.005359,0.01652,0.1492,0.04631,0.8181,0.6786,0.04743,0.0112,0.1556,0.007956,0.3093,0.1119,0.4582,0.005921,0.962,0.07341,0.006267,0.1776,0.02265,0.02515,0.07056,0.6993,0.005596,0.5767,0.04328,0.02399,0.003818,0.0284,0.2746,0.009987,0.08436,0.4828,0.2119,0.01476,0.325,0.06612,0.6605,0.005238,0.009769,0.006143,0.2184,0.2019,0.8925,0.4249,0.1978,0.07475,0.3997,0.2757,0.009487,0.8459,0.8834,0.2618,0.8397,0.01899,0.3778,0.1813,0.006202,0.008705,0.005587,0.006297,0.03363,0.5443,0.5734,0.2001,0.007368,0.05379,0.1316,0.7604,0.4799,0.04677,0.3866,0.5548,0.02247,0.5435,0.01344,0.02428,0.02088,0.09882,0.6261,0.1354,0.1615,0.1967,0.6059,0.02636,0.113,0.293,0.02218,0.2101,0.05082,0.09034,0.004266,0.00846,0.8884,0.4587,0.5957,0.0485,0.09616,0.04805,0.05566,0.5971,0.08026,0.009722,0.1691,0.06886,0.06058,0.02679,0.08718,0.8228,0.9448,0.04224,0.03718,0.9876,0.1318,0.6144,0.4738,0.1177,0.5625,0.4747,0.7924,0.6477,0.6839,0.1613,0.02291,0.5514,0.9928,0.5748,0.1322,0.5908,0.02058,0.1302,0.8797,0.1765,0.01893,0.0236,0.1026,0.187,0.6149,0.3913,0.632,0.007738,0.0291,0.003786,0.4738,0.006168,0.04437,0.05864,0.7428,0.7198,0.1443,0.04945,0.03821,0.06457,0.04684,0.1182,0.07548,0.1778,0.4069,0.005935,0.02922,0.4048,0.006827,0.09394,0.8833,0.0233,0.05242,0.3501,0.696,0.7336,0.0148,0.01233,0.004784,0.4035,0.3584,0.009674,0.004124,0.9466,0.2201,0.008215,0.5952,0.9853,0.2016,0.9263,0.4808,0.04895,0.02617,0.1481,0.005134,0.02633,0.3037,0.1831,0.4309,0.9763,0.01596,0.01695,0.1598,0.04064,0.3119,0.00764,0.3035,0.6198,0.05365,0.005444,0.03094,0.004874,0.07342,0.4916,0.4964,0.1757,0.23,0.03852,0.8019,0.06558,0.2529,0.229,0.4525,0.1757,0.004188,0.2649,0.05367,0.5679,0.1896,0.01636,0.01182,0.2863,0.07685,0.4315,0.5615,0.06121,0.3558,0.4606,0.4351,0.2713,0.1241,0.0131,0.01988,0.4662,0.564,0.6592,0.1401,0.02319,0.03267,0.02792,0.004681,0.01961,0.6515,0.176,0.004087,0.768,0.6769,0.8971,0.3042,0.04373,0.5486,0.007123,0.8209,0.03001,0.03466,0.004736,0.101,0.04136,0.8461,0.1741,0.09756,0.2272,0.1279,0.1129,0.07719,0.004557,0.0558,0.943,0.3877,0.007527,0.1411,0.1453,0.1699,0.01712,0.004299,0.142,0.01559,0.07906,0.004957,0.006705,0.4956,0.7153,0.007906,0.02104,0.7969,0.5087,0.005993,0.004138,0.7821,0.4495,0.007837,0.02892,0.01492,0.0713,0.1774,0.1204,0.007774,0.01303,0.03166,0.9359,0.4345,0.6444,0.3591,0.5135,0.6759,0.1734,0.5638,0.2737,0.04915,0.009834,0.3629,0.01171,0.09535,0.02588,0.01847,0.8322,0.01741,0.5798,0.1298,0.03655,0.02627,0.2958,0.007679,0.9654,0.1638,0.2439,0.2591,0.2753,0.005543,0.4697,0.07557,0.01544,0.05093,0.0682,0.9702,0.565,0.3698,0.006761,0.7733,0.9472,0.02879,0.1545,0.1824,0.004255,0.6429,0.00581,0.003974,0.08996,0.004334,0.3755,0.1734,0.5035,0.5441,0.03753,0.01822,0.01528,0.005368,0.06364,0.8525,0.07348,0.7679,0.1737,0.3135,0.00733,0.005516,0.9242,0.834,0.5175,0.9183,0.1622,0.1358,0.3766,0.004099,0.04152,0.5373,0.5726,0.03851,0.2507,0.6822,0.1954,0.7776,0.01219,0.005579,0.2316,0.1655,0.343,0.02706,0.5002,0.1249,0.01069,0.04221,0.9728,0.2841,0.01238,0.9407,0.1875,0.01649,0.8069,0.7269,0.03301,0.5653,0.004829,0.228,0.0246,0.8372,0.03486,0.01595,0.05746,0.006965,0.7375,0.0817,0.006072,0.4101,0.6671,0.147,0.4638,0.177,0.9155,0.005526,0.4701,0.1364,0.7619,0.03005,0.04451,0.008141,0.03171,0.1726,0.6297,0.9919,0.1711,0.5596,0.1403,0.3316,0.03568,0.0647,0.3972,0.9642,0.4614,0.005681,0.0156,0.03073,0.2604,0.05233,0.5539,0.1524,0.4032,0.00809,0.1594,0.1377,0.4673,0.6513,0.6475,0.1698,0.0176,0.3431,0.01164,0.008293,0.3998,0.02629,0.01187,0.3602,0.06396,0.03915,0.8579,0.01144,0.1733,0.01651,0.04565,0.03865,0.07737,0.05441,0.1824,0.3825,0.649,0.0124,0.00854,0.3,0.008151,0.5763,0.9559,0.05689,0.006618,0.8104,0.9187,0.02272,0.2597,0.1377,0.08094,0.5747,0.2104,0.005644,0.007238,0.5828,0.1618,0.06163,0.1089,0.07312,0.1341,0.09128,0.4426,0.004582,0.01192,0.7539,0.3205,0.7488,0.01024,0.8254,0.01436,0.1328,0.04303,0.2751,0.8013,0.1908,0.2819,0.4956,0.2779,0.7049,0.08328,0.004406,0.03682,0.3596,0.4706,0.1045,0.4124,0.2709,0.3817,0.05691,0.7227,0.9822,0.652,0.0821,0.6127,0.02424,0.05747,0.01089,0.004929,0.2428,0.907,0.03428,0.01118,0.4245,0.03234,0.3413,0.7861,0.0982,0.02633,0.4586,0.0114,0.03217,0.2132,0.004544,0.0196,0.1908,0.1222,0.5782,0.7065,0.27,0.5095,0.755,0.04441,0.004256,0.3539,0.5916,0.003974,0.01996,0.3135,0.04432,0.004365,0.01233,0.03821,0.005022,0.05333,0.1127,0.8464,0.06002,0.01852,0.3903,0.00547,0.009539,0.2922,0.02726,0.2406,0.5511,0.01407,0.1683,0.1581,0.3325,0.5489,0.006559,0.1158,0.01422,0.006653,0.38,0.01683,0.3977,0.04819,0.06518,0.06157,0.03702,0.3688,0.05087,0.04815,0.00534,0.003911,0.005761,0.02742,0.6959,0.02195,0.8663,0.1262,0.07229,0.005821,0.303,0.09262,0.03045,0.01228,0.05082,0.01827,0.6006,0.1108,0.0541,0.1653,0.7313,0.3651,0.8267,0.4947,0.1627,0.01878,0.2854,0.182,0.8444,0.3801,0.5828,0.02999,0.04003,0.1145,0.02374,0.004064,0.7721,0.3631,0.03211,0.07479,0.09879,0.4785,0.3766,0.004845,0.3177,0.03235,0.5295,0.02266,0.1112,0.7936,0.1425,0.04246,0.625,0.2417,0.6389,0.2265,0.01044,0.4866,0.8849,0.2752,0.02449,0.2803,0.01104,0.1403,0.4515,0.09935,0.6047,0.6263,0.1762,0.02277,0.6194,0.8025,0.07971,0.02425,0.9217,0.2364,0.08322,0.544,0.02717,0.1924,0.04298,0.3569,0.007731,0.07913,0.05198,0.03042,0.7212,0.4609,0.007287,0.4228,0.005656,0.7997,0.9017,0.7831,0.02736,0.09859,0.6933,0.06387,0.1686,0.443,0.01749,0.5496,0.7282,0.01086,0.06631,0.1108,0.00475,0.1808,0.02822,0.79,0.2192,0.4973,0.06782,0.7668,0.04422,0.006293,0.005095,0.5214,0.04046,0.03043,0.05019,0.04803,0.1553,0.07164,0.1402,0.09222,0.02856,0.4662,0.1056,0.08599,0.1019,0.9674,0.1161,0.9861,0.0078,0.08423,0.01756,0.1776,0.6518,0.586,0.1353,0.043,0.6372,0.8449,0.2316,0.5385,0.9711,0.02455,0.1647,0.1883,0.2344,0.08859,0.1934,0.01501,0.1206,0.3974,0.6374,0.0361,0.113,0.01116,0.2553,0.006196,0.1653,0.7015,0.01552,0.01549,0.07614,0.7196,0.112,0.2189,0.3157,0.05305,0.6921,0.7932,0.6693,0.1031,0.09345,0.1377,0.9634,0.02734,0.1596,0.1431,0.7198,0.3113,0.6118,0.03735,0.9335,0.04015,0.7111,0.1809,0.5341,0.1032,0.02467,0.3615,0.5358,0.004585,0.02934,0.5629,0.008344,0.08209,0.3257,0.07878,0.5676,0.05124,0.9454,0.2982,0.008605,0.1245,0.01836,0.01399,0.4814,0.0118,0.4538,0.2542,0.5802,0.263,0.02763,0.7012,0.7924,0.7047,0.009333,0.02091,0.4922,0.6653,0.1779,0.1102,0.6782,0.03714,0.04238,0.5608,0.9384,0.003896,0.1379,0.3706,0.1152,0.006615,0.01325,0.8481,0.03409,0.5593,0.01574,0.2725,0.4008,0.5039,0.004256,0.2834,0.3174,0.02328,0.04529,0.005118,0.01347,0.0494,0.3099,0.196,0.09821,0.8645,0.08293,0.04351,0.895,0.005138,0.3591,0.1035,0.005405,0.8402,0.575,0.1768,0.2076,0.8019,0.02935,0.2518,0.1411,0.2807,0.1357,0.105,0.2484,0.4313,0.03757,0.1265,0.07304,0.06279,0.6763,0.3193,0.005855,0.006327,0.0712,0.09383,0.04055,0.1094,0.03928,0.02982,0.9271,0.44,0.4039,0.1281,0.5041,0.6312,0.8334,0.01361,0.01505,0.4712,0.1423,0.9153,0.05196,0.3471,0.59,0.9118,0.9776,0.008577,0.005822,0.6379,0.3079,0.2436,0.02587,0.4803,0.124,0.004152,0.01082,0.6533,0.4135,0.01898,0.04887,0.6964,0.02738,0.004066,0.05765,0.08078,0.1514,0.005718,0.9517,0.1883,0.2044,0.007527,0.3648,0.2061,0.005734,0.01243,0.2454,0.5822,0.1703,0.8873,0.4382,0.04025,0.00486,0.2256,0.1225,0.02736,0.7858,0.7628,0.2407,0.724,0.6502,0.8894,0.01574,0.003952,0.3765,0.1053,0.4938,0.9377,0.8079,0.5056,0.02876,0.54,0.05293,0.03729,0.024,0.02606,0.01055,0.1451,0.3278,0.1617,0.516,0.2569,0.01461,0.3602,0.7467,0.003966,0.01672,0.4514,0.0633,0.3284,0.02629,0.9785,0.2201,0.8882,0.1933,0.5687,0.005863,0.9136,0.09751,0.1319,0.02024,0.01009,0.1703,0.0215,0.3258,0.1322,0.06441,0.4077,0.01967,0.2502,0.219,0.006306,0.02438,0.03175,0.329,0.9508,0.02797,0.5765,0.2948,0.01046,0.9827,0.1627,0.2995,0.2559,0.1649,0.233,0.8045,0.03634,0.6708,0.01717,0.01587,0.1288,0.03181,0.01564,0.5785,0.04408,0.008958,0.04469,0.2682,0.004429,0.005728,0.1028,0.137,0.3287,0.1579,0.006416,0.8687,0.4679,0.2441,0.4439,0.3462,0.4251,0.04961,0.01453,0.004993,0.004452,0.008394,0.03599,0.19,0.7203,0.995,0.02653,0.3811,0.7216,0.005986,0.3096,0.6673,0.2511,0.04614,0.007469,0.1425,0.01527,0.05485,0.008049,0.01295,0.6364,0.006425,0.007371,0.08129,0.02273,0.01427,0.02166,0.1465,0.2079,0.08526,0.9353,0.7258,0.08525,0.04716,0.2758,0.1937,0.01909,0.735,0.1187,0.05141,0.5586,0.1905,0.06288,0.5696,0.01243,0.2778,0.1268,0.4275,0.9813,0.01452,0.07851,0.05226,0.4985,0.1183,0.4118,0.005619,0.04601,0.04962,0.528,0.0513,0.2075,0.3714,0.01281,0.006231,0.007748,0.01628,0.3067,0.3328,0.9622,0.6093,0.115,0.6611,0.8658,0.9927,0.007207,0.1417,0.2849,0.2504,0.9487,0.004395,0.1922,0.009425,0.6821,0.0603,0.03102,0.3209,0.009006,0.2332,0.5319,0.2488,0.08742,0.6625,0.0412,0.06484,0.07786,0.02033,0.1234,0.553,0.3834,0.08622,0.05329,0.5157,0.007755,0.01215,0.4986,0.2296,0.9885,0.9806,0.02681,0.4567,0.4648,0.7047,0.1805,0.004687,0.165,0.6487,0.1244,0.9642,0.0553,0.1502,0.005036,0.00858,0.134,0.8359,0.0481,0.1144,0.3679,0.08928,0.009102,0.6696,0.08453,0.00416,0.1512,0.01741,0.2856,0.5704,0.004843,0.3323,0.9665,0.09972,0.1197,0.3628,0.01479,0.5367,0.1218,0.6328,0.7274,0.1468,0.1901,0.6375,0.9543,0.9002,0.09043,0.1216,0.9423,0.09392,0.01961,0.004182,0.09774,0.1848,0.5735,0.7144,0.1563,0.005206,0.008193,0.01599,0.01235,0.7742,0.01395,0.5066,0.1955,0.1054,0.9155,0.04516,0.3574,0.8521,0.1349,0.3637,0.7508,0.4956,0.08847,0.1497,0.366,0.0322,0.0203,0.117,0.2755,0.01315,0.008429,0.00544,0.09895,0.02766,0.1204,0.08468,0.1121,0.8188,0.01303,0.148,0.03861,0.07231,0.06335,0.2963,0.05566,0.2001,0.5855,0.1568,0.007789,0.06283,0.01783,0.3128,0.2281,0.06948,0.04141,0.006099,0.8946,0.6134,0.2017,0.3681,0.01087,0.2646,0.07749,0.07331,0.331,0.004043,0.5179,0.05903,0.7104,0.006007,0.4422,0.0654,0.3884,0.2359,0.01912,0.0114,0.0153,0.08439,0.004564,0.05231,0.1215,0.4203,0.07563,0.7085,0.6021,0.1305,0.0843,0.06081,0.217,0.3628,0.6288,0.0656,0.3535,0.1564,0.6199,0.04104,0.004338,0.0842,0.0183,0.3718,0.4037,0.01721,0.01489,0.0182,0.007709,0.2116,0.01008,0.7417,0.01857,0.8835,0.444,0.03972,0.6644,0.01865,0.7018,0.08595,0.3039,0.01096,0.04239,0.9116,0.1344,0.03088,0.05192,0.03262,0.5699,0.321,0.08091,0.2922,0.6061,0.0321,0.3561,0.1877,0.0286,0.3117,0.3334,0.02164,0.1705,0.5072,0.8488,0.01986,0.8206,0.7146,0.01887,0.09211,0.9628,0.1342,0.2839,0.4766,0.1287,0.04108,0.0161,0.07612,0.07962,0.1061,0.2718,0.02904,0.207,0.3431,0.1085,0.009602,0.05485,0.1361,0.2803,0.5177,0.5794,0.004382,0.8521,0.2098,0.7137,0.8852,0.08548,0.003993,0.7066,0.8757,0.8731,0.1586,0.6469,0.1035,0.144,0.1025,0.06118,0.02999,0.159,0.5803,0.2161,0.04294,0.2577,0.03271,0.102,0.06767,0.04836,0.05241,0.05652,0.6917,0.03921,0.04633,0.8656,0.2982,0.04861,0.04643,0.04981,0.575,0.1667,0.2776,0.01248,0.005718,0.09888,0.02205,0.009874,0.005639,0.6507,0.446,0.1924,0.02492,0.3709,0.3768,0.3237,0.05252,0.1134,0.1305,0.01865,0.0343,0.7366,0.03281,0.06049,0.4528,0.05049,0.008432,0.07763,0.6243,0.1309,0.1548,0.599,0.08672,0.5372,0.04754,0.6247,0.03018,0.8234,0.6269,0.03807,0.2504,0.06712,0.7686,0.3218,0.0257,0.7983,0.1072,0.7996,0.931,0.8956,0.02043,0.2777,0.4721,0.03025,0.07763,0.03452,0.3793,0.1969,0.9175,0.4011,0.8807,0.8168,0.8985,0.7267,0.09947,0.2511,0.2663,0.113,0.007996,0.1182,0.02751,0.007657,0.07294,0.4788,0.1622,0.1904,0.01654,0.2183,0.6403,0.09398,0.01235,0.01016,0.01306,0.02009,0.0746,0.06503,0.05178,0.0818,0.853,0.1047,0.0889,0.7117,0.32,0.3326,0.007409,0.03955,0.6477,0.4659,0.4843,0.5584,0.2352,0.005275,0.01389,0.419,0.8208,0.02938,0.3791,0.2755,0.05009,0.8567,0.2146,0.06383,0.0115,0.004631,0.9112,0.5311,0.193,0.05088,0.4601,0.6451,0.5774,0.8602,0.4519,0.4525,0.01697,0.6043,0.02666,0.08581,0.1803,0.2733,0.8045,0.2498,0.00692,0.7835,0.05832,0.01609,0.8963,0.8595,0.5041,0.01734,0.3041,0.04952,0.1906,0.05539,0.2393,0.0483,0.3267,0.03588,0.1802,0.09531,0.04948,0.9083,0.01949,0.07037,0.3757,0.02236,0.0182,0.3179,0.1879,0.522,0.004903,0.02275,0.01336,0.01642,0.7247,0.004485,0.109,0.08201,0.00431,0.5774,0.01508,0.9748,0.05539,0.6922,0.02916,0.6151,0.6446,0.1435,0.04442,0.8412,0.2816,0.07968,0.05509,0.7925,0.05217,0.04178,0.009865,0.02908,0.6476,0.7342,0.6865,0.4626,0.9978,0.02314,0.03879,0.4953,0.3256,0.6729,0.009582,0.1065,0.4135,0.03103,0.1162,0.6662,0.007809,0.3977,0.04663,0.01937,0.8042,0.7764,0.7896,0.1866,0.05719,0.01495,0.2077,0.3422,0.02939,0.2566,0.5104,0.06208,0.1876,0.6111,0.1177,0.01377,0.6875,0.008099,0.0147,0.04392,0.02086,0.187,0.01526,0.2433,0.003783,0.0157,0.04995,0.1747,0.02239,0.08346,0.05159,0.483,0.04772,0.01191,0.578,0.009482,0.8726,0.2169,0.09034,0.0121,0.3562,0.2673,0.6401,0.008216,0.003902,0.00544,0.4799,0.01451,0.5909,0.09875,0.05108,0.7713,0.09702,0.8329,0.005184,0.4057,0.1759,0.004216,0.007327,0.8671,0.1146,0.02531,0.237,0.009942,0.8974,0.01098,0.3555,0.1213,0.4968,0.06066,0.6997,0.01379,0.04742,0.01153,0.06525,0.09404,0.02253,0.007808,0.009782,0.003791,0.09299,0.004332,0.02456,0.02423,0.2514,0.009771,0.005359,0.01112,0.3507,0.02307,0.7483,0.08341,0.1092,0.1223,0.1198,0.0697,0.04692,0.7387,0.6865,0.4554,0.03533,0.3973,0.04982,0.455,0.2566,0.944,0.2057,0.7615,0.04789,0.01751,0.3698,0.09,0.1035,0.2744,0.05417,0.7945,0.2117,0.02554,0.1375,0.2543,0.8393,0.7866,0.02428,0.01119,0.6495,0.2611,0.006224,0.07552,0.02439,0.3417,0.1771,0.005954,0.899,0.09482,0.6394,0.1013,0.04921,0.03308,0.07596,0.003991,0.2354,0.2828,0.7696,0.1137,0.5535,0.08323,0.02448,0.01735,0.005855,0.1134,0.04188,0.5016,0.009027,0.2719,0.1231,0.03294,0.1542,0.006889,0.8438,0.2403,0.06658,0.9692,0.009248,0.5391,0.4486,0.245,0.1581,0.3748,0.102,0.5545,0.005248,0.07918,0.0114,0.07147,0.00494,0.004768,0.7779,0.1632,0.259,0.2047,0.009293,0.1452,0.4419,0.6673,0.7353,0.005011,0.04474,0.05693,0.02187,0.1905,0.03879,0.03041,0.07225,0.005389,0.09748,0.02024,0.6282,0.104,0.7517,0.415,0.1095,0.02721,0.01115,0.05489,0.0206,0.6201,0.2164,0.3663,0.04737,0.5939,0.6222,0.004887,0.2842,0.8205,0.5073,0.2778,0.06129,0.07184,0.553,0.08638,0.07196,0.1293,0.0245,0.01314,0.2842,0.04985,0.8028,0.3901,0.527,0.04816,0.03481,0.046,0.008365,0.07494,0.5061,0.09399,0.3009,0.01231,0.01013,0.0291,0.106,0.6038,0.2252,0.489,0.004368,0.01959,0.02573,0.06707,0.07923,0.1858,0.03837,0.8198,0.7707,0.003782,0.004391,0.7227,0.5412,0.01287,0.1653,0.08932,0.6128,0.06935,0.007793,0.004082,0.9637,0.02925,0.4243,0.02019,0.4317,0.2235,0.1988,0.1232,0.01359,0.4198,0.779,0.6161,0.3003,0.2352,0.8352,0.1352,0.01262,0.9194,0.5512,0.804,0.01875,0.8644,0.7974,0.4742,0.05672,0.02169,0.06717,0.3333,0.01852,0.5108,0.4813,0.08189,0.8535,0.4197,0.009284,0.04048,0.03957,0.9948,0.4242,0.114,0.004747,0.01169,0.05066,0.2747,0.03051,0.7031,0.5321,0.3792,0.1726,0.2481,0.1421,0.2478,0.02777,0.3419,0.0968,0.2493,0.01252,0.006863,0.5826,0.005227,0.7735,0.2614,0.2853,0.394,0.5698,0.008131,0.1556,0.005401,0.04699,0.09172,0.8928,0.01501,0.2159,0.9934,0.04398,0.7149,0.02198,0.1979,0.01068,0.06171,0.3836,0.03546,0.128,0.5063,0.3525,0.00553,0.1231,0.004987,0.0258,0.3601,0.201,0.5571,0.2669,0.1854,0.372,0.9299,0.4723,0.354,0.05661,0.1101,0.1254,0.41,0.01133,0.4453,0.01373,0.01155,0.005778,0.4124,0.007805,0.01064,0.02528,0.08196,0.1122,0.05245,0.01495,0.5033,0.8964,0.00817,0.6512,0.1478,0.04899,0.2042,0.0262,0.1387,0.03441,0.6278,0.291,0.05964,0.2501,0.0208,0.01538,0.1647,0.4861,0.005636,0.01285,0.1868,0.1084,0.0156,0.042,0.01729,0.9373,0.007369,0.1892,0.02243,0.01591,0.1892,0.8839,0.9566,0.03624,0.03942,0.9838,0.1088,0.2092,0.0091,0.02308,0.3189,0.4192,0.3537,0.005751,0.5483,0.5342,0.004875,0.01201,0.5526,0.006777,0.09978,0.005083,0.5684,0.09226,0.3277,0.7937,0.4041,0.515,0.02966,0.005118,0.8749,0.006688,0.04039,0.008522,0.05676,0.09999,0.006484,0.06587,0.01779,0.004806,0.8374,0.03668,0.01064,0.0927,0.24,0.8231,0.008202,0.1058,0.06417,0.6051,0.4044,0.1125,0.0242,0.08808,0.572,0.3569,0.5123,0.9018,0.02402,0.02955,0.1538,0.317,0.007405,0.3294,0.1507,0.005744,0.9786,0.2583,0.2529,0.009936,0.1171,0.7655,0.5049,0.09297,0.5726,0.02244,0.3609,0.6648,0.6063,0.004087,0.07645,0.005225,0.2816,0.1143,0.3059,0.7066,0.003973,0.1829,0.4095,0.6949,0.03179,0.4655,0.1607,0.9668,0.004817,0.03033,0.01549,0.01571,0.2203,0.03111,0.0142,0.4208,0.7784,0.04592,0.2797,0.01291,0.03175,0.6189,0.03097,0.06798,0.2798,0.6553,0.3483,0.7026,0.007735,0.01212,0.7134,0.0804,0.01293,0.7591,0.01331,0.01693,0.03609,0.1193,0.3492,0.6298,0.01046,0.06947,0.3893,0.3277,0.005063,0.01659,0.8354,0.1192,0.01698,0.003848,0.008563,0.5398,0.6923,0.2007,0.308,0.3842,0.7249,0.01639,0.01477,0.04356,0.3702,0.1175,0.1681,0.1809,0.02284,0.1129,0.01563,0.01748,0.01036,0.006663,0.07741,0.2419,0.05588,0.2192,0.01319,0.05618,0.008551,0.01396,0.5088,0.04472,0.3321,0.344,0.004033,0.01663,0.1001,0.405,0.2194,0.7679,0.1577,0.8304,0.0654,0.1325,0.1612,0.0547,0.9975,0.8859,0.07857,0.7564,0.007489,0.1191,0.04089,0.03468,0.01411,0.409,0.4948,0.39,0.1698,0.006458,0.1055,0.2838,0.1878,0.05734,0.2188,0.01458,0.08601,0.07246,0.5984,0.007617,0.7154,0.1533,0.2633,0.6213,0.4686,0.06479,0.5522,0.1234,0.01343,0.6135,0.04836,0.5651,0.1604,0.034,0.04523,0.04147,0.9543,0.003933,0.0917,0.05544,0.3311,0.269,0.07169,0.1102,0.2227,0.677,0.0153,0.004691,0.0974,0.3881,0.3286,0.9867,0.5654,0.1078,0.007539,0.3186,0.1542,0.4041,0.3326,0.2849,0.0359,0.9243,0.07977,0.1877,0.6806,0.03238,0.1682,0.00873,0.005045,0.2631,0.3303,0.04686,0.006895,0.02098,0.6172,0.1117,0.7982,0.01753,0.8308,0.09378,0.01753,0.2514,0.02245,0.007852,0.103,0.05612,0.7766,0.07081,0.01772,0.03153,0.2047,0.02791,0.02335,0.9009,0.7442,0.00426,0.5521,0.0132,0.4014,0.3523,0.9598,0.2995,0.09066,0.02055,0.08976,0.3228,0.004503,0.3975,0.07985,0.01164,0.2895,0.3097,0.03768,0.01909,0.05486,0.4122,0.09151,0.5312,0.9767,0.00671,0.5262,0.3567,0.04637,0.3521,0.004387,0.01496,0.01363,0.5825,0.01451,0.6247,0.08863,0.6001,0.3521,0.2089,0.01615,0.02327,0.1756,0.09452,0.445,0.801,0.05468,0.07796,0.758,0.01507,0.9465,0.2004,0.9082,0.02543,0.6365,0.1357,0.04929,0.1908,0.1467,0.5909,0.9949,0.7381,0.6398,0.07441,0.1917,0.00505,0.06261,0.005578,0.1596,0.2576,0.05333,0.1592,0.8516,0.5347,0.0146,0.5829,0.6965,0.6068,0.3663,0.05972,0.2652,0.07114,0.01049,0.7393,0.01028,0.1953,0.2871,0.00786,0.005839,0.009305,0.01649,0.7557,0.005469,0.008361,0.1337,0.08384,0.5876,0.3188,0.01121,0.06834,0.5331,0.1028,0.412,0.5219,0.1691,0.1801,0.009896,0.9574,0.2819,0.152,1,0.01169,0.01697,0.01255,0.009437,0.0003115,0.0002477,8.341e-06,5.141e-06,0.0002509,5.409e-05,0.0001059,0.002945,0.0001187,3.735e-05,3.628e-05,1.07e-06,0.001189,5.735e-06,0.002658,0.0005694,0.0003637,0.002181,0.0001762,6.749e-05,7.169e-05,2.504e-08,3.62e-07,0.0003729,4.767e-05,5.149e-06,2.934e-05,1.467e-05,9.827e-10,0.0006904,0.00147,1.953e-05,1.104e-07,1.317e-09,6.585e-07,0.0001718,0.00018,1.918e-08,4.1e-11,0.001577,0.000956,0.0005357,0.000463,1.868e-05,4.193e-06,4.517e-06,9.496e-09,0.003338,0.0001646,0.0001543,0.001389,0.0001323,0.0006241,9.794e-05,9.354e-05,3.982e-06,8.135e-05,2.147e-06,2.92e-05,2.065e-06,0.000437,8.171e-05,0.0004192,0.0004037,0.0003857,0.003233,3.209e-05,0.0003571,0.0001111,9.438e-08,0.002167,1.487e-08,5.895e-05,1.501e-05,0.002338,4.801e-05,0.0003521,0.0001171,6.898e-06,3.168e-06,2.663e-06,1.34e-05,2.946e-05,6.288e-06,9.361e-07,0.001107,0.0012,0.0007904,0.0009989,0.003672,5.649e-07,1.999e-06,7.848e-07,2.042e-07,3.075e-05,0.0007165,1.969e-05,0.0004917,0.0001512,6.911e-05,2.091e-06,0.001655,0.0001992,0.0007915,3.489e-05,3.296e-07,0.0008323,2.133e-05,3.391e-05,0.003058,0.002113,1.204e-05,0.0006204,6.143e-05,2.098e-07,0.0001321,0.00262,6.816e-08,1.874e-05,0.001264,1.778e-06,0.003355,0.0004885,0.0002153,2.038e-05,0.001431,4.995e-07,8.449e-05,0.000642,0.0002207,0.0006135,0.003619,0.0004671,0.00064,4.339e-06,0.0001913,0.001369,0.003095,0.0002318,0.0008984,1.463e-05,3.493e-07,2.883e-09,1.512e-07,2.087e-08,0.0001513,1.215e-07,8.131e-07,1.45e-05,7.499e-11,1.019e-06,2.434e-07,1.448e-05,0.003594,0.003116,0.0003275,0.0001646,0.00023,1.005e-07,2.573e-09,0.0002019,0.0003416,5.599e-06,0.0008341,0.0001381,0.0002192,3.64e-05,7.097e-06,0.0006614,0.002522,5.02e-07,9.719e-06,1.853e-08,3.946e-05,0.0005397,6.877e-06,0.0003314,0.0007212,0.0005935,0.003055,1.678e-05,0.0001361,1.502e-06,0.001119,4.921e-05,3.283e-05,1.265e-06,0.002283,6.697e-06,5.871e-05,0.001797,0.001674,2.406e-06,9.867e-05,6.815e-05,0.002992,0.0001398,1.187e-05,0.00046,0.0002938,0.0009127,0.000347,3.982e-06,0.0002765,9.668e-05,5.814e-05,0.0001099,0.0007138,1.184e-07,8.772e-08,3.426e-07,0.0007956,0.002641,6.732e-05,0.001939,0.003004,0.0005354,0.001378,2.316e-05,0.002446,2.433e-07,2.881e-07,5.493e-06,0.002032,6.872e-05,1.164e-05,1.519e-08,0.002977,4.866e-08,7.233e-05,2.411e-05,4.226e-05,0.001895,0.001581,0.0008671,0.002694,7.31e-06,0.0001962,4.927e-06,5.11e-05,3.093e-05,0.003741,0.001693,0.001201,0.00177,0.002065,1.153e-08,2.301e-05,0.002748,9.335e-09,3.873e-06,1.265e-05,0.0006679,9.734e-06,0.002978,4.555e-05,0.001067,0.0004605,0.001042,1.327e-06,0.001012,0.0008344,0.002365,3.929e-05,0.0001906,5.901e-09,0.0009473,1.299e-06,1.411e-05,0.000127,0.0005625,6.909e-06,1.785e-06,5.911e-05,0.003076,6.652e-08,5.631e-08,3.576e-08,0.0005084,2.646e-09,9.378e-08,1.345e-06,0.0009146,2.347e-06,0.001235,0.003469,7.595e-05,4.947e-05,0.0003752,7.226e-06,0.001235,9.092e-06,0.003198,2.379e-07,3.085e-05,1.832e-08,0.002656,3.831e-05,2.583e-06,0.001019,4.532e-05,0.0009799,2.177e-05,4.043e-05,0.001954,0.00019,0.0006067,0.00284,0.0008055,0.0004929,0.002492,0.0004315,6.138e-08,0.00136,0.0003655,3.643e-07,5.598e-07,1.018e-06,0.0005484,5.982e-05,1.601e-06,1.391e-07,0.0003099,0.001607,0.000161,0.0006678,2.737e-05,7.085e-07,1.387e-07,0.0001177,1.34e-06,0.0007604,2.873e-05,8.45e-06,5.063e-07,1.133e-05,6.862e-05,0.001171,0.001525,0.002295,0.0001227,3.881e-07,0.0004988,1.511e-07,0.002039,0.0004562,0.00285,0.0002473,7.31e-06,0.0001934,7.64e-06,6.191e-05,0.0001154,0.0001466,0.001505,4.772e-07,0.0003222,0.0001474,1.551e-07,9.135e-05,0.0008895,8.031e-10,1.942e-05,0.0002313,0.001994,2.026e-05,0.001406,0.001426,0.0001911,2.48e-05,0.0001056,0.000187,1.68e-06,0.0001598,4.688e-07,0.000177,1.258e-05,2.468e-06,2.011e-05,0.0003134,4.598e-05,0.001562,6.549e-06,0.002448,0.0002201,5.221e-05,0.0007124,0.000109,0.0009051,0.001112,0.002728,0.0006447,1.155e-05,0.0001304,0.0001658,1.46e-06,0.0001844,3.377e-07,0.0001252,1.444e-05,0.0004068,5.77e-07,7.248e-07,0.0001787,0.0007448,8.117e-07,1.036e-07,3.784e-05,0.000465,0.000344,0.001209,1.856e-05,0.0003281,0.000106,0.000338,1.164e-05,2.576e-07,0.002127,3.611e-07,0.0001881,1.929e-05,5.343e-06,0.003041,0.002834,4.488e-09,5.896e-07,4.442e-07,0.0002485,0.0004972,6.488e-06,0.0005875,8.865e-07,0.000361,1.2e-10,1.921e-08,0.0002714,8.769e-06,0.0003017,0.0002915,2.232e-05,0.0001977,0.0001483,1.39e-05,0.0001237,5.559e-08,0.0003016,0.0004835,0.001198,0.0002723,6.433e-06,1.39e-05,0.0007137,0.000388,6.252e-05,4.335e-06,0.0001196,0.00119,0.0001047,7.197e-06,0.0001558,6.953e-05,0.0004865,0.001858,0.001808,3.098e-08,0.0007056,3.033e-08,0.0005426,2.636e-05,0.001072,2.108e-05,0.0002595,3.731e-05,0.0009967,0.0002212,2.079e-05,3.928e-06,2.52e-05,5.049e-05,0.000126,6.697e-06,1.162e-06,8.164e-06,2.345e-05,3.659e-06,0.0002123,4.412e-07,1.22e-05,1.181e-05,0.0004708,1.521e-05,0.0008341,0.000118,7.609e-05,0.0005381,4.903e-05,0.003605,1.045e-05,0.0003402,1.499e-05,0.0005125,3.144e-06,6.413e-08,0.001488,0.002162,2.148e-07,7.53e-06,0.001516,0.002183,1.142e-11,0.00181,0.0007843,3.513e-05,0.002805,1.86e-09,0.001507,8.924e-05,1.429e-05,1.323e-07,0.000114,0.0001563,0.0006864,0.003673,3.561e-07,4.127e-06,7.438e-06,0.00088,0.001047,0.001633,0.0009251,0.0007415,0.0004943,6.858e-06,0.000508,0.000694,0.0003573,7.885e-08,0.0002726,0.0005891,0.0001825,6.87e-05,1.787e-05,2.295e-06,9.837e-07,0.0003222,4.468e-10,0.0004508,3.096e-06,2.217e-07,7.425e-05,0.0008908,0.000148,0.0005531,0.002264,6.619e-07,1.672e-07,0.0008303,8.328e-07,2.007e-05,0.0003373,0.0001053,0.0001172,0.001912,7.335e-06,8.771e-05,1.714e-05,0.001312,2.029e-07,0.001076,3.336e-06,3.223e-07,0.003008,0.001889,0.000797,0.0001455,0.0002718,0.000152,1.99e-06,1.564e-05,0.0001522,0.0004113,0.002938,1.363e-07,0.0004393,1.18e-05,0.0004042,0.0001592,5.334e-09,3.09e-07,1.618e-05,2.671e-06,0.001979,2.723e-05,2.981e-06,2.115e-05,0.0001931,0.0006072,1.211e-05,3.918e-07,3.435e-05,0.0001016,0.0003079,0.001717,0.0001582,0.0003571,3.408e-07,3.374e-06,1.337e-05,0.001355,0.0005874,0.0001625,1.908e-05,1.054e-07,1.603e-06,0.001229,0.00235,0.0006523,2.076e-05,0.000406,0.001568,0.0004769,2.858e-05,6.984e-06,0.00129,1.772e-05,7.941e-06,8.246e-08,4.286e-10,0.0001834,0.001371,0.0003745,1.482e-06,7.954e-07,1.294e-06,6.665e-12,0.0002878,6.443e-06,5.925e-08,0.0001961,1.217e-06,0.002407,0.001663,0.003703,3.966e-06,9.702e-05,1.355e-05,0.0003366,0.001957,0.0005762,4.667e-06,0.003301,7.323e-05,1.723e-05,1.62e-10,2.249e-08,0.001432,0.0006917,0.0003473,0.0004534,5.504e-05,3.426e-05,0.000121,0.002239,3.129e-08,0.0001007,0.0001922,0.0007912,9.106e-06,0.002524,1.44e-05,1.983e-05,0.0012,8.334e-07,0.0001348,6.49e-05,7.757e-07,2.301e-05,0.0008547,4.925e-05,0.003328,0.002538,5.369e-06,6.82e-07,0.001484,9.699e-05,0.002543,8.052e-06,2.693e-07,0.000209,0.0004289,0.001814,1.465e-05,9.925e-05,1.852e-09,2.658e-05,0.003215,6.948e-05,0.0002278,2.173e-07,5.921e-05,1.964e-08,0.002576,1.462e-05,2.834e-06,8.403e-05,0.001853,4.426e-08,2.2e-05,0.001802,0.001275,5.49e-05,4.378e-05,7.793e-05,9.23e-06,0.0001221,0.001566,2.743e-05,1.461e-07,4.359e-05,3.272e-05,0.0001604,0.0003819,0.0002669,0.0009403,2.516e-07,6.886e-07,4.046e-05,3.044e-05,3.625e-05,2.895e-06,0.0001621,1.068e-06,2.076e-06,8.392e-06,4.863e-05,3.786e-05,0.000969,6.854e-05,0.0004804,2.151e-05,0.0004896,0.001983,0.0007286,3.242e-06,0.000133,0.0001499,0.001259,1.655e-05,7.406e-06,1.357e-05,1.043e-06,0.001485,0.002137,0.0001685,0.003061,0.0003475,0.0001933,0.002228,0.0006065,0.001455,3.926e-07,1.027e-06,0.001256,0.001719,0.0005663,0.001807,4.683e-07,9.965e-05,9.068e-06,0.002327,5.053e-05,2.356e-05,3.027e-05,0.001383,8.566e-06,0.002673,0.001016,5.912e-05,1.304e-07,0.000612,3.621e-06,0.002382,0.0002689,0.0001825,1.992e-05,0.0002766,0.003389,0.0002556,0.001483,8.591e-07,6.449e-05,0.0004206,9.738e-05,0.0004939,0.000829,8.383e-08,2.929e-07,0.003114,3.347e-05,0.0001574,0.002485,0.002047,0.00358,0.0005946,0.0005094,0.001395,0.0003937,0.001969,0.001608,0.003723,0.000203,9.779e-06,0.003515,1.357e-06,1.158e-05,0.001144,3.01e-07,1.014e-05,0.0001087,0.000544,9.919e-05,0.0001063,5.198e-06,2.537e-08,2.955e-08,1.2e-06,2.584e-06,0.002189,6.778e-06,3.682e-07,0.0002343,2.929e-06,1.334e-07,1.589e-06,1.824e-07,0.0004453,0.0002776,0.0008968,6.647e-05,2.57e-07,1.945e-05,0.0009948,0.0003836,1.251e-05,2.436e-07,0.0006639,0.003284,0.0007263,0.0006062,2.118e-08,4.238e-06,0.001408,0.0001157,0.0002015,0.001854,2.502e-09,9.252e-07,1.1e-07,1.003e-07,4.602e-06,1.615e-08,0.001634,2.873e-07,0.000849,5.379e-05,0.0001162,0.001256,0.001679,0.0008559,0.002176,4.024e-06,0.003299,0.003121,2.824e-06,0.003653,0.001323,0.001137,4.967e-07,3.464e-05,0.0009861,0.0002057,0.0008311,6.28e-06,1.538e-07,1.319e-08,0.001744,0.0001352,1.528e-05,0.001639,6.538e-09,0.003467,3.272e-06,0.0004327,0.00115,8.287e-07,0.002754,6.023e-07,0.002859,0.0003661,1.234e-05,0.001676,0.001122,1.269e-08,0.002243,3.826e-06,0.0001424,4.678e-05,1.5e-05,2.148e-05,0.000486,0.0004184,3.027e-05,0.0003469,0.0002847,3.994e-05,2.7e-05,2.945e-05,0.000903,8.223e-05,3.311e-07,5.317e-08,3.541e-06,0.002818,0.001318,0.0004514,7.424e-07,0.0003398,3.086e-05,0.0007844,4.062e-05,4.37e-05,5.268e-07,7.589e-08,1.536e-05,0.0002225,0.0001177,1.715e-09,3.374e-05,8.828e-07,0.0002541,0.0001188,6.408e-07,6.773e-09,5.962e-07,0.0004386,3.705e-07,0.0002549,0.00263,3.593e-05,0.001572,0.0006399,0.002635,5.456e-07,3.305e-05,0.0007117,0.001911,4.801e-06,7.741e-05,3.984e-06,0.002021,1.994e-07,4.925e-07,0.0001345,0.002991,0.0005189,5.544e-07,3.359e-05,0.002111,0.001151,0.0001086,0.0004767,3.001e-06,0.0009683,1.032e-05,0.0002256,0.0001259,0.0001531,6.265e-06,0.000575,1.352e-07,1.732e-06,0.003657,0.002542,0.000602,0.0005463,9.542e-05,0.001753,0.002466,0.001019,0.00191,0.0007945,8.555e-05,0.001462,3.234e-06,0.001026,0.0004845,0.001266,0.0001175,1.588e-06,4.542e-05,4.634e-06,1.417e-05,0.002923,5.713e-05,0.0001498,1.235e-06,0.0003547,1.513e-06,0.001497,9.375e-06,0.0006786,7.658e-06,0.0003492,9.039e-05,3.534e-05,0.0001238,0.0009926,0.0005245,9.891e-08,0.0001342,1.444e-06,0.002576,9.682e-05,9.173e-05,7.536e-05,1.19e-05,0.001245,0.001168,0.002405,0.001406,7.444e-05,0.001144,0.0001449,6.968e-08,0.002228,0.0003315,6.035e-05,1.765e-06,6.825e-06,2.991e-06,0.002364,9.505e-05,0.000219,5.323e-06,0.0006534,0.001114,0.0001732,0.0009828,0.0001092,0.0006775,4.552e-06,0.002726,0.0004512,1.545e-09,0.0001712,9.675e-07,7.329e-06,1.893e-07,5.048e-05,4.057e-05,4.424e-05,0.001118,6.201e-05,0.001011,0.0001077,7.979e-05,0.001081,0.001474,0.002236,0.0008419,1.087e-06,1.242e-05,0.0001871,0.0003284,0.0005616,0.003314,1.323e-05,3.641e-05,0.003589,0.002868,0.002487,0.00194,1.927e-06,0.0003038,0.0001162,0.001283,0.0008503,2.118e-05,0.000441,0.0005263,8.816e-05,0.0005208,0.002858,8.549e-09,0.002589,2.859e-05,7.893e-06,0.003116,0.001545,0.0001024,0.002414,0.0003841,0.001202,0.003449,0.0001409,0.002607,5.12e-05,0.0004161,0.0002913,3.542e-05,1.678e-06,2.935e-07,0.0001248,0.0002617,0.001789,0.0002317,0.0007618,0.0005404,0.003604,4.835e-06,7.084e-05,0.0004332,0.000222,0.001519,0.0003669,0.001117,0.002088,5.281e-06,0.0001433,0.001085,1.323e-05,0.0002178,2.479e-05,6.407e-09,0.002807,0.001626,2.523e-07,0.0008873,6.885e-05,0.0009799,0.0006661,0.0001861,0.002268,6.045e-06,0.0003497,1.848e-05,0.0001413,0.002687,0.001616,0.0003898,3.685e-05,5.363e-05,7.573e-05,0.0004309,1.259e-07,0.000274,2.519e-05,0.003505,1.418e-05,1.995e-06,0.001066,0.0004395,7.454e-05,2.701e-05,1.38e-05,0.0001989,6.681e-05,1.731e-10,0.0005496,0.001266,0.0005538,7.191e-09,0.0007374,3.512e-06,0.001144,0.0003979,7.418e-05,0.0003236,0.002454,1.01e-05,0.002332,0.001477,3.72e-08,0.002346,0.001475,4.999e-06,0.002429,1.891e-05,0.00171,0.00364,9.719e-06,7.024e-06,7.277e-07,4.984e-05,0.0001901,3.78e-05,0.0001072,6.647e-05,1.059e-05,0.0007213,0.0001424,0.0003134,2.898e-06,4.024e-07,2.208e-08,0.0002957,0.0004981,0.0001552,0.0002379,0.0004735,6.021e-07,0.0003783,0.0007802,0.0007896,9.019e-06,0.0003351,4.35e-08,0.002556,2.288e-06,3.079e-05,0.00177,0.0001505,4.028e-07,5.848e-06,2.213e-05,4.038e-06,0.0006773,1.922e-06,0.001114,6.111e-09,4.092e-07,0.002094,4.878e-09,0.0006957,0.0001099,0.00272,0.0004007,0.003169,1.753e-06,0.000149,5.222e-06,0.0006911,0.0008149,6.823e-06,0.0001446,1.514e-05,1.547e-05,7.129e-07,6.182e-06,0.001853,1.387e-06,1.86e-05,0.000712,0.0004351,0.0007705,4.394e-05,1.319e-05,0.0003924,0.0004327,0.002941,1.439e-06,0.002156,1.316e-06,2.27e-05,0.000452,0.0003611,0.001182,5.51e-05,0.00265,4.995e-06,0.0002282,6.214e-06,0.0002352,3.336e-07,1.883e-07,0.001435,3.706e-07,0.0001448,0.002732,0.0008667,1.322e-05,2.041e-05,9.962e-07,0.0001555,3.245e-05,0.0003408,0.0005177,0.0002956,2.821e-08,0.001229,3.693e-05,0.003016,0.0002318,3.085e-05,1.011e-05,2.658e-07,3.778e-06,0.000494,1.715e-05,0.002705,0.0001631,0.0003451,1.769e-07,2.978e-08,8.642e-05,0.0003709,3.718e-06,2.908e-05,2.278e-05,0.001369,7.55e-08,4.745e-06,9.071e-08,0.002296,0.003254,0.0009165,2.612e-05,0.002218,0.0001533,5.08e-05,0.003718,0.001136,0.002257,0.001969,0.0004467,0.001079,6.072e-05,6.893e-07,0.0003218,8.995e-06,3.53e-05,4.906e-07,1.628e-05,0.0008896,0.0003134,0.0007272,1.366e-05,4.281e-05,3.254e-06,0.0001214,0.0001254,4.197e-07,2.963e-08,0.0005077,0.001107,0.0009512,9.95e-08,3.019e-08,0.003383,0.0002409,7.391e-06,1.662e-06,0.000133,0.001846,8.682e-06,7.085e-05,5.395e-08,6.087e-05,0.000276,9.625e-07,1.417e-05,0.0004889,1.735e-05,3.569e-06,8.11e-05,1.271e-05,1.255e-05,0.0008335,3.445e-05,1.353e-06,8.639e-05,0.001585,0.0001171,0.0009131,3.876e-05,0.0002433,0.002458,0.0006833,2.159e-06,0.002103,0.000214,2.481e-05,0.0006649,0.0001742,5.238e-07,7.034e-07,0.0001753,1.258e-05,3.113e-06,0.00353,9.983e-07,0.001251,0.0003306,1.652e-07,4.444e-05,0.0006446,2.555e-06,2.184e-05,1.024e-06,4.528e-07,2.668e-05,6.64e-10,6.163e-05,0.0003508,6.501e-06,3.298e-07,0.0004336,0.002845,0.0005081,2.405e-05,4.151e-08,0.0002749,2.515e-08,7.228e-06,4.498e-07,8.04e-05,1.948e-06,1.206e-05,0.0008185,7.198e-05,4.801e-06,0.002329,0.0003738,0.0001083,0.0009496,3.124e-10,2.201e-08,5.199e-06,1.177e-05,0.001595,0.001138,0.001356,0.0001027,0.003249,9.422e-05,0.00117,5.107e-05,7.359e-06,1.148e-06,8.77e-07,2.858e-06,4.948e-05,0.001455,2.58e-08,8.009e-07,0.000795,0.0003198,6.396e-08,8.628e-06,6.237e-05,5.156e-06,0.0004036,9.515e-06,1.27e-05,6.312e-07,8.027e-05,1.264e-05,0.0002467,0.003443,0.0006627,3.097e-05,0.0007381,0.0003829,3.574e-08,0.003732,0.003628,4.428e-05,6.336e-06,0.003294,0.002003,3.956e-07,4.651e-10,8.375e-06,0.003052,0.0006371,0.0004113,3.192e-05,1.554e-08,1.948e-07,0.0002139,3.107e-05,2.112e-06,2.449e-05,1.51e-05,1.988e-05,0.0008021,3.453e-07,0.001256,0.002767,0.002049,2.135e-06,0.000142,0.002068,0.0001219,3.934e-12,0.0004382,6.008e-05,4.736e-05,0.001288,1.66e-07,0.001013,3.274e-05,0.003582,0.002492,2.083e-05,3.114e-05,1.413e-05,2.428e-05,0.0001112,2.634e-05,1.188e-05,0.0002232,0.000556,9.235e-06,7.475e-08,6.922e-05,2.672e-06,0.0001302,6.797e-07,1.198e-05,1.667e-05,0.001938,9.225e-05,0.002028,2.171e-07,1.419e-06,0.001993,0.0004335,0.001085,6.13e-06,0.0001468,4.13e-08,8.645e-05,0.0007938,0.0001096,1.718e-06,0.002717,0.001627,6.033e-06,3.254e-08,1.148e-05,0.0009978,8.071e-05,1.656e-06,0.002243,1.528e-05,1.451e-07,0.0003985,0.003274,0.00068,2.939e-05,0.0002474,3.685e-05,0.0003645,2.952e-06,6.242e-05,0.0002398,0.001534,5.407e-05,8.732e-05,9.712e-08,0.0003063,0.0009184,1.019e-05,0.0005257,0.0002751,0.0004212,0.0003597,7.285e-06,2.197e-06,0.001302,2.086e-06,0.0001555,1.396e-08,1.96e-05,0.0005137,0.0004902,8.218e-07,1.68e-06,0.0002675,0.0003893,0.0001482,1.316e-07,1.207e-05,0.0001274,0.001414,2.544e-07,4.188e-05,0.0002159,0.0005994,0.002045,3.895e-05,2.023e-05,0.003359,1.268e-06,0.003596,6.721e-05,1.249e-05,0.0001313,0.003296,1.143e-05,0.0002931,5.024e-06,0.001991,1.436e-06,0.003617,0.0009643,0.0002668,6.355e-05,0.001146,0.0004287,0.0006914,9.259e-07,9.073e-05,2.983e-06,0.0002608,0.0008462,0.003076,1.344e-06,0.0003167,3.106e-05,0.002307,2.302e-07,2.541e-05,0.0009129,2.031e-06,0.00126,0.001803,0.0001458,0.002306,0.002708,4.235e-08,1.038e-06,4.255e-05,0.00212,0.0005201,3.569e-07,9.27e-05,0.0009992,2.429e-08,5.575e-08,8.933e-05,0.001899,4.682e-07,2.676e-07,0.0001004,0.001148,0.002539,0.0026,0.00248,0.0001239,1.053e-05,1.532e-05,8.249e-06,8.026e-05,0.0002773,0.001091,0.003045,0.001503,0.002404,3.737e-05,3.474e-05,0.002264,5.323e-05,9.456e-05,0.001885,0.000257,0.0004794,0.001754,5.45e-08,0.000558,6.613e-05,2.169e-05,0.0001357,0.0002117,2.704e-06,0.001136,0.001454,0.0002132,7.027e-06,5.067e-05,8.082e-05,2.642e-06,0.0006653,0.00246,6.101e-05,8.293e-06,1.833e-06,1.913e-06,8.353e-05,1.875e-06,9.413e-06,3.197e-07,1.291e-06,7.726e-12,0.002272,6.293e-05,0.002695,0.002714,1.363e-06,4.49e-08,0.000111,0.0007172,0.0007672,0.0002418,0.0002574,0.0001045,3.565e-07,2.966e-09,0.003397,6.95e-06,8.803e-06,6.383e-05,8.965e-07,6.92e-05,5.297e-06,7.31e-08,2.289e-07,4.882e-07,2.276e-09,7.45e-06,0.0001206,0.002075,0.001354,4.441e-07,0.001107,9.505e-05,9.449e-07,6.731e-05,2.22e-08,4.736e-06,0.0001086,3.804e-06,0.003586,3.023e-06,0.0001789,0.001963,0.00084,0.001457,0.0001503,1.85e-06,4.75e-10,7.078e-07,5.456e-06,0.0004539,3.321e-06,3.908e-06,0.00251,1.633e-05,9.004e-11,0.0005542,9.003e-05,5.08e-06,0.000166,7.686e-06,5.781e-05,0.0003888,6.519e-07,3.824e-08,6.796e-07,0.0004434,2.162e-08,7.569e-05,3.233e-05,0.0005927,0.001305,3.681e-06,2.598e-05,0.0001042,0.0007451,7.851e-05,6.328e-08,0.0006819,0.0002214,0.0006168,0.002152,7.577e-06,3.665e-05,0.0002333,5.595e-08,0.000142,1.91e-05,0.001216,0.002298,3.162e-07,6.886e-07,0.000145,0.0009365,1.182e-06,0.0003273,0.003276,0.000746,3.4e-06,0.0002849,4.281e-05,2.421e-05,0.0009461,1.52e-05,0.0007492,9.116e-08,0.0001964,0.0009353,0.0007787,0.0003087,1.778e-05,6.344e-06,0.0007574,4.892e-06,0.0001311,0.001277,0.0002621,2.384e-06,1.205e-05,4.666e-06,8.403e-08,0.0002576,1.903e-07,0.0001614,3.041e-05,1.426e-07,2.796e-06,9.116e-05,1.974e-05,3.584e-08,0.000107,1.793e-05,0.001992,9.104e-06,1.082e-09,6.339e-05,5.748e-05,0.003704,3.922e-06,4.919e-06,0.003035,8.204e-05,0.00152,9.963e-07,0.0001045,6.453e-06,0.001545,0.0001921,0.001771,2.818e-07,6.267e-07,0.0003607,9.819e-11,0.0003795,2.54e-05,0.003482,0.003299,0.002072,1.616e-05,0.0002243,0.000749,1.602e-05,1.56e-07,5.104e-07,5.072e-08,0.0005615,3.465e-07,0.0002063,4.624e-05,0.001072,2.625e-05,0.0002513,0.001561,0.0004161,2.363e-05,2.733e-06,3.359e-08,0.0001428,0.0005205,0.0005917,0.0007769,0.0002438,0.0005101,2.697e-06,5.545e-05,0.0004503,0.0001973,0.0004575,0.0001293,0.0001213,8.011e-07,0.001528,1.203e-07,6.774e-05,0.001056,0.002535,0.0002893,0.000856,0.0002647,0.002335,0.0006194,0.001876,0.0003161,0.001811,9.146e-07,1.918e-05,0.0005819,0.001762,5.96e-07,0.002462,6.485e-05,5.613e-06,4.948e-05,0.0001027,0.0004954,0.0004322,0.002271,5.569e-07,0.001003,2.037e-06,0.0002076,2.442e-06,0.0009952,0.0002808,1.472e-07,0.001922,0.003341,0.001275,0.00146,0.0009338,0.001214,1.8e-06,0.0004703,0.0008315,0.0003262,0.000267,6.186e-07,0.0001072,1.909e-05,0.003421,8.71e-05,0.0009499,0.0005967,1.117e-06,6.231e-07,2.267e-07,3.626e-07,0.0029,0.0001466,2.45e-05,0.001315,0.0001916,3.746e-06,1.678e-05,2.659e-07,4.727e-08,0.0006719,0.001859,0.0007241,1.049e-09,0.002675,0.001334,2.624e-06,1.012e-05,0.0007098,0.001286,0.0003372,0.0007119,2.14e-07,3.09e-07,0.0001509,2.88e-05,0.0005136,5.015e-05,0.0002549,6.144e-06,0.0001688,0.0001542,1.218e-08,3.465e-05,0.002687,0.0002508,0.0002698,5.049e-08,0.003159,4.942e-05,2.436e-05,0.0005182,1.472e-05,0.001942,0.0007113,4.791e-06,0.00336,0.001468,0.0005995,2.492e-07,7.47e-08,0.0002114,0.002116,0.0009623,7.443e-06,7.536e-05,0.000577,1.65e-07,0.0001229,0.0007583,2.459e-05,2.485e-05,4.995e-13,0.0001038,2.069e-08,0.002949,4.119e-05,0.00012,0.0002492,0.0001931,0.0007998,5.288e-07,0.0001603,0.001839,0.0001593,1.036e-07,9.958e-05,0.0001172,0.00354,0.0003643,2.45e-07,6.703e-07],"Contig":["Transfer.PvP01_00_1.final","Transfer.PvP01_00_1.final","Transfer.PvP01_00_1.final","Transfer.PvP01_00_1.final","Transfer.PvP01_00_1.final","Transfer.PvP01_00_1.final","Transfer.PvP01_00_1.final","Transfer.PvP01_00_1.final","Transfer.PvP01_00_1.final","Transfer.PvP01_00_1.final","Transfer.PvP01_00_1.final","Transfer.PvP01_00_1.final","Transfer.PvP01_00_1.final","Transfer.PvP01_00_1.final","Transfer.PvP01_00_1.final","Transfer.PvP01_00_1.final","Transfer.PvP01_00_1.final","Transfer.PvP01_00_2.final","Transfer.PvP01_00_2.final","Transfer.PvP01_00_2.final","Transfer.PvP01_00_2.final","Transfer.PvP01_00_2.final","Transfer.PvP01_00_3.final","Transfer.PvP01_00_3.final","Transfer.PvP01_00_3.final","Transfer.PvP01_00_3.final","Transfer.PvP01_00_3.final","Transfer.PvP01_00_3.final","Transfer.PvP01_00_3.final","Transfer.PvP01_00_3.final","Transfer.PvP01_00_4.final","Transfer.PvP01_00_4.final","Transfer.PvP01_00_4.final","Transfer.PvP01_00_4.final","Transfer.PvP01_00_4.final","Transfer.PvP01_00_4.final","Transfer.PvP01_00_4.final","Transfer.PvP01_00_5.final","Transfer.PvP01_00_5.final","Transfer.PvP01_00_5.final","Transfer.PvP01_00_6.final","Transfer.PvP01_00_6.final","Transfer.PvP01_00_6.final","Transfer.PvP01_00_6.final","Transfer.PvP01_00_7.final","Transfer.PvP01_00_7.final","Transfer.PvP01_00_7.final","Transfer.PvP01_00_8.final","Transfer.PvP01_00_8.final","Transfer.PvP01_00_8.final","Transfer.PvP01_00_8.final","Transfer.PvP01_00_8.final","Transfer.PvP01_00_8.final","Transfer.PvP01_00_8.final","Transfer.PvP01_00_8.final","Transfer.PvP01_00_8.final","Transfer.PvP01_00_9.final","Transfer.PvP01_00_9.final","Transfer.PvP01_00_9.final","Transfer.PvP01_00_10.final","Transfer.PvP01_00_10.final","Transfer.PvP01_00_10.final","Transfer.PvP01_00_10.final","Transfer.PvP01_00_10.final","Transfer.PvP01_00_11.final","Transfer.PvP01_00_11.final","Transfer.PvP01_00_11.final","Transfer.PvP01_00_11.final","Transfer.PvP01_00_11.final","Transfer.PvP01_00_11.final","Transfer.PvP01_00_11.final","Transfer.PvP01_00_11.final","Transfer.PvP01_00_11.final","Transfer.PvP01_00_11.final","Transfer.PvP01_00_11.final","Transfer.PvP01_00_11.final","Transfer.PvP01_00_12.final","Transfer.PvP01_00_12.final","Transfer.PvP01_00_13.final","Transfer.PvP01_00_13.final","Transfer.PvP01_00_14.final","Transfer.PvP01_00_14.final","Transfer.PvP01_00_14.final","Transfer.PvP01_00_14.final","Transfer.PvP01_00_14.final","Transfer.PvP01_00_15.final","Transfer.PvP01_00_15.final","Transfer.PvP01_00_15.final","Transfer.PvP01_00_15.final","Transfer.PvP01_00_15.final","Transfer.PvP01_00_16.final","Transfer.PvP01_00_16.final","Transfer.PvP01_00_17.final","Transfer.PvP01_00_17.final","Transfer.PvP01_00_17.final","Transfer.PvP01_00_17.final","Transfer.PvP01_00_17.final","Transfer.PvP01_00_17.final","Transfer.PvP01_00_18.final","Transfer.PvP01_00_18.final","Transfer.PvP01_00_18.final","Transfer.PvP01_00_19.final","Transfer.PvP01_00_22.final","Transfer.PvP01_00_22.final","Transfer.PvP01_00_22.final","Transfer.PvP01_00_22.final","Transfer.PvP01_00_22.final","Transfer.PvP01_00_23.final","Transfer.PvP01_00_23.final","Transfer.PvP01_00_24.final","Transfer.PvP01_00_25.final","Transfer.PvP01_00_26.final","Transfer.PvP01_00_26.final","Transfer.PvP01_00_26.final","Transfer.PvP01_00_26.final","Transfer.PvP01_00_29.final","Transfer.PvP01_00_29.final","Transfer.PvP01_00_29.final","Transfer.PvP01_00_30.final","Transfer.PvP01_00_30.final","Transfer.PvP01_00_31.final","Transfer.PvP01_00_31.final","Transfer.PvP01_00_31.final","Transfer.PvP01_00_32.final","Transfer.PvP01_00_34.final","Transfer.PvP01_00_37.final","Transfer.PvP01_00_38.final","Transfer.PvP01_00_38.final","Transfer.PvP01_00_39.final","Transfer.PvP01_00_41.final","Transfer.PvP01_00_42.final","Transfer.PvP01_00_42.final","Transfer.PvP01_00_45.final","Transfer.PvP01_00_45.final","Transfer.PvP01_00_48.final","Transfer.PvP01_00_48.final","Transfer.PvP01_00_48.final","Transfer.PvP01_00_51.final","Transfer.PvP01_00_52.final","Transfer.PvP01_00_53.final","Transfer.PvP01_00_55.final","Transfer.PvP01_00_57.final","Transfer.PvP01_00_61.final","Transfer.PvP01_00_61.final","Transfer.PvP01_00_63.final","Transfer.PvP01_00_64.final","Transfer.PvP01_00_64.final","Transfer.PvP01_00_66.final","Transfer.PvP01_00_67.final","Transfer.PvP01_00_67.final","Transfer.PvP01_00_67.final","Transfer.PvP01_00_69.final","Transfer.PvP01_00_71.final","Transfer.PvP01_00_74.final","Transfer.PvP01_00_77.final","Transfer.PvP01_00_77.final","Transfer.PvP01_00_79.final","Transfer.PvP01_00_81.final","Transfer.PvP01_00_82.final","Transfer.PvP01_00_83.final","Transfer.PvP01_00_86.final","Transfer.PvP01_00_88.final","Transfer.PvP01_00_89.final","Transfer.PvP01_00_89.final","Transfer.PvP01_00_90.final","Transfer.PvP01_00_90.final","Transfer.PvP01_00_104.final","Transfer.PvP01_00_108.final","Transfer.PvP01_00_119.final","Transfer.PvP01_00_124.final","Transfer.PvP01_00_129.final","Transfer.PvP01_00_138.final","Transfer.PvP01_00_143.final","Transfer.PvP01_00_150.final","Transfer.PvP01_00_152.final","Transfer.PvP01_00_157.final","Transfer.PvP01_00_158.final","Transfer.PvP01_00_180.final","Transfer.PvP01_00_203.final","Transfer.PvP01_00_208.final","Transfer.PvP01_00_209.final","Transfer.PvP01_00_217.final","Transfer.PvP01_00_221.final","Transfer.PvP01_00_223.final","Transfer.PvP01_00_233.final","Transfer.PvP01_00_300.final","PvP01_01_v1","PvP01_01_v1","PvP01_01_v1","PvP01_01_v1","PvP01_01_v1","PvP01_01_v1","PvP01_01_v1","PvP01_01_v1","PvP01_01_v1","PvP01_01_v1","PvP01_01_v1","PvP01_01_v1","PvP01_01_v1","PvP01_01_v1","PvP01_01_v1","PvP01_01_v1","PvP01_01_v1","PvP01_01_v1","PvP01_01_v1","PvP01_01_v1","PvP01_01_v1","PvP01_01_v1","PvP01_01_v1","PvP01_01_v1","PvP01_01_v1","PvP01_01_v1","PvP01_01_v1","PvP01_01_v1","PvP01_01_v1","PvP01_01_v1","PvP01_01_v1","PvP01_01_v1","PvP01_01_v1","PvP01_01_v1","PvP01_01_v1","PvP01_01_v1","PvP01_01_v1","PvP01_01_v1","PvP01_01_v1","PvP01_01_v1","PvP01_01_v1","PvP01_01_v1","PvP01_01_v1","PvP01_01_v1","PvP01_01_v1","PvP01_01_v1","PvP01_01_v1","PvP01_01_v1","PvP01_01_v1","PvP01_01_v1","PvP01_01_v1","PvP01_01_v1","PvP01_01_v1","PvP01_01_v1","PvP01_01_v1","PvP01_01_v1","PvP01_01_v1","PvP01_01_v1","PvP01_01_v1","PvP01_01_v1","PvP01_01_v1","PvP01_01_v1","PvP01_01_v1","PvP01_01_v1","PvP01_01_v1","PvP01_01_v1","PvP01_01_v1","PvP01_01_v1","PvP01_01_v1","PvP01_01_v1","PvP01_01_v1","PvP01_01_v1","PvP01_01_v1","PvP01_01_v1","PvP01_01_v1","PvP01_01_v1","PvP01_01_v1","PvP01_01_v1","PvP01_01_v1","PvP01_01_v1","PvP01_01_v1","PvP01_01_v1","PvP01_01_v1","PvP01_01_v1","PvP01_01_v1","PvP01_01_v1","PvP01_01_v1","PvP01_01_v1","PvP01_01_v1","PvP01_01_v1","PvP01_01_v1","PvP01_01_v1","PvP01_01_v1","PvP01_01_v1","PvP01_01_v1","PvP01_01_v1","PvP01_01_v1","PvP01_01_v1","PvP01_01_v1","PvP01_01_v1","PvP01_01_v1","PvP01_01_v1","PvP01_01_v1","PvP01_01_v1","PvP01_01_v1","PvP01_01_v1","PvP01_01_v1","PvP01_01_v1","PvP01_01_v1","PvP01_01_v1","PvP01_01_v1","PvP01_01_v1","PvP01_01_v1","PvP01_01_v1","PvP01_01_v1","PvP01_01_v1","PvP01_01_v1","PvP01_01_v1","PvP01_01_v1","PvP01_01_v1","PvP01_02_v1","PvP01_02_v1","PvP01_02_v1","PvP01_02_v1","PvP01_02_v1","PvP01_02_v1","PvP01_02_v1","PvP01_02_v1","PvP01_02_v1","PvP01_02_v1","PvP01_02_v1","PvP01_02_v1","PvP01_02_v1","PvP01_02_v1","PvP01_02_v1","PvP01_02_v1","PvP01_02_v1","PvP01_02_v1","PvP01_02_v1","PvP01_02_v1","PvP01_02_v1","PvP01_02_v1","PvP01_02_v1","PvP01_02_v1","PvP01_02_v1","PvP01_02_v1","PvP01_02_v1","PvP01_02_v1","PvP01_02_v1","PvP01_02_v1","PvP01_02_v1","PvP01_02_v1","PvP01_02_v1","PvP01_02_v1","PvP01_02_v1","PvP01_02_v1","PvP01_02_v1","PvP01_02_v1","PvP01_02_v1","PvP01_02_v1","PvP01_02_v1","PvP01_02_v1","PvP01_02_v1","PvP01_02_v1","PvP01_02_v1","PvP01_02_v1","PvP01_02_v1","PvP01_02_v1","PvP01_02_v1","PvP01_02_v1","PvP01_02_v1","PvP01_02_v1","PvP01_02_v1","PvP01_02_v1","PvP01_02_v1","PvP01_02_v1","PvP01_02_v1","PvP01_02_v1","PvP01_02_v1","PvP01_02_v1","PvP01_02_v1","PvP01_02_v1","PvP01_02_v1","PvP01_02_v1","PvP01_02_v1","PvP01_02_v1","PvP01_02_v1","PvP01_02_v1","PvP01_02_v1","PvP01_02_v1","PvP01_02_v1","PvP01_02_v1","PvP01_02_v1","PvP01_02_v1","PvP01_02_v1","PvP01_02_v1","PvP01_02_v1","PvP01_02_v1","PvP01_02_v1","PvP01_02_v1","PvP01_02_v1","PvP01_02_v1","PvP01_02_v1","PvP01_02_v1","PvP01_02_v1","PvP01_02_v1","PvP01_02_v1","PvP01_02_v1","PvP01_02_v1","PvP01_02_v1","PvP01_02_v1","PvP01_02_v1","PvP01_02_v1","PvP01_02_v1","PvP01_02_v1","PvP01_02_v1","PvP01_02_v1","PvP01_02_v1","PvP01_02_v1","PvP01_02_v1","PvP01_02_v1","PvP01_02_v1","PvP01_02_v1","PvP01_02_v1","PvP01_02_v1","PvP01_02_v1","PvP01_02_v1","PvP01_02_v1","PvP01_02_v1","PvP01_02_v1","PvP01_02_v1","PvP01_02_v1","PvP01_02_v1","PvP01_02_v1","PvP01_02_v1","PvP01_02_v1","PvP01_02_v1","PvP01_02_v1","PvP01_02_v1","PvP01_02_v1","PvP01_03_v1","PvP01_03_v1","PvP01_03_v1","PvP01_03_v1","PvP01_03_v1","PvP01_03_v1","PvP01_03_v1","PvP01_03_v1","PvP01_03_v1","PvP01_03_v1","PvP01_03_v1","PvP01_03_v1","PvP01_03_v1","PvP01_03_v1","PvP01_03_v1","PvP01_03_v1","PvP01_03_v1","PvP01_03_v1","PvP01_03_v1","PvP01_03_v1","PvP01_03_v1","PvP01_03_v1","PvP01_03_v1","PvP01_03_v1","PvP01_03_v1","PvP01_03_v1","PvP01_03_v1","PvP01_03_v1","PvP01_03_v1","PvP01_03_v1","PvP01_03_v1","PvP01_03_v1","PvP01_03_v1","PvP01_03_v1","PvP01_03_v1","PvP01_03_v1","PvP01_03_v1","PvP01_03_v1","PvP01_03_v1","PvP01_03_v1","PvP01_03_v1","PvP01_03_v1","PvP01_03_v1","PvP01_03_v1","PvP01_03_v1","PvP01_03_v1","PvP01_03_v1","PvP01_03_v1","PvP01_03_v1","PvP01_03_v1","PvP01_03_v1","PvP01_03_v1","PvP01_03_v1","PvP01_03_v1","PvP01_03_v1","PvP01_03_v1","PvP01_03_v1","PvP01_03_v1","PvP01_03_v1","PvP01_03_v1","PvP01_03_v1","PvP01_03_v1","PvP01_03_v1","PvP01_03_v1","PvP01_03_v1","PvP01_03_v1","PvP01_03_v1","PvP01_03_v1","PvP01_03_v1","PvP01_03_v1","PvP01_03_v1","PvP01_03_v1","PvP01_03_v1","PvP01_03_v1","PvP01_03_v1","PvP01_03_v1","PvP01_03_v1","PvP01_03_v1","PvP01_03_v1","PvP01_03_v1","PvP01_03_v1","PvP01_03_v1","PvP01_03_v1","PvP01_03_v1","PvP01_03_v1","PvP01_03_v1","PvP01_03_v1","PvP01_03_v1","PvP01_03_v1","PvP01_03_v1","PvP01_03_v1","PvP01_03_v1","PvP01_03_v1","PvP01_03_v1","PvP01_03_v1","PvP01_03_v1","PvP01_03_v1","PvP01_03_v1","PvP01_03_v1","PvP01_03_v1","PvP01_03_v1","PvP01_03_v1","PvP01_03_v1","PvP01_03_v1","PvP01_03_v1","PvP01_03_v1","PvP01_03_v1","PvP01_03_v1","PvP01_03_v1","PvP01_04_v1","PvP01_04_v1","PvP01_04_v1","PvP01_04_v1","PvP01_04_v1","PvP01_04_v1","PvP01_04_v1","PvP01_04_v1","PvP01_04_v1","PvP01_04_v1","PvP01_04_v1","PvP01_04_v1","PvP01_04_v1","PvP01_04_v1","PvP01_04_v1","PvP01_04_v1","PvP01_04_v1","PvP01_04_v1","PvP01_04_v1","PvP01_04_v1","PvP01_04_v1","PvP01_04_v1","PvP01_04_v1","PvP01_04_v1","PvP01_04_v1","PvP01_04_v1","PvP01_04_v1","PvP01_04_v1","PvP01_04_v1","PvP01_04_v1","PvP01_04_v1","PvP01_04_v1","PvP01_04_v1","PvP01_04_v1","PvP01_04_v1","PvP01_04_v1","PvP01_04_v1","PvP01_04_v1","PvP01_04_v1","PvP01_04_v1","PvP01_04_v1","PvP01_04_v1","PvP01_04_v1","PvP01_04_v1","PvP01_04_v1","PvP01_04_v1","PvP01_04_v1","PvP01_04_v1","PvP01_04_v1","PvP01_04_v1","PvP01_04_v1","PvP01_04_v1","PvP01_04_v1","PvP01_04_v1","PvP01_04_v1","PvP01_04_v1","PvP01_04_v1","PvP01_04_v1","PvP01_04_v1","PvP01_04_v1","PvP01_04_v1","PvP01_04_v1","PvP01_04_v1","PvP01_04_v1","PvP01_04_v1","PvP01_04_v1","PvP01_04_v1","PvP01_04_v1","PvP01_04_v1","PvP01_04_v1","PvP01_04_v1","PvP01_04_v1","PvP01_04_v1","PvP01_04_v1","PvP01_04_v1","PvP01_04_v1","PvP01_04_v1","PvP01_04_v1","PvP01_04_v1","PvP01_04_v1","PvP01_04_v1","PvP01_04_v1","PvP01_04_v1","PvP01_04_v1","PvP01_04_v1","PvP01_04_v1","PvP01_04_v1","PvP01_04_v1","PvP01_04_v1","PvP01_04_v1","PvP01_04_v1","PvP01_04_v1","PvP01_04_v1","PvP01_04_v1","PvP01_04_v1","PvP01_04_v1","PvP01_04_v1","PvP01_04_v1","PvP01_04_v1","PvP01_04_v1","PvP01_04_v1","PvP01_04_v1","PvP01_04_v1","PvP01_04_v1","PvP01_04_v1","PvP01_04_v1","PvP01_04_v1","PvP01_04_v1","PvP01_04_v1","PvP01_04_v1","PvP01_04_v1","PvP01_04_v1","PvP01_04_v1","PvP01_04_v1","PvP01_05_v1","PvP01_05_v1","PvP01_05_v1","PvP01_05_v1","PvP01_05_v1","PvP01_05_v1","PvP01_05_v1","PvP01_05_v1","PvP01_05_v1","PvP01_05_v1","PvP01_05_v1","PvP01_05_v1","PvP01_05_v1","PvP01_05_v1","PvP01_05_v1","PvP01_05_v1","PvP01_05_v1","PvP01_05_v1","PvP01_05_v1","PvP01_05_v1","PvP01_05_v1","PvP01_05_v1","PvP01_05_v1","PvP01_05_v1","PvP01_05_v1","PvP01_05_v1","PvP01_05_v1","PvP01_05_v1","PvP01_05_v1","PvP01_05_v1","PvP01_05_v1","PvP01_05_v1","PvP01_05_v1","PvP01_05_v1","PvP01_05_v1","PvP01_05_v1","PvP01_05_v1","PvP01_05_v1","PvP01_05_v1","PvP01_05_v1","PvP01_05_v1","PvP01_05_v1","PvP01_05_v1","PvP01_05_v1","PvP01_05_v1","PvP01_05_v1","PvP01_05_v1","PvP01_05_v1","PvP01_05_v1","PvP01_05_v1","PvP01_05_v1","PvP01_05_v1","PvP01_05_v1","PvP01_05_v1","PvP01_05_v1","PvP01_05_v1","PvP01_05_v1","PvP01_05_v1","PvP01_05_v1","PvP01_05_v1","PvP01_05_v1","PvP01_05_v1","PvP01_05_v1","PvP01_05_v1","PvP01_05_v1","PvP01_05_v1","PvP01_05_v1","PvP01_05_v1","PvP01_05_v1","PvP01_05_v1","PvP01_05_v1","PvP01_05_v1","PvP01_05_v1","PvP01_05_v1","PvP01_05_v1","PvP01_05_v1","PvP01_05_v1","PvP01_05_v1","PvP01_05_v1","PvP01_05_v1","PvP01_05_v1","PvP01_05_v1","PvP01_05_v1","PvP01_05_v1","PvP01_05_v1","PvP01_05_v1","PvP01_05_v1","PvP01_05_v1","PvP01_05_v1","PvP01_05_v1","PvP01_05_v1","PvP01_05_v1","PvP01_05_v1","PvP01_05_v1","PvP01_05_v1","PvP01_05_v1","PvP01_05_v1","PvP01_05_v1","PvP01_05_v1","PvP01_05_v1","PvP01_05_v1","PvP01_05_v1","PvP01_05_v1","PvP01_05_v1","PvP01_05_v1","PvP01_05_v1","PvP01_05_v1","PvP01_05_v1","PvP01_05_v1","PvP01_05_v1","PvP01_05_v1","PvP01_05_v1","PvP01_05_v1","PvP01_05_v1","PvP01_05_v1","PvP01_05_v1","PvP01_05_v1","PvP01_05_v1","PvP01_05_v1","PvP01_05_v1","PvP01_05_v1","PvP01_05_v1","PvP01_05_v1","PvP01_05_v1","PvP01_05_v1","PvP01_05_v1","PvP01_05_v1","PvP01_05_v1","PvP01_05_v1","PvP01_05_v1","PvP01_05_v1","PvP01_05_v1","PvP01_05_v1","PvP01_05_v1","PvP01_05_v1","PvP01_05_v1","PvP01_05_v1","PvP01_05_v1","PvP01_05_v1","PvP01_05_v1","PvP01_05_v1","PvP01_05_v1","PvP01_05_v1","PvP01_05_v1","PvP01_05_v1","PvP01_05_v1","PvP01_05_v1","PvP01_05_v1","PvP01_05_v1","PvP01_05_v1","PvP01_05_v1","PvP01_05_v1","PvP01_05_v1","PvP01_05_v1","PvP01_05_v1","PvP01_05_v1","PvP01_05_v1","PvP01_05_v1","PvP01_05_v1","PvP01_05_v1","PvP01_05_v1","PvP01_05_v1","PvP01_05_v1","PvP01_05_v1","PvP01_05_v1","PvP01_05_v1","PvP01_05_v1","PvP01_05_v1","PvP01_05_v1","PvP01_05_v1","PvP01_05_v1","PvP01_05_v1","PvP01_05_v1","PvP01_05_v1","PvP01_05_v1","PvP01_05_v1","PvP01_05_v1","PvP01_05_v1","PvP01_05_v1","PvP01_05_v1","PvP01_05_v1","PvP01_05_v1","PvP01_05_v1","PvP01_05_v1","PvP01_05_v1","PvP01_05_v1","PvP01_05_v1","PvP01_05_v1","PvP01_05_v1","PvP01_05_v1","PvP01_06_v1","PvP01_06_v1","PvP01_06_v1","PvP01_06_v1","PvP01_06_v1","PvP01_06_v1","PvP01_06_v1","PvP01_06_v1","PvP01_06_v1","PvP01_06_v1","PvP01_06_v1","PvP01_06_v1","PvP01_06_v1","PvP01_06_v1","PvP01_06_v1","PvP01_06_v1","PvP01_06_v1","PvP01_06_v1","PvP01_06_v1","PvP01_06_v1","PvP01_06_v1","PvP01_06_v1","PvP01_06_v1","PvP01_06_v1","PvP01_06_v1","PvP01_06_v1","PvP01_06_v1","PvP01_06_v1","PvP01_06_v1","PvP01_06_v1","PvP01_06_v1","PvP01_06_v1","PvP01_06_v1","PvP01_06_v1","PvP01_06_v1","PvP01_06_v1","PvP01_06_v1","PvP01_06_v1","PvP01_06_v1","PvP01_06_v1","PvP01_06_v1","PvP01_06_v1","PvP01_06_v1","PvP01_06_v1","PvP01_06_v1","PvP01_06_v1","PvP01_06_v1","PvP01_06_v1","PvP01_06_v1","PvP01_06_v1","PvP01_06_v1","PvP01_06_v1","PvP01_06_v1","PvP01_06_v1","PvP01_06_v1","PvP01_06_v1","PvP01_06_v1","PvP01_06_v1","PvP01_06_v1","PvP01_06_v1","PvP01_06_v1","PvP01_06_v1","PvP01_06_v1","PvP01_06_v1","PvP01_06_v1","PvP01_06_v1","PvP01_06_v1","PvP01_06_v1","PvP01_06_v1","PvP01_06_v1","PvP01_06_v1","PvP01_06_v1","PvP01_06_v1","PvP01_06_v1","PvP01_06_v1","PvP01_06_v1","PvP01_06_v1","PvP01_06_v1","PvP01_06_v1","PvP01_06_v1","PvP01_06_v1","PvP01_06_v1","PvP01_06_v1","PvP01_06_v1","PvP01_06_v1","PvP01_06_v1","PvP01_06_v1","PvP01_06_v1","PvP01_06_v1","PvP01_06_v1","PvP01_06_v1","PvP01_06_v1","PvP01_06_v1","PvP01_06_v1","PvP01_06_v1","PvP01_06_v1","PvP01_06_v1","PvP01_06_v1","PvP01_06_v1","PvP01_06_v1","PvP01_06_v1","PvP01_06_v1","PvP01_06_v1","PvP01_06_v1","PvP01_06_v1","PvP01_06_v1","PvP01_06_v1","PvP01_06_v1","PvP01_06_v1","PvP01_06_v1","PvP01_06_v1","PvP01_06_v1","PvP01_06_v1","PvP01_06_v1","PvP01_06_v1","PvP01_06_v1","PvP01_06_v1","PvP01_06_v1","PvP01_06_v1","PvP01_06_v1","PvP01_06_v1","PvP01_06_v1","PvP01_06_v1","PvP01_06_v1","PvP01_06_v1","PvP01_06_v1","PvP01_06_v1","PvP01_06_v1","PvP01_06_v1","PvP01_06_v1","PvP01_06_v1","PvP01_06_v1","PvP01_06_v1","PvP01_06_v1","PvP01_06_v1","PvP01_06_v1","PvP01_06_v1","PvP01_06_v1","PvP01_06_v1","PvP01_06_v1","PvP01_06_v1","PvP01_07_v1","PvP01_07_v1","PvP01_07_v1","PvP01_07_v1","PvP01_07_v1","PvP01_07_v1","PvP01_07_v1","PvP01_07_v1","PvP01_07_v1","PvP01_07_v1","PvP01_07_v1","PvP01_07_v1","PvP01_07_v1","PvP01_07_v1","PvP01_07_v1","PvP01_07_v1","PvP01_07_v1","PvP01_07_v1","PvP01_07_v1","PvP01_07_v1","PvP01_07_v1","PvP01_07_v1","PvP01_07_v1","PvP01_07_v1","PvP01_07_v1","PvP01_07_v1","PvP01_07_v1","PvP01_07_v1","PvP01_07_v1","PvP01_07_v1","PvP01_07_v1","PvP01_07_v1","PvP01_07_v1","PvP01_07_v1","PvP01_07_v1","PvP01_07_v1","PvP01_07_v1","PvP01_07_v1","PvP01_07_v1","PvP01_07_v1","PvP01_07_v1","PvP01_07_v1","PvP01_07_v1","PvP01_07_v1","PvP01_07_v1","PvP01_07_v1","PvP01_07_v1","PvP01_07_v1","PvP01_07_v1","PvP01_07_v1","PvP01_07_v1","PvP01_07_v1","PvP01_07_v1","PvP01_07_v1","PvP01_07_v1","PvP01_07_v1","PvP01_07_v1","PvP01_07_v1","PvP01_07_v1","PvP01_07_v1","PvP01_07_v1","PvP01_07_v1","PvP01_07_v1","PvP01_07_v1","PvP01_07_v1","PvP01_07_v1","PvP01_07_v1","PvP01_07_v1","PvP01_07_v1","PvP01_07_v1","PvP01_07_v1","PvP01_07_v1","PvP01_07_v1","PvP01_07_v1","PvP01_07_v1","PvP01_07_v1","PvP01_07_v1","PvP01_07_v1","PvP01_07_v1","PvP01_07_v1","PvP01_07_v1","PvP01_07_v1","PvP01_07_v1","PvP01_07_v1","PvP01_07_v1","PvP01_07_v1","PvP01_07_v1","PvP01_07_v1","PvP01_07_v1","PvP01_07_v1","PvP01_07_v1","PvP01_07_v1","PvP01_07_v1","PvP01_07_v1","PvP01_07_v1","PvP01_07_v1","PvP01_07_v1","PvP01_07_v1","PvP01_07_v1","PvP01_07_v1","PvP01_07_v1","PvP01_07_v1","PvP01_07_v1","PvP01_07_v1","PvP01_07_v1","PvP01_07_v1","PvP01_07_v1","PvP01_07_v1","PvP01_07_v1","PvP01_07_v1","PvP01_07_v1","PvP01_07_v1","PvP01_07_v1","PvP01_07_v1","PvP01_07_v1","PvP01_07_v1","PvP01_07_v1","PvP01_07_v1","PvP01_07_v1","PvP01_07_v1","PvP01_07_v1","PvP01_07_v1","PvP01_07_v1","PvP01_07_v1","PvP01_07_v1","PvP01_07_v1","PvP01_07_v1","PvP01_07_v1","PvP01_07_v1","PvP01_07_v1","PvP01_07_v1","PvP01_07_v1","PvP01_07_v1","PvP01_07_v1","PvP01_07_v1","PvP01_07_v1","PvP01_07_v1","PvP01_07_v1","PvP01_07_v1","PvP01_07_v1","PvP01_07_v1","PvP01_07_v1","PvP01_07_v1","PvP01_07_v1","PvP01_07_v1","PvP01_07_v1","PvP01_07_v1","PvP01_07_v1","PvP01_07_v1","PvP01_07_v1","PvP01_07_v1","PvP01_07_v1","PvP01_07_v1","PvP01_07_v1","PvP01_07_v1","PvP01_07_v1","PvP01_07_v1","PvP01_07_v1","PvP01_07_v1","PvP01_07_v1","PvP01_07_v1","PvP01_07_v1","PvP01_07_v1","PvP01_07_v1","PvP01_07_v1","PvP01_07_v1","PvP01_07_v1","PvP01_07_v1","PvP01_07_v1","PvP01_07_v1","PvP01_07_v1","PvP01_07_v1","PvP01_07_v1","PvP01_07_v1","PvP01_07_v1","PvP01_07_v1","PvP01_07_v1","PvP01_07_v1","PvP01_07_v1","PvP01_07_v1","PvP01_07_v1","PvP01_07_v1","PvP01_07_v1","PvP01_07_v1","PvP01_07_v1","PvP01_07_v1","PvP01_07_v1","PvP01_07_v1","PvP01_07_v1","PvP01_07_v1","PvP01_07_v1","PvP01_07_v1","PvP01_07_v1","PvP01_07_v1","PvP01_07_v1","PvP01_07_v1","PvP01_07_v1","PvP01_07_v1","PvP01_07_v1","PvP01_07_v1","PvP01_07_v1","PvP01_07_v1","PvP01_07_v1","PvP01_07_v1","PvP01_07_v1","PvP01_07_v1","PvP01_07_v1","PvP01_07_v1","PvP01_07_v1","PvP01_07_v1","PvP01_07_v1","PvP01_07_v1","PvP01_07_v1","PvP01_07_v1","PvP01_07_v1","PvP01_07_v1","PvP01_07_v1","PvP01_07_v1","PvP01_07_v1","PvP01_07_v1","PvP01_07_v1","PvP01_07_v1","PvP01_07_v1","PvP01_08_v1","PvP01_08_v1","PvP01_08_v1","PvP01_08_v1","PvP01_08_v1","PvP01_08_v1","PvP01_08_v1","PvP01_08_v1","PvP01_08_v1","PvP01_08_v1","PvP01_08_v1","PvP01_08_v1","PvP01_08_v1","PvP01_08_v1","PvP01_08_v1","PvP01_08_v1","PvP01_08_v1","PvP01_08_v1","PvP01_08_v1","PvP01_08_v1","PvP01_08_v1","PvP01_08_v1","PvP01_08_v1","PvP01_08_v1","PvP01_08_v1","PvP01_08_v1","PvP01_08_v1","PvP01_08_v1","PvP01_08_v1","PvP01_08_v1","PvP01_08_v1","PvP01_08_v1","PvP01_08_v1","PvP01_08_v1","PvP01_08_v1","PvP01_08_v1","PvP01_08_v1","PvP01_08_v1","PvP01_08_v1","PvP01_08_v1","PvP01_08_v1","PvP01_08_v1","PvP01_08_v1","PvP01_08_v1","PvP01_08_v1","PvP01_08_v1","PvP01_08_v1","PvP01_08_v1","PvP01_08_v1","PvP01_08_v1","PvP01_08_v1","PvP01_08_v1","PvP01_08_v1","PvP01_08_v1","PvP01_08_v1","PvP01_08_v1","PvP01_08_v1","PvP01_08_v1","PvP01_08_v1","PvP01_08_v1","PvP01_08_v1","PvP01_08_v1","PvP01_08_v1","PvP01_08_v1","PvP01_08_v1","PvP01_08_v1","PvP01_08_v1","PvP01_08_v1","PvP01_08_v1","PvP01_08_v1","PvP01_08_v1","PvP01_08_v1","PvP01_08_v1","PvP01_08_v1","PvP01_08_v1","PvP01_08_v1","PvP01_08_v1","PvP01_08_v1","PvP01_08_v1","PvP01_08_v1","PvP01_08_v1","PvP01_08_v1","PvP01_08_v1","PvP01_08_v1","PvP01_08_v1","PvP01_08_v1","PvP01_08_v1","PvP01_08_v1","PvP01_08_v1","PvP01_08_v1","PvP01_08_v1","PvP01_08_v1","PvP01_08_v1","PvP01_08_v1","PvP01_08_v1","PvP01_08_v1","PvP01_08_v1","PvP01_08_v1","PvP01_08_v1","PvP01_08_v1","PvP01_08_v1","PvP01_08_v1","PvP01_08_v1","PvP01_08_v1","PvP01_08_v1","PvP01_08_v1","PvP01_08_v1","PvP01_08_v1","PvP01_08_v1","PvP01_08_v1","PvP01_08_v1","PvP01_08_v1","PvP01_08_v1","PvP01_08_v1","PvP01_08_v1","PvP01_08_v1","PvP01_08_v1","PvP01_08_v1","PvP01_08_v1","PvP01_08_v1","PvP01_08_v1","PvP01_08_v1","PvP01_08_v1","PvP01_08_v1","PvP01_08_v1","PvP01_08_v1","PvP01_08_v1","PvP01_08_v1","PvP01_08_v1","PvP01_08_v1","PvP01_08_v1","PvP01_08_v1","PvP01_08_v1","PvP01_08_v1","PvP01_08_v1","PvP01_08_v1","PvP01_08_v1","PvP01_08_v1","PvP01_08_v1","PvP01_08_v1","PvP01_08_v1","PvP01_08_v1","PvP01_08_v1","PvP01_08_v1","PvP01_08_v1","PvP01_08_v1","PvP01_08_v1","PvP01_08_v1","PvP01_08_v1","PvP01_08_v1","PvP01_08_v1","PvP01_08_v1","PvP01_08_v1","PvP01_08_v1","PvP01_08_v1","PvP01_08_v1","PvP01_08_v1","PvP01_08_v1","PvP01_08_v1","PvP01_08_v1","PvP01_08_v1","PvP01_08_v1","PvP01_08_v1","PvP01_08_v1","PvP01_08_v1","PvP01_08_v1","PvP01_08_v1","PvP01_08_v1","PvP01_08_v1","PvP01_08_v1","PvP01_08_v1","PvP01_08_v1","PvP01_08_v1","PvP01_08_v1","PvP01_08_v1","PvP01_08_v1","PvP01_08_v1","PvP01_08_v1","PvP01_08_v1","PvP01_08_v1","PvP01_08_v1","PvP01_08_v1","PvP01_08_v1","PvP01_08_v1","PvP01_08_v1","PvP01_08_v1","PvP01_08_v1","PvP01_08_v1","PvP01_08_v1","PvP01_08_v1","PvP01_08_v1","PvP01_08_v1","PvP01_08_v1","PvP01_08_v1","PvP01_08_v1","PvP01_08_v1","PvP01_08_v1","PvP01_08_v1","PvP01_08_v1","PvP01_08_v1","PvP01_08_v1","PvP01_08_v1","PvP01_08_v1","PvP01_08_v1","PvP01_08_v1","PvP01_08_v1","PvP01_08_v1","PvP01_08_v1","PvP01_08_v1","PvP01_08_v1","PvP01_08_v1","PvP01_08_v1","PvP01_08_v1","PvP01_08_v1","PvP01_08_v1","PvP01_08_v1","PvP01_08_v1","PvP01_08_v1","PvP01_08_v1","PvP01_08_v1","PvP01_08_v1","PvP01_08_v1","PvP01_08_v1","PvP01_08_v1","PvP01_08_v1","PvP01_08_v1","PvP01_08_v1","PvP01_08_v1","PvP01_08_v1","PvP01_08_v1","PvP01_08_v1","PvP01_08_v1","PvP01_08_v1","PvP01_08_v1","PvP01_09_v1","PvP01_09_v1","PvP01_09_v1","PvP01_09_v1","PvP01_09_v1","PvP01_09_v1","PvP01_09_v1","PvP01_09_v1","PvP01_09_v1","PvP01_09_v1","PvP01_09_v1","PvP01_09_v1","PvP01_09_v1","PvP01_09_v1","PvP01_09_v1","PvP01_09_v1","PvP01_09_v1","PvP01_09_v1","PvP01_09_v1","PvP01_09_v1","PvP01_09_v1","PvP01_09_v1","PvP01_09_v1","PvP01_09_v1","PvP01_09_v1","PvP01_09_v1","PvP01_09_v1","PvP01_09_v1","PvP01_09_v1","PvP01_09_v1","PvP01_09_v1","PvP01_09_v1","PvP01_09_v1","PvP01_09_v1","PvP01_09_v1","PvP01_09_v1","PvP01_09_v1","PvP01_09_v1","PvP01_09_v1","PvP01_09_v1","PvP01_09_v1","PvP01_09_v1","PvP01_09_v1","PvP01_09_v1","PvP01_09_v1","PvP01_09_v1","PvP01_09_v1","PvP01_09_v1","PvP01_09_v1","PvP01_09_v1","PvP01_09_v1","PvP01_09_v1","PvP01_09_v1","PvP01_09_v1","PvP01_09_v1","PvP01_09_v1","PvP01_09_v1","PvP01_09_v1","PvP01_09_v1","PvP01_09_v1","PvP01_09_v1","PvP01_09_v1","PvP01_09_v1","PvP01_09_v1","PvP01_09_v1","PvP01_09_v1","PvP01_09_v1","PvP01_09_v1","PvP01_09_v1","PvP01_09_v1","PvP01_09_v1","PvP01_09_v1","PvP01_09_v1","PvP01_09_v1","PvP01_09_v1","PvP01_09_v1","PvP01_09_v1","PvP01_09_v1","PvP01_09_v1","PvP01_09_v1","PvP01_09_v1","PvP01_09_v1","PvP01_09_v1","PvP01_09_v1","PvP01_09_v1","PvP01_09_v1","PvP01_09_v1","PvP01_09_v1","PvP01_09_v1","PvP01_09_v1","PvP01_09_v1","PvP01_09_v1","PvP01_09_v1","PvP01_09_v1","PvP01_09_v1","PvP01_09_v1","PvP01_09_v1","PvP01_09_v1","PvP01_09_v1","PvP01_09_v1","PvP01_09_v1","PvP01_09_v1","PvP01_09_v1","PvP01_09_v1","PvP01_09_v1","PvP01_09_v1","PvP01_09_v1","PvP01_09_v1","PvP01_09_v1","PvP01_09_v1","PvP01_09_v1","PvP01_09_v1","PvP01_09_v1","PvP01_09_v1","PvP01_09_v1","PvP01_09_v1","PvP01_09_v1","PvP01_09_v1","PvP01_09_v1","PvP01_09_v1","PvP01_09_v1","PvP01_09_v1","PvP01_09_v1","PvP01_09_v1","PvP01_09_v1","PvP01_09_v1","PvP01_09_v1","PvP01_09_v1","PvP01_09_v1","PvP01_09_v1","PvP01_09_v1","PvP01_09_v1","PvP01_09_v1","PvP01_09_v1","PvP01_09_v1","PvP01_09_v1","PvP01_09_v1","PvP01_09_v1","PvP01_09_v1","PvP01_09_v1","PvP01_09_v1","PvP01_09_v1","PvP01_09_v1","PvP01_09_v1","PvP01_09_v1","PvP01_09_v1","PvP01_09_v1","PvP01_09_v1","PvP01_09_v1","PvP01_09_v1","PvP01_09_v1","PvP01_09_v1","PvP01_09_v1","PvP01_09_v1","PvP01_09_v1","PvP01_09_v1","PvP01_09_v1","PvP01_09_v1","PvP01_09_v1","PvP01_09_v1","PvP01_09_v1","PvP01_09_v1","PvP01_09_v1","PvP01_09_v1","PvP01_09_v1","PvP01_09_v1","PvP01_09_v1","PvP01_09_v1","PvP01_09_v1","PvP01_09_v1","PvP01_09_v1","PvP01_09_v1","PvP01_09_v1","PvP01_09_v1","PvP01_09_v1","PvP01_09_v1","PvP01_09_v1","PvP01_09_v1","PvP01_09_v1","PvP01_09_v1","PvP01_09_v1","PvP01_09_v1","PvP01_09_v1","PvP01_09_v1","PvP01_09_v1","PvP01_09_v1","PvP01_09_v1","PvP01_09_v1","PvP01_09_v1","PvP01_09_v1","PvP01_09_v1","PvP01_09_v1","PvP01_09_v1","PvP01_09_v1","PvP01_09_v1","PvP01_09_v1","PvP01_09_v1","PvP01_09_v1","PvP01_09_v1","PvP01_09_v1","PvP01_09_v1","PvP01_09_v1","PvP01_09_v1","PvP01_09_v1","PvP01_09_v1","PvP01_09_v1","PvP01_09_v1","PvP01_09_v1","PvP01_09_v1","PvP01_09_v1","PvP01_09_v1","PvP01_09_v1","PvP01_09_v1","PvP01_09_v1","PvP01_09_v1","PvP01_09_v1","PvP01_09_v1","PvP01_09_v1","PvP01_09_v1","PvP01_09_v1","PvP01_09_v1","PvP01_09_v1","PvP01_09_v1","PvP01_09_v1","PvP01_09_v1","PvP01_09_v1","PvP01_09_v1","PvP01_09_v1","PvP01_09_v1","PvP01_09_v1","PvP01_09_v1","PvP01_09_v1","PvP01_09_v1","PvP01_09_v1","PvP01_09_v1","PvP01_09_v1","PvP01_09_v1","PvP01_09_v1","PvP01_09_v1","PvP01_09_v1","PvP01_09_v1","PvP01_09_v1","PvP01_09_v1","PvP01_09_v1","PvP01_09_v1","PvP01_09_v1","PvP01_09_v1","PvP01_09_v1","PvP01_09_v1","PvP01_09_v1","PvP01_09_v1","PvP01_09_v1","PvP01_09_v1","PvP01_09_v1","PvP01_09_v1","PvP01_09_v1","PvP01_09_v1","PvP01_09_v1","PvP01_09_v1","PvP01_09_v1","PvP01_09_v1","PvP01_09_v1","PvP01_09_v1","PvP01_09_v1","PvP01_09_v1","PvP01_09_v1","PvP01_09_v1","PvP01_09_v1","PvP01_09_v1","PvP01_09_v1","PvP01_09_v1","PvP01_09_v1","PvP01_09_v1","PvP01_09_v1","PvP01_09_v1","PvP01_09_v1","PvP01_09_v1","PvP01_09_v1","PvP01_09_v1","PvP01_09_v1","PvP01_09_v1","PvP01_09_v1","PvP01_09_v1","PvP01_09_v1","PvP01_09_v1","PvP01_09_v1","PvP01_09_v1","PvP01_09_v1","PvP01_09_v1","PvP01_09_v1","PvP01_09_v1","PvP01_09_v1","PvP01_09_v1","PvP01_09_v1","PvP01_10_v1","PvP01_10_v1","PvP01_10_v1","PvP01_10_v1","PvP01_10_v1","PvP01_10_v1","PvP01_10_v1","PvP01_10_v1","PvP01_10_v1","PvP01_10_v1","PvP01_10_v1","PvP01_10_v1","PvP01_10_v1","PvP01_10_v1","PvP01_10_v1","PvP01_10_v1","PvP01_10_v1","PvP01_10_v1","PvP01_10_v1","PvP01_10_v1","PvP01_10_v1","PvP01_10_v1","PvP01_10_v1","PvP01_10_v1","PvP01_10_v1","PvP01_10_v1","PvP01_10_v1","PvP01_10_v1","PvP01_10_v1","PvP01_10_v1","PvP01_10_v1","PvP01_10_v1","PvP01_10_v1","PvP01_10_v1","PvP01_10_v1","PvP01_10_v1","PvP01_10_v1","PvP01_10_v1","PvP01_10_v1","PvP01_10_v1","PvP01_10_v1","PvP01_10_v1","PvP01_10_v1","PvP01_10_v1","PvP01_10_v1","PvP01_10_v1","PvP01_10_v1","PvP01_10_v1","PvP01_10_v1","PvP01_10_v1","PvP01_10_v1","PvP01_10_v1","PvP01_10_v1","PvP01_10_v1","PvP01_10_v1","PvP01_10_v1","PvP01_10_v1","PvP01_10_v1","PvP01_10_v1","PvP01_10_v1","PvP01_10_v1","PvP01_10_v1","PvP01_10_v1","PvP01_10_v1","PvP01_10_v1","PvP01_10_v1","PvP01_10_v1","PvP01_10_v1","PvP01_10_v1","PvP01_10_v1","PvP01_10_v1","PvP01_10_v1","PvP01_10_v1","PvP01_10_v1","PvP01_10_v1","PvP01_10_v1","PvP01_10_v1","PvP01_10_v1","PvP01_10_v1","PvP01_10_v1","PvP01_10_v1","PvP01_10_v1","PvP01_10_v1","PvP01_10_v1","PvP01_10_v1","PvP01_10_v1","PvP01_10_v1","PvP01_10_v1","PvP01_10_v1","PvP01_10_v1","PvP01_10_v1","PvP01_10_v1","PvP01_10_v1","PvP01_10_v1","PvP01_10_v1","PvP01_10_v1","PvP01_10_v1","PvP01_10_v1","PvP01_10_v1","PvP01_10_v1","PvP01_10_v1","PvP01_10_v1","PvP01_10_v1","PvP01_10_v1","PvP01_10_v1","PvP01_10_v1","PvP01_10_v1","PvP01_10_v1","PvP01_10_v1","PvP01_10_v1","PvP01_10_v1","PvP01_10_v1","PvP01_10_v1","PvP01_10_v1","PvP01_10_v1","PvP01_10_v1","PvP01_10_v1","PvP01_10_v1","PvP01_10_v1","PvP01_10_v1","PvP01_10_v1","PvP01_10_v1","PvP01_10_v1","PvP01_10_v1","PvP01_10_v1","PvP01_10_v1","PvP01_10_v1","PvP01_10_v1","PvP01_10_v1","PvP01_10_v1","PvP01_10_v1","PvP01_10_v1","PvP01_10_v1","PvP01_10_v1","PvP01_10_v1","PvP01_10_v1","PvP01_10_v1","PvP01_10_v1","PvP01_10_v1","PvP01_10_v1","PvP01_10_v1","PvP01_10_v1","PvP01_10_v1","PvP01_10_v1","PvP01_10_v1","PvP01_10_v1","PvP01_10_v1","PvP01_10_v1","PvP01_10_v1","PvP01_10_v1","PvP01_10_v1","PvP01_10_v1","PvP01_10_v1","PvP01_10_v1","PvP01_10_v1","PvP01_10_v1","PvP01_10_v1","PvP01_10_v1","PvP01_10_v1","PvP01_10_v1","PvP01_10_v1","PvP01_10_v1","PvP01_10_v1","PvP01_10_v1","PvP01_10_v1","PvP01_10_v1","PvP01_10_v1","PvP01_10_v1","PvP01_10_v1","PvP01_10_v1","PvP01_10_v1","PvP01_10_v1","PvP01_10_v1","PvP01_10_v1","PvP01_10_v1","PvP01_10_v1","PvP01_10_v1","PvP01_10_v1","PvP01_10_v1","PvP01_10_v1","PvP01_10_v1","PvP01_10_v1","PvP01_10_v1","PvP01_10_v1","PvP01_10_v1","PvP01_10_v1","PvP01_10_v1","PvP01_10_v1","PvP01_10_v1","PvP01_10_v1","PvP01_10_v1","PvP01_10_v1","PvP01_10_v1","PvP01_10_v1","PvP01_10_v1","PvP01_11_v1","PvP01_11_v1","PvP01_11_v1","PvP01_11_v1","PvP01_11_v1","PvP01_11_v1","PvP01_11_v1","PvP01_11_v1","PvP01_11_v1","PvP01_11_v1","PvP01_11_v1","PvP01_11_v1","PvP01_11_v1","PvP01_11_v1","PvP01_11_v1","PvP01_11_v1","PvP01_11_v1","PvP01_11_v1","PvP01_11_v1","PvP01_11_v1","PvP01_11_v1","PvP01_11_v1","PvP01_11_v1","PvP01_11_v1","PvP01_11_v1","PvP01_11_v1","PvP01_11_v1","PvP01_11_v1","PvP01_11_v1","PvP01_11_v1","PvP01_11_v1","PvP01_11_v1","PvP01_11_v1","PvP01_11_v1","PvP01_11_v1","PvP01_11_v1","PvP01_11_v1","PvP01_11_v1","PvP01_11_v1","PvP01_11_v1","PvP01_11_v1","PvP01_11_v1","PvP01_11_v1","PvP01_11_v1","PvP01_11_v1","PvP01_11_v1","PvP01_11_v1","PvP01_11_v1","PvP01_11_v1","PvP01_11_v1","PvP01_11_v1","PvP01_11_v1","PvP01_11_v1","PvP01_11_v1","PvP01_11_v1","PvP01_11_v1","PvP01_11_v1","PvP01_11_v1","PvP01_11_v1","PvP01_11_v1","PvP01_11_v1","PvP01_11_v1","PvP01_11_v1","PvP01_11_v1","PvP01_11_v1","PvP01_11_v1","PvP01_11_v1","PvP01_11_v1","PvP01_11_v1","PvP01_11_v1","PvP01_11_v1","PvP01_11_v1","PvP01_11_v1","PvP01_11_v1","PvP01_11_v1","PvP01_11_v1","PvP01_11_v1","PvP01_11_v1","PvP01_11_v1","PvP01_11_v1","PvP01_11_v1","PvP01_11_v1","PvP01_11_v1","PvP01_11_v1","PvP01_11_v1","PvP01_11_v1","PvP01_11_v1","PvP01_11_v1","PvP01_11_v1","PvP01_11_v1","PvP01_11_v1","PvP01_11_v1","PvP01_11_v1","PvP01_11_v1","PvP01_11_v1","PvP01_11_v1","PvP01_11_v1","PvP01_11_v1","PvP01_11_v1","PvP01_11_v1","PvP01_11_v1","PvP01_11_v1","PvP01_11_v1","PvP01_11_v1","PvP01_11_v1","PvP01_11_v1","PvP01_11_v1","PvP01_11_v1","PvP01_11_v1","PvP01_11_v1","PvP01_11_v1","PvP01_11_v1","PvP01_11_v1","PvP01_11_v1","PvP01_11_v1","PvP01_11_v1","PvP01_11_v1","PvP01_11_v1","PvP01_11_v1","PvP01_11_v1","PvP01_11_v1","PvP01_11_v1","PvP01_11_v1","PvP01_11_v1","PvP01_11_v1","PvP01_11_v1","PvP01_11_v1","PvP01_11_v1","PvP01_11_v1","PvP01_11_v1","PvP01_11_v1","PvP01_11_v1","PvP01_11_v1","PvP01_11_v1","PvP01_11_v1","PvP01_11_v1","PvP01_11_v1","PvP01_11_v1","PvP01_11_v1","PvP01_11_v1","PvP01_11_v1","PvP01_11_v1","PvP01_11_v1","PvP01_11_v1","PvP01_11_v1","PvP01_11_v1","PvP01_11_v1","PvP01_11_v1","PvP01_11_v1","PvP01_11_v1","PvP01_11_v1","PvP01_11_v1","PvP01_11_v1","PvP01_11_v1","PvP01_11_v1","PvP01_11_v1","PvP01_11_v1","PvP01_11_v1","PvP01_11_v1","PvP01_11_v1","PvP01_11_v1","PvP01_11_v1","PvP01_11_v1","PvP01_11_v1","PvP01_11_v1","PvP01_11_v1","PvP01_11_v1","PvP01_11_v1","PvP01_11_v1","PvP01_11_v1","PvP01_11_v1","PvP01_11_v1","PvP01_11_v1","PvP01_11_v1","PvP01_11_v1","PvP01_11_v1","PvP01_11_v1","PvP01_11_v1","PvP01_11_v1","PvP01_11_v1","PvP01_11_v1","PvP01_11_v1","PvP01_11_v1","PvP01_11_v1","PvP01_11_v1","PvP01_11_v1","PvP01_11_v1","PvP01_11_v1","PvP01_11_v1","PvP01_11_v1","PvP01_11_v1","PvP01_11_v1","PvP01_11_v1","PvP01_11_v1","PvP01_11_v1","PvP01_11_v1","PvP01_11_v1","PvP01_11_v1","PvP01_11_v1","PvP01_11_v1","PvP01_11_v1","PvP01_11_v1","PvP01_11_v1","PvP01_11_v1","PvP01_11_v1","PvP01_11_v1","PvP01_11_v1","PvP01_11_v1","PvP01_11_v1","PvP01_11_v1","PvP01_11_v1","PvP01_11_v1","PvP01_11_v1","PvP01_11_v1","PvP01_11_v1","PvP01_11_v1","PvP01_11_v1","PvP01_11_v1","PvP01_11_v1","PvP01_11_v1","PvP01_11_v1","PvP01_11_v1","PvP01_11_v1","PvP01_11_v1","PvP01_11_v1","PvP01_11_v1","PvP01_11_v1","PvP01_11_v1","PvP01_11_v1","PvP01_11_v1","PvP01_11_v1","PvP01_11_v1","PvP01_11_v1","PvP01_11_v1","PvP01_11_v1","PvP01_11_v1","PvP01_11_v1","PvP01_11_v1","PvP01_11_v1","PvP01_11_v1","PvP01_11_v1","PvP01_11_v1","PvP01_11_v1","PvP01_11_v1","PvP01_11_v1","PvP01_11_v1","PvP01_11_v1","PvP01_11_v1","PvP01_11_v1","PvP01_11_v1","PvP01_11_v1","PvP01_11_v1","PvP01_11_v1","PvP01_11_v1","PvP01_11_v1","PvP01_11_v1","PvP01_11_v1","PvP01_11_v1","PvP01_11_v1","PvP01_11_v1","PvP01_11_v1","PvP01_11_v1","PvP01_11_v1","PvP01_11_v1","PvP01_11_v1","PvP01_11_v1","PvP01_11_v1","PvP01_11_v1","PvP01_11_v1","PvP01_11_v1","PvP01_11_v1","PvP01_11_v1","PvP01_11_v1","PvP01_11_v1","PvP01_11_v1","PvP01_11_v1","PvP01_11_v1","PvP01_11_v1","PvP01_11_v1","PvP01_11_v1","PvP01_11_v1","PvP01_11_v1","PvP01_11_v1","PvP01_11_v1","PvP01_11_v1","PvP01_11_v1","PvP01_11_v1","PvP01_11_v1","PvP01_11_v1","PvP01_11_v1","PvP01_11_v1","PvP01_11_v1","PvP01_11_v1","PvP01_11_v1","PvP01_11_v1","PvP01_11_v1","PvP01_11_v1","PvP01_11_v1","PvP01_11_v1","PvP01_11_v1","PvP01_11_v1","PvP01_11_v1","PvP01_11_v1","PvP01_12_v1","PvP01_12_v1","PvP01_12_v1","PvP01_12_v1","PvP01_12_v1","PvP01_12_v1","PvP01_12_v1","PvP01_12_v1","PvP01_12_v1","PvP01_12_v1","PvP01_12_v1","PvP01_12_v1","PvP01_12_v1","PvP01_12_v1","PvP01_12_v1","PvP01_12_v1","PvP01_12_v1","PvP01_12_v1","PvP01_12_v1","PvP01_12_v1","PvP01_12_v1","PvP01_12_v1","PvP01_12_v1","PvP01_12_v1","PvP01_12_v1","PvP01_12_v1","PvP01_12_v1","PvP01_12_v1","PvP01_12_v1","PvP01_12_v1","PvP01_12_v1","PvP01_12_v1","PvP01_12_v1","PvP01_12_v1","PvP01_12_v1","PvP01_12_v1","PvP01_12_v1","PvP01_12_v1","PvP01_12_v1","PvP01_12_v1","PvP01_12_v1","PvP01_12_v1","PvP01_12_v1","PvP01_12_v1","PvP01_12_v1","PvP01_12_v1","PvP01_12_v1","PvP01_12_v1","PvP01_12_v1","PvP01_12_v1","PvP01_12_v1","PvP01_12_v1","PvP01_12_v1","PvP01_12_v1","PvP01_12_v1","PvP01_12_v1","PvP01_12_v1","PvP01_12_v1","PvP01_12_v1","PvP01_12_v1","PvP01_12_v1","PvP01_12_v1","PvP01_12_v1","PvP01_12_v1","PvP01_12_v1","PvP01_12_v1","PvP01_12_v1","PvP01_12_v1","PvP01_12_v1","PvP01_12_v1","PvP01_12_v1","PvP01_12_v1","PvP01_12_v1","PvP01_12_v1","PvP01_12_v1","PvP01_12_v1","PvP01_12_v1","PvP01_12_v1","PvP01_12_v1","PvP01_12_v1","PvP01_12_v1","PvP01_12_v1","PvP01_12_v1","PvP01_12_v1","PvP01_12_v1","PvP01_12_v1","PvP01_12_v1","PvP01_12_v1","PvP01_12_v1","PvP01_12_v1","PvP01_12_v1","PvP01_12_v1","PvP01_12_v1","PvP01_12_v1","PvP01_12_v1","PvP01_12_v1","PvP01_12_v1","PvP01_12_v1","PvP01_12_v1","PvP01_12_v1","PvP01_12_v1","PvP01_12_v1","PvP01_12_v1","PvP01_12_v1","PvP01_12_v1","PvP01_12_v1","PvP01_12_v1","PvP01_12_v1","PvP01_12_v1","PvP01_12_v1","PvP01_12_v1","PvP01_12_v1","PvP01_12_v1","PvP01_12_v1","PvP01_12_v1","PvP01_12_v1","PvP01_12_v1","PvP01_12_v1","PvP01_12_v1","PvP01_12_v1","PvP01_12_v1","PvP01_12_v1","PvP01_12_v1","PvP01_12_v1","PvP01_12_v1","PvP01_12_v1","PvP01_12_v1","PvP01_12_v1","PvP01_12_v1","PvP01_12_v1","PvP01_12_v1","PvP01_12_v1","PvP01_12_v1","PvP01_12_v1","PvP01_12_v1","PvP01_12_v1","PvP01_12_v1","PvP01_12_v1","PvP01_12_v1","PvP01_12_v1","PvP01_12_v1","PvP01_12_v1","PvP01_12_v1","PvP01_12_v1","PvP01_12_v1","PvP01_12_v1","PvP01_12_v1","PvP01_12_v1","PvP01_12_v1","PvP01_12_v1","PvP01_12_v1","PvP01_12_v1","PvP01_12_v1","PvP01_12_v1","PvP01_12_v1","PvP01_12_v1","PvP01_12_v1","PvP01_12_v1","PvP01_12_v1","PvP01_12_v1","PvP01_12_v1","PvP01_12_v1","PvP01_12_v1","PvP01_12_v1","PvP01_12_v1","PvP01_12_v1","PvP01_12_v1","PvP01_12_v1","PvP01_12_v1","PvP01_12_v1","PvP01_12_v1","PvP01_12_v1","PvP01_12_v1","PvP01_12_v1","PvP01_12_v1","PvP01_12_v1","PvP01_12_v1","PvP01_12_v1","PvP01_12_v1","PvP01_12_v1","PvP01_12_v1","PvP01_12_v1","PvP01_12_v1","PvP01_12_v1","PvP01_12_v1","PvP01_12_v1","PvP01_12_v1","PvP01_12_v1","PvP01_12_v1","PvP01_12_v1","PvP01_12_v1","PvP01_12_v1","PvP01_12_v1","PvP01_12_v1","PvP01_12_v1","PvP01_12_v1","PvP01_12_v1","PvP01_12_v1","PvP01_12_v1","PvP01_12_v1","PvP01_12_v1","PvP01_12_v1","PvP01_12_v1","PvP01_12_v1","PvP01_12_v1","PvP01_12_v1","PvP01_12_v1","PvP01_12_v1","PvP01_12_v1","PvP01_12_v1","PvP01_12_v1","PvP01_12_v1","PvP01_12_v1","PvP01_12_v1","PvP01_12_v1","PvP01_12_v1","PvP01_12_v1","PvP01_12_v1","PvP01_12_v1","PvP01_12_v1","PvP01_12_v1","PvP01_12_v1","PvP01_12_v1","PvP01_12_v1","PvP01_12_v1","PvP01_12_v1","PvP01_12_v1","PvP01_12_v1","PvP01_12_v1","PvP01_12_v1","PvP01_12_v1","PvP01_12_v1","PvP01_12_v1","PvP01_12_v1","PvP01_12_v1","PvP01_12_v1","PvP01_12_v1","PvP01_12_v1","PvP01_12_v1","PvP01_12_v1","PvP01_12_v1","PvP01_12_v1","PvP01_12_v1","PvP01_12_v1","PvP01_12_v1","PvP01_12_v1","PvP01_12_v1","PvP01_12_v1","PvP01_12_v1","PvP01_12_v1","PvP01_12_v1","PvP01_12_v1","PvP01_12_v1","PvP01_12_v1","PvP01_12_v1","PvP01_12_v1","PvP01_12_v1","PvP01_12_v1","PvP01_12_v1","PvP01_12_v1","PvP01_12_v1","PvP01_12_v1","PvP01_12_v1","PvP01_12_v1","PvP01_12_v1","PvP01_12_v1","PvP01_12_v1","PvP01_12_v1","PvP01_12_v1","PvP01_12_v1","PvP01_12_v1","PvP01_12_v1","PvP01_12_v1","PvP01_12_v1","PvP01_12_v1","PvP01_12_v1","PvP01_12_v1","PvP01_12_v1","PvP01_12_v1","PvP01_12_v1","PvP01_12_v1","PvP01_12_v1","PvP01_12_v1","PvP01_12_v1","PvP01_12_v1","PvP01_12_v1","PvP01_12_v1","PvP01_12_v1","PvP01_12_v1","PvP01_12_v1","PvP01_12_v1","PvP01_12_v1","PvP01_12_v1","PvP01_12_v1","PvP01_12_v1","PvP01_12_v1","PvP01_12_v1","PvP01_12_v1","PvP01_12_v1","PvP01_12_v1","PvP01_12_v1","PvP01_12_v1","PvP01_12_v1","PvP01_12_v1","PvP01_12_v1","PvP01_12_v1","PvP01_12_v1","PvP01_12_v1","PvP01_12_v1","PvP01_12_v1","PvP01_12_v1","PvP01_12_v1","PvP01_12_v1","PvP01_12_v1","PvP01_12_v1","PvP01_12_v1","PvP01_12_v1","PvP01_12_v1","PvP01_12_v1","PvP01_12_v1","PvP01_12_v1","PvP01_12_v1","PvP01_12_v1","PvP01_12_v1","PvP01_12_v1","PvP01_12_v1","PvP01_12_v1","PvP01_12_v1","PvP01_12_v1","PvP01_12_v1","PvP01_12_v1","PvP01_12_v1","PvP01_12_v1","PvP01_12_v1","PvP01_12_v1","PvP01_12_v1","PvP01_12_v1","PvP01_12_v1","PvP01_12_v1","PvP01_12_v1","PvP01_12_v1","PvP01_12_v1","PvP01_12_v1","PvP01_12_v1","PvP01_12_v1","PvP01_12_v1","PvP01_12_v1","PvP01_12_v1","PvP01_12_v1","PvP01_12_v1","PvP01_12_v1","PvP01_12_v1","PvP01_12_v1","PvP01_12_v1","PvP01_12_v1","PvP01_12_v1","PvP01_12_v1","PvP01_12_v1","PvP01_12_v1","PvP01_12_v1","PvP01_12_v1","PvP01_12_v1","PvP01_12_v1","PvP01_12_v1","PvP01_12_v1","PvP01_12_v1","PvP01_12_v1","PvP01_12_v1","PvP01_12_v1","PvP01_12_v1","PvP01_12_v1","PvP01_12_v1","PvP01_12_v1","PvP01_12_v1","PvP01_12_v1","PvP01_12_v1","PvP01_12_v1","PvP01_12_v1","PvP01_12_v1","PvP01_12_v1","PvP01_12_v1","PvP01_12_v1","PvP01_12_v1","PvP01_12_v1","PvP01_12_v1","PvP01_12_v1","PvP01_12_v1","PvP01_12_v1","PvP01_12_v1","PvP01_12_v1","PvP01_12_v1","PvP01_12_v1","PvP01_12_v1","PvP01_12_v1","PvP01_12_v1","PvP01_12_v1","PvP01_12_v1","PvP01_12_v1","PvP01_12_v1","PvP01_12_v1","PvP01_12_v1","PvP01_12_v1","PvP01_12_v1","PvP01_12_v1","PvP01_12_v1","PvP01_12_v1","PvP01_12_v1","PvP01_12_v1","PvP01_12_v1","PvP01_12_v1","PvP01_12_v1","PvP01_12_v1","PvP01_12_v1","PvP01_12_v1","PvP01_12_v1","PvP01_12_v1","PvP01_12_v1","PvP01_13_v1","PvP01_13_v1","PvP01_13_v1","PvP01_13_v1","PvP01_13_v1","PvP01_13_v1","PvP01_13_v1","PvP01_13_v1","PvP01_13_v1","PvP01_13_v1","PvP01_13_v1","PvP01_13_v1","PvP01_13_v1","PvP01_13_v1","PvP01_13_v1","PvP01_13_v1","PvP01_13_v1","PvP01_13_v1","PvP01_13_v1","PvP01_13_v1","PvP01_13_v1","PvP01_13_v1","PvP01_13_v1","PvP01_13_v1","PvP01_13_v1","PvP01_13_v1","PvP01_13_v1","PvP01_13_v1","PvP01_13_v1","PvP01_13_v1","PvP01_13_v1","PvP01_13_v1","PvP01_13_v1","PvP01_13_v1","PvP01_13_v1","PvP01_13_v1","PvP01_13_v1","PvP01_13_v1","PvP01_13_v1","PvP01_13_v1","PvP01_13_v1","PvP01_13_v1","PvP01_13_v1","PvP01_13_v1","PvP01_13_v1","PvP01_13_v1","PvP01_13_v1","PvP01_13_v1","PvP01_13_v1","PvP01_13_v1","PvP01_13_v1","PvP01_13_v1","PvP01_13_v1","PvP01_13_v1","PvP01_13_v1","PvP01_13_v1","PvP01_13_v1","PvP01_13_v1","PvP01_13_v1","PvP01_13_v1","PvP01_13_v1","PvP01_13_v1","PvP01_13_v1","PvP01_13_v1","PvP01_13_v1","PvP01_13_v1","PvP01_13_v1","PvP01_13_v1","PvP01_13_v1","PvP01_13_v1","PvP01_13_v1","PvP01_13_v1","PvP01_13_v1","PvP01_13_v1","PvP01_13_v1","PvP01_13_v1","PvP01_13_v1","PvP01_13_v1","PvP01_13_v1","PvP01_13_v1","PvP01_13_v1","PvP01_13_v1","PvP01_13_v1","PvP01_13_v1","PvP01_13_v1","PvP01_13_v1","PvP01_13_v1","PvP01_13_v1","PvP01_13_v1","PvP01_13_v1","PvP01_13_v1","PvP01_13_v1","PvP01_13_v1","PvP01_13_v1","PvP01_13_v1","PvP01_13_v1","PvP01_13_v1","PvP01_13_v1","PvP01_13_v1","PvP01_13_v1","PvP01_13_v1","PvP01_13_v1","PvP01_13_v1","PvP01_13_v1","PvP01_13_v1","PvP01_13_v1","PvP01_13_v1","PvP01_13_v1","PvP01_13_v1","PvP01_13_v1","PvP01_13_v1","PvP01_13_v1","PvP01_13_v1","PvP01_13_v1","PvP01_13_v1","PvP01_13_v1","PvP01_13_v1","PvP01_13_v1","PvP01_13_v1","PvP01_13_v1","PvP01_13_v1","PvP01_13_v1","PvP01_13_v1","PvP01_13_v1","PvP01_13_v1","PvP01_13_v1","PvP01_13_v1","PvP01_13_v1","PvP01_13_v1","PvP01_13_v1","PvP01_13_v1","PvP01_13_v1","PvP01_13_v1","PvP01_13_v1","PvP01_13_v1","PvP01_13_v1","PvP01_13_v1","PvP01_13_v1","PvP01_13_v1","PvP01_13_v1","PvP01_13_v1","PvP01_13_v1","PvP01_13_v1","PvP01_13_v1","PvP01_13_v1","PvP01_13_v1","PvP01_13_v1","PvP01_13_v1","PvP01_13_v1","PvP01_13_v1","PvP01_13_v1","PvP01_13_v1","PvP01_13_v1","PvP01_13_v1","PvP01_13_v1","PvP01_13_v1","PvP01_13_v1","PvP01_13_v1","PvP01_13_v1","PvP01_13_v1","PvP01_13_v1","PvP01_13_v1","PvP01_13_v1","PvP01_13_v1","PvP01_13_v1","PvP01_13_v1","PvP01_13_v1","PvP01_13_v1","PvP01_13_v1","PvP01_13_v1","PvP01_13_v1","PvP01_13_v1","PvP01_13_v1","PvP01_13_v1","PvP01_13_v1","PvP01_13_v1","PvP01_13_v1","PvP01_13_v1","PvP01_13_v1","PvP01_13_v1","PvP01_13_v1","PvP01_13_v1","PvP01_13_v1","PvP01_13_v1","PvP01_13_v1","PvP01_13_v1","PvP01_13_v1","PvP01_13_v1","PvP01_13_v1","PvP01_13_v1","PvP01_13_v1","PvP01_13_v1","PvP01_13_v1","PvP01_13_v1","PvP01_13_v1","PvP01_13_v1","PvP01_13_v1","PvP01_13_v1","PvP01_13_v1","PvP01_13_v1","PvP01_13_v1","PvP01_13_v1","PvP01_13_v1","PvP01_13_v1","PvP01_13_v1","PvP01_13_v1","PvP01_13_v1","PvP01_13_v1","PvP01_13_v1","PvP01_13_v1","PvP01_13_v1","PvP01_13_v1","PvP01_13_v1","PvP01_13_v1","PvP01_13_v1","PvP01_13_v1","PvP01_13_v1","PvP01_13_v1","PvP01_13_v1","PvP01_13_v1","PvP01_13_v1","PvP01_13_v1","PvP01_13_v1","PvP01_13_v1","PvP01_13_v1","PvP01_13_v1","PvP01_13_v1","PvP01_13_v1","PvP01_13_v1","PvP01_13_v1","PvP01_13_v1","PvP01_13_v1","PvP01_13_v1","PvP01_13_v1","PvP01_13_v1","PvP01_13_v1","PvP01_13_v1","PvP01_13_v1","PvP01_13_v1","PvP01_13_v1","PvP01_13_v1","PvP01_13_v1","PvP01_13_v1","PvP01_13_v1","PvP01_13_v1","PvP01_13_v1","PvP01_13_v1","PvP01_13_v1","PvP01_13_v1","PvP01_13_v1","PvP01_13_v1","PvP01_13_v1","PvP01_13_v1","PvP01_13_v1","PvP01_13_v1","PvP01_13_v1","PvP01_13_v1","PvP01_13_v1","PvP01_13_v1","PvP01_13_v1","PvP01_13_v1","PvP01_13_v1","PvP01_13_v1","PvP01_13_v1","PvP01_13_v1","PvP01_13_v1","PvP01_13_v1","PvP01_13_v1","PvP01_13_v1","PvP01_13_v1","PvP01_13_v1","PvP01_13_v1","PvP01_14_v1","PvP01_14_v1","PvP01_14_v1","PvP01_14_v1","PvP01_14_v1","PvP01_14_v1","PvP01_14_v1","PvP01_14_v1","PvP01_14_v1","PvP01_14_v1","PvP01_14_v1","PvP01_14_v1","PvP01_14_v1","PvP01_14_v1","PvP01_14_v1","PvP01_14_v1","PvP01_14_v1","PvP01_14_v1","PvP01_14_v1","PvP01_14_v1","PvP01_14_v1","PvP01_14_v1","PvP01_14_v1","PvP01_14_v1","PvP01_14_v1","PvP01_14_v1","PvP01_14_v1","PvP01_14_v1","PvP01_14_v1","PvP01_14_v1","PvP01_14_v1","PvP01_14_v1","PvP01_14_v1","PvP01_14_v1","PvP01_14_v1","PvP01_14_v1","PvP01_14_v1","PvP01_14_v1","PvP01_14_v1","PvP01_14_v1","PvP01_14_v1","PvP01_14_v1","PvP01_14_v1","PvP01_14_v1","PvP01_14_v1","PvP01_14_v1","PvP01_14_v1","PvP01_14_v1","PvP01_14_v1","PvP01_14_v1","PvP01_14_v1","PvP01_14_v1","PvP01_14_v1","PvP01_14_v1","PvP01_14_v1","PvP01_14_v1","PvP01_14_v1","PvP01_14_v1","PvP01_14_v1","PvP01_14_v1","PvP01_14_v1","PvP01_14_v1","PvP01_14_v1","PvP01_14_v1","PvP01_14_v1","PvP01_14_v1","PvP01_14_v1","PvP01_14_v1","PvP01_14_v1","PvP01_14_v1","PvP01_14_v1","PvP01_14_v1","PvP01_14_v1","PvP01_14_v1","PvP01_14_v1","PvP01_14_v1","PvP01_14_v1","PvP01_14_v1","PvP01_14_v1","PvP01_14_v1","PvP01_14_v1","PvP01_14_v1","PvP01_14_v1","PvP01_14_v1","PvP01_14_v1","PvP01_14_v1","PvP01_14_v1","PvP01_14_v1","PvP01_14_v1","PvP01_14_v1","PvP01_14_v1","PvP01_14_v1","PvP01_14_v1","PvP01_14_v1","PvP01_14_v1","PvP01_14_v1","PvP01_14_v1","PvP01_14_v1","PvP01_14_v1","PvP01_14_v1","PvP01_14_v1","PvP01_14_v1","PvP01_14_v1","PvP01_14_v1","PvP01_14_v1","PvP01_14_v1","PvP01_14_v1","PvP01_14_v1","PvP01_14_v1","PvP01_14_v1","PvP01_14_v1","PvP01_14_v1","PvP01_14_v1","PvP01_14_v1","PvP01_14_v1","PvP01_14_v1","PvP01_14_v1","PvP01_14_v1","PvP01_14_v1","PvP01_14_v1","PvP01_14_v1","PvP01_14_v1","PvP01_14_v1","PvP01_14_v1","PvP01_14_v1","PvP01_14_v1","PvP01_14_v1","PvP01_14_v1","PvP01_14_v1","PvP01_14_v1","PvP01_14_v1","PvP01_14_v1","PvP01_14_v1","PvP01_14_v1","PvP01_14_v1","PvP01_14_v1","PvP01_14_v1","PvP01_14_v1","PvP01_14_v1","PvP01_14_v1","PvP01_14_v1","PvP01_14_v1","PvP01_14_v1","PvP01_14_v1","PvP01_14_v1","PvP01_14_v1","PvP01_14_v1","PvP01_14_v1","PvP01_14_v1","PvP01_14_v1","PvP01_14_v1","PvP01_14_v1","PvP01_14_v1","PvP01_14_v1","PvP01_14_v1","PvP01_14_v1","PvP01_14_v1","PvP01_14_v1","PvP01_14_v1","PvP01_14_v1","PvP01_14_v1","PvP01_14_v1","PvP01_14_v1","PvP01_14_v1","PvP01_14_v1","PvP01_14_v1","PvP01_14_v1","PvP01_14_v1","PvP01_14_v1","PvP01_14_v1","PvP01_14_v1","PvP01_14_v1","PvP01_14_v1","PvP01_14_v1","PvP01_14_v1","PvP01_14_v1","PvP01_14_v1","PvP01_14_v1","PvP01_14_v1","PvP01_14_v1","PvP01_14_v1","PvP01_14_v1","PvP01_14_v1","PvP01_14_v1","PvP01_14_v1","PvP01_14_v1","PvP01_14_v1","PvP01_14_v1","PvP01_14_v1","PvP01_14_v1","PvP01_14_v1","PvP01_14_v1","PvP01_14_v1","PvP01_14_v1","PvP01_14_v1","PvP01_14_v1","PvP01_14_v1","PvP01_14_v1","PvP01_14_v1","PvP01_14_v1","PvP01_14_v1","PvP01_14_v1","PvP01_14_v1","PvP01_14_v1","PvP01_14_v1","PvP01_14_v1","PvP01_14_v1","PvP01_14_v1","PvP01_14_v1","PvP01_14_v1","PvP01_14_v1","PvP01_14_v1","PvP01_14_v1","PvP01_14_v1","PvP01_14_v1","PvP01_14_v1","PvP01_14_v1","PvP01_14_v1","PvP01_14_v1","PvP01_14_v1","PvP01_14_v1","PvP01_14_v1","PvP01_14_v1","PvP01_14_v1","PvP01_14_v1","PvP01_14_v1","PvP01_14_v1","PvP01_14_v1","PvP01_14_v1","PvP01_14_v1","PvP01_14_v1","PvP01_14_v1","PvP01_14_v1","PvP01_14_v1","PvP01_14_v1","PvP01_14_v1","PvP01_14_v1","PvP01_14_v1","PvP01_14_v1","PvP01_14_v1","PvP01_14_v1","PvP01_14_v1","PvP01_14_v1","PvP01_14_v1","PvP01_14_v1","PvP01_14_v1","PvP01_14_v1","PvP01_14_v1","PvP01_14_v1","PvP01_14_v1","PvP01_14_v1","PvP01_14_v1","PvP01_14_v1","PvP01_14_v1","PvP01_14_v1","PvP01_14_v1","PvP01_14_v1","PvP01_14_v1","PvP01_14_v1","PvP01_14_v1","PvP01_14_v1","PvP01_14_v1","PvP01_14_v1","PvP01_14_v1","PvP01_14_v1","PvP01_14_v1","PvP01_14_v1","PvP01_14_v1","PvP01_14_v1","PvP01_14_v1","PvP01_14_v1","PvP01_14_v1","PvP01_14_v1","PvP01_14_v1","PvP01_14_v1","PvP01_14_v1","PvP01_14_v1","PvP01_14_v1","PvP01_14_v1","PvP01_14_v1","PvP01_14_v1","PvP01_14_v1","PvP01_14_v1","PvP01_14_v1","PvP01_14_v1","PvP01_14_v1","PvP01_14_v1","PvP01_14_v1","PvP01_14_v1","PvP01_14_v1","PvP01_14_v1","PvP01_14_v1","PvP01_14_v1","PvP01_14_v1","PvP01_14_v1","PvP01_14_v1","PvP01_14_v1","PvP01_14_v1","PvP01_14_v1","PvP01_14_v1","PvP01_14_v1","PvP01_14_v1","PvP01_14_v1","PvP01_14_v1","PvP01_14_v1","PvP01_14_v1","PvP01_14_v1","PvP01_14_v1","PvP01_14_v1","PvP01_14_v1","PvP01_14_v1","PvP01_14_v1","PvP01_14_v1","PvP01_14_v1","PvP01_14_v1","PvP01_14_v1","PvP01_14_v1","PvP01_14_v1","PvP01_14_v1","PvP01_14_v1","PvP01_14_v1","PvP01_14_v1","PvP01_14_v1","PvP01_14_v1","PvP01_14_v1","PvP01_14_v1","PvP01_14_v1","PvP01_14_v1","PvP01_14_v1","PvP01_14_v1","PvP01_14_v1","PvP01_14_v1","PvP01_14_v1","PvP01_14_v1","PvP01_14_v1","PvP01_14_v1","PvP01_14_v1","PvP01_14_v1","PvP01_14_v1","PvP01_14_v1","PvP01_14_v1","PvP01_14_v1","PvP01_14_v1","PvP01_14_v1","PvP01_14_v1","PvP01_14_v1","PvP01_14_v1","PvP01_14_v1","PvP01_14_v1","PvP01_14_v1","PvP01_14_v1","PvP01_14_v1","PvP01_14_v1","PvP01_14_v1","PvP01_14_v1","PvP01_14_v1","PvP01_14_v1","PvP01_14_v1","PvP01_14_v1","PvP01_14_v1","PvP01_14_v1","PvP01_14_v1","PvP01_14_v1","PvP01_14_v1","PvP01_14_v1","PvP01_14_v1","PvP01_14_v1","PvP01_14_v1","PvP01_14_v1","PvP01_14_v1","PvP01_14_v1","PvP01_14_v1","PvP01_14_v1","PvP01_14_v1","PvP01_14_v1","PvP01_14_v1","PvP01_14_v1","PvP01_14_v1","PvP01_14_v1","PvP01_14_v1","PvP01_14_v1","PvP01_14_v1","PvP01_14_v1","PvP01_14_v1","PvP01_14_v1","PvP01_14_v1","PvP01_14_v1","PvP01_14_v1","PvP01_14_v1","PvP01_14_v1","PvP01_14_v1","PvP01_14_v1","PvP01_14_v1","PvP01_14_v1","PvP01_14_v1","PvP01_14_v1","PvP01_14_v1","PvP01_14_v1","PvP01_14_v1","PvP01_14_v1","PvP01_14_v1","PvP01_14_v1","PvP01_14_v1","PvP01_14_v1","PvP01_14_v1","PvP01_14_v1","PvP01_14_v1","PvP01_14_v1","PvP01_14_v1","PvP01_14_v1","PvP01_14_v1","PvP01_14_v1","PvP01_14_v1","PvP01_14_v1","PvP01_14_v1","PvP01_14_v1","PvP01_14_v1","PvP01_14_v1","PvP01_14_v1","PvP01_14_v1","PvP01_14_v1","PvP01_14_v1","PvP01_14_v1","PvP01_14_v1","PvP01_14_v1","PvP01_14_v1","PvP01_14_v1","PvP01_14_v1","PvP01_14_v1","PvP01_14_v1","PvP01_14_v1","PvP01_API_v1","PvP01_API_v1","PvP01_API_v1","PvP01_API_v1","PvP01_API_v1","Transfer.PvP01_00_1.final","Transfer.PvP01_00_1.final","Transfer.PvP01_00_1.final","Transfer.PvP01_00_1.final","Transfer.PvP01_00_1.final","Transfer.PvP01_00_2.final","Transfer.PvP01_00_2.final","Transfer.PvP01_00_8.final","Transfer.PvP01_00_10.final","Transfer.PvP01_00_11.final","Transfer.PvP01_00_11.final","Transfer.PvP01_00_11.final","Transfer.PvP01_00_11.final","Transfer.PvP01_00_11.final","Transfer.PvP01_00_11.final","Transfer.PvP01_00_11.final","Transfer.PvP01_00_16.final","Transfer.PvP01_00_17.final","Transfer.PvP01_00_26.final","Transfer.PvP01_00_61.final","Transfer.PvP01_00_89.final","Transfer.PvP01_00_161.final","Transfer.PvP01_00_199.final","Transfer.PvP01_00_227.final","PvP01_01_v1","PvP01_01_v1","PvP01_01_v1","PvP01_01_v1","PvP01_01_v1","PvP01_01_v1","PvP01_01_v1","PvP01_01_v1","PvP01_01_v1","PvP01_01_v1","PvP01_01_v1","PvP01_01_v1","PvP01_01_v1","PvP01_01_v1","PvP01_01_v1","PvP01_01_v1","PvP01_01_v1","PvP01_01_v1","PvP01_01_v1","PvP01_01_v1","PvP01_01_v1","PvP01_01_v1","PvP01_01_v1","PvP01_01_v1","PvP01_01_v1","PvP01_01_v1","PvP01_01_v1","PvP01_01_v1","PvP01_01_v1","PvP01_01_v1","PvP01_01_v1","PvP01_01_v1","PvP01_01_v1","PvP01_01_v1","PvP01_01_v1","PvP01_01_v1","PvP01_01_v1","PvP01_01_v1","PvP01_01_v1","PvP01_01_v1","PvP01_01_v1","PvP01_01_v1","PvP01_01_v1","PvP01_01_v1","PvP01_01_v1","PvP01_01_v1","PvP01_01_v1","PvP01_01_v1","PvP01_01_v1","PvP01_01_v1","PvP01_01_v1","PvP01_01_v1","PvP01_01_v1","PvP01_01_v1","PvP01_01_v1","PvP01_01_v1","PvP01_01_v1","PvP01_02_v1","PvP01_02_v1","PvP01_02_v1","PvP01_02_v1","PvP01_02_v1","PvP01_02_v1","PvP01_02_v1","PvP01_02_v1","PvP01_02_v1","PvP01_02_v1","PvP01_02_v1","PvP01_02_v1","PvP01_02_v1","PvP01_02_v1","PvP01_02_v1","PvP01_02_v1","PvP01_02_v1","PvP01_02_v1","PvP01_02_v1","PvP01_02_v1","PvP01_02_v1","PvP01_02_v1","PvP01_02_v1","PvP01_02_v1","PvP01_02_v1","PvP01_02_v1","PvP01_02_v1","PvP01_02_v1","PvP01_02_v1","PvP01_02_v1","PvP01_02_v1","PvP01_02_v1","PvP01_02_v1","PvP01_02_v1","PvP01_02_v1","PvP01_02_v1","PvP01_02_v1","PvP01_02_v1","PvP01_02_v1","PvP01_02_v1","PvP01_02_v1","PvP01_02_v1","PvP01_02_v1","PvP01_02_v1","PvP01_02_v1","PvP01_02_v1","PvP01_02_v1","PvP01_02_v1","PvP01_02_v1","PvP01_02_v1","PvP01_02_v1","PvP01_03_v1","PvP01_03_v1","PvP01_03_v1","PvP01_03_v1","PvP01_03_v1","PvP01_03_v1","PvP01_03_v1","PvP01_03_v1","PvP01_03_v1","PvP01_03_v1","PvP01_03_v1","PvP01_03_v1","PvP01_03_v1","PvP01_03_v1","PvP01_03_v1","PvP01_03_v1","PvP01_03_v1","PvP01_03_v1","PvP01_03_v1","PvP01_03_v1","PvP01_03_v1","PvP01_03_v1","PvP01_03_v1","PvP01_03_v1","PvP01_03_v1","PvP01_03_v1","PvP01_03_v1","PvP01_03_v1","PvP01_03_v1","PvP01_03_v1","PvP01_03_v1","PvP01_03_v1","PvP01_03_v1","PvP01_03_v1","PvP01_03_v1","PvP01_03_v1","PvP01_03_v1","PvP01_03_v1","PvP01_03_v1","PvP01_03_v1","PvP01_03_v1","PvP01_03_v1","PvP01_03_v1","PvP01_03_v1","PvP01_03_v1","PvP01_03_v1","PvP01_03_v1","PvP01_03_v1","PvP01_03_v1","PvP01_03_v1","PvP01_03_v1","PvP01_03_v1","PvP01_03_v1","PvP01_03_v1","PvP01_03_v1","PvP01_03_v1","PvP01_03_v1","PvP01_03_v1","PvP01_03_v1","PvP01_03_v1","PvP01_03_v1","PvP01_03_v1","PvP01_03_v1","PvP01_03_v1","PvP01_03_v1","PvP01_03_v1","PvP01_03_v1","PvP01_03_v1","PvP01_03_v1","PvP01_03_v1","PvP01_03_v1","PvP01_03_v1","PvP01_03_v1","PvP01_03_v1","PvP01_03_v1","PvP01_03_v1","PvP01_03_v1","PvP01_03_v1","PvP01_03_v1","PvP01_03_v1","PvP01_03_v1","PvP01_03_v1","PvP01_03_v1","PvP01_03_v1","PvP01_03_v1","PvP01_03_v1","PvP01_04_v1","PvP01_04_v1","PvP01_04_v1","PvP01_04_v1","PvP01_04_v1","PvP01_04_v1","PvP01_04_v1","PvP01_04_v1","PvP01_04_v1","PvP01_04_v1","PvP01_04_v1","PvP01_04_v1","PvP01_04_v1","PvP01_04_v1","PvP01_04_v1","PvP01_04_v1","PvP01_04_v1","PvP01_04_v1","PvP01_04_v1","PvP01_04_v1","PvP01_04_v1","PvP01_04_v1","PvP01_04_v1","PvP01_04_v1","PvP01_04_v1","PvP01_04_v1","PvP01_04_v1","PvP01_04_v1","PvP01_04_v1","PvP01_04_v1","PvP01_04_v1","PvP01_04_v1","PvP01_04_v1","PvP01_04_v1","PvP01_04_v1","PvP01_04_v1","PvP01_04_v1","PvP01_04_v1","PvP01_04_v1","PvP01_04_v1","PvP01_04_v1","PvP01_04_v1","PvP01_04_v1","PvP01_04_v1","PvP01_04_v1","PvP01_04_v1","PvP01_04_v1","PvP01_04_v1","PvP01_04_v1","PvP01_04_v1","PvP01_04_v1","PvP01_04_v1","PvP01_04_v1","PvP01_04_v1","PvP01_04_v1","PvP01_04_v1","PvP01_04_v1","PvP01_04_v1","PvP01_04_v1","PvP01_04_v1","PvP01_04_v1","PvP01_04_v1","PvP01_04_v1","PvP01_04_v1","PvP01_04_v1","PvP01_04_v1","PvP01_04_v1","PvP01_04_v1","PvP01_04_v1","PvP01_04_v1","PvP01_04_v1","PvP01_04_v1","PvP01_04_v1","PvP01_04_v1","PvP01_04_v1","PvP01_04_v1","PvP01_04_v1","PvP01_04_v1","PvP01_04_v1","PvP01_04_v1","PvP01_04_v1","PvP01_04_v1","PvP01_04_v1","PvP01_04_v1","PvP01_04_v1","PvP01_04_v1","PvP01_04_v1","PvP01_04_v1","PvP01_05_v1","PvP01_05_v1","PvP01_05_v1","PvP01_05_v1","PvP01_05_v1","PvP01_05_v1","PvP01_05_v1","PvP01_05_v1","PvP01_05_v1","PvP01_05_v1","PvP01_05_v1","PvP01_05_v1","PvP01_05_v1","PvP01_05_v1","PvP01_05_v1","PvP01_05_v1","PvP01_05_v1","PvP01_05_v1","PvP01_05_v1","PvP01_05_v1","PvP01_05_v1","PvP01_05_v1","PvP01_05_v1","PvP01_05_v1","PvP01_05_v1","PvP01_05_v1","PvP01_05_v1","PvP01_05_v1","PvP01_05_v1","PvP01_05_v1","PvP01_05_v1","PvP01_05_v1","PvP01_05_v1","PvP01_05_v1","PvP01_05_v1","PvP01_05_v1","PvP01_05_v1","PvP01_05_v1","PvP01_05_v1","PvP01_05_v1","PvP01_05_v1","PvP01_05_v1","PvP01_05_v1","PvP01_05_v1","PvP01_05_v1","PvP01_05_v1","PvP01_05_v1","PvP01_05_v1","PvP01_05_v1","PvP01_05_v1","PvP01_05_v1","PvP01_05_v1","PvP01_05_v1","PvP01_05_v1","PvP01_05_v1","PvP01_05_v1","PvP01_05_v1","PvP01_05_v1","PvP01_05_v1","PvP01_05_v1","PvP01_05_v1","PvP01_05_v1","PvP01_05_v1","PvP01_05_v1","PvP01_05_v1","PvP01_05_v1","PvP01_05_v1","PvP01_05_v1","PvP01_05_v1","PvP01_05_v1","PvP01_05_v1","PvP01_05_v1","PvP01_05_v1","PvP01_05_v1","PvP01_05_v1","PvP01_05_v1","PvP01_05_v1","PvP01_05_v1","PvP01_05_v1","PvP01_05_v1","PvP01_05_v1","PvP01_05_v1","PvP01_05_v1","PvP01_05_v1","PvP01_05_v1","PvP01_05_v1","PvP01_05_v1","PvP01_05_v1","PvP01_05_v1","PvP01_05_v1","PvP01_05_v1","PvP01_05_v1","PvP01_05_v1","PvP01_05_v1","PvP01_05_v1","PvP01_05_v1","PvP01_05_v1","PvP01_05_v1","PvP01_05_v1","PvP01_05_v1","PvP01_05_v1","PvP01_05_v1","PvP01_05_v1","PvP01_05_v1","PvP01_05_v1","PvP01_05_v1","PvP01_05_v1","PvP01_05_v1","PvP01_05_v1","PvP01_05_v1","PvP01_05_v1","PvP01_05_v1","PvP01_05_v1","PvP01_05_v1","PvP01_05_v1","PvP01_05_v1","PvP01_05_v1","PvP01_05_v1","PvP01_05_v1","PvP01_05_v1","PvP01_06_v1","PvP01_06_v1","PvP01_06_v1","PvP01_06_v1","PvP01_06_v1","PvP01_06_v1","PvP01_06_v1","PvP01_06_v1","PvP01_06_v1","PvP01_06_v1","PvP01_06_v1","PvP01_06_v1","PvP01_06_v1","PvP01_06_v1","PvP01_06_v1","PvP01_06_v1","PvP01_06_v1","PvP01_06_v1","PvP01_06_v1","PvP01_06_v1","PvP01_06_v1","PvP01_06_v1","PvP01_06_v1","PvP01_06_v1","PvP01_06_v1","PvP01_06_v1","PvP01_06_v1","PvP01_06_v1","PvP01_06_v1","PvP01_06_v1","PvP01_06_v1","PvP01_06_v1","PvP01_06_v1","PvP01_06_v1","PvP01_06_v1","PvP01_06_v1","PvP01_06_v1","PvP01_06_v1","PvP01_06_v1","PvP01_06_v1","PvP01_06_v1","PvP01_06_v1","PvP01_06_v1","PvP01_06_v1","PvP01_06_v1","PvP01_06_v1","PvP01_06_v1","PvP01_06_v1","PvP01_06_v1","PvP01_06_v1","PvP01_06_v1","PvP01_06_v1","PvP01_06_v1","PvP01_06_v1","PvP01_06_v1","PvP01_06_v1","PvP01_06_v1","PvP01_06_v1","PvP01_06_v1","PvP01_06_v1","PvP01_06_v1","PvP01_06_v1","PvP01_06_v1","PvP01_06_v1","PvP01_06_v1","PvP01_06_v1","PvP01_06_v1","PvP01_06_v1","PvP01_06_v1","PvP01_06_v1","PvP01_06_v1","PvP01_06_v1","PvP01_06_v1","PvP01_06_v1","PvP01_06_v1","PvP01_06_v1","PvP01_06_v1","PvP01_06_v1","PvP01_06_v1","PvP01_06_v1","PvP01_06_v1","PvP01_06_v1","PvP01_06_v1","PvP01_06_v1","PvP01_06_v1","PvP01_06_v1","PvP01_06_v1","PvP01_06_v1","PvP01_06_v1","PvP01_06_v1","PvP01_06_v1","PvP01_06_v1","PvP01_06_v1","PvP01_06_v1","PvP01_06_v1","PvP01_06_v1","PvP01_06_v1","PvP01_06_v1","PvP01_06_v1","PvP01_06_v1","PvP01_07_v1","PvP01_07_v1","PvP01_07_v1","PvP01_07_v1","PvP01_07_v1","PvP01_07_v1","PvP01_07_v1","PvP01_07_v1","PvP01_07_v1","PvP01_07_v1","PvP01_07_v1","PvP01_07_v1","PvP01_07_v1","PvP01_07_v1","PvP01_07_v1","PvP01_07_v1","PvP01_07_v1","PvP01_07_v1","PvP01_07_v1","PvP01_07_v1","PvP01_07_v1","PvP01_07_v1","PvP01_07_v1","PvP01_07_v1","PvP01_07_v1","PvP01_07_v1","PvP01_07_v1","PvP01_07_v1","PvP01_07_v1","PvP01_07_v1","PvP01_07_v1","PvP01_07_v1","PvP01_07_v1","PvP01_07_v1","PvP01_07_v1","PvP01_07_v1","PvP01_07_v1","PvP01_07_v1","PvP01_07_v1","PvP01_07_v1","PvP01_07_v1","PvP01_07_v1","PvP01_07_v1","PvP01_07_v1","PvP01_07_v1","PvP01_07_v1","PvP01_07_v1","PvP01_07_v1","PvP01_07_v1","PvP01_07_v1","PvP01_07_v1","PvP01_07_v1","PvP01_07_v1","PvP01_07_v1","PvP01_07_v1","PvP01_07_v1","PvP01_07_v1","PvP01_07_v1","PvP01_07_v1","PvP01_07_v1","PvP01_07_v1","PvP01_07_v1","PvP01_07_v1","PvP01_07_v1","PvP01_07_v1","PvP01_07_v1","PvP01_07_v1","PvP01_07_v1","PvP01_07_v1","PvP01_07_v1","PvP01_07_v1","PvP01_07_v1","PvP01_07_v1","PvP01_07_v1","PvP01_07_v1","PvP01_07_v1","PvP01_07_v1","PvP01_07_v1","PvP01_07_v1","PvP01_07_v1","PvP01_07_v1","PvP01_07_v1","PvP01_07_v1","PvP01_07_v1","PvP01_07_v1","PvP01_07_v1","PvP01_07_v1","PvP01_07_v1","PvP01_07_v1","PvP01_07_v1","PvP01_07_v1","PvP01_07_v1","PvP01_07_v1","PvP01_07_v1","PvP01_07_v1","PvP01_07_v1","PvP01_07_v1","PvP01_07_v1","PvP01_07_v1","PvP01_07_v1","PvP01_07_v1","PvP01_07_v1","PvP01_07_v1","PvP01_07_v1","PvP01_07_v1","PvP01_07_v1","PvP01_07_v1","PvP01_07_v1","PvP01_07_v1","PvP01_07_v1","PvP01_07_v1","PvP01_07_v1","PvP01_07_v1","PvP01_07_v1","PvP01_07_v1","PvP01_07_v1","PvP01_07_v1","PvP01_07_v1","PvP01_07_v1","PvP01_07_v1","PvP01_07_v1","PvP01_07_v1","PvP01_07_v1","PvP01_07_v1","PvP01_07_v1","PvP01_07_v1","PvP01_07_v1","PvP01_07_v1","PvP01_07_v1","PvP01_07_v1","PvP01_07_v1","PvP01_08_v1","PvP01_08_v1","PvP01_08_v1","PvP01_08_v1","PvP01_08_v1","PvP01_08_v1","PvP01_08_v1","PvP01_08_v1","PvP01_08_v1","PvP01_08_v1","PvP01_08_v1","PvP01_08_v1","PvP01_08_v1","PvP01_08_v1","PvP01_08_v1","PvP01_08_v1","PvP01_08_v1","PvP01_08_v1","PvP01_08_v1","PvP01_08_v1","PvP01_08_v1","PvP01_08_v1","PvP01_08_v1","PvP01_08_v1","PvP01_08_v1","PvP01_08_v1","PvP01_08_v1","PvP01_08_v1","PvP01_08_v1","PvP01_08_v1","PvP01_08_v1","PvP01_08_v1","PvP01_08_v1","PvP01_08_v1","PvP01_08_v1","PvP01_08_v1","PvP01_08_v1","PvP01_08_v1","PvP01_08_v1","PvP01_08_v1","PvP01_08_v1","PvP01_08_v1","PvP01_08_v1","PvP01_08_v1","PvP01_08_v1","PvP01_08_v1","PvP01_08_v1","PvP01_08_v1","PvP01_08_v1","PvP01_08_v1","PvP01_08_v1","PvP01_08_v1","PvP01_08_v1","PvP01_08_v1","PvP01_08_v1","PvP01_08_v1","PvP01_08_v1","PvP01_08_v1","PvP01_08_v1","PvP01_08_v1","PvP01_08_v1","PvP01_08_v1","PvP01_08_v1","PvP01_08_v1","PvP01_08_v1","PvP01_08_v1","PvP01_08_v1","PvP01_08_v1","PvP01_08_v1","PvP01_08_v1","PvP01_08_v1","PvP01_08_v1","PvP01_08_v1","PvP01_08_v1","PvP01_08_v1","PvP01_08_v1","PvP01_08_v1","PvP01_08_v1","PvP01_08_v1","PvP01_08_v1","PvP01_08_v1","PvP01_08_v1","PvP01_08_v1","PvP01_08_v1","PvP01_08_v1","PvP01_08_v1","PvP01_08_v1","PvP01_08_v1","PvP01_08_v1","PvP01_08_v1","PvP01_08_v1","PvP01_08_v1","PvP01_08_v1","PvP01_08_v1","PvP01_08_v1","PvP01_08_v1","PvP01_08_v1","PvP01_08_v1","PvP01_08_v1","PvP01_08_v1","PvP01_08_v1","PvP01_08_v1","PvP01_08_v1","PvP01_08_v1","PvP01_08_v1","PvP01_08_v1","PvP01_08_v1","PvP01_08_v1","PvP01_08_v1","PvP01_08_v1","PvP01_08_v1","PvP01_08_v1","PvP01_08_v1","PvP01_08_v1","PvP01_08_v1","PvP01_08_v1","PvP01_08_v1","PvP01_08_v1","PvP01_08_v1","PvP01_08_v1","PvP01_08_v1","PvP01_08_v1","PvP01_08_v1","PvP01_08_v1","PvP01_08_v1","PvP01_08_v1","PvP01_08_v1","PvP01_08_v1","PvP01_08_v1","PvP01_08_v1","PvP01_08_v1","PvP01_08_v1","PvP01_08_v1","PvP01_08_v1","PvP01_08_v1","PvP01_08_v1","PvP01_08_v1","PvP01_08_v1","PvP01_08_v1","PvP01_08_v1","PvP01_08_v1","PvP01_08_v1","PvP01_08_v1","PvP01_08_v1","PvP01_08_v1","PvP01_08_v1","PvP01_08_v1","PvP01_08_v1","PvP01_08_v1","PvP01_08_v1","PvP01_08_v1","PvP01_08_v1","PvP01_09_v1","PvP01_09_v1","PvP01_09_v1","PvP01_09_v1","PvP01_09_v1","PvP01_09_v1","PvP01_09_v1","PvP01_09_v1","PvP01_09_v1","PvP01_09_v1","PvP01_09_v1","PvP01_09_v1","PvP01_09_v1","PvP01_09_v1","PvP01_09_v1","PvP01_09_v1","PvP01_09_v1","PvP01_09_v1","PvP01_09_v1","PvP01_09_v1","PvP01_09_v1","PvP01_09_v1","PvP01_09_v1","PvP01_09_v1","PvP01_09_v1","PvP01_09_v1","PvP01_09_v1","PvP01_09_v1","PvP01_09_v1","PvP01_09_v1","PvP01_09_v1","PvP01_09_v1","PvP01_09_v1","PvP01_09_v1","PvP01_09_v1","PvP01_09_v1","PvP01_09_v1","PvP01_09_v1","PvP01_09_v1","PvP01_09_v1","PvP01_09_v1","PvP01_09_v1","PvP01_09_v1","PvP01_09_v1","PvP01_09_v1","PvP01_09_v1","PvP01_09_v1","PvP01_09_v1","PvP01_09_v1","PvP01_09_v1","PvP01_09_v1","PvP01_09_v1","PvP01_09_v1","PvP01_09_v1","PvP01_09_v1","PvP01_09_v1","PvP01_09_v1","PvP01_09_v1","PvP01_09_v1","PvP01_09_v1","PvP01_09_v1","PvP01_09_v1","PvP01_09_v1","PvP01_09_v1","PvP01_09_v1","PvP01_09_v1","PvP01_09_v1","PvP01_09_v1","PvP01_09_v1","PvP01_09_v1","PvP01_09_v1","PvP01_09_v1","PvP01_09_v1","PvP01_09_v1","PvP01_09_v1","PvP01_09_v1","PvP01_09_v1","PvP01_09_v1","PvP01_09_v1","PvP01_09_v1","PvP01_09_v1","PvP01_09_v1","PvP01_09_v1","PvP01_09_v1","PvP01_09_v1","PvP01_09_v1","PvP01_09_v1","PvP01_09_v1","PvP01_09_v1","PvP01_09_v1","PvP01_09_v1","PvP01_09_v1","PvP01_09_v1","PvP01_09_v1","PvP01_09_v1","PvP01_09_v1","PvP01_09_v1","PvP01_09_v1","PvP01_09_v1","PvP01_09_v1","PvP01_09_v1","PvP01_09_v1","PvP01_09_v1","PvP01_09_v1","PvP01_09_v1","PvP01_09_v1","PvP01_09_v1","PvP01_09_v1","PvP01_09_v1","PvP01_09_v1","PvP01_09_v1","PvP01_09_v1","PvP01_09_v1","PvP01_09_v1","PvP01_09_v1","PvP01_09_v1","PvP01_09_v1","PvP01_09_v1","PvP01_09_v1","PvP01_09_v1","PvP01_09_v1","PvP01_09_v1","PvP01_09_v1","PvP01_09_v1","PvP01_09_v1","PvP01_09_v1","PvP01_09_v1","PvP01_09_v1","PvP01_09_v1","PvP01_09_v1","PvP01_09_v1","PvP01_09_v1","PvP01_09_v1","PvP01_09_v1","PvP01_09_v1","PvP01_09_v1","PvP01_09_v1","PvP01_09_v1","PvP01_09_v1","PvP01_09_v1","PvP01_09_v1","PvP01_09_v1","PvP01_09_v1","PvP01_09_v1","PvP01_09_v1","PvP01_09_v1","PvP01_09_v1","PvP01_09_v1","PvP01_09_v1","PvP01_09_v1","PvP01_09_v1","PvP01_09_v1","PvP01_09_v1","PvP01_09_v1","PvP01_09_v1","PvP01_09_v1","PvP01_09_v1","PvP01_09_v1","PvP01_09_v1","PvP01_09_v1","PvP01_09_v1","PvP01_09_v1","PvP01_09_v1","PvP01_09_v1","PvP01_09_v1","PvP01_09_v1","PvP01_09_v1","PvP01_10_v1","PvP01_10_v1","PvP01_10_v1","PvP01_10_v1","PvP01_10_v1","PvP01_10_v1","PvP01_10_v1","PvP01_10_v1","PvP01_10_v1","PvP01_10_v1","PvP01_10_v1","PvP01_10_v1","PvP01_10_v1","PvP01_10_v1","PvP01_10_v1","PvP01_10_v1","PvP01_10_v1","PvP01_10_v1","PvP01_10_v1","PvP01_10_v1","PvP01_10_v1","PvP01_10_v1","PvP01_10_v1","PvP01_10_v1","PvP01_10_v1","PvP01_10_v1","PvP01_10_v1","PvP01_10_v1","PvP01_10_v1","PvP01_10_v1","PvP01_10_v1","PvP01_10_v1","PvP01_10_v1","PvP01_10_v1","PvP01_10_v1","PvP01_10_v1","PvP01_10_v1","PvP01_10_v1","PvP01_10_v1","PvP01_10_v1","PvP01_10_v1","PvP01_10_v1","PvP01_10_v1","PvP01_10_v1","PvP01_10_v1","PvP01_10_v1","PvP01_10_v1","PvP01_10_v1","PvP01_10_v1","PvP01_10_v1","PvP01_10_v1","PvP01_10_v1","PvP01_10_v1","PvP01_10_v1","PvP01_10_v1","PvP01_10_v1","PvP01_10_v1","PvP01_10_v1","PvP01_10_v1","PvP01_10_v1","PvP01_10_v1","PvP01_10_v1","PvP01_10_v1","PvP01_10_v1","PvP01_10_v1","PvP01_10_v1","PvP01_10_v1","PvP01_10_v1","PvP01_10_v1","PvP01_10_v1","PvP01_10_v1","PvP01_10_v1","PvP01_10_v1","PvP01_10_v1","PvP01_10_v1","PvP01_10_v1","PvP01_10_v1","PvP01_10_v1","PvP01_10_v1","PvP01_10_v1","PvP01_10_v1","PvP01_10_v1","PvP01_10_v1","PvP01_10_v1","PvP01_10_v1","PvP01_10_v1","PvP01_10_v1","PvP01_10_v1","PvP01_10_v1","PvP01_10_v1","PvP01_10_v1","PvP01_10_v1","PvP01_10_v1","PvP01_10_v1","PvP01_10_v1","PvP01_10_v1","PvP01_10_v1","PvP01_10_v1","PvP01_10_v1","PvP01_10_v1","PvP01_10_v1","PvP01_10_v1","PvP01_10_v1","PvP01_10_v1","PvP01_10_v1","PvP01_10_v1","PvP01_10_v1","PvP01_10_v1","PvP01_10_v1","PvP01_10_v1","PvP01_10_v1","PvP01_10_v1","PvP01_10_v1","PvP01_10_v1","PvP01_10_v1","PvP01_10_v1","PvP01_10_v1","PvP01_10_v1","PvP01_10_v1","PvP01_10_v1","PvP01_10_v1","PvP01_10_v1","PvP01_10_v1","PvP01_10_v1","PvP01_11_v1","PvP01_11_v1","PvP01_11_v1","PvP01_11_v1","PvP01_11_v1","PvP01_11_v1","PvP01_11_v1","PvP01_11_v1","PvP01_11_v1","PvP01_11_v1","PvP01_11_v1","PvP01_11_v1","PvP01_11_v1","PvP01_11_v1","PvP01_11_v1","PvP01_11_v1","PvP01_11_v1","PvP01_11_v1","PvP01_11_v1","PvP01_11_v1","PvP01_11_v1","PvP01_11_v1","PvP01_11_v1","PvP01_11_v1","PvP01_11_v1","PvP01_11_v1","PvP01_11_v1","PvP01_11_v1","PvP01_11_v1","PvP01_11_v1","PvP01_11_v1","PvP01_11_v1","PvP01_11_v1","PvP01_11_v1","PvP01_11_v1","PvP01_11_v1","PvP01_11_v1","PvP01_11_v1","PvP01_11_v1","PvP01_11_v1","PvP01_11_v1","PvP01_11_v1","PvP01_11_v1","PvP01_11_v1","PvP01_11_v1","PvP01_11_v1","PvP01_11_v1","PvP01_11_v1","PvP01_11_v1","PvP01_11_v1","PvP01_11_v1","PvP01_11_v1","PvP01_11_v1","PvP01_11_v1","PvP01_11_v1","PvP01_11_v1","PvP01_11_v1","PvP01_11_v1","PvP01_11_v1","PvP01_11_v1","PvP01_11_v1","PvP01_11_v1","PvP01_11_v1","PvP01_11_v1","PvP01_11_v1","PvP01_11_v1","PvP01_11_v1","PvP01_11_v1","PvP01_11_v1","PvP01_11_v1","PvP01_11_v1","PvP01_11_v1","PvP01_11_v1","PvP01_11_v1","PvP01_11_v1","PvP01_11_v1","PvP01_11_v1","PvP01_11_v1","PvP01_11_v1","PvP01_11_v1","PvP01_11_v1","PvP01_11_v1","PvP01_11_v1","PvP01_11_v1","PvP01_11_v1","PvP01_11_v1","PvP01_11_v1","PvP01_11_v1","PvP01_11_v1","PvP01_11_v1","PvP01_11_v1","PvP01_11_v1","PvP01_11_v1","PvP01_11_v1","PvP01_11_v1","PvP01_11_v1","PvP01_11_v1","PvP01_11_v1","PvP01_11_v1","PvP01_11_v1","PvP01_11_v1","PvP01_11_v1","PvP01_11_v1","PvP01_11_v1","PvP01_11_v1","PvP01_11_v1","PvP01_11_v1","PvP01_11_v1","PvP01_11_v1","PvP01_11_v1","PvP01_11_v1","PvP01_11_v1","PvP01_11_v1","PvP01_11_v1","PvP01_11_v1","PvP01_11_v1","PvP01_11_v1","PvP01_11_v1","PvP01_11_v1","PvP01_11_v1","PvP01_11_v1","PvP01_11_v1","PvP01_11_v1","PvP01_11_v1","PvP01_11_v1","PvP01_11_v1","PvP01_11_v1","PvP01_11_v1","PvP01_11_v1","PvP01_11_v1","PvP01_11_v1","PvP01_11_v1","PvP01_11_v1","PvP01_11_v1","PvP01_11_v1","PvP01_11_v1","PvP01_11_v1","PvP01_11_v1","PvP01_11_v1","PvP01_11_v1","PvP01_11_v1","PvP01_11_v1","PvP01_11_v1","PvP01_11_v1","PvP01_11_v1","PvP01_11_v1","PvP01_11_v1","PvP01_11_v1","PvP01_11_v1","PvP01_11_v1","PvP01_11_v1","PvP01_11_v1","PvP01_11_v1","PvP01_11_v1","PvP01_11_v1","PvP01_11_v1","PvP01_11_v1","PvP01_11_v1","PvP01_11_v1","PvP01_11_v1","PvP01_11_v1","PvP01_11_v1","PvP01_11_v1","PvP01_11_v1","PvP01_11_v1","PvP01_11_v1","PvP01_12_v1","PvP01_12_v1","PvP01_12_v1","PvP01_12_v1","PvP01_12_v1","PvP01_12_v1","PvP01_12_v1","PvP01_12_v1","PvP01_12_v1","PvP01_12_v1","PvP01_12_v1","PvP01_12_v1","PvP01_12_v1","PvP01_12_v1","PvP01_12_v1","PvP01_12_v1","PvP01_12_v1","PvP01_12_v1","PvP01_12_v1","PvP01_12_v1","PvP01_12_v1","PvP01_12_v1","PvP01_12_v1","PvP01_12_v1","PvP01_12_v1","PvP01_12_v1","PvP01_12_v1","PvP01_12_v1","PvP01_12_v1","PvP01_12_v1","PvP01_12_v1","PvP01_12_v1","PvP01_12_v1","PvP01_12_v1","PvP01_12_v1","PvP01_12_v1","PvP01_12_v1","PvP01_12_v1","PvP01_12_v1","PvP01_12_v1","PvP01_12_v1","PvP01_12_v1","PvP01_12_v1","PvP01_12_v1","PvP01_12_v1","PvP01_12_v1","PvP01_12_v1","PvP01_12_v1","PvP01_12_v1","PvP01_12_v1","PvP01_12_v1","PvP01_12_v1","PvP01_12_v1","PvP01_12_v1","PvP01_12_v1","PvP01_12_v1","PvP01_12_v1","PvP01_12_v1","PvP01_12_v1","PvP01_12_v1","PvP01_12_v1","PvP01_12_v1","PvP01_12_v1","PvP01_12_v1","PvP01_12_v1","PvP01_12_v1","PvP01_12_v1","PvP01_12_v1","PvP01_12_v1","PvP01_12_v1","PvP01_12_v1","PvP01_12_v1","PvP01_12_v1","PvP01_12_v1","PvP01_12_v1","PvP01_12_v1","PvP01_12_v1","PvP01_12_v1","PvP01_12_v1","PvP01_12_v1","PvP01_12_v1","PvP01_12_v1","PvP01_12_v1","PvP01_12_v1","PvP01_12_v1","PvP01_12_v1","PvP01_12_v1","PvP01_12_v1","PvP01_12_v1","PvP01_12_v1","PvP01_12_v1","PvP01_12_v1","PvP01_12_v1","PvP01_12_v1","PvP01_12_v1","PvP01_12_v1","PvP01_12_v1","PvP01_12_v1","PvP01_12_v1","PvP01_12_v1","PvP01_12_v1","PvP01_12_v1","PvP01_12_v1","PvP01_12_v1","PvP01_12_v1","PvP01_12_v1","PvP01_12_v1","PvP01_12_v1","PvP01_12_v1","PvP01_12_v1","PvP01_12_v1","PvP01_12_v1","PvP01_12_v1","PvP01_12_v1","PvP01_12_v1","PvP01_12_v1","PvP01_12_v1","PvP01_12_v1","PvP01_12_v1","PvP01_12_v1","PvP01_12_v1","PvP01_12_v1","PvP01_12_v1","PvP01_12_v1","PvP01_12_v1","PvP01_12_v1","PvP01_12_v1","PvP01_12_v1","PvP01_12_v1","PvP01_12_v1","PvP01_12_v1","PvP01_12_v1","PvP01_12_v1","PvP01_12_v1","PvP01_12_v1","PvP01_12_v1","PvP01_12_v1","PvP01_12_v1","PvP01_12_v1","PvP01_12_v1","PvP01_12_v1","PvP01_12_v1","PvP01_12_v1","PvP01_12_v1","PvP01_12_v1","PvP01_12_v1","PvP01_12_v1","PvP01_12_v1","PvP01_12_v1","PvP01_12_v1","PvP01_12_v1","PvP01_12_v1","PvP01_12_v1","PvP01_12_v1","PvP01_12_v1","PvP01_12_v1","PvP01_12_v1","PvP01_12_v1","PvP01_12_v1","PvP01_12_v1","PvP01_12_v1","PvP01_12_v1","PvP01_12_v1","PvP01_12_v1","PvP01_12_v1","PvP01_12_v1","PvP01_12_v1","PvP01_12_v1","PvP01_12_v1","PvP01_12_v1","PvP01_12_v1","PvP01_12_v1","PvP01_12_v1","PvP01_12_v1","PvP01_12_v1","PvP01_12_v1","PvP01_12_v1","PvP01_12_v1","PvP01_12_v1","PvP01_12_v1","PvP01_12_v1","PvP01_12_v1","PvP01_12_v1","PvP01_12_v1","PvP01_12_v1","PvP01_12_v1","PvP01_12_v1","PvP01_12_v1","PvP01_12_v1","PvP01_12_v1","PvP01_12_v1","PvP01_12_v1","PvP01_12_v1","PvP01_12_v1","PvP01_12_v1","PvP01_12_v1","PvP01_12_v1","PvP01_12_v1","PvP01_12_v1","PvP01_12_v1","PvP01_12_v1","PvP01_12_v1","PvP01_12_v1","PvP01_12_v1","PvP01_12_v1","PvP01_12_v1","PvP01_12_v1","PvP01_12_v1","PvP01_12_v1","PvP01_12_v1","PvP01_12_v1","PvP01_12_v1","PvP01_12_v1","PvP01_12_v1","PvP01_12_v1","PvP01_12_v1","PvP01_12_v1","PvP01_12_v1","PvP01_12_v1","PvP01_12_v1","PvP01_12_v1","PvP01_12_v1","PvP01_12_v1","PvP01_12_v1","PvP01_12_v1","PvP01_12_v1","PvP01_12_v1","PvP01_12_v1","PvP01_12_v1","PvP01_12_v1","PvP01_12_v1","PvP01_12_v1","PvP01_12_v1","PvP01_12_v1","PvP01_12_v1","PvP01_12_v1","PvP01_12_v1","PvP01_12_v1","PvP01_12_v1","PvP01_12_v1","PvP01_12_v1","PvP01_12_v1","PvP01_12_v1","PvP01_12_v1","PvP01_12_v1","PvP01_12_v1","PvP01_12_v1","PvP01_12_v1","PvP01_12_v1","PvP01_12_v1","PvP01_12_v1","PvP01_12_v1","PvP01_12_v1","PvP01_12_v1","PvP01_12_v1","PvP01_12_v1","PvP01_12_v1","PvP01_12_v1","PvP01_12_v1","PvP01_12_v1","PvP01_12_v1","PvP01_12_v1","PvP01_12_v1","PvP01_12_v1","PvP01_12_v1","PvP01_12_v1","PvP01_12_v1","PvP01_12_v1","PvP01_12_v1","PvP01_12_v1","PvP01_12_v1","PvP01_12_v1","PvP01_12_v1","PvP01_12_v1","PvP01_12_v1","PvP01_12_v1","PvP01_12_v1","PvP01_12_v1","PvP01_12_v1","PvP01_12_v1","PvP01_12_v1","PvP01_12_v1","PvP01_12_v1","PvP01_12_v1","PvP01_13_v1","PvP01_13_v1","PvP01_13_v1","PvP01_13_v1","PvP01_13_v1","PvP01_13_v1","PvP01_13_v1","PvP01_13_v1","PvP01_13_v1","PvP01_13_v1","PvP01_13_v1","PvP01_13_v1","PvP01_13_v1","PvP01_13_v1","PvP01_13_v1","PvP01_13_v1","PvP01_13_v1","PvP01_13_v1","PvP01_13_v1","PvP01_13_v1","PvP01_13_v1","PvP01_13_v1","PvP01_13_v1","PvP01_13_v1","PvP01_13_v1","PvP01_13_v1","PvP01_13_v1","PvP01_13_v1","PvP01_13_v1","PvP01_13_v1","PvP01_13_v1","PvP01_13_v1","PvP01_13_v1","PvP01_13_v1","PvP01_13_v1","PvP01_13_v1","PvP01_13_v1","PvP01_13_v1","PvP01_13_v1","PvP01_13_v1","PvP01_13_v1","PvP01_13_v1","PvP01_13_v1","PvP01_13_v1","PvP01_13_v1","PvP01_13_v1","PvP01_13_v1","PvP01_13_v1","PvP01_13_v1","PvP01_13_v1","PvP01_13_v1","PvP01_13_v1","PvP01_13_v1","PvP01_13_v1","PvP01_13_v1","PvP01_13_v1","PvP01_13_v1","PvP01_13_v1","PvP01_13_v1","PvP01_13_v1","PvP01_13_v1","PvP01_13_v1","PvP01_13_v1","PvP01_13_v1","PvP01_13_v1","PvP01_13_v1","PvP01_13_v1","PvP01_13_v1","PvP01_13_v1","PvP01_13_v1","PvP01_13_v1","PvP01_13_v1","PvP01_13_v1","PvP01_13_v1","PvP01_13_v1","PvP01_13_v1","PvP01_13_v1","PvP01_13_v1","PvP01_13_v1","PvP01_13_v1","PvP01_13_v1","PvP01_13_v1","PvP01_13_v1","PvP01_13_v1","PvP01_13_v1","PvP01_13_v1","PvP01_13_v1","PvP01_13_v1","PvP01_13_v1","PvP01_13_v1","PvP01_13_v1","PvP01_13_v1","PvP01_13_v1","PvP01_13_v1","PvP01_13_v1","PvP01_13_v1","PvP01_13_v1","PvP01_13_v1","PvP01_13_v1","PvP01_13_v1","PvP01_13_v1","PvP01_13_v1","PvP01_13_v1","PvP01_13_v1","PvP01_13_v1","PvP01_13_v1","PvP01_13_v1","PvP01_13_v1","PvP01_13_v1","PvP01_13_v1","PvP01_13_v1","PvP01_13_v1","PvP01_13_v1","PvP01_13_v1","PvP01_13_v1","PvP01_13_v1","PvP01_13_v1","PvP01_13_v1","PvP01_13_v1","PvP01_13_v1","PvP01_13_v1","PvP01_13_v1","PvP01_13_v1","PvP01_13_v1","PvP01_13_v1","PvP01_13_v1","PvP01_13_v1","PvP01_13_v1","PvP01_13_v1","PvP01_13_v1","PvP01_13_v1","PvP01_13_v1","PvP01_13_v1","PvP01_13_v1","PvP01_13_v1","PvP01_13_v1","PvP01_13_v1","PvP01_13_v1","PvP01_13_v1","PvP01_13_v1","PvP01_13_v1","PvP01_13_v1","PvP01_13_v1","PvP01_13_v1","PvP01_13_v1","PvP01_13_v1","PvP01_13_v1","PvP01_13_v1","PvP01_13_v1","PvP01_13_v1","PvP01_13_v1","PvP01_13_v1","PvP01_13_v1","PvP01_13_v1","PvP01_13_v1","PvP01_13_v1","PvP01_13_v1","PvP01_13_v1","PvP01_13_v1","PvP01_13_v1","PvP01_13_v1","PvP01_13_v1","PvP01_13_v1","PvP01_13_v1","PvP01_13_v1","PvP01_13_v1","PvP01_13_v1","PvP01_13_v1","PvP01_13_v1","PvP01_13_v1","PvP01_13_v1","PvP01_13_v1","PvP01_13_v1","PvP01_13_v1","PvP01_13_v1","PvP01_13_v1","PvP01_13_v1","PvP01_13_v1","PvP01_13_v1","PvP01_13_v1","PvP01_13_v1","PvP01_13_v1","PvP01_13_v1","PvP01_13_v1","PvP01_13_v1","PvP01_13_v1","PvP01_13_v1","PvP01_14_v1","PvP01_14_v1","PvP01_14_v1","PvP01_14_v1","PvP01_14_v1","PvP01_14_v1","PvP01_14_v1","PvP01_14_v1","PvP01_14_v1","PvP01_14_v1","PvP01_14_v1","PvP01_14_v1","PvP01_14_v1","PvP01_14_v1","PvP01_14_v1","PvP01_14_v1","PvP01_14_v1","PvP01_14_v1","PvP01_14_v1","PvP01_14_v1","PvP01_14_v1","PvP01_14_v1","PvP01_14_v1","PvP01_14_v1","PvP01_14_v1","PvP01_14_v1","PvP01_14_v1","PvP01_14_v1","PvP01_14_v1","PvP01_14_v1","PvP01_14_v1","PvP01_14_v1","PvP01_14_v1","PvP01_14_v1","PvP01_14_v1","PvP01_14_v1","PvP01_14_v1","PvP01_14_v1","PvP01_14_v1","PvP01_14_v1","PvP01_14_v1","PvP01_14_v1","PvP01_14_v1","PvP01_14_v1","PvP01_14_v1","PvP01_14_v1","PvP01_14_v1","PvP01_14_v1","PvP01_14_v1","PvP01_14_v1","PvP01_14_v1","PvP01_14_v1","PvP01_14_v1","PvP01_14_v1","PvP01_14_v1","PvP01_14_v1","PvP01_14_v1","PvP01_14_v1","PvP01_14_v1","PvP01_14_v1","PvP01_14_v1","PvP01_14_v1","PvP01_14_v1","PvP01_14_v1","PvP01_14_v1","PvP01_14_v1","PvP01_14_v1","PvP01_14_v1","PvP01_14_v1","PvP01_14_v1","PvP01_14_v1","PvP01_14_v1","PvP01_14_v1","PvP01_14_v1","PvP01_14_v1","PvP01_14_v1","PvP01_14_v1","PvP01_14_v1","PvP01_14_v1","PvP01_14_v1","PvP01_14_v1","PvP01_14_v1","PvP01_14_v1","PvP01_14_v1","PvP01_14_v1","PvP01_14_v1","PvP01_14_v1","PvP01_14_v1","PvP01_14_v1","PvP01_14_v1","PvP01_14_v1","PvP01_14_v1","PvP01_14_v1","PvP01_14_v1","PvP01_14_v1","PvP01_14_v1","PvP01_14_v1","PvP01_14_v1","PvP01_14_v1","PvP01_14_v1","PvP01_14_v1","PvP01_14_v1","PvP01_14_v1","PvP01_14_v1","PvP01_14_v1","PvP01_14_v1","PvP01_14_v1","PvP01_14_v1","PvP01_14_v1","PvP01_14_v1","PvP01_14_v1","PvP01_14_v1","PvP01_14_v1","PvP01_14_v1","PvP01_14_v1","PvP01_14_v1","PvP01_14_v1","PvP01_14_v1","PvP01_14_v1","PvP01_14_v1","PvP01_14_v1","PvP01_14_v1","PvP01_14_v1","PvP01_14_v1","PvP01_14_v1","PvP01_14_v1","PvP01_14_v1","PvP01_14_v1","PvP01_14_v1","PvP01_14_v1","PvP01_14_v1","PvP01_14_v1","PvP01_14_v1","PvP01_14_v1","PvP01_14_v1","PvP01_14_v1","PvP01_14_v1","PvP01_14_v1","PvP01_14_v1","PvP01_14_v1","PvP01_14_v1","PvP01_14_v1","PvP01_14_v1","PvP01_14_v1","PvP01_14_v1","PvP01_14_v1","PvP01_14_v1","PvP01_14_v1","PvP01_14_v1","PvP01_14_v1","PvP01_14_v1","PvP01_14_v1","PvP01_14_v1","PvP01_14_v1","PvP01_14_v1","PvP01_14_v1","PvP01_14_v1","PvP01_14_v1","PvP01_14_v1","PvP01_14_v1","PvP01_14_v1","PvP01_14_v1","PvP01_14_v1","PvP01_14_v1","PvP01_14_v1","PvP01_14_v1","PvP01_14_v1","PvP01_14_v1","PvP01_14_v1","PvP01_14_v1","PvP01_14_v1","PvP01_14_v1","PvP01_14_v1","PvP01_14_v1","PvP01_14_v1","PvP01_14_v1","PvP01_14_v1","PvP01_14_v1","PvP01_14_v1","PvP01_14_v1","PvP01_14_v1","PvP01_14_v1","PvP01_14_v1","PvP01_14_v1","PvP01_14_v1","PvP01_14_v1","PvP01_14_v1","PvP01_14_v1","PvP01_14_v1","PvP01_14_v1","PvP01_14_v1","PvP01_14_v1","PvP01_14_v1","PvP01_14_v1","PvP01_14_v1","PvP01_14_v1","PvP01_14_v1","PvP01_14_v1","PvP01_14_v1","PvP01_14_v1","PvP01_14_v1","PvP01_14_v1","PvP01_14_v1","PvP01_14_v1","PvP01_14_v1","PvP01_14_v1","PvP01_14_v1","PvP01_14_v1","PvP01_14_v1","PvP01_14_v1","PvP01_14_v1","PvP01_14_v1","PvP01_14_v1","PvP01_14_v1","PvP01_14_v1","PvP01_14_v1","PvP01_14_v1","PvP01_14_v1","PvP01_14_v1","PvP01_14_v1","PvP01_14_v1","PvP01_14_v1","PvP01_14_v1","PvP01_14_v1","PvP01_14_v1","PvP01_14_v1","PvP01_14_v1","PvP01_14_v1","PvP01_14_v1","PvP01_14_v1","PvP01_14_v1","PvP01_14_v1","PvP01_14_v1","PvP01_14_v1","PvP01_14_v1","PvP01_14_v1","PvP01_14_v1","PvP01_14_v1","PvP01_14_v1","PvP01_14_v1","PvP01_14_v1","PvP01_14_v1","PvP01_14_v1","PvP01_14_v1","PvP01_14_v1","PvP01_14_v1","PvP01_14_v1","PvP01_14_v1","PvP01_14_v1","PvP01_14_v1","PvP01_14_v1","PvP01_14_v1","PvP01_14_v1","PvP01_14_v1","PvP01_14_v1","PvP01_14_v1","PvP01_14_v1","PvP01_14_v1","PvP01_14_v1","PvP01_14_v1","PvP01_14_v1","PvP01_14_v1","PvP01_14_v1","PvP01_14_v1","PvP01_14_v1","PvP01_14_v1","PvP01_14_v1","PvP01_14_v1","PvP01_14_v1","PvP01_14_v1","PvP01_14_v1","PvP01_14_v1","PvP01_14_v1","PvP01_14_v1","PvP01_14_v1","PvP01_14_v1","PvP01_14_v1","PvP01_API_v1","PvP01_API_v1","PvP01_API_v1","PvP01_API_v1","PvP01_API_v1","PvP01_MIT_v1","PvP01_MIT_v1","PvP01_MIT_v1"],"Source":["EuPathDB","EuPathDB","EuPathDB","EuPathDB","EuPathDB","EuPathDB","EuPathDB","EuPathDB","EuPathDB","EuPathDB","EuPathDB","EuPathDB","EuPathDB","EuPathDB","EuPathDB","EuPathDB","EuPathDB","EuPathDB","EuPathDB","EuPathDB","EuPathDB","EuPathDB","EuPathDB","EuPathDB","EuPathDB","EuPathDB","EuPathDB","EuPathDB","EuPathDB","EuPathDB","EuPathDB","EuPathDB","EuPathDB","EuPathDB","EuPathDB","EuPathDB","EuPathDB","EuPathDB","EuPathDB","EuPathDB","EuPathDB","EuPathDB","EuPathDB","EuPathDB","EuPathDB","EuPathDB","EuPathDB","EuPathDB","EuPathDB","EuPathDB","EuPathDB","EuPathDB","EuPathDB","EuPathDB","EuPathDB","EuPathDB","EuPathDB","EuPathDB","EuPathDB","EuPathDB","EuPathDB","EuPathDB","EuPathDB","EuPathDB","EuPathDB","EuPathDB","EuPathDB","EuPathDB","EuPathDB","EuPathDB","EuPathDB","EuPathDB","EuPathDB","EuPathDB","EuPathDB","EuPathDB","EuPathDB","EuPathDB","EuPathDB","EuPathDB","EuPathDB","EuPathDB","EuPathDB","EuPathDB","EuPathDB","EuPathDB","EuPathDB","EuPathDB","EuPathDB","EuPathDB","EuPathDB","EuPathDB","EuPathDB","EuPathDB","EuPathDB","EuPathDB","EuPathDB","EuPathDB","EuPathDB","EuPathDB","EuPathDB","EuPathDB","EuPathDB","EuPathDB","EuPathDB","EuPathDB","EuPathDB","EuPathDB","EuPathDB","EuPathDB","EuPathDB","EuPathDB","EuPathDB","EuPathDB","EuPathDB","EuPathDB","EuPathDB","EuPathDB","EuPathDB","EuPathDB","EuPathDB","EuPathDB","EuPathDB","EuPathDB","EuPathDB","EuPathDB","EuPathDB","EuPathDB","EuPathDB","EuPathDB","EuPathDB","EuPathDB","EuPathDB","EuPathDB","EuPathDB","EuPathDB","EuPathDB","EuPathDB","EuPathDB","EuPathDB","EuPathDB","EuPathDB","EuPathDB","EuPathDB","EuPathDB","EuPathDB","EuPathDB","EuPathDB","EuPathDB","EuPathDB","EuPathDB","EuPathDB","EuPathDB","EuPathDB","EuPathDB","EuPathDB","EuPathDB","EuPathDB","EuPathDB","EuPathDB","EuPathDB","EuPathDB","EuPathDB","EuPathDB","EuPathDB","EuPathDB","EuPathDB","EuPathDB","EuPathDB","EuPathDB","EuPathDB","EuPathDB","EuPathDB","EuPathDB","EuPathDB","EuPathDB","EuPathDB","EuPathDB","EuPathDB","EuPathDB","EuPathDB","EuPathDB","EuPathDB","EuPathDB","EuPathDB","EuPathDB","EuPathDB","EuPathDB","EuPathDB","EuPathDB","EuPathDB","EuPathDB","EuPathDB","EuPathDB","EuPathDB","EuPathDB","EuPathDB","EuPathDB","EuPathDB","EuPathDB","EuPathDB","EuPathDB","EuPathDB","EuPathDB","EuPathDB","EuPathDB","EuPathDB","EuPathDB","EuPathDB","EuPathDB","EuPathDB","EuPathDB","EuPathDB","EuPathDB","EuPathDB","EuPathDB","EuPathDB","EuPathDB","EuPathDB","EuPathDB","EuPathDB","EuPathDB","EuPathDB","EuPathDB","EuPathDB","EuPathDB","EuPathDB","EuPathDB","EuPathDB","EuPathDB","EuPathDB","EuPathDB","EuPathDB","EuPathDB","EuPathDB","EuPathDB","EuPathDB","EuPathDB","EuPathDB","EuPathDB","EuPathDB","EuPathDB","EuPathDB","EuPathDB","EuPathDB","EuPathDB","EuPathDB","EuPathDB","EuPathDB","EuPathDB","EuPathDB","EuPathDB","EuPathDB","EuPathDB","EuPathDB","EuPathDB","EuPathDB","EuPathDB","EuPathDB","EuPathDB","EuPathDB","EuPathDB","EuPathDB","EuPathDB","EuPathDB","EuPathDB","EuPathDB","EuPathDB","EuPathDB","EuPathDB","EuPathDB","EuPathDB","EuPathDB","EuPathDB","EuPathDB","EuPathDB","EuPathDB","EuPathDB","EuPathDB","EuPathDB","EuPathDB","EuPathDB","EuPathDB","EuPathDB","EuPathDB","EuPathDB","EuPathDB","EuPathDB","EuPathDB","EuPathDB","EuPathDB","EuPathDB","EuPathDB","EuPathDB","EuPathDB","EuPathDB","EuPathDB","EuPathDB","EuPathDB","EuPathDB","EuPathDB","EuPathDB","EuPathDB","Eu
```
