## Supplementary figures and images for "Transcriptome and histone epigenome of *Plasmodium vivax* salivary-gland sporozoites point to tight regulatory control and potential mechanisms for liver-stage differentiation"

### animated-overlay.gif

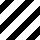

### sort_asc.png

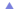

### sort_asc_disabled.png

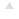

### sort_both.png

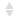

### sort_desc.png

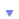

### sort_desc_disabled.png

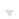

### ui-bg_flat_0_aaaaaa_40x100.png

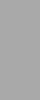

### ui-bg_flat_75_ffffff_40x100.png

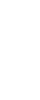

### ui-icons_2e83ff_256x240.png

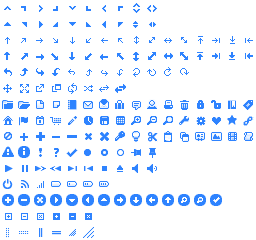

### ui-icons_222222_256x240.png

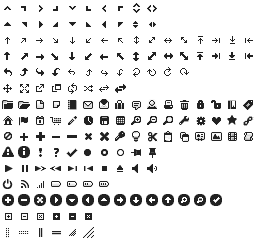

### ui-icons_454545_256x240.png

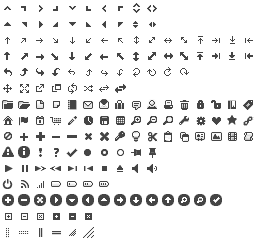

### ui-icons_888888_256x240.png

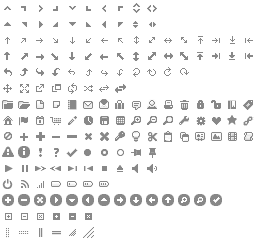

### ui-icons_cd0a0a_256x240.png

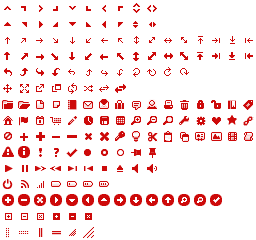
